# Systematic annotation of a complete adult male *Drosophila* nerve cord connectome reveals principles of functional organisation

**DOI:** 10.1101/2023.06.05.543407

**Authors:** Elizabeth C. Marin, Billy J. Morris, Tomke Stürner, Andrew S. Champion, Dominik Krzeminski, Griffin Badalamente, Marina Gkantia, Christopher R. Dunne, Katharina Eichler, Shin-ya Takemura, Imaan F. M. Tamimi, Siqi Fang, Sung Soo Moon, Han S. J. Cheong, Feng Li, Philipp Schlegel, Sebastian E. Ahnert, Stuart Berg, Janelia FlyEM Project Team, Gwyneth M. Card, Marta Costa, David Shepherd, Gregory S.X.E. Jefferis

## Abstract

Nervous systems function as ensembles of neurons communicating via synaptic connections, and a functional understanding of nervous systems requires extensive knowledge of their connectomes. In a companion paper (Takemura et al., 2023), we describe the acquisition of a complete fruit fly nerve cord connectome, the first for an animal that can walk or fly. Here, to efficiently navigate and to appreciate the biological significance of this connectome, we categorise and name nearly all neurons systematically and link them to the experimental literature. We employ a system of hierarchical coarse annotations and group similar neurons across the midline and across segments, then define systematic cell types for sensory neurons, intrinsic neurons, ascending neurons, and non-motor efferent neurons. Stereotyped arrays of neuroblasts generate related neuron populations called hemilineages that repeat across the segments of the nerve cord. We confirm that larval-born neurons from a given hemilineage generally express the same neurotransmitter but find that earlier born neurons often express a different one. We match over 35% of intrinsic, ascending, and non-motor efferent neurons across segments, defining serial sets which were crucial for systematic typing of motor neurons and sensory neurons. We assign a sensory modality to over 5000 sensory neurons, cluster them by connectivity, and identify serially homologous cell types and a layered organisation likely corresponding to peripheral topography. Finally, we present selected examples of sensory circuits predicated on programmatic analysis of a complete VNC connectome. Our annotations are critical for analysing the structure of descending input to the nerve cord and of motor output, both described in a third companion paper (Cheong et al., 2023). These annotations are being released as part of the neuprint.janelia.org and clio.janelia.org web applications and also serve as the basis for programmatic analysis of the connectome through dedicated tools that we describe in this paper.

## INTRODUCTION

The adult insect ventral nerve cord (**VNC**), analogous to the vertebrate spinal cord, has long been an important model system for the investigation of the sensorimotor circuits underlying key behaviours including locomotion (Bidaye et al., 2018; Cruz and Chiappe, 2023), grooming (Seeds et al., 2014), escape (Card, 2012), and reproduction (Pavlou and Goodwin, 2013). However, over a century of anatomical, physiological and functional studies necessarily focused on neurons that were easily accessible for physical and/or genetic manipulations, leaving much of the nerve cord a “black box” of undescribed interneurons that somehow connected sensory input to motor output. Recently, large-scale, systematic efforts have tackled the classification and functional analysis of these interneurons by considering related developmental populations (postembryonic neuroblast hemilineages) as anatomical and functional units (Harris et al., 2015; Shepherd et al., 2019, 2016). But these approaches omitted the larger, more diverse embryonic-born neurons and lacked the resolution needed to detect functional cell types within hemilineage populations. Acquiring a full inventory of cells and cell types and their synaptic connections - a complete connectome - would provide the necessary atlas for navigating the complexity of the adult ventral nerve cord network.

The methods and value of generating synaptic resolution connectomes have already been established by several large-scale fruit fly electron microscopy (**EM**) volumes. A pioneering project enabled the first glimpses into the anatomy and connectivity of the first instar larval CNS, including part of the VNC (Ohyama et al., 2015); later work has completed the larval brain (but not nerve cord) connectome (Winding et al., 2023). Previous adult fly connectome projects have focussed on the isolated brain (Scheffer et al., 2020; Zheng et al., 2018), now including a complete female brain connectome (Dorkenwald et al., 2023; Schlegel et al., 2023). However, recently the first female adult nerve cord connectome (**FANC**) has been partially reconstructed, focusing on the sensory and motor neurons in the prothoracic neuropils (Chen et al., 2021; Lesser et al., 2023; Phelps et al., 2021). A fully reconstructed male VNC connectome would reveal circuits for sex-shared behaviours across the CNS as well as male-specific behaviours such as courtship.

The acquisition, segmentation, and proofreading of the first ∼complete synaptic resolution connectome of an adult VNC (**MANC**, or male adult nerve cord) is described in a companion paper (Takemura et al., 2023). Another companion paper describes how the descending neurons from the brain and the motor neurons were typed and reveals detailed principles of the logic of descending control of motor circuits (Cheong et al., 2023). Here we report our efforts to make this vast data set accessible to the wider *Drosophila* neuroscience community by expert annotation of observed anatomical and inferred developmental features and the assignment of a **systematic cell typing** nomenclature based on those features to most of the neurons contributing to the VNC. This provides a comprehensive framework to understand the organisational principles of sensorimotor circuits within the VNC.

In adult flies, the VNC contains premotor circuits controlling walking and flight located in the thoracic **neuromeres**, while the abdominal neuromeres regulate and report on the gut and reproductive tracts. As such it represents the final common pathway for the majority of the nervous system’s behavioural outputs. Indeed flies without a head are still able to carry out numerous motor actions, suggesting that large parts of the motor control circuitry reside within the VNC (Yellman et al., 1997). Conversely, key sensory pathways from the legs, wings and viscera project into the VNC. While some sensory neurons ascend straight to the brain, most incoming information is processed locally.

Anatomically the VNC has both a segmental organisation along the anterior-posterior length of the nerve cord and a layered organisation along the dorso-ventral axis. The segmental organisation is directly analogous to spinal cord segments. It is inherited from the larval nervous system and is most obvious in the three thoracic neuromeres (T1, T2, T3) associated with pairs of legs in the adult (Figure 1A). The dorso-ventral axis has two main organisational features. The first is a layered arrangement in which neurons controlling the legs are located in ventral regions, while wing-control neurons reside in dorsal regions; an intermediate region contains neurons that bridge motor systems or have more complex functions (Namiki et al., 2018). The second organisational motif is the arrangement of sensory neurons in more ventral layers and motor neurons in more dorsal layers of a given region, i.e. the opposite arrangement to the mammalian spinal cord (Strausfeld, 1976). These features are visible in the gross anatomy of the VNC, resulting in the definition of named **neuropil** domains (Figure 2A) which provide a framework for describing both the regional organisation of the nerve cord and the structure of individual neurons (Court et al., 2020).

**Figure 1.**
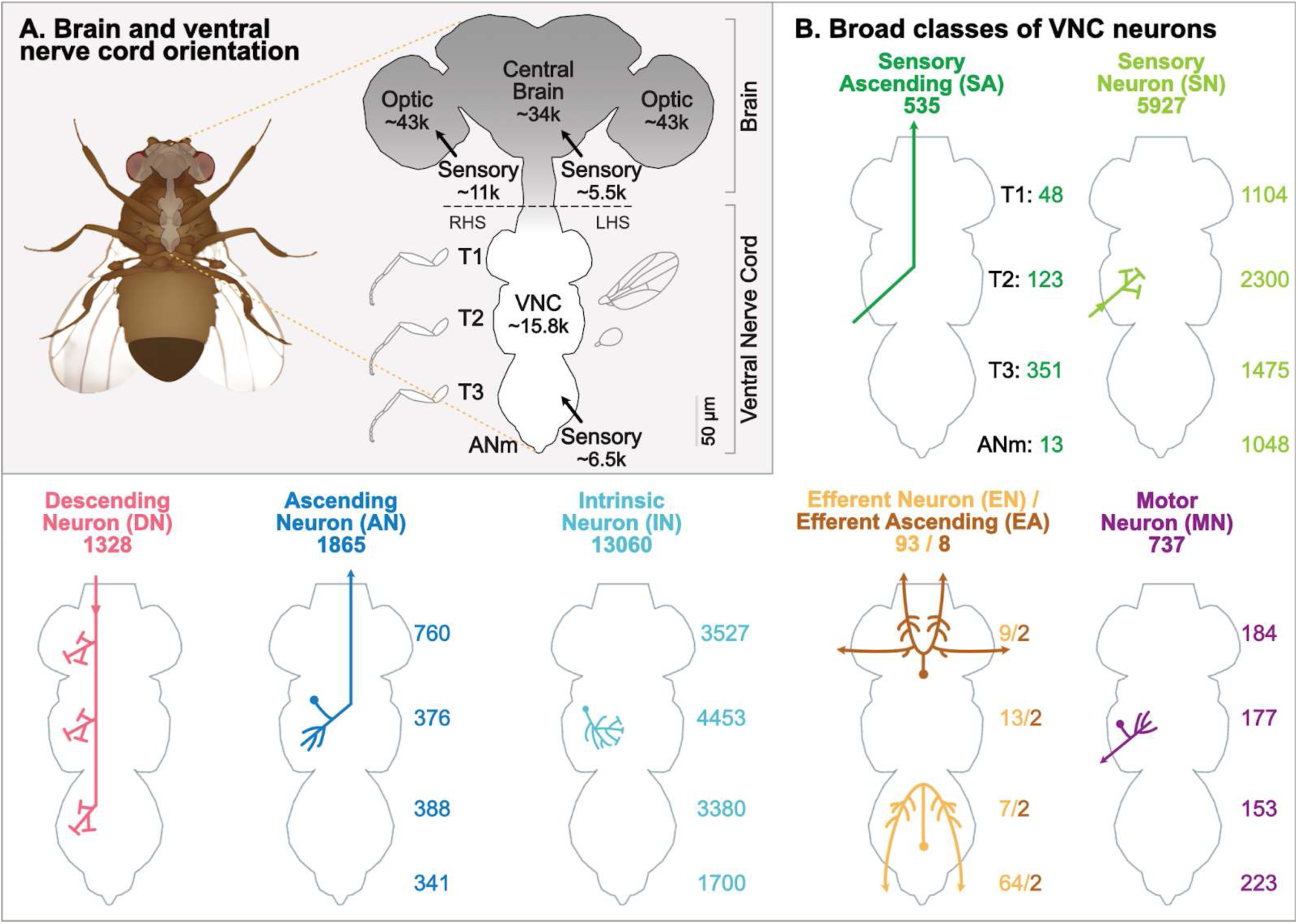
Overview of the male adult nerve cord (MANC). **A.** Orientation schematic showing a ventral view of the adult male *Drosophila* central nervous system. The ventral nerve cord (VNC) can be roughly divided into three thoracic neuromeres (T1, T2, T3) and the abdominal neuromeres (ANm). The thoracic neuromeres contain serially repeating neuropils controlling the three pairs of legs. T2 includes additional circuitry dedicated to flight. Number of neurons with cell bodies in the central brain are current estimates from FlyWire-FAFB (Schlegel et al., 2023). Cartoon legs and wings not to scale. **B.** Cartoon overview of the broad classes of neurons in the VNC including their overall and neuromere-specific neuron counts in this dataset.

**Figure 2.**
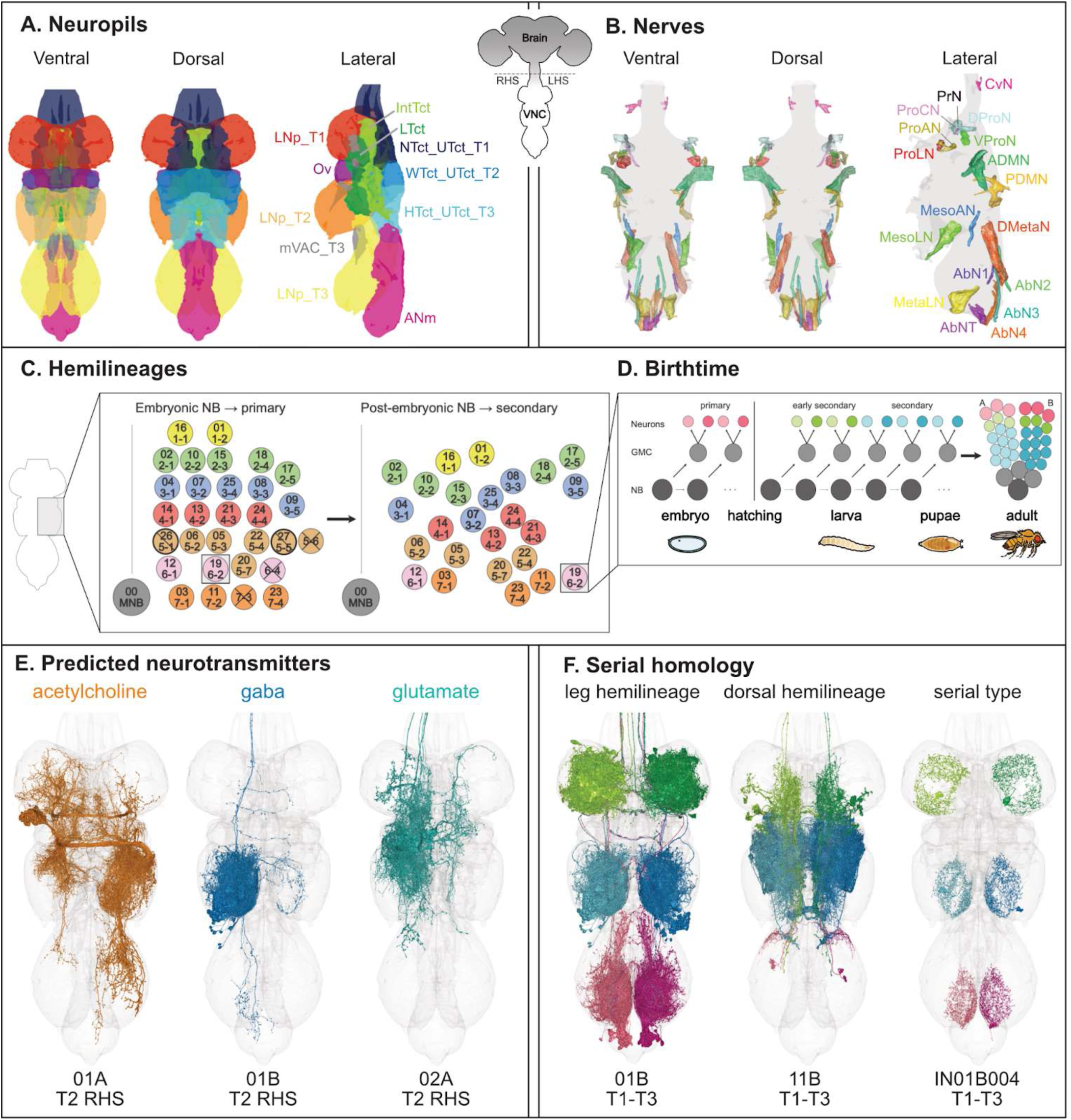
Overview of key features in MANC. **A.** Overview of the neuropils in the ventral nerve cord. The left-hand side leg neuropils are removed in the lateral view. **B.** Overview of the nerves in the ventral nerve cord. **C.** Projected cross section of half of the mesothoracic neuromere (T2) depicting the array of neuroblasts that divide to produce hemilineages in the embryo and/or postembryonically. Insect embryonic neuroblasts (e.g., 1-1) were named for their row and column of delamination (Bate, 1976; Doe and Goodman, 1985) whilst postembryonic neuroblasts (e.g., 16) were assigned sequential numbers based on their observed progeny in Drosophila (Truman et al., 2004). Embryonic and postembryonic neuroblasts were eventually matched (Birkholz et al., 2015; Lacin and Truman, 2016), but both nomenclature systems are still in use. Bolded circles denote novel matches proposed in our study. A subset of neuroblasts do not survive to produce postembryonic progeny; “/” indicates that embryonic progeny were identified in this study while “X” indicates that they were not. **D.** Overview of neuroblast (NB) division into ganglion mother cells (GMC) prior to differentiation into two hemilineages. Primary neurons are born in the embryo followed by early secondary and finally secondary neurons. Cartoon icons by DBCLS (https://togotv.dbcls.jp/en/pics.html) were identified via https://bioicons.com/?query=fly and reproduced under https://creativecommons.org/licenses/by/4.0/. **E.** Most secondary neurons in a hemilineage share the same neurotransmitter prediction (Eckstein et al., 2023; Takemura et al., 2023). **F.** Hemilineages are serially repeated along the segments of the nerve cord. This provides a basis for identifying serially homologous neurons, which are concentrated in the six leg neuropils. Note that the T1 neuropils are rotated outwards relative to the T2 and T3 neuropils. Hemilineages innervating dorsal neuropils (e.g., 11B) exhibit more variation in survival and morphology between segments.

Insect central neurons originating in the nerve cord and suboesophageal ganglia (**SEZ**) arise from stereotyped arrays of neuroblasts (neural stem cells) (Figure 2C) each generating a reproducible set of neuronal cell types. These neuroblasts are exactly reflected across the midline and repeat (with some variations) in each segment. Each neuroblast produces two distinct populations of progeny called **hemilineages**, which usually differ in gross morphology, gene expression, and neurotransmitter (Lacin et al., 2019; Truman et al., 2010) (Figure 2D-E). Members of each hemilineage may mature to function in larval and/or adult life or be eliminated soon after birth by programmed cell death, sometimes with segment-specific variation (Marin et al., 2012; Truman et al., 2010).

In holometabolous insects such as moths and flies that undergo metamorphosis, the dramatic changes in body plan and behaviour from larvae to adults necessitate significant expansion and reorganisation of the nervous system. New populations of sensory neurons are generated during development of the adult-specific cuticle and appendages such as legs and wings. Meanwhile, the larval VNC is transformed by the death or remodelling of the embryonically produced **primary** neurons and the addition of a much larger population of later born, adult-specific **secondary** neurons (Truman, 1990) (Figure 2D). Secondary neurons from the same hemilineage share grossly similar morphology (Shepherd et al., 2019; Truman et al., 2004) and appear to express the same neurotransmitter (Allen et al., 2020; Lacin et al., 2019). Furthermore, targeted activation of each postembryonic hemilineage has been associated with a specific behaviour (Harris et al., 2015), suggesting that hemilineages comprise the functional units of the VNC. However, a systematic comparison of the morphology and connectivity of the individual neurons within each hemilineage would be required to assess the conservation vs diversity of neuron structure and function.

In *Drosophila*, most sensory neurons are born peripherally from populations of cells, set aside in the embryo, that are the precursors of the adult appendages and epidermis (Bate and Arias, 1991). As these imaginal tissues grow, the sensory neurons emerge, *in situ*, in precise and reproducible patterns (Nottebohm et al., 1994; Palka et al., 1984). Axons from insect sensory neurons grow centripetally into the VNC via stereotyped **entry nerves** (Figure 2B) to form synaptic connections with their targets (Jan et al., 1985). These afferent projections are highly organised, with axons from the legs, wings, and halteres targeting distinct neuropils and axons that serve different sensory modalities terminating in different layers of neuropil (Merritt and Murphey, 1992; Murphey et al., 1989a; Tsubouchi et al., 2017). This precise ordering of the sensory system allows us to assign modality and coarse types to most sensory neurons based solely on their afferent projections.

In this study, we provide a multi-resolution catalogue of the neuronal constituents of the VNC including incoming sensory neurons, outgoing motor neurons, descending neurons from the brain and ascending neurons connecting to the head. We place particular emphasis on the ∼16,000 neurons generated in the VNC as individual neuronal hemilineages, thereby linking developmental origin to connectome structure. We also identify examples of **serial homology** - cell types that are repeated across neuromeres - that allow identification of recurring circuit motifs. This study confirms that developmental origins generate natural units of functional organisation while allowing the evolution of cell type diversity and also provide a unified way to define systematic cell types across the dataset.

In the results sections that follow, we describe the complete atlas of annotations provided by this work. We first provide an overview of the gross neuronal classes and general organisational features of the VNC dataset and how we have conducted cell typing. Then we will explore information flow within the VNC regions and how we predicted neurons with electrical and neurosecretory transmission. We then examine the general characteristics of hemilineage, birth time, and serial homology that underlie our systematic typing of neurons originating in the VNC, before dissecting each of the hemilineages annotated in this work. Finally we present the sensory neurons, their inferred modalities, the types we define within each modality by peripheral origin, and examples of specific sensorimotor circuits. The paper is structured so that the initial and concluding sections should be useful and accessible for the general reader. The intervening results sections describing each secondary hemilineage should serve as a reference for those with specific circuit interests and are ordered by hemilineage number for convenience.

## RESULTS

### Orientation and broad classes

The adult *Drosophila* central nervous system can be divided into the brain and suboesophageal ganglia in the head capsule and the thoracic and abdominal ganglia comprising the ventral nerve cord (**VNC**), linked by axons running through a narrow neck connective. The annotated male adult nerve cord (**MANC**) connectome (Takemura et al., 2023) provides the first complete and detailed inventory of neurons in the VNC as well as the neck connective.

The adult VNC can be grossly characterised as consisting of bilaterally symmetrical and segmentally repeating information processing units, with a limited number of connections to the brain and SEZ and to each other. It can be divided into three thoracic segments or **neuromeres** (**T1**/**T2**/**T3**) and abdominal neuromeres (**ANm**) consisting of segments A1-A10. T1, T2, and T3 are associated with the front legs, middle legs, and hind legs, respectively, while T2 is also associated with the wings and T3 with flight-associated balancing structures called the halteres (Figure 1A). The presence or absence of these appendages is expected to drive the number of neurons needed for information processing and motor control in each neuromere.

The MANC connectome (Takemura et al., 2023) is composed of ∼15.8K central neurons and ∼6.5K sensory neurons (Figure 1A). We first assigned every reconstructed neuron a broad **class** and corresponding abbreviation (Figure 1B). (Glia were not systematically reconstructed or typed, but 348 bodies have been annotated as class “glia”.) 1328 descending neurons (**DN**) relay information from the brain to the VNC via the neck connective. 5927 sensory neurons (**SN**) relay information through a nerve from the periphery and terminate in the VNC and 535 sensory ascending neurons (**SA**) ascend to the brain. 13,060 intrinsic neurons (**IN**) are restricted to the VNC, and 1865 ascending neurons (**AN**) originate in the VNC but extend an axon through the neck connective to the SEZ and/or brain. Efferents originate in the VNC and exit via the nerves; they can be classified either as efferent neurons (which are neuromodulatory, e.g., the octopaminergic neurons from the unpaired median neuroblast) or as motor neurons (which innervate the muscles). We annotated 93 efferent neurons (**EN**), 8 efferent ascending neurons (**EA**) that project into the neck connective, and 737 motor neurons (**MN**). These broad classes are not evenly distributed across neuromeres; for example, significantly more surviving intrinsic neurons originate in T2 than in T1 or T3, and nearly twice as many ascending neurons originate in T1 than in T2 and T3 combined.

With this basic parts list, we can make some simple but instructive comparisons between the brain and VNC. The full adult female brain (**FAFB**) is reported to contain about 133,000 cells, of which at least 115,000 are neurons (Mu et al., 2021). Recent collaborative work from our group focusing on the central brain (i.e. the brain without the massively expanded optic lobes) finds about 34,000 neurons with cell bodies in the central brain (Schlegel et al., 2023) compared with 15,763 in the VNC (this study). Although the central brain and VNC are roughly equivalent in volume, the central brain includes twice as many neurons, not including the sensory neurons from peripheral tissues (Figure 1A). VNC neurons are therefore larger on average, perhaps reflecting the need for rapid communication near the motor periphery. The central brain has a ratio of 6:1 between intrinsic and sensory neurons and 55:1 between intrinsic and motor neurons. Conversely, the VNC displays a considerably reduced ratio of 2.4:1 for intrinsic to sensory neurons and a substantially smaller ratio of 9:1 for intrinsic to motor neurons. These numbers highlight the VNC’s significance for sensory processing and as the principal centre for motor control. Nevertheless, the great majority of VNC neurons (83%) are intrinsic neurons restricted to the nerve cord that define local circuits for sensory-motor processing.

### Organisational features used for systematic typing

A common language and coordinate system aids communication of results between researchers working with different tools in different times and places. For example, descending neurons have been named based on the locations of their cell bodies in the brain rather than on the specific genetic lines used to label them (Namiki et al., 2018). We were motivated to develop a systematic nomenclature for the adult VNC because only a fraction of its cell types have been described and named in the light level literature; the principal exceptions are the descending neurons and motor neurons, both of which were typed separately (Cheong et al., 2023).

To prepare for systematic typing we sought anatomical features that could be used to subdivide the neurons – particularly the large populations of unnamed intrinsic and sensory neurons – into units likely to be relevant for function (Figures 2 and 3). Recently a systematic nomenclature was developed for the anatomy of the adult fly nerve cord, synthesising many decades of light-level experimentation in various insect species (Court et al., 2020). We based our annotation of anatomical features on this consensus framework and leveraged newly accessible features, including connectivity and predicted neurotransmitter, to contribute to classification of systematic cell types. Our systematic nomenclature doubles as concise descriptions of key features, allowing searches for neurons of interest by broad class, position, and developmental origin. Table 1 provides a complete list of annotation fields and values and demonstrates how to query for neurons with specific features at clio.janelia.org (Table 1 - video 1, Table 1 - video 2), a free public resource accessible to any user with a registered Google account.

**Figure 3.**
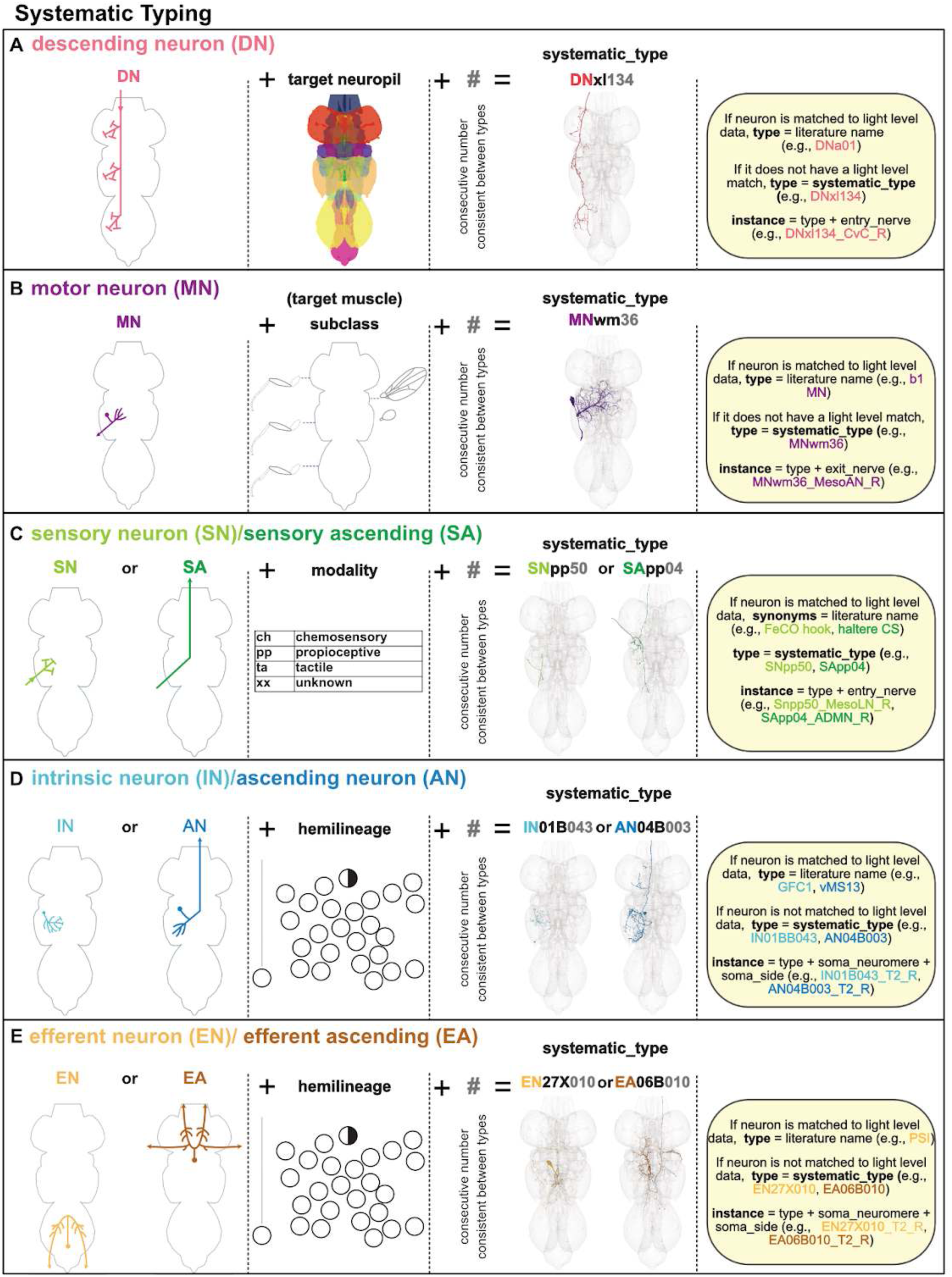
Systematic cell typing schemes for each broad cell class. **A.** Descending neurons are assigned a systematic type composed of the prefix “DN” followed by an abbreviation for the target VNC neuropil and a number consistent for members of their cell type. Their entry nerve (CvC) and root side are used to denote the instance. **B.** Motor neurons are assigned a systematic type composed of the prefix “MN” followed by the subclass (an abbreviation for the target muscle category) and a number consistent for members of their cell type. Their exit nerves are used to denote the instance. **C.** Sensory neurons are assigned a systematic type composed of the prefix “SN” (if not ascending) or “SA” (if ascending) followed by an abbreviation for their assigned modality and a number consistent for members of their connectivity cluster. Their entry nerve is used to denote the instance. **D.** Intrinsic neurons and ascending neurons are assigned a systematic type composed of the prefix “IN” or “AN” followed by their assigned hemilineage and a number consistent for members of their connectivity cluster. Their soma neuromere and side are used to denote the instance. **E.** Efferent neurons are assigned a systematic type composed of the prefix “EA” (if they ascend to the brain) or “EN” otherwise followed by their assigned hemilineage and a number consistent for members of their connectivity cluster. Their soma neuromere and side are used to denote the instance.

**Table 1.**
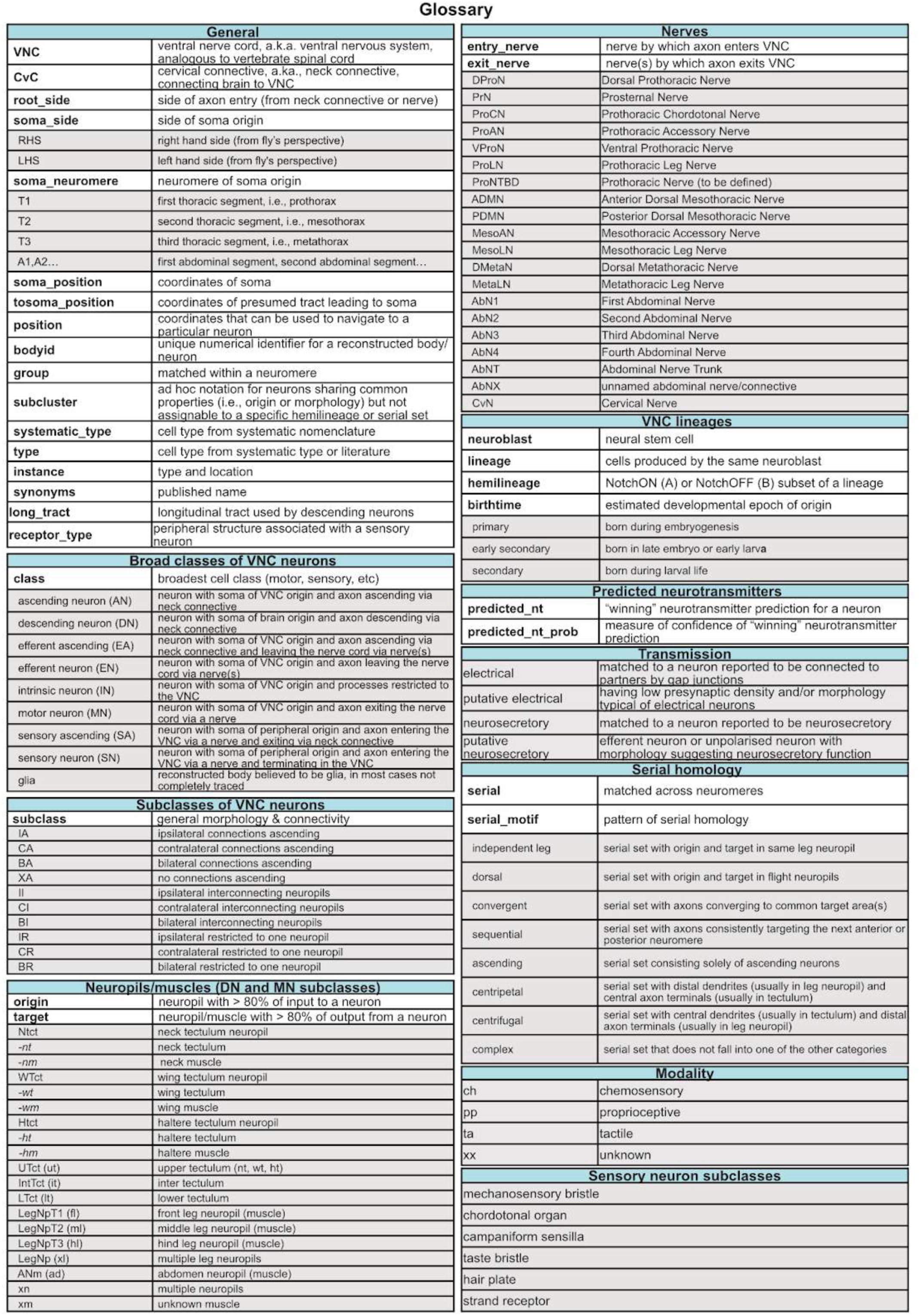
Glossary of annotation terms. Annotation field names are bolded and shown against a white background whilst the associated field values are shown against a grey background. Note that compound field names are presented in the snake_case format (e.g., entry_nerve) used in Clio Neuroglancer whereas Neuprint employs the camelCase format (e.g., entryNerve). **Table 1 - video 1. Using Clio Neuroglancer with the plain option to retrieve neurons by annotation (2:37).** https://youtu.be/EmOB8WvDHss Navigate to https://clio.janelia.org/ and select workspace “annotate” and the desired segmentation volume. On the right side click the “BODIES” tab to bring up the annotation fields. Using the plain option, type your query enclosed in curly brackets, with quotation marks around the field name, a colon, and quotations around the desired field value. Then click on the magnifying glass icon to retrieve all bodies matching your criteria. You can use the Fields checklist to hide or display particular fields for your neurons of interest. Now you can filter the list by typing into the fields. For example, let’s compare the FeCO claw neurons in the right mesothoracic **entry nerve**. Pressing the flashlight icon will display all listed neurons in random colours. I’ll remove the tissue layer and add the neuropil layer so that it’s easier to see where the neurons are positioned within the leg neuropil. To compare the two different types of FeCO claw neurons, we could select a few neurons from each **type** and assign them to a specific colour, like this. Let’s right-click on the segmentation layer, go to the Seg. section, and copy-paste the respective **bodyids** into the Neuroglancer window. This is so that they won’t be lost when we select more neurons to display. Now clear the first selection, select neurons from the other type, and assign these to a different colour. We can then click on the saved neurons to view both of them at the same time. Note that when using this method to query for a numerical field value - for example, the **bodyid, group, serial**, or **subcluster** fields - you need to enclose the number in square brackets instead of in quotation marks, like this. If we now remove the previous filters, we can view all examples of a given serial set. **Table 1 - video 2. Using Clio Neuroglancer with the structured option to retrieve neurons by annotation (2:41).** https://youtu.be/jObOKhAD7LQ Navigate to https://clio.janelia.org/ and select workspace “annotate” and the desired segmentation volume. On the right side click the “BODIES” tab and select the structured option. Type your desired annotation fields and values into the query boxes, clicking the “+” symbol to add them - for example, **hemilineage** “14A” and **serial_motif** “independent leg”. Once you’ve added your criteria, click on the magnifying glass icon to retrieve the corresponding bodies. To customise the displayed information, use the Fields checklist to select fields of interest. You can also add filters for specific characteristics. For example, to visualise neurons from a specific serial set, such as “10424” (Figure 11A), type the number into the serial field and click the flashlight icon. Now right-click on the segmentation layer, go to the Seg. section, and copy-paste the respective **bodyids** into the Neuroglancer window. Then you can colour the neurons as you like. For example, filter for **soma_neuromere** “T1”, click to select those bodyids, set their colour to green and highlight the selected neurons. You can repeat the same process for the other thoracic neuromeres, colouring T2 neurons in blue and T3 neurons in magenta. Then click on the saved list to visualise all of the neurons together. We recommend removing the “all-tissue” layer and selecting the “neuropil” layer to get a clearer view of the neuropils innervated by your neurons. Note that to search for a specific serial set directly, you could instead use the structured option by simply typing serial and the number into the query boxes.

The VNC can be divided into a set of major neuropils (Figure 2A) reflecting neuroanatomical organisation and function (Court et al., 2020). In the thorax, the six leg neuropils (**LegNp**) each consist of two sensory neuropils - the ventral association centre (**VAC**, not labelled) and medial ventral association centre (**mVAC**) - along with the intermediate neuropil (**IntNp**). The tectulum lies dorsal to the leg neuropils and is specialised for flight. It is composed of the ventralmost lower tectulum (**LTct**), the intermediate tectulum (**IntTct**), and finally the dorsalmost upper tectulum; upper tectulum can be further divided into the neck tectulum (**NTct**), wing tectulum (**WTct**), and haltere tectulum (**HTct**), proceeding from anterior to posterior. We assigned every neuron an **origin** neuropil (or peripheral structure) based on the observed or inferred locations of its postsynapses and a **target** neuropil (or peripheral structure) based on its presynapses.

The VNC has a clear segmental organisation along its anterior-posterior length, matching the major division of the insect body into thorax and abdomen and further divided into individual **neuromeres** corresponding to repeated somatic structures such as the legs in the thorax (Figure 1A). The first thoracic neuromere (**T1** or prothorax) is the most anterior and includes the NTct and prothoracic leg neuropils, LegNp(T1). The second thoracic neuromere (**T2**) or mesothorax includes the specialised wing and notum sensory neuropil called the ovoid (Ov), along with the WTct and the mesothoracic leg neuropils, LegNp(T2). The third thoracic neuromere (**T3**) or metathorax includes the HTct and the metathoracic leg neuropils, LegNp(T3). The abdominal neuromeres (**ANm**) are treated as one neuropil consisting of fused neuromeres A1-A10.

Axons enter and exit the VNC via a defined set of peripheral nerves (Court et al., 2020) (Figure 2B). The small cervical nerve (**CvN**) connects laterally to the CvC. The dorsal prothoracic nerve (**DProN**), prosternal nerve (**PrN**), prothoracic chordotonal nerve (**ProCN**), prothoracic accessory nerve (**ProAN**), ventral prothoracic nerve (**VProN**), and prothoracic leg nerve (**ProLN**) directly innervate the prothoracic neuromere (T1). The anterior dorsal mesothoracic nerve (**ADMN**), posterior dorsal mesothoracic nerve (PDMN), mesothoracic accessory nerve (**MesoAN**), and mesothoracic leg nerve (**MesoLN**) innervate the mesothoracic neuromere (T2). The dorsal metathoracic nerve (**DMetaN**) and metathoracic leg nerve (MetaLN) innervate the metathoracic neuromere (T3). Finally, the abdominal neuromeres are innervated by the bilaterally paired first, second, third, and fourth abdominal nerves (**AbN1, AbN2, AbN3, AbN4**) and by the fused abdominal nerve trunk (**AbNT**). We assigned an **entry nerve** and/or **exit nerve** to all neurons with axons entering or leaving the VNC - that is, SNs and ANs vs MNs, EN, and EAs, respectively. We also annotated the cervical connective (**CvC**), which connects the VNC to the brain.

30 paired and one unpaired neuroblast in each VNC neuromere divide embryonically to generate an initial set of **primary** neurons (Schmid et al., 1999); in T2, 23 + 1 of these also divide post-embryonically to generate adult-specific **secondary** neurons (Booker and Truman, 1987; Prokop and Technau, 1991; Truman and Bate, 1988); and two neuroblasts only divide post-embryonically (Lacin and Truman, 2016) (Figure 2D). Most (but not all) of these neuroblasts also divide postembryonically in T1 and T3, with a much smaller subset surviving in the abdomen (Truman and Bate, 1988). In most cases, the embryonic and postembryonic neuroblasts have been mapped onto each other (Birkholz et al., 2015; Lacin and Truman, 2016) (Figure 2C), but we found evidence for two new potential matches (Figure 48J,K).

Each neuroblast typically buds off a series of transient ganglion mother cells that divide once to generate a pair of post-mitotic neurons (Bate, 1976; Doe and Goodman, 1985). Through asymmetric division, one daughter neuron acquires a Notch^ON^ (A) fate while the other acquires a Notch^OFF^ (B) fate, resulting in two distinct **hemilineages** as progeny accumulate over time (Truman et al., 2010). Most or all secondary neurons from one hemilineage may be eliminated by programmed cell death in one or more neuromeres (Marin et al., 2012). When both hemilineages from one neuroblast survive, they typically have distinctive features (eg., 01A vs 01B) (Figure 2E), and their somas may be pulled far apart during development, depending on the neuropil(s) targeted by each one.

Secondary neurons from the same hemilineage are expected to share similar gross morphology (Shepherd et al., 2019; Truman et al., 2004) and express the same fast-acting neurotransmitter (Allen et al., 2020; Lacin et al., 2019), making hemilineages strong candidates as units of functional organisation in the VNC. We assigned >90% of VNC neurons (97% in thoracic neuromeres) to a hemilineage, relying primarily on soma tract entry position and gross morphology but also on several clustering methods and serially homologous cell types (see Introduction to hemilineage-based organisation and Methods). Fast-acting neurotransmitter predictions were made for every neuron in the dataset (Eckstein et al., 2023; Takemura et al., 2023), and all/most secondary neurons assigned to each hemilineage shared the same predicted neurotransmitter (**predicted nt**) (Figure 2E).

We annotated key positional features for every neuron in the dataset. For neurons originating in the VNC, we annotated the **soma position** or (**tosoma_position** if missing the expected soma), **soma neuromere** (e.g., T1, T2, etc) and the **soma side** (RHS, LHS, or midline). For neurons originating in the brain or periphery we annotated the **root side** as defined by the location of axon entry into the VNC.

In preparation for automated systematic typing, we performed expert review of neurons that appeared similar by morphology and connectivity. Neurons matched across the midline were assigned the same **group** number, whilst neurons matched across different neuromeres were assigned the same **serial** number (Figure 2F). Excluding sensory neurons and neurons from the ventral unpaired median neuroblasts, 98.7% neurons were matched across the midline (16,556 of 16,778). We identified serial homologues for 36.7% of neurons originating in the VNC; the large majority innervated the leg neuropils, whilst serial homologues innervating the dorsal neuropils were rare, consistent with segment-specific survival and specialisation.

We defined systematic types using a standardised method and format for neurons of each broad class, as described in more detail below. In each case, the **systematic type** name begins with a two-letter, class-specific prefix (e.g., DN, MN, AN) and ends with a number defined by hierarchical clustering on the basis of morphology and connectivity. To be comprehensive, the systematic type depends only on the rich feature set of the connectome itself. Each neuron was also assigned a **type**; this may be preferred by end users since it includes commonly recognised cell type names based on the light-level literature or because the systematic types defined by clustering were subsequently refined using specialised methods or external information. For example, the pheromone-responsive foreleg sensory neurons were initially defined as one systematic type but then further distinguished into subtypes based on their ipsilateral vs contralateral ascending neuron partners (Figure 64). Finally, each neuron was assigned an **instance** - that is, its type plus a suffix denoting its specific location. This distinguishes homologous neurons of the same cell type across sides and neuromeres.

We assigned **systematic type** names to all descending neurons (DN) and motor neurons (MN) based on their **target neuropils** and their **target muscles**, respectively (Figure 3A,B). Descending neurons that could be matched with confidence to light level images were assigned canonical **type** names (e.g., DNp50) and additional **synonyms** (e.g., Giant Fiber) if applicable and otherwise received the systematic type name as their type name, as described in our companion manuscript (Cheong et al., 2023). Similarly, motor neurons matched to light level images were assigned canonical type names (e.g., hg1 MN) and synonyms (e.g., vMS2) if applicable and otherwise received the systematic type name as their type name (Cheong et al., 2023). Many of the leg motor neurons are serially repeating, so we kept the same number for the systematic type, while using “fl”, “ml”, “hl” (front, <u>m</u>iddle and <u>h</u>ind leg) as the subclass (target muscle abbreviation). Each neuron was also assigned an **instance** based on its position in the volume: the entry nerve (CVC) plus root side was used for DNs, and the exit nerve (including side) was used for MNs.

The **entry nerve** was used as the instance of sensory neurons (SA if ascending to the brain, SN if not), but systematic typing was based on inferred **modality** (Figure 3C). Sensory neurons were broadly classified as *chemosensory* (responsive to chemical stimuli), *tactile* (responsive to touch), *proprioceptive* (providing information on the location and movement of the body), or of *unknown* modality. We used weighted nearest neighbour clustering to assign the **systematic types** (Figure 52 and Methods) and put the cluster number in the **subcluster** field. Many leg sensory neurons are serially repeating, so we assigned the same systematic type to all neurons that could be matched across sides and neuromeres. When possible, we annotated the sensilla type (e.g., chordotonal organ) in the **subclass** field and annotated any names from light level matching in the **synonyms** field. The assigned **type** was usually the same as the systematic type, except where additional information was used to split the systematic type into finer types (Figure 64).

Most of the remaining ∼15,000 neurons have only been described previously at light level as part of large populations deriving from the same postembryonic neuroblast (Shepherd et al., 2019) or labelled by a broad driver (Harris et al., 2015). These include the interneurons ascending to the brain (AN), intrinsic neurons restricted to the VNC (IN), neurosecretory efferent neurons exiting one or more nerves (EN), and neurosecretory efferent neurons exiting one or more nerves and ascending to the brain (EA). The **systematic type** of these neurons was based on their developmental origin, i.e., **hemilineage** (Figure 3D-E). Within each hemilineage, matching groups and serial sets (Figure 11) and predicted neurotransmitter (**predicted nt**, Figure 7D) were used to inform the clustering used to assign systematic type names and type names (Figure 3D-E, Figure 13, and Methods). The hemineuromere (**soma neuromere** and **side**) of origin was used for the instance. In most cases, the **type** was the same as the **systematic type**, although a small number of identified electrical, neurosecretory, or sexually dimorphic neurons were assigned a type based on matching to the light-level literature and also annotated in the **synonyms** field.

It is important to note that MANC annotations were improved substantially between the version of the dataset (manc:v1.0) that accompanied our initial preprints and the current version (manc:v1.2.1). In particular, hemilineage “05A” was merged into hemilineage “05B” as a minority neurotransmitter subpopulation and the tentative hemilineage designation “15A” was removed. Neurons of VNC origin and sensory neurons were also systematically retyped to emphasise serial homology across neuromeres while retaining finer resolution within neuromeres. Figures featuring type-specific information have been updated or created using current annotations and types (manc:v1.2.1). Researchers who have accessed the original dataset are strongly encouraged to refer to the annotations and type names featured in the current version of the dataset in any publications; these can be retrieved either programmatically or in Clio Neuroglancer (manc:v1.2.1) via a neuron’s conserved, unique numerical identifier, also known as the **bodyid**.

This is a living dataset, with annotations still expected to improve incrementally over time with input from the larger *Drosophila* research community. Barring major proofreading corrections, bodyids are stable and the best way to track particular neurons of interest. Future updates to annotations, including cell type matches to the light level literature, will be provided in periodic patches to Neuprint and new versions of Clio Neuroglancer.

### Neurons of VNC origin

#### Electrical and neurosecretory transmission

The MANC connectome was generated based on automated neurite segmentation and detection of chemical synapses. Therefore, classification of MANC neurons by innervated neuropil and/or synaptic partners as well as circuit analysis comes with some important caveats. Some neurons are missing synapses due to poor segmentation (typically an artefact of darkly stained profiles) or truncation (because they leave the volume or because segments could not be joined through a damaged area). Some sensory and motor neurons were challenging to reconstruct in this dataset, and in particular many neurons that enter or exit the leg nerves are missing or incomplete.

In addition, synapse prediction models were only trained to recognise chemical synapses, which in *Drosophila* can be identified by the presence of T-bars, vesicles, and postsynaptic densities. However, some neurons are known to be activated by electrical coupling via gap junctions, for example targets of the Giant Fiber neurons (Kennedy and Broadie, 2018). Innexin gap junction proteins such as ShakB are widely expressed in the VNC (Ammer et al., 2022), but gap junctions were not included in MANC segmentation or annotation. Therefore, synapse counts used for neuropil innervation or connectivity analyses were likely to be substantially underestimated for electrically coupled neurons.

To address these issues, we manually reviewed neurons with unusually low presynaptic density by volume (Figure 4A). We noted 48 matches to electrical neurons in the literature such as the peripherally synapsing interneuron (PSI) (Figure 4C) and annotated these as having electrical **transmission**. We then annotated 189 morphologically similar neurons with low presynaptic density and thick, smooth axons as having putative electrical transmission (Figure 4D). Many of these have been investigated in more detail with respect to flight circuits in our companion manuscript (Cheong et al., 2023).

**Figure 4.**
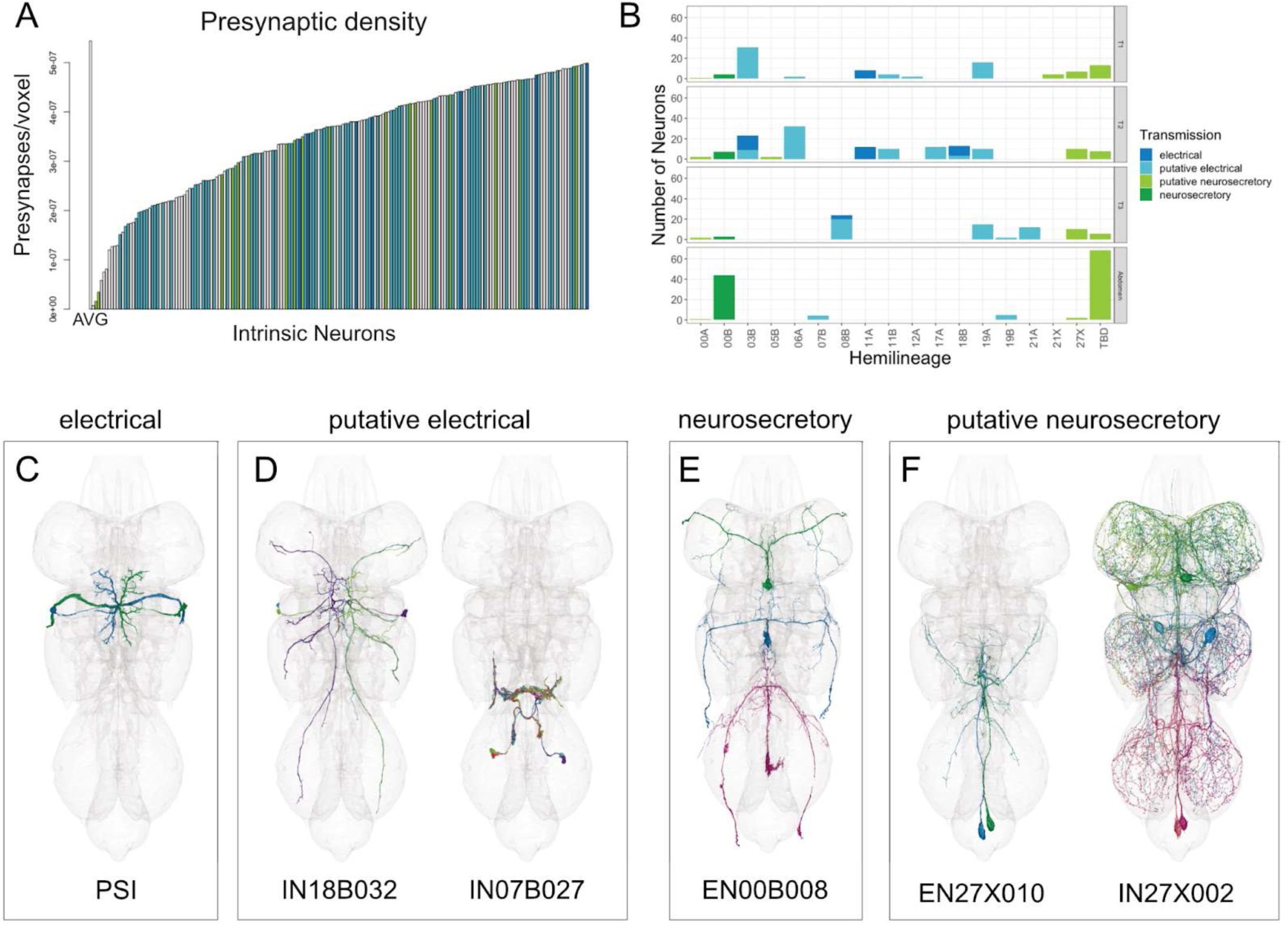
Neurons with suspected electrical or neurosecretory transmission. **A.** The 185 intrinsic neurons with the lowest presynaptic density by volume are enriched for types with electrical or neurosecretory transmission. AVG = average number of presynapses per voxel for all intrinsic neurons. **B.** Candidate electrical and neurosecretory neurons are produced by specific hemilineages. **C.** Example of an electrical neuron, PSI (group 11446). **D.** Examples of putative electrical neurons (groups 20924 and 12943). **E.** Examples of a neurosecretory neuron (serial 11170). **F.** Examples of putative neurosecretory efferent neurons (group 10985) and intrinsic neurons (serial 10083).

We also matched 58 neurons to neurosecretory cells from the literature such as the octopaminergic neurons from the median neuroblast (hemilineage 00B) (Pop et al., 2020) (Figure 4E). We annotated 39 other efferent neurons as having putative neurosecretory transmission (Figure 4F). We also annotated 152 neurons that broadly innervate the neuropil surface, with unpolarised synaptic swellings resembling beads on a string, as putative neurosecretory (Figure 4F).

These minority transmission populations were not uniformly distributed in the dataset but rather belonged to a small number of specific hemilineages, most of them innervating dorsal neuropils (Figure 4B). We were able to match one primary-only hemilineage (which we designated as 27X) to its neuroblast of origin based largely on the presence of putative neurosecretory cells (Figure 48J).

#### Information flow

We employed laterality and neuropil innervation as criteria to classify ANs and INs into distinct subclasses. The subclass nomenclature is based on these two indicators. The first letter reflects whether a neuron shows bilateral (B), ipsilateral (I), or contralateral (C) innervation of the VNC with respect to its soma side (Figure 5 - figure supplement 1A,B). The second letter indicates whether the neuron ascends to the brain (A), is restricted to a single neuropil (R), or interconnects multiple neuropils (I) (refer to Figure 5A and the Methods section for more details). Furthermore, we identified a subset of 30 ANs with fewer than 5 synapses in the VNC, which we designated as subclass XA (see Figure 5C for a corresponding image).

**Figure 5.**
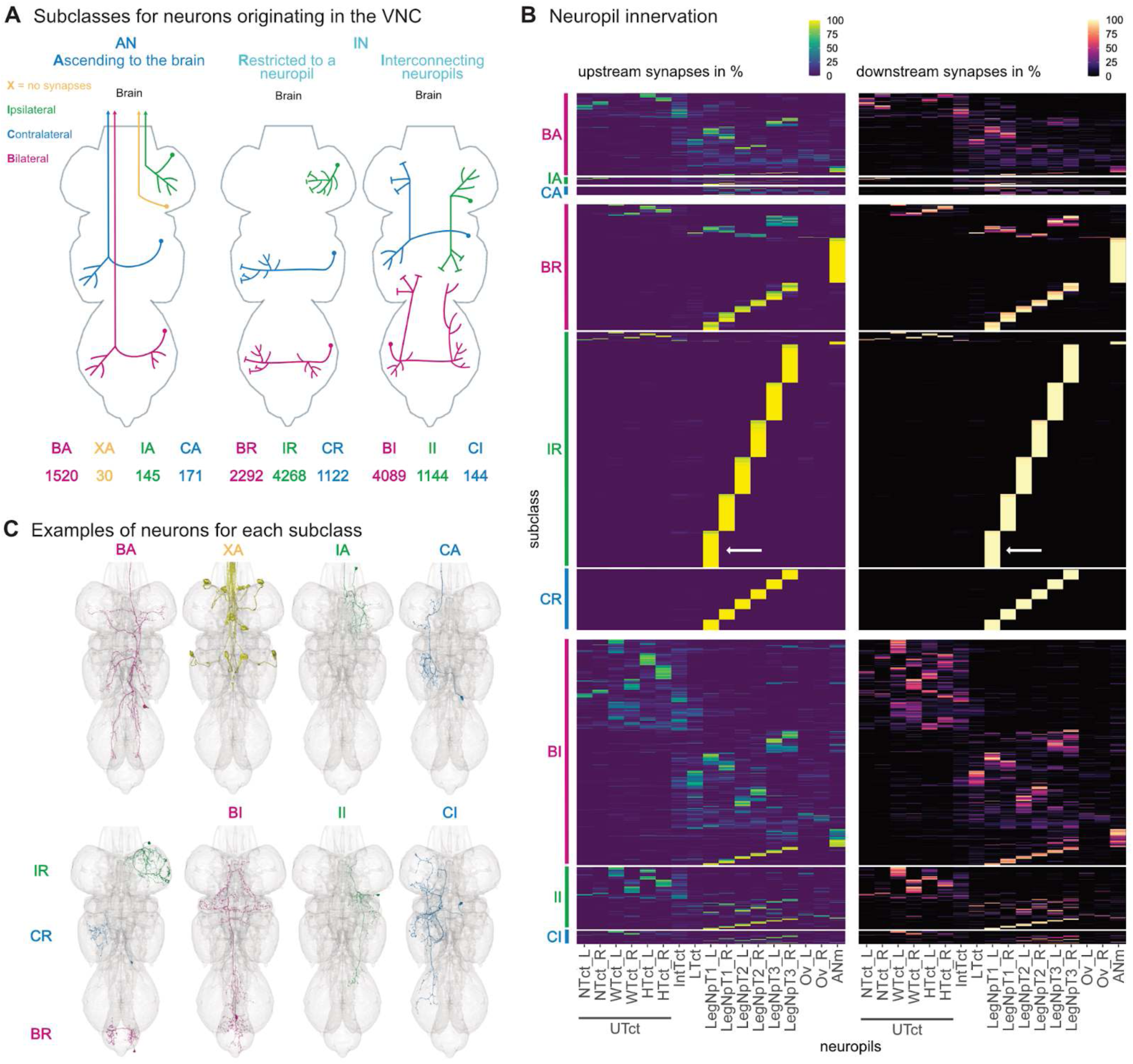
Subclasses of ANs and INs. VNC innervation of ascending neurons and intrinsic neurons. **A.** First letter of the subclass reflects if the neuronal arbour is Ipsilateral, Contralateral, Bilateral or not eXisting in the VNC. The second letter indicates if it is Ascending to the brain, Restricted to one neuropil in the VNC or Interconnecting different neuropils in the VNC. **B.** Single neuron inputs and outputs to VNC neuropils separated by subclass. Arrows pointing to a group of IR subclass neurons that receive all their input and give all their output to the front left leg neuropil. **C.** Example images for neurons of different subclasses.

**Figure 5 - figure supplement 1.**
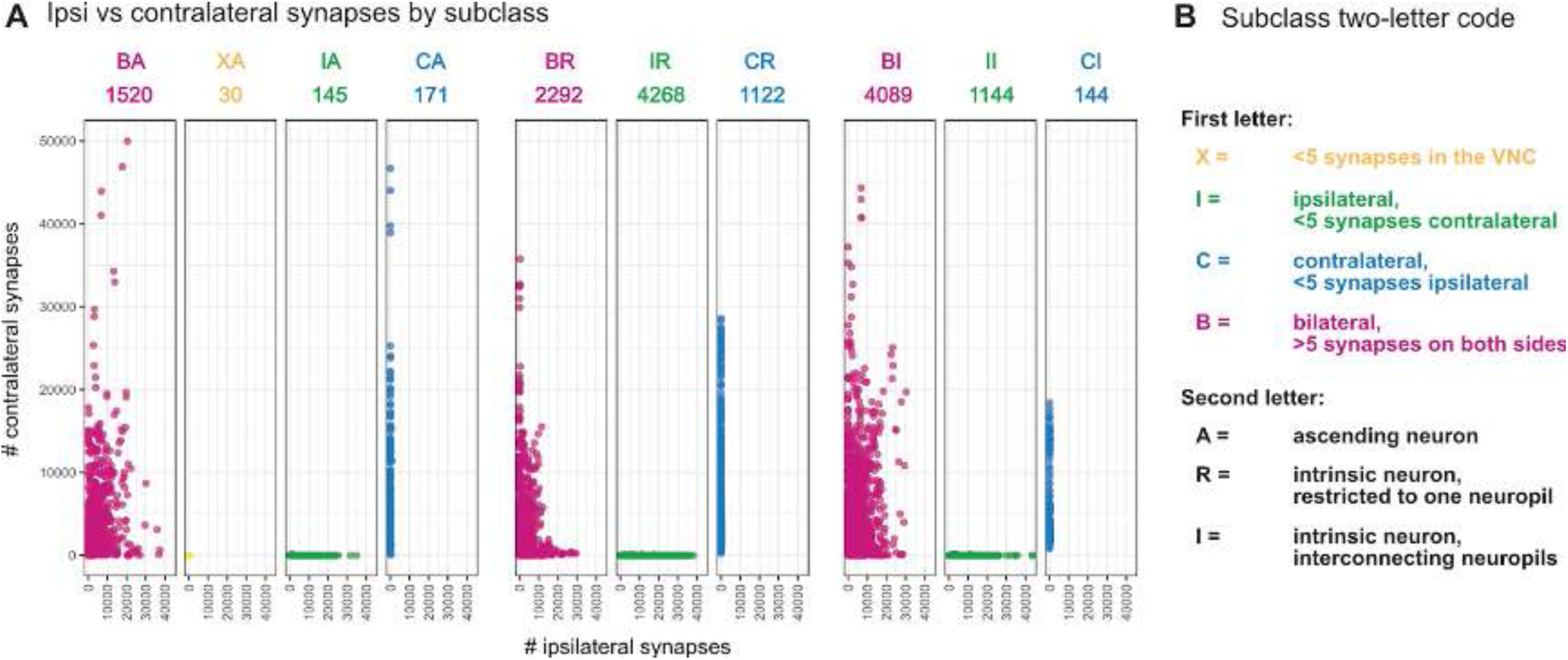
Subclass defined by ipsi and contralateral innervation of the VNC. **A.** Total number of synapses ipsilateral and contralateral for each neuron of a given subclass. Colours correspond to bilateral (magenta), no synapses (yellow), ipsilateral (green), contralateral (blue). **B.** description of the two-letter code of the subclass. The first letter refers to the laterality and the second if a neuron is ascending, restricted to one neuropil or interconnecting VNC neuropils.

The most abundant subclass among the classified neurons is IR, comprising intrinsic neurons that solely innervate a single neuropil ipsilateral to their soma location. The majority of these IR neurons target individual leg neuropils, similarly seen for the subclass CR (see Figure 5B, white arrows pointing to IR neurons that have all their input and output from the front left leg neuropil). The second largest subclass is BI, consisting of intrinsic neurons that bilaterally interconnect multiple neuropils. Upon closer examination of their neuropil innervation, this subclass could be further divided into two groups: those interconnecting the upper tectulum (a combination of NTct, WTct and HTct) and those interconnecting the leg neuropils (LegNpT1-T3) (Figure 5B).

We assigned every neuron an origin neuropil based on the observed or inferred locations of its postsynapses and a target neuropil based on its presynapses (see Methods for details). This allows us to characterise the main input and output neuropils for each individual neuron to produce a general model of information flow for ANs and INs (Figure 6A). For both classes, many neurons receive input from and provide output to the same neuropil (diagonal along the top panels in Figure 6A). Many ANs primarily innervate the T1 (front) leg neuropils, while the INs have a roughly even number of neurons innervating all six legs neuropils and the abdominal neuropil.

**Figure 6.**
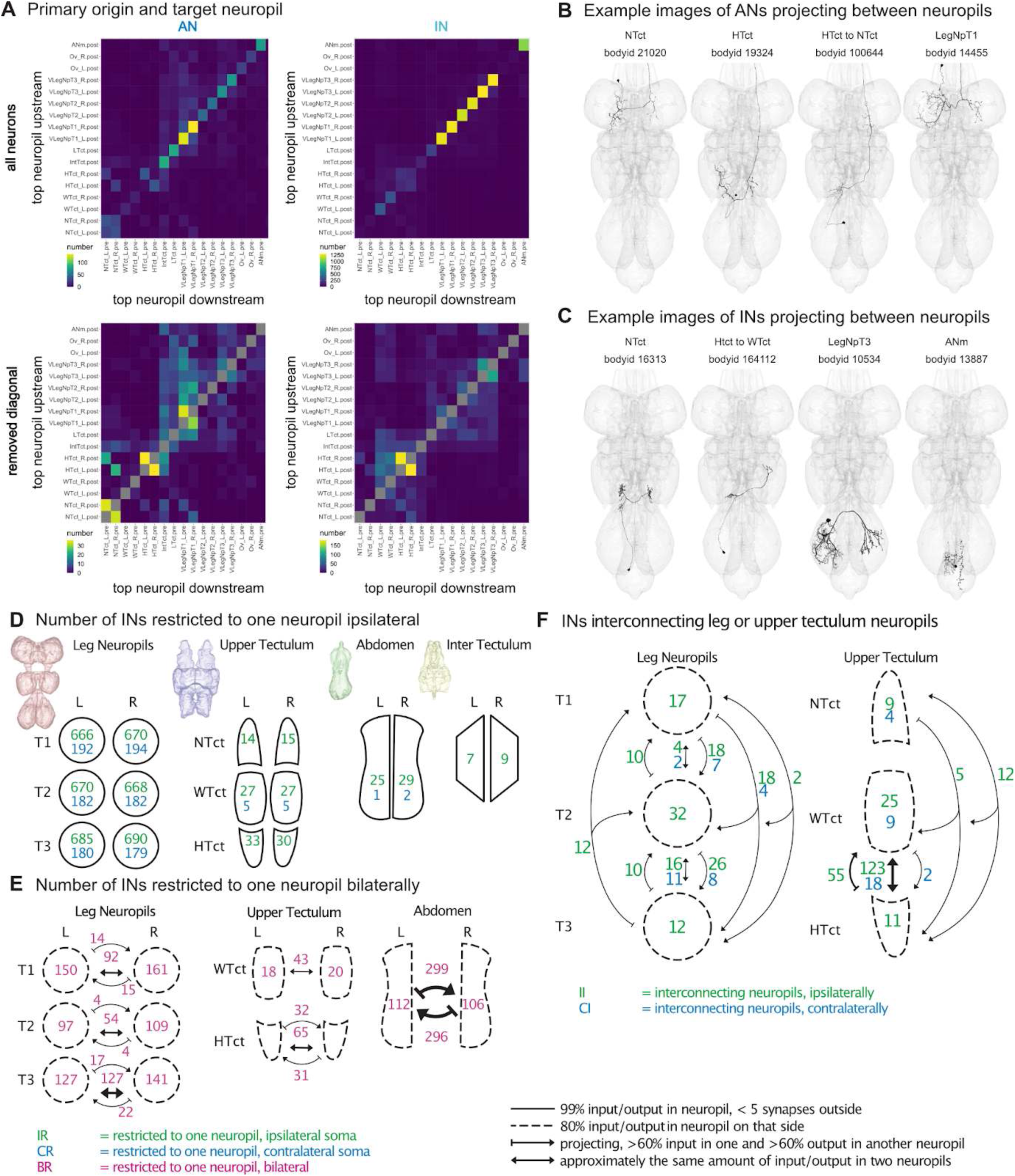
Information flow. Neurons within and projecting between neuropils. **A.** Primary origin and target of ascending (left) and intrinsic neurons (right). Top panels show all neurons and lower panels show all that do not have the same primary origin and target fields (shown in grey). **B.** Example images of ANs projecting between neuropils. **C.** Example images of INs projecting between neuropils. **D.** Illustration of the number of intrinsic neurons restricted to one neuropil ipsilaterally (green) or contralaterally (blue), neurons with origin and target in the same leg neuropil shown in the right top panel in A. **E.** Illustration of the number of neurons bilaterally restricted to one neuropil (in magenta). Numbers within the dotted neuropil schematic have 80% of their input and output in that area. Arrows indicate neurons that are projecting, having > 60% input in a different neuropil than > 60% of their output. Double arrows show neurons that connect neuropils almost evenly. **F.** Number of neurons interconnecting leg or upper tectulum neuropils on just one hemisphere. Numbers showing ipsilateral in green and contralateral numbers in blue.

After removing neurons that have the same top input and top output neuropil, we see interesting connections between the neuropils (lower panels in Figure 6A). There are four groups of connections that stand out for the ANs: bilateral connections between the NTct, HTct and LegNpT1 neuropils as well as a small group of HTct to NTct connecting ANs (Figure 6B). INs also have a high number of neurons that bilaterally connect the haltere neuropils (Figure 6C, first example). However, they also show a high number of neurons projecting from the haltere neuropils to the wing neuropils, a combination that was not seen for ANs (Figure 6C, second example). All leg neuropils show groups of IN neurons that project between the two sides, however, the largest number is seen in the hind leg neuropils (Figure 5C, third example). The abdomen neuropil mesh (ANm) is not split by hemisphere: if one compares neurons with the ANm as their top input and output neuropil, the majority project across the midline (Figure 6C, fourth example).

We used the origin and target neuropils to see how many of the IR and CR subclass innervate the leg versus other neuropils (Figure 6D). We found that 95% of IR and CR neurons are restricted to one of the leg neuropils and only a few neurons are restricted to one of the upper tectulum neuropil, abdomen or inter tectulum neuropil. Next we looked into the numbers of the BR subclass and how many of these neurons connect between the same neuropil on the two hemispheres (Figure 6E). Here as expected from Figure 6A we saw an increased bilateral connection between BR neurons of the T3 leg neuropils and the halteres. The largest connection between them was, however, in most cases not directed but with approximately the same amount of input and output on both sides. For neurons connecting the ANm however, the majority of the connections were directed, meaning with more than 60% input on one side and more than 60% output on the other side. Finally we looked at neurons connecting the upper tectulum or leg neuropils within one side (II and CI subclass) (Figure 6F). The number of neurons connecting these are few in number apart from 100 neurons that connect the WTcT and HTct in an undirected manner.

Finally, we examined the relationship between developmental hemilineage and innervated neuropils (Figure 7). Secondary hemilineages had previously been reported to innervate specific neuropils based on light-level neurite morphology (Shepherd et al., 2019); here we quantify these innervation patterns based on synapse location. We confirmed that numerous hemilineages are restricted to the leg neuropils (e.g., 13B, 03A, and 13A) and a few to the dorsal neuropils (e.g., 06A and 11B) but found that most innervate both leg and dorsal neuropils when including the primary neurons not considered in earlier studies. We also assessed the directionality for hemilineages that connect multiple neuropils (Figure 7C). For example, 01A neurons generally transmit information from the intermediate tectulum to the (contralateral) leg neuropil, whilst 10B neurons mainly transmit it from the leg neuropils to the lower tectulum (Figure 7C,F). However, it should be noted that we have not carried out axon/dendrite splits, and these synapses could engage in any of the four types of connections: dendro-dendritic, dendro-axonic, and axo-axonic in addition to the canonical axo-dendritic.

**Figure 7.**
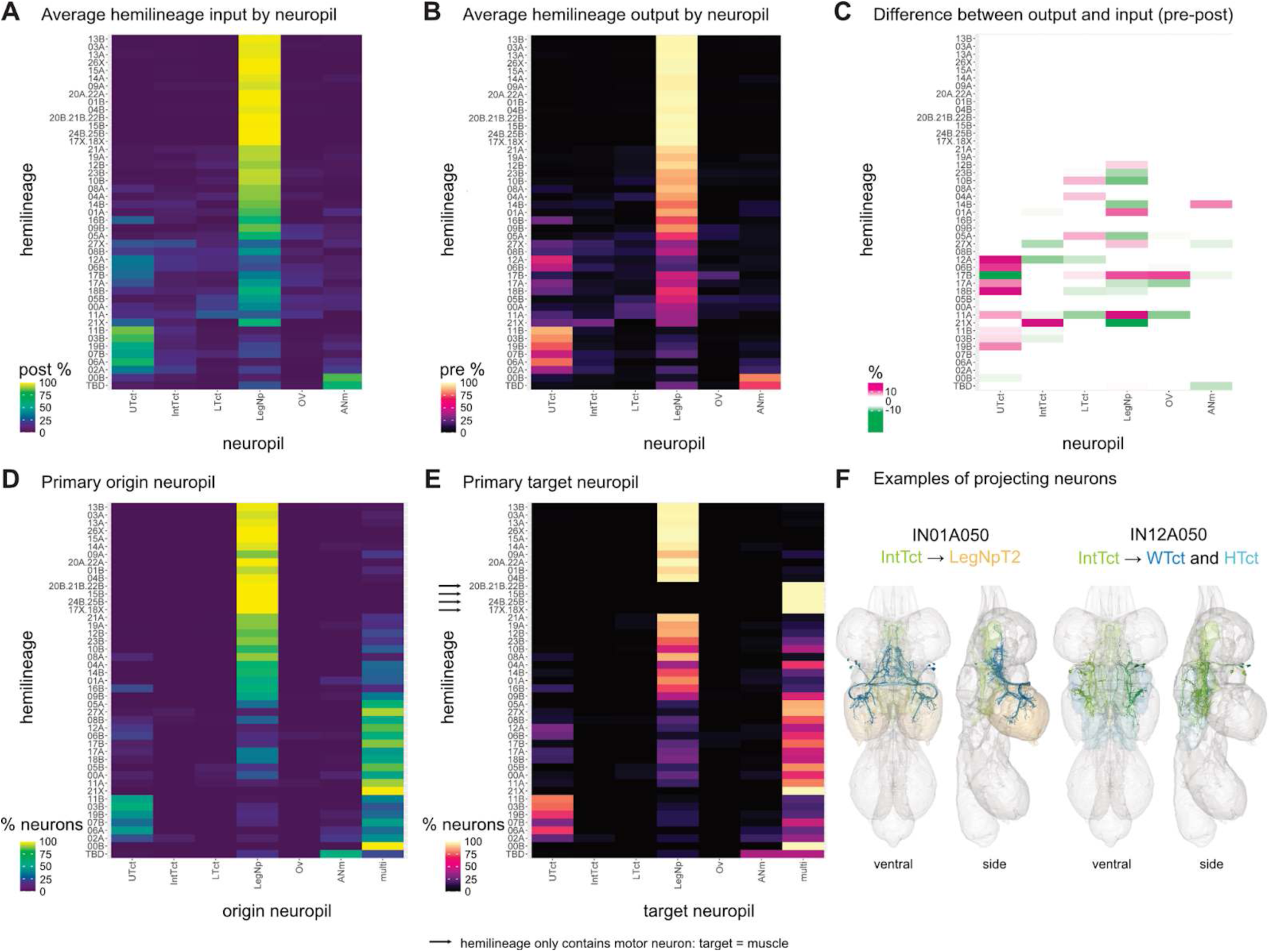
Neuropil innervation by hemilineage. **A.** The average proportion of input to each hemilineage from VNC neuropils as a percentage. **B.** The average proportion of output from each hemilineage to VNC neuropils as a percentage. **C.** The average input to neuropils subtracted from the average output to that neuropil. Magenta (positive numbers) represent more output and green (negative numbers) represent more input to that neuropil by a given hemilineage. For neuropils that do not receive any input or output see Figure 7 - figure supplement 1. **D.** INs and ANs by their primary input neuropil (origin) in the VNC. If there is more than one origin neuropil it is included in multi. **E.** INs and ANs by their primary output neuropil (target) in the VNC. If there is more than one target neuropil or a muscle target outside of the VNC it is included in multi. **F.** Examples of types that project between neuropils, partially responsible for the differences seen in **C** (origin and target neuropils are coloured). Hemilineages order on the y axis is organised by the innervation heatmap that is shown in **A** and reused for the heatmaps in **B,C,D** and **E.**

**Figure 7 - figure supplement 1.**
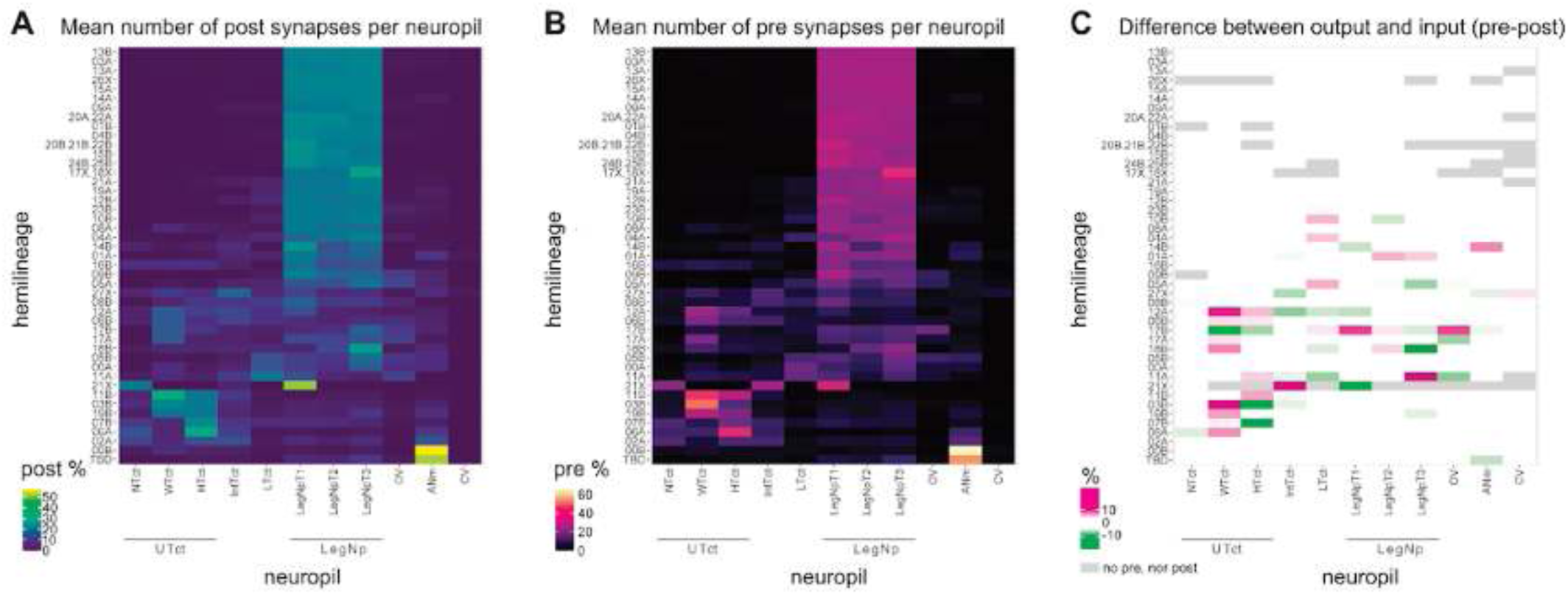
Neuropil innervation by hemilineage with differences in the leg and upper tectulum neuropils. **A.** The average proportion of input to each hemilineage from VNC neuropils as a percentage. **B.** The average proportion of output from each hemilineage to VNC neuropils as a percentage. **C.** The average input to neuropils subtracted from the average output to that neuropil. Magenta (positive numbers) represent more output and green (negative numbers) represent more input to that neuropil by a given hemilineage. Neuropils that do not receive any input or output from a given hemilineage are coloured in light grey.

#### Introduction to hemilineage-based organisation

During embryonic development, divisions of a segmentally repeated array of neuroblasts produce similar numbers of primary neurons in most neuromeres of the *Drosophila* VNC (Poulson, 1950). However, during postembryonic development, segment-specific subsets of neuroblasts generate the secondary neurons (Truman and Bate, 1988), facilitating an expansion of neuronal types in the thorax and distal abdomen for complex sensorimotor functions. In late larva, secondary hemilineages restricted to the leg neuropils are present in all three thoracic neuromeres, while those associated with dorsal (flight) neuropils are typically absent or greatly reduced in at least one (Marin et al., 2012; Truman et al., 2004).

Because **hemilineages** represent developmental and likely functional units in the VNC, we wanted them to be central to any systematic typing scheme. However, assignment of individual neurons to specific hemilineages based on light-level images of postembryonic neuroblast clones (Shepherd et al., 2019) was a complex and multifaceted process. Whilst the relative position of midline crossing is useful for distinguishing hemilineages that project contralaterally (Shepherd et al., 2016), many hemilineages are confined to ipsilateral neuropils. The locations of primary neurite bundles are also key characters (Figure 8 - figure supplement 1), but typically these remain tightly bundled during development only if all neurons target a similar location; in hemilineages with diverse cell types, they can be pulled apart.

**Figure 8 - figure supplement 1.**
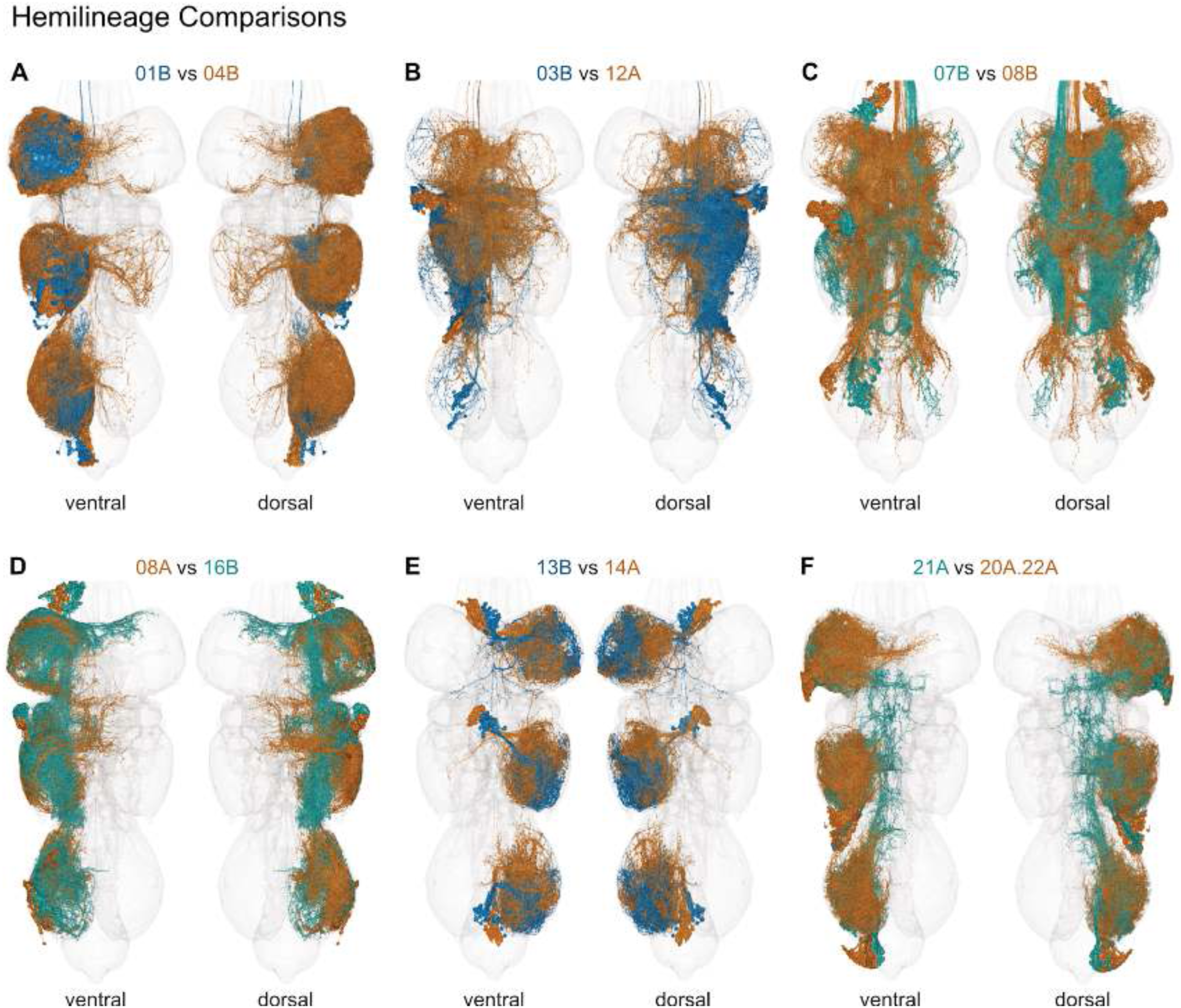
Pairwise comparisons of selected secondary hemilineages. **A.** 01B (blue, gabaergic) vs 04B (dark orange, cholinergic). **B.** 03B (blue, gabaergic) vs 12A (dark orange, cholinergic). **C.** 07B (marine, cholinergic) vs 08B (dark orange, cholinergic). **D.** 08A (dark orange, glutamatergic) vs 16B (marine, glutamatergic). **E.** 13B (blue, gabaergic) vs 14A (dark orange, glutamatergic). **F.** 20A.22A (dark orange, cholinergic) vs 21A (marine, glutamatergic).

Typically only the most common neuron types can be distinguished in neuroblast clone light-level images (Figure 8A), and if both A and B secondary hemilineages survive, it can be difficult to distinguish their respective projections. We originally seeded secondary soma tract bundles with hemilineage annotations, then mined for nearby soma tracts using a form of NBLAST (Figure 8B,C). Fast-acting neurotransmitter predictions (Eckstein et al., 2023) helped to confirm or distinguish between identifications based on morphology (Figure 8 - figure supplement 1). Left-right matching and serial set identification by cosine clustering also helped with hemilineage assignment, particularly when the primary neurite was displaced. Most primary neurons have not been associated with a specific hemilineage in the light level literature and so were difficult to assign to a hemilineage unless they shared the soma tract bundle and/or morphological features of identified secondary neurons (but many did not). Motor neurons were especially challenging; please see (Cheong et al., 2023) for more details on motor neuron identification and see below for the identification of individual hemilineages. Neurons that could not be assigned to a specific hemilineage were annotated as hemilineage “TBD”.

**Figure 8.**
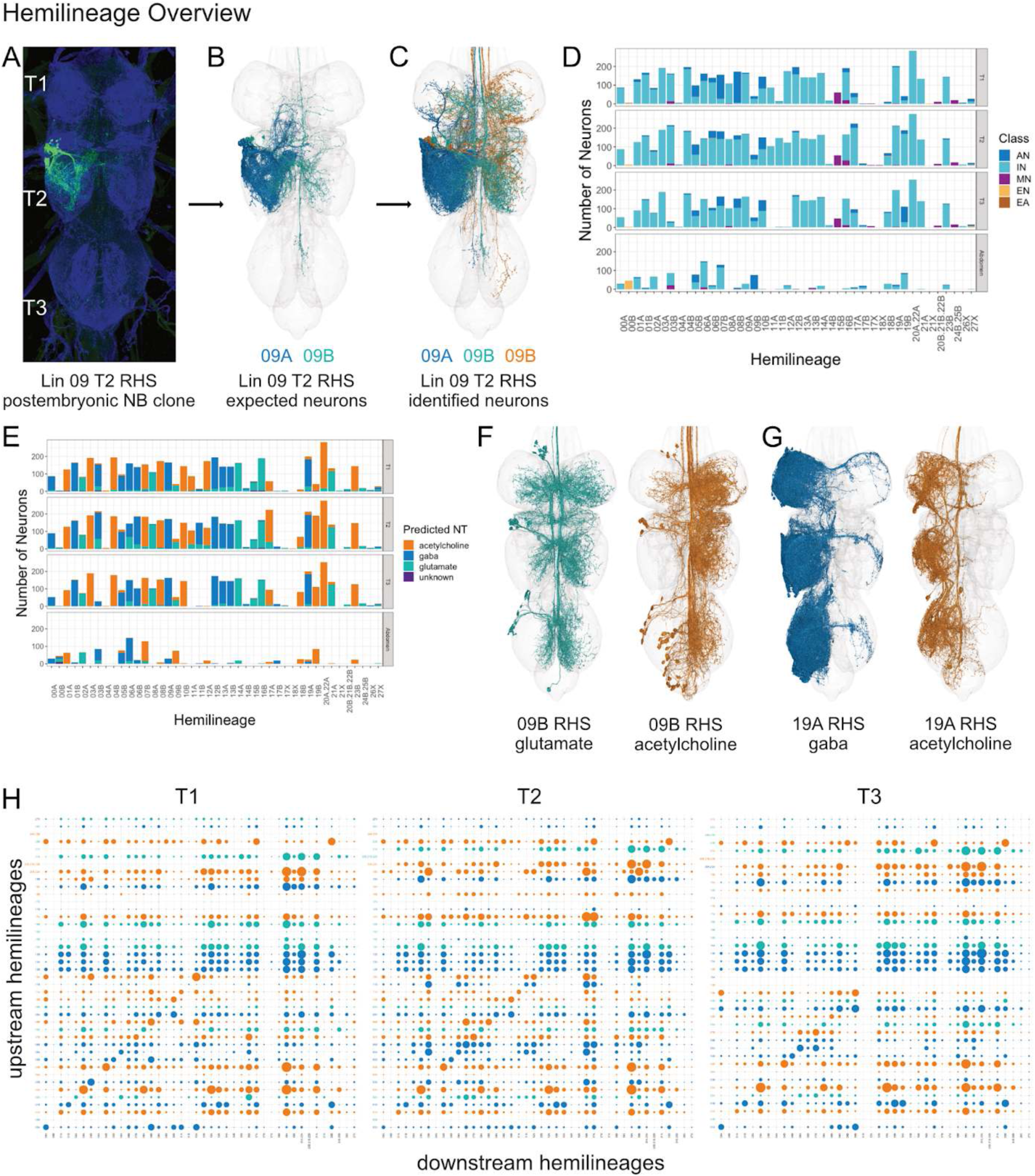
Hemilineage overview. **A.** 09A and 09B secondary neurons in a postembryonic neuroblast clone generated in T2 RHS. **B.** 09A and 09B secondary neurons in T2 RHS identified in MANC from light-level images, coloured by predicted neurotransmitter. **C.** All 09A and 09B neurons in T2 RHS identified in MANC, coloured by predicted neurotransmitter. **E.** Predicted fast-acting neurotransmitters for all VNC neurons (Eckstein et al., 2023; Takemura et al., 2023) by hemilineage and neuromere (neurons of TBD/unknown hemilineage have been omitted, and abdominal neuromeres have been pooled). **F.** Glutamatergic hemilineage 09B includes a minority cholinergic population. **G.** Gabaergic hemilineage 19A includes a minority cholinergic population. **H.** Intra-neuromere, inter-hemilineage synaptic connectivity for the three thoracic neuromeres (T1, T2, T3). Dot size correlates with number of synaptic connections, and dot colour reflects predicted neurotransmitter of upstream hemilineage (Eckstein et al., 2023; Takemura et al., 2023).

We find that broad neuron classes are not distributed homogeneously across the hemilineages (Figure 8D). For example, motor neurons are associated with a limited number of hemilineages, including 15B, 24B/25B and 20B/21B/22B as expected (Brierley et al., 2012) but also 03B and 16B, among others. Non-motor efferent neurons are almost exclusively associated with 00B and 21X. Ascending neurons are also associated with specific hemilineages and are notably more common in T1 for 05B, 07B, and 08B - likely because these hemilineages predominantly consist of intersegmental neurons that project to adjacent neuromeres, including through the neck connective to the suboesophageal ganglia. 19B is an unusual case in that most of its ascending neurons are generated in T3, perhaps for haltere input specialisation.

**Figure 8 - figure supplement 2.**
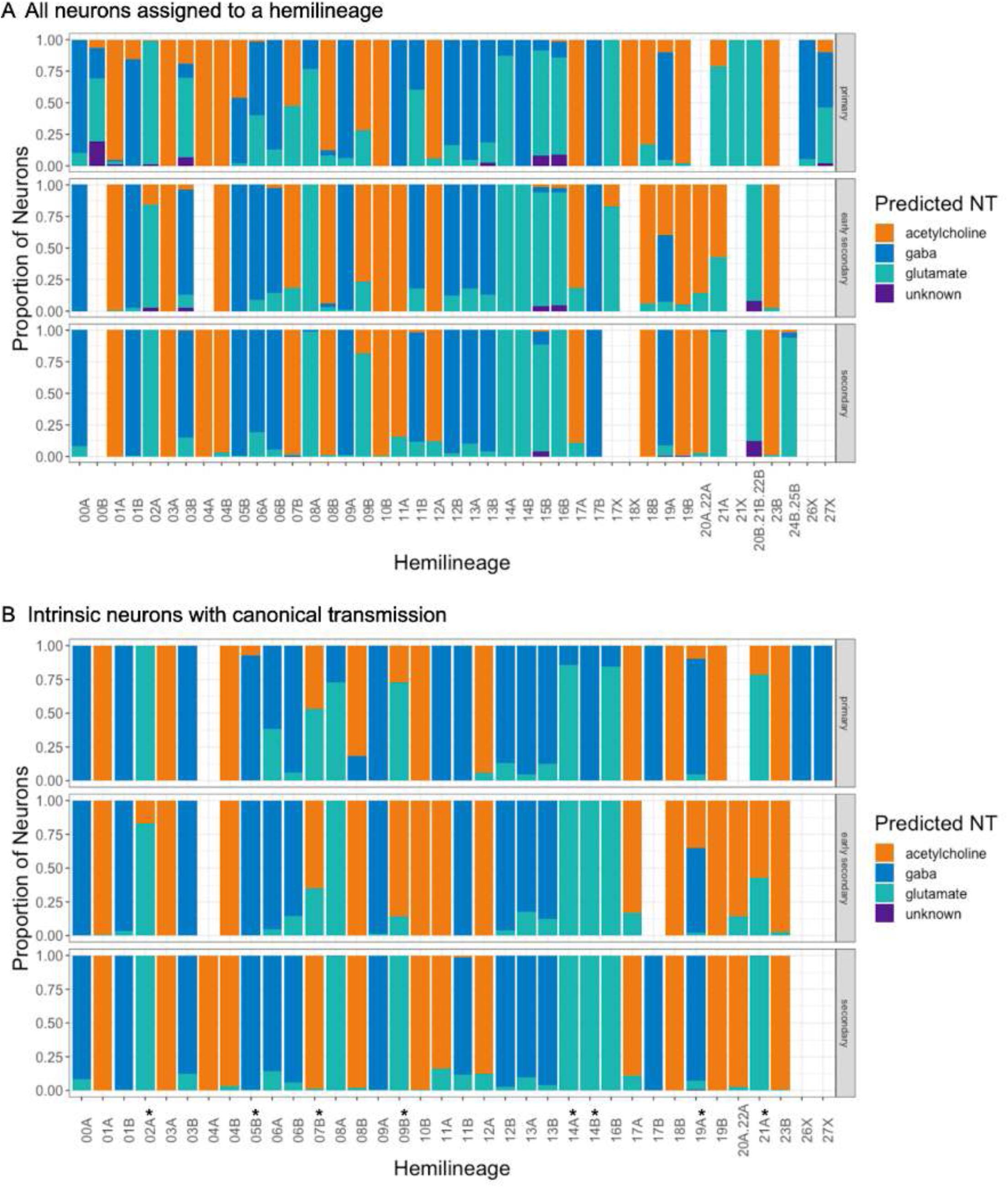
Early born neurons are more likely to express minority neurotransmitters. **A.** Proportion of fast-acting neurotransmitter expression predicted (Eckstein et al., 2023; Takemura et al., 2023) for all neurons assigned to a hemilineage, faceted by birthtime. **B.** Proportion of fast-acting neurotransmitter expression predicted for intrinsic neurons assigned to a hemilineage (with putative electrical and neurosecretory transmission excluded), faceted by birthtime. Asterisks indicate neurons with consistent expression of a minority neurotransmitter, enriched in early born neurons.

Secondary neurons belonging to the same hemilineage are expected to express the same fast-acting neurotransmitter (Lacin et al., 2019). We generated a neurotransmitter prediction for every neuron in MANC (see (Eckstein et al., 2023; Takemura et al., 2023) for details), which matched our expectations for the large majority of neurons assigned to each hemilineage (Figure 8E). There were exceptions, however - most commonly due to a paucity of chemical presynapses in the MANC volume because of class (e.g., ascending or motor neuron), transmission mode (e.g., putative electrical), or incomplete reconstruction (Figure 8 - figure supplement 2).

We also identified examples of complete, canonical cell types predicted to express a distinct neurotransmitter from the one expected. 09B is a small hemilineage that was reported to be glutamatergic but in MANC includes a substantial fraction of cholinergic neurons (Figure 8F); these appear to be early born, typically ascend through the neck connective, and have fewer ipsilateral neurites than their glutamatergic cousins. Most 19A neurons are gabaergic as expected, but we identified a population of early born, intersegmental neurons predicted to be cholinergic (Figure 8G). These two minority neurotransmitter examples are both morphologically distinct from their respective modal populations, based on NBLAST distances within vs across populations (p = 7.99 x 10^-27^ for 09B and p = 9.60 x 10^-8^ for 19A, one-sided paired t-test). They appear to represent genuine diversity in neurotransmitter expression, perhaps previously undetected because the genetic driver lines used to label specific hemilineages did not include early born neurons.

Secondary hemilineages had previously been reported to innervate specific neuropils based on neurite morphology (Shepherd et al., 2019). The members of each hemilineage typically share some aspects of gross morphology, including the neuropil(s) where they receive inputs (origin) and the neuropils where they provide outputs (target). Some hemilineages are restricted to the ipsilateral leg neuropil (e.g., 01B, 09A, 13A), and others to the contralateral leg neuropil (13B, 14A). Still others mainly receive inputs in ipsilateral neuropils and provide outputs to contralateral neuropils, but this is more common for hemilineages innervating flight or mixed neuropils (06A, 14B). For reference, we plotted all locations of postsynapses and presynapses for each annotated hemilineage by projecting the contents of all T2 neuropils onto a transverse section (Figure 9).

**Figure 8 - figure supplement 3.**
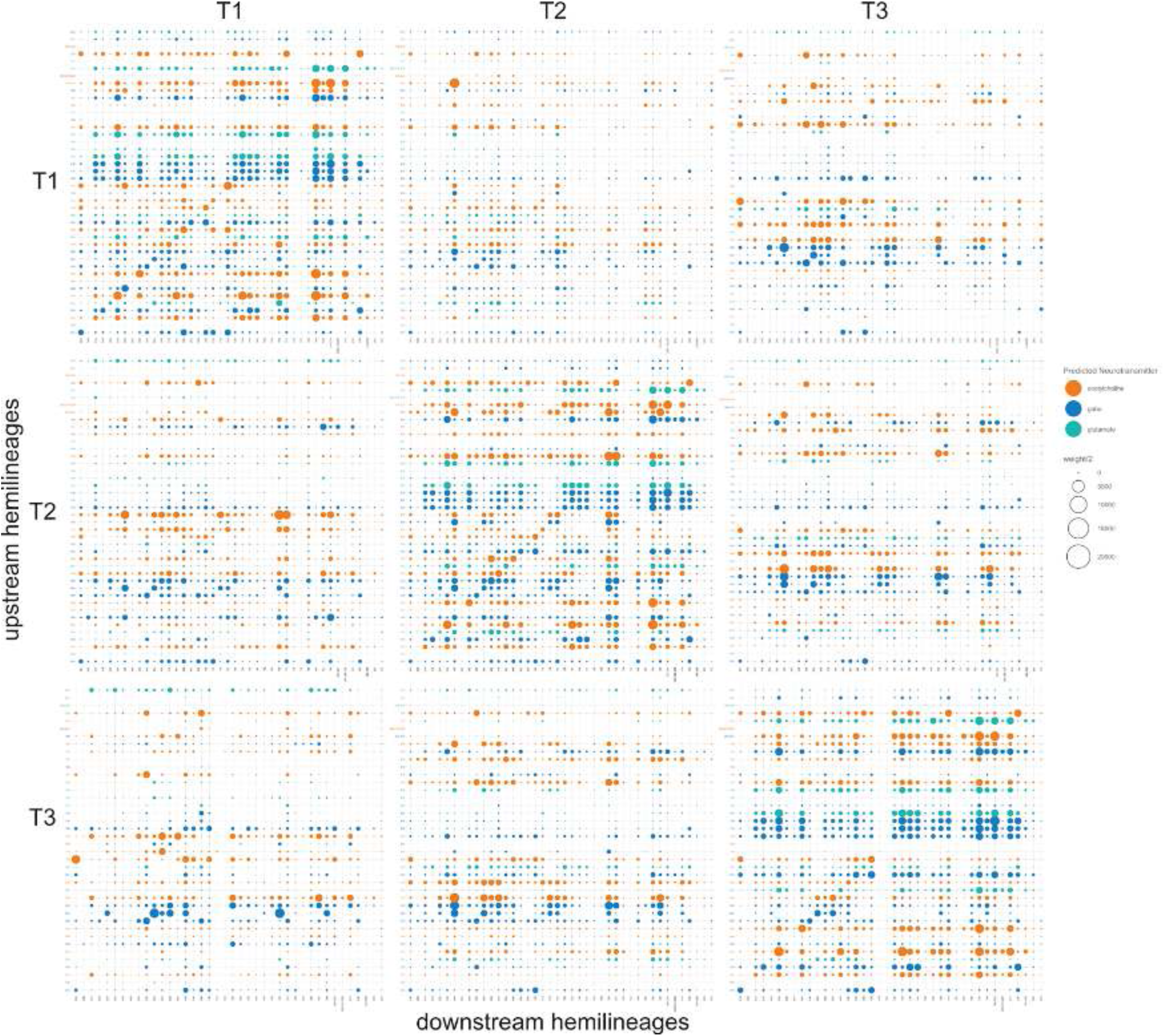
Intra-neuromere hemilineage synaptic connectivity across thoracic neuromeres (T1, T2, T3). Dot size correlates with number of synaptic connections, and dot colour reflects predicted neurotransmitter of upstream hemilineage (Eckstein et al., 2023; Takemura et al., 2023).

**Figure 9.**
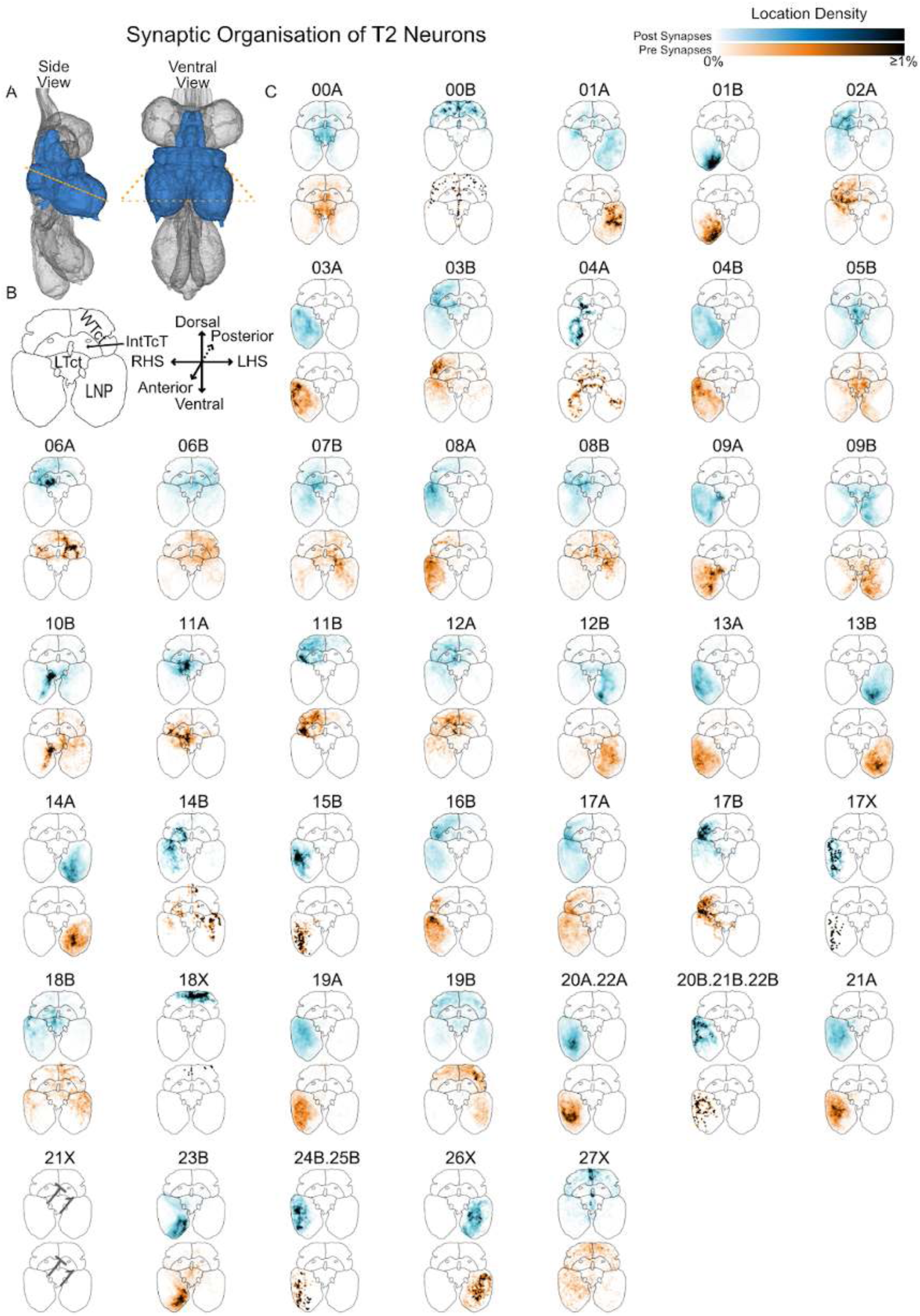
Summary of the anatomical organisation of synapses for each hemilineage originating in T2 RHS. **A.** Side and ventral views illustrating the neuropils surveyed: T2 Leg Neuropil (LNP), medial ventral association center (mVAC), ovoid (Ov), wing tectulum (WTct), lower tectulum (LTct), and Intermediate tectulum (IntTct). The orange dashed line shows the approximate location of the transverse section onto which their synapses have been projected. **B.** Template transverse section with major neuropils labelled and axes for orientation. **C.** Projected synapses for T2 RHS neurons of each hemilineage. Top (blue): postsynapses, connections to upstream neurons. Bottom (orange): presynapses, connections to downstream neurons. 21X is only observed in T1.

For the most part, we found very similar connectivity patterns between pairs of hemilineages across the thoracic neuromeres, at least for the hemilineages that innervate leg neuropils (Figure 8H). Many secondary hemilineages that innervate flight neuropils survive in T2 but are missing in T1 and/or T3. But there were some interesting differences suggesting segment-specific specialisation among flight hemilineages that survive in multiple neuromeres, particularly 06B and 19B. We detected significant across-neuromere connectivity, particularly for hemilineages primarily innervating flight neuropils such as 03B, 06A, 06B, 07B, 08B and 12A, but also for intersegmental hemilineages to leg sensory neuropils such as 05B and 10B (Figure 8 - figure supplement 3). As we have not carried out axon/dendrite splits, these synapses could engage in any of the four types of connections: dendro-dendritic, dendro-axonic, and axo-axonic in addition to the canonical axo-dendritic. This likely explains the high degree of intra-hemilineage connectivity that we observe for many hemilineages within a given neuromere (diagonals in Figure 8H).

#### Birthtime

VNC neuroblasts generate an initial set of embryonic primary neurons and post-embryonic, adult-specific secondary neurons (Booker and Truman, 1987; Prokop and Technau, 1991; Truman and Bate, 1988). Primary neurons typically mature towards the end of embryogenesis and function in the larval nervous system, although there are exceptions (Schmid et al., 1999). During metamorphosis, some primary neurons die, but others survive and mature or remodel to innervate adult targets (reviewed in (Tissot and Stocker, 2000)). These are expected to comprise <10% of the neurons in thoracic neuromeres in *Drosophila*, the remainder being secondary neurons that mature during metamorphosis (Truman and Bate, 1988). Secondary neurons belonging to the same hemilineage are expected to share similar gross morphology (Shepherd et al., 2019; Truman et al., 2004) and to express the same neurotransmitter (Allen et al., 2020; Lacin et al., 2019).

Birth order influences cell fate via a series of temporal factors expressed by neuroblasts and inherited by ganglion mother cells and their progeny (Isshiki et al., 2001; Kambadur et al., 1998). Both primary and early secondary neurons in the adult fly brain are typically larger and elaborate more extensively than late born secondary neurons from the same lineage (Lee et al., 2020). In the VNC, several neuroblasts have been reported to generate distinct cell types in a stereotyped sequence during embryogenesis (reviewed in (Doe, 2017)), but the effect of birth order on adult cell fate has not yet been examined.

We annotated all VNC-originating neurons as primary, early secondary, or (late) secondary **birthtime** based on position within the soma tract, soma size, and primary neurite diameter (Figure 10B,D; also see Methods). Neurons with unusually large, often ragged somas and thick primary neurites that run at the periphery or outside of the hemilineage soma tract bundle were designated as primary. Neurons with small, compact somas and thin primary neurites that run in the soma tract bundle were designated as secondary. Intermediate cases that could not be assigned with confidence to either category were annotated as early secondary. We annotated each hemineuromere independently before comparing within serial sets to ensure consistency. Note that our birthtime assignments are presumptive, with validation awaiting future experiments in which all primary neurons can be labeled and followed into adulthood.

**Figure 10.**
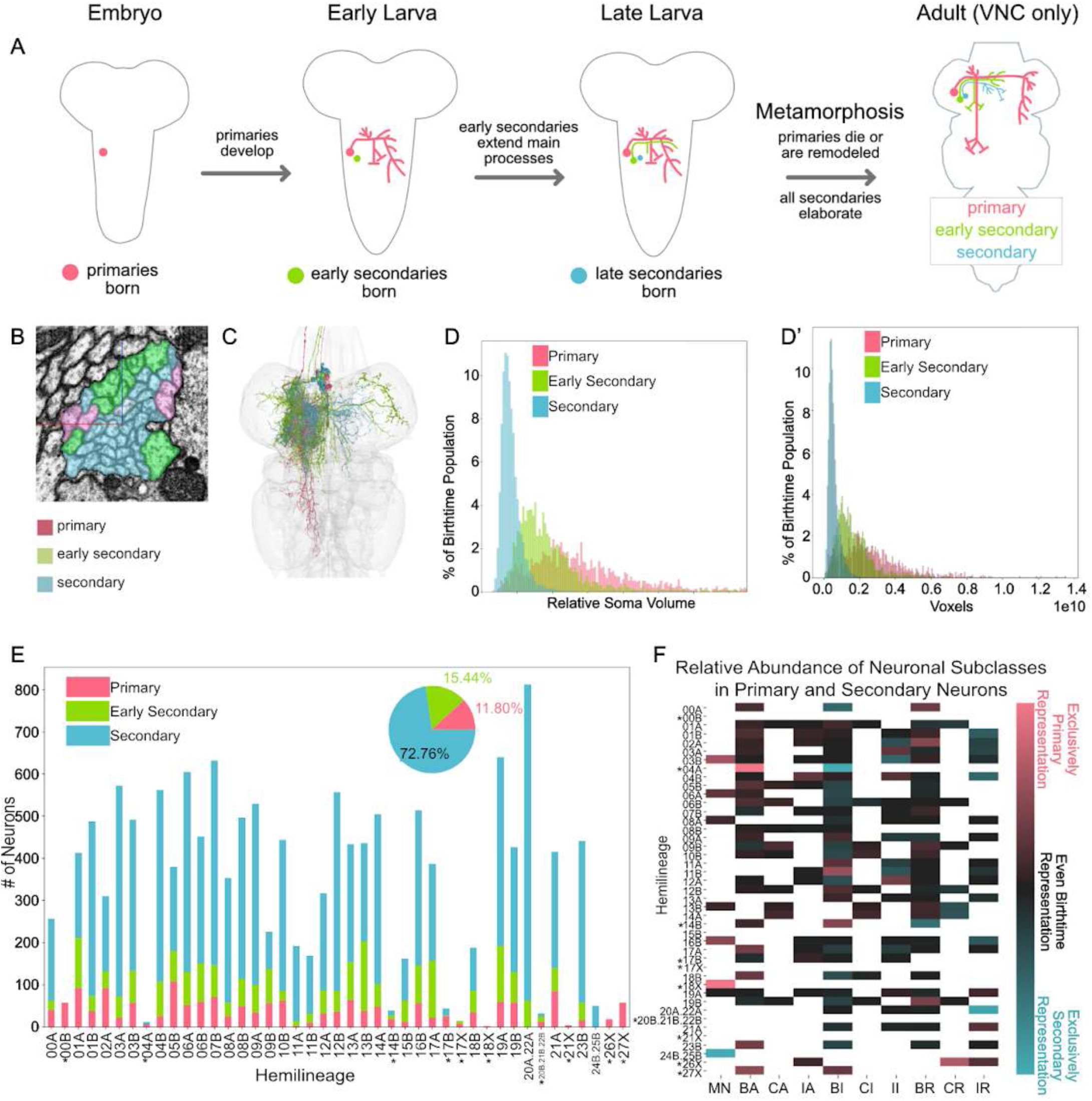
Birth order during development. **A.** Cartoon of birthtime development. Primary neurons are born during embryogenesis while secondaries are born during larval life. At metamorphosis primary neurons die or remodel while secondary neurons continue to elaborate. **B.** 02A T1 RHS primary neurites, coloured by birthtime. **C.** 02A T1 RHS neuron meshes, coloured by birthtime. **D-F.** Data is shown only for neurons originating in thoracic neuromeres; * indicates primary-only hemilineages. **D.** Neurons with larger somas were assigned to earlier birthtimes. **D’.** Primary and early secondary neurons are larger than secondary neurons. **E.** Counts of primary, early secondary, and secondary annotations for all VNC origin neurons with annotated hemilineage. Inset is frequency of birthtime amongst all annotated neurons. **F.** Comparative representation of each subclass between primary and secondary neurons within each hemilineage. Pink values have a higher percentage representation in the primary population than secondary, and vice-versa for blue, while black represents equal percentage abundance. White cells have no neurons with that subclass in that hemilineage. For more details, see **Methods**.

We annotated 11.95% of all neurons originating in the VNC as primary (Figure 10E). However, only 10.53% (1400/13,288) of thoracic neurons were annotated as primary, ∼consistent with the 10% contribution predicted in (Truman and Bate, 1988). Neurons annotated as primary or early secondary tend to be larger in volume than those designated as secondary (Figure 10D’). Primary neurons also tend to be more exuberant: they are more likely to ascend to the brain via the neck connective and to project across the midline and across neuromere boundaries (Figure 10F).

#### Serial homology and motifs

Insect neuroblasts and neurons exhibit serial homology, originally defined as “anatomical correspondence among repetitive or serial structures within a single individual” (Matsuda, 1976). Thoracic motor neurons are the best known examples of serial homology (Iles, 1976; Tyrer and Altman, 1974); however, we found that many individual interneurons could be clearly identified across multiple neuromeres. This was especially obvious for large primary neurons with discordant neurotransmitter predictions, such as the gabaergic 14A primary neurons (Figure 11A). We manually identified these initial **serial** sets (Figure 11D) and used them as seeds for an iterative graph matching process that predicted candidate sets of neurons with homologous connectivity in adjacent neuromeres (Figure 11B). These candidates were manually reviewed and annotated, generating additional seeds for the next run. Later on, we leveraged serial annotations to identify more candidates using serial cosine clustering (Figure 11C).

**Figure 11.**
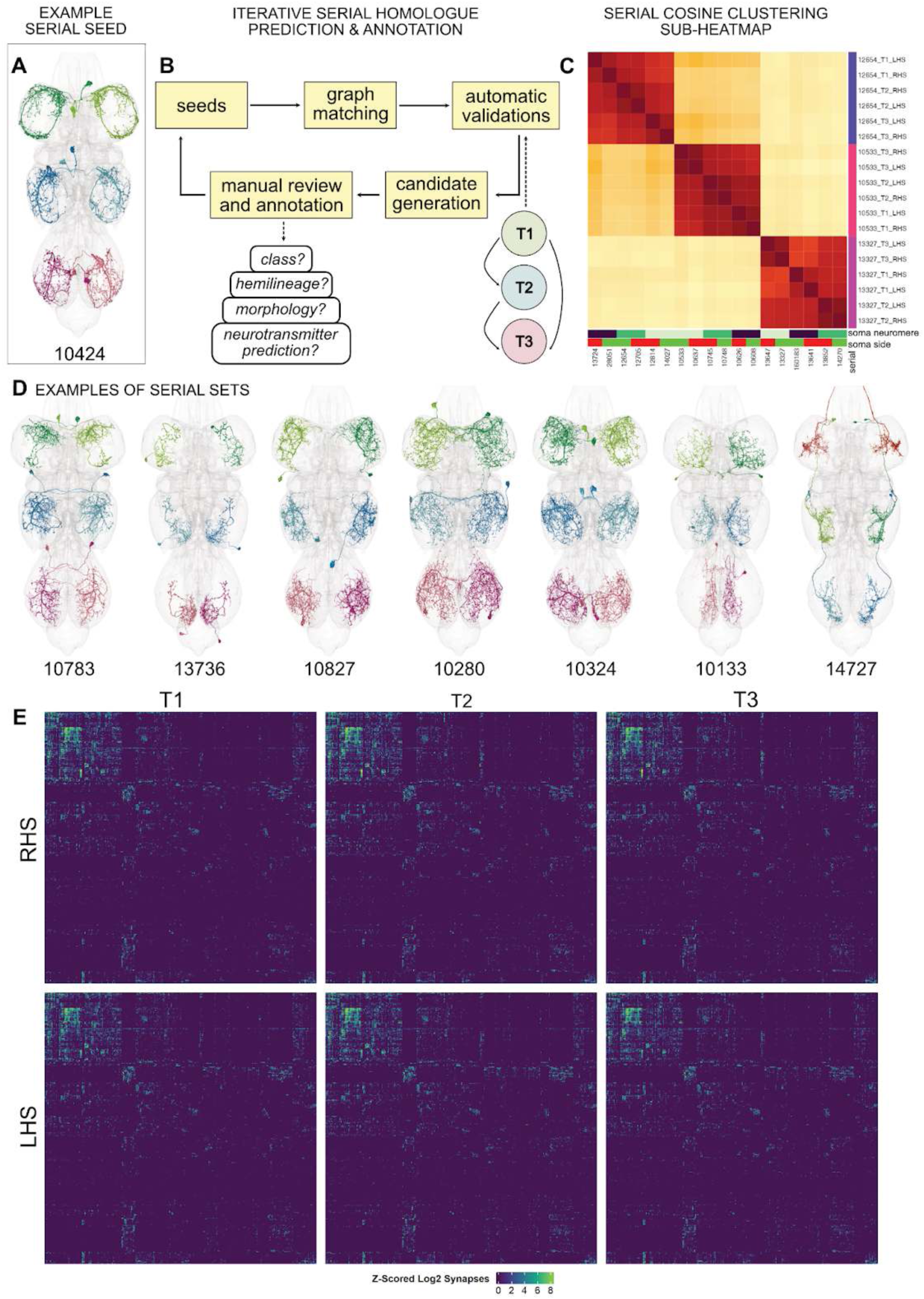
Identification of serially homologous neurons. **A.** Example of a manually identified serial set, serial 10424. **B.** Iterative serial homologue prediction and annotation procedure. Briefly, identified serial homologues were used as seeds for graph matching by **<u>F</u>**ast **<u>A</u>**pproximate **<u>Q</u>**uadratic **<u>A</u>**ssignment **<u>P</u>**roblem solver (*graspologic*) (Chung et al., 2019). Predicted matches were subjected to automated validation for consistency across sides and neuromeres, producing candidate serial matches which were manually reviewed for shared features including class, hemilineage, morphology, and predicted neurotransmitter. Those confirmed were annotated and used as seeds for the next round of predictions. Please see Methods for further details. **C.** Once an initial corpus of serial homologues was identified, clustering of serial cosine connectivity similarity scores was highly effective at revealing/confirming serially homologous types. The selected sub-heatmap shows three such groups from hemilineage 14A with a strong diagonal structure consisting of three blocks of 6 neurons, one for each side/neuromere. **D.** Selected examples of serial seeds from various hemilineages. Neuron meshes have been coloured by soma neuromere and side RHS/LHS (DN = red/dark red, T1 = green/dark green, T2 = cyan/blue, T3 = pink/magenta). **E.** Heat maps showing consistency of mutual all-by-all connectivity across hemispheres and neuromeres for thoracic serial homologues.

We were able to assign 5830 individual neurons (36.7% of neurons originating in the VNC) with similar appearance, connectivity, hemilineage, and/or predicted neurotransmitter to 968 serial sets. These sets were typically limited to one neuron per hemineuromere, but there were cases of two ∼identical neurons per hemineuromere, e.g., sets 10710 (Figure 38C) and 10894 (Figure 44C). There may be additional examples that our methods did not identify, but in most cases neurons without a serial number either are segment-specific cell types (most common in the thoracic dorsal hemilineages) or belong to a larger population of similar neurons.

The largest population (and the easiest to identify) were “independent leg” neurons that were largely restricted to a single leg neuropil (Figure 11D, Figure 12A, Figure 12 - video 1). These were typically very similar in topology and connectivity across neuromeres, albeit with differences in morphology attributable to segment-specific neuropil rotation and expansion. This category was dominated by gabaergic and glutamatergic neurons (Figure 12I), suggesting a large role for serially homologous inhibitory circuits. One striking example featured two pairs of glutamatergic descending neurons along with T1 and T2 neurons, each projecting to the next posterior neuromere (serial 14727, Figure 11D), underlining the serial homology between subesophageal and thoracic ganglia despite their physical separation during metamorphosis. As expected, most leg motor neurons were serially homologous, allowing us to propagate light level matches across thoracic neuromeres by comparing their connectivity to our annotated serial sets; please see (Cheong et al., 2023) for details.

**Figure 12.**
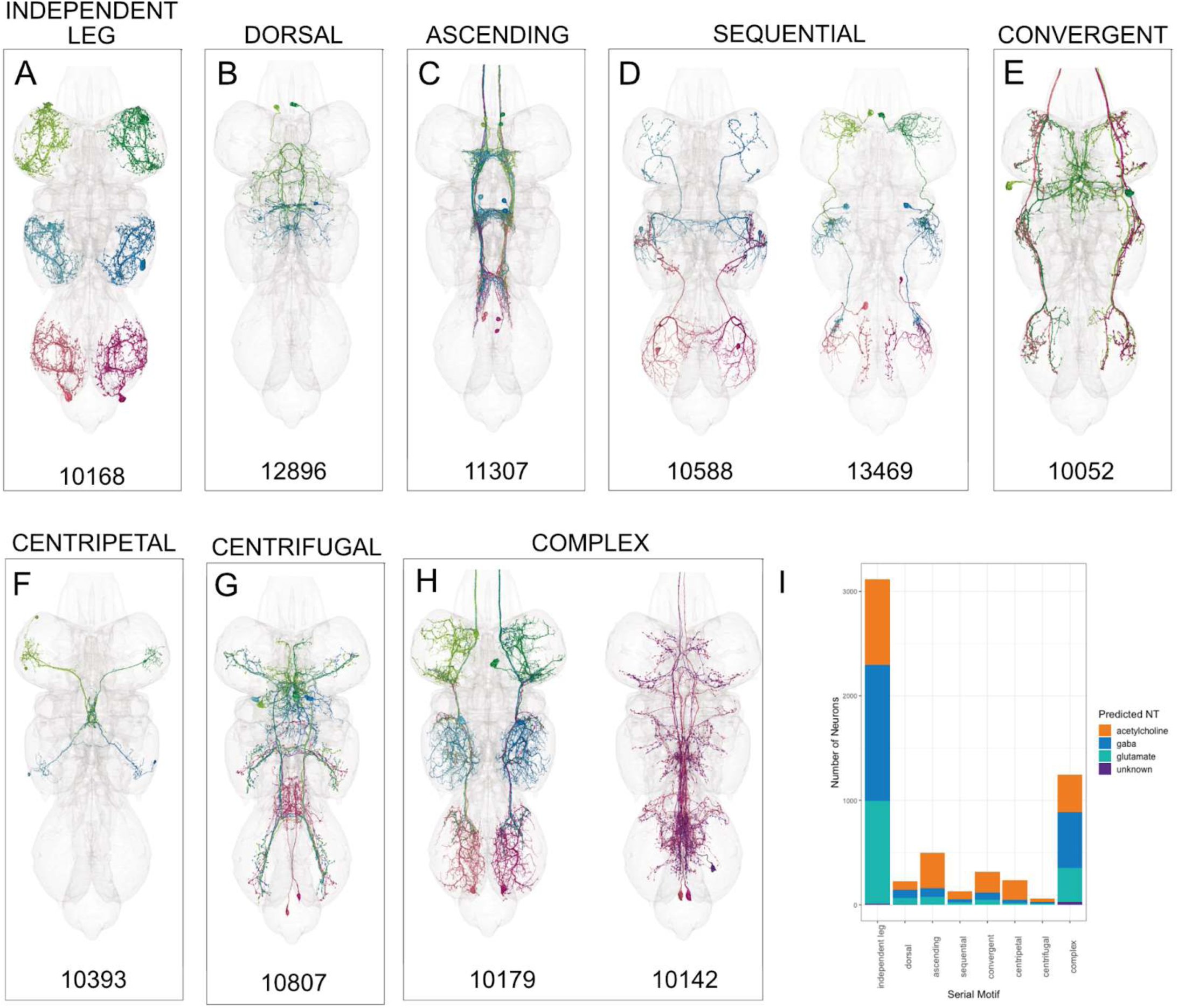
Classification of serial sets into motifs. Neuron meshes have been coloured by soma neuromere and side RHS/LHS (DN = red/dark red, T1 = green/dark green, T2 = cyan/blue, T3 = pink/magenta). **A.** Independent leg serial set 10168. **B.** Dorsal serial set 12896. **C.** Ascending serial set 11307. **D.** Sequential (ascending or descending to the next neuromere) serial sets 10588 and 13469. **E.** Convergent serial set 10052. **F.** Centripetal serial set 10393. **G.** Centrifugal serial set 10807. **H.** Complex serial set 10142. **I.** Number of neurons assigned to each serial motif, coloured by predicted neurotransmitter (Eckstein et al., 2023; Takemura et al., 2023).

However, we also identified numerous more complex **serial motifs** (Figure 12 - video 2). We assigned those that primarily innervated non-leg neuropils to a “dorsal” category (Figure 12B) and those exclusively composed of neurons ascending via the neck connective to an “ascending” category (Figure 12C). A small but interesting category were the “sequential” sets in which neurons featured dendrites in one neuromere but axons terminating in the next anterior or next posterior neuromere (Figure 12D); for a discussion of their likely role in intersegmental leg coordination, please see our accompanying manuscript (Cheong et al., 2023).

We identified numerous cases of “convergent” axons of presumed serially homologous neurons, including a few that included descending neurons (Figure 12E). We assigned those with peripheral dendrites in leg neuropils and axons converging centrally in non-leg neuropils to a special “centripetal” category (Figure 12F). Conversely, we found a very small number of “centrifugal” neurons with central, partially overlapping dendrites and axons projecting out to one or more leg neuropils (Figure 12G). The remaining serial sets with more complicated patterns of connectivity were grouped together as “complex” (Figure 12H) and form the largest motif category apart from the “independent leg” (Figure 12I).

Curiously, the proportion of neurons expected to be excitatory or inhibitory varied by serial motif (Figure 12 - figure supplement 1). Dorsal and independent leg serial sets were much more likely to be gabaergic or glutamatergic than to be cholinergic, suggesting inhibitory roles in local circuits. In contrast, the serial sets that connect and presumably coordinate between different neuropils are much more likely to be excitatory. Please see our companion manuscript (Cheong et al., 2023) for detailed examples of serial circuit analysis.

**Figure 12 - figure supplement 1.**
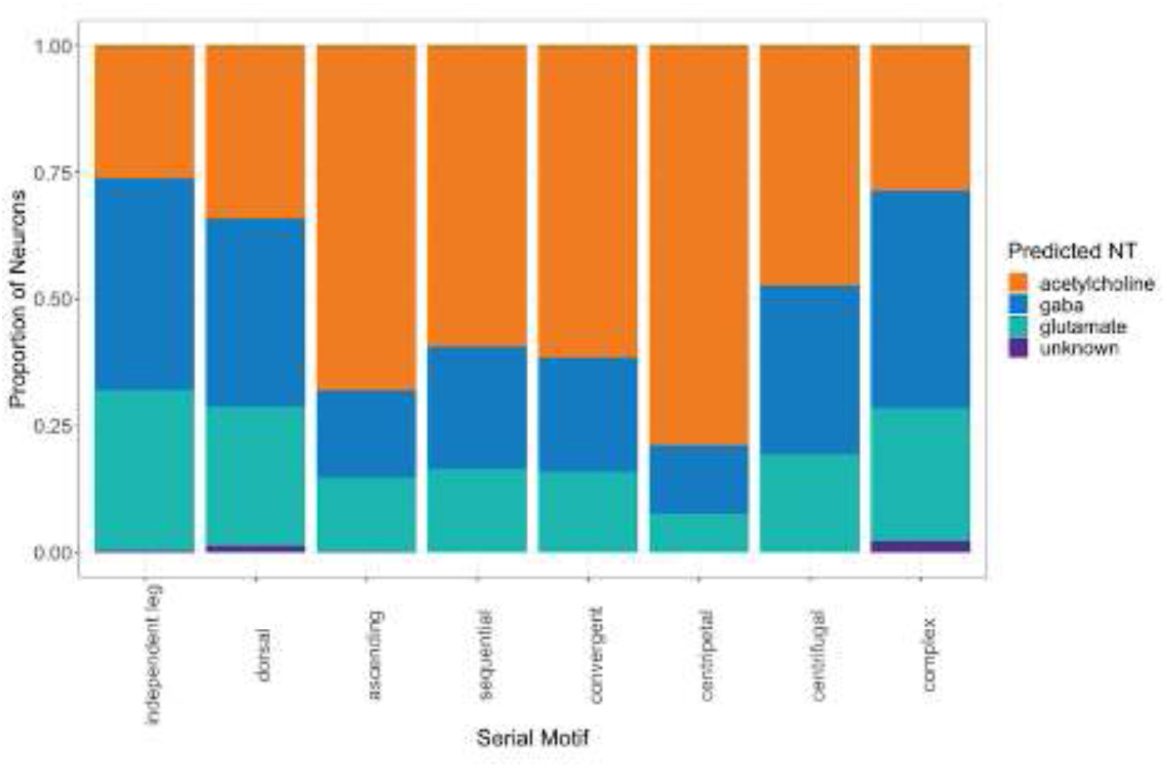
Proportions of neurons from each serial motif predicted to express a given neurotransmitter. (Eckstein et al., 2023; Takemura et al., 2023). **Figure 12 - video 1. All thoracic examples of independent leg serial sets (1:19).** https://youtu.be/O6QfQTpNjo8 Neuron meshes have been coloured by soma neuromere and side RHS/LHS (DN = red/dark red, T1 = green/dark green, T2 = cyan/blue, T3 = pink/magenta). To view this movie at a preferred speed, we recommend downloading the video and using a player with more control such as iMovie. **Figure 12 - video 2. All thoracic examples of other annotated serial motifs (0:42).** https://youtu.be/VFnhqX9Ee_g Neuron meshes have been coloured by soma neuromere and side RHS/LHS (DN = red/dark red, T1 = green/dark green, T2 = cyan/blue, T3 = pink/magenta). To view this movie at a preferred speed, we recommend downloading the video and using a player with more control such as iMovie.

Whilst these serial sets were interesting as examples of segment-specific reuse and diversification of repeated elements, they also proved to be extremely useful in the systematic typing of the VNC, including interneurons, sensory neurons (see below), and leg motor neurons (as described in our accompanying manuscript (Cheong et al., 2023)). They also informed the annotation of birthtimes and hemilineages; in particular, sets bridging the thoracic and abdominal neuromeres aided in abdominal soma neuromere and hemilineage assignments (e.g., serial 10142, Figure 12H). We also expect them to aid in assigning hemilineages to neurons in the suboesophageal ganglion once a full CNS dataset has been obtained.

#### Cell typing

We partitioned the population of intrinsic, ascending, and efferent neurons into systematic types on the basis of their synaptic connectivity, using the identified lateral and serial homologues. The aim of this typing is fourfold: first, to identify granular sets of neurons that are likely to be mutually identifiable between multiple VNCs; second, to provide units for exploratory and functional analysis, which may be broader than identified lateral and serial homologues; third, to identify seriality or other symmetries that might not be captured by the previously described approaches; and fourth, to assign a systematic nomenclature consistent with those developed for the sensory, descending, and motor neuron populations.

Because our goal in systematic typing is to identify sets of neurons that participate in similar ways in circuits homologous across the midline and across neuromeres, typing is based on synaptic connectivity rather than morphology. Consequently, types should approximate developmental homologues, but may differ, and the typing process must decide thresholds for how similar in connectivity neurons may be to belong to the same type even when finer divisions are possible. Types are systematically determined by a clustering of synaptic connectivity, which uses our annotations of laterality and seriality to inform and constrain the process at several stages. To estimate connectivity to homologous circuitry, the connectivity adjacency is symmetrized by averaging synaptic counts across annotated lateral and/or serial homologues (Figure 13A). Each annotated hemilineage is independently clustered based on how similar its neurons’ connectivity is in this symmetrised network. Clustering hemilineages independently constrains types to better approximate development homologues.

**Figure 13.**
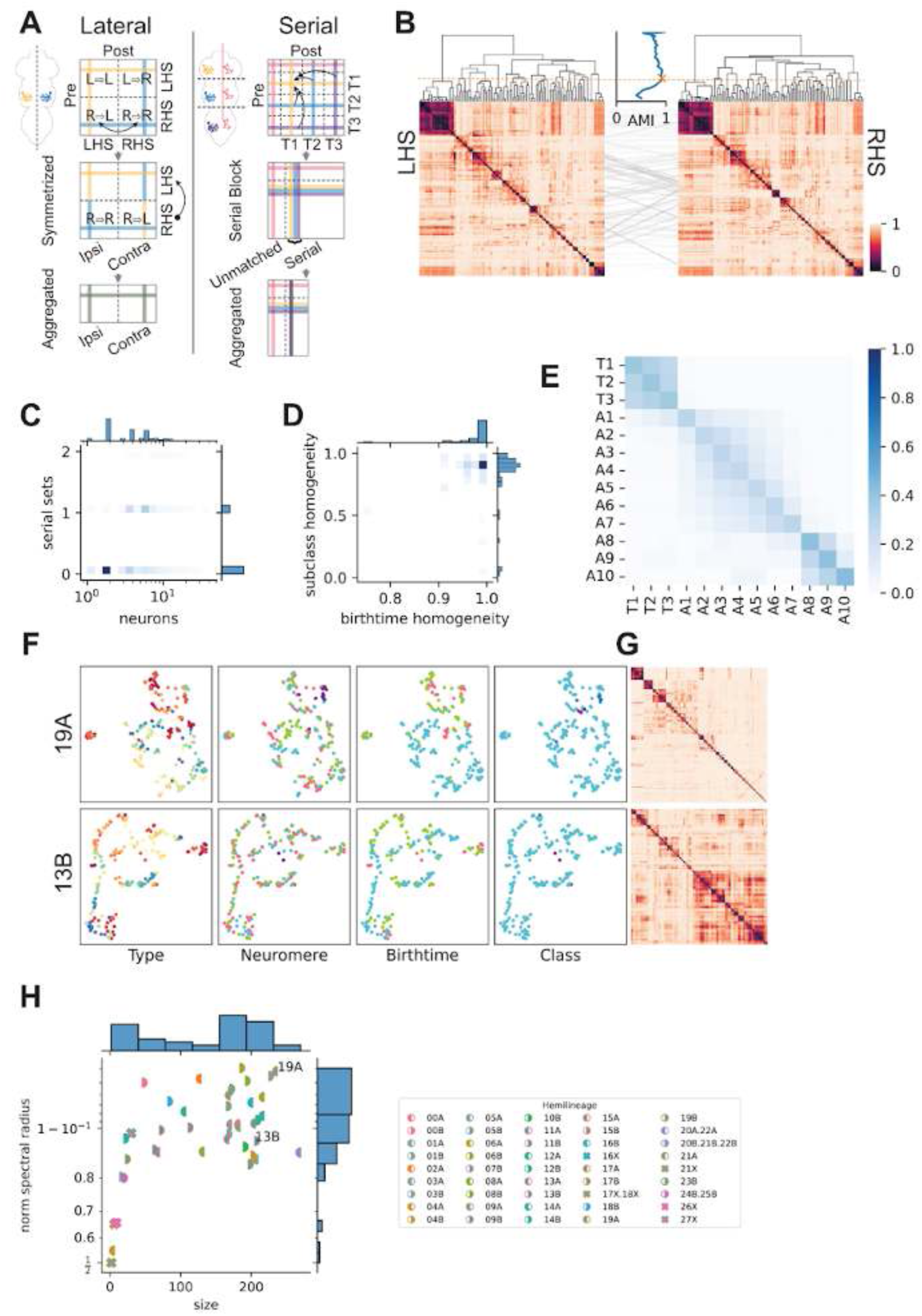
Typing of intrinsic neurons by connectivity. **A.** Schematic describing how the raw adjacency matrix of connectivity between individual neurons can be simplified by aggregating connectivity of homologous neurons; this is possible both for left-right homologues and for serial homologues (diagrammed here for 6 thoracic neurons). **B.** Example of consistency of independent connectivity clustering of each side of a hemilineage, 09B. The adjusted mutual information (AMI) between sides’ increasingly granular clusters is shown between the corresponding dendrograms for each side. Peak consistency chosen to select clusters for systematic types is indicated by a yellow cross and dashed line. Heatmaps display cosine distance between the serially aggregated connectivity for each neuron set. Lines between the heatmaps indicate neurons whose terminal position in each side’s cluster dendrogram differs, shaded by the distance of this difference. **C.** Density histogram of the number of neurons and serial sets per clustered type in all hemilineages’ intrinsic, non-motor popular. See E for colour scale. **D.** Density histogram of the birthtime and subclass homogeneity of all types in C. See E for colour scale. **E.** Type seriality, as a histogram of number of types containing any neuron in each pair of neuromeres. Values are normalised by rows, to show relative distributions when population size differs. **F.** Example comparison of the inter-hemilineage variability of intra-hemilineage connectivity similarity structure. Each panel is a UMAP embedding the hemilineage’s symmetrized, serially aggregated connectivity cosine similarity, with each point representing a lateral group. Panels are coloured by neuromere, birthtime, and class as in all other figures. Types are coloured arbitrarily for visualisation. **G.** Heatmaps of symmetrized, serially aggregated connectivity cosine distance for the hemilineages in F. **H.** Normalised spectral radius versus number of neurons for all hemilineages. Hemilineages from F and G are labelled.

To select thresholds for how similar in connectivity neurons may be to belong to the same type, clustering parameters were selected so as to produce consistent clusterings between annotated homologous populations. As an illustrative example, one goal of typing is to maximise consistency between independent clusterings of each lateral side (without lateral symmetrisation), preferring clustering parameters coinciding with a peak in consistency nearest the most granular typing (Figure 13B). Selecting parameters consistent across lateral variability is intended to make types robust to noise and error in the reconstruction and exploit lateral variability as a proxy for inter-individual variability in connectivity (Schneider-Mizell et al., 2016). While this illustrates our approach to selecting similarity thresholds for types consistent across homologous populations, because of variability of connectivity and annotations within and across hemilineages an altered, more robust strategy was used for the final clustering. In this strategy terminal type clusters for each hemilineage were chosen by selecting the clustering of the laterally symmetrised but unaggregated connectivity most consistent with lateral group annotations, then selecting the clustering of the laterally and serially aggregated connectivity most consistent with those unaggregated clusters (see Methods).

The resulting types most often consist of single pairs of laterally grouped neurons, and types containing a serial set most often contain only one (Figure 13C). Homogeneity of birthtime and subclass within each type is high (Figure 13D), though in some hemilineages broad type clusters result in types with less homogeneous birthtime and subclass. Some mixed-birthtime types may be an artefact of suboptimal systematic types or of uncertainty in annotation of early versus late secondaries, however there are cases of populations where primaries and secondaries appear to have similar connectivity consistent with our intent (e.g., IN11B004, Figure 32 - figure supplement 1). Because type clusters are based on the neuromere-serialised network determined by annotated serial homologue sets, types are highly serial, frequently containing neurons from multiple neuromeres (Figure 13E). As with other serial annotations, types are broadly structured into three block communities: the thoracic neuromeres, A1 through A7, and A8 through A10. Of these, the A8 through A10 community has the least serial frequency between its neuromeres, though it has the most seriality with neuromeres outside the block, namely A8 and A9 with T3 and the rest of the abdominal neuromeres. As expected, A1 shares many types with T3 and the other thoracic neuromeres. While type clusters often align with morphological similarity (e.g., 01A, Figure 16 – figure supplement 1C), since types are based on connectivity and not morphology they sometimes do not (e.g., 01B, Figure 17 - figure supplement 1C).

How similar neurons and types within a hemilineage are to one another varies between hemilineages. For example, hemilineage 19A has a highly block-structured connectivity similarity (Figure 13G), resulting in types containing neurons strongly similar within the type but dissimilar between types (Figure 13F). Types in these hemilineages tend to share few or no strong synaptic partners with other types. In contrast, hemilineage 13B has broad but structured connectivity similarity, which results in types that, while strongly similar within themselves, also form segments of a weaker inter-type similarity structure. That is, there are communities of types in these hemilineages that share strong synaptic partners. In the case of 13B, this larger structure is not defined by neuromere-specific communities, but is anchored by similarity to primary neuron types (Figure 13F).

To summarise how independently or similarly structured each hemilineage’s types are, a single measure of inter-type connectivity similarity was chosen. The spectral radius of the hemilineage connectivity cosine distance matrix (its largest absolute value eigenvalue, equivalently the 2-norm in this case) measures how strongly diagonal similarity is distributed (Figure 19H). Intuitively, hemilineages whose secondaries template their connectivity off of primary or early secondaries will tend to have lower spectral radii, while hemilineages composed of neurons with distinct, independent connectivity will have higher spectral radii. Hemilineages like 19A containing small type clusters with low intra-cluster variance and high inter-cluster variance have a high normalised spectral radius, while those like 13B with less distinct clusters and more inter-cluster similarity are comparatively lower. However, this single measure does not distinguish hemilineages with other patterns of connectivity structure from each other, such as large clusters with low intra-cluster variance or hemilineages composed of distinct populations with inhomogeneous similarity structure.

Ultimately, systematic types - often with members in multiple neuromeres - were assigned to every intrinsic, ascending, and efferent neuron. We chose a systematic nomenclature that emphasises three key features of these neurons: 1) their class, 2) their hemilineage of origin if identified, and 3) their average volume (because size roughly correlates with birthtime and because any other distinction would be too subjective). Neuron types within each hemilineage initially received sequential numbers, with similar neurons receiving similar numbers, but subsequent rounds of across-neuromere clustering prompted merging of some preliminary types, resulting in gaps. Neurons matched to cell types in the light level literature received distinct names in the type and/or synonyms fields.

#### Secondary Hemilineages

Segment-specific subsets of neuroblasts generate the secondary (postembryonic) neurons (Truman and Bate, 1988). Secondary hemilineages restricted to the leg neuropils are found in all three thoracic neuromeres, while those associated with dorsal (flight) neuropils (e.g., 3B, 11A, 11B, 12A, 18B) are absent or substantially reduced in at least one (Marin et al., 2012; Truman et al., 2004). Only three neuroblasts produce secondary neurons in abdominal segments A2-A7 (Birkholz et al., 2015; Truman and Bate, 1988), but 13 neuroblasts divide post-embryonically in A1, 31 in A8, 23 in A9, and 11 in A10 (Birkholz et al., 2013).

We were able to attribute 97% of neurons in the thorax and 38% of neurons in the abdomen (mainly in A1) to a specific hemilineage. The adult leg hemilineages are relatively consistent in neuron number and connectivity, while flight hemilineages tend to be more variable (Figure 7D,E). In the abdomen, the middle segments contain the fewest neurons and the highest proportion of primary neurons and serial homologues, compared to more anterior and posterior neuromeres (Figure 49B,E).

Each secondary hemilineage has been reported to exhibit a characteristic broad morphology (e.g., ipsilateral leg neuropil, bilateral flight neuropils, etc) in the adult VNC (Shepherd et al., 2019). We have summarised the locations of postsynapses and presynapses for every T2 secondary hemilineage in our dataset for reference (Figure 14). We have also created an online atlas - a neuroglancer view featuring an “average” primary neurite and backbone for every secondary hemilineage in each thoracic hemineuromere, https://clio-ng.janelia.org/#!gs://flyem-user-links/short/hemilineages.json. Hemilineages are colour-coded and can be selected for display in the segmentation layer, allowing users to compare relative positions of hemilineages of interest.

**Figure 14.**
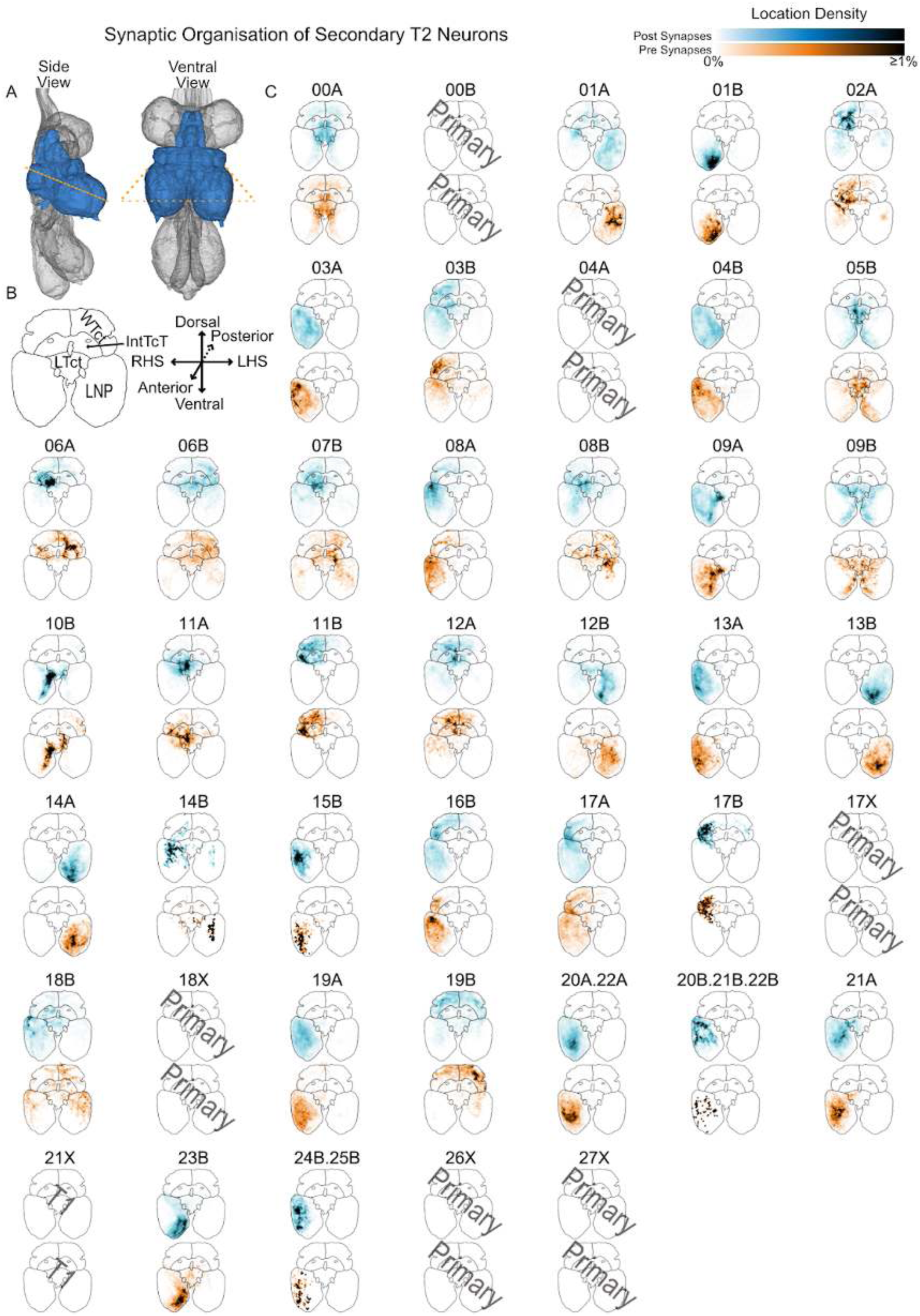
Synaptic organisation by secondary hemilineage. Summary of the anatomical organisation of synapses for secondary neurons in each hemilineage originating in T2 RHS. **A.** Side and ventral views illustrating the neuropils surveyed: T2 Leg Neuropil (LNP), medial ventral association center (mVAC), ovoid (Ov), wing tectulum (WTct), lower tectulum (LTct), and Intermediate tectulum (IntTct). The orange dashed line shows the approximate location of the transverse section onto which their synapses have been projected. **B.** Template transverse section with major neuropils labelled and axes for orientation. **C.** Projected synapses for T2 RHS neurons of each hemilineage. Top (blue): postsynapses, connections to upstream neurons. Bottom (orange): presynapses, connections to downstream neurons. 21X is only observed in T1.

Below we describe in more depth every hemilineage that produces more than one or two secondary neurons. For each of these 35 hemilineages, we show (A) the overall morphology of the secondary population, (B) representative individual neurons (as estimated by highest average NBLAST score to other members of the hemilineage), and (C) specific notable examples (which in some cases are primary). We then report (D) the locations of their connectors (postsynapses and presynapses), (E) their upstream and downstream partners by class, and (F) their upstream and downstream partners by finer subdivisions corresponding to their systematic types (secondary hemilineage, target, or sensory modality). We also provide supplementary figures showing the morphology and normalised up- and downstream connectivity of all systematic types for each hemilineage. It should be noted that this normalisation reflects the strength of connections to upstream partners but not to downstream partners (since inputs to those partners from other sources are not considered). Also, hemilineages 20A and 22A could not be distinguished based on light-level reports and have been pooled, as have hemilineages 24B and 25B.

#### Hemilineage 00A

Hemilineages 00A and 00B derive from the ventral unpaired median neuroblast (VUM or MNB) (Truman et al., 2004), which produces 3 bilaterally exiting neurosecretory cells, 6 intersegmental interneurons, and 2-6 dorsally projecting interneurons per segment in the embryo (Schmid et al., 1999). 00A neurons enter the neuropil at the midline near the posterior edge of each neuromere and project dorsally in a coherent bundle before elaborating bilaterally symmetric dendrites on both sides of the midline. In T1 and T3, secondary neurons primarily innervate the mVAC and the intermediate neuropil just dorsal to the ovoid and leg sensory neuropils, whilst in T2 they also arborise extensively in the upper tectulum (Shepherd et al., 2019). (Figure 15A,B,D). Most secondary neurons were predicted to be gabaergic, consistent with previous reports (Lacin et al., 2019). A small subpopulation was predicted to be glutamatergic (Figures 8E and 15C top), although we cannot completely exclude the possibility that these are 00B. Soma tracts of primary 00A neurons run adjacent to those of secondary neurons, although a few diverge early on to innervate ventral leg neuropil (e.g., type AN/IN00A009) (Figure 15C bottom); a small number ascend into the neck connective bilaterally (e.g., type AN00A006) (Figure 15 - figure supplement 2).

**Figure 15.**
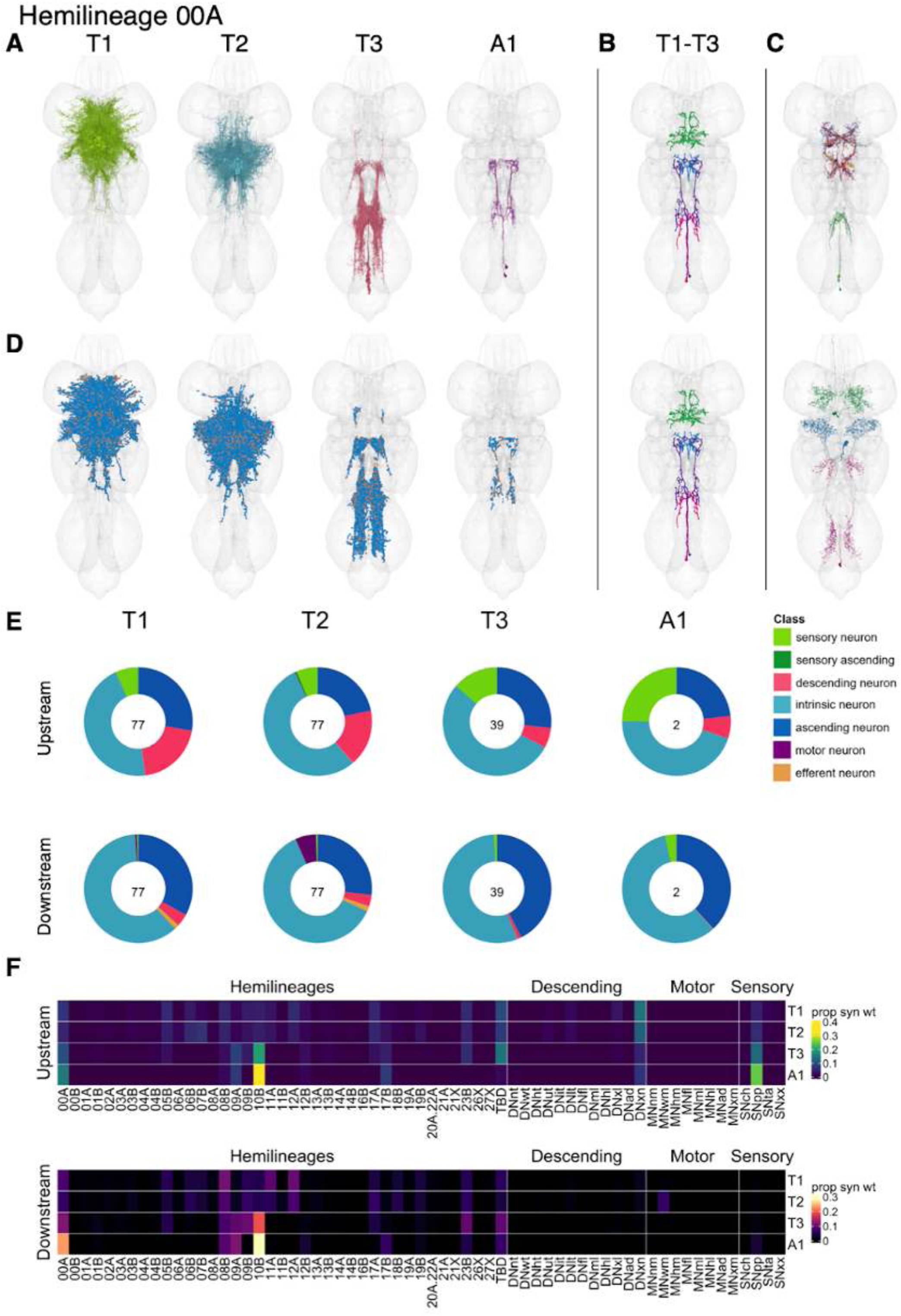
Hemilineage 00A. **A.** Meshes of all secondary neurons plotted in neuromere-specific colours. **B.** “Representative” secondary neuron skeletons plotted in neuromere-specific colours. The skeleton with the top accumulated NBLAST score among all neurons from the hemilineage in a given neuromere was used. **C.** Neuron meshes of selected examples. Top: glutamatergic subcluster 21723. Bottom: serial set 11979. **D.** Predicted synapses of secondary neurons. Blue: postsynapses; dark orange: presynapses. **E.** Proportions of connections from secondary neurons to upstream or downstream partners, normalised by neuromere and coloured by broad class. Numbers of query neurons appear in the centre. **F.** Proportions of synaptic weight from secondary neurons originating in each neuromere to upstream or downstream partners, normalised by row.

Secondary neurons survive in T1-A1 (Truman et al., 2010), with comparable numbers in neuromeres T1 and T2 but a smaller number in T3 (Figure 15E). They receive substantial input from descending neurons in T1 and T2 but more input from proprioceptive sensory neurons and hemilineages 00A and 10B in T3 and A1.They mainly inhibit target neurons of 08B, 11A, and 12A from T1-T2; 00A, 09A, 09B, 10B, and 23B from T3, and 00A, 09A, and 10B from A1 (Figure 15F).

There is considerable variability in morphology and connectivity between 00A cell types (Figure 15 - figure supplement 1-7). Specific types are dedicated to processing leg proprioceptive information from 10B and sensory neurons in the mVAC (e.g., IN00A003, IN00A005, IN00A007, IN00A011, and IN00A014) (Figure 15 - figure supplement 4-5). Other types innervate the tectulum (e.g., IN00A054), in a few cases inhibiting wing motor neurons (e.g., IN00A039 and IN00A056) (Figure 15 - figure supplement 7). Notably, IN00A015/vPR9 neurons inhibit sexually dimorphic 12A neurons in the mesothoracic triangle (Figure 15 - figure supplement 2,5) and have been implicated in normal production of courtship song (Lillvis et al., 2024; Mellert et al., 2016; von Philipsborn et al., 2011).

**Figure 15 - figure supplement 1.**
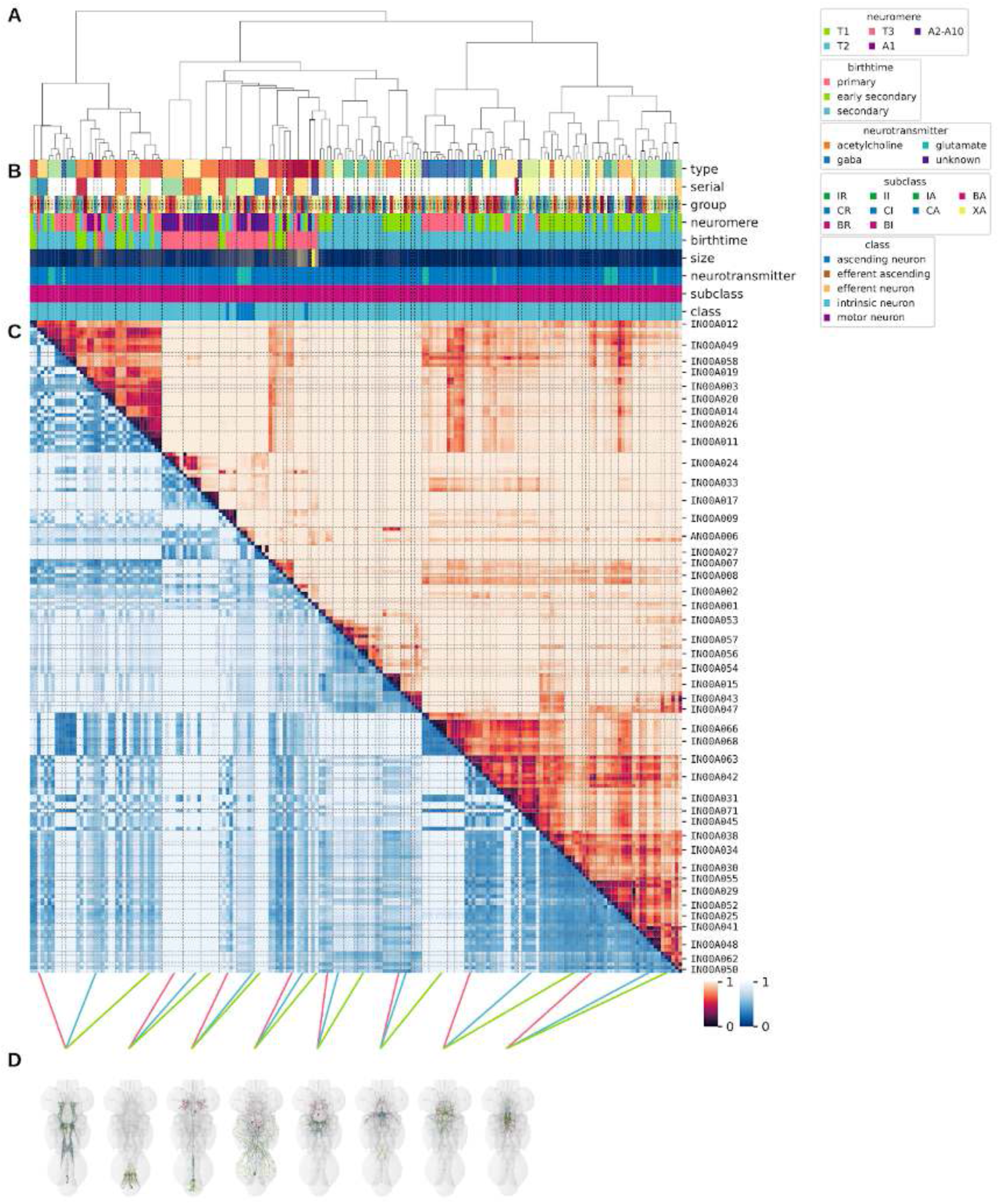
Systematic typing of hemilineage 00A. **A.** Hierarchical clustering dendrogram of hemilineage groups by laterally and serially aggregated connectivity cosine clustering. **B.** Categorical annotations of each hemilineage group, each column corresponding to the aligned leaf in A. Colours for type, serial set, and group are arbitrary for visualisation. Colours for neuromere, birthtime, neurotransmitter, subclass, and class are as in all other figures. **C.** Similarity distance heatmap for hemilineage. Cosine distance is in the upper triangle, while laterally symmetrised NBLAST distance is in the lower triangle. Systematic type names of some types are labelled. **D.** Morphologically representative groups from dendrogram subtrees. Each group, indicated by colour and line connecting to its column in B and C, is the most morphologically representative group (medoid of NBLAST distance) from a subtree of A. The subtrees (flat clusters) are equal height cuts of A determined to yield the number of groups per plot and plots in D.

**Figure 15 - figure supplement 2.**
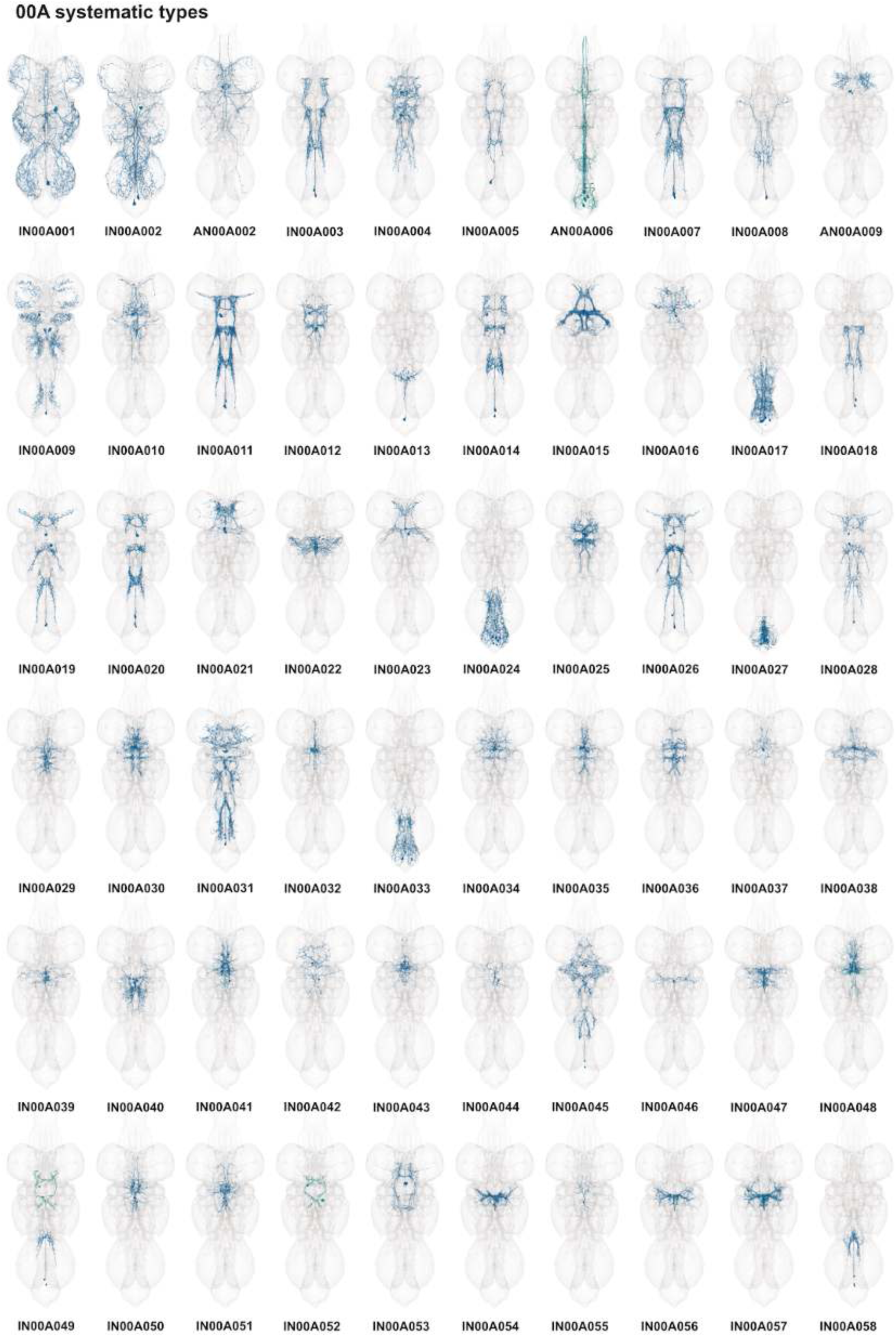
Systematic types of hemilineage 00A. Systematic types have been arranged in numerical order, with neurons of the same type that belong to distinct classes (e.g., intrinsic neuron vs ascending neuron) plotted separately but placed adjacent to each other. Individual neuron meshes have been coloured based on their predicted neurotransmitters: dark orange = acetylcholine, blue = gaba, marine = glutamate, dark purple = unknown.

**Figure 15 - figure supplement 3.**
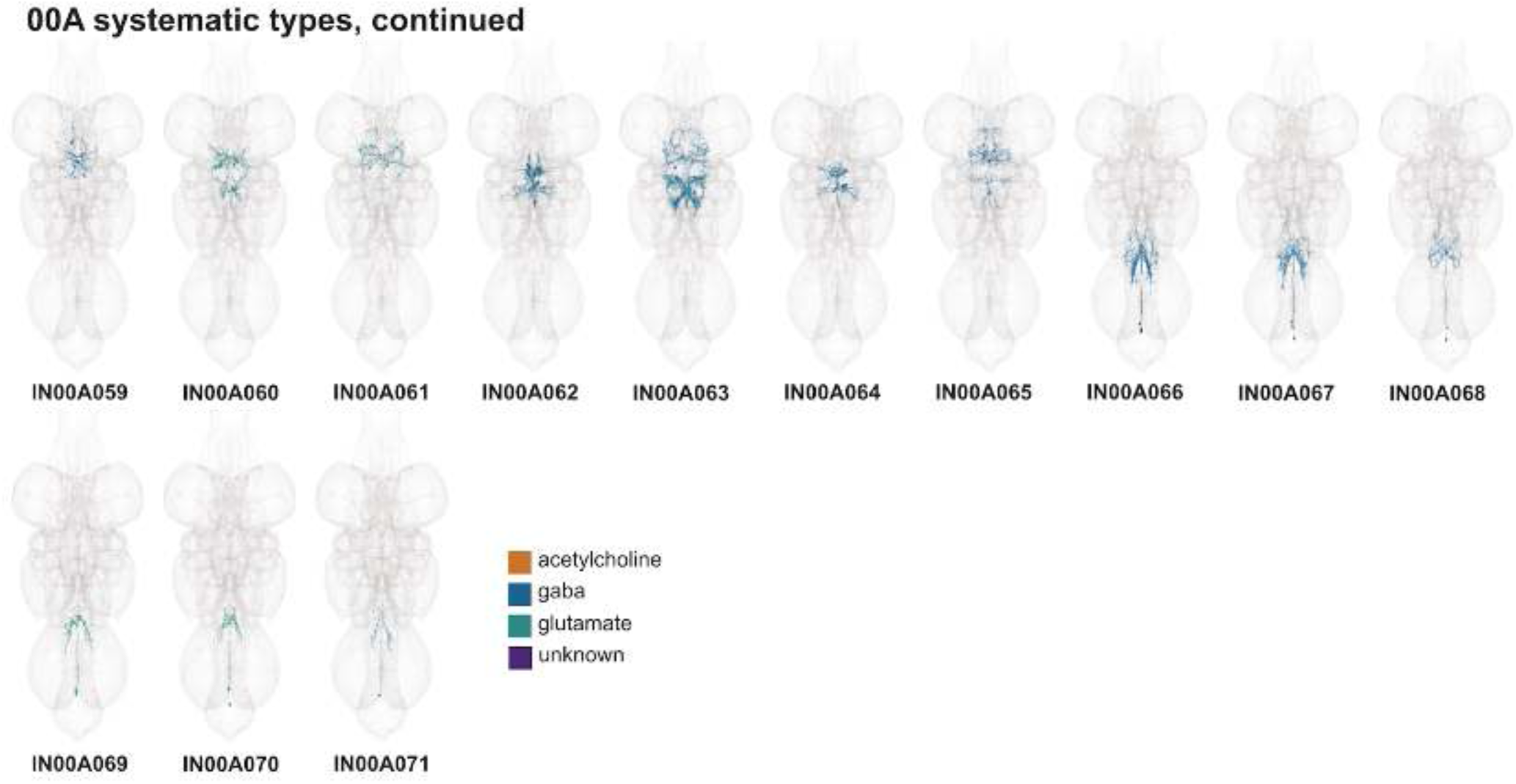
Systematic types of hemilineage 00A, continued. Systematic types have been arranged in numerical order, with neurons of the same type that belong to distinct classes (e.g., intrinsic neuron vs ascending neuron) plotted separately but placed adjacent to each other. Individual neuron meshes have been coloured based on predicted neurotransmitter: dark orange = acetylcholine, blue = gaba, marine = glutamate, dark purple = unknown.

**Figure 15 - figure supplement 4.**
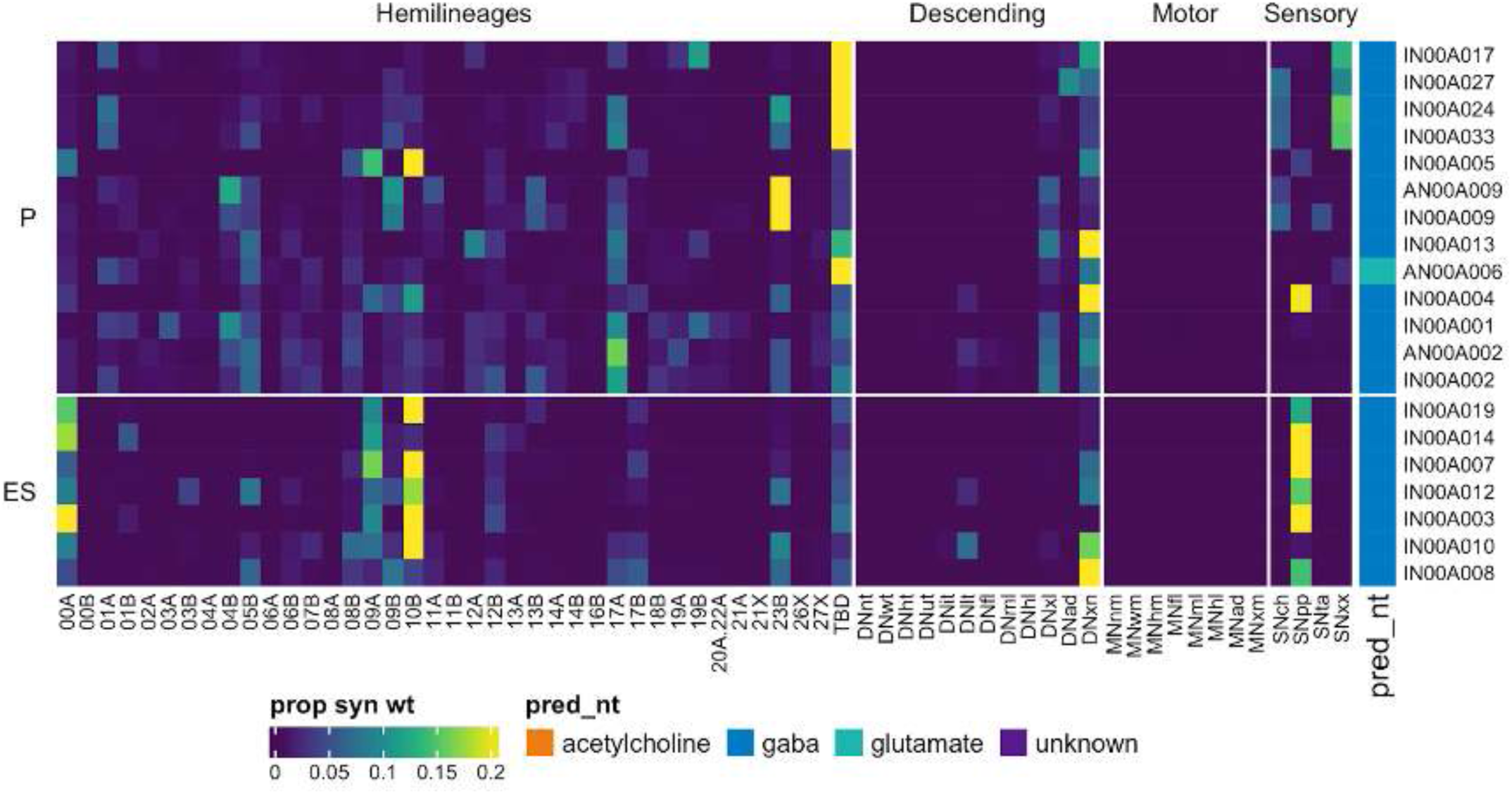
Connectivity to upstream partners by 00A primary and early secondary systematic types. Proportions of synaptic weight to systematic types from upstream partners, normalised by row. 00A neurons have been clustered within each assigned birthtime window (P = primary, ES = early secondary, S = secondary) based on both upstream and downstream connectivity to hemilineages, descending neuron subclasses, motor neuron subclasses, and sensory neuron modalities. Annotation bar is coloured by the most common predicted neurotransmitter for the neurons of each type.

**Figure 15 - figure supplement 5.**
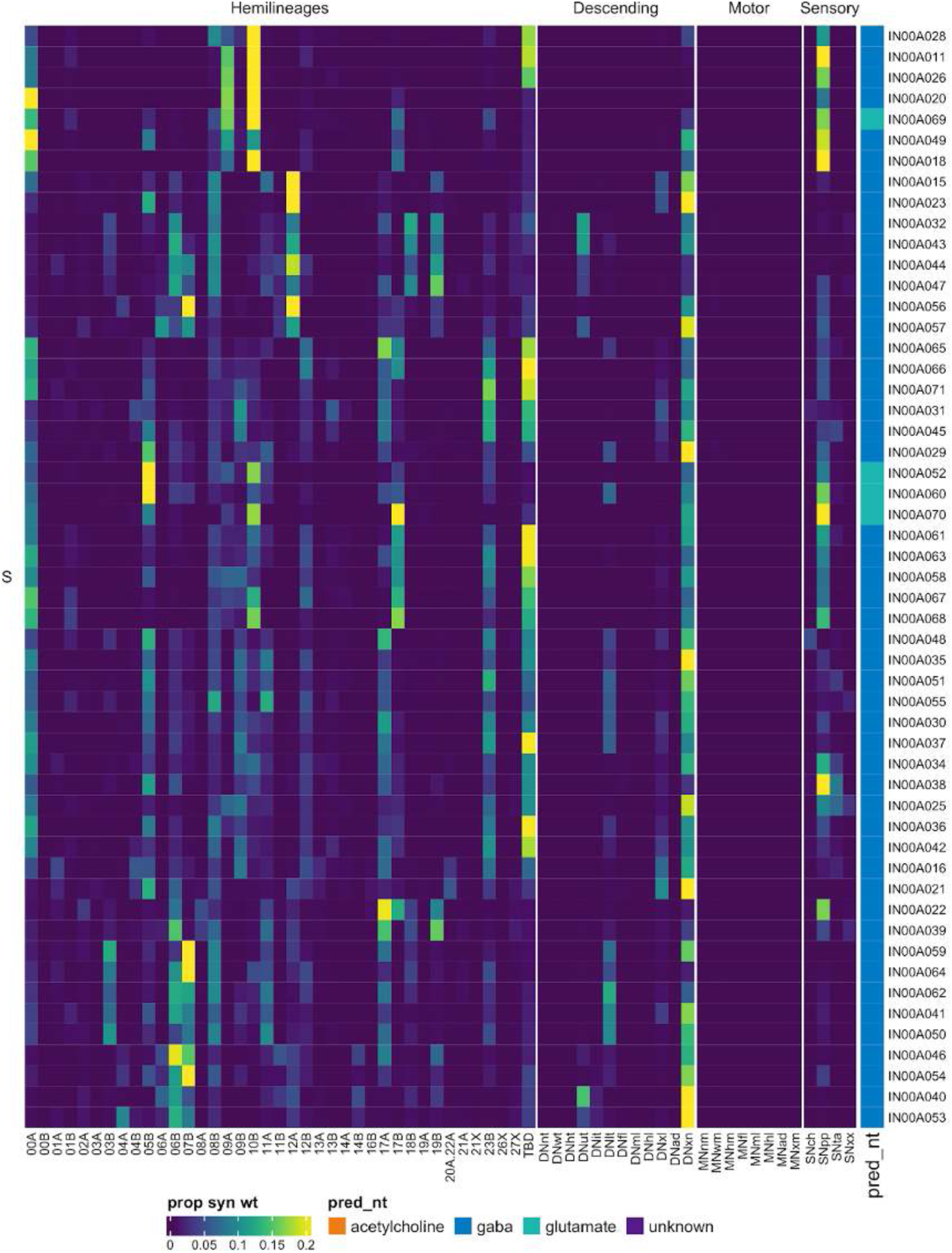
Connectivity to upstream partners by 00A secondary systematic types. Proportions of synaptic weight to systematic types from upstream partners, normalised by row. 00A neurons have been clustered within each assigned birthtime window (P = primary, ES = early secondary, S = secondary) based on both upstream and downstream connectivity to hemilineages, descending neuron subclasses, motor neuron subclasses, and sensory neuron modalities. The annotation bar is coloured by the most common predicted neurotransmitter within each type.

**Figure 15 - figure supplement 6.**
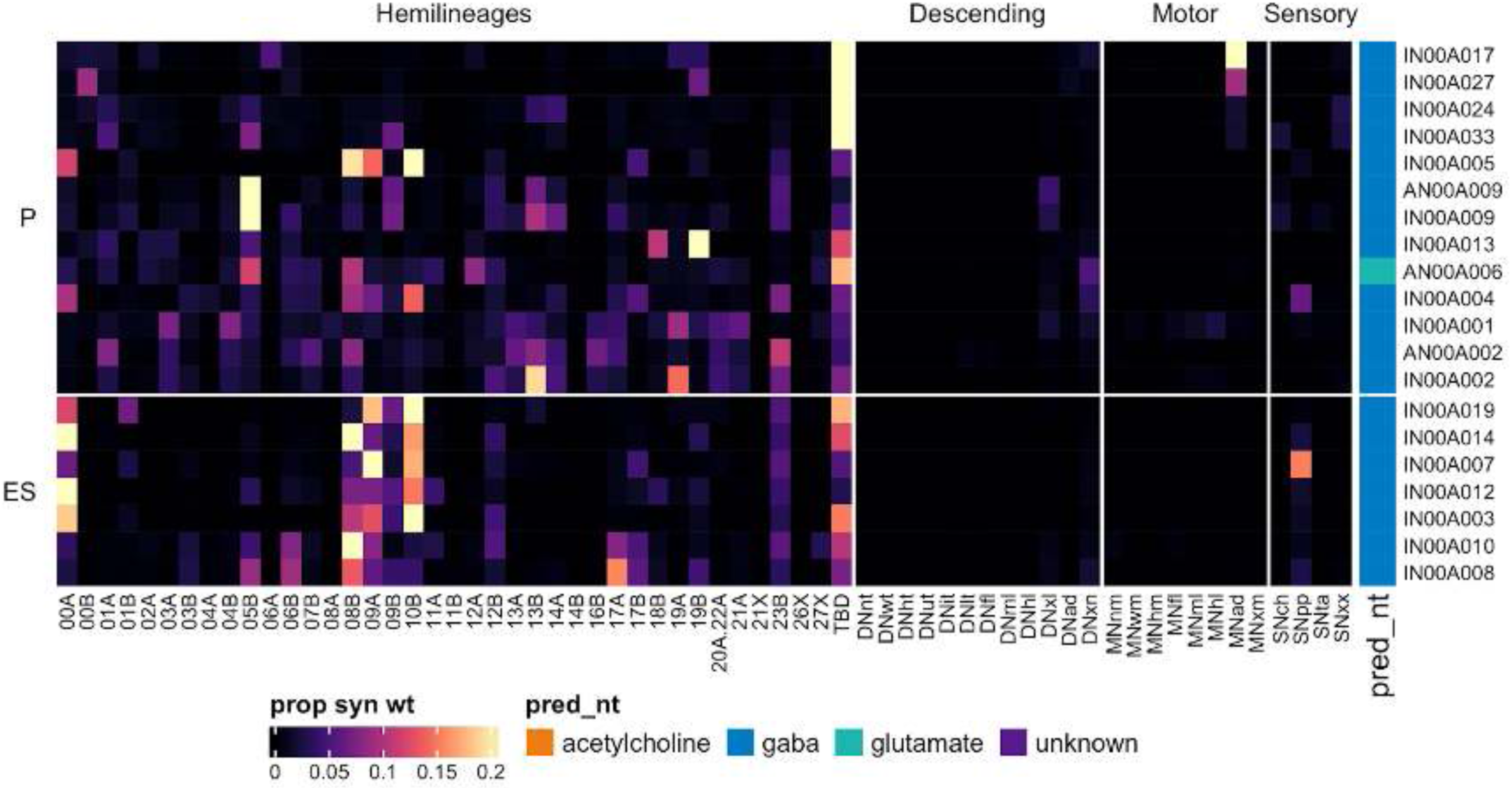
Connectivity to downstream partners by 00A primary and early secondary systematic types. Proportions of synaptic weight from systematic types to downstream partners, normalised by row. 00A neurons have been clustered within each assigned birthtime window (P = primary, ES = early secondary, S = secondary) based on both upstream and downstream connectivity to hemilineages, descending neuron subclasses, motor neuron subclasses, and sensory neuron modalities. The annotation bar is coloured by the most common predicted neurotransmitter within each type.

**Figure 15 - figure supplement 7.**
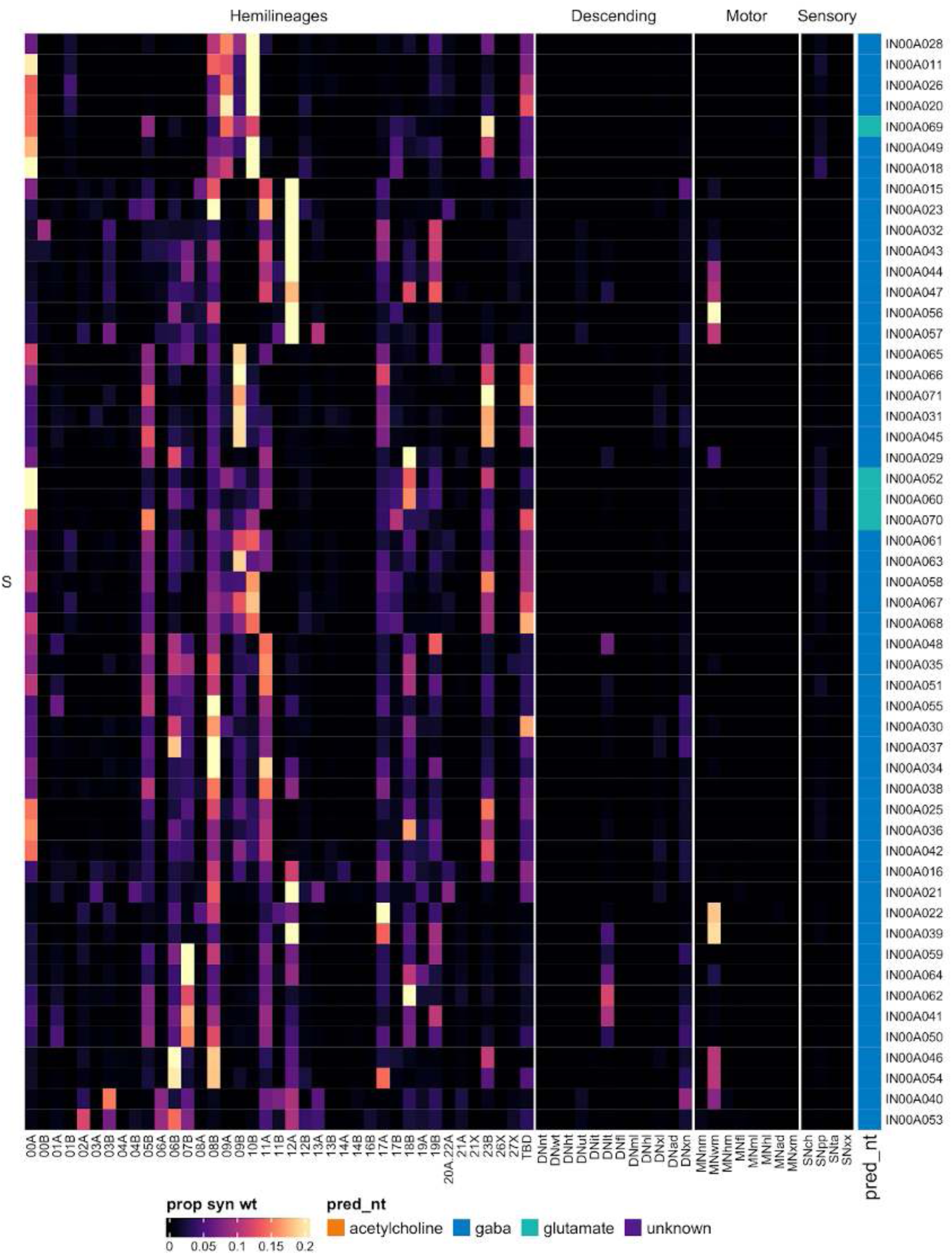
Connectivity to downstream partners by 00A secondary systematic types. Proportions of synaptic weight from systematic types to downstream partners, normalised by row. 00A neurons have been clustered within each assigned birthtime window (P = primary, ES = early secondary, S = secondary) based on both upstream and downstream connectivity to hemilineages, descending neuron subclasses, motor neuron subclasses, and sensory neuron modalities. The annotation bar is coloured by the most common predicted neurotransmitter for the neurons of each type.

#### Hemilineage 01A

Hemilineages 01A and 01B derive from NB1-2 at the anterior edge of each neuromere (Truman et al., 2004) that generates 2-4 MNs in T2, 1 MN in T3, 3-4 intersegmental neurons, and 10-15 local interneurons in the embryo (Schmid et al., 1999). Neurons of hemilineage 01A enter the neuropil from the anterior edge of each neuromere and project their axons across the ventral surface to curve around in the contralateral leg neuropil in a characteristic “crook” (Figure 16A,B) (Truman et al., 2004) before forming the terminal axon branches (Shepherd et al., 2019, 2016). A subset is intersegmental, with dendrites innervating a leg neuropil in the neuromere of origin and axons projecting to a more posterior neuropil (e.g., Figure 16C top). We also identified 01A neurons that enter the neuropil more medially to cross the midline in the aV and exclusively innervate the contralateral leg neuropil (Figure 16C bottom); unusually, these types (IN01A032 and IN01A039) are strongly connected to 01B (Figure 16 - figure supplement 4).

**Figure 16.**
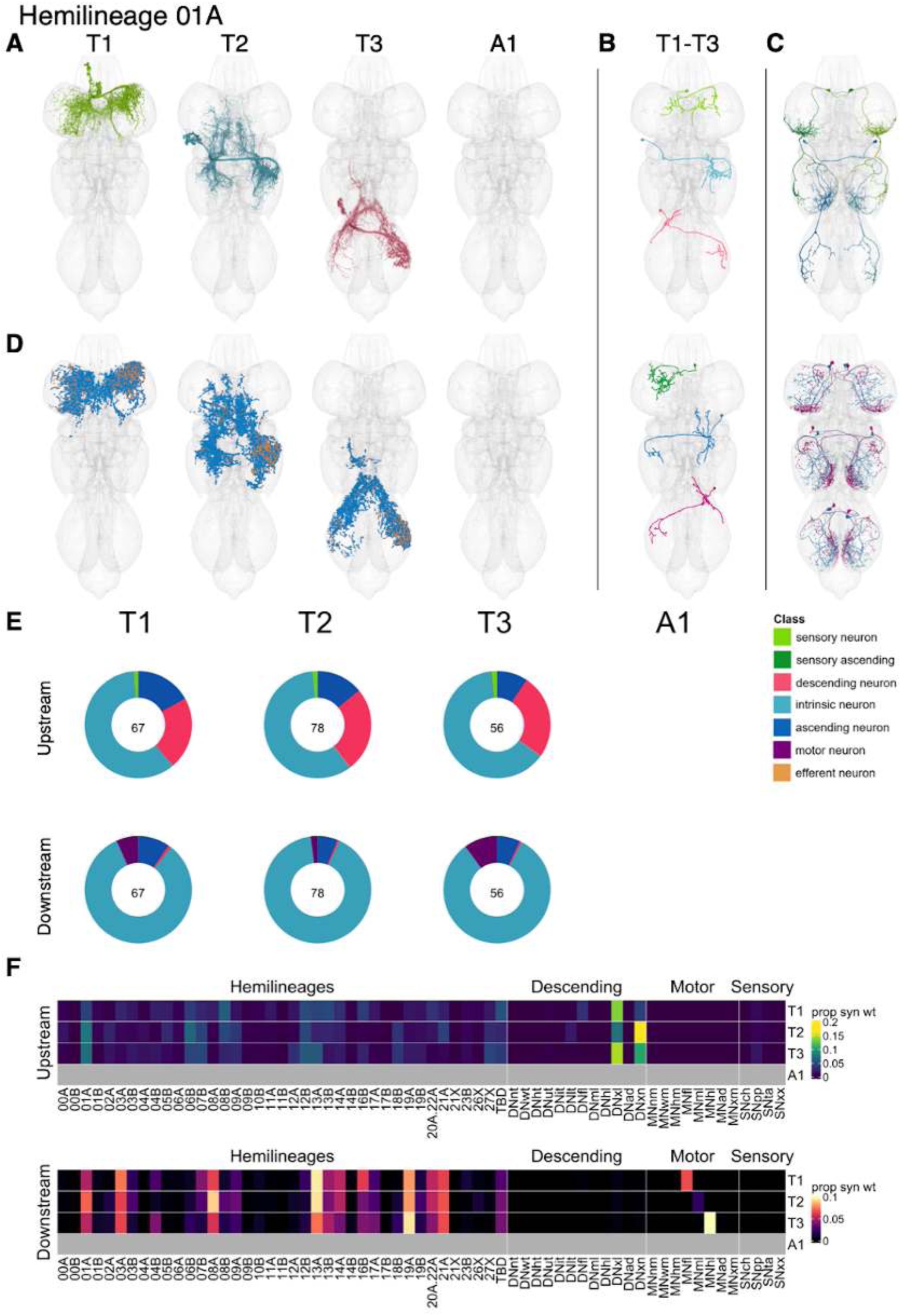
Hemilineage 01A. **A.** Meshes of all RHS secondary neurons plotted in hemineuromere-specific colours. **B.** “Representative” secondary neuron skeletons plotted in neuromere-specific colours. The skeleton with the top accumulated NBLAST score among all neurons from the hemilineage in a given hemineuromere was used. **C.** Neuron meshes of selected examples. Top: sequential serial set 10039. Bottom: contralateral independent leg serial sets 12457 and 13010. **D.** Predicted synapses of RHS secondary neurons. Blue: postsynapses; dark orange: presynapses. **E.** Proportions of connections from secondary neurons to upstream or downstream partners, normalised by neuromere and coloured by broad class. Numbers of query neurons appear in the centre. **F.** Proportions of synaptic weight from secondary neurons originating in each neuromere to upstream or downstream partners, normalised by row.

Secondary 01A neurons survive in T1-T3 with comparable numbers of neurons in all three neuromeres (Figure 16E). They receive much of their input from descending neurons, particularly those innervating multiple leg neuropils (Figure 16E,F). They are predicted to be cholinergic as reported (Figure 8E) (Lacin et al., 2019) and activate target neurons from many leg hemilineages, notably 01A, 03A, 08A, 13A, 19A, and 21A in T1-T3, 16B in T1, and leg MNs in T1 and T3 (Figure 16F). Activation of 01A secondary neurons bilaterally results in erratic forward locomotion interrupted by bouts of grooming (Harris et al., 2015), consistent with a role in exciting the contralateral T1 and T3 legs.

Several early-born 01A types receive substantial input from tactile sensory neurons (e.g., IN01A012 and IN01A036) (Figure 16 - figure supplement 4). A few types activate wing motor neurons (e.g., IN01A017 and IN01A020) (Figure 16 - figure supplement 6). Unusually, AN01A055 and AN01A086 immediately ascend through the neck connective without forming many synapses (Figure 16 - figure supplement 2-3).

**Figure 16 - figure supplement 1.**
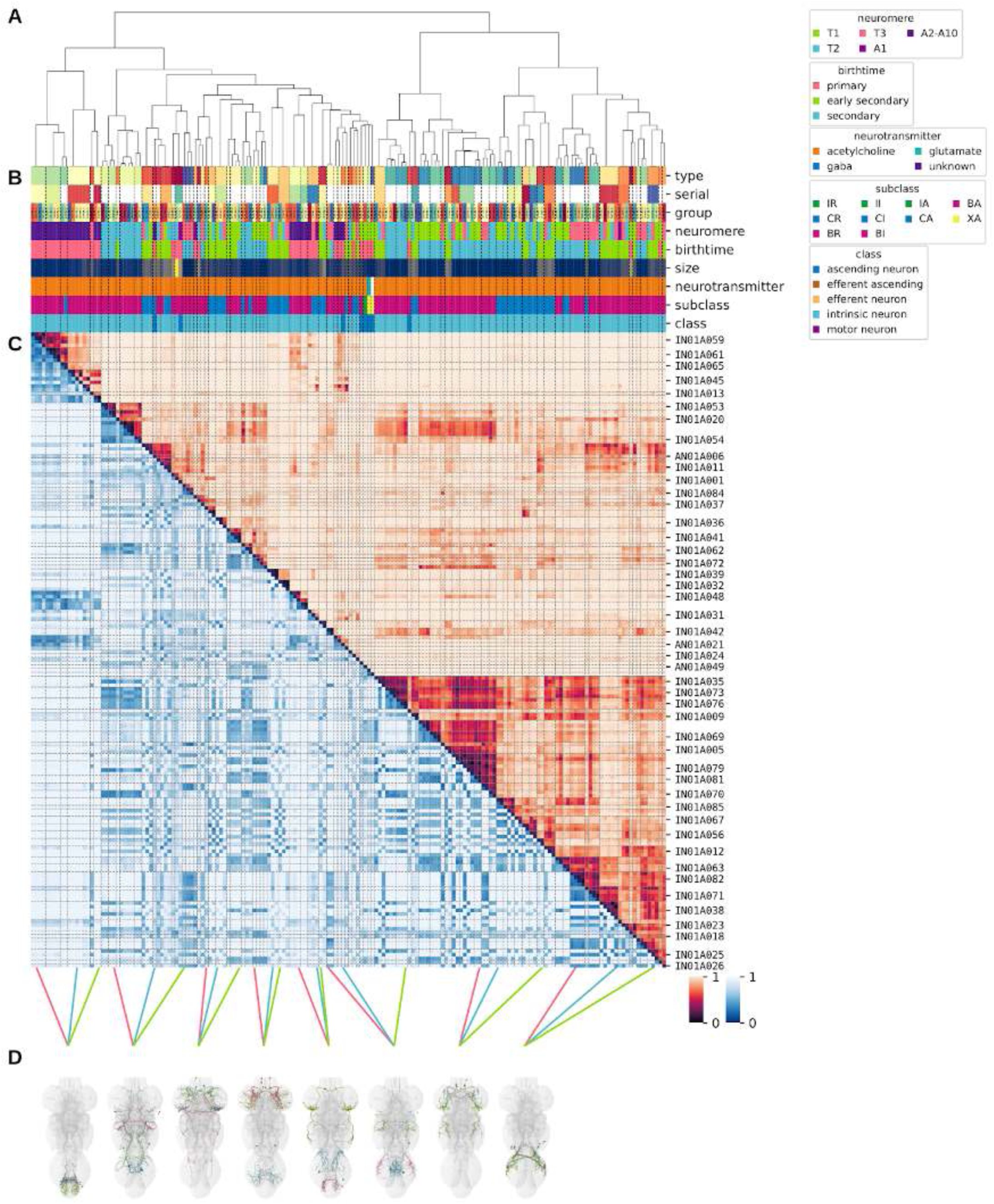
Systematic typing of hemilineage 01A. **A.** Hierarchical clustering dendrogram of hemilineage groups by laterally and serially aggregated connectivity cosine clustering. **B.** Categorical annotations of each hemilineage group, each column corresponding to the aligned leaf in A. Colours for type, serial set, and group are arbitrary for visualisation. Colours for neuromere, birthtime, neurotransmitter, subclass, and class are as in all other figures. **C.** Similarity distance heatmap for hemilineage. Cosine distance is in the upper triangle, while laterally symmetrised NBLAST distance is in the lower triangle. Systematic type names of some types are labelled. **D.** Morphologically representative groups from dendrogram subtrees. Each group, indicated by colour and line connecting to its column in B and C, is the most morphologically representative group (medoid of NBLAST distance) from a subtree of A. The subtrees (flat clusters) are equal height cuts of A determined to yield the number of groups per plot and plots in D.

**Figure 16 - figure supplement 2.**
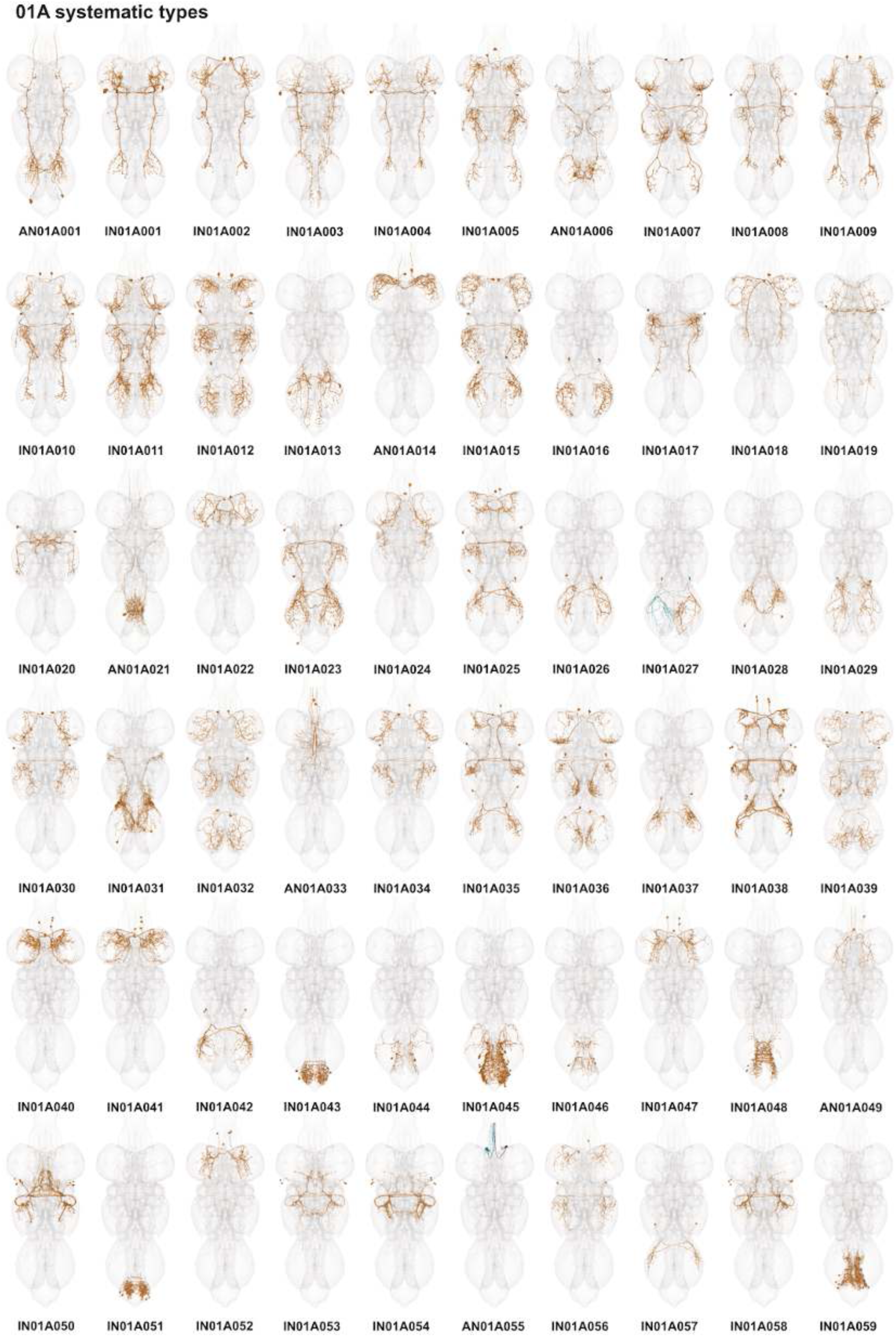
Systematic types of hemilineage 01A. Systematic types have been arranged in numerical order, with neurons of the same type that belong to distinct classes (e.g., intrinsic neuron vs ascending neuron) plotted separately but placed adjacent to each other. Individual neuron meshes have been coloured based on predicted neurotransmitter: dark orange = acetylcholine, blue = gaba, marine = glutamate, dark purple = unknown.

**Figure 16 - figure supplement 3.**
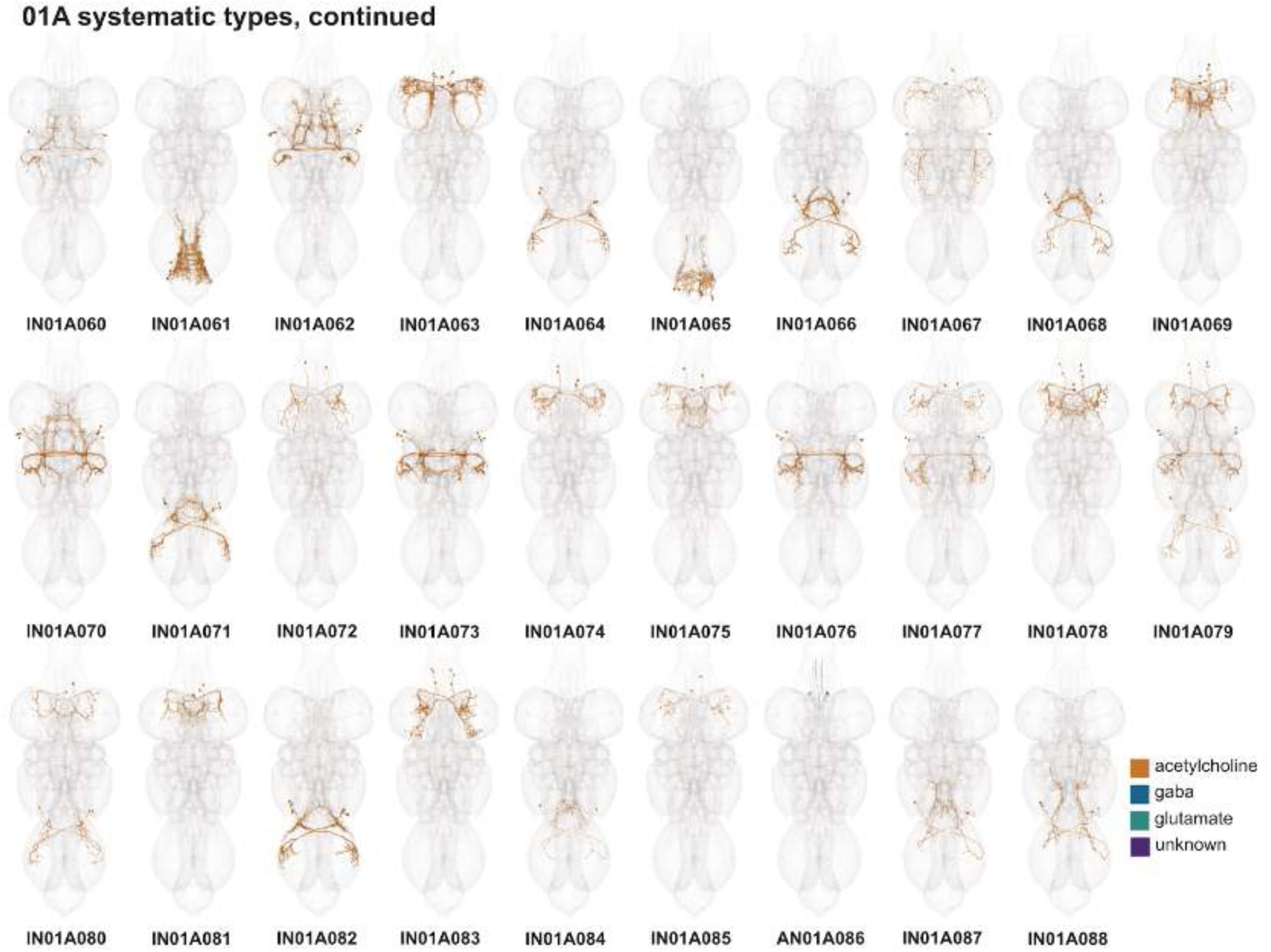
Systematic types of hemilineage 01A, continued. Systematic types have been arranged in numerical order, with neurons of the same type that belong to distinct classes (e.g., intrinsic neuron vs ascending neuron) plotted separately but placed adjacent to each other. Individual neuron meshes have been coloured based on predicted neurotransmitter: dark orange = acetylcholine, blue = gaba, marine = glutamate, dark purple = unknown.

**Figure 16 - figure supplement 4.**
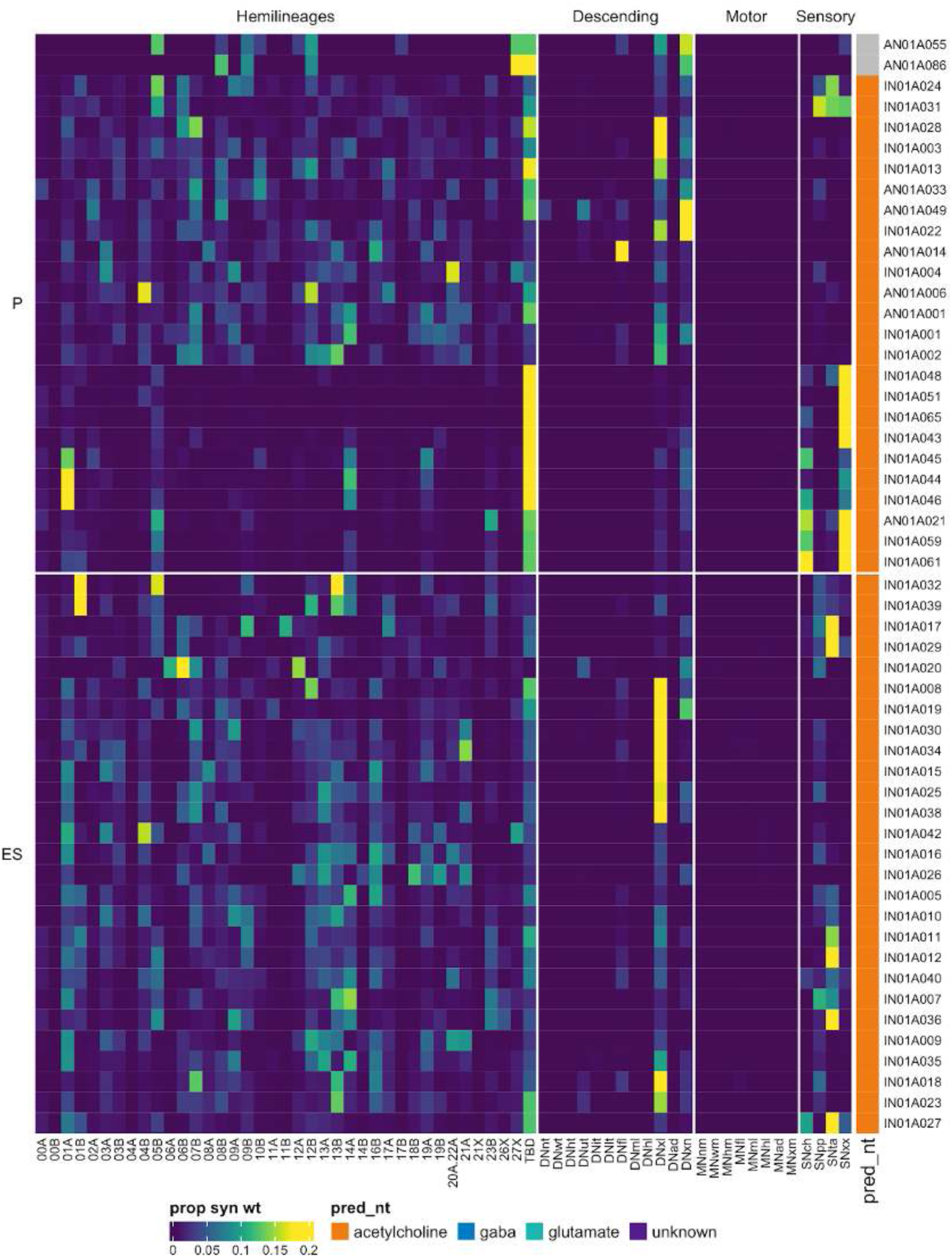
Connectivity to upstream partners by 01A primary and early secondary systematic types. Proportions of synaptic weight to systematic types from upstream partners, normalised by row. 01A neurons have been clustered within each assigned birthtime window (P = primary, ES = early secondary, S = secondary) based on both upstream and downstream connectivity to hemilineages, descending neuron subclasses, motor neuron subclasses, and sensory neuron modalities. Annotation bar is coloured by the most common predicted neurotransmitter for the neurons of each type.

**Figure 16 - figure supplement 5.**
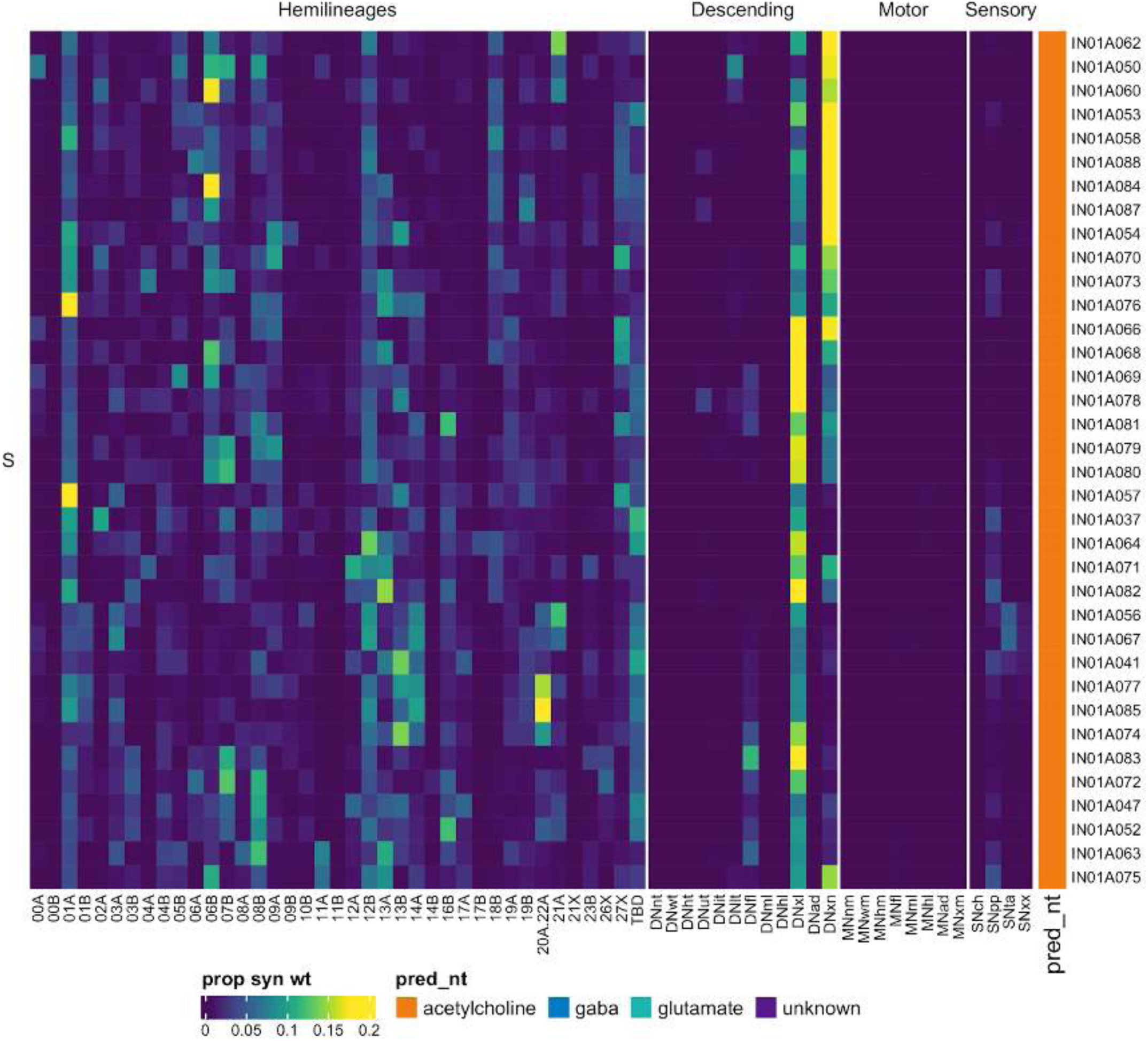
Connectivity to upstream partners by 01A secondary systematic types. Proportions of synaptic weight to systematic types from upstream partners, normalised by row. 01A neurons have been clustered within each assigned birthtime window (P = primary, ES = early secondary, S = secondary) based on both upstream and downstream connectivity to hemilineages, descending neuron subclasses, motor neuron subclasses, and sensory neuron modalities. The annotation bar is coloured by the most common predicted neurotransmitter within each type.

**Figure 16 - figure supplement 6.**
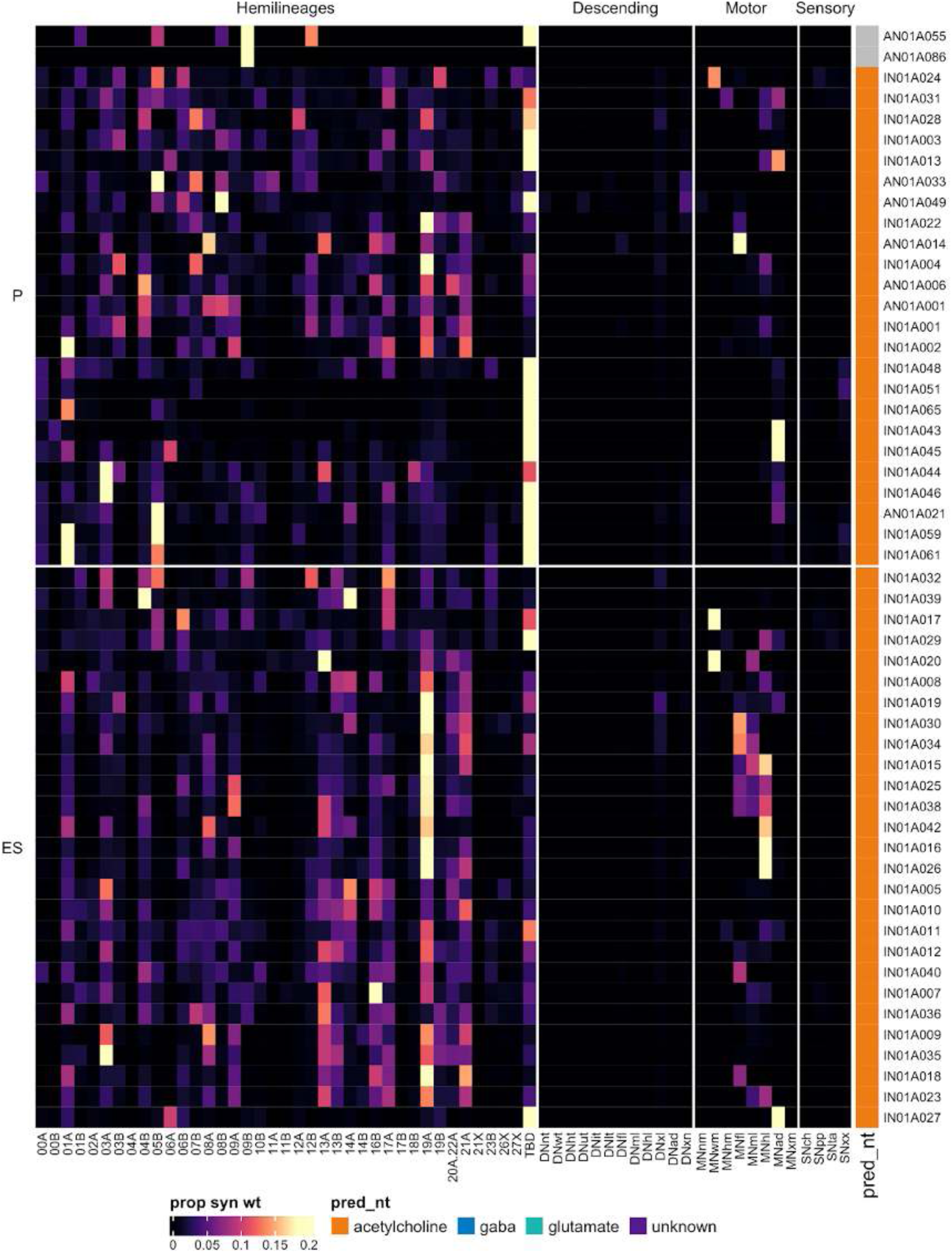
Connectivity to downstream partners by 01A primary and early secondary systematic types. Proportions of synaptic weight from systematic types to downstream partners, normalised by row. 01A neurons have been clustered within each assigned birthtime window (P = primary, ES = early secondary, S = secondary) based on both upstream and downstream connectivity to hemilineages, descending neuron subclasses, motor neuron subclasses, and sensory neuron modalities. The annotation bar is coloured by the most common predicted neurotransmitter within each type.

**Figure 16 - figure supplement 7.**
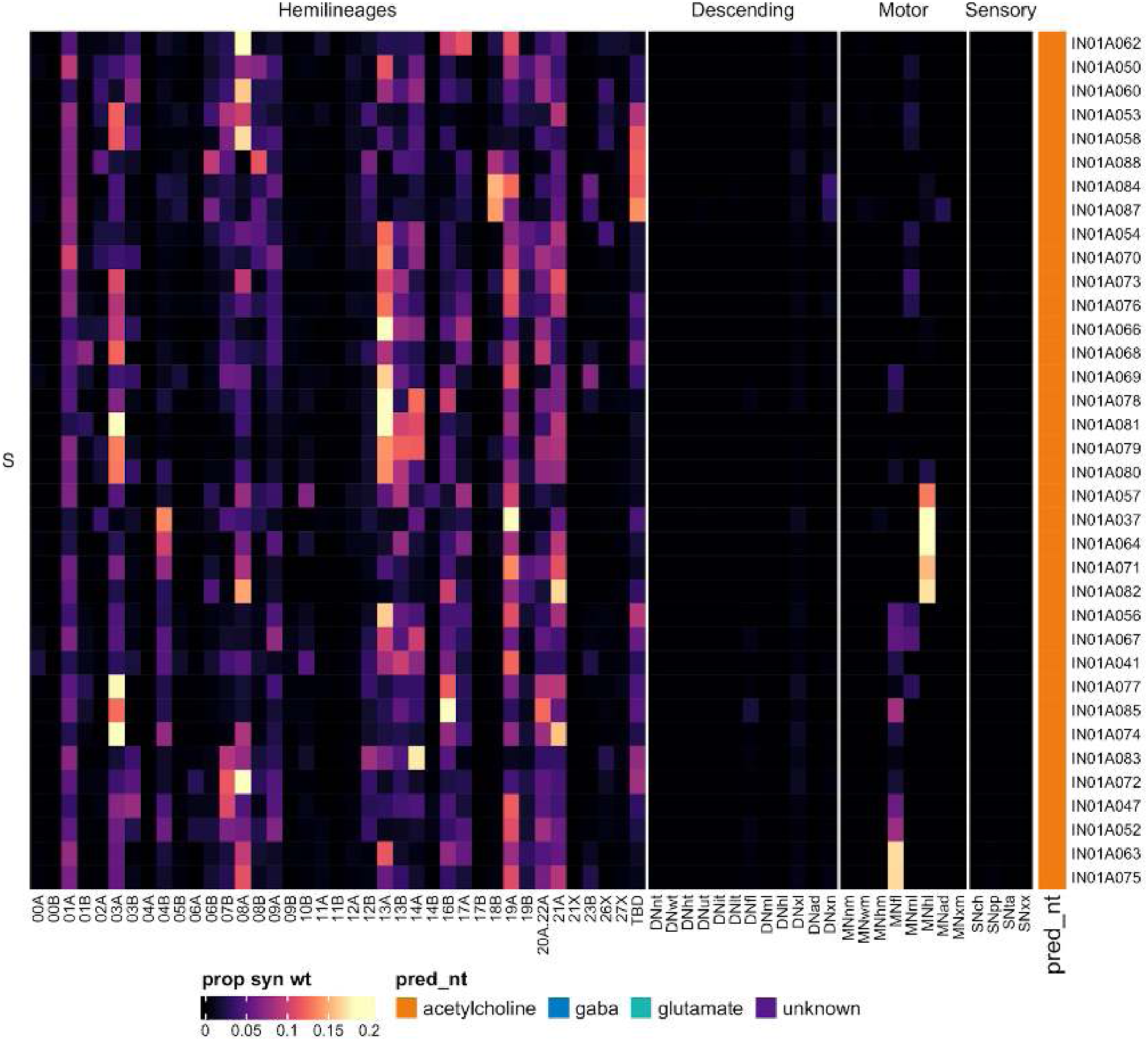
Connectivity to downstream partners by 01A secondary systematic types. Proportions of synaptic weight from systematic types to downstream partners, normalised by row. 01A neurons have been clustered within each assigned birthtime window (P = primary, ES = early secondary, S = secondary) based on both upstream and downstream connectivity to hemilineages, descending neuron subclasses, motor neuron subclasses, and sensory neuron modalities. The annotation bar is coloured by the most common predicted neurotransmitter for the neurons of each type.

#### Hemilineage 01B

Neurons from hemilineage 01B innervate the ipsilateral leg sensory neuropil of the next anterior neuromere and migrate (or are dragged) towards it during development. Therefore, secondary 01B neurons survive in T2-A1 but innervate T1-T3 (Shepherd et al., 2019); to simplify connectivity analyses, we opted to assign the soma neuromere based on innervation rather than true developmental origin. 01B primary neurites only converge as they enter the ventral leg neuropil, and their dendrites are largely restricted to the leg sensory association neuropils - the VAC (e.g., Figure 17C bottom) or the mVAC (e.g., Figure 17C top). There were several primary types, particularly in T3, that we found challenging to assign to 01B vs 04B; the latter overlaps with 01B but generally enters the neuropil more medially.

**Figure 17.**
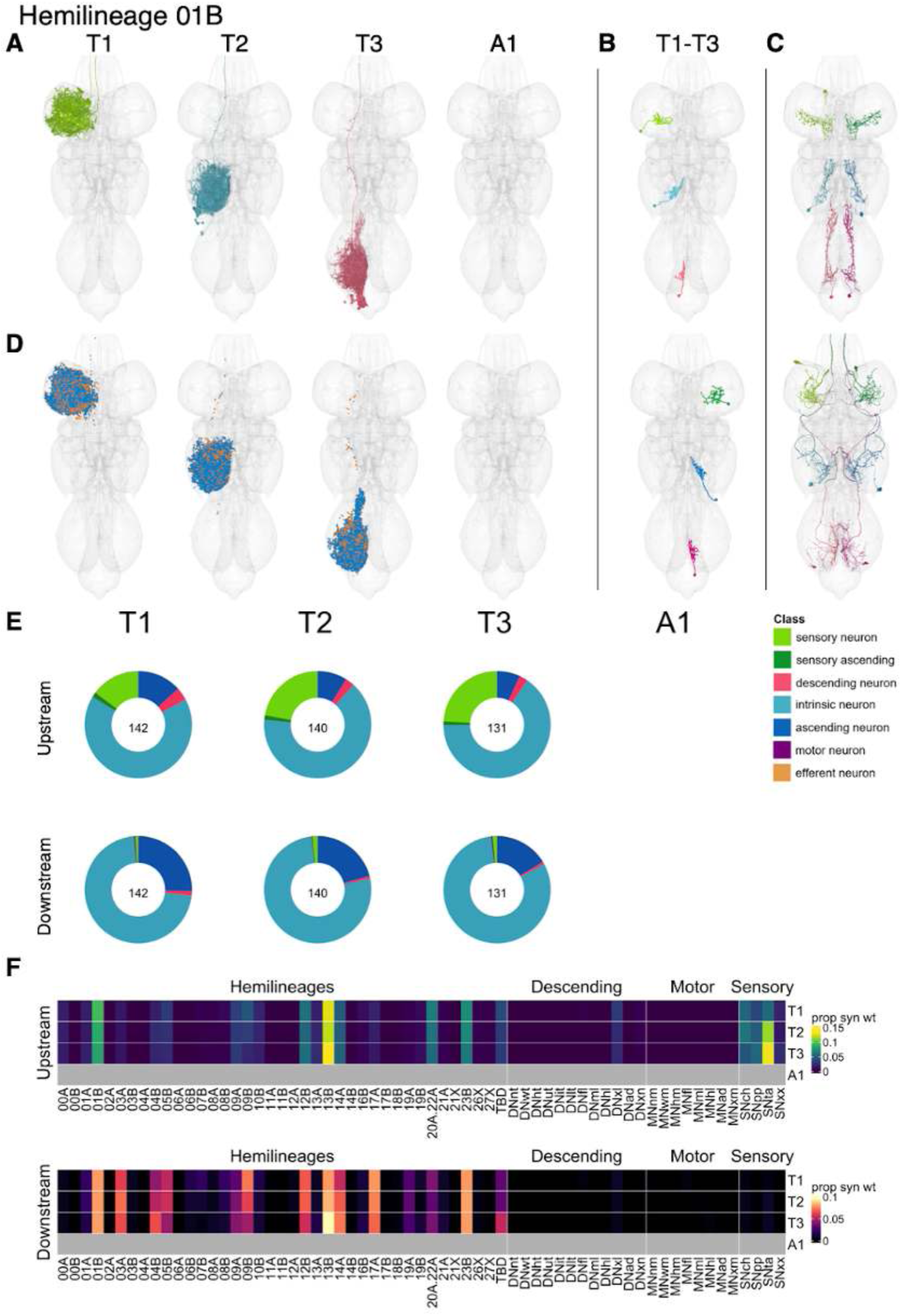
Hemilineage 01B. **A.** Meshes of all RHS secondary neurons plotted in hemineuromere-specific colours. **B.** “Representative” secondary neuron skeletons plotted in hemineuromere-specific colours. The skeleton with the top accumulated NBLAST score among all neurons from the hemilineage in a given hemineuromere was used. **C.** Neuron meshes of selected examples. Top: independent leg serial set 12532. Bottom: cholinergic ascending serial set 12577. **D.** Predicted synapses of RHS secondary neurons. Blue: postsynapses; dark orange: presynapses. **E.** Proportions of connections from secondary neurons to upstream or downstream partners, normalised by neuromere and coloured by broad class. Numbers of query neurons appear in the centre. **F.** Proportions of synaptic weight from secondary neurons originating in each neuromere to upstream or downstream partners, normalised by row.

There are comparable numbers of 01B secondary neurons innervating all three thoracic neuromeres (Figure 17E). They receive input from chemosensory, proprioceptive, and notably tactile sensory neurons and from leg hemilineages 01B, 12B, 14A, 20A/22A, 23B and especially 13B (Figure 17F). They are predicted to be gabergic as reported (Figure 8E) (Lacin et al., 2019) and inhibit the same hemilineages mentioned above plus 03A, 04B, 05B, 09B, and especially 17A (Figure 17F). No functional studies have been published for secondary 01B neurons.

Most 01B neurons are restricted to a single ipsilateral hemineuromere, but we identified five ascending types, one of which is cholinergic (AN01B004) (Figure 17C bottom, Figure 17 - figure supplement 2). Specific types can be dedicated to chemosensory (e.g., IN01B075), proprioceptive (e.g., IN01B007), or tactile (e.g., IN01B020-21) information or can process multiple modalities (e.g., IN01B039) (Figure 17 - figure supplement 4-5). A subset of chemosensory and proprioceptive 01B types are downstream of descending neurons targeting all legs (Figure 17 - figure supplement 4-5). Notably, a few 01B types inhibit chemosensory (e.g., IN01B080) or tactile sensory neurons (e.g., AN/IN01B002) (Figure 17 - figure supplement 7-8).

**Figure 17 - figure supplement 1.**
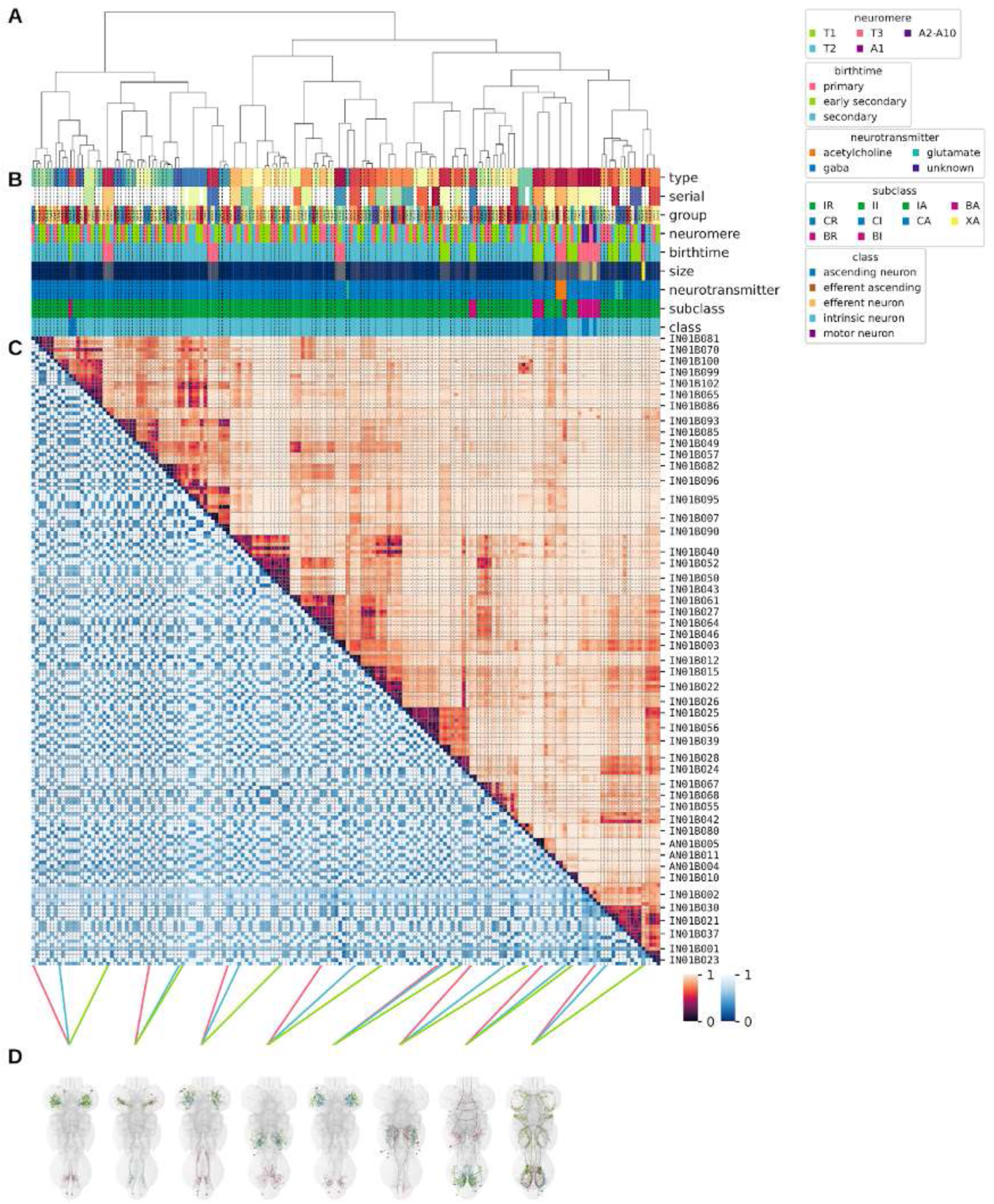
Systematic typing of hemilineage 01B. **A.** Hierarchical clustering dendrogram of hemilineage groups by laterally and serially aggregated connectivity cosine clustering. **B.** Categorical annotations of each hemilineage group, each column corresponding to the aligned leaf in A. Colours for type, serial set, and group are arbitrary for visualisation. Colours for neuromere, birthtime, neurotransmitter, subclass, and class are as in all other figures. **C.** Similarity distance heatmap for hemilineage. Cosine distance is in the upper triangle, while laterally symmetrised NBLAST distance is in the lower triangle. Systematic type names of some types are labelled. **D.** Morphologically representative groups from dendrogram subtrees. Each group, indicated by colour and line connecting to its column in B and C, is the most morphologically representative group (medoid of NBLAST distance) from a subtree of A. The subtrees (flat clusters) are equal height cuts of A determined to yield the number of groups per plot and plots in D.

**Figure 17 - figure supplement 2.**
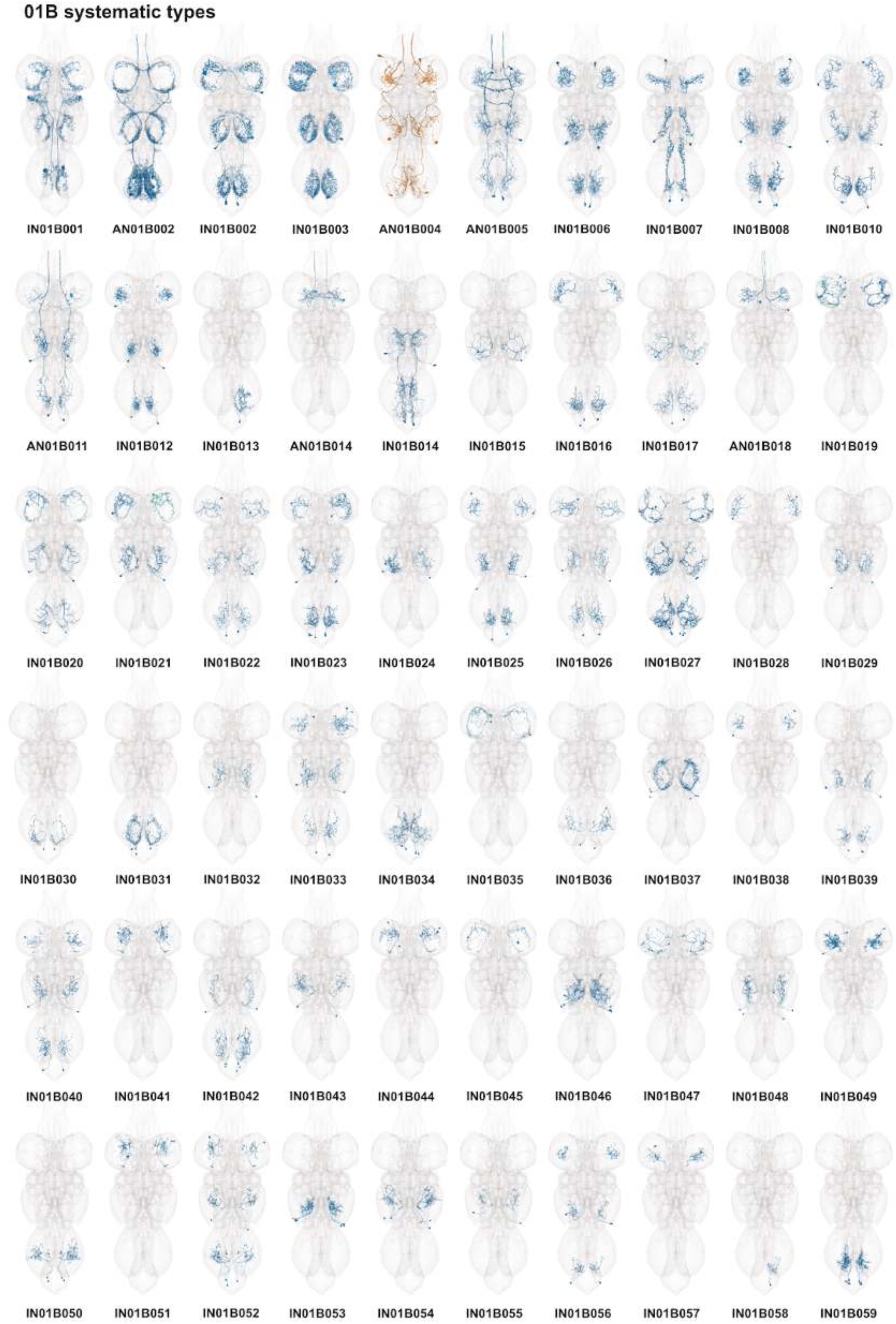
Systematic types of hemilineage 01B. Systematic types have been arranged in numerical order, with neurons of the same type that belong to distinct classes (e.g., intrinsic neuron vs ascending neuron) plotted separately but placed adjacent to each other. Individual neuron meshes have been coloured based on predicted neurotransmitter: dark orange = acetylcholine, blue = gaba, marine = glutamate, dark purple = unknown.

**Figure 17 - figure supplement 3.**
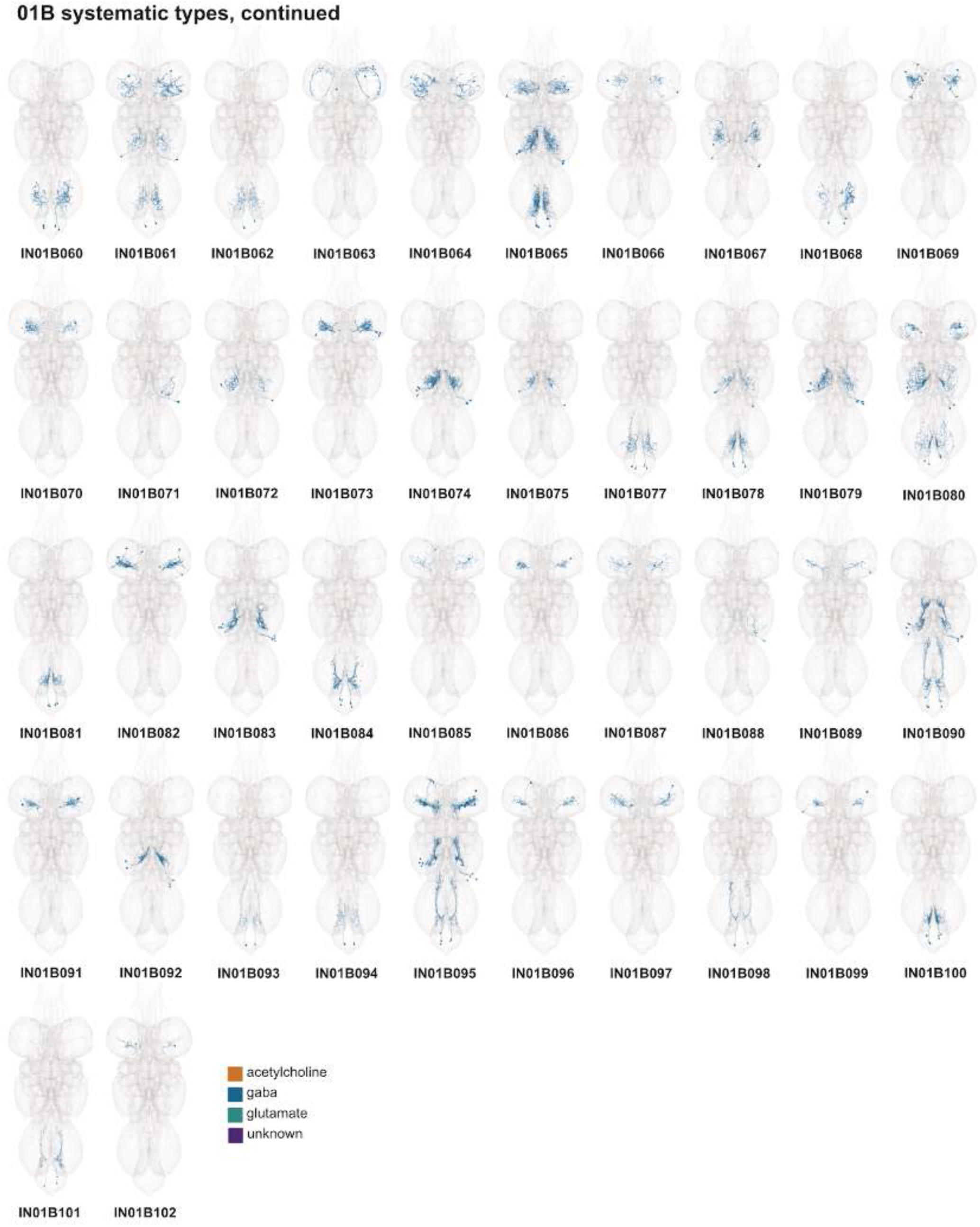
Systematic types of hemilineage 01B, continued. Systematic types have been arranged in numerical order, with neurons of the same type that belong to distinct classes (e.g., intrinsic neuron vs ascending neuron) plotted separately but placed adjacent to each other. Individual neuron meshes have been coloured based on predicted neurotransmitter: dark orange = acetylcholine, blue = gaba, marine = glutamate, dark purple = unknown.

**Figure 17 - figure supplement 4.**
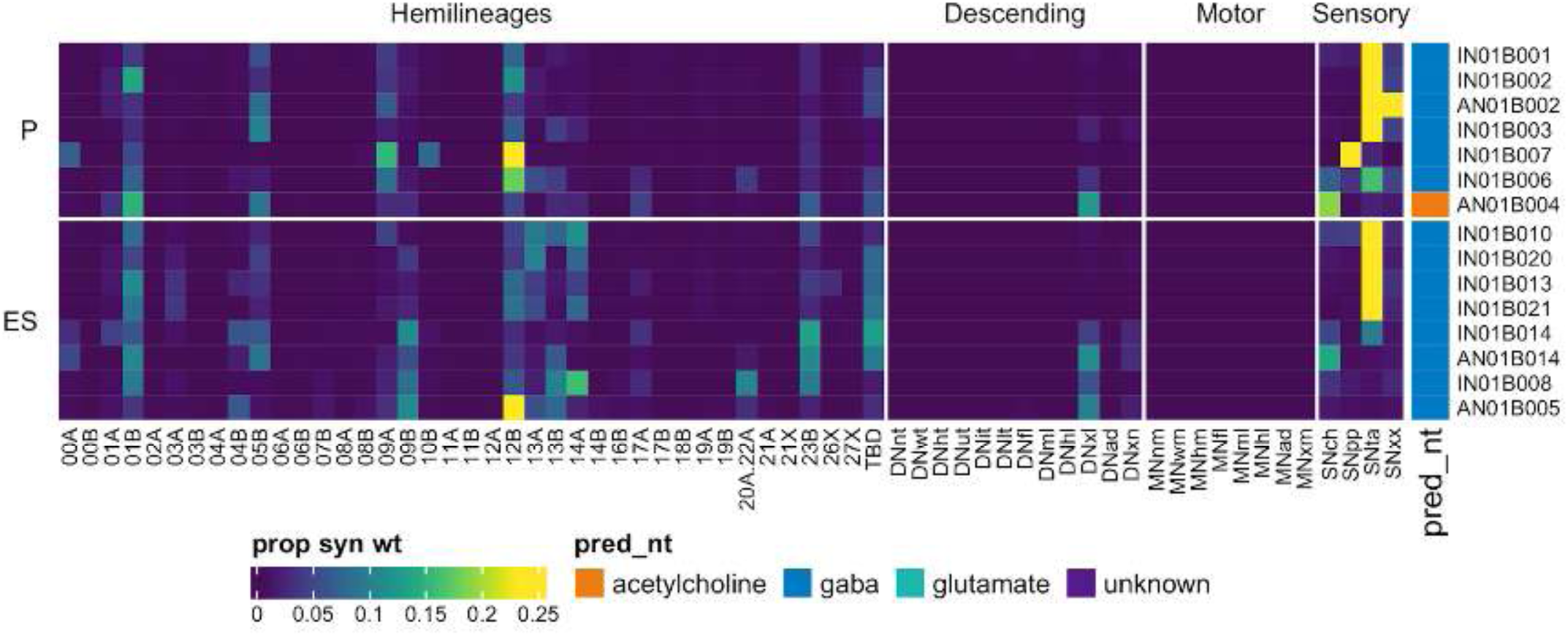
Connectivity to upstream partners by 01B primary and early secondary systematic types. Proportions of synaptic weight to systematic types from upstream partners, normalised by row. 01B neurons have been clustered within each assigned birthtime window (P = primary, ES = early secondary, S = secondary) based on both upstream and downstream connectivity to hemilineages, descending neuron subclasses, motor neuron subclasses, and sensory neuron modalities. Annotation bar is coloured by the most common predicted neurotransmitter for the neurons of each type.

**Figure 17 - figure supplement 5a.**
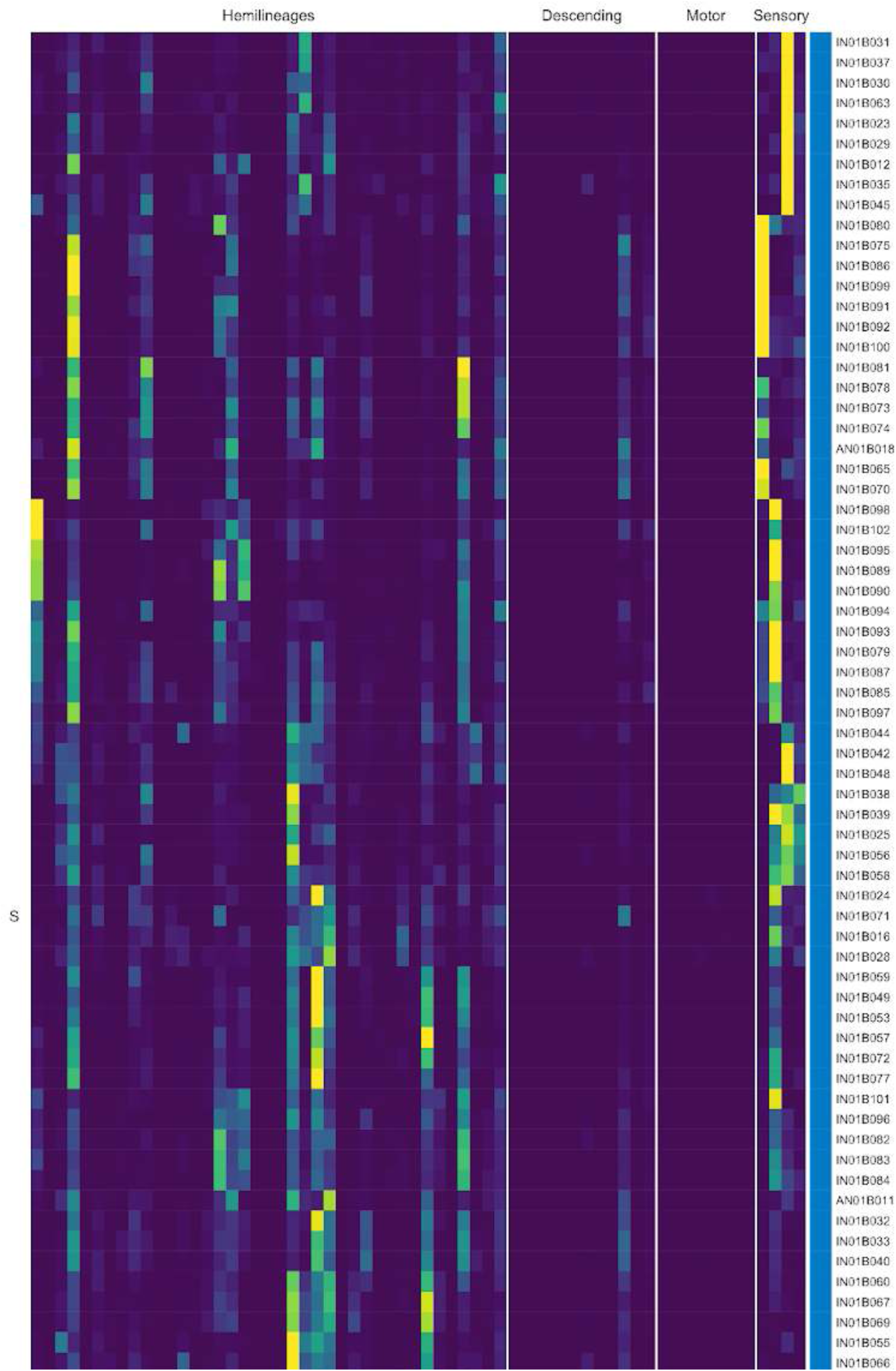
Connectivity to upstream partners by 01B secondary systematic types. Proportions of synaptic weight to systematic types from upstream partners, normalised by row. 01B neurons have been clustered within each assigned birthtime window (P = primary, ES = early secondary, S = secondary) based on both upstream and downstream connectivity to hemilineages, descending neuron subclasses, motor neuron subclasses, and sensory neuron modalities. The annotation bar is coloured by the most common predicted neurotransmitter within each type.

**Figure 17 - figure supplement 5b.**
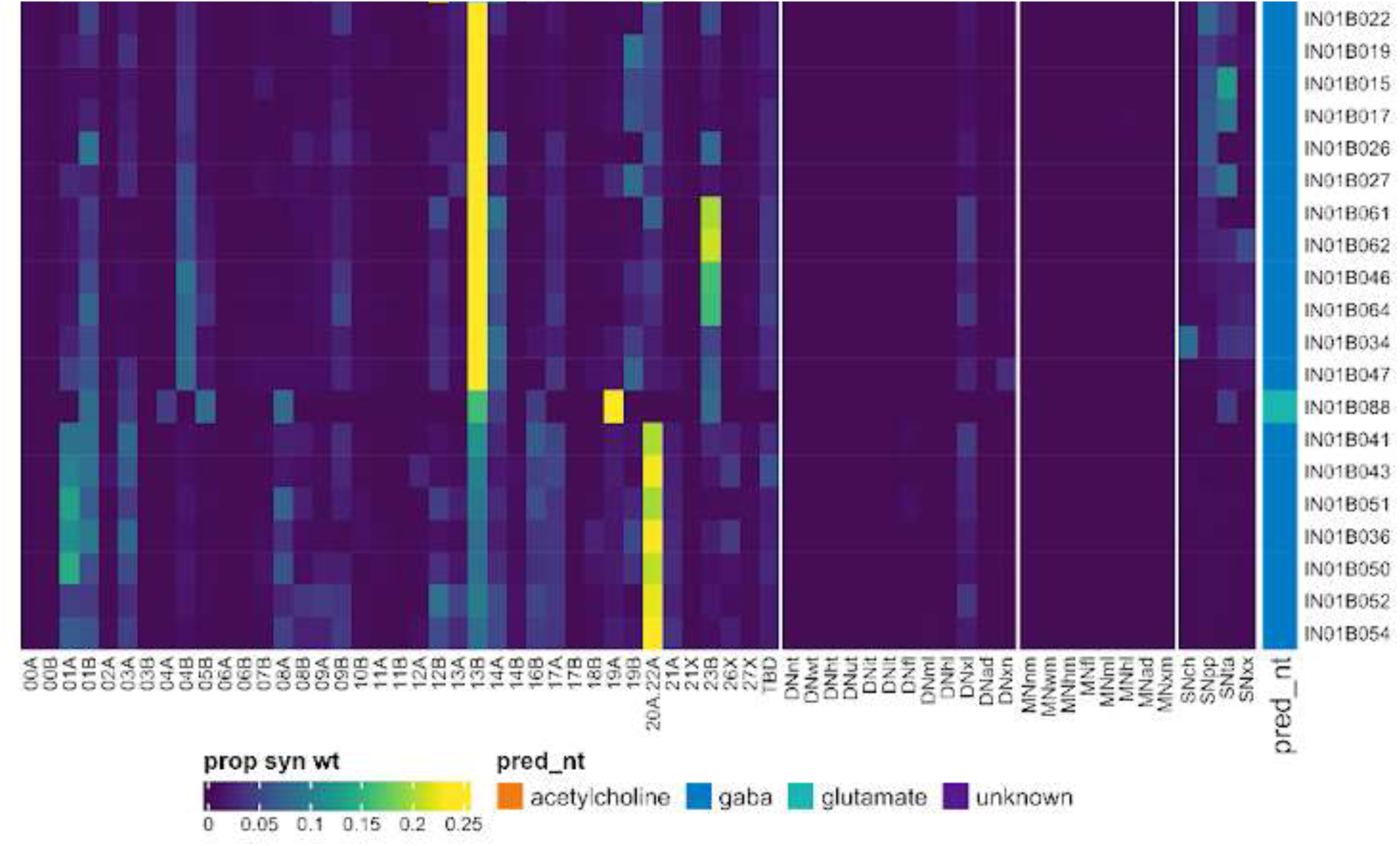
Connectivity to upstream partners by 01B secondary systematic types, continued. Proportions of synaptic weight to systematic types from upstream partners, normalised by row. 01B neurons have been clustered within each assigned birthtime window (P = primary, ES = early secondary, S = secondary) based on both upstream and downstream connectivity to hemilineages, descending neuron subclasses, motor neuron subclasses, and sensory neuron modalities. The annotation bar is coloured by the most common predicted neurotransmitter within each type.

**Figure 17 - figure supplement 6.**
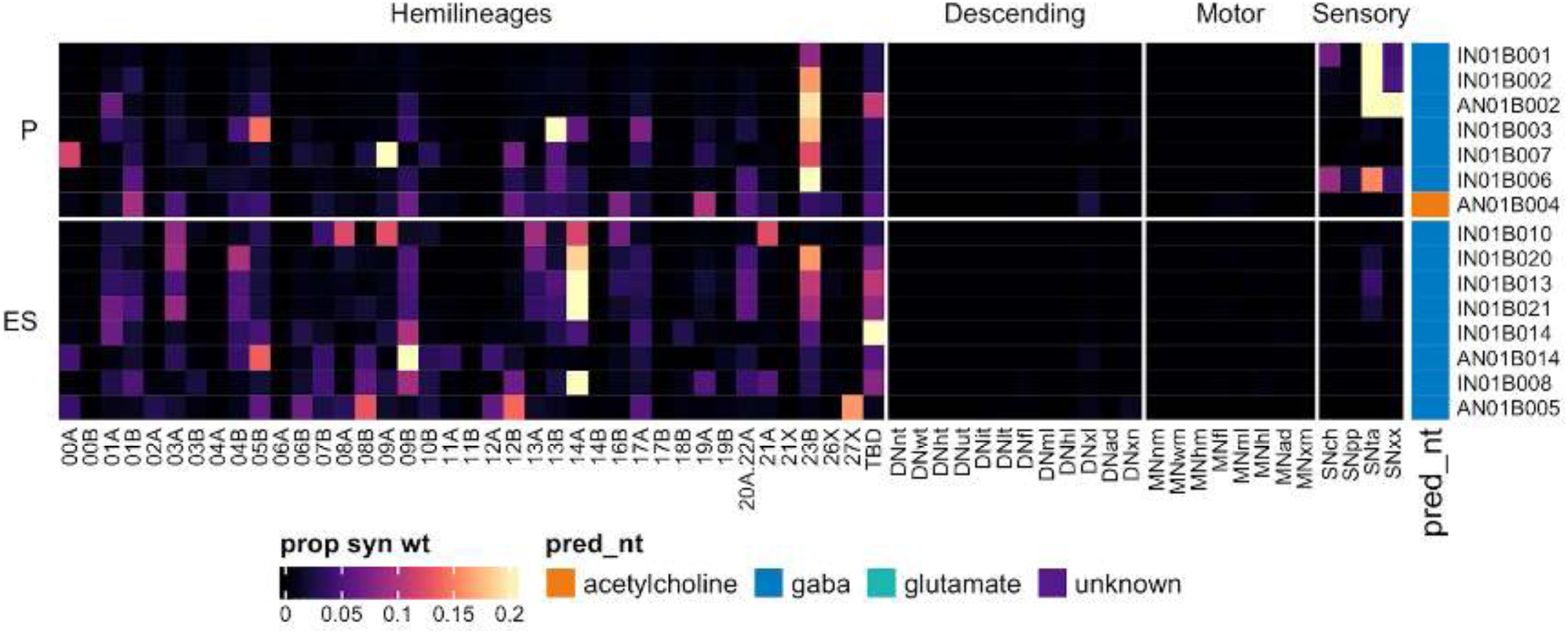
Connectivity to downstream partners by 01B primary and early secondary systematic types. Proportions of synaptic weight from systematic types to downstream partners, normalised by row. 01B neurons have been clustered within each assigned birthtime window (P = primary, ES = early secondary, S = secondary) based on both upstream and downstream connectivity to hemilineages, descending neuron subclasses, motor neuron subclasses, and sensory neuron modalities. The annotation bar is coloured by the most common predicted neurotransmitter within each type.

**Figure 17 - figure supplement 7a.**
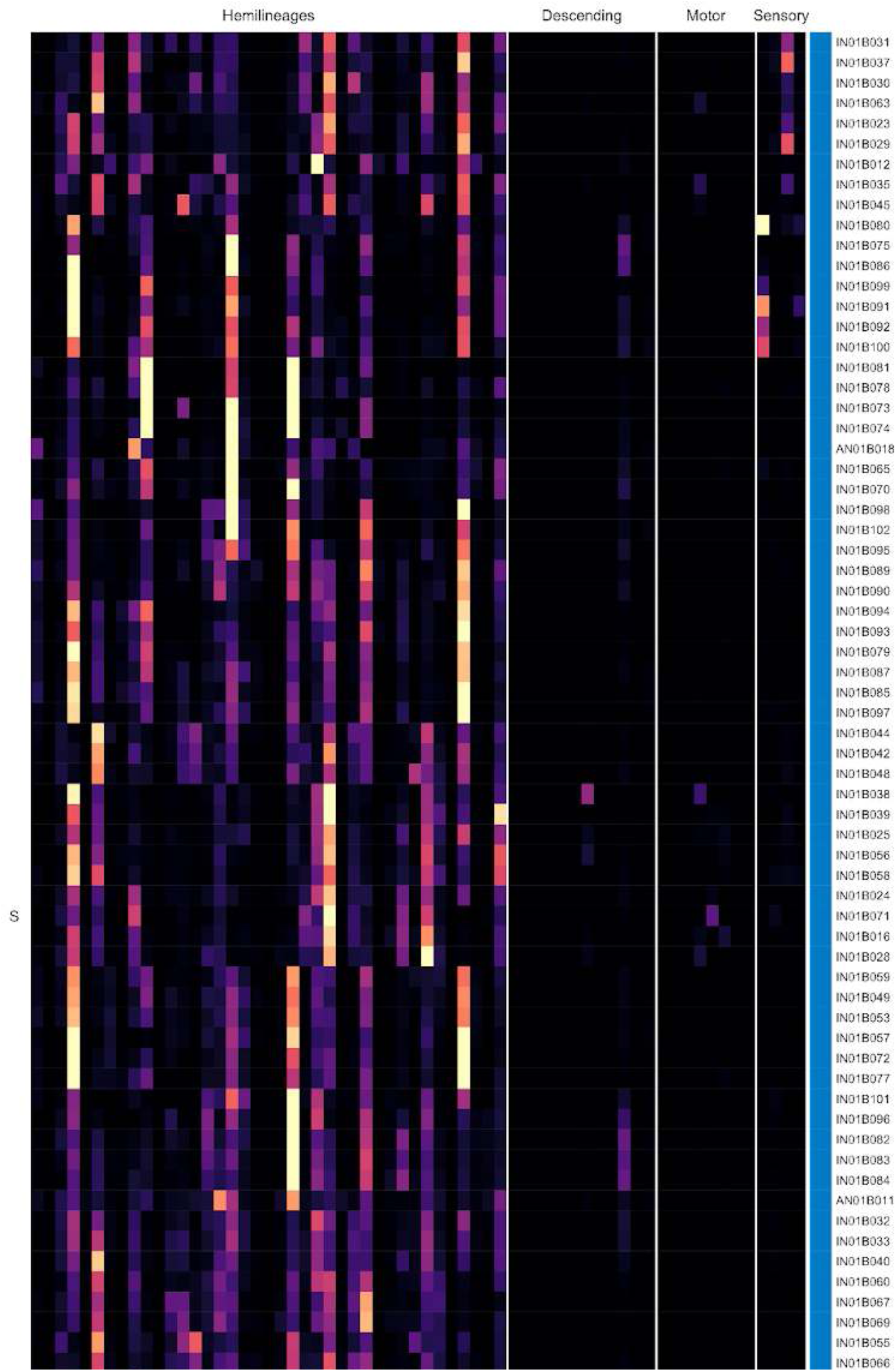
Connectivity to downstream partners by 01B secondary systematic types. Proportions of synaptic weight from systematic types to downstream partners, normalised by row. 01B neurons have been clustered within each assigned birthtime window (P = primary, ES = early secondary, S = secondary) based on both upstream and downstream connectivity to hemilineages, descending neuron subclasses, motor neuron subclasses, and sensory neuron modalities. The annotation bar is coloured by the most common predicted neurotransmitter for the neurons of each type.

**Figure 17 - figure supplement 7b.**
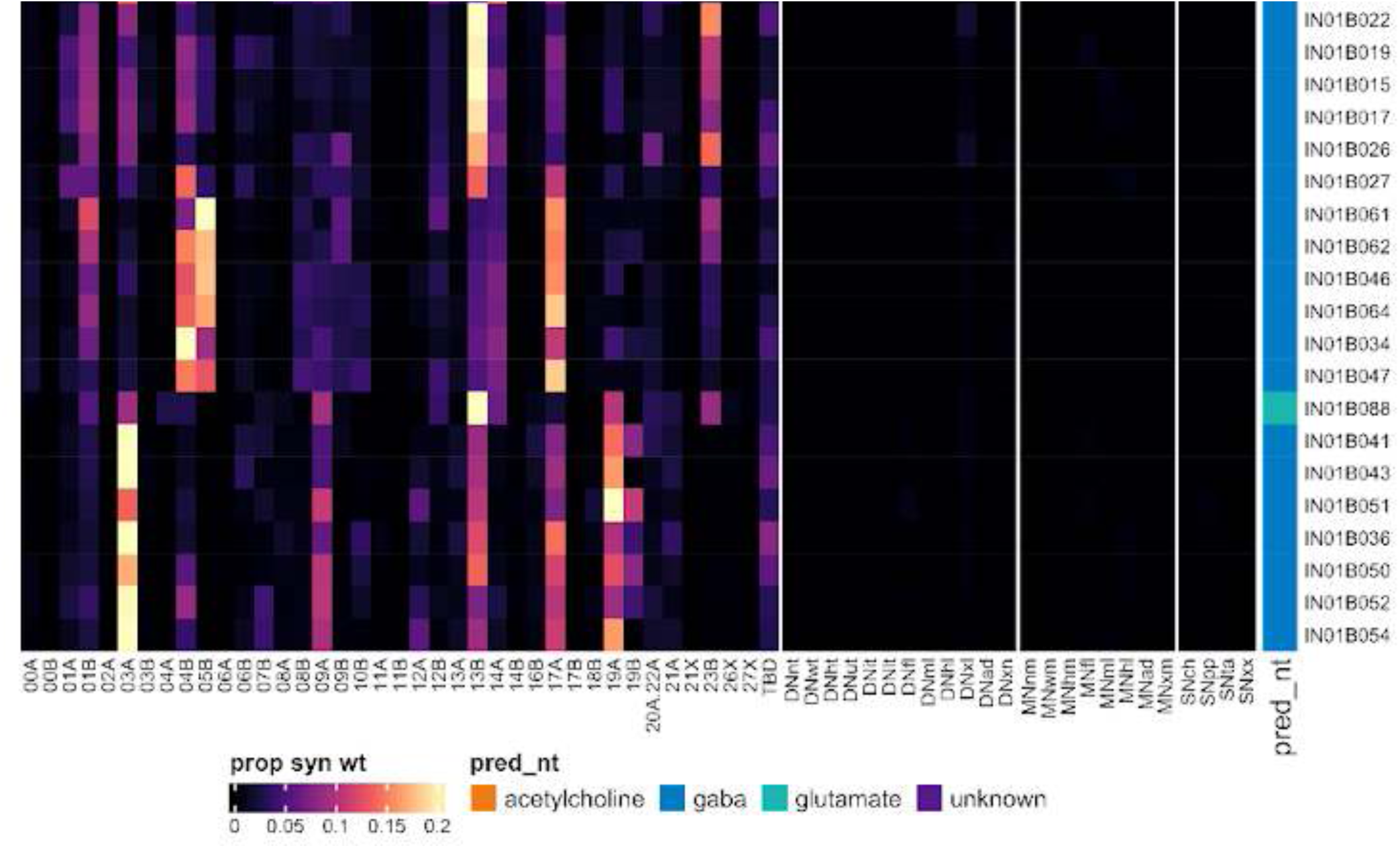
Connectivity to downstream partners by 01B secondary systematic types, continued. Proportions of synaptic weight from systematic types to downstream partners, normalised by row. 01B neurons have been clustered within each assigned birthtime window (P = primary, ES = early secondary, S = secondary) based on both upstream and downstream connectivity to hemilineages, descending neuron subclasses, motor neuron subclasses, and sensory neuron modalities. The annotation bar is coloured by the most common predicted neurotransmitter for the neurons of each type.

#### Hemilineage 02A

02A is the only surviving secondary hemilineage of anterior neuroblast NB2-1 (Truman et al., 2004), which produces 15 local interneurons in the embryo (Schmid et al., 1999). 02A neurons enter the neuropil at the anterior margin of each neuromere close to the midline and project dorsally into the tectulum, turning laterally to arborise in ipsilateral flight neuropil, with some having a small contralateral projection (Figure 18A) (Shepherd et al., 2019). In this data set, 02A neurons in T2 exhibit a striking developmental defect: the majority of the RHS neurons are shifted laterally while a few remain close to the midline but are shifted anteriorly; however, both populations exhibit RHS-appropriate morphology and connectivity (Figure 18 - figure supplement 1).

Whilst most thoracic 02A neurons were glutamatergic and restricted to the ipsilateral tectulum as expected (Lacin et al., 2019), we identified 9 types that ascend through the neck connective, two of which cross the midline to ascend on the contralateral side (AN02A005 and AN02A009) (Figure 18 - figure supplement 3). A few types innervate the tectulum bilaterally (e.g., IN02A008) (Figure 18 - figure supplement 3), and an unusual cholinergic type projects to the contralateral leg neuropil (IN02A015) (Figure 18C bottom). We also identified serial sets of intersegmental neurons that either converge in an anterior neuromere or ascend sequentially to the next anterior neuromere (e.g., IN02A003) (Figure 18C top). We have annotated all of these types as 02A, although it’s possible that some early born neurons are 02B, which might be expected to share the main projection but extend further laterally (Truman et al., 2010).

**Figure 18.**
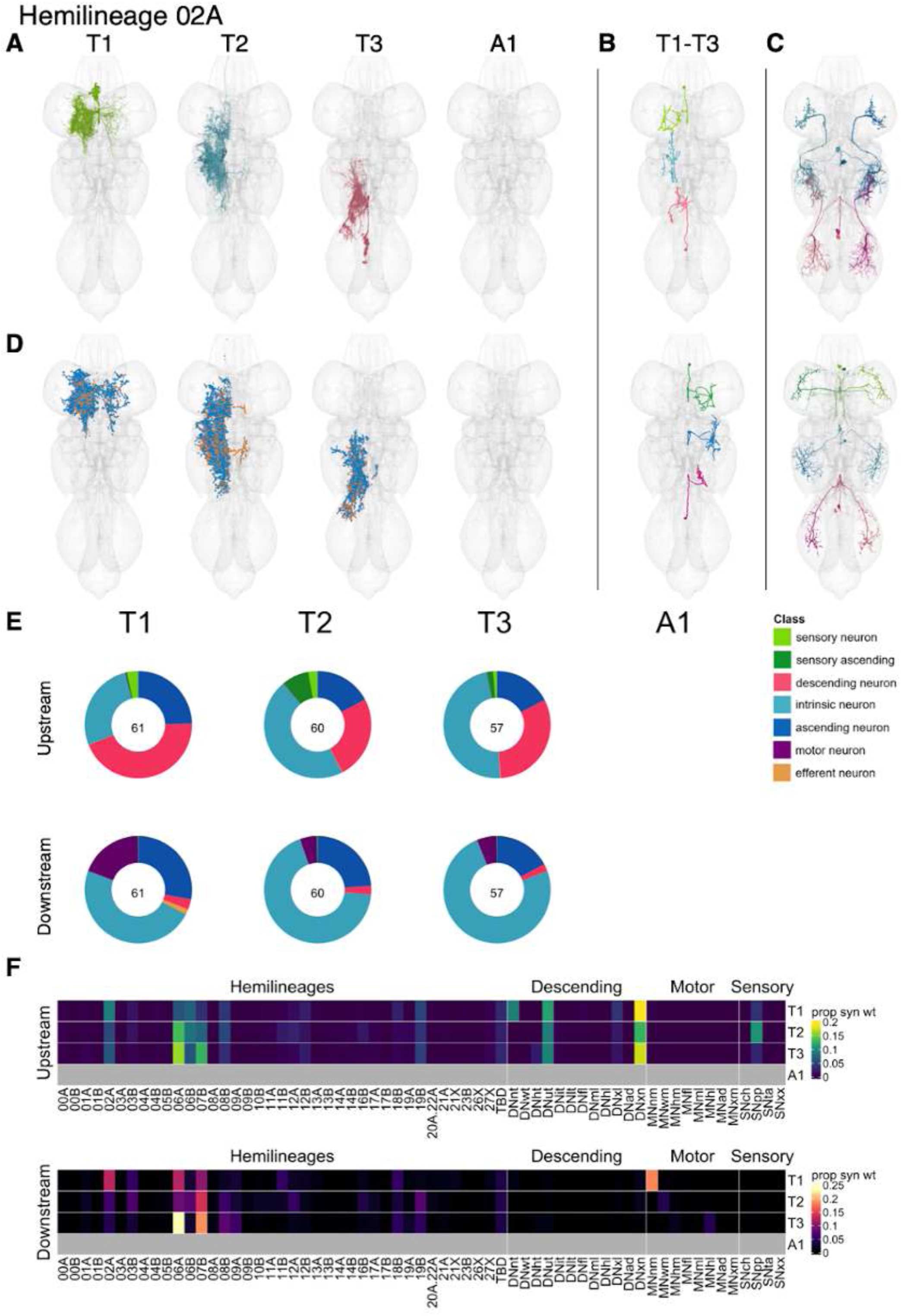
Hemilineage 02A. **A.** Meshes of all RHS secondary neurons plotted in neuromere-specific colours. **B.** “Representative” secondary neuron skeletons plotted in hemineuromere-specific colours. The skeleton with the top accumulated NBLAST score among all neurons from the hemilineage in a given hemineuromere was used. **C.** Neuron meshes of selected examples. Top: sequential serial set 10193. Bottom: cholinergic, contralateral independent leg serial set 13011. **D.** Predicted synapses of RHS secondary neurons. Blue: postsynapses; dark orange: presynapses. **E.** Proportions of connections from secondary neurons to upstream or downstream partners, normalised by neuromere and coloured by broad class. Numbers of query neurons appear in the centre. **F.** Proportions of synaptic weight from secondary neurons originating in each neuromere to upstream or downstream partners, normalised by row.

Comparable numbers of 02A secondary neurons survive in all three thoracic neuromeres (Figure 18E). They receive inputs mostly from descending neurons innervating the tectulum or multiple neuropils as well as from hemilineages 06A (T2 and T3) and 07B (T3) and proprioceptive sensory neurons, particularly in T2 (Figure 18F). They are predicted to be glutamatergic (Figure 8E) as expected (LACIN) and mainly target hemilineages 06A and 07B; in T1 they strongly target 02A neurons and neck motor neurons (Figure 18F). There are also 02A types targeting wing motor neurons (e.g., IN02A042) and hind leg motor neurons (e.g., IN02A035) (Figure 18 - figure supplement 8). Activation of 02A secondary neurons bilaterally drives high frequency wing flapping, occasionally interrupted by a bout of wing grooming (Harris et al., 2015).

**Figure 18 - figure supplement 1.**
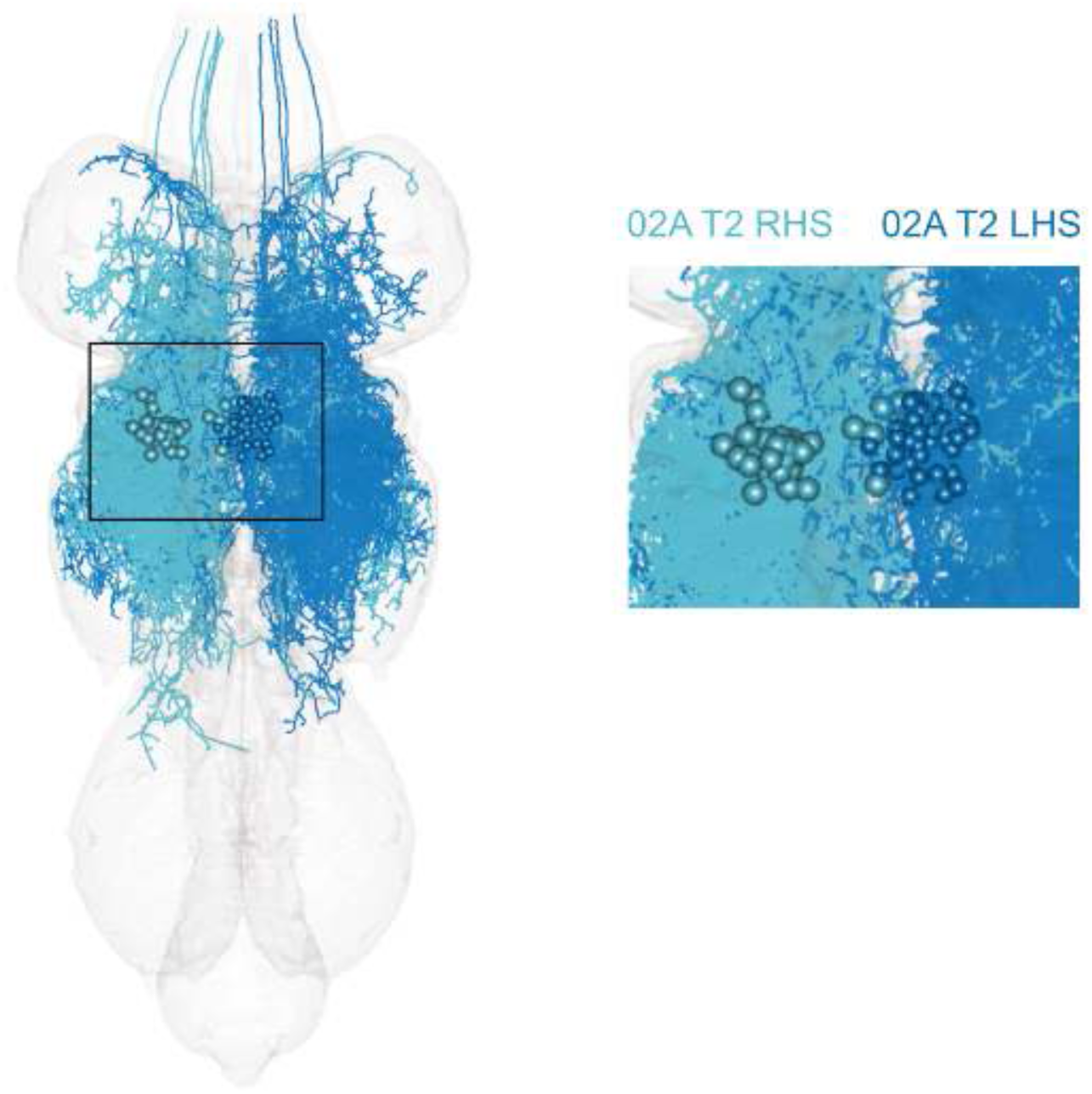
Developmental aberration in MANC of 02A neurons in RHS T2. Most 02A RHS somas (large, cyan) have been displaced laterally from their expected position adjacent to the midline (inset), but their gross dendritic and axonal morphologies appear normal when compared to their LHS counterparts (small, blue).

**Figure 18 - figure supplement 2.**
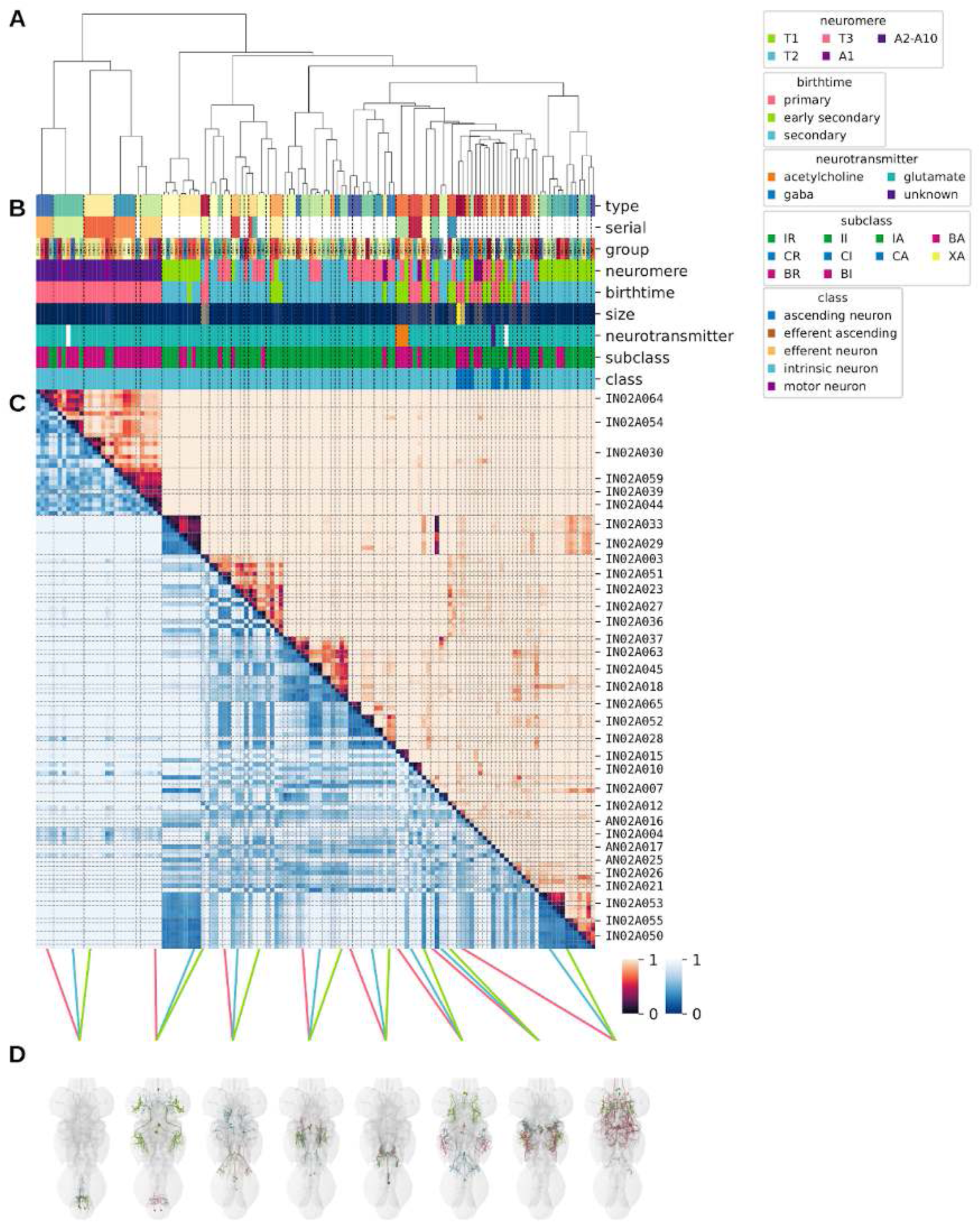
Systematic typing of hemilineage 02A. **A.** Hierarchical clustering dendrogram of hemilineage groups by laterally and serially aggregated connectivity cosine clustering. **B.** Categorical annotations of each hemilineage group, each column corresponding to the aligned leaf in A. Colours for type, serial set, and group are arbitrary for visualisation. Colours for neuromere, birthtime, neurotransmitter, subclass, and class are as in all other figures. **C.** Similarity distance heatmap for hemilineage. Cosine distance is in the upper triangle, while laterally symmetrised NBLAST distance is in the lower triangle. Systematic type names of some types are labelled. **D.** Morphologically representative groups from dendrogram subtrees. Each group, indicated by colour and line connecting to its column in B and C, is the most morphologically representative group (medoid of NBLAST distance) from a subtree of A. The subtrees (flat clusters) are equal height cuts of A determined to yield the number of groups per plot and plots in D.

**Figure 18 - figure supplement 3.**
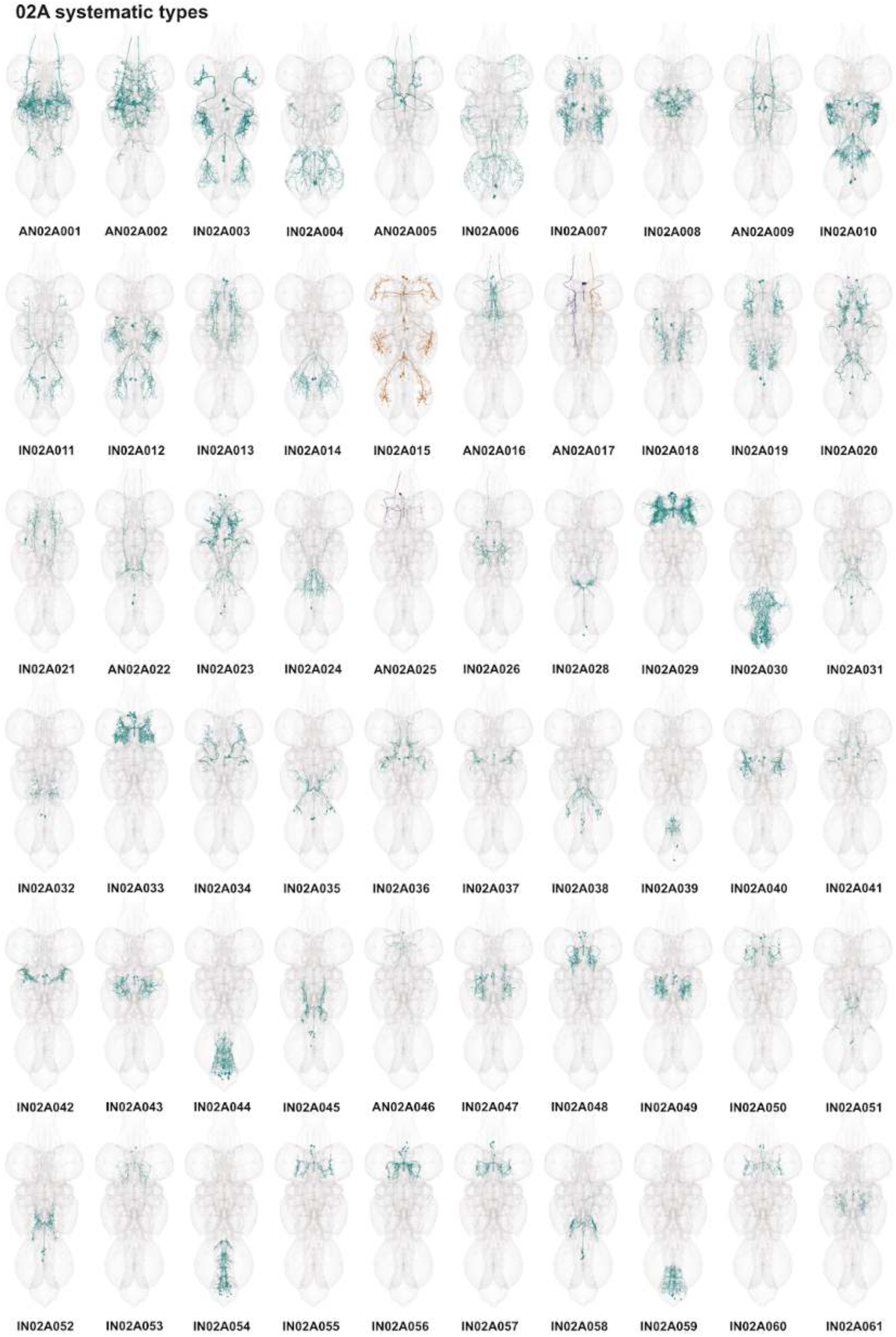
Systematic types of hemilineage 02A. Systematic types have been arranged in numerical order, with neurons of the same type that belong to distinct classes (e.g., intrinsic neuron vs ascending neuron) plotted separately but placed adjacent to each other. Individual neuron meshes have been coloured based on predicted neurotransmitter: dark orange = acetylcholine, blue = gaba, marine = glutamate, dark purple = unknown.

**Figure 18 - figure supplement 4.**
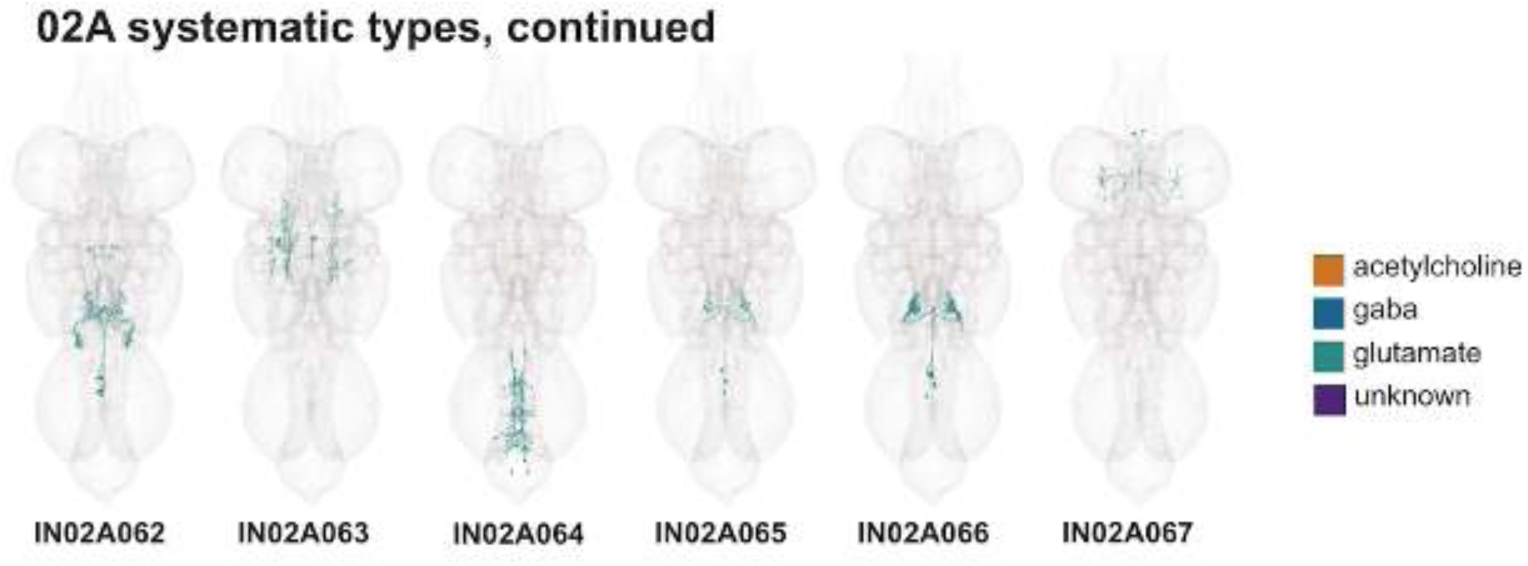
Systematic types of hemilineage 02A, continued. Systematic types have been arranged in numerical order, with neurons of the same type that belong to distinct classes (e.g., intrinsic neuron vs ascending neuron) plotted separately but placed adjacent to each other. Individual neuron meshes have been coloured based on predicted neurotransmitter: dark orange = acetylcholine, blue = gaba, marine = glutamate, dark purple = unknown.

**Figure 18 - figure supplement 5.**
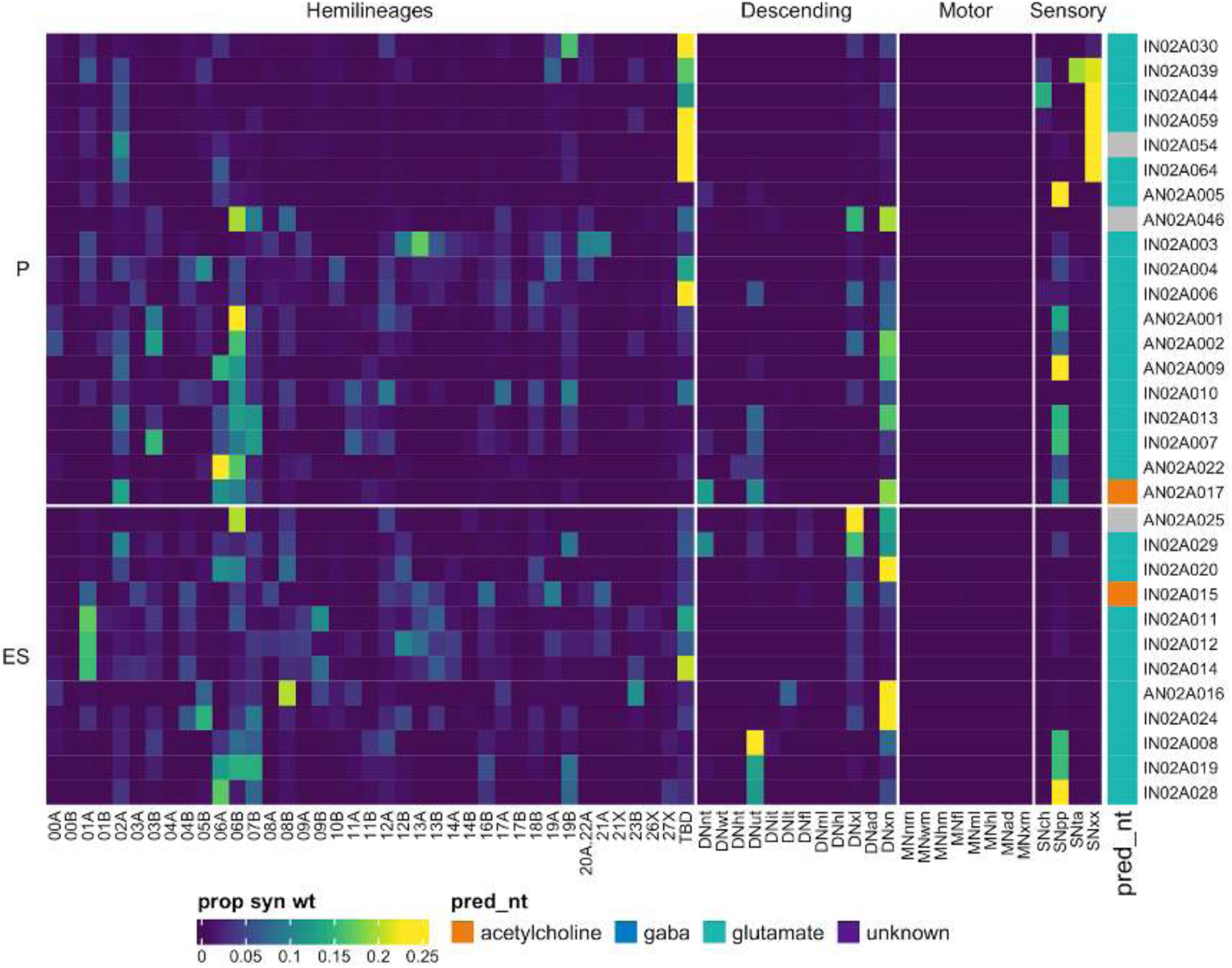
Connectivity to upstream partners by 02A primary and early secondary systematic types. Proportions of synaptic weight to systematic types from upstream partners, normalised by row. 02A neurons have been clustered within each assigned birthtime window (P = primary, ES = early secondary, S = secondary) based on both upstream and downstream connectivity to hemilineages, descending neuron subclasses, motor neuron subclasses, and sensory neuron modalities. Annotation bar is coloured by the most common predicted neurotransmitter for the neurons of each type.

**Figure 18 - figure supplement 6.**
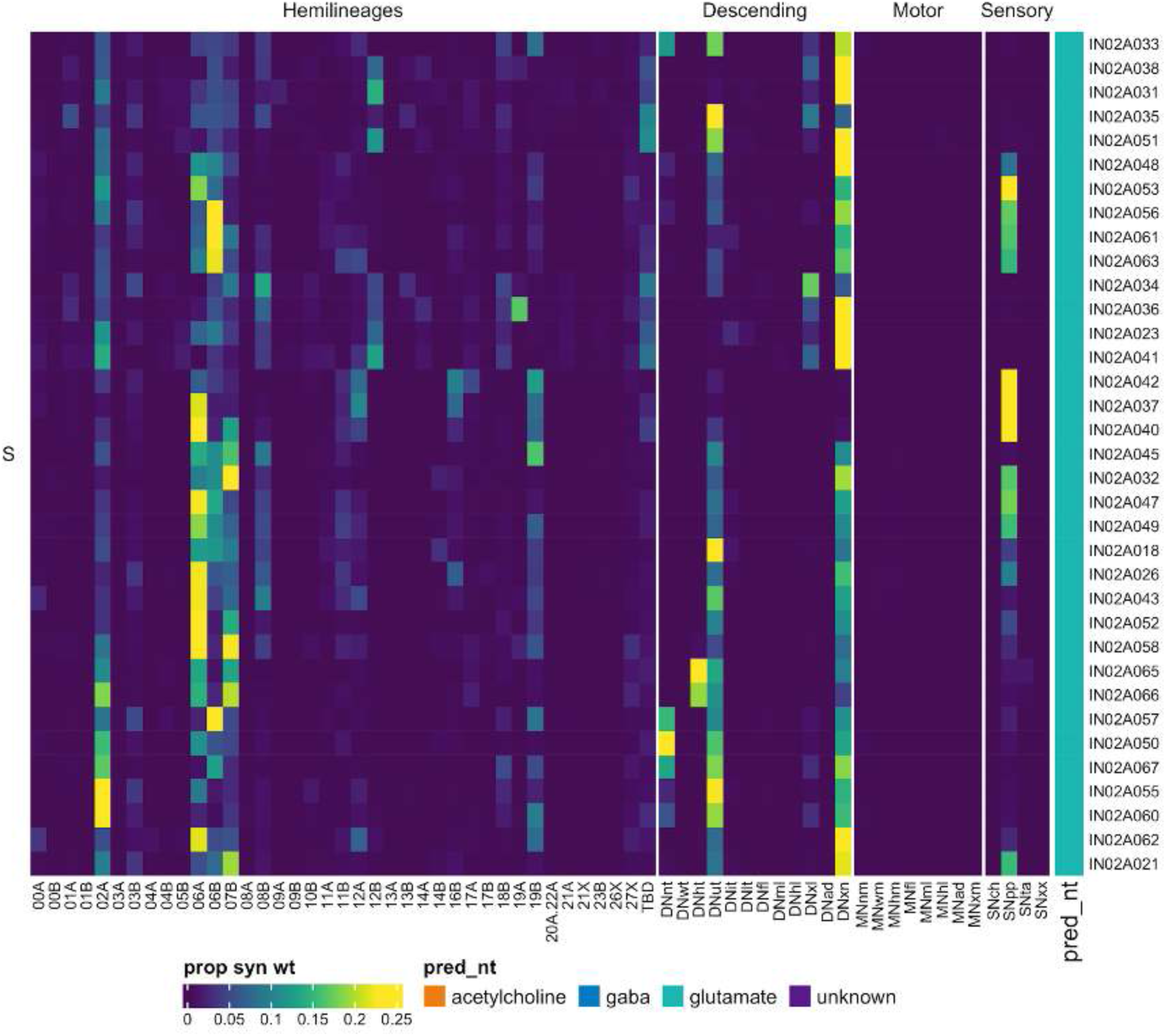
Connectivity to upstream partners by 02A secondary systematic types. Proportions of synaptic weight to systematic types from upstream partners, normalised by row. 02A neurons have been clustered within each assigned birthtime window (P = primary, ES = early secondary, S = secondary) based on both upstream and downstream connectivity to hemilineages, descending neuron subclasses, motor neuron subclasses, and sensory neuron modalities. The annotation bar is coloured by the most common predicted neurotransmitter within each type.

**Figure 18 - figure supplement 7.**
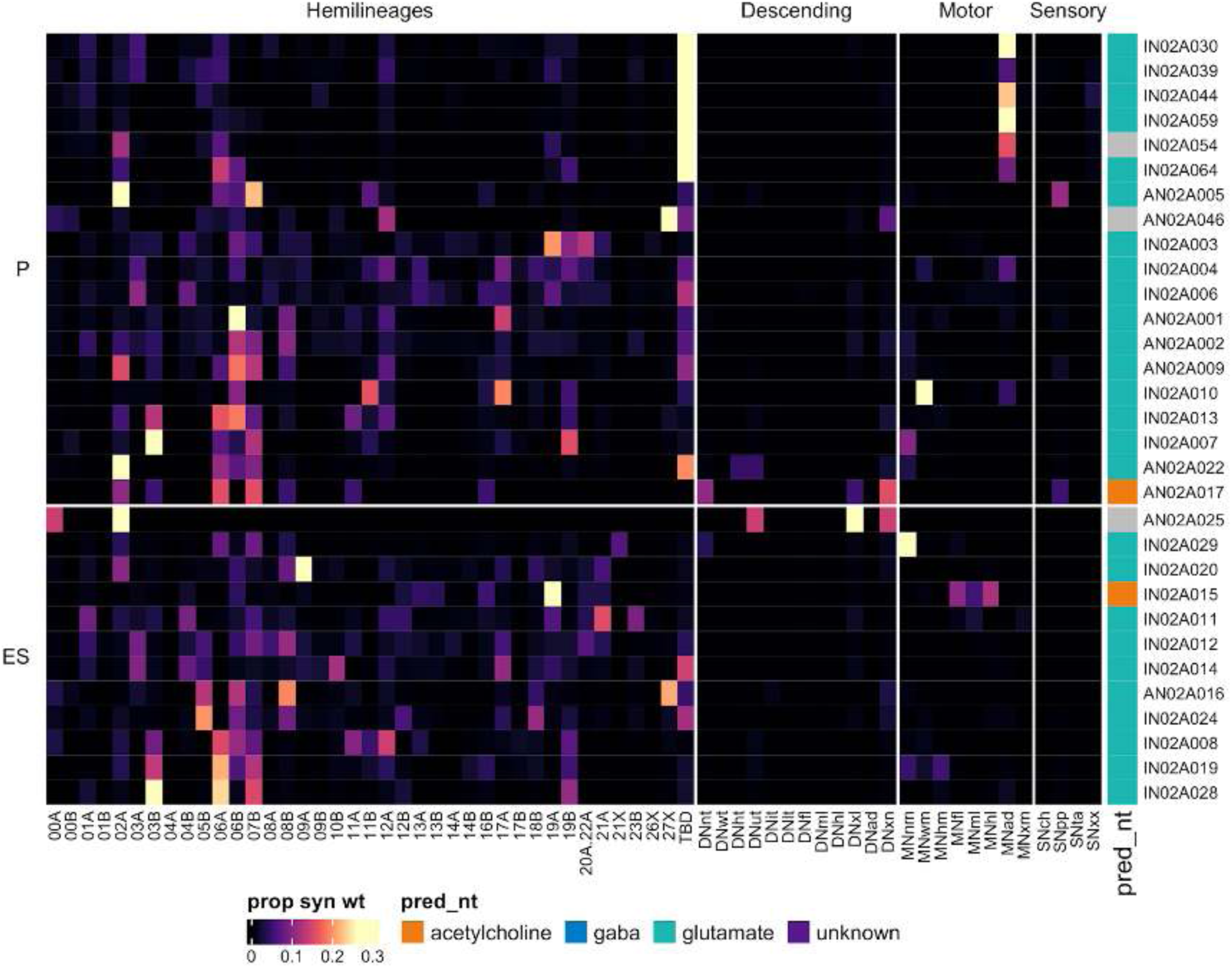
Connectivity to downstream partners by 02A primary and early secondary systematic types. Proportions of synaptic weight from systematic types to downstream partners, normalised by row. 02A neurons have been clustered within each assigned birthtime window (P = primary, ES = early secondary, S = secondary) based on both upstream and downstream connectivity to hemilineages, descending neuron subclasses, motor neuron subclasses, and sensory neuron modalities. The annotation bar is coloured by the most common predicted neurotransmitter within each type.

**Figure 18 - figure supplement 8.**
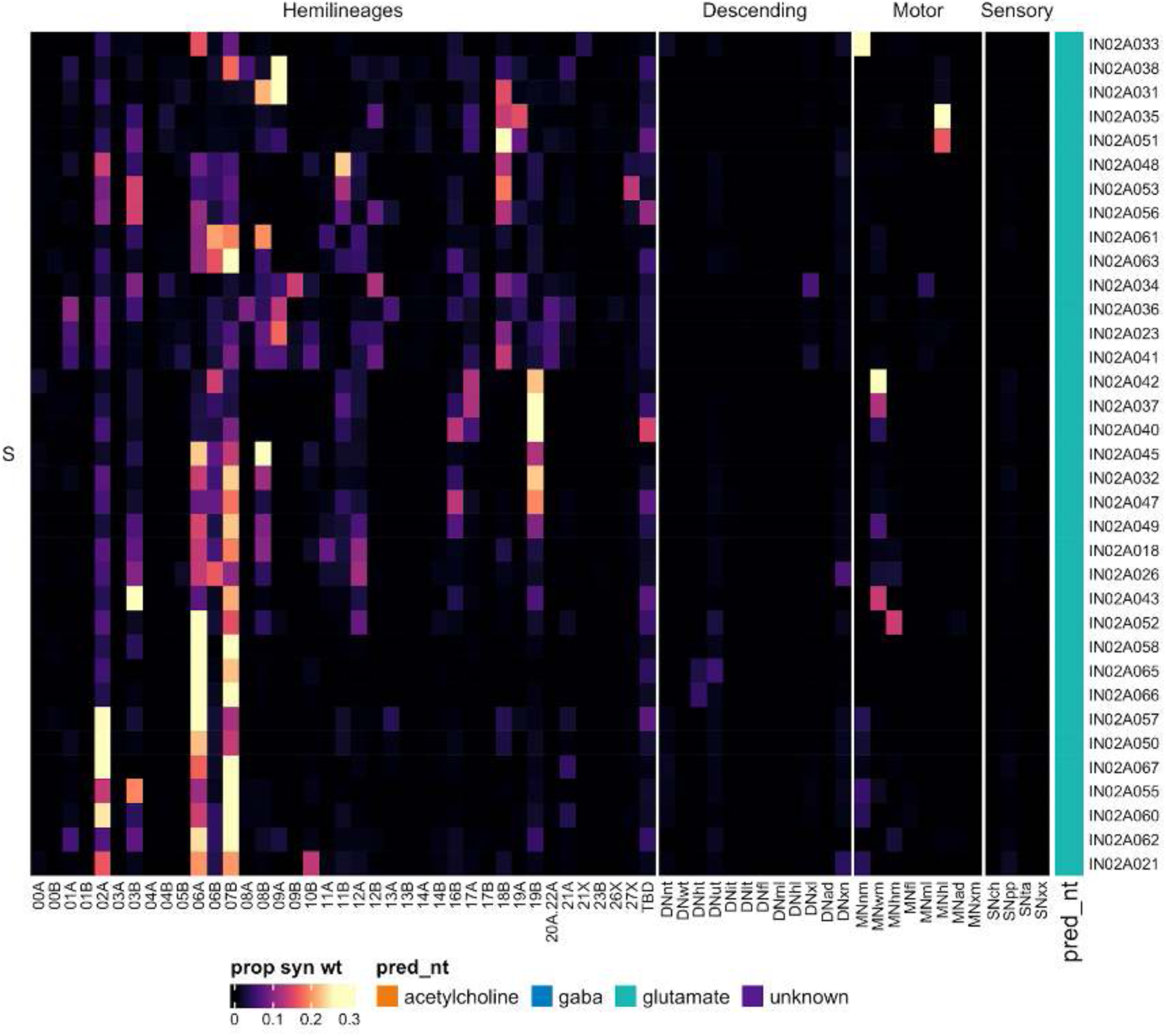
Connectivity to downstream partners by 02A secondary systematic types. Proportions of synaptic weight from systematic types to downstream partners, normalised by row. 02A neurons have been clustered within each assigned birthtime window (P = primary, ES = early secondary, S = secondary) based on both upstream and downstream connectivity to hemilineages, descending neuron subclasses, motor neuron subclasses, and sensory neuron modalities. The annotation bar is coloured by the most common predicted neurotransmitter for the neurons of each type.

#### Hemilineage 03A

Hemilineages 03A and 03B derive from posterior neuroblast NB7-1 (Truman et al., 2004), which generates 4-6 “U” MNs and ∼30 local interneurons in the embryo (Schmid et al., 1999). 03A is one of several posterior cholinergic secondary hemilineages that innervate the ipsilateral leg neuropil (Lacin et al., 2019). It was distinguished from 20A and 22A by two key features. First, 03A somas are interspersed with 03B somas and their primary neurites run in the same tract before 03B neurons diverge to project to dorsal neuropil. Second, 03A neurites wrap around the surface of the ipsilateral leg neuropil, with minimal innervation of the interior (Shepherd et al., 2019).

03A neurons enter the neuropil at the posterior margin of the neuropil, projecting ventrally to arborise extensively in the ipsilateral leg neuropil (Figure 19A), with a subset in T1 and T2 projecting posteriorly into the adjacent neuromere (e.g., subcluster 15430) (Figure 19C bottom). We also identified 03A neurons with axons ascending or descending to the adjacent neuromere (e.g., IN03A001) (Figure 19C top). There are two early born 03A types in T1 that ascend via the neck connective (AN03A002 and AN03A008) (Figure 19 - figure supplement 2); the first is an unusually strong target of descending neurons (Figure 19 - figure supplement 3).

**Figure 19.**
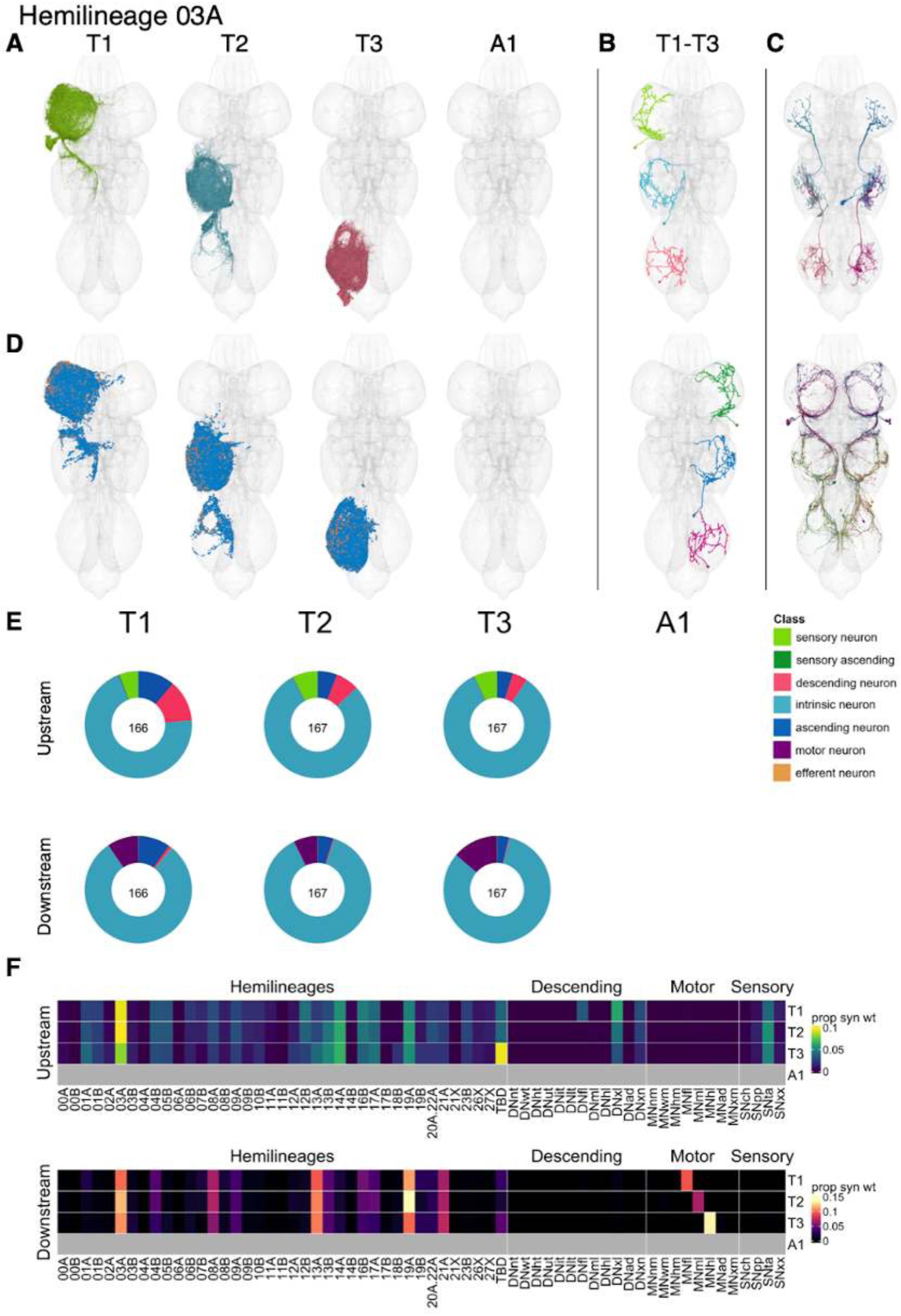
Hemilineage 03A. **A.** Meshes of all RHS secondary neurons plotted in neuromere-specific colours. **B.** “Representative” secondary neuron skeletons plotted in hemineuromere-specific colours. The skeleton with the top accumulated NBLAST score among all neurons from the hemilineage in a given hemineuromere was used. **C.** Neuron meshes of selected examples. Top: sequential serial set 10262. Bottom: sequential subcluster 15430. **D.** Predicted synapses of RHS secondary neurons. Blue: postsynapses; dark orange: presynapses. **E.** Proportions of connections from secondary neurons to upstream or downstream partners, normalised by neuromere and coloured by broad class. Numbers of query neurons appear in the centre. **F.** Proportions of synaptic weight from secondary neurons originating in each neuromere to upstream or downstream partners, normalised by row.

Comparable numbers of 03A secondary neurons survive in all three thoracic neuromeres (Figure 19E). Secondary 03A neurons receive inputs mostly from leg hemilineages 03A, 14A, 16B, 17A, and 19A and from descending neurons, particularly those innervating multiple leg neuropils (Figure 19F). A subset of secondary 03A types receive significant input from tactile sensory neurons (e.g., IN03A092-097) (Figure 19 - figure supplement 3). Secondary 03A neurons are predicted to be cholinergic (Figure 8E) and output primarily onto leg hemilineages 03A, 08A, 13A, 19A, and 21A and onto leg motor neurons (Figure 19F). We also identified two early born types that strongly target neck or wing motor neurons (AN03A002 and IN03A011, respectively) (Figure 19 - figure supplement 7).

**Figure 19 - figure supplement 1.**
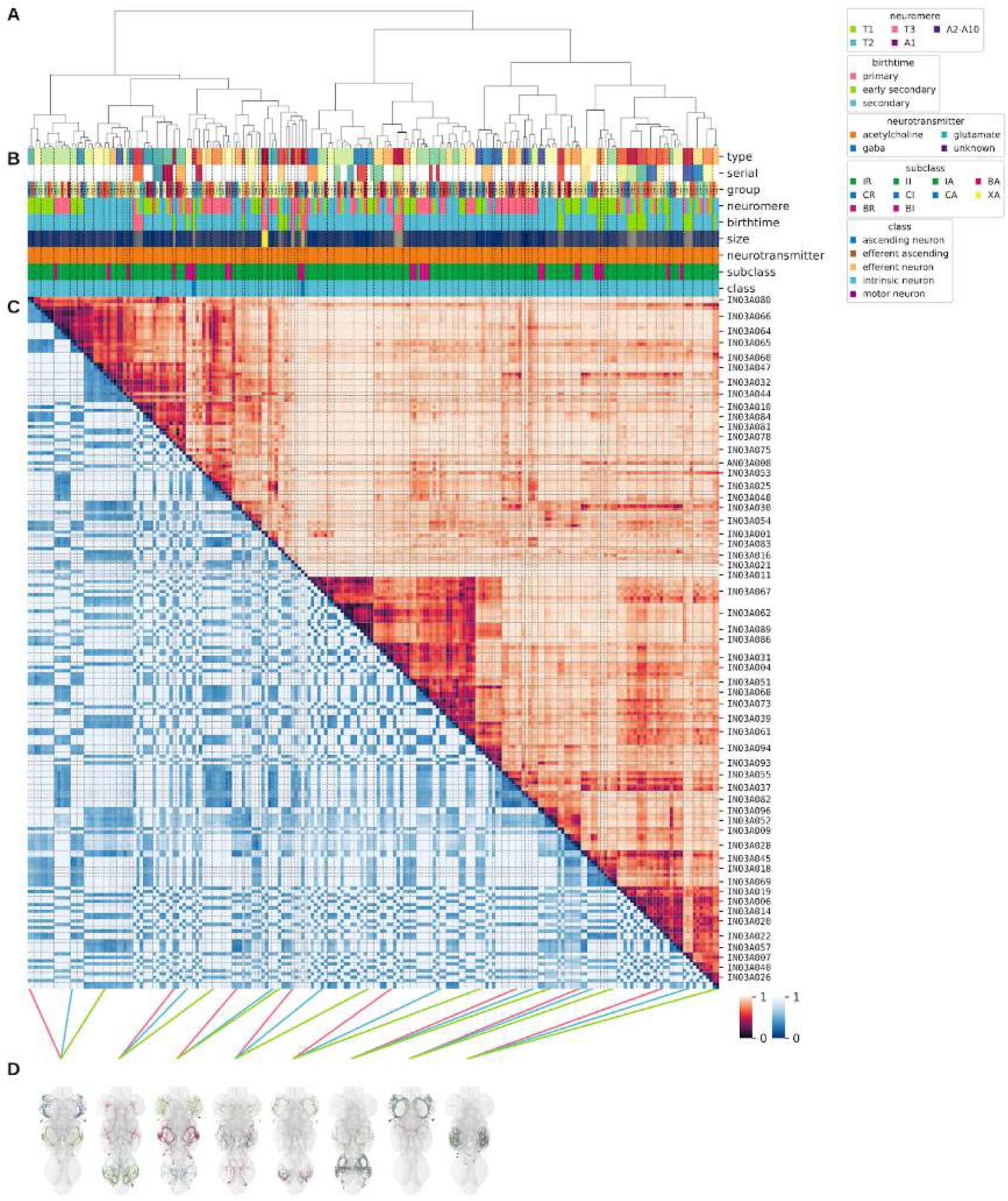
Systematic typing of hemilineage 03A. **A.** Hierarchical clustering dendrogram of hemilineage groups by laterally and serially aggregated connectivity cosine clustering. **B.** Categorical annotations of each hemilineage group, each column corresponding to the aligned leaf in A. Colours for type, serial set, and group are arbitrary for visualisation. Colours for neuromere, birthtime, neurotransmitter, subclass, and class are as in all other figures. **C.** Similarity distance heatmap for hemilineage. Cosine distance is in the upper triangle, while laterally symmetrised NBLAST distance is in the lower triangle. Systematic type names of some types are labelled. **D.** Morphologically representative groups from dendrogram subtrees. Each group, indicated by colour and line connecting to its column in B and C, is the most morphologically representative group (medoid of NBLAST distance) from a subtree of A. The subtrees (flat clusters) are equal height cuts of A determined to yield the number of groups per plot and plots in D.

**Figure 19 - figure supplement 2.**
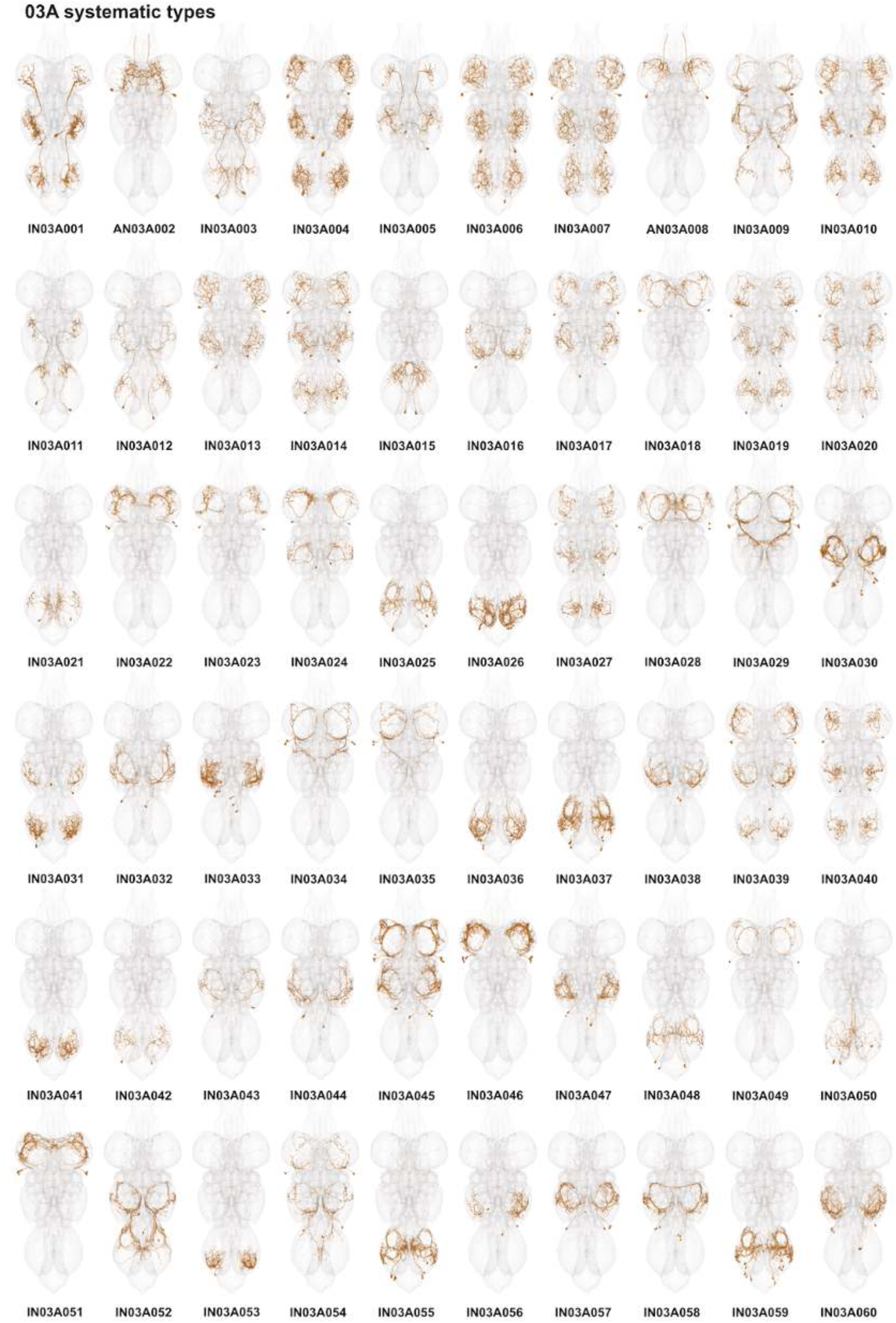
Systematic types of hemilineage 03A. Systematic types have been arranged in numerical order, with neurons of the same type that belong to distinct classes (e.g., intrinsic neuron vs ascending neuron) plotted separately but placed adjacent to each other. Individual neuron meshes have been coloured based on predicted neurotransmitter: dark orange = acetylcholine, blue = gaba, marine = glutamate, dark purple = unknown.

**Figure 19 - figure supplement 3.**
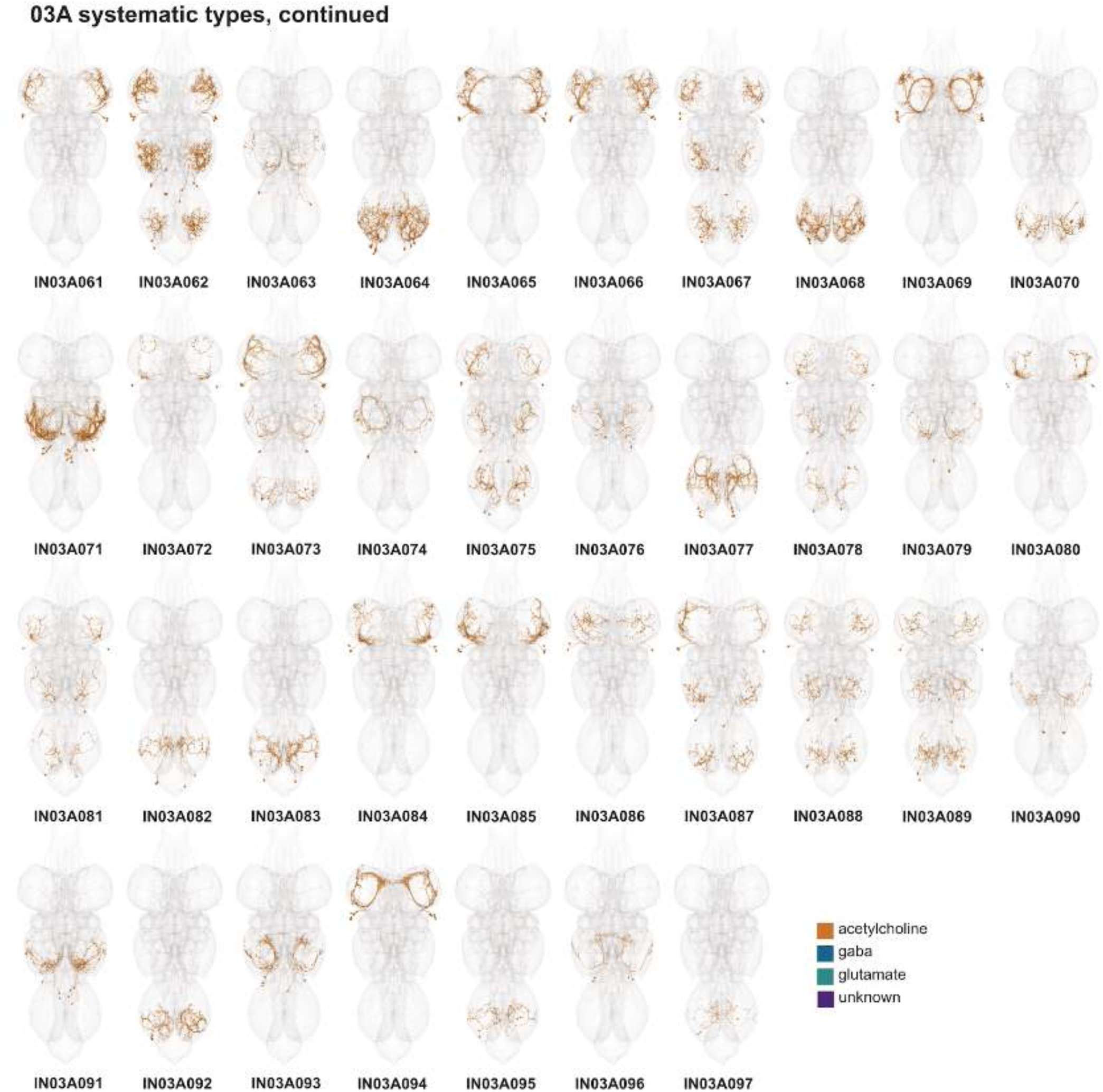
Systematic types of hemilineage 03A, continued. Systematic types have been arranged in numerical order, with neurons of the same type that belong to distinct classes (e.g., intrinsic neuron vs ascending neuron) plotted separately but placed adjacent to each other. Individual neuron meshes have been coloured based on predicted neurotransmitter: dark orange = acetylcholine, blue = gaba, marine = glutamate, dark purple = unknown.

**Figure 19 - figure supplement 4.**
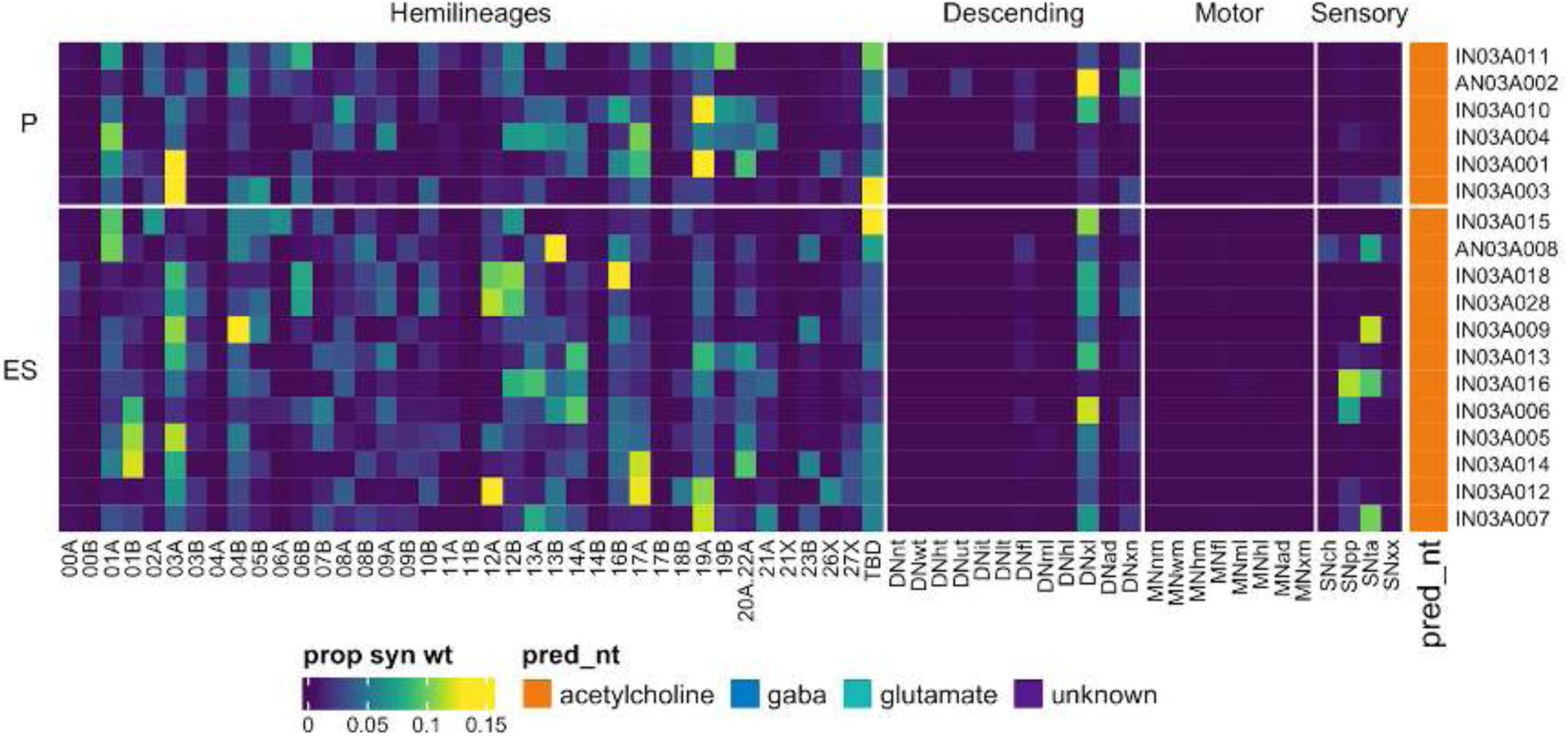
Connectivity to upstream partners by 03A primary and early secondary systematic types. Proportions of synaptic weight to systematic types from upstream partners, normalised by row. 03A neurons have been clustered within each assigned birthtime window (P = primary, ES = early secondary, S = secondary) based on both upstream and downstream connectivity to hemilineages, descending neuron subclasses, motor neuron subclasses, and sensory neuron modalities. Annotation bar is coloured by the most common predicted neurotransmitter for the neurons of each type.

**Figure 19 - figure supplement 5.**
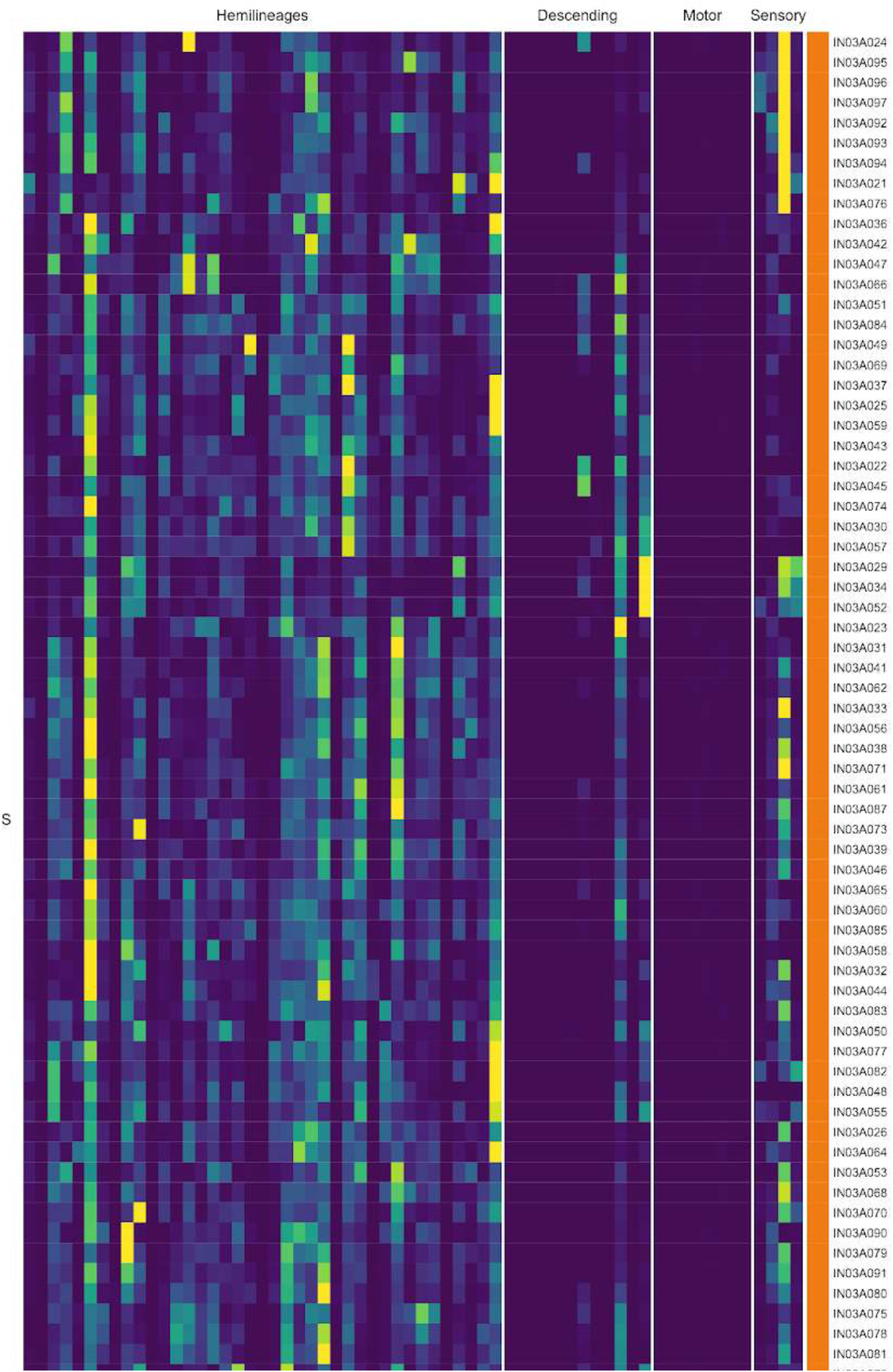
Connectivity to upstream partners by 03A secondary systematic types. Proportions of synaptic weight to systematic types from upstream partners, normalised by row. 03A neurons have been clustered within each assigned birthtime window (P = primary, ES = early secondary, S = secondary) based on both upstream and downstream connectivity to hemilineages, descending neuron subclasses, motor neuron subclasses, and sensory neuron modalities. The annotation bar is coloured by the most common predicted neurotransmitter within each type.

**Figure 19 - figure supplement 6.**
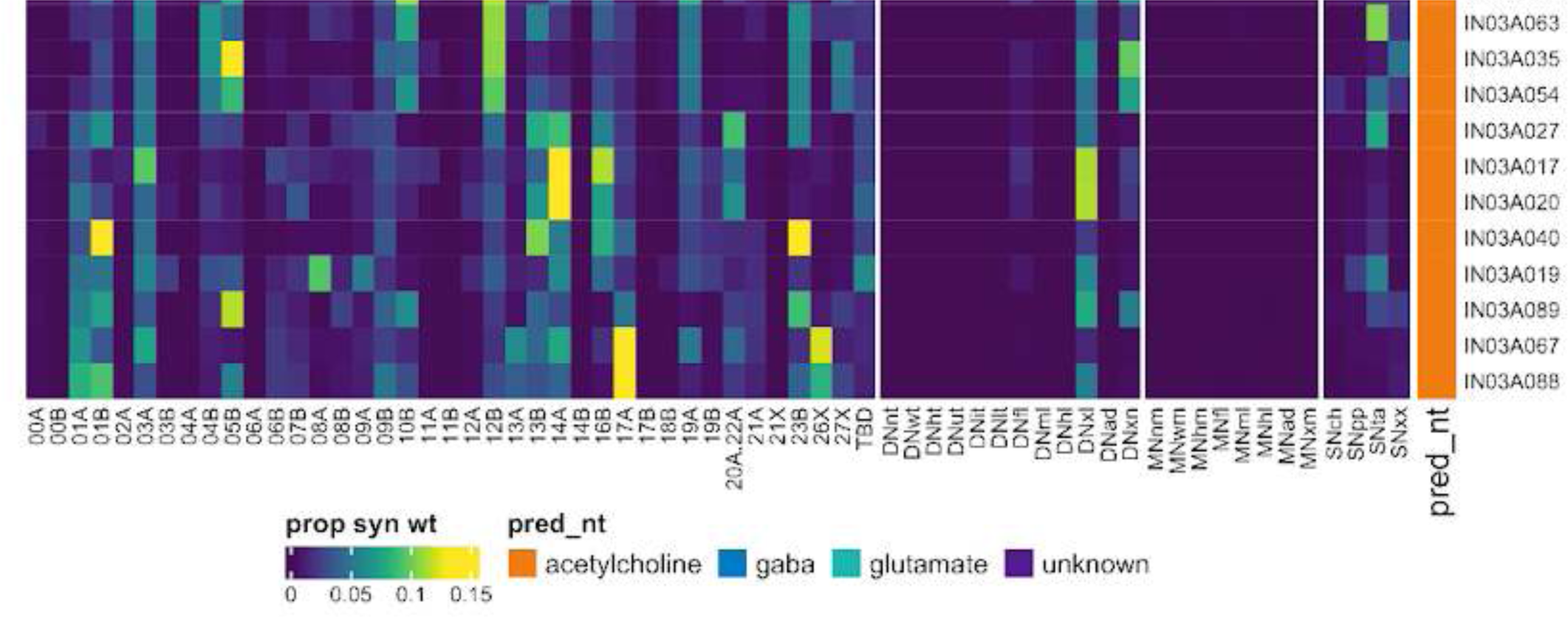
Connectivity to upstream partners by 03A secondary systematic types, continued. Proportions of synaptic weight to systematic types from upstream partners, normalised by row. 03A neurons have been clustered within each assigned birthtime window (P = primary, ES = early secondary, S = secondary) based on both upstream and downstream connectivity to hemilineages, descending neuron subclasses, motor neuron subclasses, and sensory neuron modalities. The annotation bar is coloured by the most common predicted neurotransmitter within each type.

**Figure 19 - figure supplement 7.**
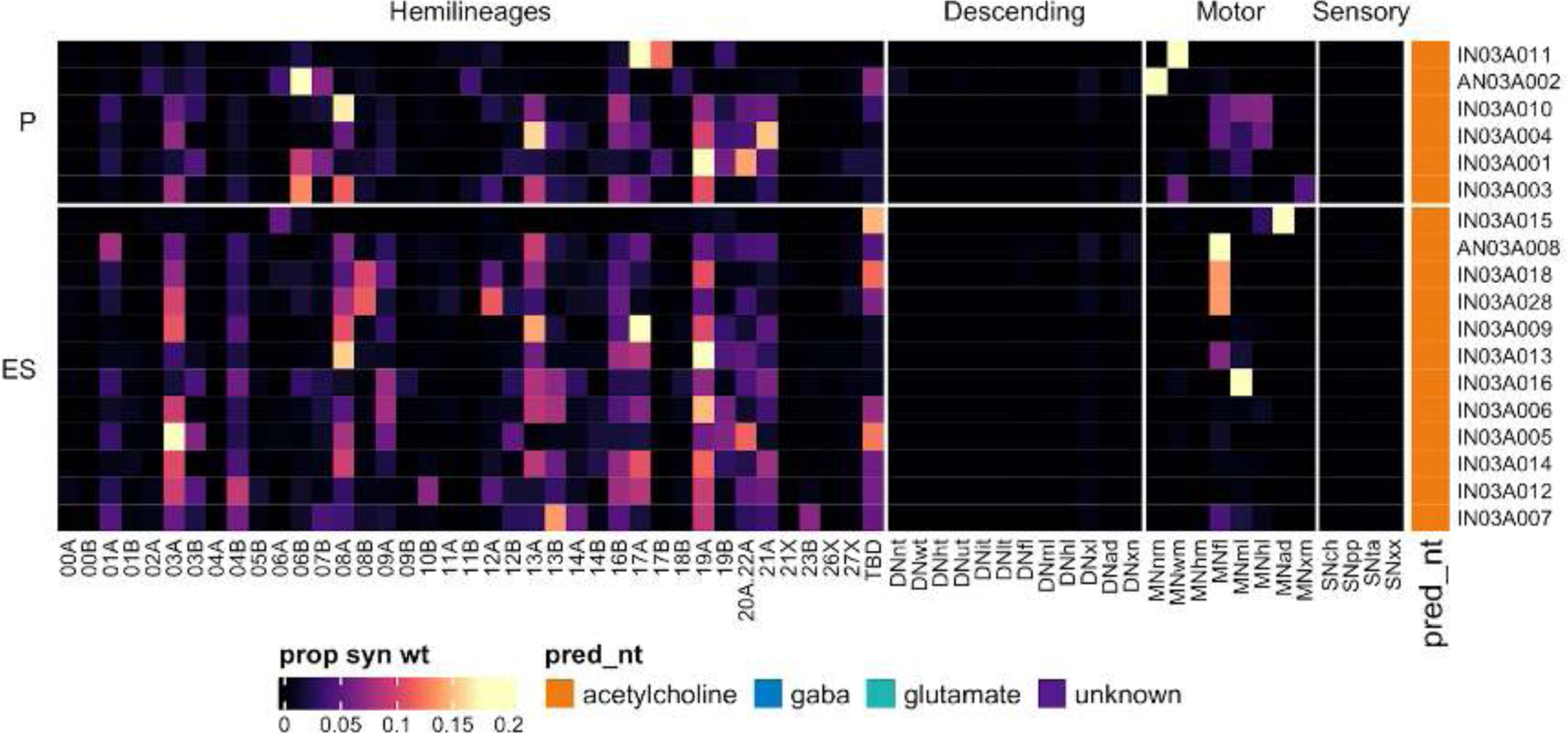
Connectivity to downstream partners by 03A primary and early secondary systematic types. Proportions of synaptic weight from systematic types to downstream partners, normalised by row. 03A neurons have been clustered within each assigned birthtime window (P = primary, ES = early secondary, S = secondary) based on both upstream and downstream connectivity to hemilineages, descending neuron subclasses, motor neuron subclasses, and sensory neuron modalities. The annotation bar is coloured by the most common predicted neurotransmitter within each type.

**Figure 19 - figure supplement 8.**
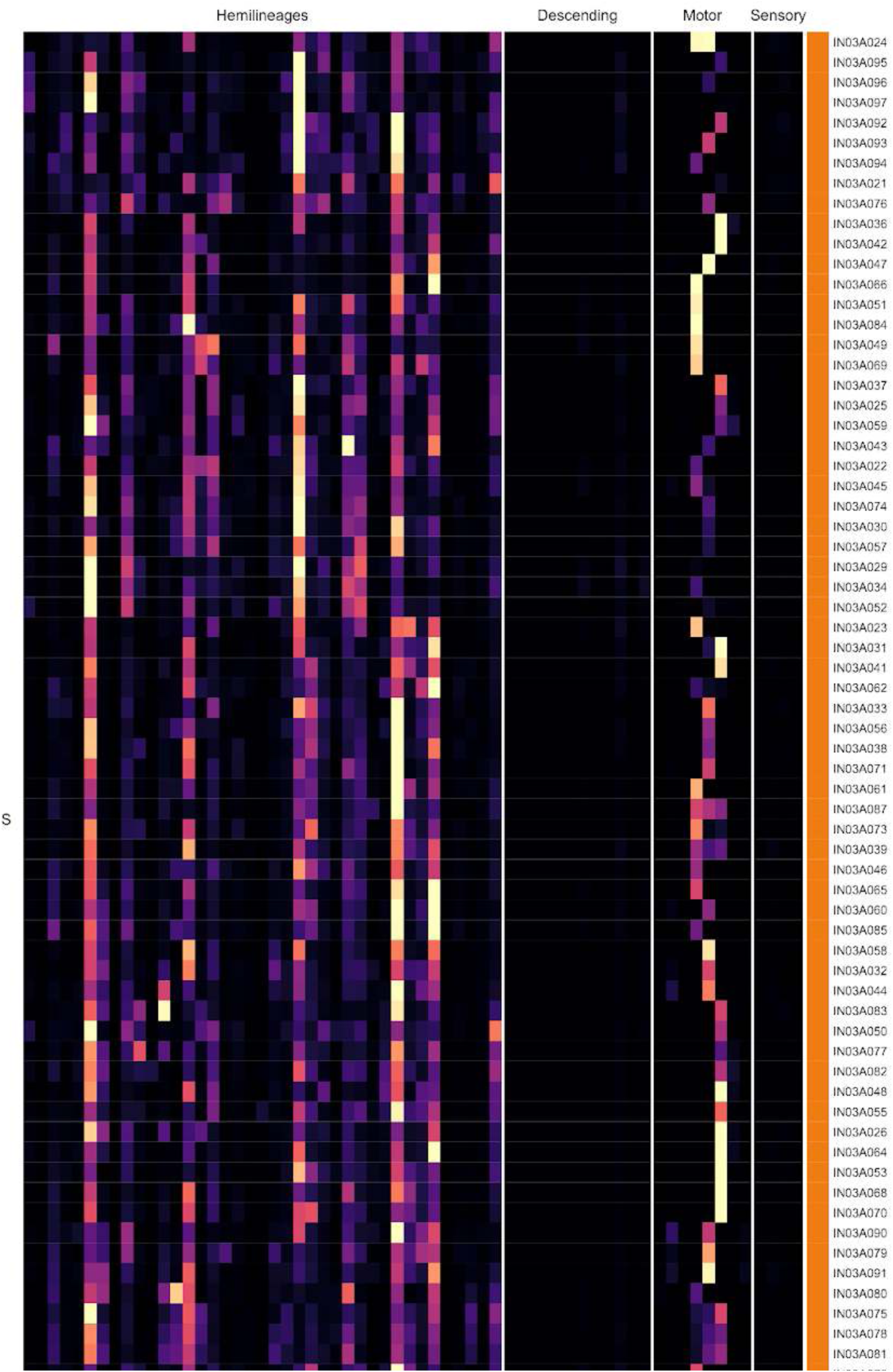
Connectivity to downstream partners by 03A secondary systematic types. Proportions of synaptic weight from systematic types to downstream partners, normalised by row. 03A neurons have been clustered within each assigned birthtime window (P = primary, ES = early secondary, S = secondary) based on both upstream and downstream connectivity to hemilineages, descending neuron subclasses, motor neuron subclasses, and sensory neuron modalities. The annotation bar is coloured by the most common predicted neurotransmitter for the neurons of each type.

**Figure 19 - figure supplement 9.**
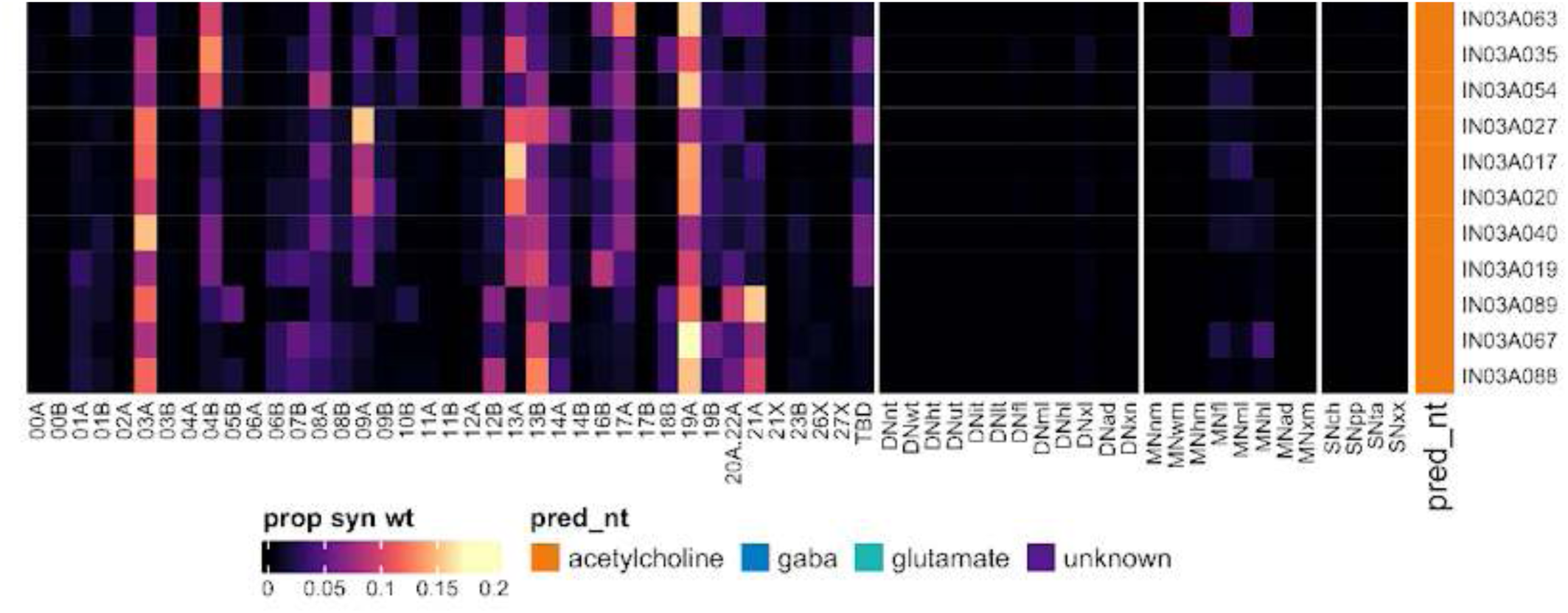
Connectivity to downstream partners by 03A secondary systematic types, continued. Proportions of synaptic weight from systematic types to downstream partners, normalised by row. 03A neurons have been clustered within each assigned birthtime window (P = primary, ES = early secondary, S = secondary) based on both upstream and downstream connectivity to hemilineages, descending neuron subclasses, motor neuron subclasses, and sensory neuron modalities. The annotation bar is coloured by the most common predicted neurotransmitter for the neurons of each type.

#### Hemilineage 03B

03B neurons derive from posterior neuroblast NB7-1 (Truman et al., 2004), which generates 4-6 “U” MNs and ∼30 local interneurons in the embryo (Schmid et al., 1999). Four “U” MNs survive metamorphosis to become the MN1-MN4 flight neurons in the adult (e.g., Figure 20C top) (Consoulas et al., 2002; Jacobs et al., 2000; Lacin et al., 2020). Neurons of hemilineage 03B enter the neuropil at the posterior margin of the neuropil with 03A and project dorsally to the tectulum, with arborisation from T1 and T2 tending to converge onto the ipsilateral T2 neuropil, T3 arborisations largely restricted to the haltere tectulum, and the A1 neurons projecting anteriorly to arborise in the ipsilateral wing tectulum in T2 (Figure 20A). We found that 03B primary neurites overlapped with those of 12A in T1, making it more difficult to distinguish them; neurotransmitter prediction was used as a defining character in these cases. Also, a surprising number of 03B neurons in T1 and T2 were predicted to be glutamatergic, some but not all of which featured thick, simple axons with an unusually low density of presynapses, suggesting they may be electrically coupled to partners via gap junctions (Figures 6B and 20C bottom).

**Figure 20.**
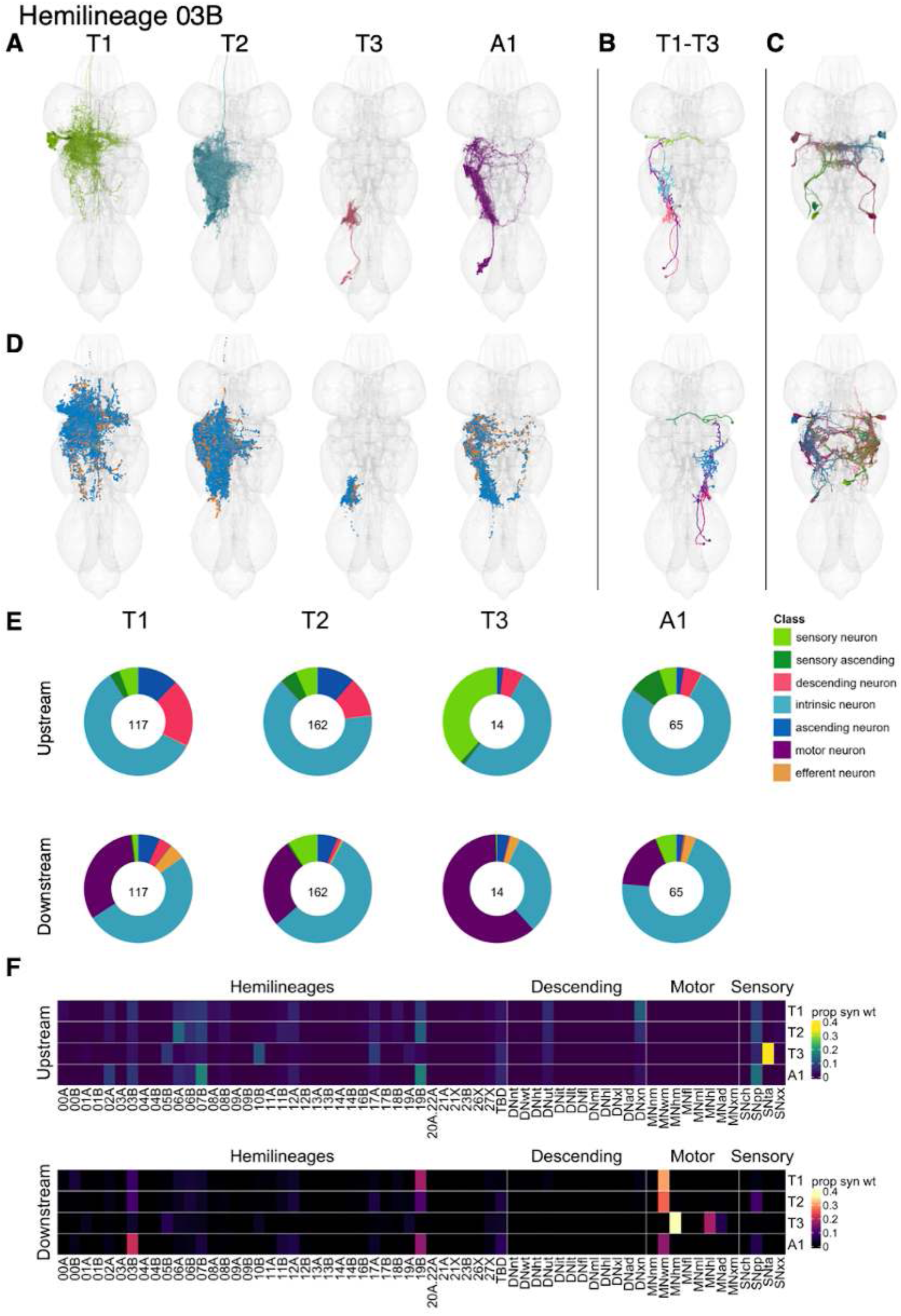
Hemilineage 03B. **A.** Meshes of all RHS secondary neurons plotted in neuromere-specific colours. **B.** “Representative” secondary neuron skeletons plotted in hemineuromere-specific colours. The skeleton with the top accumulated NBLAST score among all neurons from the hemilineage in a given hemineuromere was used. **C.** Neuron meshes of selected examples. Top: motor neuron serial set 11663 (DVMn). Bottom: putative electrical subcluster 10042. **D.** Predicted synapses of RHS secondary neurons. Blue: postsynapses; dark orange: presynapses. **E.** Proportions of connections from secondary neurons to upstream or downstream partners, normalised by neuromere and coloured by broad class. Numbers of query neurons appear in the centre. **F.** Proportions of synaptic weight from secondary neurons originating in each neuromere to upstream or downstream partners, normalised by row.

Secondary 03B neurons survive in T1-A1 (Marin et al., 2012), but the number of neurons in each neuromere varies, with a substantial reduction in T3 (Shepherd et al., 2019) (Figure 20E). 03B secondary neurons receive broadly similar inputs in T1, T2 and A1: from proprioceptive sensory neurons, hemilineages 06A, 07B and 19B, and descending neurons innervating the upper tectulum (Figure 20F). They are predicted to be gabaergic (Figure 8E) and output primarily onto wing motor neurons (Figure 20E), proprioceptive sensory neurons, and hemilineage 19B (Figure 20F). In contrast, the neurons in T3 receive tactile sensory inputs from the haltere knob and output onto haltere and hind leg motor neurons (Figure 20F), suggesting a specialised role.

Whilst most 03B neurons are restricted to the tectulum, a subset of types innervate the intermediate neuropil at the junction of leg neuropil and tectulum (e.g., IN03B015-16, which are strongly downstream of descending neurons) (Figure 6 - figure supplement 2,4). Bilateral activation of 03B secondary neurons results in repetitive, poorly coordinated leg movements, occasional grooming bouts, wing flicking, and wing scissoring movements (Harris et al., 2015).

**Figure 20 - figure supplement 1.**
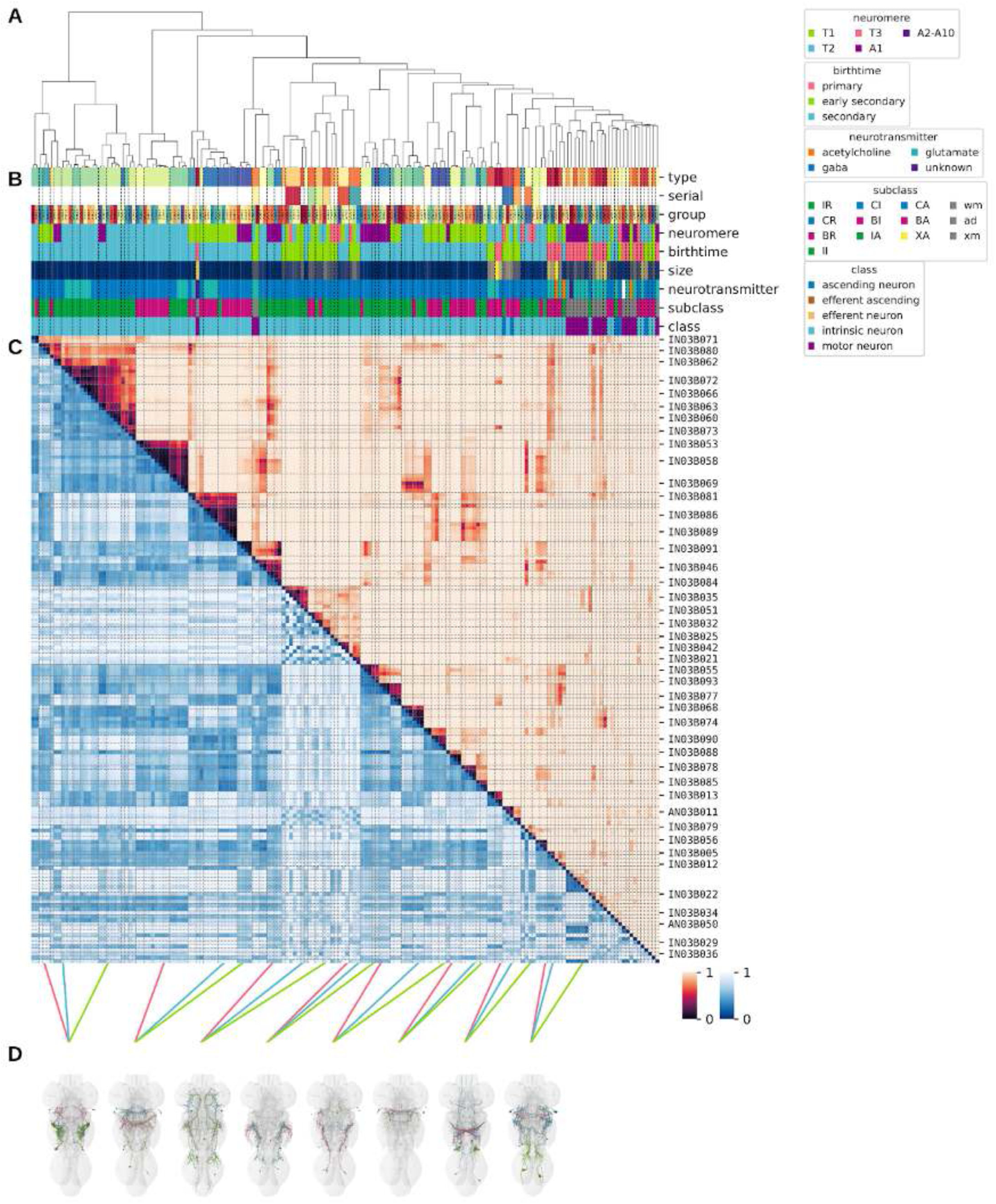
Systematic typing of hemilineage 03B. Types for motor neurons were assigned separately as outlined in our accompanying manuscript (Cheong et al., 2023). **A.** Hierarchical clustering dendrogram of hemilineage groups by laterally and serially aggregated connectivity cosine clustering. **B.** Categorical annotations of each hemilineage group, each column corresponding to the aligned leaf in A. Colours for type, serial set, and group are arbitrary for visualisation. Colours for neuromere, birthtime, neurotransmitter, subclass, and class are as in all other figures. **C.** Similarity distance heatmap for hemilineage. Cosine distance is in the upper triangle, while laterally symmetrised NBLAST distance is in the lower triangle. Systematic type names of some types are labelled. **D.** Morphologically representative groups from dendrogram subtrees. Each group, indicated by colour and line connecting to its column in B and C, is the most morphologically representative group (medoid of NBLAST distance) from a subtree of A. The subtrees (flat clusters) are equal height cuts of A determined to yield the number of groups per plot and plots in D.

**Figure 20 - figure supplement 2.**
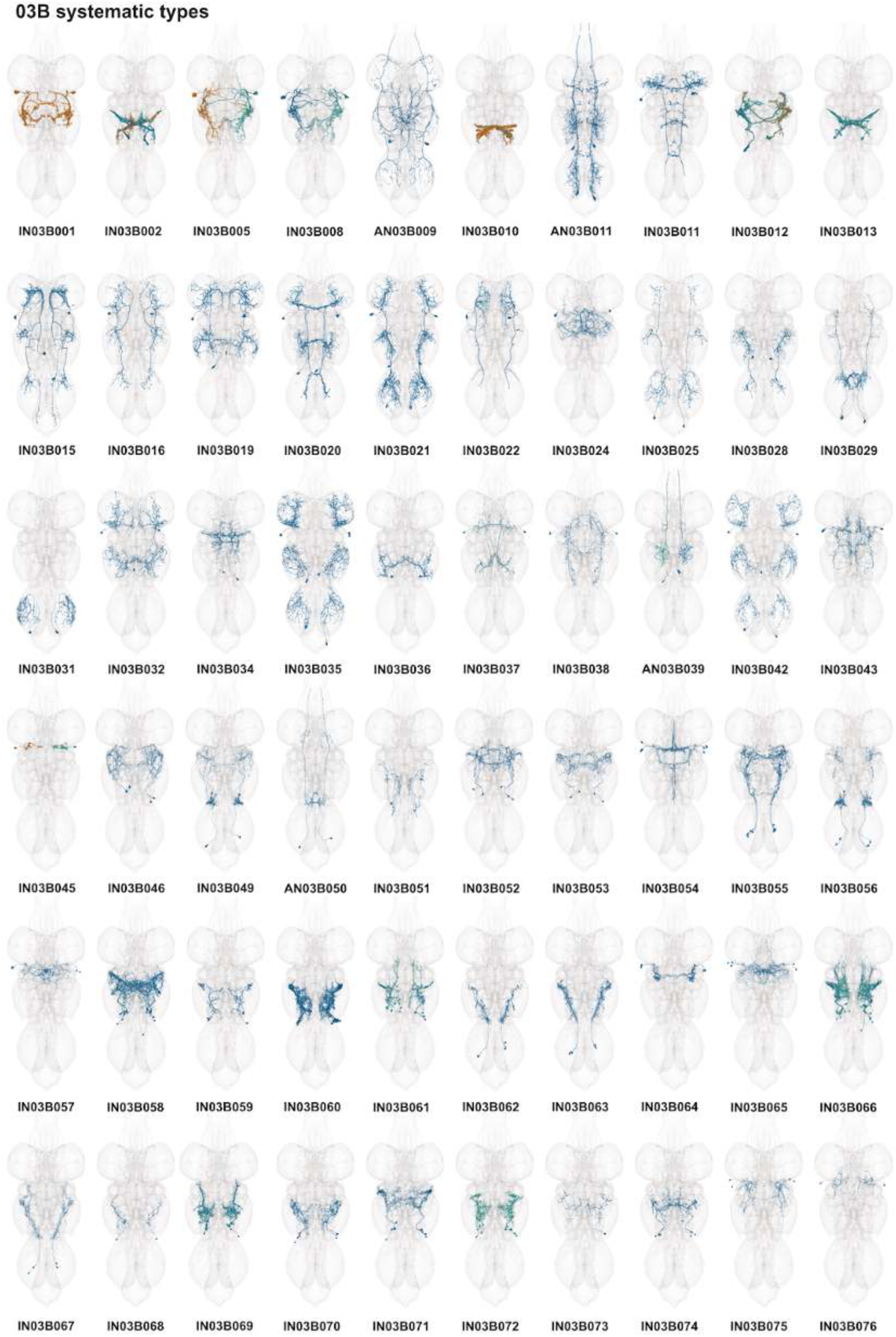
Systematic types of hemilineage 03B. Systematic types have been arranged in numerical order, with neurons of the same type that belong to distinct classes (e.g., intrinsic neuron vs ascending neuron) plotted separately but placed adjacent to each other. Individual neuron meshes have been coloured based on predicted neurotransmitter: dark orange = acetylcholine, blue = gaba, marine = glutamate, dark purple = unknown. Motor neurons (typed separately in (Cheong et al., 2023)) have been plotted by serial set if identified in multiple neuromeres and by systematic type if not. Individual motor neuron meshes have been coloured based on soma neuromere: dark green = T1, blue = T2, purple = A1.

**Figure 20 - figure supplement 3.**
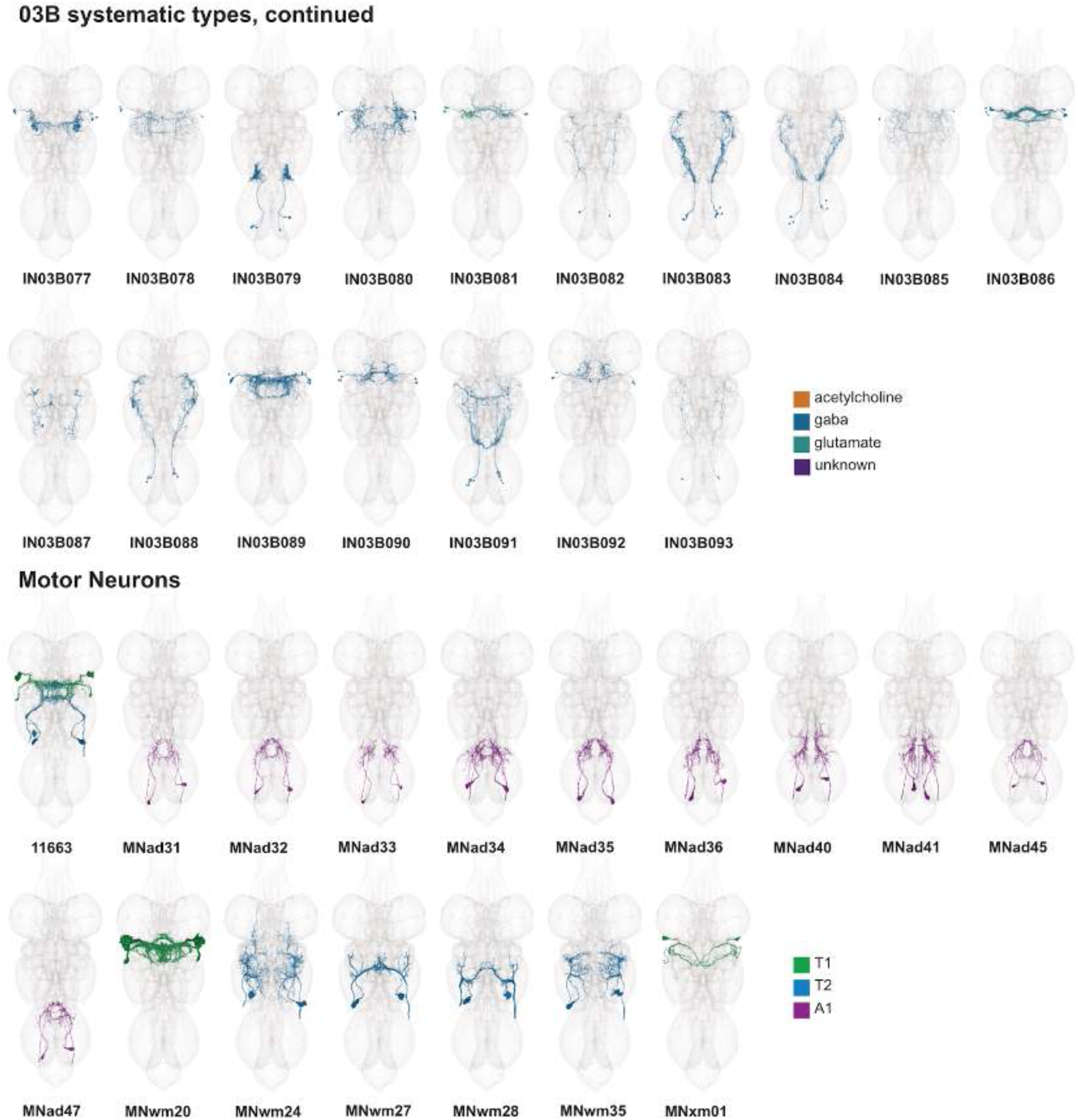
Systematic types of hemilineage 03B, continued. Systematic types have been arranged in numerical order, with neurons of the same type that belong to distinct classes (e.g., intrinsic neuron vs ascending neuron) plotted separately but placed adjacent to each other. Individual neuron meshes have been coloured based on predicted neurotransmitter: dark orange = acetylcholine, blue = gaba, marine = glutamate, dark purple = unknown. Motor neurons (typed separately in (Cheong et al., 2023)) have been plotted by serial set if identified in multiple neuromeres and by systematic type if not. Individual motor neuron meshes have been coloured based on soma neuromere: dark green = T1, blue = T2, purple = A1.

**Figure 20 - figure supplement 4.**
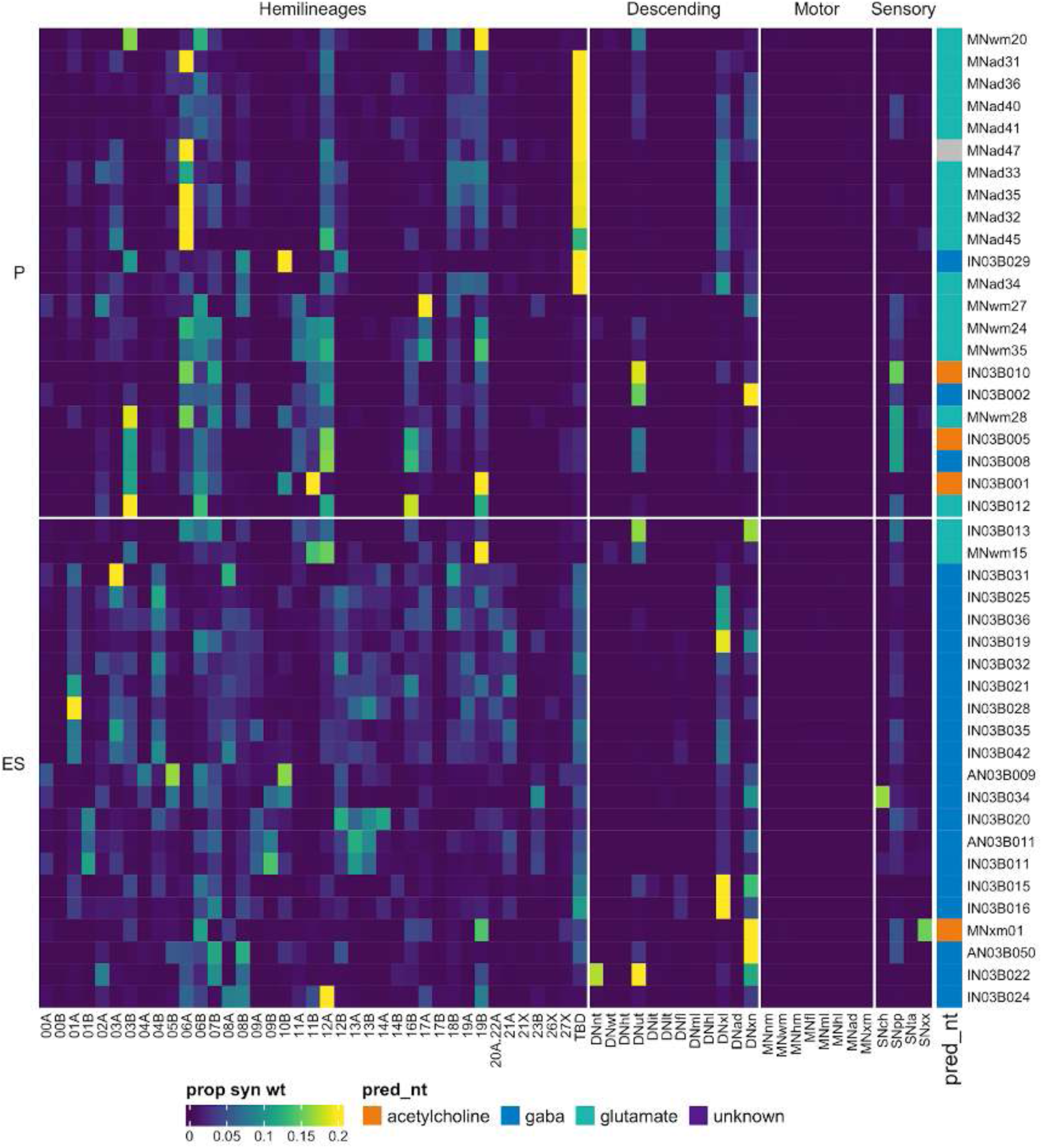
Connectivity to upstream partners by 03B primary and early secondary systematic types. Proportions of synaptic weight to systematic types from upstream partners, normalised by row. 03B neurons have been clustered within each assigned birthtime window (P = primary, ES = early secondary, S = secondary) based on both upstream and downstream connectivity to hemilineages, descending neuron subclasses, motor neuron subclasses, and sensory neuron modalities. Annotation bar is coloured by the most common predicted neurotransmitter for the neurons of each type.

**Figure 20 - figure supplement 5.**
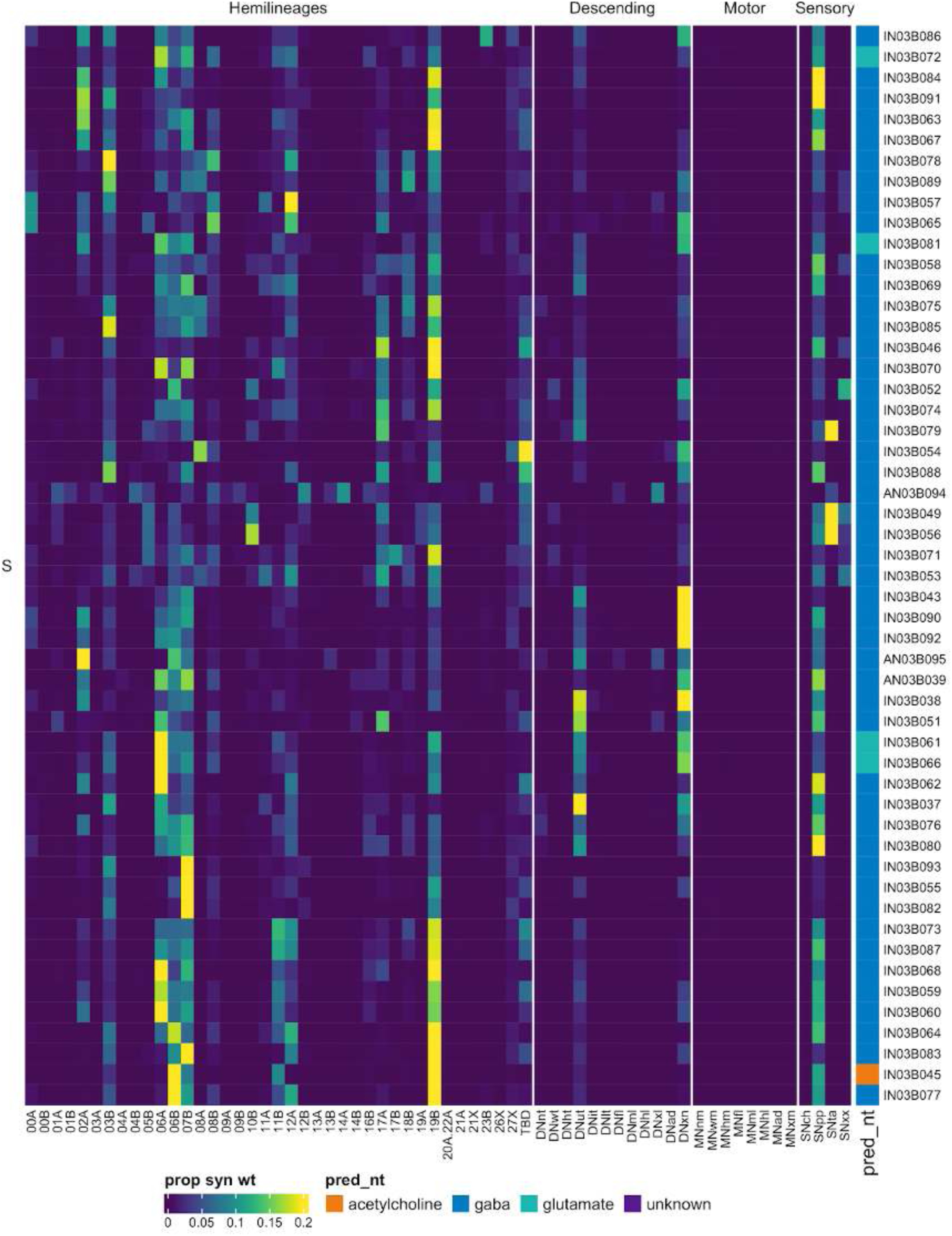
Connectivity to upstream partners by 03B secondary systematic types. Proportions of synaptic weight to systematic types from upstream partners, normalised by row. 03B neurons have been clustered within each assigned birthtime window (P = primary, ES = early secondary, S = secondary) based on both upstream and downstream connectivity to hemilineages, descending neuron subclasses, motor neuron subclasses, and sensory neuron modalities. The annotation bar is coloured by the most common predicted neurotransmitter within each type.

**Figure 20 - figure supplement 6.**
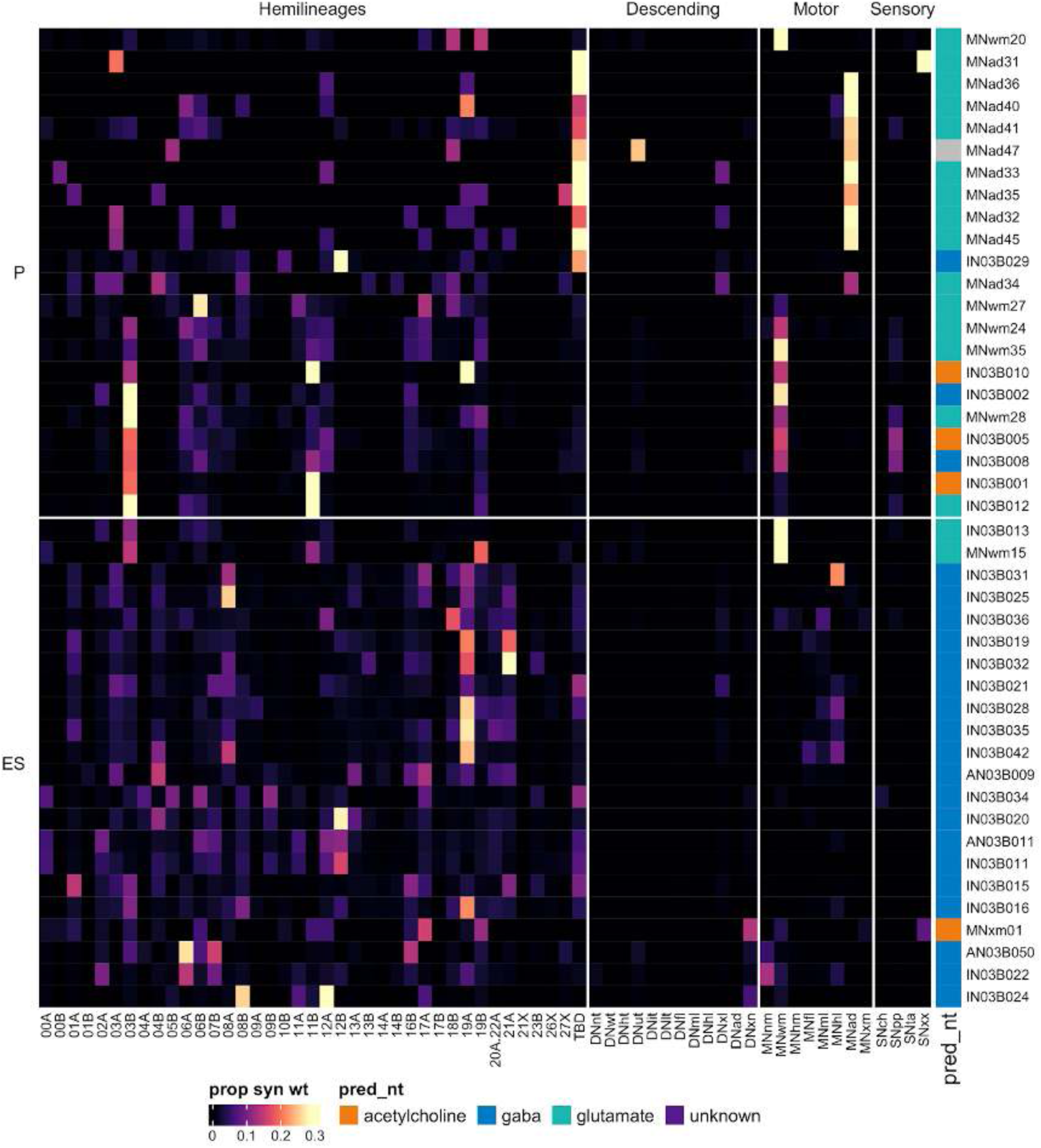
Connectivity to downstream partners by 03B primary and early secondary systematic types. Proportions of synaptic weight from systematic types to downstream partners, normalised by row. 03B neurons have been clustered within each assigned birthtime window (P = primary, ES = early secondary, S = secondary) based on both upstream and downstream connectivity to hemilineages, descending neuron subclasses, motor neuron subclasses, and sensory neuron modalities. The annotation bar is coloured by the most common predicted neurotransmitter within each type.

**Figure 20 - figure supplement 7.**
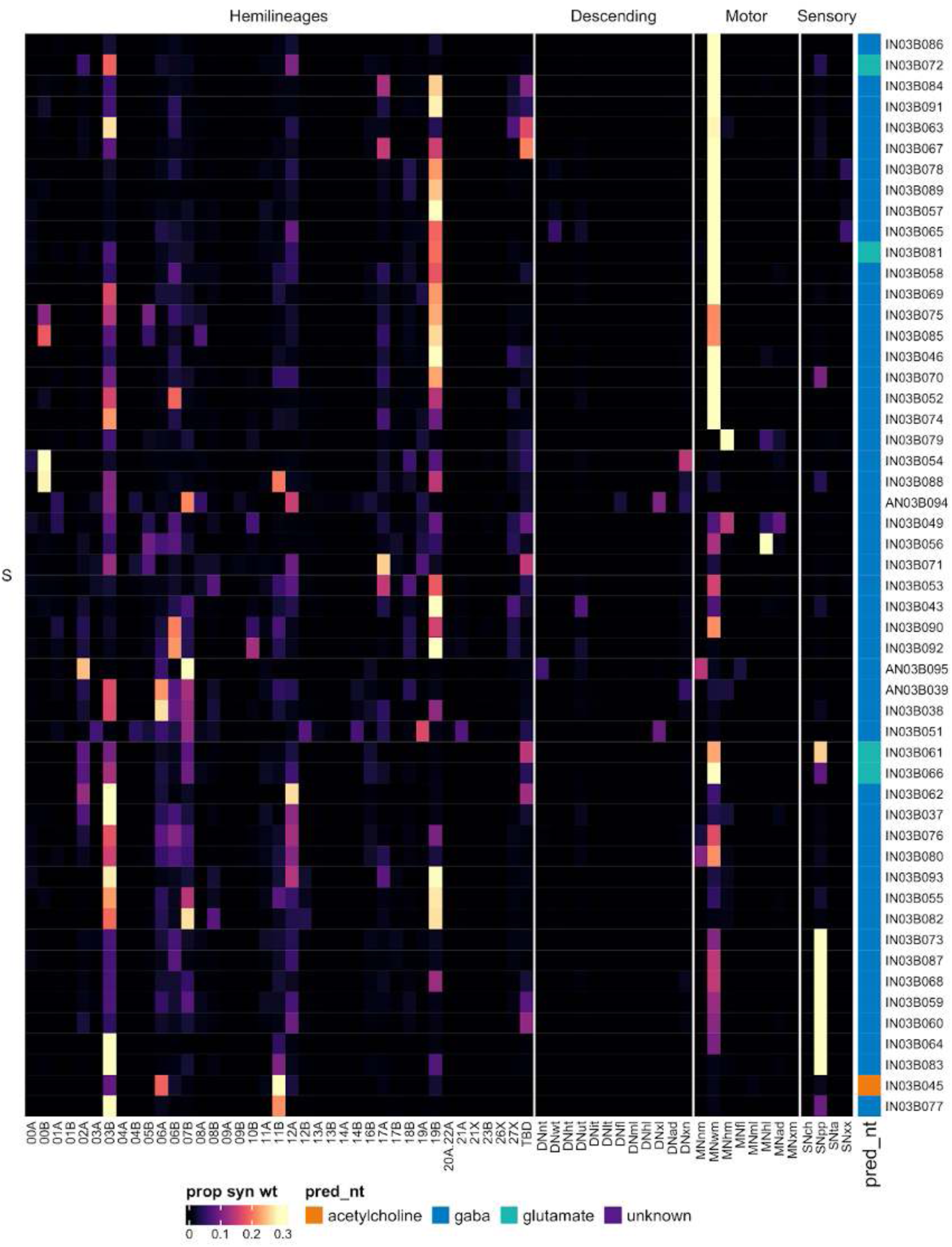
Connectivity to downstream partners by 03B secondary systematic types. Proportions of synaptic weight from systematic types to downstream partners, normalised by row. 03B neurons have been clustered within each assigned birthtime window (P = primary, ES = early secondary, S = secondary) based on both upstream and downstream connectivity to hemilineages, descending neuron subclasses, motor neuron subclasses, and sensory neuron modalities. The annotation bar is coloured by the most common predicted neurotransmitter for the neurons of each type.

#### Hemilineage 04B

04B is the main secondary hemilineage (Truman et al., 2010) from medial neuroblast NB3-1 (Truman et al., 2004), which generates 4 contralateral MNs, 3-6 intersegmental interneurons, and a variable number of local interneurons in the embryo (Schmid et al., 1999). 04B is a relatively large hemilineage (Figure 8D), and its neurons do not enter the neuropil in a single coherent bundle but rather project from the ventral midline in multiple clusters. Most 04B secondary neurons innervate the ipsilateral leg neuropil diffusely, especially the surface, but a small subset exhibit intersegmental projections (Shepherd et al., 2019). 04B neurons arborise extensively in the ipsilateral leg neuropil, sometimes spreading across the midline (Figure 21A). Five types of early born 04B neurons ascend via the neck connective (e.g., AN04B003) (Figure 21C top), but most 04B neurons are restricted to their neuromere of origin (e.g., IN04B008) (Figure 21C bottom).

Secondary 04B neurons survive in T1-T3 with comparable numbers of neurons in all thoracic neuromeres (Figure 21E). They mainly receive inputs from hemilineages 01A, 04B, 12B, 13A, and 13B, from descending neurons that innervate leg neuropils, and from tactile sensory neurons (Figure 21F), although a few types also receive chemosensory input (e.g., IN04B005 and IN04B079) (Figure 21 - figure supplement 4-5). They are predicted to be cholinergic (Figure 8E) as expected (Lacin et al., 2019) and output onto leg motor neurons and leg hemilineages, notably 04B, 19A, and 21A (Figure 21F). No functional studies have been published for secondary 04B neurons.

**Figure 21.**
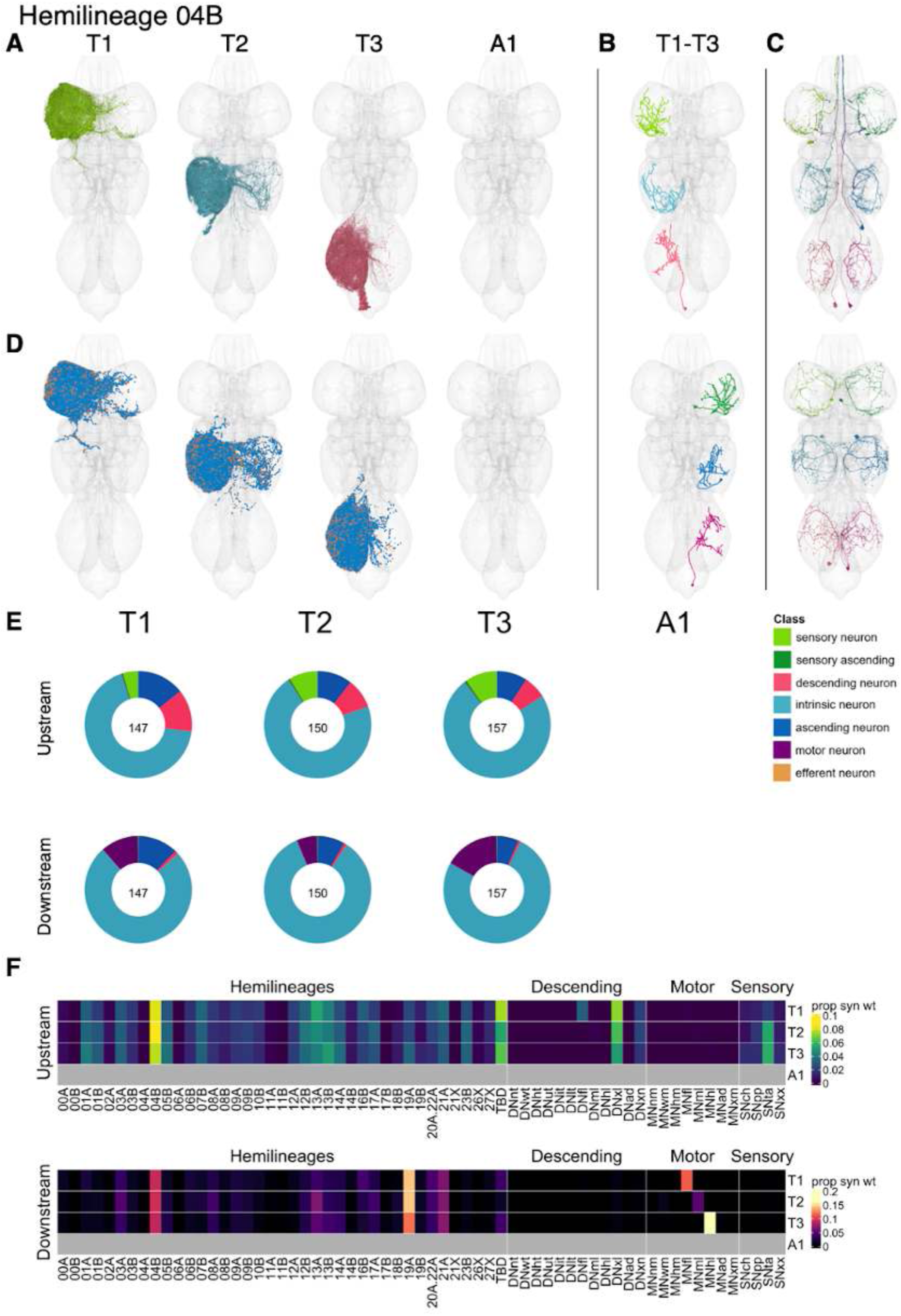
Hemilineage 04B. **A.** Meshes of all RHS secondary neurons plotted in neuromere-specific colours. **B.** “Representative” secondary neuron skeletons plotted in hemineuromere-specific colours. The skeleton with the top accumulated NBLAST score among all neurons from the hemilineage in a given hemineuromere was used. **C.** Neuron meshes of selected examples. Top: ascending serial set 10394. Bottom: independent leg serial set 11945. **D.** Predicted synapses of RHS secondary neurons. Blue: postsynapses; dark orange: presynapses. **E.** Proportions of connections from secondary neurons to upstream or downstream partners, normalised by neuromere and coloured by broad class. Numbers of query neurons appear in the centre. **F.** Proportions of synaptic weight from secondary neurons originating in each neuromere to upstream or downstream partners, normalised by row.

**Figure 21 - figure supplement 1.**
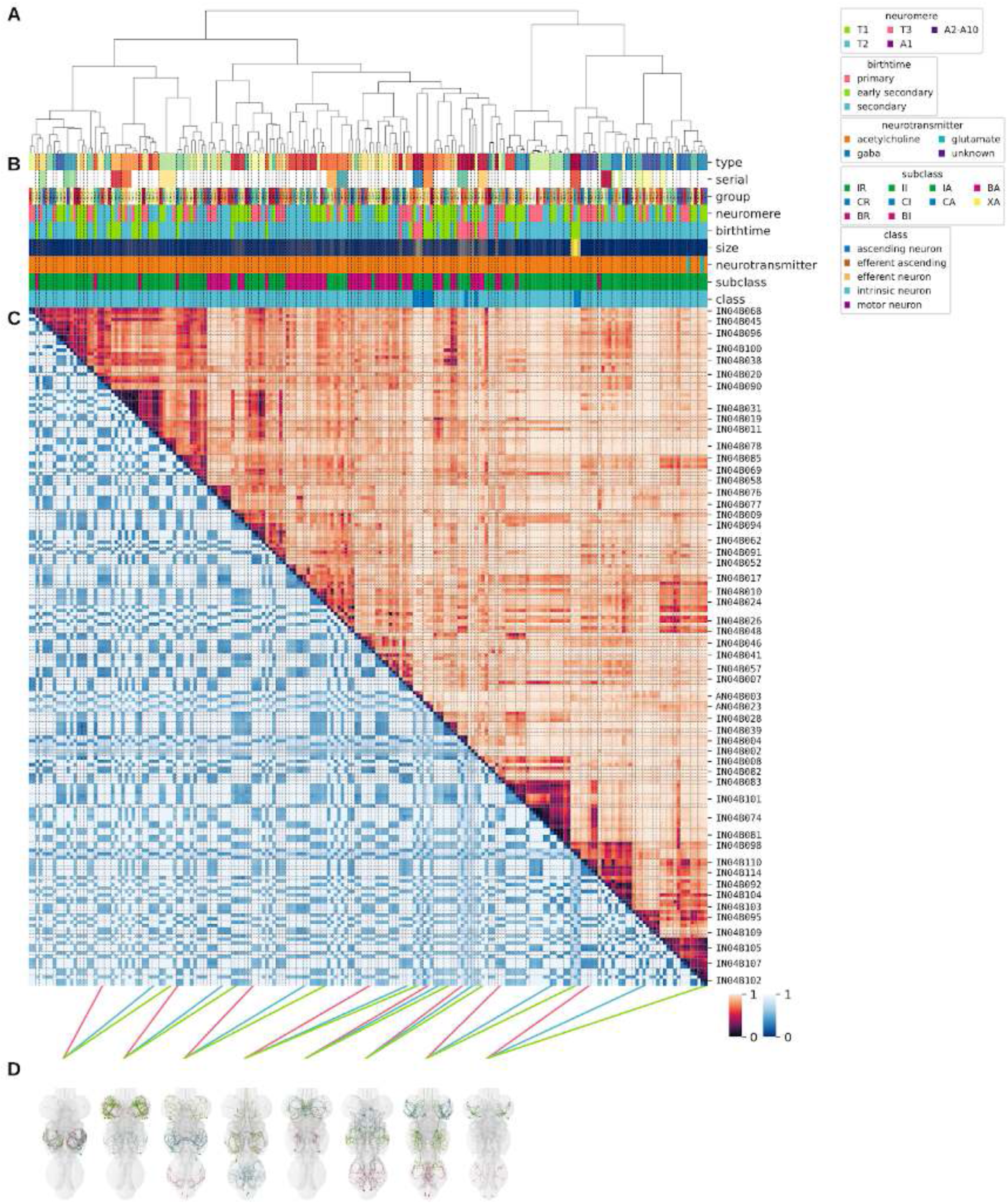
Systematic typing of hemilineage 04B. **A.** Hierarchical clustering dendrogram of hemilineage groups by laterally and serially aggregated connectivity cosine clustering. **B.** Categorical annotations of each hemilineage group, each column corresponding to the aligned leaf in A. Colours for type, serial set, and group are arbitrary for visualisation. Colours for neuromere, birthtime, neurotransmitter, subclass, and class are as in all other figures. **C.** Similarity distance heatmap for hemilineage. Cosine distance is in the upper triangle, while laterally symmetrised NBLAST distance is in the lower triangle. Systematic type names of some types are labelled. **D.** Morphologically representative groups from dendrogram subtrees. Each group, indicated by colour and line connecting to its column in B and C, is the most morphologically representative group (medoid of NBLAST distance) from a subtree of A. The subtrees (flat clusters) are equal height cuts of A determined to yield the number of groups per plot and plots in D.

**Figure 21 - figure supplement 2.**
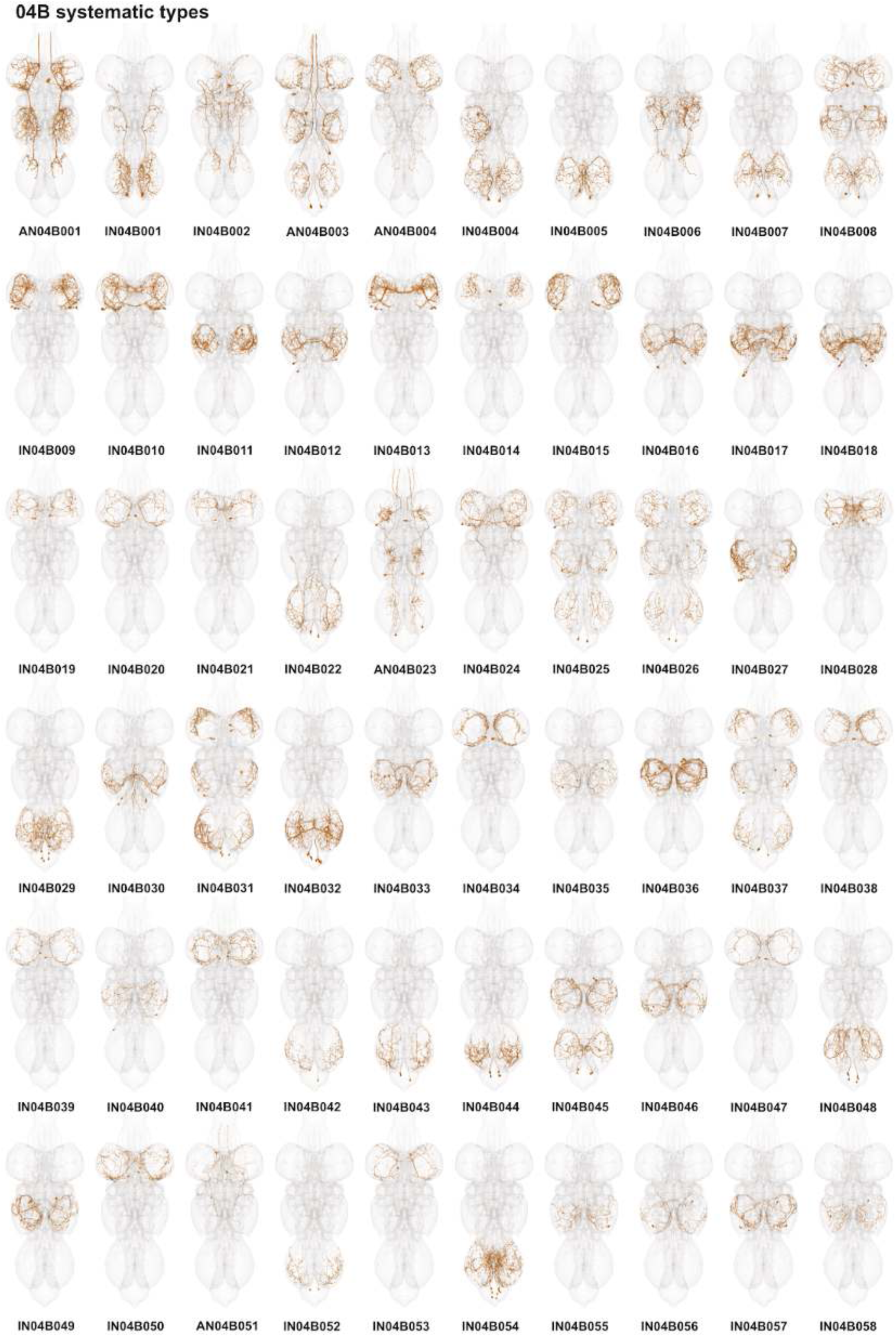
Systematic types of hemilineage 04B. Systematic types have been arranged in numerical order, with neurons of the same type that belong to distinct classes (e.g., intrinsic neuron vs ascending neuron) plotted separately but placed adjacent to each other. Individual neuron meshes have been coloured based on predicted neurotransmitter: dark orange = acetylcholine, blue = gaba, marine = glutamate, dark purple = unknown.

**Figure 21 - figure supplement 3.**
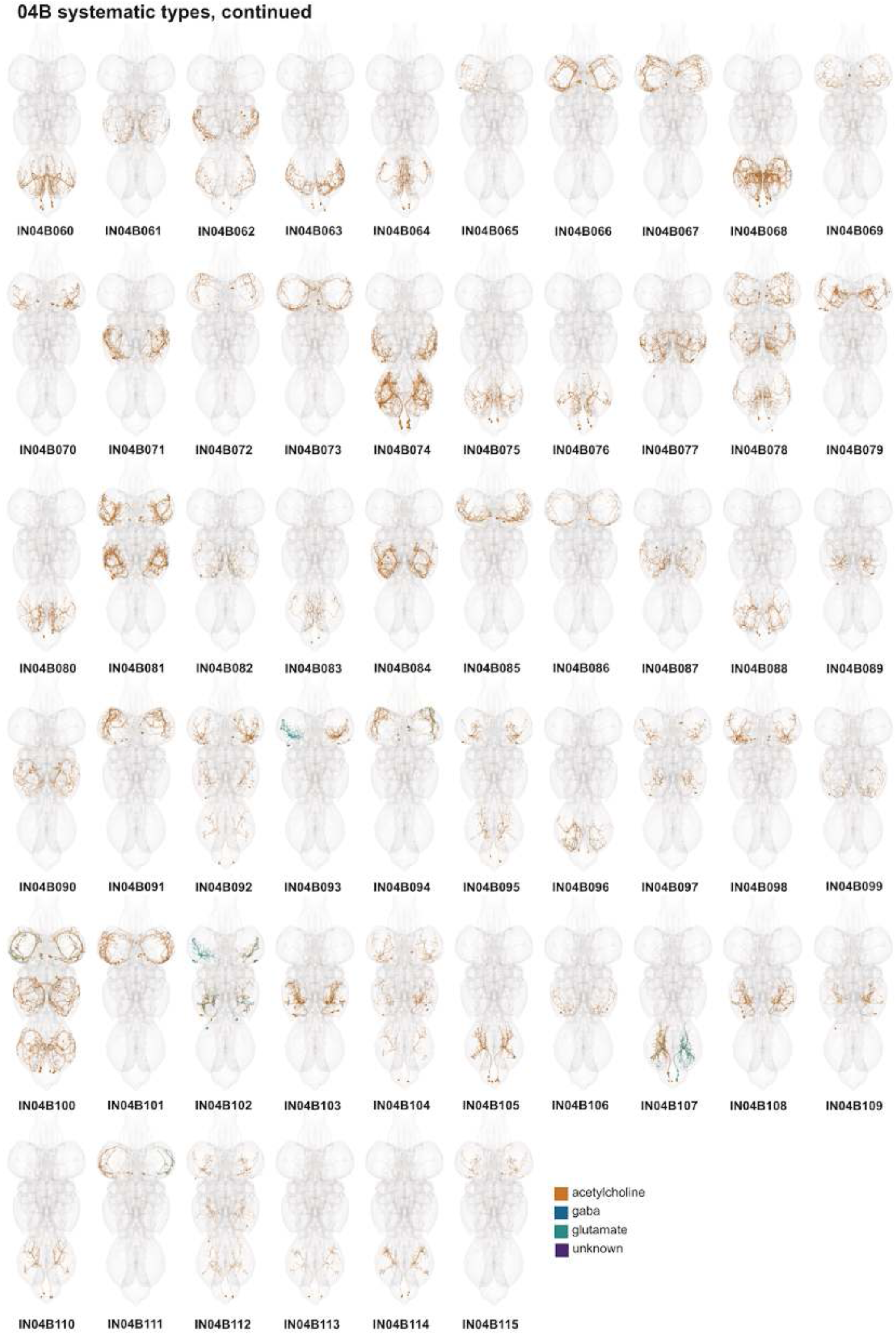
Systematic types of hemilineage 04B, continued. Systematic types have been arranged in numerical order, with neurons of the same type that belong to distinct classes (e.g., intrinsic neuron vs ascending neuron) plotted separately but placed adjacent to each other. Individual neuron meshes have been coloured based on predicted neurotransmitter: dark orange = acetylcholine, blue = gaba, marine = glutamate, dark purple = unknown.

**Figure 21 - figure supplement 4.**
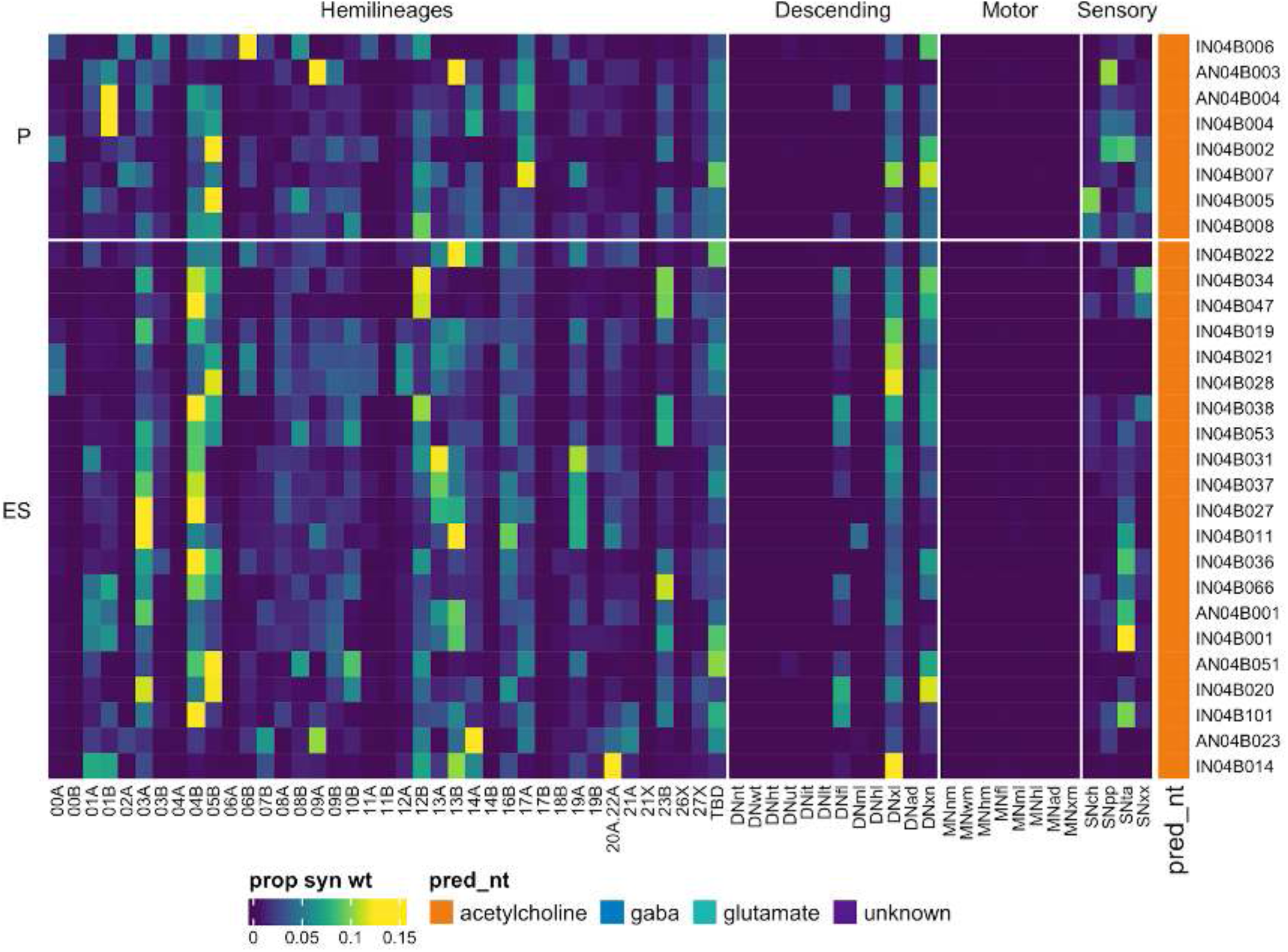
Connectivity to upstream partners by 04B primary and early secondary systematic types. Proportions of synaptic weight to systematic types from upstream partners, normalised by row. 04B neurons have been clustered within each assigned birthtime window (P = primary, ES = early secondary, S = secondary) based on both upstream and downstream connectivity to hemilineages, descending neuron subclasses, motor neuron subclasses, and sensory neuron modalities. Annotation bar is coloured by the most common predicted neurotransmitter for the neurons of each type.

**Figure 21 - figure supplement 5.**
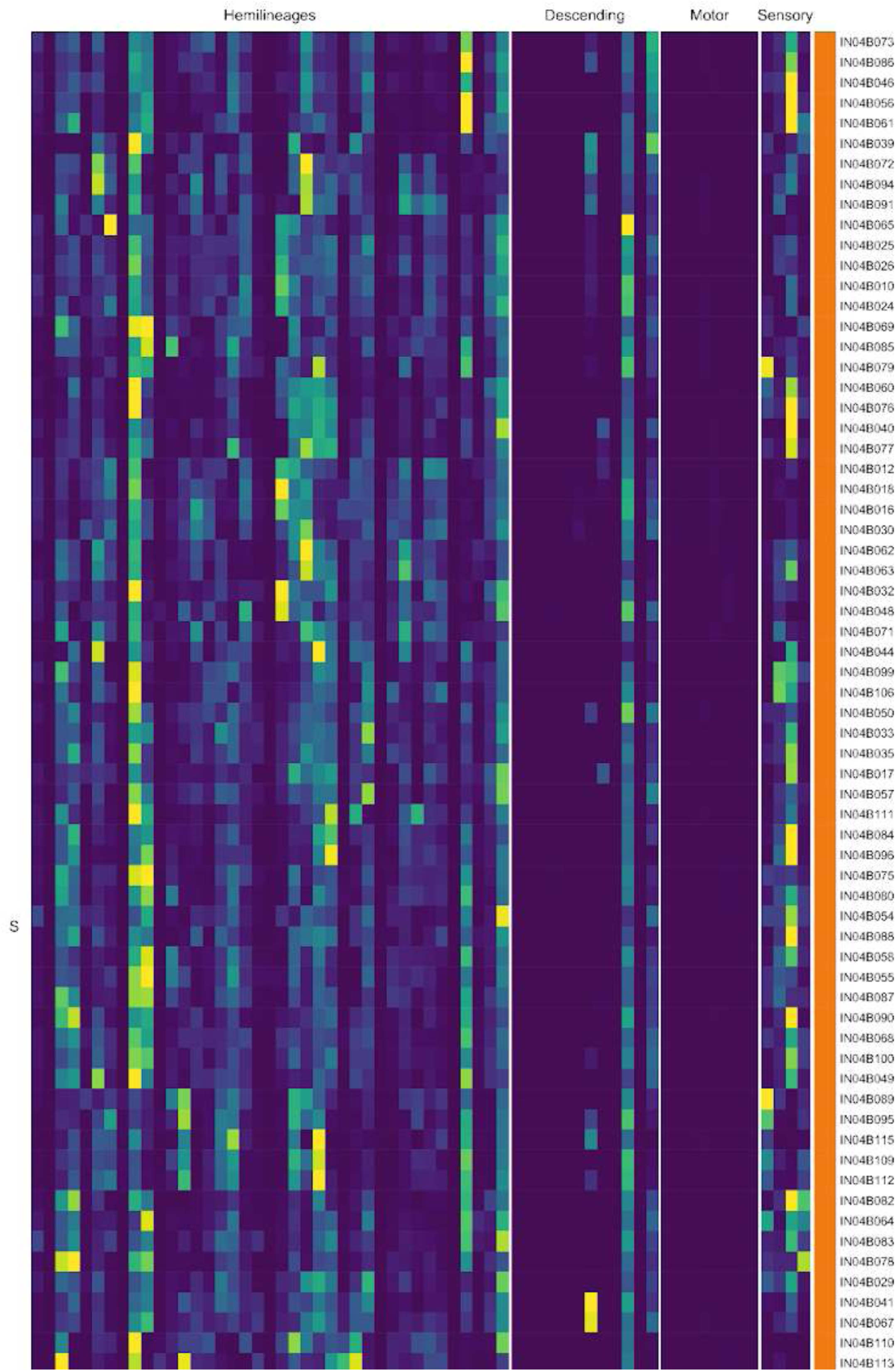
Connectivity to upstream partners by 04B secondary systematic types. Proportions of synaptic weight to systematic types from upstream partners, normalised by row. 04B neurons have been clustered within each assigned birthtime window (P = primary, ES = early secondary, S = secondary) based on both upstream and downstream connectivity to hemilineages, descending neuron subclasses, motor neuron subclasses, and sensory neuron modalities. The annotation bar is coloured by the most common predicted neurotransmitter within each type.

**Figure 21 - figure supplement 6.**
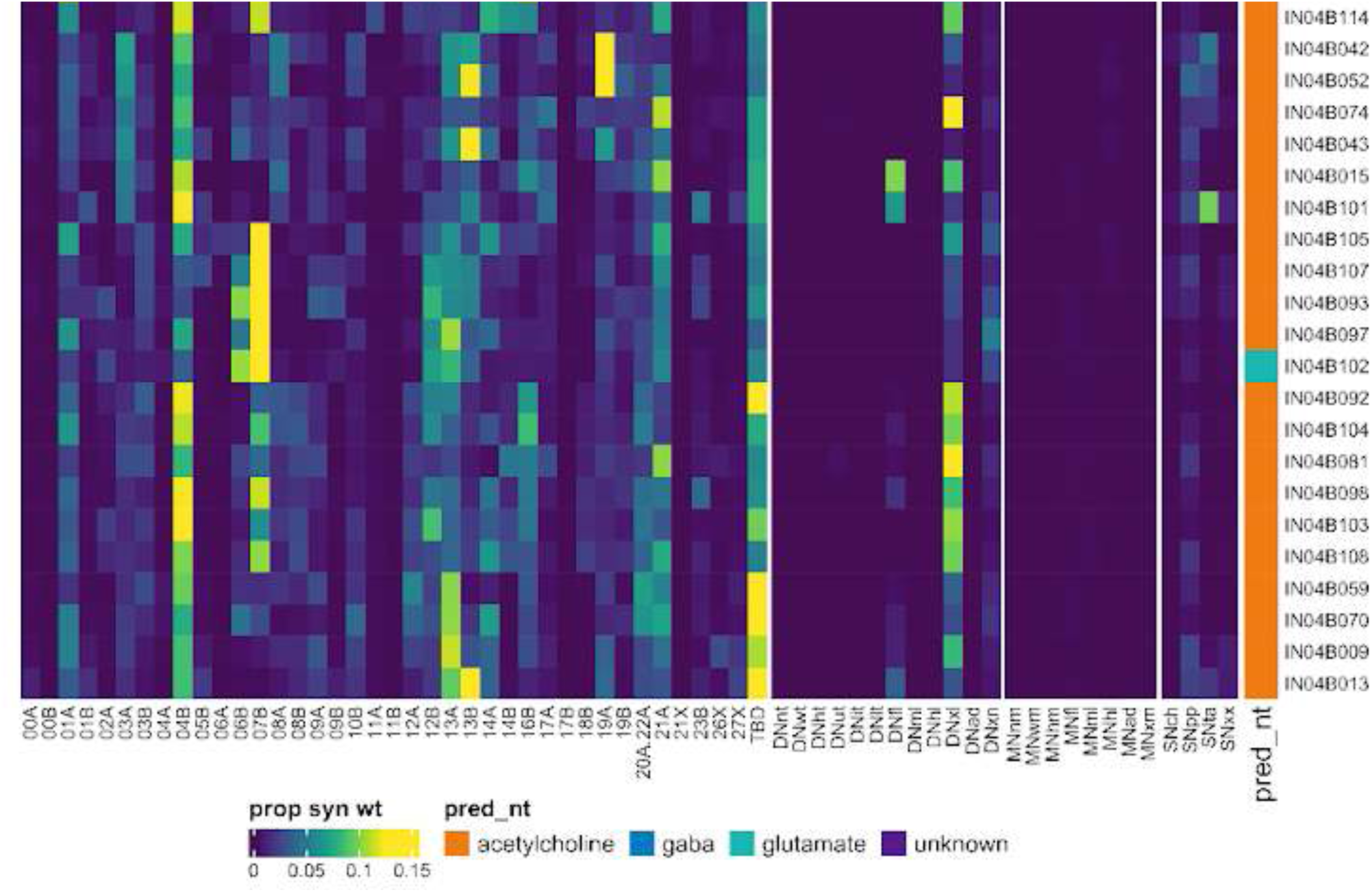
Connectivity to upstream partners by 04B secondary systematic types, continued. Proportions of synaptic weight to systematic types from upstream partners, normalised by row. 04B neurons have been clustered within each assigned birthtime window (P = primary, ES = early secondary, S = secondary) based on both upstream and downstream connectivity to hemilineages, descending neuron subclasses, motor neuron subclasses, and sensory neuron modalities. The annotation bar is coloured by the most common predicted neurotransmitter within each type.

**Figure 21 - figure supplement 7.**
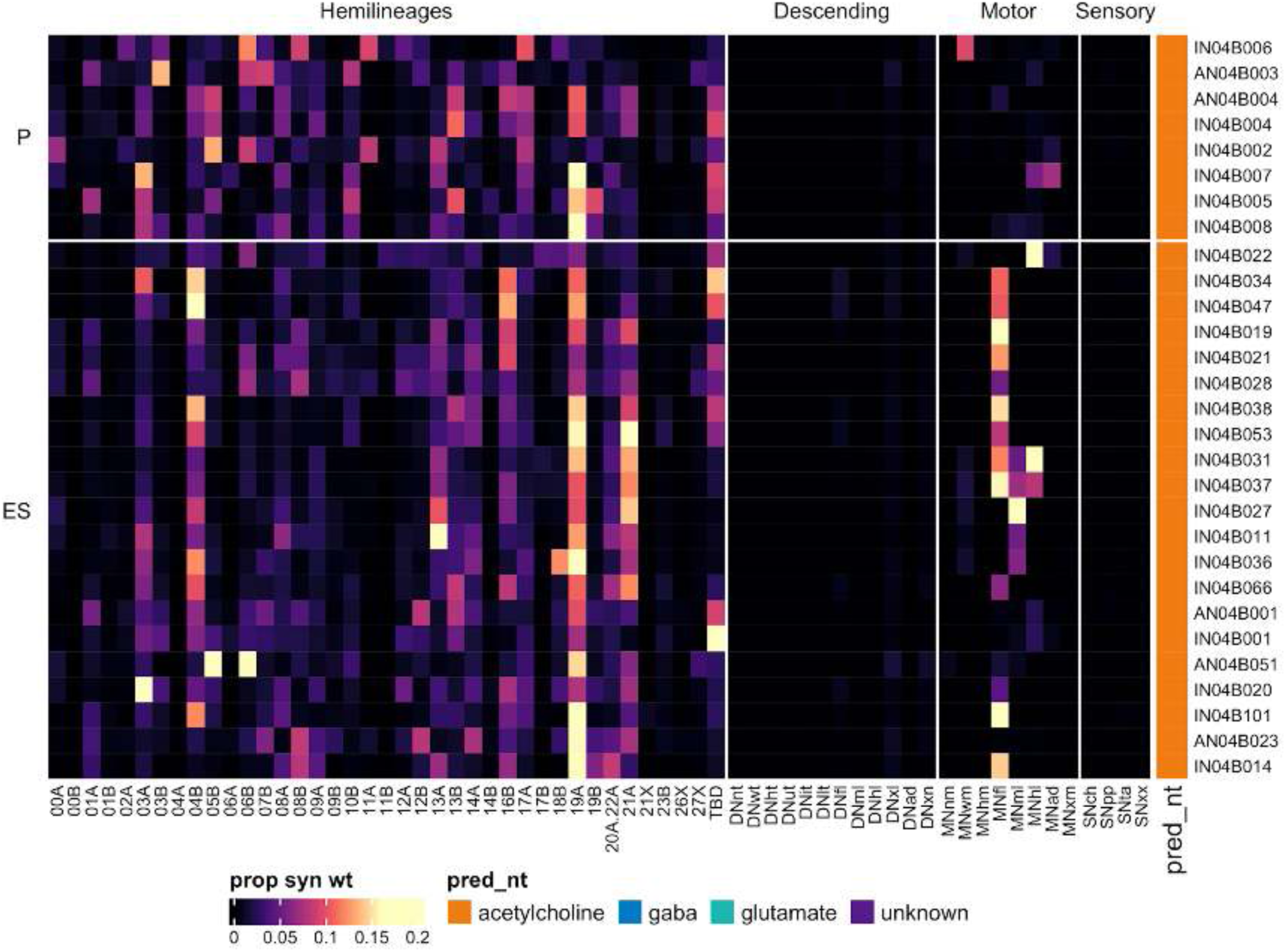
Connectivity to downstream partners by 04B primary and early secondary systematic types. Proportions of synaptic weight from systematic types to downstream partners, normalised by row. 04B neurons have been clustered within each assigned birthtime window (P = primary, ES = early secondary, S = secondary) based on both upstream and downstream connectivity to hemilineages, descending neuron subclasses, motor neuron subclasses, and sensory neuron modalities. The annotation bar is coloured by the most common predicted neurotransmitter within each type.

**Figure 21 - figure supplement 8.**
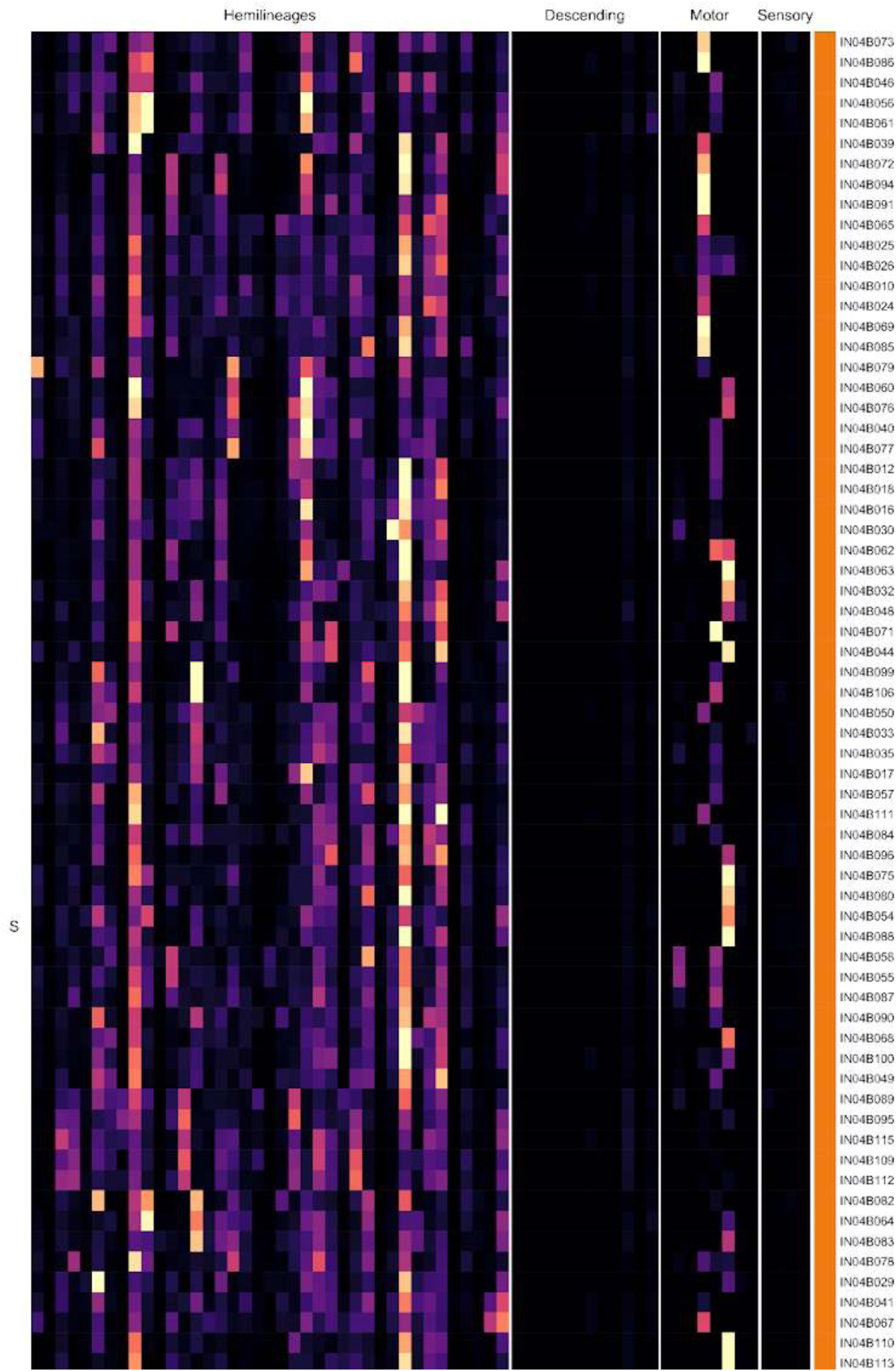
Connectivity to downstream partners by 04B secondary systematic types. Proportions of synaptic weight from systematic types to downstream partners, normalised by row. 04B neurons have been clustered within each assigned birthtime window (P = primary, ES = early secondary, S = secondary) based on both upstream and downstream connectivity to hemilineages, descending neuron subclasses, motor neuron subclasses, and sensory neuron modalities. The annotation bar is coloured by the most common predicted neurotransmitter for the neurons of each type.

**Figure 21 - figure supplement 9.**
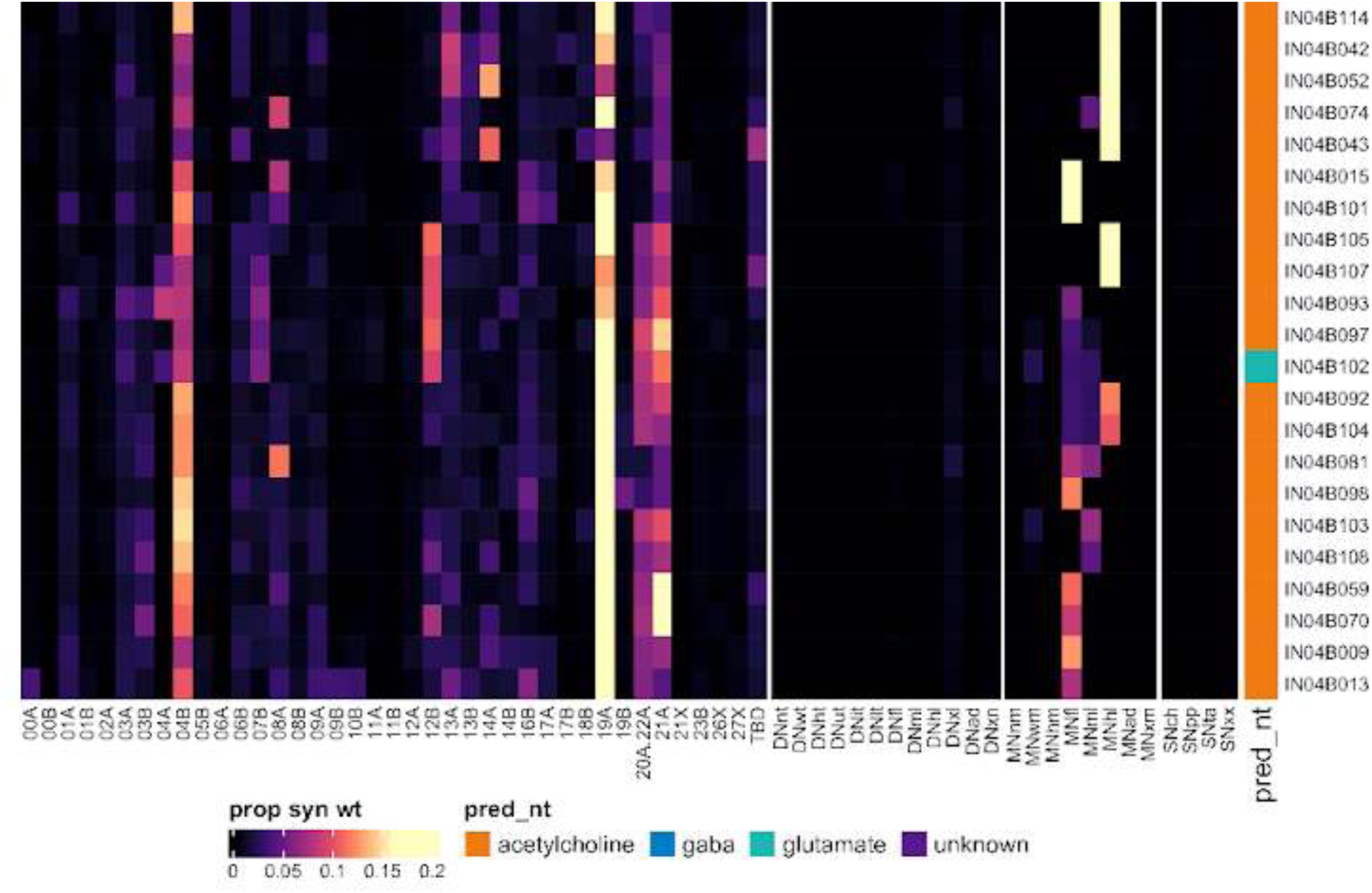
Connectivity to downstream partners by 04B secondary systematic types, continued. Proportions of synaptic weight from systematic types to downstream partners, normalised by row. 04B neurons have been clustered within each assigned birthtime window (P = primary, ES = early secondary, S = secondary) based on both upstream and downstream connectivity to hemilineages, descending neuron subclasses, motor neuron subclasses, and sensory neuron modalities. The annotation bar is coloured by the most common predicted neurotransmitter for the neurons of each type.

#### Hemilineage 05B

05B is the only surviving secondary hemilineage (Truman et al., 2010) from posterior neuroblast NB5-3 (Birkholz et al., 2015; Lacin and Truman, 2016), which generates a single ipsilateral MN in T1 as well as 12-15 intersegmental interneurons and 12-15 local interneurons per segment in the embryo (Schmid et al., 1999). In the adult, the primary neurites of 05B secondary neurons enter the neuropil near the posterior edge of each neuromere near the midline, cross in the posterior commissure (ventral to 06B in T1-T2) (Shepherd et al., 2016), and run anteriorly and/or posteriorly in a ventral tract to innervate the primary sensory neuropil of up to all six leg neuropils and the ovoid (Shepherd et al., 2019) (Figure 22A). Many 05B types ascend via the neck connective (e.g., AN05B006) (Figure 22 - figure supplement 2). 05B neurons innervate both ventral leg and flight neuropils, with some receiving inputs in the VAC (e.g., Figure 22C bottom) while others are restricted to the tectulum (e.g., Figure 22C top). 05B secondary neurons are reported to be gabaergic (Lacin et al., 2019); cholinergic neurons with similar morphology were annotated as 05A.

**Figure 22.**
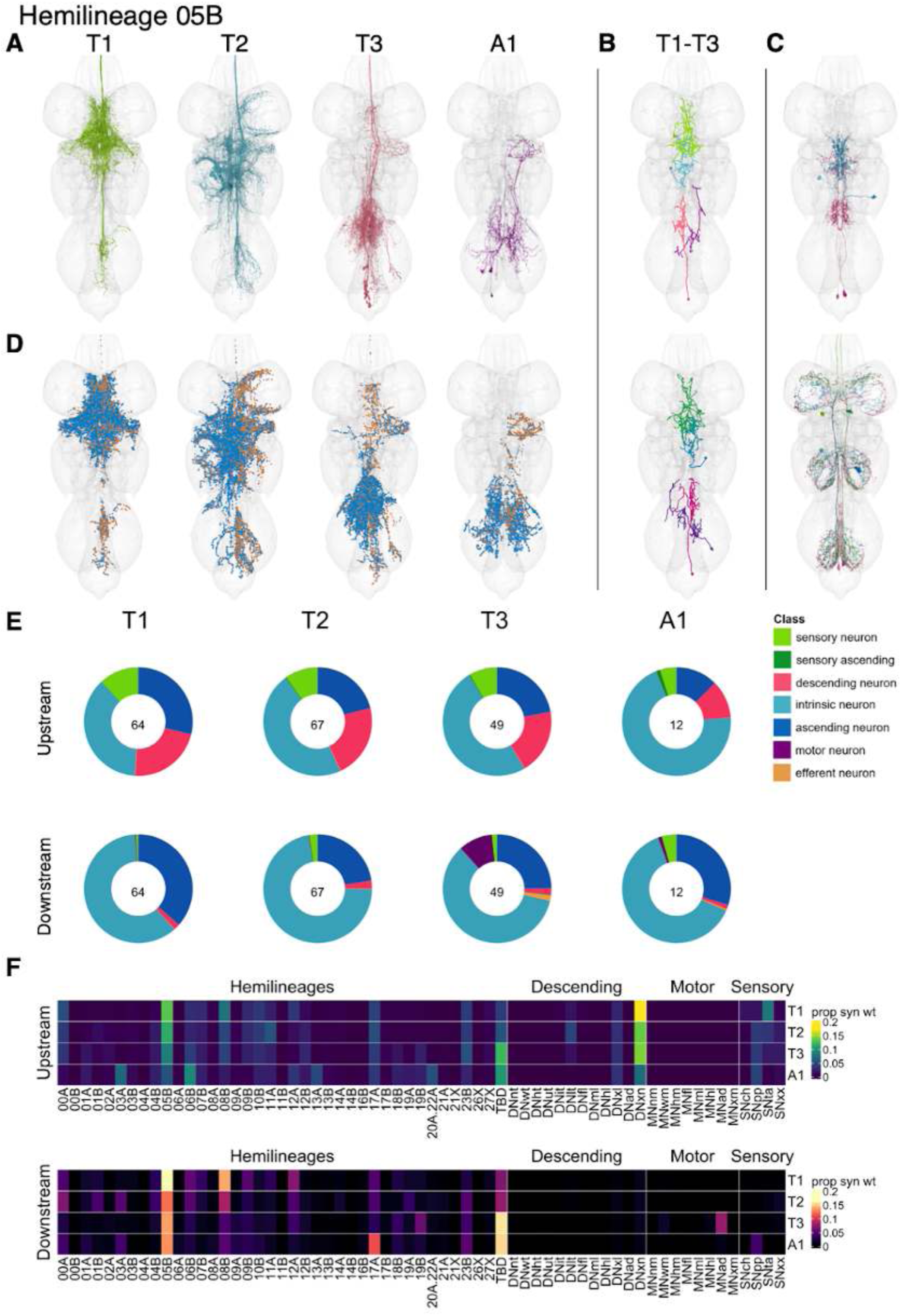
Hemilineage 05B. **A.** Meshes of all RHS secondary neurons plotted in neuromere-specific colours. **B.** “Representative” secondary neuron skeletons plotted in hemineuromere-specific colours. The skeleton with the top accumulated NBLAST score among all neurons from the hemilineage in a given hemineuromere was used. **C.** Neuron meshes of selected examples. Top: dorsal serial set 12983. Bottom: complex serial set 11757. **D.** Predicted synapses of RHS secondary neurons. Blue: postsynapses; dark orange: presynapses. **E.** Proportions of connections from secondary neurons to upstream or downstream partners, normalised by neuromere and coloured by broad class. Numbers of query neurons appear in the centre. **F.** Proportions of synaptic weight from secondary neurons originating in each neuromere to upstream or downstream partners, normalised by row.

Secondary 05B neurons survive in T1-T3 with comparable neuron numbers in T1 and T2 and slightly fewer in T3 (Figure 22E). Their inputs are primarily from descending interneurons that innervate multiple neuropils, with lesser inputs from 05B neurons and sensory neurons (Figure 22F). The most common sensory input is tactile, but there are several 05B types dedicated to proprioceptive input (e.g., IN05B001 and IN05B016) as well as a few dedicated to chemosensory input, notably IN05B011 and AN05B23 (Figure 22 - figure supplement 4). 05B secondary neurons inhibit neurons from a range of hemilineages including 08B (particularly in T1 and T2), abdominal motor neurons in T3, 17A in A1, and 05B in all neuromeres (Figure 22E,F). Bilateral activation of 05B secondary neurons results in limb repositioning/splaying that brings the fly closer to the ground (Harris et al., 2015).

**Figure 22 - figure supplement 1.**
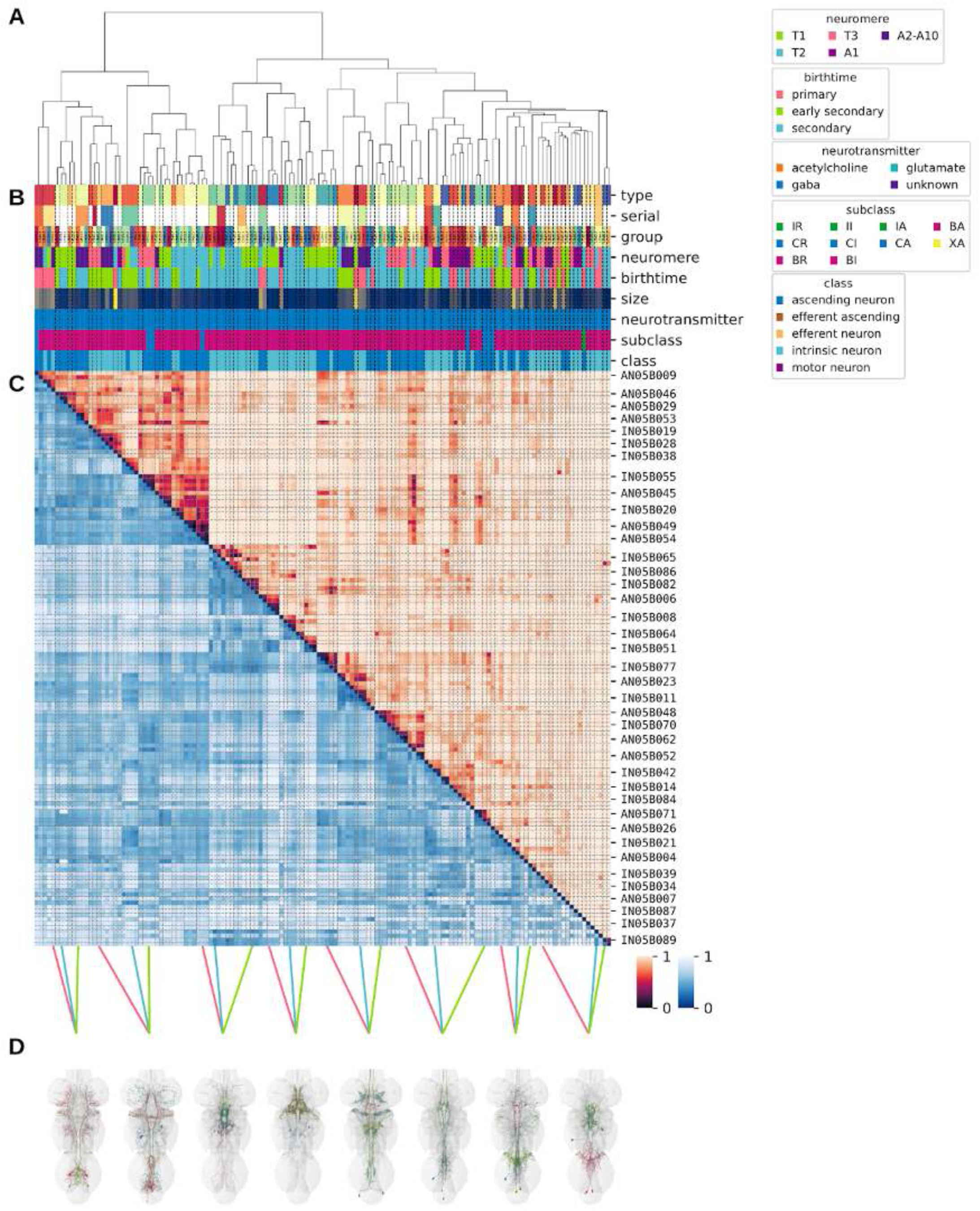
Systematic typing of hemilineage 05B. **A.** Hierarchical clustering dendrogram of hemilineage groups by laterally and serially aggregated connectivity cosine clustering. **B.** Categorical annotations of each hemilineage group, each column corresponding to the aligned leaf in A. Colours for type, serial set, and group are arbitrary for visualisation. Colours for neuromere, birthtime, neurotransmitter, subclass, and class are as in all other figures. **C.** Similarity distance heatmap for hemilineage. Cosine distance is in the upper triangle, while laterally symmetrised NBLAST distance is in the lower triangle. Systematic type names of some types are labelled. **D.** Morphologically representative groups from dendrogram subtrees. Each group, indicated by colour and line connecting to its column in B and C, is the most morphologically representative group (medoid of NBLAST distance) from a subtree of A. The subtrees (flat clusters) are equal height cuts of A determined to yield the number of groups per plot and plots in D.

**Figure 22 - figure supplement 2.**
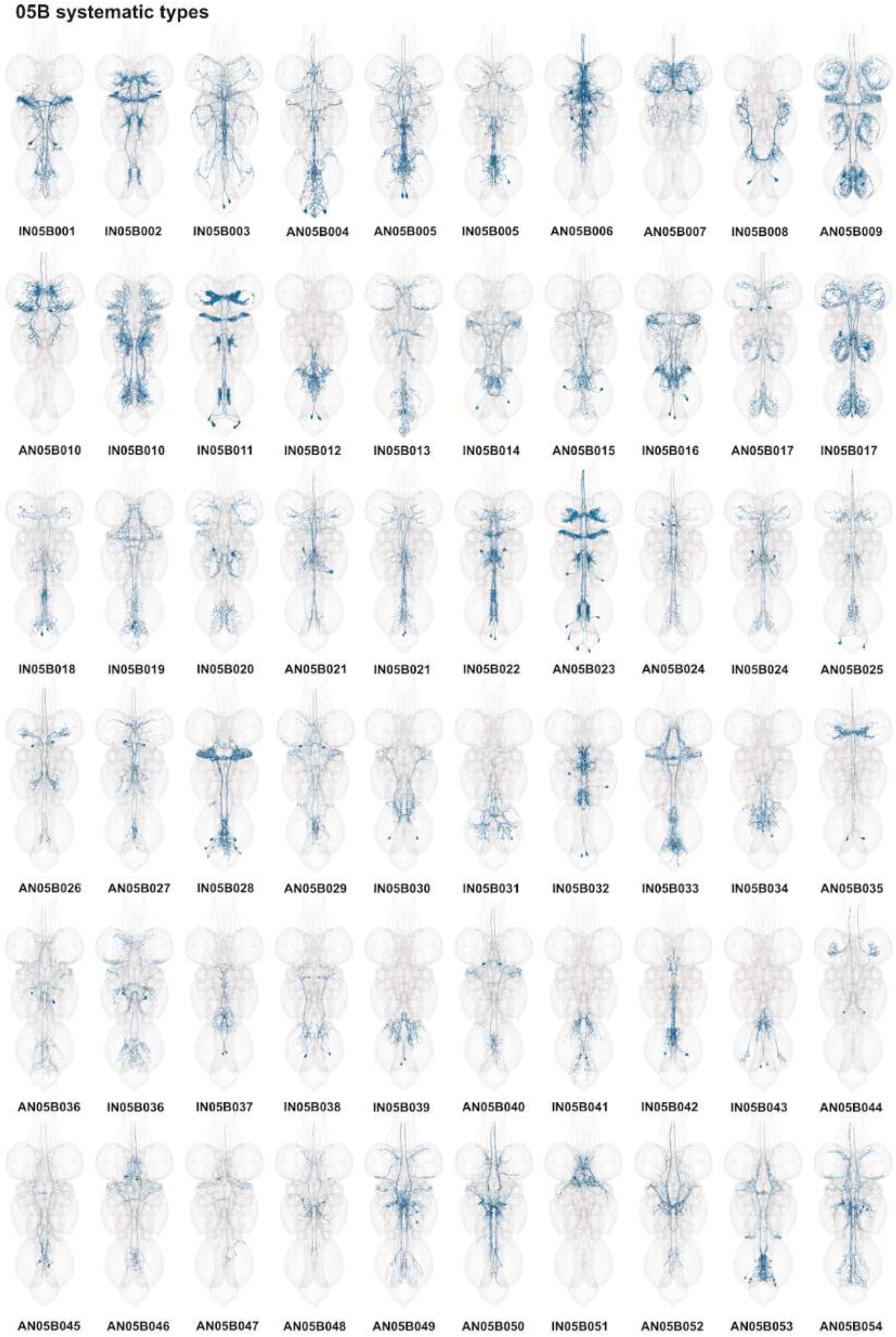
Systematic types of hemilineage 05B. Systematic types have been arranged in numerical order, with neurons of the same type that belong to distinct classes (e.g., intrinsic neuron vs ascending neuron) plotted separately but placed adjacent to each other. Individual neuron meshes have been coloured based on predicted neurotransmitter: dark orange = acetylcholine, blue = gaba, marine = glutamate, dark purple = unknown.

**Figure 22 - figure supplement 3.**
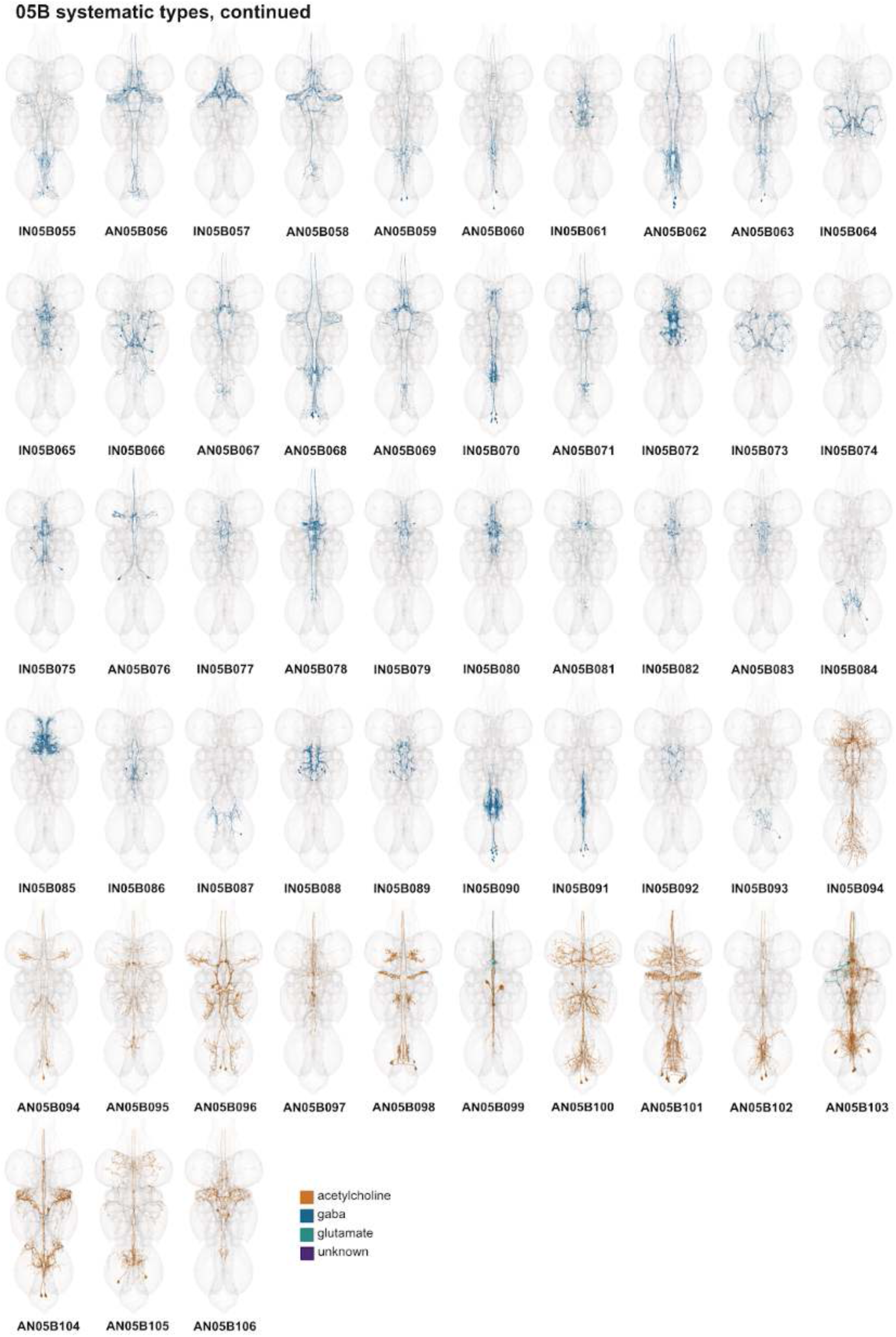
Systematic types of hemilineage 05B, continued. Systematic types have been arranged in numerical order, with neurons of the same type that belong to distinct classes (e.g., intrinsic neuron vs ascending neuron) plotted separately but placed adjacent to each other. Individual neuron meshes have been coloured based on predicted neurotransmitter: dark orange = acetylcholine, blue = gaba, marine = glutamate, dark purple = unknown.

**Figure 22 - figure supplement 4.**
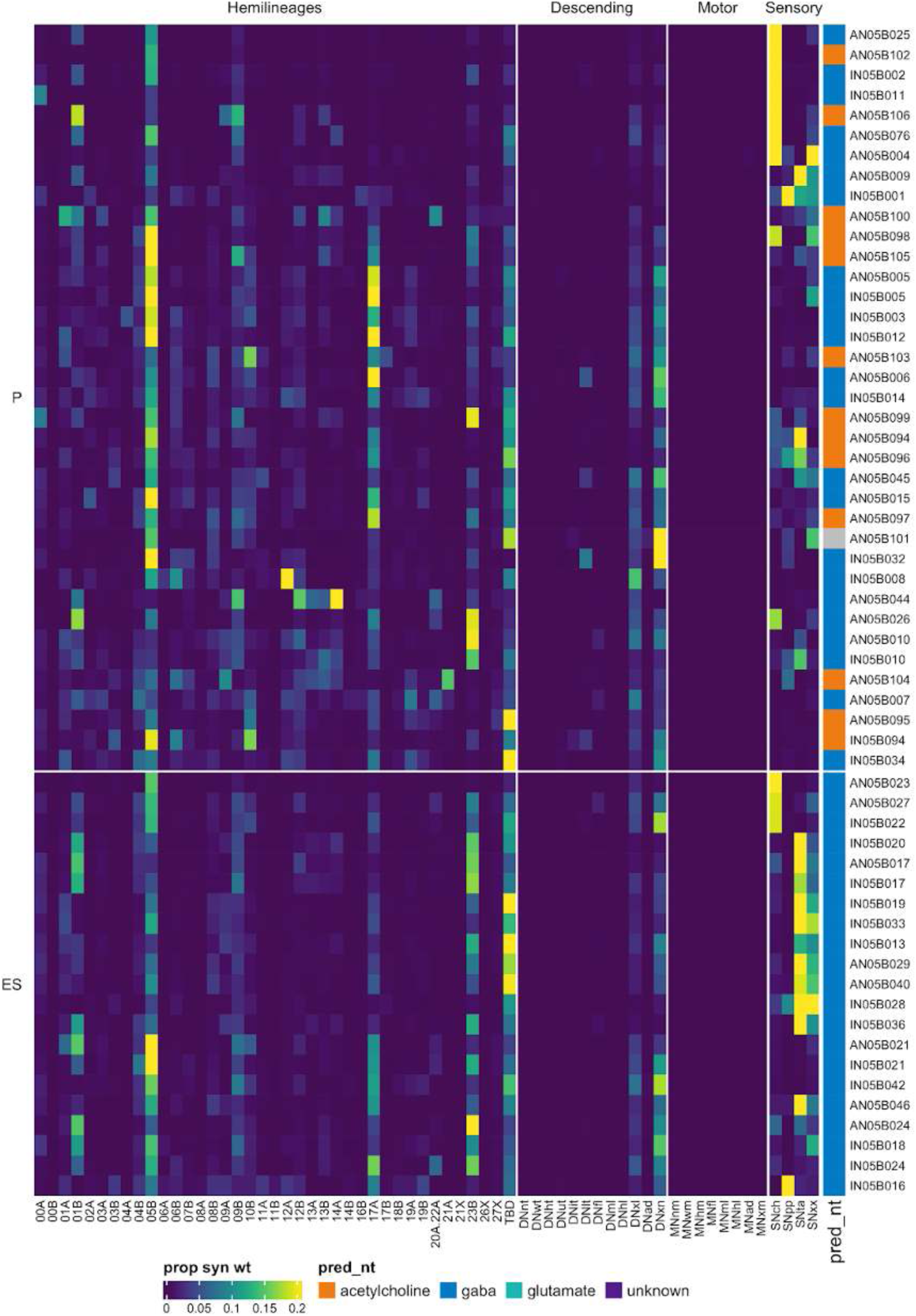
Connectivity to upstream partners by 05B primary and early secondary systematic types. Proportions of synaptic weight to systematic types from upstream partners, normalised by row. 05B neurons have been clustered within each assigned birthtime window (P = primary, ES = early secondary, S = secondary) based on both upstream and downstream connectivity to hemilineages, descending neuron subclasses, motor neuron subclasses, and sensory neuron modalities. Annotation bar is coloured by the most common predicted neurotransmitter for the neurons of each type.

**Figure 22 - figure supplement 5.**
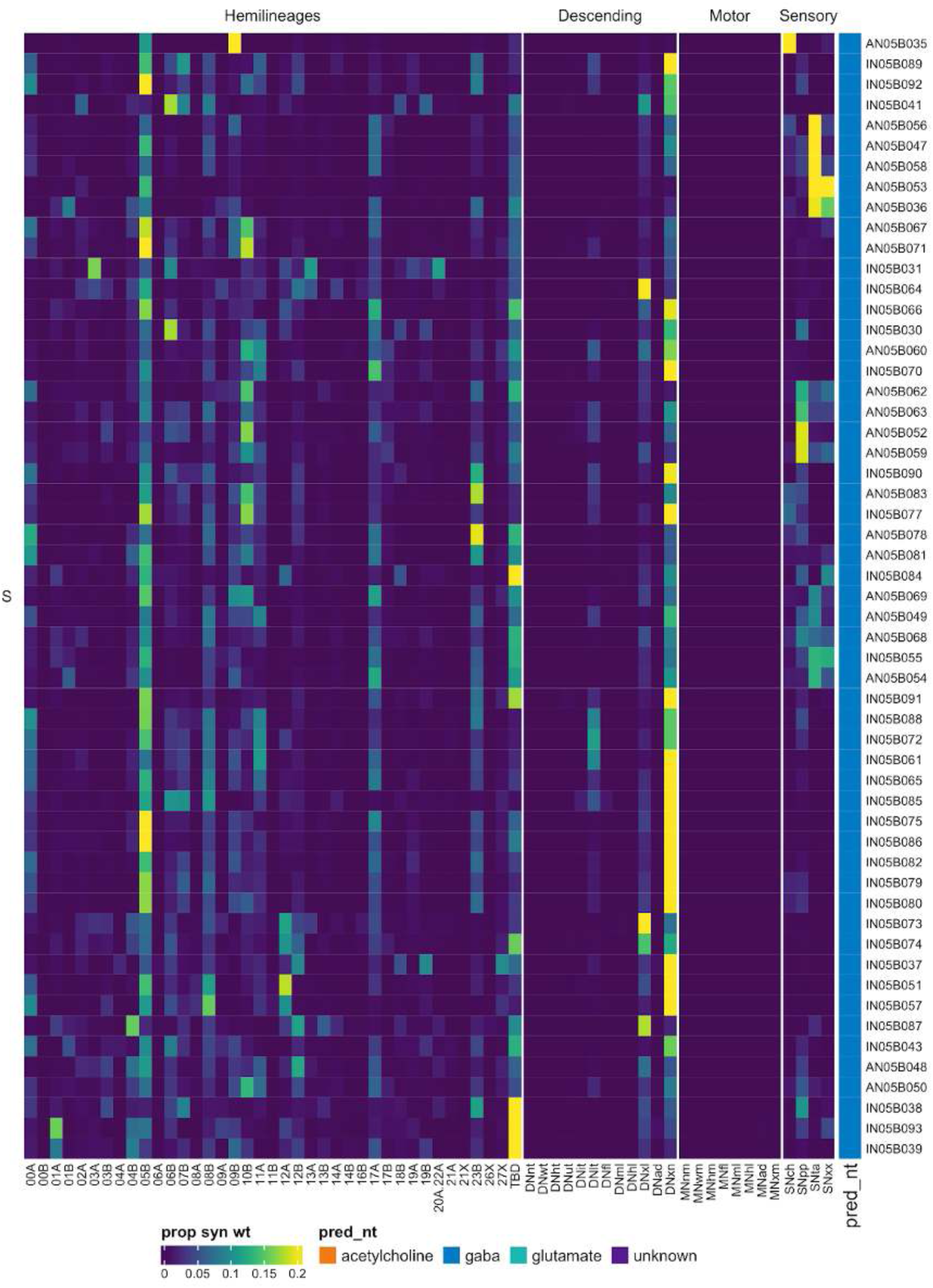
Connectivity to upstream partners by 05B secondary systematic types. Proportions of synaptic weight to systematic types from upstream partners, normalised by row. 05B neurons have been clustered within each assigned birthtime window (P = primary, ES = early secondary, S = secondary) based on both upstream and downstream connectivity to hemilineages, descending neuron subclasses, motor neuron subclasses, and sensory neuron modalities. The annotation bar is coloured by the most common predicted neurotransmitter within each type.

**Figure 22 - figure supplement 6.**
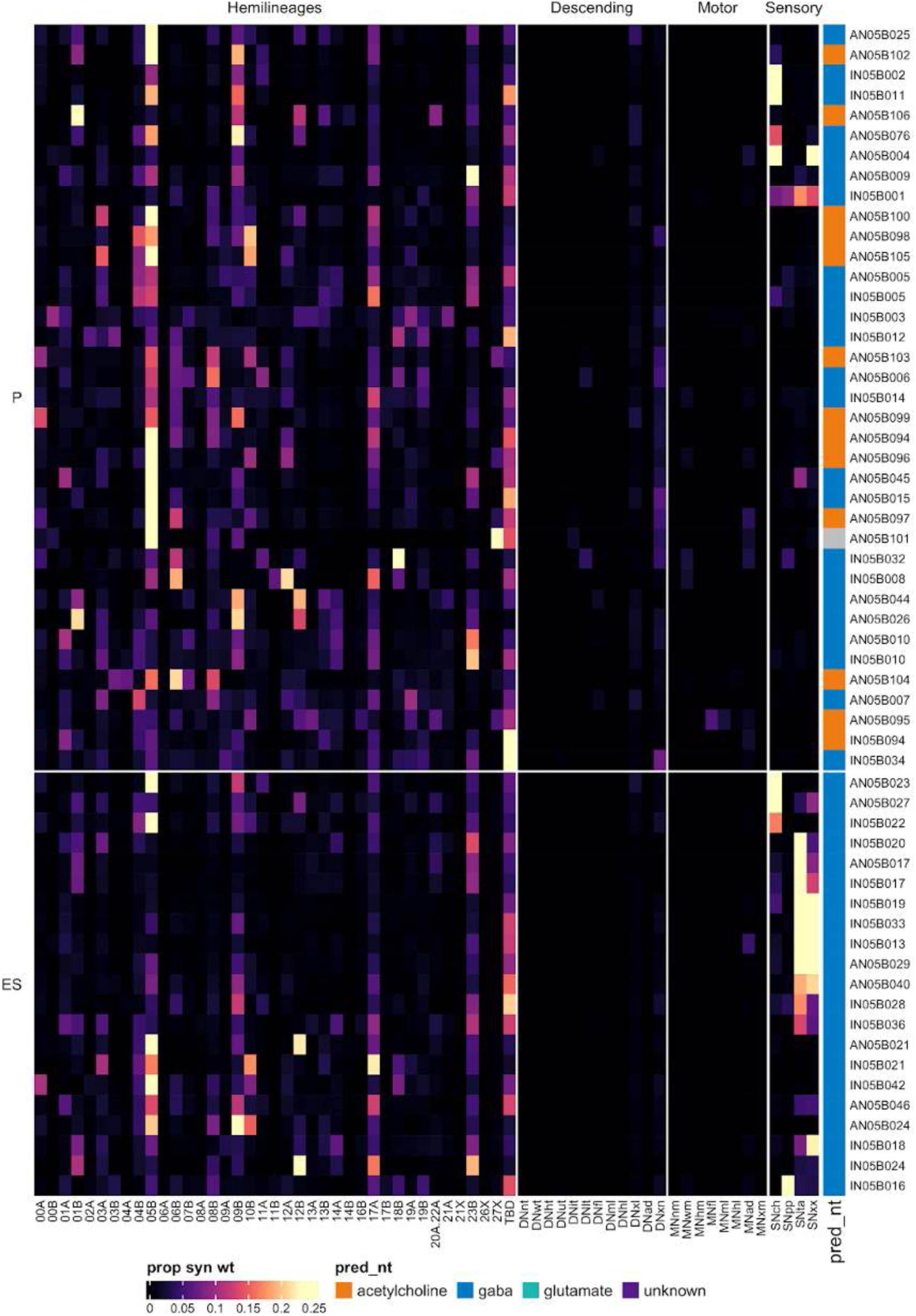
Connectivity to downstream partners by 05B primary and early secondary systematic types. Proportions of synaptic weight from systematic types to downstream partners, normalised by row. 05B neurons have been clustered within each assigned birthtime window (P = primary, ES = early secondary, S = secondary) based on both upstream and downstream connectivity to hemilineages, descending neuron subclasses, motor neuron subclasses, and sensory neuron modalities. The annotation bar is coloured by the most common predicted neurotransmitter within each type.

**Figure 22 - figure supplement 7.**
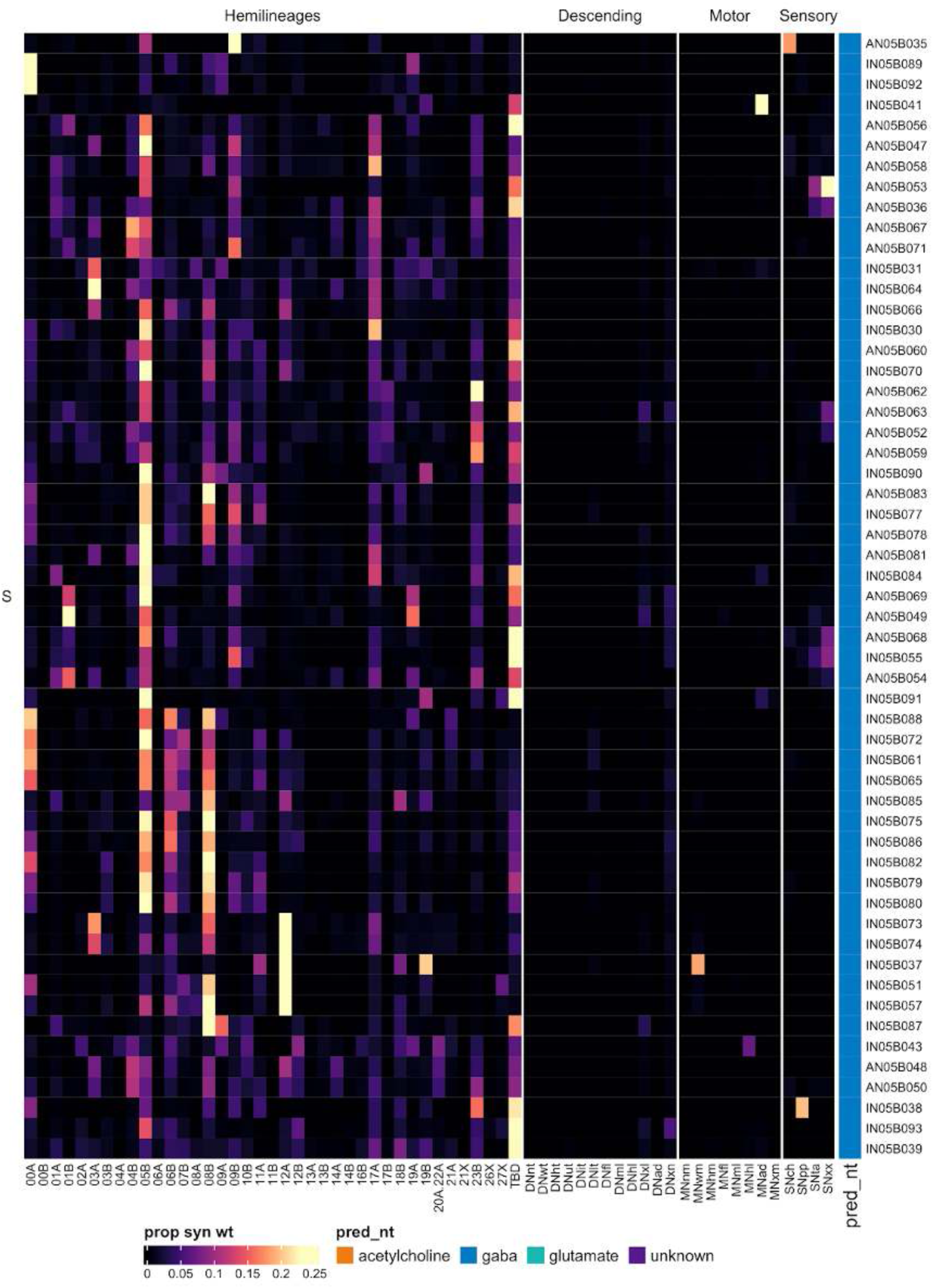
Connectivity to downstream partners by 05B secondary systematic types. Proportions of synaptic weight from systematic types to downstream partners, normalised by row. 05B neurons have been clustered within each assigned birthtime window (P = primary, ES = early secondary, S = secondary) based on both upstream and downstream connectivity to hemilineages, descending neuron subclasses, motor neuron subclasses, and sensory neuron modalities. The annotation bar is coloured by the most common predicted neurotransmitter for the neurons of each type.

#### Hemilineage 06A

Hemilineages 06A and 06B derive from posterior medial neuroblast NB5-2 (Birkholz et al., 2015; Lacin and Truman, 2016), which generates one contralateral MN, 4-8 intersegmental neurons, and ∼30 local interneurons in the embryo (Schmid et al., 1999). Their primary neurites separate into two distinct hemilineages during metamorphosis (Harris et al., 2015). 06A neurons have a very characteristic gross morphology, entering near the posterior edge of the neuromere and projecting very dorsally before crossing the midline. They feature dendrites in ipsilateral flight neuropils in the neuromere of origin, i.e. T1 in neck tectulum, T2 in wing tectulum and T3 in haltere tectulum (Shepherd et al., 2019), but they project to contralateral flight neuropils and are typically intersegmental, with T2 being a neuromere of convergence; A1 neurons innervate the ipsilateral haltere tectulum in T3 and ascend into contralateral T1 (Figure 23A).

**Figure 23.**
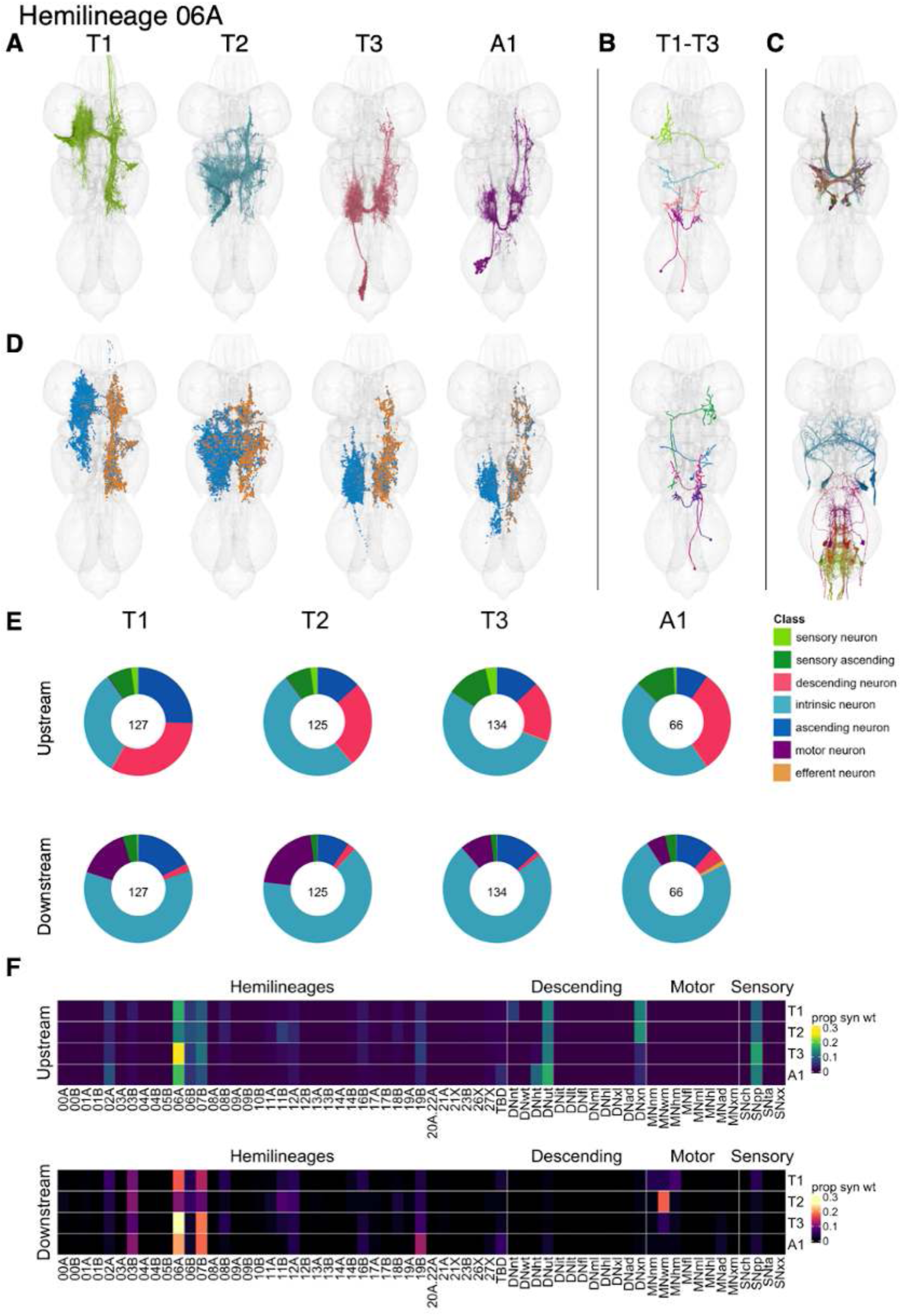
Hemilineage 06A. **A.** Meshes of all RHS secondary neurons plotted in neuromere-specific colours. **B.** “Representative” secondary neuron skeletons plotted in hemineuromere-specific colours. The skeleton with the top accumulated NBLAST score among all neurons from the hemilineage in a given hemineuromere was used. **C.** Neuron meshes of selected examples. Top: putative electrical subcluster 12112. Bottom: motor neuron subcluster 10204. **D.** Predicted synapses of RHS secondary neurons. Blue: postsynapses; dark orange: presynapses. **E.** Proportions of connections from secondary neurons to upstream or downstream partners, normalised by neuromere and coloured by broad class. Numbers of query neurons appear in the centre. **F.** Proportions of synaptic weight from secondary neurons originating in each neuromere to upstream or downstream partners, normalised by row.

We identified a single motor neuron in T2 - A5 that shares the primary neurite tract of 06A (Figure 23C bottom). 06A neurons were reported to be gabaergic (Lacin et al., 2019), but many 06A neurons in T2 are predicted to be glutamatergic, some but not all of which featured thick, simple axons with an unusually low density of presynapses (e.g., subcluster 12112) (Figure 23C top), suggesting that they might be electrically coupled to partners via gap junctions; please see our companion manuscript for details (Cheong et al., 2023).

Secondary 06A neurons survive in T1-A1 (Marin et al., 2012), with comparable neuron numbers in T1-T3 and slightly fewer in A1 (Figure 23E). Their inputs are from descending neurons in the tectulum, proprioceptive sensory neurons, and hemilineages 06A and 07B (Figure 23F). A few types feature especially strong descending control (e.g., IN06A056 and IN06A121), while others receive a greater proportion of sensory input (e.g, IN06A052 and IN06A071) (Figure 23 - figure supplement 6-7). 06A secondary neurons output onto hemilineages 03B, 06A, 07B, and 19B with segmental variation (Figure 23F) and strongly onto wing motor neurons in T2 (Figure 23E). Early born type AN06A016 in T1 strongly targets neck motor neurons (Figure 23 - figure supplement 8). Bilateral activation of 06A secondary neurons results in erratic leg movements with occasional wing flicking and high frequency flapping in a partially spread position (Harris et al., 2015); perhaps 06A inhibition of flight circuits tends to disinhibit leg movement.

**Figure 23 - figure supplement 1.**
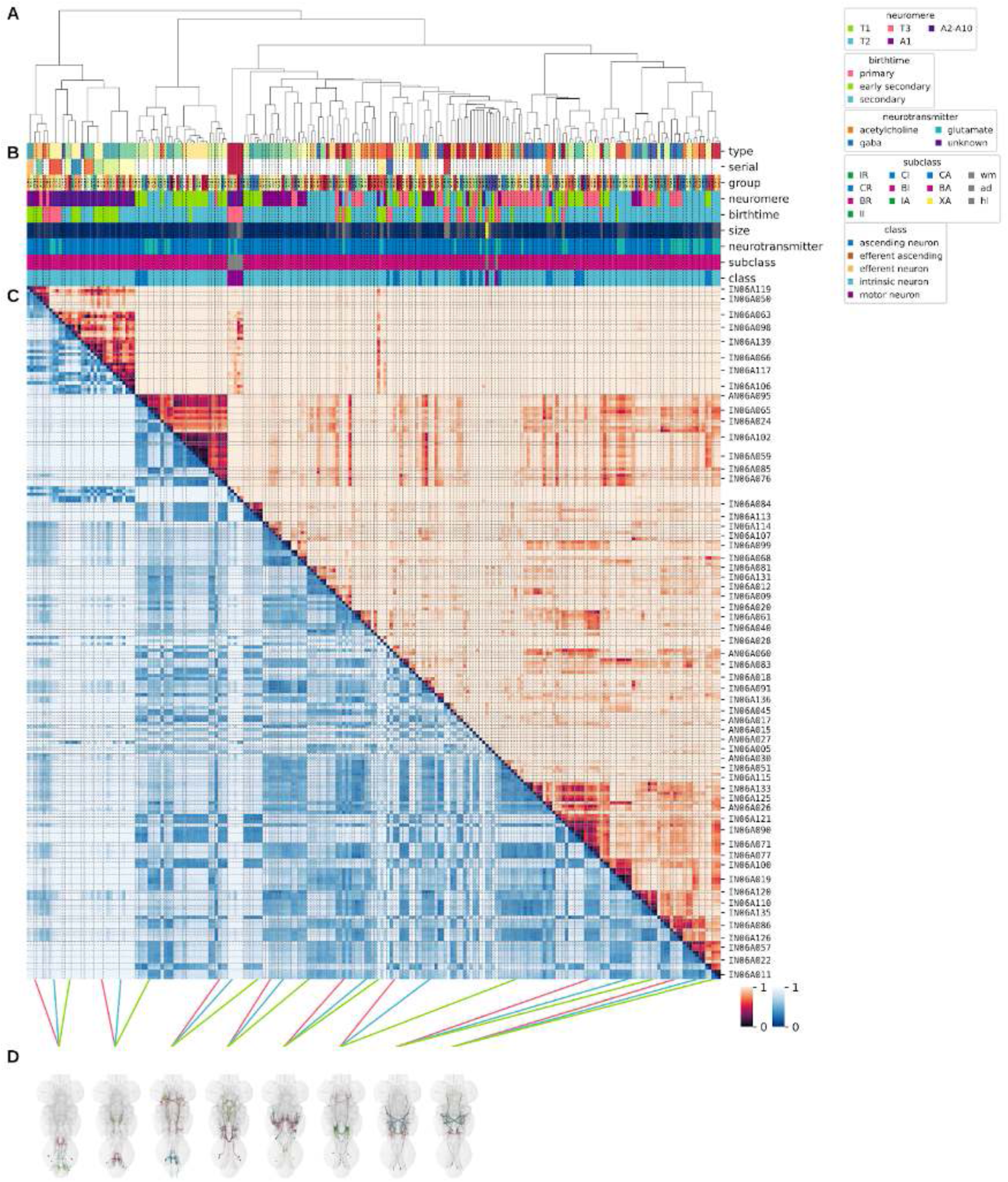
Systematic typing of hemilineage 06A. Types for motor neurons were assigned separately as outlined in our accompanying manuscript (Cheong et al., 2023). **A.** Hierarchical clustering dendrogram of hemilineage groups by laterally and serially aggregated connectivity cosine clustering. **B.** Categorical annotations of each hemilineage group, each column corresponding to the aligned leaf in A. Colours for type, serial set, and group are arbitrary for visualisation. Colours for neuromere, birthtime, neurotransmitter, subclass, and class are as in all other figures. **C.** Similarity distance heatmap for hemilineage. Cosine distance is in the upper triangle, while laterally symmetrised NBLAST distance is in the lower triangle. Systematic type names of some types are labelled. **D.** Morphologically representative groups from dendrogram subtrees. Each group, indicated by colour and line connecting to its column in B and C, is the most morphologically representative group (medoid of NBLAST distance) from a subtree of A. The subtrees (flat clusters) are equal height cuts of A determined to yield the number of groups per plot and plots in D.

**Figure 23 - figure supplement 2.**
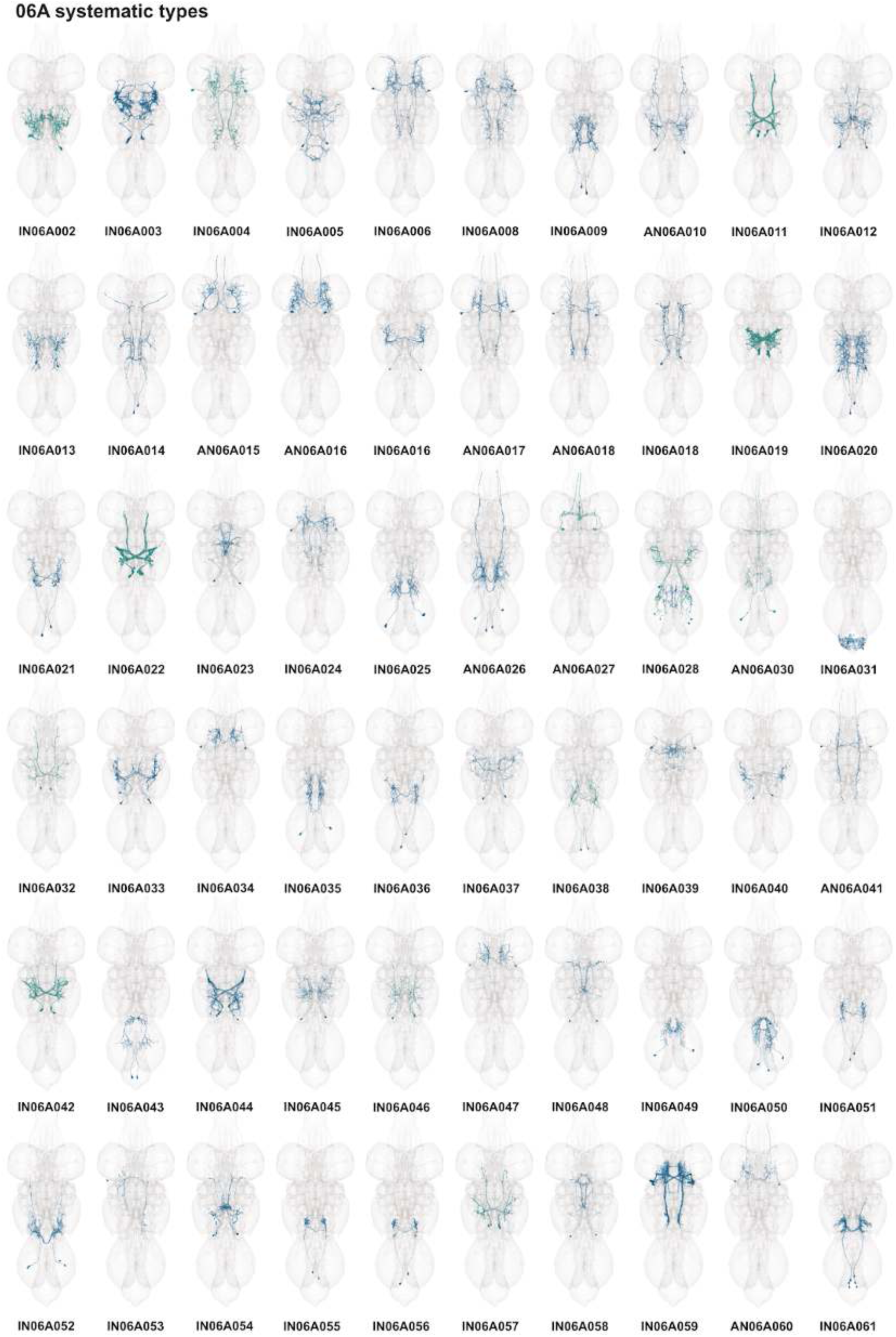
Systematic types of hemilineage 06A. Systematic types have been arranged in numerical order, with neurons of the same type that belong to distinct classes (e.g., intrinsic neuron vs ascending neuron) plotted separately but placed adjacent to each other. Individual neuron meshes have been coloured based on predicted neurotransmitter: dark orange = acetylcholine, blue = gaba, marine = glutamate, dark purple = unknown.

**Figure 23 - figure supplement 3.**
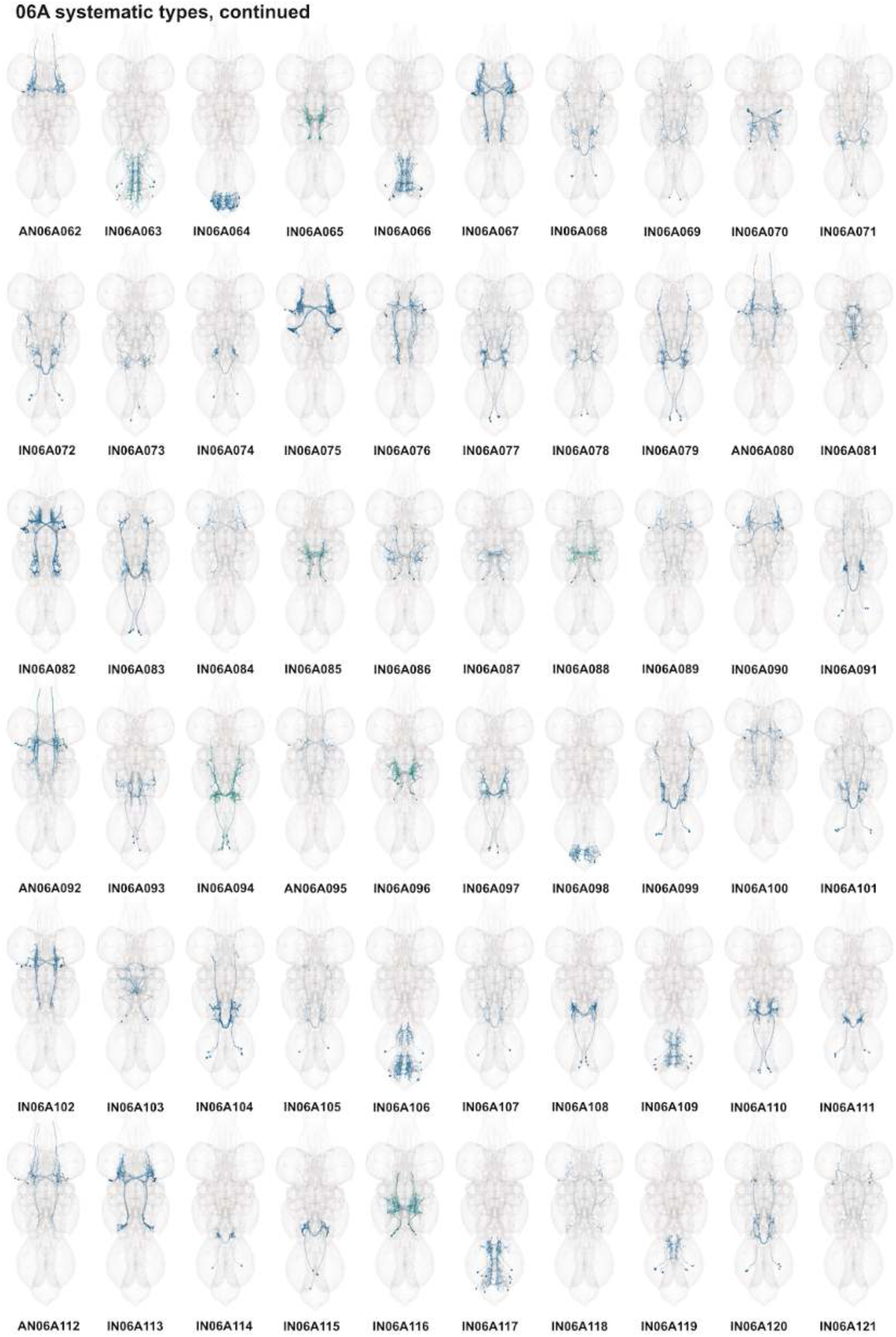
Systematic types of hemilineage 06A, continued. Systematic types have been arranged in numerical order, with neurons of the same type that belong to distinct classes (e.g., intrinsic neuron vs ascending neuron) plotted separately but placed adjacent to each other. Individual neuron meshes have been coloured based on predicted neurotransmitter: dark orange = acetylcholine, blue = gaba, marine = glutamate, dark purple = unknown.

**Figure 23 - figure supplement 4.**
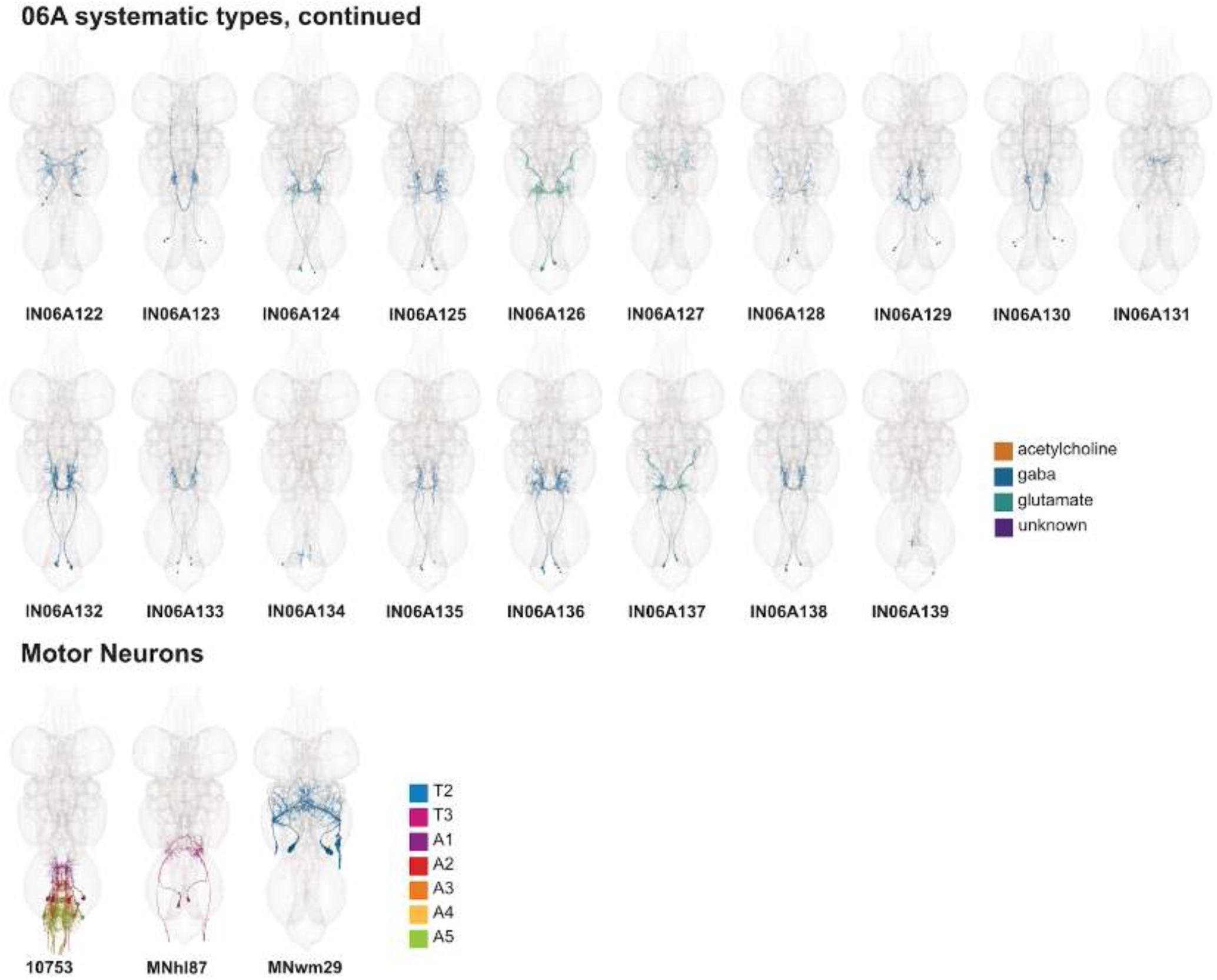
Systematic types of hemilineage 06A, continued. Systematic types have been arranged in numerical order, with neurons of the same type that belong to distinct classes (e.g., intrinsic neuron vs ascending neuron) plotted separately but placed adjacent to each other. Individual neuron meshes have been coloured based on predicted neurotransmitter: dark orange = acetylcholine, blue = gaba, marine = glutamate, dark purple = unknown. Motor neurons (typed separately in (Cheong et al., 2023)) have been plotted by serial set if identified in multiple neuromeres and by systematic type if not. Individual motor neuron meshes have been coloured based on soma neuromere: blue = T2, magenta = T3, purple = A1, red = A2, dark orange = A3, dark yellow = A4, green = A5.

**Figure 23 - figure supplement 5.**
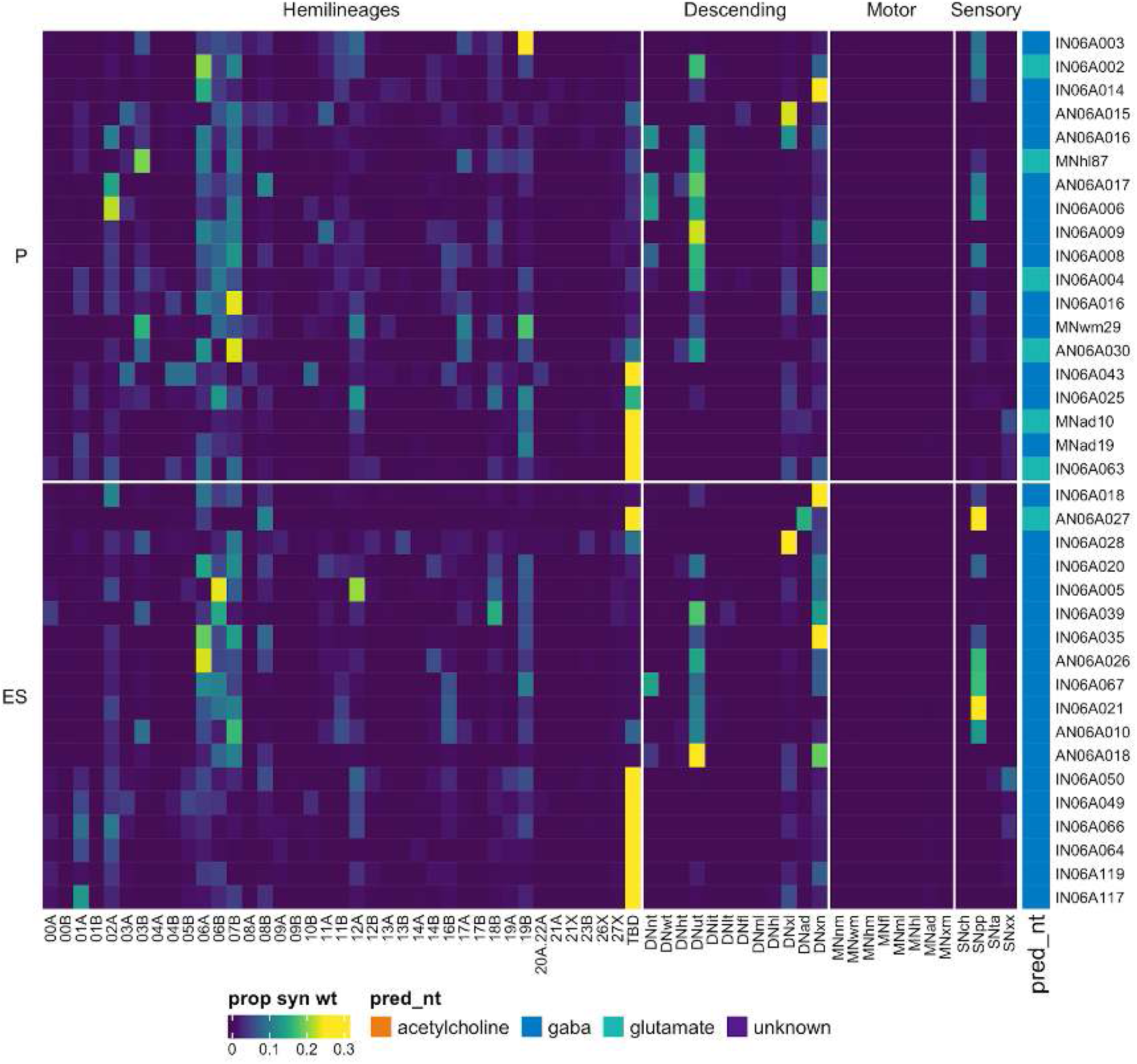
Connectivity to upstream partners by 06A primary and early secondary systematic types. Proportions of synaptic weight to systematic types from upstream partners, normalised by row. 06A neurons have been clustered within each assigned birthtime window (P = primary, ES = early secondary, S = secondary) based on both upstream and downstream connectivity to hemilineages, descending neuron subclasses, motor neuron subclasses, and sensory neuron modalities. Annotation bar is coloured by the most common predicted neurotransmitter for the neurons of each type.

**Figure 23 - figure supplement 6.**
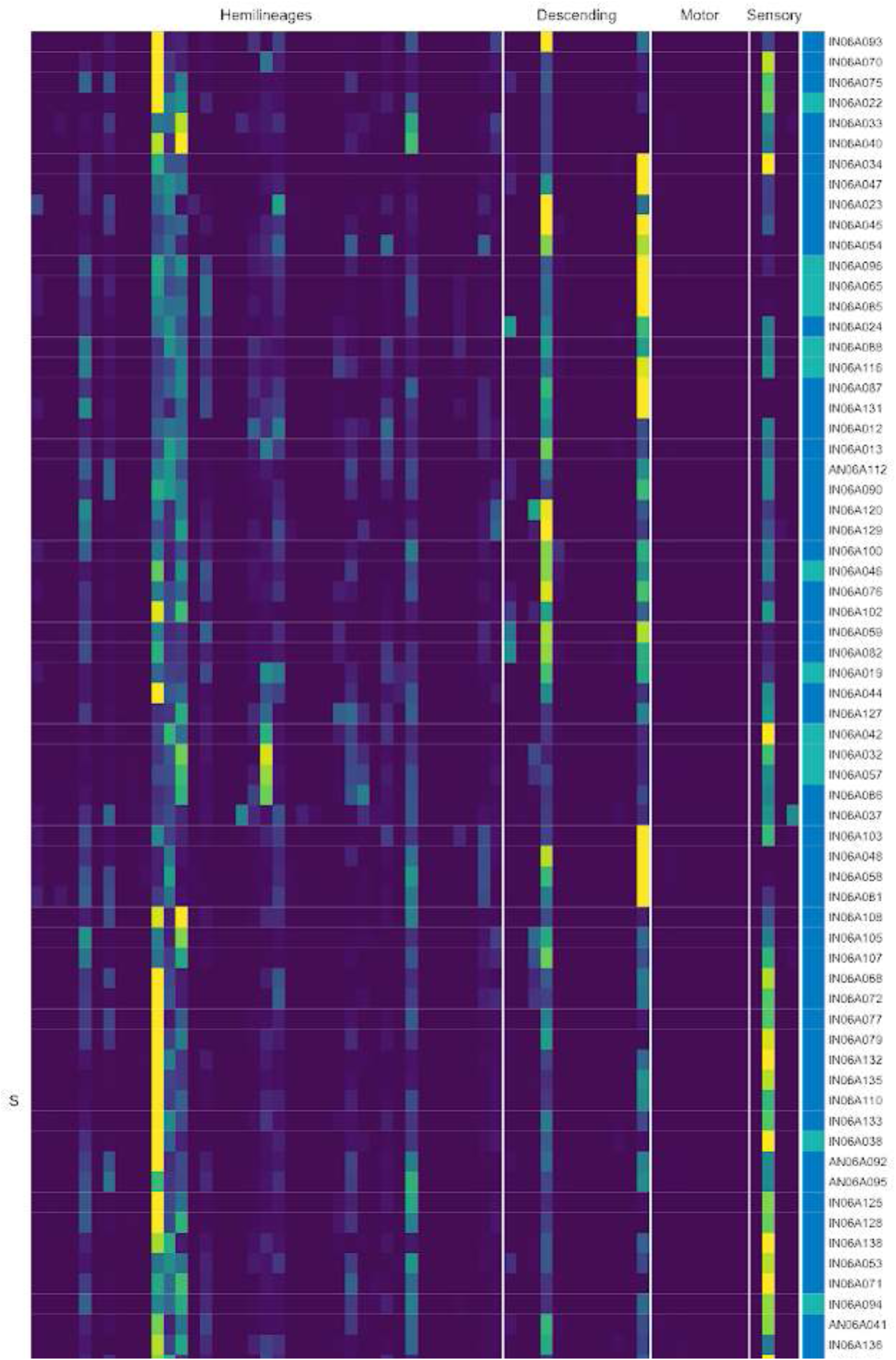
Connectivity to upstream partners by 06A secondary systematic types. Proportions of synaptic weight to systematic types from upstream partners, normalised by row. 06A neurons have been clustered within each assigned birthtime window (P = primary, ES = early secondary, S = secondary) based on both upstream and downstream connectivity to hemilineages, descending neuron subclasses, motor neuron subclasses, and sensory neuron modalities. The annotation bar is coloured by the most common predicted neurotransmitter within each type.

**Figure 23 - figure supplement 7.**
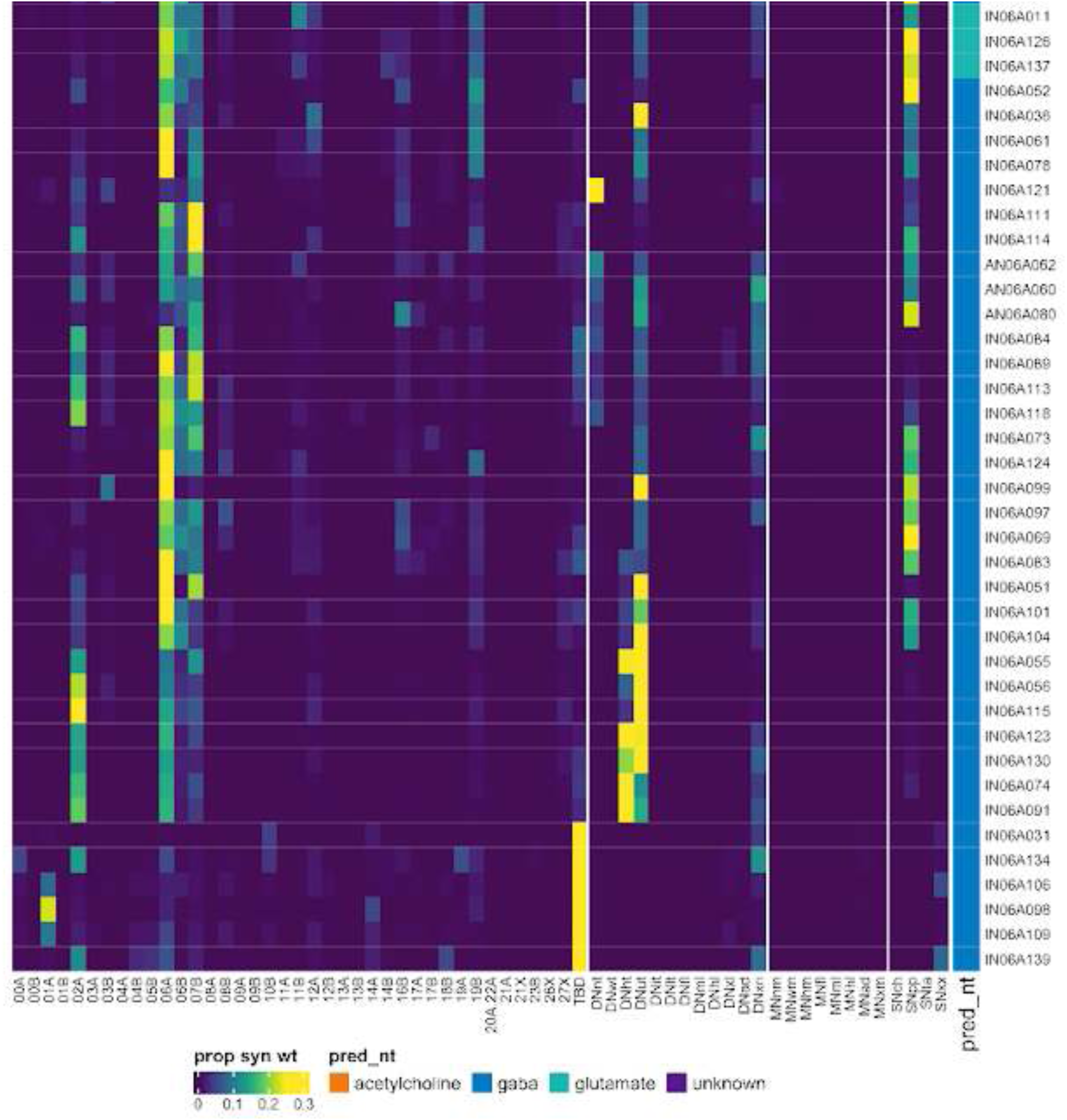
Connectivity to upstream partners by 06A secondary systematic types, continued. Proportions of synaptic weight to systematic types from upstream partners, normalised by row. 06A neurons have been clustered within each assigned birthtime window (P = primary, ES = early secondary, S = secondary) based on both upstream and downstream connectivity to hemilineages, descending neuron subclasses, motor neuron subclasses, and sensory neuron modalities. The annotation bar is coloured by the most common predicted neurotransmitter within each type.

**Figure 23 - figure supplement 8.**
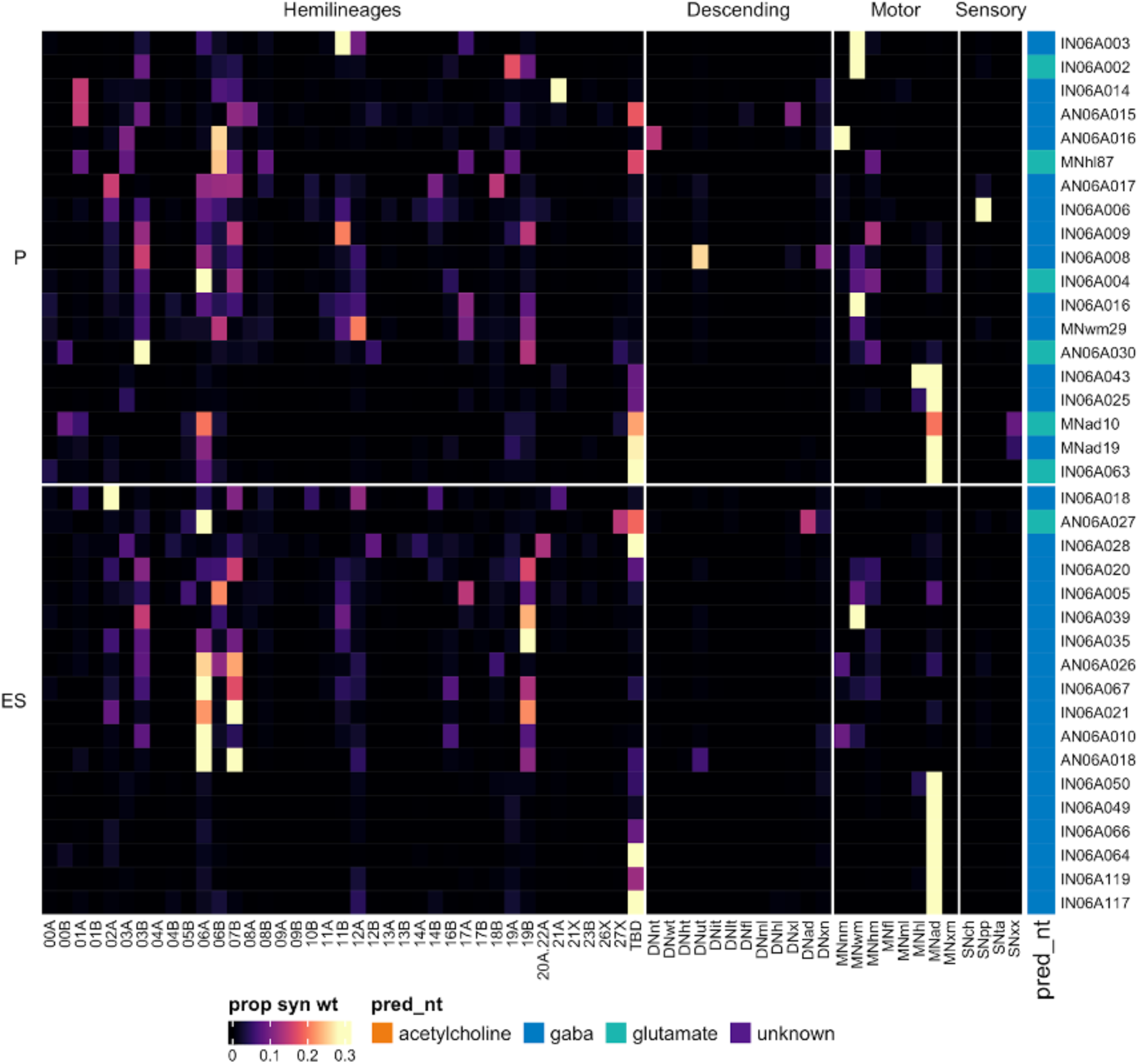
Connectivity to downstream partners by 06A primary and early secondary systematic types. Proportions of synaptic weight from systematic types to downstream partners, normalised by row. 06A neurons have been clustered within each assigned birthtime window (P = primary, ES = early secondary, S = secondary) based on both upstream and downstream connectivity to hemilineages, descending neuron subclasses, motor neuron subclasses, and sensory neuron modalities. The annotation bar is coloured by the most common predicted neurotransmitter within each type.

**Figure 23 - figure supplement 9.**
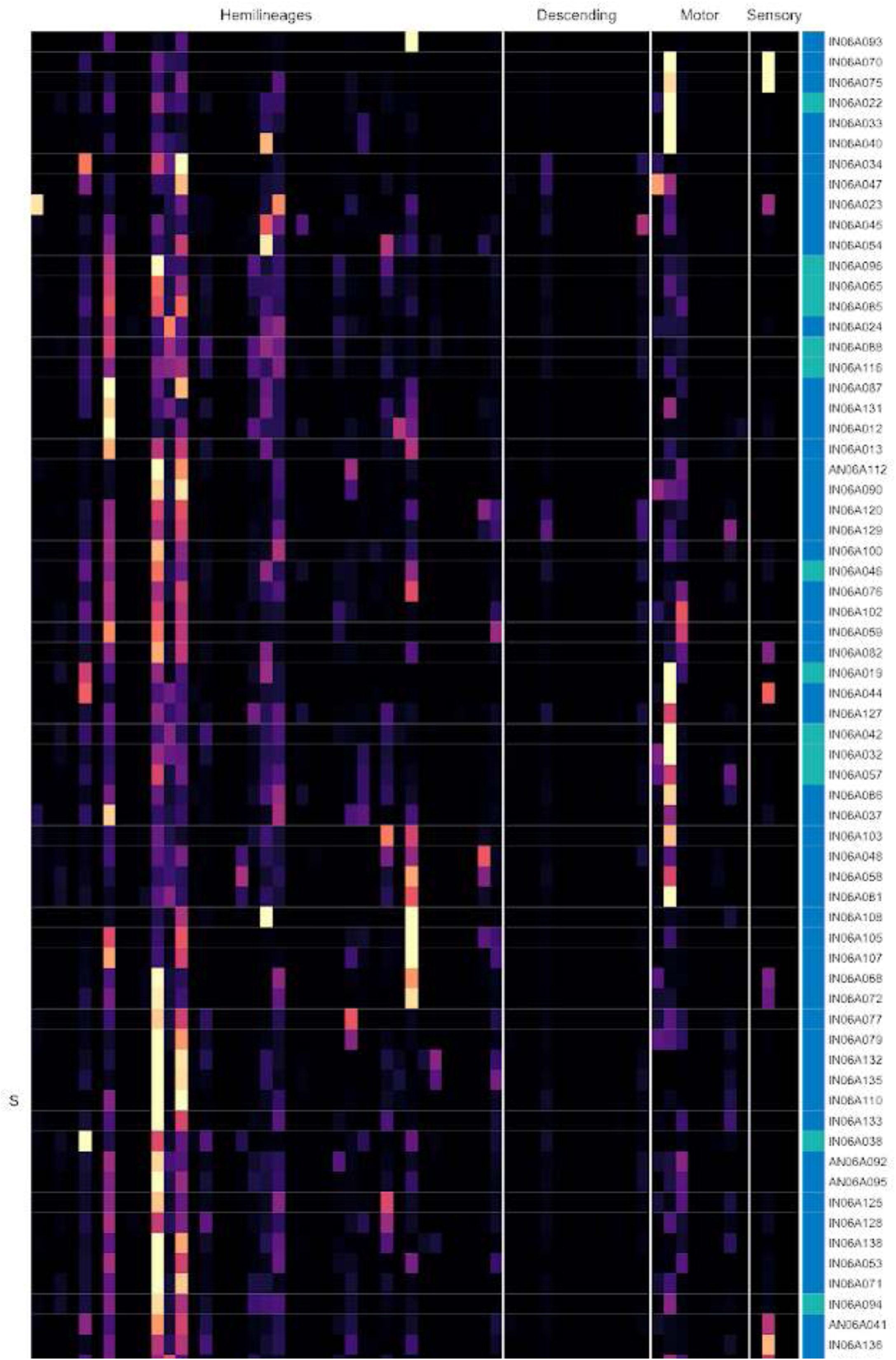
Connectivity to downstream partners by 06A secondary systematic types. Proportions of synaptic weight from systematic types to downstream partners, normalised by row. 06A neurons have been clustered within each assigned birthtime window (P = primary, ES = early secondary, S = secondary) based on both upstream and downstream connectivity to hemilineages, descending neuron subclasses, motor neuron subclasses, and sensory neuron modalities. The annotation bar is coloured by the most common predicted neurotransmitter for the neurons of each type.

**Figure 23 - figure supplement 10.**
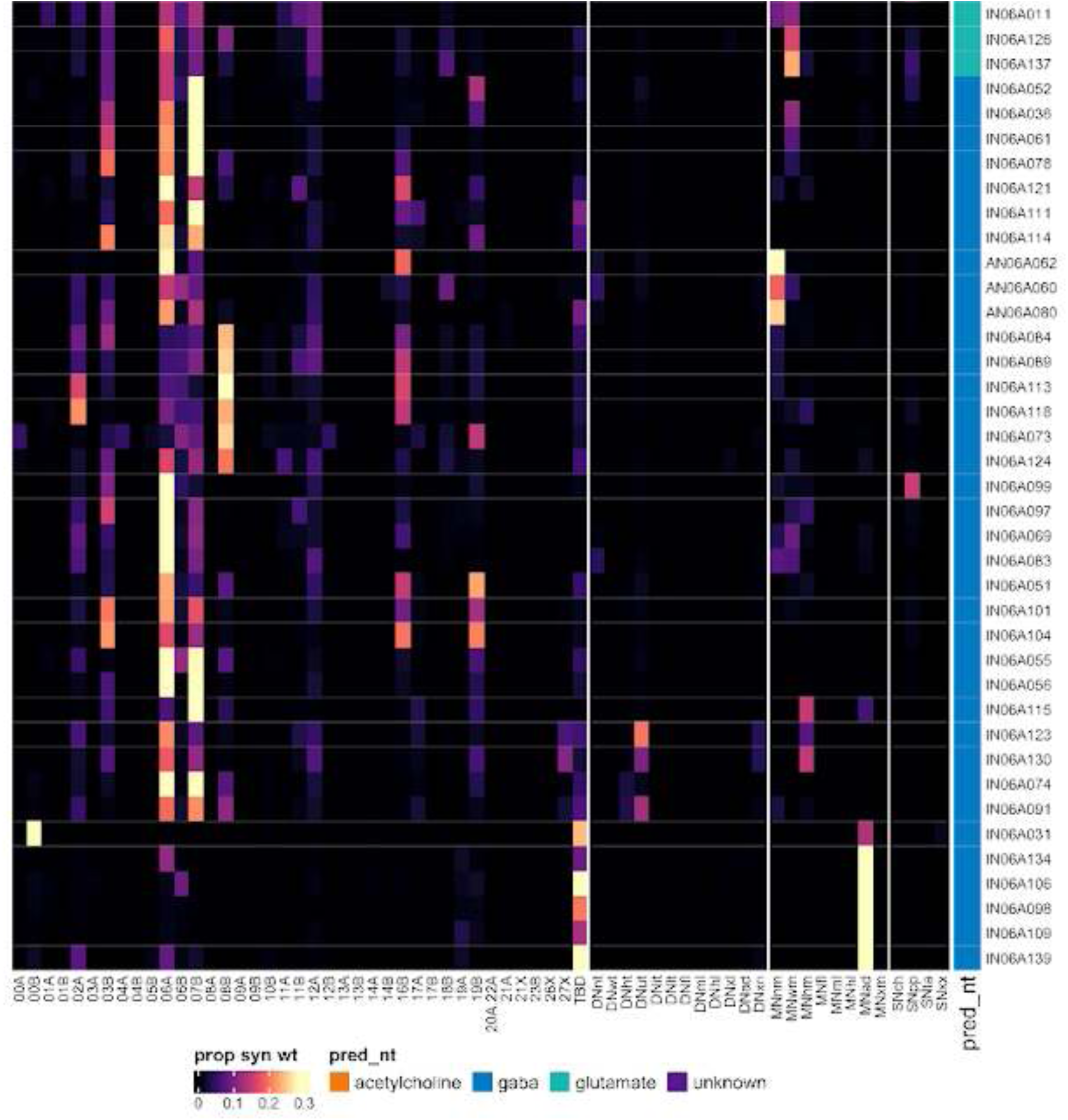
Connectivity to downstream partners by 06A secondary systematic types, continued. Proportions of synaptic weight from systematic types to downstream partners, normalised by row. 06A neurons have been clustered within each assigned birthtime window (P = primary, ES = early secondary, S = secondary) based on both upstream and downstream connectivity to hemilineages, descending neuron subclasses, motor neuron subclasses, and sensory neuron modalities. The annotation bar is coloured by the most common predicted neurotransmitter for the neurons of each type.

#### Hemilineage 06B

Neurons from hemilineage 06B enter the neuropil in the posterior of the neuromere, medially to the 06A neurons, and project dorso-medially to cross the midline in the posterior intermediate commissure, anterior to the 12B neurons (Shepherd et al., 2016). They arborise extensively both sides of the midline in the ventral tectulum and the majority are intersegmental, with the mass of the arborisation from all three neuromeres concentrated in T2 (Figure 24A). Based on their primary neurites, the intersegmental neurons forming varicose projections in lateral leg neuropils (Shepherd et al., 2019) likely belong to 06B rather than 06A (e.g., IN06B035) (Figure 24C bottom). Like 05B, 06B neurons innervate both leg and flight neuropils and are especially prominent in the ovoid (Shepherd et al., 2019), and many types ascend via the neck connective (e.g., AN06B002) (Figure 24C top).

**Figure 24.**
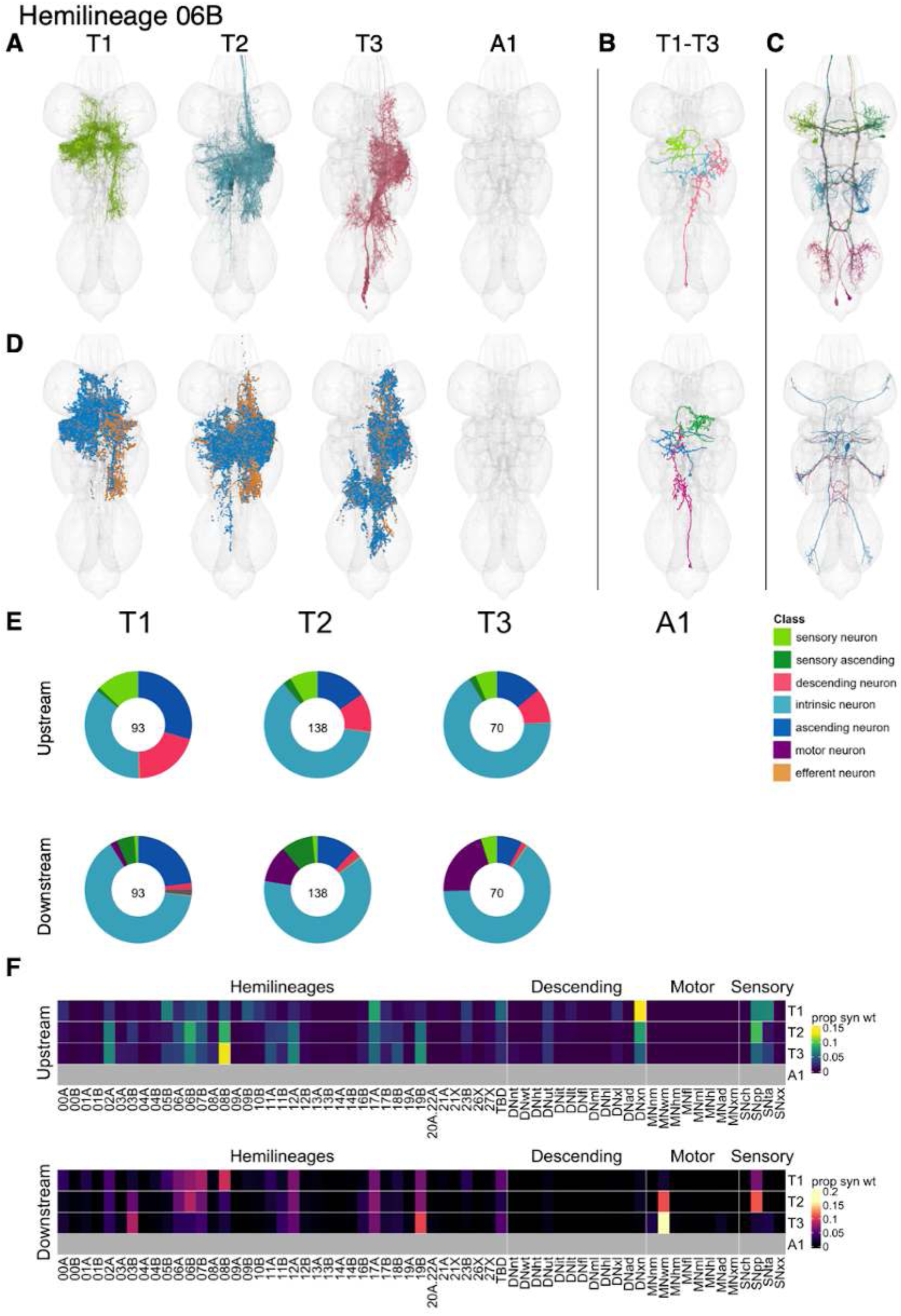
Hemilineage 06B. **A.** Meshes of all RHS secondary neurons plotted in neuromere-specific colours. **B.** “Representative” secondary neuron skeletons plotted in hemineuromere-specific colours. The skeleton with the top accumulated NBLAST score among all neurons from the hemilineage in a given hemineuromere was used. **C.** Neuron meshes of selected examples. Top: ascending serial set 10257. Bottom: centrifugal serial set 11261. **D.** Predicted synapses of RHS secondary neurons. Blue: postsynapses; dark orange: presynapses. **E.** Proportions of connections from secondary neurons to upstream or downstream partners, normalised by neuromere and coloured by broad class. Numbers of query neurons appear in the centre. **F.** Proportions of synaptic weight from secondary neurons originating in each neuromere to upstream or downstream partners, normalised by row.

Secondary 06B neurons survive in all thoracic neuromeres but with many more neurons in T2 than T1 or T3 (Figure 24E). Their inputs come mainly from descending neurons, tactile and proprioceptive sensory neurons, and numerous hemilineages including 02A, 06B, 08B, 12A, 17A, and 19B, with some segmental variation (Figure 24F). Specific 06B types can be dedicated to proprioceptive (e.g., AN/IN06B14), tactile (e.g., IN06B016) or both (e.g., IN06B078) (Figure 24 - figure supplement 4-5). 06B secondary neurons are predicted to be gabaergic as expected (Lacin et al., 2019) and can inhibit wing motor neurons (Figure 24E), proprioceptive sensory neurons, and hemilineages 03B, 06A, 06B, 07B, 08B, 12A, 17A, and 19B with some segmental variation (Figure 24F). AN/IN06B040 neurons target neck motor neurons (Figure 24 - figure supplement 6-7). Bilateral activation of 06B neurons elicits leg-related movements without substantial forward locomotion along with occasional wing grooming bouts (Harris et al., 2015).

**Figure 24 - figure supplement 1.**
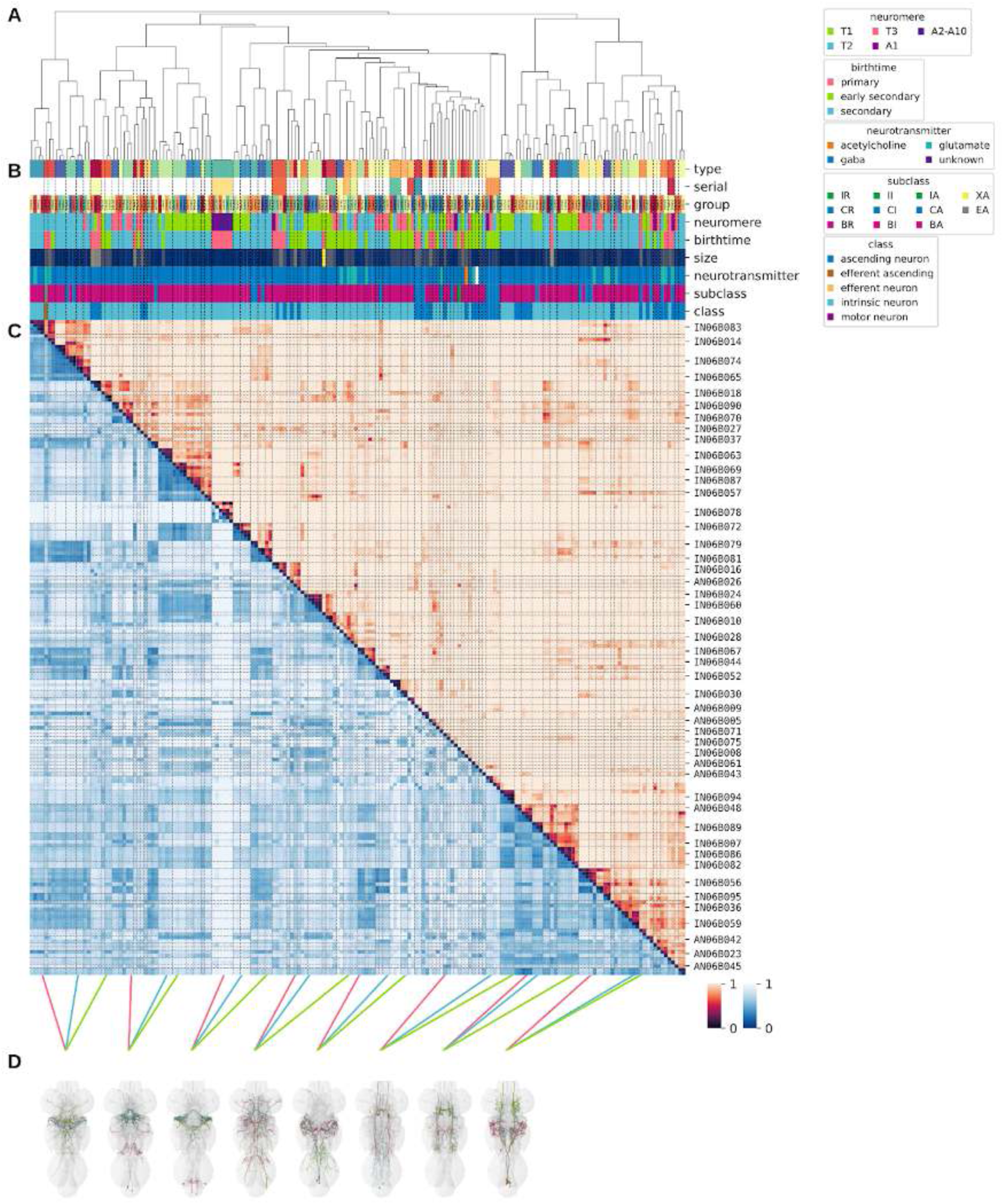
Systematic typing of hemilineage 06B. **A.** Hierarchical clustering dendrogram of hemilineage groups by laterally and serially aggregated connectivity cosine clustering. **B.** Categorical annotations of each hemilineage group, each column corresponding to the aligned leaf in A. Colours for type, serial set, and group are arbitrary for visualisation. Colours for neuromere, birthtime, neurotransmitter, subclass, and class are as in all other figures. **C.** Similarity distance heatmap for hemilineage. Cosine distance is in the upper triangle, while laterally symmetrised NBLAST distance is in the lower triangle. Systematic type names of some types are labelled. **D.** Morphologically representative groups from dendrogram subtrees. Each group, indicated by colour and line connecting to its column in B and C, is the most morphologically representative group (medoid of NBLAST distance) from a subtree of A. The subtrees (flat clusters) are equal height cuts of A determined to yield the number of groups per plot and plots in D.

**Figure 24 - figure supplement 2.**
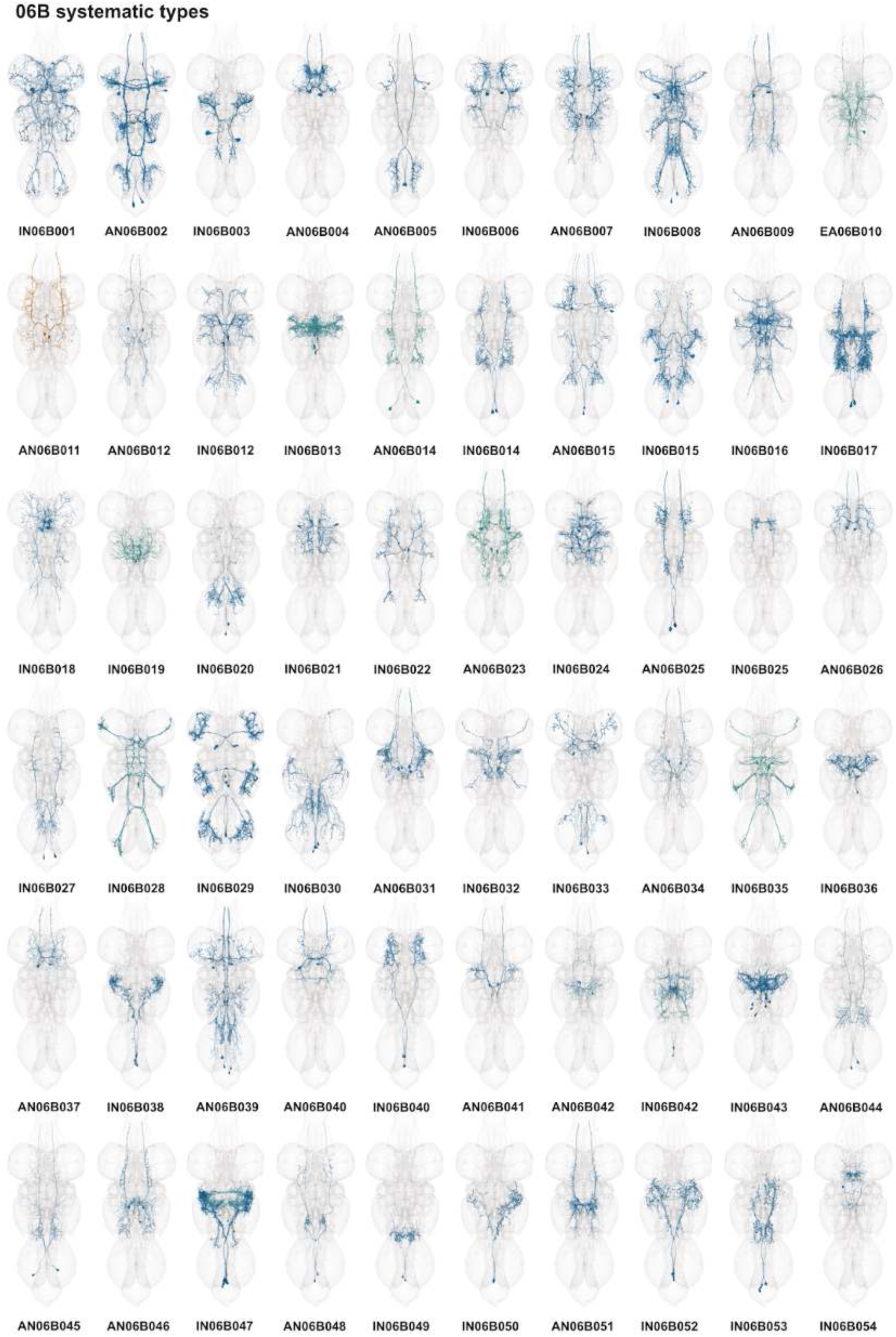
Systematic types of hemilineage 06B. Systematic types have been arranged in numerical order, with neurons of the same type that belong to distinct classes (e.g., intrinsic neuron vs ascending neuron) plotted separately but placed adjacent to each other. Individual neuron meshes have been coloured based on predicted neurotransmitter: dark orange = acetylcholine, blue = gaba, marine = glutamate, dark purple = unknown.

**Figure 24 - figure supplement 3.**
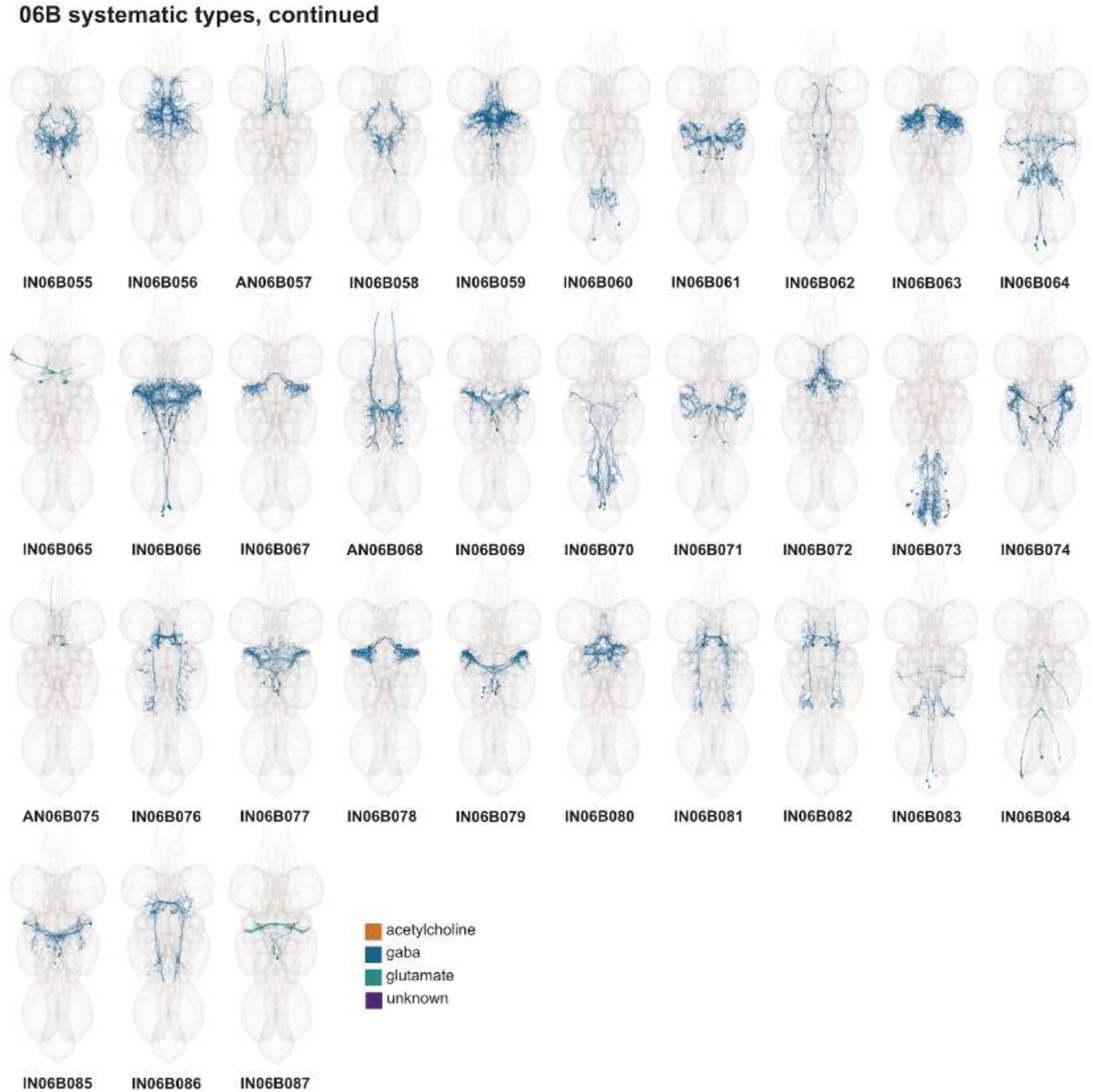
Systematic types of hemilineage 06B, continued. Systematic types have been arranged in numerical order, with neurons of the same type that belong to distinct classes (e.g., intrinsic neuron vs ascending neuron) plotted separately but placed adjacent to each other. Individual neuron meshes have been coloured based on predicted neurotransmitter: dark orange = acetylcholine, blue = gaba, marine = glutamate, dark purple = unknown.

**Figure 24 - figure supplement 4.**
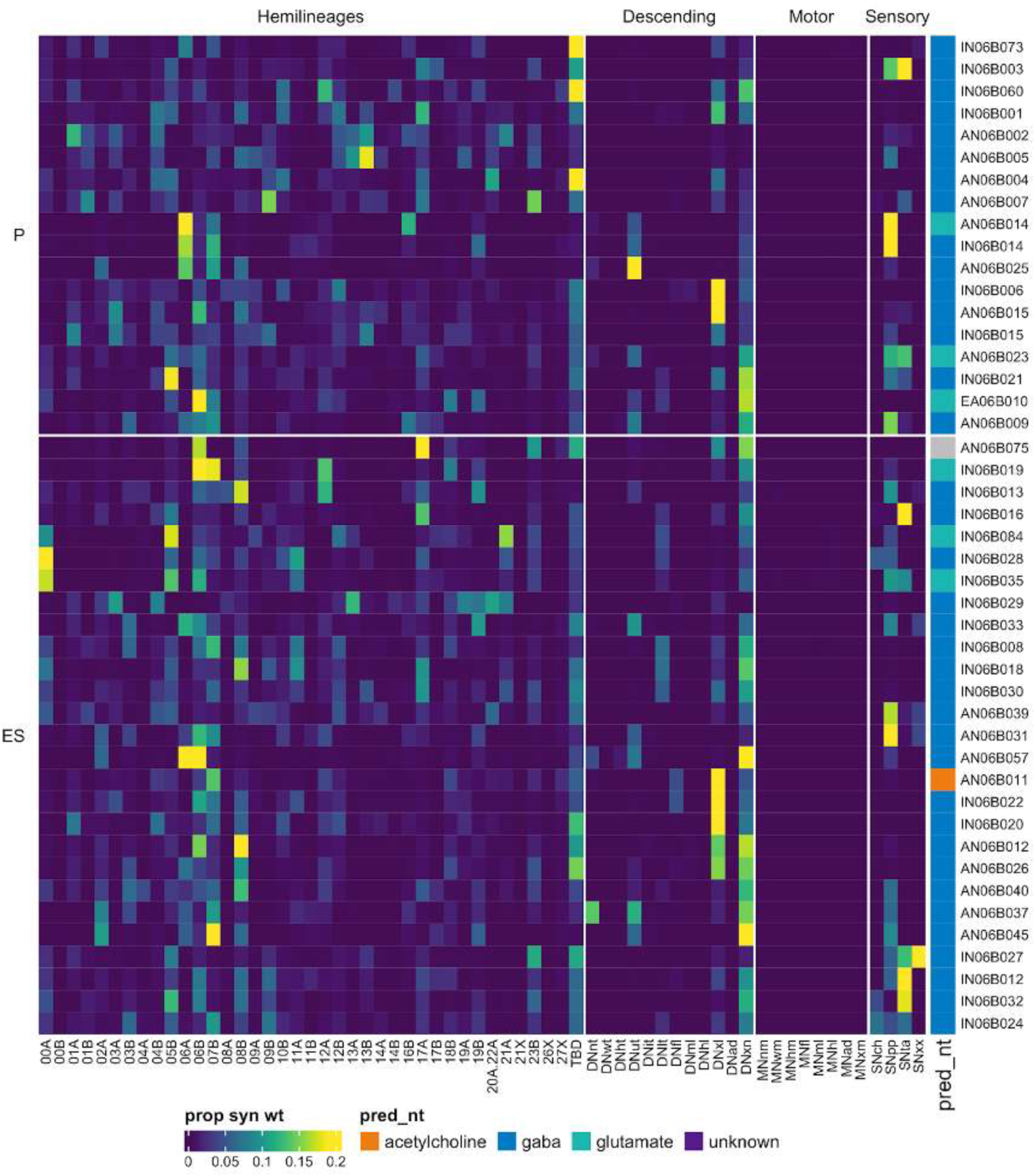
Connectivity to upstream partners by 06B primary and early secondary systematic types. Proportions of synaptic weight to systematic types from upstream partners, normalised by row. 06B neurons have been clustered within each assigned birthtime window (P = primary, ES = early secondary, S = secondary) based on both upstream and downstream connectivity to hemilineages, descending neuron subclasses, motor neuron subclasses, and sensory neuron modalities. Annotation bar is coloured by the most common predicted neurotransmitter for the neurons of each type.

**Figure 24 - figure supplement 5.**
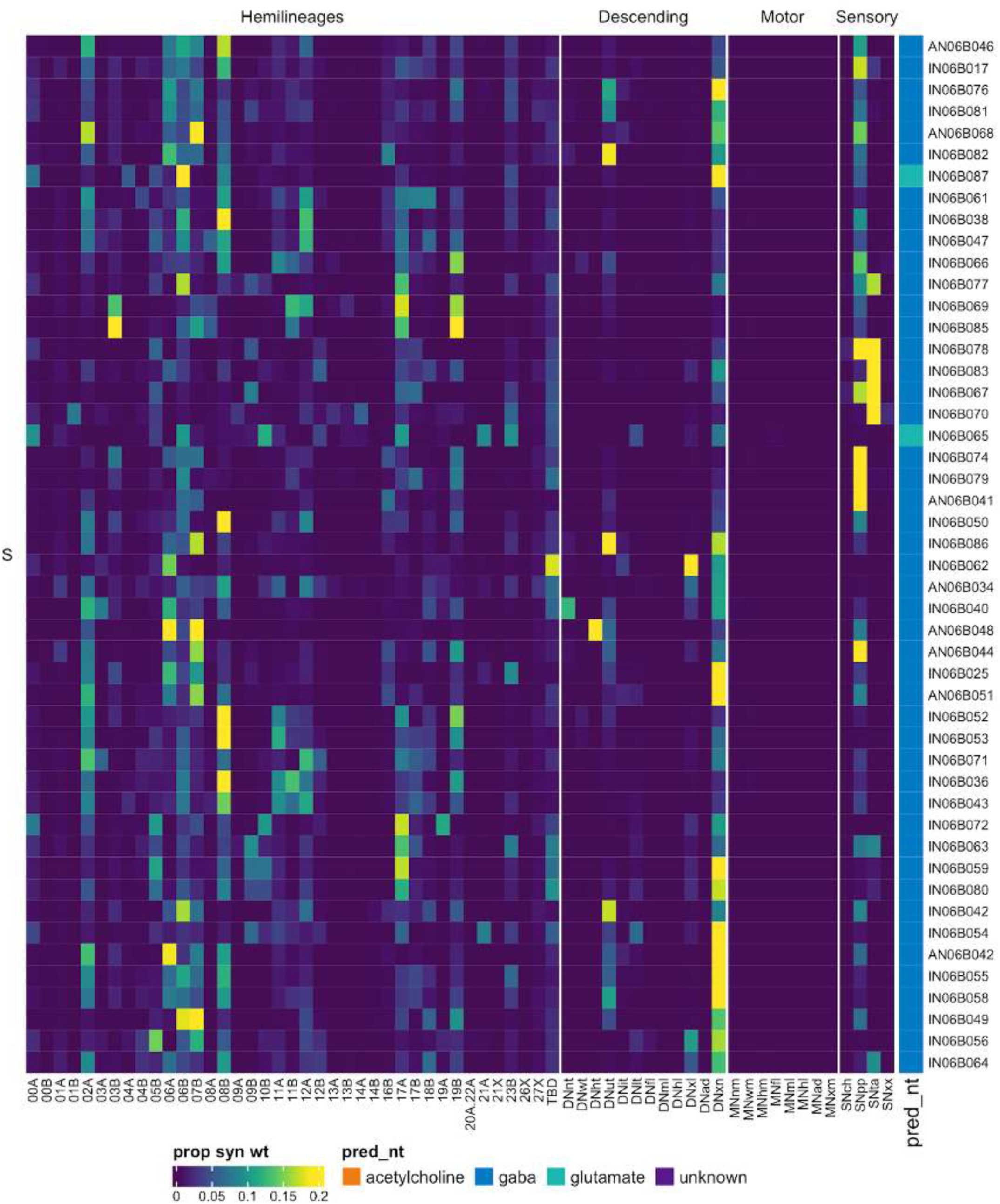
Connectivity to upstream partners by 06B secondary systematic types. Proportions of synaptic weight to systematic types from upstream partners, normalised by row. 06B neurons have been clustered within each assigned birthtime window (P = primary, ES = early secondary, S = secondary) based on both upstream and downstream connectivity to hemilineages, descending neuron subclasses, motor neuron subclasses, and sensory neuron modalities. The annotation bar is coloured by the most common predicted neurotransmitter within each type.

**Figure 24 - figure supplement 6.**
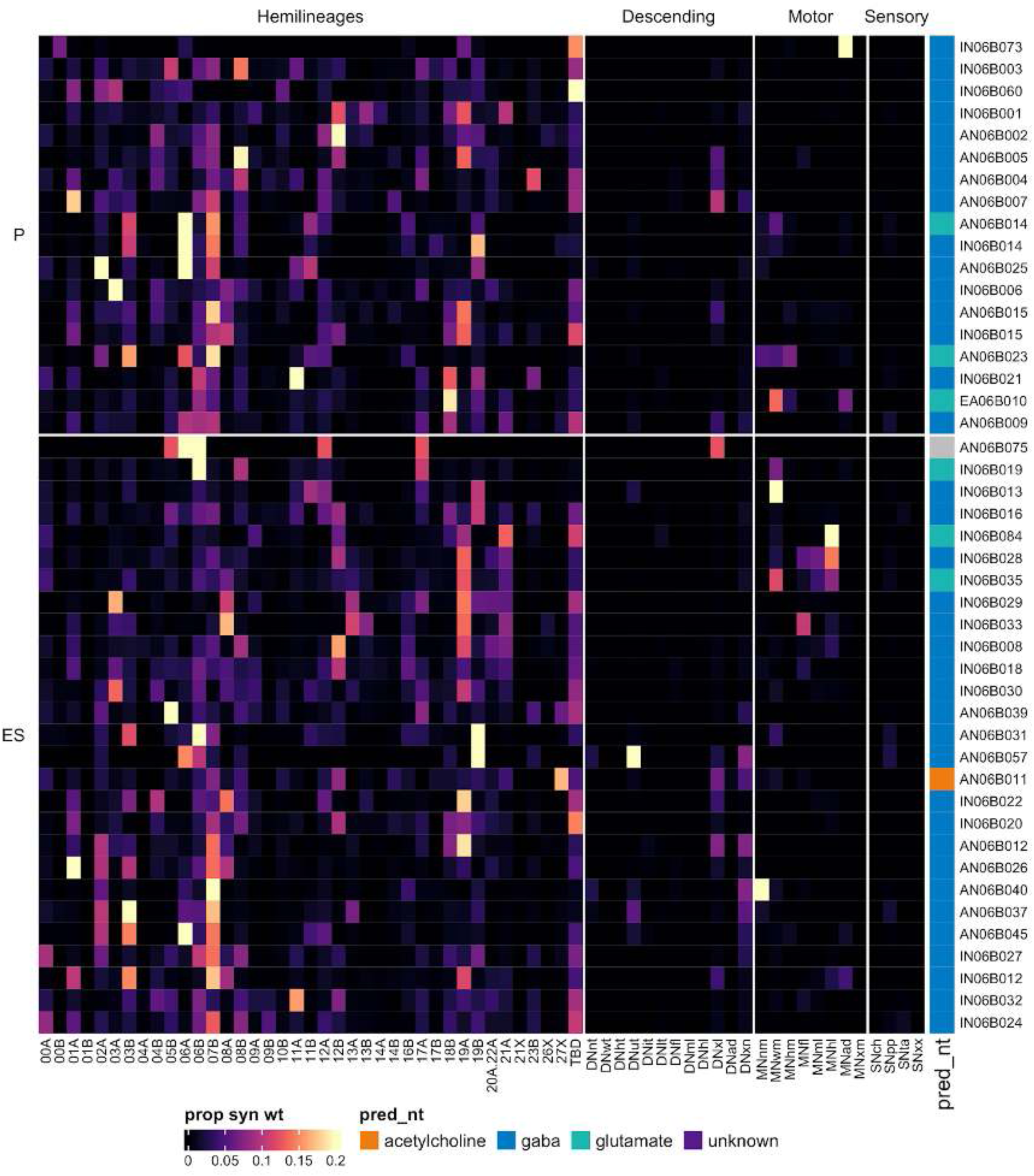
Connectivity to downstream partners by 06B primary and early secondary systematic types. Proportions of synaptic weight from systematic types to downstream partners, normalised by row. 06B neurons have been clustered within each assigned birthtime window (P = primary, ES = early secondary, S = secondary) based on both upstream and downstream connectivity to hemilineages, descending neuron subclasses, motor neuron subclasses, and sensory neuron modalities. The annotation bar is coloured by the most common predicted neurotransmitter within each type.

**Figure 24 - figure supplement 7.**
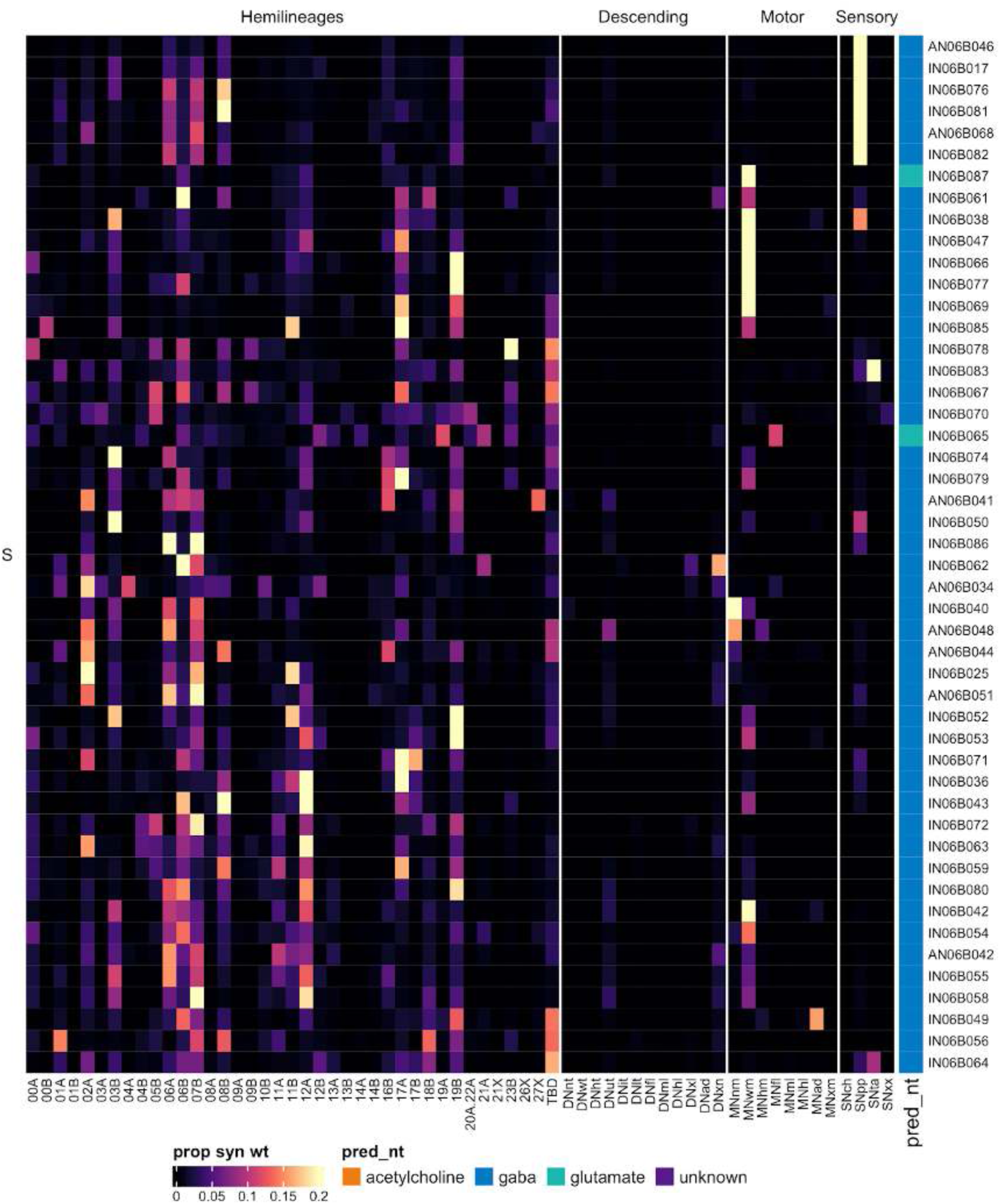
Connectivity to downstream partners by 06B secondary systematic types. Proportions of synaptic weight from systematic types to downstream partners, normalised by row. 06B neurons have been clustered within each assigned birthtime window (P = primary, ES = early secondary, S = secondary) based on both upstream and downstream connectivity to hemilineages, descending neuron subclasses, motor neuron subclasses, and sensory neuron modalities. The annotation bar is coloured by the most common predicted neurotransmitter for the neurons of each type.

#### Hemilineage 07B

Hemilineage 07B is believed to derive from NB3-2 (Lacin and Truman, 2016) (but see also (Birkholz et al., 2015; Truman et al., 2004)), which generates 6 ipsilateral MNs and 10-16 contralateral interneurons in the embryo (Schmid et al., 1999). 07B secondary neurons are found in T1-A1 in the adult (Truman et al., 2010, 2004) and are reported to be cholinergic (Lacin et al., 2019). 07B neurons enter the neuropil near 08B in the anterior of the neuromeres, projecting dorsomedially to cross the midline posterior to 08B (Shepherd et al., 2019, 2016). We found that 07B primary neurites enter the neuropil lateral to those of 08B in T1 but medial to 8B in T2-T3 (although with some overlap in T3). We were not able to distinguish 07A cells from the rest of the leg motor neurons.

07B secondary neurons arborise in the tectulum on both sides of the midline but more profusely ipsilaterally (Shepherd et al., 2019, 2016). The T1, T3 and A1 hemilineages ascend into the contralateral cervical connective. The T2 hemilineage likewise extends contralateral axons anteriorly into the connective but also posteriorly into T3 (Figure 25A). A subset of secondary neurons in T2 cross the midline before terminating in all three contralateral leg neuropils (subcluster 14004) (Figure 25C bottom) (Shepherd et al., 2019). Whilst the majority of 07B neurons in our dataset are predicted to be cholinergic as expected (Lacin et al., 2019), some are consistently predicted to be glutamatergic (e.g., IN07B007) (Figure 25C top) and are significantly different in morphology from the cholinergic population (p = 1.13 x 10^-2^).

**Figure 25.**
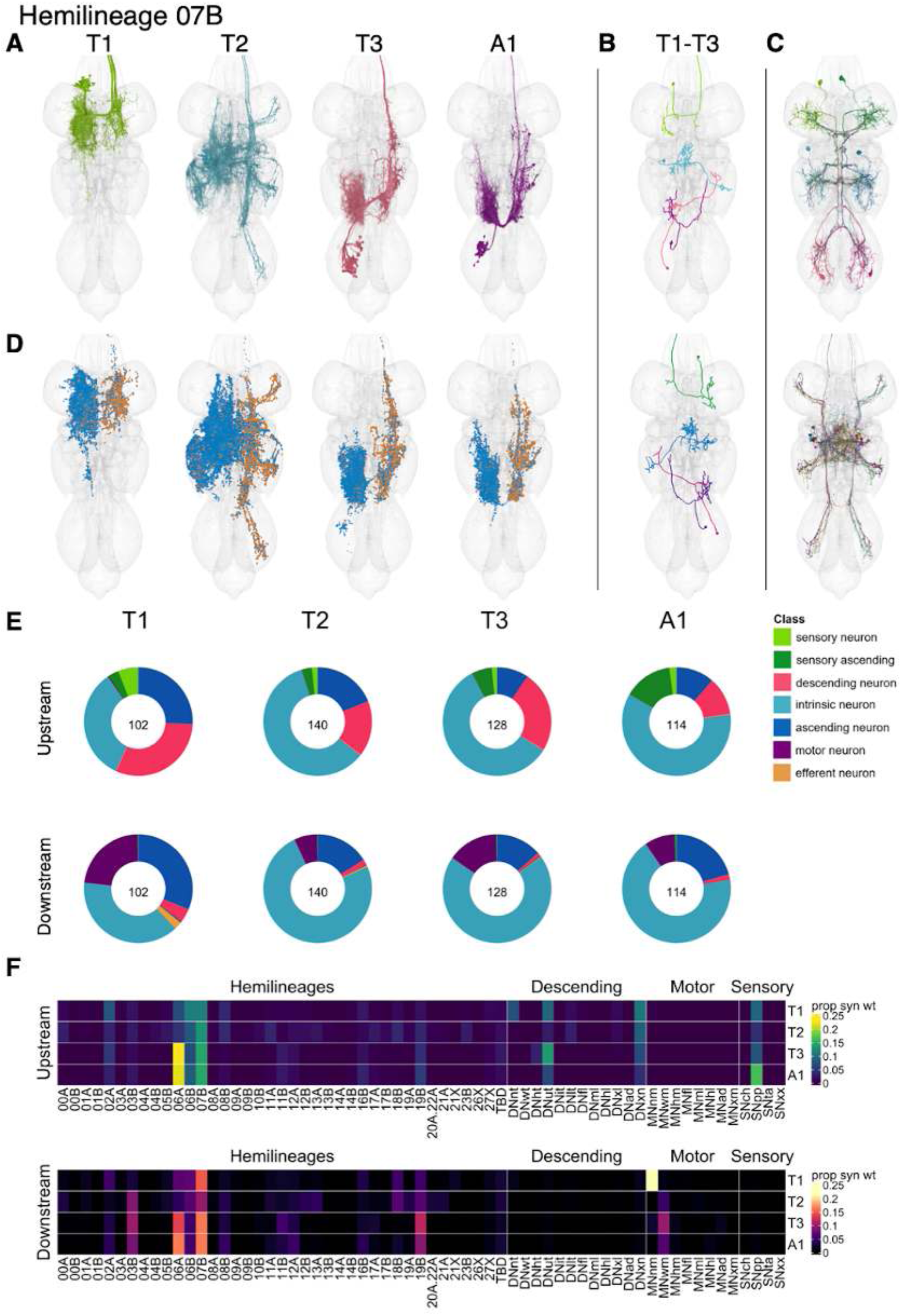
Hemilineage 07B. **A.** Meshes of all RHS secondary neurons plotted in neuromere-specific colours. **B.** “Representative” secondary neuron skeletons plotted in hemineuromere-specific colours. The skeleton with the top accumulated NBLAST score among all neurons from the hemilineage in a given hemineuromere was used. **C.** Neuron meshes of selected examples. Top: glutamatergic convergent serial set 10306. Bottom: centrifugal subcluster 14004. **D.** Predicted synapses of secondary neurons. Blue: postsynapses; dark orange: presynapses. **E.** Proportions of connections from RHS secondary neurons to upstream or downstream partners, normalised by neuromere and coloured by broad class. Numbers of query neurons appear in the centre. **F.** Proportions of synaptic weight from secondary neurons originating in each neuromere to upstream or downstream partners, normalised by row.

Secondary 07B neurons survive in T1-A1, with T2 having the largest population (Figure 25E). Their inputs come from descending neurons, proprioceptive sensory neurons, and flight-related hemilineages, especially 06A, 06B, and 07B. Some types are under unusually strong levels of descending control (e.g., IN07B023 and IN07B064) whilst others receive predominantly sensory inputs (e.g., AN07B069) (Figure 25 - figure supplement 4-6). Secondary 07B neurons target hemilineages 03B, 06A, 07B, and 19B (Figure 25F) and neck and wing motor neurons (Figure 25E). One type, IN07B090, targets hind leg motor neurons (Figure 25 - figure supplement 8). Bilateral activation of 07B secondary neurons results in spontaneous grooming with occasional wing flicking, ending in a characteristic takeoff sequence (wings raised, mesothoracic legs extended in a jump, wing depression and flapping) (Harris et al., 2015).

**Figure 25 - figure supplement 1.**
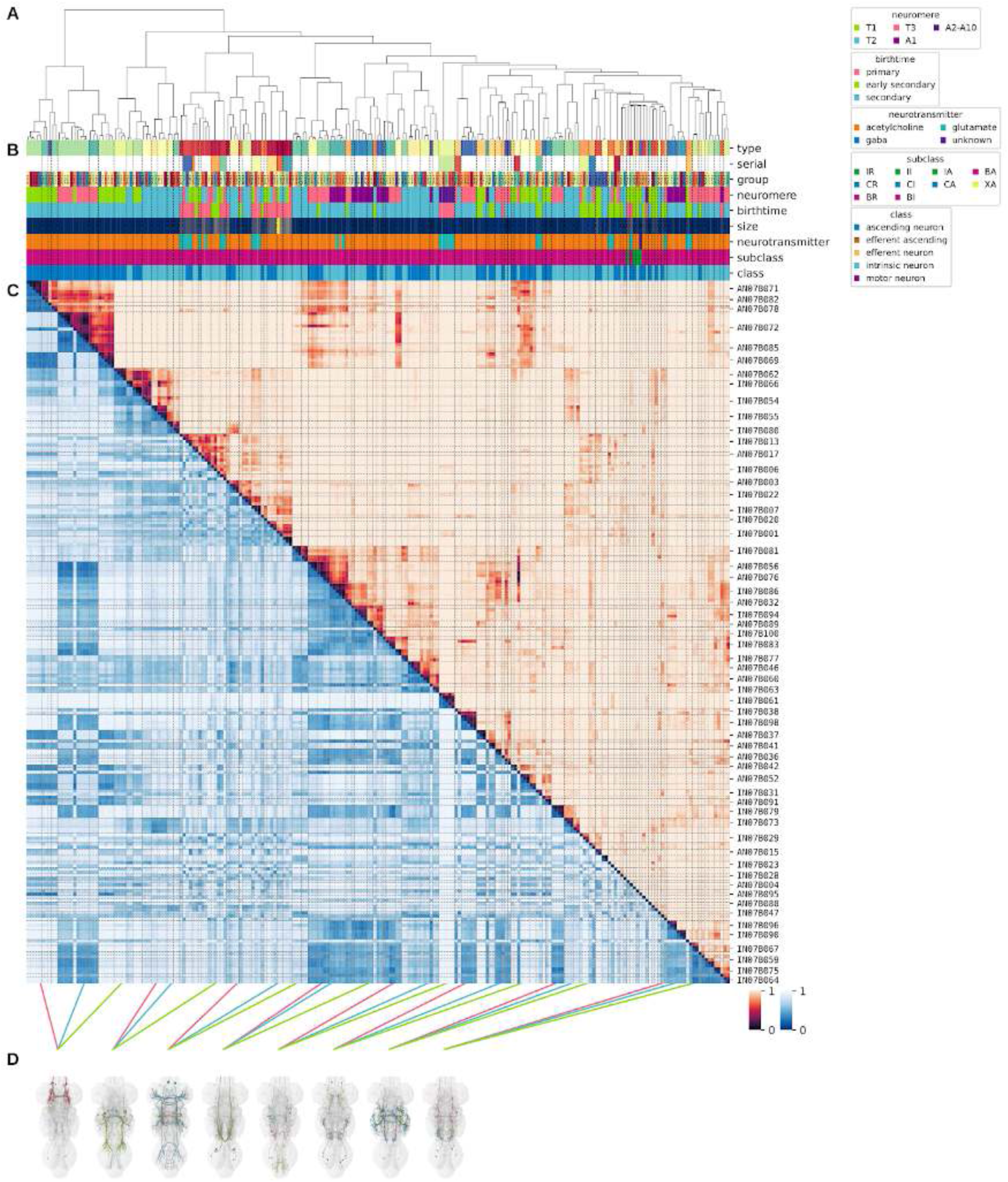
Systematic typing of hemilineage 07B. **A.** Hierarchical clustering dendrogram of hemilineage groups by laterally and serially aggregated connectivity cosine clustering. **B.** Categorical annotations of each hemilineage group, each column corresponding to the aligned leaf in A. Colours for type, serial set, and group are arbitrary for visualisation. Colours for neuromere, birthtime, neurotransmitter, subclass, and class are as in all other figures. **C.** Similarity distance heatmap for hemilineage. Cosine distance is in the upper triangle, while laterally symmetrised NBLAST distance is in the lower triangle. Systematic type names of some types are labelled. **D.** Morphologically representative groups from dendrogram subtrees. Each group, indicated by colour and line connecting to its column in B and C, is the most morphologically representative group (medoid of NBLAST distance) from a subtree of A. The subtrees (flat clusters) are equal height cuts of A determined to yield the number of groups per plot and plots in D.

**Figure 25 - figure supplement 2.**
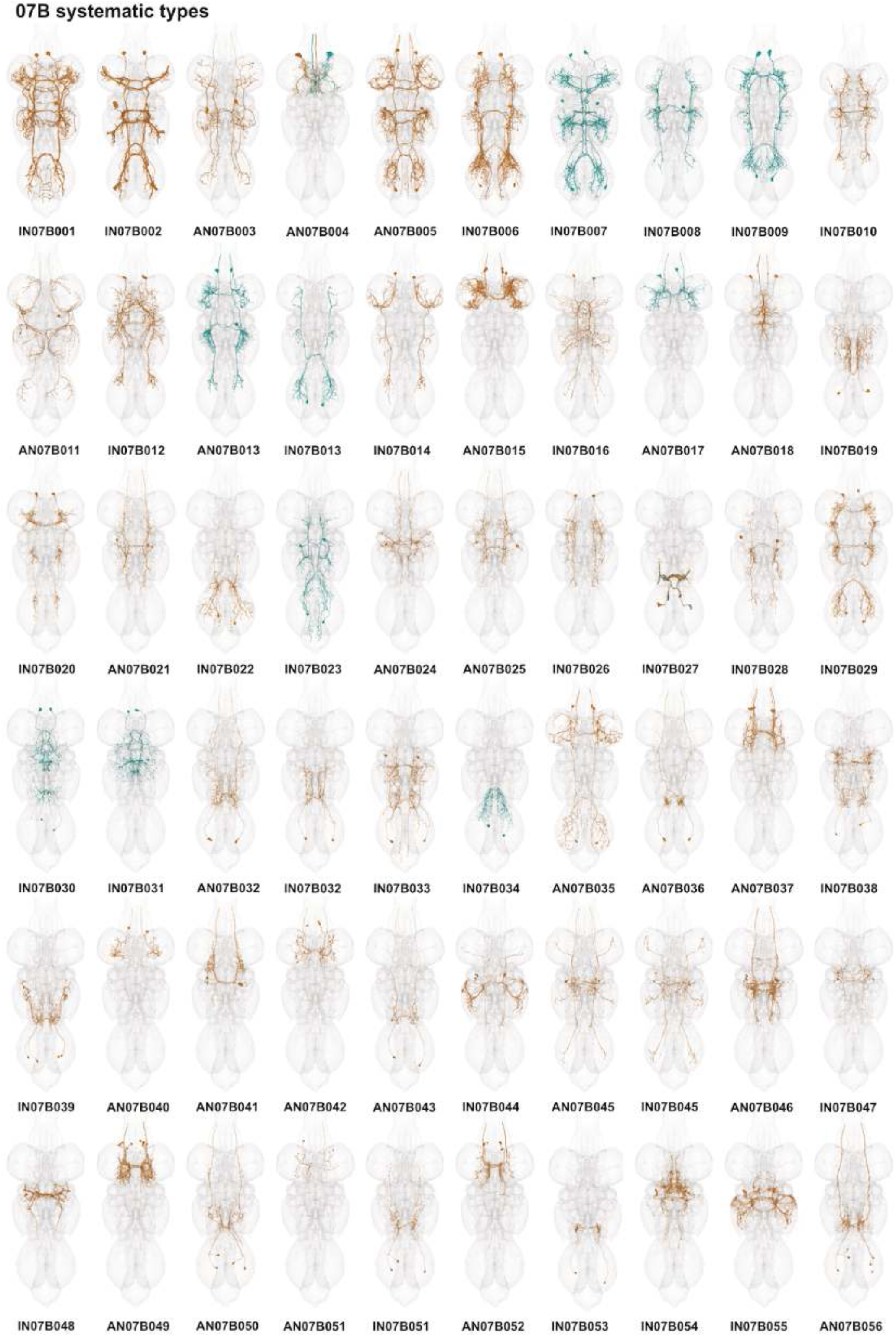
Systematic types of hemilineage 07B. Systematic types have been arranged in numerical order, with neurons of the same type that belong to distinct classes (e.g., intrinsic neuron vs ascending neuron) plotted separately but placed adjacent to each other. Individual neuron meshes have been coloured based on predicted neurotransmitter: dark orange = acetylcholine, blue = gaba, marine = glutamate, dark purple = unknown.

**Figure 25 - figure supplement 3.**
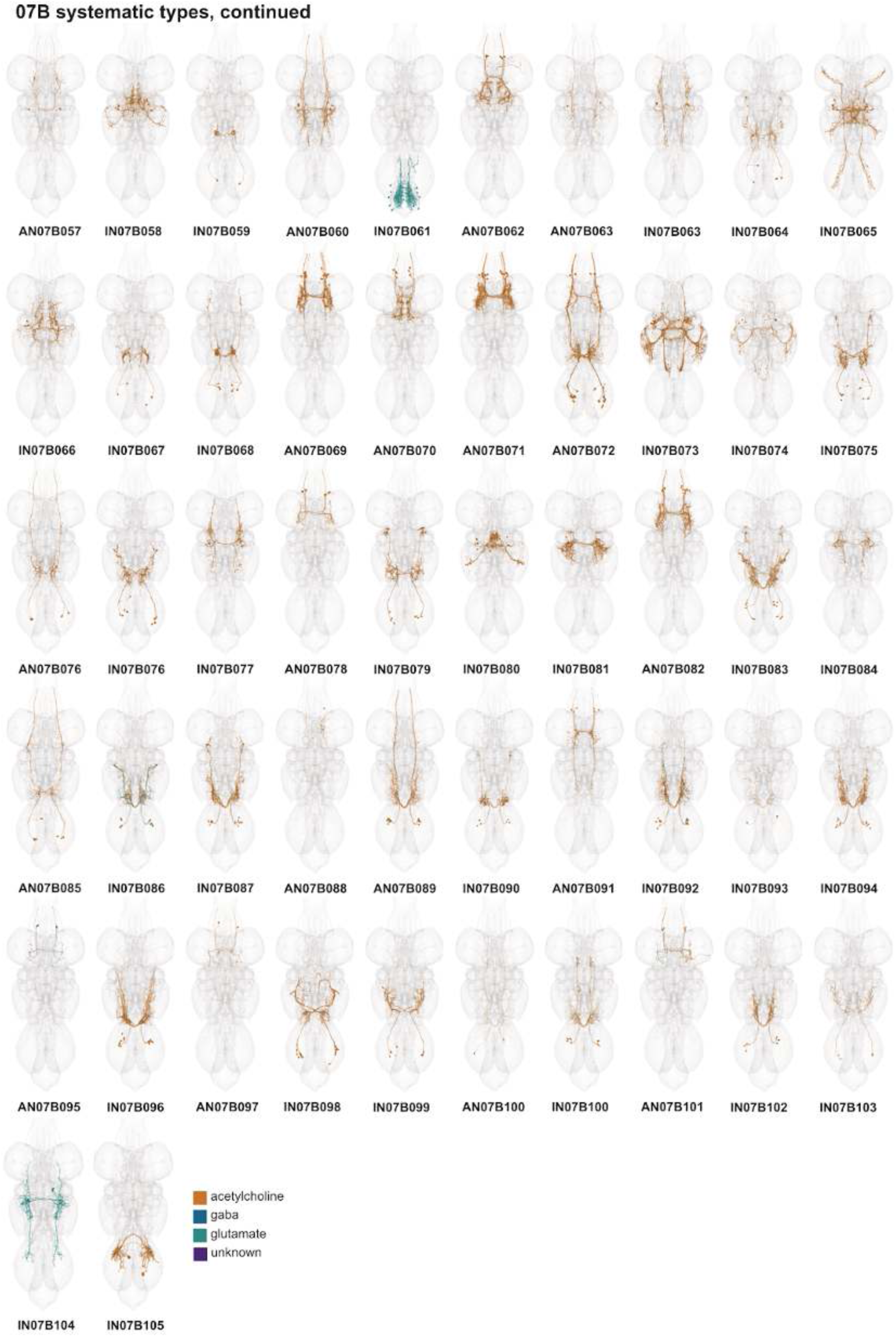
Systematic types of hemilineage 07B, continued. Systematic types have been arranged in numerical order, with neurons of the same type that belong to distinct classes (e.g., intrinsic neuron vs ascending neuron) plotted separately but placed adjacent to each other. Individual neuron meshes have been coloured based on predicted neurotransmitter: dark orange = acetylcholine, blue = gaba, marine = glutamate, dark purple = unknown.

**Figure 25 - figure supplement 4.**
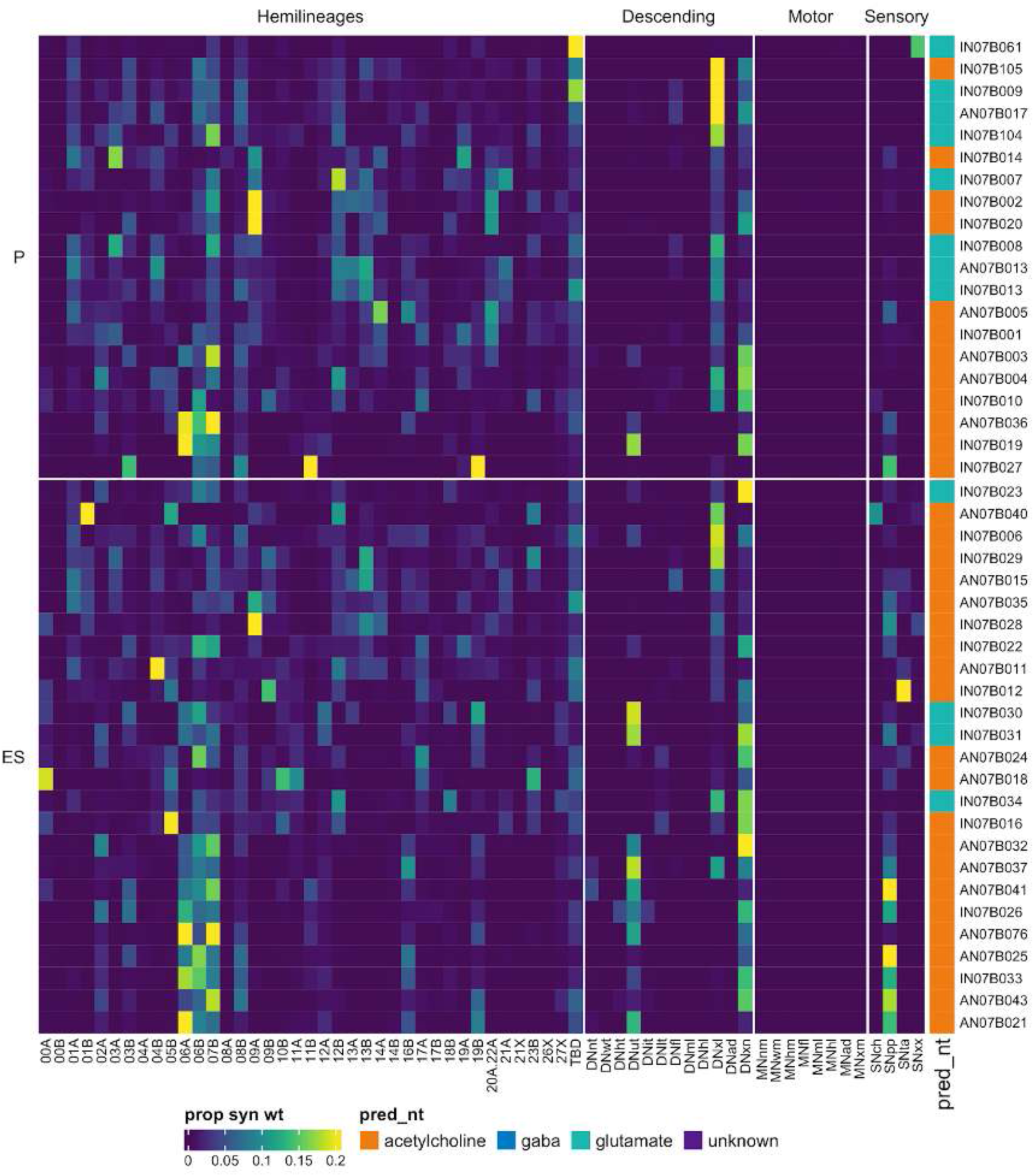
Connectivity to upstream partners by 07B primary and early secondary systematic types. Proportions of synaptic weight to systematic types from upstream partners, normalised by row. 07B neurons have been clustered within each assigned birthtime window (P = primary, ES = early secondary, S = secondary) based on both upstream and downstream connectivity to hemilineages, descending neuron subclasses, motor neuron subclasses, and sensory neuron modalities. Annotation bar is coloured by the most common predicted neurotransmitter for the neurons of each type.

**Figure 25 - figure supplement 5.**
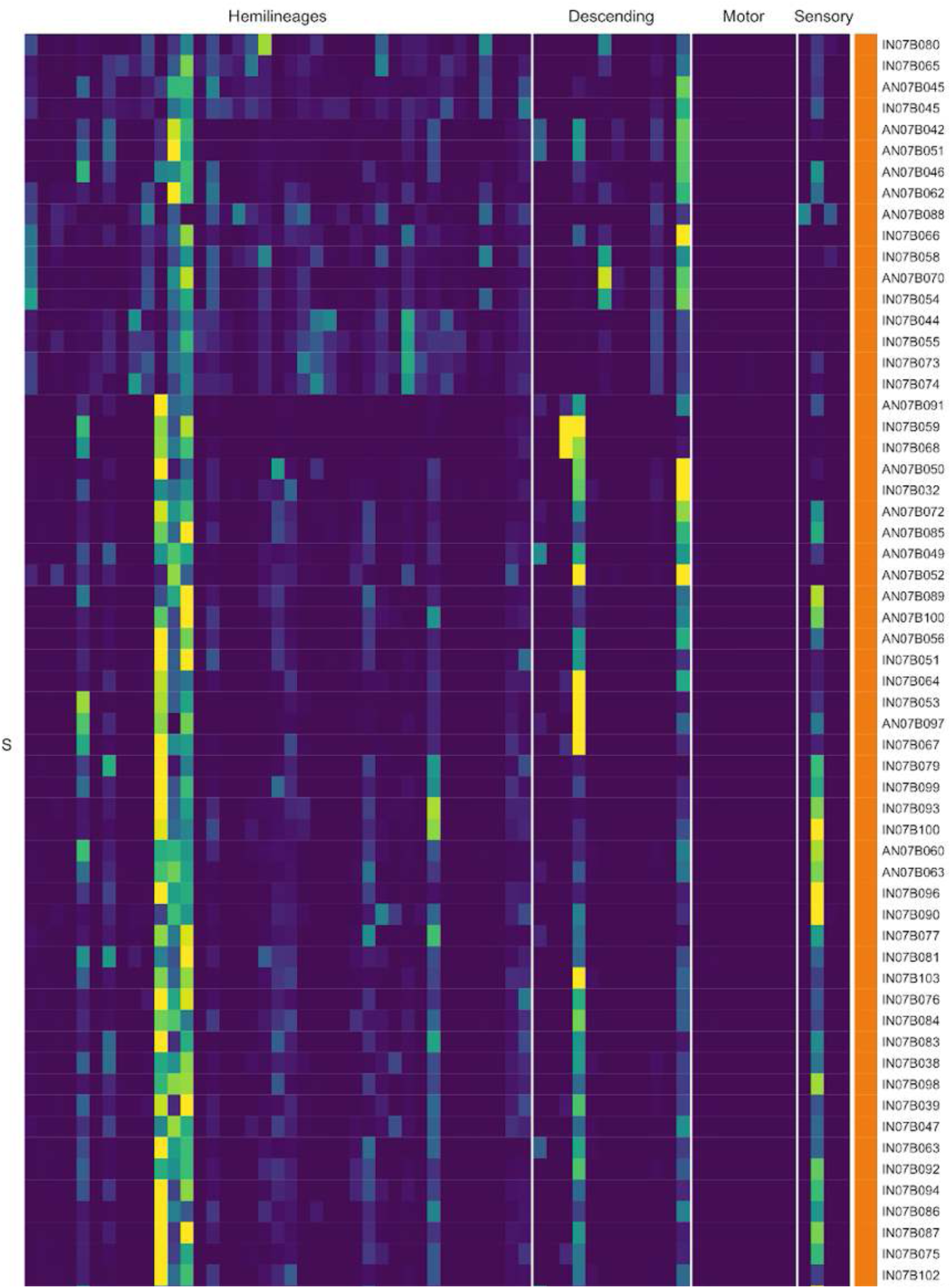
Connectivity to upstream partners by 07B secondary systematic types. Proportions of synaptic weight to systematic types from upstream partners, normalised by row. 07B neurons have been clustered within each assigned birthtime window (P = primary, ES = early secondary, S = secondary) based on both upstream and downstream connectivity to hemilineages, descending neuron subclasses, motor neuron subclasses, and sensory neuron modalities. The annotation bar is coloured by the most common predicted neurotransmitter within each type.

**Figure 25 - figure supplement 6.**
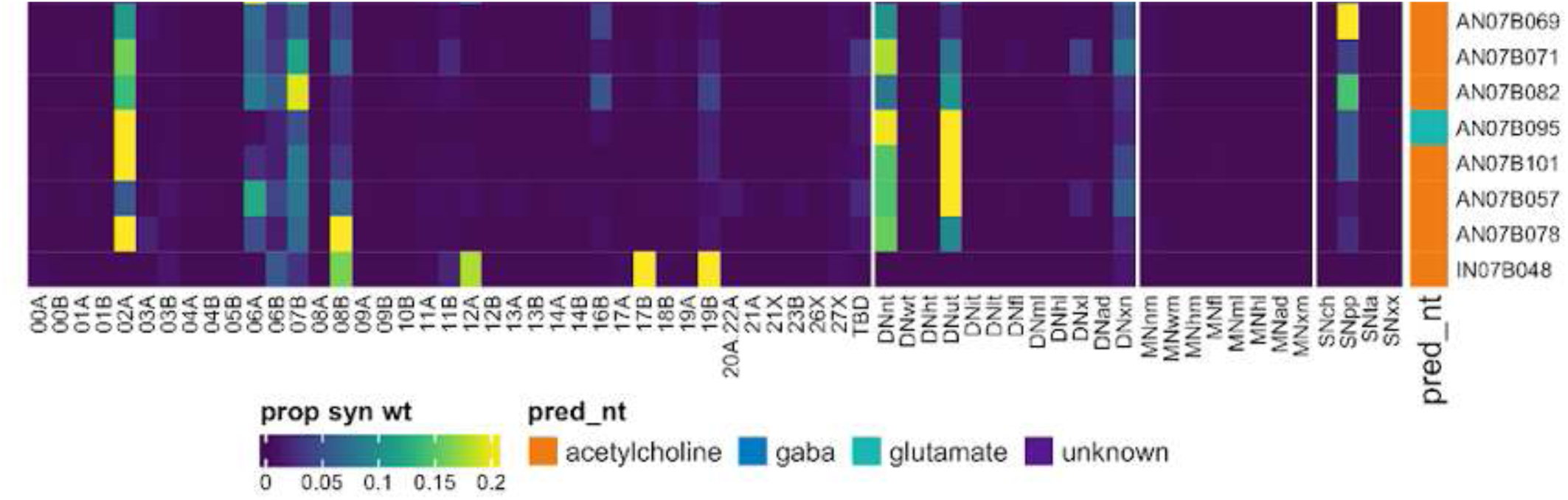
Connectivity to upstream partners by 07B secondary systematic types, continued. Proportions of synaptic weight to systematic types from upstream partners, normalised by row. 07B neurons have been clustered within each assigned birthtime window (P = primary, ES = early secondary, S = secondary) based on both upstream and downstream connectivity to hemilineages, descending neuron subclasses, motor neuron subclasses, and sensory neuron modalities. The annotation bar is coloured by the most common predicted neurotransmitter within each type.

**Figure 25 - figure supplement 7.**
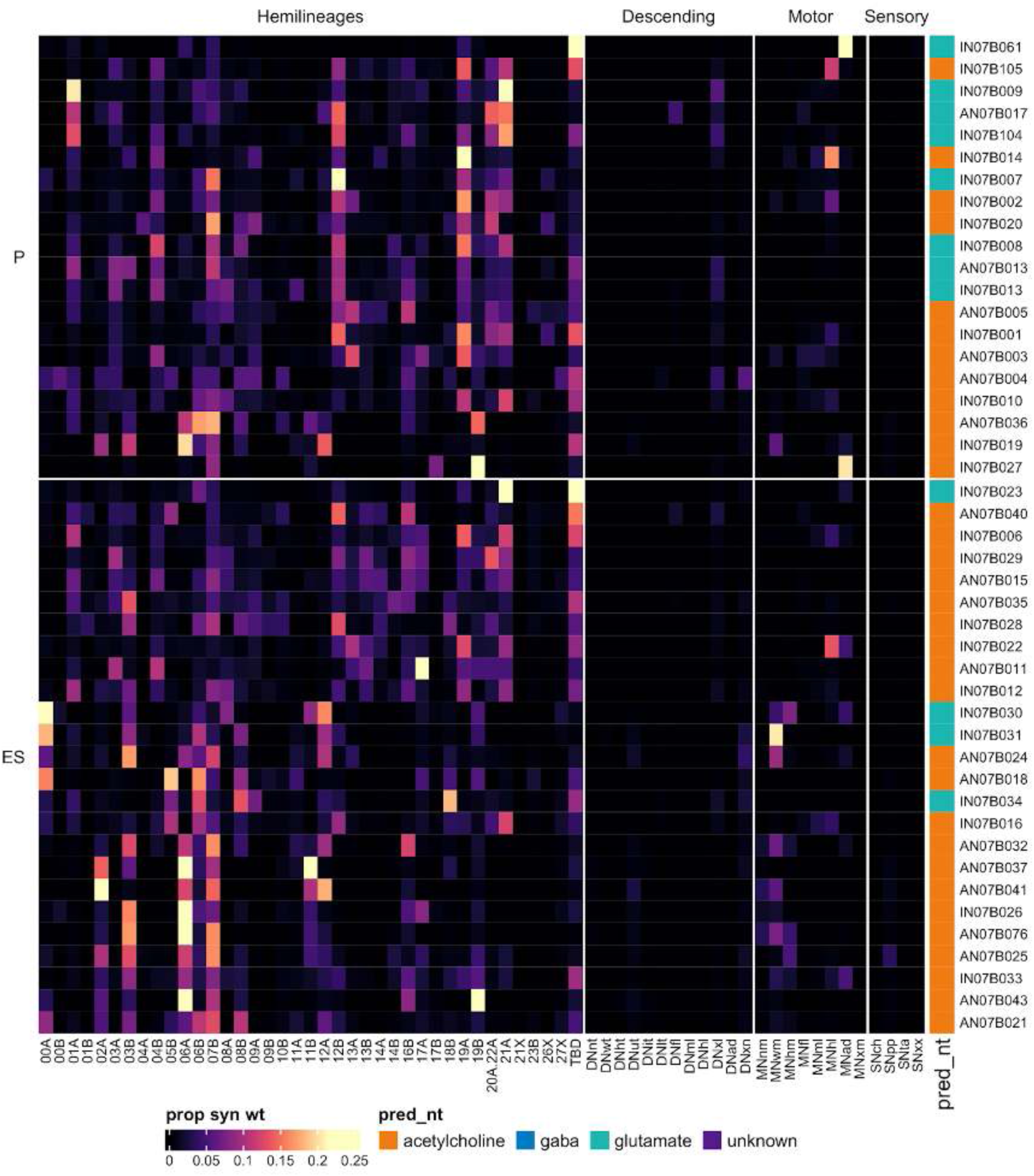
Connectivity to downstream partners by 07B primary and early secondary systematic types. Proportions of synaptic weight from systematic types to downstream partners, normalised by row. 07B neurons have been clustered within each assigned birthtime window (P = primary, ES = early secondary, S = secondary) based on both upstream and downstream connectivity to hemilineages, descending neuron subclasses, motor neuron subclasses, and sensory neuron modalities. The annotation bar is coloured by the most common predicted neurotransmitter within each type.

**Figure 25 - figure supplement 8.**
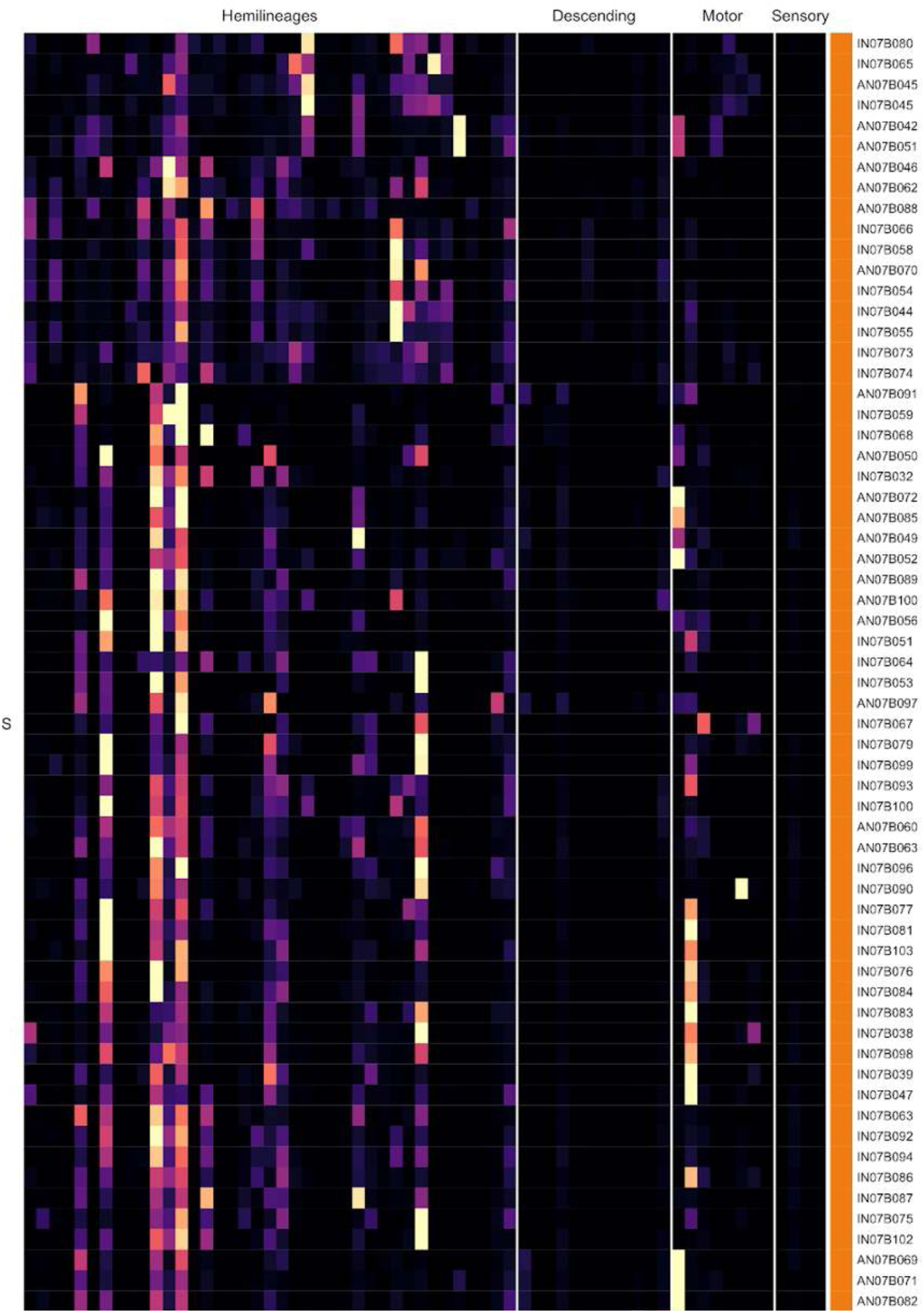
Connectivity to downstream partners by 07B secondary systematic types. Proportions of synaptic weight from systematic types to downstream partners, normalised by row. 07B neurons have been clustered within each assigned birthtime window (P = primary, ES = early secondary, S = secondary) based on both upstream and downstream connectivity to hemilineages, descending neuron subclasses, motor neuron subclasses, and sensory neuron modalities. The annotation bar is coloured by the most common predicted neurotransmitter for the neurons of each type.

**Figure 25 - figure supplement 9.**
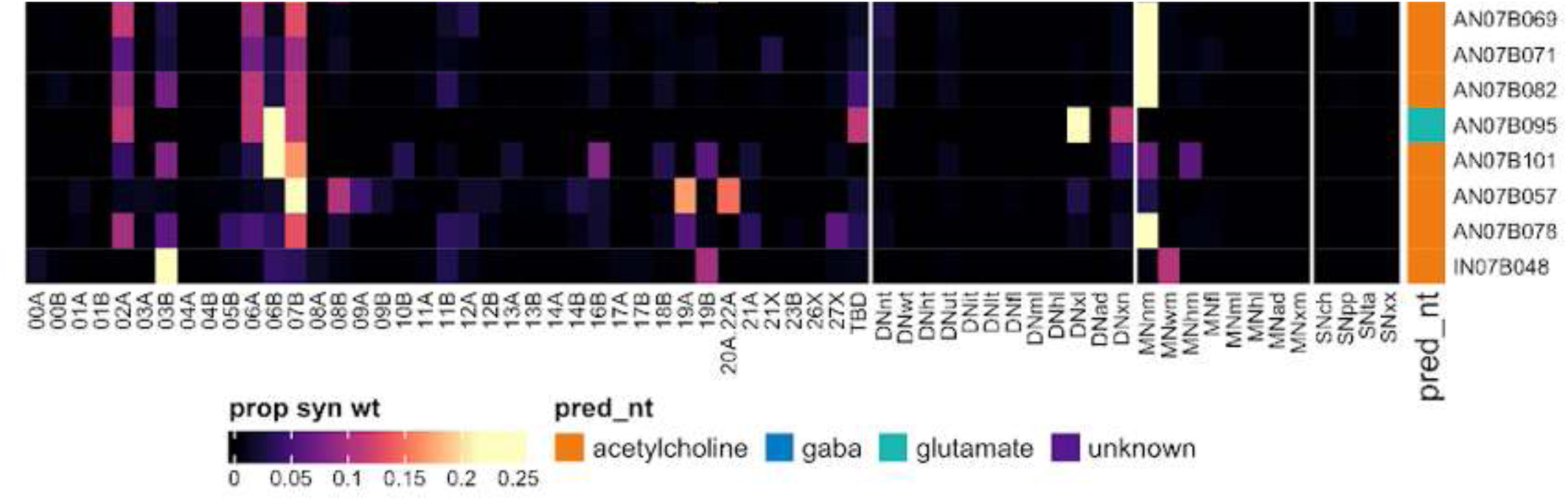
Connectivity to downstream partners by 07B secondary systematic types, continued. Proportions of synaptic weight from systematic types to downstream partners, normalised by row. 07B neurons have been clustered within each assigned birthtime window (P = primary, ES = early secondary, S = secondary) based on both upstream and downstream connectivity to hemilineages, descending neuron subclasses, motor neuron subclasses, and sensory neuron modalities. The annotation bar is coloured by the most common predicted neurotransmitter for the neurons of each type.

#### Hemilineage 08A

Hemilineages 08A and 08B are believed to derive from NB3-3 (Lacin and Truman, 2016) (but see also (Birkholz et al., 2015)), which produces 1 MN in abdominal segments plus 3 intersegmental neurons and 4-10 local interneurons in the embryo (Schmid et al., 1999). This lineage also appears to produce one thoracic motor neuron postembryonically (Truman et al., 2004). In the larva, lineage 08 is lateral to lineage 16 in all three thoracic segments (Truman et al., 2004), but in the adult, 08A and 16B primary neurites enter the neuropil close together (Shepherd et al., 2016), both are glutamatergic (Lacin et al., 2019), and both are largely restricted to the ipsilateral leg neuropil (Shepherd et al., 2019). Diagnostic dorsal projections of 08A in T2 in MANC (Figure 26C bottom) were identified from Fru+ neuroblast clones (Cachero et al., 2010) (E Dona and S Cachero, personal communication) and used to assign all neurons sharing the soma tract. We identified one primary and one secondary motor neuron that innervate a similar domain (Figure 26C top). It should be noted that while we are confident that the 08A vs 16B secondary neurons have been assigned correctly, their primary neurons are not closely associated with them and could have been misassigned. We also assigned one associated gabaergic serial set as IN08A006 (Figure 26 - figure supplement 2).

**Figure 26.**
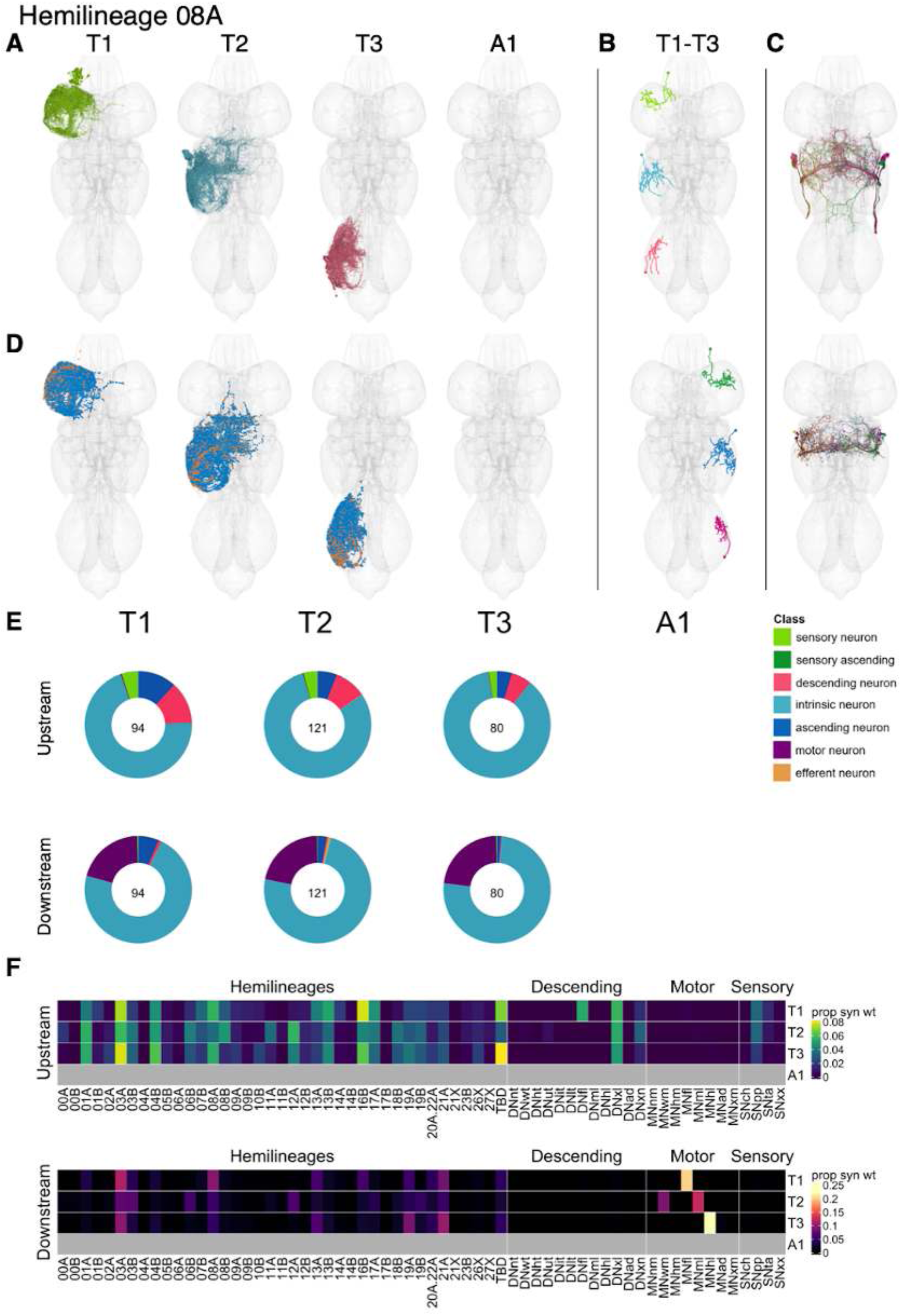
Hemilineage 08A. **A.** Meshes of all RHS secondary neurons plotted in neuromere-specific colours. **B.** “Representative” secondary neuron skeletons plotted in hemineuromere-specific colours. The skeleton with the top accumulated NBLAST score among all neurons from the hemilineage in a given hemineuromere was used. **C.** Neuron meshes of selected examples. Top: motor neuron groups 10178 and 10694. Bottom: dorsal subcluster 16785. **D.** Predicted synapses of RHS secondary neurons. Blue: postsynapses; dark orange: presynapses. **E.** Proportions of connections from secondary neurons to upstream or downstream partners, normalised by neuromere and coloured by broad class. Numbers of query neurons appear in the centre. **F.** Proportions of synaptic weight from secondary neurons originating in each neuromere to upstream or downstream partners, normalised by row.

Secondary 08A neurons survive in all thoracic neuromeres (Truman et al., 2004), with comparable numbers in T1 and T3 but a greater number in T2 (Figure 26E). Their inputs come from descending neurons that innervate front leg or multiple leg neuropils and from proprioceptive sensory neurons and numerous hemilineages including 01A, 03A, 04B, 08A, 08B, 14B, 16B and 17A (Figure 26F). A few 08A types receive tactile sensory inputs (IN08A012, IN08A017, IN08A021, IN08A041) (Figure 26 - figure supplement 3 - 4). Secondary 08A neurons are predicted to be glutamatergic (Figure 8E) and output onto leg motor neurons in their respective neuromeres, and also onto hemilineages 03A, 08A, 19A, and 21A with some segmental variation (Figure 26F). The Fru+ dorsal types IN08A018 and IN08A020 (aka vMS11) target wing motor neurons (Figure 26 - figure supplement 6), consistent with their role in wing extension during courtship song (von Philipsborn et al., 2011). Bilateral activation of 08A secondary neurons does not drive any noticeable change in behaviour in the isolated fly aside from leg repositioning (Harris et al., 2015).

**Figure 26 - figure supplement 1.**
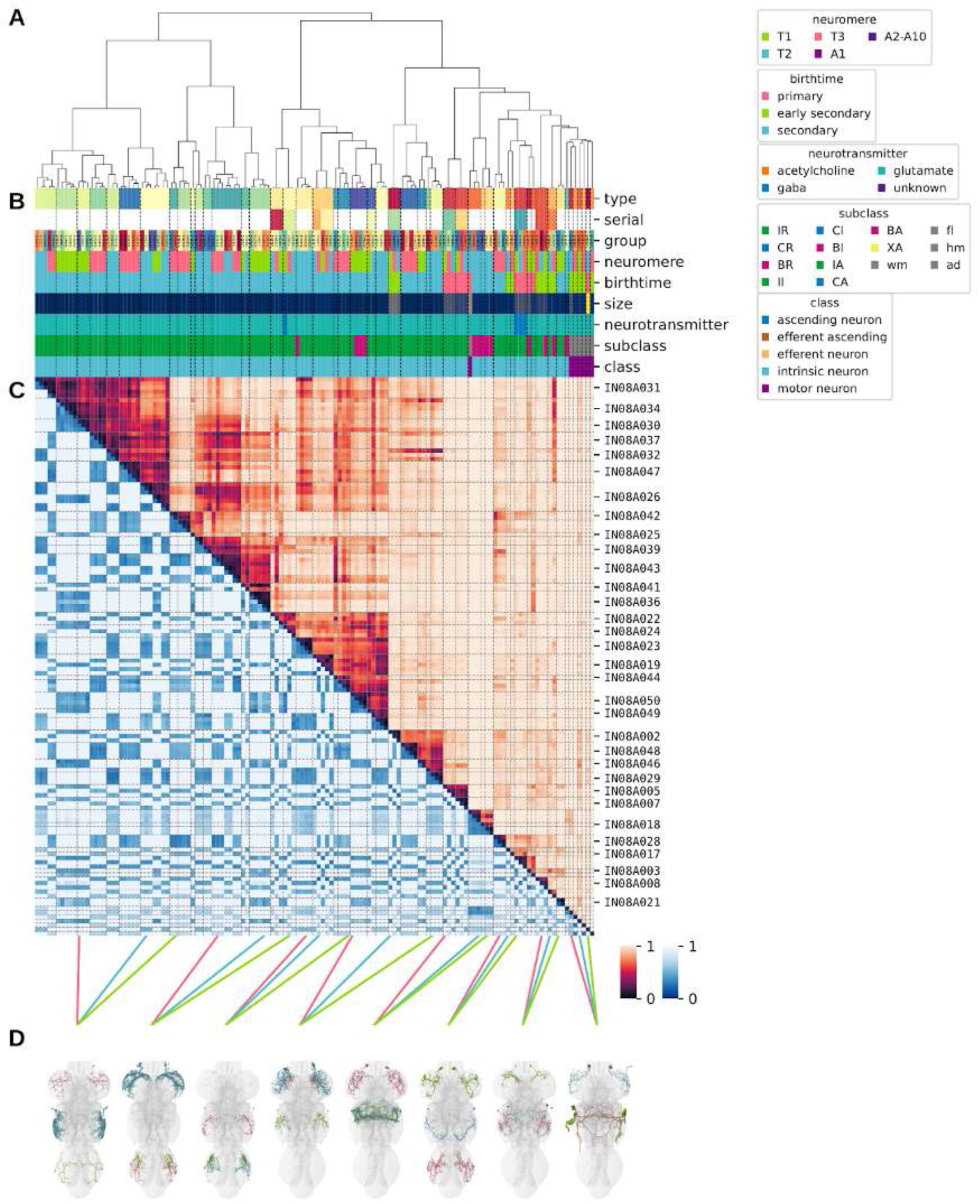
Systematic typing of hemilineage 08A. Types for motor neurons were assigned separately as outlined in our accompanying manuscript (Cheong et al., 2023). **A.** Hierarchical clustering dendrogram of hemilineage groups by laterally and serially aggregated connectivity cosine clustering. **B.** Categorical annotations of each hemilineage group, each column corresponding to the aligned leaf in A. Colours for type, serial set, and group are arbitrary for visualisation. Colours for neuromere, birthtime, neurotransmitter, subclass, and class are as in all other figures. **C.** Similarity distance heatmap for hemilineage. Cosine distance is in the upper triangle, while laterally symmetrised NBLAST distance is in the lower triangle. Systematic type names of some types are labelled. **D.** Morphologically representative groups from dendrogram subtrees. Each group, indicated by colour and line connecting to its column in B and C, is the most morphologically representative group (medoid of NBLAST distance) from a subtree of A. The subtrees (flat clusters) are equal height cuts of A determined to yield the number of groups per plot and plots in D.

**Figure 26 - figure supplement 2.**
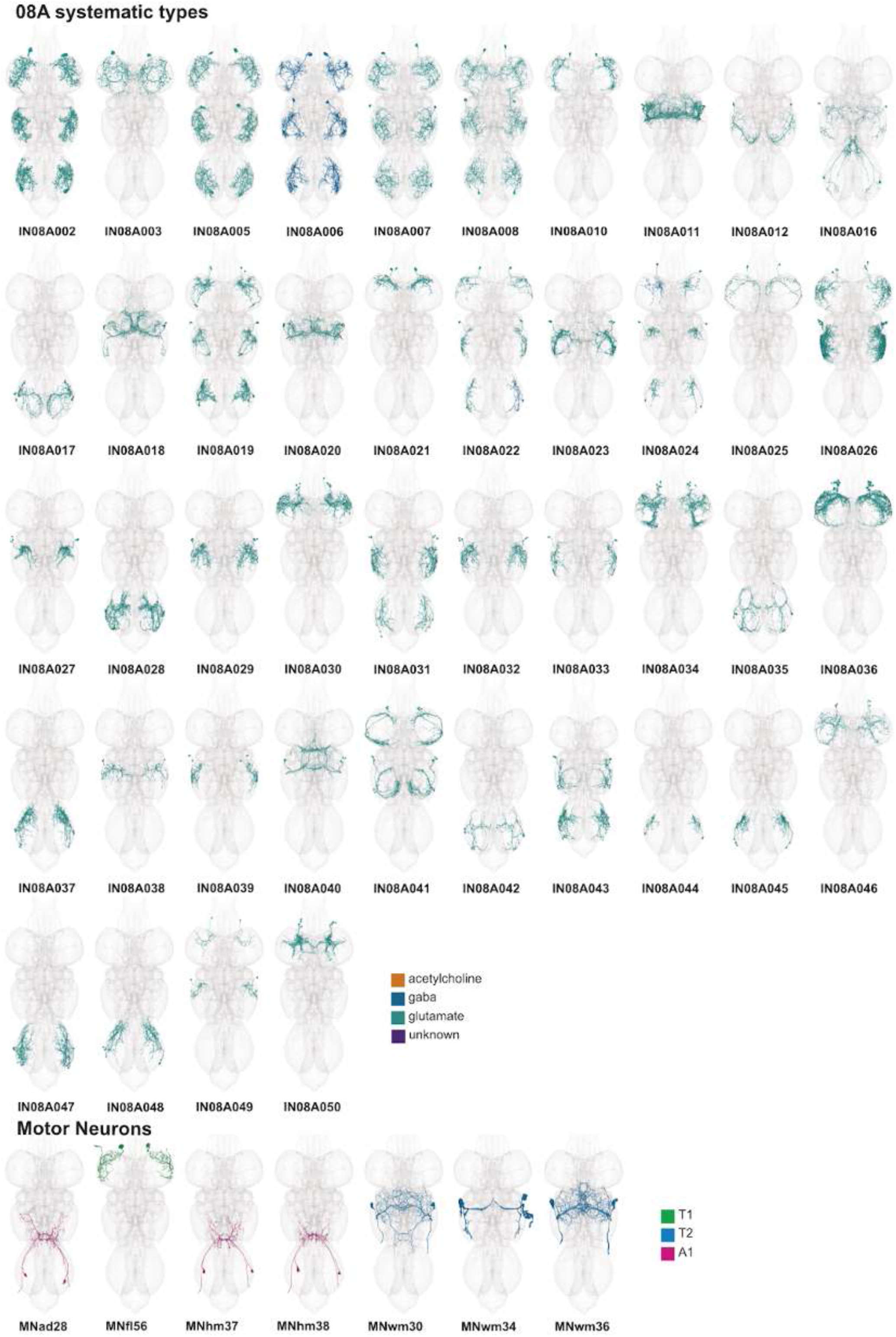
Systematic types of hemilineage 08A. Systematic types have been arranged in numerical order, with neurons of the same type that belong to distinct classes (e.g., intrinsic neuron vs ascending neuron) plotted separately but placed adjacent to each other. Individual neuron meshes have been coloured based on predicted neurotransmitter: dark orange = acetylcholine, blue = gaba, marine = glutamate, dark purple = unknown. Motor neurons (typed separately in (Cheong et al., 2023)) have been plotted by serial set if identified in multiple neuromeres and by systematic type if not. Individual motor neuron meshes have been coloured based on soma neuromere: dark green=T1, blue=T2, purple=A1.

**Figure 26 - figure supplement 3.**
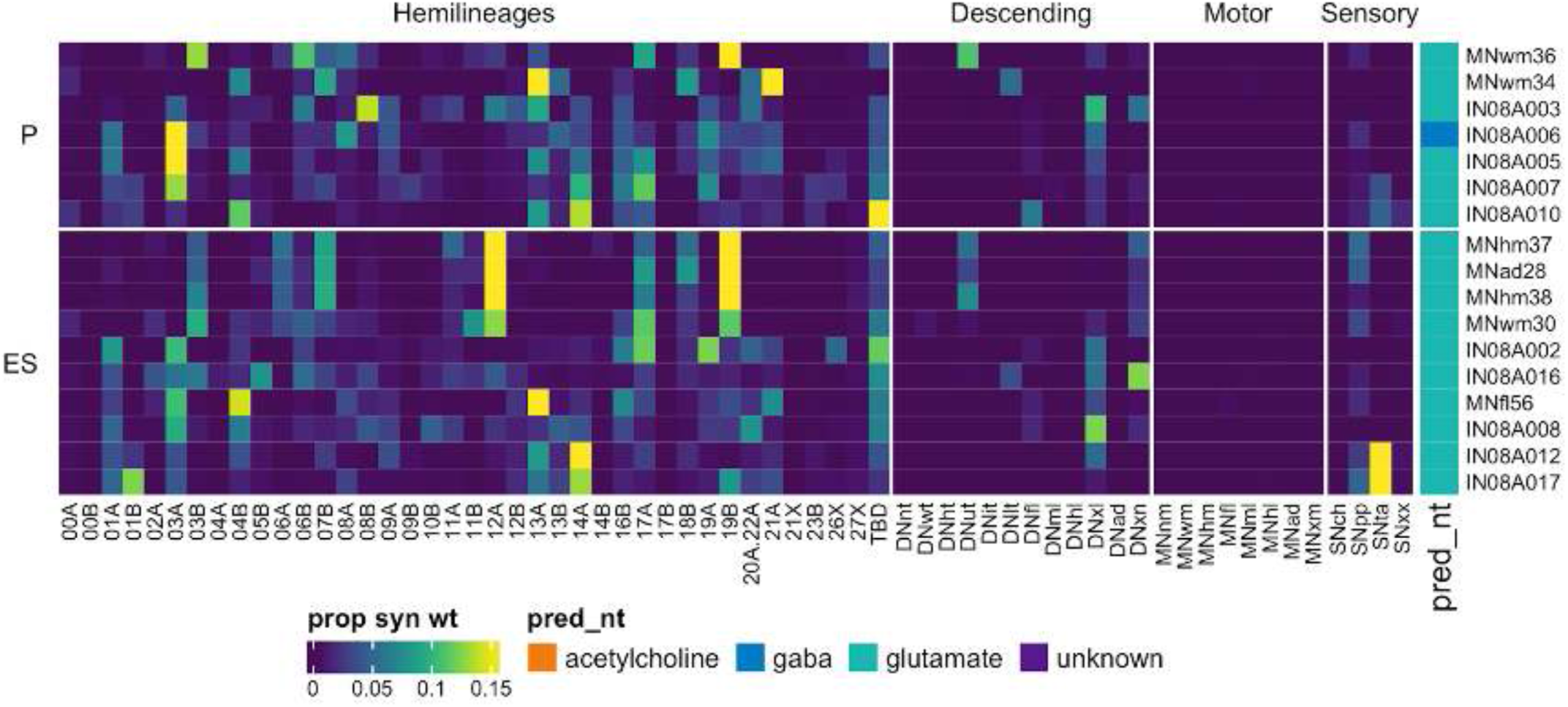
Connectivity to upstream partners by 08A primary and early secondary systematic types. Proportions of synaptic weight to systematic types from upstream partners, normalised by row. 08A neurons have been clustered within each assigned birthtime window (P = primary, ES = early secondary, S = secondary) based on both upstream and downstream connectivity to hemilineages, descending neuron subclasses, motor neuron subclasses, and sensory neuron modalities. Annotation bar is coloured by the most common predicted neurotransmitter for the neurons of each type.

**Figure 26 - figure supplement 4.**
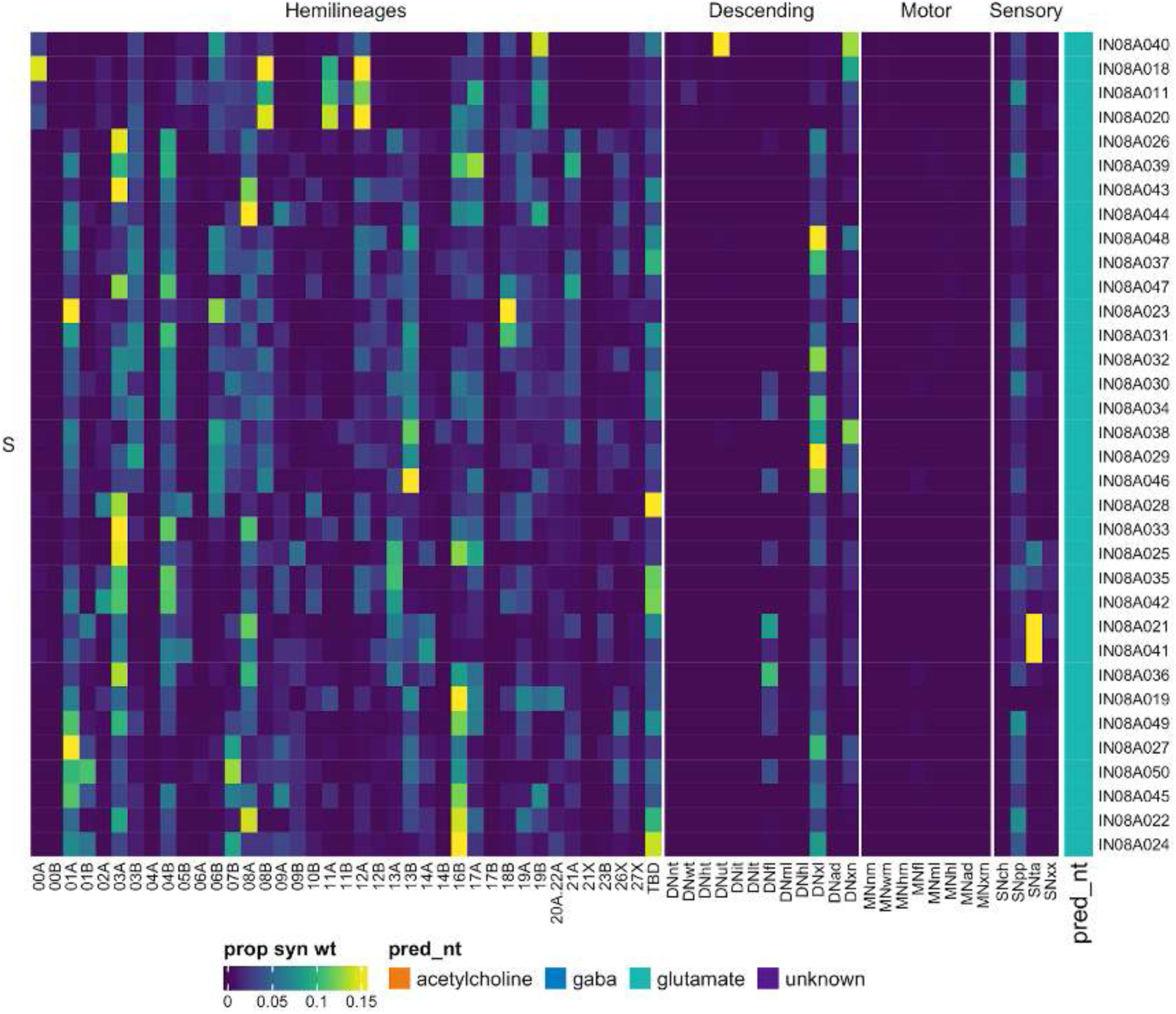
Connectivity to upstream partners by 08A secondary systematic types. Proportions of synaptic weight to systematic types from upstream partners, normalised by row. 08A neurons have been clustered within each assigned birthtime window (P = primary, ES = early secondary, S = secondary) based on both upstream and downstream connectivity to hemilineages, descending neuron subclasses, motor neuron subclasses, and sensory neuron modalities. Annotation bar is coloured by the most common predicted neurotransmitter for the neurons of each type.

**Figure 26 - figure supplement 5.**
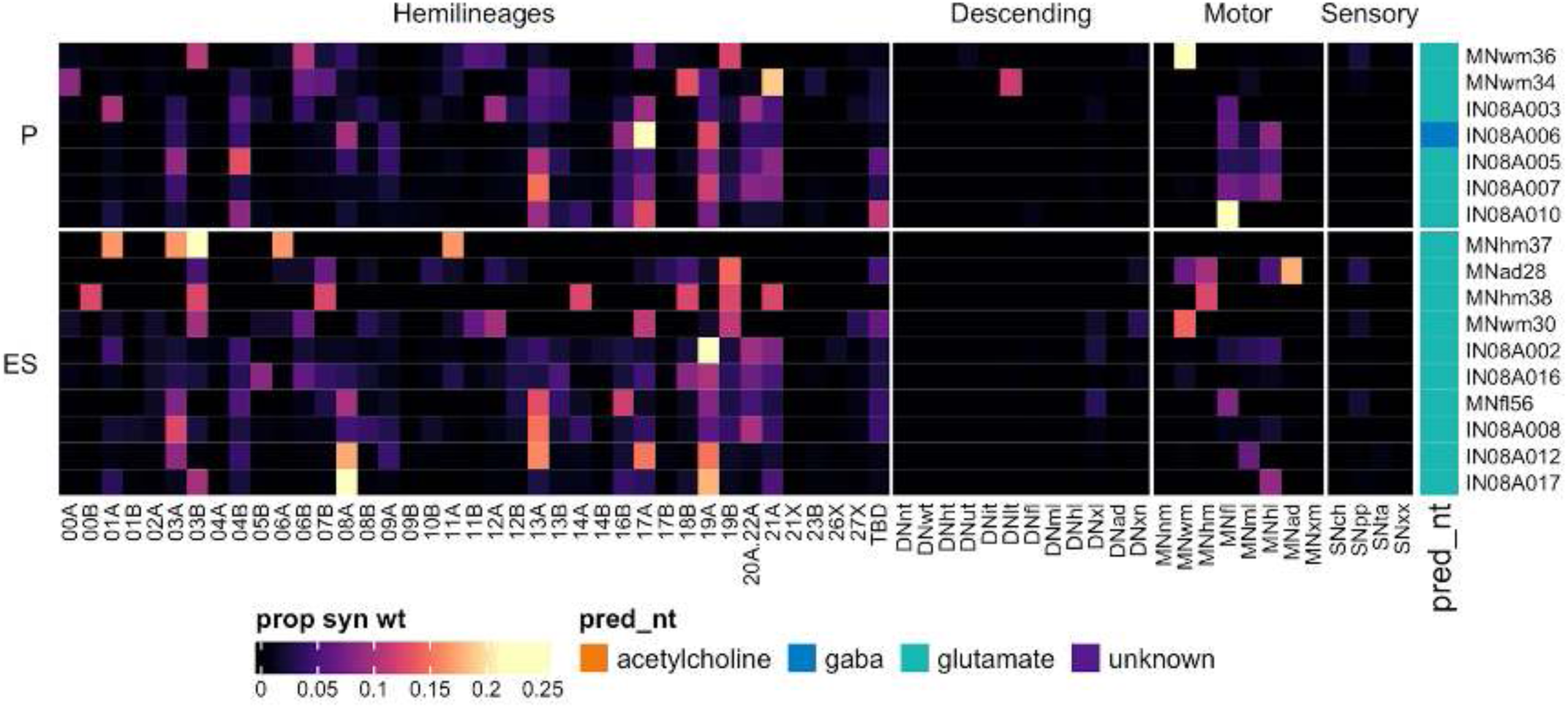
Connectivity to downstream partners by 08A primary and early secondary systematic types. Proportions of synaptic weight from systematic types to downstream partners, normalised by row. 08A neurons have been clustered within each assigned birthtime window (P = primary, ES = early secondary, S = secondary) based on both upstream and downstream connectivity to hemilineages, descending neuron subclasses, motor neuron subclasses, and sensory neuron modalities. The annotation bar is coloured by the most common predicted neurotransmitter within each type.

**Figure 26 - figure supplement 6.**
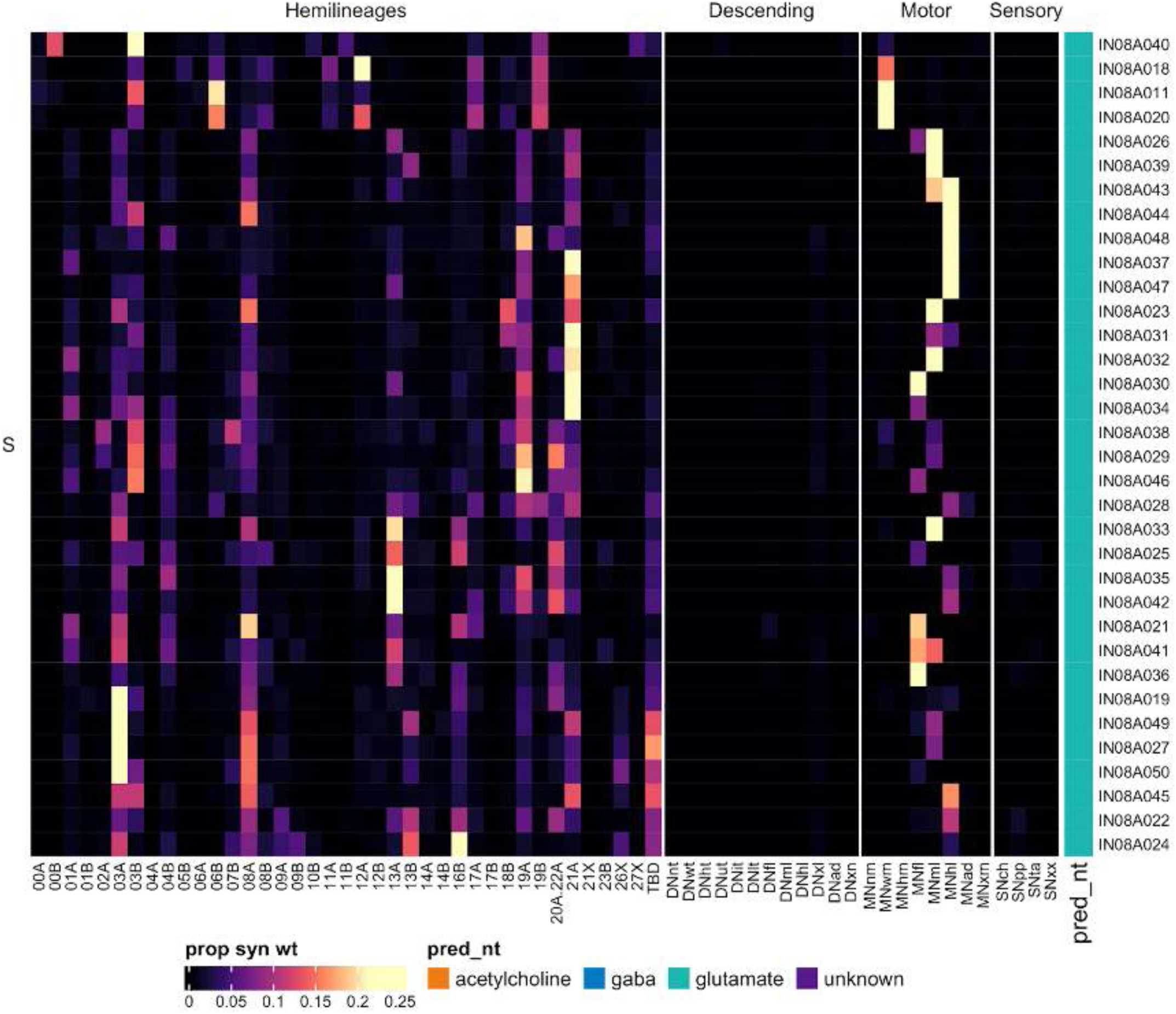
Connectivity to downstream partners by 08A secondary systematic types. Proportions of synaptic weight from systematic types to downstream partners, normalised by row. 08A neurons have been clustered within each assigned birthtime window (P = primary, ES = early secondary, S = secondary) based on both upstream and downstream connectivity to hemilineages, descending neuron subclasses, motor neuron subclasses, and sensory neuron modalities. The annotation bar is coloured by the most common predicted neurotransmitter within each type.

#### Hemilineage 08B

The primary neurites of 08B neurons enter the neuropil in the anterior of the neuromeres, medial to those of 08A (Shepherd et al., 2016). The 08B population crosses the midline in the anterior intermediate commissure and is among the most diverse hemilineages, with segment-specific morphologies innervating dorsal neuropils (Shepherd et al., 2019). We found that 08B primary neurites enter the neuropil medial to those of 07B in T1 but lateral to 07B in T2-T3 (although with some overlap in T3). They project dorsomedially to cross the midline and arborise both sides of the midline in the tectulum (Figure 27A). Secondary 08B neurons survive in T1-T3 with comparable neuron numbers in all neuromeres but T3 having a smaller population (Figure 27E).

**Figure 27.**
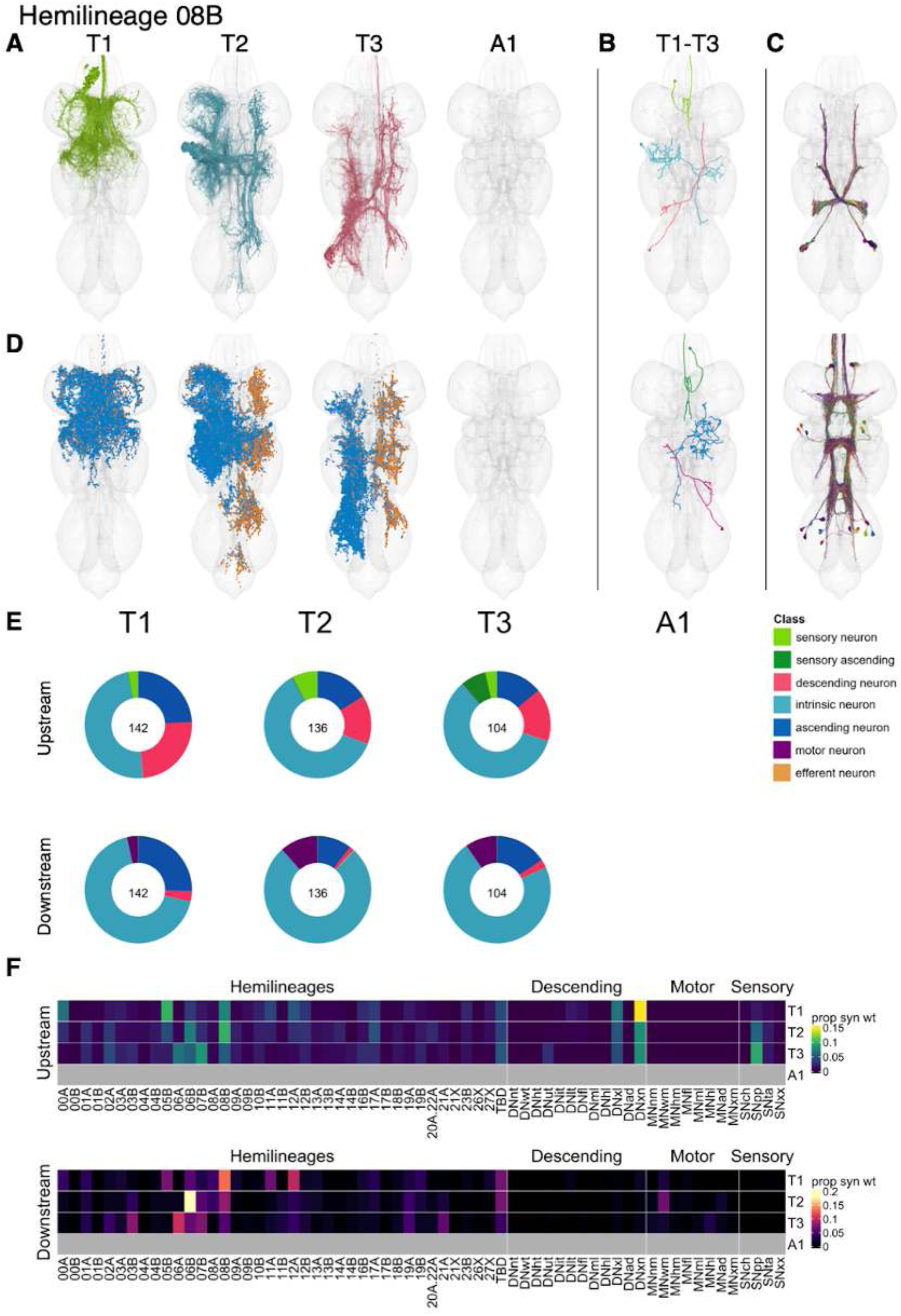
Hemilineage 08B. **A.** Meshes of all RHS secondary neurons plotted in neuromere-specific colours. **B.** “Representative” secondary neuron skeletons plotted in hemineuromere-specific colours. The skeleton with the top accumulated NBLAST score among all neurons from the hemilineage in a given hemineuromere was used. **C.** Neuron meshes of selected examples. Top: electrical n-CHINs and putative electrical cHINs, subcluster 10598. Bottom: intersegmental mVAC population, subcluster 10997. **D.** Predicted synapses of RHS secondary neurons. Blue: postsynapses; dark orange: presynapses. **E.** Proportions of connections from secondary neurons to upstream or downstream partners, normalised by neuromere and coloured by broad class. Numbers of query neurons appear in the centre. **F.** Proportions of synaptic weight from secondary neurons originating in each neuromere to upstream or downstream partners, normalised by row.

Their inputs come mainly from descending neurons that innervate multiple neuropils, proprioceptive sensory neurons in T2 and especially T3, and from hemilineages 00A, 05B, and 08B in T1, 06B and 08B in T2, and 06A, 06B, and 07B in T3. Their outputs also vary by neuromere: in T1 to hemilineages 08B and 12A, in T2 mostly to 06B, 08B, and wing motor neurons, and in T3 to 03B, 06A, and 07B (Figure 27F). 08B secondary neurons are mainly cholinergic (Figure 8E), as expected (Lacin et al., 2019). No functional studies have been published for secondary 08B neurons.

Hemilineage 08B is unusually diverse. We identified a subpopulation of ascending 08B neurons with dendrites restricted to the mVAC (Figure 27C bottom); specific types receive direct input from proprioceptive sensory neurons (AN08B011 and AN08B018), while others receive it mainly from inhibitory hemilineages 00A and 09A and excitatory hemilineage 10B (AN08B024, AN08B025, and AN08B028). 08B produces putative electrical cHINs (contralateral haltere interneurons) that receive input from haltere afferents (Strausfeld and Seyan, 1985) (Figure 27C top); we distinguished three distinct types: IN08B036, IN08B070, and IN08B008 (Figure 27 - figure supplement 2-3), the last being n-cHINs projecting up to the neck tectulum (Trimarchi and Murphey, 1997). We identified several sexually dimorphic 08B types including IN08B002/dPR1 (Lillvis et al., 2024; von Philipsborn et al., 2011) or vPR1 (Yu et al., 2010), which is strongly downstream of descending neurons and upstream of 12A neurons, and IN08B040/vMS4 (Yu et al., 2010), which receives substantial descending input and targets 13A and 19A (Figure 27 - figure supplement 2,4-5,7-8). We can also distinguish several connectivity subtypes of vPR13 (AN08B031, AN08B043, AN08B059, and AN08B069) (Figure 27 - figure supplement 2-3,6,9), which triggers courtship song when experimentally activated but is not required for the core song circuit (Lillvis et al., 2024).

**Figure 27 - figure supplement 1.**
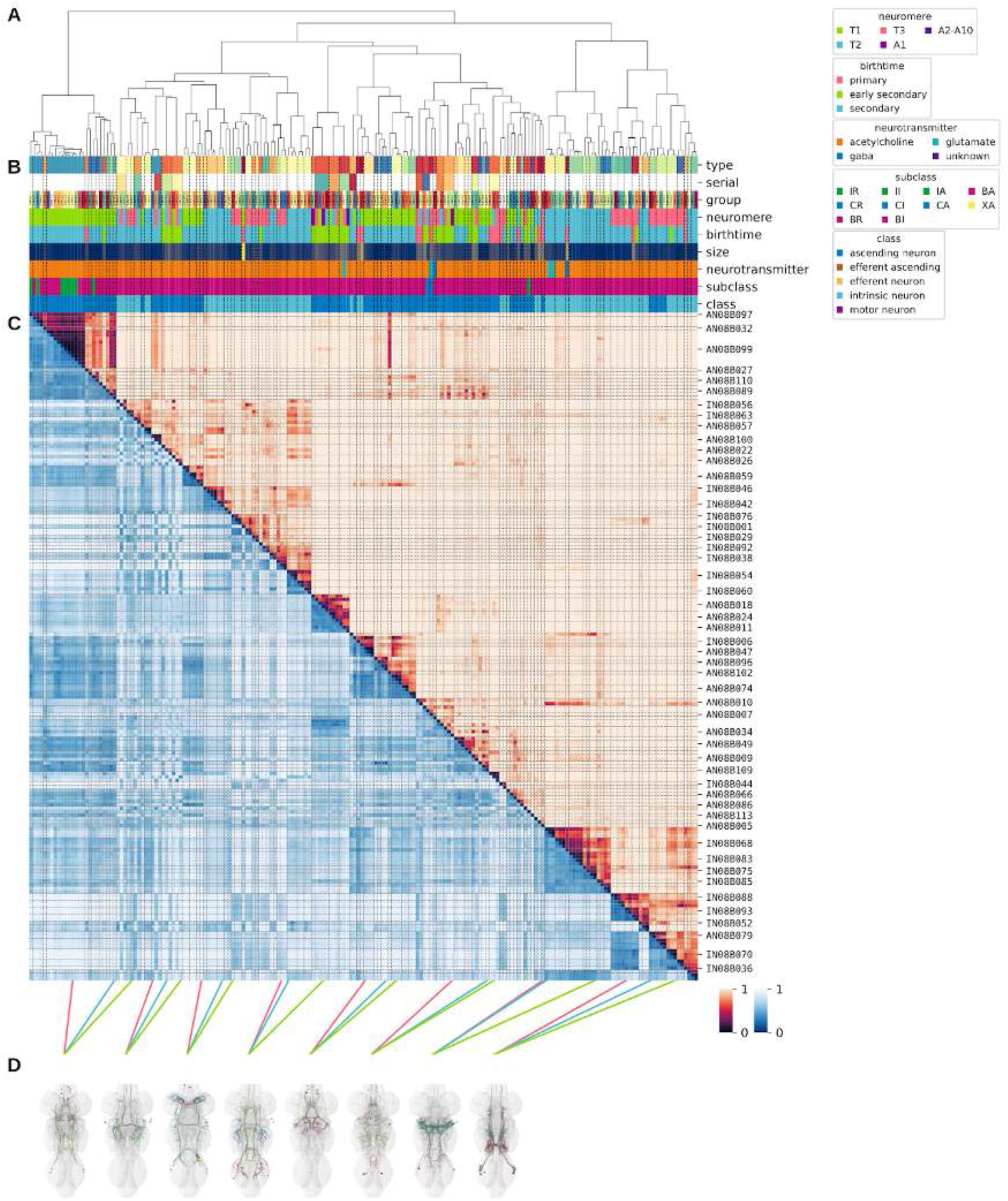
Systematic typing of hemilineage 08B. **A.** Hierarchical clustering dendrogram of hemilineage groups by laterally and serially aggregated connectivity cosine clustering. **B.** Categorical annotations of each hemilineage group, each column corresponding to the aligned leaf in A. Colours for type, serial set, and group are arbitrary for visualisation. Colours for neuromere, birthtime, neurotransmitter, subclass, and class are as in all other figures. **C.** Similarity distance heatmap for hemilineage. Cosine distance is in the upper triangle, while laterally symmetrised NBLAST distance is in the lower triangle. Systematic type names of some types are labelled. **D.** Morphologically representative groups from dendrogram subtrees. Each group, indicated by colour and line connecting to its column in B and C, is the most morphologically representative group (medoid of NBLAST distance) from a subtree of A. The subtrees (flat clusters) are equal height cuts of A determined to yield the number of groups per plot and plots in D.

**Figure 27 - figure supplement 2.**
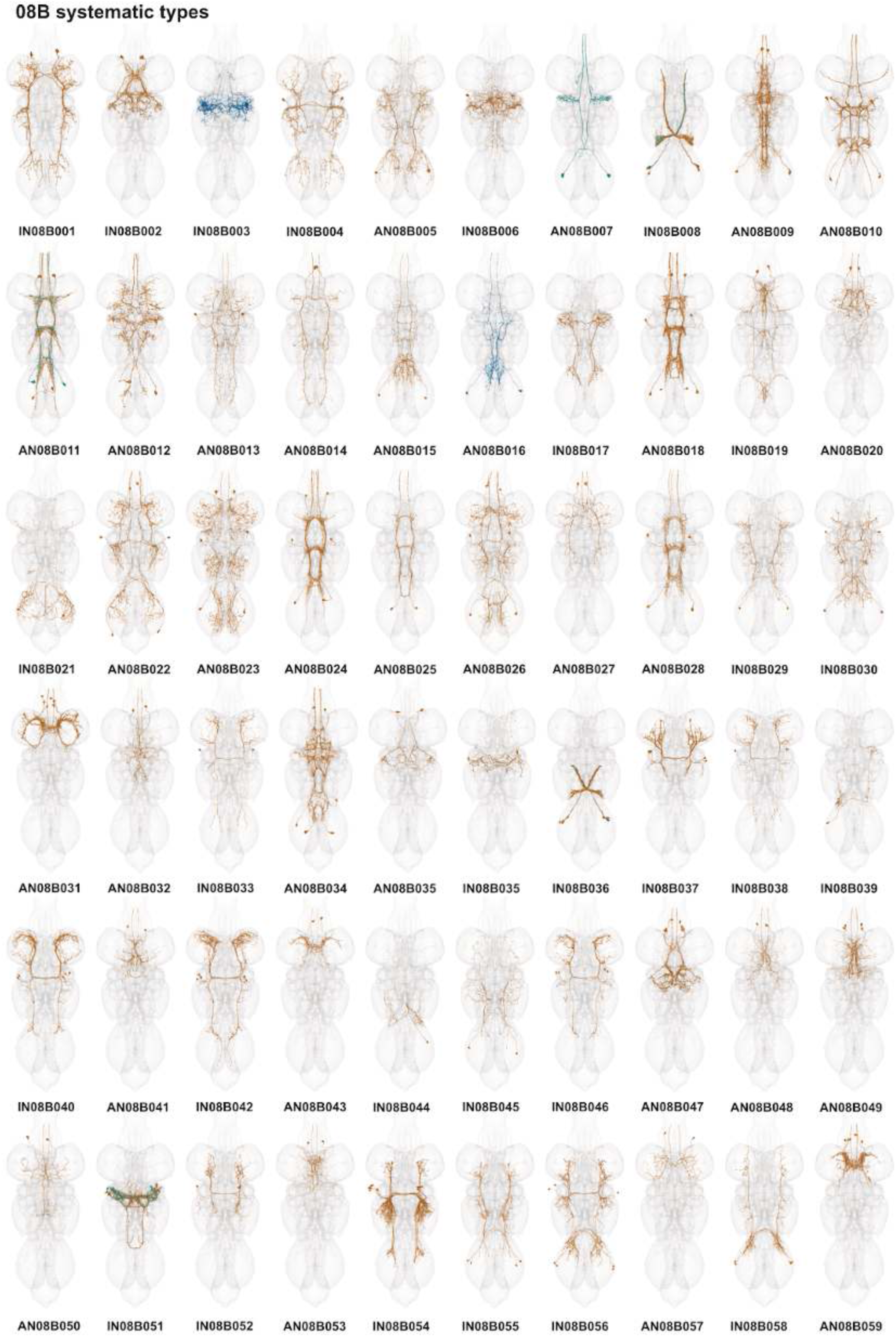
Systematic types of hemilineage 08B. Systematic types have been arranged in numerical order, with neurons of the same type that belong to distinct classes (e.g., intrinsic neuron vs ascending neuron) plotted separately but placed adjacent to each other. Individual neuron meshes have been coloured based on predicted neurotransmitter: dark orange = acetylcholine, blue = gaba, marine = glutamate, dark purple = unknown.

**Figure 27 - figure supplement 3.**
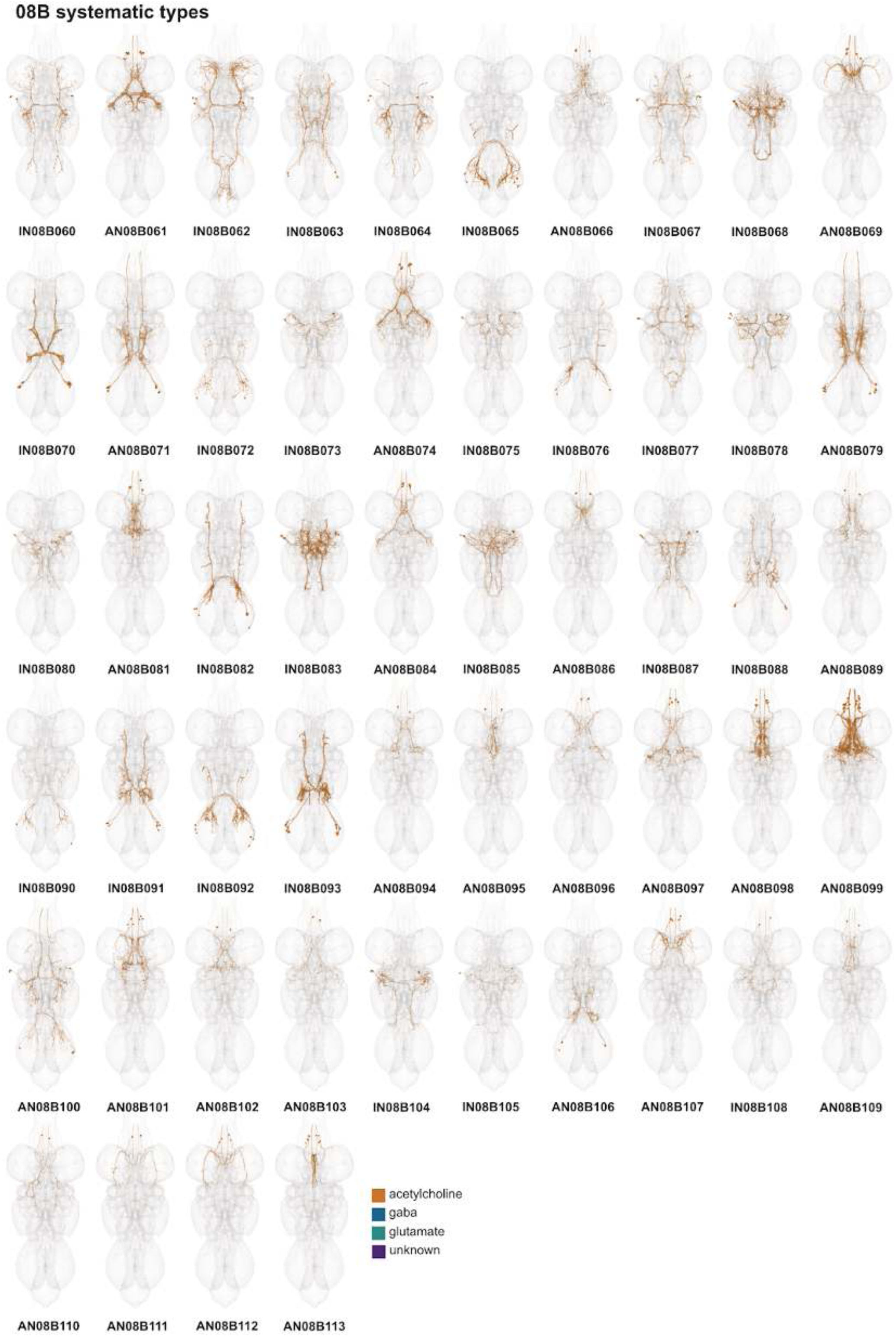
Systematic types of hemilineage 08B, continued. Systematic types have been arranged in numerical order, with neurons of the same type that belong to distinct classes (e.g., intrinsic neuron vs ascending neuron) plotted separately but placed adjacent to each other. Individual neuron meshes have been coloured based on predicted neurotransmitter: dark orange = acetylcholine, blue = gaba, marine = glutamate, dark purple = unknown.

**Figure 27 - figure supplement 4.**
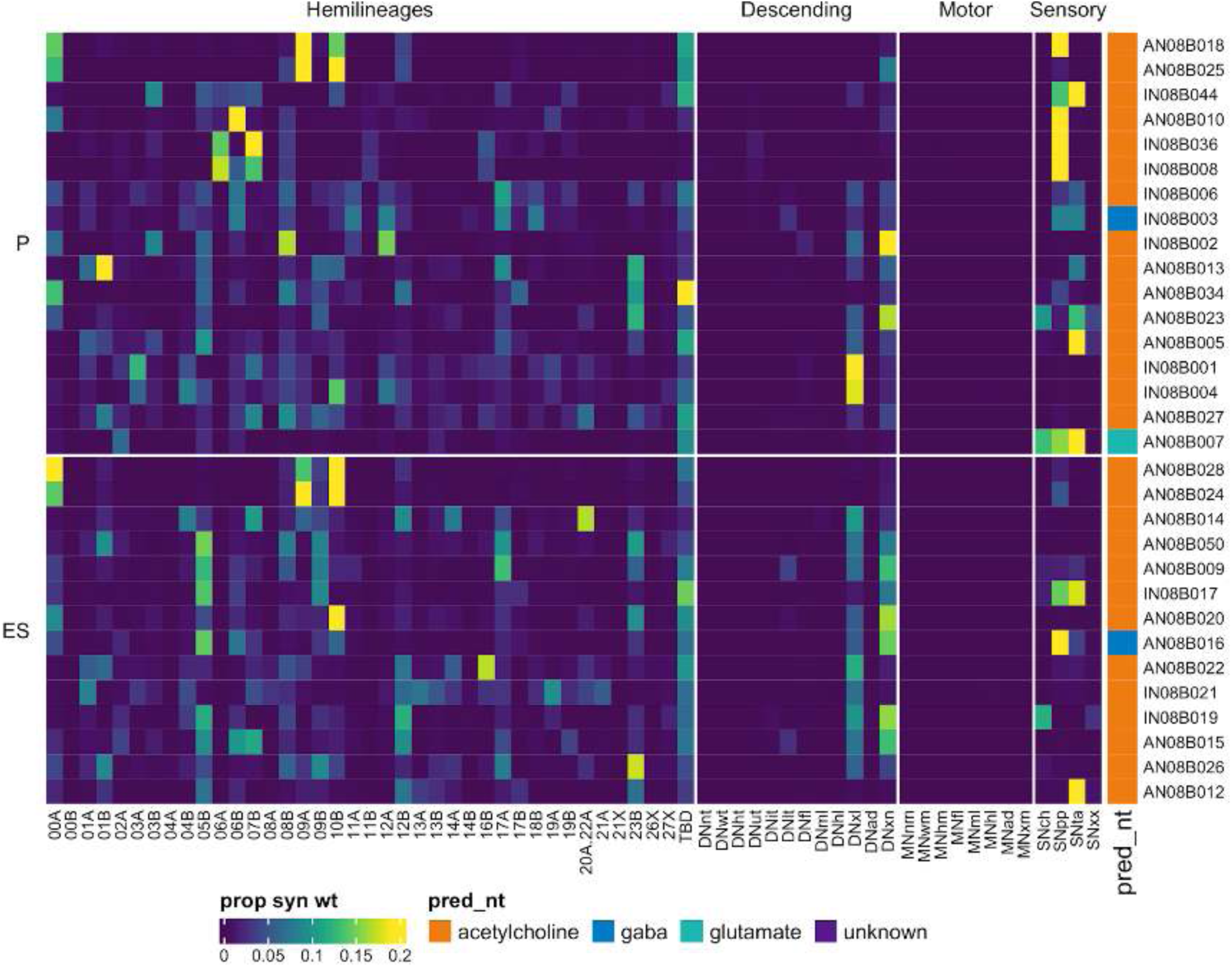
Connectivity to upstream partners by 08B primary and early secondary systematic types. Proportions of synaptic weight to systematic types from upstream partners, normalised by row. 08B neurons have been clustered within each assigned birthtime window (P = primary, ES = early secondary, S = secondary) based on both upstream and downstream connectivity to hemilineages, descending neuron subclasses, motor neuron subclasses, and sensory neuron modalities. Annotation bar is coloured by the most common predicted neurotransmitter for the neurons of each type.

**Figure 27 - figure supplement 5.**
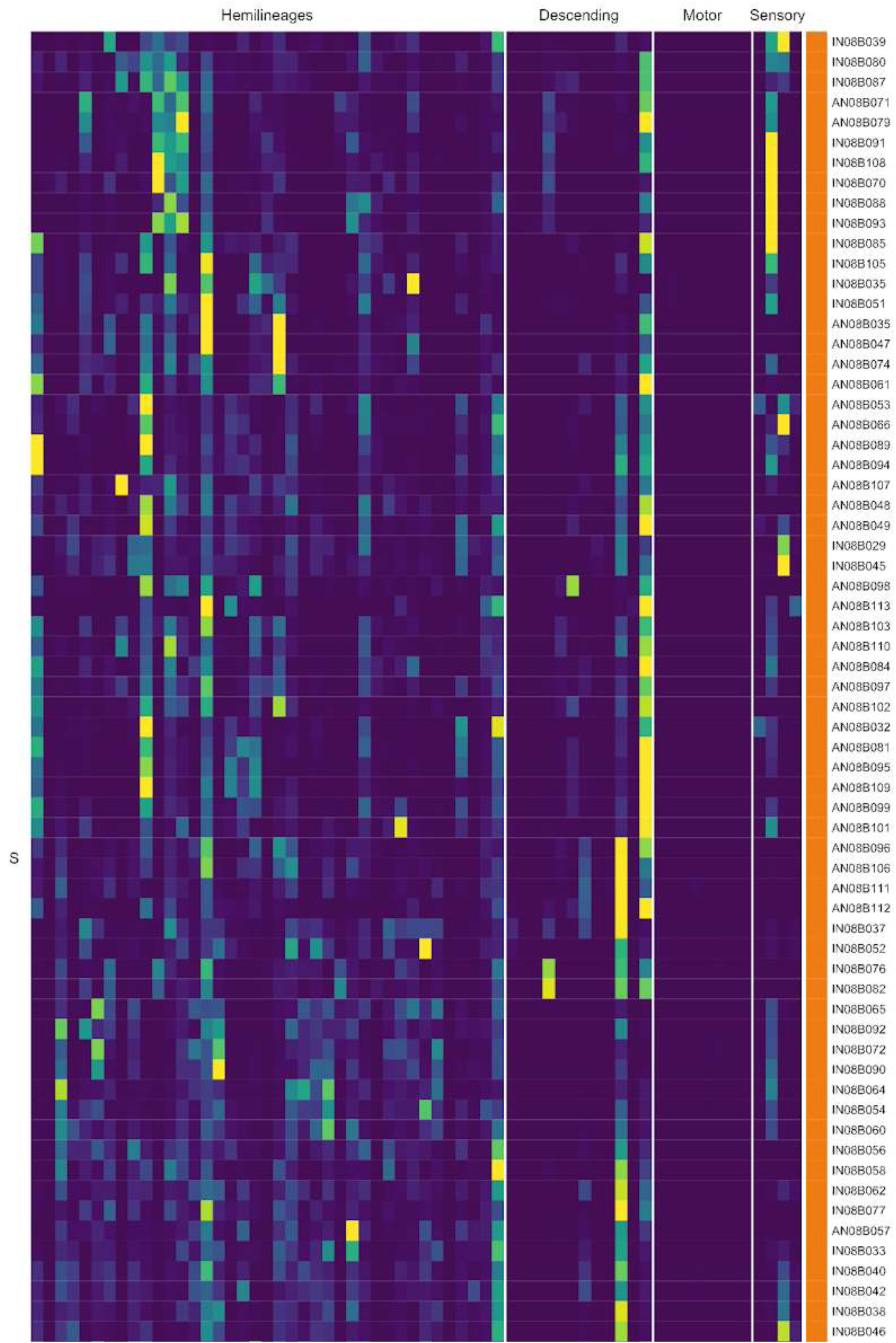
Connectivity to upstream partners by 08B secondary systematic types. Proportions of synaptic weight to systematic types from upstream partners, normalised by row. 08B neurons have been clustered within each assigned birthtime window (P = primary, ES = early secondary, S = secondary) based on both upstream and downstream connectivity to hemilineages, descending neuron subclasses, motor neuron subclasses, and sensory neuron modalities. The annotation bar is coloured by the most common predicted neurotransmitter within each type.

**Figure 27 - figure supplement 6.**
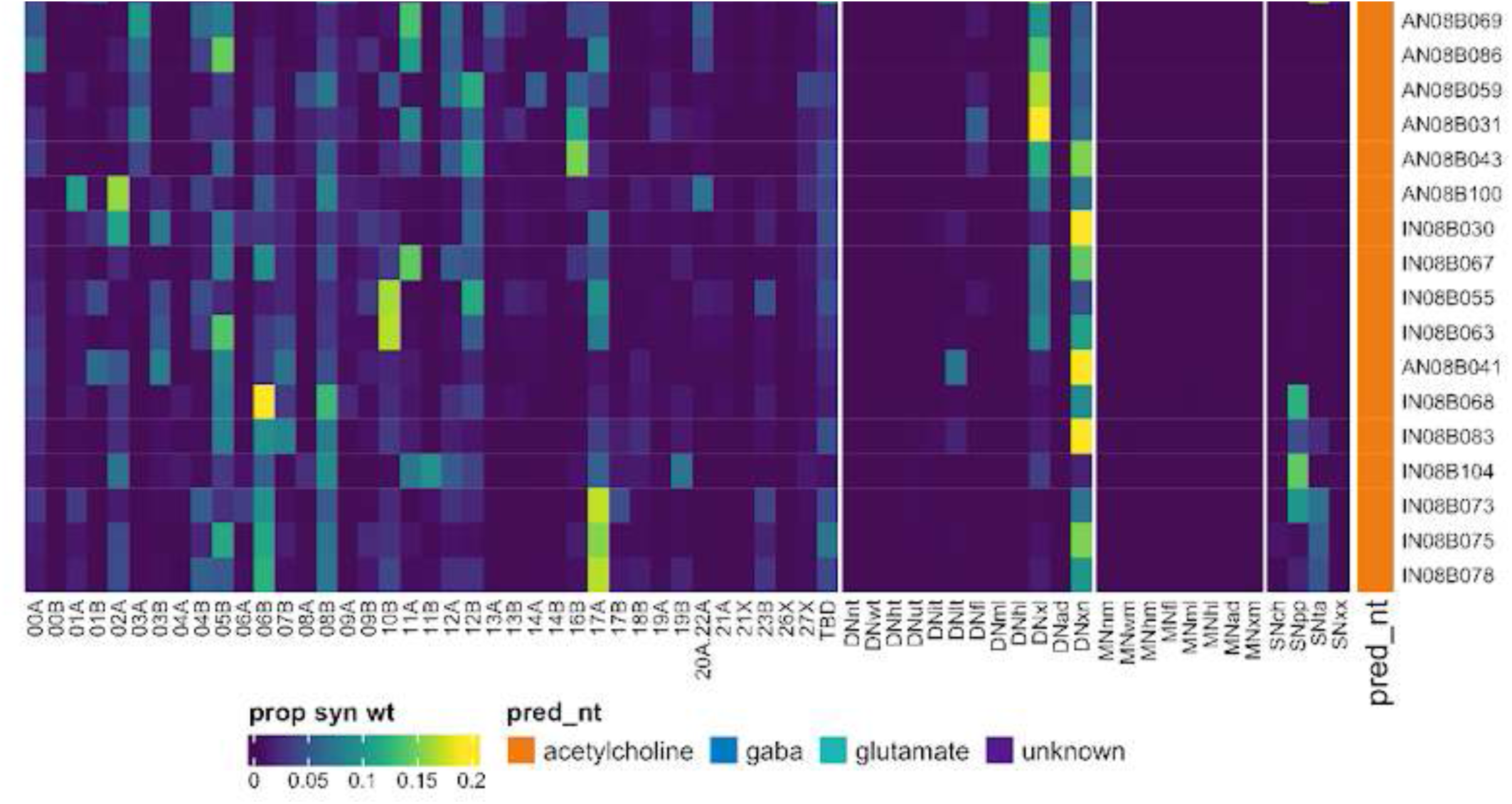
Connectivity to upstream partners by 08B secondary systematic types, continued. Proportions of synaptic weight to systematic types from upstream partners, normalised by row. 08B neurons have been clustered within each assigned birthtime window (P = primary, ES = early secondary, S = secondary) based on both upstream and downstream connectivity to hemilineages, descending neuron subclasses, motor neuron subclasses, and sensory neuron modalities. The annotation bar is coloured by the most common predicted neurotransmitter within each type.

**Figure 27 - figure supplement 7.**
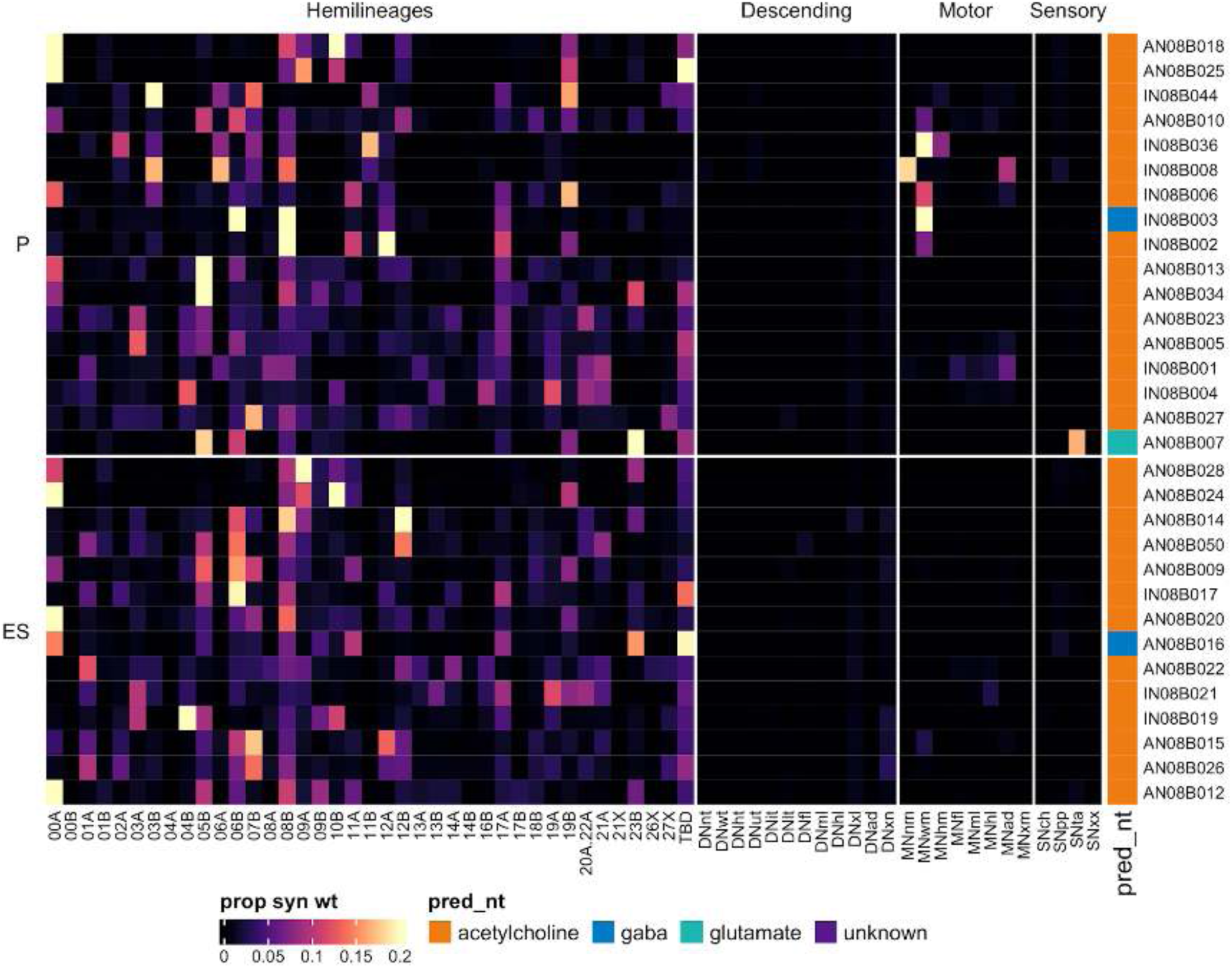
Connectivity to downstream partners by 08B primary and early secondary systematic types. Proportions of synaptic weight from systematic types to downstream partners, normalised by row. 08B neurons have been clustered within each assigned birthtime window (P = primary, ES = early secondary, S = secondary) based on both upstream and downstream connectivity to hemilineages, descending neuron subclasses, motor neuron subclasses, and sensory neuron modalities. The annotation bar is coloured by the most common predicted neurotransmitter within each type.

**Figure 27 - figure supplement 8.**
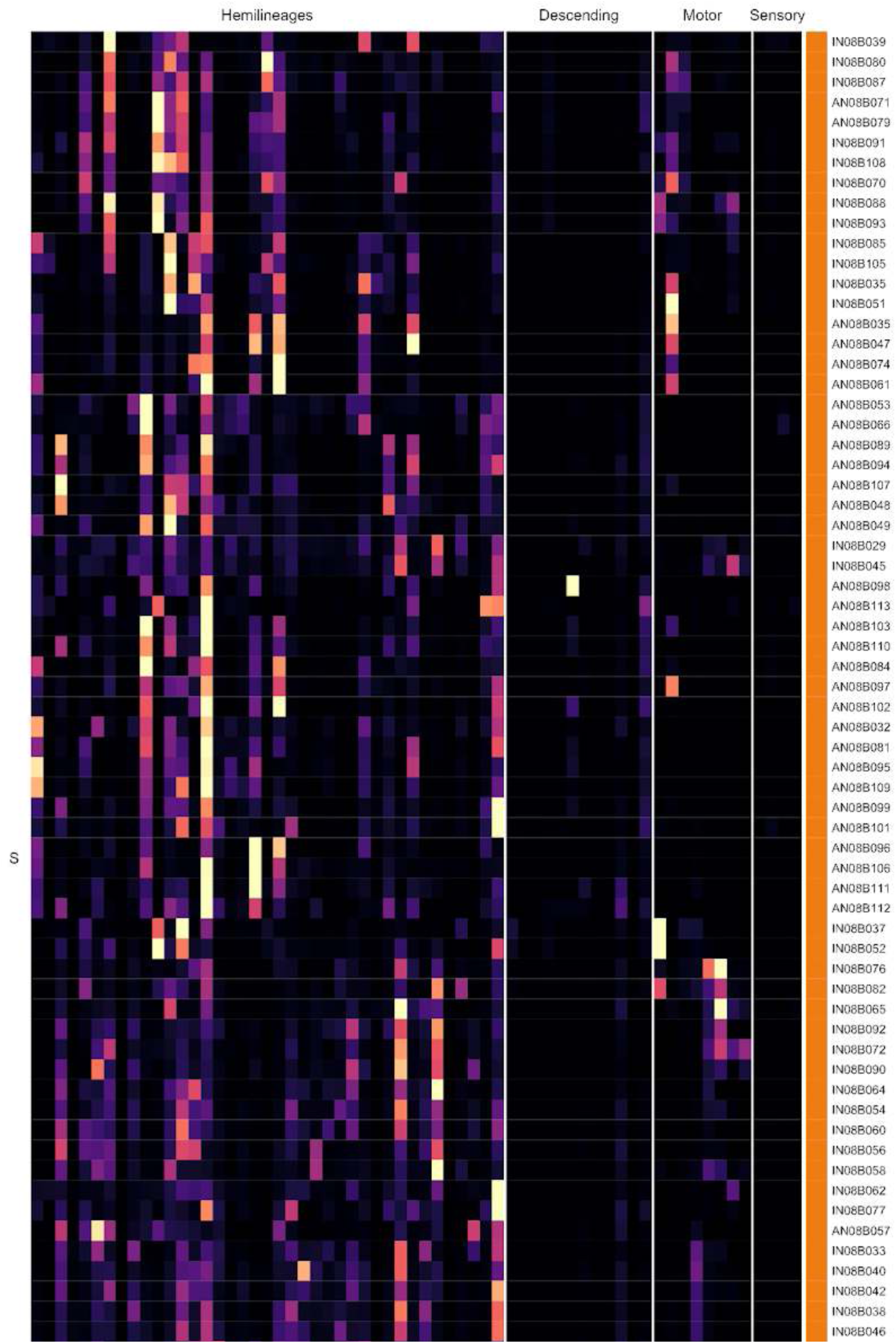
Connectivity to downstream partners by 08B secondary systematic types. Proportions of synaptic weight from systematic types to downstream partners, normalised by row. 08B neurons have been clustered within each assigned birthtime window (P = primary, ES = early secondary, S = secondary) based on both upstream and downstream connectivity to hemilineages, descending neuron subclasses, motor neuron subclasses, and sensory neuron modalities. The annotation bar is coloured by the most common predicted neurotransmitter for the neurons of each type.

**Figure 27 - figure supplement 9.**
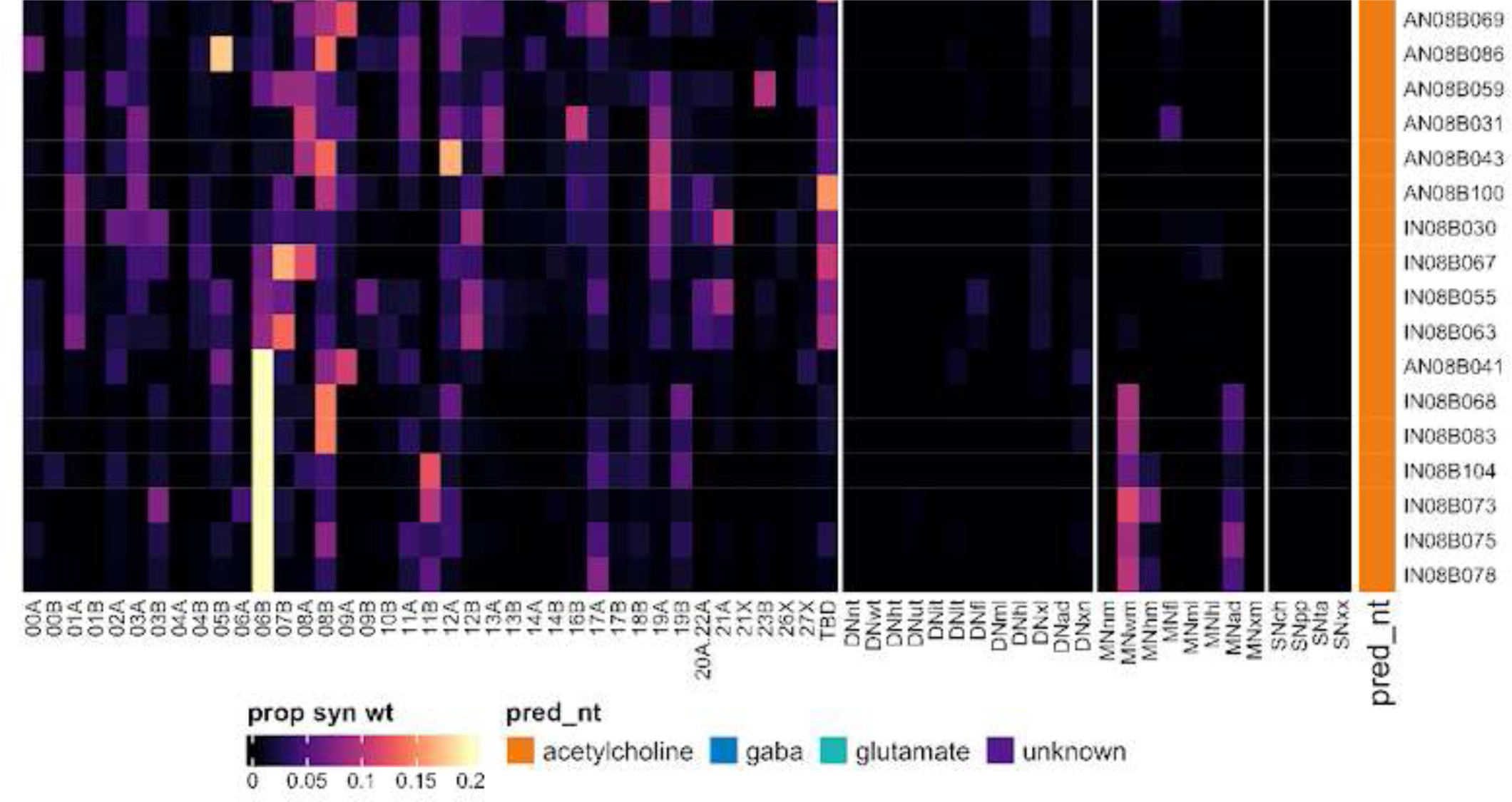
Connectivity to downstream partners by 08B secondary systematic types, continued. Proportions of synaptic weight from systematic types to downstream partners, normalised by row. 08B neurons have been clustered within each assigned birthtime window (P = primary, ES = early secondary, S = secondary) based on both upstream and downstream connectivity to hemilineages, descending neuron subclasses, motor neuron subclasses, and sensory neuron modalities. The annotation bar is coloured by the most common predicted neurotransmitter for the neurons of each type.

#### Hemilineage 09A

Hemilineages 09A and 09B derive from anterior dorsal neuroblast NB3-5 (Lacin and Truman, 2016), which generates an MN in T1 only as well as 12-15 intersegmental neurons and 12-15 local interneurons in the embryo (Schmid et al., 1999). Both are preserved in T1-T3 as well as (at reduced numbers) in abdominal segments (Truman et al., 2004). The projections of 09A secondary neurons are largely restricted to the ipsilateral leg neuropil (Shepherd et al., 2019), although we find a small subset that ascends to the next neuromere (e.g., Figure 28C top). The distinctive “curl” of its axons observed in the larva (Truman et al., 2004) is preserved as a landmark feature of many secondary neurons in the adult, but there is another population that innervates the ipsilateral mVAC (Shepherd et al., 2019) (e.g., Figure 28C bottom). 09A neurons in A1 are intersegmental, innervating the ipsilateral mVAC in all thoracic segments (Figure 28A).

**Figure 28.**
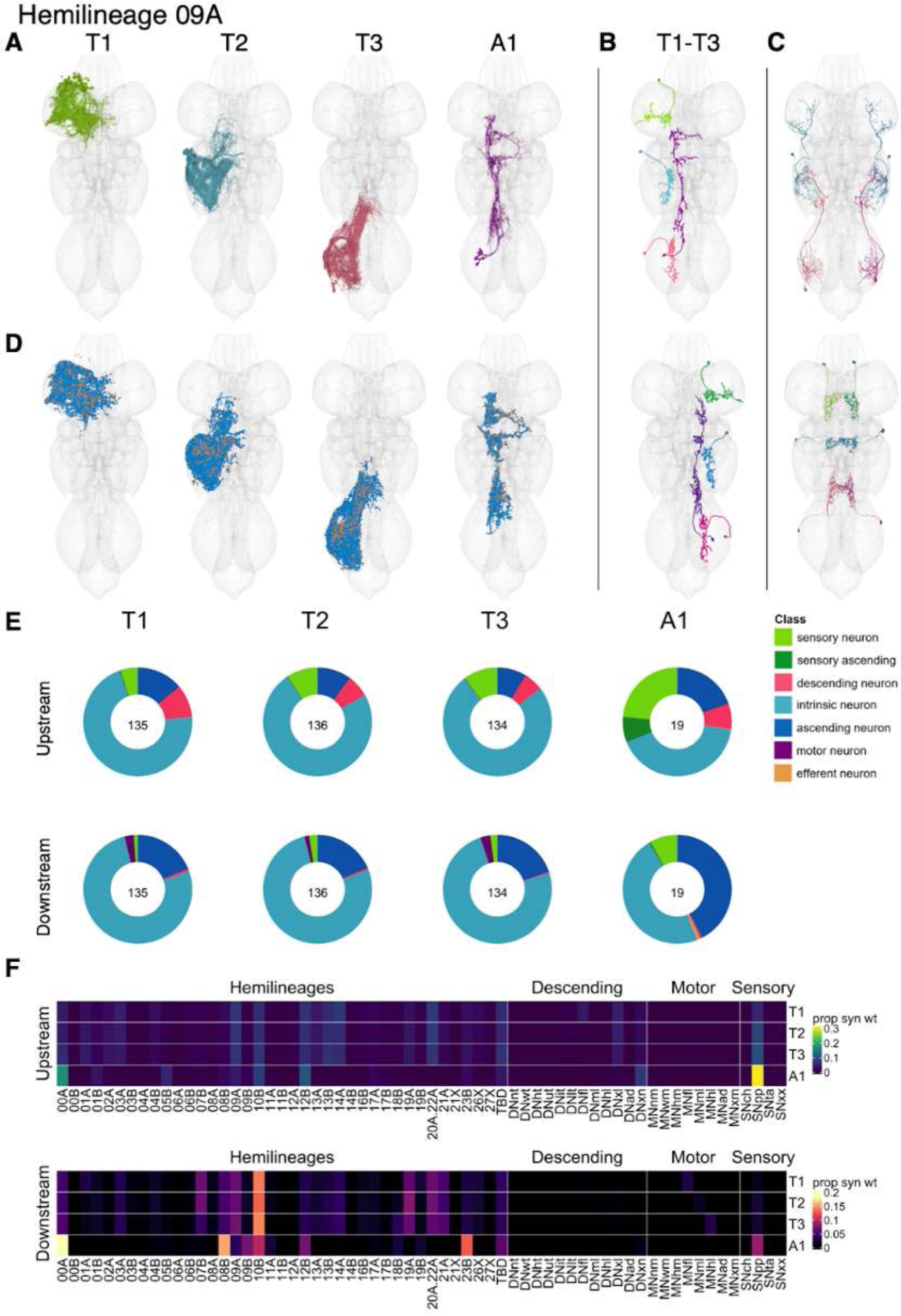
Hemilineage 09A. **A.** Meshes of all RHS secondary neurons plotted in neuromere-specific colours. **B.** “Representative” secondary neuron skeletons plotted in hemineuromere-specific colours. The skeleton with the top accumulated NBLAST score among all neurons from the hemilineage in a given hemineuromere was used. **C.** Neuron meshes of selected examples. Top: sequential serial set 11194. Bottom: mVAC-restricted independent leg serial set 14731. **D.** Predicted synapses of RHS secondary neurons. Blue: postsynapses; dark orange: presynapses. **E.** Proportions of connections from secondary neurons to upstream or downstream partners, normalised by neuromere and coloured by broad class. Numbers of query neurons appear in the centre. **F.** Proportions of synaptic weight from secondary neurons originating in each neuromere to upstream or downstream partners, normalised by row.

We identified 09A secondary neurons in ∼identical numbers in the three thoracic neuromeres, with a much smaller population in A1 (Figure 28E). They receive inputs from multiple hemilineages including 09A, 10B, 12B, and 20A.22A and also from proprioceptive neurons, particularly the A1 neurons, which also receive significant input from 00A. They are predicted to be gabaergic as expected (Lacin et al., 2019), and neurons from thoracic neuromeres primarily inhibit 10B, 09A, 19A, and 20A.22A whilst those from A1 inhibit 00A, 08B, 10B, and 23B and proprioceptive neurons (Figure 28F). Bilateral activation of 09A neurons results in a subtle repositioning/splaying out of legs with occasional bouts of spontaneous grooming (Harris et al., 2015), consistent with a role in mediating leg proprioception.

Although the 09A population as a whole does not receive a high proportion of direct input from descending neurons, several specific types do (e.g., IN09A012, IN09A045, and IN09A085) (Figure 28 - figure supplement 4-5). Whilst numerous 09A types receive strong proprioceptive input in the mVAC, several inhibit proprioceptive sensory neurons instead (e.g., IN09A012, IN09A025, IN09A026, IN09A030, IN09A046). We also identified several types that inhibit leg motor neurons (e.g., IN09A002, IN09A057, IN09A071, and IN09A079) (Figure 28 - figure supplement 7-9).

**Figure 28 - figure supplement 1.**
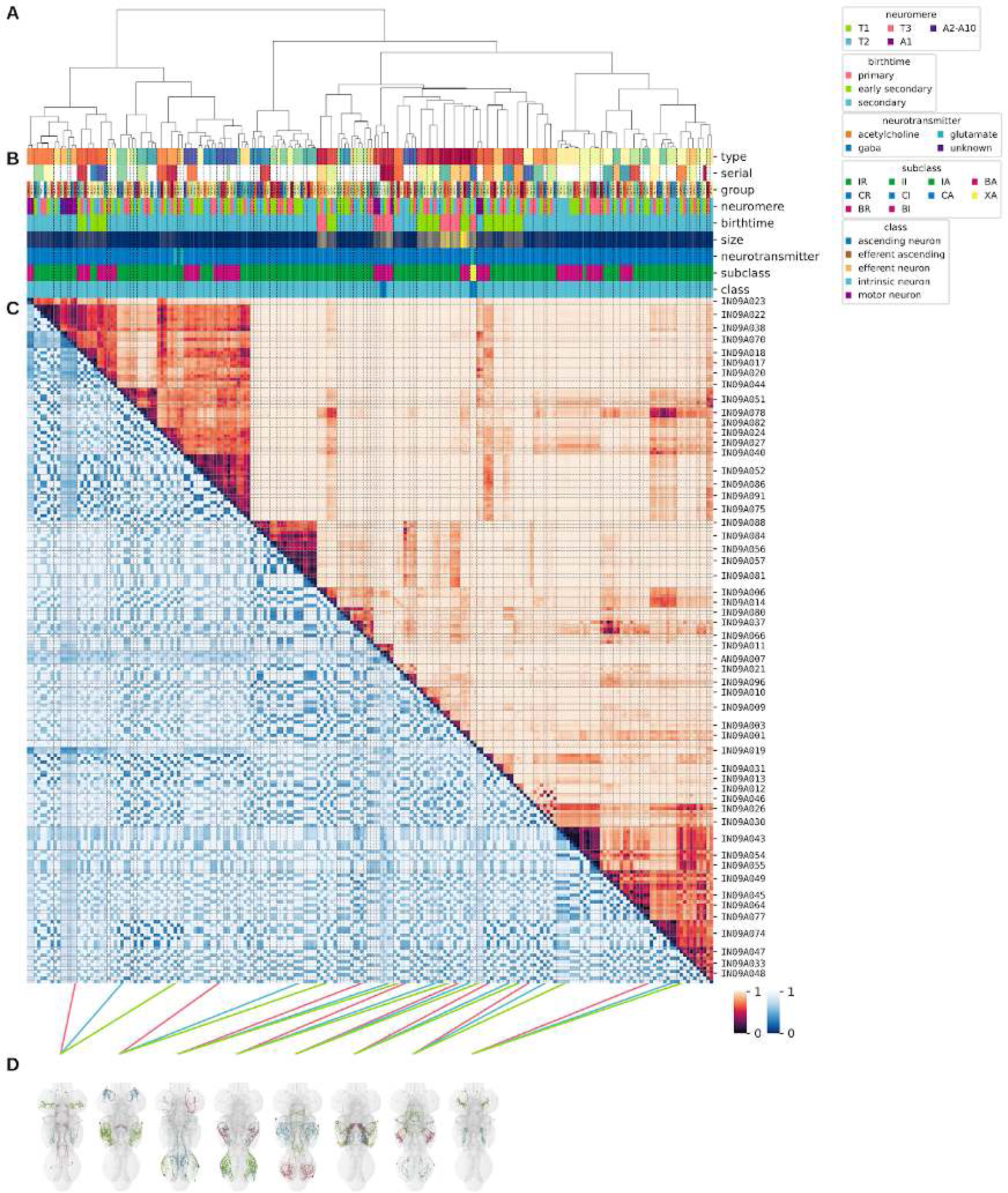
Systematic typing of hemilineage 09A. **A.** Hierarchical clustering dendrogram of hemilineage groups by laterally and serially aggregated connectivity cosine clustering. **B.** Categorical annotations of each hemilineage group, each column corresponding to the aligned leaf in A. Colours for type, serial set, and group are arbitrary for visualisation. Colours for neuromere, birthtime, neurotransmitter, subclass, and class are as in all other figures. **C.** Similarity distance heatmap for hemilineage. Cosine distance is in the upper triangle, while laterally symmetrised NBLAST distance is in the lower triangle. Systematic type names of some types are labelled. **D.** Morphologically representative groups from dendrogram subtrees. Each group, indicated by colour and line connecting to its column in B and C, is the most morphologically representative group (medoid of NBLAST distance) from a subtree of A. The subtrees (flat clusters) are equal height cuts of A determined to yield the number of groups per plot and plots in D.

**Figure 28 - figure supplement 2.**
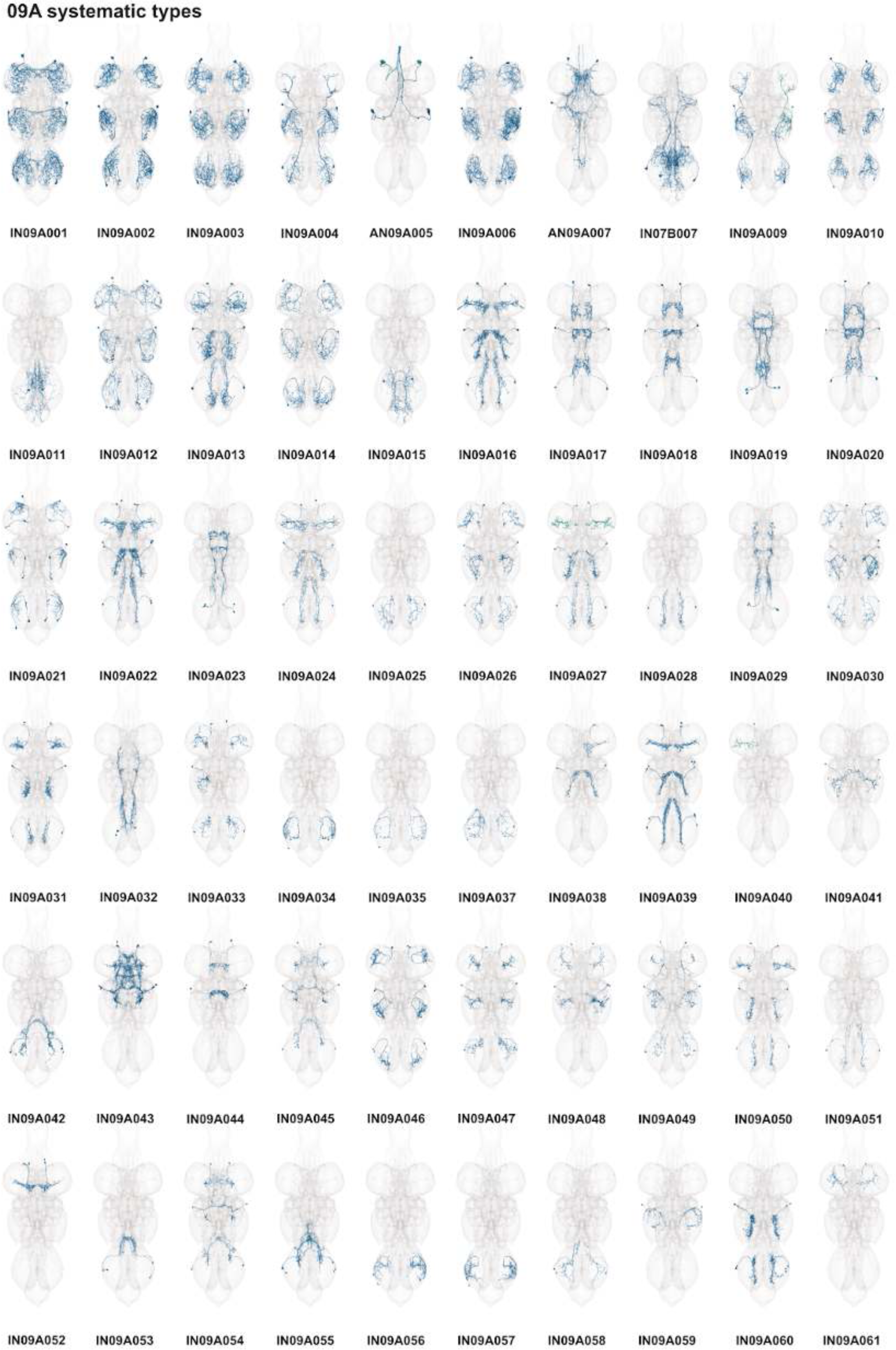
Systematic types of hemilineage 09A. Systematic types have been arranged in numerical order, with neurons of the same type that belong to distinct classes (e.g., intrinsic neuron vs ascending neuron) plotted separately but placed adjacent to each other. Individual neuron meshes have been coloured based on predicted neurotransmitter: dark orange = acetylcholine, blue = gaba, marine = glutamate, dark purple = unknown.

**Figure 28 - figure supplement 3.**
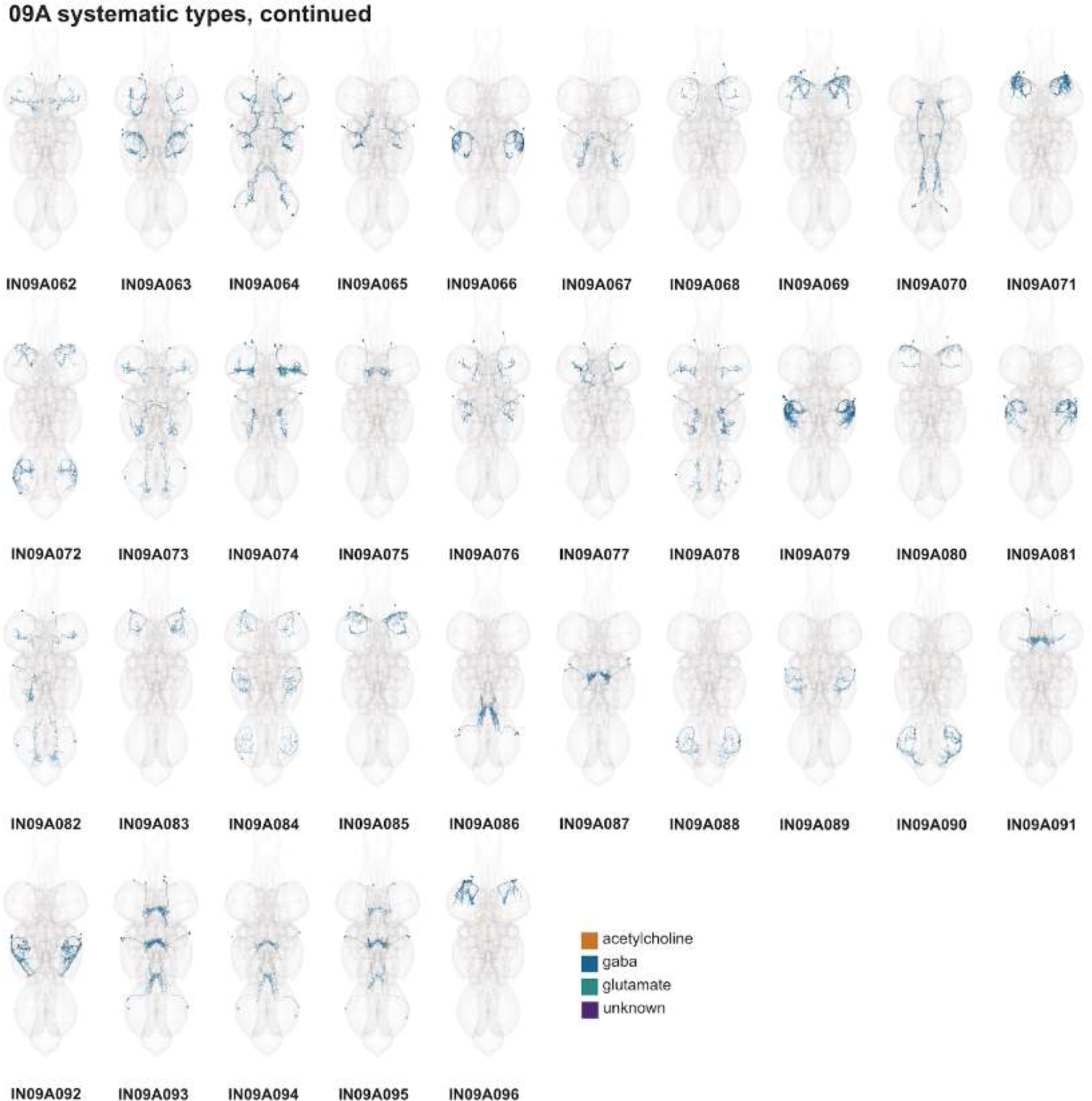
Systematic types of hemilineage 09A, continued. Systematic types have been arranged in numerical order, with neurons of the same type that belong to distinct classes (e.g., intrinsic neuron vs ascending neuron) plotted separately but placed adjacent to each other. Individual neuron meshes have been coloured based on predicted neurotransmitter: dark orange = acetylcholine, blue = gaba, marine = glutamate, dark purple = unknown.

**Figure 28 - figure supplement 4.**
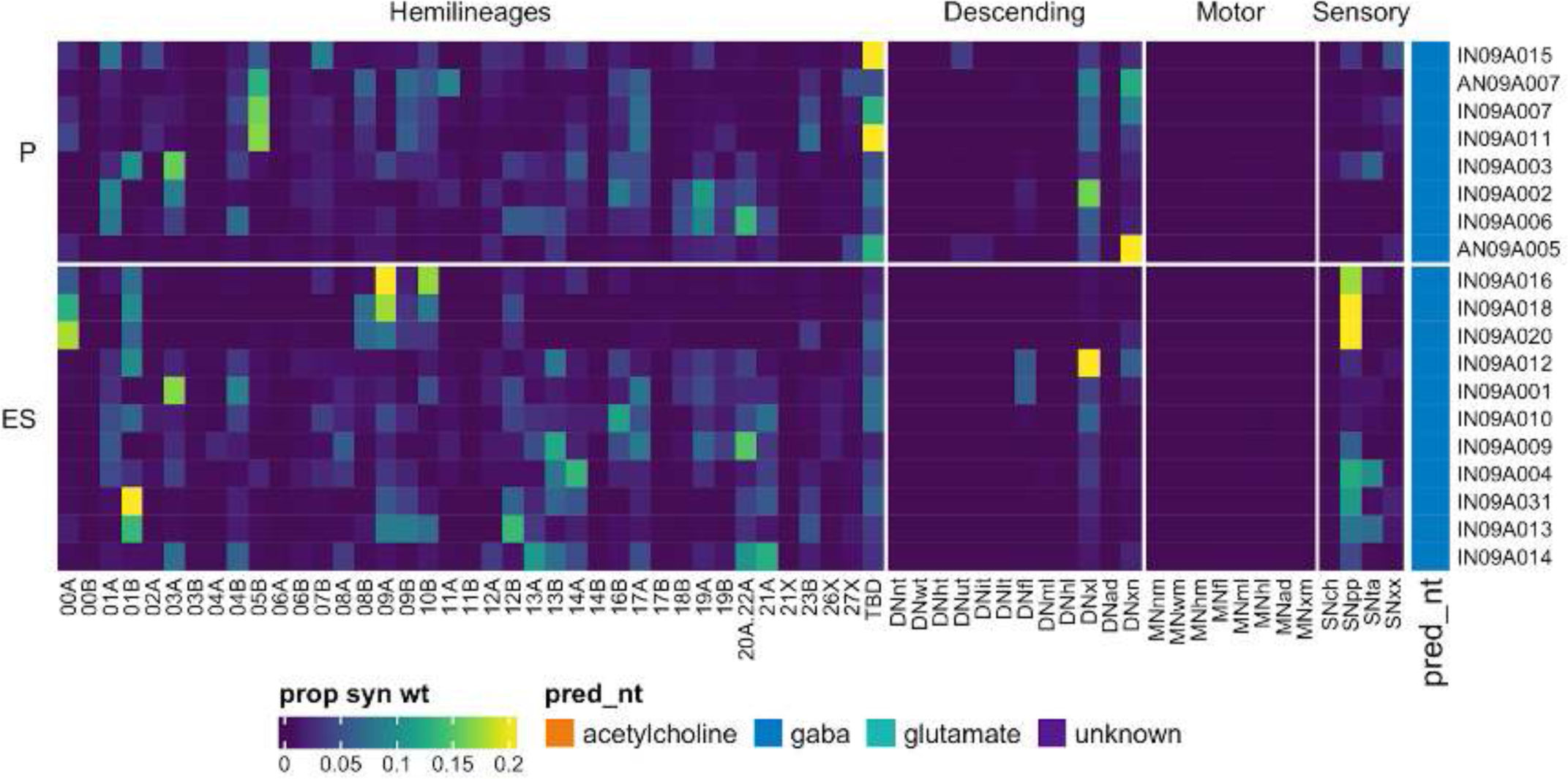
Connectivity to upstream partners by 09A primary and early secondary systematic types. Proportions of synaptic weight to systematic types from upstream partners, normalised by row. 09A neurons have been clustered within each assigned birthtime window (P = primary, ES = early secondary, S = secondary) based on both upstream and downstream connectivity to hemilineages, descending neuron subclasses, motor neuron subclasses, and sensory neuron modalities. Annotation bar is coloured by the most common predicted neurotransmitter for the neurons of each type.

**Figure 28 - figure supplement 5.**
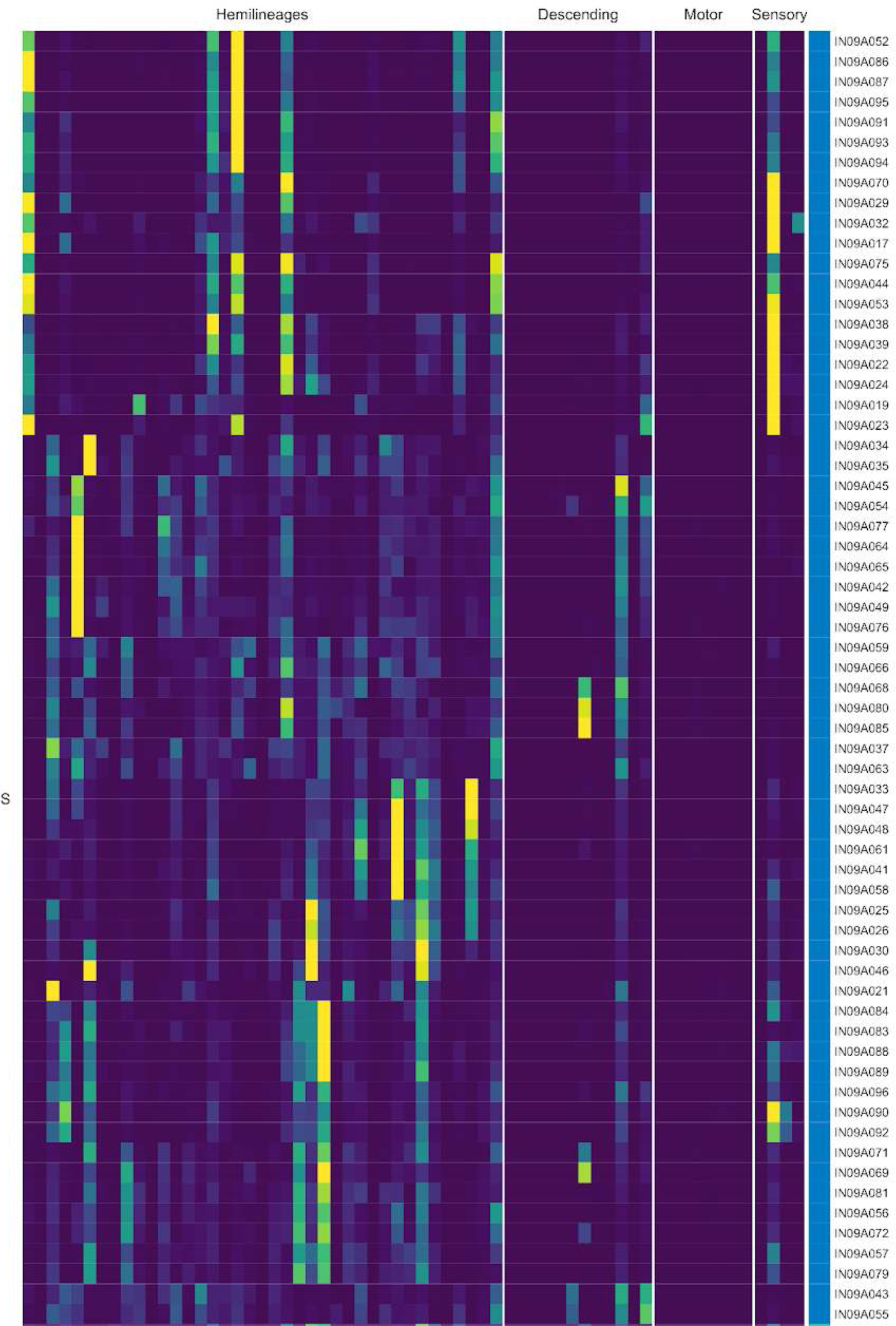
Connectivity to upstream partners by 09A secondary systematic types. Proportions of synaptic weight to systematic types from upstream partners, normalised by row. 09A neurons have been clustered within each assigned birthtime window (P = primary, ES = early secondary, S = secondary) based on both upstream and downstream connectivity to hemilineages, descending neuron subclasses, motor neuron subclasses, and sensory neuron modalities. The annotation bar is coloured by the most common predicted neurotransmitter within each type.

**Figure 28 - figure supplement 6.**
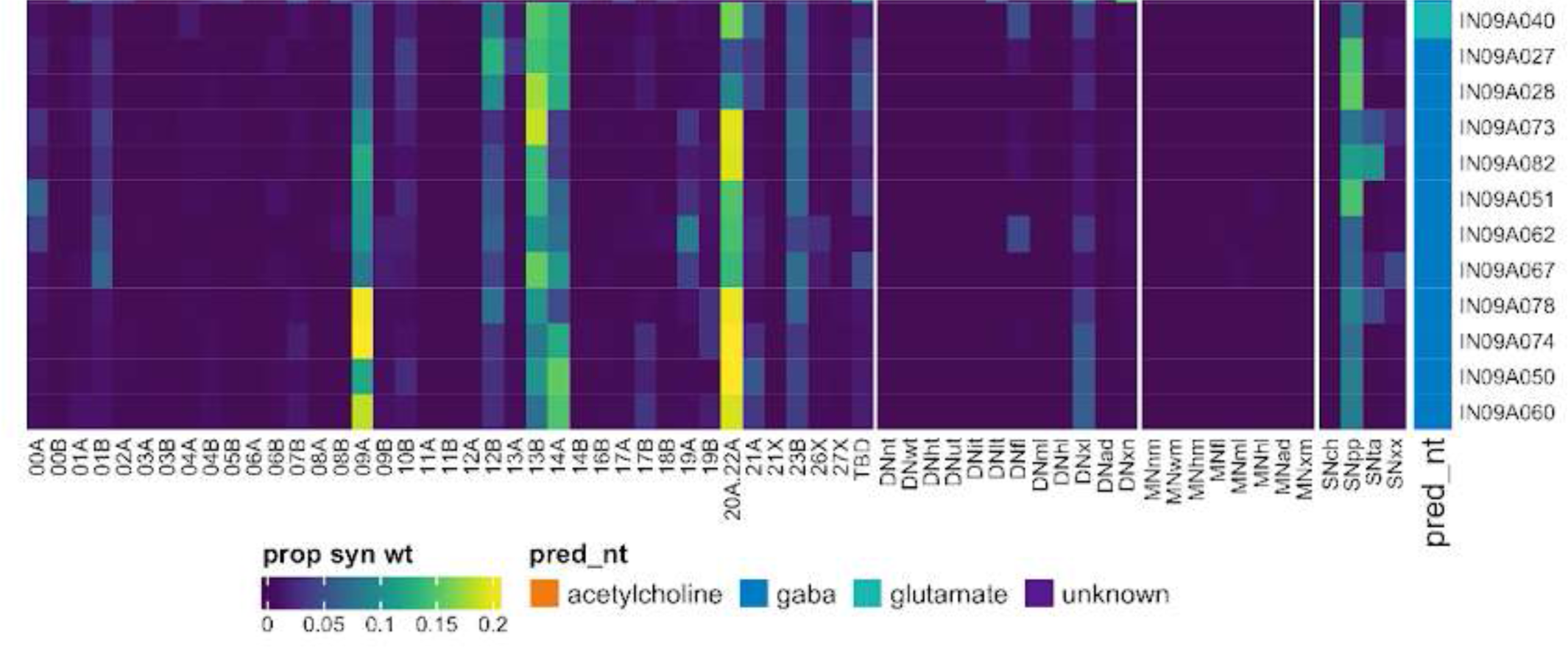
Connectivity to upstream partners by 09A secondary systematic types, continued. Proportions of synaptic weight to systematic types from upstream partners, normalised by row. 09A neurons have been clustered within each assigned birthtime window (P = primary, ES = early secondary, S = secondary) based on both upstream and downstream connectivity to hemilineages, descending neuron subclasses, motor neuron subclasses, and sensory neuron modalities. The annotation bar is coloured by the most common predicted neurotransmitter within each type.

**Figure 28 - figure supplement 7.**
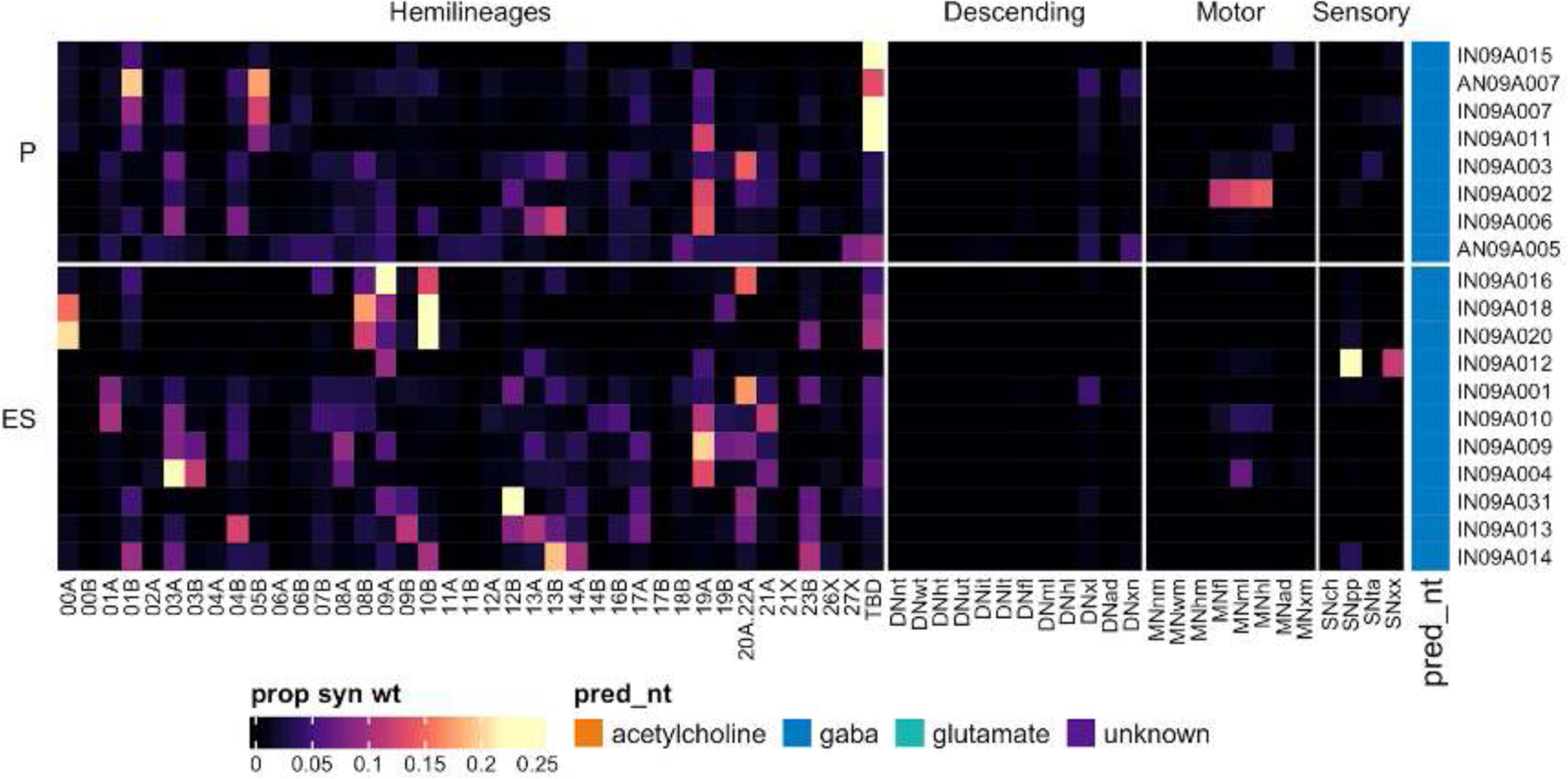
Connectivity to downstream partners by 09A primary and early secondary systematic types. Proportions of synaptic weight from systematic types to downstream partners, normalised by row. 09A neurons have been clustered within each assigned birthtime window (P = primary, ES = early secondary, S = secondary) based on both upstream and downstream connectivity to hemilineages, descending neuron subclasses, motor neuron subclasses, and sensory neuron modalities. The annotation bar is coloured by the most common predicted neurotransmitter within each type.

**Figure 28 - figure supplement 8.**
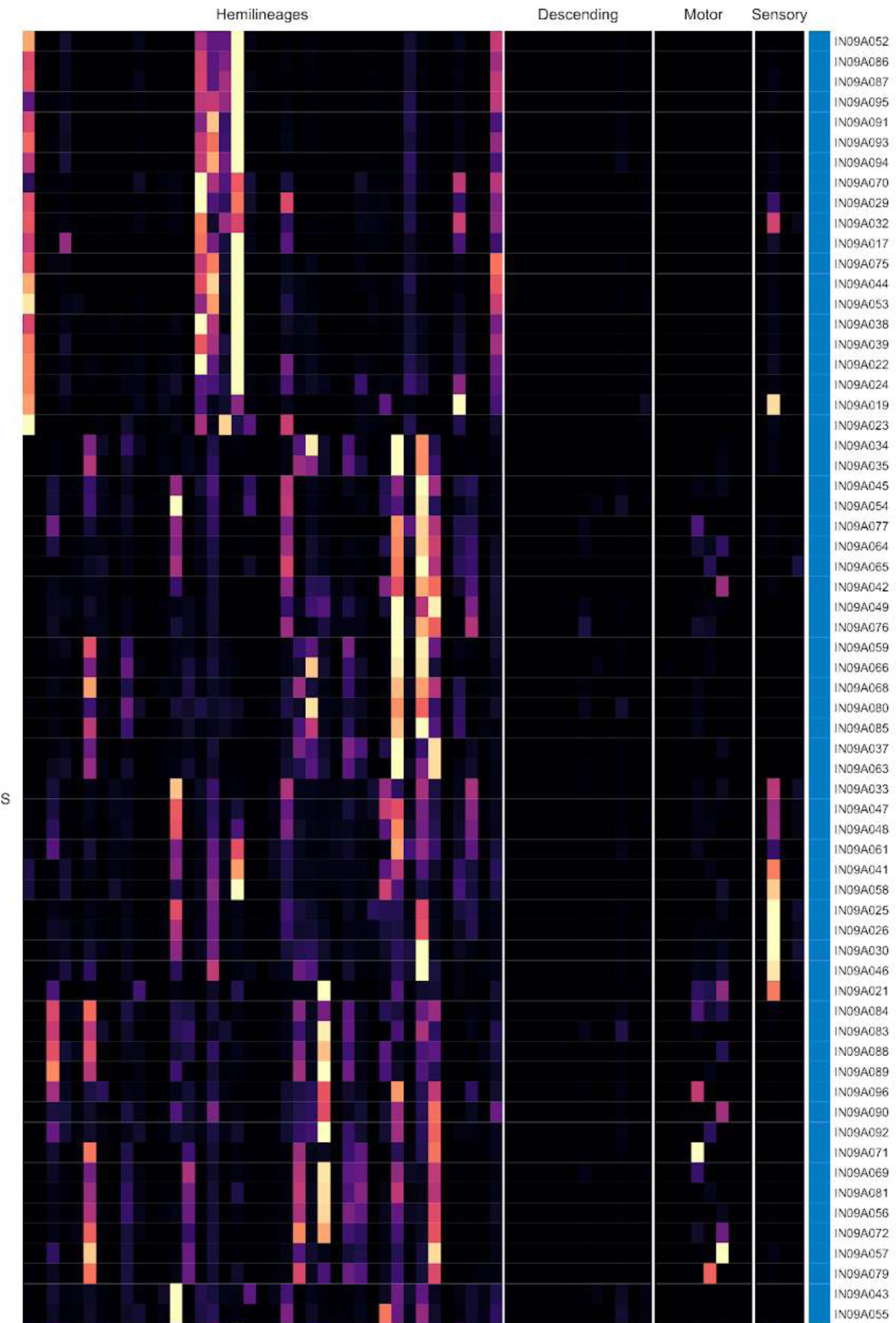
Connectivity to downstream partners by 09A secondary systematic types. Proportions of synaptic weight from systematic types to downstream partners, normalised by row. 09A neurons have been clustered within each assigned birthtime window (P = primary, ES = early secondary, S = secondary) based on both upstream and downstream connectivity to hemilineages, descending neuron subclasses, motor neuron subclasses, and sensory neuron modalities. The annotation bar is coloured by the most common predicted neurotransmitter for the neurons of each type.

**Figure 28 - figure supplement 9.**
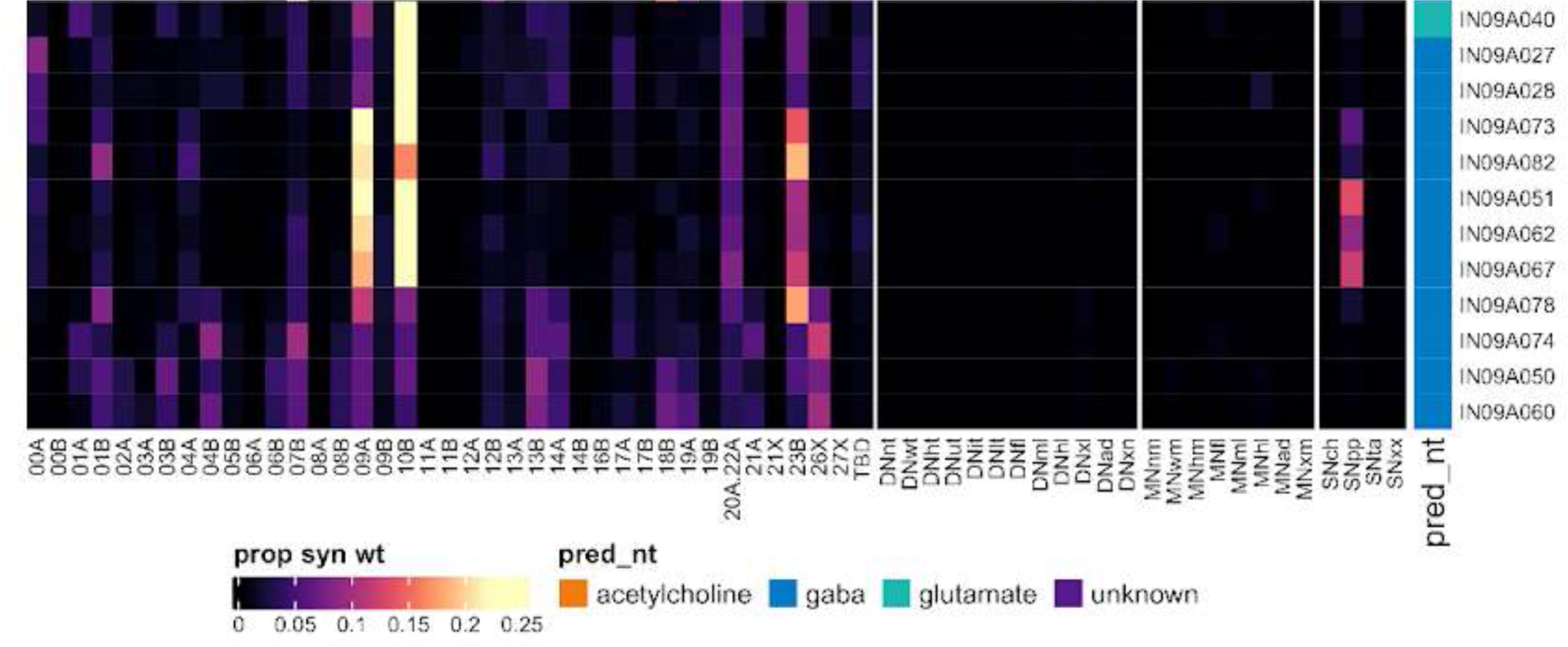
Connectivity to downstream partners by 09A secondary systematic types, continued. Proportions of synaptic weight from systematic types to downstream partners, normalised by row. 09A neurons have been clustered within each assigned birthtime window (P = primary, ES = early secondary, S = secondary) based on both upstream and downstream connectivity to hemilineages, descending neuron subclasses, motor neuron subclasses, and sensory neuron modalities. The annotation bar is coloured by the most common predicted neurotransmitter for the neurons of each type.

#### Hemilineage 09B

Hemilineages 09A and 09B derive from anterior dorsal neuroblast NB3-5 (Lacin and Truman, 2016), which generates an MN in T1 only as well as 12-15 intersegmental neurons and 12-15 local interneurons in the embryo (Schmid et al., 1999). Both are preserved in T1-T3 as well as (at reduced numbers) in abdominal segments (Truman et al., 2004). 09B is a small hemilineage that enters the neuropil with 09A; its neurons cross the midline near the ventral surface, generally between 01A and 14A, and elaborate in the leg and/or wing sensory neuropil on either side (Shepherd et al., 2019). Most 09B neurons project to the brain and/or other neuromeres (Figure 29A). The majority have bilateral dendrites (e.g., AN/IN09B006) Figure 29C top), but a subpopulation only innervates contralateral neuropils (e.g., IN09B022) (Figure 29C bottom).

**Figure 29.**
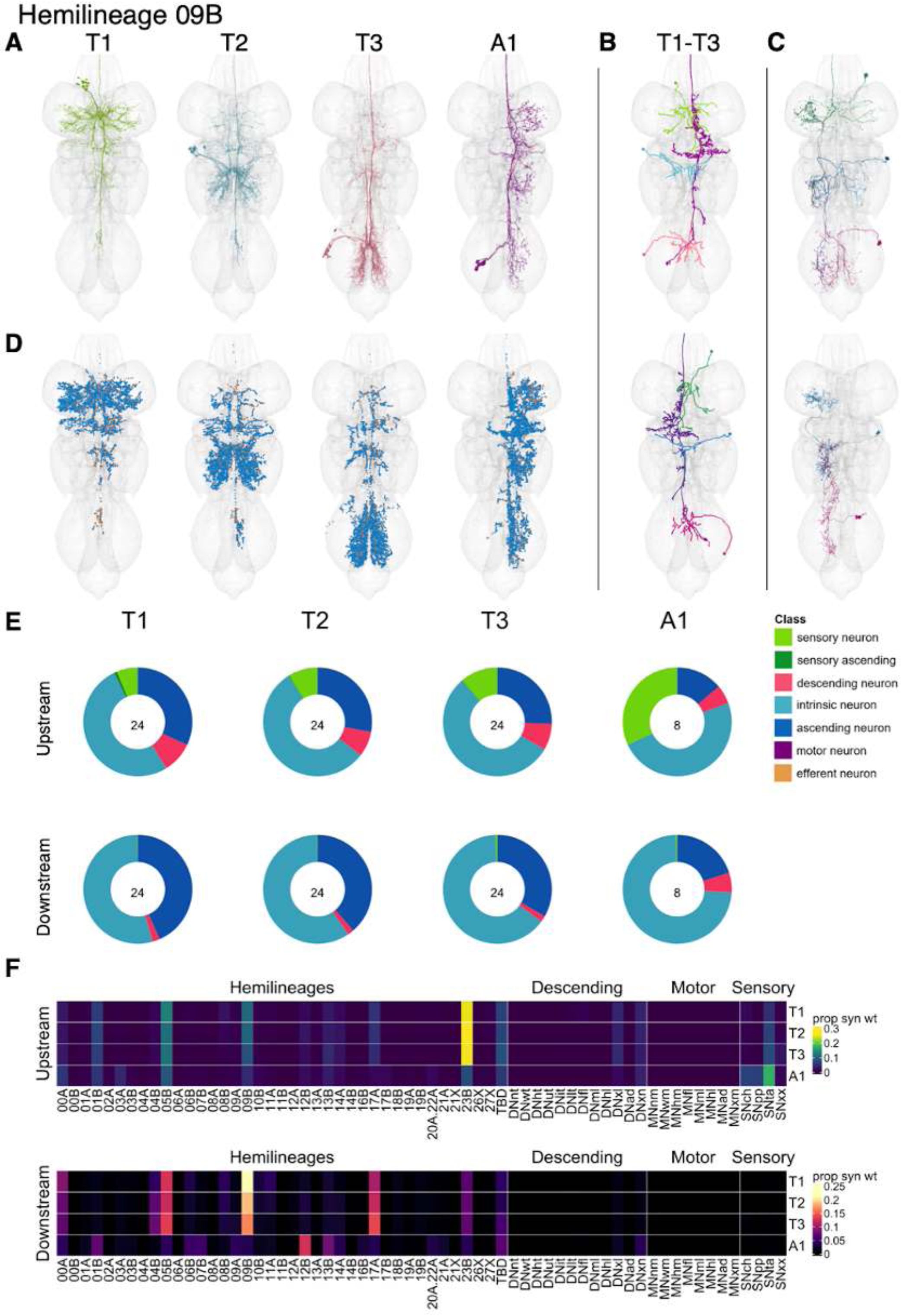
Hemilineage 09B. **A.** Meshes of all RHS secondary neurons plotted in neuromere-specific colours. **B.** “Representative” secondary neuron skeletons plotted in hemineuromere-specific colours. The skeleton with the top accumulated NBLAST score among all neurons from the hemilineage in a given hemineuromere was used. **C.** Neuron meshes of selected examples. Top: LHS neurons from bilateral, complex serial set 11491. Bottom: LHS neurons from contralateral, sequential serial set 17610. **D.** Predicted synapses of RHS secondary neurons. Blue: postsynapses; dark orange: presynapses. **E.** Proportions of connections from secondary neurons to upstream or downstream partners, normalised by neuromere and coloured by broad class. Numbers of query neurons appear in the centre. **F.** Proportions of synaptic weight from secondary neurons originating in each neuromere to upstream or downstream partners, normalised by row.

09B secondary neurons survive in identical numbers in each thoracic neuromere. They were reported to be glutamatergic (Lacin et al., 2019), but we identified two subpopulations of 09B neurons in thoracic neuromeres, one predicted to be glutamatergic and the other cholinergic (Figure 8F, Figure 29 - figure supplement 2-3), confirmed by gene expression studies (S Cachero and E Dona, personal communication). Apart from reduced innervation of the ipsilateral leg neuropil, we did not discern any obvious differences in morphology or in connectivity to hemilineages, descending neuron or motor neuron subclasses, or sensory neuron modalities between glutamatergic and cholinergic 09B neurons (Figure 8F, Figure 29 - figure supplement 2-7). For example, both glutamatergic and cholinergic 09B types (e.g., AN09B017 vs AN09B028) innervate the ventral area of T1 thought to receive pheromonal input (Figure 29 - figure supplement 2).

In T1-T3, secondary 09B neurons primarily receive inputs from hemilineages 01B, 05B, 09B, and especially 23B but also from tactile sensory neurons and descending neurons to multiple legs and neuromeres. In A1, they primarily receive inputs from tactile neurons, but also from chemosensory and proprioceptive sensory neurons and hemilineages 01B and 05B. Individual 09B types are typically restricted to one sensory modality (e.g., AN09B007 and AN09B033) but can be mixed (e.g., AN09B029 and AN09B032) (Figure 29 - figure supplement 4-5). Secondary 09B neurons target hemilineages 00A, 04B, 05B, 17A, 23B, and especially 09B in T1-T3, and the A1 secondary neurons target hemilineages 01B, 12B, and and 13B (Figure 29F). No functional studies have been published for secondary 09B neurons.

**Figure 29 - figure supplement 1.**
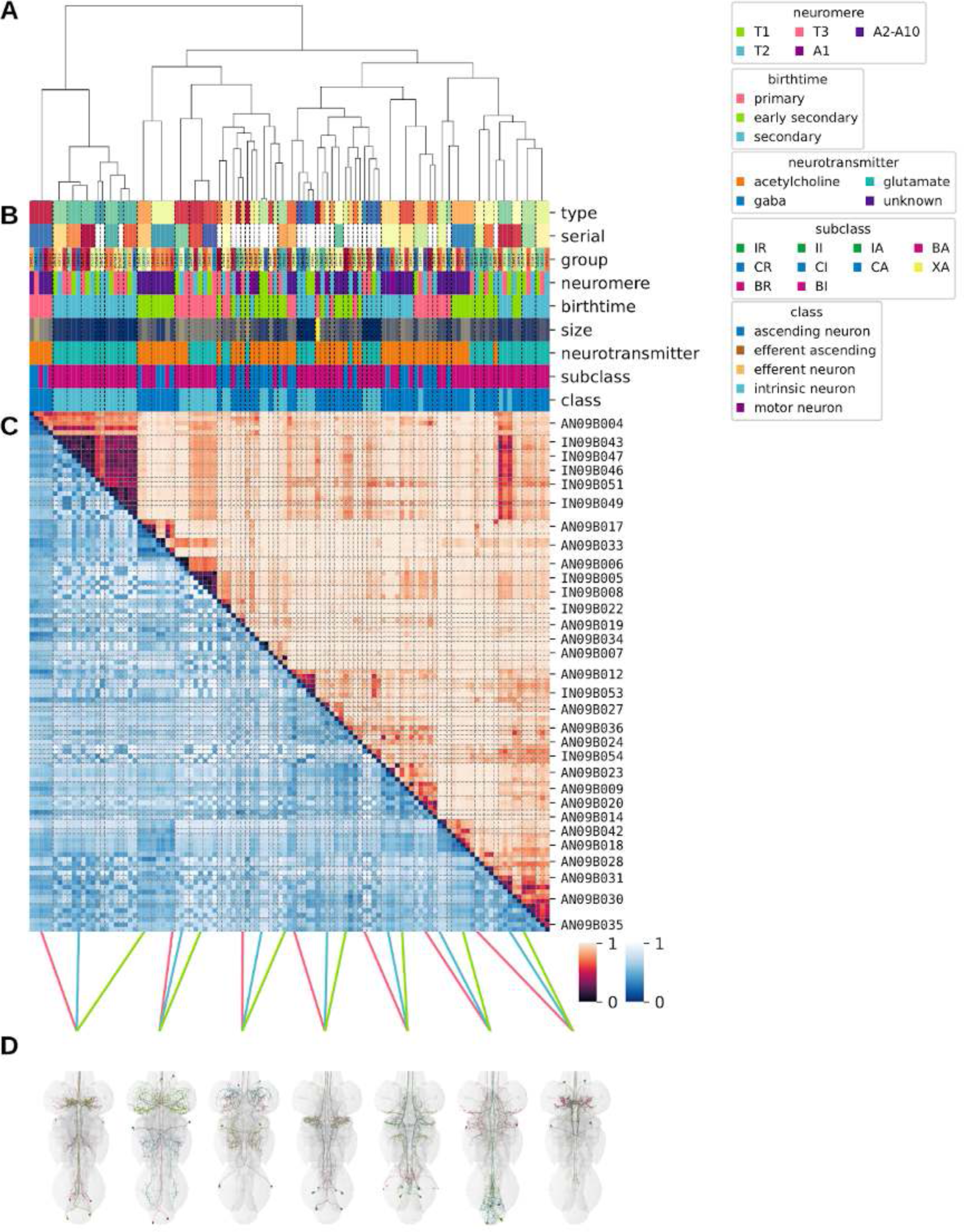
Systematic typing of hemilineage 09B. **A.** Hierarchical clustering dendrogram of hemilineage groups by laterally and serially aggregated connectivity cosine clustering. **B.** Categorical annotations of each hemilineage group, each column corresponding to the aligned leaf in A. Colours for type, serial set, and group are arbitrary for visualisation. Colours for neuromere, birthtime, neurotransmitter, subclass, and class are as in all other figures. **C.** Similarity distance heatmap for hemilineage. Cosine distance is in the upper triangle, while laterally symmetrised NBLAST distance is in the lower triangle. Systematic type names of some types are labelled. **D.** Morphologically representative groups from dendrogram subtrees. Each group, indicated by colour and line connecting to its column in B and C, is the most morphologically representative group (medoid of NBLAST distance) from a subtree of A. The subtrees (flat clusters) are equal height cuts of A determined to yield the number of groups per plot and plots in D.

**Figure 29 - figure supplement 2.**
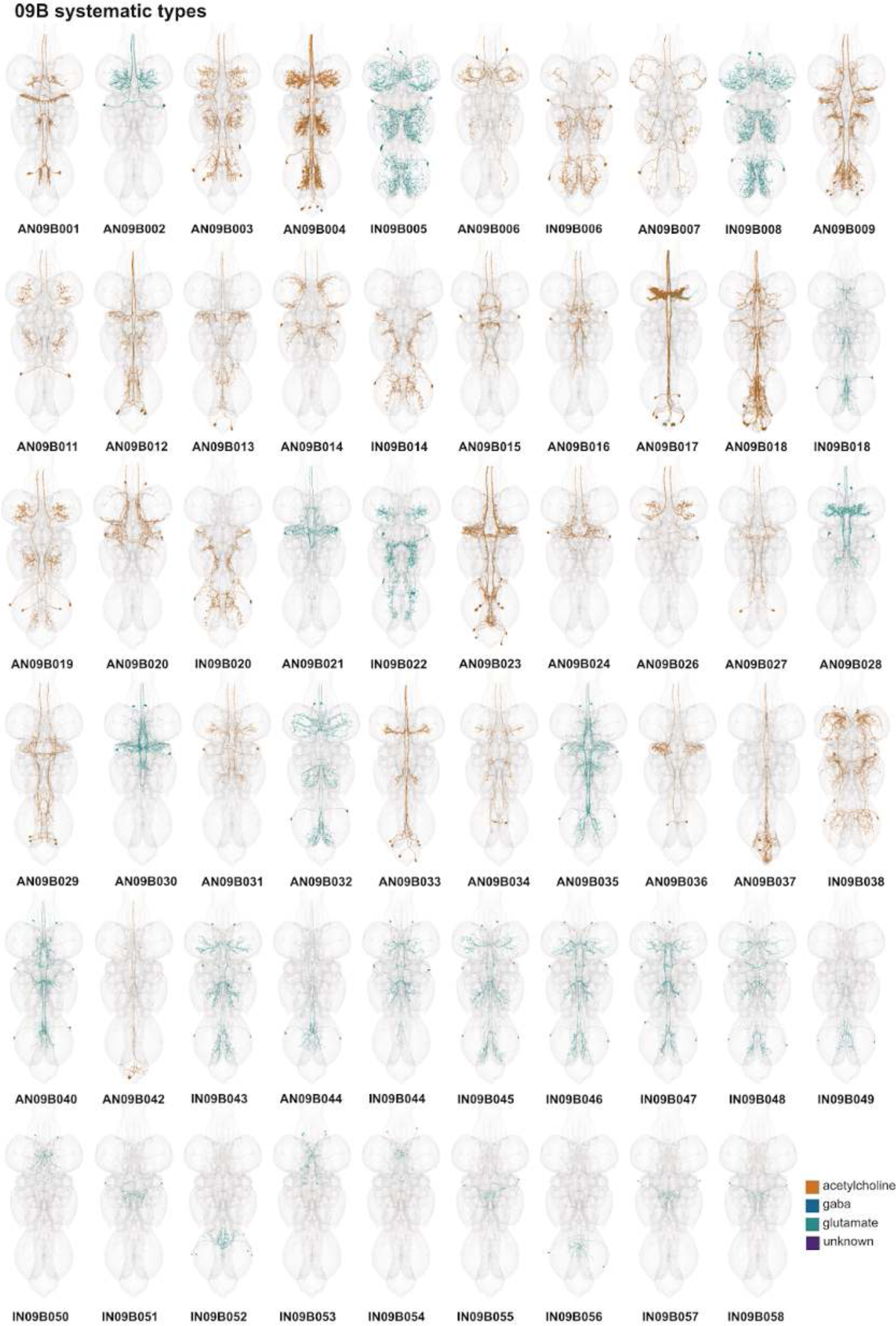
Systematic types of hemilineage 09B. Systematic types have been arranged in numerical order, with neurons of the same type that belong to distinct classes (e.g., intrinsic neuron vs ascending neuron) plotted separately but placed adjacent to each other. Individual neuron meshes have been coloured based on predicted neurotransmitter: dark orange = acetylcholine, blue = gaba, marine = glutamate, dark purple = unknown.

**Figure 29 - figure supplement 3.**
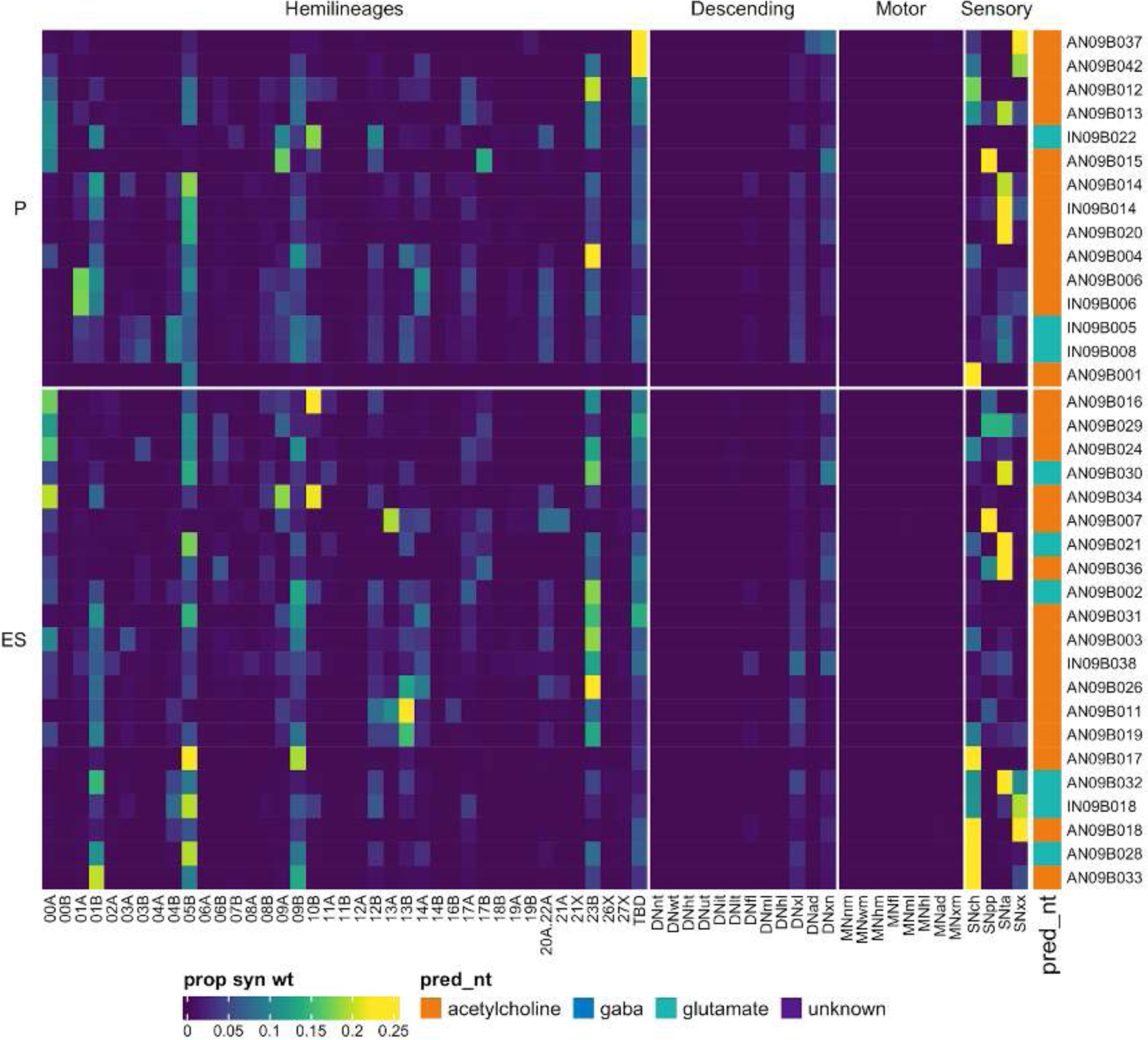
Connectivity to upstream partners by 09B primary and early secondary systematic types. Proportions of synaptic weight to systematic types from upstream partners, normalised by row. 09B neurons have been clustered within each assigned birthtime window (P = primary, ES = early secondary, S = secondary) based on both upstream and downstream connectivity to hemilineages, descending neuron subclasses, motor neuron subclasses, and sensory neuron modalities. Annotation bar is coloured by the most common predicted neurotransmitter for the neurons of each type.

**Figure 29 - figure supplement 4.**
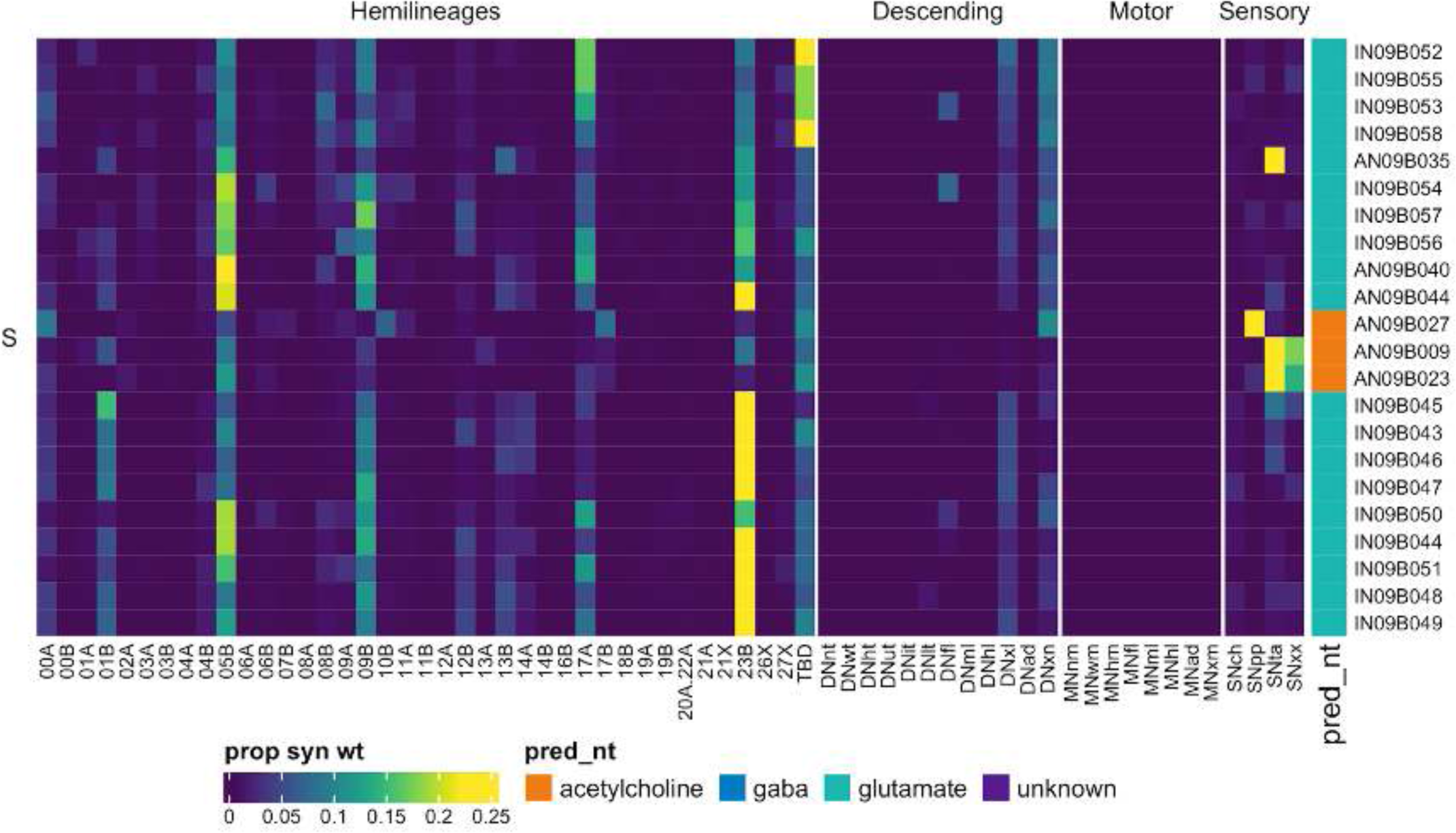
Connectivity to upstream partners by 09B secondary systematic types. Proportions of synaptic weight to systematic types from upstream partners, normalised by row. 09B neurons have been clustered within each assigned birthtime window (P = primary, ES = early secondary, S = secondary) based on both upstream and downstream connectivity to hemilineages, descending neuron subclasses, motor neuron subclasses, and sensory neuron modalities. The annotation bar is coloured by the most common predicted neurotransmitter within each type.

**Figure 29 - figure supplement 5.**
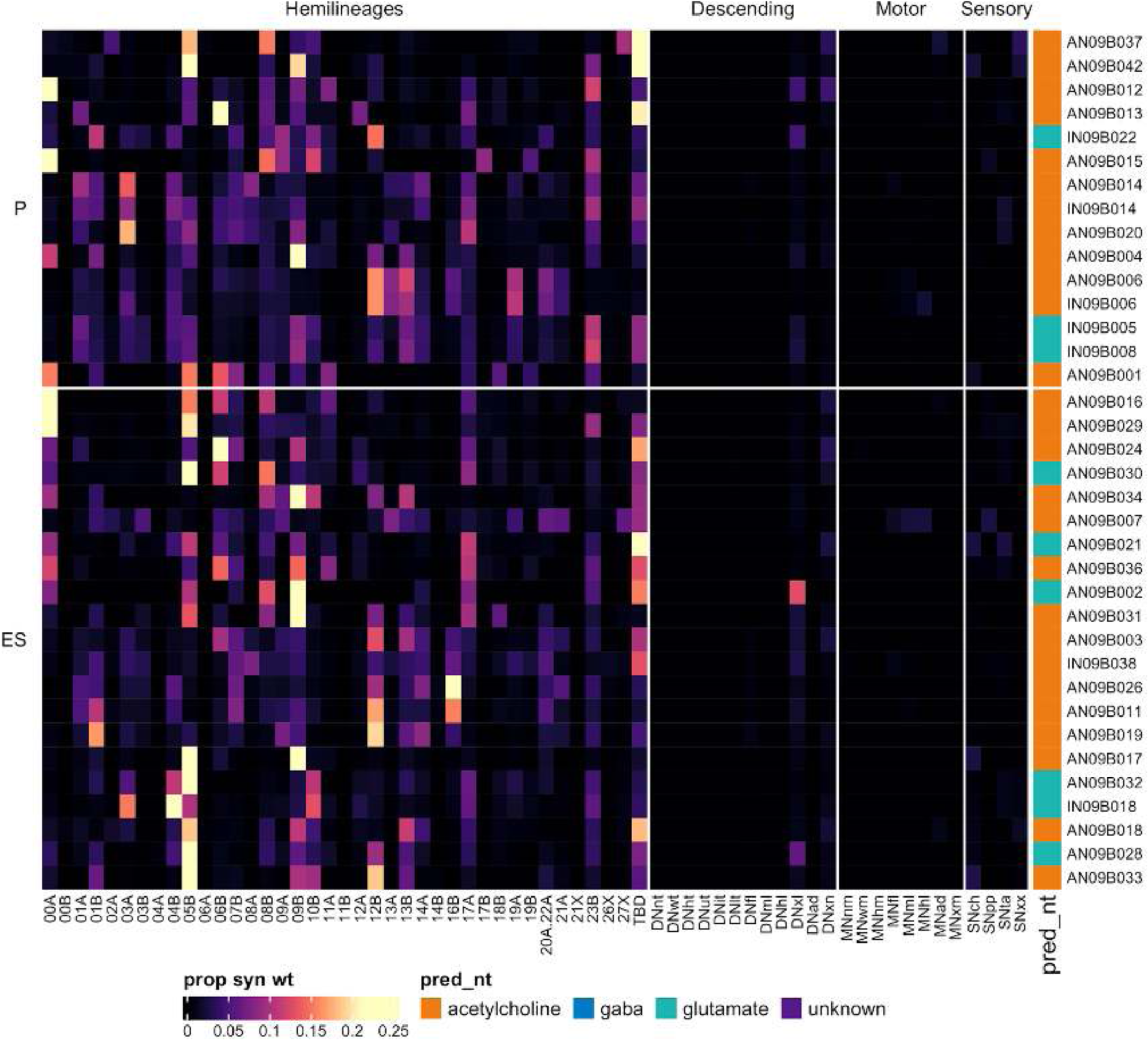
Connectivity to downstream partners by 09B primary and early secondary systematic types. Proportions of synaptic weight from systematic types to downstream partners, normalised by row. 09B neurons have been clustered within each assigned birthtime window (P = primary, ES = early secondary, S = secondary) based on both upstream and downstream connectivity to hemilineages, descending neuron subclasses, motor neuron subclasses, and sensory neuron modalities. The annotation bar is coloured by the most common predicted neurotransmitter within each type.

**Figure 29 - figure supplement 6.**
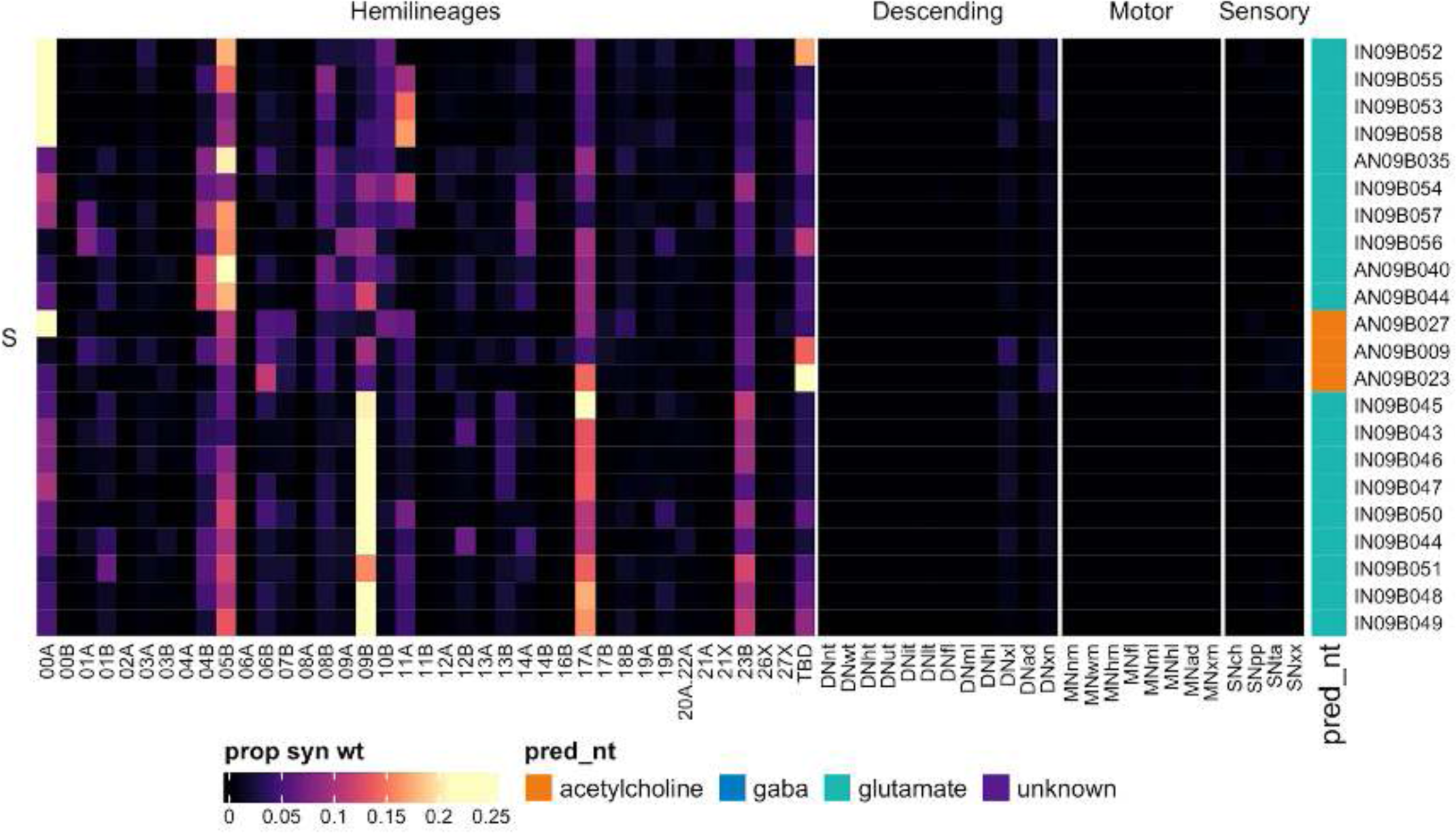
Connectivity to downstream partners by 09B secondary systematic types. Proportions of synaptic weight from systematic types to downstream partners, normalised by row. 09B neurons have been clustered within each assigned birthtime window (P = primary, ES = early secondary, S = secondary) based on both upstream and downstream connectivity to hemilineages, descending neuron subclasses, motor neuron subclasses, and sensory neuron modalities. The annotation bar is coloured by the most common predicted neurotransmitter for the neurons of each type.

#### Hemilineage 10B

10B is the only surviving secondary hemilineage (Truman et al., 2010) of anterior medial neuroblast NB2-2 (Birkholz et al., 2015; Lacin and Truman, 2016), which generates 2-3 MNs, 2-4 intersegmental interneurons, and 15 local interneurons in the embryo (Schmid et al., 1999). In the adult, 10B secondary neurons enter the anterior neuromere, typically innervating the ipsilateral mVAC (Shepherd et al., 2019) before crossing the midline quite ventrally in the anterior intermediate commissure, just anterior to the orthogonally projecting 2A primary neurites (Shepherd et al., 2016)). They are typically intersegmental, extending their axons anteriorly and/or posteriorly, and are mainly cholinergic (Lacin et al., 2019).

10B secondary neurons are restricted to thoracic segments (Truman et al., 2004), and we identified roughly similar numbers of neurons in each neuromere (Figure 30E). Whilst specific secondary types can be distinguished, they are generally fairly similar to each other in morphology and connectivity, with three main populations (Figure 30 - figure supplement 2-7). Early born 10B neurons cross in the same commissure but are much more diverse (e.g., Figure 30C top, bottom).

**Figure 30.**
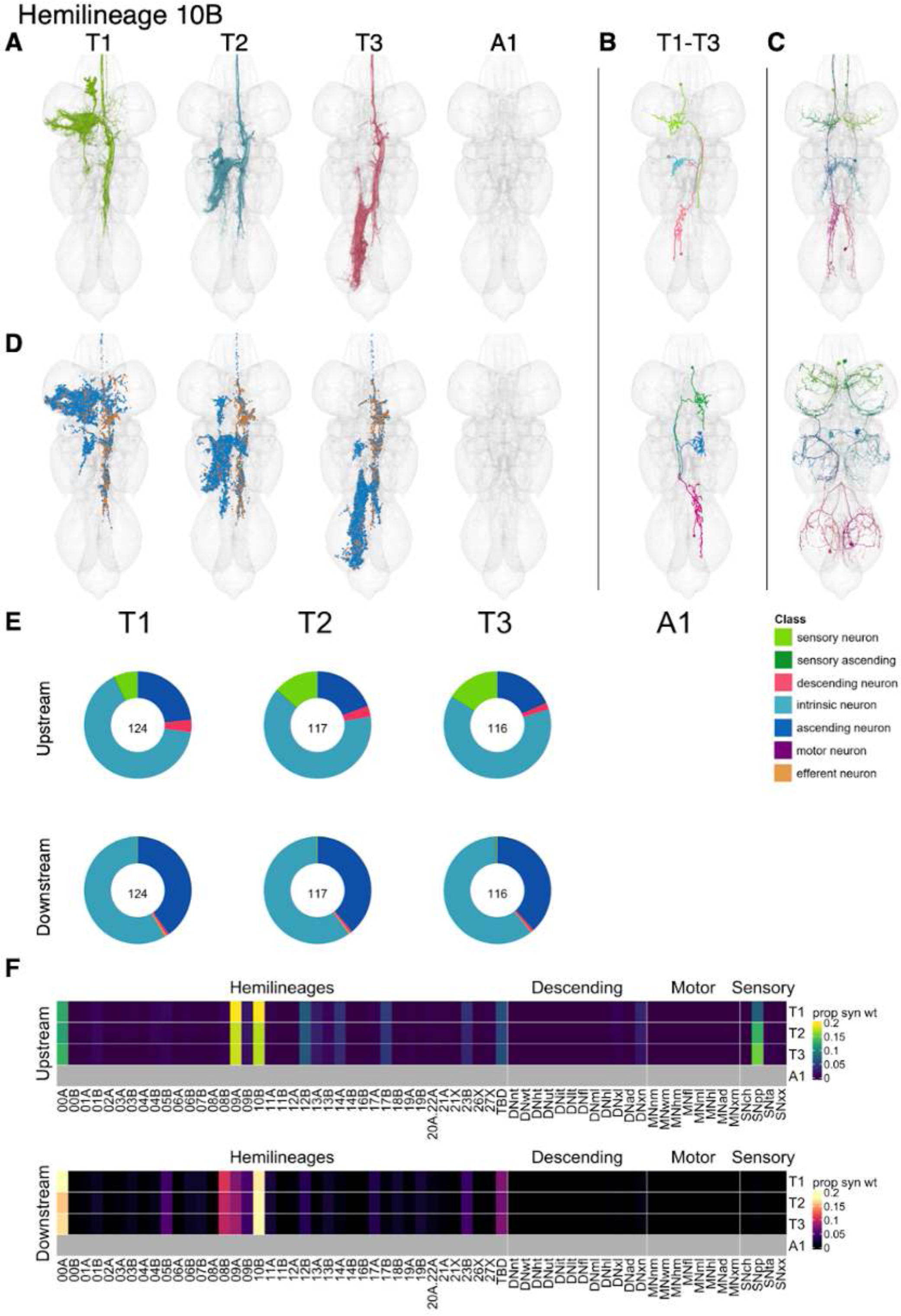
Hemilineage 10B. **A.** Meshes of all RHS secondary neurons plotted in neuromere-specific colours. **B.** “Representative” secondary neuron skeletons plotted in hemineuromere-specific colours. The skeleton with the top accumulated NBLAST score among all neurons from the hemilineage in a given hemineuromere was used. **C.** Neuron meshes of selected examples. Top: ascending serial set 15289. Bottom: complex serial set 11781. Bottom: LHS neurons from contralateral, sequential serial set 17610. **D.** Predicted synapses of RHS secondary neurons. Blue: postsynapses; dark orange: presynapses. **E.** Proportions of connections from secondary neurons to upstream or downstream partners, normalised by neuromere and coloured by broad class. Numbers of query neurons appear in the centre. **F.** Proportions of synaptic weight from secondary neurons originating in each neuromere to upstream or downstream partners, normalised by row.

Secondary 10B neurons receive inputs primarily from proprioceptive sensory neurons and from hemilineages 00A, 09A, and 10B, and they primarily activate 00A, 08B, 09A, and especially other 10B neurons (Figure 30F). These features support a role in leg coordination. Indeed, bilateral activation of 10B secondary neurons results in somewhat erratic leg movements, with occasional pivoting or backwards walking, sometimes accompanied by wing flicking and wing buzzing (Harris et al., 2015). A small number of early born 10B types target wing and abdominal motor neurons (e.g., IN10B006, IN10B011, IN10B023) (Figure 30 - figure supplement 6).

**Figure 30 - figure supplement 1.**
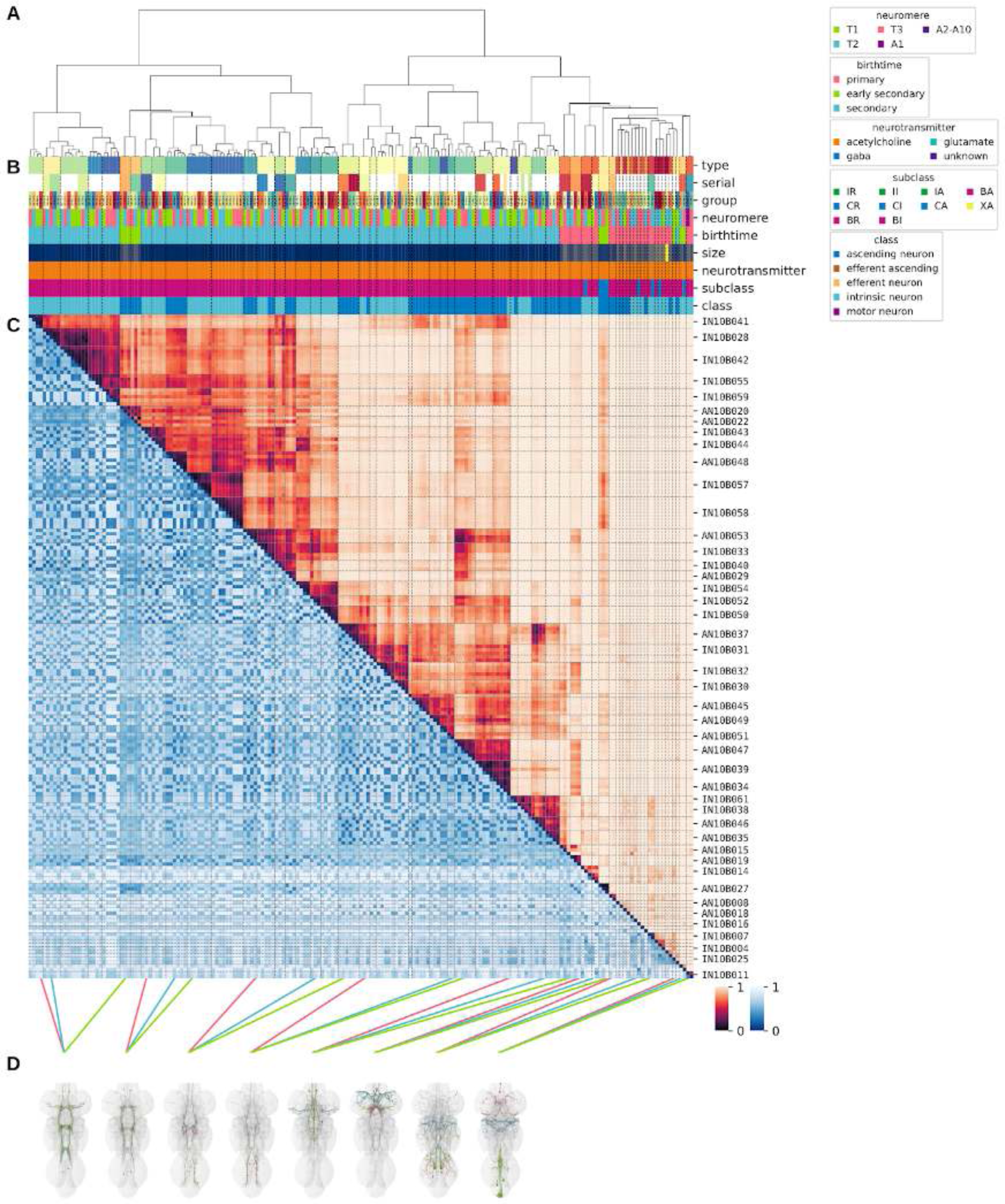
Systematic typing of hemilineage 10B. **A.** Hierarchical clustering dendrogram of hemilineage groups by laterally and serially aggregated connectivity cosine clustering. **B.** Categorical annotations of each hemilineage group, each column corresponding to the aligned leaf in A. Colours for type, serial set, and group are arbitrary for visualisation. Colours for neuromere, birthtime, neurotransmitter, subclass, and class are as in all other figures. **C.** Similarity distance heatmap for hemilineage. Cosine distance is in the upper triangle, while laterally symmetrised NBLAST distance is in the lower triangle. Systematic type names of some types are labelled. **D.** Morphologically representative groups from dendrogram subtrees. Each group, indicated by colour and line connecting to its column in B and C, is the most morphologically representative group (medoid of NBLAST distance) from a subtree of A. The subtrees (flat clusters) are equal height cuts of A determined to yield the number of groups per plot and plots in D.

**Figure 30 - figure supplement 2.**
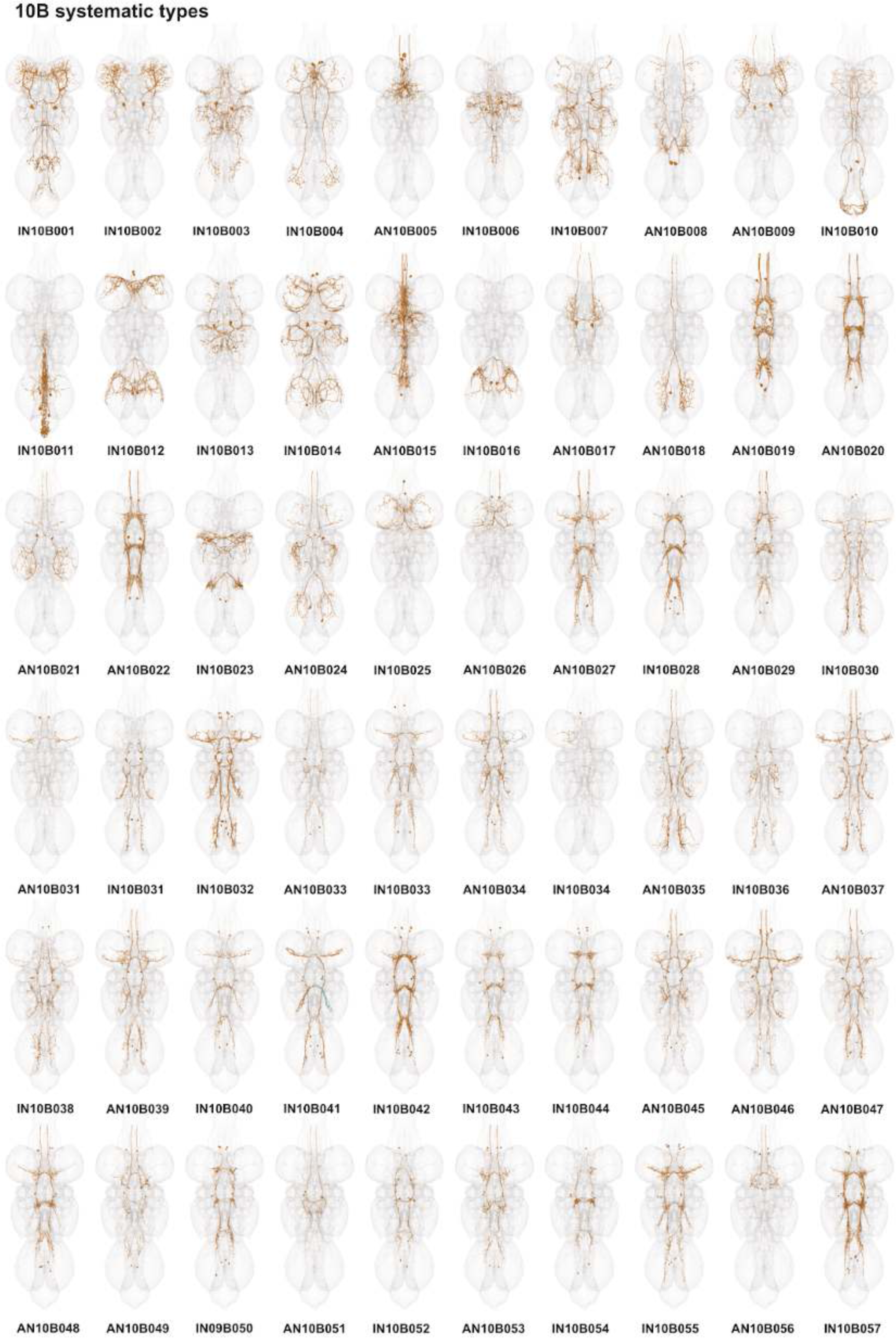
Systematic types of hemilineage 10B. Systematic types have been arranged in numerical order, with neurons of the same type that belong to distinct classes (e.g., intrinsic neuron vs ascending neuron) plotted separately but placed adjacent to each other. Individual neuron meshes have been coloured based on predicted neurotransmitter: dark orange = acetylcholine, blue = gaba, marine = glutamate, dark purple = unknown.

**Figure 30 - figure supplement 3.**
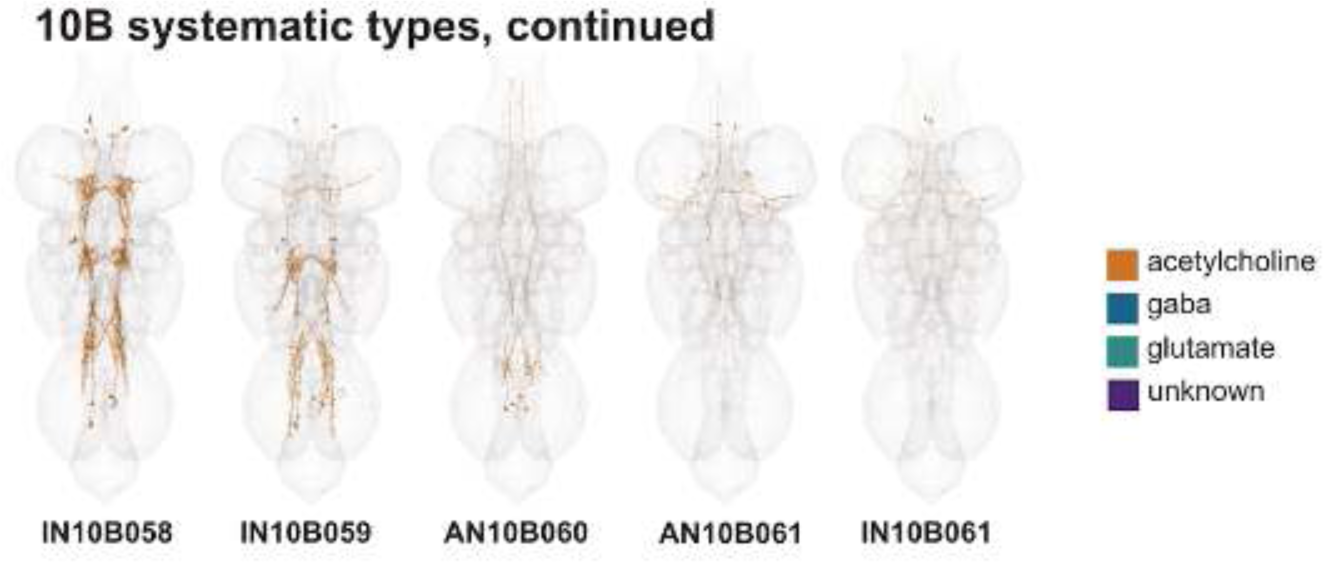
Systematic types of hemilineage 10B, continued. Systematic types have been arranged in numerical order, with neurons of the same type that belong to distinct classes (e.g., intrinsic neuron vs ascending neuron) plotted separately but placed adjacent to each other. Individual neuron meshes have been coloured based on predicted neurotransmitter: dark orange = acetylcholine, blue = gaba, marine = glutamate, dark purple = unknown.

**Figure 30 - figure supplement 4.**
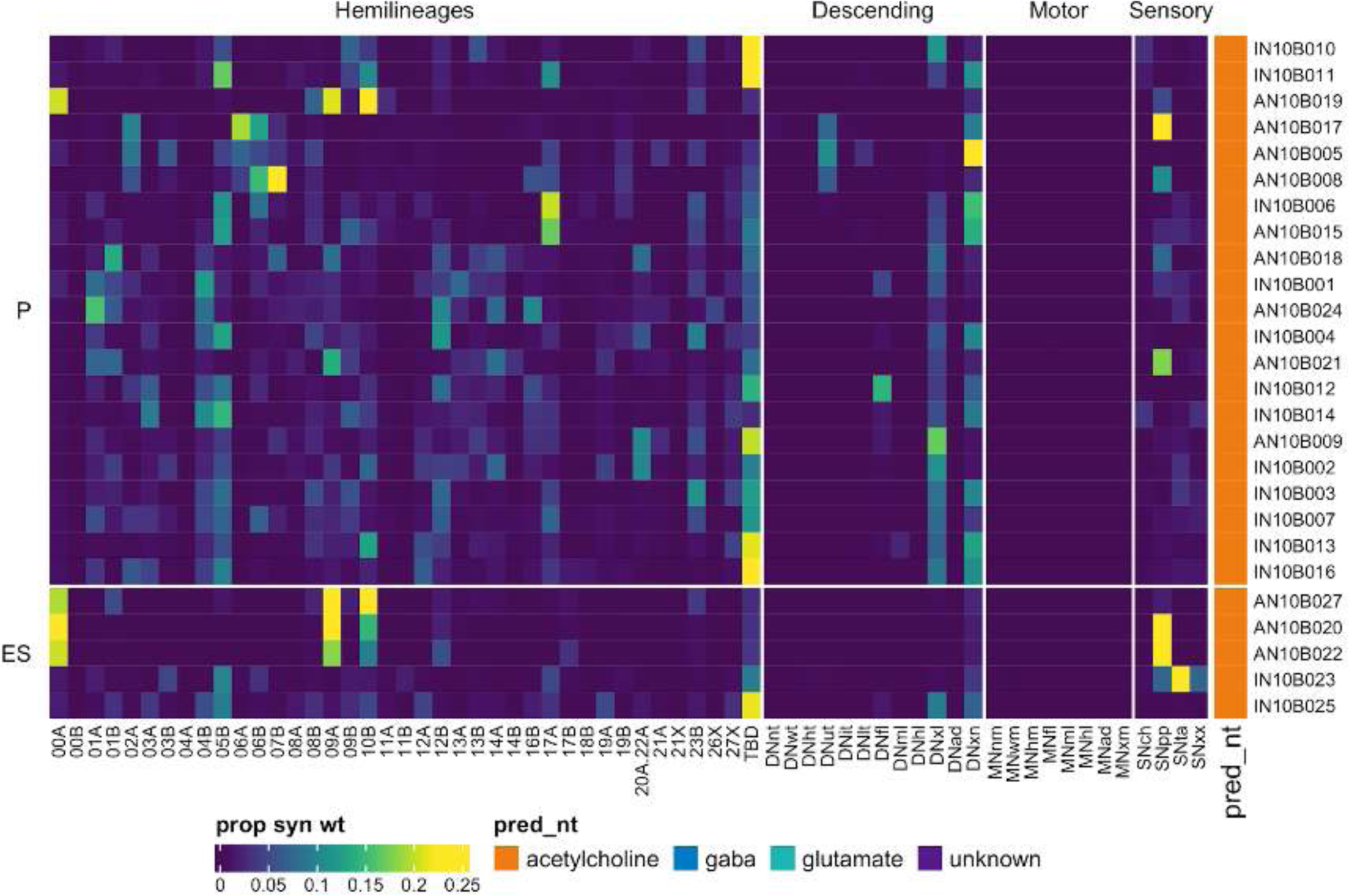
Connectivity to upstream partners by 10B primary and early secondary systematic types. Proportions of synaptic weight to systematic types from upstream partners, normalised by row. 10B neurons have been clustered within each assigned birthtime window (P = primary, ES = early secondary, S = secondary) based on both upstream and downstream connectivity to hemilineages, descending neuron subclasses, motor neuron subclasses, and sensory neuron modalities. Annotation bar is coloured by the most common predicted neurotransmitter for the neurons of each type.

**Figure 30 - figure supplement 5.**
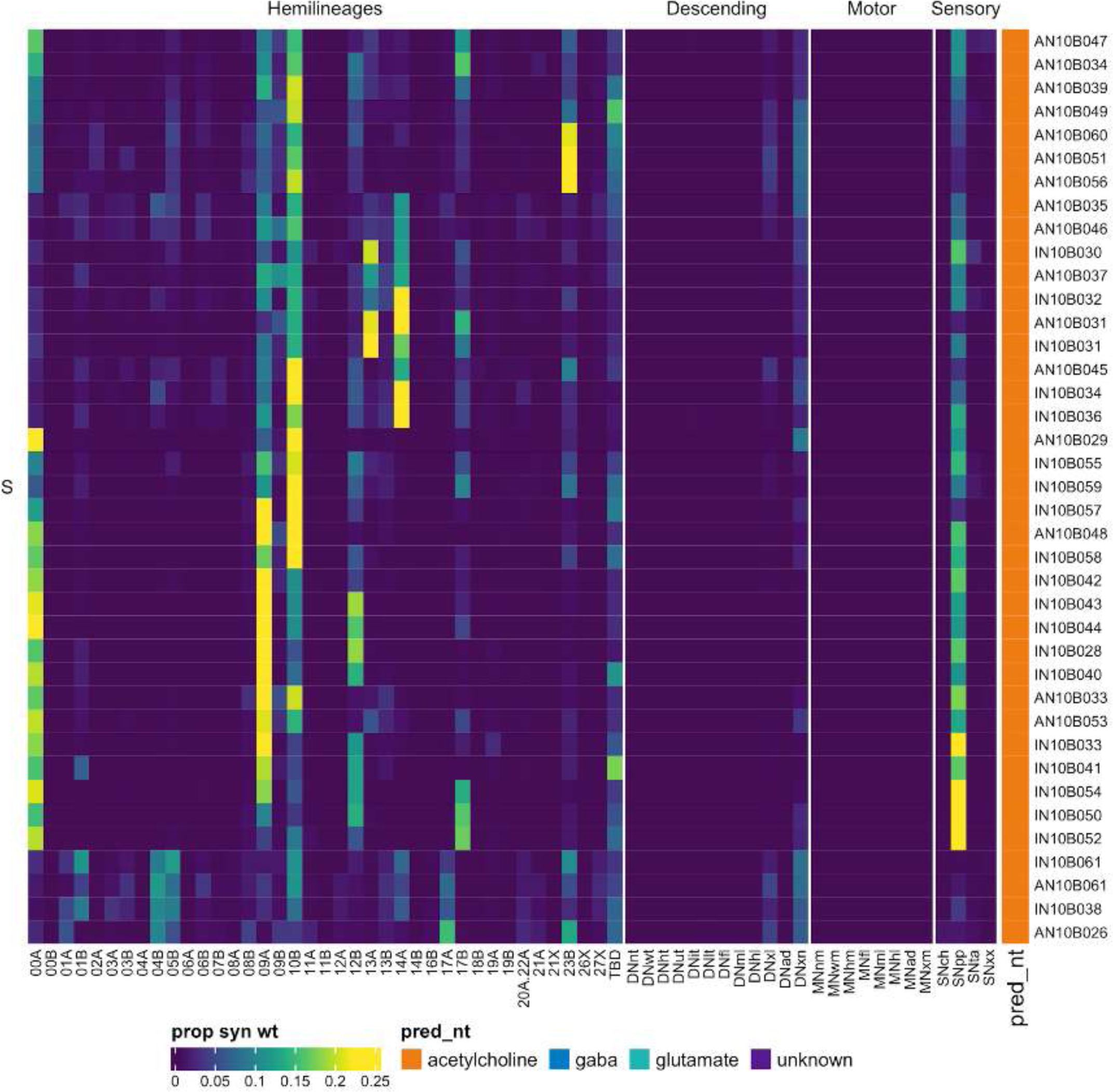
Connectivity to upstream partners by 10B secondary systematic types. Proportions of synaptic weight to systematic types from upstream partners, normalised by row. 10B neurons have been clustered within each assigned birthtime window (P = primary, ES = early secondary, S = secondary) based on both upstream and downstream connectivity to hemilineages, descending neuron subclasses, motor neuron subclasses, and sensory neuron modalities. The annotation bar is coloured by the most common predicted neurotransmitter within each type.

**Figure 30 - figure supplement 6.**
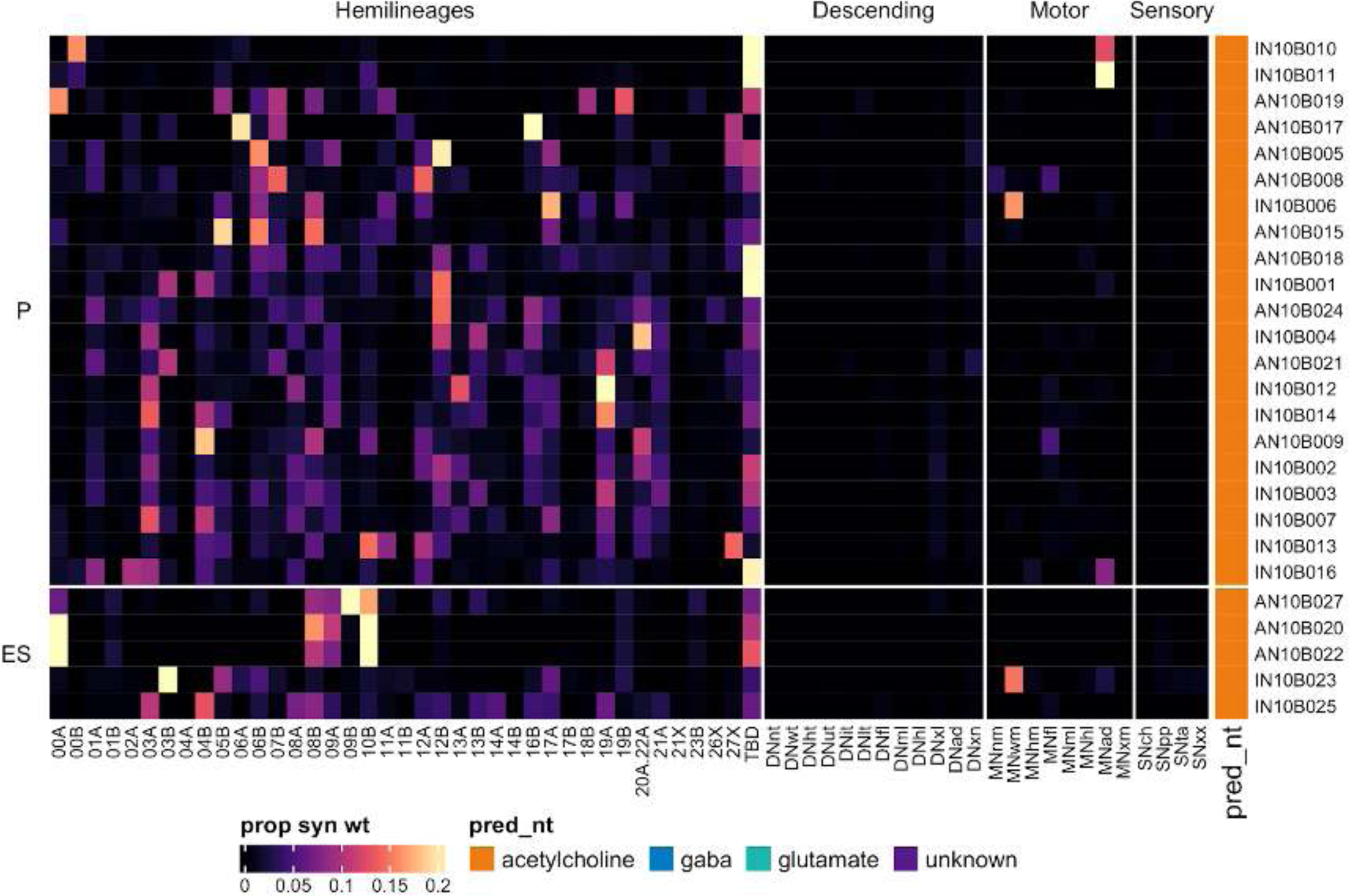
Connectivity to downstream partners by 10B primary and early secondary systematic types. Proportions of synaptic weight from systematic types to downstream partners, normalised by row. 10B neurons have been clustered within each assigned birthtime window (P = primary, ES = early secondary, S = secondary) based on both upstream and downstream connectivity to hemilineages, descending neuron subclasses, motor neuron subclasses, and sensory neuron modalities. The annotation bar is coloured by the most common predicted neurotransmitter within each type.

**Figure 30 - figure supplement 7.**
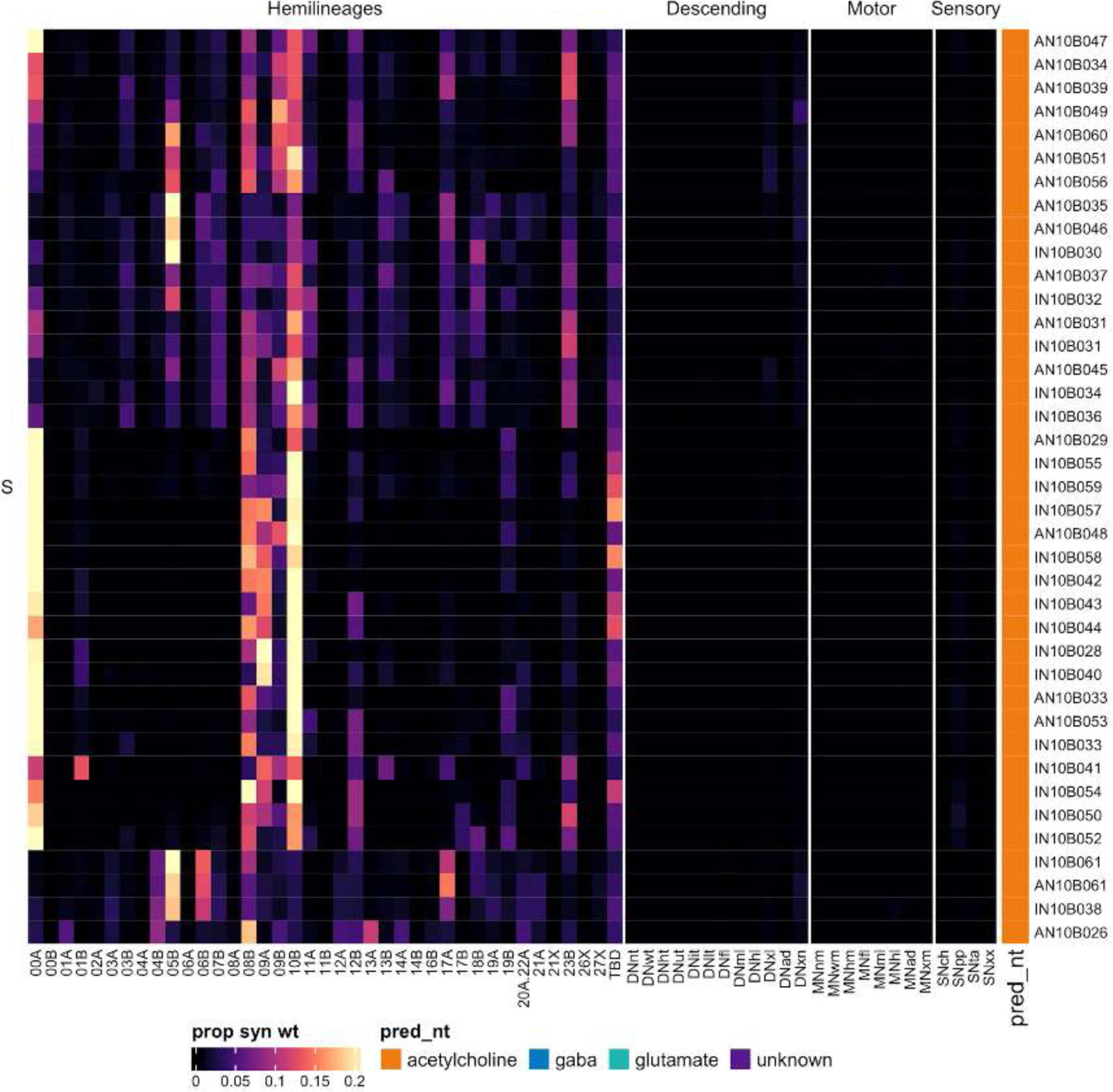
Connectivity to downstream partners by 10B secondary systematic types. Proportions of synaptic weight from systematic types to downstream partners, normalised by row. 10B neurons have been clustered within each assigned birthtime window (P = primary, ES = early secondary, S = secondary) based on both upstream and downstream connectivity to hemilineages, descending neuron subclasses, motor neuron subclasses, and sensory neuron modalities. The annotation bar is coloured by the most common predicted neurotransmitter for the neurons of each type.

#### Hemilineage 11A

Hemilineages 11A and 11B are believed to derive from posterior dorsal neuroblast 7-2 (Lacin and Truman, 2016) (but see also (Birkholz et al., 2015)), which produces ∼10 intersegmental interneurons and 5 local interneurons in thoracic segments of the embryo (Schmid et al., 1999). We note here that the morphological descriptions of 11A vs 11B have been reversed in the adult lineages atlas (Shepherd et al., 2019) when compared to neuroblast clones in the late larva (Truman et al., 2010). Hemilineage 11A secondary neurons only survive in T1 and T2. They enter the posterior neuromere with 19B and 23B, projecting ventrally and medially, curving around the tectulum to innervate the dorsal leg neuropil, ventral tectulum, and mechanosensory neuropil of the ovoid (Shepherd et al., 2019).

Secondary 11A neurons receive significant input from chemosensory neurons in T1 and T2 and from proprioceptive and tactile sensory neurons in T2; they also receive it from numerous hemilineages including 00A, 05B, 06B, 08B, 11A, and 23B and from descending neurons targeting the lower tectulum or multiple legs or neuropils (Figure 31F). Most of the types receiving a large proportion of sensory information appear to integrate multiple modalities; IN11A022 in particular is a strong target of wing chemosensory and tactile neurons (Figure 31 - figure supplement 2-3).

**Figure 31.**
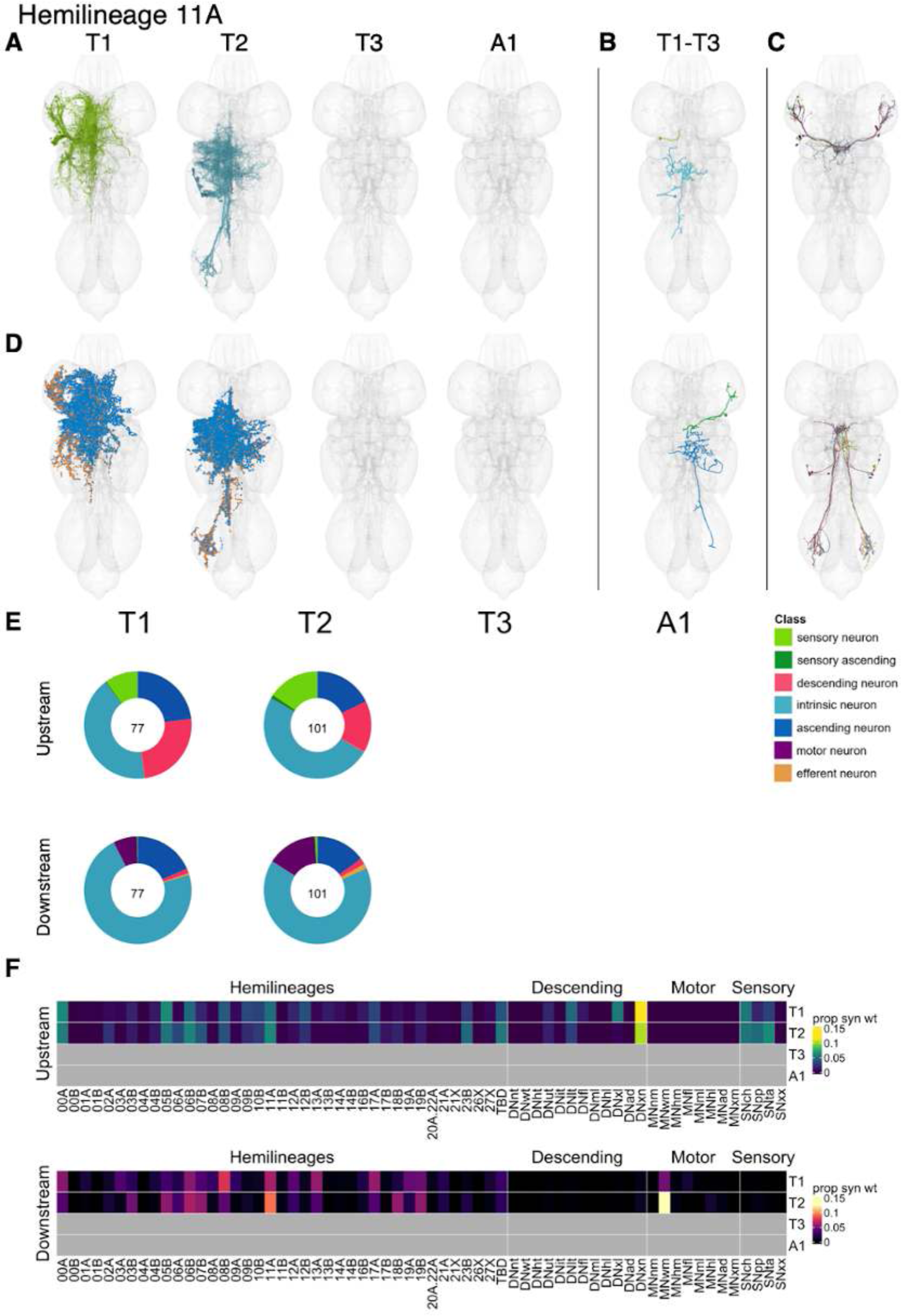
Hemilineage 11A. **A.** Meshes of all RHS secondary neurons plotted in neuromere-specific colours. **B.** “Representative” secondary neuron skeletons plotted in hemineuromere-specific colours. The skeleton with the top accumulated NBLAST score among all neurons from the hemilineage in a given hemineuromere was used. **C.** Neuron meshes of selected examples. Top: electrical GFC4, subcluster 23460. Bottom: electrical GFC3, subcluster 17383. **D.** Predicted synapses of RHS secondary neurons. Blue: postsynapses; dark orange: presynapses. **E.** Proportions of connections from secondary neurons to upstream or downstream partners, normalised by neuromere and coloured by broad class. Numbers of query neurons appear in the centre. **F.** Proportions of synaptic weight from secondary neurons originating in each neuromere to upstream or downstream partners, normalised by row.

11A neurons are predicted to be cholinergic as expected (Lacin et al., 2019) with a few exceptions and primarily activate 00A, 08B, 13A, and 17A in T1 and 03B, 05B, 06B, 07B, 18B, 19B, and especially 11A in T2 as well as wing motor neurons (Figure 31F). We matched a few specific 11A types (Figure 31C top, bottom) to neurons electrically coupled to the Giant Fiber (Kennedy and Broadie, 2018); these target front and/or hind leg motor neurons (Figure 31 - figure supplement 2,4) and are likely involved in takeoff behaviour. These results are consistent with experimental bilateral activation of 11A and 11B secondary neurons, which evokes takeoff behaviour with wing flapping prior to the jump (Harris et al., 2015).

**Figure 31 - figure supplement 1.**
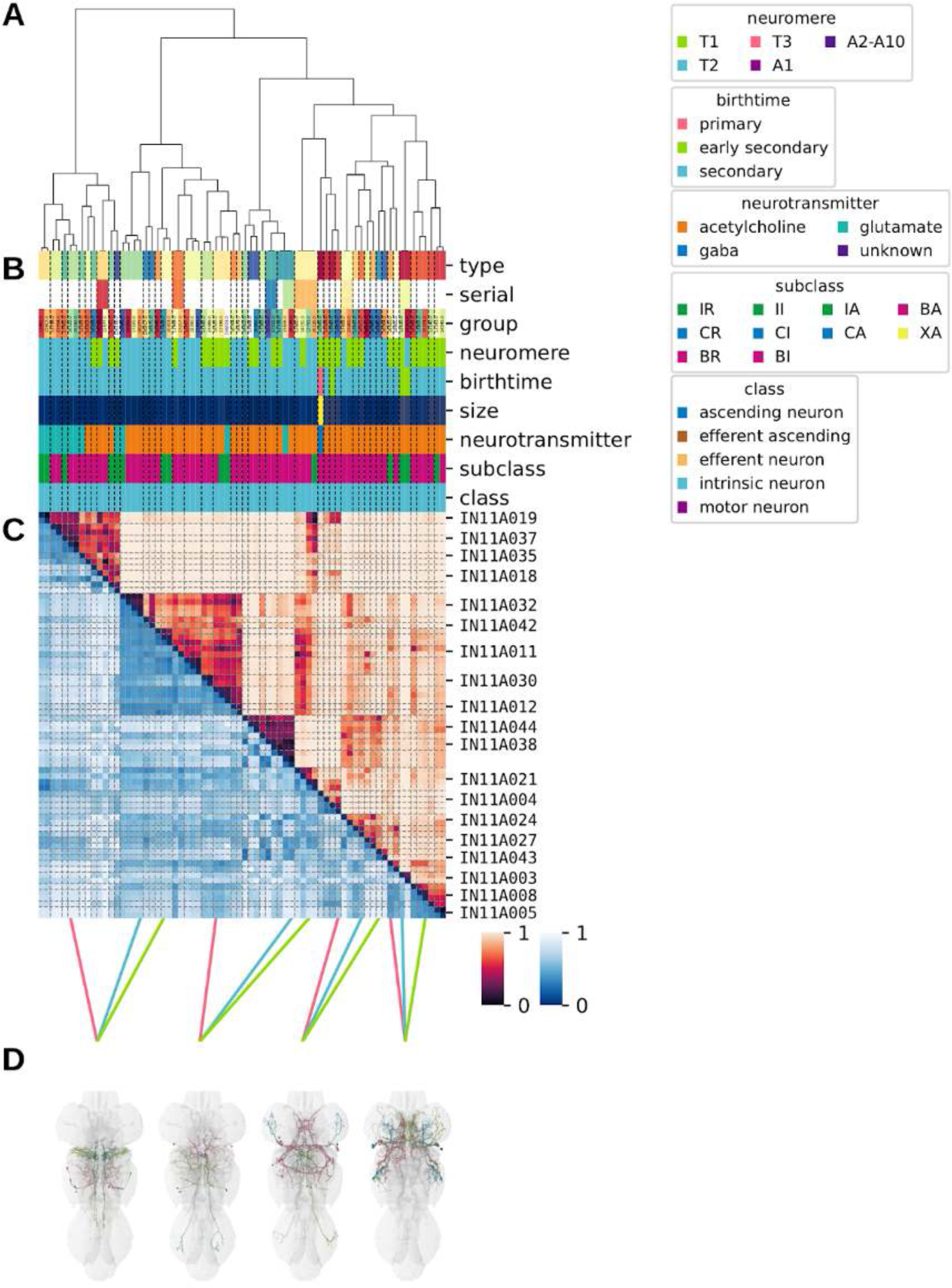
Systematic typing of hemilineage 11A. **A.** Hierarchical clustering dendrogram of hemilineage groups by laterally and serially aggregated connectivity cosine clustering. **B.** Categorical annotations of each hemilineage group, each column corresponding to the aligned leaf in A. Colours for type, serial set, and group are arbitrary for visualisation. Colours for neuromere, birthtime, neurotransmitter, subclass, and class are as in all other figures. **C.** Similarity distance heatmap for hemilineage. Cosine distance is in the upper triangle, while laterally symmetrised NBLAST distance is in the lower triangle. Systematic type names of some types are labelled. **D.** Morphologically representative groups from dendrogram subtrees. Each group, indicated by colour and line connecting to its column in B and C, is the most morphologically representative group (medoid of NBLAST distance) from a subtree of A. The subtrees (flat clusters) are equal height cuts of A determined to yield the number of groups per plot and plots in D.

**Figure 31 - figure supplement 2.**
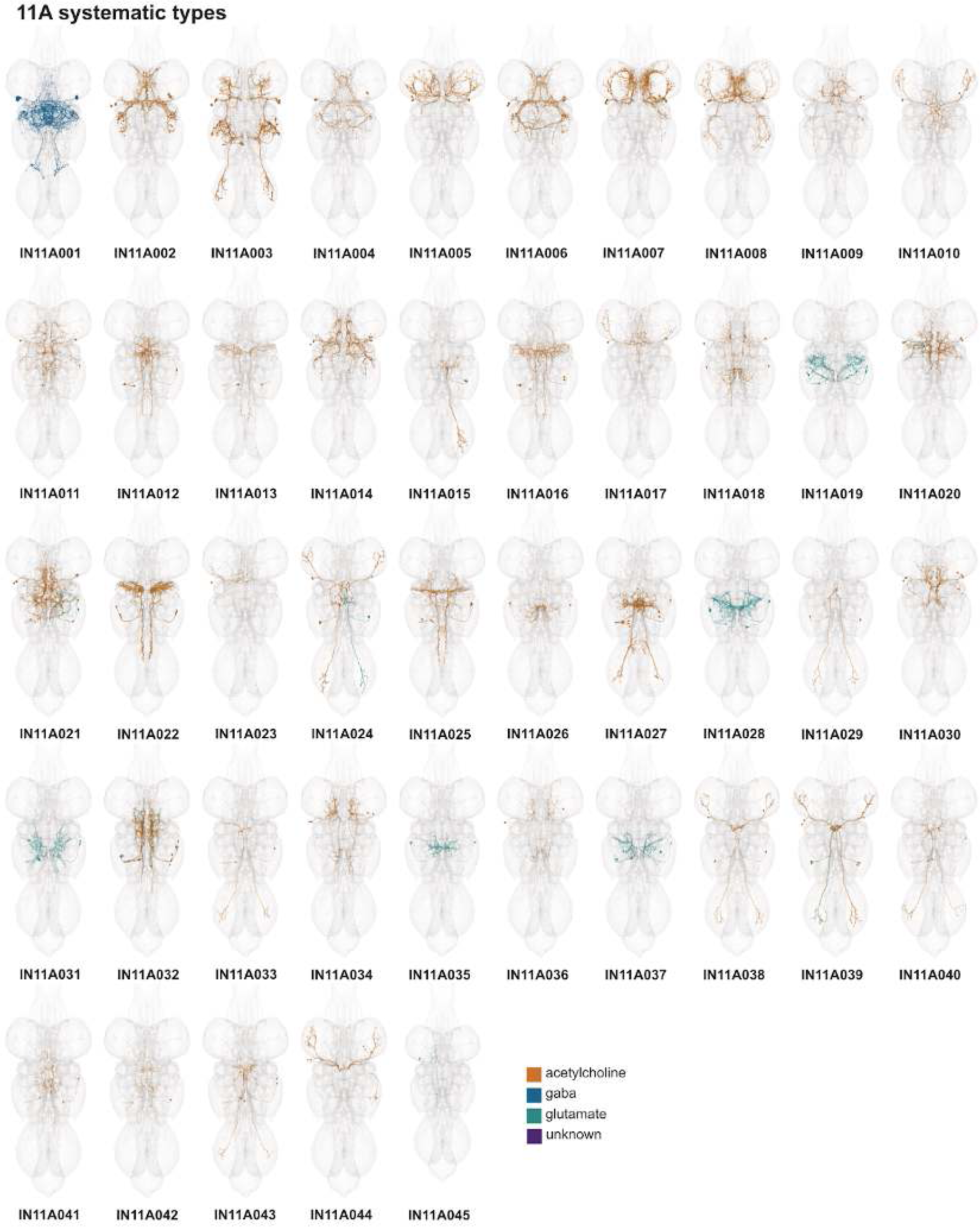
Systematic types of hemilineage 11A. Systematic types have been arranged in numerical order, with neurons of the same type that belong to distinct classes (e.g., intrinsic neuron vs ascending neuron) plotted separately but placed adjacent to each other. Individual neuron meshes have been coloured based on predicted neurotransmitter: dark orange = acetylcholine, blue = gaba, marine = glutamate, dark purple = unknown.

**Figure 31 - figure supplement 3.**
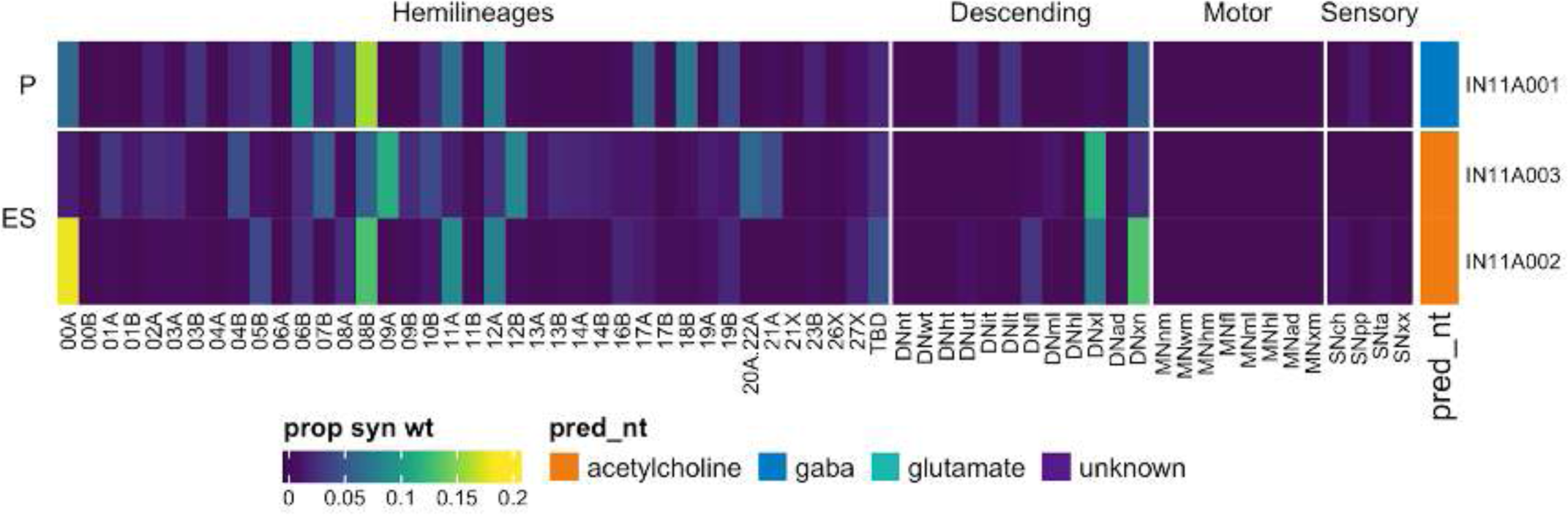
Connectivity to upstream partners by 11A primary and early secondary systematic types. Proportions of synaptic weight to systematic types from upstream partners, normalised by row. 11A neurons have been clustered within each assigned birthtime window (P = primary, ES = early secondary, S = secondary) based on both upstream and downstream connectivity to hemilineages, descending neuron subclasses, motor neuron subclasses, and sensory neuron modalities. Annotation bar is coloured by the most common predicted neurotransmitter for the neurons of each type.

**Figure 31 - figure supplement 4.**
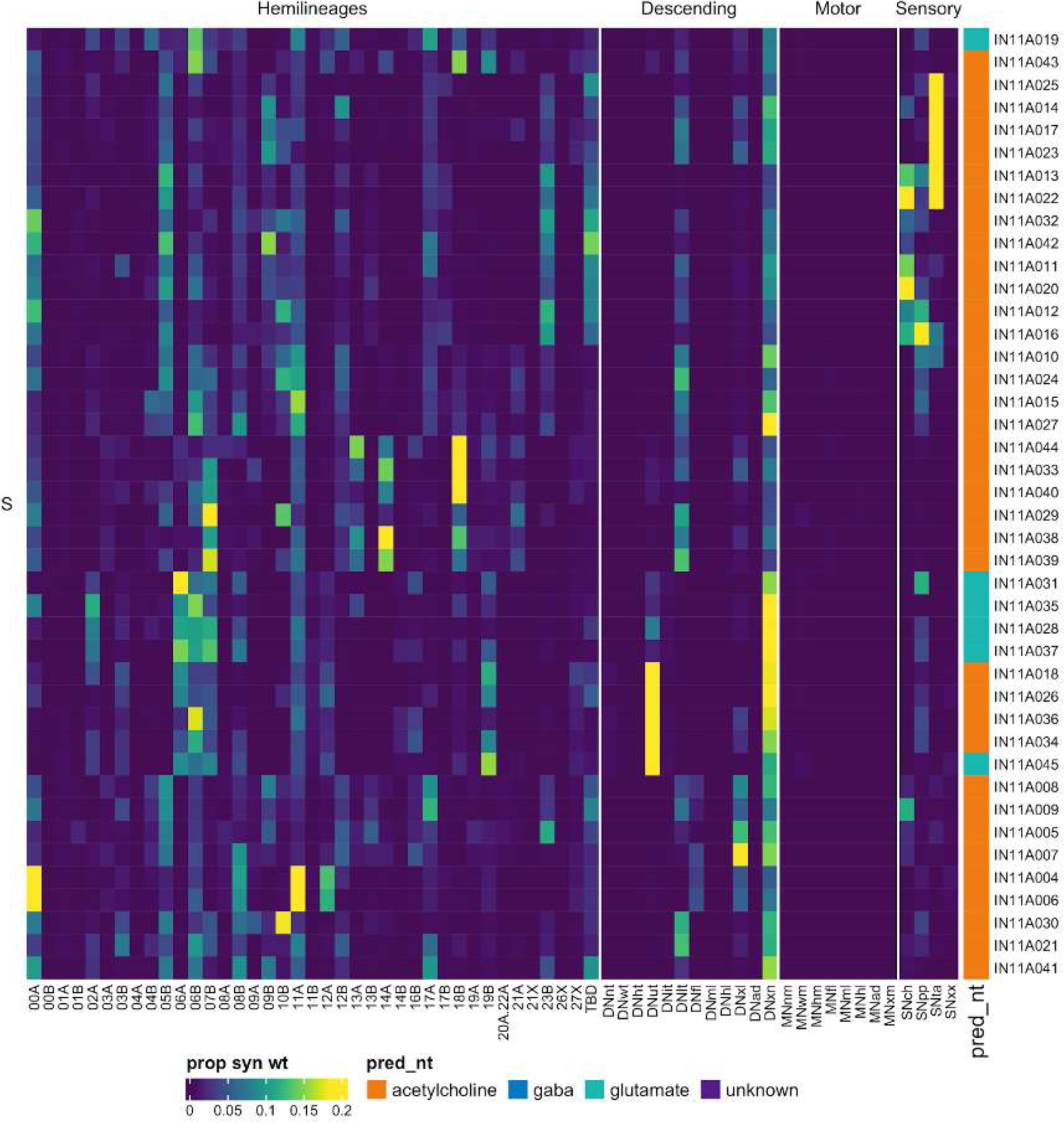
Connectivity to upstream partners by 11A secondary systematic types. Proportions of synaptic weight to systematic types from upstream partners, normalised by row. 11A neurons have been clustered within each assigned birthtime window (P = primary, ES = early secondary, S = secondary) based on both upstream and downstream connectivity to hemilineages, descending neuron subclasses, motor neuron subclasses, and sensory neuron modalities. Annotation bar is coloured by the most common predicted neurotransmitter for the neurons of each type.

**Figure 31 - figure supplement 5.**
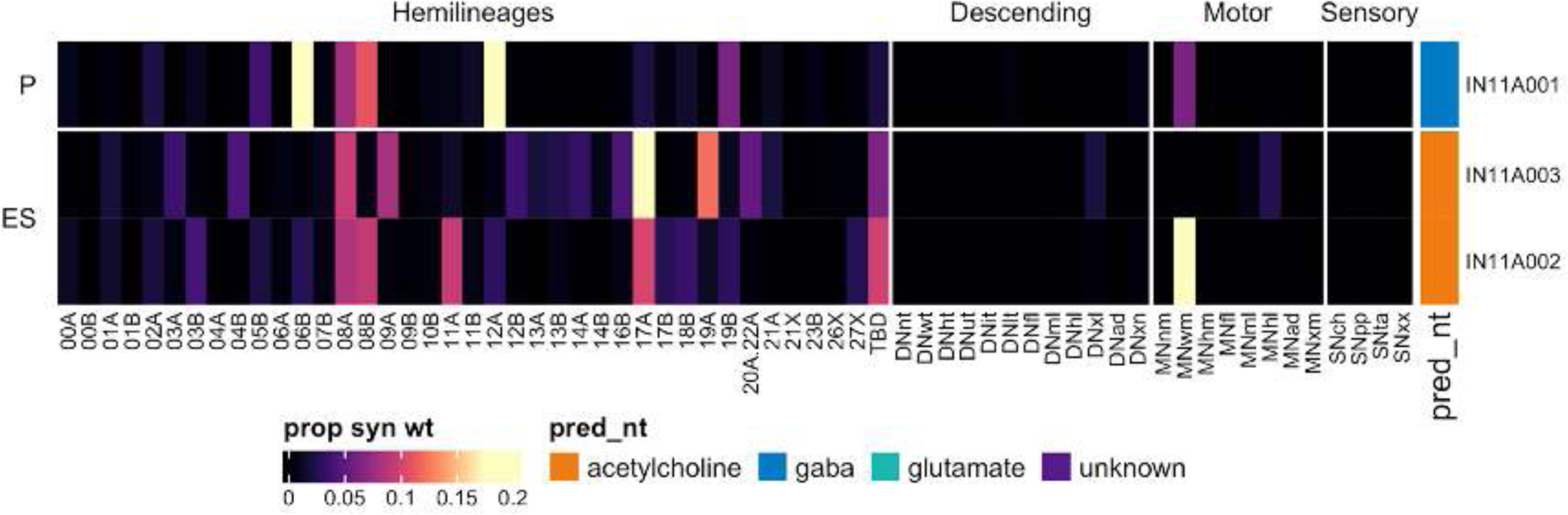
Connectivity to downstream partners by 11A primary and early secondary systematic types. Proportions of synaptic weight from systematic types to downstream partners, normalised by row. 11A neurons have been clustered within each assigned birthtime window (P = primary, ES = early secondary, S = secondary) based on both upstream and downstream connectivity to hemilineages, descending neuron subclasses, motor neuron subclasses, and sensory neuron modalities. The annotation bar is coloured by the most common predicted neurotransmitter within each type.

**Figure 31 - figure supplement 6.**
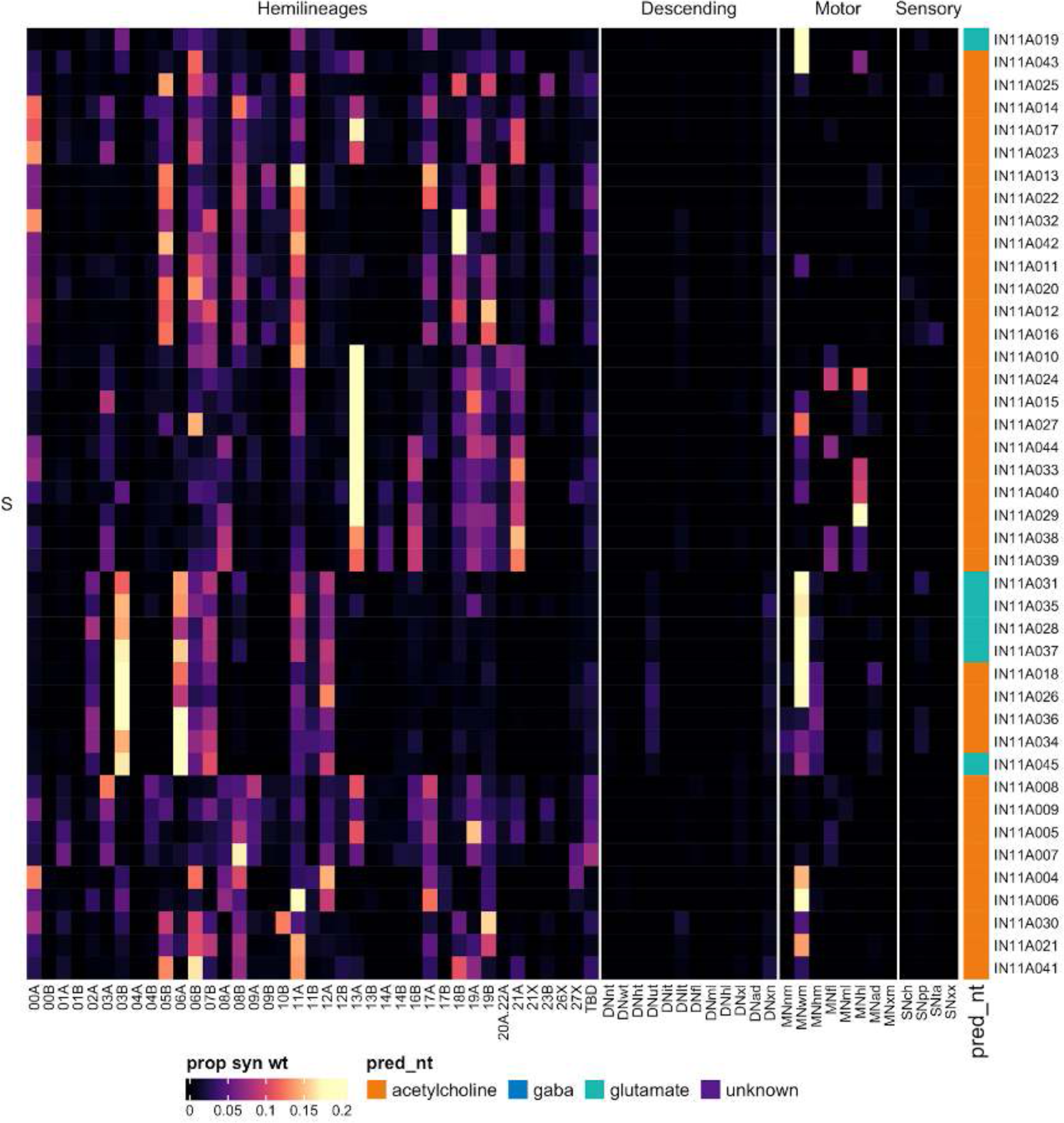
Connectivity to downstream partners by 11A secondary systematic types. Proportions of synaptic weight from systematic types to downstream partners, normalised by row. 11A neurons have been clustered within each assigned birthtime window (P = primary, ES = early secondary, S = secondary) based on both upstream and downstream connectivity to hemilineages, descending neuron subclasses, motor neuron subclasses, and sensory neuron modalities. The annotation bar is coloured by the most common predicted neurotransmitter within each type.

#### Hemilineage 11B

Hemilineage 11B is the most extreme of the segment-specific dorsal hemilineages, with a large population of secondary neurons surviving only in T2 (Marin et al., 2012); we identified seven 11B neurons in T1 and two in T3, consistent with immortalisation experiments (H Lacin, personal communication). 11B secondary neurons feature a sharp bend in their primary neurites as they turn to innervate the dorsal tectulum (Shepherd et al., 2019). They are mainly gabaergic (Lacin et al., 2019), but we identified a subset of predicted glutamatergic neurons with unusually thick axons and low presynaptic density (e.g., Figure 32C top) which we suggest are electrically coupled to flight neuropil partners via gap junctions (for details please see (Cheong et al., 2023)). IN11B004 is one example from the 54 systematic types containing neurons that have been assigned to different birthtimes (Figure 32C bottom).

**Figure 32.**
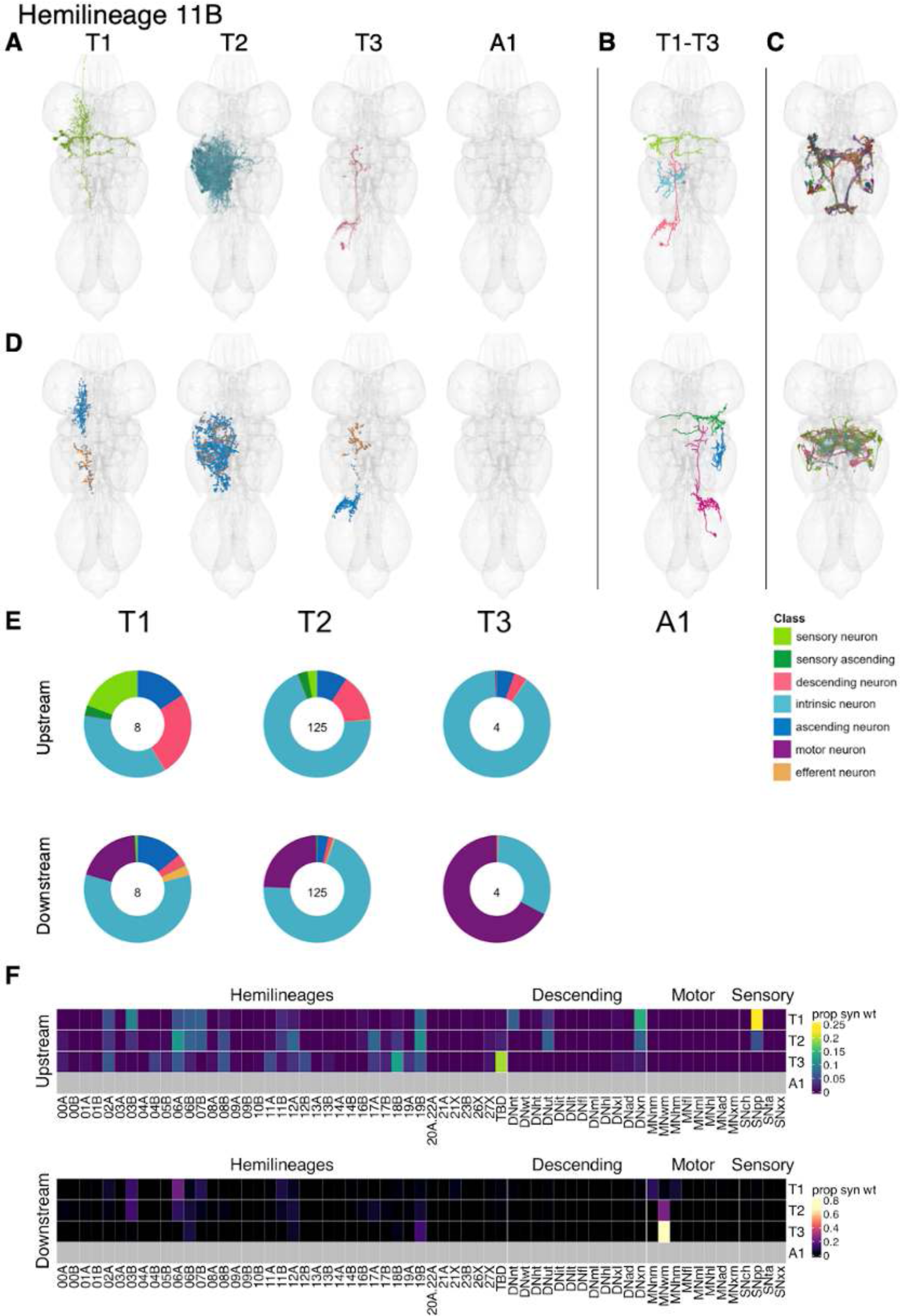
Hemilineage 11B. **A.** Meshes of all RHS secondary neurons plotted in neuromere-specific colours. **B.** “Representative” secondary neuron skeletons plotted in hemineuromere-specific colours. The skeleton with the top accumulated NBLAST score among all neurons from the hemilineage in a given hemineuromere was used. **C.** Neuron meshes of selected examples. Top: putative electrical subcluster 10326. Bottom: systematic type IN11B004 containing primary (pink), early secondary (green), and secondary (cyan) neurons. **D.** Predicted synapses of RHS secondary neurons. Blue: postsynapses; dark orange: presynapses. **E.** Proportions of connections from secondary neurons to upstream or downstream partners, normalised by neuromere and coloured by broad class. Numbers of query neurons appear in the centre. **F.** Proportions of synaptic weight from secondary neurons originating in each neuromere to upstream or downstream partners, normalised by row.

Secondary 11B neurons in T1 and T2 receive most inputs from hemilineages 06B, 07B, 08B, 17A, 19B, and especially 06A as well as from descending neurons targeting the upper tectulum or multiple neuropils and from proprioceptive sensory neurons (Figure 32F); those in T3 have a distinct connectivity profile and in particular receive most input from hemilineage 18B. Primary types IN11B009, AN11B008, and AN11B012 receive an especially large proportion of proprioceptive input (Figure 32 - figure supplement 3). Secondary neurons are predicted to be gabaergic and inhibit wing motor neurons as well as hemilineages 03B, 06B, 12A, 17A, 19B, and especially 06A (Figure 32F). Bilateral activation of 11A and 11B secondary neurons evokes takeoff behaviour but with wing flapping prior to the jump (Harris et al., 2015). There are also two primary types (AN11B008 and AN11B012) that inhibit neck motor neurons (Figure 32 - figure supplement 5).

**Figure 32 - figure supplement 1.**
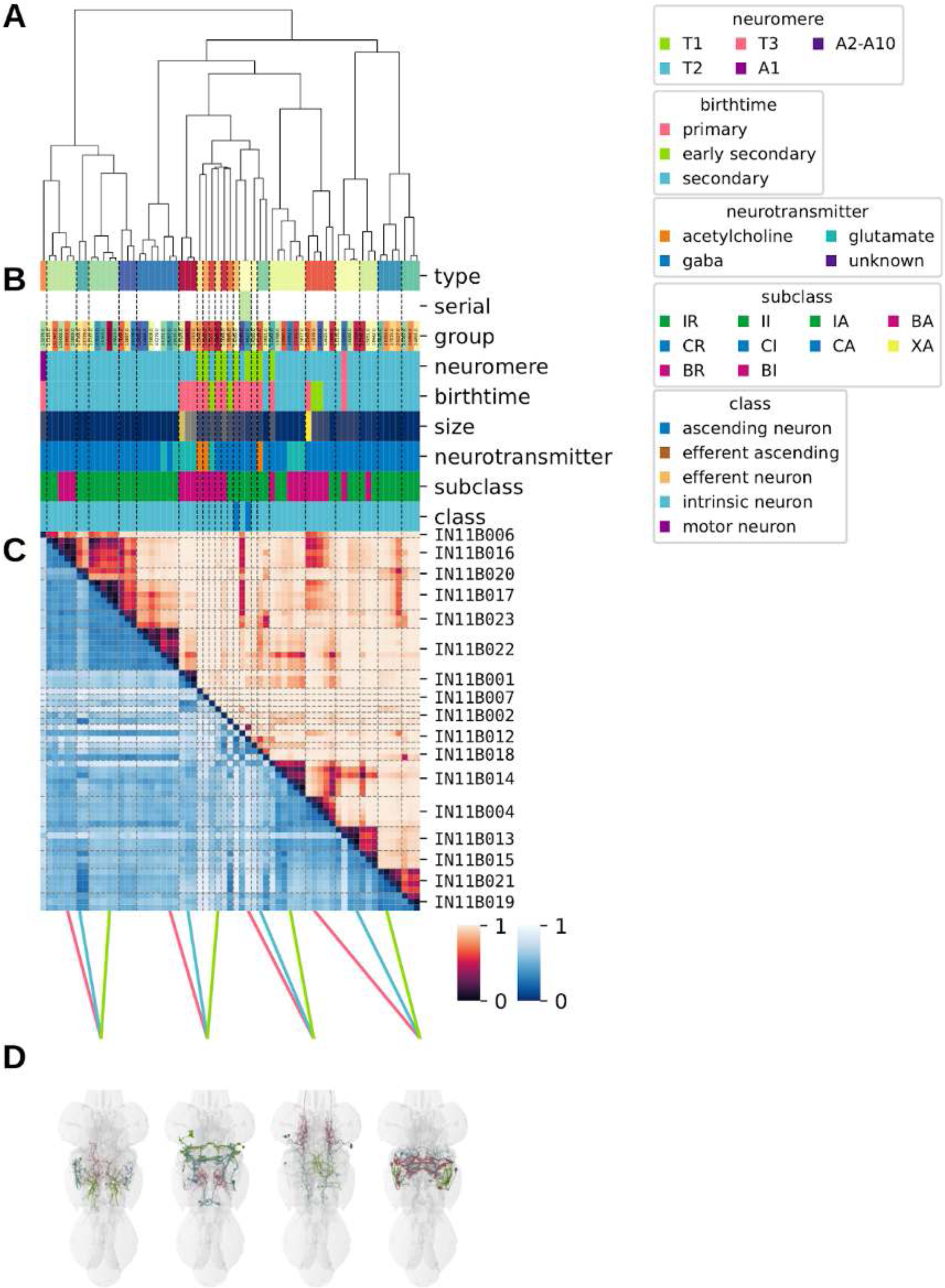
Systematic typing of hemilineage 11B. **A.** Hierarchical clustering dendrogram of hemilineage groups by laterally and serially aggregated connectivity cosine clustering. **B.** Categorical annotations of each hemilineage group, each column corresponding to the aligned leaf in A. Colours for type, serial set, and group are arbitrary for visualisation. Colours for neuromere, birthtime, neurotransmitter, subclass, and class are as in all other figures. **C.** Similarity distance heatmap for hemilineage. Cosine distance is in the upper triangle, while laterally symmetrised NBLAST distance is in the lower triangle. Systematic type names of some types are labelled. **D.** Morphologically representative groups from dendrogram subtrees. Each group, indicated by colour and line connecting to its column in B and C, is the most morphologically representative group (medoid of NBLAST distance) from a subtree of A. The subtrees (flat clusters) are equal height cuts of A determined to yield the number of groups per plot and plots in D.

**Figure 32 - figure supplement 2.**
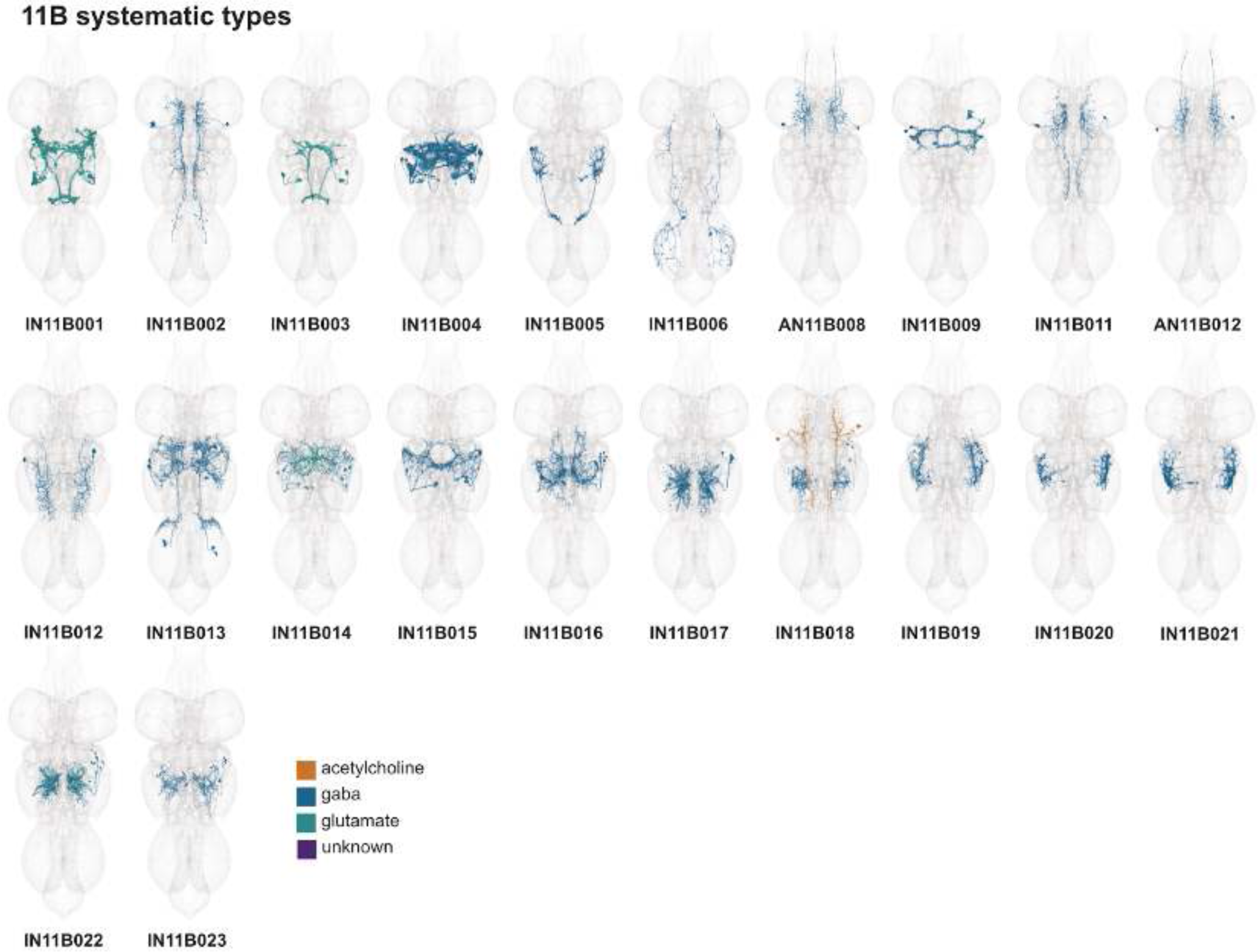
Systematic types of hemilineage 11B. Systematic types have been arranged in numerical order, with neurons of the same type that belong to distinct classes (e.g., intrinsic neuron vs ascending neuron) plotted separately but placed adjacent to each other. Individual neuron meshes have been coloured based on predicted neurotransmitter: dark orange = acetylcholine, blue = gaba, marine = glutamate, dark purple = unknown.

**Figure 32 - figure supplement 3.**
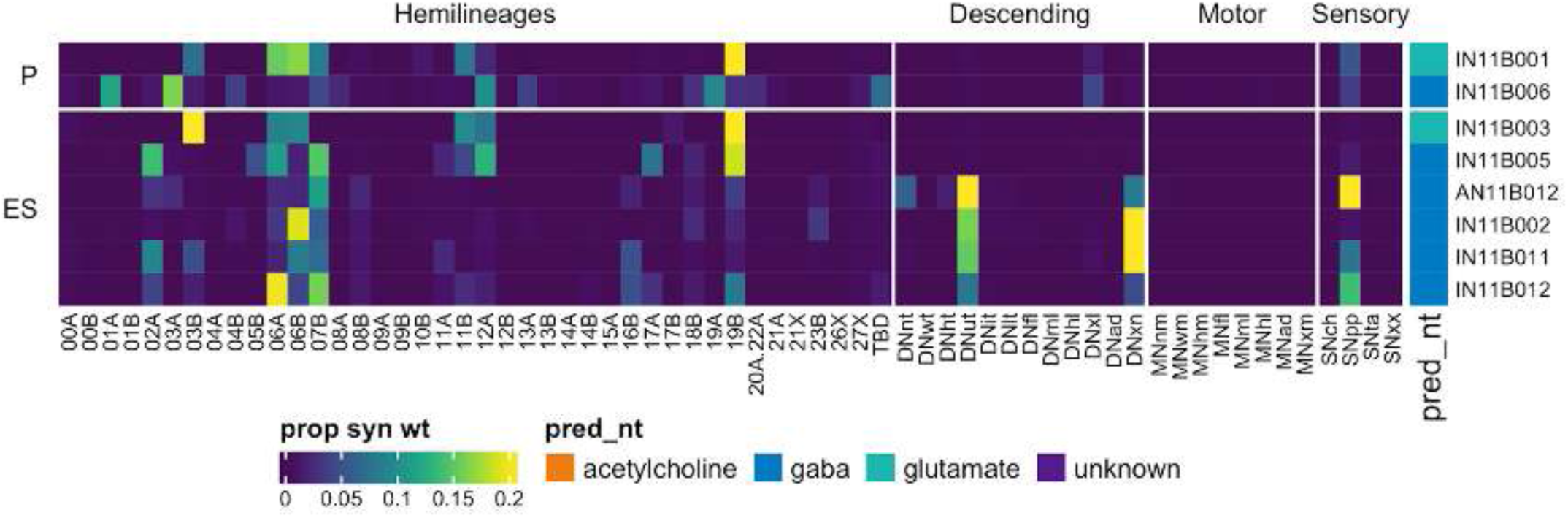
Connectivity to upstream partners by 11B primary and early secondary systematic types. Proportions of synaptic weight to systematic types from upstream partners, normalised by row. 11B neurons have been clustered within each assigned birthtime window (P = primary, ES = early secondary, S = secondary) based on both upstream and downstream connectivity to hemilineages, descending neuron subclasses, motor neuron subclasses, and sensory neuron modalities. Annotation bar is coloured by the most common predicted neurotransmitter for the neurons of each type.

**Figure 32 - figure supplement 4.**
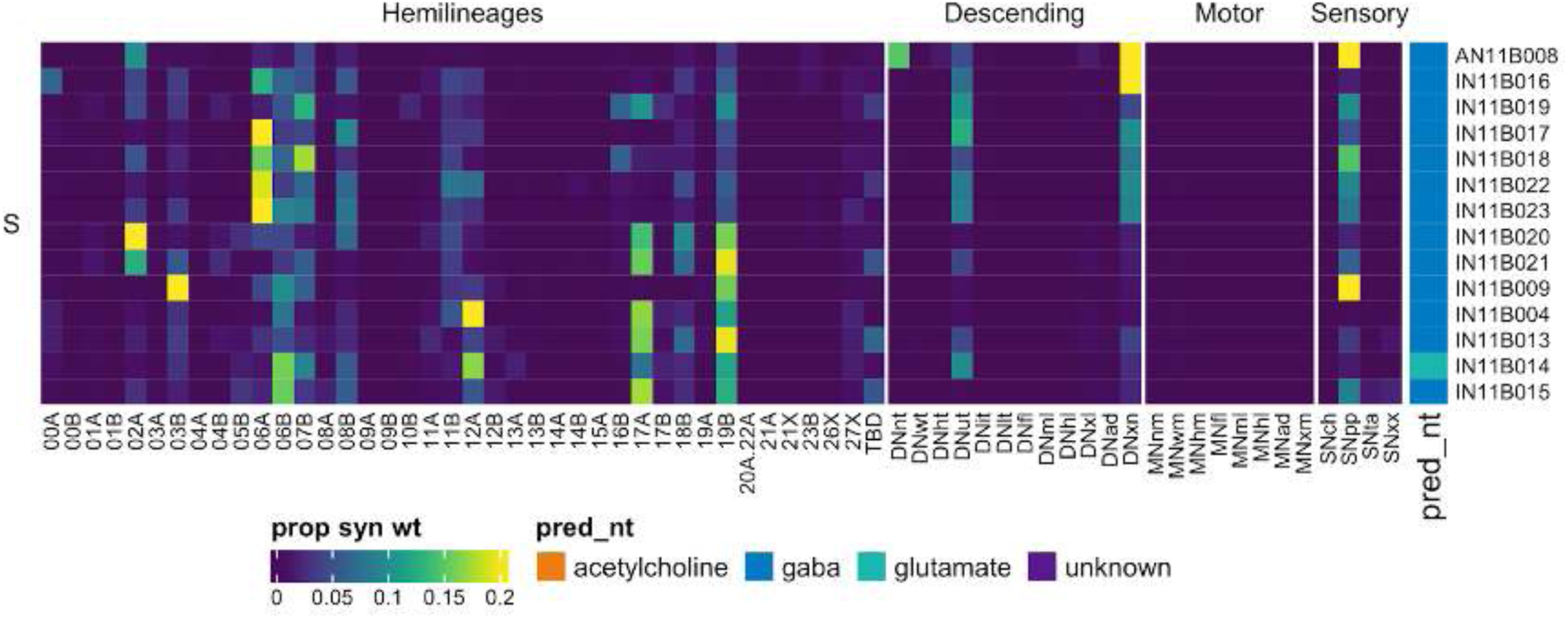
Connectivity to upstream partners by 11B secondary systematic types. Proportions of synaptic weight to systematic types from upstream partners, normalised by row. 11B neurons have been clustered within each assigned birthtime window (P = primary, ES = early secondary, S = secondary) based on both upstream and downstream connectivity to hemilineages, descending neuron subclasses, motor neuron subclasses, and sensory neuron modalities. Annotation bar is coloured by the most common predicted neurotransmitter for the neurons of each type.

**Figure 32 - figure supplement 5.**
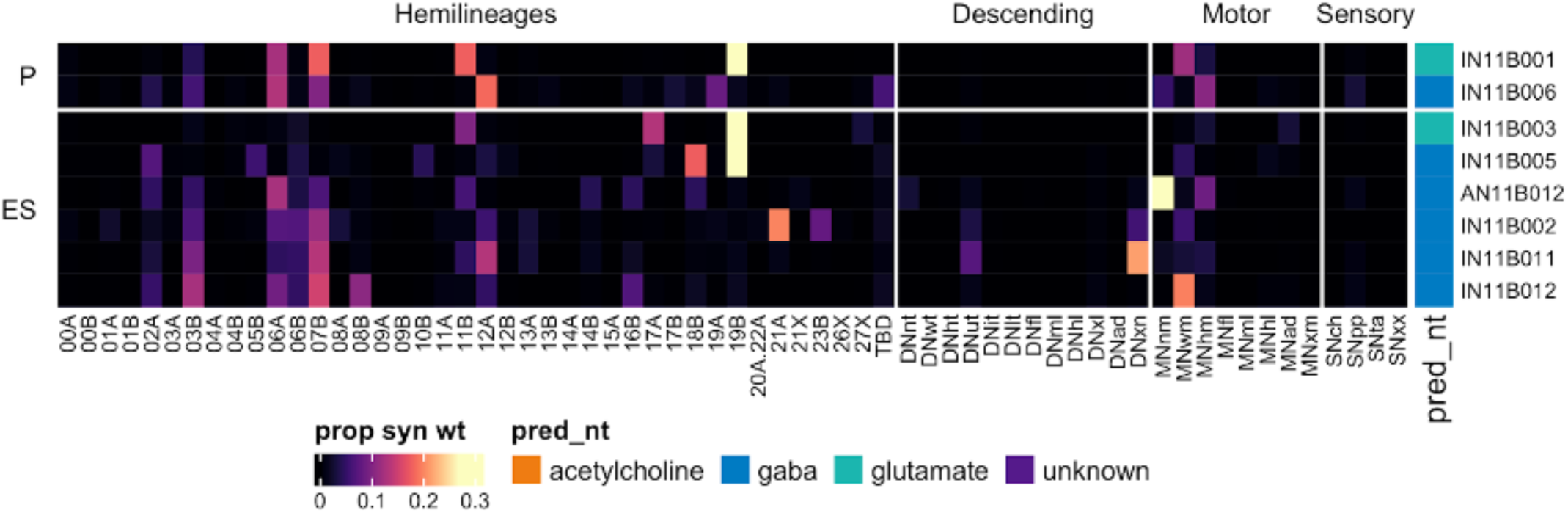
Connectivity to downstream partners by 11B primary and early secondary systematic types. Proportions of synaptic weight from systematic types to downstream partners, normalised by row. 11B neurons have been clustered within each assigned birthtime window (P = primary, ES = early secondary, S = secondary) based on both upstream and downstream connectivity to hemilineages, descending neuron subclasses, motor neuron subclasses, and sensory neuron modalities. The annotation bar is coloured by the most common predicted neurotransmitter within each type.

**Figure 32 - figure supplement 6.**
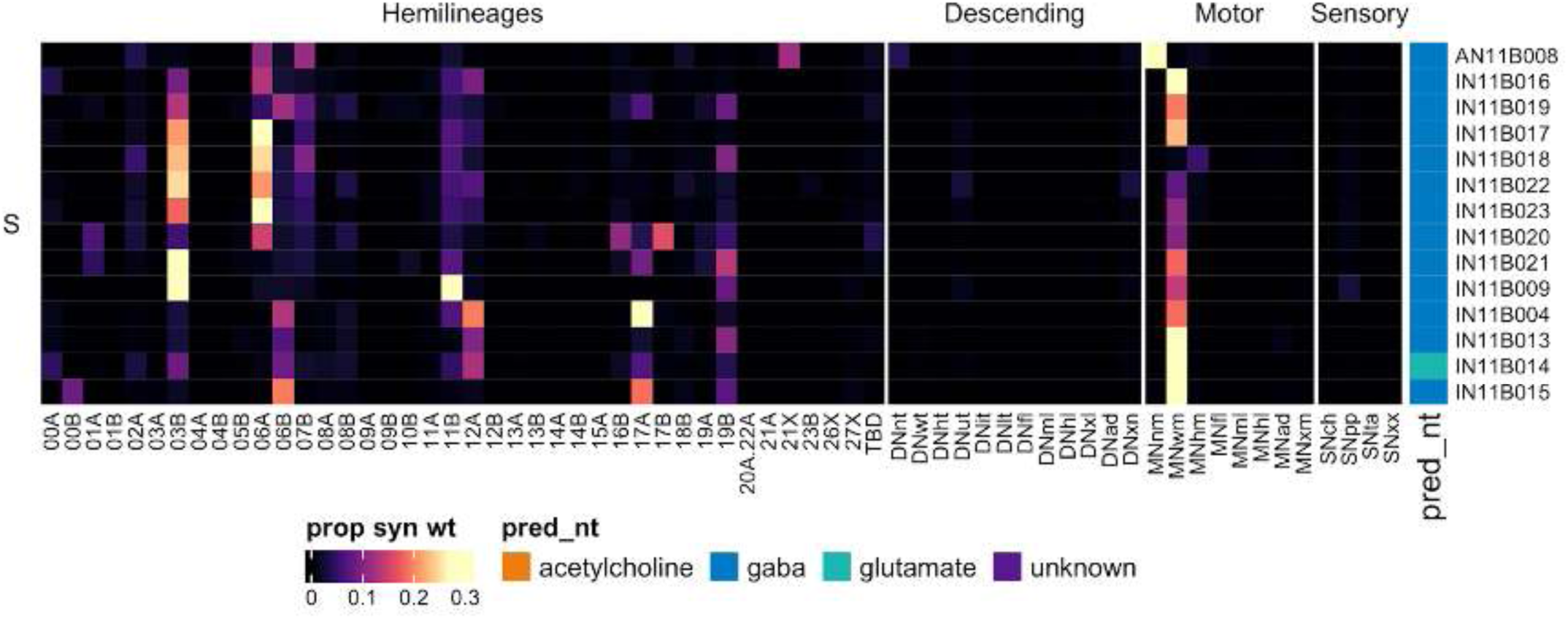
Connectivity to downstream partners by 11B secondary systematic types. Proportions of synaptic weight from systematic types to downstream partners, normalised by row. 11B neurons have been clustered within each assigned birthtime window (P = primary, ES = early secondary, S = secondary) based on both upstream and downstream connectivity to hemilineages, descending neuron subclasses, motor neuron subclasses, and sensory neuron modalities. The annotation bar is coloured by the most common predicted neurotransmitter within each type.

#### Hemilineage 12A

Hemilineages 12A and 12B derive from posterior ventral neuroblast NB6-1 (Truman et al., 2004), which generates 1 MN (in T1 only), 3-5 intersegmental interneurons, and ∼20 local interneurons in the embryo (Schmid et al., 1999). The primary neurites of 12A secondary neurons enter in the posterior of the neuromere, lateral to those of 12B (Shepherd et al., 2016), before projecting to the tectulum. 12A neurons exhibit a segment-specific pattern of survival consistent with a role in wing control, with populations in T1 and T2 but not in T3 (Figure 33A,E), although a small population survives in A1 (Marin et al., 2012). Consequently, most 12A serial sets we identified only had members in T1 and T2 (e.g., Figure 33C bottom); however, we did find one serial set in T1, T2, and A1 (Figure 33C top). In T1, 12A neurons could be difficult to separate from 03A and 03B.

**Figure 33.**
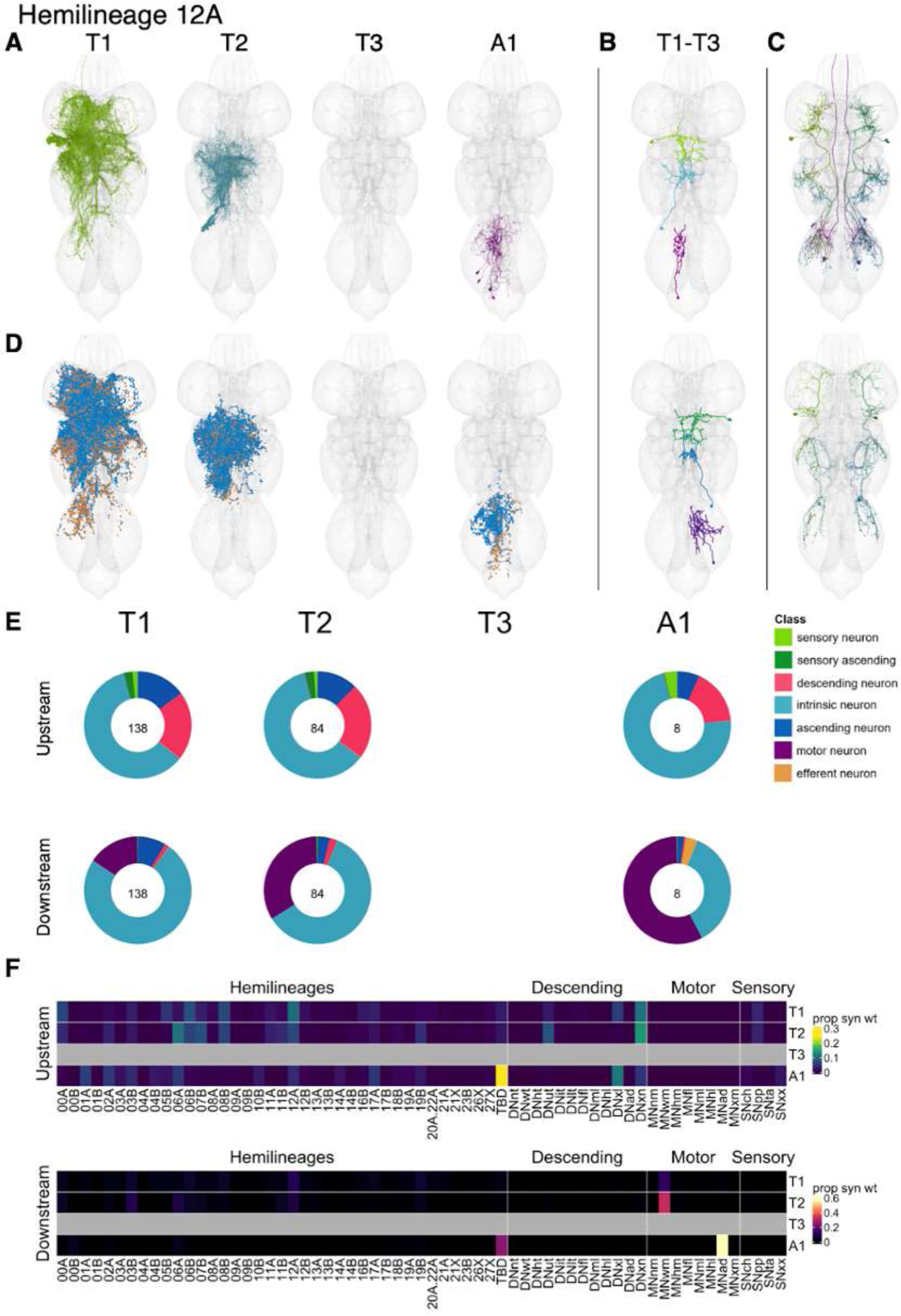
Hemilineage 12A. **A.** Meshes of all RHS secondary neurons plotted in neuromere-specific colours. **B.** “Representative” secondary neuron skeletons plotted in hemineuromere-specific colours. The skeleton with the top accumulated NBLAST score among all neurons from the hemilineage in a given hemineuromere was used. **C.** Neuron meshes of selected examples. Top: convergent serial set 10554. Bottom: convergent serial set 11769. **D.** Predicted synapses of RHS secondary neurons. Blue: postsynapses; dark orange: presynapses. **E.** Proportions of connections from secondary neurons to upstream or downstream partners, normalised by neuromere and coloured by broad class. Numbers of query neurons appear in the centre. **F.** Proportions of synaptic weight from secondary neurons originating in each neuromere to upstream or downstream partners, normalised by row.

Much of the input to 12A secondary neurons is from descending neurons to the upper tectulum, to multiple legs, or to multiple neuropils, but they also receive input from proprioceptive sensory neurons and from flight hemilineages 00A, 06B, and 08B, and 12A in T1-T2 and additionally from 05B in T1 and from 07B and especially 06A in T2 (Figure 33F). A few 12A types receive sensory input of mixed modalities, including IN12A005 and IN12A006 (Figure 33 - figure supplement 4). Secondary 12A neurons are predicted to be cholinergic, as expected (Lacin et al., 2019), and activate wing and abdominal motor neurons. Bilateral activation of 12A secondary neurons results in a sequence of behaviours: walking, wing flicking, and lateral extension and vibration of one wing (akin to male courtship song) (Harris et al., 2015).

Prothoracic 12A neurons include several sexually dimorphic populations targeting wing motor neurons and exhibiting a characteristic “A-Frame”-shaped arborisation when viewed from dorsal called the mesothoracic triangle (e.g., IN12A020/TN1a, IN12A022/vPR6, and IN12A028/TN1a) (Figure 33 - figure supplement 2, 6-7) that have been implicated in courtship song (Lillvis et al., 2024; Mellert et al., 2016; Shirangi et al., 2016; von Philipsborn et al., 2011; Yu et al., 2010). Interestingly, IN12A014 and IN12A023 have both been matched morphologically to song circuit type vMS12 (Lillvis et al., 2024), but only IN12A023 is directly upstream of wing motor neurons (Figure 33 - figure supplement 2,6). We also identified several connectivity subtypes (IN12A031, IN12A038, and IN12A045) of TN1c (Figure 33 - figure supplement 2,5,7), which affects sine song frequency and amplitude (Lillvis et al., 2024; Shirangi et al., 2016).

**Figure 33 - figure supplement 1.**
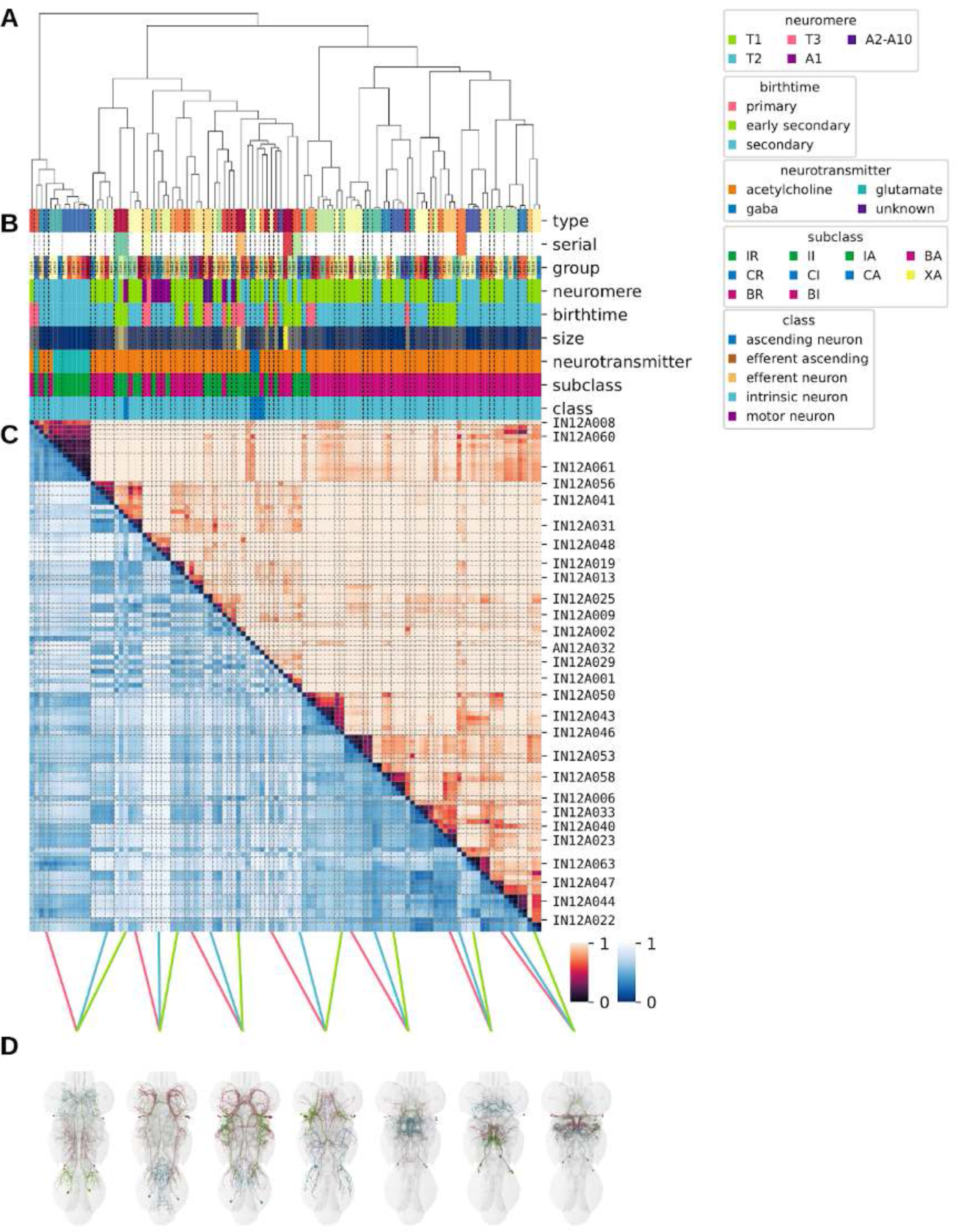
Systematic typing of hemilineage 12A. **A.** Hierarchical clustering dendrogram of hemilineage groups by laterally and serially aggregated connectivity cosine clustering. **B.** Categorical annotations of each hemilineage group, each column corresponding to the aligned leaf in A. Colours for type, serial set, and group are arbitrary for visualisation. Colours for neuromere, birthtime, neurotransmitter, subclass, and class are as in all other figures. **C.** Similarity distance heatmap for hemilineage. Cosine distance is in the upper triangle, while laterally symmetrised NBLAST distance is in the lower triangle. Systematic type names of some types are labelled. **D.** Morphologically representative groups from dendrogram subtrees. Each group, indicated by colour and line connecting to its column in B and C, is the most morphologically representative group (medoid of NBLAST distance) from a subtree of A. The subtrees (flat clusters) are equal height cuts of A determined to yield the number of groups per plot and plots in D.

**Figure 33 - figure supplement 2.**
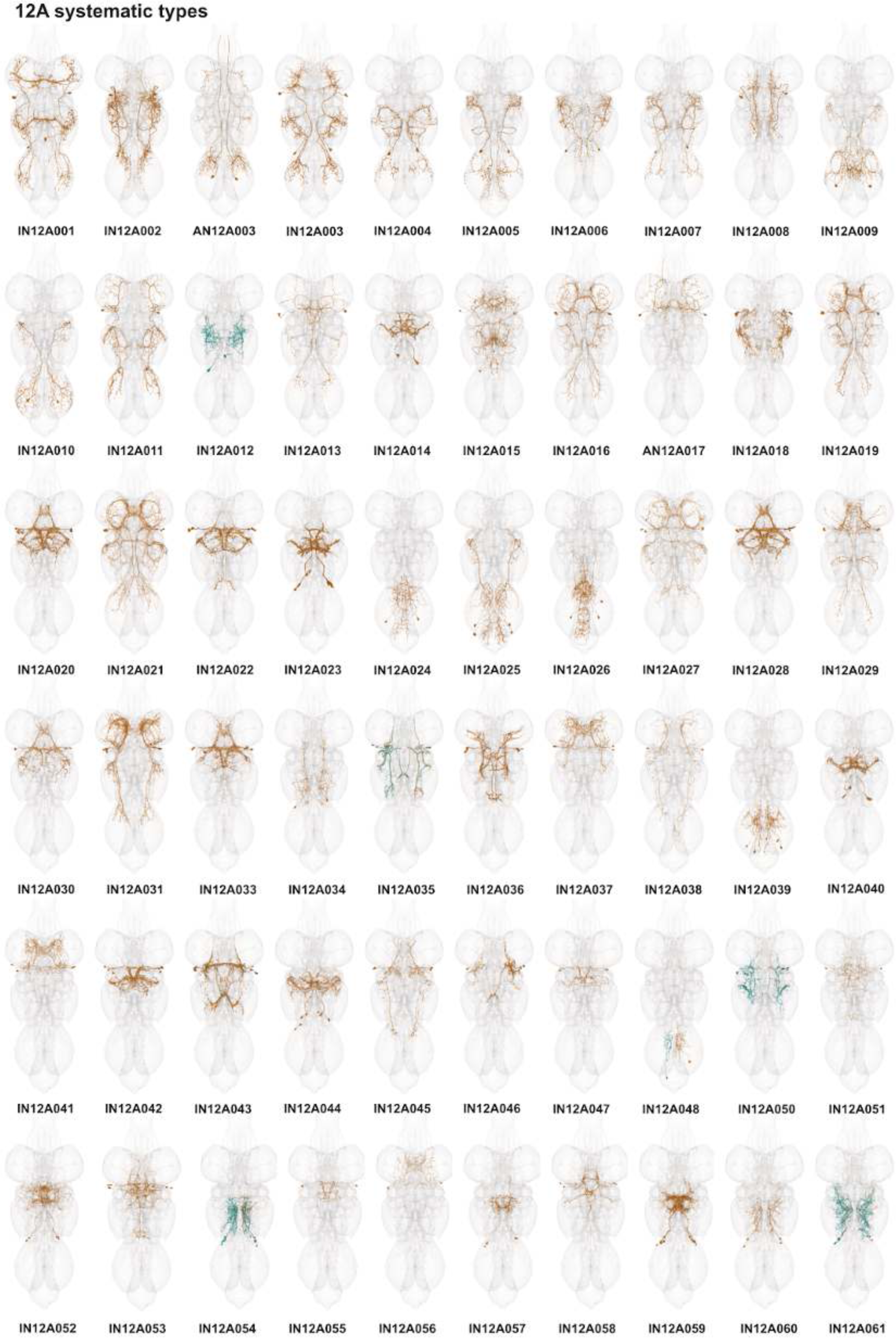
Systematic types of hemilineage 12A. Systematic types have been arranged in numerical order, with neurons of the same type that belong to distinct classes (e.g., intrinsic neuron vs ascending neuron) plotted separately but placed adjacent to each other. Individual neuron meshes have been coloured based on predicted neurotransmitter: dark orange = acetylcholine, blue = gaba, marine = glutamate, dark purple = unknown.

**Figure 33 - figure supplement 3.**
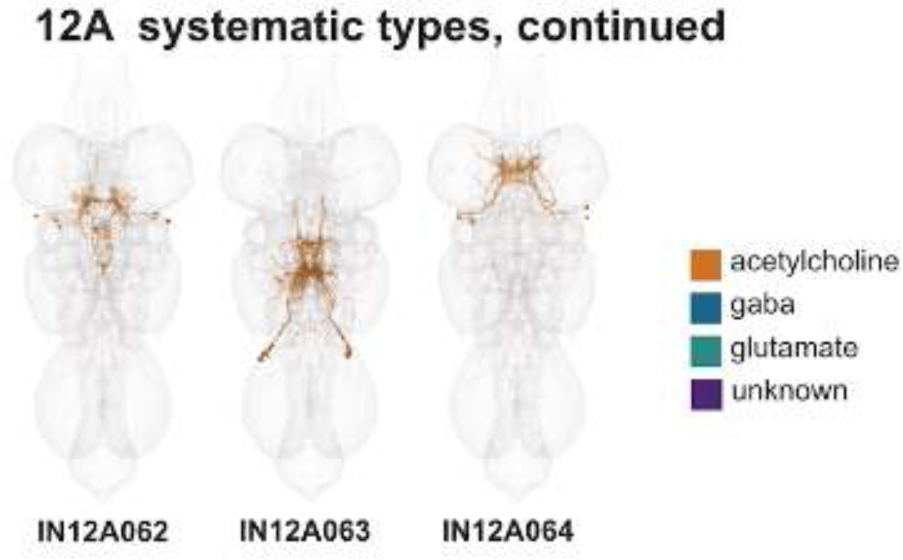
Systematic types of hemilineage 12A, continued. Systematic types have been arranged in numerical order, with neurons of the same type that belong to distinct classes (e.g., intrinsic neuron vs ascending neuron) plotted separately but placed adjacent to each other. Individual neuron meshes have been coloured based on predicted neurotransmitter: dark orange = acetylcholine, blue = gaba, marine = glutamate, dark purple = unknown.

**Figure 33 - figure supplement 4.**
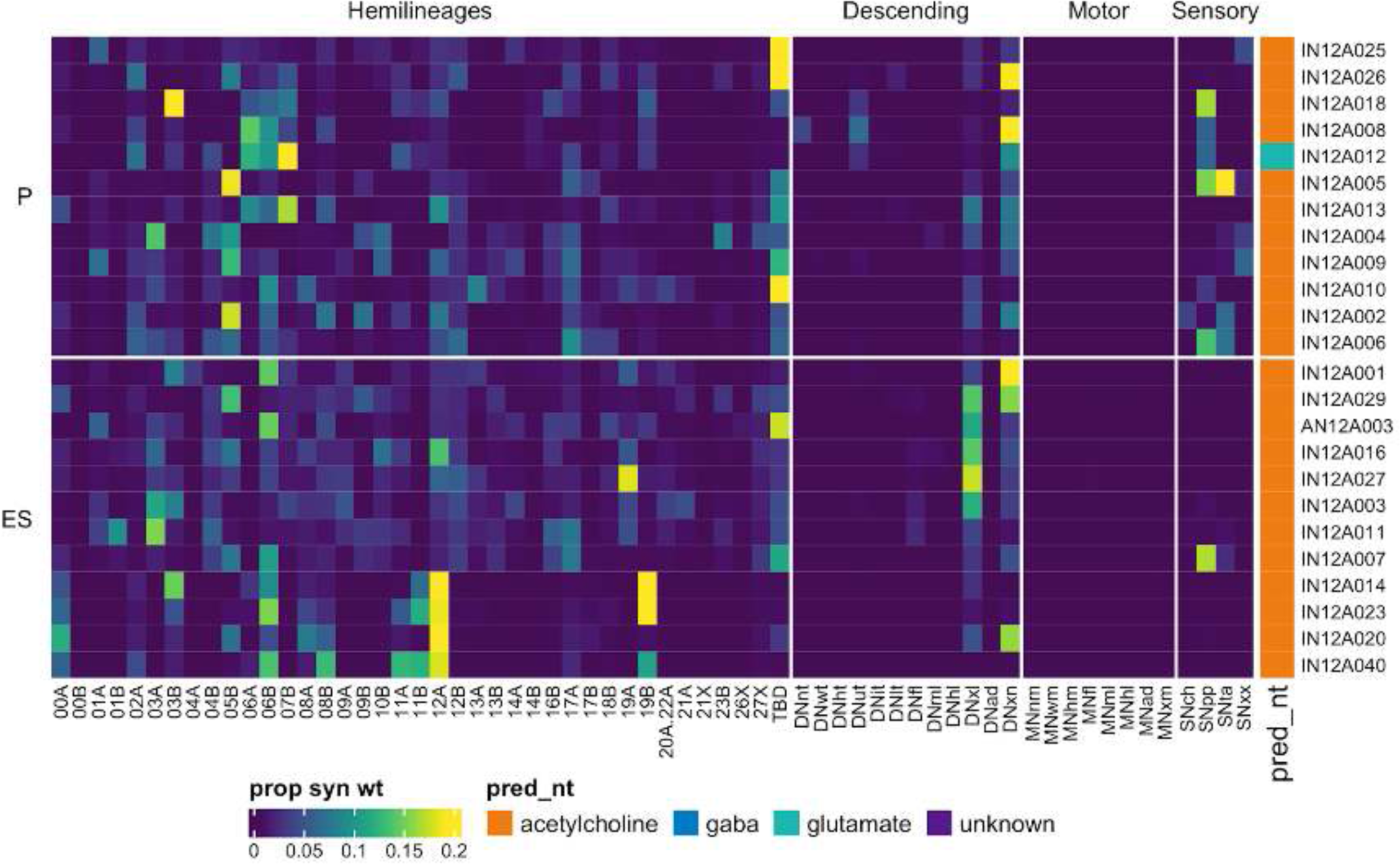
Connectivity to upstream partners by 12A primary and early secondary systematic types. Proportions of synaptic weight to systematic types from upstream partners, normalised by row. 12A neurons have been clustered within each assigned birthtime window (P = primary, ES = early secondary, S = secondary) based on both upstream and downstream connectivity to hemilineages, descending neuron subclasses, motor neuron subclasses, and sensory neuron modalities. Annotation bar is coloured by the most common predicted neurotransmitter for the neurons of each type.

**Figure 33 - figure supplement 5.**
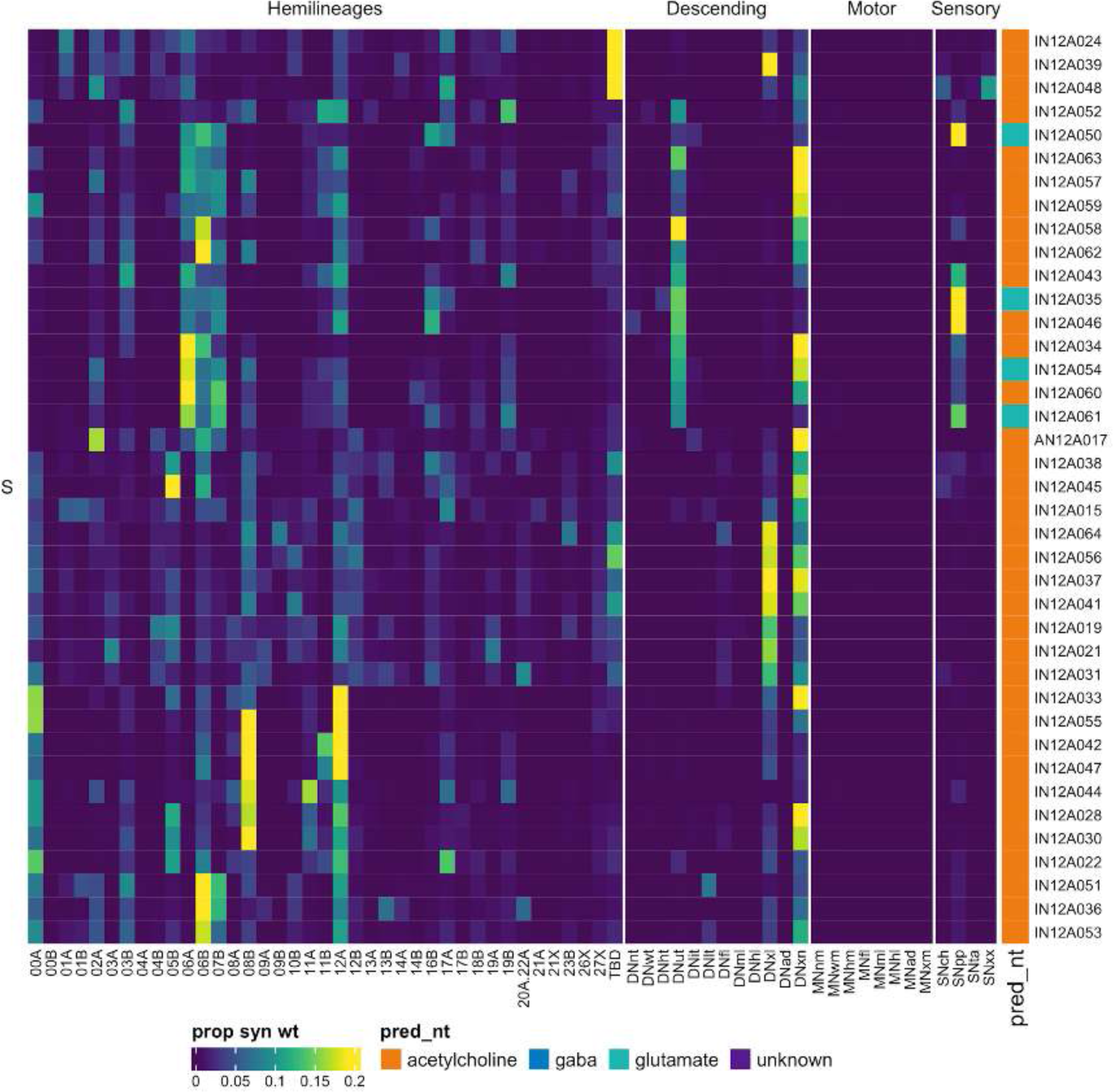
Connectivity to upstream partners by 12A secondary systematic types. Proportions of synaptic weight to systematic types from upstream partners, normalised by row. 12A neurons have been clustered within each assigned birthtime window (P = primary, ES = early secondary, S = secondary) based on both upstream and downstream connectivity to hemilineages, descending neuron subclasses, motor neuron subclasses, and sensory neuron modalities. The annotation bar is coloured by the most common predicted neurotransmitter within each type.

**Figure 33 - figure supplement 6.**
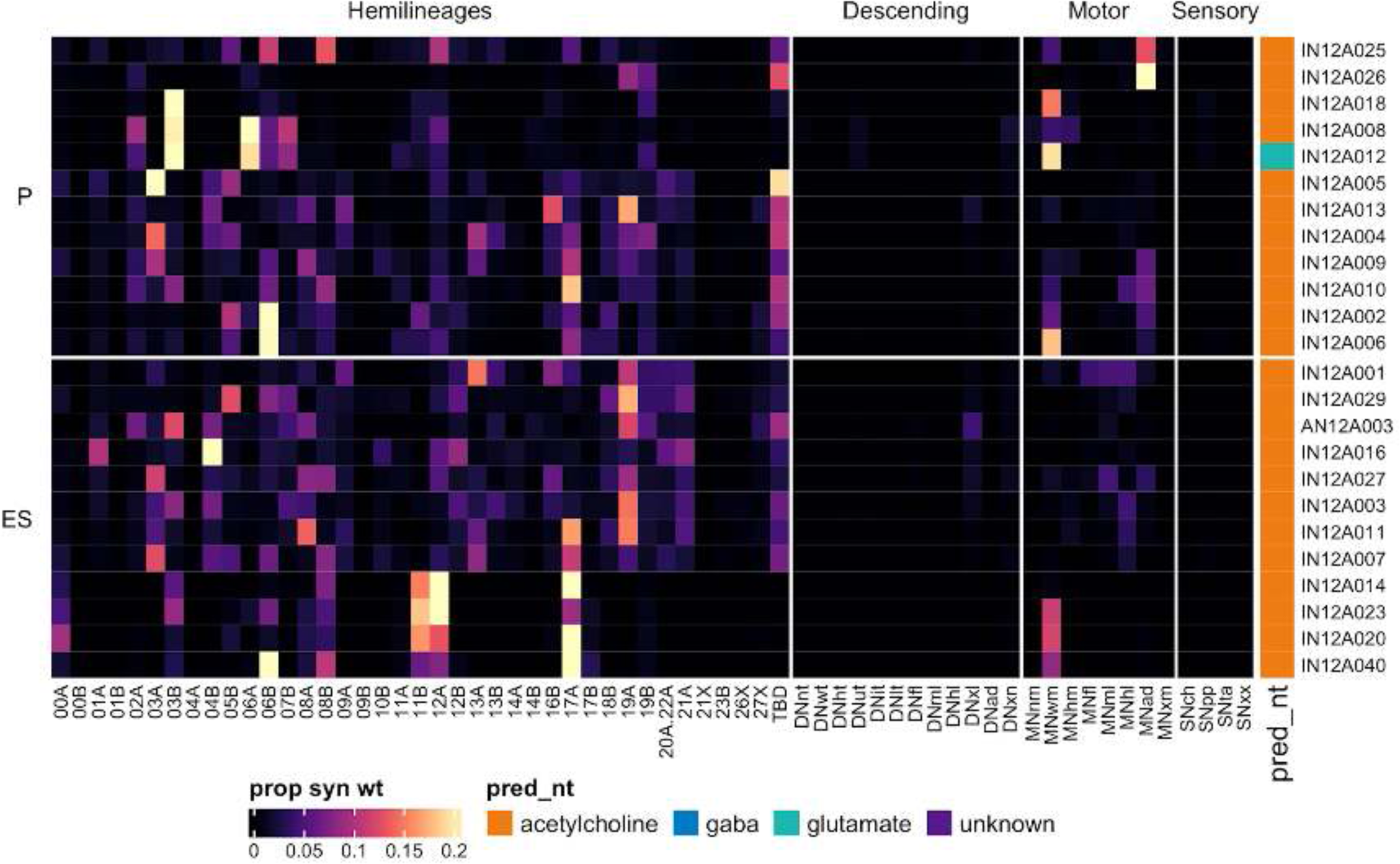
Connectivity to downstream partners by 12A primary and early secondary systematic types. Proportions of synaptic weight from systematic types to downstream partners, normalised by row. 12A neurons have been clustered within each assigned birthtime window (P = primary, ES = early secondary, S = secondary) based on both upstream and downstream connectivity to hemilineages, descending neuron subclasses, motor neuron subclasses, and sensory neuron modalities. The annotation bar is coloured by the most common predicted neurotransmitter within each type.

**Figure 33 - figure supplement 7.**
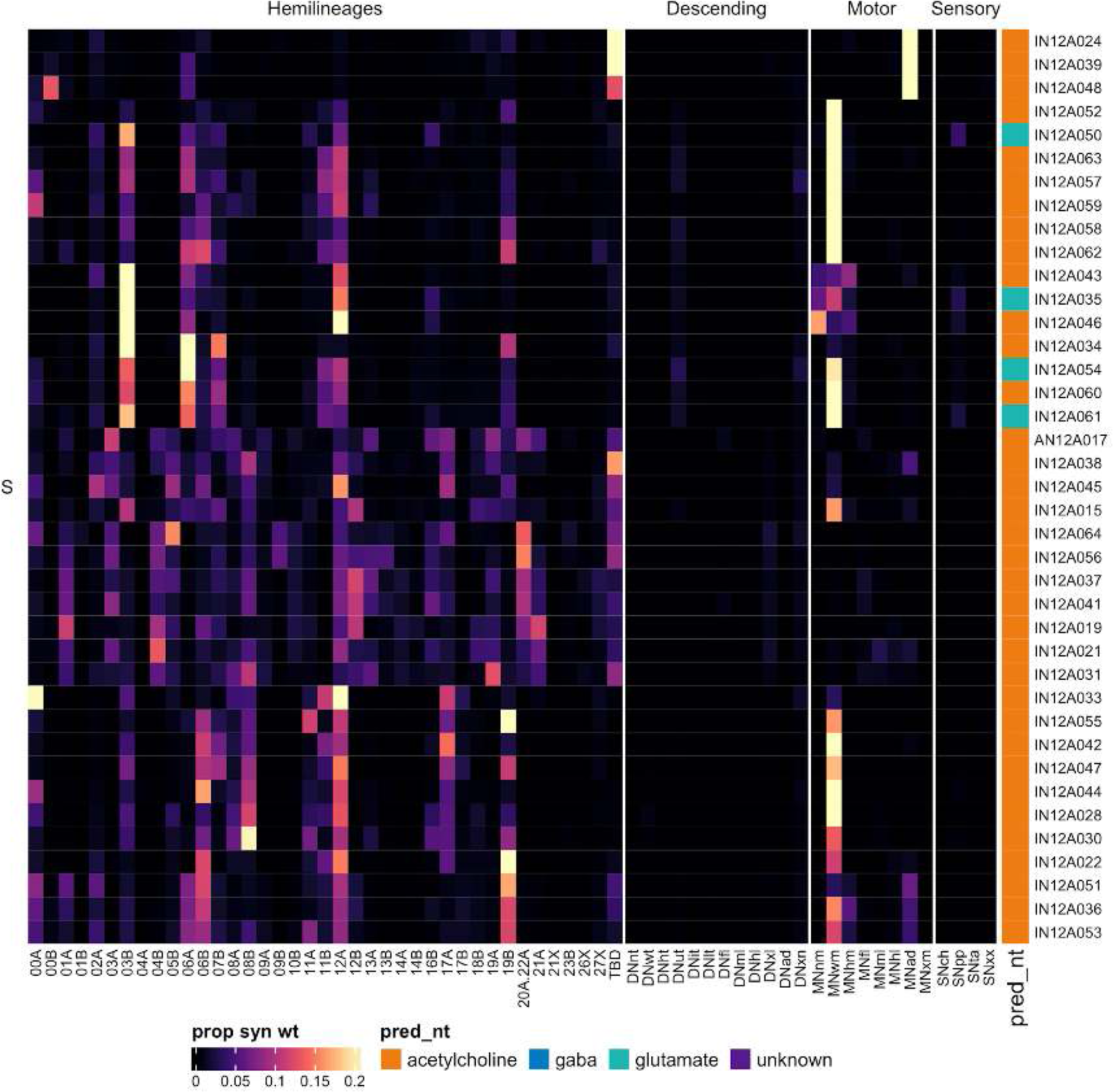
Connectivity to downstream partners by 12A secondary systematic types. Proportions of synaptic weight from systematic types to downstream partners, normalised by row. 12A neurons have been clustered within each assigned birthtime window (P = primary, ES = early secondary, S = secondary) based on both upstream and downstream connectivity to hemilineages, descending neuron subclasses, motor neuron subclasses, and sensory neuron modalities. The annotation bar is coloured by the most common predicted neurotransmitter for the neurons of each type.

#### Hemilineage 12B

12B neurons appear to be superficially similar in survival and appearance across all thoracic segments (Shepherd et al., 2019; Truman et al., 2004) (Figure 34E). They enter the posterior of the neuromere and typically cross the midline in the posterior intermediate commissure (posterior to 06B) (Shepherd et al., 2016) to innervate the ventral tectulum and contralateral leg neuropil (e.g., Figure 34C top). We identified a minority intersegmental population that innervates the mVACs (e.g., Figure 34C bottom); these types (AN/IN12B004 and AN12B006) receive direct proprioceptive input and inhibit 09A and 10B (Figure 34 - figure supplement 4,7).

**Figure 34.**
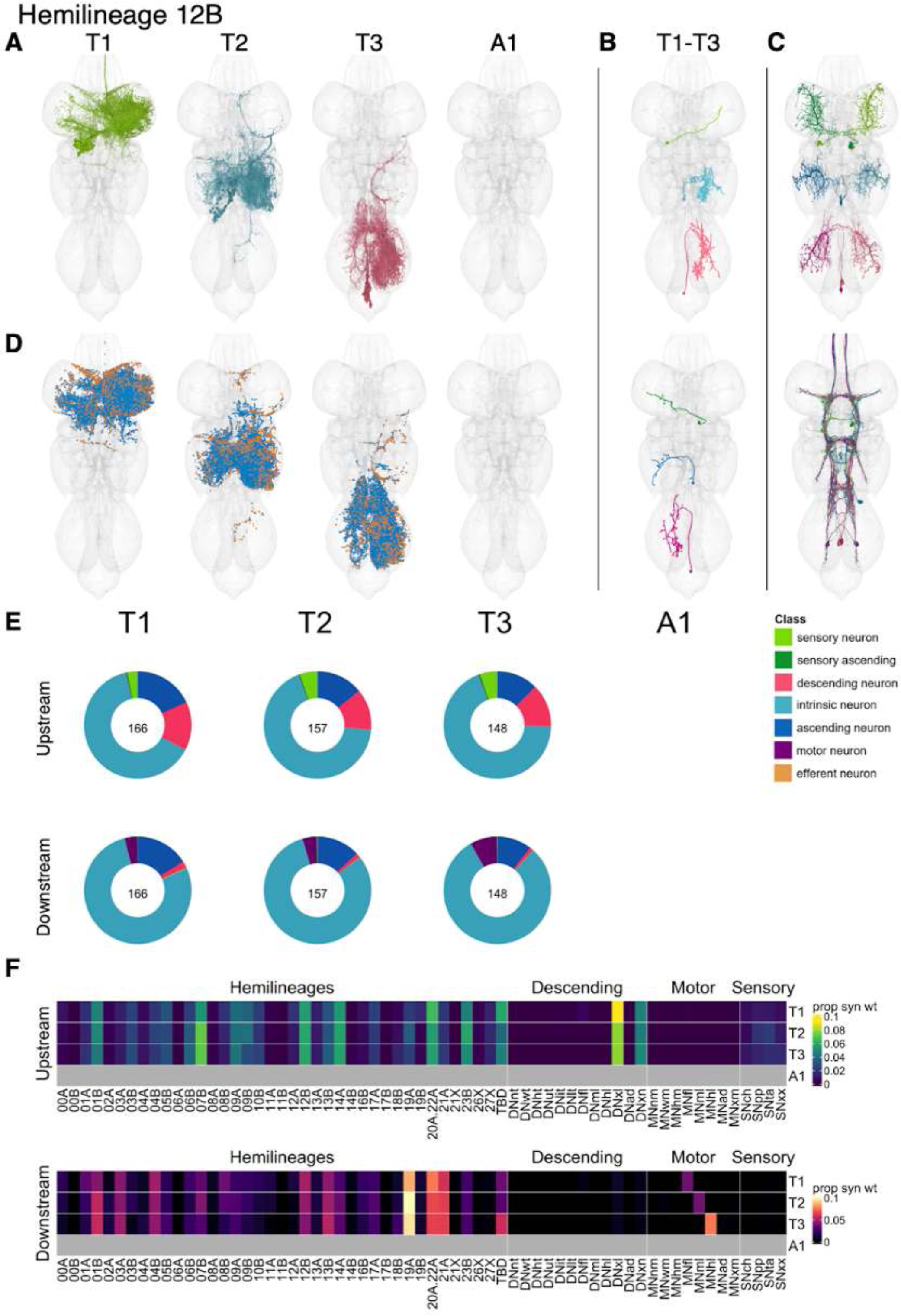
Hemilineage 12B. **A.** Meshes of all RHS secondary neurons plotted in neuromere-specific colours. **B.** “Representative” secondary neuron skeletons plotted in hemineuromere-specific colours. The skeleton with the top accumulated NBLAST score among all neurons from the hemilineage in a given hemineuromere was used. **C.** Neuron meshes of selected examples. Top: independent leg serial 10370. Bottom: ascending mVAC serial 10208. **D.** Predicted synapses of RHS secondary neurons. Blue: postsynapses; dark orange: presynapses. **E.** Proportions of connections from secondary neurons to upstream or downstream partners, normalised by neuromere and coloured by broad class. Numbers of query neurons appear in the centre. **F.** Proportions of synaptic weight from secondary neurons originating in each neuromere to upstream or downstream partners, normalised by row.

12B secondary neurons in T1-T3 receive inputs from descending neurons to multiple legs and from hemilineages 01B, 07B, 09A, 12B, 14A, 20A/22A, and 23B. They are mainly predicted to be gabaergic, as expected (Lacin et al., 2019), and inhibit leg motor neurons and hemilineages 19A, 20A/22A, and 21A, and to a lesser extent 01B, 03A, 04B, 12B, and 13B (Figure 34F). Bilateral activation of 12B secondary neurons sometimes results in variable movements of T1 legs with extreme extension of the T2 and T3 legs (Harris et al., 2015).

Although 12B neurons do not receive much sensory input as a population, a few early born types receive direct proprioceptive (e.g., A/IN12B004) or tactile (e.g., A/IN12B011) input (Figure 34 - figure supplement 4). And whilst 12B neurons generally target leg motor neurons, one primary type (IN12B015) inhibits wing motor neurons instead (Figure 34 - figure supplement 7).

**Figure 34 - figure supplement 1.**
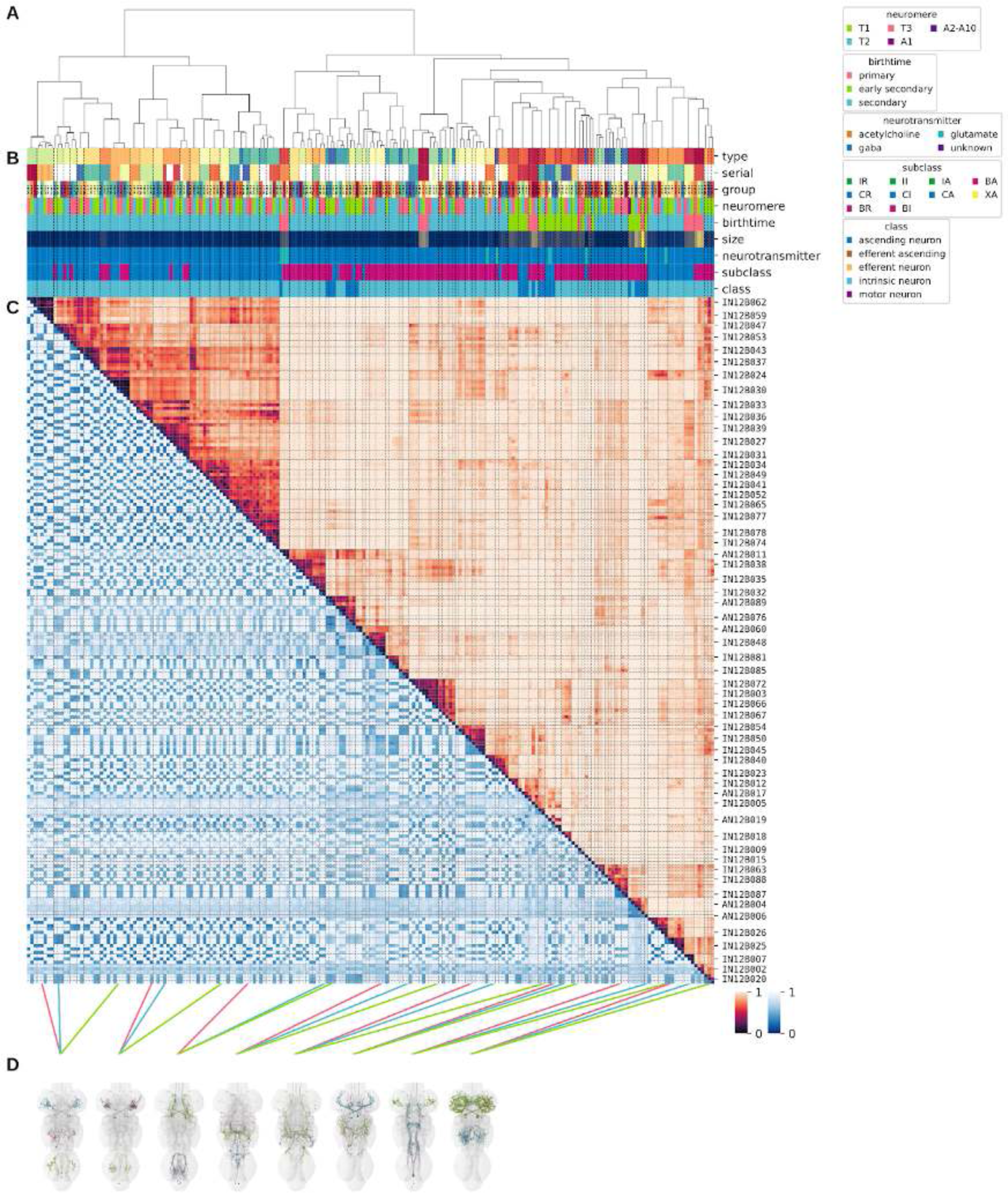
Systematic typing of hemilineage 12B. **A.** Hierarchical clustering dendrogram of hemilineage groups by laterally and serially aggregated connectivity cosine clustering. **B.** Categorical annotations of each hemilineage group, each column corresponding to the aligned leaf in A. Colours for type, serial set, and group are arbitrary for visualisation. Colours for neuromere, birthtime, neurotransmitter, subclass, and class are as in all other figures. **C.** Similarity distance heatmap for hemilineage. Cosine distance is in the upper triangle, while laterally symmetrised NBLAST distance is in the lower triangle. Systematic type names of some types are labelled. **D.** Morphologically representative groups from dendrogram subtrees. Each group, indicated by colour and line connecting to its column in B and C, is the most morphologically representative group (medoid of NBLAST distance) from a subtree of A. The subtrees (flat clusters) are equal height cuts of A determined to yield the number of groups per plot and plots in D.

**Figure 34 - figure supplement 2.**
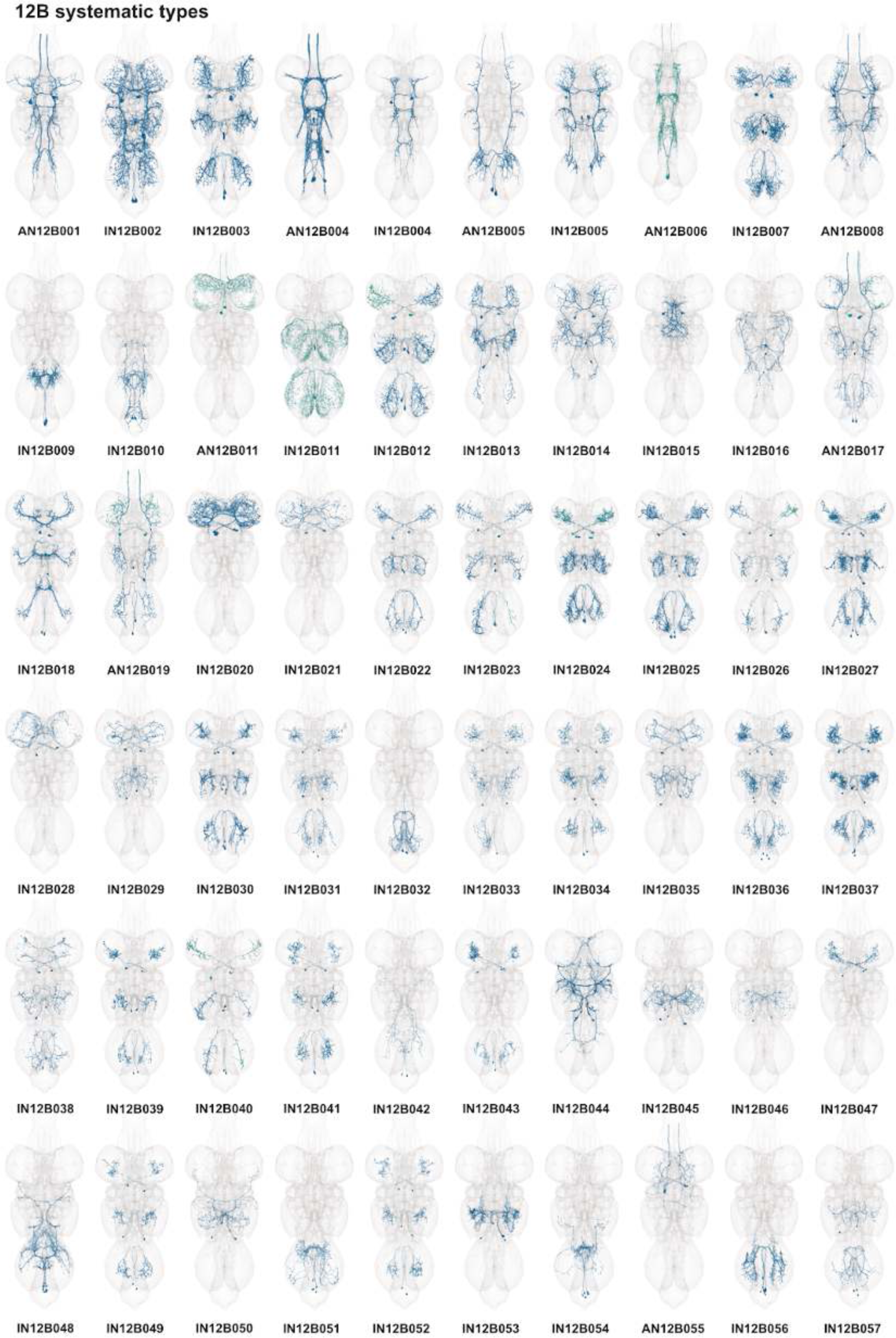
Systematic types of hemilineage 12B. Systematic types have been arranged in numerical order, with neurons of the same type that belong to distinct classes (e.g., intrinsic neuron vs ascending neuron) plotted separately but placed adjacent to each other. Individual neuron meshes have been coloured based on predicted neurotransmitter: dark orange = acetylcholine, blue = gaba, marine = glutamate, dark purple = unknown.

**Figure 34 - figure supplement 3.**
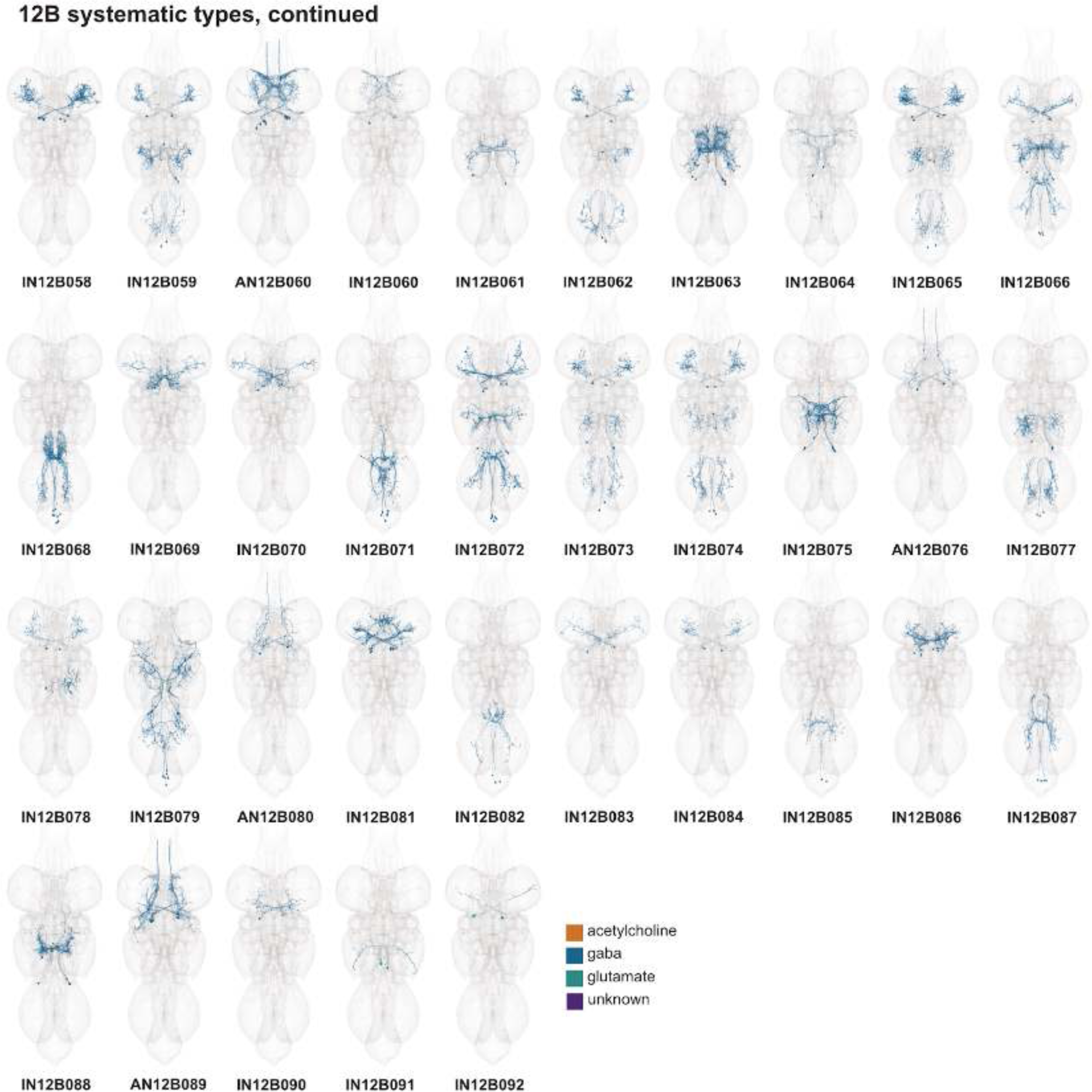
Systematic types of hemilineage 12B, continued. Systematic types have been arranged in numerical order, with neurons of the same type that belong to distinct classes (e.g., intrinsic neuron vs ascending neuron) plotted separately but placed adjacent to each other. Individual neuron meshes have been coloured based on predicted neurotransmitter: dark orange = acetylcholine, blue = gaba, marine = glutamate, dark purple = unknown.

**Figure 34 - figure supplement 4.**
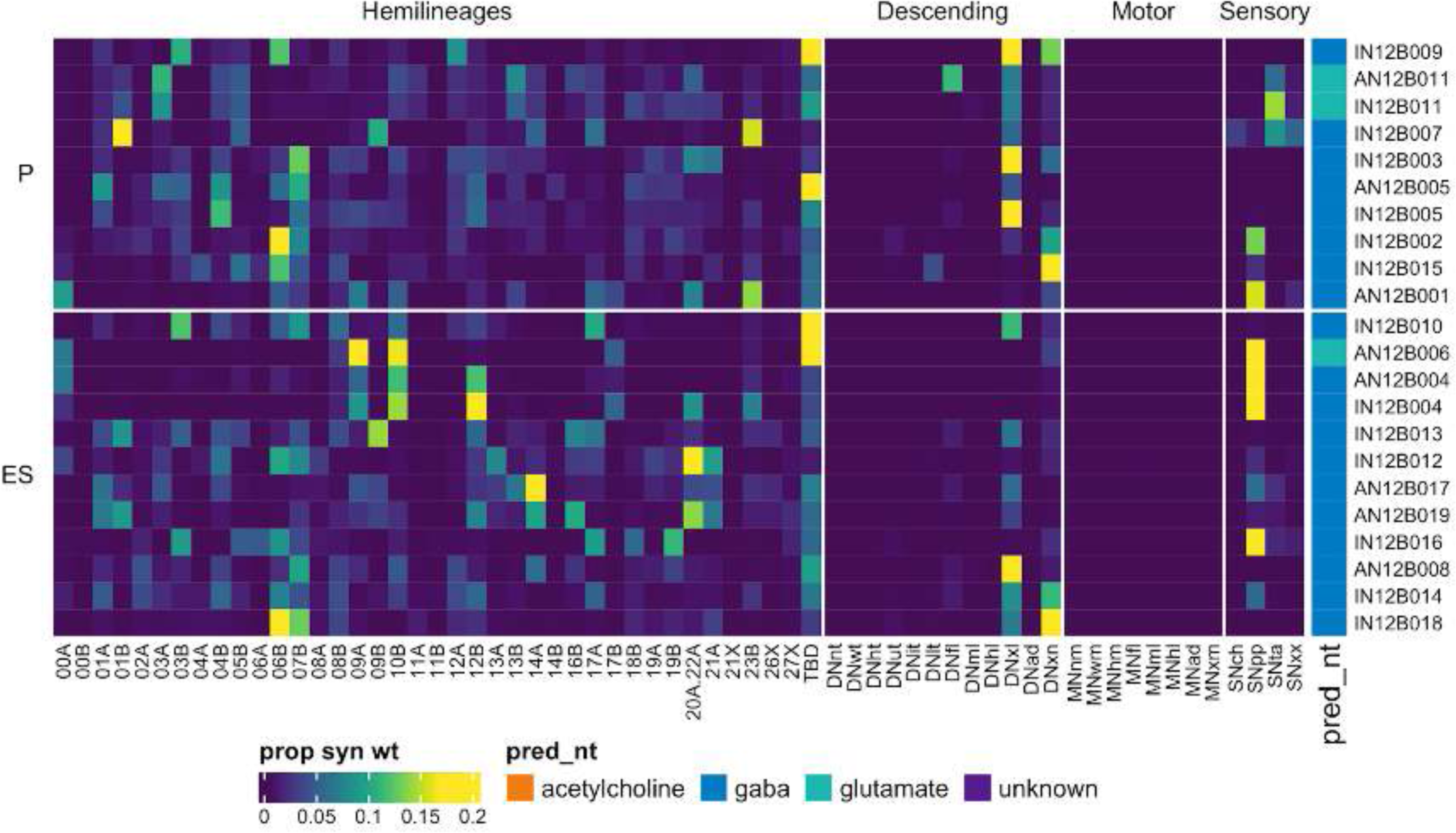
Connectivity to upstream partners by 12B primary and early secondary systematic types. Proportions of synaptic weight to systematic types from upstream partners, normalised by row. 12B neurons have been clustered within each assigned birthtime window (P = primary, ES = early secondary, S = secondary) based on both upstream and downstream connectivity to hemilineages, descending neuron subclasses, motor neuron subclasses, and sensory neuron modalities. Annotation bar is coloured by the most common predicted neurotransmitter for the neurons of each type.

**Figure 34 - figure supplement 5.**
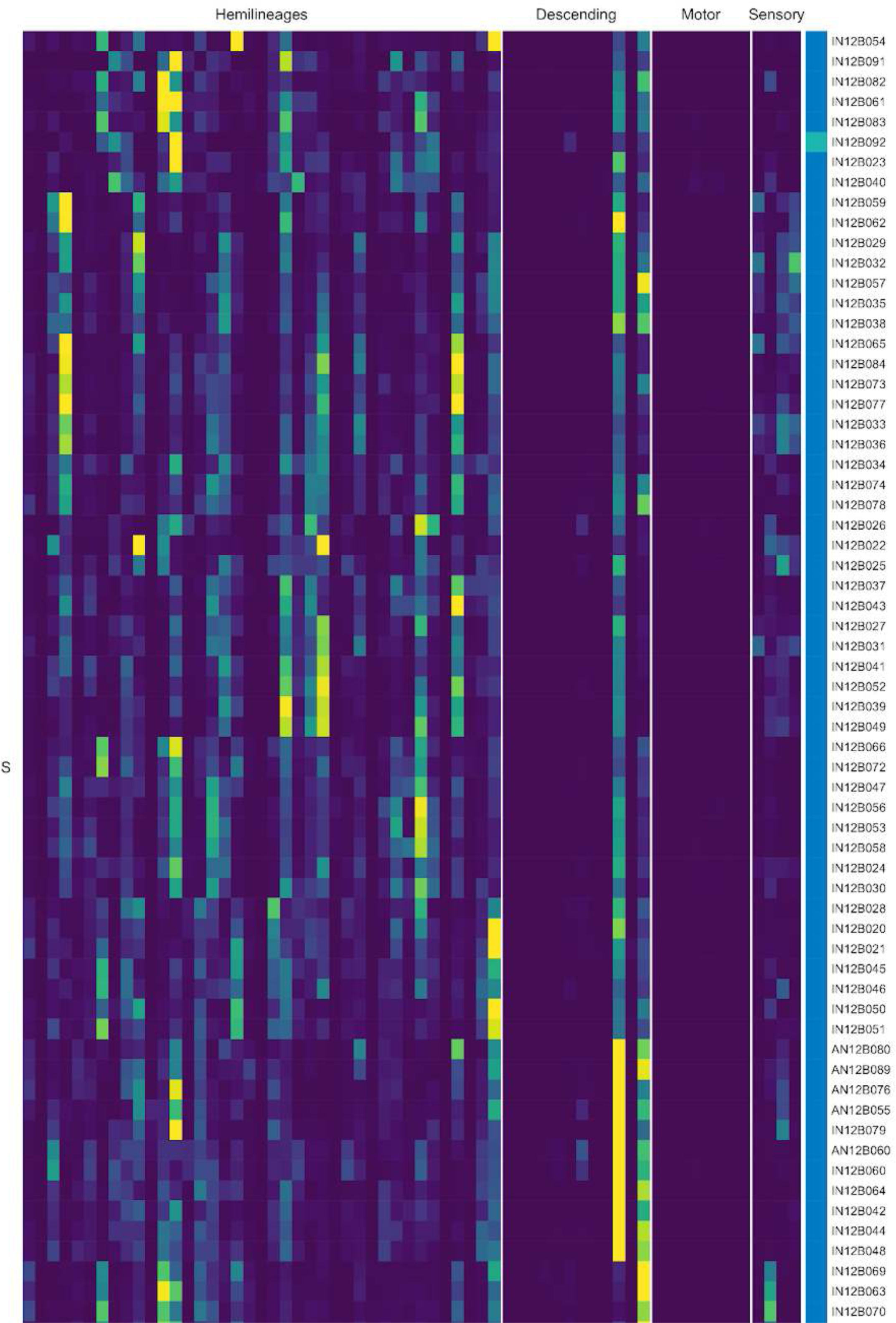
Connectivity to upstream partners by 12B secondary systematic types. Proportions of synaptic weight to systematic types from upstream partners, normalised by row. 12B neurons have been clustered within each assigned birthtime window (P = primary, ES = early secondary, S = secondary) based on both upstream and downstream connectivity to hemilineages, descending neuron subclasses, motor neuron subclasses, and sensory neuron modalities. The annotation bar is coloured by the most common predicted neurotransmitter within each type.

**Figure 34 - figure supplement 6.**
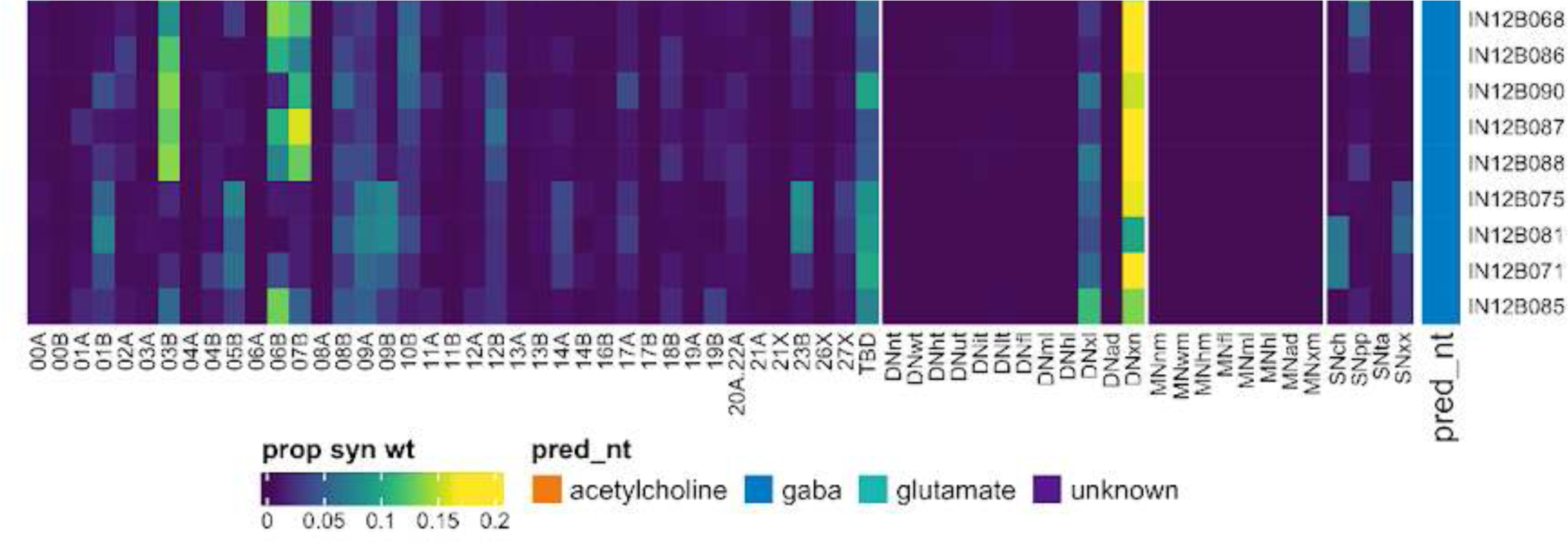
Connectivity to upstream partners by 12B secondary systematic types, continued. Proportions of synaptic weight to systematic types from upstream partners, normalised by row. 12B neurons have been clustered within each assigned birthtime window (P = primary, ES = early secondary, S = secondary) based on both upstream and downstream connectivity to hemilineages, descending neuron subclasses, motor neuron subclasses, and sensory neuron modalities. The annotation bar is coloured by the most common predicted neurotransmitter within each type.

**Figure 34 - figure supplement 7.**
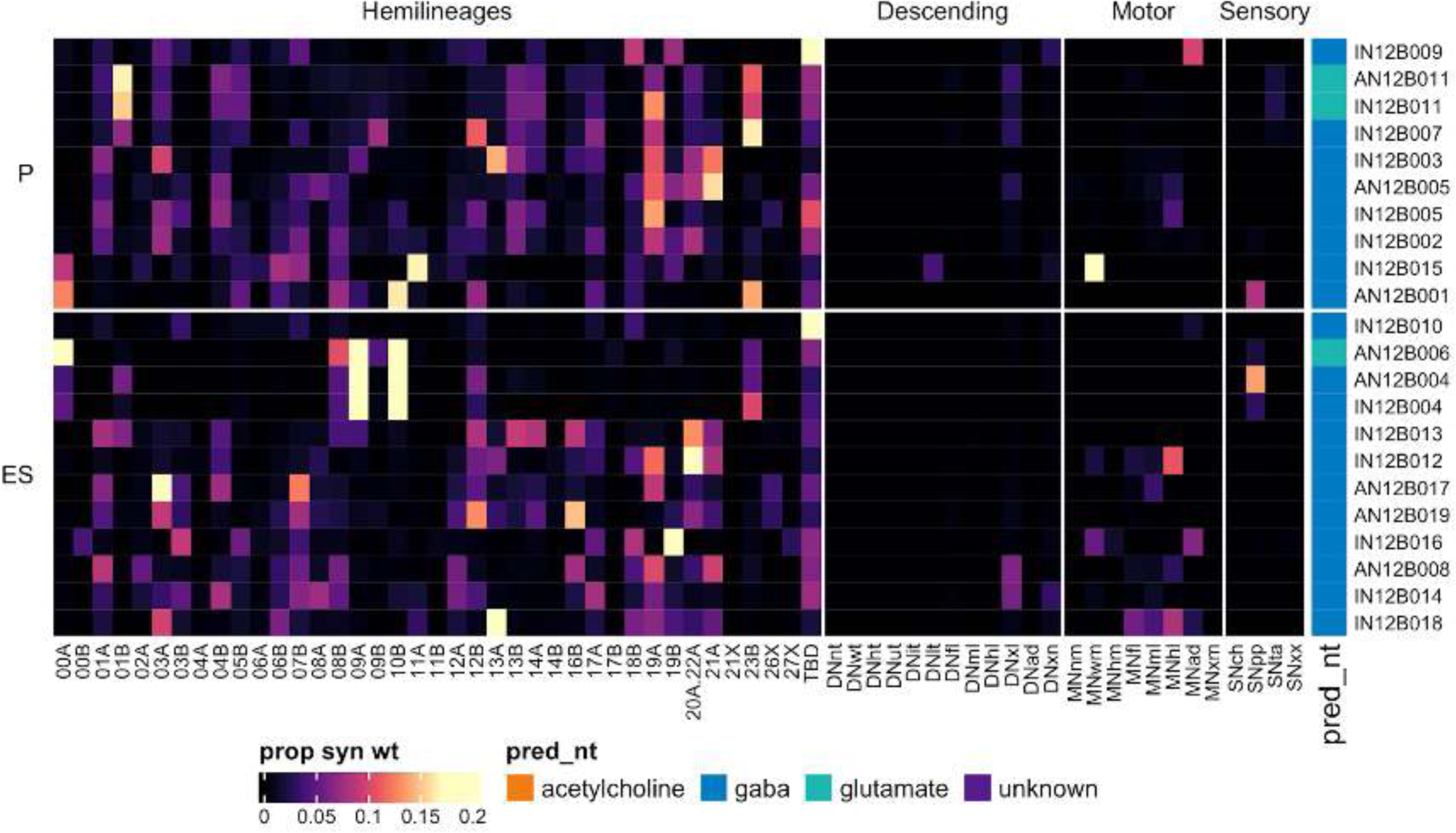
Connectivity to downstream partners by 12B primary and early secondary systematic types. Proportions of synaptic weight from systematic types to downstream partners, normalised by row. 12B neurons have been clustered within each assigned birthtime window (P = primary, ES = early secondary, S = secondary) based on both upstream and downstream connectivity to hemilineages, descending neuron subclasses, motor neuron subclasses, and sensory neuron modalities. The annotation bar is coloured by the most common predicted neurotransmitter within each type.

**Figure 34 - figure supplement 8.**
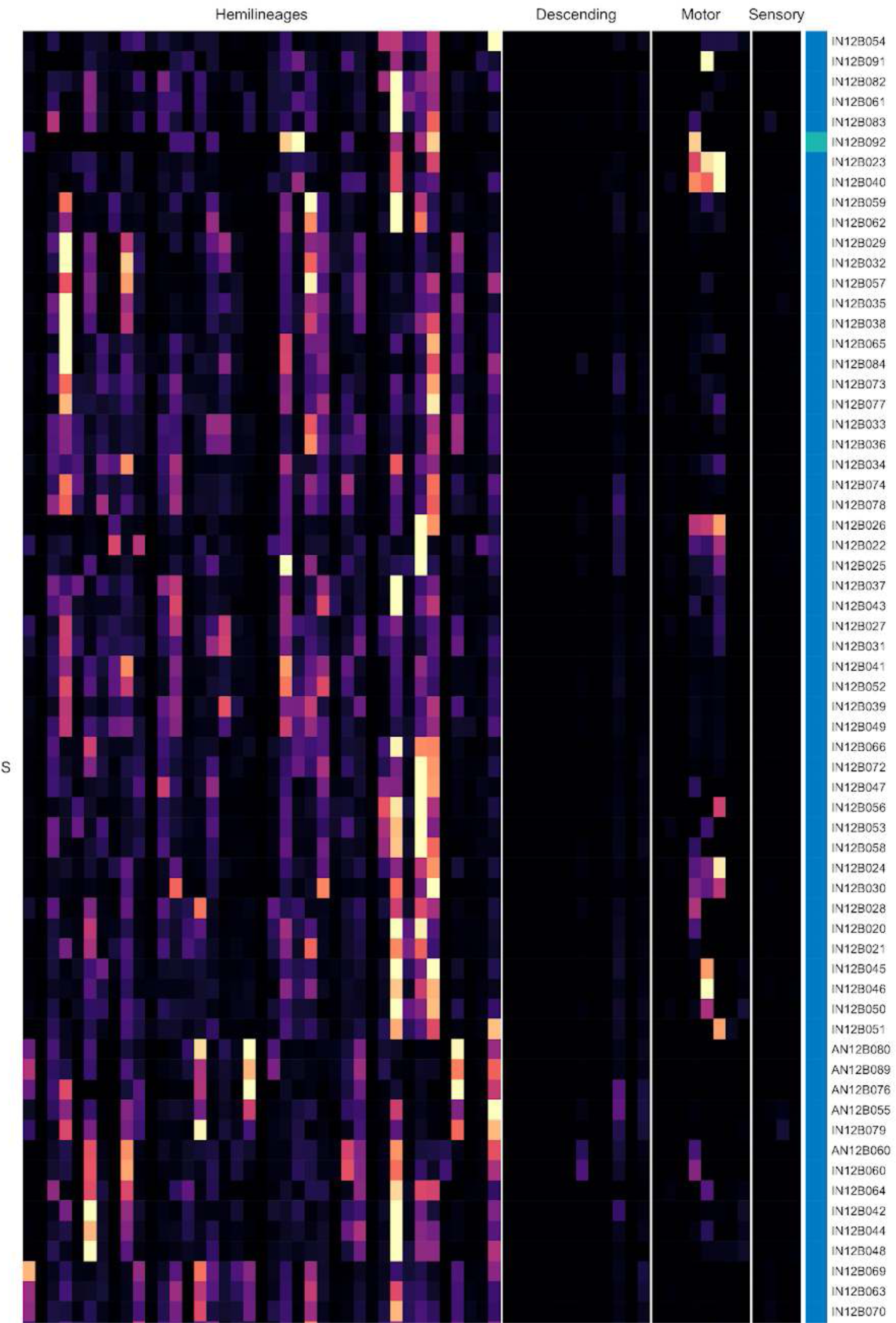
Connectivity to downstream partners by 12B secondary systematic types. Proportions of synaptic weight from systematic types to downstream partners, normalised by row. 12B neurons have been clustered within each assigned birthtime window (P = primary, ES = early secondary, S = secondary) based on both upstream and downstream connectivity to hemilineages, descending neuron subclasses, motor neuron subclasses, and sensory neuron modalities. The annotation bar is coloured by the most common predicted neurotransmitter for the neurons of each type.

**Figure 34 - figure supplement 9.**
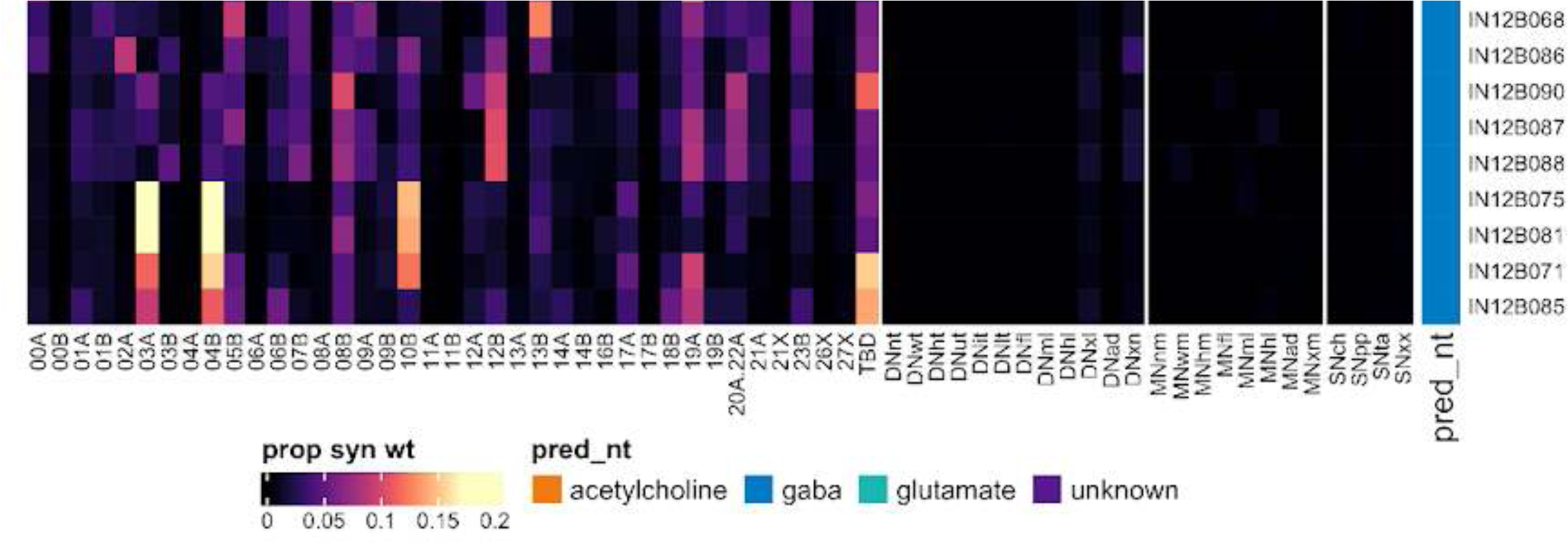
Connectivity to downstream partners by 12B secondary systematic types, continued. Proportions of synaptic weight from systematic types to downstream partners, normalised by row. 12B neurons have been clustered within each assigned birthtime window (P = primary, ES = early secondary, S = secondary) based on both upstream and downstream connectivity to hemilineages, descending neuron subclasses, motor neuron subclasses, and sensory neuron modalities. The annotation bar is coloured by the most common predicted neurotransmitter for the neurons of each type.

#### Hemilineage 13A

Hemilineages 13A and 13B are believed to derive from neuroblast NB4-2 (Lacin and Truman, 2016) (but see also (Birkholz et al., 2015; Truman et al., 2004)), which produces 4 MNs and ∼20 local interneurons in the embryo (Schmid et al., 1999). 13A secondary neurons survive in roughly comparable numbers across the thoracic neuromeres (Figure 35E). Their primary neurites spread over the VNC surface and only converge upon neuropil entry, and their processes are largely restricted to the ipsilateral leg neuropil of origin (Figure 35A,C bottom). We distinguished 13A neurons from other anterior ipsilateral leg neuropil hemilineages by their characteristic subpopulation of neurons projecting medially and dorsally (Shepherd et al., 2019).

**Figure 35.**
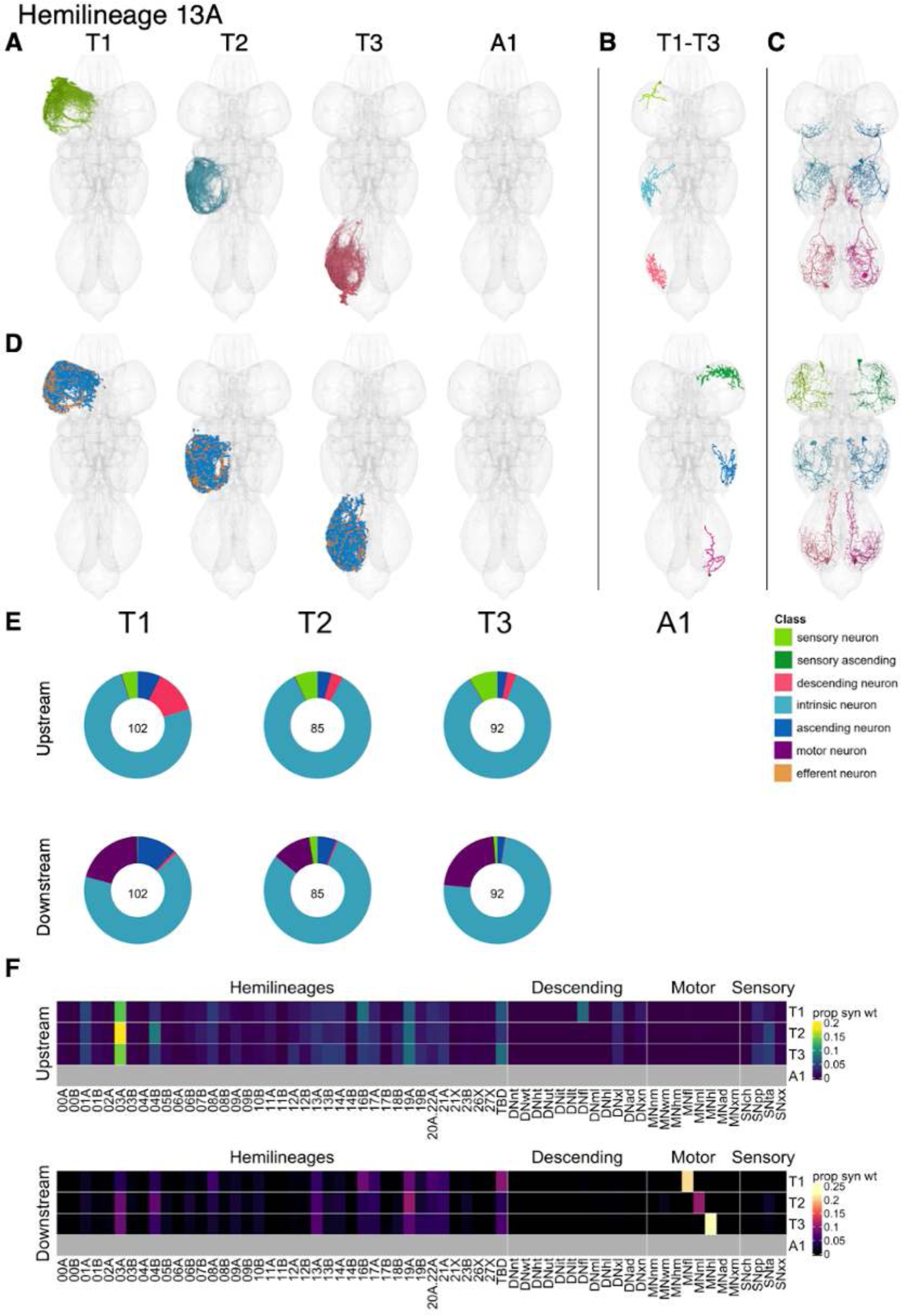
Hemilineage 13A. **A.** Meshes of all RHS secondary neurons plotted in neuromere-specific colours. **B.** “Representative” secondary neuron skeletons plotted in hemineuromere-specific colours. The skeleton with the top accumulated NBLAST score among all neurons from the hemilineage in a given hemineuromere was used. **C.** Neuron meshes of selected examples. Top: sequential serial set 10655. Bottom: independent leg serial set 11077. **D.** Predicted synapses of RHS secondary neurons. Blue: postsynapses; dark orange: presynapses. **E.** Proportions of connections from secondary neurons to upstream or downstream partners, normalised by neuromere and coloured by broad class. Numbers of query neurons appear in the centre. **F.** Proportions of synaptic weight from secondary neurons originating in each neuromere to upstream or downstream partners, normalised by row.

13A neurons receive input predominantly from hemilineage 03A as well as from 04B, 16B, and 19A, from descending neurons targeting the front leg or all leg neuropils and from proprioceptive and tactile sensory neurons (Figure 35F). They are predicted to be gabaergic, as expected (Lacin et al., 2019), and provide outputs mainly to 16B in T1 and to 03A, 04B, 13A, and 19A in T2-T3. They also inhibit front and hind leg motor neurons and to a lesser extent middle leg motor neurons. No functional studies have been published for secondary 13A neurons.

Most 13A types target leg motor neurons, but we found several types that inhibit wing motor neurons instead (e.g., IN13A013 and IN13A022) (Figure 35 - figure supplement 6). We also identified one early born sequential serial type, IN13A004 (Figure 35C top), that ascends to the next anterior neuromere and clusters by connectivity with type IN13A007, receiving direct tactile inputs and strongly targeting neurons from hemilineage 23B (Figure 35 - figure supplement 4-6).

**Figure 35 - figure supplement 1.**
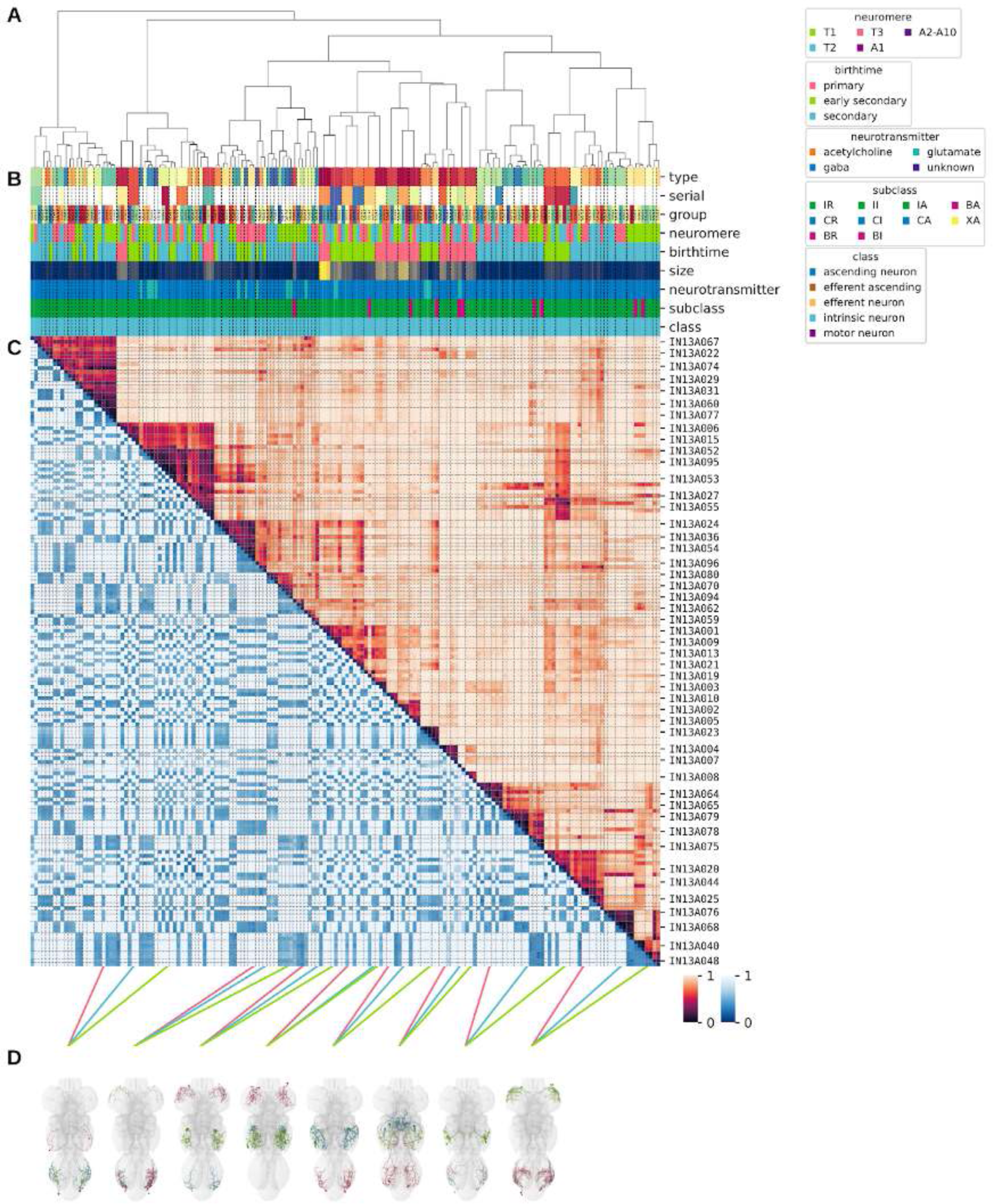
Systematic typing of hemilineage 13A. **A.** Hierarchical clustering dendrogram of hemilineage groups by laterally and serially aggregated connectivity cosine clustering. **B.** Categorical annotations of each hemilineage group, each column corresponding to the aligned leaf in A. Colours for type, serial set, and group are arbitrary for visualisation. Colours for neuromere, birthtime, neurotransmitter, subclass, and class are as in all other figures. **C.** Similarity distance heatmap for hemilineage. Cosine distance is in the upper triangle, while laterally symmetrised NBLAST distance is in the lower triangle. Systematic type names of some types are labelled. **D.** Morphologically representative groups from dendrogram subtrees. Each group, indicated by colour and line connecting to its column in B and C, is the most morphologically representative group (medoid of NBLAST distance) from a subtree of A. The subtrees (flat clusters) are equal height cuts of A determined to yield the number of groups per plot and plots in D.

**Figure 35 - figure supplement 2.**
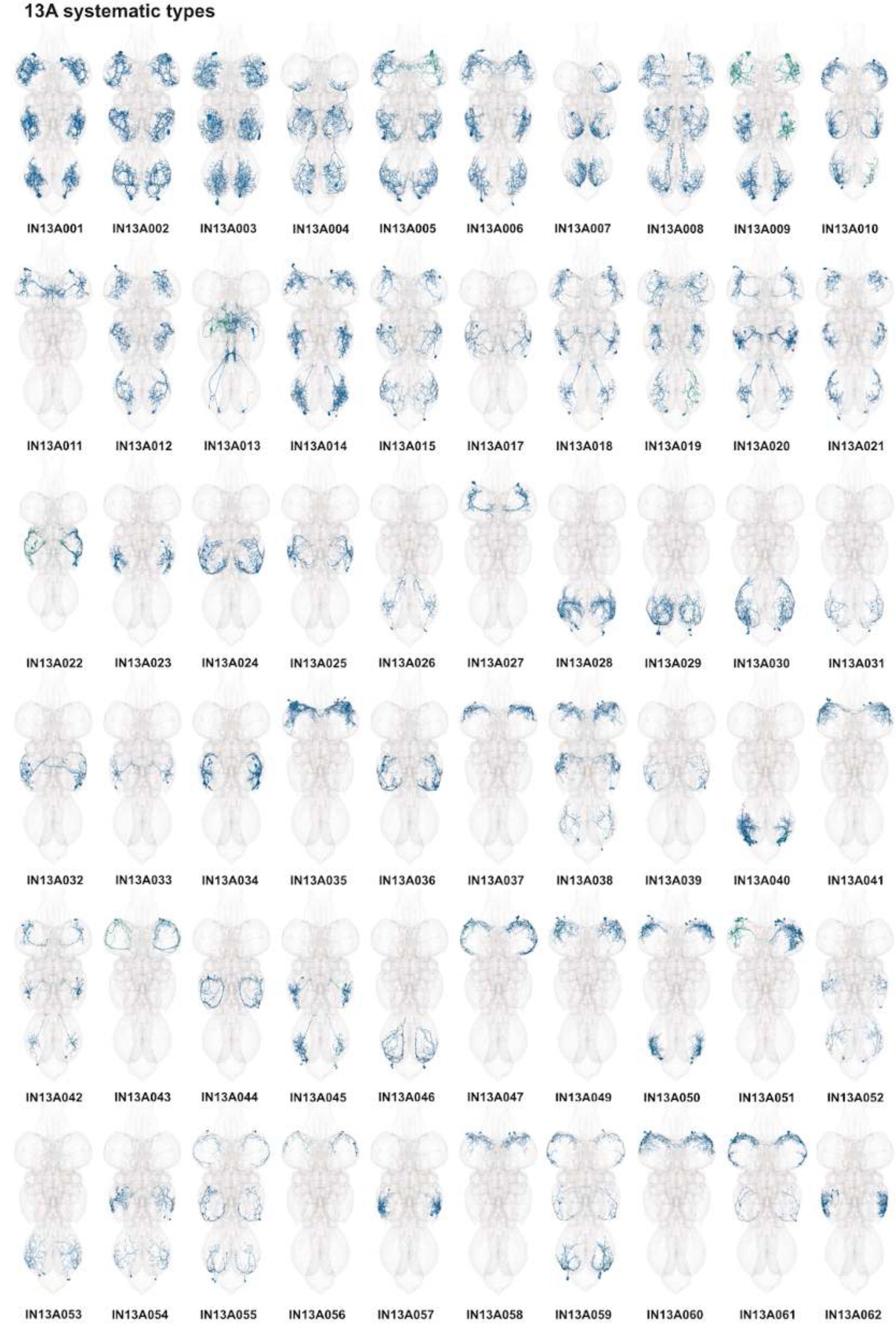
Systematic types of hemilineage 13A. Systematic types have been arranged in numerical order, with neurons of the same type that belong to distinct classes (e.g., intrinsic neuron vs ascending neuron) plotted separately but placed adjacent to each other. Individual neuron meshes have been coloured based on predicted neurotransmitter: dark orange = acetylcholine, blue = gaba, marine = glutamate, dark purple = unknown.

**Figure 35 - figure supplement 3.**
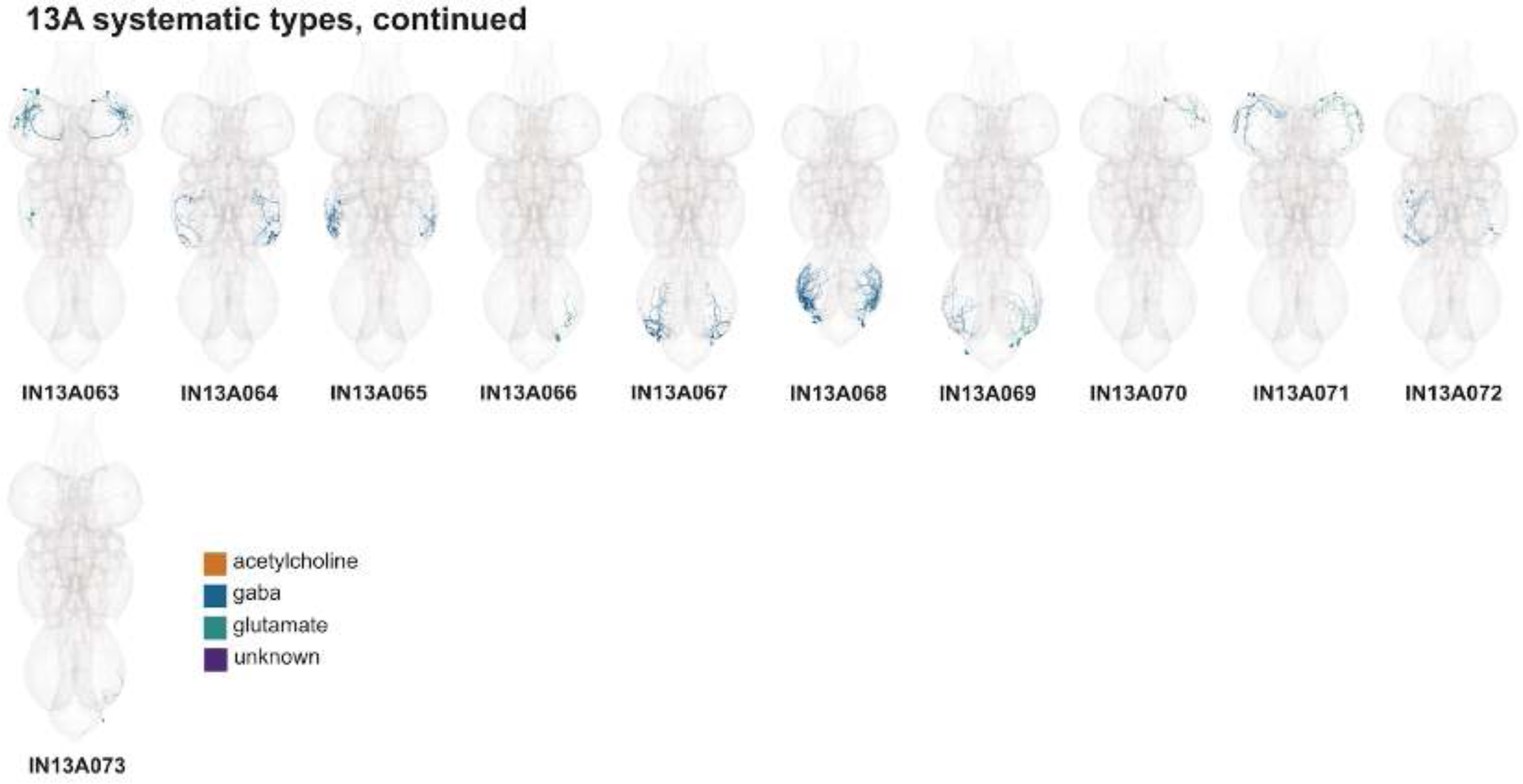
Systematic types of hemilineage 13A, continued. Systematic types have been arranged in numerical order, with neurons of the same type that belong to distinct classes (e.g., intrinsic neuron vs ascending neuron) plotted separately but placed adjacent to each other. Individual neuron meshes have been coloured based on predicted neurotransmitter: dark orange = acetylcholine, blue = gaba, marine = glutamate, dark purple = unknown.

**Figure 35 - figure supplement 4.**
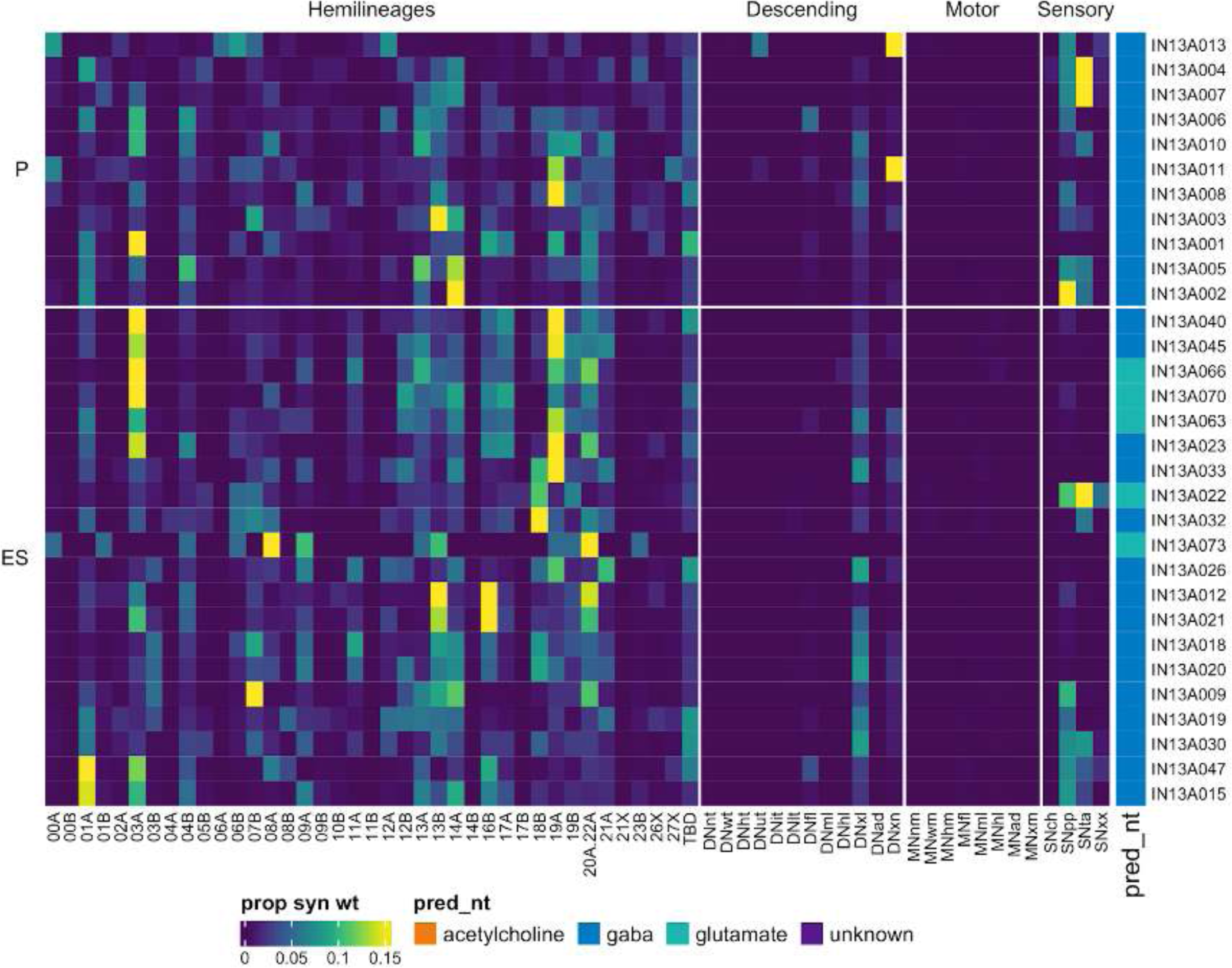
Connectivity to upstream partners by 13A primary and early secondary systematic types. Proportions of synaptic weight to systematic types from upstream partners, normalised by row. 13A neurons have been clustered within each assigned birthtime window (P = primary, ES = early secondary, S = secondary) based on both upstream and downstream connectivity to hemilineages, descending neuron subclasses, motor neuron subclasses, and sensory neuron modalities. Annotation bar is coloured by the most common predicted neurotransmitter for the neurons of each type.

**Figure 35 - figure supplement 5.**
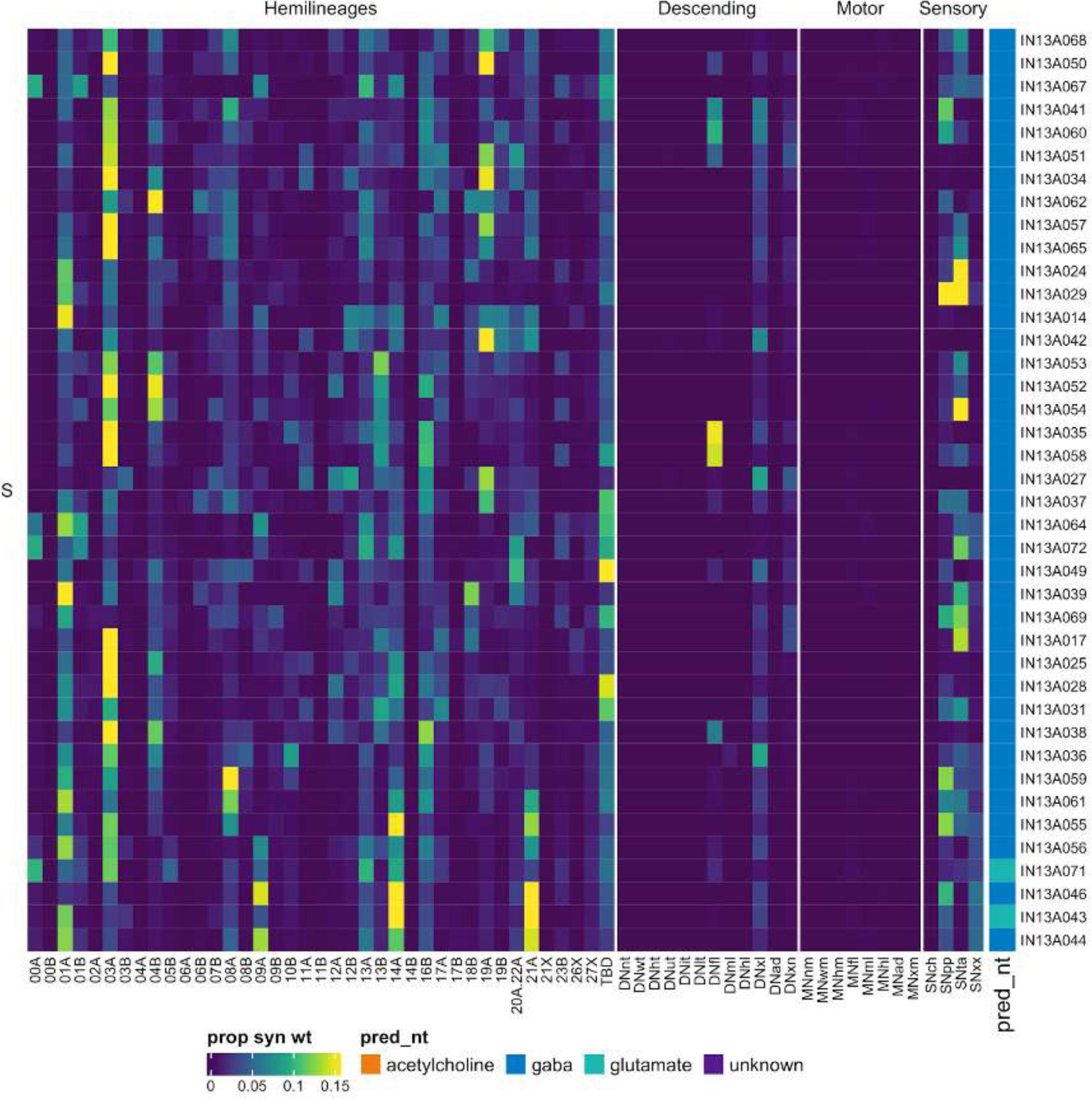
Connectivity to upstream partners by 13A secondary systematic types. Proportions of synaptic weight to systematic types from upstream partners, normalised by row. 13A neurons have been clustered within each assigned birthtime window (P = primary, ES = early secondary, S = secondary) based on both upstream and downstream connectivity to hemilineages, descending neuron subclasses, motor neuron subclasses, and sensory neuron modalities. The annotation bar is coloured by the most common predicted neurotransmitter within each type.

**Figure 35 - figure supplement 6.**
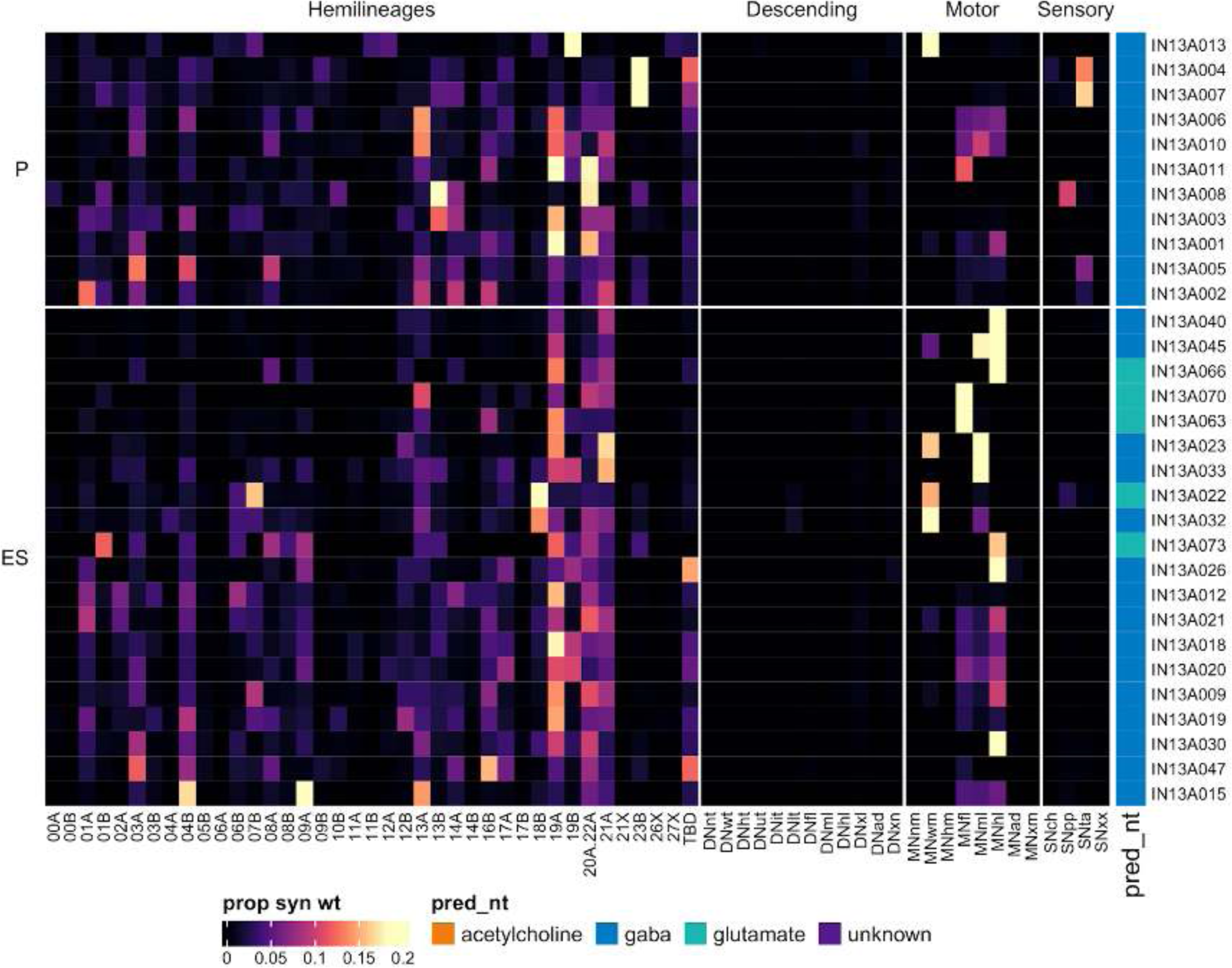
Connectivity to downstream partners by 13A primary and early secondary systematic types. Proportions of synaptic weight from systematic types to downstream partners, normalised by row. 13A neurons have been clustered within each assigned birthtime window (P = primary, ES = early secondary, S = secondary) based on both upstream and downstream connectivity to hemilineages, descending neuron subclasses, motor neuron subclasses, and sensory neuron modalities. The annotation bar is coloured by the most common predicted neurotransmitter within each type.

**Figure 35 - figure supplement 7.**
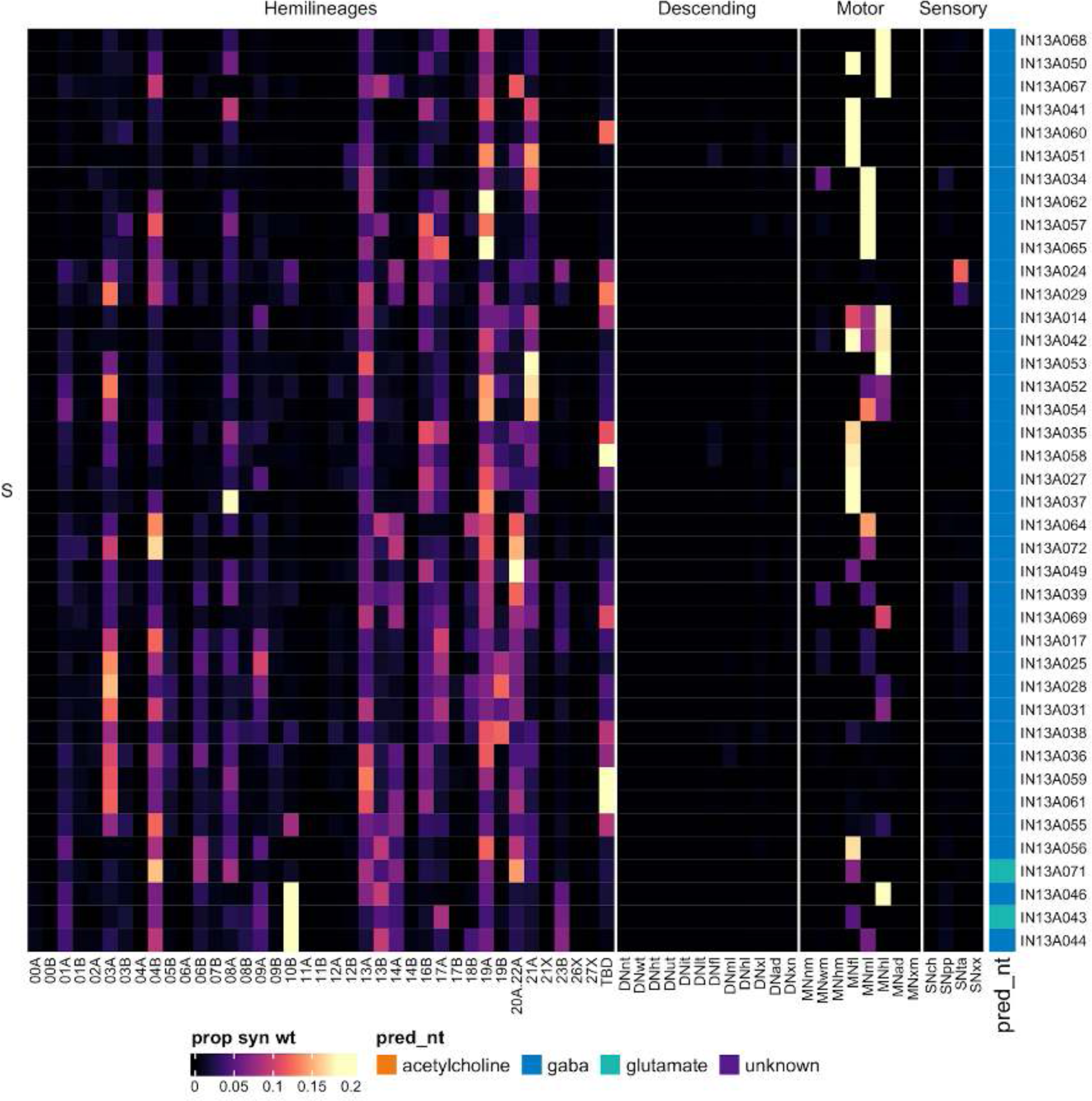
Connectivity to downstream partners by 13A secondary systematic types. Proportions of synaptic weight from systematic types to downstream partners, normalised by row. 13A neurons have been clustered within each assigned birthtime window (P = primary, ES = early secondary, S = secondary) based on both upstream and downstream connectivity to hemilineages, descending neuron subclasses, motor neuron subclasses, and sensory neuron modalities. The annotation bar is coloured by the most common predicted neurotransmitter for the neurons of each type.

#### Hemilineage 13B

Neurons from hemilineage 13B survive in very similar numbers across the thoracic neuromeres (Figure 36E) and cross the midline very ventrally, just posterior to 14A (Figure 8 - figure supplement 1E) (Shepherd et al., 2016; Truman et al., 2004). Secondary 13B neurons innervate the contralateral leg neuropil, and their somas are sometimes pulled across the midline (Shepherd et al., 2019). We opted to use the soma side of origin (as inferred by morphology) for the instance. Almost all 13B neurons are restricted to the contralateral leg neuropil of the same neuromere (e.g., Figure 36C bottom), although we did identify one primary type that spreads into dorsal neuropil (Figure 36C top).

**Figure 36.**
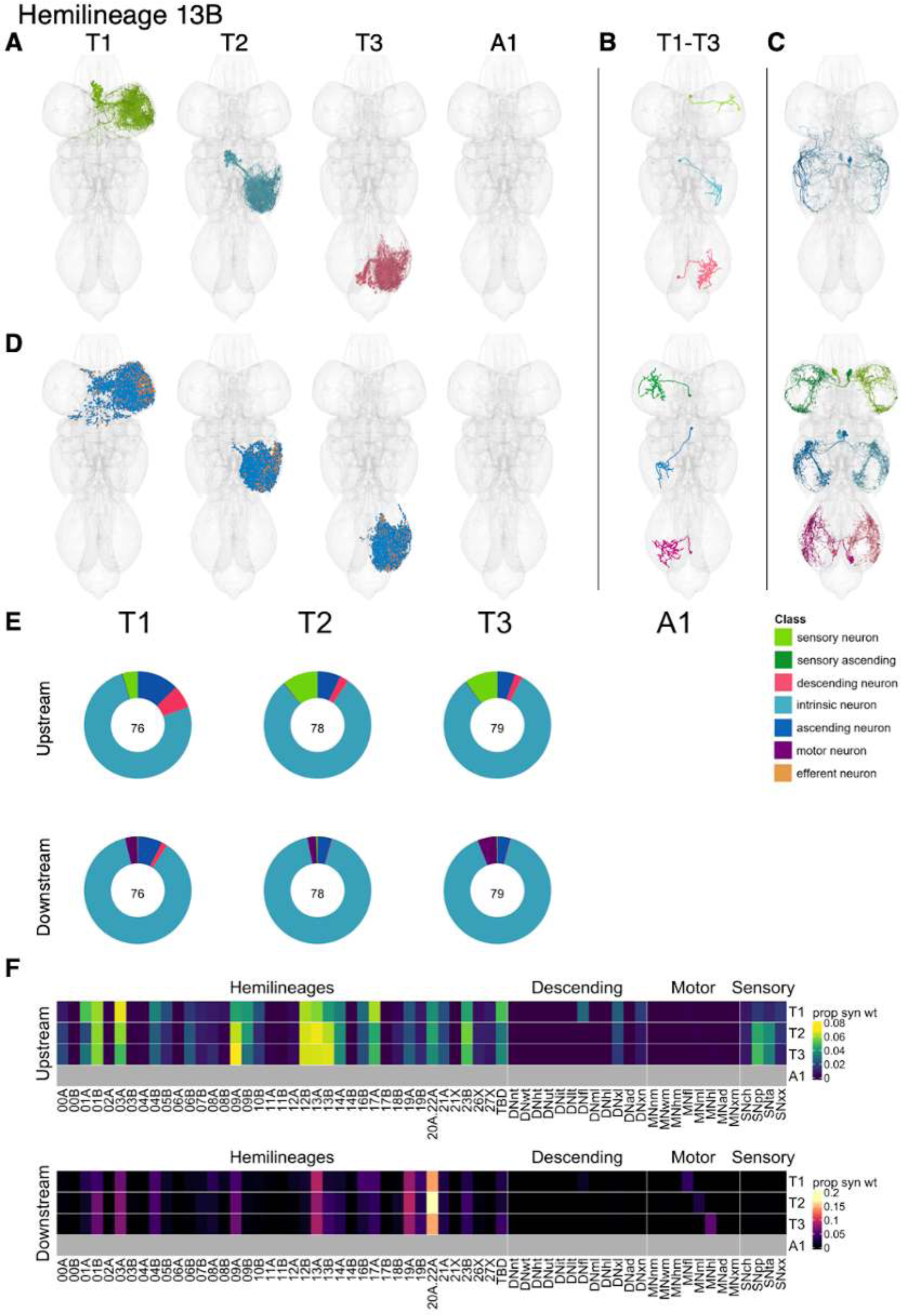
Hemilineage 13B. **A.** Meshes of all RHS secondary neurons plotted in neuromere-specific colours. **B.** “Representative” secondary neuron skeletons plotted in hemineuromere-specific colours. The skeleton with the top accumulated NBLAST score among all neurons from the hemilineage in a given hemineuromere was used. **C.** Neuron meshes of selected examples. Top: group 11111. Bottom: independent leg serial 10116. **D.** Predicted synapses of RHS secondary neurons. Blue: postsynapses; dark orange: presynapses. **E.** Proportions of connections from secondary neurons to upstream or downstream partners, normalised by neuromere and coloured by broad class. Numbers of query neurons appear in the centre. **F.** Proportions of synaptic weight from secondary neurons originating in each neuromere to upstream or downstream partners, normalised by row.

Bilateral activation of 13B secondary neurons results in a tonic postural change with progressive sideways leg extension (Harris et al., 2015). 13B secondary neurons receive significant inputs from proprioceptive and tactile sensory neurons and descending neurons targeting the front leg or multiple leg neuropils, as well as from numerous leg hemilineages, especially 01B, 03A, 09A, 12B, 13A, 13B, 14A, 20A/22A, and 23B, along with 17A (Figure 37F). They are predicted to be gabaergic, as expected (Lacin et al., 2019), and provide output primarily to 20A/22A and to a lesser extent to 01B, 03A, 09A, 13A, and 19A. Several 13B types (e.g., IN13B012 and IN13B048) inhibit leg motor neurons (Figure 36 - figure supplement 7-8); IN13B006 mainly targets hind leg MNs and is under strong descending control. A single primary type, IN13B008, inhibits wing motor neurons instead (Figure 36 - figure supplement 6).

**Figure 36 - figure supplement 1.**
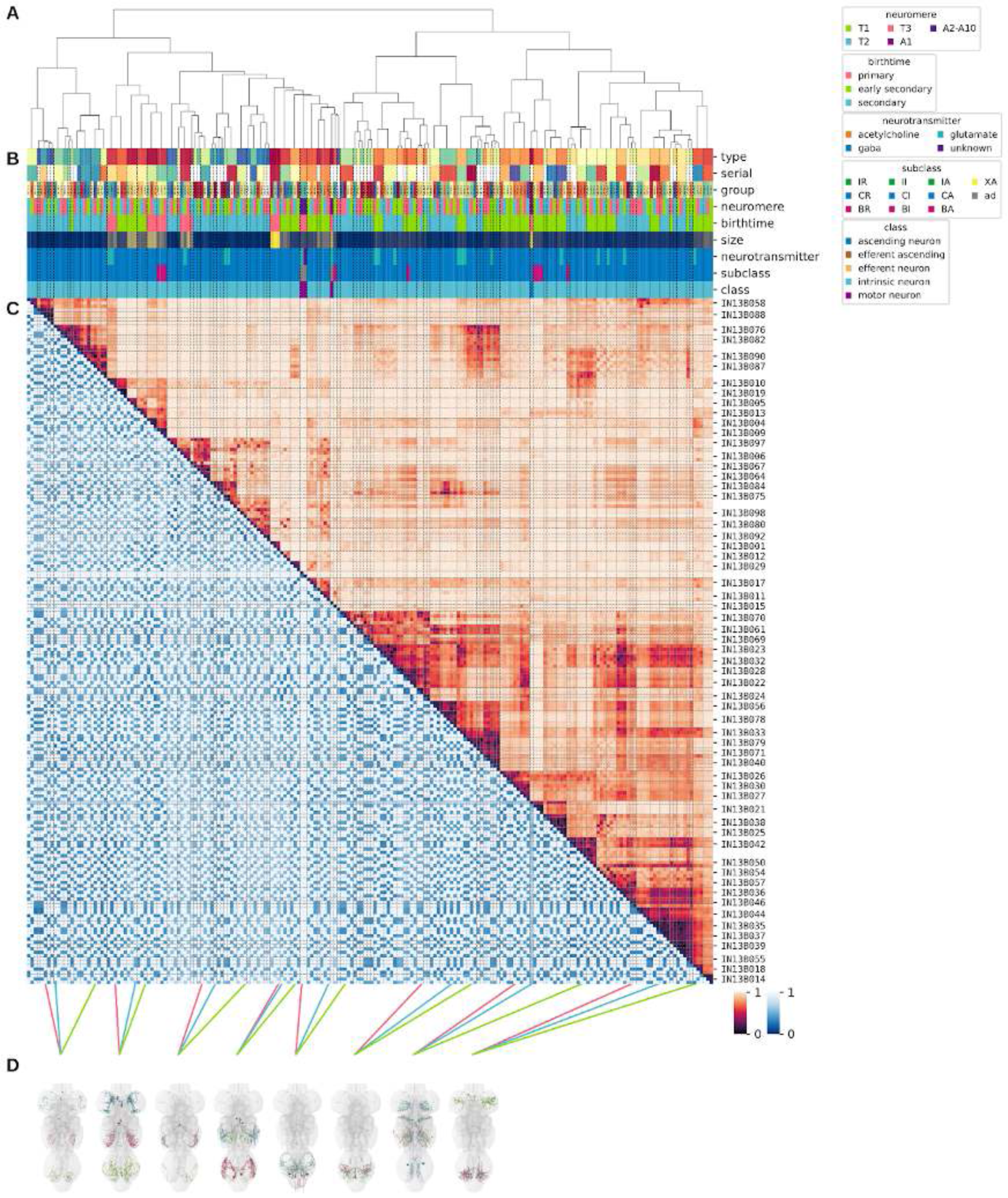
Systematic typing of hemilineage 13B. **A.** Hierarchical clustering dendrogram of hemilineage groups by laterally and serially aggregated connectivity cosine clustering. **B.** Categorical annotations of each hemilineage group, each column corresponding to the aligned leaf in A. Colours for type, serial set, and group are arbitrary for visualisation. Colours for neuromere, birthtime, neurotransmitter, subclass, and class are as in all other figures. **C.** Similarity distance heatmap for hemilineage. Cosine distance is in the upper triangle, while laterally symmetrised NBLAST distance is in the lower triangle. Systematic type names of some types are labelled. **D.** Morphologically representative groups from dendrogram subtrees. Each group, indicated by colour and line connecting to its column in B and C, is the most morphologically representative group (medoid of NBLAST distance) from a subtree of A. The subtrees (flat clusters) are equal height cuts of A determined to yield the number of groups per plot and plots in D.

**Figure 36 - figure supplement 2.**
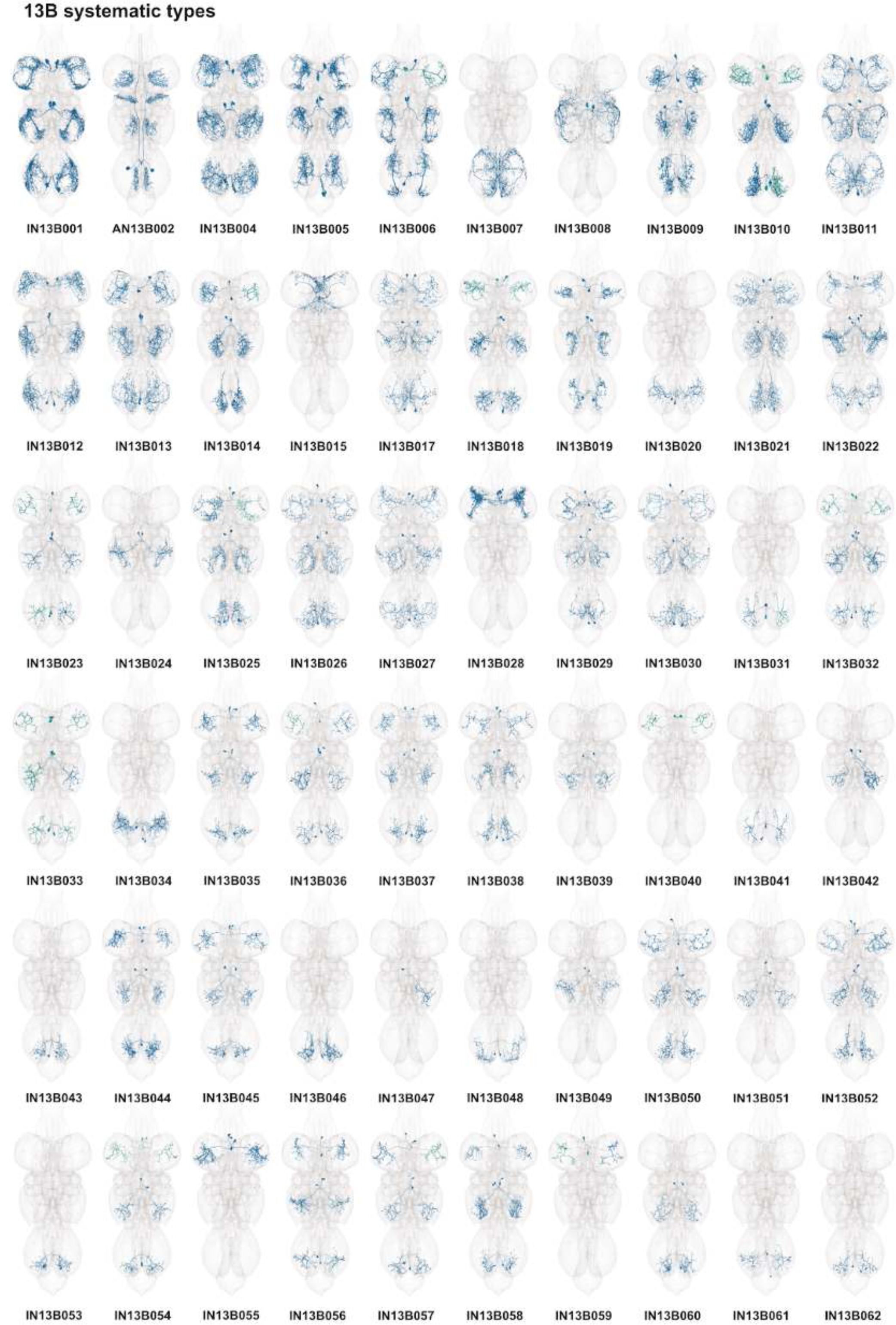
Systematic types of hemilineage 13B. Systematic types have been arranged in numerical order, with neurons of the same type that belong to distinct classes (e.g., intrinsic neuron vs ascending neuron) plotted separately but placed adjacent to each other. Individual neuron meshes have been coloured based on predicted neurotransmitter: dark orange = acetylcholine, blue = gaba, marine = glutamate, dark purple = unknown.

**Figure 36 - figure supplement 3.**
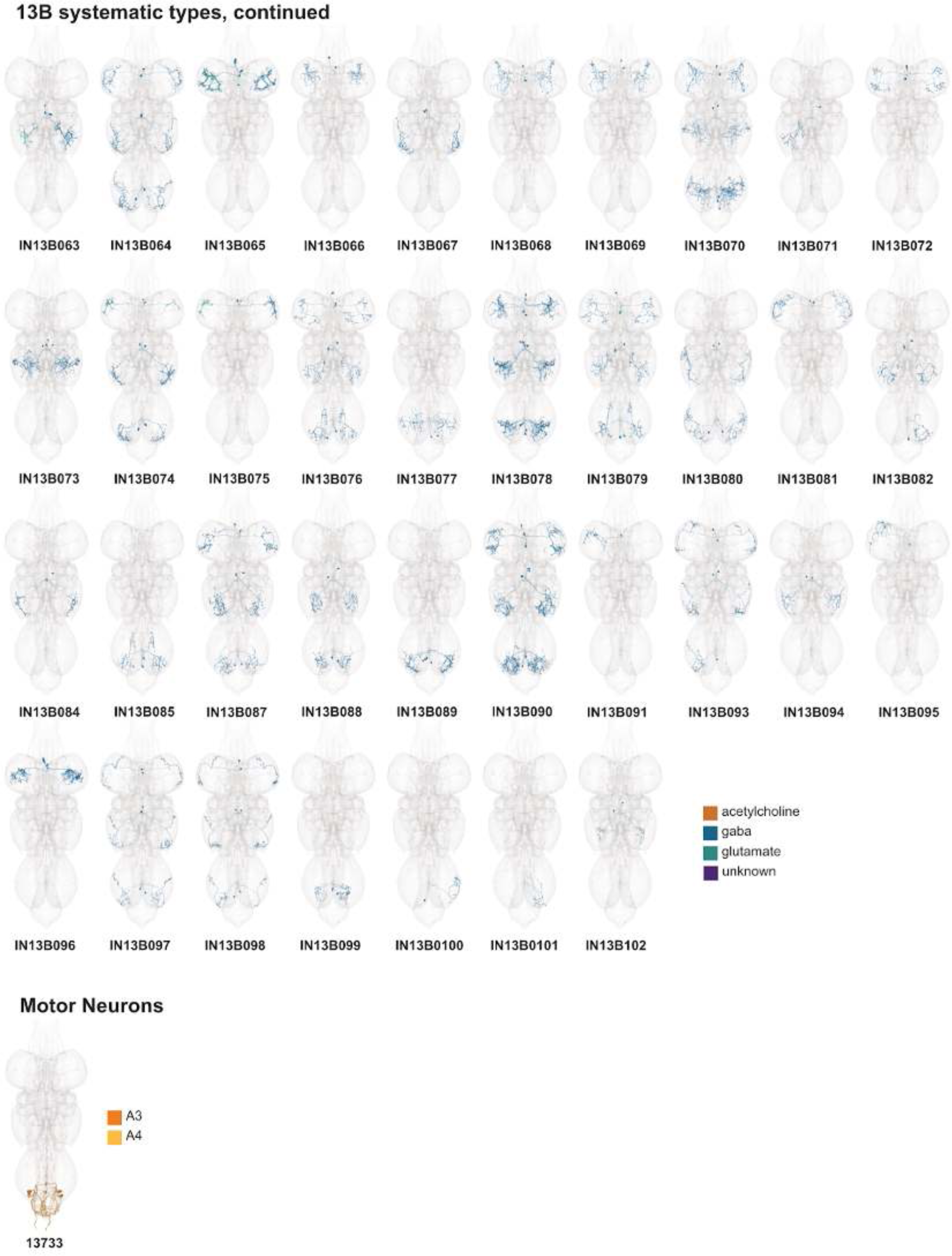
Systematic types of hemilineage 13B, continued. Systematic types have been arranged in numerical order, with neurons of the same type that belong to distinct classes (e.g., intrinsic neuron vs ascending neuron) plotted separately but placed adjacent to each other. Individual neuron meshes have been coloured based on predicted neurotransmitter: dark orange = acetylcholine, blue = gaba, marine = glutamate, dark purple = unknown. Motor neurons (typed separately in (Cheong et al., 2023)) have been plotted by serial set if identified in multiple neuromeres and by systematic type if not. Individual motor neuron meshes have been coloured based on soma neuromere: dark orange = A3, dark yellow = A4.

**Figure 36 - figure supplement 4.**
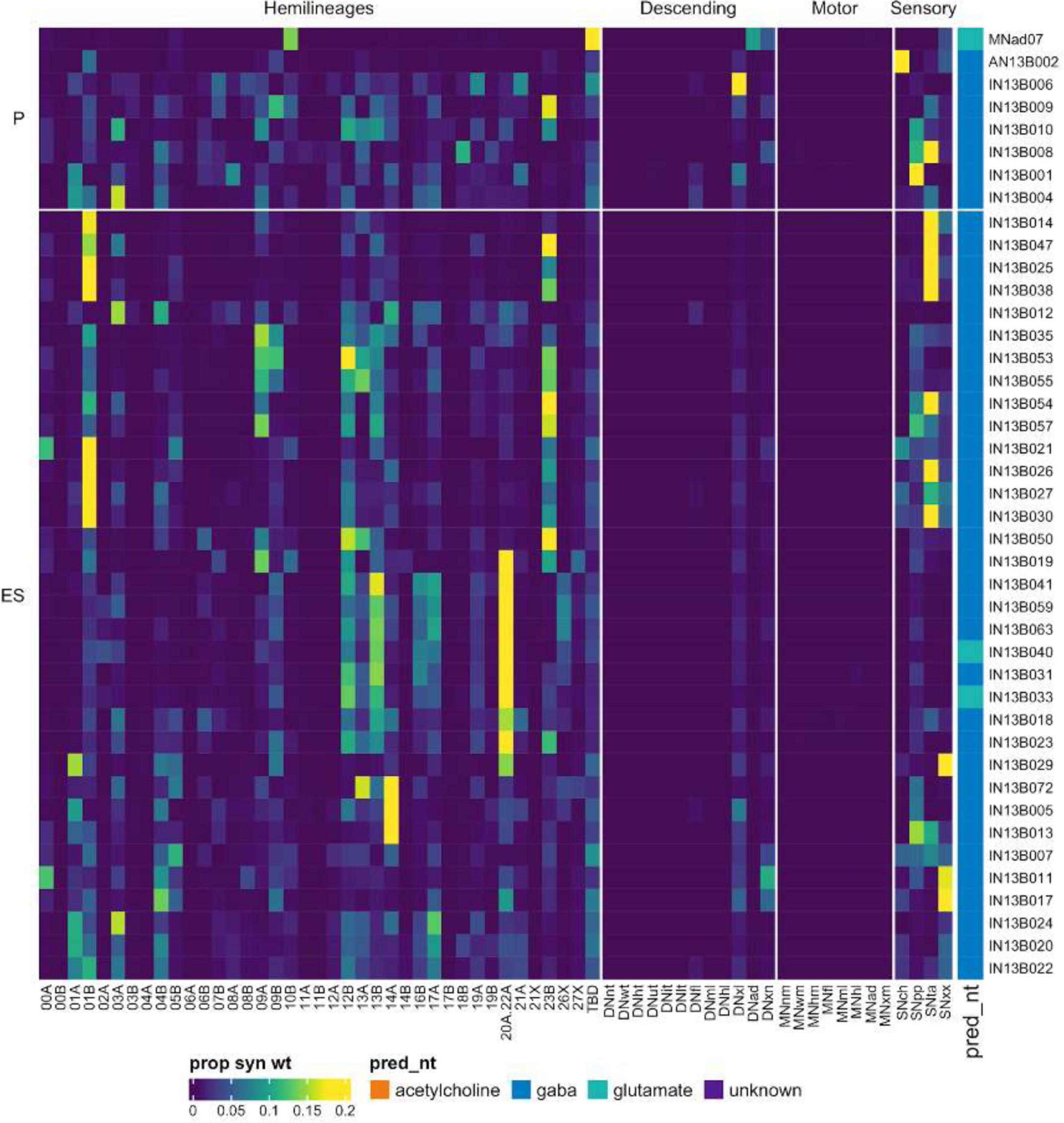
Connectivity to upstream partners by 13B primary and early secondary systematic types. Proportions of synaptic weight to systematic types from upstream partners, normalised by row. 13B neurons have been clustered within each assigned birthtime window (P = primary, ES = early secondary, S = secondary) based on both upstream and downstream connectivity to hemilineages, descending neuron subclasses, motor neuron subclasses, and sensory neuron modalities. Annotation bar is coloured by the most common predicted neurotransmitter for the neurons of each type.

**Figure 36 - figure supplement 5.**
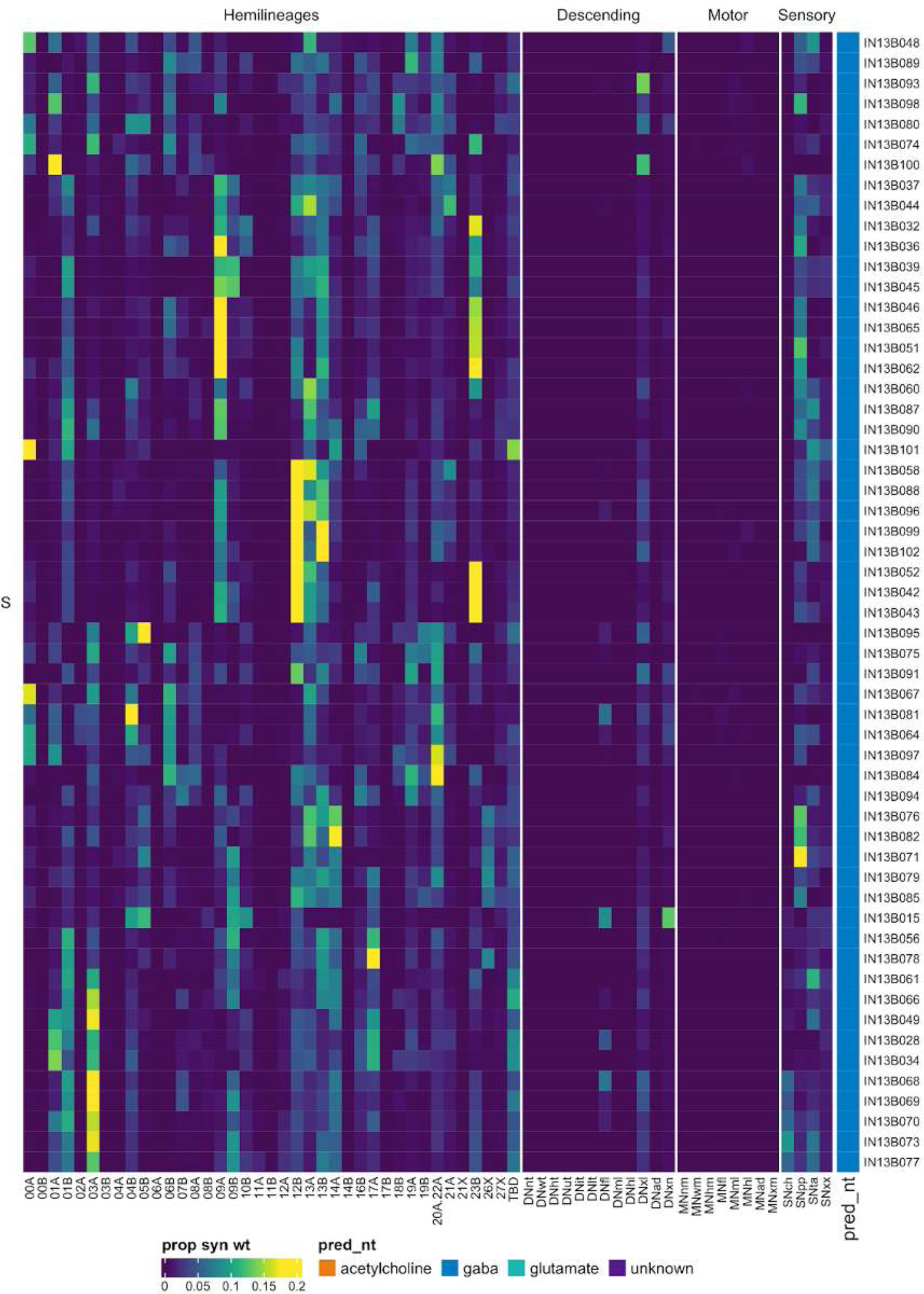
Connectivity to upstream partners by 13B secondary systematic types. Proportions of synaptic weight to systematic types from upstream partners, normalised by row. 13B neurons have been clustered within each assigned birthtime window (P = primary, ES = early secondary, S = secondary) based on both upstream and downstream connectivity to hemilineages, descending neuron subclasses, motor neuron subclasses, and sensory neuron modalities. The annotation bar is coloured by the most common predicted neurotransmitter within each type.

**Figure 36 - figure supplement 6.**
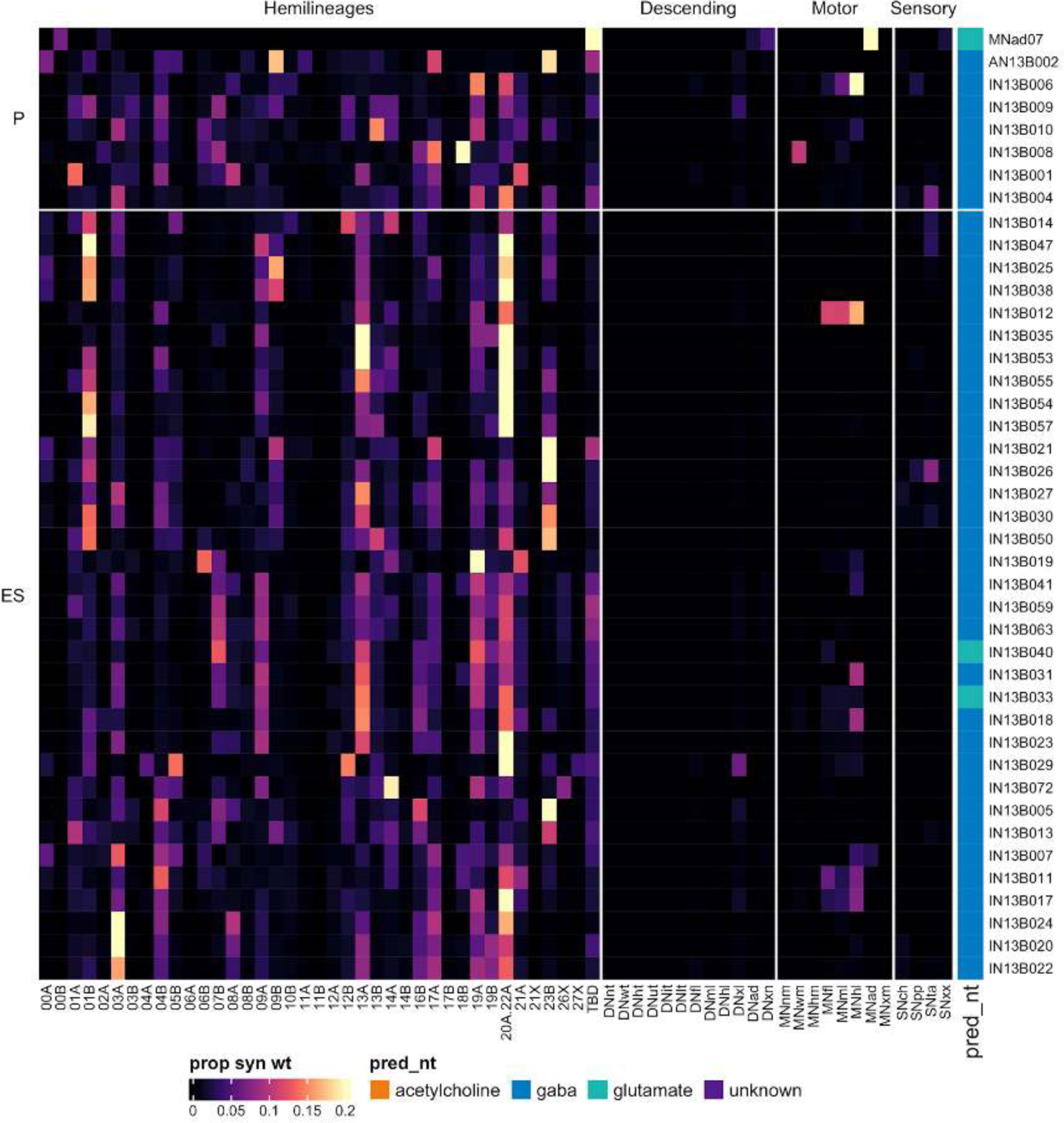
Connectivity to downstream partners by 13B primary and early secondary systematic types. Proportions of synaptic weight from systematic types to downstream partners, normalised by row. 13B neurons have been clustered within each assigned birthtime window (P = primary, ES = early secondary, S = secondary) based on both upstream and downstream connectivity to hemilineages, descending neuron subclasses, motor neuron subclasses, and sensory neuron modalities. The annotation bar is coloured by the most common predicted neurotransmitter within each type.

**Figure 36 - figure supplement 7.**
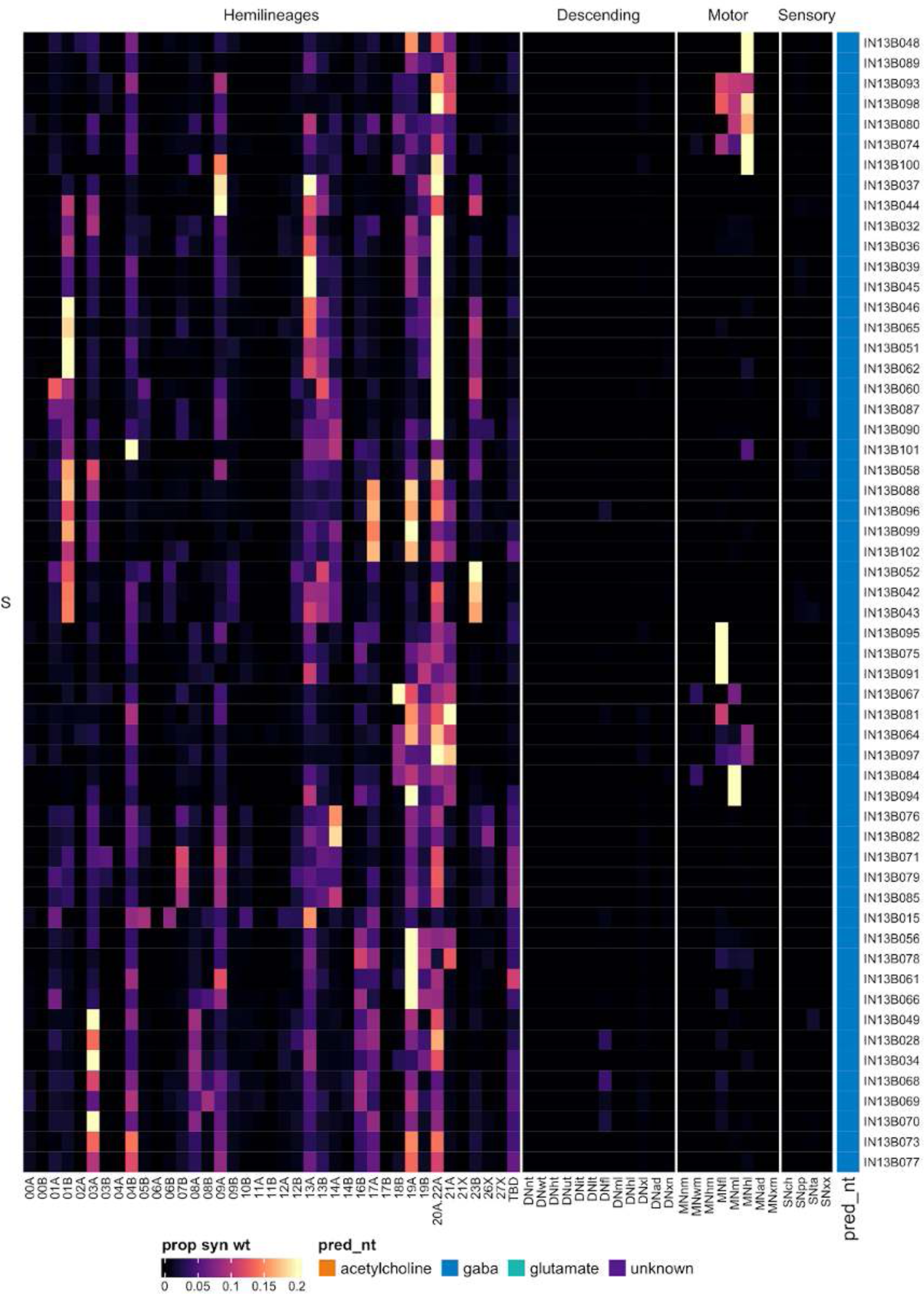
Connectivity to downstream partners by 13B secondary systematic types. Proportions of synaptic weight from systematic types to downstream partners, normalised by row. 13B neurons have been clustered within each assigned birthtime window (P = primary, ES = early secondary, S = secondary) based on both upstream and downstream connectivity to hemilineages, descending neuron subclasses, motor neuron subclasses, and sensory neuron modalities. The annotation bar is coloured by the most common predicted neurotransmitter for the neurons of each type.

#### Hemilineage 14A

Hemilineages 14A and 14B derive from medial neuroblast NB4-1 (Birkholz et al., 2015; Lacin and Truman, 2016), which generates the motor neuron of the transverse nerve, ∼8 intersegmental interneurons, and 20-24 local interneurons in the embryo (Schmid et al., 1999). Hemilineage 14A secondary neurons survive in ∼identical numbers across the thoracic neuromeres and cross the midline very ventrally, just anterior to 13B (Shepherd et al., 2016; Truman et al., 2004) (Figure 37A,E). 14A secondary neurons innervate the contralateral leg neuropil, and their somas are sometimes pulled across the midline (Shepherd et al., 2019). We opted to use the soma side of origin (as inferred by morphology) for the instance. Most 14A neurons are restricted to the contralateral leg neuropil of the same neuromere (Figure 37C top), but we identified one ascending serial set of primary neurons (Figure 37C bottom).

**Figure 37.**
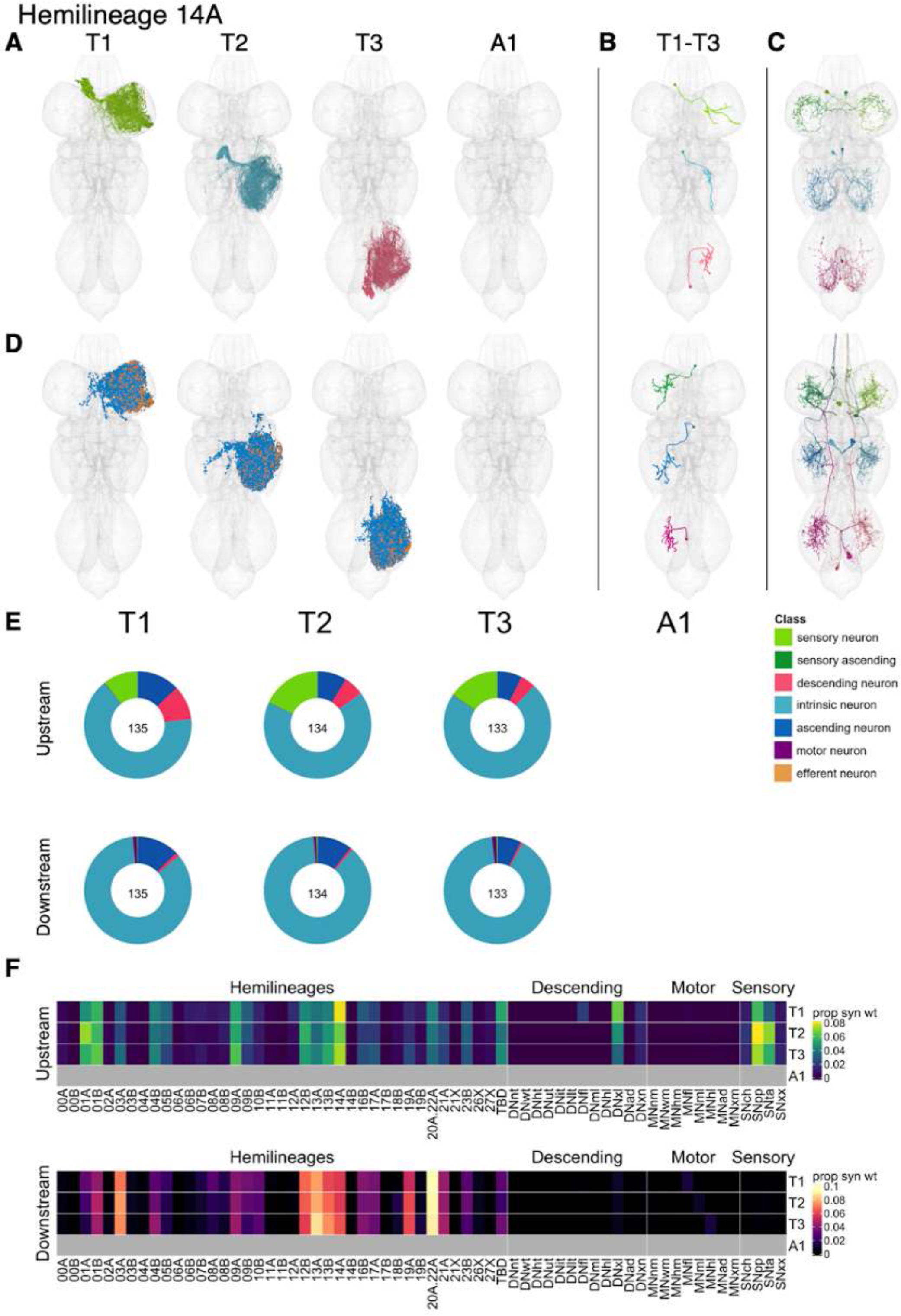
Hemilineage 14A. **A.** Meshes of all RHS secondary neurons plotted in neuromere-specific colours. **B.** “Representative” secondary neuron skeletons plotted in hemineuromere-specific colours. The skeleton with the top accumulated NBLAST score among all neurons from the hemilineage in a given hemineuromere was used. **C.** Neuron meshes of selected examples. Top: independent leg serial set 10019. Bottom: ascending serial set 10411. **D.** Predicted synapses of RHS secondary neurons. Blue: postsynapses; dark orange: presynapses. **E.** Proportions of connections from secondary neurons to upstream or downstream partners, normalised by neuromere and coloured by broad class. Numbers of query neurons appear in the centre. **F.** Proportions of synaptic weight from secondary neurons originating in each neuromere to upstream or downstream partners, normalised by row.

14A neurons receive significant inputs from proprioceptive and tactile sensory neurons and descending neurons targeting multiple leg neuropils, as well as from numerous leg hemilineages, especially 01A, 01B, 09A, 12B, 13B, and 14A (Figure 37F). 14A secondary neurons are predicted to be glutamatergic, as expected (Lacin et al., 2019), and provide output to hemilineages 20A/22A and to a lesser extent to 03A, 12B, 13A, 13B, 14A, and 19A. However, we identified a distinctive primary serial set of gabergic neurons (IN14A001) that receives input from 04B, 13A, and proprioceptive neurons and inhibits neurons from 13B and 21A. We also identified a single 14A ascending type (AN14A003); these neurons receive input primarily from 12B and 21A and provide it to 07B and 19A (Figure 38 - figure supplement 2,5,8). No functional studies have been published for secondary 14A neurons.

**Figure 37 - figure supplement 1.**
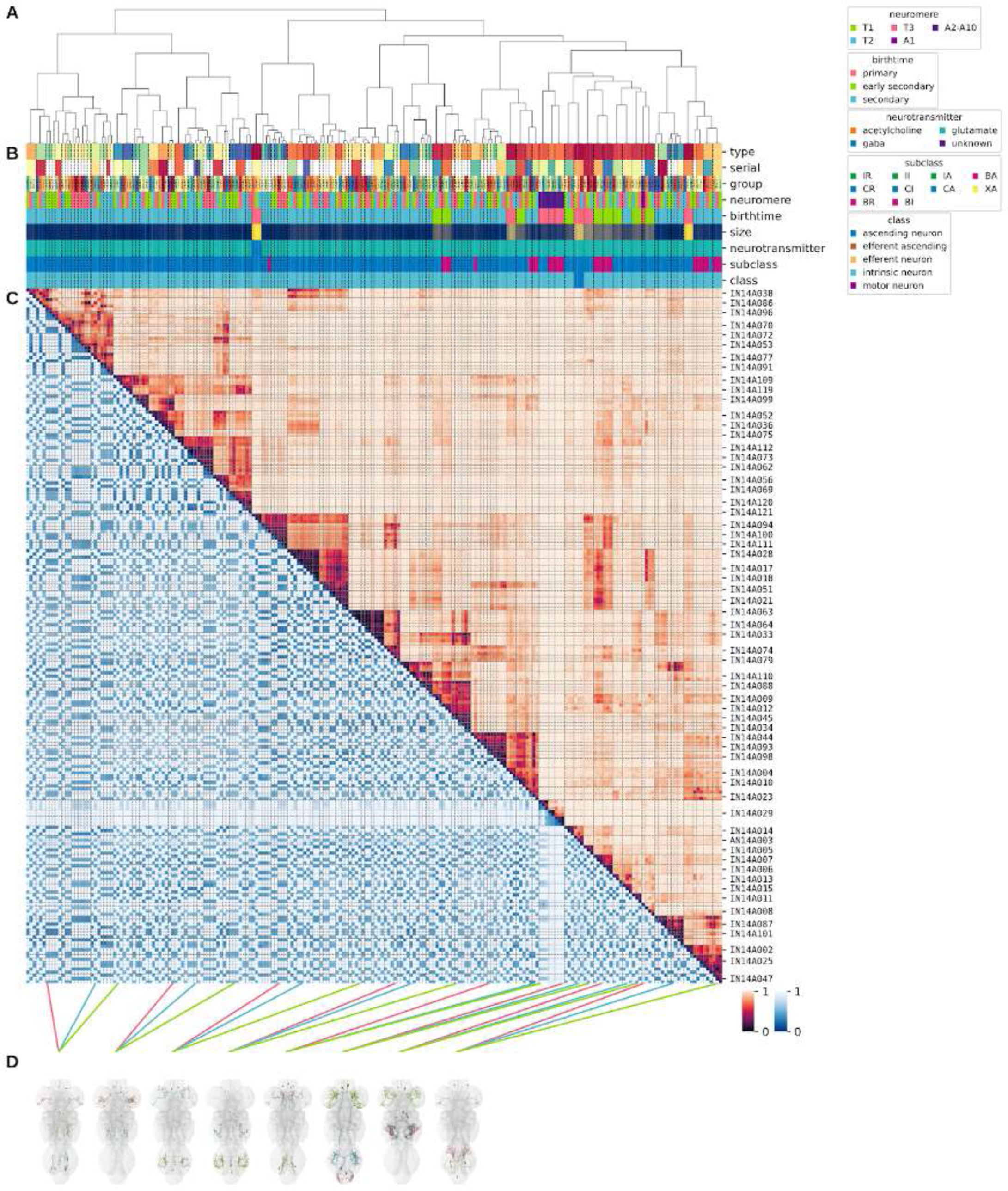
Systematic typing of hemilineage 14A. **A.** Hierarchical clustering dendrogram of hemilineage groups by laterally and serially aggregated connectivity cosine clustering. **B.** Categorical annotations of each hemilineage group, each column corresponding to the aligned leaf in A. Colours for type, serial set, and group are arbitrary for visualisation. Colours for neuromere, birthtime, neurotransmitter, subclass, and class are as in all other figures. **C.** Similarity distance heatmap for hemilineage. Cosine distance is in the upper triangle, while laterally symmetrised NBLAST distance is in the lower triangle. Systematic type names of some types are labelled. **D.** Morphologically representative groups from dendrogram subtrees. Each group, indicated by colour and line connecting to its column in B and C, is the most morphologically representative group (medoid of NBLAST distance) from a subtree of A. The subtrees (flat clusters) are equal height cuts of A determined to yield the number of groups per plot and plots in D.

**Figure 37 - figure supplement 2.**
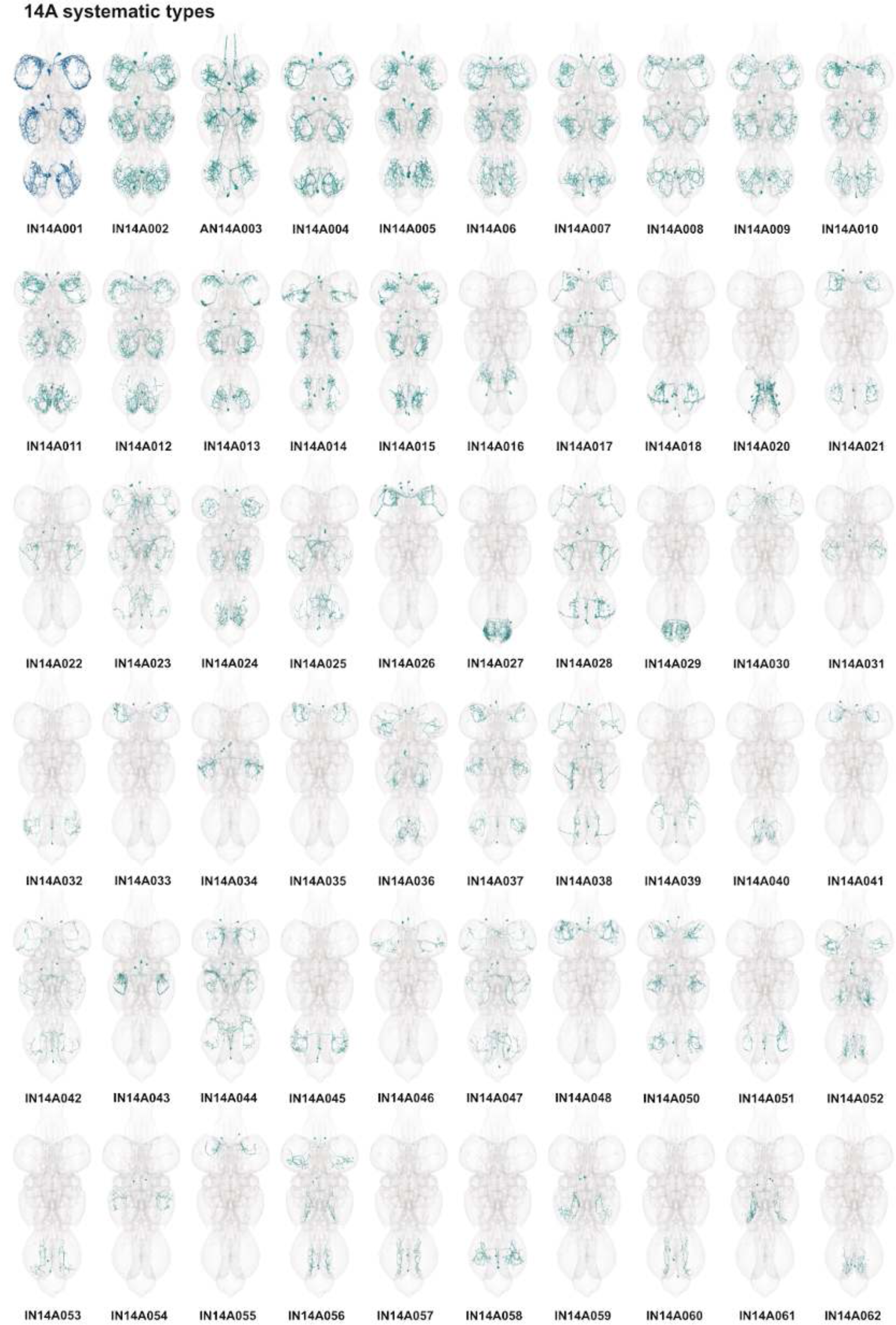
Systematic types of hemilineage 14A. Systematic types have been arranged in numerical order, with neurons of the same type that belong to distinct classes (e.g., intrinsic neuron vs ascending neuron) plotted separately but placed adjacent to each other. Individual neuron meshes have been coloured based on predicted neurotransmitter: dark orange = acetylcholine, blue = gaba, marine = glutamate, dark purple = unknown.

**Figure 37 - figure supplement 3.**
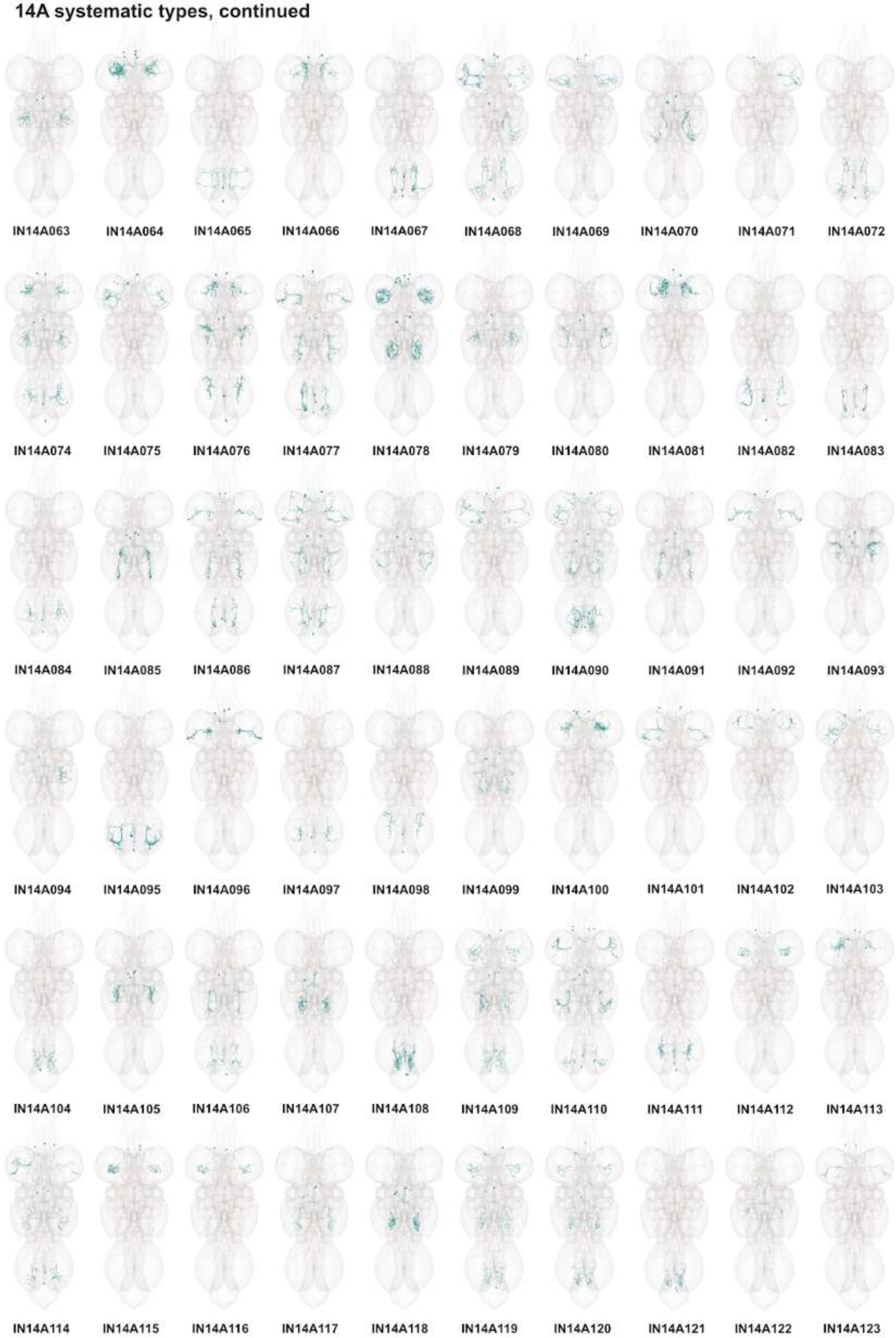
Systematic types of hemilineage 14A, continued. Systematic types have been arranged in numerical order, with neurons of the same type that belong to distinct classes (e.g., intrinsic neuron vs ascending neuron) plotted separately but placed adjacent to each other. Individual neuron meshes have been coloured based on predicted neurotransmitter: dark orange = acetylcholine, blue = gaba, marine = glutamate, dark purple = unknown.

**Figure 37 - figure supplement 4.**
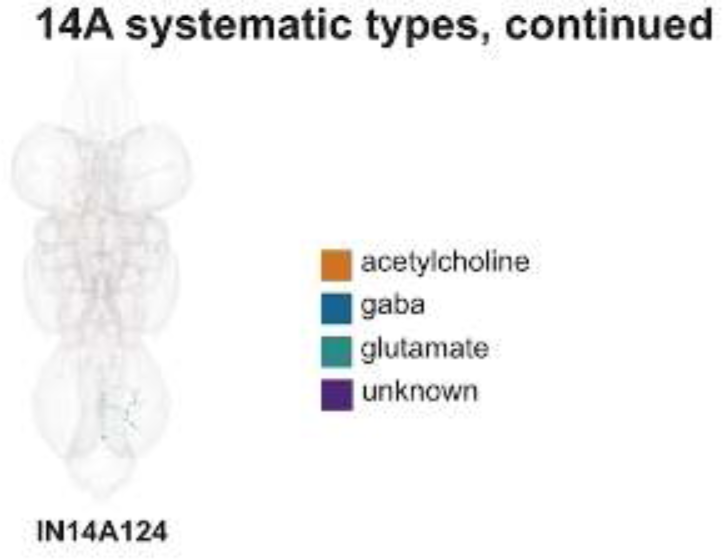
Systematic types of hemilineage 14A, continued. Systematic types have been arranged in numerical order, with neurons of the same type that belong to distinct classes (e.g., intrinsic neuron vs ascending neuron) plotted separately but placed adjacent to each other. Individual neuron meshes have been coloured based on predicted neurotransmitter: dark orange = acetylcholine, blue = gaba, marine = glutamate, dark purple = unknown.

**Figure 37 - figure supplement 5.**
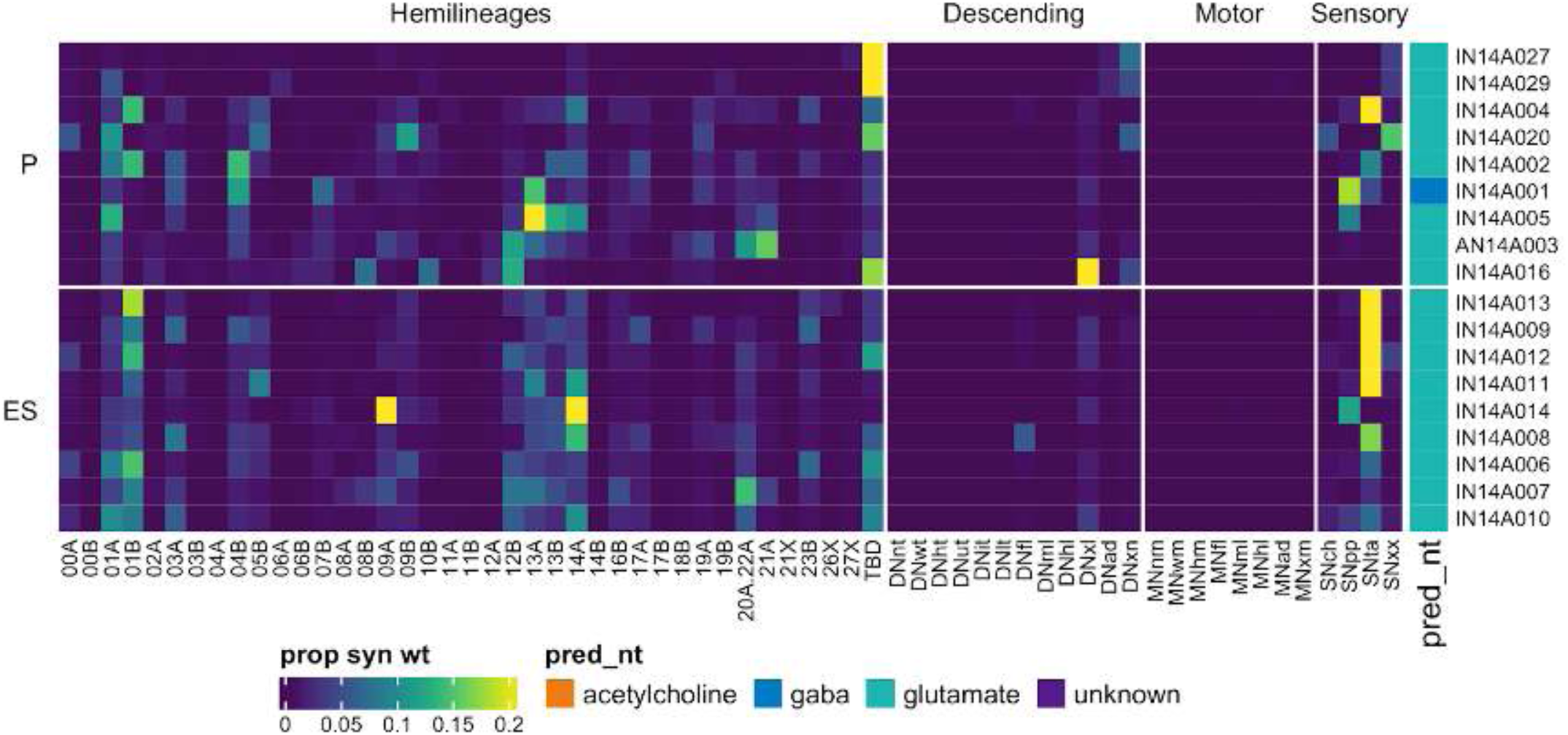
Connectivity to upstream partners by 14A primary and early secondary systematic types. Proportions of synaptic weight to systematic types from upstream partners, normalised by row. 14A neurons have been clustered within each assigned birthtime window (P = primary, ES = early secondary, S = secondary) based on both upstream and downstream connectivity to hemilineages, descending neuron subclasses, motor neuron subclasses, and sensory neuron modalities. Annotation bar is coloured by the most common predicted neurotransmitter for the neurons of each type.

**Figure 37 - figure supplement 6.**
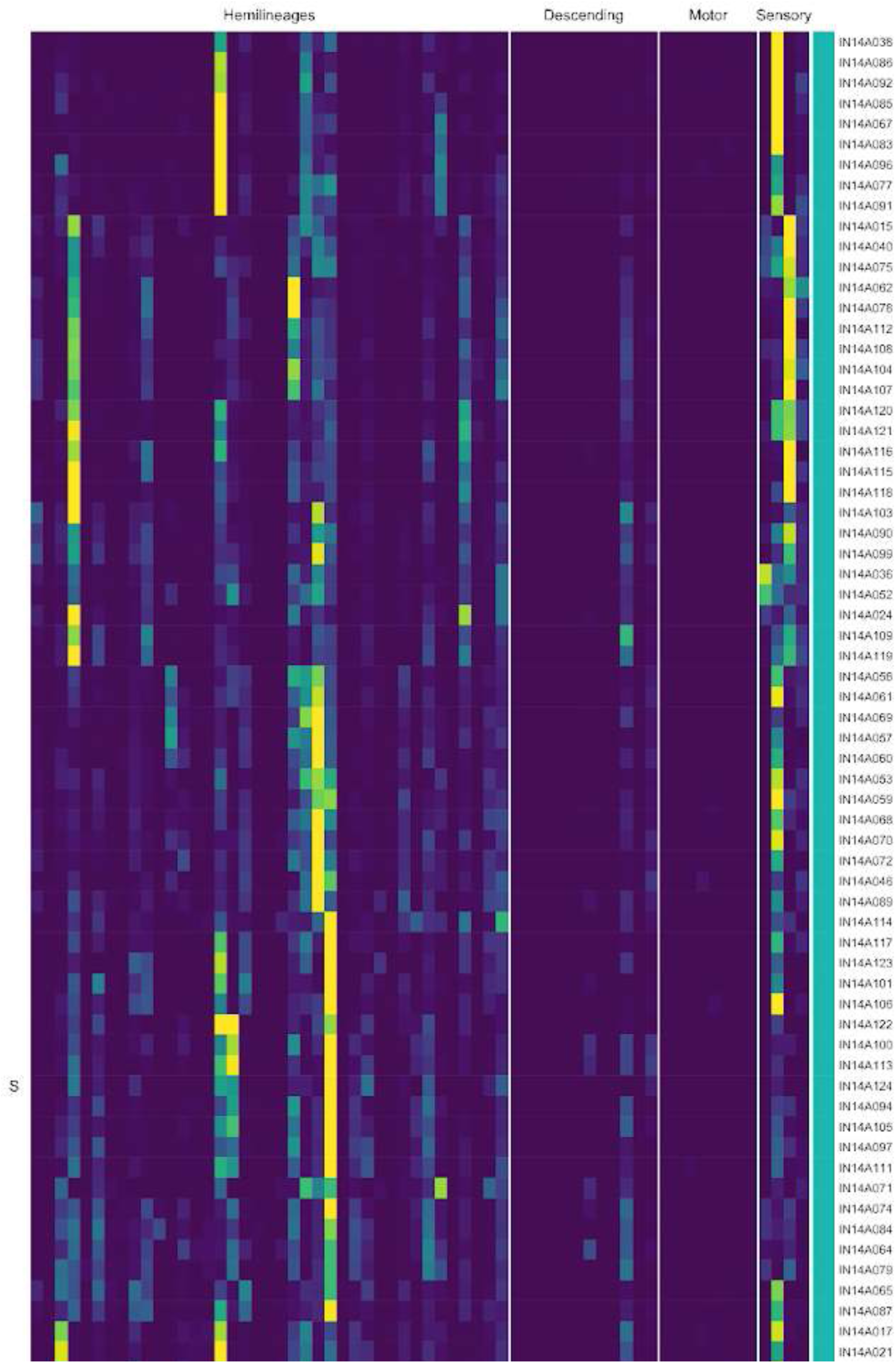
Connectivity to upstream partners by 14A secondary systematic types. Proportions of synaptic weight to systematic types from upstream partners, normalised by row. 14A neurons have been clustered within each assigned birthtime window (P = primary, ES = early secondary, S = secondary) based on both upstream and downstream connectivity to hemilineages, descending neuron subclasses, motor neuron subclasses, and sensory neuron modalities. The annotation bar is coloured by the most common predicted neurotransmitter within each type.

**Figure 37 - figure supplement 7.**
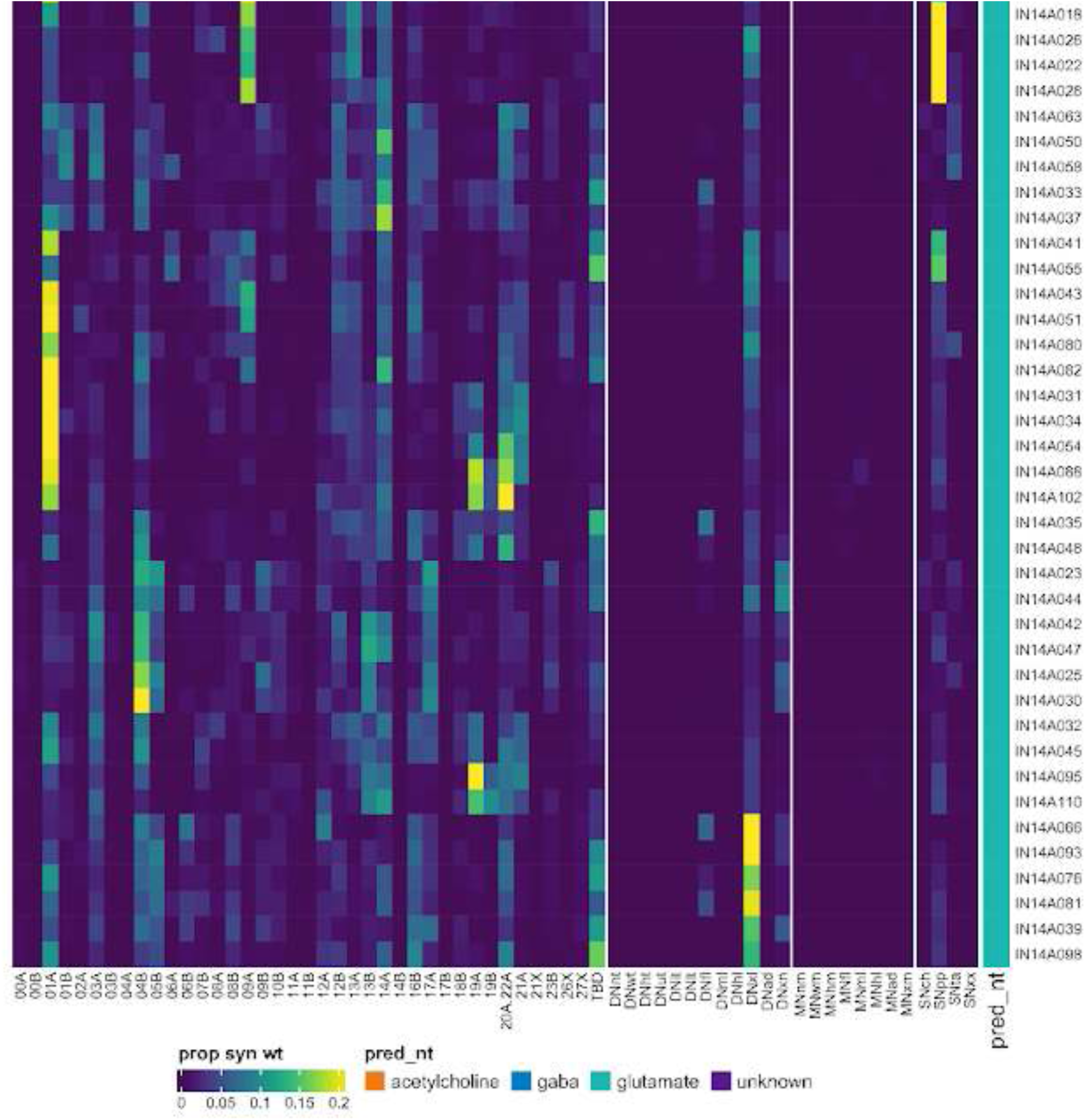
Connectivity to upstream partners by 14A secondary systematic types, continued. Proportions of synaptic weight to systematic types from upstream partners, normalised by row. 14A neurons have been clustered within each assigned birthtime window (P = primary, ES = early secondary, S = secondary) based on both upstream and downstream connectivity to hemilineages, descending neuron subclasses, motor neuron subclasses, and sensory neuron modalities. The annotation bar is coloured by the most common predicted neurotransmitter within each type.

**Figure 37 - figure supplement 8.**
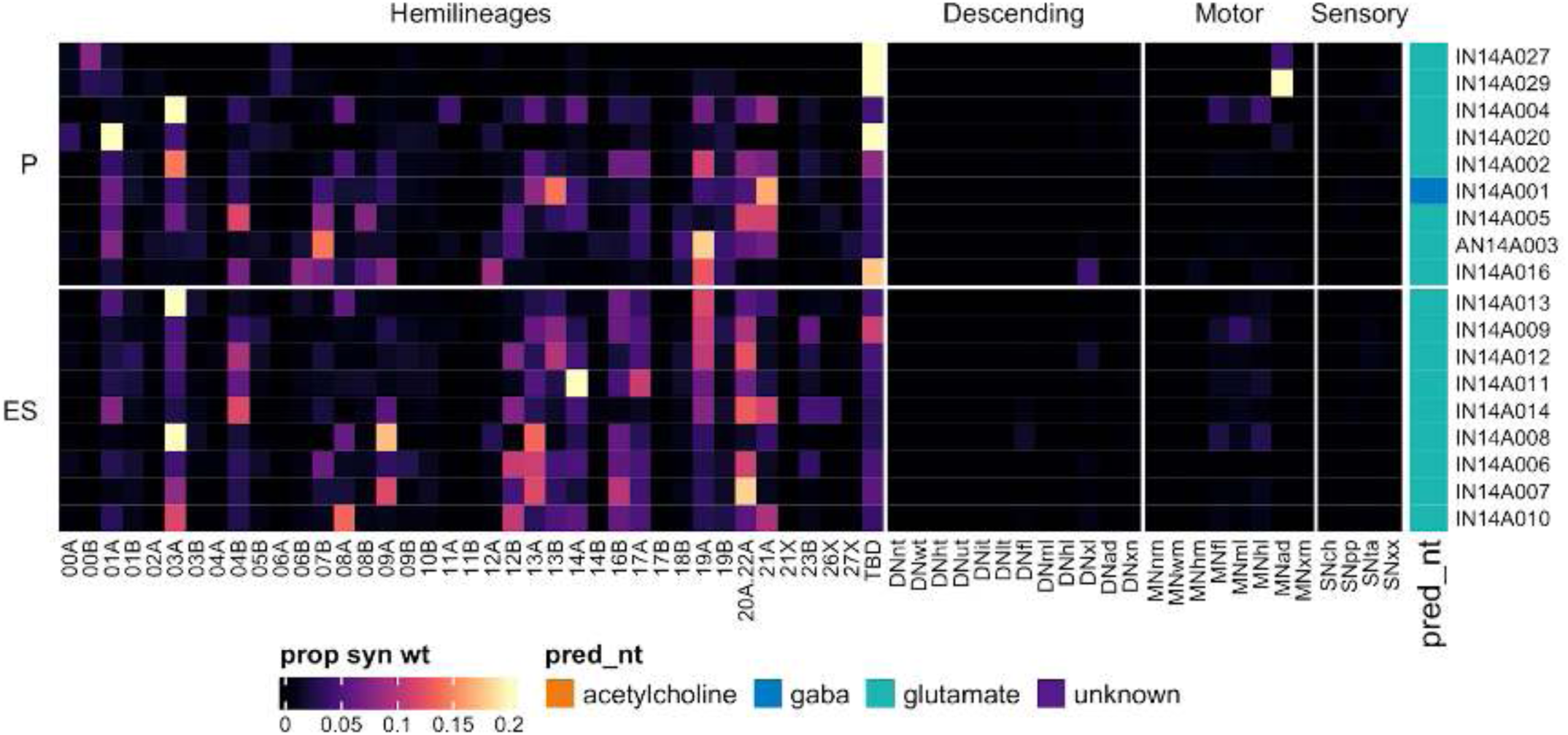
Connectivity to downstream partners by 14A primary and early secondary systematic types. Proportions of synaptic weight from systematic types to downstream partners, normalised by row. 14A neurons have been clustered within each assigned birthtime window (P = primary, ES = early secondary, S = secondary) based on both upstream and downstream connectivity to hemilineages, descending neuron subclasses, motor neuron subclasses, and sensory neuron modalities. The annotation bar is coloured by the most common predicted neurotransmitter within each type.

**Figure 37 - figure supplement 9.**
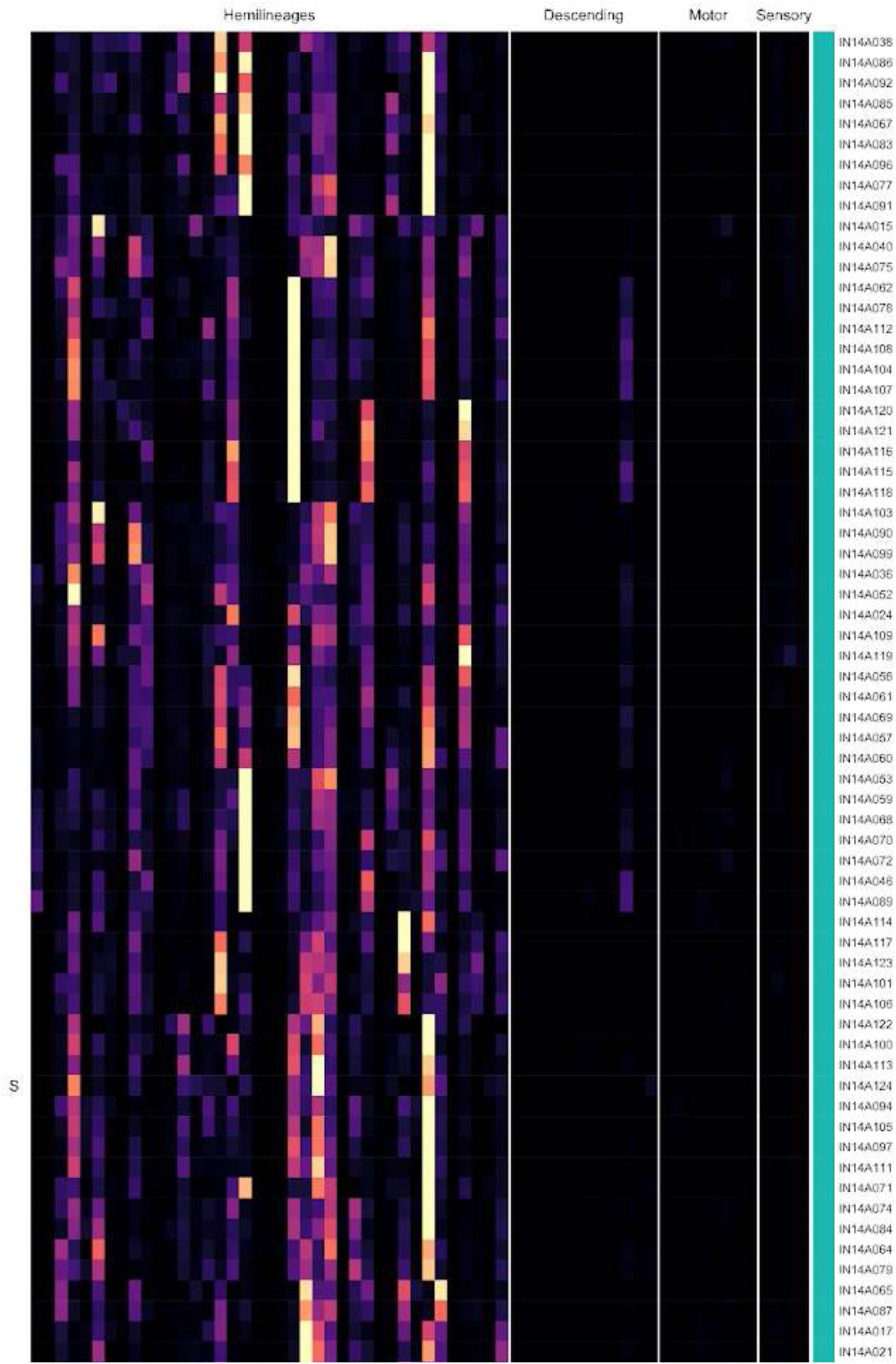
Connectivity to downstream partners by 14A secondary systematic types. Proportions of synaptic weight from systematic types to downstream partners, normalised by row. 14A neurons have been clustered within each assigned birthtime window (P = primary, ES = early secondary, S = secondary) based on both upstream and downstream connectivity to hemilineages, descending neuron subclasses, motor neuron subclasses, and sensory neuron modalities. The annotation bar is coloured by the most common predicted neurotransmitter for the neurons of each type.

**Figure 37 - figure supplement 10.**
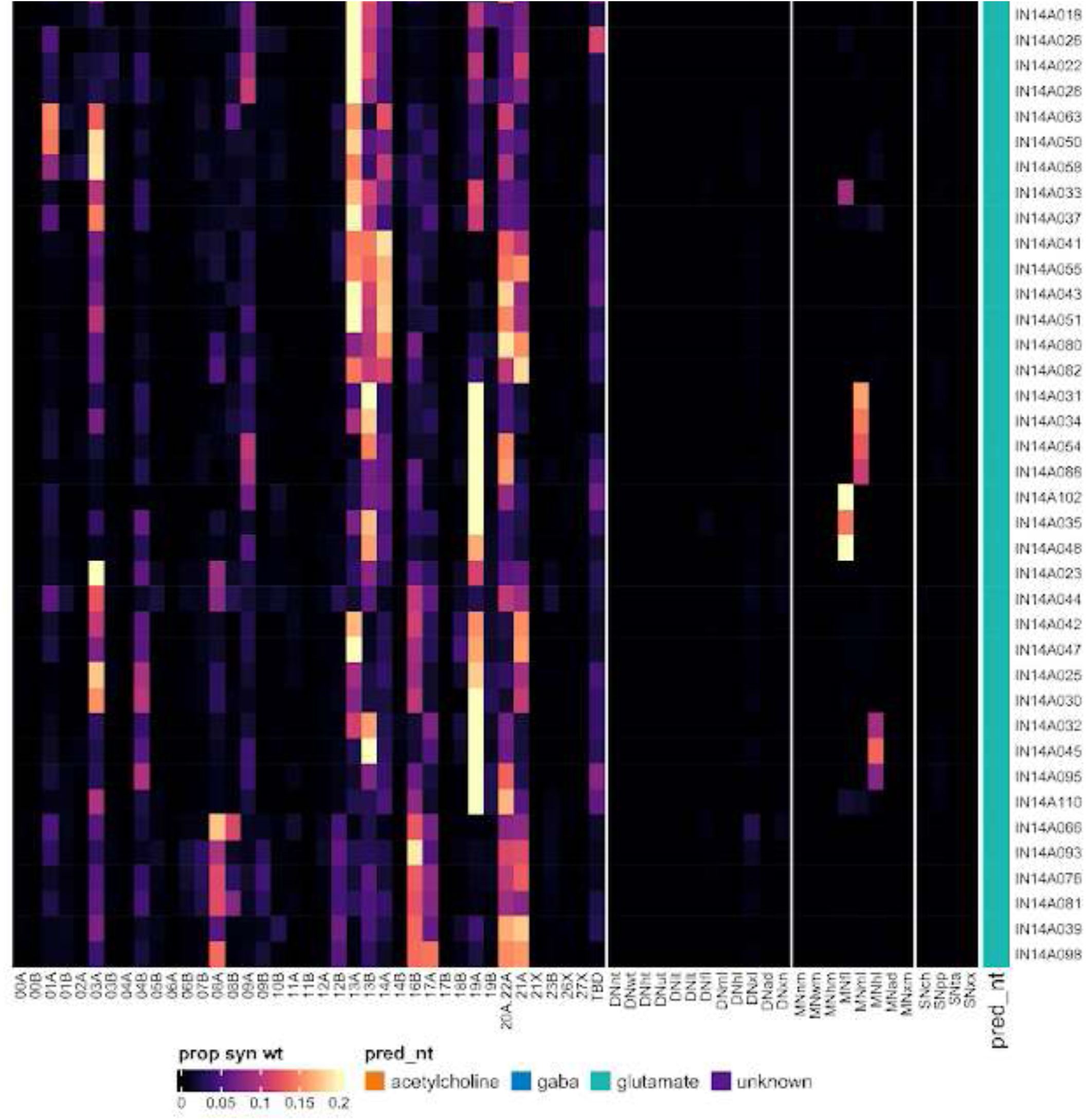
Connectivity to downstream partners by 14A secondary systematic types, continued. Proportions of synaptic weight from systematic types to downstream partners, normalised by row. 14A neurons have been clustered within each assigned birthtime window (P = primary, ES = early secondary, S = secondary) based on both upstream and downstream connectivity to hemilineages, descending neuron subclasses, motor neuron subclasses, and sensory neuron modalities. The annotation bar is coloured by the most common predicted neurotransmitter for the neurons of each type.

#### Hemilineage 15B

Hemilineage 15B is believed to derive from anterior neuroblast NB2-3 (Lacin and Truman, 2016) (but see also (Birkholz et al., 2015)), which produces 3 putative leg MNs and 3-5 local interneurons in the embryo (Schmid et al., 1999). 15B secondary neurons are glutamatergic (Lacin et al., 2019) motor neurons found only in T1-T3 (Truman et al., 2004). Individual 15B (aka Lin A) neuron dendritic morphologies have been characterised at light level (Baek and Mann, 2009; Brierley et al., 2012).

Many leg motor neurons in the MANC dataset are poorly reconstructed due to nerve damage and/or staining artefacts, and several hemilineages producing leg motor neurons enter the anterior neuropil together, making them very difficult to differentiate. As described in detail in our companion manuscript (Cheong et al., 2023), we matched motor neurons in T1 by morphology to cells annotated in the FANC dataset (Lesser et al., 2023; Phelps et al., 2021) and then identified their serial homologues in T2 and T3 using morphology and connectivity. We then annotated hemilineage 15B neurons based on their inferred target muscles in the femur and tibia (Baek and Mann, 2009; Brierley et al., 2012). Not all 15B neurons could be assigned to serial sets or even to left-right groups, likely due to reconstruction quality (Figure 38 - figure supplement 2).

We identified 15B secondary leg motor neurons in all three thoracic neuromeres, but more in T1 and fewer in T3 than in T2 (Figure 38A,E). 15B secondary neurons receive most input from leg hemilineages 03A, 04B, 09A, 12B, 13A, 13B, and especially 20A/22A and 21A and also from descending neurons to multiple legs (Figure 38F). However, there are distinct populations with respect to these inputs - for example, types MNf/m/hl30 and 40 (tibia flexor MNs) receive significantly more input from 03A and less input from 06B than do types MNf/m/hl15 and 32 (ltm-2 and ltm MNs) (Figure 38 - figure supplement 3).

**Figure 38.**
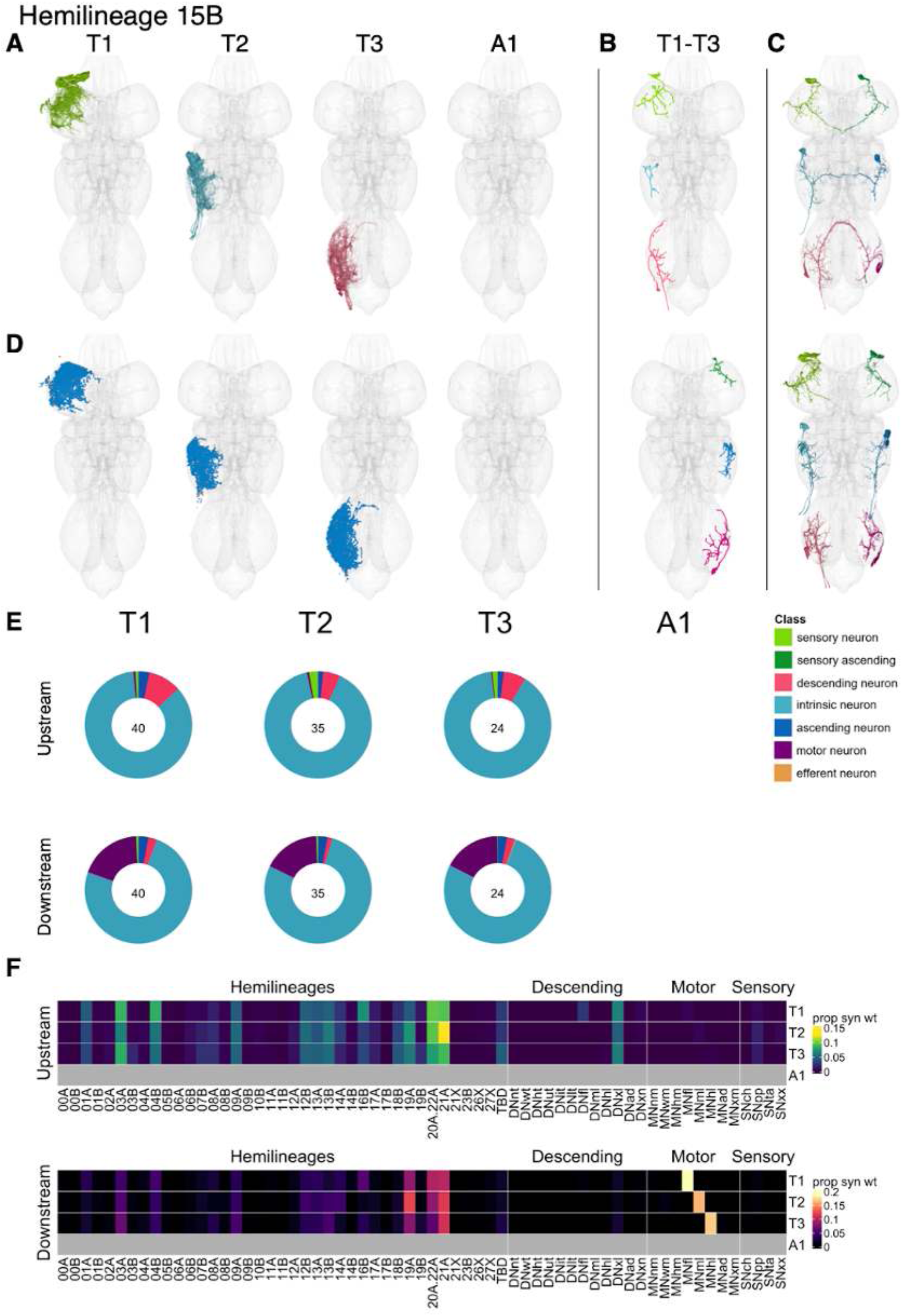
Hemilineage 15B. **A.** Meshes of all RHS secondary neurons plotted in neuromere-specific colours. **B.** “Representative” secondary neuron skeletons plotted in hemineuromere-specific colours. The skeleton with the top accumulated NBLAST score among all neurons from the hemilineage in a given hemineuromere was used. **C.** Neuron meshes of selected examples. Top: ltm1-tibia MN independent leg serial set 10811. Bottom: Ti flexor MN independent leg serial set 10710. **D.** Predicted synapses of RHS secondary neurons. Blue: postsynapses; dark orange: presynapses. **E.** Proportions of connections from secondary neurons to upstream or downstream partners, normalised by neuromere and coloured by broad class. Numbers of query neurons appear in the centre. **F.** Proportions of synaptic weight from secondary neurons originating in each neuromere to upstream or downstream partners, normalised by row.

**Figure 38 - figure supplement 1.**
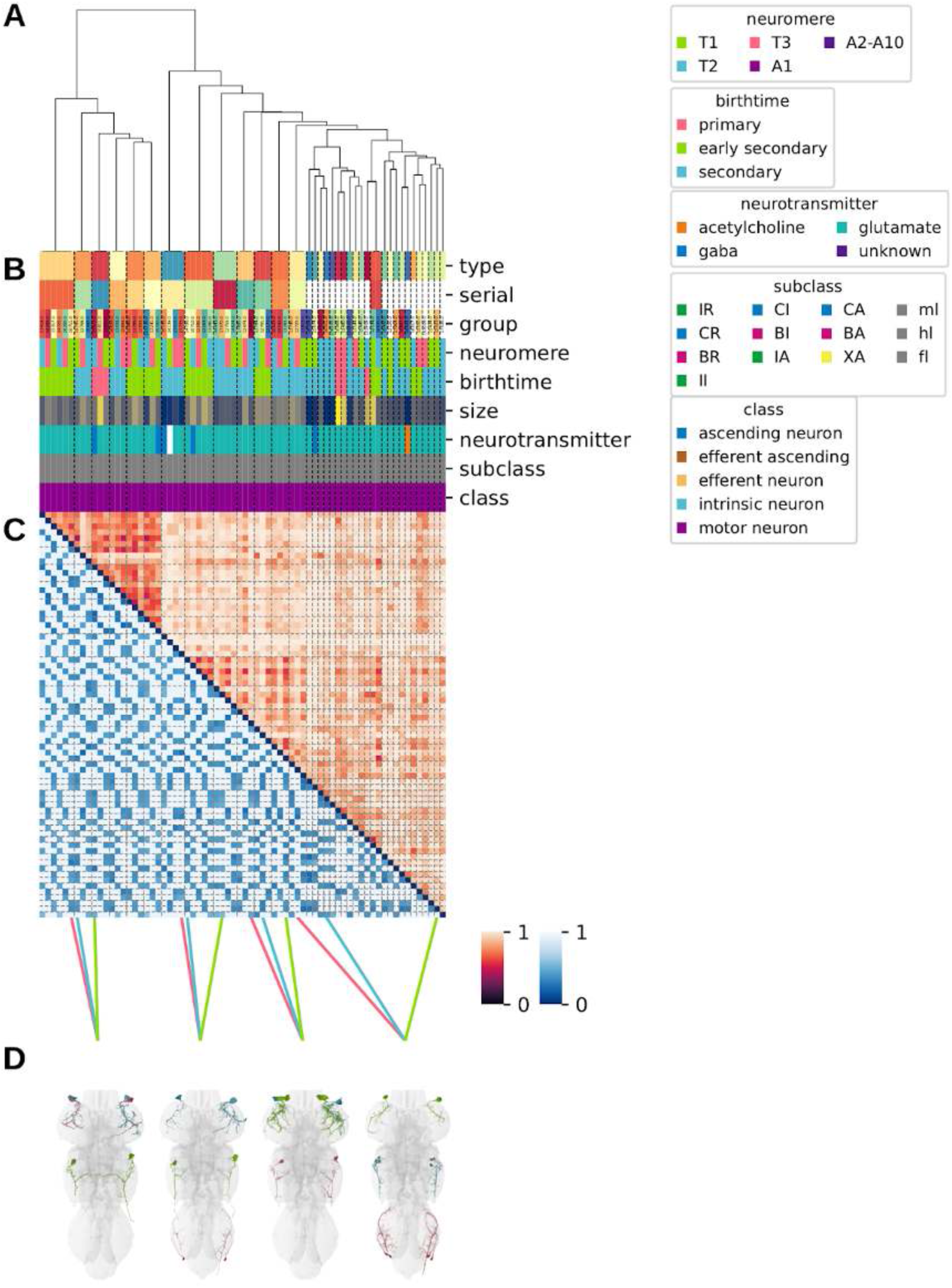
Systematic typing of hemilineage 15B. Types for motor neurons were assigned separately as outlined in our accompanying manuscript (Cheong et al., 2023). **A.** Hierarchical clustering dendrogram of hemilineage groups by laterally and serially aggregated connectivity cosine clustering. **B.** Categorical annotations of each hemilineage group, each column corresponding to the aligned leaf in A. Colours for cluster, serial set, and group are arbitrary for visualisation. Colours for neuromere, birthtime, neurotransmitter, subclass, and class are as in all other figures. **C.** Similarity distance heatmap for hemilineage. Cosine distance is in the upper triangle, while laterally symmetrised NBLAST distance is in the lower triangle. Systematic type names of some types are labelled. **D.** Morphologically representative groups from dendrogram subtrees. Each group, indicated by colour and line connecting to its column in B and C, is the most morphologically representative group (medoid of NBLAST distance) from a subtree of A. The subtrees (flat clusters) are equal height cuts of A determined to yield the number of groups per plot and plots in D.

**Figure 38 - figure supplement 2.**
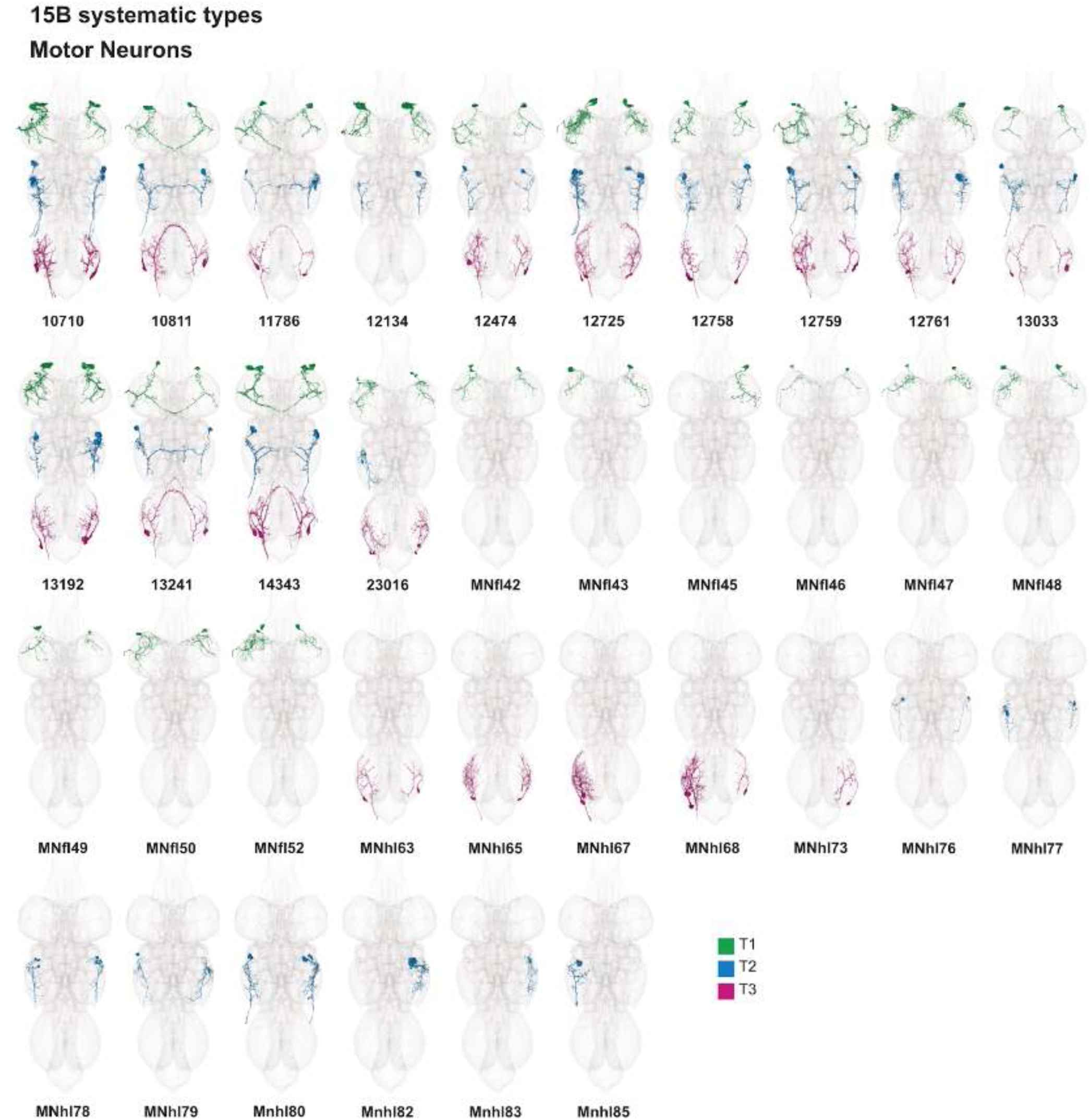
Systematic types of hemilineage 15B. Motor neurons (typed separately in (Cheong et al., 2023)) have been plotted by serial set if identified in multiple neuromeres and by systematic type if not. Individual motor neuron meshes have been coloured based on soma neuromere: dark green = T1, blue = T2, magenta = T3.

**Figure 38 - figure supplement 3.**
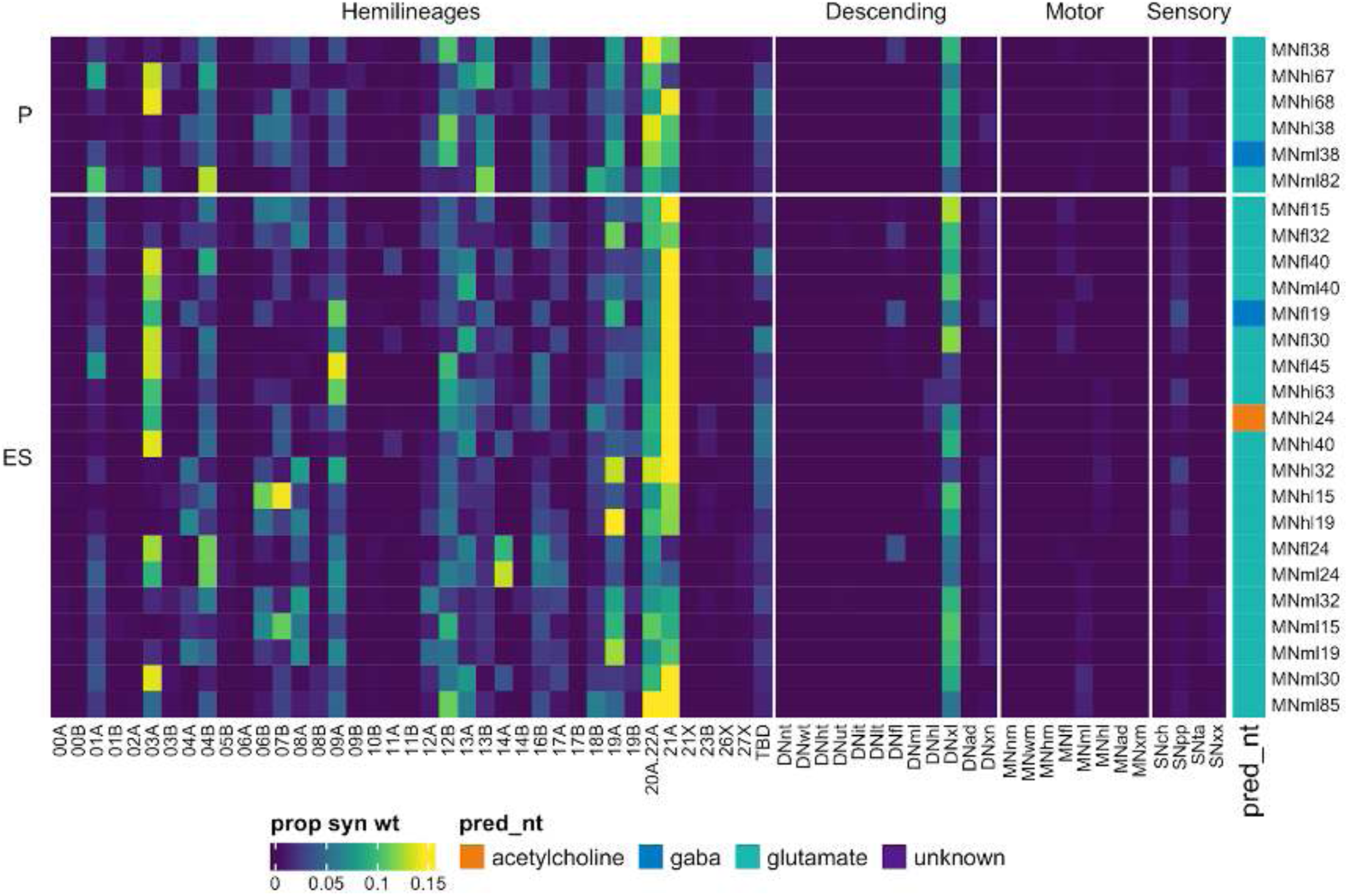
Connectivity to upstream partners by 15B primary and early secondary systematic types. Proportions of synaptic weight to systematic types from upstream partners, normalised by row. 15B neurons have been clustered within each assigned birthtime window (P = primary, ES = early secondary, S = secondary) based on both upstream and downstream connectivity to hemilineages, descending neuron subclasses, motor neuron subclasses, and sensory neuron modalities. Annotation bar is coloured by the most common predicted neurotransmitter for the neurons of each type.

**Figure 38 - figure supplement 4.**
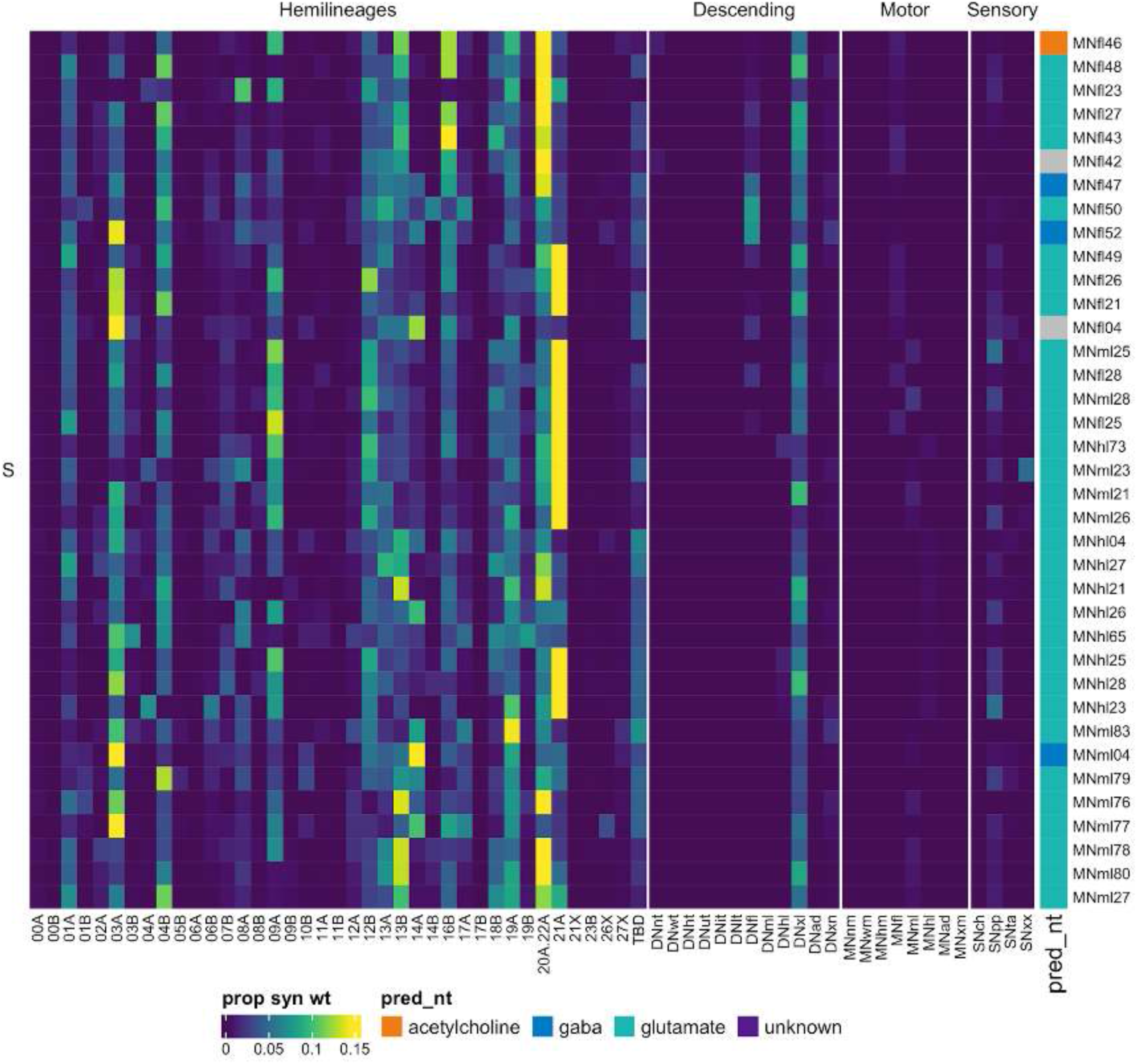
Connectivity to upstream partners by 15B secondary systematic types. Proportions of synaptic weight to systematic types from upstream partners, normalised by row. 15B neurons have been clustered within each assigned birthtime window (P = primary, ES = early secondary, S = secondary) based on both upstream and downstream connectivity to hemilineages, descending neuron subclasses, motor neuron subclasses, and sensory neuron modalities. The annotation bar is coloured by the most common predicted neurotransmitter within each type.

**Figure 38 - figure supplement 5.**
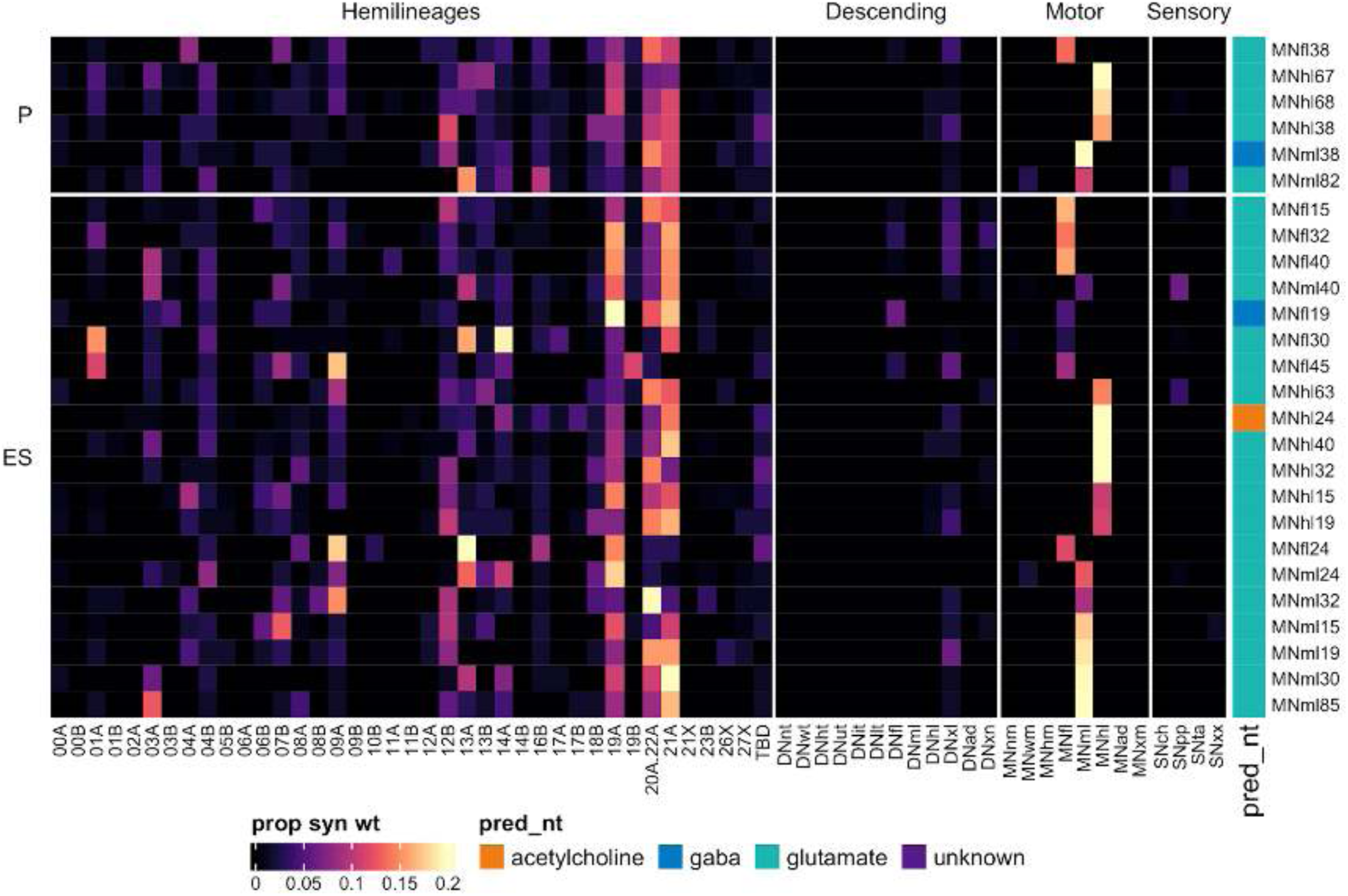
Connectivity to downstream partners by 15B primary and early secondary systematic types. Proportions of synaptic weight from systematic types to downstream partners, normalised by row. 15B neurons have been clustered within each assigned birthtime window (P = primary, ES = early secondary, S = secondary) based on both upstream and downstream connectivity to hemilineages, descending neuron subclasses, motor neuron subclasses, and sensory neuron modalities. The annotation bar is coloured by the most common predicted neurotransmitter within each type.

**Figure 38 - figure supplement 6.**
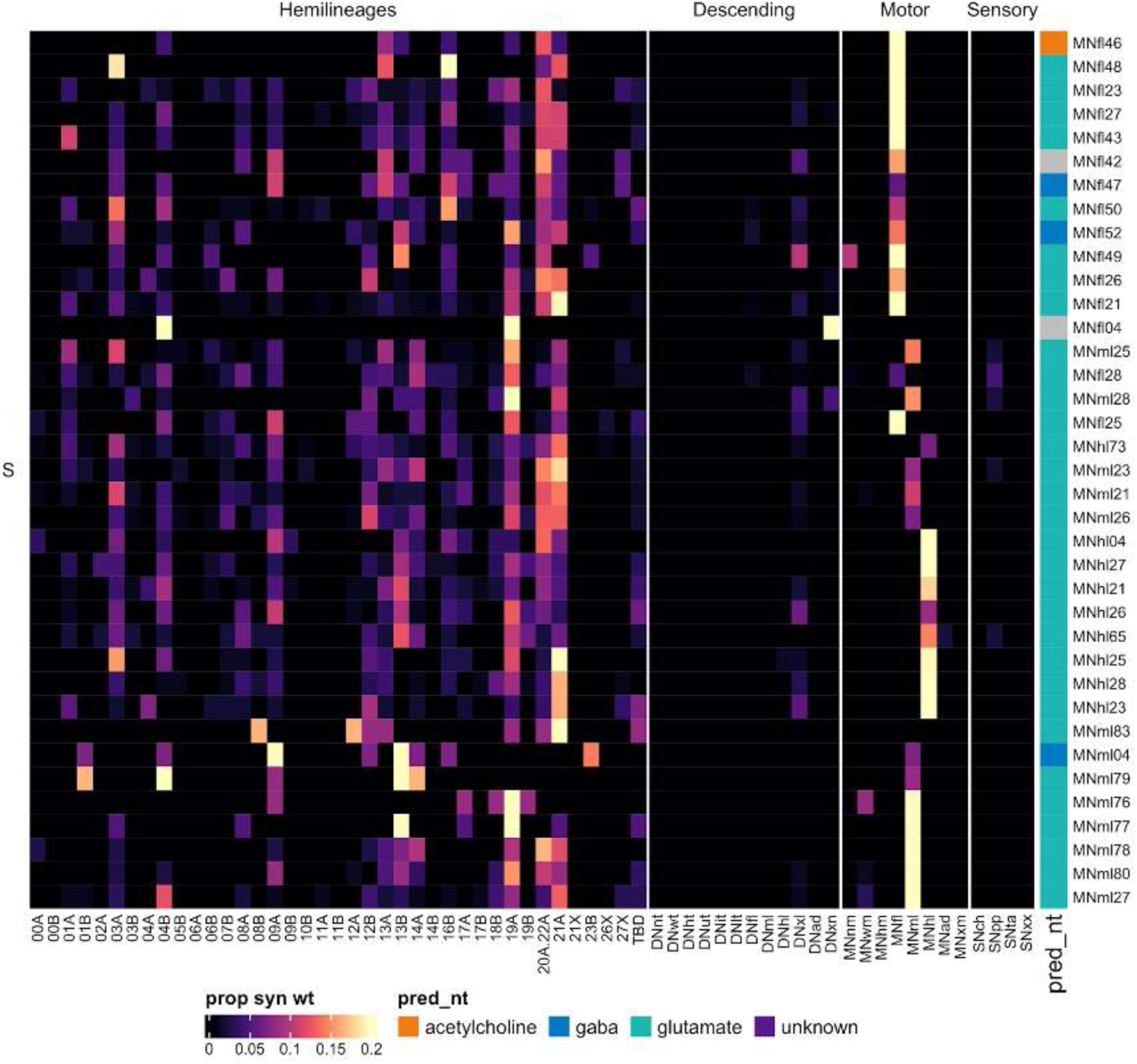
Connectivity to downstream partners by 15B secondary systematic types. Proportions of synaptic weight from systematic types to downstream partners, normalised by row. 15B neurons have been clustered within each assigned birthtime window (P = primary, ES = early secondary, S = secondary) based on both upstream and downstream connectivity to hemilineages, descending neuron subclasses, motor neuron subclasses, and sensory neuron modalities. The annotation bar is coloured by the most common predicted neurotransmitter for the neurons of each type.

#### Hemilineage 16B

Hemilineage 16B derives from anterior neuroblast NB1-1 (Birkholz et al., 2015; Lacin et al., 2019), which generates 1 or 2 motor neurons, 1 thoracic intersegmental interneuron, and 6-10 local interneurons in the embryo (Schmid et al., 1999). In the larva, lineage 16 is medial to lineage 08 in all three thoracic segments (Truman et al., 2004), but as metamorphosis progresses, 08A neurons and a subpopulation of 16B neurons are pulled laterally towards their target ipsilateral leg neuropil. In the adult, 08A and 16B primary neurites enter the neuropil close together (Shepherd et al., 2016). Most 16B neurons are restricted to their ipsilateral hemineuromere, but we identified a sequentially descending serial set (type IN16B037) and a pair of neurons that innervate both neck and wing tectulum (IN16B046) (Figure 39 - figure supplement 2).

We initially had difficulty distinguishing between the glutamatergic hemilineages 08A and 16B in MANC but assigned neurons with prominent T1 ventral dendrites crossing the midline to 16B (Figure 39C top), based on light-level neuroblast clones (Shepherd et al., 2019). We extended this annotation to neurons entering in the same soma tract bundle. Serial homology then allowed us to assign the corresponding secondary hemilineages in T2 and T3. However, we could not be confident about our 08A vs 16B primary neuron assignments because they did not enter with the secondary neurons.

**Figure 39.**
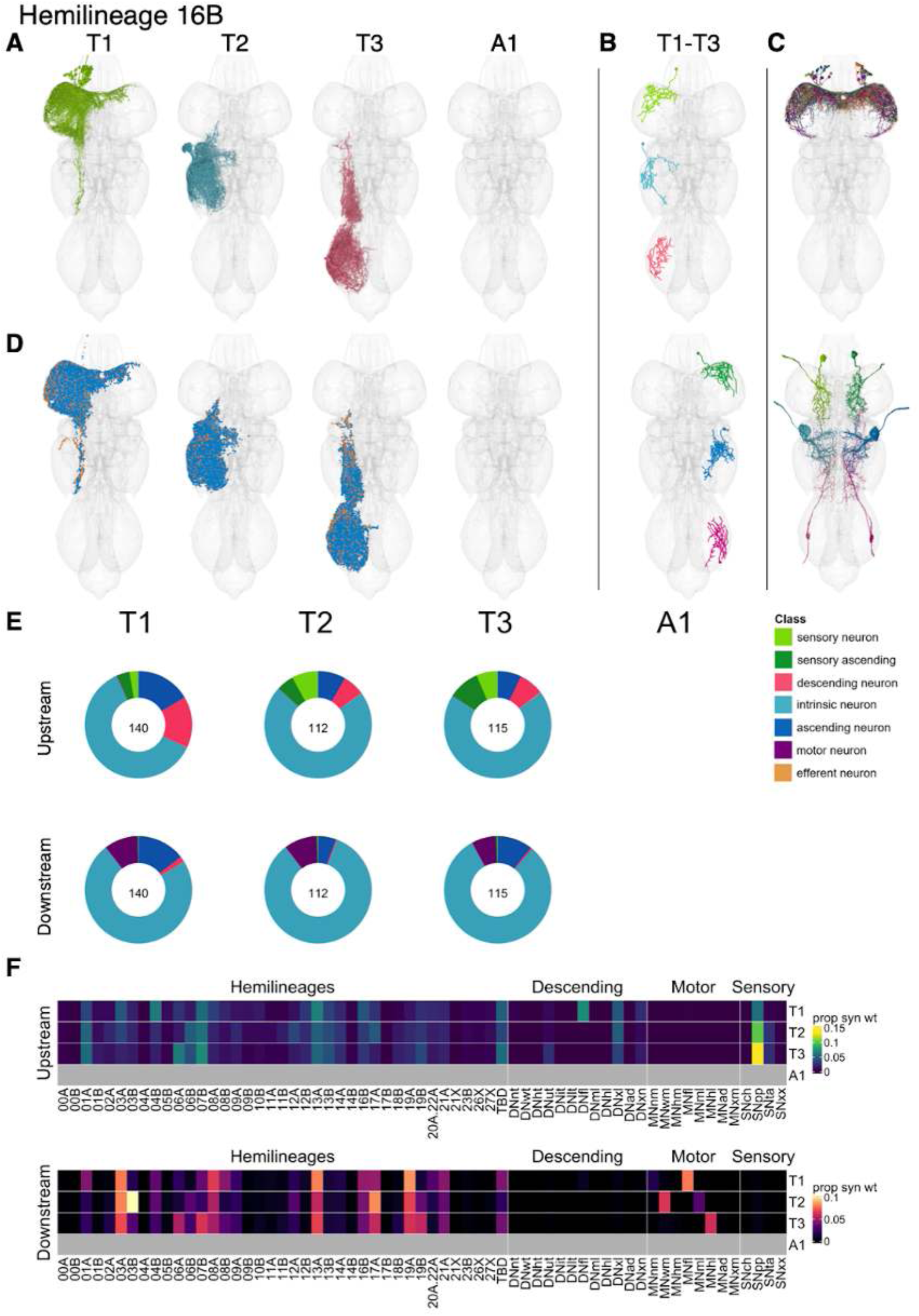
Hemilineage 16B. **A.** Meshes of all RHS secondary neurons plotted in neuromere-specific colours. **B.** “Representative” secondary neuron skeletons plotted in hemineuromere-specific colours. The skeleton with the top accumulated NBLAST score among all neurons from the hemilineage in a given hemineuromere was used. **C.** Neuron meshes of selected examples. Top: T1 BR subclass. Bottom: motor neuron dorsal serial set 10011. **D.** Predicted synapses of RHS secondary neurons. Blue: postsynapses; dark orange: presynapses. **E.** Proportions of connections from secondary neurons to upstream or downstream partners, normalised by neuromere and coloured by broad class. Numbers of query neurons appear in the centre. **F.** Proportions of synaptic weight from secondary neurons originating in each neuromere to upstream or downstream partners, normalised by row.

We also identified a glutamatergic subpopulation entering the neuropil near 08A, projecting straight dorsally, and including morphologically similar ascending neurons (T1 only), intrinsic neurons, and motor neurons that innervate neck and/or wing tectulum and exit via DProN, ADMN, and AbN1 in T1, T2, and T3 respectively (e.g., Figure 39C bottom). The motor neurons are not recovered in postembryonic clones (Haluk Lacin, personal communication), so we suspect that they are embryonic-born but remain immature until metamorphosis. Most of the T2 motor neurons linked to courtship song and flight (O’Sullivan et al., 2018) belong to this population. The intrinsic neurons are visible in postembryonic lineage 16 clones (Shepherd et al., 2019); they are Hb9+ and Glu+ and are absent in *numb* loss of function clones (Lacin et al., 2014), indicating that they belong to 16B rather than 16A. We have therefore tentatively annotated this entire dorsal glutamatergic subpopulation as 16B.

16B neurons receive strong input from proprioceptive sensory neurons and input from descending neurons targeting the front leg neuropils or targeting multiple leg neuropils and from hemilineages including 01A, 04B, 06A, 07B, and 13A, with some segmental variation. The 16B types split into clear leg vs dorsal categories by upstream connectivity, with dorsal types (e.g., IN16B046-48) receiving high levels of input from proprioceptive neurons and dorsal hemilineages 02A, 06A, 06B, and 07B while leg types (e.g., IN16B045, IN16B061) receive input from a wider range of sources (Figure 39 - figure supplement 6). Secondary 16B neurons are predicted to be glutamatergic as expected (Lacin et al., 2019) and mainly target hemilineages 03A, 03B (in T2), 08A, 13A, 17A, and 19A in thoracic neuromeres and wing or leg motor neurons (Figure 39F). A few types target neck motor neurons (e.g., AN16B081 and IN16B100) (Figure 39 - figure supplement 9). No functional studies have been published for secondary 16B neurons.

**Figure 39 - figure supplement 1.**
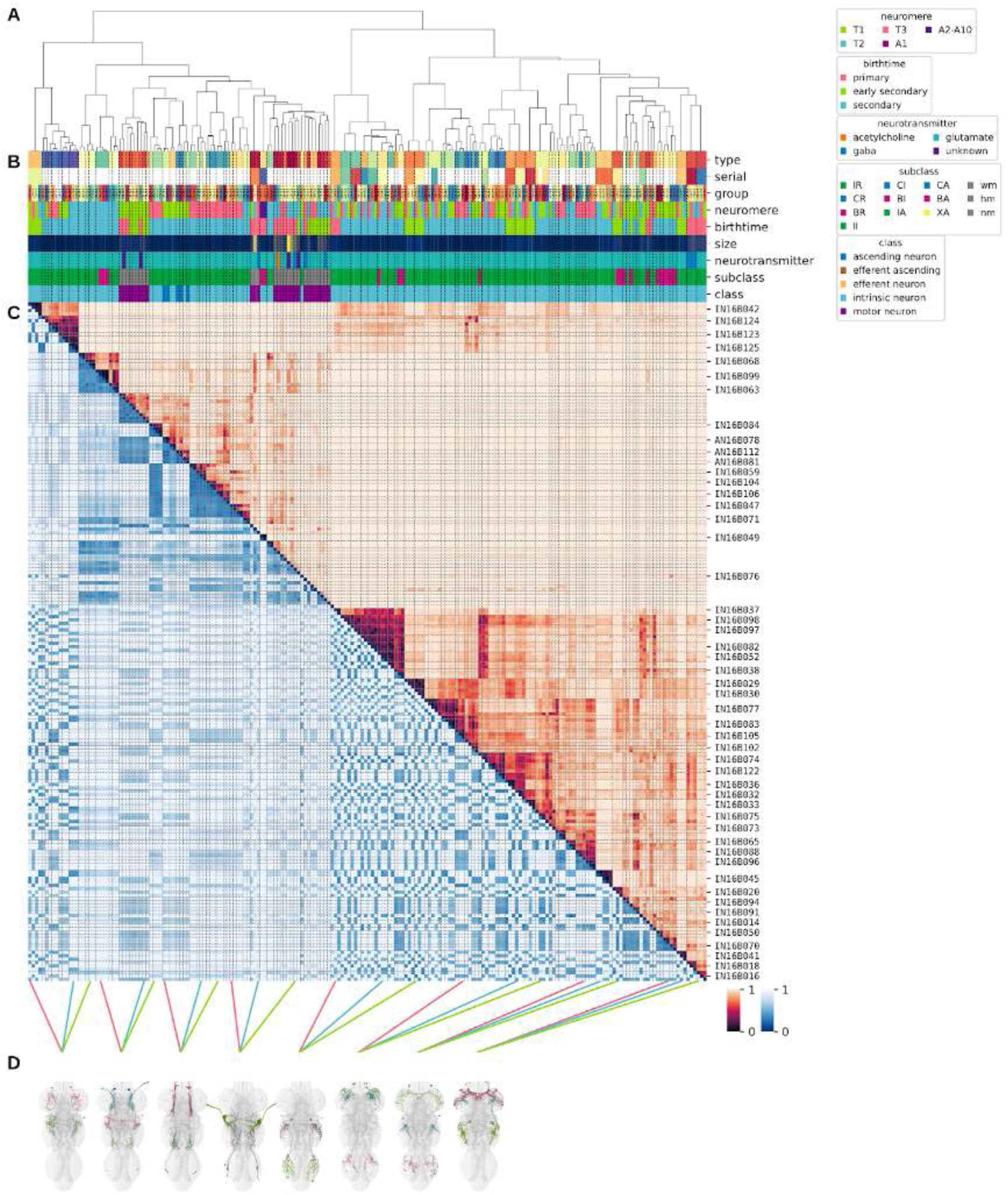
Systematic typing of hemilineage 16B. **A.** Hierarchical clustering dendrogram of hemilineage groups by laterally and serially aggregated connectivity cosine clustering. **B.** Categorical annotations of each hemilineage group, each column corresponding to the aligned leaf in A. Colours for type, serial set, and group are arbitrary for visualisation. Colours for neuromere, birthtime, neurotransmitter, subclass, and class are as in all other figures. **C.** Similarity distance heatmap for hemilineage. Cosine distance is in the upper triangle, while laterally symmetrised NBLAST distance is in the lower triangle. Systematic type names of some types are labelled. **D.** Morphologically representative groups from dendrogram subtrees. Each group, indicated by colour and line connecting to its column in B and C, is the most morphologically representative group (medoid of NBLAST distance) from a subtree of A. The subtrees (flat clusters) are equal height cuts of A determined to yield the number of groups per plot and plots in D.

**Figure 39 - figure supplement 2.**
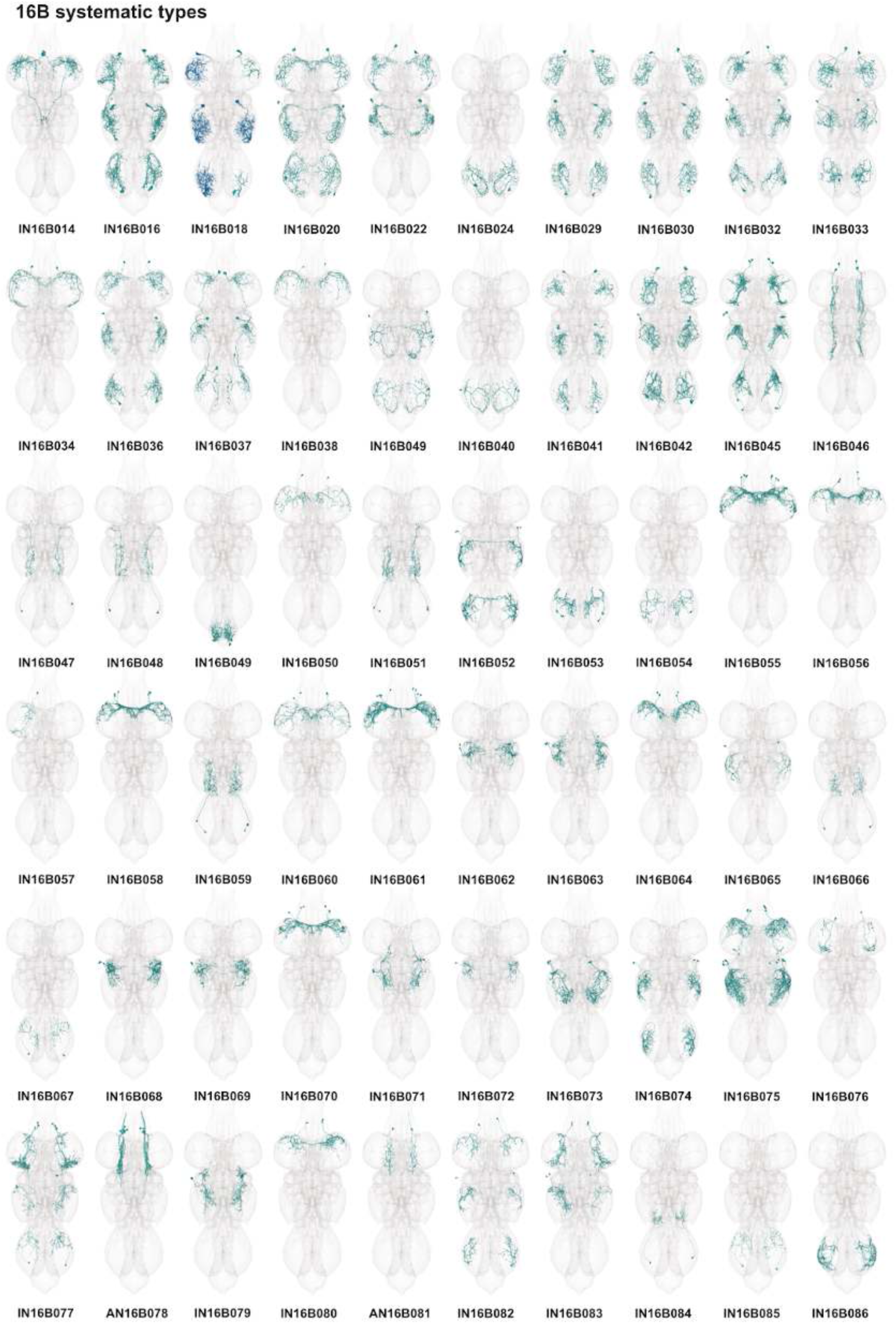
Systematic types of hemilineage 16B. Systematic types have been arranged in numerical order, with neurons of the same type that belong to distinct classes (e.g., intrinsic neuron vs ascending neuron) plotted separately but placed adjacent to each other. Individual neuron meshes have been coloured based on predicted neurotransmitter: dark orange = acetylcholine, blue = gaba, marine = glutamate, dark purple = unknown.

**Figure 39 - figure supplement 3.**
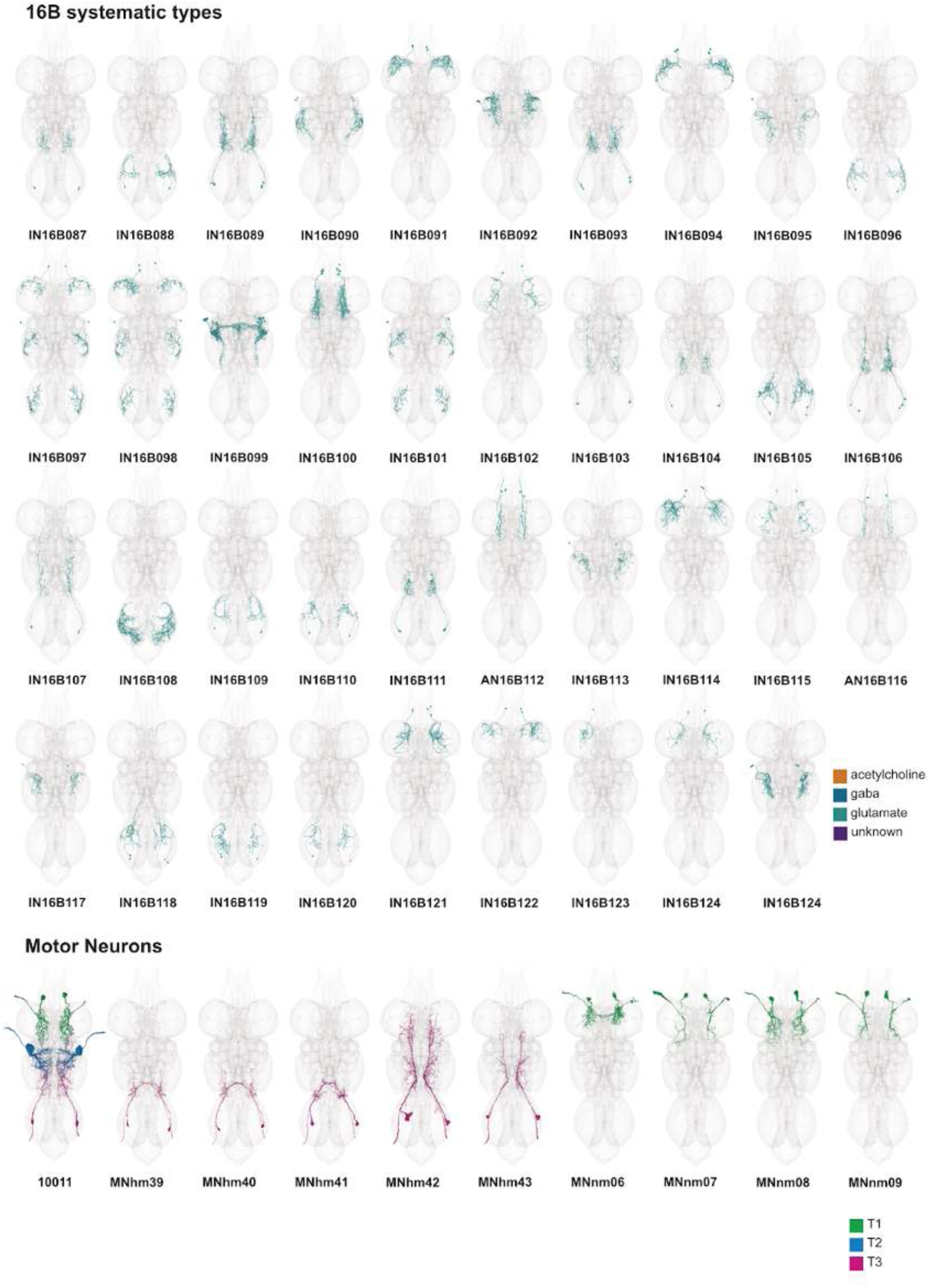
Systematic types of hemilineage 16B, continued. Systematic types have been arranged in numerical order, with neurons of the same type that belong to distinct classes (e.g., intrinsic neuron vs ascending neuron) plotted separately but placed adjacent to each other. Individual neuron meshes have been coloured based on predicted neurotransmitter: dark orange = acetylcholine, blue = gaba, marine = glutamate, dark purple = unknown. Motor neurons (typed separately in (Cheong et al., 2023)) have been plotted by serial set if identified in multiple neuromeres and by systematic type if not. Individual motor neuron meshes have been coloured based on soma neuromere: dark green = T1, blue = T2, magenta = T3.

**Figure 39 - figure supplement 4.**
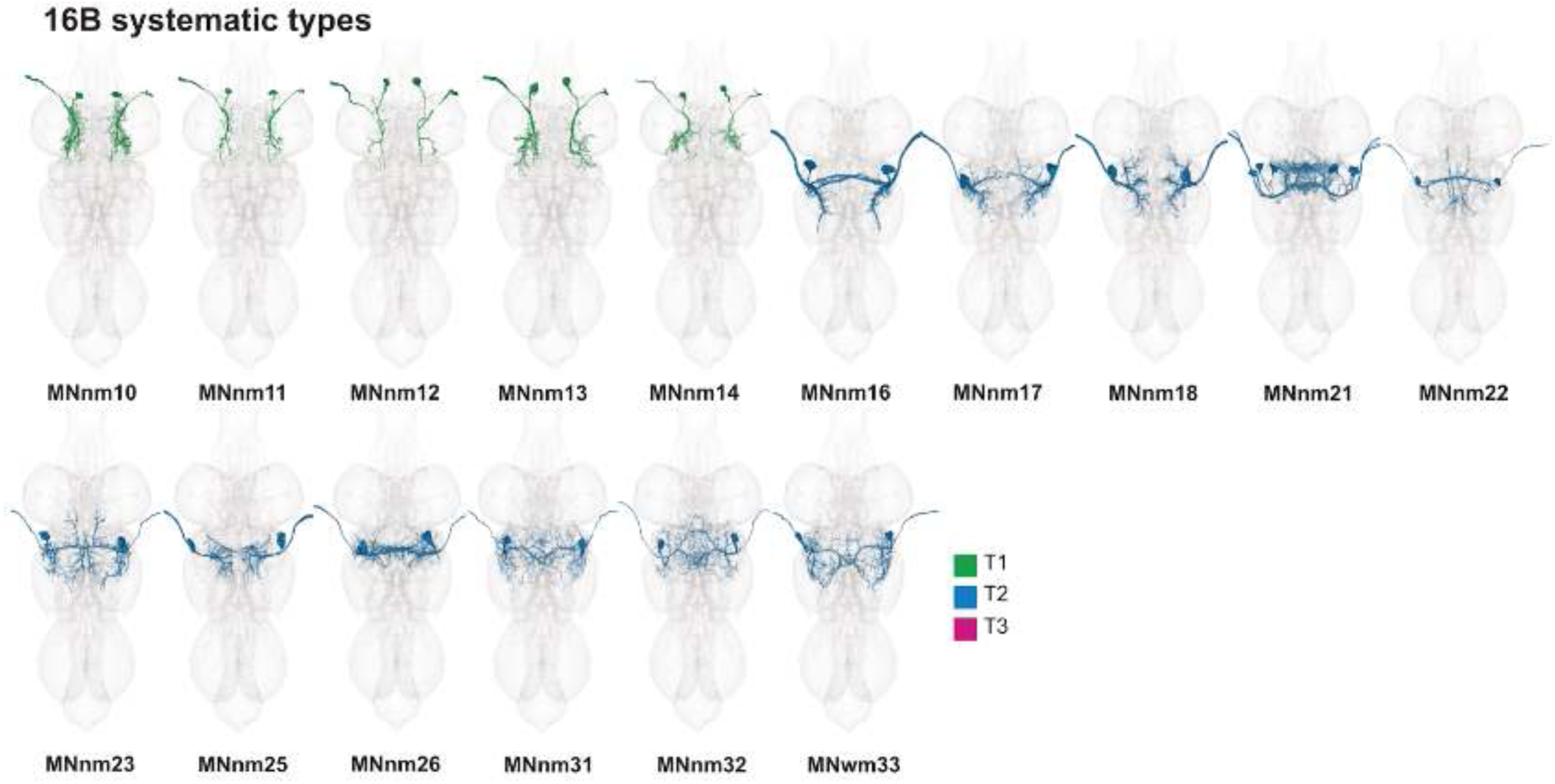
Systematic types of hemilineage 16B, continued. Systematic types have been arranged in numerical order, with neurons of the same type that belong to distinct classes (e.g., intrinsic neuron vs ascending neuron) plotted separately but placed adjacent to each other. Individual neuron meshes have been coloured based on predicted neurotransmitter: dark orange = acetylcholine, blue = gaba, marine = glutamate, dark purple = unknown. Motor neurons (typed separately in (Cheong et al., 2023)) have been plotted by serial set if identified in multiple neuromeres and by systematic type if not. Individual motor neuron meshes have been coloured based on soma neuromere: dark green = T1, blue = T2, magenta = T3.

**Figure 39 - figure supplement 5.**
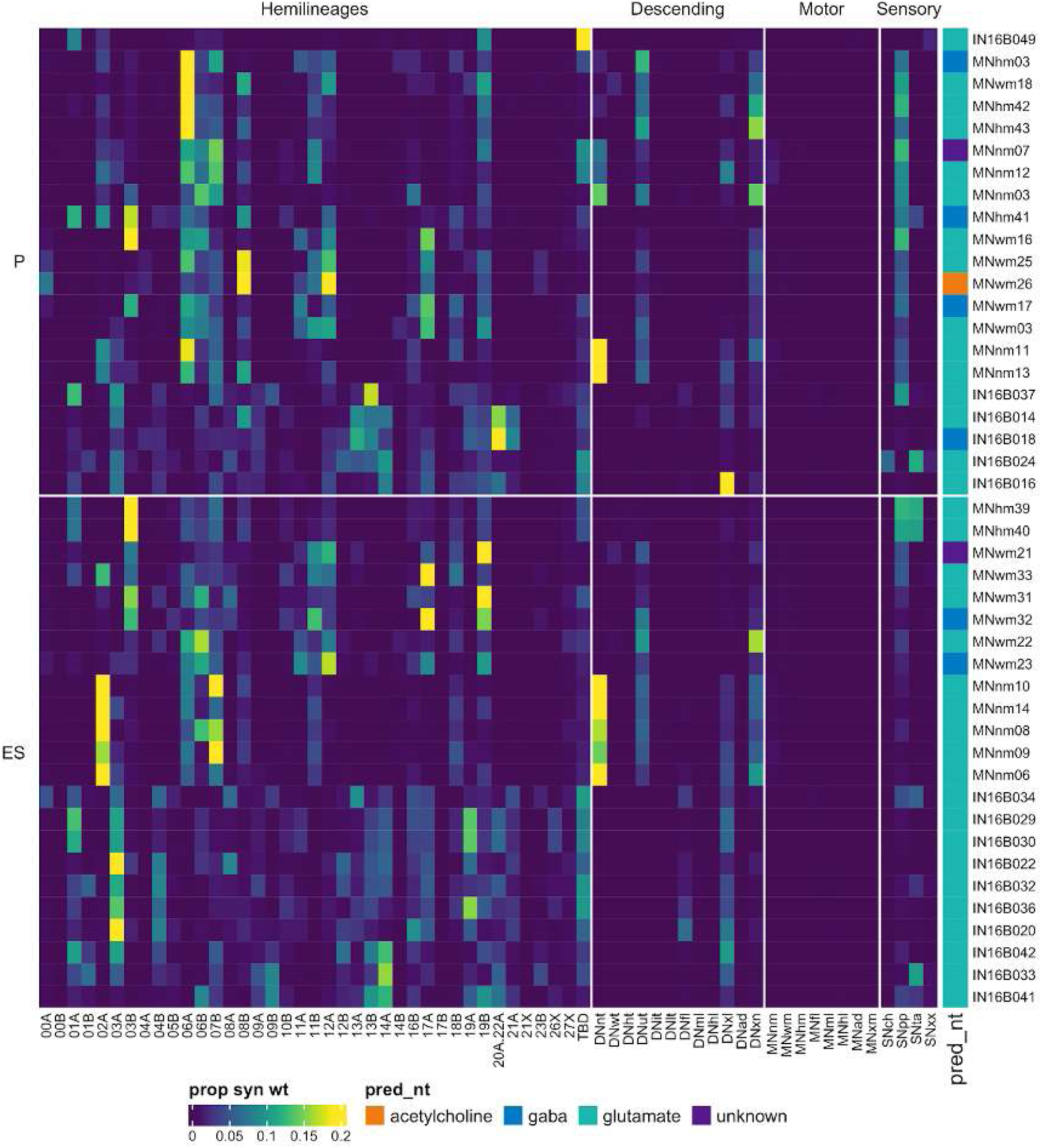
Connectivity to upstream partners by 16B primary and early secondary systematic types. Proportions of synaptic weight to systematic types from upstream partners, normalised by row. 16B neurons have been clustered within each assigned birthtime window (P = primary, ES = early secondary, S = secondary) based on both upstream and downstream connectivity to hemilineages, descending neuron subclasses, motor neuron subclasses, and sensory neuron modalities. Annotation bar is coloured by the most common predicted neurotransmitter for the neurons of each type.

**Figure 39 - figure supplement 6.**
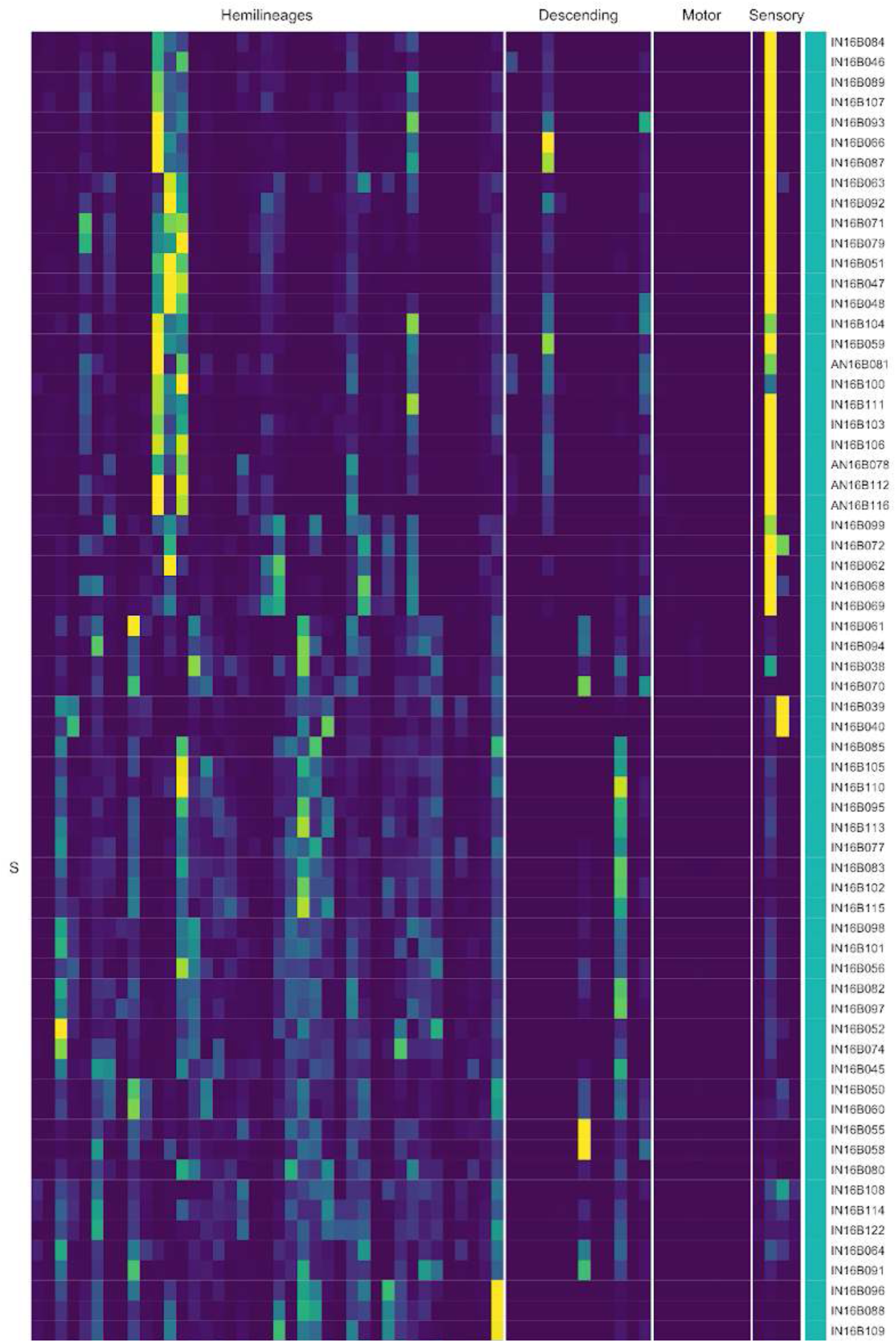
Connectivity to upstream partners by 16B secondary systematic types. Proportions of synaptic weight to systematic types from upstream partners, normalised by row. 16B neurons have been clustered within each assigned birthtime window (P = primary, ES = early secondary, S = secondary) based on both upstream and downstream connectivity to hemilineages, descending neuron subclasses, motor neuron subclasses, and sensory neuron modalities. The annotation bar is coloured by the most common predicted neurotransmitter within each type.

**Figure 39 - figure supplement 7.**
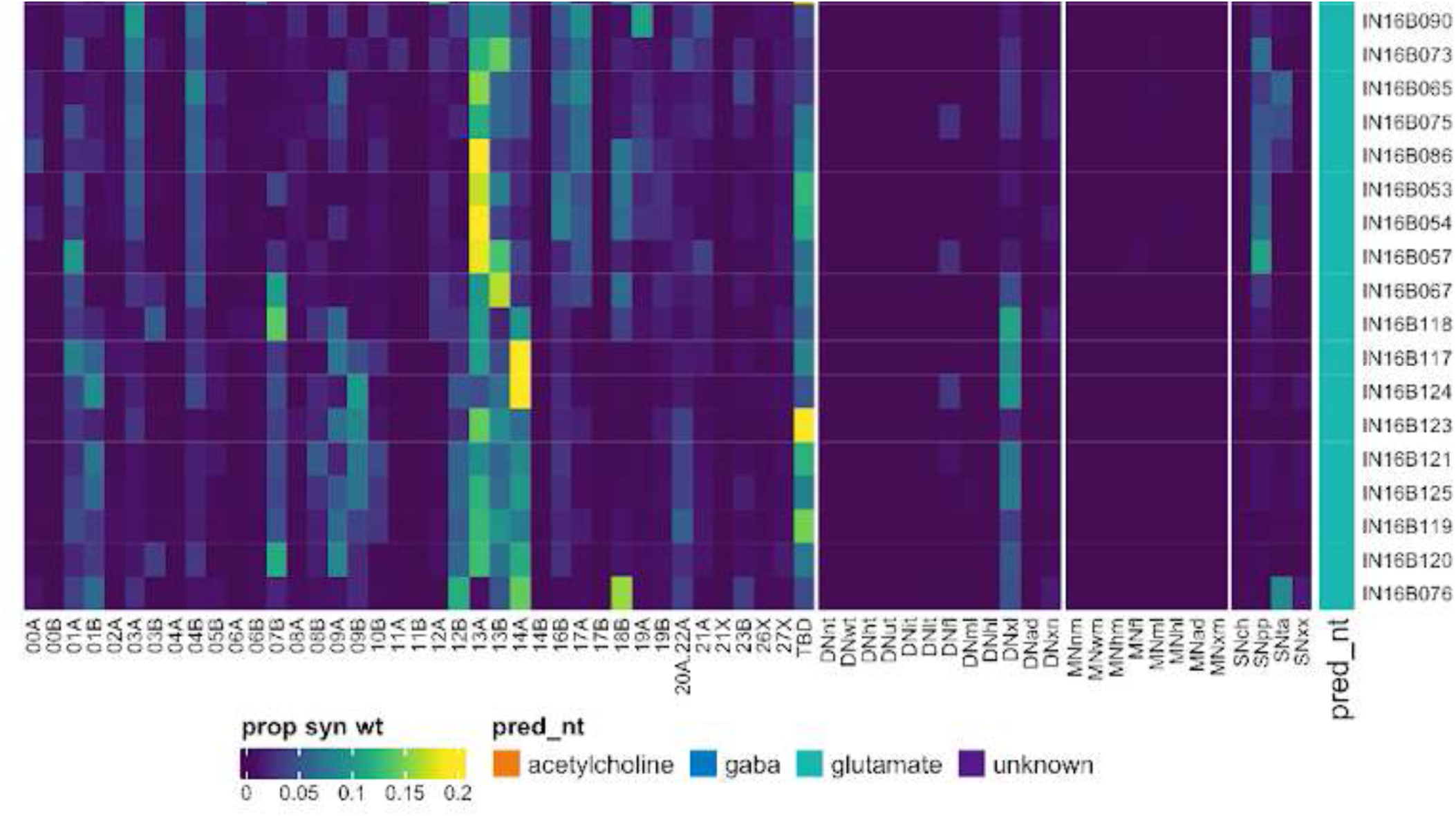
Connectivity to upstream partners by 16B secondary systematic types, continued. Proportions of synaptic weight to systematic types from upstream partners, normalised by row. 16B neurons have been clustered within each assigned birthtime window (P = primary, ES = early secondary, S = secondary) based on both upstream and downstream connectivity to hemilineages, descending neuron subclasses, motor neuron subclasses, and sensory neuron modalities. The annotation bar is coloured by the most common predicted neurotransmitter within each type.

**Figure 39 - figure supplement 8.**
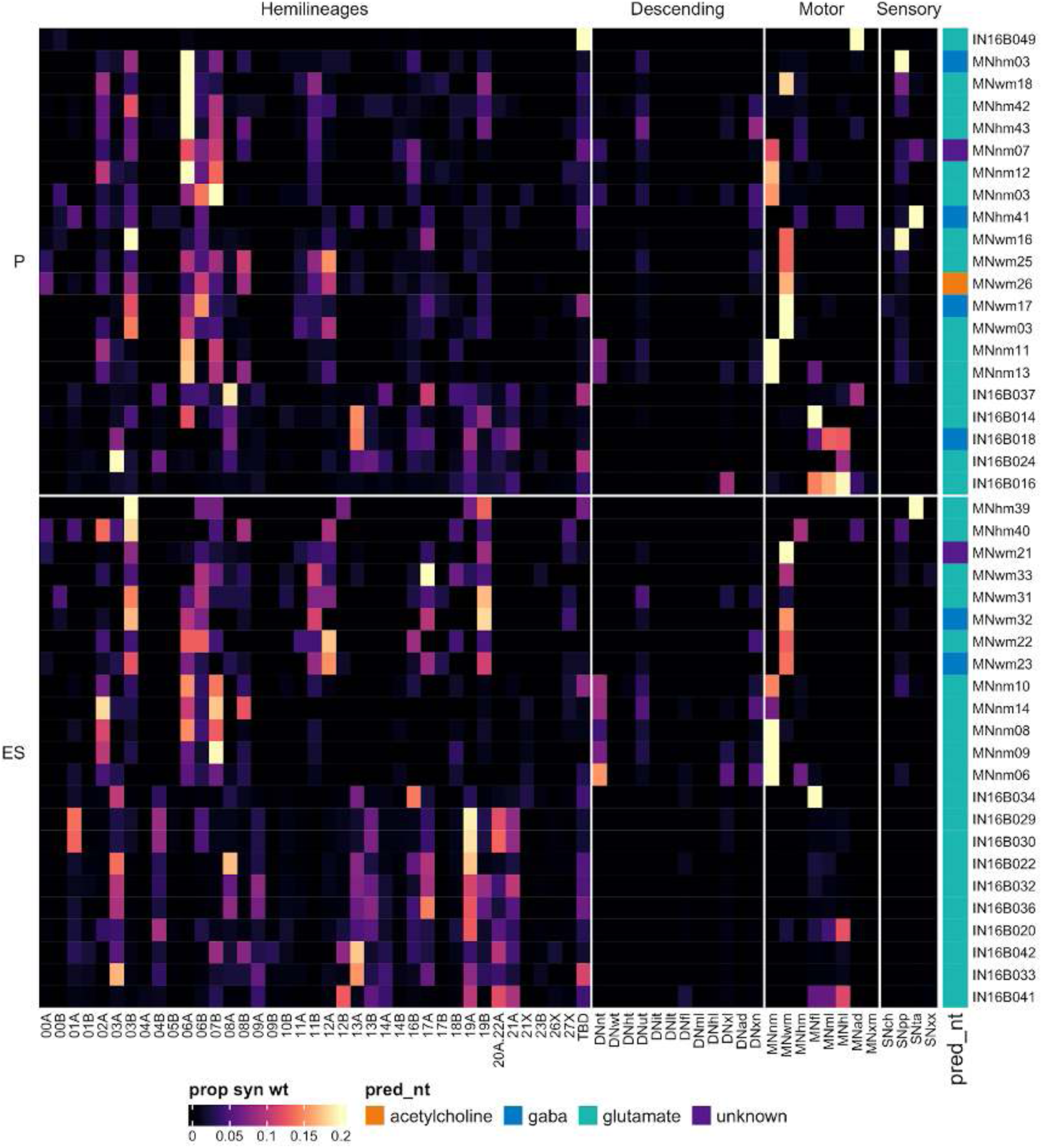
Connectivity to downstream partners by 16B primary and early secondary systematic types. Proportions of synaptic weight from systematic types to downstream partners, normalised by row. 16B neurons have been clustered within each assigned birthtime window (P = primary, ES = early secondary, S = secondary) based on both upstream and downstream connectivity to hemilineages, descending neuron subclasses, motor neuron subclasses, and sensory neuron modalities. The annotation bar is coloured by the most common predicted neurotransmitter within each type.

**Figure 39 - figure supplement 9.**
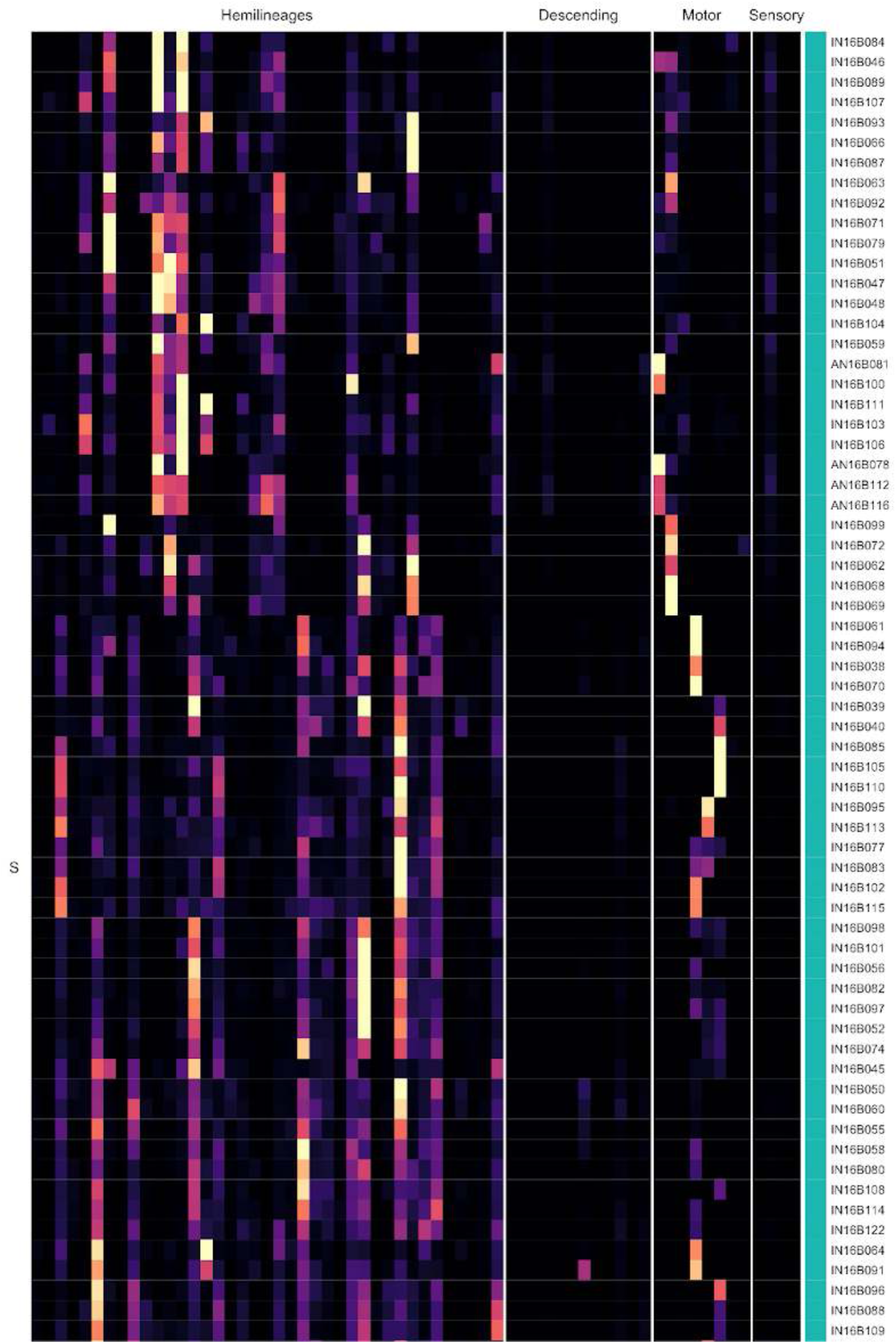
Connectivity to downstream partners by 16B econdary systematic types. Proportions of synaptic weight from systematic types to downstream partners, normalised by row. 16B neurons have been clustered within each assigned birthtime window (P = primary, ES = early secondary, S = secondary) based on both upstream and downstream connectivity to hemilineages, descending neuron subclasses, motor neuron subclasses, and sensory neuron modalities. The annotation bar is coloured by the most common predicted neurotransmitter for the neurons of each type.

**Figure 39 - figure supplement 10.**
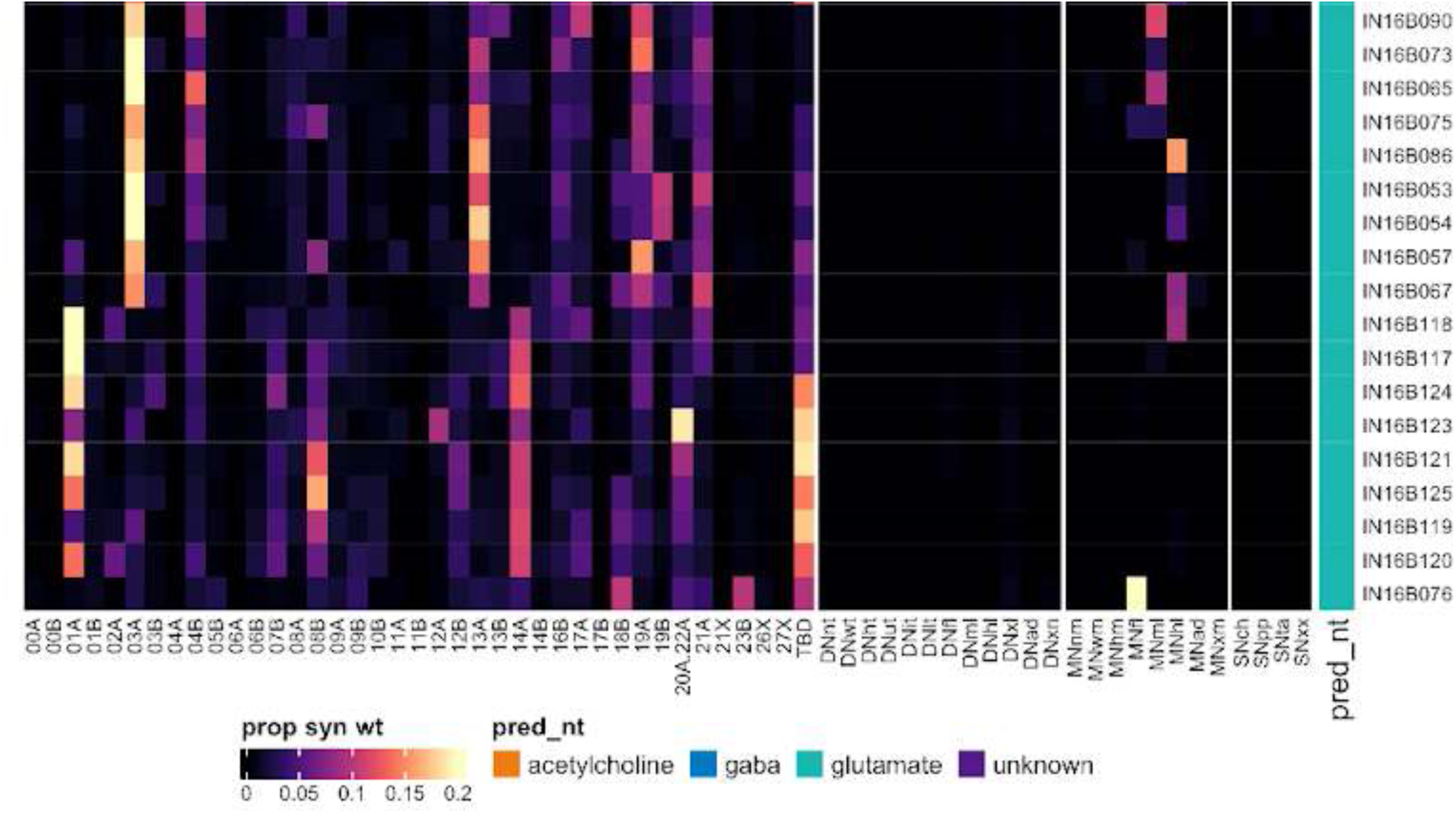
Connectivity to downstream partners by 16B secondary systematic types, continued. Proportions of synaptic weight from systematic types to downstream partners, normalised by row. 16B neurons have been clustered within each assigned birthtime window (P = primary, ES = early secondary, S = secondary) based on both upstream and downstream connectivity to hemilineages, descending neuron subclasses, motor neuron subclasses, and sensory neuron modalities. The annotation bar is coloured by the most common predicted neurotransmitter for the neurons of each type.

#### Hemilineage 17A

Hemilineage 17A derives from anterior dorsal neuroblast NB2-5 (Birkholz et al., 2015; Lacin and Truman, 2016), which generates at least one ipsilateral motor neuron, 4-6 intersegmental interneurons, and 6-8 local interneurons in the embryo (Schmid et al., 1999; Schmidt et al., 1997). 17A secondary neurons have been reported in T2-A1 (Truman et al., 2004) and T1 (Birkholz et al., 2015). We identified 17A neurons in T1-A1, although their numbers were greatly reduced in T1 and A1 and expanded in T2 (Figure 40A,E). They enter the anterior neuromere near 18B and are likewise cholinergic as predicted (Lacin et al., 2019) but remain largely restricted to ipsilateral neuropils, except for some bilateral T2 and T3 types that project to the upper tectulum (Figure 40C bottom). We identified the small 17A population in A1 based on serial homology with thoracic neurons (e.g., Figure 40C top).

**Figure 40.**
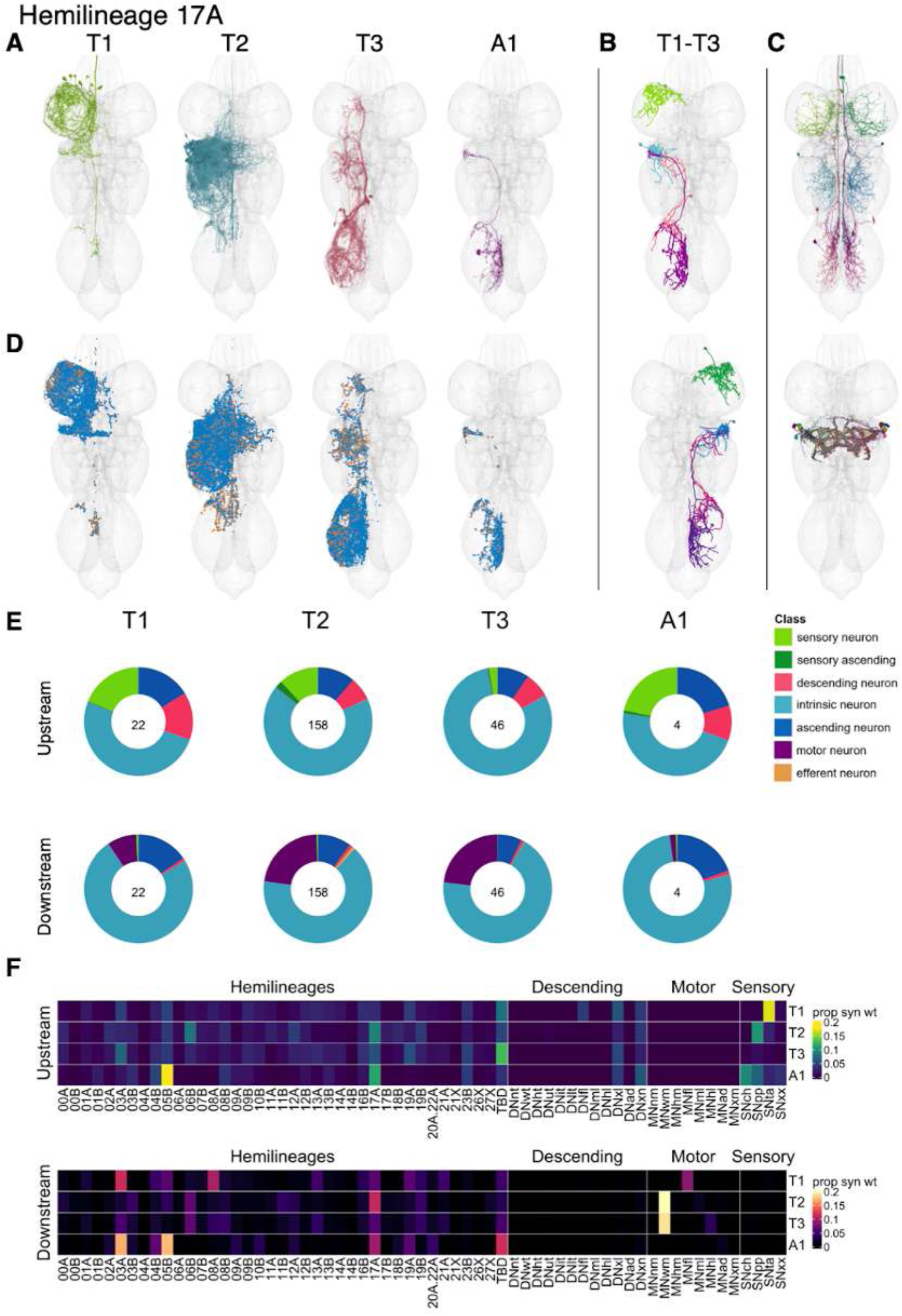
Hemilineage 17A. **A.** Meshes of all RHS secondary neurons plotted in neuromere-specific colours. **B.** “Representative” secondary neuron skeletons plotted in hemineuromere-specific colours. The skeleton with the top accumulated NBLAST score among all neurons from the hemilineage in a given hemineuromere was used. **C.** Neuron meshes of selected examples. Top: ascending serial set 11036 (T1-A1). Bottom: putative electrical subcluster 10629/dMS2. **D.** Predicted synapses of RHS secondary neurons. Blue: postsynapses; dark orange: presynapses. **E.** Proportions of connections from secondary neurons to upstream or downstream partners, normalised by neuromere and coloured by broad class. Numbers of query neurons appear in the centre. **F.** Proportions of synaptic weight from secondary neurons originating in each neuromere to upstream or downstream partners, normalised by row.

17A secondary neurons are mainly predicted to be cholinergic (Figure 8E) as expected (Lacin et al., 2019), but we also identified populations consistently predicted to be glutamatergic, with low presynaptic density, which may be electrically coupled to partners via gap junctions; among them are the Fru+ dMS2 neurons (Figure 40C bottom) involved in courtship song (Lillvis et al., 2024; Yu et al., 2010). Gabaergic neurons that appeared to originate from the same lineage were tentatively assigned to hemilineage 17B (see “Primary” hemilineages, below).

17A neurons are morphologically diverse, with T2-specific types innervating flight neuropil but serially recurring types innervating leg neuropil (Figure 40 - figure supplement 2-4). No functional studies have been published for secondary 17A neurons. They receive diverse inputs, notably from tactile sensory neurons in T1, proprioceptive sensory neurons in T2 and A1, and chemosensory neurons in A1. Most types receive mixed chemosensory, proprioceptive, and/or tactile input, although a few are specific for one modality (Figure 40 - figure supplement 5-7). The postsynaptic partners of 17A secondary neurons include 03A and 08A in T1, 17A in T2, wing motor neurons in T2 and T3, 03A, 05B, and 17A in A1, and leg motor neurons in all thoracic neuromeres but especially T1 (Figure 40E,F). The putative electrical dMS2 types (IN17A05, IN17A06, IN17A010, and IN17A021) receive most of their input from hemilineages 11B and 12A and target 11B and wing motor neurons (Figure 40 - figure supplement 5, 8).

**Figure 40 - figure supplement 1.**
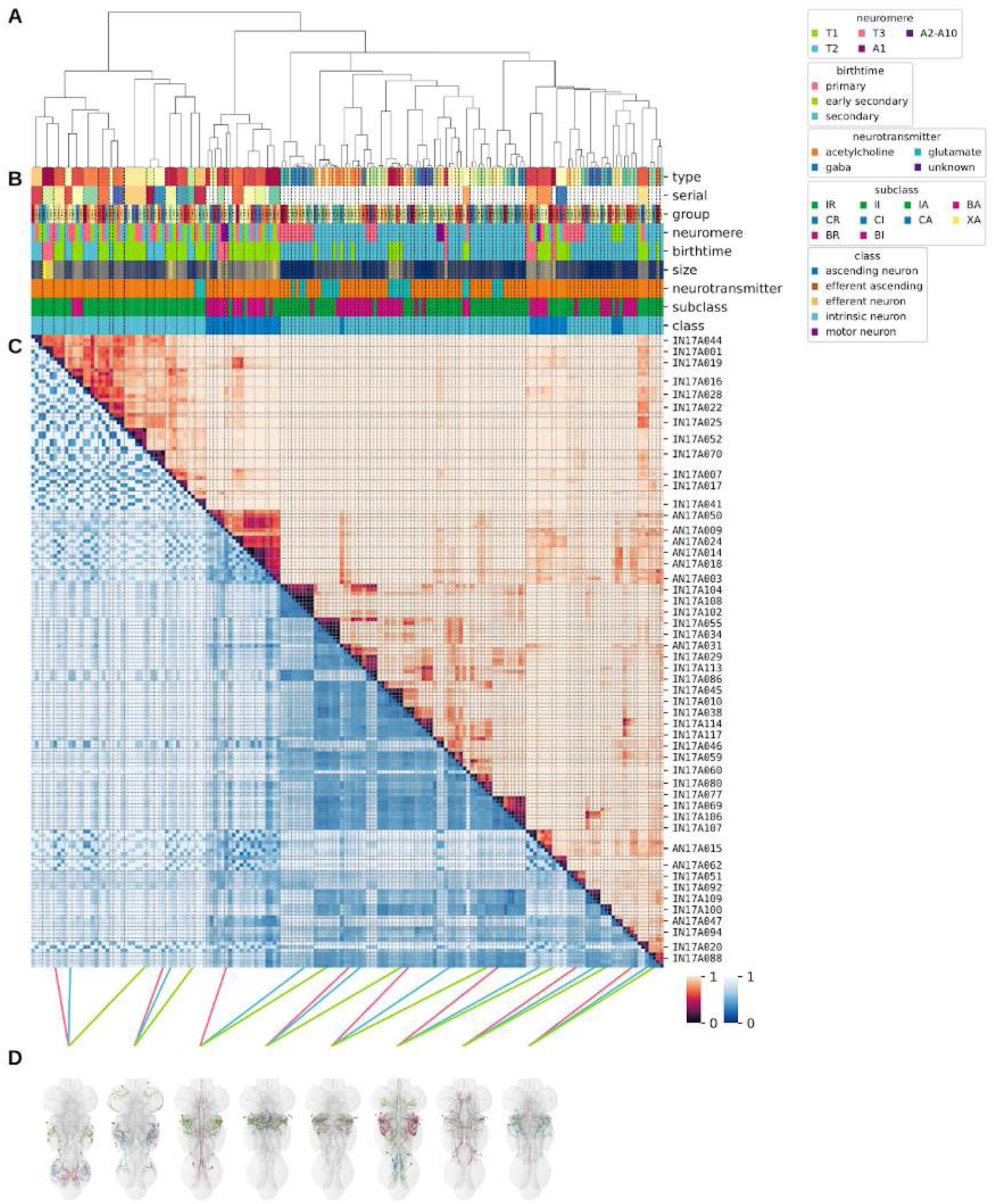
Systematic typing of hemilineage 17A. **A.** Hierarchical clustering dendrogram of hemilineage groups by laterally and serially aggregated connectivity cosine clustering. **B.** Categorical annotations of each hemilineage group, each column corresponding to the aligned leaf in A. Colours for type, serial set, and group are arbitrary for visualisation. Colours for neuromere, birthtime, neurotransmitter, subclass, and class are as in all other figures. **C.** Similarity distance heatmap for hemilineage. Cosine distance is in the upper triangle, while laterally symmetrised NBLAST distance is in the lower triangle. Systematic type names of some types are labelled. **D.** Morphologically representative groups from dendrogram subtrees. Each group, indicated by colour and line connecting to its column in B and C, is the most morphologically representative group (medoid of NBLAST distance) from a subtree of A. The subtrees (flat clusters) are equal height cuts of A determined to yield the number of groups per plot and plots in D.

**Figure 40 - figure supplement 2.**
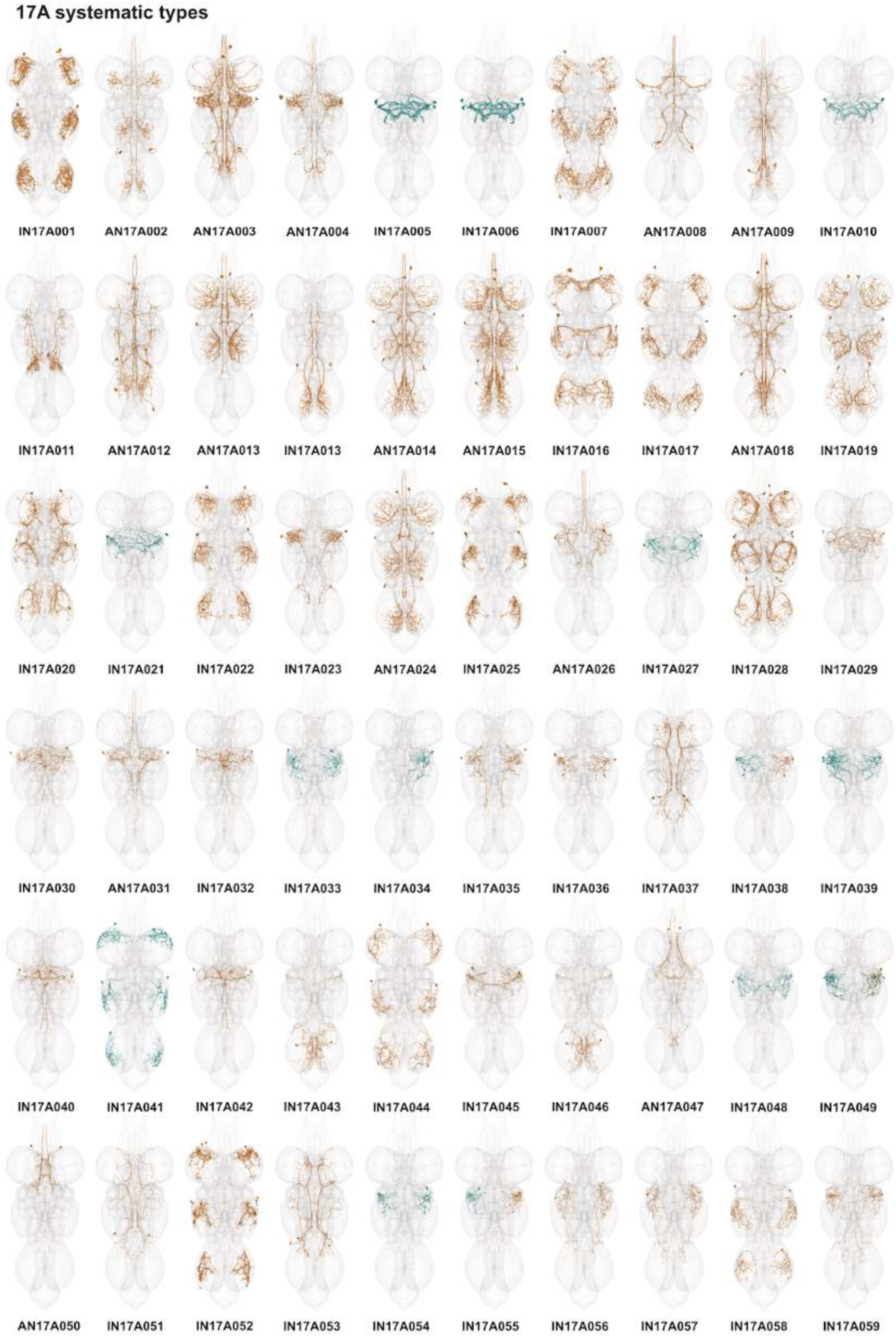
Systematic types of hemilineage 17A. Systematic types have been arranged in numerical order, with neurons of the same type that belong to distinct classes (e.g., intrinsic neuron vs ascending neuron) plotted separately but placed adjacent to each other. Individual neuron meshes have been coloured based on predicted neurotransmitter: dark orange = acetylcholine, blue = gaba, marine = glutamate, dark purple = unknown.

**Figure 40 - figure supplement 3.**
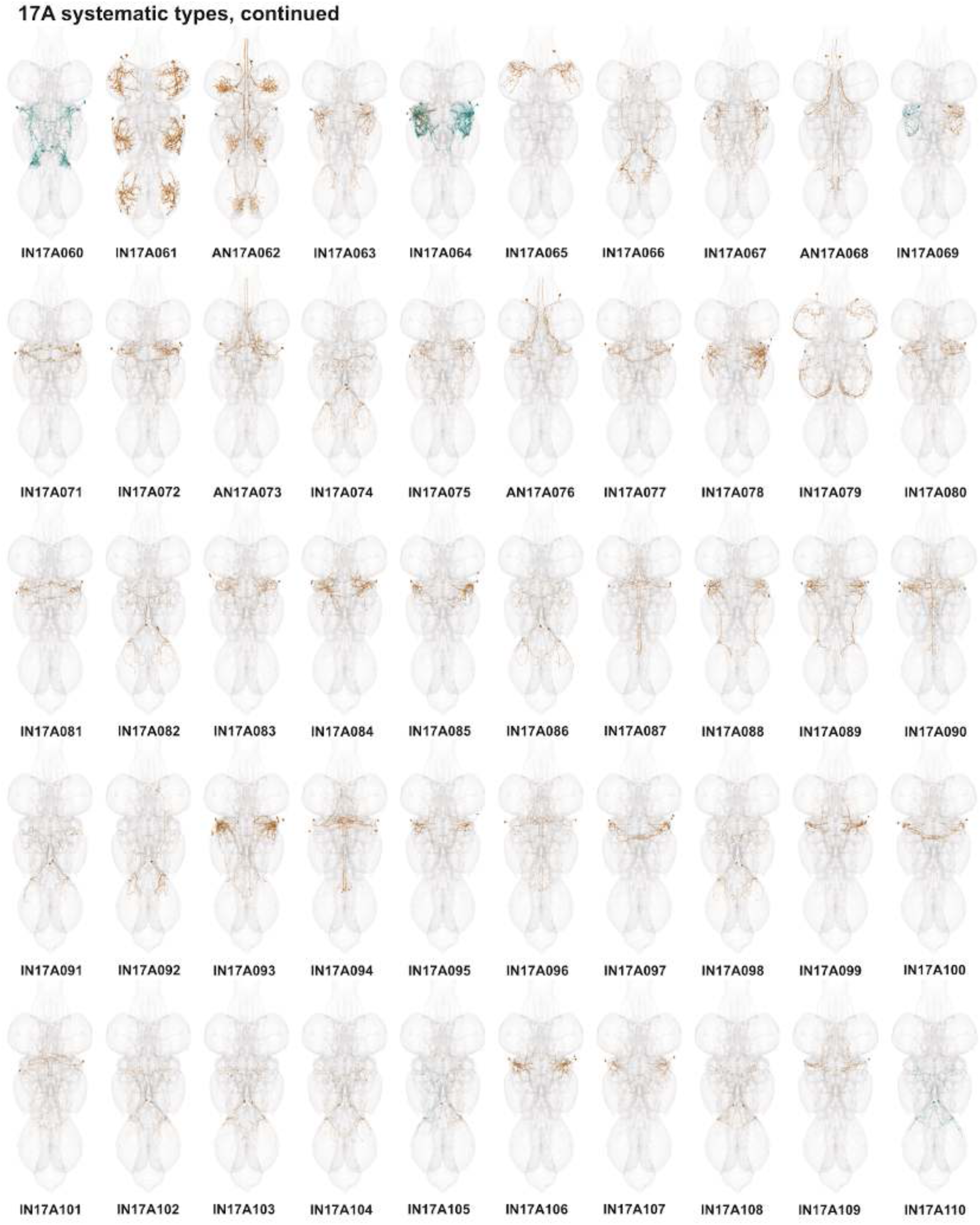
Systematic types of hemilineage 17A, continued. Systematic types have been arranged in numerical order, with neurons of the same type that belong to distinct classes (e.g., intrinsic neuron vs ascending neuron) plotted separately but placed adjacent to each other. Individual neuron meshes have been coloured based on predicted neurotransmitter: dark orange = acetylcholine, blue = gaba, marine = glutamate, dark purple = unknown.

**Figure 40 - figure supplement 4.**
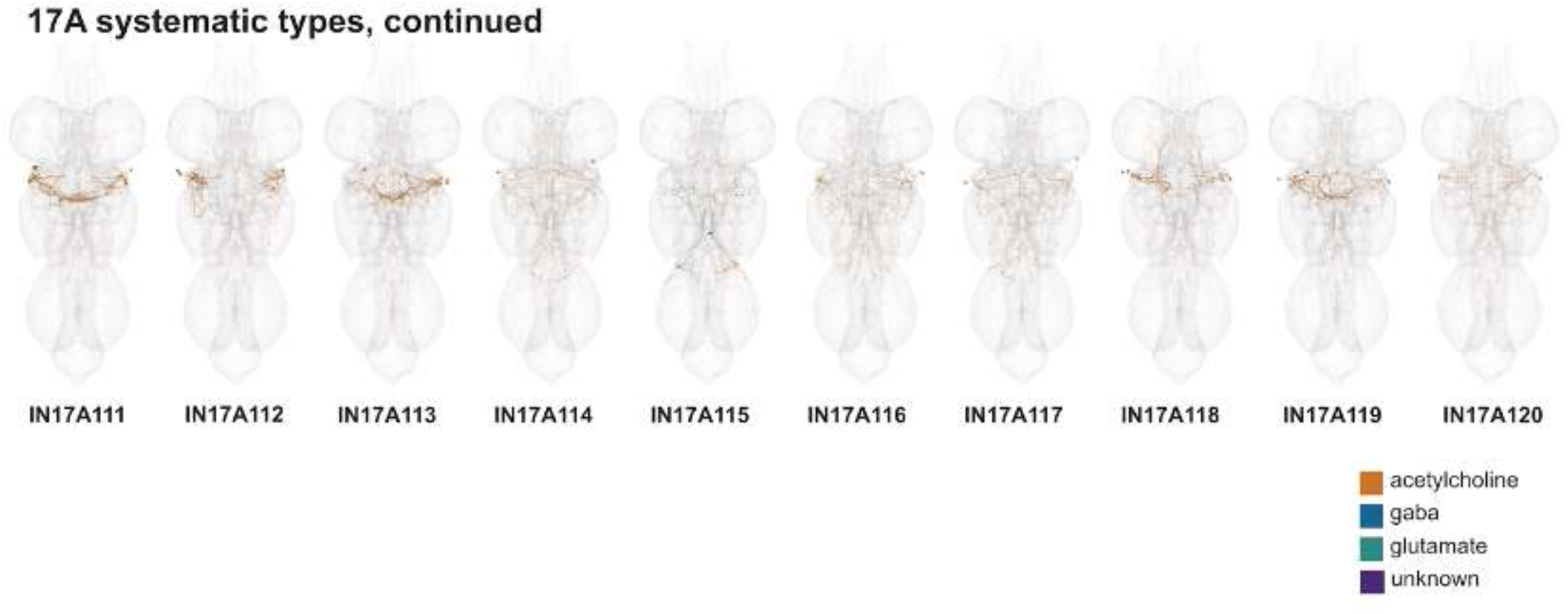
Systematic types of hemilineage 17A, continued. Systematic types have been arranged in numerical order, with neurons of the same type that belong to distinct classes (e.g., intrinsic neuron vs ascending neuron) plotted separately but placed adjacent to each other. Individual neuron meshes have been coloured based on predicted neurotransmitter: dark orange = acetylcholine, blue = gaba, marine = glutamate, dark purple = unknown.

**Figure 40 - figure supplement 5.**
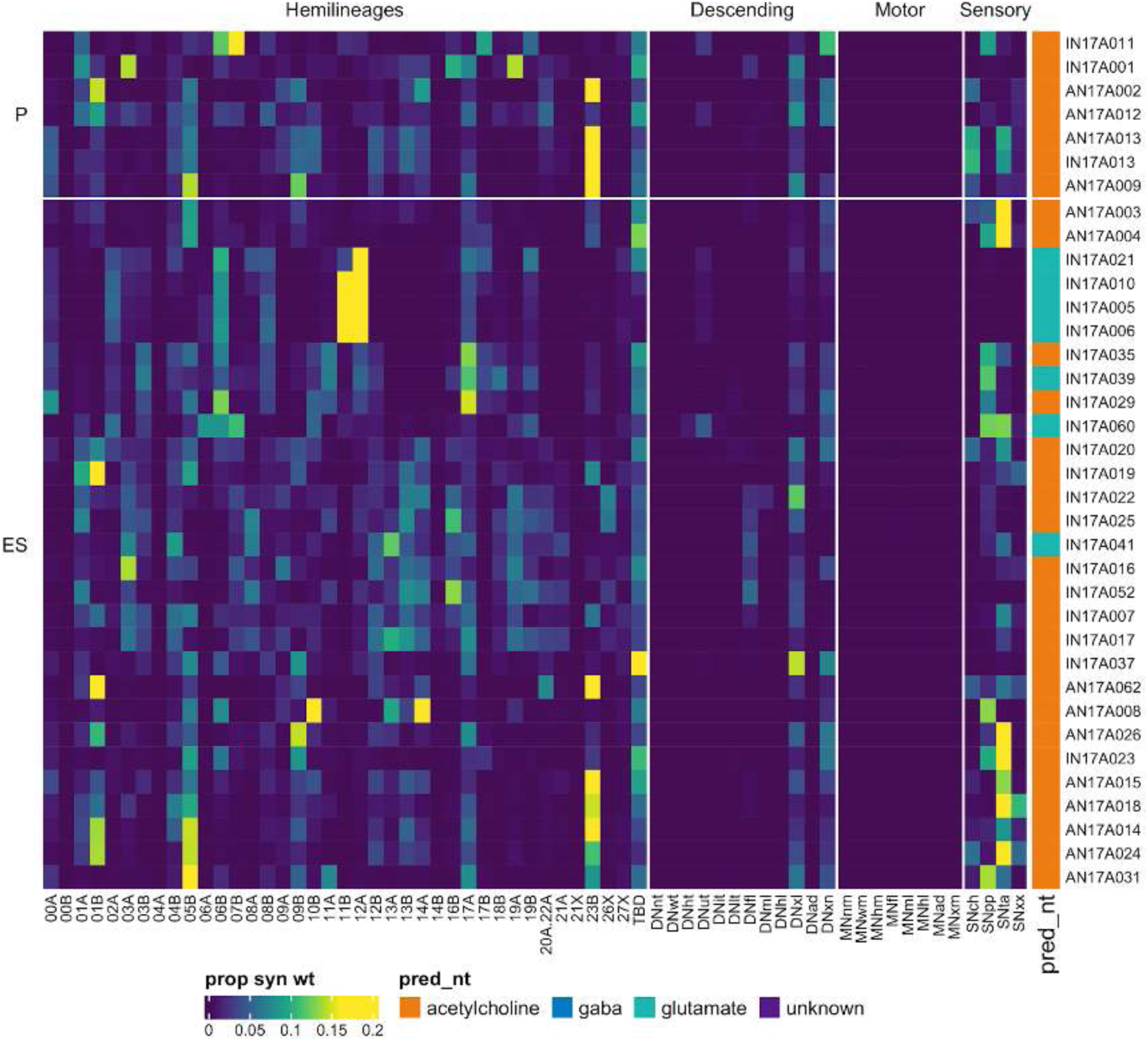
Connectivity to upstream partners by 17A primary and early secondary systematic types. Proportions of synaptic weight to systematic types from upstream partners, normalised by row. 17A neurons have been clustered within each assigned birthtime window (P = primary, ES = early secondary, S = secondary) based on both upstream and downstream connectivity to hemilineages, descending neuron subclasses, motor neuron subclasses, and sensory neuron modalities. Annotation bar is coloured by the most common predicted neurotransmitter for the neurons of each type.

**Figure 40 - figure supplement 6.**
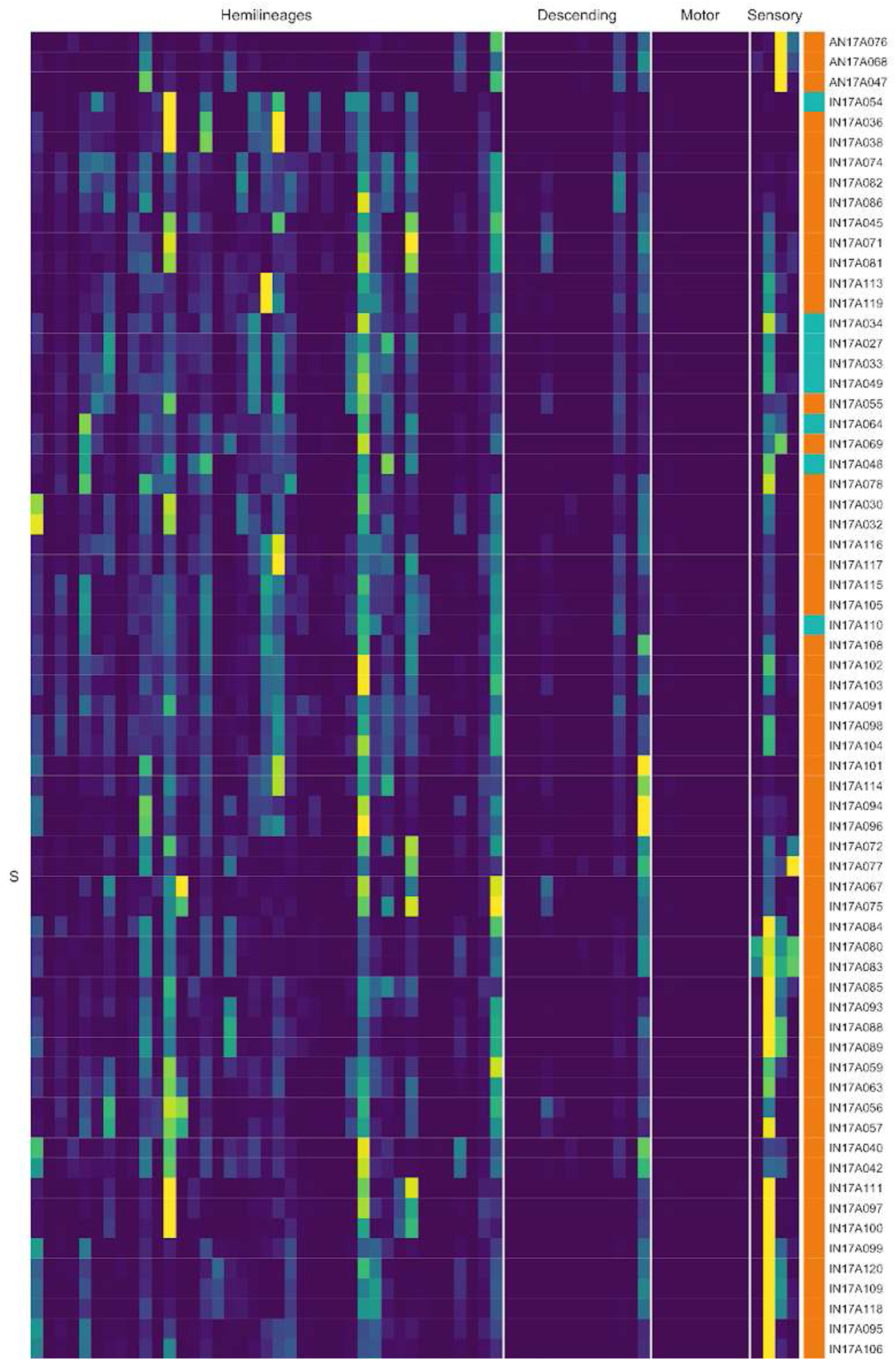
Connectivity to upstream partners by 17A secondary systematic types. Proportions of synaptic weight to systematic types from upstream partners, normalised by row. 17A neurons have been clustered within each assigned birthtime window (P = primary, ES = early secondary, S = secondary) based on both upstream and downstream connectivity to hemilineages, descending neuron subclasses, motor neuron subclasses, and sensory neuron modalities. The annotation bar is coloured by the most common predicted neurotransmitter within each type.

**Figure 40 - figure supplement 7.**
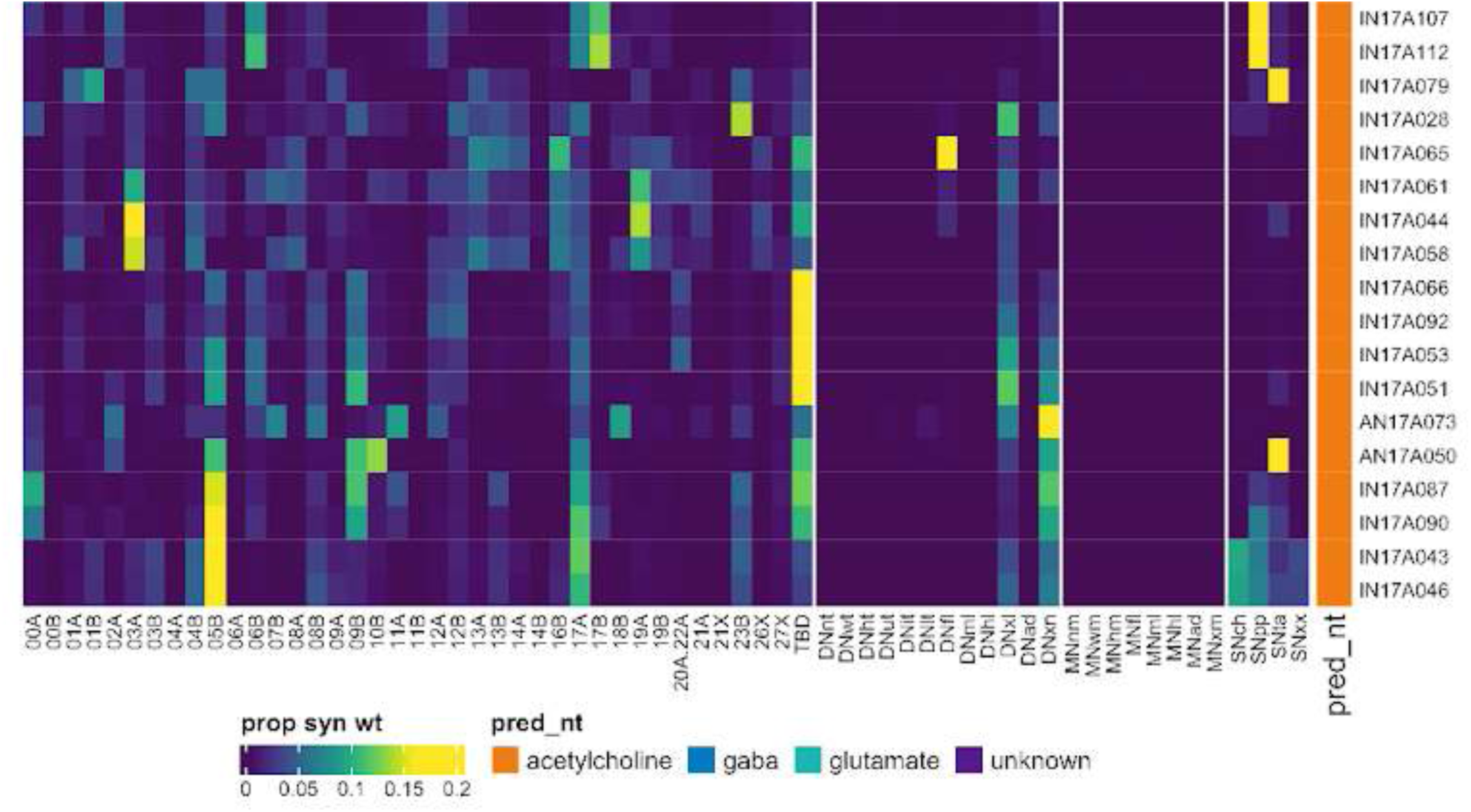
Connectivity to upstream partners by 17A secondary systematic types, continued. Proportions of synaptic weight to systematic types from upstream partners, normalised by row. 17A neurons have been clustered within each assigned birthtime window (P = primary, ES = early secondary, S = secondary) based on both upstream and downstream connectivity to hemilineages, descending neuron subclasses, motor neuron subclasses, and sensory neuron modalities. The annotation bar is coloured by the most common predicted neurotransmitter within each type.

**Figure 40 - figure supplement 8.**
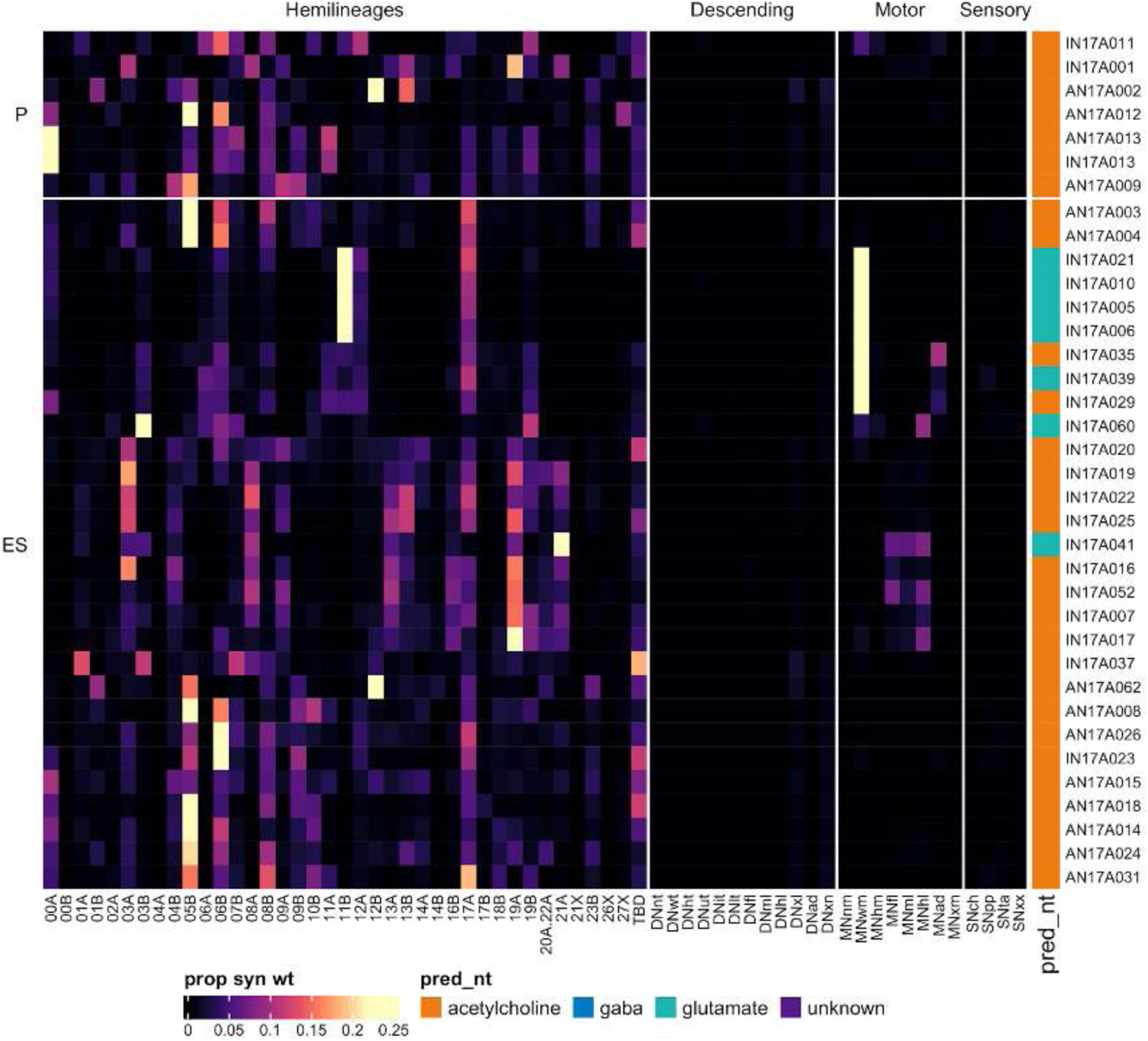
Connectivity to downstream partners by 17A primary and early secondary systematic types. Proportions of synaptic weight from systematic types to downstream partners, normalised by row. 17A neurons have been clustered within each assigned birthtime window (P = primary, ES = early secondary, S = secondary) based on both upstream and downstream connectivity to hemilineages, descending neuron subclasses, motor neuron subclasses, and sensory neuron modalities. The annotation bar is coloured by the most common predicted neurotransmitter within each type.

**Figure 40 - figure supplement 9.**
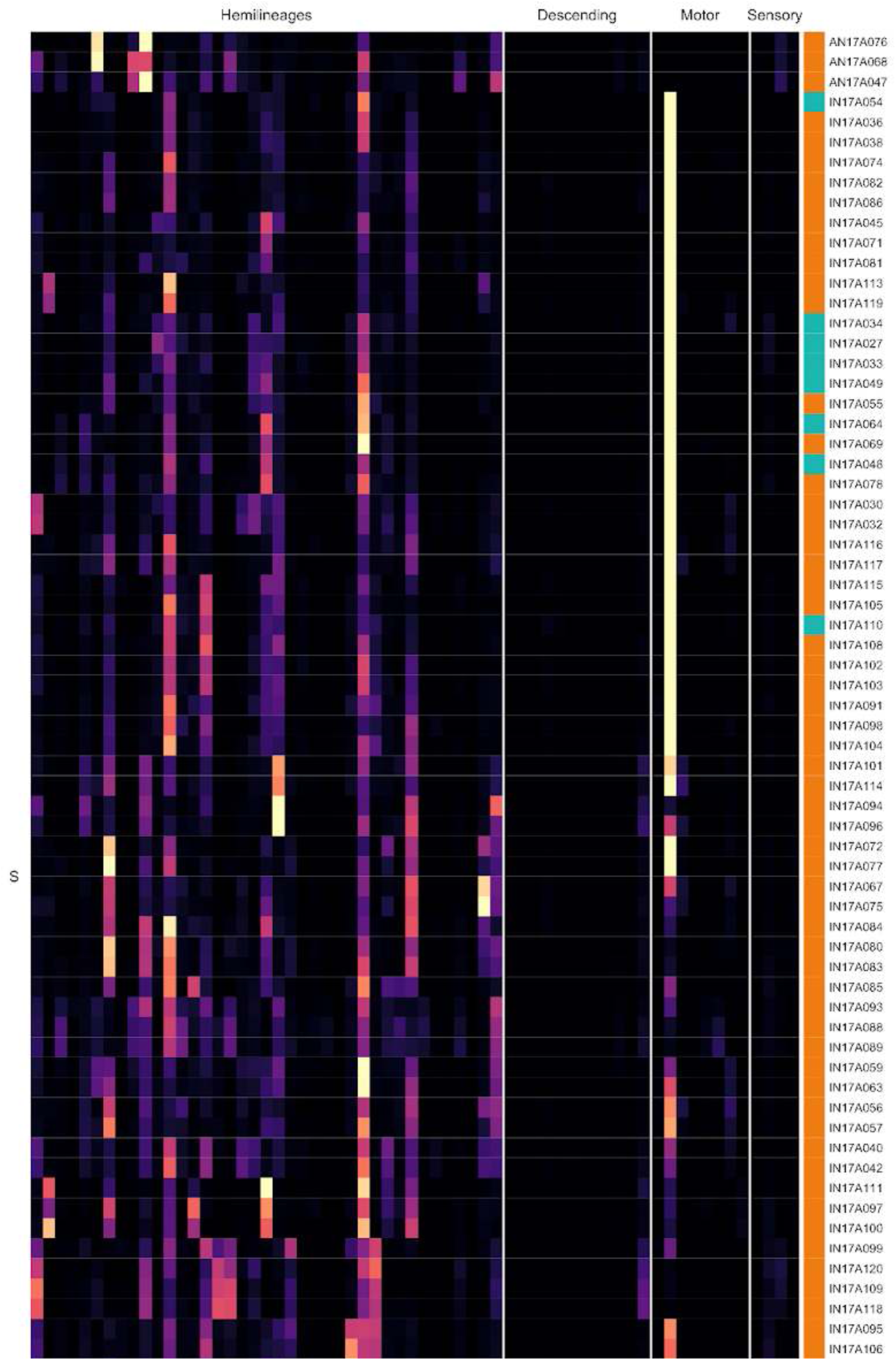
Connectivity to downstream partners by 17A secondary systematic types. Proportions of synaptic weight from systematic types to downstream partners, normalised by row. 17A neurons have been clustered within each assigned birthtime window (P = primary, ES = early secondary, S = secondary) based on both upstream and downstream connectivity to hemilineages, descending neuron subclasses, motor neuron subclasses, and sensory neuron modalities. The annotation bar is coloured by the most common predicted neurotransmitter for the neurons of each type.

**Figure 40 - figure supplement 10.**
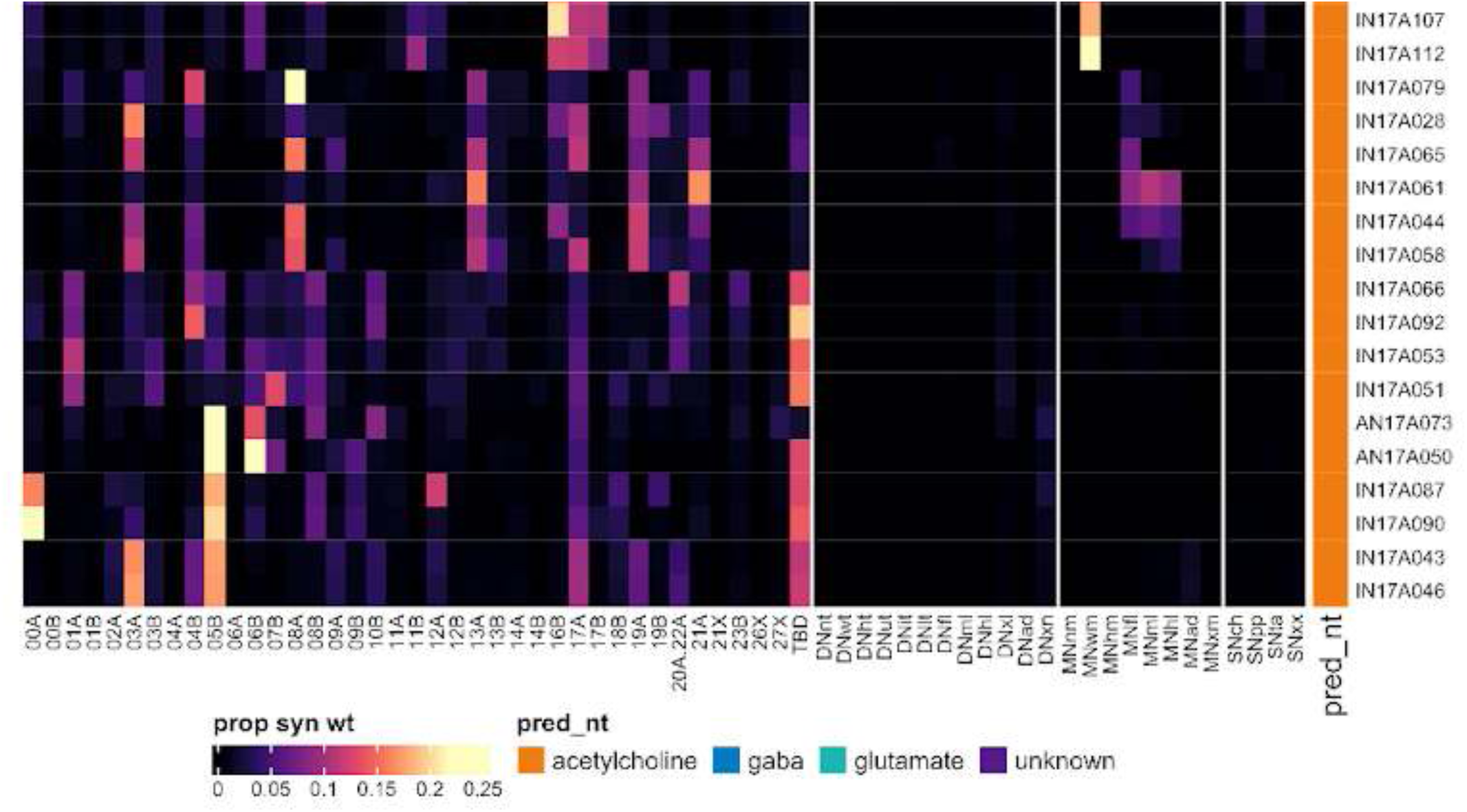
Connectivity to downstream partners by 17A secondary systematic types, continued. Proportions of synaptic weight from systematic types to downstream partners, normalised by row. 17A neurons have been clustered within each assigned birthtime window (P = primary, ES = early secondary, S = secondary) based on both upstream and downstream connectivity to hemilineages, descending neuron subclasses, motor neuron subclasses, and sensory neuron modalities. The annotation bar is coloured by the most common predicted neurotransmitter for the neurons of each type.

#### Hemilineage 18B

Hemilineage 18B is believed to derive from anterior dorsal neuroblast NB2-4 (Lacin and Truman, 2016) (but see also (Birkholz et al., 2015)), which generates one motor neuron and 7-10 local interneurons in the embryo (Schmid et al., 1999). The development of the motor neuron is arrested during larval life, and it becomes flight motor neuron MN5 in the adult (Lacin et al., 2020); we assigned this as hemilineage 18X. The 18B secondary neurons survive in T2-A1 only and enter the anterior dorsal neuromere near 17A, crossing the midline in the anterior intermediate commissure (Truman et al., 2004) to innervate the tectulum and dorsolateral leg neuropils (Harris et al., 2015) (Figure 41A). 18B secondary neurons are mainly cholinergic (Lacin et al., 2019), but we also identified a small subset predicted glutamatergic (Figure 41C top, bottom) that we matched to neurons reported to be electrically coupled to the Giant Fiber descending neurons (Kennedy and Broadie, 2018).

**Figure 41.**
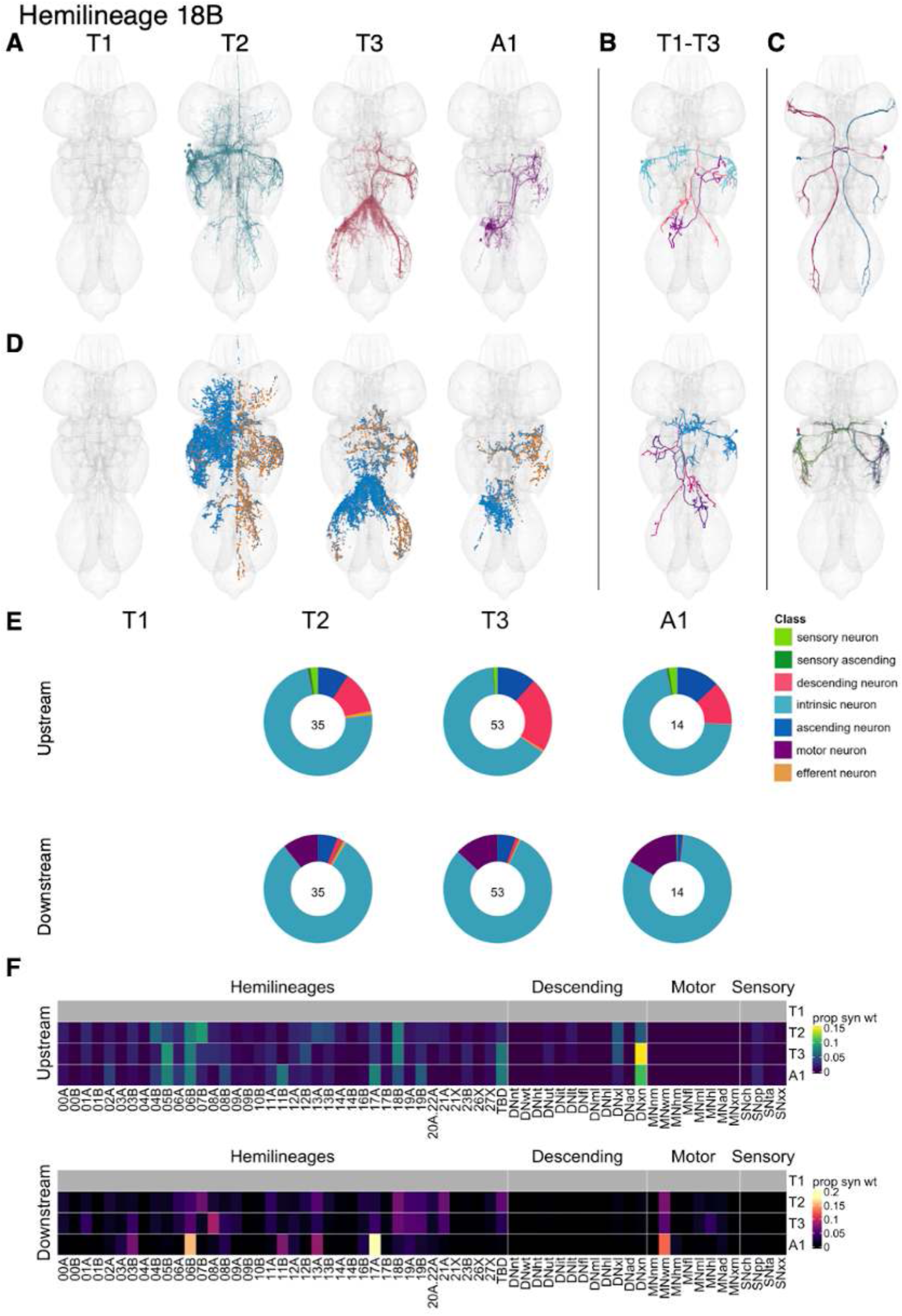
Hemilineage 18B. **A.** Meshes of all RHS secondary neurons plotted in neuromere-specific colours. **B.** “Representative” secondary neuron skeletons plotted in hemineuromere-specific colours. The skeleton with the top accumulated NBLAST score among all neurons from the hemilineage in a given hemineuromere was used. **C.** Neuron meshes of selected examples. Top: GFC1, electrical group 10228. Bottom: GFC2, electrical subcluster 13645. **D.** Predicted synapses of RHS secondary neurons. Blue: postsynapses; dark orange: presynapses. **E.** Proportions of connections from secondary neurons to upstream or downstream partners, normalised by neuromere and coloured by broad class. Numbers of query neurons appear in the centre. **F.** Proportions of synaptic weight from secondary neurons originating in each neuromere to upstream or downstream partners, normalised by row.

Bilateral activation of 18B secondary neurons initiates walking, sometimes accompanied by partially folded wing movements, eventually followed by a jump and wing flapping (Harris et al., 2015). 18B inputs vary from neuromere to neuromere but include 05B, 06B, 11B, 17A, 18B, and 19B with segmental variation, and descending neurons throughout. Their postsynaptic partners also vary but are predominantly wing motor neurons in T2-A1 and 03B, 06B, 08A, 11B, 13A, and especially 17A, again with segmental variation (Figure 41F). A few 18B types receive substantial input from proprioceptive (e.g., IN18B026) or tactile (e.g., IN18B012) sensory neurons (Figure 41 - figure supplement 3-4). Several types activate neck (AN18B023), leg (e.g., IN18B006), or abdominal (IN18B033) motor neurons (Figure 41 - figure supplement 5-6).

**Figure 41 - figure supplement 1.**
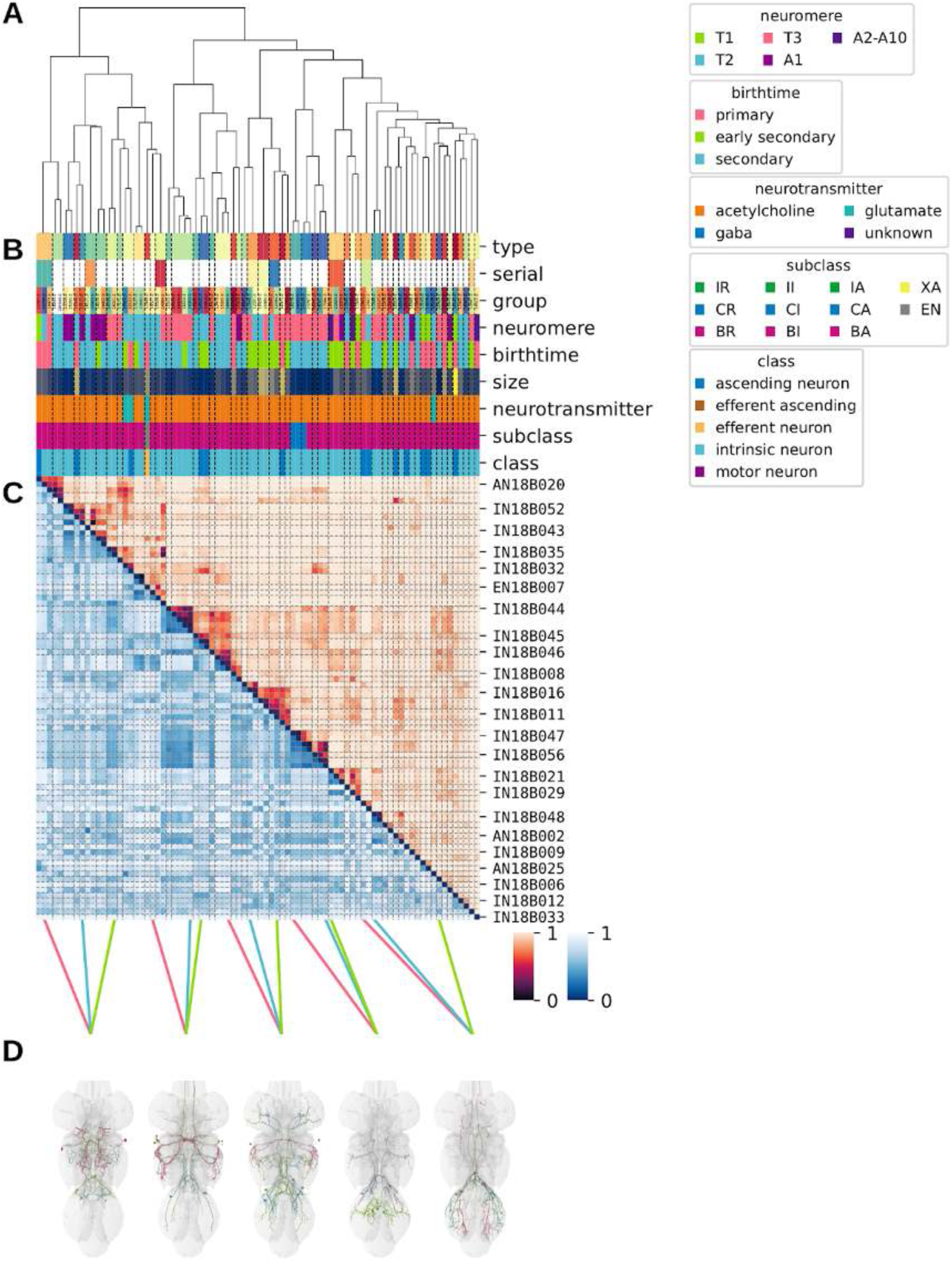
Systematic typing of hemilineage 18B. **A.** Hierarchical clustering dendrogram of hemilineage groups by laterally and serially aggregated connectivity cosine clustering. **B.** Categorical annotations of each hemilineage group, each column corresponding to the aligned leaf in A. Colours for type, serial set, and group are arbitrary for visualisation. Colours for neuromere, birthtime, neurotransmitter, subclass, and class are as in all other figures. **C.** Similarity distance heatmap for hemilineage. Cosine distance is in the upper triangle, while laterally symmetrised NBLAST distance is in the lower triangle. Systematic type names of some types are labelled. **D.** Morphologically representative groups from dendrogram subtrees. Each group, indicated by colour and line connecting to its column in B and C, is the most morphologically representative group (medoid of NBLAST distance) from a subtree of A. The subtrees (flat clusters) are equal height cuts of A determined to yield the number of groups per plot and plots in D.

**Figure 41 - figure supplement 2.**
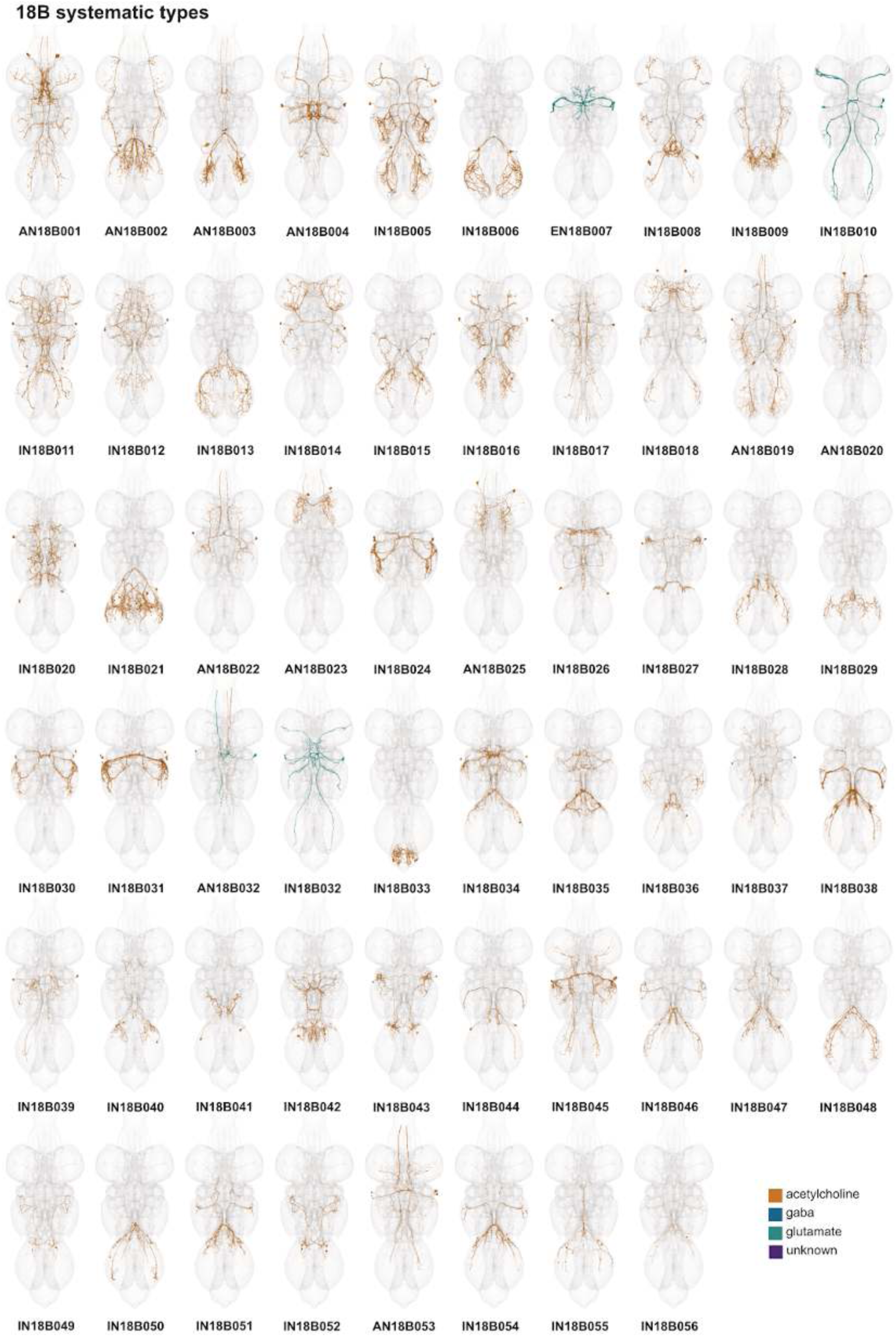
Systematic types of hemilineage 18B. Systematic types have been arranged in numerical order, with neurons of the same type that belong to distinct classes (e.g., intrinsic neuron vs ascending neuron) plotted separately but placed adjacent to each other. Individual neuron meshes have been coloured based on predicted neurotransmitter: dark orange = acetylcholine, blue = gaba, marine = glutamate, dark purple = unknown.

**Figure 41 - figure supplement 3.**
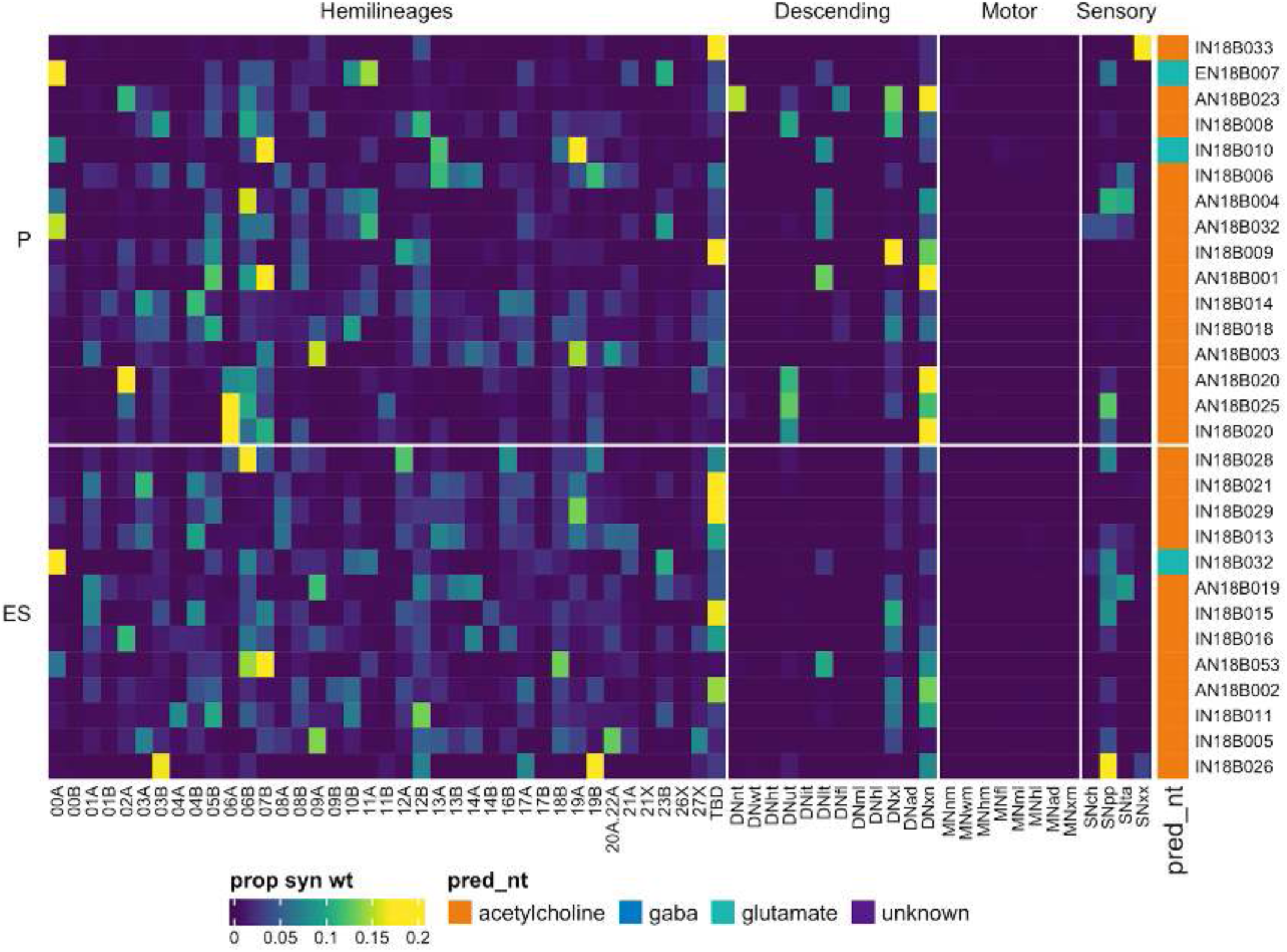
Connectivity to upstream partners by 18B primary and early secondary systematic types. Proportions of synaptic weight to systematic types from upstream partners, normalised by row. 18B neurons have been clustered within each assigned birthtime window (P = primary, ES = early secondary, S = secondary) based on both upstream and downstream connectivity to hemilineages, descending neuron subclasses, motor neuron subclasses, and sensory neuron modalities. Annotation bar is coloured by the most common predicted neurotransmitter for the neurons of each type.

**Figure 41 - figure supplement 4.**
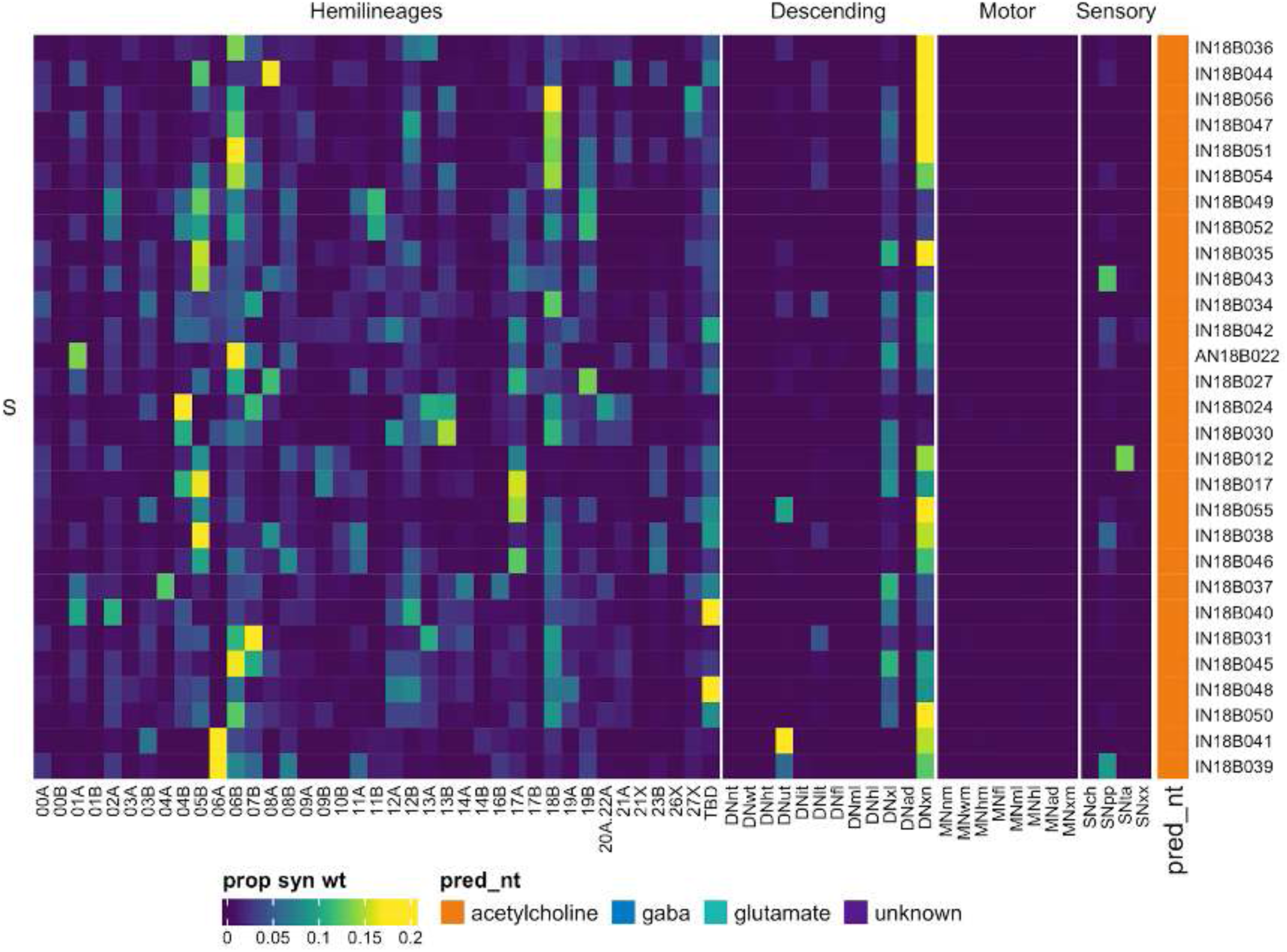
Connectivity to upstream partners by 18B secondary systematic types. Proportions of synaptic weight to systematic types from upstream partners, normalised by row. 18B neurons have been clustered within each assigned birthtime window (P = primary, ES = early secondary, S = secondary) based on both upstream and downstream connectivity to hemilineages, descending neuron subclasses, motor neuron subclasses, and sensory neuron modalities. Annotation bar is coloured by the most common predicted neurotransmitter for the neurons of each type.

**Figure 41 - figure supplement 5.**
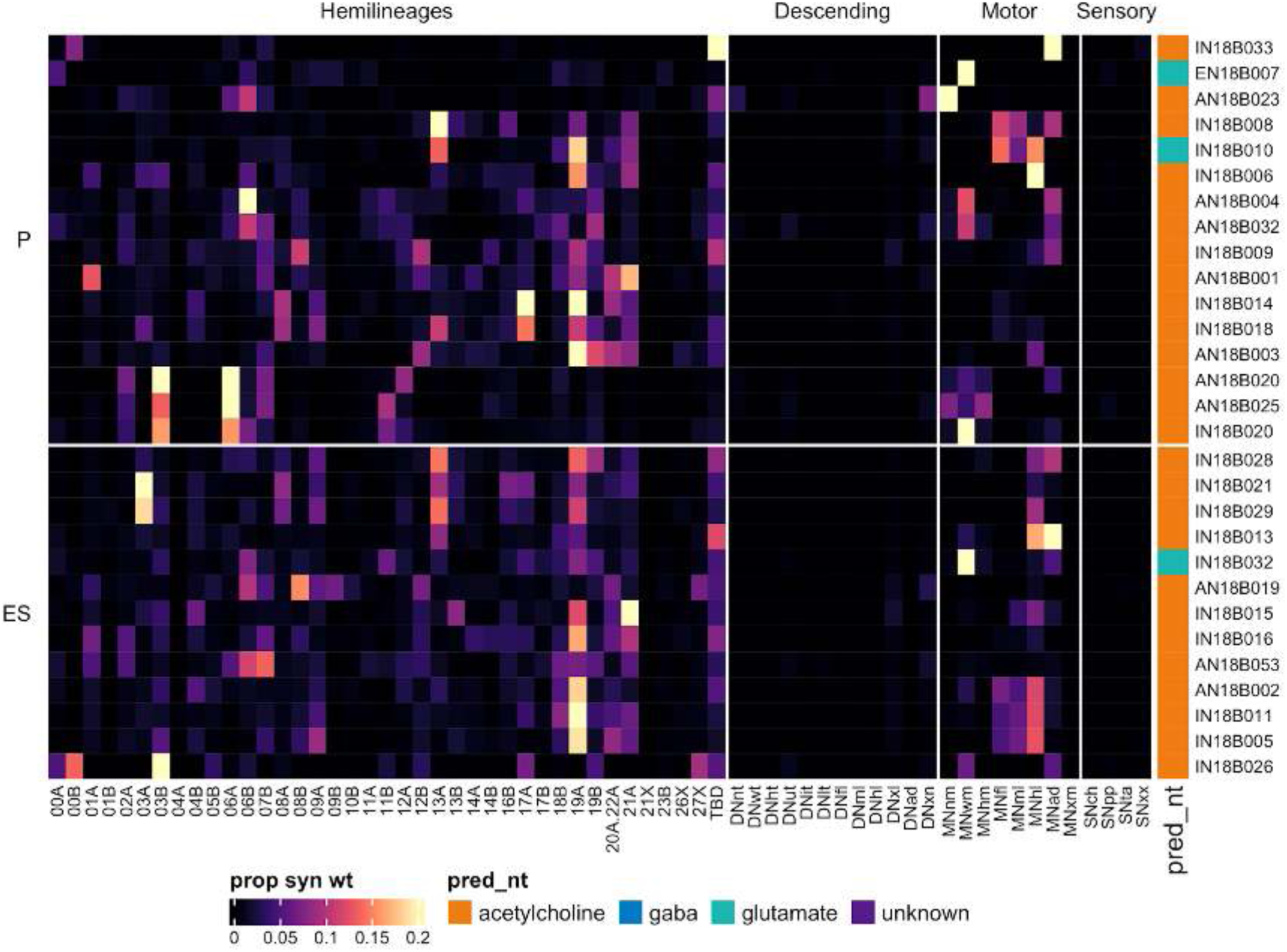
Connectivity to downstream partners by 18B primary and early secondary systematic types. Proportions of synaptic weight from systematic types to downstream partners, normalised by row. 18B neurons have been clustered within each assigned birthtime window (P = primary, ES = early secondary, S = secondary) based on both upstream and downstream connectivity to hemilineages, descending neuron subclasses, motor neuron subclasses, and sensory neuron modalities. The annotation bar is coloured by the most common predicted neurotransmitter within each type.

**Figure 41 - figure supplement 6.**
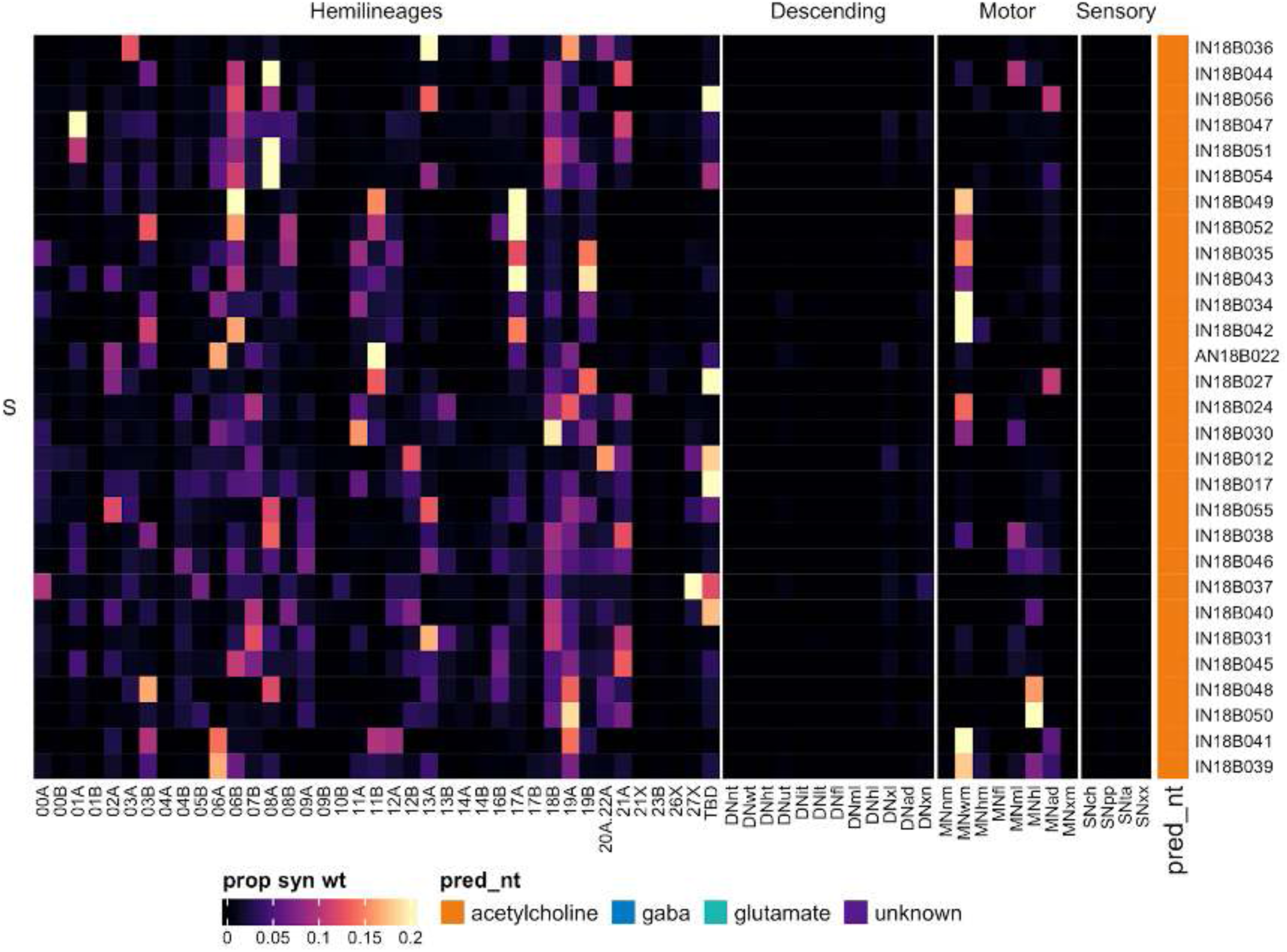
Connectivity to downstream partners by 18B secondary systematic types. Proportions of synaptic weight from systematic types to downstream partners, normalised by row. 18B neurons have been clustered within each assigned birthtime window (P = primary, ES = early secondary, S = secondary) based on both upstream and downstream connectivity to hemilineages, descending neuron subclasses, motor neuron subclasses, and sensory neuron modalities. The annotation bar is coloured by the most common predicted neurotransmitter within each type.

#### Hemilineage 19A

Hemilineages 19A and 19B derive from posterior dorsal neuroblast NB6-2 (Birkholz et al., 2015; Lacin and Truman, 2016), which generates 3-4 intersegmental interneurons and ∼20 local interneurons in the embryo (Schmid et al., 1999). Both secondary hemilineages survive in all thoracic neuromeres, but the number of 19B in T1 is greatly reduced (Truman et al., 2004). The 19A secondary hemilineage is mainly restricted to ipsilateral leg neuropil (Harris et al., 2015) but also includes subtypes that cross the midline in a small anterior and ventral tract to wrap around the contralateral leg neuropil.

We identified 19A secondary neurons in similar numbers in all three thoracic neuromeres (Figure 42E). They enter the posterior of each neuromere quite dorsally and mainly innervate the ipsilateral leg neuropil (Figure 42C), although a subpopulation crosses the midline just anterior to the leg sensory neuropils (Figure 42C). Hemilineage 19A secondary neurons are typically gabaergic (Lacin et al., 2019), but we also identified an early born cholinergic population that descend or ascend to adjacent neuromeres (Figure 7G).

**Figure 42.**
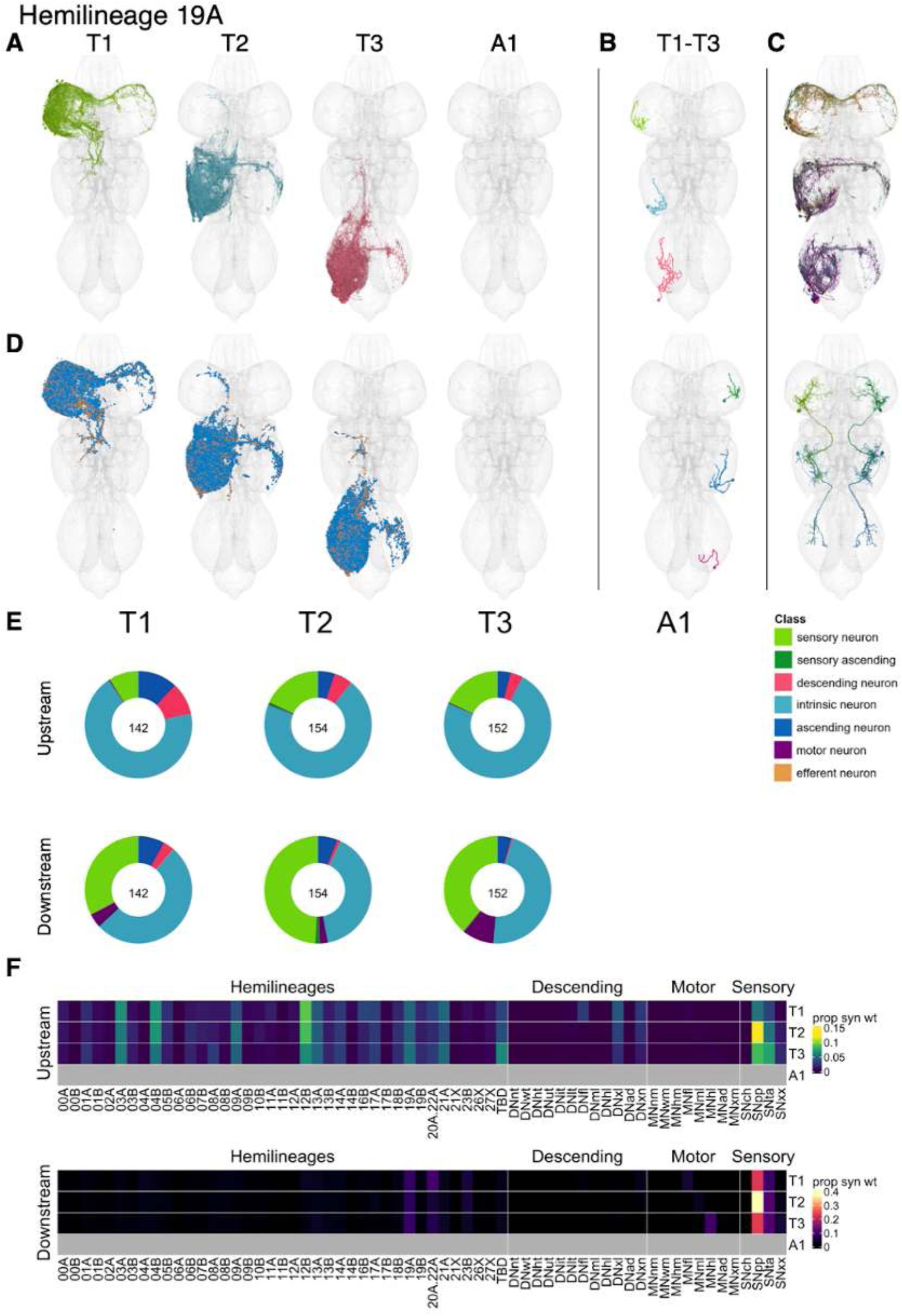
Hemilineage 19A. **A.** Meshes of all RHS secondary neurons plotted in neuromere-specific colours. **B.** “Representative” secondary neuron skeletons plotted in hemineuromere-specific colours. The skeleton with the top accumulated NBLAST score among all neurons from the hemilineage in a given hemineuromere was used. **C.** Neuron meshes of selected examples. Top: cholinergic sequential serial set 10173. Bottom: independent leg serial set 14080. **D.** Predicted synapses of RHS secondary neurons. Blue: postsynapses; dark orange: presynapses. **E.** Proportions of connections from secondary neurons to upstream or downstream partners, normalised by neuromere and coloured by broad class. Numbers of query neurons appear in the centre. **F.** Ratios of connections from secondary neurons originating in each neuromere to upstream or downstream partners, normalised by row.

19A secondary neurons receive considerable input from 03A, 04B, 12B, and both proprioceptive and tactile sensory neurons. Unusually, their main downstream targets are sensory neurons, particularly proprioceptive neurons and especially in T2 (Figure 42F). Bilateral activation of 19A results in incessant waving of T2 legs without movement of T1 or T3 legs (Harris et al., 2015).

**Figure 42 - figure supplement 1.**
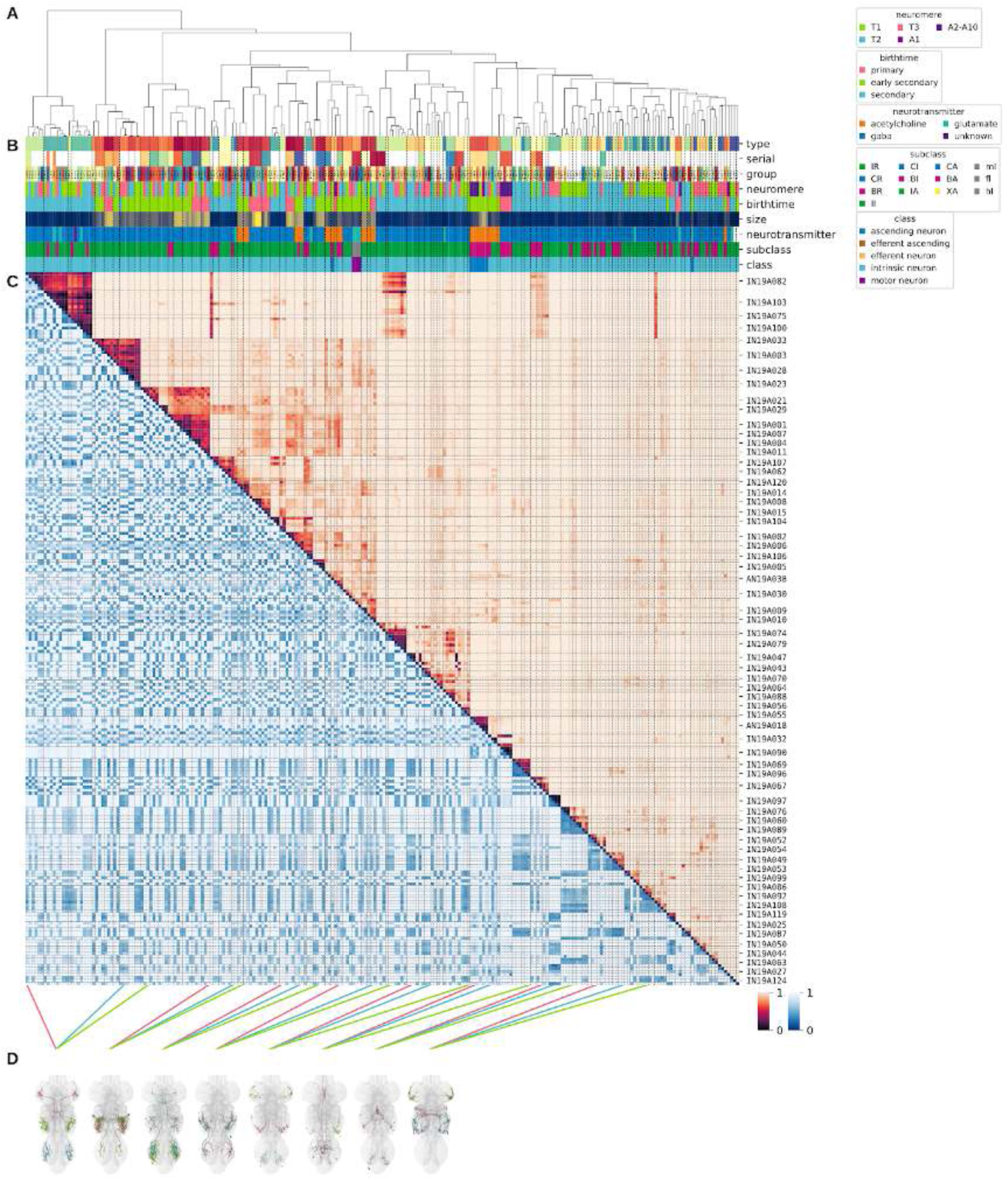
Systematic typing of hemilineage 19A. Types for motor neurons were assigned separately as outlined in our accompanying manuscript (Cheong et al., 2023). **A.** Hierarchical clustering dendrogram of hemilineage groups by laterally and serially aggregated connectivity cosine clustering. **B.** Categorical annotations of each hemilineage group, each column corresponding to the aligned leaf in A. Colours for type, serial set, and group are arbitrary for visualisation. Colours for neuromere, birthtime, neurotransmitter, subclass, and class are as in all other figures. **C.** Similarity distance heatmap for hemilineage. Cosine distance is in the upper triangle, while laterally symmetrised NBLAST distance is in the lower triangle. Systematic type names of some types are labelled. **D.** Morphologically representative groups from dendrogram subtrees. Each group, indicated by colour and line connecting to its column in B and C, is the most morphologically representative group (medoid of NBLAST distance) from a subtree of A. The subtrees (flat clusters) are equal height cuts of A determined to yield the number of groups per plot and plots in D.

**Figure 42 - figure supplement 2.**
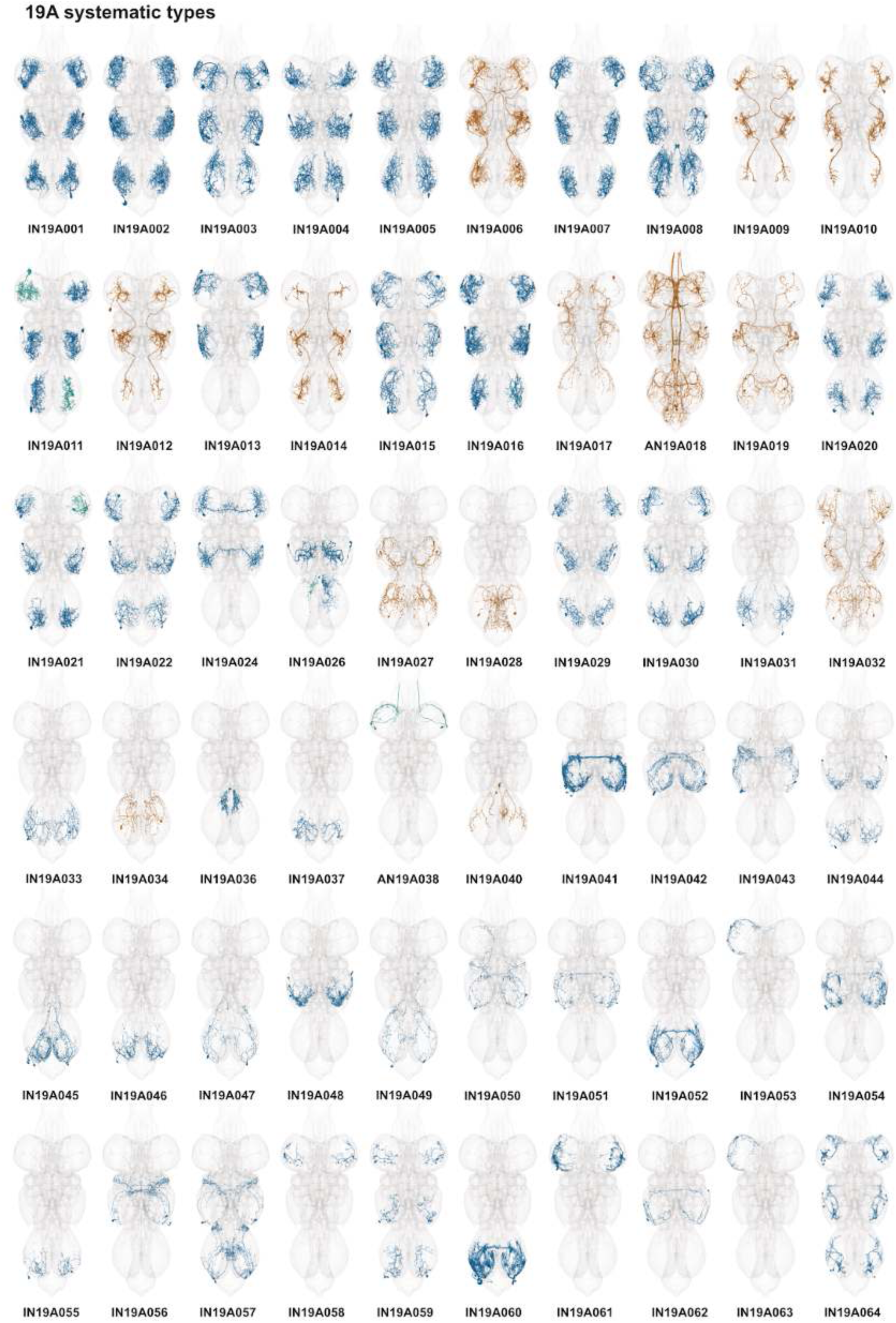
Systematic types of hemilineage 19A. Systematic types have been arranged in numerical order, with neurons of the same type that belong to distinct classes (e.g., intrinsic neuron vs ascending neuron) plotted separately but placed adjacent to each other. Individual neuron meshes have been coloured based on predicted neurotransmitter: dark orange = acetylcholine, blue = gaba, marine = glutamate, dark purple = unknown.

**Figure 42 - figure supplement 3.**
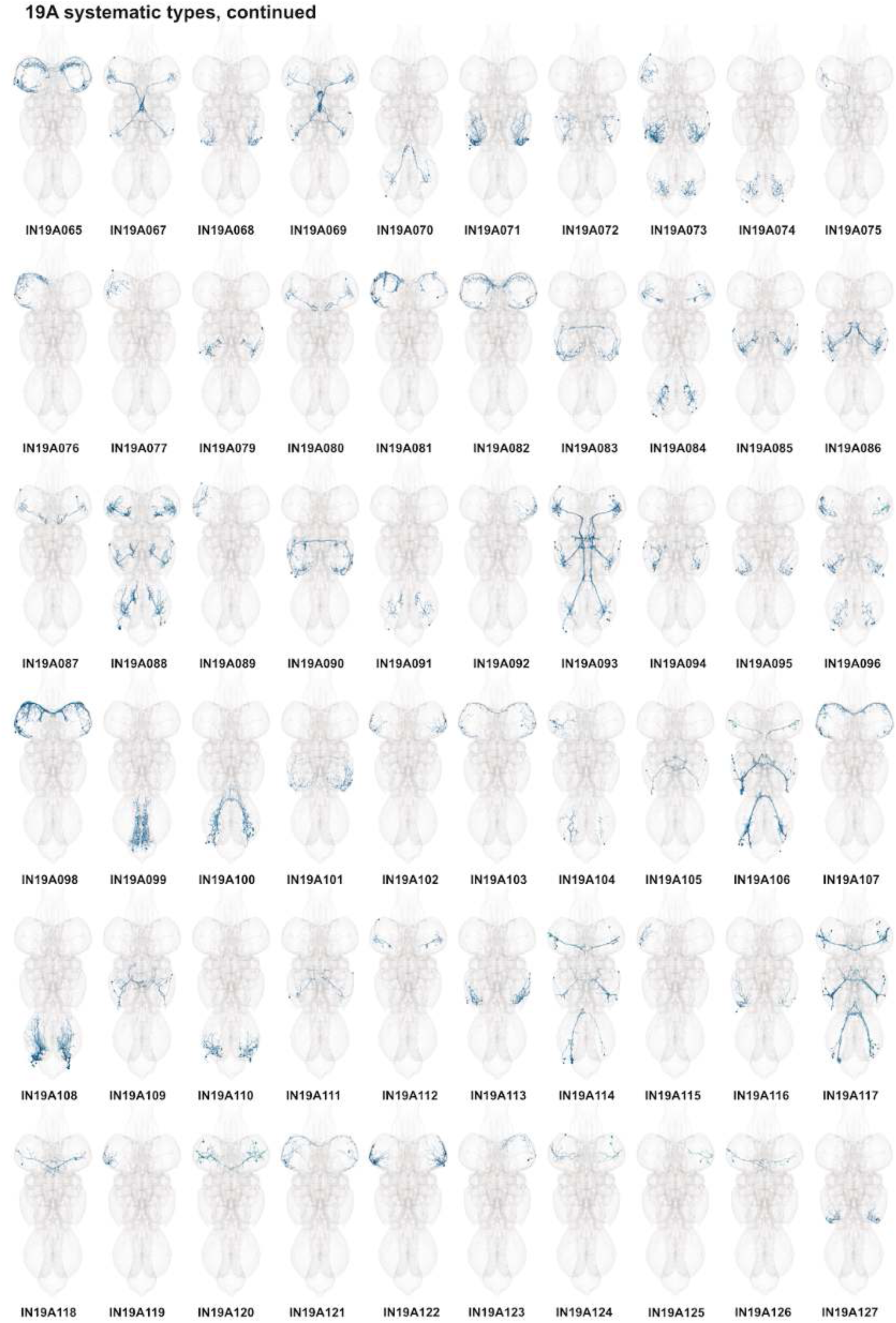
Systematic types of hemilineage 19A, continued. Systematic types have been arranged in numerical order, with neurons of the same type that belong to distinct classes (e.g., intrinsic neuron vs ascending neuron) plotted separately but placed adjacent to each other. Individual neuron meshes have been coloured based on predicted neurotransmitter: dark orange = acetylcholine, blue = gaba, marine = glutamate, dark purple = unknown.

**Figure 42 - figure supplement 4.**
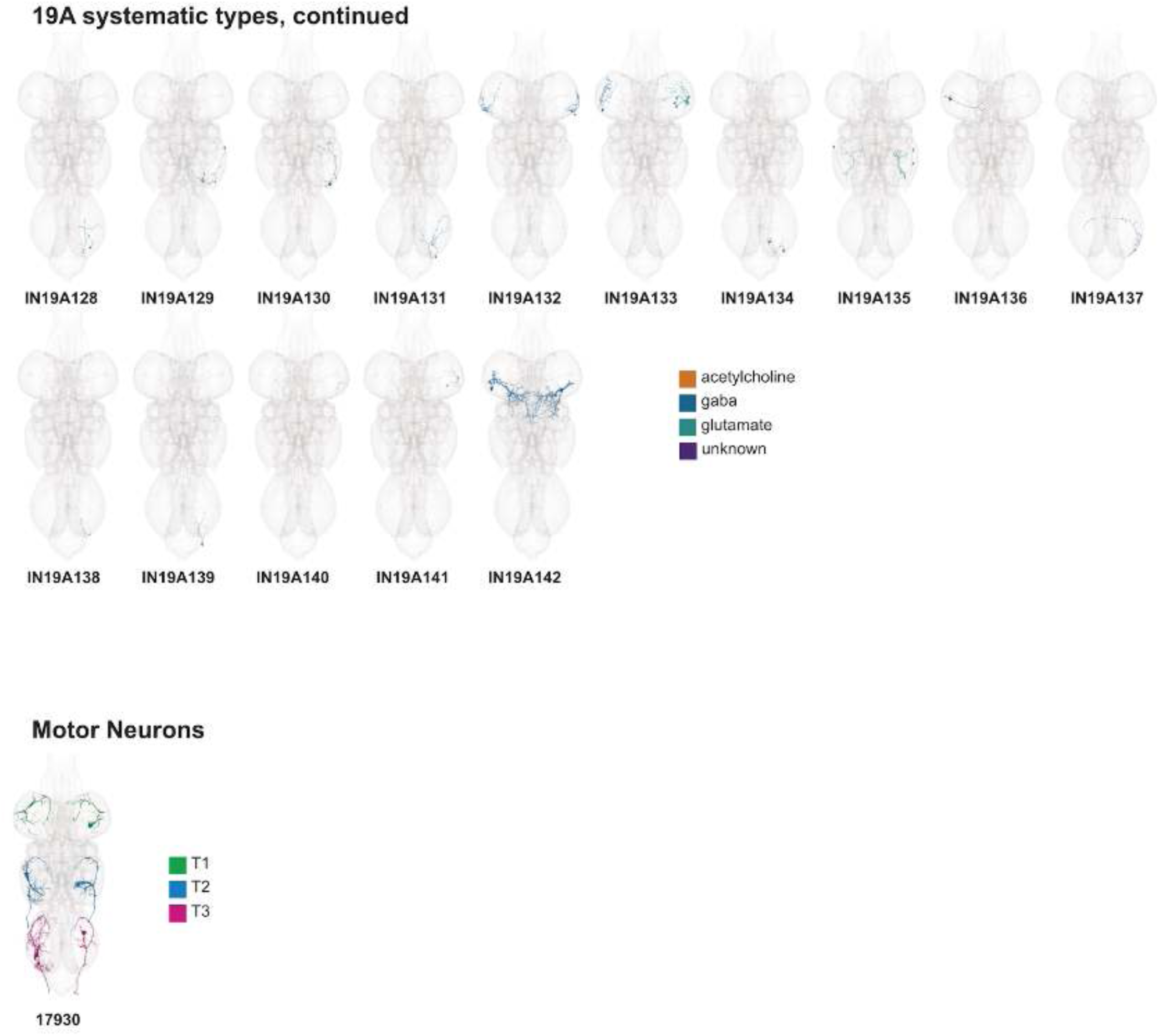
Systematic types of hemilineage 19A, continued. Systematic types have been arranged in numerical order, with neurons of the same type that belong to distinct classes (e.g., intrinsic neuron vs ascending neuron) plotted separately but placed adjacent to each other. Individual neuron meshes have been coloured based on predicted neurotransmitter: dark orange = acetylcholine, blue = gaba, marine = glutamate, dark purple = unknown. Motor neurons (typed separately in (Cheong et al., 2023)) have been plotted by serial set if identified in multiple neuromeres and by systematic type if not. Individual motor neuron meshes have been coloured based on soma neuromere: dark green = T1, blue = T2, magenta = T3.

**Figure 42 - figure supplement 5.**
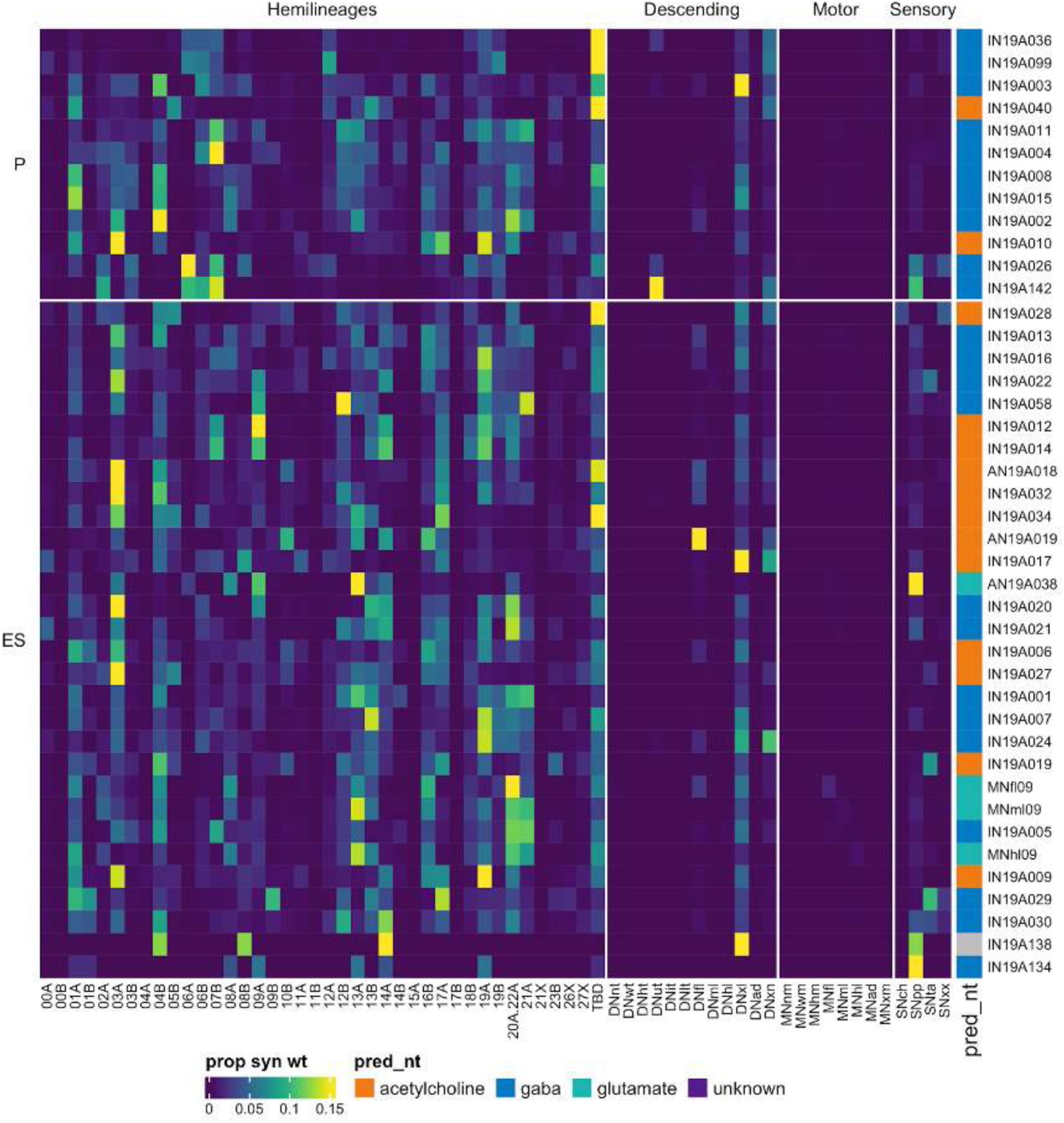
Connectivity to upstream partners by 19A primary and early secondary systematic types. Proportions of synaptic weight to systematic types from upstream partners, normalised by row. 19A neurons have been clustered within each assigned birthtime window (P = primary, ES = early secondary, S = secondary) based on both upstream and downstream connectivity to hemilineages, descending neuron subclasses, motor neuron subclasses, and sensory neuron modalities. Annotation bar is coloured by the most common predicted neurotransmitter for the neurons of each type.

**Figure 42 - figure supplement 6.**
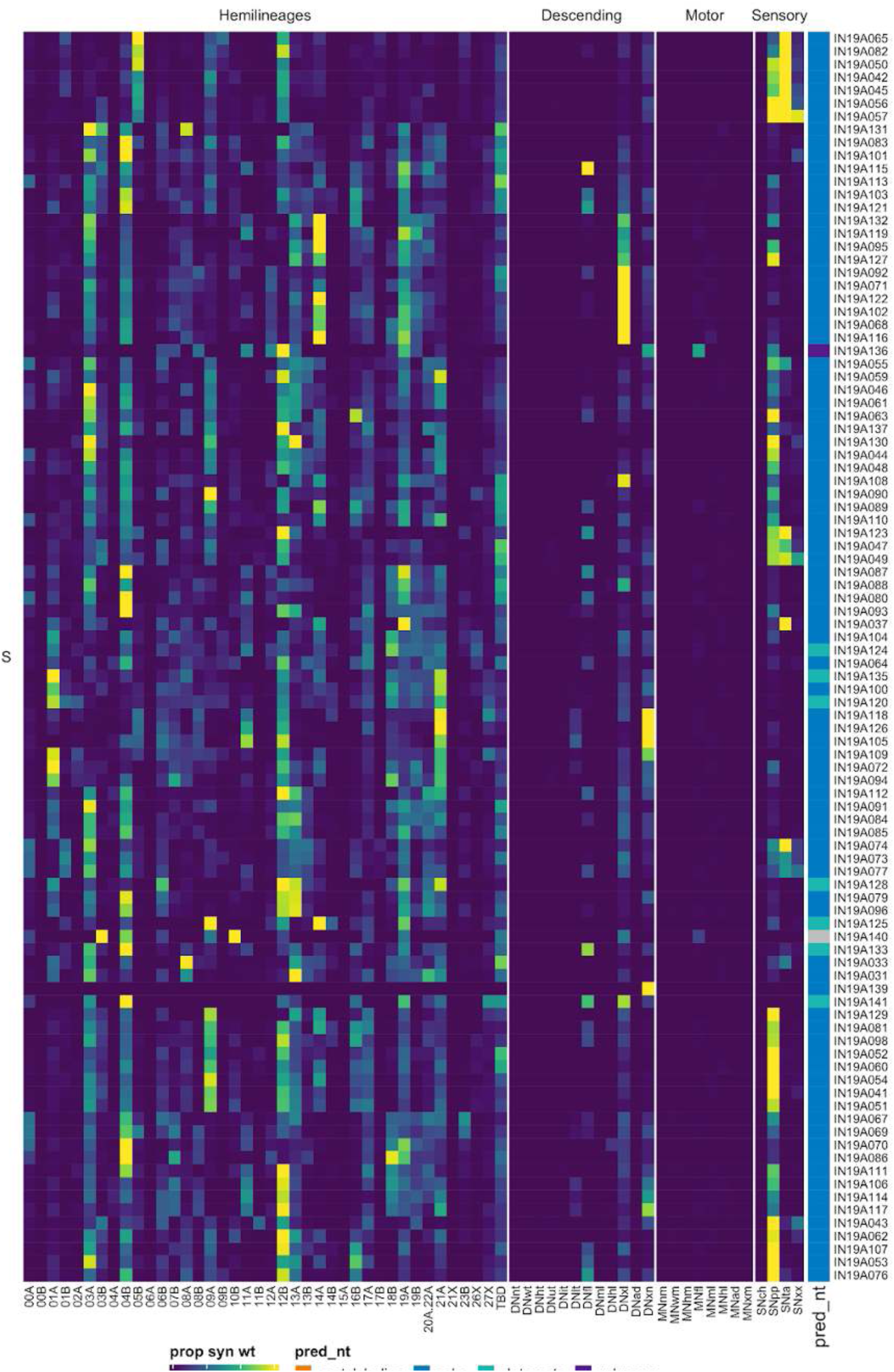
Connectivity to upstream partners by 19A secondary systematic types. Proportions of synaptic weight to systematic types from upstream partners, normalised by row. 19A neurons have been clustered within each assigned birthtime window (P = primary, ES = early secondary, S = secondary) based on both upstream and downstream connectivity to hemilineages, descending neuron subclasses, motor neuron subclasses, and sensory neuron modalities. The annotation bar is coloured by the most common predicted neurotransmitter within each type.

**Figure 42 - figure supplement 7.**
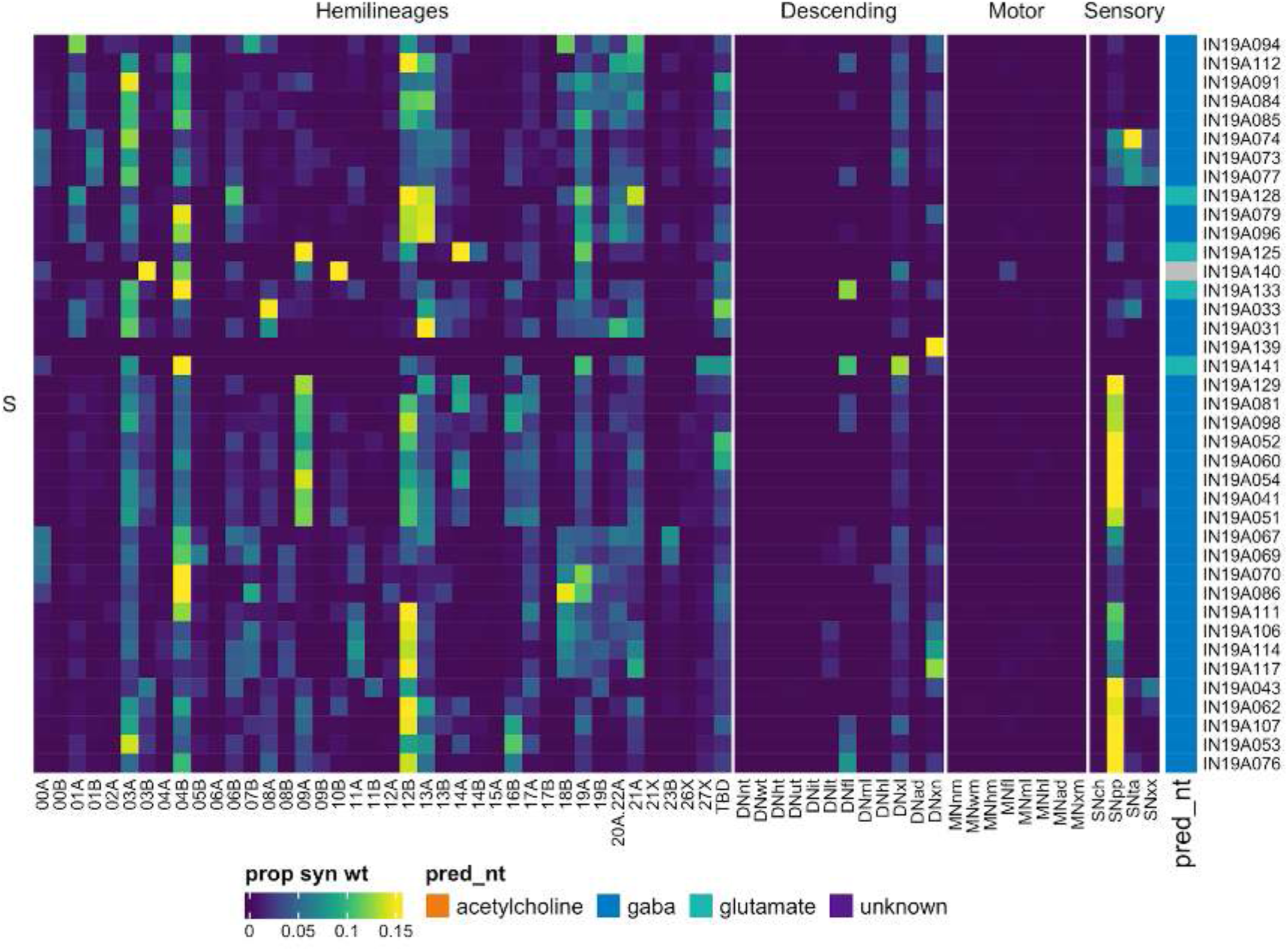
Connectivity to upstream partners by 19A secondary systematic types, continued. Proportions of synaptic weight to systematic types from upstream partners, normalised by row. 19A neurons have been clustered within each assigned birthtime window (P = primary, ES = early secondary, S = secondary) based on both upstream and downstream connectivity to hemilineages, descending neuron subclasses, motor neuron subclasses, and sensory neuron modalities. The annotation bar is coloured by the most common predicted neurotransmitter within each type.

**Figure 42 - figure supplement 8.**
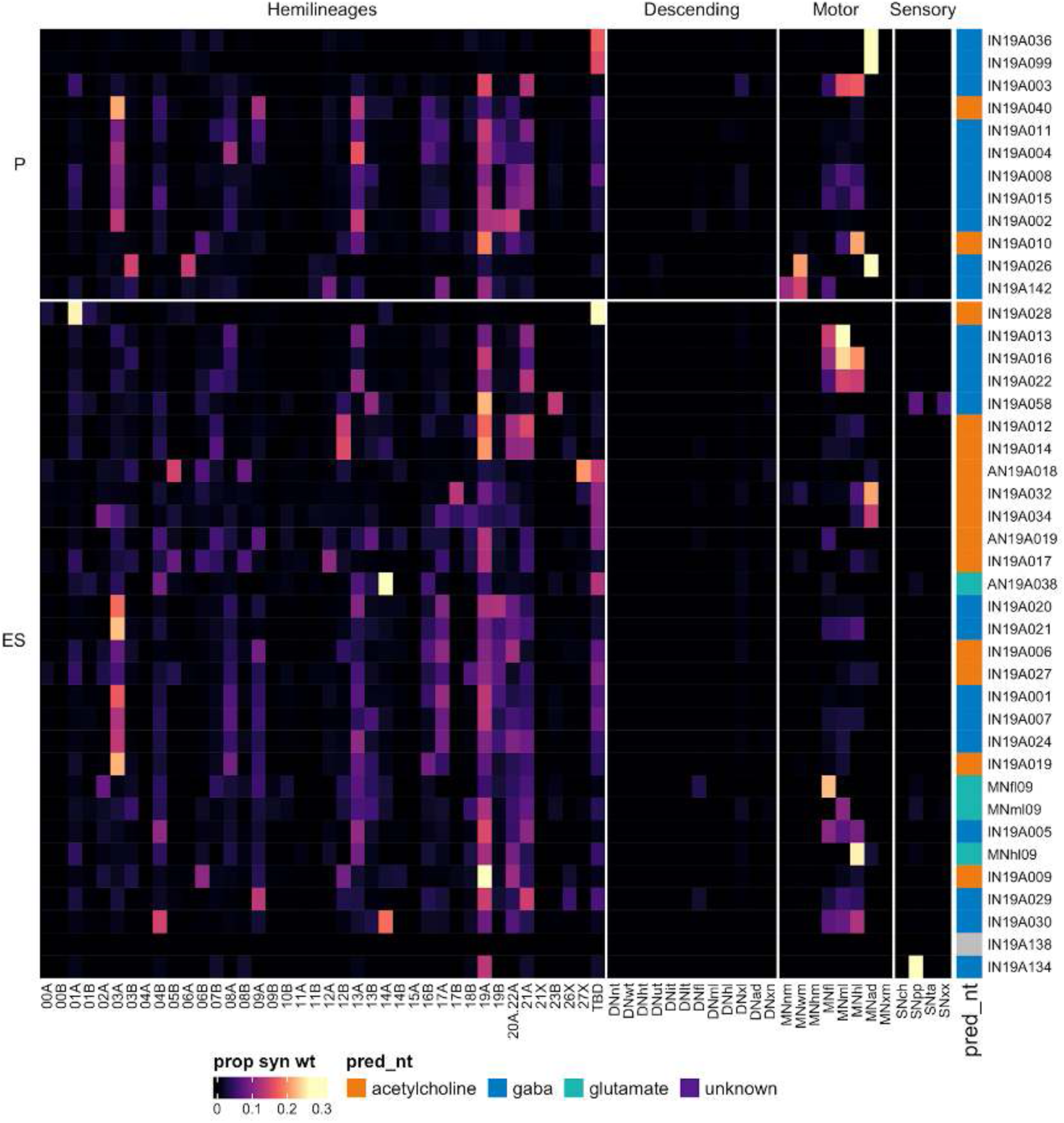
Connectivity to downstream partners by 19A primary and early secondary systematic types. Proportions of synaptic weight from systematic types to downstream partners, normalised by row. 19A neurons have been clustered within each assigned birthtime window (P = primary, ES = early secondary, S = secondary) based on both upstream and downstream connectivity to hemilineages, descending neuron subclasses, motor neuron subclasses, and sensory neuron modalities. The annotation bar is coloured by the most common predicted neurotransmitter within each type.

**Figure 42 - figure supplement 9.**
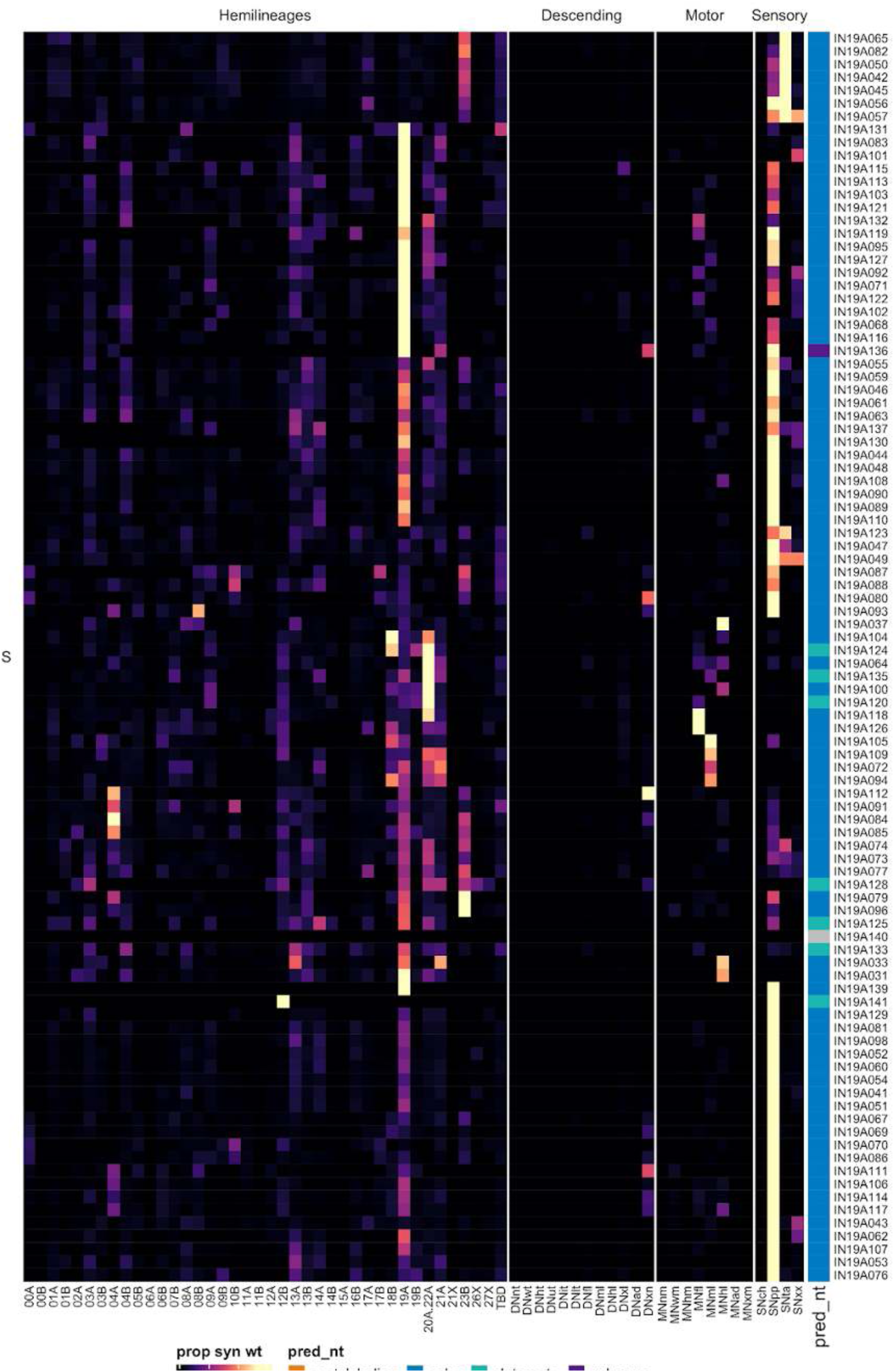
Connectivity to downstream partners by 19A secondary systematic types. Proportions of synaptic weight from systematic types to downstream partners, normalised by row. 19A neurons have been clustered within each assigned birthtime window (P = primary, ES = early secondary, S = secondary) based on both upstream and downstream connectivity to hemilineages, descending neuron subclasses, motor neuron subclasses, and sensory neuron modalities. The annotation bar is coloured by the most common predicted neurotransmitter for the neurons of each type.

**Figure 42 - figure supplement 10.**
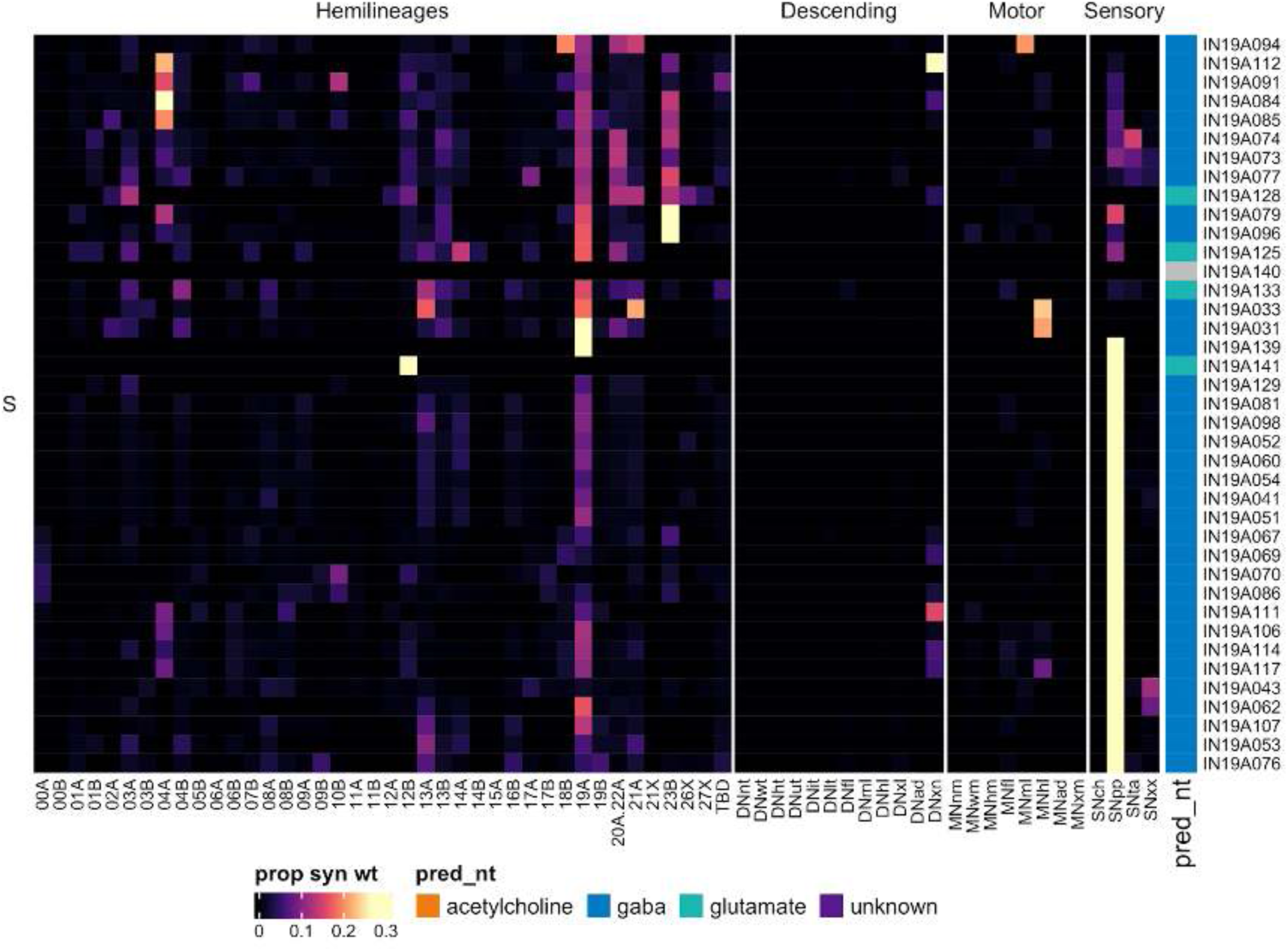
Connectivity to downstream partners by 19A secondary systematic types, continued. Proportions of synaptic weight from systematic types to downstream partners, normalised by row. 19A neurons have been clustered within each assigned birthtime window (P = primary, ES = early secondary, S = secondary) based on both upstream and downstream connectivity to hemilineages, descending neuron subclasses, motor neuron subclasses, and sensory neuron modalities. The annotation bar is coloured by the most common predicted neurotransmitter for the neurons of each type.

#### Hemilineage 19B

19B secondary neurons are cholinergic (Lacin et al., 2019), share a large, complex primary neurite tract with 11A and 23B (Shepherd et al., 2016), and generally cross the midline in the posterior intermediate commissure (Truman et al., 2004) to elaborate in contralateral dorsal tectulum. They are found in T1-A1 (Truman et al., 2004), but their numbers and connectivity vary dramatically, likely reflecting segment-specific specialisations for flight (Figure 43A,E,F). They enter the neuropil with 11A and 23B near the posterior edge of the neuromere and generally cross the midline in an intermediate commissure to innervate the contralateral tectulum (Figure 43A). We also assigned a small population of cholinergic cells to 19B that cross much more dorsally (Figure 43C bottom). The secondary population includes ascending neurons in T1 and T3 but not in T2 (Figure 43A); however, there are ascending primary neurons in T2 (e.g., Figure 43C top).

**Figure 43.**
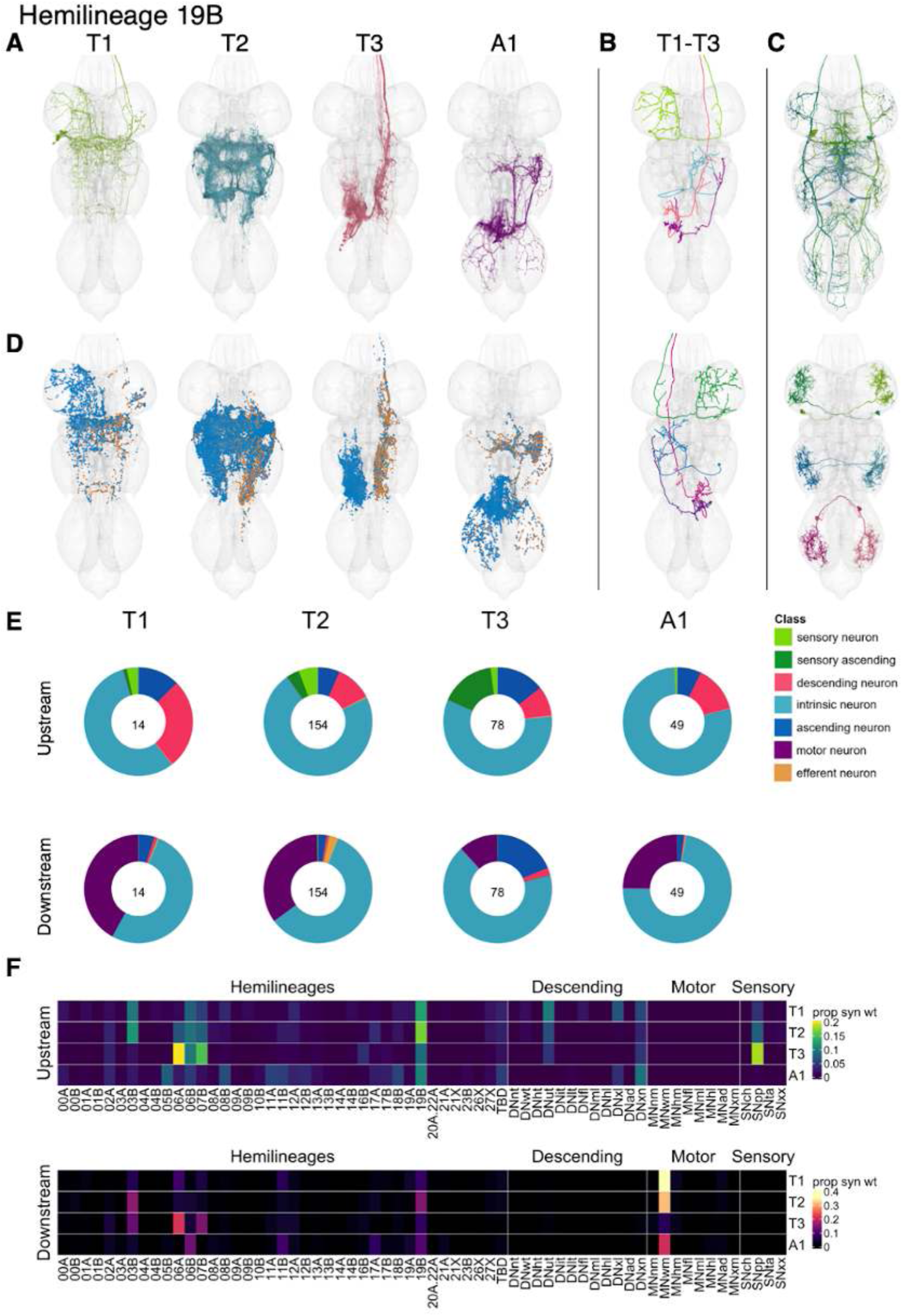
Hemilineage 19B. **A.** Meshes of all RHS secondary neurons plotted in neuromere-specific colours. **B.** “Representative” secondary neuron skeletons plotted in hemineuromere-specific colours. The skeleton with the top accumulated NBLAST score among all neurons from the hemilineage in a given hemineuromere was used. **C.** Neuron meshes of selected examples. Top: primary ascending serial set 10016. Bottom: independent leg serial set 10715. **D.** Predicted synapses of RHS secondary neurons. Blue: postsynapses; dark orange: presynapses. **E.** Proportions of connections from secondary neurons to upstream or downstream partners, normalised by neuromere and coloured by broad class. Numbers of query neurons appear in the centre. **F.** Proportions of synaptic weight from secondary neurons originating in each neuromere to upstream or downstream partners, normalised by row.

In T1 and T2, 19B secondary neurons receive input mainly from 03B, 06B, 19B, and descending neurons, while in T3 they receive it mainly from 06A, 06B, and 07B and proprioceptive sensory neurons (Figure 43F). The A1 neurons mainly have distinct upstream and downstream partners but were identified via serial minority subtypes IN19B013 and IN19B050 (Figure 43 - figure supplement 2). A subset of 19B types receive significant input from tactile sensory neurons (e.g., AN19B032 and IN19B033) (Figure 43 - figure supplement 4).

19B neurons in T1 activate wing motor neurons, those in T2 are upstream of 03B and 19B as well, those in T3 are upstream of 03B, 06A, and 07B instead, and those in A1 ascend to the tectulum and are upstream of 06B, 11B, and 19B and wing motor neurons (Figure 43F). A few 19B types target neck (e.g., AN19B025 and AN19B044), leg (e.g., IN19B02 and IN19B030), or abdominal motor neurons (e.g., IN19B016 and IN19B078) (Figure 43 - figure supplement 7-9). No functional studies have been published for secondary 19B neurons. However, AN019B007 (dMS9), which is strongly upstream of 12A neurons ( 2,8), is essential for pulse song production during courtship (Lillvis et al., 2024).

**Figure 43 - figure supplement 1.**
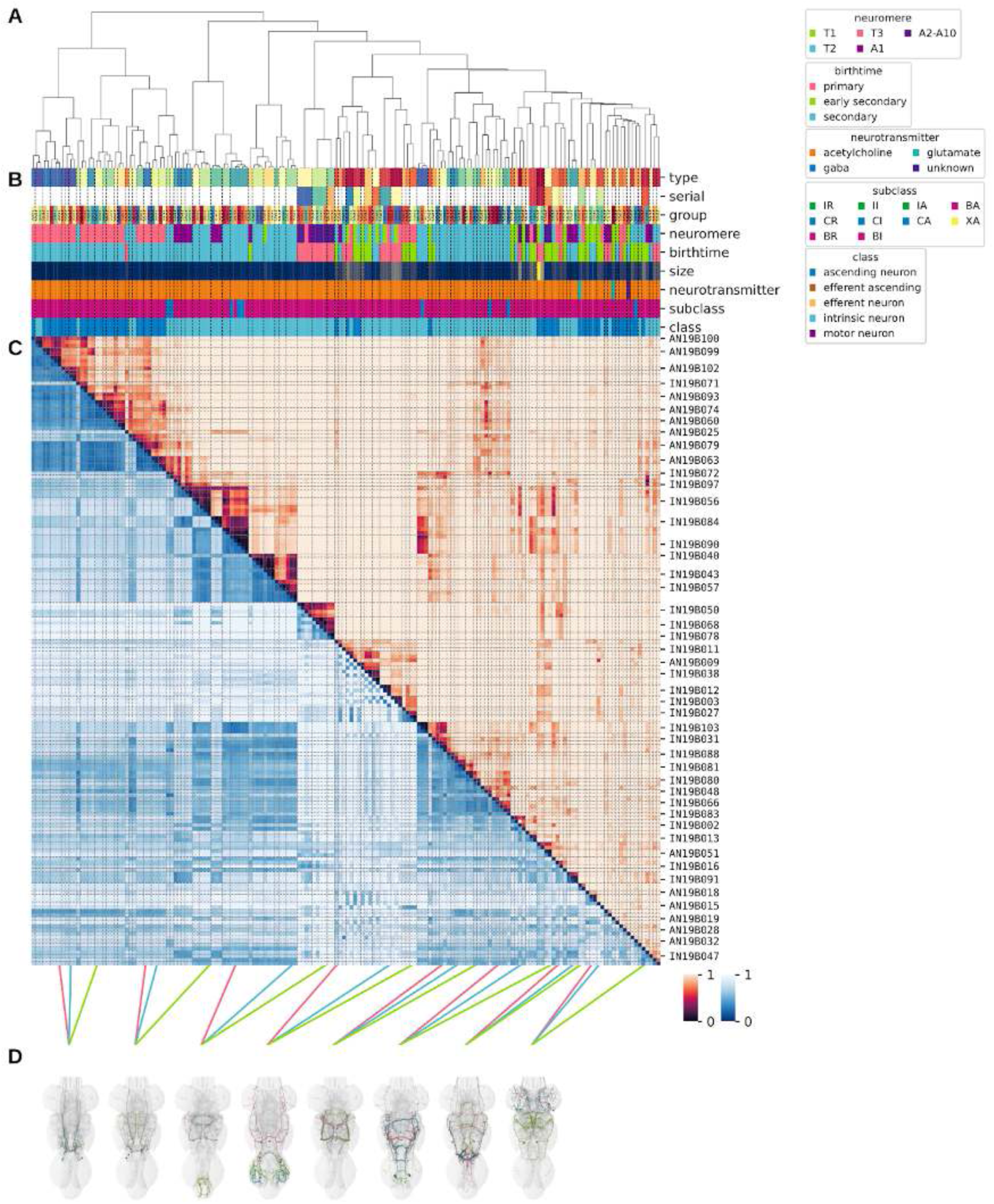
Systematic typing of hemilineage 19B. **A.** Hierarchical clustering dendrogram of hemilineage groups by laterally and serially aggregated connectivity cosine clustering. **B.** Categorical annotations of each hemilineage group, each column corresponding to the aligned leaf in A. Colours for type, serial set, and group are arbitrary for visualisation. Colours for neuromere, birthtime, neurotransmitter, subclass, and class are as in all other figures. **C.** Similarity distance heatmap for hemilineage. Cosine distance is in the upper triangle, while laterally symmetrised NBLAST distance is in the lower triangle. Systematic type names of some types are labelled. **D.** Morphologically representative groups from dendrogram subtrees. Each group, indicated by colour and line connecting to its column in B and C, is the most morphologically representative group (medoid of NBLAST distance) from a subtree of A. The subtrees (flat clusters) are equal height cuts of A determined to yield the number of groups per plot and plots in D.

**Figure 43 - figure supplement 2.**
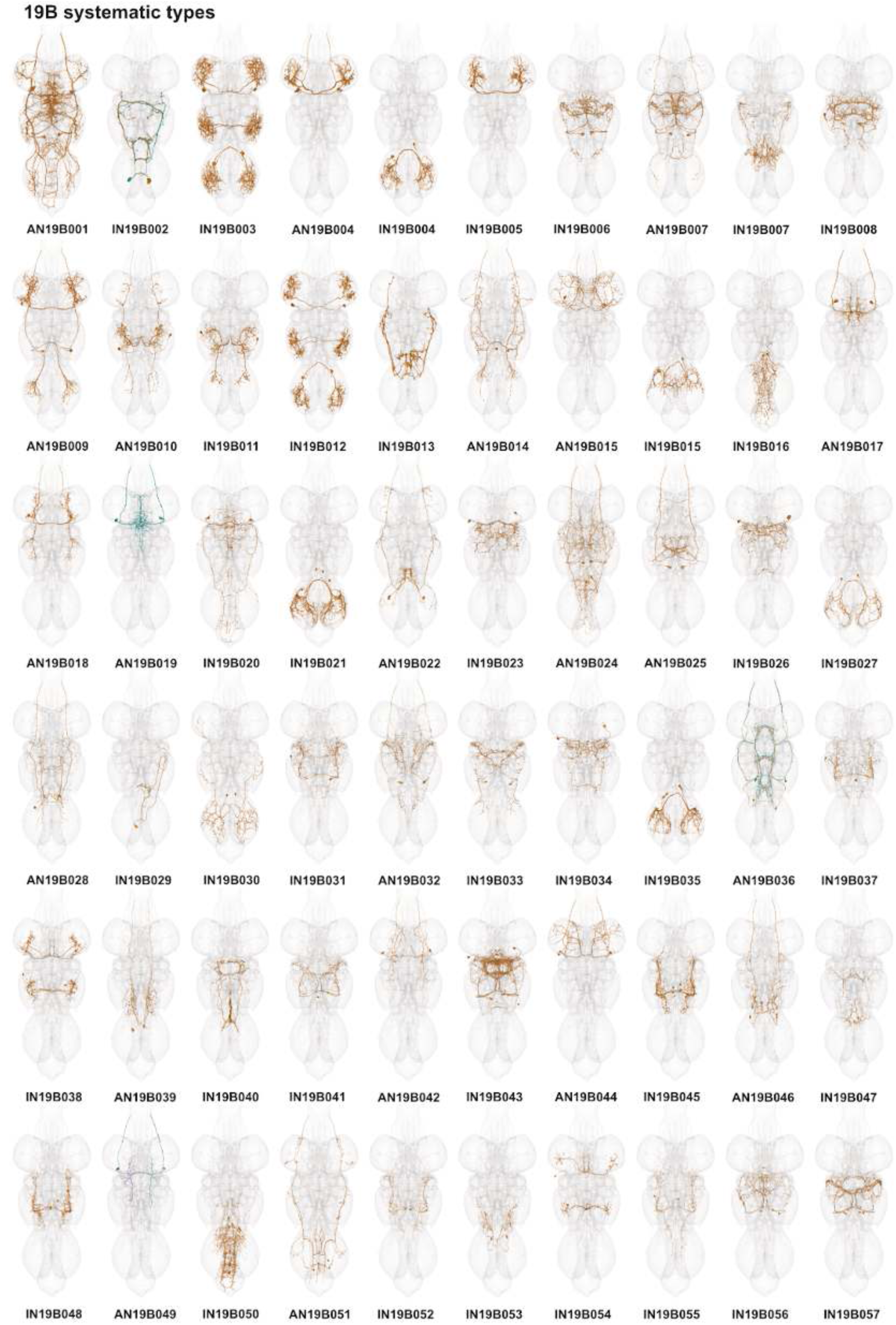
Systematic types of hemilineage 19B. Systematic types have been arranged in numerical order, with neurons of the same type that belong to distinct classes (e.g., intrinsic neuron vs ascending neuron) plotted separately but placed adjacent to each other. Individual neuron meshes have been coloured based on predicted neurotransmitter: dark orange = acetylcholine, blue = gaba, marine = glutamate, dark purple = unknown.

**Figure 43 - figure supplement 3.**
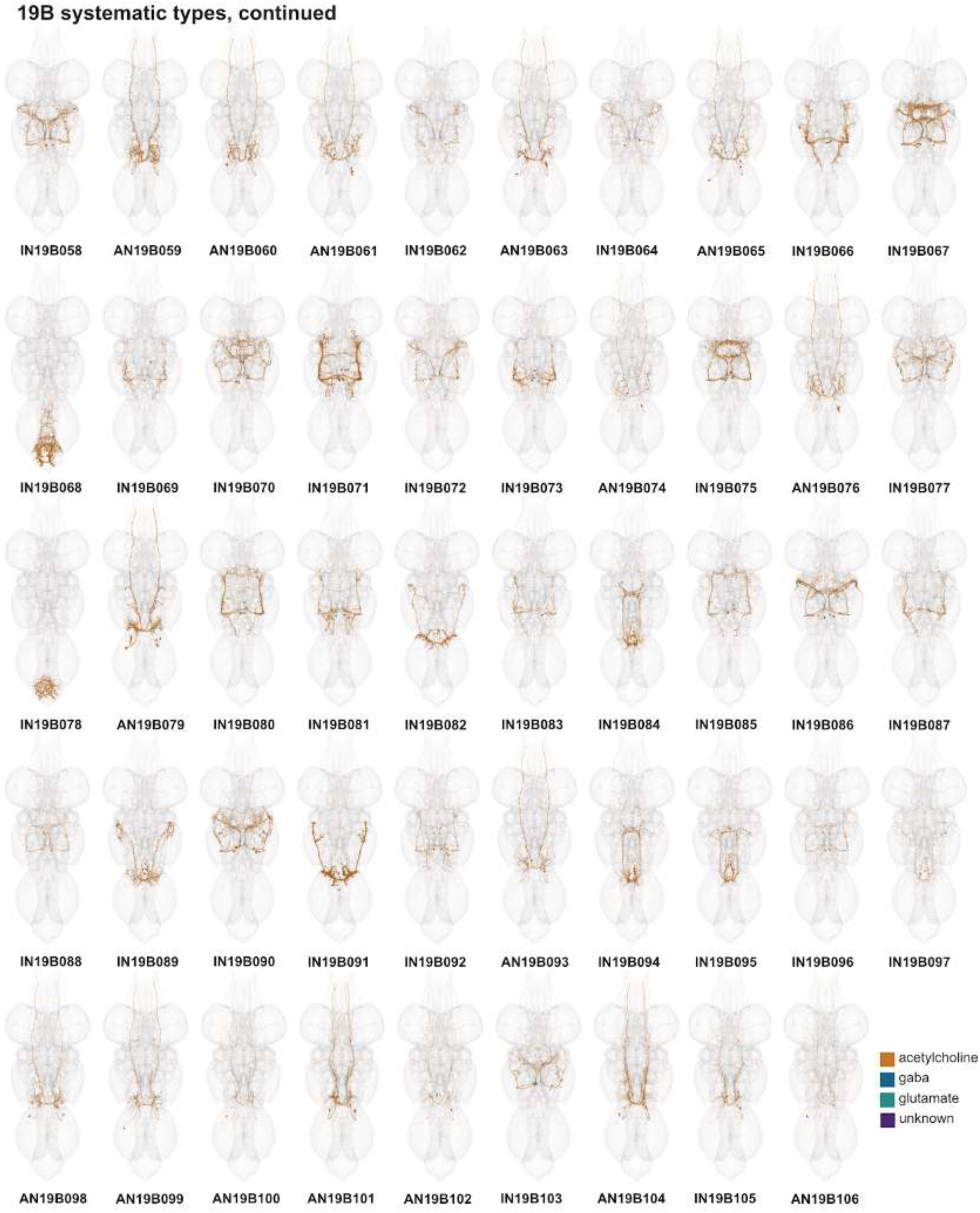
Systematic types of hemilineage 19B, continued. Systematic types have been arranged in numerical order, with neurons of the same type that belong to distinct classes (e.g., intrinsic neuron vs ascending neuron) plotted separately but placed adjacent to each other. Individual neuron meshes have been coloured based on predicted neurotransmitter: dark orange = acetylcholine, blue = gaba, marine = glutamate, dark purple = unknown.

**Figure 43 - figure supplement 4.**
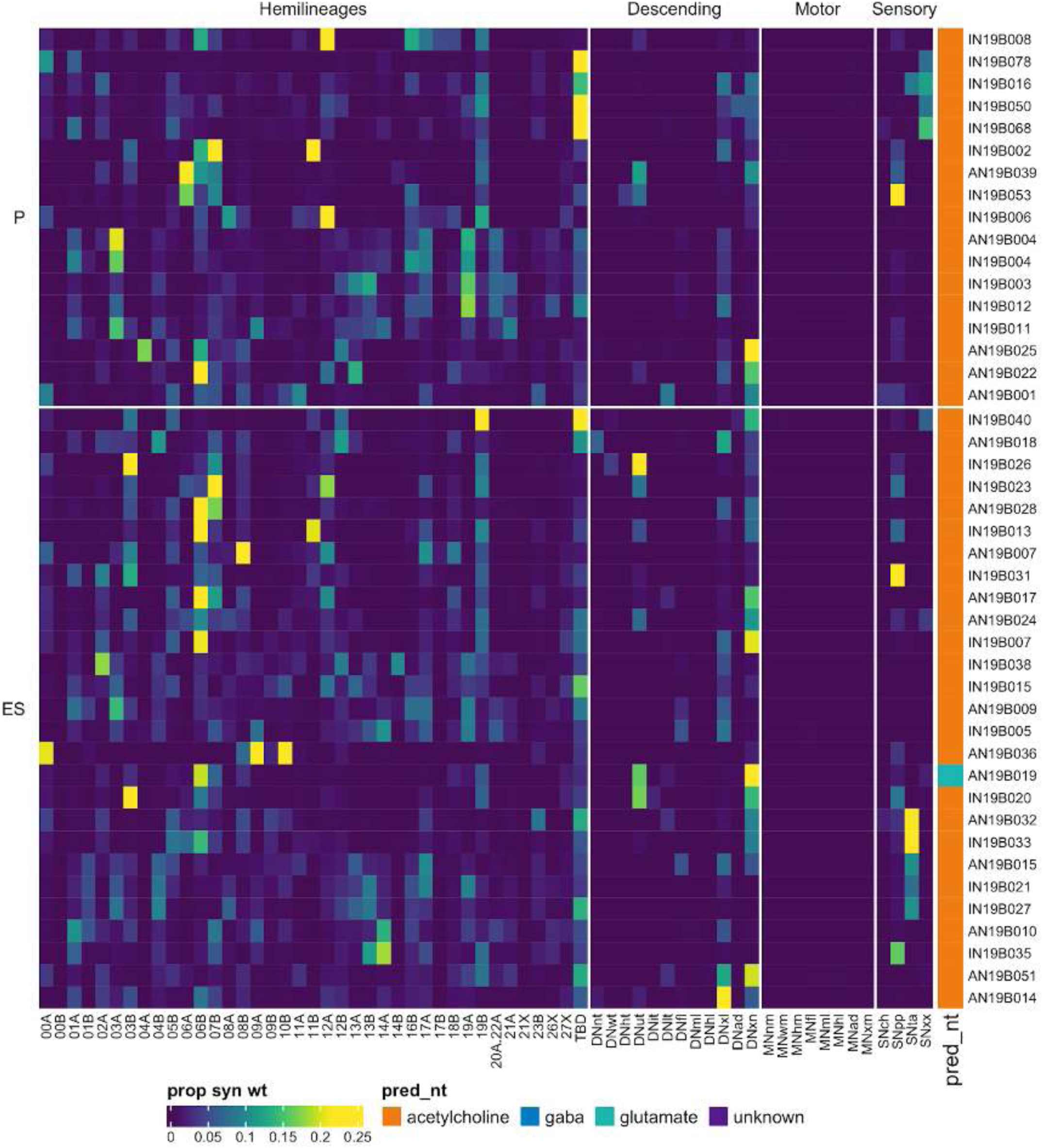
Connectivity to upstream partners by 19B primary and early secondary systematic types. Proportions of synaptic weight to systematic types from upstream partners, normalised by row. 19B neurons have been clustered within each assigned birthtime window (P = primary, ES = early secondary, S = secondary) based on both upstream and downstream connectivity to hemilineages, descending neuron subclasses, motor neuron subclasses, and sensory neuron modalities. Annotation bar is coloured by the most common predicted neurotransmitter for the neurons of each type.

**Figure 43 - figure supplement 5.**
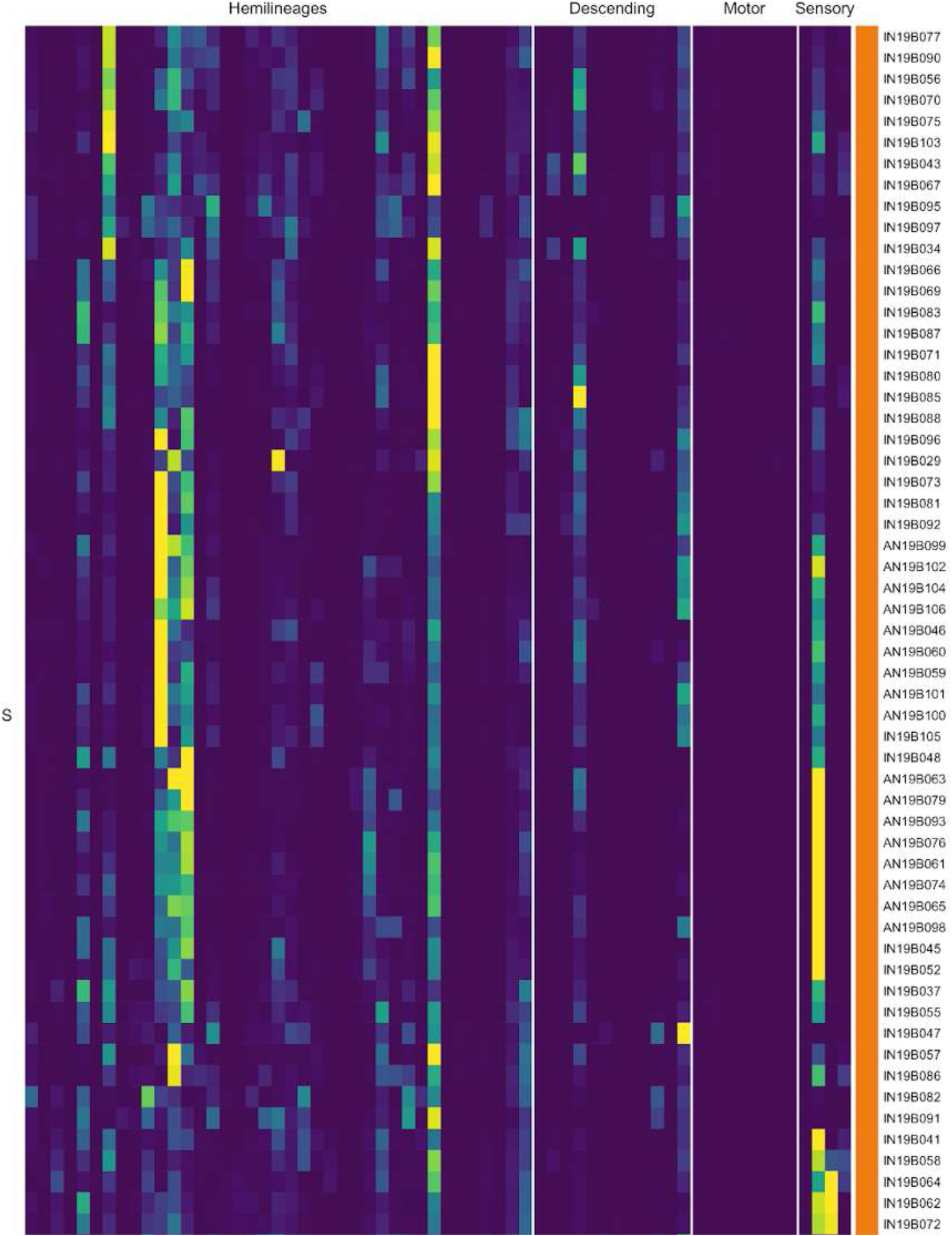
Connectivity to upstream partners by 19B secondary systematic types. Proportions of synaptic weight to systematic types from upstream partners, normalised by row. 19B neurons have been clustered within each assigned birthtime window (P = primary, ES = early secondary, S = secondary) based on both upstream and downstream connectivity to hemilineages, descending neuron subclasses, motor neuron subclasses, and sensory neuron modalities. The annotation bar is coloured by the most common predicted neurotransmitter within each type.

**Figure 43 - figure supplement 6.**
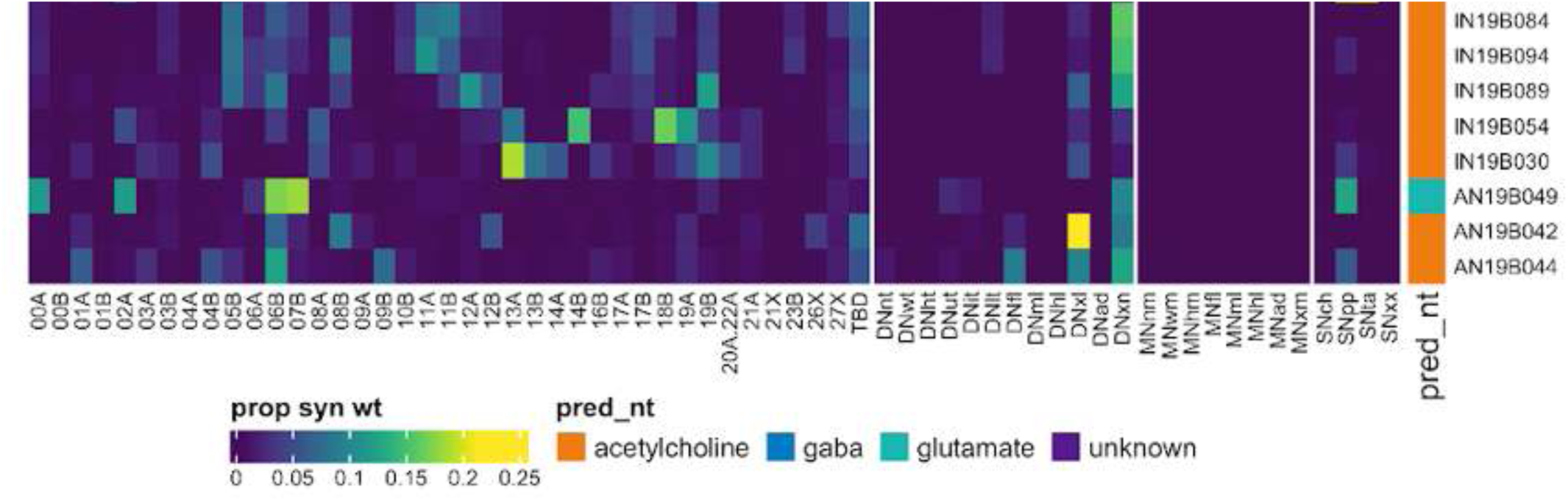
Connectivity to upstream partners by 19B secondary systematic types, continued. Proportions of synaptic weight to systematic types from upstream partners, normalised by row. 19B neurons have been clustered within each assigned birthtime window (P = primary, ES = early secondary, S = secondary) based on both upstream and downstream connectivity to hemilineages, descending neuron subclasses, motor neuron subclasses, and sensory neuron modalities. The annotation bar is coloured by the most common predicted neurotransmitter within each type.

**Figure 43 - figure supplement 7.**
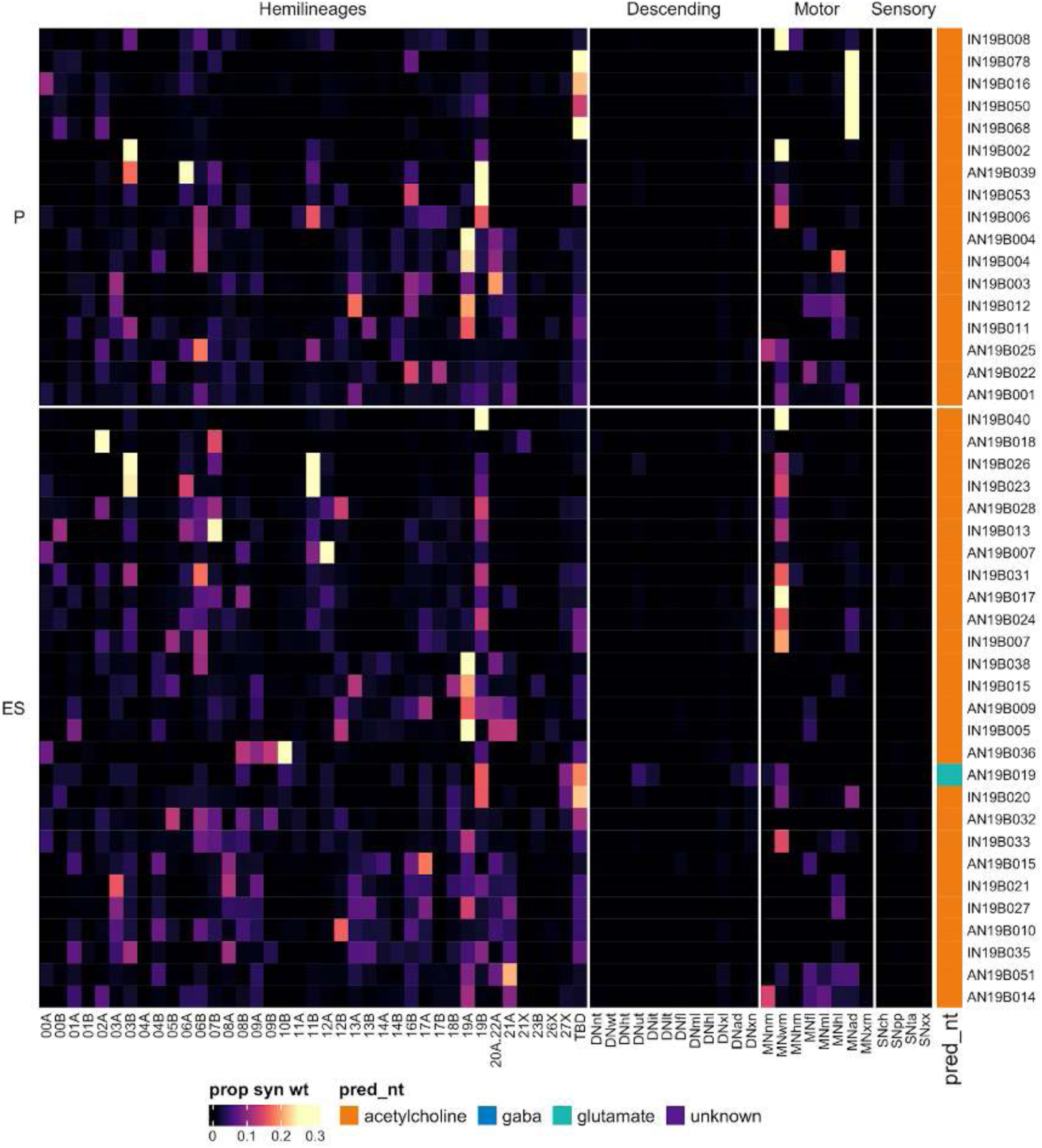
Connectivity to downstream partners by 19B primary and early secondary systematic types. Proportions of synaptic weight from systematic types to downstream partners, normalised by row. 19B neurons have been clustered within each assigned birthtime window (P = primary, ES = early secondary, S = secondary) based on both upstream and downstream connectivity to hemilineages, descending neuron subclasses, motor neuron subclasses, and sensory neuron modalities. The annotation bar is coloured by the most common predicted neurotransmitter within each type.

**Figure 43 - figure supplement 8.**
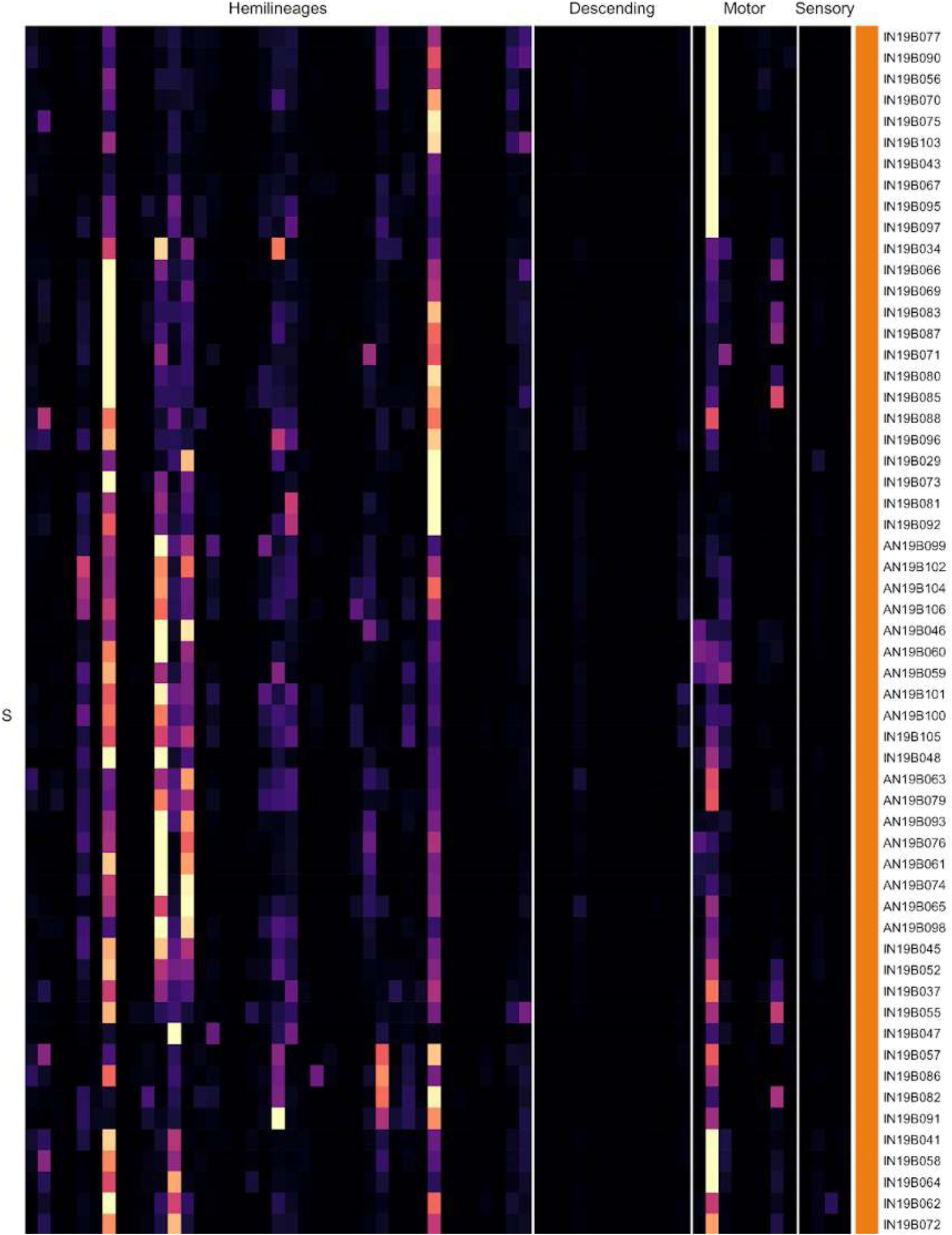
Connectivity to downstream partners by 19B secondary systematic types. Proportions of synaptic weight from systematic types to downstream partners, normalised by row. 19B neurons have been clustered within each assigned birthtime window (P = primary, ES = early secondary, S = secondary) based on both upstream and downstream connectivity to hemilineages, descending neuron subclasses, motor neuron subclasses, and sensory neuron modalities. The annotation bar is coloured by the most common predicted neurotransmitter for the neurons of each type.

**Figure 43 - figure supplement 9.**
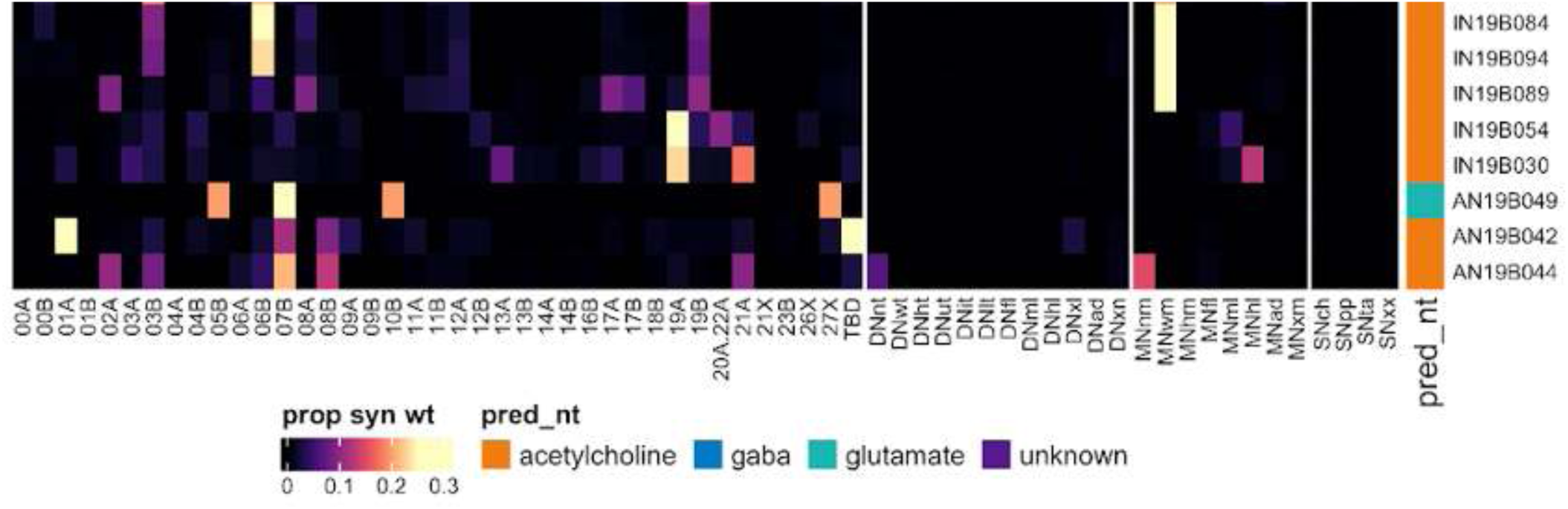
Connectivity to downstream partners by 19B secondary systematic types, continued. Proportions of synaptic weight from systematic types to downstream partners, normalised by row. 19B neurons have been clustered within each assigned birthtime window (P = primary, ES = early secondary, S = secondary) based on both upstream and downstream connectivity to hemilineages, descending neuron subclasses, motor neuron subclasses, and sensory neuron modalities. The annotation bar is coloured by the most common predicted neurotransmitter for the neurons of each type.

#### Hemilineages 20A and 22A

Hemilineages 20A, 21A, and 22A are posterior lateral hemilineages that innervate the six ipsilateral leg neuropils, with all three entering the neuropil in approximately the same location. In late larva they can be distinguished by separate primary neurite bundles (Truman et al., 2004). Both 20A and 22A have been ascribed to NB5-4 (Birkholz et al., 2015), which generates 2-4 efferent cells in the embryo and 2-4 local interneurons in the abdomen (Schmid et al., 1999). However, more recently 20A was mapped to a novel neuroblast, NB5-7, which does not produce any embryonic progeny (Lacin and Truman, 2016).

Both 20A and 22A are expected to be cholinergic (Lacin et al., 2019). We identified two largely separable but overlapping cholinergic populations in T1-T3 that we annotated collectively as “20A.22A” (Figure 44A). Left-right groups and serial sets sometimes contained two morphologically identical neurons entering the leg neuropil in distinct soma tracts (e.g., Figure 45C top), suggesting that some cell types have been preserved in both hemilineages since the hypothetical ancestral duplication of a common neuroblast (Lacin and Truman, 2016). Most 20A/22A neurons innervate the ipsilateral neuropil, but there is a prominent bilateral population in T1 (Figure 45C bottom).

**Figure 44.**
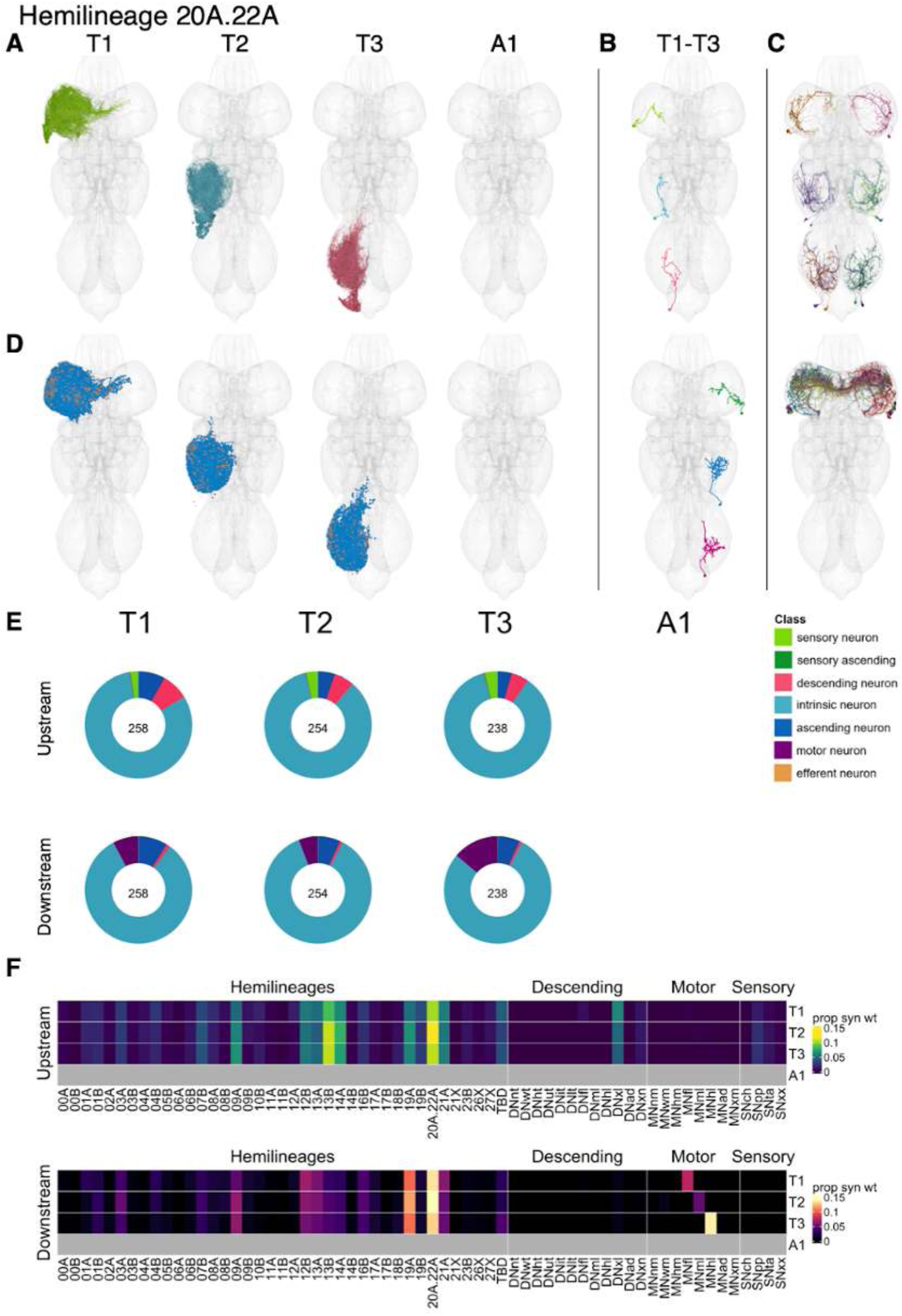
Hemilineages 20A and 22A. **A.** Meshes of all RHS secondary neurons plotted in neuromere-specific colours. **B.** “Representative” secondary neuron skeletons plotted in hemineuromere-specific colours. The skeleton with the top accumulated NBLAST score among all neurons from the hemilineage in a given hemineuromere was used. **C.** Neuron meshes of selected examples. Top: independent leg serial set 10894. Bottom: T1 bilateral subcluster 11636. **D.** Predicted synapses of RHS secondary neurons. Blue: postsynapses; dark orange: presynapses. **E.** Proportions of connections from secondary neurons to upstream or downstream partners, normalised by neuromere and coloured by broad class. Numbers of query neurons appear in the centre. **F.** Proportions of synaptic weight from secondary neurons originating in each neuromere to upstream or downstream partners, normalised by row.

The secondary 20A/22A neurons receive input from hemilineages including 09A, 12B, 13A, 13B, and 14A, 19A, 20A/22A, and 21A as well as from descending neurons targeting multiple legs and from proprioceptive sensory neurons (Figure 45F), although some early born types receive tactile inputs (e.g., IN20A.22A006 and IN20A.22A007) (Figure 44 - figure supplement 4). Secondary 20A/22A neurons activate hemilineages 09A, 12B, 13A, 19A, and 20A/22A and leg motor neurons. However, this is type-specific, with some types (e.g., IN20A.22A005 and IN20A.22A028) directly targeting motor neurons and others (e.g., IN20A.22A077 and IN20A.22A092) strongly targeting 12B or other interneurons instead. Bilateral activation of 20A/22A neurons results in extension/splaying of the legs with occasional grooming bouts (Harris et al., 2015).

**Figure 44 - figure supplement 1.**
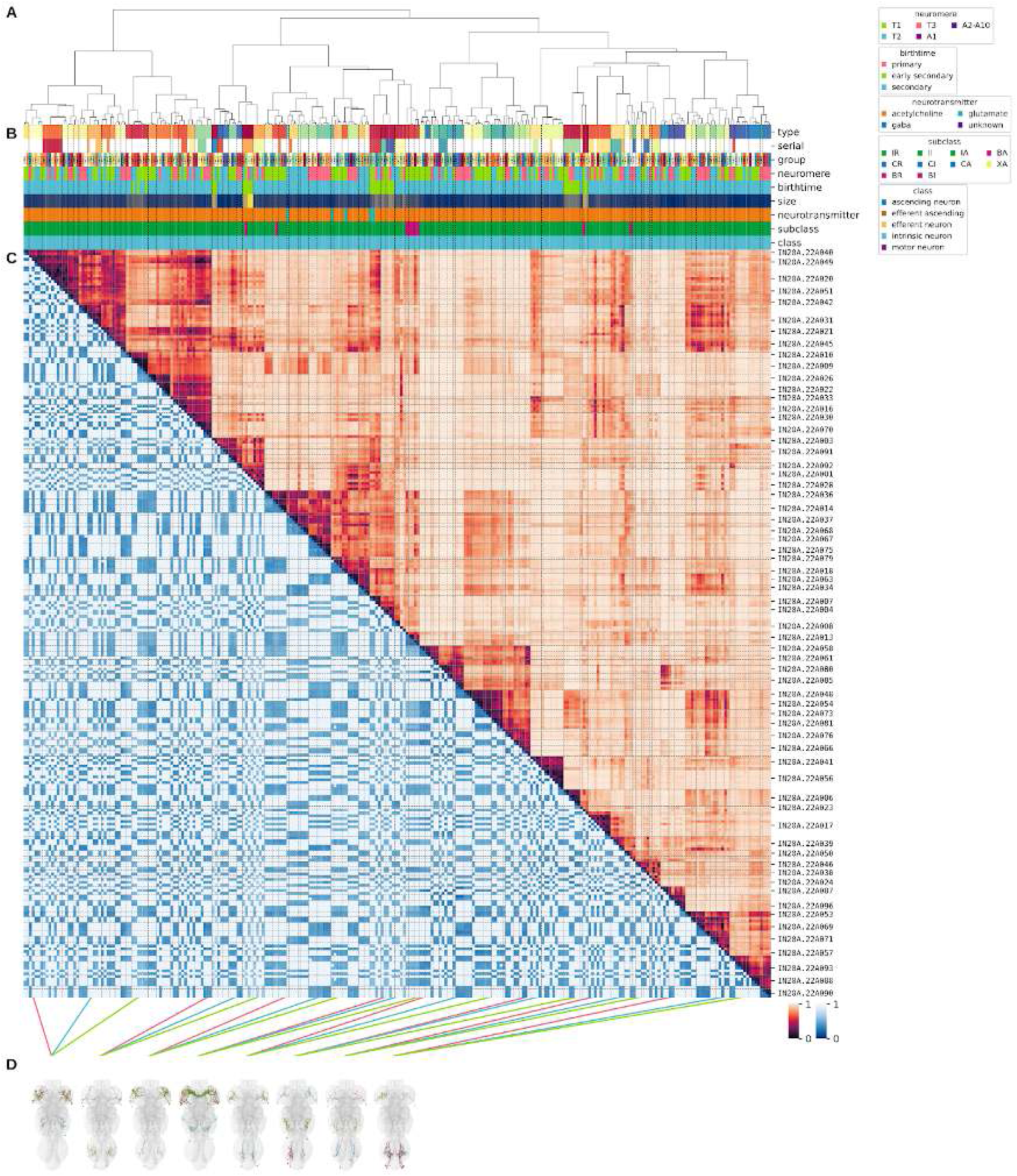
Systematic typing of hemilineage 20A and 22A. **A.** Hierarchical clustering dendrogram of hemilineage groups by laterally and serially aggregated connectivity cosine clustering. **B.** Categorical annotations of each hemilineage group, each column corresponding to the aligned leaf in A. Colours for type, serial set, and group are arbitrary for visualisation. Colours for neuromere, birthtime, neurotransmitter, subclass, and class are as in all other figures. **C.** Similarity distance heatmap for hemilineage. Cosine distance is in the upper triangle, while laterally symmetrised NBLAST distance is in the lower triangle. Systematic type names of some types are labelled. **D.** Morphologically representative groups from dendrogram subtrees. Each group, indicated by colour and line connecting to its column in B and C, is the most morphologically representative group (medoid of NBLAST distance) from a subtree of A. The subtrees (flat clusters) are equal height cuts of A determined to yield the number of groups per plot and plots in D.

**Figure 44 - figure supplement 2.**
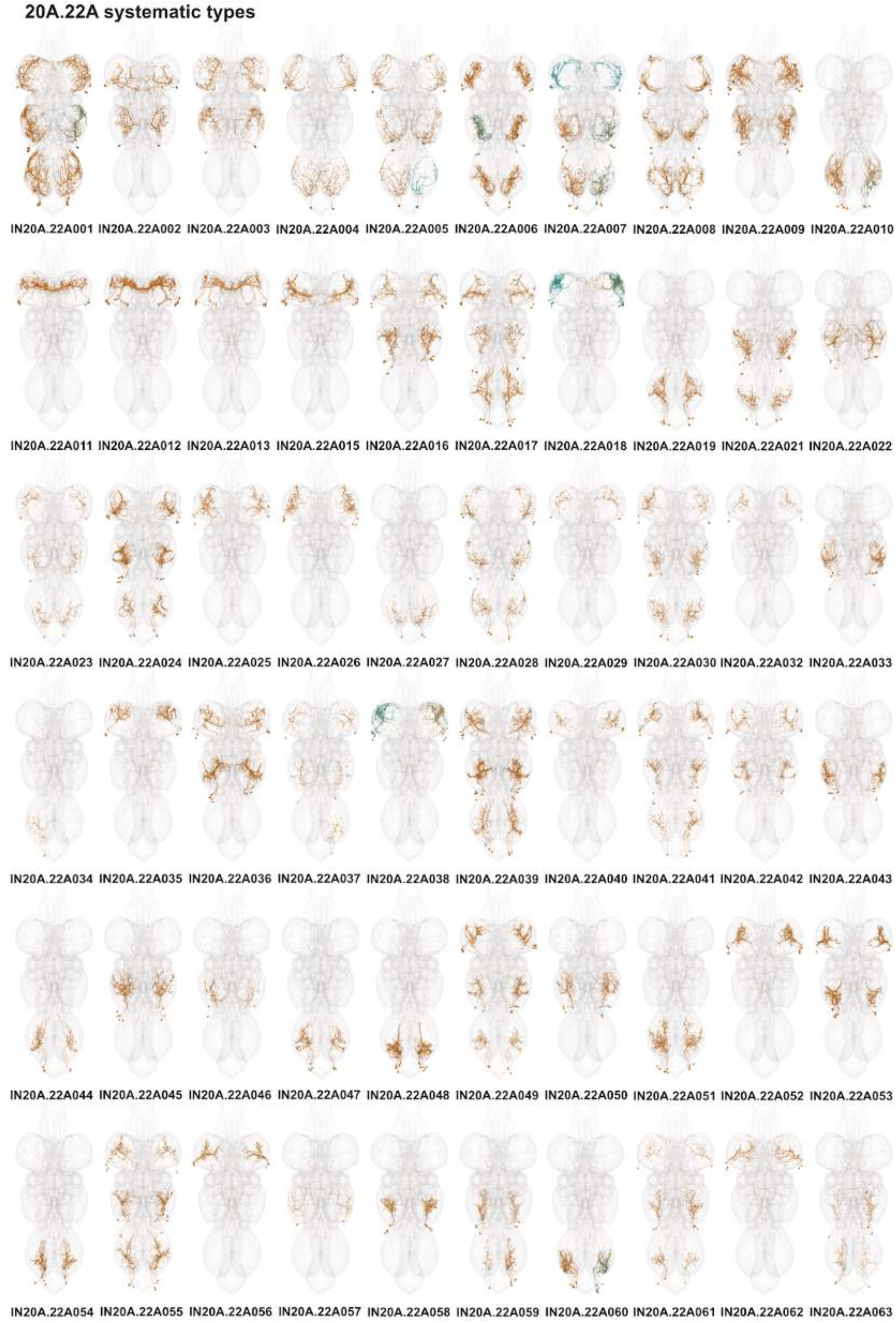
Systematic types of hemilineages 20A and 22A. Systematic types have been arranged in numerical order, with neurons of the same type that belong to distinct classes (e.g., intrinsic neuron vs ascending neuron) plotted separately but placed adjacent to each other. Individual neuron meshes have been coloured based on predicted neurotransmitter: dark orange = acetylcholine, blue = gaba, marine = glutamate, dark purple = unknown.

**Figure 44 - figure supplement 3.**
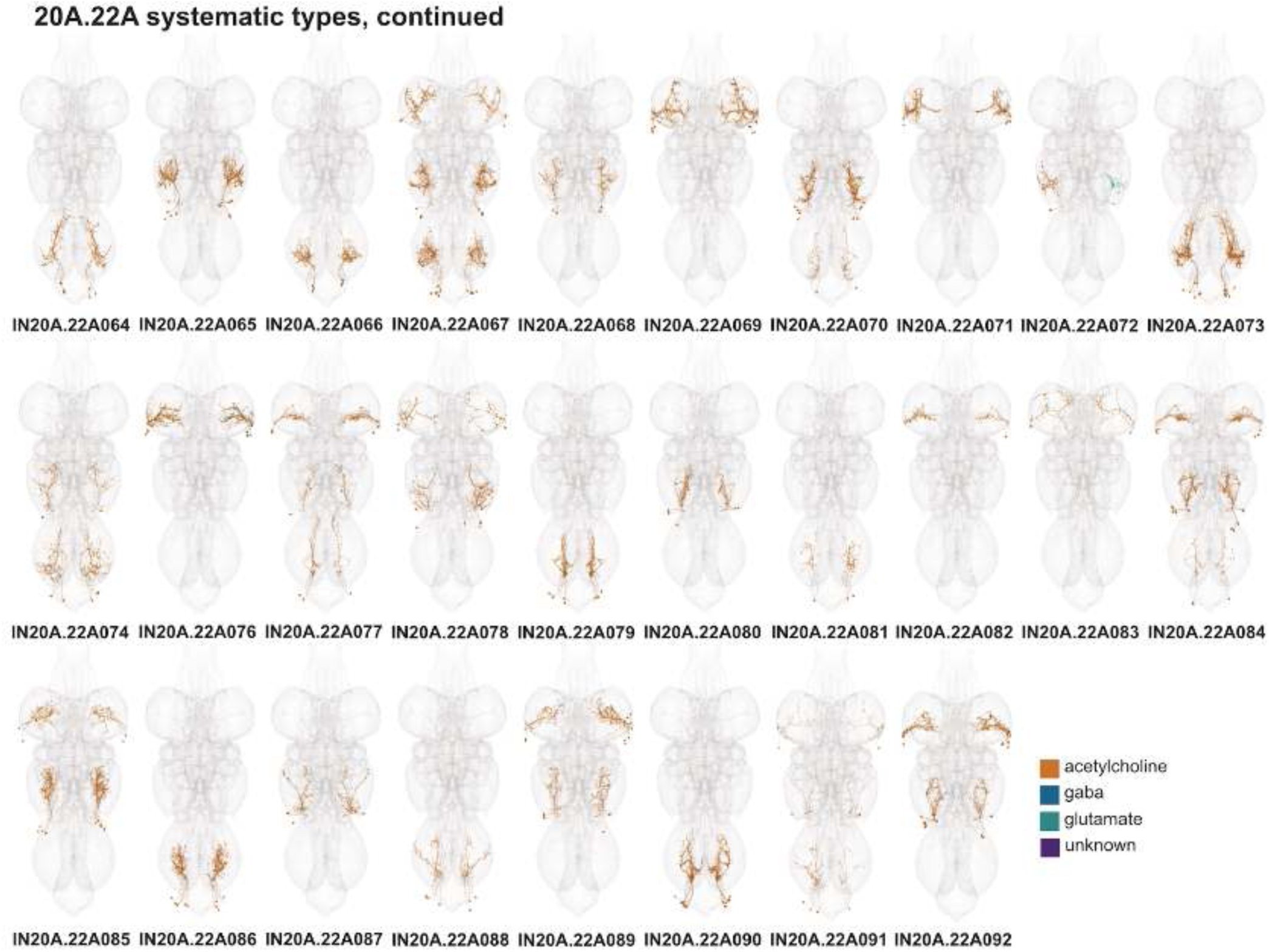
Systematic types of hemilineages 20A and 22A, continued. Systematic types have been arranged in numerical order, with neurons of the same type that belong to distinct classes (e.g., intrinsic neuron vs ascending neuron) plotted separately but placed adjacent to each other. Individual neuron meshes have been coloured based on predicted neurotransmitter: dark orange = acetylcholine, blue = gaba, marine = glutamate, dark purple = unknown.

**Figure 44 - figure supplement 4.**
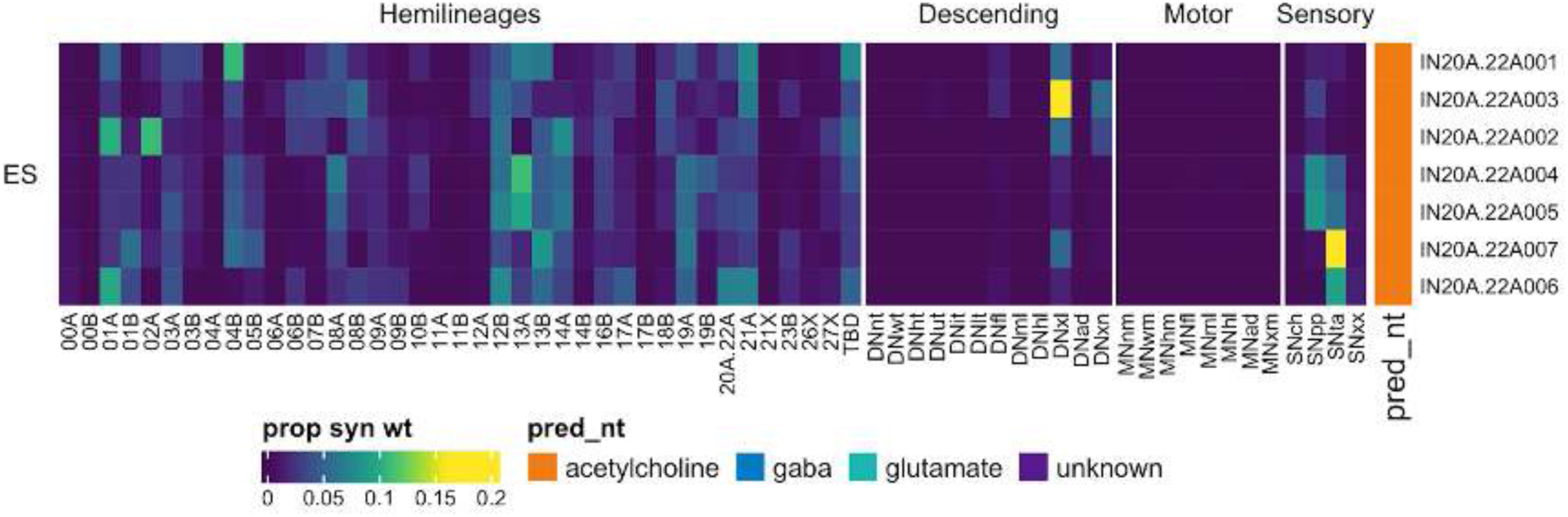
Connectivity to upstream partners by 20A.22A early secondary systematic types. Proportions of synaptic weight to systematic types from upstream partners, normalised by row. 20A.22A neurons have been clustered within each assigned birthtime window (P = primary, ES = early secondary, S = secondary) based on both upstream and downstream connectivity to hemilineages, descending neuron subclasses, motor neuron subclasses, and sensory neuron modalities. Annotation bar is coloured by the most common predicted neurotransmitter for the neurons of each type.

**Figure 44 - figure supplement 5.**
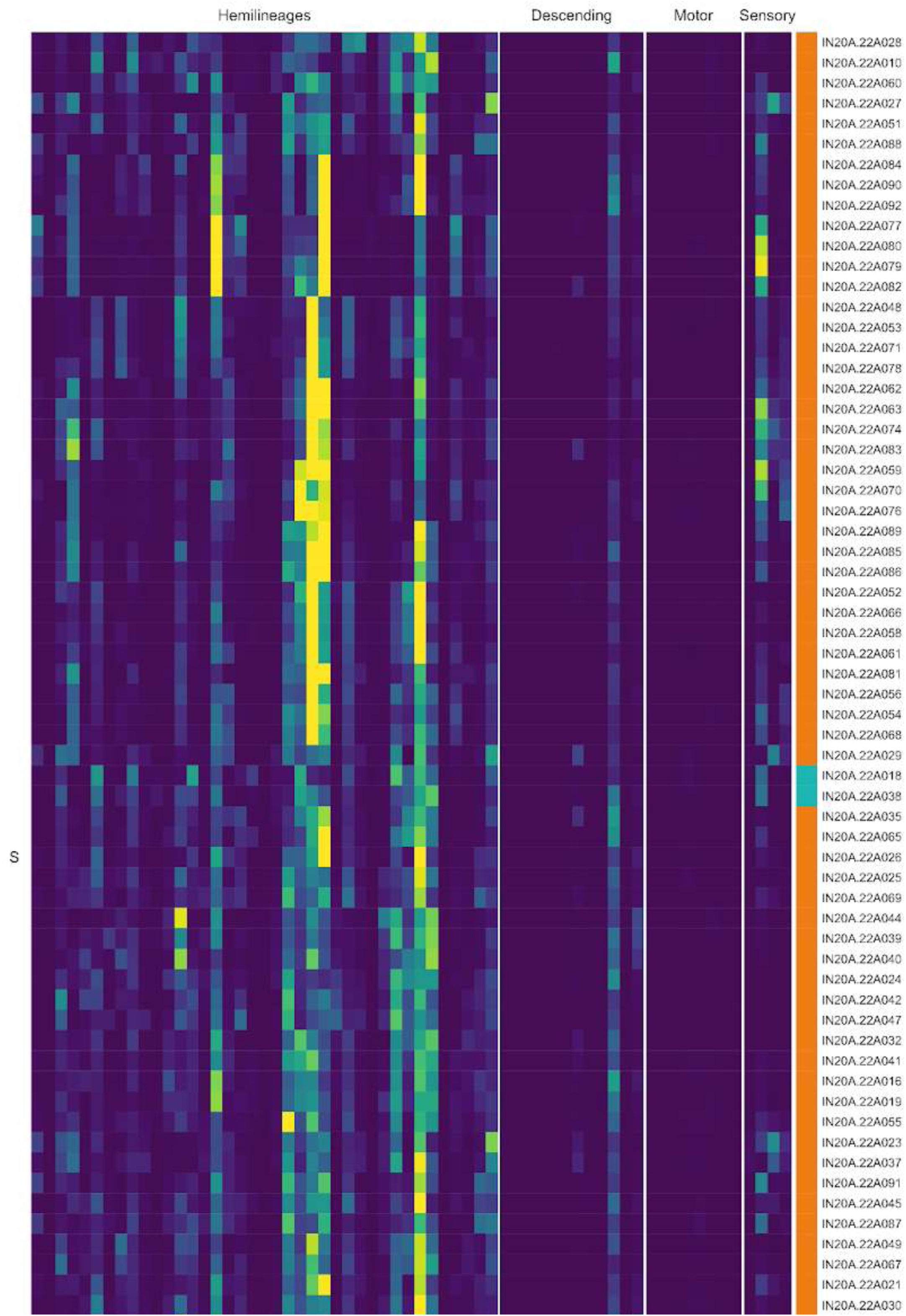
Connectivity to upstream partners by 20A.22A secondary systematic types. Proportions of synaptic weight to systematic types from upstream partners, normalised by row. 20A.22A neurons have been clustered within each assigned birthtime window (P = primary, ES = early secondary, S = secondary) based on both upstream and downstream connectivity to hemilineages, descending neuron subclasses, motor neuron subclasses, and sensory neuron modalities. The annotation bar is coloured by the most common predicted neurotransmitter within each type.

**Figure 44 - figure supplement 6.**
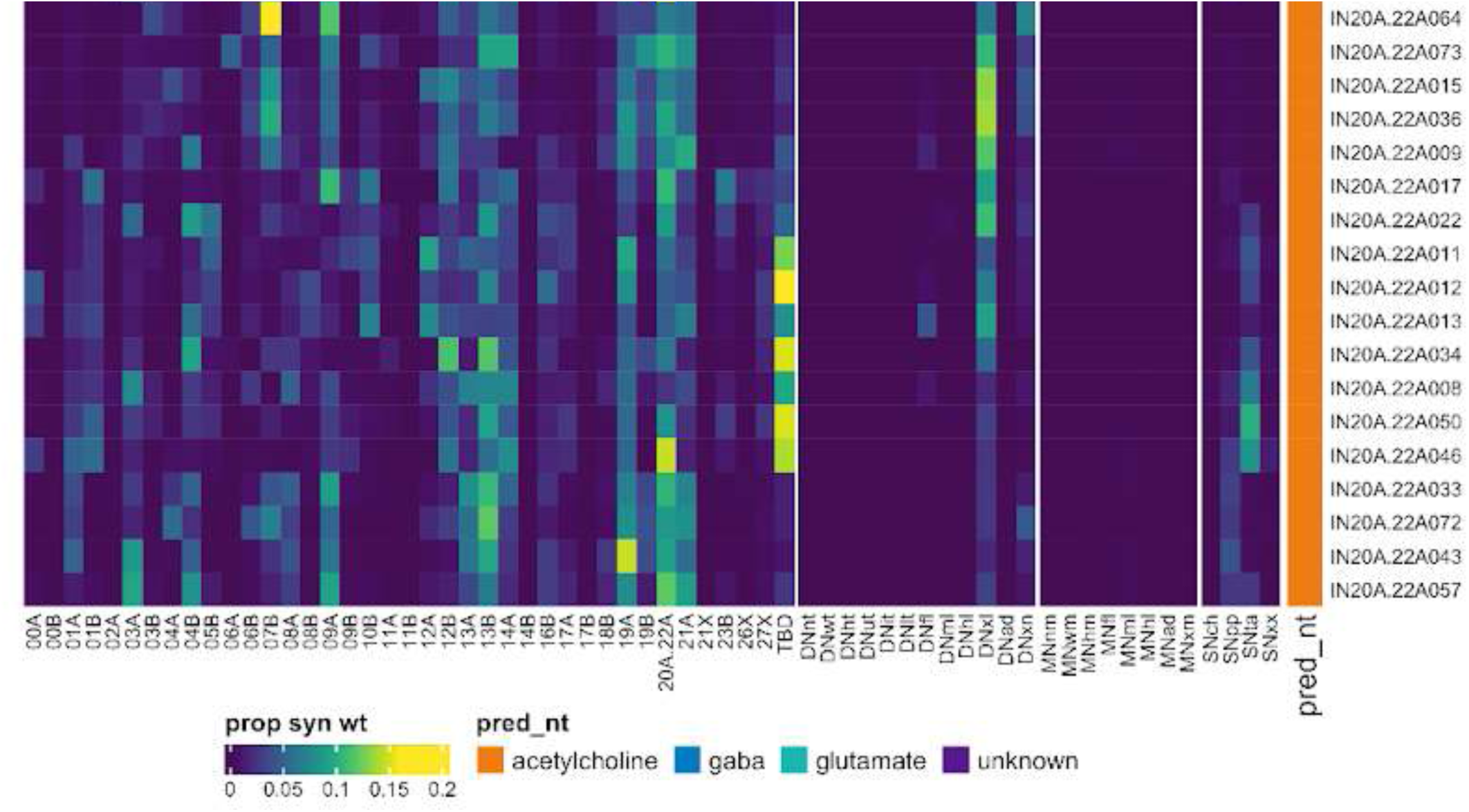
Connectivity to upstream partners by 20A.22A secondary systematic types, continued. Proportions of synaptic weight to systematic types from upstream partners, normalised by row. 20A.22A neurons have been clustered within each assigned birthtime window (P = primary, ES = early secondary, S = secondary) based on both upstream and downstream connectivity to hemilineages, descending neuron subclasses, motor neuron subclasses, and sensory neuron modalities. The annotation bar is coloured by the most common predicted neurotransmitter within each type.

**Figure 44 - figure supplement 7.**
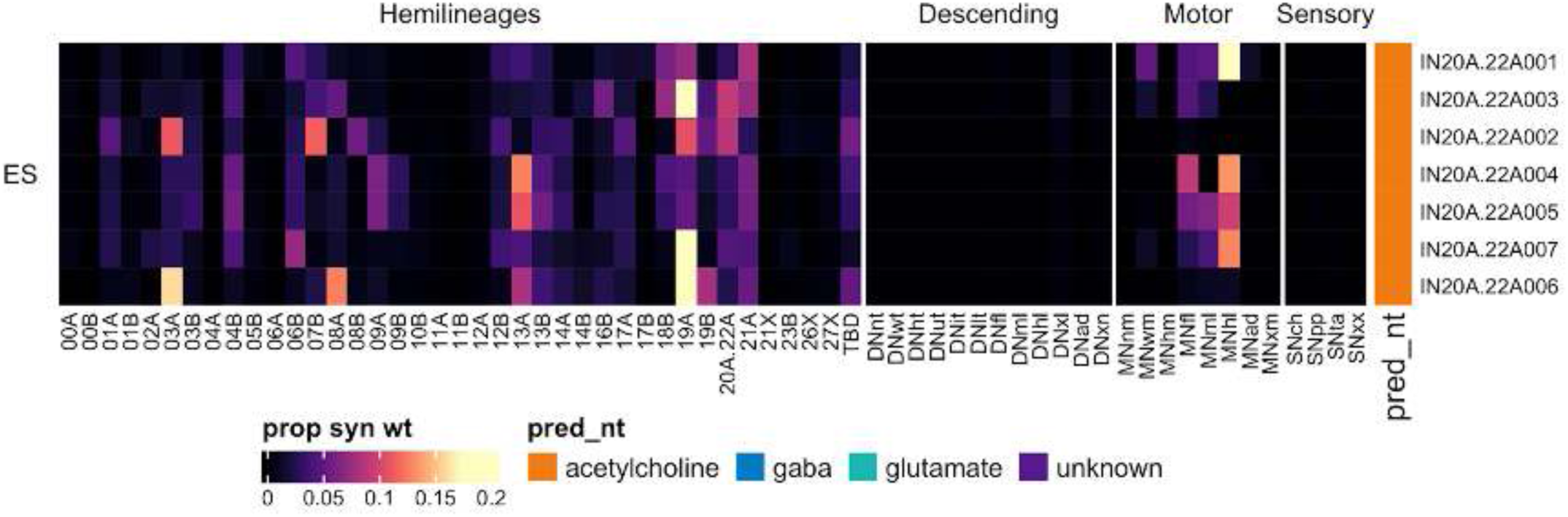
Connectivity to downstream partners by 20A.22A early secondary systematic types. Proportions of synaptic weight from systematic types to downstream partners, normalised by row. 20A.22A neurons have been clustered within each assigned birthtime window (P = primary, ES = early secondary, S = secondary) based on both upstream and downstream connectivity to hemilineages, descending neuron subclasses, motor neuron subclasses, and sensory neuron modalities. The annotation bar is coloured by the most common predicted neurotransmitter within each type.

**Figure 44 - figure supplement 8.**
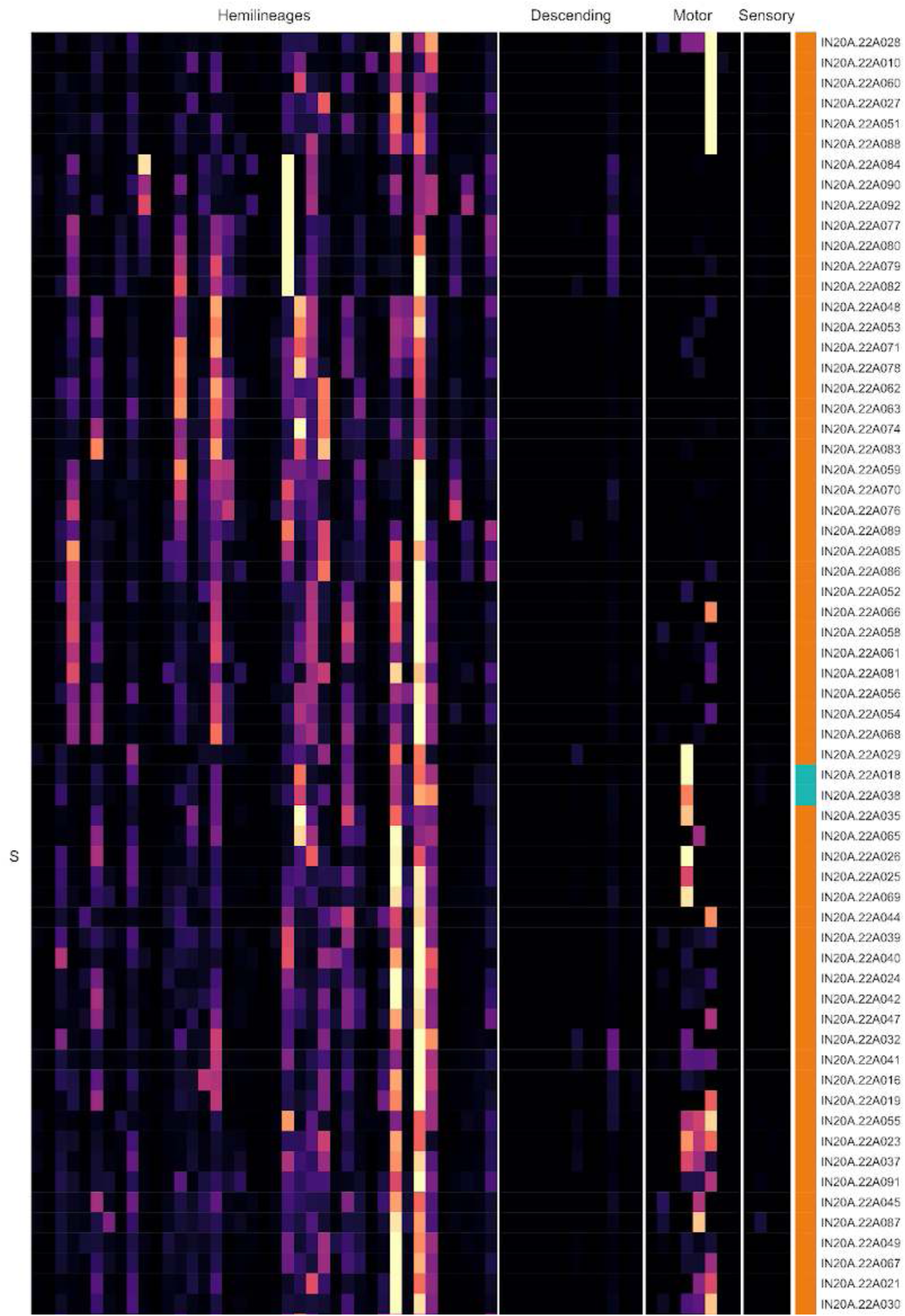
Connectivity to downstream partners by 20A.22A secondary systematic types. Proportions of synaptic weight from systematic types to downstream partners, normalised by row. 20A.22A neurons have been clustered within each assigned birthtime window (P = primary, ES = early secondary, S = secondary) based on both upstream and downstream connectivity to hemilineages, descending neuron subclasses, motor neuron subclasses, and sensory neuron modalities. The annotation bar is coloured by the most common predicted neurotransmitter for the neurons of each type.

**Figure 44 - figure supplement 9.**
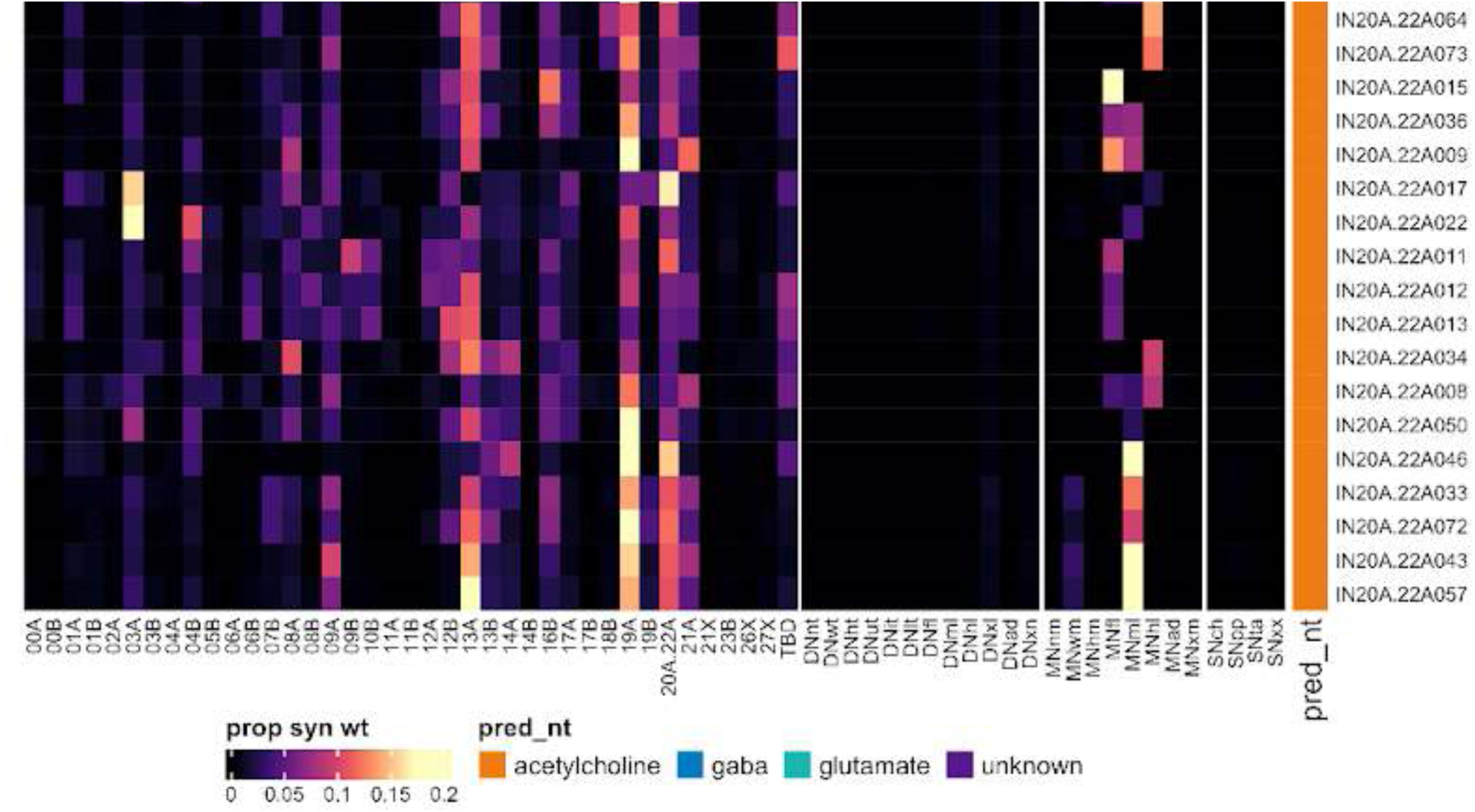
Connectivity to downstream partners by 20A.22A secondary systematic types, continued. Proportions of synaptic weight from systematic types to downstream partners, normalised by row. 20A.22A neurons have been clustered within each assigned birthtime window (P = primary, ES = early secondary, S = secondary) based on both upstream and downstream connectivity to hemilineages, descending neuron subclasses, motor neuron subclasses, and sensory neuron modalities. The annotation bar is coloured by the most common predicted neurotransmitter for the neurons of each type.

#### Hemilineage 21A

Hemilineages 20A, 21A, and 22A are posterior lateral hemilineages that innervate the six ipsilateral leg neuropils, with all three entering the neuropil in approximately the same location. Hemilineage 21A derives from neuroblast NB4-3 (Birkholz et al., 2015; Lacin and Truman, 2016), which generates 2-3 neurosecretory cells and 10-12 local interneurons in the embryo (Schmid et al., 1999). 21A neurons are mainly glutamatergic (Lacin et al., 2019) and include a distinctive neuronal subtype with a “T-junction” morphology that innervates the mVAC (Shepherd et al., 2019). They are mainly restricted to the ipsilateral leg neuropils but also project to the tectulum.

Secondary 21A neurons are found in similar numbers in all thoracic neuromeres (Figure 45E). We identified them from a diagnostic subpopulation that splits to diverge orthogonally to the primary neurite (Figure 45C top). Whilst most 21A neurons were predicted to be glutamatergic as expected, an early born population was predicted to be cholinergic (Figure 45C bottom). As mentioned above, no 20A or 22A primary neurons were expected in the thorax, so we tentatively assigned these cells to 21A.

**Figure 45.**
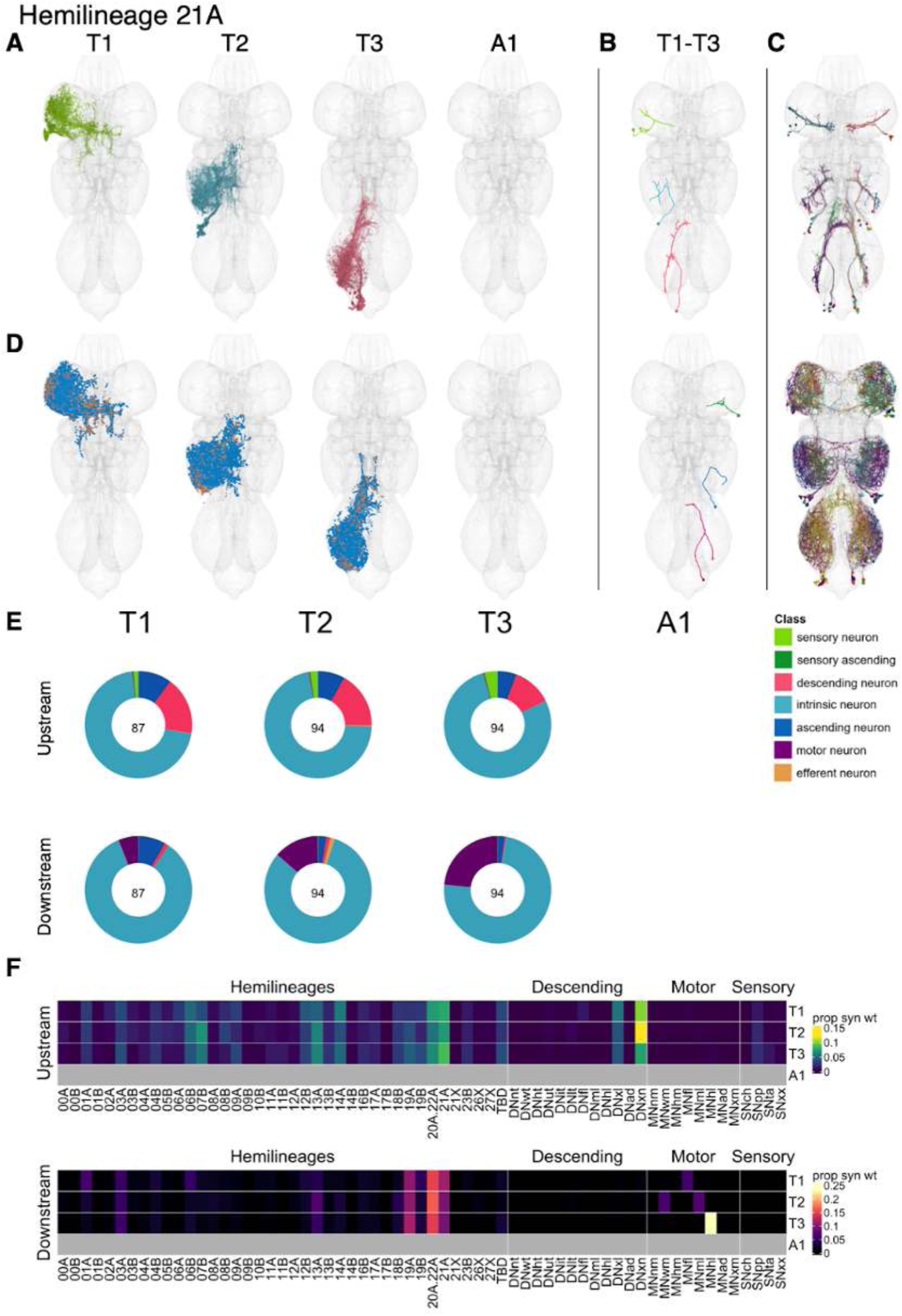
Hemilineage 21A. **A.** Meshes of all RHS secondary neurons plotted in neuromere-specific colours. **B.** “Representative” secondary neuron skeletons plotted in hemineuromere-specific colours. The skeleton with the top accumulated NBLAST score among all neurons from the hemilineage in a given hemineuromere was used. **C.** Neuron meshes of selected examples. Top: independent leg subcluster 23565. Bottom: cholinergic population. **D.** Predicted synapses of RHS secondary neurons. Blue: postsynapses; dark orange: presynapses. **E.** Proportions of connections from secondary neurons to upstream or downstream partners, normalised by neuromere and coloured by broad class. Numbers of query neurons appear in the centre. **F.** Proportions of synaptic weight from secondary neurons originating in each neuromere to upstream or downstream partners, normalised by row.

21A secondary neurons receive input mainly from 07B, 13A, 20A/22A, and 21A and from descending neurons that target the legs or the VNC more broadly (Figure 45F). They primarily target 19A, 20A/22A, 21A, and leg motor neurons, especially in T3 (Figure 45F). A subset of dorsal secondary types (e.g., IN21A025-027) are under especially strong descending control and target wing motor neurons instead (Figure 45 - figure supplement 5,8). Bilateral activation of 21A secondary neurons evokes incessant, uncoordinated leg movements without locomotion (Harris et al., 2015). There was no obvious difference in connectivity between glutamatergic and cholinergic early born types (Figure 45 - figure supplement 4,7).

**Figure 45 - figure supplement 1.**
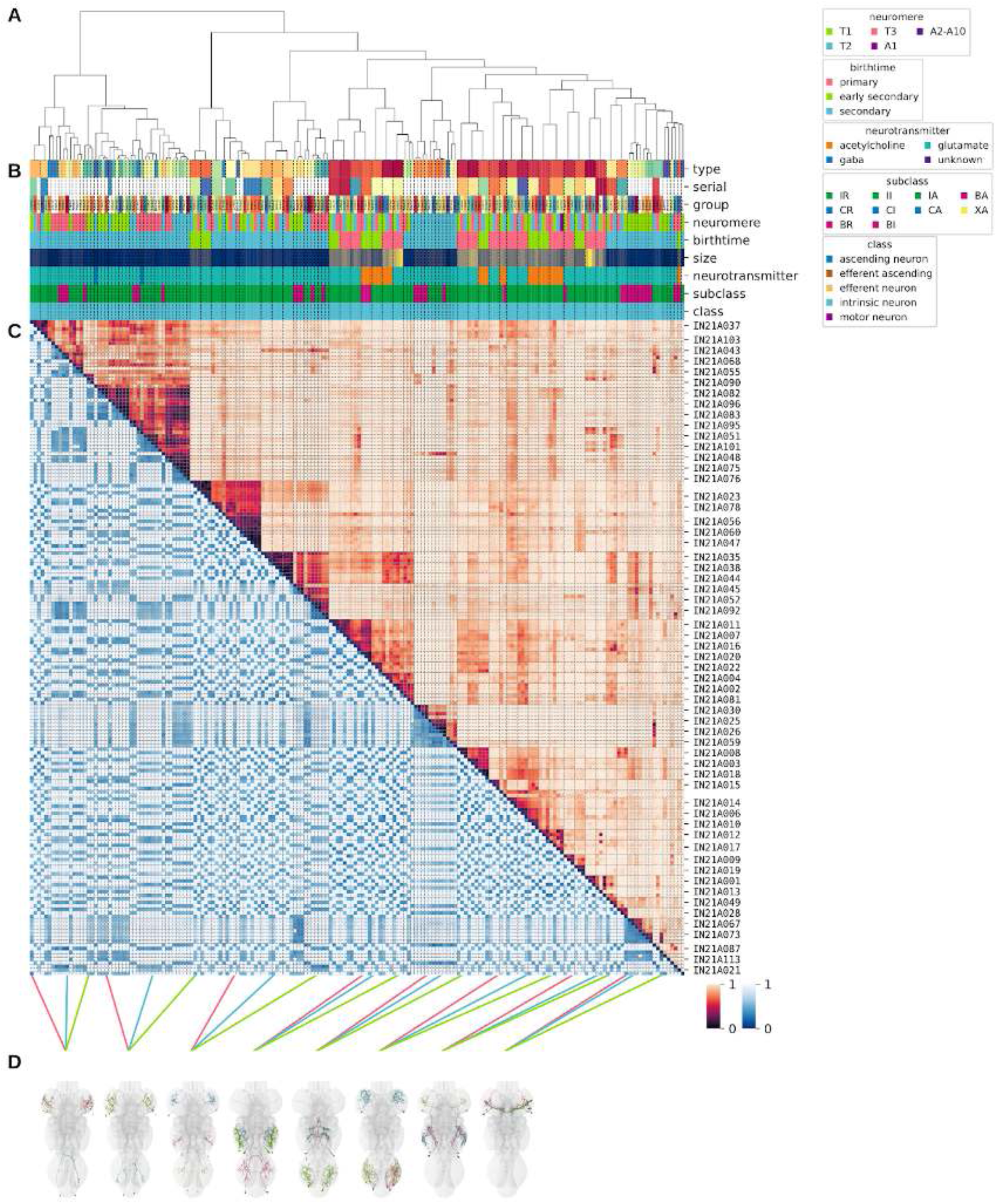
Systematic typing of hemilineage 21A. **A.** Hierarchical clustering dendrogram of hemilineage groups by laterally and serially aggregated connectivity cosine clustering. **B.** Categorical annotations of each hemilineage group, each column corresponding to the aligned leaf in A. Colours for type, serial set, and group are arbitrary for visualisation. Colours for neuromere, birthtime, neurotransmitter, subclass, and class are as in all other figures. **C.** Similarity distance heatmap for hemilineage. Cosine distance is in the upper triangle, while laterally symmetrised NBLAST distance is in the lower triangle. Systematic type names of some types are labelled. **D.** Morphologically representative groups from dendrogram subtrees. Each group, indicated by colour and line connecting to its column in B and C, is the most morphologically representative group (medoid of NBLAST distance) from a subtree of A. The subtrees (flat clusters) are equal height cuts of A determined to yield the number of groups per plot and plots in D.

**Figure 45 - figure supplement 2.**
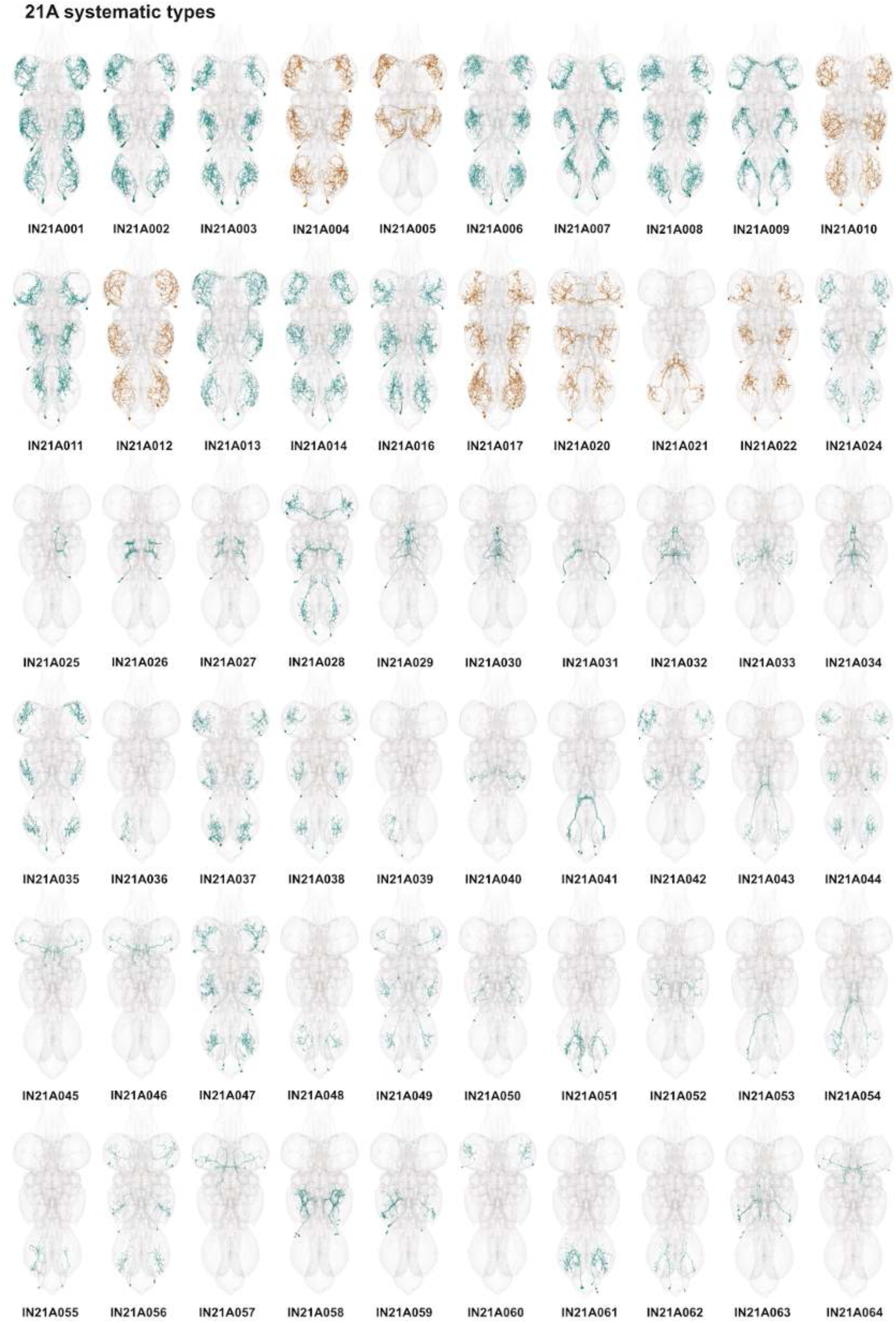
Systematic types of hemilineage 21A. Systematic types have been arranged in numerical order, with neurons of the same type that belong to distinct classes (e.g., intrinsic neuron vs ascending neuron) plotted separately but placed adjacent to each other. Individual neuron meshes have been coloured based on predicted neurotransmitter: dark orange = acetylcholine, blue = gaba, marine = glutamate, dark purple = unknown.

**Figure 45 - figure supplement 3.**
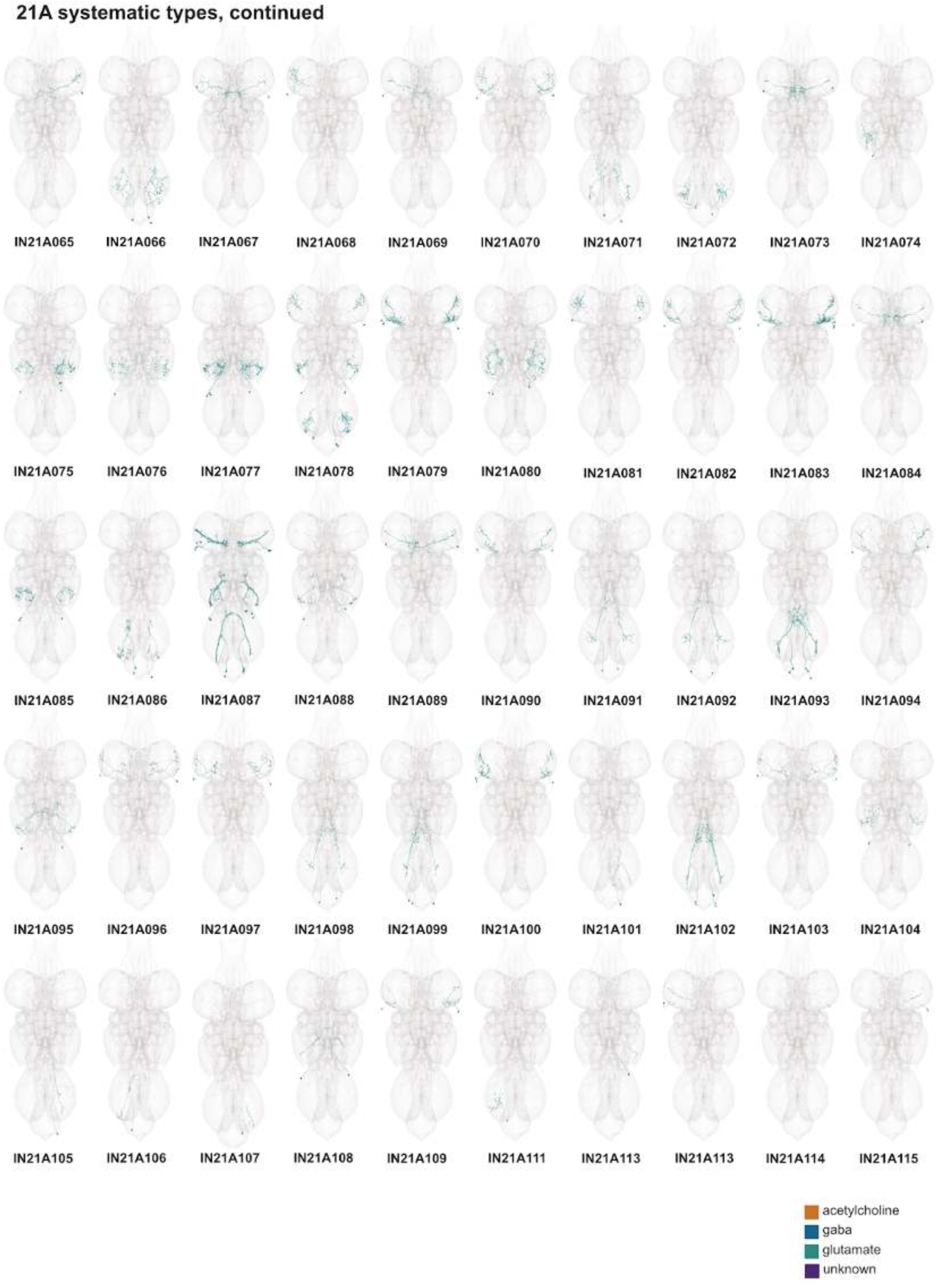
Systematic types of hemilineage 21A, continued. Systematic types have been arranged in numerical order, with neurons of the same type that belong to distinct classes (e.g., intrinsic neuron vs ascending neuron) plotted separately but placed adjacent to each other. Individual neuron meshes have been coloured based on predicted neurotransmitter: dark orange = acetylcholine, blue = gaba, marine = glutamate, dark purple = unknown.

**Figure 45 - figure supplement 4.**
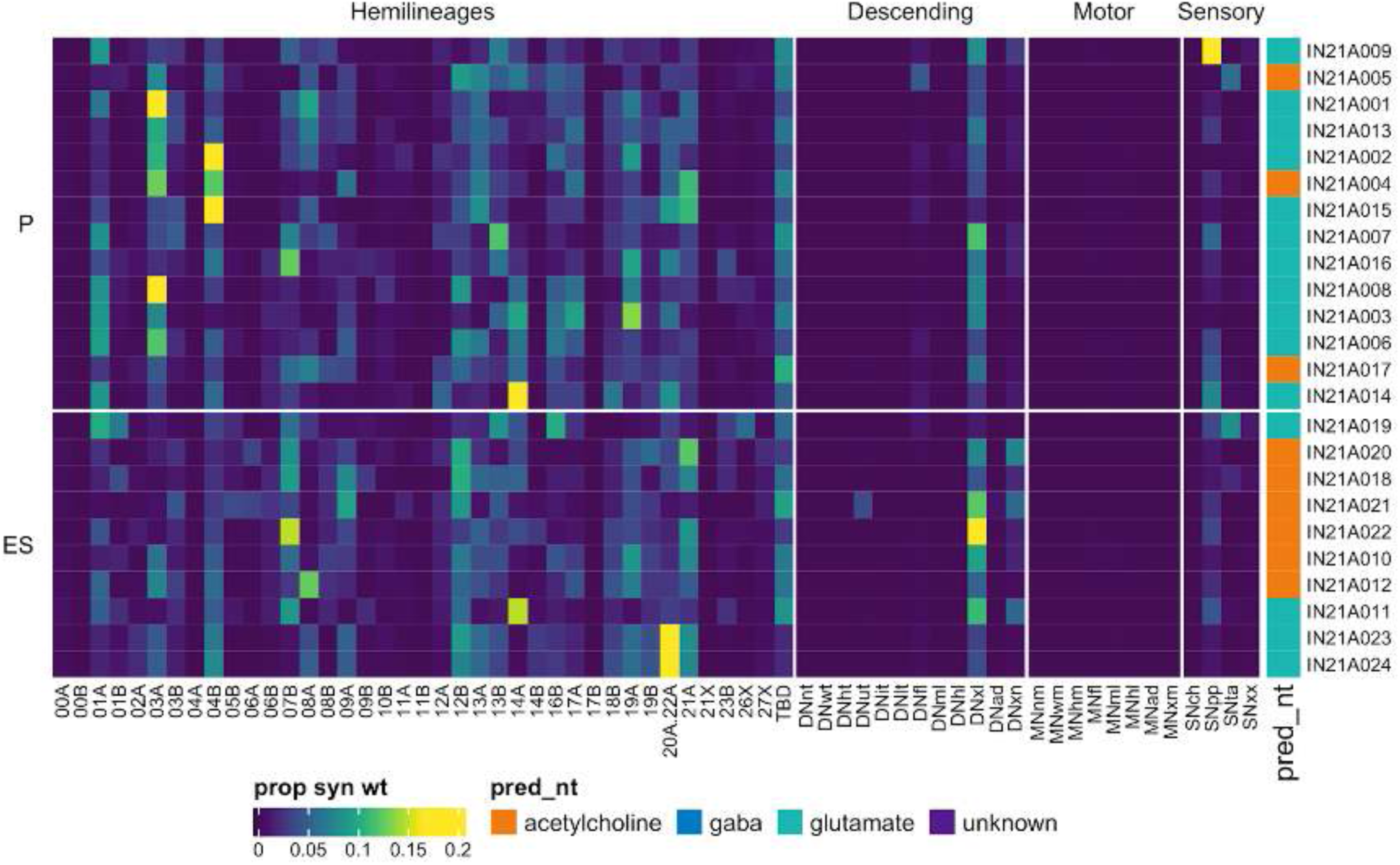
Connectivity to upstream partners by 21A primary and early secondary systematic types. Proportions of synaptic weight to systematic types from upstream partners, normalised by row. 21A neurons have been clustered within each assigned birthtime window (P = primary, ES = early secondary, S = secondary) based on both upstream and downstream connectivity to hemilineages, descending neuron subclasses, motor neuron subclasses, and sensory neuron modalities. Annotation bar is coloured by the most common predicted neurotransmitter for the neurons of each type.

**Figure 45 - figure supplement 5.**
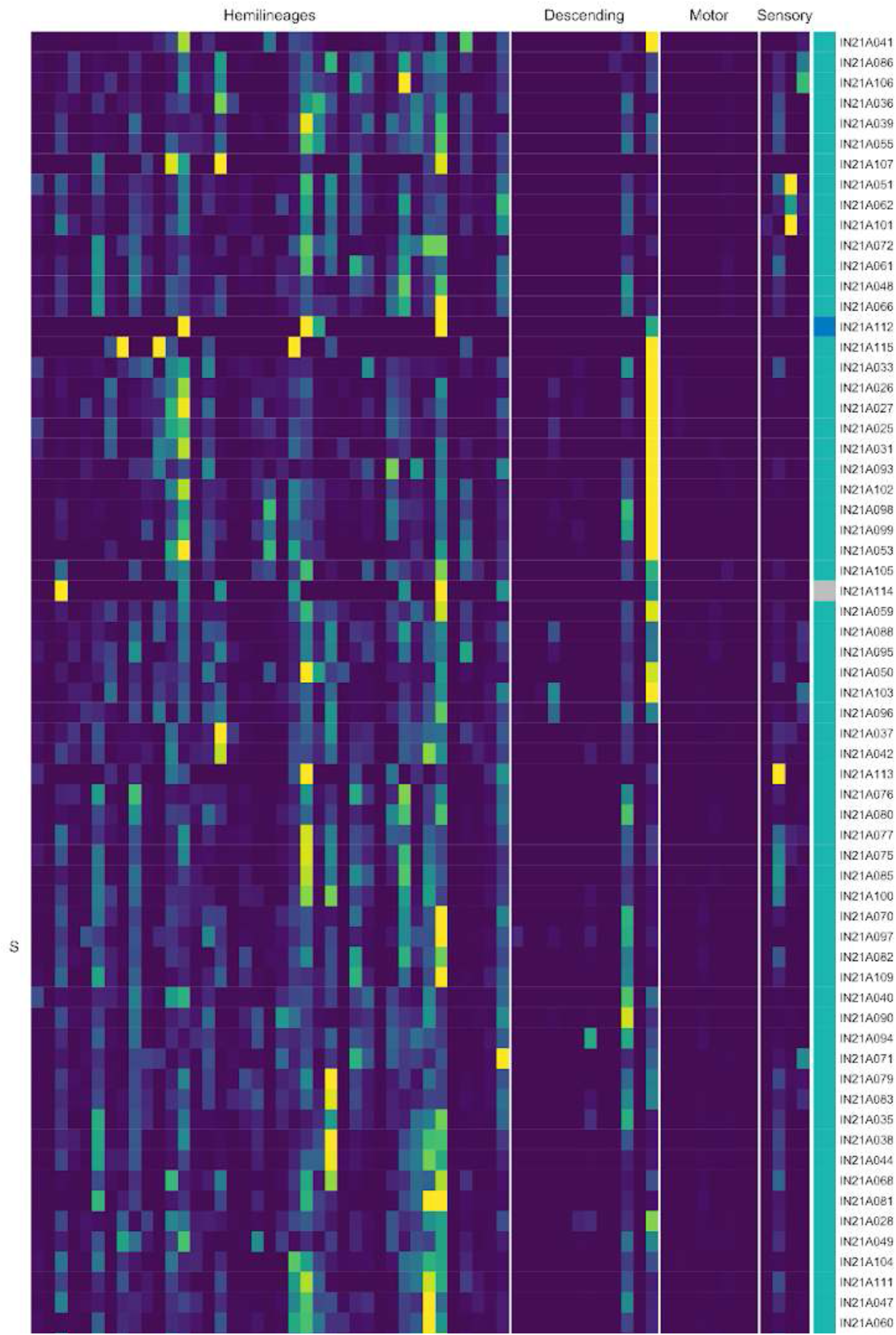
Connectivity to upstream partners by 21A secondary systematic types. Proportions of synaptic weight to systematic types from upstream partners, normalised by row. 21A neurons have been clustered within each assigned birthtime window (P = primary, ES = early secondary, S = secondary) based on both upstream and downstream connectivity to hemilineages, descending neuron subclasses, motor neuron subclasses, and sensory neuron modalities. The annotation bar is coloured by the most common predicted neurotransmitter within each type.

**Figure 45 - figure supplement 6.**
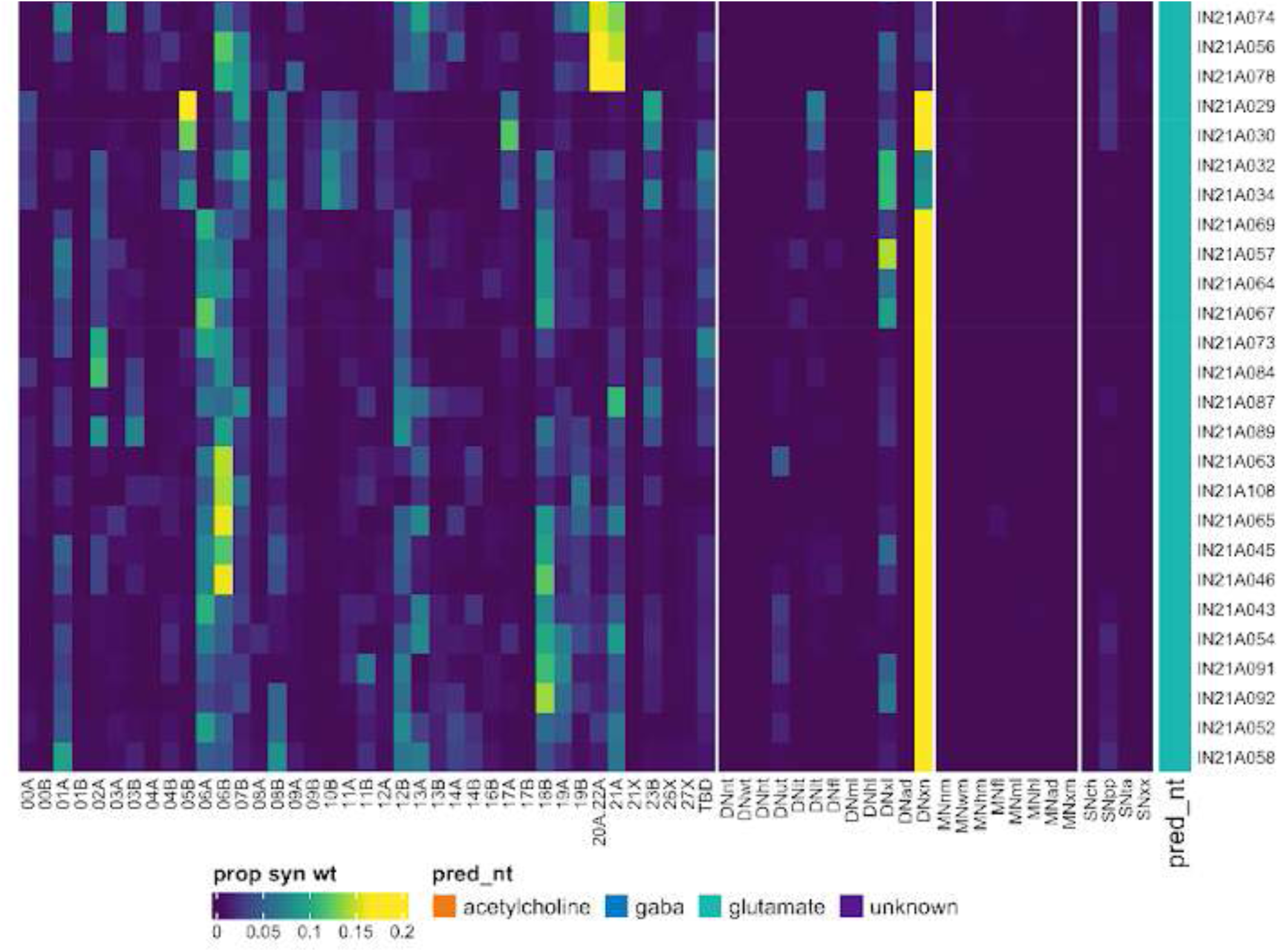
Connectivity to upstream partners by 21A secondary systematic types, continued. Proportions of synaptic weight to systematic types from upstream partners, normalised by row. 21A neurons have been clustered within each assigned birthtime window (P = primary, ES = early secondary, S = secondary) based on both upstream and downstream connectivity to hemilineages, descending neuron subclasses, motor neuron subclasses, and sensory neuron modalities. The annotation bar is coloured by the most common predicted neurotransmitter within each type.

**Figure 45 - figure supplement 7.**
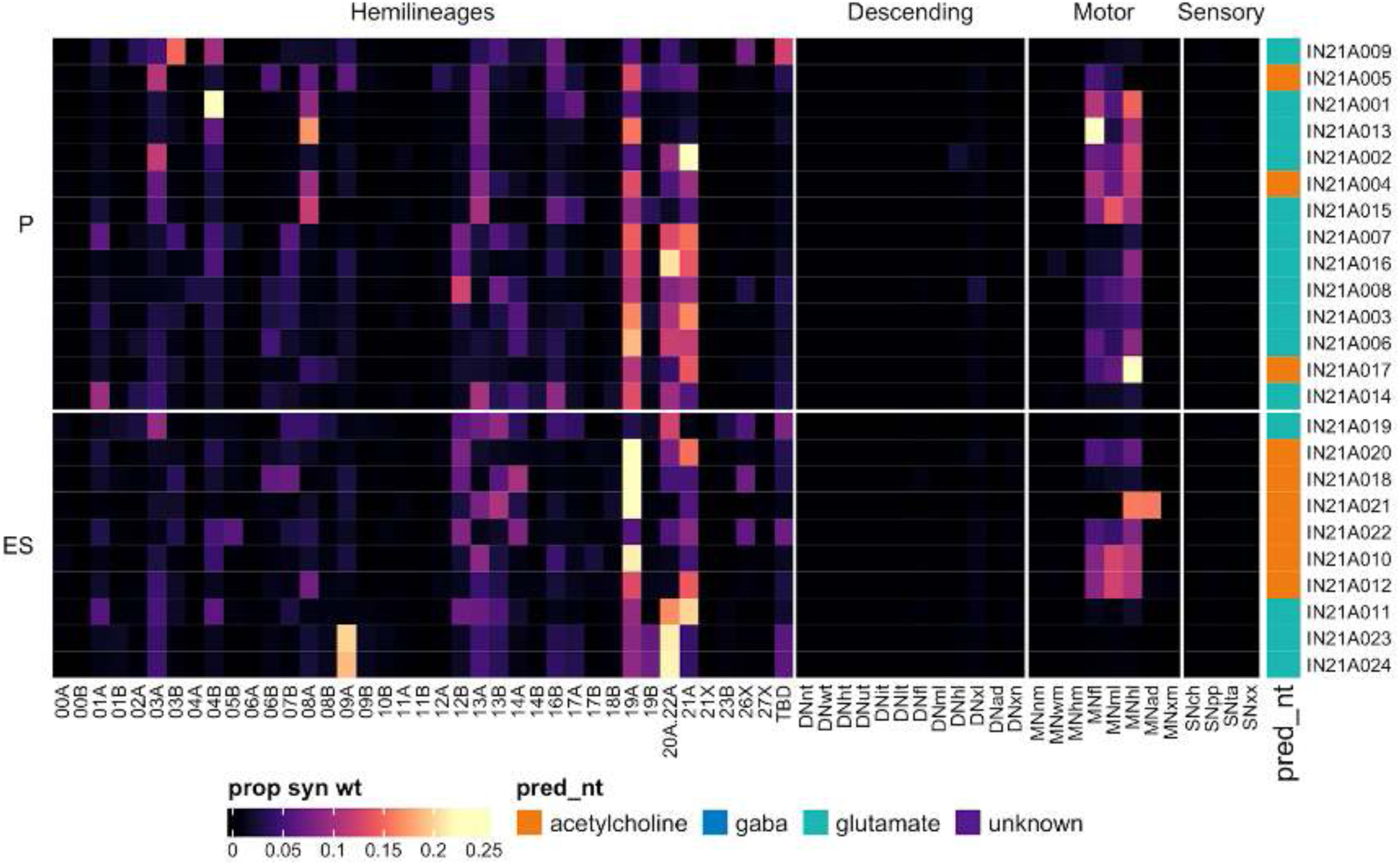
Connectivity to downstream partners by 21A primary and early secondary systematic types. Proportions of synaptic weight from systematic types to downstream partners, normalised by row. 21A neurons have been clustered within each assigned birthtime window (P = primary, ES = early secondary, S = secondary) based on both upstream and downstream connectivity to hemilineages, descending neuron subclasses, motor neuron subclasses, and sensory neuron modalities. The annotation bar is coloured by the most common predicted neurotransmitter within each type.

**Figure 45 - figure supplement 8.**
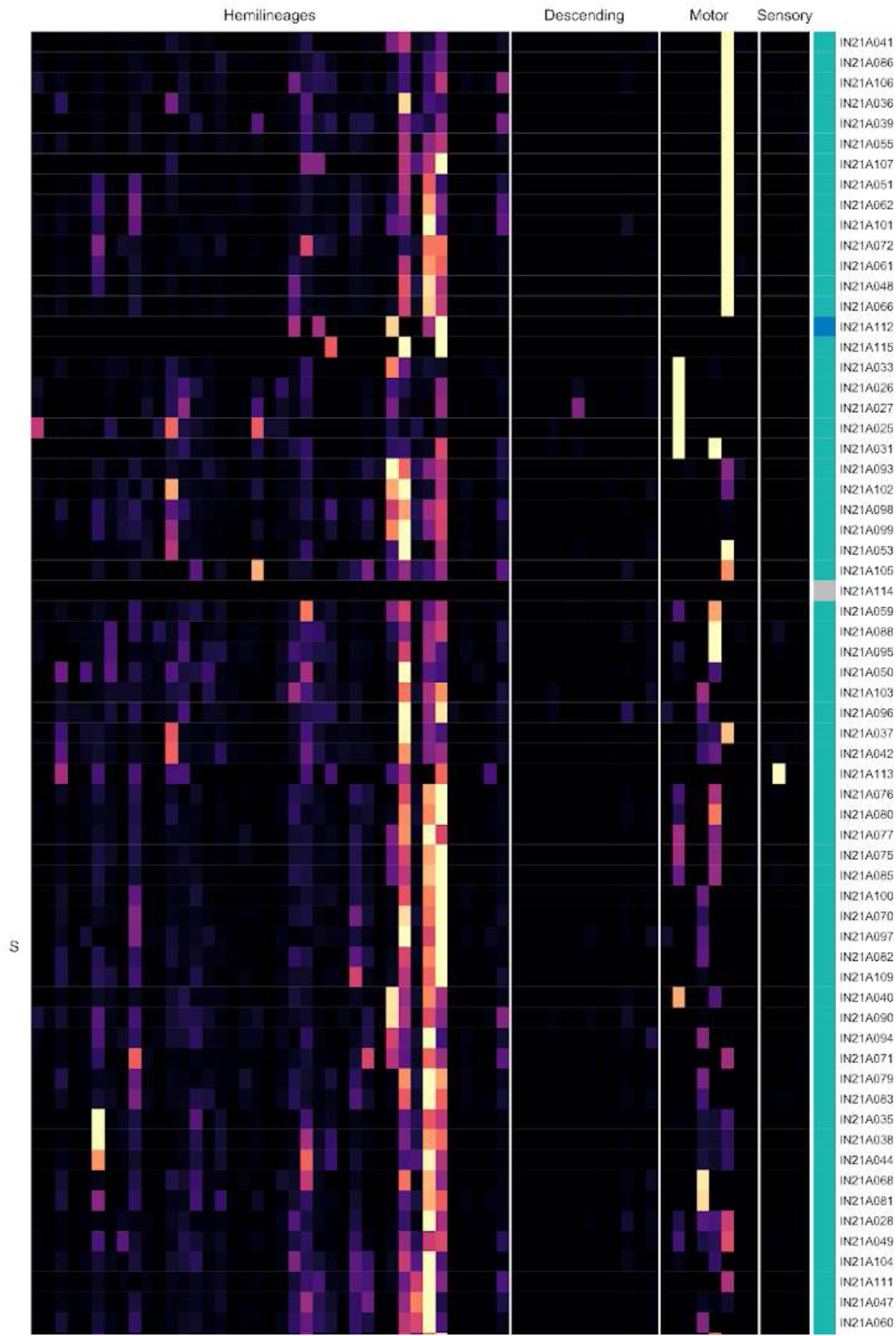
Connectivity to downstream partners by 21A secondary systematic types. Proportions of synaptic weight from systematic types to downstream partners, normalised by row. 21A neurons have been clustered within each assigned birthtime window (P = primary, ES = early secondary, S = secondary) based on both upstream and downstream connectivity to hemilineages, descending neuron subclasses, motor neuron subclasses, and sensory neuron modalities. The annotation bar is coloured by the most common predicted neurotransmitter for the neurons of each type.

**Figure 45 - figure supplement 9.**
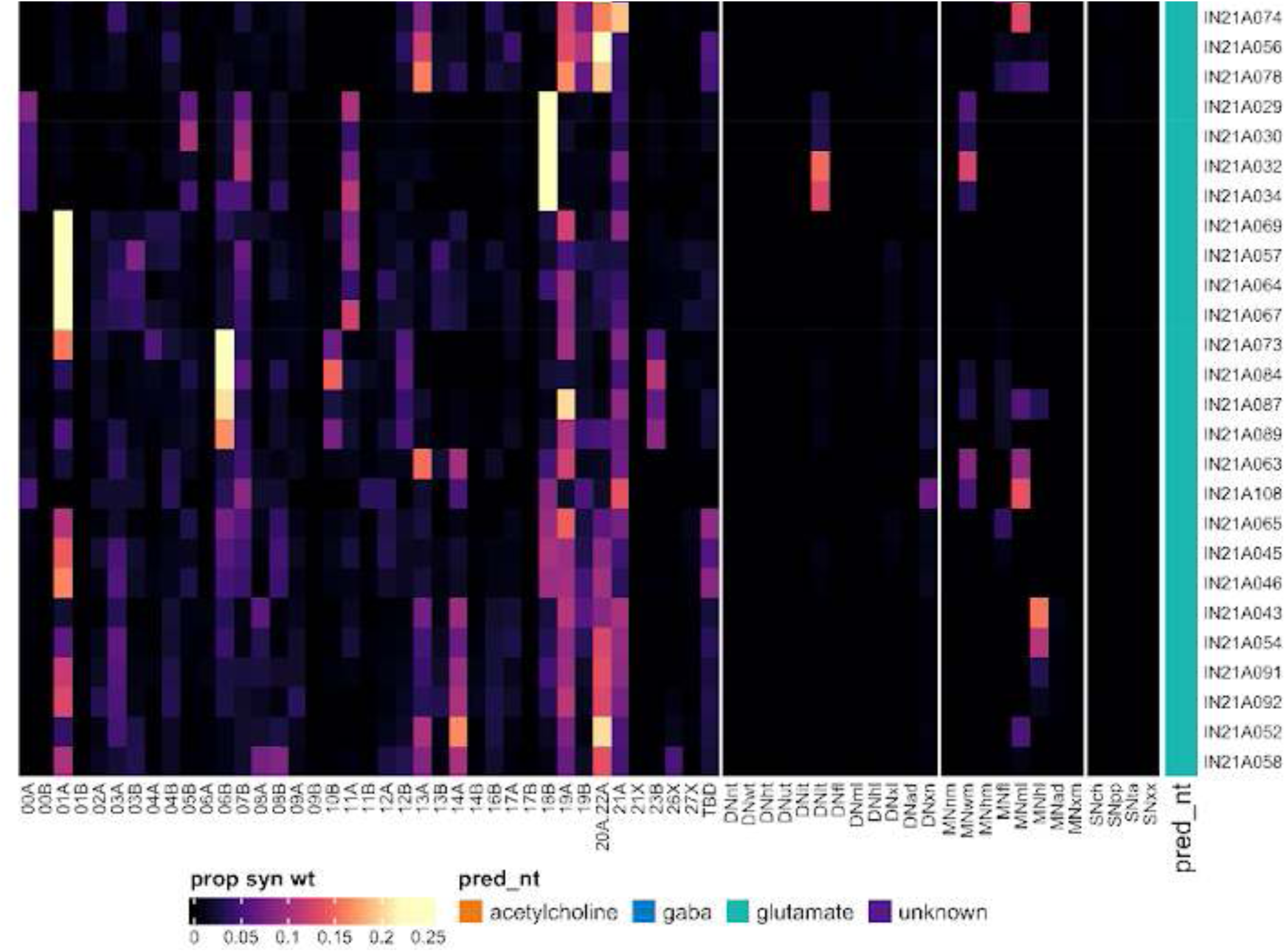
Connectivity to downstream partners by 21A secondary systematic types, continued. Proportions of synaptic weight from systematic types to downstream partners, normalised by row. 21A neurons have been clustered within each assigned birthtime window (P = primary, ES = early secondary, S = secondary) based on both upstream and downstream connectivity to hemilineages, descending neuron subclasses, motor neuron subclasses, and sensory neuron modalities. The annotation bar is coloured by the most common predicted neurotransmitter for the neurons of each type.

#### Hemilineage 23B

Hemilineage 23B derives from posterior dorsal neuroblast NB7-4 (Birkholz et al., 2015; Lacin and Truman, 2016), which generates 8-15 thoracic interneurons and 5-6 glia in the embryo (Schmid et al., 1999). 23B secondary neurons are mainly cholinergic (Lacin et al., 2019) and share a large, complex primary neurite tract with 11A and 19B (Shepherd et al., 2016). The 23B primary neurites bend towards the midline as they project ventrally to innervate the ipsilateral sensory neuropil, resulting in a characteristic “hourglass” morphology when viewed in a transverse section (Shepherd et al., 2019). The A1 neurons were identified via several serial minority subtypes (e.g., Figure 46C).

**Figure 46.**
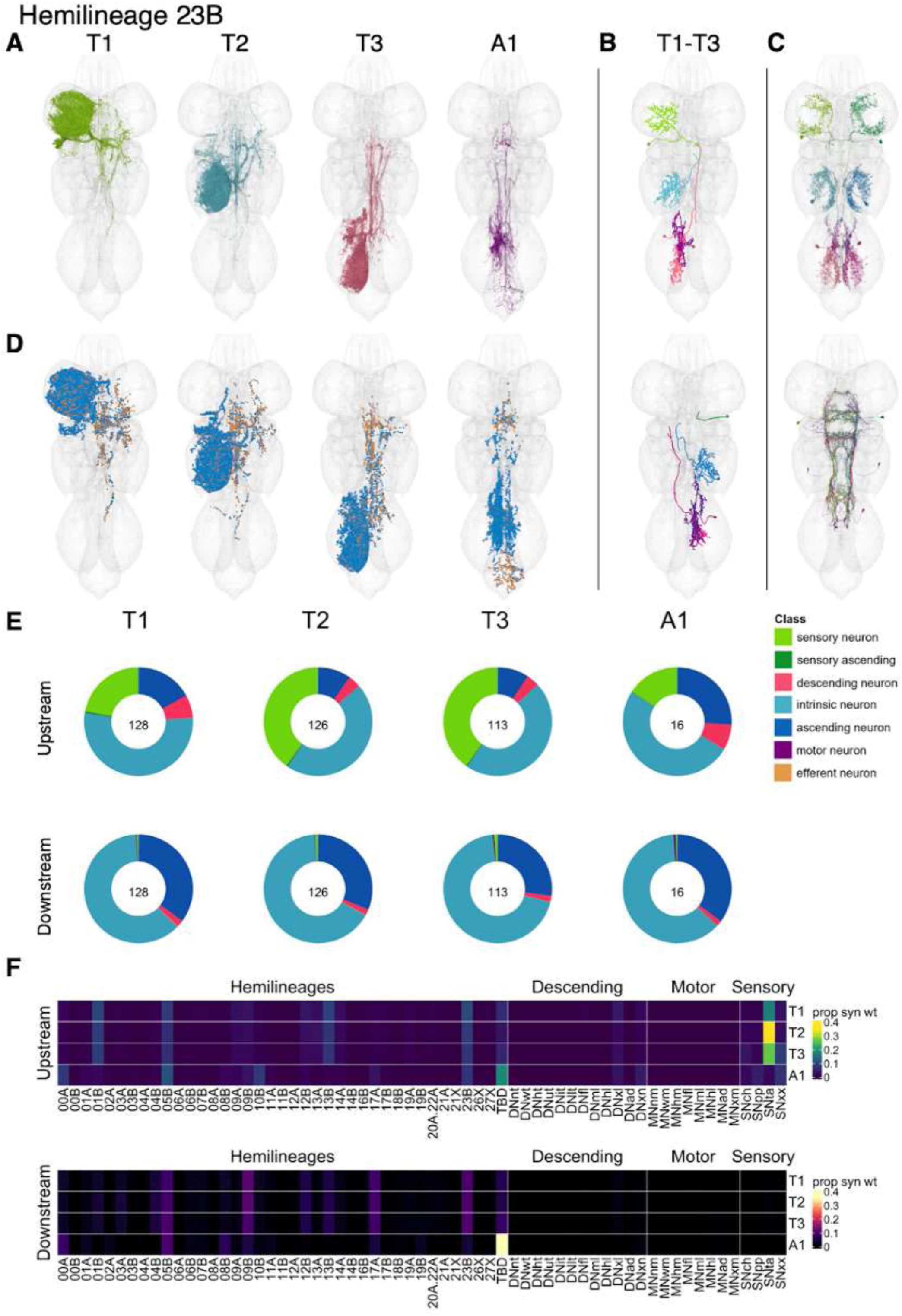
Hemilineage 23B. **A.** Meshes of all RHS secondary neurons plotted in neuromere-specific colours. **B.** “Representative” secondary neuron skeletons plotted in hemineuromere-specific colours. The skeleton with the top accumulated NBLAST score among all neurons from the hemilineage in a given hemineuromere was used. **C.** Neuron meshes of selected examples. Top: independent leg serial 12339 (T1-A1). Bottom: intersegmental complex serial 11371 (T1-A1). **D.** Predicted synapses of RHS secondary neurons. Blue: postsynapses; dark orange: presynapses. **E.** Proportions of connections from secondary neurons to upstream or downstream partners, normalised by neuromere and coloured by broad class. Numbers of query neurons appear in the centre. **F.** Proportions of synaptic weight from secondary neurons originating in each neuromere to upstream or downstream partners, normalised by row.

We identified 23B neurons in T1-A1, although the number in A1 is much reduced (Figure 46E). Most innervate the ipsilateral leg sensory neuropil, then project across the midline and almost immediately ascend or descend (Figure 46A,C top). However, there are also minority types with distinct morphology and connectivity (e.g., Figure 46C bottom).

Bilateral activation of 23B secondary neurons results in intersegmental limb movements with poor intralimb coordination (Harris et al., 2015). 23B secondary neurons in T1-T3 receive most input from tactile sensory neurons and from multiple hemilineages including 01B, 05B, 13B, and 23B; they activate neurons from hemilineages 05B, 09B, 17A, and 23B. The A1 neurons are distinct in morphology and connectivity, mainly receiving input from 05B, 10B, and 23B and synapsing onto 00A, 05B, 08B, 09B, and neurons of unknown hemilineage; they were validated using several serial minority types (e.g., IN23B008 and IN23B009) (Figure 46 - figure supplement 2).

We found that some specific 23B types receive chemosensory information (e.g., AN23B002 and AN23B010) and others proprioceptive information (e.g., IN23B008 and IN23B024) (Figure 46 - figure supplement 4). Moreover, several early born types activate wing (AN23B002), hind leg (IN23B001), or abdominal motor neurons (IN23B016) (Figure 46 - figure supplement 7).

**Figure 46 - figure supplement 1.**
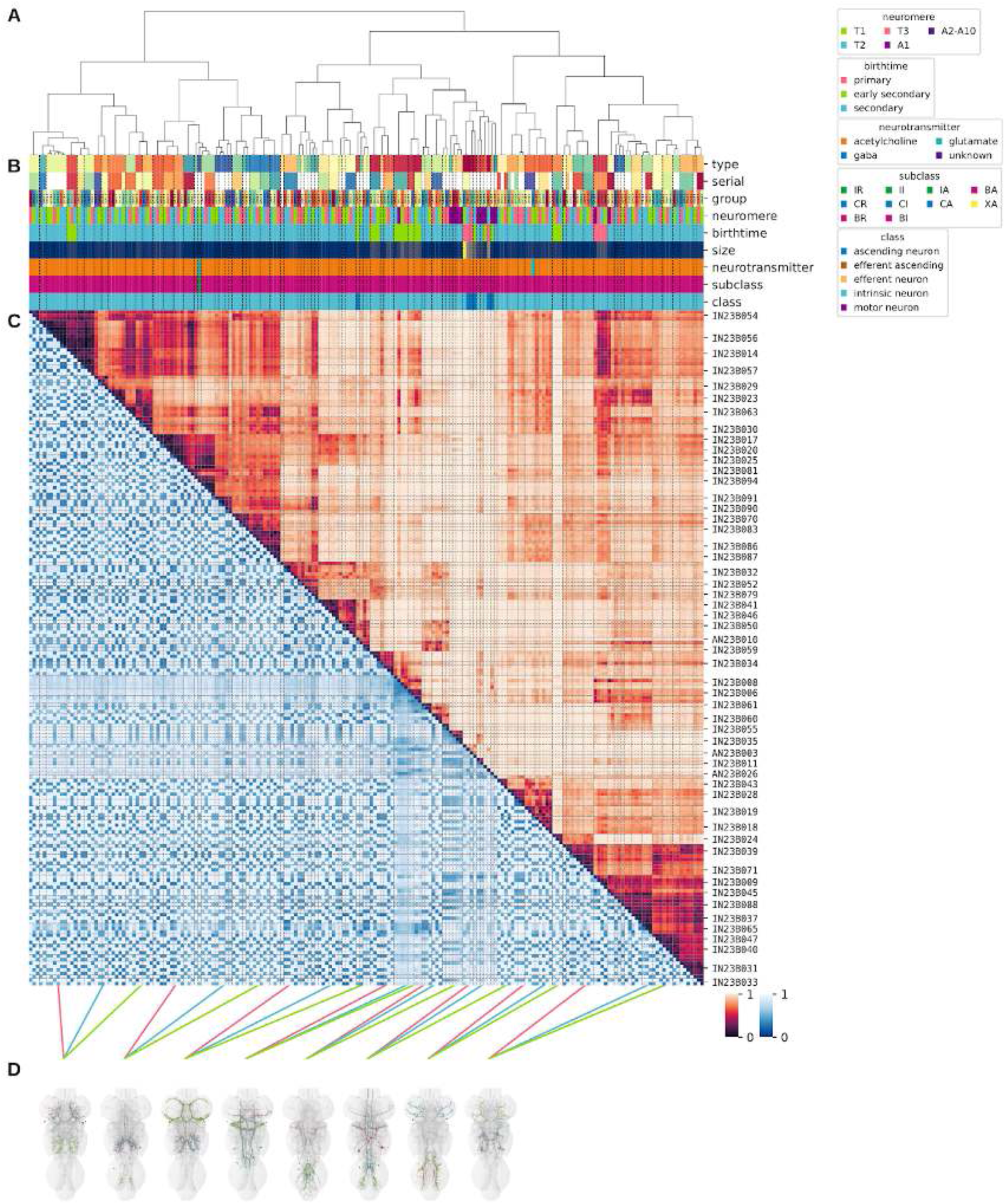
Systematic typing of hemilineage 23B. **A.** Hierarchical clustering dendrogram of hemilineage groups by laterally and serially aggregated connectivity cosine clustering. **B.** Categorical annotations of each hemilineage group, each column corresponding to the aligned leaf in A. Colours for type, serial set, and group are arbitrary for visualisation. Colours for neuromere, birthtime, neurotransmitter, subclass, and class are as in all other figures. **C.** Similarity distance heatmap for hemilineage. Cosine distance is in the upper triangle, while laterally symmetrised NBLAST distance is in the lower triangle. Systematic type names of some types are labelled. **D.** Morphologically representative groups from dendrogram subtrees. Each group, indicated by colour and line connecting to its column in B and C, is the most morphologically representative group (medoid of NBLAST distance) from a subtree of A. The subtrees (flat clusters) are equal height cuts of A determined to yield the number of groups per plot and plots in D.

**Figure 46 - figure supplement 2.**
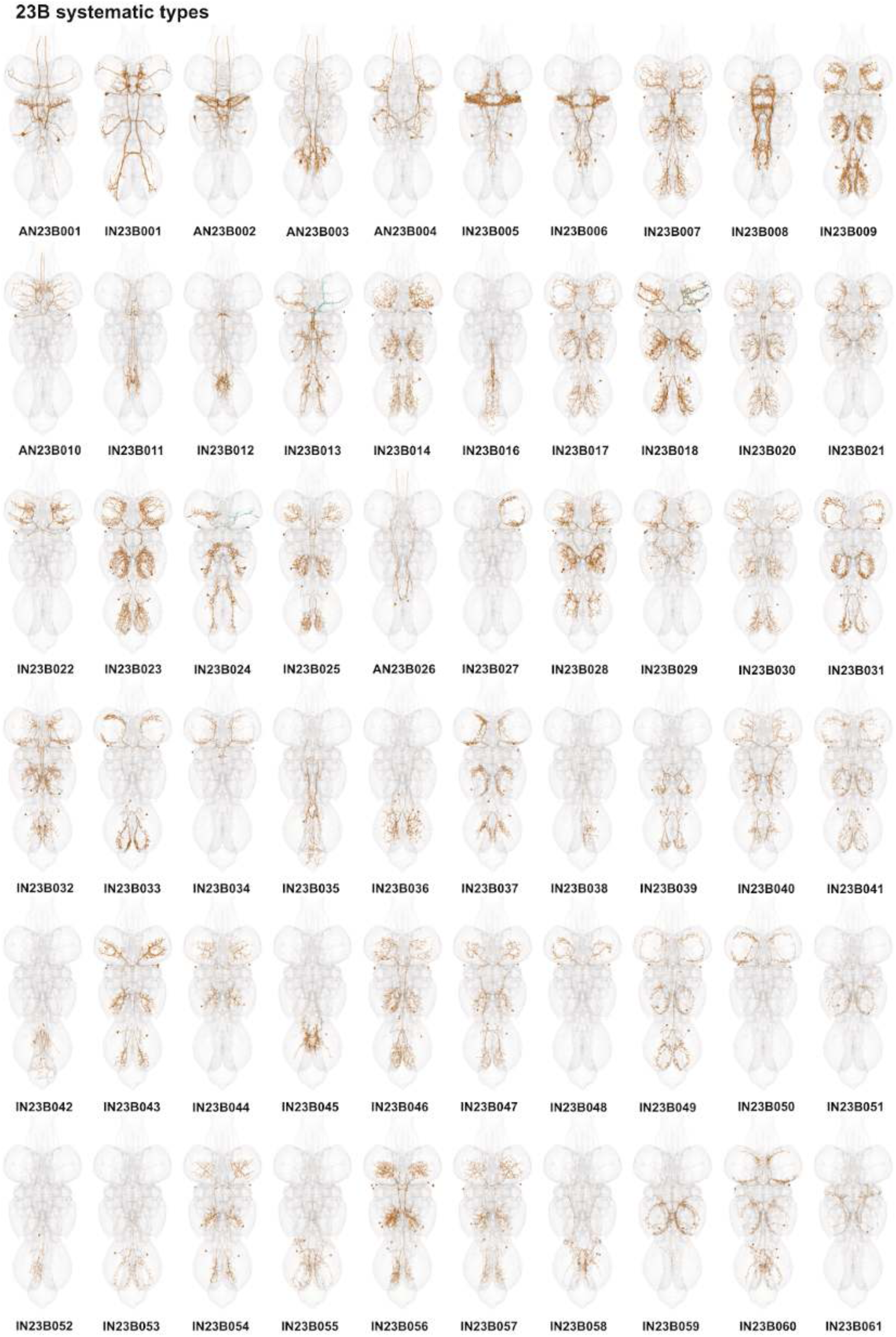
Systematic types of hemilineage 23B. Systematic types have been arranged in numerical order, with neurons of the same type that belong to distinct classes (e.g., intrinsic neuron vs ascending neuron) plotted separately but placed adjacent to each other. Individual neuron meshes have been coloured based on predicted neurotransmitter: dark orange = acetylcholine, blue = gaba, marine = glutamate, dark purple = unknown.

**Figure 46 - figure supplement 3.**
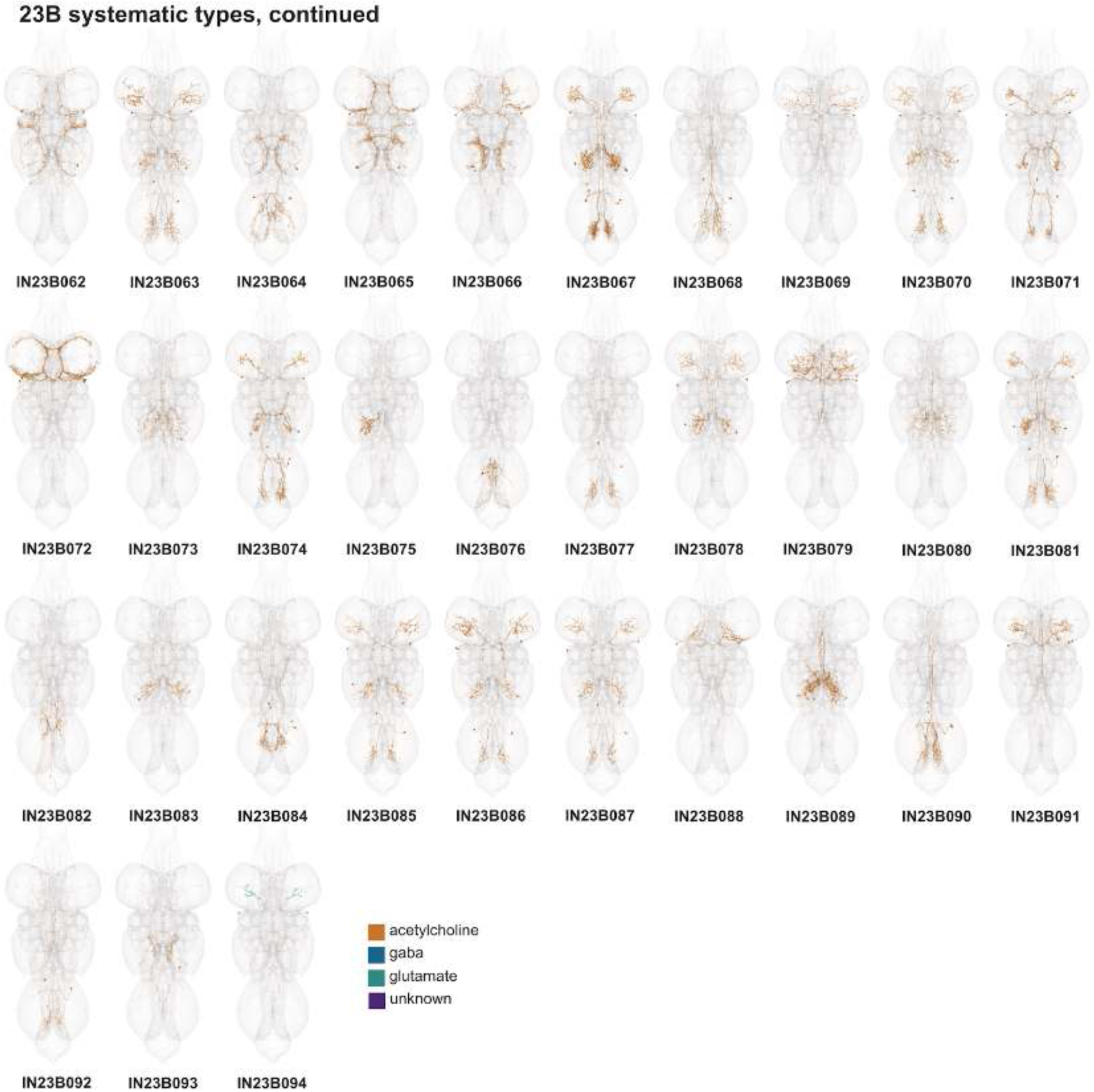
Systematic types of hemilineage 23B, continued. Systematic types have been arranged in numerical order, with neurons of the same type that belong to distinct classes (e.g., intrinsic neuron vs ascending neuron) plotted separately but placed adjacent to each other. Individual neuron meshes have been coloured based on predicted neurotransmitter: dark orange = acetylcholine, blue = gaba, marine = glutamate, dark purple = unknown.

**Figure 46 - figure supplement 4.**
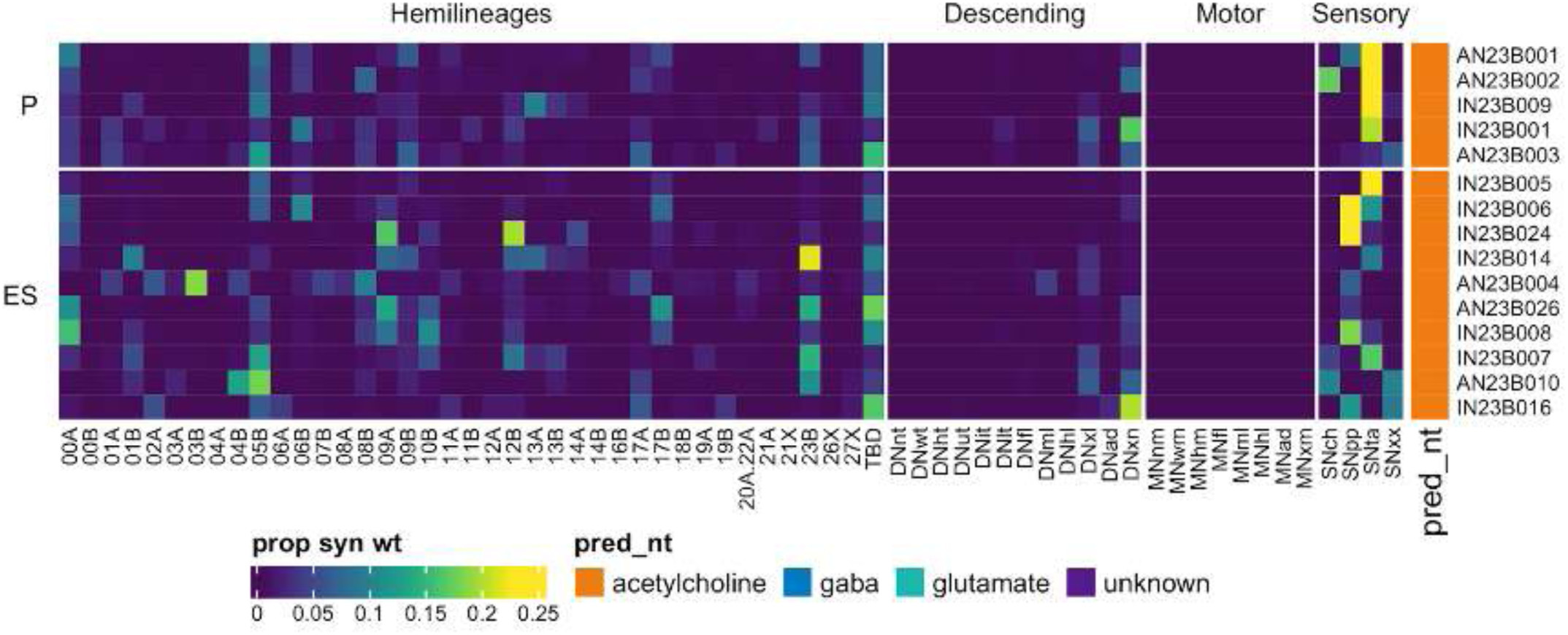
Connectivity to upstream partners by 23B primary and early secondary systematic types. Proportions of synaptic weight to systematic types from upstream partners, normalised by row. 23B neurons have been clustered within each assigned birthtime window (P = primary, ES = early secondary, S = secondary) based on both upstream and downstream connectivity to hemilineages, descending neuron subclasses, motor neuron subclasses, and sensory neuron modalities. Annotation bar is coloured by the most common predicted neurotransmitter for the neurons of each type.

**Figure 46 - figure supplement 5.**
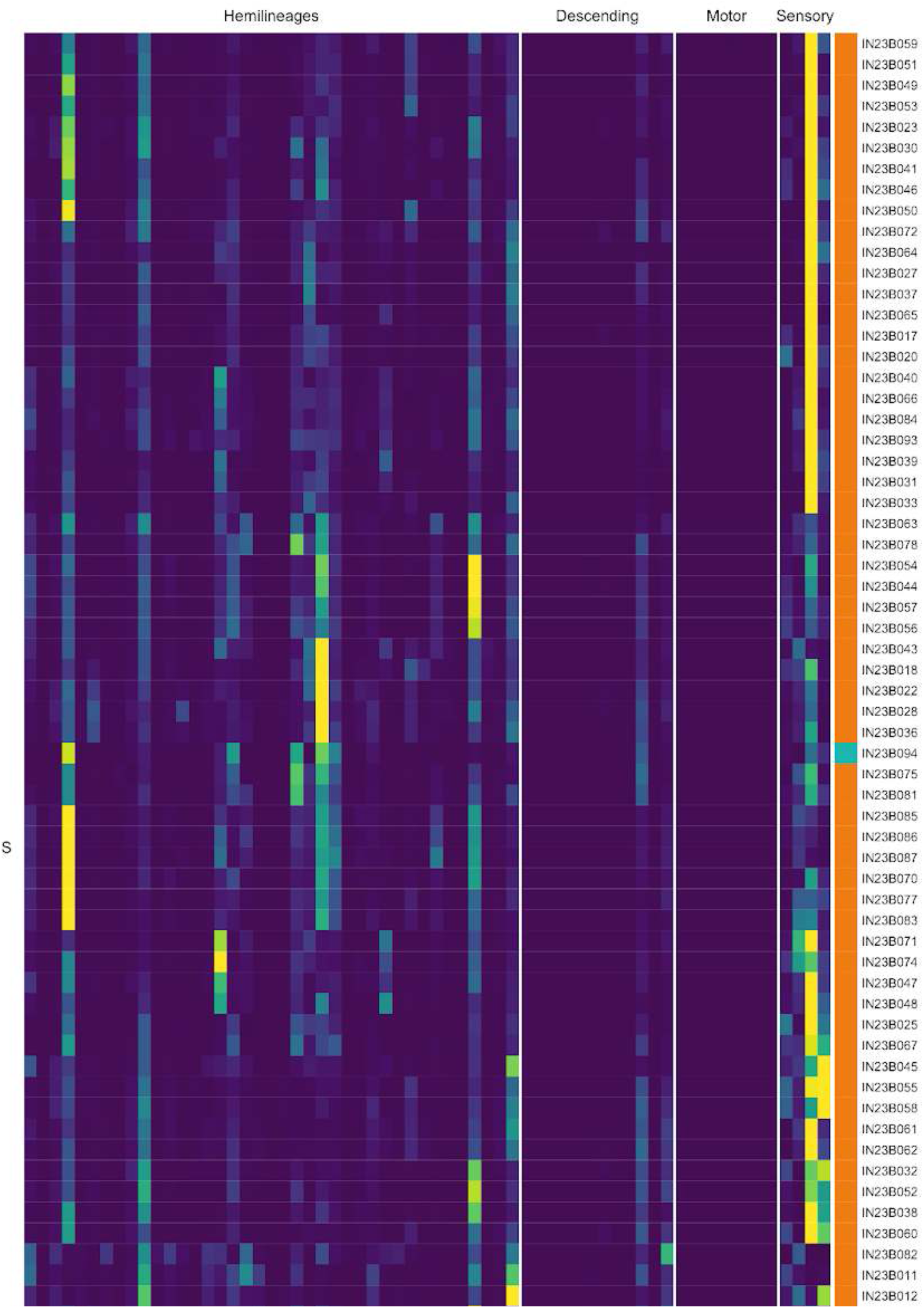
Connectivity to upstream partners by 23B secondary systematic types. Proportions of synaptic weight to systematic types from upstream partners, normalised by row. 23B neurons have been clustered within each assigned birthtime window (P = primary, ES = early secondary, S = secondary) based on both upstream and downstream connectivity to hemilineages, descending neuron subclasses, motor neuron subclasses, and sensory neuron modalities. The annotation bar is coloured by the most common predicted neurotransmitter within each type.

**Figure 46 - figure supplement 6.**
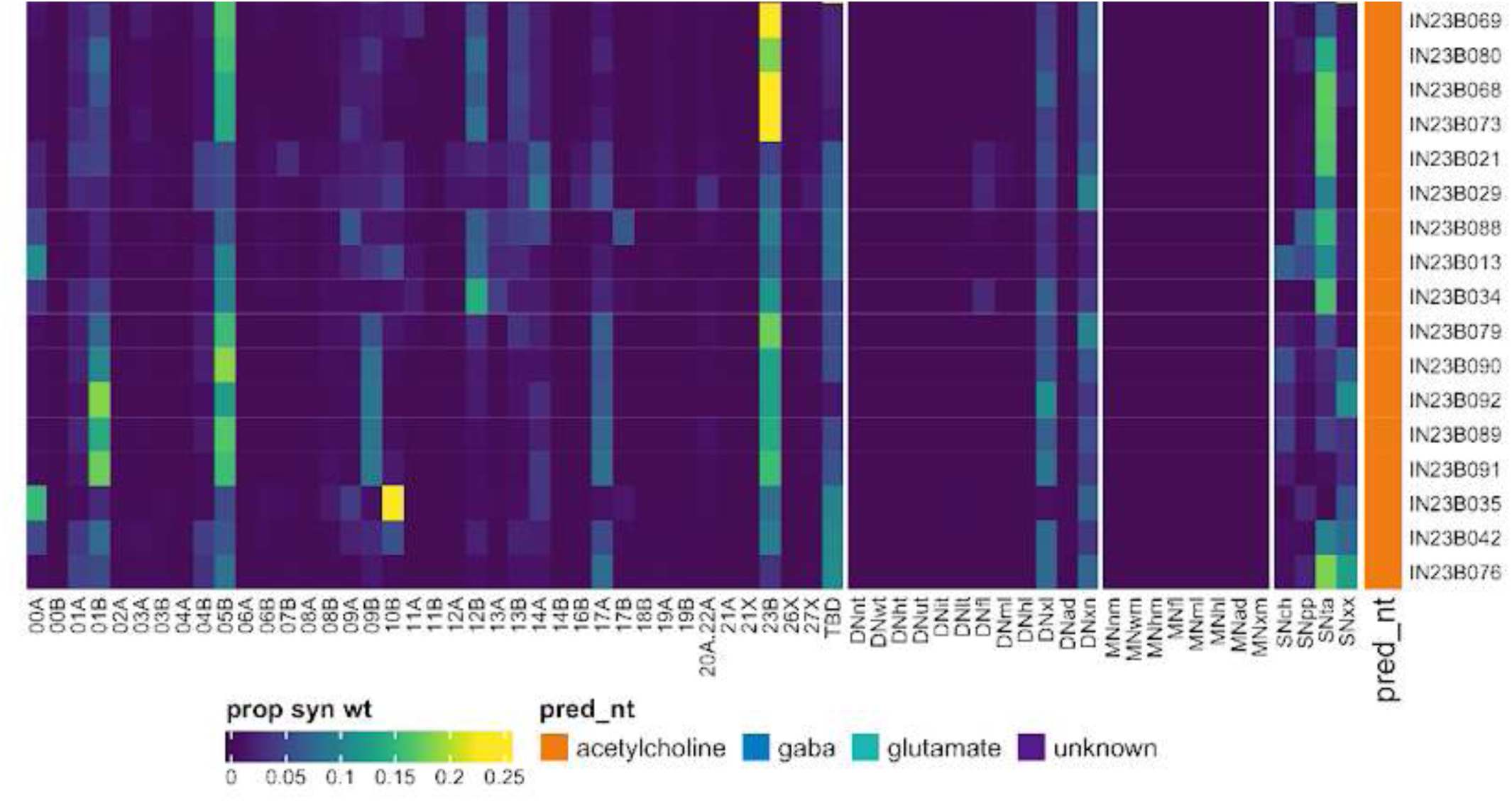
Connectivity to upstream partners by 23B secondary systematic types, continued. Proportions of synaptic weight to systematic types from upstream partners, normalised by row. 23B neurons have been clustered within each assigned birthtime window (P = primary, ES = early secondary, S = secondary) based on both upstream and downstream connectivity to hemilineages, descending neuron subclasses, motor neuron subclasses, and sensory neuron modalities. The annotation bar is coloured by the most common predicted neurotransmitter within each type.

**Figure 46 - figure supplement 7.**
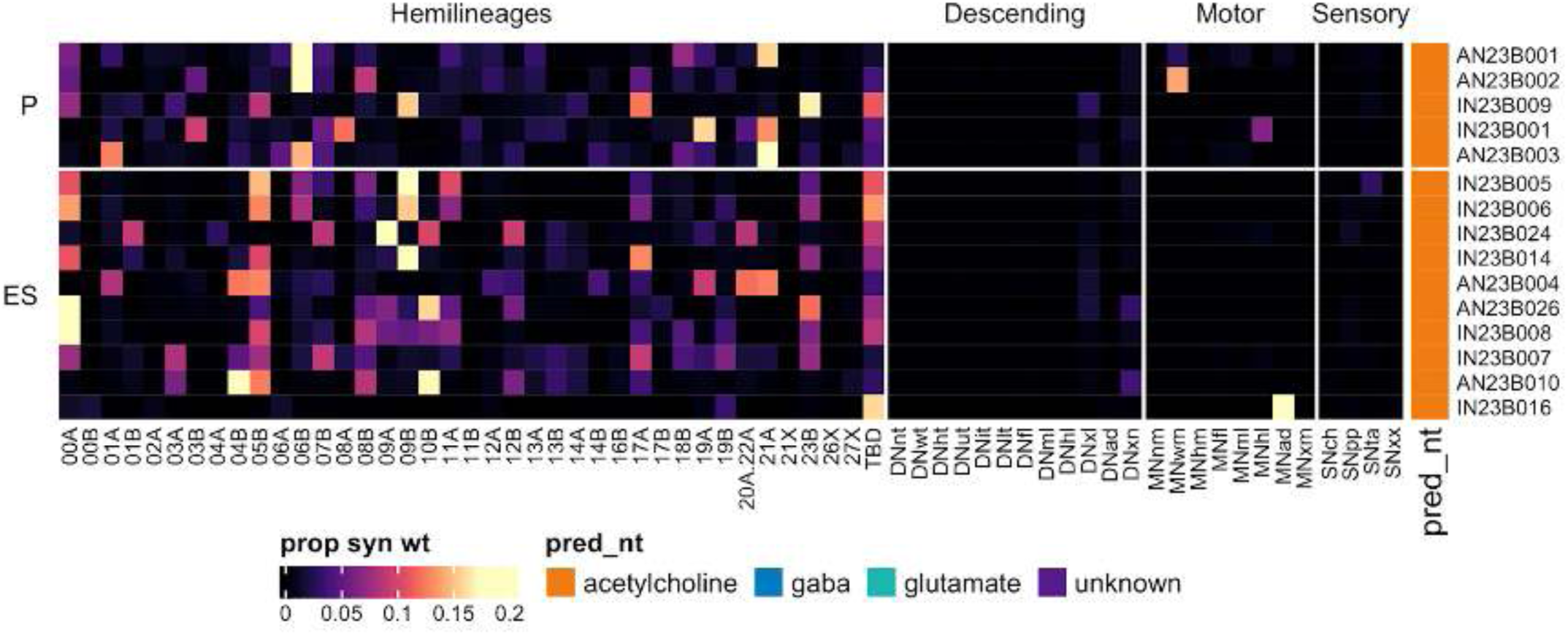
Connectivity to downstream partners by 23B primary and early secondary systematic types. Proportions of synaptic weight from systematic types to downstream partners, normalised by row. 23B neurons have been clustered within each assigned birthtime window (P = primary, ES = early secondary, S = secondary) based on both upstream and downstream connectivity to hemilineages, descending neuron subclasses, motor neuron subclasses, and sensory neuron modalities. The annotation bar is coloured by the most common predicted neurotransmitter within each type.

**Figure 46 - figure supplement 8.**
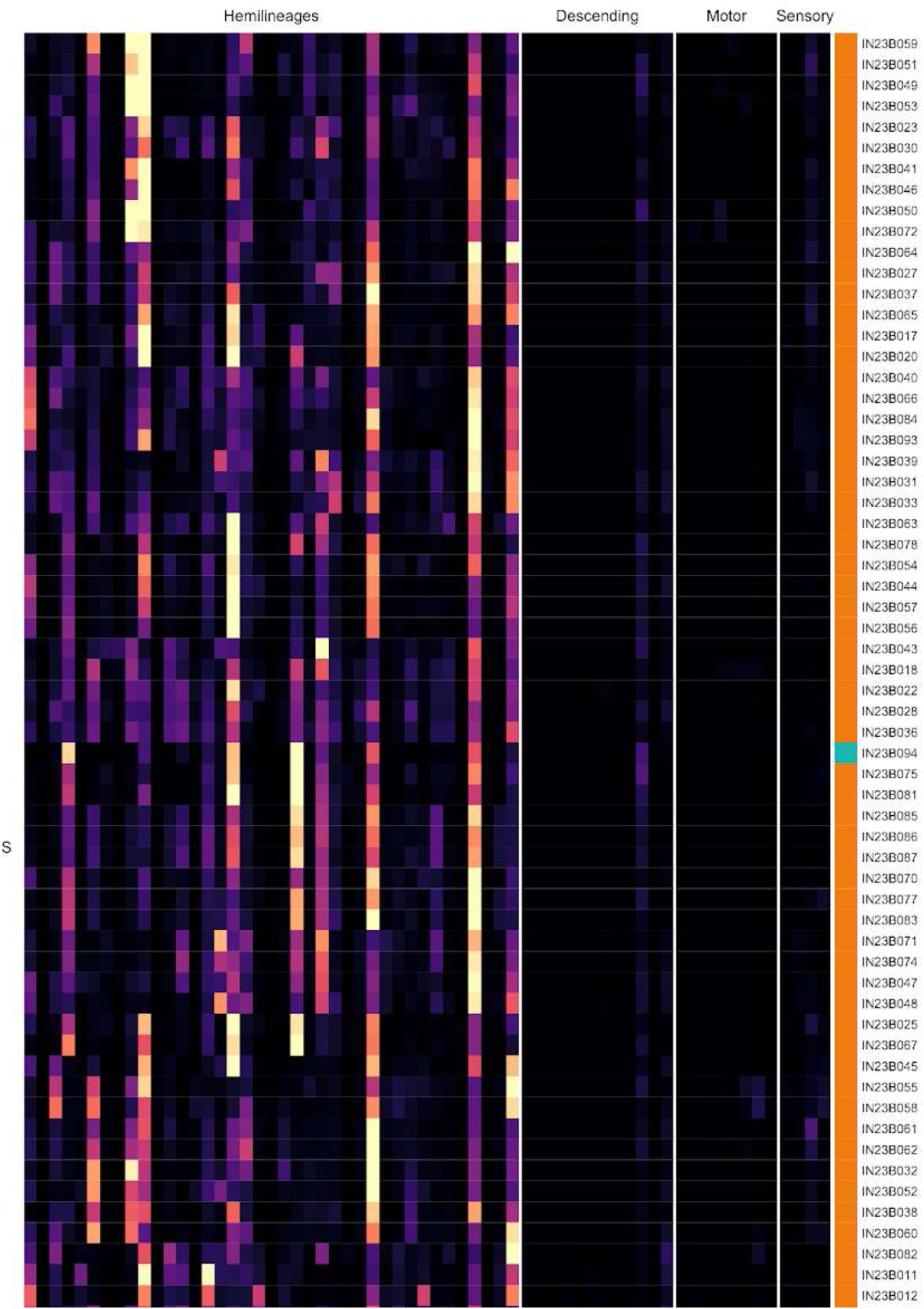
Connectivity to downstream partners by 23B secondary systematic types. Proportions of synaptic weight from systematic types to downstream partners, normalised by row. 23B neurons have been clustered within each assigned birthtime window (P = primary, ES = early secondary, S = secondary) based on both upstream and downstream connectivity to hemilineages, descending neuron subclasses, motor neuron subclasses, and sensory neuron modalities. The annotation bar is coloured by the most common predicted neurotransmitter for the neurons of each type.

**Figure 46 - figure supplement 9.**
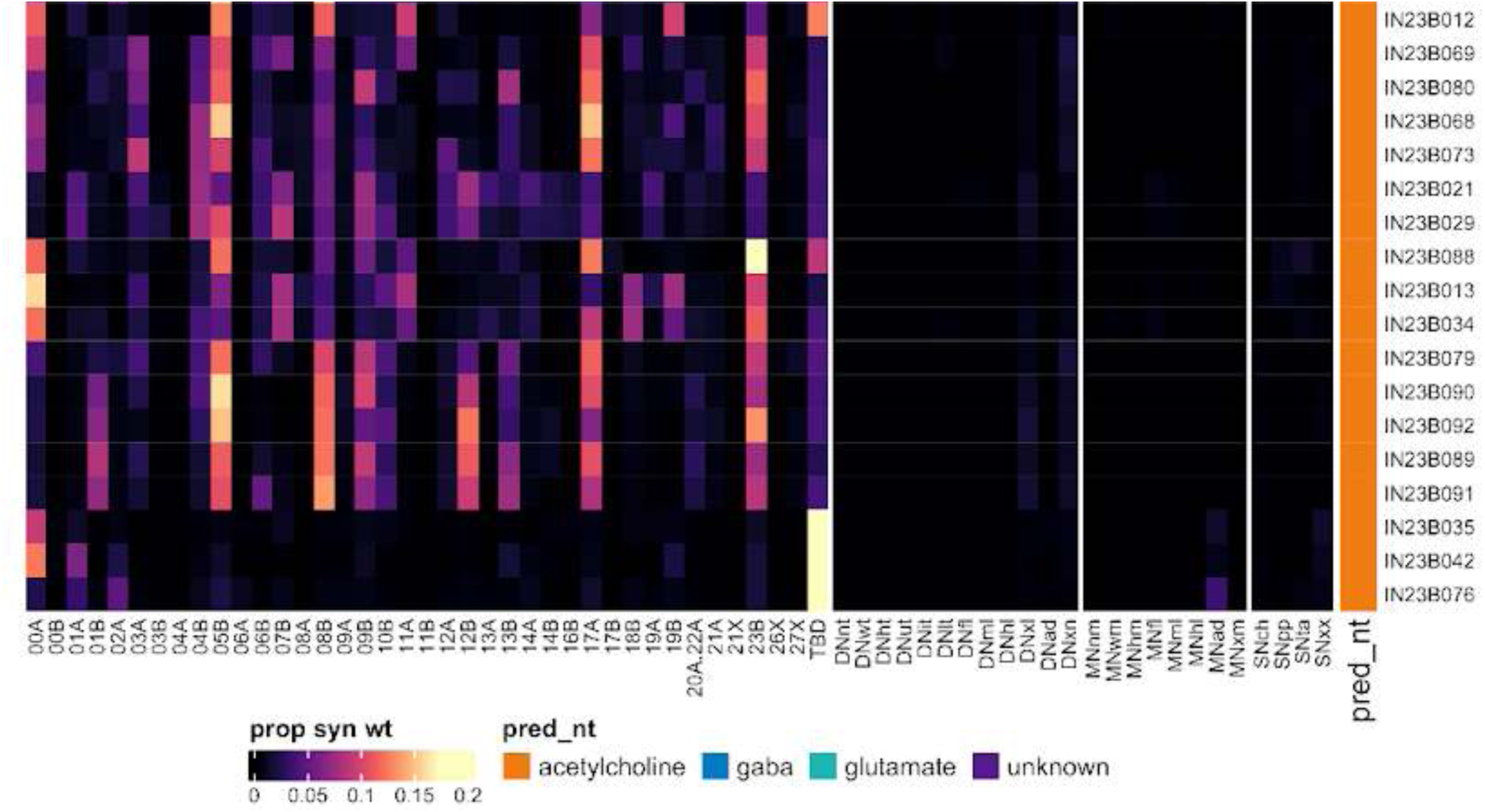
Connectivity to downstream partners by 23B secondary systematic types, continued. Proportions of synaptic weight from systematic types to downstream partners, normalised by row. 23B neurons have been clustered within each assigned birthtime window (P = primary, ES = early secondary, S = secondary) based on both upstream and downstream connectivity to hemilineages, descending neuron subclasses, motor neuron subclasses, and sensory neuron modalities. The annotation bar is coloured by the most common predicted neurotransmitter for the neurons of each type.

#### Hemilineages 24B and 25B

Hemilineage 24B is believed to derive from NB4-4 (Birkholz et al., 2015; Lacin and Truman, 2016), which generates 1 MN, two intersegmental interneurons, and 8-15 local interneurons in the embryo (Schmid et al., 1999). 24B is present in thoracic neuromeres, produces only leg motor neurons postembryonically (Brown and Truman, 2009), and has been reported to be glutamatergic (Lacin et al., 2019). The dendrites of its secondary motor neurons are confined to lateral ipsilateral leg neuropil, and their axons target muscles in the coxa, trochanter, and femur (Baek and Mann, 2009; Brierley et al., 2012). However, NB4-4 is virtually indistinguishable from adjacent NB3-4, which is believed to correspond to lineage 25 (Lacin and Truman, 2016) (but see also (Birkholz et al., 2015)). And 25 is also a small thoracic postembryonic lineage predicted to generate leg motor neurons (Lacin and Truman, 2016).

As reported in our companion manuscript (Cheong et al., 2023), we matched motor neurons in T1 by morphology to cells annotated in the FANC dataset (Lesser et al., 2023) and then identified their serial homologues in T2 and T3 using morphology and connectivity. We then annotated 24B MNs based on their inferred target muscles in the coxa, trochanter, and femur (Baek and Mann, 2009; Brierley et al., 2012). Given that 25 had not yet been identified/reported at the time that leg motor neurons were mapped to their target muscles, we strongly suspect that some of the MNs previously ascribed to 24B actually belong to 25.

In MANC, “24B” or “25” MN types exit the ProLN, ProAN, or VProN in T1 (and their serial homologues also segregate into distinct populations in the MesoLN and MetaLN) (Figure 47C bottom), consistent with reports that the 7 LinB neurons’ axons enter the prothoracic leg at three different points (Enriquez et al., 2015). As we cannot distinguish between 24B and 25, we annotated all of these neurons as “24B.25B” and assigned them to serial sets (e.g., Figure 47C top) but anticipate that further light-level work will be required to assign them correctly to two or possibly even three distinct small hemilineages.

**Figure 47.**
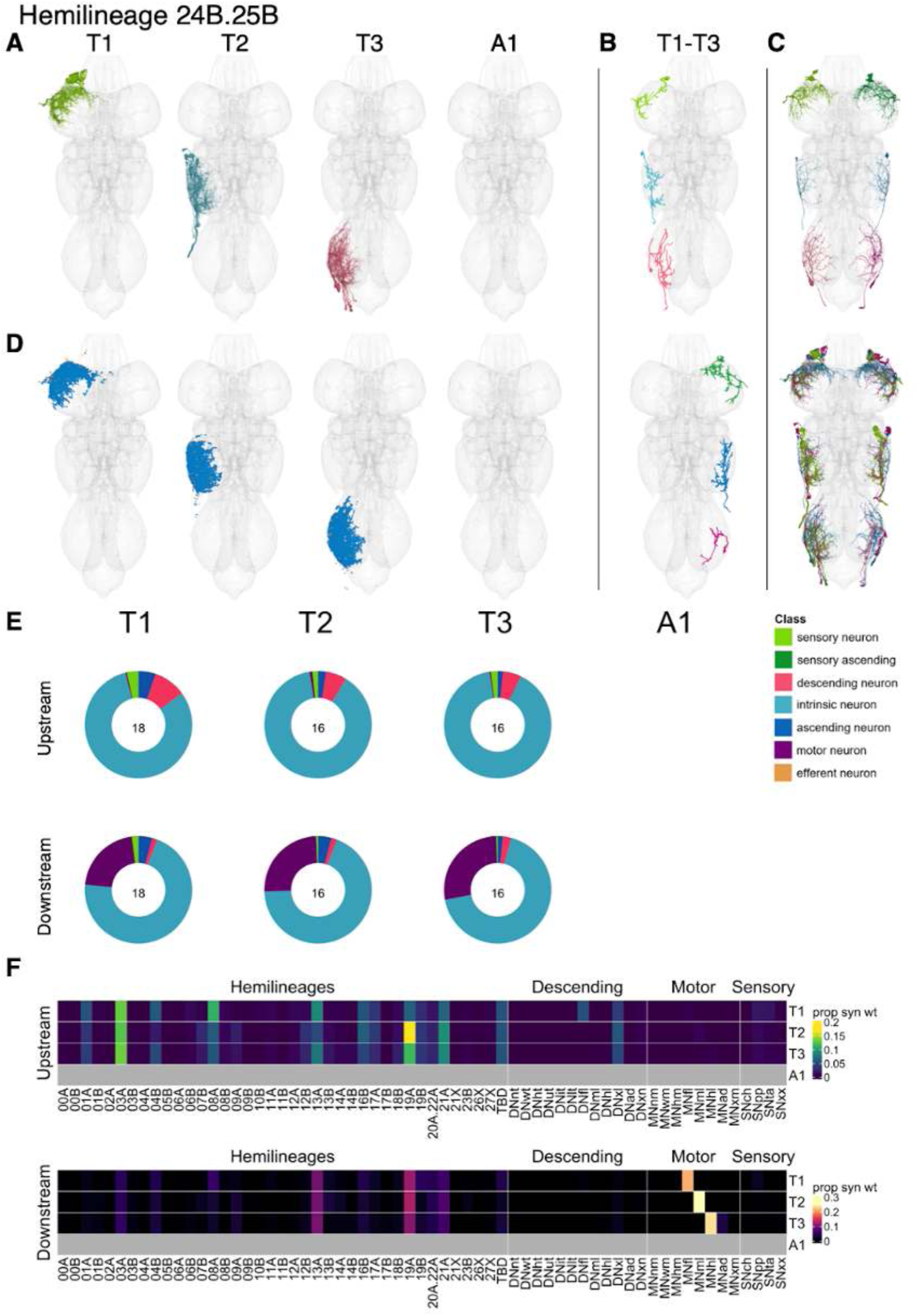
Hemilineages 24B and 25B. **A.** Meshes of all RHS secondary neurons plotted in neuromere-specific colours. **B.** “Representative” secondary neuron skeletons plotted in hemineuromere-specific colours. The skeleton with the top accumulated NBLAST score among all neurons from the hemilineage in a given hemineuromere was used. **C.** Neuron meshes of selected examples. Top: Fe reductor MN independent leg serial 11226 (T1-T3). Bottom: Serial populations exiting T1 via ProLN (blue), VProN (magenta), and ProAN (green) (T1-T3). **D.** Predicted synapses of RHS secondary neurons. Blue: postsynapses; dark orange: presynapses. **E.** Proportions of connections from secondary neurons to upstream or downstream partners, normalised by neuromere and coloured by broad class. Numbers of query neurons appear in the centre. **F.** Proportions of synaptic weight from secondary neurons originating in each neuromere to upstream or downstream partners, normalised by row.

Bilateral stimulation of 24B neurons produces repetitive leg movements (Harris et al., 2015). The 24B/25B secondary neurons receive input mainly from hemilineages 03A, 08A, 13A, 19A, and 21A as well as from descending neurons targeting leg neuropils (Figure 47F). They provide most output to 13A and 19A and to leg motor neurons as expected. Clustering by connectivity to hemilineages, descending neuron subclasses, and sensory modalities did not separate the 24B/25B population into distinct subpopulations that might represent different hemilineages (Figure 47 - figure supplement 3-4).

**Figure 47 - figure supplement 1.**
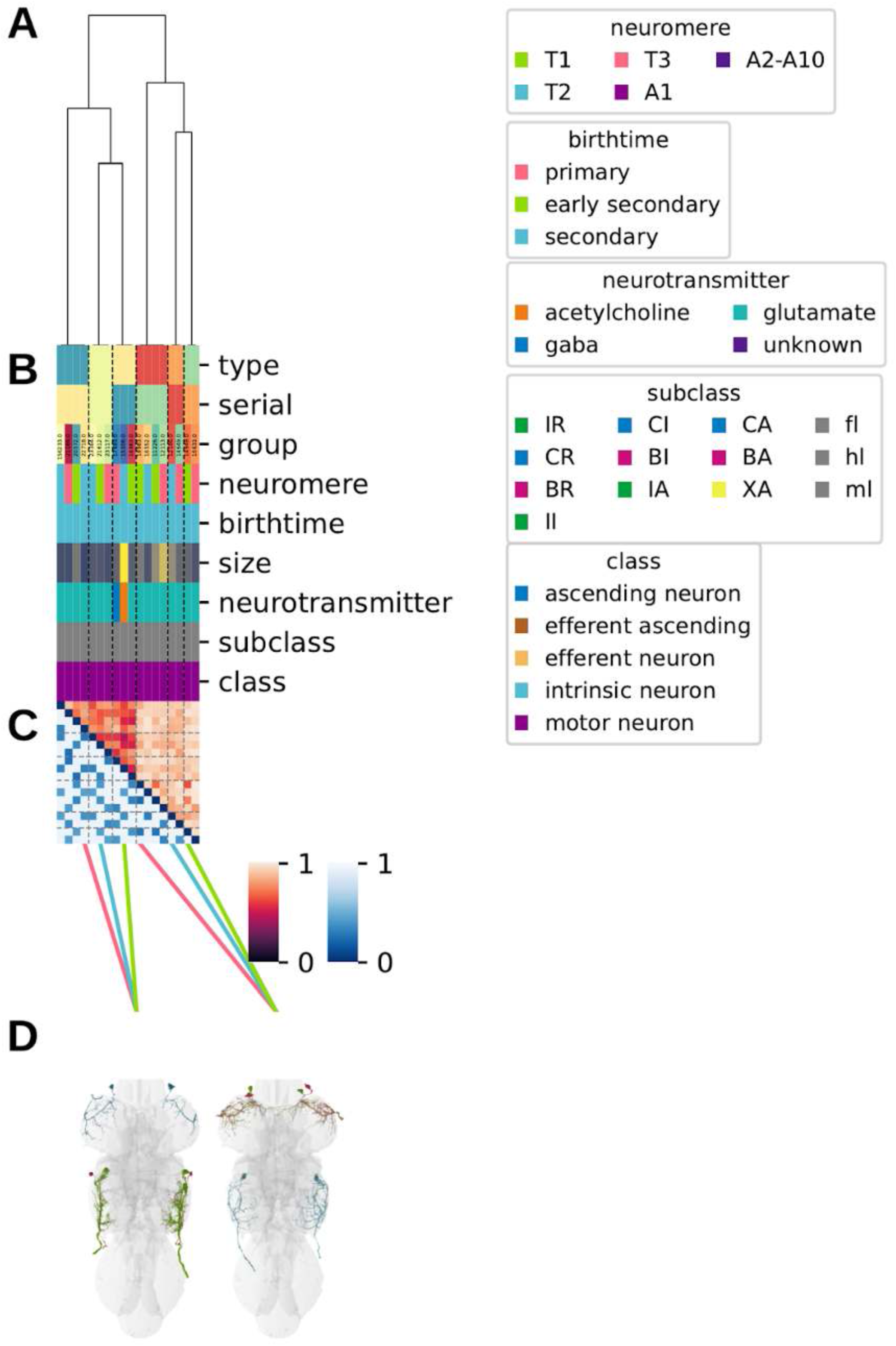
Systematic typing of hemilineages 24B and 25B. Types for motor neurons were assigned separately as outlined in our accompanying manuscript (Cheong et al., 2023)**. A.** Hierarchical clustering dendrogram of hemilineage groups by laterally and serially aggregated connectivity cosine clustering. **B.** Categorical annotations of each hemilineage group, each column corresponding to the aligned leaf in A. Colours for cluster, serial set, and group are arbitrary for visualisation. Colours for neuromere, birthtime, neurotransmitter, subclass, and class are as in all other figures. **C.** Similarity distance heatmap for hemilineage. Cosine distance is in the upper triangle, while laterally symmetrised NBLAST distance is in the lower triangle. Systematic type names of some types are labelled. **D.** Morphologically representative groups from dendrogram subtrees. Each group, indicated by colour and line connecting to its column in B and C, is the most morphologically representative group (medoid of NBLAST distance) from a subtree of A. The subtrees (flat clusters) are equal height cuts of A determined to yield the number of groups per plot and plots in D.

**Figure 47 - figure supplement 2.**
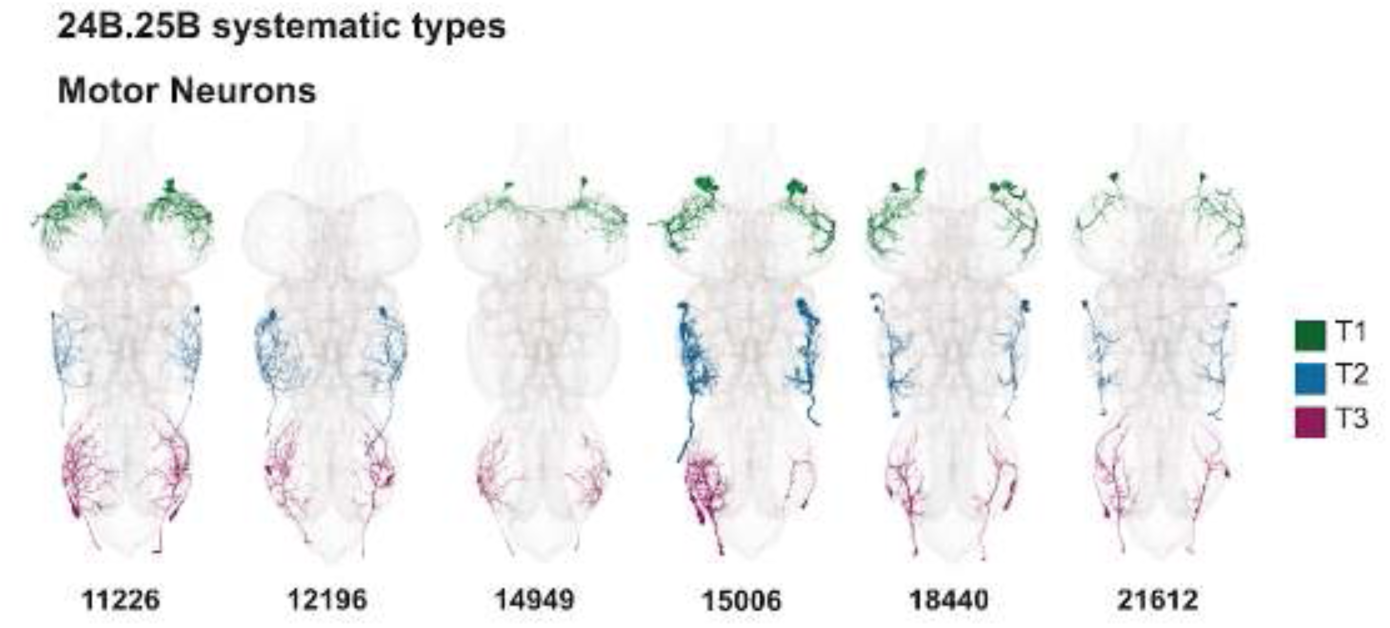
Systematic types of hemilineages 24B and 25B. Motor neurons (typed separately in (Cheong et al., 2023)) have been plotted by serial set if identified in multiple neuromeres and by systematic type if not. Individual motor neuron meshes have been coloured based on soma neuromere: dark green = T1, blue = T2, magenta = T3.

**Figure 47 - figure supplement 3.**
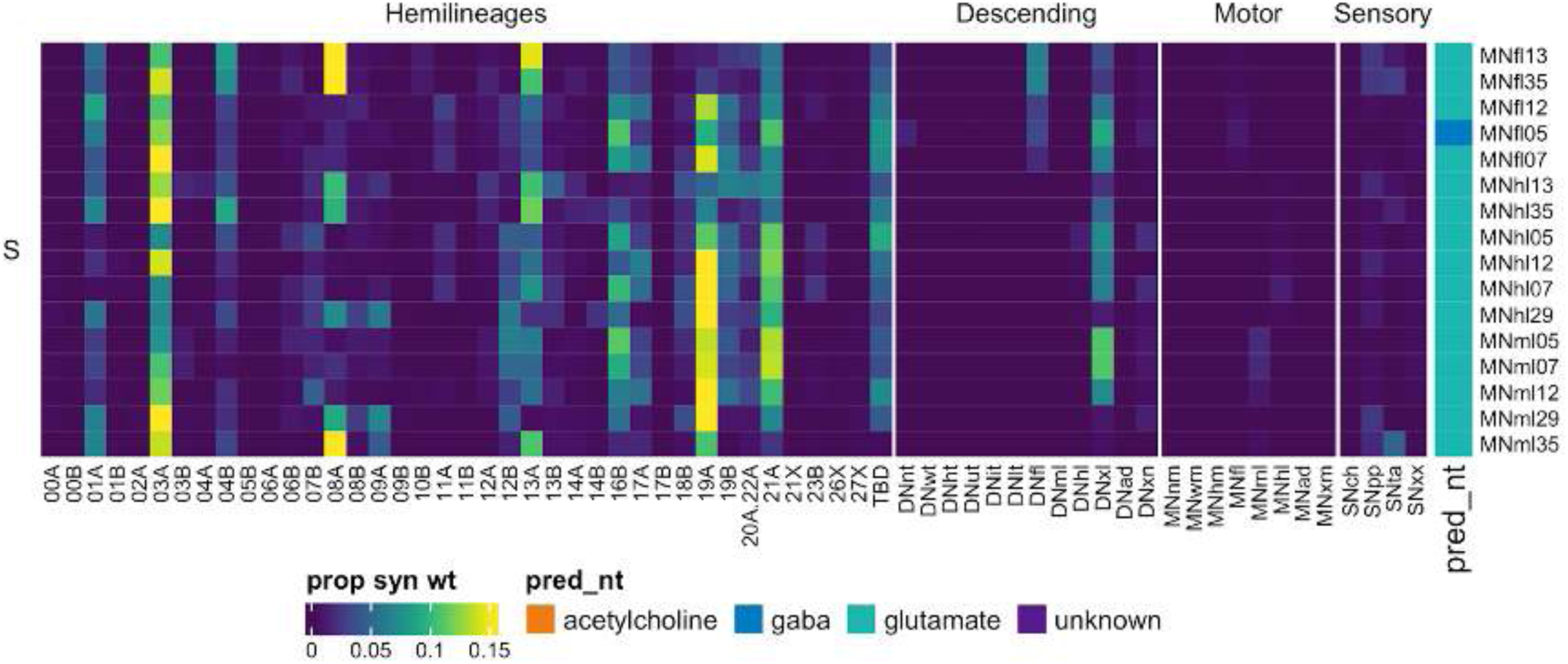
Connectivity to upstream partners by 24B.25B secondary systematic types. Proportions of synaptic weight to systematic types from upstream partners, normalised by row. 24B.25B neurons have been clustered within each assigned birthtime window (P = primary, ES = early secondary, S = secondary) based on both upstream and downstream connectivity to hemilineages, descending neuron subclasses, motor neuron subclasses, and sensory neuron modalities. Annotation bar is coloured by the most common predicted neurotransmitter for the neurons of each type.

**Figure 47 - figure supplement 4.**
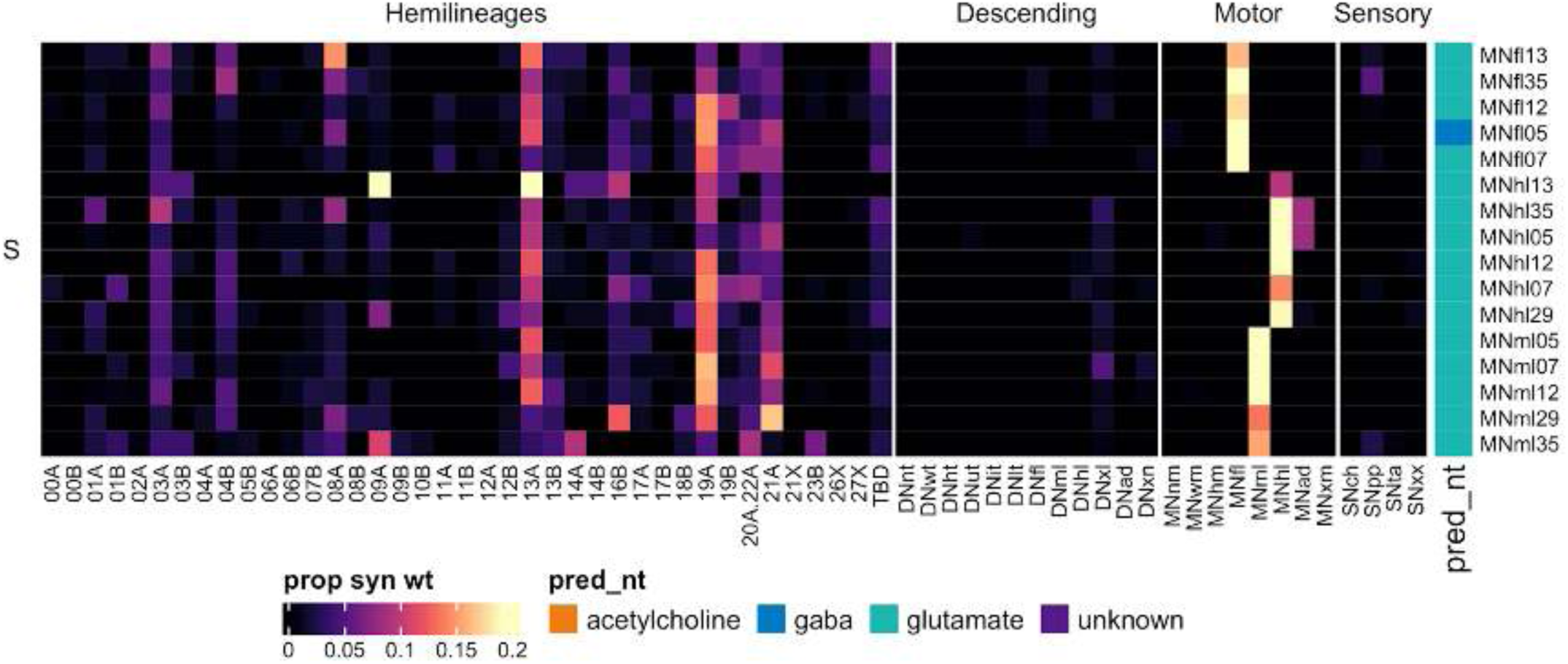
Connectivity to downstream partners by 24B.25B secondary systematic types. Proportions of synaptic weight from systematic types to downstream partners, normalised by row. 24B.25B neurons have been clustered within each assigned birthtime window (P = primary, ES = early secondary, S = secondary) based on both upstream and downstream connectivity to hemilineages, descending neuron subclasses, motor neuron subclasses, and sensory neuron modalities. The annotation bar is coloured by the most common predicted neurotransmitter within each type.

“Primary” hemilineages

Several hemilineages do not include a significant secondary population, either because the parent neuroblast is eliminated after generating neurons in the embryo or because all/most of the secondary neurons die soon after birth (Truman et al., 2010). The morphologies of 00B, 04A, 14B, and 20B/21B/22B had been previously predicted and/or reported at light level. We tentatively assigned a neuroblast of origin to 5 populations of early-born neurons that are not associated with, or appear significantly distinct from, an identifiable secondary hemilineage (Figure 48). In some cases, light-level data regarding Notch expression was unavailable, so we could not confidently assign them to “A” vs “B” and instead opted for “X”. For reference, we plotted the postsynapses and presynapses for all primary neurons in T2 neuropils by annotated hemilineage (Figure 48 - figure supplement 1).

**Figure 48.**
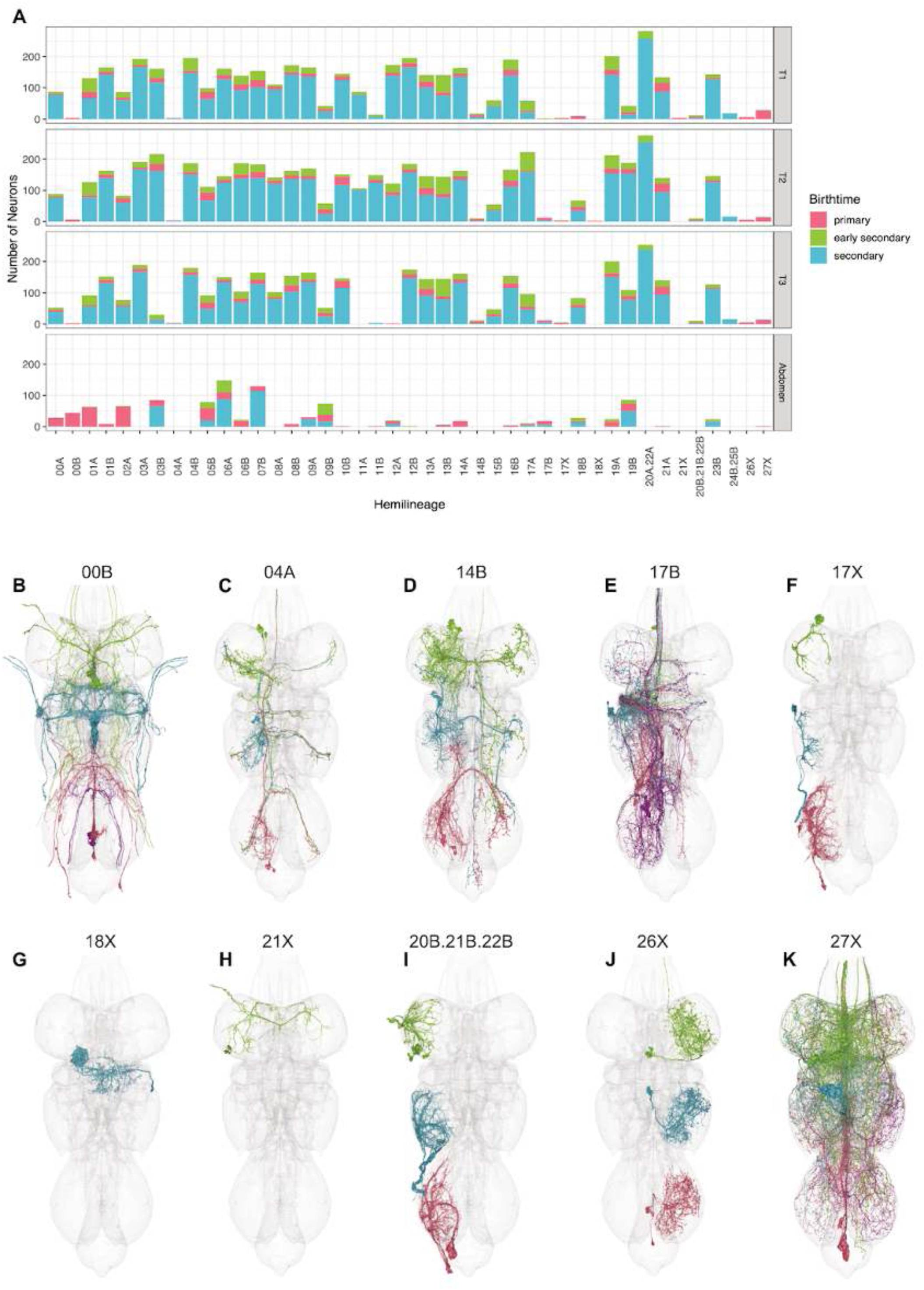
Hemilineages with few or no associated secondary neurons. **A.** Estimated birthtime by hemilineage and soma neuromere (neurons of unknown hemilineage have been omitted and all abdominal neuromeres have been combined). **B.** Octopaminergic hemilineage 00B. **C.** Cholinergic hemilineage 04A. **D.** Gabaergic/glutamatergic hemilineage 14B. **E.** Gabaergic hemilineage “17B”. **F.** Motor neurons from hemilineage “17X”. **G.** Motor neurons from hemilineage “18X”. **H.** Hemilineage “21X”. **I.** Motor neurons from hemilineages 20B, 21B, or 22B. **J.** Gabergic hemilineage “26X”. **K.** Putative neurosecretory hemilineage “27X”. Neuron meshes are coloured by soma neuromere and side (green: T1 RHS, cyan: T2 RHS, pink: T3 RHS, purple: A1 RHS).

00B neurons (also called mVUMs) are octopaminergic efferents, with small clusters of primary neurons found in each segment in adults (Monastirioti et al., 1995; Pop et al., 2020). Each neuron has bilaterally symmetric dendrites and an axon that splits to exit the VNC in homologous nerves on both sides (Figure 48A). 00B neurons receive most input from 03B and 19B in T2, from 05B and 08B in T3, and from descending neurons to multiple neuromeres (Figure 48 - figure supplement 2).

There are one or two contralaterally projecting 04A secondary neurons (Truman et al., 2010) which we identified as a pair of serially homologous, cholinergic, intersegmental neurons closely associated with 04B neurons in the adult and crossing the midline in an intermediate commissure (Figure 48B). They receive inputs from multiple hemilineages, especially 06B and 12B, and from descending neurons to multiple neuropils and chemosensory, proprioceptive, and tactile sensory neurons. They provide outputs to hemilineages 08B, 12B, and especially 20A/22A (Figure 48 - figure supplement 2).

14B secondary neurons project dorsally to cross the midline in the anterior dorsal commissure (Truman et al., 2010) and innervate both ipsilateral and contralateral leg neuropil (Shepherd et al., 2019). We identified a serially homologous set of glutamatergic secondary neurons matching these characteristics as 14B. They share a primary neurite tract with the cholinergic hemilineage 07B and with additional gabaergic and/or glutamatergic neurons that we also assigned to 14B (Figure 48D). We find that 14B neurons receive inputs from numerous hemilineages, descending neurons to the upper tectulum and legs, and proprioceptive sensory neurons whilst providing output to 20A/22A, 21A, and especially 19A and to wing and hind leg motor neurons (Figure 48 - figure supplement 2).

Anterior dorsal neuroblasts NB2-5/17 and NB2-4/18 each generate one motor neuron in the embryo (Schmid et al., 1999). We identified two serially homologous sets of motor neurons with anterior soma tracts that we initially annotated as 17X.18X but have since changed to 17X (Figure 48F). These sets exit through different nerves in T3 and so might belong to two different hemilineages, but we do not have light level data to determine which one comes from 17 or to know whether they express activated Notch during development. The two “17X” leg motor neuron types in T1-T3 receive inputs from 04B, 13A, and especially 20A/22A and from descending neurons to leg neuropils (Figure 48 - figure supplement 3).

We identified a single large, anterodorsal, bilaterally projecting motor neuron in T2 as the primary flight motor neuron MN5 produced by lineage 18 (Lacin et al., 2020). We annotated its hemilineage as “18X” since we do not know whether it expresses activated Notch (Figure 48H). This motor neuron receives strong input from hemilineage 19B and to a lesser extent from hemilineages 03B and 06B (Figure 48 - figure supplement 4).

Lineage 20 was reported to include two secondary motor neurons while lineage 22 included a single secondary motor neuron, all of which are thought to originate with the “B” hemilineage (Truman et al., 2010). These MNs have not been distinguished at light level from those of 21B or each other in the adult (Brierley et al., 2012). In MANC, all of these MNs enter the neuropil together and exit via the ProAN in T1, the MesoLN in T2, and the MetaLN in T3, so differential origins cannot be inferred. We therefore opted to designate all of these posterior MNs as “20B.21B.22B.” The leg motor neurons from hemilineages 20B, 21B, and/or 22B in T1-T3 (Figure 48I) receive most input from 03A, 08A, and 19A (Figure 48 - figure supplement 3).

Hemilineages 21A and 21B derive from neuroblast NB4-3 (Birkholz et al., 2015; Lacin and Truman, 2016), which generates 2-3 neurosecretory cells and 10-12 local interneurons in the embryo (Schmid et al., 1999). We have annotated a pair of bilateral efferent neurons associated with 21A primary neurites in T1 as “21X”, believing them to be the NB4-3 NSCs (Figure 48G). They mainly receive inputs from 02A and 07B and from descending neurons to front leg and multiple leg neuropils (Figure 48 - figure supplement 4).

In addition, we associated several cell populations with specific neuroblasts based on cell types described in the late embryo (Schmid et al., 1999). First, we identified a set of neurons that entered the neuropil with the 09/17/18 anterodorsal triad but were predicted to be gabaergic and had a characteristic morphology, with the primary neurite turning dorsally upon entry then projecting straight ventrally before ascending (Figure 48F). These neurons are quite distinct from 09A (the only gabaergic secondary hemilineage in the vicinity) in both morphology and connectivity and so could only belong to 17A, 17B, or 18A. We have tentatively assigned them as 17B because NB2-5/17 (but not NB2-4/18) has been reported to generate intersegmental neurons in the embryo (however they might be a gabaergic subpopulation of 17A). They receive most input from proprioceptive and tactile sensory neurons and mainly inhibit 09B and 11A in T1, 06B and 17A in T2, 10B in T3, 23B in A1, and proprioceptive and tactile sensory neurons (Figure 48 - figure supplement 3).

We identified a small population of gabaergic primary neurons that seem likely to derive from the same neuroblast but do not align with any previously reported secondary hemilineage. The neurons appear to originate at a posterior and medial position but project in a ventral commissure to target the contralateral leg neuropil, often resulting in T1 somas being dragged across the midline. Based on this soma position and neurotransmitter prediction, we suggest that these neurons might originate from embryonic NB5-1 and have designated them as hemilineage “26X” (Figure 48J). They receive input in the contralateral leg neuropil from 07B, 09A, 14A, 20A/22A, and 21A, descending neurons, and proprioceptive and tactile sensory neurons. They primarily inhibit 03A, 08A, 09A, 13A, 17A, 19A, and 19B (Figure 48 - figure supplement 4).

Finally, we identified a distinctive population of primary neurons in T1-A1 that appear to originate from the same neuroblast but do not align with any previously reported secondary hemilineage (Figure 48K). The neurons enter the neuropil near 5B and 6B but project very dorsally and nearly converge with their contralateral homologues before crossing the midline. As this primary population is enriched in putative neurosecretory cells, we suggest that it might originate from embryonic NB5-5, which is located at approximately the same anteroposterior position as NB5-2 (6B) and NB5-3 (5B), although NB5-6 is another possibility (H Lacin, personal communication). These “27X” neurons mainly receive input from descending neurons to upper tectulum or multiple neuropils and provide output to most hemilineages, descending neurons to multiple legs, and wing motor neurons (Figure 48 - figure supplement 4).

**Figure 48 - figure supplement 1.**
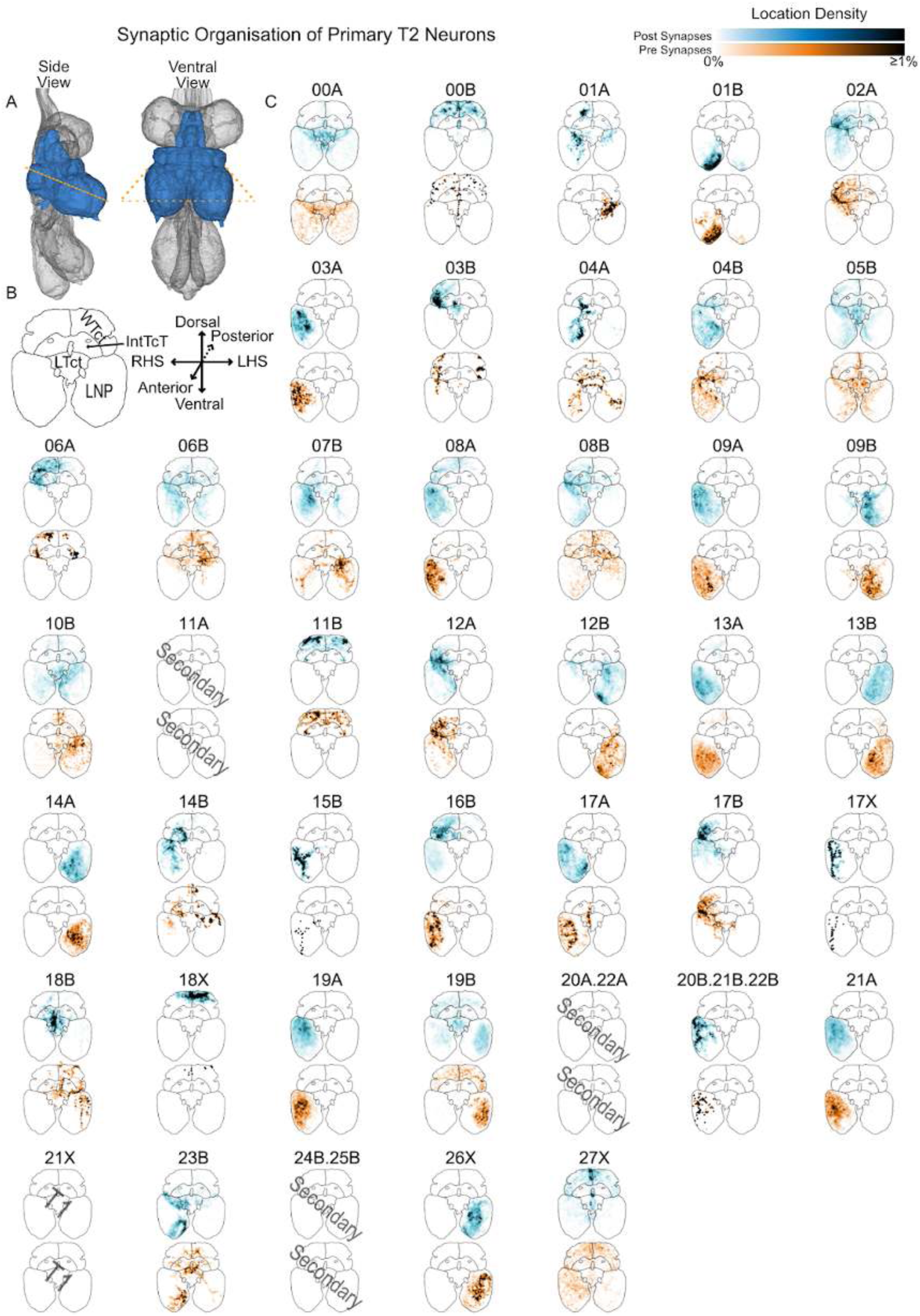
Summary of the anatomical organisation of synapses for primary neurons in each hemilineage originating in T2 RHS. **A.** Side and ventral views illustrating the neuropils surveyed: T2 Leg Neuropil (LNP), medial ventral association center (mVAC), ovoid (Ov), wing tectulum (WTct), lower tectulum (LTct), and Intermediate tectulum (IntTct). The orange dashed line shows the approximate location of the transverse section onto which their synapses have been projected. **B.** Template transverse section with major neuropils labelled and axes for orientation. **C.** Projected synapses for T2 RHS neurons of each hemilineage. Top (blue): postsynapses, connections to upstream neurons. Bottom (orange): presynapses, connections to downstream neurons. 21X is only observed in T1.

**Figure 48 - figure supplement 2.**
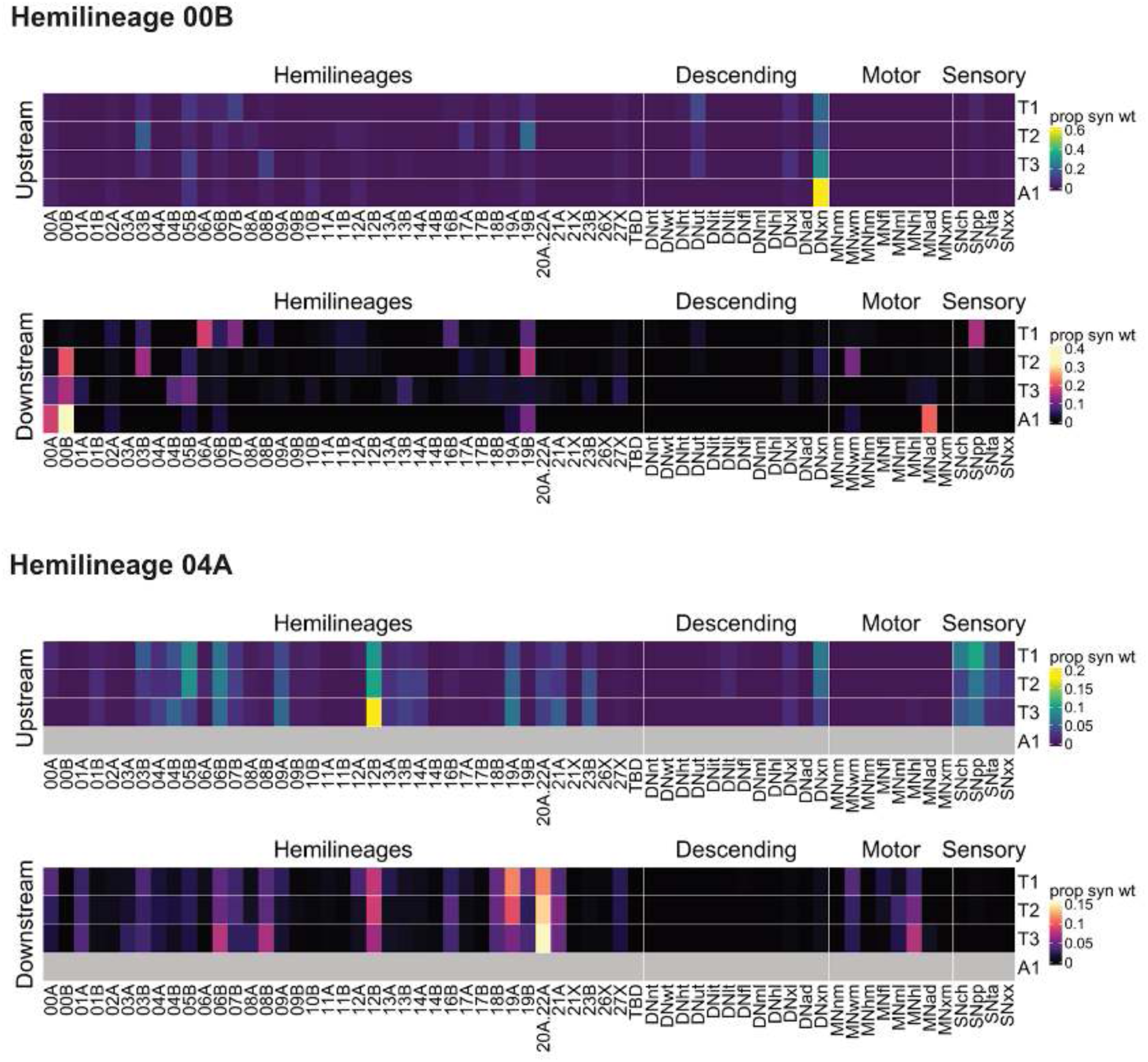
Connectivity heat maps for primary hemilineages 00B and 04A. Ratios of connections from neurons originating in each neuromere to upstream or downstream partners, normalised by row.

**Figure 48 - figure supplement 3.**
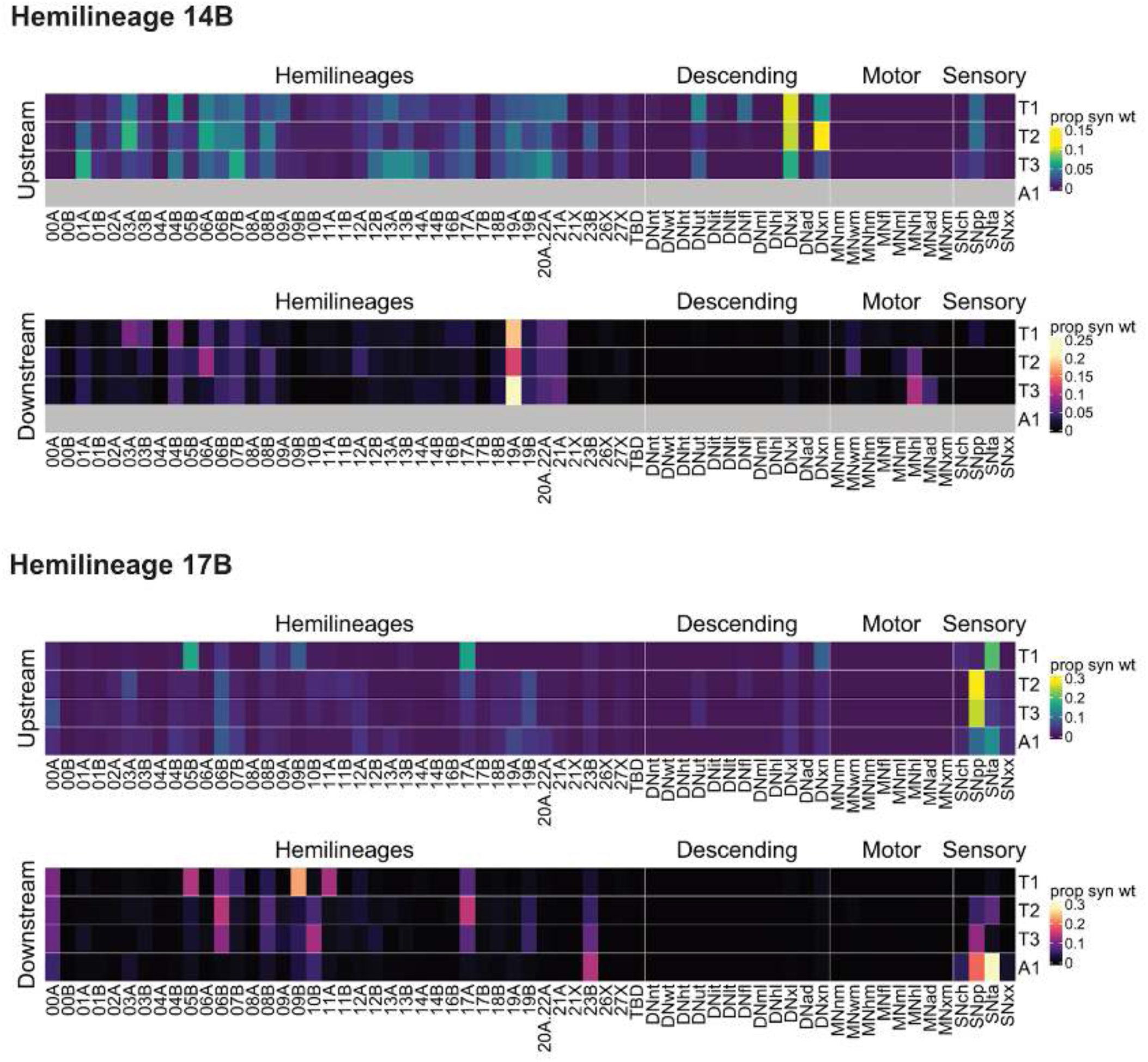
Connectivity heat maps for primary hemilineages 14B and 17B. Ratios of connections from neurons originating in each neuromere to upstream or downstream partners, normalised by row.

**Figure 48 - figure supplement 4.**
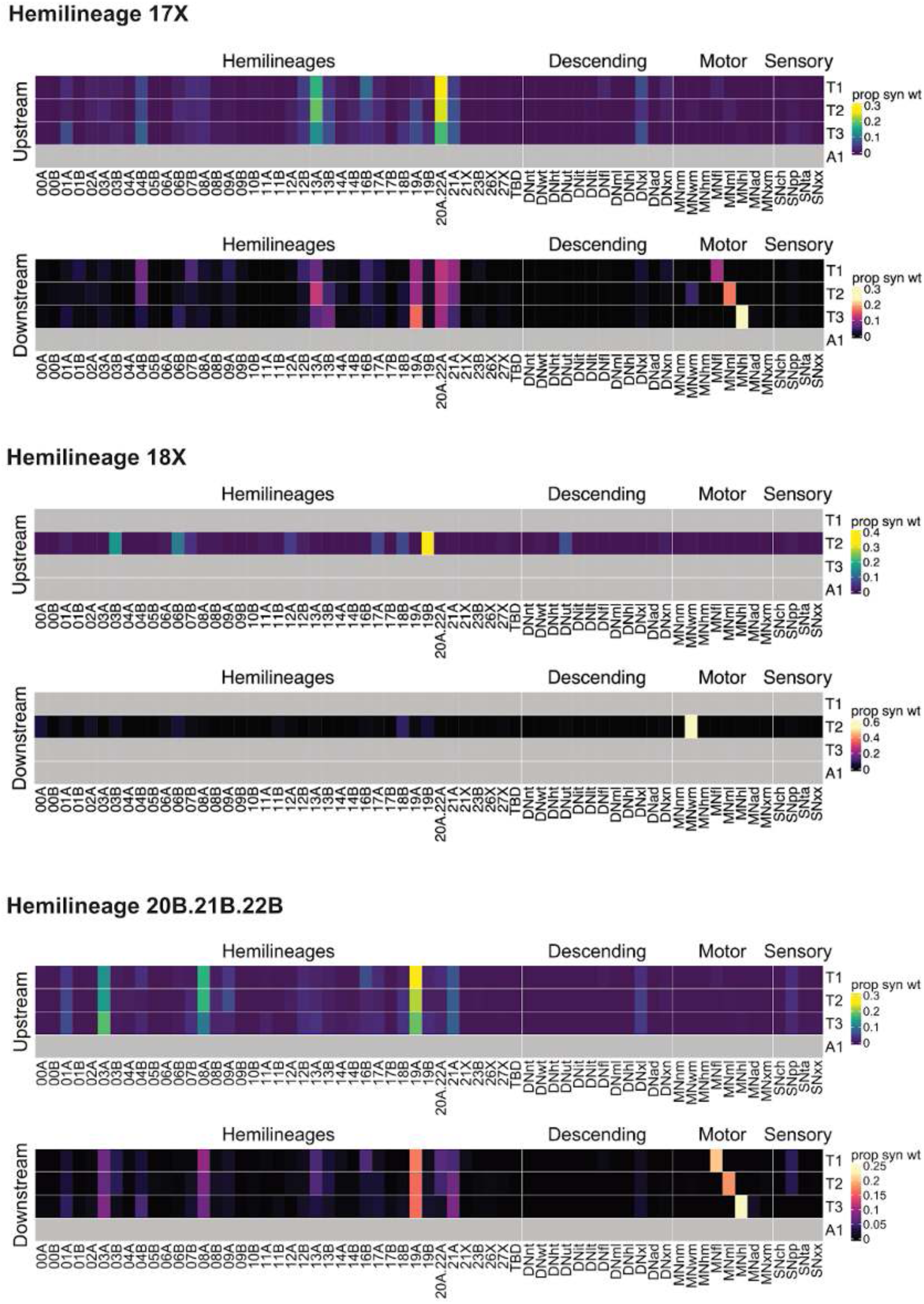
Connectivity heat maps for primary hemilineages 17X.18X and 20B.21B.22B. Ratios of connections from neurons originating in each neuromere to upstream or downstream partners, normalised by row.

**Figure 48 - figure supplement 5.**
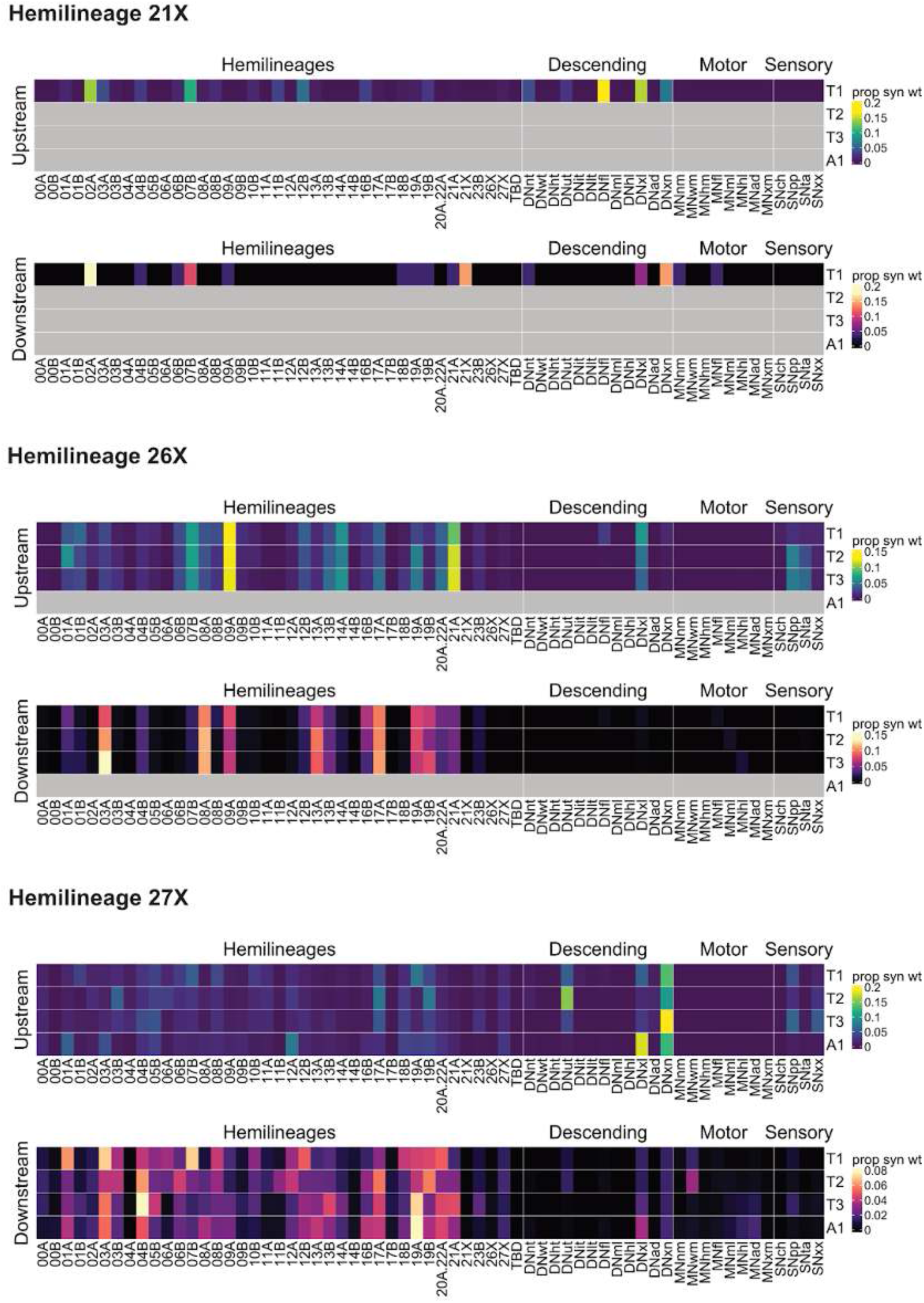
Connectivity heat maps for primary hemilineages 21X, 26X and 27X. Ratios of connections from neurons originating in each neuromere to upstream or downstream partners, normalised by row.

**Figure 48 - figure supplement 6.**
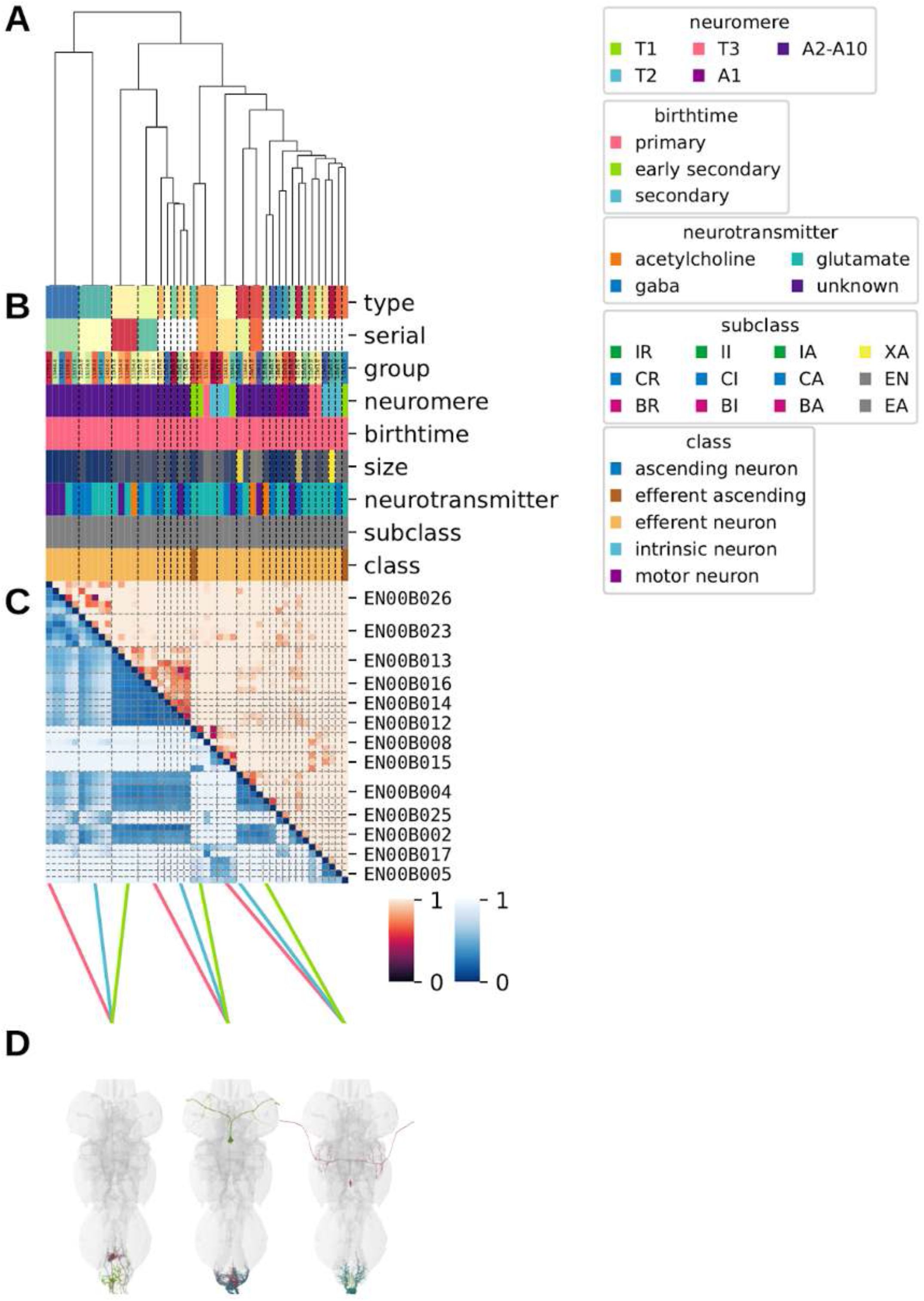
Systematic typing of hemilineage 00B. **A.** Hierarchical clustering dendrogram of hemilineage groups by laterally and serially aggregated connectivity cosine clustering. **B.** Categorical annotations of each hemilineage group, each column corresponding to the aligned leaf in A. Colours for type, serial set, and group are arbitrary for visualisation. Colours for neuromere, birthtime, neurotransmitter, subclass, and class are as in all other figures. **C.** Similarity distance heatmap for hemilineage. Cosine distance is in the upper triangle, while laterally symmetrised NBLAST distance is in the lower triangle. Systematic type names of some types are labelled. **D.** Morphologically representative groups from dendrogram subtrees. Each group, indicated by colour and line connecting to its column in B and C, is the most morphologically representative group (medoid of NBLAST distance) from a subtree of A. The subtrees (flat clusters) are equal height cuts of A determined to yield the number of groups per plot and plots in D.

**Figure 48 - figure supplement 7.**
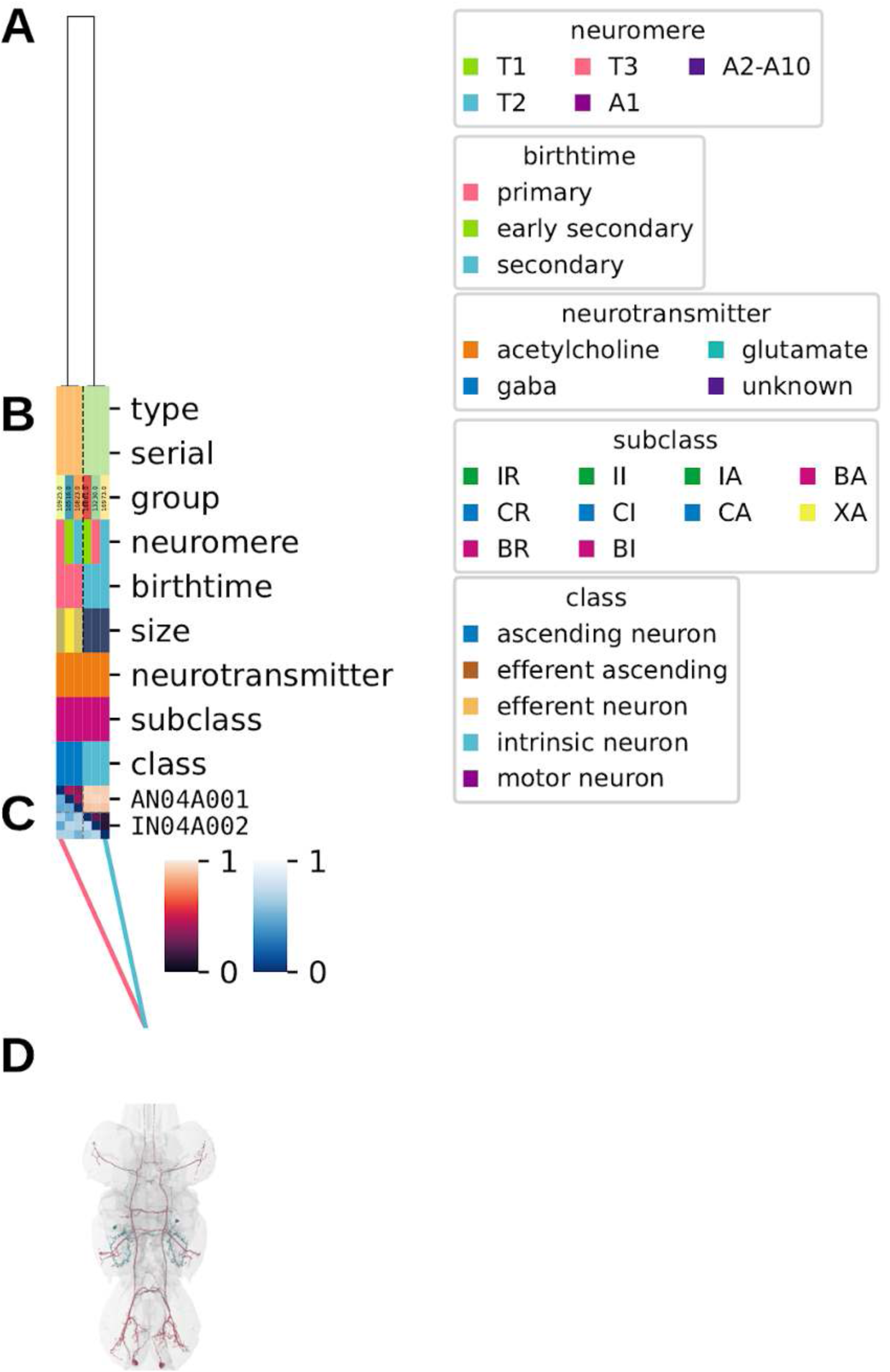
Systematic typing of hemilineage 04A. **A.** Hierarchical clustering dendrogram of hemilineage groups by laterally and serially aggregated connectivity cosine clustering. **B.** Categorical annotations of each hemilineage group, each column corresponding to the aligned leaf in A. Colours for type, serial set, and group are arbitrary for visualisation. Colours for neuromere, birthtime, neurotransmitter, subclass, and class are as in all other figures. **C.** Similarity distance heatmap for hemilineage. Cosine distance is in the upper triangle, while laterally symmetrised NBLAST distance is in the lower triangle. Systematic type names of some types are labelled. **D.** Morphologically representative groups from dendrogram subtrees. Each group, indicated by colour and line connecting to its column in B and C, is the most morphologically representative group (medoid of NBLAST distance) from a subtree of A. The subtrees (flat clusters) are equal height cuts of A determined to yield the number of groups per plot and plots in D.

**Figure 48 - figure supplement 8.**
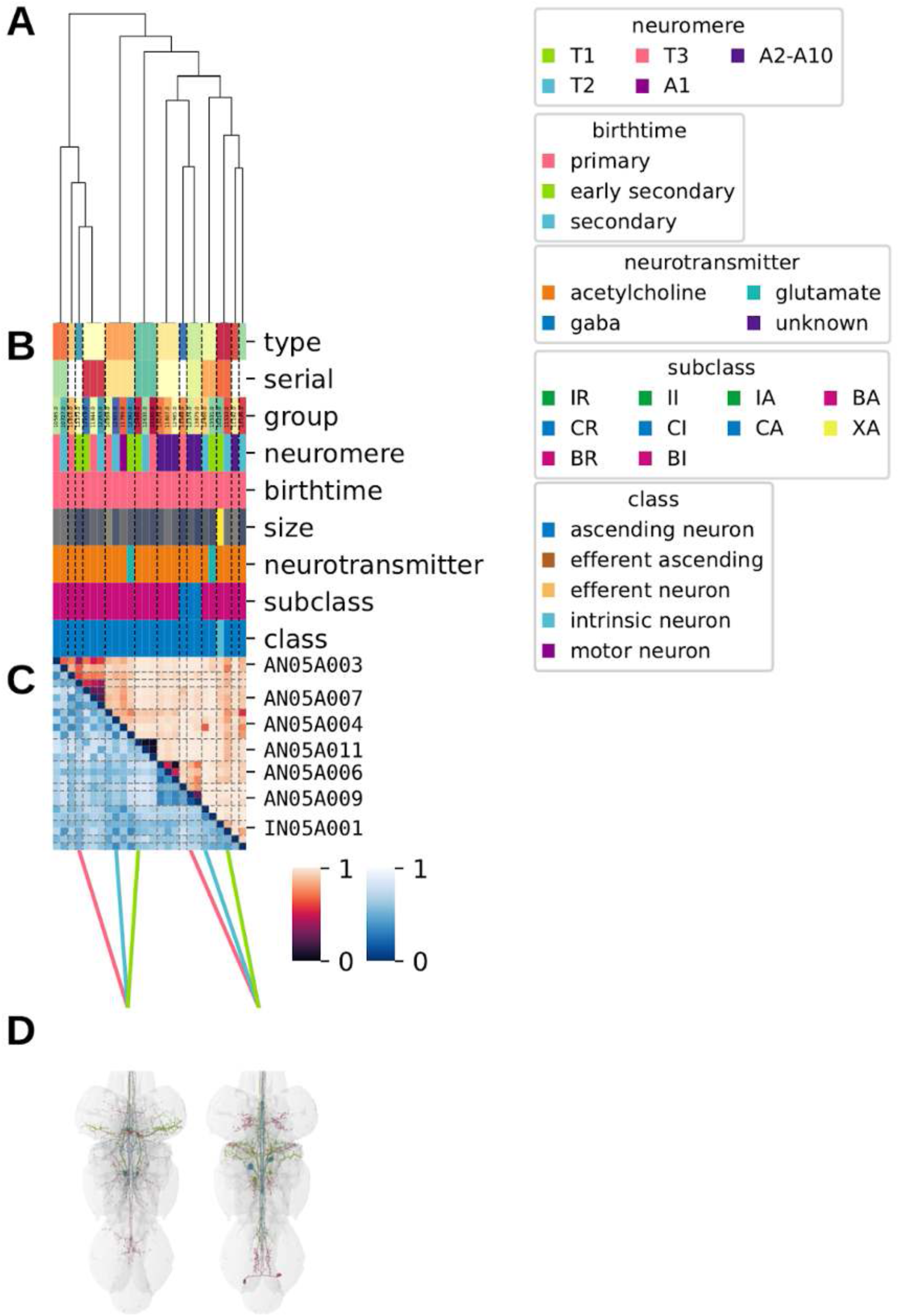
Systematic typing of hemilineage 05A. **A.** Hierarchical clustering dendrogram of hemilineage groups by laterally and serially aggregated connectivity cosine clustering. **B.** Categorical annotations of each hemilineage group, each column corresponding to the aligned leaf in A. Colours for type, serial set, and group are arbitrary for visualisation. Colours for neuromere, birthtime, neurotransmitter, subclass, and class are as in all other figures. **C.** Similarity distance heatmap for hemilineage. Cosine distance is in the upper triangle, while laterally symmetrised NBLAST distance is in the lower triangle. Systematic type names of some types are labelled. **D.** Morphologically representative groups from dendrogram subtrees. Each group, indicated by colour and line connecting to its column in B and C, is the most morphologically representative group (medoid of NBLAST distance) from a subtree of A. The subtrees (flat clusters) are equal height cuts of A determined to yield the number of groups per plot and plots in D.

**Figure 48 - figure supplement 9.**
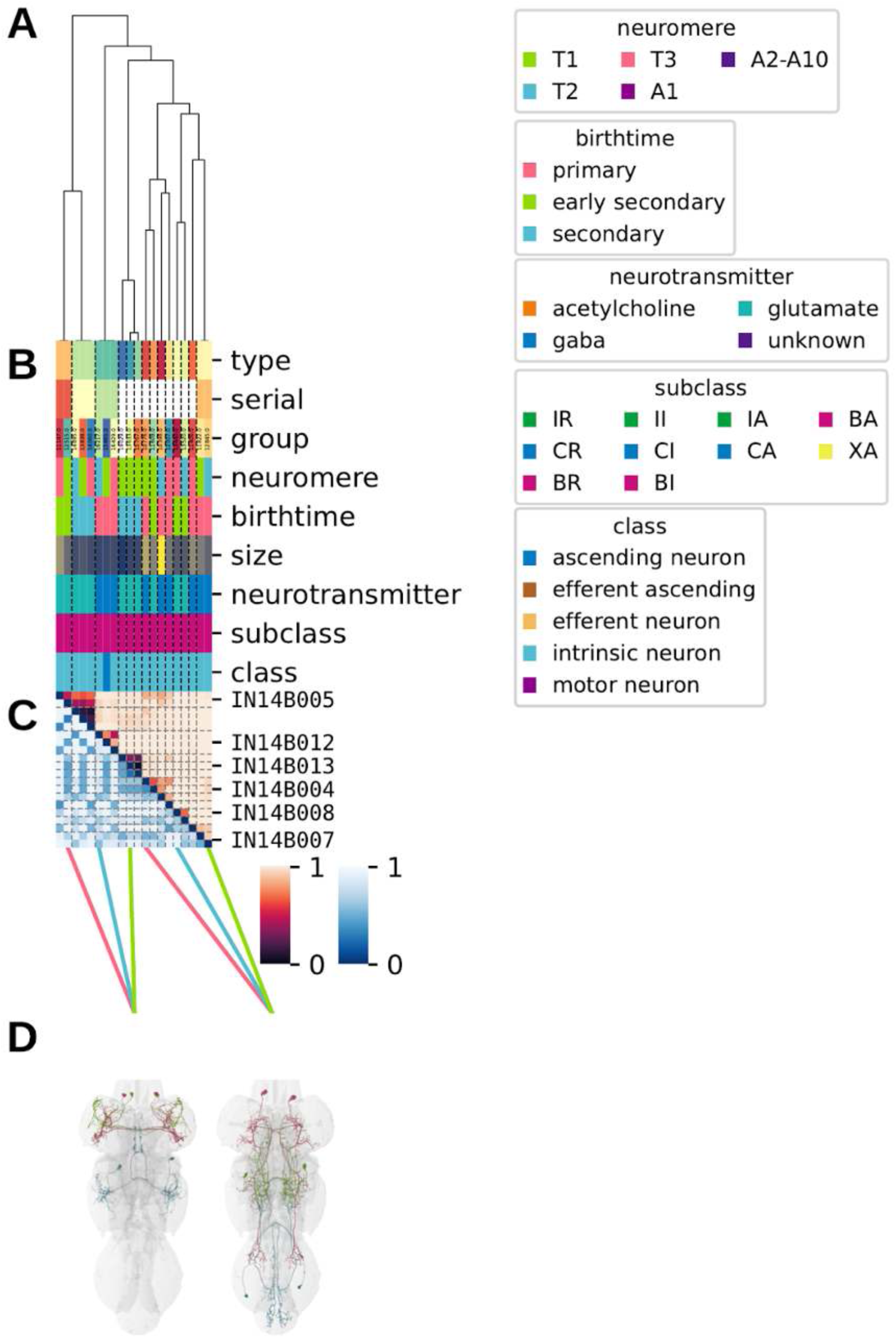
Systematic typing of hemilineage 14B. **A.** Hierarchical clustering dendrogram of hemilineage groups by laterally and serially aggregated connectivity cosine clustering. **B.** Categorical annotations of each hemilineage group, each column corresponding to the aligned leaf in A. Colours for type, serial set, and group are arbitrary for visualisation. Colours for neuromere, birthtime, neurotransmitter, subclass, and class are as in all other figures. **C.** Similarity distance heatmap for hemilineage. Cosine distance is in the upper triangle, while laterally symmetrised NBLAST distance is in the lower triangle. Systematic type names of some types are labelled. **D.** Morphologically representative groups from dendrogram subtrees. Each group, indicated by colour and line connecting to its column in B and C, is the most morphologically representative group (medoid of NBLAST distance) from a subtree of A. The subtrees (flat clusters) are equal height cuts of A determined to yield the number of groups per plot and plots in D.

**Figure 48 - figure supplement 10.**
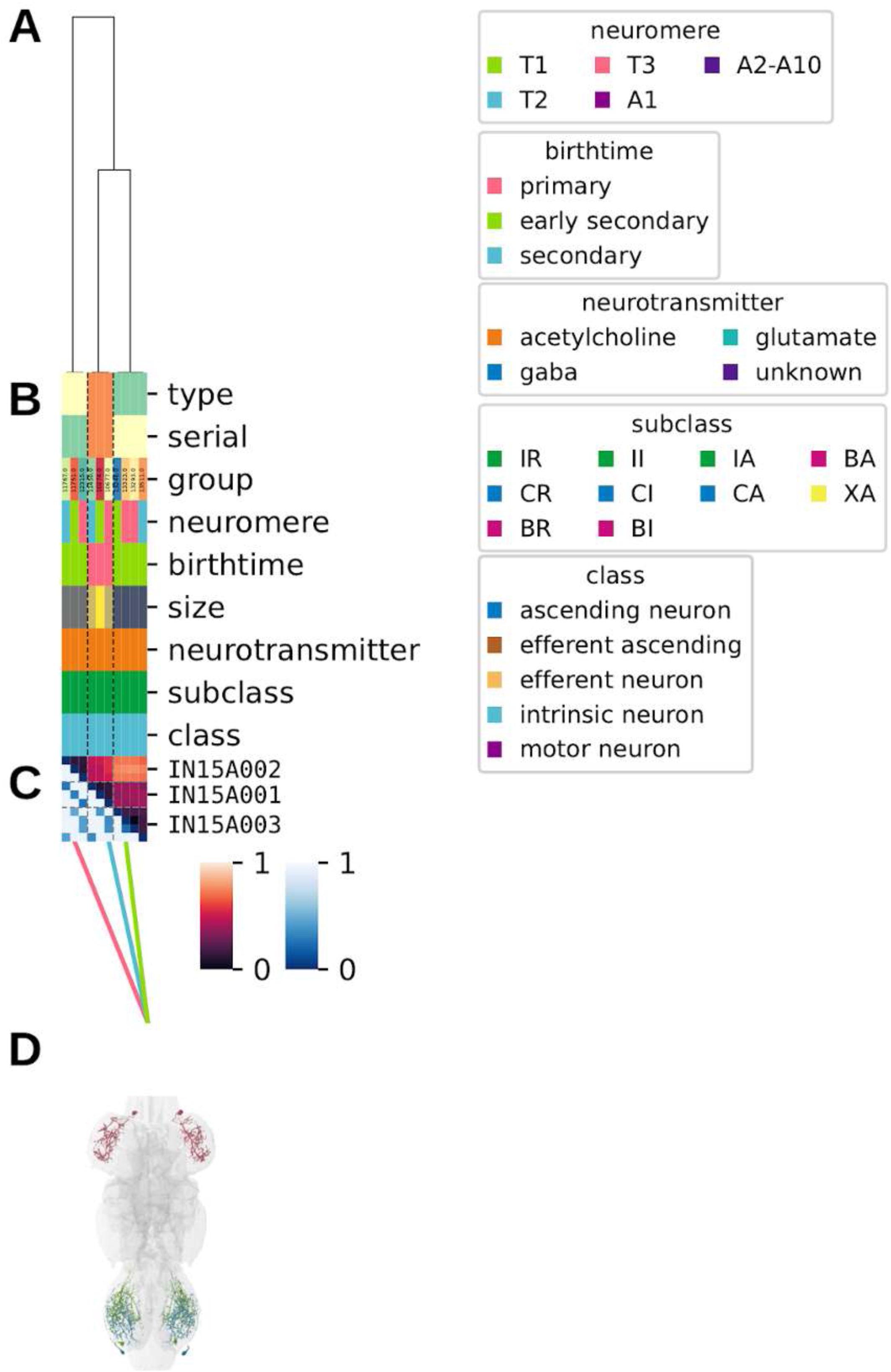
Systematic typing of hemilineage 15A. **A.** Hierarchical clustering dendrogram of hemilineage groups by laterally and serially aggregated connectivity cosine clustering. **B.** Categorical annotations of each hemilineage group, each column corresponding to the aligned leaf in A. Colours for type, serial set, and group are arbitrary for visualisation. Colours for neuromere, birthtime, neurotransmitter, subclass, and class are as in all other figures. **C.** Similarity distance heatmap for hemilineage. Cosine distance is in the upper triangle, while laterally symmetrised NBLAST distance is in the lower triangle. Systematic type names of some types are labelled. **D.** Morphologically representative groups from dendrogram subtrees. Each group, indicated by colour and line connecting to its column in B and C, is the most morphologically representative group (medoid of NBLAST distance) from a subtree of A. The subtrees (flat clusters) are equal height cuts of A determined to yield the number of groups per plot and plots in D.

**Figure 48 - figure supplement 11.**
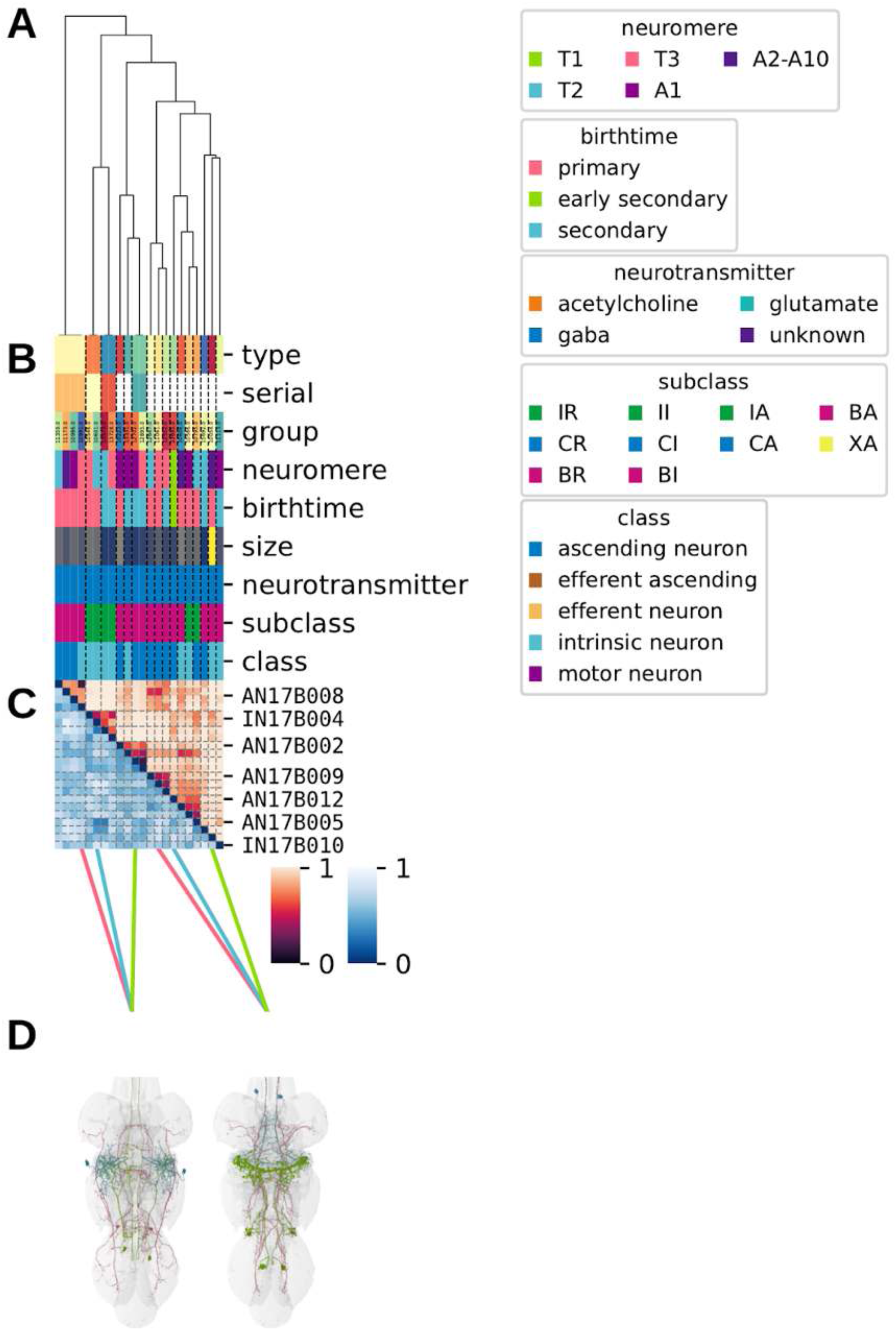
Systematic typing of hemilineage 17B. **A.** Hierarchical clustering dendrogram of hemilineage groups by laterally and serially aggregated connectivity cosine clustering. **B.** Categorical annotations of each hemilineage group, each column corresponding to the aligned leaf in A. Colours for type, serial set, and group are arbitrary for visualisation. Colours for neuromere, birthtime, neurotransmitter, subclass, and class are as in all other figures. **C.** Similarity distance heatmap for hemilineage. Cosine distance is in the upper triangle, while laterally symmetrised NBLAST distance is in the lower triangle. Systematic type names of some types are labelled. **D.** Morphologically representative groups from dendrogram subtrees. Each group, indicated by colour and line connecting to its column in B and C, is the most morphologically representative group (medoid of NBLAST distance) from a subtree of A. The subtrees (flat clusters) are equal height cuts of A determined to yield the number of groups per plot and plots in D.

**Figure 48 - figure supplement 12.**
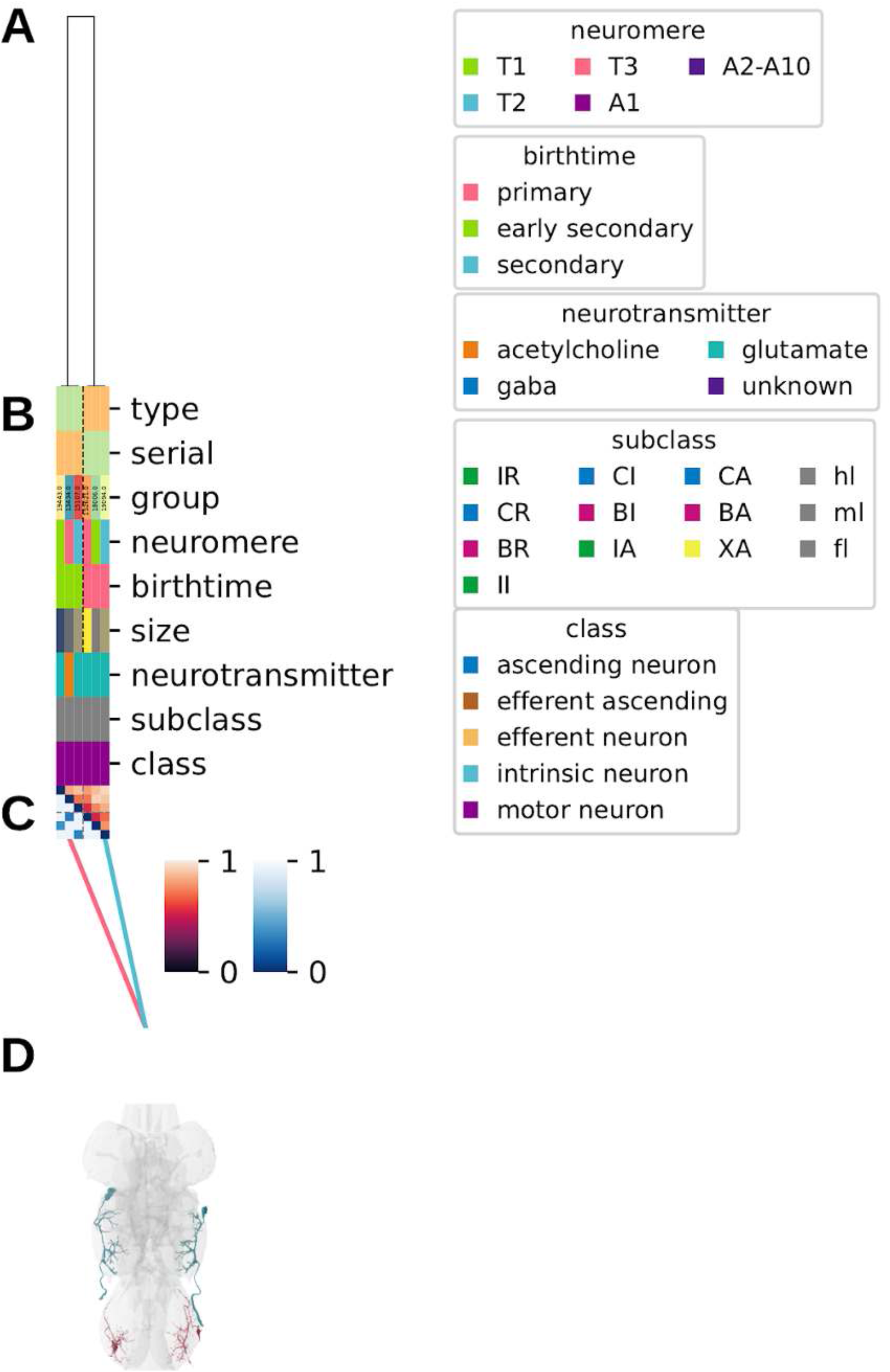
Systematic typing of hemilineage 17X and 18X. Types for motor neurons were assigned separately as outlined in our accompanying manuscript (Cheong et al., 2023). **A.** Hierarchical clustering dendrogram of hemilineage groups by laterally and serially aggregated connectivity cosine clustering. **B.** Categorical annotations of each hemilineage group, each column corresponding to the aligned leaf in A. Colours for cluster, serial set, and group are arbitrary for visualisation. Colours for neuromere, birthtime, neurotransmitter, subclass, and class are as in all other figures. **C.** Similarity distance heatmap for hemilineage. Cosine distance is in the upper triangle, while laterally symmetrised NBLAST distance is in the lower triangle. Systematic type names of some types are labelled. **D.** Morphologically representative groups from dendrogram subtrees. Each group, indicated by colour and line connecting to its column in B and C, is the most morphologically representative group (medoid of NBLAST distance) from a subtree of A. The subtrees (flat clusters) are equal height cuts of A determined to yield the number of groups per plot and plots in D.

**Figure 48 - figure supplement 13.**
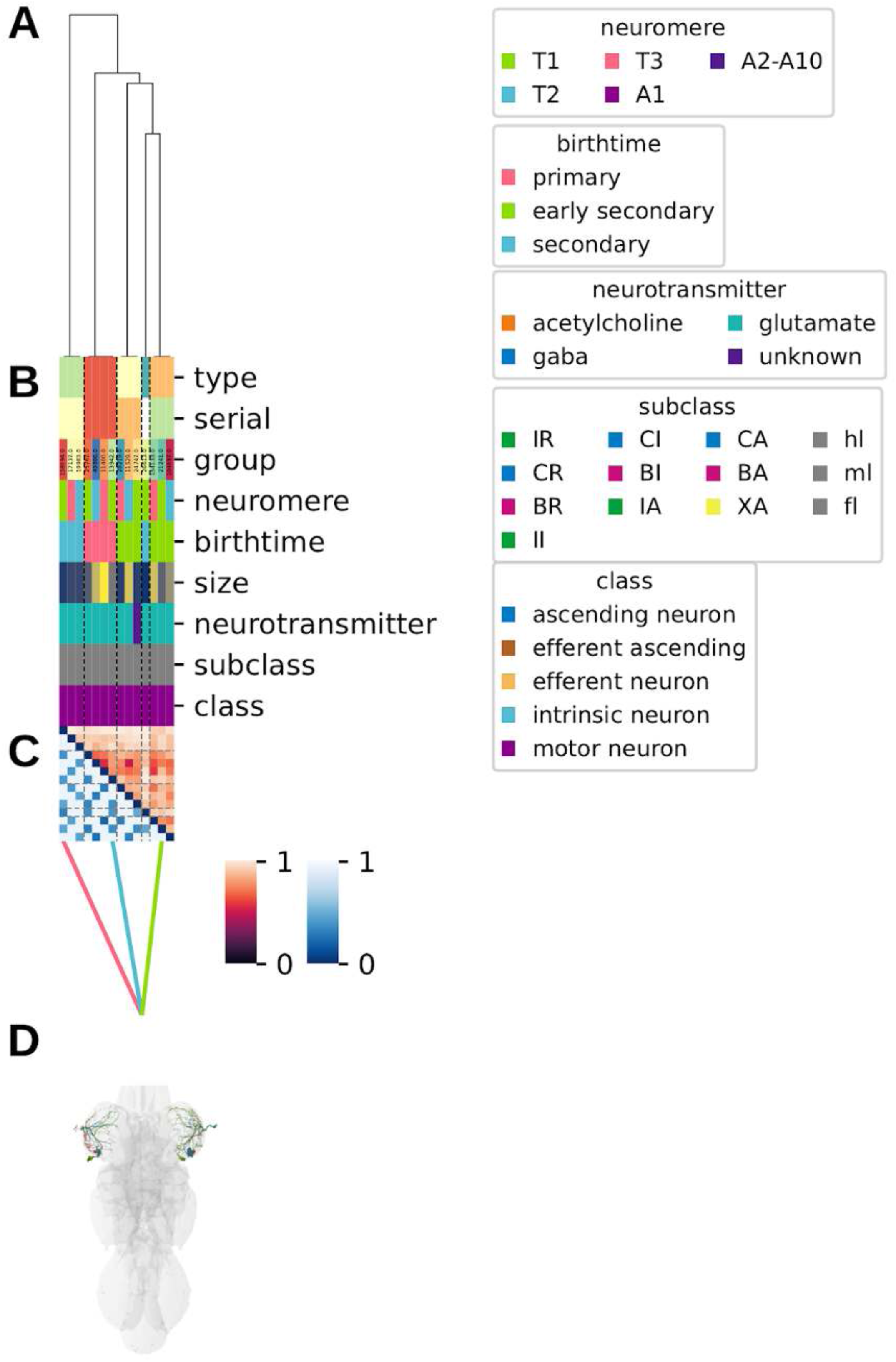
Systematic typing of hemilineage 20B, 21B, and 22B. Types for motor neurons were assigned separately as outlined in our accompanying manuscript (Cheong et al., 2023). **A.** Hierarchical clustering dendrogram of hemilineage groups by laterally and serially aggregated connectivity cosine clustering. **B.** Categorical annotations of each hemilineage group, each column corresponding to the aligned leaf in A. Colours for cluster, serial set, and group are arbitrary for visualisation. Colours for neuromere, birthtime, neurotransmitter, subclass, and class are as in all other figures. **C.** Similarity distance heatmap for hemilineage. Cosine distance is in the upper triangle, while laterally symmetrised NBLAST distance is in the lower triangle. Systematic type names of some types are labelled. **D.** Morphologically representative groups from dendrogram subtrees. Each group, indicated by colour and line connecting to its column in B and C, is the most morphologically representative group (medoid of NBLAST distance) from a subtree of A. The subtrees (flat clusters) are equal height cuts of A determined to yield the number of groups per plot and plots in D.

**Figure 48 - figure supplement 14.**
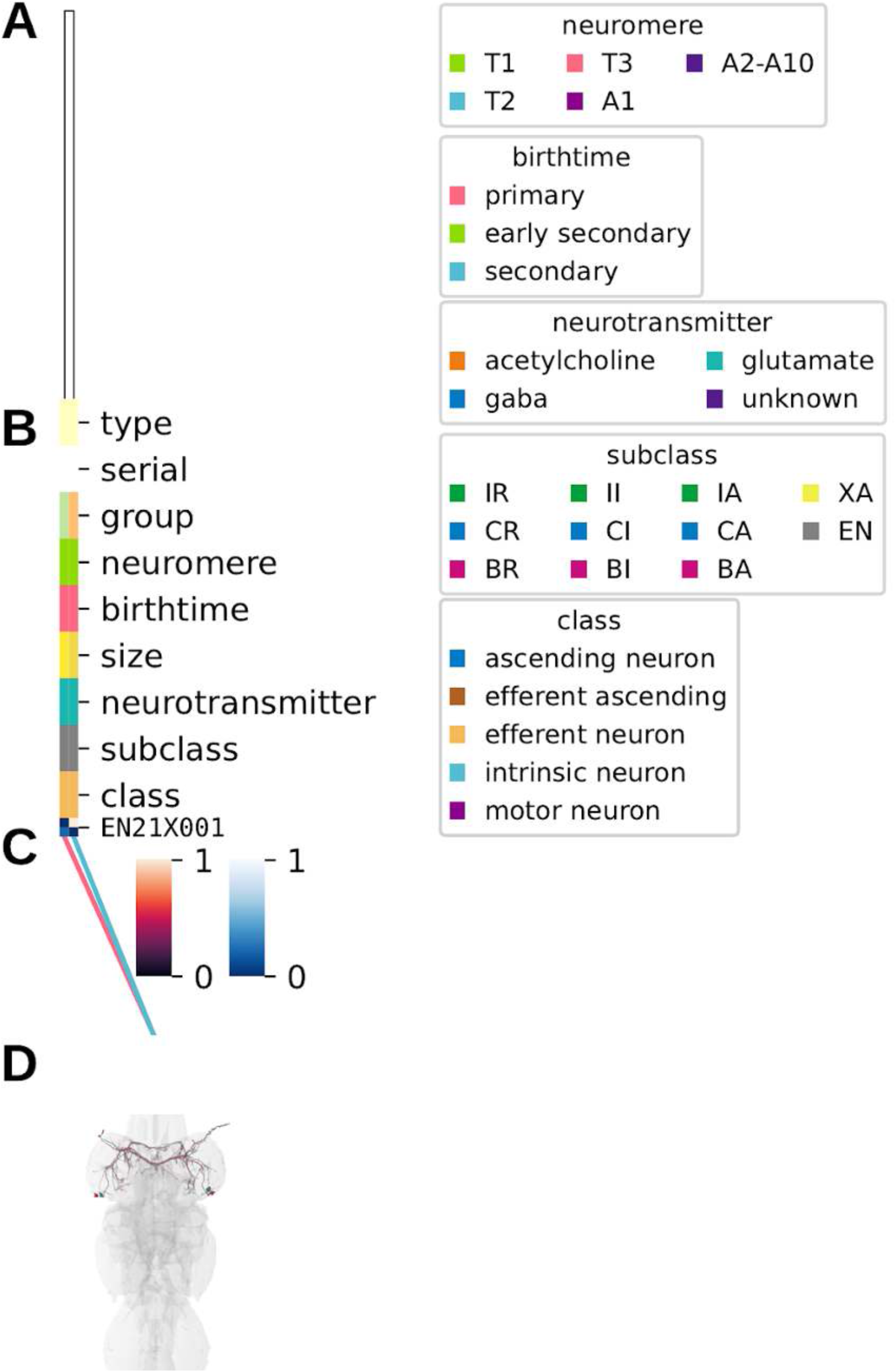
Systematic typing of hemilineage 21X. **A.** Hierarchical clustering dendrogram of hemilineage groups by laterally and serially aggregated connectivity cosine clustering. **B.** Categorical annotations of each hemilineage group, each column corresponding to the aligned leaf in A. Colours for type, serial set, and group are arbitrary for visualisation. Colours for neuromere, birthtime, neurotransmitter, subclass, and class are as in all other figures. **C.** Similarity distance heatmap for hemilineage. Cosine distance is in the upper triangle, while laterally symmetrised NBLAST distance is in the lower triangle. Systematic type names of some types are labelled. **D.** Morphologically representative groups from dendrogram subtrees. Each group, indicated by colour and line connecting to its column in B and C, is the most morphologically representative group (medoid of NBLAST distance) from a subtree of A. The subtrees (flat clusters) are equal height cuts of A determined to yield the number of groups per plot and plots in D.

**Figure 48 - figure supplement 15.**
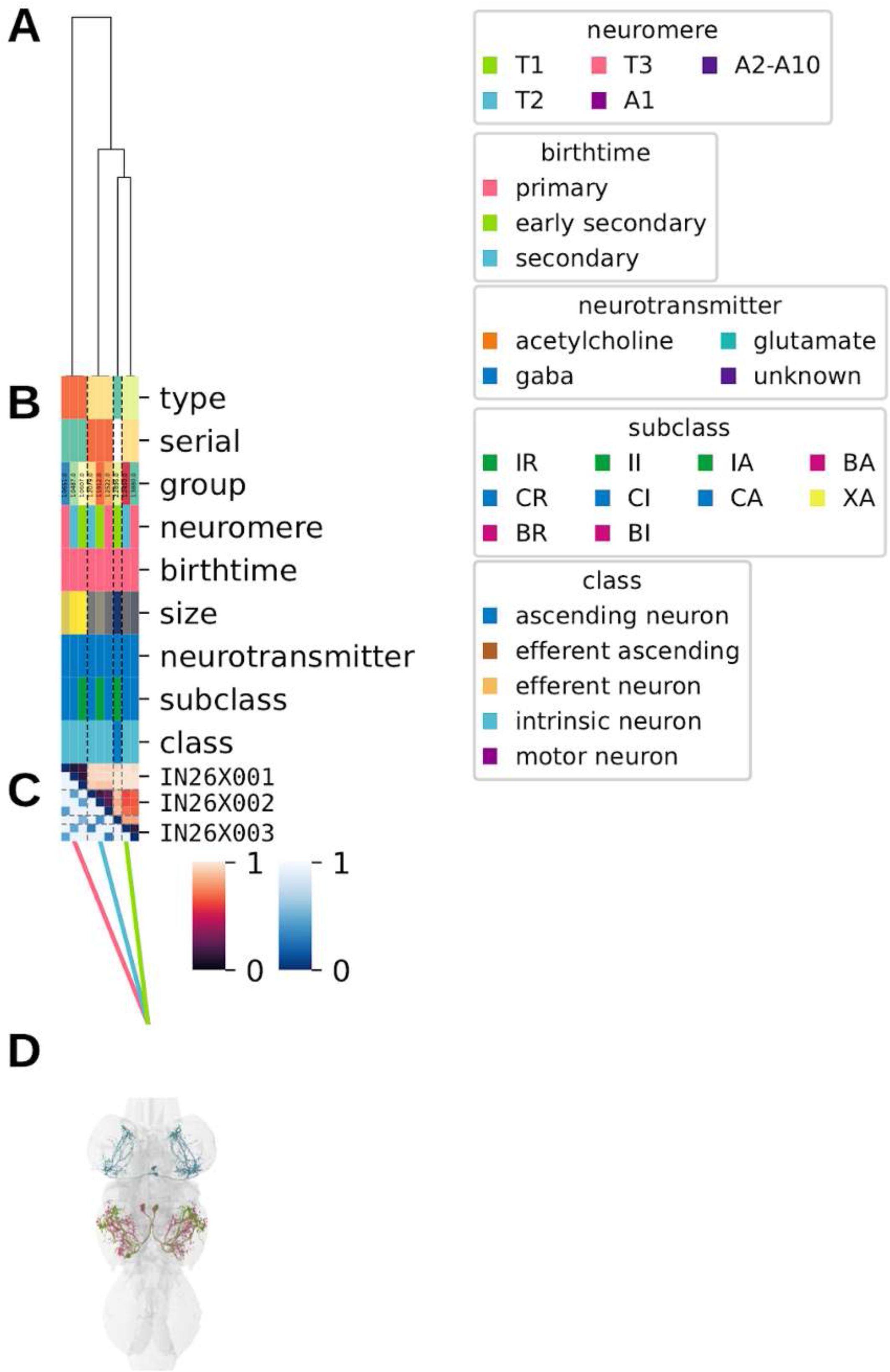
Systematic typing of hemilineage 26X. **A.** Hierarchical clustering dendrogram of hemilineage groups by laterally and serially aggregated connectivity cosine clustering. **B.** Categorical annotations of each hemilineage group, each column corresponding to the aligned leaf in A. Colours for type, serial set, and group are arbitrary for visualisation. Colours for neuromere, birthtime, neurotransmitter, subclass, and class are as in all other figures. **C.** Similarity distance heatmap for hemilineage. Cosine distance is in the upper triangle, while laterally symmetrised NBLAST distance is in the lower triangle. Systematic type names of some types are labelled. **D.** Morphologically representative groups from dendrogram subtrees. Each group, indicated by colour and line connecting to its column in B and C, is the most morphologically representative group (medoid of NBLAST distance) from a subtree of A. The subtrees (flat clusters) are equal height cuts of A determined to yield the number of groups per plot and plots in D.

**Figure 48 - figure supplement 16.**
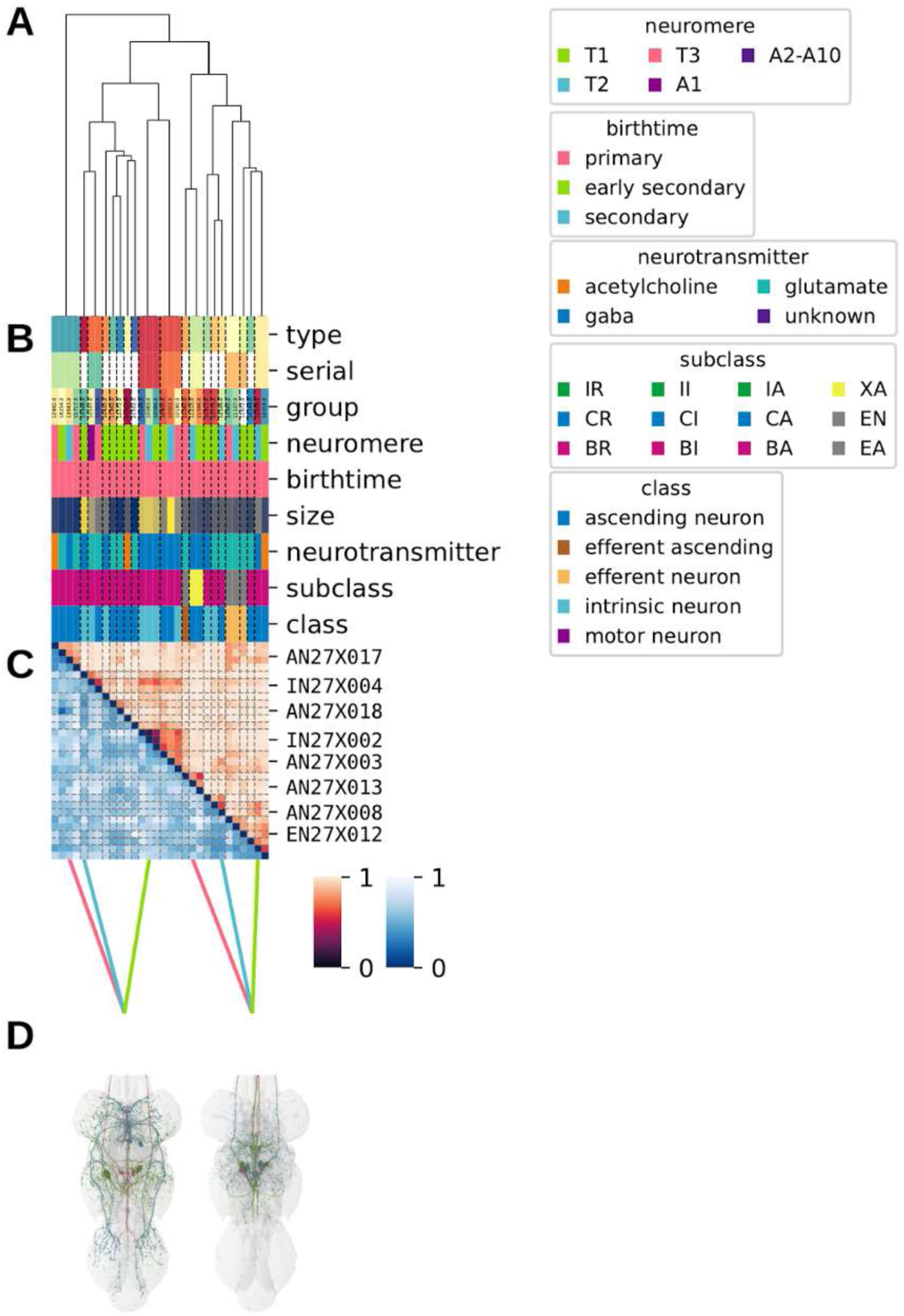
Systematic typing of hemilineage 27X. **A.** Hierarchical clustering dendrogram of hemilineage groups by laterally and serially aggregated connectivity cosine clustering. **B.** Categorical annotations of each hemilineage group, each column corresponding to the aligned leaf in A. Colours for type, serial set, and group are arbitrary for visualisation. Colours for neuromere, birthtime, neurotransmitter, subclass, and class are as in all other figures. **C.** Similarity distance heatmap for hemilineage. Cosine distance is in the upper triangle, while laterally symmetrised NBLAST distance is in the lower triangle. Systematic type names of some types are labelled. **D.** Morphologically representative groups from dendrogram subtrees. Each group, indicated by colour and line connecting to its column in B and C, is the most morphologically representative group (medoid of NBLAST distance) from a subtree of A. The subtrees (flat clusters) are equal height cuts of A determined to yield the number of groups per plot and plots in D.

**Figure 48 - figure supplement 17.**
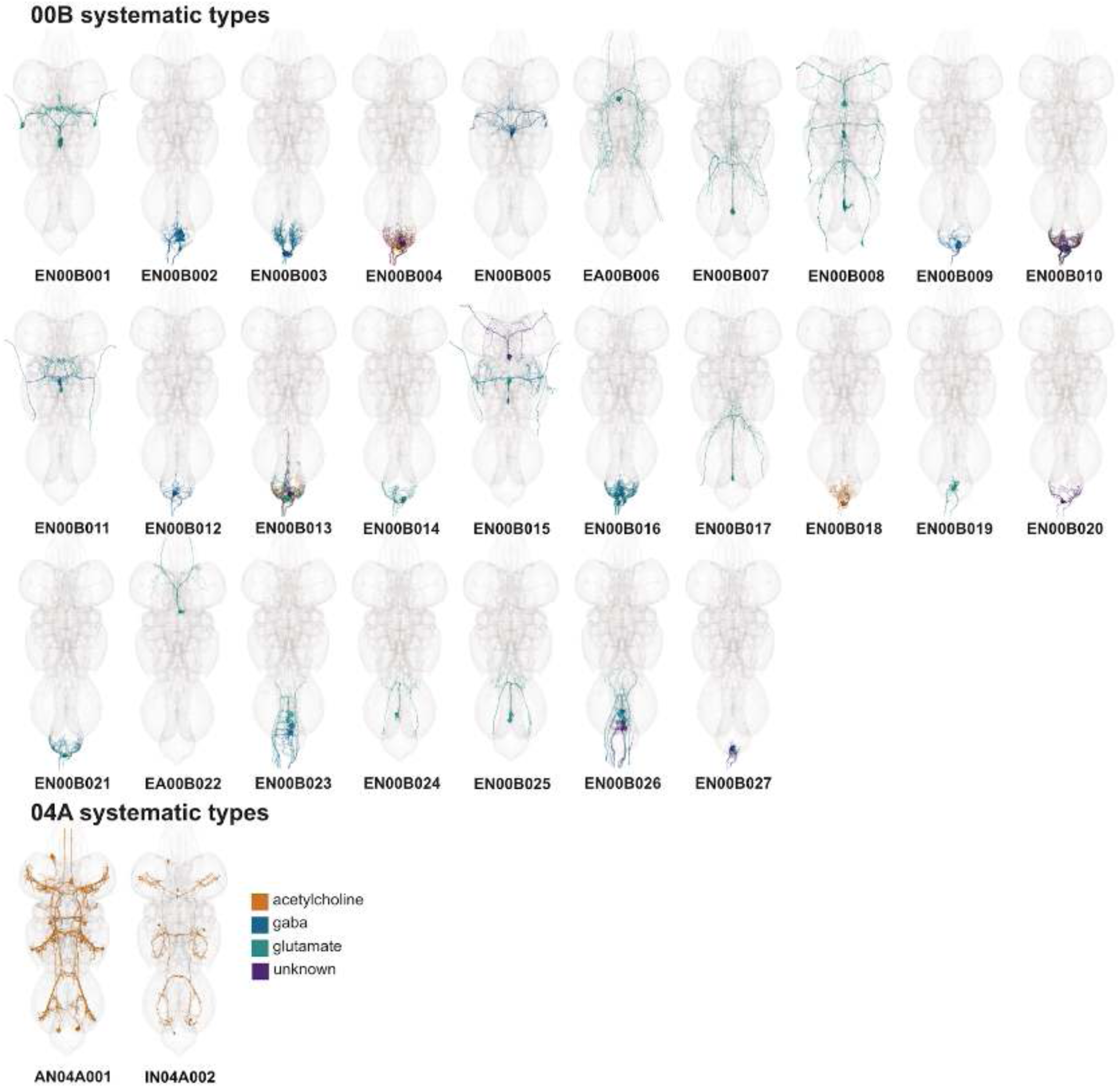
Systematic types of hemilineages 00B and 04A. Systematic types have been arranged in numerical order, with neurons of the same type that belong to distinct classes (e.g., intrinsic neuron vs ascending neuron) plotted separately but placed adjacent to each other. Individual neuron meshes have been coloured based on their predicted neurotransmitters: dark orange = acetylcholine, blue = gaba, marine = glutamate, dark purple = unknown.

**Figure 48 - figure supplement 18.**
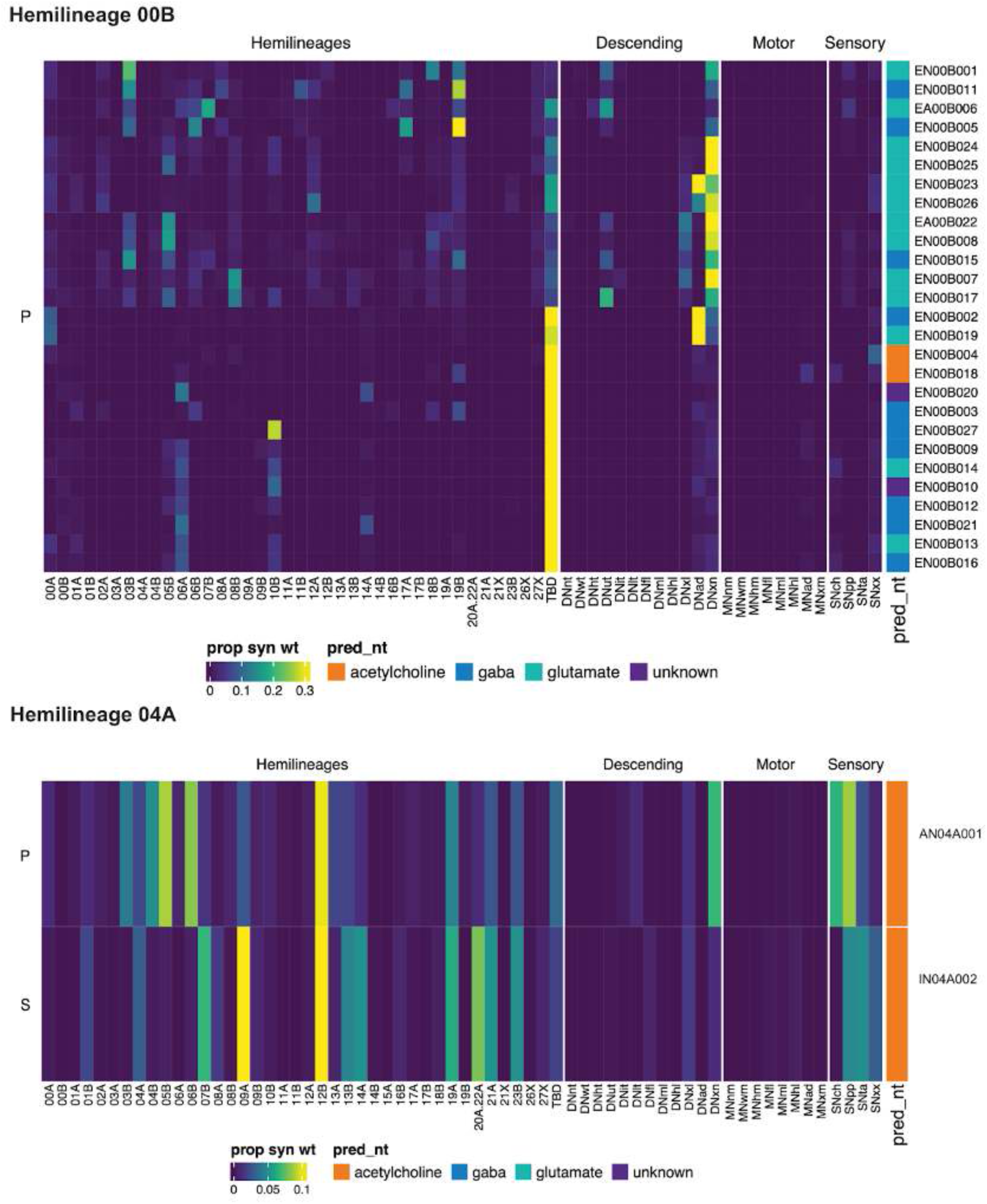
Connectivity to upstream partners by 00B and 04A systematic types. Proportions of synaptic weight to systematic types from upstream partners, normalised by row. 00B and 04A neurons have been clustered within each assigned birthtime window (P = primary, ES = early secondary, S = secondary) based on both upstream and downstream connectivity to hemilineages, descending neuron subclasses, motor neuron subclasses, and sensory neuron modalities. Annotation bar is coloured by the most common predicted neurotransmitter for the neurons of each type.

**Figure 48 - figure supplement 19.**
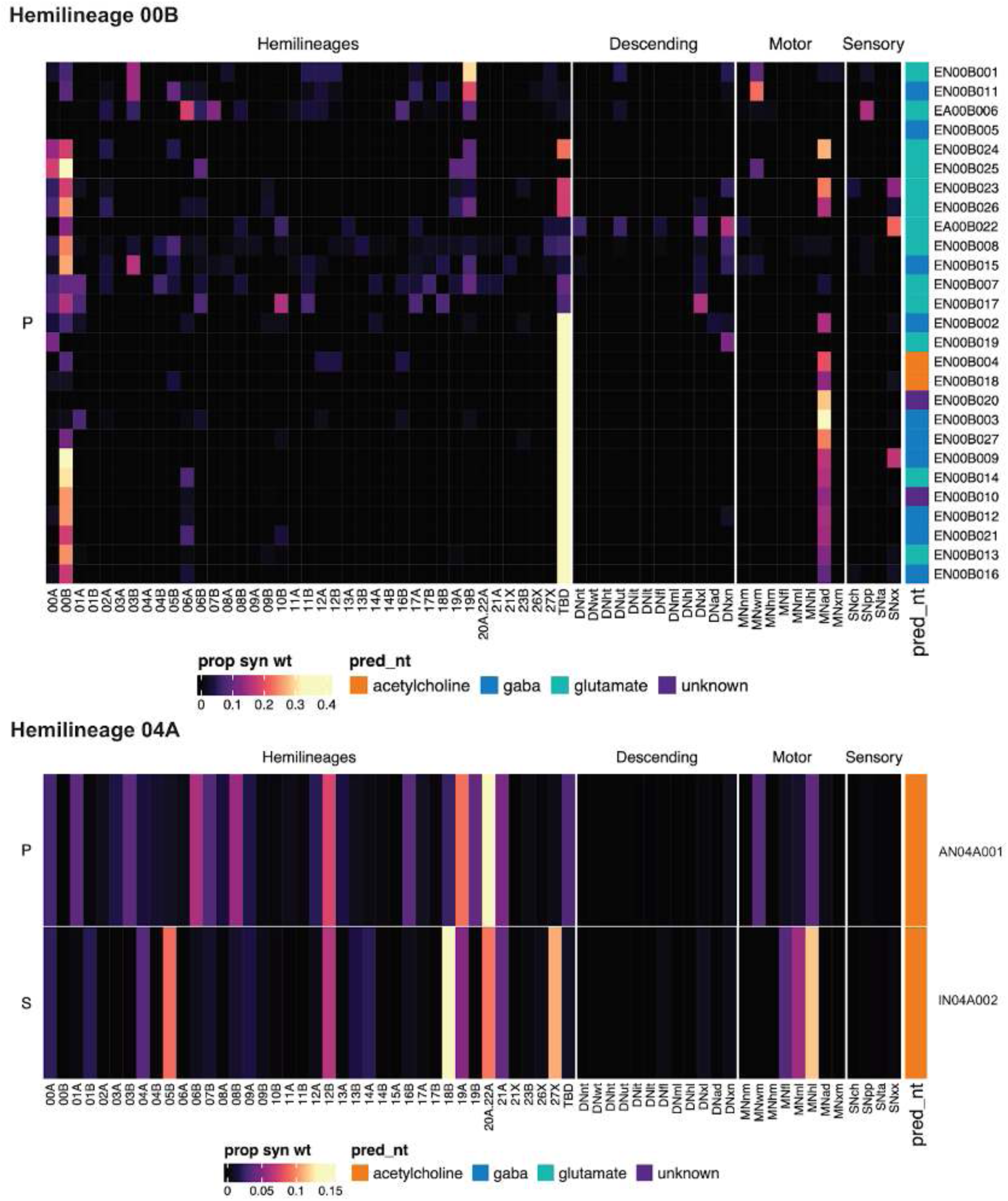
Connectivity to downstream partners by 00B and 04A systematic types. Proportions of synaptic weight from systematic types to downstream partners, normalised by row. 00B and 04A neurons have been clustered within each assigned birthtime window (P = primary, ES = early secondary, S = secondary) based on both upstream and downstream connectivity to hemilineages, descending neuron subclasses, motor neuron subclasses, and sensory neuron modalities. The annotation bar is coloured by the most common predicted neurotransmitter within each type.

**Figure 48 - figure supplement 20.**
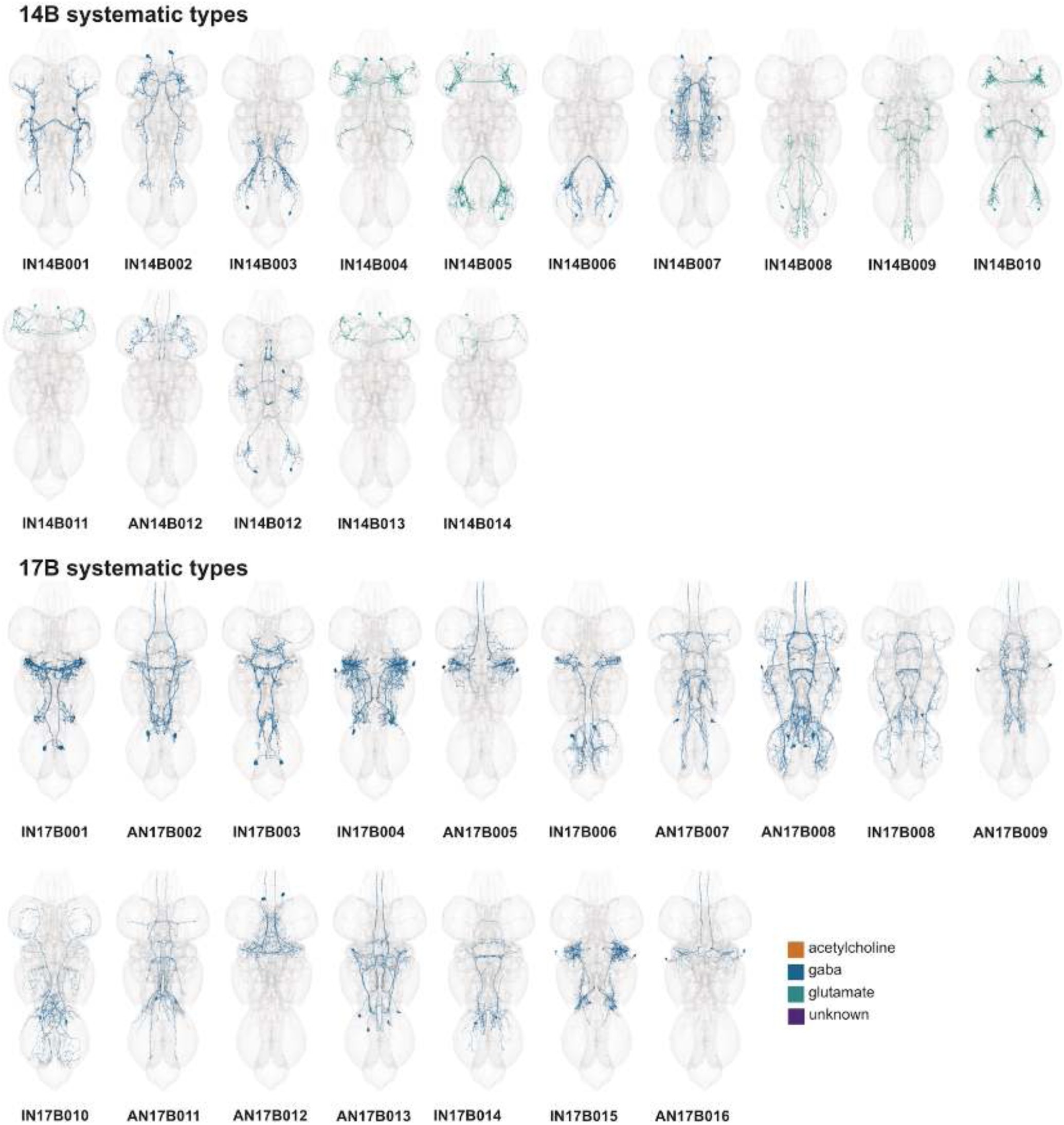
Systematic types of hemilineages 14B and 17B. Systematic types have been arranged in numerical order, with neurons of the same type that belong to distinct classes (e.g., intrinsic neuron vs ascending neuron) plotted separately but placed adjacent to each other. Individual neuron meshes have been coloured based on their predicted neurotransmitters: dark orange = acetylcholine, blue = gaba, marine = glutamate, dark purple = unknown.

**Figure 48 - figure supplement 21.**
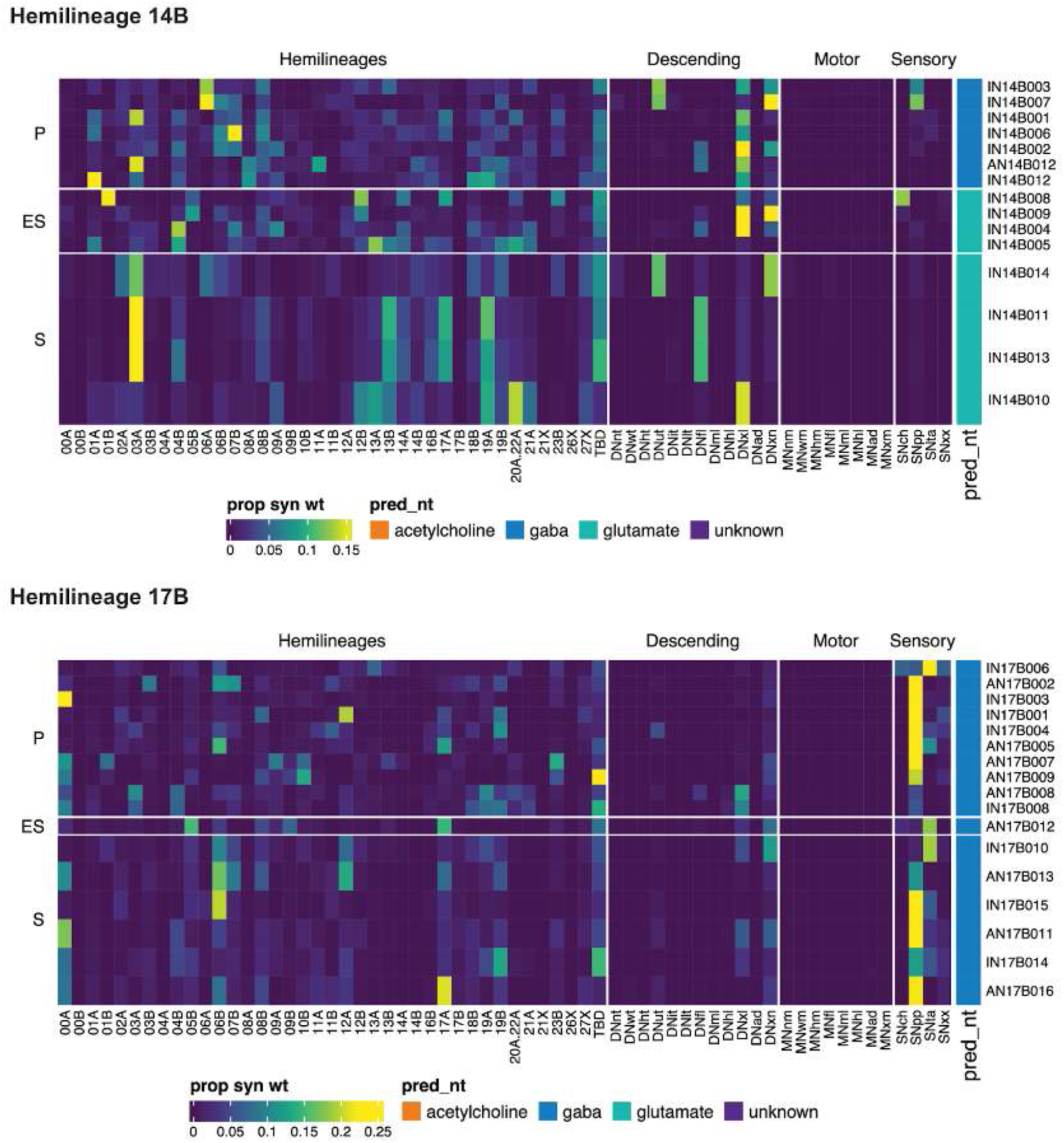
Connectivity to upstream partners by 14B and 17B systematic types. Proportions of synaptic weight to systematic types from upstream partners, normalised by row. 14B and 17B have been clustered within each assigned birthtime window (P = primary, ES = early secondary, S = secondary) based on both upstream and downstream connectivity to hemilineages, descending neuron subclasses, motor neuron subclasses, and sensory neuron modalities. Annotation bar is coloured by the most common predicted neurotransmitter for the neurons of each type.

**Figure 48 - figure supplement 22.**
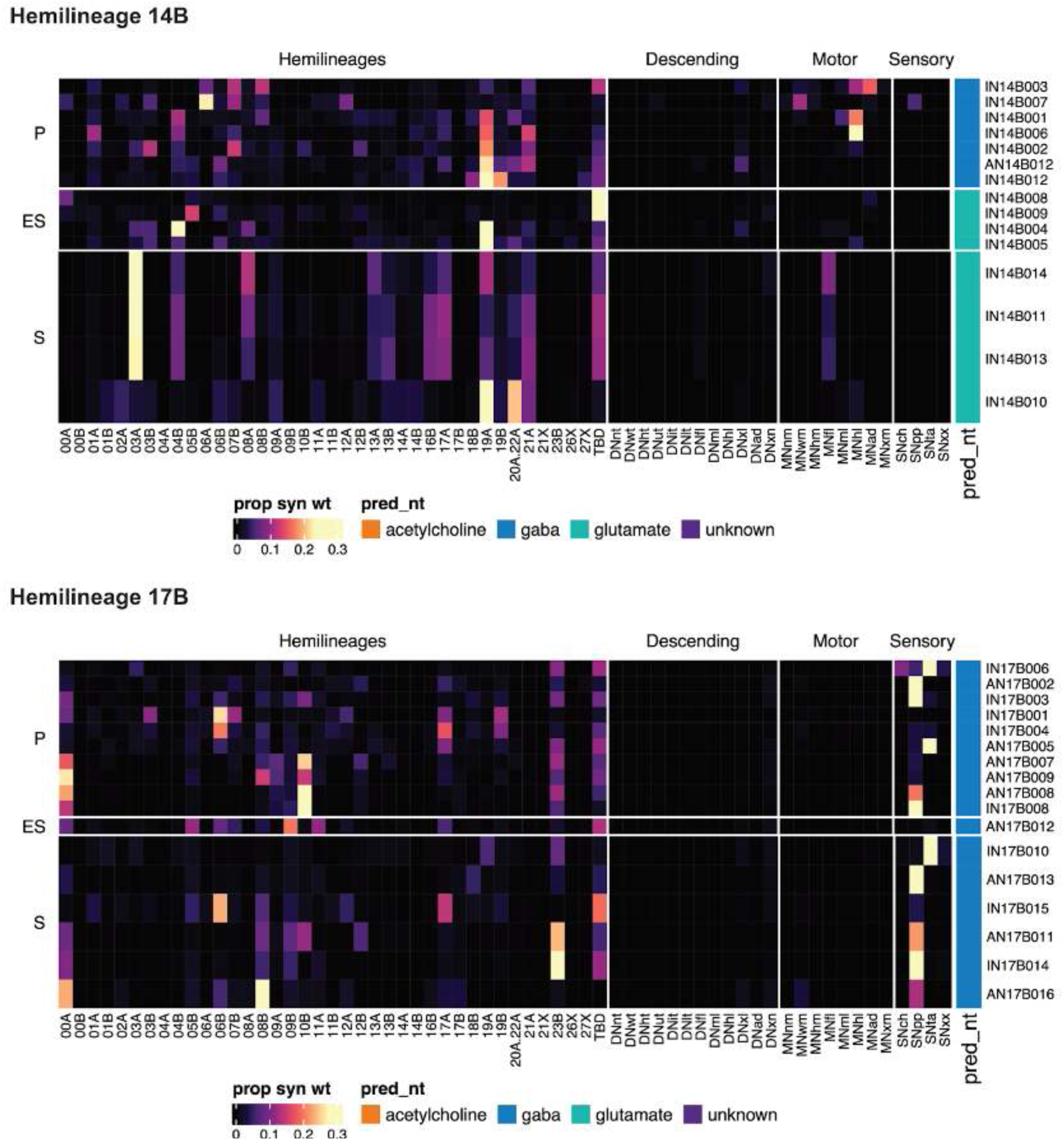
Connectivity to downstream partners by 14B and 17B systematic types. Proportions of synaptic weight from systematic types to downstream partners, normalised by row. 14B and 17B neurons have been clustered within each assigned birthtime window (P = primary, ES = early secondary, S = secondary) based on both upstream and downstream connectivity to hemilineages, descending neuron subclasses, motor neuron subclasses, and sensory neuron modalities. The annotation bar is coloured by the most common predicted neurotransmitter within each type.

**Figure 48 - figure supplement 23.**
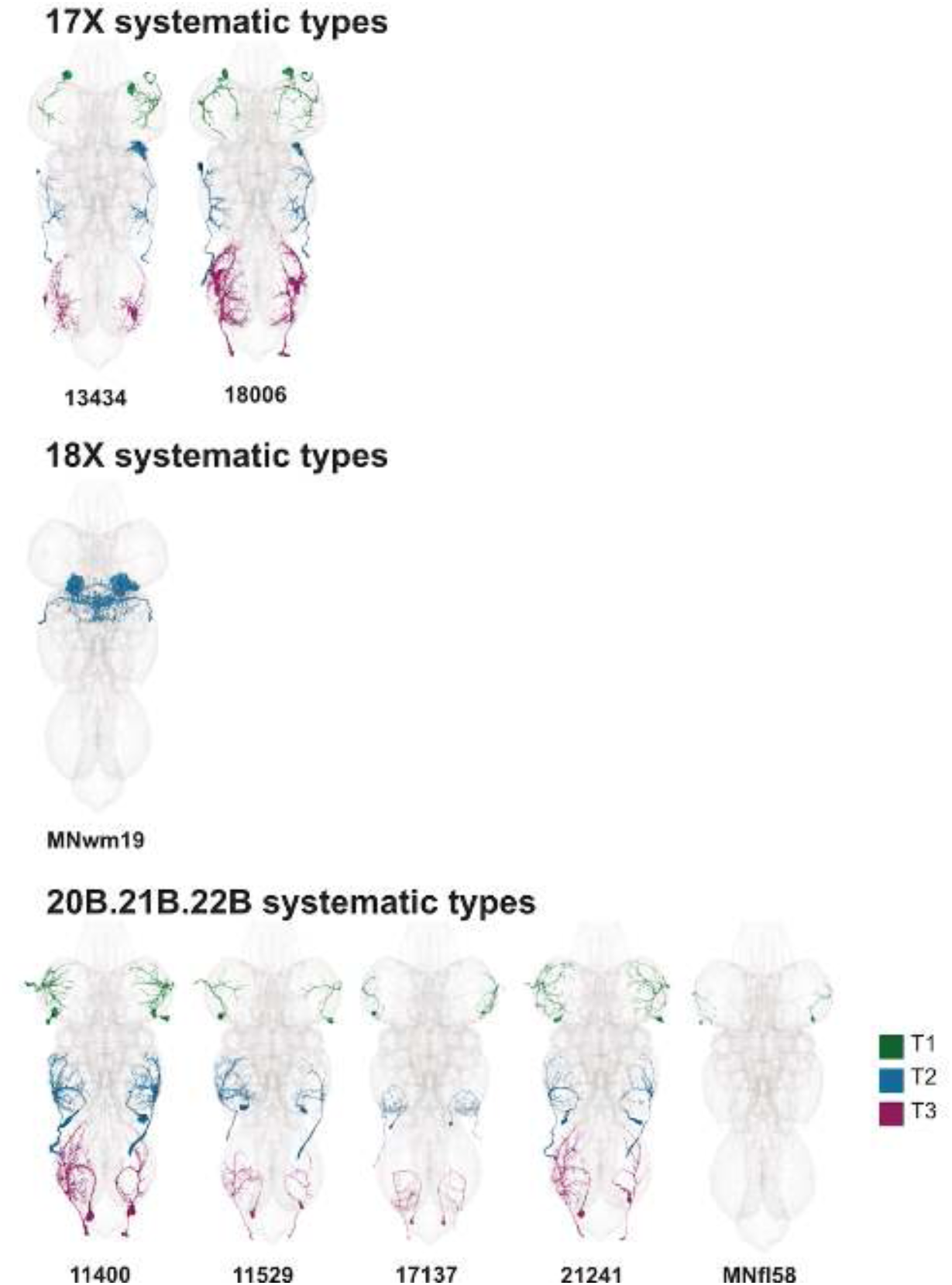
Systematic types of hemilineages 17X, 18X, and 20B.21B.22B. Motor neurons (typed separately in (Cheong et al., 2023)) have been plotted by serial set if identified in multiple neuromeres and by systematic type if not. Individual motor neuron meshes have been coloured based on soma neuromere: dark green = T1, blue = T2, magenta = T3.

**Figure 48 - figure supplement 24.**
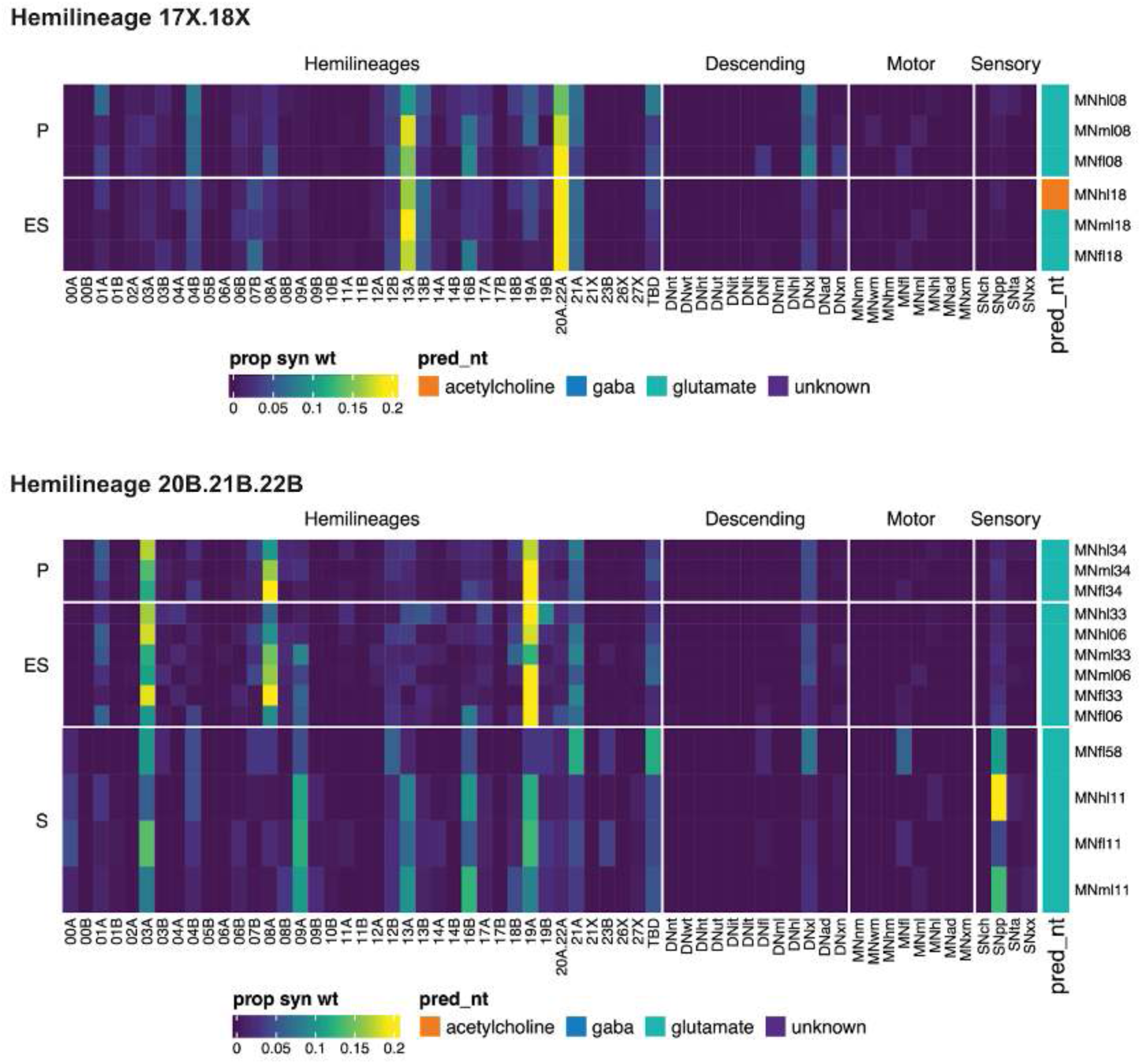
Connectivity to upstream partners by 17X, 18X, and 20B.21B.22B systematic types. Proportions of synaptic weight to systematic types from upstream partners, normalised by row. 17X, 18X, and 20B.21B.22B neurons have been clustered within each assigned birthtime window (P = primary, ES = early secondary, S = secondary) based on both upstream and downstream connectivity to hemilineages, descending neuron subclasses, motor neuron subclasses, and sensory neuron modalities. Annotation bar is coloured by the most common predicted neurotransmitter for the neurons of each type.

**Figure 48 - figure supplement 25.**
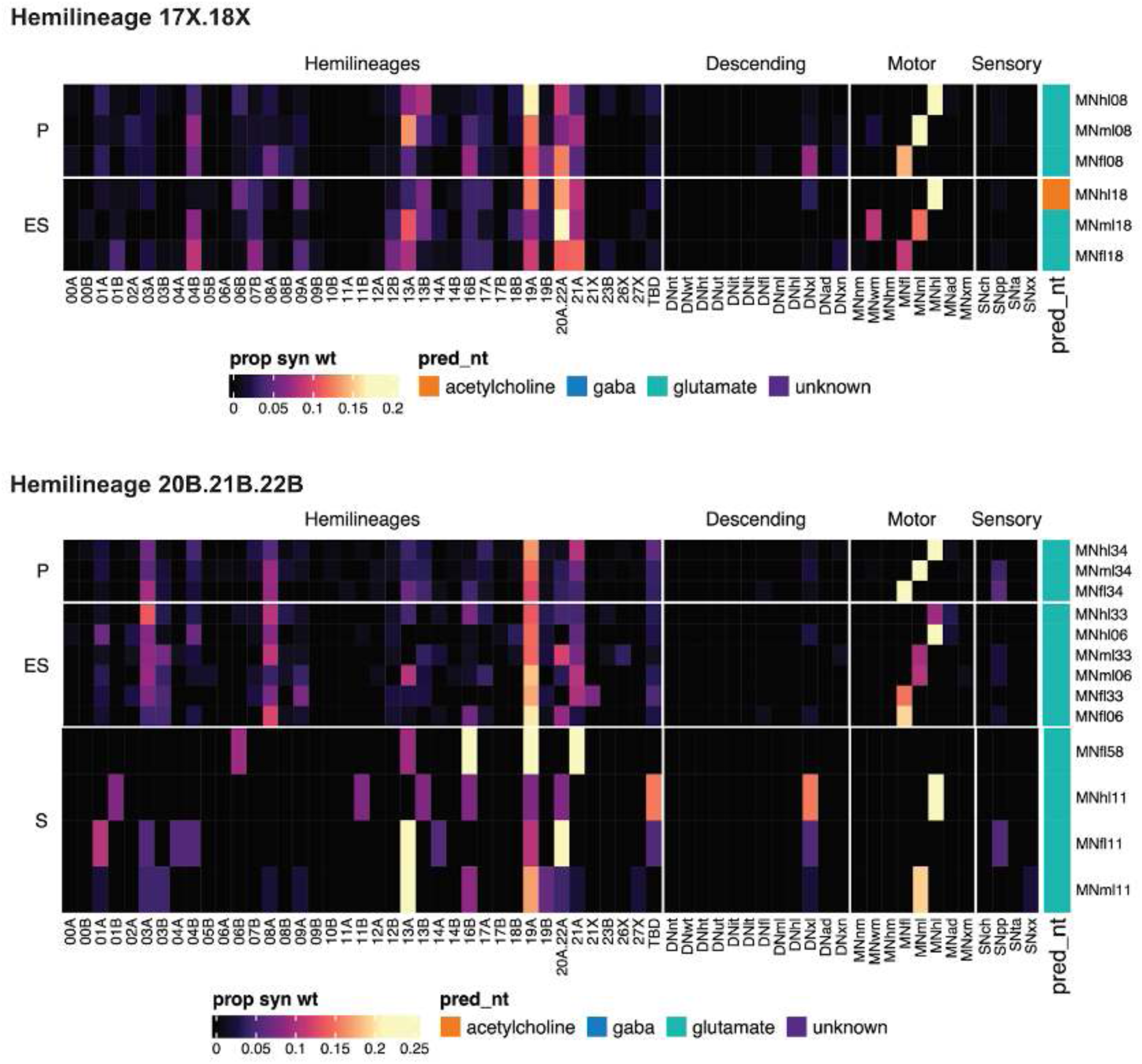
Connectivity to downstream partners by 17X, 18X, and 20B.21B.22B systematic types. Proportions of synaptic weight from systematic types to downstream partners, normalised by row. 17X, 18X, and 20B.21B.22B neurons have been clustered within each assigned birthtime window (P = primary, ES = early secondary, S = secondary) based on both upstream and downstream connectivity to hemilineages, descending neuron subclasses, motor neuron subclasses, and sensory neuron modalities. The annotation bar is coloured by the most common predicted neurotransmitter within each type.

**Figure 48 - figure supplement 26.**
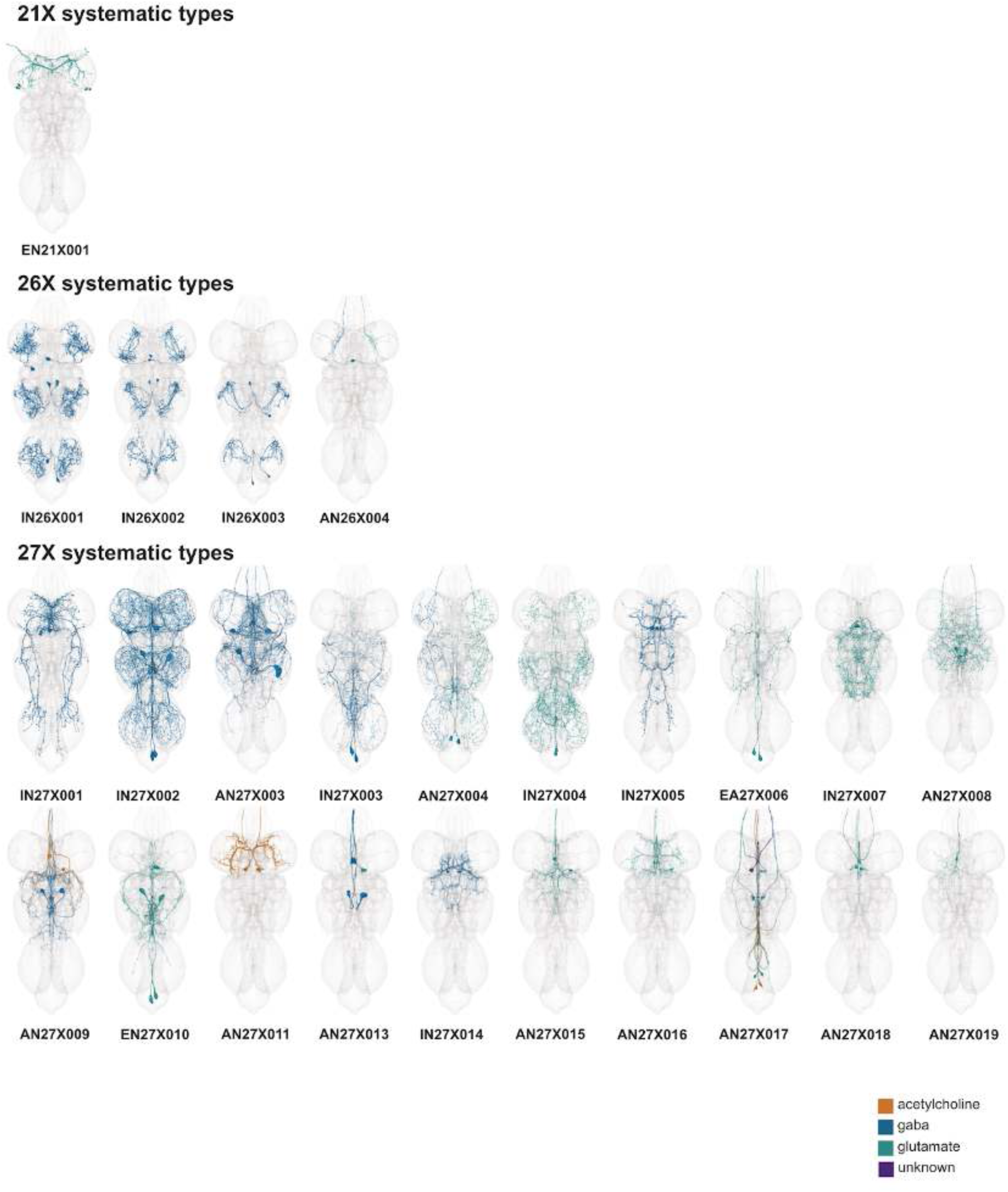
Systematic types of hemilineages 21X, 26X, and 27X. Systematic types have been arranged in numerical order, with neurons of the same type that belong to distinct classes (e.g., intrinsic neuron vs ascending neuron) plotted separately but placed adjacent to each other. Individual neuron meshes have been coloured based on their predicted neurotransmitters: dark orange = acetylcholine, blue = gaba, marine = glutamate, dark purple = unknown.

**Figure 48 - figure supplement 27.**
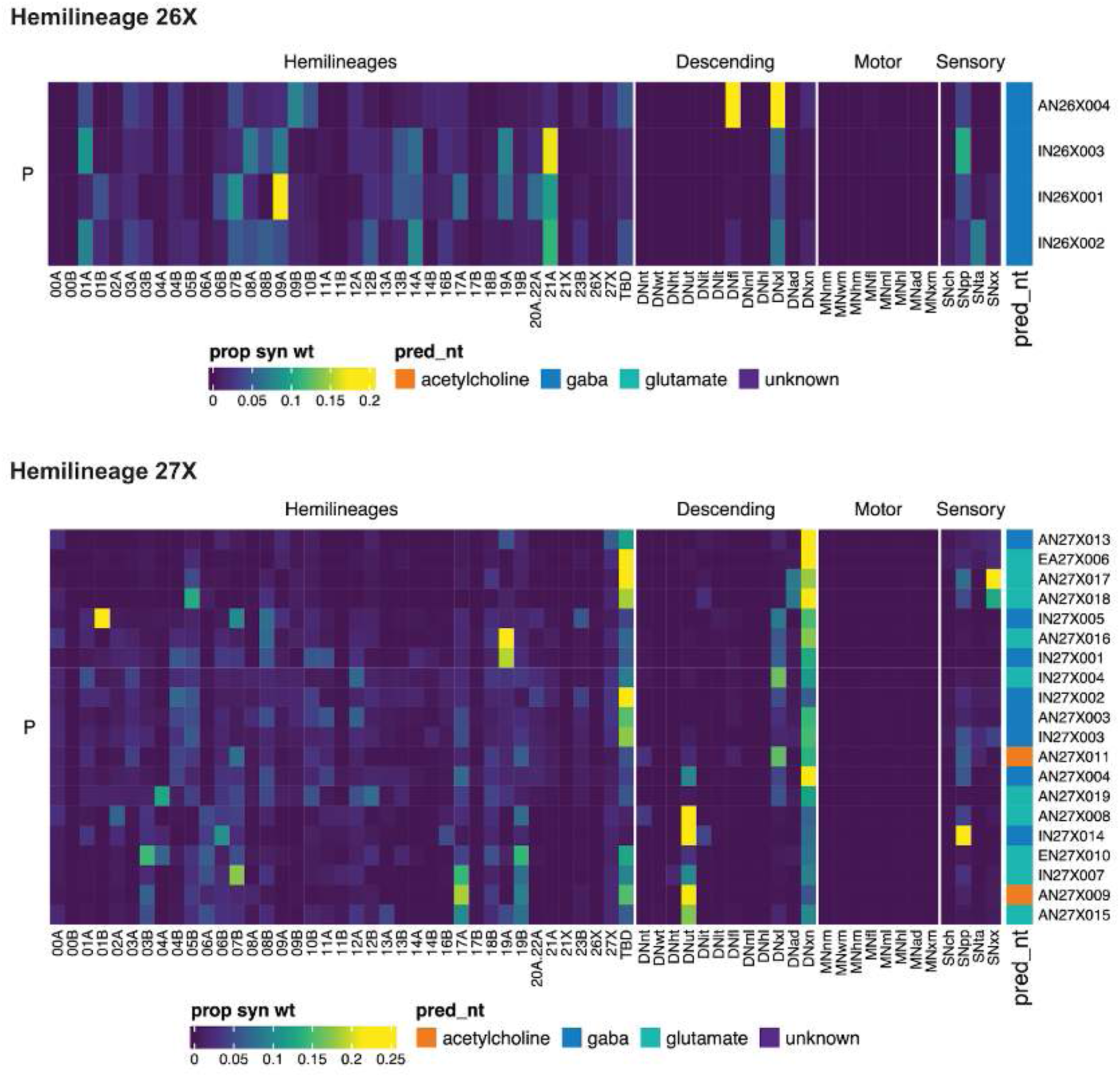
Connectivity to upstream partners by 26X and 27X systematic types. Proportions of synaptic weight to systematic types from upstream partners, normalised by row. 26X and 27X neurons have been clustered within each assigned birthtime window (P = primary, ES = early secondary, S = secondary) based on both upstream and downstream connectivity to hemilineages, descending neuron subclasses, motor neuron subclasses, and sensory neuron modalities. Annotation bar is coloured by the most common predicted neurotransmitter for the neurons of each type.

**Figure 48 - figure supplement 28.**
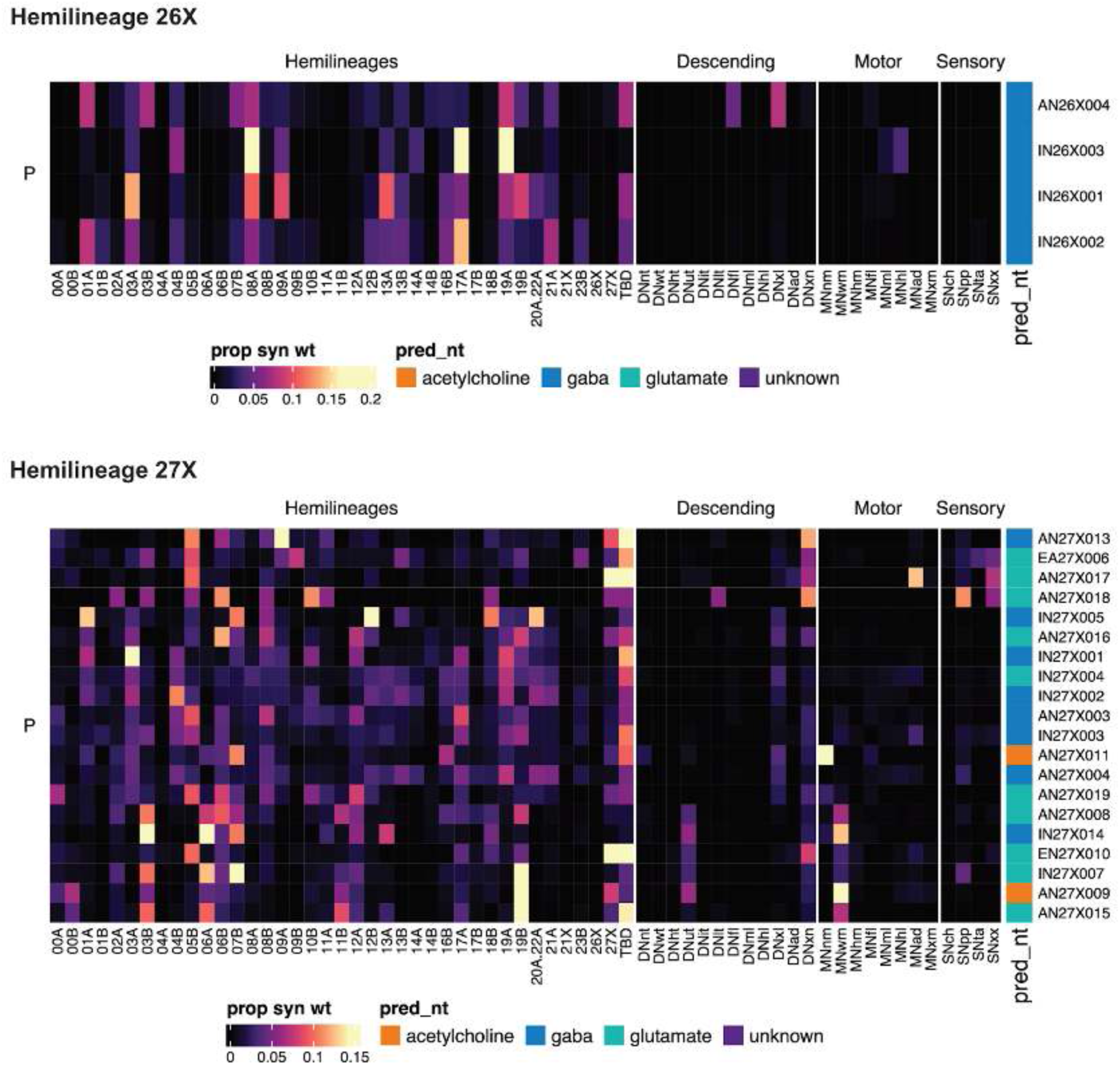
Connectivity to downstream partners by 26X and 27X systematic types. Proportions of synaptic weight from systematic types to downstream partners, normalised by row. 26X and 27X neurons have been clustered within each assigned birthtime window (P = primary, ES = early secondary, S = secondary) based on both upstream and downstream connectivity to hemilineages, descending neuron subclasses, motor neuron subclasses, and sensory neuron modalities. The annotation bar is coloured by the most common predicted neurotransmitter within each type.

#### Organisation of abdominal ganglia

The tremendous expansion of the thoracic neuromeres by secondary neurons has been documented extensively at light level. In contrast, little has been reported regarding the morphology or expected connectivity of neurons in the adult abdominal ganglia. This paucity of light-level data, in combination with the lack of clear segmental boundaries and the increase in dorsal relative to ventral neuropil, made the assignment of soma neuromere much more challenging than in the thorax, with significant refinement dependent on identification of serial sets.

Most abdominal neuromeres are expected to consist of a relatively high proportion of primary neurons that have persisted and remodeled during metamorphosis. Only three neuroblasts - NB3-5 (lin 09), NB5-2 (lin 06), and NB3-5 (lin 05) - produce secondary neurons in segments A3-A7, plus NB6-2 (lin 19) in A2 (Birkholz et al., 2015; Truman and Bate, 1988). In contrast, 13 neuroblasts (lin 00, 01, 03, 05, 06, 07, 09, 10, 12, 17, 18, 19, 23) divide postembryonically in A1 (Birkholz et al., 2015; Truman et al., 2004), 31 in A8, 23 in A9, and 11 in A10 (Birkholz et al., 2013).

We first examined general patterns of connectivity in the abdominal neuromeres. Whilst 888 abdominal neurons are restricted to the boundaries of the ANm (Figure 49A), the vast majority of these project across the midline, connecting both sides (Figure 5B). We identified 571 neurons that strongly link the abdomen with the thoracic neuropils and 245 that solely innervate thoracic neuropils (Figure 49A) - together comprising nearly half of all abdominal neurons. Most of these neurons originate in A1 (Figure 49B), suggesting that many A1 secondary neuron populations survive because they have been co-opted into processing required for adult-specific appendages. The majority of neurons in A3 - A8 are restricted to the abdomen, although a substantial proportion ascend to the thorax and/or higher centres. In contrast, a very small proportion of neurons in A8-A10 span both thorax and abdomen (Figure 49 - figure supplement 1). In summary, the reported expansion of the secondary neuron population at either end of the abdominal neuromeres likely reflects the more complex functions associated with thoracic appendages or genitalia.

**Figure 49.**
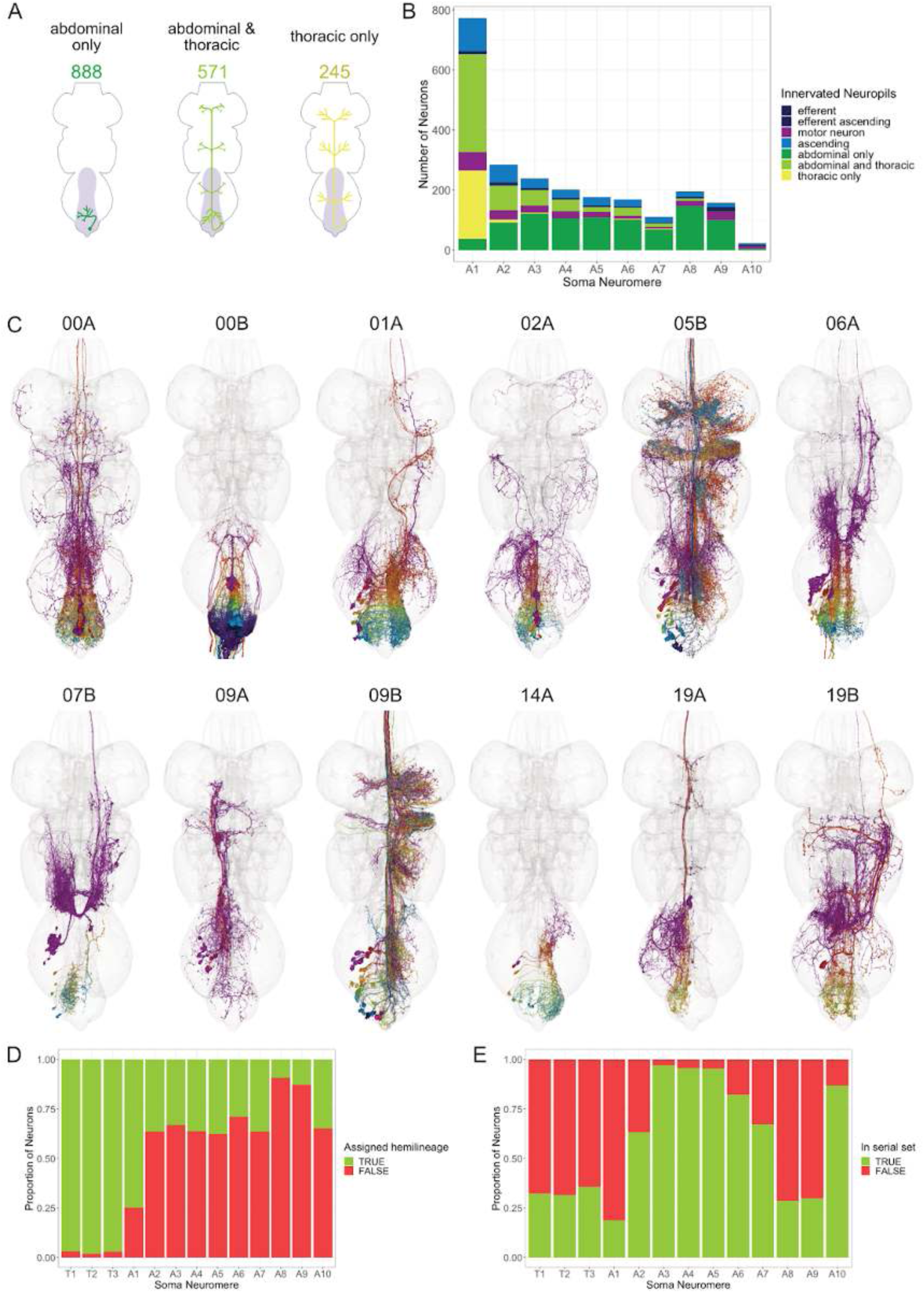
Organisation of abdominal ganglia. **A.** Cartoon schematic depicting VNC innervation by neurons originating in abdominal neuromeres. Abdominal neurons have no more than 5 synapses outside of the ANm, abdominal & thoracic neurons have at least 5% of synapses in the ANm and more than 5 synapses outside of the ANm, and thoracic neurons have less than 5% of synapses in the ANm. **B.** Quantification of abdominal neuron VNC innervation categories by soma neuromere. **C.** Examples of hemilineages identified in abdominal neuromeres (midline or RHS soma side only). Neuron meshes are coloured by soma neuromere: purple = A1, red = A2, dark orange = A3, dark yellow = A4, green = A5, cyan = A6, blue = A7, navy = A8, dark purple = A9, magenta = A10. **D.** Proportion of neurons assigned to a hemilineage for each soma neuromere. **E.** Proportion of neurons assigned to a serial set for each soma neuromere.

We were able to assign 97.0% of thoracic neurons to a hemilineage (Figure 49D). There was not enough light level data available to assign the majority of abdominal neurons to a hemilineage. However, we were able to assign 47.7% of them (78.6% of those originating in A1), based on serial homology to thoracic neurons and predicted neurotransmitter expression (Figure 49C-D). We found that hemilineages 00A, 01B, 03B, 05B, 06A, 07B, 09A, 09B, 12A, 17A, 18B, 19B, and 23B in A1 contribute populations of secondary neurons innervating thoracic neuropils (Figures 15, 16, 20, 22-23, 25, 28-29, 33, 40-41, 43, 46, 49C). (Note: 01B neurons originating in A1 ascend to innervate T3 leg neuropils and have been annotated and analysed elsewhere in this manuscript as “T3” but counted as A1 here.)

Posterior to A1, we find that most neurons are segmentally repeated primary neurons that participate in local circuits. Whilst only 29.3% of neurons in A2-A10 could be assigned to a hemilineage, the majority could be matched to serial homologues in other neuromeres; the proportion was highest for neurons in A3-A5 (Figure 49E). Overall we matched 54% of abdominal neurons to serial homologues, compared with 33% of thoracic neurons, reflecting the segment-specific specialisation of adult neuron populations in the thorax. Most of the unmatched abdominal neurons either originate in A1 and innervate the thoracic neuropils or originate in A8-A9 and innervate the posterior abdomen, presumably functioning in excretory and reproductive circuits - again reflecting segment-specific expansion and specialisation.

**Figure 49 - figure supplement 1.**
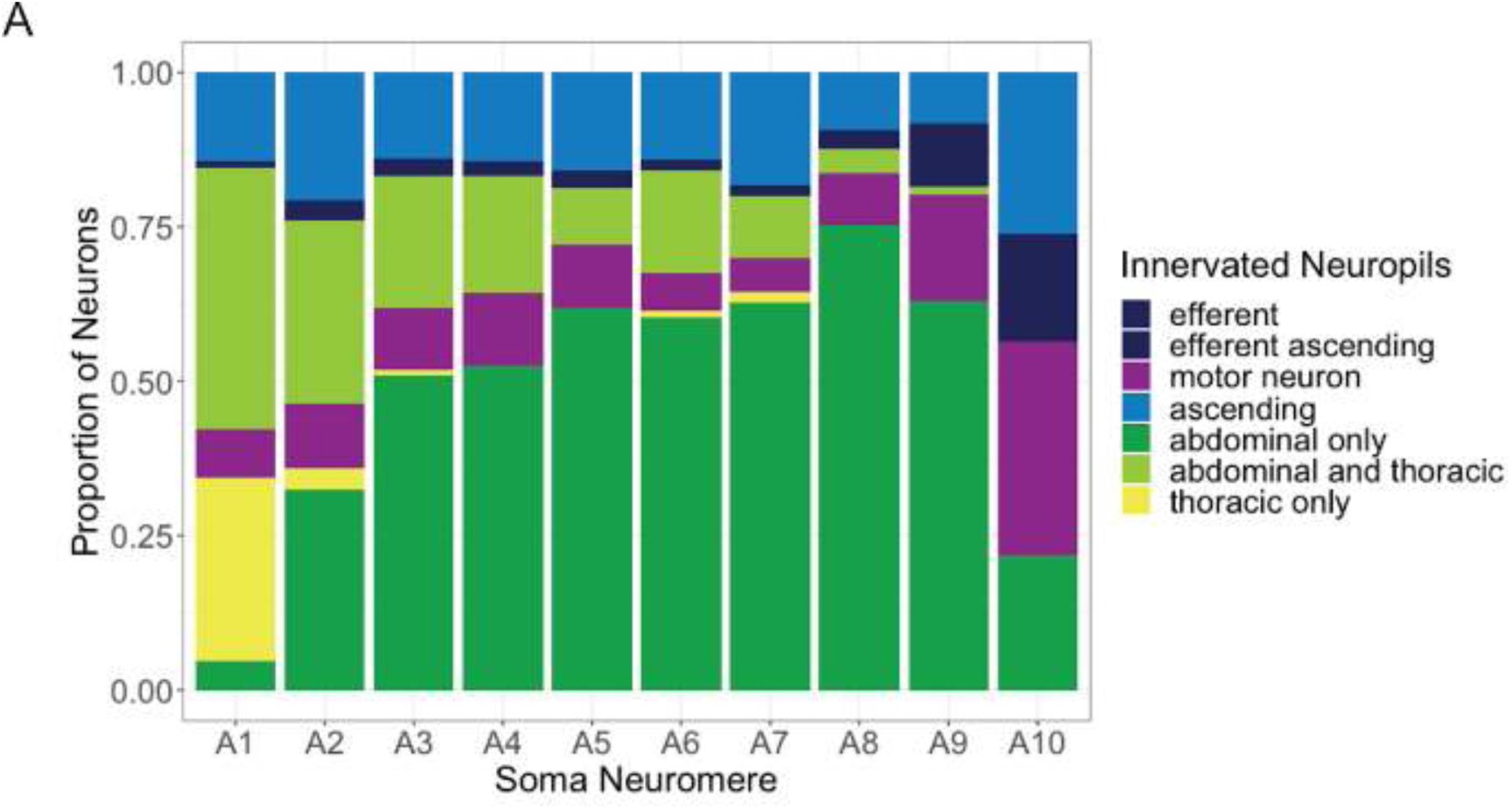
Proportion of neurons originating in each abdominal neuromere that innervate the abdominal (dark green), thoracic (yellow), or abdominal and thoracic neuromeres (light green).

#### Hemilineage diversity

The neurons in the VNC are responsible for a diverse and flexible set of actions, and because those neurons come from a limited number of hemilineages, there must be extensive variety across the hemilineages. Secondary neurons show considerable morphological variety across most hemilineages (Figure 50A), and their roles in the VNC network reflect this diversity. The diversity of different neuron subclasses within a hemilineage is variable: some hemilineages consist exclusively of motor neurons, a few are almost entirely restricted to the ipsilateral hemineuromere (IR), and some are more evenly distributed between the subclasses (Figure 50B). Hemilineages with a higher proportion of IR neurons tend to retain received signals in their local neuropils for longer than those with lower proportion of IR neurons (Figure 50C). On average, signals sent to neurons restricted to the ipsilateral or contralateral neuropil (IR or CR) tend to require 3 more hops (intermediary neurons) before exiting the original neuropil than those sent to intersegmental neurons (BI or II) (data not shown). Hemilineages with more distributed occurrences of subclasses may have more bimodal distributions of intra-neuropil depth (e.g. 09A) while others with more concentrated subclass occurrences are more unimodal (e.g. 20A/22A, 13B).

**Figure 50.**
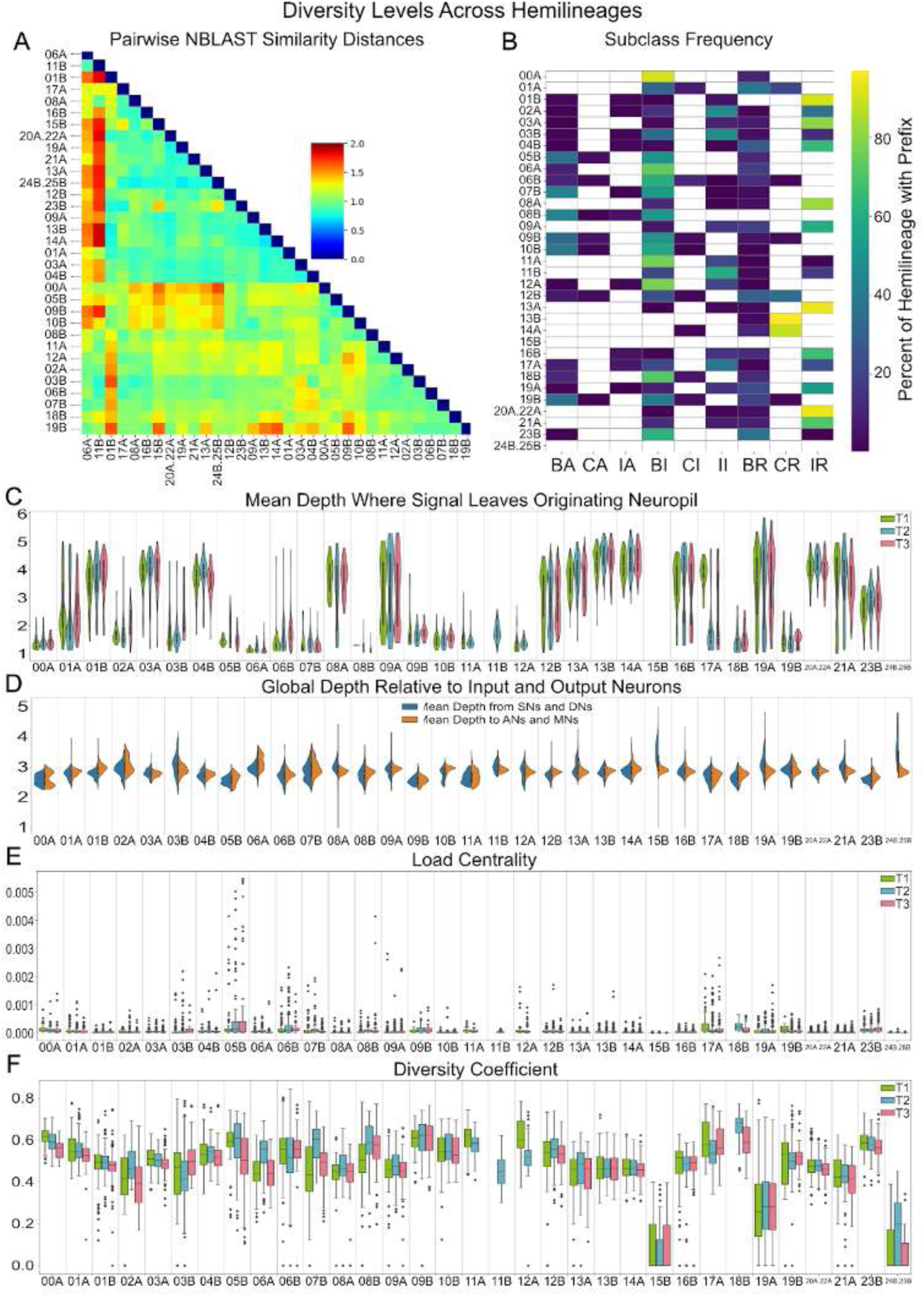
Diversity levels by hemilineage. Data is shown for secondary neurons in T1, T2, and T3 with a hemilineage assignment. **A.** Morphologic similarity between hemilineages (only considering secondary neurons). NBLAST (Costa et al., 2016) score was calculated as the mean of all distance scores (from 0 to 2) between each neuron in the query target and its most similar match in the target hemilineage. **B.** Percentage of neurons in each hemilineage annotated with each respective subclass. White cell indicates no neurons annotated with respective prefix within hemilineage. Hemilineages 15B and and 24B.25B are motor neuron only hemilineages. **C.** Depth at which a signal leaves the originating neuropil, or else reaches a motor or ascending neuron. Higher values indicate a signal is likely to spend more time in its respective neuropil before exiting. **D.** Mean global shortest path distance to and from all output and input neurons, respectively. Calculated via a breadth-first search. **E.** Load centrality (Newman, 2001) distributions of all secondary neurons in each hemilineage, subdivided by neuromere. Load centrality is the fraction of all shortest paths between each pair of neurons that go through a given node. **F.** Diversity coefficient (inspired by (Eagle et al., 2010), except with weights to hemilineage-in-a-hemineuromere communities instead of individual nodes) distributions for all secondary neurons in each hemilineage, subdivided by neuromere. Diversity coefficient was calculated via BCT (Rubinov and Sporns, 2010). Data shown here were measured via iGraph (Csárdi and Nepusz, 2006) and NetworkX (Hagberg et al., 2008) graph implementations of the VNC.

There is also significant variety in the position that hemilineages take within the global structure of the VNC network, even amongst just secondary neurons (Figure 50D). Hemilineages closer to the edges of the graph (e.g. 05B, 09B) tend to have higher values for an important measure of network centrality called load centrality, which is the fraction of all shortest paths between each pair of neurons that go through a given node (Newman, 2001) (Figure 50E). These hemilineages also tend to have lower intra-neuropil depths, but this trend does not always hold (e.g. 16B, 04B). The position that individual hemilineages take in global and local circuits can vary across neuromeres. Hemilineage 10B, for instance, has a flat distribution amongst subclasses, and it has a bimodal distribution of depths from inputs but a relatively uniform distance to outputs (Figure 50D). The neurons of 09A have a relatively unimodal location in the global network, but are split into two different locations within the neuropil, with about half the neurons retaining signals for 2 to 3 more synapses than the other half.

The network position of individual hemilineages across neuromeres is not entirely consistent either. In T1, signals received by hemilineage 17A tend to stay in the local neuropil 3 hops deeper than in T2 or T3 (Figure 50C). For hemilineage 07B, the global location varies significantly within T1, but is more limited in other neuromeres, with T2 being closer to both inputs and outputs than T3 (data not shown). These network differences are often reflected morphologically. For example, 07B is split into two morphologically similar groups in T3, is incredibly morphologically diverse in T2, and in T1 about half the neurons are in one large group of similar neurons while the rest are relatively dissimilar (data not shown). In summary, across the VNC, postembryonic hemilineages show high levels of diversity in both morphology and circuit role and function.

Position and role in the VNC network also differ by birthtime. Despite there being far fewer primary neurons than secondary neurons, primary neurons are more centrally positioned within the VNC network (Figure 51A). Primary neurons are markedly more likely to be involved in the shortest path between pairs of neurons, an important measure of centrality called load centrality (Newman, 2001). Conversely, primary neurons are less insular than their secondary counterparts, connecting more abundantly and evenly with other hemilineages across the VNC (Figure 51A’). Their centrality in the network is reflected in their overrepresentation in the rich-club relative to their population (Figure 51B). In a network, a rich-club is a set of high-centrality nodes, here measured by load centrality, that are more strongly interconnected than other neurons (Colizza et al., 2006). Secondary neurons also tend to be further from input and output neurons (Figure 51C), and signals received by them take longer to leave their originating neuropil (Figure 51D). Overall, earlier born neurons seem to play more central and global roles than their later born counterparts.

**Figure 51.**
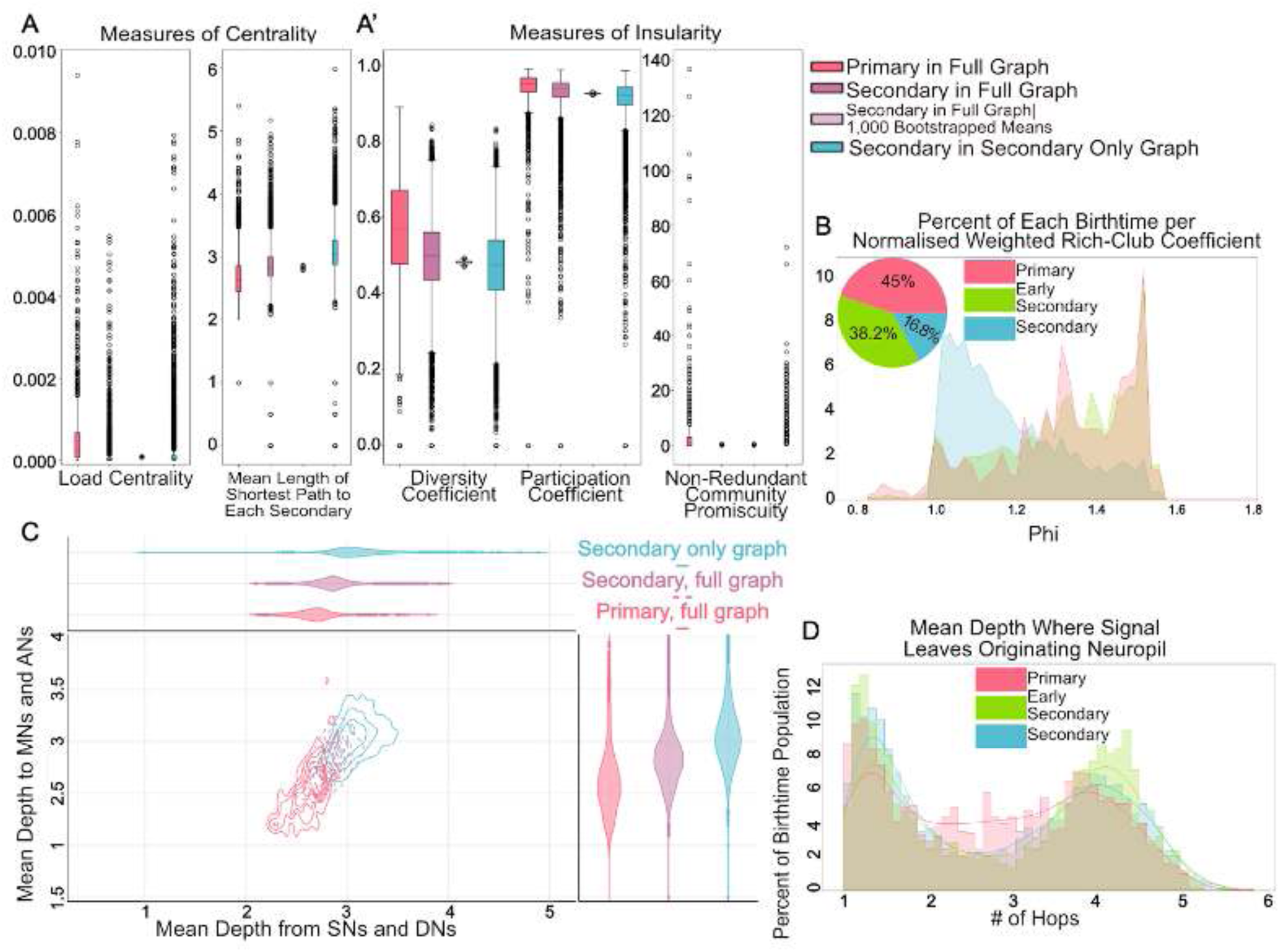
Roles in the VNC network differ by birthtime. **A-D**. Data is shown only for VNC-origin neurons in T1, T2 and T3. **A-A’.** Key measures of centrality and insularity within the network, compared between primary and secondary neurons in the full graph, as well as in a network excluding primary intrinsic neurons. One community is defined as all neurons of a hemilineage with a soma in a hemineuromere (e.g. all neurons identified as hemilineage 19A that have somas in the left side of T2 would be one community). **A.** Key measures of centrality. **A’.** Key measures of community insularity and diversity. Higher values suggest a higher variety of connection to communities. Non-redundant community promiscuity is a measure of how essential an individual neuron is in connecting its community to others in the graph. Primary neurons are uniquely responsible for connecting their respective communities to 1 other community (on median). **B.** Normalised weighted rich-club coefficient across birthtimes. Inset shows the percentage occurrence of the three birthtimes in the rich-club. **C.** Contour plot of depth from and to input neurons (all SNs and DNs) and output neurons (all MNs and ANs), respectively, calculated via a breadth-first search. **D.** Distribution of mean depth where a received signal leaves the neuropil at which it originated. Higher values indicate a signal is likely to spend more time in its respective neuropil before exiting. Data shown here were measured via iGraph (Csardi et al., 2006) and NetworkX (Hagberg et al., 2008) graph implementations of the VNC, based on either the full dataset or a subset of the dataset without primary intrinsic neurons (referred to as “secondary only graph” in the figure).

#### Sensory Neurons

Except for a small number of embryonically produced sensory neurons (Tix et al., 1989; Williams and Shepherd, 1999) adult *Drosophila* sensory systems are generated during postembryonic life. In contrast to central neurons, sensory neurons are born peripherally from cell populations set aside in the embryo that are the precursors of the adult appendages and epidermis (Bate and Arias, 1991). As these imaginal tissues grow, the sensory neurons emerge, *in situ*, in precise and stereotyped patterns (Nottebohm et al., 1994; Palka et al., 1984). Axons from sensory neurons grow into the VNC via specific nerves to develop synaptic connections with their targets (Jan et al., 1985).

In common with other insects, the sensory projections in the *Drosophila* VNC neuropil are highly organised. Sensory neurons from different classes of sensilla and serving different sensory modalities project into different regions of neuropil to establish connections (Merritt and Murphey, 1992; Murphey et al., 1989a); (Jan et al., 1985). Within these modality-specific regions, a more intricate order becomes apparent. Tactile afferents, for example, form a somatotopic map of the body surface (Murphey et al., 1989b), and chordotonal neurons encoding different features of limb movement, projecting to specific regions of the proprioceptive layers (Mamiya et al., 2018). This systematic arrangement facilitates the identification of sensory neuron modalities based on their afferent projections within the VNC.

#### Systematic typing of sensory neurons

Whilst light level images have been published for many sensory neurons, these typically label either large populations (making it difficult to distinguish between distinct types) or randomly labelled single cells (which may not represent all distinct types). Moreover, many sensory neurons in MANC were difficult to reconstruct completely due to dark staining and/or physical damage, particularly in the leg nerves. Consequently, our emphasis shifted towards developing a robust classification system for reliable type assignment, prioritising reliability over fine distinctions between cell types.

We used a variety of features for the systematic classification of sensory neurons. First, we annotated the peripheral nerve by which each neuron entered the VNC and pooled neurons according to peripheral origin; for example, neurons from the ProLN_R, DProN_R, VProN_R, and ProAN_R were all inferred to originate in the right front leg. We then clustered the neurons of each origin with respect to synaptic input and output partners (considering both individual neuron partners and aggregated hemilineages), morphological distances (using NBLAST), and graph traversal metric (Figure 52A). We integrated these various features using a weighted nearest neighbour (WNN) approach (Hao et al., 2021) and identified neuron clusters within each nerve (Figure 52Bi; see Methods for more details).

**Figure 52.**
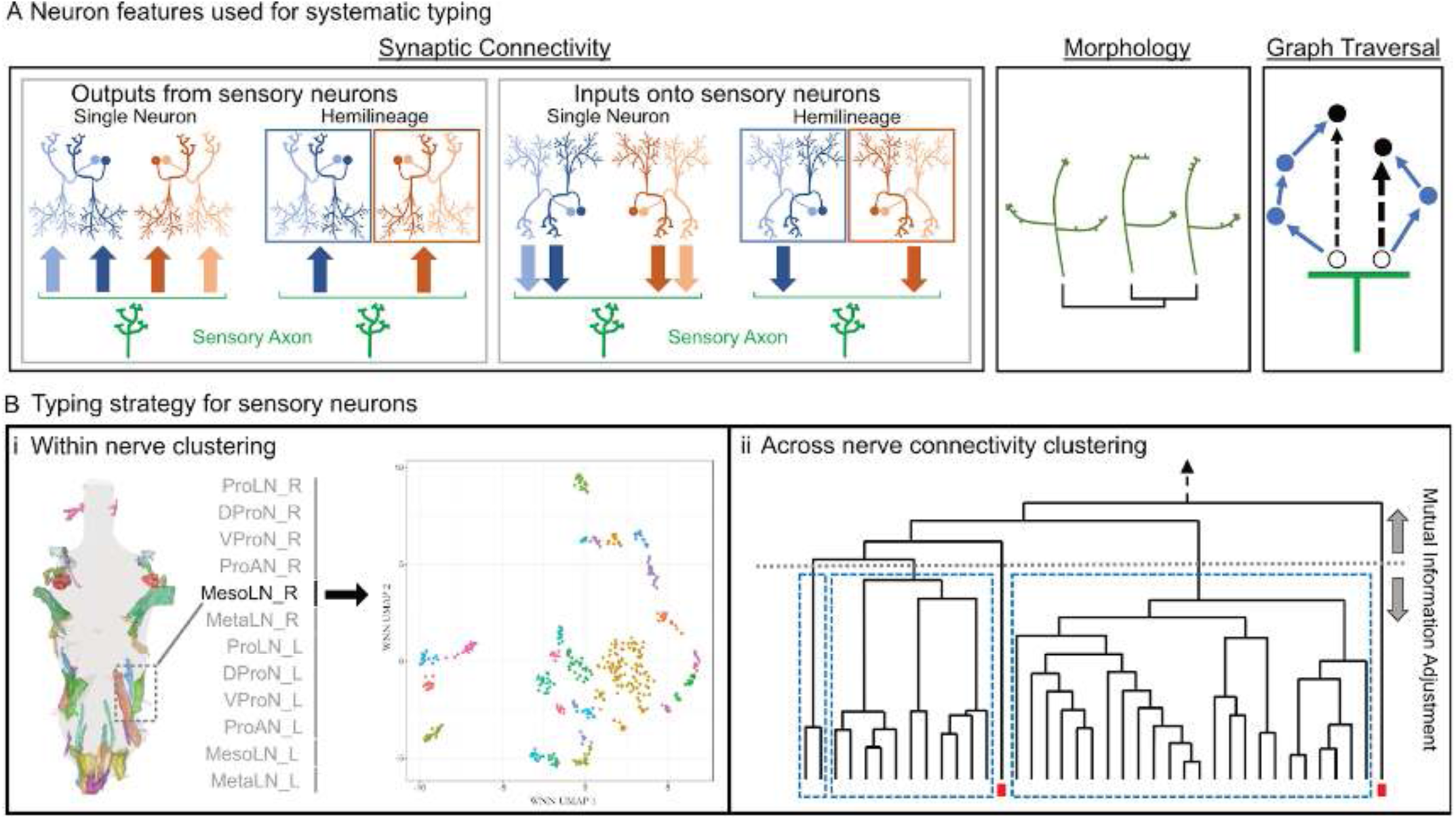
Systematic typing of reconstructed sensory neurons. **A.** Neuron features utilised for sensory neuron typing with a weighted nearest neighbour (WNN) approach included synaptic connectivity (synaptic outputs to individual neurons, synaptic outputs aggregated by hemilineage, synaptic inputs to individual neurons, synaptic inputs aggregated by hemilineage), morphology (inter-sensory neuron NBLAST scores), and graph measures (graph traversal distances). **B.** Depiction of typing process. **i.** Sensory neurons were initially clustered for each peripheral origin individually and individual nerve clusters were detected using HDBSCAN from a UMAP representation of the WNN output (Campello et al., 2013). **ii.** These clusterings were then utilised to inform a cut height of pan-region cosine clustering of synaptic connectivity onto serial neuron groupings.

In order to associate homologous cell populations across nerves, we pooled neurons from each of the three main peripheral regions - leg, dorsal, and abdomen - and used the cosine distances of their outputs to VNC neurons in annotated serial sets to generate a hierarchical clustering of these neurons. From this, we then defined serially homologous cell types by determining a cut height with which to separate neurons into clusters through partial maximisation of the mutual information of individual nerve clusterings derived previously (Figure 52Bii). These connectivity clusters (annotated in the **subcluster** field) were then manually reviewed to remove obvious outliers and to separate morphologically distinct subpopulations.

Final sensory cell types were assigned a prefix indicating whether they ascend to the neck connective (SA) or not (SN) and an abbreviation based on their inferred modality - “ch” for chemosensory, “pp” for proprioceptive, “ta” for tactile, or “xx” for unknown. Each distinct type within a modality was given a unique number; neurons that could not be classified as a specific type were assigned “xx” as their number. We see high intratype connectivity for sensory neurons (Figure 52 - figure supplement 1), as expected based on previous connectomics work (e.g., on the antennal lobe thermosensory neurons (Marin et al., 2020)). Significant intertype connectivity is also evident (Figure 52 - figure supplement 1,2), and this could be ascribed either to a continuous spectrum rather than discrete types, to functionally related types, or to noise level connectivity due to types targeting similar regions in space.

**Figure 52 - figure supplement 1.**
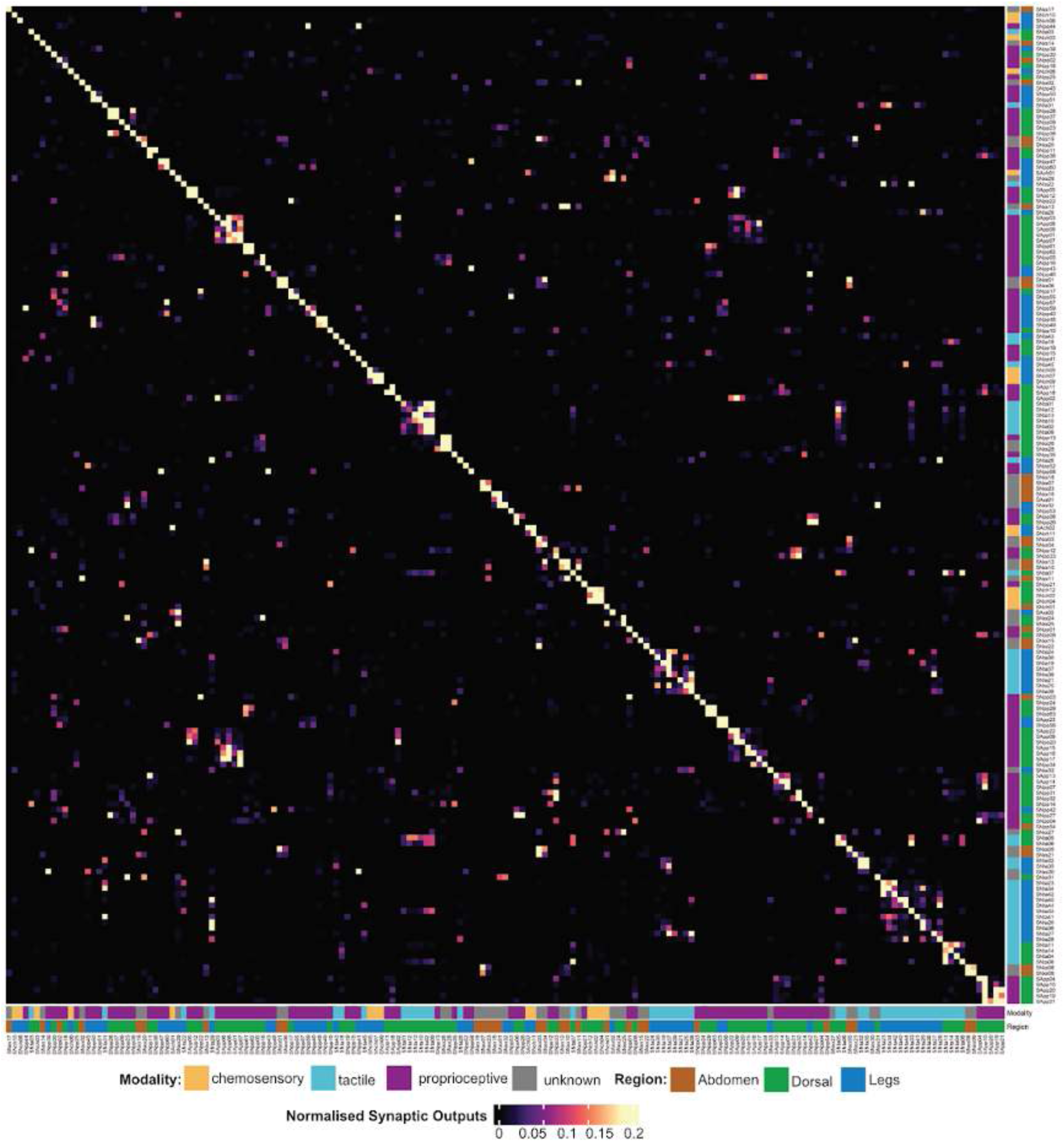
Sensory cell type interconnectivity. Annotation bars indicate modality and peripheral region for each type. Rows are the upstream systematic type, with columns being the downstream systematic type.

**Figure 52 - figure supplement 2.**
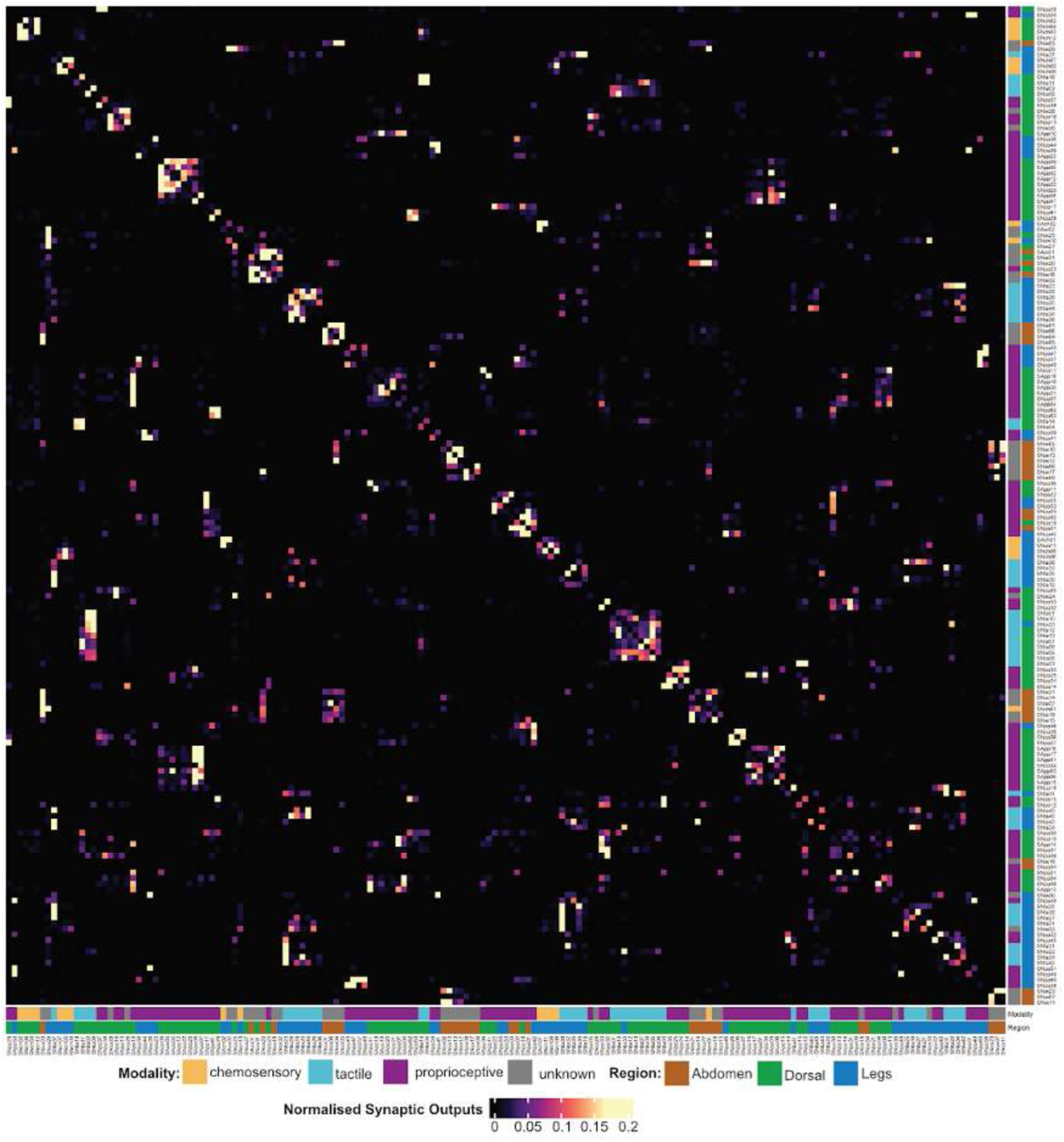
Sensory cell type interconnectivity, excluding intra-type connectivity. Annotation bars indicate modality and peripheral region for each type. Rows are the upstream systematic type, with columns being the downstream systematic type.

#### Organisation by Modality

Within the VNC, sensory neurons target specific regions of neuropil depending upon the modality of the information they convey (Merritt and Murphey, 1992; Murphey et al., 1989a; Tsubouchi et al., 2017). Chemosensory neurons, primarily from gustatory sensilla (taste), terminate in the ventral most layer of the VAC, followed by tactile neurons from external sensilla (bristle contact), then several classes of proprioceptive neurons: stretch receptors (stretch, tension), chordotonal organs (tension, vibration), and campaniform sensilla (deformation). Utilising this knowledge together with light-level images from the literature (see Methods), we classified 1211 neurons as **chemosensory** (responsive to chemical stimuli), 1347 as **proprioceptive** (providing information as to the location and movement of the body), and 2497 as **tactile** (responsive to touch) (Figure 53A-B). We were unable to assign a further 1422 neurons to any of those three modalities; these were annotated as modality “unknown”, but a subpopulation was later identified as heat nociceptive (Figure 55).

**Figure 53.**
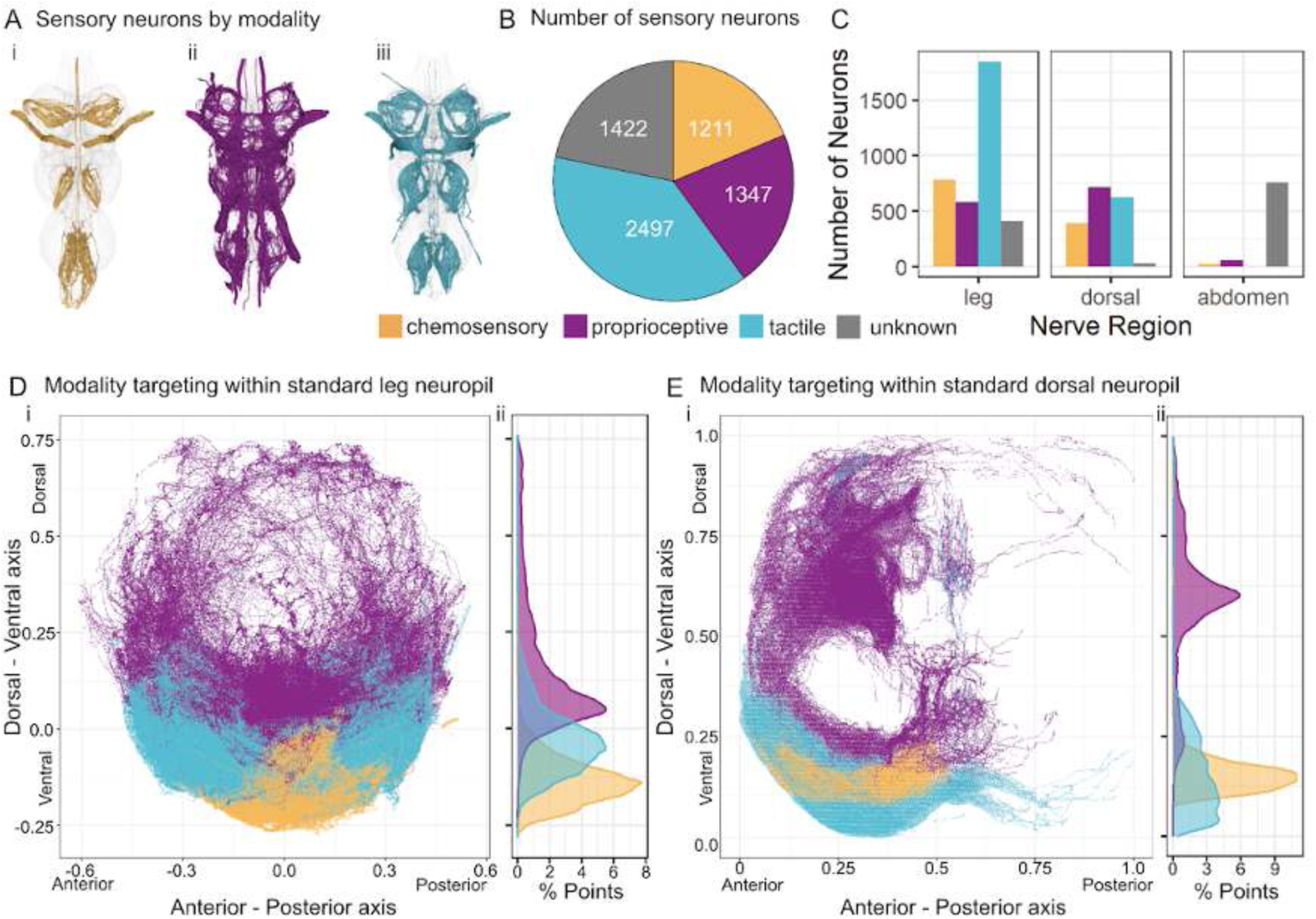
Overview of sensory neurons by modality. **A.** Plotted meshes of i. chemosensory neurons, ii. proprioceptive neurons, and iii. tactile neurons. **B.** Quantification of sensory neurons by assigned modality. C. Quantification of sensory neurons by modality, faceted by peripheral origin. **D.** i. Standard leg neuropil depiction, see Methods, of the regions innervated by sensory neurons of each modality. ii. Histogram showing the relative distribution of neuron innervation through the dorsal ventral axis. **E.** i. Standard dorsal neuropil depiction, see methods, of the regions innervated by sensory neurons of each modality. ii. Histogram showing the relative distribution of neuron innervation through the dorsal ventral axis.

The relative proportions of the different modalities vary according to the peripheral region (Figure 53C). For example, sensory neurons entering via the leg nerves have a far higher proportion of tactile neurons, whereas sensory neurons entering via the dorsal nerves represent the three modalities more equally. The sensory neurons entering via the abdominal nerves account for the majority of the neurons of unknown modality, due to a limited published literature for abdominal sensory neurons in the adult.

We created a standard leg neuropil cross-section in order to visualise neurons from all six legs in a single, consistent space (see Methods); this shows the relative placement of the afferent projections of the different sensory modalities in modality-specific layers (Figure 53D). The leg chemosensory neurons project to the central, ventral-most neuropil, tactile neurons spread over the adjacent ventral neuropil much more broadly, and the proprioceptive neurons arborise in the more dorsal regions of leg neuropil but with significant variations dependent on the specific type of proprioceptive organ (see below).

A comparable projection for the dorsal neuropil (Fig 53E) reveals a slightly different organisation, with the chemosensory neurons forming a coherent core of afferent axons within the tactile projections from the notum and wing, rather than being layered directly ventral to the tactile neurons as in the leg neuropils. These patterns in leg and dorsal neuropil are consistent with the layering previously reported from light level microscopy (Tsubouchi et al., 2017).

#### Downstream Sensory Targets

Many central neurons in the VNC are involved in the primary processing of sensory information, receiving direct synaptic input from sensory neurons. Using a threshold of 30% of total connectivity received from sensory neurons (Figure 54 - figure supplement 1), based on studies of sensory inputs to projection neurons in the antennal lobe (Schlegel et al., 2021), we estimate that ∼20% of neurons in the VNC (excluding sensory neurons) could be considered second order neurons involved primarily in sensory processing (Figure 54B). However, we find that the percentage of sensory processing neurons varies by class: 20% of INs, >25% of ANs, and <15% of DNs (Figure 54B). We also note that these are lower limits, given that we were not able to reconstruct all sensory neurons, particularly in the leg neuropils.

**Figure 54.**
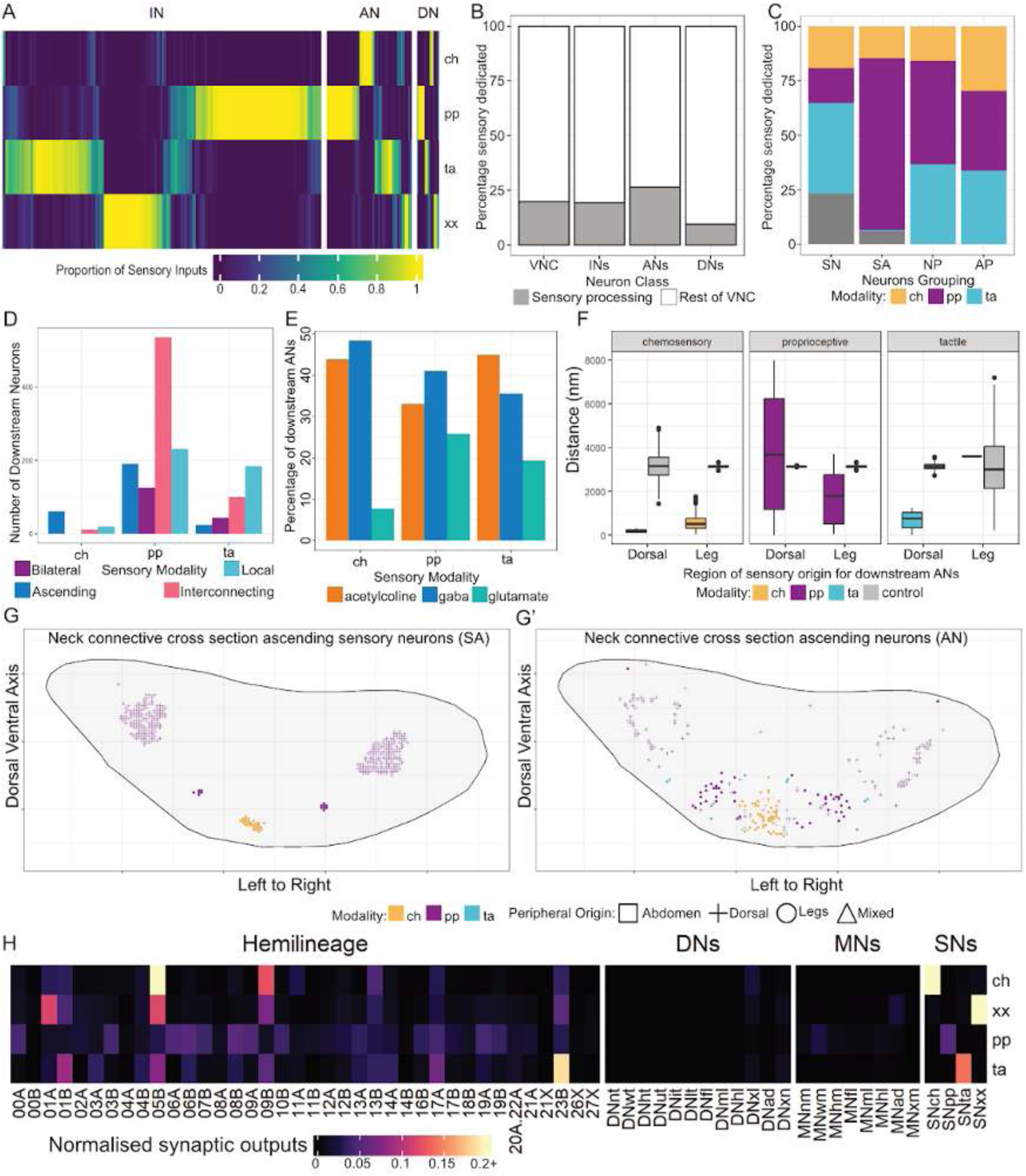
Overview of sensory neuron targets by modality. **A.** Proportion of inputs from sensory modalities to neurons implicated in direct sensory processing. **B.** Percentage of neurons from each class identified as directly sensory processing. VNC neurons = all VNC neurons (excluding sensory neurons), INs = intrinsic neurons, ANs = ascending neurons, and DNs = descending neurons. **C.** Proportion of neurons assigned to each modality for SN, sensory neurons, SA, sensory ascending neurons, NP, non ascending modality specific partners, AP, ascending modality specific partners. **D.** Number of downstream partners of each modality aggregated by expanded subclass: ascending (ANs), bilateral (connecting both hemispheres of the VNC), interconnecting (projecting across multiple neuromeres within a single hemisphere), and local (restricted to the hemi-neuromere associate to the entry nerve). **E.** Percentage breakdown for each modality of predicted neurotransmitter for ascending neurons implicated in direct sensory processing. **F.** Distance within a neck connective plane between neurons of the same region and modality vs. a randomised control of all ascending neurons. **G.** Cross-section of the neck connective displaying the positioning of ascending sensory neurons. Coloured by modality. **G’.** Cross-section of the neck connective displaying the positioning of ascending partners of sensory neurons. Coloured by modality. **H.**Heatmap showing normalised sensory neuron modality connectivity to hemilineage, DN groups, MN groups, and SN modalities in the VNC. Connectivity to neurons of undetermined hemilineage (TBD) have been omitted.

**Figure 54 - figure supplement 1.**
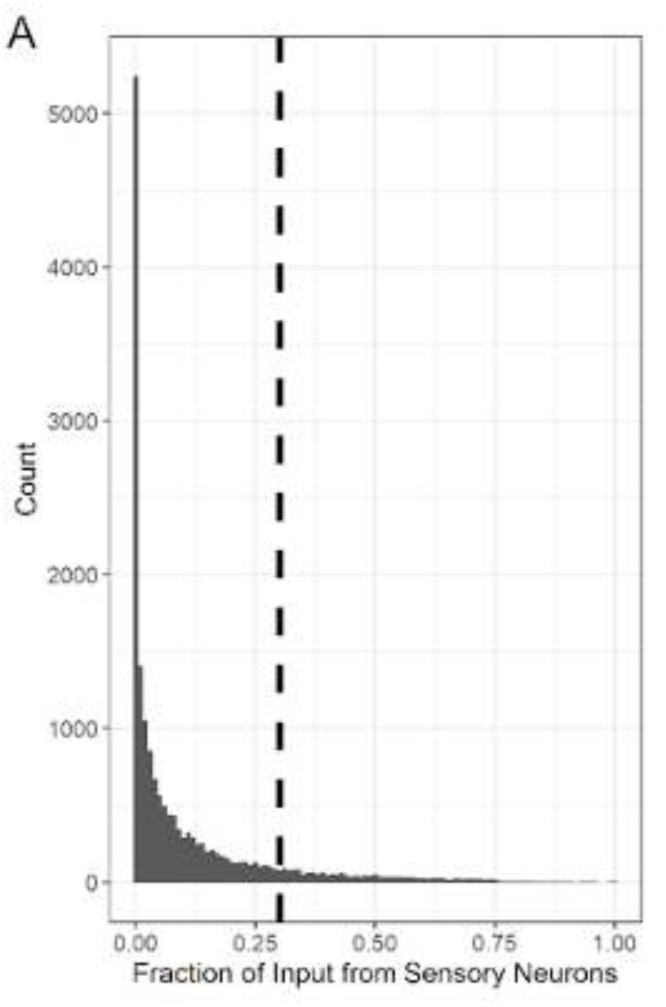
Sensory neuron targets. Histogram of proportion of inputs to VNC neurons from sensory neurons. Dashed line represents the cutoff (0.30) used for classifying neurons as implicated in direct sensory processing.

**Figure 54 - figure supplement 2.**
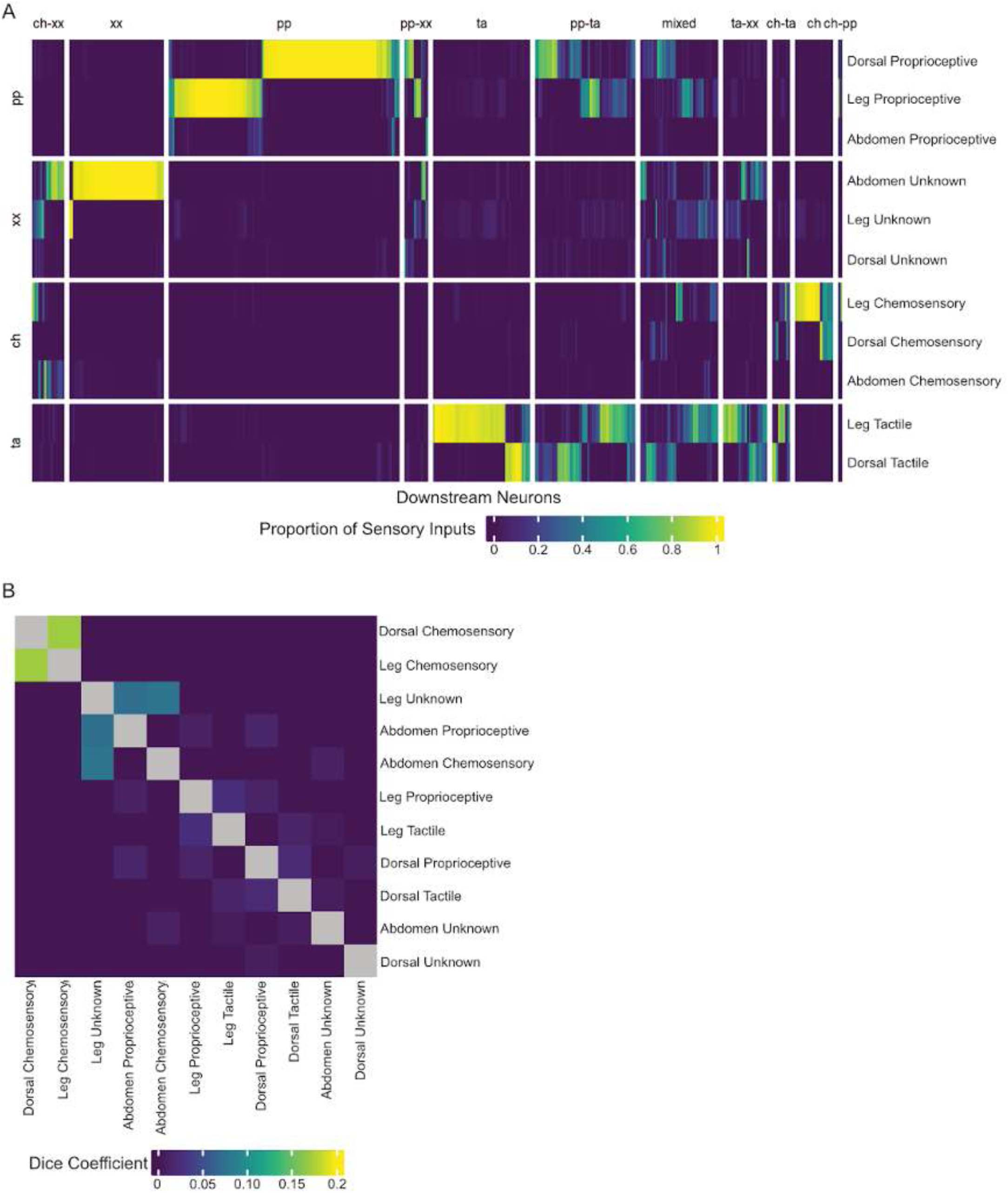
Sensory neuron targets by modality. **A.** Heatmap showing relative specificity of sensory associated downstream neurons with each modality within each region, normalised by total inputs for each downstream neuron. **B.** Dice coefficient for downstream modality within each region.

**Figure 54 - figure supplement 3.**
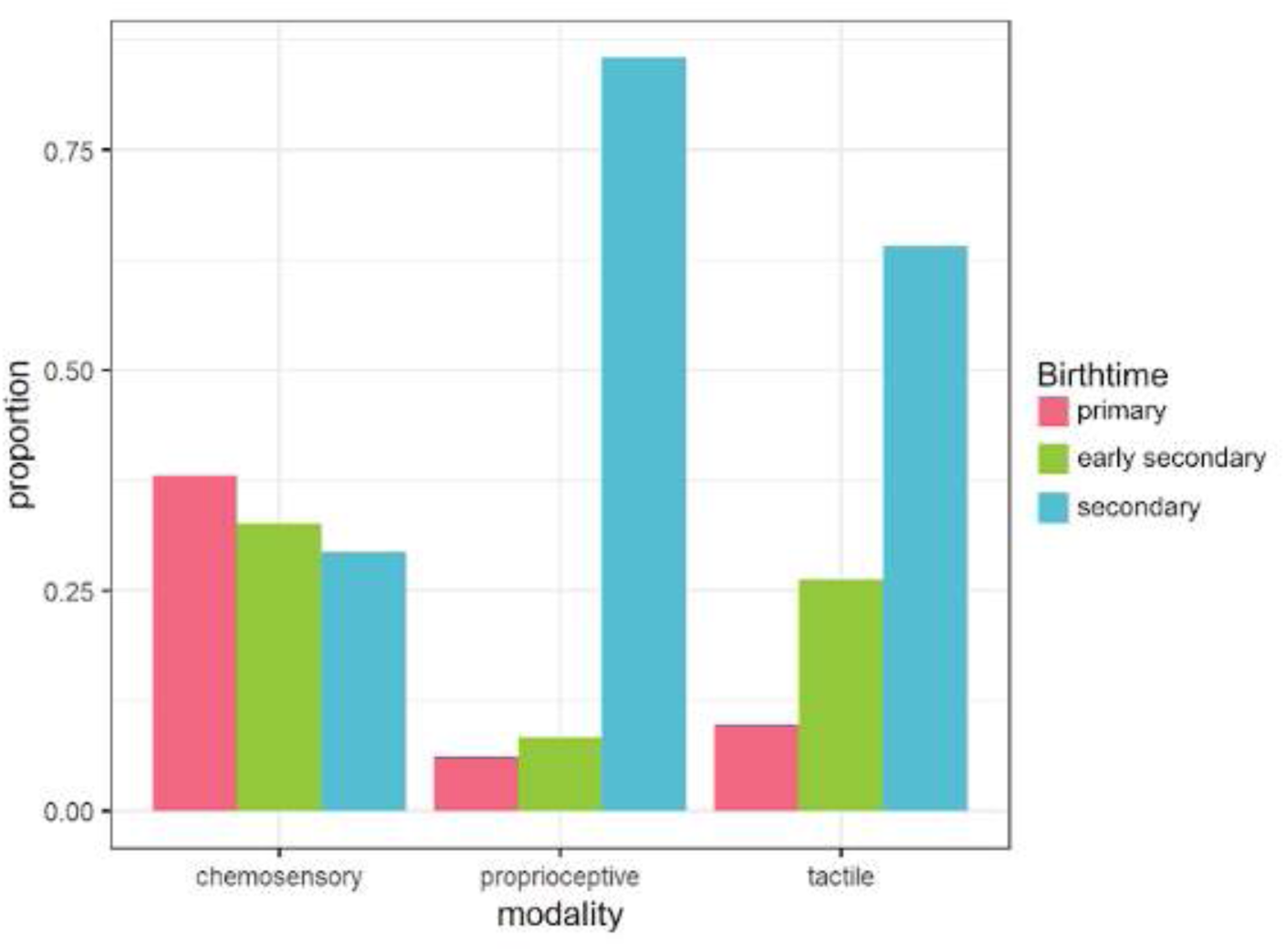
Proportion of downstream ascending neurons of each birthtime associated with each modality. Pink = primary, green = early secondary and cyan = secondary.

**Figure 54 - figure supplement 4.**
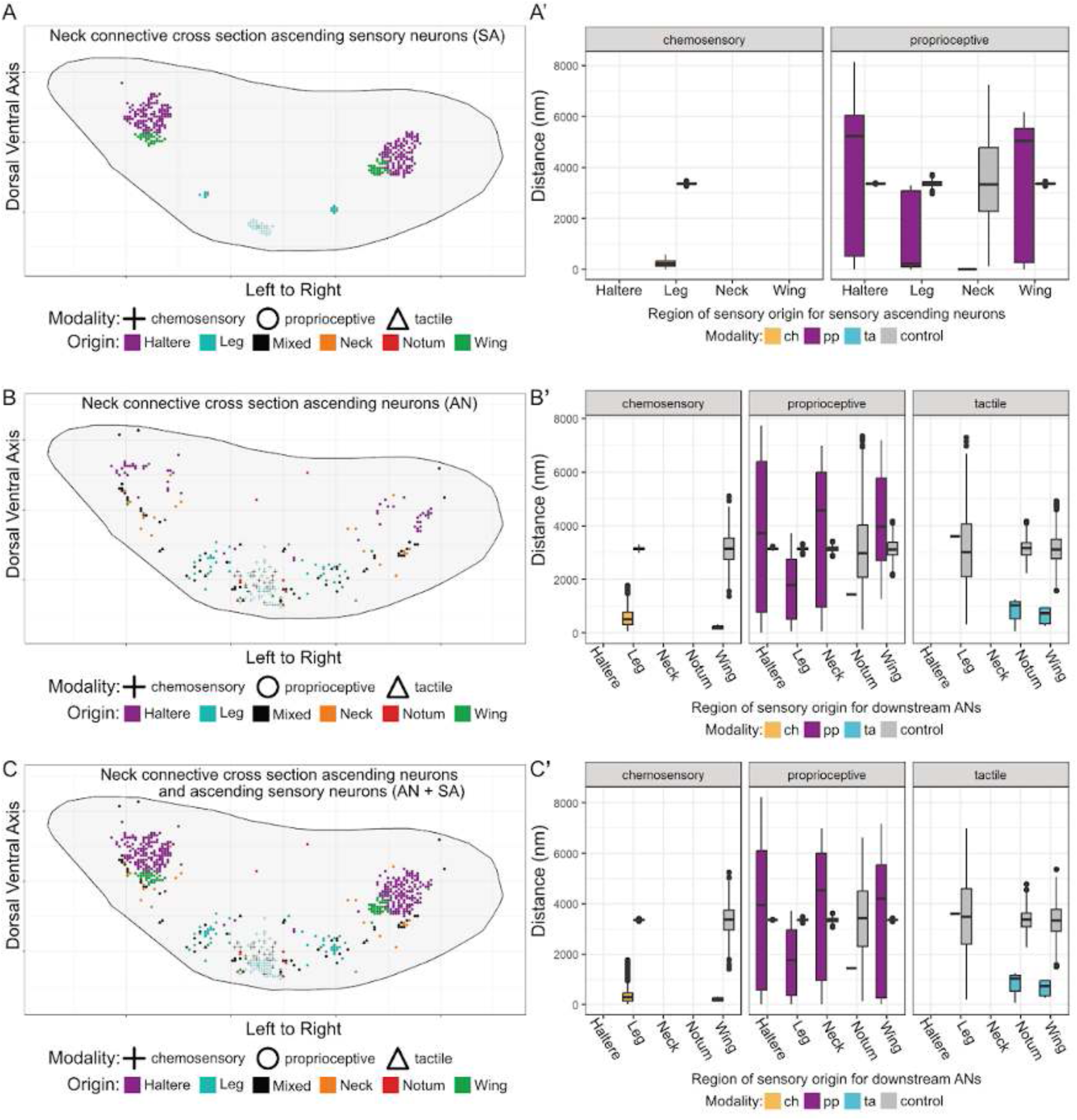
Targeting of sensory ascending neurons and ascending neurons within the neck connective. **A.** Sensory ascending neuron targeting within the neck connective, coloured by region of peripheral origin. **A’.** Distance within the plane between neurons of the same region and modality vs. a randomised control of all sensory ascending neurons. **B.** Ascending neuron targeting within the neck connective, coloured by region of peripheral origin. **B’.** Distance within the plane between neurons of the same region and modality vs. a randomised control of all ascending neurons. **C.** Sensory ascending neuron and ascending neuron targeting within the neck connective, coloured by region of peripheral origin. **C’.** Distance within the plane between neurons of the same region and modality vs. a randomised control of all sensory ascending neurons and ascending neurons.

We find that 70.4% of the second order neurons can be considered to be specific to a single sensory modality (excluding unknown modality) (Figure 54A). However, the rest integrate two or more modalities, most commonly proprioceptive and tactile (Figure 54 - figure supplement 2). All second order sensory processing neurons have been annotated in the **modality** field with the associated one or two modalities or as “mixed” if appearing to integrate all three. We also find that, whilst most second order neurons receive input only from one region (leg, dorsal, or abdominal), the chemosensory inputs from the wings are integrated with a subset of those from the legs (Figure 54 - figure supplement 2).

We next looked at distributions of modality across sensory neurons and their downstream partners. Whilst most sensory neurons are local (SN), terminating within the VNC, a small proportion (535/6461) ascend to the SEZ and brain (SA). These different classes of sensory neuron also show differences in modality dominance, with local sensory neurons being predominantly tactile whereas the sensory ascending neurons are predominantly proprioceptive (Figure 54C). Despite this difference, the downstream non-ascending partners (NP) of both types of sensory neuron in the VNC feature equal representation of tactile and proprioceptive information and a lower proportion of chemosensory inputs from local sensory neurons (Figure 54C). In contrast, there is greater equality in the number of downstream ascending neurons (AP) devoted to each modality.

Furthermore, different sensory modalities are processed by second order neurons from different birthtimes (Figure 54 - figure supplement 3). Chemosensory neurons target a much higher proportion of early born partners than either tactile or proprioceptive sensory neurons, which may reflect conservation of information content and/or valence across life stages. In contrast, tactile and especially proprioceptive sensory neurons have higher proportions of secondary neuron targets, possibly related to the development of novel appendages for the adult with their peripheral sensory structures.

Further breakdown of the downstream partner classes into coarse subclasses reveals additional diversity in downstream neuron targets (Figure 54D). The primary downstream subclass of proprioceptive neurons are neurons that interconnect thoracic neuromeres, suggesting a role in segmental coordination, but there are substantial bilateral and ascending populations as well. In contrast, chemosensory downstream targets are far more likely to be ascending, with few local or interconnecting examples, suggesting that the majority of chemosensory processing occurs in the brain. In contrast, tactile target neurons are typically local, with additional interconnecting and bilateral populations but with the lowest proportion of ascending partners, supporting a role in reflexive rather than directed behaviours.

Downstream ascending neurons of each modality are predicted to express roughly the same proportions of acetylcholine (excitatory) vs gaba (inhibitory), with slight biases towards gaba for chemosensory and proprioceptive and towards acetylcholine for tactile (Figure 53E). Chemosensory ascending neurons are notably less likely to express glutamate; however, the effects of glutamate depends on receptor expression in downstream partners, which cannot be determined here.

Ascending sensory information appears to be organised by modality, with ascending sensory neurons of each modality retaining close association with the downstream ascending partners of that modality as they pass through the neck connective (Figure 54F). The pattern of organisation seen in the synapses of each modality in the leg neuropil (Figure 53D) appears to be largely maintained by ascending sensory and partner neurons’ axons in the neck connective, with chemosensory axons near the ventral midline, tactile axons adjacent, and proprioceptive axons more dorsal (Figure 54G-G’). These ascending sensory and partner neurons’ axons are also organised by peripheral origin, with neurons transmitting information from the leg and notum ascending in the ventromedial neck connective and neurons transmitting it from the neck, wing, and haltere ascending in progressively more dorsal positions (Figure 54 - figure supplement 4). It will be interesting in future to see whether and how the organisation of descending neurons reflects the sensory information they carry.

Finally, we see modality-specific patterns of connectivity when we break down second order neurons by hemilineage or subclass (Figure 54G). For example, chemosensory neurons mainly target hemilineages 05B and 09B, whilst tactile neurons target 01B, 05B, 09B, and especially 23B. Proprioceptive neurons exhibit a broader pattern of connectivity, possibly due to a higher number of discrete peripheral origins. Neurons of unknown modality have a distinct profile of hemilineage connectivity, likely because 1) few neurons in the abdomen (where the majority of unknown sensory neurons originate, Figure 53C) have been assigned to a hemilineage, 2) true connectivity is poorly represented due to incomplete reconstruction, and 3) multiple modalities are represented. Sensory neurons also exhibit a very high degree of intra-modality and low degree of inter-modality connectivity, presumably due to the aforementioned intratype connectivity, as was previously observed in the antennal lobe connectome (Marin et al., 2020).

#### Leg Sensory Types

Sensory neurons from the front legs enter the prothoracic neuromere via four different nerves (ProLN, DProN, VProN, and ProAN), whilst those from the middle and hind legs enter the mesothoracic via the MesoLN and the metathoracic via the MetaLN, respectively (Figure 55A). As described above (Figure 52), neurons from each leg-associated nerve were clustered and then associated across leg nerves by connectivity to previously defined serial sets in order to define serial leg sensory types. Degradation and dark staining resulted in significant variance in the number and quality of sensory neurons recovered in each nerve: the fewest were recovered from the front legs, whilst in general more neurons were recovered on the right side than on the left (Figure 55B). Nevertheless, we successfully reconstructed the majority of leg sensory neurons, surpassing previous efforts (Phelps et al., 2021), and identified 63 distinct cell types (9 chemosensory, 22 proprioceptive, 27 tactile, and 5 of unknown modality). Here we highlight only a subset of these types, but reconstructions and connectivity heatmaps are provided for all (Figure 55 - figure supplement 1-4).

Broadly speaking, leg sensory neurons mainly target intrinsic neurons (∼67%), followed by ascending neurons (∼22%) and each other (∼9%), with other classes receiving 1% or fewer of leg sensory outputs (Figure 55C). This is a significantly smaller proportion of ascending neuron targets than observed for dorsal sensory neurons, perhaps suggesting greater involvement in reflexive sensorimotor circuits that do not require communication with the brain. Leg sensory neurons of each modality tend to target neurons of specific hemilineages: chemosensory neurons mainly target 05B, tactile neurons mainly target 01B and 23B, and proprioceptive neurons have more diverse targets but show a preference for 09A (Figure 55E).

**Figure 55.**
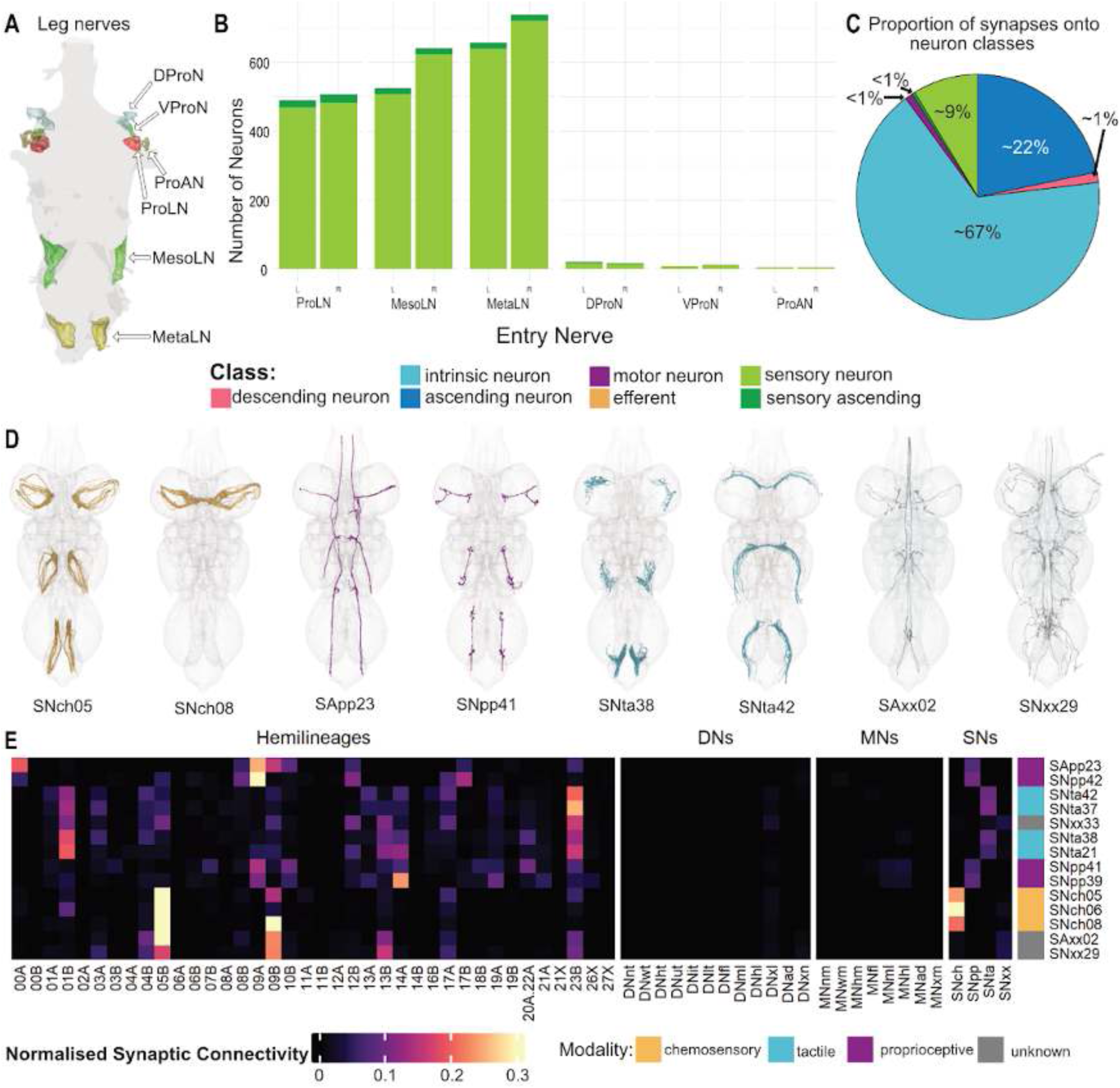
Leg sensory neuron systematic types. **A.** Anatomy of leg nerves (ventral view). **B.** Numbers of sensory neurons from each leg nerve. **C.** Proportions of downstream partners of leg sensory neurons, coloured by class. **D.** Plotted meshes of exemplar leg sensory types, coloured by assigned modality. **E.** Heatmap showing normalised sensory neuron cluster connectivity to hemilineages and other neuron subclasses in the VNC. Annotation bar displays annotated modality of component sensory neurons within the sensory neuron cluster.

**Figure 55 - figure supplement 1.**
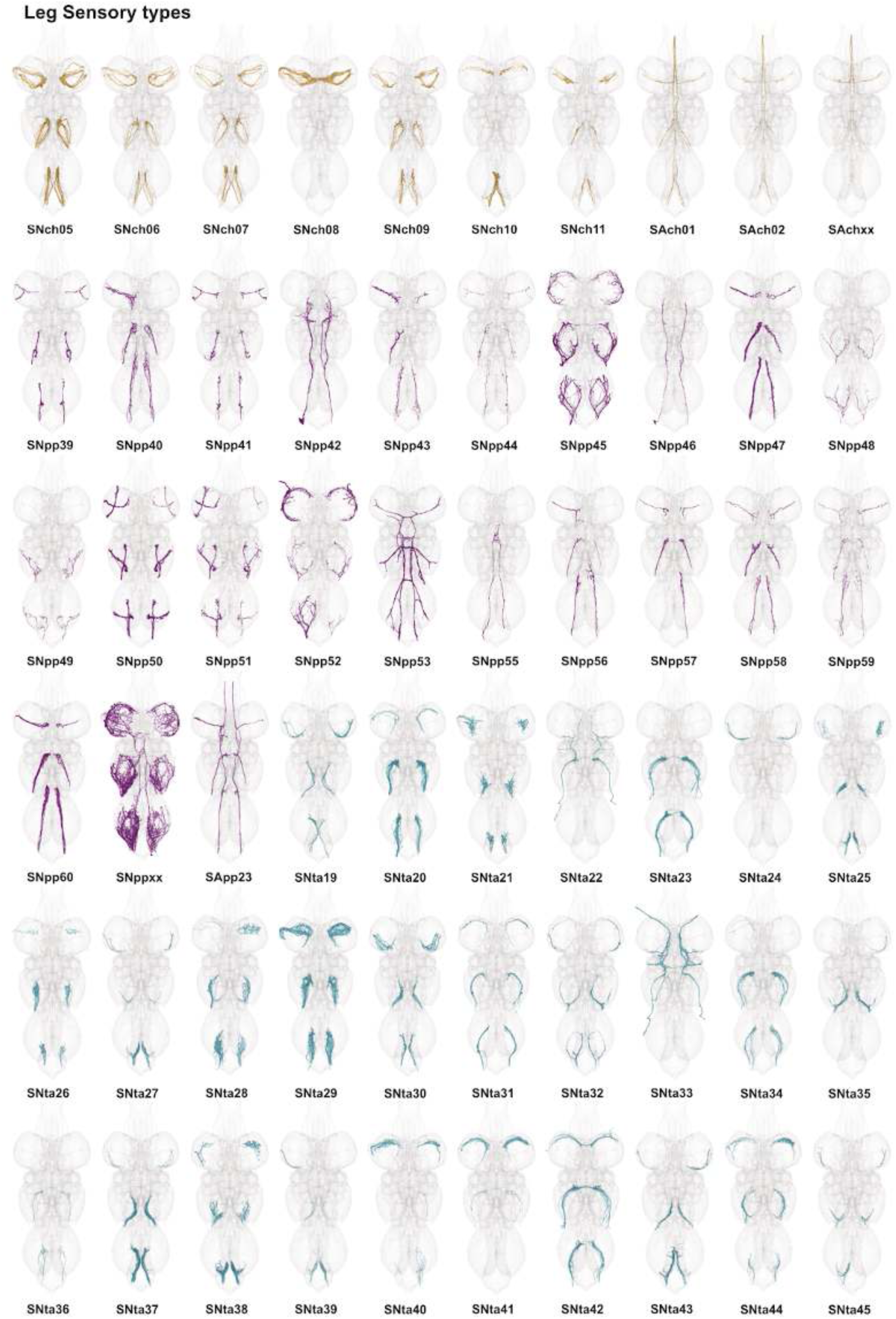
Leg sensory neuron types. Plotted meshes of neurons that comprise each systematic type from leg nerves, coloured by modality: orange = chemosensory, purple = proprioceptive, cyan = tactile, grey = unknown.

**Figure 55 - figure supplement 2.**
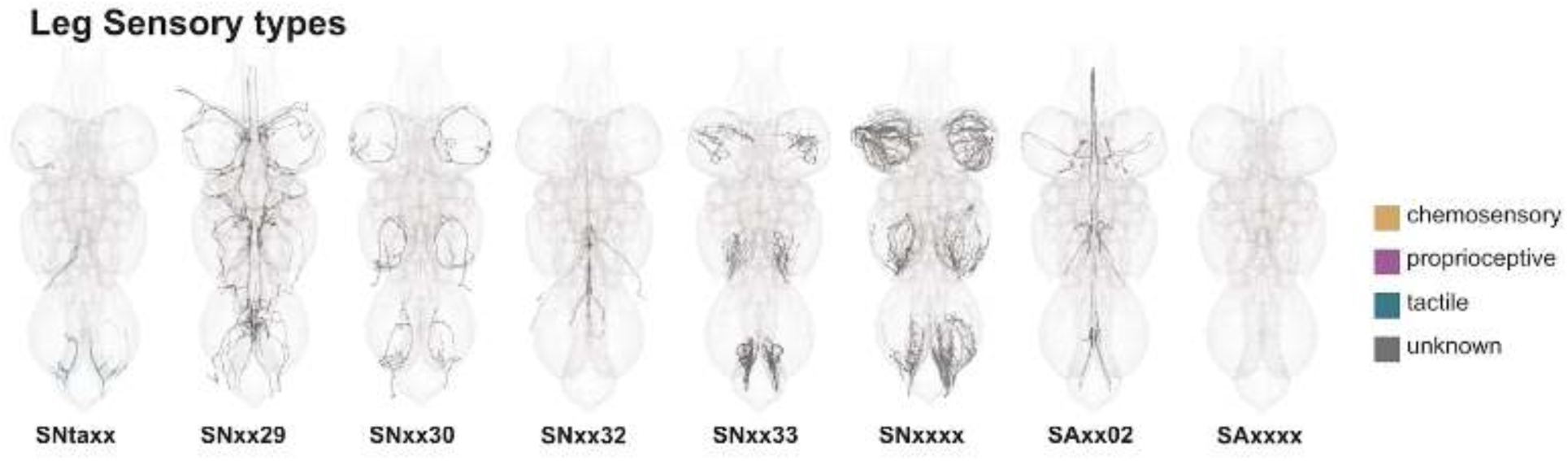
Leg sensory neuron types, continued. Plotted meshes of neurons that comprise each systematic type from leg nerves, coloured by modality: orange = chemosensory, purple = proprioceptive, cyan = tactile, grey = unknown.

**Figure 55 - figure supplement 3.**
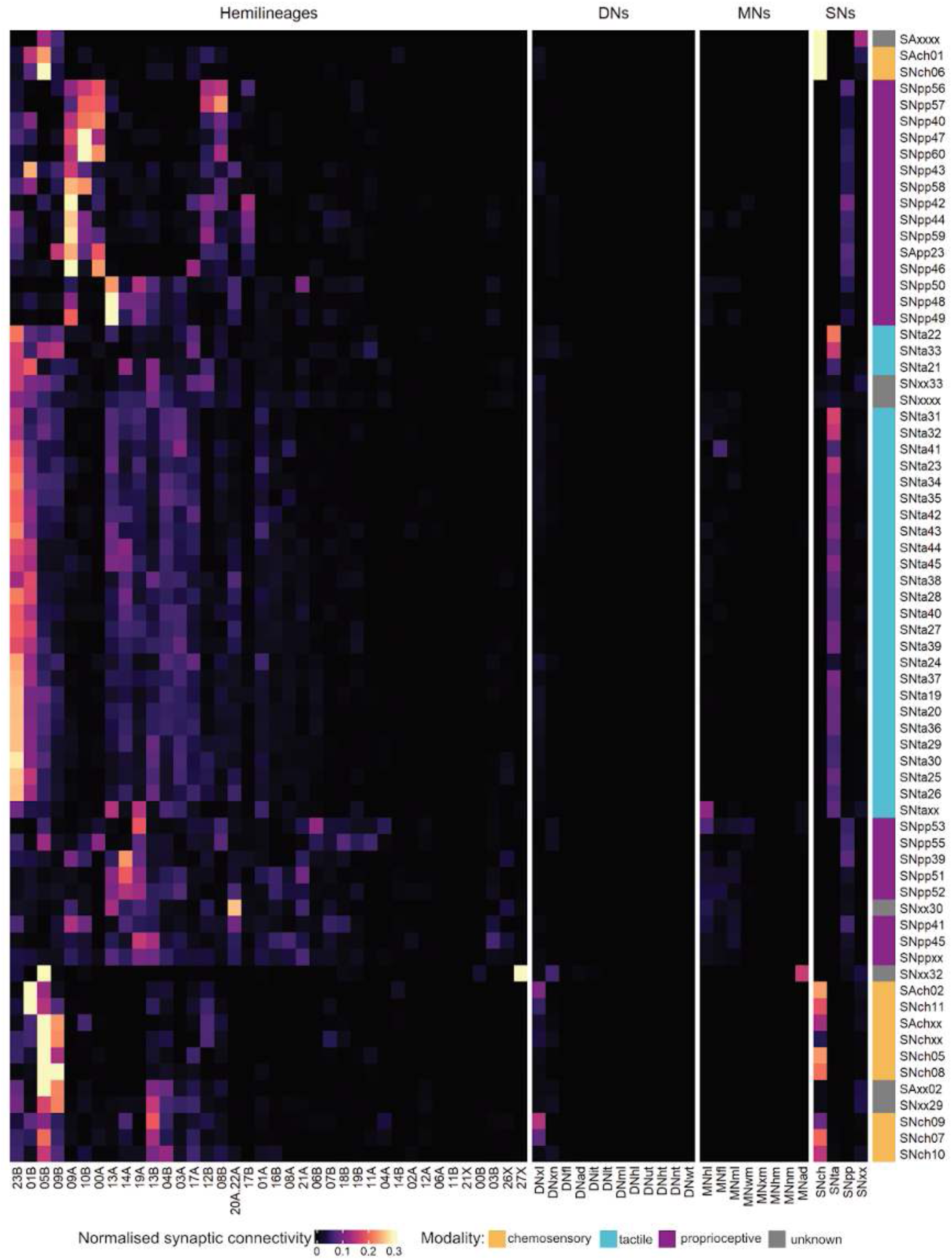
Leg sensory neuron type output connectivity by hemilineage. Right annotation bar is coloured by modality: orange = chemosensory, purple = proprioceptive, cyan = tactile, grey = unknown. Rows and columns are ordered based on hierarchical clustering of data.

**Figure 55 - figure supplement 4.**
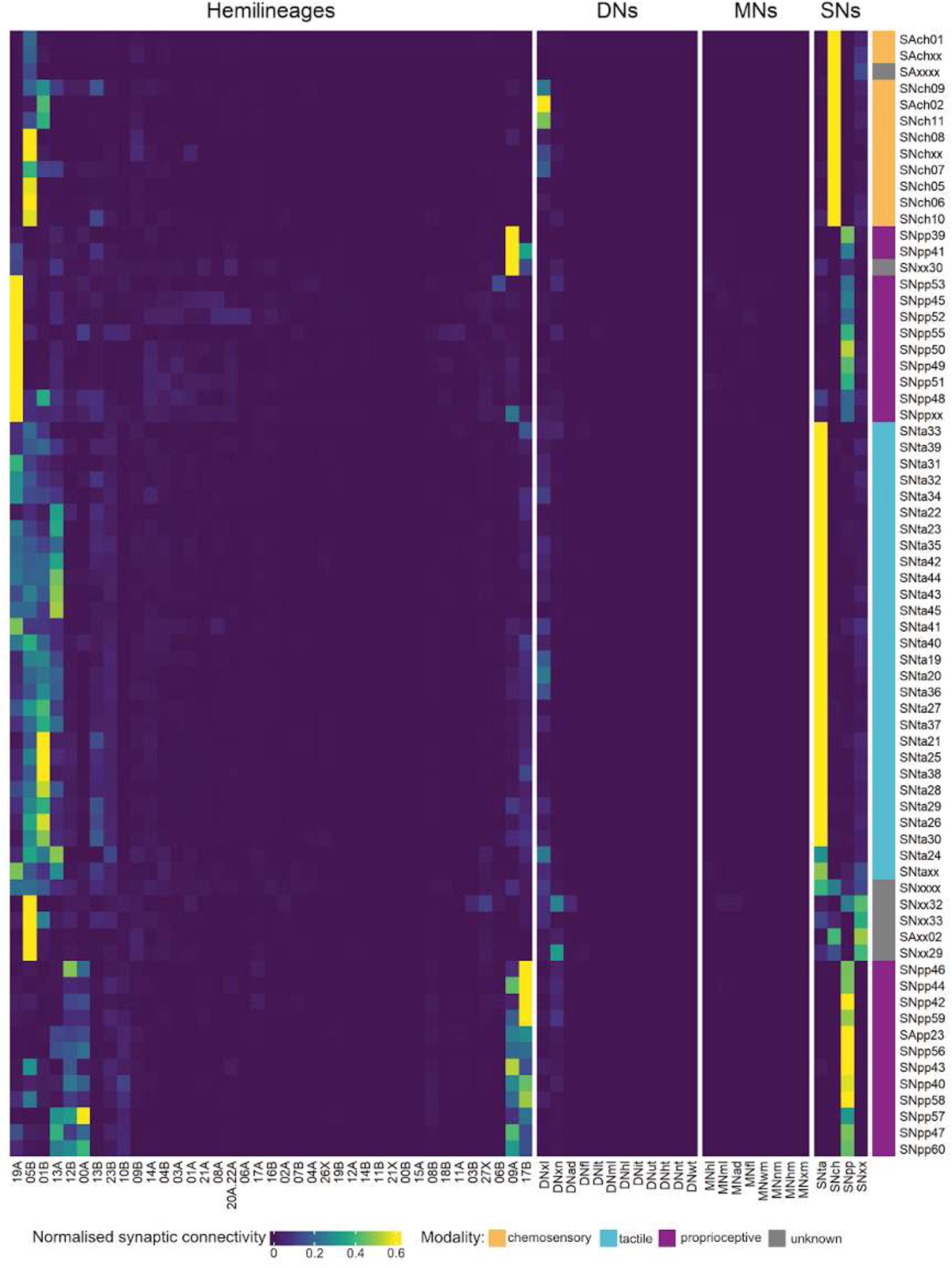
Leg sensory neuron type input connectivity by hemilineage. Right annotation bar is coloured by modality: orange = chemosensory, purple = proprioceptive, cyan = tactile, grey = unknown. Rows and columns are ordered based on hierarchical clustering of data.

**Figure 55 - figure supplement 5.**
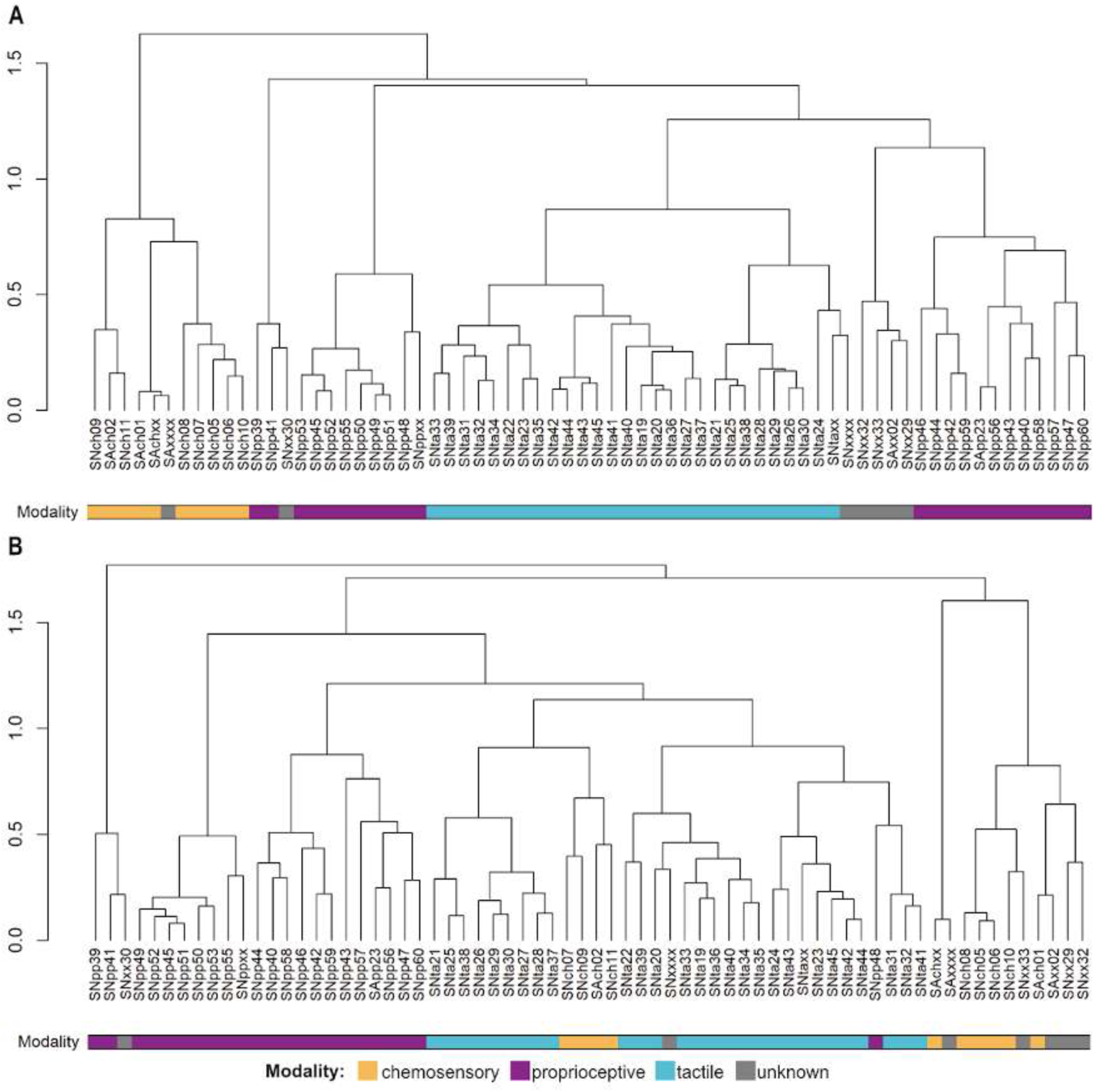
Leg sensory neuron type dendrograms showing combined inputs and outputs clustering. **A.** Dendrogram showing combined clustering with intra-type connectivity included. **B.** Dendrogram showing combined clustering with intra-type connectivity excluded. Bottom bar is coloured by modality: orange = chemosensory, purple = proprioceptive, cyan = tactile, grey = unknown.

##### Chemosensory leg sensory types

Four leg chemosensory types had been previously distinguished based solely on gustatory receptor expression (Hopkins et al., 2023), but more types are expected based on the observed diversity of responses to tastants. Here we separated the chemosensory neurons into 9 distinguishable leg sensory neuron types. 7 of these types originate in all three pairs of legs (e.g., SNch05) whilst two are specific either to T1 (SNch08) or to T1 and T3 (SNch10) (Figure 55 - figure supplement 1). We have matched SNch08 (Figure 55D) by morphology to a male-specific, fru+ cell type representing pheromone information (Yu et al., 2010).

We can identify three main clusters of chemosensory types based on their downstream connectivity (Figure 55 - figure supplement 3). The first is composed of SNch11 and SAch02, which target hemilineage 01B more strongly and have been identified as sweet (or appetitive) gustatory neurons from their characteristic morphology (Koh et al., 2014). A closely associated second cluster consists of types SNch05, SNch06, and SNch01 plus the male-specific type SNch08. The third cluster targets distinct downstream partners (particularly hemilineage 13B), consists of types SNch10, SNch07, and SNch09; and is closely associated with SNxx29 (Figure 55D), the ppk+, Gr28b.d+ neurons carrying heat nociceptive information (Khuong et al., 2019; Kwon et al., 2014) (S Rumpf, personal communication). Downstream connectivity therefore suggests that the first two clusters might represent appetitive valence while the third represents aversive valence. However, these groupings are not strictly maintained with respect to inputs to these types: the pheromone- and sweet-responsive types receive distinct inputs from inhibitory hemilineage 01B and DNxl neurons whilst the other types receive input mainly from inhibitory hemilineage 05B (Figure 55 - figure supplement 4). This may reflect different contexts and/or organismal states in which specific chemosensory information is suppressed or filtered.

##### Proprioceptive leg sensory types

We detect 22 proprioceptive leg sensory types (Figure 55 - figure supplement 1), of which 19 can be suggested to correspond to known cell types or functional classes, the majority to chordotonal organ types (see Figures 59 - 61 for more details). Numbers varied widely between hemineuromeres for some types (e.g., SNpp51 and SNpp52), most likely because proprioceptive neurons were susceptible to degradation and poor segmentation and there was regional bias in their reconstruction despite focused efforts. For similar reasons, ∼25% of proprioceptive neurons could not be assigned to a specific type and were annotated as SNppxx.

Proprioceptive leg sensory types exhibit a higher degree of inter-type variation in connectivity to downstream hemilineages than do chemosensory or tactile types (Figure 55 - figure supplement 3). Connectivity clustering suggests four main groupings of proprioceptive neurons: 1) 5 types with strong connectivity to hemilineages 00A and 10B; 2) types strongly downstream of inhibitory hemilineage 09A including SNpp39 and SNpp41 as well as SNxx30, which might therefore also be proprioceptive (Figure 55 - figure supplement 4); 3) a highly variable group with outputs to hemilineages 13A, 13B, 14A, 19A, and (weakly) leg motor neurons, and 4) two types with sparse connectivity biased towards dorsal hemilineages 00A, 06B, and/or 08B, suggesting a role in coordination between leg and other body movements.

##### Tactile leg sensory types

Classifying tactile sensory neurons into discrete types posed particular challenges. In part, this may be because the pattern of tactile axon terminals in the VNC is thought to represent a somatotopic map of the leg, which naturally lends itself to gradations rather than sharp boundaries. But poor segmentation quality also plays a part, as evidenced by the variation in number of recovered neurons of each type across sides and neuromeres. Due to technical difficulties, we cannot conclude that the absence of some tactile types in specific neuromeres (Figure 55 - figure supplement 1) is biologically relevant.

We observe three broad morphological categories of tactile neurons: intersegmental (SNta22 and SNta33), bilateral (SNta42) (Figure 55D), and local (constituting the vast majority of types and typically paired as anterior and posterior instances, please see Figure 62 for details). All types share ∼similar outputs, mainly onto hemilineages 01B and 23B (Figure 55 - figure supplement 3). SNta41 is a notable exception for its direct connectivity to motor neurons, mainly in the prothorax where most examples of this type were reconstructed.

##### Unknown leg sensory types

We identified five types of leg sensory neurons that we could not initially associate with any particular modality. First, SNxx29 has been identified as the ppk+, Gr28b.d-expressing heat nociceptive leg neurons (Khuong et al., 2019; Kwon et al., 2014). Intriguingly, the connectivity pattern of this type overlaps with that of both chemosensory and tactile types, suggesting that it may convey positional information as well as valence (Figure 55 - figure supplement 3). SAxx02 is highly associated with SNxx29 and exhibits distinctive morphology, but we have not been able to match it to a known type, although it may also express Gr28b.d (Kwon et al., 2014). SNxx30 is also distinctive, appears once in each hemineuromere, and is most likely proprioceptive. One SNxx32 neuron enters from each of the mesothoracic leg nerves to innervate the midline; this type exhibits an unusual pattern of connectivity, notably to hemilineage 27X and abdominal motor neurons. Finally, SNxx33 was originally classified as chemosensory but has a pattern of output connectivity more similar to that of tactile types. However, through other connectivity measures this type co-clusters with chemosensory neurons (Figure 55 - figure supplement 5B), which may suggest functional association. We hypothesise that SNxx33 neurons may represent the single mechanosensory cell within each taste bristle (Nayak and Singh, 1983).

#### Dorsal Sensory Types

Sensory neurons from the neck, ventral prothorax, notum, wings, and halteres enter the VNC via the PrN, ProCN, ADMN, PDMN, and DMetaN, respectively (Figure 56A). In clustering the dorsal sensory neurons, we grouped those from the prothoracic nerves PrN and ProCN but treated each of the other nerves independently. The quality of reconstruction in the ADMN, PDMN, and DMetaN was significantly higher than for the leg nerves, allowing any numerical asymmetry to be interpreted as biological variation rather than due to technical limitations (Figure 56B). Notably, a higher proportion of dorsal than leg sensory neurons ascend via the neck connective, consistent with the need for rapid communication between the wings and especially the halteres and the brain during flight. Ascending, sensory, and motor neurons also comprise a higher proportion of direct targets from dorsal sensory neurons as compared to leg neurons (Figure 56C).

**Figure 56.**
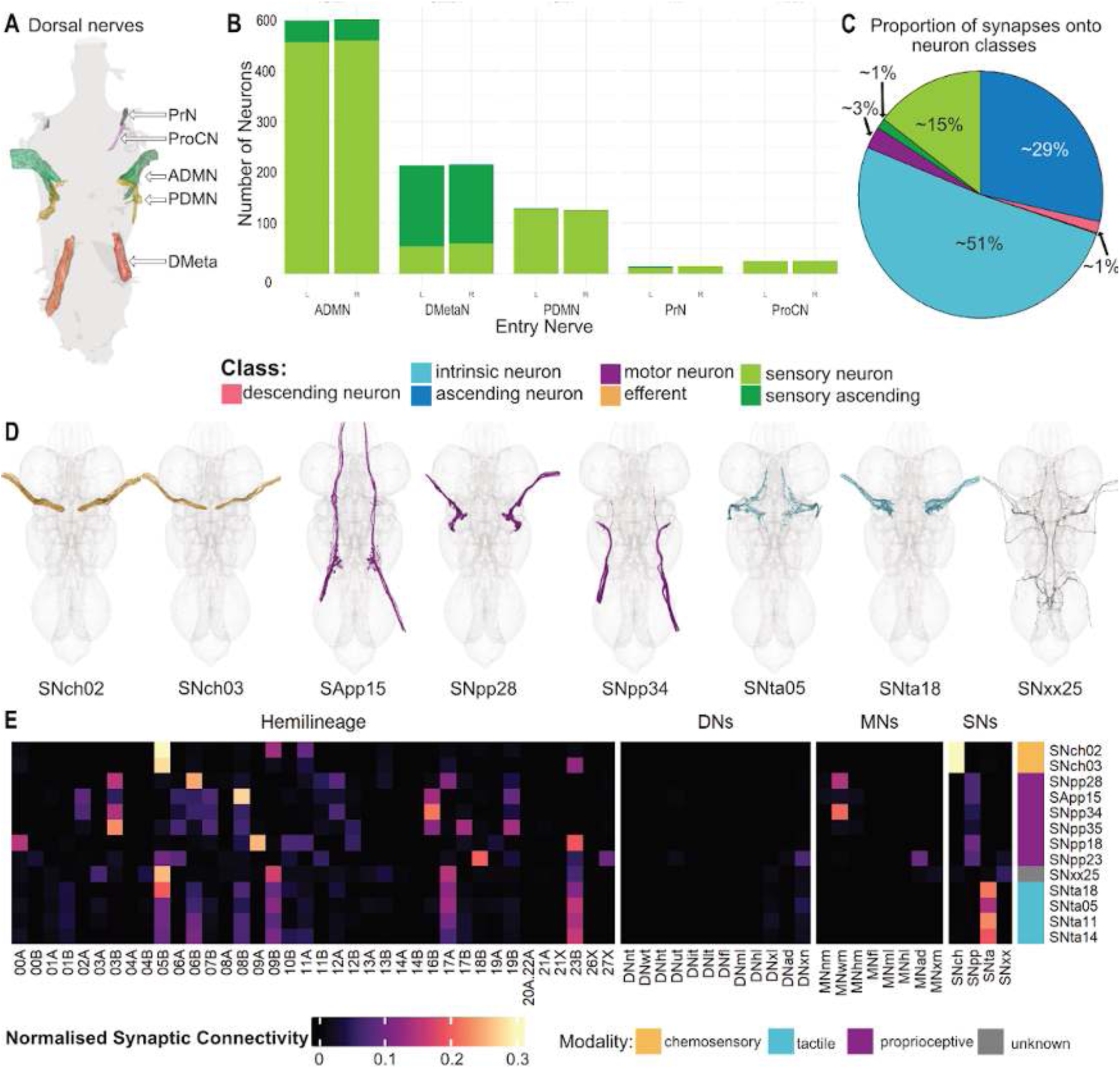
Dorsal sensory neuron systematic types. **A.** Anatomy of dorsal nerves (dorsal view). **B.** Numbers of sensory neurons from each dorsal nerve. **C.** Proportions of downstream partners of dorsal sensory neurons, coloured by class. **D.** Plotted meshes of exemplar dorsal types, coloured by assigned modality: orange = chemosensory, purple = proprioceptive, cyan = tactile, grey = unknown. **E.** Heatmap showing normalised sensory neuron cluster connectivity to hemilineages in the VNC. Annotation bar displays annotated modality of component sensory neurons within the sensory neuron cluster; orange = chemosensory, purple = proprioceptive, cyan = tactile, grey = unknown.

We define 85 dorsal sensory types: 4 chemosensory, 60 proprioceptive, 15 tactile, and 6 of unknown modality. It should be noted that many of the ascending sensory neurons from the halteres form few chemical presynapses in the VNC and that we could not distinguish electrical connections (Fayyazuddin and Dickinson, 1996), so these types might be refined if and when gap junction expression data and/or a full CNS connectome become available. Neurons of each modality tend to target characteristic hemilineages and other cell subclasses, with chemosensory types strongly connecting to 05B, tactile types targeting 05B, 06B, 08B, 09B, 17A, and 23B, and proprioceptive types exhibiting more variation in hemilineage targets and sometimes connecting directly to motor neurons (Figure 56E).

**Figure 56 - figure supplement 1.**
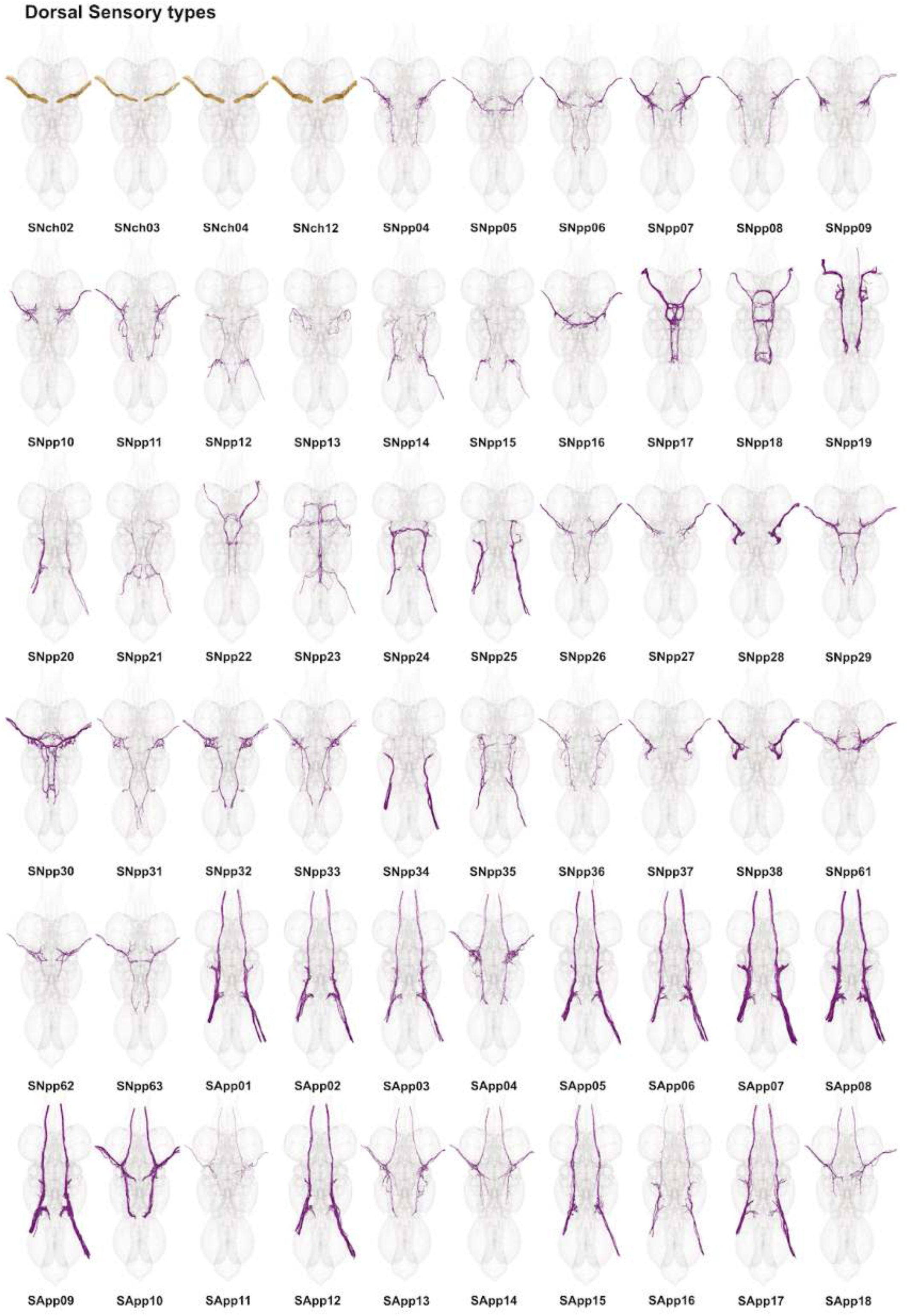
Dorsal sensory neuron types. Plotted meshes of neurons that comprise each systematic type from dorsal nerves, coloured by modality: orange = chemosensory, purple = proprioceptive, cyan = tactile, grey = unknown.

**Figure 56 - figure supplement 2.**
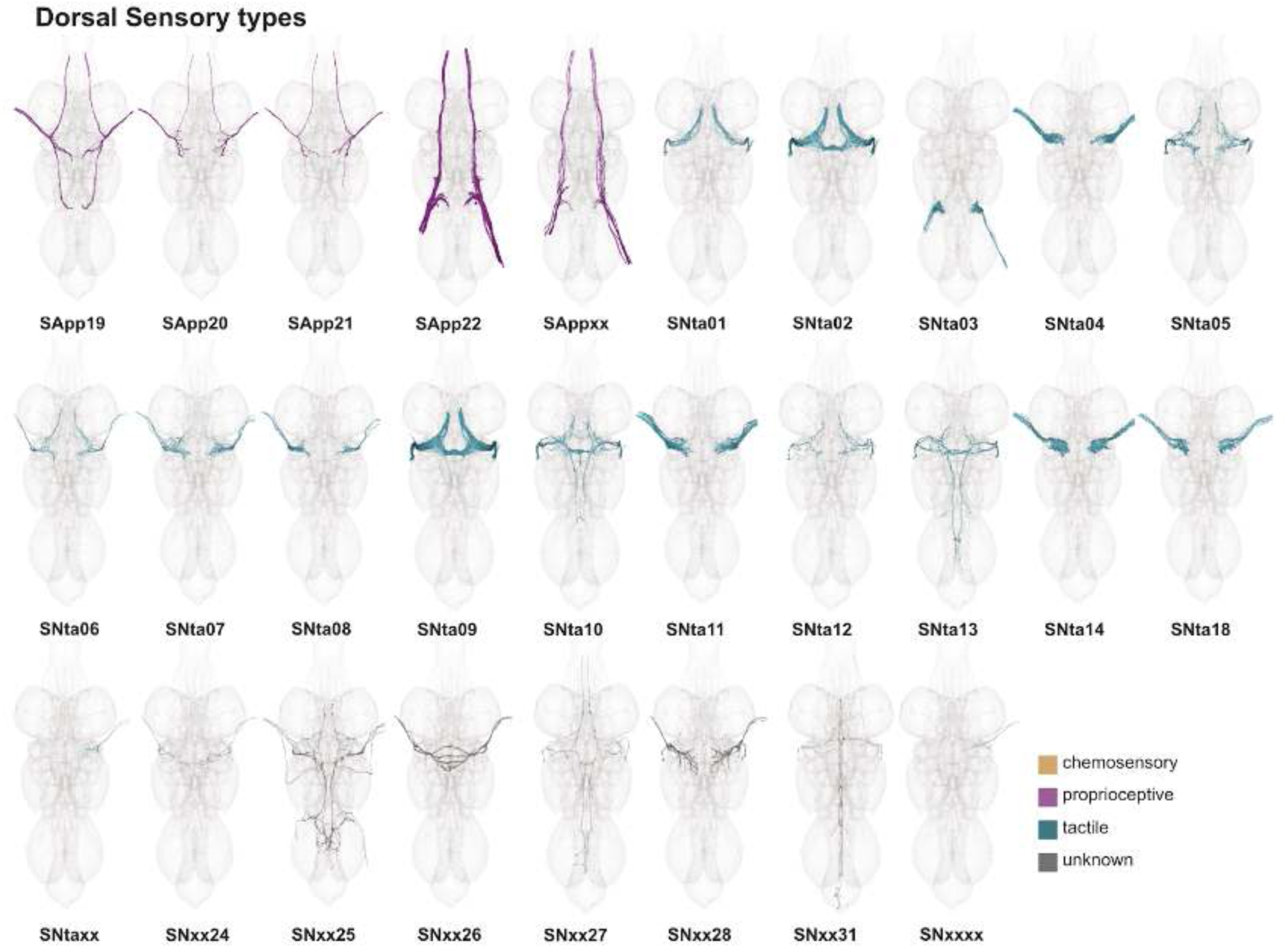
Dorsal sensory neuron types, continued. Plotted meshes of neurons that comprise each systematic type from dorsal nerves, coloured by modality: orange = chemosensory, purple = proprioceptive, cyan = tactile, grey = unknown.

**Figure 56 - figure supplement 3.**
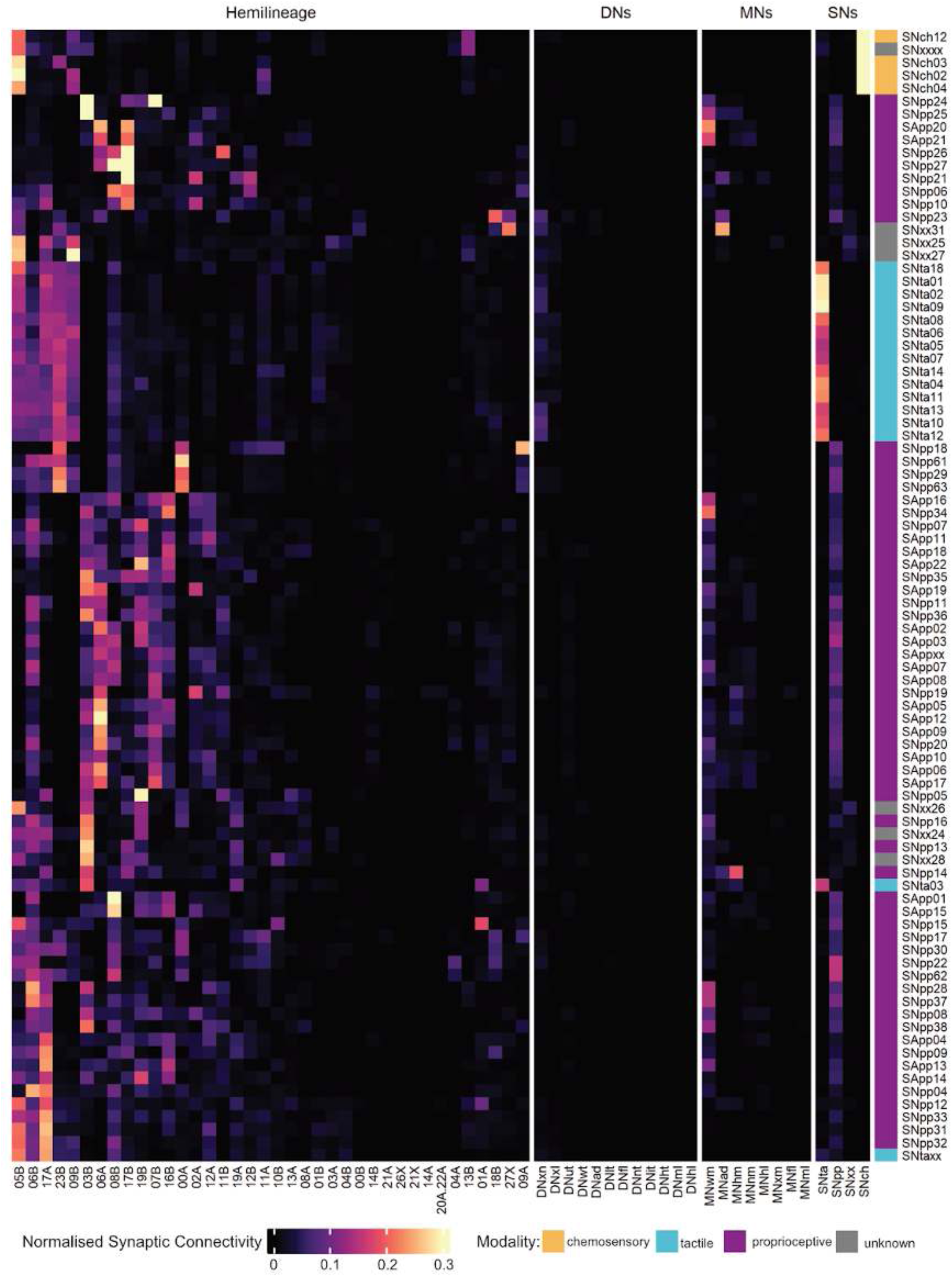
Dorsal sensory neuron type output connectivity by hemilineage. Right annotation bar is coloured by modality: orange = chemosensory, purple = proprioceptive, cyan = tactile, grey = unknown. Rows and columns are ordered based on hierarchical clustering of data.

**Figure 56 - figure supplement 4.**
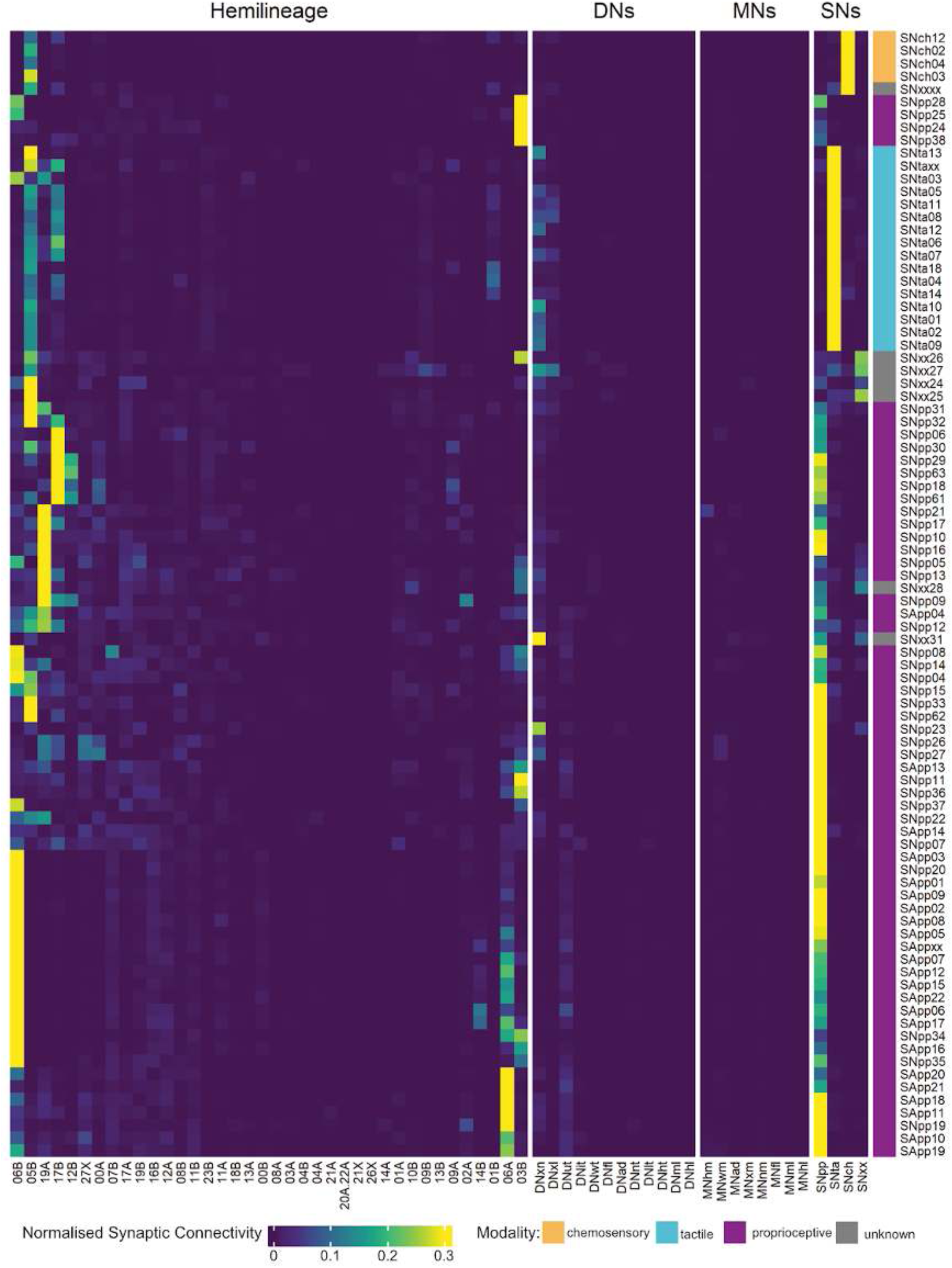
Dorsal sensory neuron type input connectivity by hemilineage. Right annotation bar is coloured by modality: orange = chemosensory, purple = proprioceptive, cyan = tactile, grey = unknown. Rows and columns are ordered based on hierarchical clustering of data.

**Figure 56 - figure supplement 5.**
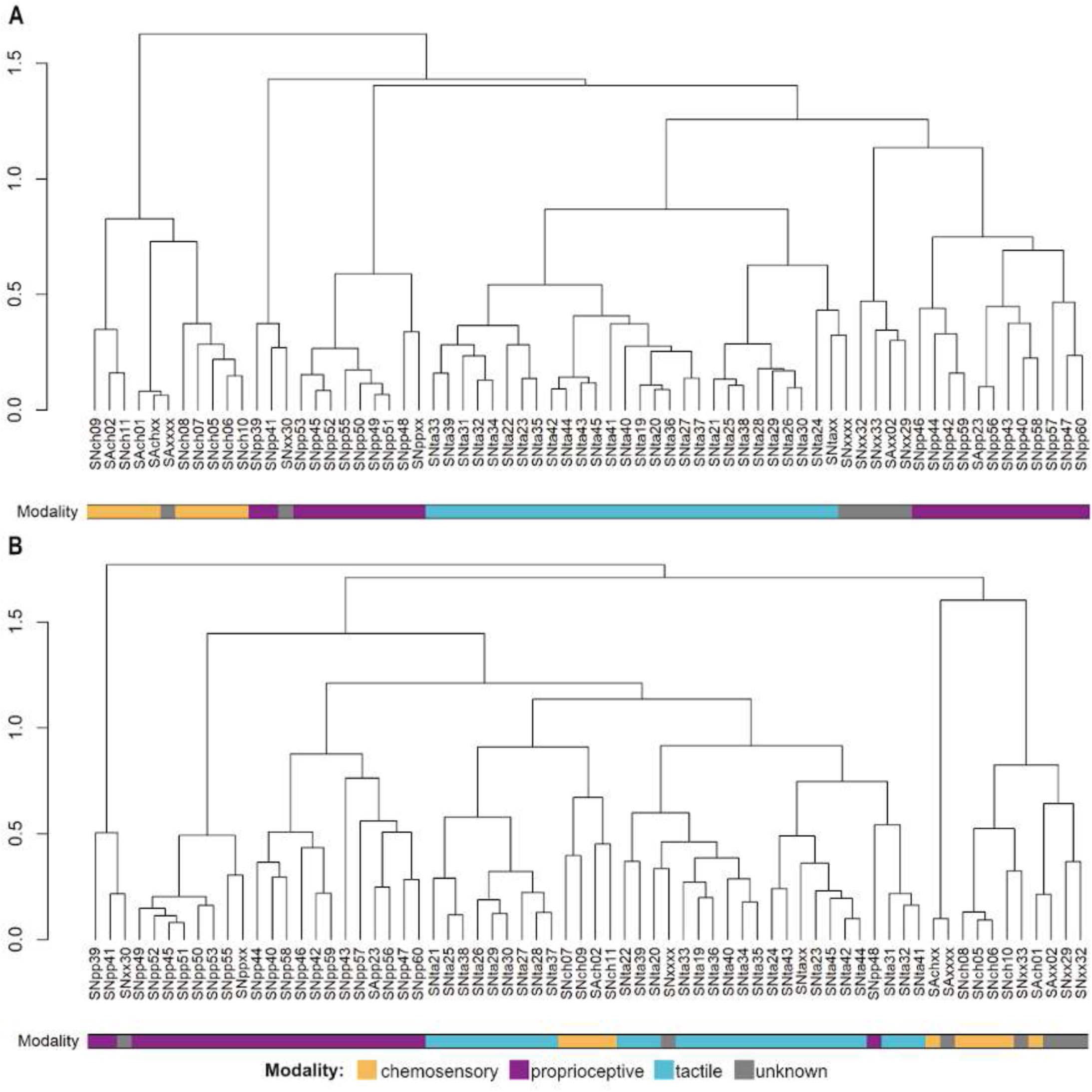
Dorsal sensory neuron type dendrograms showing combined inputs and outputs clustering. **A.** Dendrogram showing combined clustering with intra-type connectivity included. **B.** Dendrogram showing combined clustering with intra-type connectivity excluded. Bottom bar is coloured by modality: orange = chemosensory, purple = proprioceptive, cyan = tactile, grey = unknown.

##### Chemosensory dorsal sensory neurons

All identified chemosensory dorsal sensory neurons originate in the wings (Figure 56 - figure supplement 1). When considering either upstream or downstream connectivity, they all cluster together along with SNxxxx, untyped neurons that therefore may also be chemosensory (Figure 56 - figure supplement 2-4). 05B is the principal hemilineage upstream as well as downstream of all chemosensory dorsal neurons. However, we observe type-specific connectivity onto 13B, 23B, and 11A. We were not able to associate any of these types with a particular valence based on comparisons to light-level morphology as they are all very similar.

##### Proprioceptive dorsal sensory neurons

The majority of dorsal sensory types are proprioceptive (Figure 56 - figure supplement 1-2). Many types ascend via the neck connective, primarily from DMetaN. We observe significant diversity in their downstream targets, with hemilineages 00A, 03B, 05B, 06A, 06B, 17A, and 23B strongly downstream of subsets of types. Notably, numerous proprioceptive types directly target motor neurons, e.g., SNpp23, SNpp28, and SNpp34 (Figure 56E, Figure 56 - figure supplement 3). Most proprioceptive types are also strongly downstream of gabaergic hemilineages 05B, 06A, 06B, 17B, and 19A (Figure 56E, Figure 56 - figure supplement 4). These connectivity patterns are consistent with roles in rapid posture adjustment.

##### Tactile dorsal sensory neurons

All sensory neurons from the notum (via the PDMN) are tactile, as are a subset of wing types and a single haltere type from the knob hairs (Figure 56 - figure supplement 2). Dorsal tactile types feature a broad, consistent pattern of connectivity onto hemilineages 05B, 09B, 17A, and 23B, with 06A also targeted by roughly half of the types (Figure 56 - figure supplement 3). They broadly receive input from gabaergic hemilineages 05B and in some cases 17B (Figure 56 - figure supplement 4). SNta03, the haltere type, exhibits an atypically strong connection onto hemilineage 03B in T3 and clusters with proprioceptive neurons based on output connectivity. However, it clusters solidly with other tactile types based on input connectivity, supporting its modality assignment and suggesting that its output connectivity is due to its unique peripheral origin.

#### Abdominal Sensory Types

We reconstructed sensory neurons from four abdominal nerves: AbN2, AbN3, AbN4, and AbNT (Figure 57A); one afferent from AbN1 was expected (Smith and Shepherd, 1996) but not recovered. Approximately equal numbers of neurons were identified on the left and right, with some slight asymmetry in AbN2 (Figure 57B). Abdominal sensory neuron targets included a higher proportion of sensory neurons and lower proportion of ascending neurons than those of either leg or dorsal sensory neurons (Figure 57C), suggesting roles in local sensorimotor circuits.

**Figure 57.**
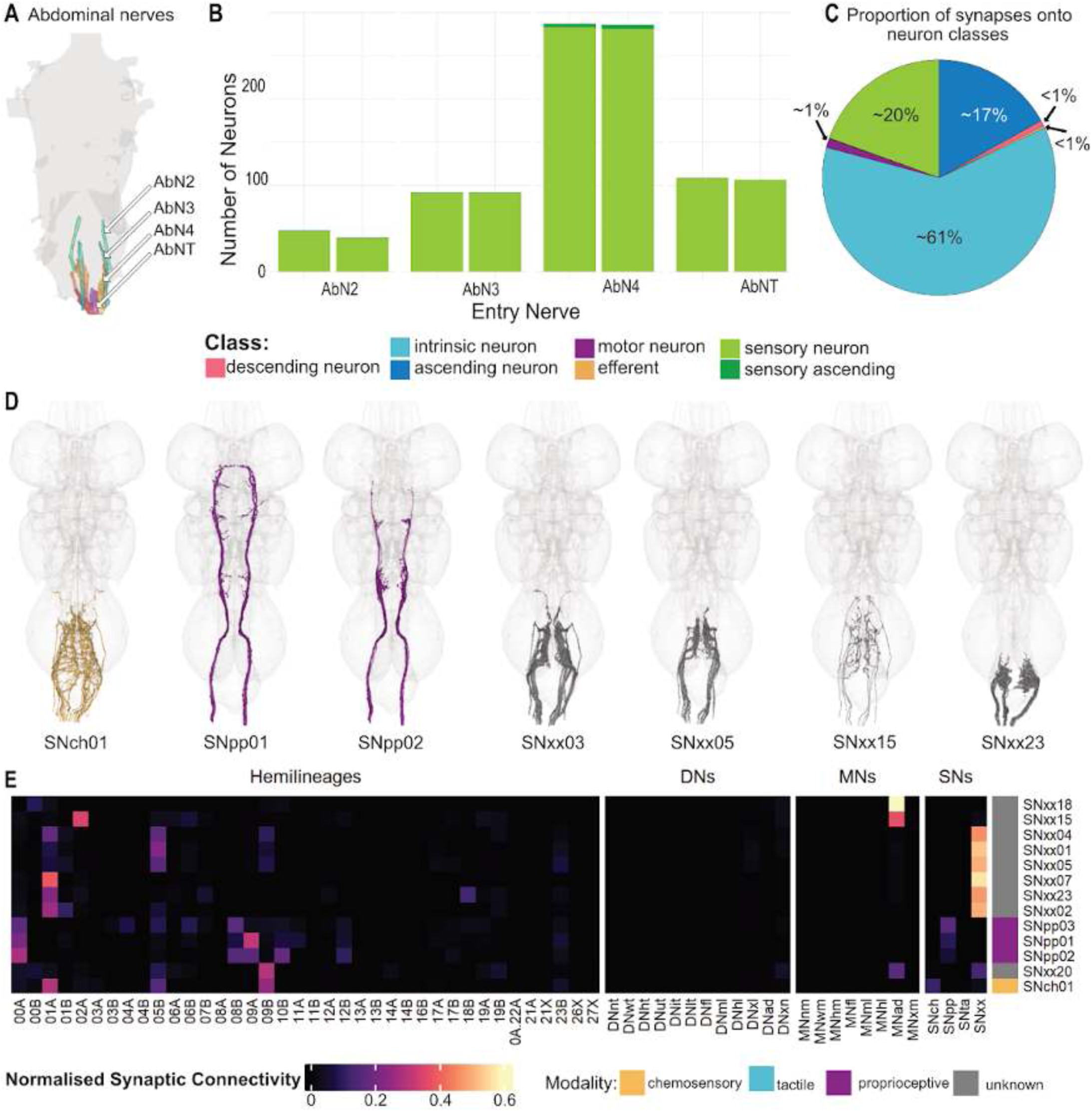
Abdominal sensory neuron systematic types. **A.** Anatomy of abdominal nerves (dorsal view). **B.** Numbers of sensory neurons from each abdominal nerve. **C.** Proportions of downstream partners of abdominal sensory neurons, coloured by class. **D.** Plotted meshes of exemplar abdominal types, coloured by assigned modality. **E.** Heatmap showing normalised sensory neuron cluster connectivity to hemilineages in the VNC. Annotation bar displays annotated modality of component sensory neurons within the sensory neuron cluster; orange = chemosensory, purple = proprioceptive, cyan = tactile, grey = unknown.

We identify 30 sensory cell types that enter the VNC via abdominal nerves, with only a single type (SAxx01) ascending via the neck connective (Figure 57 - figure supplement 1). The sparseness of hemilineage assignments and published sources for the adult abdomen made it difficult to define discrete abdominal sensory types and to assign them to a modality. We identified only a single candidate chemosensory type - SNch01, which likely corresponds to ppk+ nociceptive neurons that include the c4da (class IV dendritic arborisation neurons) (S Rumpf, personal communication) - and three proprioceptive types (e.g., SNpp01 and SNpp02) (Figure 57D) from Wheeler’s organ (Smith and Shepherd, 1996). However, the high degree of seriality in abdominal segments facilitated association of abdominal types across multiple neuromeres. Cells entering through the AbNT were less likely to be associated with cells from the other nerves, possibly because A8 and A9 include more secondary neurons with a lower proportion of hemilineages and serial sets identified (Figure 49D, E). As previously noted, this segment-specific expansion likely underlies specialisation for excretory and reproductive functions.

Clustering the abdominal types by downstream or upstream connectivity suggests some additional modality assignments, but we could not be confident enough to annotate them. For example, types SAxxxx, SNxx22, and SNxxxx cluster with SNch01 by their downstream connectivity to hemilineages (especially to 01A and 05B) but more closely with SNpp03 by their upstream connectivity (especially to 19A) (Figure 57 - figure supplement 2 - 3). Conversely, SNxx02, SNxx07, SNxx10, SNxx11, SNxx12, SNxx13, SNxx14, and SNxx23 cluster together and next to SNpp01-03 based on their downstream connectivity (especially to 01A), but their upstream connectivity is more variable, placing only SNxx07 next to SNpp01-02. Based on our earlier analyses of dorsal sensory types, direct output to motor neurons might suggest a proprioceptive role for SAxx01, SNxx08, SNxx09, SNxx15, SNxx16m SNxx18, SNxx19, and SNxx20 (Figure 57 - figure supplement 2); however, leg sensory types targeting motor neurons were more likely to be tactile. SNxx18 is particularly striking; it is a single unpaired neuron that enters the AbNT and targets the posterior tip of the abdomen bilaterally.

**Figure 57 - figure supplement 1.**
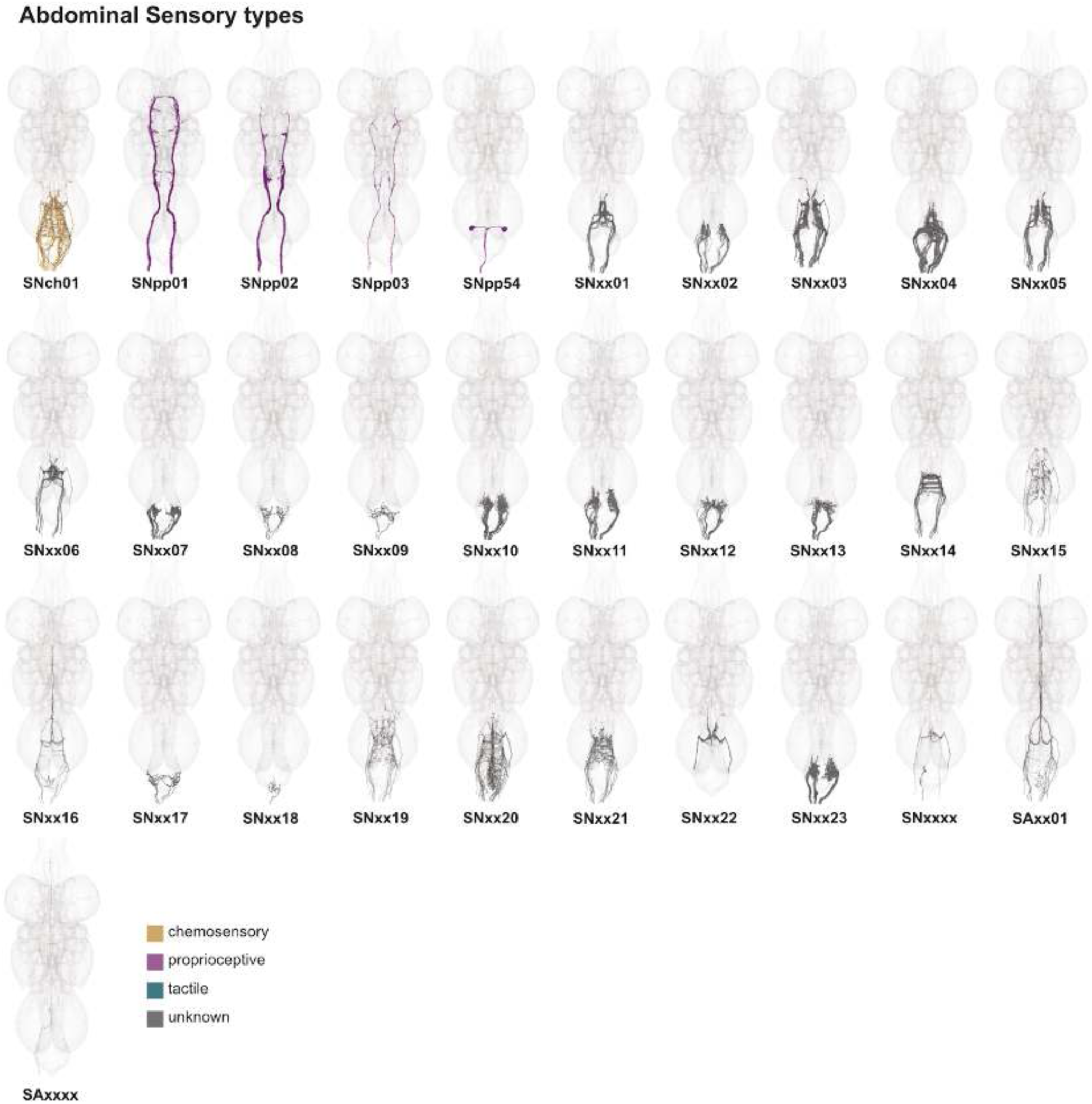
Abdominal sensory neuron types. Plotted meshes of neurons that comprise each systematic type from abdominal nerves, coloured by modality: orange = chemosensory, purple = proprioceptive, cyan = tactile, grey = unknown.

**Figure 57 - figure supplement 2.**
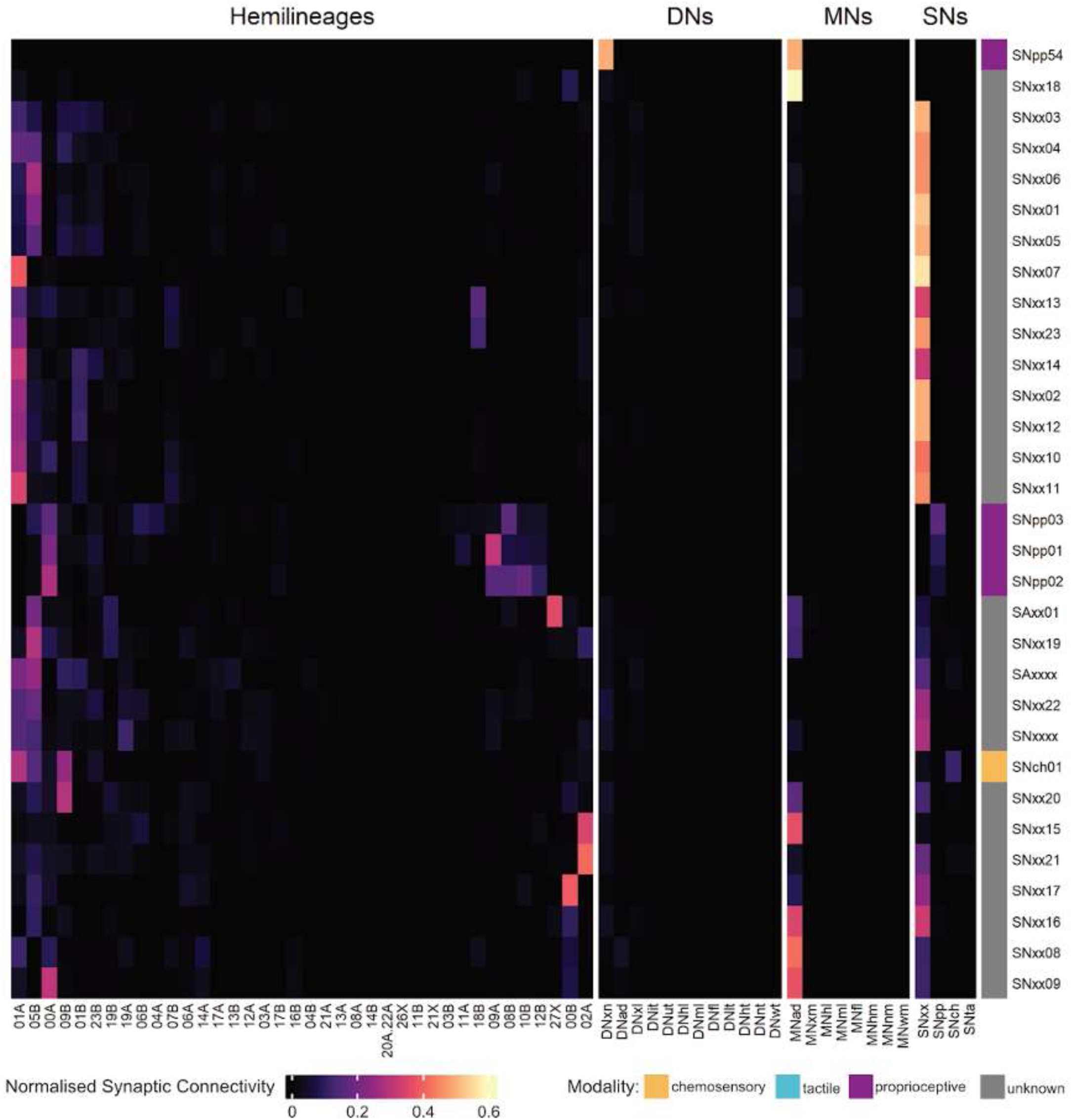
Abdominal sensory neuron type output connectivity by hemilineage. Right annotation bar is coloured by modality: orange = chemosensory, purple = proprioceptive, cyan = tactile, grey = unknown. Rows and columns are ordered based on hierarchical clustering of data.

**Figure 57 - figure supplement 3.**
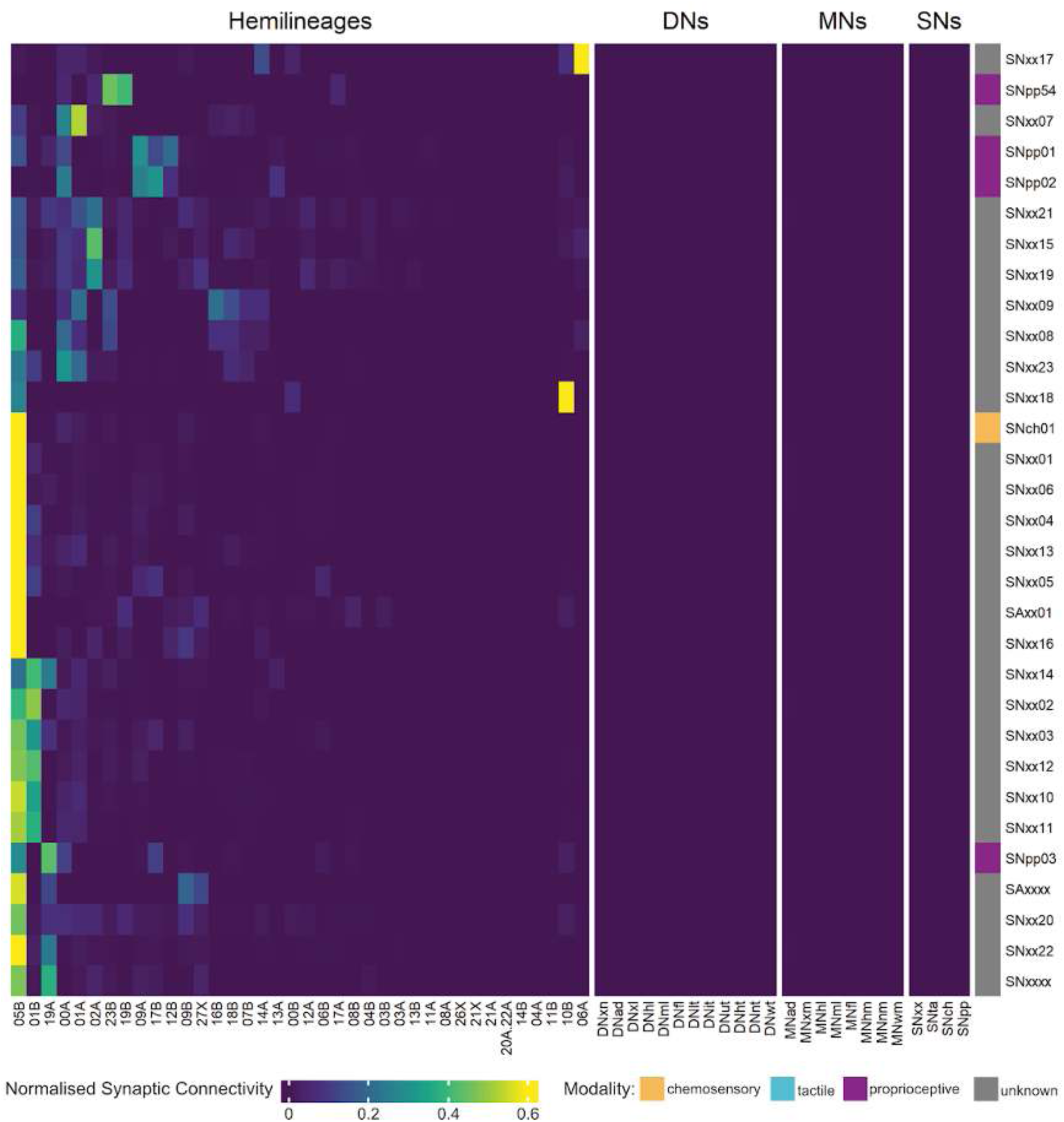
Abdominal sensory neuron type input connectivity by hemilineage. Right annotation bar is coloured by modality: orange = chemosensory, purple = proprioceptive, cyan = tactile, grey = unknown. Rows and columns are ordered based on hierarchical clustering of data.

**Figure 57 - figure supplement 4.**
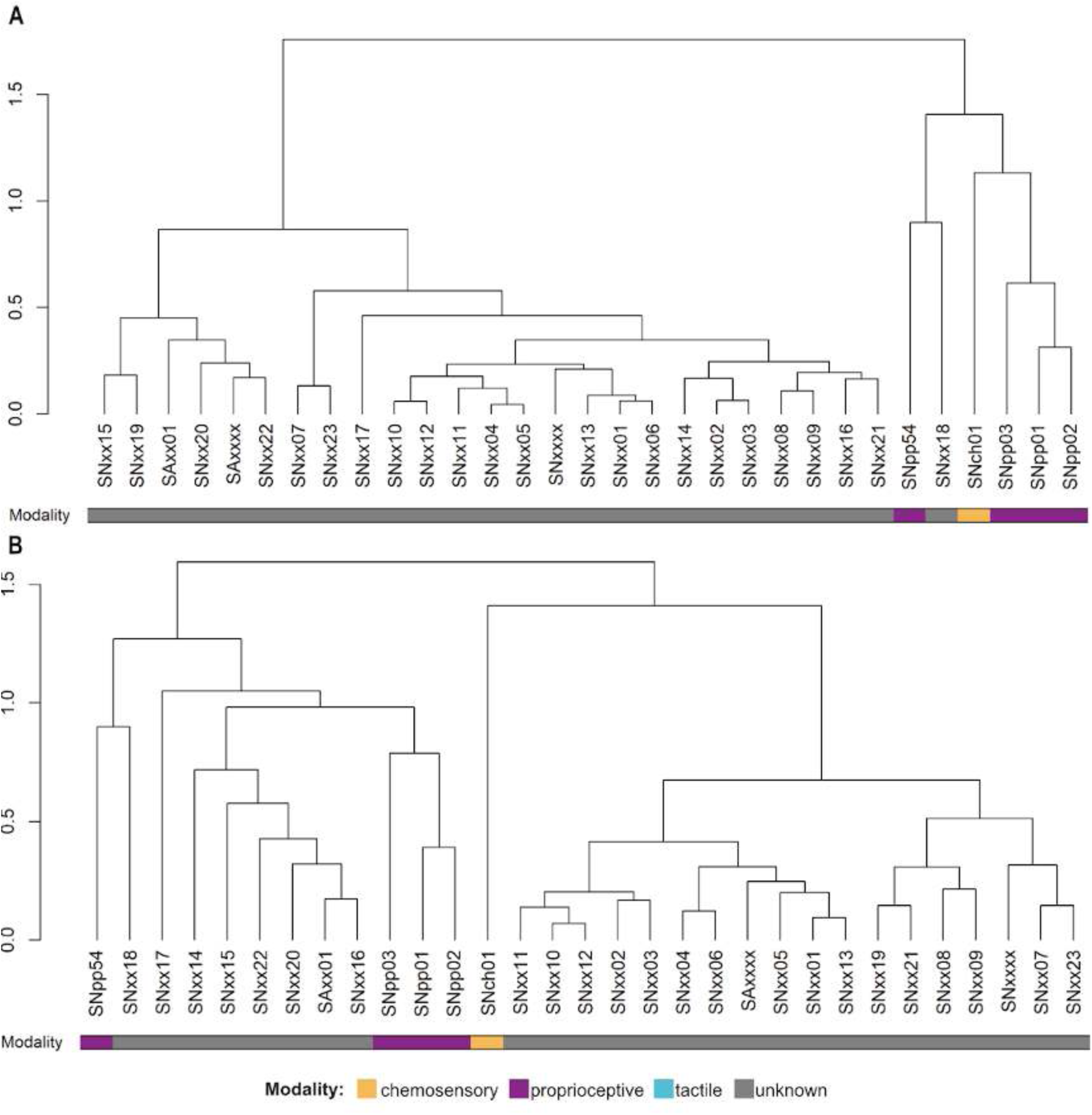
Abdominal sensory neuron type dendrograms showing combined inputs and outputs clustering. **A.** Dendrogram showing combined clustering with intra-type connectivity included. **B.** Dendrogram showing combined clustering with intra-type connectivity excluded. Bottom bar is coloured by modality: orange = chemosensory, purple = proprioceptive, cyan = tactile, grey = unknown.

#### Inter-regional associations between sensory types

We suspected that there could be sensory types sharing similar functions across two or more of the three main regions (leg, dorsal, and abdominal). To address this, we used cosine clustering of outputs to serial neurons across the entire sensory neuron population (Figure 58 top, Figure 58 - figure supplement 1), rather than within each region as we had done originally to generate the types. We do not claim that the associations detected here represent genuine cross-regional types, although this may turn out to be true for at least a subset of cases; rather, we suggest that co-clustering implies functional association and may also indicate modality.

**Figure 58.**
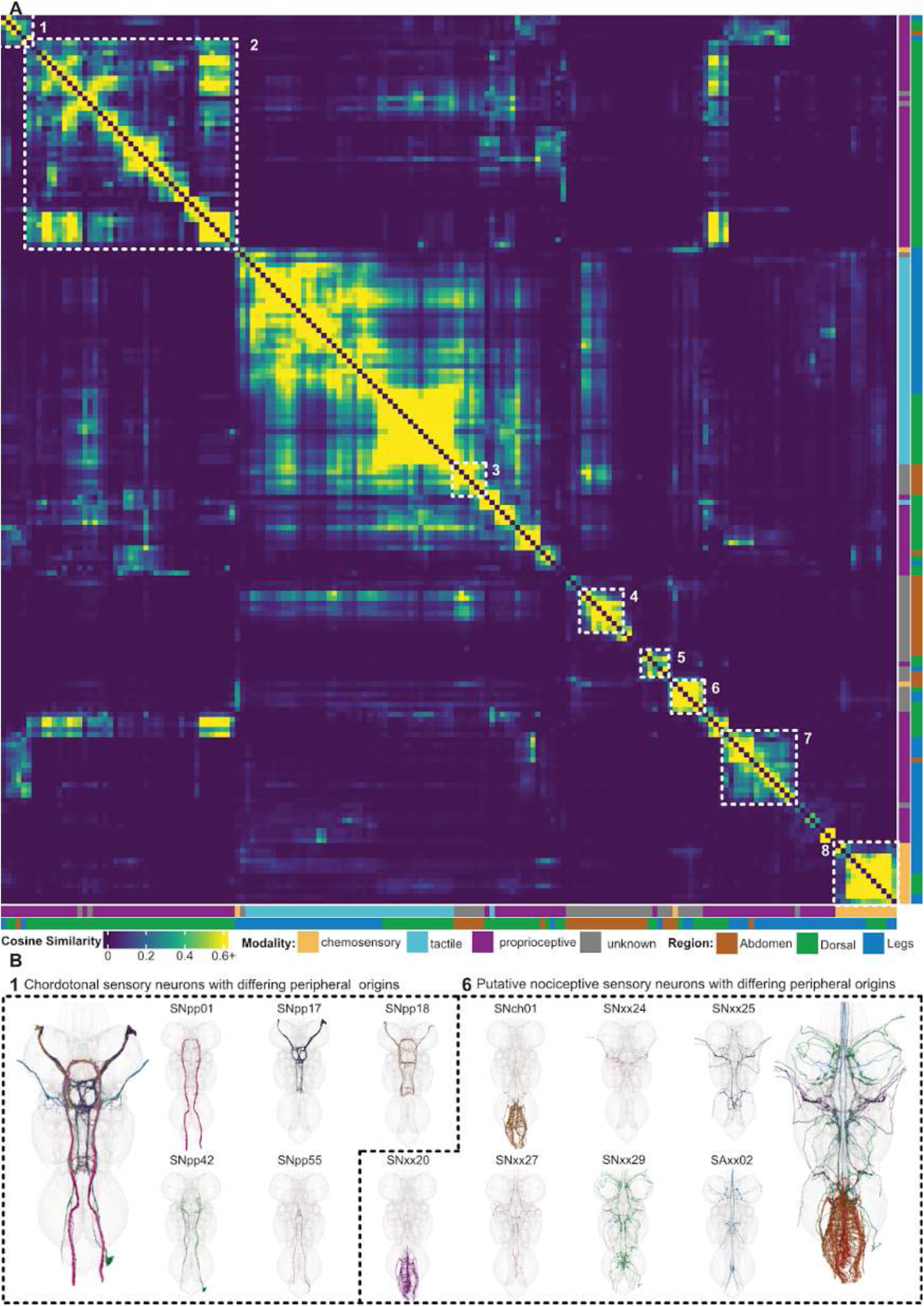
Inter-regional associations between sensory cell types. Top. Heatmap showing cosine similarity between the outputs of sensory cell types on serial neuron groups. Coloured annotation bars indicate the assigned modality of the sensory cell type and its region of peripheral origin, inner and outer bars respectively. Heatmap row and column ordering generated through travelling sales problem optimisation. **Bottom.** Exemplar cell type clusters indicated by dashed line boxes 1 and 6 in heatmap.

**Figure 58 - figure supplement 1.**
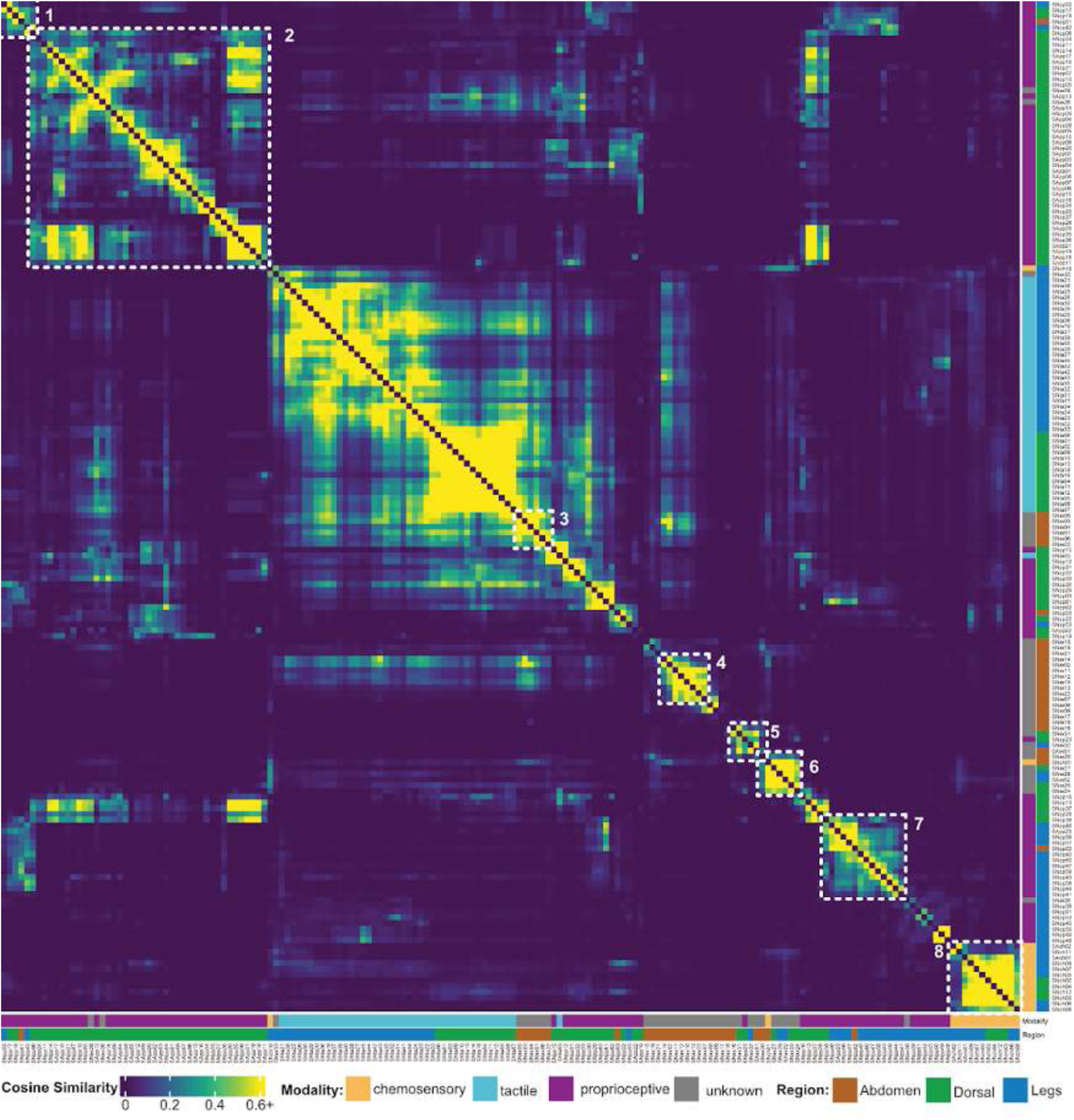
Cosine similarity between the outputs of sensory cell types (labelled) on serial neuron group. Coloured annotation bars indicate the assigned modality of the sensory cell type and its region of peripheral origin, inner and outer bars respectively. Heatmap row and column ordering generated through travelling sales problem optimisation.

**Figure 58 - table 1.**
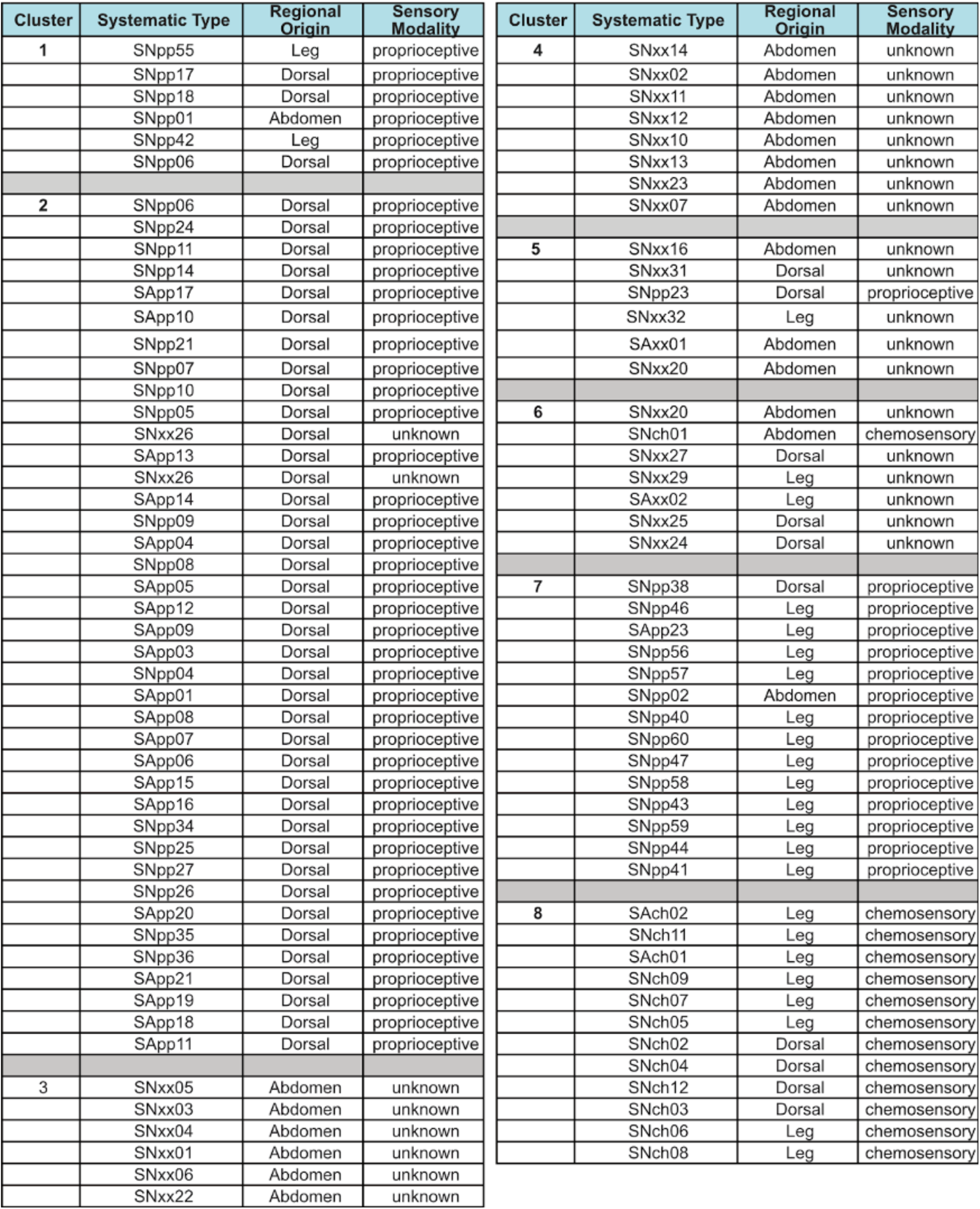
Highlighted cosine similarity clusters from Figure 58A. Cluster numbers, systematic types, regional origins, and annotated sensory modalities are provided.

Although we were able to assign a modality to very few cell types in the abdomen in isolation based on the available light-level literature, cosine clustering across regions suggests modalities for many more. For example, types SNxx01-06 (Figure 58 top, cluster 3) co-cluster with almost all of the tactile types from the leg and dorsal nerves, implying that they are likely to be tactile as well. And the utility of this approach is not limited to the abdomen: dorsal types SNxx26 and SNxx28 (Figure 58 top, cluster 2) are likely to be proprioceptive because they co-cluster with identified campaniform sensilla.

In cases where co-clustering does not provide additional information regarding modality, it can still suggest functional connections. For example, the types in one proprioceptive cluster spanning all three regions (Figure 58 bottom, cluster 1) is distinct in connectivity and shows a high degree of inter-type overlap in downstream partners. Indeed, three of these types (SNpp01, SNpp18, and SNpp42) are morphologically similar intersegmental neurons (Figure 58 bottom, cluster 1) that might represent a single cross-regional chordotonal organ type.

Finally, based on this analysis we have identified one strong candidate for a cross-regional type (Figure 58 top, cluster 6). This cluster includes SNch01 (the abdominal ppk+ nociceptive neurons that include c4da, the class IV dendritic arborisation neurons) and SNxx29 (heat nociceptive ppk+ leg neurons that express thermoreceptor Gr28b.d (Khuong et al., 2019; Kwon et al., 2014)) as well as several other types of unknown modality from the legs, wings, and/or halteres. These types share similar terminal morphology as well as serial connectivity (Figure 58 bottom, cluster 6) and may all represent aversive stimuli.

#### Proprioceptive neuron types

Classically defined proprioceptors are mechanosensory neurons located within muscles, tendons, and joints that provide information on body position and movement. Three main subclasses of proprioceptors have been identified in insects (reviewed in (Tuthill and Azim, 2018)). Chordotonal organs contain populations of mechanosensory neurons connected to muscles by tendons and encode position and velocity during joint displacement. Campaniform sensilla are small dome-like structures, usually clustered close to joints, that are sensitive to cuticular strain and detect mechanical load on limbs as resistance to muscle contraction. Hair plates are tightly packed arrays of sensory hairs positioned on the cuticle so as to be deflected when the joint reaches a particular position and may anticipate phase changes in walking such as the swing to stance transition.

##### Chordotonal organ types

We identified 24 distinct connectivity types of chordotonal sensory neurons: 13 from the femoral chordotonal organ (FeCO), three from Wheeler’s organ, three from the prosternal chordotonal organ (pCO), and 5 unclassified types (Figure 59A). Two of the FeCO types morphologically resemble the “hooks,” two the “claws,” and 9 the “clubs” previously reported in the light-level literature (Mamiya et al., 2018). Many chordotonal types are strongly upstream of neurons from hemilineage 09A, with one FeCO claw type targeting 13A and one FeCO hook type targeting 14A (and not the morphologically similar hemilineage 13B as was previously reported (Agrawal et al., 2020)).

**Figure 59.**
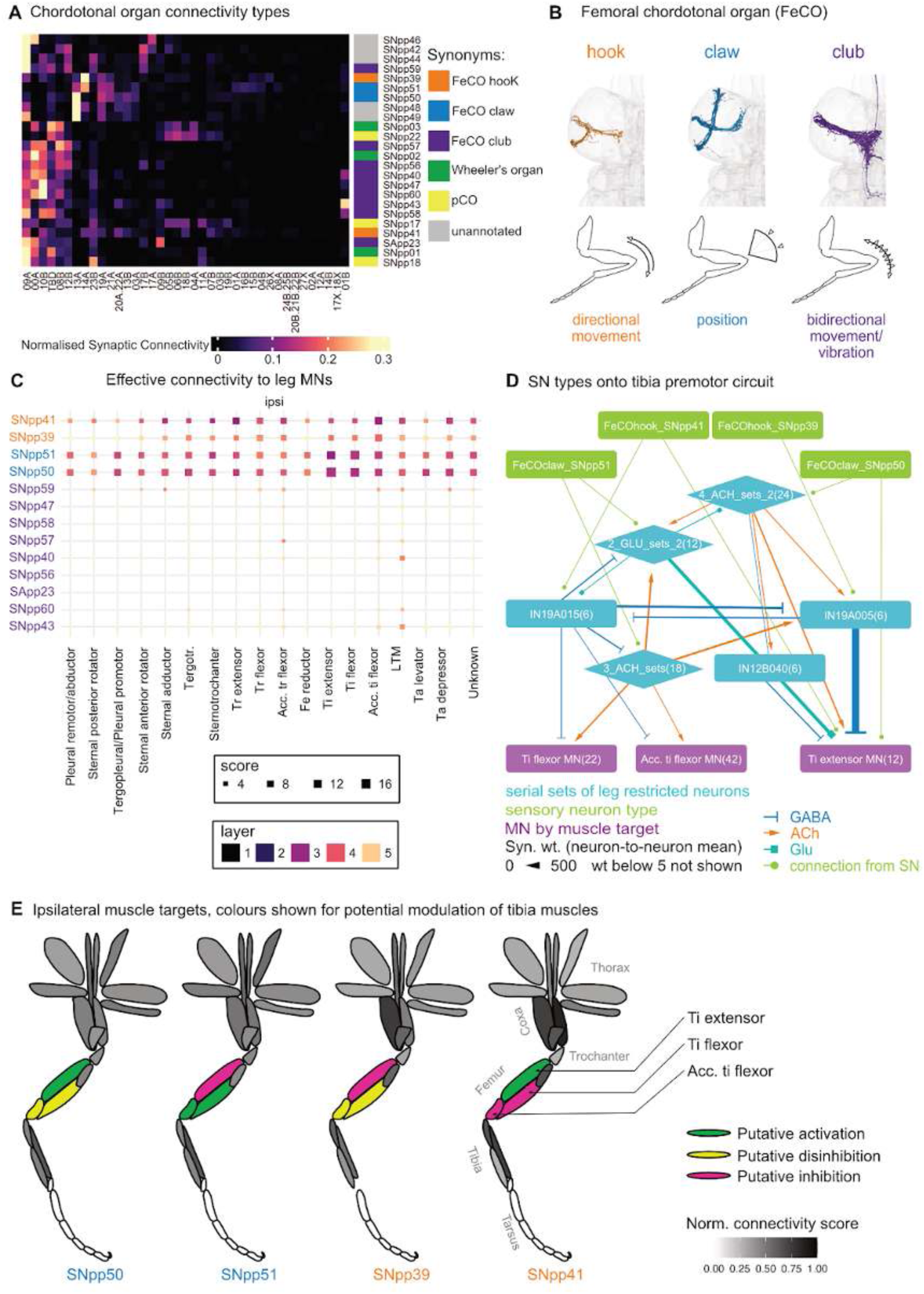
Chordotonal organ connectivity. **A.** Chordotonal organ types clustered by connectivity to hemilineages. **B.** Neuron meshes for categories of femoral chordotonal organ (FeCO hook, FeCO claw, and FeCO club) from entry nerve ProLN_R. Cartoons depict leg movements detected by each neuron (after (Mamiya et al., 2018)). **C.** Effective connectivity from FeCO types to ipsilateral leg motor neuron targets. **D.** Relationship of FeCO hook and claw types to the tibia premotor circuit. Edges are coloured by predicted neurotransmitter. Premotor neuron types in D were grouped by similar connectivity to legMNs: IN17A061, INXXX466, IN19B003 (4_ACH_sets_2), IN08A005, IN08A007 (2_GLU_sets_2) and IN03A004, IN21A004, IN20A.22A023 (3_ACH_sets). The number in brackets indicates the number of neurons included in each node. **E.** Ipsilateral leg muscle targets for FeCO types, coloured by excitation (green), disinhibition (yellow), and inhibition (magenta).

We chose to focus on the defined FeCO types for further analysis because they constitute an active area of research and their morphology and connectivity have been well-described at light level. Briefly, the femoral chordotonal organ is composed of ∼152 neurons located in the proximal femur and attached by tendons to the tibia (Kuan et al., 2020). FeCO neurons have been divided into three main categories based on their response profiles and morphologies within the VNC (Figure 59B). Hooks respond to directional movement of the tibia, claws to the position of the tibia relative to the femur, and clubs to bidirectional movement/vibration (Mamiya et al., 2018). Two distinct subpopulations of hook neurons have been found to respond to extension vs flexion and to target different downstream partner hemilineages (Chen et al., 2021).

With the complete VNC connectome at our disposal, we sought to place these FeCO types in the context of motor circuits. First we examined the effective connectivity of FeCO types to motor neurons using the approach developed in our companion paper (Cheong et al., 2023). Briefly, we calculated the percent of the total input to the postsynaptic neuron for every edge in the VNC connectome. We then calculated the matrix-vector-product of the input percent matrix for neurons of interest for five downstream layers. We normalised by the non-zero mean of all scores in each layer, for comparisons across the layers. To calculate effective connectivity strength for each given SN-MN-pair, we picked the highest score of the five layer matrices and recorded the layer in which it occurred.

Using this method to calculate effective connectivity to leg motor neurons, we find that the FeCO club types are not appreciably connected to any motor neurons but that the claw and hook types exhibit indirect connectivity to the ipsilateral accessory tibia flexor MN and to the ipsilateral tibia extensor MN and tibia flexor MN, respectively (Figure 59C). They are also weakly upstream of the TTM and STTMn, which control the tergotrochanter muscle in the middle legs during escape/rapid take-off (Figure 59 - figure supplement 1). Notably, they are not effectively connected to contralateral motor neurons (Figure 59 - figure supplement 1A-B), suggesting that they mainly participate in local leg reflexive actions.

Our effective connectivity calculations do not take the sign of connection into account, but this can sometimes be determined by a closer look at specific circuits. We therefore decided to investigate the roles of the FeCO hook and claw types in the tibia premotor circuit identified in our companion paper (Cheong et al., 2023). Hook type SNpp41 is predicted to inhibit the tibia flexor MN and accessory tibia flexor MN via gabaergic serial type IN19A015 while activating the tibia extensor MN (Figure 59D). Conversely, hook type SNpp39 would inhibit the tibia extensor via gabaergic serial type IN19A005, which also inhibits IN19A015, disinhibiting the tibia flexor MN and accessory tibia flexor MN. Assuming that glutamate is inhibitory in this circuit, claw type SNpp51 would activate the tibia flexor MN and accessory tibia flexor MN while inhibiting the tibia extensor MN via two glutamatergic serial types; conversely, claw type SNpp50 would activate the tibia extensor MN directly and via two cholinergic serial types but weakly disinhibit the tibia flexor and accessory tibia flexor MNs after a time delay.

In summary, our reconstructed circuit predicts that the two FeCO hook types should exert opposing effects on the tibia extensor vs the flexor and accessory tibia flexor muscles (Figure 59E), consistent with their observed responses to directional tibia movement (Chen et al., 2021). We also identify two distinct claw types, as yet unreported in the light-level literature, that would likewise be expected to exert opposing effects on the same muscles.

**Figure 59 - figure supplement 1.**
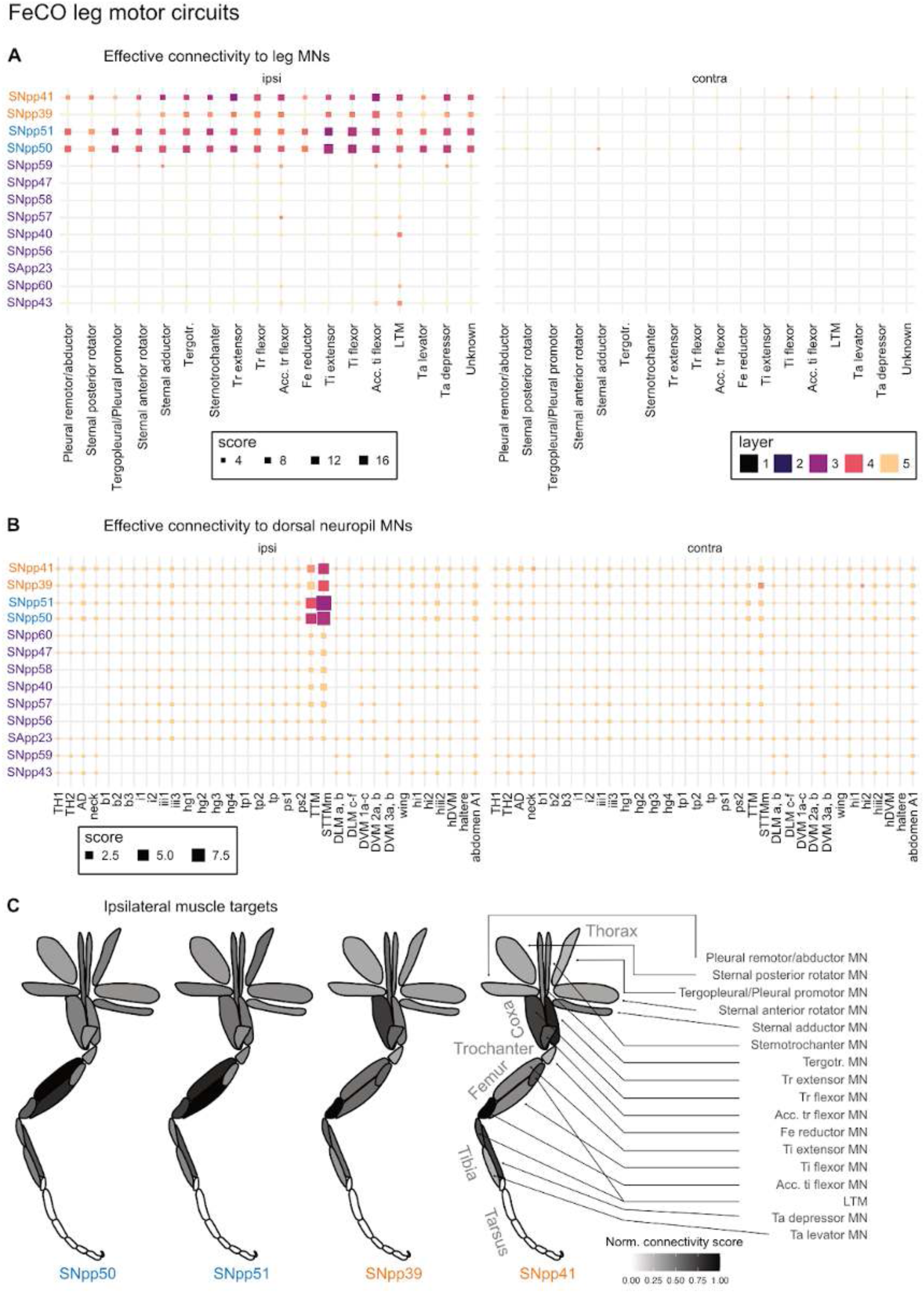
FeCO leg motor circuits. **A.** Effective connectivity to leg motor neuron targets. **B.** Effective connectivity to dorsal motor neuron targets. **C.** Ipsilateral leg muscle targets for FeCO types.

##### Campaniform sensilla types

Approximately 1200 campaniform sensilla (CS) are distributed over the legs, wings, halteres, and antennae of the fly (Gnatzy et al., 1987). We identified 46 CS types: 21 from the wings, 6 from the base of the wings, 14 from the halteres, one serial type from the legs, and one from the antenna. These types target a broad array of dorsal and mixed neuropil hemilineages - 00A, 03B, 06A, 06B, 07B, 08B, 16B, 17A, 17B, 19A, 19B, and 23B - in a type-specific manner (Figure 60A). We also expected to identify leg types with axons morphologically similar to those of hair plate neurons, but unfortunately we were unable to distinguish between them in our unclassified proprioceptive population (SNppxx) (A Büschges, personal communication).

**Figure 60.**
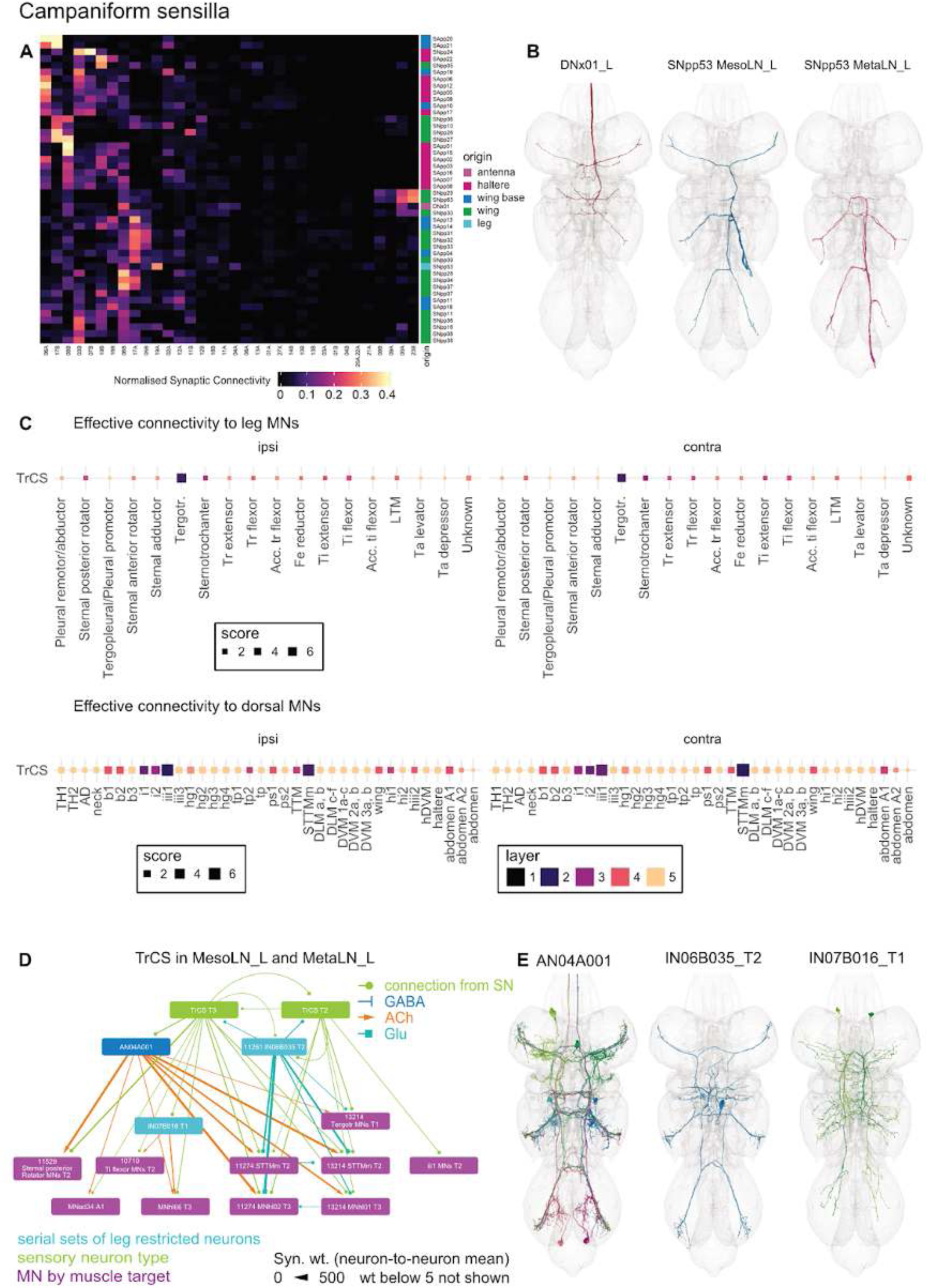
Campaniform sensilla connectivity. **A.** Campaniform sensilla types clustered by connectivity to hemilineages. **B.** Plotted meshes of bilateral TrCS from MesoLN_L vs MetaLN_L. **C.** Effective connectivity of TrCS to leg motor neuron targets vs dorsal motor neuron targets. **D.** Relationship of TrCS (MesoLN and MetaLN only) to standard leg premotor circuit. **E.** Plotted meshes of non-motor elements in the TrCS circuit shown in D.

We focus on the trochanter campaniform sensilla (TrCS), also referred to as bilateral campaniform sensilla, as they could be matched reliably to the light level literature. Interestingly, the antennal CS (DNx01) descends to the VNC and shares a similar pattern of connectivity with the TrCS (Figure 60A,B). We identified two candidate TrCS neurons (type SNpp53 in our clustering) from each leg nerve except for the left prothoracic, consistent with previous reports (Phelps et al., 2021). Unfortunately, the TrCS neurons proved especially difficult to reconstruct in MANC, with dark staining that obscured neuronal profiles and resulted in small fragments and frequent false merges. We therefore focused the rest of our analysis on the four best reconstructed examples: the pairs from the MesoLN_L and the MetaLN_L (Figure 60B).

Consistent with their bilateral morphology, we find that the TrCS neurons associate with ipsilateral and contralateral motor neuron homologues with a similar pattern of effective connectivity. They contribute only weakly to leg motor circuits, with strong association limited to the Tergotrochanter motor neurons, both ipsi- and contralateral to their nerve entry point (Figure 60C top row). Additionally the TrCS neurons associate with a diverse suite of dorsal motor neurons, with strong connections with iii1 and the STTMm motor neurons and weaker ones to i1 and i2 MNs (Figure 60C bottom row). The Tergotrochanter muscle sits in the thorax, similar to the TTM muscle targeted by the STTMm motor neuron, and both by morphology in the VNC and by connectivity we have placed them in two serial sets (13214 and 11274) with motor neurons of unknown motor output in T3. We postulate that a circuit (Figure 60D) including two glutamatergic IN types and a serially repeating cholinergic AN (Figure 60E) might play a role in the posture changes due to mechanical load applied and sensed by the TrCS proprioceptors.

##### Hair plate types

Hair plates can be found at most leg joints in flies and have been shown in larger insects to provide sensory feedback to motor neurons that control walking (Wong and Pearson, 1976). Hair plate axon projections from the legs are morphologically quite similar to those of some classes of campaniform sensilla, making them difficult to match reliably to light level (A Büschges, personal communication). We annotated two leg hair plate types, SNpp45 and SNpp52 (Figure 61A), but there may be more in our unclassified proprioceptive population (SNppxx). The leg hair plate neurons we identified vary widely in their quality of reconstruction and are unevenly represented across the nerves, implying that some examples either were not recovered or were not identified.

**Figure 61.**
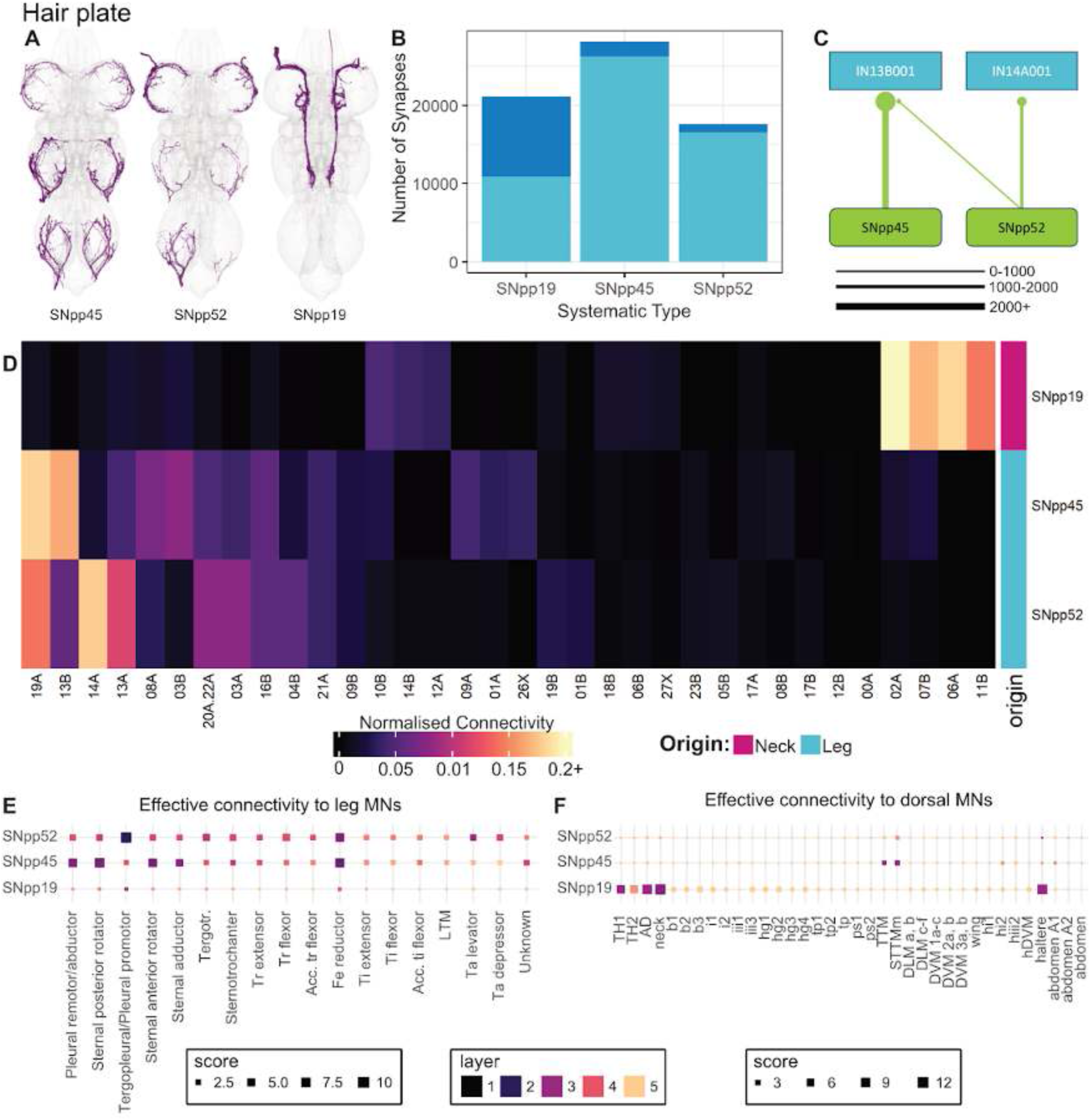
Hair plate connectivity. **A.** Plotted meshes of hair plate types. **B.** Number of synaptic outputs of hair plate types onto intrinsic neurons (cyan) and ascending neurons (blue). **C.** Diagram displaying the principal downstream connections of leg hairplate types. **D.** Hair plate types clustered by connectivity to hemilineages. **E.** Effective connectivity to ipsilateral leg motor neuron targets. **F.** Effective connectivity to ipsilateral dorsal motor neuron targets. Layer legend is shared between E and F.

The prosternal organ consists of two fused hair plates on the ventral surface of the neck and encodes head rotations along all three axes, contributing to compensatory head movements (Paulk and Gilbert, 2006). We identified one neck hair plate type, SNpp19, which makes roughly 50% of its outputs onto ascending neurons (Figure 61B). In contrast, the leg types appear to be much more heavily involved in local circuitry.

Our three hair plate types differ in their connectivity to downstream hemilineages, with the neck type strongly targeting dorsal hemilineages 02A, 07B, 06A, and 11B, whereas the leg types target leg hemilineages (Figure 61D). The leg types share strong connectivity to 19A, but they have distinctive connections to other hemilineages, with SNpp45 strongly targeting 13B whilst SNpp52 targets 14A and 13A. This differential connectivity to hemilineages is reflected in each type’s strongest respective partners, primary serial type IN13B001 for SNpp45 vs IN14A001 for SNpp52 (Figure 61C).

The functional split between the neck and leg types is maintained in their effective connectivity to motor neurons (Figure 61E). The two leg types share relatively strong indirect connectivity to the femur reductor MN and are weakly upstream of many other ipsilateral motor neurons (Figure 61 - figure supplement 1). However, SNpp45 neurons have greater effective connectivity to the plural remotor/abductor, sternal posterior and anterior rotators, and sternal abductor MNs, whereas SNpp52 neurons are strongly upstream of the tergopleural/pleural promotor MNs. These differences suggest distinct functional roles, with SNpp45 potentially more involved in walking speed changes or walking modes and SNpp52 more involved in posture adjustments. We note here that SNpp52 could possibly be a campaniform sensilla rather than hair plate type, as the morphologies are similar and the effective motor connectivity would also be consistent. The neck type, SNpp19, features strong effective connectivity to both ipsi- and contralateral TH1, TH2, AD, neck, and haltere motor neurons (Figure 61 - figure supplement 1) even though it is restricted to one hemisphere. This suggests a role in bilateral coordination of the neck and halteres in response to neck posture during flight.

**Figure 61 - figure supplement 1.**
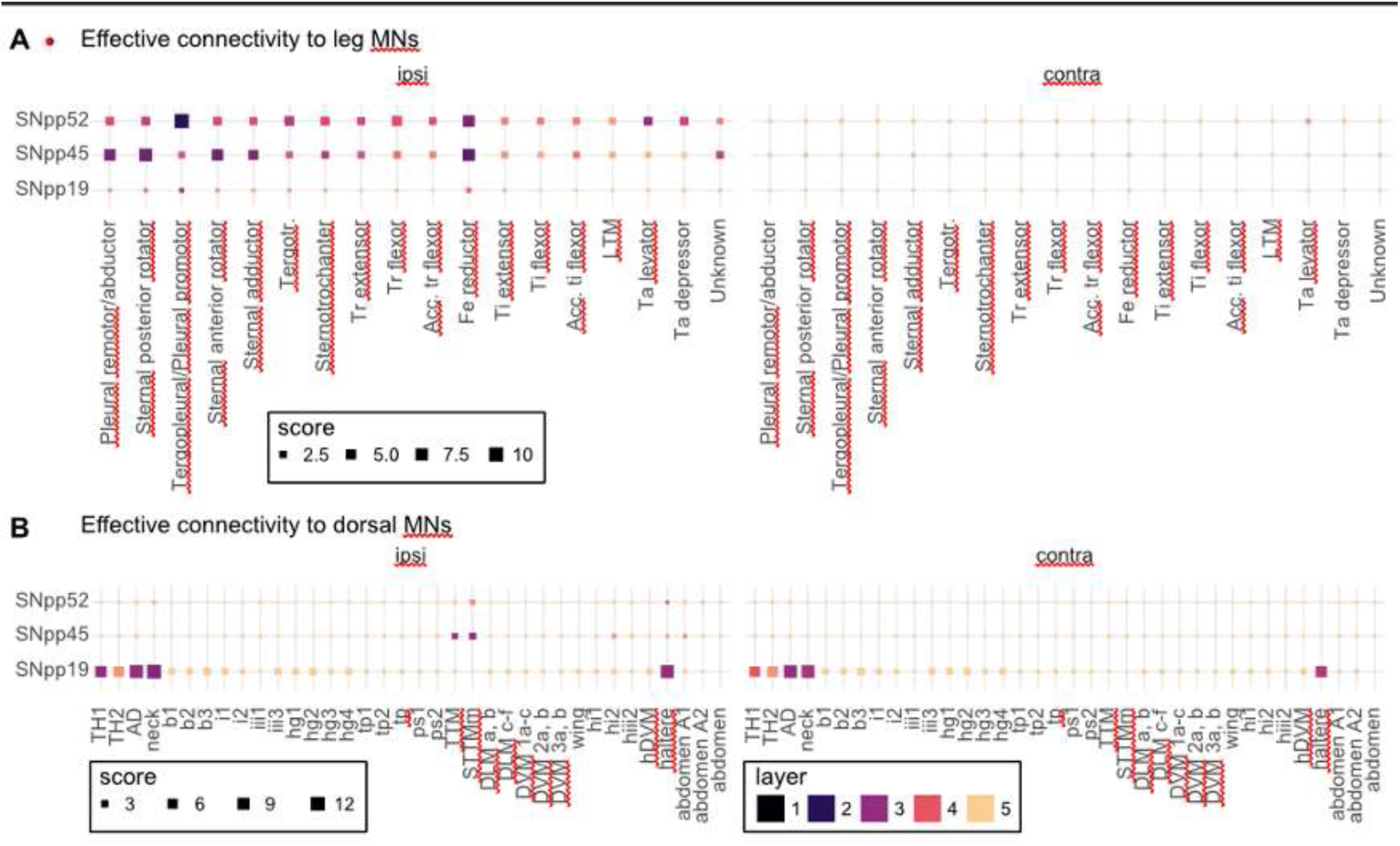
Hair plate effective connectivity. **A.** Effective connectivity of selected hair plate types to ipsilateral and contralateral leg motor neurons. **B.** Effective connectivity of selected hair plate types to ipsilateral and contralateral dorsal motor neurons.

#### Tactile neuron types

Tactile hairs, referred to as “bristles” in the *Drosophila* literature, are the primary detectors of externally generated mechanical forces (reviewed in (Tuthill and Wilson, 2016a)). An adult fruit fly has several thousand bristles distributed over the cuticle: large macrochaetes in stereotyped positions and small microchaetes distributed with more variability but still in regularly spaced rows. Anatomical studies in various insect species have shown that bristle axons in the CNS are systematically organised depending on mechanoreceptor location, physiological tuning, and receptor type. For example, axon terminals of neurons from sensilla on distal leg segments terminate in the most central/ventral part of the leg neuropil, whilst those from proximal segments terminate in more peripheral rings (Tsubouchi et al., 2017).

Here we identify 45 tactile sensory neuron types from the wing (4), wing margin (6), haltere (1), legs (27), and notum (7) (Figure 62A). We first compared the downstream connectivity of tactile neurons from each region. Haltere knob hair sensilla neurons exhibit strong, specific connections to hemilineages 03B, 01A, and 19A (Figure 62Bi). Tactile neurons from all other regions share strong connectivity to hemilineage 23B. However, we also observe region-specific patterns of downstream connectivity to hemilineages, with leg types targeting 01B and wing and notum types connecting more broadly to 17A, 09B, and 05B. We see more striking differences between the wing margin, wing, and notum populations when examining connections to single cell types (Figure 62Bii). Intriguingly, we find that wing, wing margin, and particularly notum tactile neurons directly target a descending neuron, DNxn166.

**Figure 62.**
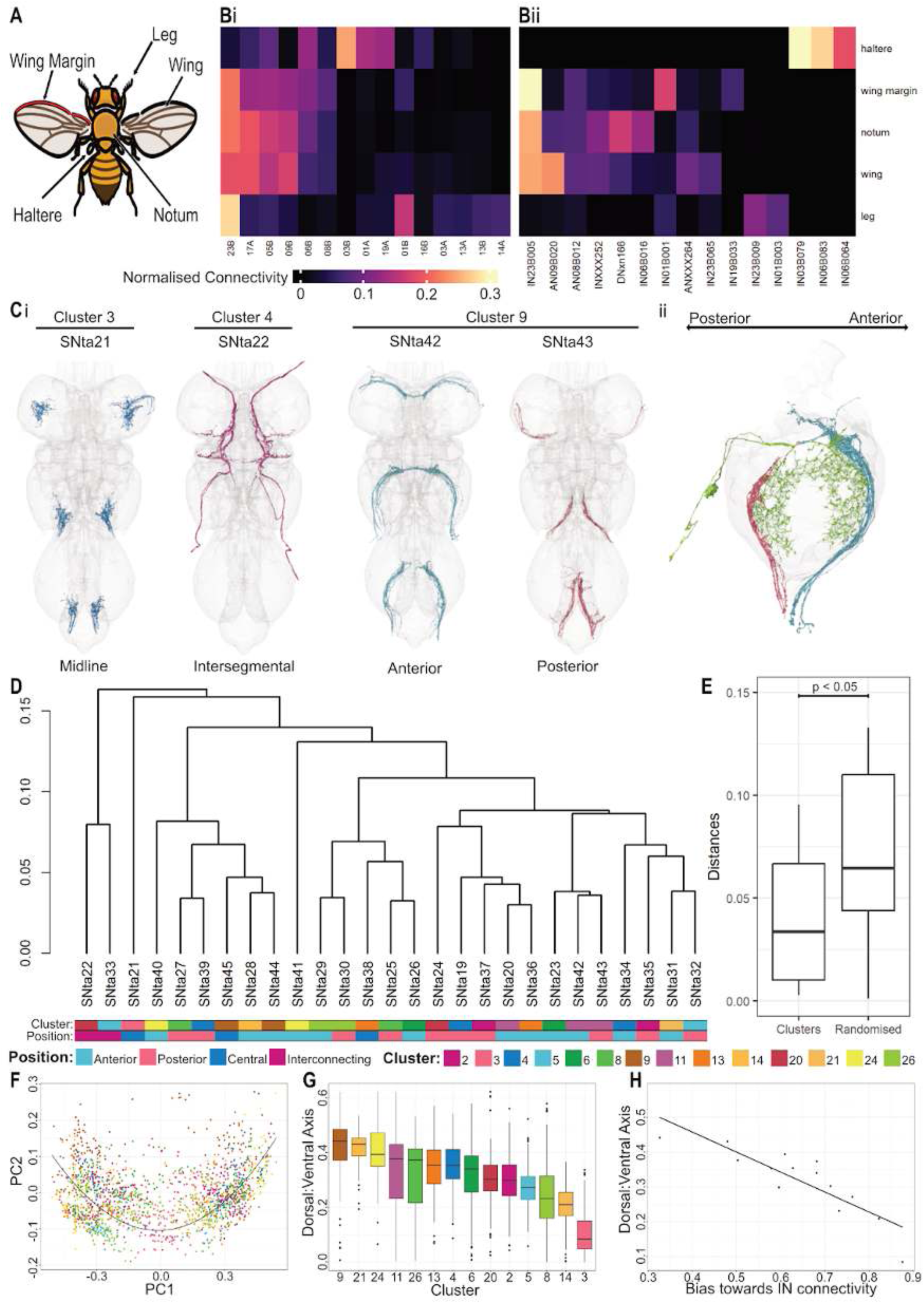
Tactile neuron types. **A.** Principal regions of origin for tactile sensory neurons within the VNC. **B.** Heatmap showing downstream connectivity of tactile neurons to i. hemilineages, and ii. strongly associated downstream partner types, downstream partners shown receive a minimum of 5% of outputs from at least one regional group. **C.** i. Neuron plots showing type representatives from different positions and clusters. ii. Neuron plot showing overlap between SNta42 and SNta43 with a strong downstream partner (bodyid: 12405). **D.** Downstream connectivity of leg tactile sensory types. Row annotation displays relative position within the neuropil and the cluster from which these types were assigned (see methods). **E.** Quantification of intra-cluster distances from clustering outputs onto ascending and intrinsic neurons vs. a randomised control (p = 0.036). **F.** Relative positions of clusters within the standard leg neuropil. Colours are consistent to those indicated in C. **G.** Patterning of tactile clusters across the dorsal ventral axis. **H.** Relative bias towards intrinsic neuron connectivity for each cluster, broken down by birthtime.

As part of our cell typing process, we distinguished between morphological categories of tactile neurons; for example, the leg tactile neurons can be separated into four categories based on their projections in the leg neuropil: midline, intersegmental, anterior, and posterior (Figure 62C). Previously, anterior projections have been shown to map to the anterior compartment of the leg and posterior projections to the posterior compartment (Murphey et al., 1989b). But we have observed that strong downstream partners of tactile neurons generally innervate both the anterior and posterior ventral neuropil (example in Figure 62C). Furthermore, we split many connectivity-based cosine clusters into anterior vs posterior types *post facto* based on their projections, suggesting that morphologically distinct types can share a consistent connectivity profile. We quantified this by performing hierarchical clustering with respect to strong downstream partners (Figure 62 - figure supplement 1), focusing on secondary partners (Figure 62D). This suggested an association between cell types within the same cluster that was significant when compared to randomised control (Figure 62E).

Therefore, when we sought to identify a topographic tactile map within the VNC, we used our original connectivity clusters rather than the final morphological types. To compare innervation patterns across all legs, we repurposed our standard leg neuropil, fitting a curve to the positions of each tactile and chemosensory neuron to access the patterning (including the chemosensory neurons qualitatively improved curve fit) (Figure 62F). We assigned each tactile neuron its closest point along the curve, considering this to be its position along the adjusted dorsoventral axis (Figure 62 supplementary 2). A comparison of this metric across clusters reveals an overlapping tiling of the dorsoventral axis (Figure 62G). Unexpectedly, we find that dorsoventral position correlates with a preference for intrinsic vs ascending neuron targets (Pearson coefficient = 0.86), with more ventral clusters showing a stronger bias towards intrinsic neurons (Figure 62H). Taken together with light-level studies (Tsubouchi et al., 2017), this suggests that information from the distal parts of the leg tend to be processed locally within the VNC, whereas information from the more proximal parts are more likely to be communicated to the brain.

**Figure 62 - figure supplement 1.**
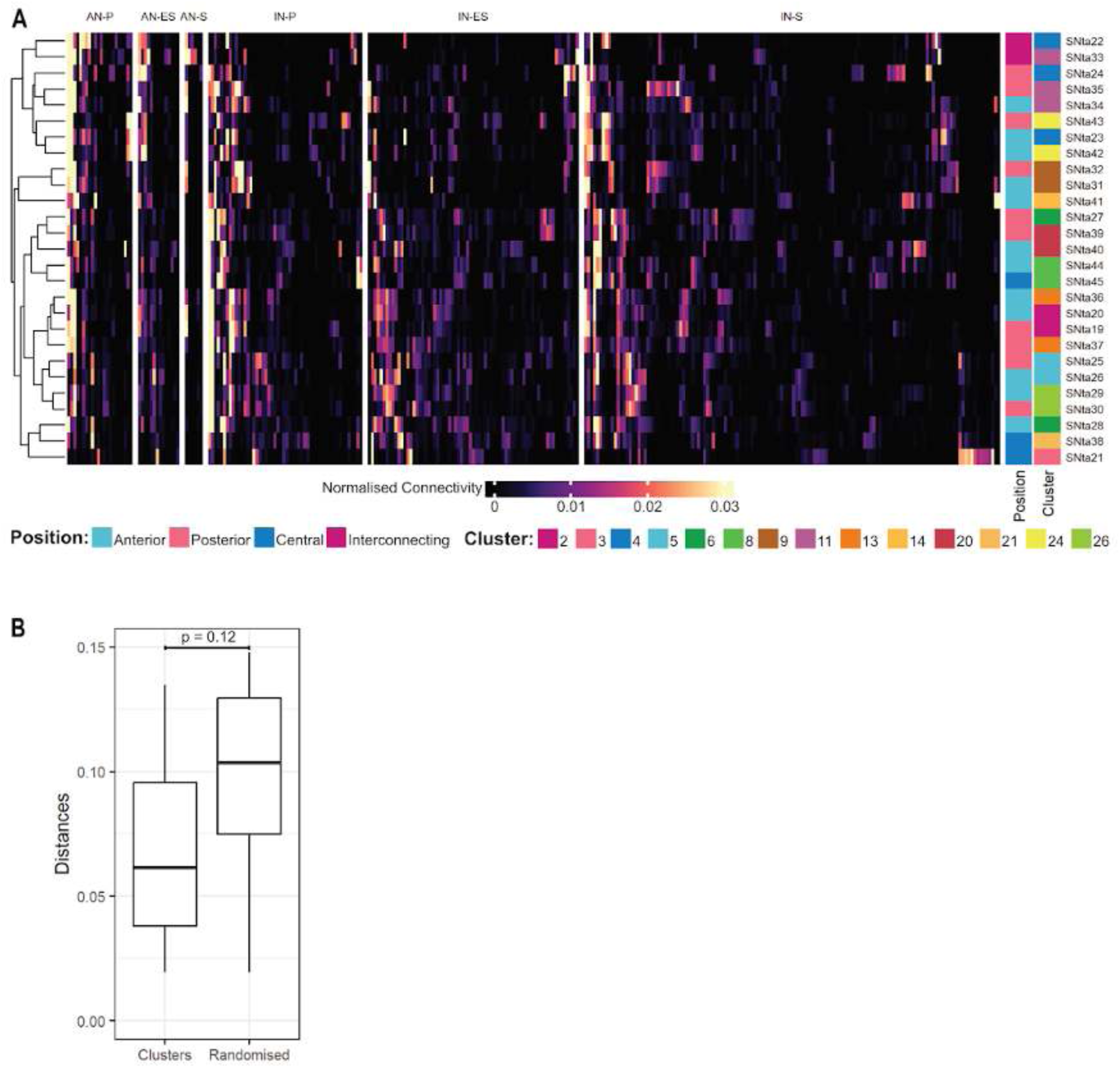
Tactile neuron types. **A.** Connectivity clustering of tactile types across all birthtimes (AN-P = ascending neurons, primary; AN-ES = ascending neurons, early secondary; AN-S = ascending neurons, secondary; IN-P = intrinsic neurons, primary; IN-ES = intrinsic neurons, early secondary; IN-S = intrinsic neurons, secondary). **B.** Quantification of intra-cluster distances from clustering outputs onto ascending and intrinsic neurons across all birthtimes vs. a randomised control (p = 0.12).

**Figure 62 - figure supplement 2.**
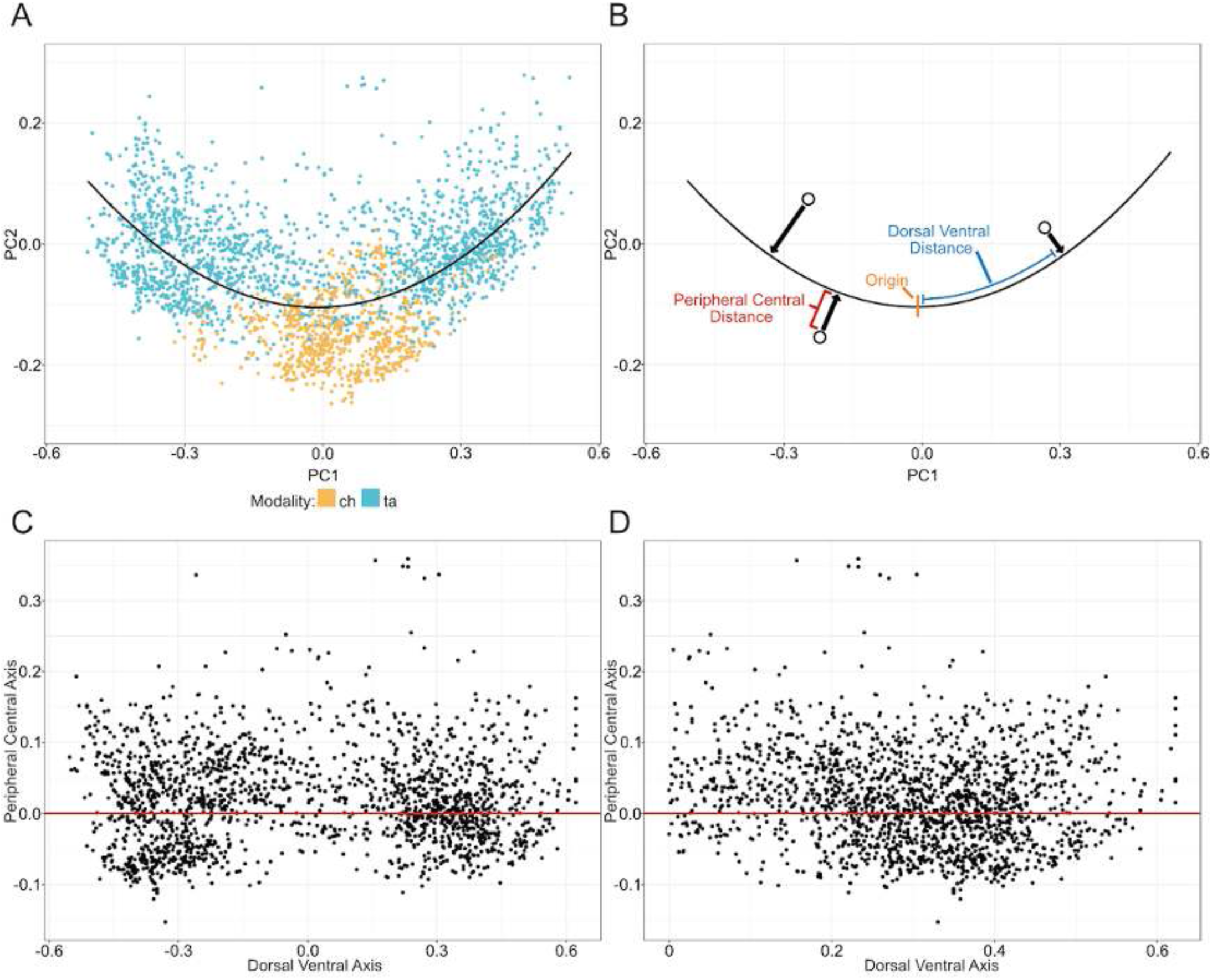
Adjusted dorsal-ventral axis methods. **A.** Positioning within the standard leg neuropil of tactile (ta) and chemosensory (ch) leg-originating sensory neurons. Line (black) indicates the line of best fit through the data points. **B.** Demonstration of calculation of adjusted axes. Dorsal ventral distance is the distance along the line of best fit between the nearest point on the line for each neuron to the minimum point on the line (origin). The peripheral central distance is the distance from each datapoint to the line of best fit. **C** Replotted data points according to newly calculated axes. **D** Replotted data points showing the absolute value of the dorsal ventral axis used for data analysis.

#### Chemosensory neuron types

Besides the well-characterised chemosensory neurons on head structures that project to the antennal lobe and suboesophageal ganglion, flies possess chemosensory neurons on the tarsae and tibia of all legs and on the anterior wing margin (reviewed in (Stocker, 1994)). Taste neurons from the legs project to the ipsilateral VAC of the associated neuromere, with a small subset ascending directly to the brain, whereas those from the wing margin project to the ovoid. Expression patterns of all known gustatory receptor isoforms have elucidated the axon projection patterns of specific populations of leg and wing chemosensory neurons in the VNC (Kwon et al., 2014); however, in most cases these are too similar to distinguish from morphology alone.

In MANC we distinguished 14 chemosensory systematic types, two of which ascend directly to the brain with few or no connections in the VNC. Four types originate in the wing, nine in the legs, and one in the abdomen. The abdominal type (SNch01) repeats in each abdominal segment and is very likely to include the abdominal ppk+ nociceptive class IV da cells (S Rumpf, personal communication). We have been able to match three of the remaining types to functional populations in the published literature. As mentioned previously, SNch08 (Figure 55D) is a male-specific, pheromone-responsive population from the sex comb on the front leg. We have also identified SAch02 and SNch11 as sweet-responsive leg gustatory neurons based on their characteristic axon projections (Kwon et al., 2014).

The remaining populations were not well enough distinguished in light-level images to match against our types. We can, however, discern distinct projection patterns of connectivity-defined cell types in MANC neuropils. In the VAC, there is clear lamination of many leg types including SNch05, SNch06, SNch07, and SNch09 (Figure 63A). Afferents from the wings show a different pattern: two of the four types form discrete microglomeruli whilst the other two overlap significantly, suggesting that they either convey related information or are two subsets of the same overall type (Figure 63B).

**Figure 63.**
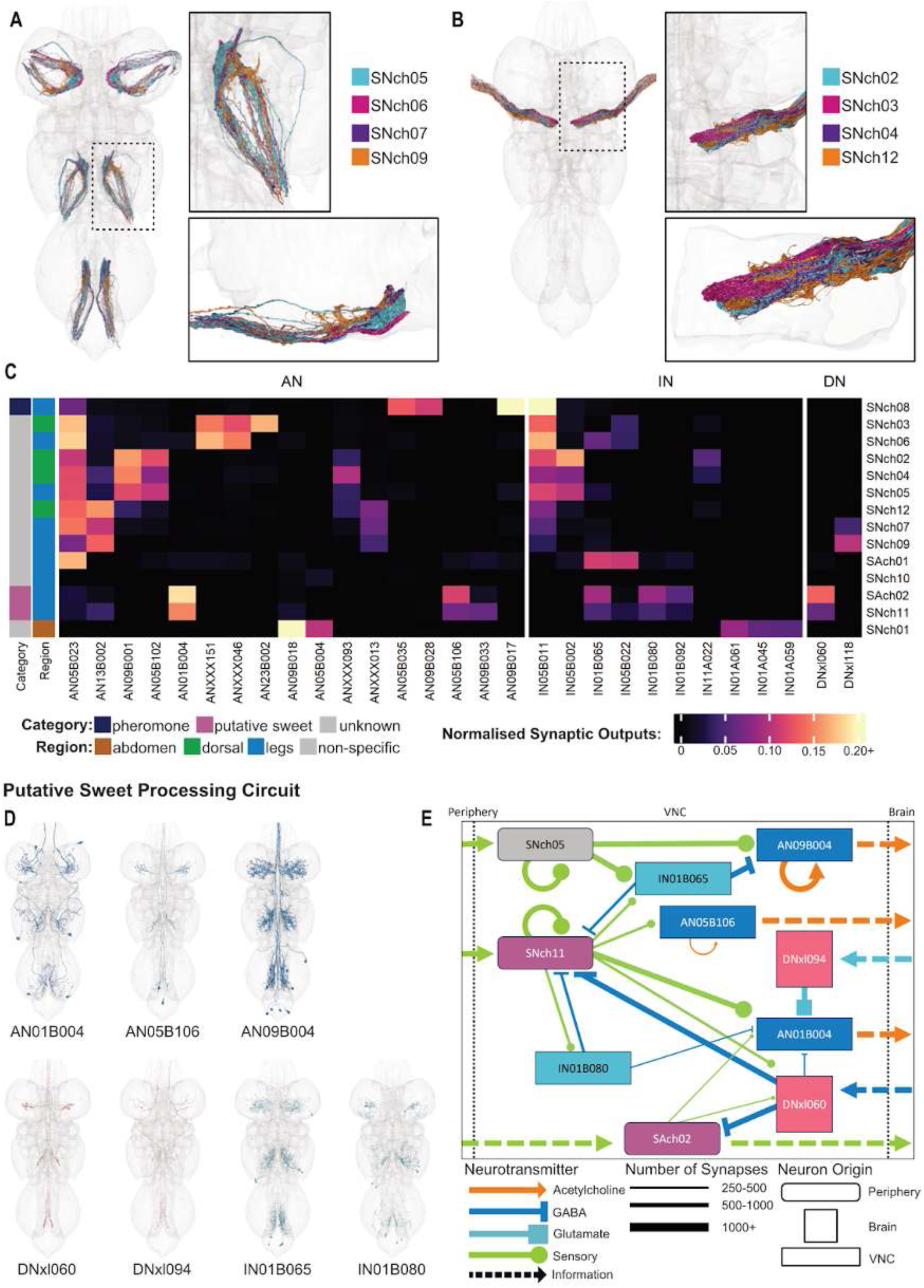
Chemosensory neurons and a putative sweet taste circuit. **A.** Chemosensory neurons originating in the legs, coloured by type. i. All types plotted in symmetrised volume, ventral view. ii. Close-up of neuropil in dotted rectangle, ventral view. iii. Rotation of selected neuropil to reveal type-specific layers. **B.** Chemosensory neurons originating in the wings, coloured by type. i. All types plotted in symmetrised volume, ventral view. ii. Close-up of neuropil in dotted rectangle, ventral view. iii. Rotation of selected neuropil to reveal type-specific layers. **C.** Heatmap showing downstream connectivity to chemosensory implicated neuronal types. Row annotations indicate the region of origin and the tastant category assigned to the sensory neuron type. **D.** Plotted meshes of strongly connected downstream partners of putative sweet taste neurons. **E.** Putative sweet taste circuit with specific downstream partners.

We explored the downstream connectivity of our chemosensory types with respect to dedicated chemosensory target neurons (Figure 63C). SNch01 does not share downstream partners with any other type; its top ascending partners AN09B018 and AN05B004 presumably alert the brain to noxious chemical stimuli, while its top intrinsic partners are all from hemilineage 01A and might be expected to initiate reflexive abdominal withdrawal. The male-specific pheromonal type (SNch08) also has some unique downstream partners (AN05B35, AN09B017, and AN09B028) that likely convey signal-specific information to the brain. Our two putative sweet types share many downstream partners including AN01B004, AN05B109, IN05B065, IN05B080, and DNxl118, suggesting that they may elicit similar responses.

Of the remaining chemosensory types, we can detect three main clusters: 1) SNch03 and SNch06,which have strong connections onto ANXXX046, ANXXX151, and AN23B002; 2) SNch02, SNch04, and SNch05, which have strong connections to IN05B002, AN09B001, and AN05B102; and 3) SNch07, SNch09, and SNch12, which connect strongly to ANXXX013 and AN13B002. We note that the types belonging to each of these connectivity groups originate in more than one peripheral region. This suggests that the downstream neurons from each cluster may be integrating cues with the same functional relevance from different appendages.

##### Putative sweet taste processing circuit

Finally, we investigated the local circuitry of our putative sweet types and their downstream neurons, uncovering a highly parallel local circuit (Figure 63D-E) which suggests that peripheral sweet information is integrated in several distinct pathways. First, SAch02 ascends directly. Second, information from SNch11 ascends in an independent, parallel pathway via AN05B106. Both sweet types target DNxl060, which in turn inhibits both sweet populations. Contextually embedded ascending information is conveyed through AN01B004, which receives local connections from both sweet sensory types as well as sweet intrinsic (IN01B080) and descending (DNxl060) partners and is also strongly downstream of a second descending neuron, DNxl094. IN01B080 also feeds back locally onto SNch11. We also observe inhibitory connections from sweet sensitive types onto other chemosensory pathways: SNch11 activates IN01B065, which inhibits ascending neurons downstream of SNch05.

##### Pheromone-responsive chemosensory types

Male flies’ forelegs feature additional chemosensory bristles containing neurons that express the sex determination gene *fruitless* (Demir and Dickson, 2005; Manoli et al., 2005) and have sexually dimorphic axonal projection patterns in the VNC (Possidente and Murphey, 1989). These ∼31.5 neurons express ppk23 and fall into at least two functional classes (Thistle et al., 2012; Toda et al., 2012): a subset also express ppk25 and respond to female pheromones including 7,11 heptacosadiene, whilst others are ppk25^-^ and respond to male pheromones that inhibit courtship (Kallman et al., 2015; Starostina et al., 2012; Vijayan et al., 2014). These male- and female-responsive subpopulations are very similar in appearance.

We initially identified one systematic type of foreleg chemosensory neurons, SNch08, as a morphological match to the ppk23^+^ pheromone-responsive population. However, closer examination revealed that SNch08 could be further subdivided into at least four subtypes by morphology and connectivity: two unilateral subtypes, SNch13 and SNch14, and two bilateral subtypes, SNch15 and SNch16 (Figure 64A-B). Unilateral type SNch14 is strongly upstream ipsilaterally of the PPN1 (group 12714, AN05B023 in A2) (Figure 64 - figure supplement 1 and Figure 22 - figure supplement 2) and upstream of group 12383 (Figure 64A,C), both of which have been reported to be activated by female pheromones (Clowney et al., 2015; Kallman et al., 2015), but we cannot associate the other types with any specific chemosensory cues at this time.

**Figure 64.**
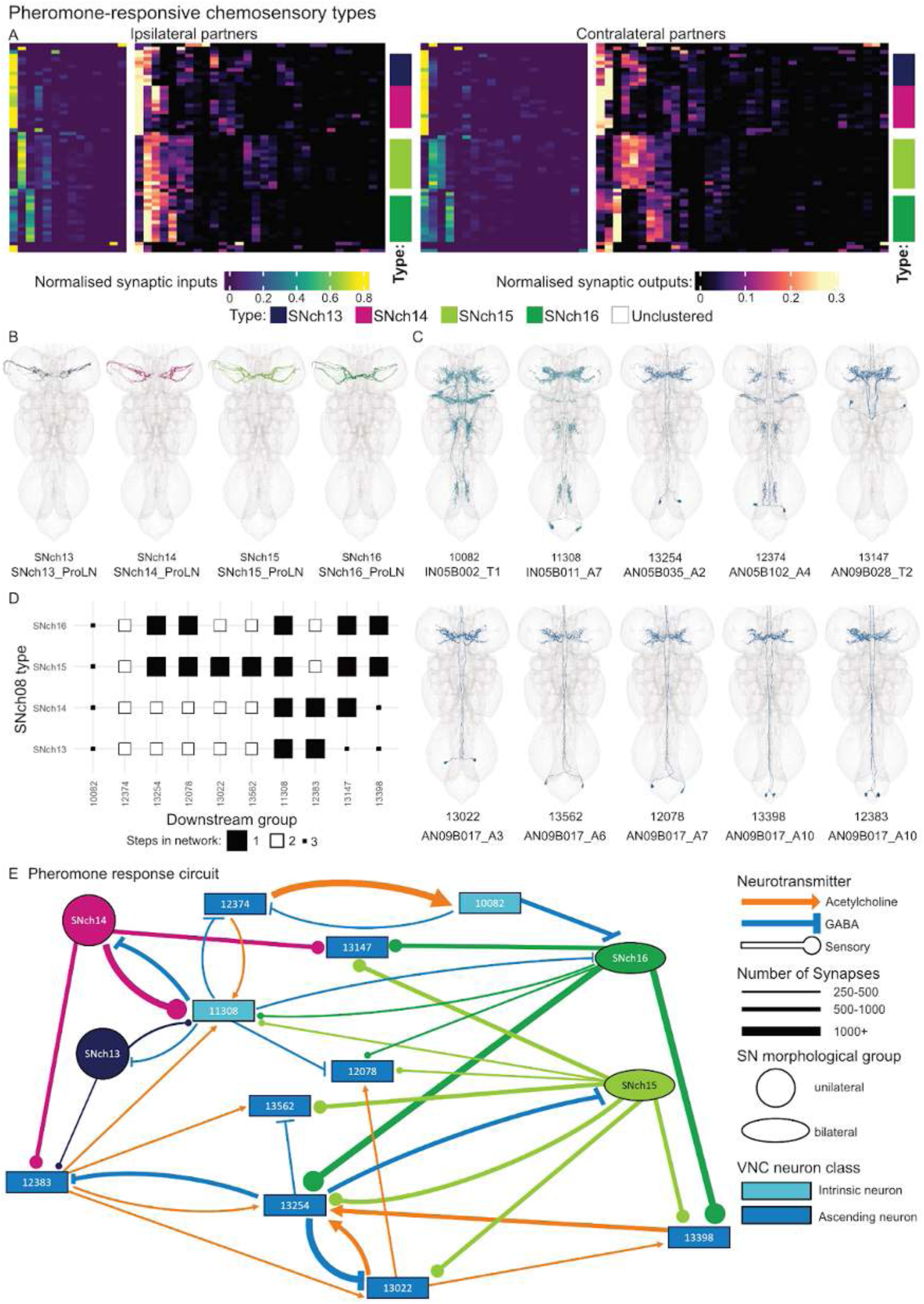
Pheromone-responsive chemosensory types. **A.** Heatmaps showing individual SNch08 neurons, clustered by connectivity to ascending neuron types. Row order was determined by hierarchical clustering with respect to both inputs and outputs (see Figure 64 - figure supplement 1) and maintained in all plots. **B.** Plotted meshes of four SNch08 subpopulations/types. **C.** Plotted meshes of VNC neurons strongly downstream of pheromone-responsive type SNch08. **D.** Minimum layer plot. **E.** Putative pheromone processing circuit with selected downstream partners. Nodes are coloured by type for sensory neurons and by class for target neurons (cyan = intrinsic neuron, blue = ascending neuron). Edges are coloured by type for sensory neurons and by predicted neurotransmitter for target neurons (dark orange = acetylcholine, blue = GABA).

Most of the strongest partners of these putative pheromone-responsive chemosensory types are morphologically similar ascending neurons, with a characteristic bowtie-shaped dendritic elaboration in the T1 leg neuropils, making them challenging to identify with certainty at light level (Figure 64C). All belong to hemilineage 05B or 09B, and at least some are likely to be serial homologues; however, it is difficult to determine whether targeting the SNch08 domain is a conserved feature of serial homologues across neuromeres or a convergent one between specific members of distinct serial sets. In addition, each within-neuromere group features distinct connectivity in the VNC. We have therefore focused on groups instead of serial systematic types for our analysis.

We find that pheromonal cues are likely to be extensively processed in the VNC before they are relayed to the brain (Figure 64E). The exception is ascending neuron group 13147, which integrates signals from types SNch14, SNch15, and SNch16 (but not SNch13). The putative 7,11 heptacosadiene-responsive type SNch14 also targets ascending neuron group 12383, which is expected to activate 13254, a gabaergic ascending neuron that is downstream of all four sensory types and of cholinergic targets 13022 and 13398 and which in turn inhibits 13022 as well as SNch15 and its target 13562. SNch14 also targets intrinsic neuron group 10082, which is activated by all four types and inhibits them and several of their targets, possibly damping down the pheromone response or sharpening the timing of pheromone perception. Understanding the biological significance of this network awaits functional experiments targeting specific types as well as elucidation of their ascending neuron-mediated circuits in the brain. It will also be very interesting to compare these pheromone-responsive circuits with the analogous circuits in female datasets such as FANC (Phelps et al., 2021).

**Figure 64 - figure supplement 1.**
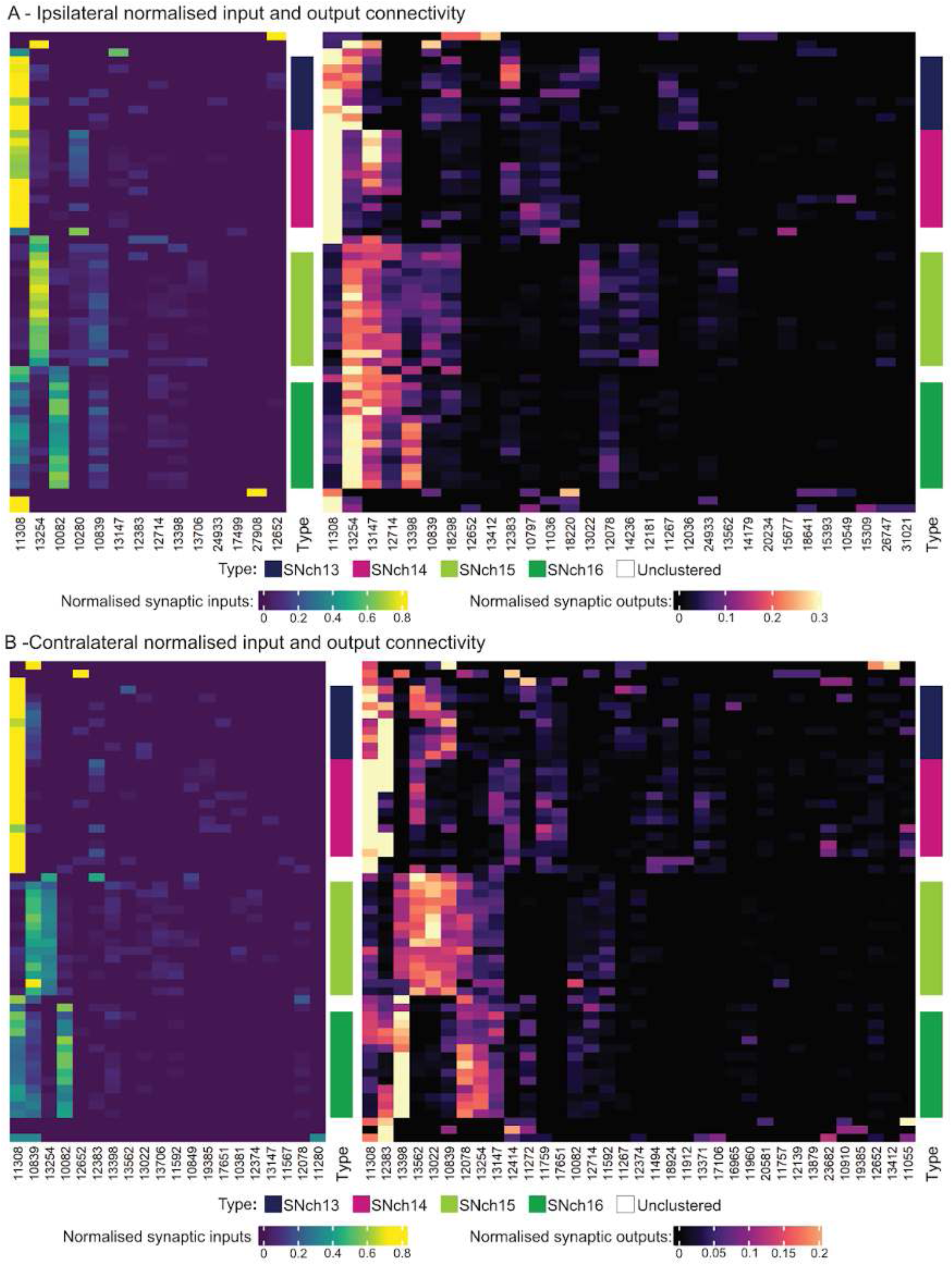
Pheromone response circuit. **A.** Ipsilateral normalised input and output connectivity between four SNch08 subtypes and their strongest downstream targets. **B.** Contralateral normalised input and output connectivity between four SNch08 subtypes and their strongest downstream targets.

**Figure 64 - figure supplement 2.**
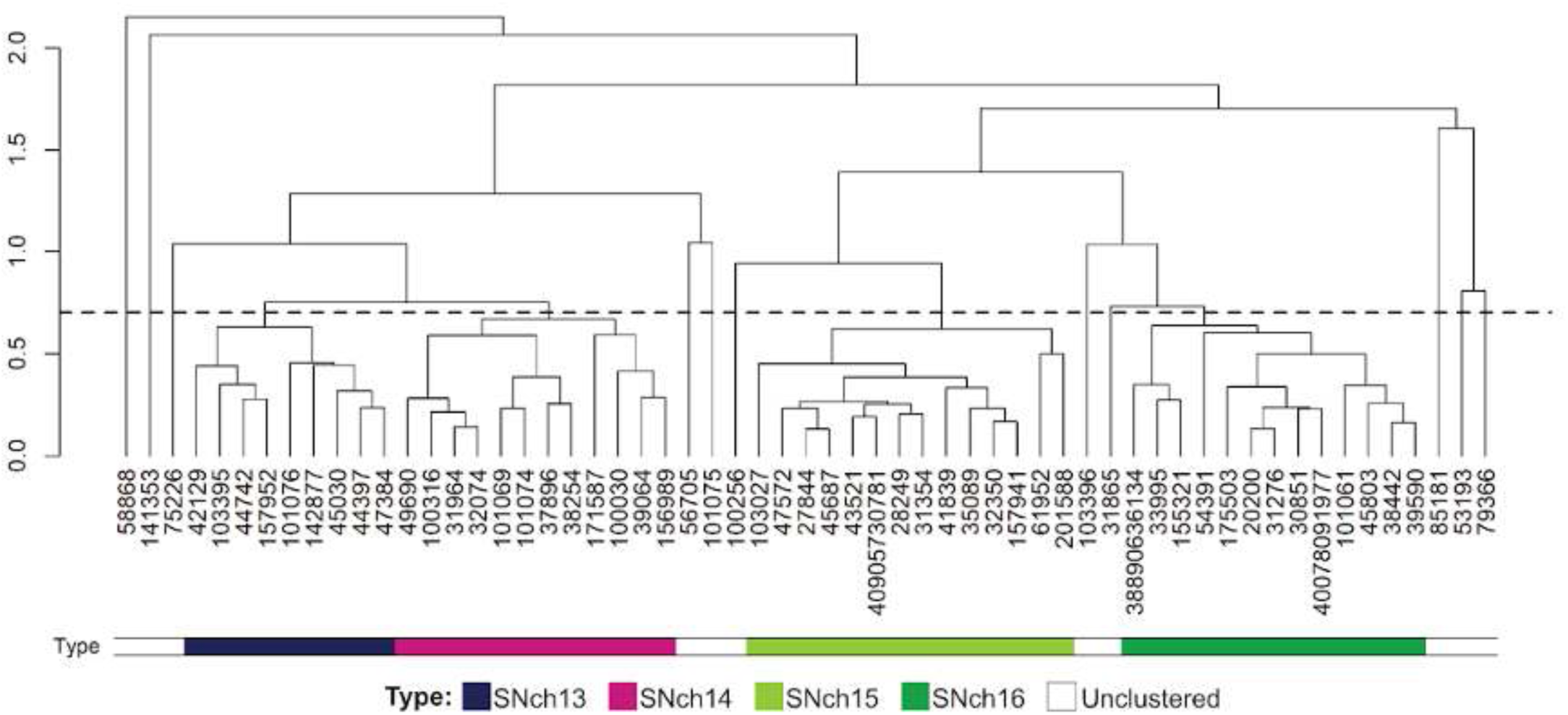
Pheromone response circuit. Dendrogram used for hierarchical clustering of SNch08 neurons. The dashed line represents the cut height used to define the four main clusters/subtypes.

## DISCUSSION

### Annotation and systematic classification

The ventral nerve cord is the seat of most insect behaviours and a long-standing model system for sensorimotor circuits. Recent synaptic resolution VNC connectome projects (Lesser et al., 2023; Ohyama et al., 2015; Phelps et al., 2021) have taken the first steps towards fully understanding its composition and organisation. In this study we have described the functional organisation of the first complete male adult nerve cord connectome, MANC (Takemura et al., 2023). We have compiled a ∼complete inventory of contributing neurons, annotated key anatomical features, developed a systematic cell typing and nomenclature for ease of identification and analysis, and presented preliminary results on selected sensory circuits.

We have reconstructed and classified 13,060 VNC intrinsic neurons, 185 ascending neurons, 1328 descending neurons, 737 motor neurons, ∼100 other efferent neurons, and nearly 6500 sensory neurons (Figure 1). Altogether, we found 4483 neurons originating in T1, 5080 in T2, and 3988 in T3 (after correcting for 01B), which is remarkably close to the 4100 to 5000 neurons per thoracic neuromere estimated from postembryonic neuroblast division rates (Truman and Bate, 1988). In total we annotated 15,763 neurons originating in the thoracic and abdominal segments. For comparison, the first instar larval VNC would be expected to include approximately 300 neurons per segment (Poulson, 1950), although the most posterior neuromeres feature fewer neuroblasts; the first instar total of 3000 - 3900 neurons is therefore four to fivefold less than the adult. We annotated 1400 out of 13,288 thoracic neurons as primary (correcting for 01B in A1) or 10.5%. This is consistent with the 10% contribution predicted in (Truman and Bate, 1988). However, it exceeds the upper limit of estimates for the number of neurons in the three first instar thoracic segments, which range from 900 (Poulson, 1950) to 1026 (Winding et al., 2023). Although there is some cell death during metamorphosis, it seems likely that most primary neurons persist until adulthood, but as their numbers have only been estimated in the literature, firm conclusions await tabulation in a complete larval VNC data set.

Our extensive annotation (Figure 2, Table 1) allowed us to go far beyond mere cell counts and formed the basis of our systematic typing and nomenclature (Figure 3) that should prove extremely valuable in leveraging this connectome for light-level developmental and functional studies. We have assigned a developmental hemilineage to 88% of the 15762 VNC neurons (97% of the 13429 originating in thoracic neuromeres) (Figure 8) and a soma side, soma neuromere, and estimated birthtime (Figure 10) to each one. We have classified their input (origin) and output (target) neuropils and assigned them a subclass reflecting their innervation pattern (Figure 5). We have also annotated neurons with low presynaptic density that are likely to be electrical or neurosecretory (Figure 4). These annotated features, together with neurotransmitter predictions (Takemura et al., 2023), will allow dataset users to focus queries for particular neurons of interest. We did not attempt to identify every published image and cell type in the literature in our data set. Rather, as predicted coding regions had to be investigated as candidate genes in the original *Drosophila* genome project, future work by experts in specific areas will be required to match most VNC cell types to light level in order to bridge connectivity and functional analyses.

The ventral nerve cord has long been understood to consist of segmentally repeating ganglia generated in large part by repeated neuroblast arrays during embryonic and larval life. We were able to identify serial homologues in other neuromeres for over a third of VNC neurons (excluding descending and sensory) and to leverage these serial sets in numerous ways (Figure 64). Recognising the same neurons across neuromeres via similar morphology and connectivity helped us to assign individual examples to the correct hemilineage, soma neuromere, and/or birthtime, particularly in the abdomen (Figure 49). Moreover, our annotated serial sets were critical for classifying the remaining VNC neurons into segmentally repeating cell types (Figure 13), for typing leg motor neurons in T2 and T3 based on T1 identifications (Cheong et al., 2023), and for developing a systematic classification for sensory neuron cell types (Figure 52). We also used serial connectivity to detect functionally related neurons from distinct peripheral origins (Figure 58).

### Hemilineages as morphological and functional units

Earlier studies of adult VNC organisation have focused on neuroblast hemilineages as developmental, morphological, and functional units (Harris et al., 2015; Lacin et al., 2019; Shepherd et al., 2019, 2016; Truman et al., 2010, 2004). This work provided an invaluable framework for our classification of neurons and characterisation of circuit elements in MANC. However, many minority cell types could not be distinguished in light-level postembryonic neuroblast clones (Shepherd et al., 2019) and so have not yet been captured in driver lines or tested for function (Harris et al., 2015). Only with this synapse-resolution connectome can we assess the hemilineage as a functional unit, by estimating the diversity of morphology and connectivity of neurons from each hemilineage and by locating each neuron within the VNC network.

Based on light level studies performed at lineage or hemilineage resolution, neurons within a given hemilineage are expected to share numerous characteristics including gross morphology (Shepherd et al., 2019), general function (Harris et al., 2015), and fast-acting neurotransmitter (Lacin et al., 2019). We broadly confirm these expectations, although we find that several hemilineages, notably 09B and 19A, include neurons expressing distinct neurotransmitters (Figure 8). Specialised cell types such as electrical or neurosecretory neurons appear repeatedly in a small number of hemilineages (Figure 4). Also, specific hemilineages such as 05B, 08A, 08B, 09B, 12A and 17A are enriched for identified sexually dimorphic neurons, mainly involved in the pheromone-sensing or courtship song circuits (Figure 26C, Figure 27 - figure supplement 2-9, Figure 29 - figure supplement 2, Figure 33 - figure supplement 2-7, Figure 40C, Figure 64).

Annotating the estimated birthtime of each neuron based on soma tract morphology and location allowed us to revisit the predicted secondary hemilineage neuron network first posited on the basis of immature neuron morphology in the late larva (Truman et al., 2004). Whilst our observed T2 secondary network largely validates these predictions, several sets of hemilineages predicted to connect based on common commissures in the larva are not significant synaptic partners (e.g., 07B/08B/10B, 13B/14A, and 05B/06B/12B) (Figure 66A). Conversely, our work reveals many more strong connections between members of different hemilineages than could have been anticipated - a testament to the power of whole-volume connectomics (Figure 66B). We also added 23B, the 24B/25B motor neurons, and sensory neurons to this network. Finally, we have predicted neurotransmitters for every secondary hemilineage (Lacin et al., 2019; Takemura et al., 2023), refining our understanding of how they might function in such a network. Notably, we observe a high level of intra-hemilineage communication for specific hemilineages, particularly 03A, 04B, 07B, 08B, 17A, 19B, 20A/22A, and 23B (Figure 65 - figure supplement 1) - all cholinergic - suggesting the potential for a local feedforward mechanism.

**Figure 65.**
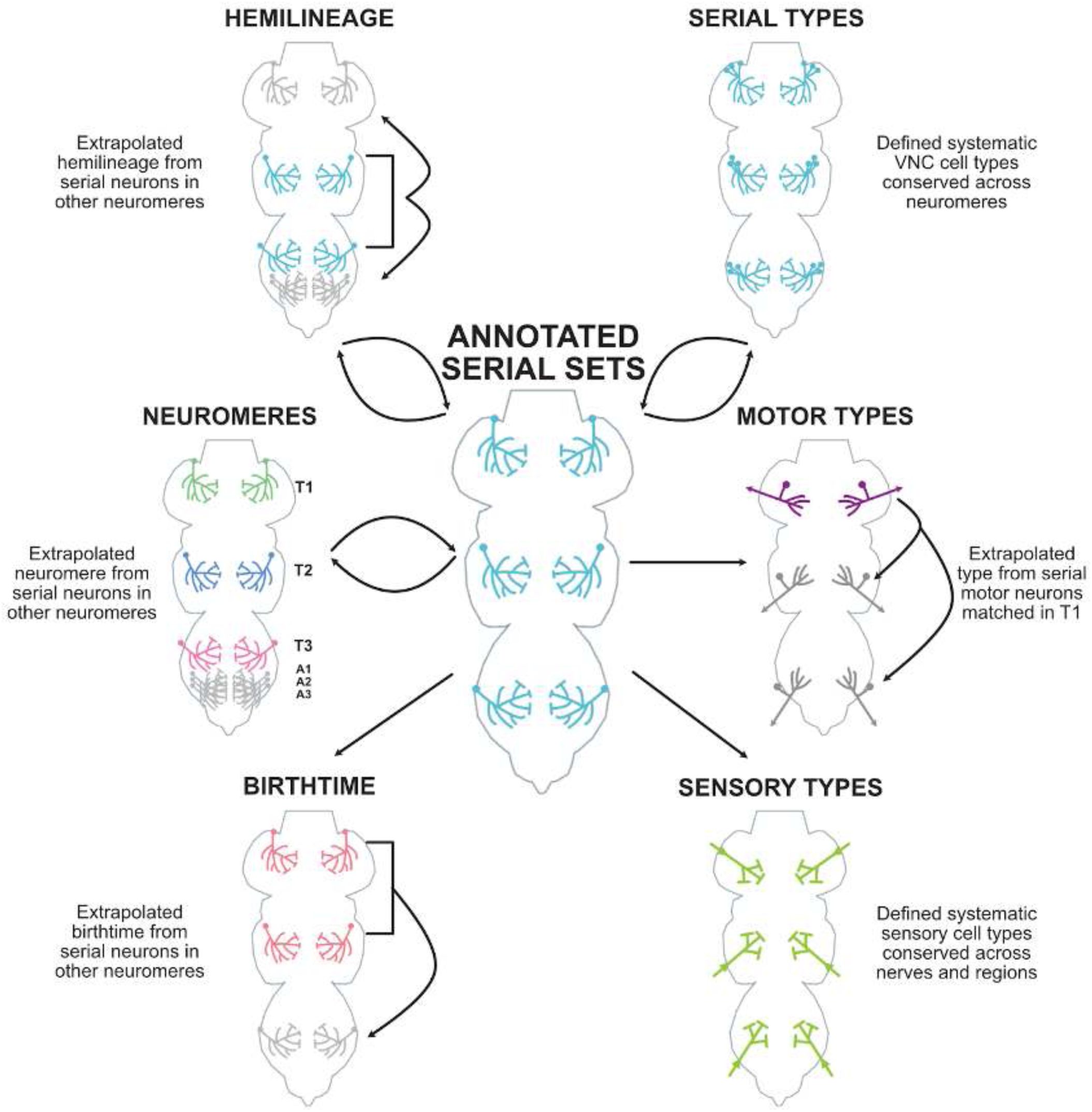
Uses of serial homology for annotation and systematic cell typing in MANC. The manual and programmatic identification of individual neurons that repeat in multiple neuromeres - referred to as serial sets (centre, cyan) - contributed to many aspects of the MANC annotation process: 1. Assignment of unknown neurons (grey) to the correct **hemilineage** based on serial homology to known neurons (cyan), especially in T1 or the abdomen. 2. Assignment of neurons to the correct left-right partner and/or soma **neuromere** in the abdomen (grey) based on serial homology to neurons in thoracic neuromeres (green = T1, blue = T2, magenta = T3). 3. Extrapolation of **birthtime** assignments in T1 and T2 (pink = primary) to inform those in T3 (grey), where somas are typically smaller. 4. Generation of systematic **serial types** (cyan) for neurons originating in the VNC. 5. Identification of leg **motor neuron types** in T2 and T3 (grey) based on serial homology to light level-matched cells in T1 (purple) (Cheong et al., 2023). 6. Generation of systematic **sensory cell types** (green) across nerves and peripheral regions.

**Figure 66.**
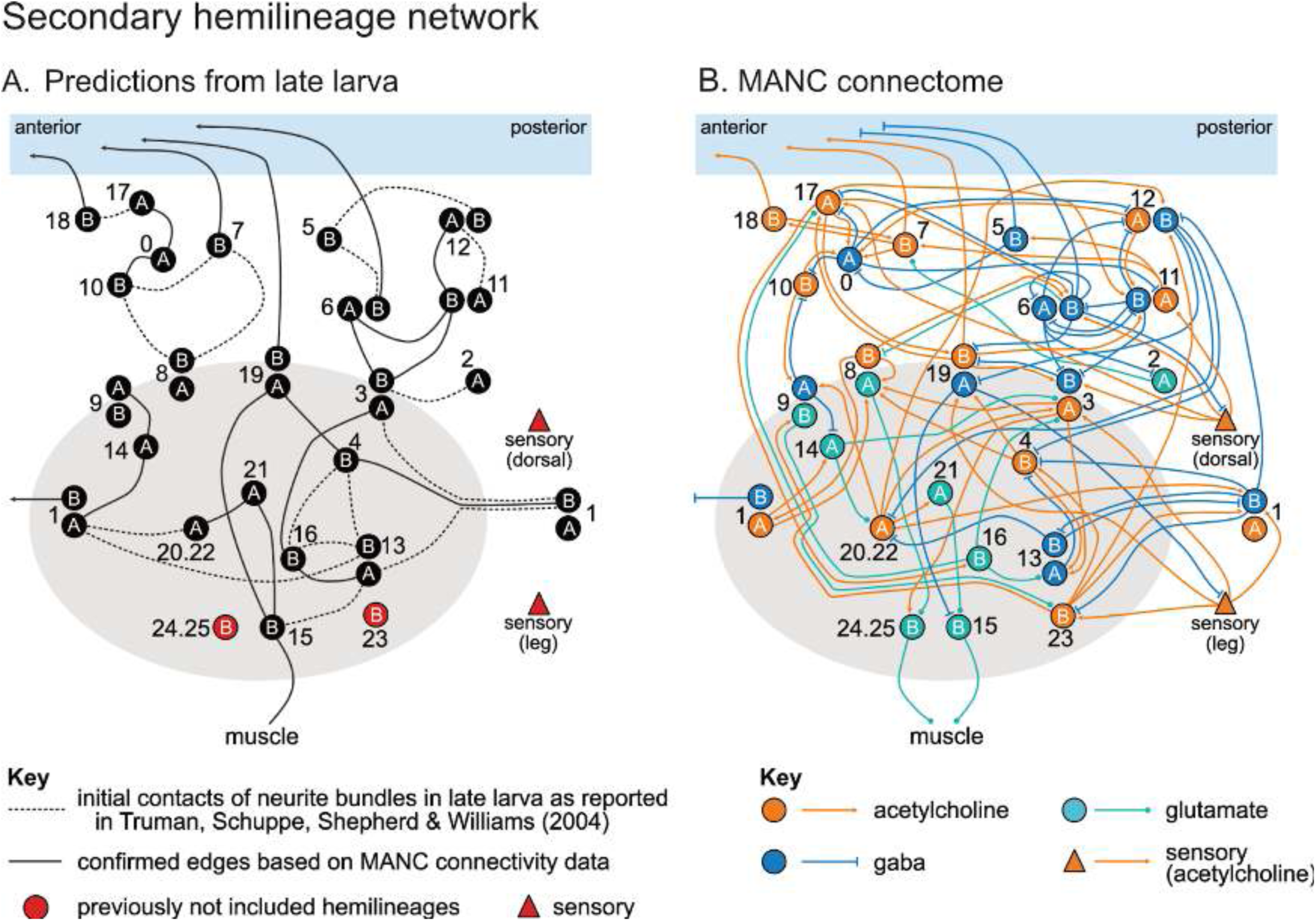
Secondary hemilineage network. Schematic summary of secondary hemilineage-to-hemilineage connectivity showing the top partners and consensus neurotransmitter prediction per hemilineage in the adult male T2 neuromere (B) compared against previously reported predictions from late larva (A). Within hemilineage connections are excluded from the schematic. **A.** The lines between hemilineages represent initial contacts of neurite bundles made in late larva as reported by Truman et. al., 2004. Bundles within the grey circle were hypothesised to terminate in the ventrolateral (leg) neuropil while those in blue stripe hypothesised to project in dorsal tracts. Solid black lines represent edges between hemilineages confirmed in the adult male based on a threshold of 5% of input or being in the top 4 partners. Thresholds were considered from the perspective of both hemilineage (i.e., hemilineage A receives at least 5% of its input from hemilineage B and hemilineage B provides at least 5% of its output to hemilinage A. **B.** All confirmed edges in (A) plus all newly reported hemilineage connections that are in the top 4 partners per hemilineage (considered from both hemilineage’s perspective). Neurotransmitter identity is based on the upstream hemilinage’s consensus neurotransmitter prediction.

However, we show that not all hemilineages can be considered to be morphological and functional units to the same degree. A few, such as 13B, 14A, and 20A/22A appear to be nearly homogeneous with respect to morphology, connectivity, and position within the VNC network, including across neuromeres (e.g., Figure 36 - figure supplement 1, Figure 37 - figure supplement 1, Figure 44 - figure supplement 1). However, many hemilineages are much more diverse (Figure 50). Positions in both local and global circuits differ across neuromeres for several hemilineages, suggesting different functions in different local circuits, as well as distinct functions in the global circuit of the VNC. Even when not identifiable by intra-hemilineage morphological disparity, these local circuit distinctions are identifiable in typing assignments (see supplemental materials for Figures 15-48). Hemilineages that have greater differences in network positions across neuromeres generally have less equal distributions of neuromeres in each type (i.e. they have lower Shannon Entropy in their distributions of counts of neuromere identity per type) (data not shown).

Early born neurons tend to be much more diverse, form many more connections, and likely play unique roles in the adult VNC network (Figure 51). However, very few have been previously characterised in the adult VNC; we are the first to report primary neuron morphologies and connectivity at scale (Figure 48, Figure 48 - figure supplement 1). Notably, primary neurons generally exhibit less intra-hemilineage and more inter-hemilineage connectivity than secondary neurons (Figure 65 - figure supplement 2), suggesting that they serve to connect functional populations rather than contribute to them. Primary neurons could be considered the most primitive cell types that define the ancestral bauplan of the CNS, with secondary neurons representing the post-embryonic expansion of particular types that slot into the existing network.

**Figure 66 - figure supplement 1.**
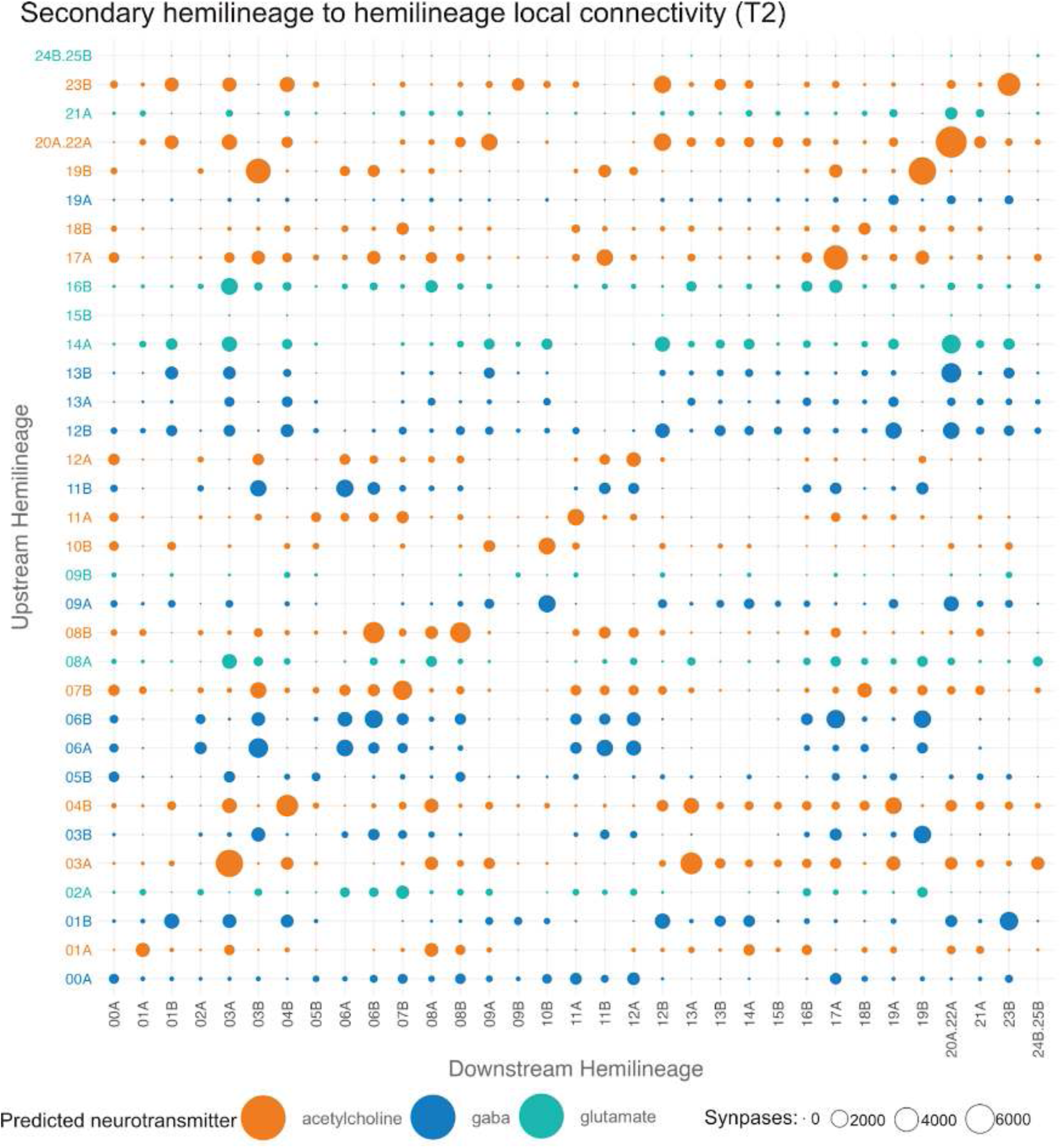
T2 secondary hemilineage to hemilineage connectivity. Bubble plot showing secondary hemilineage to hemilineage connectivity restricted to T2. Colour of bubble is based on the consensus neurotransmitter prediction for the given upstream hemilineage. Size of each bubble is proportional to the synaptic weight between hemilineages.

**Figure 66 - figure supplement 2.**
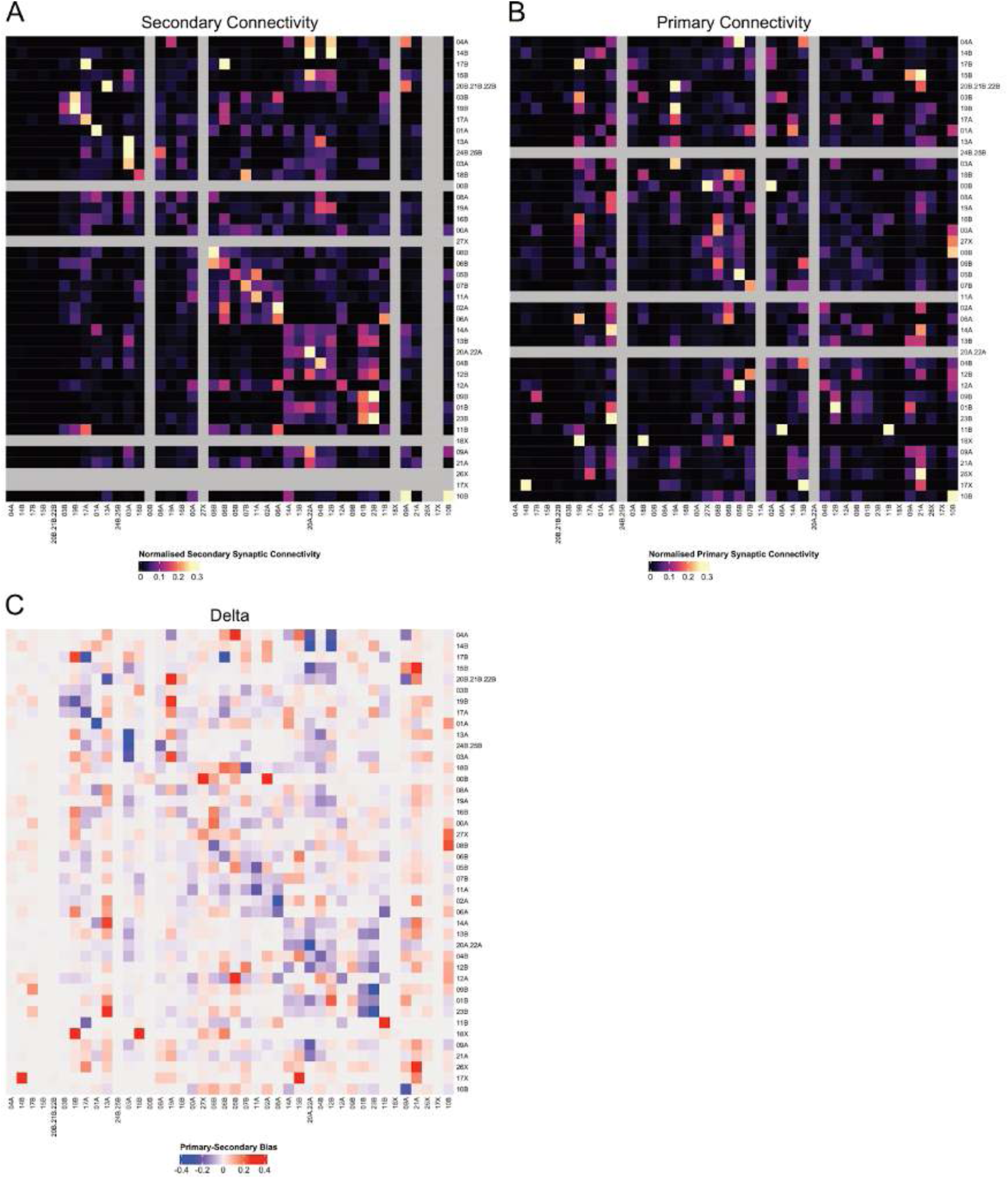
T2 hemilineage to hemilineage connectivity. **A.** Clustering of secondary hemilineages by secondary hemilineage to secondary hemilineage synaptic connectivity (grey cells represent primary-only hemilineages). **B.** Heatmap of primary hemilineages by primary hemilineage to primary hemilineage synaptic connectivity (grey cells represent secondary-only hemilineages). Ordering enforced to be consistent with A. **C.** Bias in connectivity to hemilineages by birthtime, calculated by subtracting the normalised primary connectivity from the normalised secondary connectivity. In cases where a hemilineage is primary or secondary only, a normalised connectivity score of 0 is assigned to the respective matrix. Ordering enforced to be consistent with A.

### Circuits for coordination of leg movements

*Drosophila* possess intricate leg circuits that enable them to navigate their environment with remarkable agility. While studies have successfully mapped the proprioceptive sensory input and motor output to individual leg neuropils (Azevedo et al., 2020; Brierley et al., 2012; Kuan et al., 2020; Tuthill and Wilson, 2016b), the INs that integrate these functions remain largely unknown. In our research, we have undertaken a significant effort to reconstruct and identify leg sensory neurons and (in a companion paper, (Cheong et al., 2023)) the leg motor neurons of the MANC dataset. Additionally, we have conducted a thorough annotation effort for the INs, encompassing cell typing, hemilineage annotation, birthtime, and innervation patterns, which enabled a comprehensive description of this neuron class.

This detailed understanding has provided us with the first complete grasp of the number of neurons involved in the leg circuits, both downstream of SNs and upstream of MNs (Figure 67). Notably, we have discovered that over 950 INs are concentrated within each of the six leg neuropils, accounting for 40% of all INs in the VNC (see also Figures 5 and 6). It is not surprising that the intrinsic leg circuits exhibit such complexity, considering their role in controlling 18 leg muscles per leg and coordinating various behaviours like turning during walking (more details in (Cheong et al., 2023)). For coordination between the leg neuropils specifically, we find an additional 930 neurons, with the largest subgroup connecting the left and right T3 leg neuropils. The uneven distribution of these bilateral connections (92+29 neurons for T1, 54+8 for T2, and 127+39 for T3, Figure 6E) indicates their role in the distinct coordination required for the various legs. The reduced neuron count for T2 coordination can be justified by its absence in the direct leg premotor circuit, as observed in the study by Cheong et al., suggesting that bilateral T2 coordination may not be essential for behaviour such as walking. Notably the prominence of additional neurons connecting T1 and T3 leg neuropils aligns with the specific behaviours associated with these legs that include simultaneous as well as alternating leg movements. For the T3 legs this includes grooming the abdomen, wing and thorax (Zhang et al., 2020).

**Figure 67.**
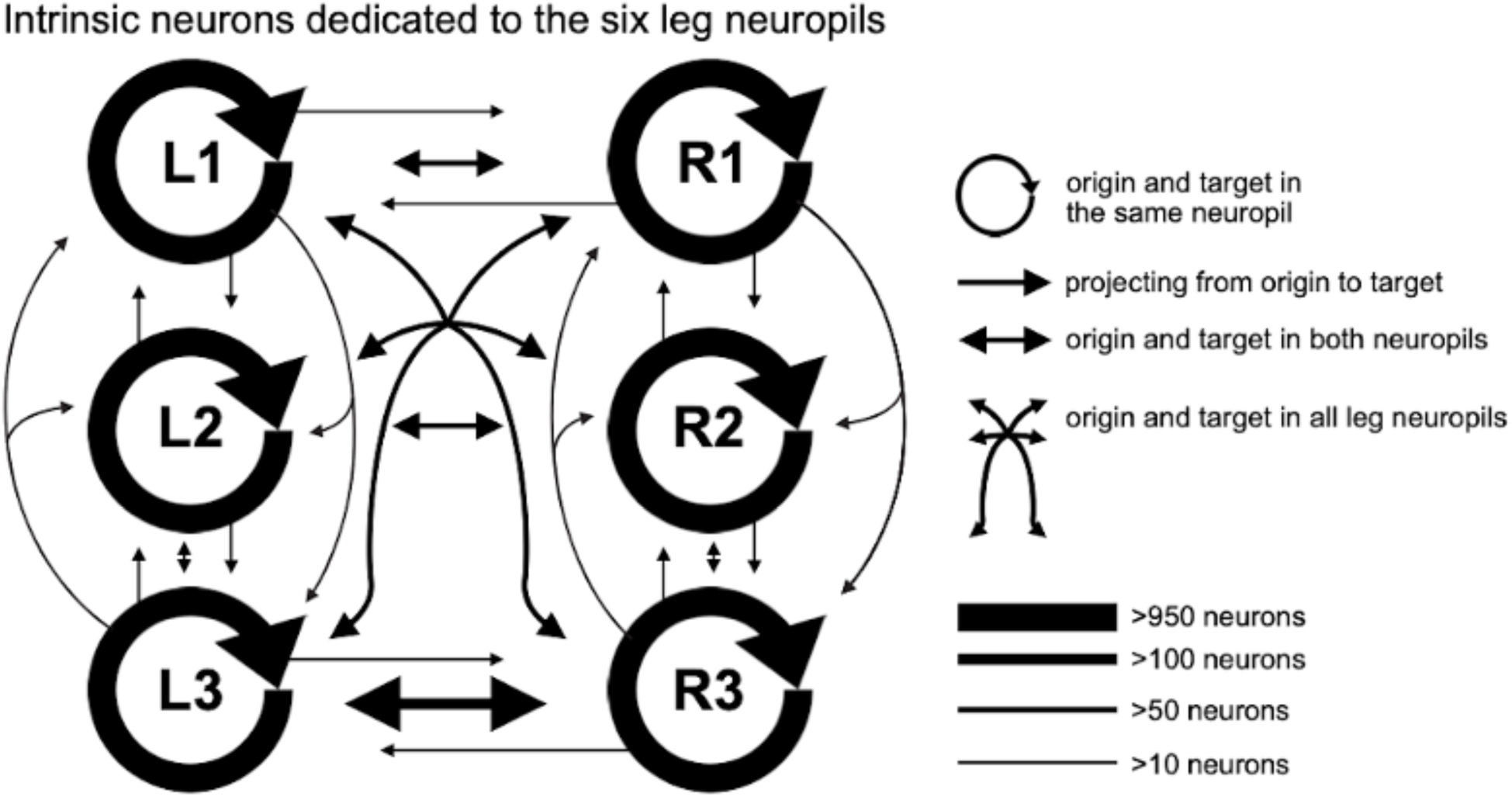
Leg circuit schematic. 6960 INs (>50%) have both origin and target specifically in the leg neuropil. Shown are the numbers of connections of these neurons, in the form of an arrow. Arrow is only shown if >10 neurons have the same origin and target. The great majority of intrinsic neurons of the leg have their origin and target within a single leg neuropil, over 950 neurons per leg neuropil. The number of neurons with the same leg connectivity pattern between legs ipsilaterally are all below 50. The most numerous bilateral connections between legs are in the T3 leg neuropils (>100).

One might have expected a comparable number of neurons within other single neuropils of the VNC. However, the maximum number of neurons dedicated to any other single neuropil is 60 neurons, found within each of the wing neuropils (Figure 6D). The INs and ANs in the upper tectulum have a different organisation, with a relatively large amount of diagonal connections between the haltere and wing neuropils or the halteres and neck neuropil, respectively (Figure 6A). We assume that because the wings of the fly act to offset haltere movement and slight asymmetries during flight can cause large and rapid effects (Deora et al., 2015), this behaviour requires a strong diagonal coordination between them, a type of connection which is rarely found in the leg neuropils.

### Sensory processing in the VNC

The VNC can be considered to be dedicated to sensorimotor processing, as opposed to the higher functions carried out by the brain such as navigation and learning and memory. Consistent with this view, sensory neurons make up more than 27% of the neurons contributing to the adult male VNC neuropils (Figure 1). We also find that roughly 20% of neurons originating in the VNC receive at least 30% of their input directly from sensory neurons and so can be regarded as second order neurons involved primarily in sensory processing (Figure 54B). In contrast, only ∼4% of neurons in the brain (whether including or excluding the optic lobes) meet these criteria (Schlegel et al., 2023).

Sensory neuron axons project into the adult VNC from specific nerves and target particular neuropils in defined patterns, based on their modality and tissue of origin. However, matching reconstructed sensory axons to specific light-level cell types based on morphology and anatomy alone proved to be extremely challenging. In particular, whilst the axons of some subsets of leg sensory neurons such as the FeCO had already been well characterised in the literature (Chen et al., 2021; Mamiya et al., 2018), many had not been reported at high enough resolution, including most of the tactile neurons, gustatory neurons, and unilateral campaniform sensilla. We therefore developed a method for the systematic cell typing of 6462 sensory neurons in MANC that leveraged connectivity to and from single cells, hemilineages, and serial sets as well as position in the network graph (Figure 52). We then assigned modality at the cell type level using morphological comparisons to light level data, which reproduced the modality-specific layering in sensory neuropils previously reported (Figure 53) (Tsubouchi et al., 2017).

**Figure 68.**
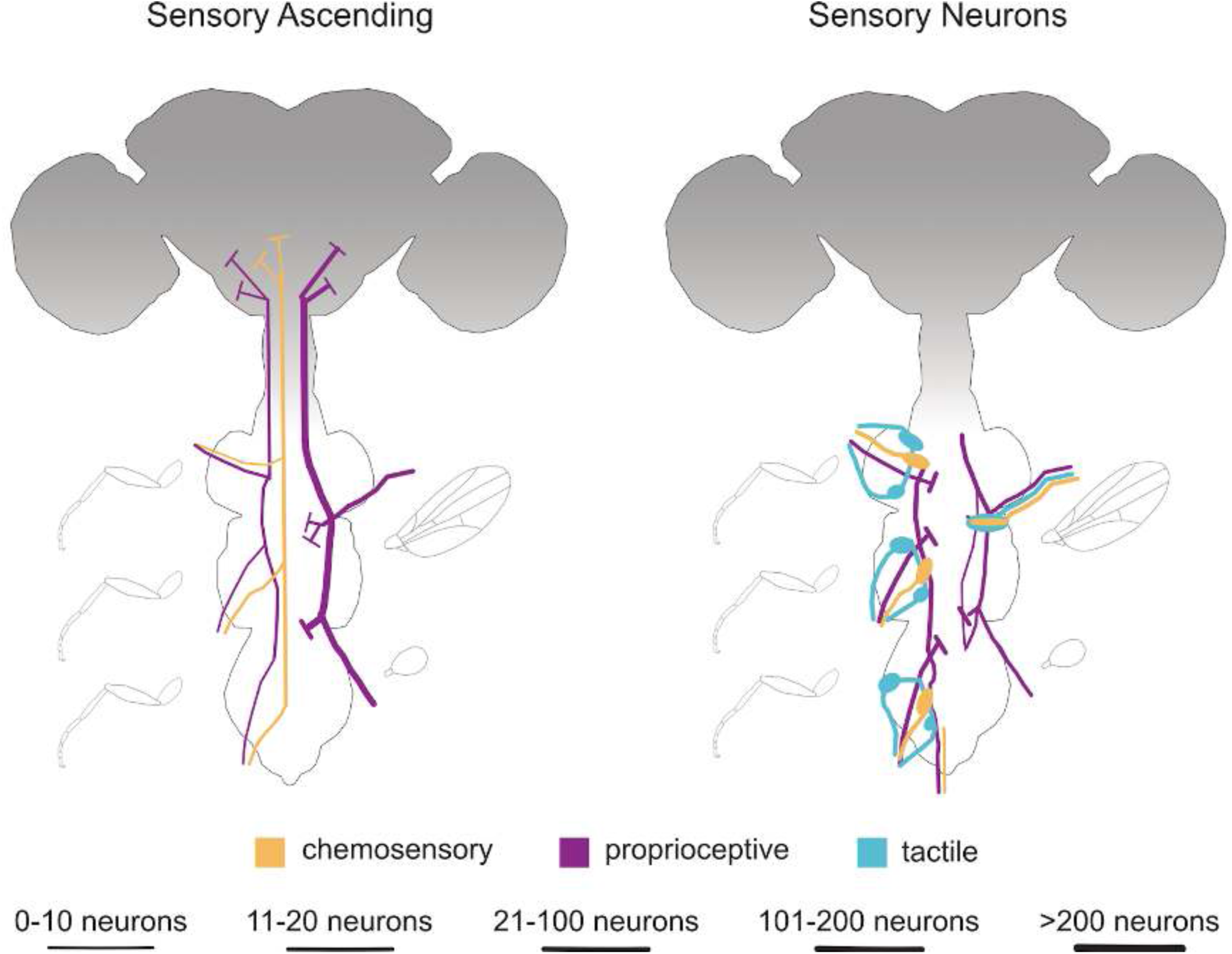
Sensory flow schematic. Global summary of MANC sensory neurons and their projections. Depicted origins are prothoracic leg (plus ventral prothorax), mesothoracic leg, metathoracic leg, wing (including wing margin), haltere, and abdomen. Line widths reflect higher neuron number when compared across sides and line colours reflect modality. Neurons of unknown modality have been omitted.

Here we summarise the projections of sensory neurons in the VNC by modality (Figure 67). Only 535 (8.28%) ascend directly through the cervical connective, the majority of which are campaniform sensilla from the wings and halteres, with a smaller number of proprioceptive and chemosensory neurons from the legs. No tactile neurons ascend directly to the head, but they comprise more than 38% of the sensory neurons terminating in the nerve cord; moreover, about a third of strongly connected downstream ascending neurons carry tactile information (Figure 54C), implying that it reaches the brain after one layer of local integration and/or processing. We also show that there are robust relationships between sensory neurons of each modality and specific hemilineages; for example, tactile neurons largely target hemilineage 23B, whilst chemosensory neurons target 05B and 09B (Figure 54G).

When we break each modality down into cell types, we uncover novel features of sensory organisation in the VNC. One of our most striking findings was that sensory types originating in distinct peripheral regions (legs vs wings/halteres vs abdomen) can share connectivity to serial partners, implying that they are functionally related and in certain cases might even represent the same cell type (Figure 58). We also found that chemosensory information from leg sensory types is organised locally into at least four discrete microglomeruli in the VAC (Figure 63A), whilst chemosensory information from wing chemosensory types targets three microglomeruli in the ovoid but with overlap between two types (Figure 63B). Unexpectedly, tactile information from the anterior and posterior leg compartments appears to be largely integrated by second order neurons in the VNC (Figure 62D), suggesting that locating the source of a stimulus along the proximodistal axis of the leg might be more important for eliciting an appropriate behavioural response than locating it on one side of the leg vs the other.

We also identify novel connectivity types within previously reported light-level morphological types and place them in the context of sensorimotor networks. For example, we have found two distinct types of FeCO hooks and two types of FeCO claws and have estimated their effective connectivity to leg motor neurons to predict opposing effects on the tibia extensor vs the flexor and accessory tibia flexor muscles (Figure 59), consistent with their observed physiological responses (Chen et al., 2021). We also identified two types of leg hair plates with differential effective connectivity to leg motor neurons (Figure 61) and proposed a sweet taste processing circuit (Figure 63). Notably, connectivity analysis predicts at least four distinct pheromone-sensitive sensory types on the foreleg and a processing network within the VNC that includes numerous morphologically similar target cell types not yet distinguished at light level (Figure 64).

Finally, we have used cosine similarity clustering between the outputs of sensory neurons to serial neuron groups to associate sensory types across the three main peripheral regions (leg, dorsal, and abdomen) (Figure 58). This method has suggested likely modalities for several unidentified types in the abdomen and elsewhere. It has also suggested functional connections between types of the same modality and has identified one strong candidate for a cross-regional type representing aversive stimuli.

In closing, we would like to emphasise that the generation, annotation, and preliminary analysis of the MANC connectome derived not only on the coordinated efforts of dedicated teams at multiple institutions but also on the decades of experimental studies that preceded it. Moreover, its ultimate utility depends on how widely it is leveraged in the future experimental and computational work of the entire neuroscience community. We have only revealed the tip of the iceberg in this report, with a wealth of opportunities now available in this publicly available dataset for forthcoming connectomic analyses that will feed into testable functional hypotheses.

This is a dynamic dataset, with potential for the improvement of annotations with every iteration. Whilst we do not anticipate substantial changes to the systematic types on par with those we implemented between MANC v1.0 and v1.2.1, we expect the type and synonyms fields to be updated over time as experts worldwide identify their neurons of interest. We encourage readers to contact the corresponding authors by email with suggested annotation changes based on matching to light level cell types.

Finally, we acknowledge that MANC is the isolated VNC connectome of a single adult male specimen. We cannot yet determine how representative it is, nor can we reconstruct the circuits that span the brain and nerve cord. Having more connectomes and more complete connectomes will allow us to assess stereotypy vs developmental variability, to quantify and characterise sexually dimorphic cell types and circuits, and to define sensorimotor circuits underlying coordination between the body and the brain.

## METHODS

### Systematic types for neurons originating in the VNC

We have given **systematic type** names to all neurons in the MANC data set using a prefix based on their broad class (intrinsic neuron, IN; ascending neuron, AN; efferent non-ascending, EN; efferent ascending, EA; motor neuron, MN; descending neuron, DN; sensory non-ascending, SN; sensory ascending, SA). Motor neurons and descending neurons were typed separately as described in our companion manuscript (Cheong et al., 2023). Reconstructed bodies that entered the volume via a nerve were classified as sensory neurons, assigned to an entry nerve, and clustered within that nerve (details below). The remaining neurons were assigned to a developmental hemilineage where possible and clustered within that hemilineage. Serially repeating cell types generally have identical systematic type names; in some cases, ascending neurons and intrinsic neurons were judged to be the same type and so have the same hemilineage and number but differing class prefixes.

### Subclass of ANs and INs

ANs and INs were given a two letter **subclass** defined by their innervation pattern in the VNC. The first letter represents the laterality of their dendritic and axonic arbour in the dataset. To this end the total number of pre and post synapses was scored in both hemispheres of the VNC for all ANs and INs. If a neuron had more than 5 synapses in total on both sides it was classified as a neuron with bilateral innervation and given the first letter B. If it has less than 5 synapses on one of the two hemispheres of the VNC the neuron either receives the letter I for ipsilateral innervation or C for contralateral innervation with respect to its soma location. The second letter of the subclass defines if a neuron is ascending, A, restricted to a single neuropil, R, or interconnecting neuropils, I. Restricted to a given neuropil was defined by having no less than 5 synapses outside of a neuropil mesh. In cases where different subclasses were given to neurons of the same group, the average across the group was used to make the assignment. There were 30 ANs that had less than 5 synapses in the entire VNC, so we gave them the subclass XA.

### Hemilineage assignments

Each hemilineage is generated by a specific neuroblast during development; in most cases the primary neurites of the secondary neurons converge to enter the neuropil at the same point and/or bundle together before branching. Secondary neurons were assigned to hemilineages by comparison with light microscopy images of neuroblast clones (Shepherd et al., 2019) and sparse driver lines (Harris et al., 2015; Shepherd et al., 2016). Key diagnostic features included the relative positions of soma tract entry points and backbones, particularly at midline crossing, as well as neuropils innervated and gross morphology. We describe these features in more detail for each hemilineage in Results.

Initially, expert annotators seeded the thoracic soma tract bundles of manually identified secondary neurons and ported those hemilineage and soma neuromere annotations to Clio Neuroglancer. Later we used computational methods to assign morphologically similar bodies with somas to the appropriate hemilineage.

We calculated NBLAST on cell body fibres to assign scores of similarity between the neurons (Costa et al., 2016). Next, hierarchical Ward clustering was performed on all-by-all NBLAST scores transformed to a distance space. 12 major clusters were selected arbitrarily, after analysing a clustering dendrogram structure. Next, for each of the 12 clusters, a subsequent clustering procedure was done with a cut-off point resulting in *k*_opt_ subclusters. In each of the resulting subclusters, a candidate hemilineage assignment was proposed based on the existing preannotated neurons for a given list of hemilineages. If a subcluster did not contain any previously annotated neurons, or the neurons within the cluster did not have consistent labels (at least 50%), the subcluster was left without any candidate assignment.

The number of subclusters *k*_opt_ for each of 12 clusters was selected automatically based on the following criterion:

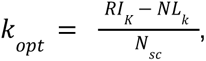

where *RI_K_* is a Rand Index of previously annotated labels and generated candidates, and *NL_K_* is a number of unlabelled subclusters as a result of a split *k*; and *N_sc_* is a total number of neurons within a subcluster. The resulting candidates for hemilineage labels were manually verified by expert annotators. The procedure was repeated iteratively with confirmed candidates used as priors.

Several hemilineages proved challenging to differentiate from proximal hemilineages due to similar soma tract entry points to the neuropil and overlapping neuron morphologies. To assist in hemilineage assignment, neurons from these hemilineages were clustered using the Weighted Nearest Neighbour (WNN) workflow from the Seurat package (Hao et al., 2021). Briefly, WNN is a form of multimodality integration developed for single-cell genomics workflows, whereby each modality is integrated with a weighting calculated for each individual cell based on its ability to predict similarity of to its nearest neighbours across each separate modality. The modalities utilised for this analysis were single cell synaptic connectivity, hemilineage aggregated synaptic connectivity, whole neuron NBLAST scores, and cell body fibre NBLAST scores. Community detection was performed on the integrated data using the Louvain algorithm. These communities were then used to assist in hemilineage assignment, alongside UMAP visualisations of the integrated data.

At a later stage of annotation, hemilineage assignments were refined using existing automatic and manual assignments to predict hemilineages for many neurons not yet assigned. Hemilineage labels for unassigned neurons were predicted with label spreading semi-supervised classification (Zhou et al. 2003), using an equally-weighted mixture of features from laterally-symmetrised cosine connectivity similarity, whole neuron NBLAST distances, and Jenson-Shannon distance between neurotransmitter prediction probabilities. The classifier and 10-nearest-neighbours kernel were fit to existing hemilineage assignments, then inferences for unassigned neurons were manually reviewed by descending confidence.

### Birthtime assignment and analysis

We sought to classify the VNC neurons into the three birthtime categories: “primary”, “early secondary” and “secondary”. For this, we used the following morphological features estimated from neurons segmentation after the skeletonization procedure: soma size, average spine skeleton radius, average total skeleton radius, spine length, total arbours length, the proportion of pre- to post-synaptic connections, and relative cell-body fibre radius within hemilineage. To simplify the problem, we considered only binary labels of “primary”, and “secondary” neurons. In total, we used 946 examples as our training set, spanning across the whole VNC. Our final classification combined predictions from 51 independently trained support vector machine classifiers with radial-basis kernel and cost factor *C* = 10. Next, we used a permutation-based variable importance measure to assess what features have the biggest discriminatory power (Biecek, 2018). Our analysis suggested that 5 features (soma size, average spine radius, average total radius, spine length, total arbours length) correspond to 80% of variance, and only those were subsequently used to train an ensemble of classifiers. We performed a 10-fold cross-validation to verify the robustness of our classification and achieved an F1 score of 58%. The neurons with top final classification scores (as sorted by a measure of confidence — normalised length of margin vectors) were manually inspected by annotators. It should be noted that we annotated 430-496 primary neurons per thoracic neuromere, a significant overestimate when compared to reported numbers of ∼300 per neuromere but resulting in comparable percentages of primary neurons (Poulson, 1950).

#### Network analysis of birthtimes

The majority of the comparative birthtime analysis (Figure 9) was done on a directed network constructed in Python, with individual neurons as nodes and edge weights as number of synapses. The network was constructed in iGraph (Csardi et al., 2006) and NetworkX (Hagberg et al., 2008), and was limited to edges with 3 more total synapses. All neurons with a class that is not one of ‘TBD’, ‘Obsolete’, ‘Unknown’, or ‘Glia’ were included, and all neurons with abdominal soma locations were excluded. Due to the difficulty of assigning accurate birthtime annotations to abdominal neurons, excluding them allows for more accurate analysis, and the analysis in T1-T3 is likely applicable to the abdominal neuromeres. The secondary only graph produced for Figures 3D and 3F were made by excluding all intrinsic neurons labelled primary. Ascending primary neurons and motor neurons were preserved in the graph, but their data is excluded from the figures.

#### Measures of centrality

Load centrality (Figure 3D’) was measured via the load_centrality() function in NetworkX, which is based on (Goh et al., 2011). Load centrality of a node is the fraction of all shortest paths between each pair of nodes in the network that passes through that node. Edge weights were ignored, and the scores shown were normalized by the total number of possible connections in the graph. A higher load centrality suggests higher centrality. The mean length of shortest paths to each secondary neuron (Figure 3D’) was calculated exactly as named: for each neuron in the respective graph, we found the shortest distance between it and each secondary neuron. The mean of that distribution is the data shown. A lower value here suggests average faster global influence on the circuit

#### Measures of insularity

These are measures of how diverse a neuron interacts with communities across the VNC. Here, we defined one community as one hemilineage-within-a-hemineuromere (e.g. 06B T1 Left, 19A T3 Right, etc.). Diversity coefficient is calculated as the Shannon Entropy of the distribution of a node’s connections to all communities divided by the log of the number of communities (inspired by (Eagle et al., 2010)). The diversity coefficient is 1 minus the sum of the squares of the fraction of a node’s total degree that is associated with each community (introduced in (Guimerà and Nunes Amaral, 2005)). The implementation of the diversity coefficient, and inspiration for the implementation of the participation coefficient, is implemented via BCT (Rubinov and Sporns, 2010). Non-redundant community promiscuity is the count of the total number of communities that are connected to a node’s community solely because of a node. A node X belonging to community Y has a score of n if, by removing X from the graph, Y is directly connected to n less communities. It is related to, but not identical to, the linchpin measure in (Nemesure et al., 2021).

#### Rich club assignment and analysis

Rich club assignment was produced using the full graph discussed above. Instead of using degree as the parameter, we used load centrality. This was done to ensure that there was no bias towards neurons involved in more circuits. It was also done to ensure that it was neither too big to be meaningless, or so small that it has exclusively primary neurons. To make up for the load centrality distribution being heavily right-skewed, the parameter used was log_10_ of the load centrality. With these adjustments in mind, the rich club coefficient (phi) generated was directed and weighted, and was calculated using the normalized, general form of the metric in (Alstott et al., 2014). From the distribution of log_10_ of load centralities, we made 1,000 bins, from minimum to maximum value in the network (nodes with 0 load centrality were assigned a value below the otherwise minimum value). At each bin, we removed all nodes with lower log_10_ load centrality than the bin and recorded the full weight of the remaining network. To normalise, we performed the same calculation on 1,000 identical graphs with shuffled edge weights. The rich club coefficient per each bin is the weight of the original graph divided by the mean of the 1,000 shuffled graphs for that bin. The rich club was chosen based on the bin with the highest phi value, such that all neurons with a greater log_10_ load centrality were assigned to the rich club. No rich club analysis was done on the secondary only graph.

#### Global depth measurements

The global depth birthtime comparison was done in a straightforward manner. Using networks constructed as described above, a breadth-first search (BFS) was done from each node. For each VNC-origin neuron, the mean BFS distance to each MN or AN was recorded. Similarly, the mean BFS distance from each SN and DN to each VNC-origin neuron. For the secondary only graph, the ANs, MNs, DNs, and SNs were not removed, so no further adjustments were needed.

#### Neuropil depth measurements (synapse traversal)

This metric did not utilise the constructed networks, and did not remove abdominal born neurons, and all synapses between annotated neurons was preserved. A table containing the body IDs of the upstream and downstream partners of each synapse, as well as the neuropil that the synapse is located in, was used. One run of the traversal simulation goes like this:

1. For the original neuron, A_O_, randomly select a synapse where AO is the downstream partner of the synapse. This is the origin synapse, and its associated neuropil, N_o_ is the origin neuropil. The starting total distance is 1. Let A be A_O_.
2. Choose a random synapse where A is the upstream partner. If the neuropil associated with this synapse, N, is the same as N_o_, then add 1 to the total distance and go to step 3. If N is not the same as N_o_, then record the total distance and exit the traversal.
3. The downstream neuron of the synapse chosen in step 2. is now A. If A is an AN or an MN, record the total distance and exit the traversal. Otherwise, repeat step 2 until the traversal exits, or until the total distance is greater than 10, at which point discard data and start a new traversal with the original neuron A_O_.

This was repeated 100,000 times per VNC-origin neuron, and the mean value recorded.

### Origin and target

Neurons belonging to the classes DN, IN, AN, SN and MN were ascribed an origin and target in accordance with the neuropils they innervate within the VNC. The percentage of post-synapses or pre-synapses in each neuropil was scored for origin and target respectively. Neurons with over 80% of their post-synapses or pre-synapses concentrated in a single neuropil were considered to have that particular neuropil as their origin or target. In instances where a neuropil is divided into left or right hemispheres, an underscore L or R denoted the respective hemisphere (i.e. *LegNpT1_L*). If the cumulative percentage of two neuropils surpassed the 80% threshold and both neuropils contributed at least 5%, both neuropils were included with a full stop (.) between them. Thus, a neuron receiving input from the front leg neuropil and the neck tectulum on the left hemisphere of the VNC would have *LegNpT1_L.NTct_L* as its origin. A substantial proportion of neurons innervate a combination of leg neuropils (LegNpT1-T3) or upper tectulum neuropils (NTct, WTct, HTct). Consequently, we also assessed neurons based on their innervation within these two combined neuropil groups, assigning them the origin or target LegNp or UTct, respectively. Anything that did not fall into one of these three categories was given *multi* as origin or target, which corresponds to 16% of the VNC neurons.

Exceptions were made for different classes of neurons, as their origin or target lie outside of the VNC. DNs were systematically assigned the brain as their origin, MNs were designated their respective identified muscle as target, and SNs were assigned their appendage as origin. ANs target the brain; however, the majority possess axons with numerous synapses in the VNC. Consequently, we decided to represent the target within the VNC for this neuron class. ANs of the XA subclass have both their origin and target annotated as brain, as they have less than 5 synapses within the VNC.

Primary origin or target, used in Figure 5 for ANs and INs, is defined as the neuropil with the highest percentage of post-synapses or pre-synapses, respectively. For all neurons that do not have multi as an origin or target this top neuropil upstream or downstream is the first neuropil in origin and target.

### Group and serial homology assignment

Similar neurons within each neuromere were assigned to groups as described in our companion paper (Takemura et al., 2023). For the identification of serially homologous neurons, we used a semi-automated iterative procedure. We started with a small pool of homologous neurons defined by expert curation based on manual assessment of morphological and connection similarity. We then obtained connectivity information for upstream and downstream partners of this initial set, where the neurons from specific VNC segments were first matched based on their connectivity patterns. We employed a “seeded” version of the Fast Approximate Quadratic Assignment solver (Fishkind et al., 2019), which aims to minimise the number of edge disagreements between the two graphs. The “candidates” were automatically screened based on consistency between the matches on the sides and among neuromeres. The remaining groups of “candidates” were manually revised, and the procedure kept repeating with confirmed matches as the new seeds. This approach allowed us to group 2725 neurons into 450 serially homologous groups across both thoracic and abdominal segments. Next, we also used serial cosine clustering to identify more candidates with similar connectivity within each annotated hemilineage.

For INs restricted to the 6 leg neuropils (subclass IR, CR and BR) we took an additional approach with hemilineage connectivity vectors. To construct a connectivity vector for a given target neuron, we group connections by available hemilineage labels divided into ipsilateral and contralateral (with respect to the target), further split with regard to incoming and outgoing edges from the target. We normalise the in- and out- contributions separately, so that the in- and out-vectors both sum to 1. The connectivity to hemilineages allows us to distinguish and cluster neurons of the same serial set. We perform hierarchical clustering of these vectors using the Euclidean metric and weighted linkage method in the scipy.cluster.hierarchy.linkage implementation. The resulting dendrogram provides a hierarchical structure of the clustering that can be extracted. We consider all possible clusters and evaluate them using a metric that rewards clusters that are evenly distributed across the leg neuropils. The clusters are then ranked and filtered to produce the proposed serial groups for proofreading. Below we give the details of the filtering procedure. Firstly, we implement a ‘balance’ score B to measure how balanced the number of neurons in each of the leg neuropils is for a given cluster. This score B is Shannon’s entropy normalised by the maximum possible entropy value:

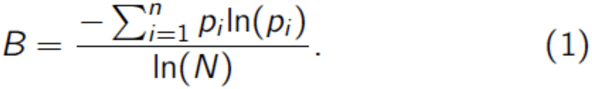

To calculate the leg neuropil balance for a given cluster, n is the number of leg neuropils represented in the cluster, and pi gives the proportion of neurons in each leg neuropil. *N* is the total number of leg neuropils so ln(6) is the largest entropy value when the cluster is most balanced - in this case, pi = 1/6 for all i gives the score B = 1. The same formula above can be used to measure the balance of other indicators such as the proportion of neurons with somas on the left or right sides. The left-right balance score is useful for ensuring that clusters are symmetric without penalising them for missing neurons in some leg neuropils.

Initially, all clusters are sorted by the leg neuropil balance score. We then discard a cluster if it overlaps (i.e. has common neurons) with a more balanced cluster. This procedure will retain the most balanced clusters across the leg neuropils but discards the clusters that do not span all regions but are still useful components of serial homologues. The simple filtering did well to find balanced clusters of multiples of six in the leg interneurons. For the bilateral neurons, there were few clusters that were perfectly balanced and the clusters that were highly balanced were too large to meaningfully extract serial homologues. To search for useful groupings of size four or multiples of two, we implement a ranking based on clustered height in the dendrogram (priority to tighter clusters), left-right balance score, size (priority to larger clusters), and leg neuropil balance score. These ranks are aggregated by taking the mean to give an overall ranking, which we go through and discard any cluster that has overlaps with a higher-ranked cluster. By performing the above procedures for the leg interneurons and bilateral neurons, we can retrieve clusters of multiples of six, as well as those with four-fold symmetry across two neuromeres. This method proposed clusters which when proofread allowed us to assign or correct 103 serial sets.

### Systematic cell typing and nomenclature

Intrinsic, ascending, and efferent neurons were clustered into systematic types on the basis of their synaptic connectivity, considering both lateral and serial symmetries. Connectivity adjacency between all neurons with an annotated class was considered. Initial systematic typing was performed based on the annotations in the manc:v1.0 dataset release and was repeated on v1.1 (available via programmatic query), following extensive annotation curation and seeking to refine within-neuromere types and maximise across-neuromere types. Systematic typing supplemental figures represent the state of annotations and systematic types on v1.1. Additional adjustments to annotations and to systematic type, type, and instance values were performed manually prior to the manc:v1.2.1 dataset release.

To aggregate connectivity across lateral symmetry, the columns of adjacency matrix blocks corresponding to right-hand soma side neurons for neurons in annotated lateral groups were permuted to match their left-hand homologues (Figure 13A). Rows and columns for each group were then aggregated as sums (as cosine similarity is used, mean aggregation is equivalent). Midline and ungrouped adjacency blocks were unaffected. To aggregate connectivity across serial symmetry, the columns of the adjacency matrix corresponding to each serial set were sum aggregated. Columns not in an annotated serial set were unaffected. Distances for clustering were computed as cosine distance for this whole-network adjacency matrix with either or both aggregations applied, as described below.

Type clusterings of the cosine distance matrix were then performed with Ward’s method independently for each hemilineage. Serial sets were constrained to be in a leaf when initialising the cluster linkage. Two clusterings were then performed to select flat type clusterings: first, of the symmetrised, unaggregated connectivity distance to maximise consistency with annotated lateral groups; second, of the laterally aggregated connectivity distance while maximising consistency with the first, intermediate clustering. For the first, intermediate flat clustering, at each distance in the symmetrized clustering linkage where an agglomeration was made consensus was measured as the adjusted mutual information (Vinh et al., 2009) (within the intersection set of annotated lateral groups) between the clustering and lateral group annotations. The symmetrized clustering threshold was chosen as the distance corresponding to the peak consensus metric nearest the dendrogram leaves. While this clustering selects for consistency with group annotations, it does not guarantee it. To generate flat type clusters consistent with lateral group annotations such that none are split, clustering was performed on the aggregated connectivity distance in the same manner. In this second clustering, terminal clusters were selected to maximise consistency with the intermediate cluster labels, aggregated by mode, on the basis of adjusted rand index (Vinh et al., 2009). Different selection criteria are used in each clustering because the first is seeking consistency with group annotations we believe to be correlated with, but not identical to, desirable type partitioning, while the second is simply trying to best recapitulate that intermediate clustering subject to the constraints of lateral group annotations.

In the special cases of the midline hemilineages 00A and 00B, for lateral consensus soma side annotations were augmented with a bootstrapped side assignment based on connectivity. At each iteration, unassigned neurons were assigned a side preference based on their total synaptic connectivity to all neurons with a side assignment (either annotated or already bootstrapped). Then the ratio of left- to right-side maximal presynaptic preference for each yet-to-be-assigned group was considered, and for those within 10% of balanced preference (meaning groups with a nearly equal number of member neurons preferring each side) the preferences were taken as side assignments. This process was repeated until no further groups were assigned, resulting in the assignment of 85 neurons, sufficient for lateral consistency analysis. These assignments were only for lateral consistency analysis and no side annotations were made.

For per-hemilineage type nomenclature, flat type clusters resulting from this process were assigned numeric serial identifiers. Serial identifiers are the descending ordering of hemilineages’ type clusters by mean neuron size. Preliminary work showed this ordering to be highly reliable between sides, and it depends only on neuron segmentation, not annotation or synapse or other predictions. Motor neurons were included in the lateral consistency and symmetrized clustering process, as they may be important to hemilineage connectivity similarity structure, but were not assigned a type based on the result.

### Sensory neuron reconstruction and classification

In general, reconstructed bodies that 1) appeared to enter the volume via a nerve and 2) lacked a soma were classified as sensory neurons; those that also appeared to leave via the CvC were classified as ascending sensory neurons. However, many neurons highly similar in morphology and connectivity could not be followed all the way to the nerve due to staining artefacts and possible degeneration, particularly in the leg neuropils, and these were also classified as sensory neurons. Also, a small number of sensory neuron somas located at the base of the nerves were preserved in the volume.

Initially bodies with no class assignment were compared to previously reconstructed sensory neurons using NBLAST (Costa et al., 2016). The resultant score matrix was then rearranged to produce a ranked spreadsheet of candidates, progressing from high score to low score, which were manually reviewed and either extended into sensory neurons or discarded. Secondarily, we selected classless bodies that were presynaptic to neurons downstream of previously annotated sensory neurons. These classless bodies were similarly manually reviewed and processed.

Following reconstruction, we compared the morphologies of sensory neurons to light level data to classify them to known types. However, due to limitations in the segmentation quality, reconstruction status, and underlying light level data we were unable to assign the majority of sensory neurons to known types. We therefore proceeded to cluster the neurons of each nerve independently using Weighted Nearest Neighbour clustering (Hao et al., 2021). For this analysis we used a range of synaptic connectivity, morphological and network traversal features, including synaptic outputs to single neurons, synaptic inputs from single neurons, synaptic outputs to neurons aggregated by hemilineage, synaptic inputs from neurons aggregated by hemilineage, NBLAST scores between sensory neurons, and mean graph traversal distances from sensory neurons to all neurons within the VNC. Individual nerve clusters were then detected using HDBSCAN from a UMAP representation of the WNN output (Campello et al., 2013). We then looked to sterilise our SN types. Sensory neurons were aggregated into regional groups based on their entry nerve: dorsal nerves, leg nerves, and abdominal nerves. Dendrograms were then generated for each of these regional groups using the cosine distance of their outputs to serial neurons, from which clusters were determined to be below an optimal cut height, taken as that which maximised the adjusted mutual information across each of the component independent nerve clusterings. All final clusters were manually reviewed by an expert annotator and assigned as systematic types. The first round of sensory neuron systematic types, released in our preprint, were generated using mancv1.0 annotations. The systematic types in this manuscript were generated from mancv1.1 annotations. The sensory neuron analysis in this paper is based on manc1.2 annotations.

The sensory afferent projections in the VNC are highly ordered and organised, projecting to specific neuropil regions according to their sensory modality (proprioceptive, tactile or chemosensory) (Tsubouchi et al., 2017). We used these structured innervation patterns, alongside literature references where available, to assign the sensory modality to each of our systematic types. Final systematic type values were derived from consecutive numbering within each modality.

### Sensory Neuron Annotation

Using detailed knowledge of the sensory afferent arrays we used key diagnostic features such as entry nerve, neuropils innervated and morphology to further assign sensory neurons into subclasses that reflect the sensory structures from which each neuron originates.

#### Proprioceptive Neurons

Proprioceptive neurons in *Drosophila* do not fall into a single morphological class and are derived from a mix of external receptor types (Campaniform Sensilla and Hair plates) and internal stretch receptors (e.g femoral chordotonal organ) (Merritt and Murphey, 1992). Each receptor type produces a characteristic afferent projection, and it is possible to identify each sensory neuron type from its projections within the VNC.

#### Leg Proprioceptive neurons

##### FeCO

The FeCO has a complex sensory afferent projection reflecting the diverse functional properties of its neurons. All FeCO afferents enter the VNC via their respective leg nerve but segregate into 3 functionally distinct projections known as the “Club”, “Claw” and “Hook” (Mamiya et al., 2018). The club neurons form a coherent medially projecting club shaped projection restricted to ipsilateral mVAC. The claw has a characteristic three-pointed projection in ipsilateral LegNp. The hook is the smallest of the FeCO classes and forms a projection similar to the club but with a single bifurcating process close to the entry point to the leg neuropil.

##### Hair plates

Hair plates are small, tightly packed clusters of short hairs, with each hair innervated by a single neuron. They are positioned at cuticular articulations and are deflected by joint movement. Afferents from HPs enter the VNC via the respective leg neuropil and bifurcate on entry to project around the anterior and posterior margins of the ipsilateral leg neuropil (Merritt and Murphey, 1992). There is anatomical similarity between hair plate sensory afferents and afferents from some leg campaniform sensilla (Merritt and Murphey, 1992), and it is not possible to fully distinguish these.

##### Campaniform Sensilla

Campaniform sensilla are small cuticular domes which detect tension in the surrounding cuticle and are typically grouped in fields in regions of high strain (Dinges et al., 2021). The projections of afferents from leg campaniform sensilla are diverse, and some are uniquely identifiable (Merritt and Murphey, 1992). Not all leg campaniform sensilla afferents have been described, and it was not possible to annotate them all reliably. Moreover, these sensory populations suffered most severely from degradation and therefore are underrepresented in our dataset.

#### Dorsal Proprioceptive neurons

##### Prothorax

###### Prosternal organ/Hair plates

The prosternal organ is a hair plate located on the anterior wall of the thorax immediately under the neck (Power, 1948). Sensory afferents from the hair plates enter the VNC via the prosternal nerve, a thin nerve that enters the VNC just medial to the dorsal prothoracic nerve (Court et al., 2020). On entry to the prothoracic neuromere, the hair plate afferents project medially around the anterior margin of the ipsilateral leg neuropil before turning dorsally to arborise in ipsilateral NTct and then extending posteriorly to arborise further in the ipsilateral HTct (Smith and Shepherd, 1996).

###### Prothoracic Chordotonal Nerve (ProCN)

The prothoracic chordotonal organ is located at the anterior/lateral margin of the VNC and connects to the VNC via the ProCN which is a short, thick connection between each prothoracic chordotonal (pCO) sense organ and the first thoracic neuromere (Power, 1948). The pCO afferents project posterior-medially from their entry to arborise in the dorsal most part of the ipsilateral ovoid and lower tectulum before projecting further posteriorly to arborise in the anteriormost region of the MetaLNp (Smith and Shepherd, 1996); (Stölting et al., 2007).

##### Mesothorax

###### Wing blade campaniform sensilla

The Drosophila wing has 8 Campaniform Sensilla found at specific locations on the anterior wing blade (Dickinson and Palka, 1987; Smith and Shepherd, 1996). The sensory afferent from these sensilla enter the VNC via the ADMN and arborise in the lateral regions of the ipsilateral ovoid. The axons then project posteriorly following either a medial or a lateral tract that run just ventral to the LTct to terminate with arborisations in the MetaLNp and the anterior ANm (Dickinson and Palka, 1987; Smith and Shepherd, 1996).

###### Proximal wing hinge campaniform sensilla

The proximal wing blade has three clusters of campaniform sensilla (Smith and Shepherd, 1996). The sensory afferents from these sensilla enter the VNC via the ipsilateral ADMN and form a complex but characteristic projection that extends anteriorly through the CvC to the brain and posteriorly to produce terminal arborisations in ipsilateral HTct. The largest arborisations are, however, in the ipsilateral WTct (Smith and Shepherd, 1996).

##### Metathorax

###### Halteres

The halteres have a large number of campaniform sensilla, organised into fields at the base of each haltere. The sensory afferents from the campaniform fields enter the VNC via the ipsilateral DMetaN and form a robust anterior-directed projection that ascends through the CvC to the SEZ and brain (Smith and Shepherd, 1996). The major arborisation in the VNC is within ipsilateral HTct with minor arborisations in the ipsilateral WTct and NTct. Afferents from different fields have different projection profiles within this major projection and have been described in detail at light level (Chan and Dickinson, 1996).

##### Tactile Neurons

The body surface of the fly is covered with bristles, each innervated by a single neuron that is activated by mechanical deflection. In *Drosophila* there are two types of tactile bristles, the larger macrochaetes and the smaller, more numerous microchaetes.

##### Leg Tactile Neurons

Leg bristles are all microchaetes, and their associated sensory neurons enter the VNC via their respective LegN to form a dense topographic projection in the ipsilateral, ventral legNp. These projections have two components, an anterior-lateral group projection from bristles on anterior leg and a posterior-medial projection from bristles on posterior leg (Murphey et al., 1989b).

##### Wing and Notum Tactile neurons

Wing bristles are all microchaetes distributed across the wing blade and anterior margin. The neurons from these bristles enter the VNC via the ipsilateral ADMN and terminate in ipsilateral ovoid with a small number projecting anteriorly into the ProNm. The tactile afferents enter the VNC in close association with wing chemosensory afferents loosely enveloping the more densely packed chemosensory neurons (Tsubouchi et al., 2017).

The notum features both micro- and macrochaetes, the microchaetes being far more numerous and evenly distributed. The macrochaetes are fewer in number, lie at highly conserved positions, and can be uniquely identified by position and sensory projections ((Ghysen, 1980; Usui-Ishihara and Simpson, 2005). The sensory neuron associated with each macrochaete has a unique projection in the VNC, and these have been described in detail (Ghysen, 1980) and are readily identified. The macrochaete sensory axons have large diameter axons and enter the VNC by the PDMN and run to the ventral most ovoid before projecting anteriorly into the ProNm and posteriorly, with a small number extending as far as the MesoNm and MetaNm. Macrochaete axons project across the midline in the posterior ovoid to produce a small contralateral projection that mirrors the ipsilateral projection.

Microchaete neurons are not individually identifiable and form a coherent and recognisable projection within the CNS. The microchaete neurons are smaller in diameter but share features with macrochaete projections, entering the VNC via the PDMN to run to the ventral most ovoid where they project anteriorly into the ProNm and cross the midline in posterior ovoid to form a short contralateral projection. Unlike the macrochaetes, none of the microchaetes have a posterior projection(Usui-Ishihara and Simpson, 2005).

##### Chemosensory Neurons

###### Leg Chemosensory Neurons

Leg chemosensory sensory neurons are associated with sensilla on the tarsus and tibia (Ling et al., 2014). Each sensillum is innervated by four chemosensory neurons, each with a different sensitivity (Sweet, high salt, low salt and water)(Kwon et al., 2014). Leg chemosensory afferents enter the VNC via the respective leg nerve and project medially to terminate ipsilaterally in the ventral most LegNp adjacent to the midline (Murphey et al, 1989b). Within this projection there are nine projection types, six of which also project anteriorly to the SEZ (Kwon et al., 2014). There is also a single mechanosensory neuron associated with each chemosensory bristle; the projections of these neurons are indistinguishable from the tactile bristle afferents(Murphey et al., 1989a).

##### Wing chemosensory Neurons

The wing of *Drosophila* has approximately 40 chemosensory sensilla running along its anterior margin(He et al., 2019). The afferents from these bristles enter the VNC via the relevant ADMN to terminate in ipsilateral ovoid. The chemosensory afferents project in close association with the wing tactile afferents, but form a more densely packed projection and terminate more medially than the tactile afferents (Kallman et al., 2015).

### Standard Neuropil Generation

As part of the sensory neuron analysis we generated a standard neuropil transform in order to visualise neurons in a single, consistent space. This is necessary as neuropils, particularly the leg neuropils, are variable in shape making consistent comparison difficult. In order to do this, we first generated a PCA based on the leg neuropil and mVAC meshes, alongside points from the skeletons of a focal neuron population (FeCO clubs), for each separate leg neuropil. This enabled the transformation of innervating sensory neurons into a single space. However, neurons from separate leg neuropils had differing rotational alignments in this space. We then conducted a second PCA of the transformed sensory neurons for each nerve, which aligned the principle axes across the leg neuropils. Lastly we z-scored within each transformed leg neuropil sensory neuron population, giving our final standard neuropil alignment. This process was repeated for the dorsal standard neuropil, with the first PCA transform performed on the OV.

### Programmatic Tools Analysis

The neuprint website (neuprint.janelia.org) provides a graphical front end to explore the VNC connectome by selecting the VNC dataset from the main dropdown menu (Plaza et al., 2022). Authentication via is required but this is available to anyone with a Google account. Most connectome scale analysis was carried out using the natverse suite of tools (Bates et al., 2020) (see natverse.org) written in the R statistical programming language (www.r-project.org). Connectome data was queried via the neuprint API (Plaza et al., 2022) using the neuprintr package (natverse.org/neuprintr) (Schlegel et al., 2021). A specialised open source natverse package, malevnc (documented at natverse.org/malevnc and downloadable from github.com/natverse/malevnc) was written to facilitate connectome annotations and analysis. Annotations were initially written programmatically using the clio API or interactively via the clio front end (clio.janelia.org, select **annotate** from the first drop down menu and then the **VNC** dataset) (Takemura et al., 2023) and then synchronised with neuprint where they can now be queried.

## Supporting information

Higher resolution figures

## ACKNOWLEDGEMENTS

We would like to thank Joseph Hsu and Tyler Paterson for assistance with proofreading, Sebastian Cachero and Erika Dona for LM-EM matching that helped to elucidate the 08A vs 16B hemilineage assignments, Haluk Lacin and Sebastian Rumpf for sharing unpublished light level data, and James W. Truman for discussions on nomenclature and access to unpublished light level data. We are very grateful to Julie Simpson, Troy Shirangi, Sebastian Rumpf, Haluk Lacin, and members of the Tuthill Lab, especially Tony Azevedo and Ellen Lesser, for extensive comments on the manuscript that informed revisions prior to submission for publication.

This work was principally funded by Wellcome Trust collaborative awards (220343/Z/20/Z and 221300/Z/20/Z to GSXEJ with GMC, GM Rubin, S Waddell and M Landgraf) for proofreading, annotation and analysis of MANC neurons. Additional funding came from the Howard Hughes Medical Institute at Janelia to GMC and SB; Neuronex2 (NSF 2014862, MRC MC_EX_MR/T046279/1) award and core support from the MRC (MC-U105188491) to GSXEJ; and a Walter-Benjamin-Fellowship from Deutsche Forschungsgemeinschaft to TS (STU 793/2-1).

## REFERENCES

Agrawal S, Dickinson ES, Sustar A, Gurung P, Shepherd D, Truman JW, Tuthill JC. 2020. Central processing of leg proprioception in. Elife 9. doi:10.7554/eLife.60299

Allen AM, Neville MC, Birtles S, Croset V, Treiber CD, Waddell S, Goodwin SF. 2020. A single-cell transcriptomic atlas of the adult ventral nerve cord. Elife 9. doi:10.7554/eLife.54074

Alstott J, Panzarasa P, Rubinov M, Bullmore ET, Vértes PE. 2014. A unifying framework for measuring weighted rich clubs. Sci Rep 4:7258.

Ammer G, Vieira RM, Fendl S, Borst A. 2022. Anatomical distribution and functional roles of electrical synapses in Drosophila. Curr Biol 32:2022–2036.e4.

Azevedo AW, Dickinson ES, Gurung P, Venkatasubramanian L, Mann RS, Tuthill JC. 2020. A size principle for recruitment of leg motor neurons. Elife 9. doi:10.7554/eLife.56754

Baek M, Mann RS. 2009. Lineage and birth date specify motor neuron targeting and dendritic architecture in adult Drosophila. J Neurosci 29:6904–6916.

Bate CM. 1976. Embryogenesis of an insect nervous system I. A map of the thoracic and abdominal neuroblasts in *Locusta migratoria*. Development. doi:10.1242/dev.35.1.107

Bate M, Arias AM. 1991. The embryonic origin of imaginal discs in Drosophila. Development 112:755–761.

Bates AS, Manton JD, Jagannathan SR, Costa M, Schlegel P, Rohlfing T, Jefferis GS. 2020. The natverse, a versatile toolbox for combining and analysing neuroanatomical data. doi:10.7554/eLife.53350

Bidaye SS, Bockemühl T, Büschges A. 2018. Six-legged walking in insects: how CPGs, peripheral feedback, and descending signals generate coordinated and adaptive motor rhythms. J Neurophysiol 119:459–475.

Biecek P. 2018. DALEX: explainers for complex predictive models in R. J Mach Learn Res 19:3245–3249.

Birkholz O, Rickert C, Berger C, Urbach R, Technau GM. 2013. Neuroblast pattern and identity in the Drosophila tail region and role of doublesex in the survival of sex-specific precursors. Development 140:1830–1842.

Birkholz O, Rickert C, Nowak J, Coban IC, Technau GM. 2015. Bridging the gap between postembryonic cell lineages and identified embryonic neuroblasts in the ventral nerve cord of Drosophila melanogaster. Biol Open 4:420–434.

Booker R, Truman JW. 1987. Postembryonic neurogenesis in the CNS of the tobacco hornworm, Manduca sexta. I. Neuroblast arrays and the fate of their progeny during metamorphosis. J Comp Neurol 255:548–559.

Brierley DJ, Rathore K, VijayRaghavan K, Williams DW. 2012. Developmental origins and architecture of Drosophila leg motoneurons. J Comp Neurol 520:1629–1649.

Brown HLD, Truman JW. 2009. Fine-tuning of secondary arbor development: the effects of the ecdysone receptor on the adult neuronal lineages of the Drosophila thoracic CNS. Development 136:3247–3256.

Cachero S, Ostrovsky AD, Yu JY, Dickson BJ, Jefferis GSXE. 2010. Sexual dimorphism in the fly brain. Curr Biol 20:1589–1601.

Campello RJGB, Moulavi D, Sander J. 2013. Density-Based Clustering Based on Hierarchical Density EstimatesAdvances in Knowledge Discovery and Data Mining. Springer Berlin Heidelberg. pp. 160–172.

Card GM. 2012. Escape behaviors in insects. Curr Opin Neurobiol 22:180–186.

Chan WP, Dickinson MH. 1996. Position-specific central projections of mechanosensory neurons on the haltere of the blow fly, Calliphora vicina. J Comp Neurol 369:405–418.

Chen C, Agrawal S, Mark B, Mamiya A, Sustar A, Phelps JS, Lee W-CA, Dickson BJ, Card GM, Tuthill JC. 2021. Functional architecture of neural circuits for leg proprioception in Drosophila. Curr Biol 31:5163–5175.e7.

Cheong HSJ, Eichler K, Stuerner T, Asinof SK, Champion AS, Marin EC, Oram TB, Sumathipala M, Venkatasubramanian L, Namiki S, Siwanowicz I, Costa M, Berg S, Jefferis GSX, Card GM. 2023. Transforming descending input into behavior: The organization of premotor circuits in the Drosophila Male Adult Nerve Cord connectome. bioRxiv. doi:10.1101/2023.06.07.543976

Chung J, Pedigo BD, Bridgeford EW, Varjavand BK, Helm HS, Vogelstein JT. 2019. GraSPy: Graph Statistics in Python. J Mach Learn Res 20:1–7.

Clowney EJ, Iguchi S, Bussell JJ, Scheer E, Ruta V. 2015. Multimodal Chemosensory Circuits Controlling Male Courtship in Drosophila. Neuron 87:1036–1049.

Colizza V, Flammini A, Serrano MA, Vespignani A. 2006. Detecting rich-club ordering in complex networks. Nat Phys 2:110–115.

Consoulas C, Restifo LL, Levine RB. 2002. Dendritic remodeling and growth of motoneurons during metamorphosis of Drosophila melanogaster. J Neurosci 22:4906–4917.

Costa M, Manton JD, Ostrovsky AD, Prohaska S, Jefferis GSXE. 2016. NBLAST: Rapid, Sensitive Comparison of Neuronal Structure and Construction of Neuron Family Databases. Neuron 91:293–311.

Court R, Namiki S, Armstrong JD, Börner J, Card G, Costa M, Dickinson M, Duch C, Korff W, Mann R, Merritt D, Murphey RK, Seeds AM, Shirangi T, Simpson JH, Truman JW, Tuthill JC, Williams DW, Shepherd D. 2020. A Systematic Nomenclature for the Drosophila Ventral Nerve Cord. Neuron 107:1071–1079.e2.

Cruz TL, Chiappe ME. 2023. Multilevel visuomotor control of locomotion in Drosophila. Curr Opin Neurobiol 82:102774.

Csardi G, Nepusz T, Others. 2006. The igraph software package for complex network research. InterJournal, complex systems 1695:1–9.

Demir E, Dickson BJ. 2005. fruitless splicing specifies male courtship behavior in Drosophila. Cell 121:785–794.

Deora T, Singh AK, Sane SP. 2015. Biomechanical basis of wing and haltere coordination in flies. Proc Natl Acad Sci U S A 112:1481–1486.

Dickinson MH, Palka J. 1987. Physiological properties, time of development, and central projection are correlated in the wing mechanoreceptors of Drosophila. J Neurosci 7:4201–4208.

Dinges GF, Chockley AS, Bockemühl T, Ito K, Blanke A, Büschges A. 2021. Location and arrangement of campaniform sensilla in Drosophila melanogaster. J Comp Neurol 529:905–925.

Doe CQ. 2017. Temporal Patterning in the Drosophila CNS. Annu Rev Cell Dev Biol 33:219–240.

Doe CQ, Goodman CS. 1985. Early events in insect neurogenesis. I. Development and segmental differences in the pattern of neuronal precursor cells. Dev Biol 111:193–205.

Dorkenwald S, Matsliah A, Sterling AR, Schlegel P, Yu S-C, McKellar CE, Lin A, Costa M, Eichler K, Yin Y, Silversmith W, Schneider-Mizell C, Jordan CS, Brittain D, Halageri A, Kuehner K, Ogedengbe O, Morey R, Gager J, Kruk K, Perlman E, Yang R, Deutsch D, Bland D, Sorek M, Lu R, Macrina T, Lee K, Bae JA, Mu S, Nehoran B, Mitchell E, Popovych S, Wu J, Jia Z, Castro M, Kemnitz N, Ih D, Bates AS, Eckstein N, Funke J, Collman F, Bock DD, Jefferis GSXE, Seung HS, Murthy M, FlyWire Consortium. 2023. Neuronal wiring diagram of an adult brain. bioRxiv. doi:10.1101/2023.06.27.546656

Eagle N, Macy M, Claxton R. 2010. Network diversity and economic development. Science 328:1029–1031.

Eckstein N, Bates AS, Du M, Hartenstein V. 2023. Neurotransmitter classification from electron microscopy images at synaptic sites in Drosophila. BioRxiv.

Enriquez J, Venkatasubramanian L, Baek M, Peterson M, Aghayeva U, Mann RS. 2015. Specification of individual adult motor neuron morphologies by combinatorial transcription factor codes. Neuron 86:955–970.

Fayyazuddin A, Dickinson MH. 1996. Haltere afferents provide direct, electrotonic input to a steering motor neuron in the blowfly, Calliphora. J Neurosci 16:5225–5232.

Fishkind DE, Adali S, Patsolic HG, Meng L, Singh D, Lyzinski V, Priebe CE. 2019. Seeded graph matching. Pattern Recognition. doi:10.1016/j.patcog.2018.09.014

Ghysen A. 1980. The projection of sensory neurons in the central nervous system of Drosophila: choice of the appropriate pathway. Dev Biol 78:521–541.

Gnatzy W, Grünert U, Bender M. 1987. Campaniform sensilla of Calliphora vicina (Insecta, Diptera). Zoomorphology 106:312–319.

Goh KI, Kahng B, Kim D. 2011. Universal behavior of load distribution in scale-free networks. The Structure and Dynamics.

Guimerà R, Nunes Amaral LA. 2005. Functional cartography of complex metabolic networks. Nature 433:895–900.

Hagberg A, Swart P, S Chult D. 2008. Exploring network structure, dynamics, and function using networkx (No. LA-UR-08-05495; LA-UR-08-5495). Los Alamos National Lab. (LANL), Los Alamos, NM (United States).

Hao Y, Hao S, Andersen-Nissen E, Mauck WM 3rd, Zheng S, Butler A, Lee MJ, Wilk AJ, Darby C, Zager M, Hoffman P, Stoeckius M, Papalexi E, Mimitou EP, Jain J, Srivastava A, Stuart T, Fleming LM, Yeung B, Rogers AJ, McElrath JM, Blish CA, Gottardo R, Smibert P, Satija R. 2021. Integrated analysis of multimodal single-cell data. Cell 184:3573–3587.e29.

Harris RM, Pfeiffer BD, Rubin GM, Truman JW. 2015. Neuron hemilineages provide the functional ground plan for the Drosophila ventral nervous system. Elife 4. doi:10.7554/eLife.04493

He Z, Luo Y, Shang X, Sun JS, Carlson JR. 2019. Chemosensory sensilla of the Drosophila wing express a candidate ionotropic pheromone receptor. PLoS Biol 17:e2006619.

Hopkins BR, Barmina O, Kopp A. 2023. A single-cell atlas of the sexually dimorphic Drosophila foreleg and its sensory organs during development. PLoS Biol 21:e3002148.

Iles JF. 1976. Organization of motoneurones in the prothoracic ganglion of the cockroach Periplaneta americana (L.). Philos Trans R Soc Lond B Biol Sci 276:205–219.

Isshiki T, Pearson B, Holbrook S, Doe CQ. 2001. Drosophila neuroblasts sequentially express transcription factors which specify the temporal identity of their neuronal progeny. Cell 106:511–521.

Jacobs K, Todman MG, Allen MJ, Davies JA, Bacon JP. 2000. Synaptogenesis in the giant-fibre system of Drosophila: interaction of the giant fibre and its major motorneuronal target. Development 127:5203–5212.

Jan YN, Ghysen A, Christoph I, Barbel S, Jan LY. 1985. Formation of neuronal pathways in the imaginal discs of Drosophila melanogaster. J Neurosci 5:2453–2464.

Kallman BR, Kim H, Scott K. 2015. Excitation and inhibition onto central courtship neurons biases Drosophila mate choice. Elife 4:e11188.

Kambadur R, Koizumi K, Stivers C, Nagle J, Poole SJ, Odenwald WF. 1998. Regulation of POU genes by castor and hunchback establishes layered compartments in the Drosophila CNS. Genes Dev 12:246–260.

Kennedy T, Broadie K. 2018. Newly Identified Electrically Coupled Neurons Support Development of the Giant Fiber Model Circuit. eNeuro 5. doi:10.1523/ENEURO.0346-18.2018

Khuong TM, Wang Q-P, Manion J, Oyston LJ, Lau M-T, Towler H, Lin YQ, Neely GG. 2019. Nerve injury drives a heightened state of vigilance and neuropathic sensitization in. Sci Adv 5:eaaw4099.

Koh T-W, He Z, Gorur-Shandilya S, Menuz K, Larter NK, Stewart S, Carlson JR. 2014. The Drosophila IR20a clade of ionotropic receptors are candidate taste and pheromone receptors. Neuron 83:850–865.

Kuan AT, Phelps JS, Thomas LA, Nguyen TM, Han J, Chen C-L, Azevedo AW, Tuthill JC, Funke J, Cloetens P, Pacureanu A, Lee W-CA. 2020. Dense neuronal reconstruction through X-ray holographic nano-tomography. Nat Neurosci 23:1637–1643.

Kwon JY, Dahanukar A, Weiss LA, Carlson JR. 2014. A map of taste neuron projections in the Drosophila CNS. J Biosci 39:565–574.

Lacin H, Chen H-M, Long X, Singer RH, Lee T, Truman JW. 2019. Neurotransmitter identity is acquired in a lineage-restricted manner in the CNS. Elife 8. doi:10.7554/eLife.43701

Lacin H, Truman JW. 2016. Lineage mapping identifies molecular and architectural similarities between the larval and adult Drosophila central nervous system. Elife 5:e13399.

Lacin H, Williamson WR, Card GM, Skeath JB, Truman JW. 2020. Unc-4 acts to promote neuronal identity and development of the take-off circuit in the Drosophila CNS. Elife 9. doi:10.7554/eLife.55007

Lacin H, Zhu Y, Wilson BA, Skeath JB. 2014. Transcription factor expression uniquely identifies most postembryonic neuronal lineages in the Drosophila thoracic central nervous system. Development 141:1011–1021.

Lee Y-J, Yang C-P, Miyares RL, Huang Y-F, He Y, Ren Q, Chen H-M, Kawase T, Ito M, Otsuna H, Sugino K, Aso Y, Ito K, Lee T. 2020. Conservation and divergence of related neuronal lineages in the Drosophila central brain. eLife. doi:10.7554/elife.53518

Lesser E, Azevedo AW, Phelps JS, Elabbady L, Cook AP, Mark B, Kuroda S, Sustar A, Moussa AJ, Dallmann CJ, Agrawal S, Lee S-YJ, Pratt BG, Skutt-Kakari K, Gerhard S, Lu R, Kemnitz N, Lee K, Halageri A, Castro M, Ih D, Gager J, Tammam M, Dorkenwald S, Collman FC, Schneider-Mizell CM, Brittain D, Jordan CS, Sebastian Seung H, Macrina T, Dickinson MH, Lee W-CA, Tuthill JC. 2023. Synaptic architecture of leg and wing motor control networks in Drosophila. bioRxiv. doi:10.1101/2023.05.30.542725

Lillvis JL, Wang K, Shiozaki HM, Xu M, Stern DL, Dickson BJ. 2024. Nested neural circuits generate distinct acoustic signals during Drosophila courtship. Curr Biol. doi:10.1016/j.cub.2024.01.015

Ling F, Dahanukar A, Weiss LA, Kwon JY, Carlson JR. 2014. The molecular and cellular basis of taste coding in the legs of Drosophila. J Neurosci 34:7148–7164.

Mamiya A, Gurung P, Tuthill JC. 2018. Neural Coding of Leg Proprioception in Drosophila. Neuron 100:636–650.e6.

Manoli DS, Foss M, Villella A, Taylor BJ, Hall JC, Baker BS. 2005. Male-specific fruitless specifies the neural substrates of Drosophila courtship behaviour. Nature 436:395–400.

Marin EC, Büld L, Theiss M, Sarkissian T, Roberts RJV, Turnbull R, Tamimi IFM, Pleijzier MW, Laursen WJ, Drummond N, Schlegel P, Bates AS, Li F, Landgraf M, Costa M, Bock DD, Garrity PA, Jefferis GSXE. 2020. Connectomics Analysis Reveals First-, Second-, and Third-Order Thermosensory and Hygrosensory Neurons in the Adult Drosophila Brain. Curr Biol 30:3167–3182.e4.

Marin EC, Dry KE, Alaimo DR, Rudd KT, Cillo AR, Clenshaw ME, Negre N, White KP, Truman JW. 2012. Ultrabithorax confers spatial identity in a context-specific manner in the Drosophila postembryonic ventral nervous system. Neural Dev 7:31.

Matsuda R. 1976. Morphology and Evolution of the Insect Abdomen. Oxford: Pergamon Press.

Mellert DJ, Williamson WR, Shirangi TR, Card GM, Truman JW. 2016. Genetic and Environmental Control of Neurodevelopmental Robustness in Drosophila. PLoS One 11:e0155957.

Merritt DJ, Murphey RK. 1992. Projections of leg proprioceptors within the CNS of the fly Phormia in relation to the generalized insect ganglion. J Comp Neurol 322:16–34.

Monastirioti M, Gorczyca M, Rapus J, Eckert M, White K, Budnik V. 1995. Octopamine immunoreactivity in the fruit fly Drosophila melanogaster. J Comp Neurol 356:275–287.

Murphey RK, Possidente D, Pollack G, Merritt DJ. 1989a. Modality-specific axonal projections in the CNS of the flies Phormia and Drosophila. J Comp Neurol 290:185–200.

Murphey RK, Possidente DR, Vandervorst P, Ghysen A. 1989b. Compartments and the topography of leg afferent projections in Drosophila. J Neurosci 9:3209–3217.

Mu S, Yu S-C, Turner NL, McKellar CE, Dorkenwald S, Collman F, Koolman S, Moore M, Morejohn S, Silverman B, Willie K, Willie R, Bland D, Burke A, Ashwood Z, Luther K, Castro M, Ogedengbe O, Silversmith W, Wu J, Halageri A, Macrina T, Kemnitz N, Murthy M, Sebastian Seung H. 2021. 3D reconstruction of cell nuclei in a full Drosophila brain. bioRxiv. doi:10.1101/2021.11.04.467197

Namiki S, Dickinson MH, Wong AM, Korff W, Card GM. 2018. The functional organization of descending sensory-motor pathways in. Elife 7. doi:10.7554/eLife.34272

Nayak SV, Singh RN. 1983. Sensilla on the tarsal segments and mouthparts of adult Drosophila melanogaster meigen (Diptera : Drosophilidae). Int J Insect Morphol Embryol 12:273–291.

Nemesure MD, Schwedhelm TM, Sacerdote S, O’Malley AJ, Rozema LR, Moen EL. 2021. A measure of local uniqueness to identify linchpins in a social network with node attributes. Appl Netw Sci 6. doi:10.1007/s41109-021-00400-8

Newman ME. 2001. Scientific collaboration networks. II. Shortest paths, weighted networks, and centrality. Phys Rev E Stat Nonlin Soft Matter Phys 64:016132.

Nottebohm E, Ramaekers A, Dambly-Chaudière C, Ghysen A. 1994. The leg of Drosophila as a model system for the analysis of neuronal diversity. J Physiol Paris 88:141–151.

Ohyama T, Schneider-Mizell CM, Fetter RD, Aleman JV, Franconville R, Rivera-Alba M, Mensh BD, Branson KM, Simpson JH, Truman JW, Cardona A, Zlatic M. 2015. A multilevel multimodal circuit enhances action selection in Drosophila. Nature 520:633–639.

O’Sullivan A, Lindsay T, Prudnikova A, Erdi B, Dickinson M, von Philipsborn AC. 2018. Multifunctional Wing Motor Control of Song and Flight. Current Biology. doi:10.1016/j.cub.2018.06.038

Palka J, Schubiger M, Murray MA. 1984. Peripheral Neurogenesis in “Drosophila.” Bioscience 34:318–321.

Paulk A, Gilbert C. 2006. Proprioceptive encoding of head position in the black soldier fly, Hermetia illucens (L.) (Stratiomyidae). J Exp Biol 209:3913–3924.

Pavlou HJ, Goodwin SF. 2013. Courtship behavior in Drosophila melanogaster: towards a “courtship connectome.” Curr Opin Neurobiol 23:76–83.

Phelps JS, Hildebrand DGC, Graham BJ, Kuan AT, Thomas LA, Nguyen TM, Buhmann J, Azevedo AW, Sustar A, Agrawal S, Liu M, Shanny BL, Funke J, Tuthill JC, Lee W-CA. 2021. Reconstruction of motor control circuits in adult Drosophila using automated transmission electron microscopy. Cell 184:759–774.e18.

Plaza SM, Clements J, Dolafi T, Umayam L, Neubarth NN, Scheffer LK, Berg S. 2022. neuPrint: An open access tool for EM connectomics. Front Neuroinform 16. doi:10.3389/fninf.2022.896292

Pop S, Chen C-L, Sproston CJ, Kondo S, Ramdya P, Williams DW. 2020. Extensive and diverse patterns of cell death sculpt neural networks in insects. Elife 9. doi:10.7554/eLife.59566

Possidente DR, Murphey RK. 1989. Genetic control of sexually dimorphic axon morphology in Drosophila sensory neurons. Dev Biol 132:448–457.

Poulson. 1950. Histogenesis, organogenesis, and differentiation in the embryo of Drosophila melanogaster Meigen In: Demerec M, editor. Biology of Drosophila. New York: Wiley. pp. 168–274.

Power ME. 1948. The thoracico-abdominal nervous system of an adult insect, Drosophila melanogaster. J Comp Neurol 88:347–409.

Prokop A, Technau GM. 1991. The origin of postembryonic neuroblasts in the ventral nerve cord of Drosophila melanogaster. Development 111:79–88.

Rubinov M, Sporns O. 2010. Complex network measures of brain connectivity: uses and interpretations. Neuroimage 52:1059–1069.

Scheffer LK, Xu CS, Januszewski M, Lu Z, Takemura S-Y, Hayworth KJ, Huang GB, Shinomiya K, Maitlin-Shepard J, Berg S, Clements J, Hubbard PM, Katz WT, Umayam L, Zhao T, Ackerman D, Blakely T, Bogovic J, Dolafi T, Kainmueller D, Kawase T, Khairy KA, Leavitt L, Li PH, Lindsey L, Neubarth N, Olbris DJ, Otsuna H, Trautman ET, Ito M, Bates AS, Goldammer J, Wolff T, Svirskas R, Schlegel P, Neace E, Knecht CJ, Alvarado CX, Bailey DA, Ballinger S, Borycz JA, Canino BS, Cheatham N, Cook M, Dreher M, Duclos O, Eubanks B, Fairbanks K, Finley S, Forknall N, Francis A, Hopkins GP, Joyce EM, Kim S, Kirk NA, Kovalyak J, Lauchie SA, Lohff A, Maldonado C, Manley EA, McLin S, Mooney C, Ndama M, Ogundeyi O, Okeoma N, Ordish C, Padilla N, Patrick CM, Paterson T, Phillips EE, Phillips EM, Rampally N, Ribeiro C, Robertson MK, Rymer JT, Ryan SM, Sammons M, Scott AK, Scott AL, Shinomiya A, Smith C, Smith K, Smith NL, Sobeski MA, Suleiman A, Swift J, Takemura S, Talebi I, Tarnogorska D, Tenshaw E, Tokhi T, Walsh JJ, Yang T, Horne JA, Li F, Parekh R, Rivlin PK, Jayaraman V, Costa M, Jefferis GS, Ito K, Saalfeld S, George R, Meinertzhagen IA, Rubin GM, Hess HF, Jain V, Plaza SM. 2020. A connectome and analysis of the adult central brain. Elife 9. doi:10.7554/eLife.57443

Schlegel P, Bates AS, Stürner T, Jagannathan SR, Drummond N, Hsu J, Serratosa Capdevila L, Javier A, Marin EC, Barth-Maron A, Tamimi IF, Li F, Rubin GM, Plaza SM, Costa M, Jefferis GSXE. 2021. Information flow, cell types and stereotypy in a full olfactory connectome. Elife 10. doi:10.7554/eLife.66018

Schlegel P, Yin Y, Bates AS, Dorkenwald S, Eichler K, Brooks P, Han DS, Gkantia M, Dos Santos M, Munnelly EJ, Badalamente G, Capdevila LS, Sane VA, Pleijzier MW, Tamimi IFM, Dunne CR, Salgarella I, Javier A, Fang S, Perlman E, Kazimiers T, Jagannathan SR, Matsliah A, Sterling AR, Yu S-C, McKellar CE, FlyWire Consortium, Costa M, Seung HS, Murthy M, Hartenstein V, Bock DD, Jefferis GSXE. 2023. Whole-brain annotation and multi-connectome cell typing quantifies circuit stereotypy in. bioRxiv. doi:10.1101/2023.06.27.546055

Schmid A, Chiba A, Doe CQ. 1999. Clonal analysis of Drosophila embryonic neuroblasts: neural cell types, axon projections and muscle targets. Development 126:4653–4689.

Schmidt H, Rickert C, Bossing T, Vef O, Urban J, Technau GM. 1997. The embryonic central nervous system lineages of Drosophila melanogaster. II. Neuroblast lineages derived from the dorsal part of the neuroectoderm. Dev Biol 189:186–204.

Schneider-Mizell CM, Gerhard S, Longair M, Kazimiers T, Li F, Zwart MF, Champion A, Midgley FM, Fetter RD, Saalfeld S, Cardona A. 2016. Quantitative neuroanatomy for connectomics in Drosophila. Elife 5. doi:10.7554/eLife.12059

Seeds AM, Ravbar P, Chung P, Hampel S, Midgley FM Jr, Mensh BD, Simpson JH. 2014. A suppression hierarchy among competing motor programs drives sequential grooming in Drosophila. Elife 3:e02951.

Shepherd D, Harris R, Williams DW, Truman JW. 2016. Postembryonic lineages of the Drosophila ventral nervous system: Neuroglian expression reveals the adult hemilineage associated fiber tracts in the adult thoracic neuromeres. J Comp Neurol 524:2677–2695.

Shepherd D, Sahota V, Court R, Williams DW, Truman JW. 2019. Developmental organization of central neurons in the adult Drosophila ventral nervous system. J Comp Neurol 527:2573–2598.

Shirangi TR, Wong AM, Truman JW, Stern DL. 2016. Doublesex Regulates the Connectivity of a Neural Circuit Controlling Drosophila Male Courtship Song. Dev Cell 37:533–544.

Smith SA, Shepherd D. 1996. Central afferent projections of proprioceptive sensory neurons in Drosophila revealed with the enhancer-trap technique. J Comp Neurol 364:311–323.

Starostina E, Liu T, Vijayan V, Zheng Z, Siwicki KK, Pikielny CW. 2012. A Drosophila DEG/ENaC subunit functions specifically in gustatory neurons required for male courtship behavior. J Neurosci 32:4665–4674.

Stocker RF. 1994. The organization of the chemosensory system in Drosophila melanogaster: a review. Cell Tissue Res 275:3–26.

Stölting H, Stumpner A, Lakes-Harlan R. 2007. Morphology and physiology of the prosternal chordotonal organ of the sarcophagid fly Sarcophaga bullata (Parker). J Insect Physiol 53:444–454.

Strausfeld NJ. 1976. Atlas of an Insect Brain.

Strausfeld NJ, Seyan HS. 1985. Convergence of visual, haltere, and prosternal inputs at neck motor neurons of Calliphora erythrocephala. Cell Tissue Res 240:601–615.

Takemura S-Y, Huang GB, Januszewski M, Lu Z, Marin EC, Preibisch S, Xu CS, Champion A, Cheong HSJ, Costa M, Eichler K, Knecht CJ, Li F, Morris BJ, Schlegel P, Stuerner T, Badalamente G, Bogovic J, Brooks P, Canino BS, Clements J, Cook M, Dunne CR, Fang S, Finley-May S, Funke J, George R, Gkantia M, Harrington K, Hayworth KJ, Hazlett A, Hopkins GP, Hsu J, Hubbard PM, Javier A, Kainmueller D, Katz WT, Korff W, Kovalyak J, Krzeminski D, Lauchie SA, Manley EA, Mooney C, Neace E, Nichols M, Okeoma N, Ordish C, Parekh R, Paterson T, Phillips EM, Plaza SM, Rivlin PK, Saalfeld S, Shepherd D, Smith C, Takemura S, Talebi I, Tamimi I, Tiscareno C, Trautman ET, Umayam L, Walsh JJ, Yang T, Zhao T, Hess HF, Rubin GM, Scheffer LK, Card G, Jefferis GSXE, Berg S. 2023. A Connectome of the Male Drosophila Ventral Nerve Cord. bioRxiv.

Thistle R, Cameron P, Ghorayshi A, Dennison L, Scott K. 2012. Contact chemoreceptors mediate male-male repulsion and male-female attraction during Drosophila courtship. Cell 149:1140–1151.

Tissot M, Stocker RF. 2000. Metamorphosis in drosophila and other insects: the fate of neurons throughout the stages. Prog Neurobiol 62:89–111.

Tix S, Bate M, Technau GM. 1989. Pre-existing neuronal pathways in the leg imaginal discs of Drosophila. Development 107:855–862.

Toda H, Zhao X, Dickson BJ. 2012. The Drosophila female aphrodisiac pheromone activates ppk23(+) sensory neurons to elicit male courtship behavior. Cell Rep 1:599–607.

Trimarchi JR, Murphey RK. 1997. The shaking-B2 mutation disrupts electrical synapses in a flight circuit in adult Drosophila. J Neurosci 17:4700–4710.

Truman JW. 1990. Metamorphosis of the central nervous system of Drosophila. J Neurobiol 21:1072–1084.

Truman JW, Bate M. 1988. Spatial and temporal patterns of neurogenesis in the central nervous system of Drosophila melanogaster. Dev Biol 125:145–157.

Truman JW, Moats W, Altman J, Marin EC, Williams DW. 2010. Role of Notch signaling in establishing the hemilineages of secondary neurons in Drosophila melanogaster. Development 137:53–61.

Truman JW, Schuppe H, Shepherd D, Williams DW. 2004. Developmental architecture of adult-specific lineages in the ventral CNS of *Drosophila*. Development. doi:10.1242/dev.01371

Tsubouchi A, Yano T, Yokoyama TK, Murtin C, Otsuna H, Ito K. 2017. Topological and modality-specific representation of somatosensory information in the fly brain. Science 358:615–623.

Tuthill JC, Azim E. 2018. Proprioception. Curr Biol 28:R194–R203.

Tuthill JC, Wilson RI. 2016a. Mechanosensation and Adaptive Motor Control in Insects. Curr Biol 26:R1022–R1038.

Tuthill JC, Wilson RI. 2016b. Parallel Transformation of Tactile Signals in Central Circuits of Drosophila. Cell 164:1046–1059.

Tyrer NM, Altman JS. 1974. Motor and sensory flight neurones in a locust demonstrated using cobalt chloride. J Comp Neurol 157:117–138.

Usui-Ishihara A, Simpson P. 2005. Differences in sensory projections between macro- and microchaetes in Drosophilid flies. Dev Biol 277:170–183.

Vijayan V, Thistle R, Liu T, Starostina E, Pikielny CW. 2014. Drosophila pheromone-sensing neurons expressing the ppk25 ion channel subunit stimulate male courtship and female receptivity. PLoS Genet 10:e1004238.

Vinh NX, Epps J, Bailey J. 2009. Information theoretic measures for clusterings comparisonProceedings of the 26th Annual International Conference on Machine Learning. Presented at the ICML ‘09: The 26th Annual International Conference on Machine Learning held in conjunction with the 2007 International Conference on Inductive Logic Programming. New York, NY, USA: ACM. doi:10.1145/1553374.1553511

von Philipsborn AC, Liu T, Yu JY, Masser C, Bidaye SS, Dickson BJ. 2011. Neuronal control of Drosophila courtship song. Neuron 69:509–522.

Williams DW, Shepherd D. 1999. Persistent larval sensory neurons in adult Drosophila melanogaster. J Neurobiol 39:275–286.

Winding M, Pedigo BD, Barnes CL, Patsolic HG, Park Y, Kazimiers T, Fushiki A, Andrade IV, Khandelwal A, Valdes-Aleman J, Li F, Randel N, Barsotti E, Correia A, Fetter RD, Hartenstein V, Priebe CE, Vogelstein JT, Cardona A, Zlatic M. 2023. The connectome of an insect brain. Science 379:eadd9330.

Wong RK, Pearson KG. 1976. Properties of the trochanteral hair plate and its function in the control of walking in the cockroach. J Exp Biol 64:233–249.

Yellman C, Tao H, He B, Hirsh J. 1997. Conserved and sexually dimorphic behavioral responses to biogenic amines in decapitated Drosophila. Proc Natl Acad Sci U S A 94:4131–4136.

Yu JY, Kanai MI, Demir E, Jefferis GSXE, Dickson BJ. 2010. Cellular organization of the neural circuit that drives Drosophila courtship behavior. Curr Biol 20:1602–1614.

Zhang N, Guo L, Simpson JH. 2020. Spatial Comparisons of Mechanosensory Information Govern the Grooming Sequence in Drosophila. Curr Biol 30:3697–3698.

Zheng Z, Lauritzen JS, Perlman E, Robinson CG, Nichols M, Milkie D, Torrens O, Price J, Fisher CB, Sharifi N, Calle-Schuler SA, Kmecova L, Ali IJ, Karsh B, Trautman ET, Bogovic JA, Hanslovsky P, Jefferis GSXE, Kazhdan M, Khairy K, Saalfeld S, Fetter RD, Bock DD. 2018. A Complete Electron Microscopy Volume of the Brain of Adult Drosophila melanogaster. Cell 174:730–743.e22.

