## Supplementary material for "Systematic annotation of a complete adult male *Drosophila* nerve cord connectome reveals principles of functional organisation": Higher resolution figures

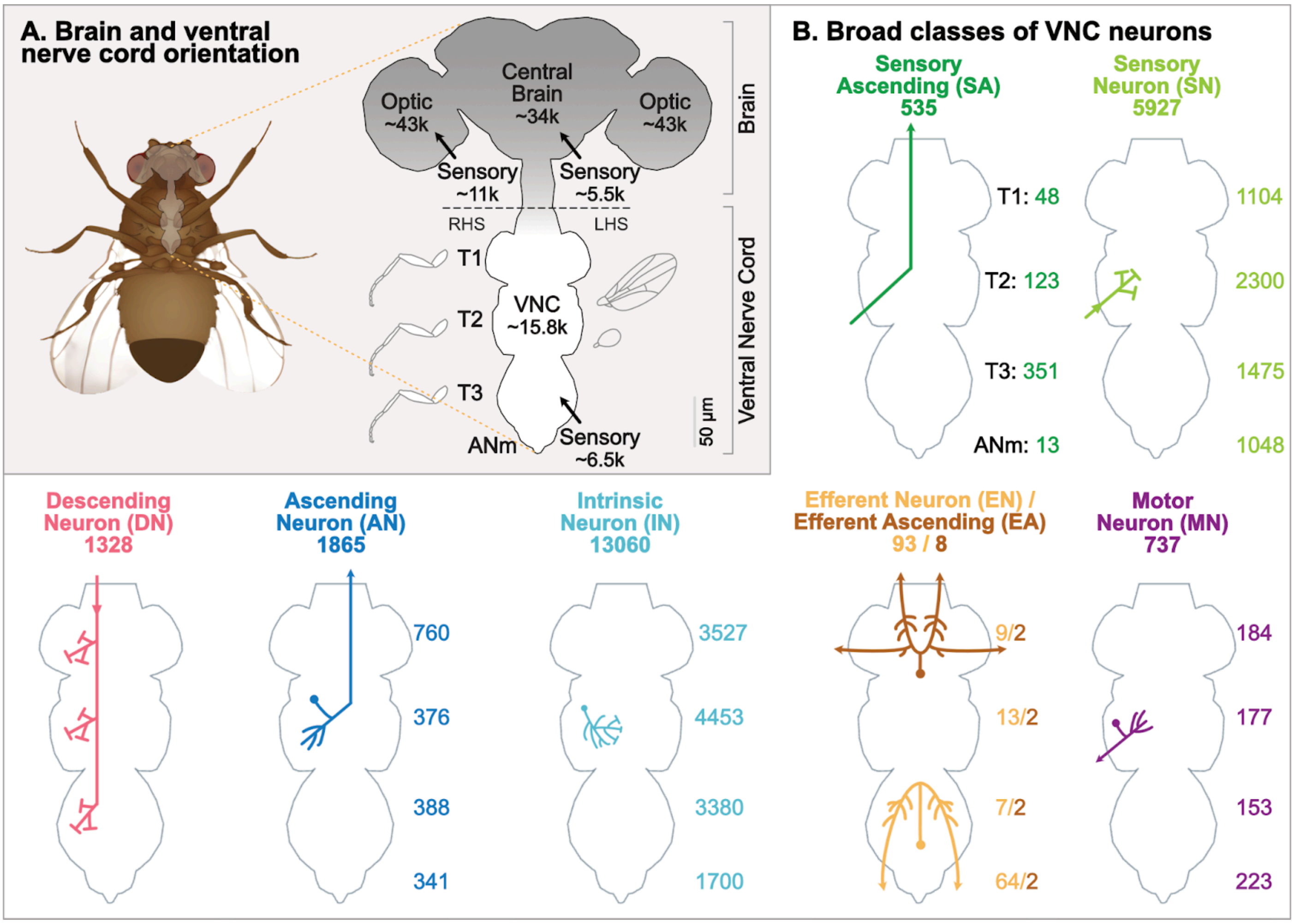

**Figure 1. Overview of the male adult nerve cord (MANC). A.** Orientation schematic showing a ventral view of the adult male *Drosophila* central nervous system. The ventral nerve cord (VNC) can be roughly divided into three thoracic neuromeres (T1, T2, T3) and the abdominal neuromeres (ANm). The thoracic neuromeres contain serially repeating neuropils controlling the three pairs of legs. T2 includes additional circuitry dedicated to flight. Number of neurons with cell bodies in the central brain are current estimates from FlyWire-FAFB (Schlegel et al., 2023). Cartoon legs and wings not to scale. **B.** Cartoon overview of the broad classes of neurons in the VNC including their overall and neuromere-specific neuron counts in this dataset.

### Glossary

#### General

| | |
|---|---|
| **VNC** | ventral nerve cord, a.k.a. ventral nervous system, analogous to vertebrate spinal cord |
| **CvC** | cervical connective, a.ka., neck connective, connecting brain to VNC |
| **root_side** | side of axon entry (from neck connective or nerve) |
| **soma_side** | side of soma origin |
| RHS | right hand side (from fly's perspective) |
| LHS | left hand side (from fly's perspective) |
| **soma_neuromere** | neuromere of soma origin |
| T1 | first thoracic segment, i.e., prothorax |
| T2 | second thoracic segment, i.e., mesothorax |
| T3 | third thoracic segment, i.e., metathorax |
| A1,A2… | first abdominal segment, second abdominal segment… |
| **soma_position** | coordinates of soma |
| **tosoma_position** | coordinates of presumed tract leading to soma |
| **position** | coordinates that can be used to navigate to a particular neuron |
| **bodyid** | unique numerical identifier for a reconstructed body/ neuron |
| **group** | matched within a neuromere |
| **subcluster** | ad hoc notation for neurons sharing common properties (i.e., origin or morphology) but not assignable to a specific hemilineage or serial set |
| **systematic_type** | cell type from systematic nomenclature |
| **type** | cell type from systematic type or literature |
| **instance** | type and location |
| **synonyms** | published name |
| **long_tract** | longitudinal tract used by descending neurons |
| **receptor_type** | peripheral structure associated with a sensory neuron |

#### Broad classes of VNC neurons

| | |
|---|---|
| **class** | broadest cell class (motor, sensory, etc) |
| ascending neuron (AN) | neuron with soma of VNC origin and axon ascending via neck connective |
| descending neuron (DN) | neuron with soma of brain origin and axon descending via neck connective |
| efferent ascending (EA) | neuron with soma of VNC origin and axon ascending via neck connective and leaving the nerve cord via nerve(s) |
| efferent neuron (EN) | neuron with soma of VNC origin and axon leaving the nerve cord via nerve(s) |
| intrinsic neuron (IN) | neuron with soma of VNC origin and processes restricted to the VNC |
| motor neuron (MN) | neuron with soma of VNC origin and axon exiting the nerve cord via a nerve |
| sensory ascending (SA) | neuron with soma of peripheral origin and axon entering the VNC via a nerve and exiting via neck connective |
| sensory neuron (SN) | neuron with soma of peripheral origin and axon entering the VNC via a nerve and terminating in the VNC |
| glia | reconstructed body believed to be glia, in most cases not completely traced |

#### Subclasses of VNC neurons

| | |
|---|---|
| **subclass** | general morphology & connectivity |
| IA | ipsilateral connections ascending |
| CA | contralateral connections ascending |
| BA | bilateral connections ascending |
| XA | no connections ascending |
| II | ipsilateral interconnecting neuropils |
| CI | contralateral interconnecting neuropils |
| BI | bilateral interconnecting neuropils |
| IR | ipsilateral restricted to one neuropil |
| CR | contralateral restricted to one neuropil |
| BR | bilateral restricted to one neuropil |

#### Neuropils/muscles (DN and MN subclasses)

| | |
|---|---|
| **origin** | neuropil with > 80% of input to a neuron |
| **target** | neuropil/muscle with > 80% of output from a neuron |
| Ntct | neck tectulum neuropil |
| *-nt* | neck tectulum |
| *-nm* | neck muscle |
| WTct | wing tectulum neuropil |
| *-wt* | wing tectulum |
| *-wm* | wing muscle |
| Htct | haltere tectulum neuropil |
| *-ht* | haltere tectulum |
| *-hm* | haltere muscle |
| UTct (ut) | upper tectulum (nt, wt, ht) |
| IntTct (it) | inter tectulum |
| LTct (lt) | lower tectulum |
| LegNpT1 (fl) | front leg neuropil (muscle) |
| LegNpT2 (ml) | middle leg neuropil (muscle) |
| LegNpT3 (hl) | hind leg neuropil (muscle) |
| LegNp (xl) | multiple leg neuropils |
| ANm (ad) | abdomen neuropil (muscle) |
| xn | multiple neuropils |
| xm | unknown muscle |

#### Nerves

| | |
|---|---|
| **entry_nerve** | nerve by which axon enters VNC |
| **exit_nerve** | nerve(s) by which axon exits VNC |
| DProN | Dorsal Prothoracic Nerve |
| PrN | Prosternal Nerve |
| ProCN | Prothoracic Chordotonal Nerve |
| ProAN | Prothoracic Accessory Nerve |
| VProN | Ventral Prothoracic Nerve |
| ProLN | Prothoracic Leg Nerve |
| ProNTBD | Prothoracic Nerve (to be defined) |
| ADMN | Anterior Dorsal Mesothoracic Nerve |
| PDMN | Posterior Dorsal Mesothoracic Nerve |
| MesoAN | Mesothoracic Accessory Nerve |
| MesoLN | Mesothoracic Leg Nerve |
| DMetaN | Dorsal Metathoracic Nerve |
| MetaLN | Metathoracic Leg Nerve |
| AbN1 | First Abdominal Nerve |
| AbN2 | Second Abdominal Nerve |
| AbN3 | Third Abdominal Nerve |
| AbN4 | Fourth Abdominal Nerve |
| AbNT | Abdominal Nerve Trunk |
| AbNX | unnamed abdominal nerve/connective |
| CvN | Cervical Nerve |

#### VNC lineages

| | |
|---|---|
| **neuroblast** | neural stem cell |
| **lineage** | cells produced by the same neuroblast |
| **hemilineage** | NotchON (A) or NotchOFF (B) subset of a lineage |
| **birthtime** | estimated developmental epoch of origin |
| primary | born during embryogenesis |
| early secondary | born in late embryo or early larva |
| secondary | born during larval life |

#### Predicted neurotransmitters

| | |
|---|---|
| **predicted_nt** | "winning" neurotransmitter prediction for a neuron |
| **predicted_nt_prob** | measure of confidence of "winning" neurotransmitter prediction |

#### Transmission

| | |
|---|---|
| electrical | matched to a neuron reported to be connected to partners by gap junctions |
| putative electrical | having low presynaptic density and/or morphology typical of electrical neurons |
| neurosecretory | matched to a neuron reported to be neurosecretory |
| putative neurosecretory | efferent neuron or unpolarised neuron with morphology suggesting neurosecretory function |

#### Serial homology

| | |
|---|---|
| **serial** | matched across neuromeres |
| **serial_motif** | pattern of serial homology |
| independent leg | serial set with origin and target in same leg neuropil |
| dorsal | serial set with origin and target in flight neuropils |
| convergent | serial set with axons converging to common target area(s) |
| sequential | serial set with axons consistently targeting the next anterior or posterior neuromere |
| ascending | serial set consisting solely of ascending neurons |
| centripetal | serial set with distal dendrites (usually in leg neuropil) and central axon terminals (usually in tectulum) |
| centrifugal | serial set with central dendrites (usually in tectulum) and distal axon terminals (usually in leg neuropil) |
| complex | serial set that does not fall into one of the other categories |

#### Modality

| | |
|---|---|
| ch | chemosensory |
| pp | proprioceptive |
| ta | tactile |
| xx | unknown |

#### Sensory neuron subclasses

| |
|---|
| mechanosensory bristle |
| chordotonal organ |
| campaniform sensilla |
| taste bristle |
| hair plate |
| strand receptor |

**Table 1. Glossary of annotation terms.** Annotation field names are bolded and shown against a white background whilst the associated field values are shown against a grey background. Note that compound field names are presented in the snake_case format (e.g., entry_nerve) used in Clio Neuroglancer whereas Neuprint employs the camelCase format (e.g., entryNerve).

**Table 1 - video 1. Using Clio Neuroglancer with the plain option to retrieve neurons by annotation (2:37).** https://youtu.be/EmOB8WvDHss Navigate to https://clio.janelia.org/ and select workspace "annotate" and the desired segmentation volume. On the right side click the "BODIES" tab to bring up the annotation fields. Using the plain option, type your query enclosed in curly brackets, with quotation marks around the field name, a colon, and quotations around the desired field value. Then click on the magnifying glass icon to retrieve all bodies matching your criteria. You can use the Fields checklist to hide or display particular fields for your neurons of interest. Now you can filter the list by typing into the fields. For example, let's compare the FeCO claw neurons in the right mesothoracic **entry nerve**. Pressing the flashlight icon will display all listed neurons in random colours. I'll remove the tissue layer and add the neuropil layer so that it's easier to see where the neurons are positioned within the leg neuropil. To compare the two different types of FeCO claw neurons, we could select a few neurons from each **type** and assign them to a specific colour, like this. Let's right-click on the segmentation layer, go to the Seg. section, and copy-paste the respective **bodyids** into the Neuroglancer window. This is so that they won't be lost when we select more neurons to display. Now clear the first selection, select neurons from the other type, and assign these to a different colour. We can then click on the saved neurons to view both of them at the same time. Note that when using this method to query for a numerical field value - for example, the **bodyid, group, serial**, or **subcluster** fields - you need to enclose the number in square brackets instead of in quotation marks, like this. If we now remove the previous filters, we can view all examples of a given serial set.

**Table 1 - video 2. Using Clio Neuroglancer with the structured option to retrieve neurons by annotation (2:41).** https://youtu.be/jObOKhAD7LQ Navigate to https://clio.janelia.org/ and select workspace "annotate" and the desired segmentation volume. On the right side click the "BODIES" tab and select the structured option. Type your desired annotation fields and values into the query boxes, clicking the "+" symbol to add them - for example, **hemilineage** "14A" and **serial_motif** "independent leg". Once you've added your criteria, click on the magnifying glass icon to retrieve the corresponding bodies. To customise the displayed information, use the Fields checklist to select fields of interest. You can also add filters for specific characteristics. For example, to visualise neurons from a specific serial set, such as "10424" (Figure 11A), type the number into the serial field and click the flashlight icon. Now right-click on the segmentation layer, go to the Seg. section, and copy-paste the respective **bodyids** into the Neuroglancer window. Then you can colour the neurons as you like. For example, filter for **soma_neuromere** "T1", click to select those bodyids, set their colour to green and highlight the selected neurons. You can repeat the same process for the other thoracic neuromeres, colouring T2 neurons in blue and T3 neurons in magenta. Then click on the saved list to visualise all of the neurons together. We recommend removing the "all-tissue" layer and selecting the "neuropil" layer to get a clearer view of the neuropils innervated by your neurons. Note that to search for a specific serial set directly, you could instead use the structured option by simply typing serial and the number into the query boxes.

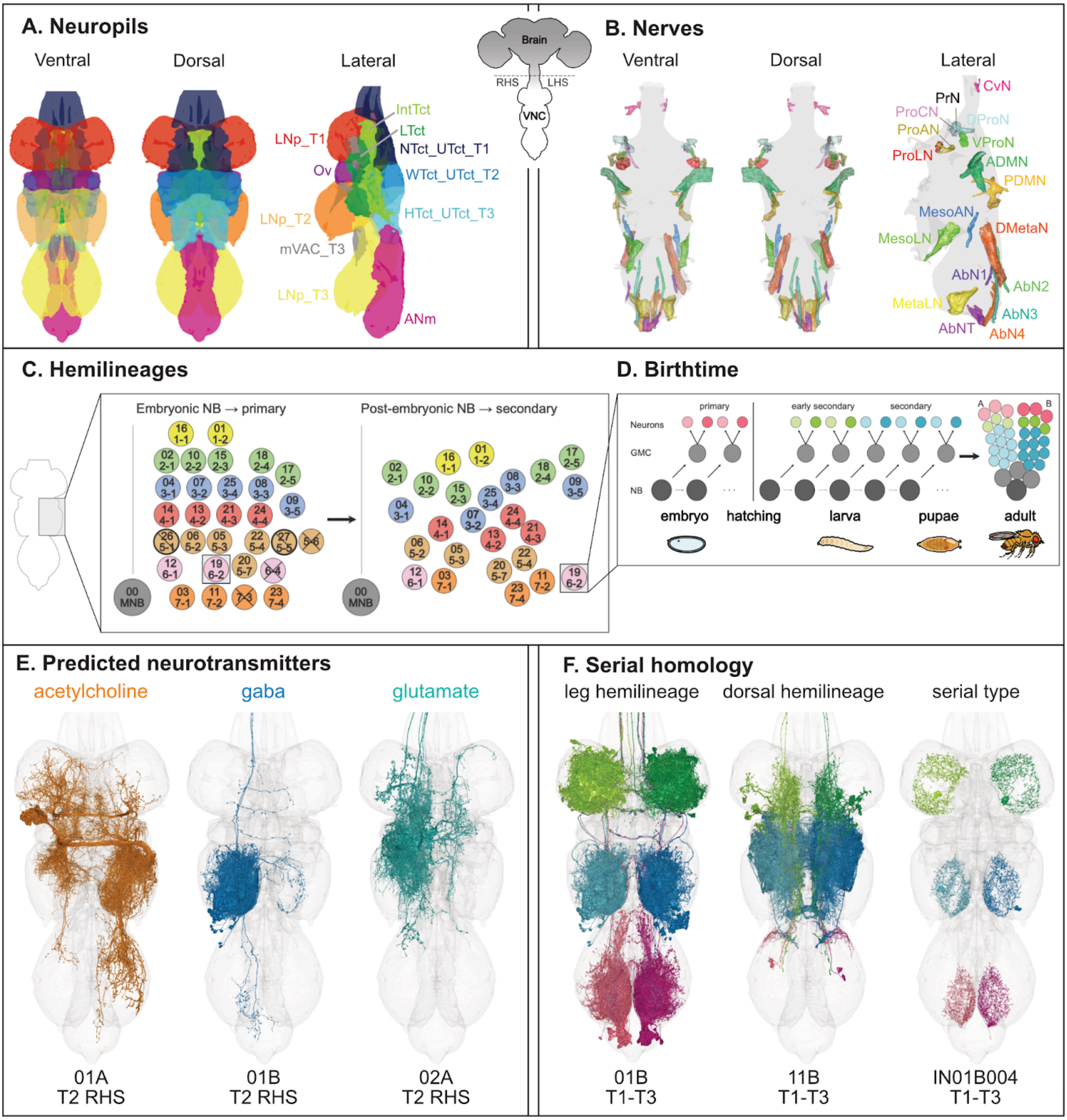
A. Neuropils
Ventral
Dorsal
Lateral
Brain
RHS
LHS
VNC
IntTct
LTct
LNp_T1
NTct_UTct_T1
Ov
WTct_UTct_T2
LNp_T2
HTct_UTct_T3
mVAC_T3
LNp_T3
ANm
B. Nerves
Ventral
Dorsal
Lateral
CvN
PrN
ProCN
DProN
ProAN
VProN
ProLN
ADMN
PDMN
MesoAN
DMetaN
MesoLN
AbN1
AbN2
MetaLN
AbN3
AbNT
AbN4
C. Hemilineages
Embryonic NB → primary
Post-embryonic NB → secondary
00 MNB
D. Birthtime
primary
early secondary
secondary
Neurons
GMC
NB
embryo
hatching
larva
pupae
adult
E. Predicted neurotransmitters
acetylcholine
gaba
glutamate
01A T2 RHS
01B T2 RHS
02A T2 RHS
F. Serial homology
leg hemilineage
dorsal hemilineage
serial type
01B T1-T3
11B T1-T3
IN01B004 T1-T3

**Figure 2. Overview of key features in MANC. A.** Overview of the neuropils in the ventral nerve cord. The left-hand side leg neuropils are removed in the lateral view. **B.** Overview of the nerves in the ventral nerve cord. **C.** Projected cross section of half of the mesothoracic neuromere (T2) depicting the array of neuroblasts that divide to produce hemilineages in the embryo and/or postembryonically. Insect embryonic neuroblasts (e.g., 1-1) were named for their row and column of delamination (Bate, 1976; Doe and Goodman, 1985) whilst postembryonic neuroblasts (e.g., 16) were assigned sequential numbers based on their observed progeny in Drosophila (Truman et al., 2004). Embryonic and postembryonic neuroblasts were eventually matched (Birkholz et al., 2015; Lacin and Truman, 2016), but both nomenclature systems are still in use. Bolded circles denote novel matches proposed in our study. A subset of neuroblasts do not survive to produce postembryonic progeny; "/" indicates that embryonic progeny were identified in this study while "X" indicates that they were not. **D.** Overview of neuroblast (NB) division into ganglion mother cells (GMC) prior to differentiation into two hemilineages. Primary neurons are born in the embryo followed by early secondary and finally secondary neurons. Cartoon icons by DBCLS (https://togotv.dbcls.jp/en/pics.html) were identified via https://bioicons.com/?query=fly and reproduced under https://creativecommons.org/licenses/by/4.0/. **E.** Most secondary neurons in a hemilineage share the same neurotransmitter prediction (Eckstein et al., 2023; Takemura et al., 2023). **F.** Hemilineages are serially repeated along the segments of the nerve cord. This provides a basis for identifying serially homologous neurons, which are concentrated in the six leg neuropils. Note that the T1 neuropils are rotated outwards relative to the T2 and T3 neuropils. Hemilineages innervating dorsal neuropils (e.g., 11B) exhibit more variation in survival and morphology between segments.

#### Systematic Typing

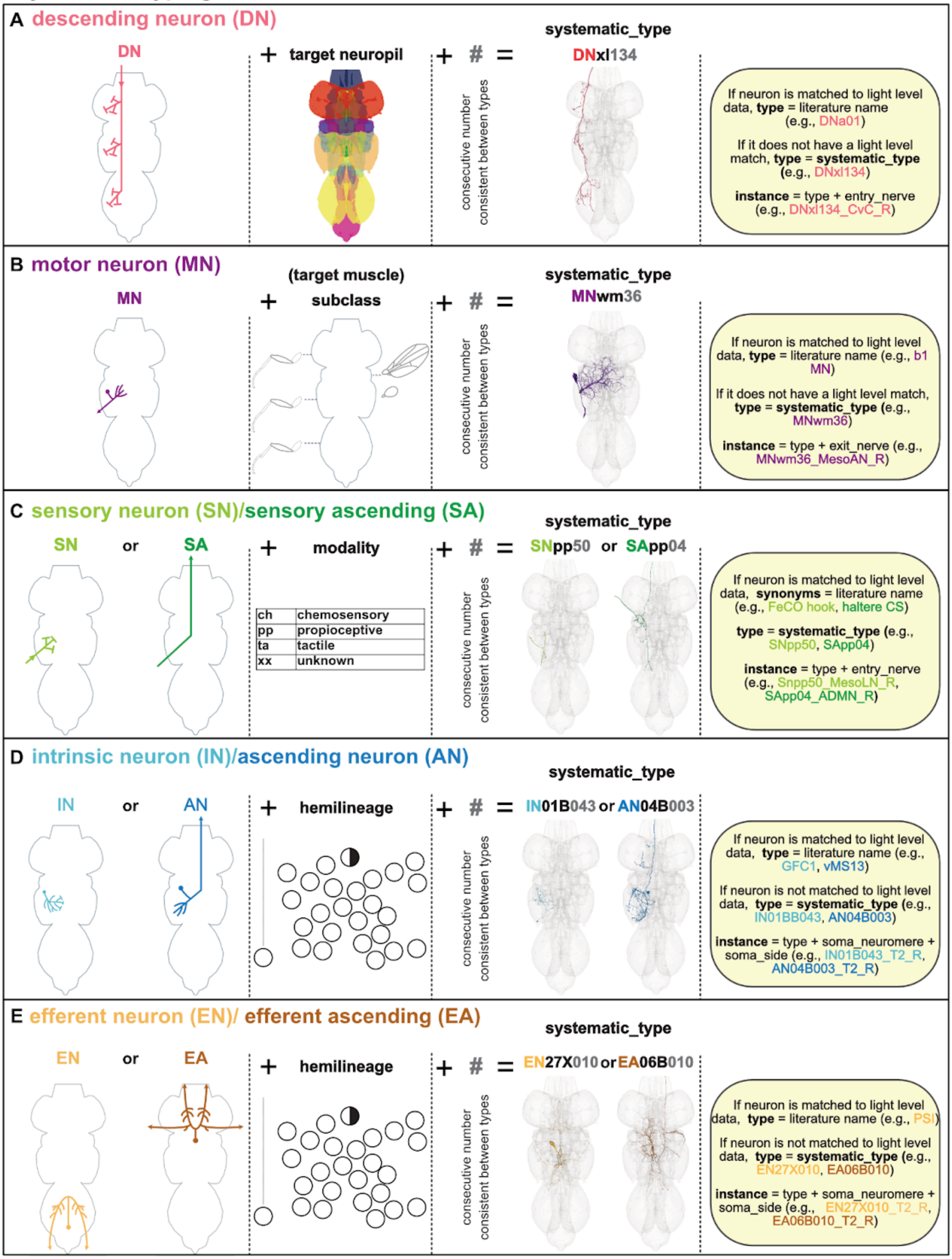

**Figure 3. Systematic cell typing schemes for each broad cell class. A.** Descending neurons are assigned a systematic type composed of the prefix "DN" followed by an abbreviation for the target VNC neuropil and a number consistent for members of their cell type. Their entry nerve (CvC) and root side are used to denote the instance. **B.** Motor neurons are assigned a systematic type composed of the prefix "MN" followed by the subclass (an abbreviation for the target muscle category) and a number consistent for members of their cell type. Their exit nerves are used to denote the instance. **C.** Sensory neurons are assigned a systematic type composed of the prefix "SN" (if not ascending) or "SA" (if ascending) followed by an abbreviation for their assigned modality and a number consistent for members of their connectivity cluster. Their entry nerve is used to denote the instance. **D.** Intrinsic neurons and ascending neurons are assigned a systematic type composed of the prefix "IN" or "AN" followed by their assigned hemilineage and a number consistent for members of their connectivity cluster. Their soma neuromere and side are used to denote the instance. **E.** Efferent neurons are assigned a systematic type composed of the prefix "EA" (if they ascend to the brain) or "EN" otherwise followed by their assigned hemilineage and a number consistent for members of their connectivity cluster. Their soma neuromere and side are used to denote the instance.

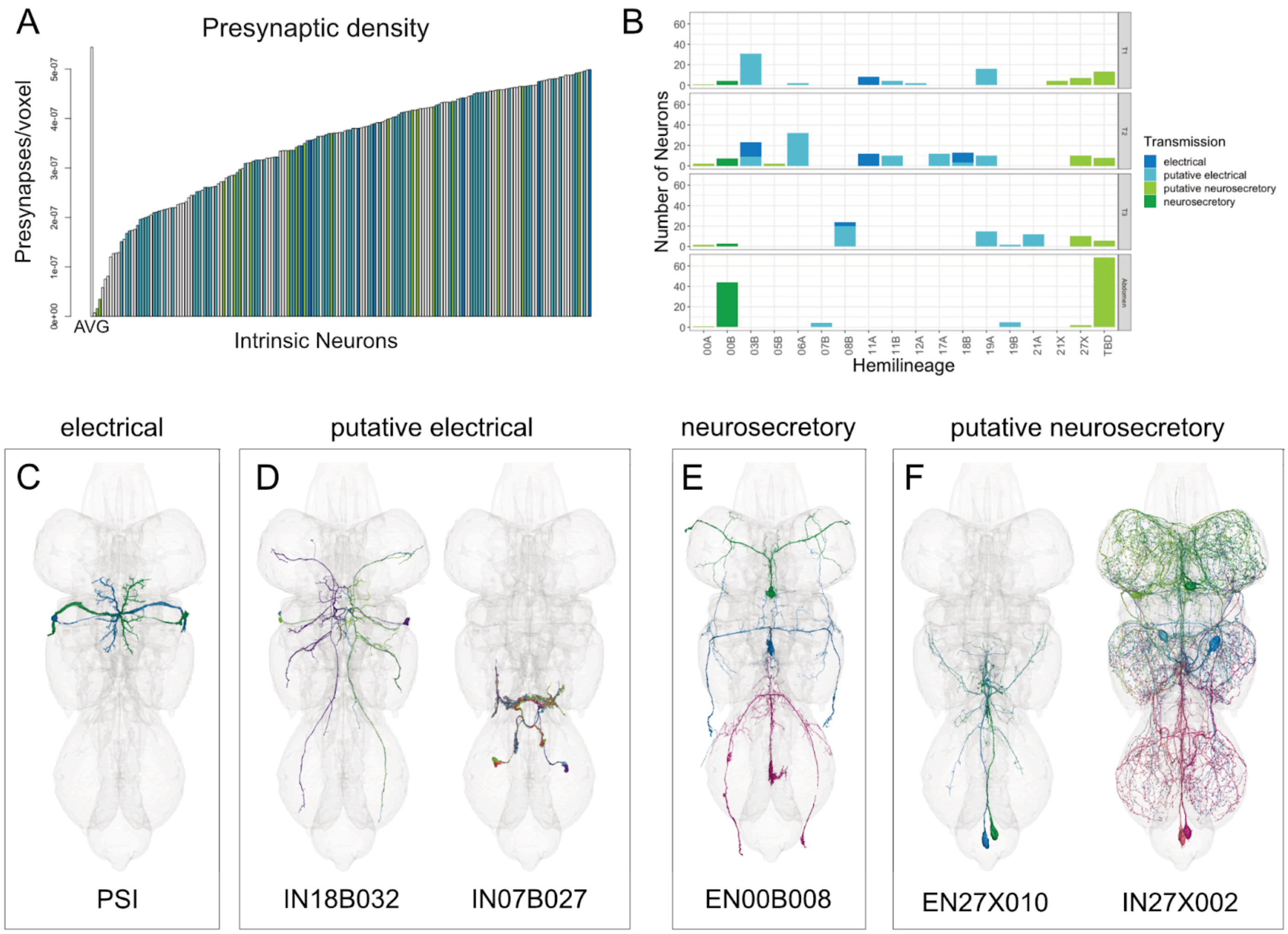

**Figure 4. Neurons with suspected electrical or neurosecretory transmission. A.** The 185 intrinsic neurons with the lowest presynaptic density by volume are enriched for types with electrical or neurosecretory transmission. AVG = average number of presynapses per voxel for all intrinsic neurons. **B.** Candidate electrical and neurosecretory neurons are produced by specific hemilineages. **C.** Example of an electrical neuron, PSI (group 11446). **D.** Examples of putative electrical neurons (groups 20924 and 12943). **E.** Examples of a neurosecretory neuron (serial 11170). **F.** Examples of putative neurosecretory efferent neurons (group 10985) and intrinsic neurons (serial 10083).

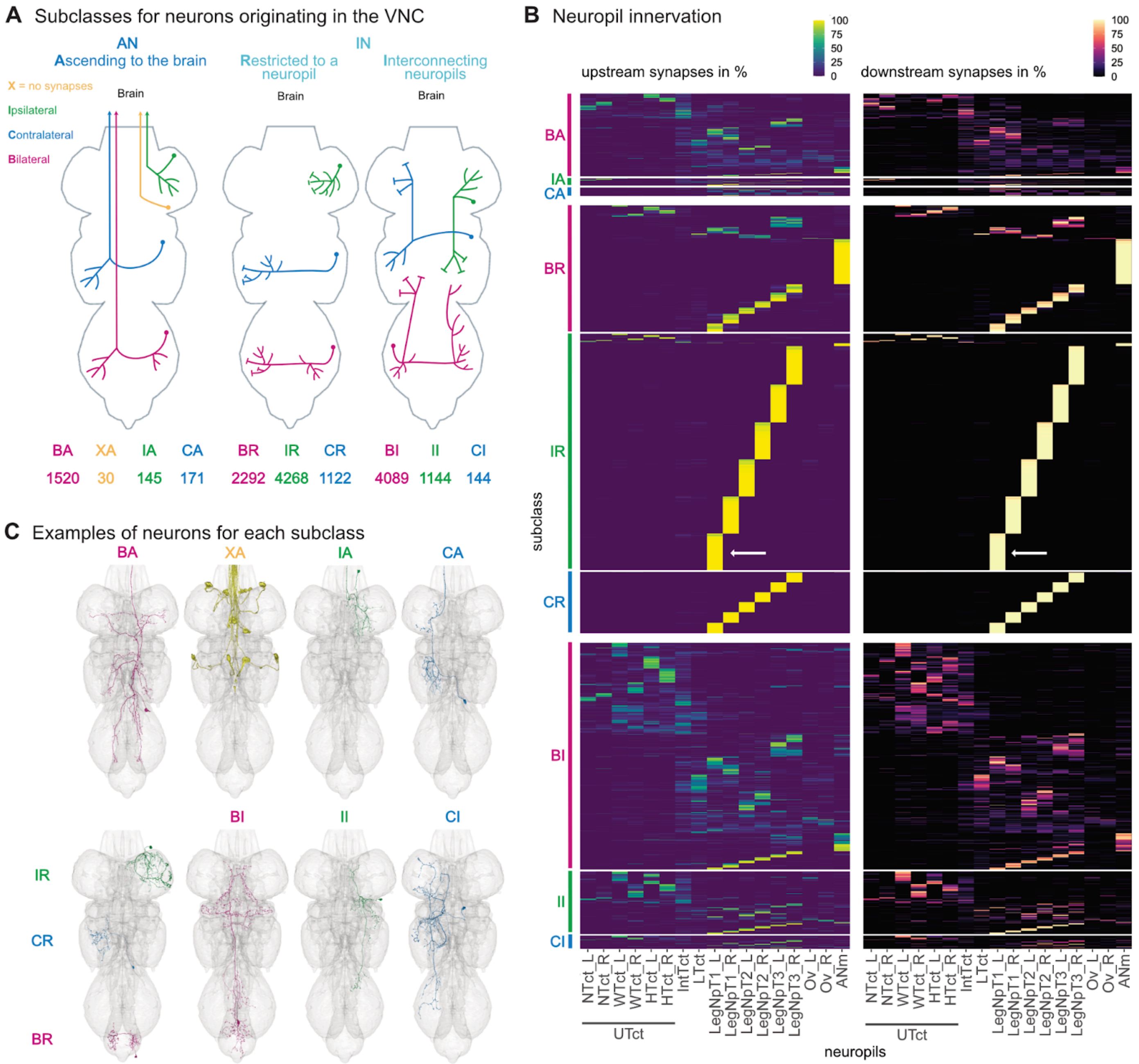

**Figure 5. Subclasses of ANs and INs.** VNC innervation of ascending neurons and intrinsic neurons. **A.** First letter of the subclass reflects if the neuronal arbour is Ipsilateral, Contralateral, Bilateral or not eXisting in the VNC. The second letter indicates if it is Ascending to the brain, Restricted to one neuropil in the VNC or Interconnecting different neuropils in the VNC. **B.** Single neuron inputs and outputs to VNC neuropils separated by subclass. Arrows pointing to a group of IR subclass neurons that receive all their input and give all their output to the front left leg neuropil. **C.** Example images for neurons of different subclasses.

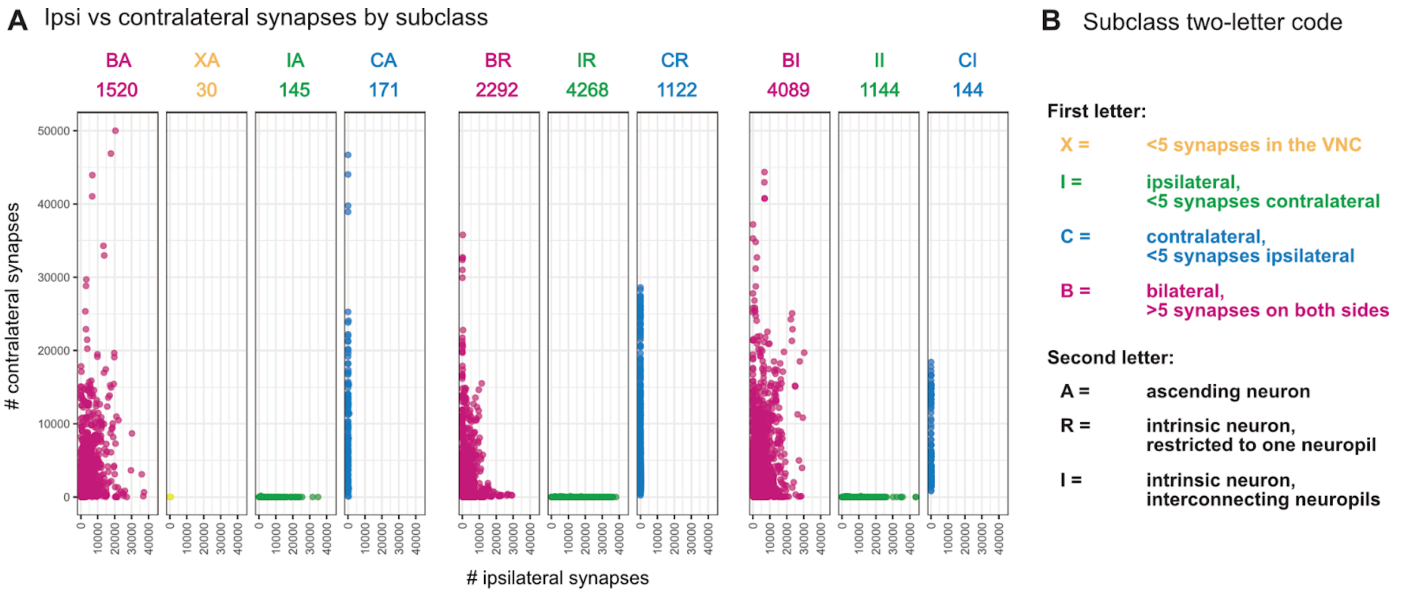

**Figure 5 - figure supplement 1. Subclass defined by ipsi and contralateral innervation of the VNC. A.** Total number of synapses ipsilateral and contralateral for each neuron of a given subclass. Colours correspond to bilateral (magenta), no synapses (yellow), ipsilateral (green), contralateral (blue). **B.** description of the two-letter code of the subclass. The first letter refers to the laterality and the second if a neuron is ascending, restricted to one neuropil or interconnecting VNC neuropils.

The most abundant subclass among the classified neurons is IR, comprising intrinsic neurons that solely innervate a single neuropil ipsilateral to their soma location. The majority of these IR neurons target individual leg neuropils, similarly seen for the subclass CR (see Figure 5B, white arrows pointing to IR neurons that have all their input and output from the front left leg neuropil). The second largest subclass is BI, consisting of intrinsic neurons that bilaterally interconnect multiple neuropils. Upon closer examination of their neuropil innervation, this subclass could be further divided into two groups: those interconnecting the upper tectulum (a combination of NTct, WTct and HTct) and those interconnecting the leg neuropils (LegNpT1-T3) (Figure 5B).

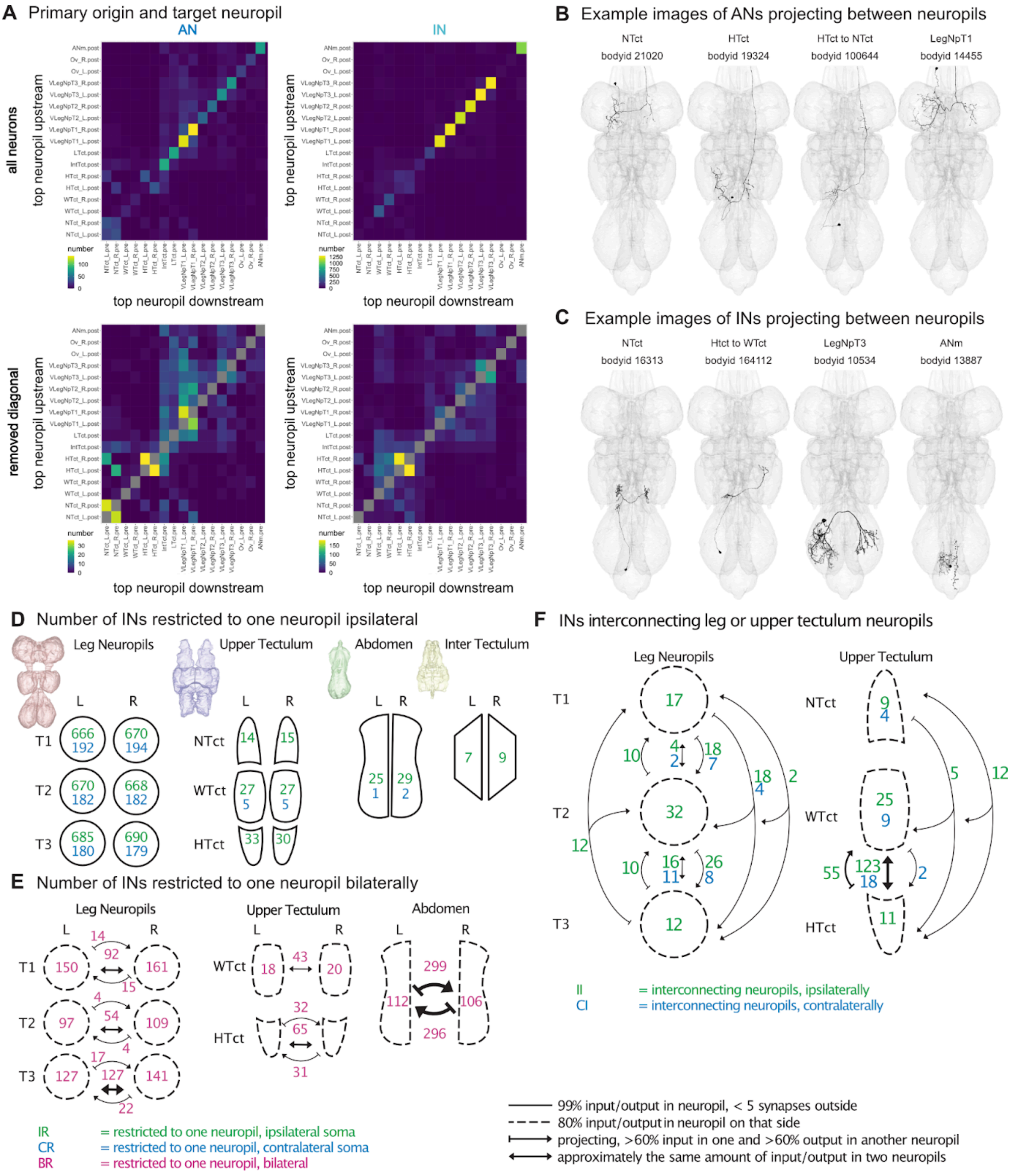

A Primary origin and target neuropil
AN
IN
all neurons
removed diagonal
top neuropil upstream
top neuropil downstream
B Example images of ANs projecting between neuropils
NTct bodyid 21020
HTct bodyid 19324
HTct to NTct bodyid 100644
LegNpT1 bodyid 14455
C Example images of INs projecting between neuropils
NTct bodyid 16313
Htct to WTct bodyid 164112
LegNpT3 bodyid 10534
ANm bodyid 13887
D Number of INs restricted to one neuropil ipsilateral
Leg Neuropils
Upper Tectulum
Abdomen
Inter Tectulum
E Number of INs restricted to one neuropil bilaterally
F INs interconnecting leg or upper tectulum neuropils
IR = restricted to one neuropil, ipsilateral soma
CR = restricted to one neuropil, contralateral soma
BR = restricted to one neuropil, bilateral
II = interconnecting neuropils, ipsilaterally
CI = interconnecting neuropils, contralaterally
99% input/output in neuropil, < 5 synapses outside
80% input/output in neuropil on that side
projecting, >60% input in one and >60% output in another neuropil
approximately the same amount of input/output in two neuropils

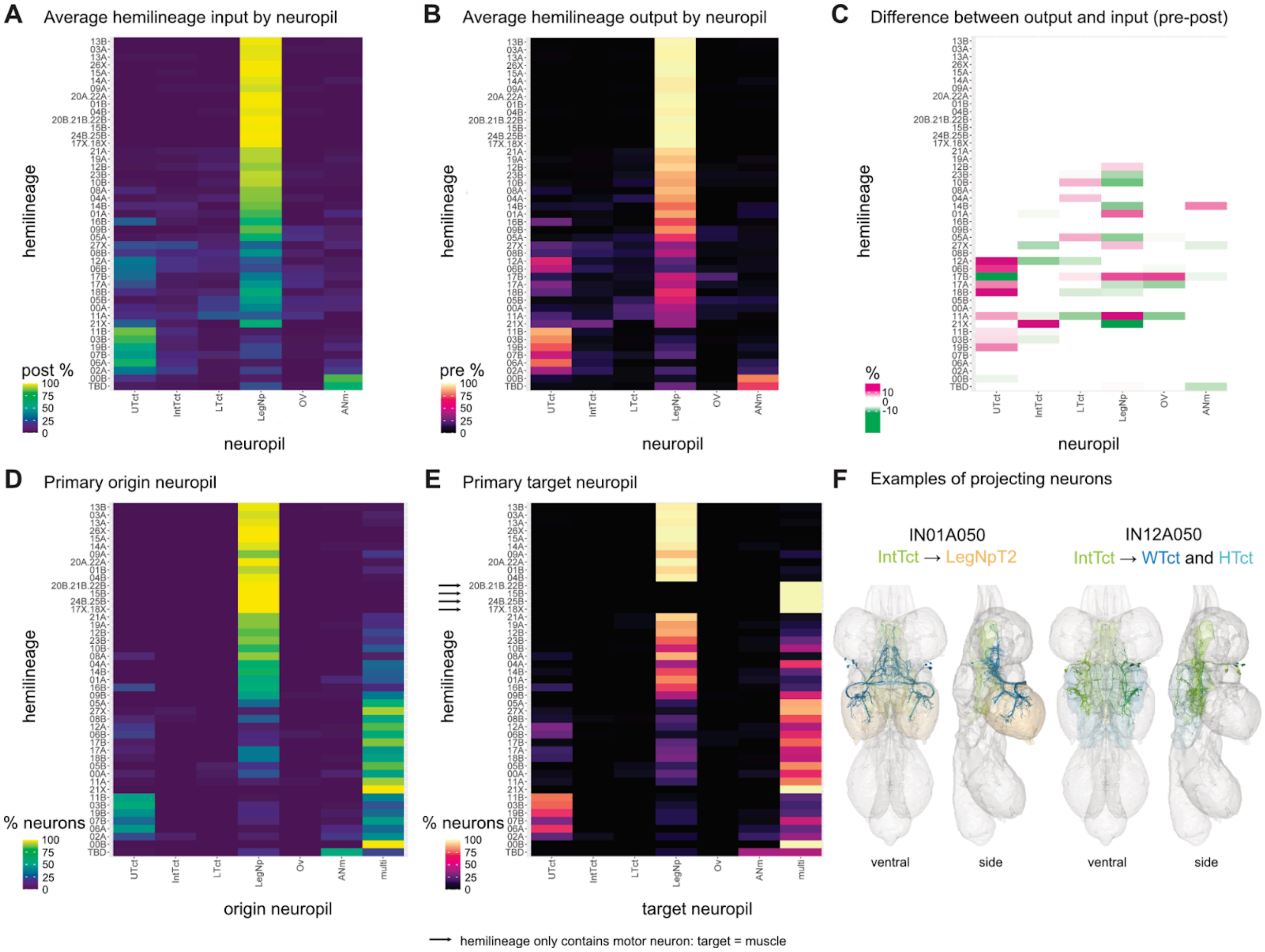

**Figure 7. Neuropil innervation by hemilineage. A.** The average proportion of input to each hemilineage from VNC neuropils as a percentage. **B.** The average proportion of output from each hemilineage to VNC neuropils as a percentage. **C.** The average input to neuropils subtracted from the average output to that neuropil. Magenta (positive numbers) represent more output and green (negative numbers) represent more input to that neuropil by a given hemilineage. For neuropils that do not receive any input or output see Figure 7 - figure supplement 1. **D.** INs and ANs by their primary input neuropil (origin) in the VNC. If there is more than one origin neuropil it is included in multi. **E.** INs and ANs by their primary output neuropil (target) in the VNC. If there is more than one target neuropil or a muscle target outside of the VNC it is included in multi. **F.** Examples of types that project between neuropils, partially responsible for the differences seen in **C** (origin and target neuropils are coloured). Hemilineages order on the y axis is organised by the innervation heatmap that is shown in **A** and reused for the heatmaps in **B,C,D** and **E.**

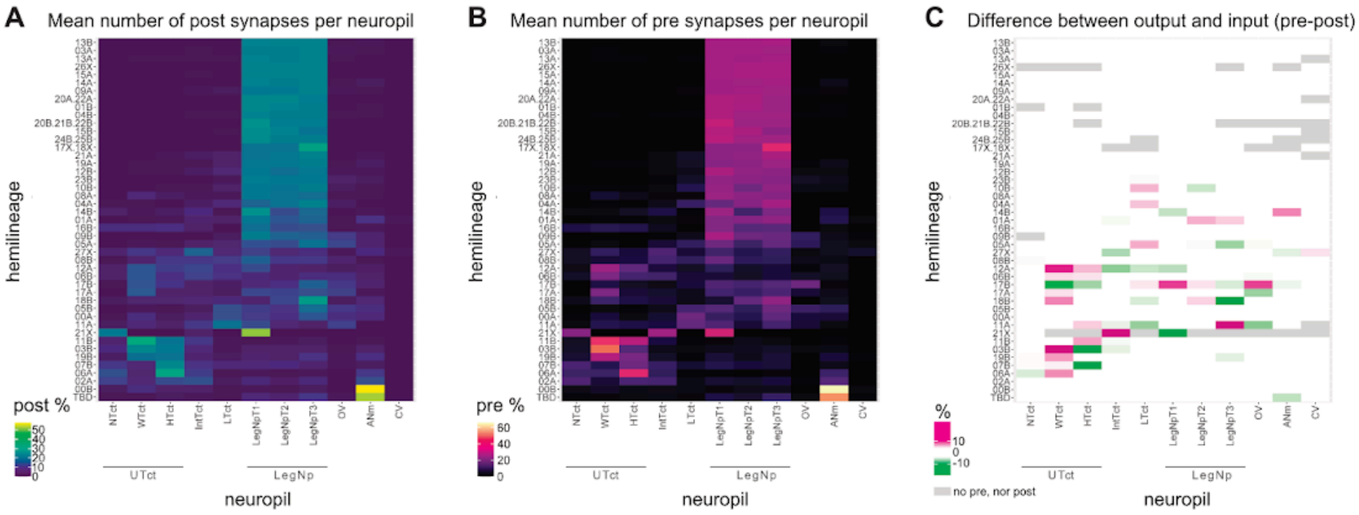

**Figure 7 - figure supplement 1. Neuropil innervation by hemilineage with differences in the leg and upper tectulum neuropils.** **A.** The average proportion of input to each hemilineage from VNC neuropils as a percentage. **B.** The average proportion of output from each hemilineage to VNC neuropils as a percentage. **C.** The average input to neuropils subtracted from the average output to that neuropil. Magenta (positive numbers) represent more output and green (negative numbers) represent more input to that neuropil by a given hemilineage. Neuropils that do not receive any input or output from a given hemilineage are coloured in light grey.

#### Introduction to hemilineage-based organisation

During embryonic development, divisions of a segmentally repeated array of neuroblasts produce similar numbers of primary neurons in most neuromeres of the *Drosophila* VNC (Poulson, 1950). However, during postembryonic development, segment-specific subsets of neuroblasts generate the secondary neurons (Truman and Bate, 1988), facilitating an expansion of neuronal types in the thorax and distal abdomen for complex sensorimotor functions. In late larva, secondary hemilineages restricted to the leg neuropils are present in all three thoracic neuromeres, while those associated with dorsal (flight) neuropils are typically absent or greatly reduced in at least one (Marin et al., 2012; Truman et al., 2004).

Because **hemilineages** represent developmental and likely functional units in the VNC, we wanted them to be central to any systematic typing scheme. However, assignment of individual neurons to specific hemilineages based on light-level images of postembryonic neuroblast clones (Shepherd et al., 2019) was a complex and multifaceted process. Whilst the relative position of midline crossing is useful for distinguishing hemilineages that project contralaterally (Shepherd et al., 2016), many hemilineages are confined to ipsilateral neuropils. The locations of primary neurite bundles are also key characters (Figure 8 - figure supplement 1), but typically these remain tightly bundled during development only if all neurons target a similar location; in hemilineages with diverse cell types, they can be pulled apart.

#### Hemilineage Overview

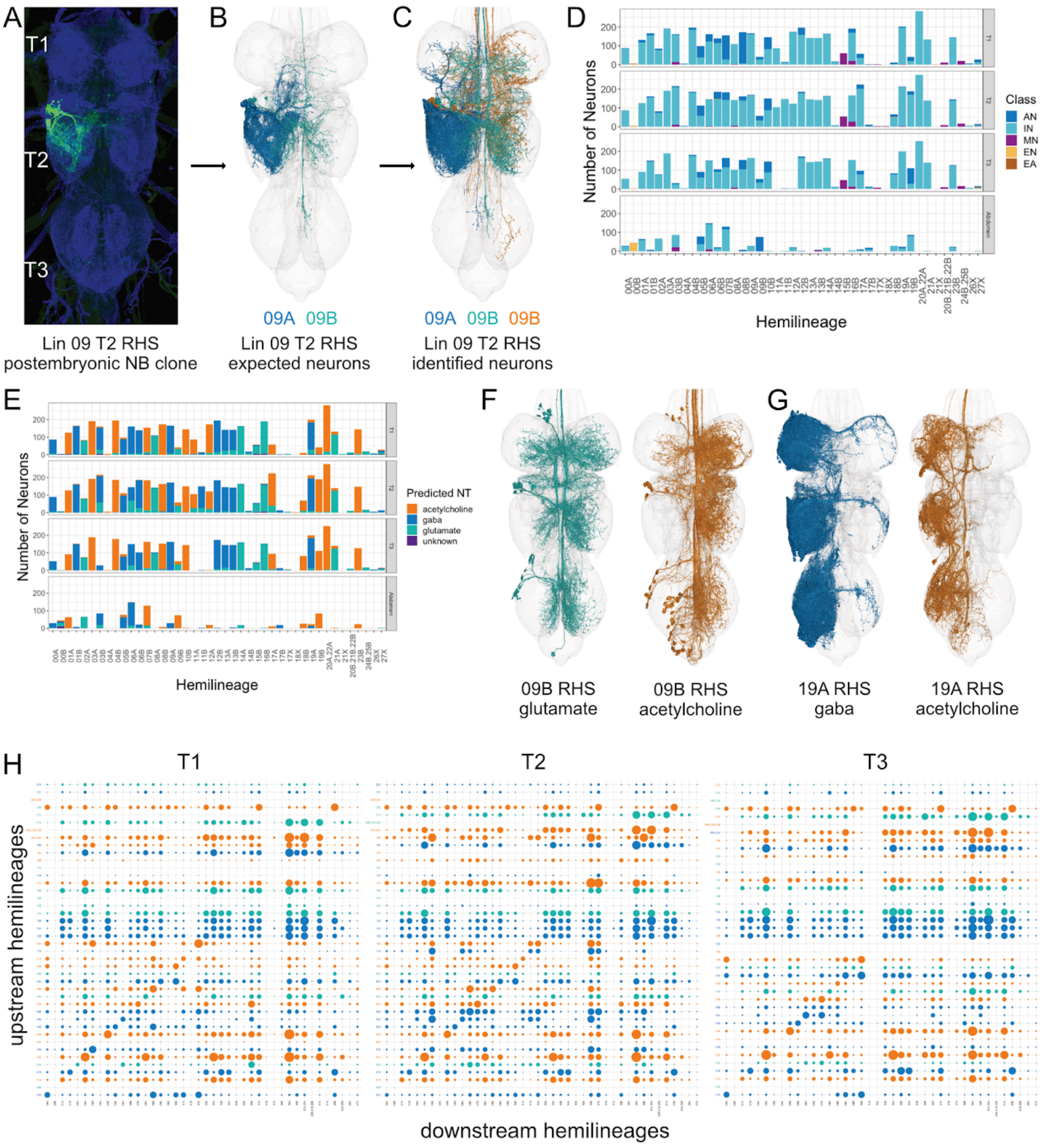

**Figure 8. Hemilineage overview. A.** 09A and 09B secondary neurons in a postembryonic neuroblast clone generated in T2 RHS. **B.** 09A and 09B secondary neurons in T2 RHS identified in MANC from light-level images, coloured by predicted neurotransmitter. **C.** All 09A and 09B neurons in T2 RHS identified in MANC, coloured by predicted neurotransmitter. **E.** Predicted fast-acting neurotransmitters for all VNC neurons (Eckstein et al., 2023; Takemura et al., 2023) by hemilineage and neuromere (neurons of TBD/unknown hemilineage have been omitted, and abdominal neuromeres have been pooled). **F.** Glutamatergic hemilineage 09B includes a minority cholinergic population. **G.** Gabaergic hemilineage 19A includes a minority cholinergic population. **H.** Intra-neuromere, inter-hemilineage synaptic connectivity for the three thoracic neuromeres (T1, T2, T3). Dot size correlates with number of synaptic connections, and dot colour reflects predicted neurotransmitter of upstream hemilineage (Eckstein et al., 2023; Takemura et al., 2023).

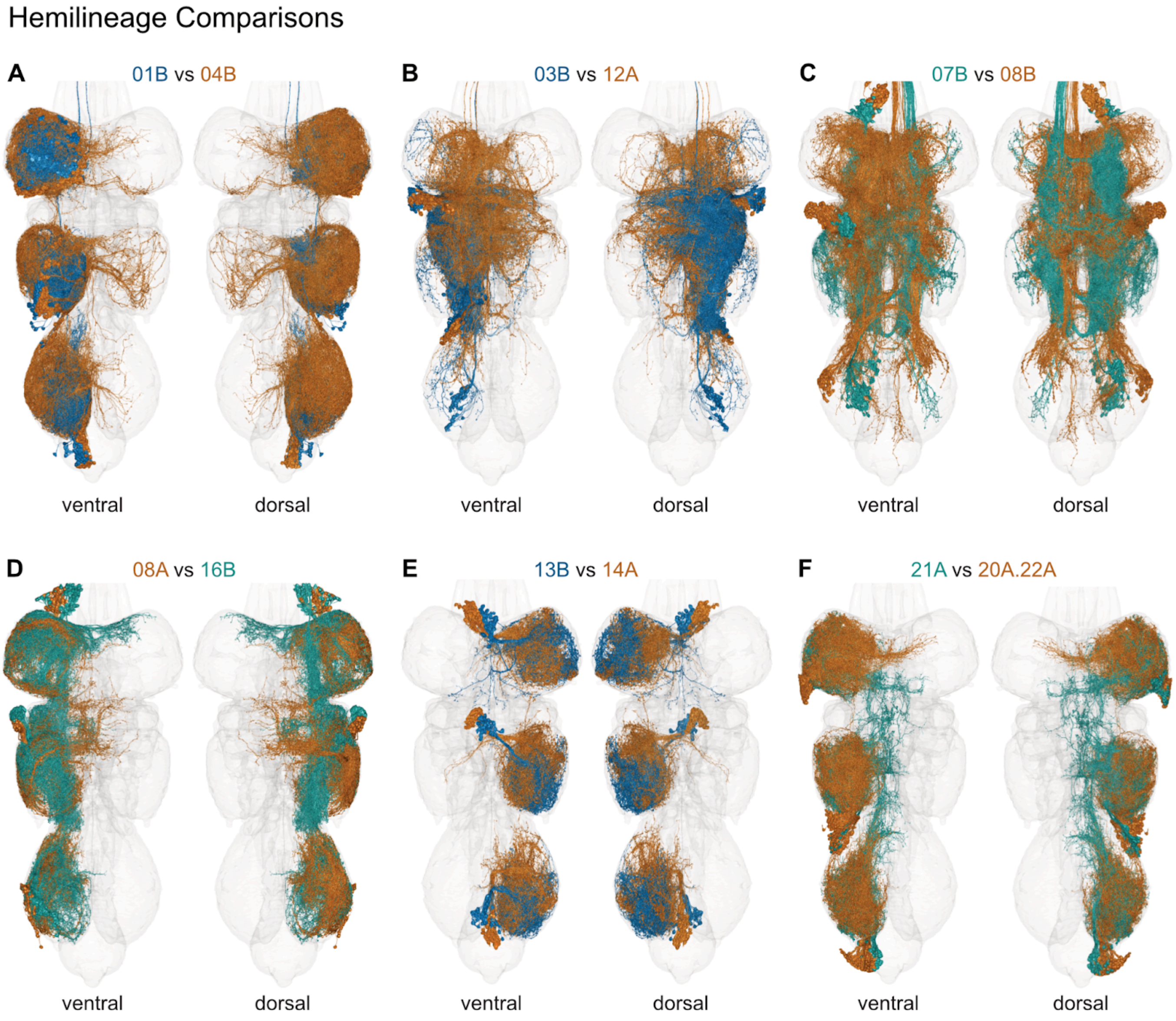

**Figure 8 - figure supplement 1. Pairwise comparisons of selected secondary hemilineages. A.** 01B (blue, gabaergic) vs 04B (dark orange, cholinergic). **B.** 03B (blue, gabaergic) vs 12A (dark orange, cholinergic). **C.** 07B (marine, cholinergic) vs 08B (dark orange, cholinergic). **D.** 08A (dark orange, glutamatergic) vs 16B (marine, glutamatergic). **E.** 13B (blue, gabaergic) vs 14A (dark orange, glutamatergic). **F.** 20A.22A (dark orange, cholinergic) vs 21A (marine, glutamatergic).

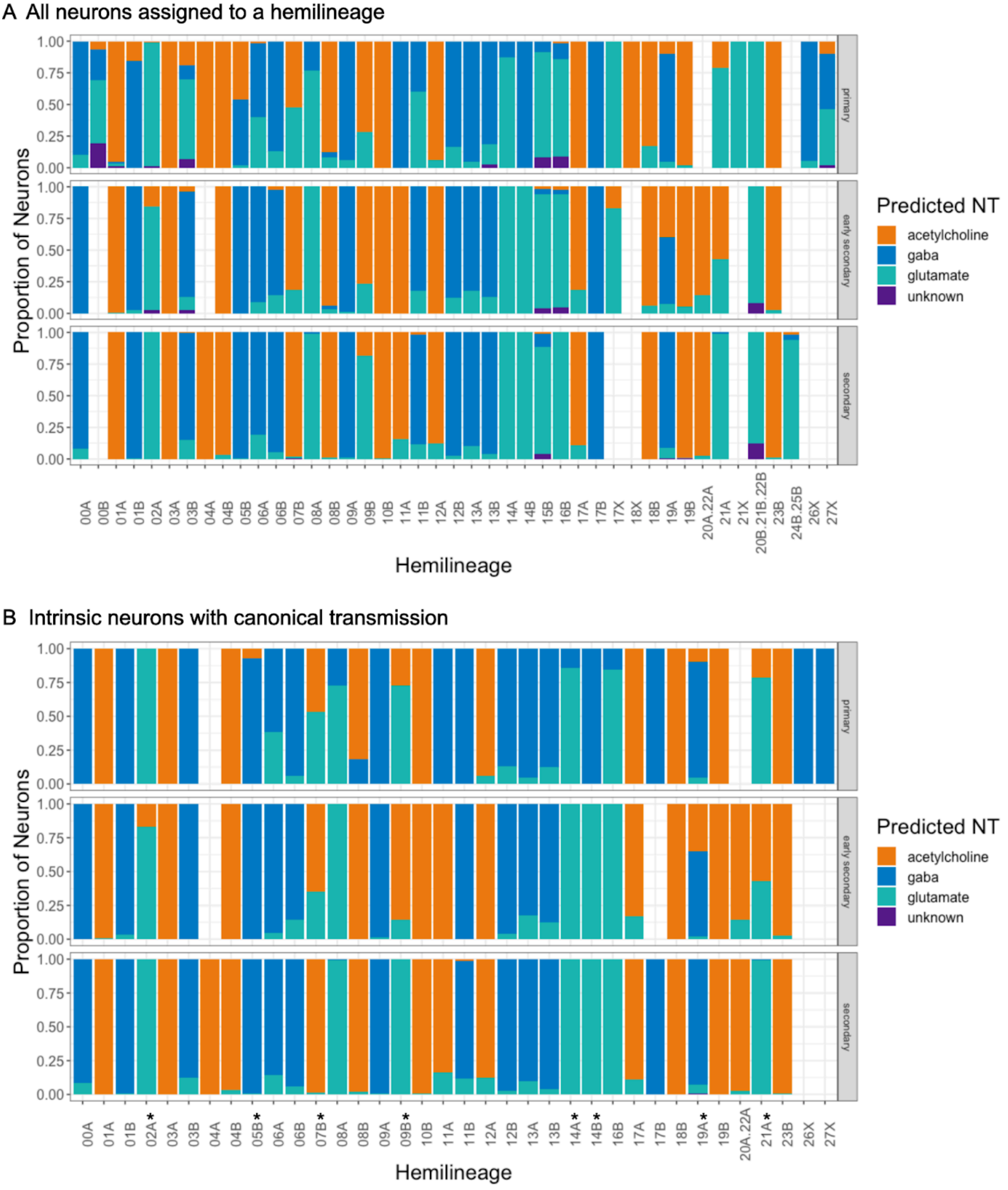

**Figure 8 - figure supplement 2. Early born neurons are more likely to express minority neurotransmitters. A.** Proportion of fast-acting neurotransmitter expression predicted (Eckstein et al., 2023; Takemura et al., 2023) for all neurons assigned to a hemilineage, faceted by birthtime. **B.** Proportion of fast-acting neurotransmitter expression predicted for intrinsic neurons assigned to a hemilineage (with putative electrical and neurosecretory transmission excluded), faceted by birthtime. Asterisks indicate neurons with consistent expression of a minority neurotransmitter, enriched in early born neurons.

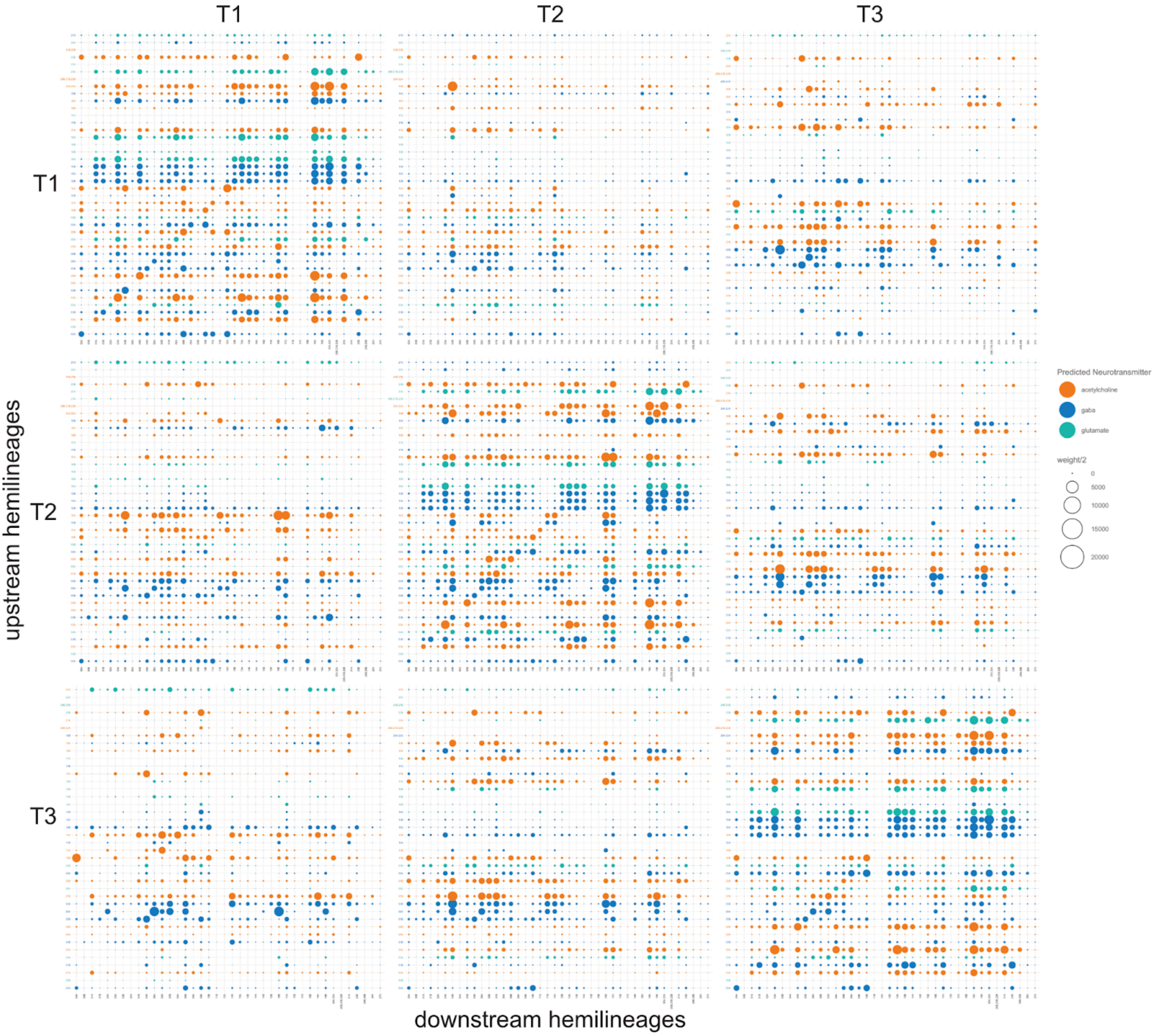

**Figure 8 - figure supplement 3. Intra-neuromere hemilineage synaptic connectivity across thoracic neuromeres (T1, T2, T3).** Dot size correlates with number of synaptic connections, and dot colour reflects predicted neurotransmitter of upstream hemilineage (Eckstein et al., 2023; Takemura et al., 2023).

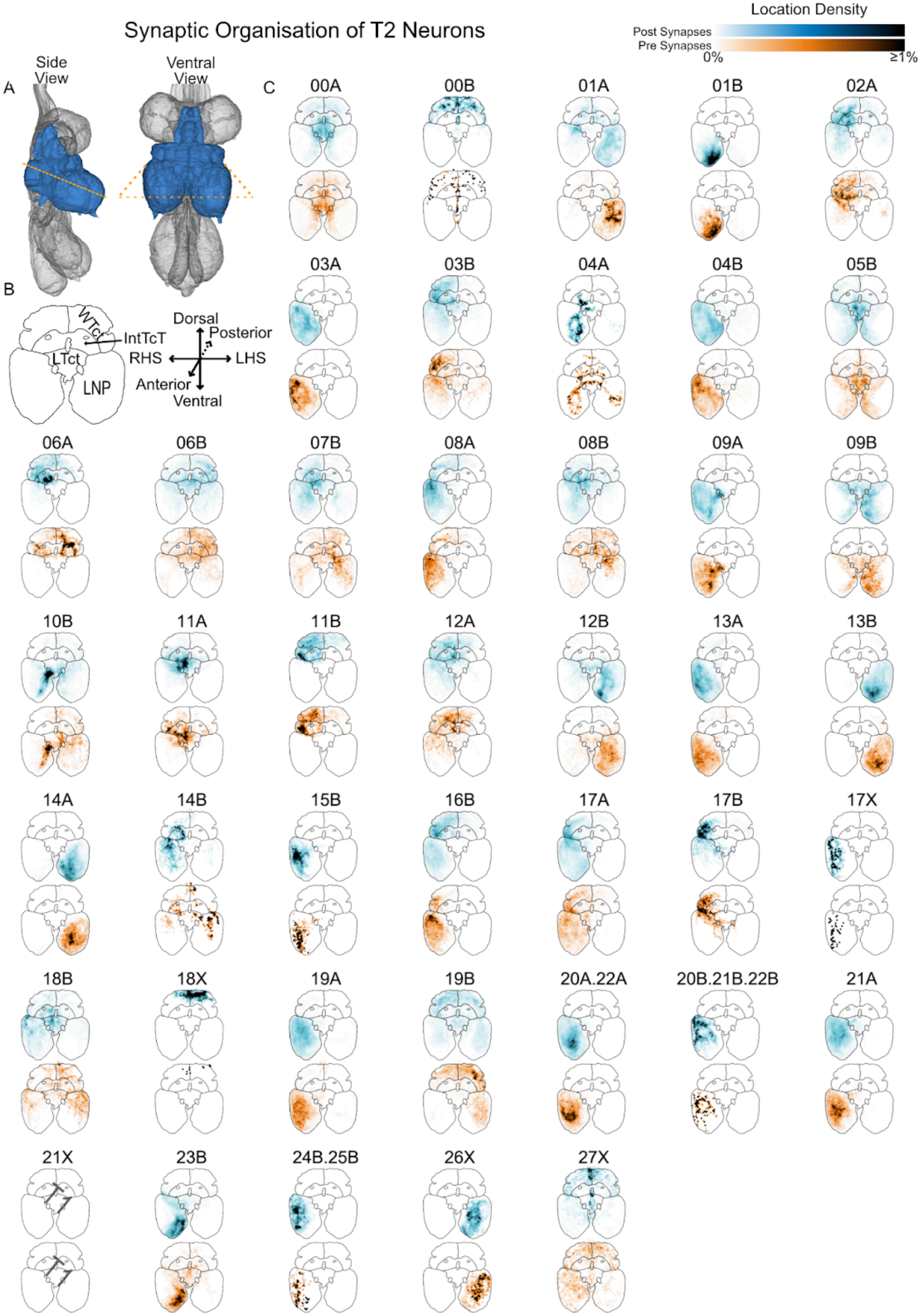
Synaptic Organisation of T2 Neurons
Location Density
Post Synapses
Pre Synapses
0%
≥1%
A
Side View
Ventral View
B
WTct
IntTcT
LTct
LNP
Dorsal
Posterior
RHS
LHS
Anterior
Ventral
C
00A
00B
01A
01B
02A
03A
03B
04A
04B
05B
06A
06B
07B
08A
08B
09A
09B
10B
11A
11B
12A
12B
13A
13B
14A
14B
15B
16B
17A
17B
17X
18B
18X
19A
19B
20A.22A
20B.21B.22B
21A
21X
23B
24B.25B
26X
27X
T1

**Figure 9. Summary of the anatomical organisation of synapses for each hemilineage originating in T2 RHS. A.** Side and ventral views illustrating the neuropils surveyed: T2 Leg Neuropil (LNP), medial ventral association center (mVAC), ovoid (Ov), wing tectulum (WTct), lower tectulum (LTct), and Intermediate tectulum (IntTct). The orange dashed line shows the approximate location of the transverse section onto which their synapses have been projected. **B.** Template transverse section with major neuropils labelled and axes for orientation. **C.** Projected synapses for T2 RHS neurons of each hemilineage. Top (blue): postsynapses, connections to upstream neurons. Bottom (orange): presynapses, connections to downstream neurons. 21X is only observed in T1.

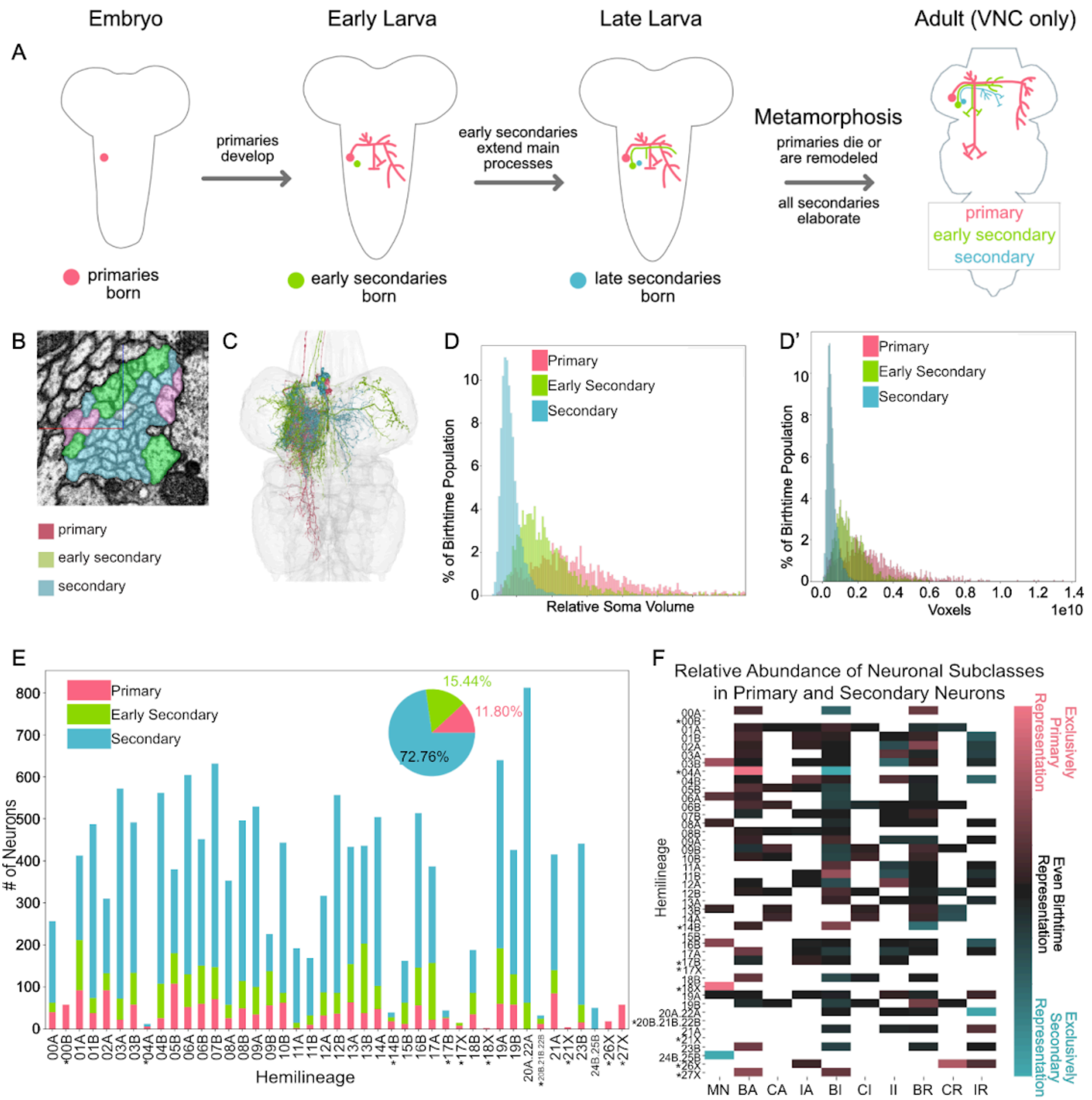

**Figure 10. Birth order during development. A.** Cartoon of birthtime development. Primary neurons are born during embryogenesis while secondaries are born during larval life. At metamorphosis primary neurons die or remodel while secondary neurons continue to elaborate. **B.** 02A T1 RHS primary neurites, coloured by birthtime. **C.** 02A T1 RHS neuron meshes, coloured by birthtime. **D-F.** Data is shown only for neurons originating in thoracic neuromeres; * indicates primary-only hemilineages. **D.** Neurons with larger somas were assigned to earlier birthtimes. **D'.** Primary and early secondary neurons are larger than secondary neurons. **E.** Counts of primary, early secondary, and secondary annotations for all VNC origin neurons with annotated hemilineage. Inset is frequency of birthtime amongst all annotated neurons. **F.** Comparative representation of each subclass between primary and secondary neurons within each hemilineage. Pink values have a higher percentage representation in the primary population than secondary, and vice-versa for blue, while black represents equal percentage abundance. White cells have no neurons with that subclass in that hemilineage. For more details, see **Methods**.

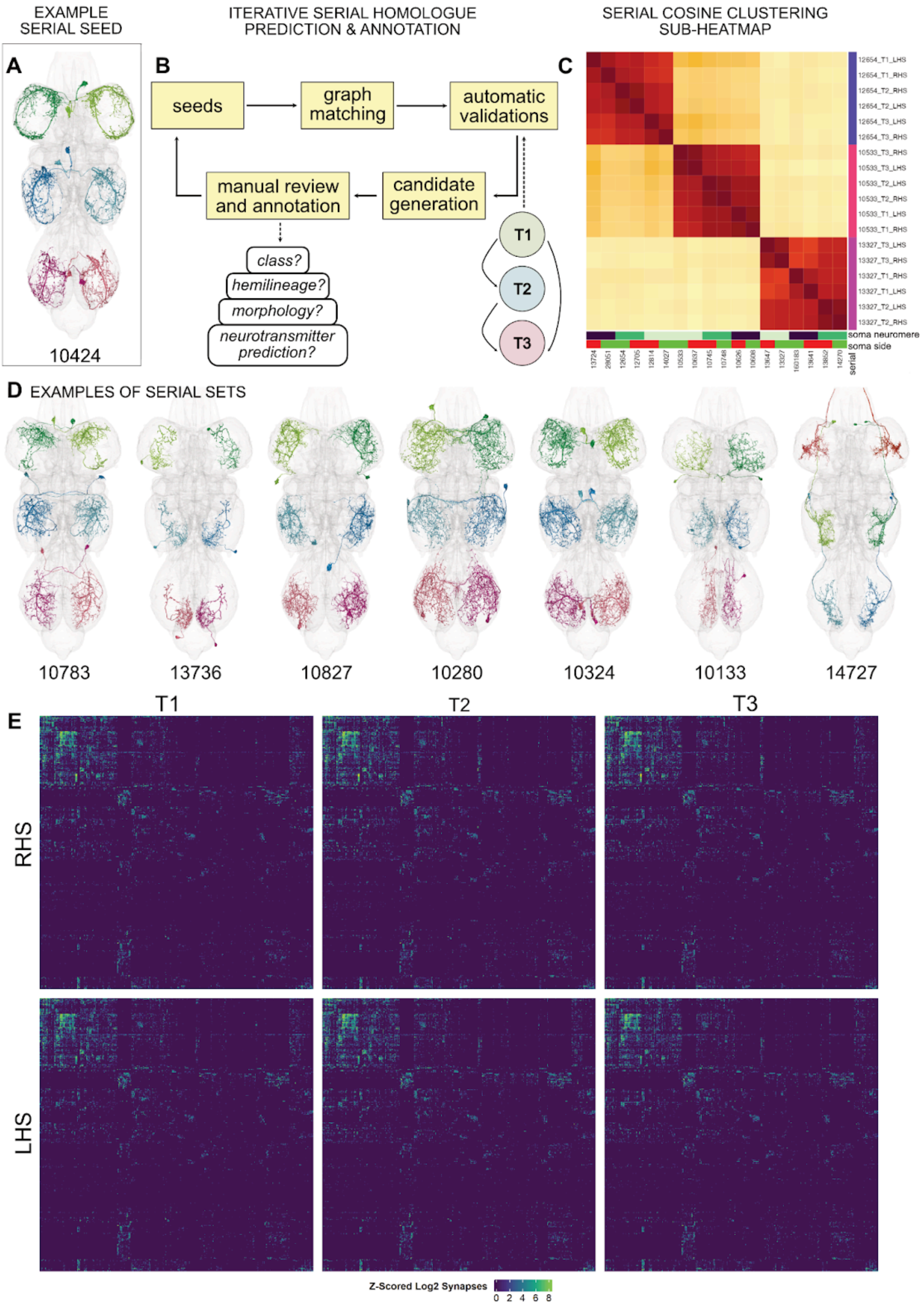

EXAMPLE SERIAL SEED
A
10424
ITERATIVE SERIAL HOMOLOGUE PREDICTION & ANNOTATION
B
seeds
graph matching
automatic validations
candidate generation
manual review and annotation
class?
hemilineage?
morphology?
neurotransmitter prediction?
T1
T2
T3
SERIAL COSINE CLUSTERING SUB-HEATMAP
C
12654_T1_LHS
12654_T1_RHS
12654_T2_RHS
12654_T2_LHS
12654_T3_LHS
12654_T3_RHS
10533_T3_RHS
10533_T3_LHS
10533_T2_LHS
10533_T2_RHS
10533_T1_LHS
10533_T1_RHS
13327_T3_LHS
13327_T3_RHS
13327_T1_RHS
13327_T1_LHS
13327_T2_LHS
13327_T2_RHS
soma neuromere
soma side
serial
13724
28051
12654
12705
12814
14027
10533
10637
10745
10748
10626
10608
13647
13327
160183
13641
13852
14270
D EXAMPLES OF SERIAL SETS
10783
13736
10827
10280
10324
10133
14727
E
T1
T2
T3
RHS
LHS
Z-Scored Log2 Synapses
0 2 4 6 8

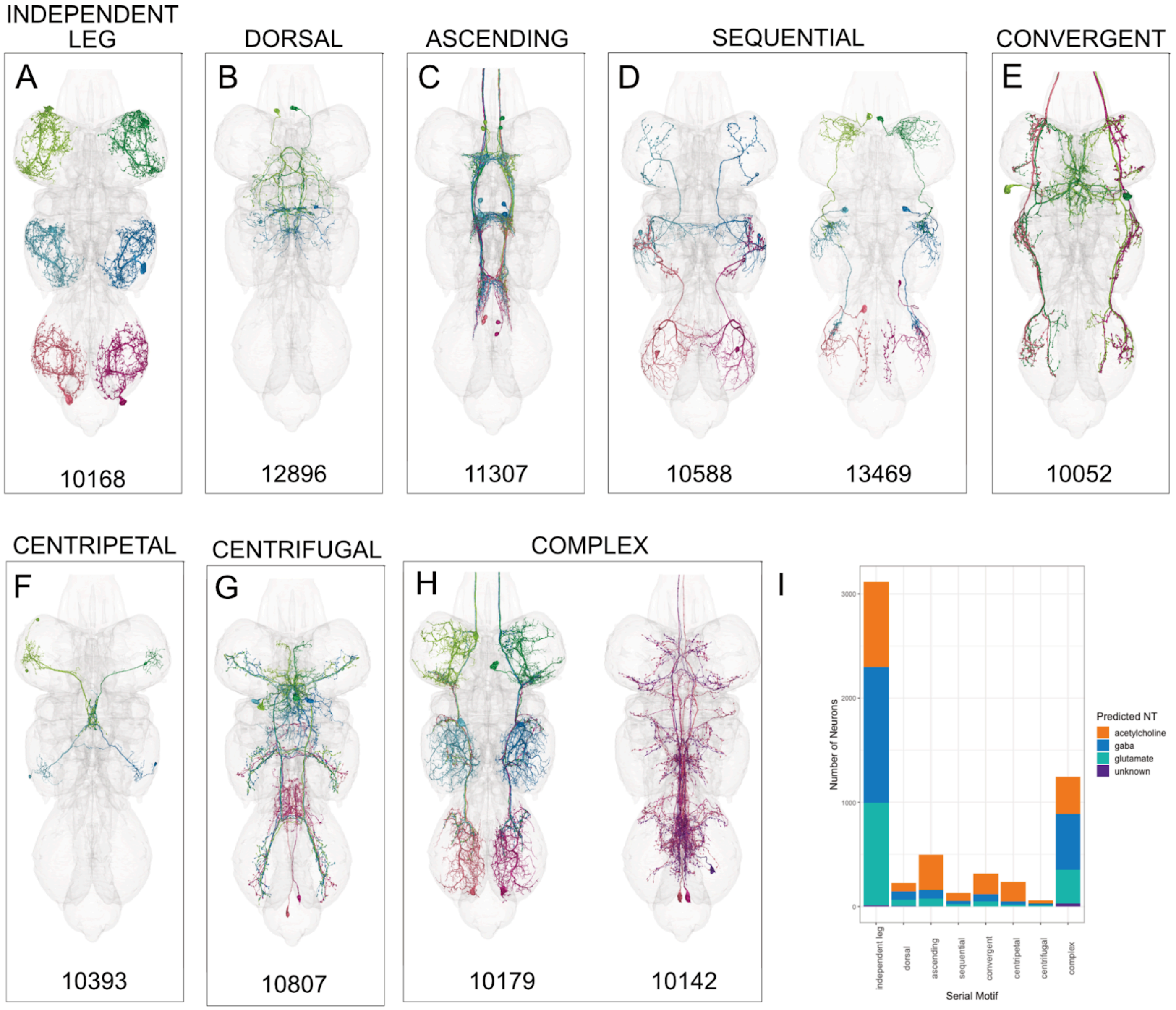

**Figure 12. Classification of serial sets into motifs.** Neuron meshes have been coloured by soma neuromere and side RHS/LHS (DN = red/dark red, T1 = green/dark green, T2 = cyan/blue, T3 = pink/magenta). **A.** Independent leg serial set 10168. **B.** Dorsal serial set 12896. **C.** Ascending serial set 11307. **D.** Sequential (ascending or descending to the next neuromere) serial sets 10588 and 13469. **E.** Convergent serial set 10052. **F.** Centripetal serial set 10393. **G.** Centrifugal serial set 10807. **H.** Complex serial set 10142. **I.** Number of neurons assigned to each serial motif, coloured by predicted neurotransmitter (Eckstein et al., 2023; Takemura et al., 2023).

.

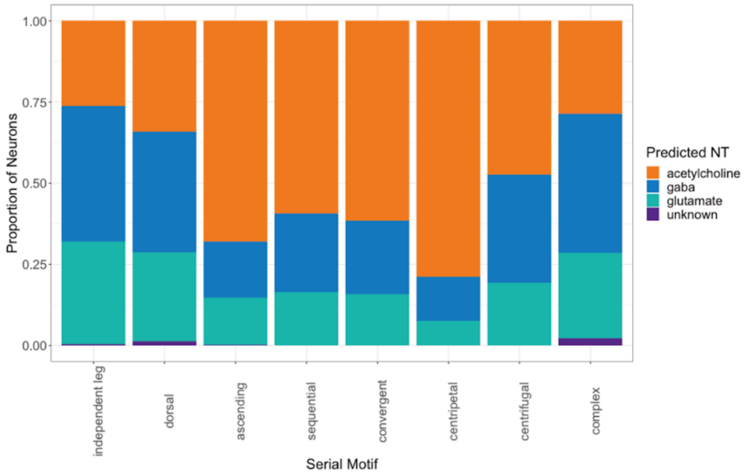

**Figure 12 - figure supplement 1. Proportions of neurons from each serial motif predicted to express a given neurotransmitter** (Eckstein et al., 2023; Takemura et al., 2023).

**Figure 12 - video 1. All thoracic examples of independent leg serial sets (1:19).** https://youtu.be/O6QfQTpNjo8 Neuron meshes have been coloured by soma neuromere and side RHS/LHS (DN = red/dark red, T1 = green/dark green, T2 = cyan/blue, T3 = pink/magenta). To view this movie at a preferred speed, we recommend downloading the video and using a player with more control such as iMovie.

**Figure 12 - video 2. All thoracic examples of other annotated serial motifs (0:42).** https://youtu.be/VFnhqX9Ee_g Neuron meshes have been coloured by soma neuromere and side RHS/LHS (DN = red/dark red, T1 = green/dark green, T2 = cyan/blue, T3 = pink/magenta). To view this movie at a preferred speed, we recommend downloading the video and using a player with more control such as iMovie.

Whilst these serial sets were interesting as examples of segment-specific reuse and diversification of repeated elements, they also proved to be extremely useful in the systematic typing of the VNC, including interneurons, sensory neurons (see below), and leg motor neurons (as described in our accompanying manuscript (Cheong et al., 2023)). They also informed the annotation of birthtimes and hemilineages; in particular, sets bridging the thoracic and abdominal neuromeres aided in abdominal soma neuromere and hemilineage assignments (e.g., serial 10142, Figure 12H). We also expect them to aid in assigning hemilineages to neurons in the suboesophageal ganglion once a full CNS dataset has been obtained.

#### Cell typing

We partitioned the population of intrinsic, ascending, and efferent neurons into systematic types on the basis of their synaptic connectivity, using the identified lateral and serial homologues. The aim of this typing is fourfold: first, to identify granular sets of neurons that are likely to be mutually identifiable between multiple VNCs; second, to provide units for exploratory and functional analysis, which may be broader than identified lateral and serial homologues; third, to identify seriality or other symmetries that might not be captured by the previously described approaches; and fourth, to assign a systematic nomenclature consistent with those developed for the sensory, descending, and motor neuron populations.

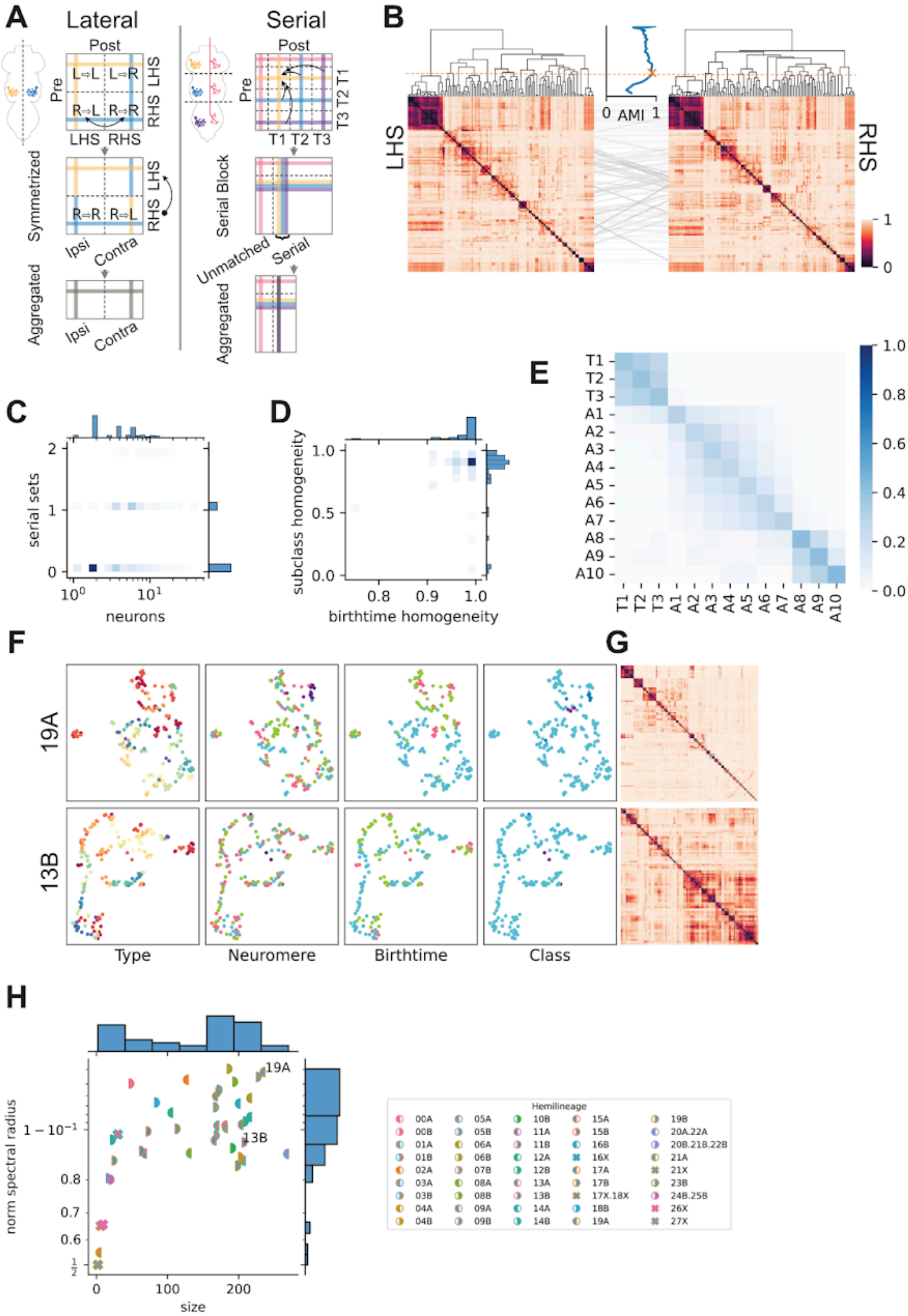
A
Lateral
Post
Pre
L⇨L
L⇨R
R⇨L
R⇨R
LHS
RHS
Symmetrized
Ipsi
Contra
Aggregated
Serial
T1 T2 T3
Serial Block
Unmatched
B
AMI
LHS
RHS
C
serial sets
neurons
D
subclass homogeneity
birthtime homogeneity
E
T1
T2
T3
A1
A2
A3
A4
A5
A6
A7
A8
A9
A10
F
19A
13B
Type
Neuromere
Birthtime
Class
G
H
norm spectral radius
size
Hemilineage
00A
00B
01A
01B
02A
03A
03B
04A
04B
05A
05B
06A
06B
07B
08A
08B
09A
09B
10B
11A
11B
12A
12B
13A
13B
14A
14B
15A
15B
16B
16X
17A
17B
17X.18X
18B
19A
19B
20A.22A
20B.21B.22B
21A
21X
23B
24B.25B
26X
27X

**Figure 13. Typing of intrinsic neurons by connectivity. A.** Schematic describing how the raw adjacency matrix of connectivity between individual neurons can be simplified by aggregating connectivity of homologous neurons; this is possible both for left-right homologues and for serial homologues (diagrammed here for 6 thoracic neurons). **B.** Example of consistency of independent connectivity clustering of each side of a hemilineage, 09B. The adjusted mutual information (AMI) between sides' increasingly granular clusters is shown between the corresponding dendrograms for each side. Peak consistency chosen to select clusters for systematic types is indicated by a yellow cross and dashed line. Heatmaps display cosine distance between the serially aggregated connectivity for each neuron set. Lines between the heatmaps indicate neurons whose terminal position in each side's cluster dendrogram differs, shaded by the distance of this difference. **C.** Density histogram of the number of neurons and serial sets per clustered type in all hemilineages' intrinsic, non-motor popular. See E for colour scale. **D.** Density histogram of the birthtime and subclass homogeneity of all types in C. See E for colour scale. **E.** Type seriality, as a histogram of number of types containing any neuron in each pair of neuromeres. Values are normalised by rows, to show relative distributions when population size differs. **F.** Example comparison of the inter-hemilineage variability of intra-hemilineage connectivity similarity structure. Each panel is a UMAP embedding the hemilineage's symmetrized, serially aggregated connectivity cosine similarity, with each point representing a lateral group. Panels are coloured by neuromere, birthtime, and class as in all other figures. Types are coloured arbitrarily for visualisation. **G.** Heatmaps of symmetrized, serially aggregated connectivity cosine distance for the hemilineages in F. **H.** Normalised spectral radius versus number of neurons for all hemilineages. Hemilineages from F and G are labelled.

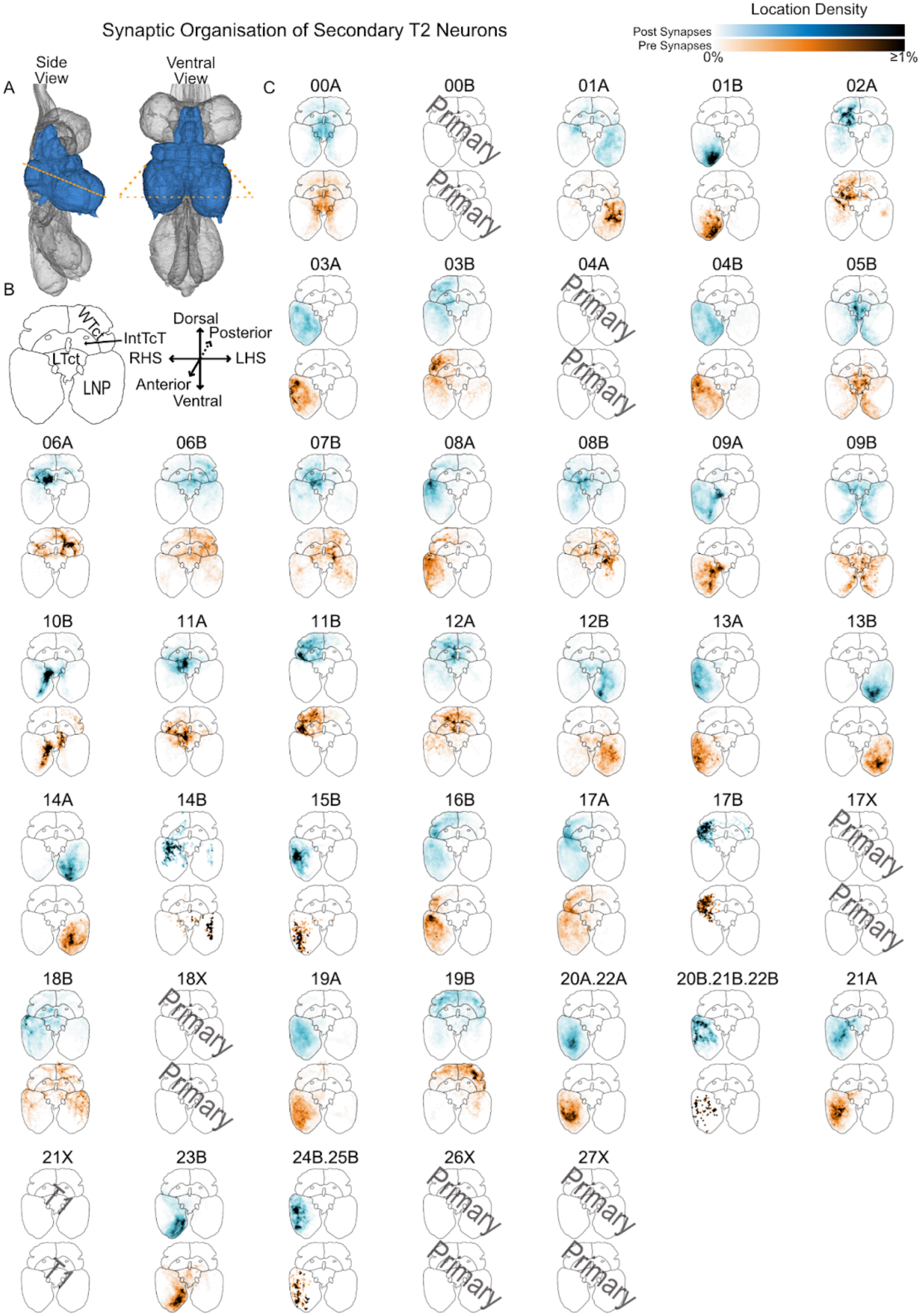

Synaptic Organisation of Secondary T2 Neurons
Location Density
Post Synapses
Pre Synapses
0%
≥1%
A
Side View
Ventral View
B
WTct
IntTcT
LTct
LNP
Dorsal
Posterior
RHS
LHS
Anterior
Ventral
C
00A
00B
01A
01B
02A
03A
03B
04A
04B
05B
06A
06B
07B
08A
08B
09A
09B
10B
11A
11B
12A
12B
13A
13B
14A
14B
15B
16B
17A
17B
17X
18B
18X
19A
19B
20A.22A
20B.21B.22B
21A
21X
23B
24B.25B
26X
27X
Primary
T1

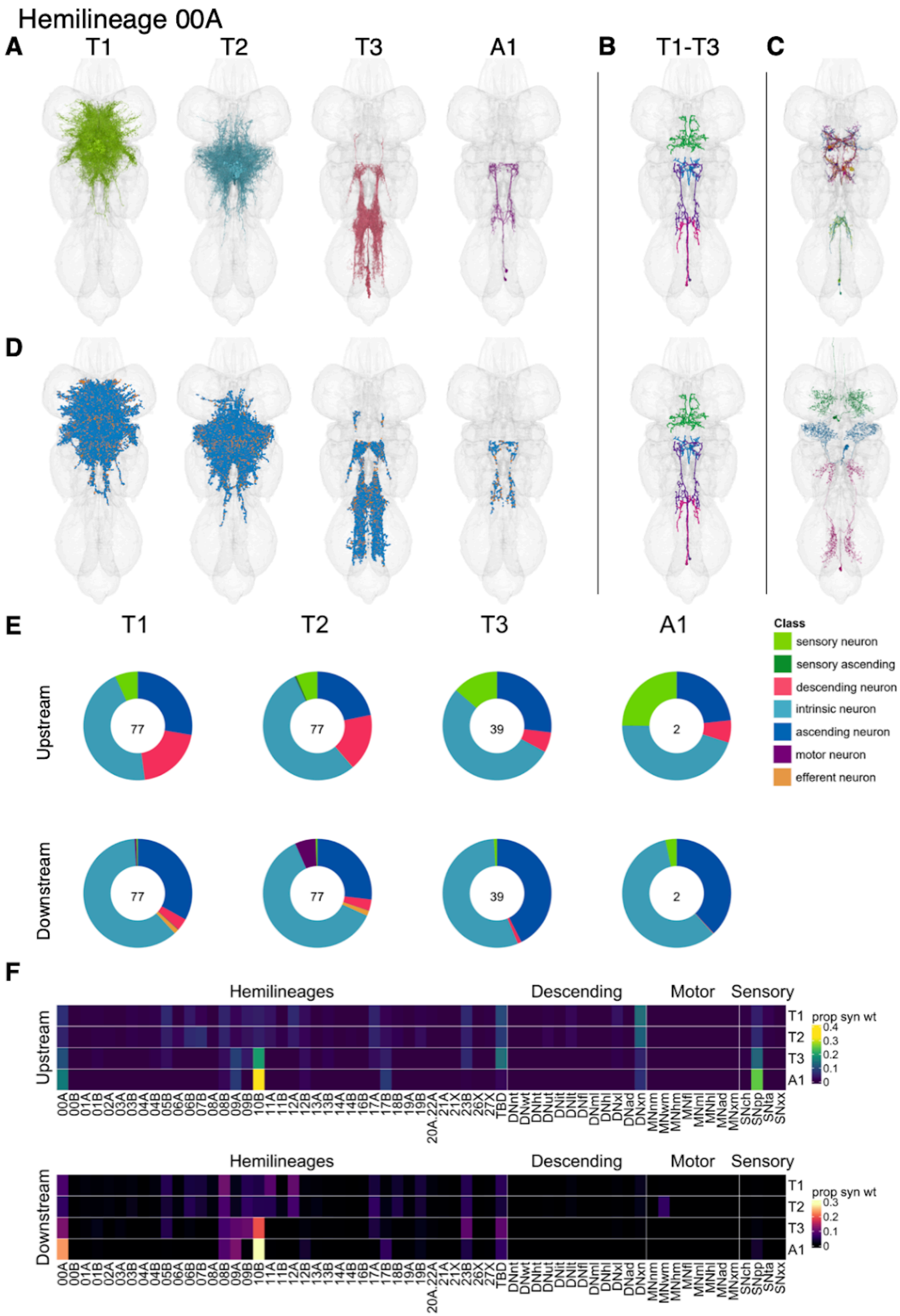

Hemilineage 00A
A
T1
T2
T3
A1
B
T1-T3
C
D
E
T1
T2
T3
A1
Upstream
Downstream
77
77
39
2
Class
sensory neuron
sensory ascending
descending neuron
intrinsic neuron
ascending neuron
motor neuron
efferent neuron
F
Hemilineages
Descending
Motor
Sensory
Upstream
Downstream
prop syn wt
00A 00B 01A 01B 02A 03A 03B 04A 04B 05B 06A 06B 07B 08A 08B 09A 09B 10B 11A 11B 12A 12B 13A 13B 14A 14B 16B 17A 17B 18B 19A 19B 20A.22A 21X 23B 26X 27X TBD DNnt DNwt DNht DNut DNlt DNfl DNml DNhl DNxl DNad DNxn MNnm MNwm MNhm MNfl MNml MNhl MNad MNxm SNch SNpp SNta SNxx

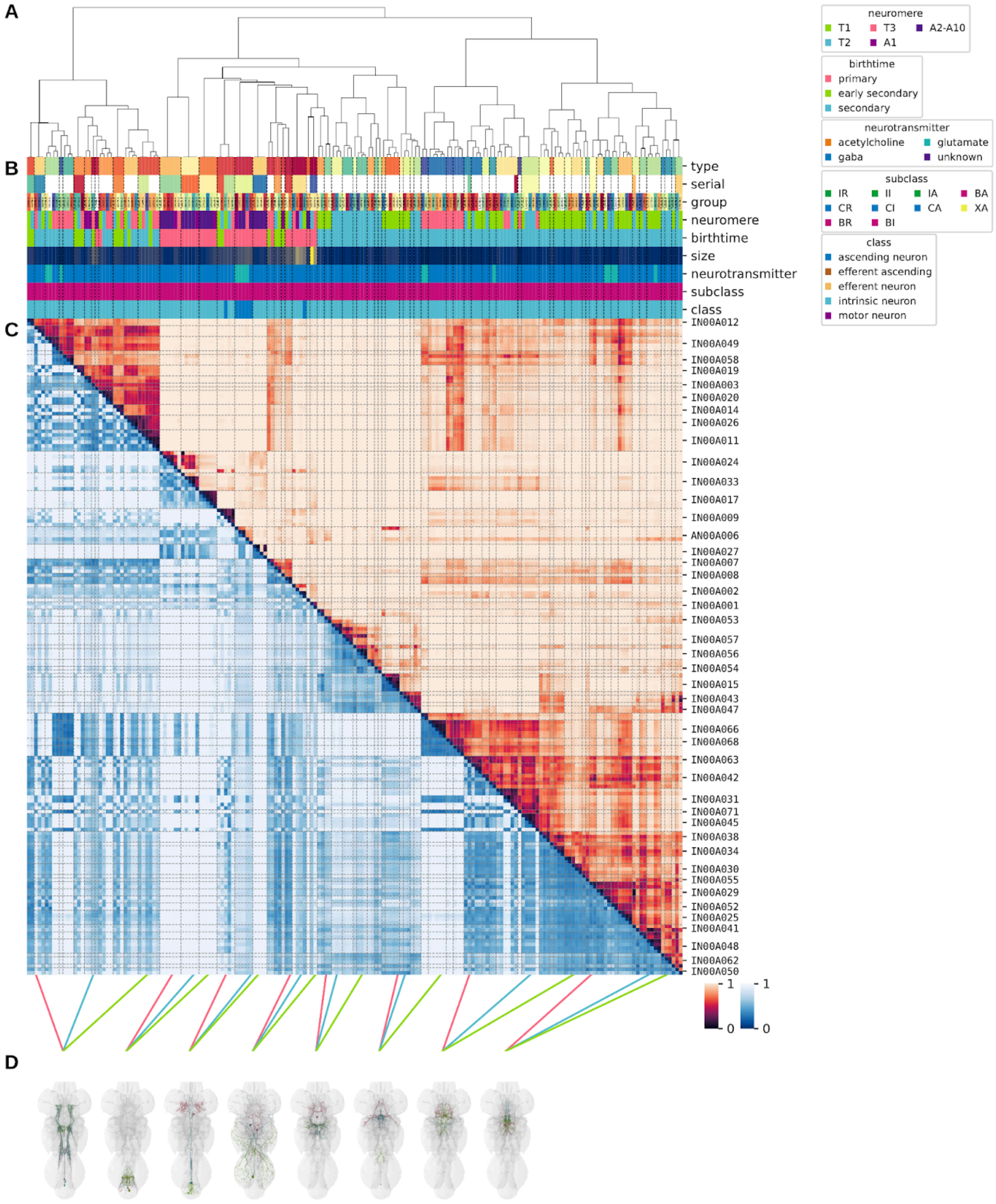

**Figure 15 - figure supplement 1. Systematic typing of hemilineage 00A. A.** Hierarchical clustering dendrogram of hemilineage groups by laterally and serially aggregated connectivity cosine clustering. **B.** Categorical annotations of each hemilineage group, each column corresponding to the aligned leaf in A. Colours for type, serial set, and group are arbitrary for visualisation. Colours for neuromere, birthtime, neurotransmitter, subclass, and class are as in all other figures. **C.** Similarity distance heatmap for hemilineage. Cosine distance is in the upper triangle, while laterally symmetrised NBLAST distance is in the lower triangle. Systematic type names of some types are labelled. **D.** Morphologically representative groups from dendrogram subtrees. Each group, indicated by colour and line connecting to its column in B and C, is the most morphologically representative group (medoid of NBLAST distance)

from a subtree of A. The subtrees (flat clusters) are equal height cuts of A determined to yield the number of groups per plot and plots in D.

#### 00A systematic types

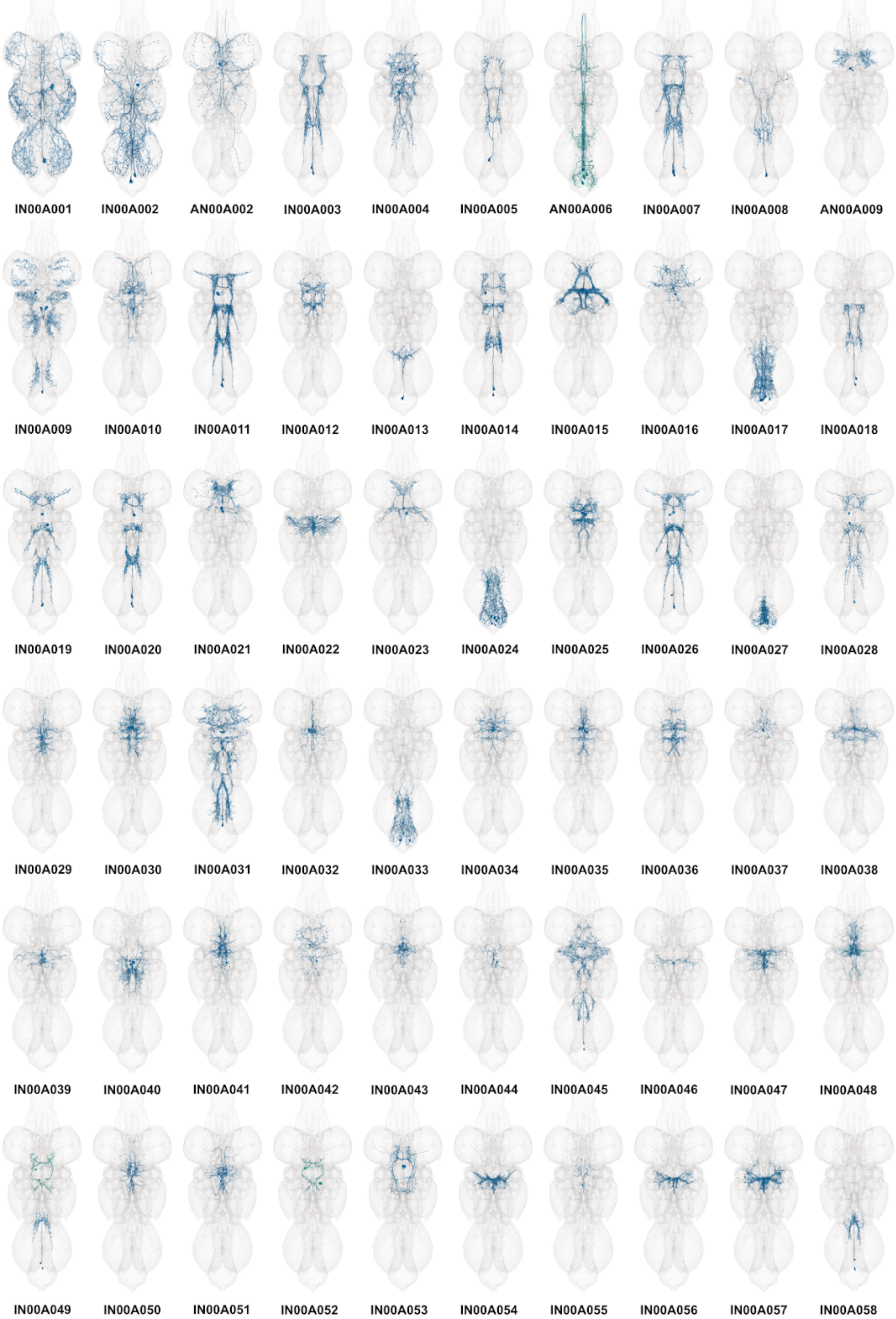

**Figure 15 - figure supplement 2. Systematic types of hemilineage 00A.** Systematic types have been arranged in numerical order, with neurons of the same type that belong to distinct classes (e.g., intrinsic neuron vs ascending neuron) plotted separately but placed adjacent to each other. Individual neuron meshes have been coloured based on their predicted neurotransmitters: dark orange = acetylcholine, blue = gaba, marine = glutamate, dark purple = unknown.

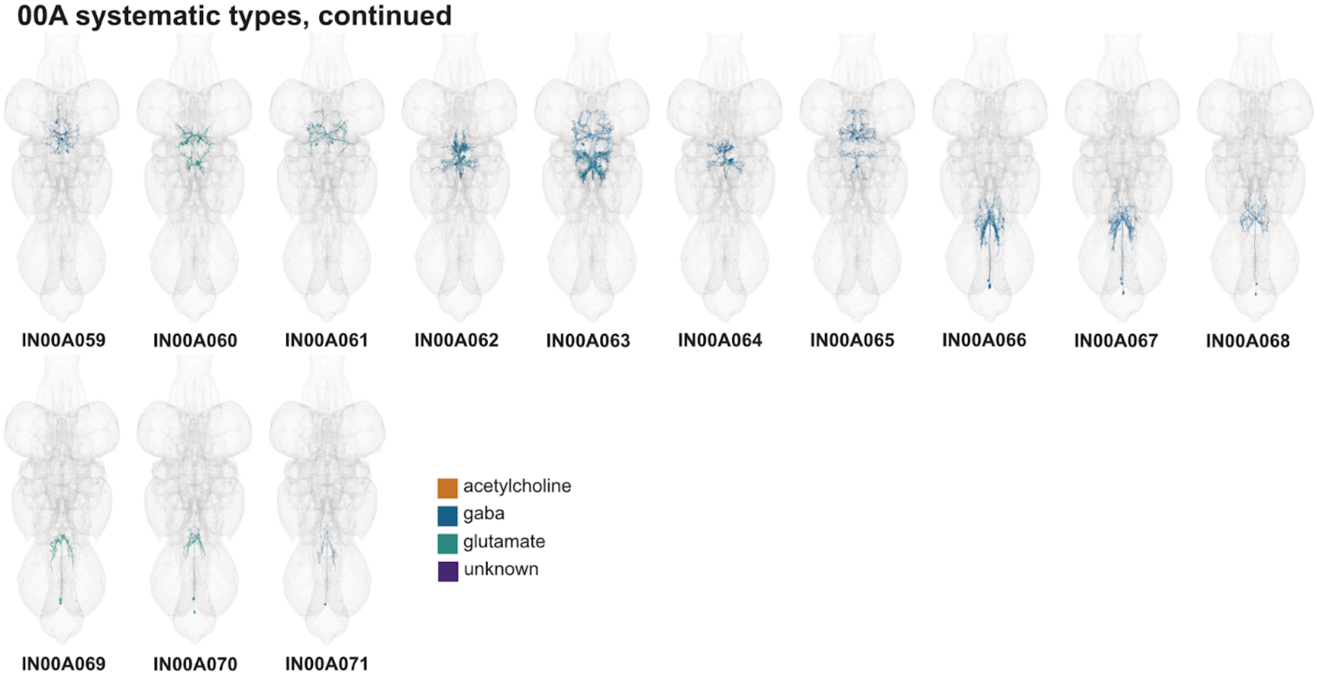

**Figure 15 - figure supplement 3. Systematic types of hemilineage 00A, continued.** Systematic types have been arranged in numerical order, with neurons of the same type that belong to distinct classes (e.g., intrinsic neuron vs ascending neuron) plotted separately but placed adjacent to each other. Individual neuron meshes have been coloured based on predicted neurotransmitter: dark orange = acetylcholine, blue = gaba, marine = glutamate, dark purple = unknown.

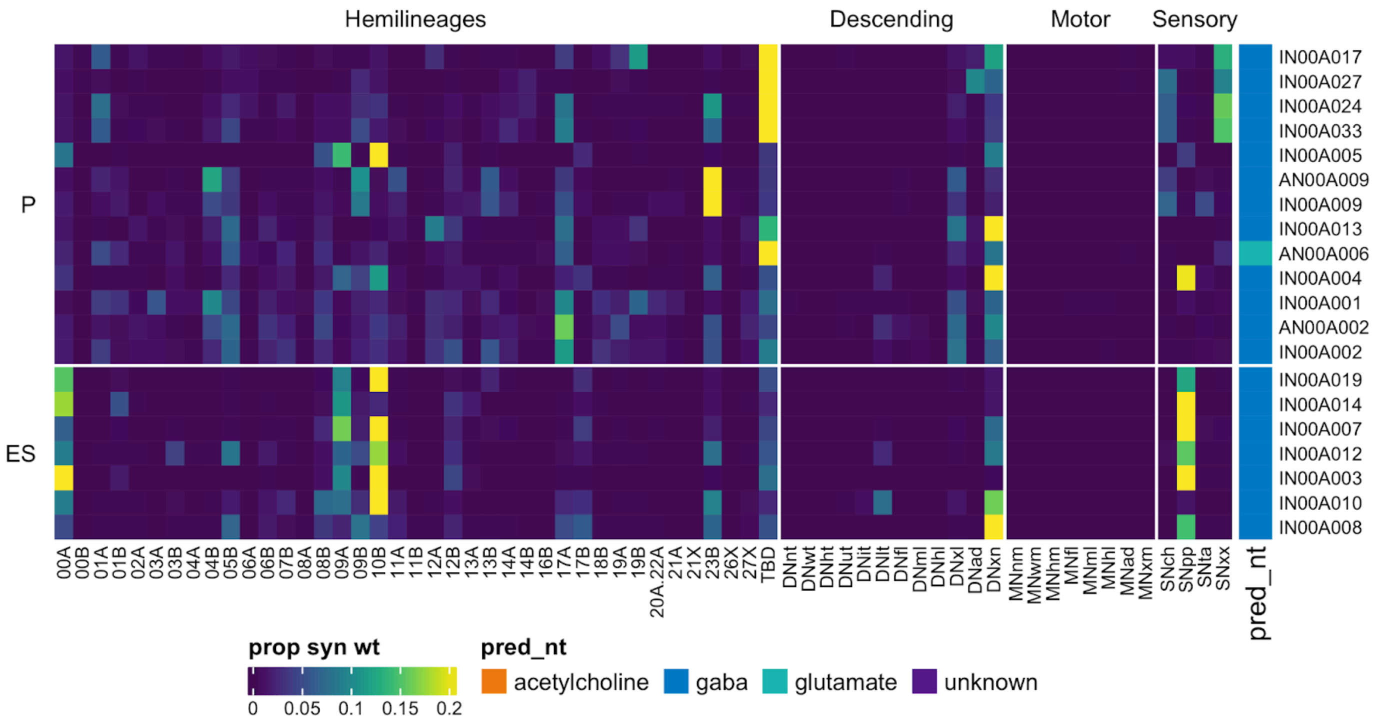

**Figure 15 - figure supplement 4. Connectivity to upstream partners by 00A primary and early secondary systematic types.** Proportions of synaptic weight to systematic types from upstream partners, normalised by row. 00A neurons have been clustered within each assigned birthtime window (P = primary, ES = early secondary, S = secondary) based on both upstream and downstream connectivity to hemilineages, descending neuron subclasses, motor neuron subclasses, and sensory neuron modalities. Annotation bar is coloured by the most common predicted neurotransmitter for the neurons of each type.

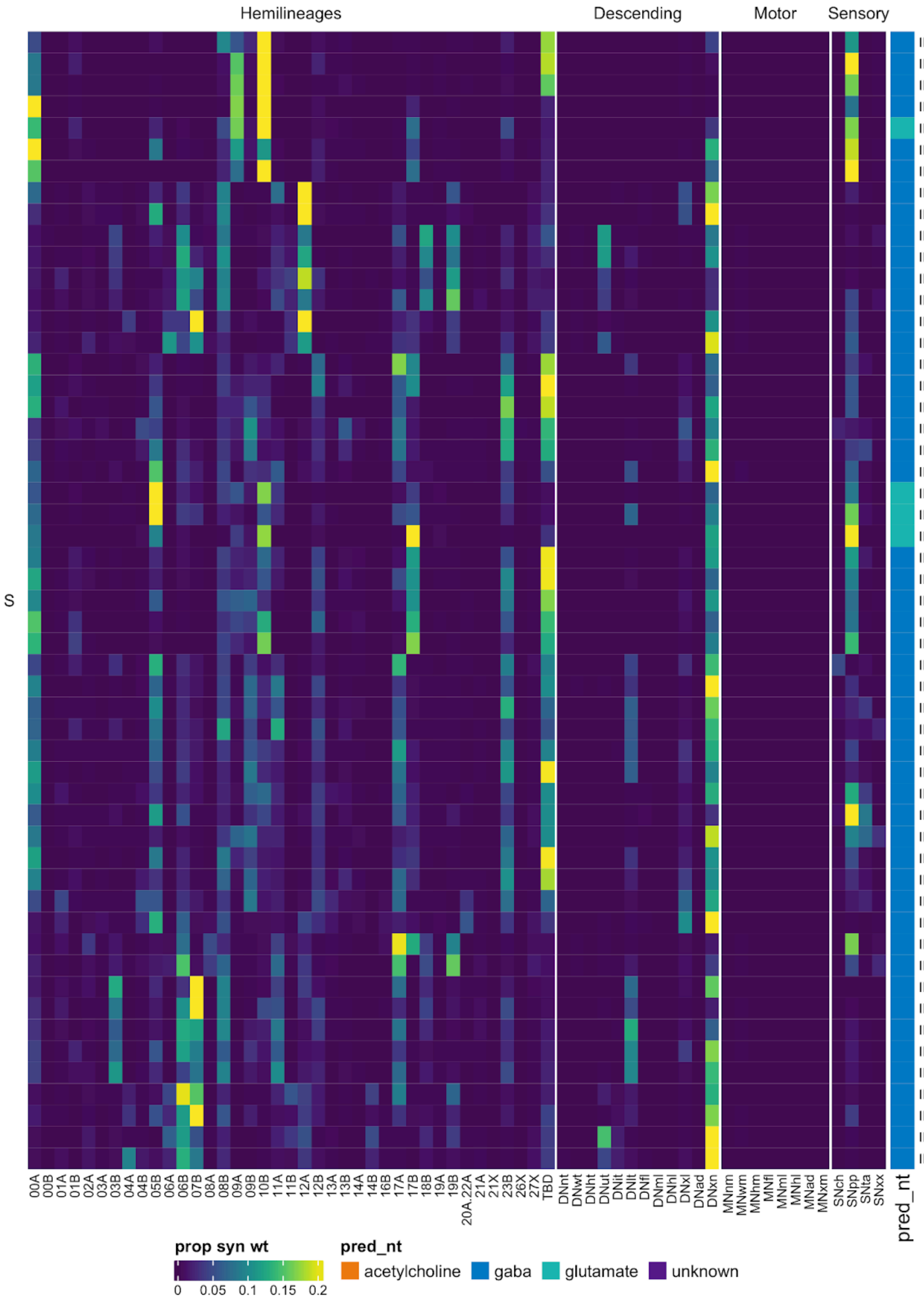

**Figure 15 - figure supplement 5. Connectivity to upstream partners by 00A secondary systematic types.** Proportions of synaptic weight to systematic types from upstream partners, normalised by row. 00A neurons have been clustered within each assigned birthtime window (P = primary, ES = early secondary, S = secondary) based on both upstream and downstream connectivity to hemilineages, descending neuron subclasses, motor neuron subclasses, and sensory neuron modalities. The annotation bar is coloured by the most common predicted neurotransmitter within each type.

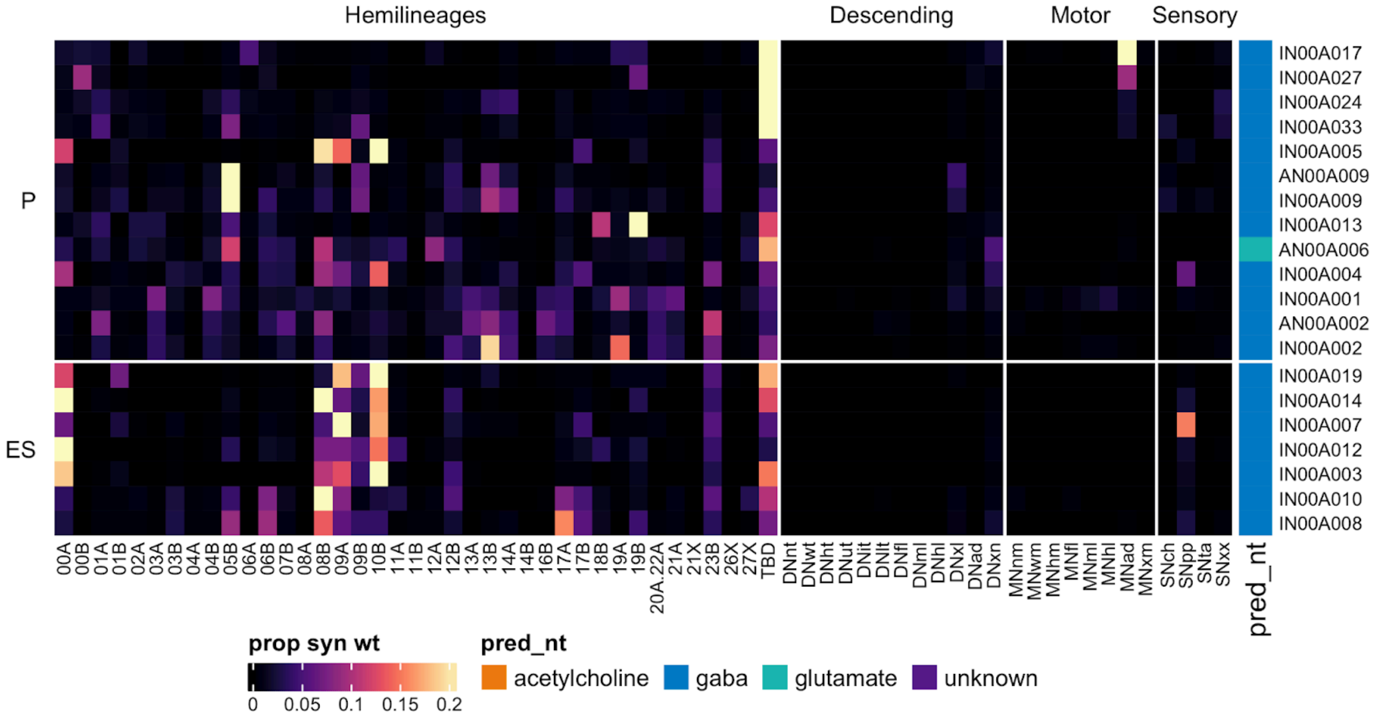

**Figure 15 - figure supplement 6. Connectivity to downstream partners by 00A primary and early secondary systematic types.** Proportions of synaptic weight from systematic types to downstream partners, normalised by row. 00A neurons have been clustered within each assigned birthtime window (P = primary, ES = early secondary, S = secondary) based on both upstream and downstream connectivity to hemilineages, descending neuron subclasses, motor neuron subclasses, and sensory neuron modalities. The annotation bar is coloured by the most common predicted neurotransmitter within each type.

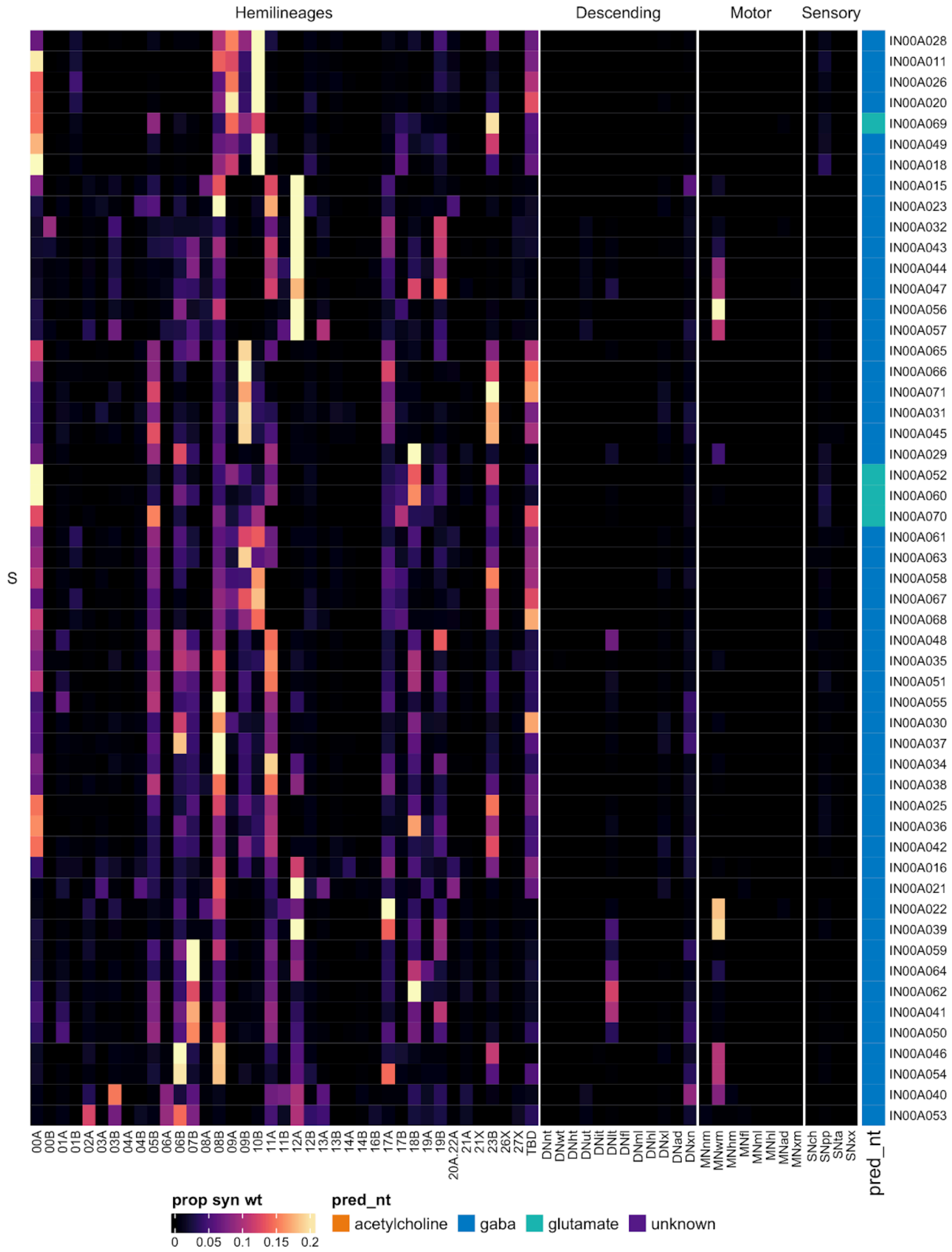

**Figure 15 - figure supplement 7. Connectivity to downstream partners by 00A secondary systematic types.** Proportions of synaptic weight from systematic types to downstream partners, normalised by row. 00A neurons have been clustered within each assigned birthtime window (P = primary, ES = early secondary, S = secondary) based on both upstream and downstream connectivity to hemilineages, descending neuron subclasses, motor neuron subclasses, and sensory neuron modalities. The annotation bar is coloured by the most common predicted

neurotransmitter for the neurons of each type.

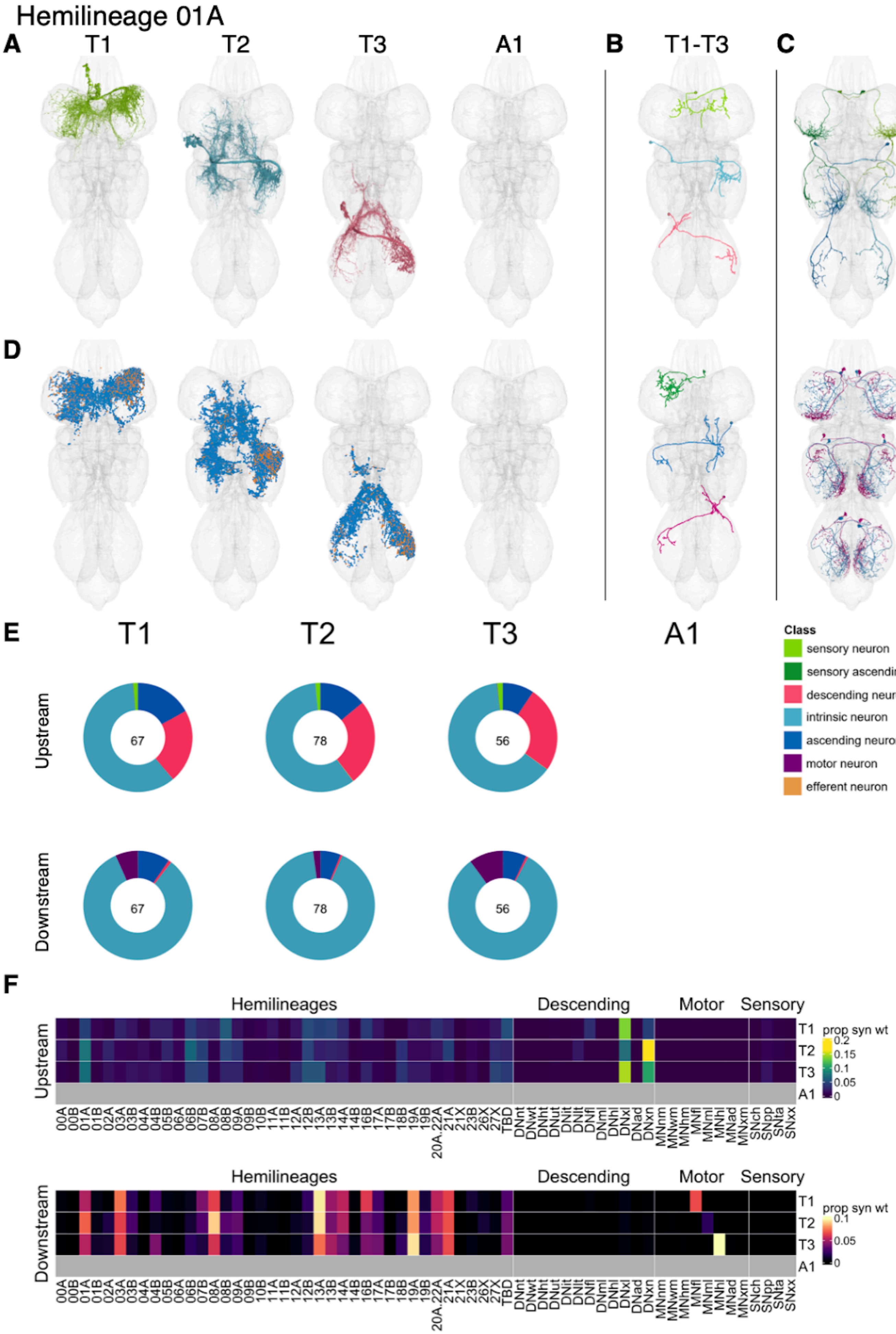
Hemilineage 01A
A
T1
T2
T3
A1
B
T1-T3
C
D
E
T1
T2
T3
A1
Upstream
Downstream
67
78
56
Class
sensory neuron
sensory ascending
descending neuron
intrinsic neuron
ascending neuron
motor neuron
efferent neuron
F
Hemilineages
Descending
Motor
Sensory
prop syn wt
00A 00B 01A 01B 02A 03A 03B 04A 04B 05B 06A 06B 07B 08A 08B 09A 09B 10B 11A 11B 12A 12B 13A 13B 14A 14B 16B 17A 17B 18B 19A 19B 20A.22A 21A 21X 23B 26X 27X TBD
DNnt DNwt DNht DNut DNit DNfl DNml DNhl DNxl DNad DNxn
MNnm MNwm MNhm MNfl MNml MNhl MNad MNxm
SNch SNpp SNta SNxx

**Figure 16. Hemilineage 01A. A.** Meshes of all RHS secondary neurons plotted in hemineuromere-specific colours. **B.** “Representative” secondary neuron skeletons plotted in neuromere-specific colours. The skeleton with the top accumulated NBLAST score among all neurons from the hemilineage in a given hemineuromere was used. **C.** Neuron meshes of selected examples. Top: sequential serial set 10039. Bottom: contralateral independent leg serial sets 12457 and 13010. **D.** Predicted synapses of RHS secondary neurons. Blue: postsynapses; dark orange: presynapses. **E.** Proportions of connections from secondary neurons to upstream or downstream partners, normalised by neuromere and coloured by broad class. Numbers of query neurons appear in the centre. **F.** Proportions of synaptic weight from secondary neurons originating in each neuromere to upstream or downstream partners, normalised by row.

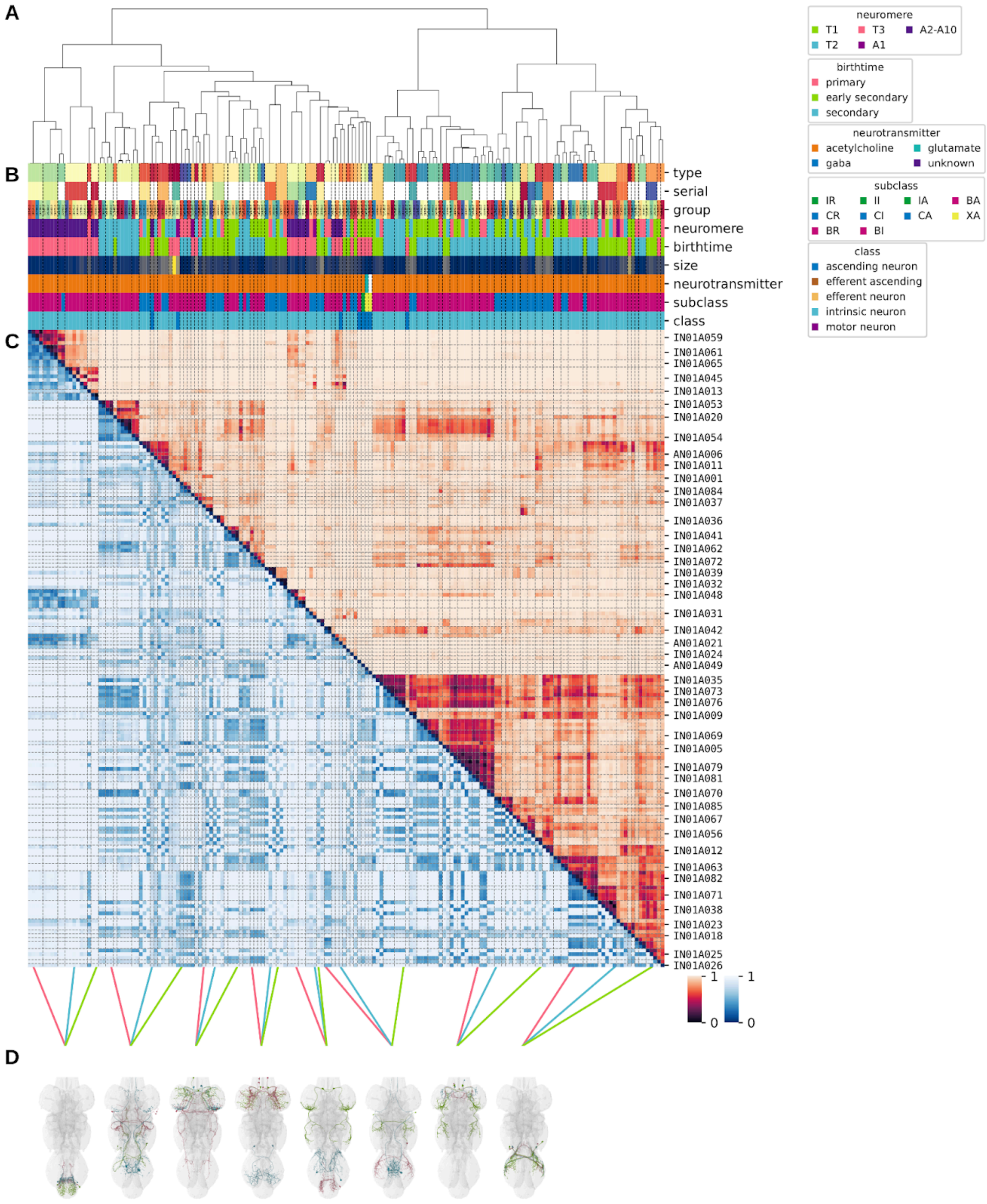

**Figure 16 - figure supplement 1. Systematic typing of hemilineage 01A. A.** Hierarchical clustering dendrogram of hemilineage groups by laterally and serially aggregated connectivity cosine clustering. **B.** Categorical annotations of each hemilineage group, each column corresponding to the aligned leaf in A. Colours for type, serial set, and group are arbitrary for visualisation. Colours for neuromere, birthtime, neurotransmitter, subclass, and class are as in all other figures. **C.** Similarity distance heatmap for hemilineage. Cosine distance is in the upper triangle, while laterally symmetrised NBLAST distance is in the lower triangle. Systematic type names of some types are labelled. **D.** Morphologically representative groups from dendrogram subtrees. Each group, indicated by colour and line

connecting to its column in B and C, is the most morphologically representative group (medoid of NBLAST distance) from a subtree of A. The subtrees (flat clusters) are equal height cuts of A determined to yield the number of groups per plot and plots in D.

#### 01A systematic types

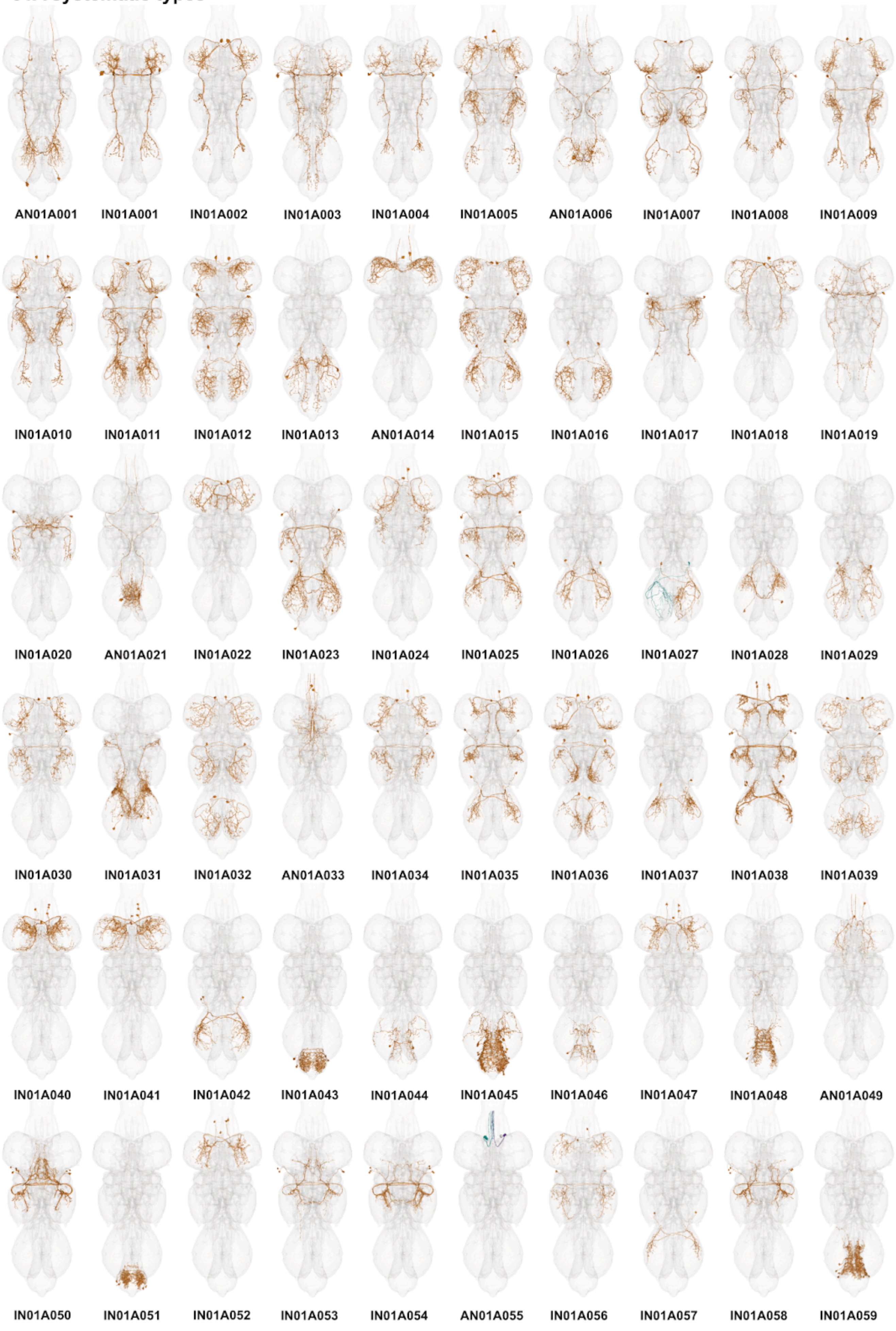

**Figure 16 - figure supplement 2. Systematic types of hemilineage 01A.** Systematic types have been arranged in numerical order, with neurons of the same type that belong to distinct classes (e.g., intrinsic neuron vs ascending neuron) plotted separately but placed adjacent to each other. Individual neuron meshes have been coloured based on predicted neurotransmitter: dark orange = acetylcholine, blue = gaba, marine = glutamate, dark purple = unknown.

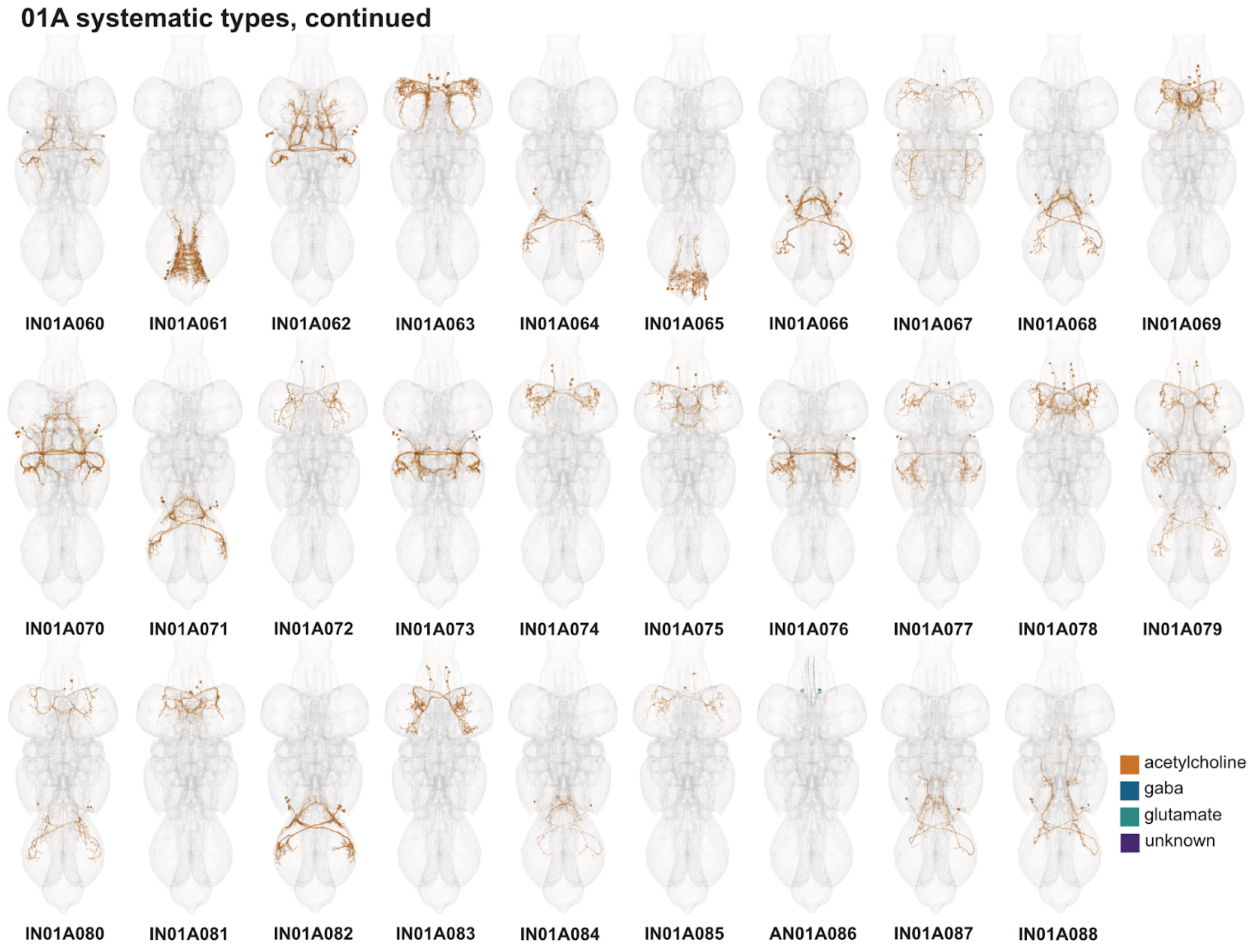

**Figure 16 - figure supplement 3.** Systematic types of hemilineage 01A, continued. Systematic types have been arranged in numerical order, with neurons of the same type that belong to distinct classes (e.g., intrinsic neuron vs ascending neuron) plotted separately but placed adjacent to each other. Individual neuron meshes have been coloured based on predicted neurotransmitter: dark orange = acetylcholine, blue = gaba, marine = glutamate, dark purple = unknown.

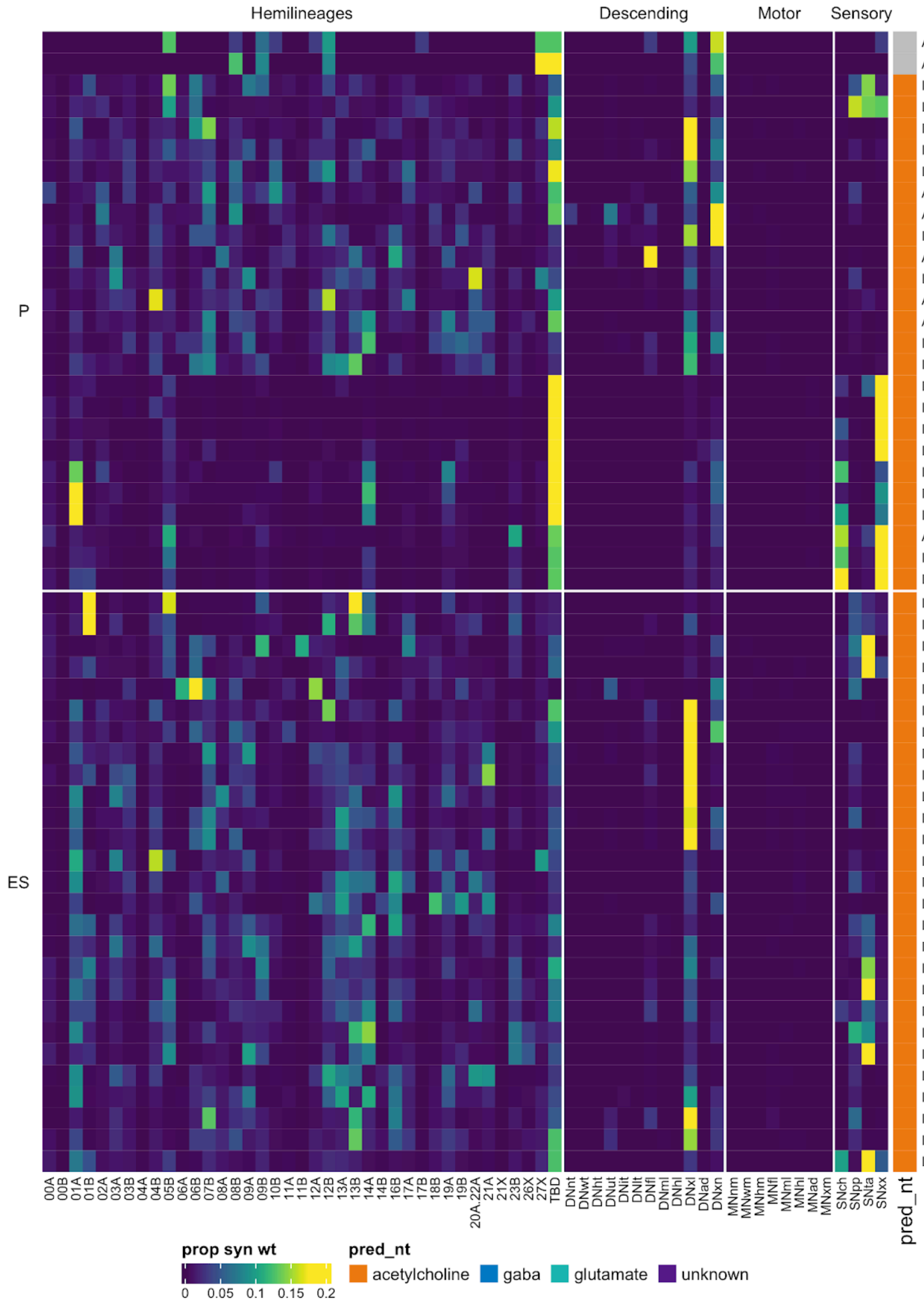

**Figure 16 - figure supplement 4. Connectivity to upstream partners by 01A primary and early secondary systematic types.** Proportions of synaptic weight to systematic types from upstream partners, normalised by row. 01A neurons have been clustered within each assigned birthtime window (P = primary, ES = early secondary, S = secondary) based on both upstream and downstream connectivity to hemilineages, descending neuron subclasses, motor neuron subclasses, and sensory neuron modalities. Annotation bar is coloured by the most common predicted

neurotransmitter for the neurons of each type.

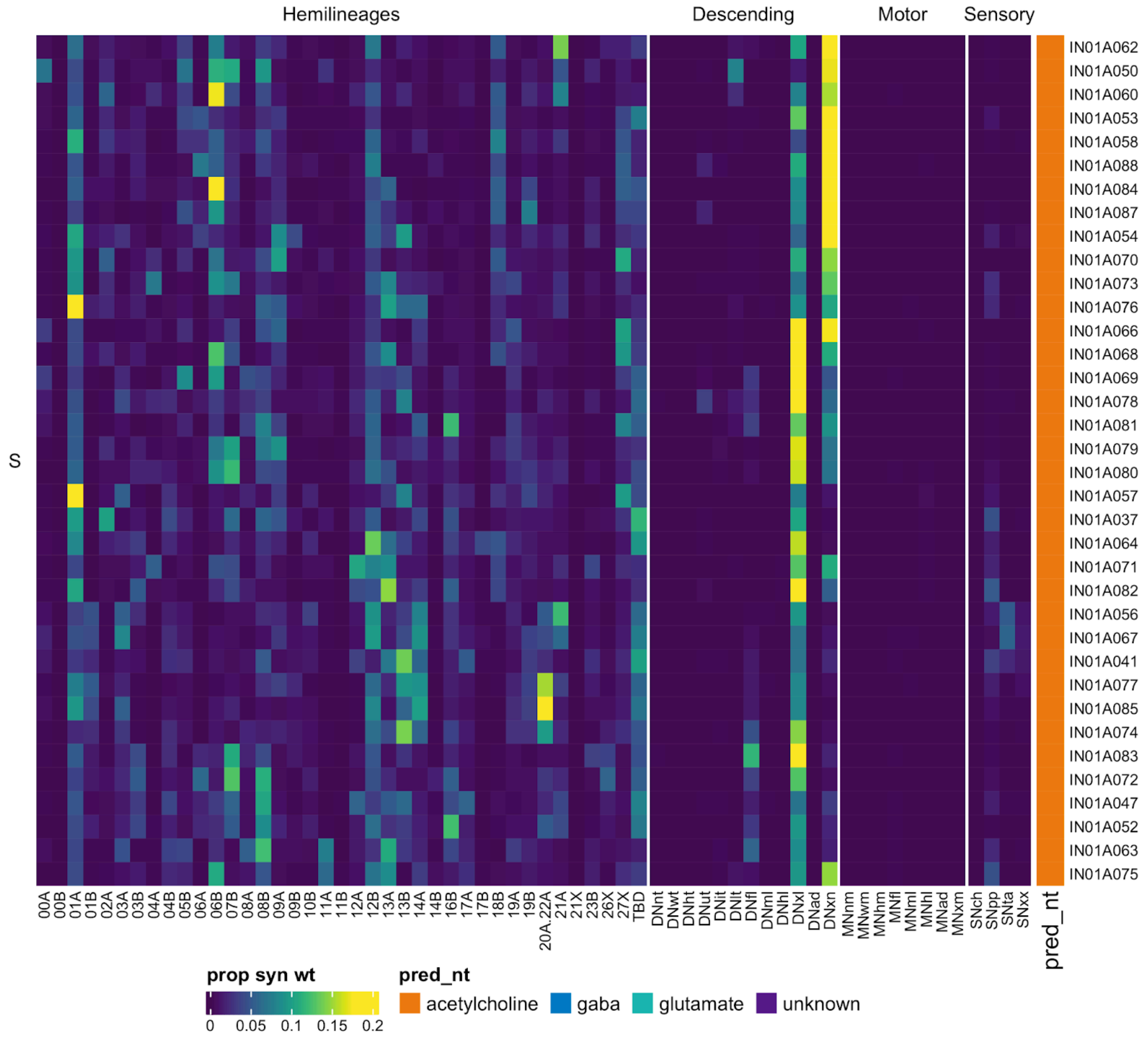

**Figure 16 - figure supplement 5. Connectivity to upstream partners by 01A secondary systematic types.** Proportions of synaptic weight to systematic types from upstream partners, normalised by row. 01A neurons have been clustered within each assigned birthtime window (P = primary, ES = early secondary, S = secondary) based on both upstream and downstream connectivity to hemilineages, descending neuron subclasses, motor neuron subclasses, and sensory neuron modalities. The annotation bar is coloured by the most common predicted neurotransmitter within each type.

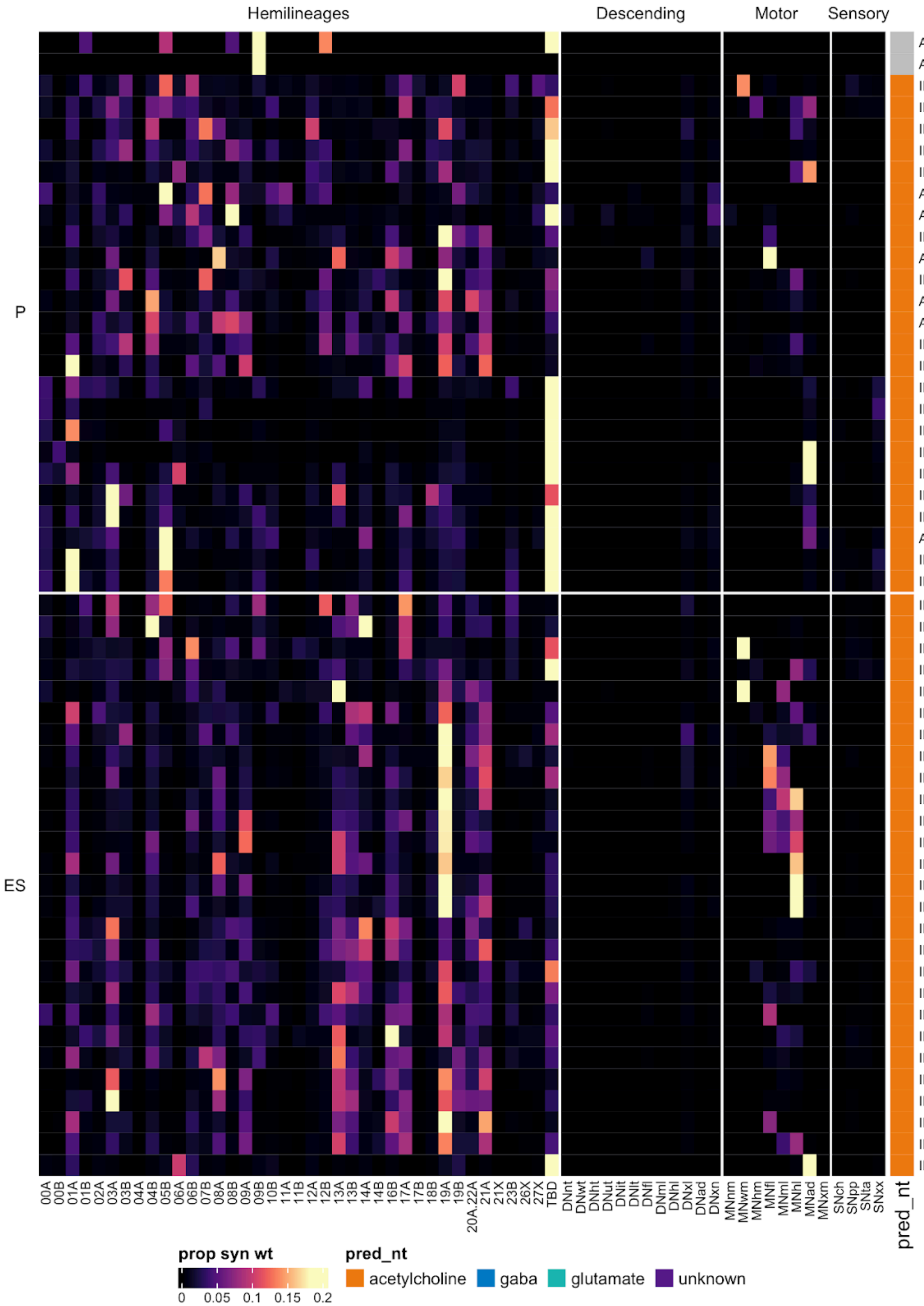

**Figure 16 - figure supplement 6. Connectivity to downstream partners by 01A primary and early secondary systematic types.** Proportions of synaptic weight from systematic types to downstream partners, normalised by row. 01A neurons have been clustered within each assigned birthtime window (P = primary, ES = early secondary, S =

secondary) based on both upstream and downstream connectivity to hemilineages, descending neuron subclasses, motor neuron subclasses, and sensory neuron modalities. The annotation bar is coloured by the most common predicted neurotransmitter within each type.

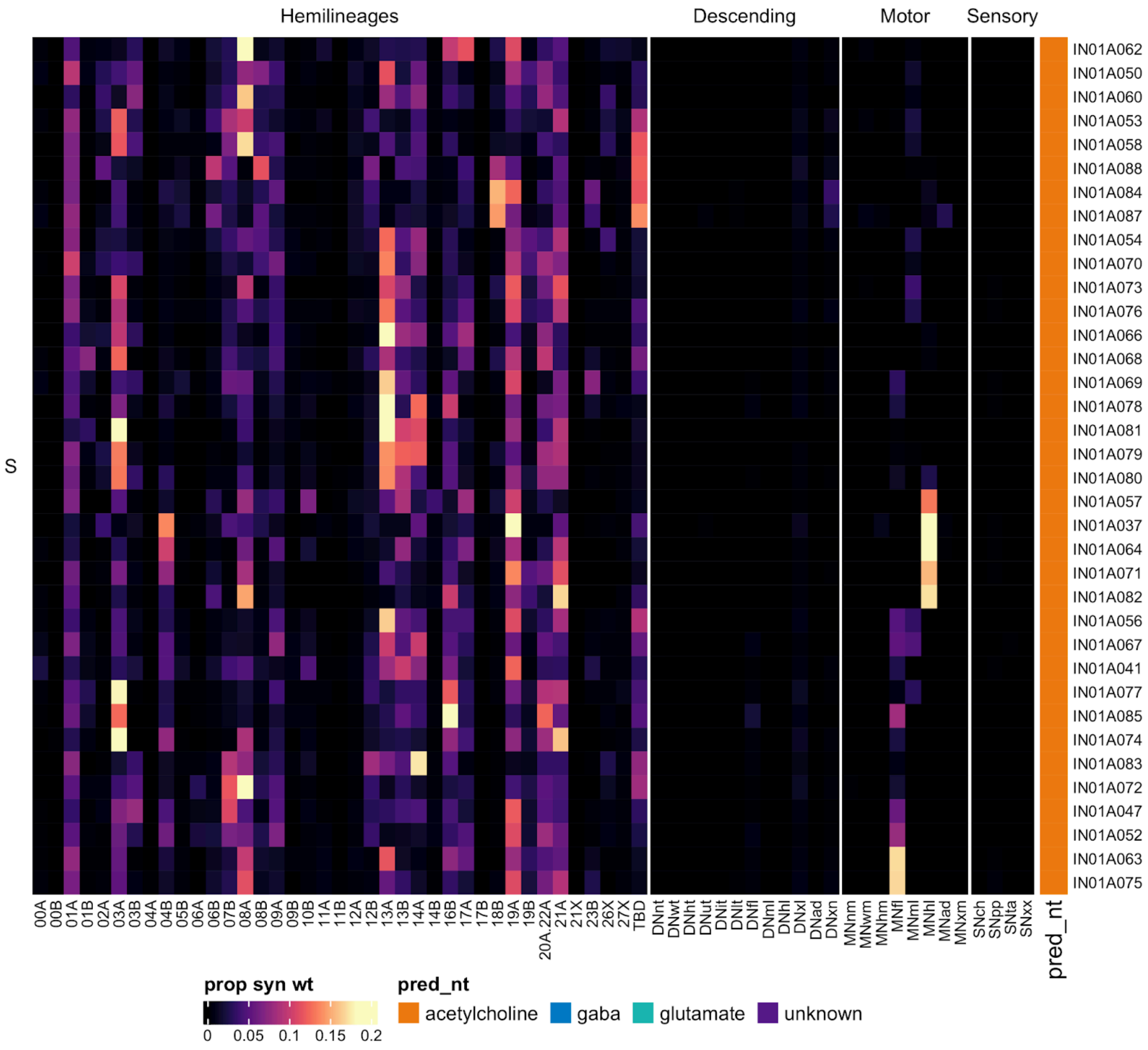

**Figure 16 - figure supplement 7.Connectivity to downstream partners by 01A secondary systematic types.** Proportions of synaptic weight from systematic types to downstream partners, normalised by row. 01A neurons have been clustered within each assigned birthtime window (P = primary, ES = early secondary, S = secondary) based on both upstream and downstream connectivity to hemilineages, descending neuron subclasses, motor neuron subclasses, and sensory neuron modalities. The annotation bar is coloured by the most common predicted neurotransmitter for the neurons of each type.

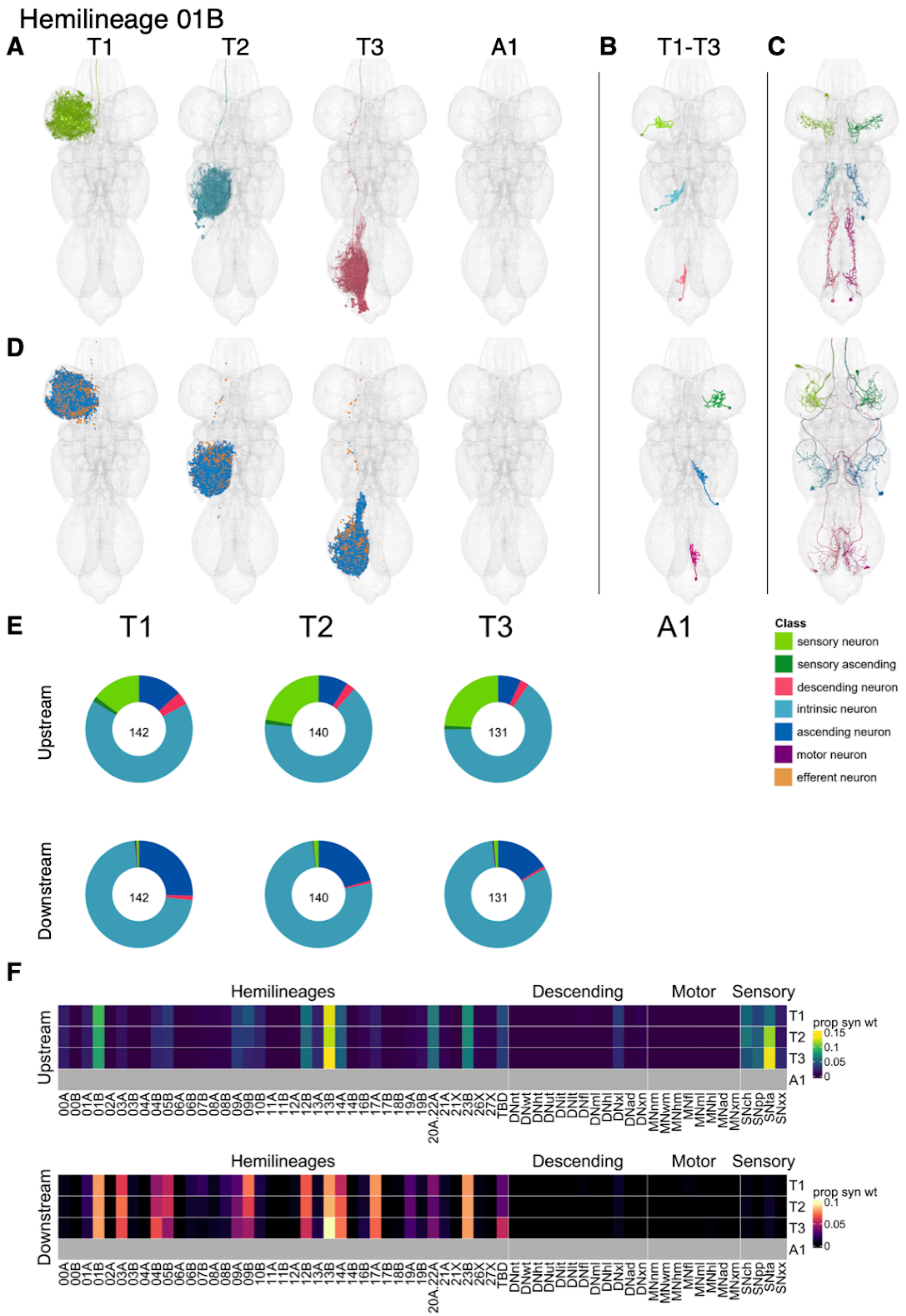

Hemilineage 01B
A
T1
T2
T3
A1
B
T1-T3
C
D
E
T1
T2
T3
A1
Upstream
142
140
131
Downstream
142
140
131
Class
sensory neuron
sensory ascending
descending neuron
intrinsic neuron
ascending neuron
motor neuron
efferent neuron
F
Hemilineages
Descending
Motor
Sensory
Upstream
T1
T2
T3
A1
prop syn wt
0.15
0.1
0.05
0
00A 00B 01A 01B 02A 03A 03B 04A 04B 05B 06A 06B 07B 08A 08B 09A 09B 10B 11A 11B 12A 12B 13A 13B 14A 14B 16B 17A 17B 18B 19A 19B 20A.22A 21A 21X 23B 26X 27X TBD DNnt DNwt DNht DNut DNlt DNfl DNml DNhl DNxl DNad DNxn MNnm MNwm MNhm MNfl MNml MNhl MNad MNxm SNch SNpp SNta SNxx
Hemilineages
Descending
Motor
Sensory
Downstream
T1
T2
T3
A1
prop syn wt
0.1
0.05
0
00A 00B 01A 01B 02A 03A 03B 04A 04B 05B 06A 06B 07B 08A 08B 09A 09B 10B 11A 11B 12A 12B 13A 13B 14A 14B 16B 17A 17B 18B 19A 19B 20A.22A 21A 21X 23B 26X 27X TBD DNnt DNwt DNht DNut DNlt DNfl DNml DNhl DNxl DNad DNxn MNnm MNwm MNhm MNfl MNml MNhl MNad MNxm SNch SNpp SNta SNxx

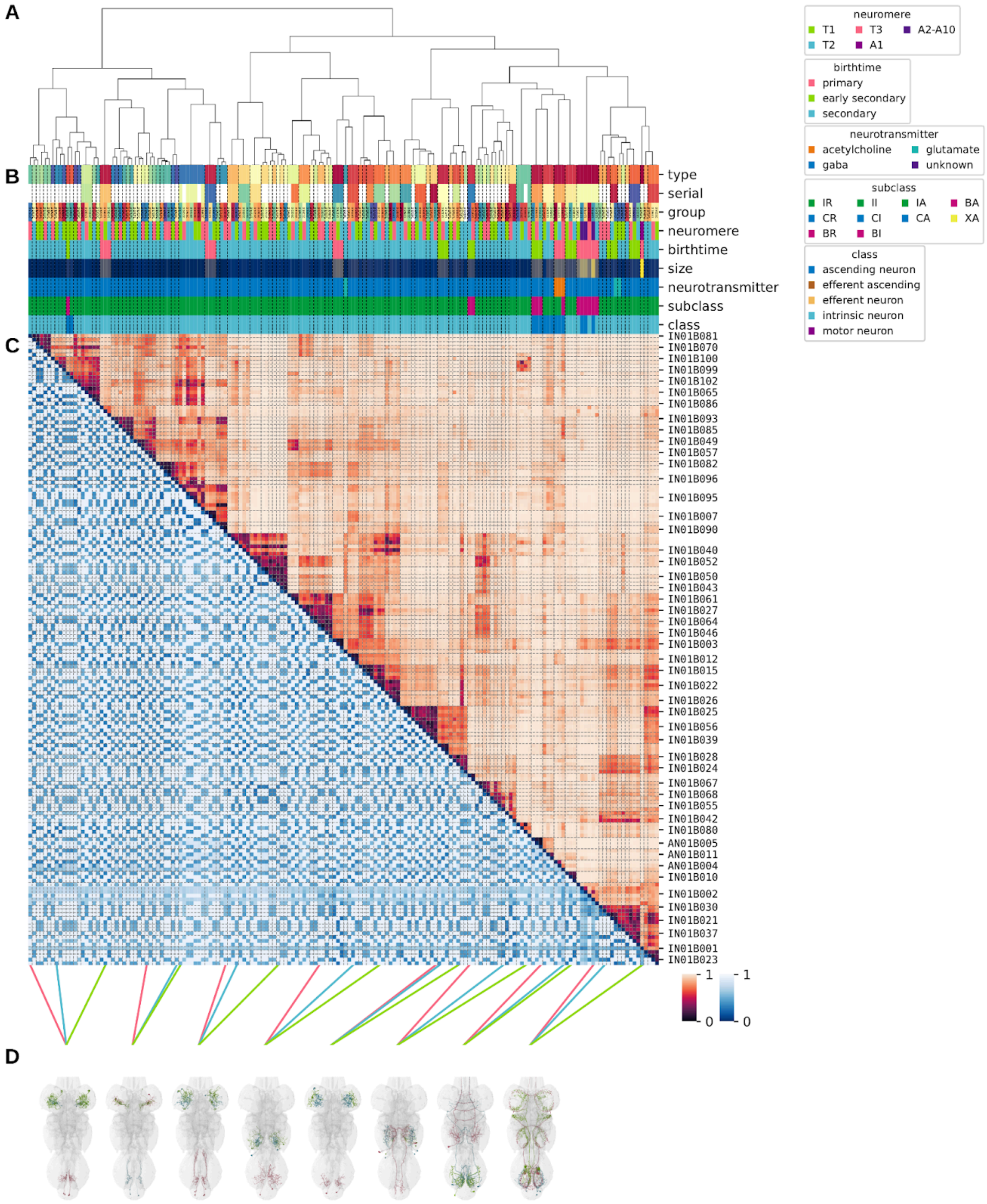

**Figure 17 - figure supplement 1. Systematic typing of hemilineage 01B. A.** Hierarchical clustering dendrogram of hemilineage groups by laterally and serially aggregated connectivity cosine clustering. **B.** Categorical annotations of each hemilineage group, each column corresponding to the aligned leaf in A. Colours for type, serial set, and group are arbitrary for visualisation. Colours for neuromere, birthtime, neurotransmitter, subclass, and class are as in all other figures. **C.** Similarity distance heatmap for hemilineage. Cosine distance is in the upper triangle, while laterally symmetrised NBLAST distance is in the lower triangle. Systematic type names of some types are labelled. **D.** Morphologically representative groups from dendrogram subtrees. Each group, indicated by colour and line

connecting to its column in B and C, is the most morphologically representative group (medoid of NBLAST distance) from a subtree of A. The subtrees (flat clusters) are equal height cuts of A determined to yield the number of groups per plot and plots in D.

#### 01B systematic types

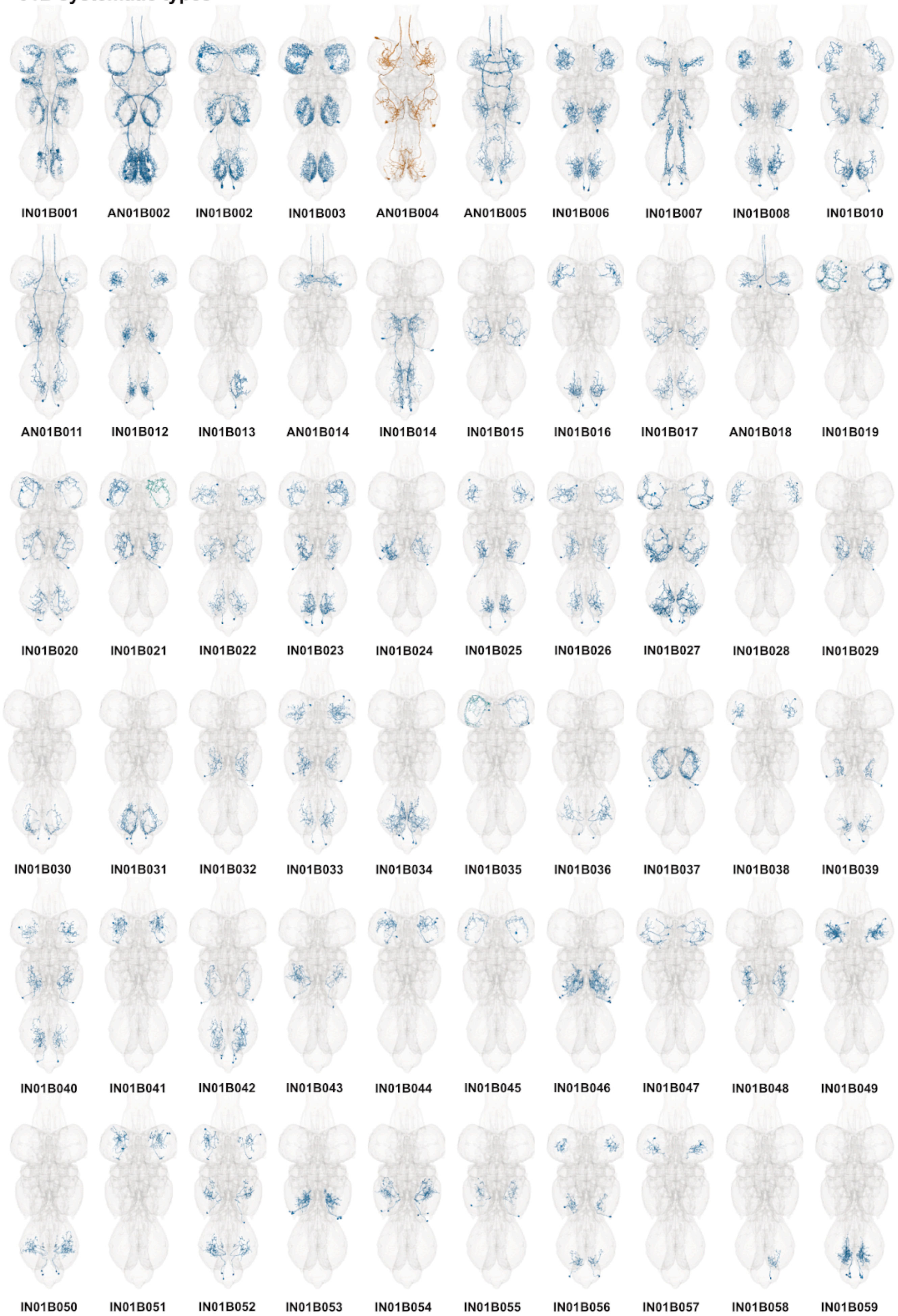

**Figure 17 - figure supplement 2. Systematic types of hemilineage 01B.** Systematic types have been arranged in numerical order, with neurons of the same type that belong to distinct classes (e.g., intrinsic neuron vs ascending neuron) plotted separately but placed adjacent to each other. Individual neuron meshes have been coloured based on predicted neurotransmitter: dark orange = acetylcholine, blue = gaba, marine = glutamate, dark purple = unknown.

#### 01B systematic types, continued

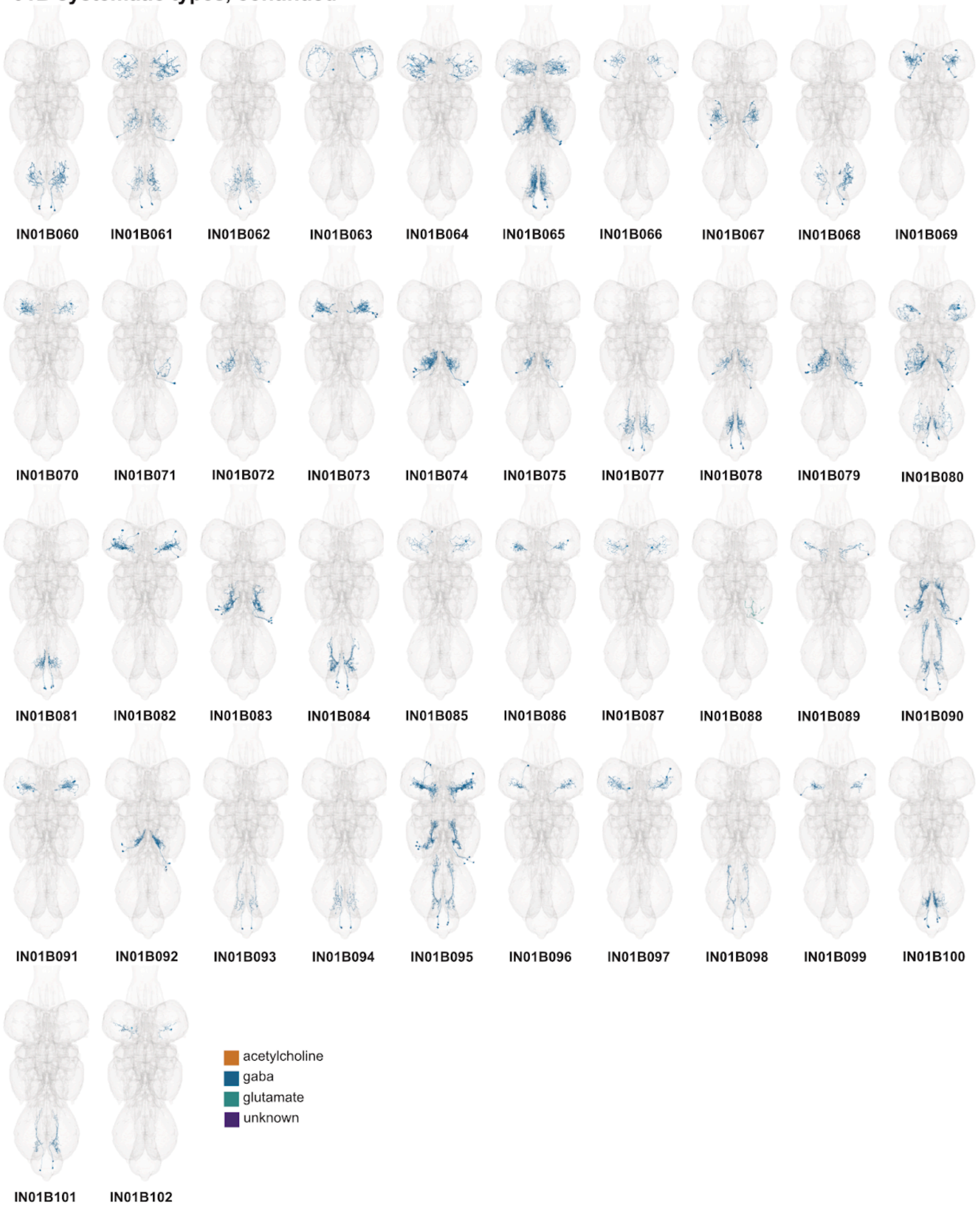

**Figure 17 - figure supplement 3. Systematic types of hemilineage 01B, continued.** Systematic types have been arranged in numerical order, with neurons of the same type that belong to distinct classes (e.g., intrinsic neuron vs ascending neuron) plotted separately but placed adjacent to each other. Individual neuron meshes have been coloured based on predicted neurotransmitter: dark orange = acetylcholine, blue = gaba, marine = glutamate, dark purple = unknown.

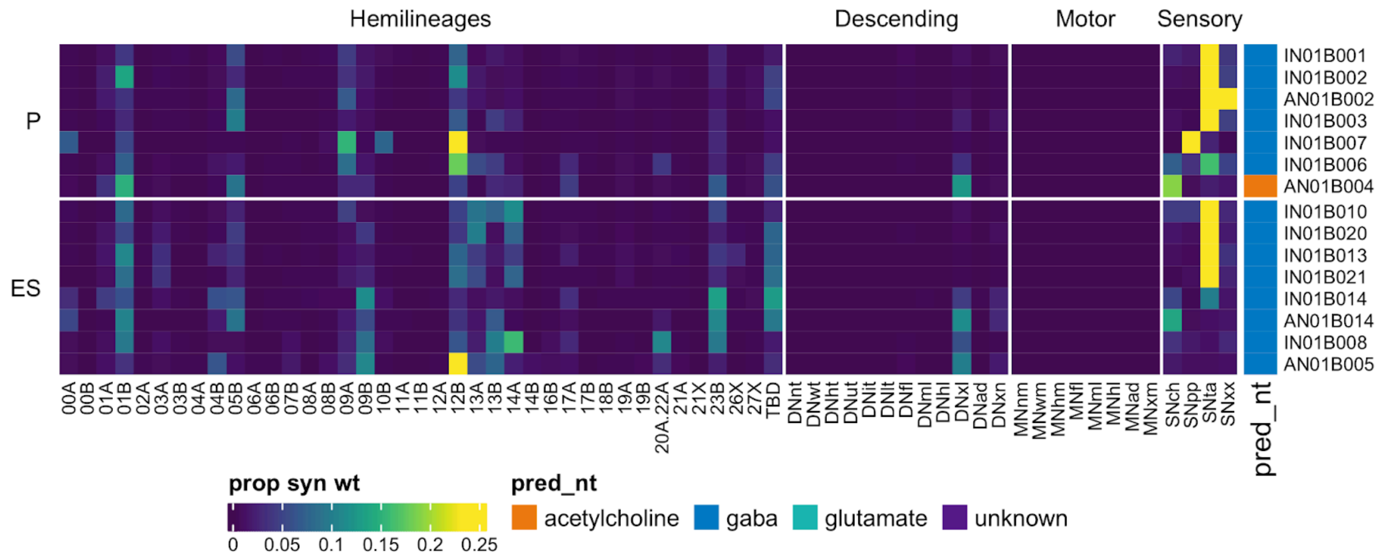

**Figure 17 - figure supplement 4. Connectivity to upstream partners by 01B primary and early secondary systematic types.** Proportions of synaptic weight to systematic types from upstream partners, normalised by row. 01B neurons have been clustered within each assigned birthtime window (P = primary, ES = early secondary, S = secondary) based on both upstream and downstream connectivity to hemilineages, descending neuron subclasses, motor neuron subclasses, and sensory neuron modalities. Annotation bar is coloured by the most common predicted neurotransmitter for the neurons of each type.

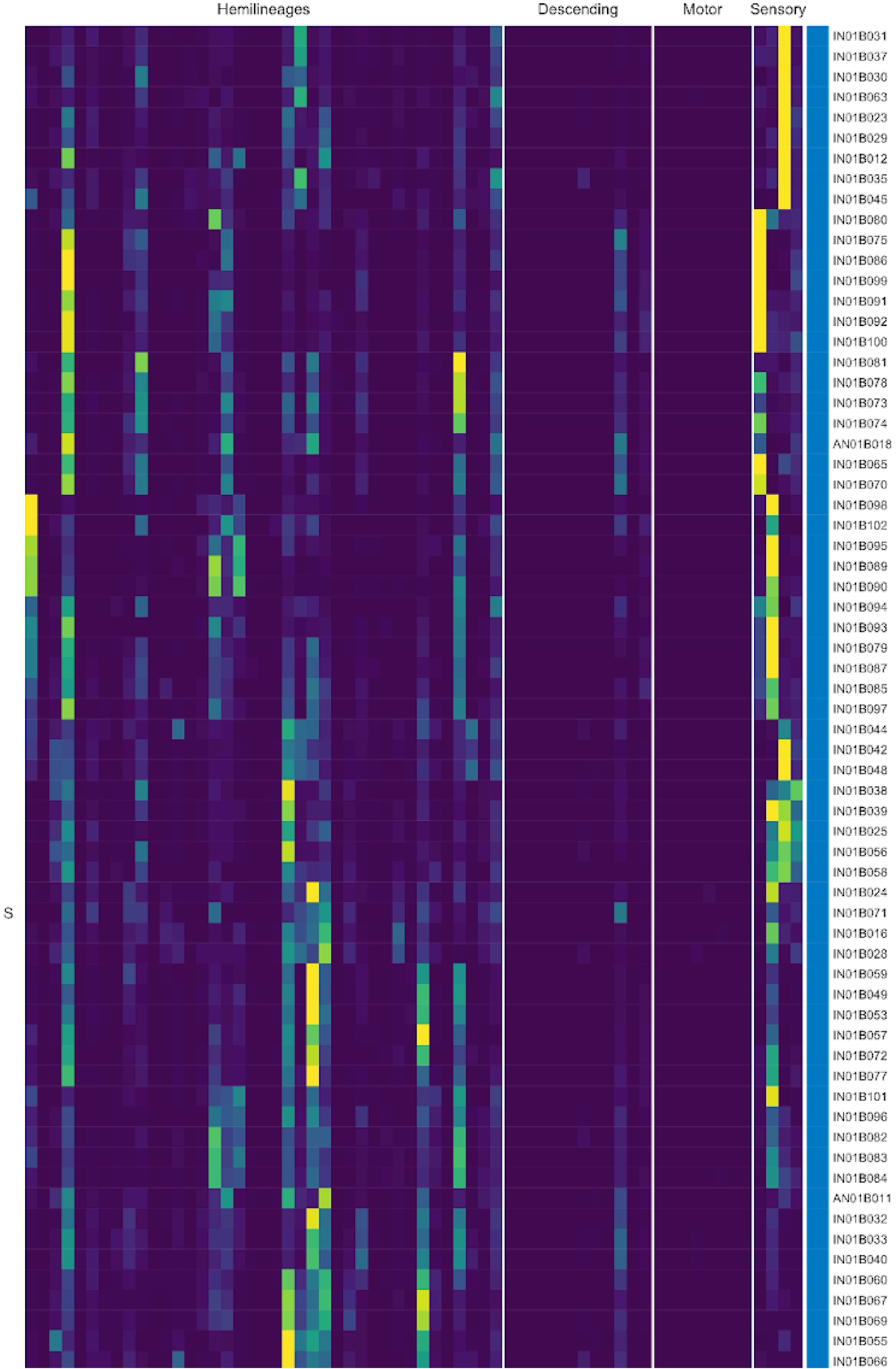

Hemilineages
Descending
Motor
Sensory
S
IN01B031
IN01B037
IN01B030
IN01B063
IN01B023
IN01B029
IN01B012
IN01B035
IN01B045
IN01B080
IN01B075
IN01B086
IN01B099
IN01B091
IN01B092
IN01B100
IN01B081
IN01B078
IN01B073
IN01B074
AN01B018
IN01B065
IN01B070
IN01B098
IN01B102
IN01B095
IN01B089
IN01B090
IN01B094
IN01B093
IN01B079
IN01B087
IN01B085
IN01B097
IN01B044
IN01B042
IN01B048
IN01B038
IN01B039
IN01B025
IN01B056
IN01B058
IN01B024
IN01B071
IN01B016
IN01B028
IN01B059
IN01B049
IN01B053
IN01B057
IN01B072
IN01B077
IN01B101
IN01B096
IN01B082
IN01B083
IN01B084
AN01B011
IN01B032
IN01B033
IN01B040
IN01B060
IN01B067
IN01B069
IN01B055
IN01B066

**Figure 17 - figure supplement 5a. Connectivity to upstream partners by 01B secondary systematic types.** Proportions of synaptic weight to systematic types from upstream partners, normalised by row. 01B neurons have been clustered within each assigned birthtime window (P = primary, ES = early secondary, S = secondary) based on both upstream and downstream connectivity to hemilineages, descending neuron subclasses, motor neuron subclasses, and sensory neuron modalities. The annotation bar is coloured by the most common predicted neurotransmitter within each type.

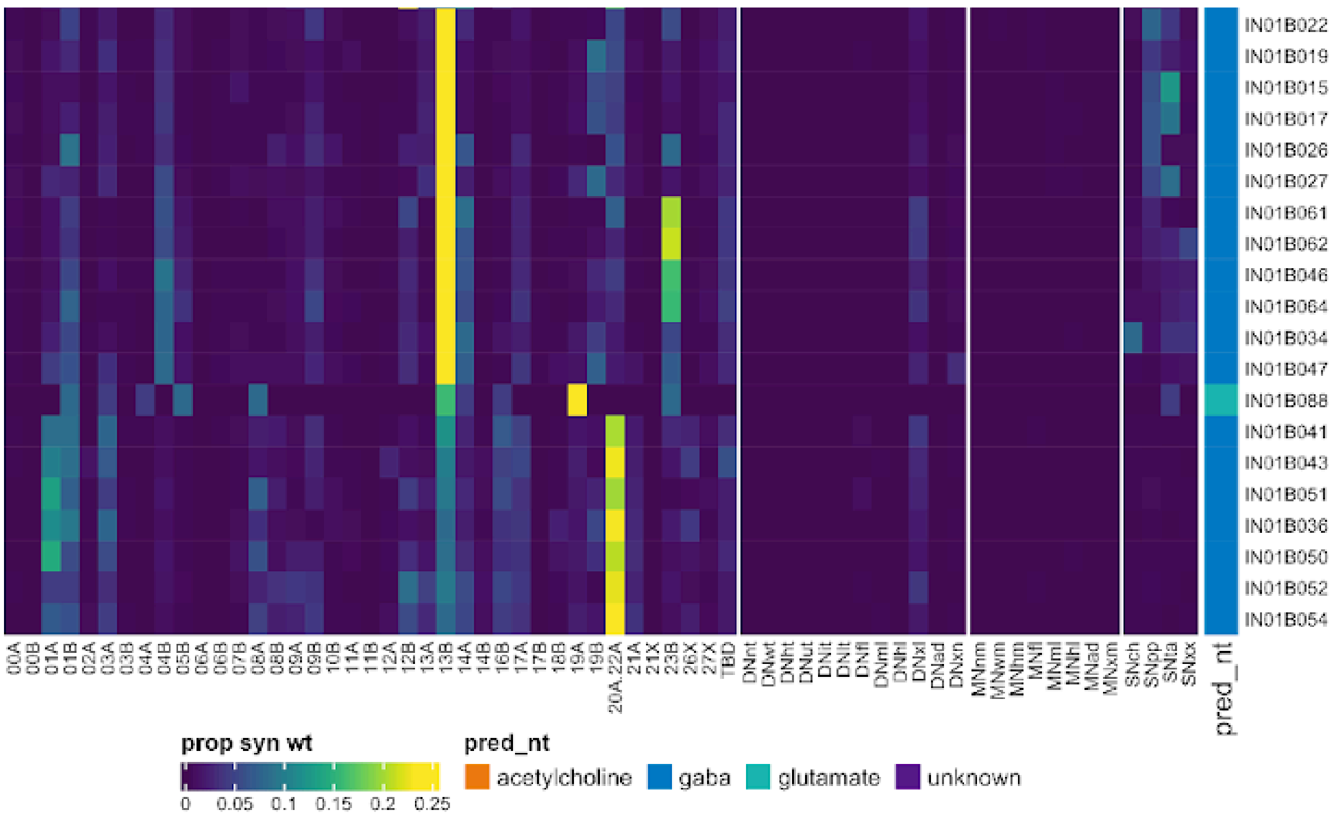

**Figure 17 - figure supplement 5b. Connectivity to upstream partners by 01B secondary systematic types, continued.** Proportions of synaptic weight to systematic types from upstream partners, normalised by row. 01B neurons have been clustered within each assigned birthtime window (P = primary, ES = early secondary, S = secondary) based on both upstream and downstream connectivity to hemilineages, descending neuron subclasses, motor neuron subclasses, and sensory neuron modalities. The annotation bar is coloured by the most common predicted neurotransmitter within each type.

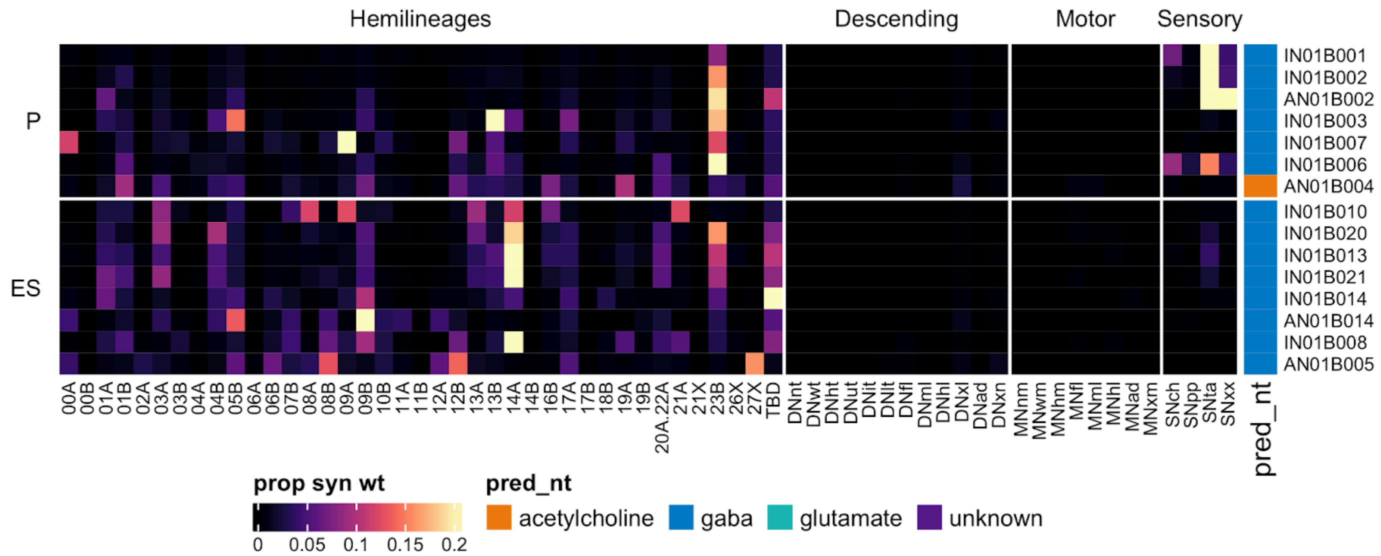

**Figure 17 - figure supplement 6. Connectivity to downstream partners by 01B primary and early secondary systematic types.** Proportions of synaptic weight from systematic types to downstream partners, normalised by row. 01B neurons have been clustered within each assigned birthtime window (P = primary, ES = early secondary, S = secondary) based on both upstream and downstream connectivity to hemilineages, descending neuron subclasses, motor neuron subclasses, and sensory neuron modalities. The annotation bar is coloured by the most common predicted neurotransmitter within each type.

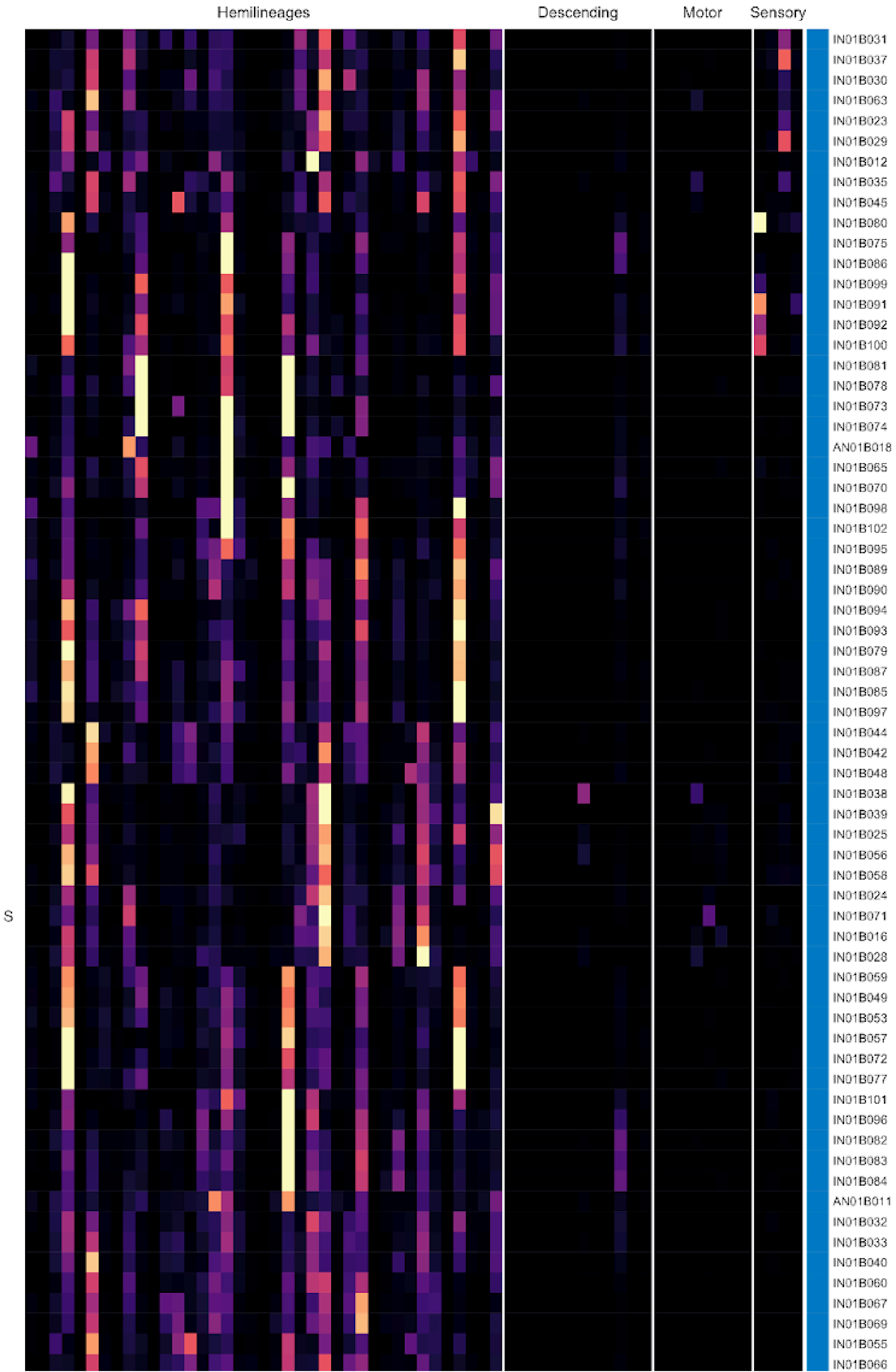
Hemilineages
Descending
Motor
Sensory
S
IN01B031
IN01B037
IN01B030
IN01B063
IN01B023
IN01B029
IN01B012
IN01B035
IN01B045
IN01B080
IN01B075
IN01B086
IN01B099
IN01B091
IN01B092
IN01B100
IN01B081
IN01B078
IN01B073
IN01B074
AN01B018
IN01B065
IN01B070
IN01B098
IN01B102
IN01B095
IN01B089
IN01B090
IN01B094
IN01B093
IN01B079
IN01B087
IN01B085
IN01B097
IN01B044
IN01B042
IN01B048
IN01B038
IN01B039
IN01B025
IN01B056
IN01B058
IN01B024
IN01B071
IN01B016
IN01B028
IN01B059
IN01B049
IN01B053
IN01B057
IN01B072
IN01B077
IN01B101
IN01B096
IN01B082
IN01B083
IN01B084
AN01B011
IN01B032
IN01B033
IN01B040
IN01B060
IN01B067
IN01B069
IN01B055
IN01B066

**Figure 17 - figure supplement 7a. Connectivity to downstream partners by 01B secondary systematic types.** Proportions of synaptic weight from systematic types to downstream partners, normalised by row. 01B neurons have been clustered within each assigned birthtime window (P = primary, ES = early secondary, S = secondary) based on both upstream and downstream connectivity to hemilineages, descending neuron subclasses, motor neuron subclasses, and sensory neuron modalities. The annotation bar is coloured by the most common predicted neurotransmitter for the neurons of each type.

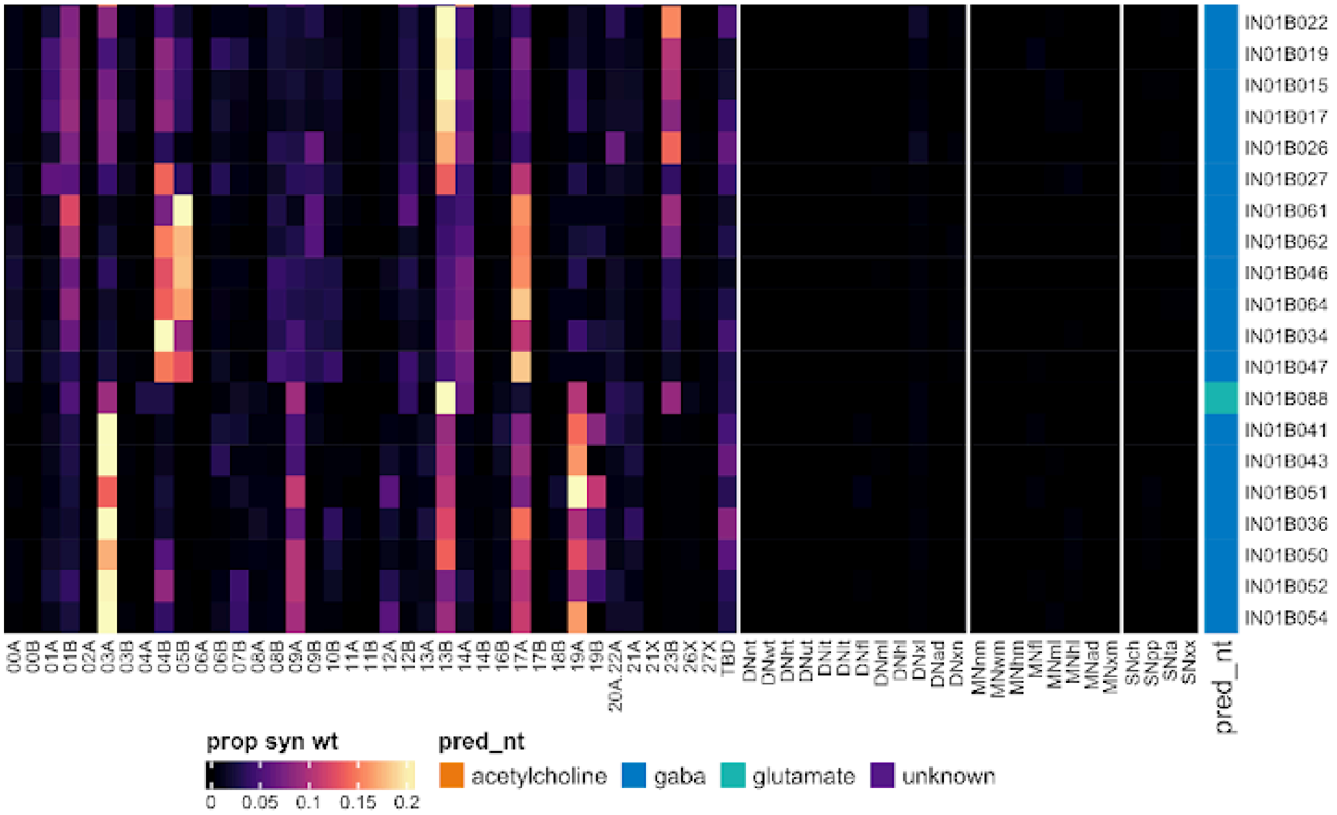

**Figure 17 - figure supplement 7b. Connectivity to downstream partners by 01B secondary systematic types, continued.** Proportions of synaptic weight from systematic types to downstream partners, normalised by row. 01B neurons have been clustered within each assigned birthtime window (P = primary, ES = early secondary, S = secondary) based on both upstream and downstream connectivity to hemilineages, descending neuron subclasses, motor neuron subclasses, and sensory neuron modalities. The annotation bar is coloured by the most common predicted neurotransmitter for the neurons of each type.

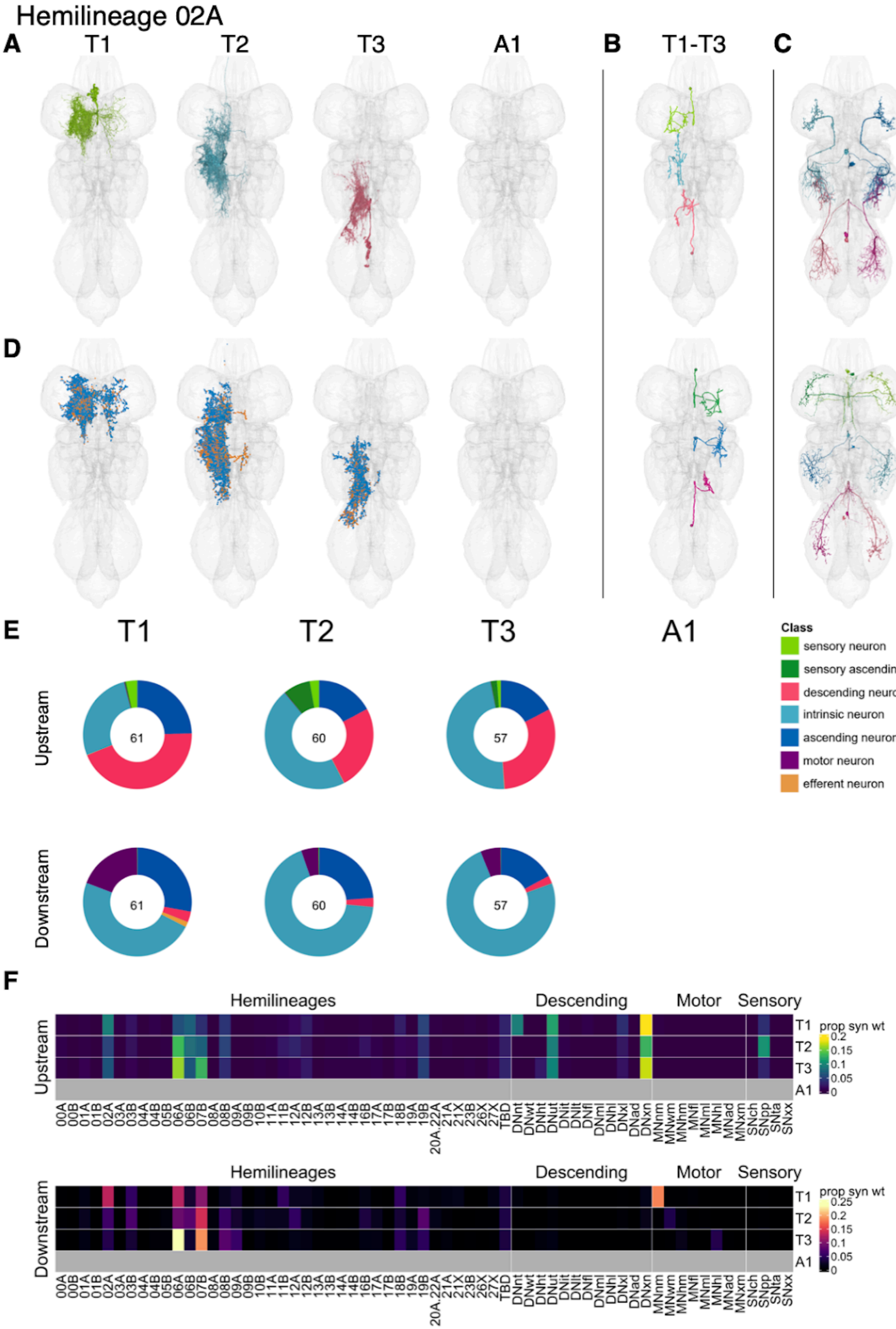

Hemilineage 02A
A
T1
T2
T3
A1
B
T1-T3
C
D
E
T1
T2
T3
A1
Upstream
61
60
57
Downstream
61
60
57
Class
sensory neuron
sensory ascending
descending neuron
intrinsic neuron
ascending neuron
motor neuron
efferent neuron
F
Hemilineages
Descending
Motor
Sensory
Upstream
T1
T2
T3
A1
prop syn wt
0.2
0.15
0.1
0.05
0
00A 00B 01A 01B 02A 03A 03B 04A 04B 05B 06A 06B 07B 08A 08B 09A 09B 10B 11A 11B 12A 12B 13A 13B 14A 14B 16B 17A 17B 18B 19A 19B 20A.22A 21A 21X 23B 26X 27X TBD DNnt DNwt DNht DNut DNit DNlt DNfl DNml DNhl DNxl DNad DNxn MNnm MNwm MNhm MNfl MNml MNhl MNad MNxm SNch SNpp SNta SNxx
Hemilineages
Descending
Motor
Sensory
Downstream
T1
T2
T3
A1
prop syn wt
0.25
0.2
0.15
0.1
0.05
0
00A 00B 01A 01B 02A 03A 03B 04A 04B 05B 06A 06B 07B 08A 08B 09A 09B 10B 11A 11B 12A 12B 13A 13B 14A 14B 16B 17A 17B 18B 19A 19B 20A.22A 21A 21X 23B 26X 27X TBD DNnt DNwt DNht DNut DNit DNlt DNfl DNml DNhl DNxl DNad DNxn MNnm MNwm MNhm MNfl MNml MNhl MNad MNxm SNch SNpp SNta SNxx

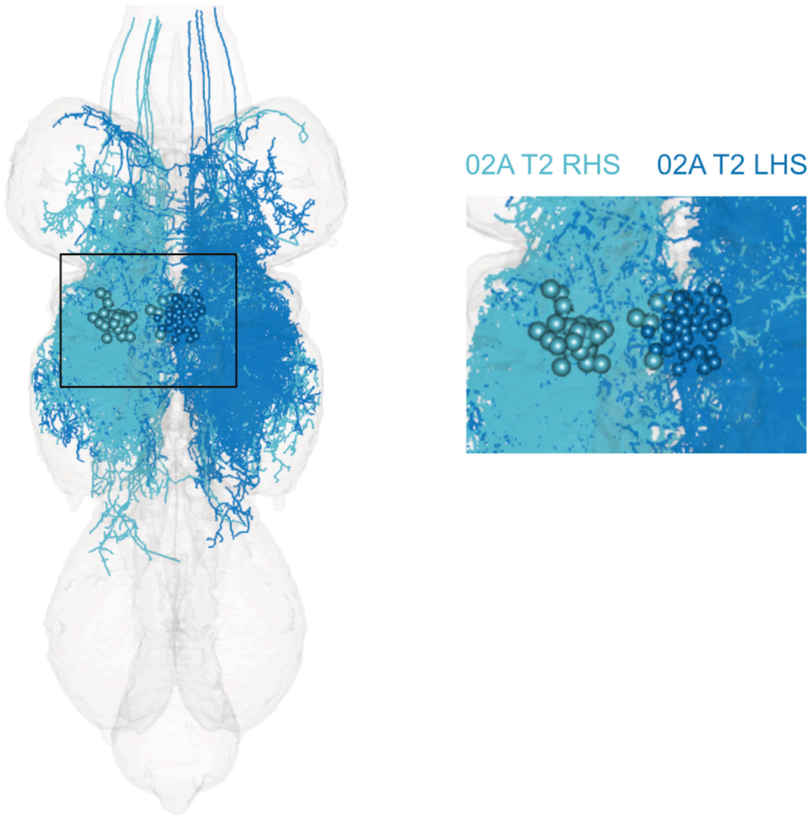

**Figure 18 - figure supplement 1. Developmental aberration in MANC of 02A neurons in RHS T2.** Most 02A RHS somas (large, cyan) have been displaced laterally from their expected position adjacent to the midline (inset), but their gross dendritic and axonal morphologies appear normal when compared to their LHS counterparts (small, blue).

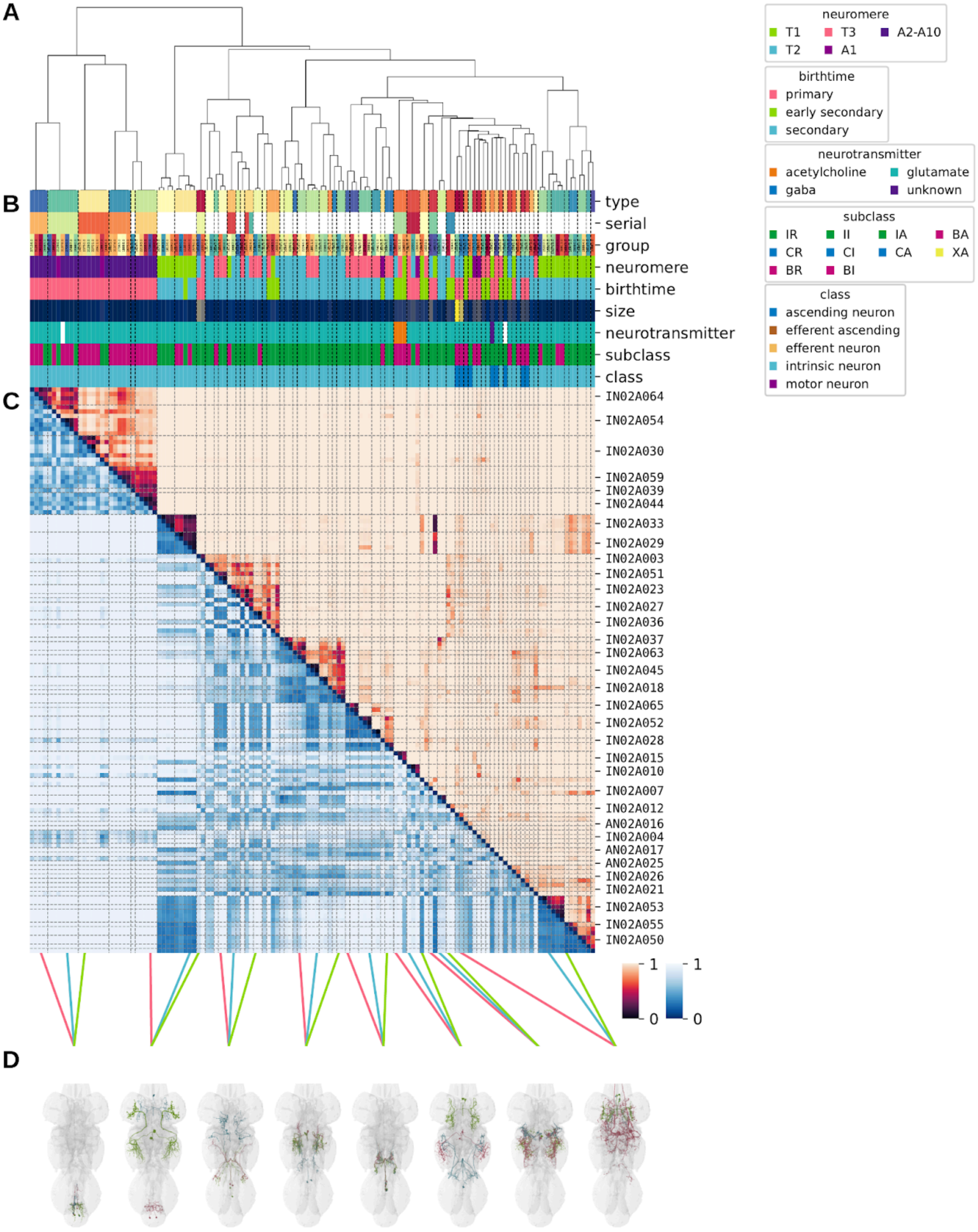

**Figure 18 - figure supplement 2. Systematic typing of hemilineage 02A. A.** Hierarchical clustering dendrogram of hemilineage groups by laterally and serially aggregated connectivity cosine clustering. **B.** Categorical annotations of each hemilineage group, each column corresponding to the aligned leaf in A. Colours for type, serial set, and group are arbitrary for visualisation. Colours for neuromere, birthtime, neurotransmitter, subclass, and class are as in all other figures. **C.** Similarity distance heatmap for hemilineage. Cosine distance is in the upper triangle, while laterally symmetrised NBLAST distance is in the lower triangle. Systematic type names of some types are labelled. **D.**

Morphologically representative groups from dendrogram subtrees. Each group, indicated by colour and line connecting to its column in B and C, is the most morphologically representative group (medoid of NBLAST distance) from a subtree of A. The subtrees (flat clusters) are equal height cuts of A determined to yield the number of groups per plot and plots in D.

#### 02A systematic types

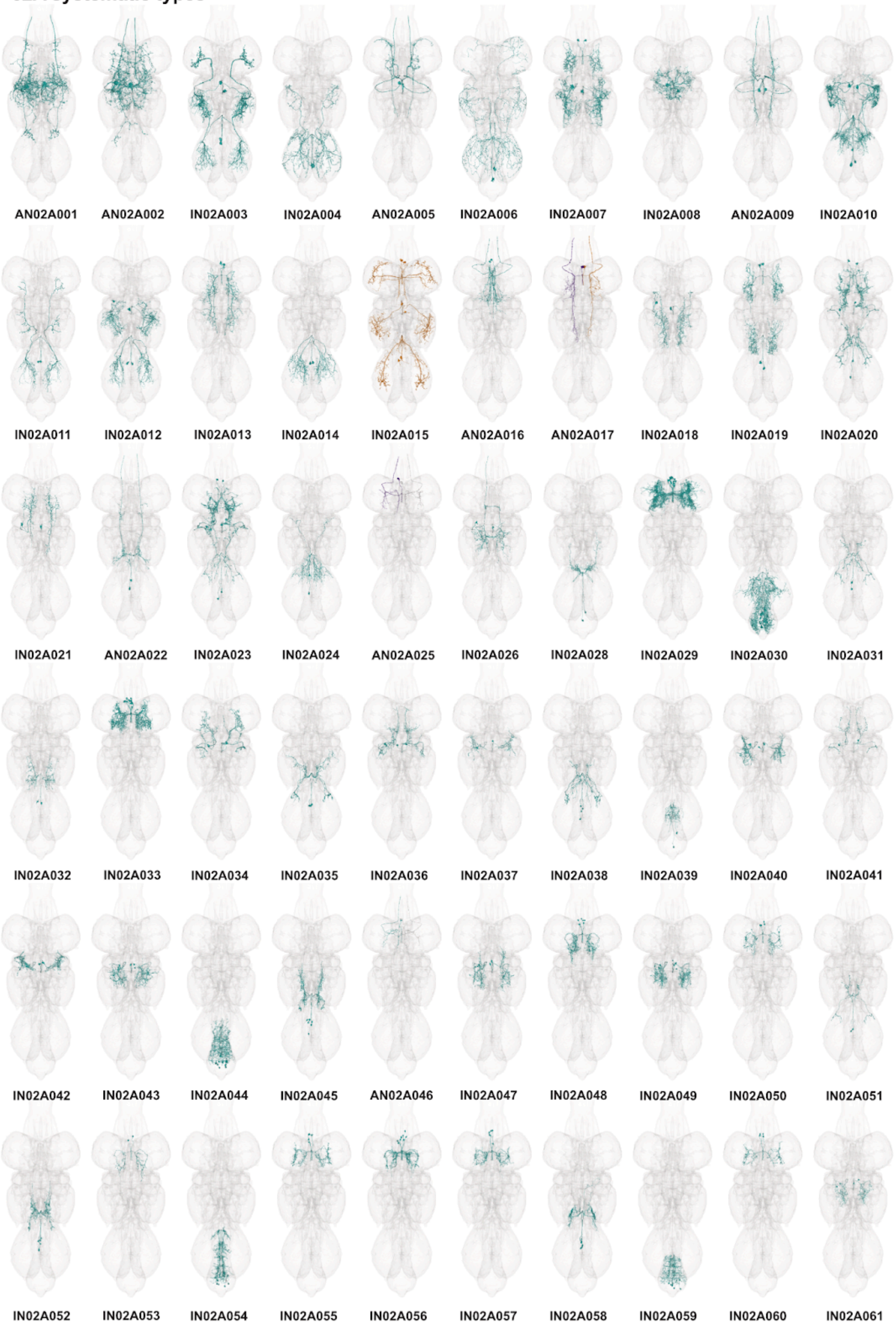

**Figure 18 - figure supplement 3. Systematic types of hemilineage 02A.** Systematic types have been arranged in numerical order, with neurons of the same type that belong to distinct classes (e.g., intrinsic neuron vs ascending neuron) plotted separately but placed adjacent to each other. Individual neuron meshes have been coloured based on predicted neurotransmitter: dark orange = acetylcholine, blue = gaba, marine = glutamate, dark purple = unknown.

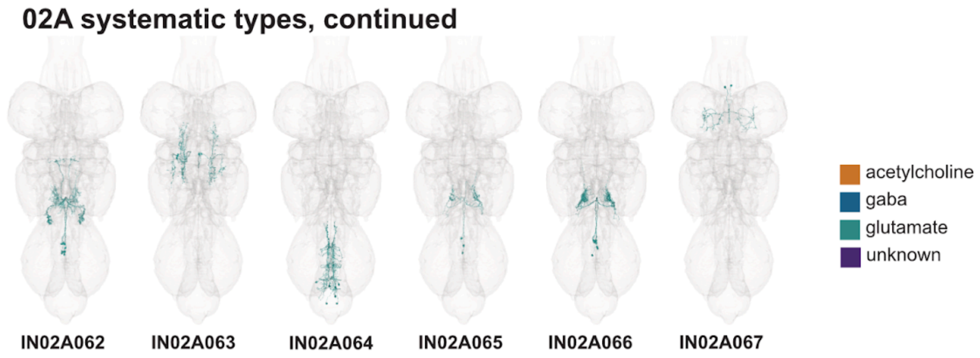

**Figure 18 - figure supplement 4. Systematic types of hemilineage 02A, continued.** Systematic types have been arranged in numerical order, with neurons of the same type that belong to distinct classes (e.g., intrinsic neuron vs ascending neuron) plotted separately but placed adjacent to each other. Individual neuron meshes have been coloured based on predicted neurotransmitter: dark orange = acetylcholine, blue = gaba, marine = glutamate, dark purple = unknown.

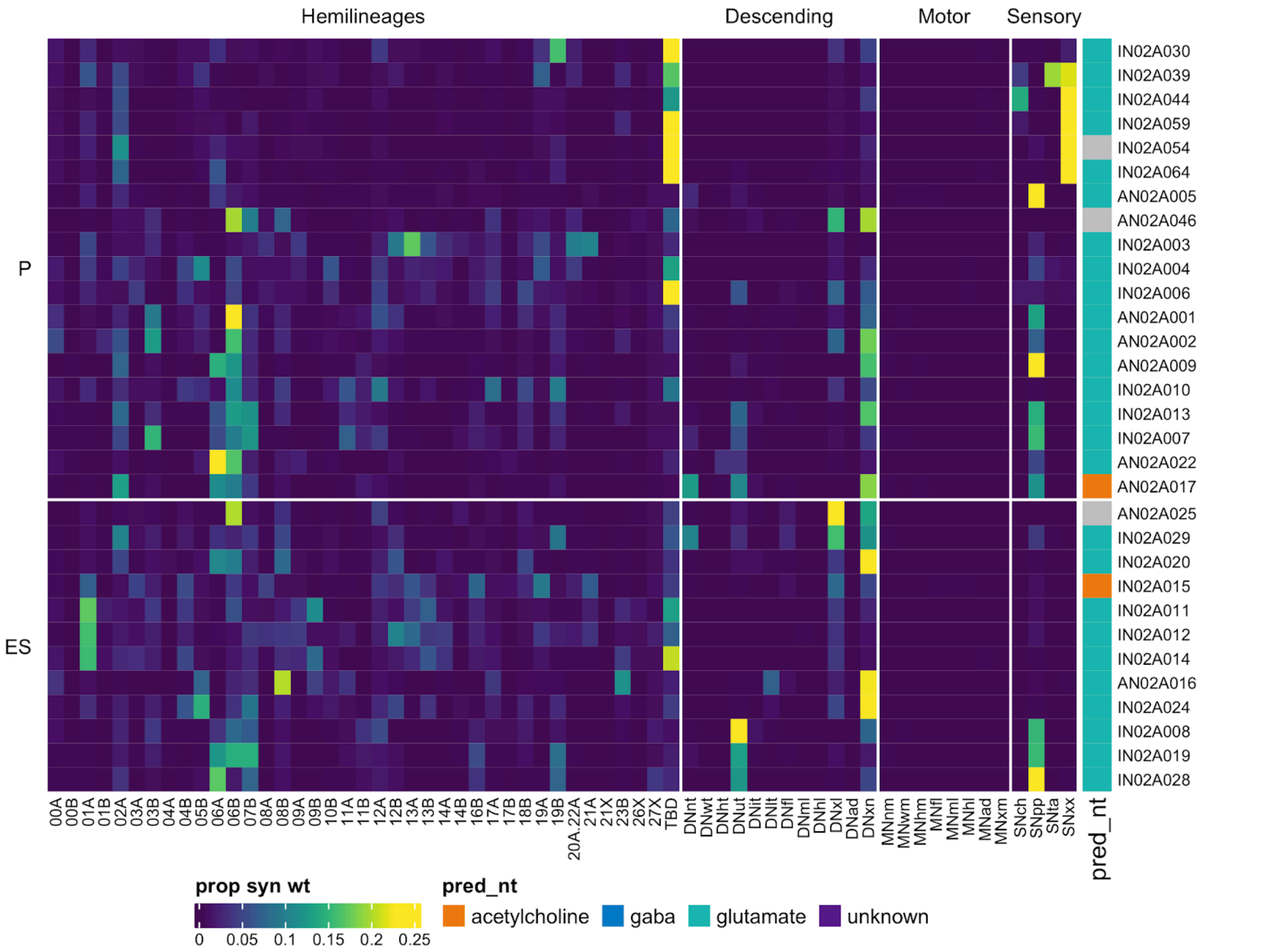

**Figure 18 - figure supplement 5. Connectivity to upstream partners by 02A primary and early secondary systematic types.** Proportions of synaptic weight to systematic types from upstream partners, normalised by row. 02A neurons have been clustered within each assigned birthtime window (P = primary, ES = early secondary, S = secondary) based on both upstream and downstream connectivity to hemilineages, descending neuron subclasses, motor neuron subclasses, and sensory neuron modalities. Annotation bar is coloured by the most common predicted neurotransmitter for the neurons of each type.

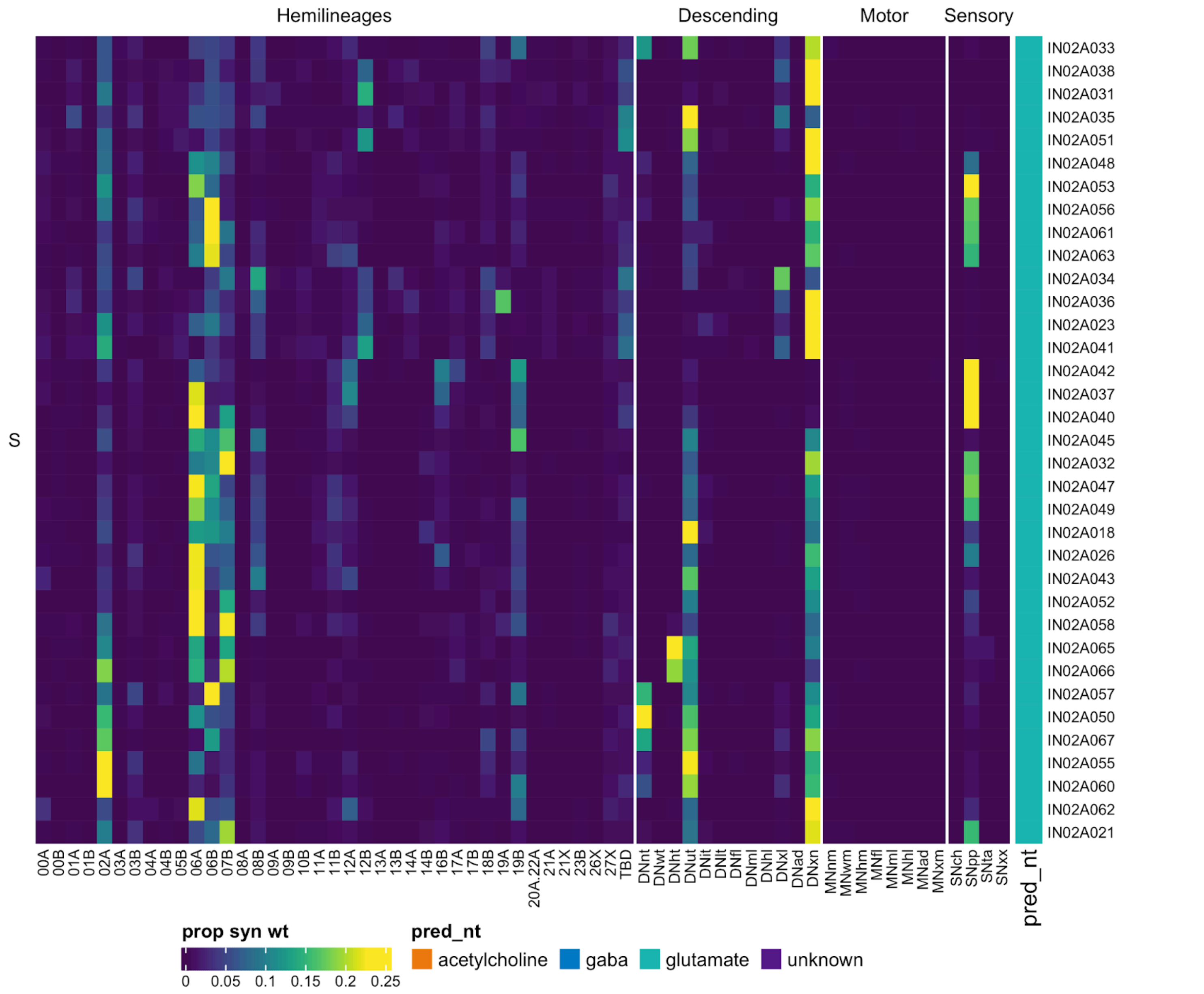

**Figure 18 - figure supplement 6. Connectivity to upstream partners by 02A secondary systematic types.** Proportions of synaptic weight to systematic types from upstream partners, normalised by row. 02A neurons have been clustered within each assigned birthtime window (P = primary, ES = early secondary, S = secondary) based on both upstream and downstream connectivity to hemilineages, descending neuron subclasses, motor neuron subclasses, and sensory neuron modalities. The annotation bar is coloured by the most common predicted neurotransmitter within each type.

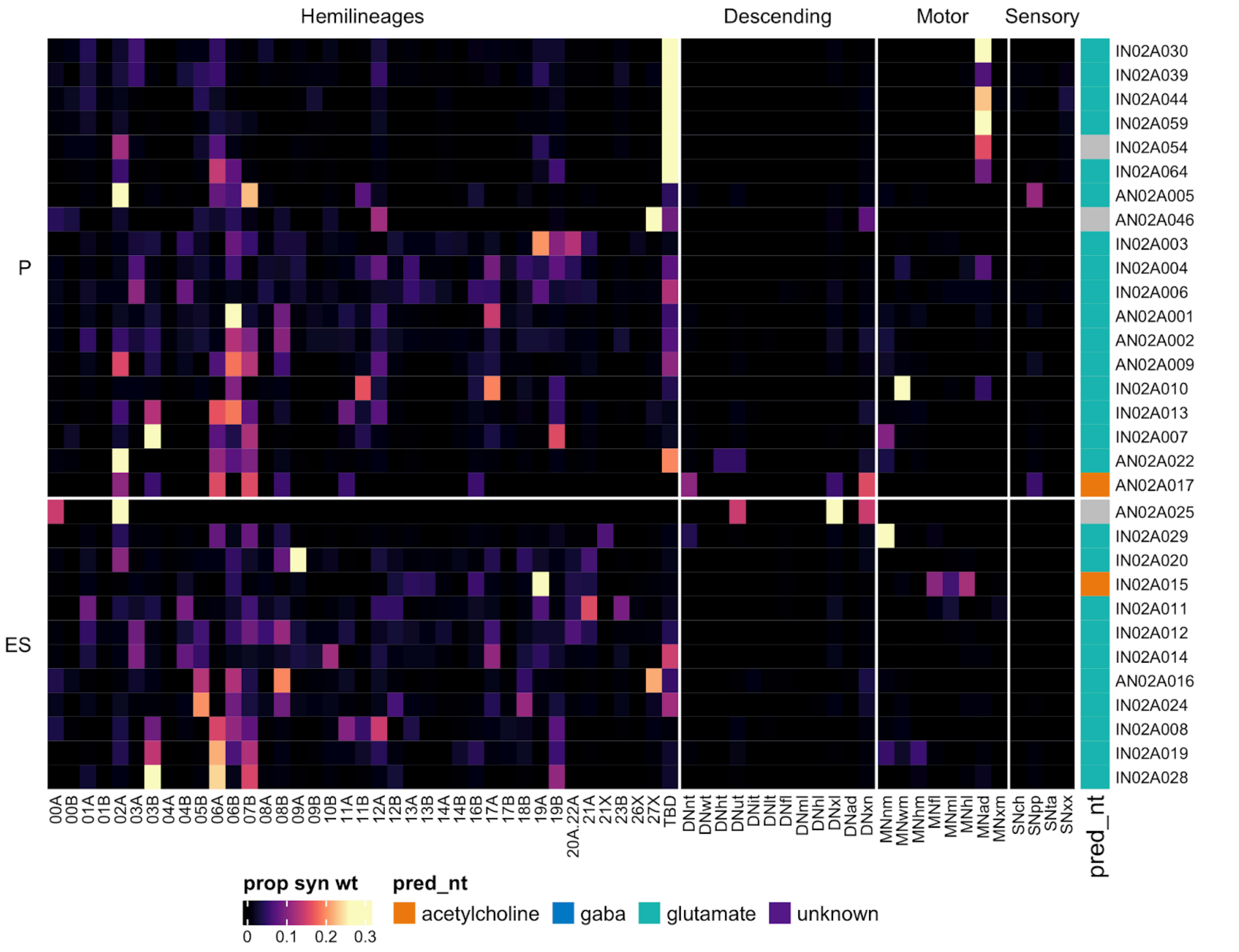

**Figure 18 - figure supplement 7. Connectivity to downstream partners by 02A primary and early secondary systematic types.** Proportions of synaptic weight from systematic types to downstream partners, normalised by row. 02A neurons have been clustered within each assigned birthtime window (P = primary, ES = early secondary, S = secondary) based on both upstream and downstream connectivity to hemilineages, descending neuron subclasses, motor neuron subclasses, and sensory neuron modalities. The annotation bar is coloured by the most common predicted neurotransmitter within each type.

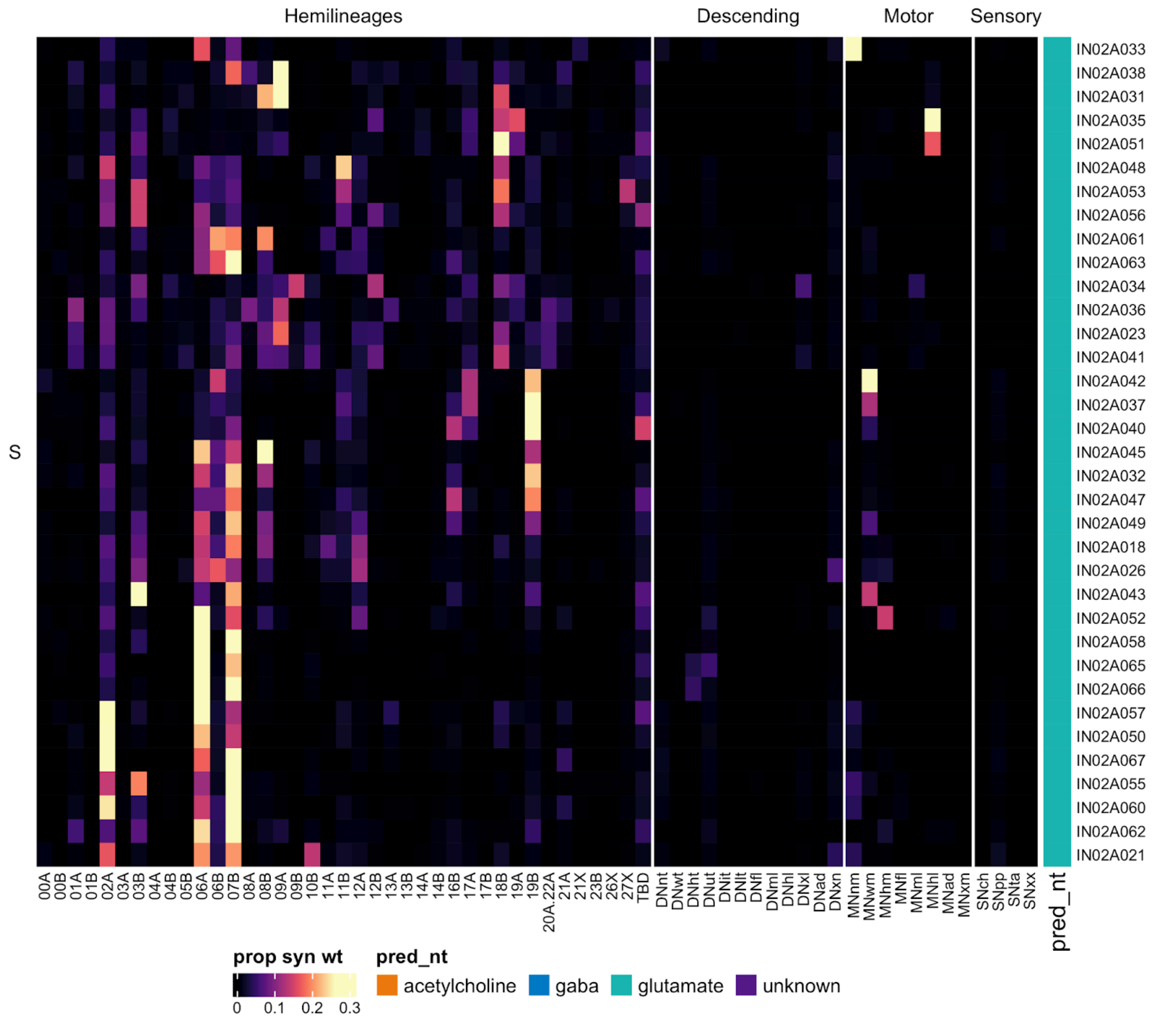

**Figure 18 - figure supplement 8.Connectivity to downstream partners by 02A secondary systematic types.** Proportions of synaptic weight from systematic types to downstream partners, normalised by row. 02A neurons have been clustered within each assigned birthtime window (P = primary, ES = early secondary, S = secondary) based on both upstream and downstream connectivity to hemilineages, descending neuron subclasses, motor neuron subclasses, and sensory neuron modalities. The annotation bar is coloured by the most common predicted neurotransmitter for the neurons of each type.

### Hemilineage 03A

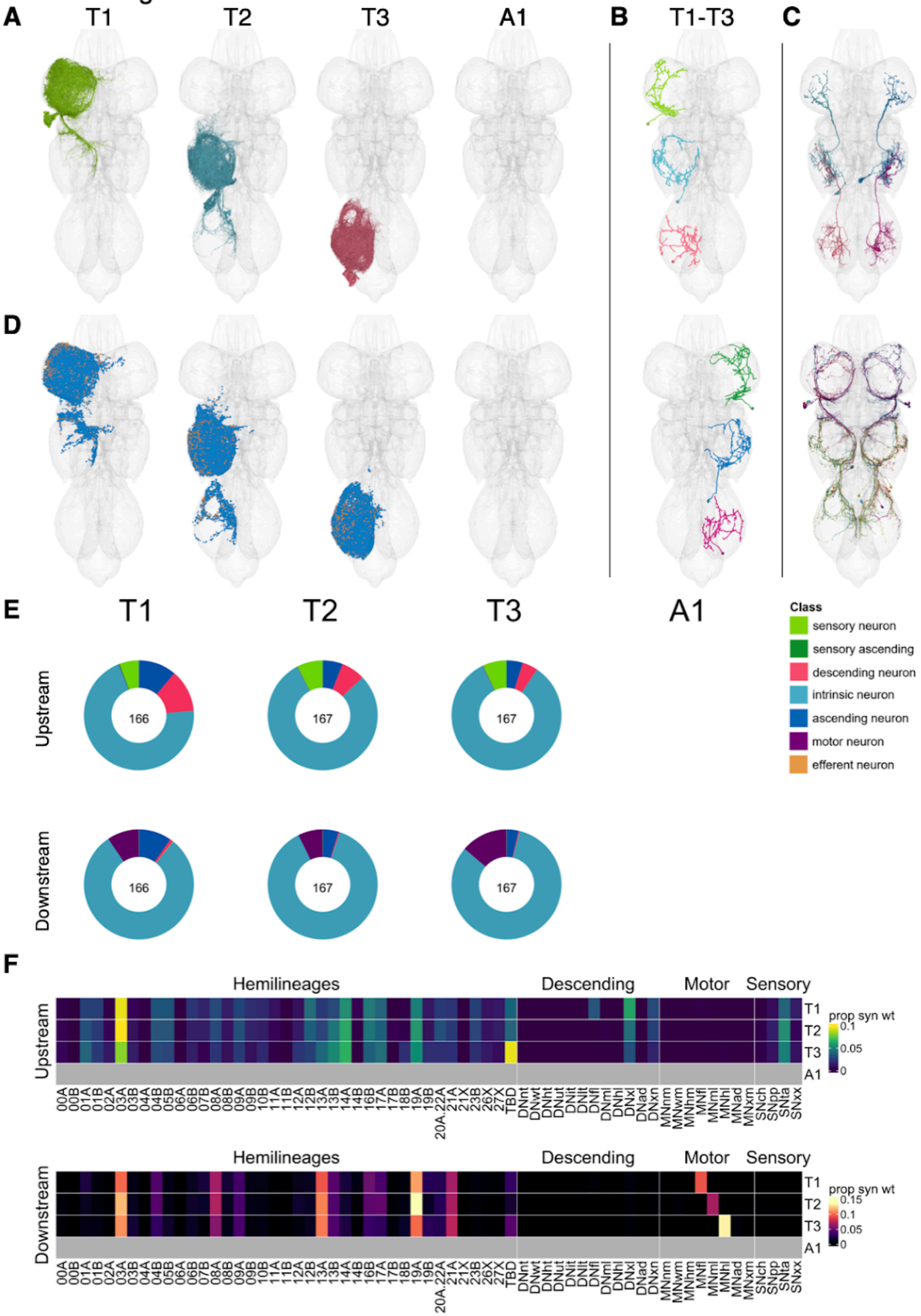

**Figure 19. Hemilineage 03A. A.** Meshes of all RHS secondary neurons plotted in neuromere-specific colours. **B.** “Representative” secondary neuron skeletons plotted in hemineuromere-specific colours. The skeleton with the top accumulated NBLAST score among all neurons from the hemilineage in a given hemineuromere was used. **C.** Neuron meshes of selected examples. Top: sequential serial set 10262. Bottom: sequential subcluster 15430. **D.** Predicted synapses of RHS secondary neurons. Blue: postsynapses; dark orange: presynapses. **E.** Proportions of connections from secondary neurons to upstream or downstream partners, normalised by neuromere and coloured by broad class. Numbers of query neurons appear in the centre. **F.** Proportions of synaptic weight from secondary neurons originating in each neuromere to upstream or downstream partners, normalised by row.

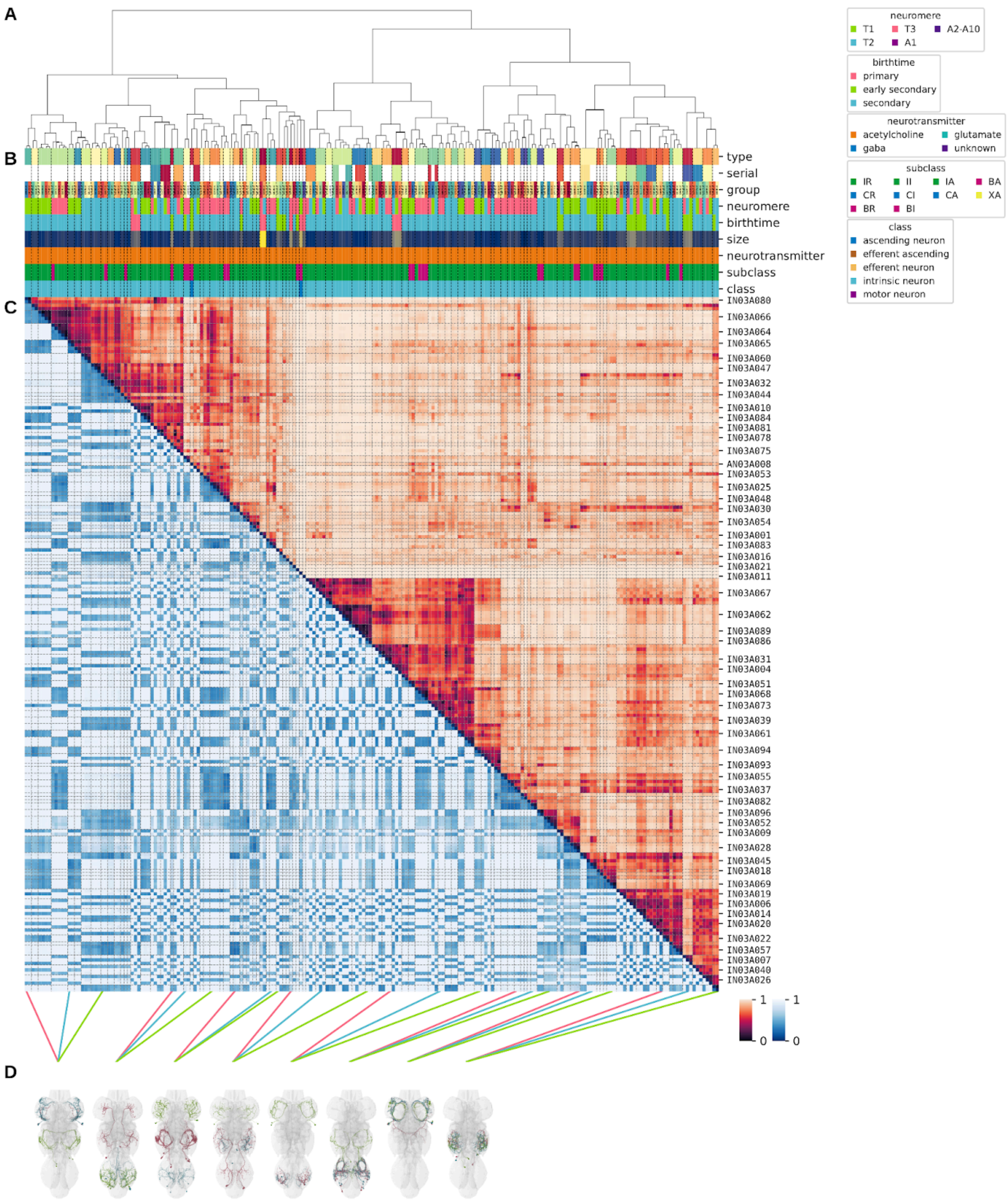

**Figure 19 - figure supplement 1. Systematic typing of hemilineage 03A. A.** Hierarchical clustering dendrogram of hemilineage groups by laterally and serially aggregated connectivity cosine clustering. **B.** Categorical annotations of each hemilineage group, each column corresponding to the aligned leaf in A. Colours for type, serial set, and group are arbitrary for visualisation. Colours for neuromere, birthtime, neurotransmitter, subclass, and class are as in all other figures. **C.** Similarity distance heatmap for hemilineage. Cosine distance is in the upper triangle, while laterally symmetrised NBLAST distance is in the lower triangle. Systematic type names of some types are labelled. **D.** Morphologically representative groups from dendrogram subtrees. Each group, indicated by colour and line connecting to its column in B and C, is the most morphologically representative group (medoid of NBLAST distance) from a subtree of A. The subtrees (flat clusters) are equal height cuts of A determined to yield the number of groups

per plot and plots in D.

#### 03A systematic types

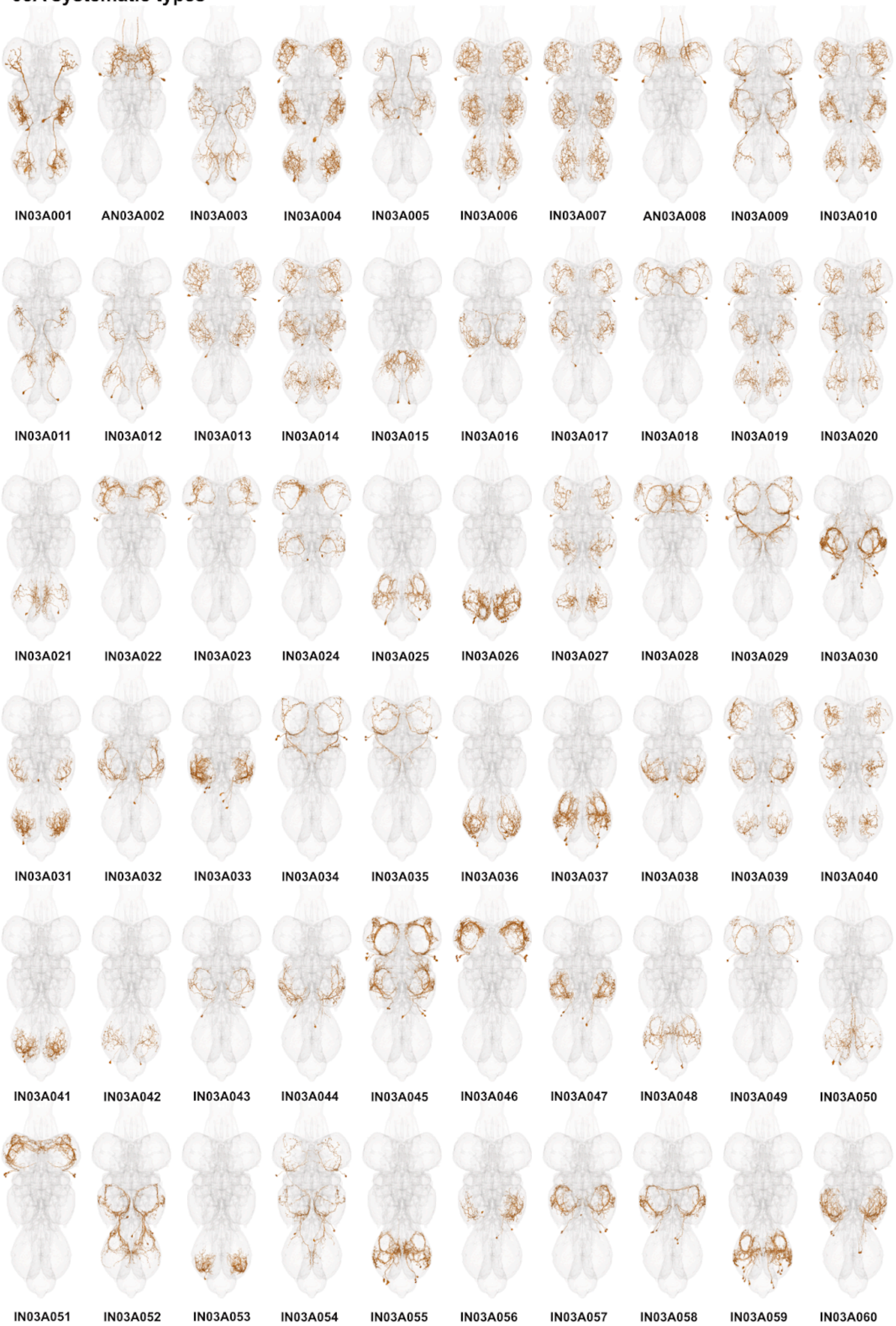

**Figure 19 - figure supplement 2. Systematic types of hemilineage 03A.** Systematic types have been arranged in numerical order, with neurons of the same type that belong to distinct classes (e.g., intrinsic neuron vs ascending neuron) plotted separately but placed adjacent to each other. Individual neuron meshes have been coloured based on predicted neurotransmitter: dark orange = acetylcholine, blue = gaba, marine = glutamate, dark purple = unknown.

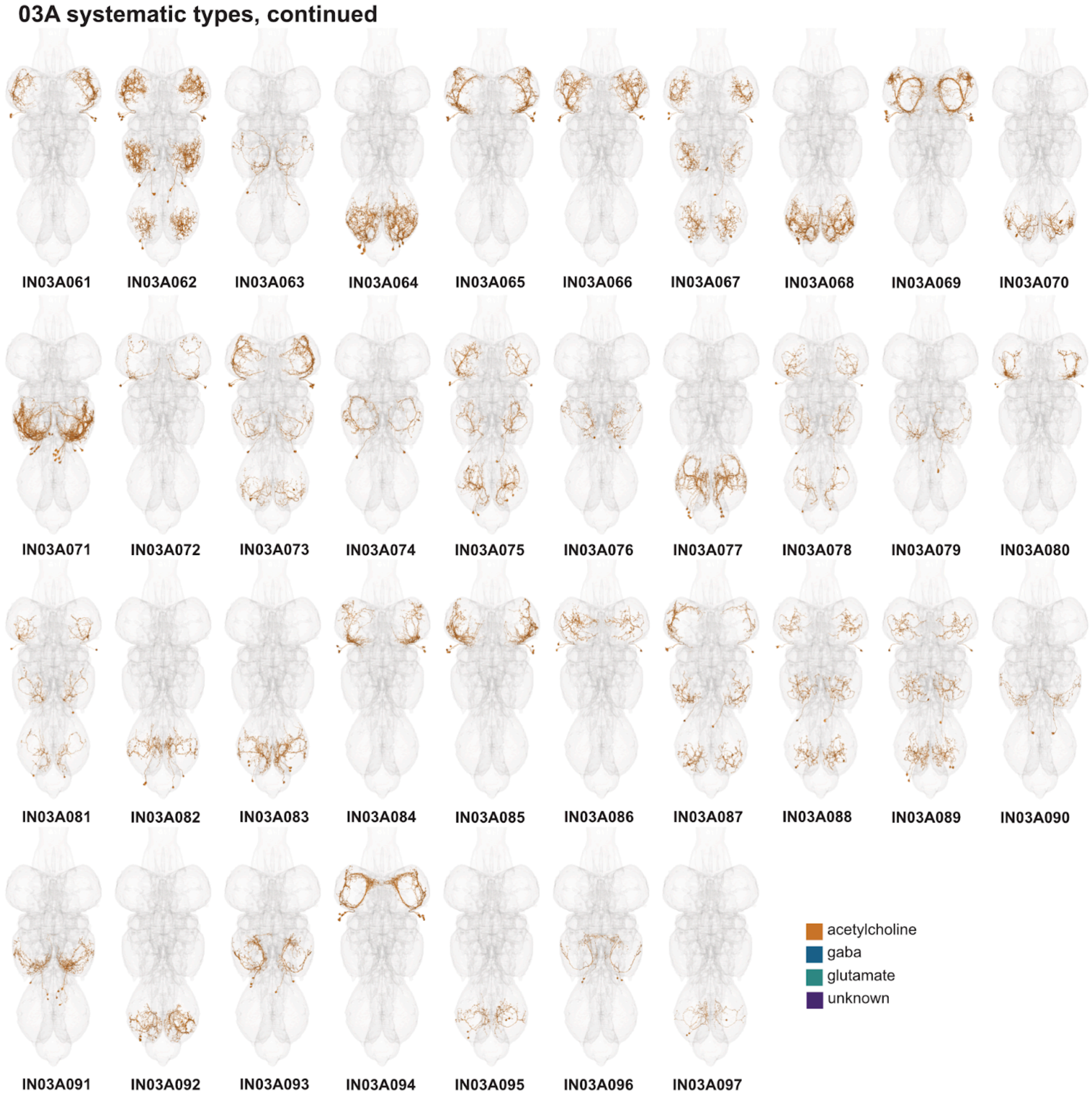

**Figure 19 - figure supplement 3. Systematic types of hemilineage 03A, continued.** Systematic types have been arranged in numerical order, with neurons of the same type that belong to distinct classes (e.g., intrinsic neuron vs ascending neuron) plotted separately but placed adjacent to each other. Individual neuron meshes have been coloured based on predicted neurotransmitter: dark orange = acetylcholine, blue = gaba, marine = glutamate, dark purple = unknown.

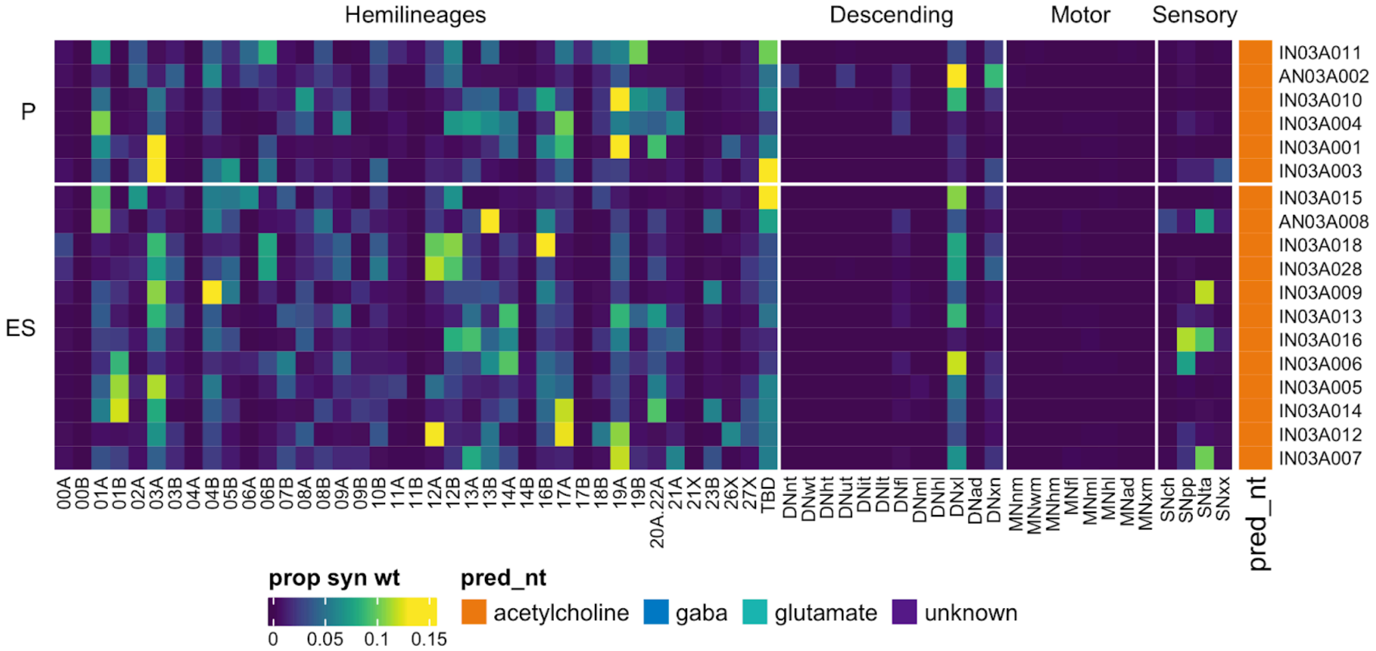

**Figure 19 - figure supplement 4. Connectivity to upstream partners by 03A primary and early secondary systematic types.** Proportions of synaptic weight to systematic types from upstream partners, normalised by row. 03A neurons have been clustered within each assigned birthtime window (P = primary, ES = early secondary, S = secondary) based on both upstream and downstream connectivity to hemilineages, descending neuron subclasses, motor neuron subclasses, and sensory neuron modalities. Annotation bar is coloured by the most common predicted neurotransmitter for the neurons of each type.

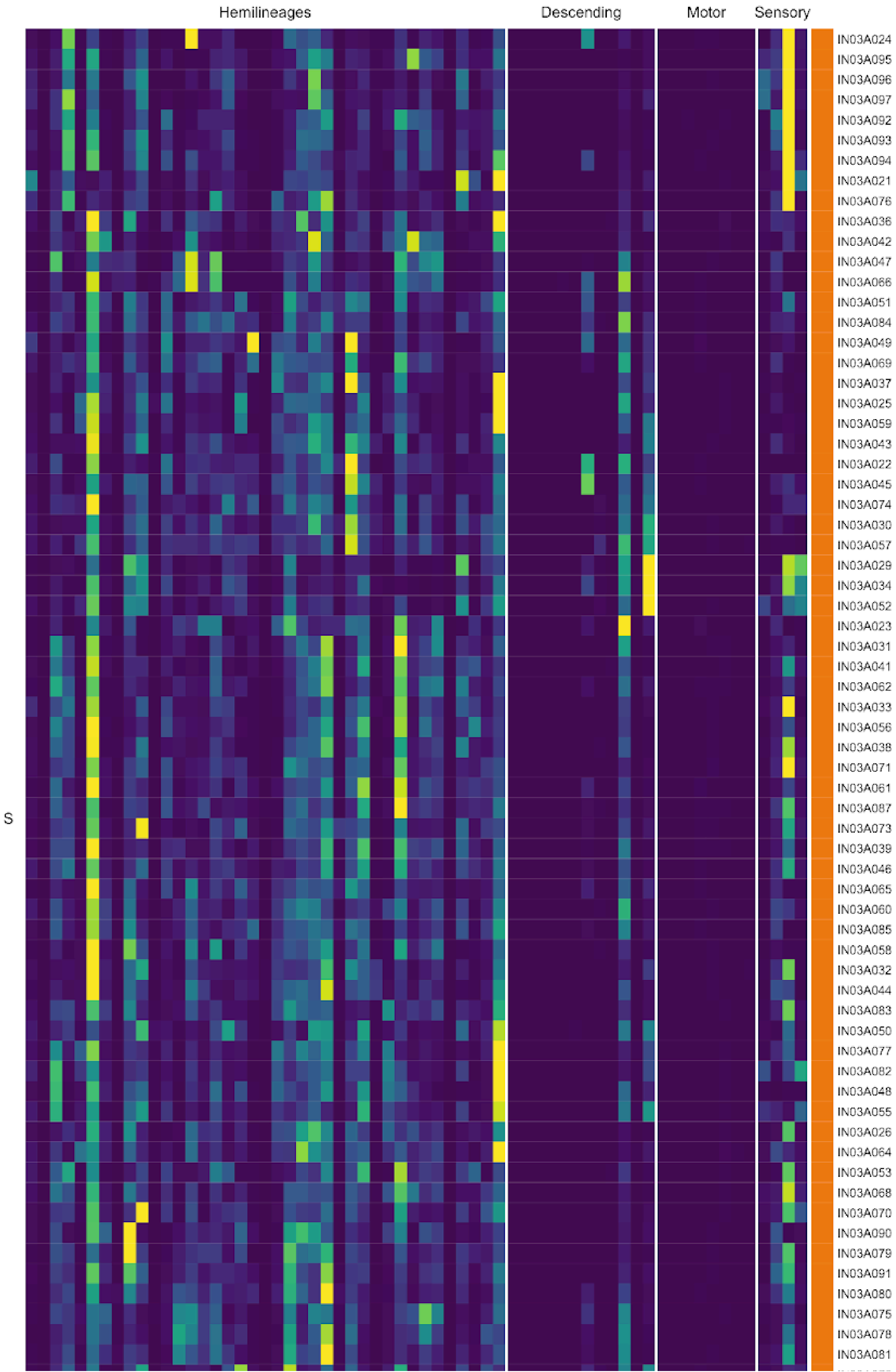
Hemilineages
Descending
Motor
Sensory
S
IN03A024
IN03A095
IN03A096
IN03A097
IN03A092
IN03A093
IN03A094
IN03A021
IN03A076
IN03A036
IN03A042
IN03A047
IN03A066
IN03A051
IN03A084
IN03A049
IN03A069
IN03A037
IN03A025
IN03A059
IN03A043
IN03A022
IN03A045
IN03A074
IN03A030
IN03A057
IN03A029
IN03A034
IN03A052
IN03A023
IN03A031
IN03A041
IN03A062
IN03A033
IN03A056
IN03A038
IN03A071
IN03A061
IN03A087
IN03A073
IN03A039
IN03A046
IN03A065
IN03A060
IN03A085
IN03A058
IN03A032
IN03A044
IN03A083
IN03A050
IN03A077
IN03A082
IN03A048
IN03A055
IN03A026
IN03A064
IN03A053
IN03A068
IN03A070
IN03A090
IN03A079
IN03A091
IN03A080
IN03A075
IN03A078
IN03A081

**Figure 19 - figure supplement 5. Connectivity to upstream partners by 03A secondary systematic types.** Proportions of synaptic weight to systematic types from upstream partners, normalised by row. 03A neurons have been clustered within each assigned birthtime window (P = primary, ES = early secondary, S = secondary) based on both upstream and downstream connectivity to hemilineages, descending neuron subclasses, motor neuron subclasses, and sensory neuron modalities. The annotation bar is coloured by the most common predicted neurotransmitter within each type.

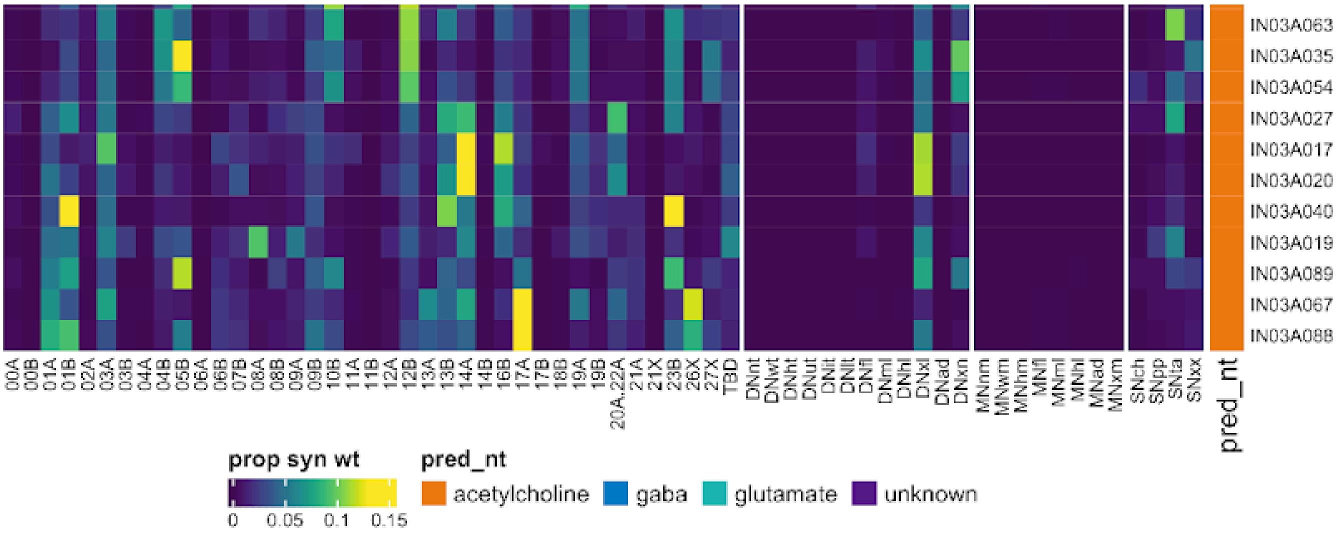

**Figure 19 - figure supplement 6. Connectivity to upstream partners by 03A secondary systematic types, continued.** Proportions of synaptic weight to systematic types from upstream partners, normalised by row. 03A neurons have been clustered within each assigned birthtime window (P = primary, ES = early secondary, S = secondary) based on both upstream and downstream connectivity to hemilineages, descending neuron subclasses, motor neuron subclasses, and sensory neuron modalities. The annotation bar is coloured by the most common predicted neurotransmitter within each type.

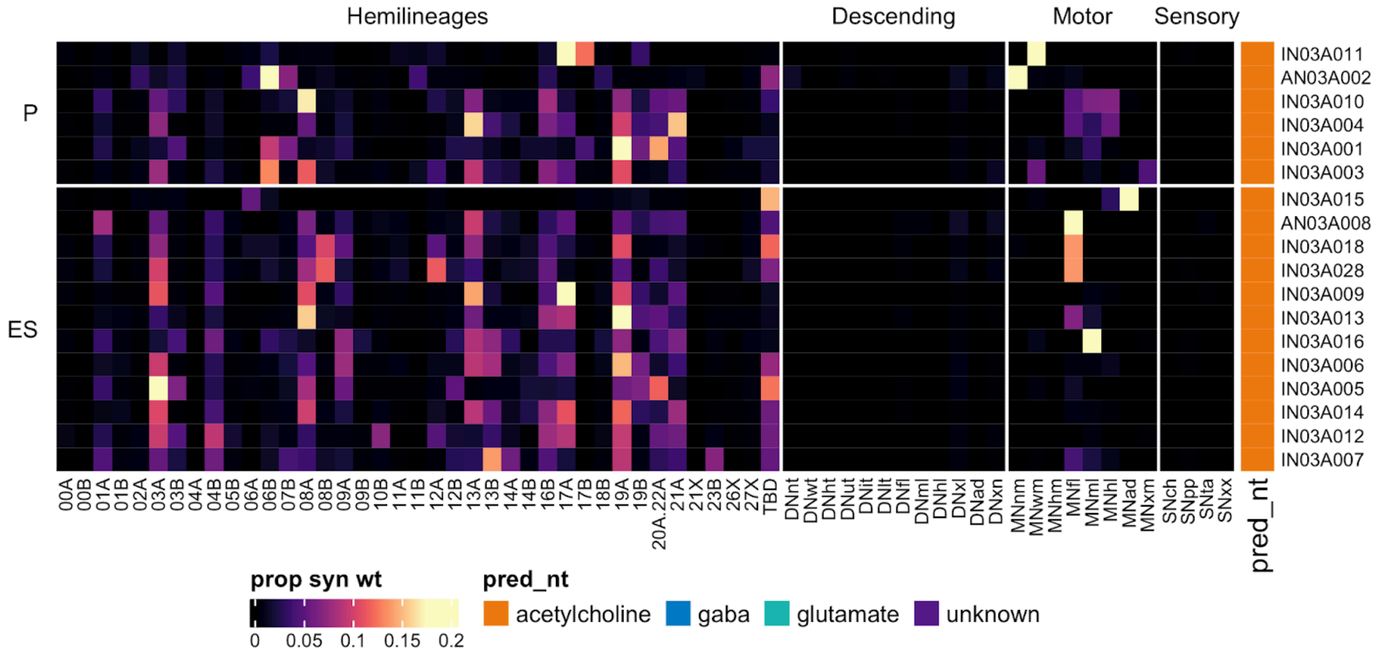

**Figure 19 - figure supplement 7. Connectivity to downstream partners by 03A primary and early secondary systematic types.** Proportions of synaptic weight from systematic types to downstream partners, normalised by row. 03A neurons have been clustered within each assigned birthtime window (P = primary, ES = early secondary, S = secondary) based on both upstream and downstream connectivity to hemilineages, descending neuron subclasses, motor neuron subclasses, and sensory neuron modalities. The annotation bar is coloured by the most common predicted neurotransmitter within each type.

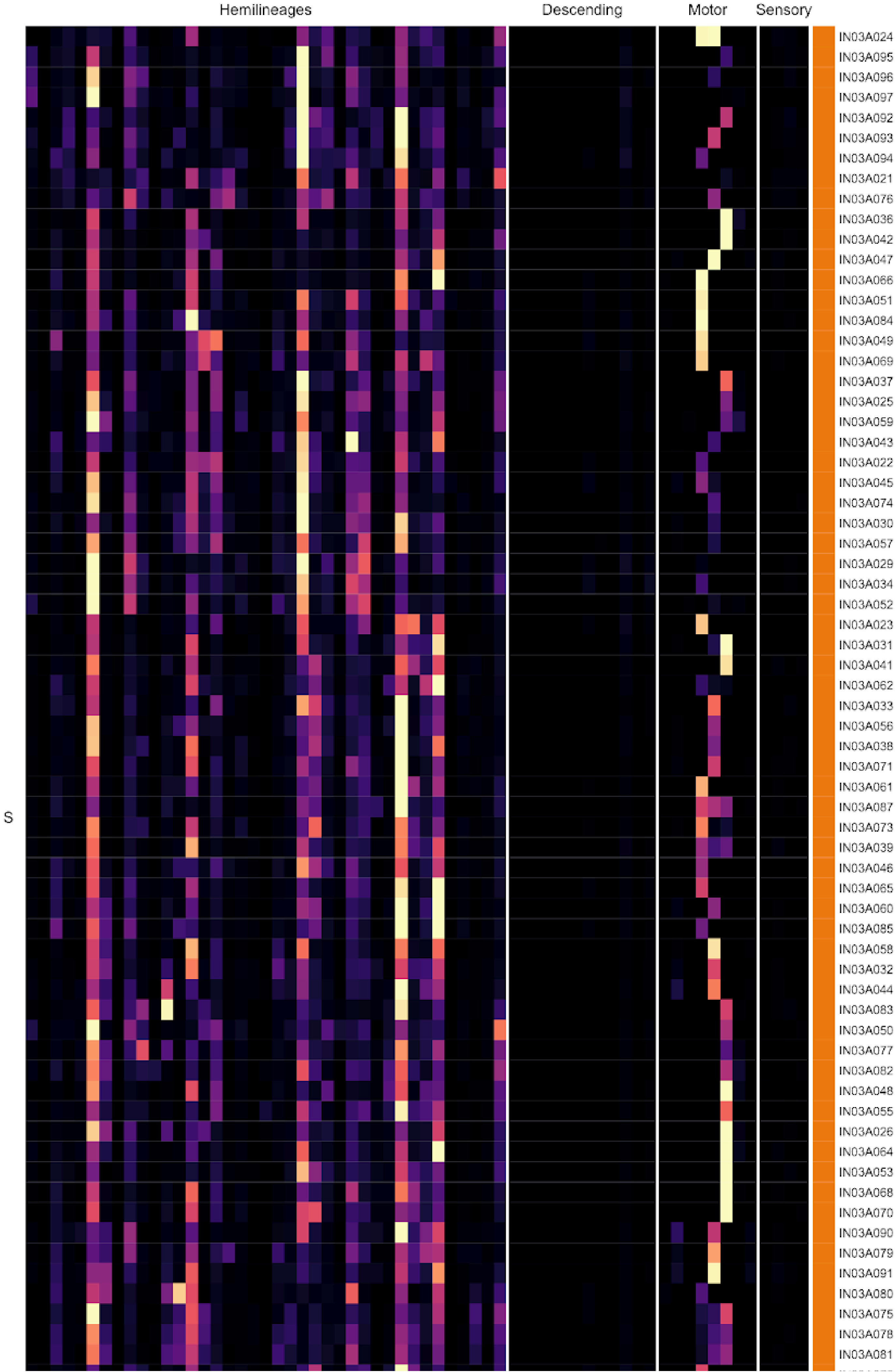
Hemilineages
Descending
Motor
Sensory
S
IN03A024
IN03A095
IN03A096
IN03A097
IN03A092
IN03A093
IN03A094
IN03A021
IN03A076
IN03A036
IN03A042
IN03A047
IN03A066
IN03A051
IN03A084
IN03A049
IN03A069
IN03A037
IN03A025
IN03A059
IN03A043
IN03A022
IN03A045
IN03A074
IN03A030
IN03A057
IN03A029
IN03A034
IN03A052
IN03A023
IN03A031
IN03A041
IN03A062
IN03A033
IN03A056
IN03A038
IN03A071
IN03A061
IN03A087
IN03A073
IN03A039
IN03A046
IN03A065
IN03A060
IN03A085
IN03A058
IN03A032
IN03A044
IN03A083
IN03A050
IN03A077
IN03A082
IN03A048
IN03A055
IN03A026
IN03A064
IN03A053
IN03A068
IN03A070
IN03A090
IN03A079
IN03A091
IN03A080
IN03A075
IN03A078
IN03A081

**Figure 19 - figure supplement 8. Connectivity to downstream partners by 03A secondary systematic types.** Proportions of synaptic weight from systematic types to downstream partners, normalised by row. 03A neurons have been clustered within each assigned birthtime window (P = primary, ES = early secondary, S = secondary) based on both upstream and downstream connectivity to hemilineages, descending neuron subclasses, motor neuron subclasses, and sensory neuron modalities. The annotation bar is coloured by the most common predicted neurotransmitter for the neurons of each type.

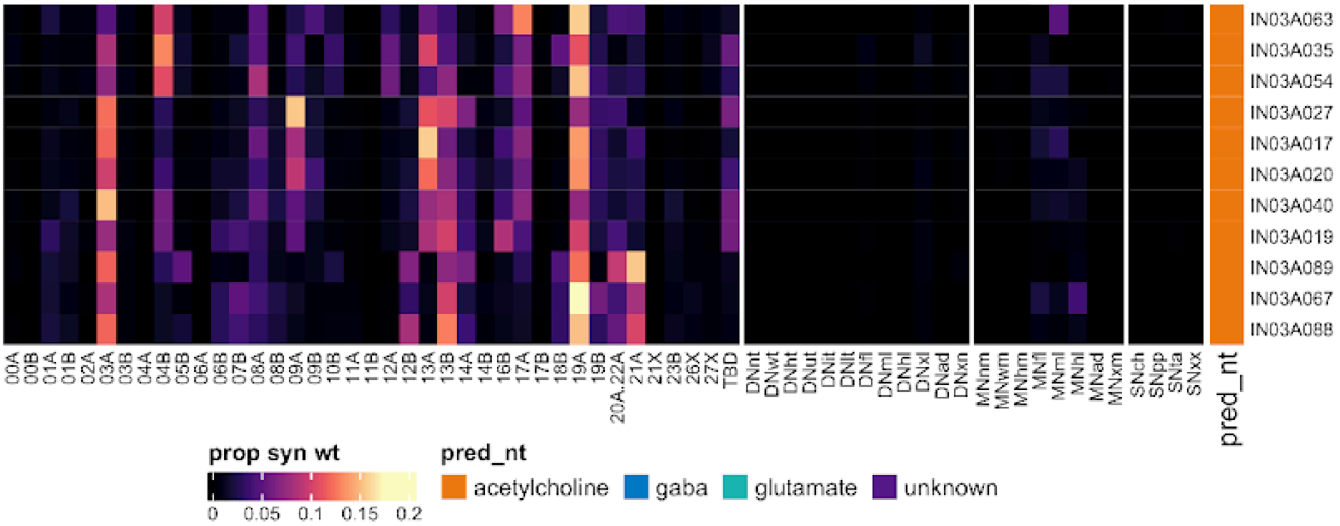

**Figure 19 - figure supplement 9. Connectivity to downstream partners by 03A secondary systematic types, continued.** Proportions of synaptic weight from systematic types to downstream partners, normalised by row. 03A neurons have been clustered within each assigned birthtime window (P = primary, ES = early secondary, S = secondary) based on both upstream and downstream connectivity to hemilineages, descending neuron subclasses, motor neuron subclasses, and sensory neuron modalities. The annotation bar is coloured by the most common predicted neurotransmitter for the neurons of each type.

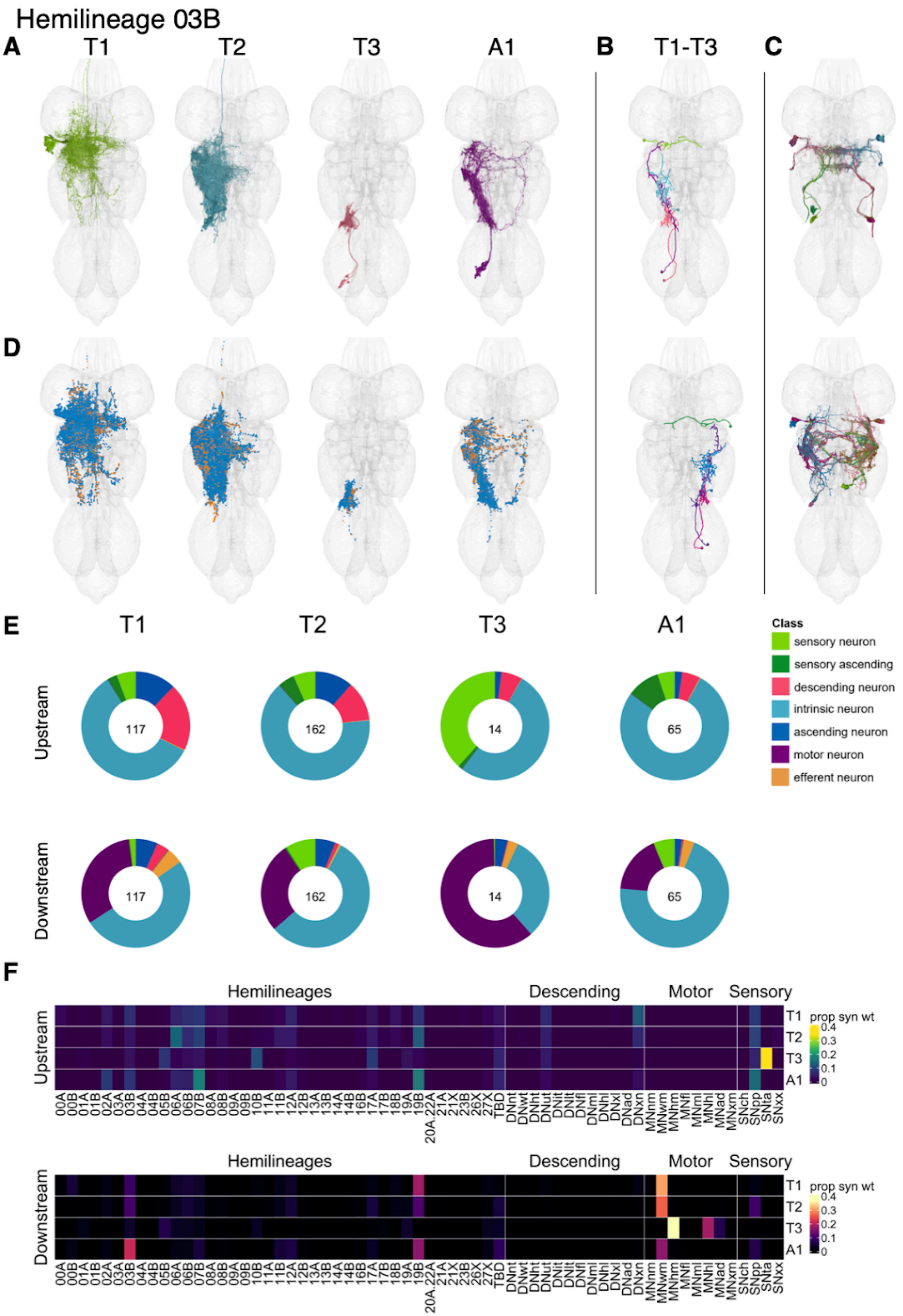

Hemilineage 03B
A
T1
T2
T3
A1
B
T1-T3
C
D
E
T1
T2
T3
A1
Upstream
Downstream
117
162
14
65
Class
sensory neuron
sensory ascending
descending neuron
intrinsic neuron
ascending neuron
motor neuron
efferent neuron
F
Hemilineages
Descending
Motor
Sensory
prop syn wt
0.4
0.3
0.2
0.1
0
00A 00B 01A 01B 02A 03A 03B 04A 04B 05B 06A 06B 07B 08A 08B 09A 09B 10B 11A 11B 12A 12B 13A 13B 14A 14B 16B 17A 17B 18B 19A 19B 20A.22A 21A 21X 23B 26X 27X TBD DNnt DNwt DNht DNut DNlt DNfl DNml DNhl DNxl DNad DNxn MNnm MNwm MNhm MNfl MNml MNhl MNad MNxm SNch SNpp SNta SNxx

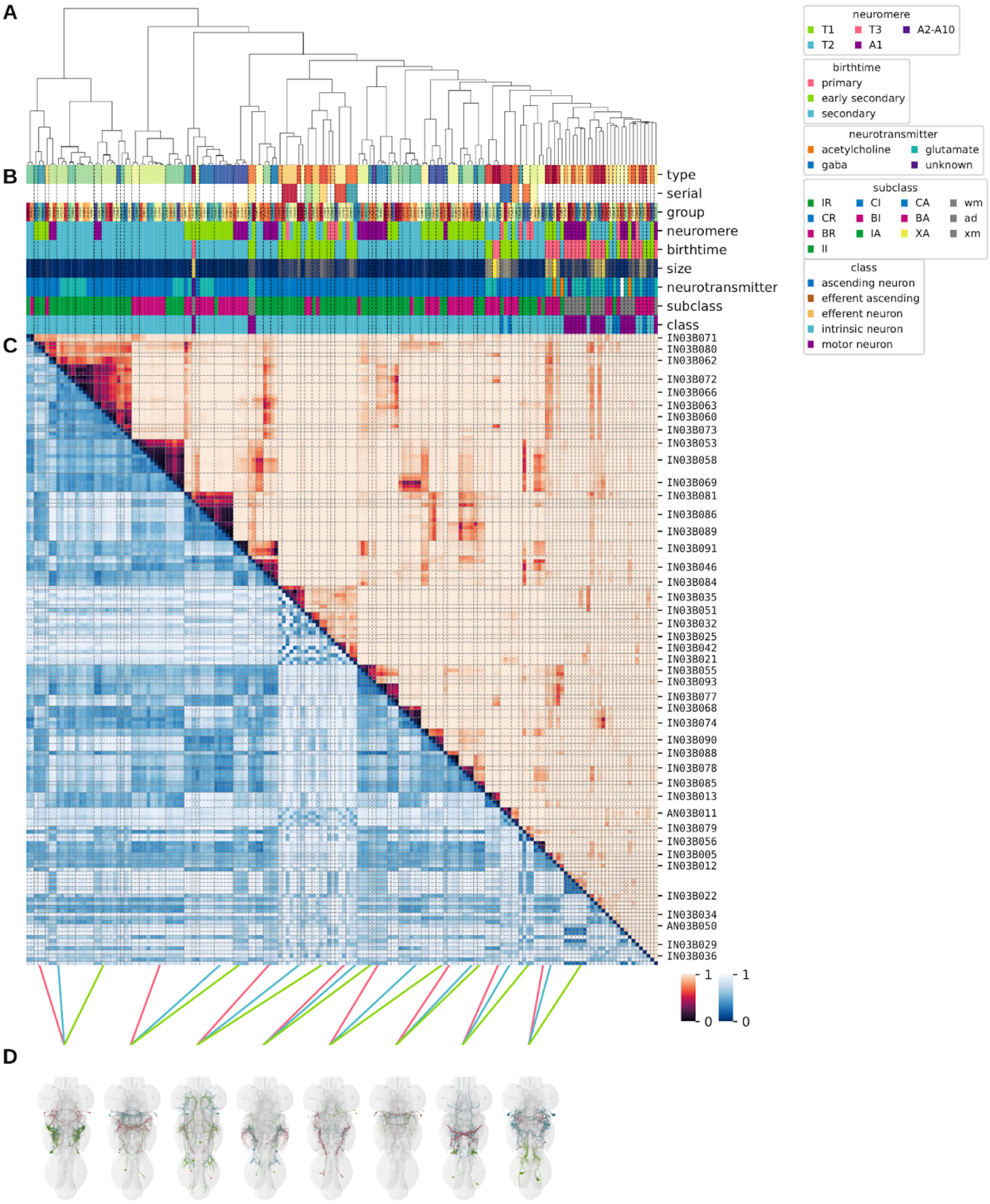

**Figure 20 - figure supplement 1. Systematic typing of hemilineage 03B.** Types for motor neurons were assigned separately as outlined in our accompanying manuscript (Cheong et al., 2023). **A.** Hierarchical clustering dendrogram of hemilineage groups by laterally and serially aggregated connectivity cosine clustering. **B.** Categorical annotations of each hemilineage group, each column corresponding to the aligned leaf in A. Colours for type, serial set, and group are arbitrary for visualisation. Colours for neuromere, birthtime, neurotransmitter, subclass, and class are as in all other figures. **C.** Similarity distance heatmap for hemilineage. Cosine distance is in the upper triangle, while laterally symmetrised NBLAST distance is in the lower triangle. Systematic type names of some types are labelled. **D.**

Morphologically representative groups from dendrogram subtrees. Each group, indicated by colour and line connecting to its column in B and C, is the most morphologically representative group (medoid of NBLAST distance) from a subtree of A. The subtrees (flat clusters) are equal height cuts of A determined to yield the number of groups per plot and plots in D.

#### 03B systematic types

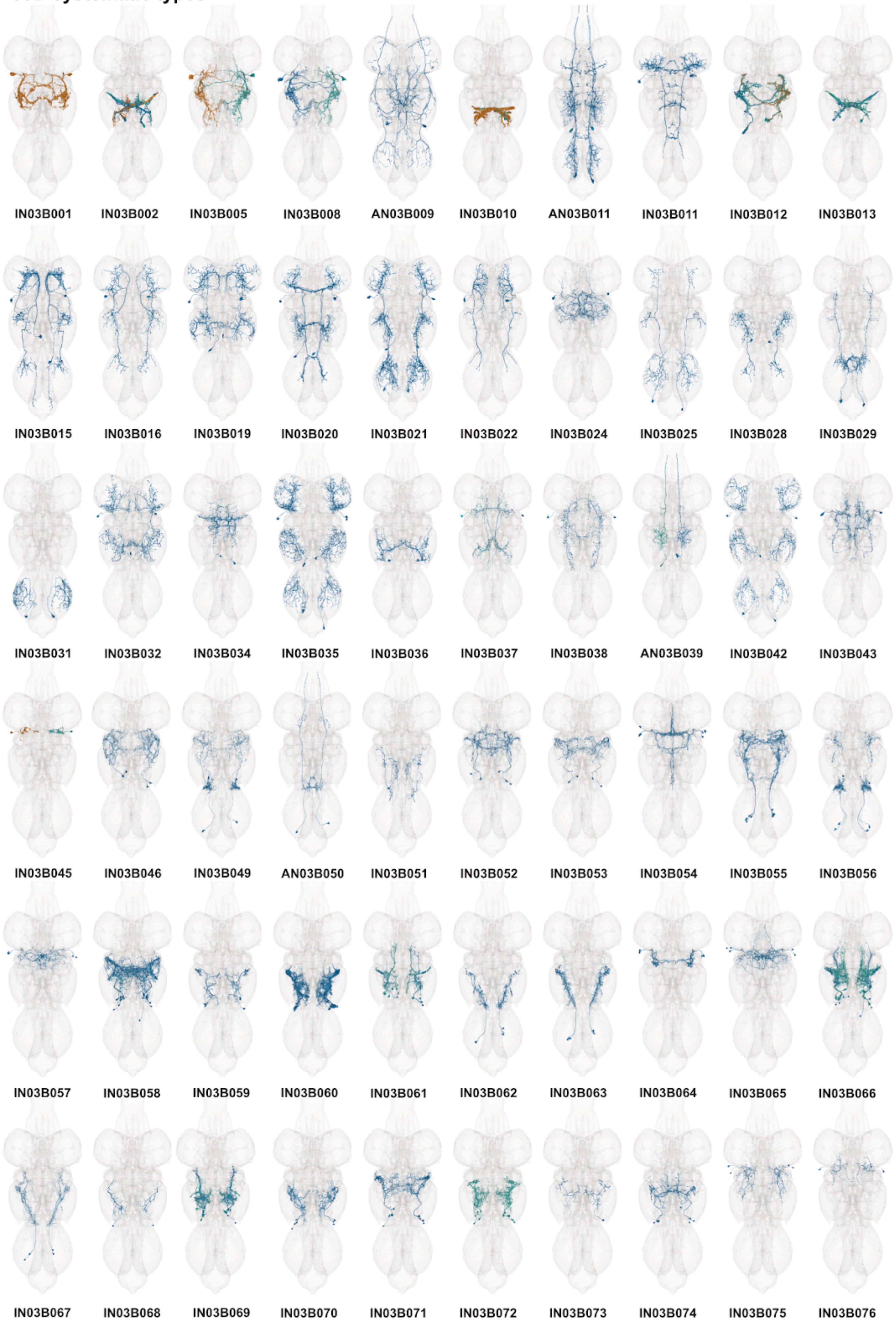

**Figure 20 - figure supplement 2. Systematic types of hemilineage 03B.** Systematic types have been arranged in numerical order, with neurons of the same type that belong to distinct classes (e.g., intrinsic neuron vs ascending neuron) plotted separately but placed adjacent to each other. Individual neuron meshes have been coloured based on predicted neurotransmitter: dark orange = acetylcholine, blue = gaba, marine = glutamate, dark purple = unknown. Motor neurons (typed separately in (Cheong et al., 2023)) have been plotted by serial set if identified in multiple neuromeres and by systematic type if not. Individual motor neuron meshes have been coloured based on soma neuromere: dark green = T1, blue = T2, purple = A1.

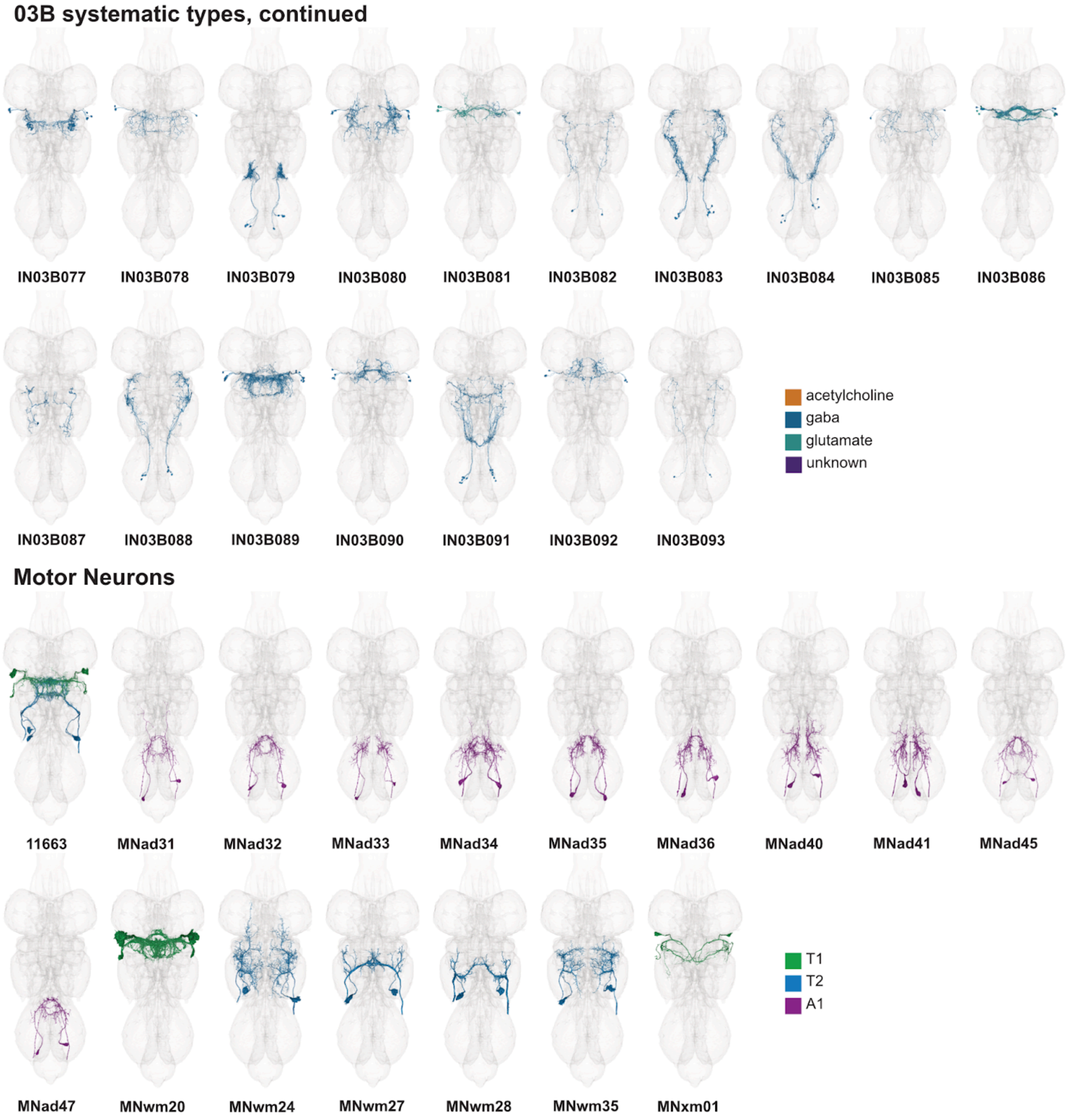

**Figure 20 - figure supplement 3. Systematic types of hemilineage 03B, continued.** Systematic types have been arranged in numerical order, with neurons of the same type that belong to distinct classes (e.g., intrinsic neuron vs ascending neuron) plotted separately but placed adjacent to each other. Individual neuron meshes have been coloured based on predicted neurotransmitter: dark orange = acetylcholine, blue = gaba, marine = glutamate, dark purple = unknown. Motor neurons (typed separately in (Cheong et al., 2023)) have been plotted by serial set if identified in multiple neuromeres and by systematic type if not. Individual motor neuron meshes have been coloured based on soma neuromere: dark green = T1, blue = T2, purple = A1.

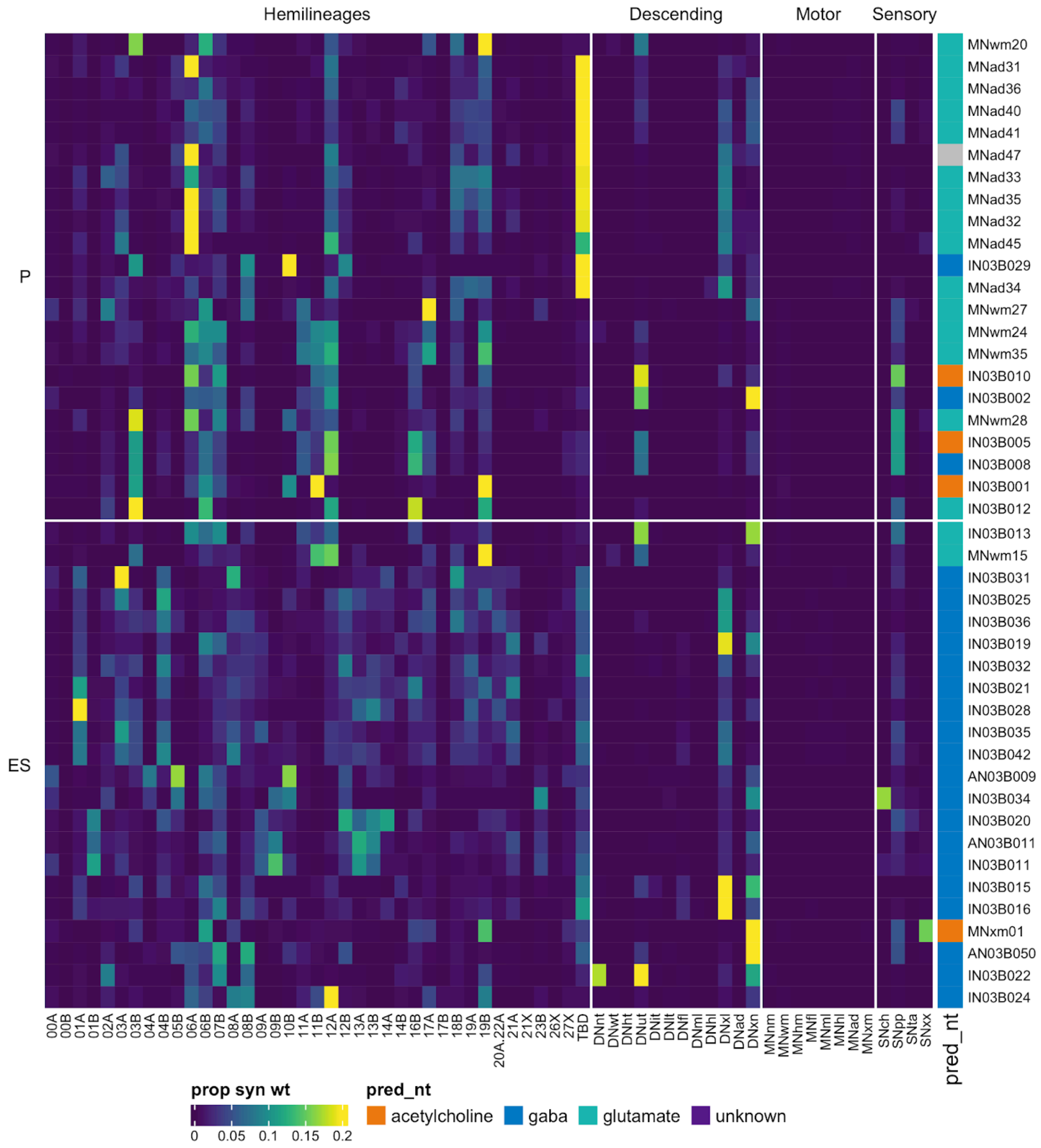

**Figure 20 - figure supplement 4. Connectivity to upstream partners by 03B primary and early secondary systematic types.** Proportions of synaptic weight to systematic types from upstream partners, normalised by row. 03B neurons have been clustered within each assigned birthtime window (P = primary, ES = early secondary, S = secondary) based on both upstream and downstream connectivity to hemilineages, descending neuron subclasses, motor neuron subclasses, and sensory neuron modalities. Annotation bar is coloured by the most common predicted neurotransmitter for the neurons of each type.

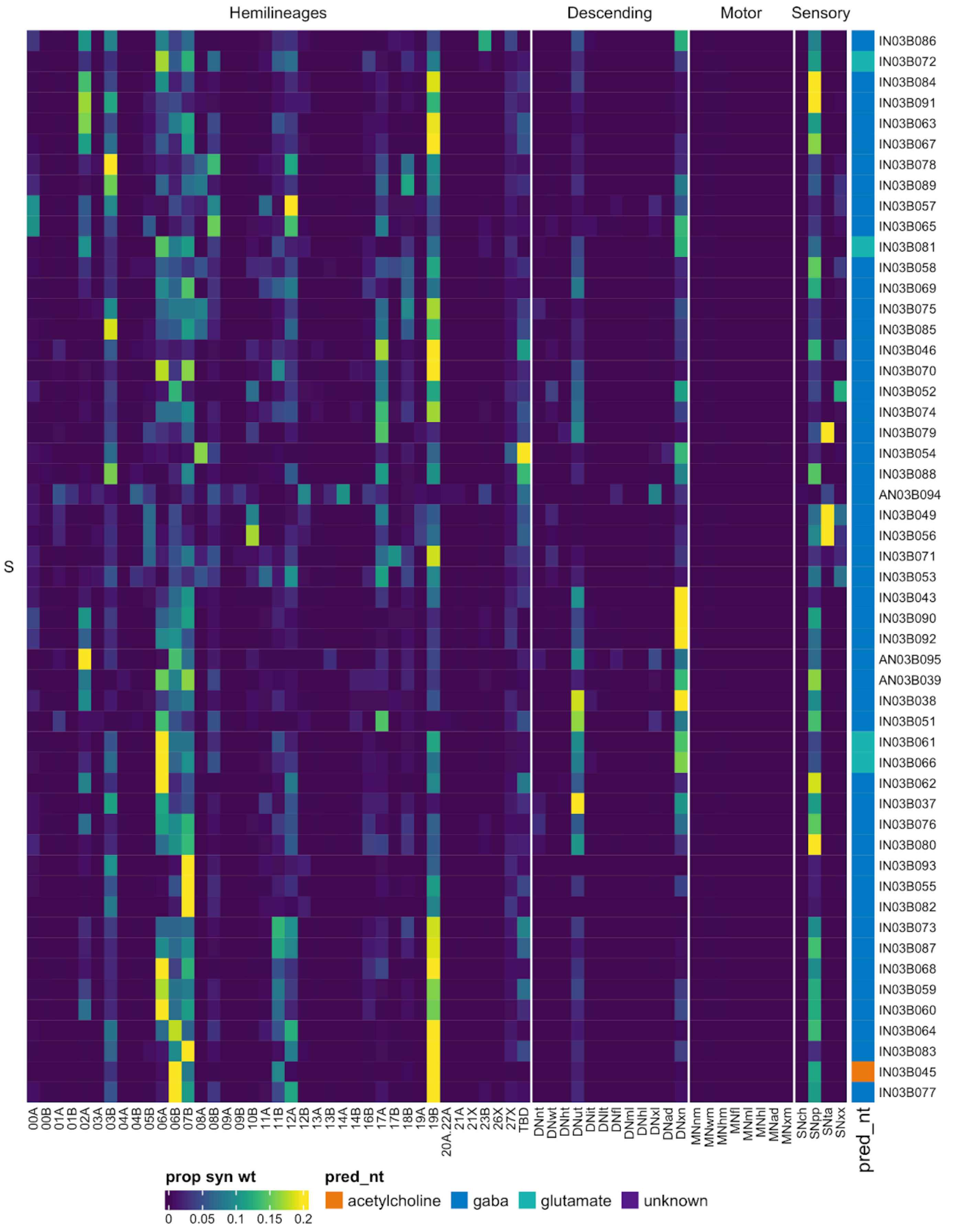

**Figure 20 - figure supplement 5. Connectivity to upstream partners by 03B secondary systematic types.** Proportions of synaptic weight to systematic types from upstream partners, normalised by row. 03B neurons have been clustered within each assigned birthtime window (P = primary, ES = early secondary, S = secondary) based on both upstream and downstream connectivity to hemilineages, descending neuron subclasses, motor neuron subclasses, and sensory neuron modalities. The annotation bar is coloured by the most common predicted neurotransmitter within each type.

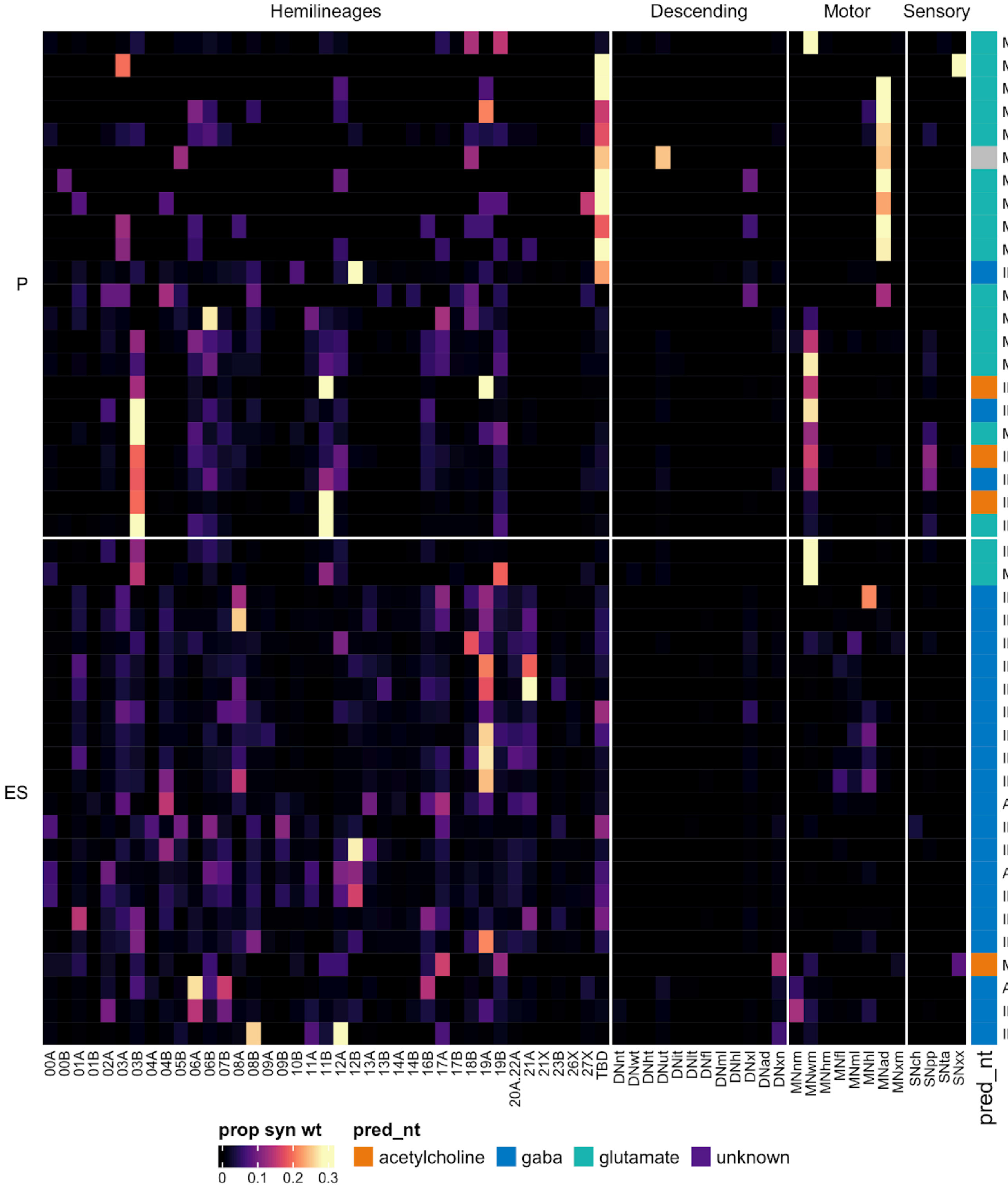

**Figure 20 - figure supplement 6. Connectivity to downstream partners by 03B primary and early secondary systematic types.** Proportions of synaptic weight from systematic types to downstream partners, normalised by row. 03B neurons have been clustered within each assigned birthtime window (P = primary, ES = early secondary, S = secondary) based on both upstream and downstream connectivity to hemilineages, descending neuron subclasses, motor neuron subclasses, and sensory neuron modalities. The annotation bar is coloured by the most common predicted neurotransmitter within each type.

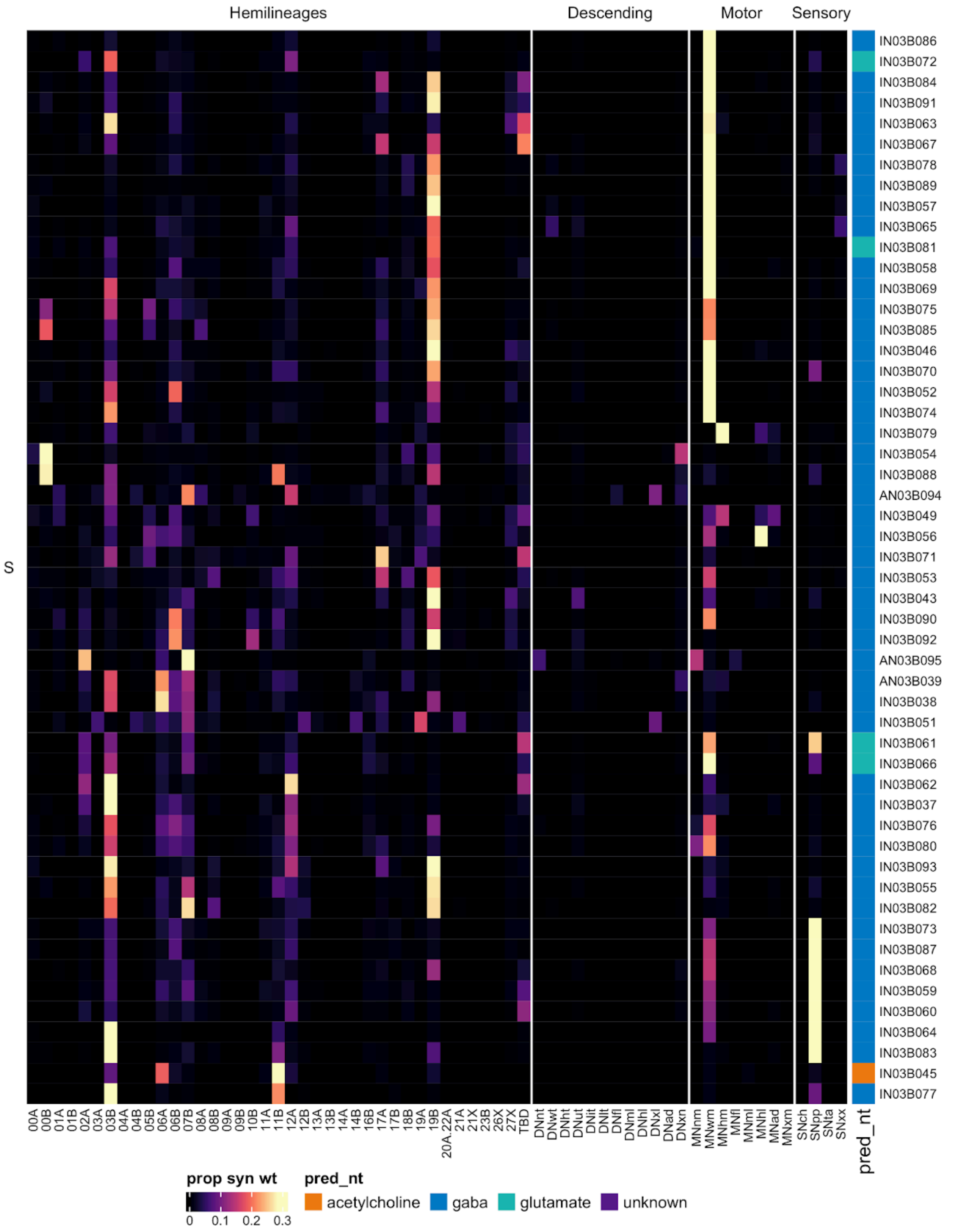

**Figure 20 - figure supplement 7. Connectivity to downstream partners by 03B secondary systematic types.** Proportions of synaptic weight from systematic types to downstream partners, normalised by row. 03B neurons have been clustered within each assigned birthtime window (P = primary, ES = early secondary, S = secondary) based on both upstream and downstream connectivity to hemilineages, descending neuron subclasses, motor neuron subclasses, and sensory neuron modalities. The annotation bar is coloured by the most common predicted

neurotransmitter for the neurons of each type.

Hemilineage 04B

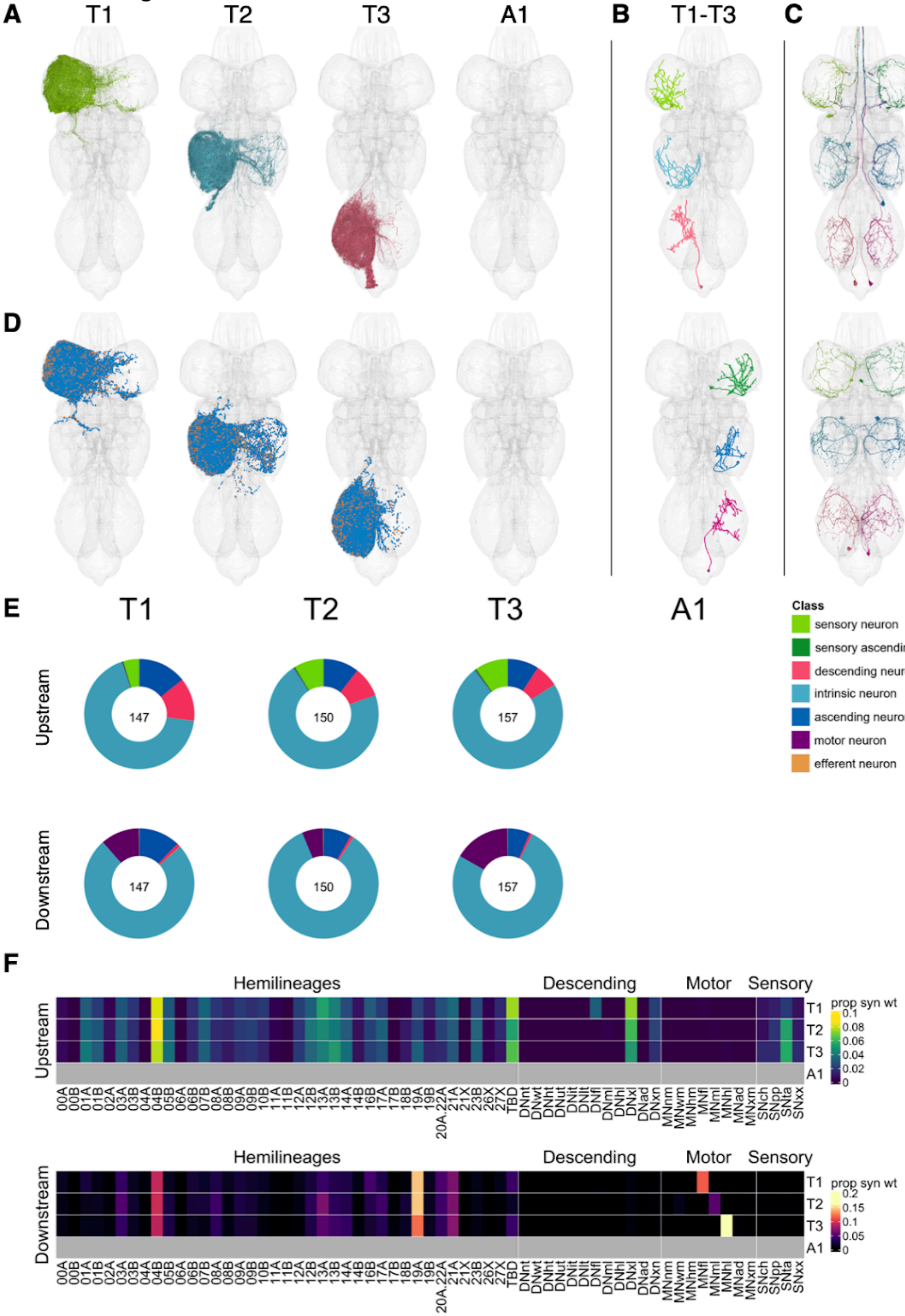

**Figure 21. Hemilineage 04B. A.** Meshes of all RHS secondary neurons plotted in neuromere-specific colours. **B.** "Representative" secondary neuron skeletons plotted in hemineuromere-specific colours. The skeleton with the top accumulated NBLAST score among all neurons from the hemilineage in a given hemineuromere was used. **C.** Neuron meshes of selected examples. Top: ascending serial set 10394. Bottom: independent leg serial set 11945. **D.** Predicted synapses of RHS secondary neurons. Blue: postsynapses; dark orange: presynapses. **E.** Proportions of connections from secondary neurons to upstream or downstream partners, normalised by neuromere and coloured by broad class. Numbers of query neurons appear in the centre. **F.** Proportions of synaptic weight from secondary neurons originating in each neuromere to upstream or downstream partners, normalised by row.

#### Hemilineage 04B

04B is the main secondary hemilineage (Truman et al., 2010) from medial neuroblast NB3-1 (Truman et al., 2004), which generates 4 contralateral MNs, 3-6 intersegmental interneurons, and a variable number of local interneurons in the embryo (Schmid et al., 1999). 04B is a relatively large hemilineage (Figure 8D), and its neurons do not enter the neuropil in a single coherent bundle but rather project from the ventral midline in multiple clusters. Most 04B secondary neurons innervate the ipsilateral leg neuropil diffusely, especially the surface, but a small subset exhibit intersegmental projections (Shepherd et al., 2019). 04B neurons arborise extensively in the ipsilateral leg neuropil, sometimes spreading across the midline (Figure 21A). Five types of early born 04B neurons ascend via the neck connective (e.g., AN04B003) (Figure 21C top), but most 04B neurons are restricted to their neuromere of origin (e.g., IN04B008) (Figure 21C bottom).

Secondary 04B neurons survive in T1-T3 with comparable numbers of neurons in all thoracic neuromeres (Figure 21E). They mainly receive inputs from hemilineages 01A, 04B, 12B, 13A, and 13B, from descending neurons that innervate leg neuropils, and from tactile sensory neurons (Figure 21F), although a few types also receive chemosensory input (e.g., IN04B005 and IN04B079) (Figure 21 - figure supplement 4-5). They are predicted to be cholinergic (Figure 8E) as expected (Lacin et al., 2019) and output onto leg motor neurons and leg hemilineages, notably 04B, 19A, and 21A (Figure 21F). No functional studies have been published for secondary 04B neurons.

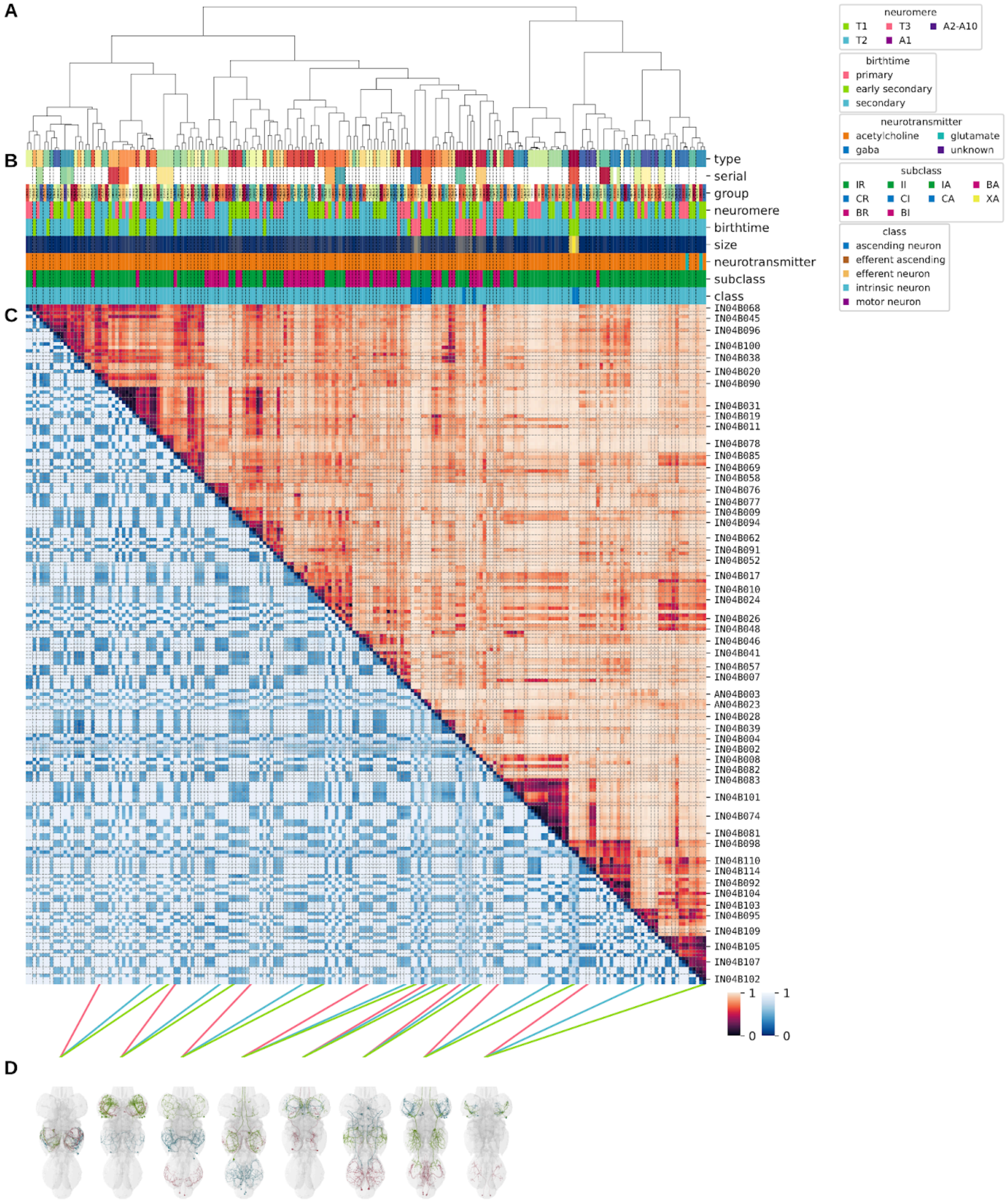

**Figure 21 - figure supplement 1. Systematic typing of hemilineage 04B. A.** Hierarchical clustering dendrogram of hemilineage groups by laterally and serially aggregated connectivity cosine clustering. **B.** Categorical annotations of each hemilineage group, each column corresponding to the aligned leaf in A. Colours for type, serial set, and group are arbitrary for visualisation. Colours for neuromere, birthtime, neurotransmitter, subclass, and class are as in all other figures. **C.** Similarity distance heatmap for hemilineage. Cosine distance is in the upper triangle, while laterally symmetrised NBLAST distance is in the lower triangle. Systematic type names of some types are labelled. **D.** Morphologically representative groups from dendrogram subtrees. Each group, indicated by colour and line connecting to its column in B and C, is the most morphologically representative group (medoid of NBLAST distance) from a subtree of A. The subtrees (flat clusters) are equal height cuts of A determined to yield the number of groups

per plot and plots in D.

#### 04B systematic types

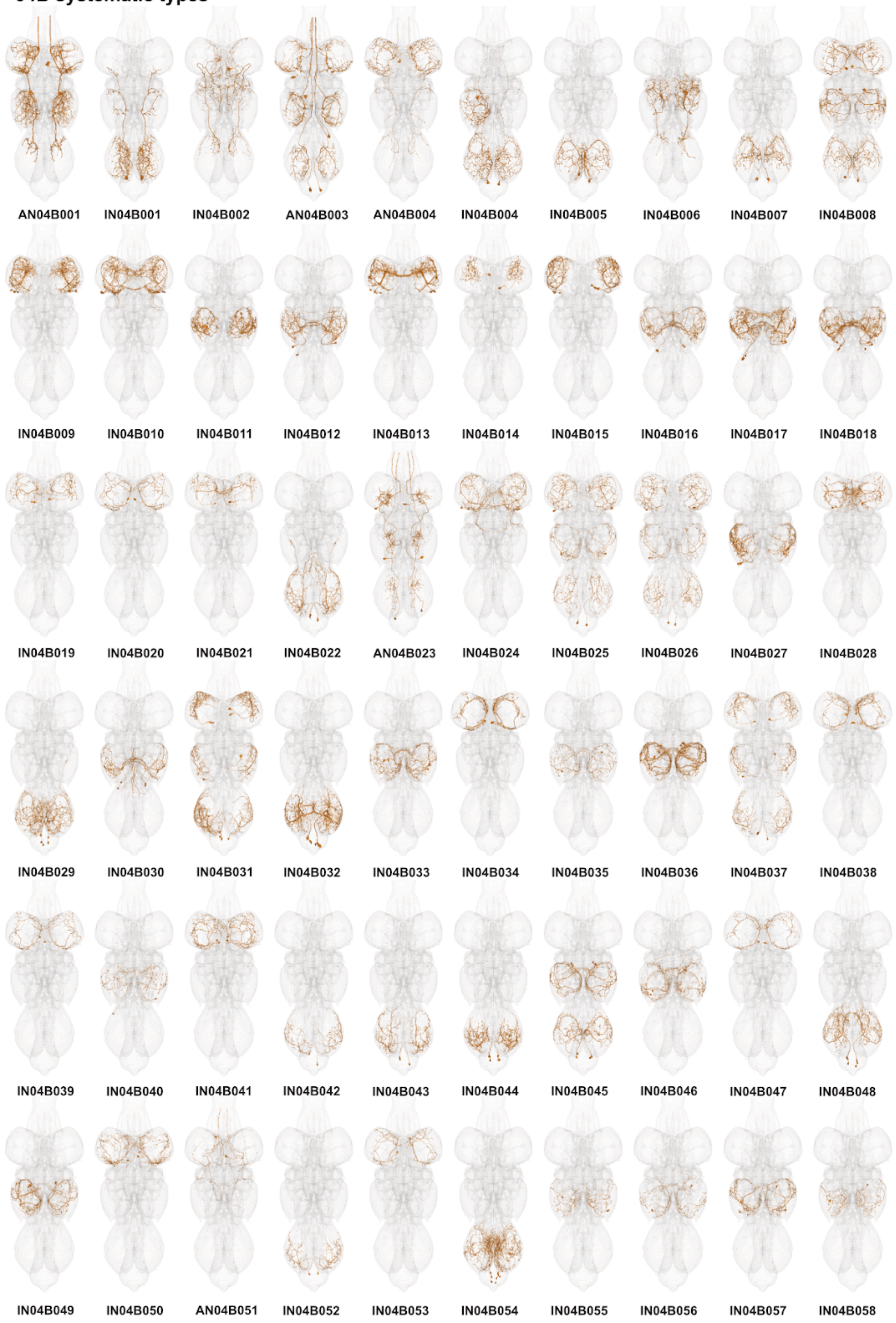

**Figure 21 - figure supplement 2. Systematic types of hemilineage 04B.** Systematic types have been arranged in numerical order, with neurons of the same type that belong to distinct classes (e.g., intrinsic neuron vs ascending neuron) plotted separately but placed adjacent to each other. Individual neuron meshes have been coloured based on predicted neurotransmitter: dark orange = acetylcholine, blue = gaba, marine = glutamate, dark purple = unknown.

### 04B systematic types, continued

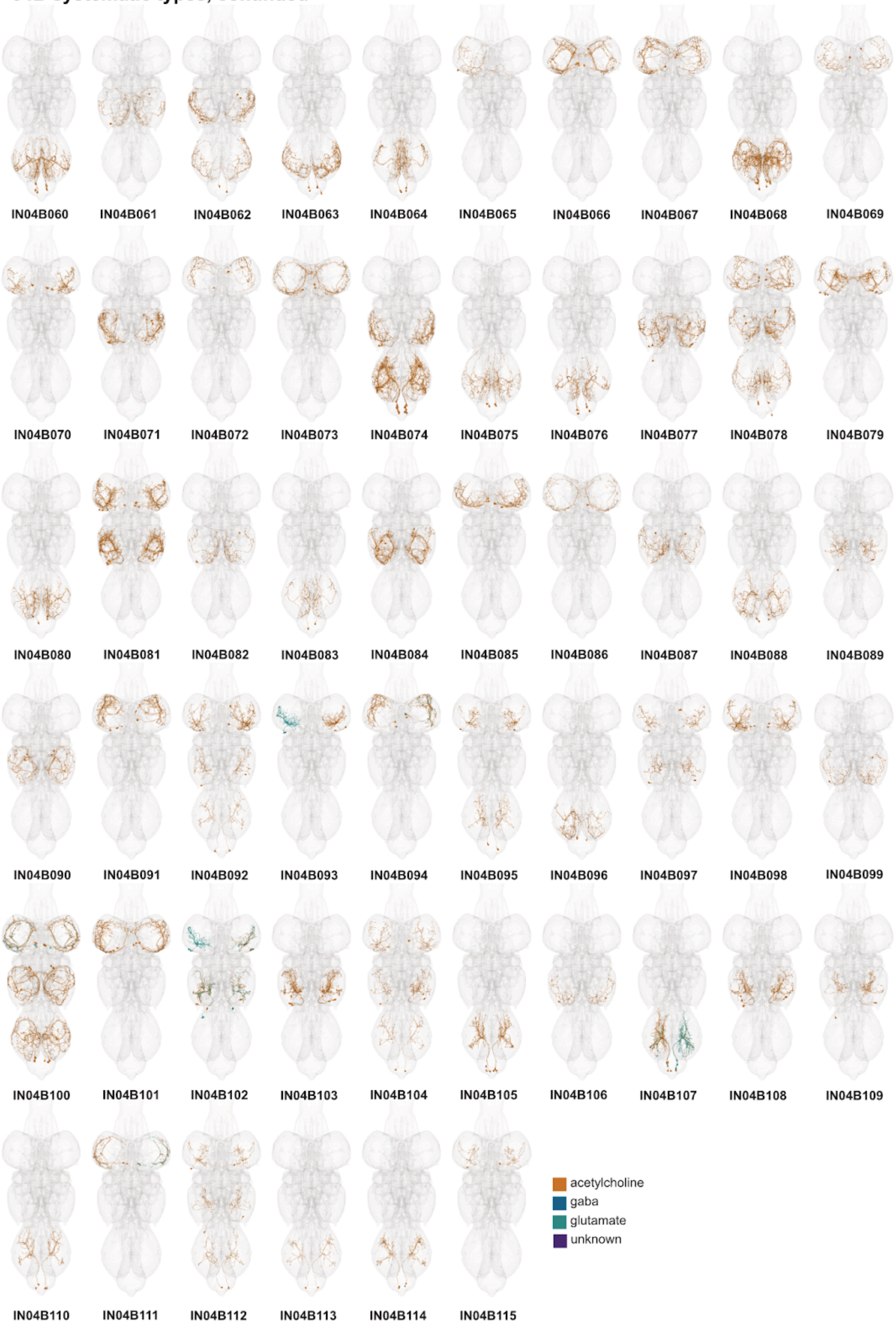

**Figure 21 - figure supplement 3. Systematic types of hemilineage 04B, continued.** Systematic types have been arranged in numerical order, with neurons of the same type that belong to distinct classes (e.g., intrinsic neuron vs ascending neuron) plotted separately but placed adjacent to each other. Individual neuron meshes have been coloured based on predicted neurotransmitter: dark orange = acetylcholine, blue = gaba, marine = glutamate, dark purple = unknown.

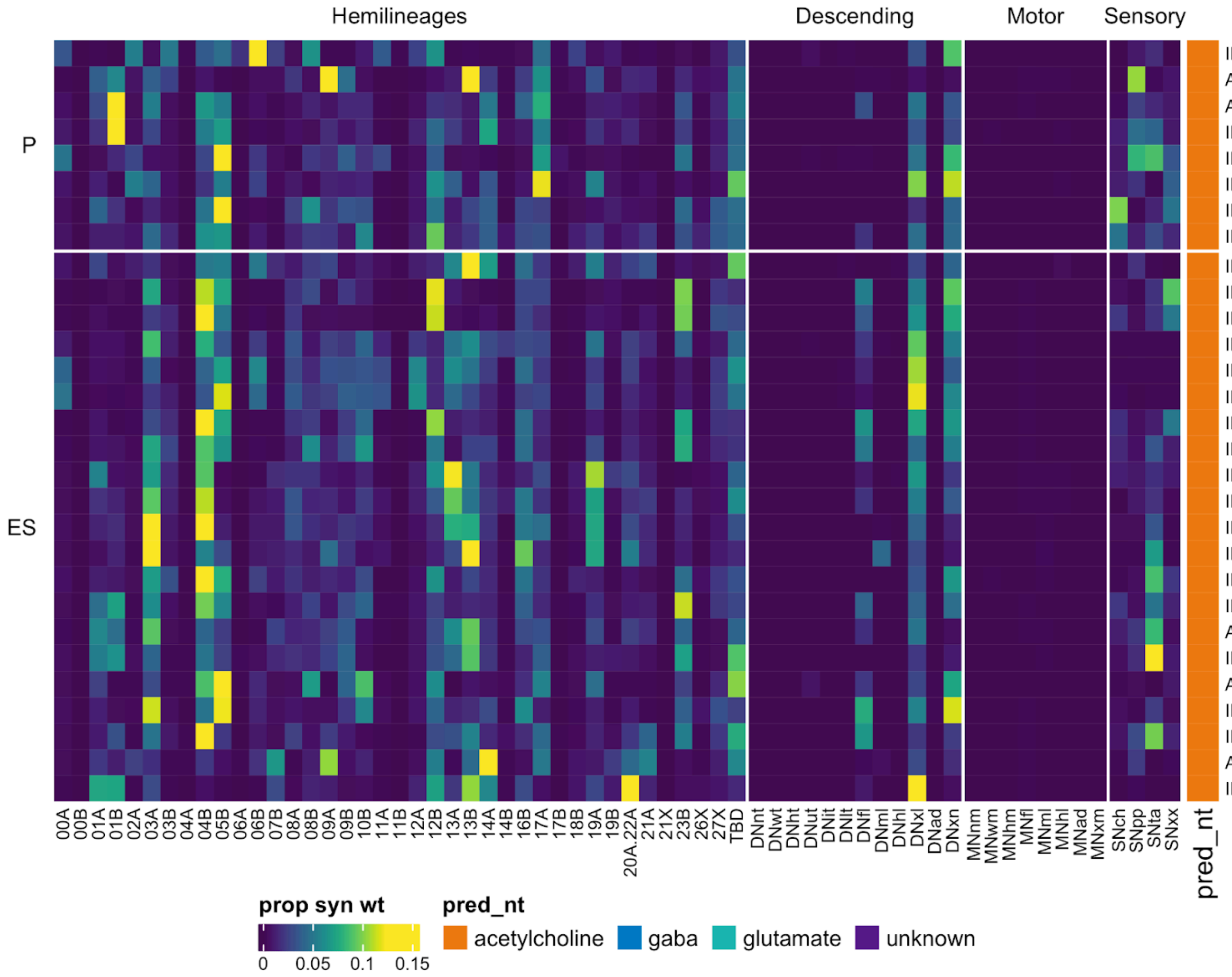

**Figure 21 - figure supplement 4. Connectivity to upstream partners by 04B primary and early secondary systematic types.** Proportions of synaptic weight to systematic types from upstream partners, normalised by row. 04B neurons have been clustered within each assigned birthtime window (P = primary, ES = early secondary, S = secondary) based on both upstream and downstream connectivity to hemilineages, descending neuron subclasses, motor neuron subclasses, and sensory neuron modalities. Annotation bar is coloured by the most common predicted neurotransmitter for the neurons of each type.

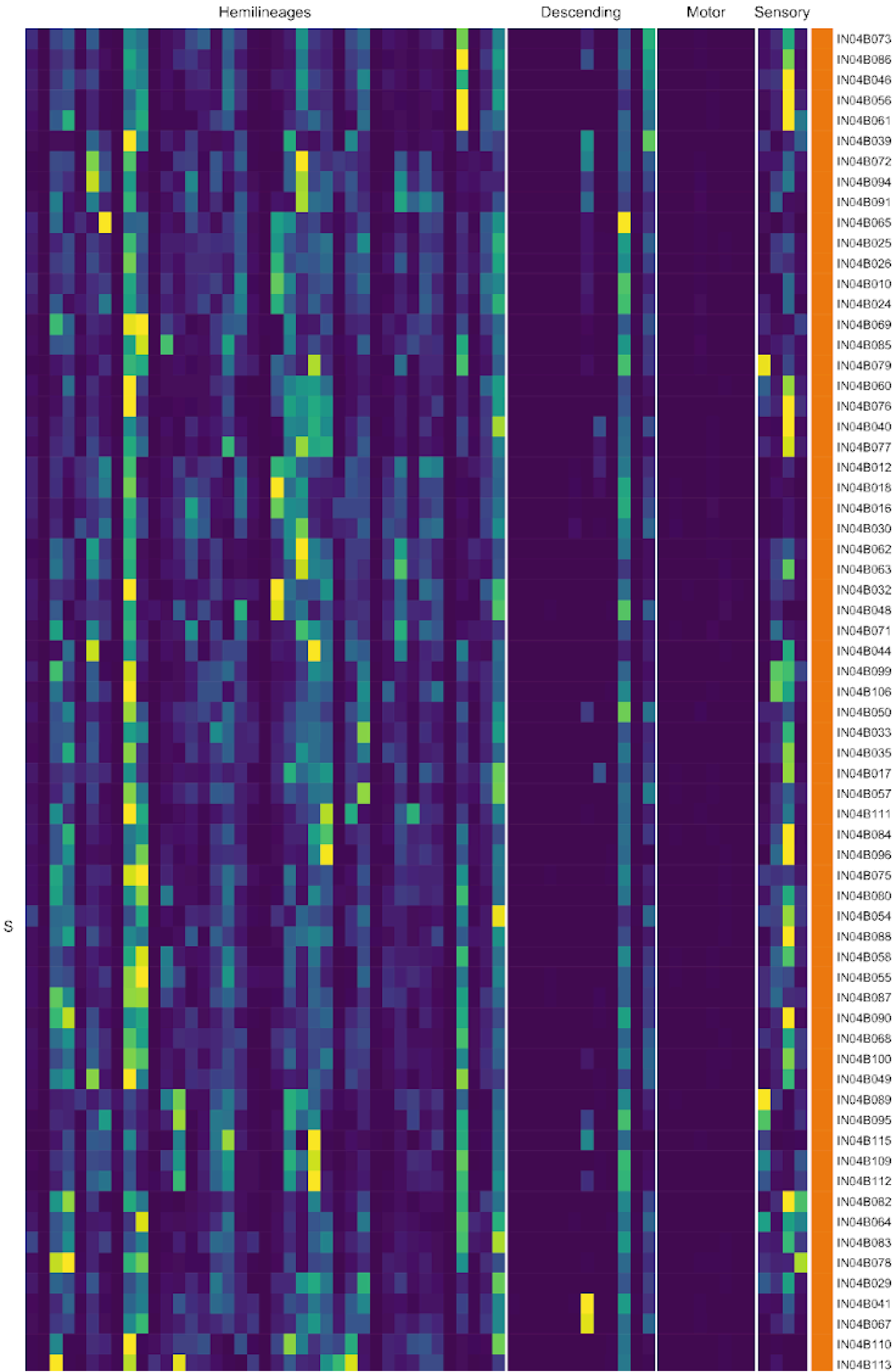
Hemilineages
Descending
Motor
Sensory
S
IN04B073
IN04B086
IN04B046
IN04B056
IN04B061
IN04B039
IN04B072
IN04B094
IN04B091
IN04B065
IN04B025
IN04B026
IN04B010
IN04B024
IN04B069
IN04B085
IN04B079
IN04B060
IN04B076
IN04B040
IN04B077
IN04B012
IN04B018
IN04B016
IN04B030
IN04B062
IN04B063
IN04B032
IN04B048
IN04B071
IN04B044
IN04B099
IN04B106
IN04B050
IN04B033
IN04B035
IN04B017
IN04B057
IN04B111
IN04B084
IN04B096
IN04B075
IN04B080
IN04B054
IN04B088
IN04B058
IN04B055
IN04B087
IN04B090
IN04B068
IN04B100
IN04B049
IN04B089
IN04B095
IN04B115
IN04B109
IN04B112
IN04B082
IN04B064
IN04B083
IN04B078
IN04B029
IN04B041
IN04B067
IN04B110
IN04B113

**Figure 21 - figure supplement 5. Connectivity to upstream partners by 04B secondary systematic types.** Proportions of synaptic weight to systematic types from upstream partners, normalised by row. 04B neurons have been clustered within each assigned birthtime window (P = primary, ES = early secondary, S = secondary) based on both upstream and downstream connectivity to hemilineages, descending neuron subclasses, motor neuron subclasses, and sensory neuron modalities. The annotation bar is coloured by the most common predicted neurotransmitter within each type.

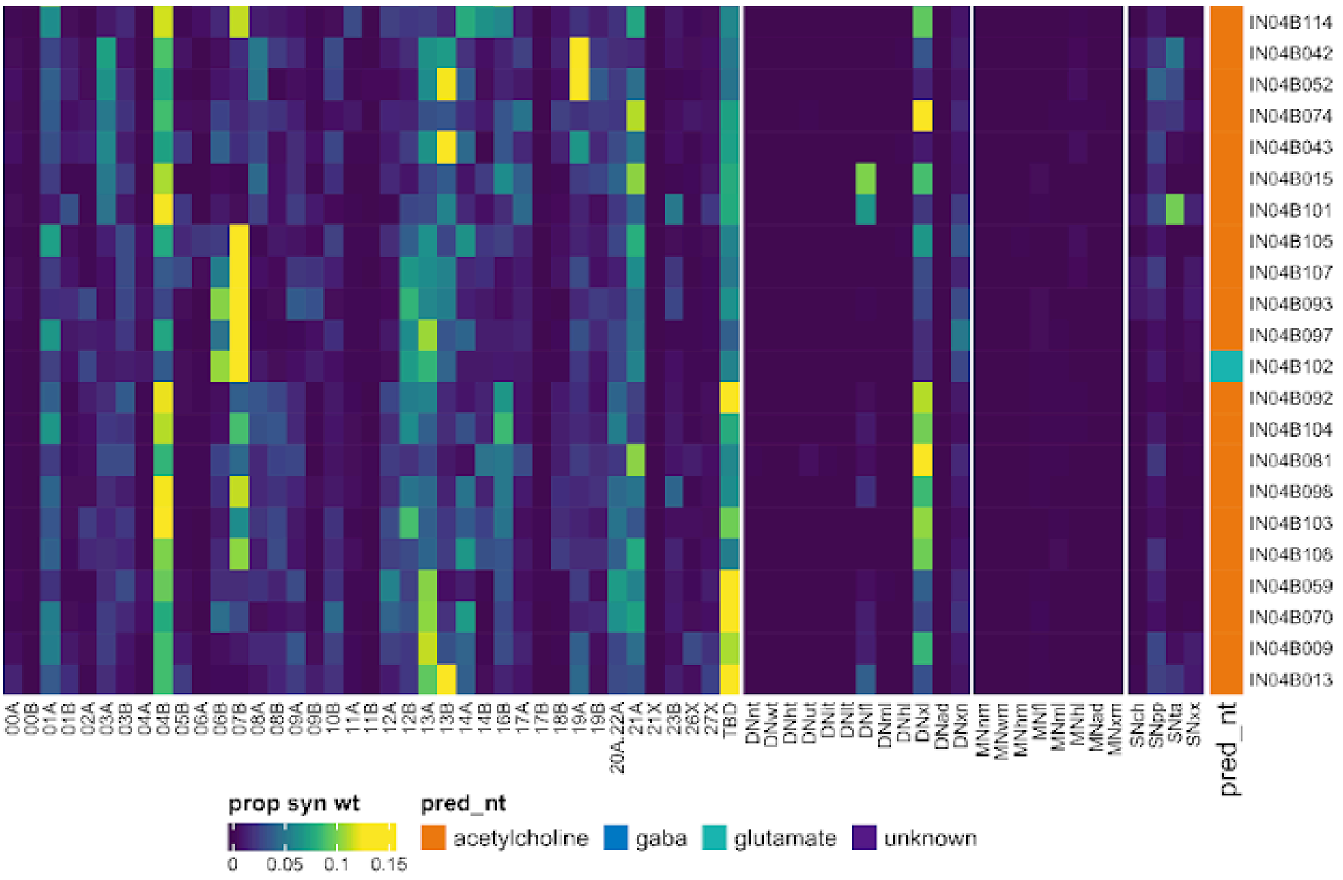

**Figure 21 - figure supplement 6. Connectivity to upstream partners by 04B secondary systematic types, continued.** Proportions of synaptic weight to systematic types from upstream partners, normalised by row. 04B neurons have been clustered within each assigned birthtime window (P = primary, ES = early secondary, S = secondary) based on both upstream and downstream connectivity to hemilineages, descending neuron subclasses, motor neuron subclasses, and sensory neuron modalities. The annotation bar is coloured by the most common predicted neurotransmitter within each type.

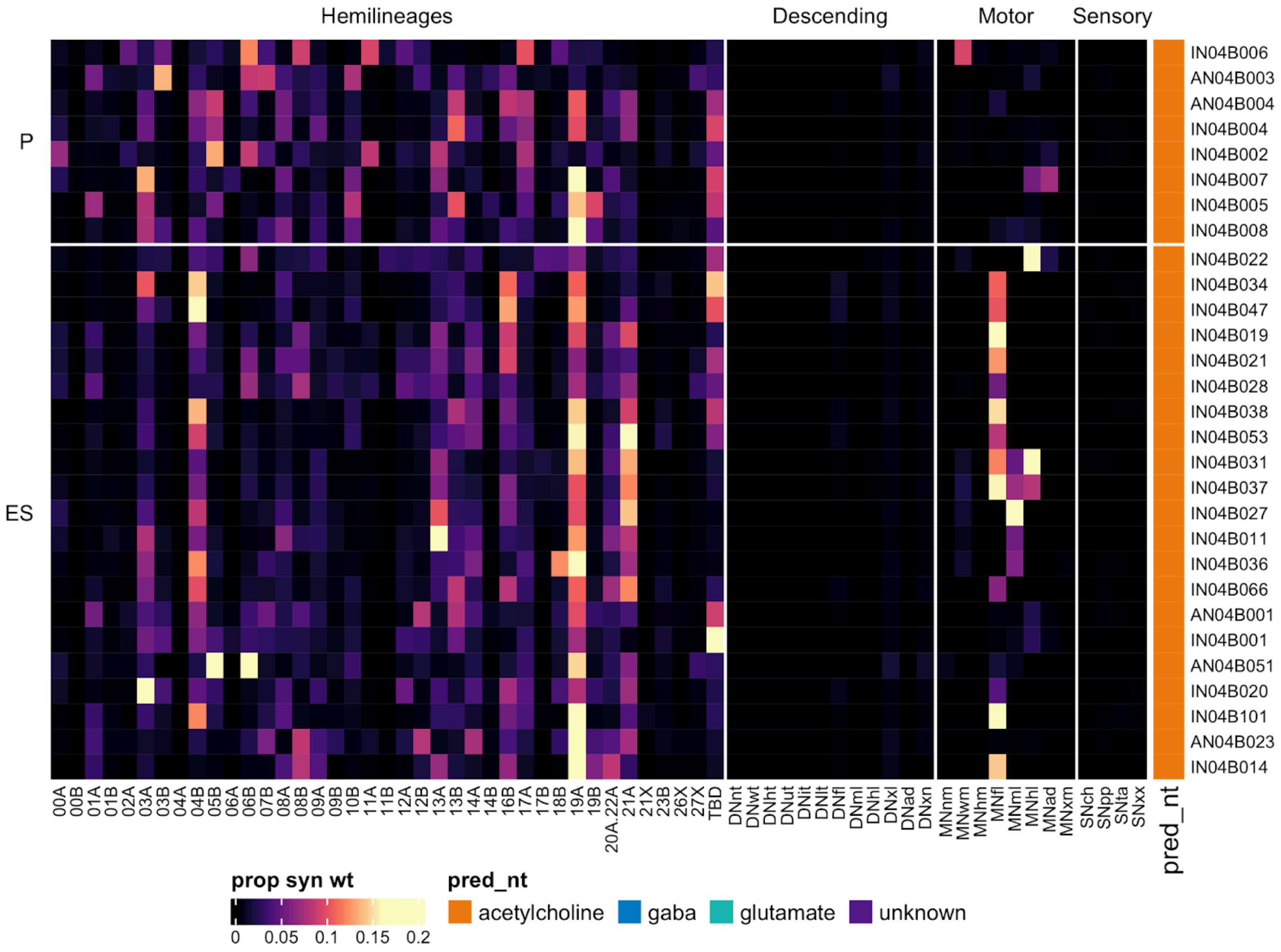

**Figure 21 - figure supplement 7. Connectivity to downstream partners by 04B primary and early secondary systematic types.** Proportions of synaptic weight from systematic types to downstream partners, normalised by row. 04B neurons have been clustered within each assigned birthtime window (P = primary, ES = early secondary, S = secondary) based on both upstream and downstream connectivity to hemilineages, descending neuron subclasses, motor neuron subclasses, and sensory neuron modalities. The annotation bar is coloured by the most common predicted neurotransmitter within each type.

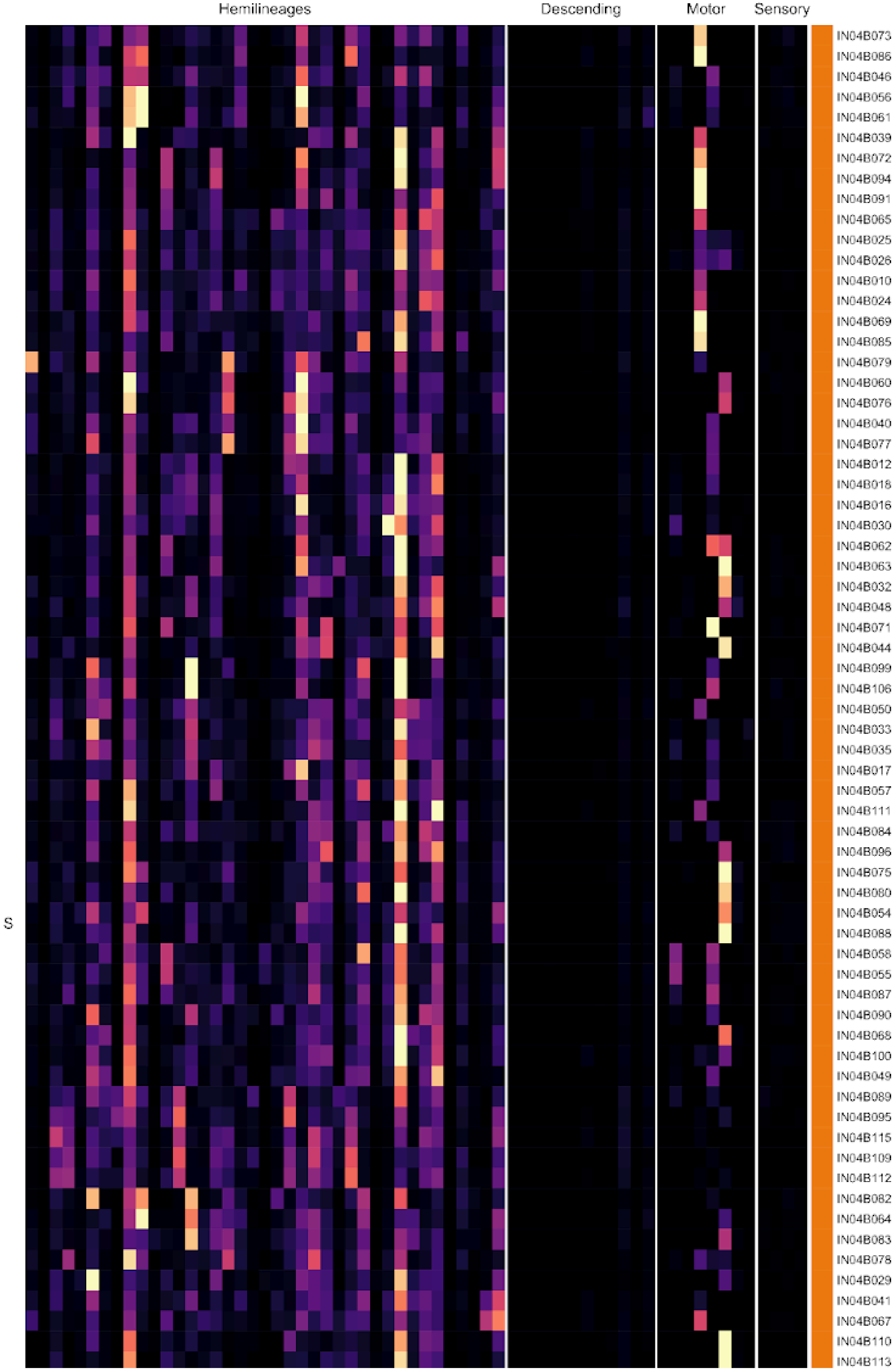
Hemilineages
Descending
Motor
Sensory
S
IN04B073
IN04B086
IN04B046
IN04B056
IN04B061
IN04B039
IN04B072
IN04B094
IN04B091
IN04B065
IN04B025
IN04B026
IN04B010
IN04B024
IN04B069
IN04B085
IN04B079
IN04B060
IN04B076
IN04B040
IN04B077
IN04B012
IN04B018
IN04B016
IN04B030
IN04B062
IN04B063
IN04B032
IN04B048
IN04B071
IN04B044
IN04B099
IN04B106
IN04B050
IN04B033
IN04B035
IN04B017
IN04B057
IN04B111
IN04B084
IN04B096
IN04B075
IN04B080
IN04B054
IN04B088
IN04B058
IN04B055
IN04B087
IN04B090
IN04B068
IN04B100
IN04B049
IN04B089
IN04B095
IN04B115
IN04B109
IN04B112
IN04B082
IN04B064
IN04B083
IN04B078
IN04B029
IN04B041
IN04B067
IN04B110
IN04B113

**Figure 21 - figure supplement 8. Connectivity to downstream partners by 04B secondary systematic types.** Proportions of synaptic weight from systematic types to downstream partners, normalised by row. 04B neurons have been clustered within each assigned birthtime window (P = primary, ES = early secondary, S = secondary) based on both upstream and downstream connectivity to hemilineages, descending neuron subclasses, motor neuron subclasses, and sensory neuron modalities. The annotation bar is coloured by the most common predicted neurotransmitter for the neurons of each type.

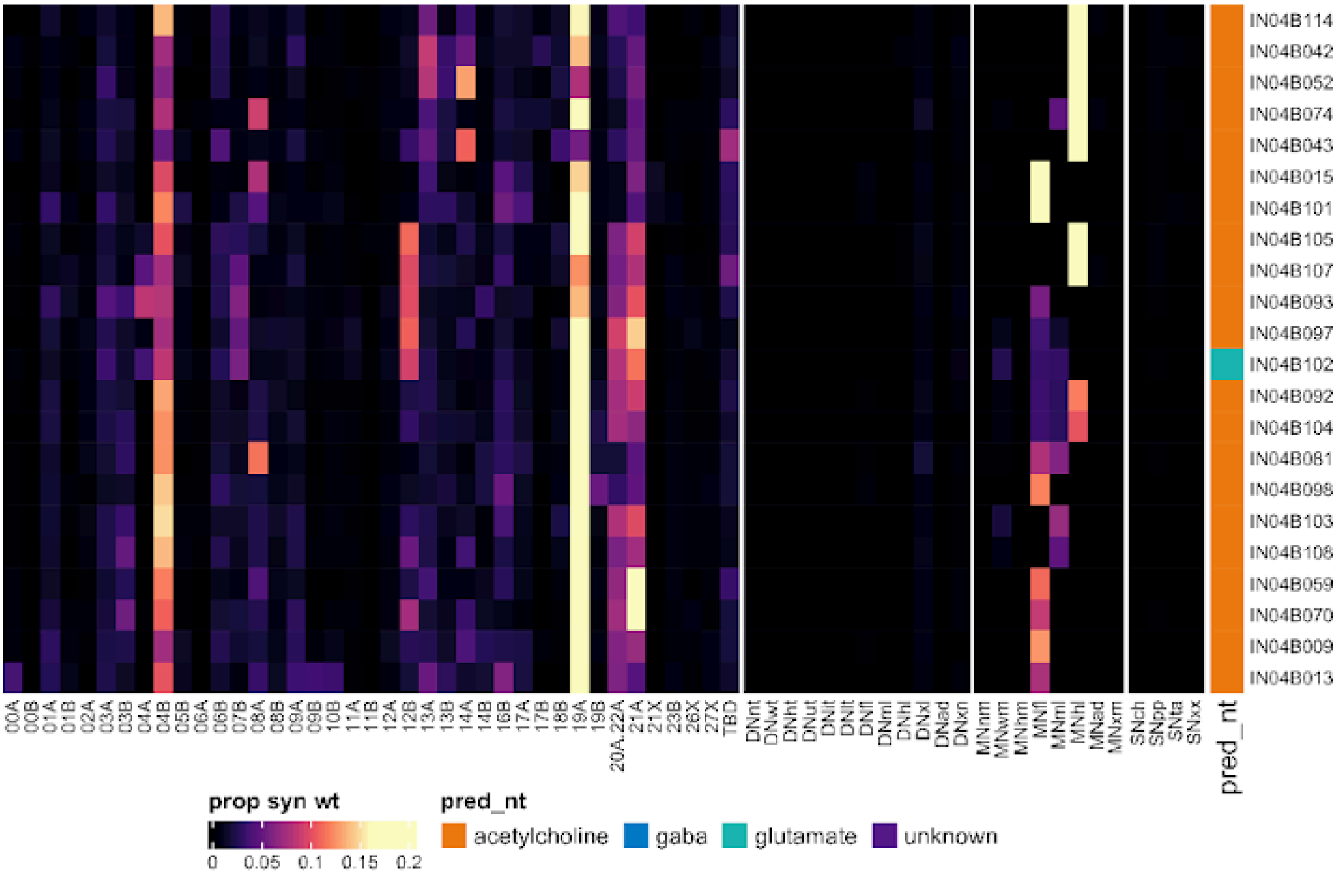

**Figure 21 - figure supplement 9.Connectivity to downstream partners by 04B secondary systematic types, continued.** Proportions of synaptic weight from systematic types to downstream partners, normalised by row. 04B neurons have been clustered within each assigned birthtime window (P = primary, ES = early secondary, S = secondary) based on both upstream and downstream connectivity to hemilineages, descending neuron subclasses, motor neuron subclasses, and sensory neuron modalities. The annotation bar is coloured by the most common predicted neurotransmitter for the neurons of each type.

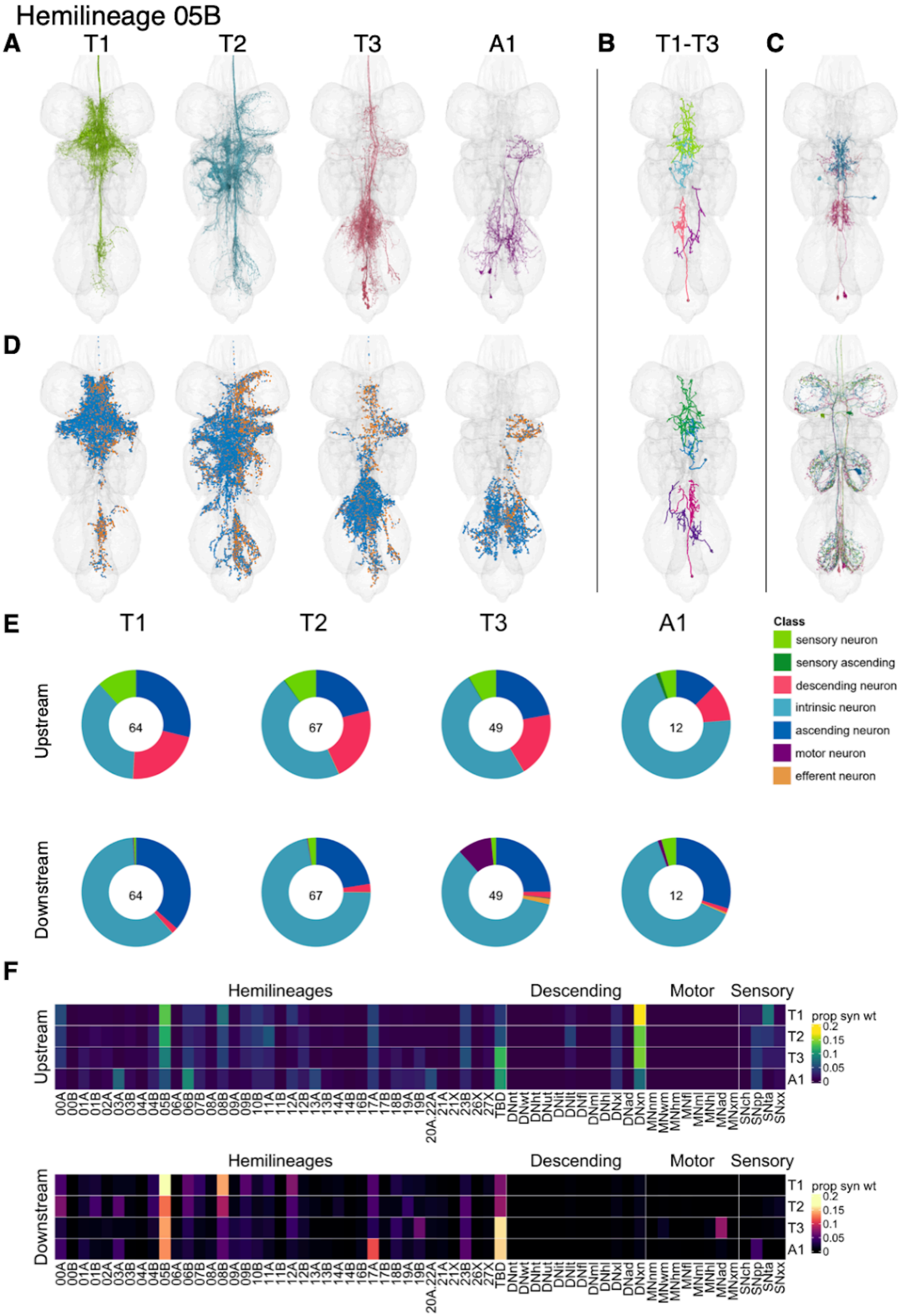

Hemilineage 05B
A
T1
T2
T3
A1
B
T1-T3
C
D
E
T1
T2
T3
A1
Upstream
64
67
49
12
Downstream
64
67
49
12
Class
sensory neuron
sensory ascending
descending neuron
intrinsic neuron
ascending neuron
motor neuron
efferent neuron
F
Hemilineages
Descending
Motor
Sensory
Upstream
T1
T2
T3
A1
prop syn wt
0.2
0.15
0.1
0.05
0
00A 00B 01A 01B 02A 03A 03B 04A 04B 05B 06A 06B 07B 08A 08B 09A 09B 10B 11A 11B 12A 12B 13A 13B 14A 14B 16B 17A 17B 18B 19A 19B 20A.22A 21A 21X 23B 26X 27X TBD DNnt DNwt DNht DNut DNit DNlt DNfl DNml DNhl DNxl DNad DNxn MNnm MNwm MNhm MNfl MNml MNhl MNad MNxm SNch SNpp SNta SNxx
Hemilineages
Descending
Motor
Sensory
Downstream
T1
T2
T3
A1
prop syn wt
0.2
0.15
0.1
0.05
0
00A 00B 01A 01B 02A 03A 03B 04A 04B 05B 06A 06B 07B 08A 08B 09A 09B 10B 11A 11B 12A 12B 13A 13B 14A 14B 16B 17A 17B 18B 19A 19B 20A.22A 21A 21X 23B 26X 27X TBD DNnt DNwt DNht DNut DNit DNlt DNfl DNml DNhl DNxl DNad DNxn MNnm MNwm MNhm MNfl MNml MNhl MNad MNxm SNch SNpp SNta SNxx

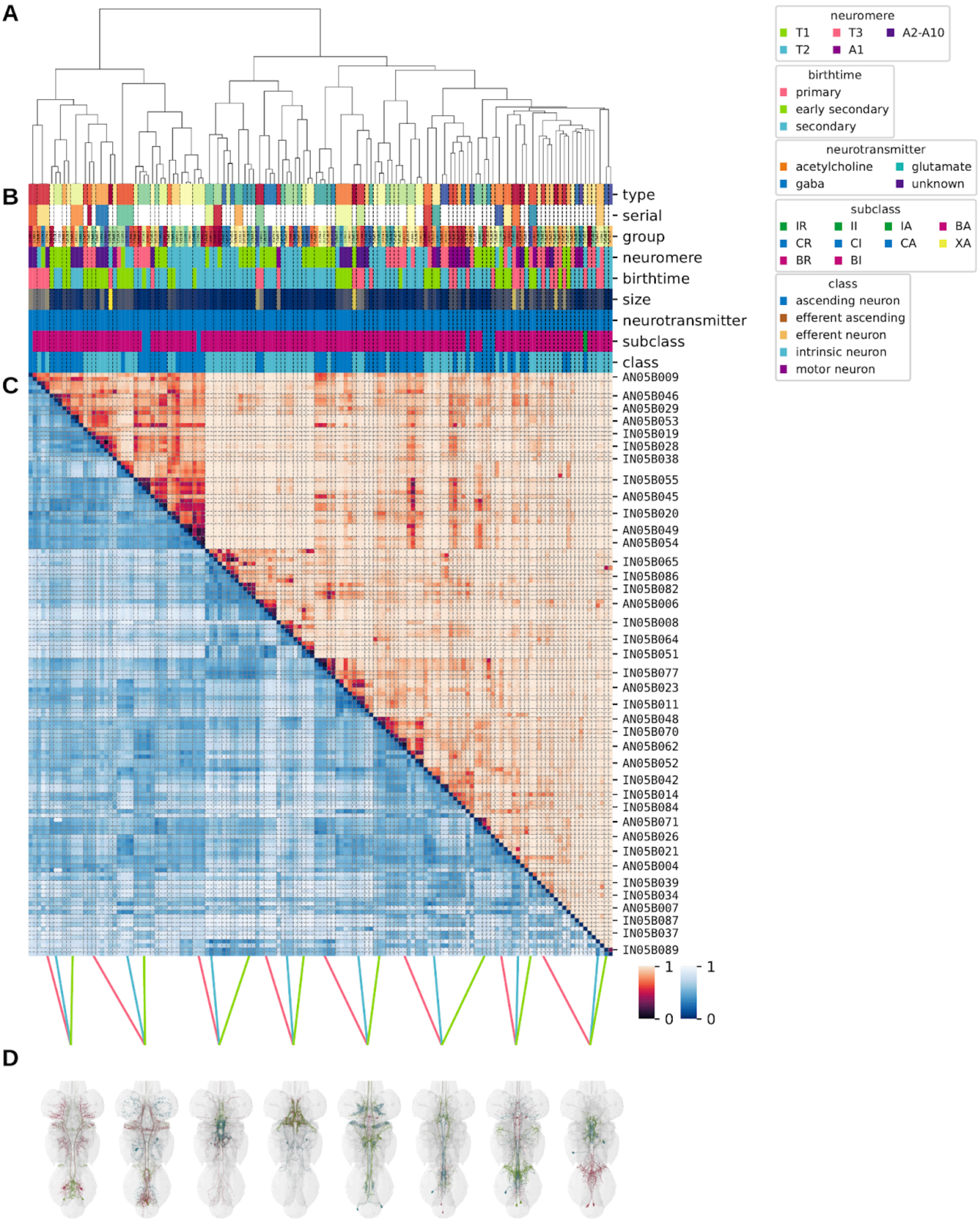

**Figure 22 - figure supplement 1. Systematic typing of hemilineage 05B. A.** Hierarchical clustering dendrogram of hemilineage groups by laterally and serially aggregated connectivity cosine clustering. **B.** Categorical annotations of each hemilineage group, each column corresponding to the aligned leaf in A. Colours for type, serial set, and group are arbitrary for visualisation. Colours for neuromere, birthtime, neurotransmitter, subclass, and class are as in all other figures. **C.** Similarity distance heatmap for hemilineage. Cosine distance is in the upper triangle, while laterally symmetrised NBLAST distance is in the lower triangle. Systematic type names of some types are labelled. **D.**

Morphologically representative groups from dendrogram subtrees. Each group, indicated by colour and line connecting to its column in B and C, is the most morphologically representative group (medoid of NBLAST distance) from a subtree of A. The subtrees (flat clusters) are equal height cuts of A determined to yield the number of groups per plot and plots in D.

**05B systematic types**

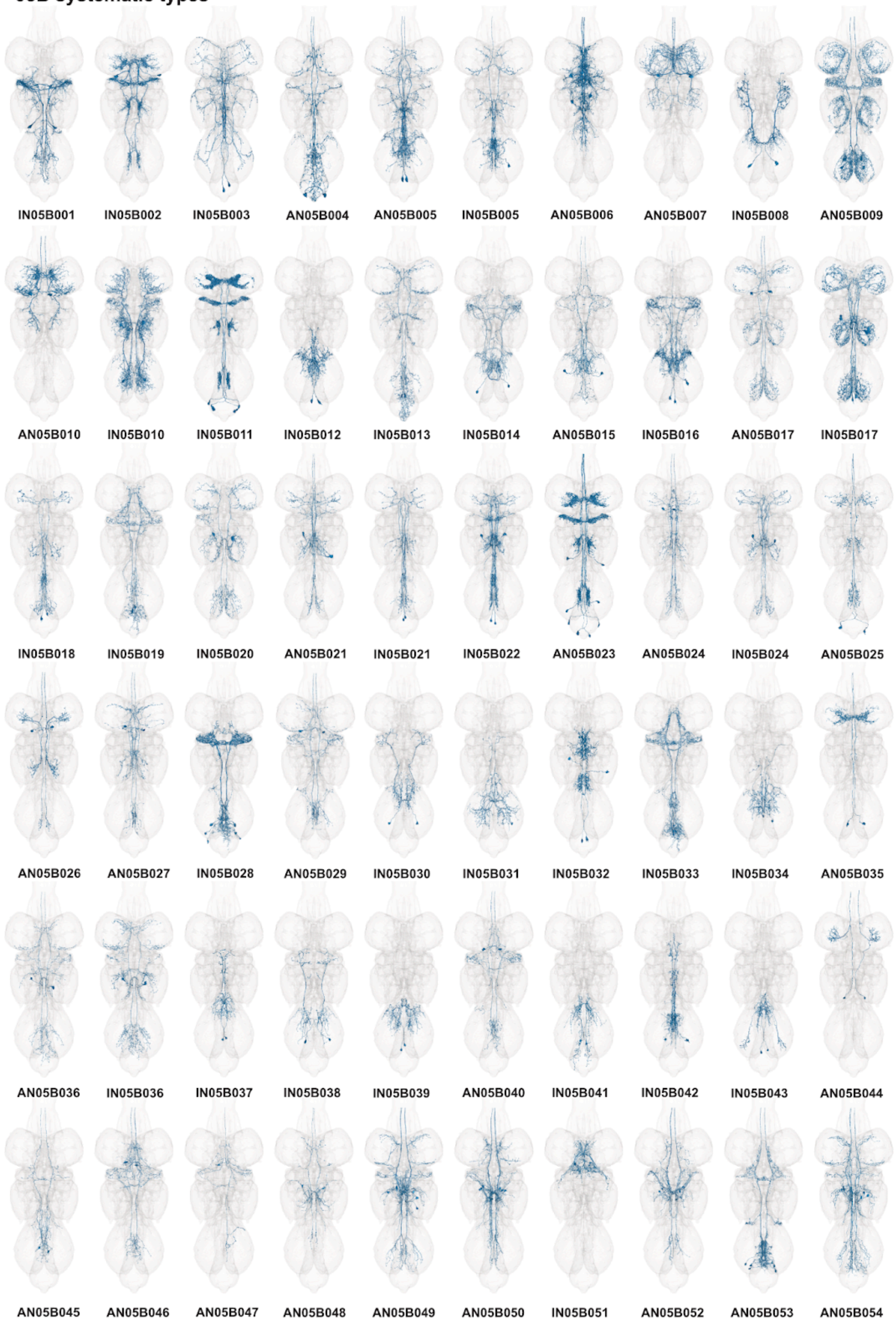

**Figure 22 - figure supplement 2. Systematic types of hemilineage 05B.** Systematic types have been arranged in numerical order, with neurons of the same type that belong to distinct classes (e.g., intrinsic neuron vs ascending neuron) plotted separately but placed adjacent to each other. Individual neuron meshes have been coloured based on predicted neurotransmitter: dark orange = acetylcholine, blue = gaba, marine = glutamate, dark purple = unknown.

#### 05B systematic types, continued

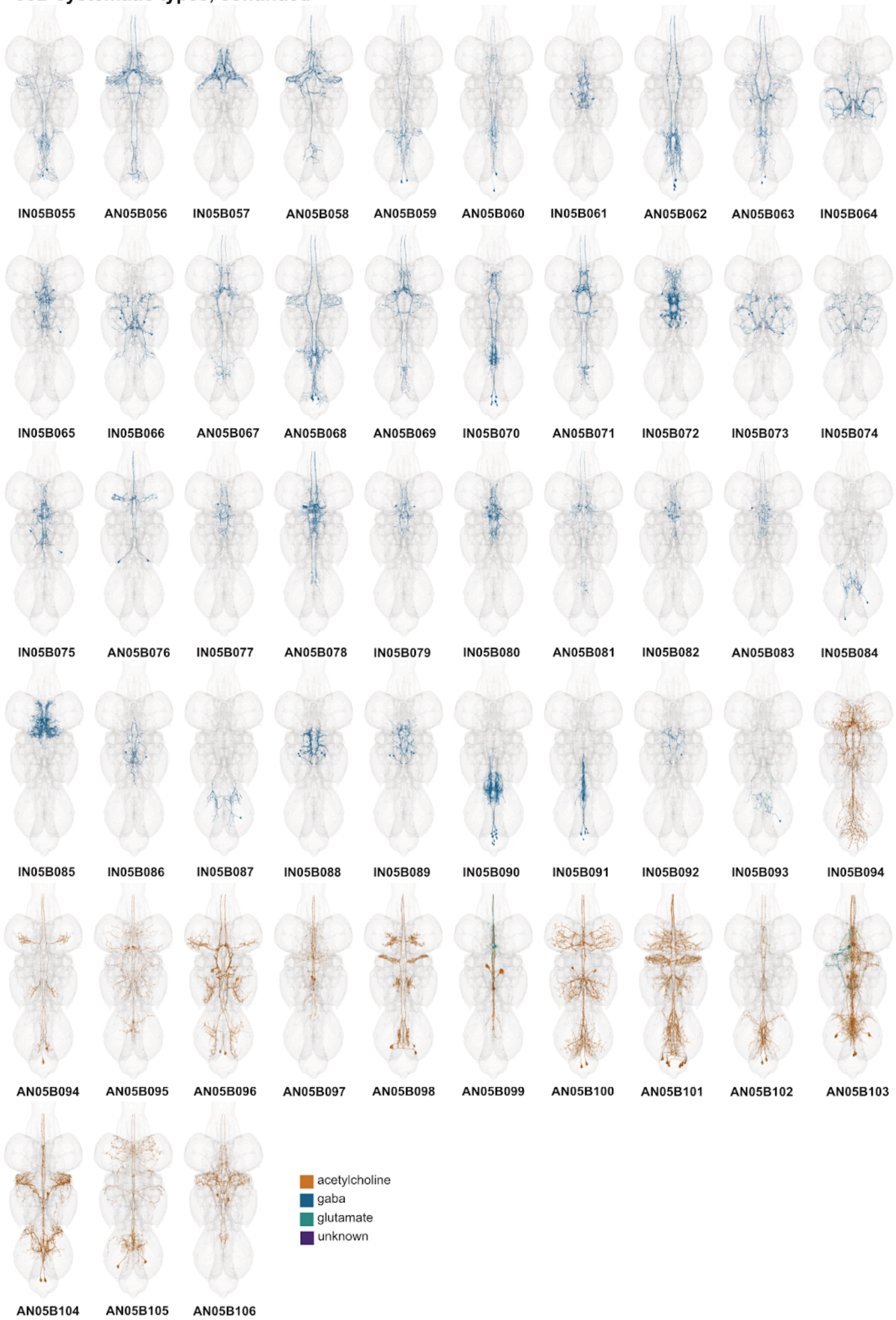

**Figure 22 - figure supplement 3. Systematic types of hemilineage 05B, continued.** Systematic types have been arranged in numerical order, with neurons of the same type that belong to distinct classes (e.g., intrinsic neuron vs ascending neuron) plotted separately but placed adjacent to each other. Individual neuron meshes have been coloured based on predicted neurotransmitter: dark orange = acetylcholine, blue = gaba, marine = glutamate, dark purple = unknown.

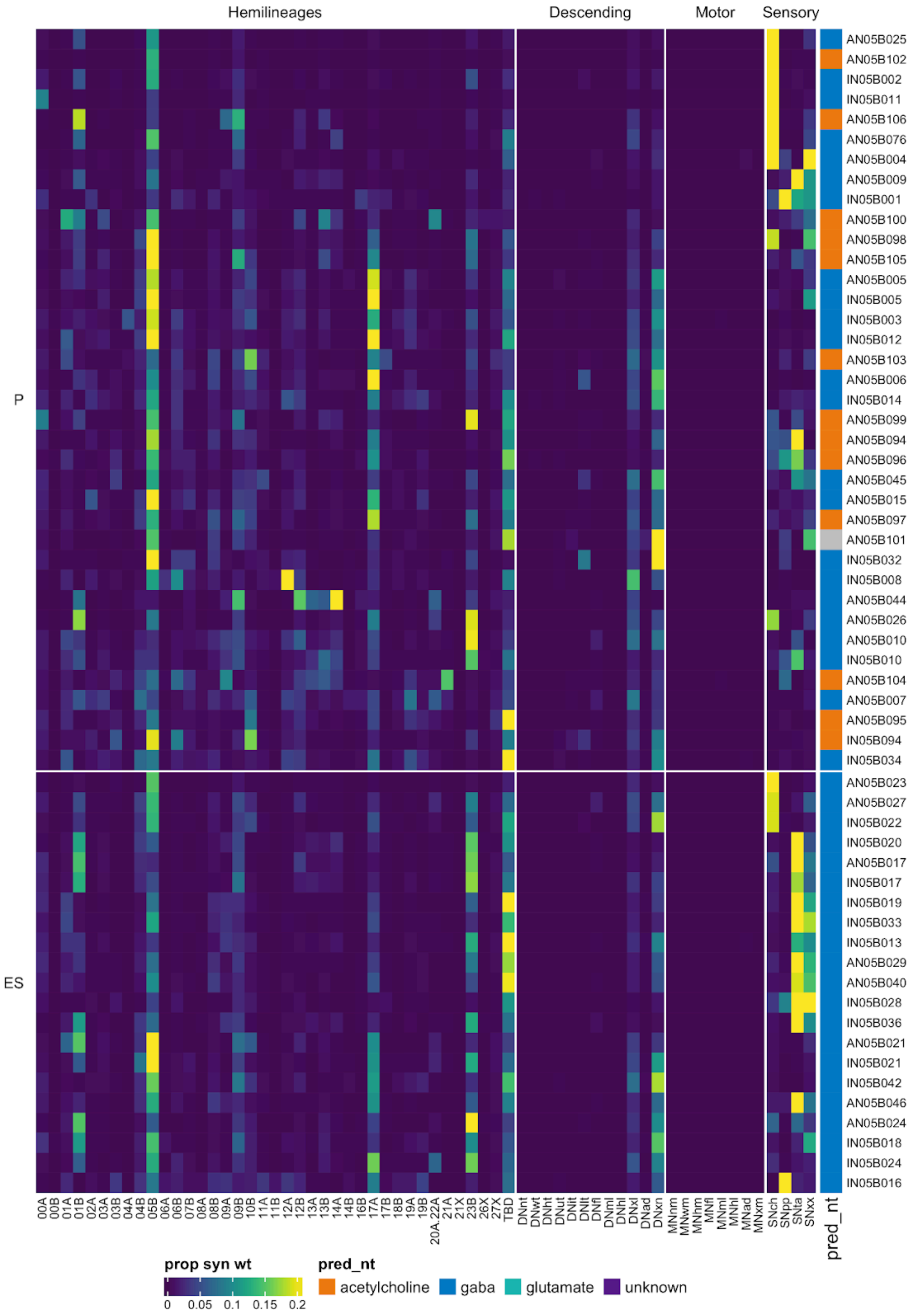

Hemilineages
Descending
Motor
Sensory
P
ES
AN05B025
AN05B102
IN05B002
IN05B011
AN05B106
AN05B076
AN05B004
AN05B009
IN05B001
AN05B100
AN05B098
AN05B105
AN05B005
IN05B005
IN05B003
IN05B012
AN05B103
AN05B006
IN05B014
AN05B099
AN05B094
AN05B096
AN05B045
AN05B015
AN05B097
AN05B101
IN05B032
IN05B008
AN05B044
AN05B026
AN05B010
IN05B010
AN05B104
AN05B007
AN05B095
IN05B094
IN05B034
AN05B023
AN05B027
IN05B022
IN05B020
AN05B017
IN05B017
IN05B019
IN05B033
IN05B013
AN05B029
AN05B040
IN05B028
IN05B036
AN05B021
IN05B021
IN05B042
AN05B046
AN05B024
IN05B018
IN05B024
IN05B016
00A 00B 01A 01B 02A 03A 03B 04A 04B 05B 06A 06B 07B 08A 08B 09A 09B 10B 11A 11B 12A 12B 13A 13B 14A 14B 16B 17A 17B 18B 19A 19B 20A.22A 21A 21X 23B 26X 27X TBD
DNnt DNwt DNht DNut DNit DNlt DNfl DNml DNhl DNxl DNad DNxn
MNnm MNwm MNhm MNfl MNml MNhl MNad MNxm
SNch SNpp SNta SNxx
pred_nt
prop syn wt
0 0.05 0.1 0.15 0.2
pred_nt
acetylcholine
gaba
glutamate
unknown

**Figure 22 - figure supplement 4. Connectivity to upstream partners by 05B primary and early secondary systematic types.** Proportions of synaptic weight to systematic types from upstream partners, normalised by row. 05B neurons have been clustered within each assigned birthtime window (P = primary, ES = early secondary, S = secondary) based on both upstream and downstream connectivity to hemilineages, descending neuron subclasses, motor neuron subclasses, and sensory neuron modalities. Annotation bar is coloured by the most common predicted neurotransmitter for the neurons of each type.

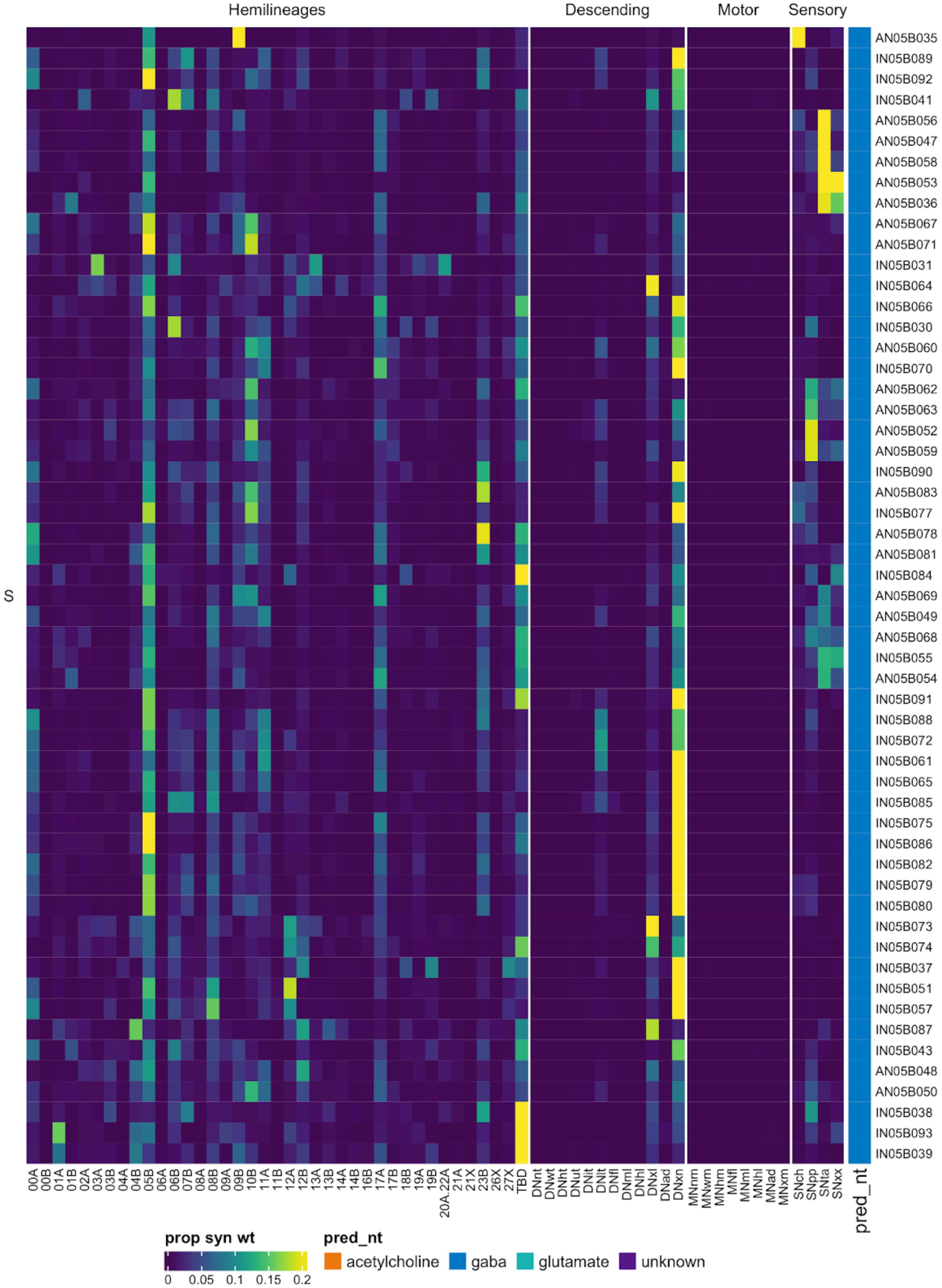
Hemilineages
Descending
Motor
Sensory
S
AN05B035
IN05B089
IN05B092
IN05B041
AN05B056
AN05B047
AN05B058
AN05B053
AN05B036
AN05B067
AN05B071
IN05B031
IN05B064
AN05B066
IN05B030
AN05B060
IN05B070
AN05B062
AN05B063
AN05B052
AN05B059
IN05B090
AN05B083
IN05B077
AN05B078
AN05B081
IN05B084
AN05B069
AN05B049
AN05B068
IN05B055
AN05B054
IN05B091
IN05B088
IN05B072
IN05B061
IN05B065
IN05B085
IN05B075
IN05B086
IN05B082
IN05B079
IN05B080
IN05B073
IN05B074
IN05B037
IN05B051
IN05B057
IN05B087
IN05B043
AN05B048
AN05B050
IN05B038
IN05B093
IN05B039
00A 00B 01A 01B 02A 03A 03B 04A 04B 05B 06A 06B 07B 08A 08B 09A 09B 10B 11A 11B 12A 12B 13A 13B 14A 14B 16B 17A 17B 18B 19A 19B 20A.22A 21X 23B 26X 27X TBD
DNnt DNwt DNht DNut DNlt DNft DNfl DNml DNhl DNxl DNad DNxn
MNnm MNwm MNhm MNfl MNml MNhl MNad MNxm
SNch SNpp SNta SNxx
pred_nt
prop syn wt
0 0.05 0.1 0.15 0.2
pred_nt
acetylcholine
gaba
glutamate
unknown

**Figure 22 - figure supplement 5. Connectivity to upstream partners by 05B secondary systematic types.** Proportions of synaptic weight to systematic types from upstream partners, normalised by row. 05B neurons have been clustered within each assigned birthtime window (P = primary, ES = early secondary, S = secondary) based on both upstream and downstream connectivity to hemilineages, descending neuron subclasses, motor neuron subclasses, and sensory neuron modalities. The annotation bar is coloured by the most common predicted neurotransmitter within each type.

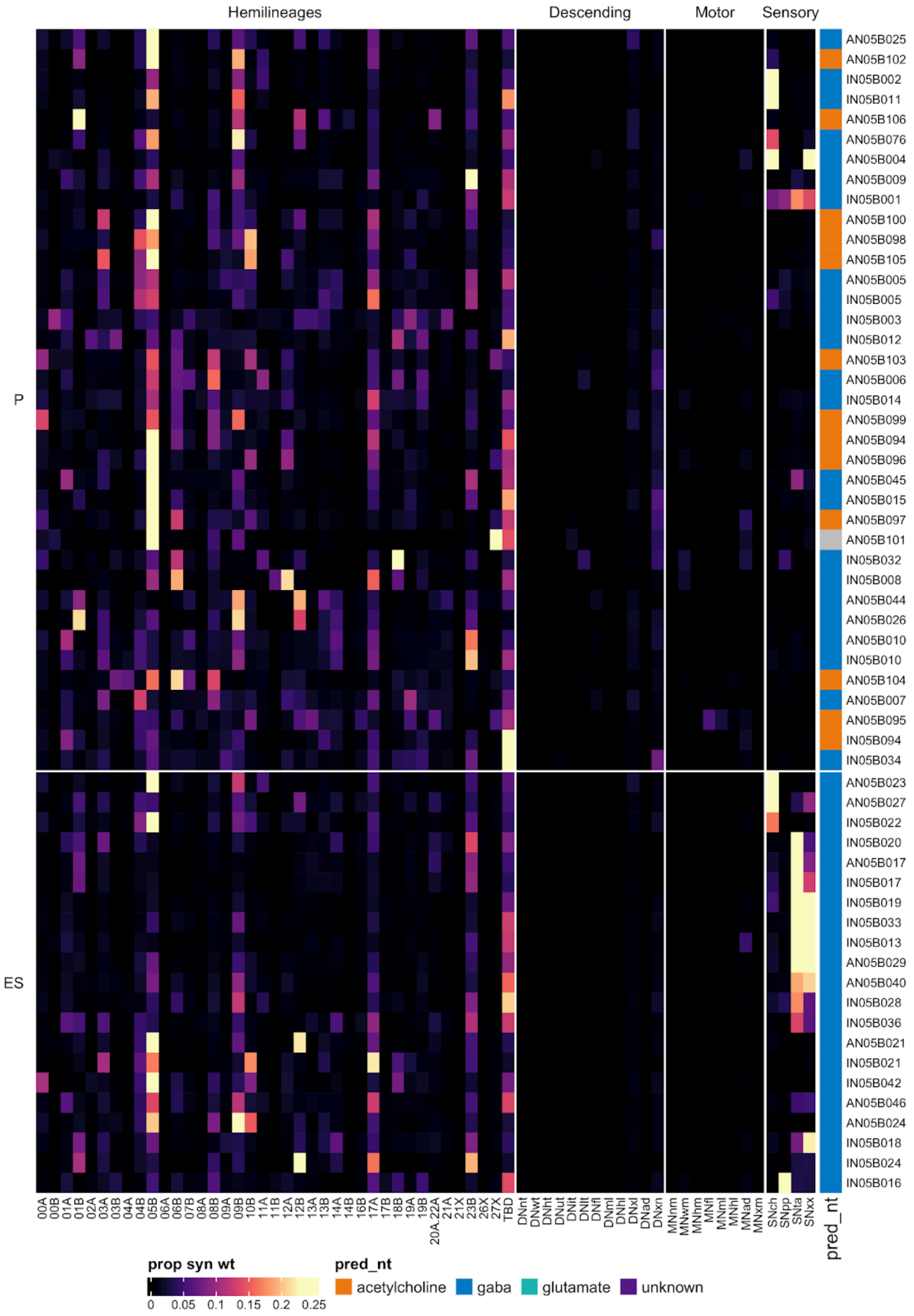
Hemilineages
Descending
Motor
Sensory
P
ES
AN05B025
AN05B102
IN05B002
IN05B011
AN05B106
AN05B076
AN05B004
AN05B009
IN05B001
AN05B100
AN05B098
AN05B105
AN05B005
IN05B005
IN05B003
IN05B012
AN05B103
AN05B006
IN05B014
AN05B099
AN05B094
AN05B096
AN05B045
AN05B015
AN05B097
AN05B101
IN05B032
IN05B008
AN05B044
AN05B026
AN05B010
IN05B010
AN05B104
AN05B007
AN05B095
IN05B094
IN05B034
AN05B023
AN05B027
IN05B022
IN05B020
AN05B017
IN05B017
IN05B019
IN05B033
IN05B013
AN05B029
AN05B040
IN05B028
IN05B036
AN05B021
IN05B021
IN05B042
AN05B046
AN05B024
IN05B018
IN05B024
IN05B016
00A 00B 01A 01B 02A 03A 03B 04A 04B 05B 06A 06B 07B 08A 08B 09A 09B 10B 11A 11B 12A 12B 13A 13B 14A 14B 16B 17A 17B 18B 19A 19B 20A.22A 21A 21X 23B 26X 27X TBD
DNnt DNwt DNht DNut DNlt DNfl DNml DNhl DNxl DNad DNxn
MNnm MNwm MNhm MNfl MNml MNhl MNad MNxm
SNch SNpp SNta SNxx
pred_nt
prop syn wt
0 0.05 0.1 0.15 0.2 0.25
pred_nt
acetylcholine
gaba
glutamate
unknown

**Figure 22 - figure supplement 6. Connectivity to downstream partners by 05B primary and early secondary systematic types.** Proportions of synaptic weight from systematic types to downstream partners, normalised by row. 05B neurons have been clustered within each assigned birthtime window (P = primary, ES = early secondary, S = secondary) based on both upstream and downstream connectivity to hemilineages, descending neuron subclasses, motor neuron subclasses, and sensory neuron modalities. The annotation bar is coloured by the most common predicted neurotransmitter within each type.

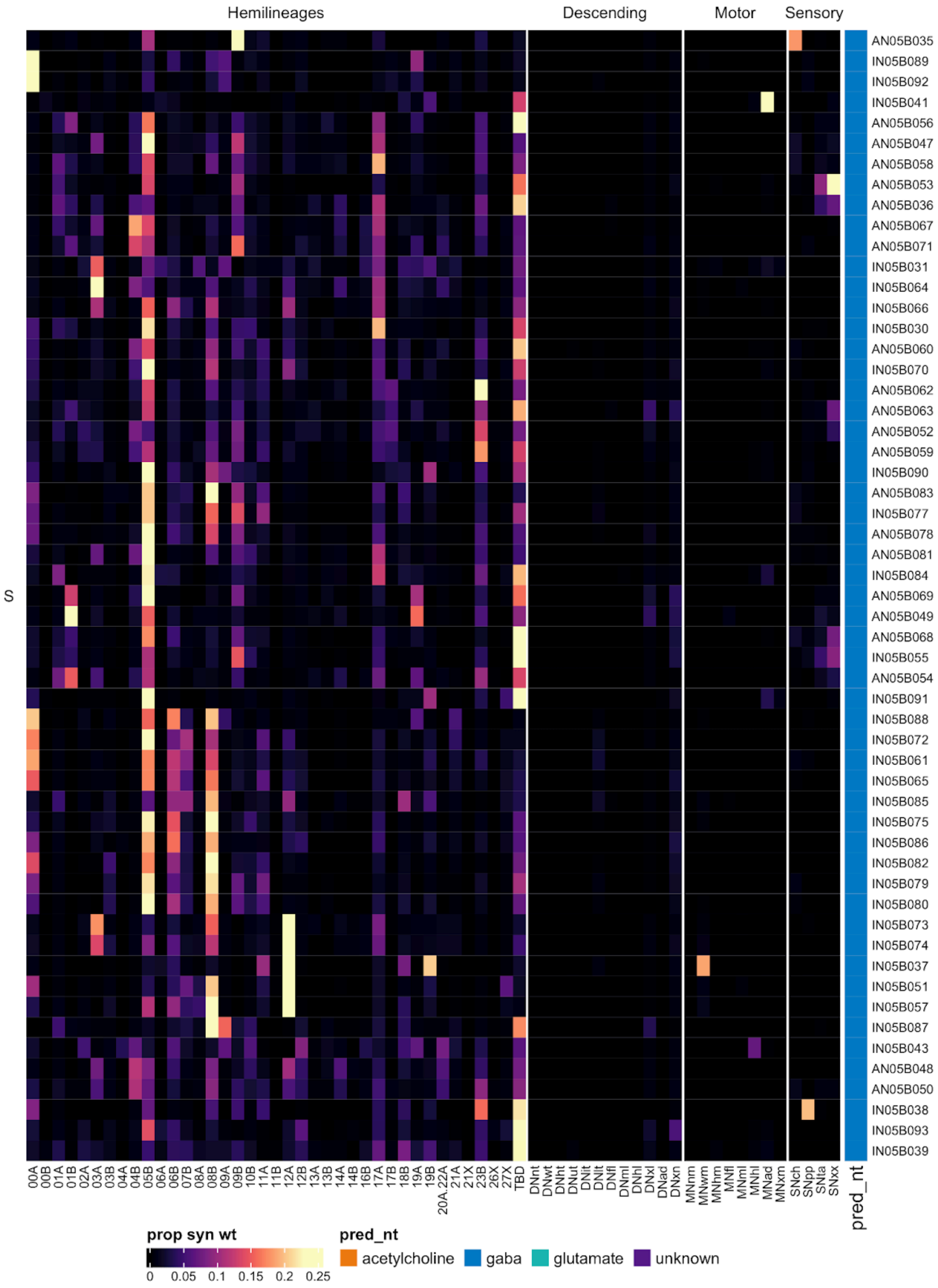
Hemilineages
Descending
Motor
Sensory
S
AN05B035
IN05B089
IN05B092
IN05B041
AN05B056
AN05B047
AN05B058
AN05B053
AN05B036
AN05B067
AN05B071
IN05B031
IN05B064
IN05B066
IN05B030
AN05B060
IN05B070
AN05B062
AN05B063
AN05B052
AN05B059
IN05B090
AN05B083
IN05B077
AN05B078
AN05B081
IN05B084
AN05B069
AN05B049
AN05B068
IN05B055
AN05B054
IN05B091
IN05B088
IN05B072
IN05B061
IN05B065
IN05B085
IN05B075
IN05B086
IN05B082
IN05B079
IN05B080
IN05B073
IN05B074
IN05B037
IN05B051
IN05B057
IN05B087
IN05B043
AN05B048
AN05B050
IN05B038
IN05B093
IN05B039
00A 00B 01A 01B 02A 03A 03B 04A 04B 05B 06A 06B 07B 08A 08B 09A 09B 10B 11A 11B 12A 12B 13A 13B 14A 14B 16B 17A 17B 18B 19A 19B 20A.22A 21A 21X 23B 26X 27X TBD
DNnt DNwt DNht DNut DNlt DNfl DNml DNhl DNxl DNad DNxn
MNnm MNwm MNhm MNfl MNml MNhl MNad MNxm
SNch SNpp SNta SNxx
pred_nt
prop syn wt
0 0.05 0.1 0.15 0.2 0.25
pred_nt
acetylcholine
gaba
glutamate
unknown

**Figure 22 - figure supplement 7. Connectivity to downstream partners by 05B secondary systematic types.** Proportions of synaptic weight from systematic types to downstream partners, normalised by row. 05B neurons have been clustered within each assigned birthtime window (P = primary, ES = early secondary, S = secondary) based on both upstream and downstream connectivity to hemilineages, descending neuron subclasses, motor neuron subclasses, and sensory neuron modalities. The annotation bar is coloured by the most common predicted neurotransmitter for the neurons of each type.

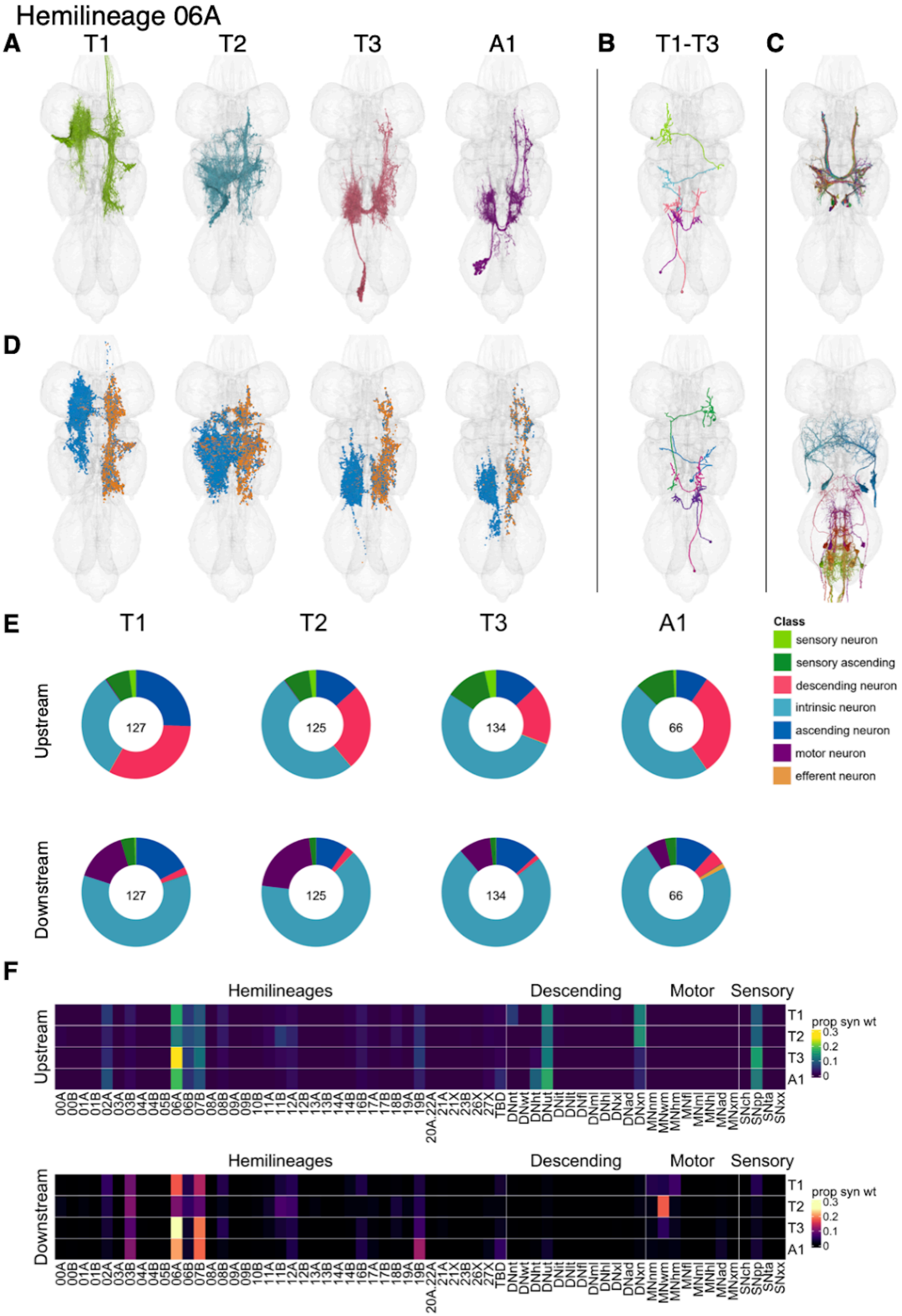

Hemilineage 06A
A
T1
T2
T3
A1
B
T1-T3
C
D
E
T1
T2
T3
A1
Upstream
127
125
134
66
Downstream
127
125
134
66
Class
sensory neuron
sensory ascending
descending neuron
intrinsic neuron
ascending neuron
motor neuron
efferent neuron
F
Hemilineages
Descending
Motor
Sensory
Upstream
Downstream
T1
T2
T3
A1
prop syn wt
0.3
0.2
0.1
0
00A
00B
01A
01B
02A
03A
03B
04A
04B
05B
06A
06B
07B
08A
08B
09A
09B
10B
11A
11B
12A
12B
13A
13B
14A
14B
16B
17A
17B
18B
19A
19B
20A.22A
21A
21X
23B
26X
27X
TBD
DNnt
DNwt
DNht
DNut
DNit
DNlt
DNfl
DNml
DNhl
DNxl
DNad
DNxn
MNnm
MNwm
MNhm
MNfl
MNml
MNhl
MNad
MNxm
SNch
SNpp
SNta
SNxx

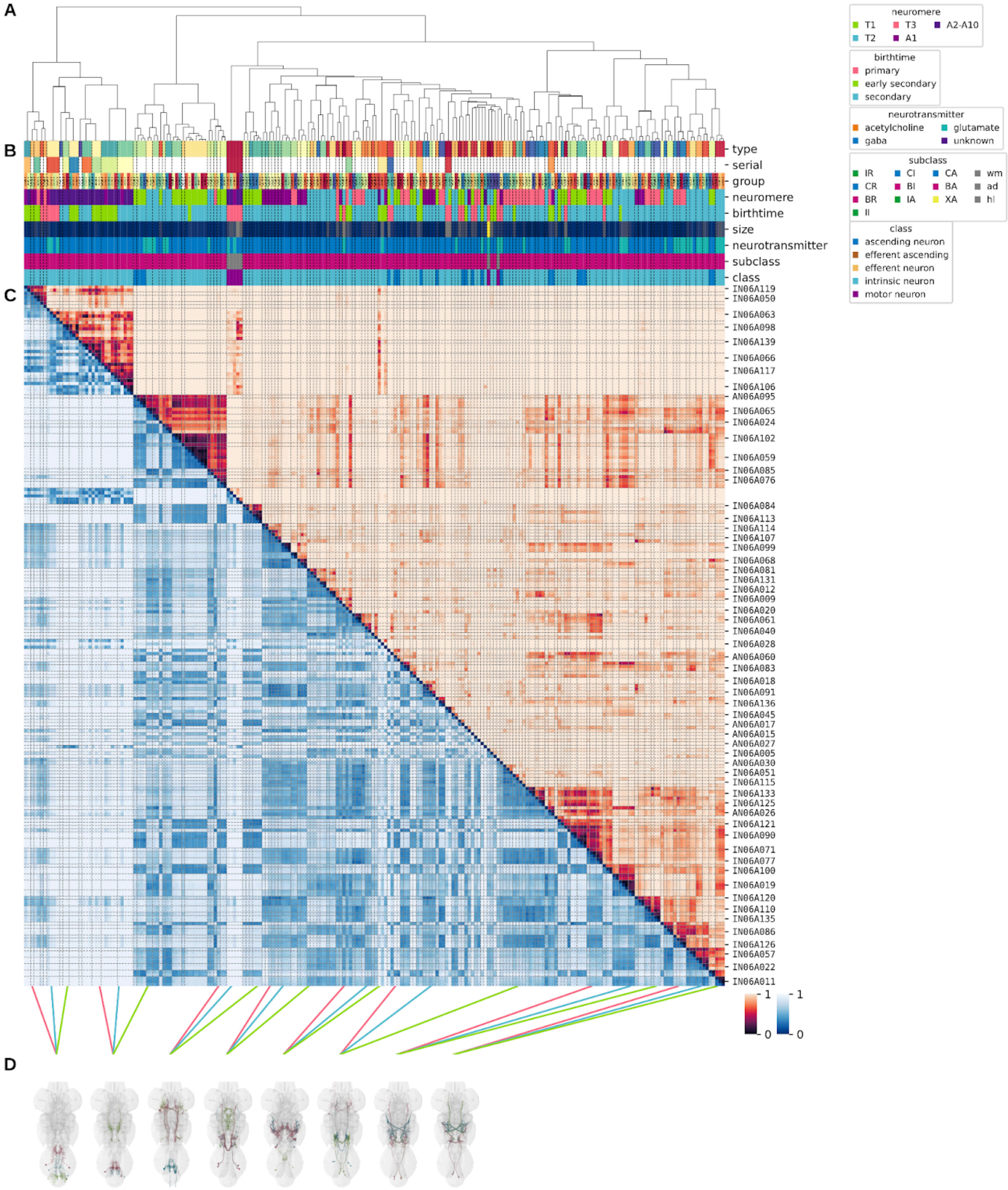

**Figure 23. Hemilineage 06A. A.** Meshes of all RHS secondary neurons plotted in neuromere-specific colours. **B.** “Representative” secondary neuron skeletons plotted in hemineuromere-specific colours. The skeleton with the top accumulated NBLAST score among all neurons from the hemilineage in a given hemineuromere was used. **C.** Neuron meshes of selected examples. Top: putative electrical subcluster 12112. Bottom: motor neuron subcluster 10204. **D.** Predicted synapses of RHS secondary neurons. Blue: postsynapses; dark orange: presynapses. **E.** Proportions of connections from secondary neurons to upstream or downstream partners, normalised by neuromere and coloured by broad class. Numbers of query neurons appear in the centre. **F.** Proportions of synaptic weight from secondary neurons originating in each neuromere to upstream or downstream partners, normalised by row.

**Figure 23 - figure supplement 1. Systematic typing of hemilineage 06A.** Types for motor neurons were assigned separately as outlined in our accompanying manuscript (Cheong et al., 2023). **A.** Hierarchical clustering dendrogram of hemilineage groups by laterally and serially aggregated connectivity cosine clustering. **B.** Categorical annotations of each hemilineage group, each column corresponding to the aligned leaf in A. Colours for type, serial set, and group are arbitrary for visualisation. Colours for neuromere, birthtime, neurotransmitter, subclass, and class are as in all other figures. **C.** Similarity distance heatmap for hemilineage. Cosine distance is in the upper triangle, while laterally symmetrised NBLAST distance is in the lower triangle. Systematic type names of some types are labelled. **D.** Morphologically representative groups from dendrogram subtrees. Each group, indicated by colour and line connecting to its column in B and C, is the most morphologically representative group (medoid of NBLAST distance) from a subtree of A. The subtrees (flat clusters) are equal height cuts of A determined to yield the number of groups per plot and plots in D.

**06A systematic types**

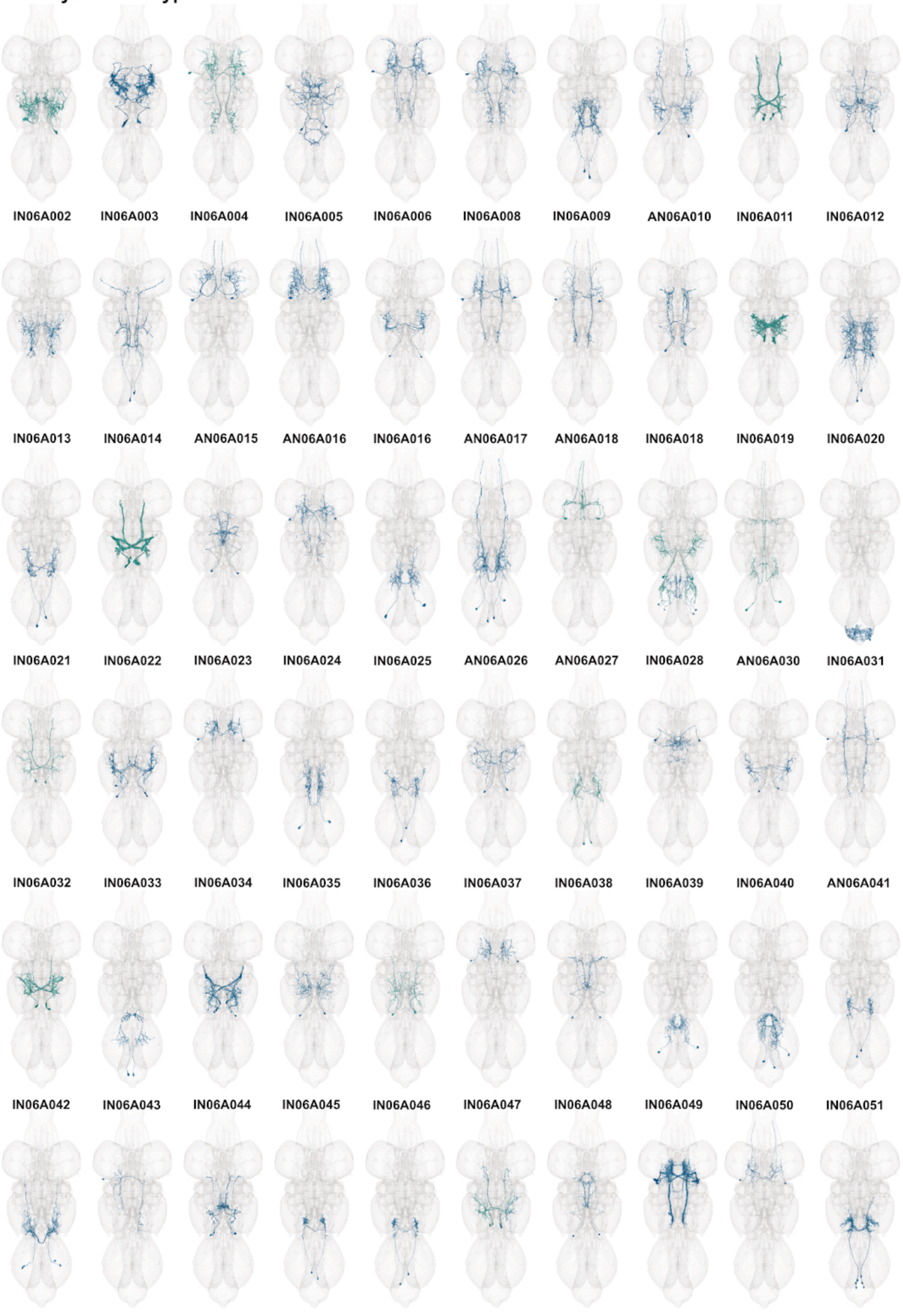

**Figure 23 - figure supplement 2. Systematic types of hemilineage 06A.** Systematic types have been arranged in numerical order, with neurons of the same type that belong to distinct classes (e.g., intrinsic neuron vs ascending neuron) plotted separately but placed adjacent to each other. Individual neuron meshes have been coloured based on predicted neurotransmitter: dark orange = acetylcholine, blue = gaba, marine = glutamate, dark purple = unknown.

#### 06A systematic types, continued

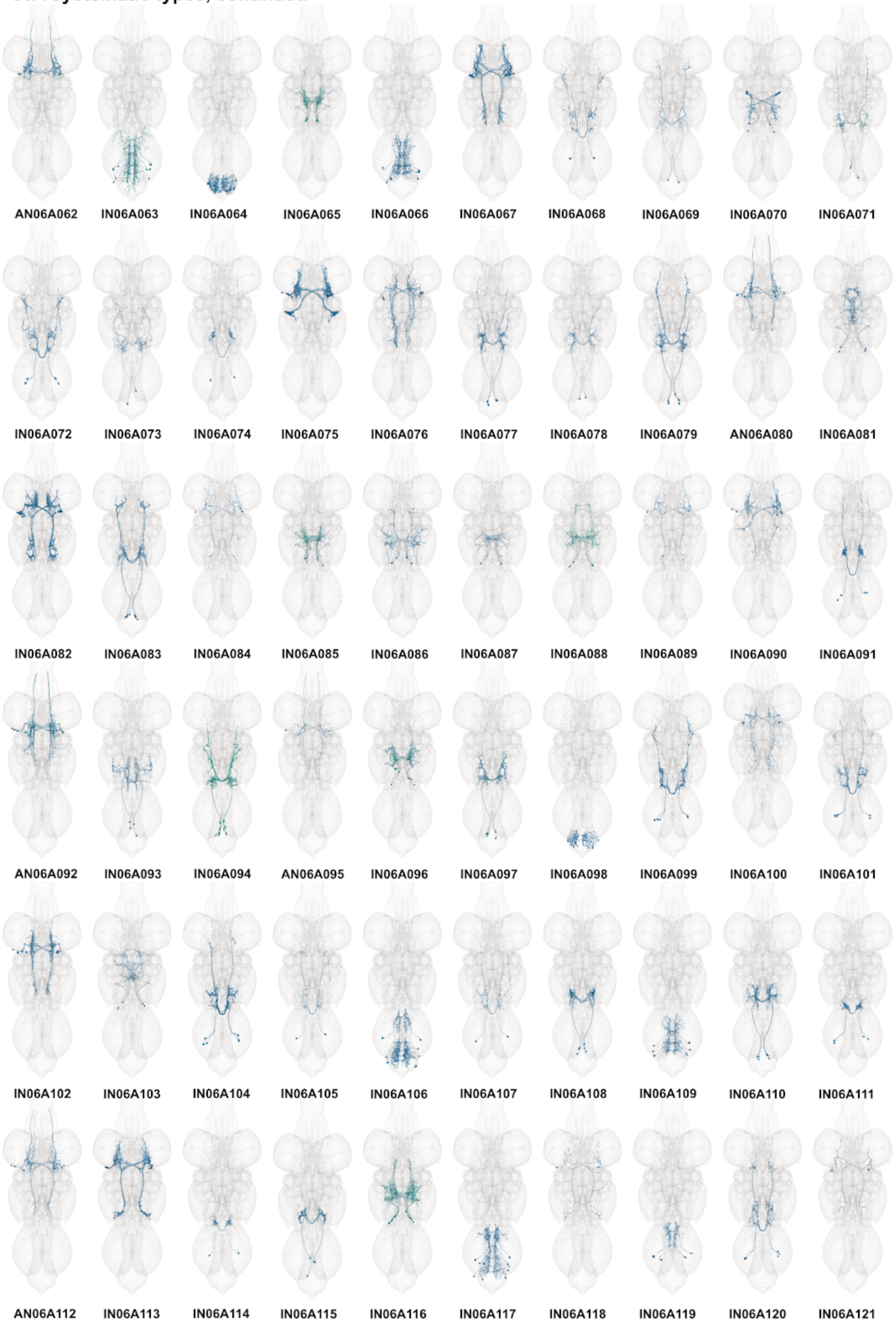

**Figure 23 - figure supplement 3. Systematic types of hemilineage 06A, continued.** Systematic types have been arranged in numerical order, with neurons of the same type that belong to distinct classes (e.g., intrinsic neuron vs ascending neuron) plotted separately but placed adjacent to each other. Individual neuron meshes have been coloured based on predicted neurotransmitter: dark orange = acetylcholine, blue = gaba, marine = glutamate, dark purple = unknown.

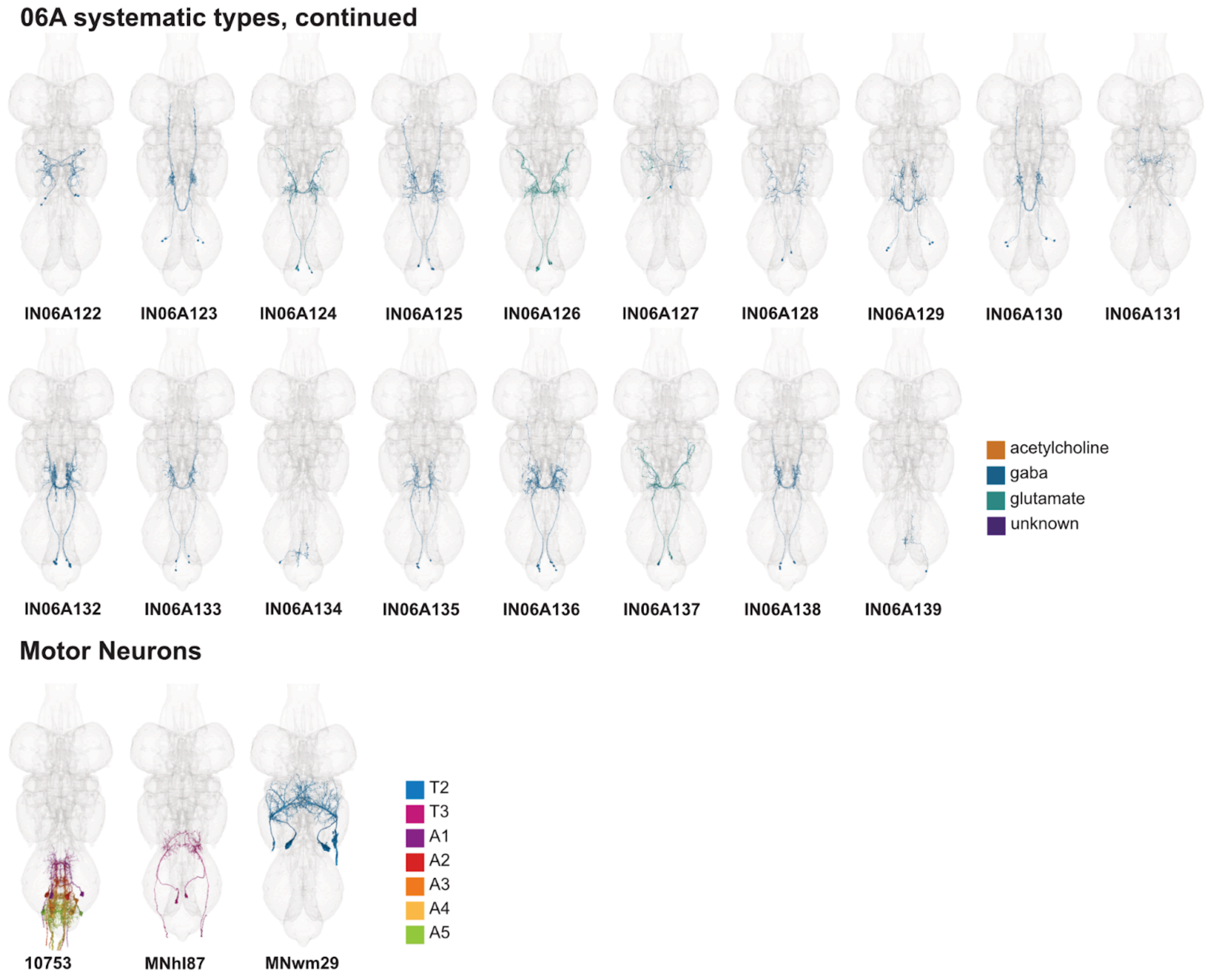

**Figure 23 - figure supplement 4. Systematic types of hemilineage 06A, continued.** Systematic types have been arranged in numerical order, with neurons of the same type that belong to distinct classes (e.g., intrinsic neuron vs ascending neuron) plotted separately but placed adjacent to each other. Individual neuron meshes have been coloured based on predicted neurotransmitter: dark orange = acetylcholine, blue = gaba, marine = glutamate, dark purple = unknown. Motor neurons (typed separately in (Cheong et al., 2023)) have been plotted by serial set if identified in multiple neuromeres and by systematic type if not. Individual motor neuron meshes have been coloured based on soma neuromere: blue = T2, magenta = T3, purple = A1, red = A2, dark orange = A3, dark yellow = A4, green = A5.

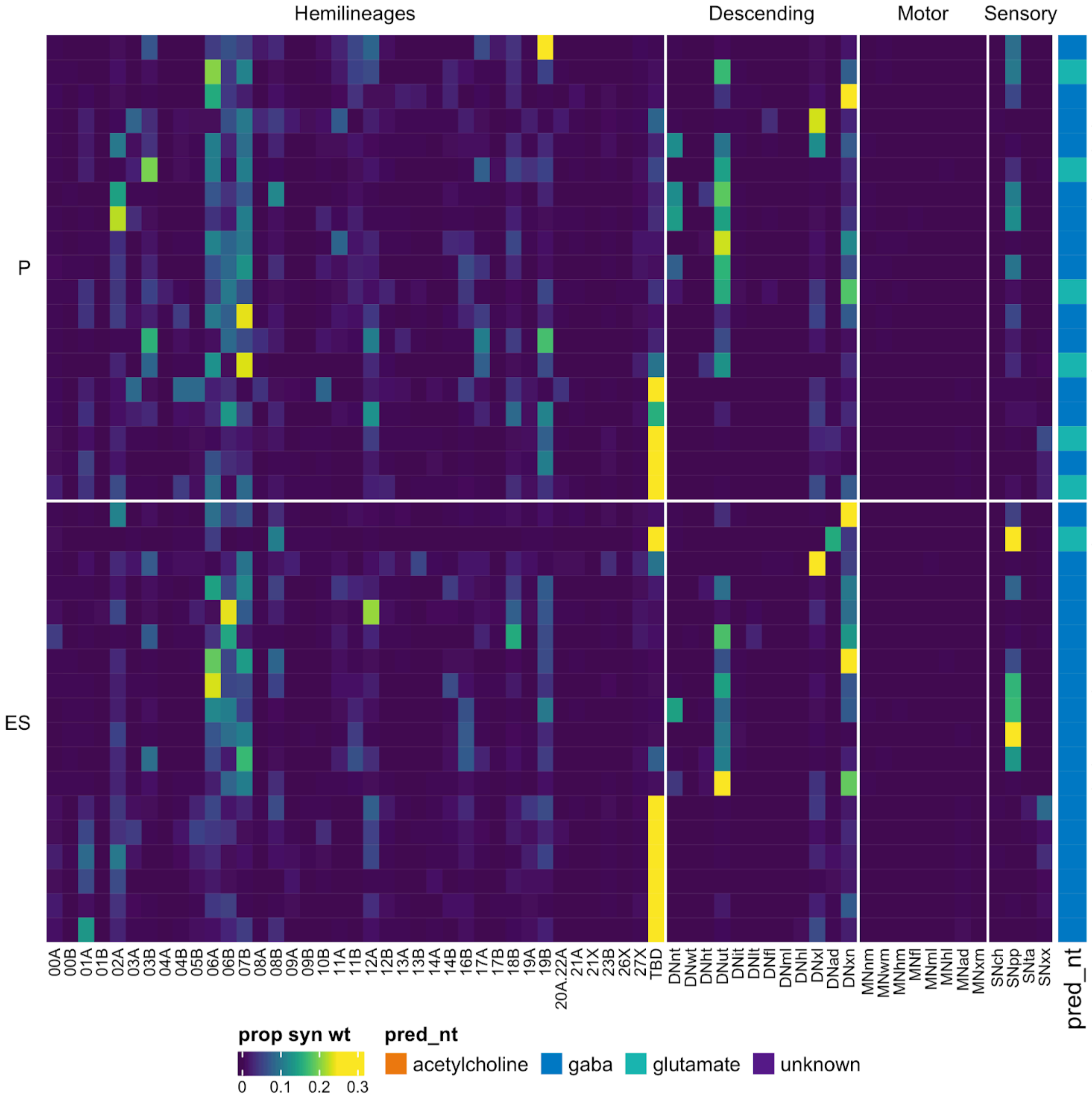

**Figure 23 - figure supplement 5. Connectivity to upstream partners by 06A primary and early secondary systematic types.** Proportions of synaptic weight to systematic types from upstream partners, normalised by row. 06A neurons have been clustered within each assigned birthtime window (P = primary, ES = early secondary, S = secondary) based on both upstream and downstream connectivity to hemilineages, descending neuron subclasses, motor neuron subclasses, and sensory neuron modalities. Annotation bar is coloured by the most common predicted neurotransmitter for the neurons of each type.

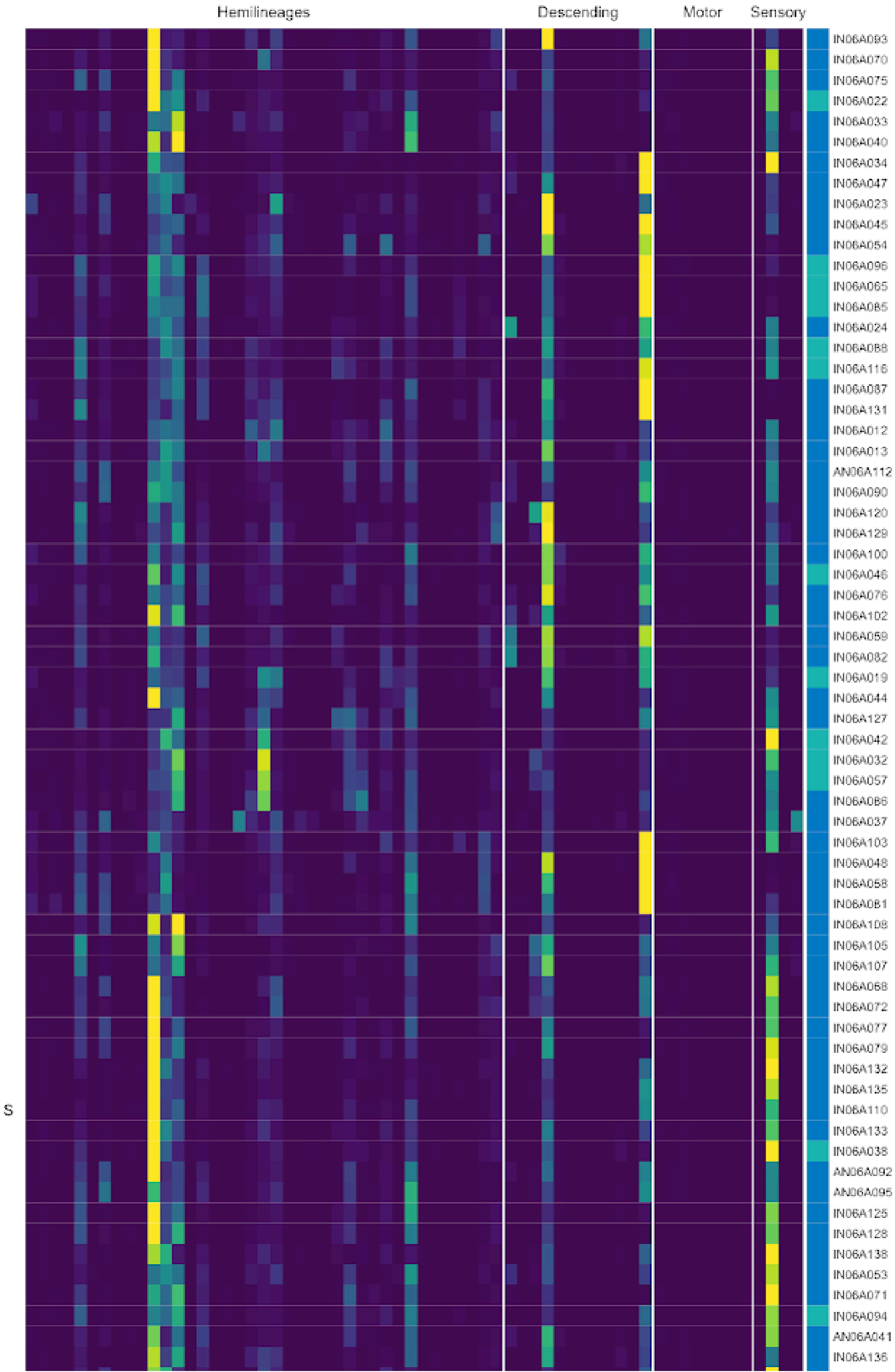
Hemilineages
Descending
Motor
Sensory
S
IN06A093
IN06A070
IN06A075
IN06A022
IN06A033
IN06A040
IN06A034
IN06A047
IN06A023
IN06A045
IN06A054
IN06A096
IN06A065
IN06A085
IN06A024
IN06A088
IN06A116
IN06A087
IN06A131
IN06A012
IN06A013
AN06A112
IN06A090
IN06A120
IN06A129
IN06A100
IN06A046
IN06A076
IN06A102
IN06A059
IN06A082
IN06A019
IN06A044
IN06A127
IN06A042
IN06A032
IN06A057
IN06A086
IN06A037
IN06A103
IN06A048
IN06A058
IN06A081
IN06A108
IN06A105
IN06A107
IN06A068
IN06A072
IN06A077
IN06A079
IN06A132
IN06A135
IN06A110
IN06A133
IN06A038
AN06A092
AN06A095
IN06A125
IN06A128
IN06A138
IN06A053
IN06A071
IN06A094
AN06A041
IN06A136

**Figure 23 - figure supplement 6. Connectivity to upstream partners by 06A secondary systematic types.** Proportions of synaptic weight to systematic types from upstream partners, normalised by row. 06A neurons have been clustered within each assigned birthtime window (P = primary, ES = early secondary, S = secondary) based on both upstream and downstream connectivity to hemilineages, descending neuron subclasses, motor neuron subclasses, and sensory neuron modalities. The annotation bar is coloured by the most common predicted neurotransmitter within each type.

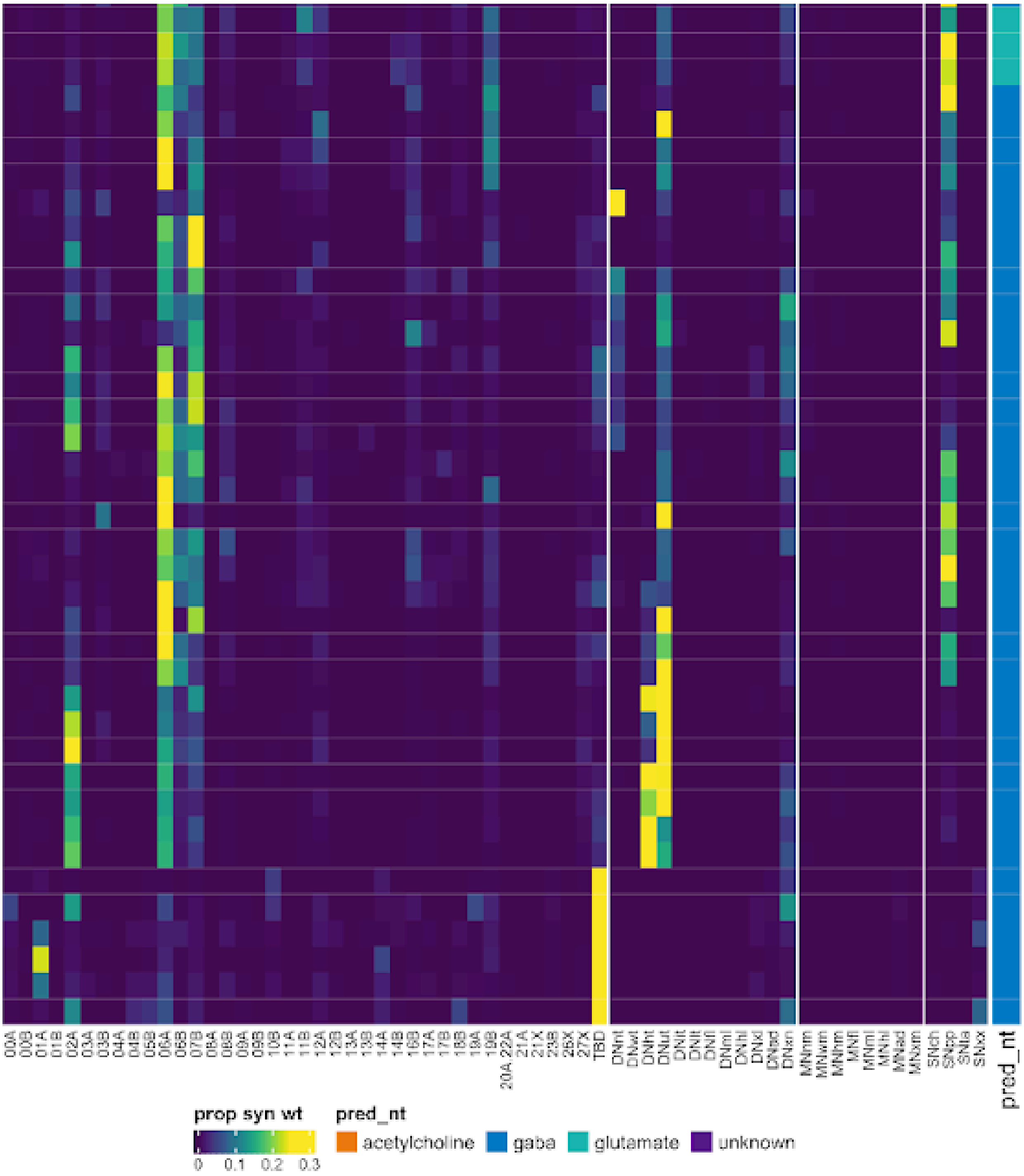

**Figure 23 - figure supplement 7. Connectivity to upstream partners by 06A secondary systematic types, continued.** Proportions of synaptic weight to systematic types from upstream partners, normalised by row. 06A neurons have been clustered within each assigned birthtime window (P = primary, ES = early secondary, S = secondary) based on both upstream and downstream connectivity to hemilineages, descending neuron subclasses, motor neuron subclasses, and sensory neuron modalities. The annotation bar is coloured by the most common predicted neurotransmitter within each type.

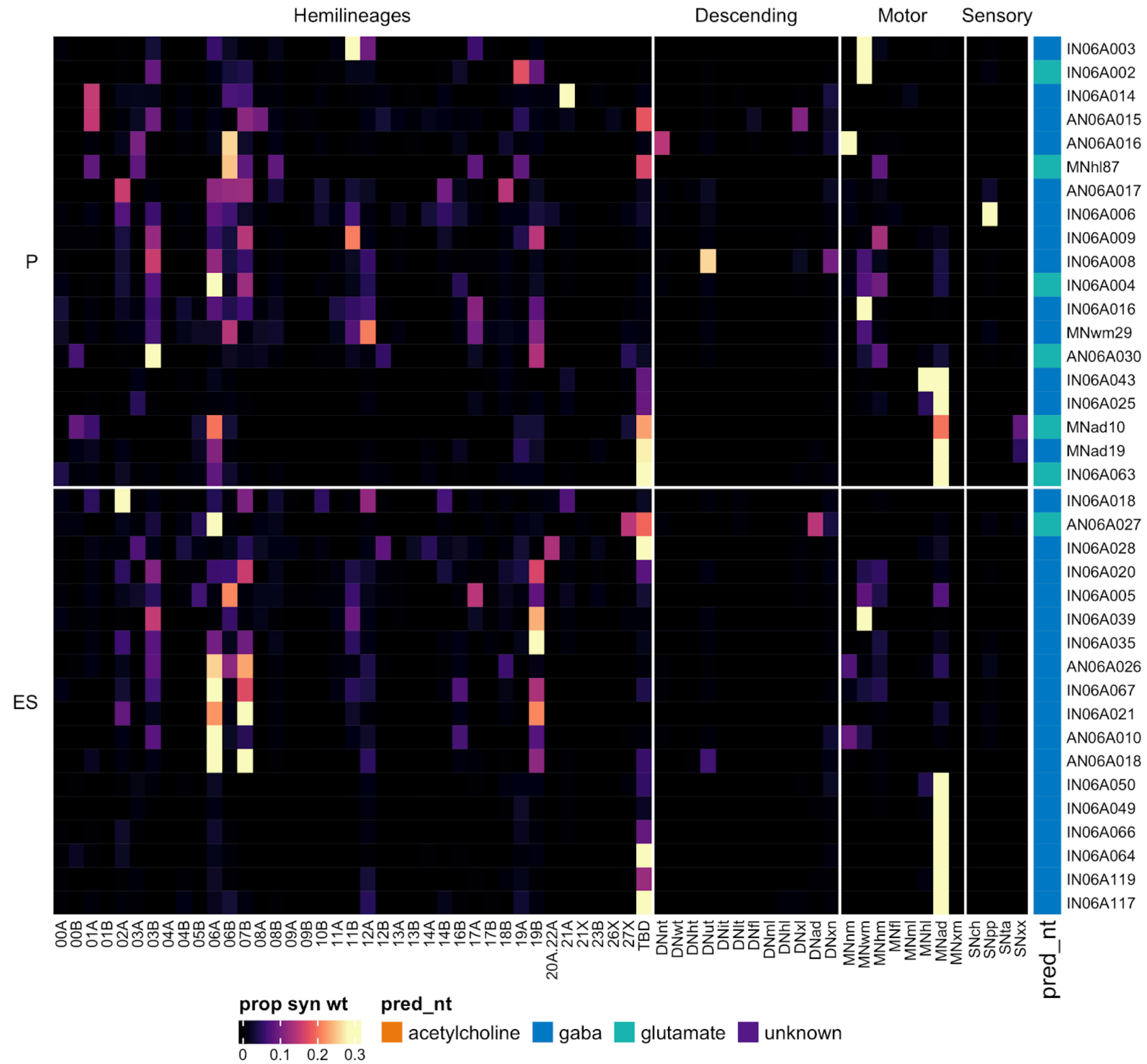

**Figure 23 - figure supplement 8. Connectivity to downstream partners by 06A primary and early secondary systematic types.** Proportions of synaptic weight from systematic types to downstream partners, normalised by row. 06A neurons have been clustered within each assigned birthtime window (P = primary, ES = early secondary, S = secondary) based on both upstream and downstream connectivity to hemilineages, descending neuron subclasses, motor neuron subclasses, and sensory neuron modalities. The annotation bar is coloured by the most common predicted neurotransmitter within each type.

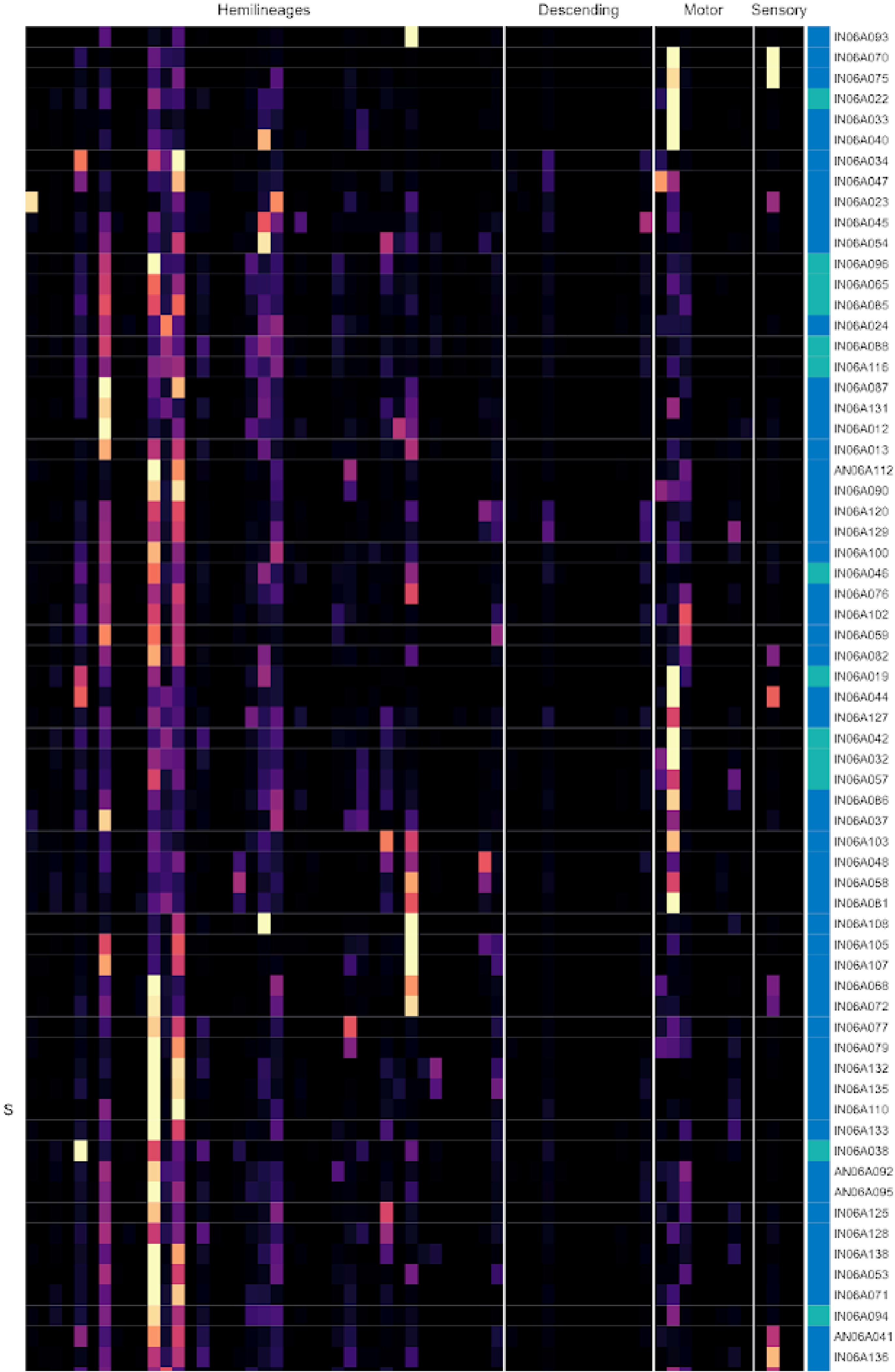
Hemilineages
Descending
Motor
Sensory
S
IN06A093
IN06A070
IN06A075
IN06A022
IN06A033
IN06A040
IN06A034
IN06A047
IN06A023
IN06A045
IN06A054
IN06A096
IN06A065
IN06A085
IN06A024
IN06A088
IN06A116
IN06A087
IN06A131
IN06A012
IN06A013
AN06A112
IN06A090
IN06A120
IN06A129
IN06A100
IN06A046
IN06A076
IN06A102
IN06A059
IN06A082
IN06A019
IN06A044
IN06A127
IN06A042
IN06A032
IN06A057
IN06A086
IN06A037
IN06A103
IN06A048
IN06A058
IN06A081
IN06A108
IN06A105
IN06A107
IN06A068
IN06A072
IN06A077
IN06A079
IN06A132
IN06A135
IN06A110
IN06A133
IN06A038
AN06A092
AN06A095
IN06A125
IN06A128
IN06A138
IN06A053
IN06A071
IN06A094
AN06A041
IN06A136

**Figure 23 - figure supplement 9. Connectivity to downstream partners by 06A secondary systematic types.** Proportions of synaptic weight from systematic types to downstream partners, normalised by row. 06A neurons have been clustered within each assigned birthtime window (P = primary, ES = early secondary, S = secondary) based on both upstream and downstream connectivity to hemilineages, descending neuron subclasses, motor neuron subclasses, and sensory neuron modalities. The annotation bar is coloured by the most common predicted neurotransmitter for the neurons of each type.

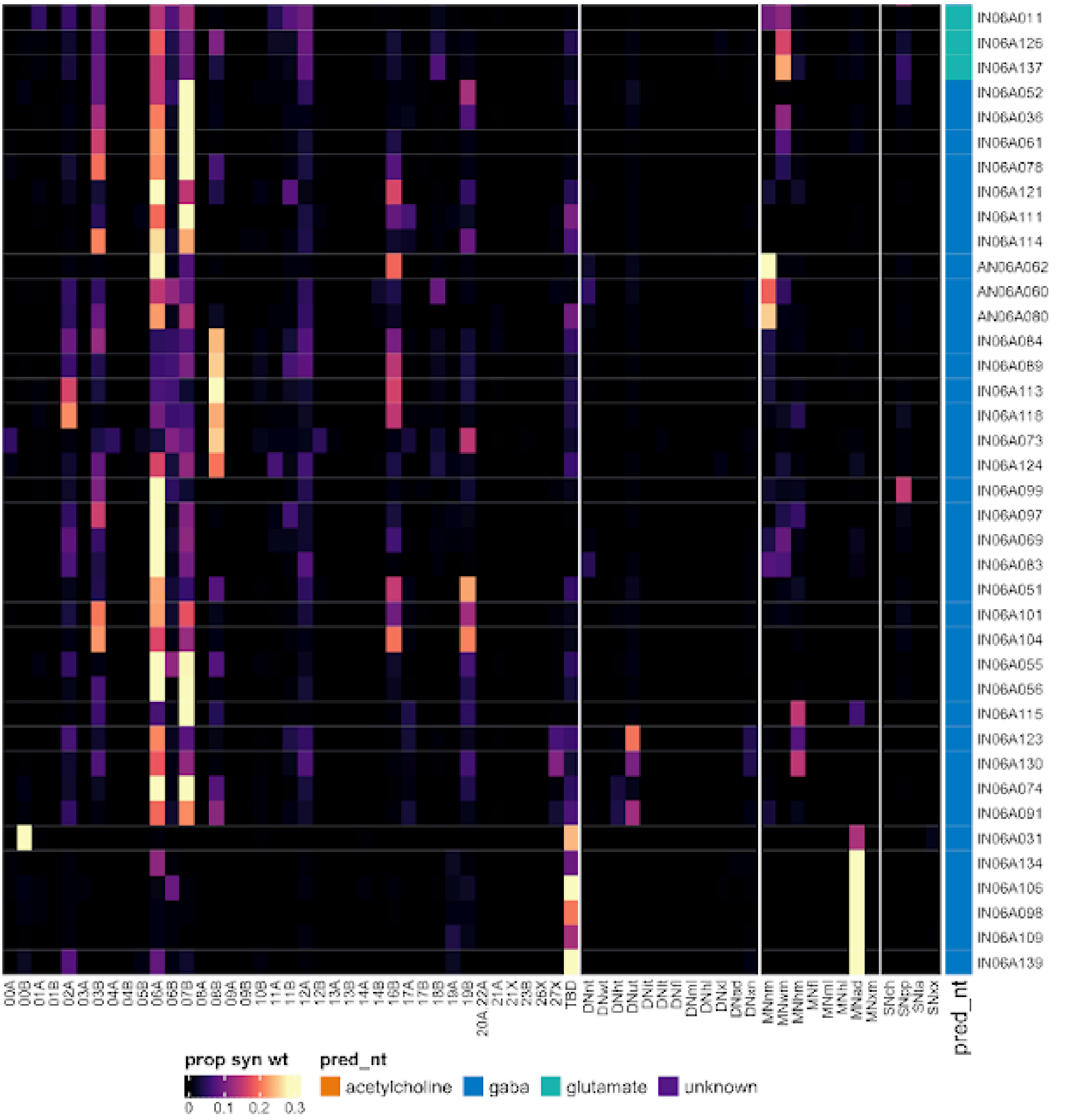

**Figure 23 - figure supplement 10. Connectivity to downstream partners by 06A secondary systematic types, continued.** Proportions of synaptic weight from systematic types to downstream partners, normalised by row. 06A neurons have been clustered within each assigned birthtime window (P = primary, ES = early secondary, S = secondary) based on both upstream and downstream connectivity to hemilineages, descending neuron subclasses, motor neuron subclasses, and sensory neuron modalities. The annotation bar is coloured by the most common predicted neurotransmitter for the neurons of each type.

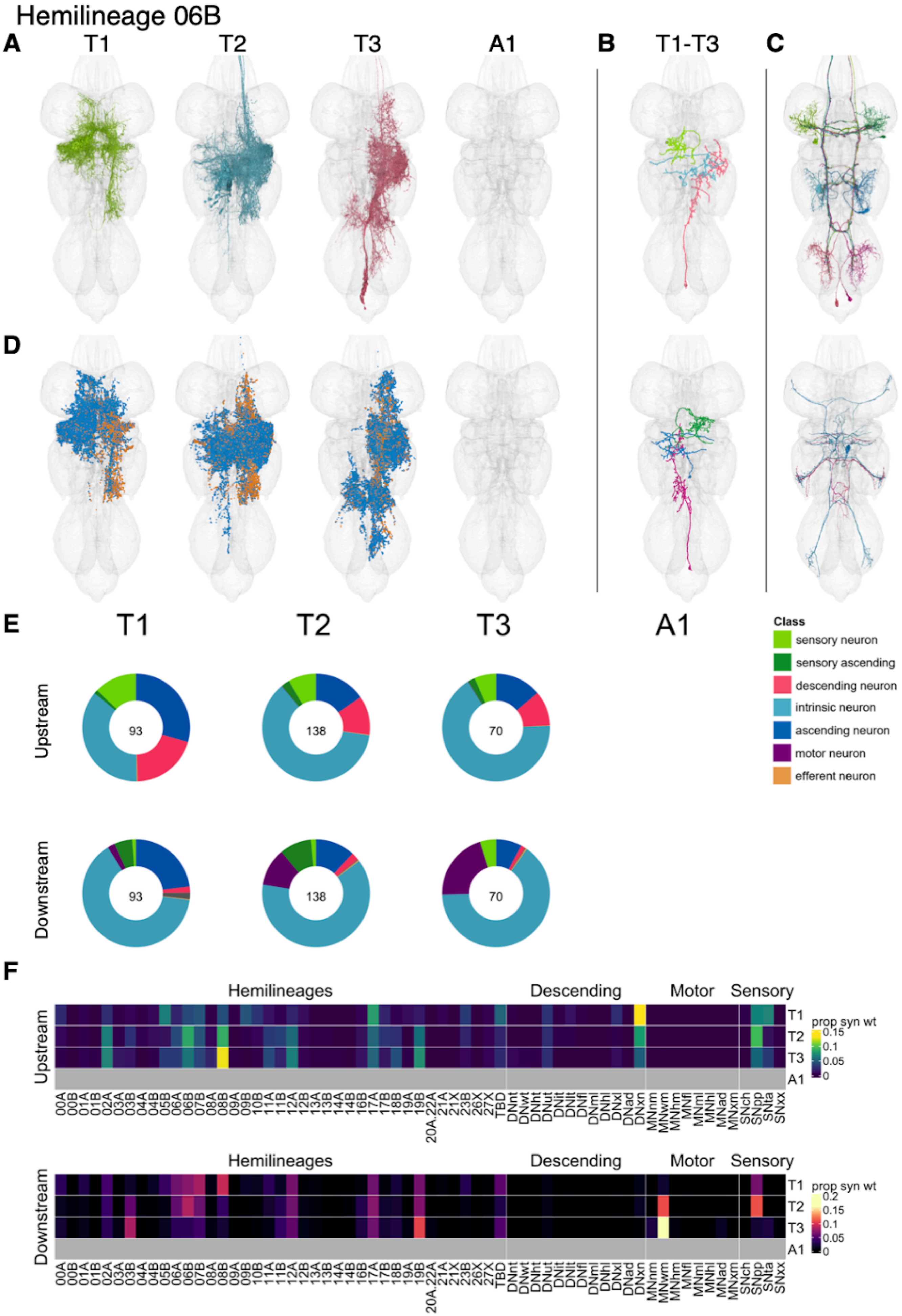
Hemilineage 06B
A
T1
T2
T3
A1
B
T1-T3
C
D
E
T1
T2
T3
A1
Upstream
Downstream
93
138
70
Class
sensory neuron
sensory ascending
descending neuron
intrinsic neuron
ascending neuron
motor neuron
efferent neuron
F
Hemilineages
Descending
Motor
Sensory
prop syn wt
0.15
0.1
0.05
0
0.2
T1
T2
T3
A1
00A 00B 01A 01B 02A 03A 03B 04A 04B 05B 06A 06B 07B 08A 08B 09A 09B 10B 11A 11B 12A 12B 13A 13B 14A 14B 16B 17A 17B 18B 19A 19B 20A.22A 21A 21X 23B 26X 27X TBD DNnt DNwt DNht DNut DNit DNfl DNml DNhl DNxl DNad DNxn MNnm MNwm MNhm MNfl MNml MNhl MNad MNxm SNch SNpp SNta SNxx

**Figure 24. Hemilineage 06B. A.** Meshes of all RHS secondary neurons plotted in neuromere-specific colours. **B.** “Representative” secondary neuron skeletons plotted in hemineuromere-specific colours. The skeleton with the top accumulated NBLAST score among all neurons from the hemilineage in a given hemineuromere was used. **C.** Neuron meshes of selected examples. Top: ascending serial set 10257. Bottom: centrifugal serial set 11261. **D.** Predicted synapses of RHS secondary neurons. Blue: postsynapses; dark orange: presynapses. **E.** Proportions of connections from secondary neurons to upstream or downstream partners, normalised by neuromere and coloured by broad class. Numbers of query neurons appear in the centre. **F.** Proportions of synaptic weight from secondary neurons originating in each neuromere to upstream or downstream partners, normalised by row.

#### Hemilineage 06B

Neurons from hemilineage 06B enter the neuropil in the posterior of the neuromere, medially to the 06A neurons, and project dorso-medially to cross the midline in the posterior intermediate commissure, anterior to the 12B neurons (Shepherd et al., 2016). They arborise extensively both sides of the midline in the ventral tectulum and the majority are intersegmental, with the mass of the arborisation from all three neuromeres concentrated in T2 (Figure 24A). Based on their primary neurites, the intersegmental neurons forming varicose projections in lateral leg neuropils (Shepherd et al., 2019) likely belong to 06B rather than 06A (e.g., IN06B035) (Figure 24C bottom). Like 05B, 06B neurons innervate both leg and flight neuropils and are especially prominent in the ovoid (Shepherd et al., 2019), and many types ascend via the neck connective (e.g., AN06B002) (Figure 24C top).

Secondary 06B neurons survive in all thoracic neuromeres but with many more neurons in T2 than T1 or T3 (Figure 24E). Their inputs come mainly from descending neurons, tactile and proprioceptive sensory neurons, and numerous hemilineages including 02A, 06B, 08B, 12A, 17A, and 19B, with some segmental variation (Figure 24F). Specific 06B types can be dedicated to proprioceptive (e.g., AN/IN06B14), tactile (e.g., IN06B016) or both (e.g., IN06B078) (Figure 24 - figure supplement 4-5). 06B secondary neurons are predicted to be gabaergic as expected (Lacin et al., 2019) and can inhibit wing motor neurons (Figure 24E), proprioceptive sensory neurons, and hemilineages 03B, 06A, 06B, 07B, 08B, 12A, 17A, and 19B with some segmental variation (Figure 24F). AN/IN06B040 neurons target neck motor neurons (Figure 24 - figure supplement 6-7). Bilateral activation of 06B neurons elicits leg-related movements without substantial forward locomotion along with occasional wing grooming bouts (Harris et al., 2015).

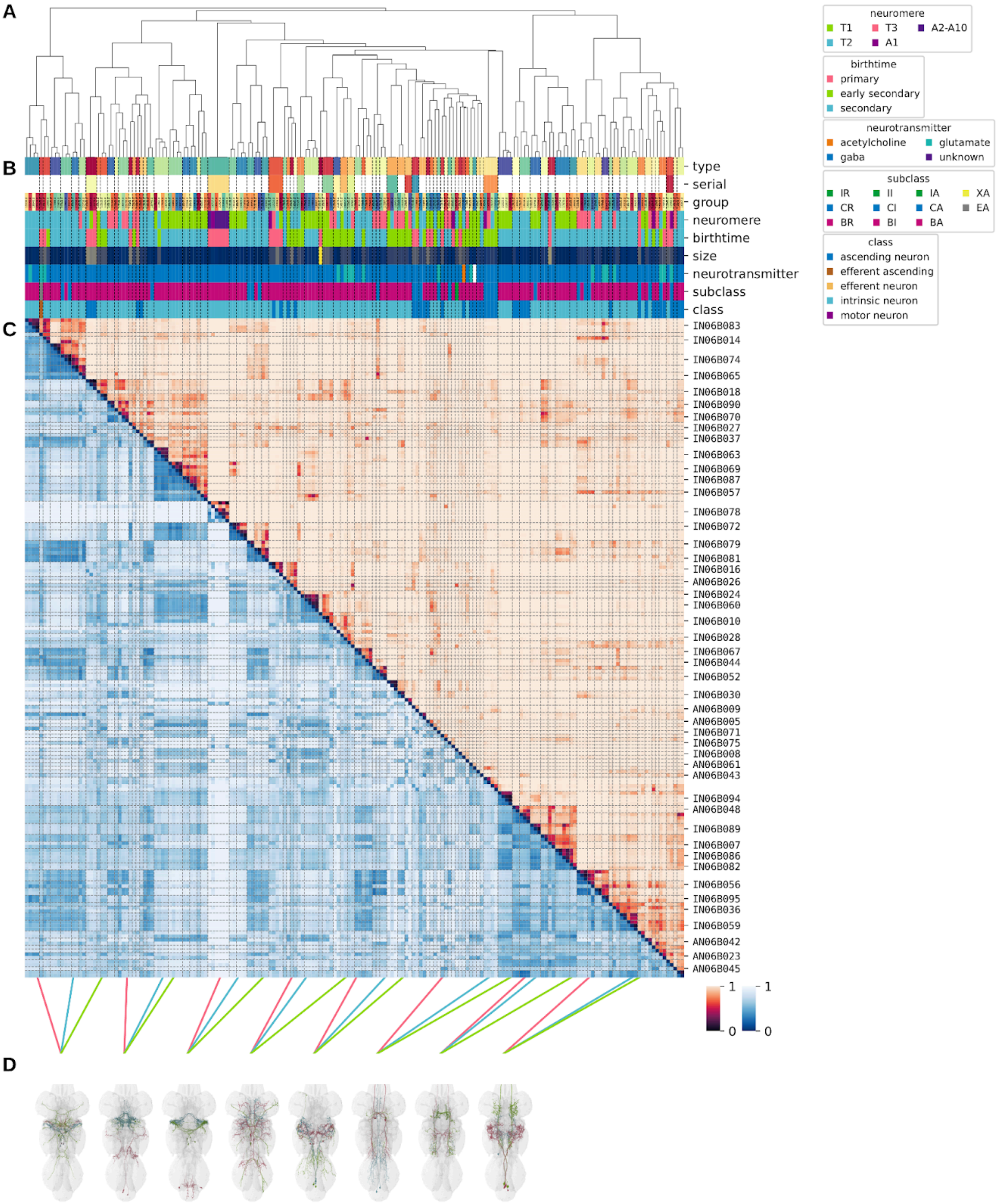

**Figure 24 - figure supplement 1. Systematic typing of hemilineage 06B. A.** Hierarchical clustering dendrogram of hemilineage groups by laterally and serially aggregated connectivity cosine clustering. **B.** Categorical annotations of each hemilineage group, each column corresponding to the aligned leaf in A. Colours for type, serial set, and group are arbitrary for visualisation. Colours for neuromere, birthtime, neurotransmitter, subclass, and class are as in all other figures. **C.** Similarity distance heatmap for hemilineage. Cosine distance is in the upper triangle, while laterally symmetrised NBLAST distance is in the lower triangle. Systematic type names of some types are labelled. **D.** Morphologically representative groups from dendrogram subtrees. Each group, indicated by colour and line connecting to its column in B and C, is the most morphologically representative group (medoid of NBLAST distance)

from a subtree of A. The subtrees (flat clusters) are equal height cuts of A determined to yield the number of groups per plot and plots in D.

#### 06B systematic types

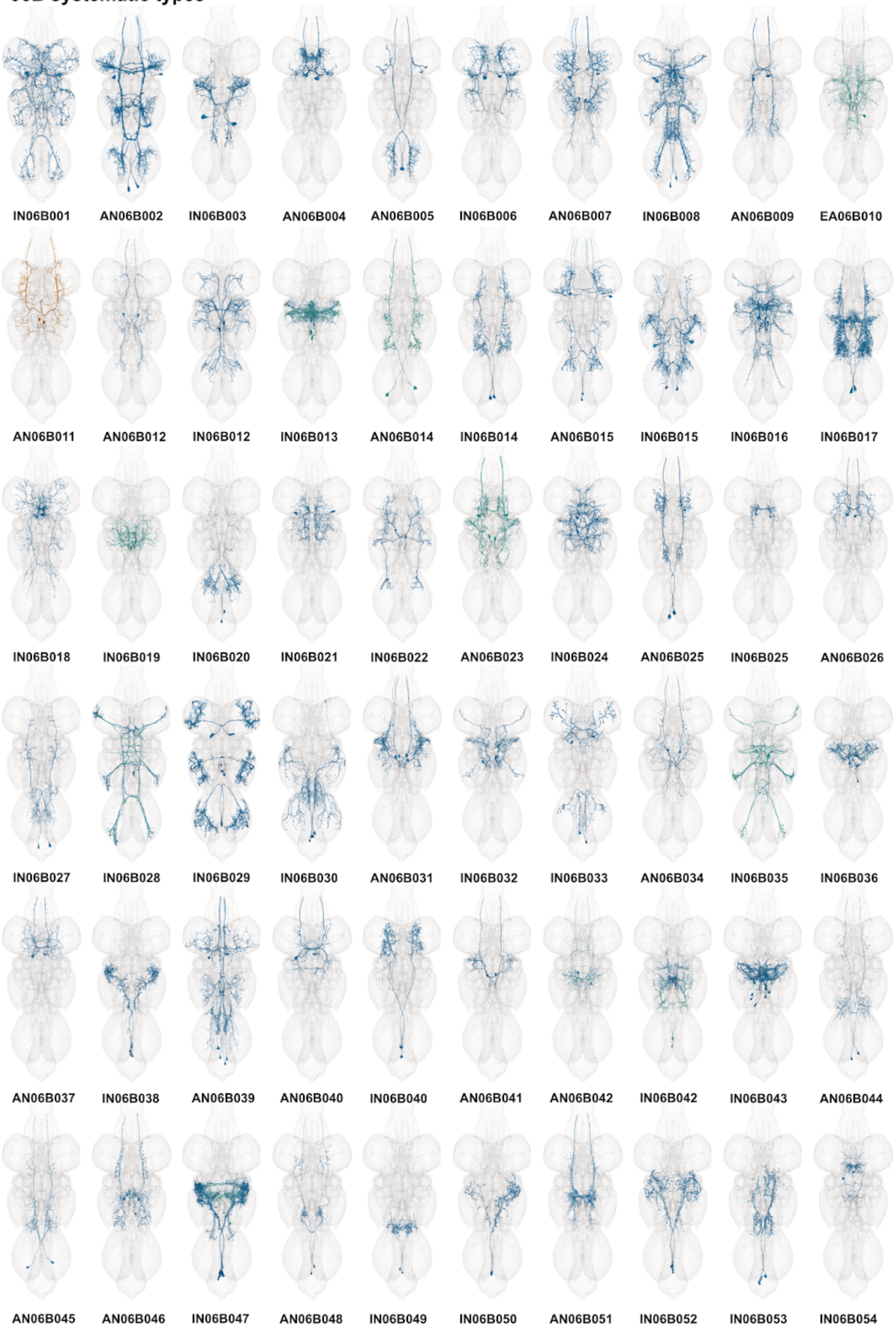

**Figure 24 - figure supplement 2. Systematic types of hemilineage 06B.** Systematic types have been arranged in numerical order, with neurons of the same type that belong to distinct classes (e.g., intrinsic neuron vs ascending neuron) plotted separately but placed adjacent to each other. Individual neuron meshes have been coloured based on predicted neurotransmitter: dark orange = acetylcholine, blue = gaba, marine = glutamate, dark purple = unknown.

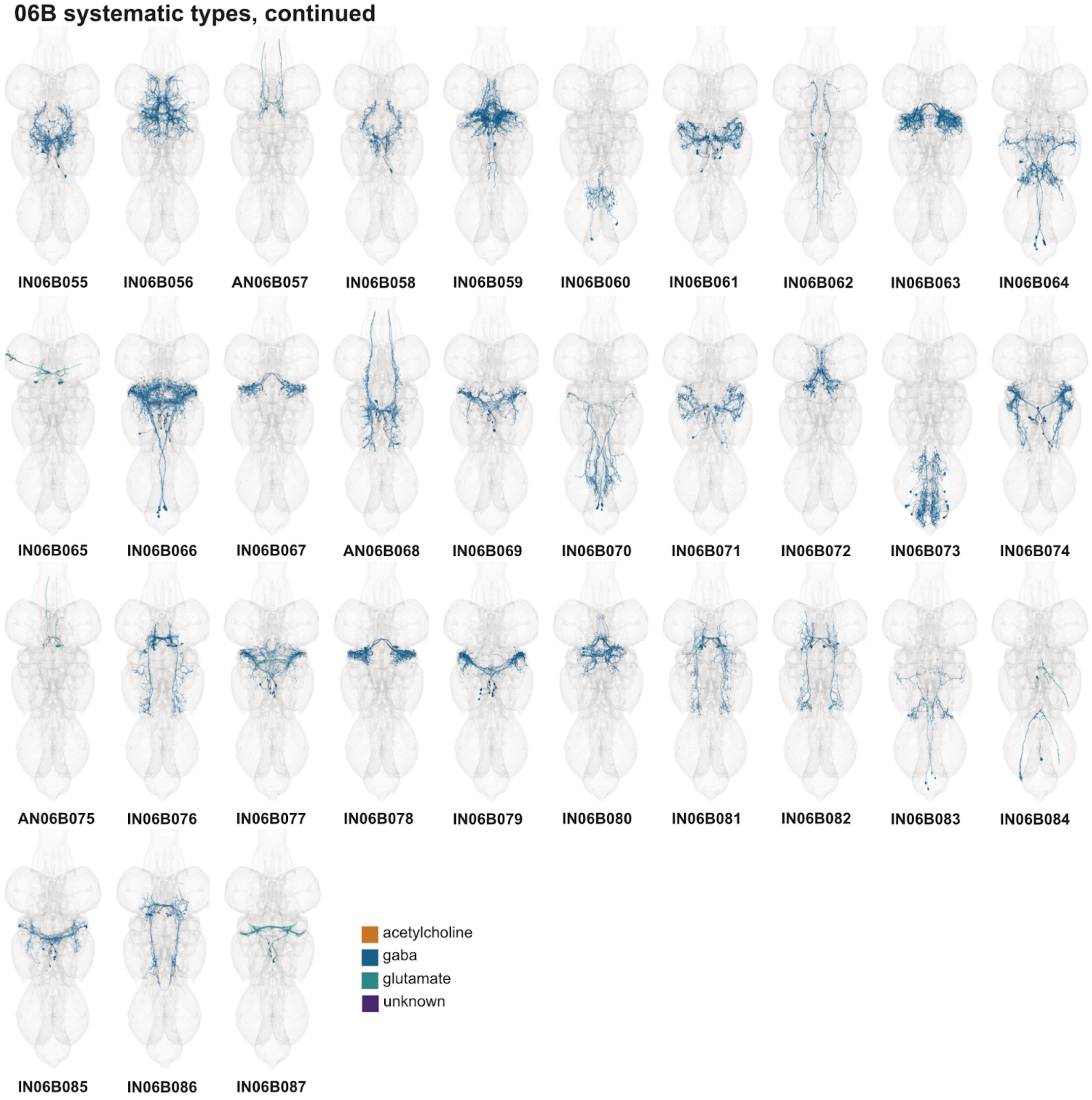

**Figure 24 - figure supplement 3. Systematic types of hemilineage 06B, continued.** Systematic types have been arranged in numerical order, with neurons of the same type that belong to distinct classes (e.g., intrinsic neuron vs ascending neuron) plotted separately but placed adjacent to each other. Individual neuron meshes have been coloured based on predicted neurotransmitter: dark orange = acetylcholine, blue = gaba, marine = glutamate, dark purple = unknown.

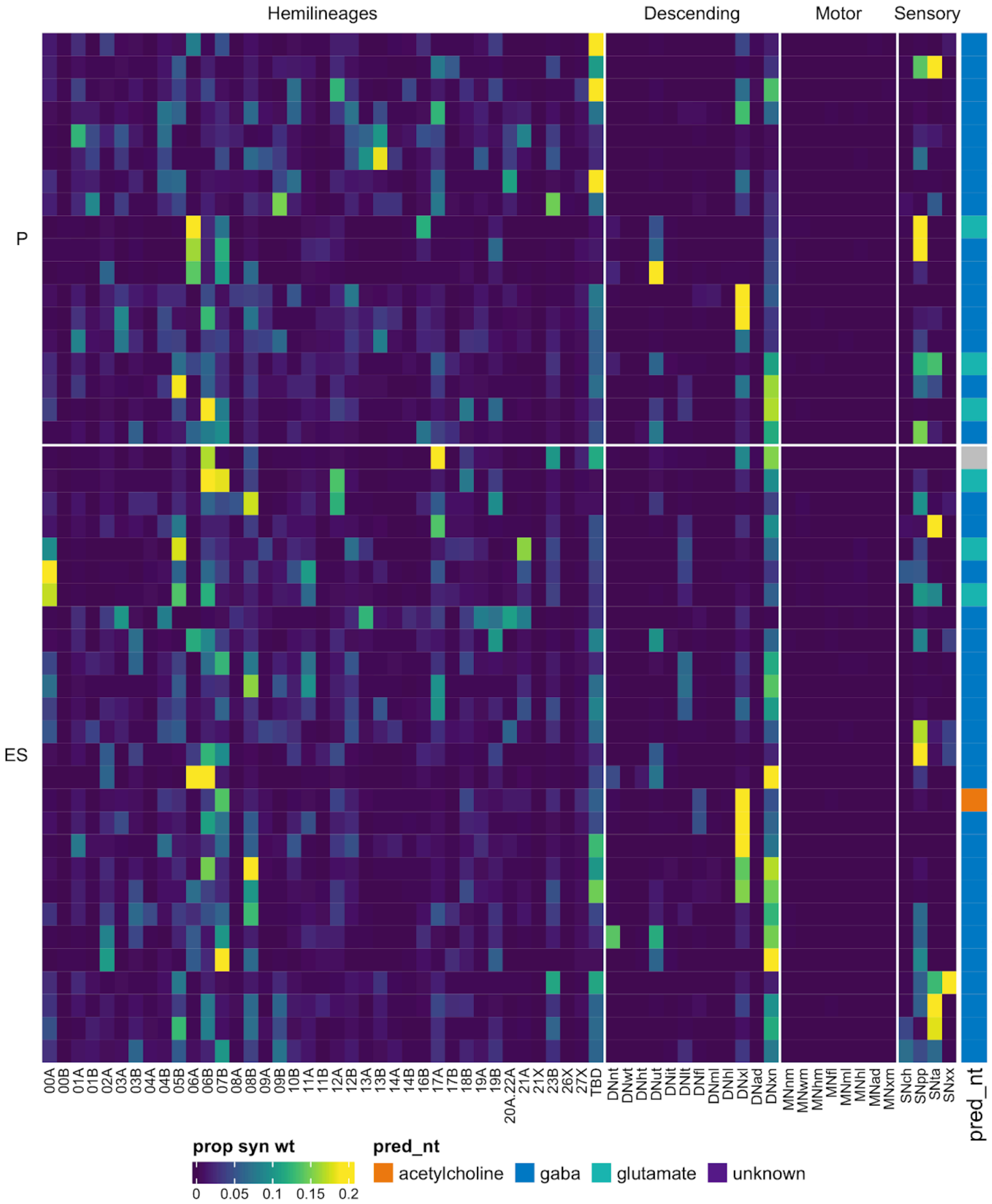

**Figure 24 - figure supplement 4. Connectivity to upstream partners by 06B primary and early secondary systematic types.** Proportions of synaptic weight to systematic types from upstream partners, normalised by row. 06B neurons have been clustered within each assigned birthtime window (P = primary, ES = early secondary, S = secondary) based on both upstream and downstream connectivity to hemilineages, descending neuron subclasses, motor neuron subclasses, and sensory neuron modalities. Annotation bar is coloured by the most common predicted neurotransmitter for the neurons of each type.

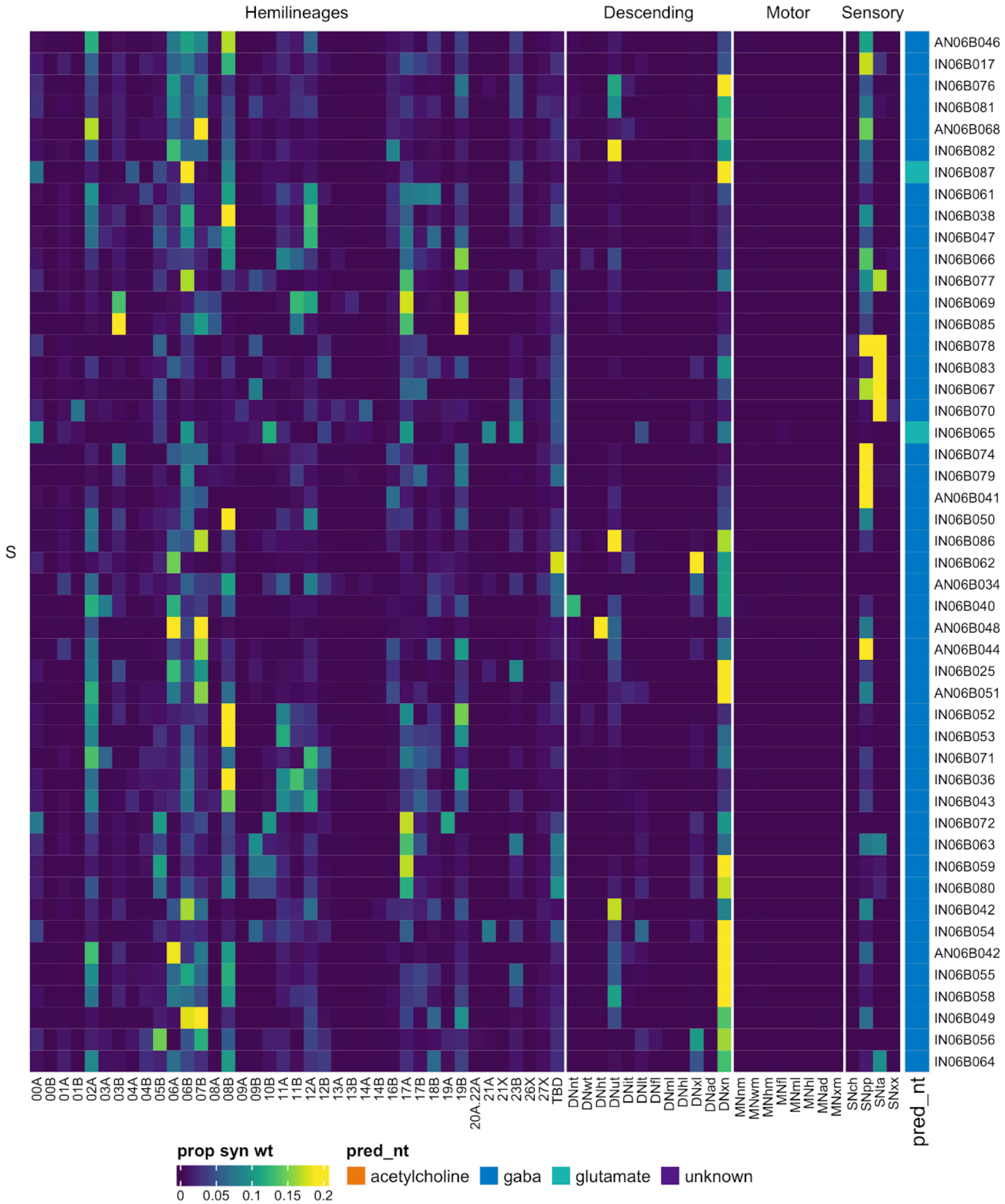

**Figure 24 - figure supplement 5. Connectivity to upstream partners by 06B secondary systematic types.** Proportions of synaptic weight to systematic types from upstream partners, normalised by row. 06B neurons have been clustered within each assigned birthtime window (P = primary, ES = early secondary, S = secondary) based on both upstream and downstream connectivity to hemilineages, descending neuron subclasses, motor neuron subclasses, and sensory neuron modalities. The annotation bar is coloured by the most common predicted neurotransmitter within each type.

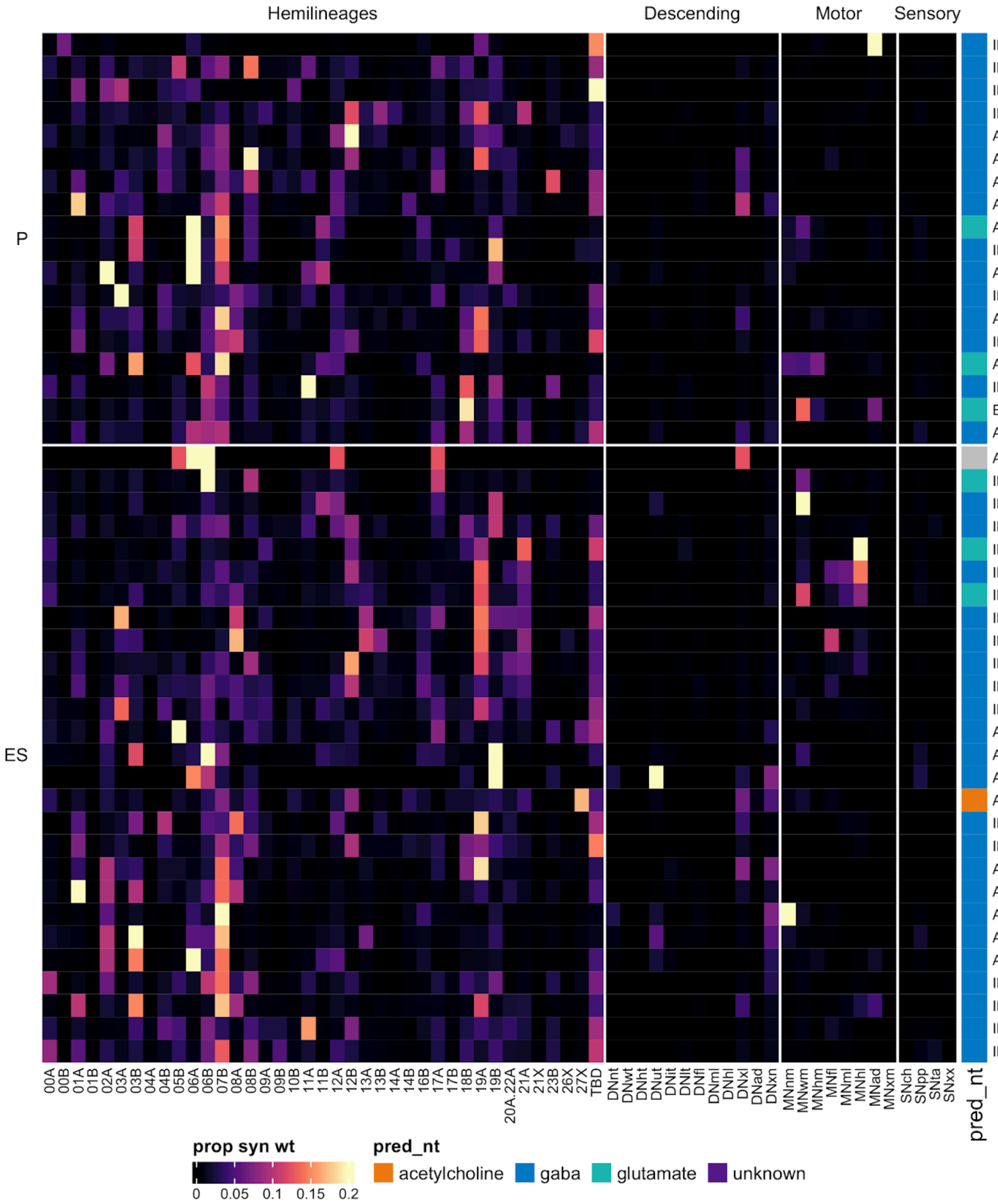

**Figure 24 - figure supplement 6. Connectivity to downstream partners by 06B primary and early secondary systematic types.** Proportions of synaptic weight from systematic types to downstream partners, normalised by row. 06B neurons have been clustered within each assigned birthtime window (P = primary, ES = early secondary, S = secondary) based on both upstream and downstream connectivity to hemilineages, descending neuron subclasses, motor neuron subclasses, and sensory neuron modalities. The annotation bar is coloured by the most common predicted neurotransmitter within each type.

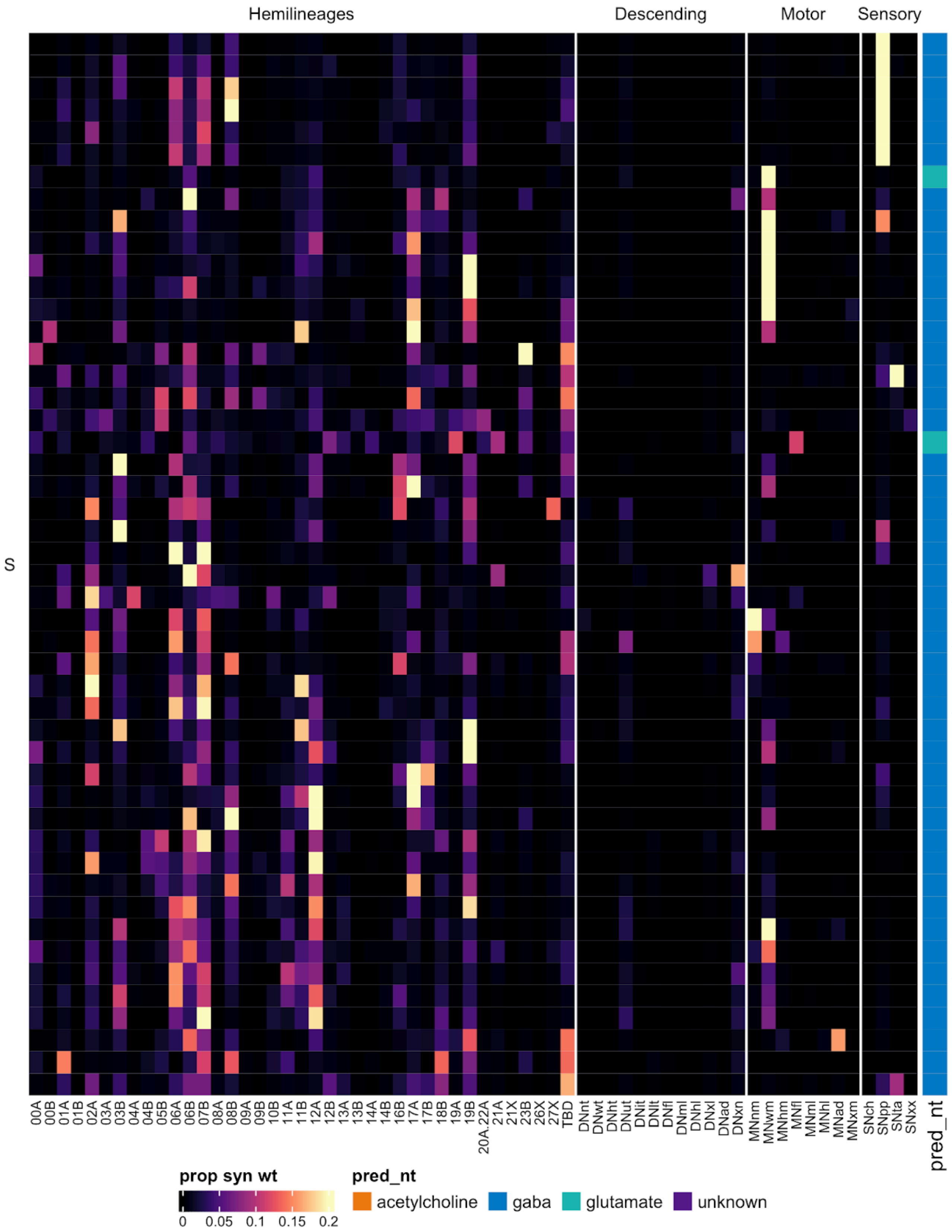

**Figure 24 - figure supplement 7. Connectivity to downstream partners by 06B secondary systematic types.** Proportions of synaptic weight from systematic types to downstream partners, normalised by row. 06B neurons have been clustered within each assigned birthtime window (P = primary, ES = early secondary, S = secondary) based on both upstream and downstream connectivity to hemilineages, descending neuron subclasses, motor neuron subclasses, and sensory neuron modalities. The annotation bar is coloured by the most common predicted neurotransmitter for the neurons of each type.

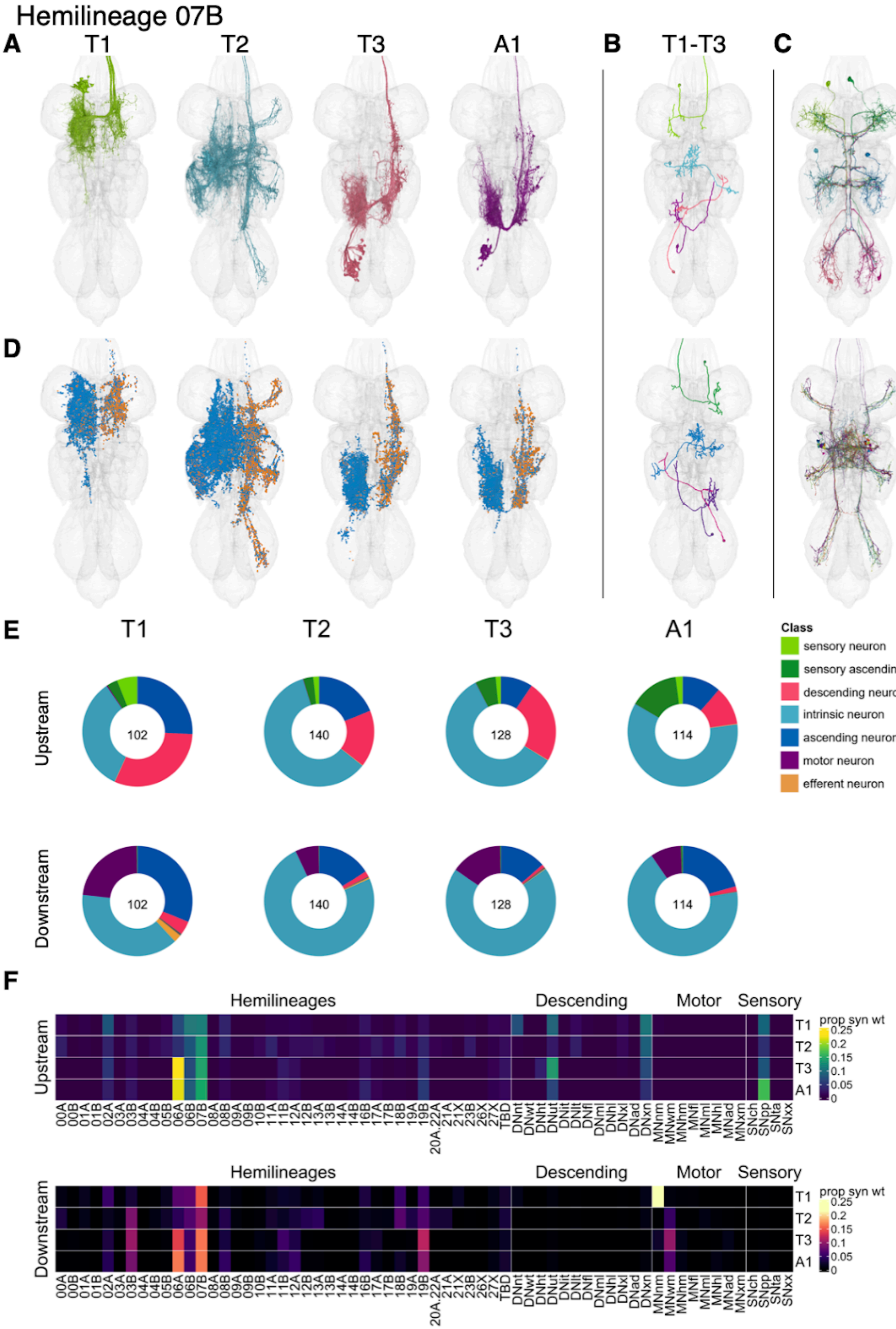

Hemilineage 07B
A
T1
T2
T3
A1
B
T1-T3
C
D
E
T1
T2
T3
A1
Upstream
Downstream
102
140
128
114
Class
sensory neuron
sensory ascending
descending neuron
intrinsic neuron
ascending neuron
motor neuron
efferent neuron
F
Hemilineages
Descending
Motor
Sensory
prop syn wt
0.25
0.2
0.15
0.1
0.05
0
T1
T2
T3
A1
00A 00B 01A 01B 02A 03A 03B 04A 04B 05B 06A 06B 07B 08A 08B 09A 09B 10B 11A 11B 12A 12B 13A 13B 14A 14B 16B 17A 17B 18B 19A 19B 20A.22A 21A 21X 23B 26X 27X TBD DNnt DNwt DNht DNut DNit DNlt DNfl DNml DNhl DNxl DNad DNxn MNnm MNwm MNhm MNfl MNml MNhl MNad MNxm SNch SNpp SNta SNxx

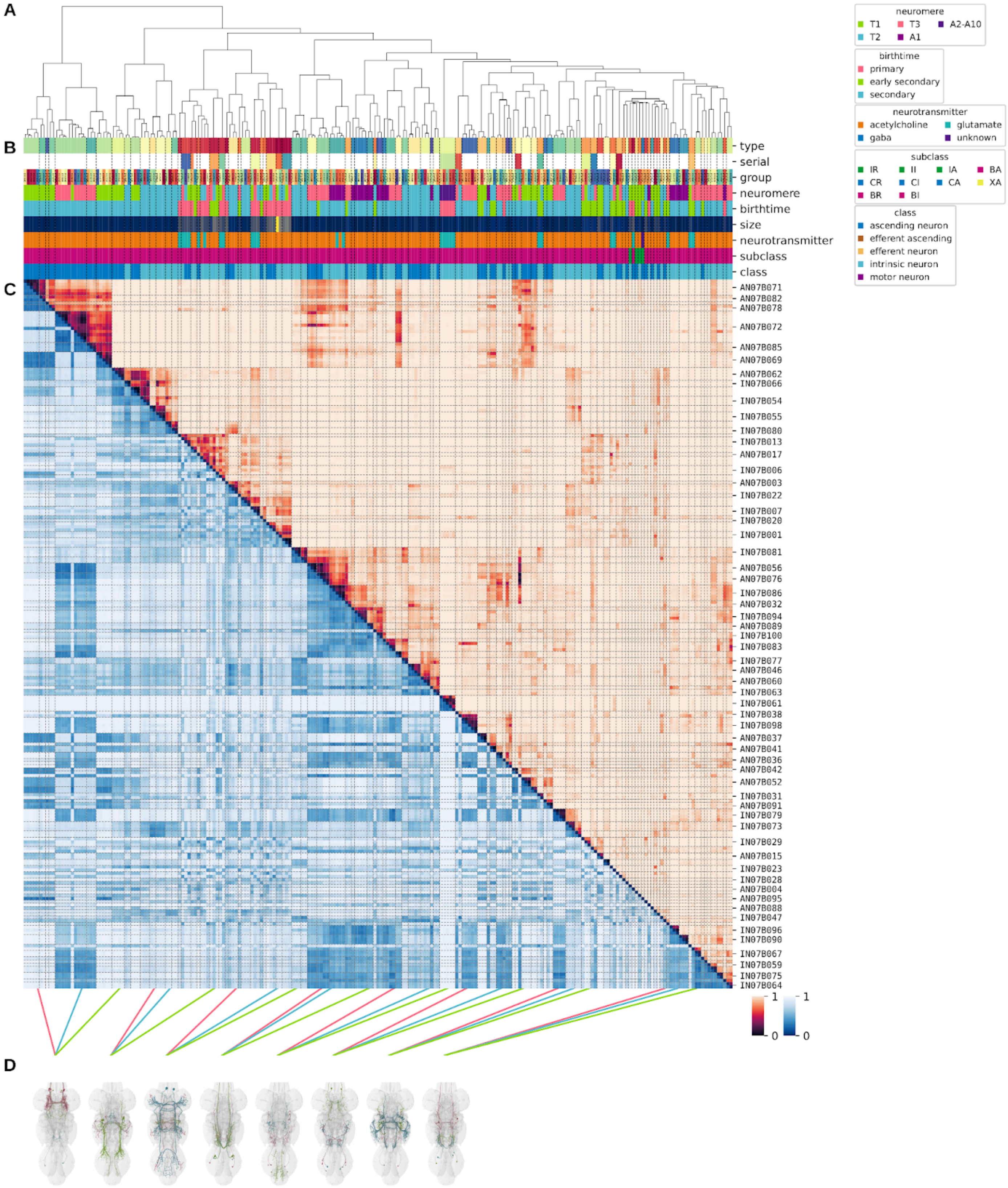

**Figure 25 - figure supplement 1. Systematic typing of hemilineage 07B. A.** Hierarchical clustering dendrogram of hemilineage groups by laterally and serially aggregated connectivity cosine clustering. **B.** Categorical annotations of each hemilineage group, each column corresponding to the aligned leaf in A. Colours for type, serial set, and group are arbitrary for visualisation. Colours for neuromere, birthtime, neurotransmitter, subclass, and class are as in all other figures. **C.** Similarity distance heatmap for hemilineage. Cosine distance is in the upper triangle, while laterally symmetrised NBLAST distance is in the lower triangle. Systematic type names of some types are labelled. **D.** Morphologically representative groups from dendrogram subtrees. Each group, indicated by colour and line connecting to its column in B and C, is the most morphologically representative group (medoid of NBLAST distance) from a subtree of A. The subtrees (flat clusters) are equal height cuts of A determined to yield the number of groups

per plot and plots in D.

#### 07B systematic types

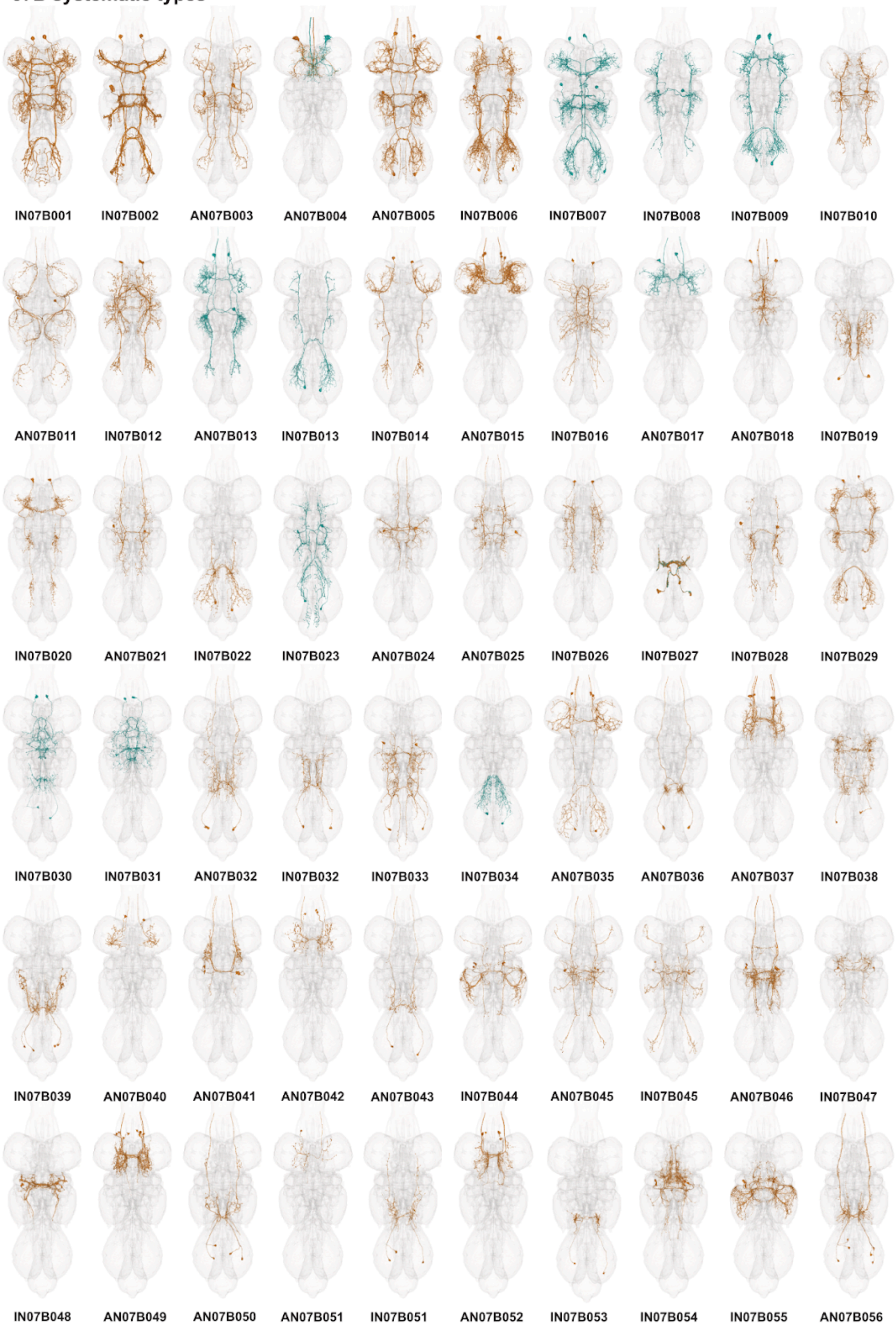

**Figure 25 - figure supplement 2. Systematic types of hemilineage 07B.** Systematic types have been arranged in numerical order, with neurons of the same type that belong to distinct classes (e.g., intrinsic neuron vs ascending neuron) plotted separately but placed adjacent to each other. Individual neuron meshes have been coloured based on predicted neurotransmitter: dark orange = acetylcholine, blue = gaba, marine = glutamate, dark purple = unknown.

#### 07B systematic types, continued

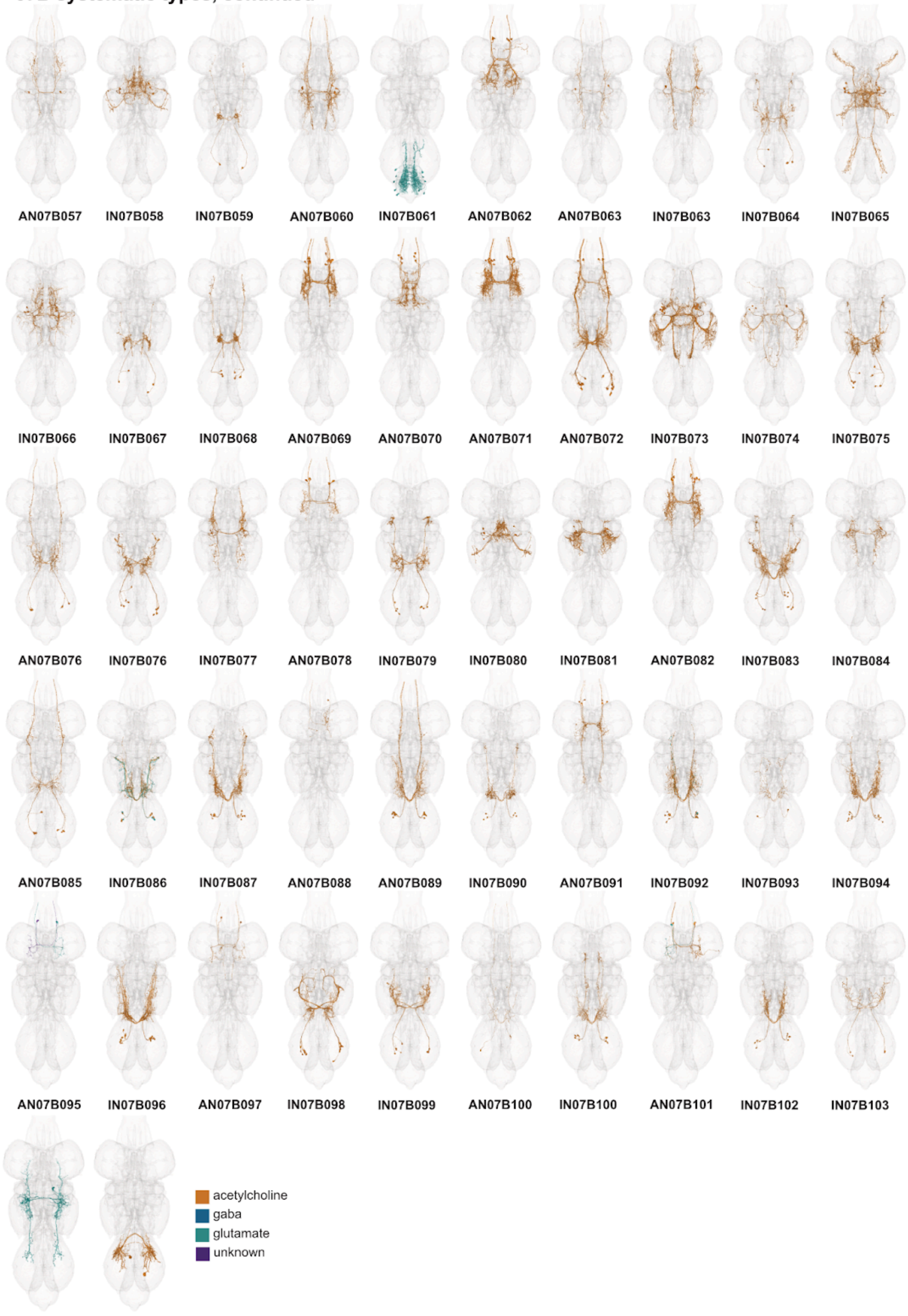

**Figure 25 - figure supplement 3. Systematic types of hemilineage 07B, continued.** Systematic types have been arranged in numerical order, with neurons of the same type that belong to distinct classes (e.g., intrinsic neuron vs ascending neuron) plotted separately but placed adjacent to each other. Individual neuron meshes have been coloured based on predicted neurotransmitter: dark orange = acetylcholine, blue = gaba, marine = glutamate, dark purple = unknown.

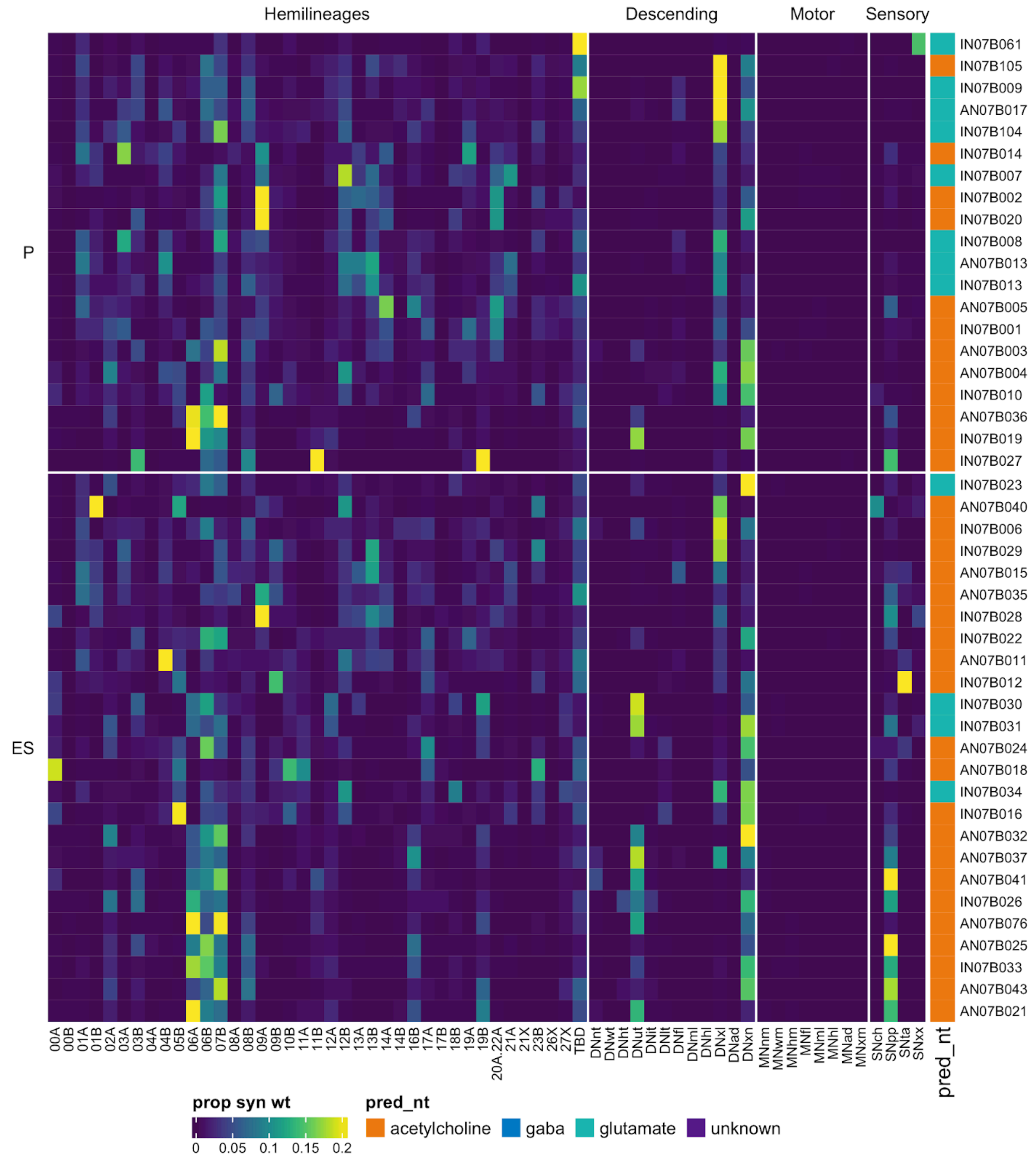

**Figure 25 - figure supplement 4. Connectivity to upstream partners by 07B primary and early secondary systematic types.** Proportions of synaptic weight to systematic types from upstream partners, normalised by row. 07B neurons have been clustered within each assigned birthtime window (P = primary, ES = early secondary, S = secondary) based on both upstream and downstream connectivity to hemilineages, descending neuron subclasses, motor neuron subclasses, and sensory neuron modalities. Annotation bar is coloured by the most common predicted neurotransmitter for the neurons of each type.

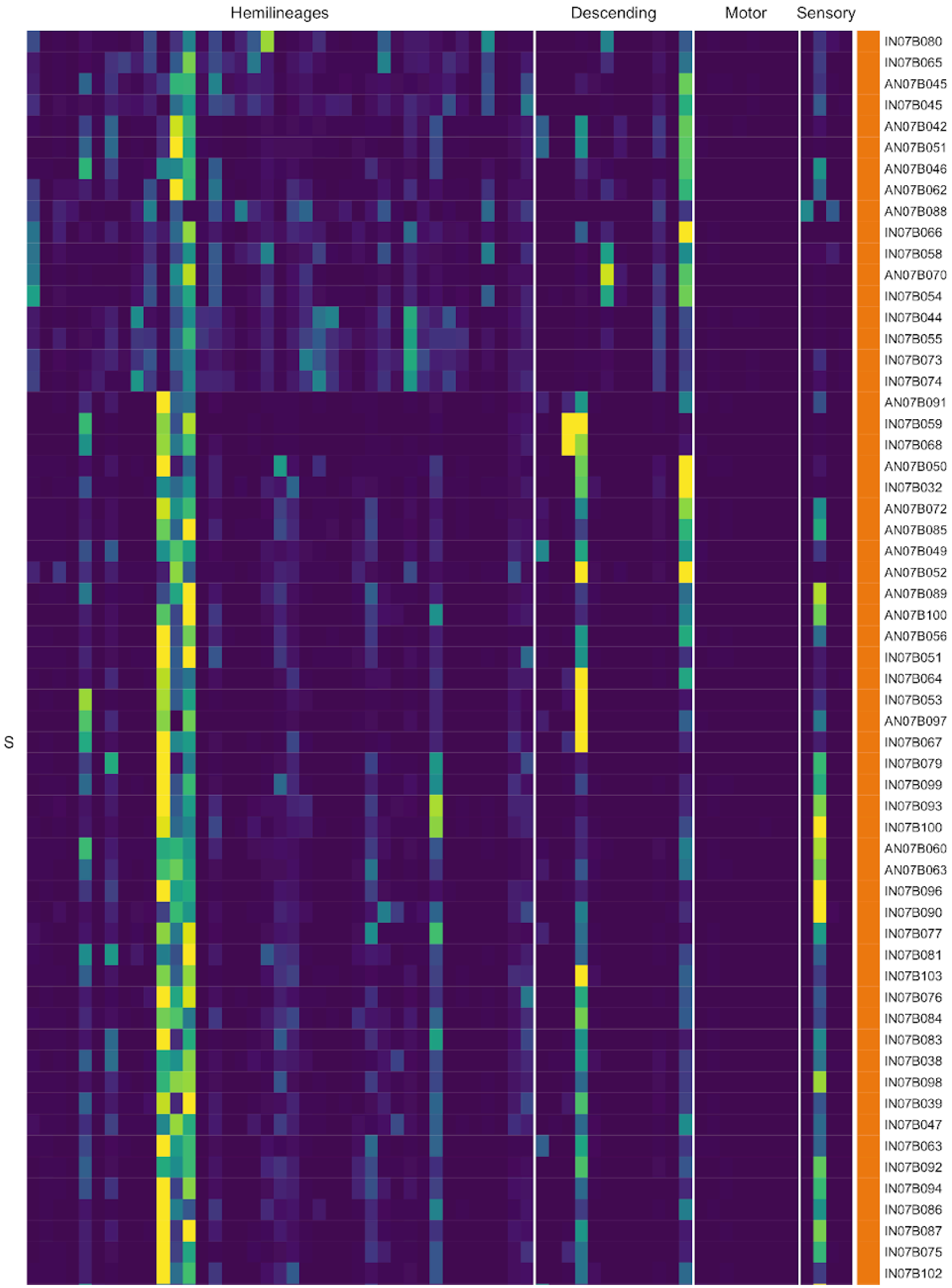
Hemilineages
Descending
Motor
Sensory
S
IN07B080
IN07B065
AN07B045
IN07B045
AN07B042
AN07B051
AN07B046
AN07B062
AN07B088
IN07B066
IN07B058
AN07B070
IN07B054
IN07B044
IN07B055
IN07B073
IN07B074
AN07B091
IN07B059
IN07B068
AN07B050
IN07B032
AN07B072
AN07B085
AN07B049
AN07B052
AN07B089
AN07B100
AN07B056
IN07B051
IN07B064
IN07B053
AN07B097
IN07B067
IN07B079
IN07B099
IN07B093
IN07B100
AN07B060
AN07B063
IN07B096
IN07B090
IN07B077
IN07B081
IN07B103
IN07B076
IN07B084
IN07B083
IN07B038
IN07B098
IN07B039
IN07B047
IN07B063
IN07B092
IN07B094
IN07B086
IN07B087
IN07B075
IN07B102

**Figure 25 - figure supplement 5. Connectivity to upstream partners by 07B secondary systematic types.** Proportions of synaptic weight to systematic types from upstream partners, normalised by row. 07B neurons have been clustered within each assigned birthtime window (P = primary, ES = early secondary, S = secondary) based on both upstream and downstream connectivity to hemilineages, descending neuron subclasses, motor neuron subclasses, and sensory neuron modalities. The annotation bar is coloured by the most common predicted neurotransmitter within each type.

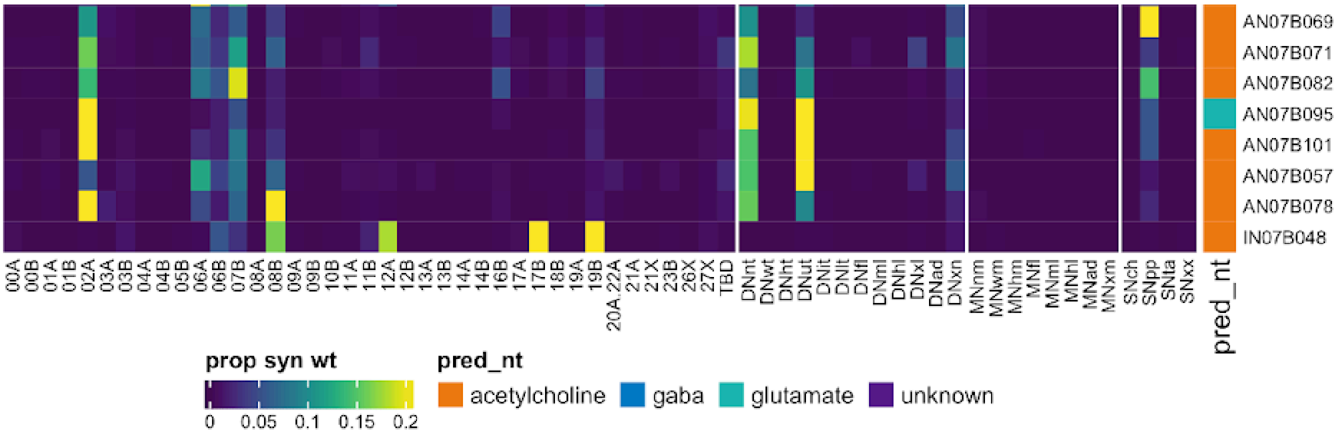

**Figure 25 - figure supplement 6. Connectivity to upstream partners by 07B secondary systematic types, continued.** Proportions of synaptic weight to systematic types from upstream partners, normalised by row. 07B neurons have been clustered within each assigned birthtime window (P = primary, ES = early secondary, S = secondary) based on both upstream and downstream connectivity to hemilineages, descending neuron subclasses, motor neuron subclasses, and sensory neuron modalities. The annotation bar is coloured by the most common predicted neurotransmitter within each type.

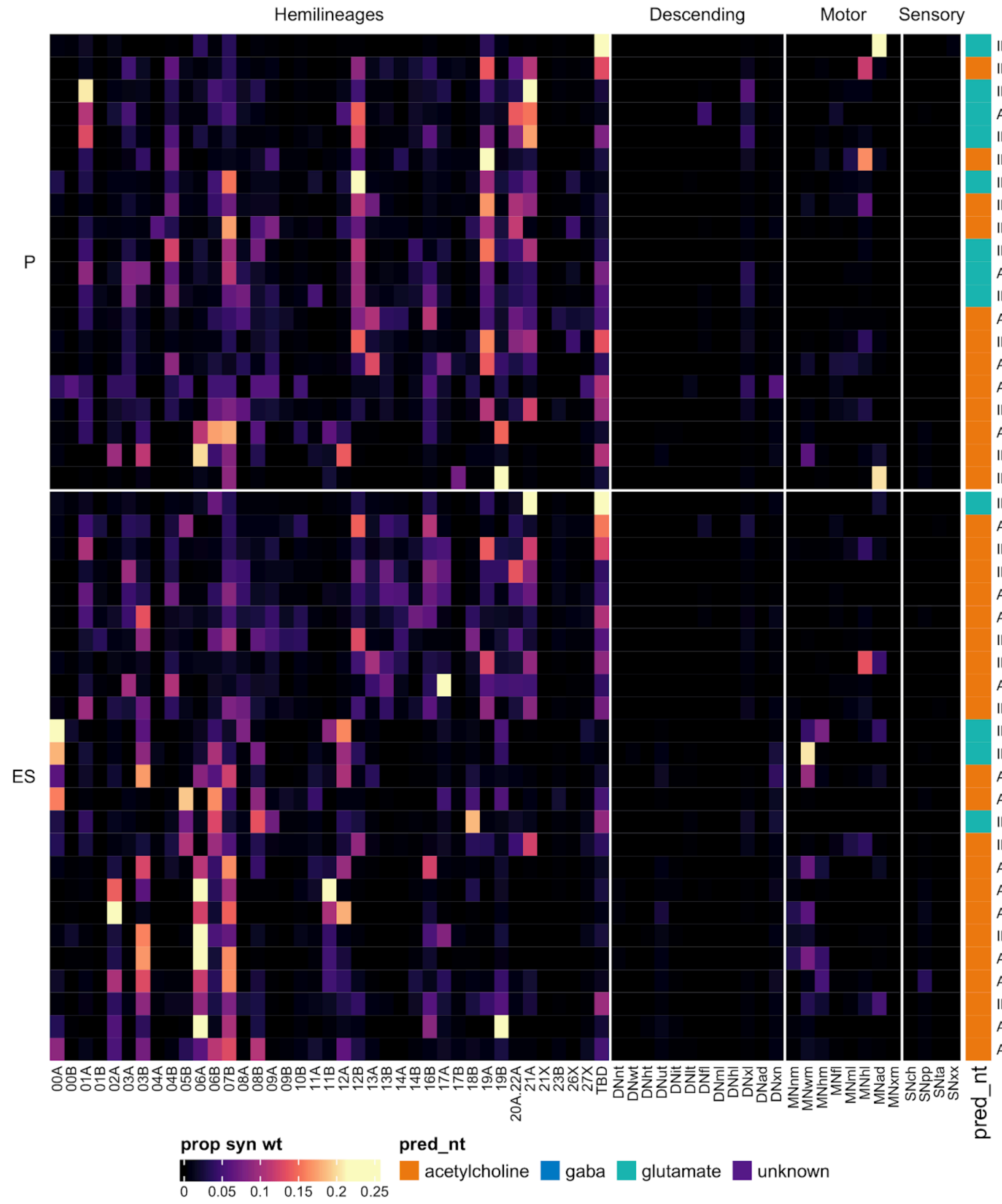

**Figure 25 - figure supplement 7. Connectivity to downstream partners by 07B primary and early secondary systematic types.** Proportions of synaptic weight from systematic types to downstream partners, normalised by row. 07B neurons have been clustered within each assigned birthtime window (P = primary, ES = early secondary, S = secondary) based on both upstream and downstream connectivity to hemilineages, descending neuron subclasses, motor neuron subclasses, and sensory neuron modalities. The annotation bar is coloured by the most common predicted neurotransmitter within each type.

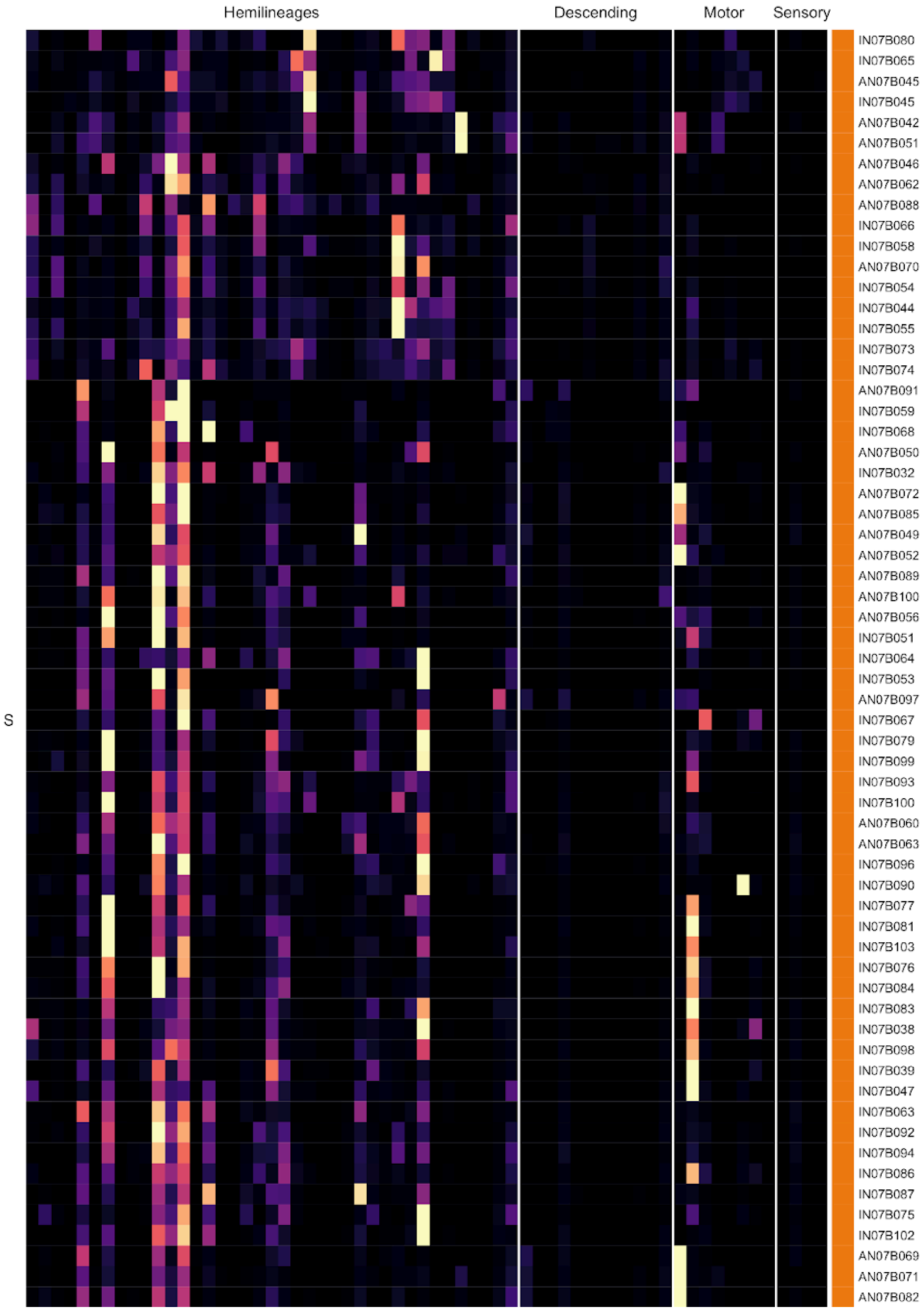

Hemilineages
Descending
Motor
Sensory
S
IN07B080
IN07B065
AN07B045
IN07B045
AN07B042
AN07B051
AN07B046
AN07B062
AN07B088
IN07B066
IN07B058
AN07B070
IN07B054
IN07B044
IN07B055
IN07B073
IN07B074
AN07B091
IN07B059
IN07B068
AN07B050
IN07B032
AN07B072
AN07B085
AN07B049
AN07B052
AN07B089
AN07B100
AN07B056
IN07B051
IN07B064
IN07B053
AN07B097
IN07B067
IN07B079
IN07B099
IN07B093
IN07B100
AN07B060
AN07B063
IN07B096
IN07B090
IN07B077
IN07B081
IN07B103
IN07B076
IN07B084
IN07B083
IN07B038
IN07B098
IN07B039
IN07B047
IN07B063
IN07B092
IN07B094
IN07B086
IN07B087
IN07B075
IN07B102
AN07B069
AN07B071
AN07B082

**Figure 25 - figure supplement 8. Connectivity to downstream partners by 07B secondary systematic types.** Proportions of synaptic weight from systematic types to downstream partners, normalised by row. 07B neurons have been clustered within each assigned birthtime window (P = primary, ES = early secondary, S = secondary) based on both upstream and downstream connectivity to hemilineages, descending neuron subclasses, motor neuron subclasses, and sensory neuron modalities. The annotation bar is coloured by the most common predicted neurotransmitter for the neurons of each type.

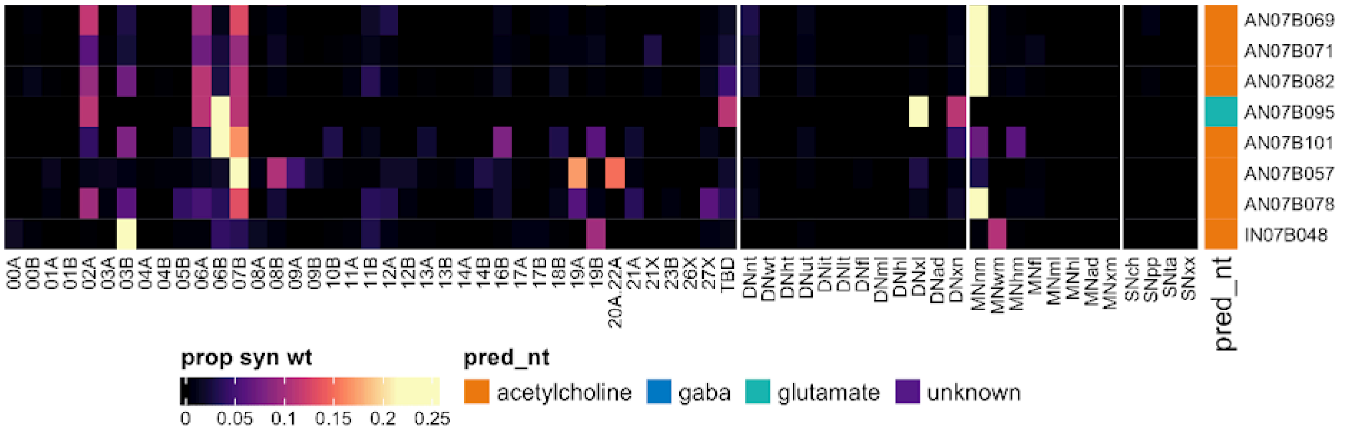

**Figure 25 - figure supplement 9. Connectivity to downstream partners by 07B secondary systematic types, continued.** Proportions of synaptic weight from systematic types to downstream partners, normalised by row. 07B neurons have been clustered within each assigned birthtime window (P = primary, ES = early secondary, S = secondary) based on both upstream and downstream connectivity to hemilineages, descending neuron subclasses, motor neuron subclasses, and sensory neuron modalities. The annotation bar is coloured by the most common predicted neurotransmitter for the neurons of each type.

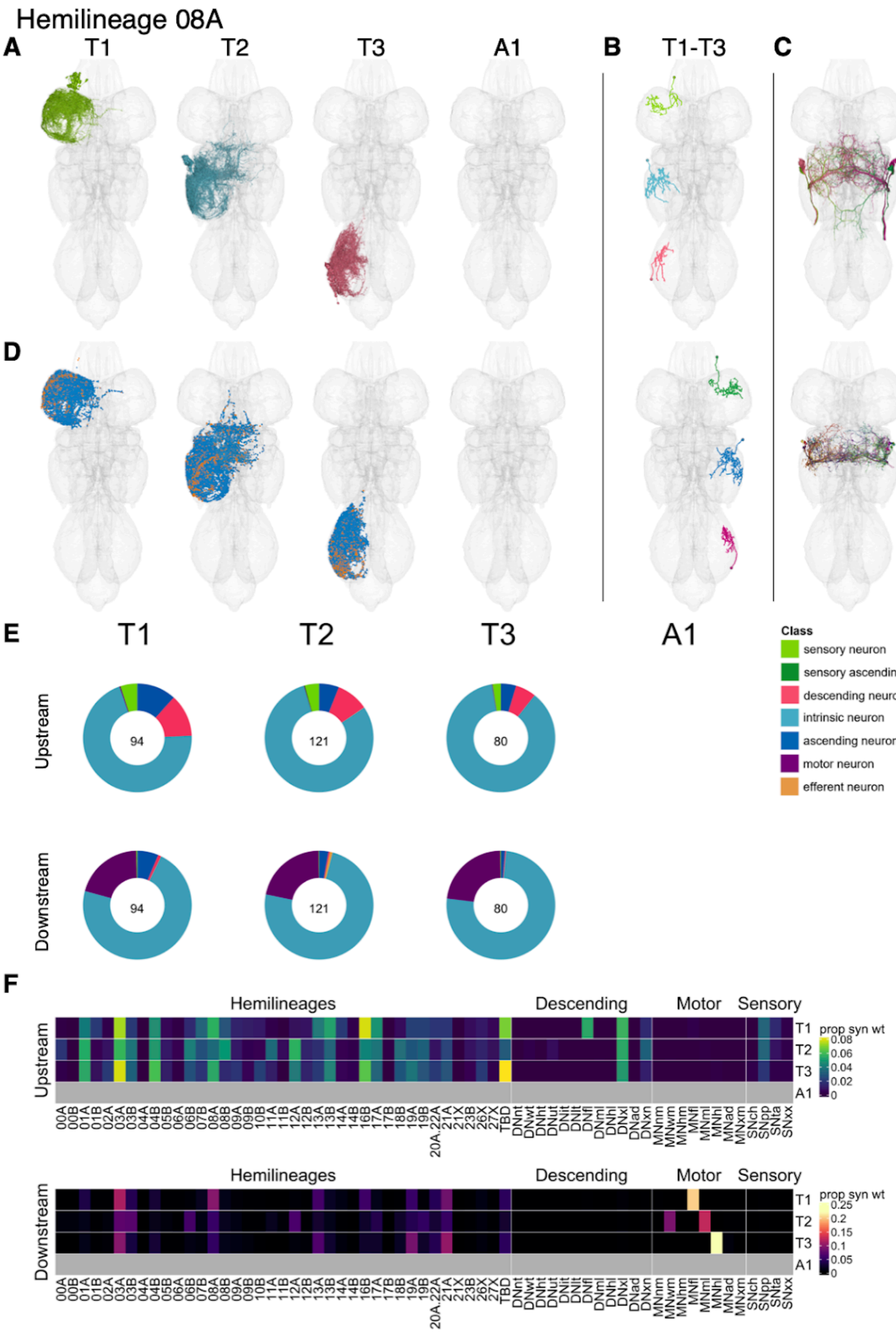

Hemilineage 08A
A
T1
T2
T3
A1
B
T1-T3
C
D
E
T1
T2
T3
A1
Upstream
Downstream
94
121
80
94
121
80
Class
sensory neuron
sensory ascending
descending neuron
intrinsic neuron
ascending neuron
motor neuron
efferent neuron
F
Hemilineages
Descending
Motor
Sensory
Upstream
Downstream
prop syn wt
T1
T2
T3
A1

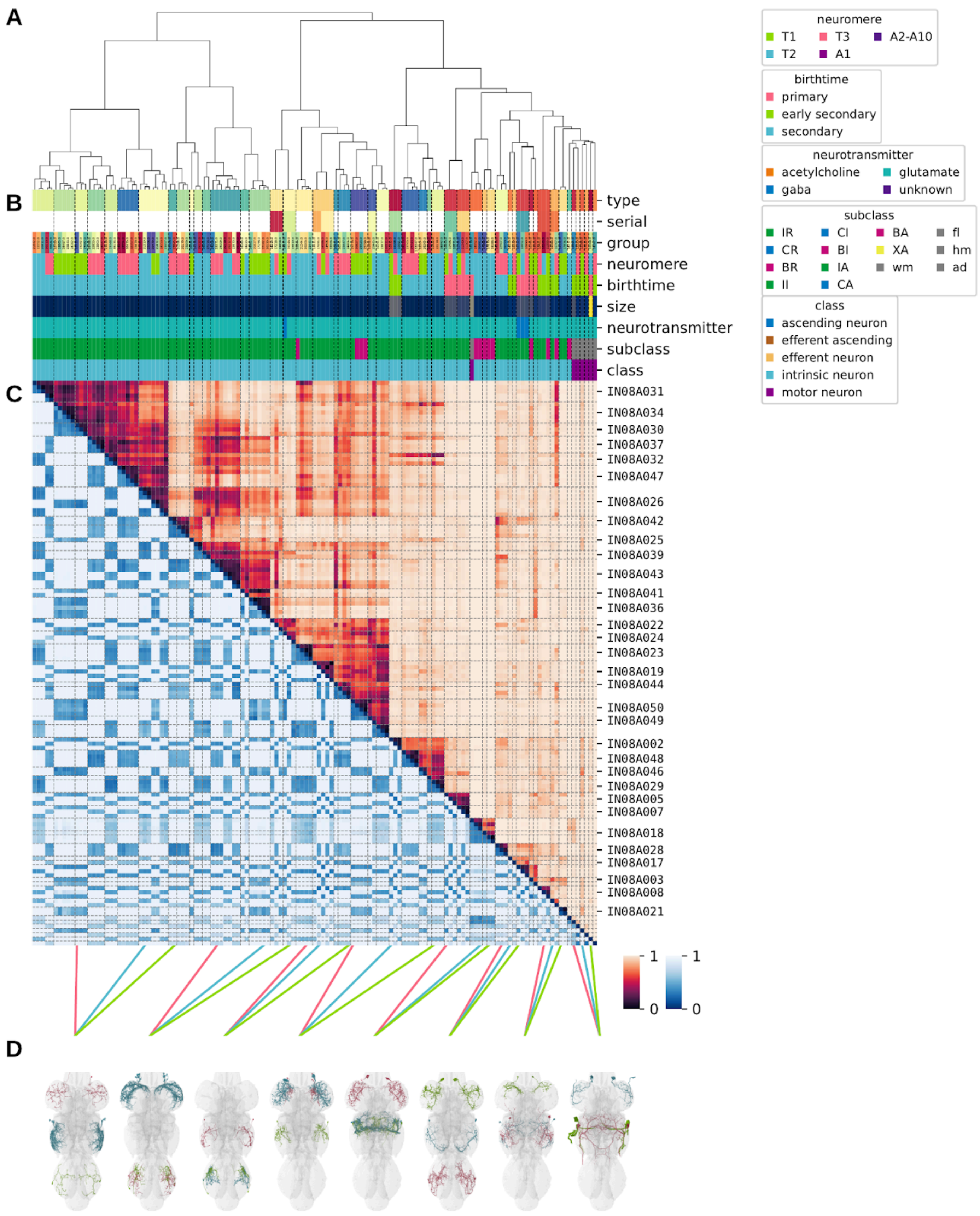

**Figure 26 - figure supplement 1. Systematic typing of hemilineage 08A.** Types for motor neurons were assigned separately as outlined in our accompanying manuscript (Cheong et al., 2023). **A.** Hierarchical clustering dendrogram of hemilineage groups by laterally and serially aggregated connectivity cosine clustering. **B.** Categorical annotations of each hemilineage group, each column corresponding to the aligned leaf in A. Colours for type, serial set, and group are arbitrary for visualisation. Colours for neuromere, birthtime, neurotransmitter, subclass, and class are as in all other figures. **C.** Similarity distance heatmap for hemilineage. Cosine distance is in the upper triangle, while laterally

symmetrised NBLAST distance is in the lower triangle. Systematic type names of some types are labelled. **D.** Morphologically representative groups from dendrogram subtrees. Each group, indicated by colour and line connecting to its column in B and C, is the most morphologically representative group (medoid of NBLAST distance) from a subtree of A. The subtrees (flat clusters) are equal height cuts of A determined to yield the number of groups per plot and plots in D.

#### 08A systematic types

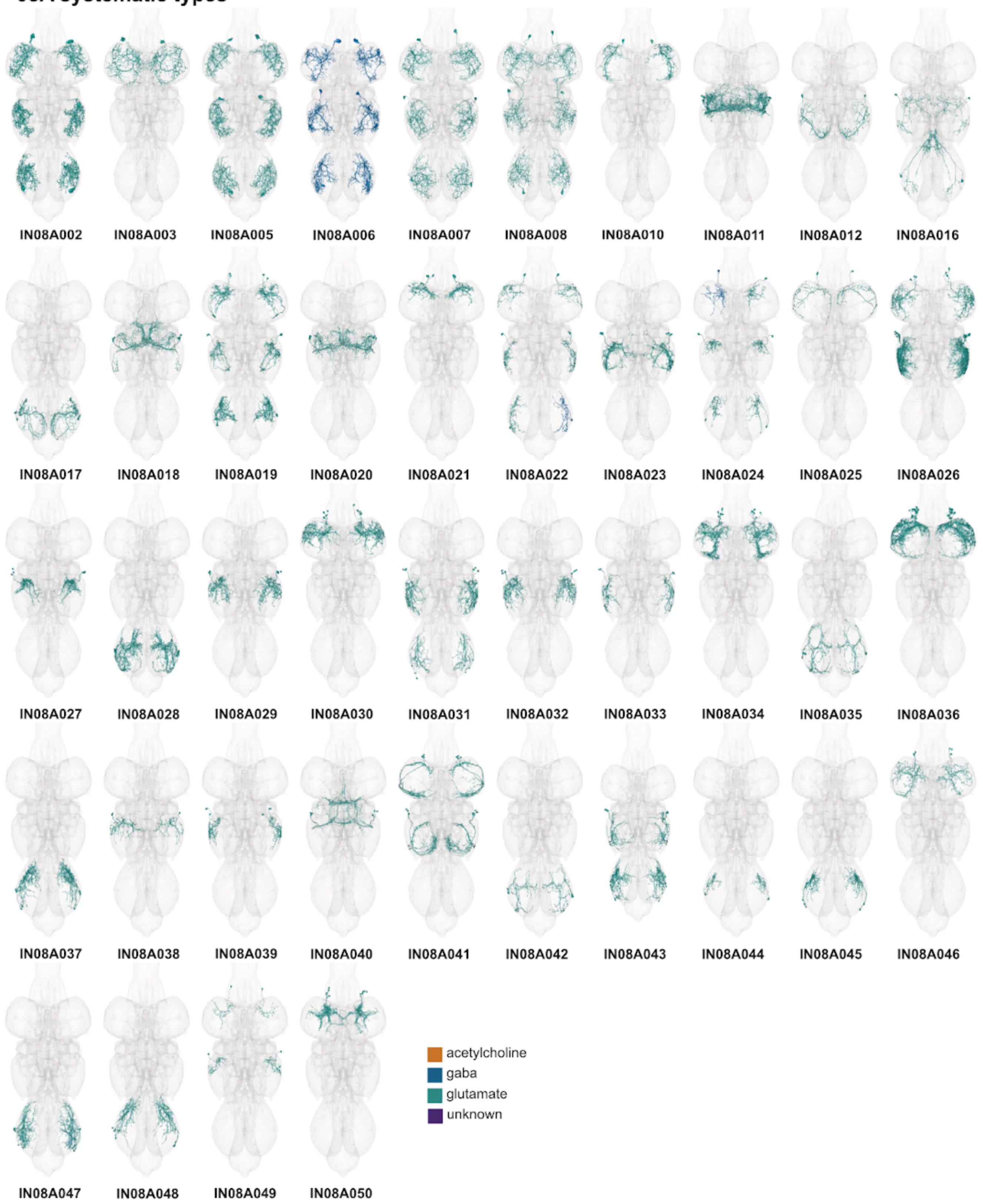

#### Motor Neurons

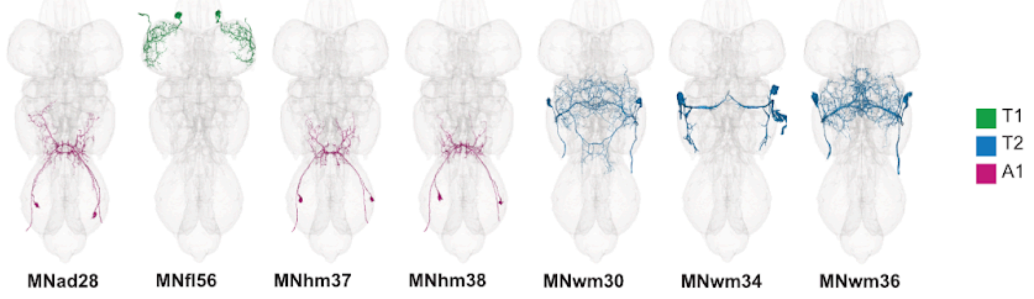

**Figure 26 - figure supplement 2. Systematic types of hemilineage 08A.** Systematic types have been arranged in numerical order, with neurons of the same type that belong to distinct classes (e.g., intrinsic neuron vs ascending neuron) plotted separately but placed adjacent to each other. Individual neuron meshes have been coloured based on predicted neurotransmitter: dark orange = acetylcholine, blue = gaba, marine = glutamate, dark purple = unknown. Motor neurons (typed separately in (Cheong et al., 2023)) have been plotted by serial set if identified in multiple neuromeres and by systematic type if not. Individual motor neuron meshes have been coloured based on soma neuromere: dark green=T1, blue=T2, purple=A1.

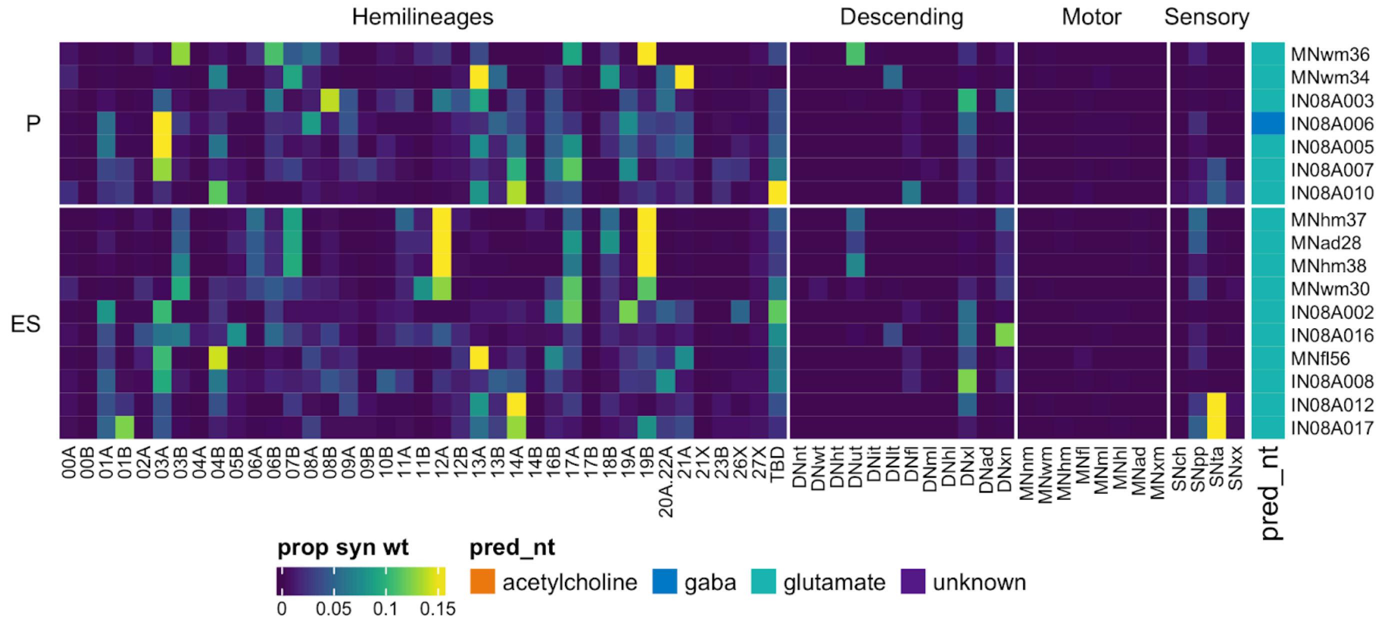

**Figure 26 - figure supplement 3. Connectivity to upstream partners by 08A primary and early secondary systematic types.** Proportions of synaptic weight to systematic types from upstream partners, normalised by row. 08A neurons have been clustered within each assigned birthtime window (P = primary, ES = early secondary, S = secondary) based on both upstream and downstream connectivity to hemilineages, descending neuron subclasses, motor neuron subclasses, and sensory neuron modalities. Annotation bar is coloured by the most common predicted neurotransmitter for the neurons of each type.

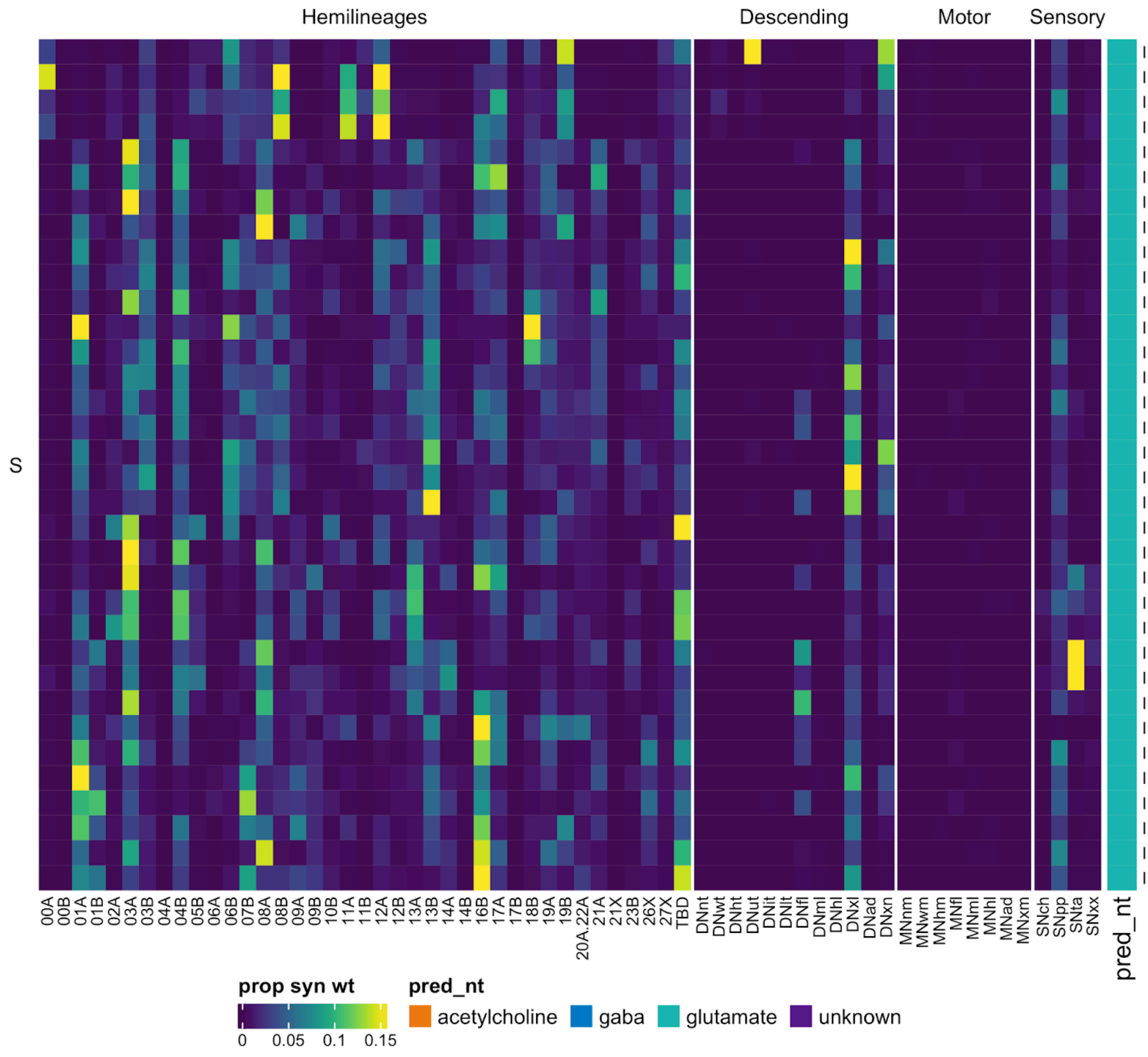

**Figure 26 - figure supplement 4. Connectivity to upstream partners by 08A secondary systematic types.** Proportions of synaptic weight to systematic types from upstream partners, normalised by row. 08A neurons have been clustered within each assigned birthtime window (P = primary, ES = early secondary, S = secondary) based on both upstream and downstream connectivity to hemilineages, descending neuron subclasses, motor neuron subclasses, and sensory neuron modalities. Annotation bar is coloured by the most common predicted neurotransmitter for the neurons of each type.

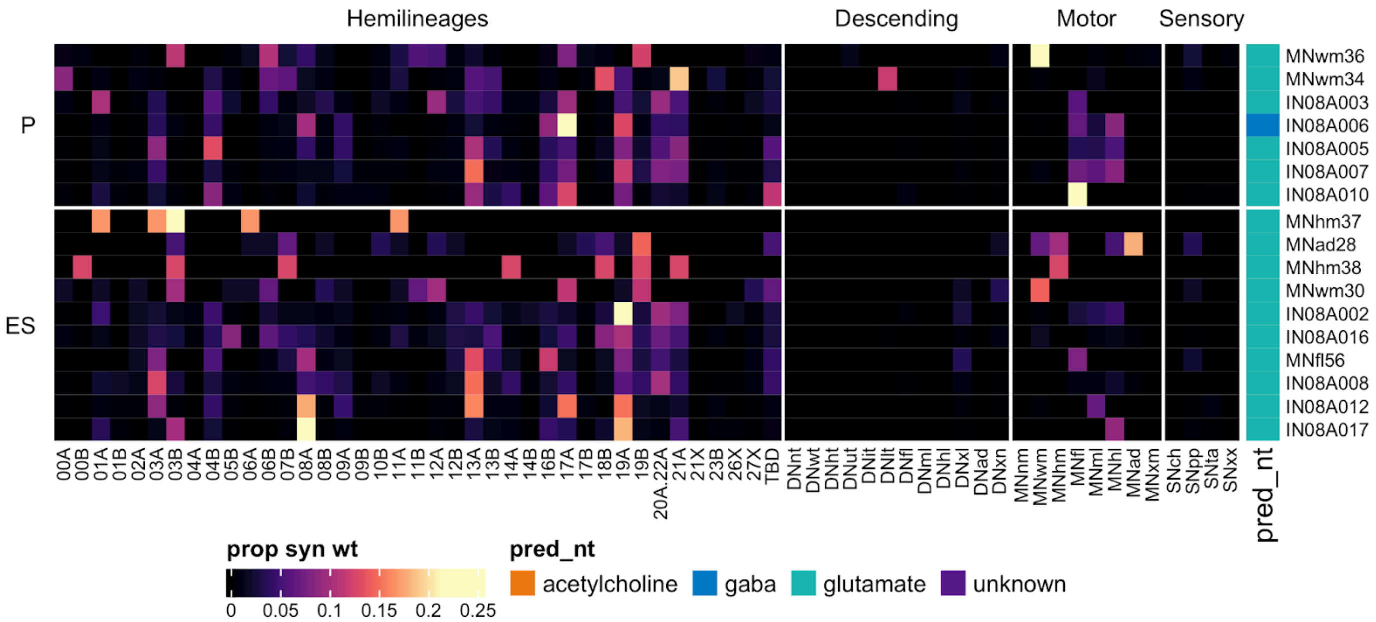

**Figure 26 - figure supplement 5. Connectivity to downstream partners by 08A primary and early secondary systematic types.** Proportions of synaptic weight from systematic types to downstream partners, normalised by row. 08A neurons have been clustered within each assigned birthtime window (P = primary, ES = early secondary, S = secondary) based on both upstream and downstream connectivity to hemilineages, descending neuron subclasses, motor neuron subclasses, and sensory neuron modalities. The annotation bar is coloured by the most common predicted neurotransmitter within each type.

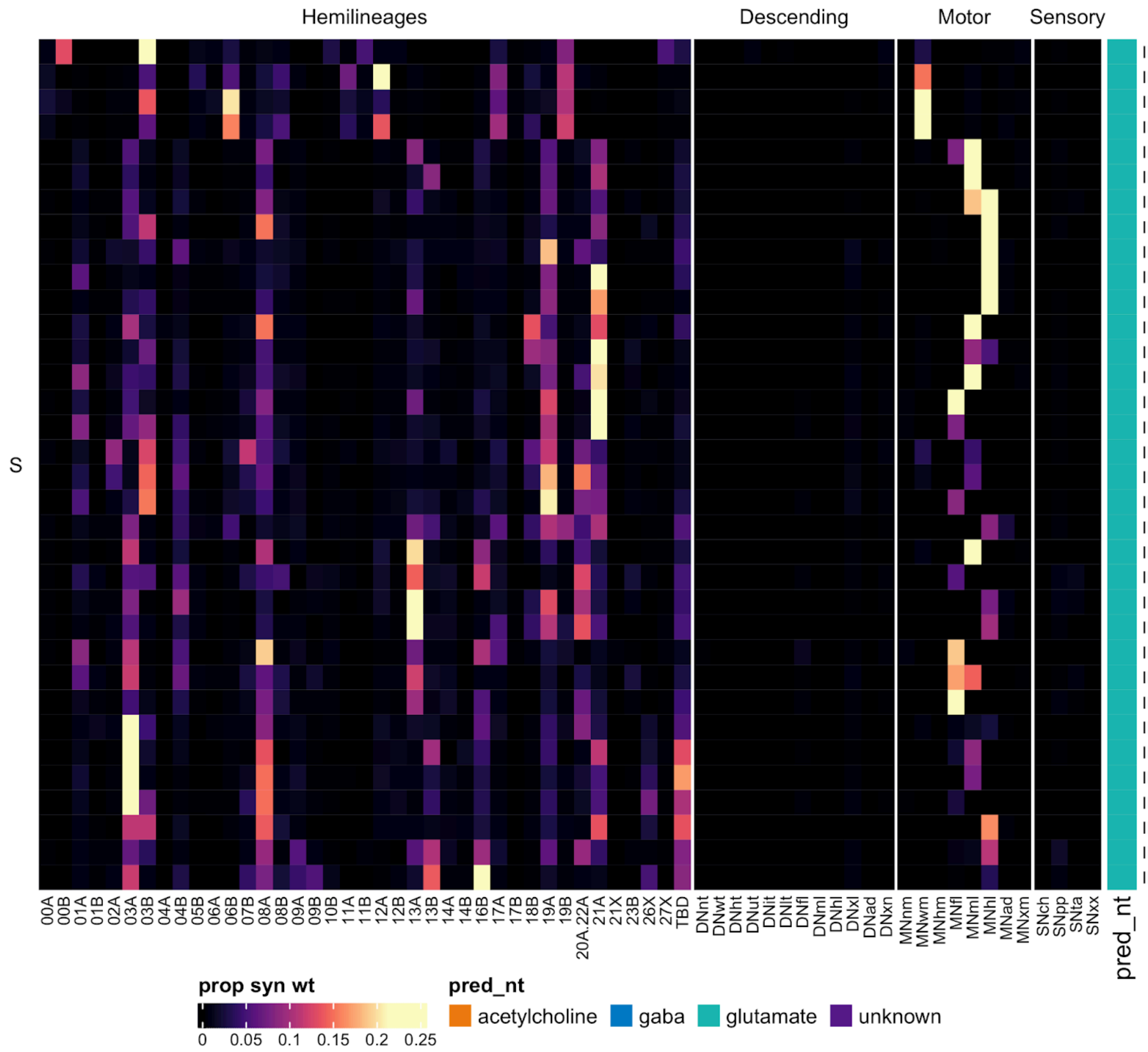

**Figure 26 - figure supplement 6. Connectivity to downstream partners by 08A secondary systematic types.** Proportions of synaptic weight from systematic types to downstream partners, normalised by row. 08A neurons have been clustered within each assigned birthtime window (P = primary, ES = early secondary, S = secondary) based on both upstream and downstream connectivity to hemilineages, descending neuron subclasses, motor neuron subclasses, and sensory neuron modalities. The annotation bar is coloured by the most common predicted neurotransmitter within each type.

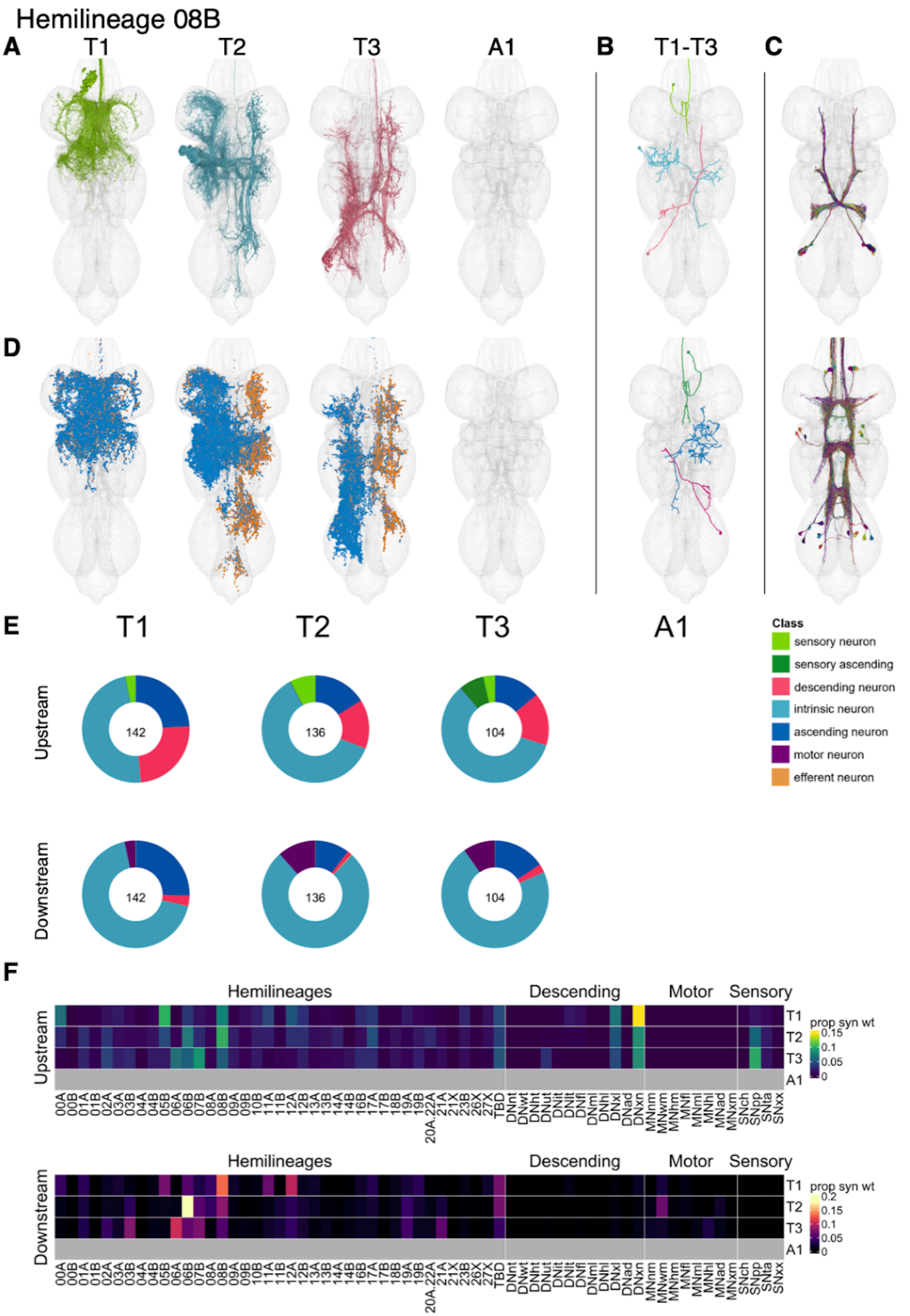
Hemilineage 08B
A
T1
T2
T3
A1
B
T1-T3
C
D
E
T1
T2
T3
A1
Upstream
142
136
104
Downstream
142
136
104
Class
sensory neuron
sensory ascending
descending neuron
intrinsic neuron
ascending neuron
motor neuron
efferent neuron
F
Hemilineages
Descending
Motor
Sensory
Upstream
T1
T2
T3
A1
prop syn wt
0.15
0.1
0.05
0
00A 00B 01A 01B 02A 03A 03B 04A 04B 05B 06A 06B 07B 08A 08B 09A 09B 10B 11A 11B 12A 12B 13A 13B 14A 14B 16B 17A 17B 18B 19A 19B 20A.22A 21A 21X 23B 26X 27X TBD DNnt DNwt DNht DNut DNlt DNfl DNml DNhl DNxl DNad DNxn MNnm MNwm MNhm MNfl MNml MNhl MNad MNxm SNch SNpp SNta SNxx
Hemilineages
Descending
Motor
Sensory
Downstream
T1
T2
T3
A1
prop syn wt
0.2
0.15
0.1
0.05
0
00A 00B 01A 01B 02A 03A 03B 04A 04B 05B 06A 06B 07B 08A 08B 09A 09B 10B 11A 11B 12A 12B 13A 13B 14A 14B 16B 17A 17B 18B 19A 19B 20A.22A 21A 21X 23B 26X 27X TBD DNnt DNwt DNht DNut DNlt DNfl DNml DNhl DNxl DNad DNxn MNnm MNwm MNhm MNfl MNml MNhl MNad MNxm SNch SNpp SNta SNxx

**Figure 27. Hemilineage 08B. A.** Meshes of all RHS secondary neurons plotted in neuromere-specific colours. **B.** "Representative" secondary neuron skeletons plotted in hemineuromere-specific colours. The skeleton with the top accumulated NBLAST score among all neurons from the hemilineage in a given hemineuromere was used. **C.** Neuron meshes of selected examples. Top: electrical n-CHINs and putative electrical cHINs, subcluster 10598. Bottom: intersegmental mVAC population, subcluster 10997. **D.** Predicted synapses of RHS secondary neurons. Blue: postsynapses; dark orange: presynapses. **E.** Proportions of connections from secondary neurons to upstream or downstream partners, normalised by neuromere and coloured by broad class. Numbers of query neurons appear in the centre. **F.** Proportions of synaptic weight from secondary neurons originating in each neuromere to upstream or downstream partners, normalised by row.

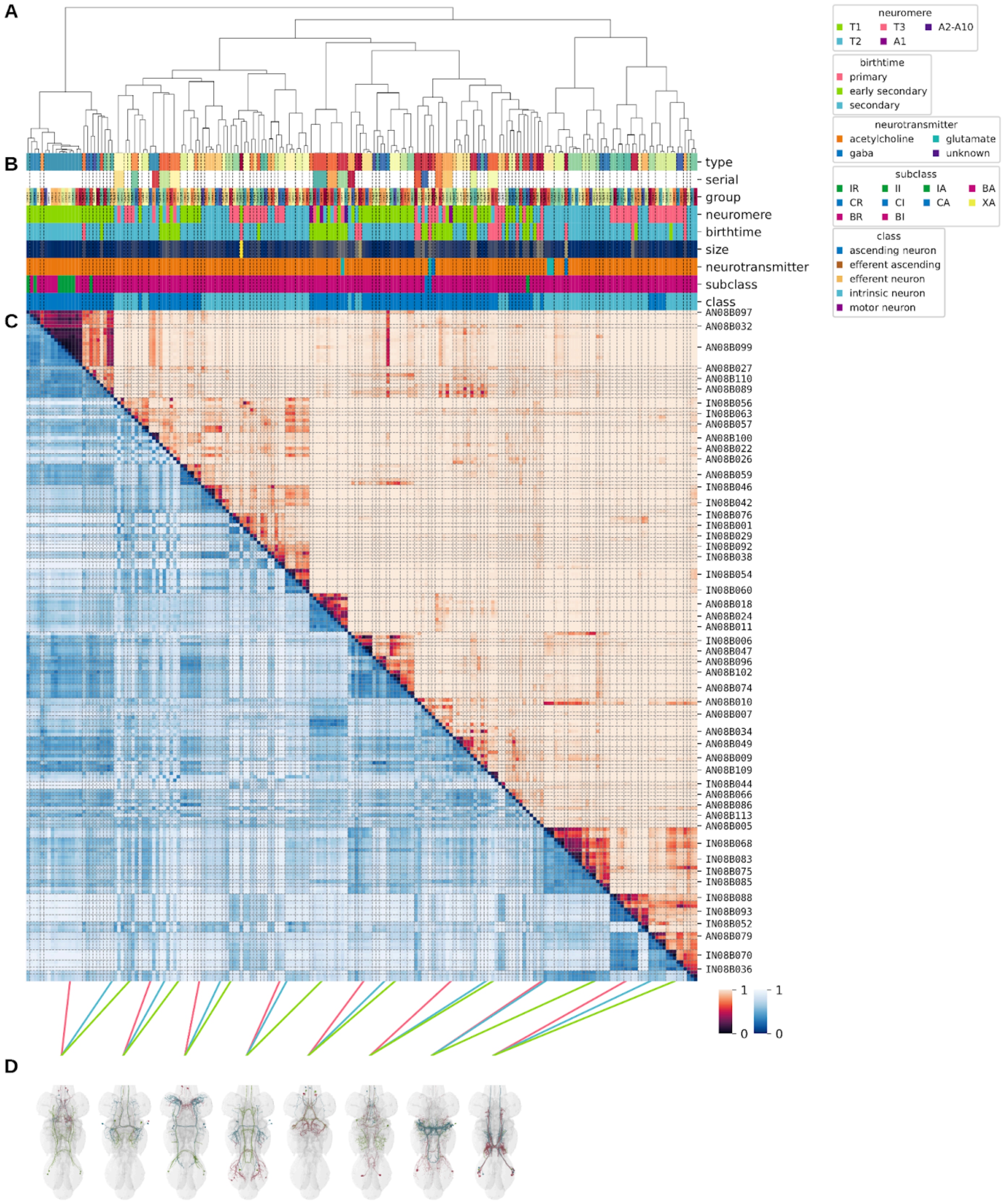

**Figure 27 - figure supplement 1. Systematic typing of hemilineage 08B. A.** Hierarchical clustering dendrogram of hemilineage groups by laterally and serially aggregated connectivity cosine clustering. **B.** Categorical annotations of each hemilineage group, each column corresponding to the aligned leaf in A. Colours for type, serial set, and group are arbitrary for visualisation. Colours for neuromere, birthtime, neurotransmitter, subclass, and class are as in all other figures. **C.** Similarity distance heatmap for hemilineage. Cosine distance is in the upper triangle, while laterally symmetrised NBLAST distance is in the lower triangle. Systematic type names of some types are labelled. **D.** Morphologically representative groups from dendrogram subtrees. Each group, indicated by colour and line connecting to its column in B and C, is the most morphologically representative group (medoid of NBLAST distance)

from a subtree of A. The subtrees (flat clusters) are equal height cuts of A determined to yield the number of groups per plot and plots in D.

#### 08B systematic types

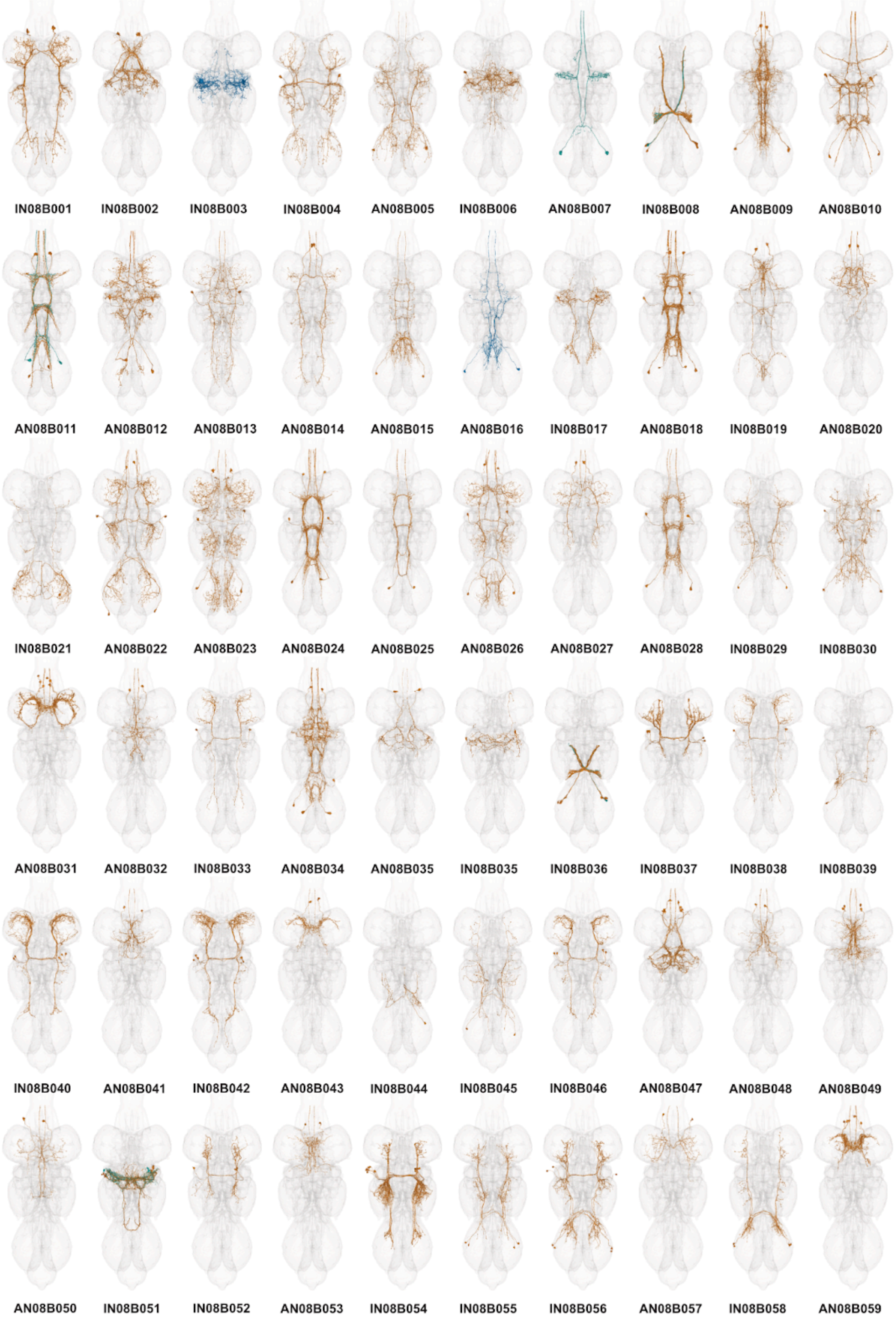

**Figure 27 - figure supplement 2. Systematic types of hemilineage 08B.** Systematic types have been arranged in numerical order, with neurons of the same type that belong to distinct classes (e.g., intrinsic neuron vs ascending neuron) plotted separately but placed adjacent to each other. Individual neuron meshes have been coloured based on predicted neurotransmitter: dark orange = acetylcholine, blue = gaba, marine = glutamate, dark purple = unknown.

#### 08B systematic types

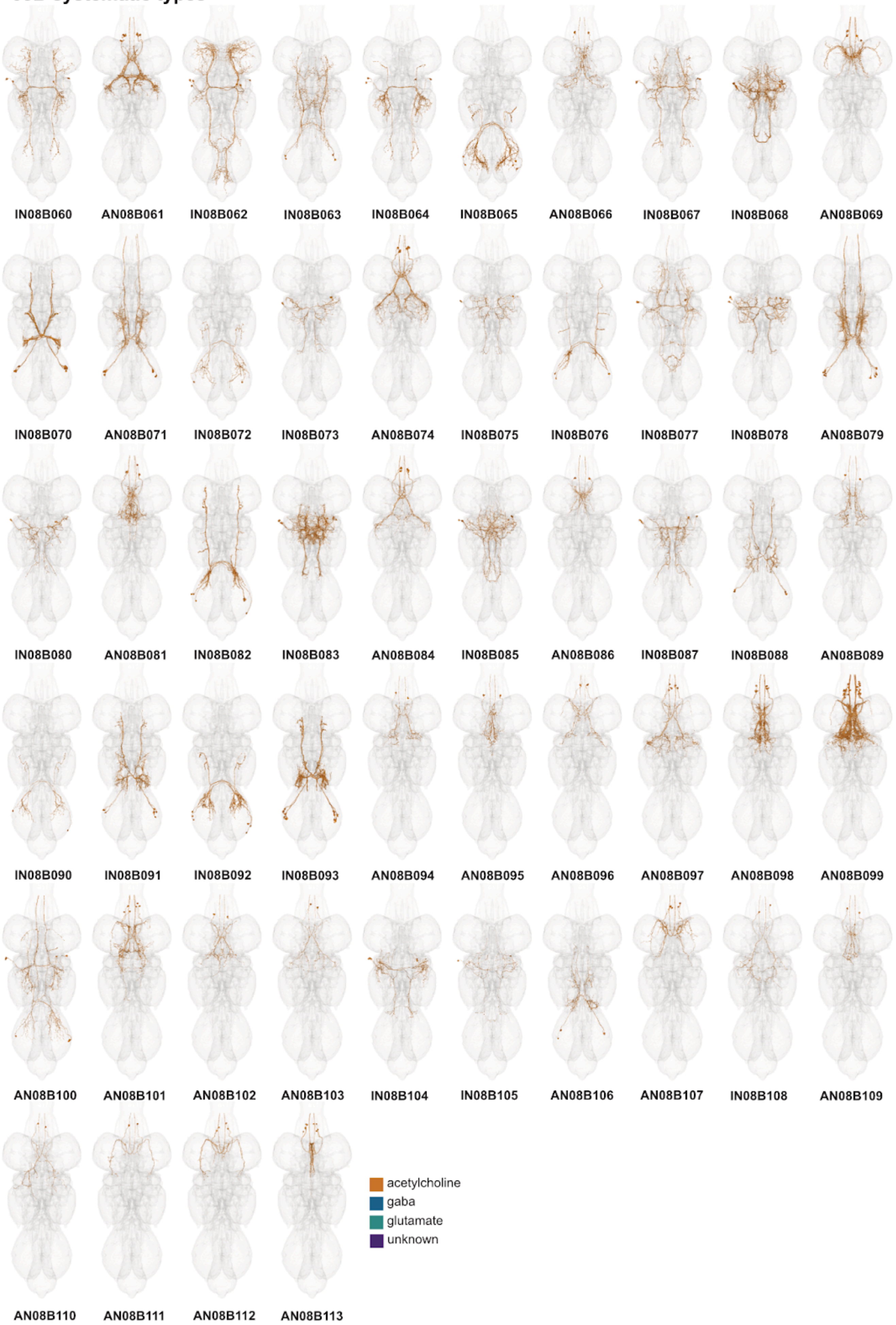

**Figure 27 - figure supplement 3. Systematic types of hemilineage 08B, continued.** Systematic types have been arranged in numerical order, with neurons of the same type that belong to distinct classes (e.g., intrinsic neuron vs ascending neuron) plotted separately but placed adjacent to each other. Individual neuron meshes have been coloured based on predicted neurotransmitter: dark orange = acetylcholine, blue = gaba, marine = glutamate, dark purple = unknown.

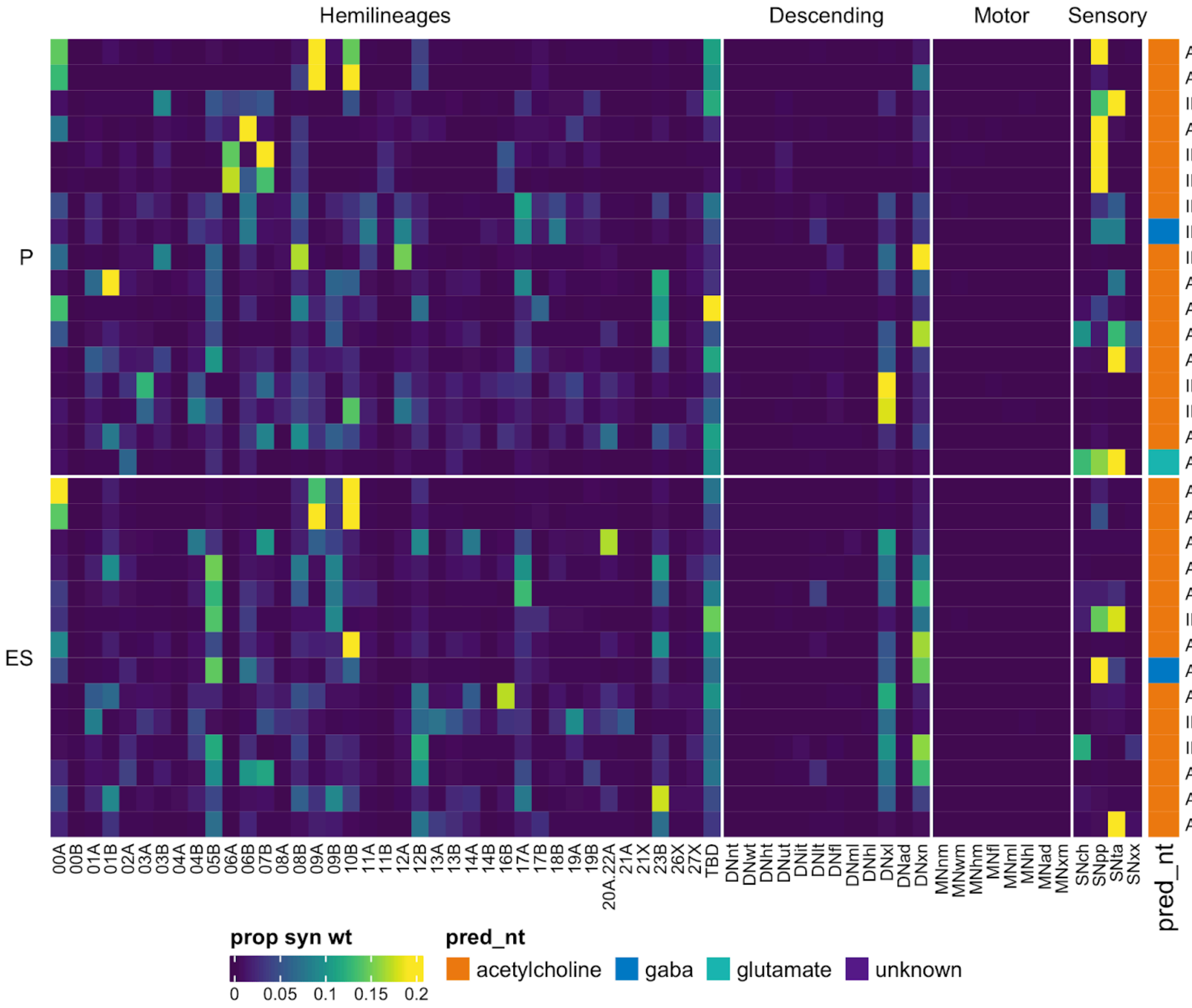

**Figure 27 - figure supplement 4. Connectivity to upstream partners by 08B primary and early secondary systematic types.** Proportions of synaptic weight to systematic types from upstream partners, normalised by row. 08B neurons have been clustered within each assigned birthtime window (P = primary, ES = early secondary, S = secondary) based on both upstream and downstream connectivity to hemilineages, descending neuron subclasses, motor neuron subclasses, and sensory neuron modalities. Annotation bar is coloured by the most common predicted neurotransmitter for the neurons of each type.

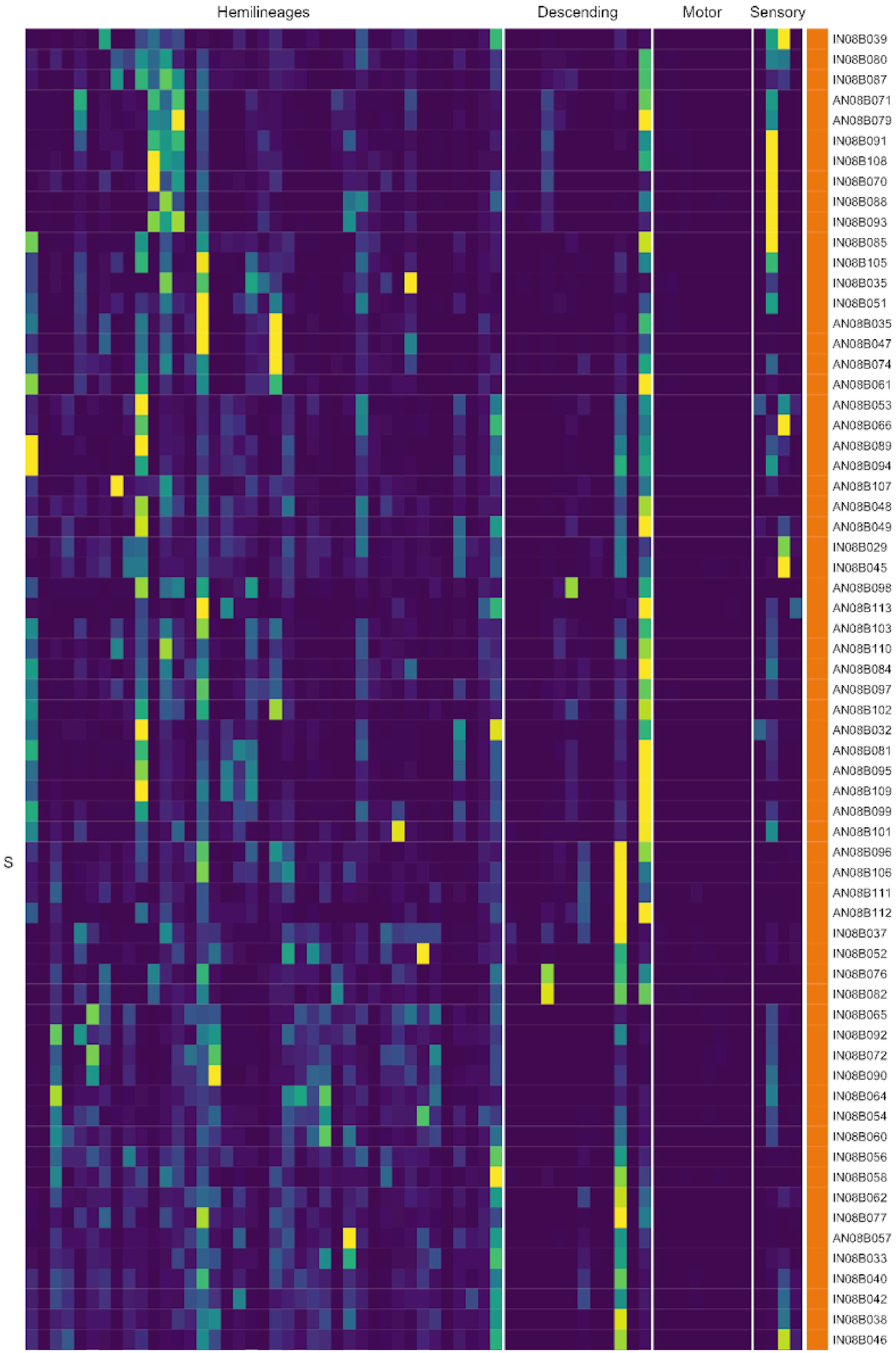
Hemilineages
Descending
Motor
Sensory
S
IN08B039
IN08B080
IN08B087
AN08B071
AN08B079
IN08B091
IN08B108
IN08B070
IN08B088
IN08B093
IN08B085
IN08B105
IN08B035
IN08B051
AN08B035
AN08B047
AN08B074
AN08B061
AN08B053
AN08B066
AN08B089
AN08B094
AN08B107
AN08B048
AN08B049
IN08B029
IN08B045
AN08B098
AN08B113
AN08B103
AN08B110
AN08B084
AN08B097
AN08B102
AN08B032
AN08B081
AN08B095
AN08B109
AN08B099
AN08B101
AN08B096
AN08B106
AN08B111
AN08B112
IN08B037
IN08B052
IN08B076
IN08B082
IN08B065
IN08B092
IN08B072
IN08B090
IN08B064
IN08B054
IN08B060
IN08B056
IN08B058
IN08B062
IN08B077
AN08B057
IN08B033
IN08B040
IN08B042
IN08B038
IN08B046

**Figure 27 - figure supplement 5. Connectivity to upstream partners by 08B secondary systematic types.** Proportions of synaptic weight to systematic types from upstream partners, normalised by row. 08B neurons have been clustered within each assigned birthtime window (P = primary, ES = early secondary, S = secondary) based on both upstream and downstream connectivity to hemilineages, descending neuron subclasses, motor neuron subclasses, and sensory neuron modalities. The annotation bar is coloured by the most common predicted neurotransmitter within each type.

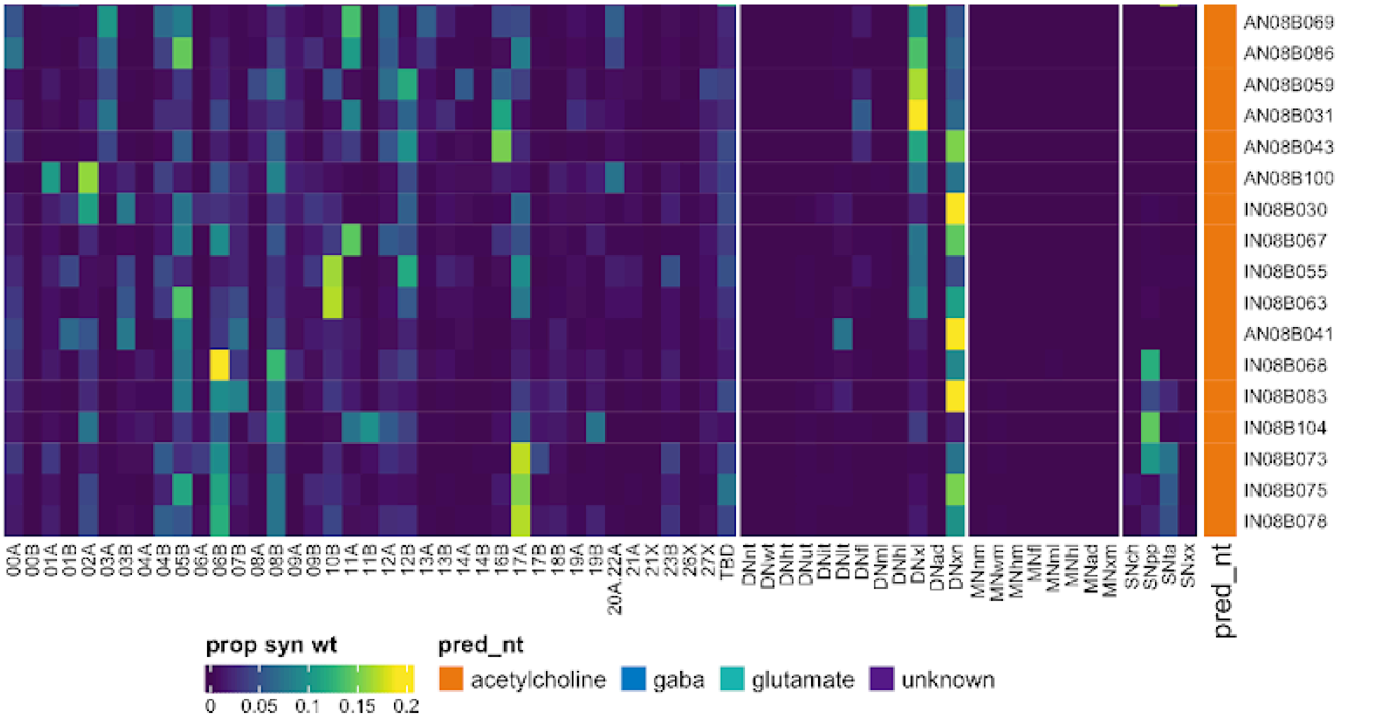

**Figure 27 - figure supplement 6. Connectivity to upstream partners by 08B secondary systematic types, continued.** Proportions of synaptic weight to systematic types from upstream partners, normalised by row. 08B neurons have been clustered within each assigned birthtime window (P = primary, ES = early secondary, S = secondary) based on both upstream and downstream connectivity to hemilineages, descending neuron subclasses, motor neuron subclasses, and sensory neuron modalities. The annotation bar is coloured by the most common predicted neurotransmitter within each type.

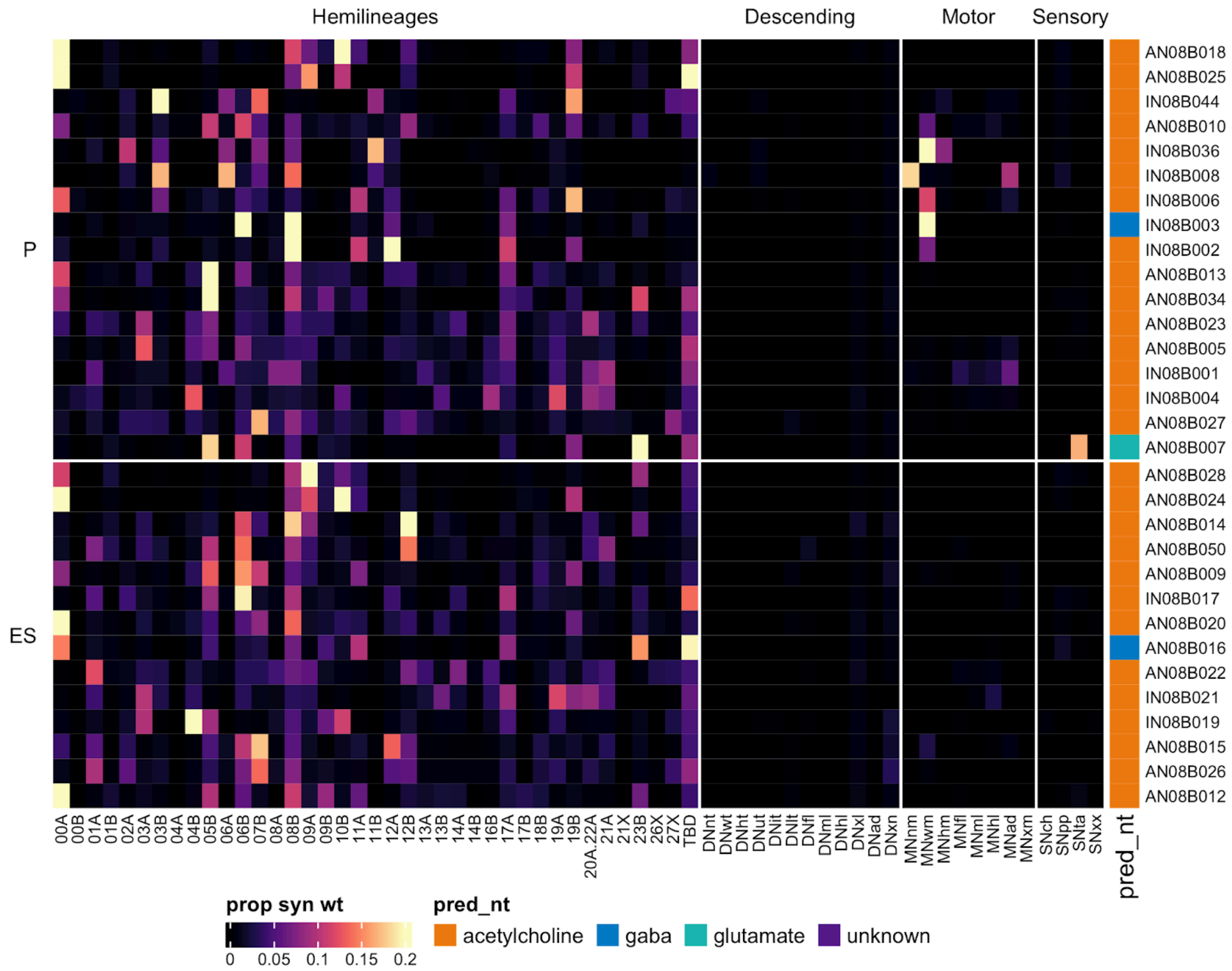

**Figure 27 - figure supplement 7. Connectivity to downstream partners by 08B primary and early secondary systematic types.** Proportions of synaptic weight from systematic types to downstream partners, normalised by row. 08B neurons have been clustered within each assigned birthtime window (P = primary, ES = early secondary, S = secondary) based on both upstream and downstream connectivity to hemilineages, descending neuron subclasses, motor neuron subclasses, and sensory neuron modalities. The annotation bar is coloured by the most common predicted neurotransmitter within each type.

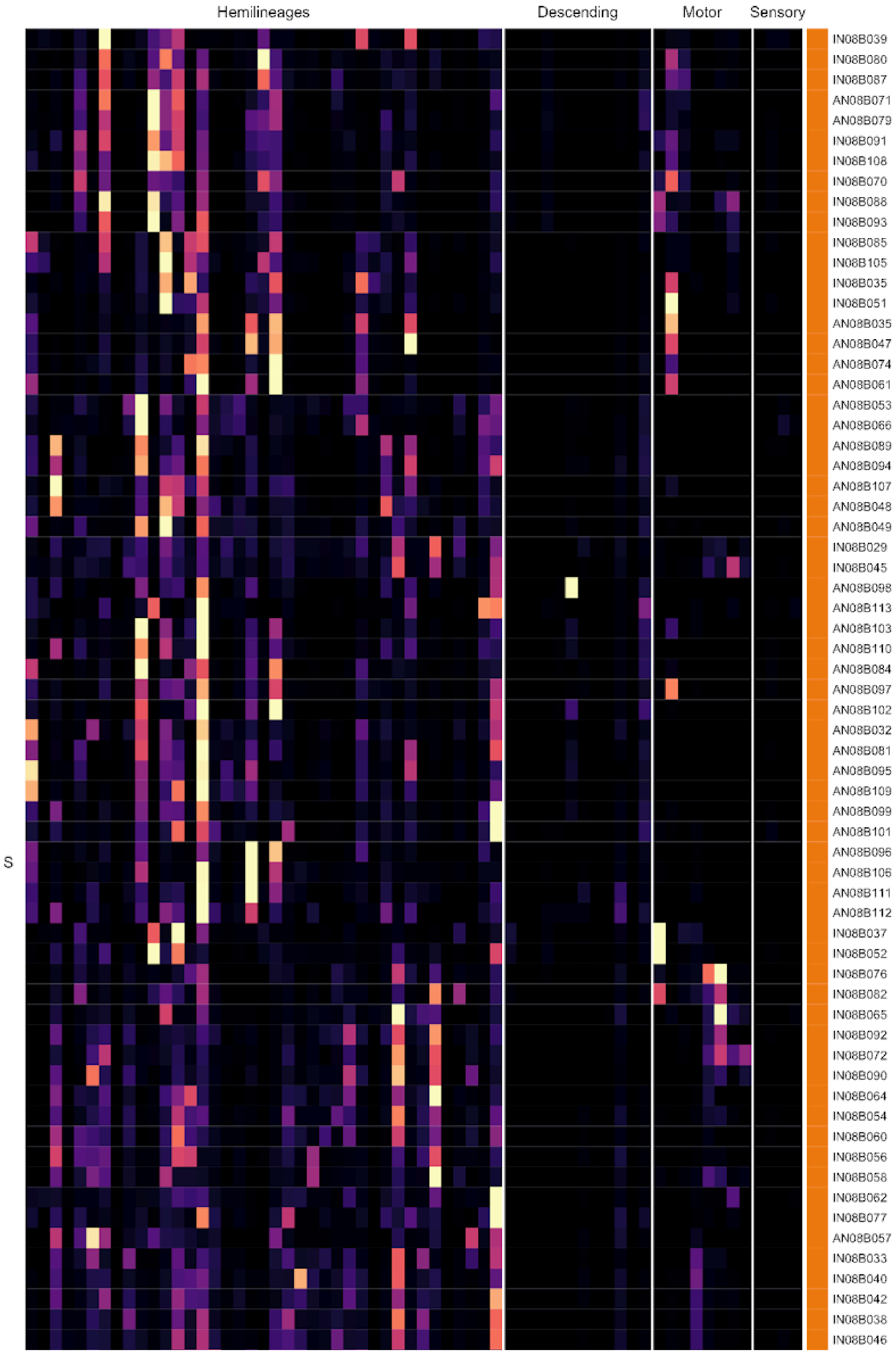

Hemilineages
Descending
Motor
Sensory
S
IN08B039
IN08B080
IN08B087
AN08B071
AN08B079
IN08B091
IN08B108
IN08B070
IN08B088
IN08B093
IN08B085
IN08B105
IN08B035
IN08B051
AN08B035
AN08B047
AN08B074
AN08B061
AN08B053
AN08B066
AN08B089
AN08B094
AN08B107
AN08B048
AN08B049
IN08B029
IN08B045
AN08B098
AN08B113
AN08B103
AN08B110
AN08B084
AN08B097
AN08B102
AN08B032
AN08B081
AN08B095
AN08B109
AN08B099
AN08B101
AN08B096
AN08B106
AN08B111
AN08B112
IN08B037
IN08B052
IN08B076
IN08B082
IN08B065
IN08B092
IN08B072
IN08B090
IN08B064
IN08B054
IN08B060
IN08B056
IN08B058
IN08B062
IN08B077
AN08B057
IN08B033
IN08B040
IN08B042
IN08B038
IN08B046

**Figure 27 - figure supplement 8. Connectivity to downstream partners by 08B secondary systematic types.** Proportions of synaptic weight from systematic types to downstream partners, normalised by row. 08B neurons have been clustered within each assigned birthtime window (P = primary, ES = early secondary, S = secondary) based on both upstream and downstream connectivity to hemilineages, descending neuron subclasses, motor neuron subclasses, and sensory neuron modalities. The annotation bar is coloured by the most common predicted neurotransmitter for the neurons of each type.

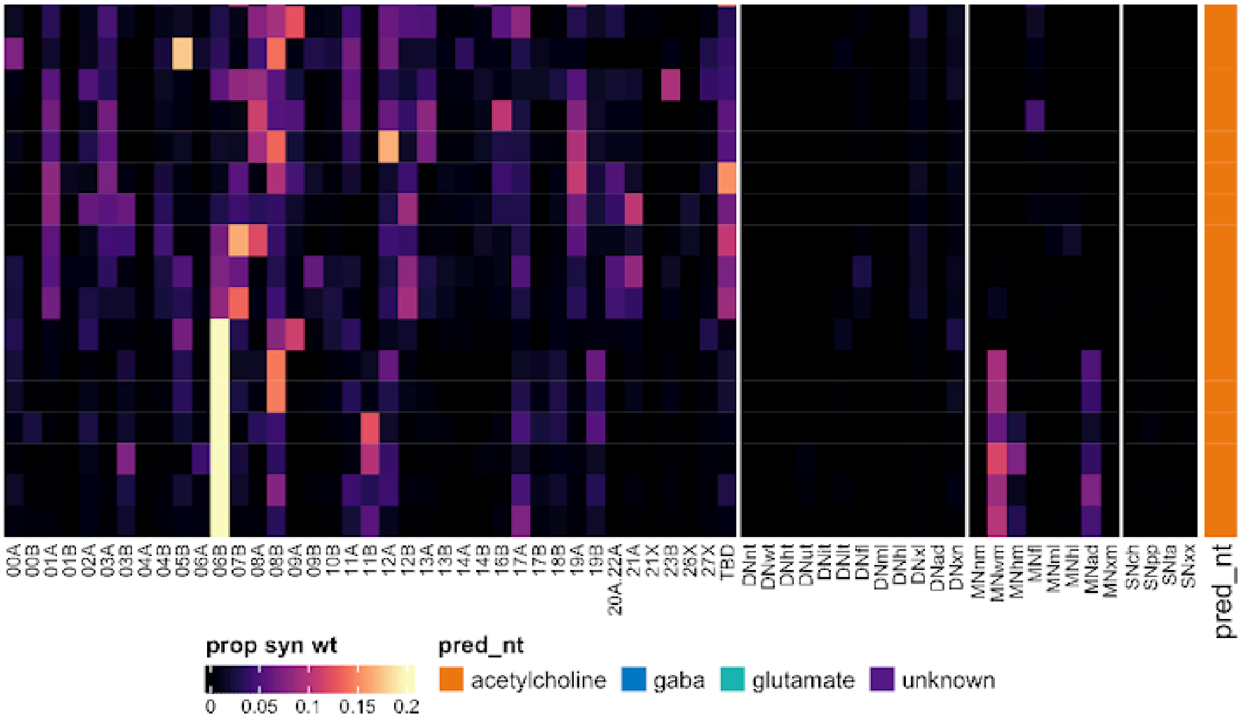

**Figure 27 - figure supplement 9. Connectivity to downstream partners by 08B secondary systematic types, continued.** Proportions of synaptic weight from systematic types to downstream partners, normalised by row. 08B neurons have been clustered within each assigned birthtime window (P = primary, ES = early secondary, S = secondary) based on both upstream and downstream connectivity to hemilineages, descending neuron subclasses, motor neuron subclasses, and sensory neuron modalities. The annotation bar is coloured by the most common predicted neurotransmitter for the neurons of each type.

Hemilineage 09A

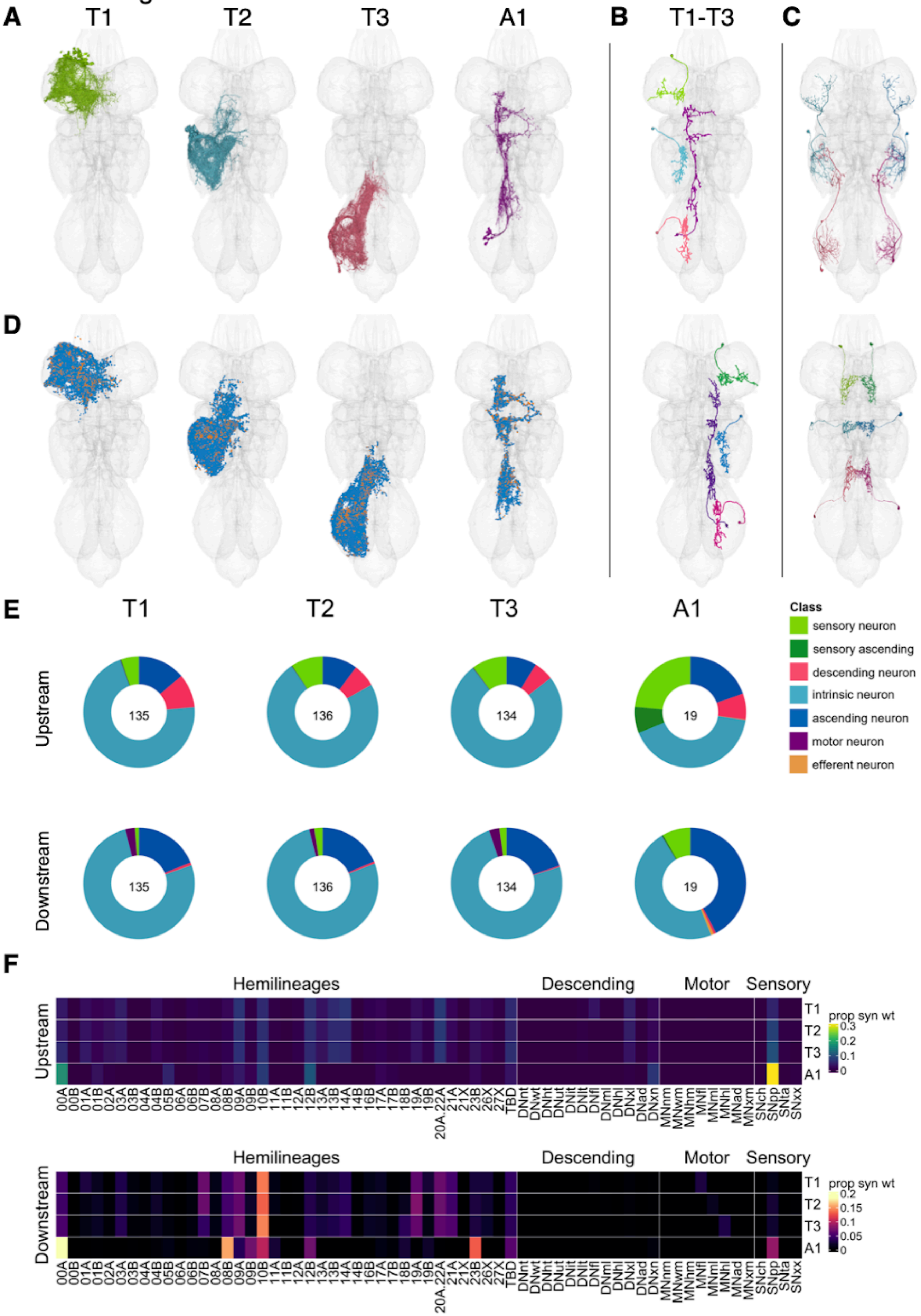

**Figure 28. Hemilineage 09A. A.** Meshes of all RHS secondary neurons plotted in neuromere-specific colours. **B.** "Representative" secondary neuron skeletons plotted in hemineuromere-specific colours. The skeleton with the top accumulated NBLAST score among all neurons from the hemilineage in a given hemineuromere was used. **C.** Neuron meshes of selected examples. Top: sequential serial set 11194. Bottom: mVAC-restricted independent leg serial set 14731. **D.** Predicted synapses of RHS secondary neurons. Blue: postsynapses; dark orange: presynapses. **E.** Proportions of connections from secondary neurons to upstream or downstream partners, normalised by neuromere and coloured by broad class. Numbers of query neurons appear in the centre. **F.** Proportions of synaptic weight from secondary neurons originating in each neuromere to upstream or downstream partners, normalised by row.

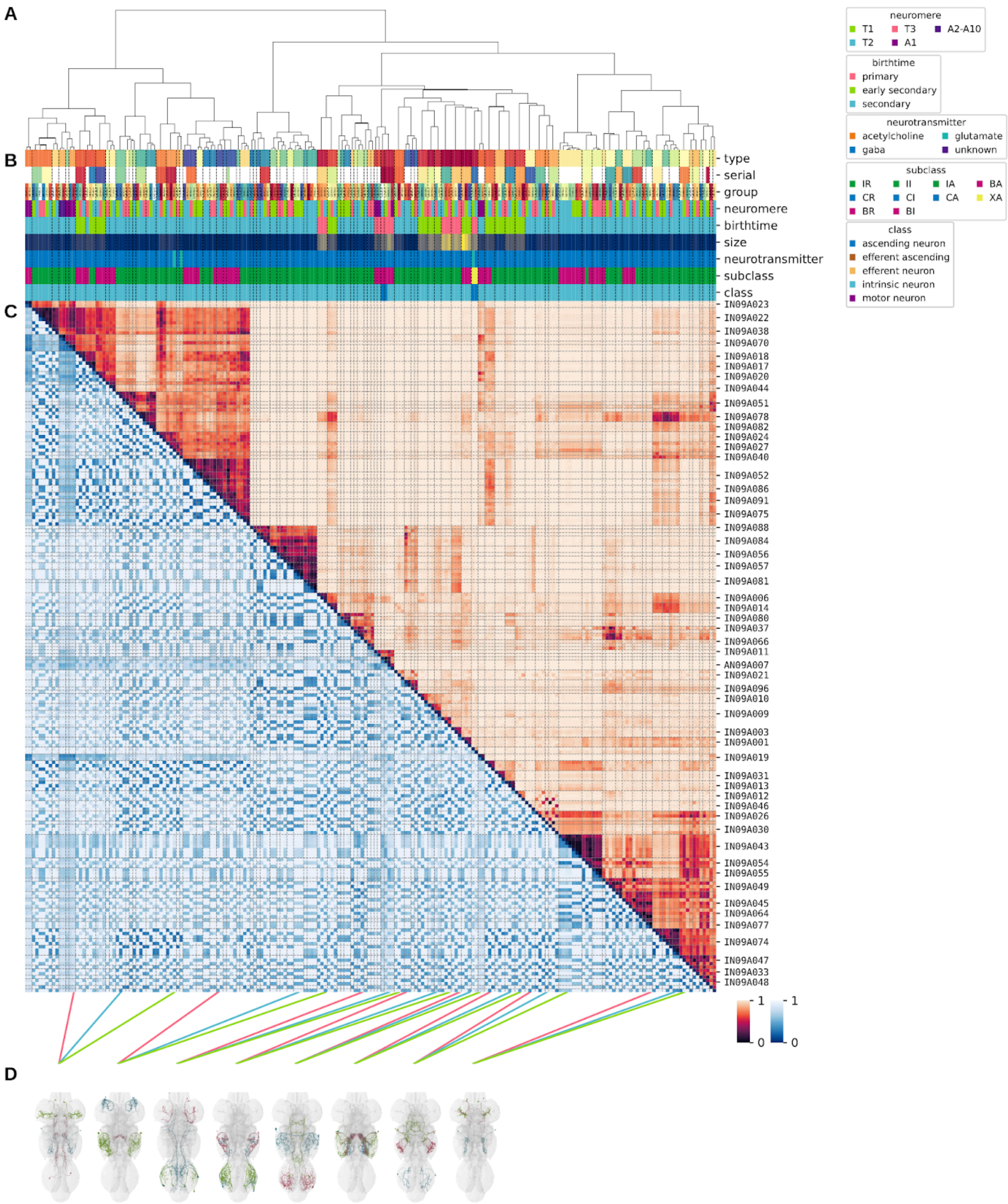

**Figure 28 - figure supplement 1. Systematic typing of hemilineage 09A. A.** Hierarchical clustering dendrogram of hemilineage groups by laterally and serially aggregated connectivity cosine clustering. **B.** Categorical annotations of each hemilineage group, each column corresponding to the aligned leaf in A. Colours for type, serial set, and group are arbitrary for visualisation. Colours for neuromere, birthtime, neurotransmitter, subclass, and class are as in all other figures. **C.** Similarity distance heatmap for hemilineage. Cosine distance is in the upper triangle, while laterally symmetrised NBLAST distance is in the lower triangle. Systematic type names of some types are labelled. **D.** Morphologically representative groups from dendrogram subtrees. Each group, indicated by colour and line connecting to its column in B and C, is the most morphologically representative group (medoid of NBLAST distance)

from a subtree of A. The subtrees (flat clusters) are equal height cuts of A determined to yield the number of groups per plot and plots in D.

#### 09A systematic types

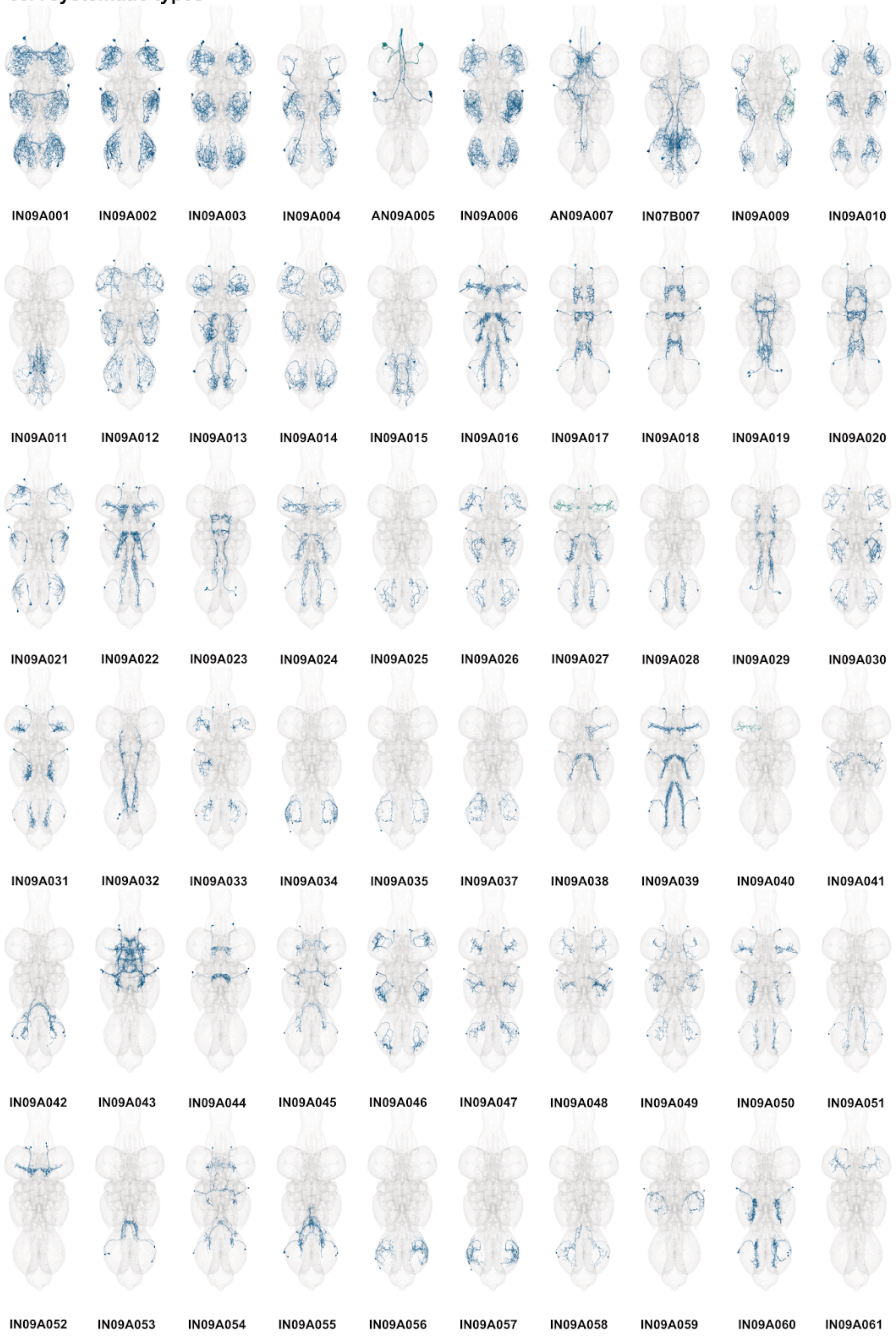

**Figure 28 - figure supplement 2. Systematic types of hemilineage 09A.** Systematic types have been arranged in numerical order, with neurons of the same type that belong to distinct classes (e.g., intrinsic neuron vs ascending neuron) plotted separately but placed adjacent to each other. Individual neuron meshes have been coloured based on predicted neurotransmitter: dark orange = acetylcholine, blue = gaba, marine = glutamate, dark purple = unknown.

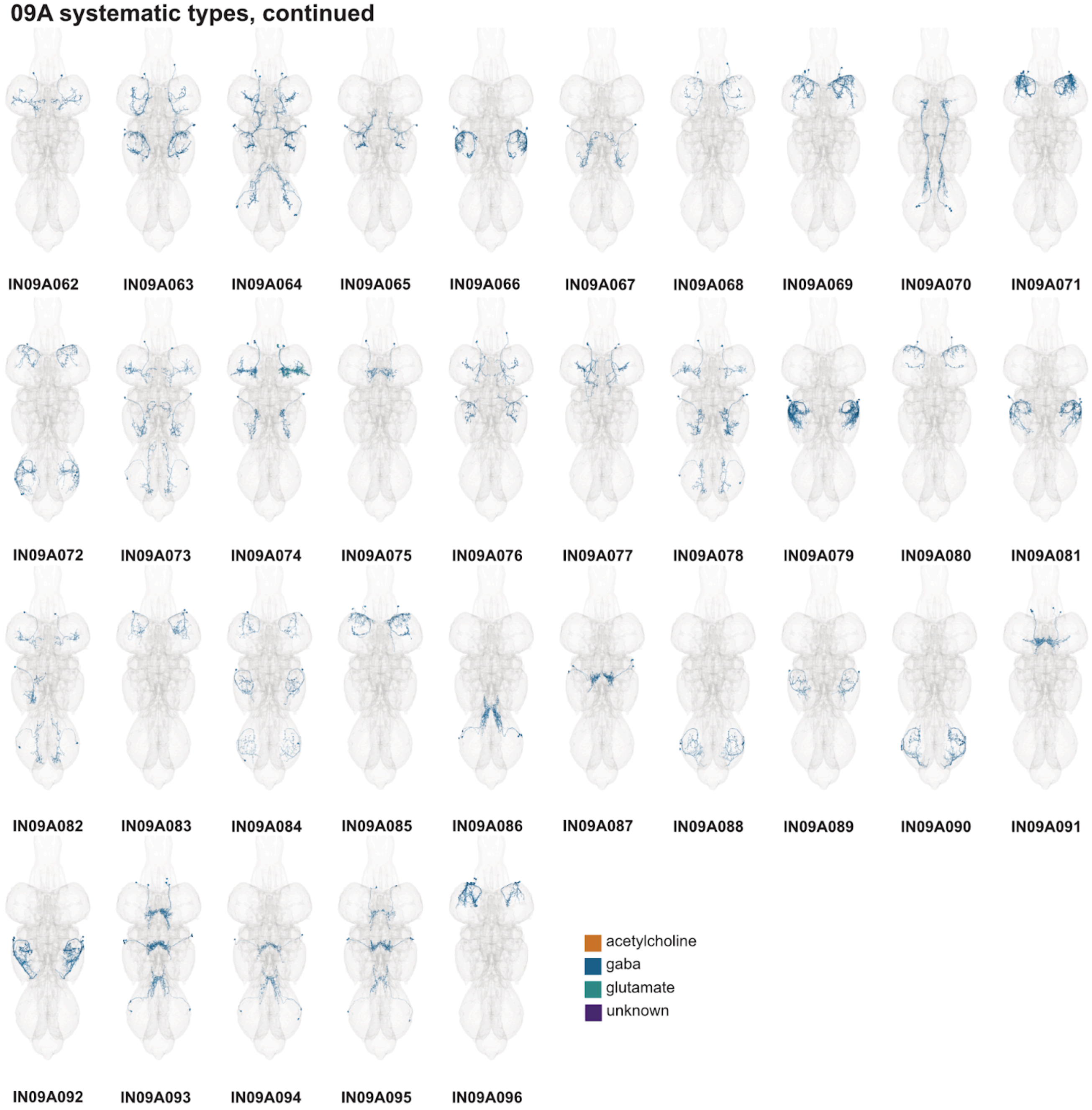

**Figure 28 - figure supplement 3. Systematic types of hemilineage 09A, continued.** Systematic types have been arranged in numerical order, with neurons of the same type that belong to distinct classes (e.g., intrinsic neuron vs ascending neuron) plotted separately but placed adjacent to each other. Individual neuron meshes have been coloured based on predicted neurotransmitter: dark orange = acetylcholine, blue = gaba, marine = glutamate, dark purple = unknown.

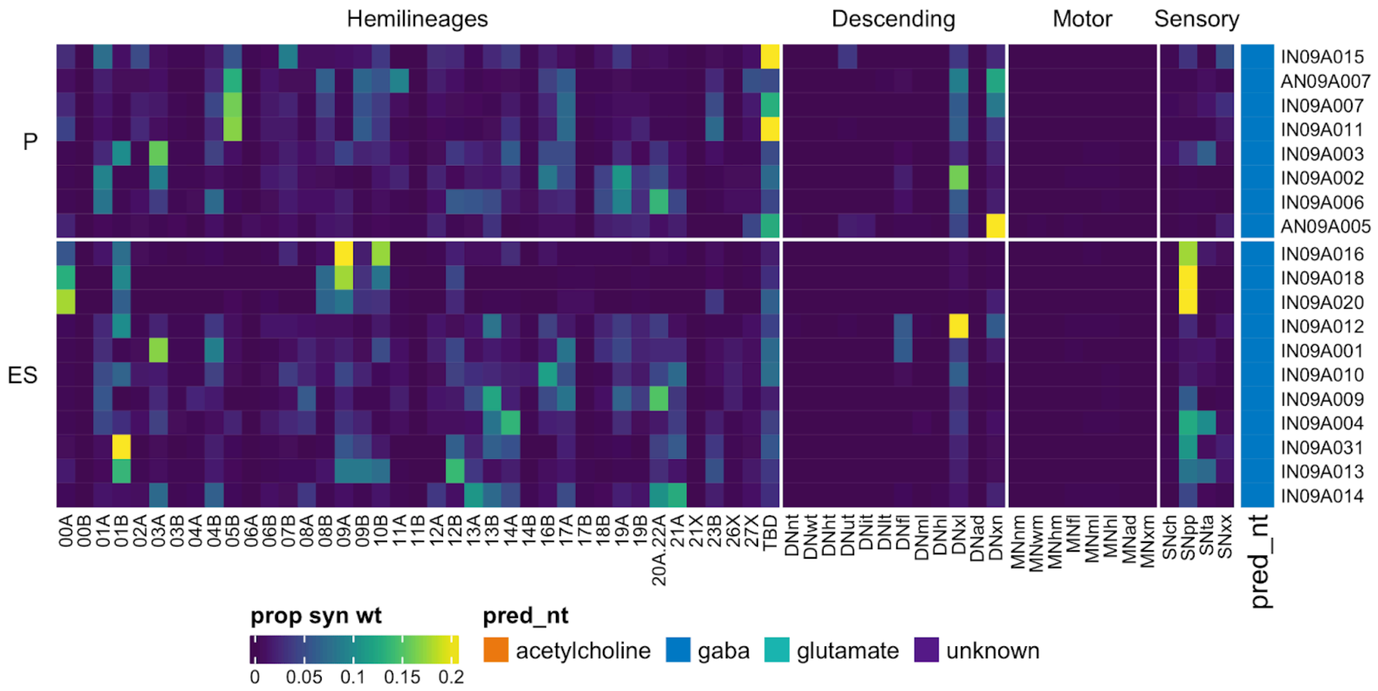

**Figure 28 - figure supplement 4. Connectivity to upstream partners by 09A primary and early secondary systematic types.** Proportions of synaptic weight to systematic types from upstream partners, normalised by row. 09A neurons have been clustered within each assigned birthtime window (P = primary, ES = early secondary, S = secondary) based on both upstream and downstream connectivity to hemilineages, descending neuron subclasses, motor neuron subclasses, and sensory neuron modalities. Annotation bar is coloured by the most common predicted neurotransmitter for the neurons of each type.

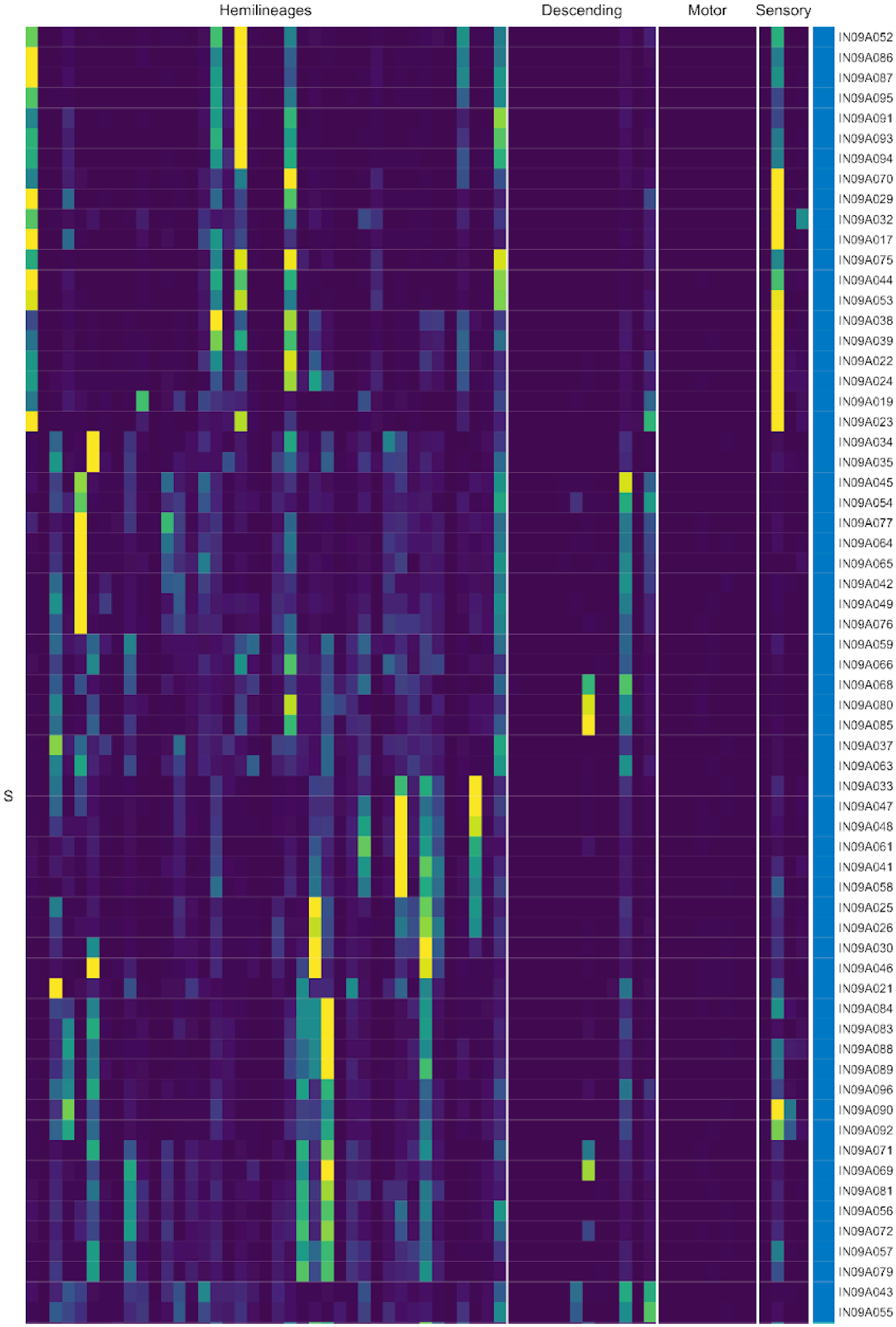
Hemilineages
Descending
Motor
Sensory
S
IN09A052
IN09A086
IN09A087
IN09A095
IN09A091
IN09A093
IN09A094
IN09A070
IN09A029
IN09A032
IN09A017
IN09A075
IN09A044
IN09A053
IN09A038
IN09A039
IN09A022
IN09A024
IN09A019
IN09A023
IN09A034
IN09A035
IN09A045
IN09A054
IN09A077
IN09A064
IN09A065
IN09A042
IN09A049
IN09A076
IN09A059
IN09A066
IN09A068
IN09A080
IN09A085
IN09A037
IN09A063
IN09A033
IN09A047
IN09A048
IN09A061
IN09A041
IN09A058
IN09A025
IN09A026
IN09A030
IN09A046
IN09A021
IN09A084
IN09A083
IN09A088
IN09A089
IN09A096
IN09A090
IN09A092
IN09A071
IN09A069
IN09A081
IN09A056
IN09A072
IN09A057
IN09A079
IN09A043
IN09A055

**Figure 28 - figure supplement 5. Connectivity to upstream partners by 09A secondary systematic types.** Proportions of synaptic weight to systematic types from upstream partners, normalised by row. 09A neurons have been clustered within each assigned birthtime window (P = primary, ES = early secondary, S = secondary) based on both upstream and downstream connectivity to hemilineages, descending neuron subclasses, motor neuron subclasses, and sensory neuron modalities. The annotation bar is coloured by the most common predicted neurotransmitter within each type.

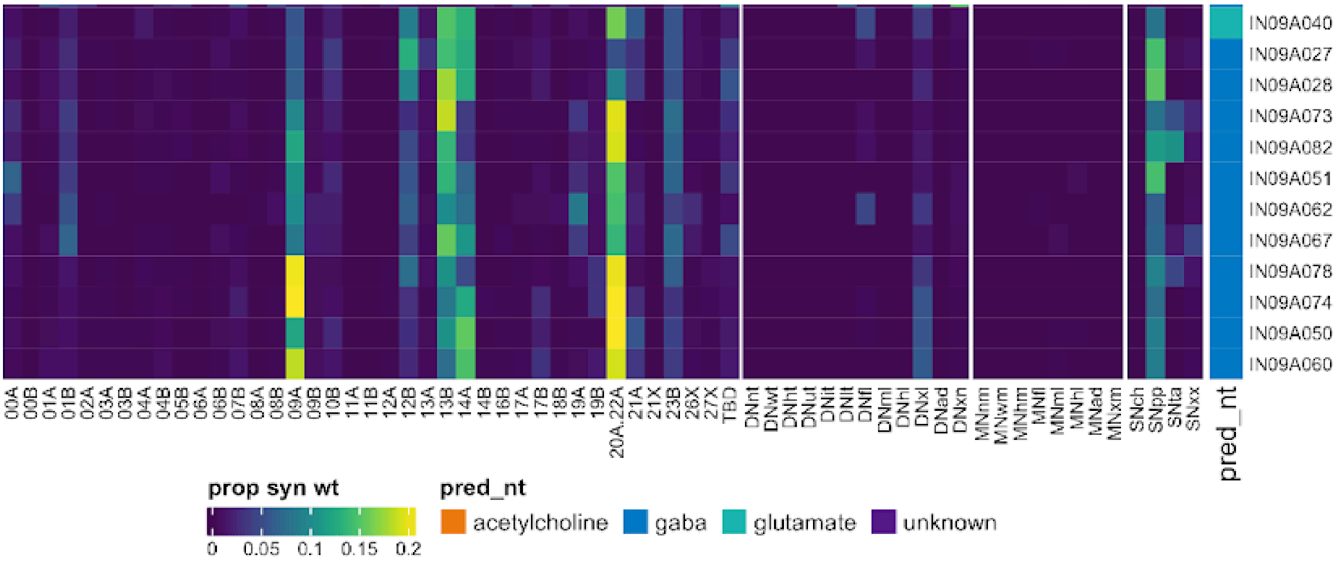

**Figure 28 - figure supplement 6. Connectivity to upstream partners by 09A secondary systematic types, continued.** Proportions of synaptic weight to systematic types from upstream partners, normalised by row. 09A neurons have been clustered within each assigned birthtime window (P = primary, ES = early secondary, S = secondary) based on both upstream and downstream connectivity to hemilineages, descending neuron subclasses, motor neuron subclasses, and sensory neuron modalities. The annotation bar is coloured by the most common predicted neurotransmitter within each type.

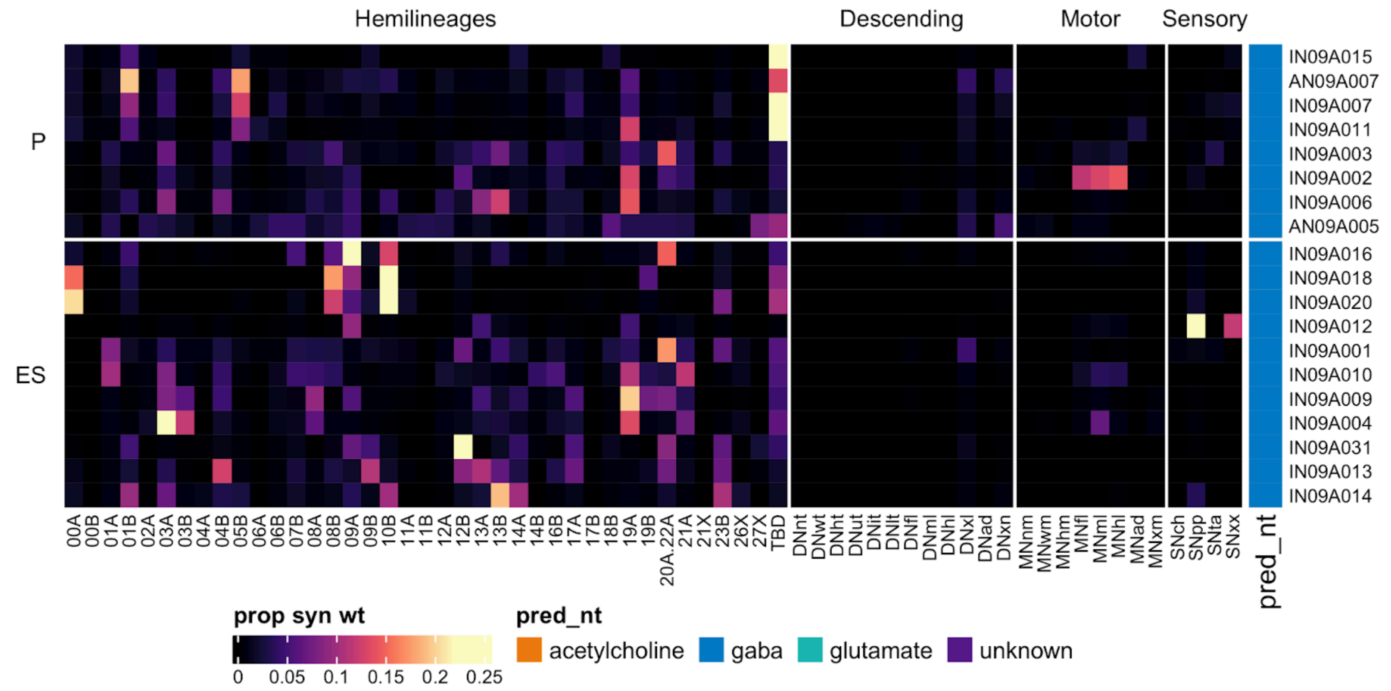

**Figure 28 - figure supplement 7. Connectivity to downstream partners by 09A primary and early secondary systematic types.** Proportions of synaptic weight from systematic types to downstream partners, normalised by row. 09A neurons have been clustered within each assigned birthtime window (P = primary, ES = early secondary, S = secondary) based on both upstream and downstream connectivity to hemilineages, descending neuron subclasses, motor neuron subclasses, and sensory neuron modalities. The annotation bar is coloured by the most common predicted neurotransmitter within each type.

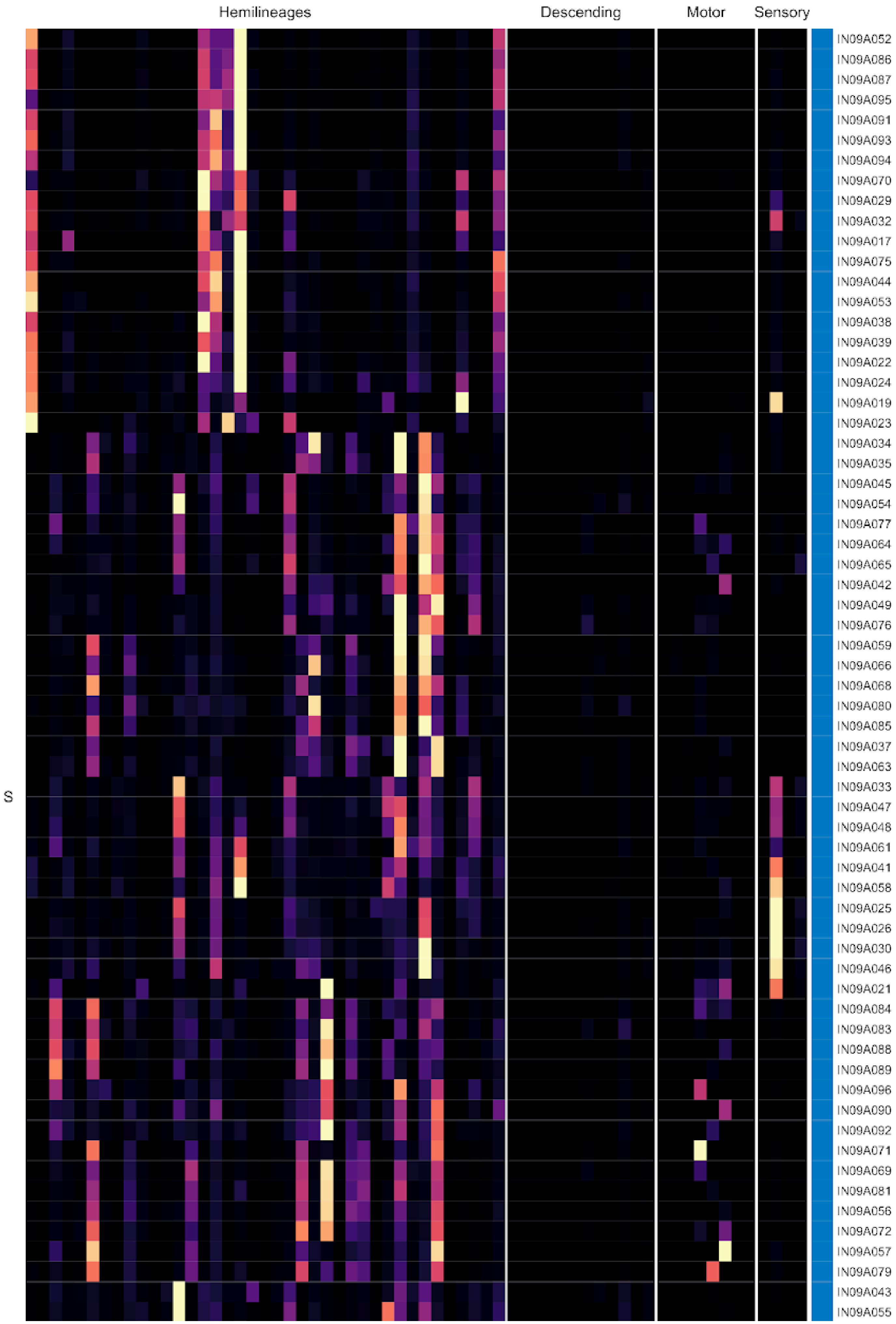

Hemilineages
Descending
Motor
Sensory
S
IN09A052
IN09A086
IN09A087
IN09A095
IN09A091
IN09A093
IN09A094
IN09A070
IN09A029
IN09A032
IN09A017
IN09A075
IN09A044
IN09A053
IN09A038
IN09A039
IN09A022
IN09A024
IN09A019
IN09A023
IN09A034
IN09A035
IN09A045
IN09A054
IN09A077
IN09A064
IN09A065
IN09A042
IN09A049
IN09A076
IN09A059
IN09A066
IN09A068
IN09A080
IN09A085
IN09A037
IN09A063
IN09A033
IN09A047
IN09A048
IN09A061
IN09A041
IN09A058
IN09A025
IN09A026
IN09A030
IN09A046
IN09A021
IN09A084
IN09A083
IN09A088
IN09A089
IN09A096
IN09A090
IN09A092
IN09A071
IN09A069
IN09A081
IN09A056
IN09A072
IN09A057
IN09A079
IN09A043
IN09A055

**Figure 28 - figure supplement 8. Connectivity to downstream partners by 09A secondary systematic types.** Proportions of synaptic weight from systematic types to downstream partners, normalised by row. 09A neurons have been clustered within each assigned birthtime window (P = primary, ES = early secondary, S = secondary) based on both upstream and downstream connectivity to hemilineages, descending neuron subclasses, motor neuron subclasses, and sensory neuron modalities. The annotation bar is coloured by the most common predicted neurotransmitter for the neurons of each type.

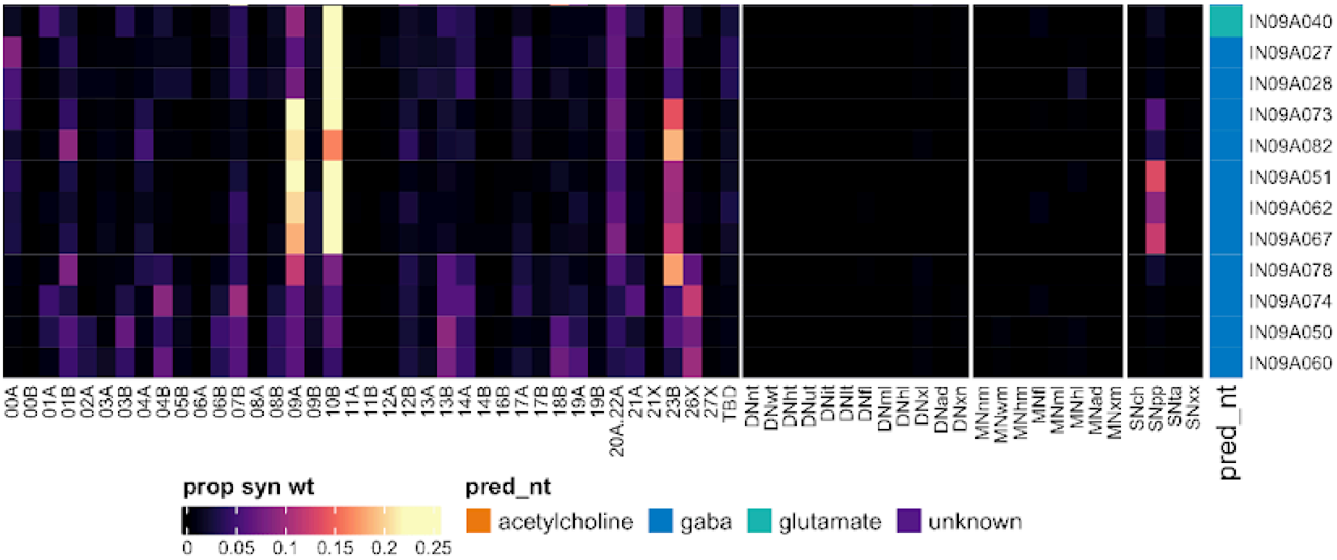

**Figure 28 - figure supplement 9. Connectivity to downstream partners by 09A secondary systematic types, continued.** Proportions of synaptic weight from systematic types to downstream partners, normalised by row. 09A neurons have been clustered within each assigned birthtime window (P = primary, ES = early secondary, S = secondary) based on both upstream and downstream connectivity to hemilineages, descending neuron subclasses, motor neuron subclasses, and sensory neuron modalities. The annotation bar is coloured by the most common predicted neurotransmitter for the neurons of each type.

Hemilineage 09B

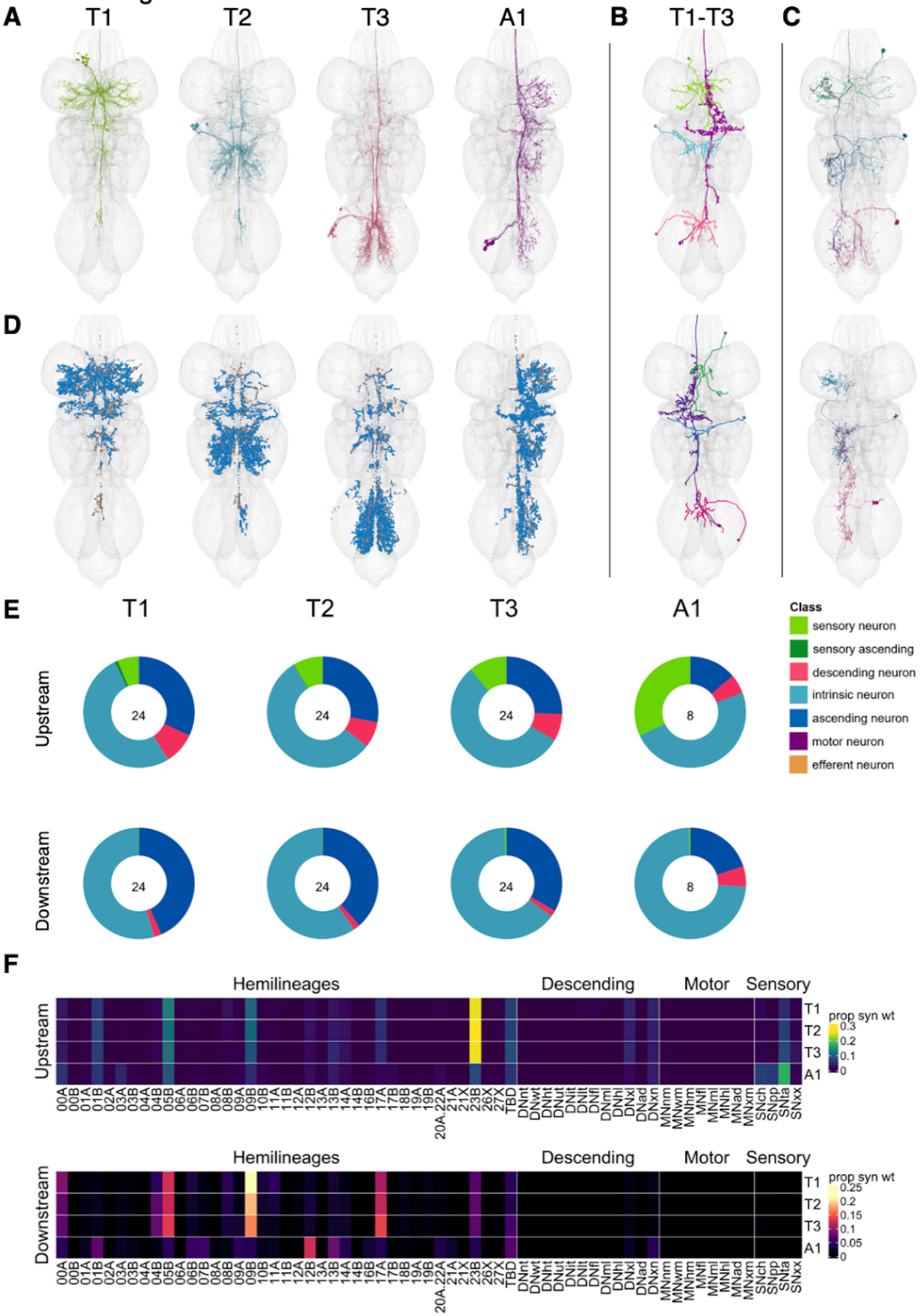

**Figure 29. Hemilineage 09B. A.** Meshes of all RHS secondary neurons plotted in neuromere-specific colours. **B.** “Representative” secondary neuron skeletons plotted in hemineuromere-specific colours. The skeleton with the top accumulated NBLAST score among all neurons from the hemilineage in a given hemineuromere was used. **C.** Neuron meshes of selected examples. Top: LHS neurons from bilateral, complex serial set 11491. Bottom: LHS neurons from contralateral, sequential serial set 17610. **D.** Predicted synapses of RHS secondary neurons. Blue: postsynapses; dark orange: presynapses. **E.** Proportions of connections from secondary neurons to upstream or downstream partners, normalised by neuromere and coloured by broad class. Numbers of query neurons appear in the centre. **F.** Proportions of synaptic weight from secondary neurons originating in each neuromere to upstream or downstream partners, normalised by row.

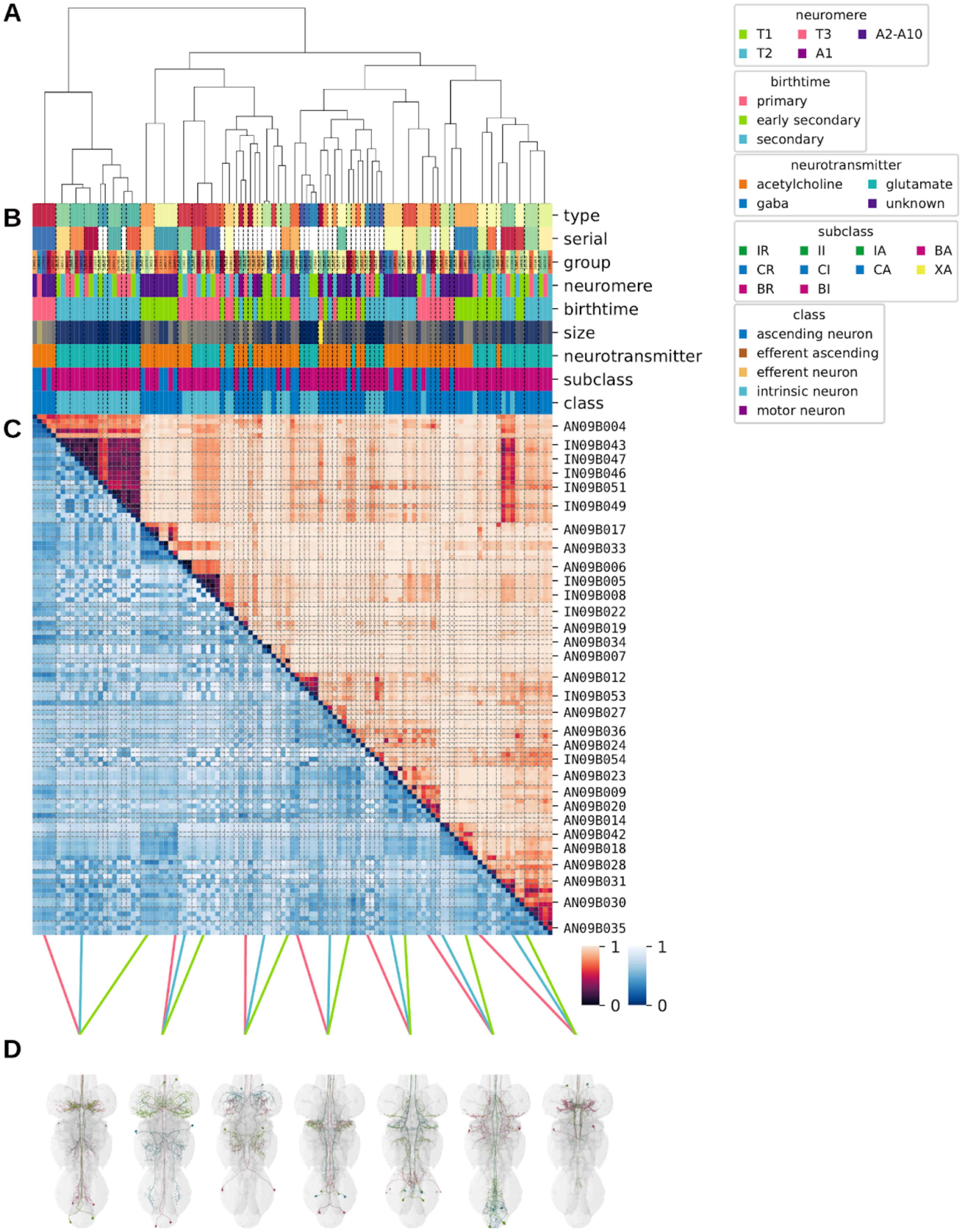

**Figure 29 - figure supplement 1. Systematic typing of hemilineage 09B. A.** Hierarchical clustering dendrogram of hemilineage groups by laterally and serially aggregated connectivity cosine clustering. **B.** Categorical annotations of each hemilineage group, each column corresponding to the aligned leaf in A. Colours for type, serial set, and group are arbitrary for visualisation. Colours for neuromere, birthtime, neurotransmitter, subclass, and class are as in all other figures. **C.** Similarity distance heatmap for hemilineage. Cosine distance is in the upper triangle, while laterally

symmetrised NBLAST distance is in the lower triangle. Systematic type names of some types are labelled. **D.** Morphologically representative groups from dendrogram subtrees. Each group, indicated by colour and line connecting to its column in B and C, is the most morphologically representative group (medoid of NBLAST distance) from a subtree of A. The subtrees (flat clusters) are equal height cuts of A determined to yield the number of groups per plot and plots in D.

### 09B systematic types

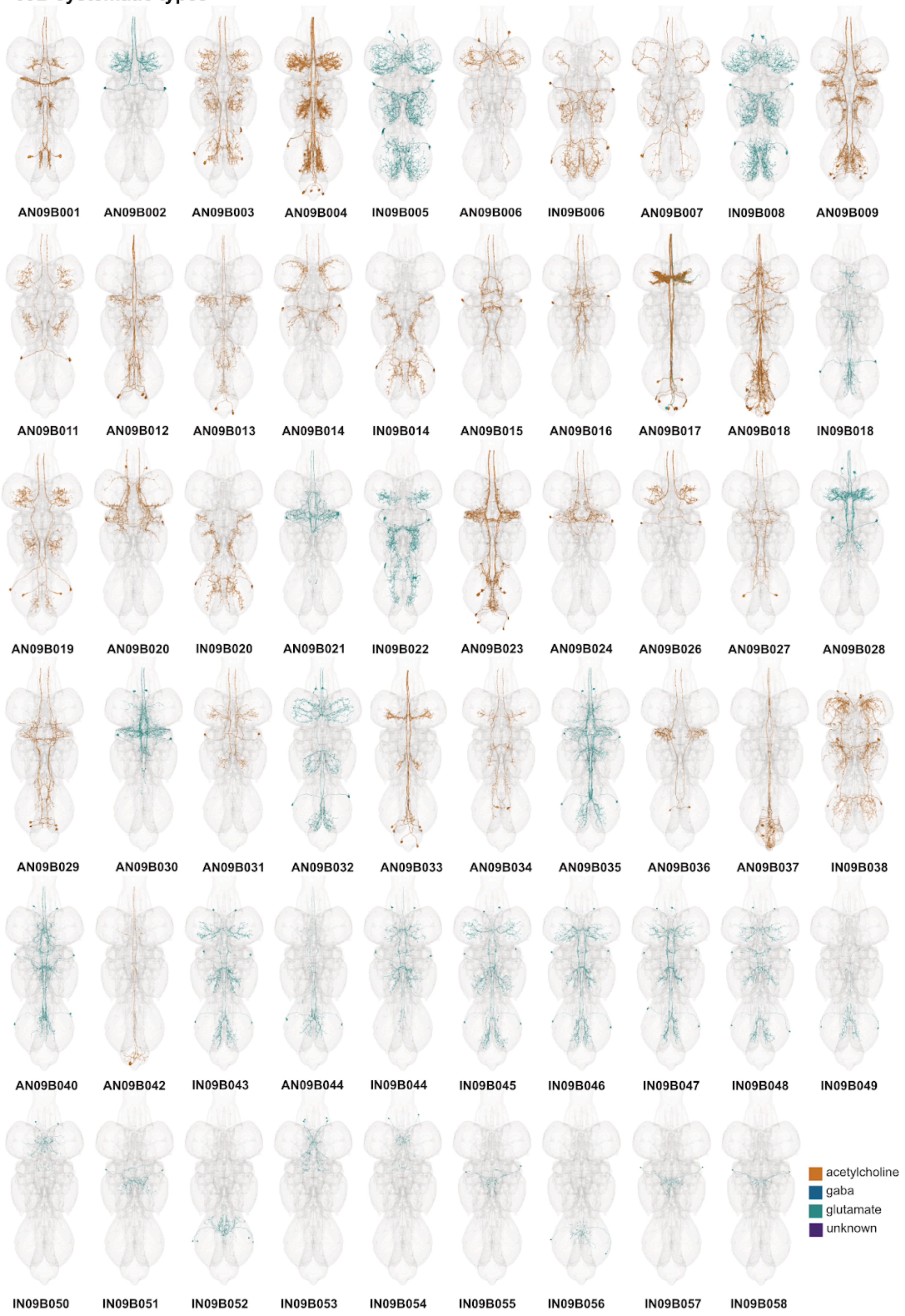

**Figure 29 - figure supplement 2. Systematic types of hemilineage 09B.** Systematic types have been arranged in numerical order, with neurons of the same type that belong to distinct classes (e.g., intrinsic neuron vs ascending neuron) plotted separately but placed adjacent to each other. Individual neuron meshes have been coloured based on predicted neurotransmitter: dark orange = acetylcholine, blue = gaba, marine = glutamate, dark purple = unknown.

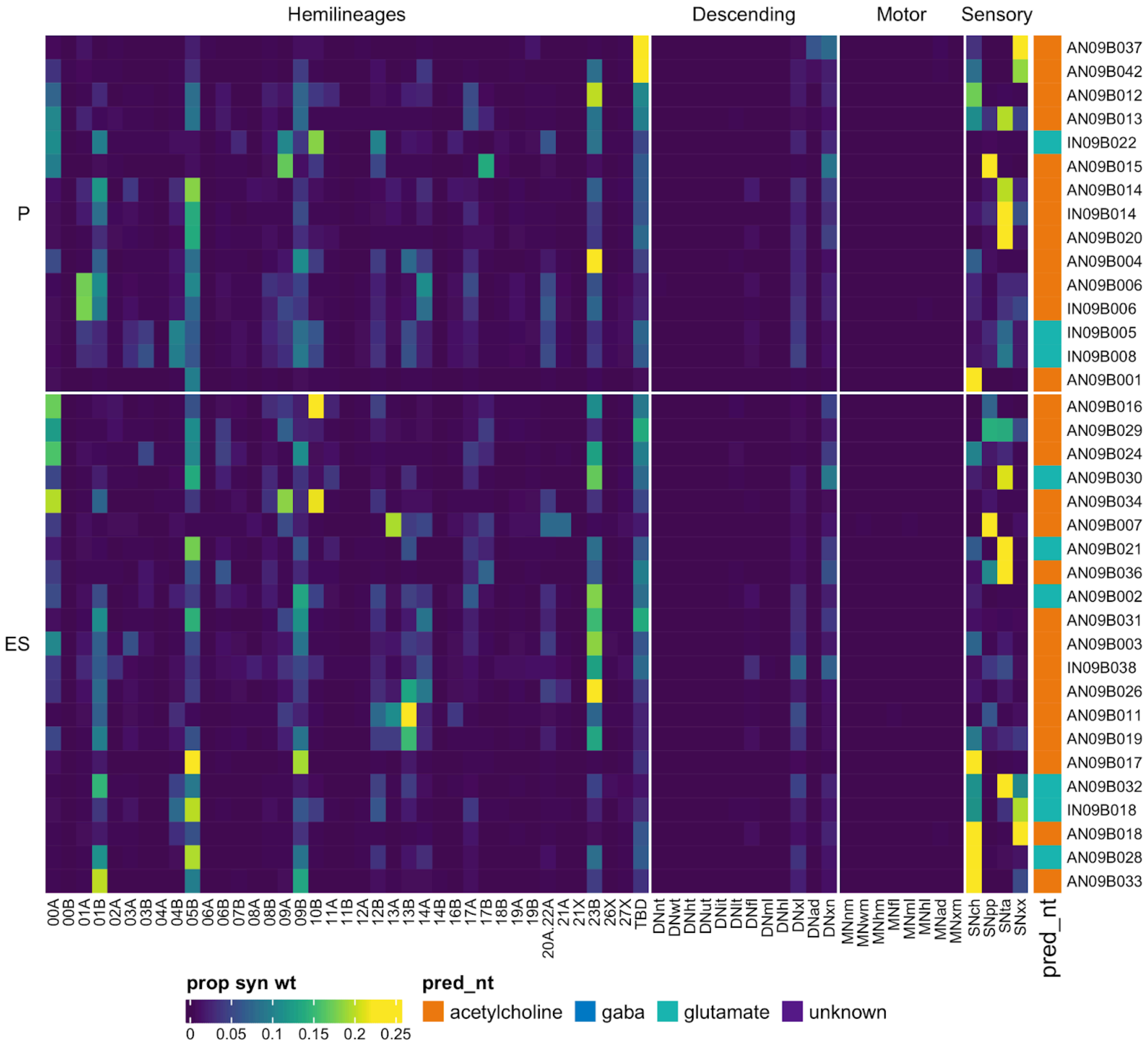

**Figure 29 - figure supplement 3. Connectivity to upstream partners by 09B primary and early secondary systematic types.** Proportions of synaptic weight to systematic types from upstream partners, normalised by row. 09B neurons have been clustered within each assigned birthtime window (P = primary, ES = early secondary, S = secondary) based on both upstream and downstream connectivity to hemilineages, descending neuron subclasses, motor neuron subclasses, and sensory neuron modalities. Annotation bar is coloured by the most common predicted neurotransmitter for the neurons of each type.

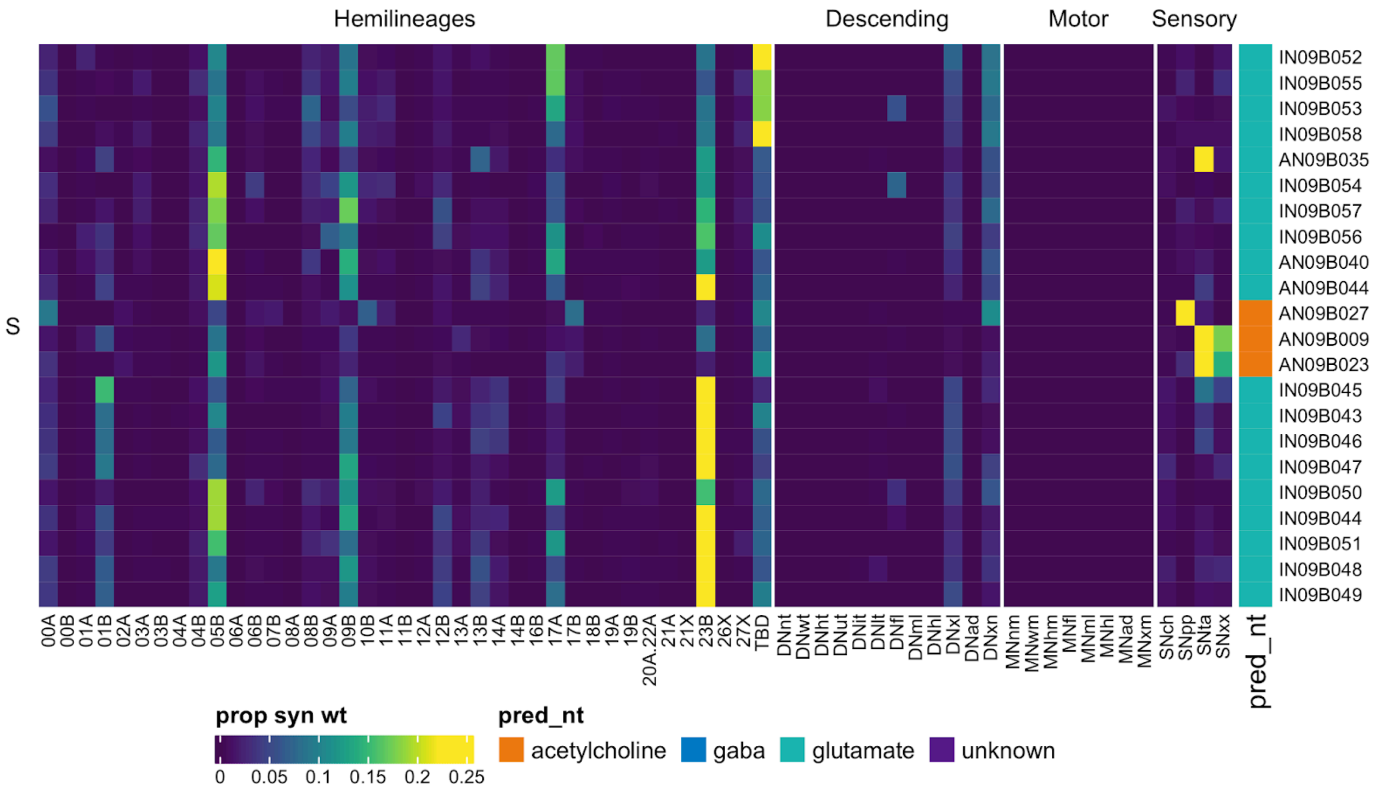

**Figure 29 - figure supplement 4. Connectivity to upstream partners by 09B secondary systematic types.** Proportions of synaptic weight to systematic types from upstream partners, normalised by row. 09B neurons have been clustered within each assigned birthtime window (P = primary, ES = early secondary, S = secondary) based on both upstream and downstream connectivity to hemilineages, descending neuron subclasses, motor neuron subclasses, and sensory neuron modalities. The annotation bar is coloured by the most common predicted neurotransmitter within each type.

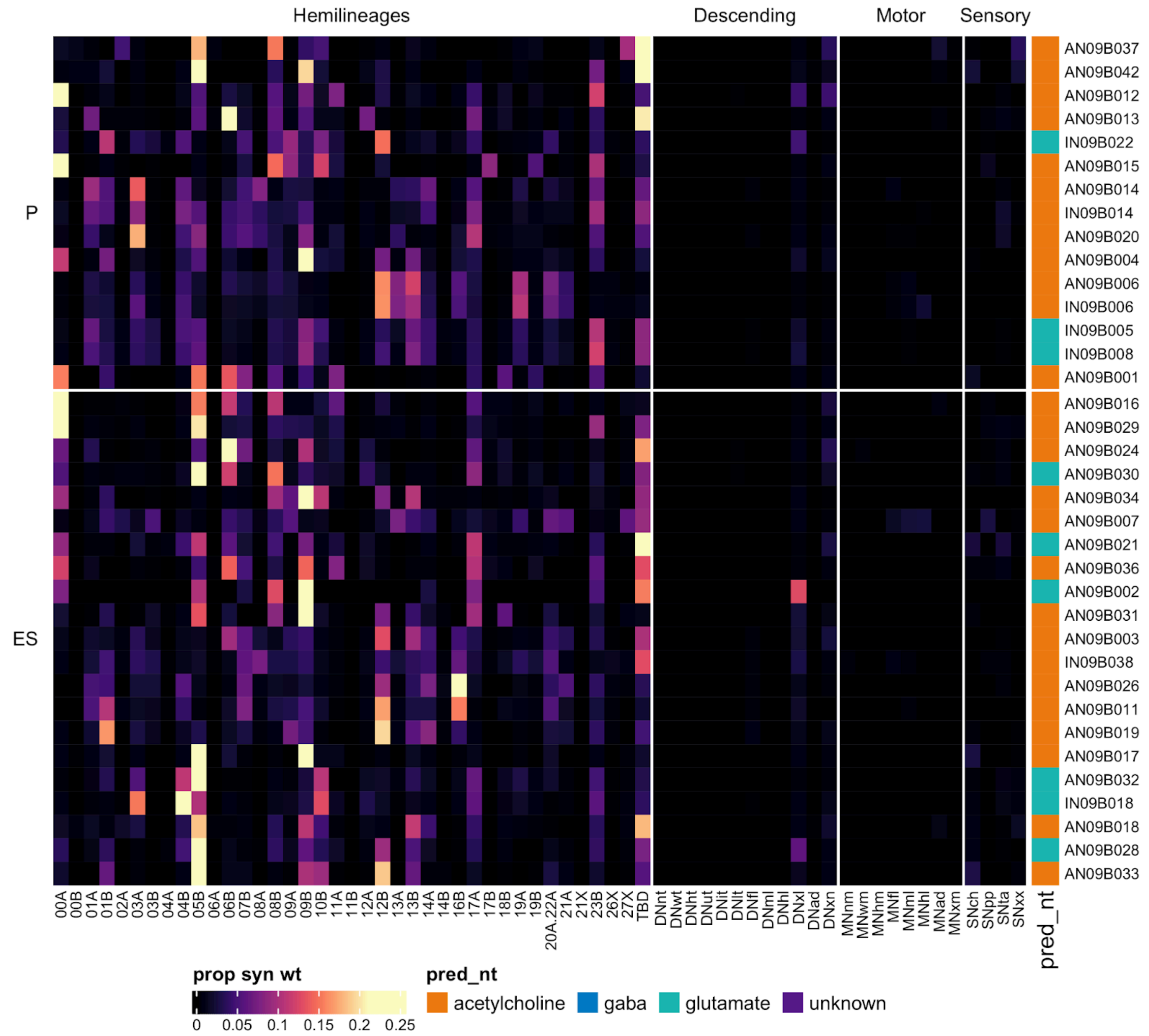

**Figure 29 - figure supplement 5. Connectivity to downstream partners by 09B primary and early secondary systematic types.** Proportions of synaptic weight from systematic types to downstream partners, normalised by row. 09B neurons have been clustered within each assigned birthtime window (P = primary, ES = early secondary, S = secondary) based on both upstream and downstream connectivity to hemilineages, descending neuron subclasses, motor neuron subclasses, and sensory neuron modalities. The annotation bar is coloured by the most common predicted neurotransmitter within each type.

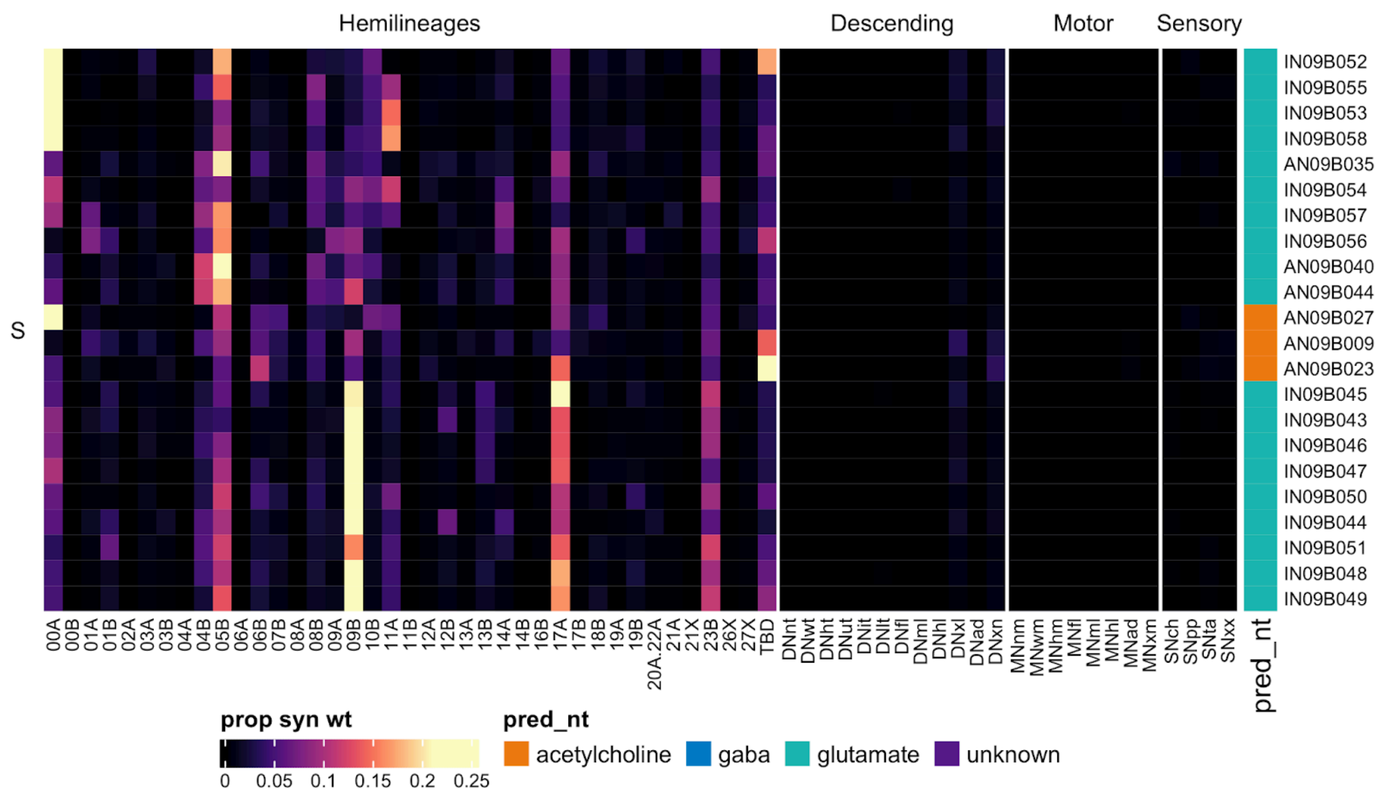

**Figure 29 - figure supplement 6. Connectivity to downstream partners by 09B secondary systematic types.** Proportions of synaptic weight from systematic types to downstream partners, normalised by row. 09B neurons have been clustered within each assigned birthtime window (P = primary, ES = early secondary, S = secondary) based on both upstream and downstream connectivity to hemilineages, descending neuron subclasses, motor neuron subclasses, and sensory neuron modalities. The annotation bar is coloured by the most common predicted neurotransmitter for the neurons of each type.

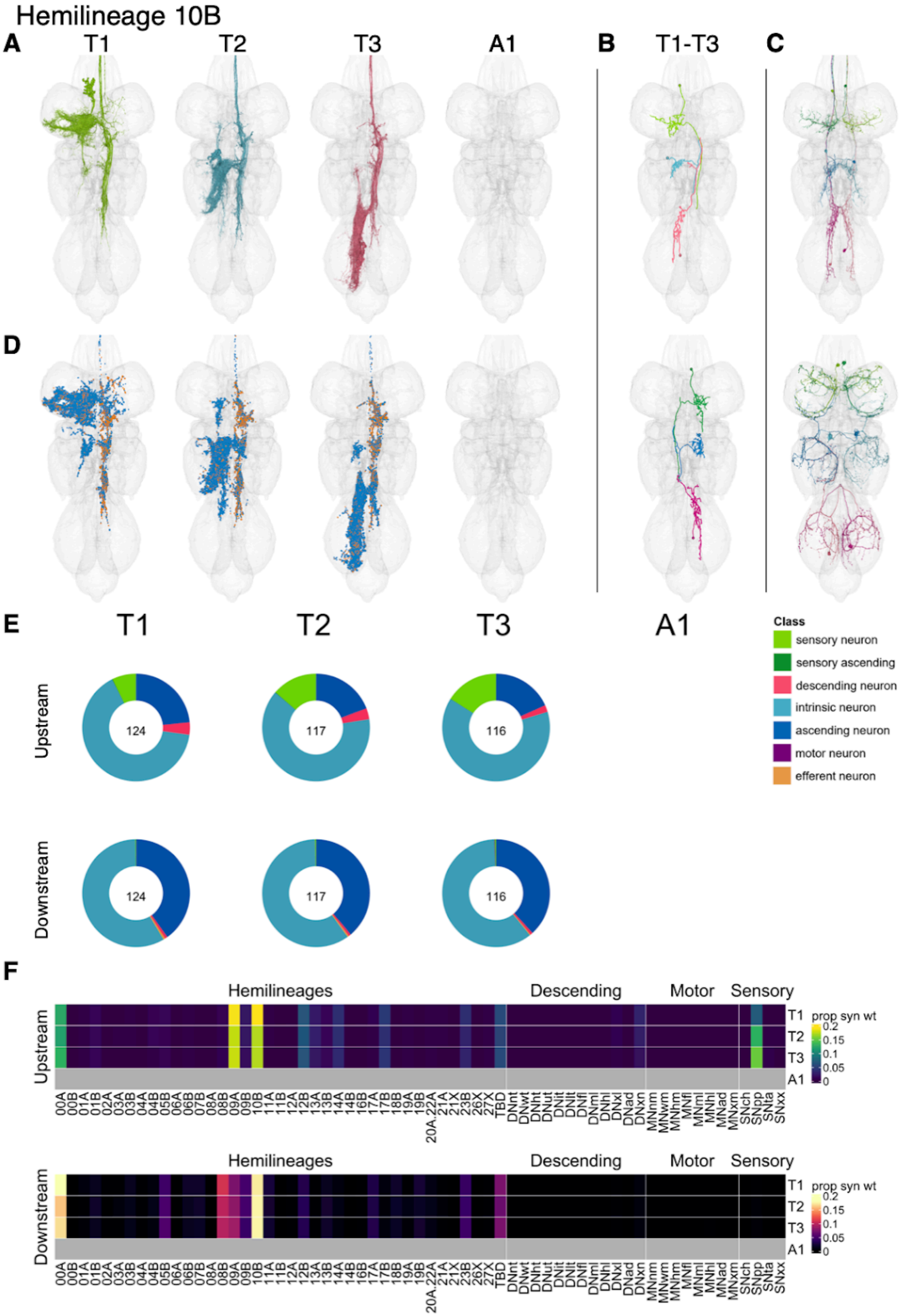
Hemilineage 10B
A
T1
T2
T3
A1
B
T1-T3
C
D
E
T1
T2
T3
A1
Upstream
124
117
116
Downstream
124
117
116
Class
sensory neuron
sensory ascending
descending neuron
intrinsic neuron
ascending neuron
motor neuron
efferent neuron
F
Hemilineages
Descending
Motor
Sensory
Upstream
T1
T2
T3
A1
prop syn wt
0.2
0.15
0.1
0.05
0
00A 00B 01A 01B 02A 03A 03B 04A 04B 05B 06A 06B 07B 08A 08B 09A 09B 10B 11A 11B 12A 12B 13A 13B 14A 14B 16B 17A 17B 18B 19A 19B 20A.22A 21A 21X 23B 26X 27X TBD DNnt DNwt DNht DNut DNit DNlt DNfl DNml DNhl DNxl DNad DNxn MNnm MNwm MNhm MNfl MNml MNhl MNad MNxm SNch SNpp SNta SNxx
Hemilineages
Descending
Motor
Sensory
Downstream
T1
T2
T3
A1
prop syn wt
0.2
0.15
0.1
0.05
0
00A 00B 01A 01B 02A 03A 03B 04A 04B 05B 06A 06B 07B 08A 08B 09A 09B 10B 11A 11B 12A 12B 13A 13B 14A 14B 16B 17A 17B 18B 19A 19B 20A.22A 21A 21X 23B 26X 27X TBD DNnt DNwt DNht DNut DNit DNlt DNfl DNml DNhl DNxl DNad DNxn MNnm MNwm MNhm MNfl MNml MNhl MNad MNxm SNch SNpp SNta SNxx

**Figure 30. Hemilineage 10B. A.** Meshes of all RHS secondary neurons plotted in neuromere-specific colours. **B.** “Representative” secondary neuron skeletons plotted in hemineuromere-specific colours. The skeleton with the top accumulated NBLAST score among all neurons from the hemilineage in a given hemineuromere was used. **C.** Neuron meshes of selected examples. Top: ascending serial set 15289. Bottom: complex serial set 11781. Bottom: LHS neurons from contralateral, sequential serial set 17610. **D.** Predicted synapses of RHS secondary neurons. Blue: postsynapses; dark orange: presynapses. **E.** Proportions of connections from secondary neurons to upstream or downstream partners, normalised by neuromere and coloured by broad class. Numbers of query neurons appear in the centre. **F.** Proportions of synaptic weight from secondary neurons originating in each neuromere to upstream or downstream partners, normalised by row.

#### Hemilineage 10B

10B is the only surviving secondary hemilineage (Truman et al., 2010) of anterior medial neuroblast NB2-2 (Birkholz et al., 2015; Lacin and Truman, 2016), which generates 2-3 MNs, 2-4 intersegmental interneurons, and 15 local interneurons in the embryo (Schmid et al., 1999). In the adult, 10B secondary neurons enter the anterior neuromere, typically innervating the ipsilateral mVAC (Shepherd et al., 2019) before crossing the midline quite ventrally in the anterior intermediate commissure, just anterior to the orthogonally projecting 2A primary neurites (Shepherd et al., 2016)). They are typically intersegmental, extending their axons anteriorly and/or posteriorly, and are mainly cholinergic (Lacin et al., 2019).

10B secondary neurons are restricted to thoracic segments (Truman et al., 2004), and we identified roughly similar numbers of neurons in each neuromere (Figure 30E). Whilst specific secondary types can be distinguished, they are generally fairly similar to each other in morphology and connectivity, with three main populations (Figure 30 - figure supplement 2-7). Early born 10B neurons cross in the same commissure but are much more diverse (e.g., Figure 30C top, bottom).

Secondary 10B neurons receive inputs primarily from proprioceptive sensory neurons and from hemilineages 00A, 09A, and 10B, and they primarily activate 00A, 08B, 09A, and especially other 10B neurons (Figure 30F). These features support a role in leg coordination. Indeed, bilateral activation of 10B secondary neurons results in somewhat erratic leg movements, with occasional pivoting or backwards walking, sometimes accompanied by wing flicking and wing buzzing (Harris et al., 2015). A small number of early born 10B types target wing and abdominal motor neurons (e.g., IN10B006, IN10B011, IN10B023) (Figure 30 - figure supplement 6).

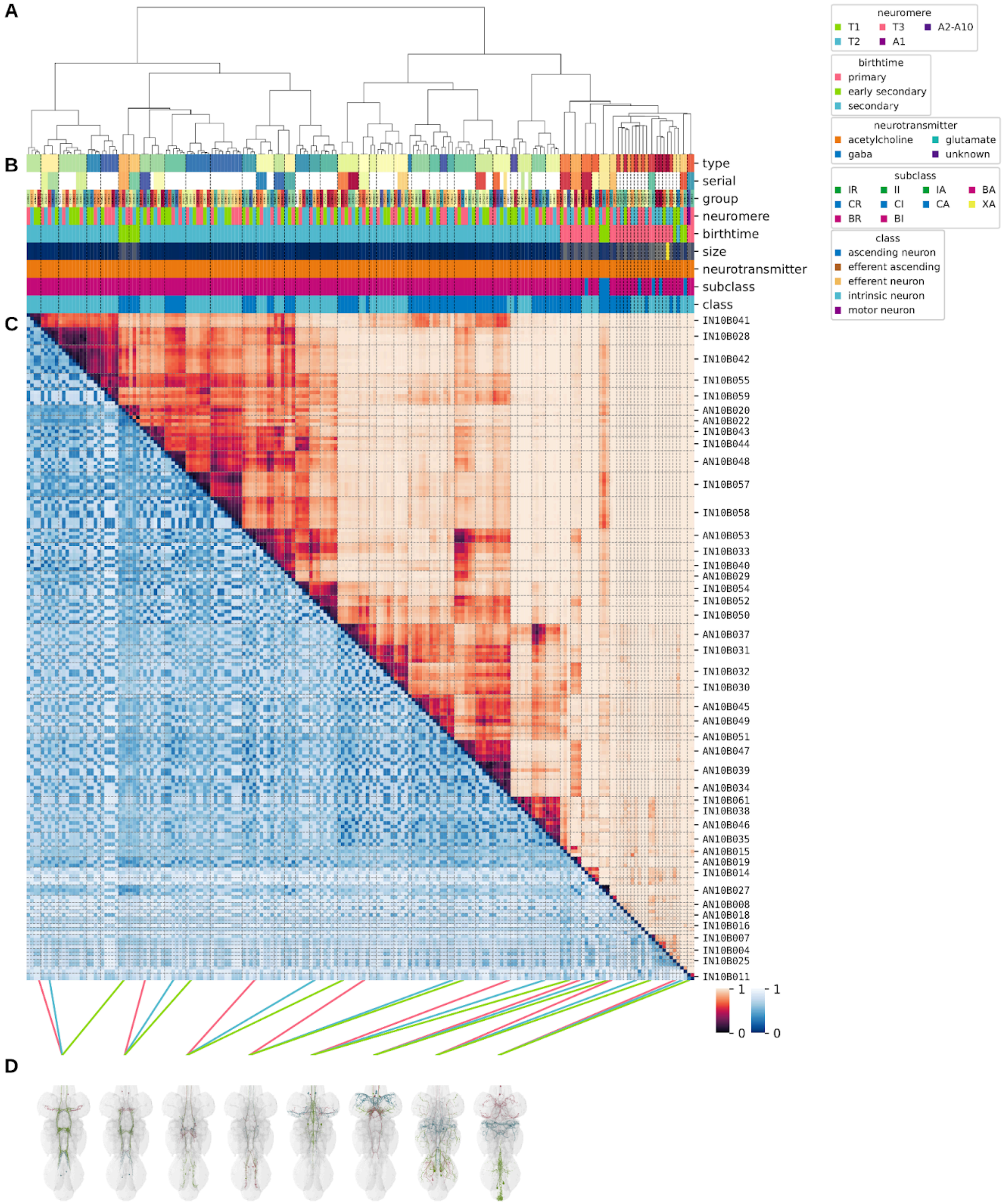

**Figure 30 - figure supplement 1. Systematic typing of hemilineage 10B. A.** Hierarchical clustering dendrogram of hemilineage groups by laterally and serially aggregated connectivity cosine clustering. **B.** Categorical annotations of each hemilineage group, each column corresponding to the aligned leaf in A. Colours for type, serial set, and group are arbitrary for visualisation. Colours for neuromere, birthtime, neurotransmitter, subclass, and class are as in all other figures. **C.** Similarity distance heatmap for hemilineage. Cosine distance is in the upper triangle, while laterally symmetrised NBLAST distance is in the lower triangle. Systematic type names of some types are labelled. **D.** Morphologically representative groups from dendrogram subtrees. Each group, indicated by colour and line connecting to its column in B and C, is the most morphologically representative group (medoid of NBLAST distance)

from a subtree of A. The subtrees (flat clusters) are equal height cuts of A determined to yield the number of groups per plot and plots in D.

#### 10B systematic types

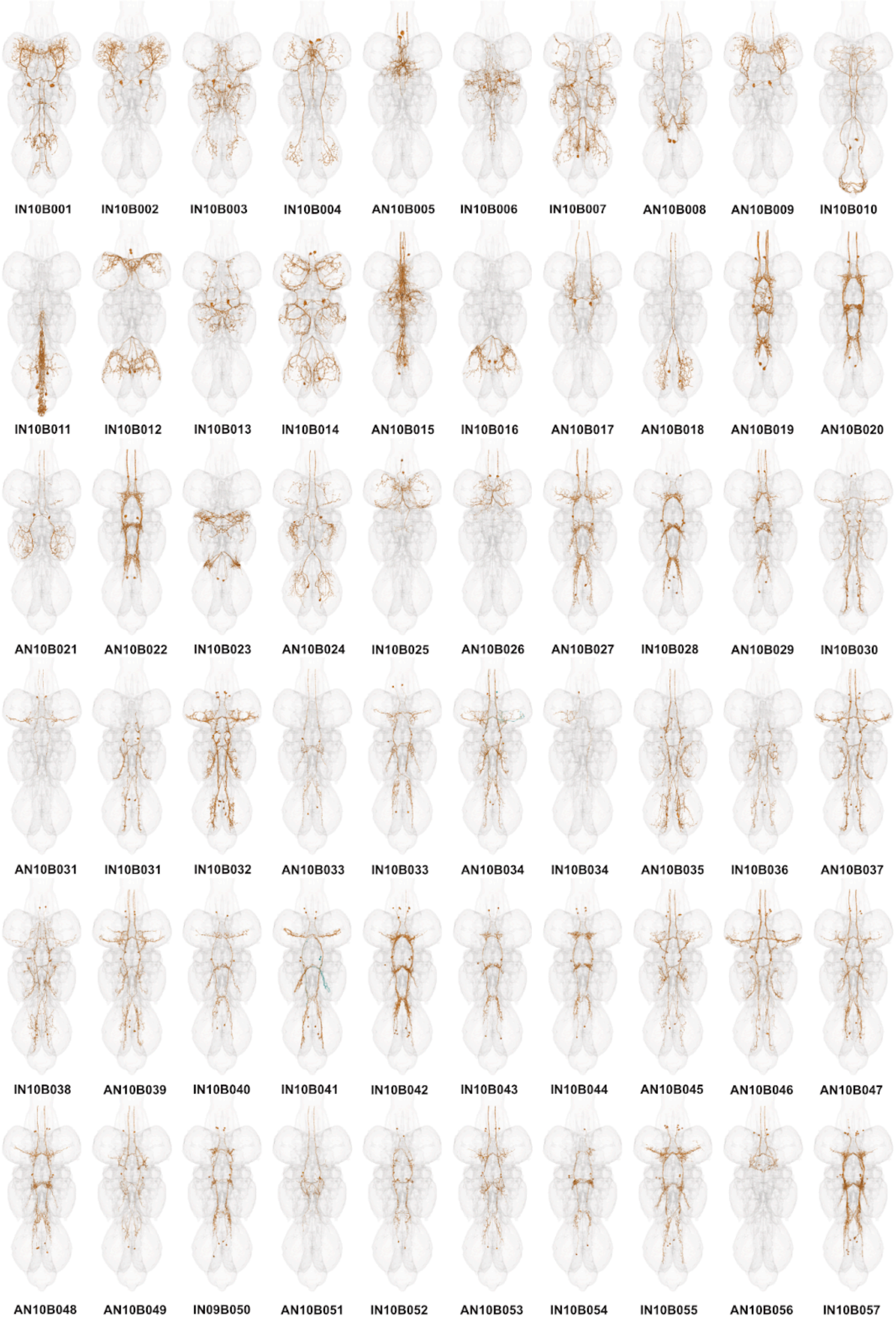

**Figure 30 - figure supplement 2. Systematic types of hemilineage 10B.** Systematic types have been arranged in numerical order, with neurons of the same type that belong to distinct classes (e.g., intrinsic neuron vs ascending neuron) plotted separately but placed adjacent to each other. Individual neuron meshes have been coloured based on predicted neurotransmitter: dark orange = acetylcholine, blue = gaba, marine = glutamate, dark purple = unknown.

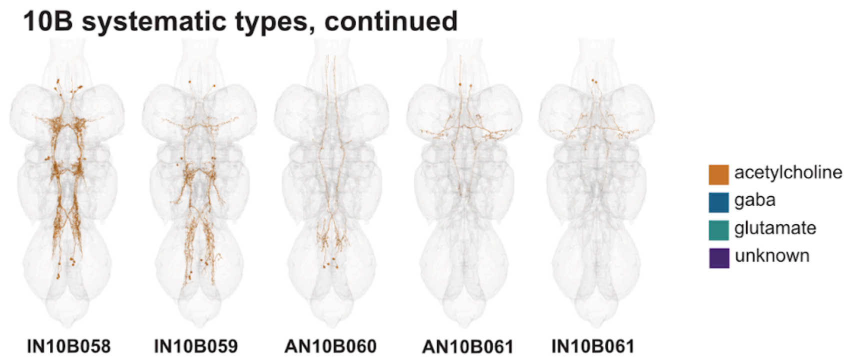

**Figure 30 - figure supplement 3. Systematic types of hemilineage 10B, continued.** Systematic types have been arranged in numerical order, with neurons of the same type that belong to distinct classes (e.g., intrinsic neuron vs ascending neuron) plotted separately but placed adjacent to each other. Individual neuron meshes have been coloured based on predicted neurotransmitter: dark orange = acetylcholine, blue = gaba, marine = glutamate, dark purple = unknown.

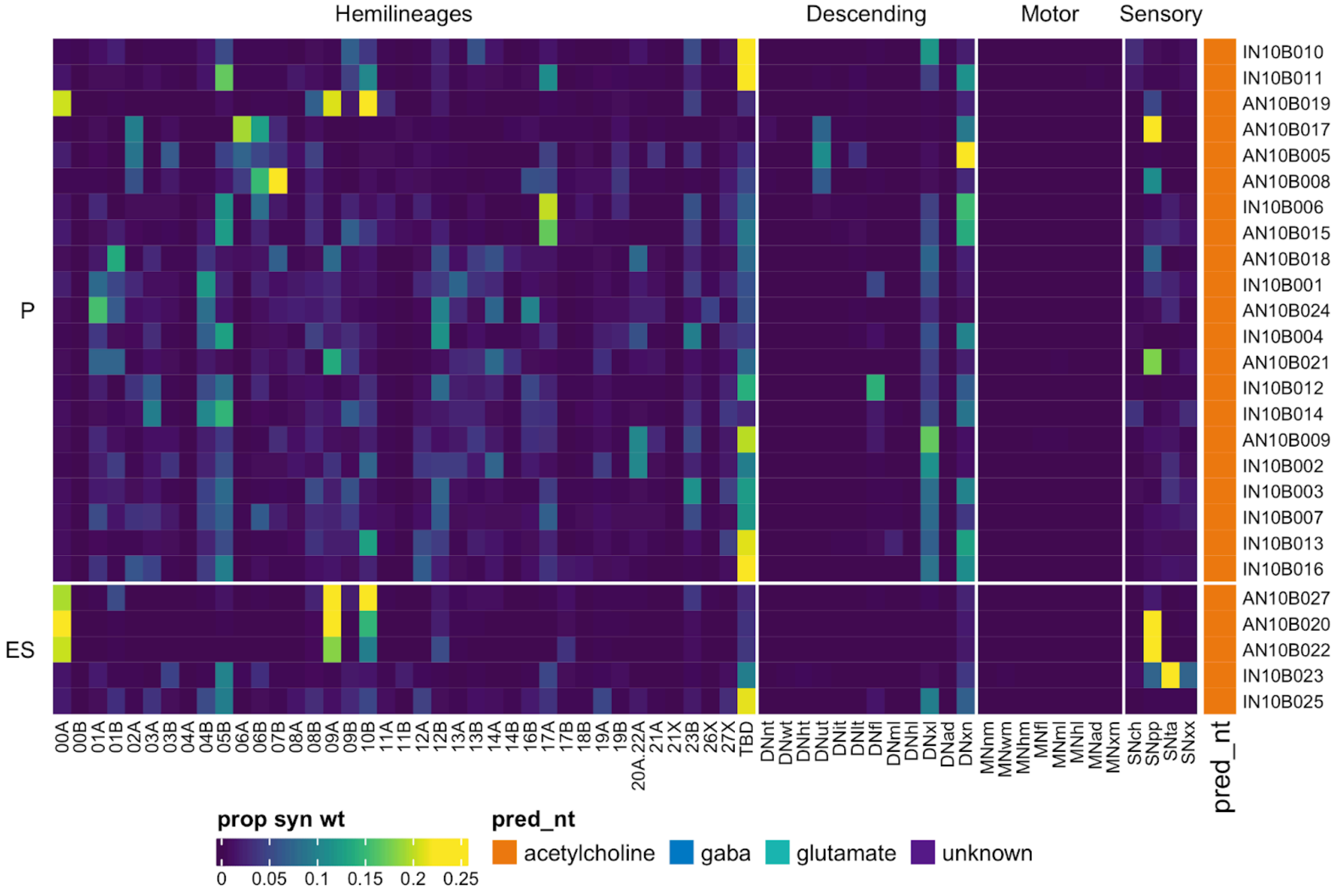

**Figure 30 - figure supplement 4. Connectivity to upstream partners by 10B primary and early secondary systematic types.** Proportions of synaptic weight to systematic types from upstream partners, normalised by row. 10B neurons have been clustered within each assigned birthtime window (P = primary, ES = early secondary, S = secondary) based on both upstream and downstream connectivity to hemilineages, descending neuron subclasses, motor neuron subclasses, and sensory neuron modalities. Annotation bar is coloured by the most common predicted neurotransmitter for the neurons of each type.

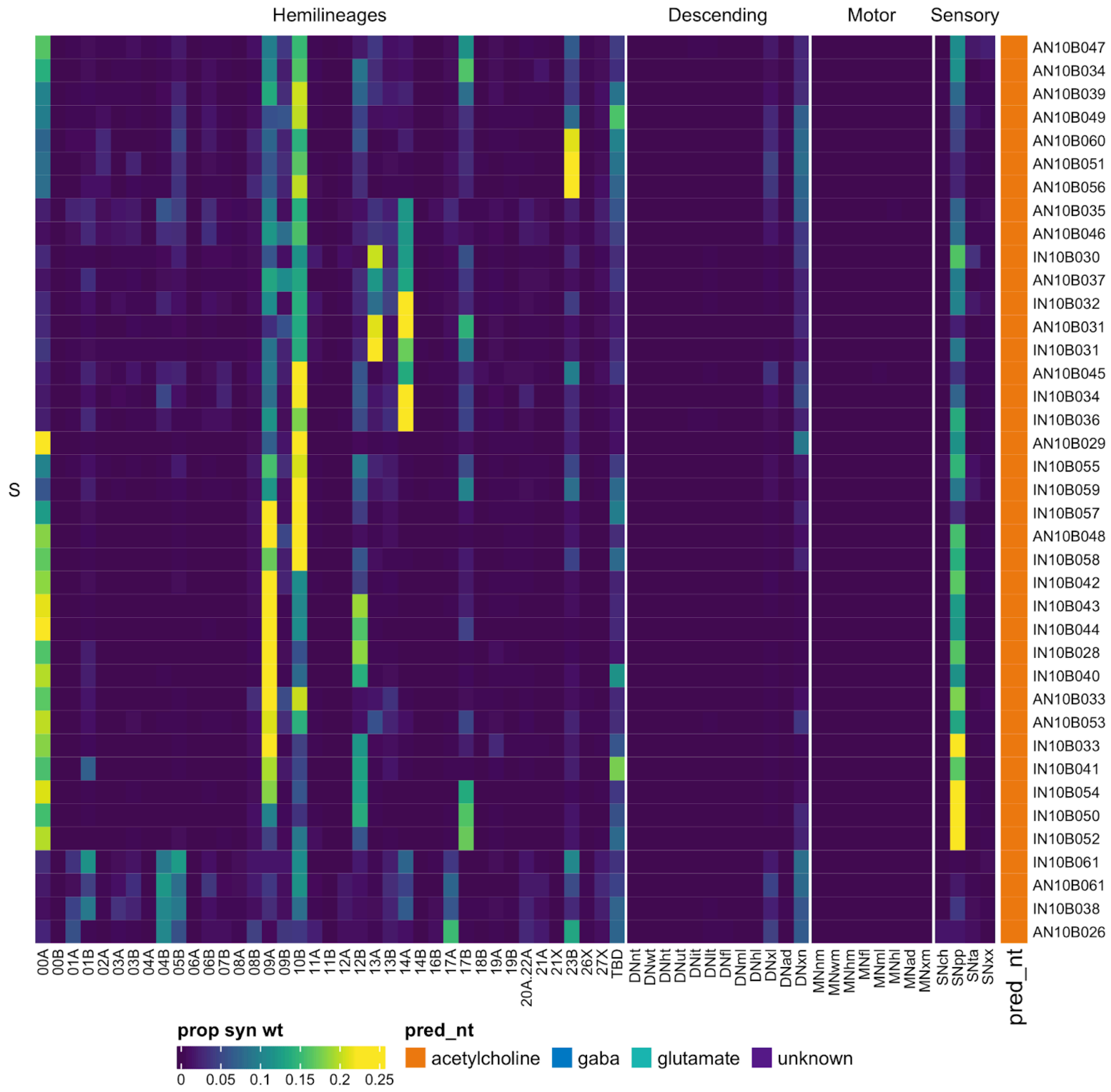

**Figure 30 - figure supplement 5. Connectivity to upstream partners by 10B secondary systematic types.** Proportions of synaptic weight to systematic types from upstream partners, normalised by row. 10B neurons have been clustered within each assigned birthtime window (P = primary, ES = early secondary, S = secondary) based on both upstream and downstream connectivity to hemilineages, descending neuron subclasses, motor neuron subclasses, and sensory neuron modalities. The annotation bar is coloured by the most common predicted neurotransmitter within each type.

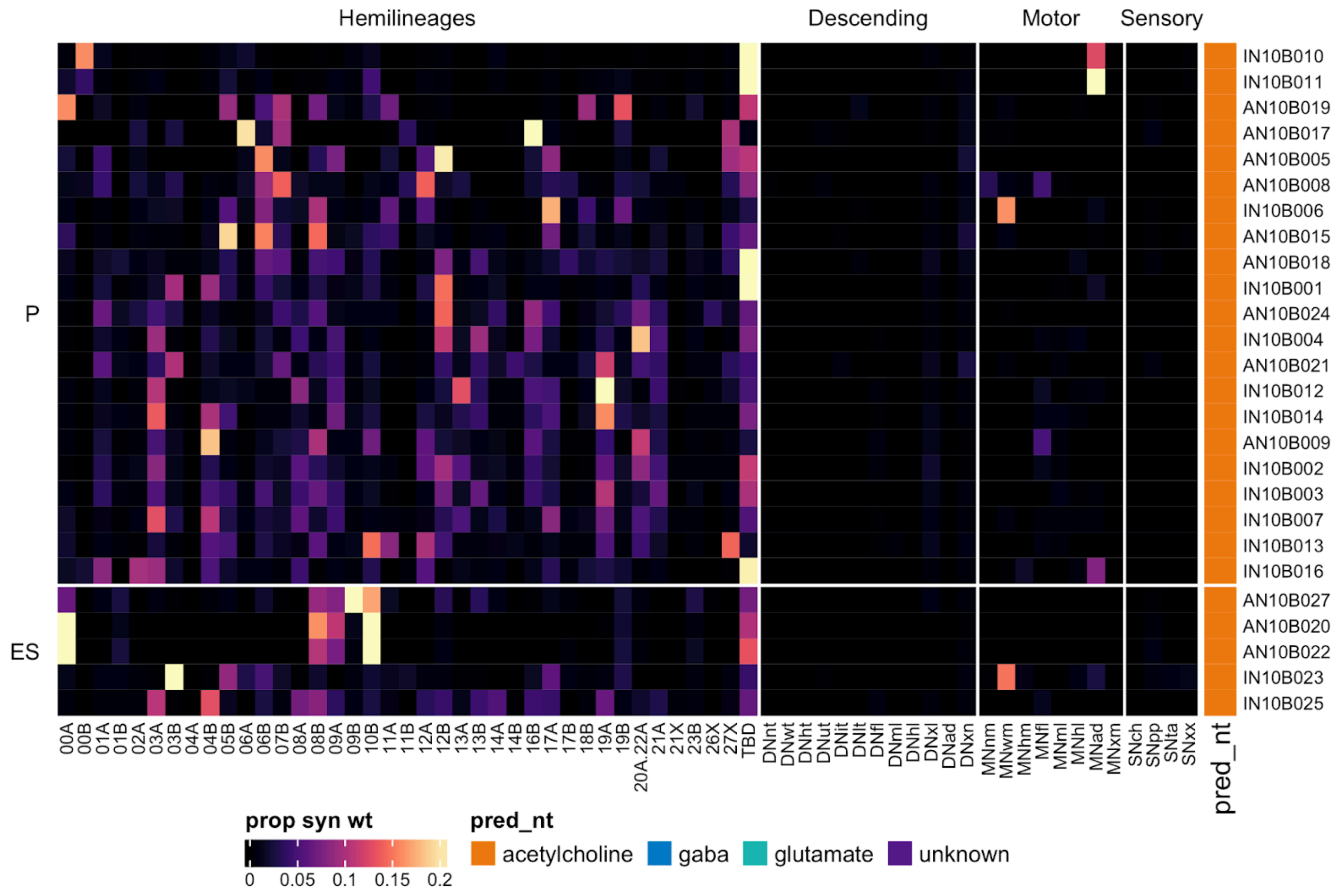

**Figure 30 - figure supplement 6. Connectivity to downstream partners by 10B primary and early secondary systematic types.** Proportions of synaptic weight from systematic types to downstream partners, normalised by row. 10B neurons have been clustered within each assigned birthtime window (P = primary, ES = early secondary, S = secondary) based on both upstream and downstream connectivity to hemilineages, descending neuron subclasses, motor neuron subclasses, and sensory neuron modalities. The annotation bar is coloured by the most common predicted neurotransmitter within each type.

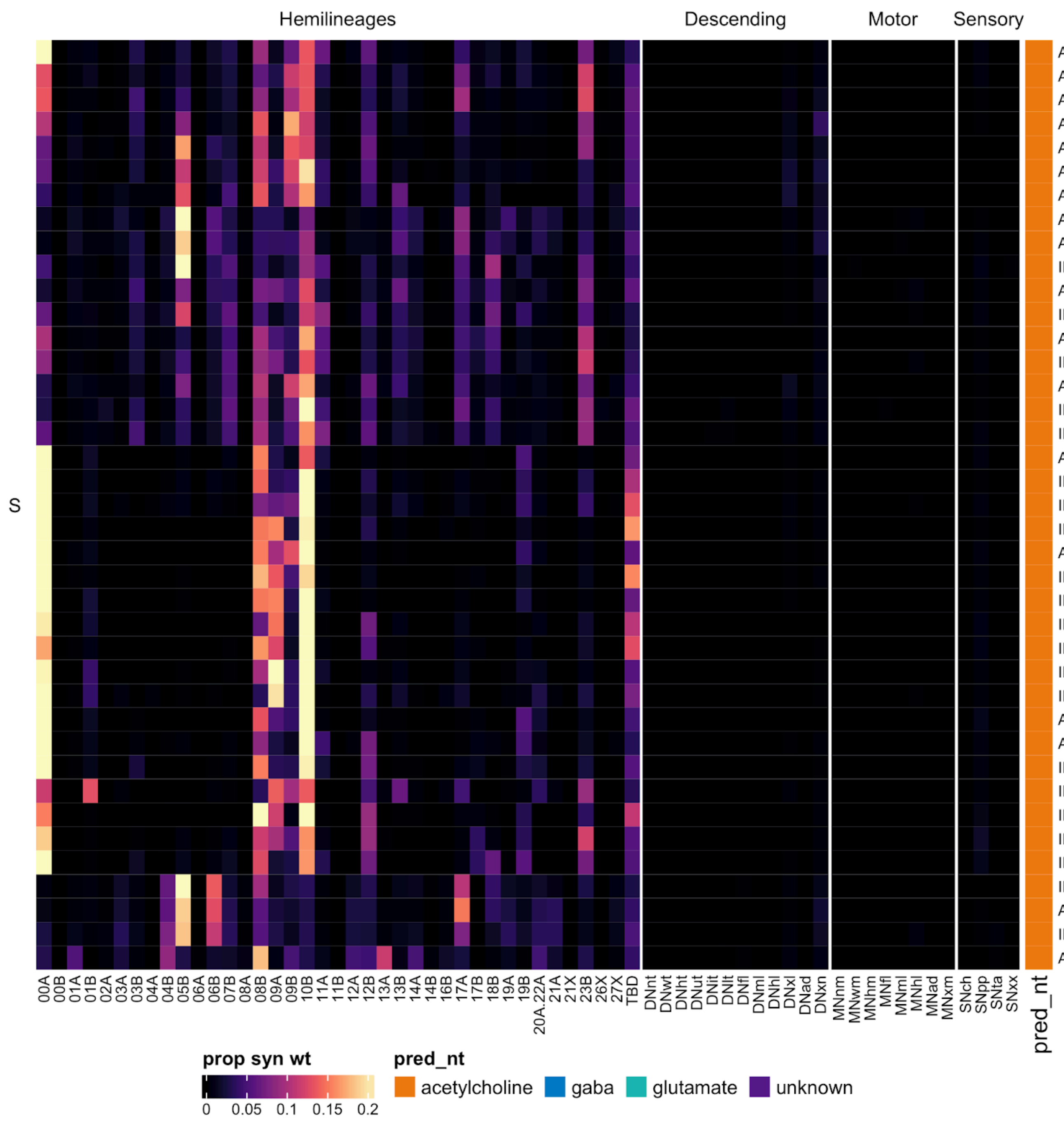

**Figure 30 - figure supplement 7. Connectivity to downstream partners by 10B secondary systematic types.** Proportions of synaptic weight from systematic types to downstream partners, normalised by row. 10B neurons have been clustered within each assigned birthtime window (P = primary, ES = early secondary, S = secondary) based on both upstream and downstream connectivity to hemilineages, descending neuron subclasses, motor neuron subclasses, and sensory neuron modalities. The annotation bar is coloured by the most common predicted neurotransmitter for the neurons of each type.

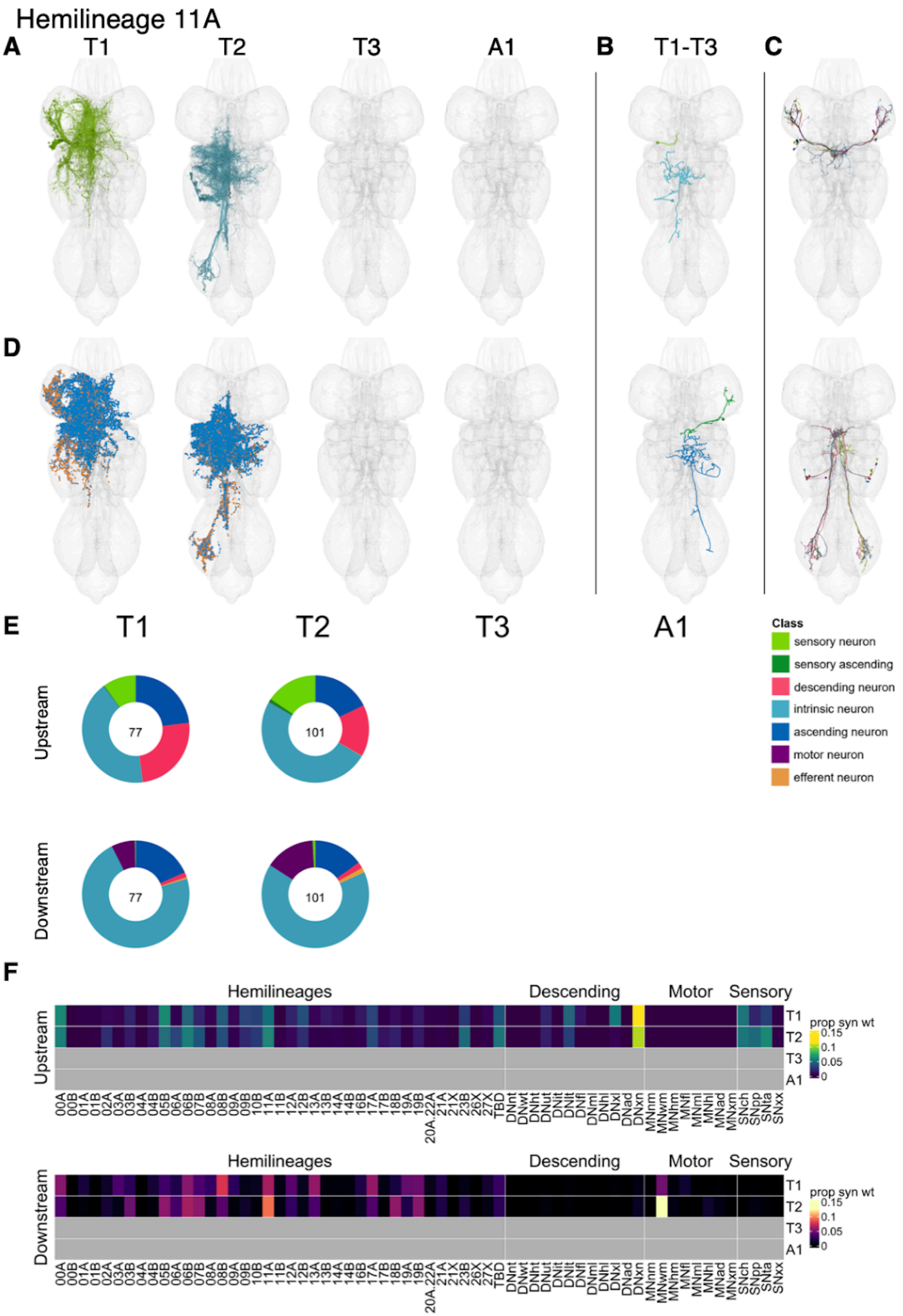
Hemilineage 11A
A
T1
T2
T3
A1
B
T1-T3
C
D
E
T1
T2
T3
A1
Upstream
Downstream
77
101
Class
sensory neuron
sensory ascending
descending neuron
intrinsic neuron
ascending neuron
motor neuron
efferent neuron
F
Hemilineages
Descending
Motor
Sensory
prop syn wt
0.15
0.1
0.05
0
00A 00B 01A 01B 02A 03A 03B 04A 04B 05B 06A 06B 07B 08A 08B 09A 09B 10B 11A 11B 12A 12B 13A 13B 14A 14B 16B 17A 17B 18B 19A 19B 20A.22A 21A 21X 23B 26X 27X TBD
DNnt DNwt DNht DNut DNit DNlt DNfl DNml DNhl DNxl DNad DNxn
MNnm MNwm MNhm MNfl MNml MNhl MNad MNxm
SNch SNpp SNta SNxx

**Figure 31. Hemilineage 11A. A.** Meshes of all RHS secondary neurons plotted in neuromere-specific colours. **B.** “Representative” secondary neuron skeletons plotted in hemineuromere-specific colours. The skeleton with the top accumulated NBLAST score among all neurons from the hemilineage in a given hemineuromere was used. **C.** Neuron meshes of selected examples. Top: electrical GFC4, subcluster 23460. Bottom: electrical GFC3, subcluster 17383. **D.** Predicted synapses of RHS secondary neurons. Blue: postsynapses; dark orange: presynapses. **E.** Proportions of connections from secondary neurons to upstream or downstream partners, normalised by neuromere and coloured by broad class. Numbers of query neurons appear in the centre. **F.** Proportions of synaptic weight from secondary neurons originating in each neuromere to upstream or downstream partners, normalised by row.

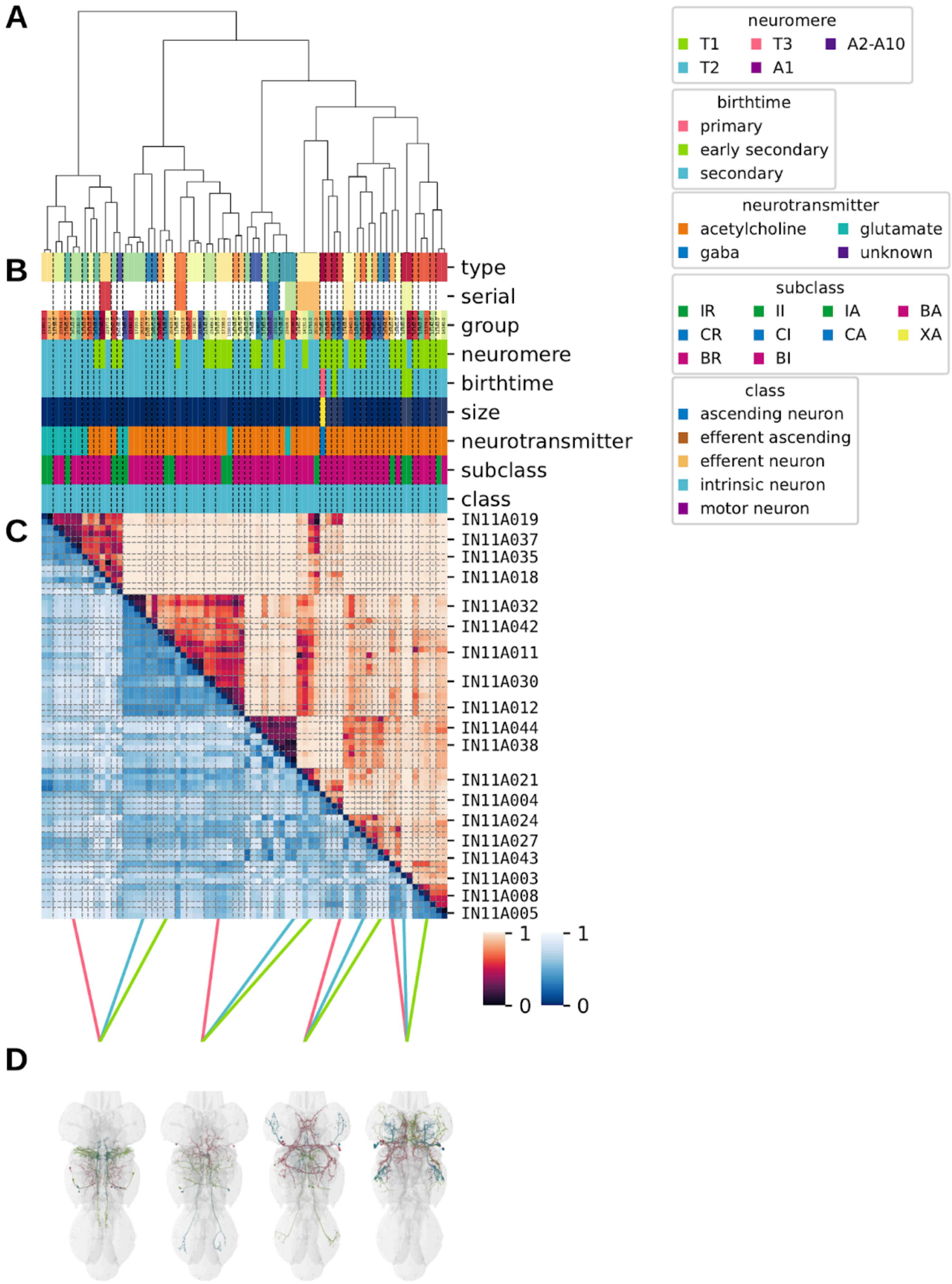

**Figure 31 - figure supplement 1. Systematic typing of hemilineage 11A. A.** Hierarchical clustering dendrogram of hemilineage groups by laterally and serially aggregated connectivity cosine clustering. **B.** Categorical annotations of each hemilineage group, each column corresponding to the aligned leaf in A. Colours for type, serial set, and group are arbitrary for visualisation. Colours for neuromere, birthtime, neurotransmitter, subclass, and class are as in all other figures. **C.** Similarity distance heatmap for hemilineage. Cosine distance is in the upper triangle, while laterally symmetrised NBLAST distance is in the lower triangle. Systematic type names of some types are labelled. **D.** Morphologically representative groups from dendrogram subtrees. Each group, indicated by colour and line connecting to its column in B and C, is the most morphologically representative group (medoid of NBLAST distance) from a subtree of A. The subtrees (flat clusters) are equal height cuts of A determined to yield the number of groups per plot and plots in D.

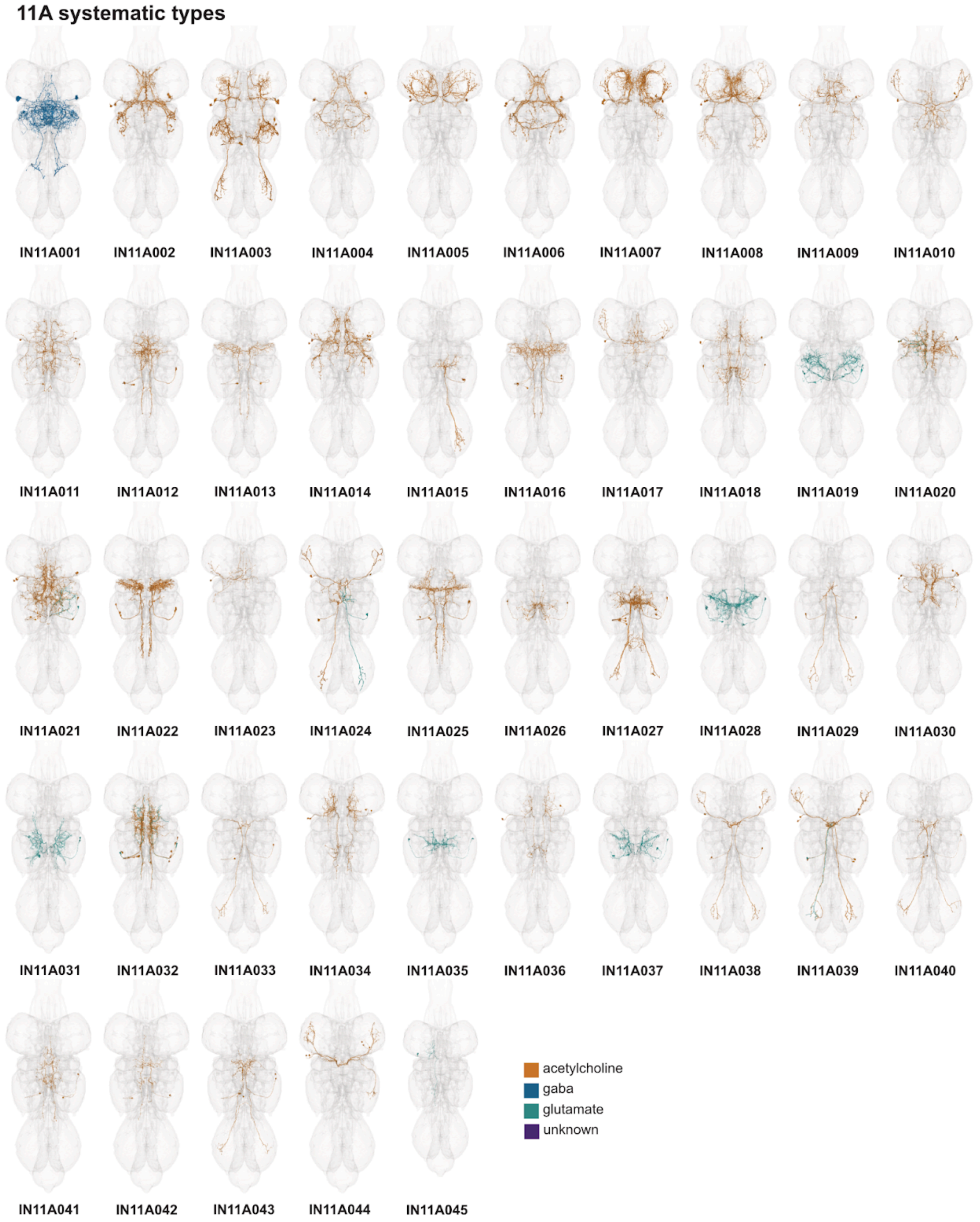

**Figure 31 - figure supplement 2. Systematic types of hemilineage 11A.** Systematic types have been arranged in numerical order, with neurons of the same type that belong to distinct classes (e.g., intrinsic neuron vs ascending neuron) plotted separately but placed adjacent to each other. Individual neuron meshes have been coloured based on predicted neurotransmitter: dark orange = acetylcholine, blue = gaba, marine = glutamate, dark purple = unknown.

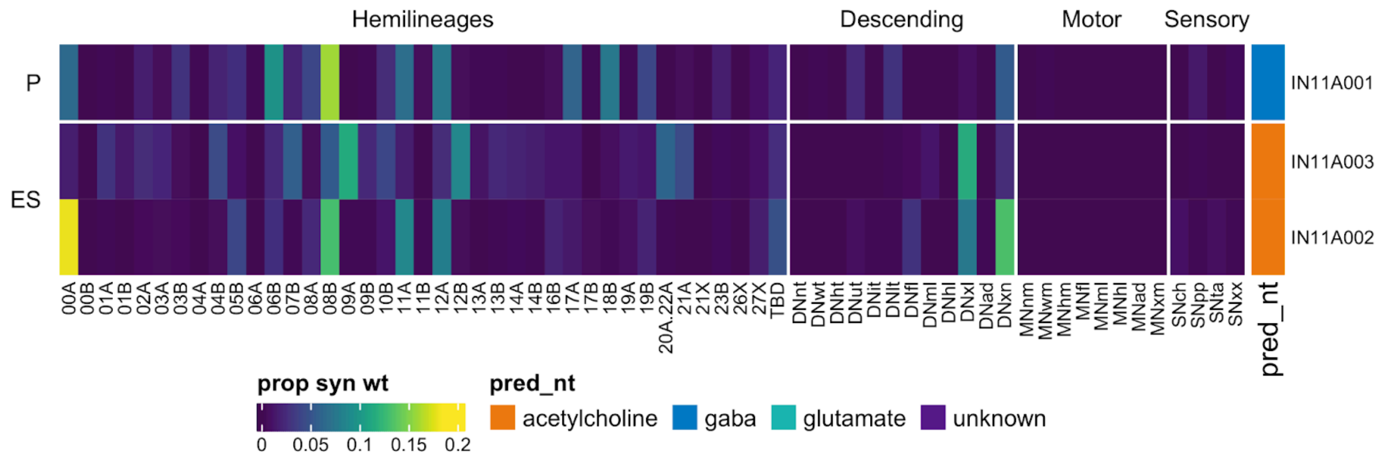

**Figure 31 - figure supplement 3. Connectivity to upstream partners by 11A primary and early secondary systematic types.** Proportions of synaptic weight to systematic types from upstream partners, normalised by row. 11A neurons have been clustered within each assigned birthtime window (P = primary, ES = early secondary, S = secondary) based on both upstream and downstream connectivity to hemilineages, descending neuron subclasses, motor neuron subclasses, and sensory neuron modalities. Annotation bar is coloured by the most common predicted neurotransmitter for the neurons of each type.

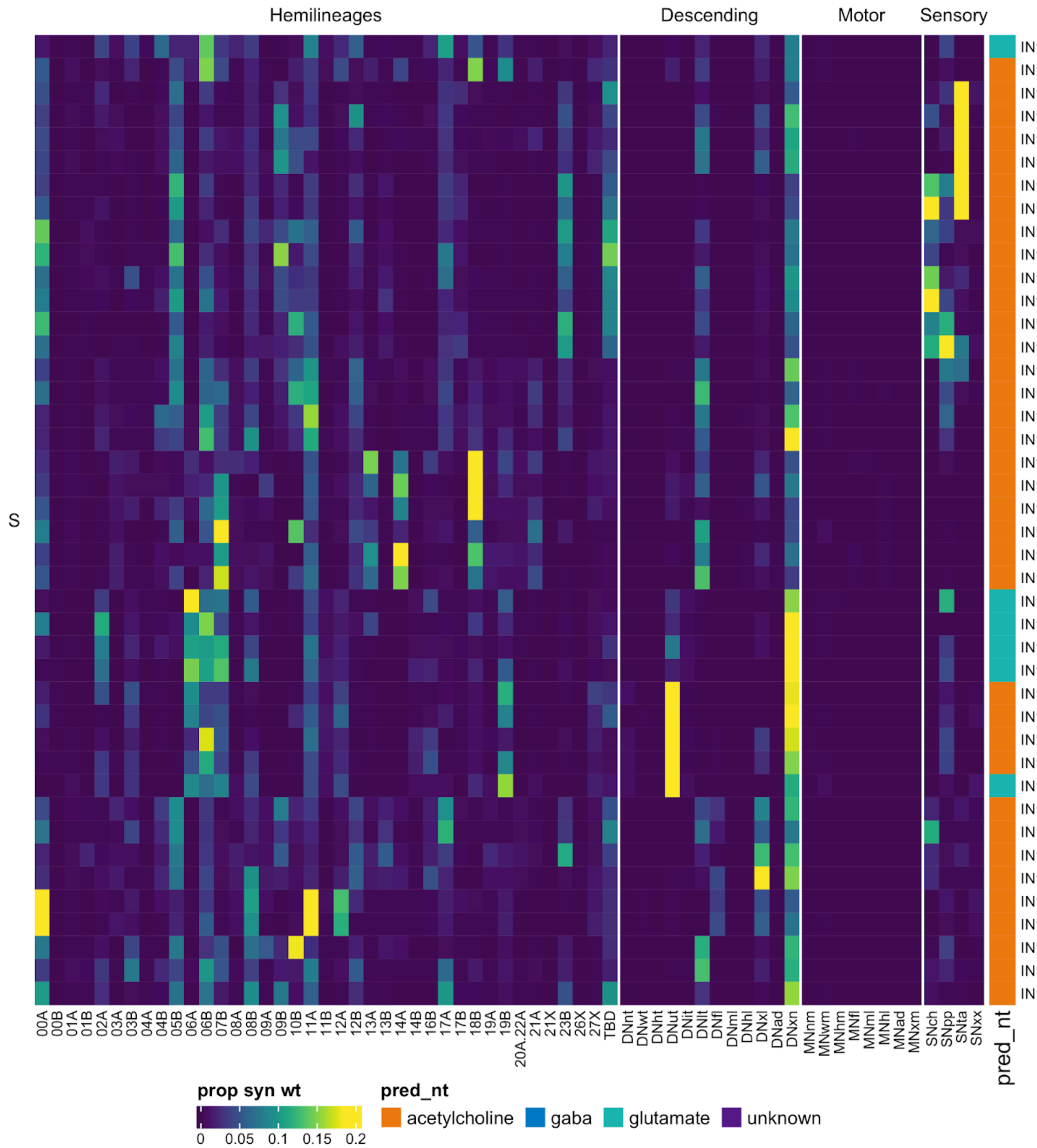

**Figure 31 - figure supplement 4. Connectivity to upstream partners by 11A secondary systematic types.** Proportions of synaptic weight to systematic types from upstream partners, normalised by row. 11A neurons have been clustered within each assigned birthtime window (P = primary, ES = early secondary, S = secondary) based on both upstream and downstream connectivity to hemilineages, descending neuron subclasses, motor neuron subclasses, and sensory neuron modalities. Annotation bar is coloured by the most common predicted neurotransmitter for the neurons of each type.

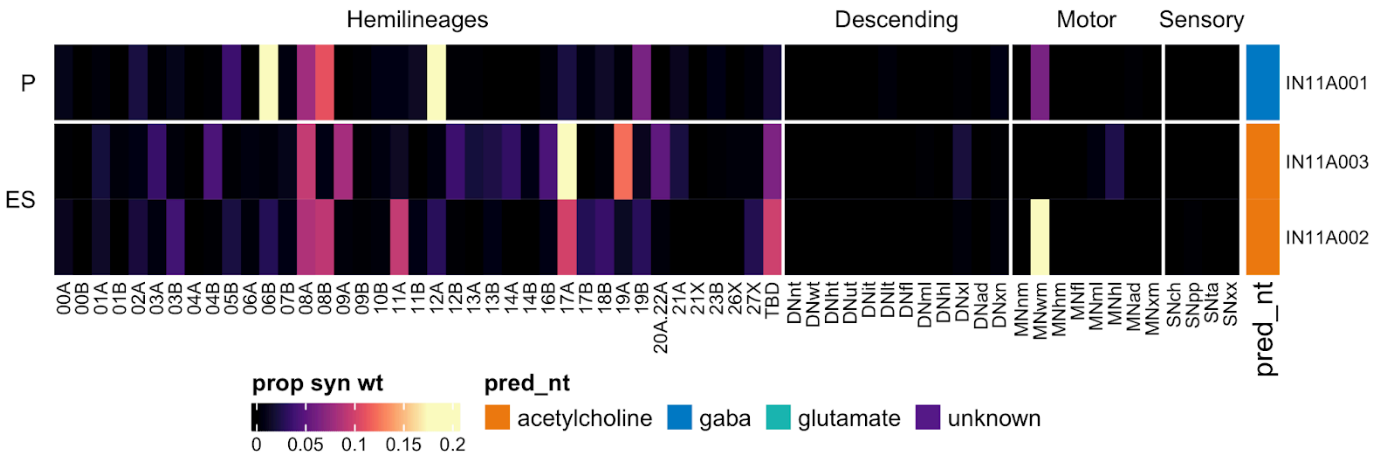

**Figure 31 - figure supplement 5. Connectivity to downstream partners by 11A primary and early secondary systematic types.** Proportions of synaptic weight from systematic types to downstream partners, normalised by row. 11A neurons have been clustered within each assigned birthtime window (P = primary, ES = early secondary, S = secondary) based on both upstream and downstream connectivity to hemilineages, descending neuron subclasses, motor neuron subclasses, and sensory neuron modalities. The annotation bar is coloured by the most common predicted neurotransmitter within each type.

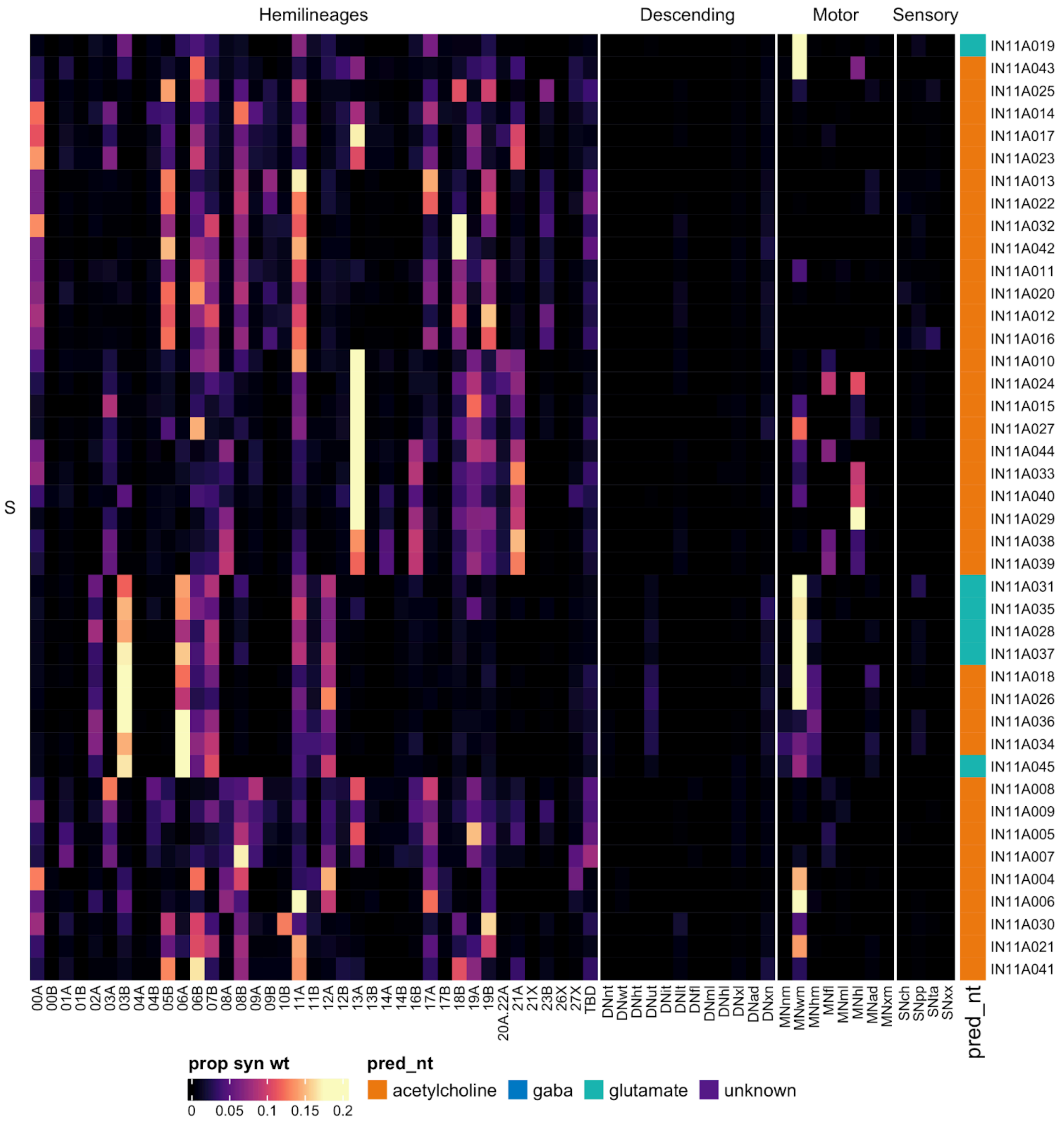

**Figure 31 - figure supplement 6. Connectivity to downstream partners by 11A secondary systematic types.** Proportions of synaptic weight from systematic types to downstream partners, normalised by row. 11A neurons have been clustered within each assigned birthtime window (P = primary, ES = early secondary, S = secondary) based on both upstream and downstream connectivity to hemilineages, descending neuron subclasses, motor neuron subclasses, and sensory neuron modalities. The annotation bar is coloured by the most common predicted neurotransmitter within each type.

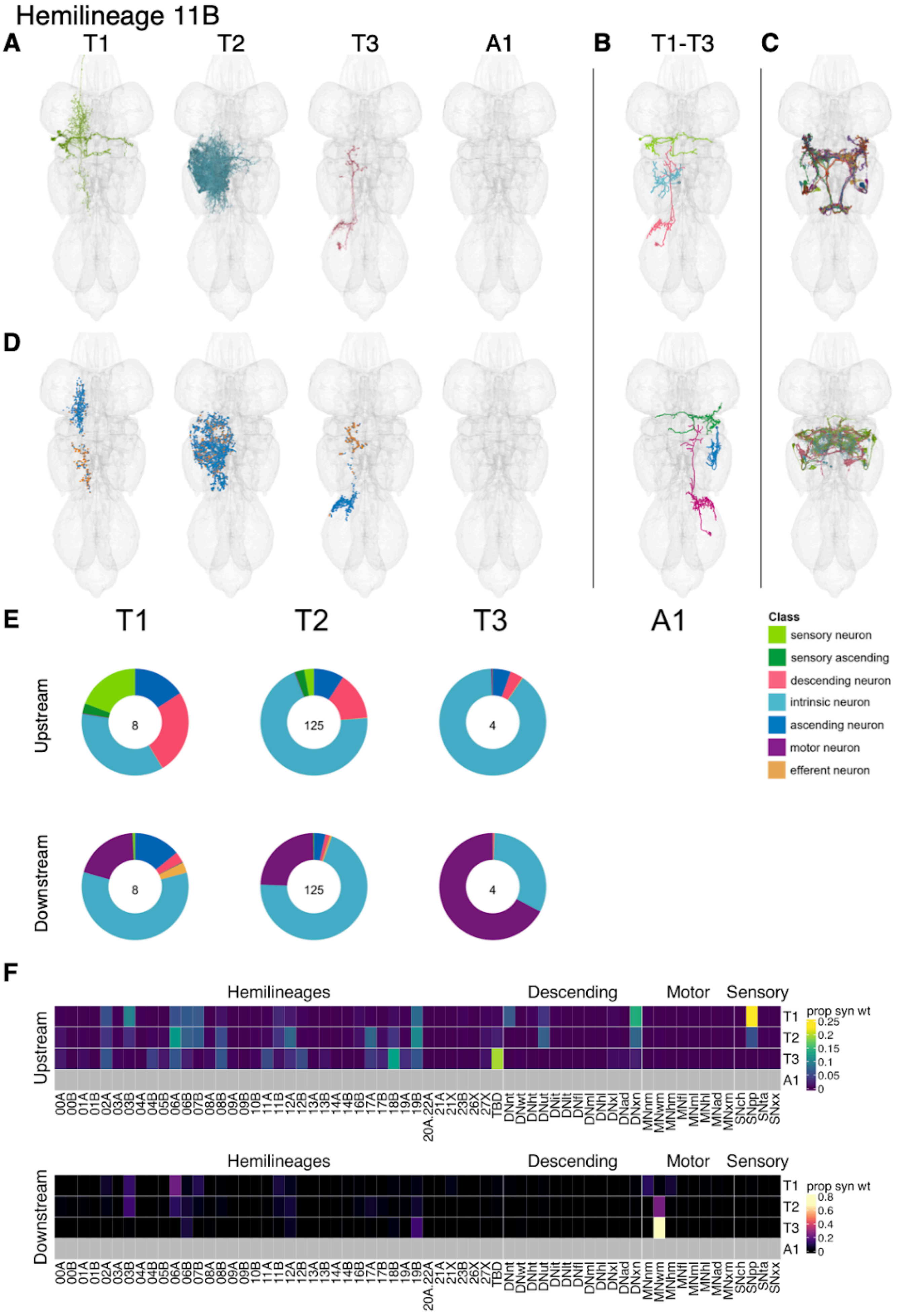

Hemilineage 11B
A
T1
T2
T3
A1
B
T1-T3
C
D
E
T1
T2
T3
A1
Upstream
Downstream
8
125
4
Class
sensory neuron
sensory ascending
descending neuron
intrinsic neuron
ascending neuron
motor neuron
efferent neuron
F
Hemilineages
Descending
Motor
Sensory
prop syn wt

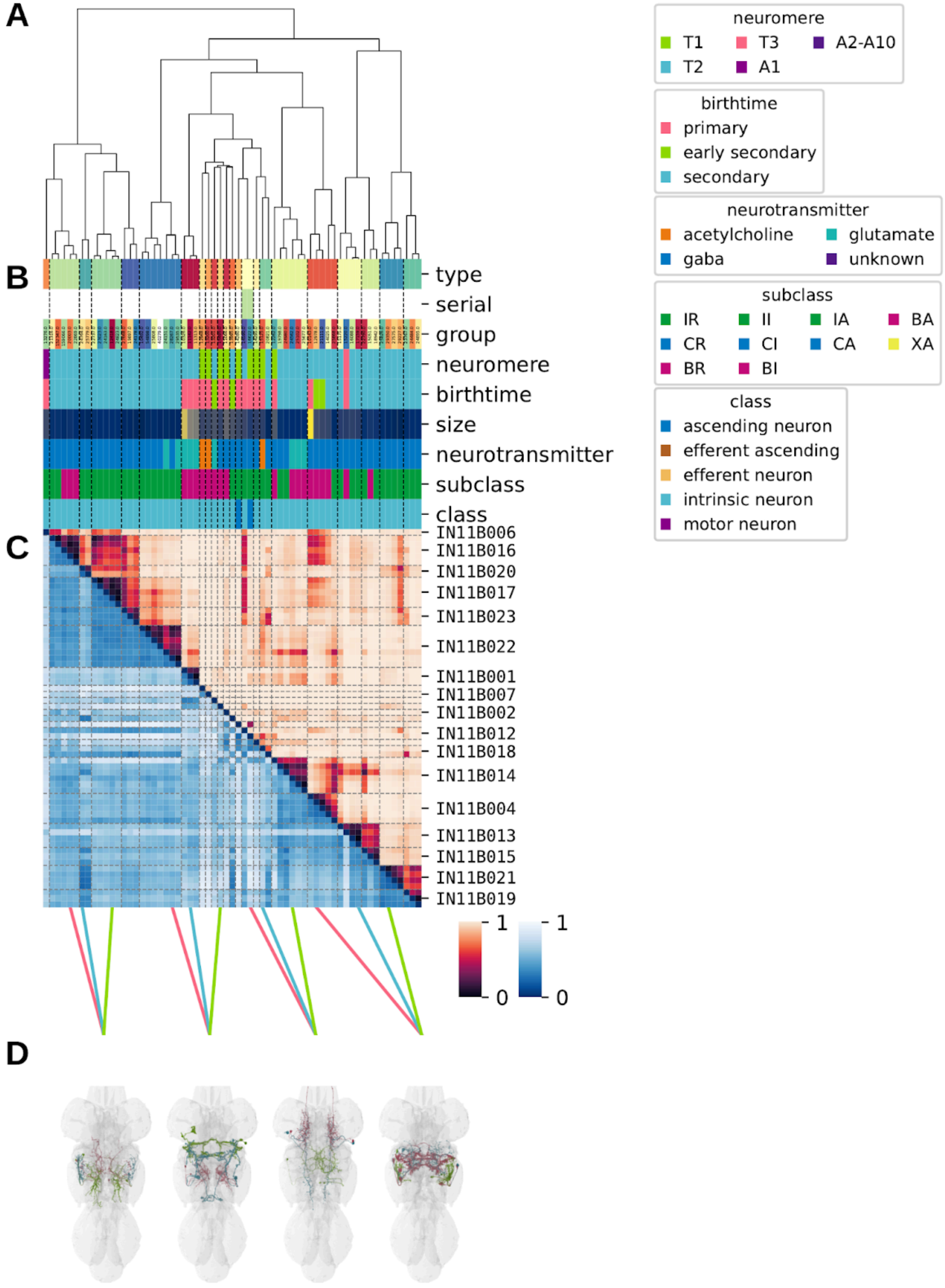

**Figure 32 - figure supplement 1. Systematic typing of hemilineage 11B. A.** Hierarchical clustering dendrogram of hemilineage groups by laterally and serially aggregated connectivity cosine clustering. **B.** Categorical annotations of each hemilineage group, each column corresponding to the aligned leaf in A. Colours for type, serial set, and group are arbitrary for visualisation. Colours for neuromere, birthtime, neurotransmitter, subclass, and class are as in all other figures. **C.** Similarity distance heatmap for hemilineage. Cosine distance is in the upper triangle, while laterally symmetrised NBLAST distance is in the lower triangle. Systematic type names of some types are labelled. **D.** Morphologically representative groups from dendrogram subtrees. Each group, indicated by colour and line connecting to its column in B and C, is the most morphologically representative group (medoid of NBLAST distance) from a subtree of A. The subtrees (flat clusters) are equal height cuts of A determined to yield the number of groups per plot and plots in D.

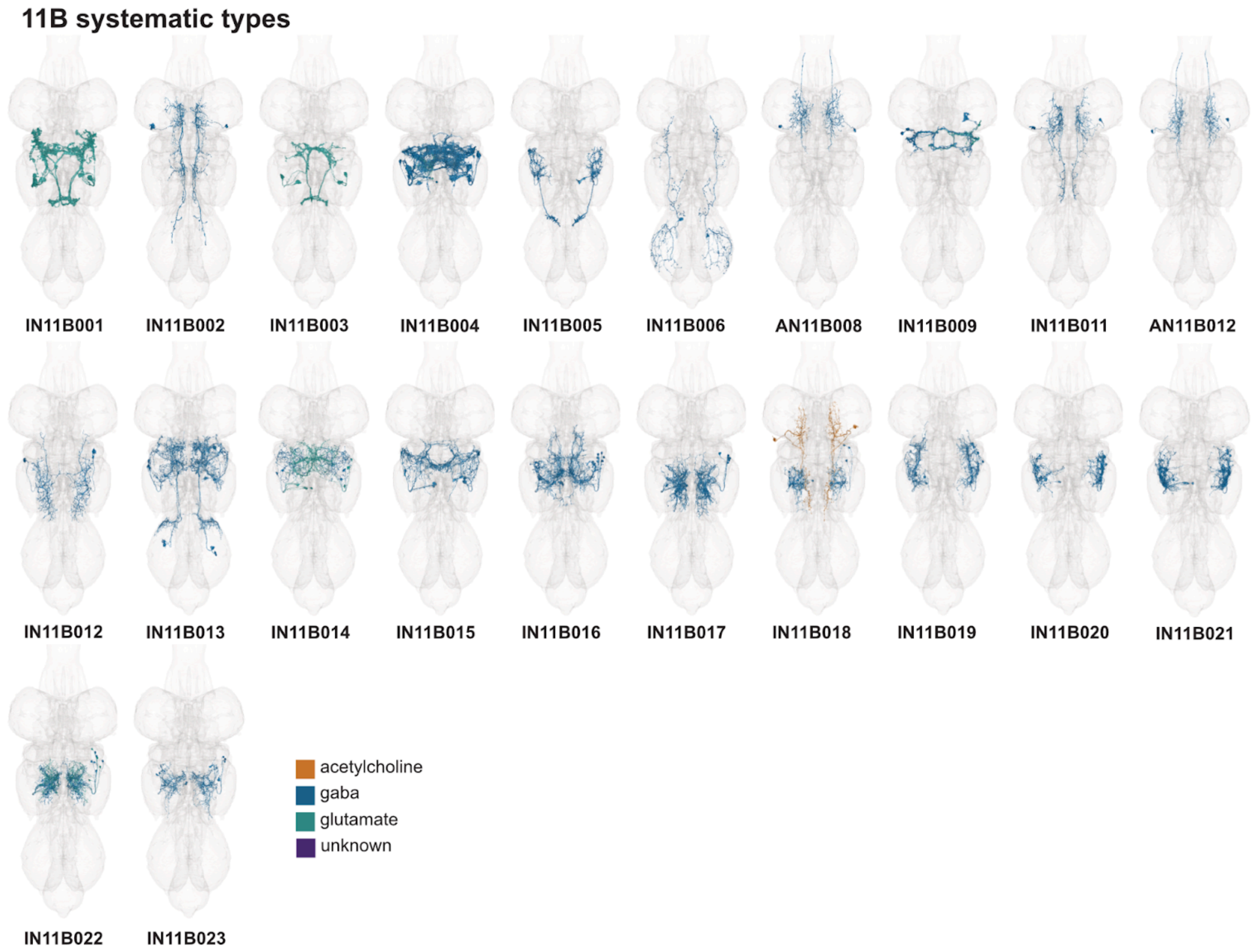

**Figure 32 - figure supplement 2. Systematic types of hemilineage 11B.** Systematic types have been arranged in numerical order, with neurons of the same type that belong to distinct classes (e.g., intrinsic neuron vs ascending neuron) plotted separately but placed adjacent to each other. Individual neuron meshes have been coloured based on predicted neurotransmitter: dark orange = acetylcholine, blue = gaba, marine = glutamate, dark purple = unknown.

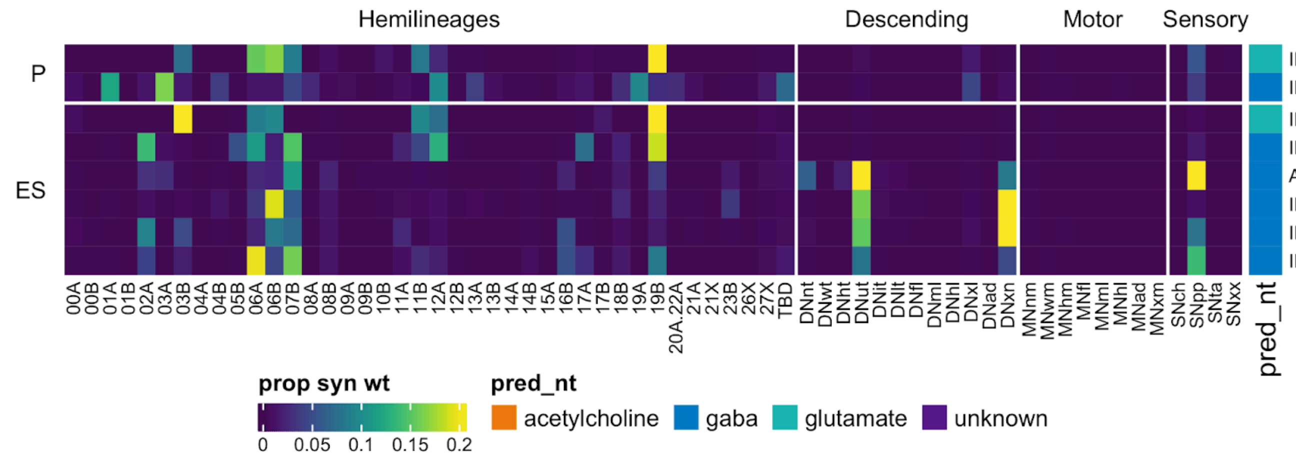

**Figure 32 - figure supplement 3. Connectivity to upstream partners by 11B primary and early secondary systematic types.** Proportions of synaptic weight to systematic types from upstream partners, normalised by row. 11B neurons have been clustered within each assigned birthtime window (P = primary, ES = early secondary, S = secondary) based on both upstream and downstream connectivity to hemilineages, descending neuron subclasses, motor neuron subclasses, and sensory neuron modalities. Annotation bar is coloured by the most common predicted neurotransmitter for the neurons of each type.

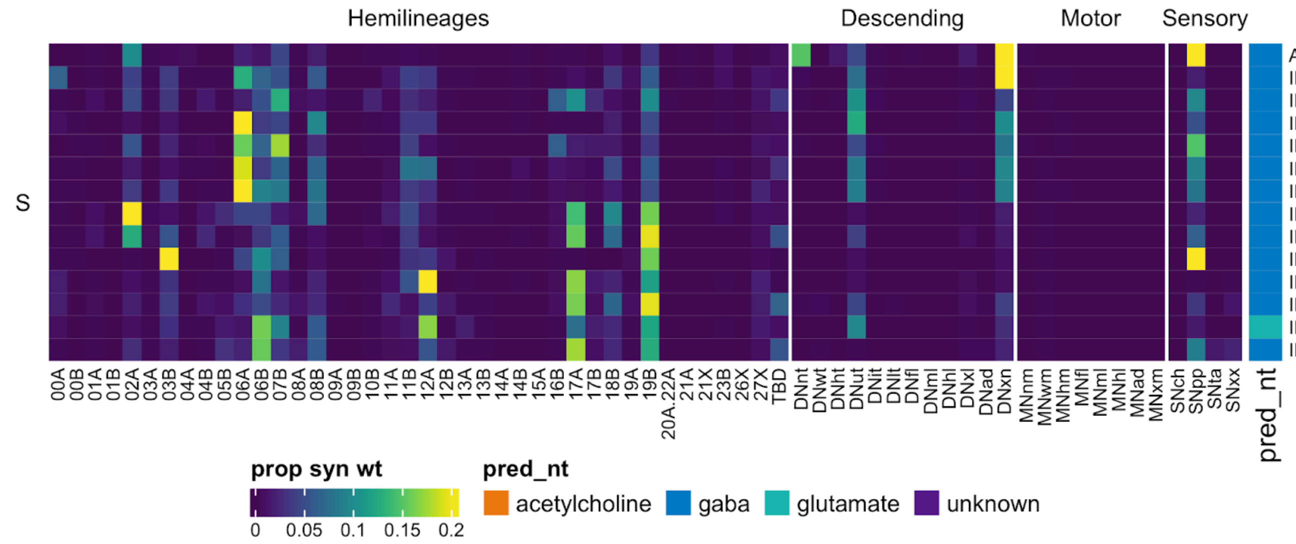

**Figure 32 - figure supplement 4. Connectivity to upstream partners by 11B secondary systematic types.** Proportions of synaptic weight to systematic types from upstream partners, normalised by row. 11B neurons have been clustered within each assigned birthtime window (P = primary, ES = early secondary, S = secondary) based on both upstream and downstream connectivity to hemilineages, descending neuron subclasses, motor neuron subclasses, and sensory neuron modalities. Annotation bar is coloured by the most common predicted neurotransmitter for the neurons of each type.

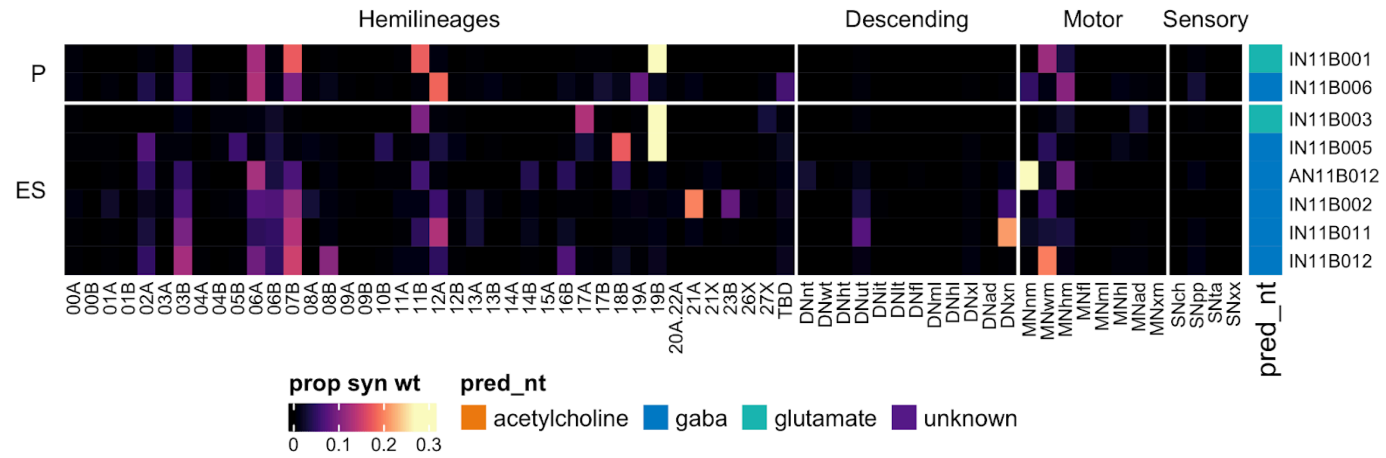

**Figure 32 - figure supplement 5. Connectivity to downstream partners by 11B primary and early secondary systematic types.** Proportions of synaptic weight from systematic types to downstream partners, normalised by row. 11B neurons have been clustered within each assigned birthtime window (P = primary, ES = early secondary, S = secondary) based on both upstream and downstream connectivity to hemilineages, descending neuron subclasses, motor neuron subclasses, and sensory neuron modalities. The annotation bar is coloured by the most common predicted neurotransmitter within each type.

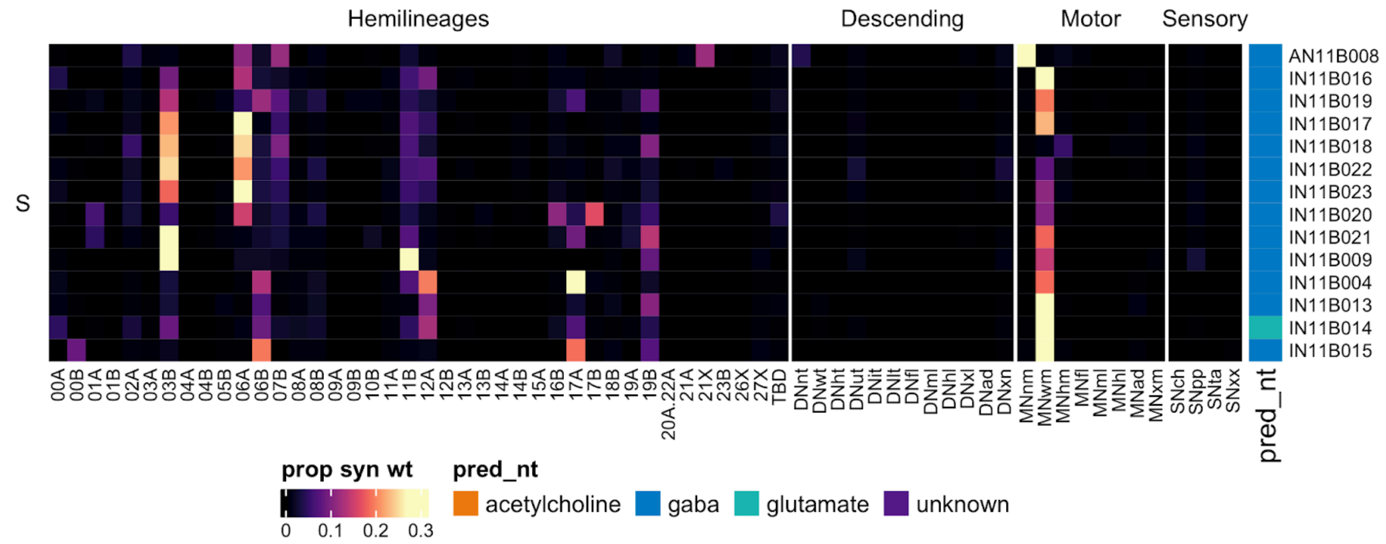

**Figure 32 - figure supplement 6. Connectivity to downstream partners by 11B secondary systematic types.** Proportions of synaptic weight from systematic types to downstream partners, normalised by row. 11B neurons have been clustered within each assigned birthtime window (P = primary, ES = early secondary, S = secondary) based on both upstream and downstream connectivity to hemilineages, descending neuron subclasses, motor neuron subclasses, and sensory neuron modalities. The annotation bar is coloured by the most common predicted neurotransmitter within each type.

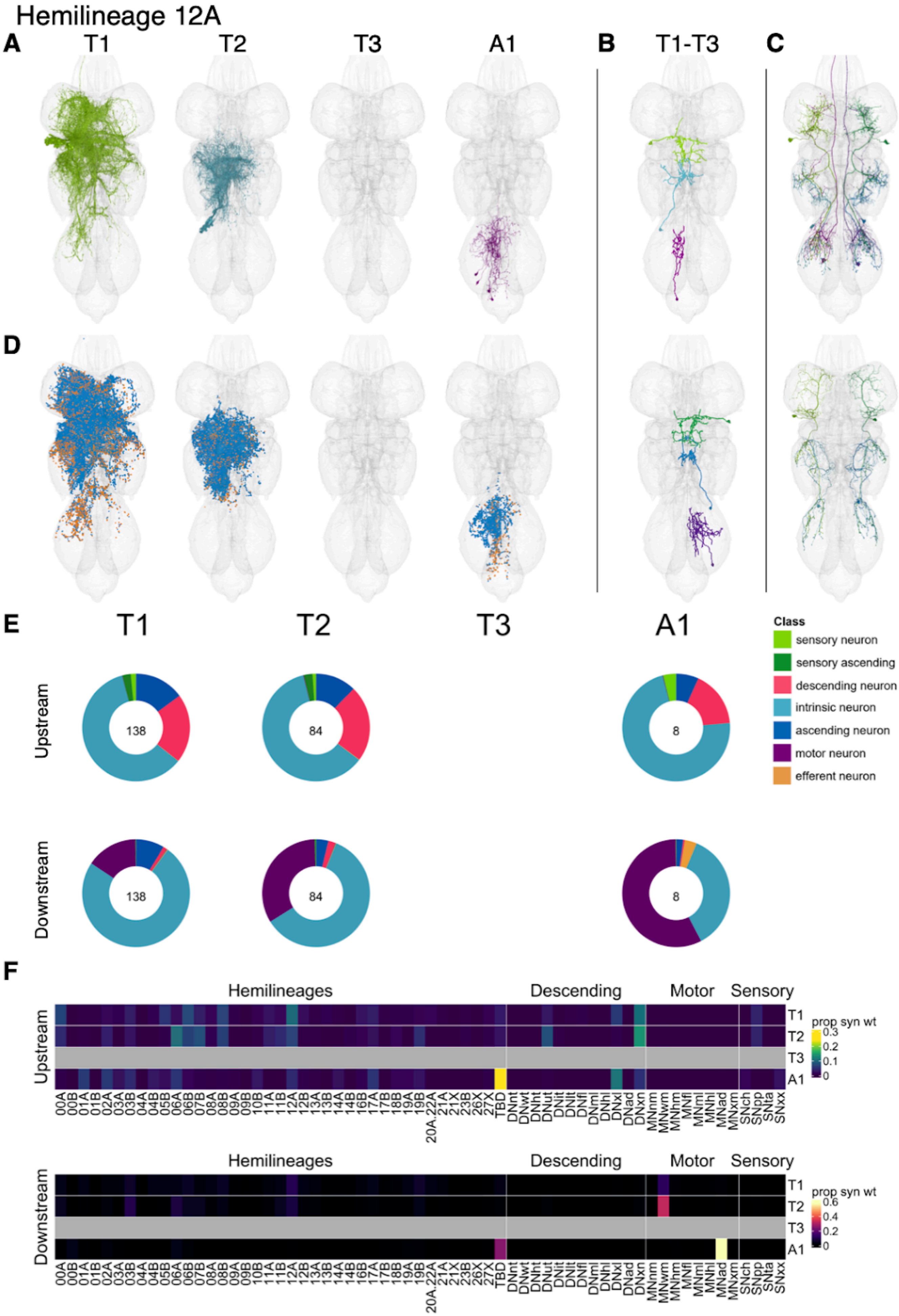

Hemilineage 12A
A
T1
T2
T3
A1
B
T1-T3
C
D
E
Upstream
Downstream
138
84
8
Class
sensory neuron
sensory ascending
descending neuron
intrinsic neuron
ascending neuron
motor neuron
efferent neuron
F
Hemilineages
Descending
Motor
Sensory
prop syn wt
0.3
0.2
0.1
0
0.6
0.4
0.2
00A
00B
01A
01B
02A
03A
03B
04A
04B
05B
06A
06B
07B
08A
08B
09A
09B
10B
11A
11B
12A
12B
13A
13B
14A
14B
16B
17A
17B
18B
19A
19B
20A.22A
21A
21X
23B
26X
27X
TBD
DNnt
DNwt
DNht
DNut
DNit
DNlt
DNfl
DNml
DNhl
DNxl
DNad
DNxn
MNnm
MNwm
MNhm
MNfl
MNml
MNhl
MNad
MNxm
SNch
SNpp
SNta
SNxx

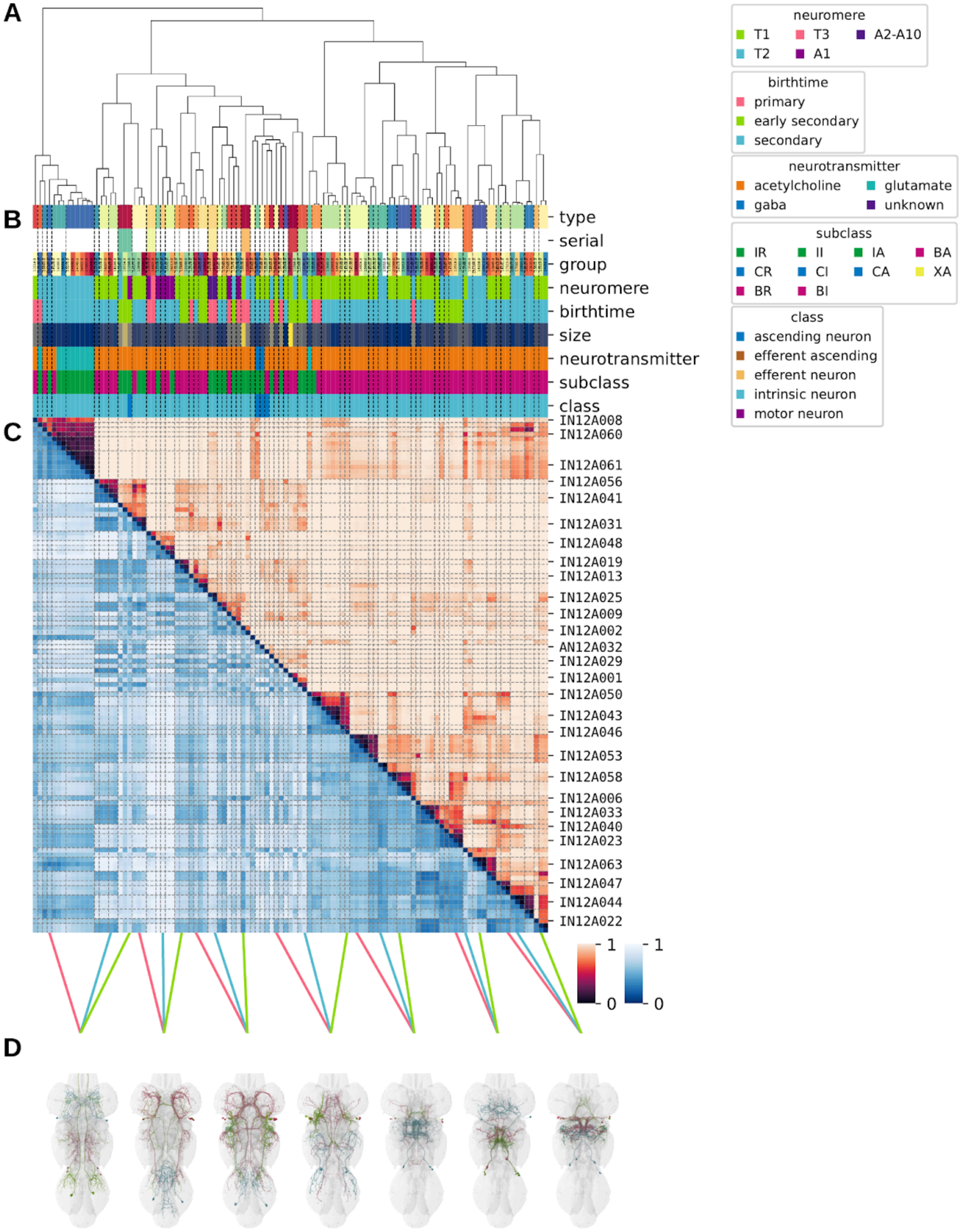

**Figure 33 - figure supplement 1. Systematic typing of hemilineage 12A. A.** Hierarchical clustering dendrogram of hemilineage groups by laterally and serially aggregated connectivity cosine clustering. **B.** Categorical annotations of each hemilineage group, each column corresponding to the aligned leaf in A. Colours for type, serial set, and group are arbitrary for visualisation. Colours for neuromere, birthtime, neurotransmitter, subclass, and class are as in all other figures. **C.** Similarity distance heatmap for hemilineage. Cosine distance is in the upper triangle, while laterally

symmetrised NBLAST distance is in the lower triangle. Systematic type names of some types are labelled. **D.** Morphologically representative groups from dendrogram subtrees. Each group, indicated by colour and line connecting to its column in B and C, is the most morphologically representative group (medoid of NBLAST distance) from a subtree of A. The subtrees (flat clusters) are equal height cuts of A determined to yield the number of groups per plot and plots in D.

#### 12A systematic types

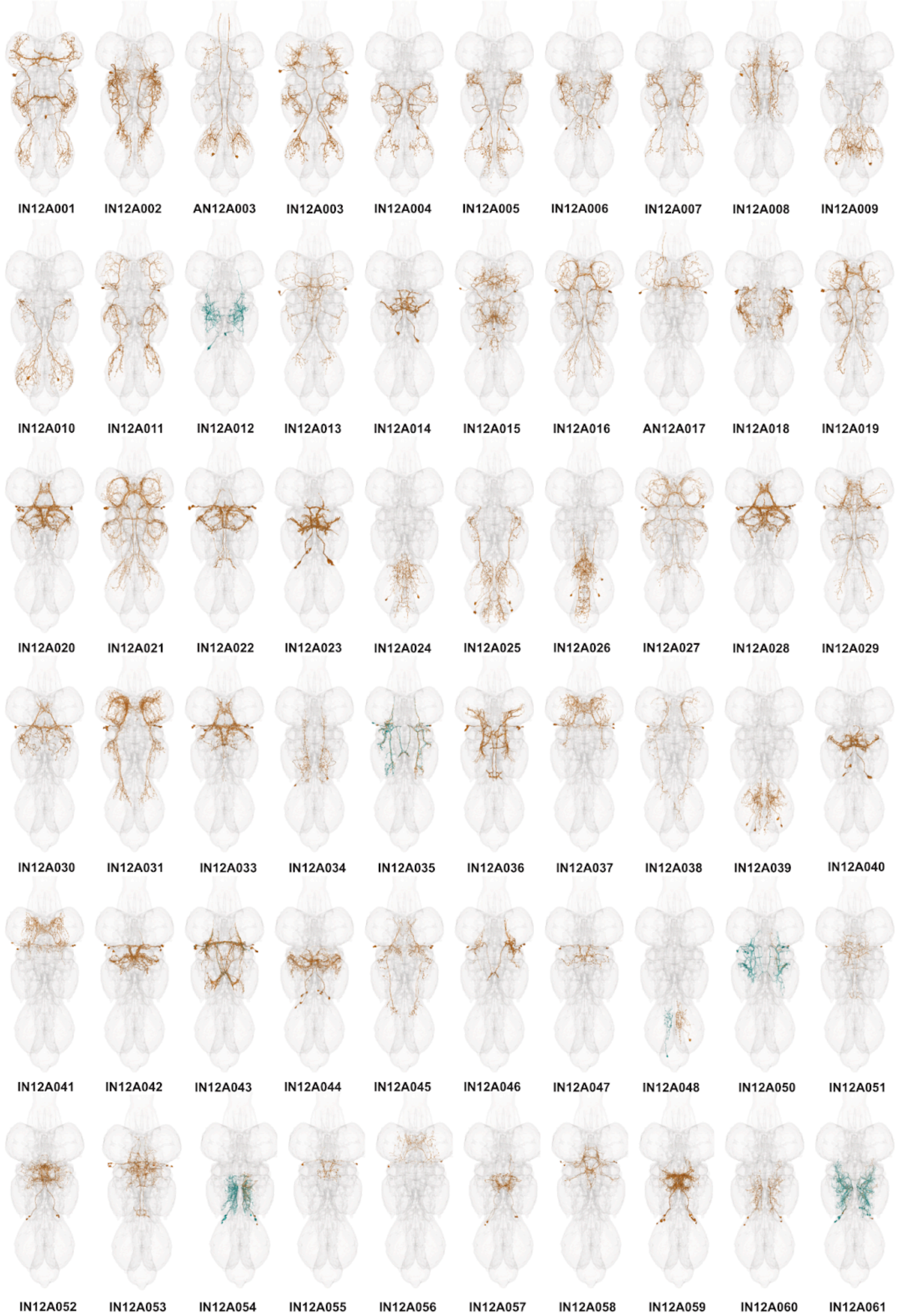

**Figure 33 - figure supplement 2. Systematic types of hemilineage 12A.** Systematic types have been arranged in numerical order, with neurons of the same type that belong to distinct classes (e.g., intrinsic neuron vs ascending neuron) plotted separately but placed adjacent to each other. Individual neuron meshes have been coloured based on predicted neurotransmitter: dark orange = acetylcholine, blue = gaba, marine = glutamate, dark purple = unknown.

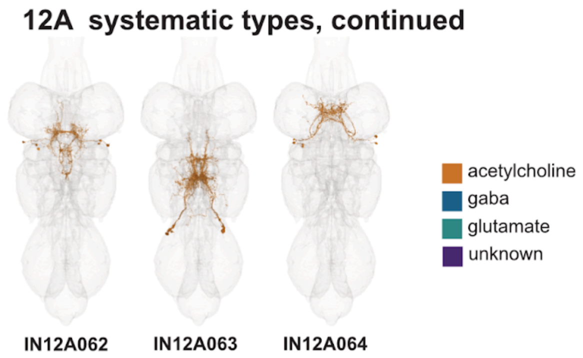

**Figure 33 - figure supplement 3. Systematic types of hemilineage 12A, continued.** Systematic types have been arranged in numerical order, with neurons of the same type that belong to distinct classes (e.g., intrinsic neuron vs ascending neuron) plotted separately but placed adjacent to each other. Individual neuron meshes have been coloured based on predicted neurotransmitter: dark orange = acetylcholine, blue = gaba, marine = glutamate, dark purple = unknown.

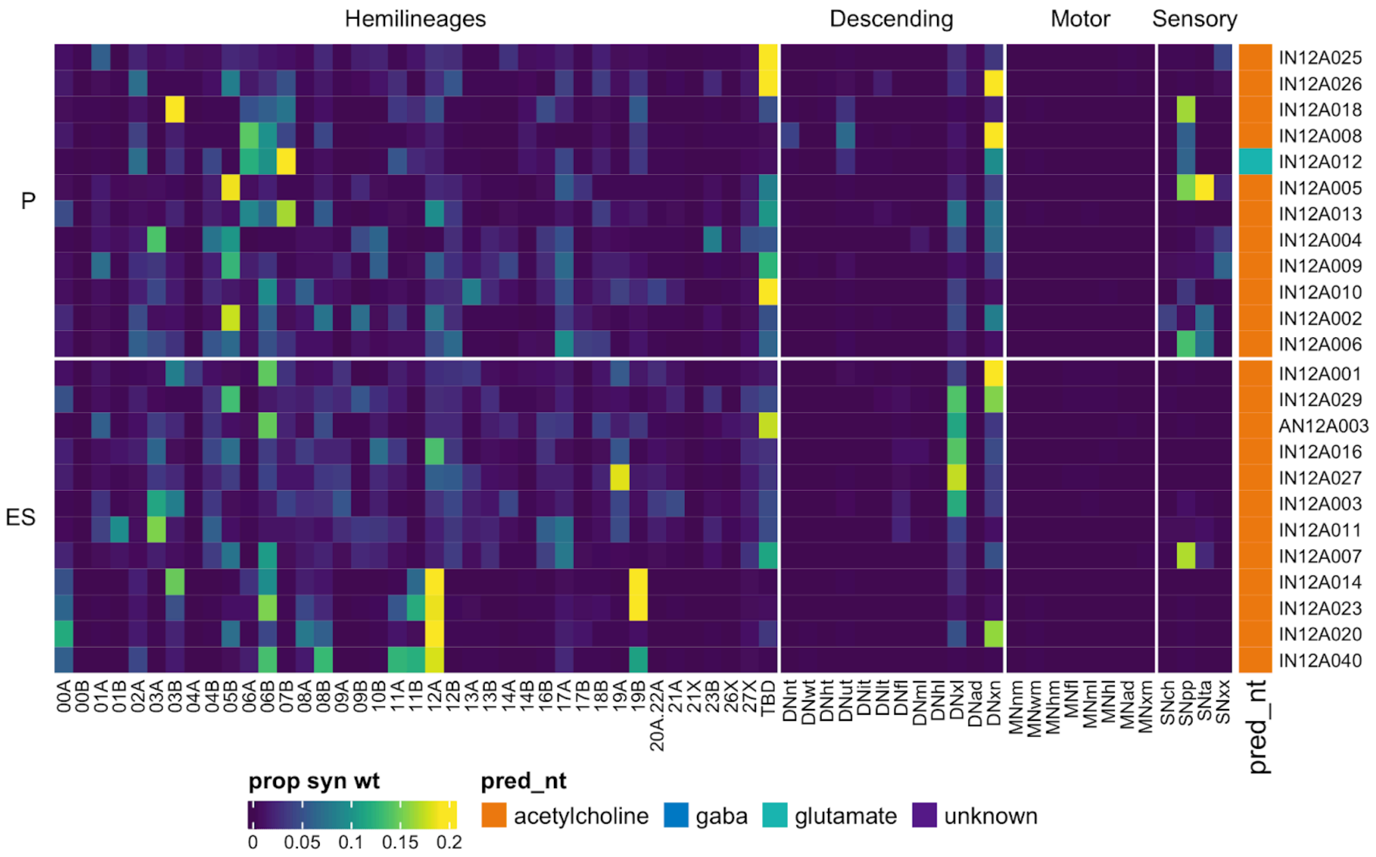

**Figure 33 - figure supplement 4. Connectivity to upstream partners by 12A primary and early secondary systematic types.** Proportions of synaptic weight to systematic types from upstream partners, normalised by row. 12A neurons have been clustered within each assigned birthtime window (P = primary, ES = early secondary, S = secondary) based on both upstream and downstream connectivity to hemilineages, descending neuron subclasses, motor neuron subclasses, and sensory neuron modalities. Annotation bar is coloured by the most common predicted neurotransmitter for the neurons of each type.

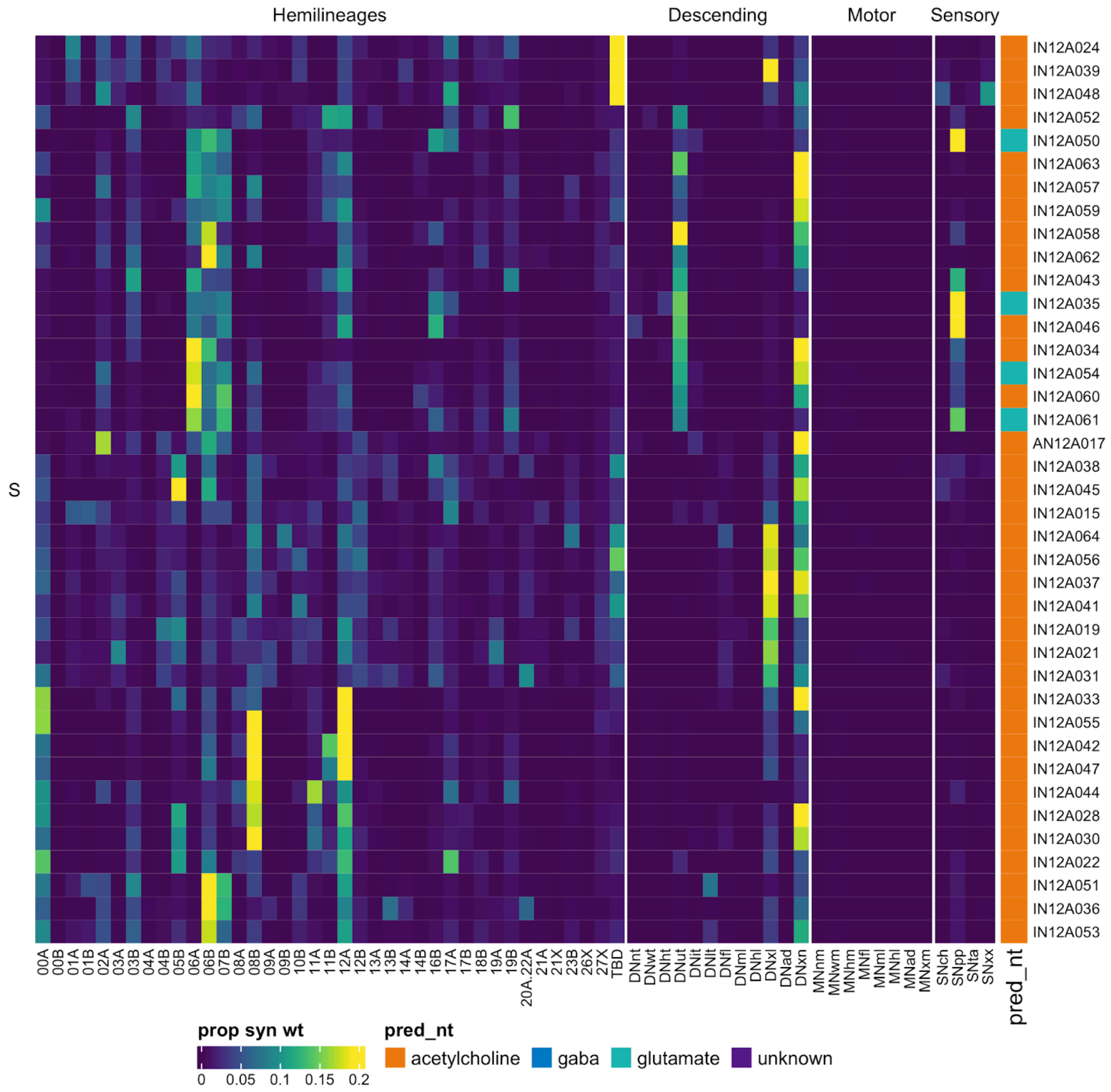

**Figure 33 - figure supplement 5. Connectivity to upstream partners by 12A secondary systematic types.** Proportions of synaptic weight to systematic types from upstream partners, normalised by row. 12A neurons have been clustered within each assigned birthtime window (P = primary, ES = early secondary, S = secondary) based on both upstream and downstream connectivity to hemilineages, descending neuron subclasses, motor neuron subclasses, and sensory neuron modalities. The annotation bar is coloured by the most common predicted neurotransmitter within each type.

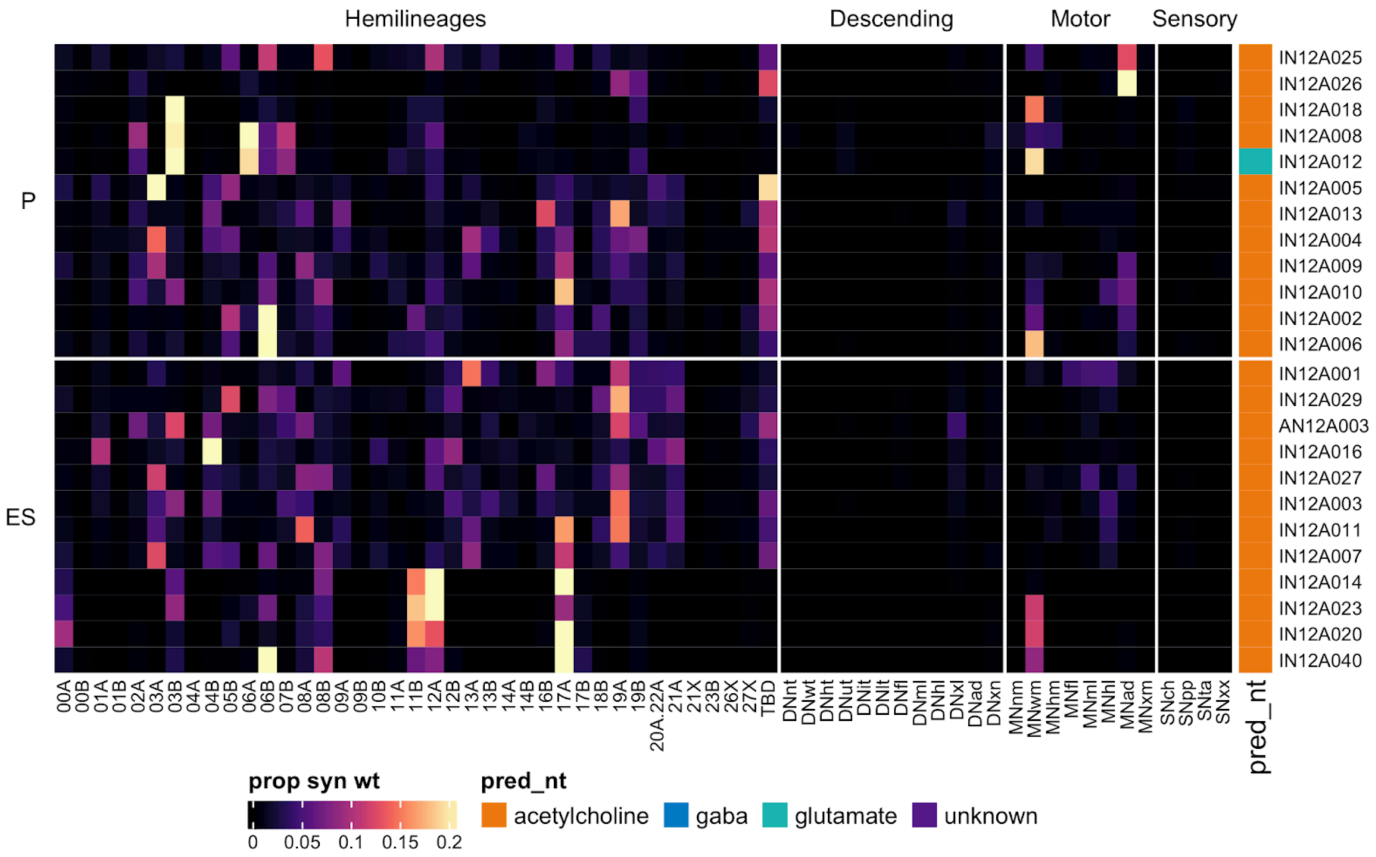

**Figure 33 - figure supplement 6. Connectivity to downstream partners by 12A primary and early secondary systematic types.** Proportions of synaptic weight from systematic types to downstream partners, normalised by row. 12A neurons have been clustered within each assigned birthtime window (P = primary, ES = early secondary, S = secondary) based on both upstream and downstream connectivity to hemilineages, descending neuron subclasses, motor neuron subclasses, and sensory neuron modalities. The annotation bar is coloured by the most common predicted neurotransmitter within each type.

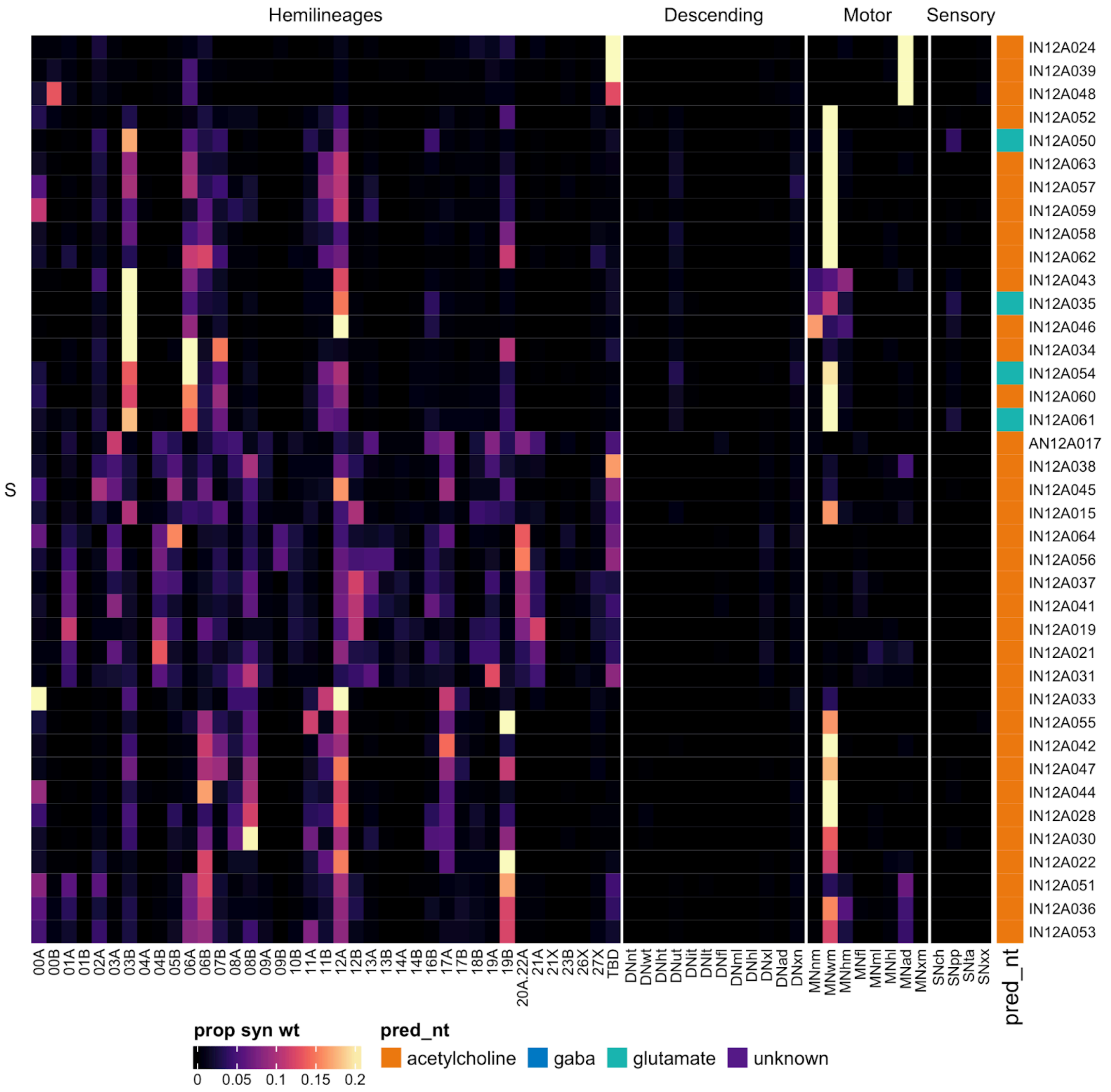

**Figure 33 - figure supplement 7. Connectivity to downstream partners by 12A secondary systematic types.** Proportions of synaptic weight from systematic types to downstream partners, normalised by row. 12A neurons have been clustered within each assigned birthtime window (P = primary, ES = early secondary, S = secondary) based on both upstream and downstream connectivity to hemilineages, descending neuron subclasses, motor neuron subclasses, and sensory neuron modalities. The annotation bar is coloured by the most common predicted neurotransmitter for the neurons of each type.

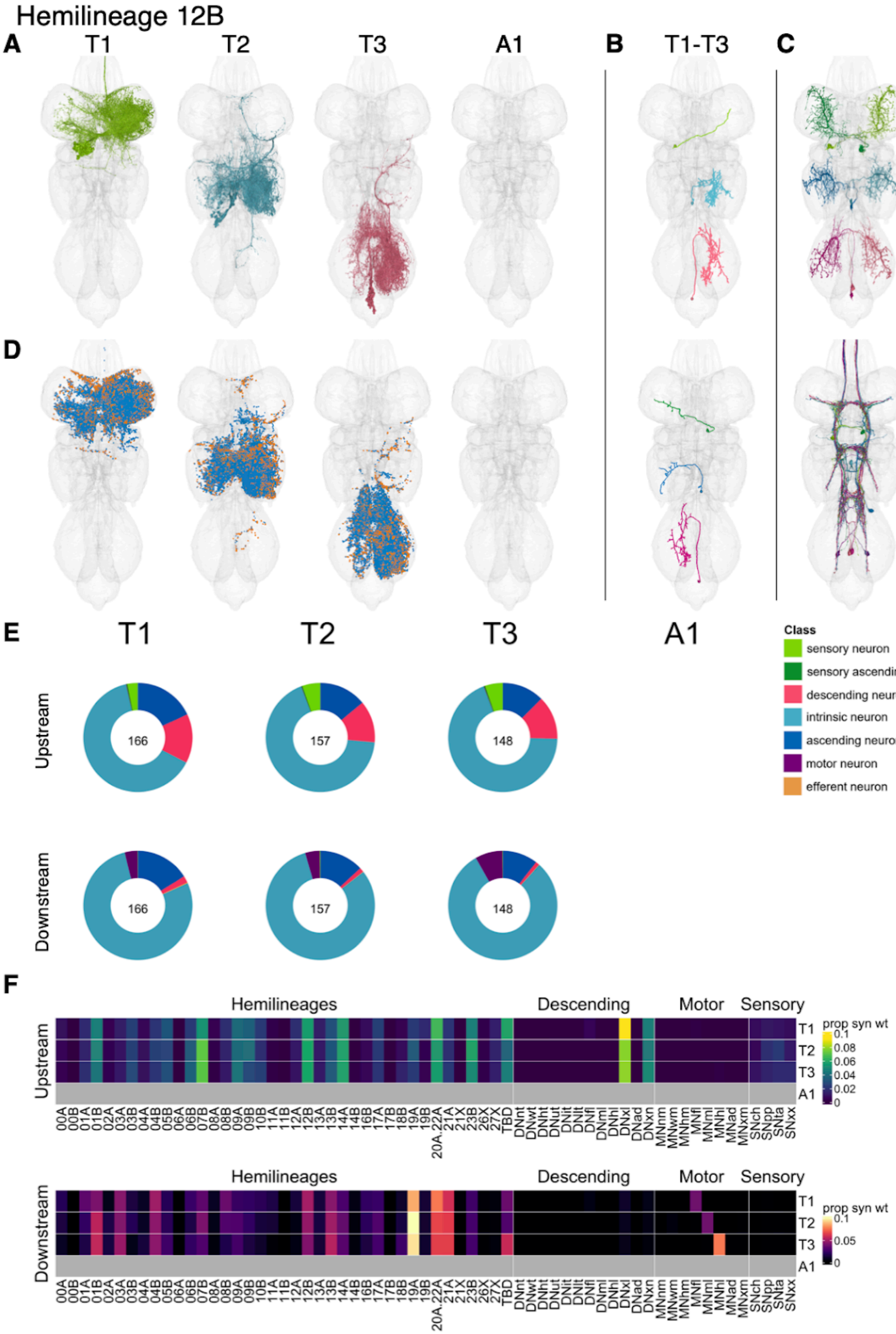

Hemilineage 12B
A
T1
T2
T3
A1
B
T1-T3
C
D
E
T1
T2
T3
A1
Upstream
Downstream
166
157
148
166
157
148
Class
sensory neuron
sensory ascending
descending neuron
intrinsic neuron
ascending neuron
motor neuron
efferent neuron
F
Hemilineages
Descending
Motor
Sensory
Upstream
Downstream
T1
T2
T3
A1
prop syn wt
0.1
0.08
0.06
0.04
0.02
0
0.05
00A 00B 01A 01B 02A 03A 03B 04A 04B 05B 06A 06B 07B 08A 08B 09A 09B 10B 11A 11B 12A 12B 13A 13B 14A 14B 16B 17A 17B 18B 19A 19B 20A.22A 21A 21X 23B 26X 27X TBD
DNnt DNwt DNht DNut DNit DNlt DNfl DNml DNhl DNxl DNad DNxn
MNnm MNwm MNhm MNfl MNml MNhl MNad MNxm
SNch SNpp SNta SNxx

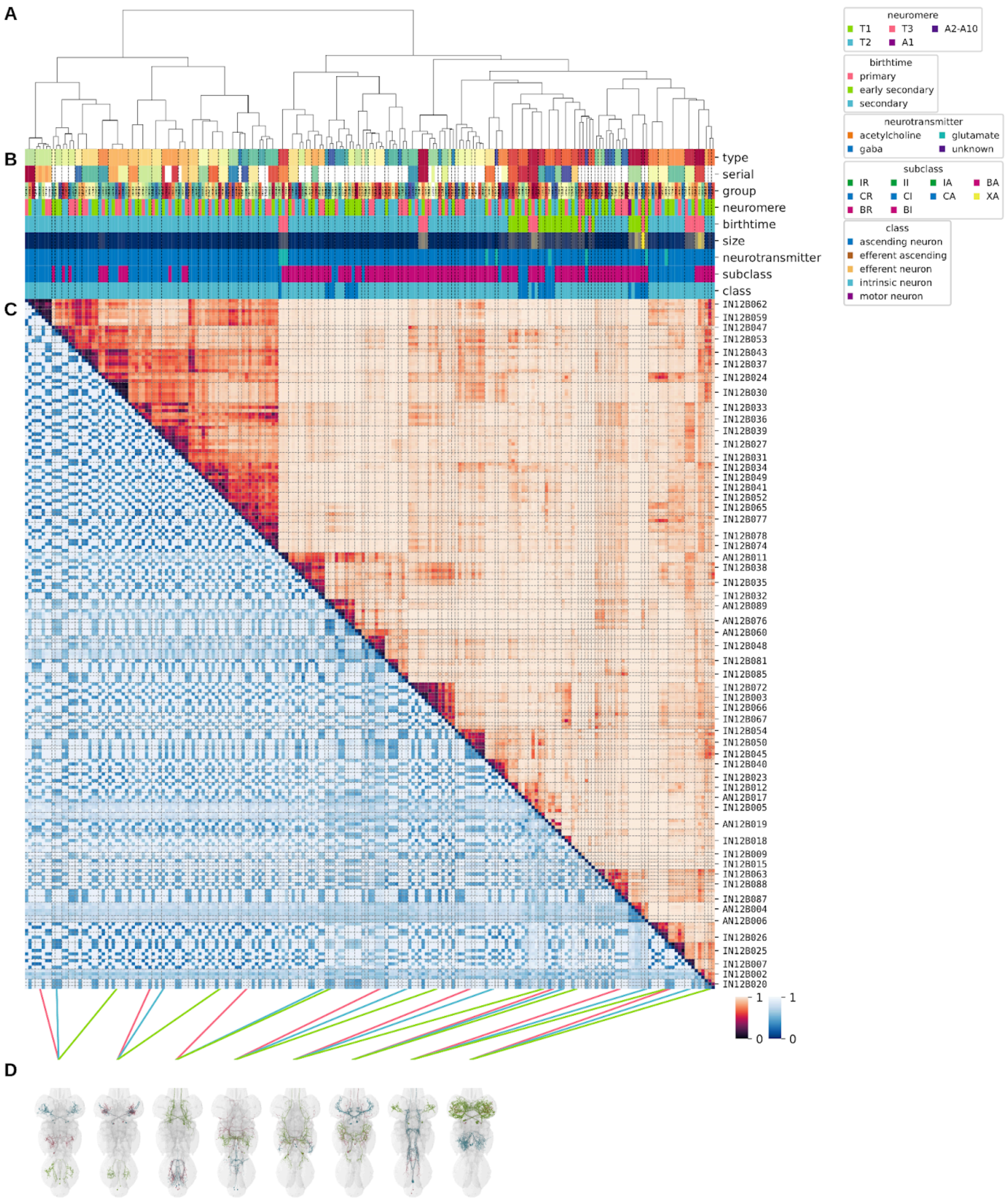

**Figure 34 - figure supplement 1. Systematic typing of hemilineage 12B. A.** Hierarchical clustering dendrogram of hemilineage groups by laterally and serially aggregated connectivity cosine clustering. **B.** Categorical annotations of each hemilineage group, each column corresponding to the aligned leaf in A. Colours for type, serial set, and group are arbitrary for visualisation. Colours for neuromere, birthtime, neurotransmitter, subclass, and class are as in all other figures. **C.** Similarity distance heatmap for hemilineage. Cosine distance is in the upper triangle, while laterally symmetrised NBLAST distance is in the lower triangle. Systematic type names of some types are labelled. **D.** Morphologically representative groups from dendrogram subtrees. Each group, indicated by colour and line connecting to its column in B and C, is the most morphologically representative group (medoid of NBLAST distance)

from a subtree of A. The subtrees (flat clusters) are equal height cuts of A determined to yield the number of groups per plot and plots in D.

### 12B systematic types

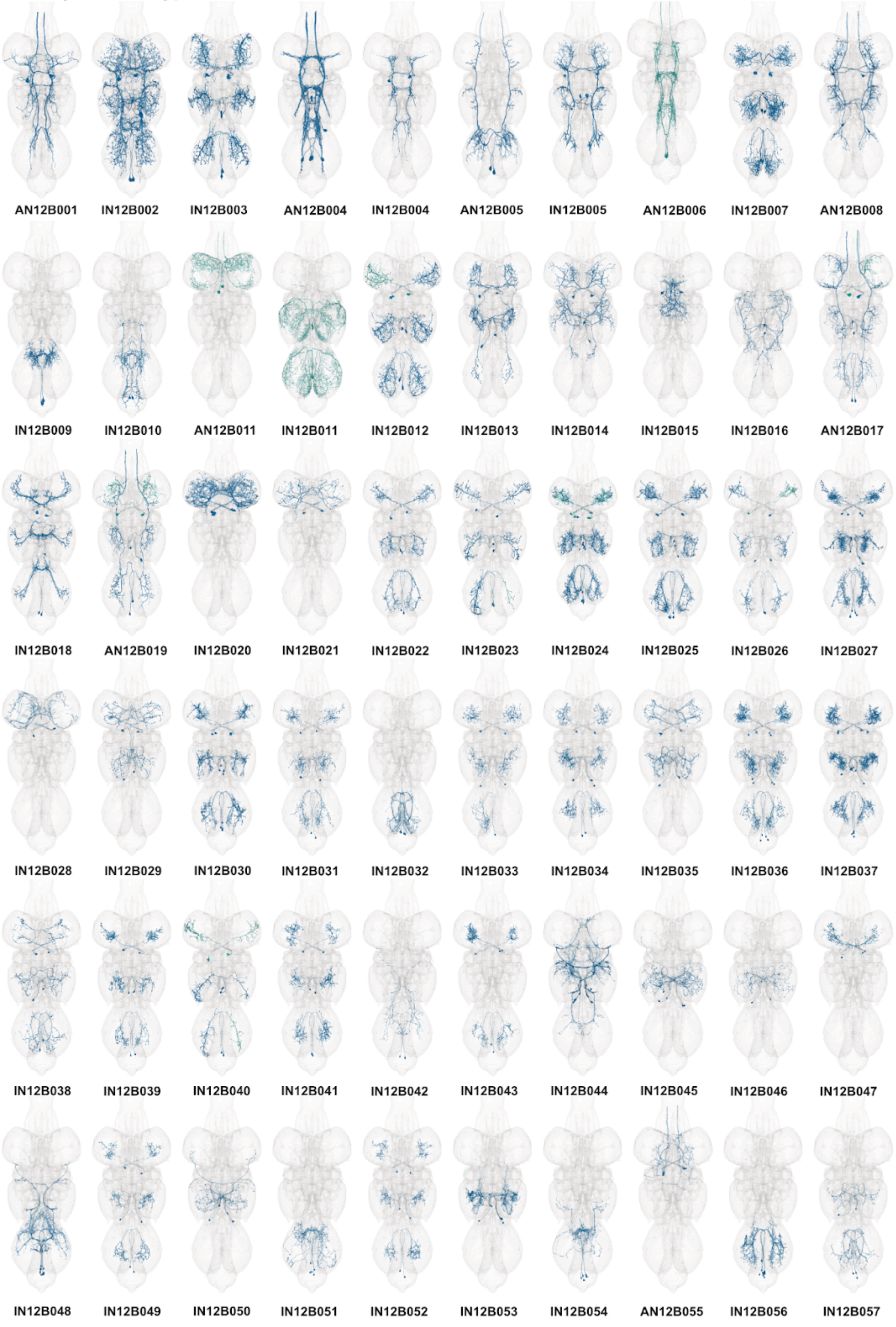

**Figure 34 - figure supplement 2. Systematic types of hemilineage 12B.** Systematic types have been arranged in numerical order, with neurons of the same type that belong to distinct classes (e.g., intrinsic neuron vs ascending neuron) plotted separately but placed adjacent to each other. Individual neuron meshes have been coloured based on predicted neurotransmitter: dark orange = acetylcholine, blue = gaba, marine = glutamate, dark purple = unknown.

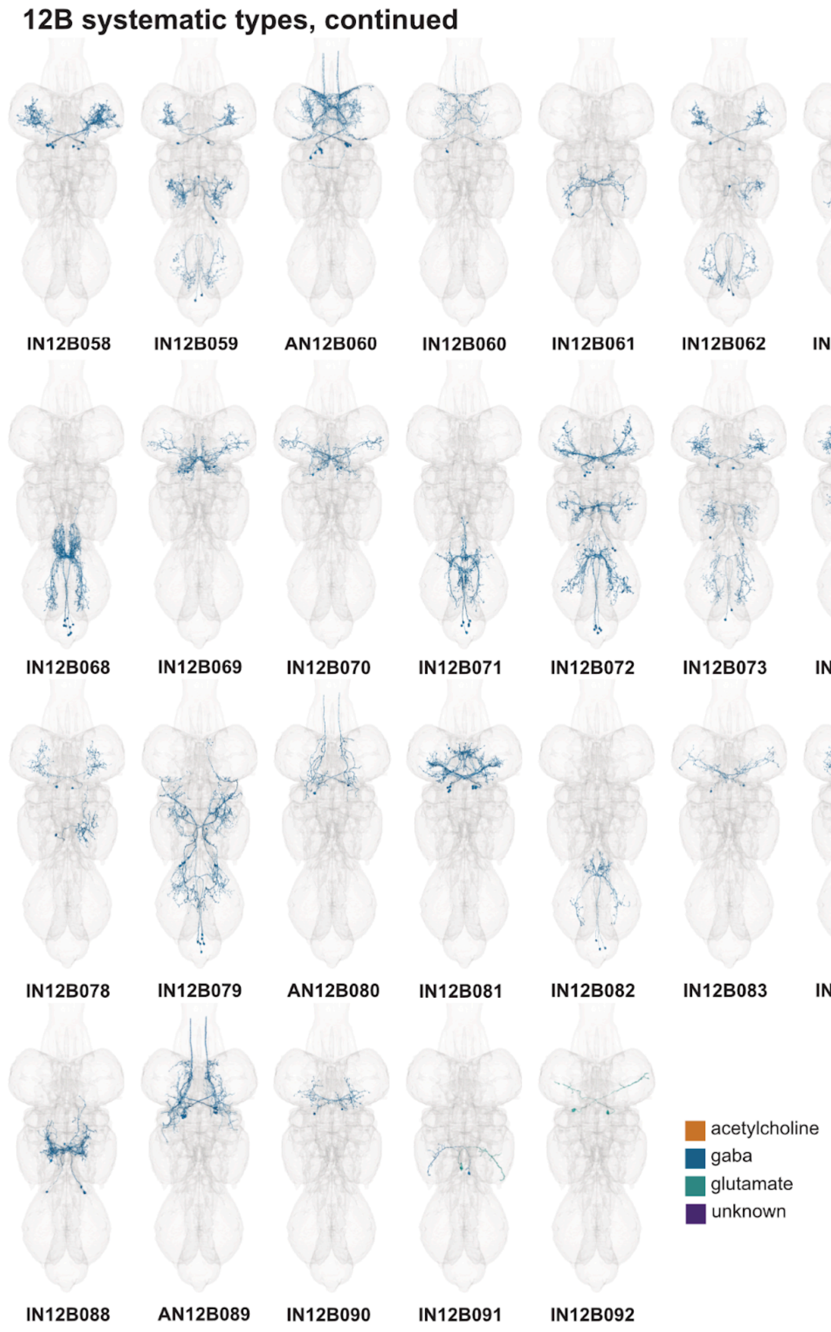

**Figure 34 - figure supplement 3. Systematic types of hemilineage 12B, continued.** Systematic types have been arranged in numerical order, with neurons of the same type that belong to distinct classes (e.g., intrinsic neuron vs ascending neuron) plotted separately but placed adjacent to each other. Individual neuron meshes have been coloured based on predicted neurotransmitter: dark orange = acetylcholine, blue = gaba, marine = glutamate, dark purple = unknown.

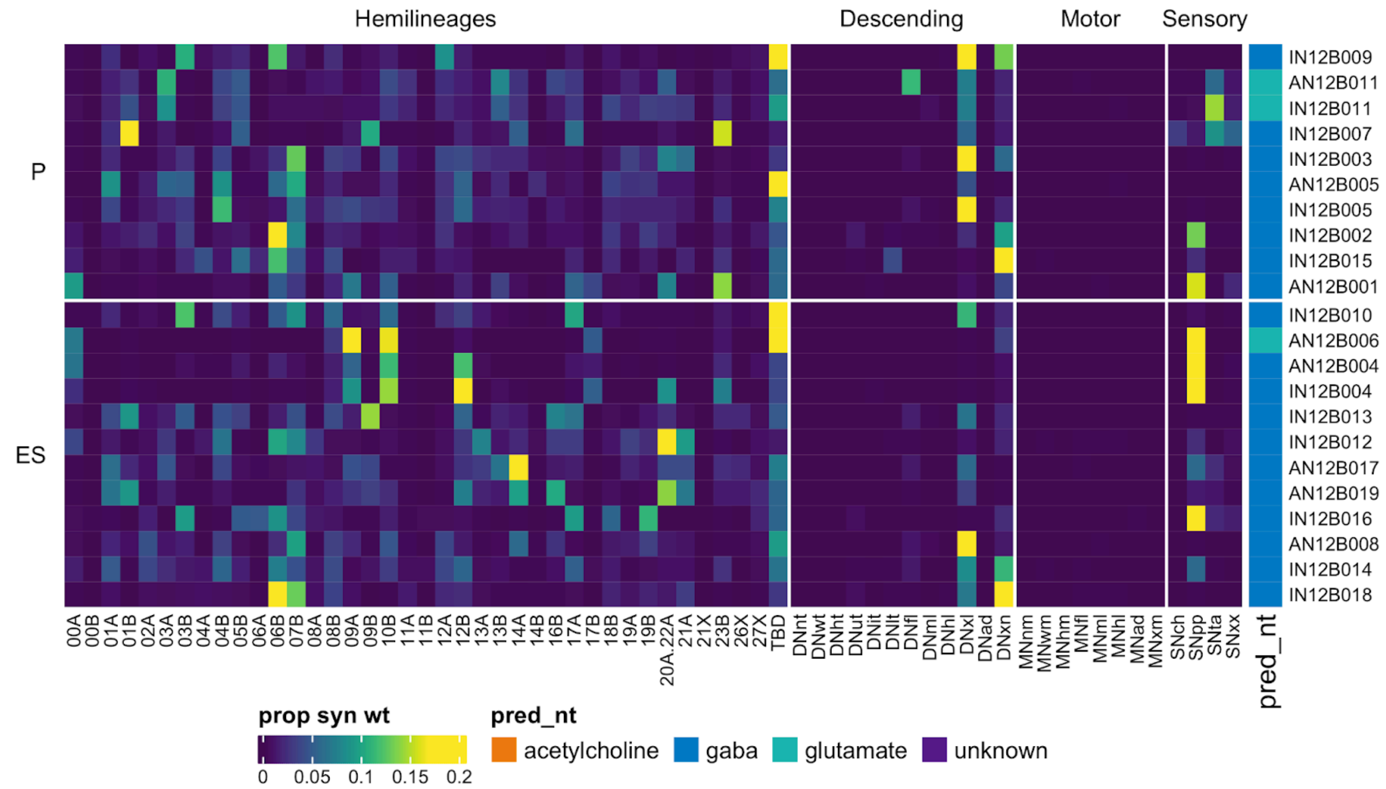

**Figure 34 - figure supplement 4. Connectivity to upstream partners by 12B primary and early secondary systematic types.** Proportions of synaptic weight to systematic types from upstream partners, normalised by row. 12B neurons have been clustered within each assigned birthtime window (P = primary, ES = early secondary, S = secondary) based on both upstream and downstream connectivity to hemilineages, descending neuron subclasses, motor neuron subclasses, and sensory neuron modalities. Annotation bar is coloured by the most common predicted neurotransmitter for the neurons of each type.

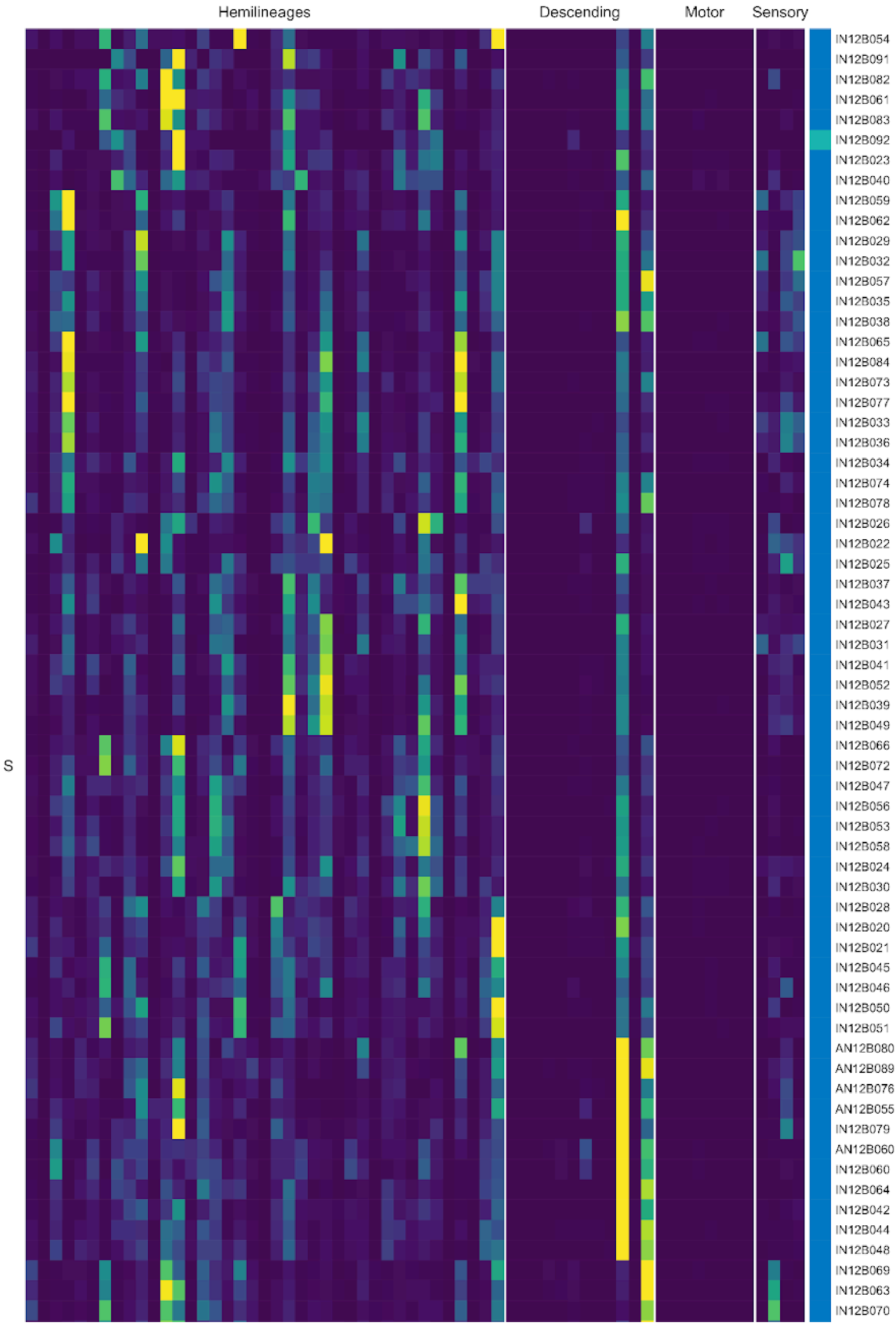

Hemilineages
Descending
Motor
Sensory
S
IN12B054
IN12B091
IN12B082
IN12B061
IN12B083
IN12B092
IN12B023
IN12B040
IN12B059
IN12B062
IN12B029
IN12B032
IN12B057
IN12B035
IN12B038
IN12B065
IN12B084
IN12B073
IN12B077
IN12B033
IN12B036
IN12B034
IN12B074
IN12B078
IN12B026
IN12B022
IN12B025
IN12B037
IN12B043
IN12B027
IN12B031
IN12B041
IN12B052
IN12B039
IN12B049
IN12B066
IN12B072
IN12B047
IN12B056
IN12B053
IN12B058
IN12B024
IN12B030
IN12B028
IN12B020
IN12B021
IN12B045
IN12B046
IN12B050
IN12B051
AN12B080
AN12B089
AN12B076
AN12B055
IN12B079
AN12B060
IN12B060
IN12B064
IN12B042
IN12B044
IN12B048
IN12B069
IN12B063
IN12B070

**Figure 34 - figure supplement 5. Connectivity to upstream partners by 12B secondary systematic types.** Proportions of synaptic weight to systematic types from upstream partners, normalised by row. 12B neurons have been clustered within each assigned birthtime window (P = primary, ES = early secondary, S = secondary) based on both upstream and downstream connectivity to hemilineages, descending neuron subclasses, motor neuron subclasses, and sensory neuron modalities. The annotation bar is coloured by the most common predicted neurotransmitter within each type.

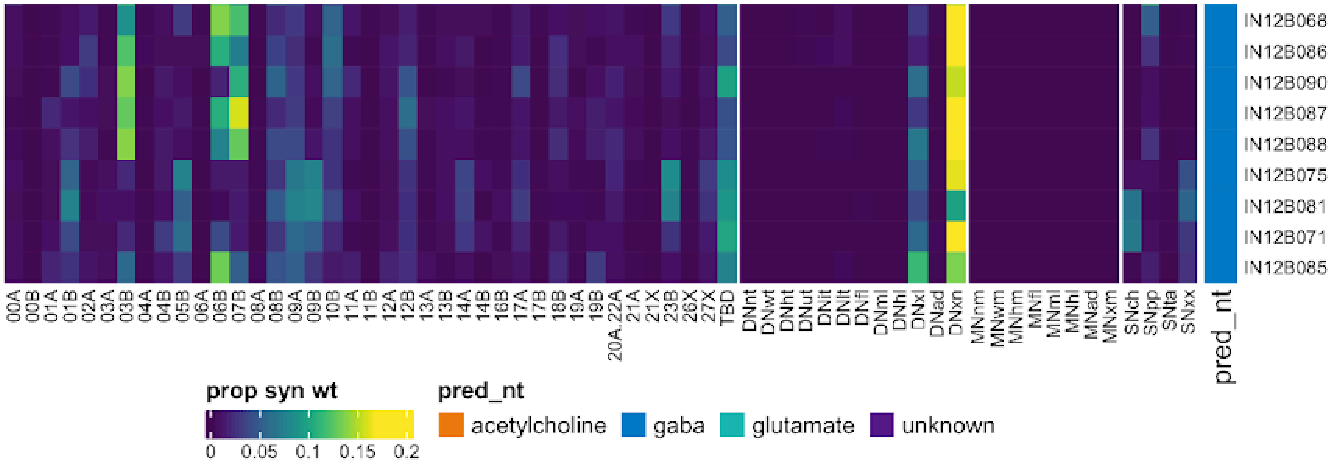

**Figure 34 - figure supplement 6. Connectivity to upstream partners by 12B secondary systematic types, continued.** Proportions of synaptic weight to systematic types from upstream partners, normalised by row. 12B neurons have been clustered within each assigned birthtime window (P = primary, ES = early secondary, S = secondary) based on both upstream and downstream connectivity to hemilineages, descending neuron subclasses, motor neuron subclasses, and sensory neuron modalities. The annotation bar is coloured by the most common predicted neurotransmitter within each type.

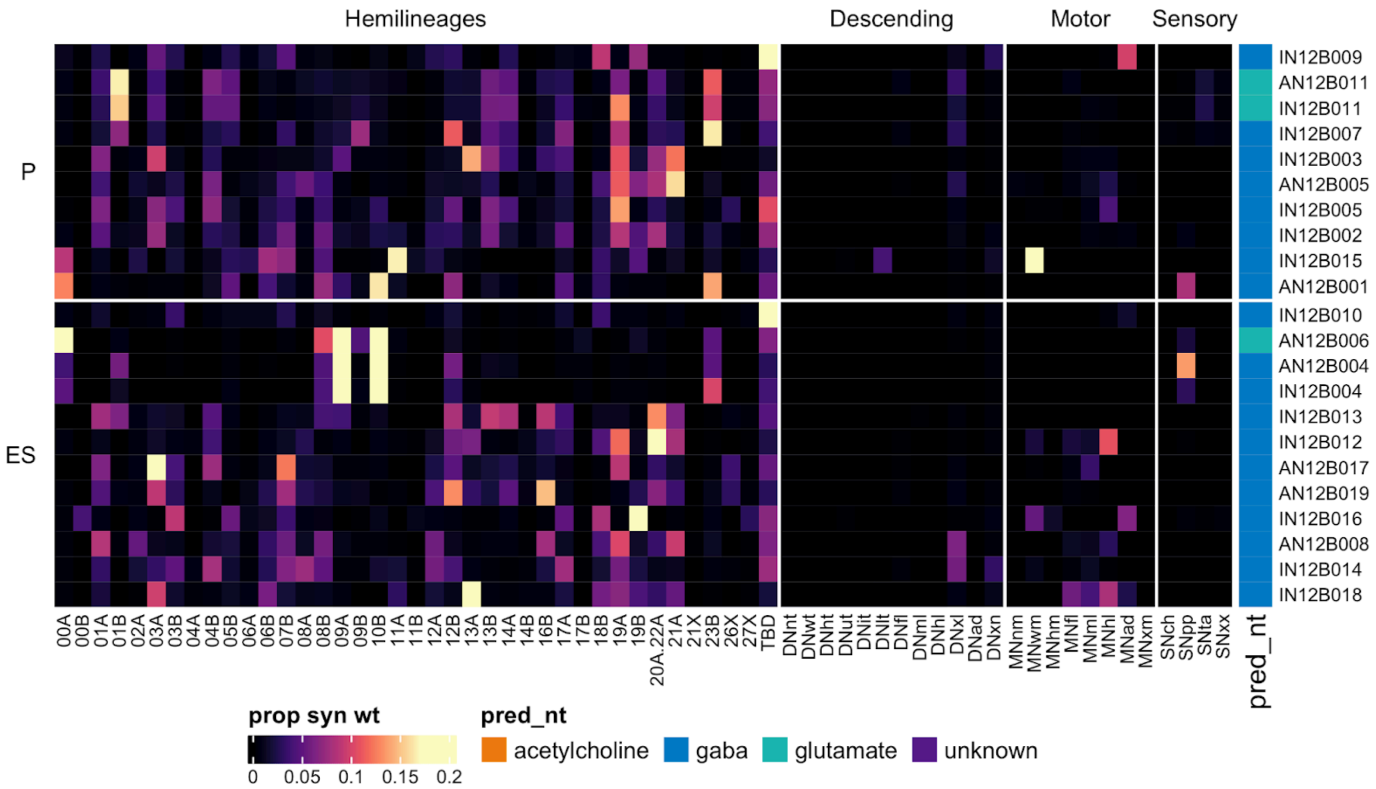

**Figure 34 - figure supplement 7. Connectivity to downstream partners by 12B primary and early secondary systematic types.** Proportions of synaptic weight from systematic types to downstream partners, normalised by row. 12B neurons have been clustered within each assigned birthtime window (P = primary, ES = early secondary, S = secondary) based on both upstream and downstream connectivity to hemilineages, descending neuron subclasses, motor neuron subclasses, and sensory neuron modalities. The annotation bar is coloured by the most common predicted neurotransmitter within each type.

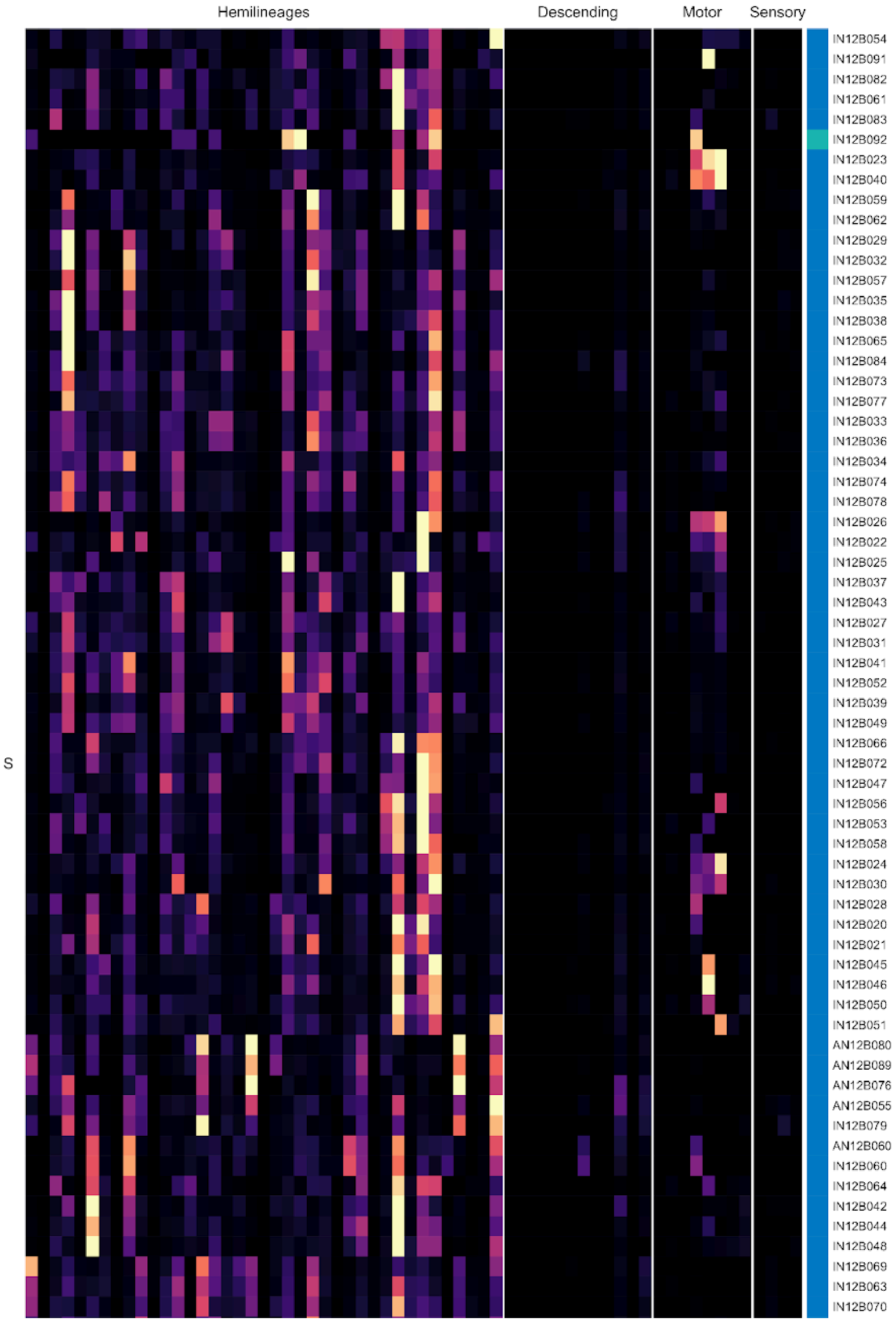

Hemilineages
Descending
Motor
Sensory
S
IN12B054
IN12B091
IN12B082
IN12B061
IN12B083
IN12B092
IN12B023
IN12B040
IN12B059
IN12B062
IN12B029
IN12B032
IN12B057
IN12B035
IN12B038
IN12B065
IN12B084
IN12B073
IN12B077
IN12B033
IN12B036
IN12B034
IN12B074
IN12B078
IN12B026
IN12B022
IN12B025
IN12B037
IN12B043
IN12B027
IN12B031
IN12B041
IN12B052
IN12B039
IN12B049
IN12B066
IN12B072
IN12B047
IN12B056
IN12B053
IN12B058
IN12B024
IN12B030
IN12B028
IN12B020
IN12B021
IN12B045
IN12B046
IN12B050
IN12B051
AN12B080
AN12B089
AN12B076
AN12B055
IN12B079
AN12B060
IN12B060
IN12B064
IN12B042
IN12B044
IN12B048
IN12B069
IN12B063
IN12B070

**Figure 34 - figure supplement 8. Connectivity to downstream partners by 12B secondary systematic types.** Proportions of synaptic weight from systematic types to downstream partners, normalised by row. 12B neurons have been clustered within each assigned birthtime window (P = primary, ES = early secondary, S = secondary) based on both upstream and downstream connectivity to hemilineages, descending neuron subclasses, motor neuron subclasses, and sensory neuron modalities. The annotation bar is coloured by the most common predicted neurotransmitter for the neurons of each type.

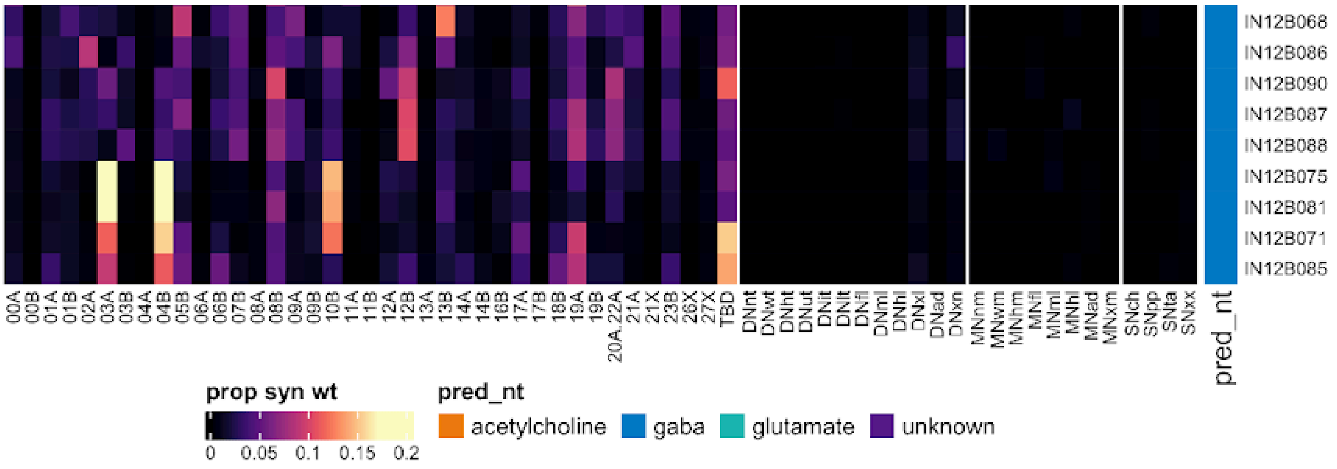

**Figure 34 - figure supplement 9. Connectivity to downstream partners by 12B secondary systematic types, continued.** Proportions of synaptic weight from systematic types to downstream partners, normalised by row. 12B neurons have been clustered within each assigned birthtime window (P = primary, ES = early secondary, S = secondary) based on both upstream and downstream connectivity to hemilineages, descending neuron subclasses, motor neuron subclasses, and sensory neuron modalities. The annotation bar is coloured by the most common predicted neurotransmitter for the neurons of each type.

Hemilineage 13A

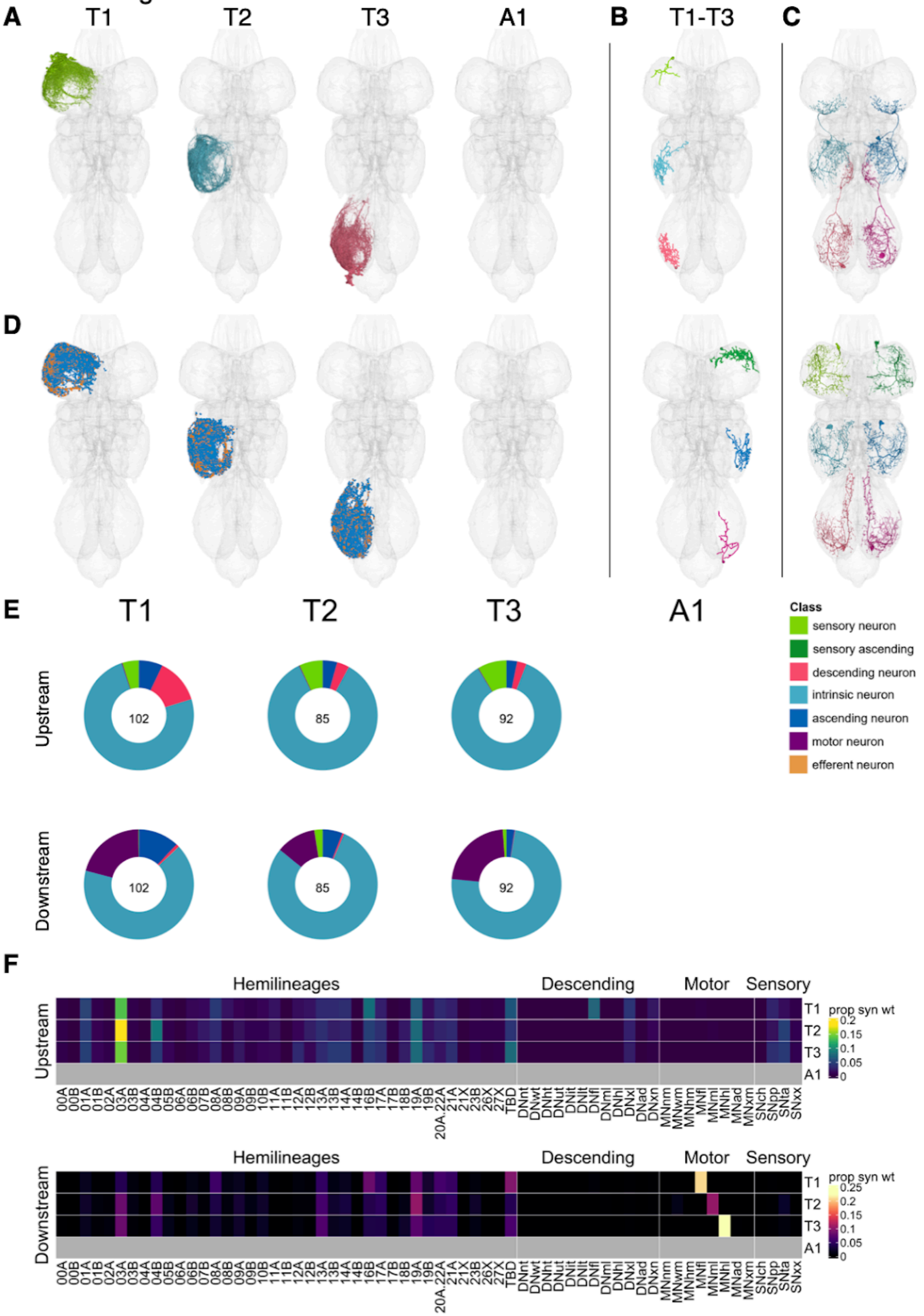

**Figure 35. Hemilineage 13A. A.** Meshes of all RHS secondary neurons plotted in neuromere-specific colours. **B.** “Representative” secondary neuron skeletons plotted in hemineuromere-specific colours. The skeleton with the top accumulated NBLAST score among all neurons from the hemilineage in a given hemineuromere was used. **C.** Neuron meshes of selected examples. Top: sequential serial set 10655. Bottom: independent leg serial set 11077. **D.** Predicted synapses of RHS secondary neurons. Blue: postsynapses; dark orange: presynapses. **E.** Proportions of connections from secondary neurons to upstream or downstream partners, normalised by neuromere and coloured by broad class. Numbers of query neurons appear in the centre. **F.** Proportions of synaptic weight from secondary neurons originating in each neuromere to upstream or downstream partners, normalised by row.

#### Hemilineage 13A

Hemilineages 13A and 13B are believed to derive from neuroblast NB4-2 (Lacin and Truman, 2016) (but see also (Birkholz et al., 2015; Truman et al., 2004)), which produces 4 MNs and ~20 local interneurons in the embryo (Schmid et al., 1999). 13A secondary neurons survive in roughly comparable numbers across the thoracic neuromeres (Figure 35E). Their primary neurites spread over the VNC surface and only converge upon neuropil entry, and their processes are largely restricted to the ipsilateral leg neuropil of origin (Figure 35A,C bottom). We distinguished 13A neurons from other anterior ipsilateral leg neuropil hemilineages by their characteristic subpopulation of neurons projecting medially and dorsally (Shepherd et al., 2019).

13A neurons receive input predominantly from hemilineage 03A as well as from 04B, 16B, and 19A, from descending neurons targeting the front leg or all leg neuropils and from proprioceptive and tactile sensory neurons (Figure 35F). They are predicted to be gabaergic, as expected (Lacin et al., 2019), and provide outputs mainly to 16B in T1 and to 03A, 04B, 13A, and 19A in T2-T3. They also inhibit front and hind leg motor neurons and to a lesser extent middle leg motor neurons. No functional studies have been published for secondary 13A neurons.

Most 13A types target leg motor neurons, but we found several types that inhibit wing motor neurons instead (e.g., IN13A013 and IN13A022) (Figure 35 - figure supplement 6). We also identified one early born sequential serial type, IN13A004 (Figure 35C top), that ascends to the next anterior neuromere and clusters by connectivity with type IN13A007, receiving direct tactile inputs and strongly targeting neurons from hemilineage 23B (Figure 35 - figure supplement 4-6).

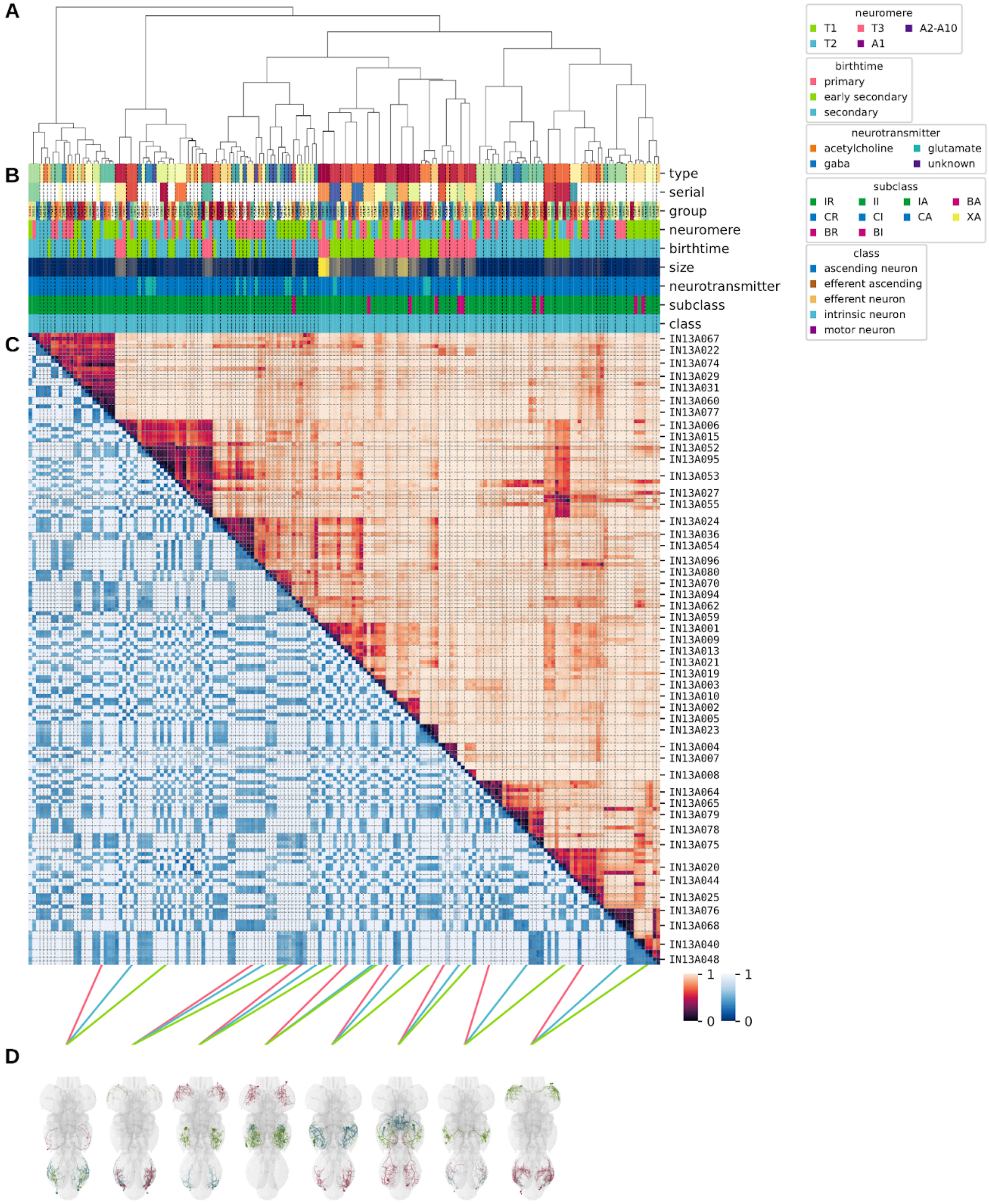

**Figure 35 - figure supplement 1. Systematic typing of hemilineage 13A. A.** Hierarchical clustering dendrogram of hemilineage groups by laterally and serially aggregated connectivity cosine clustering. **B.** Categorical annotations of each hemilineage group, each column corresponding to the aligned leaf in A. Colours for type, serial set, and group are arbitrary for visualisation. Colours for neuromere, birthtime, neurotransmitter, subclass, and class are as in all other figures. **C.** Similarity distance heatmap for hemilineage. Cosine distance is in the upper triangle, while laterally symmetrised NBLAST distance is in the lower triangle. Systematic type names of some types are labelled. **D.** Morphologically representative groups from dendrogram subtrees. Each group, indicated by colour and line

connecting to its column in B and C, is the most morphologically representative group (medoid of NBLAST distance) from a subtree of A. The subtrees (flat clusters) are equal height cuts of A determined to yield the number of groups per plot and plots in D.

#### 13A systematic types

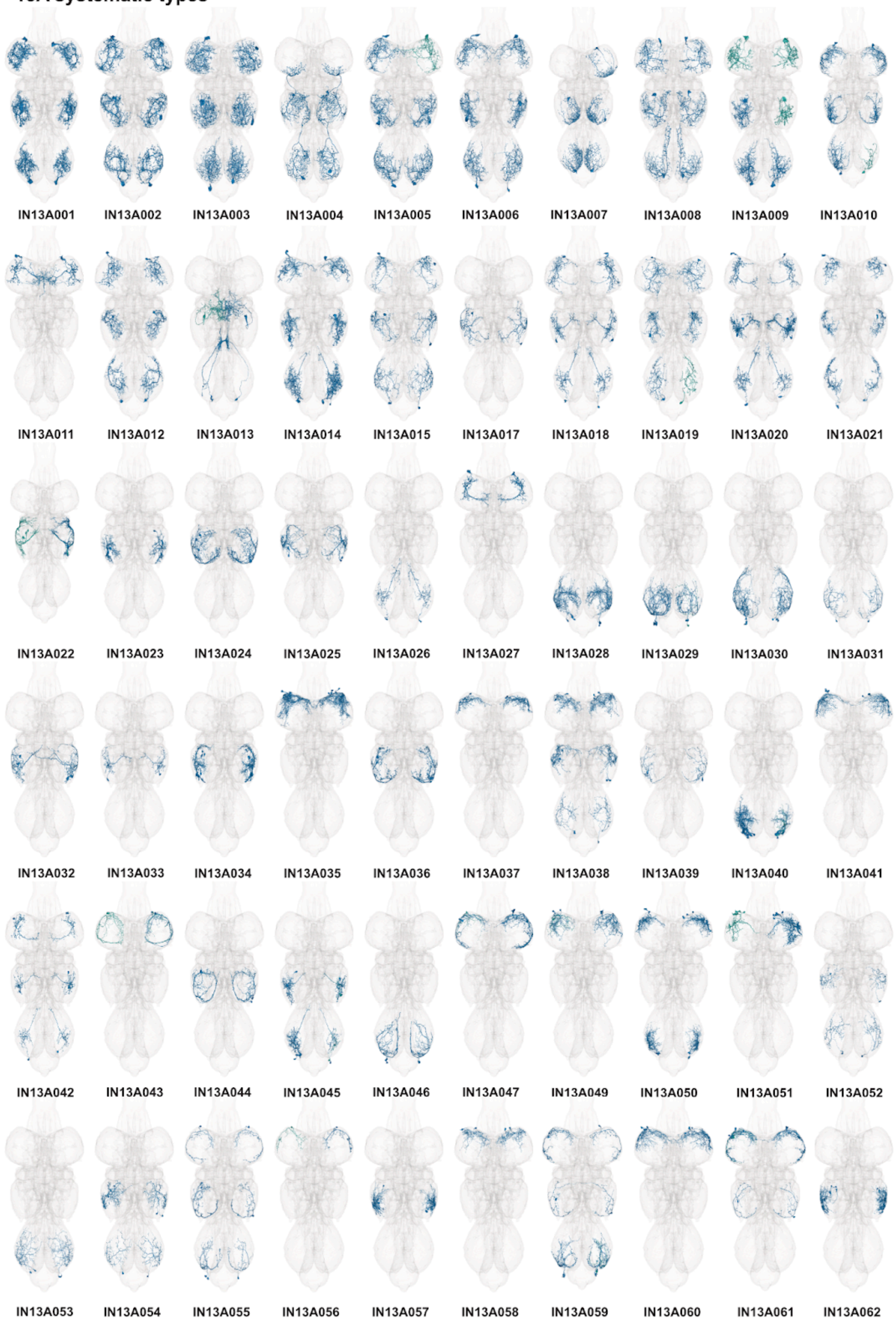

**Figure 35 - figure supplement 2. Systematic types of hemilineage 13A.** Systematic types have been arranged in numerical order, with neurons of the same type that belong to distinct classes (e.g., intrinsic neuron vs ascending neuron) plotted separately but placed adjacent to each other. Individual neuron meshes have been coloured based on predicted neurotransmitter: dark orange = acetylcholine, blue = gaba, marine = glutamate, dark purple = unknown.

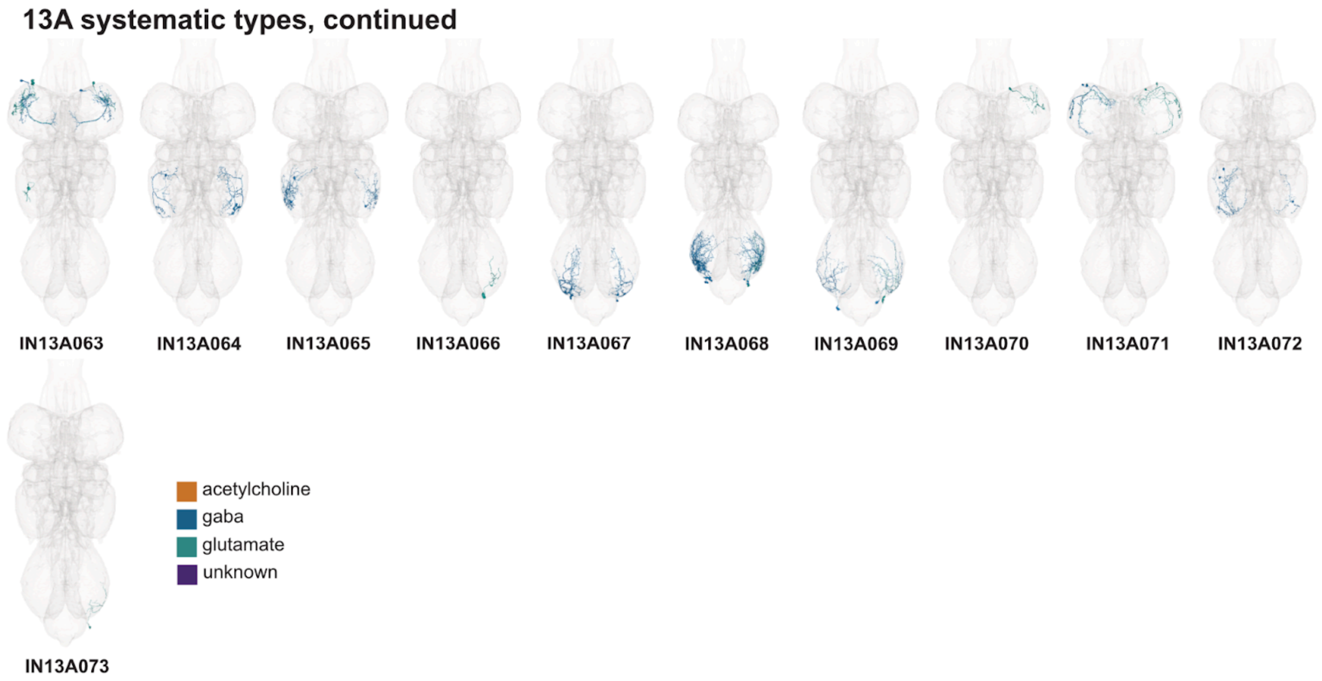

**Figure 35 - figure supplement 3. Systematic types of hemilineage 13A, continued.** Systematic types have been arranged in numerical order, with neurons of the same type that belong to distinct classes (e.g., intrinsic neuron vs ascending neuron) plotted separately but placed adjacent to each other. Individual neuron meshes have been coloured based on predicted neurotransmitter: dark orange = acetylcholine, blue = gaba, marine = glutamate, dark purple = unknown.

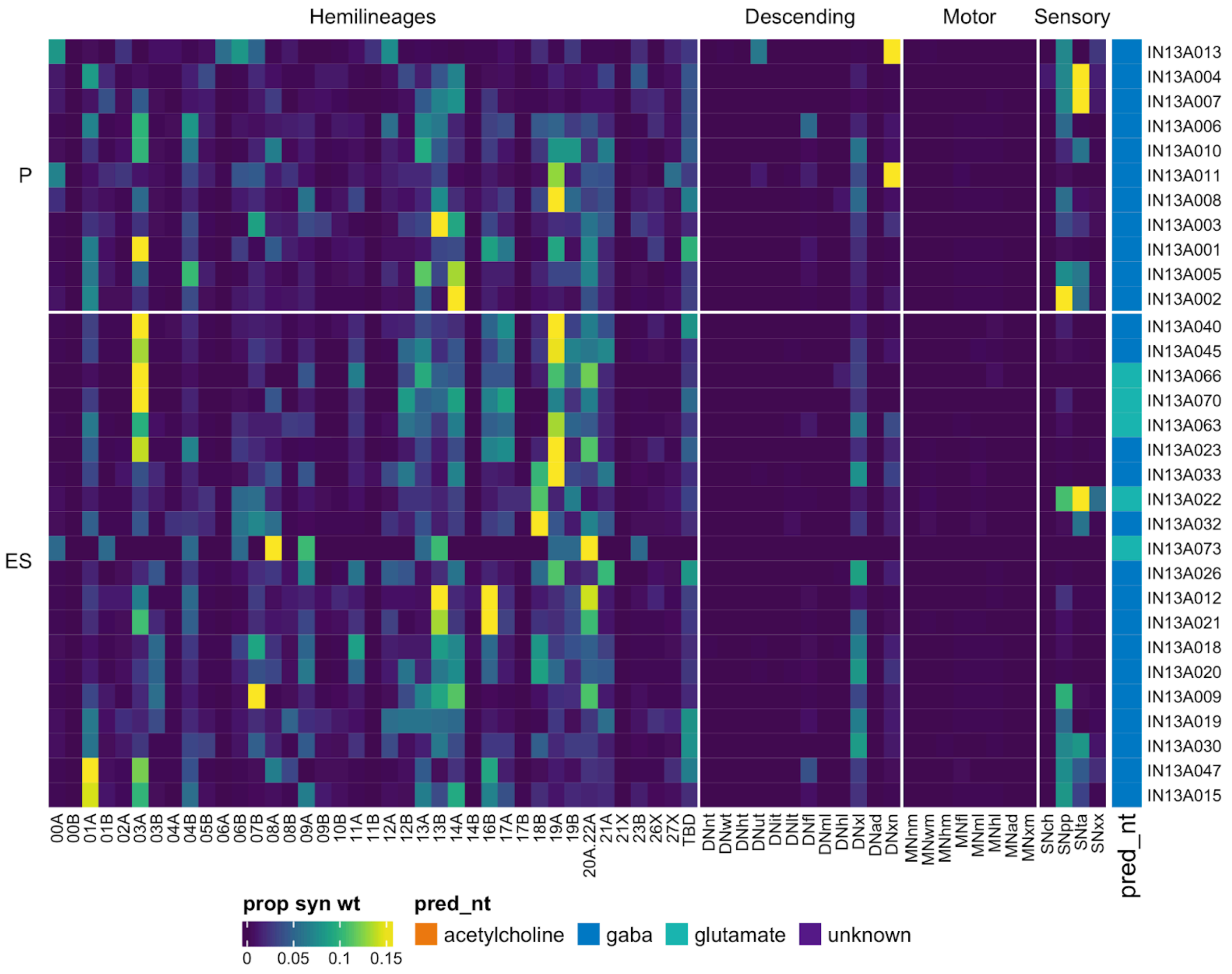

**Figure 35 - figure supplement 4. Connectivity to upstream partners by 13A primary and early secondary systematic types.** Proportions of synaptic weight to systematic types from upstream partners, normalised by row. 13A neurons have been clustered within each assigned birthtime window (P = primary, ES = early secondary, S = secondary) based on both upstream and downstream connectivity to hemilineages, descending neuron subclasses, motor neuron subclasses, and sensory neuron modalities. Annotation bar is coloured by the most common predicted neurotransmitter for the neurons of each type.

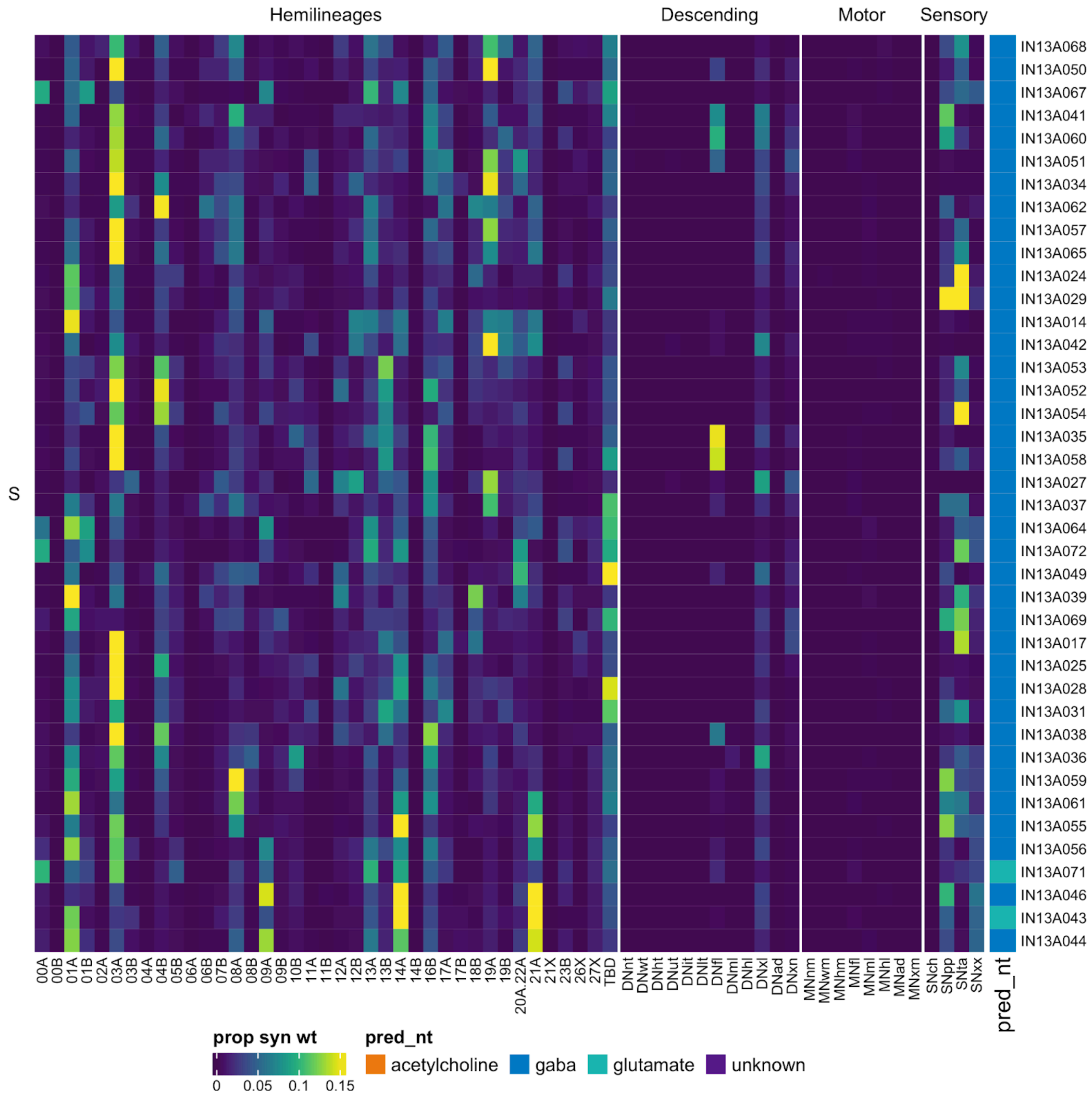

**Figure 35 - figure supplement 5. Connectivity to upstream partners by 13A secondary systematic types.** Proportions of synaptic weight to systematic types from upstream partners, normalised by row. 13A neurons have been clustered within each assigned birthtime window (P = primary, ES = early secondary, S = secondary) based on both upstream and downstream connectivity to hemilineages, descending neuron subclasses, motor neuron subclasses, and sensory neuron modalities. The annotation bar is coloured by the most common predicted neurotransmitter within each type.

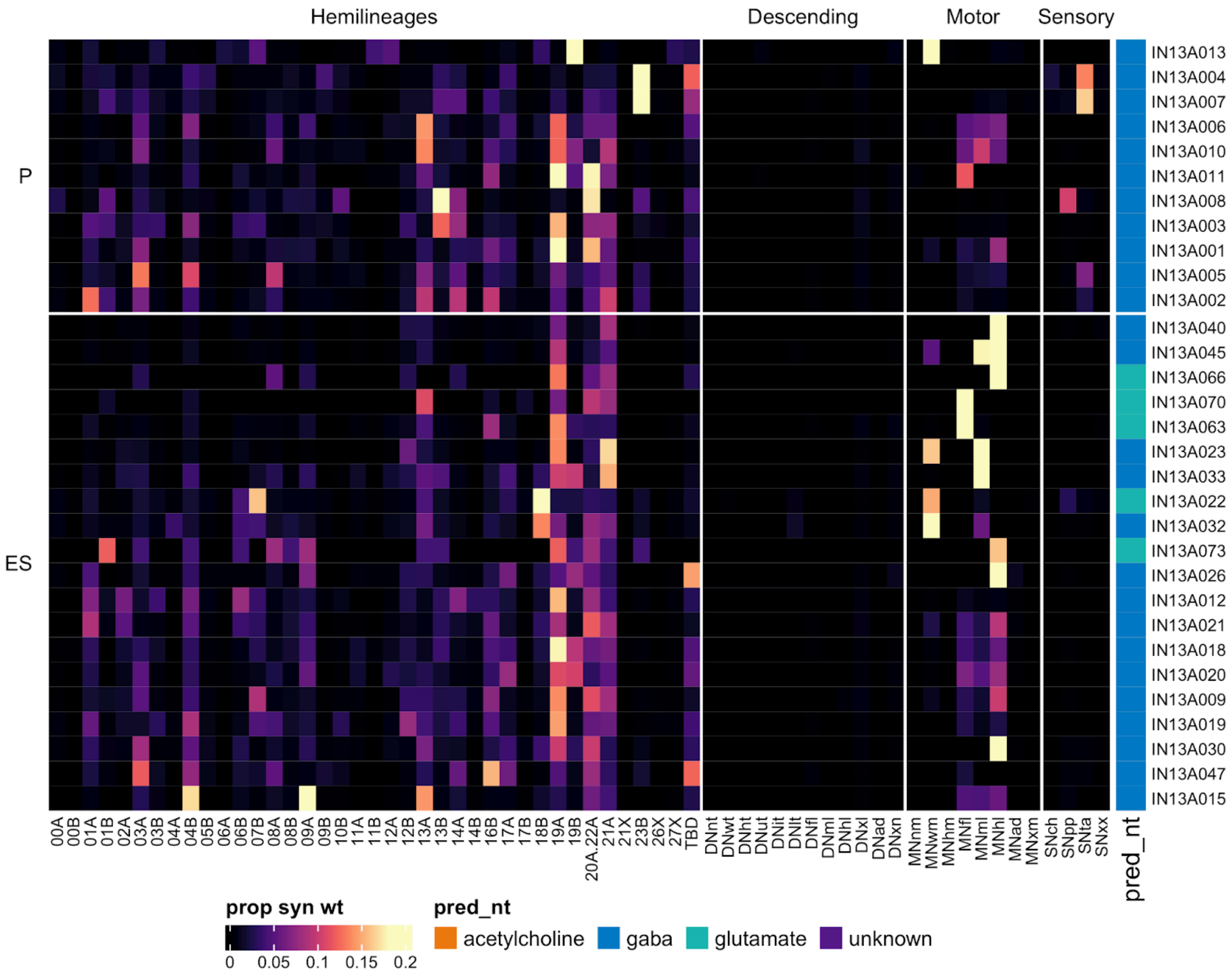

**Figure 35 - figure supplement 6. Connectivity to downstream partners by 13A primary and early secondary systematic types.** Proportions of synaptic weight from systematic types to downstream partners, normalised by row. 13A neurons have been clustered within each assigned birthtime window (P = primary, ES = early secondary, S = secondary) based on both upstream and downstream connectivity to hemilineages, descending neuron subclasses, motor neuron subclasses, and sensory neuron modalities. The annotation bar is coloured by the most common predicted neurotransmitter within each type.

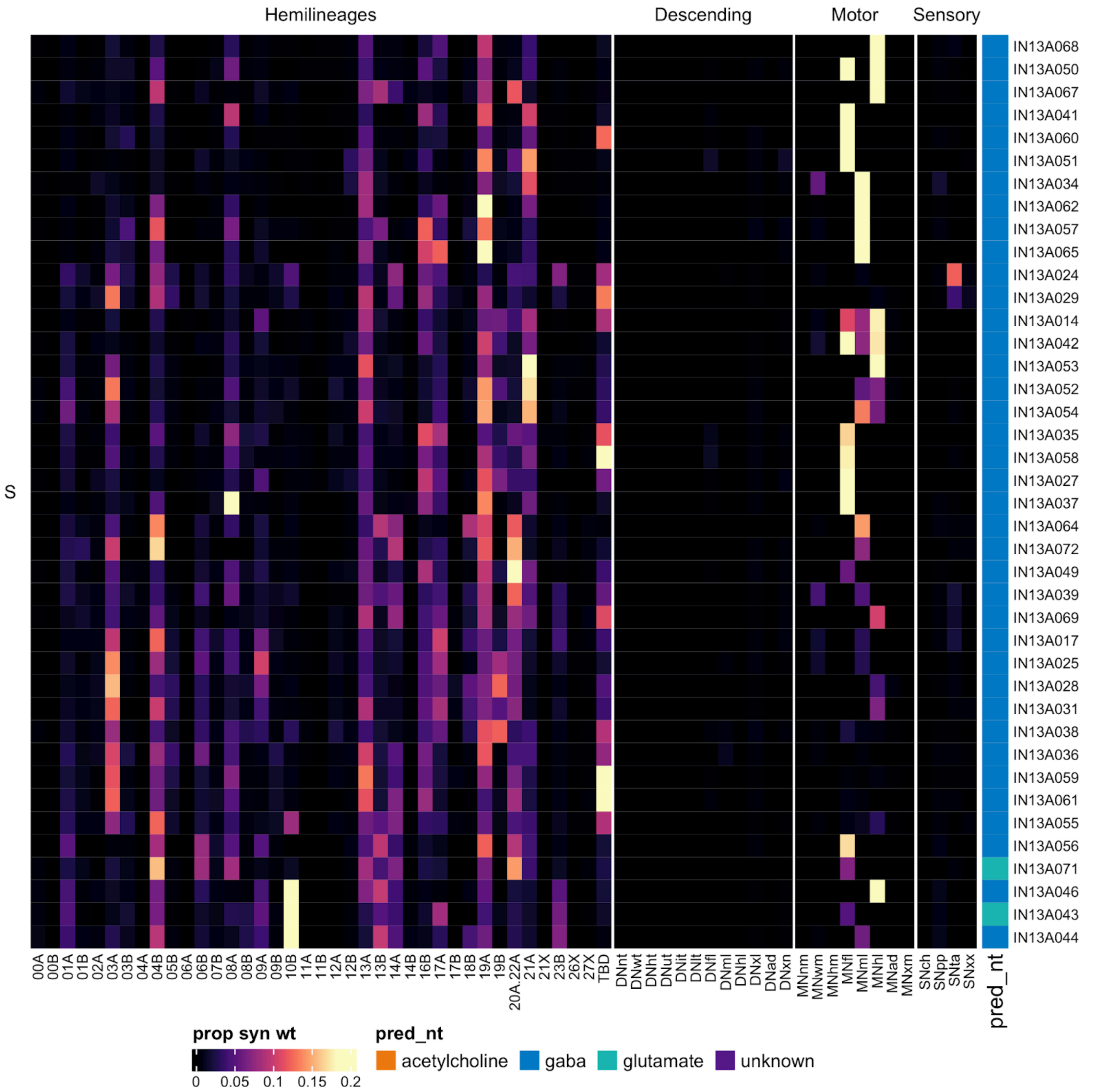

**Figure 35 - figure supplement 7. Connectivity to downstream partners by 13A secondary systematic types.** Proportions of synaptic weight from systematic types to downstream partners, normalised by row. 13A neurons have been clustered within each assigned birthtime window (P = primary, ES = early secondary, S = secondary) based on both upstream and downstream connectivity to hemilineages, descending neuron subclasses, motor neuron subclasses, and sensory neuron modalities. The annotation bar is coloured by the most common predicted neurotransmitter for the neurons of each type.

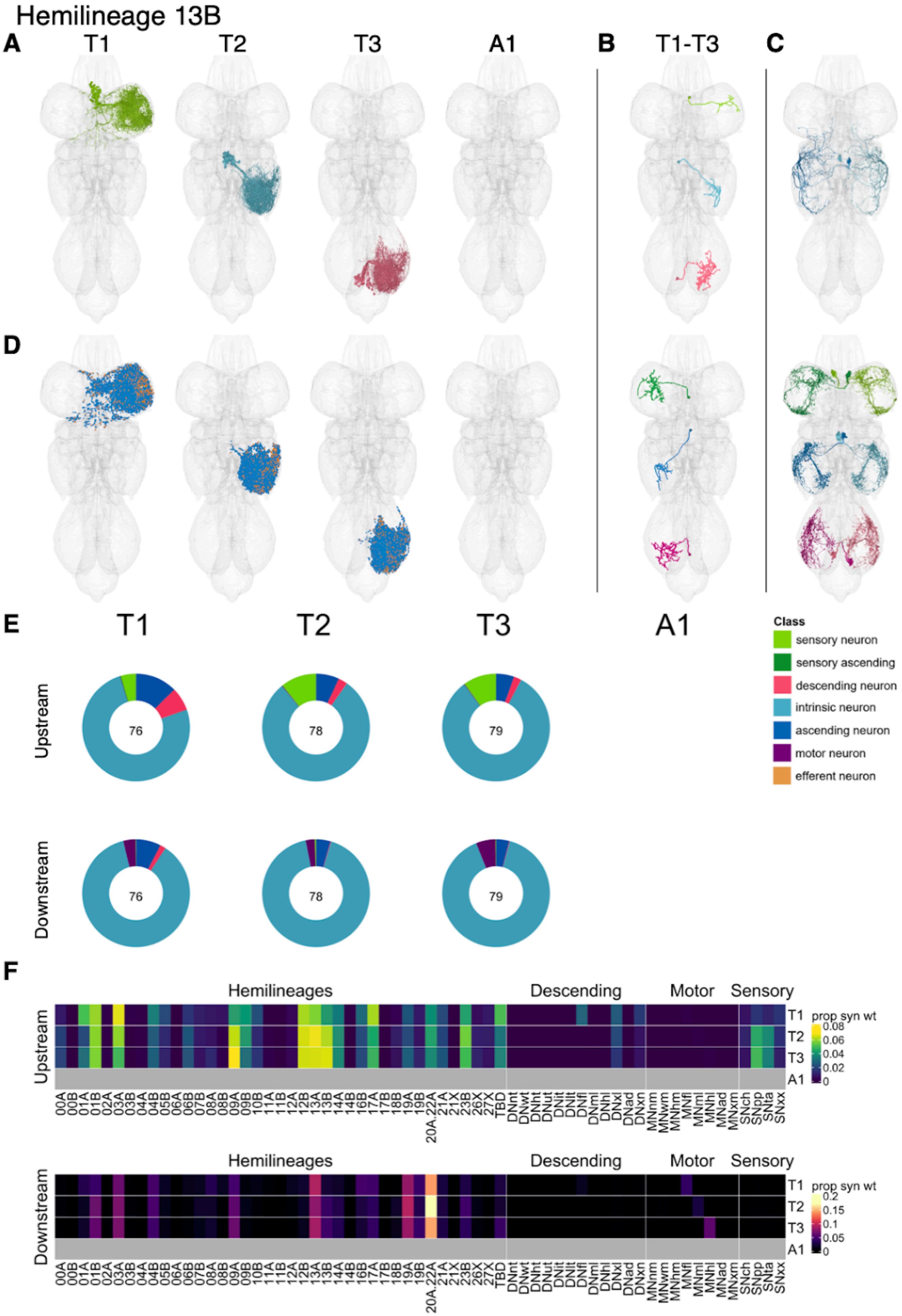

Hemilineage 13B
A
T1
T2
T3
A1
B
T1-T3
C
D
E
T1
T2
T3
A1
Upstream
Downstream
76
78
79
Class
sensory neuron
sensory ascending
descending neuron
intrinsic neuron
ascending neuron
motor neuron
efferent neuron
F
Hemilineages
Descending
Motor
Sensory
prop syn wt

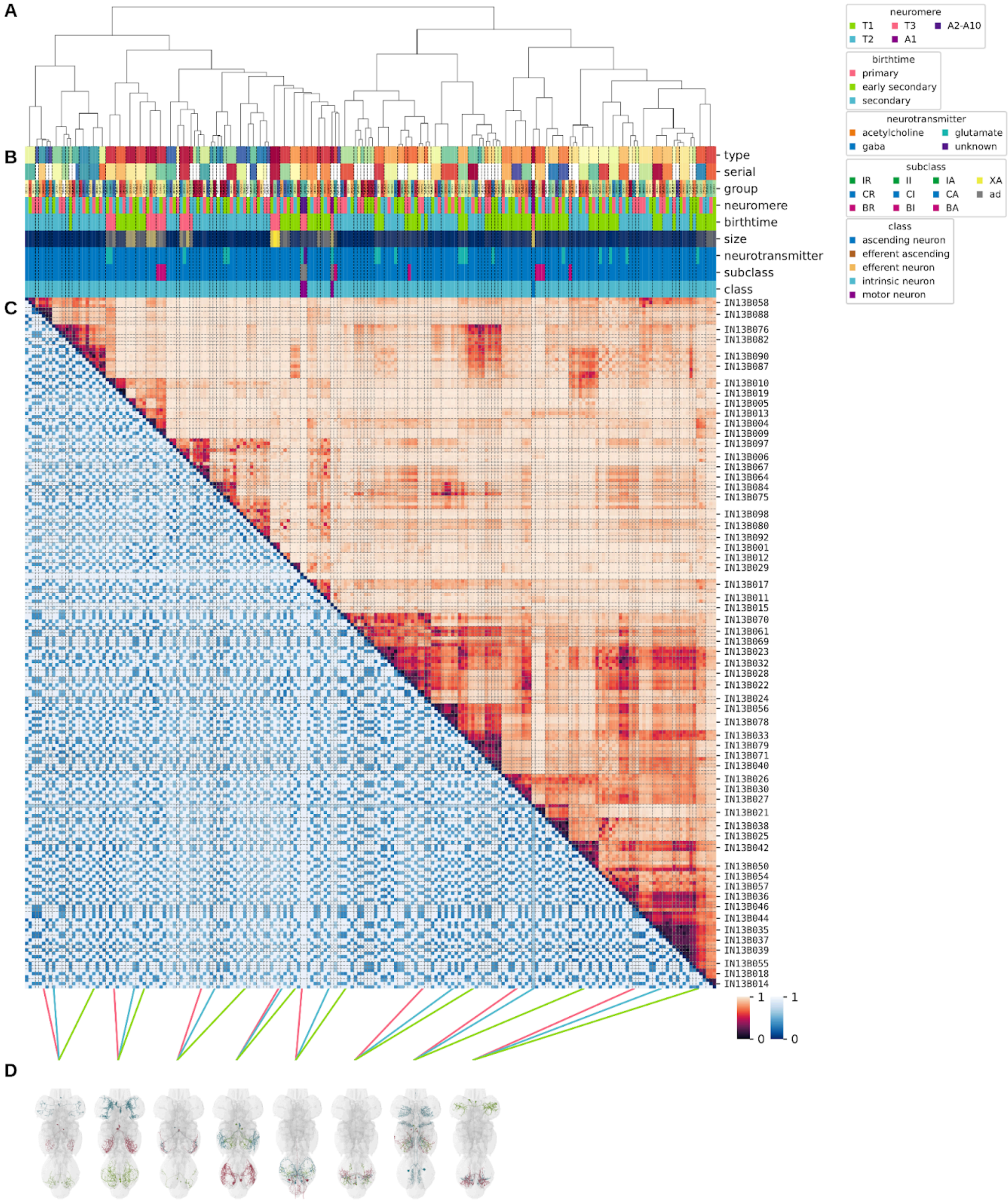

**Figure 36 - figure supplement 1. Systematic typing of hemilineage 13B. A.** Hierarchical clustering dendrogram of hemilineage groups by laterally and serially aggregated connectivity cosine clustering. **B.** Categorical annotations of each hemilineage group, each column corresponding to the aligned leaf in A. Colours for type, serial set, and group are arbitrary for visualisation. Colours for neuromere, birthtime, neurotransmitter, subclass, and class are as in all other figures. **C.** Similarity distance heatmap for hemilineage. Cosine distance is in the upper triangle, while laterally symmetrised NBLAST distance is in the lower triangle. Systematic type names of some types are labelled. **D.** Morphologically representative groups from dendrogram subtrees. Each group, indicated by colour and line connecting to its column in B and C, is the most morphologically representative group (medoid of NBLAST distance)

from a subtree of A. The subtrees (flat clusters) are equal height cuts of A determined to yield the number of groups per plot and plots in D.

#### 13B systematic types

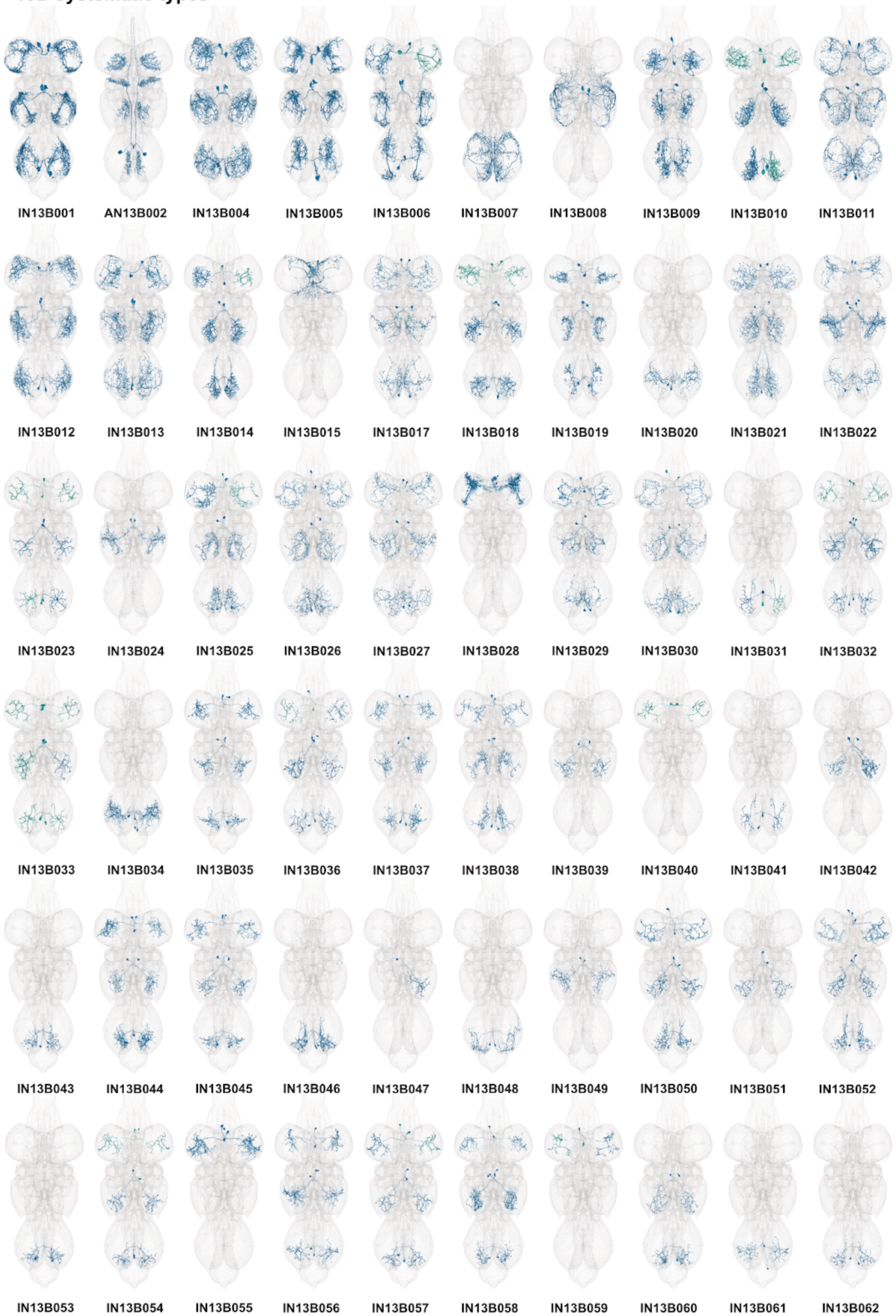

**Figure 36 - figure supplement 2. Systematic types of hemilineage 13B.** Systematic types have been arranged in numerical order, with neurons of the same type that belong to distinct classes (e.g., intrinsic neuron vs ascending neuron) plotted separately but placed adjacent to each other. Individual neuron meshes have been coloured based on predicted neurotransmitter: dark orange = acetylcholine, blue = gaba, marine = glutamate, dark purple = unknown.

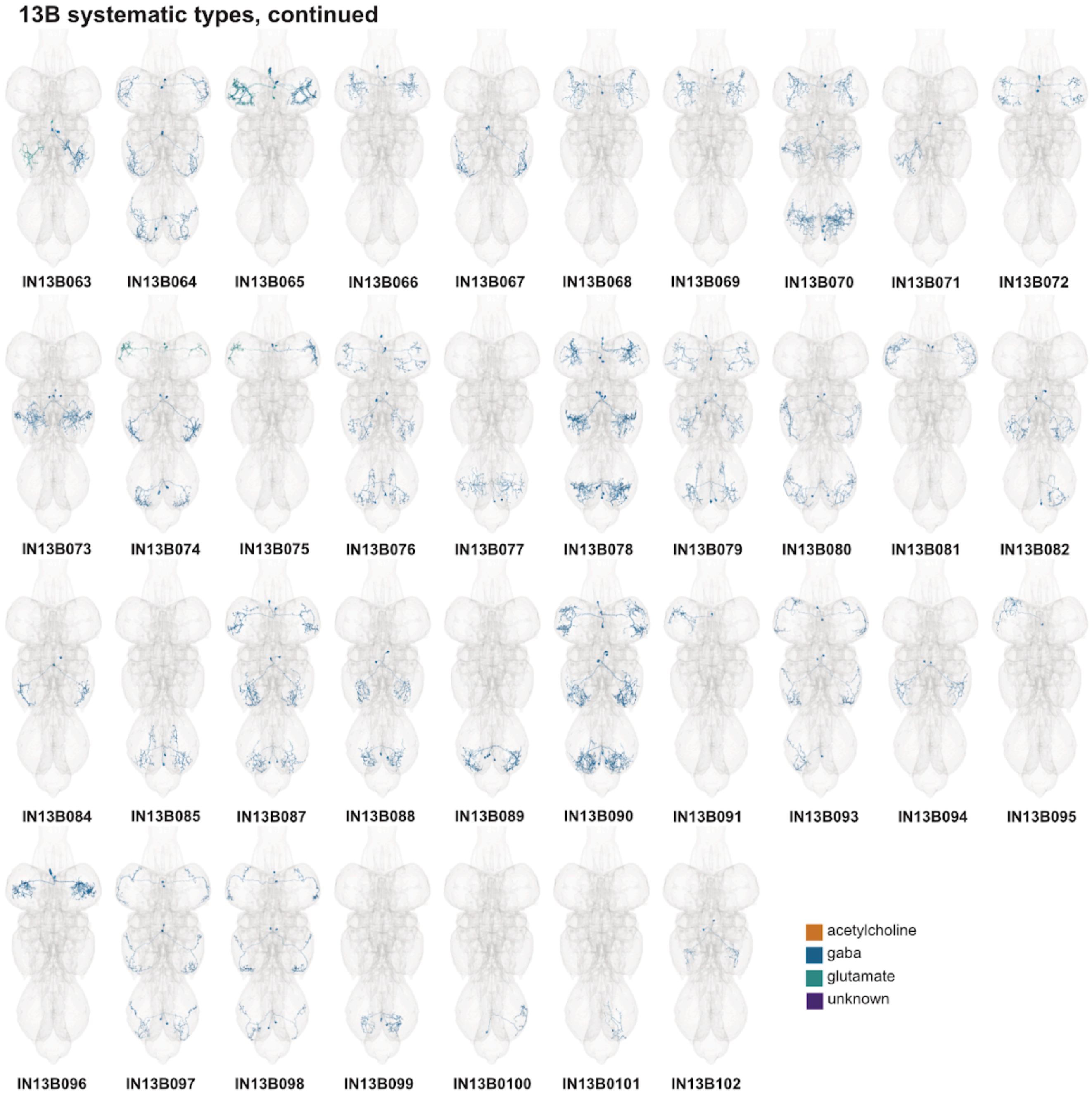

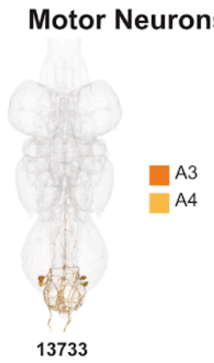

**Figure 36 - figure supplement 3. Systematic types of hemilineage 13B, continued.** Systematic types have been arranged in numerical order, with neurons of the same type that belong to distinct classes (e.g., intrinsic neuron vs ascending neuron) plotted separately but placed adjacent to each other. Individual neuron meshes have been coloured based on predicted neurotransmitter: dark orange = acetylcholine, blue = gaba, marine = glutamate, dark purple = unknown. Motor neurons (typed separately in (Cheong et al., 2023)) have been plotted by serial set if identified in multiple neuromeres and by systematic type if not. Individual motor neuron meshes have been coloured based on soma neuromere: dark orange = A3, dark yellow = A4.

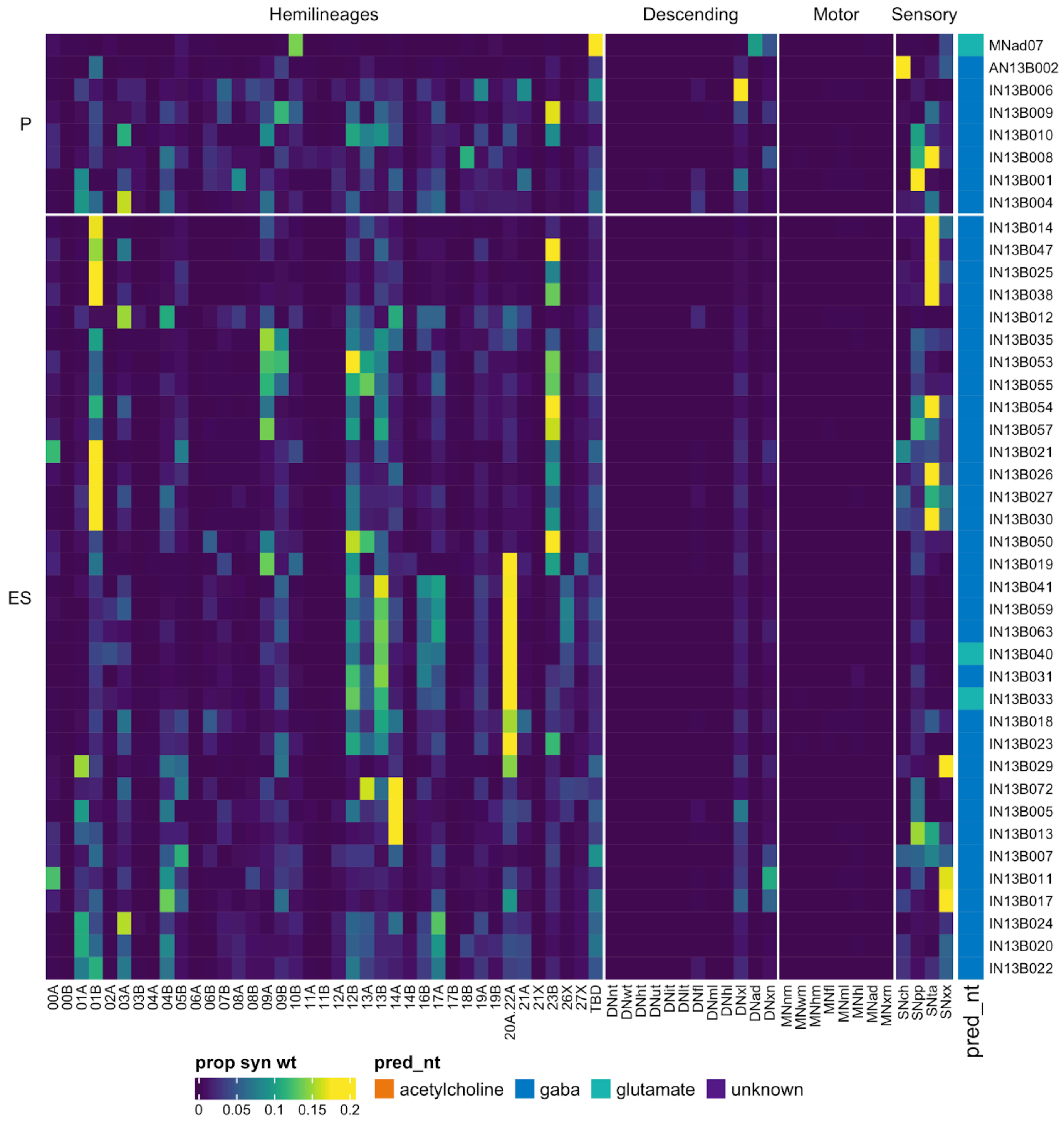

**Figure 36 - figure supplement 4. Connectivity to upstream partners by 13B primary and early secondary systematic types.** Proportions of synaptic weight to systematic types from upstream partners, normalised by row. 13B neurons have been clustered within each assigned birthtime window (P = primary, ES = early secondary, S = secondary) based on both upstream and downstream connectivity to hemilineages, descending neuron subclasses, motor neuron subclasses, and sensory neuron modalities. Annotation bar is coloured by the most common predicted neurotransmitter for the neurons of each type.

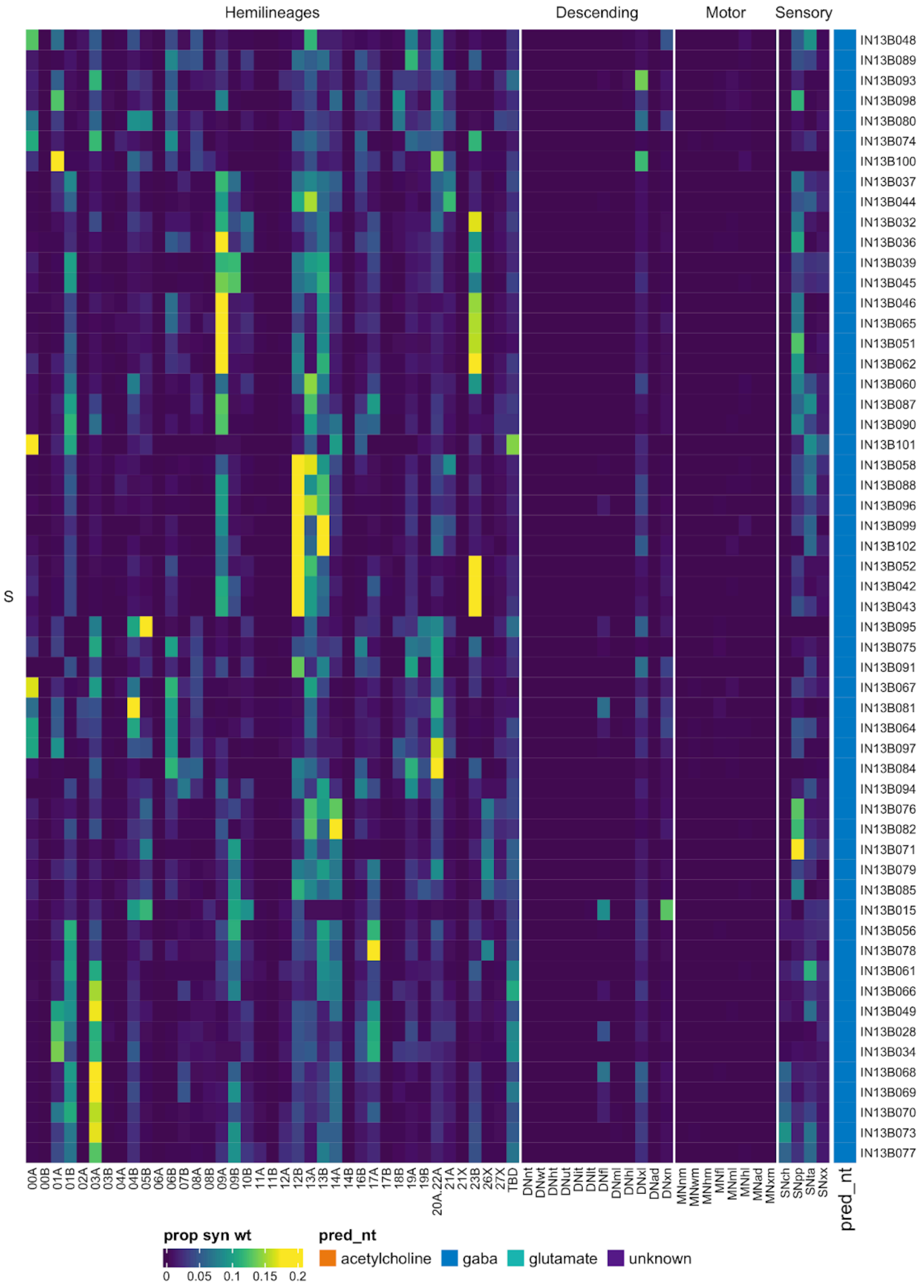

Hemilineages
Descending
Motor
Sensory
S
IN13B048
IN13B089
IN13B093
IN13B098
IN13B080
IN13B074
IN13B100
IN13B037
IN13B044
IN13B032
IN13B036
IN13B039
IN13B045
IN13B046
IN13B065
IN13B051
IN13B062
IN13B060
IN13B087
IN13B090
IN13B101
IN13B058
IN13B088
IN13B096
IN13B099
IN13B102
IN13B052
IN13B042
IN13B043
IN13B095
IN13B075
IN13B091
IN13B067
IN13B081
IN13B064
IN13B097
IN13B084
IN13B094
IN13B076
IN13B082
IN13B071
IN13B079
IN13B085
IN13B015
IN13B056
IN13B078
IN13B061
IN13B066
IN13B049
IN13B028
IN13B034
IN13B068
IN13B069
IN13B070
IN13B073
IN13B077
00A
00B
01A
01B
02A
03A
03B
04A
04B
05B
06A
06B
07B
08A
08B
09A
09B
10B
11A
11B
12A
12B
13A
13B
14A
14B
16B
17A
17B
18B
19A
19B
20A.22A
21A
21X
23B
26X
27X
TBD
DNnt
DNwt
DNht
DNut
DNit
DNlt
DNfl
DNml
DNhl
DNxl
DNad
DNxn
MNnm
MNwm
MNhm
MNfl
MNml
MNhl
MNad
MNxm
SNch
SNpp
SNta
SNxx
pred_nt
prop syn wt
0
0.05
0.1
0.15
0.2
pred_nt
acetylcholine
gaba
glutamate
unknown

**Figure 36 - figure supplement 5. Connectivity to upstream partners by 13B secondary systematic types.** Proportions of synaptic weight to systematic types from upstream partners, normalised by row. 13B neurons have been clustered within each assigned birthtime window (P = primary, ES = early secondary, S = secondary) based on both upstream and downstream connectivity to hemilineages, descending neuron subclasses, motor neuron subclasses, and sensory neuron modalities. The annotation bar is coloured by the most common predicted neurotransmitter within each type.

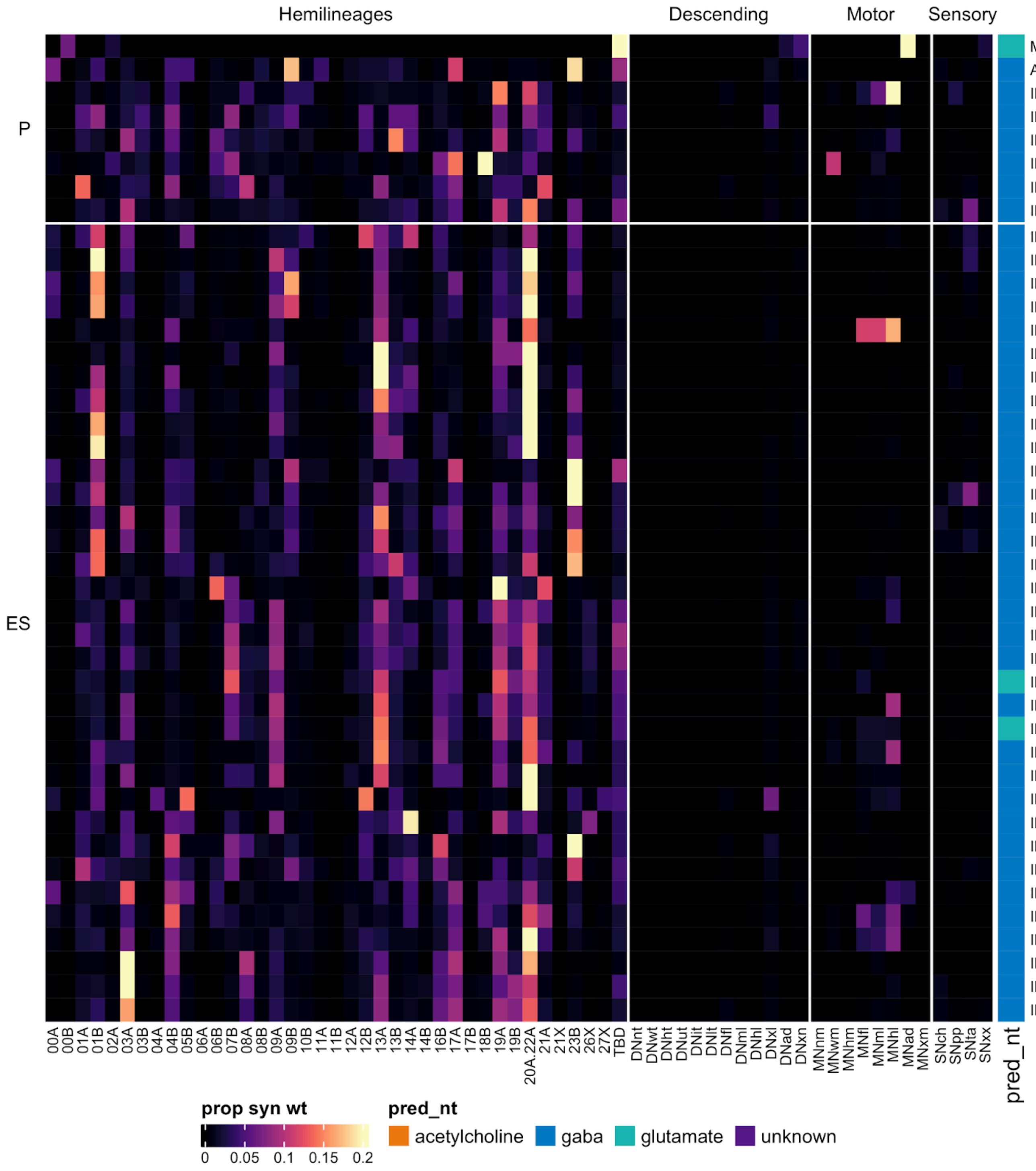

**Figure 36 - figure supplement 6. Connectivity to downstream partners by 13B primary and early secondary systematic types.** Proportions of synaptic weight from systematic types to downstream partners, normalised by row. 13B neurons have been clustered within each assigned birthtime window (P = primary, ES = early secondary, S = secondary) based on both upstream and downstream connectivity to hemilineages, descending neuron subclasses, motor neuron subclasses, and sensory neuron modalities. The annotation bar is coloured by the most common predicted neurotransmitter within each type.

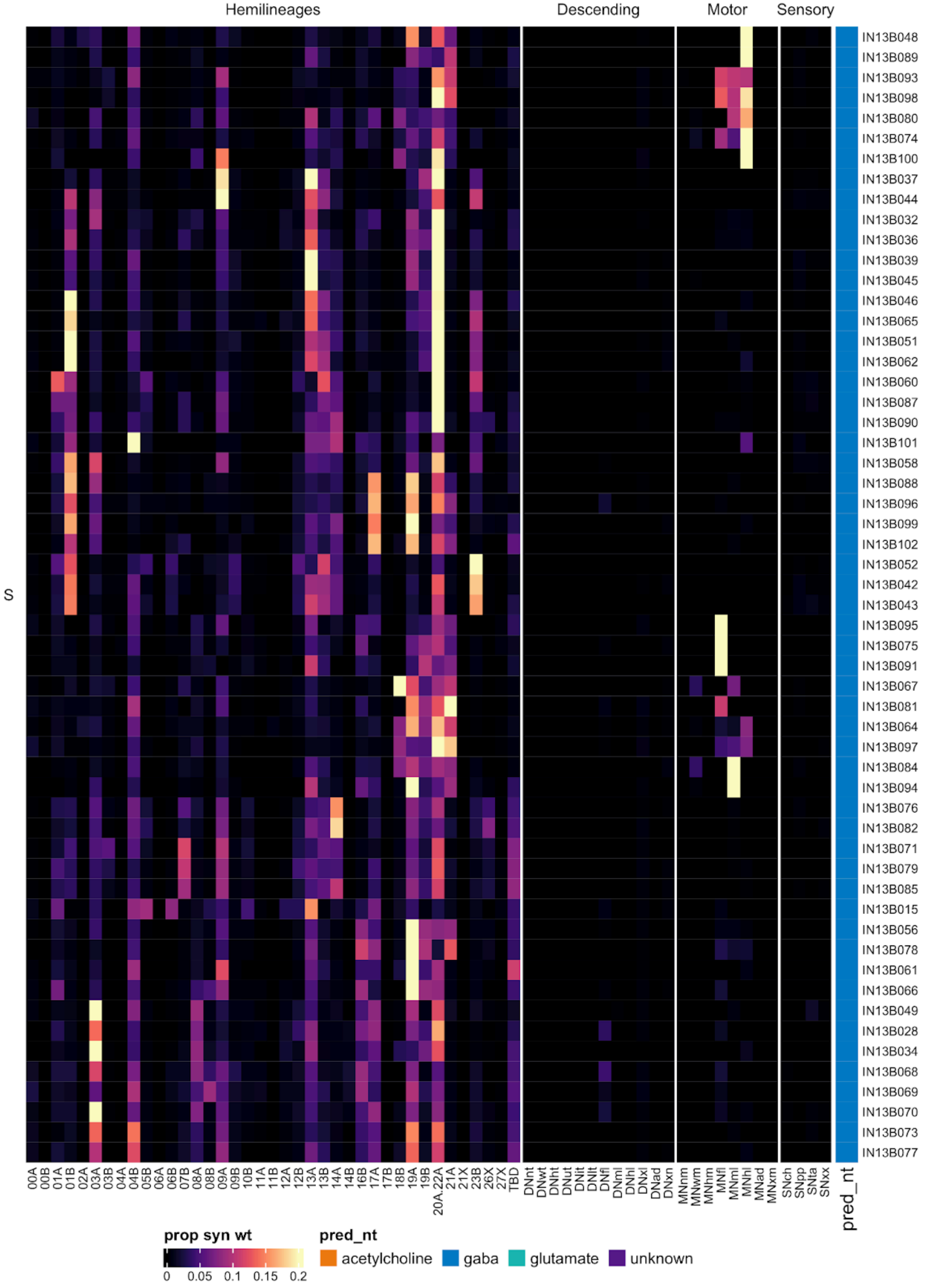
Hemilineages
Descending
Motor
Sensory
S
IN13B048
IN13B089
IN13B093
IN13B098
IN13B080
IN13B074
IN13B100
IN13B037
IN13B044
IN13B032
IN13B036
IN13B039
IN13B045
IN13B046
IN13B065
IN13B051
IN13B062
IN13B060
IN13B087
IN13B090
IN13B101
IN13B058
IN13B088
IN13B096
IN13B099
IN13B102
IN13B052
IN13B042
IN13B043
IN13B095
IN13B075
IN13B091
IN13B067
IN13B081
IN13B064
IN13B097
IN13B084
IN13B094
IN13B076
IN13B082
IN13B071
IN13B079
IN13B085
IN13B015
IN13B056
IN13B078
IN13B061
IN13B066
IN13B049
IN13B028
IN13B034
IN13B068
IN13B069
IN13B070
IN13B073
IN13B077
00A 00B 01A 01B 02A 03A 03B 04A 04B 05B 06A 06B 07B 08A 08B 09A 09B 10B 11A 11B 12A 12B 13A 13B 14A 14B 16B 17A 17B 18B 19A 19B 20A.22A 21A 21X 23B 26X 27X TBD
DNnt DNwt DNht DNut DNit DNlt DNfl DNml DNhl DNxl DNad DNxn
MNnm MNwm MNhm MNfl MNml MNhl MNad MNxm
SNch SNpp SNta SNxx
pred_nt
prop syn wt
0 0.05 0.1 0.15 0.2
pred_nt
acetylcholine
gaba
glutamate
unknown

**Figure 36 - figure supplement 7. Connectivity to downstream partners by 13B secondary systematic types.** Proportions of synaptic weight from systematic types to downstream partners, normalised by row. 13B neurons have been clustered within each assigned birthtime window (P = primary, ES = early secondary, S = secondary) based on both upstream and downstream connectivity to hemilineages, descending neuron subclasses, motor neuron subclasses, and sensory neuron modalities. The annotation bar is coloured by the most common predicted neurotransmitter for the neurons of each type.

Hemilineage 14A

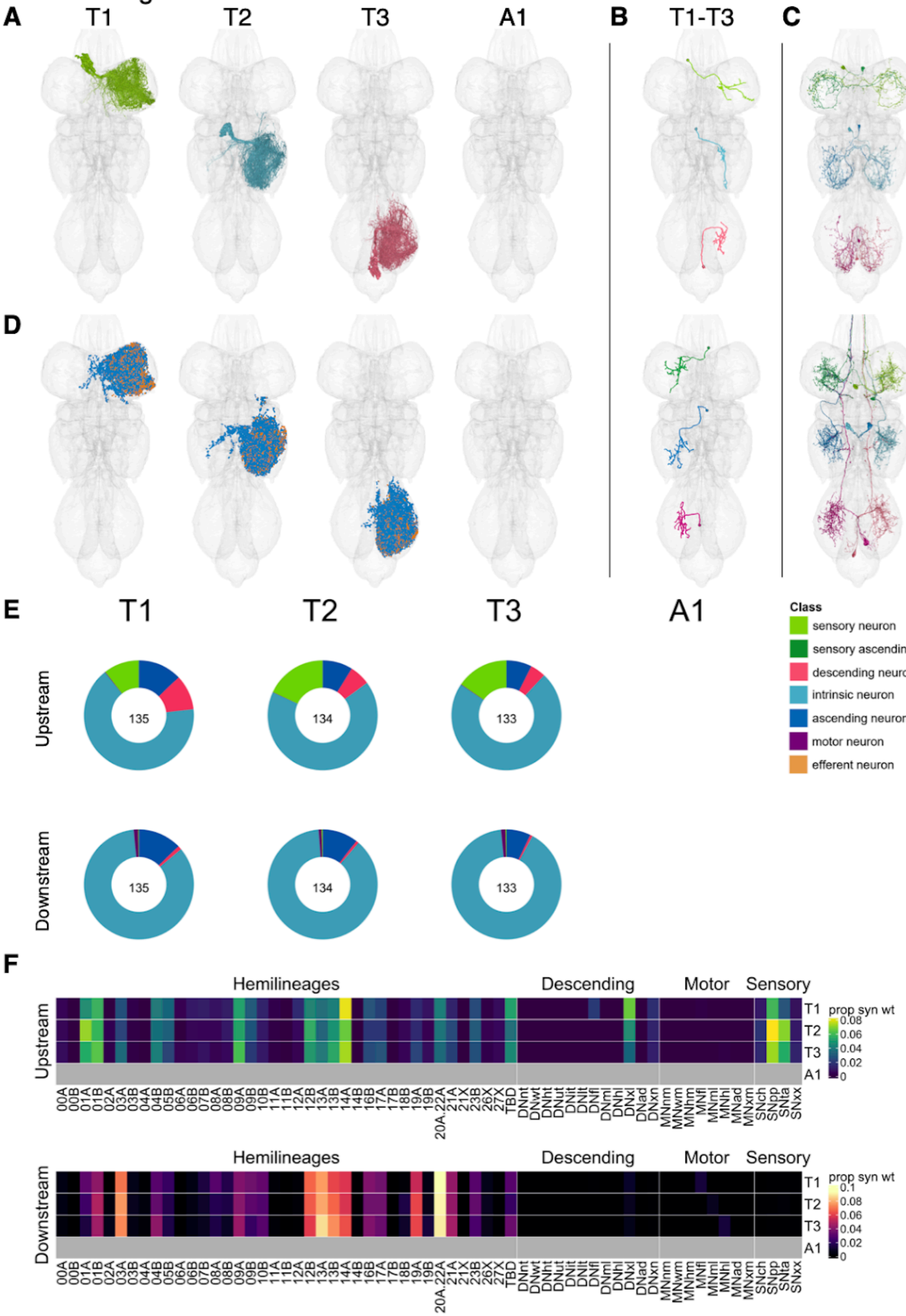

**Figure 37. Hemilineage 14A. A.** Meshes of all RHS secondary neurons plotted in neuromere-specific colours. **B.** “Representative” secondary neuron skeletons plotted in hemineuromere-specific colours. The skeleton with the top accumulated NBLAST score among all neurons from the hemilineage in a given hemineuromere was used. **C.** Neuron meshes of selected examples. Top: independent leg serial set 10019. Bottom: ascending serial set 10411. **D.** Predicted synapses of RHS secondary neurons. Blue: postsynapses; dark orange: presynapses. **E.** Proportions of connections from secondary neurons to upstream or downstream partners, normalised by neuromere and coloured by broad class. Numbers of query neurons appear in the centre. **F.** Proportions of synaptic weight from secondary neurons originating in each neuromere to upstream or downstream partners, normalised by row.

#### Hemilineage 14A

Hemilineages 14A and 14B derive from medial neuroblast NB4-1 (Birkholz et al., 2015; Lacin and Truman, 2016), which generates the motor neuron of the transverse nerve, ~8 intersegmental interneurons, and 20-24 local interneurons in the embryo (Schmid et al., 1999). Hemilineage 14A secondary neurons survive in ~identical numbers across the thoracic neuromeres and cross the midline very ventrally, just anterior to 13B (Shepherd et al., 2016; Truman et al., 2004) (Figure 37A,E). 14A secondary neurons innervate the contralateral leg neuropil, and their somas are sometimes pulled across the midline (Shepherd et al., 2019). We opted to use the soma side of origin (as inferred by morphology) for the instance. Most 14A neurons are restricted to the contralateral leg neuropil of the same neuromere (Figure 37C top), but we identified one ascending serial set of primary neurons (Figure 37C bottom).

14A neurons receive significant inputs from proprioceptive and tactile sensory neurons and descending neurons targeting multiple leg neuropils, as well as from numerous leg hemilineages, especially 01A, 01B, 09A, 12B, 13B, and 14A (Figure 37F). 14A secondary neurons are predicted to be glutamatergic, as expected (Lacin et al., 2019), and provide output to hemilineages 20A/22A and to a lesser extent to 03A, 12B, 13A, 13B, 14A, and 19A. However, we identified a distinctive primary serial set of gabergic neurons (IN14A001) that receives input from 04B, 13A, and proprioceptive neurons and inhibits neurons from 13B and 21A. We also identified a single 14A ascending type (AN14A003); these neurons receive input primarily from 12B and 21A and provide it to 07B and 19A (Figure 38 - figure supplement 2,5,8). No functional studies have been published for secondary 14A neurons.

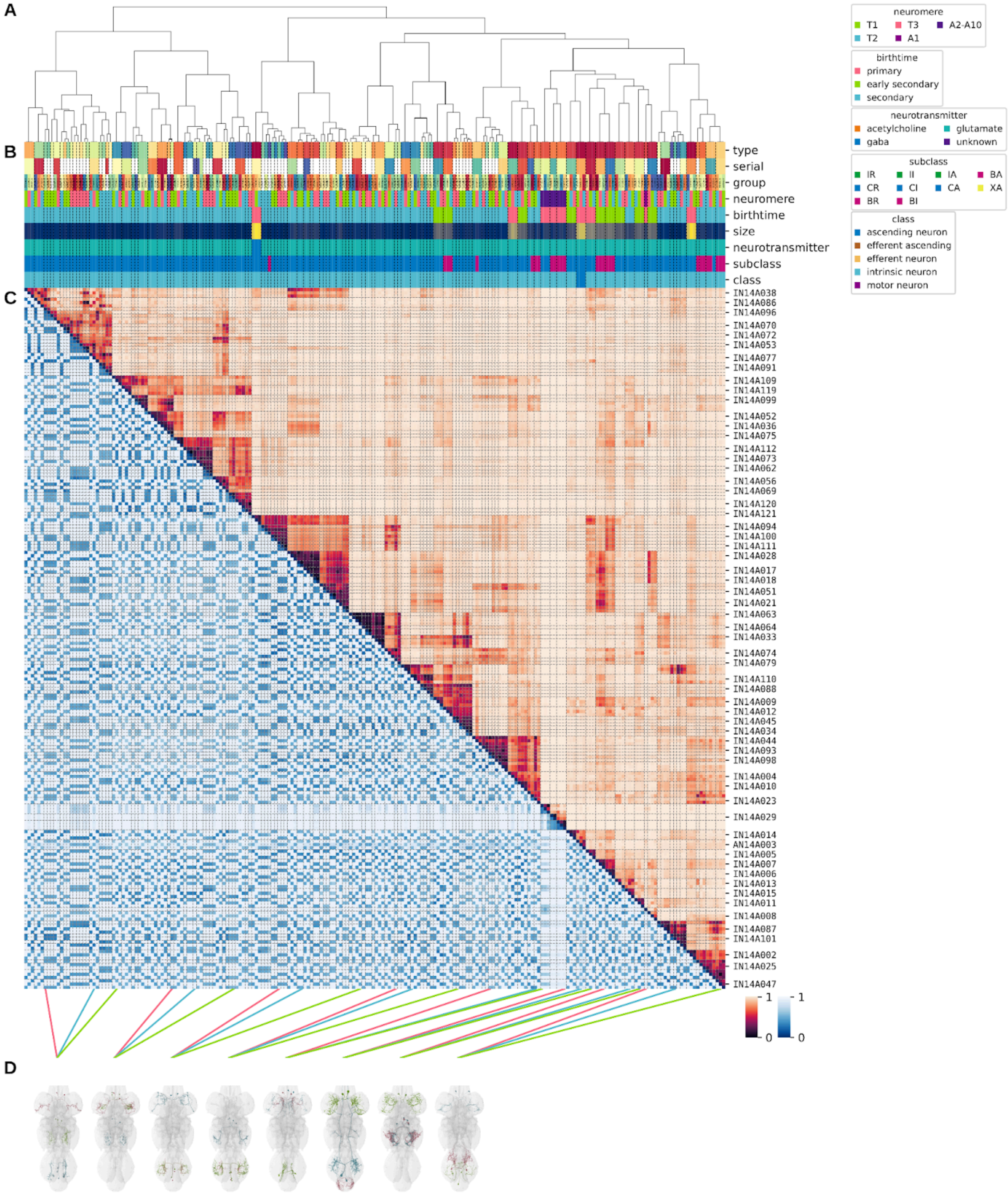

**Figure 37 - figure supplement 1. Systematic typing of hemilineage 14A. A.** Hierarchical clustering dendrogram of hemilineage groups by laterally and serially aggregated connectivity cosine clustering. **B.** Categorical annotations of each hemilineage group, each column corresponding to the aligned leaf in A. Colours for type, serial set, and group are arbitrary for visualisation. Colours for neuromere, birthtime, neurotransmitter, subclass, and class are as in all other figures. **C.** Similarity distance heatmap for hemilineage. Cosine distance is in the upper triangle, while laterally symmetrised NBLAST distance is in the lower triangle. Systematic type names of some types are labelled. **D.** Morphologically representative groups from dendrogram subtrees. Each group, indicated by colour and line connecting to its column in B and C, is the most morphologically representative group (medoid of NBLAST distance) from a subtree of A. The subtrees (flat clusters) are equal height cuts of A determined to yield the number of groups

per plot and plots in D.

#### 14A systematic types

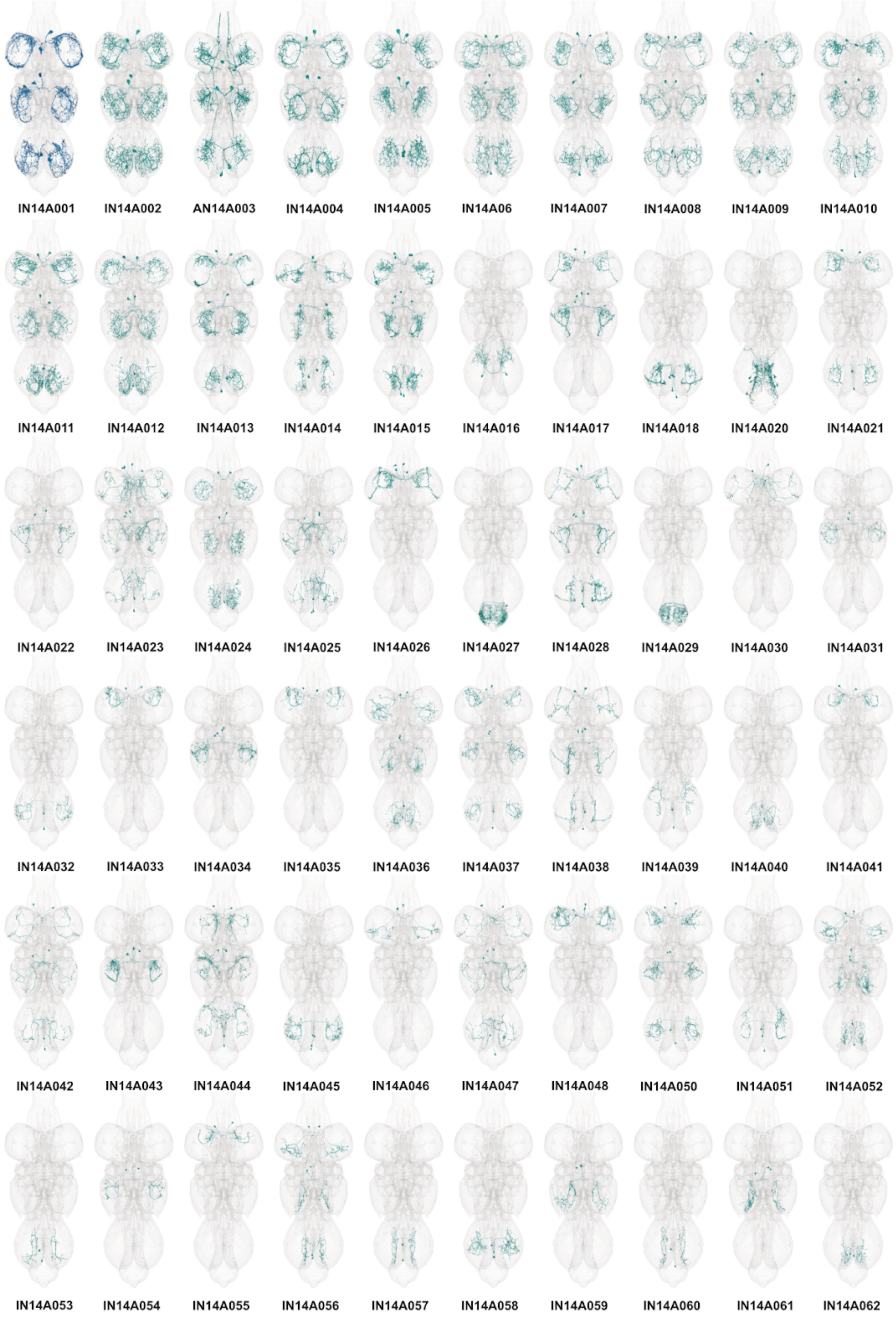

**Figure 37 - figure supplement 2. Systematic types of hemilineage 14A.** Systematic types have been arranged in numerical order, with neurons of the same type that belong to distinct classes (e.g., intrinsic neuron vs ascending neuron) plotted separately but placed adjacent to each other. Individual neuron meshes have been coloured based on predicted neurotransmitter: dark orange = acetylcholine, blue = gaba, marine = glutamate, dark purple = unknown.

#### 14A systematic types, continued

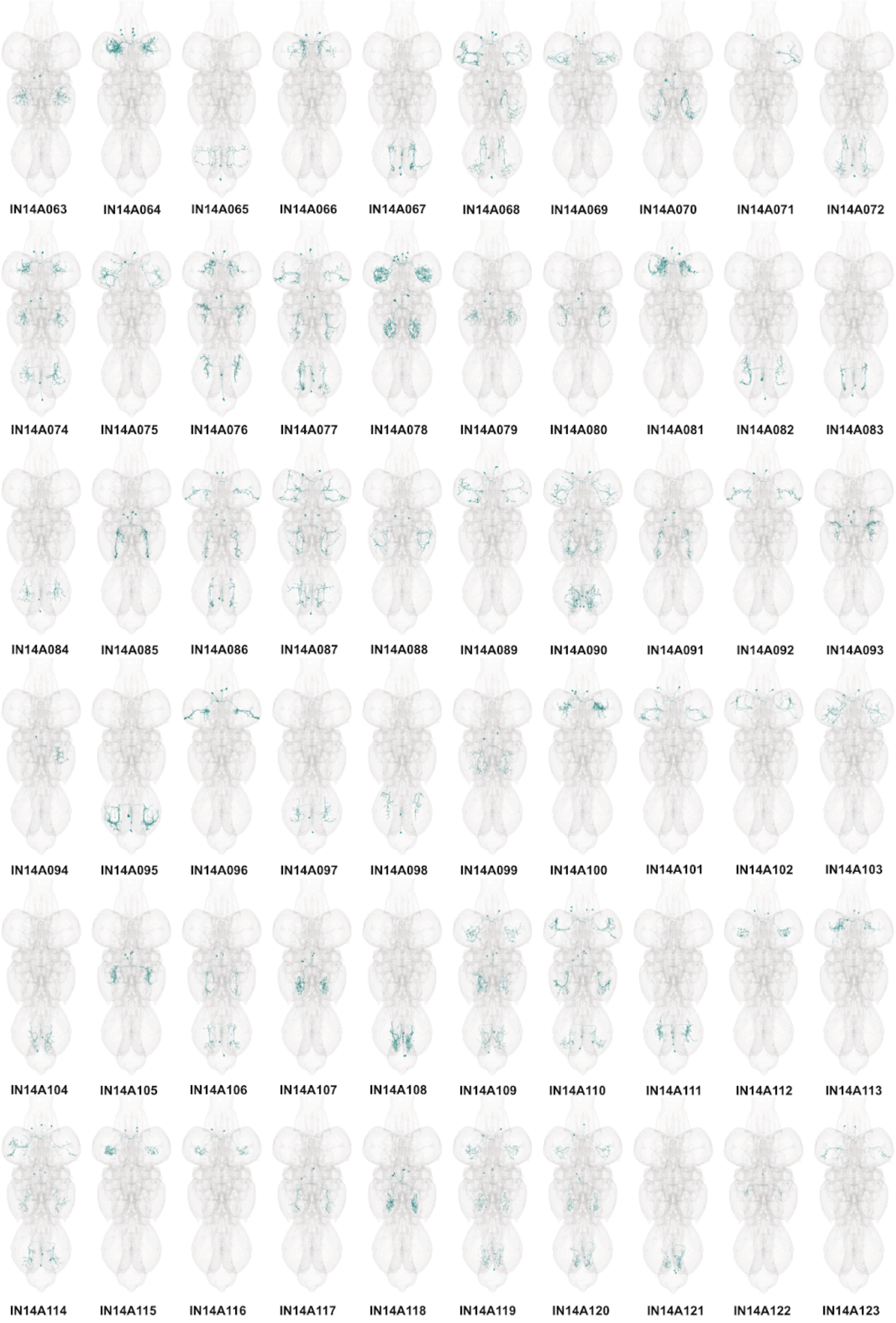

**Figure 37 - figure supplement 3. Systematic types of hemilineage 14A, continued.** Systematic types have been arranged in numerical order, with neurons of the same type that belong to distinct classes (e.g., intrinsic neuron vs ascending neuron) plotted separately but placed adjacent to each other. Individual neuron meshes have been coloured based on predicted neurotransmitter: dark orange = acetylcholine, blue = gaba, marine = glutamate, dark purple = unknown.

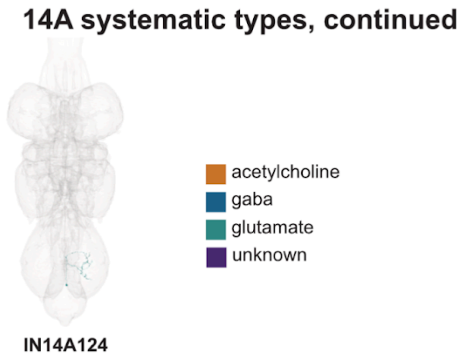

**Figure 37 - figure supplement 4. Systematic types of hemilineage 14A, continued.** Systematic types have been arranged in numerical order, with neurons of the same type that belong to distinct classes (e.g., intrinsic neuron vs ascending neuron) plotted separately but placed adjacent to each other. Individual neuron meshes have been coloured based on predicted neurotransmitter: dark orange = acetylcholine, blue = gaba, marine = glutamate, dark purple = unknown.

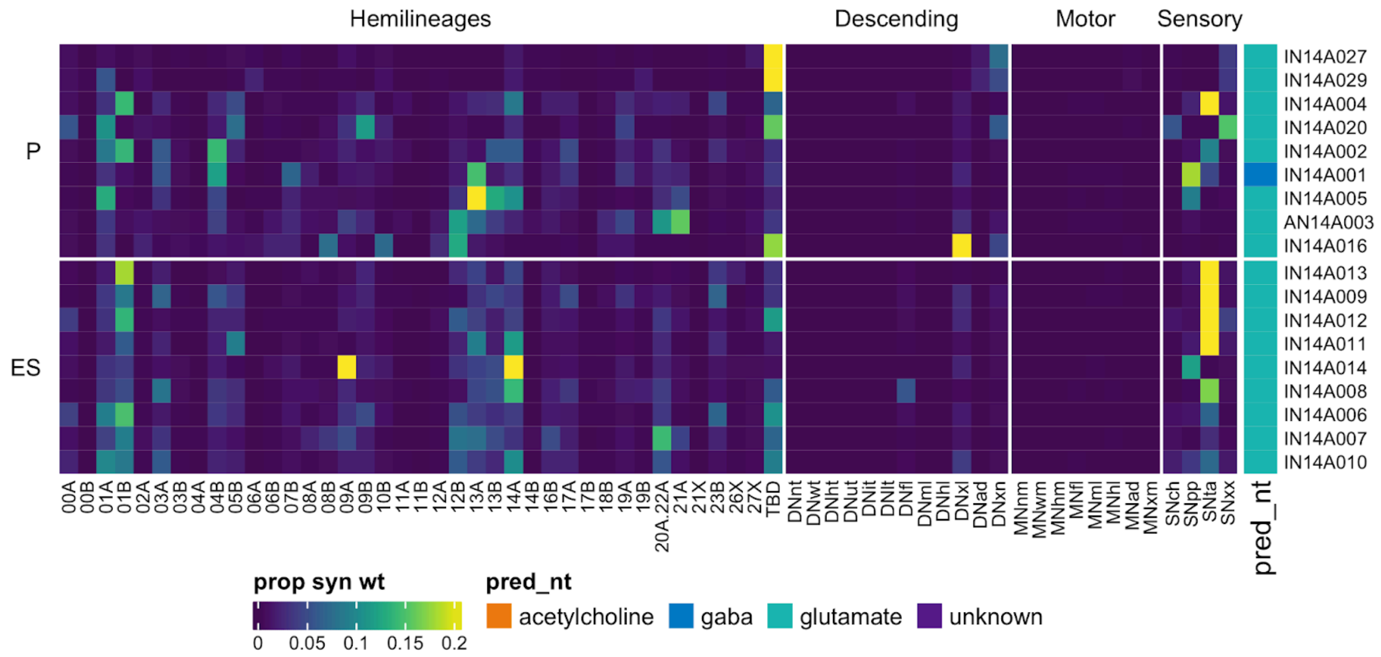

**Figure 37 - figure supplement 5. Connectivity to upstream partners by 14A primary and early secondary systematic types.** Proportions of synaptic weight to systematic types from upstream partners, normalised by row. 14A neurons have been clustered within each assigned birthtime window (P = primary, ES = early secondary, S = secondary) based on both upstream and downstream connectivity to hemilineages, descending neuron subclasses, motor neuron subclasses, and sensory neuron modalities. Annotation bar is coloured by the most common predicted neurotransmitter for the neurons of each type.

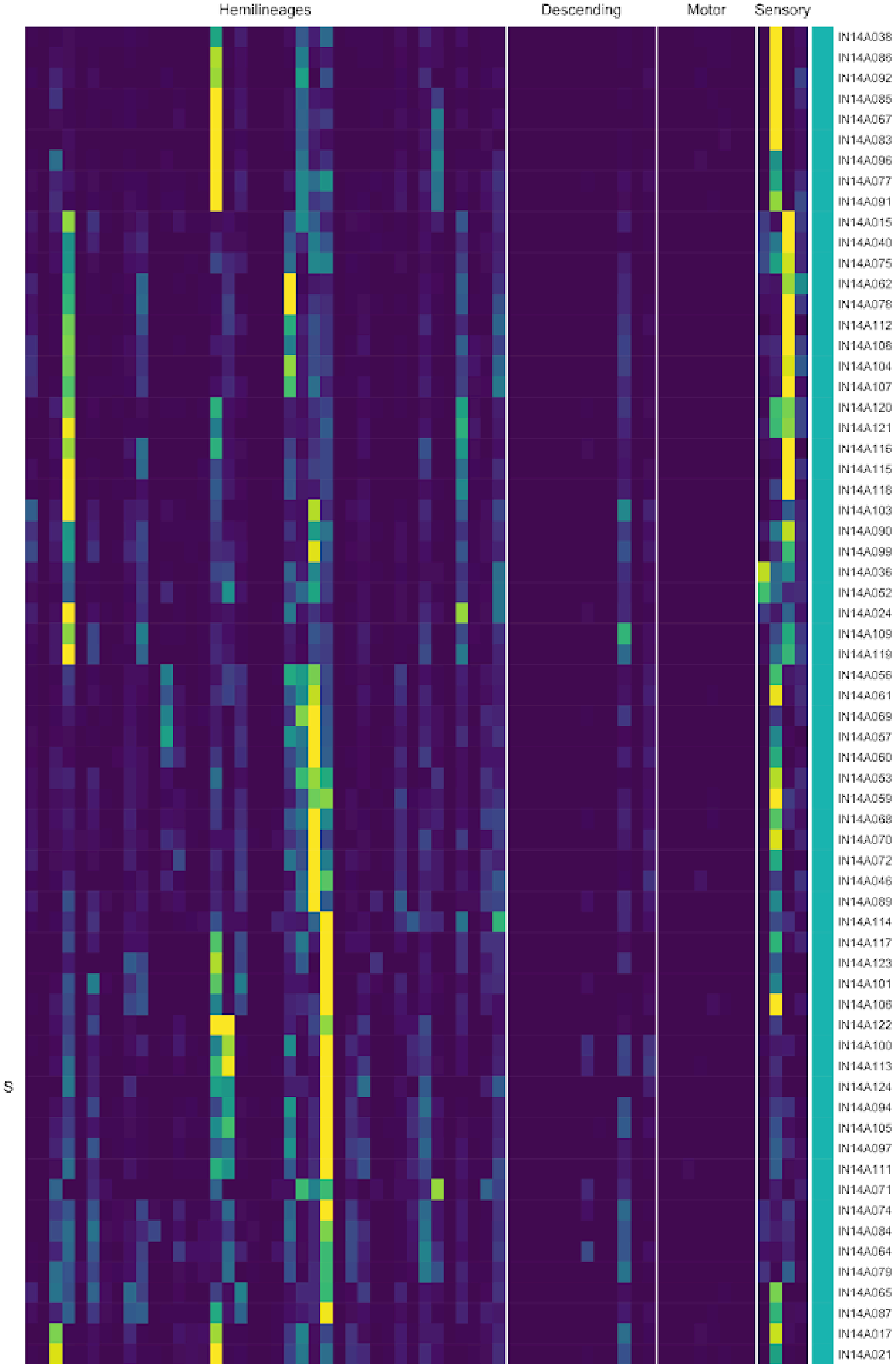

Hemilineages
Descending
Motor
Sensory
S
IN14A038
IN14A086
IN14A092
IN14A085
IN14A067
IN14A083
IN14A096
IN14A077
IN14A091
IN14A015
IN14A040
IN14A075
IN14A062
IN14A078
IN14A112
IN14A108
IN14A104
IN14A107
IN14A120
IN14A121
IN14A116
IN14A115
IN14A118
IN14A103
IN14A090
IN14A099
IN14A036
IN14A052
IN14A024
IN14A109
IN14A119
IN14A056
IN14A061
IN14A069
IN14A057
IN14A060
IN14A053
IN14A059
IN14A068
IN14A070
IN14A072
IN14A046
IN14A089
IN14A114
IN14A117
IN14A123
IN14A101
IN14A106
IN14A122
IN14A100
IN14A113
IN14A124
IN14A094
IN14A105
IN14A097
IN14A111
IN14A071
IN14A074
IN14A084
IN14A064
IN14A079
IN14A065
IN14A087
IN14A017
IN14A021

**Figure 37 - figure supplement 6. Connectivity to upstream partners by 14A secondary systematic types.** Proportions of synaptic weight to systematic types from upstream partners, normalised by row. 14A neurons have been clustered within each assigned birthtime window (P = primary, ES = early secondary, S = secondary) based on both upstream and downstream connectivity to hemilineages, descending neuron subclasses, motor neuron subclasses, and sensory neuron modalities. The annotation bar is coloured by the most common predicted neurotransmitter within each type.

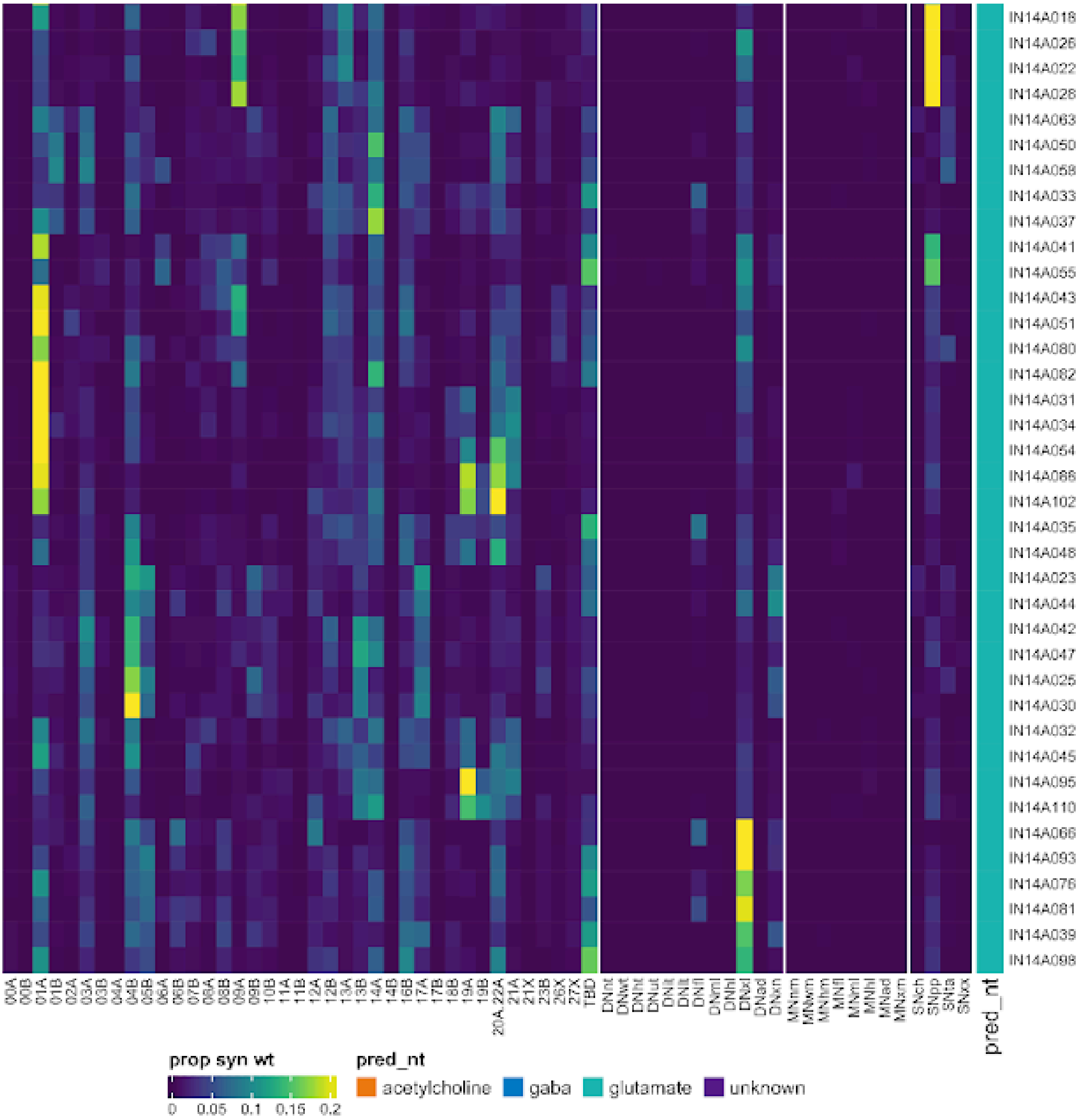

**Figure 37 - figure supplement 7. Connectivity to upstream partners by 14A secondary systematic types, continued.** Proportions of synaptic weight to systematic types from upstream partners, normalised by row. 14A neurons have been clustered within each assigned birthtime window (P = primary, ES = early secondary, S = secondary) based on both upstream and downstream connectivity to hemilineages, descending neuron subclasses, motor neuron subclasses, and sensory neuron modalities. The annotation bar is coloured by the most common predicted neurotransmitter within each type.

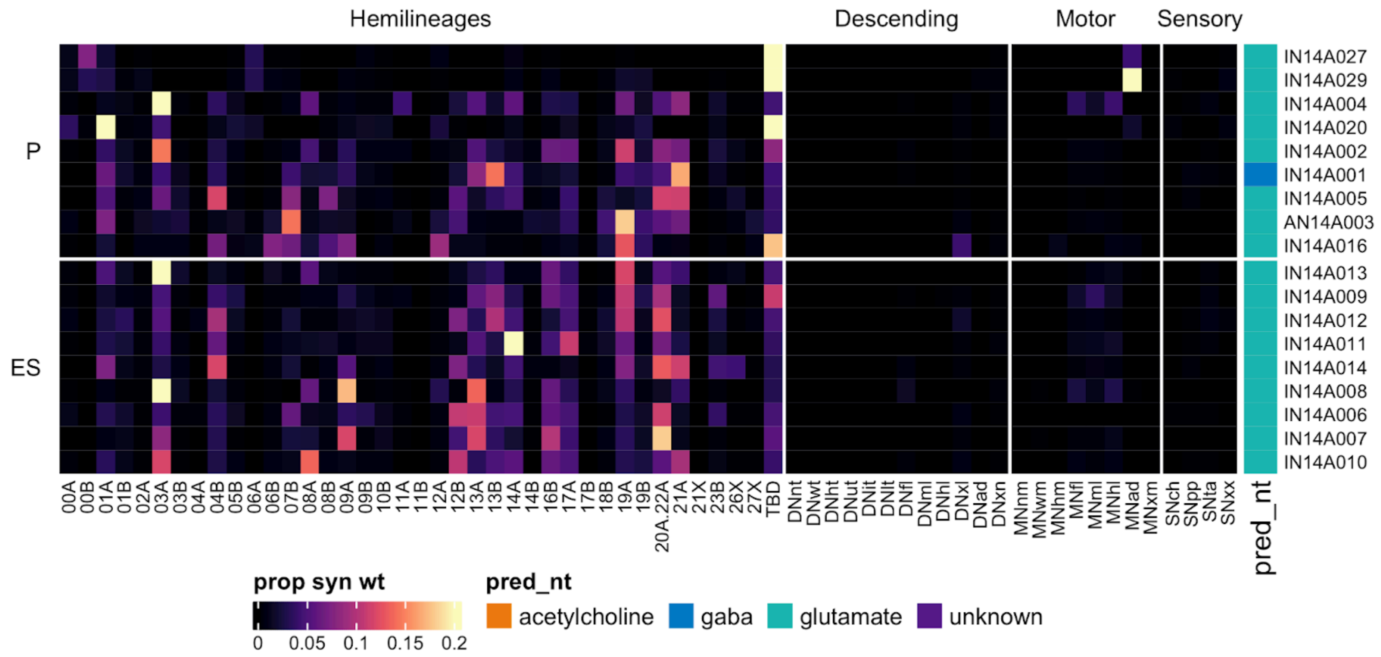

**Figure 37 - figure supplement 8. Connectivity to downstream partners by 14A primary and early secondary systematic types.** Proportions of synaptic weight from systematic types to downstream partners, normalised by row. 14A neurons have been clustered within each assigned birthtime window (P = primary, ES = early secondary, S = secondary) based on both upstream and downstream connectivity to hemilineages, descending neuron subclasses, motor neuron subclasses, and sensory neuron modalities. The annotation bar is coloured by the most common predicted neurotransmitter within each type.

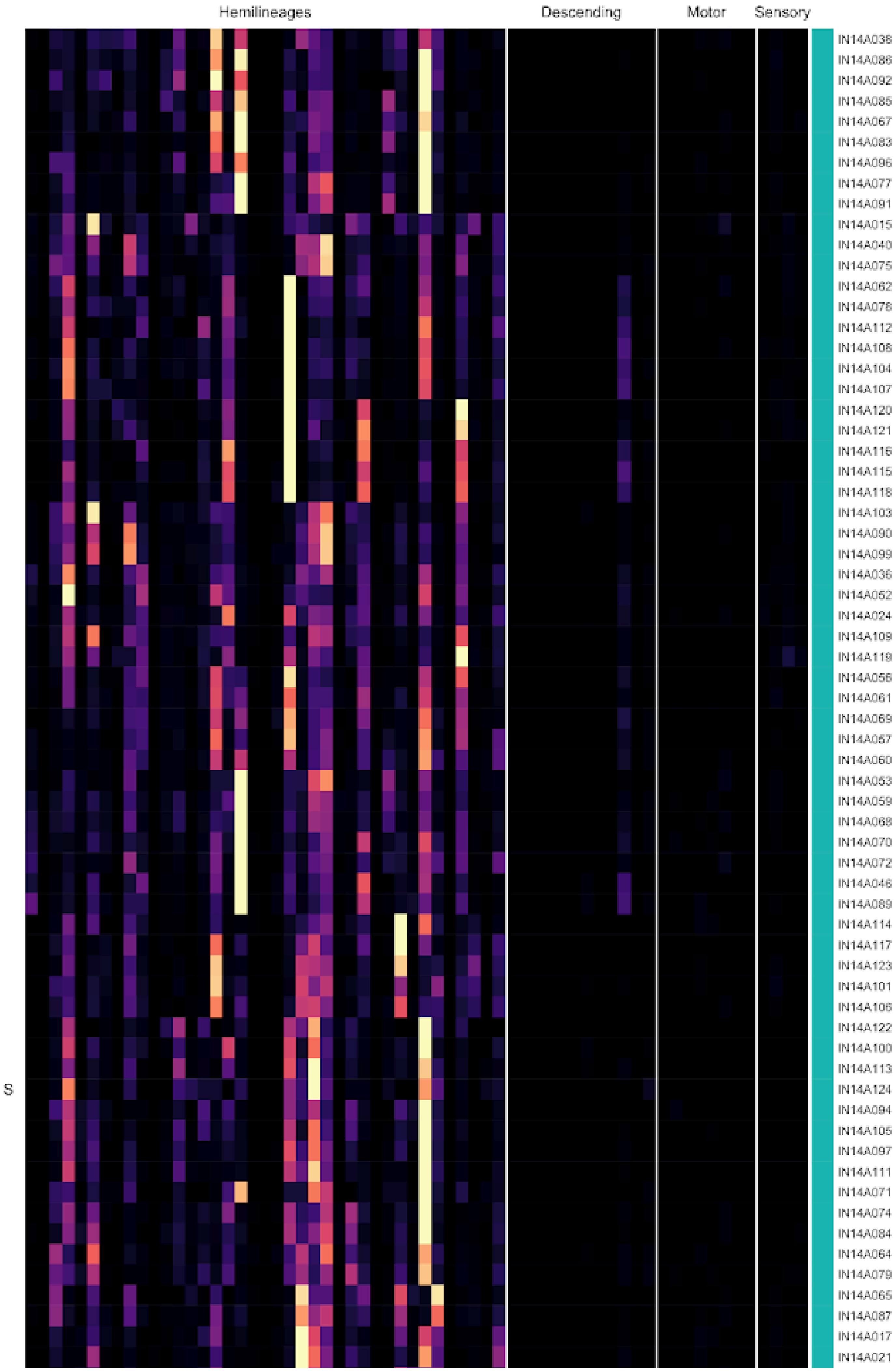
Hemilineages
Descending
Motor
Sensory
S
IN14A038
IN14A086
IN14A092
IN14A085
IN14A067
IN14A083
IN14A096
IN14A077
IN14A091
IN14A015
IN14A040
IN14A075
IN14A062
IN14A078
IN14A112
IN14A108
IN14A104
IN14A107
IN14A120
IN14A121
IN14A116
IN14A115
IN14A118
IN14A103
IN14A090
IN14A099
IN14A036
IN14A052
IN14A024
IN14A109
IN14A119
IN14A056
IN14A061
IN14A069
IN14A057
IN14A060
IN14A053
IN14A059
IN14A068
IN14A070
IN14A072
IN14A046
IN14A089
IN14A114
IN14A117
IN14A123
IN14A101
IN14A106
IN14A122
IN14A100
IN14A113
IN14A124
IN14A094
IN14A105
IN14A097
IN14A111
IN14A071
IN14A074
IN14A084
IN14A064
IN14A079
IN14A065
IN14A087
IN14A017
IN14A021

**Figure 37 - figure supplement 9. Connectivity to downstream partners by 14A secondary systematic types.** Proportions of synaptic weight from systematic types to downstream partners, normalised by row. 14A neurons have been clustered within each assigned birthtime window (P = primary, ES = early secondary, S = secondary) based on both upstream and downstream connectivity to hemilineages, descending neuron subclasses, motor neuron subclasses, and sensory neuron modalities. The annotation bar is coloured by the most common predicted neurotransmitter for the neurons of each type.

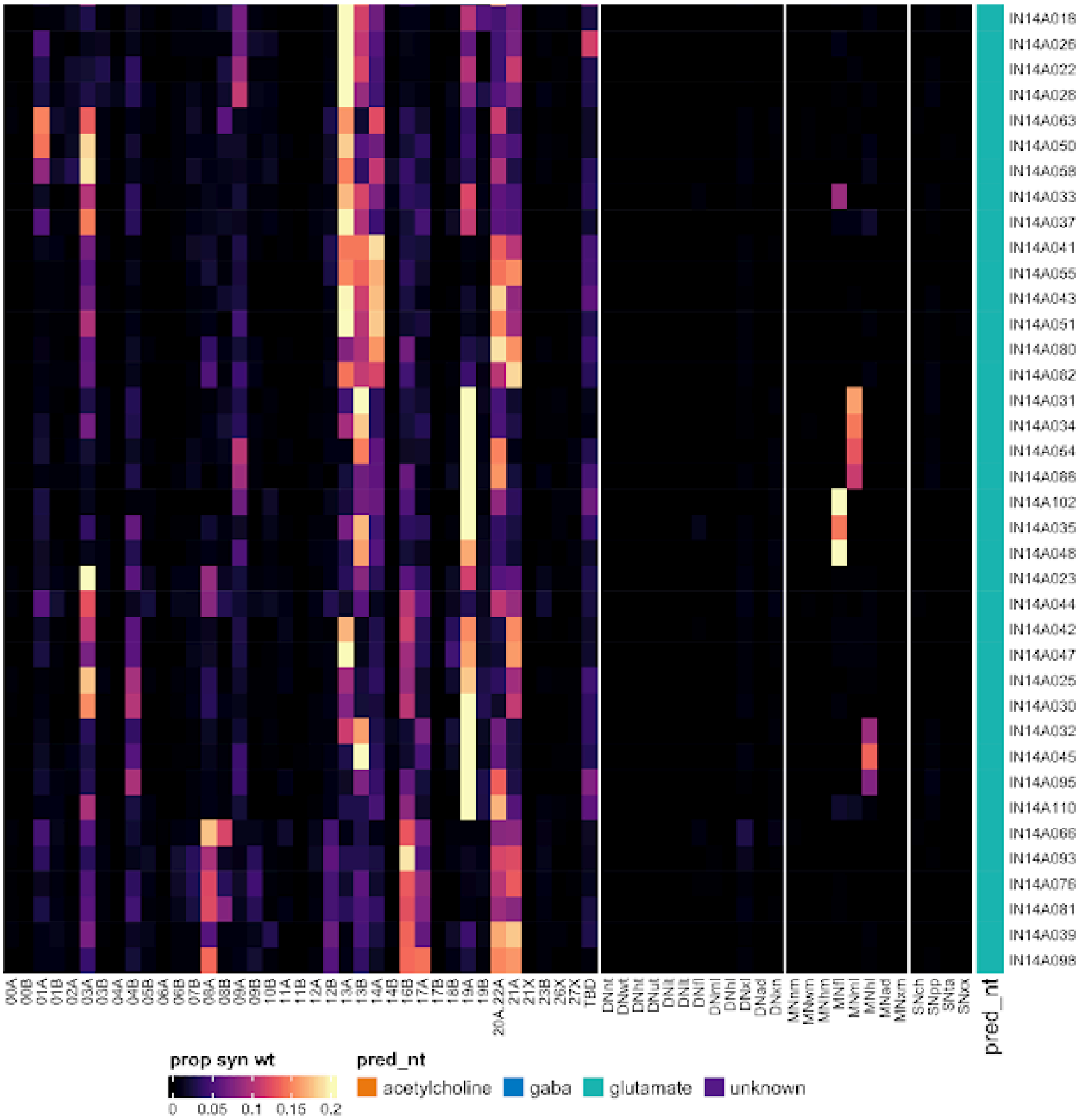

**Figure 37 - figure supplement 10. Connectivity to downstream partners by 14A secondary systematic types, continued.** Proportions of synaptic weight from systematic types to downstream partners, normalised by row. 14A neurons have been clustered within each assigned birthtime window (P = primary, ES = early secondary, S = secondary) based on both upstream and downstream connectivity to hemilineages, descending neuron subclasses, motor neuron subclasses, and sensory neuron modalities. The annotation bar is coloured by the most common predicted neurotransmitter for the neurons of each type.

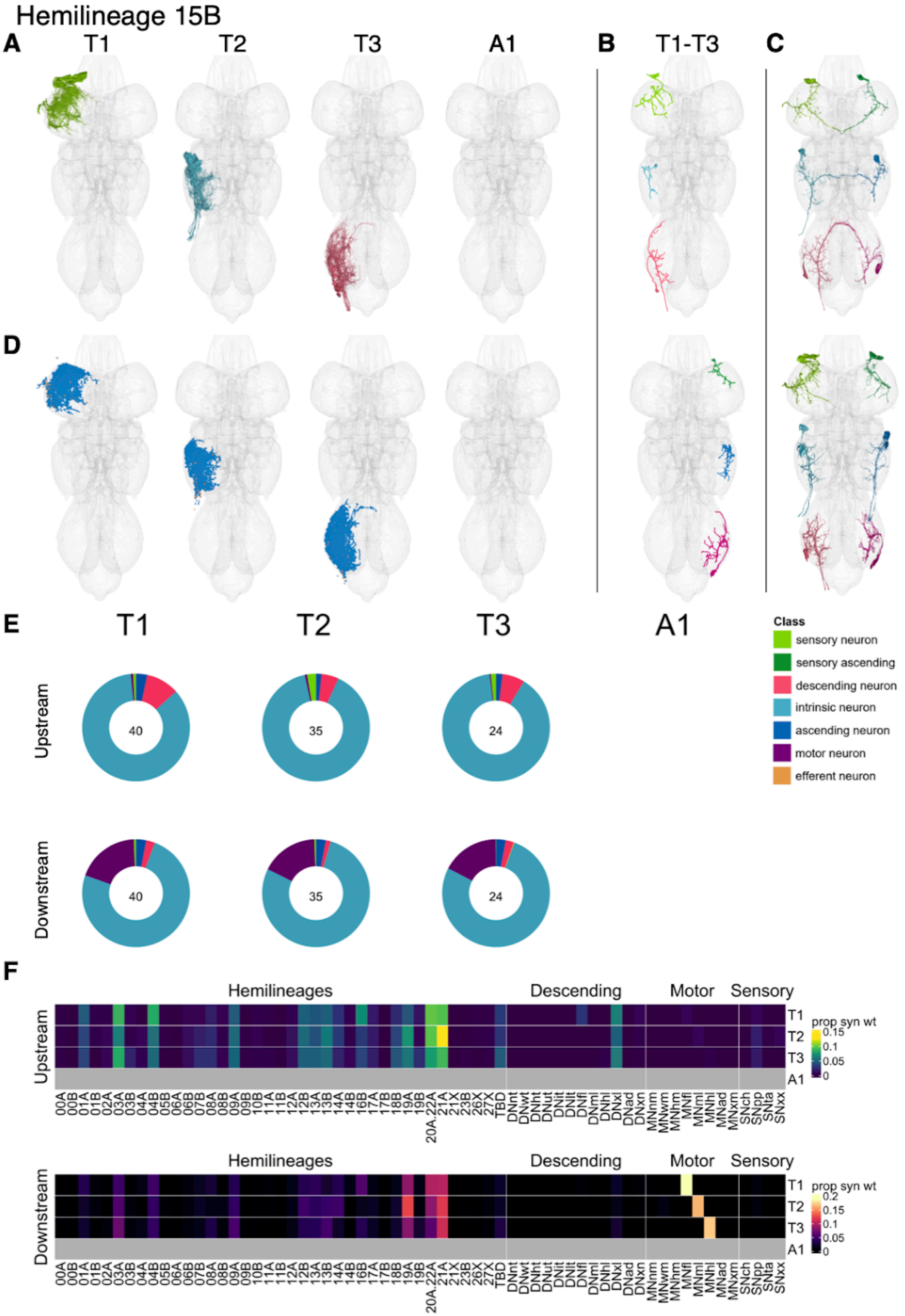

Hemilineage 15B
A
T1
T2
T3
A1
B
T1-T3
C
D
E
T1
T2
T3
A1
Upstream
Downstream
40
35
24
40
35
24
Class
sensory neuron
sensory ascending
descending neuron
intrinsic neuron
ascending neuron
motor neuron
efferent neuron
F
Hemilineages
Descending
Motor
Sensory
Upstream
Downstream
T1
T2
T3
A1
prop syn wt
00A 00B 01A 01B 02A 03A 03B 04A 04B 05B 06A 06B 07B 08A 08B 09A 09B 10B 11A 11B 12A 12B 13A 13B 14A 14B 16B 17A 17B 18B 19A 19B 20A.22A 21A 21X 23B 26X 27X TBD DNnt DNwt DNht DNut DNlt DNfl DNml DNhl DNxl DNad DNxn MNnm MNwm MNhm MNfl MNml MNhl MNad MNxm SNch SNpp SNta SNxx

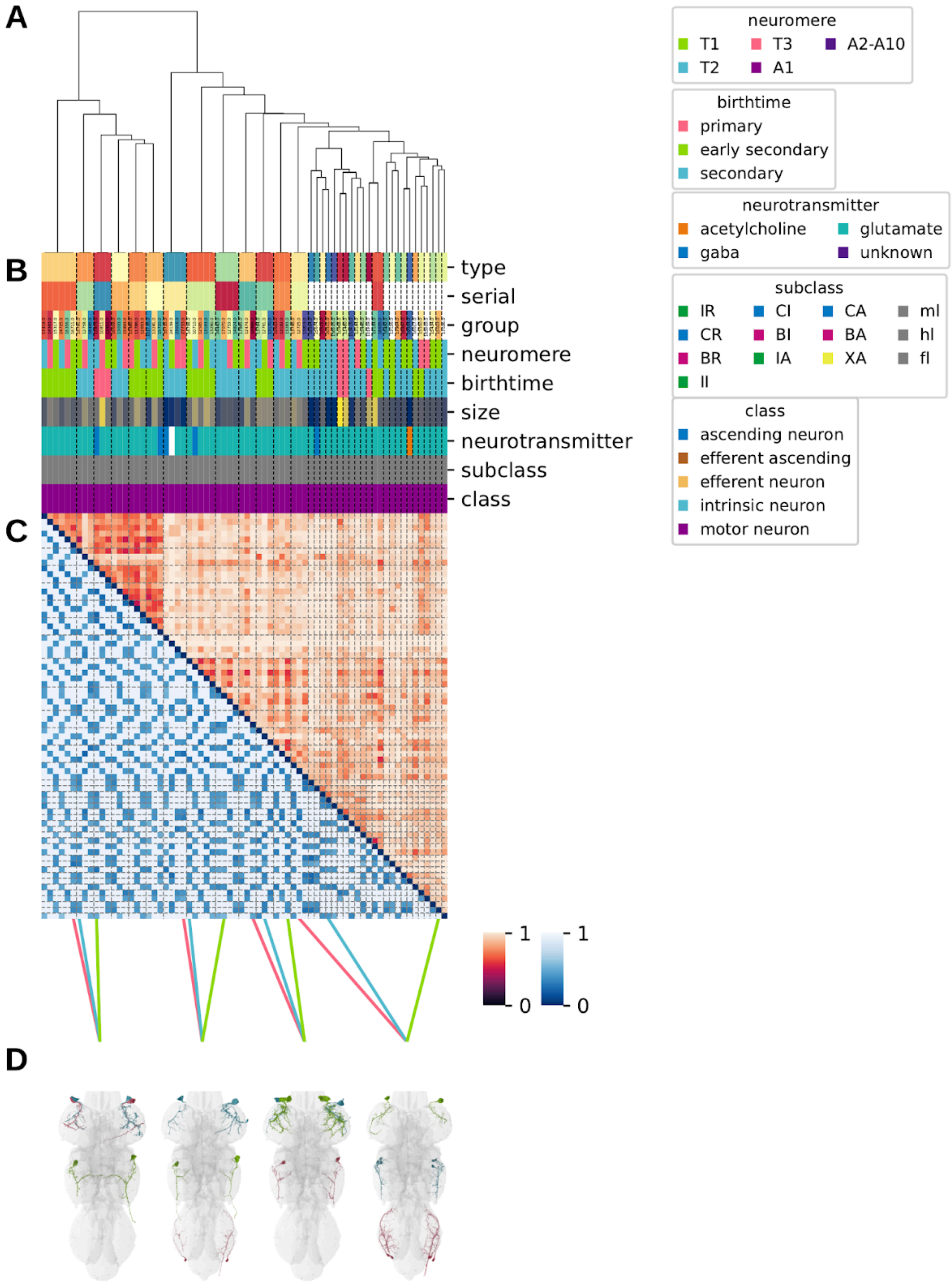

**Figure 38 - figure supplement 1. Systematic typing of hemilineage 15B.** Types for motor neurons were assigned separately as outlined in our accompanying manuscript (Cheong et al., 2023). **A.** Hierarchical clustering dendrogram of hemilineage groups by laterally and serially aggregated connectivity cosine clustering. **B.** Categorical annotations of each hemilineage group, each column corresponding to the aligned leaf in A. Colours for cluster, serial set, and group are arbitrary for visualisation. Colours for neuromere, birthtime, neurotransmitter, subclass, and class are as in all other figures. **C.** Similarity distance heatmap for hemilineage. Cosine distance is in the upper triangle, while laterally symmetrised NBLAST distance is in the lower triangle. Systematic type names of some types are labelled. **D.** Morphologically representative groups from dendrogram subtrees. Each group, indicated by colour and line connecting to its column in B and C, is the most morphologically representative group (medoid of NBLAST distance) from a subtree of A. The subtrees (flat clusters) are equal height cuts of A determined to yield the number of groups per plot and plots in D.

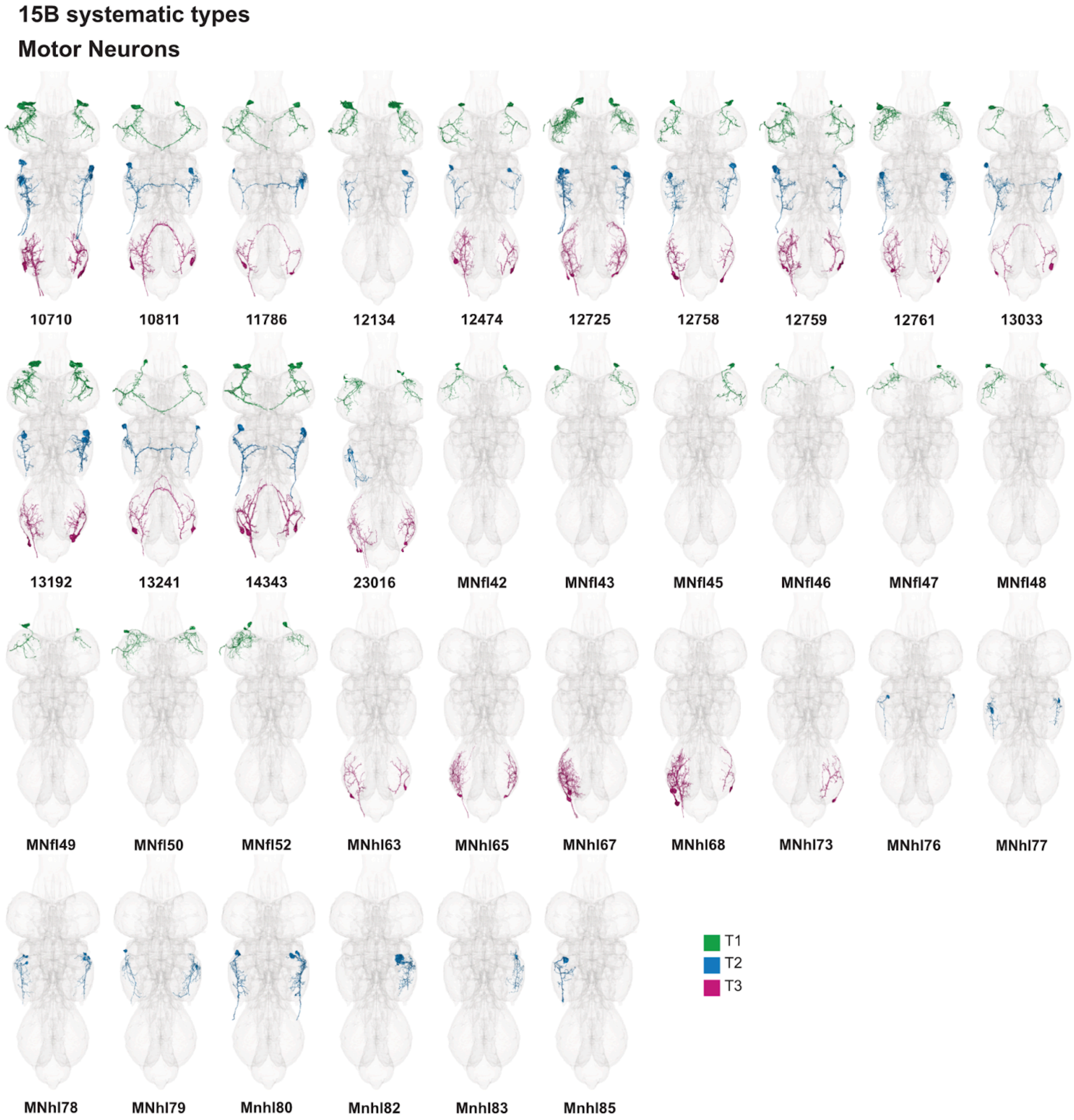

**Figure 38 - figure supplement 2. Systematic types of hemilineage 15B.** Motor neurons (typed separately in (Cheong et al., 2023)) have been plotted by serial set if identified in multiple neuromeres and by systematic type if not. Individual motor neuron meshes have been coloured based on soma neuromere: dark green = T1, blue = T2, magenta = T3.

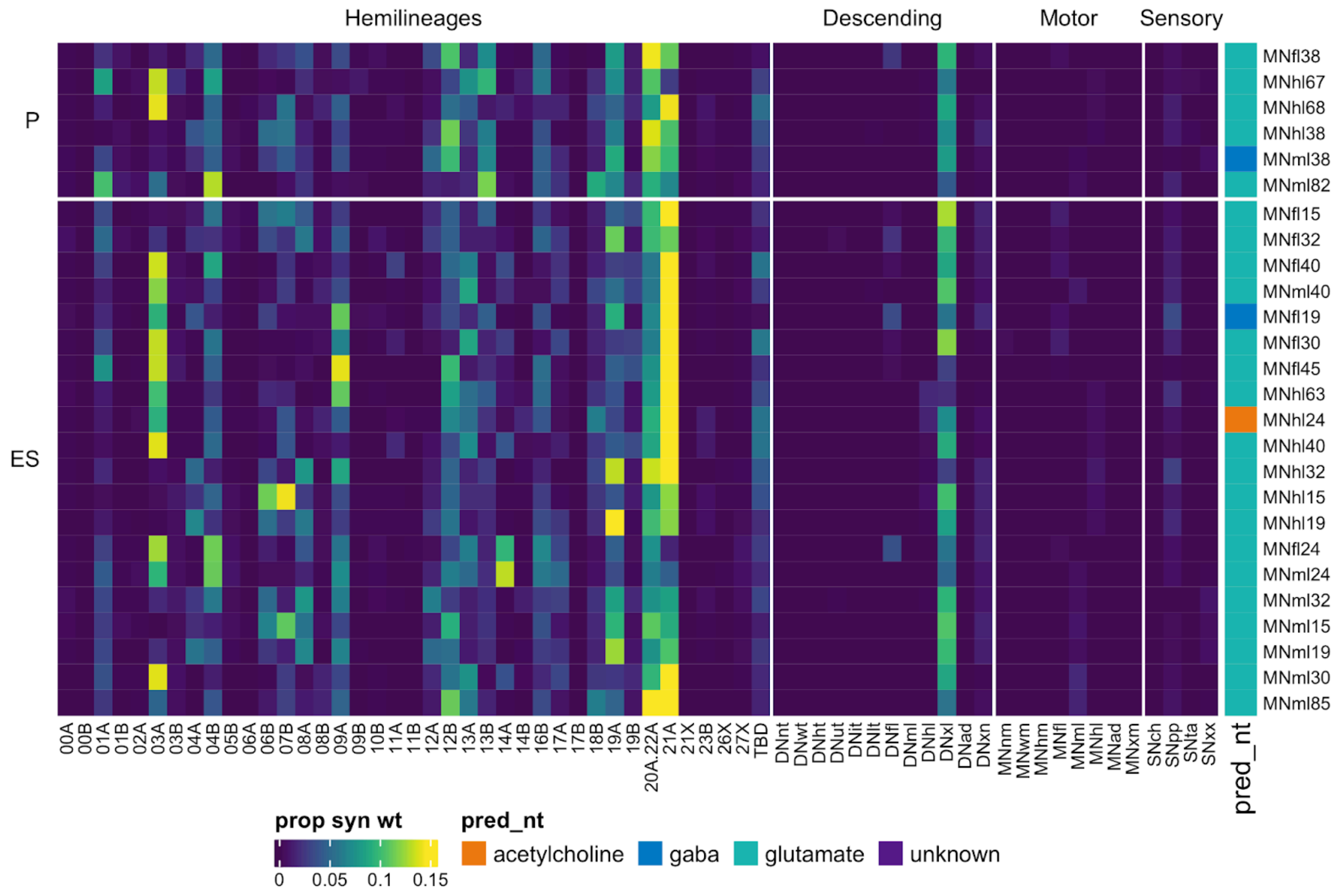

**Figure 38 - figure supplement 3. Connectivity to upstream partners by 15B primary and early secondary systematic types.** Proportions of synaptic weight to systematic types from upstream partners, normalised by row. 15B neurons have been clustered within each assigned birthtime window (P = primary, ES = early secondary, S = secondary) based on both upstream and downstream connectivity to hemilineages, descending neuron subclasses, motor neuron subclasses, and sensory neuron modalities. Annotation bar is coloured by the most common predicted neurotransmitter for the neurons of each type.

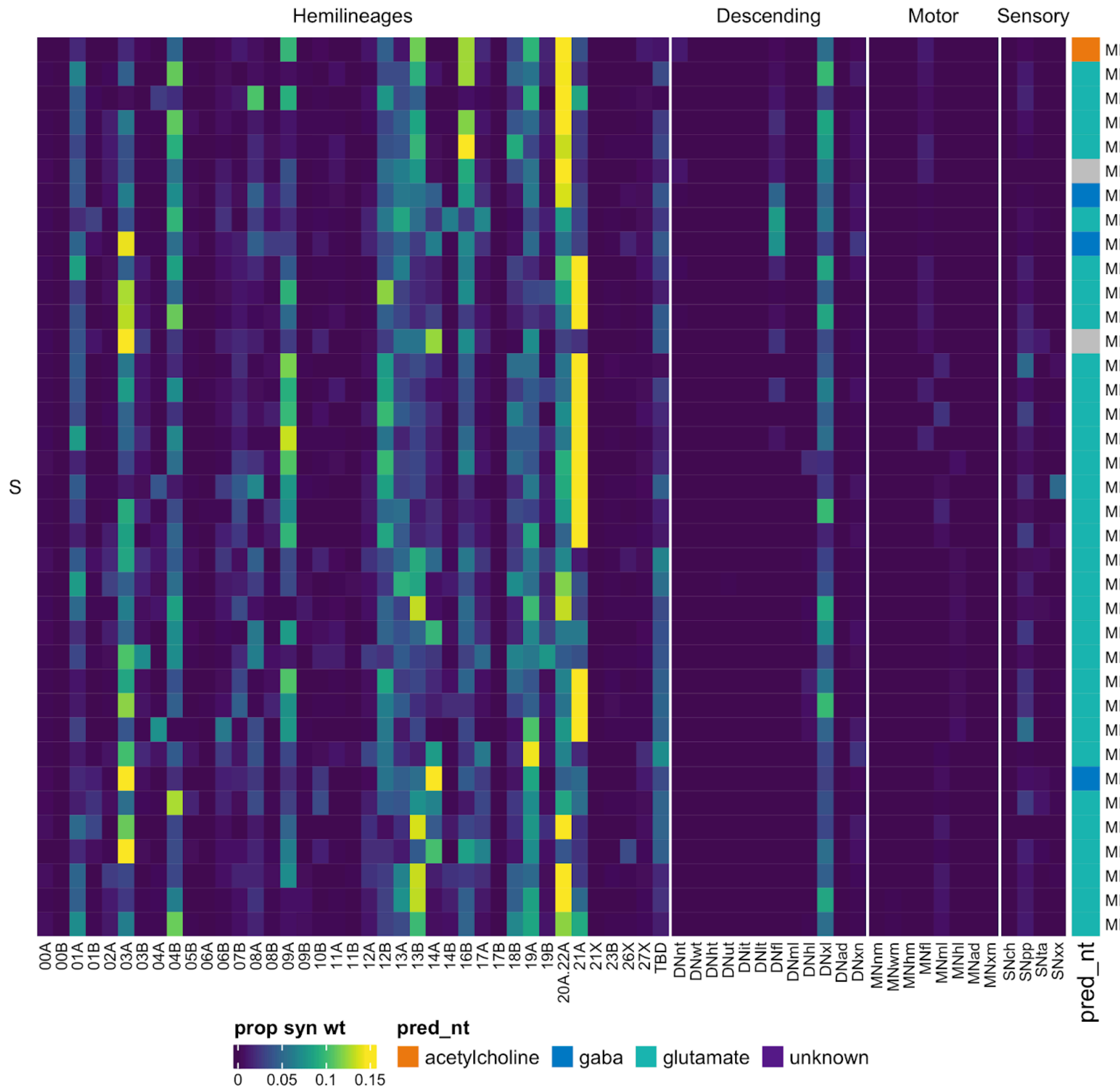

**Figure 38 - figure supplement 4. Connectivity to upstream partners by 15B secondary systematic types.** Proportions of synaptic weight to systematic types from upstream partners, normalised by row. 15B neurons have been clustered within each assigned birthtime window (P = primary, ES = early secondary, S = secondary) based on both upstream and downstream connectivity to hemilineages, descending neuron subclasses, motor neuron subclasses, and sensory neuron modalities. The annotation bar is coloured by the most common predicted neurotransmitter within each type.

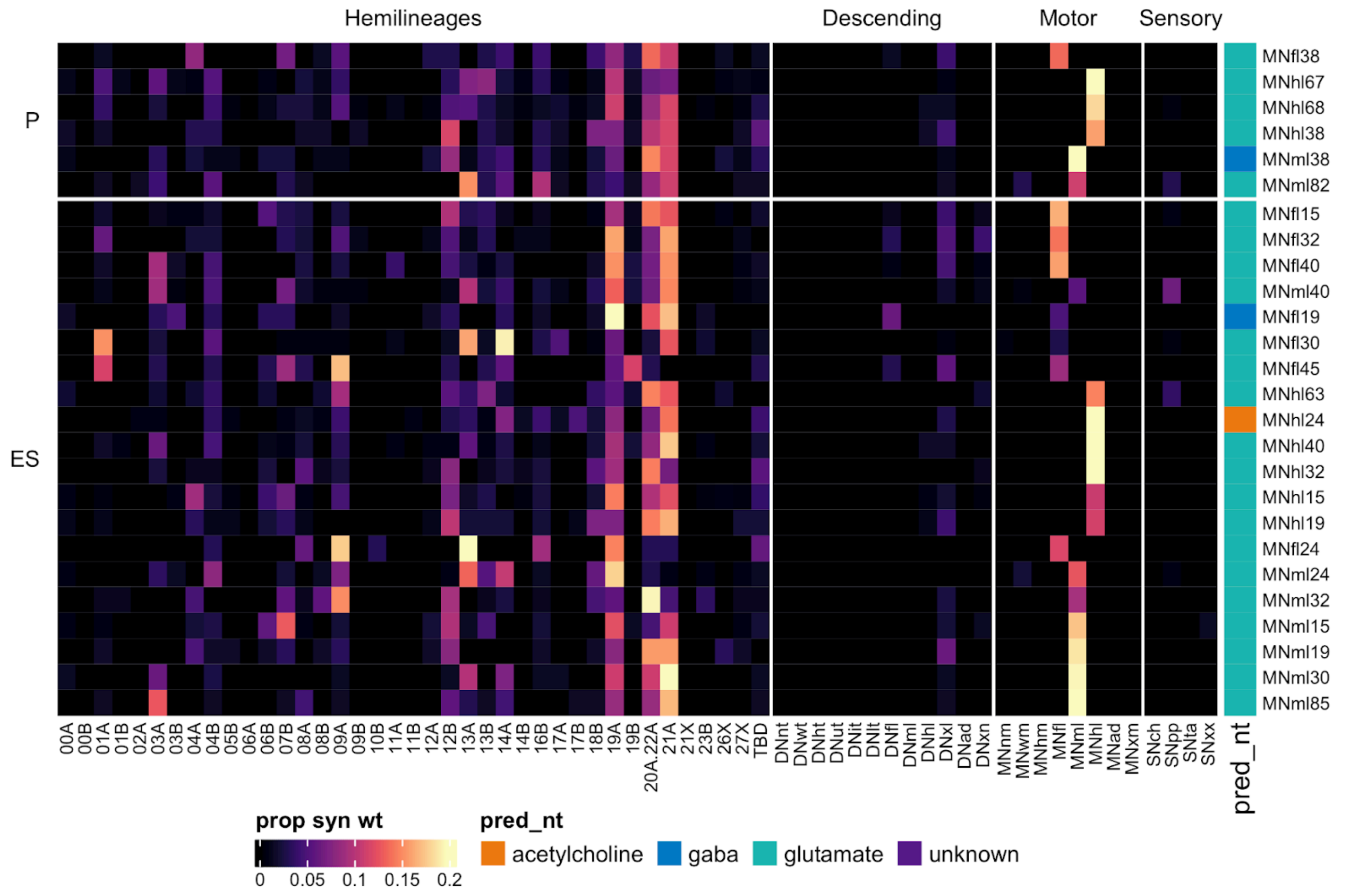

**Figure 38 - figure supplement 5. Connectivity to downstream partners by 15B primary and early secondary systematic types.** Proportions of synaptic weight from systematic types to downstream partners, normalised by row. 15B neurons have been clustered within each assigned birthtime window (P = primary, ES = early secondary, S = secondary) based on both upstream and downstream connectivity to hemilineages, descending neuron subclasses, motor neuron subclasses, and sensory neuron modalities. The annotation bar is coloured by the most common predicted neurotransmitter within each type.

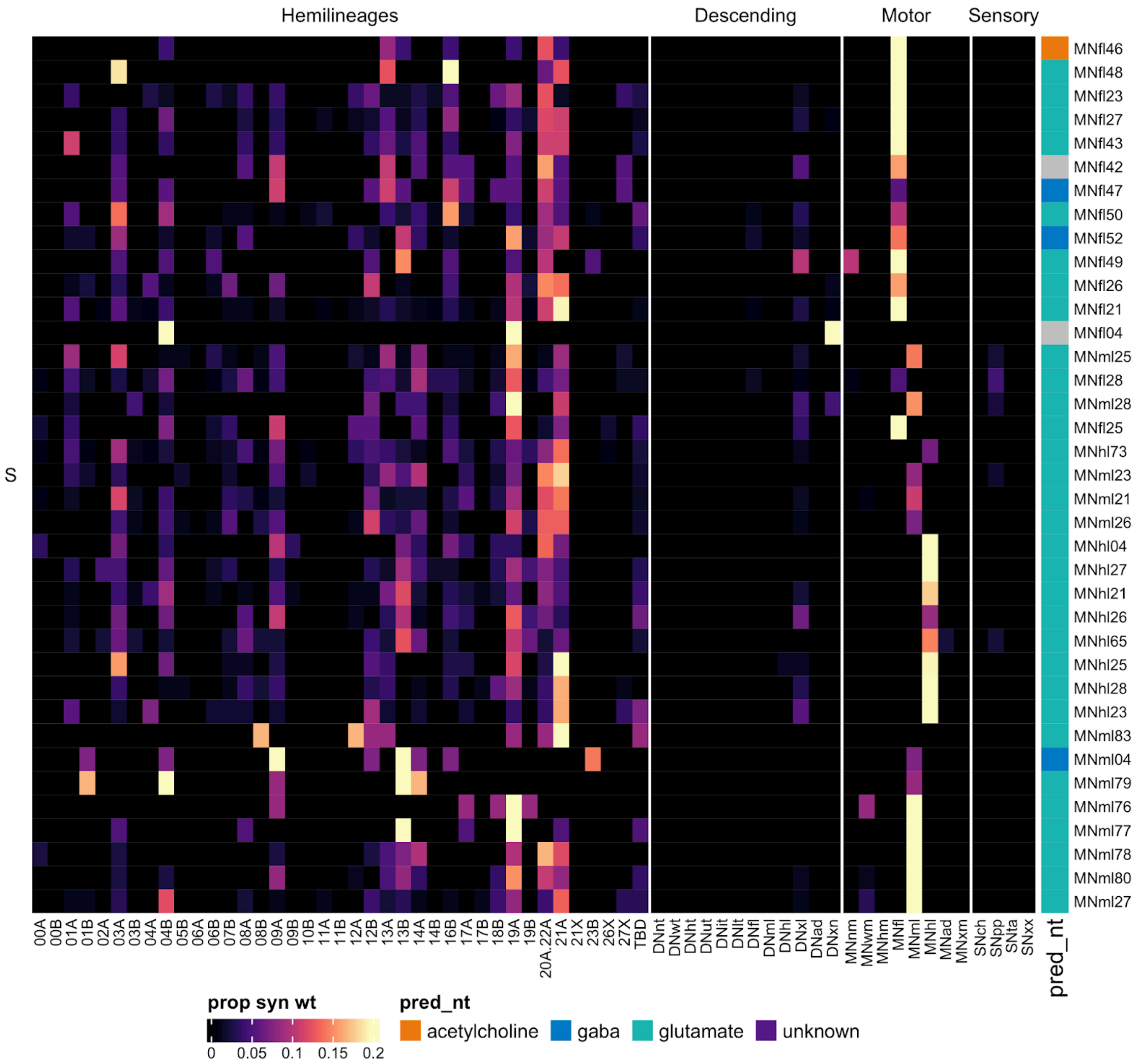

**Figure 38 - figure supplement 6. Connectivity to downstream partners by 15B secondary systematic types.** Proportions of synaptic weight from systematic types to downstream partners, normalised by row. 15B neurons have been clustered within each assigned birthtime window (P = primary, ES = early secondary, S = secondary) based on both upstream and downstream connectivity to hemilineages, descending neuron subclasses, motor neuron subclasses, and sensory neuron modalities. The annotation bar is coloured by the most common predicted neurotransmitter for the neurons of each type.

Hemilineage 16B

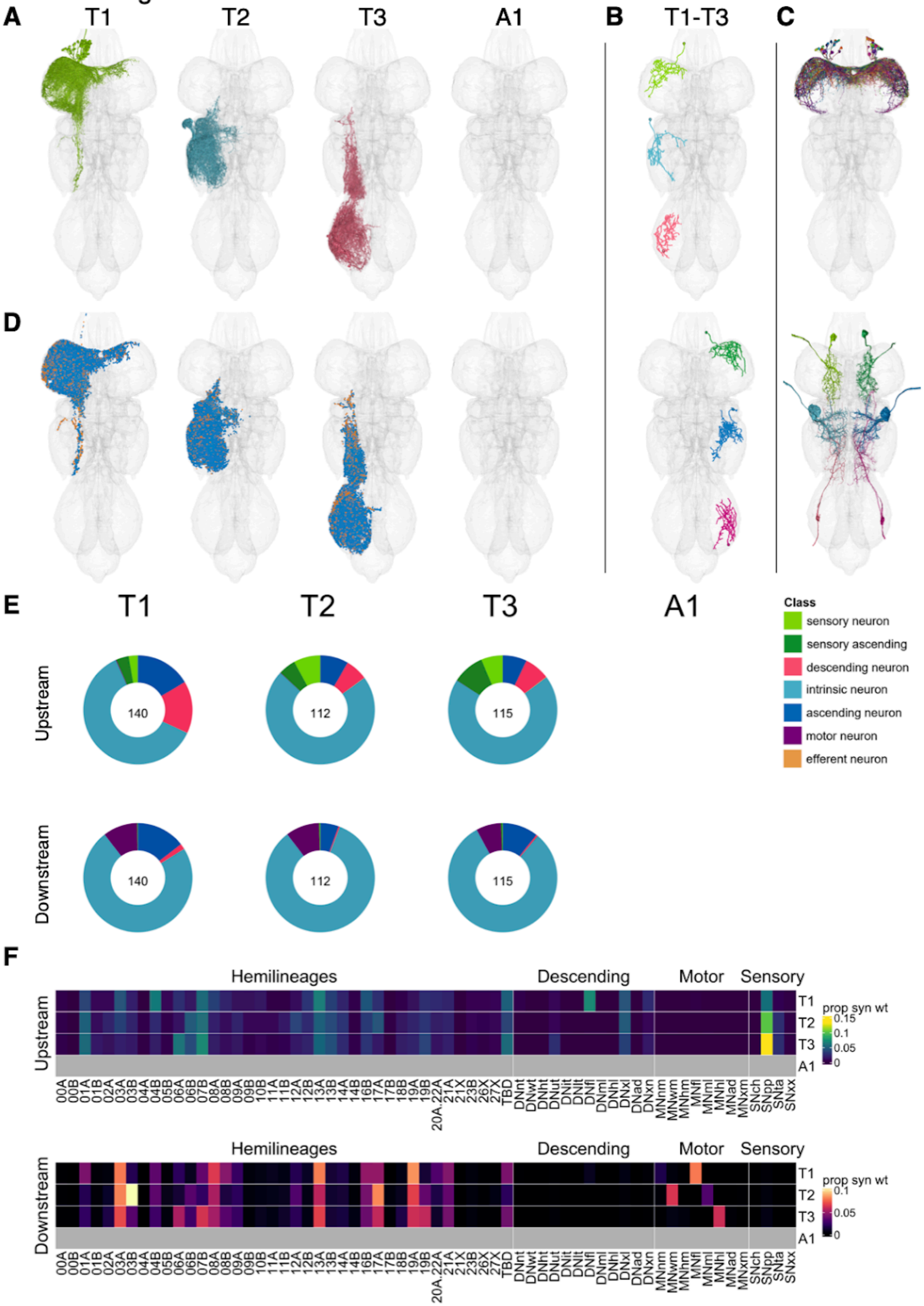

**Figure 39. Hemilineage 16B. A.** Meshes of all RHS secondary neurons plotted in neuromere-specific colours. **B.** "Representative" secondary neuron skeletons plotted in hemineuromere-specific colours. The skeleton with the top accumulated NBLAST score among all neurons from the hemilineage in a given hemineuromere was used. **C.** Neuron meshes of selected examples. Top: T1 BR subclass. Bottom: motor neuron dorsal serial set 10011. **D.** Predicted synapses of RHS secondary neurons. Blue: postsynapses; dark orange: presynapses. **E.** Proportions of connections from secondary neurons to upstream or downstream partners, normalised by neuromere and coloured by broad class. Numbers of query neurons appear in the centre. **F.** Proportions of synaptic weight from secondary neurons originating in each neuromere to upstream or downstream partners, normalised by row.

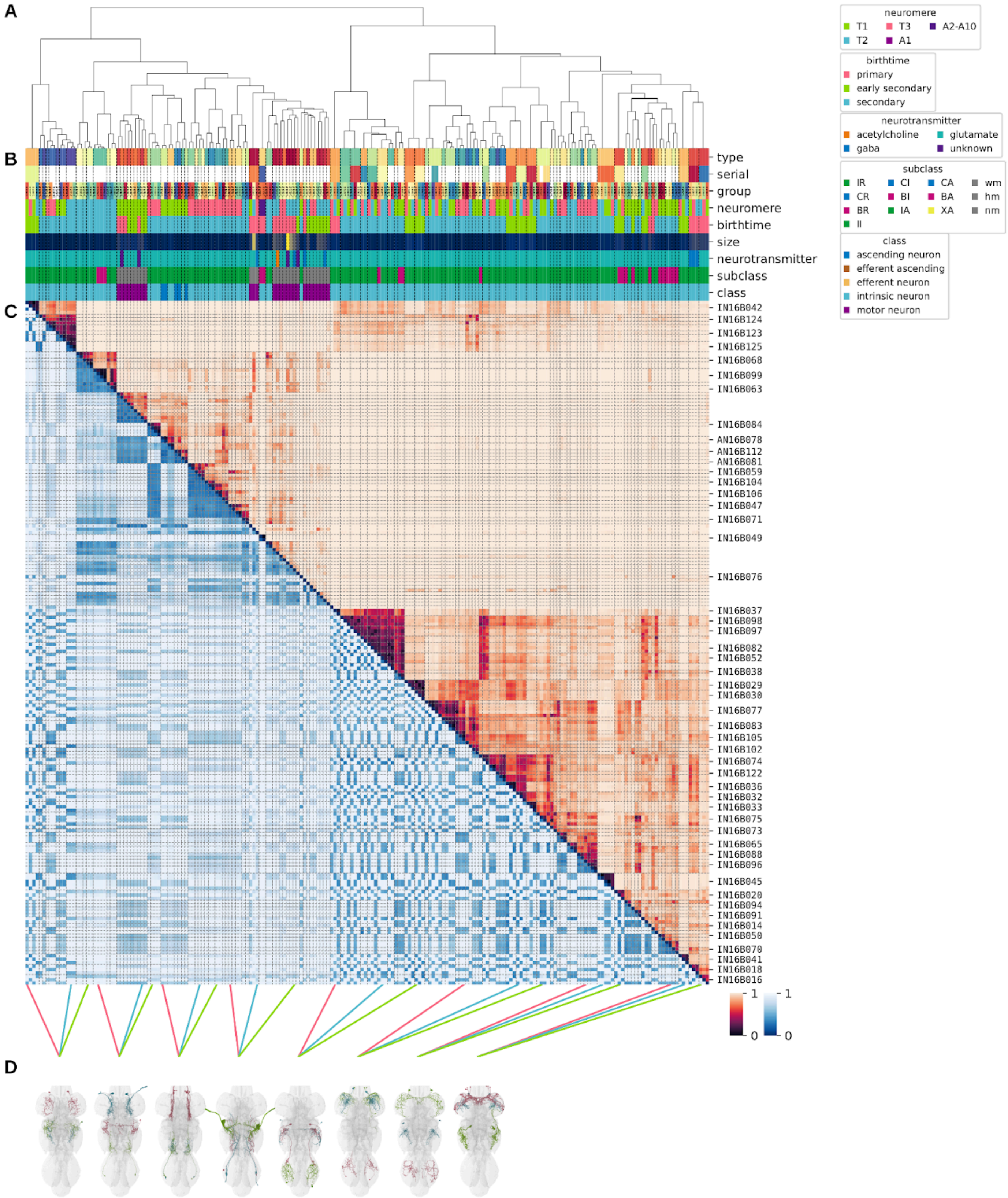

**Figure 39 - figure supplement 1. Systematic typing of hemilineage 16B. A.** Hierarchical clustering dendrogram of hemilineage groups by laterally and serially aggregated connectivity cosine clustering. **B.** Categorical annotations of each hemilineage group, each column corresponding to the aligned leaf in A. Colours for type, serial set, and group are arbitrary for visualisation. Colours for neuromere, birthtime, neurotransmitter, subclass, and class are as in all other figures. **C.** Similarity distance heatmap for hemilineage. Cosine distance is in the upper triangle, while laterally symmetrised NBLAST distance is in the lower triangle. Systematic type names of some types are labelled. **D.** Morphologically representative groups from dendrogram subtrees. Each group, indicated by colour and line connecting to its column in B and C, is the most morphologically representative group (medoid of NBLAST distance) from a subtree of A. The subtrees (flat clusters) are equal height cuts of A determined to yield the number of groups

per plot and plots in D.

#### 16B systematic types

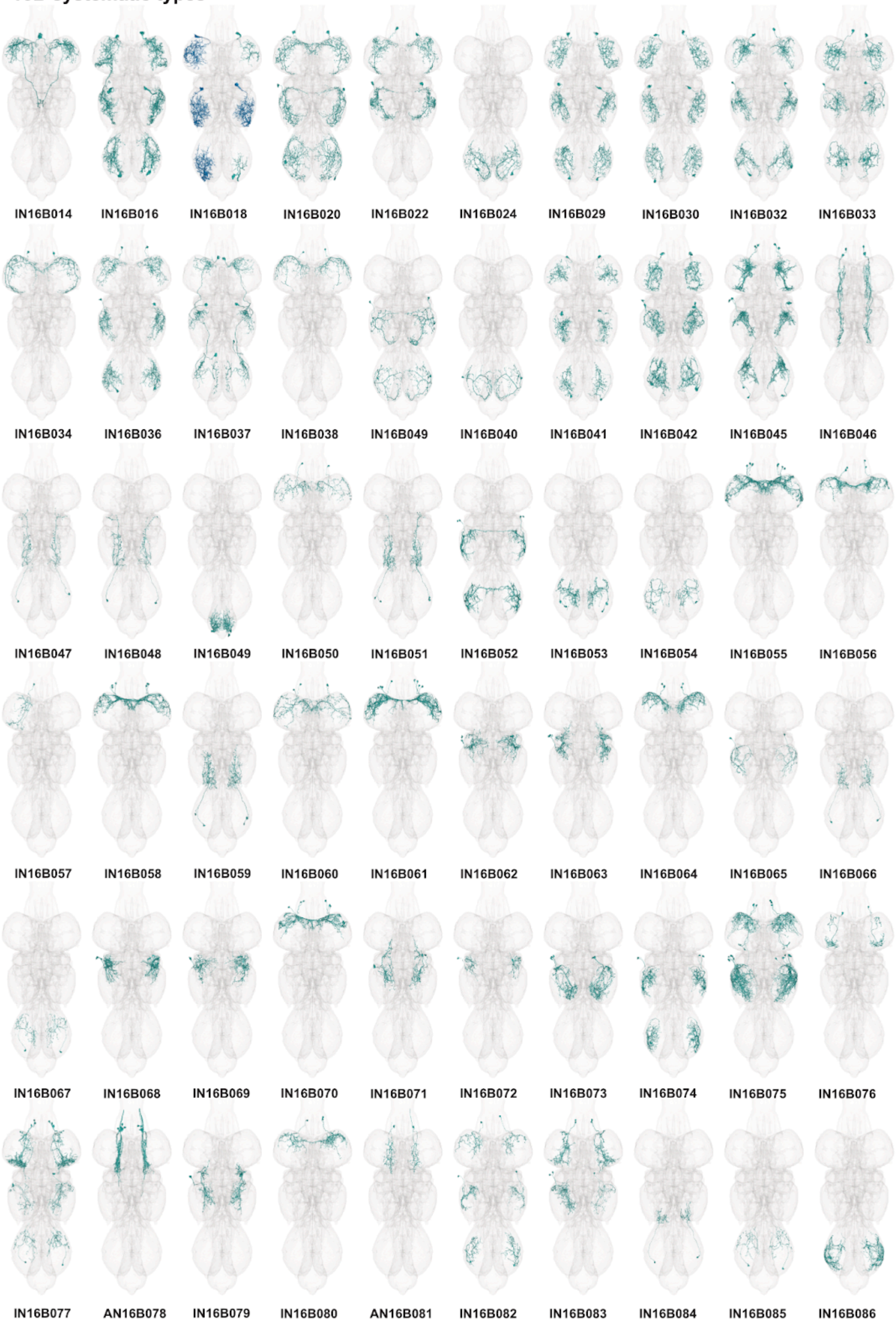

**Figure 39 - figure supplement 2. Systematic types of hemilineage 16B.** Systematic types have been arranged in numerical order, with neurons of the same type that belong to distinct classes (e.g., intrinsic neuron vs ascending neuron) plotted separately but placed adjacent to each other. Individual neuron meshes have been coloured based on predicted neurotransmitter: dark orange = acetylcholine, blue = gaba, marine = glutamate, dark purple = unknown.

#### 16B systematic types

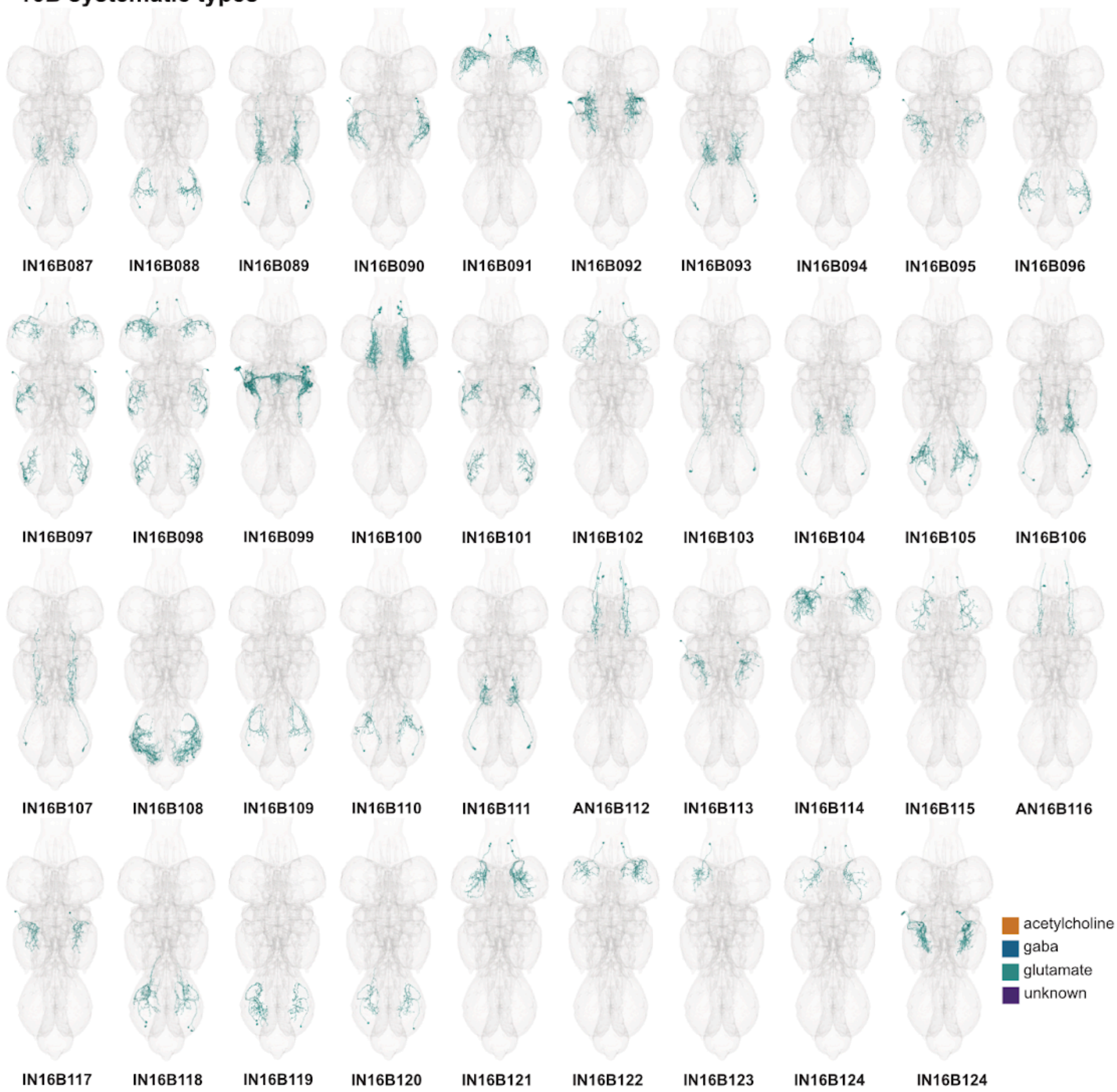

#### Motor Neurons

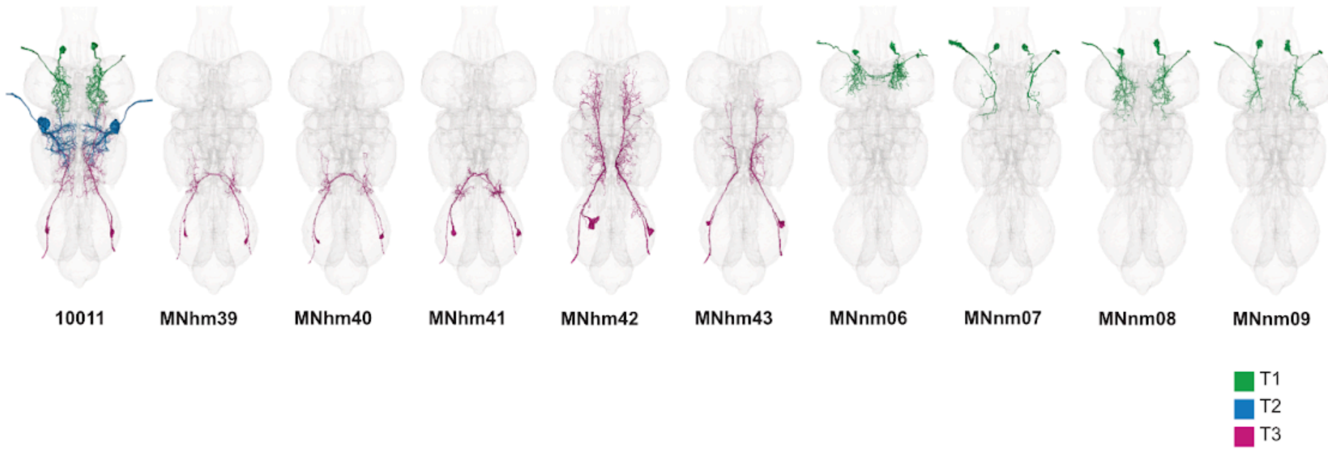

**Figure 39 - figure supplement 3. Systematic types of hemilineage 16B, continued.** Systematic types have been arranged in numerical order, with neurons of the same type that belong to distinct classes (e.g., intrinsic neuron vs ascending neuron) plotted separately but placed adjacent to each other. Individual neuron meshes have been coloured based on predicted neurotransmitter: dark orange = acetylcholine, blue = gaba, marine = glutamate, dark purple = unknown. Motor neurons (typed separately in (Cheong et al., 2023)) have been plotted by serial set if identified in multiple neuromeres and by systematic type if not. Individual motor neuron meshes have been coloured based on soma neuromere: dark green = T1, blue = T2, magenta = T3.

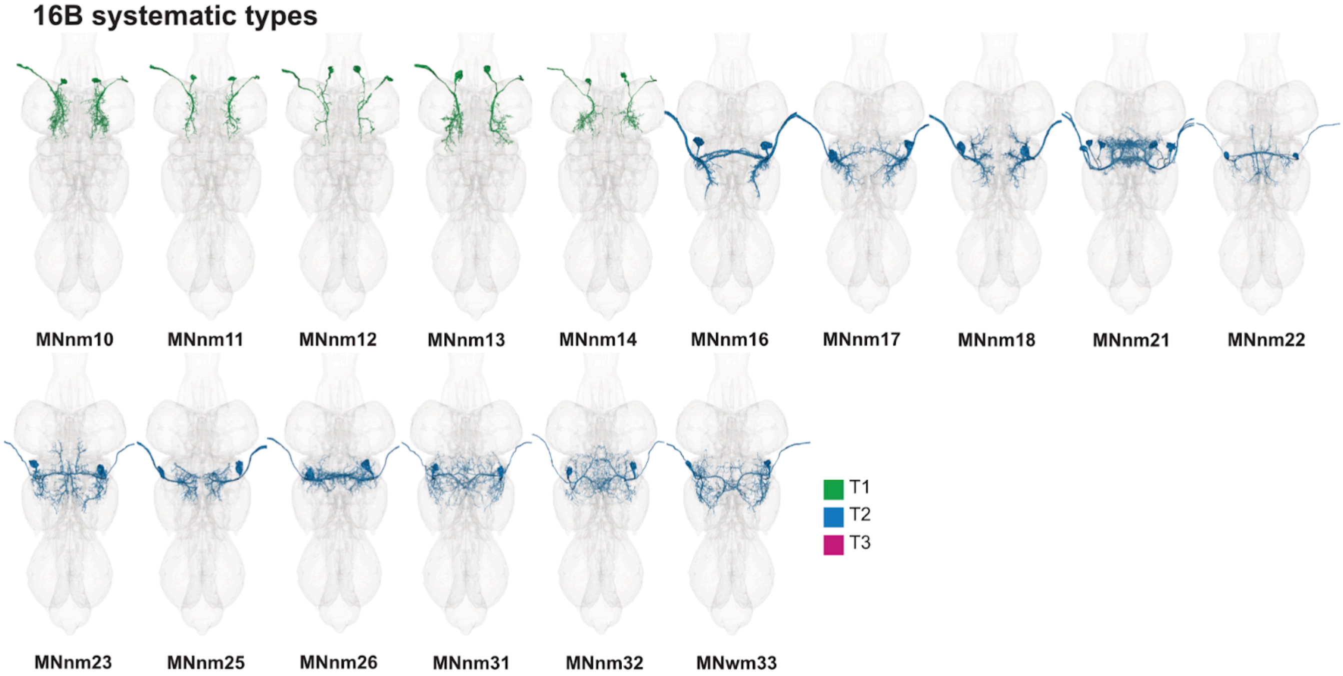

**Figure 39 - figure supplement 4. Systematic types of hemilineage 16B, continued.** Systematic types have been arranged in numerical order, with neurons of the same type that belong to distinct classes (e.g., intrinsic neuron vs ascending neuron) plotted separately but placed adjacent to each other. Individual neuron meshes have been coloured based on predicted neurotransmitter: dark orange = acetylcholine, blue = gaba, marine = glutamate, dark purple = unknown. Motor neurons (typed separately in (Cheong et al., 2023)) have been plotted by serial set if identified in multiple neuromeres and by systematic type if not. Individual motor neuron meshes have been coloured based on soma neuromere: dark green = T1, blue = T2, magenta = T3.

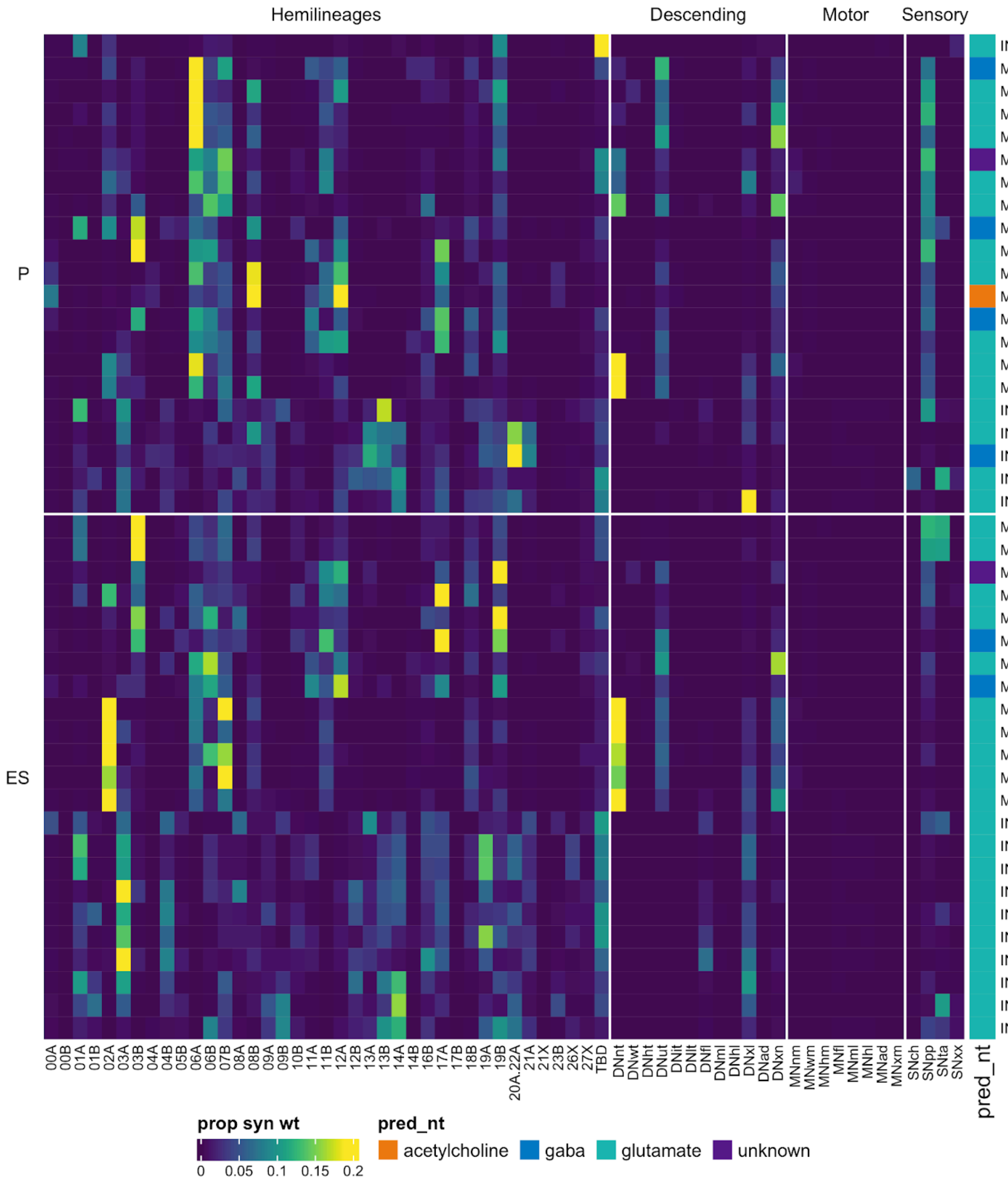

**Figure 39 - figure supplement 5. Connectivity to upstream partners by 16B primary and early secondary systematic types.** Proportions of synaptic weight to systematic types from upstream partners, normalised by row. 16B neurons have been clustered within each assigned birthtime window (P = primary, ES = early secondary, S = secondary) based on both upstream and downstream connectivity to hemilineages, descending neuron subclasses, motor neuron subclasses, and sensory neuron modalities. Annotation bar is coloured by the most common predicted neurotransmitter for the neurons of each type.

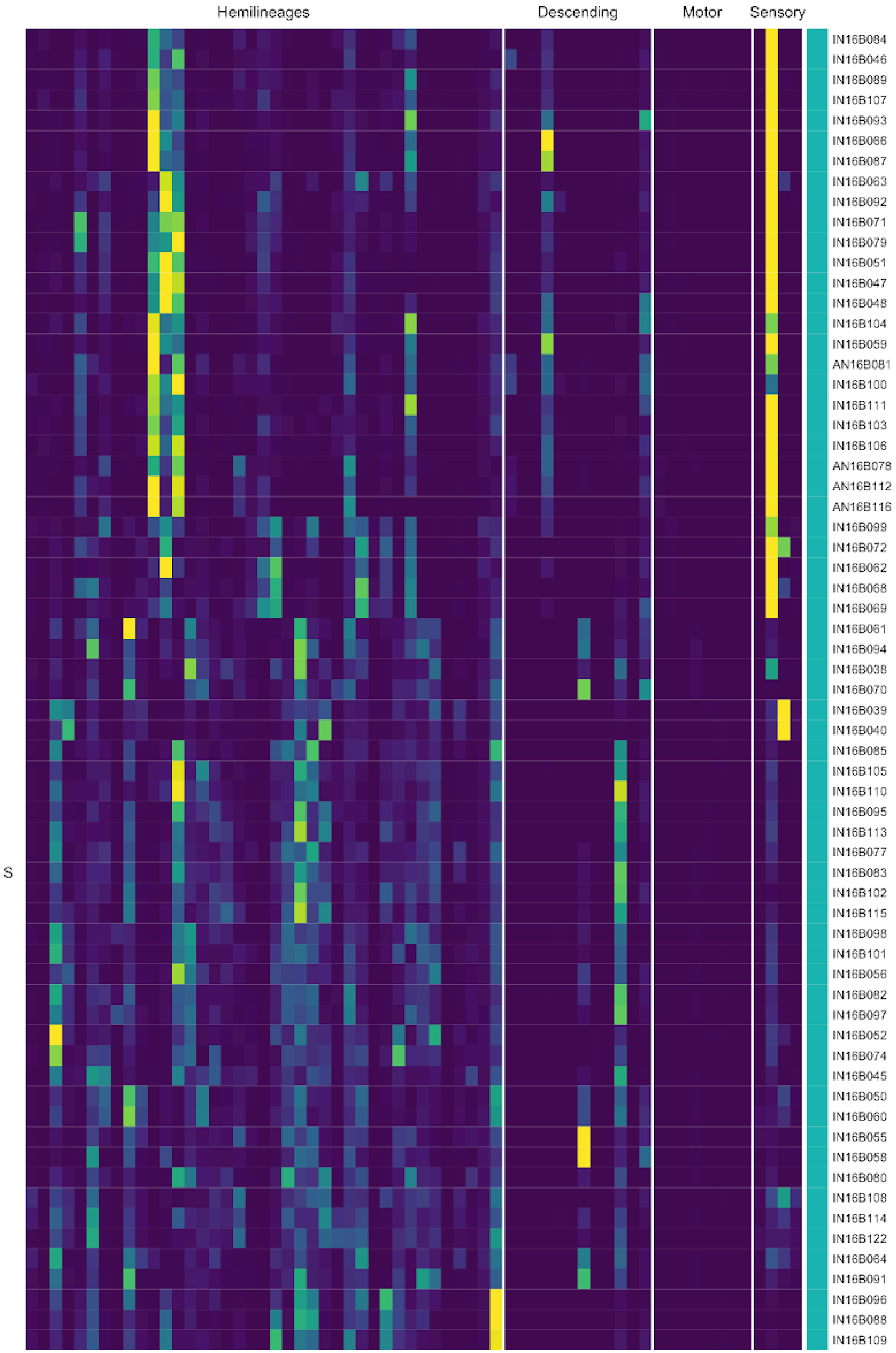
Hemilineages
Descending
Motor
Sensory
S
IN16B084
IN16B046
IN16B089
IN16B107
IN16B093
IN16B066
IN16B087
IN16B063
IN16B092
IN16B071
IN16B079
IN16B051
IN16B047
IN16B048
IN16B104
IN16B059
AN16B081
IN16B100
IN16B111
IN16B103
IN16B106
AN16B078
AN16B112
AN16B116
IN16B099
IN16B072
IN16B062
IN16B068
IN16B069
IN16B061
IN16B094
IN16B038
IN16B070
IN16B039
IN16B040
IN16B085
IN16B105
IN16B110
IN16B095
IN16B113
IN16B077
IN16B083
IN16B102
IN16B115
IN16B098
IN16B101
IN16B056
IN16B082
IN16B097
IN16B052
IN16B074
IN16B045
IN16B050
IN16B060
IN16B055
IN16B058
IN16B080
IN16B108
IN16B114
IN16B122
IN16B064
IN16B091
IN16B096
IN16B088
IN16B109

**Figure 39 - figure supplement 6. Connectivity to upstream partners by 16B secondary systematic types.** Proportions of synaptic weight to systematic types from upstream partners, normalised by row. 16B neurons have been clustered within each assigned birthtime window (P = primary, ES = early secondary, S = secondary) based on both upstream and downstream connectivity to hemilineages, descending neuron subclasses, motor neuron subclasses, and sensory neuron modalities. The annotation bar is coloured by the most common predicted neurotransmitter within each type.

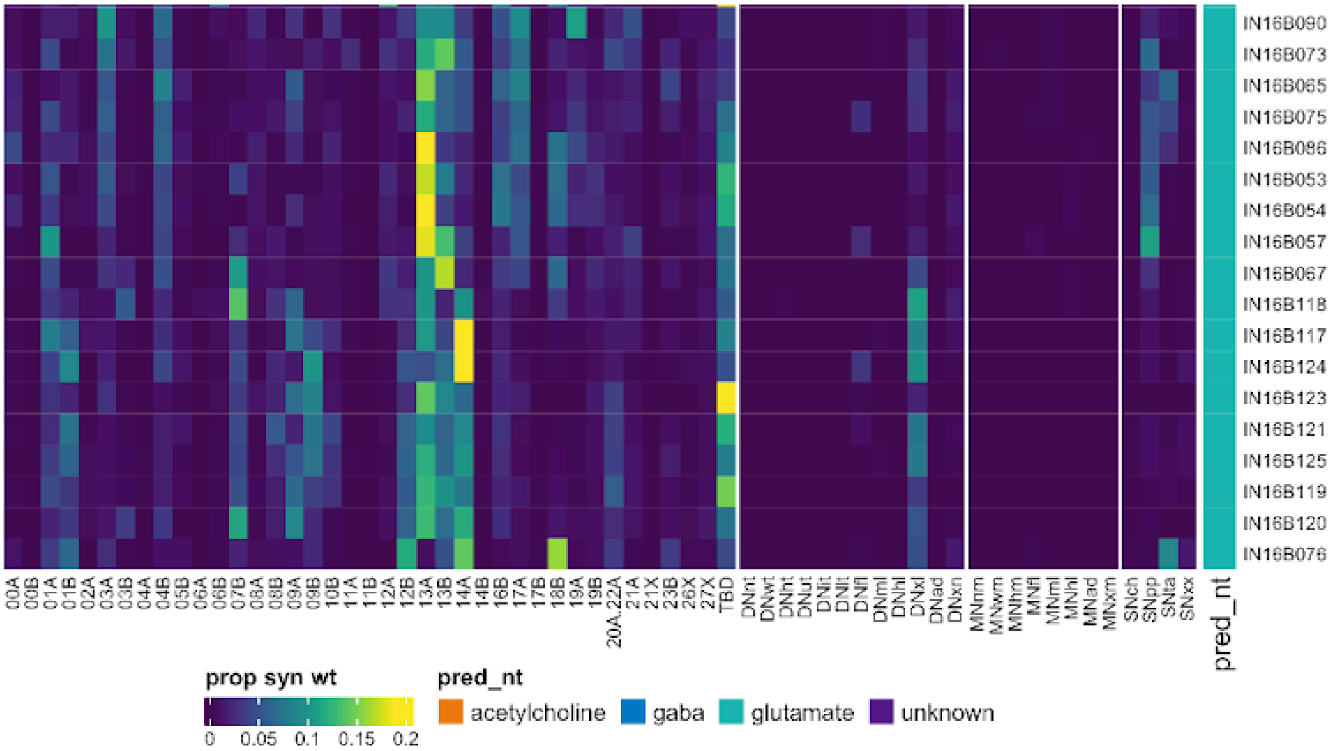

**Figure 39 - figure supplement 7. Connectivity to upstream partners by 16B secondary systematic types, continued.** Proportions of synaptic weight to systematic types from upstream partners, normalised by row. 16B neurons have been clustered within each assigned birthtime window (P = primary, ES = early secondary, S = secondary) based on both upstream and downstream connectivity to hemilineages, descending neuron subclasses, motor neuron subclasses, and sensory neuron modalities. The annotation bar is coloured by the most common predicted neurotransmitter within each type.

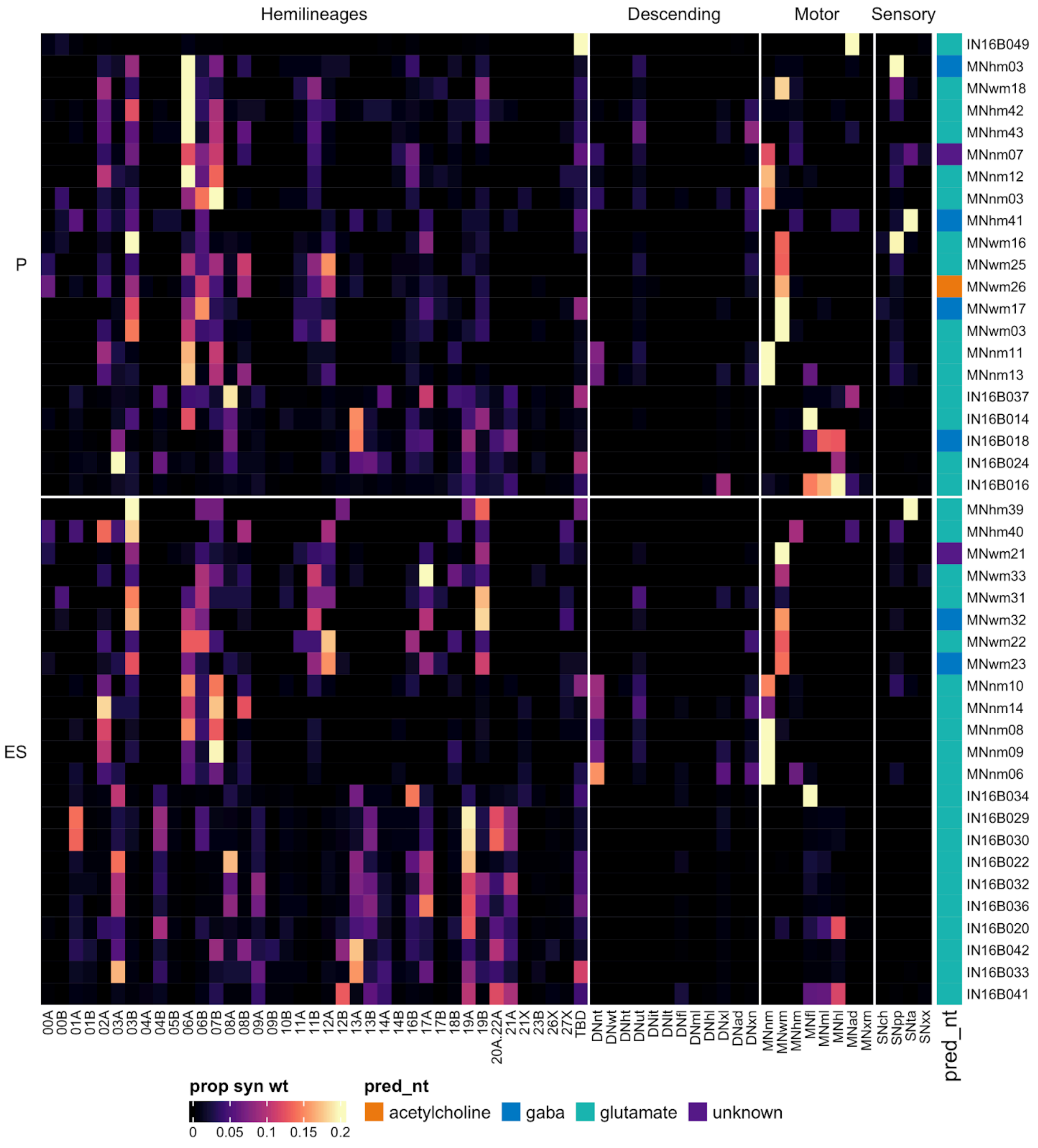

**Figure 39 - figure supplement 8. Connectivity to downstream partners by 16B primary and early secondary systematic types.** Proportions of synaptic weight from systematic types to downstream partners, normalised by row. 16B neurons have been clustered within each assigned birthtime window (P = primary, ES = early secondary, S = secondary) based on both upstream and downstream connectivity to hemilineages, descending neuron subclasses, motor neuron subclasses, and sensory neuron modalities. The annotation bar is coloured by the most common predicted neurotransmitter within each type.

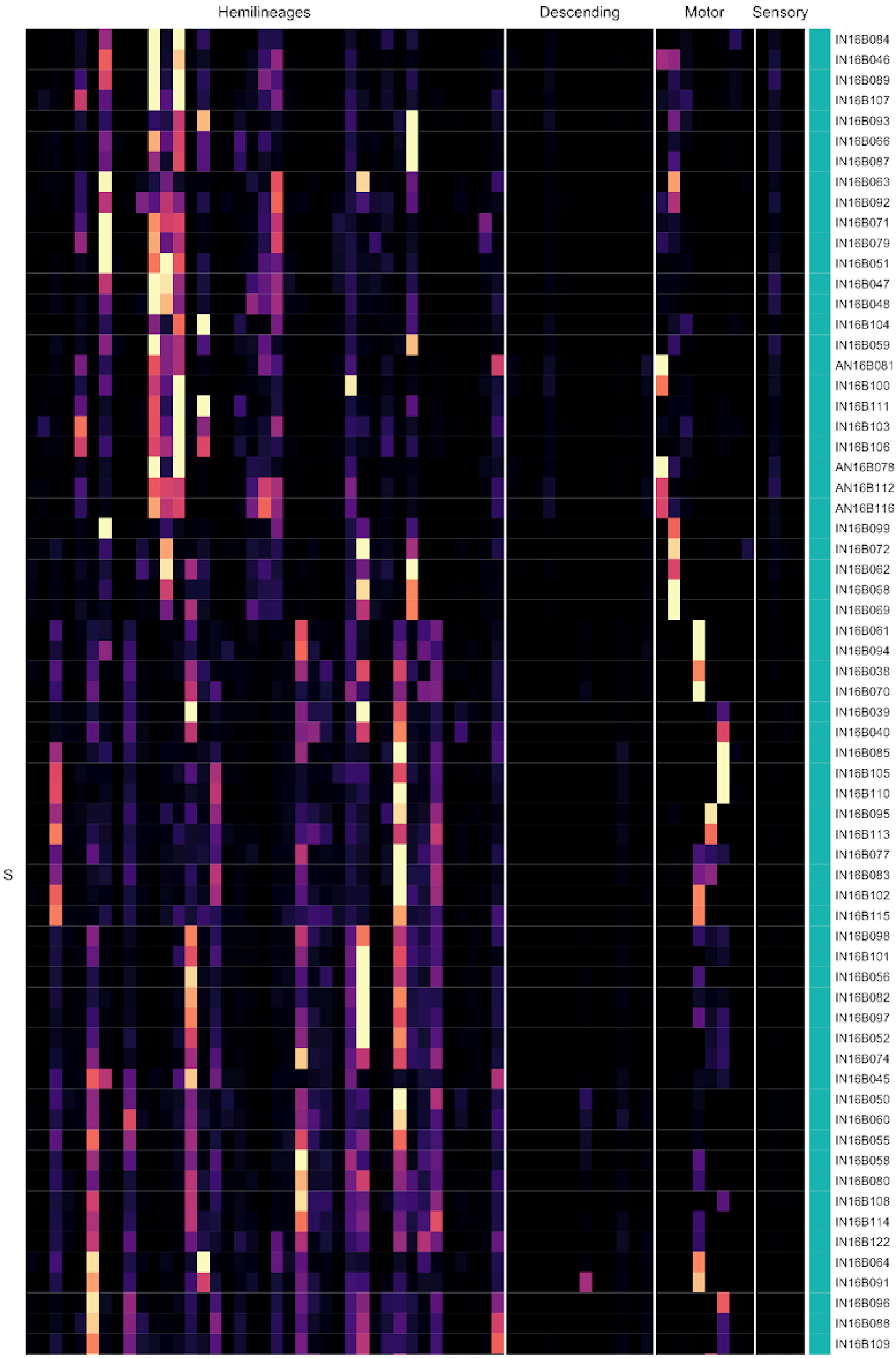
Hemilineages
Descending
Motor
Sensory
S
IN16B084
IN16B046
IN16B089
IN16B107
IN16B093
IN16B066
IN16B087
IN16B063
IN16B092
IN16B071
IN16B079
IN16B051
IN16B047
IN16B048
IN16B104
IN16B059
AN16B081
IN16B100
IN16B111
IN16B103
IN16B106
AN16B078
AN16B112
AN16B116
IN16B099
IN16B072
IN16B062
IN16B068
IN16B069
IN16B061
IN16B094
IN16B038
IN16B070
IN16B039
IN16B040
IN16B085
IN16B105
IN16B110
IN16B095
IN16B113
IN16B077
IN16B083
IN16B102
IN16B115
IN16B098
IN16B101
IN16B056
IN16B082
IN16B097
IN16B052
IN16B074
IN16B045
IN16B050
IN16B060
IN16B055
IN16B058
IN16B080
IN16B108
IN16B114
IN16B122
IN16B064
IN16B091
IN16B096
IN16B088
IN16B109

**Figure 39 - figure supplement 9. Connectivity to downstream partners by 16B econdary systematic types.** Proportions of synaptic weight from systematic types to downstream partners, normalised by row. 16B neurons have been clustered within each assigned birthtime window (P = primary, ES = early secondary, S = secondary) based on both upstream and downstream connectivity to hemilineages, descending neuron subclasses, motor neuron subclasses, and sensory neuron modalities. The annotation bar is coloured by the most common predicted neurotransmitter for the neurons of each type.

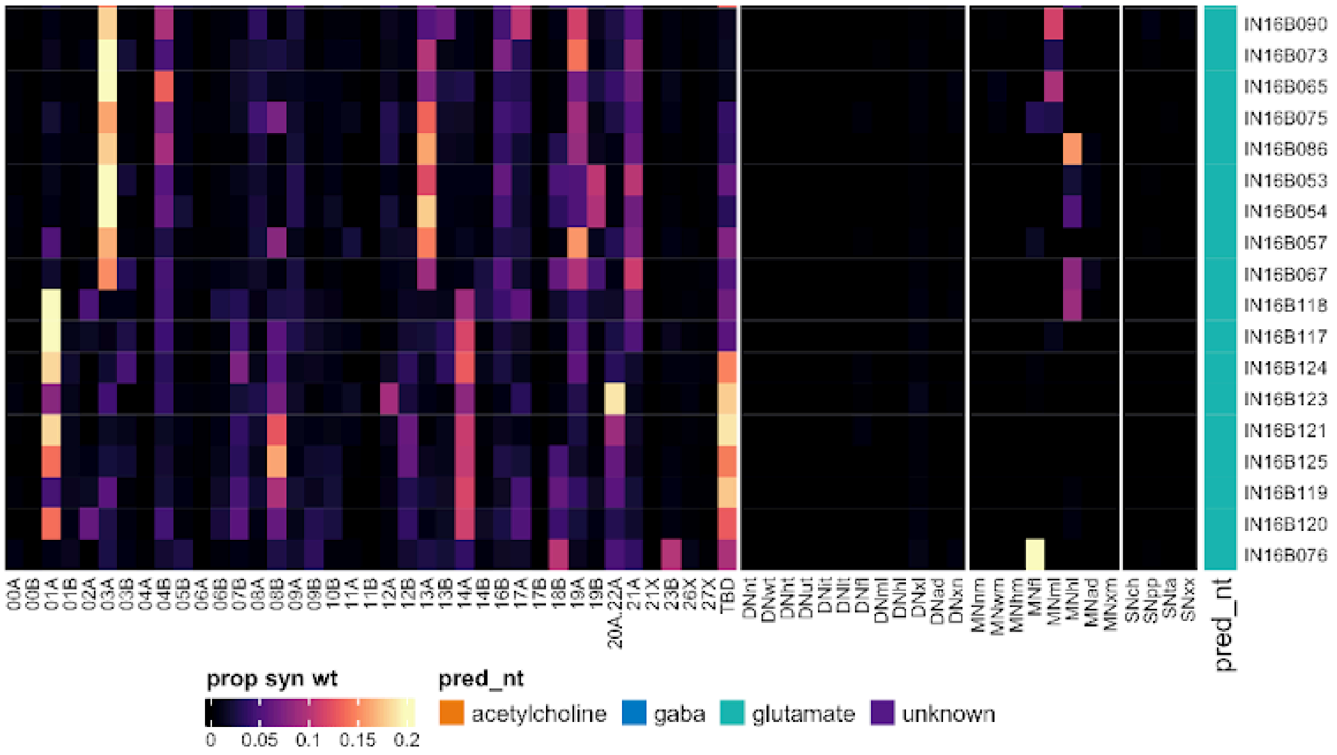

**Figure 39 - figure supplement 10. Connectivity to downstream partners by 16B secondary systematic types, continued.** Proportions of synaptic weight from systematic types to downstream partners, normalised by row. 16B neurons have been clustered within each assigned birthtime window (P = primary, ES = early secondary, S = secondary) based on both upstream and downstream connectivity to hemilineages, descending neuron subclasses, motor neuron subclasses, and sensory neuron modalities. The annotation bar is coloured by the most common predicted neurotransmitter for the neurons of each type.

Hemilineage 17A

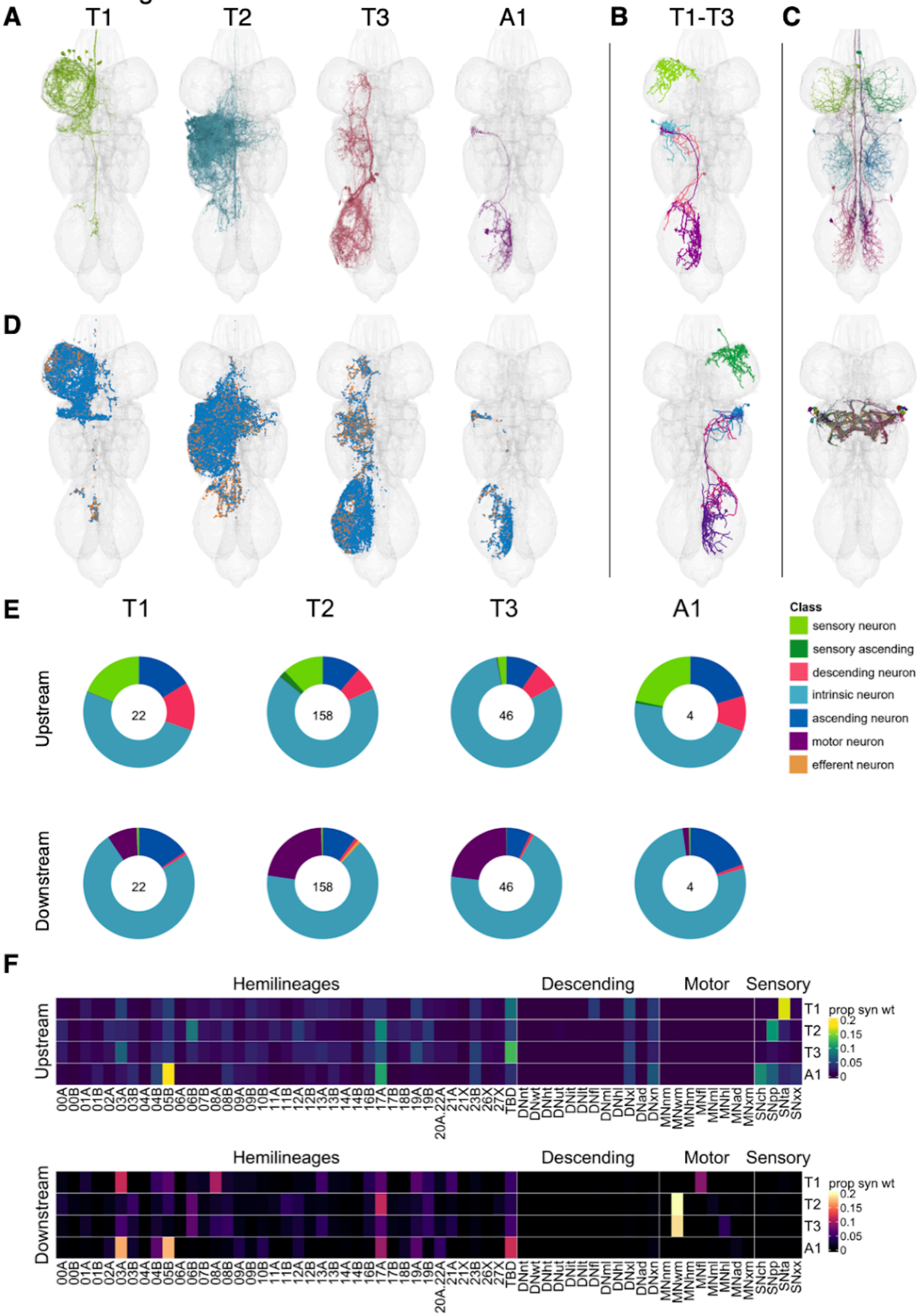

**Figure 40. Hemilineage 17A. A.** Meshes of all RHS secondary neurons plotted in neuromere-specific colours. **B.** "Representative" secondary neuron skeletons plotted in hemineuromere-specific colours. The skeleton with the top accumulated NBLAST score among all neurons from the hemilineage in a given hemineuromere was used. **C.** Neuron meshes of selected examples. Top: ascending serial set 11036 (T1-A1). Bottom: putative electrical subcluster 10629/dMS2. **D.** Predicted synapses of RHS secondary neurons. Blue: postsynapses; dark orange: presynapses. **E.** Proportions of connections from secondary neurons to upstream or downstream partners, normalised by neuromere and coloured by broad class. Numbers of query neurons appear in the centre. **F.** Proportions of synaptic weight from secondary neurons originating in each neuromere to upstream or downstream partners, normalised by row.

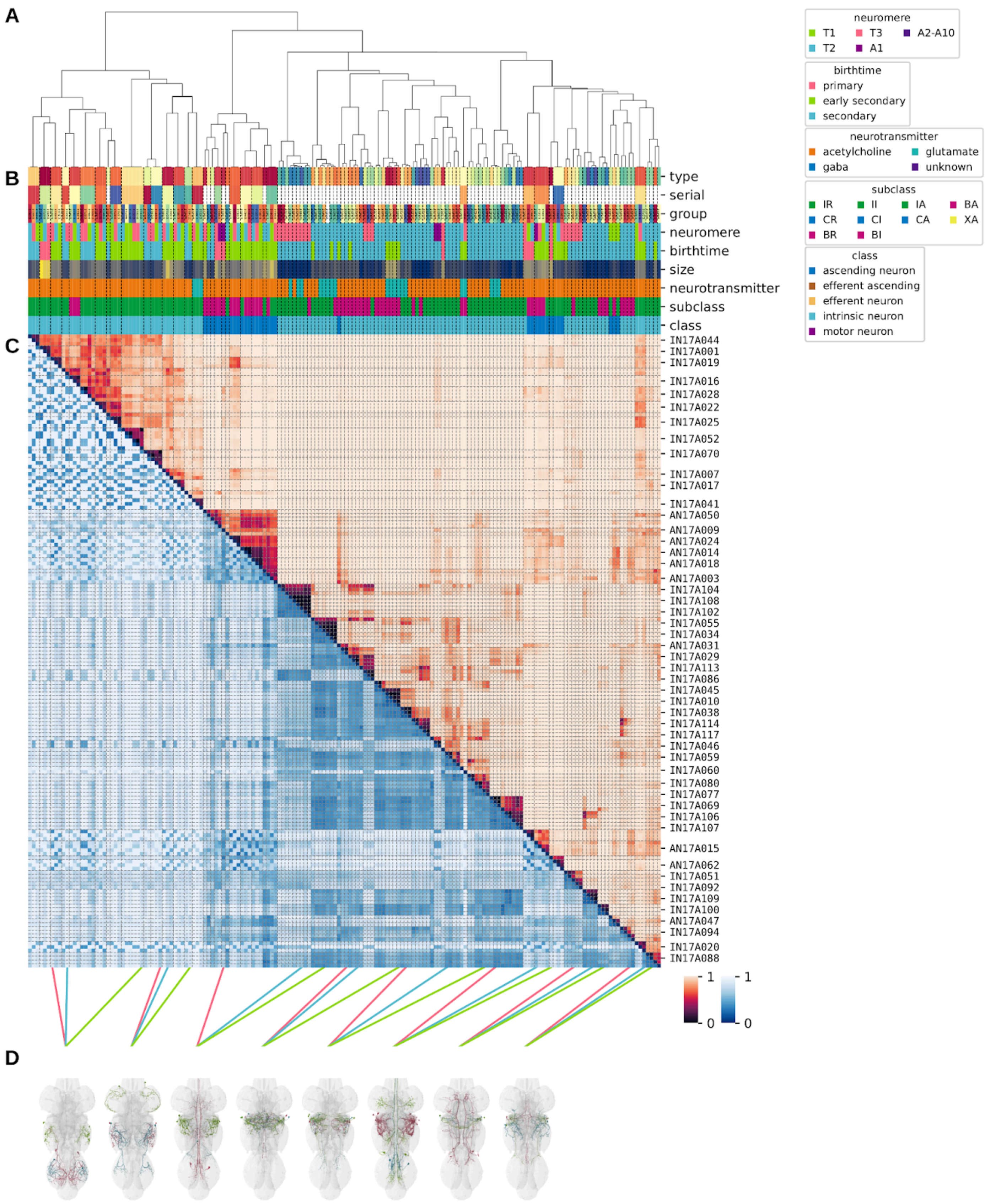

**Figure 40 - figure supplement 1. Systematic typing of hemilineage 17A. A.** Hierarchical clustering dendrogram of hemilineage groups by laterally and serially aggregated connectivity cosine clustering. **B.** Categorical annotations of each hemilineage group, each column corresponding to the aligned leaf in A. Colours for type, serial set, and group are arbitrary for visualisation. Colours for neuromere, birthtime, neurotransmitter, subclass, and class are as in all other figures. **C.** Similarity distance heatmap for hemilineage. Cosine distance is in the upper triangle, while laterally symmetrised NBLAST distance is in the lower triangle. Systematic type names of some types are labelled. **D.** Morphologically representative groups from dendrogram subtrees. Each group, indicated by colour and line

connecting to its column in B and C, is the most morphologically representative group (medoid of NBLAST distance) from a subtree of A. The subtrees (flat clusters) are equal height cuts of A determined to yield the number of groups per plot and plots in D.

#### 17A systematic types

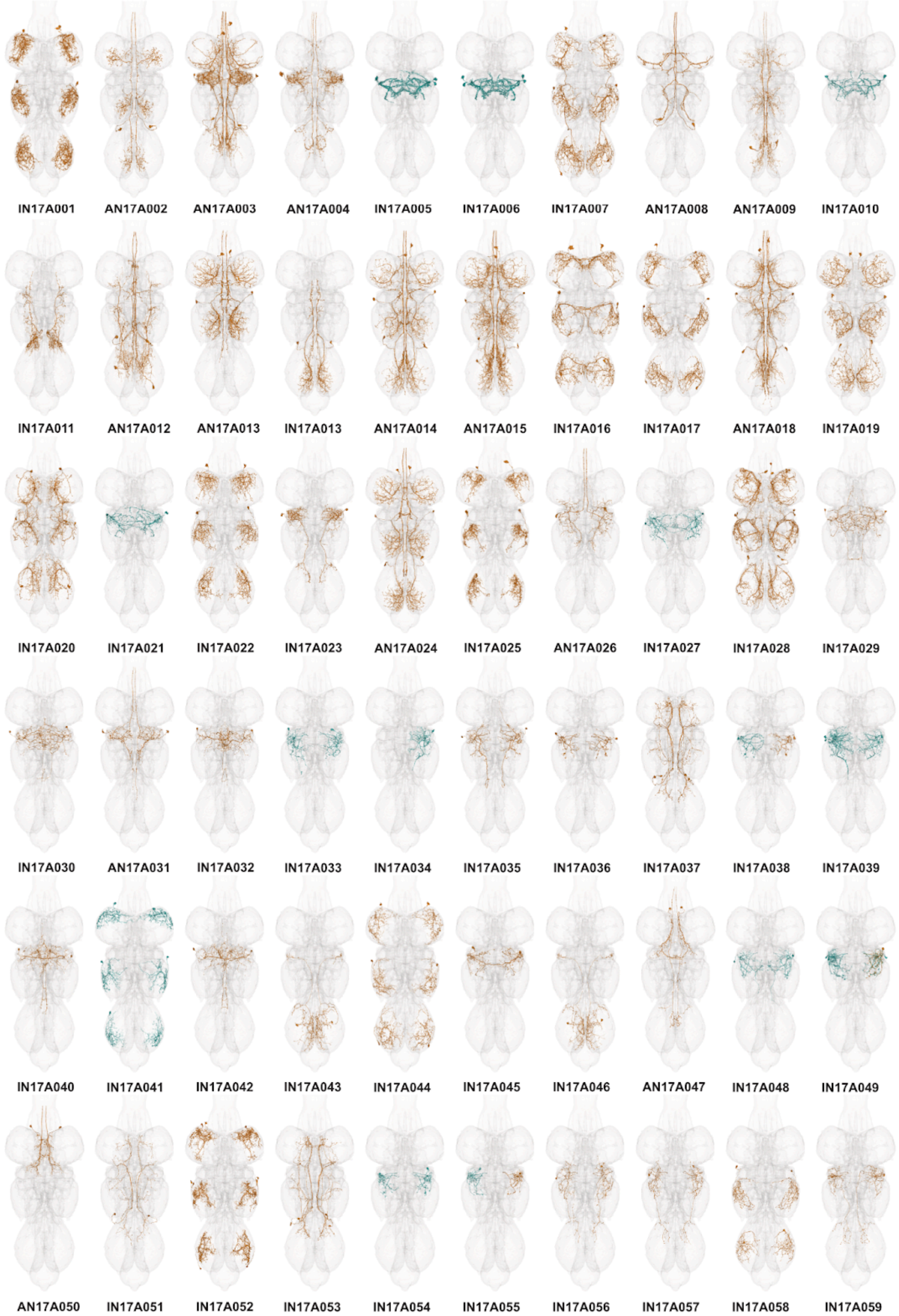

**Figure 40 - figure supplement 2. Systematic types of hemilineage 17A.** Systematic types have been arranged in numerical order, with neurons of the same type that belong to distinct classes (e.g., intrinsic neuron vs ascending neuron) plotted separately but placed adjacent to each other. Individual neuron meshes have been coloured based on predicted neurotransmitter: dark orange = acetylcholine, blue = gaba, marine = glutamate, dark purple = unknown.

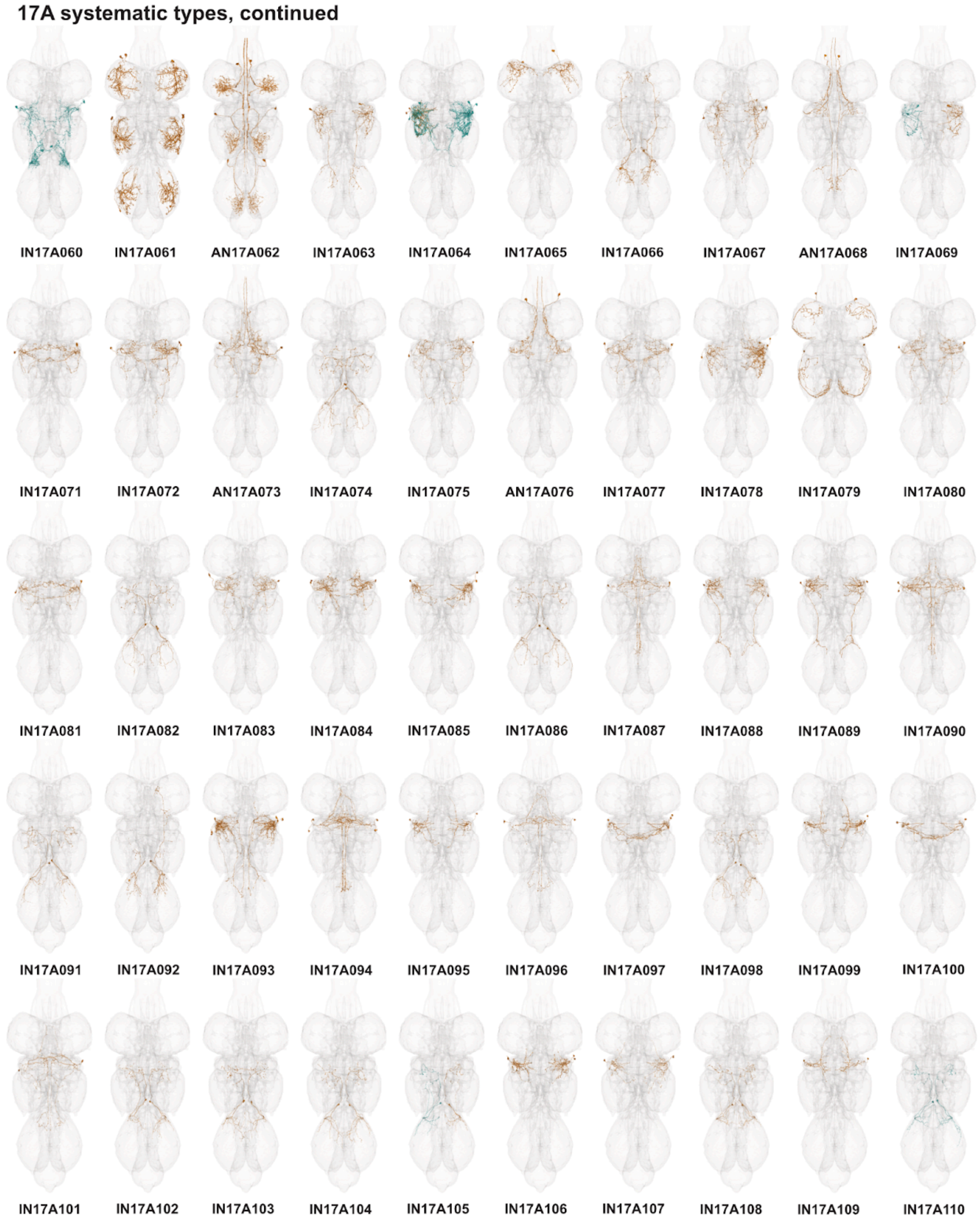

**Figure 40 - figure supplement 3. Systematic types of hemilineage 17A, continued.** Systematic types have been arranged in numerical order, with neurons of the same type that belong to distinct classes (e.g., intrinsic neuron vs ascending neuron) plotted separately but placed adjacent to each other. Individual neuron meshes have been coloured based on predicted neurotransmitter: dark orange = acetylcholine, blue = gaba, marine = glutamate, dark purple = unknown.

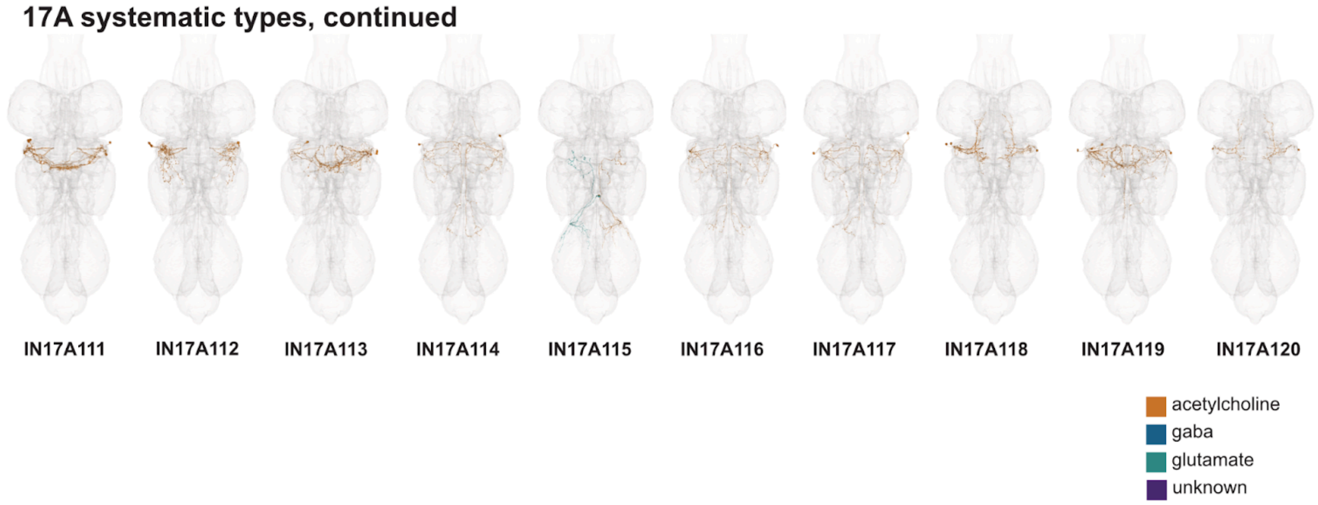

**Figure 40 - figure supplement 4. Systematic types of hemilineage 17A, continued.** Systematic types have been arranged in numerical order, with neurons of the same type that belong to distinct classes (e.g., intrinsic neuron vs ascending neuron) plotted separately but placed adjacent to each other. Individual neuron meshes have been coloured based on predicted neurotransmitter: dark orange = acetylcholine, blue = gaba, marine = glutamate, dark purple = unknown.

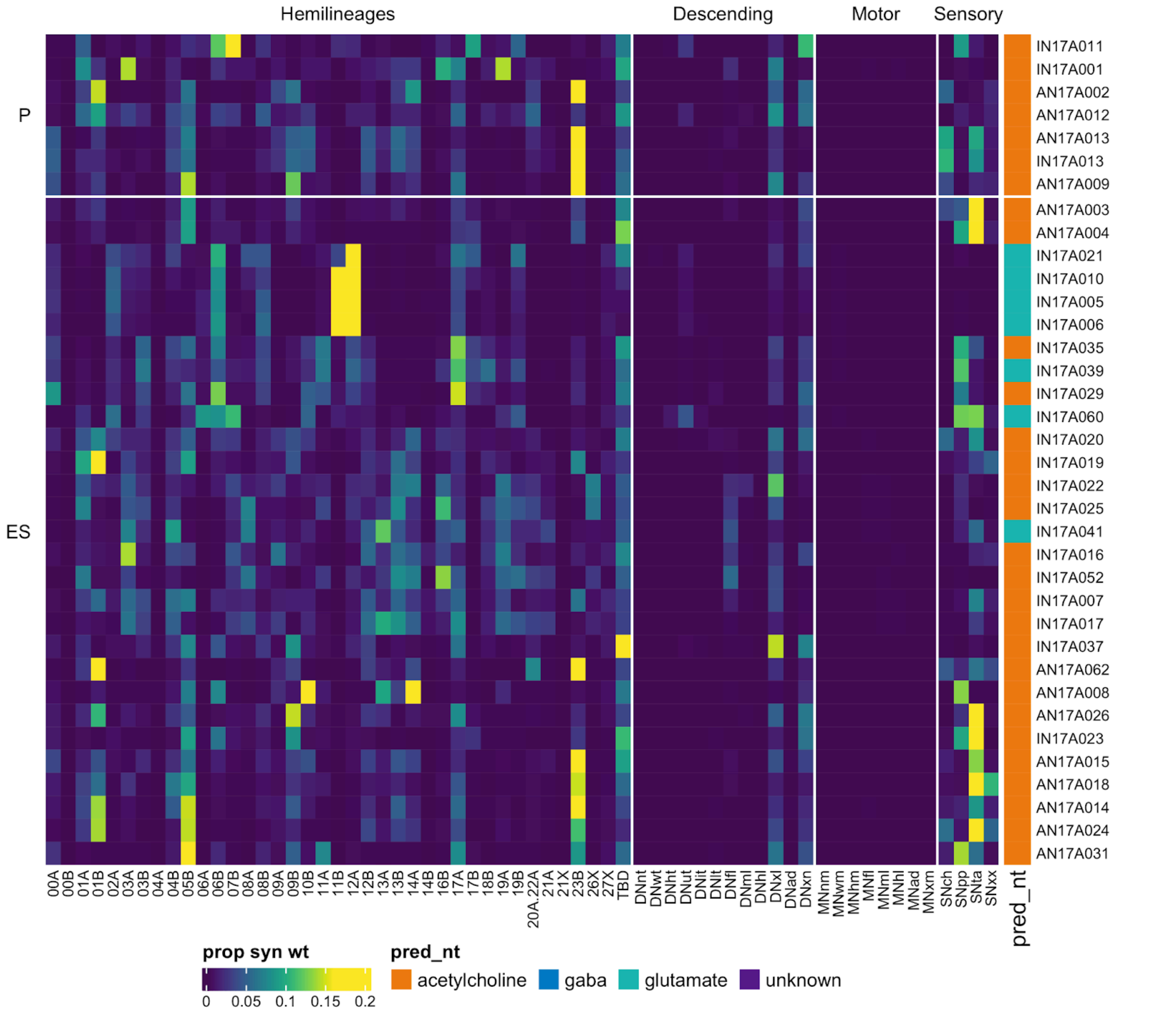

**Figure 40 - figure supplement 5. Connectivity to upstream partners by 17A primary and early secondary systematic types.** Proportions of synaptic weight to systematic types from upstream partners, normalised by row. 17A neurons have been clustered within each assigned birthtime window (P = primary, ES = early secondary, S = secondary) based on both upstream and downstream connectivity to hemilineages, descending neuron subclasses, motor neuron subclasses, and sensory neuron modalities. Annotation bar is coloured by the most common predicted neurotransmitter for the neurons of each type.

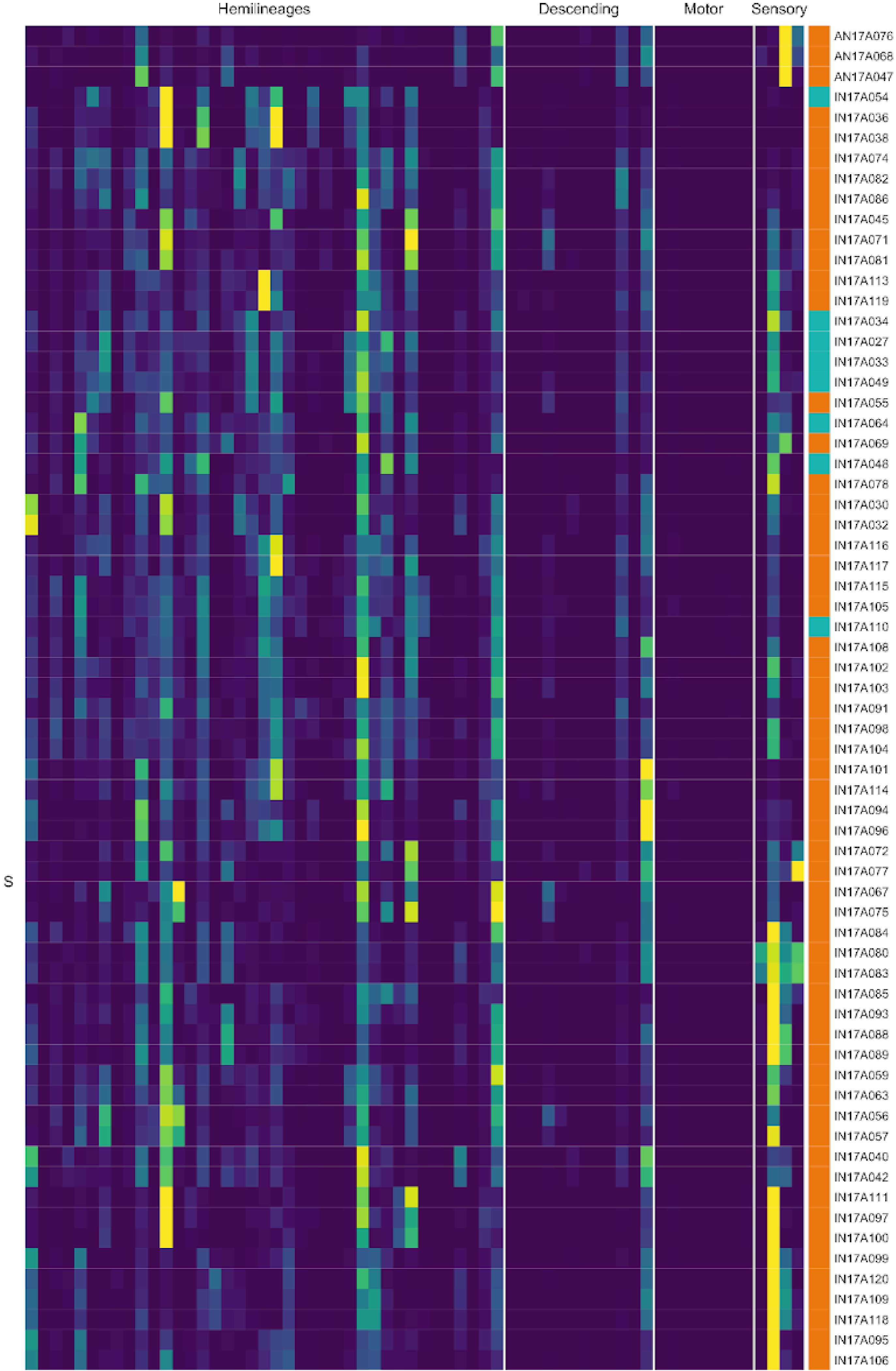
Hemilineages
Descending
Motor
Sensory
S
AN17A076
AN17A068
AN17A047
IN17A054
IN17A036
IN17A038
IN17A074
IN17A082
IN17A086
IN17A045
IN17A071
IN17A081
IN17A113
IN17A119
IN17A034
IN17A027
IN17A033
IN17A049
IN17A055
IN17A064
IN17A069
IN17A048
IN17A078
IN17A030
IN17A032
IN17A116
IN17A117
IN17A115
IN17A105
IN17A110
IN17A108
IN17A102
IN17A103
IN17A091
IN17A098
IN17A104
IN17A101
IN17A114
IN17A094
IN17A096
IN17A072
IN17A077
IN17A067
IN17A075
IN17A084
IN17A080
IN17A083
IN17A085
IN17A093
IN17A088
IN17A089
IN17A059
IN17A063
IN17A056
IN17A057
IN17A040
IN17A042
IN17A111
IN17A097
IN17A100
IN17A099
IN17A120
IN17A109
IN17A118
IN17A095
IN17A106

**Figure 40 - figure supplement 6. Connectivity to upstream partners by 17A secondary systematic types.** Proportions of synaptic weight to systematic types from upstream partners, normalised by row. 17A neurons have been clustered within each assigned birthtime window (P = primary, ES = early secondary, S = secondary) based on both upstream and downstream connectivity to hemilineages, descending neuron subclasses, motor neuron subclasses, and sensory neuron modalities. The annotation bar is coloured by the most common predicted neurotransmitter within each type.

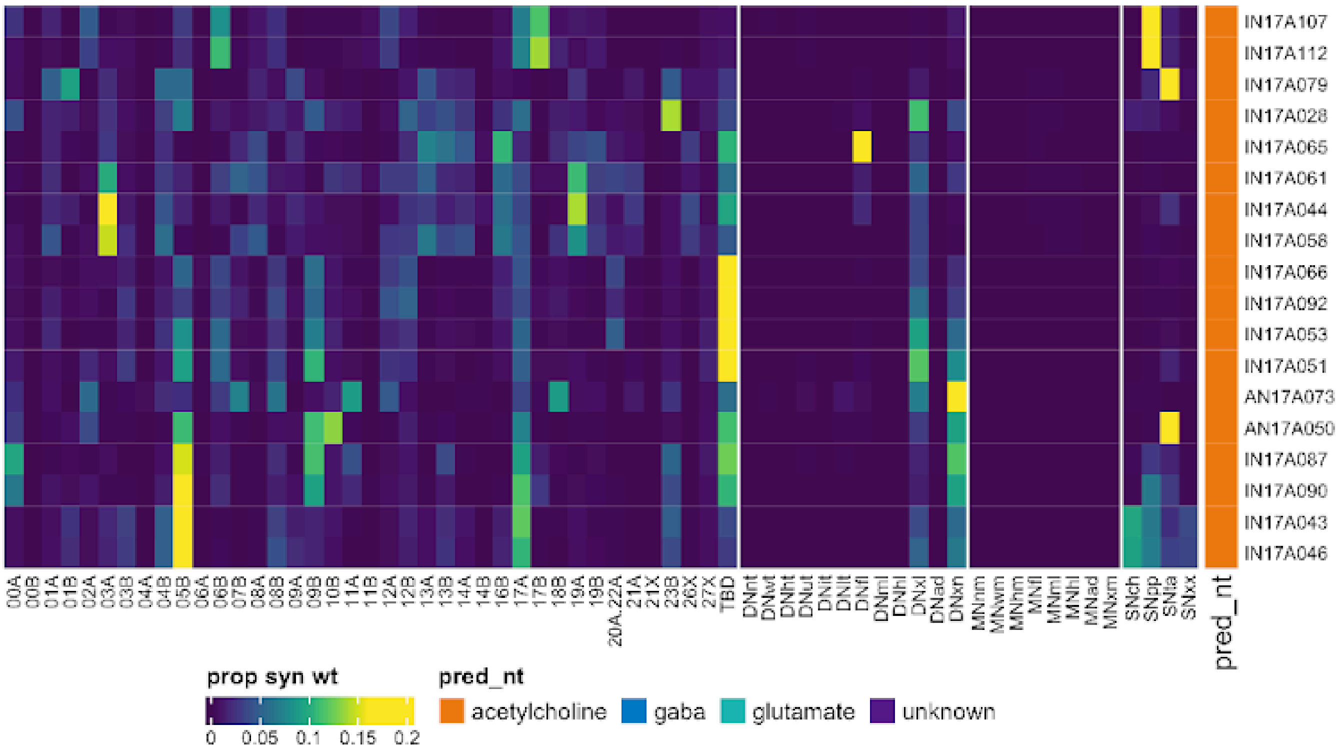

**Figure 40 - figure supplement 7. Connectivity to upstream partners by 17A secondary systematic types, continued.** Proportions of synaptic weight to systematic types from upstream partners, normalised by row. 17A neurons have been clustered within each assigned birthtime window (P = primary, ES = early secondary, S = secondary) based on both upstream and downstream connectivity to hemilineages, descending neuron subclasses, motor neuron subclasses, and sensory neuron modalities. The annotation bar is coloured by the most common predicted neurotransmitter within each type.

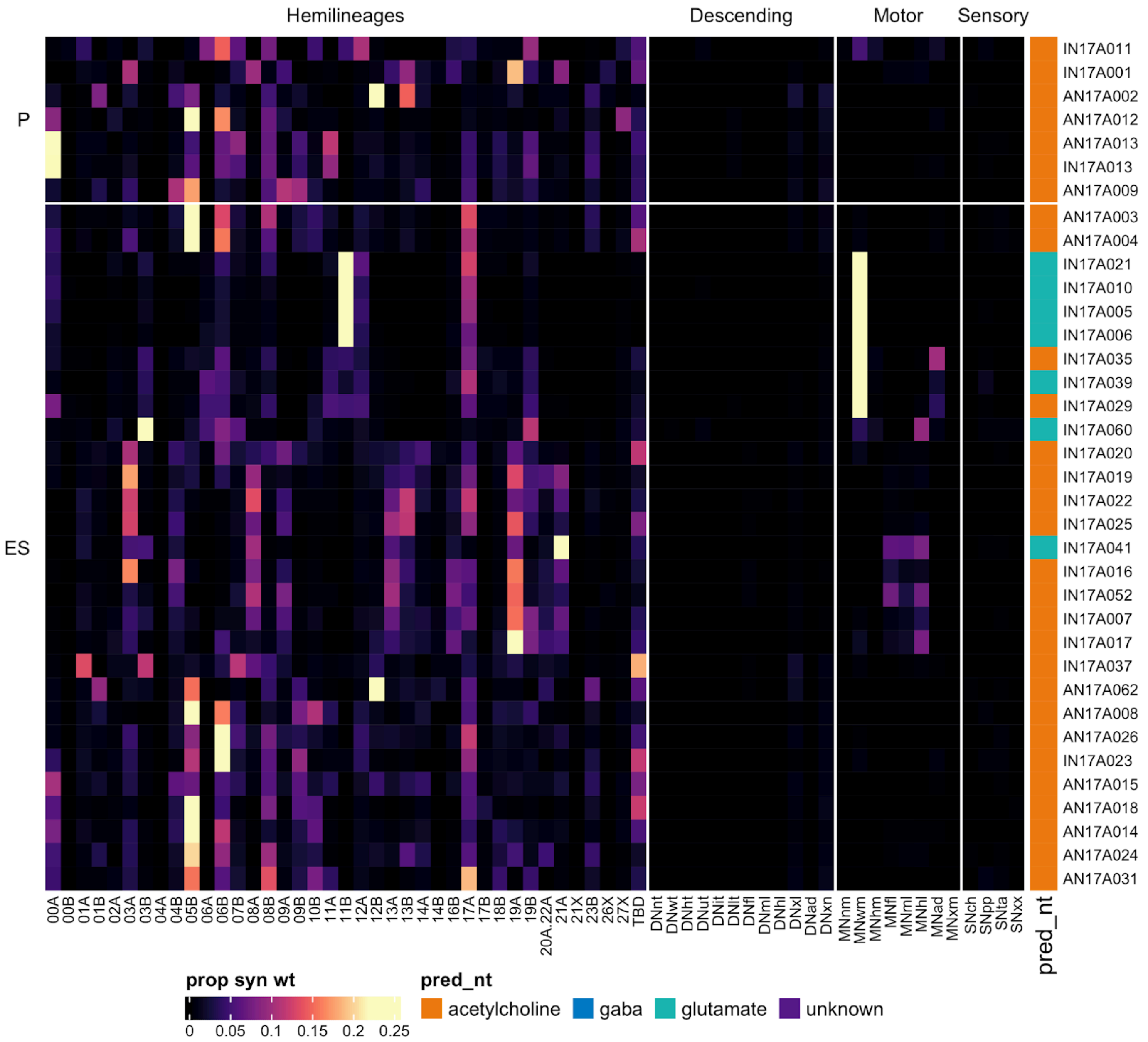

**Figure 40 - figure supplement 8. Connectivity to downstream partners by 17A primary and early secondary systematic types.** Proportions of synaptic weight from systematic types to downstream partners, normalised by row. 17A neurons have been clustered within each assigned birthtime window (P = primary, ES = early secondary, S = secondary) based on both upstream and downstream connectivity to hemilineages, descending neuron subclasses, motor neuron subclasses, and sensory neuron modalities. The annotation bar is coloured by the most common predicted neurotransmitter within each type.

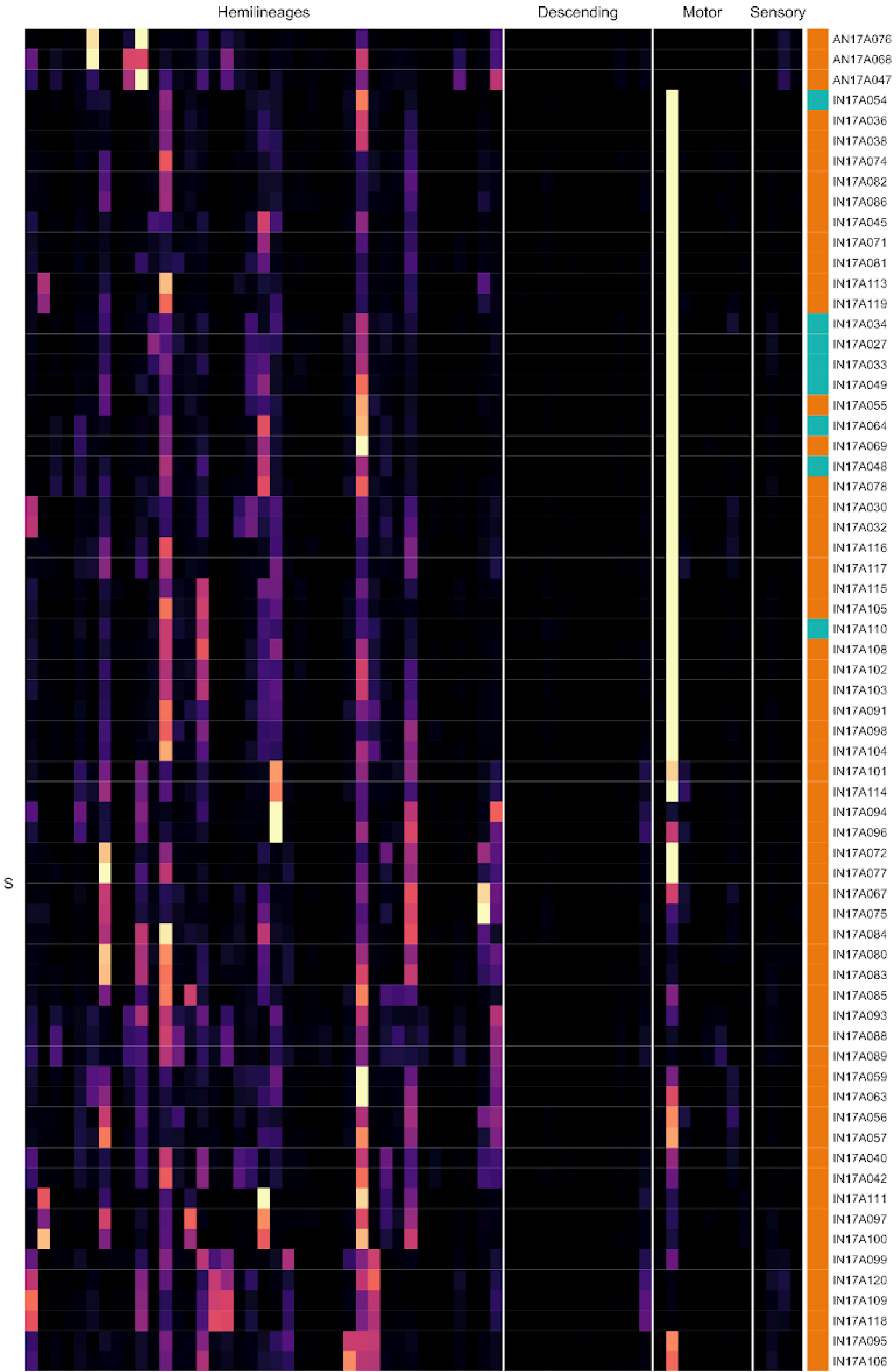

Hemilineages
Descending
Motor
Sensory
S
AN17A076
AN17A068
AN17A047
IN17A054
IN17A036
IN17A038
IN17A074
IN17A082
IN17A086
IN17A045
IN17A071
IN17A081
IN17A113
IN17A119
IN17A034
IN17A027
IN17A033
IN17A049
IN17A055
IN17A064
IN17A069
IN17A048
IN17A078
IN17A030
IN17A032
IN17A116
IN17A117
IN17A115
IN17A105
IN17A110
IN17A108
IN17A102
IN17A103
IN17A091
IN17A098
IN17A104
IN17A101
IN17A114
IN17A094
IN17A096
IN17A072
IN17A077
IN17A067
IN17A075
IN17A084
IN17A080
IN17A083
IN17A085
IN17A093
IN17A088
IN17A089
IN17A059
IN17A063
IN17A056
IN17A057
IN17A040
IN17A042
IN17A111
IN17A097
IN17A100
IN17A099
IN17A120
IN17A109
IN17A118
IN17A095
IN17A106

**Figure 40 - figure supplement 9. Connectivity to downstream partners by 17A secondary systematic types.** Proportions of synaptic weight from systematic types to downstream partners, normalised by row. 17A neurons have been clustered within each assigned birthtime window (P = primary, ES = early secondary, S = secondary) based on both upstream and downstream connectivity to hemilineages, descending neuron subclasses, motor neuron subclasses, and sensory neuron modalities. The annotation bar is coloured by the most common predicted neurotransmitter for the neurons of each type.

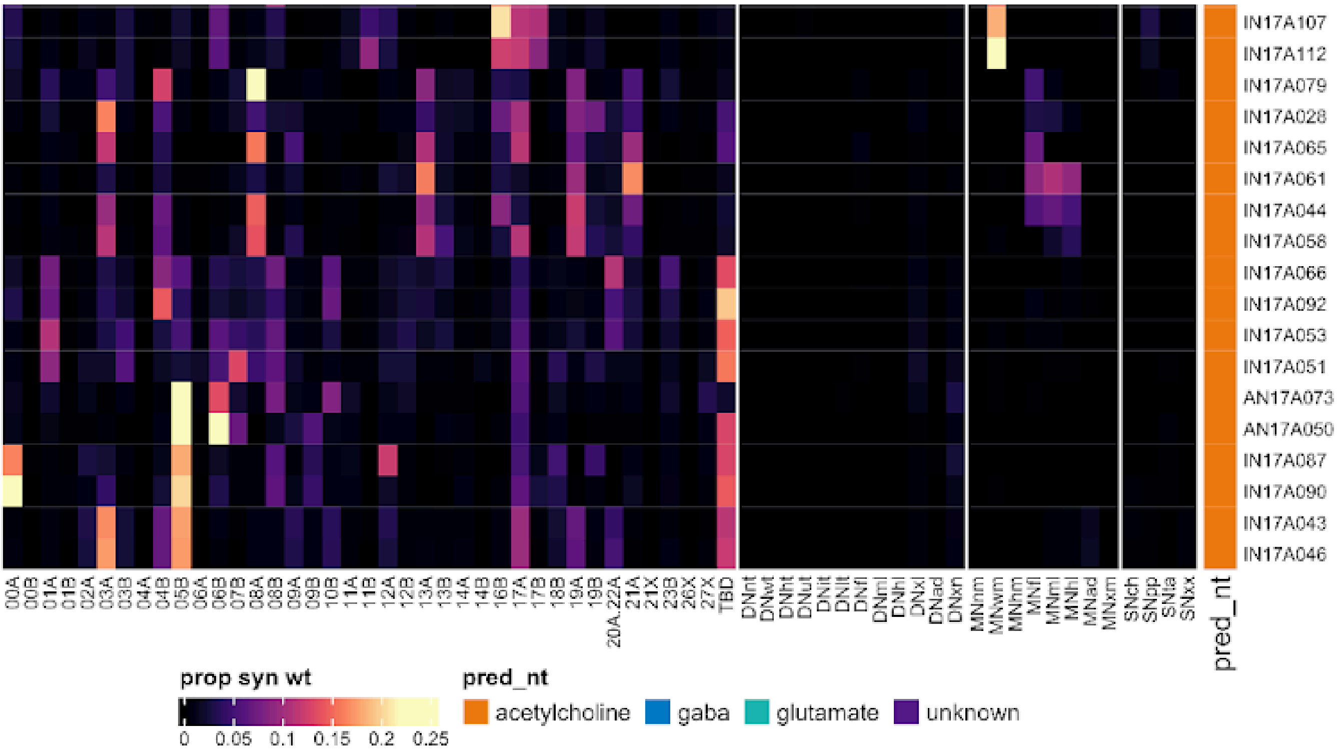

**Figure 40 - figure supplement 10. Connectivity to downstream partners by 17A secondary systematic types, continued.** Proportions of synaptic weight from systematic types to downstream partners, normalised by row. 17A neurons have been clustered within each assigned birthtime window (P = primary, ES = early secondary, S = secondary) based on both upstream and downstream connectivity to hemilineages, descending neuron subclasses, motor neuron subclasses, and sensory neuron modalities. The annotation bar is coloured by the most common predicted neurotransmitter for the neurons of each type.

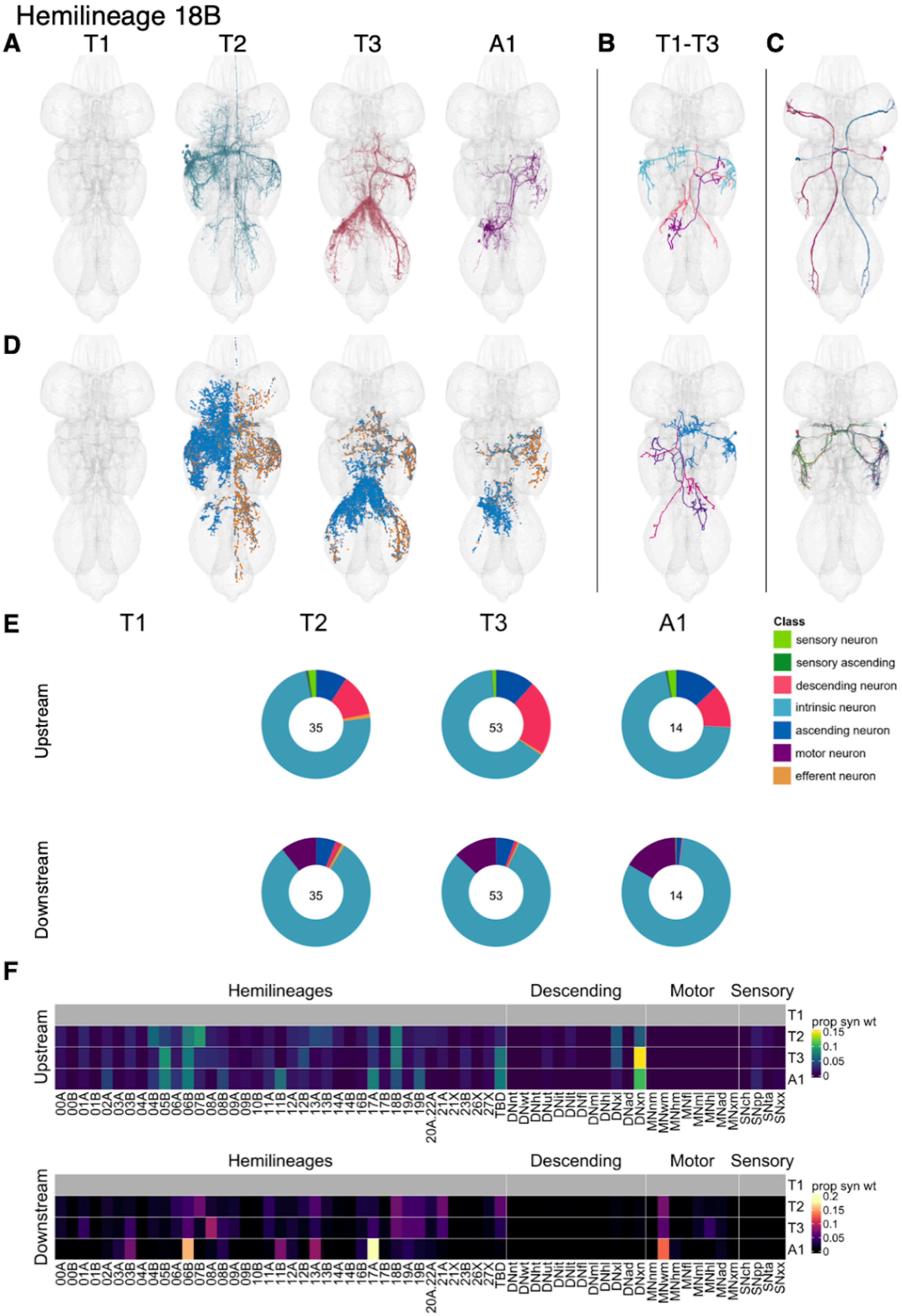

Hemilineage 18B
A
T1
T2
T3
A1
B
T1-T3
C
D
E
T1
T2
T3
A1
Upstream
Downstream
35
53
14
35
53
14
Class
sensory neuron
sensory ascending
descending neuron
intrinsic neuron
ascending neuron
motor neuron
efferent neuron
F
Hemilineages
Descending
Motor
Sensory
Upstream
Downstream
T1
T2
T3
A1
prop syn wt
0.15
0.1
0.05
0
0.2
0.15
0.1
0.05
0
00A 00B 01A 01B 02A 03A 03B 04A 04B 05B 06A 06B 07B 08A 08B 09A 09B 10B 11A 11B 12A 12B 13A 13B 14A 14B 16B 17A 17B 18B 19A 19B 20A.22A 21A 21X 23B 26X 27X TBD
DNnt DNwt DNht DNut DNit DNlt DNfl DNml DNnl DNhl DNxl DNad DNxn
MNnm MNwm MNhm MNfl MNml MNhl MNad MNxm
SNch SNpp SNta SNxx

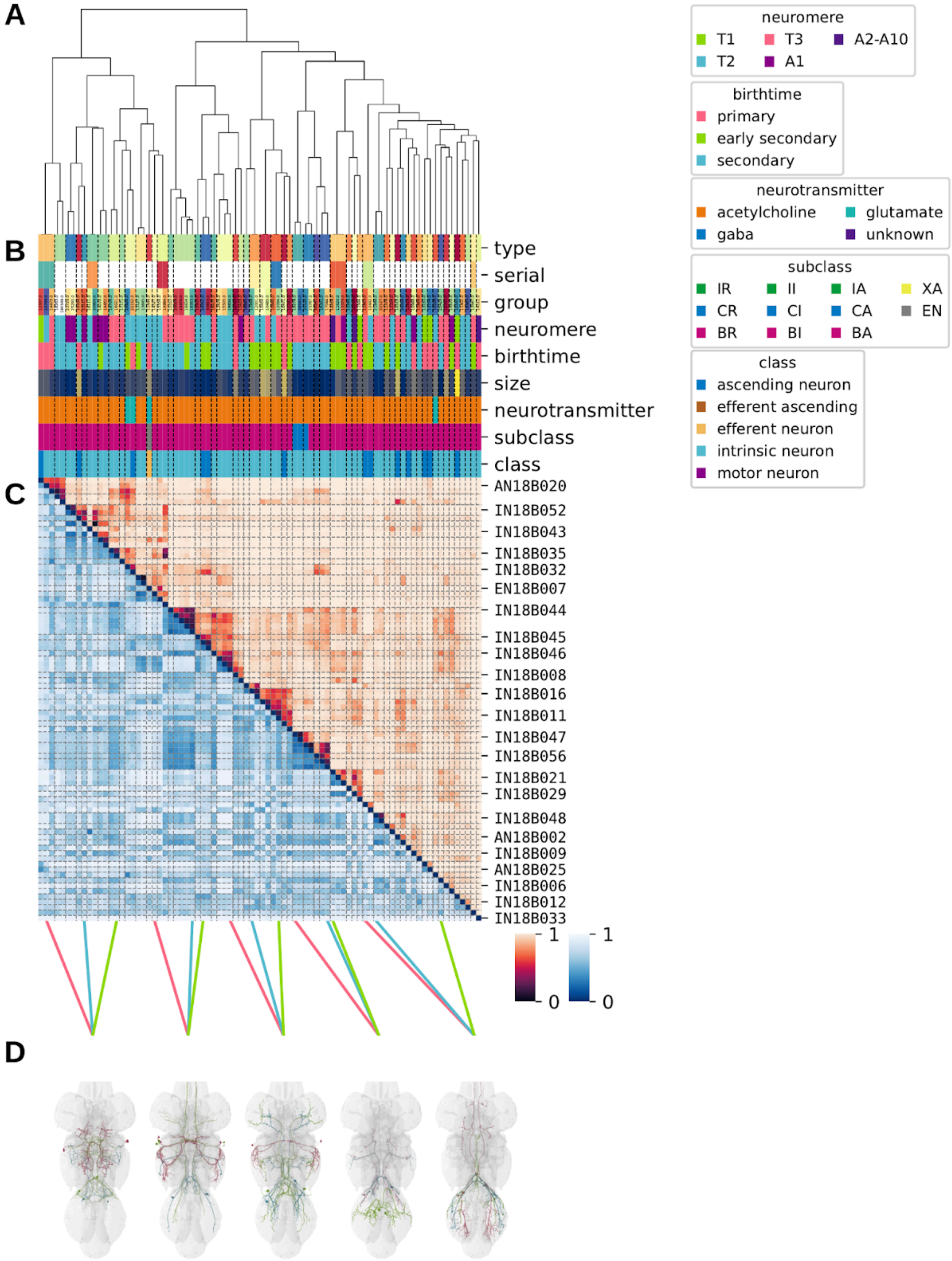

**Figure 41 - figure supplement 1. Systematic typing of hemilineage 18B. A.** Hierarchical clustering dendrogram of hemilineage groups by laterally and serially aggregated connectivity cosine clustering. **B.** Categorical annotations of each hemilineage group, each column corresponding to the aligned leaf in A. Colours for type, serial set, and group are arbitrary for visualisation. Colours for neuromere, birthtime, neurotransmitter, subclass, and class are as in all other figures. **C.** Similarity distance heatmap for hemilineage. Cosine distance is in the upper triangle, while laterally symmetrised NBLAST distance is in the lower triangle. Systematic type names of some types are labelled. **D.** Morphologically representative groups from dendrogram subtrees. Each group, indicated by colour and line connecting to its column in B and C, is the most morphologically representative group (medoid of NBLAST distance) from a subtree of A. The subtrees (flat clusters) are equal height cuts of A determined to yield the number of groups per plot and plots in D.

#### 18B systematic types

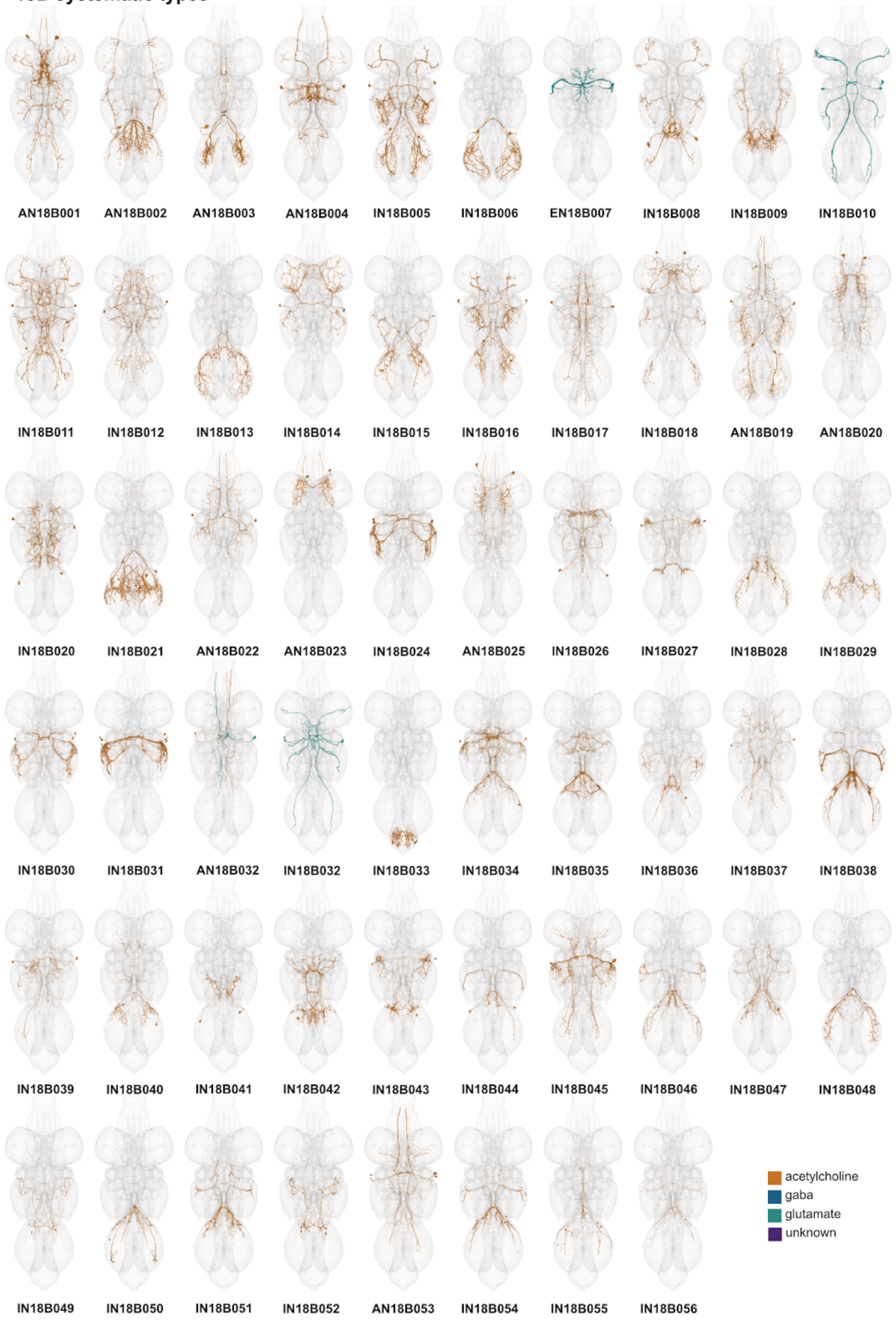

**Figure 41 - figure supplement 2. Systematic types of hemilineage 18B.** Systematic types have been arranged in numerical order, with neurons of the same type that belong to distinct classes (e.g., intrinsic neuron vs ascending neuron) plotted separately but placed adjacent to each other. Individual neuron meshes have been coloured based on predicted neurotransmitter: dark orange = acetylcholine, blue = gaba, marine = glutamate, dark purple = unknown.

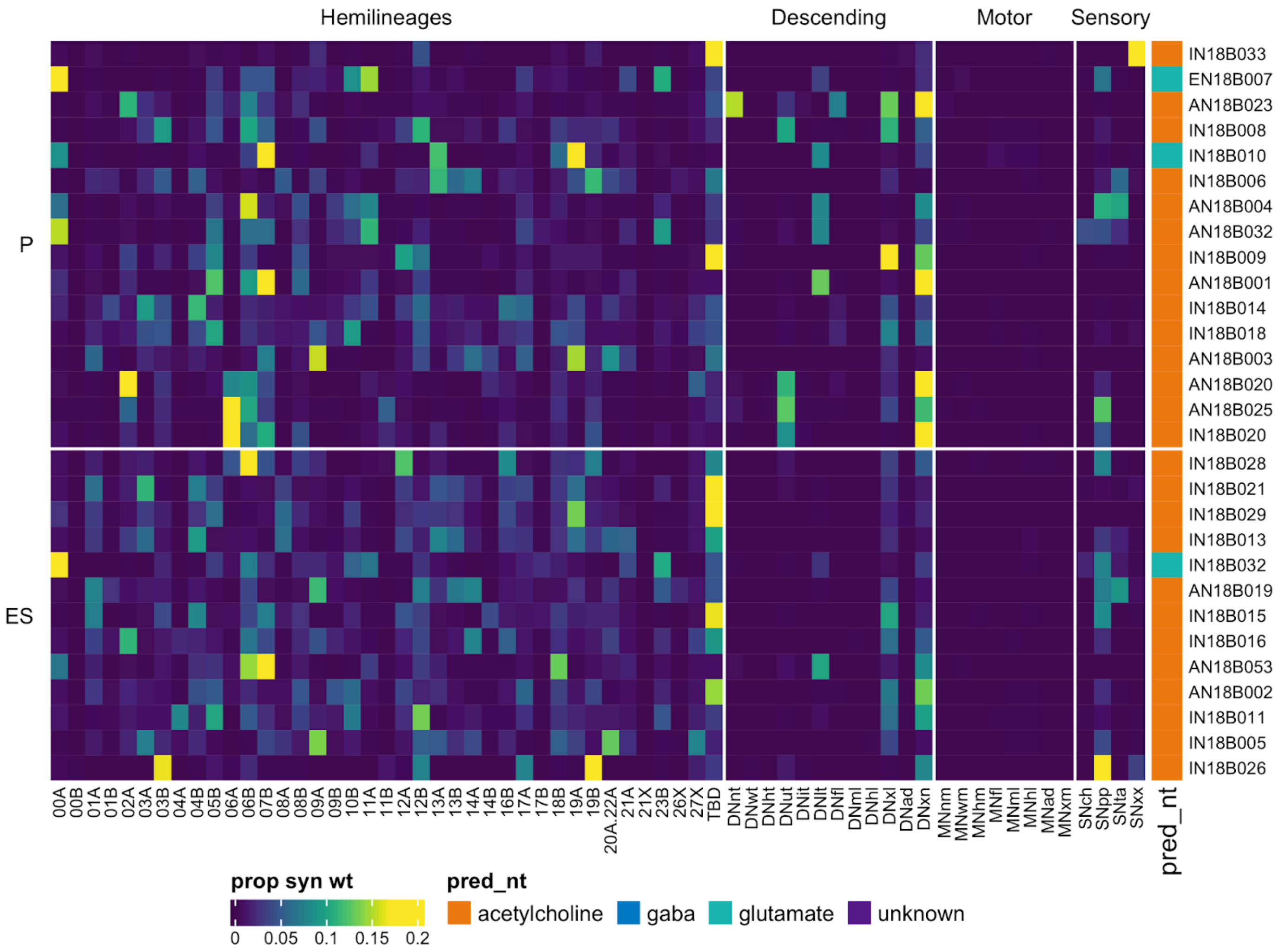

**Figure 41 - figure supplement 3. Connectivity to upstream partners by 18B primary and early secondary systematic types.** Proportions of synaptic weight to systematic types from upstream partners, normalised by row. 18B neurons have been clustered within each assigned birthtime window (P = primary, ES = early secondary, S = secondary) based on both upstream and downstream connectivity to hemilineages, descending neuron subclasses, motor neuron subclasses, and sensory neuron modalities. Annotation bar is coloured by the most common predicted neurotransmitter for the neurons of each type.

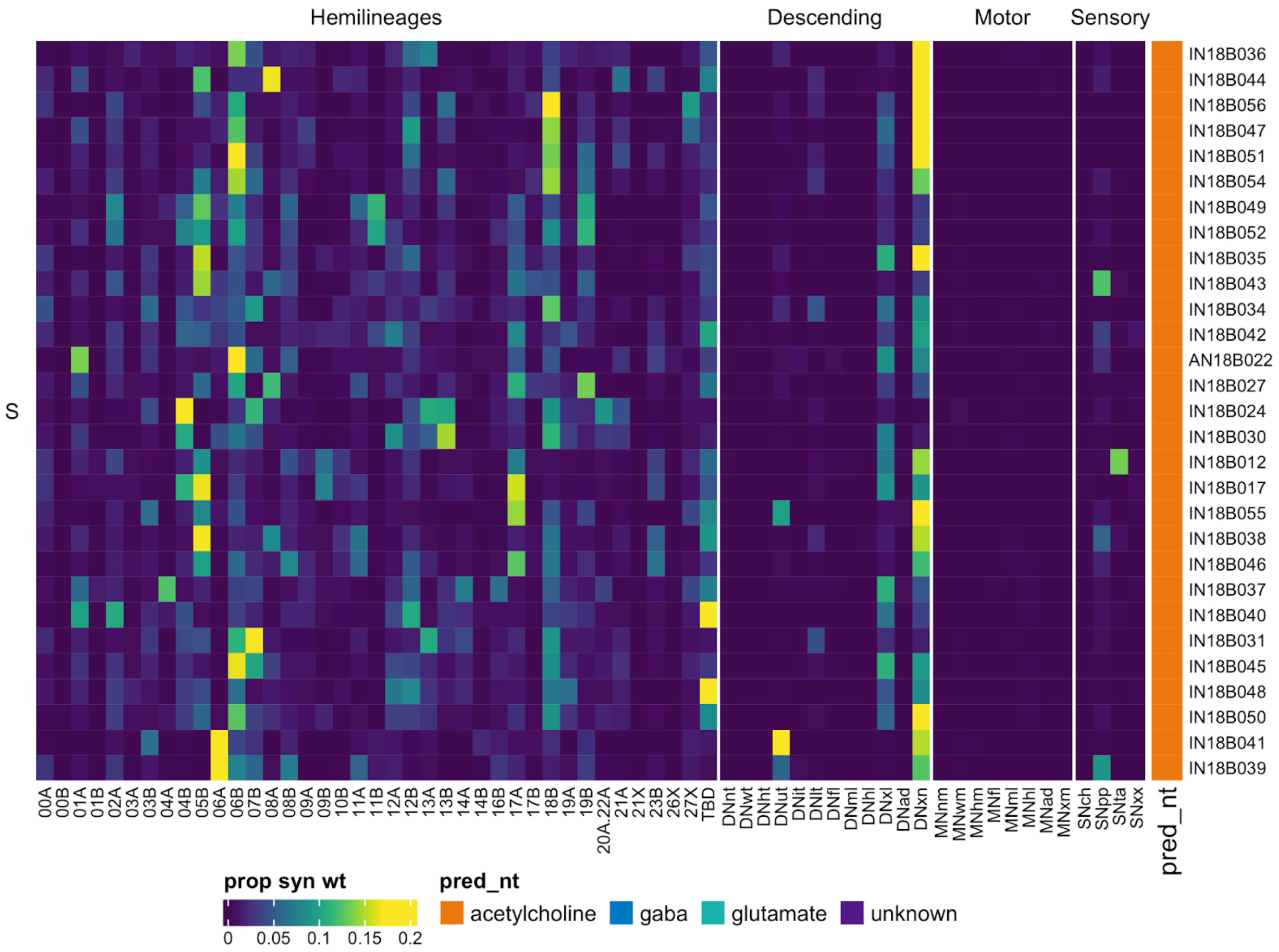

**Figure 41 - figure supplement 4. Connectivity to upstream partners by 18B secondary systematic types.** Proportions of synaptic weight to systematic types from upstream partners, normalised by row. 18B neurons have been clustered within each assigned birthtime window (P = primary, ES = early secondary, S = secondary) based on both upstream and downstream connectivity to hemilineages, descending neuron subclasses, motor neuron subclasses, and sensory neuron modalities. Annotation bar is coloured by the most common predicted neurotransmitter for the neurons of each type.

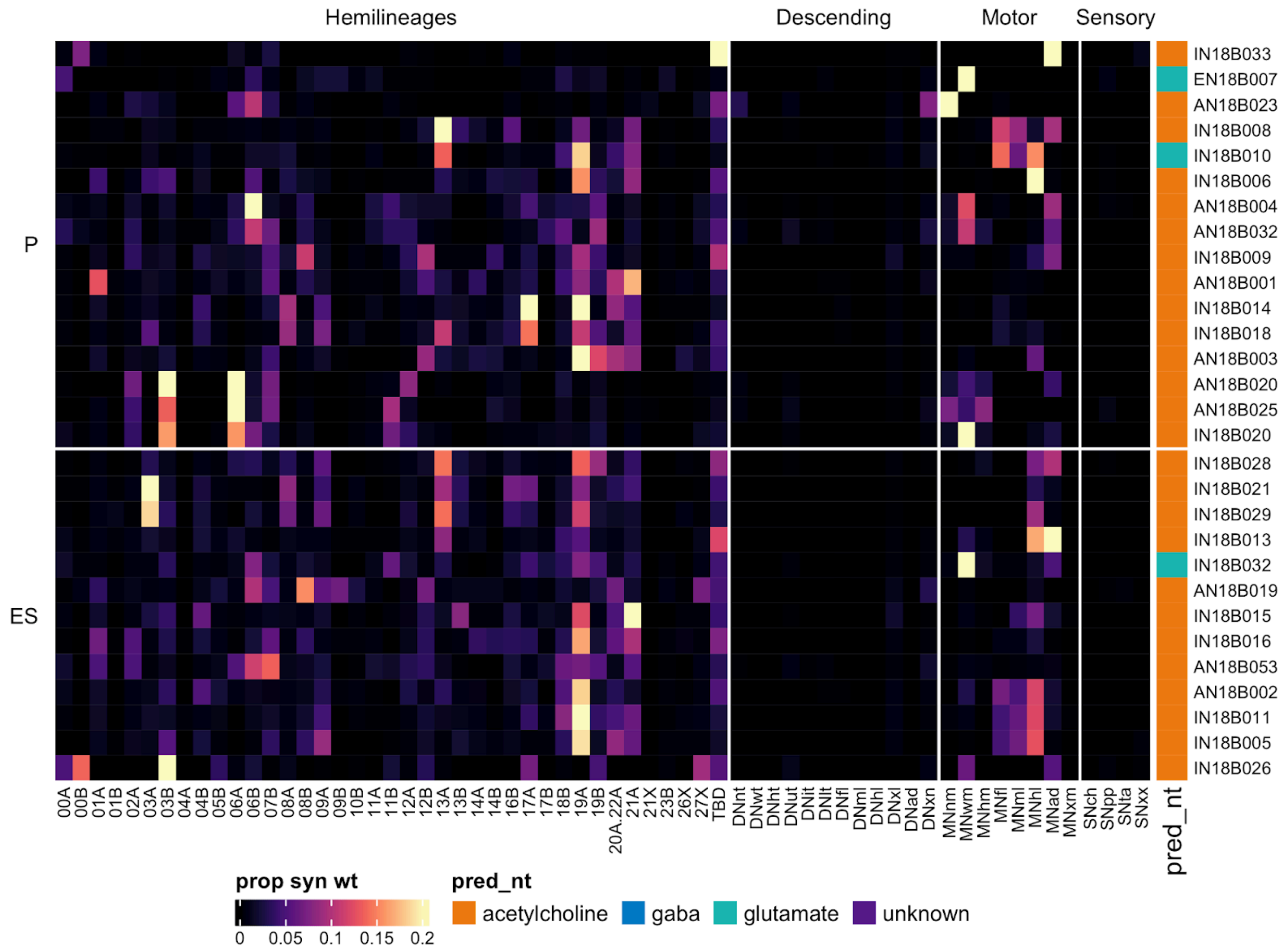

**Figure 41 - figure supplement 5. Connectivity to downstream partners by 18B primary and early secondary systematic types.** Proportions of synaptic weight from systematic types to downstream partners, normalised by row. 18B neurons have been clustered within each assigned birthtime window (P = primary, ES = early secondary, S = secondary) based on both upstream and downstream connectivity to hemilineages, descending neuron subclasses, motor neuron subclasses, and sensory neuron modalities. The annotation bar is coloured by the most common predicted neurotransmitter within each type.

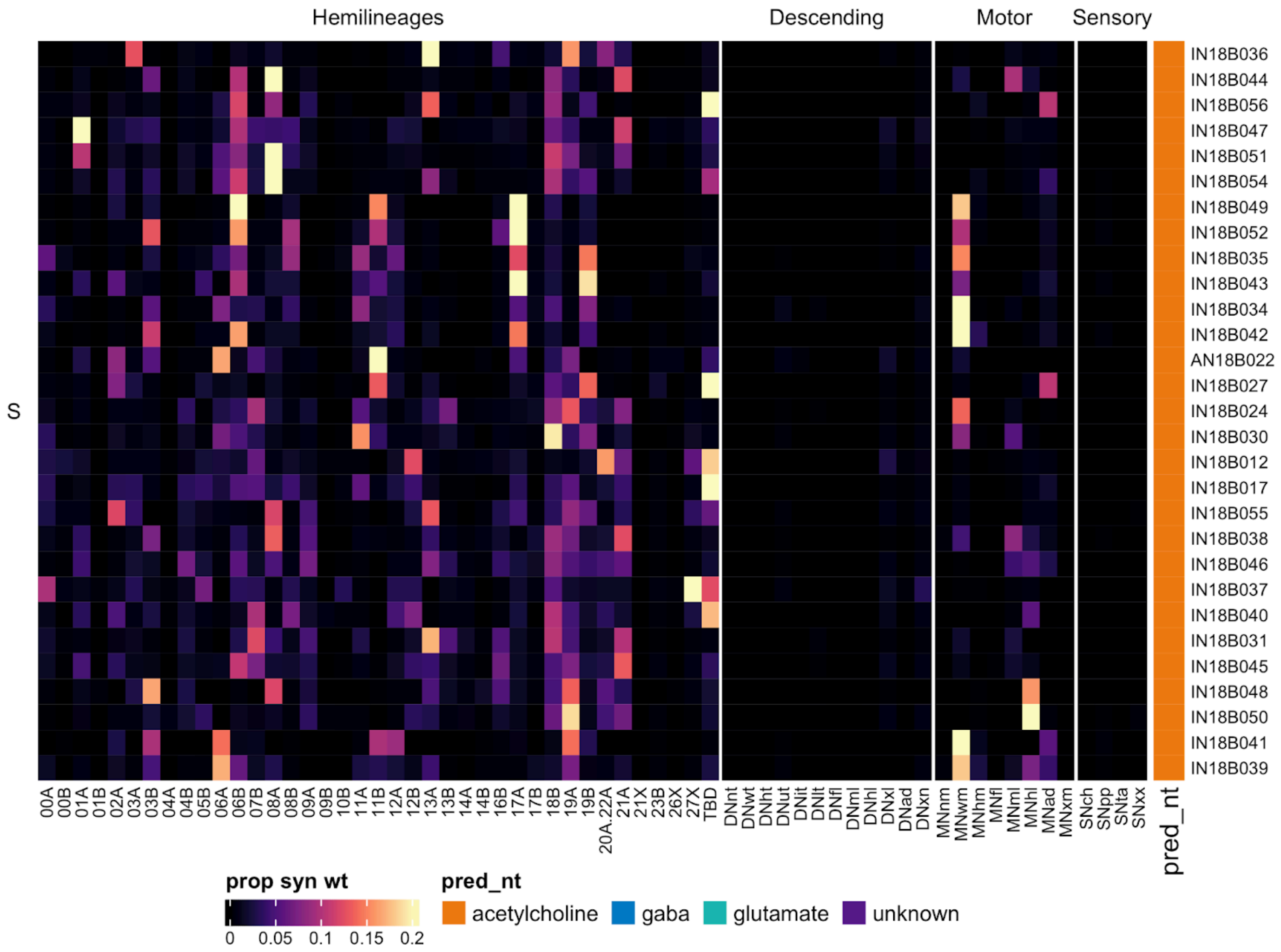

**Figure 41 - figure supplement 6. Connectivity to downstream partners by 18B secondary systematic types.** Proportions of synaptic weight from systematic types to downstream partners, normalised by row. 18B neurons have been clustered within each assigned birthtime window (P = primary, ES = early secondary, S = secondary) based on both upstream and downstream connectivity to hemilineages, descending neuron subclasses, motor neuron subclasses, and sensory neuron modalities. The annotation bar is coloured by the most common predicted neurotransmitter within each type.

Hemilineage 19A

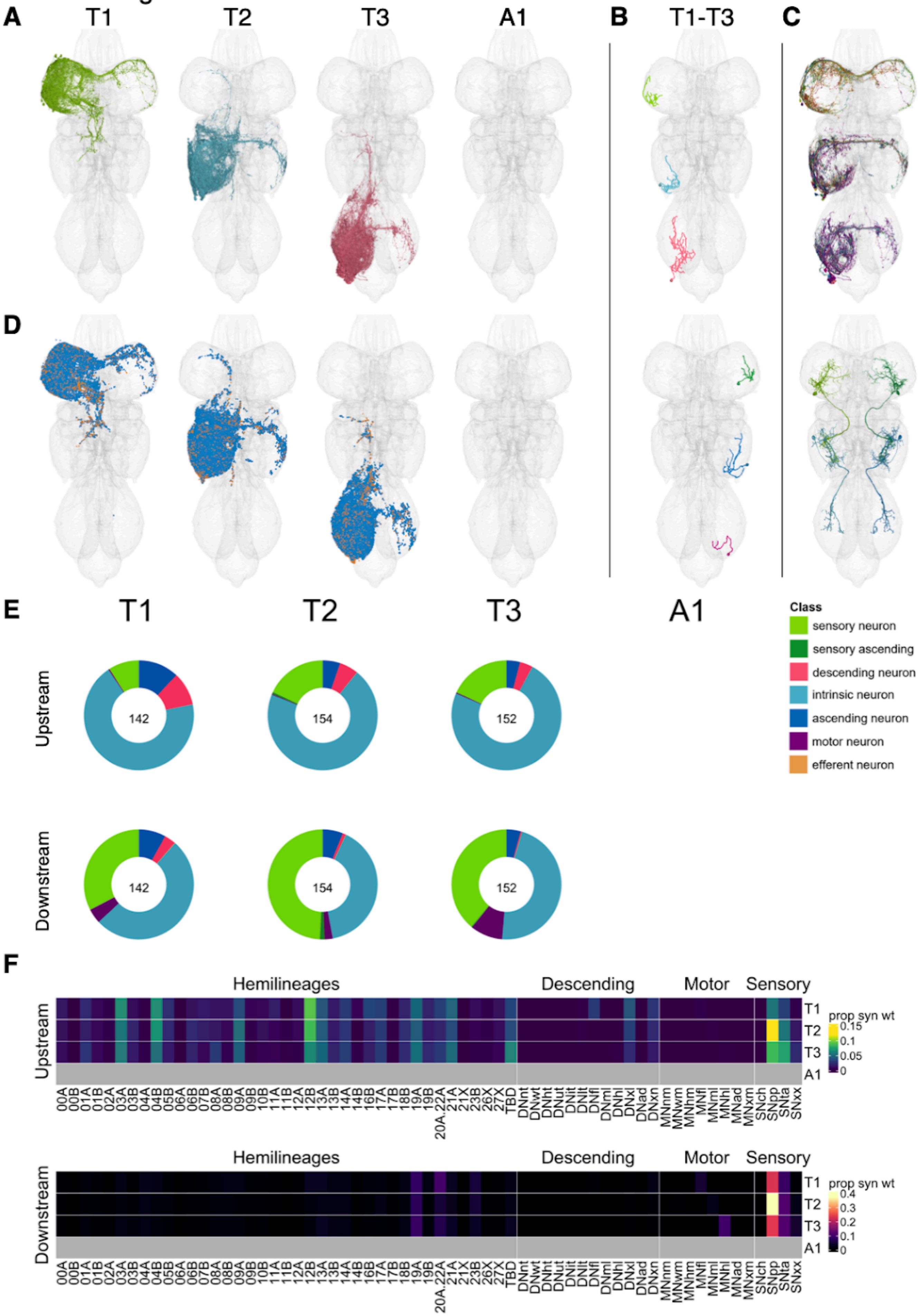

**Figure 42. Hemilineage 19A. A.** Meshes of all RHS secondary neurons plotted in neuromere-specific colours. **B.** "Representative" secondary neuron skeletons plotted in hemineuromere-specific colours. The skeleton with the top accumulated NBLAST score among all neurons from the hemilineage in a given hemineuromere was used. **C.** Neuron meshes of selected examples. Top: cholinergic sequential serial set 10173. Bottom: independent leg serial set 14080. **D.** Predicted synapses of RHS secondary neurons. Blue: postsynapses; dark orange: presynapses. **E.** Proportions of connections from secondary neurons to upstream or downstream partners, normalised by neuromere and coloured by broad class. Numbers of query neurons appear in the centre. **F.** Ratios of connections from secondary neurons originating in each neuromere to upstream or downstream partners, normalised by row.

#### Hemilineage 19A

Hemilineages 19A and 19B derive from posterior dorsal neuroblast NB6-2 (Birkholz et al., 2015; Lacin and Truman, 2016), which generates 3-4 intersegmental interneurons and ~20 local interneurons in the embryo (Schmid et al., 1999). Both secondary hemilineages survive in all thoracic neuromeres, but the number of 19B in T1 is greatly reduced (Truman et al., 2004). The 19A secondary hemilineage is mainly restricted to ipsilateral leg neuropil (Harris et al., 2015) but also includes subtypes that cross the midline in a small anterior and ventral tract to wrap around the contralateral leg neuropil.

We identified 19A secondary neurons in similar numbers in all three thoracic neuromeres (Figure 42E). They enter the posterior of each neuromere quite dorsally and mainly innervate the ipsilateral leg neuropil (Figure 42C), although a subpopulation crosses the midline just anterior to the leg sensory neuropils (Figure 42C). Hemilineage 19A secondary neurons are typically gabaergic (Lacin et al., 2019), but we also identified an early born cholinergic population that descend or ascend to adjacent neuromeres (Figure 7G).

19A secondary neurons receive considerable input from 03A, 04B, 12B, and both proprioceptive and tactile sensory neurons. Unusually, their main downstream targets are sensory neurons, particularly proprioceptive neurons and especially in T2 (Figure 42F). Bilateral activation of 19A results in incessant waving of T2 legs without movement of T1 or T3 legs (Harris et al., 2015).

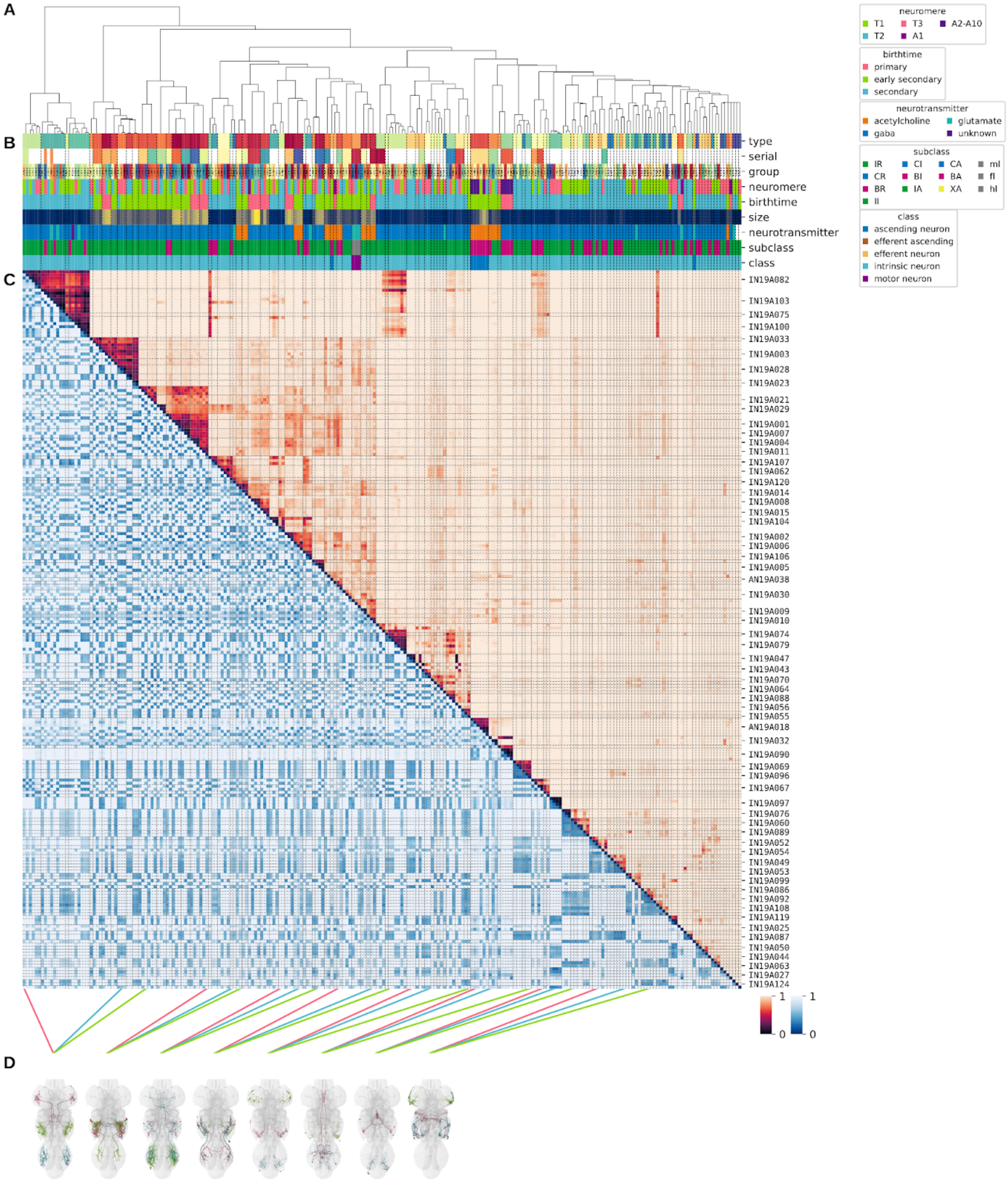

**Figure 42 - figure supplement 1. Systematic typing of hemilineage 19A.** Types for motor neurons were assigned separately as outlined in our accompanying manuscript (Cheong et al., 2023). **A.** Hierarchical clustering dendrogram of hemilineage groups by laterally and serially aggregated connectivity cosine clustering. **B.** Categorical annotations of each hemilineage group, each column corresponding to the aligned leaf in A. Colours for type, serial set, and group are arbitrary for visualisation. Colours for neuromere, birthtime, neurotransmitter, subclass, and class are as in all other figures. **C.** Similarity distance heatmap for hemilineage. Cosine distance is in the upper triangle, while laterally symmetrised NBLAST distance is in the lower triangle. Systematic type names of some types are labelled. **D.** Morphologically representative groups from dendrogram subtrees. Each group, indicated by colour and line connecting to its column in B and C, is the most morphologically representative group (medoid of NBLAST distance)

from a subtree of A. The subtrees (flat clusters) are equal height cuts of A determined to yield the number of groups per plot and plots in D.

#### 19A systematic types

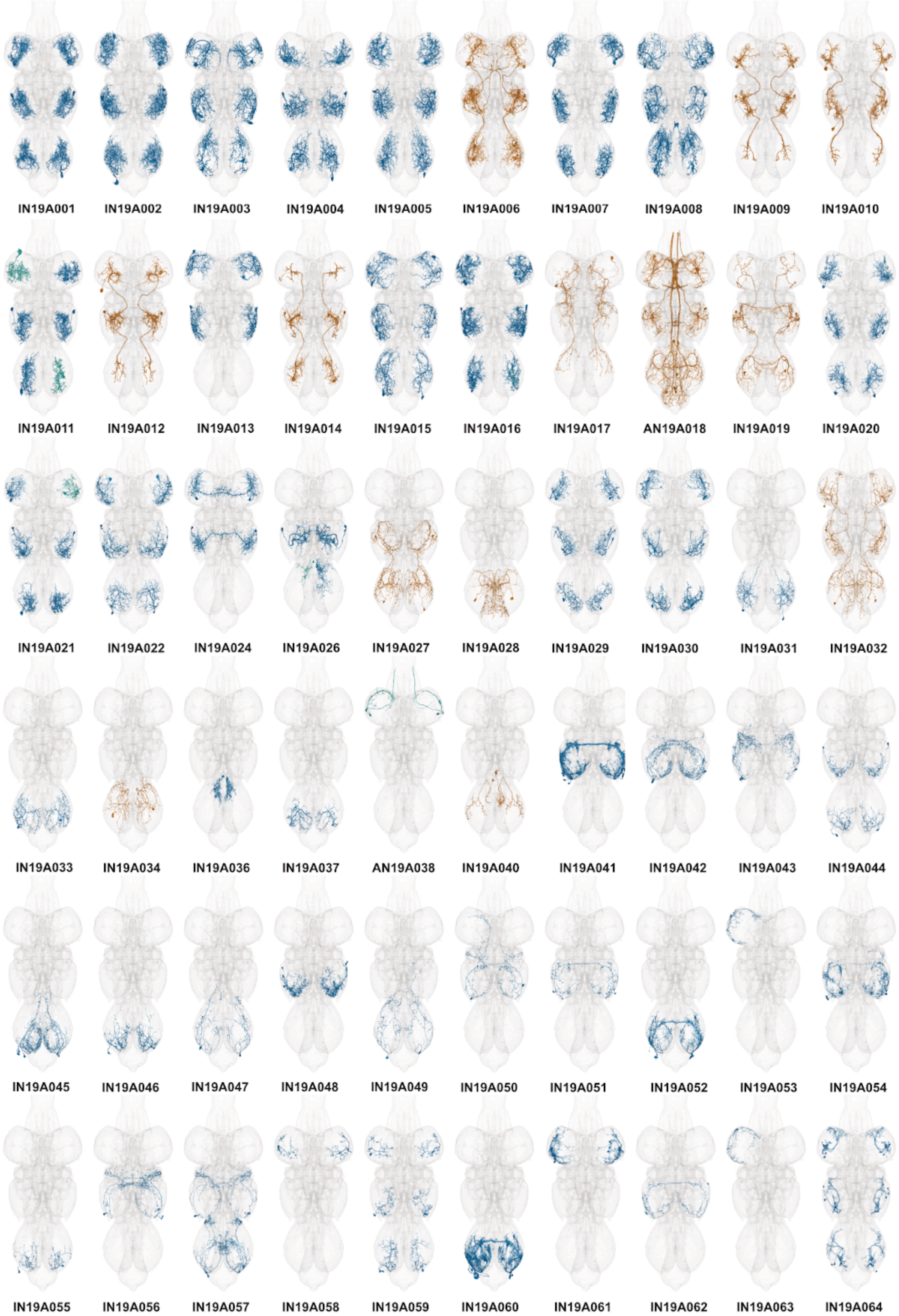

**Figure 42 - figure supplement 2. Systematic types of hemilineage 19A.** Systematic types have been arranged in numerical order, with neurons of the same type that belong to distinct classes (e.g., intrinsic neuron vs ascending neuron) plotted separately but placed adjacent to each other. Individual neuron meshes have been coloured based on predicted neurotransmitter: dark orange = acetylcholine, blue = gaba, marine = glutamate, dark purple = unknown.

#### 19A systematic types, continued

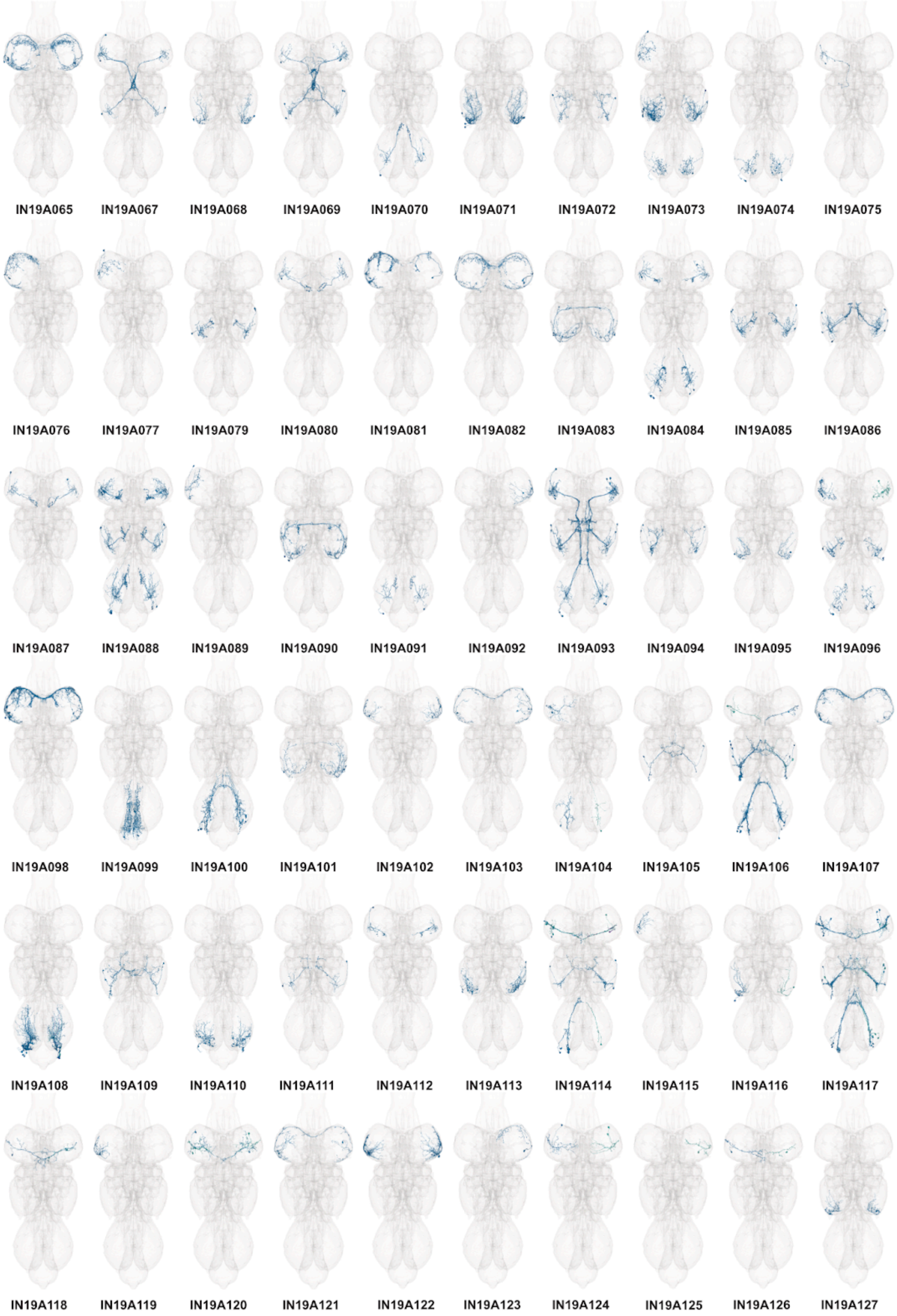

**Figure 42 - figure supplement 3. Systematic types of hemilineage 19A, continued.** Systematic types have been arranged in numerical order, with neurons of the same type that belong to distinct classes (e.g., intrinsic neuron vs ascending neuron) plotted separately but placed adjacent to each other. Individual neuron meshes have been coloured based on predicted neurotransmitter: dark orange = acetylcholine, blue = gaba, marine = glutamate, dark purple = unknown.

**19A systematic types, continued**

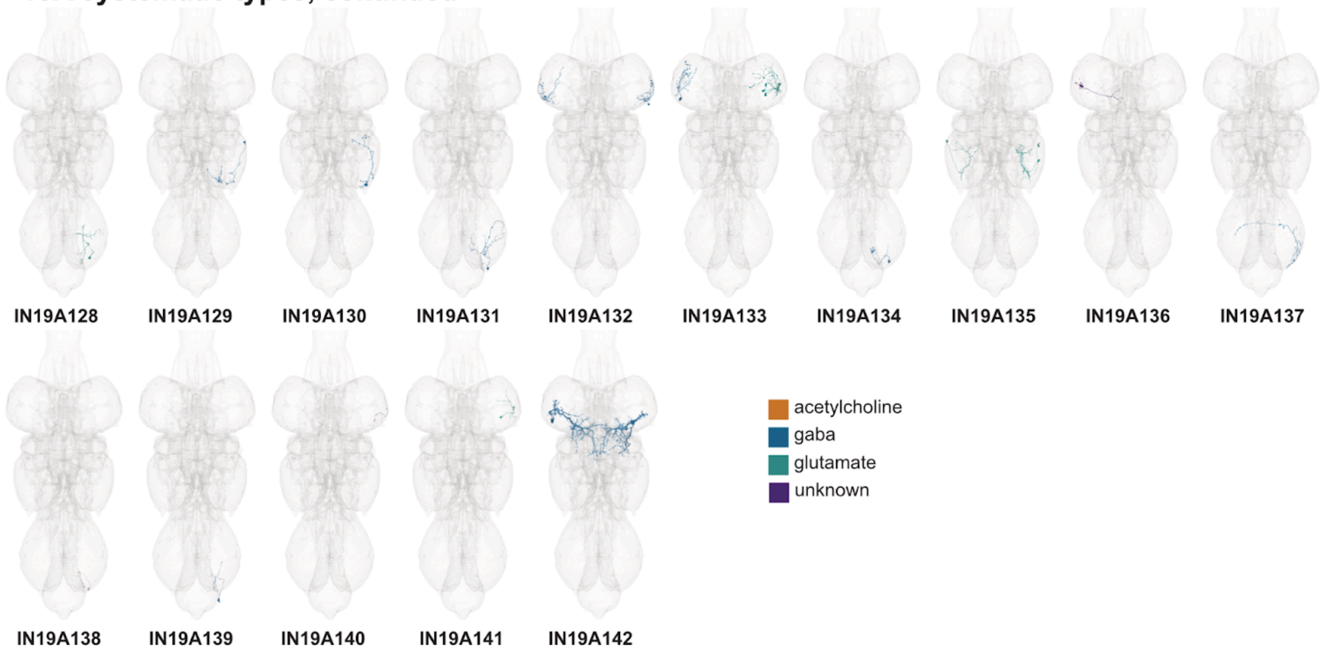

**Motor Neurons**

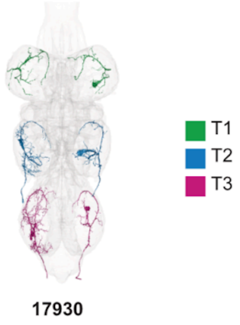

**Figure 42 - figure supplement 4. Systematic types of hemilineage 19A, continued.** Systematic types have been arranged in numerical order, with neurons of the same type that belong to distinct classes (e.g., intrinsic neuron vs ascending neuron) plotted separately but placed adjacent to each other. Individual neuron meshes have been coloured based on predicted neurotransmitter: dark orange = acetylcholine, blue = gaba, marine = glutamate, dark purple = unknown. Motor neurons (typed separately in (Cheong et al., 2023)) have been plotted by serial set if identified in multiple neuromeres and by systematic type if not. Individual motor neuron meshes have been coloured based on soma neuromere: dark green = T1, blue = T2, magenta = T3.

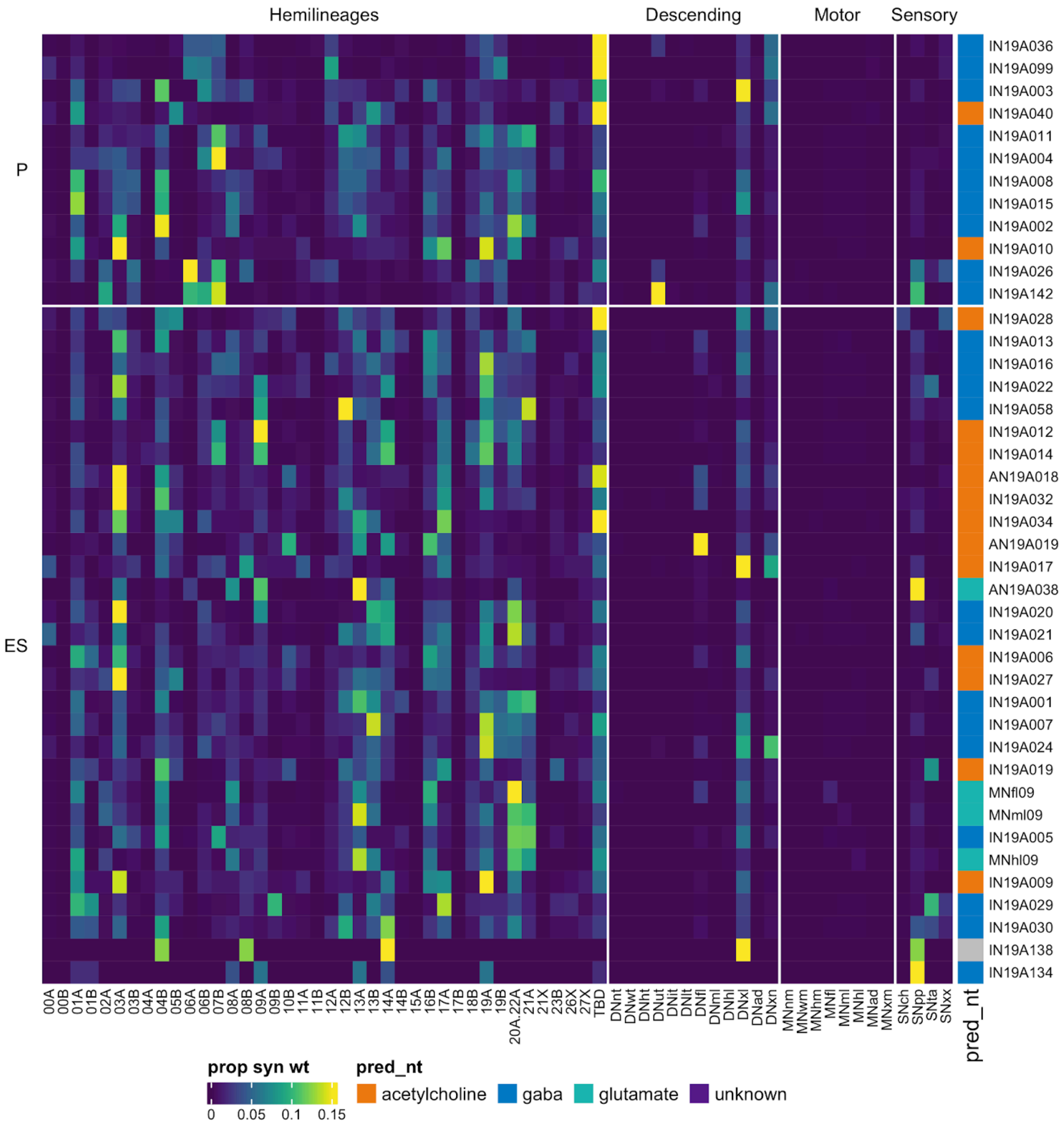

**Figure 42 - figure supplement 5. Connectivity to upstream partners by 19A primary and early secondary systematic types.** Proportions of synaptic weight to systematic types from upstream partners, normalised by row. 19A neurons have been clustered within each assigned birthtime window (P = primary, ES = early secondary, S = secondary) based on both upstream and downstream connectivity to hemilineages, descending neuron subclasses, motor neuron subclasses, and sensory neuron modalities. Annotation bar is coloured by the most common predicted neurotransmitter for the neurons of each type.

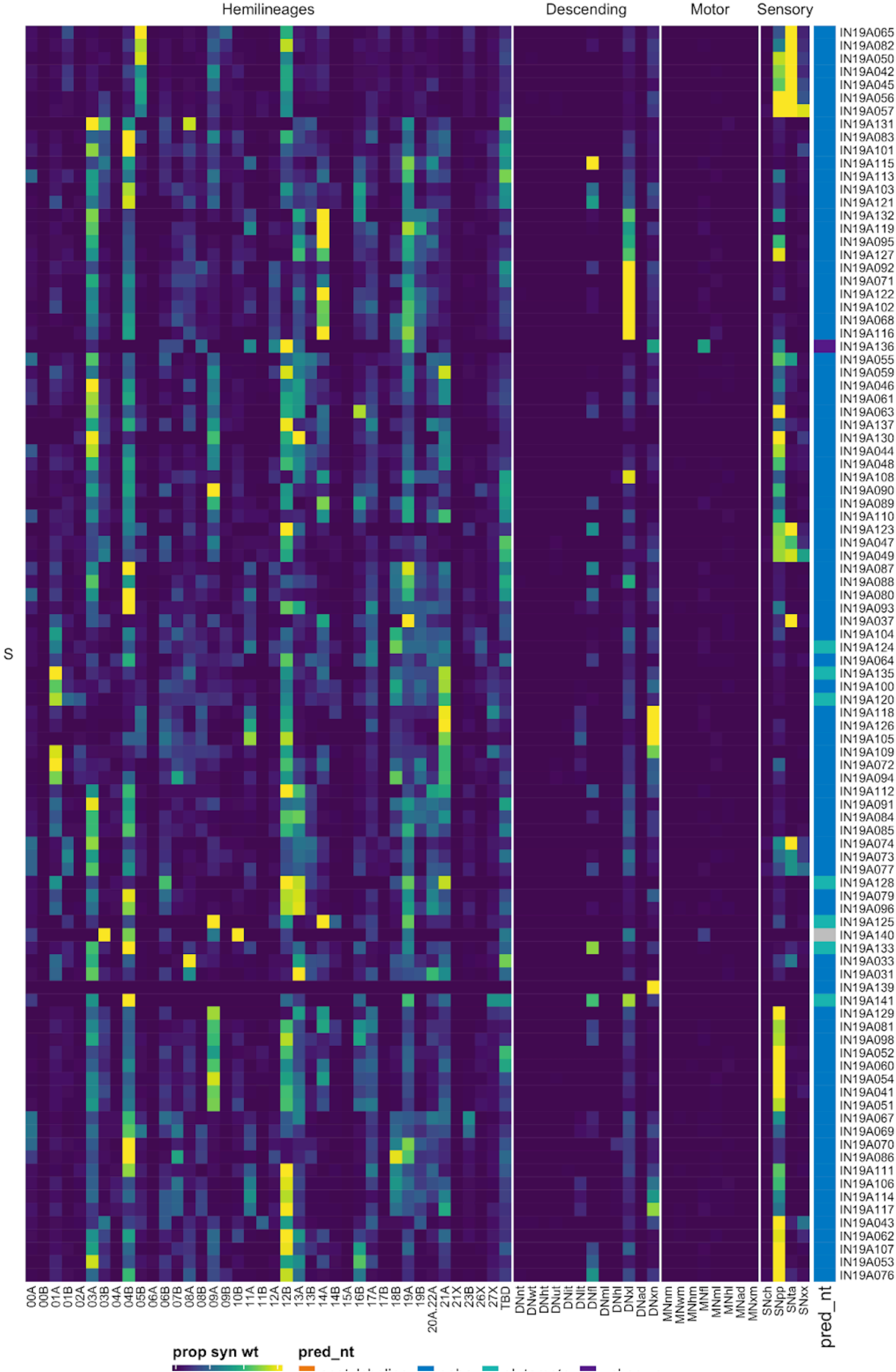

Hemilineages
Descending
Motor
Sensory
S
IN19A065
IN19A082
IN19A050
IN19A042
IN19A045
IN19A056
IN19A057
IN19A131
IN19A083
IN19A101
IN19A115
IN19A113
IN19A103
IN19A121
IN19A132
IN19A119
IN19A095
IN19A127
IN19A092
IN19A071
IN19A122
IN19A102
IN19A068
IN19A116
IN19A136
IN19A055
IN19A059
IN19A046
IN19A061
IN19A063
IN19A137
IN19A130
IN19A044
IN19A048
IN19A108
IN19A090
IN19A089
IN19A110
IN19A123
IN19A047
IN19A049
IN19A087
IN19A088
IN19A080
IN19A093
IN19A037
IN19A104
IN19A124
IN19A064
IN19A135
IN19A100
IN19A120
IN19A118
IN19A126
IN19A105
IN19A109
IN19A072
IN19A094
IN19A112
IN19A091
IN19A084
IN19A085
IN19A074
IN19A073
IN19A077
IN19A128
IN19A079
IN19A096
IN19A125
IN19A140
IN19A133
IN19A033
IN19A031
IN19A139
IN19A141
IN19A129
IN19A081
IN19A098
IN19A052
IN19A060
IN19A054
IN19A041
IN19A051
IN19A067
IN19A069
IN19A070
IN19A086
IN19A111
IN19A106
IN19A114
IN19A117
IN19A043
IN19A062
IN19A107
IN19A053
IN19A076
00A 00B 01A 01B 02A 03A 03B 04A 04B 05B 06A 06B 07B 08A 08B 09A 09B 10B 11A 11B 12A 12B 13A 13B 14A 14B 15A 16B 17A 17B 18B 19A 19B 20A.22A 21A 21X 23B 26X 27X TBD
DNnt DNwt DNht DNut DNlt DNfl DNml DNhl DNxl DNad DNxn
MNnm MNwm MNhm MNfl MNml MNhl MNad MNxm
SNch SNpp SNta SNxx
pred_nt
prop syn wt
pred_nt

**Figure 42 - figure supplement 6. Connectivity to upstream partners by 19A secondary systematic types.** Proportions of synaptic weight to systematic types from upstream partners, normalised by row. 19A neurons have been clustered within each assigned birthtime window (P = primary, ES = early secondary, S = secondary) based on both upstream and downstream connectivity to hemilineages, descending neuron subclasses, motor neuron subclasses, and sensory neuron modalities. The annotation bar is coloured by the most common predicted neurotransmitter within each type.

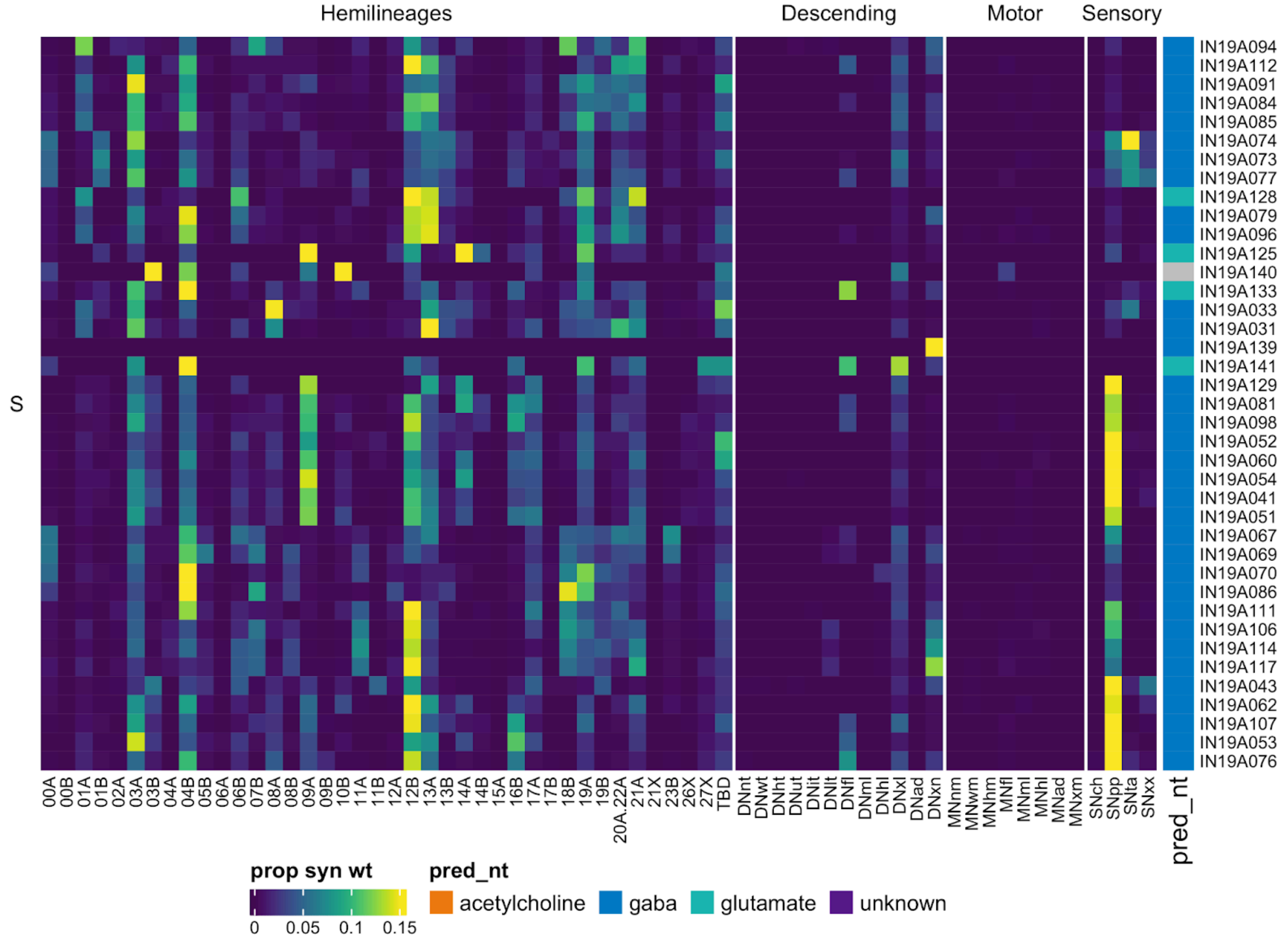

**Figure 42 - figure supplement 7. Connectivity to upstream partners by 19A secondary systematic types, continued.** Proportions of synaptic weight to systematic types from upstream partners, normalised by row. 19A neurons have been clustered within each assigned birthtime window (P = primary, ES = early secondary, S = secondary) based on both upstream and downstream connectivity to hemilineages, descending neuron subclasses, motor neuron subclasses, and sensory neuron modalities. The annotation bar is coloured by the most common predicted neurotransmitter within each type.

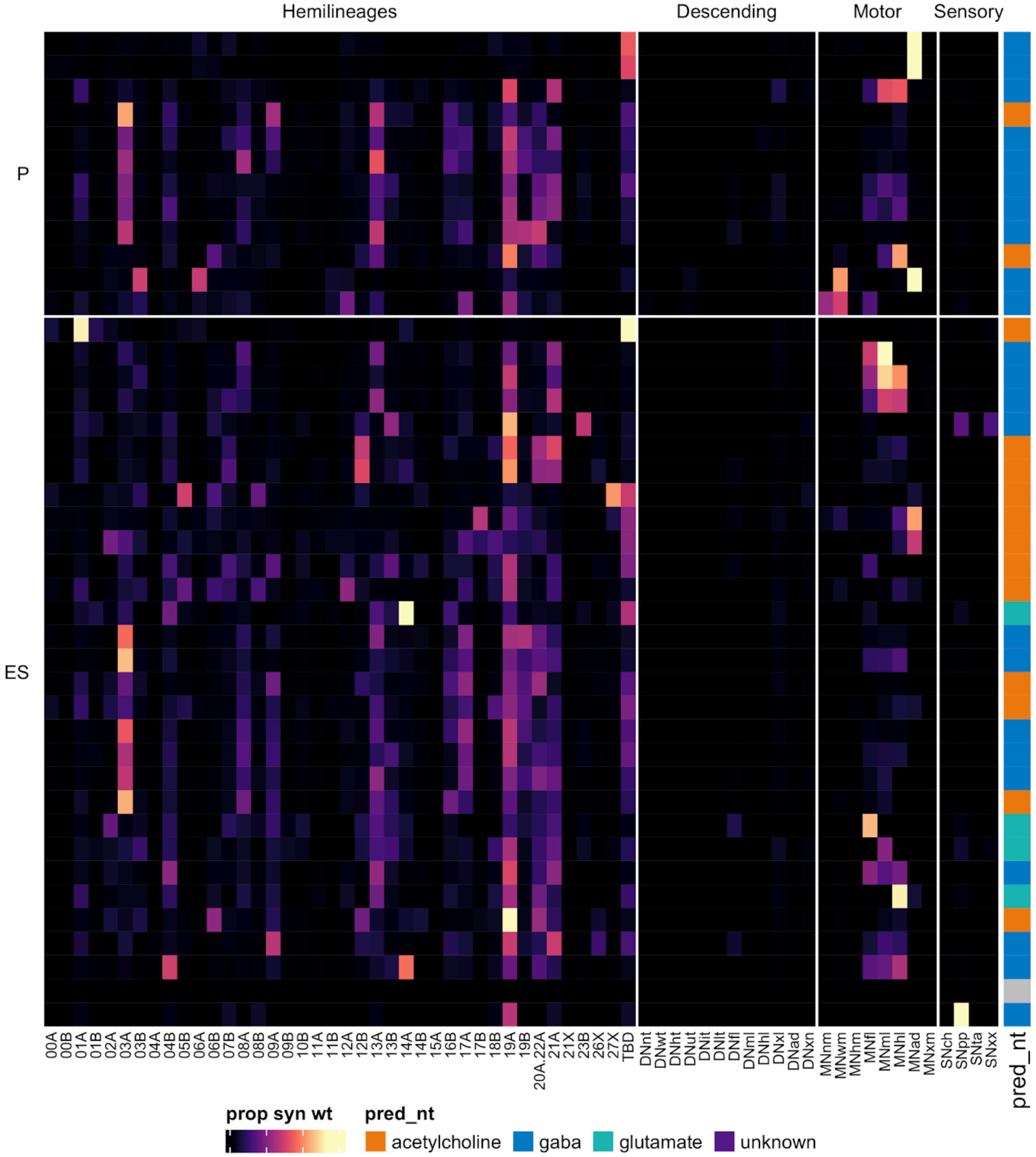

**Figure 42 - figure supplement 8. Connectivity to downstream partners by 19A primary and early secondary systematic types.** Proportions of synaptic weight from systematic types to downstream partners, normalised by row. 19A neurons have been clustered within each assigned birthtime window (P = primary, ES = early secondary, S = secondary) based on both upstream and downstream connectivity to hemilineages, descending neuron subclasses, motor neuron subclasses, and sensory neuron modalities. The annotation bar is coloured by the most common predicted neurotransmitter within each type.

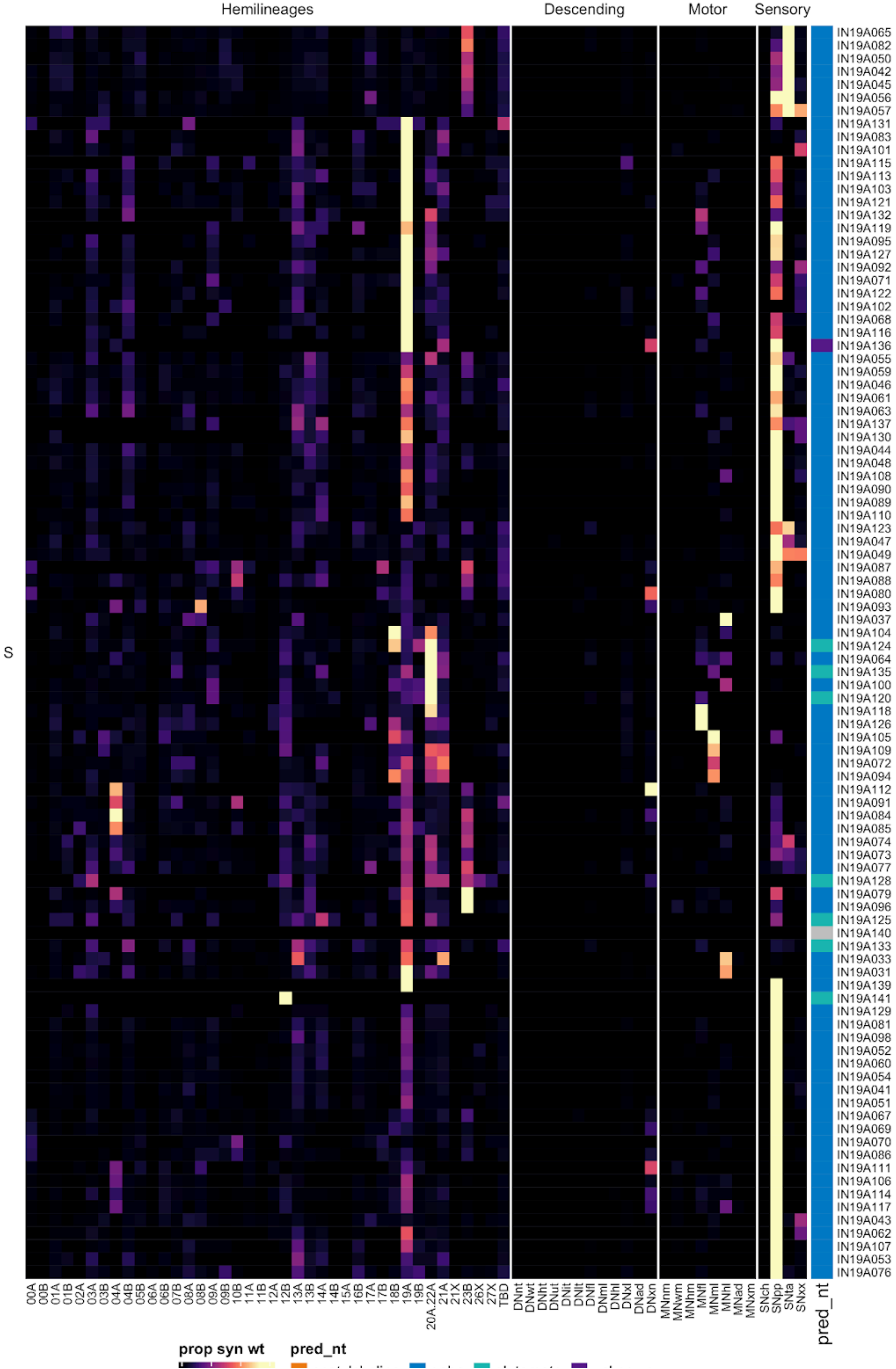

Hemilineages
Descending
Motor
Sensory
S
IN19A065
IN19A082
IN19A050
IN19A042
IN19A045
IN19A056
IN19A057
IN19A131
IN19A083
IN19A101
IN19A115
IN19A113
IN19A103
IN19A121
IN19A132
IN19A119
IN19A095
IN19A127
IN19A092
IN19A071
IN19A122
IN19A102
IN19A068
IN19A116
IN19A136
IN19A055
IN19A059
IN19A046
IN19A061
IN19A063
IN19A137
IN19A130
IN19A044
IN19A048
IN19A108
IN19A090
IN19A089
IN19A110
IN19A123
IN19A047
IN19A049
IN19A087
IN19A088
IN19A080
IN19A093
IN19A037
IN19A104
IN19A124
IN19A064
IN19A135
IN19A100
IN19A120
IN19A118
IN19A126
IN19A105
IN19A109
IN19A072
IN19A094
IN19A112
IN19A091
IN19A084
IN19A085
IN19A074
IN19A073
IN19A077
IN19A128
IN19A079
IN19A096
IN19A125
IN19A140
IN19A133
IN19A033
IN19A031
IN19A139
IN19A141
IN19A129
IN19A081
IN19A098
IN19A052
IN19A060
IN19A054
IN19A041
IN19A051
IN19A067
IN19A069
IN19A070
IN19A086
IN19A111
IN19A106
IN19A114
IN19A117
IN19A043
IN19A062
IN19A107
IN19A053
IN19A076
00A 00B 01A 01B 02A 03A 03B 04A 04B 05B 06A 06B 07B 08A 08B 09A 09B 10B 11A 11B 12A 12B 13A 13B 14A 14B 15A 16B 17A 17B 18B 19A 19B 20A.22A 21A 21X 23B 26X 27X TBD
DNnt DNwt DNht DNut DNit DNlt DNfl DNml DNhl DNxl DNad DNxn
MNnm MNwm MNhm MNfl MNml MNhl MNad MNxm
SNch SNpp SNta SNxx
pred_nt
prop syn wt
pred_nt

**Figure 42 - figure supplement 9. Connectivity to downstream partners by 19A secondary systematic types.** Proportions of synaptic weight from systematic types to downstream partners, normalised by row. 19A neurons have been clustered within each assigned birthtime window (P = primary, ES = early secondary, S = secondary) based on both upstream and downstream connectivity to hemilineages, descending neuron subclasses, motor neuron subclasses, and sensory neuron modalities. The annotation bar is coloured by the most common predicted neurotransmitter for the neurons of each type.

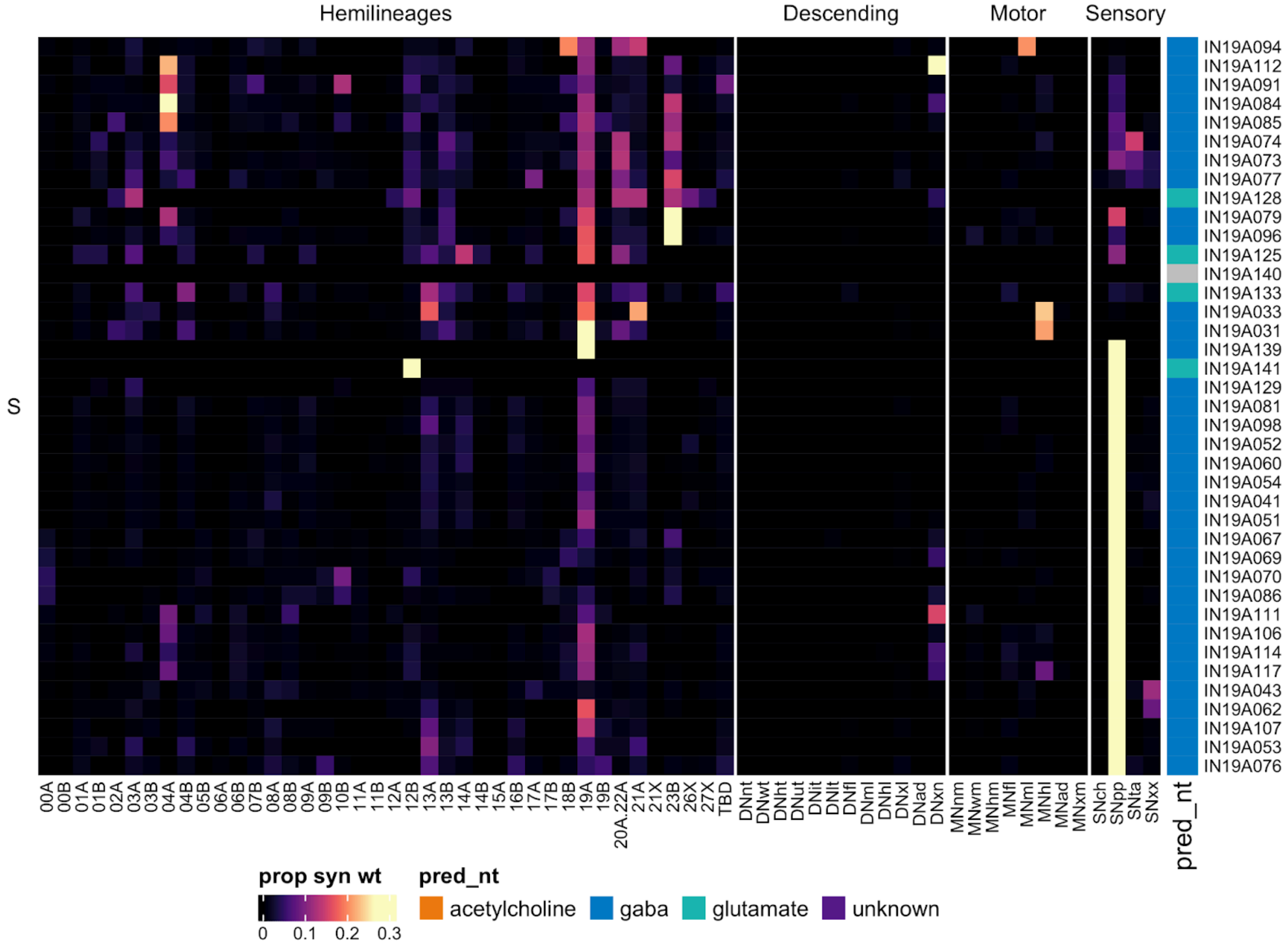

**Figure 42 - figure supplement 10. Connectivity to downstream partners by 19A secondary systematic types, continued.** Proportions of synaptic weight from systematic types to downstream partners, normalised by row. 19A neurons have been clustered within each assigned birthtime window (P = primary, ES = early secondary, S = secondary) based on both upstream and downstream connectivity to hemilineages, descending neuron subclasses, motor neuron subclasses, and sensory neuron modalities. The annotation bar is coloured by the most common predicted neurotransmitter for the neurons of each type.

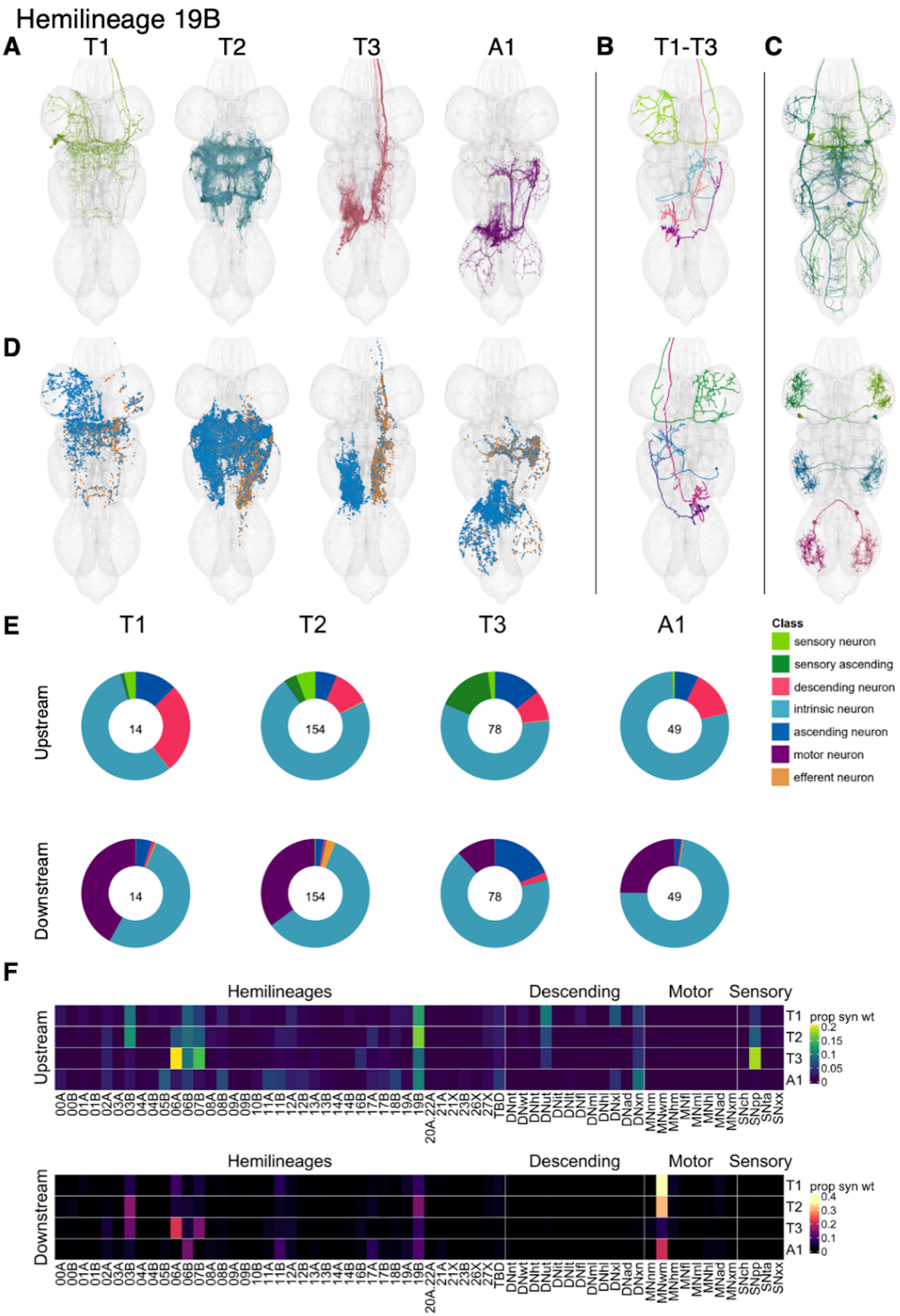

Hemilineage 19B
A
T1
T2
T3
A1
B
T1-T3
C
D
E
T1
T2
T3
A1
Upstream
Downstream
14
154
78
49
Class
sensory neuron
sensory ascending
descending neuron
intrinsic neuron
ascending neuron
motor neuron
efferent neuron
F
Hemilineages
Descending
Motor
Sensory
prop syn wt
00A 00B 01A 01B 02A 03A 03B 04A 04B 05B 06A 06B 07B 08A 08B 09A 09B 10B 11A 11B 12A 12B 13A 13B 14A 14B 16B 17A 17B 18B 19A 19B 20A.22A 21A 21X 23B 26X 27X TBD DNnt DNwt DNht DNut DNit DNlt DNfl DNml DNhl DNxl DNad DNxn MNnm MNwm MNhm MNfl MNml MNhl MNad MNxm SNch SNpp SNta SNxx
0.2 0.15 0.1 0.05 0
0.4 0.3 0.2 0.1 0

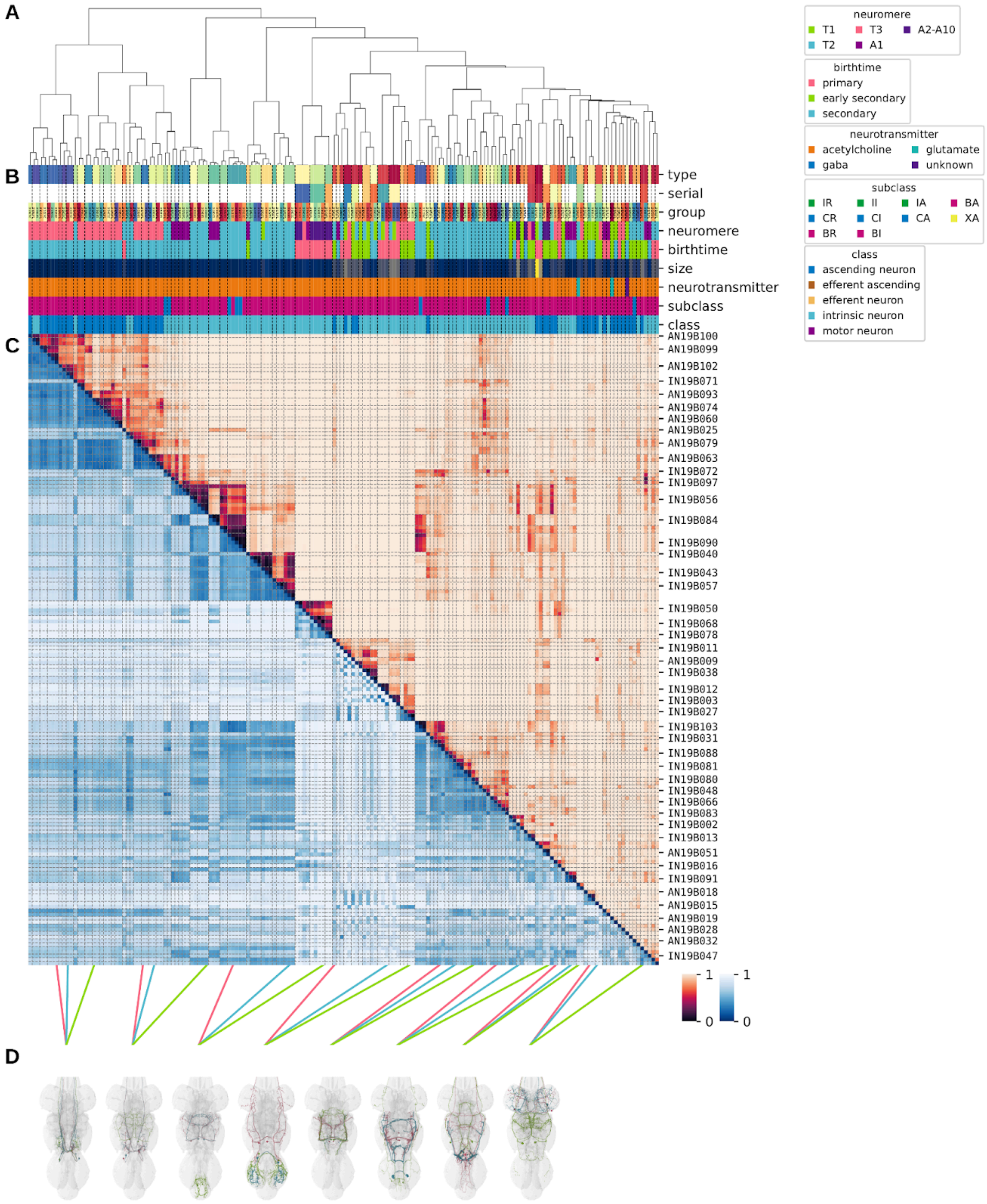

**Figure 43 - figure supplement 1. Systematic typing of hemilineage 19B. A.** Hierarchical clustering dendrogram of hemilineage groups by laterally and serially aggregated connectivity cosine clustering. **B.** Categorical annotations of each hemilineage group, each column corresponding to the aligned leaf in A. Colours for type, serial set, and group are arbitrary for visualisation. Colours for neuromere, birthtime, neurotransmitter, subclass, and class are as in all other figures. **C.** Similarity distance heatmap for hemilineage. Cosine distance is in the upper triangle, while laterally symmetrised NBLAST distance is in the lower triangle. Systematic type names of some types are labelled. **D.** Morphologically representative groups from dendrogram subtrees. Each group, indicated by colour and line

connecting to its column in B and C, is the most morphologically representative group (medoid of NBLAST distance) from a subtree of A. The subtrees (flat clusters) are equal height cuts of A determined to yield the number of groups per plot and plots in D.

**19B systematic types**

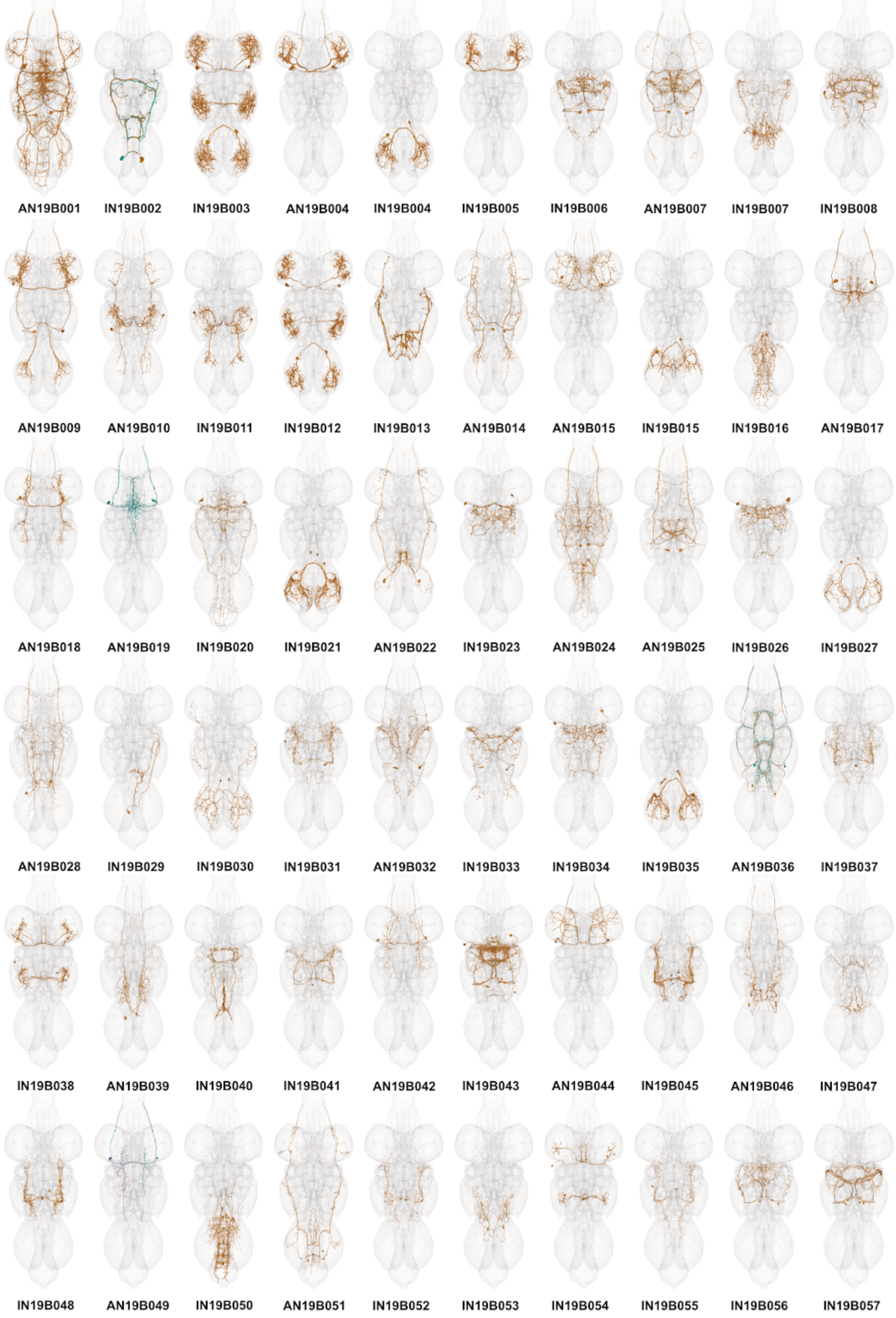

**Figure 43 - figure supplement 2. Systematic types of hemilineage 19B.** Systematic types have been arranged in numerical order, with neurons of the same type that belong to distinct classes (e.g., intrinsic neuron vs ascending neuron) plotted separately but placed adjacent to each other. Individual neuron meshes have been coloured based on predicted neurotransmitter: dark orange = acetylcholine, blue = gaba, marine = glutamate, dark purple = unknown.

#### 19B systematic types, continued

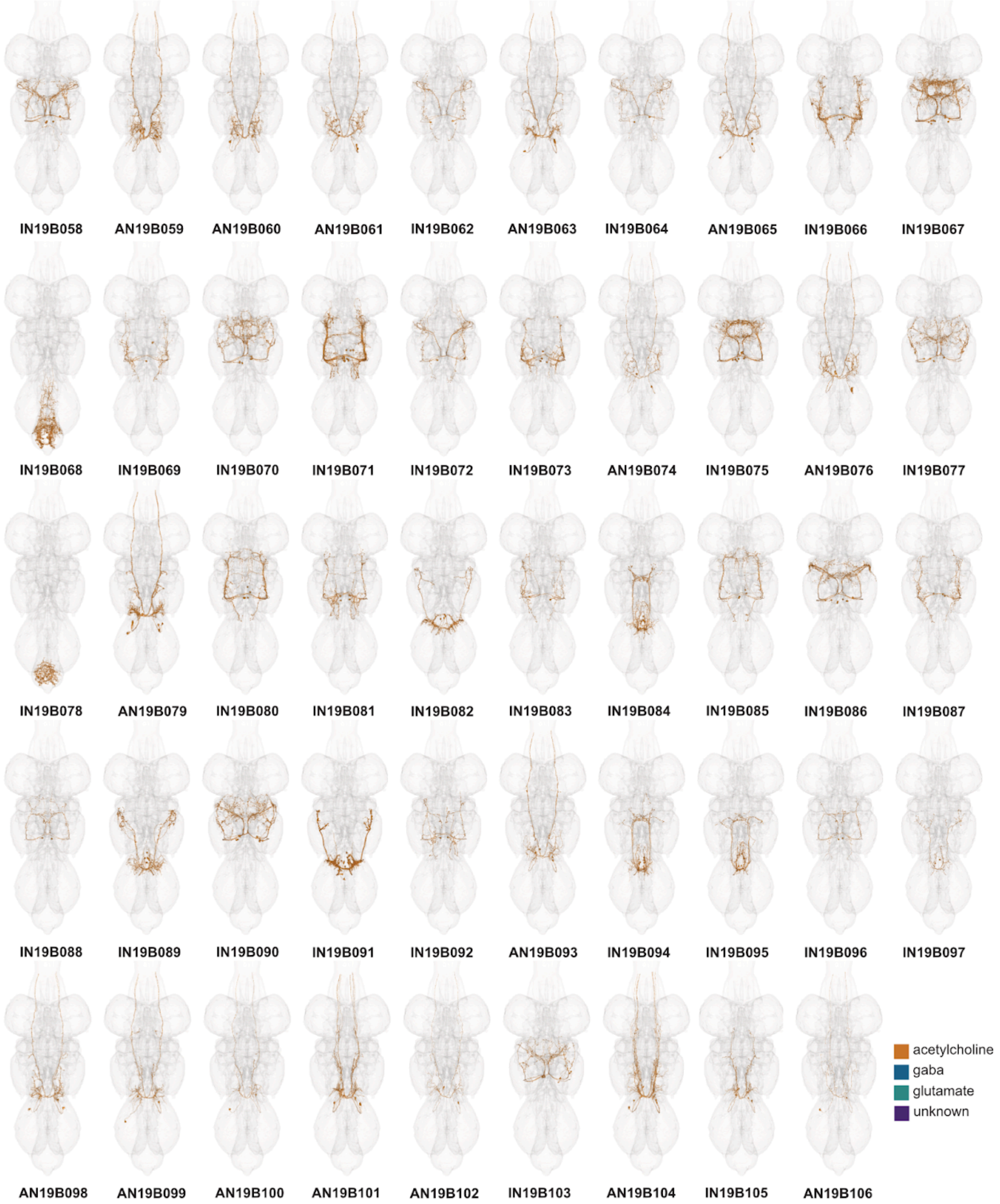

**Figure 43 - figure supplement 3. Systematic types of hemilineage 19B, continued.** Systematic types have been arranged in numerical order, with neurons of the same type that belong to distinct classes (e.g., intrinsic neuron vs ascending neuron) plotted separately but placed adjacent to each other. Individual neuron meshes have been coloured based on predicted neurotransmitter: dark orange = acetylcholine, blue = gaba, marine = glutamate, dark purple = unknown.

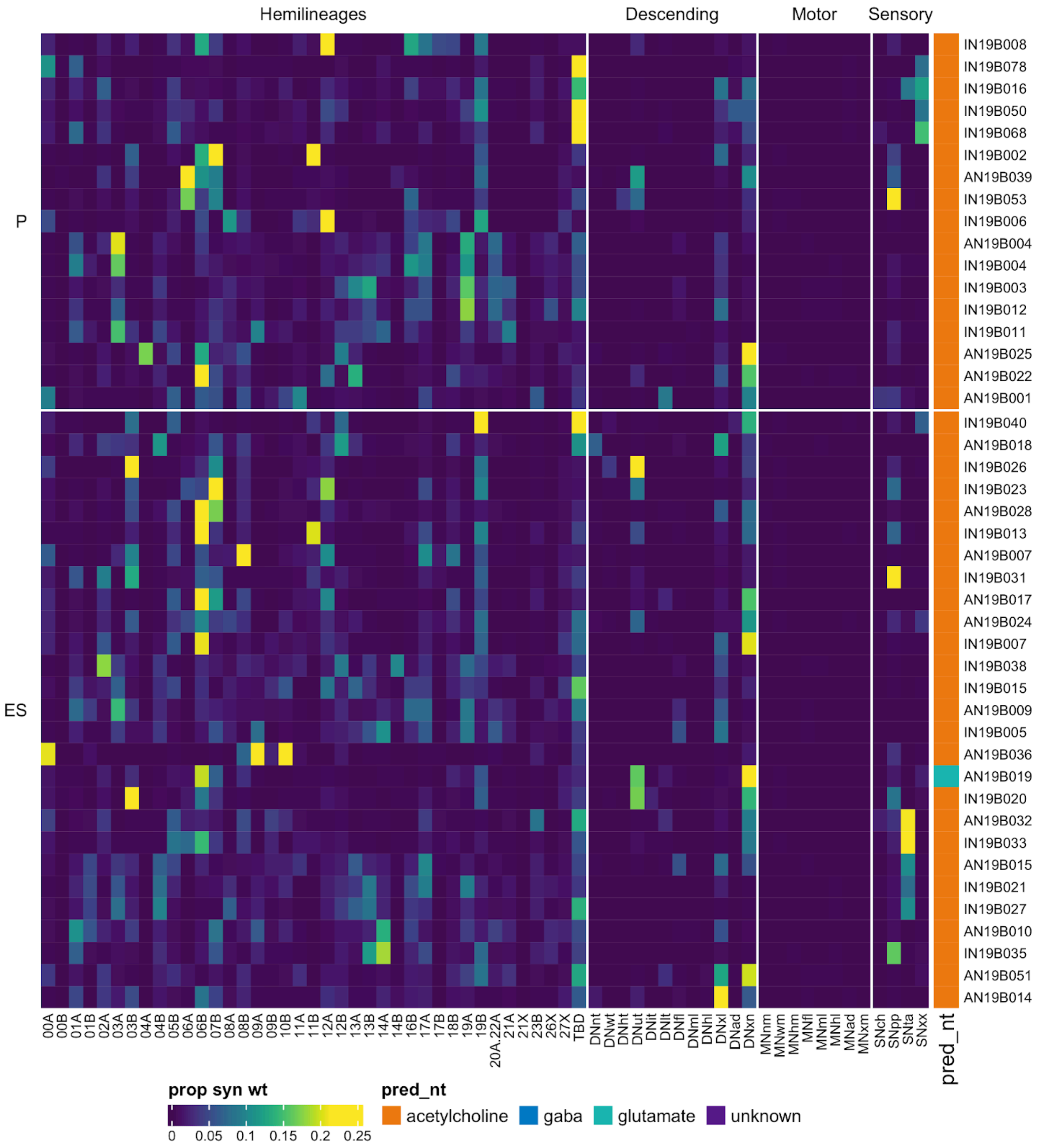

**Figure 43 - figure supplement 4. Connectivity to upstream partners by 19B primary and early secondary systematic types.** Proportions of synaptic weight to systematic types from upstream partners, normalised by row. 19B neurons have been clustered within each assigned birthtime window (P = primary, ES = early secondary, S = secondary) based on both upstream and downstream connectivity to hemilineages, descending neuron subclasses, motor neuron subclasses, and sensory neuron modalities. Annotation bar is coloured by the most common predicted neurotransmitter for the neurons of each type.

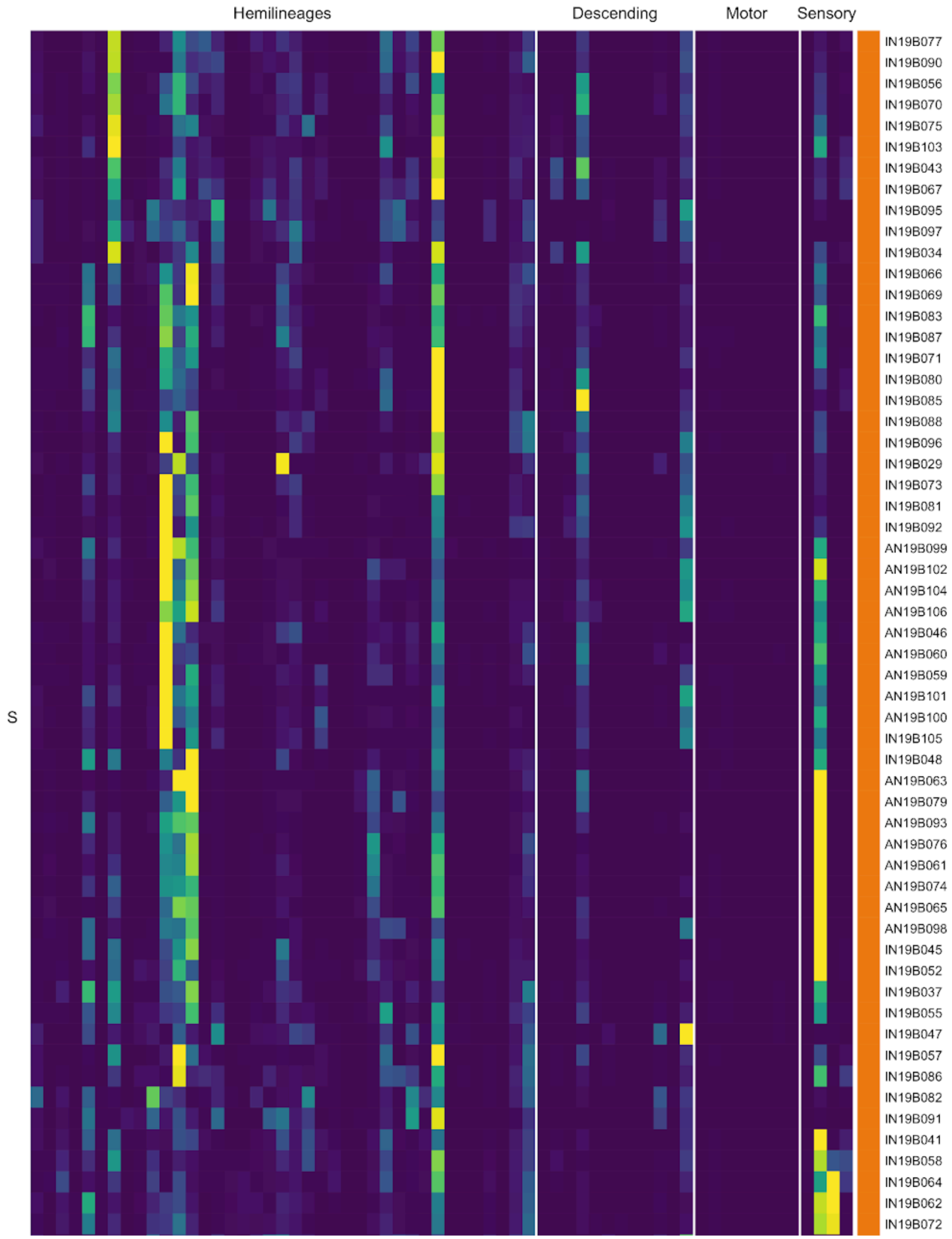

**Figure 43 - figure supplement 5. Connectivity to upstream partners by 19B secondary systematic types.** Proportions of synaptic weight to systematic types from upstream partners, normalised by row. 19B neurons have been clustered within each assigned birthtime window (P = primary, ES = early secondary, S = secondary) based on both upstream and downstream connectivity to hemilineages, descending neuron subclasses, motor neuron subclasses, and sensory neuron modalities. The annotation bar is coloured by the most common predicted neurotransmitter within each type.

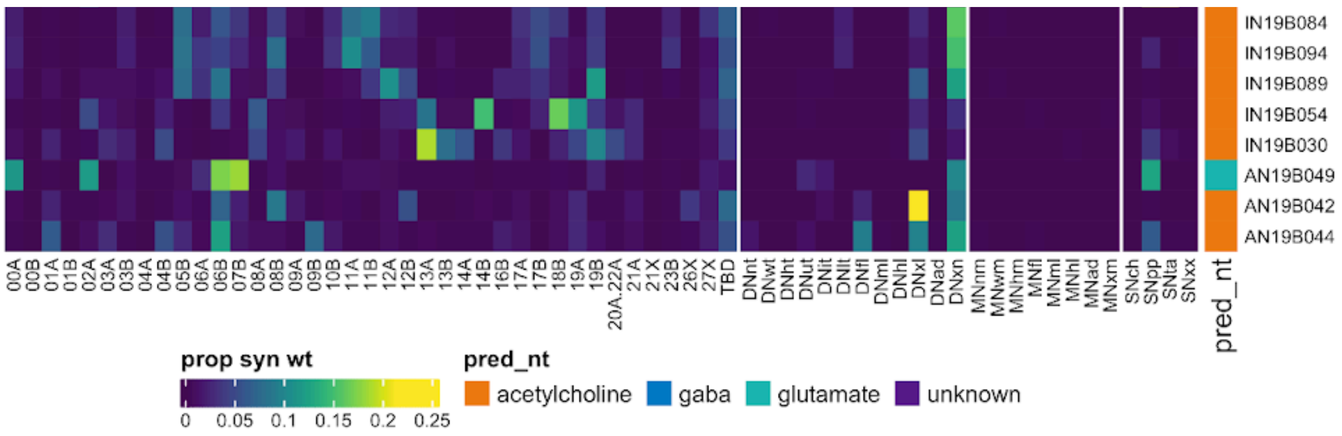

**Figure 43 - figure supplement 6. Connectivity to upstream partners by 19B secondary systematic types, continued.** Proportions of synaptic weight to systematic types from upstream partners, normalised by row. 19B neurons have been clustered within each assigned birthtime window (P = primary, ES = early secondary, S = secondary) based on both upstream and downstream connectivity to hemilineages, descending neuron subclasses, motor neuron subclasses, and sensory neuron modalities. The annotation bar is coloured by the most common predicted neurotransmitter within each type.

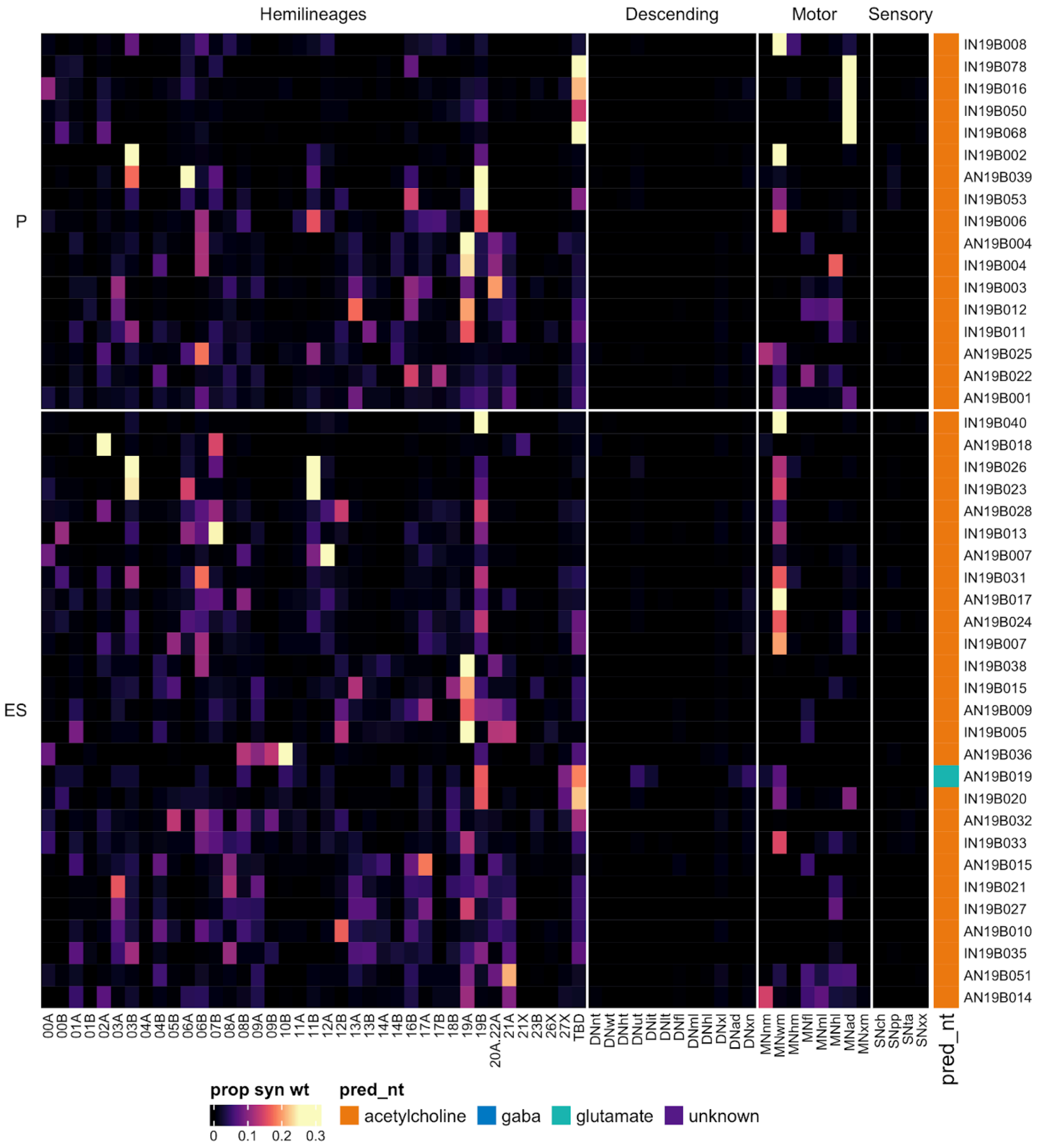

**Figure 43 - figure supplement 7. Connectivity to downstream partners by 19B primary and early secondary systematic types.** Proportions of synaptic weight from systematic types to downstream partners, normalised by row. 19B neurons have been clustered within each assigned birthtime window (P = primary, ES = early secondary, S = secondary) based on both upstream and downstream connectivity to hemilineages, descending neuron subclasses, motor neuron subclasses, and sensory neuron modalities. The annotation bar is coloured by the most common predicted neurotransmitter within each type.

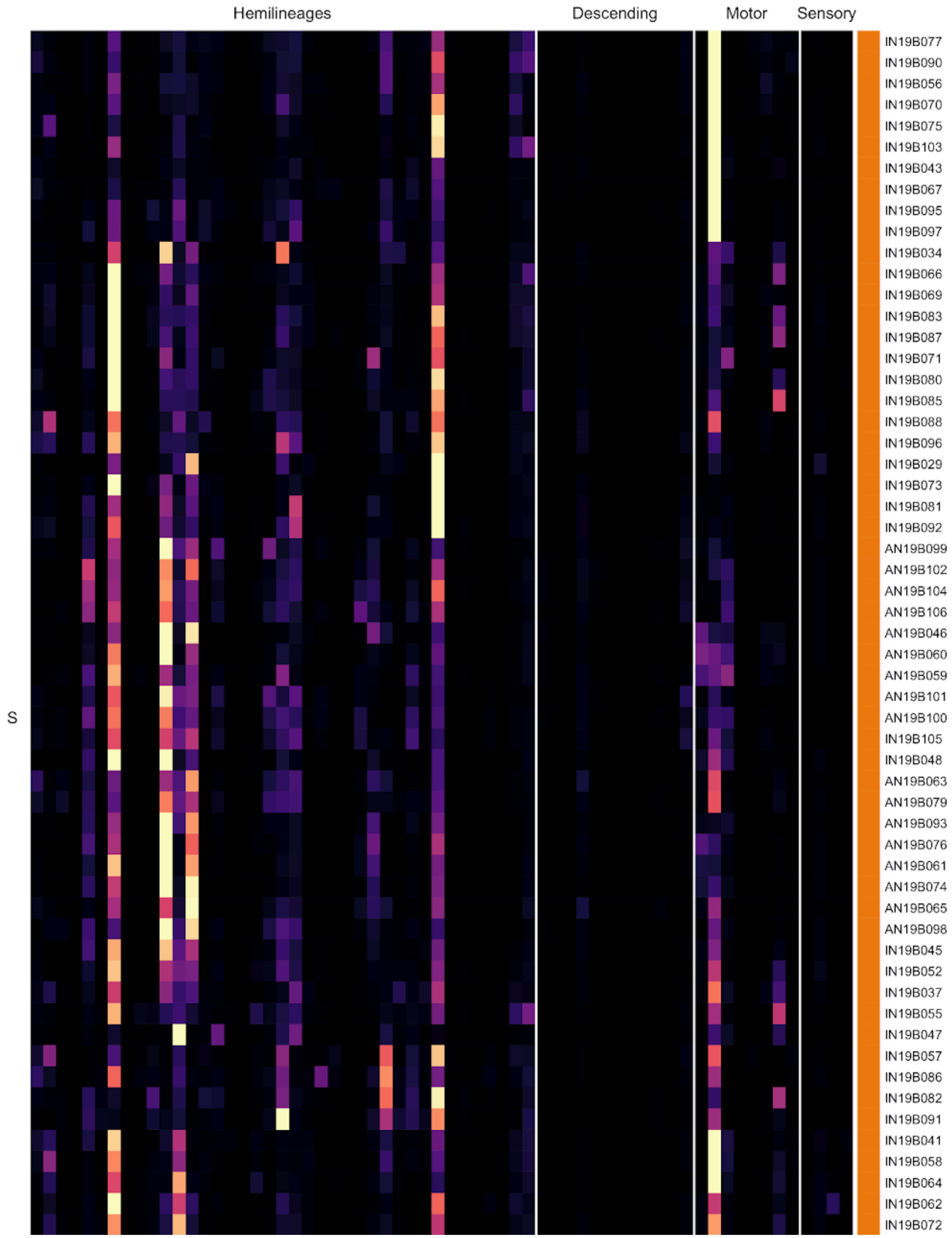

**Figure 43 - figure supplement 8. Connectivity to downstream partners by 19B secondary systematic types.** Proportions of synaptic weight from systematic types to downstream partners, normalised by row. 19B neurons have been clustered within each assigned birthtime window (P = primary, ES = early secondary, S = secondary) based on both upstream and downstream connectivity to hemilineages, descending neuron subclasses, motor neuron subclasses, and sensory neuron modalities. The annotation bar is coloured by the most common predicted neurotransmitter for the neurons of each type.

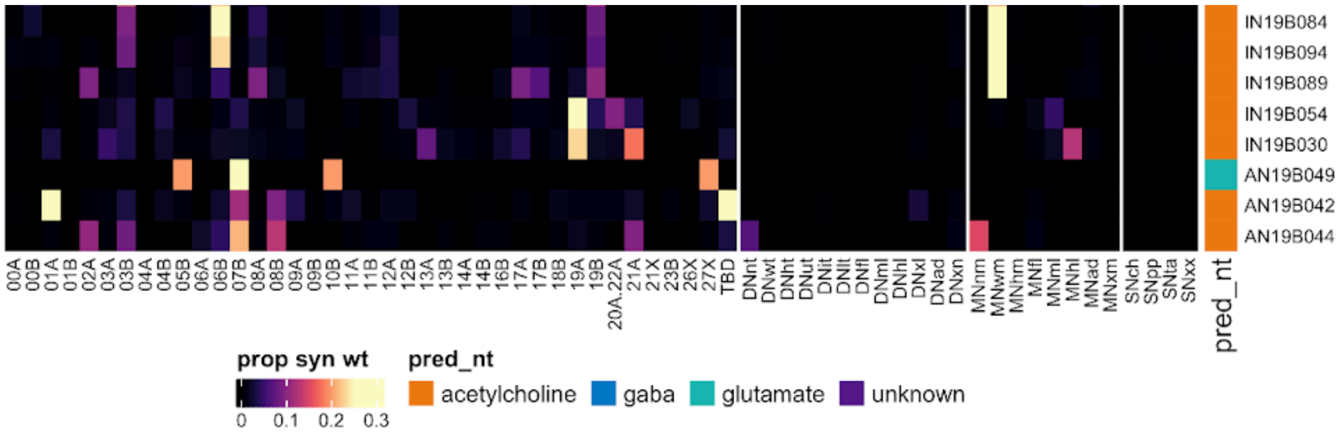

**Figure 43 - figure supplement 9. Connectivity to downstream partners by 19B secondary systematic types, continued.** Proportions of synaptic weight from systematic types to downstream partners, normalised by row. 19B neurons have been clustered within each assigned birthtime window (P = primary, ES = early secondary, S = secondary) based on both upstream and downstream connectivity to hemilineages, descending neuron subclasses, motor neuron subclasses, and sensory neuron modalities. The annotation bar is coloured by the most common predicted neurotransmitter for the neurons of each type.

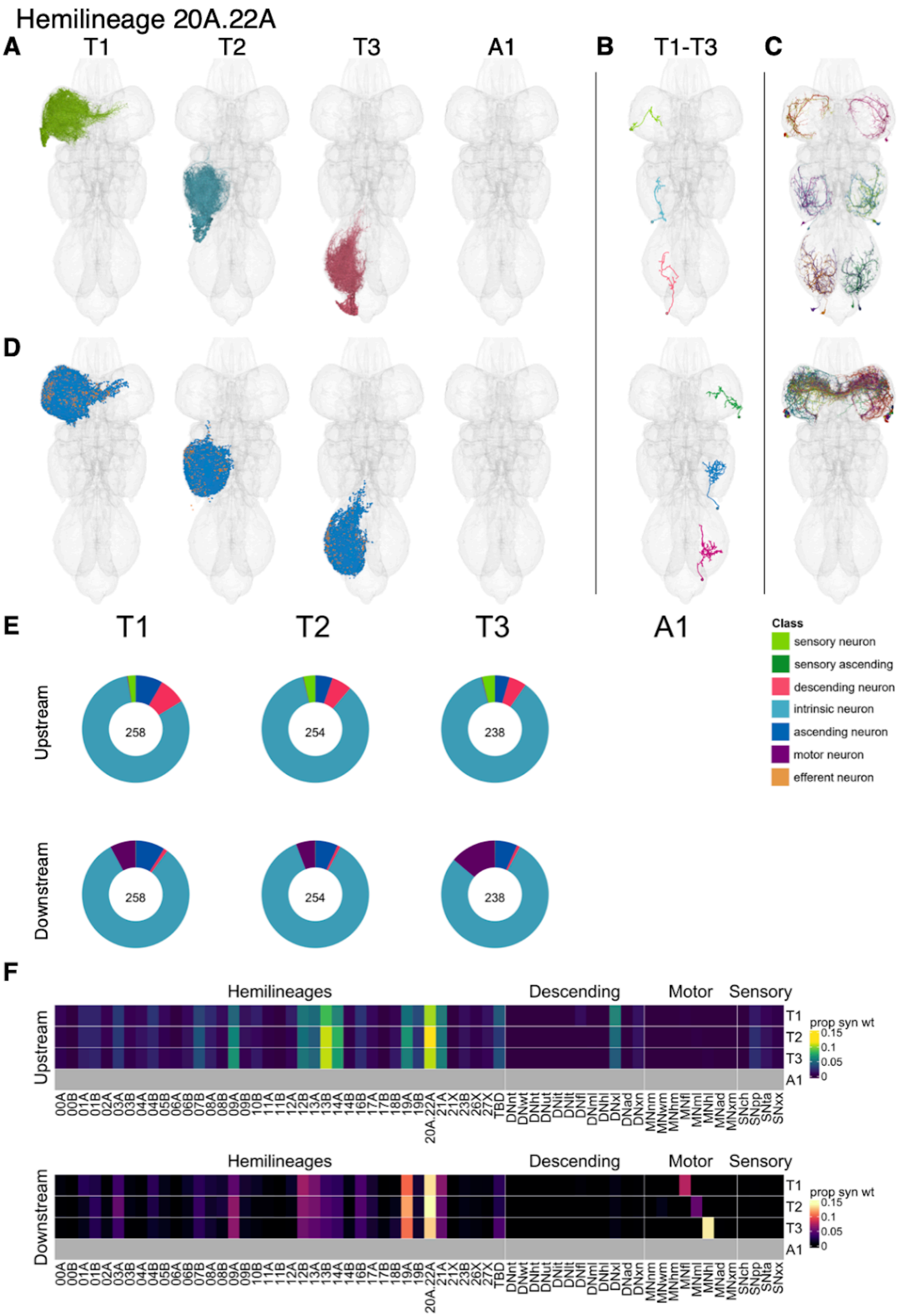

Hemilineage 20A.22A
A
T1
T2
T3
A1
B
T1-T3
C
D
E
T1
T2
T3
A1
Upstream
Downstream
258
254
238
Class
sensory neuron
sensory ascending
descending neuron
intrinsic neuron
ascending neuron
motor neuron
efferent neuron
F
Hemilineages
Descending
Motor
Sensory
prop syn wt
0.15
0.1
0.05
0
00A 00B 01A 01B 02A 03A 03B 04A 04B 05B 06A 06B 07B 08A 08B 09A 09B 10B 11A 11B 12A 12B 13A 13B 14A 14B 16B 17A 17B 18B 19A 19B 20A.22A 21A 21X 23B 26X 27X TBD DNnt DNwt DNht DNut DNlt DNfl DNml DNhl DNxl DNad DNxn MNnm MNwm MNhm MNfl MNml MNhl MNad MNxm SNch SNpp SNta SNxx

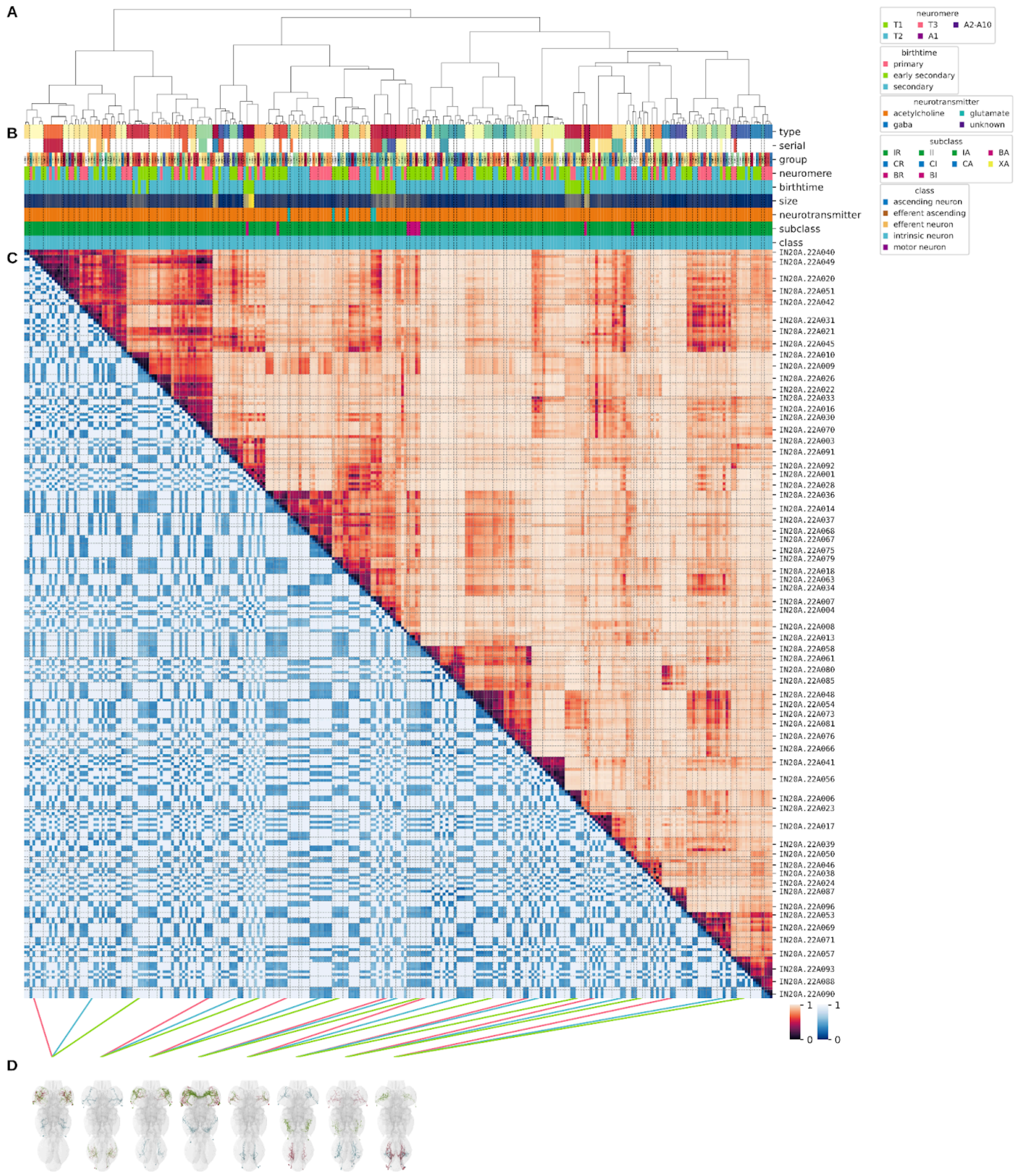

**Figure 44 - figure supplement 1. Systematic typing of hemilineage 20A and 22A. A.** Hierarchical clustering dendrogram of hemilineage groups by laterally and serially aggregated connectivity cosine clustering. **B.** Categorical annotations of each hemilineage group, each column corresponding to the aligned leaf in A. Colours for type, serial set, and group are arbitrary for visualisation. Colours for neuromere, birthtime, neurotransmitter, subclass, and class are as in all other figures. **C.** Similarity distance heatmap for hemilineage. Cosine distance is in the upper triangle, while laterally symmetrised NBLAST distance is in the lower triangle. Systematic type names of some types are labelled. **D.** Morphologically representative groups from dendrogram subtrees. Each group, indicated by colour and line connecting to its column in B and C, is the most morphologically representative group (medoid of NBLAST distance) from a subtree of A. The subtrees (flat clusters) are equal height cuts of A determined to yield the number of groups per plot and plots in D.

#### 20A.22A systematic types

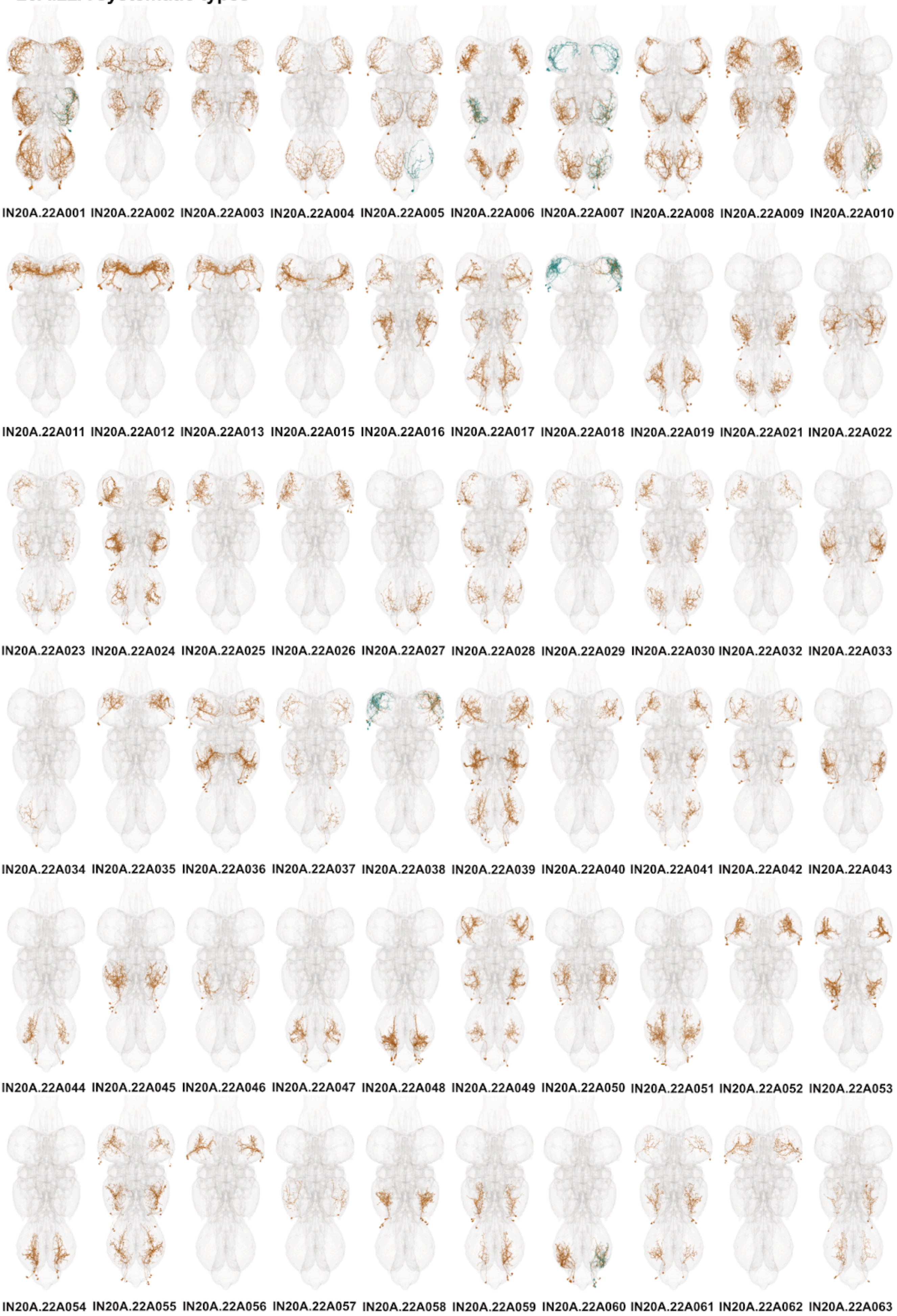

**Figure 44 - figure supplement 2. Systematic types of hemilineages 20A and 22A.** Systematic types have been arranged in numerical order, with neurons of the same type that belong to distinct classes (e.g., intrinsic neuron vs ascending neuron) plotted separately but placed adjacent to each other. Individual neuron meshes have been coloured based on predicted neurotransmitter: dark orange = acetylcholine, blue = gaba, marine = glutamate, dark purple = unknown.

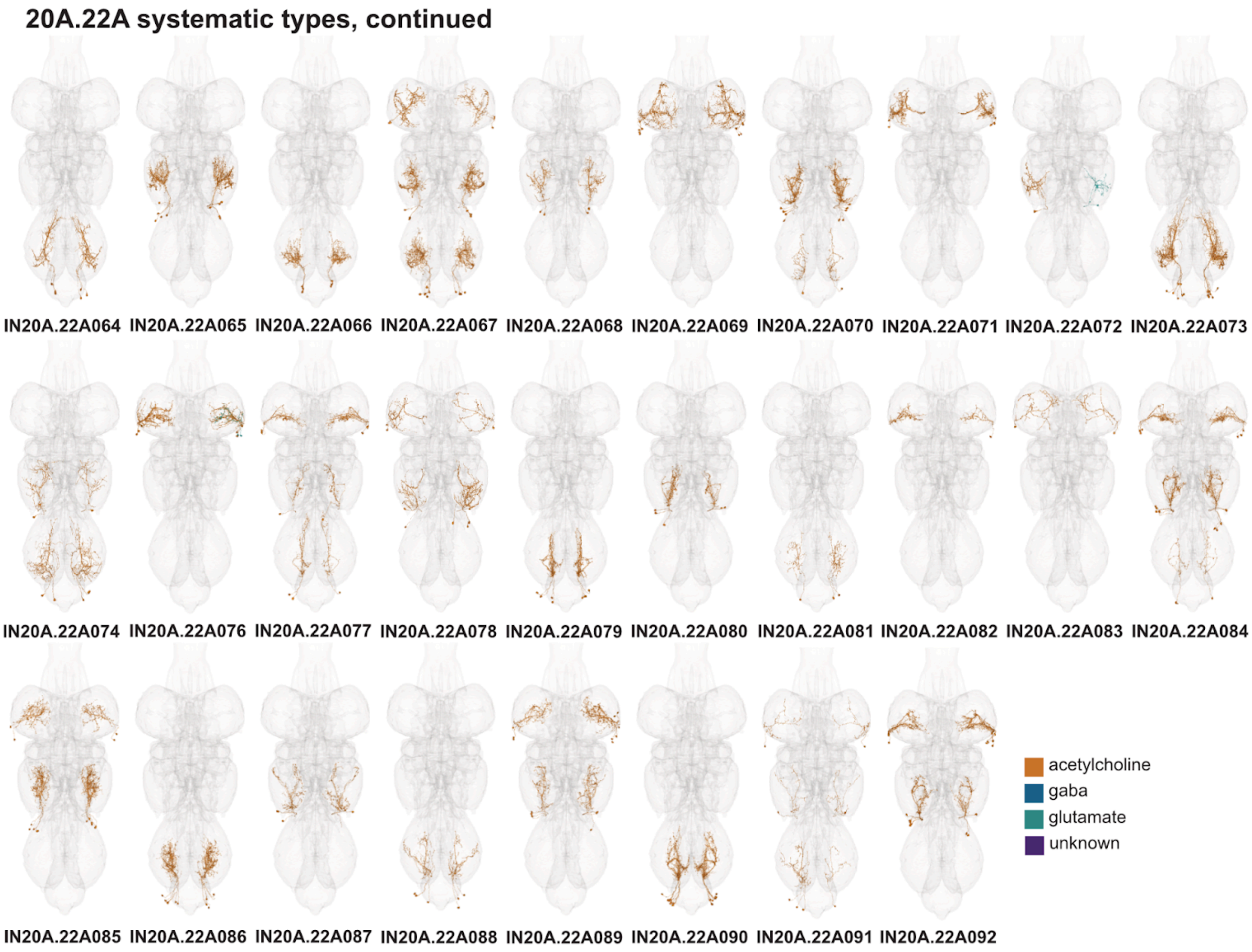

**Figure 44 - figure supplement 3. Systematic types of hemilineages 20A and 22A, continued.** Systematic types have been arranged in numerical order, with neurons of the same type that belong to distinct classes (e.g., intrinsic neuron vs ascending neuron) plotted separately but placed adjacent to each other. Individual neuron meshes have been coloured based on predicted neurotransmitter: dark orange = acetylcholine, blue = gaba, marine = glutamate, dark purple = unknown.

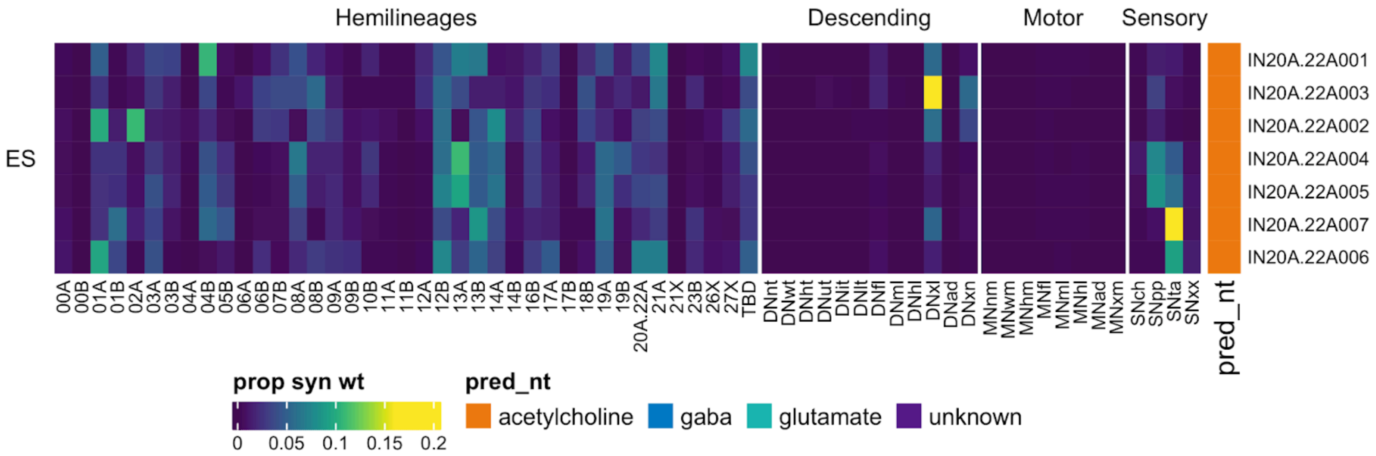

**Figure 44 - figure supplement 4. Connectivity to upstream partners by 20A.22A early secondary systematic types.** Proportions of synaptic weight to systematic types from upstream partners, normalised by row. 20A.22A neurons have been clustered within each assigned birthtime window (P = primary, ES = early secondary, S = secondary) based on both upstream and downstream connectivity to hemilineages, descending neuron subclasses, motor neuron subclasses, and sensory neuron modalities. Annotation bar is coloured by the most common predicted neurotransmitter for the neurons of each type.

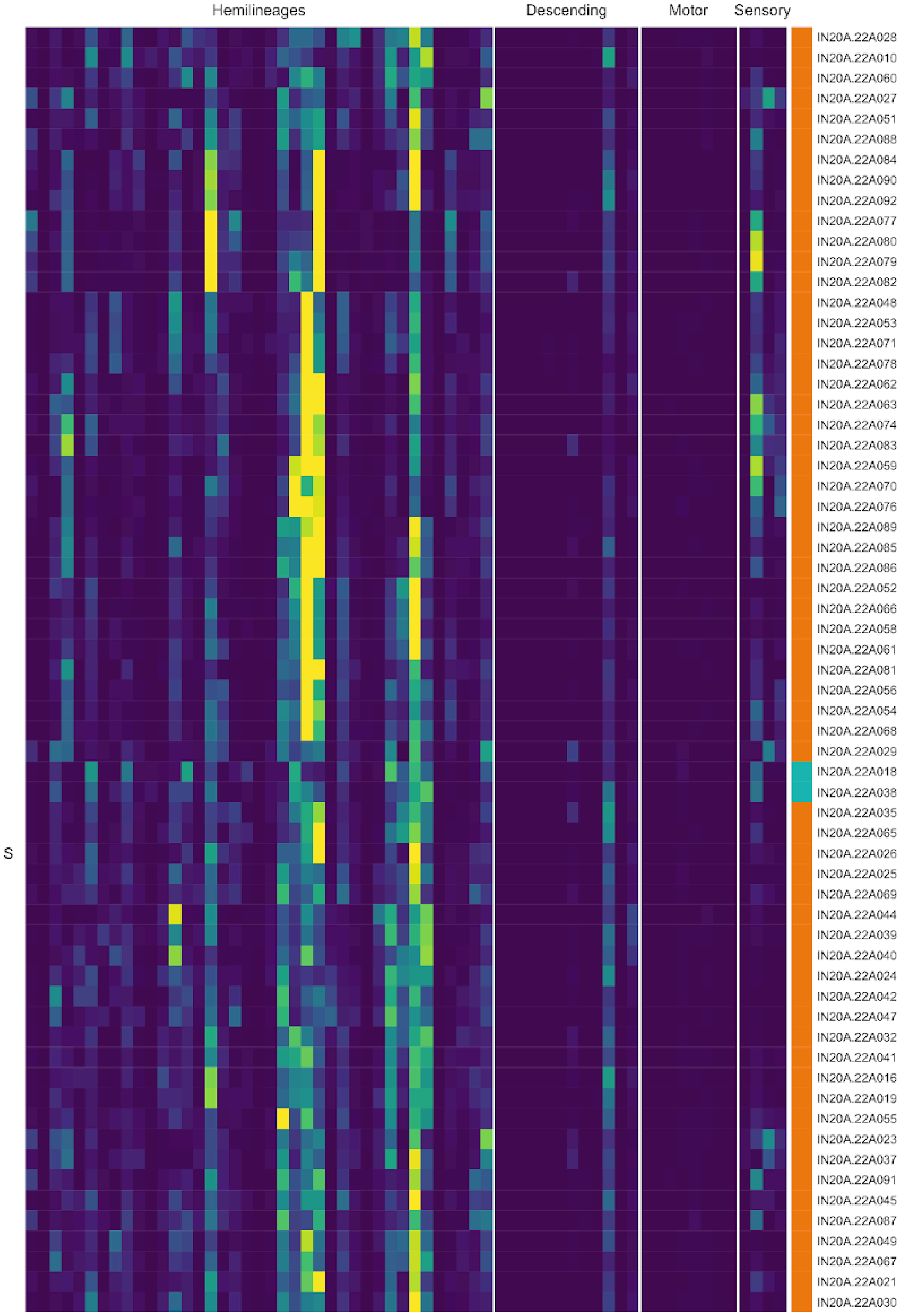
Hemilineages
Descending
Motor
Sensory
S
IN20A.22A028
IN20A.22A010
IN20A.22A060
IN20A.22A027
IN20A.22A051
IN20A.22A088
IN20A.22A084
IN20A.22A090
IN20A.22A092
IN20A.22A077
IN20A.22A080
IN20A.22A079
IN20A.22A082
IN20A.22A048
IN20A.22A053
IN20A.22A071
IN20A.22A078
IN20A.22A062
IN20A.22A063
IN20A.22A074
IN20A.22A083
IN20A.22A059
IN20A.22A070
IN20A.22A076
IN20A.22A089
IN20A.22A085
IN20A.22A086
IN20A.22A052
IN20A.22A066
IN20A.22A058
IN20A.22A061
IN20A.22A081
IN20A.22A056
IN20A.22A054
IN20A.22A068
IN20A.22A029
IN20A.22A018
IN20A.22A038
IN20A.22A035
IN20A.22A065
IN20A.22A026
IN20A.22A025
IN20A.22A069
IN20A.22A044
IN20A.22A039
IN20A.22A040
IN20A.22A024
IN20A.22A042
IN20A.22A047
IN20A.22A032
IN20A.22A041
IN20A.22A016
IN20A.22A019
IN20A.22A055
IN20A.22A023
IN20A.22A037
IN20A.22A091
IN20A.22A045
IN20A.22A087
IN20A.22A049
IN20A.22A067
IN20A.22A021
IN20A.22A030

**Figure 44 - figure supplement 5. Connectivity to upstream partners by 20A.22A secondary systematic types.** Proportions of synaptic weight to systematic types from upstream partners, normalised by row. 20A.22A neurons have been clustered within each assigned birthtime window (P = primary, ES = early secondary, S = secondary) based on both upstream and downstream connectivity to hemilineages, descending neuron subclasses, motor neuron subclasses, and sensory neuron modalities. The annotation bar is coloured by the most common predicted neurotransmitter within each type.

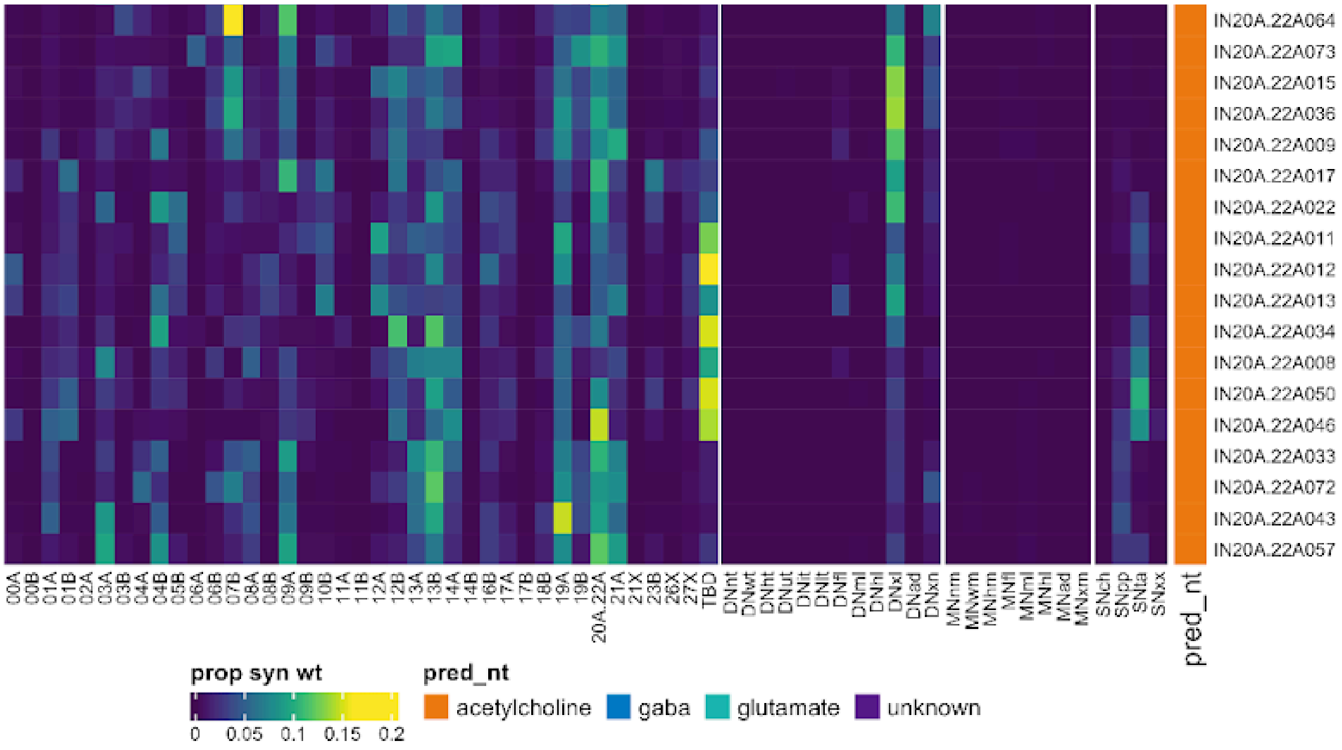

**Figure 44 - figure supplement 6.Connectivity to upstream partners by 20A.22A secondary systematic types, continued.** Proportions of synaptic weight to systematic types from upstream partners, normalised by row. 20A.22A neurons have been clustered within each assigned birthtime window (P = primary, ES = early secondary, S = secondary) based on both upstream and downstream connectivity to hemilineages, descending neuron subclasses, motor neuron subclasses, and sensory neuron modalities. The annotation bar is coloured by the most common predicted neurotransmitter within each type.

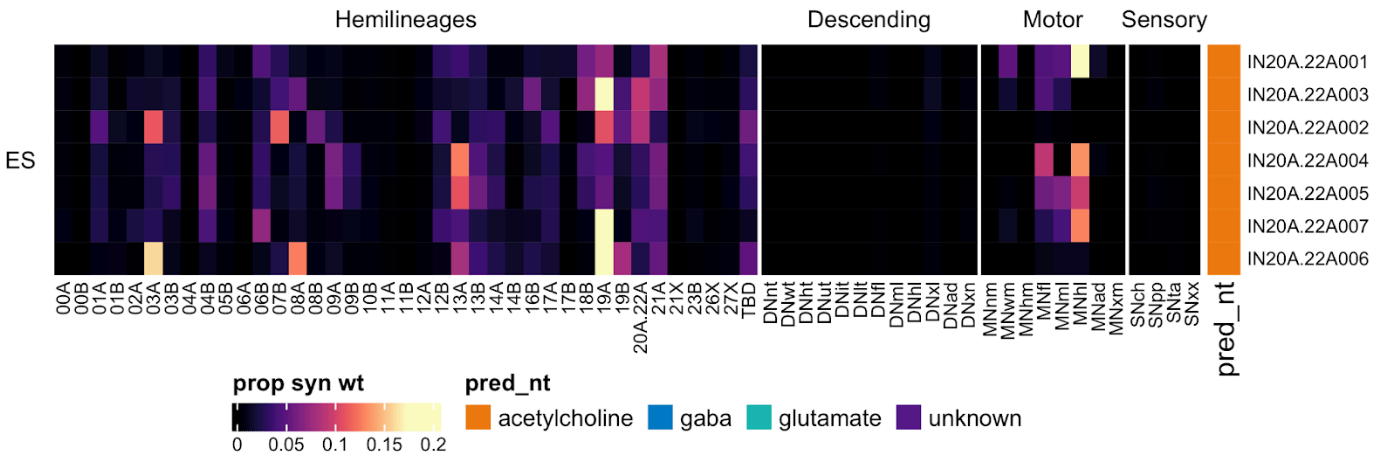

**Figure 44 - figure supplement 7. Connectivity to downstream partners by 20A.22A early secondary systematic types.** Proportions of synaptic weight from systematic types to downstream partners, normalised by row. 20A.22A neurons have been clustered within each assigned birthtime window (P = primary, ES = early secondary, S = secondary) based on both upstream and downstream connectivity to hemilineages, descending neuron subclasses, motor neuron subclasses, and sensory neuron modalities. The annotation bar is coloured by the most common predicted neurotransmitter within each type.

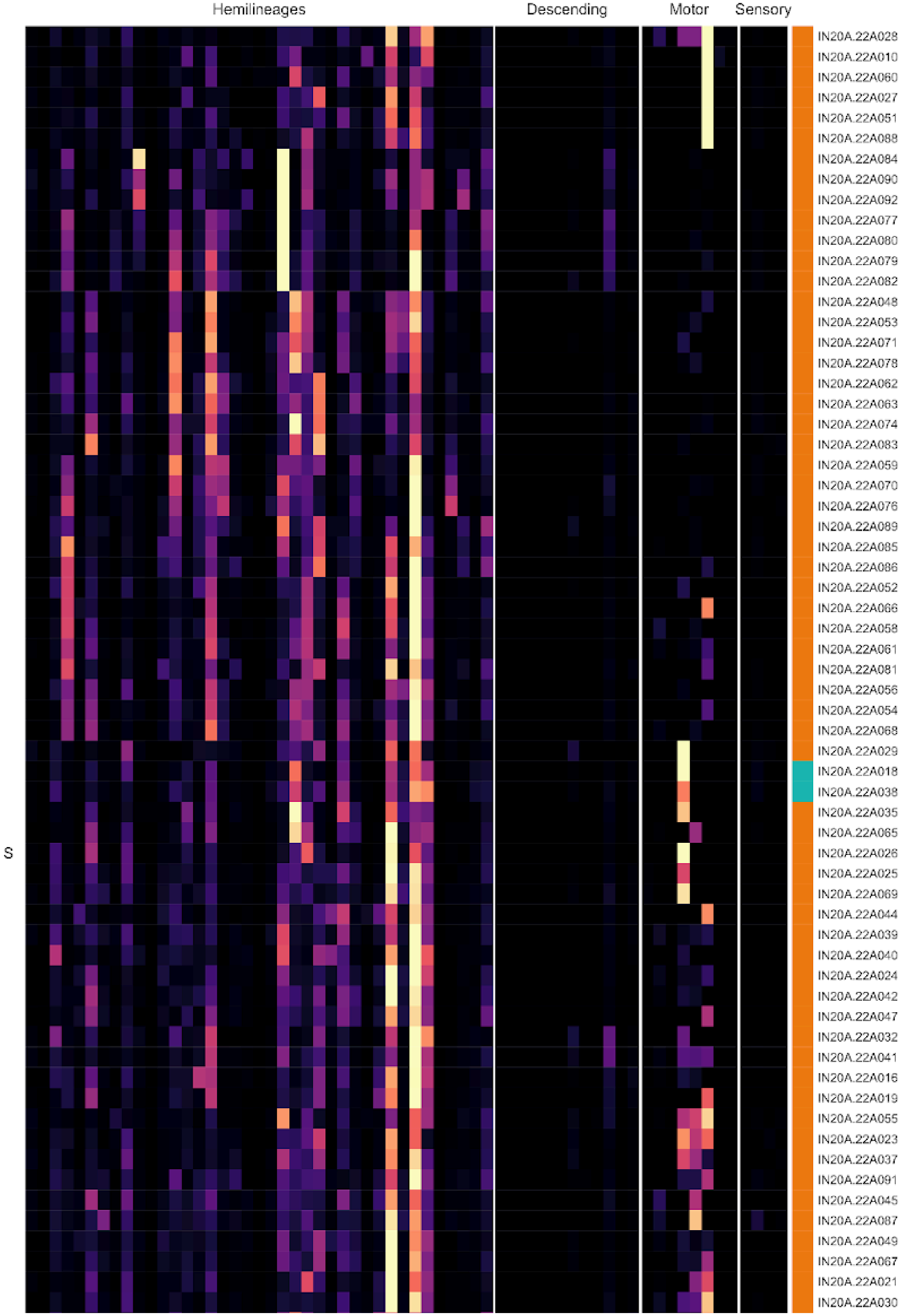

Hemilineages
Descending
Motor
Sensory
S
IN20A.22A028
IN20A.22A010
IN20A.22A060
IN20A.22A027
IN20A.22A051
IN20A.22A088
IN20A.22A084
IN20A.22A090
IN20A.22A092
IN20A.22A077
IN20A.22A080
IN20A.22A079
IN20A.22A082
IN20A.22A048
IN20A.22A053
IN20A.22A071
IN20A.22A078
IN20A.22A062
IN20A.22A063
IN20A.22A074
IN20A.22A083
IN20A.22A059
IN20A.22A070
IN20A.22A076
IN20A.22A089
IN20A.22A085
IN20A.22A086
IN20A.22A052
IN20A.22A066
IN20A.22A058
IN20A.22A061
IN20A.22A081
IN20A.22A056
IN20A.22A054
IN20A.22A068
IN20A.22A029
IN20A.22A018
IN20A.22A038
IN20A.22A035
IN20A.22A065
IN20A.22A026
IN20A.22A025
IN20A.22A069
IN20A.22A044
IN20A.22A039
IN20A.22A040
IN20A.22A024
IN20A.22A042
IN20A.22A047
IN20A.22A032
IN20A.22A041
IN20A.22A016
IN20A.22A019
IN20A.22A055
IN20A.22A023
IN20A.22A037
IN20A.22A091
IN20A.22A045
IN20A.22A087
IN20A.22A049
IN20A.22A067
IN20A.22A021
IN20A.22A030

**Figure 44 - figure supplement 8. Connectivity to downstream partners by 20A.22A secondary systematic types.** Proportions of synaptic weight from systematic types to downstream partners, normalised by row. 20A.22A neurons have been clustered within each assigned birthtime window (P = primary, ES = early secondary, S = secondary) based on both upstream and downstream connectivity to hemilineages, descending neuron subclasses, motor neuron subclasses, and sensory neuron modalities. The annotation bar is coloured by the most common predicted neurotransmitter for the neurons of each type.

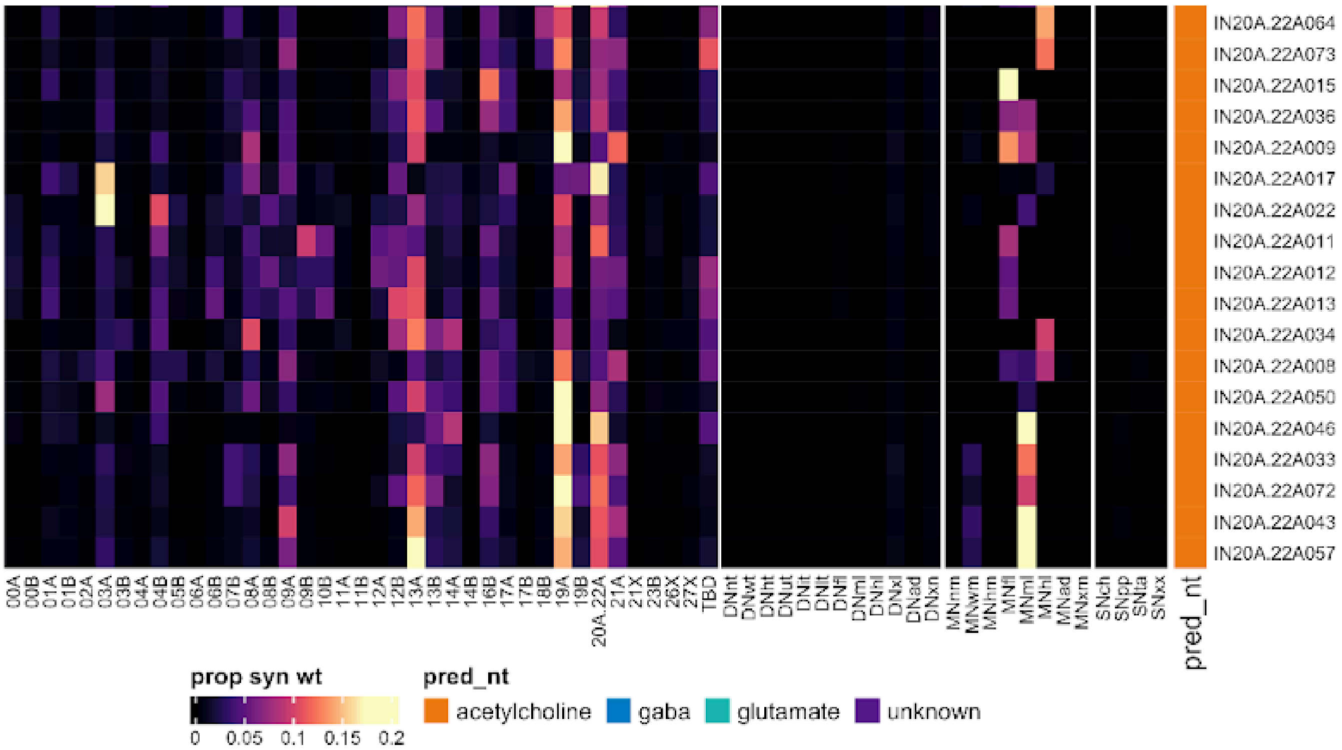

**Figure 44 - figure supplement 9. Connectivity to downstream partners by 20A.22A secondary systematic types, continued.** Proportions of synaptic weight from systematic types to downstream partners, normalised by row. 20A.22A neurons have been clustered within each assigned birthtime window (P = primary, ES = early secondary, S = secondary) based on both upstream and downstream connectivity to hemilineages, descending neuron subclasses, motor neuron subclasses, and sensory neuron modalities. The annotation bar is coloured by the most common predicted neurotransmitter for the neurons of each type.

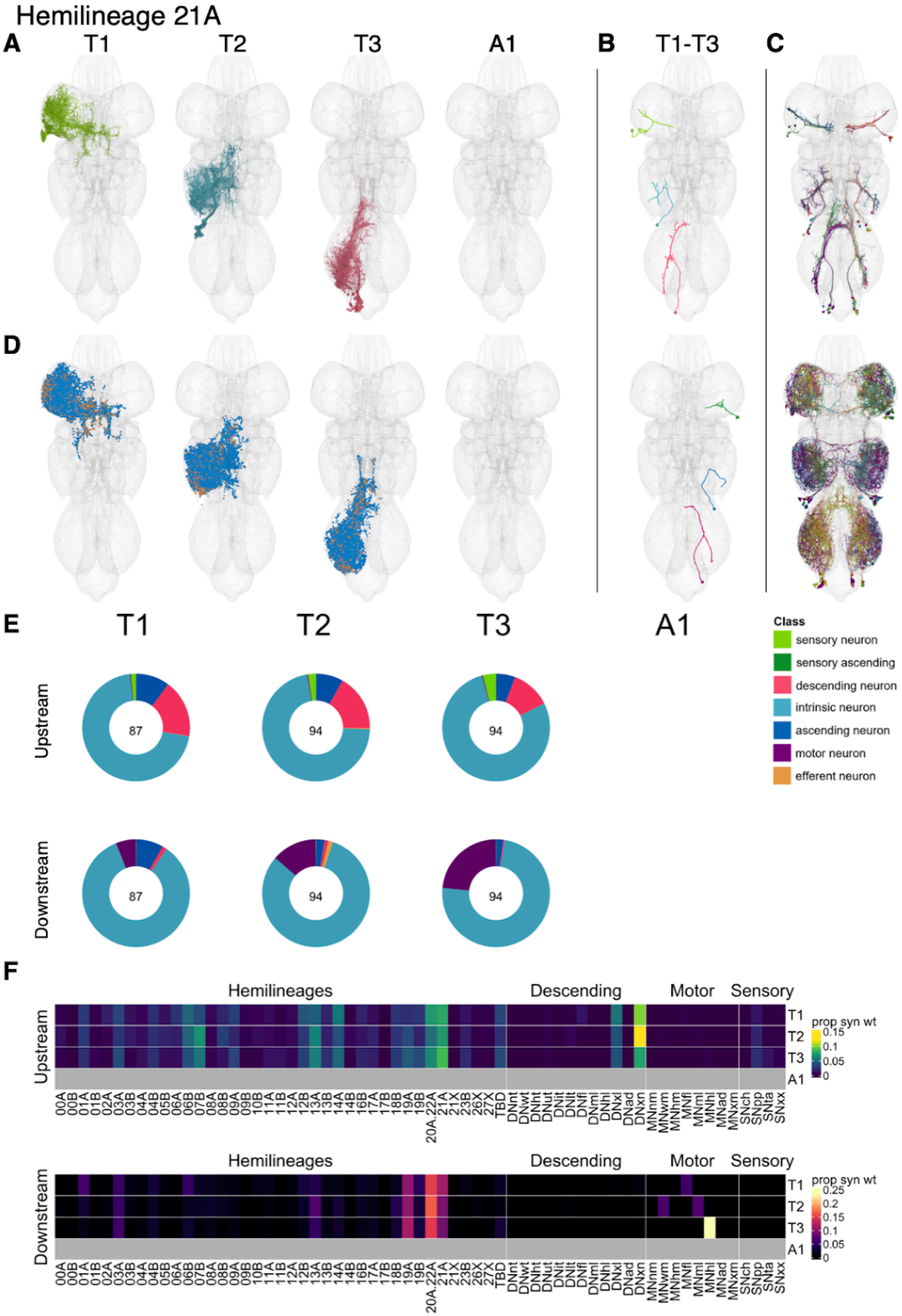
Hemilineage 21A
A
T1
T2
T3
A1
B
T1-T3
C
D
E
T1
T2
T3
A1
Upstream
Downstream
87
94
94
87
94
94
Class
sensory neuron
sensory ascending
descending neuron
intrinsic neuron
ascending neuron
motor neuron
efferent neuron
F
Hemilineages
Descending
Motor
Sensory
prop syn wt
0.15
0.1
0.05
0
0.25
0.2
0.15
0.1
0.05
0
T1
T2
T3
A1
00A 00B 01A 01B 02A 03A 03B 04A 04B 05B 06A 06B 07B 08A 08B 09A 09B 10B 11A 11B 12A 12B 13A 13B 14A 14B 16B 17A 17B 18B 19A 19B 20A.22A 21A 21X 23B 26X 27X TBD DNnt DNwt DNht DNut DNit DNlt DNfl DNml DNhl DNxl DNad DNxn MNnm MNwm MNhm MNfl MNml MNhl MNad MNxm SNch SNpp SNta SNxx

**Figure 45. Hemilineage 21A. A.** Meshes of all RHS secondary neurons plotted in neuromere-specific colours. **B.** "Representative" secondary neuron skeletons plotted in hemineuromere-specific colours. The skeleton with the top accumulated NBLAST score among all neurons from the hemilineage in a given hemineuromere was used. **C.** Neuron meshes of selected examples. Top: independent leg subcluster 23565. Bottom: cholinergic population. **D.** Predicted synapses of RHS secondary neurons. Blue: postsynapses; dark orange: presynapses. **E.** Proportions of connections from secondary neurons to upstream or downstream partners, normalised by neuromere and coloured by broad class. Numbers of query neurons appear in the centre. **F.** Proportions of synaptic weight from secondary neurons originating in each neuromere to upstream or downstream partners, normalised by row.

#### Hemilineage 21A

Hemilineages 20A, 21A, and 22A are posterior lateral hemilineages that innervate the six ipsilateral leg neuropils, with all three entering the neuropil in approximately the same location. Hemilineage 21A derives from neuroblast NB4-3 (Birkholz et al., 2015; Lacin and Truman, 2016), which generates 2-3 neurosecretory cells and 10-12 local interneurons in the embryo (Schmid et al., 1999). 21A neurons are mainly glutamatergic (Lacin et al., 2019) and include a distinctive neuronal subtype with a "T-junction" morphology that innervates the mVAC (Shepherd et al., 2019). They are mainly restricted to the ipsilateral leg neuropils but also project to the tectulum.

Secondary 21A neurons are found in similar numbers in all thoracic neuromeres (Figure 45E). We identified them from a diagnostic subpopulation that splits to diverge orthogonally to the primary neurite (Figure 45C top). Whilst most 21A neurons were predicted to be glutamatergic as expected, an early born population was predicted to be cholinergic (Figure 45C bottom). As mentioned above, no 20A or 22A primary neurons were expected in the thorax, so we tentatively assigned these cells to 21A.

21A secondary neurons receive input mainly from 07B, 13A, 20A/22A, and 21A and from descending neurons that target the legs or the VNC more broadly (Figure 45F). They primarily target 19A, 20A/22A, 21A, and leg motor neurons, especially in T3 (Figure 45F). A subset of dorsal secondary types (e.g., IN21A025-027) are under especially strong descending control and target wing motor neurons instead (Figure 45 - figure supplement 5,8). Bilateral activation of 21A secondary neurons evokes incessant, uncoordinated leg movements without locomotion (Harris et al., 2015). There was no obvious difference in connectivity between glutamatergic and cholinergic early born types (Figure 45 - figure supplement 4,7).

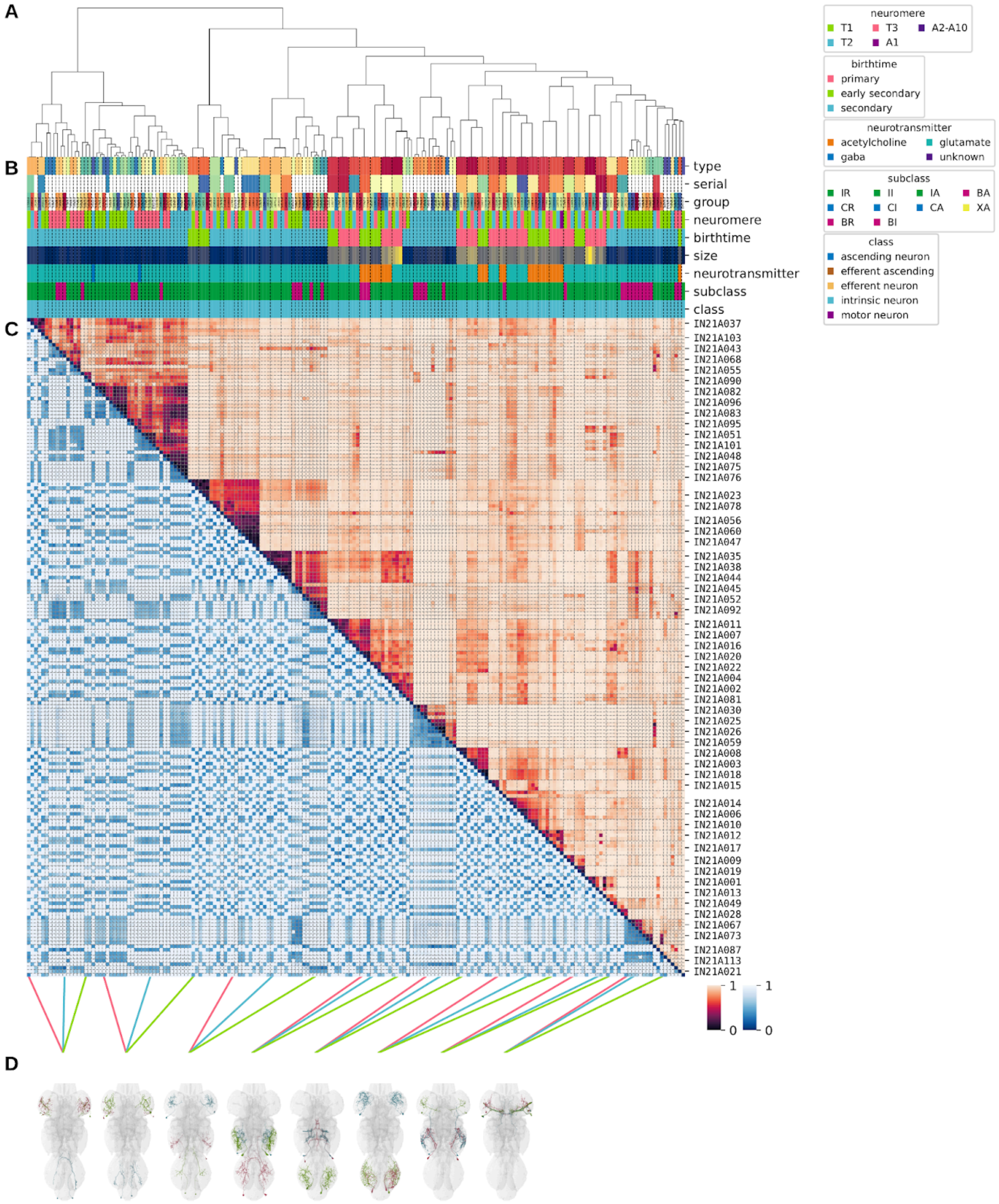

**Figure 45 - figure supplement 1. Systematic typing of hemilineage 21A. A.** Hierarchical clustering dendrogram of hemilineage groups by laterally and serially aggregated connectivity cosine clustering. **B.** Categorical annotations of each hemilineage group, each column corresponding to the aligned leaf in A. Colours for type, serial set, and group are arbitrary for visualisation. Colours for neuromere, birthtime, neurotransmitter, subclass, and class are as in all other figures. **C.** Similarity distance heatmap for hemilineage. Cosine distance is in the upper triangle, while laterally symmetrised NBLAST distance is in the lower triangle. Systematic type names of some types are labelled. **D.** Morphologically representative groups from dendrogram subtrees. Each group, indicated by colour and line connecting to its column in B and C, is the most morphologically representative group (medoid of NBLAST distance) from a subtree of A. The subtrees (flat clusters) are equal height cuts of A determined to yield the number of groups

per plot and plots in D.

#### 21A systematic types

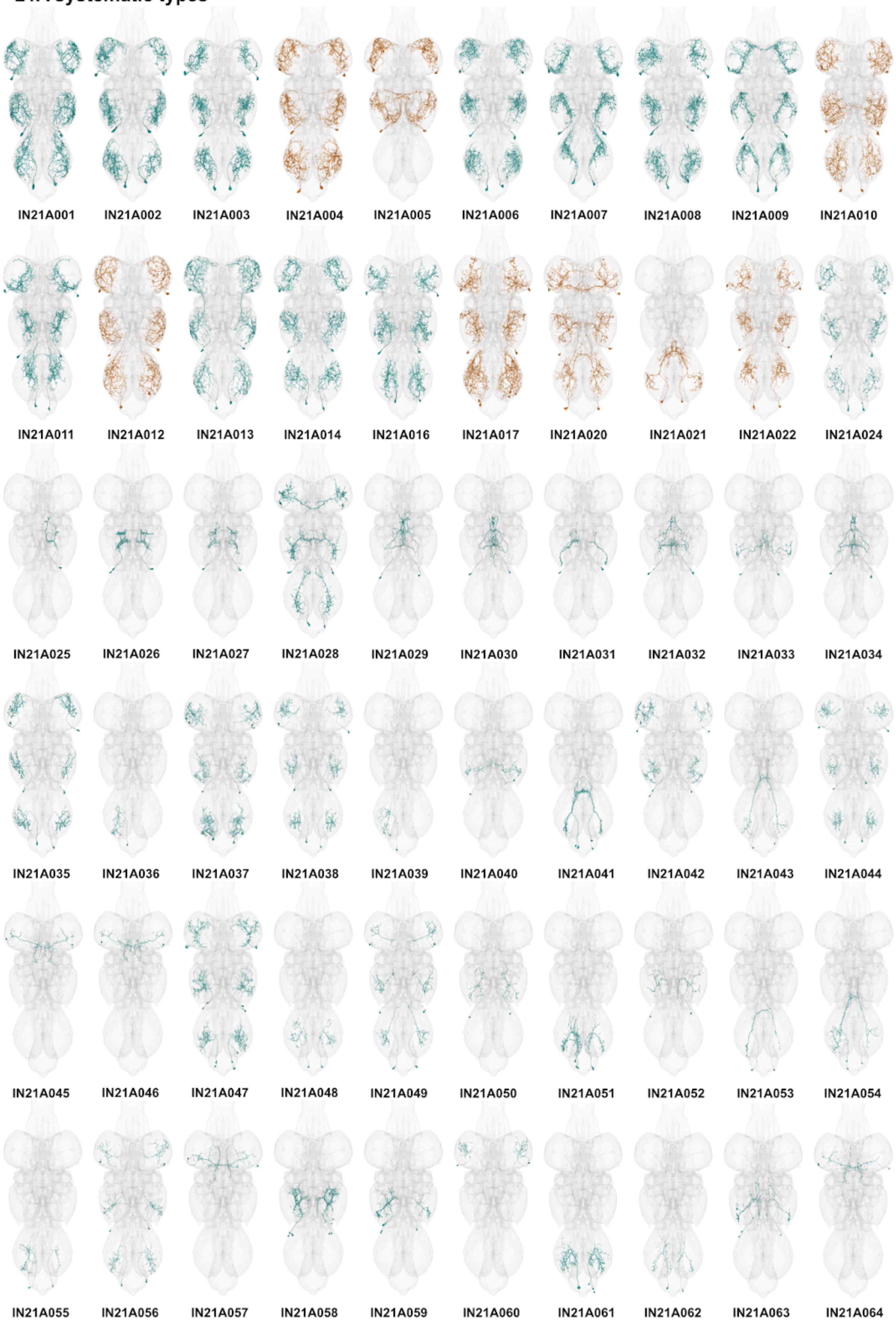

**Figure 45 - figure supplement 2. Systematic types of hemilineage 21A.** Systematic types have been arranged in numerical order, with neurons of the same type that belong to distinct classes (e.g., intrinsic neuron vs ascending neuron) plotted separately but placed adjacent to each other. Individual neuron meshes have been coloured based on predicted neurotransmitter: dark orange = acetylcholine, blue = gaba, marine = glutamate, dark purple = unknown.

#### 21A systematic types, continued

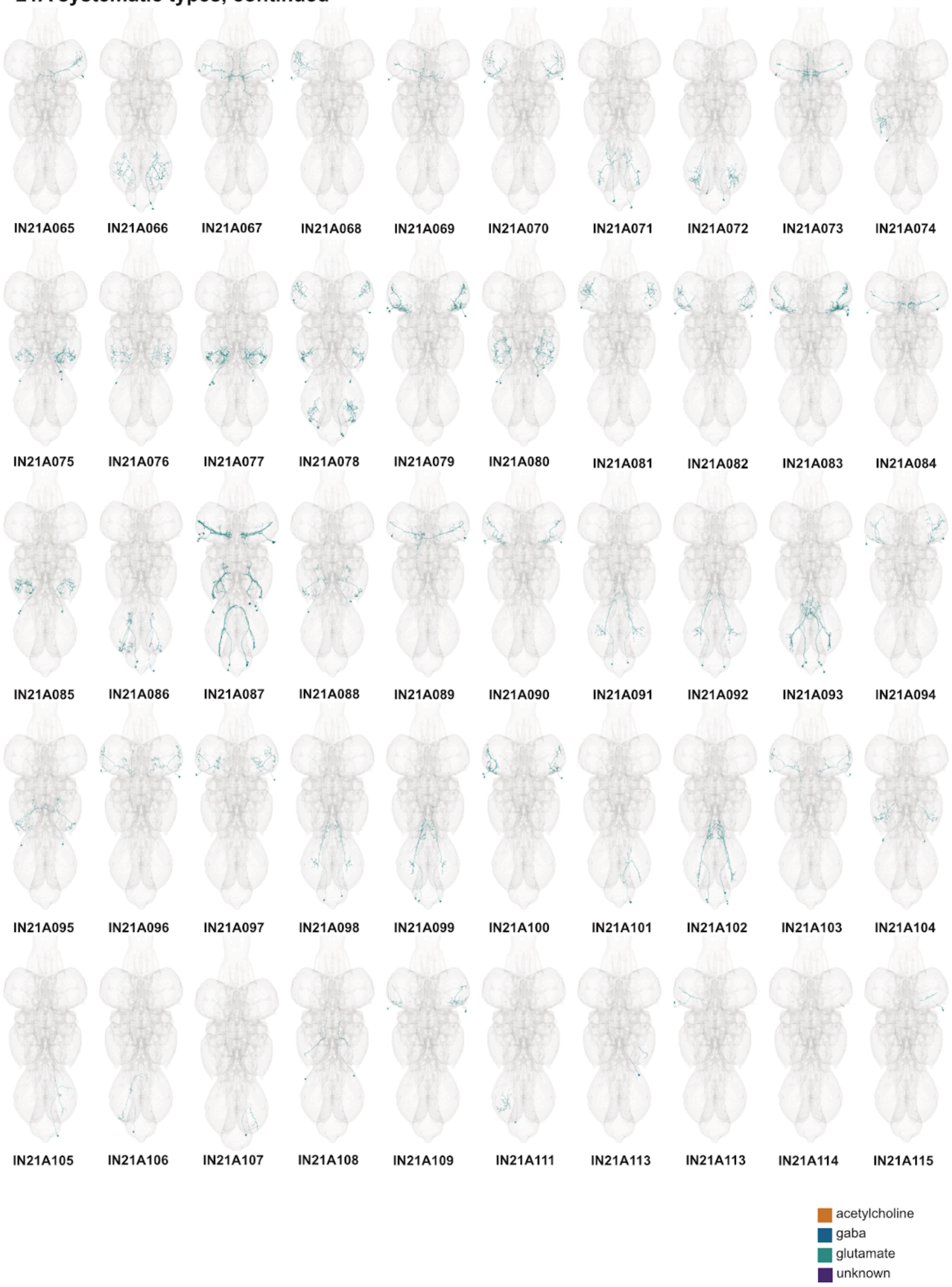

**Figure 45 - figure supplement 3. Systematic types of hemilineage 21A, continued.** Systematic types have been arranged in numerical order, with neurons of the same type that belong to distinct classes (e.g., intrinsic neuron vs ascending neuron) plotted separately but placed adjacent to each other. Individual neuron meshes have been coloured based on predicted neurotransmitter: dark orange = acetylcholine, blue = gaba, marine = glutamate, dark purple = unknown.

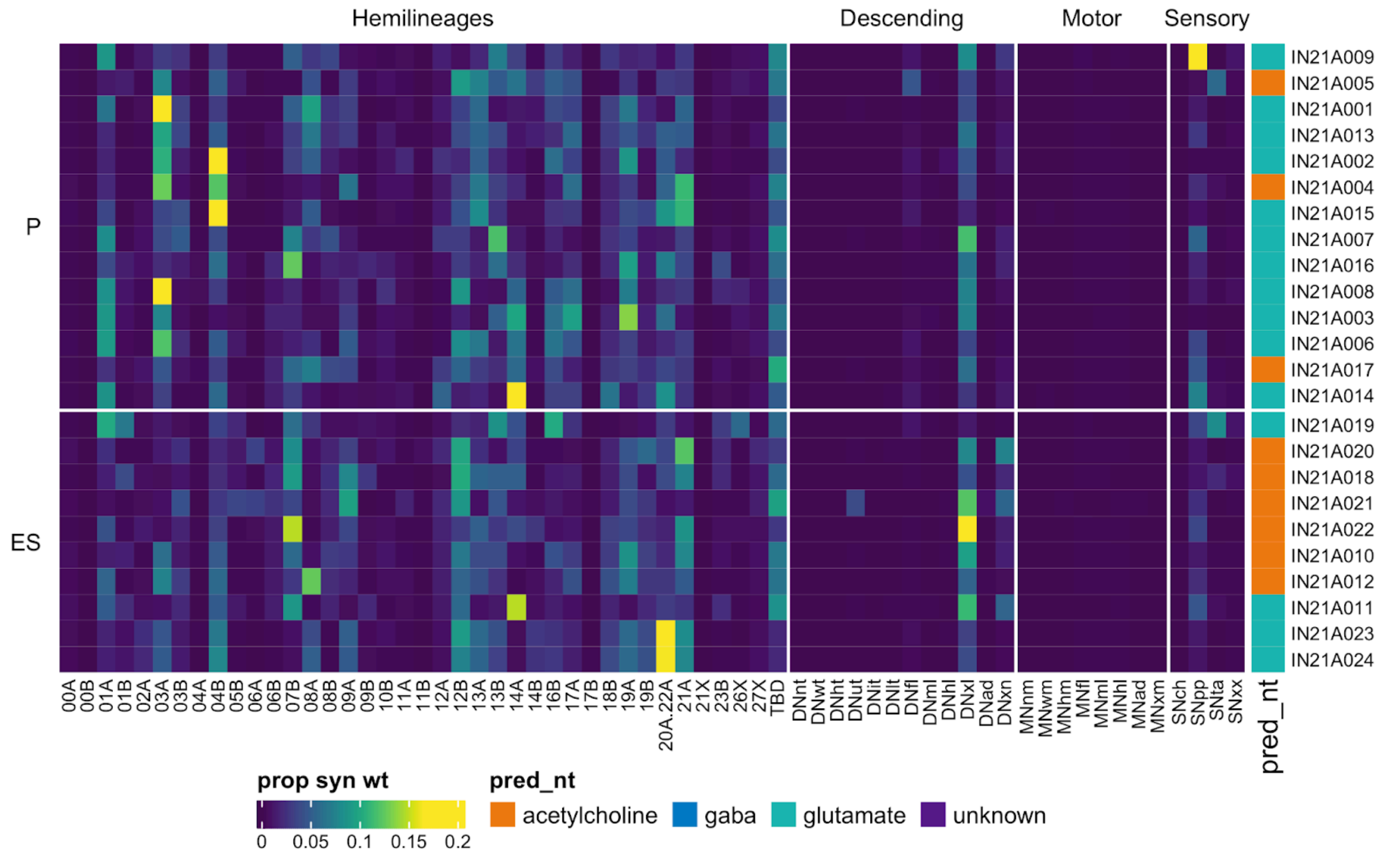

**Figure 45 - figure supplement 4. Connectivity to upstream partners by 21A primary and early secondary systematic types.** Proportions of synaptic weight to systematic types from upstream partners, normalised by row. 21A neurons have been clustered within each assigned birthtime window (P = primary, ES = early secondary, S = secondary) based on both upstream and downstream connectivity to hemilineages, descending neuron subclasses, motor neuron subclasses, and sensory neuron modalities. Annotation bar is coloured by the most common predicted neurotransmitter for the neurons of each type.

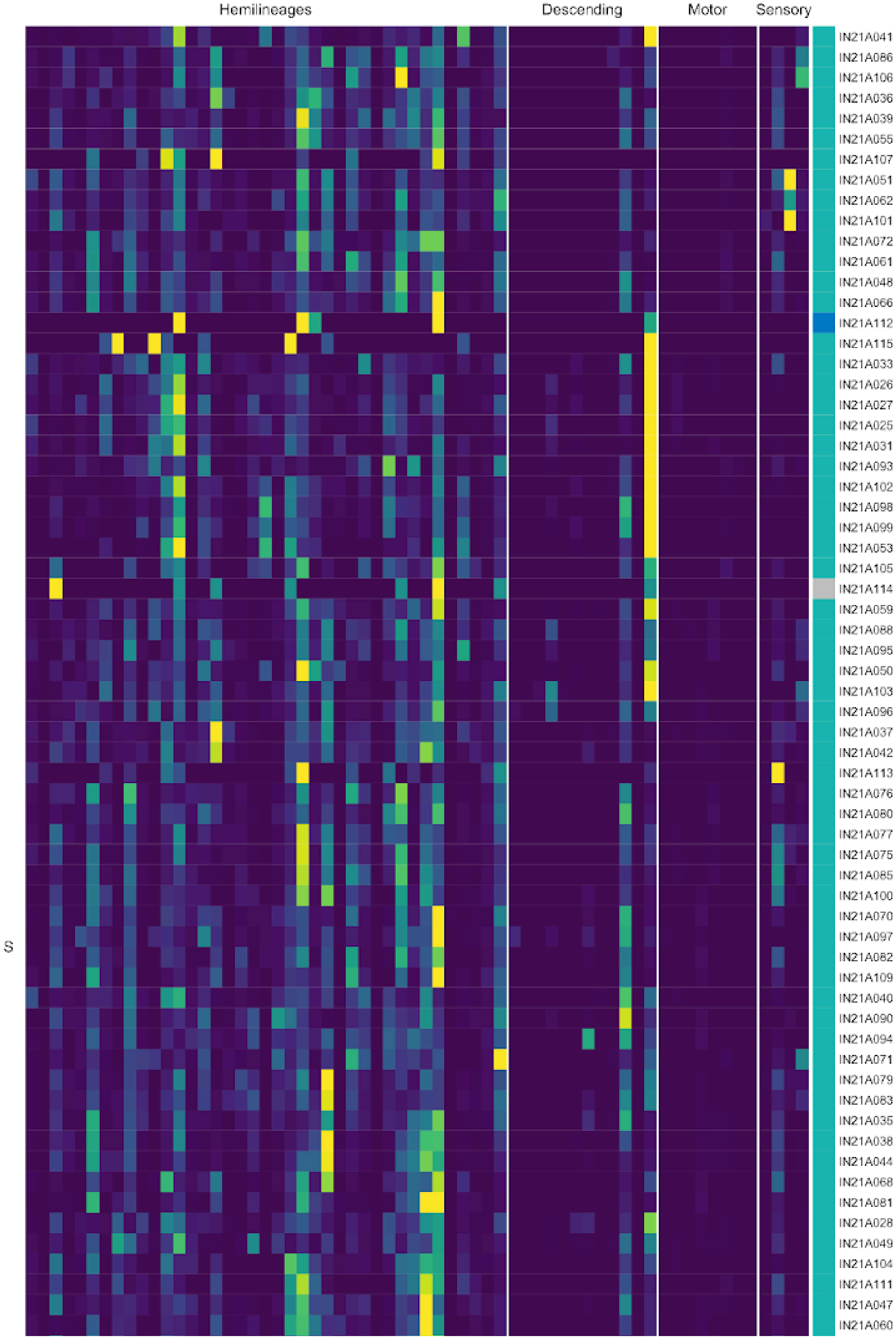
Hemilineages
Descending
Motor
Sensory
S
IN21A041
IN21A086
IN21A106
IN21A036
IN21A039
IN21A055
IN21A107
IN21A051
IN21A062
IN21A101
IN21A072
IN21A061
IN21A048
IN21A066
IN21A112
IN21A115
IN21A033
IN21A026
IN21A027
IN21A025
IN21A031
IN21A093
IN21A102
IN21A098
IN21A099
IN21A053
IN21A105
IN21A114
IN21A059
IN21A088
IN21A095
IN21A050
IN21A103
IN21A096
IN21A037
IN21A042
IN21A113
IN21A076
IN21A080
IN21A077
IN21A075
IN21A085
IN21A100
IN21A070
IN21A097
IN21A082
IN21A109
IN21A040
IN21A090
IN21A094
IN21A071
IN21A079
IN21A083
IN21A035
IN21A038
IN21A044
IN21A068
IN21A081
IN21A028
IN21A049
IN21A104
IN21A111
IN21A047
IN21A060

**Figure 45 - figure supplement 5. Connectivity to upstream partners by 21A secondary systematic types.** Proportions of synaptic weight to systematic types from upstream partners, normalised by row. 21A neurons have been clustered within each assigned birthtime window (P = primary, ES = early secondary, S = secondary) based on both upstream and downstream connectivity to hemilineages, descending neuron subclasses, motor neuron subclasses, and sensory neuron modalities. The annotation bar is coloured by the most common predicted neurotransmitter within each type.

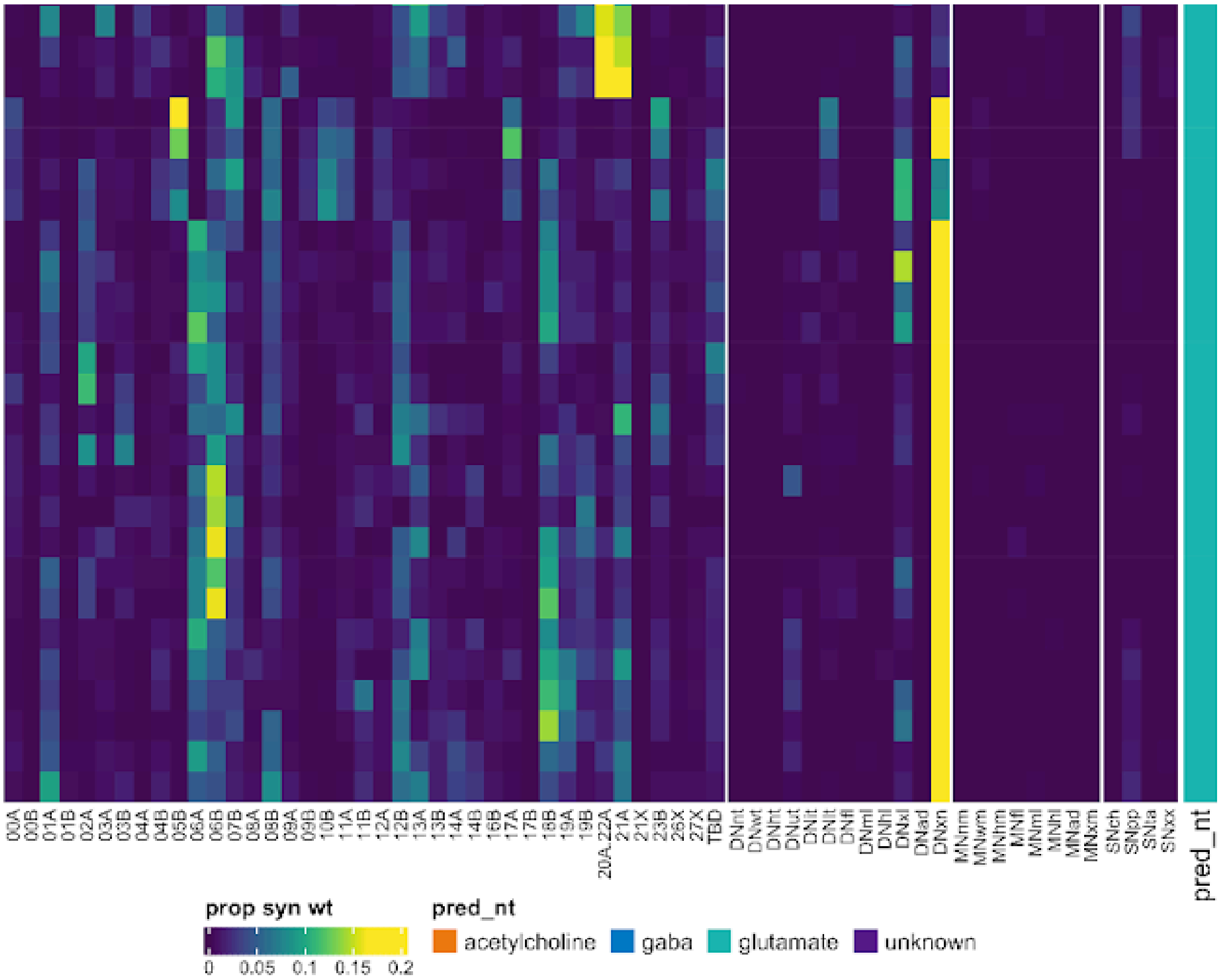

**Figure 45 - figure supplement 6. Connectivity to upstream partners by 21A secondary systematic types, continued.** Proportions of synaptic weight to systematic types from upstream partners, normalised by row. 21A neurons have been clustered within each assigned birthtime window (P = primary, ES = early secondary, S = secondary) based on both upstream and downstream connectivity to hemilineages, descending neuron subclasses, motor neuron subclasses, and sensory neuron modalities. The annotation bar is coloured by the most common predicted neurotransmitter within each type.

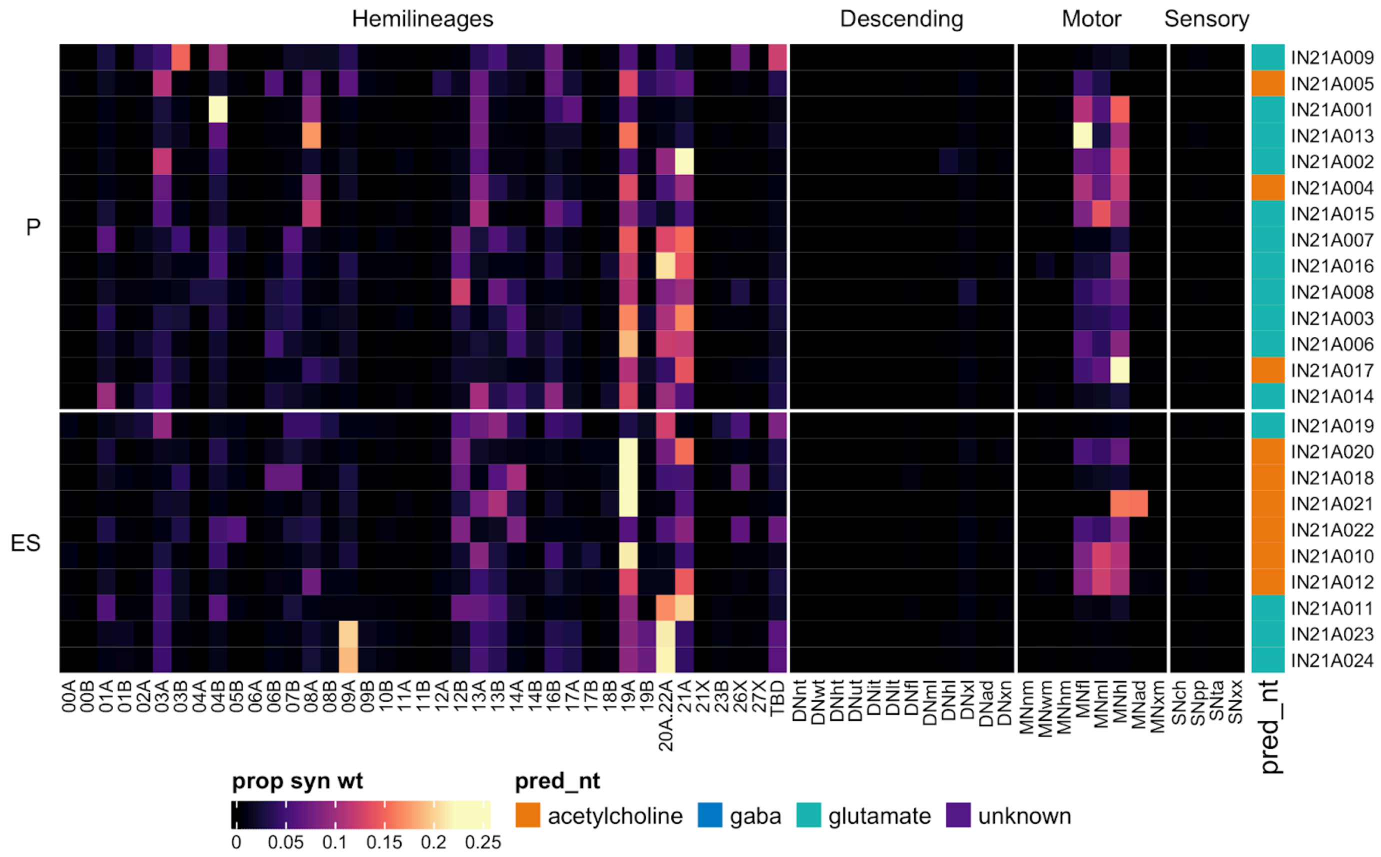

**Figure 45 - figure supplement 7. Connectivity to downstream partners by 21A primary and early secondary systematic types.** Proportions of synaptic weight from systematic types to downstream partners, normalised by row. 21A neurons have been clustered within each assigned birthtime window (P = primary, ES = early secondary, S = secondary) based on both upstream and downstream connectivity to hemilineages, descending neuron subclasses, motor neuron subclasses, and sensory neuron modalities. The annotation bar is coloured by the most common predicted neurotransmitter within each type.

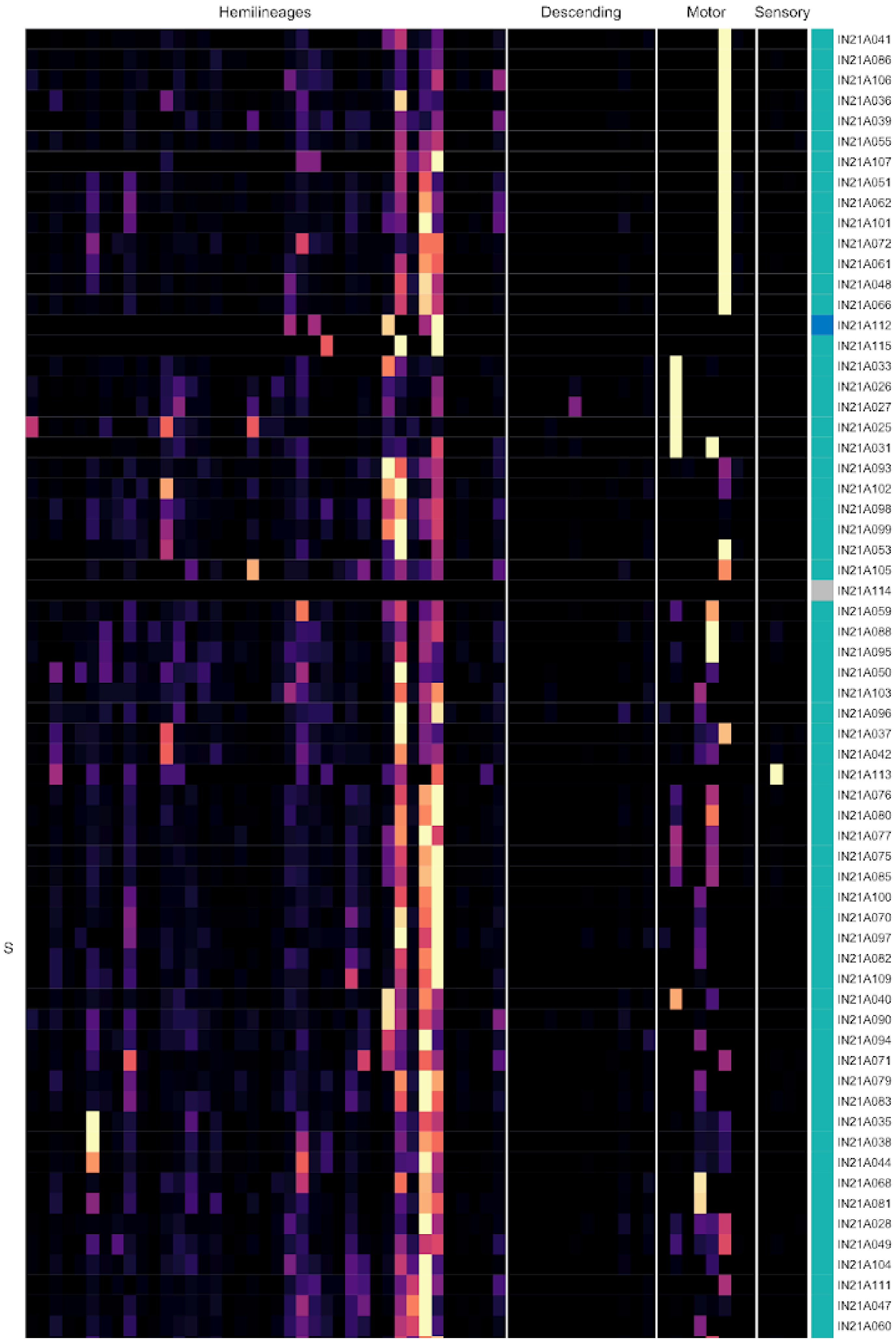
Hemilineages
Descending
Motor
Sensory
S
IN21A041
IN21A086
IN21A106
IN21A036
IN21A039
IN21A055
IN21A107
IN21A051
IN21A062
IN21A101
IN21A072
IN21A061
IN21A048
IN21A066
IN21A112
IN21A115
IN21A033
IN21A026
IN21A027
IN21A025
IN21A031
IN21A093
IN21A102
IN21A098
IN21A099
IN21A053
IN21A105
IN21A114
IN21A059
IN21A088
IN21A095
IN21A050
IN21A103
IN21A096
IN21A037
IN21A042
IN21A113
IN21A076
IN21A080
IN21A077
IN21A075
IN21A085
IN21A100
IN21A070
IN21A097
IN21A082
IN21A109
IN21A040
IN21A090
IN21A094
IN21A071
IN21A079
IN21A083
IN21A035
IN21A038
IN21A044
IN21A068
IN21A081
IN21A028
IN21A049
IN21A104
IN21A111
IN21A047
IN21A060

**Figure 45 - figure supplement 8. Connectivity to downstream partners by 21A secondary systematic types.** Proportions of synaptic weight from systematic types to downstream partners, normalised by row. 21A neurons have been clustered within each assigned birthtime window (P = primary, ES = early secondary, S = secondary) based on both upstream and downstream connectivity to hemilineages, descending neuron subclasses, motor neuron subclasses, and sensory neuron modalities. The annotation bar is coloured by the most common predicted neurotransmitter for the neurons of each type.

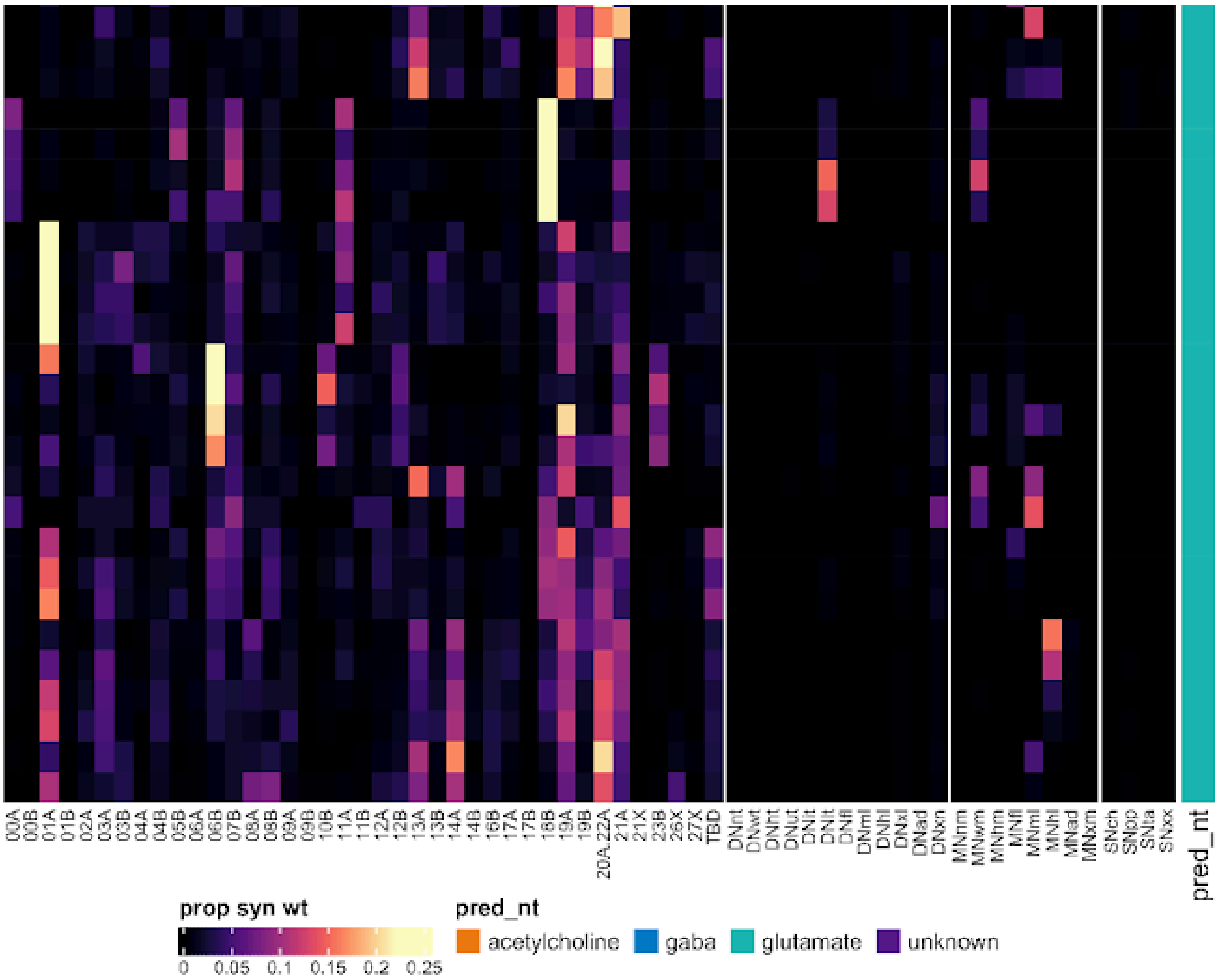

**Figure 45 - figure supplement 9. Connectivity to downstream partners by 21A secondary systematic types, continued.** Proportions of synaptic weight from systematic types to downstream partners, normalised by row. 21A neurons have been clustered within each assigned birthtime window (P = primary, ES = early secondary, S = secondary) based on both upstream and downstream connectivity to hemilineages, descending neuron subclasses, motor neuron subclasses, and sensory neuron modalities. The annotation bar is coloured by the most common predicted neurotransmitter for the neurons of each type.

#### Hemilineage 23B

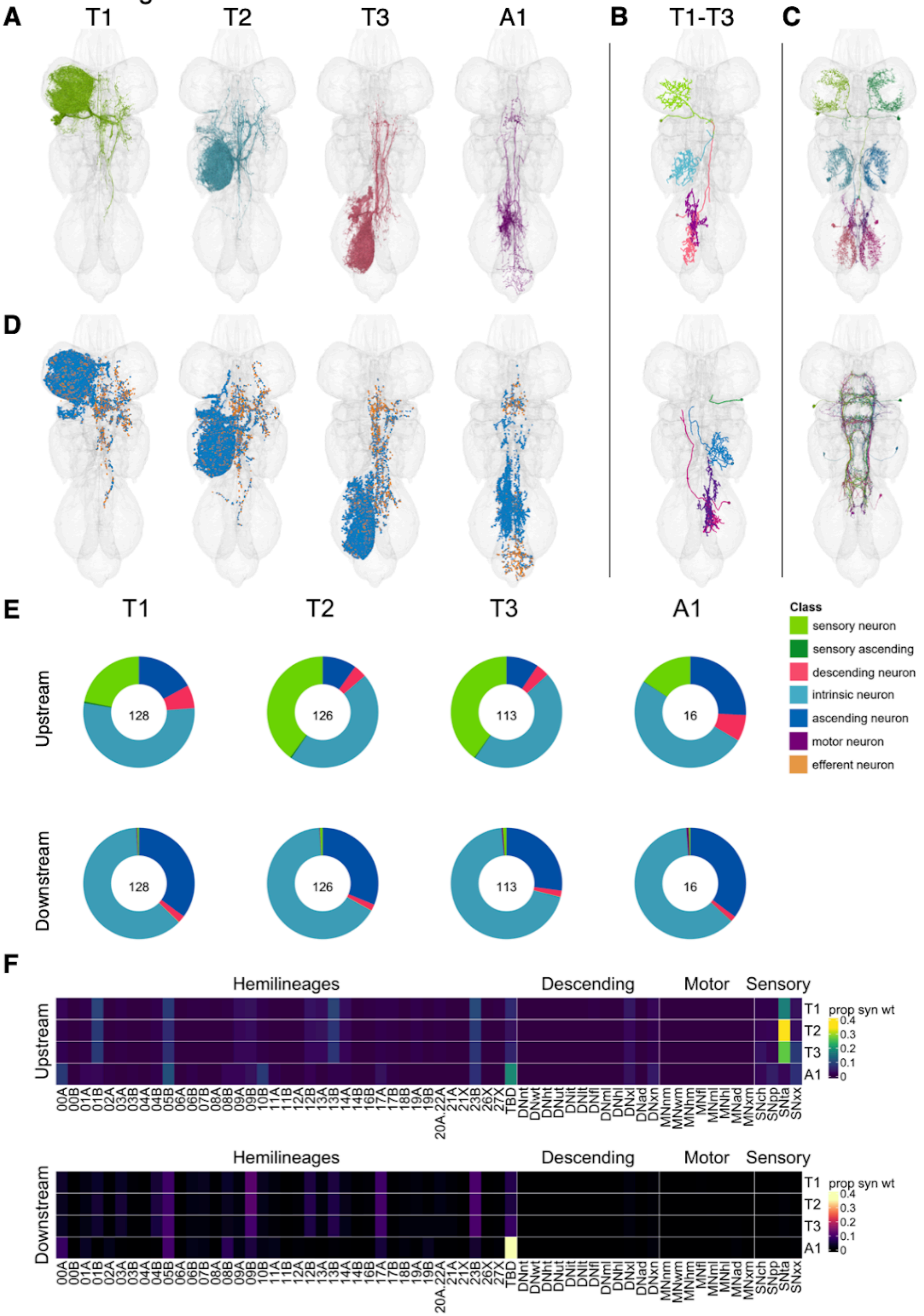

**Figure 46. Hemilineage 23B. A.** Meshes of all RHS secondary neurons plotted in neuromere-specific colours. **B.** “Representative” secondary neuron skeletons plotted in hemineuromere-specific colours. The skeleton with the top accumulated NBLAST score among all neurons from the hemilineage in a given hemineuromere was used. **C.** Neuron meshes of selected examples. Top: independent leg serial 12339 (T1-A1). Bottom: intersegmental complex serial 11371 (T1-A1). **D.** Predicted synapses of RHS secondary neurons. Blue: postsynapses; dark orange: presynapses. **E.** Proportions of connections from secondary neurons to upstream or downstream partners, normalised by neuromere and coloured by broad class. Numbers of query neurons appear in the centre. **F.** Proportions of synaptic weight from secondary neurons originating in each neuromere to upstream or downstream partners, normalised by row.

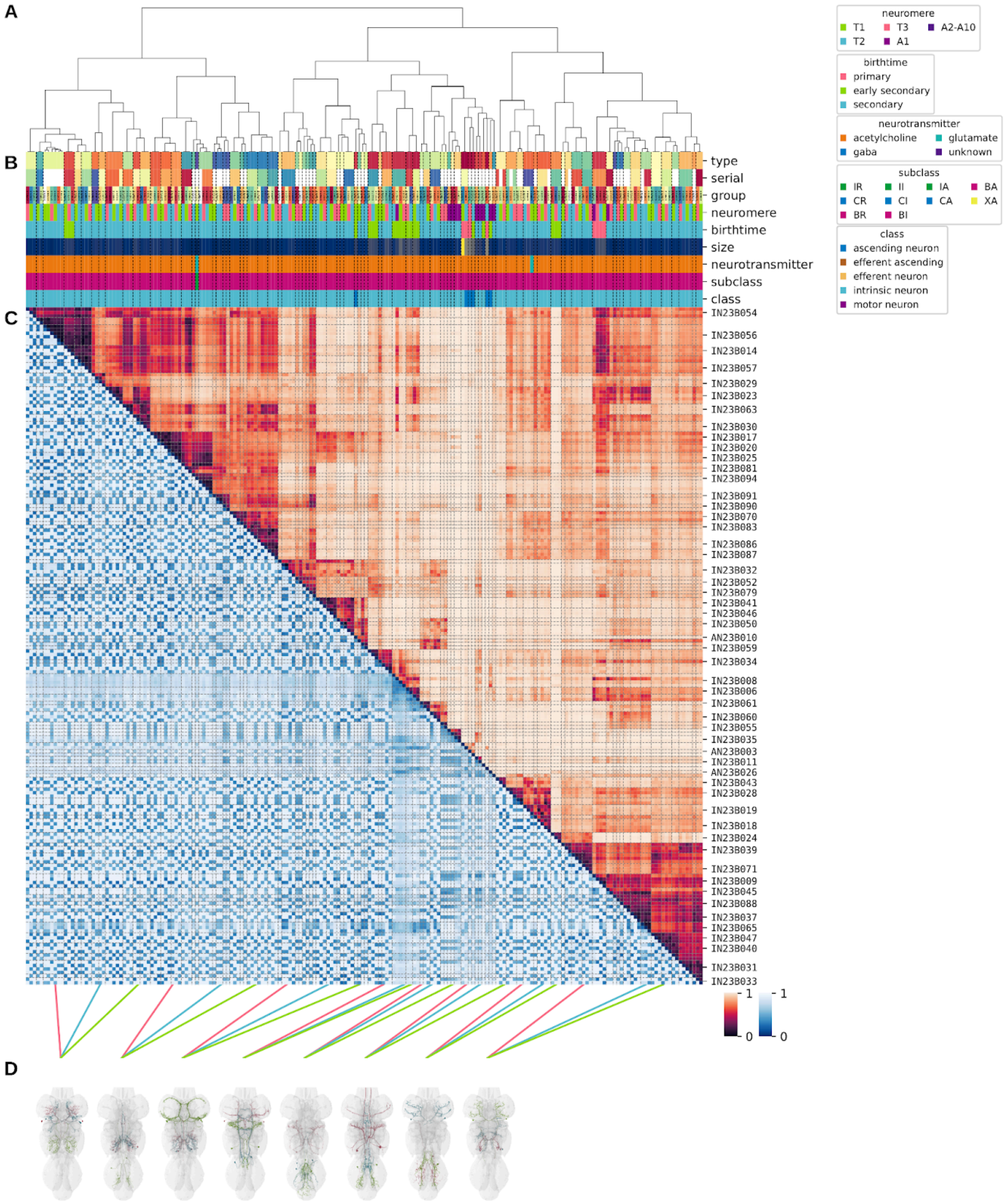

**Figure 46 - figure supplement 1. Systematic typing of hemilineage 23B. A.** Hierarchical clustering dendrogram of hemilineage groups by laterally and serially aggregated connectivity cosine clustering. **B.** Categorical annotations of each hemilineage group, each column corresponding to the aligned leaf in A. Colours for type, serial set, and group are arbitrary for visualisation. Colours for neuromere, birthtime, neurotransmitter, subclass, and class are as in all other figures. **C.** Similarity distance heatmap for hemilineage. Cosine distance is in the upper triangle, while laterally symmetrised NBLAST distance is in the lower triangle. Systematic type names of some types are labelled. **D.** Morphologically representative groups from dendrogram subtrees. Each group, indicated by colour and line connecting to its column in B and C, is the most morphologically representative group (medoid of NBLAST distance) from a subtree of A. The subtrees (flat clusters) are equal height cuts of A determined to yield the number of groups

per plot and plots in D.

#### 23B systematic types

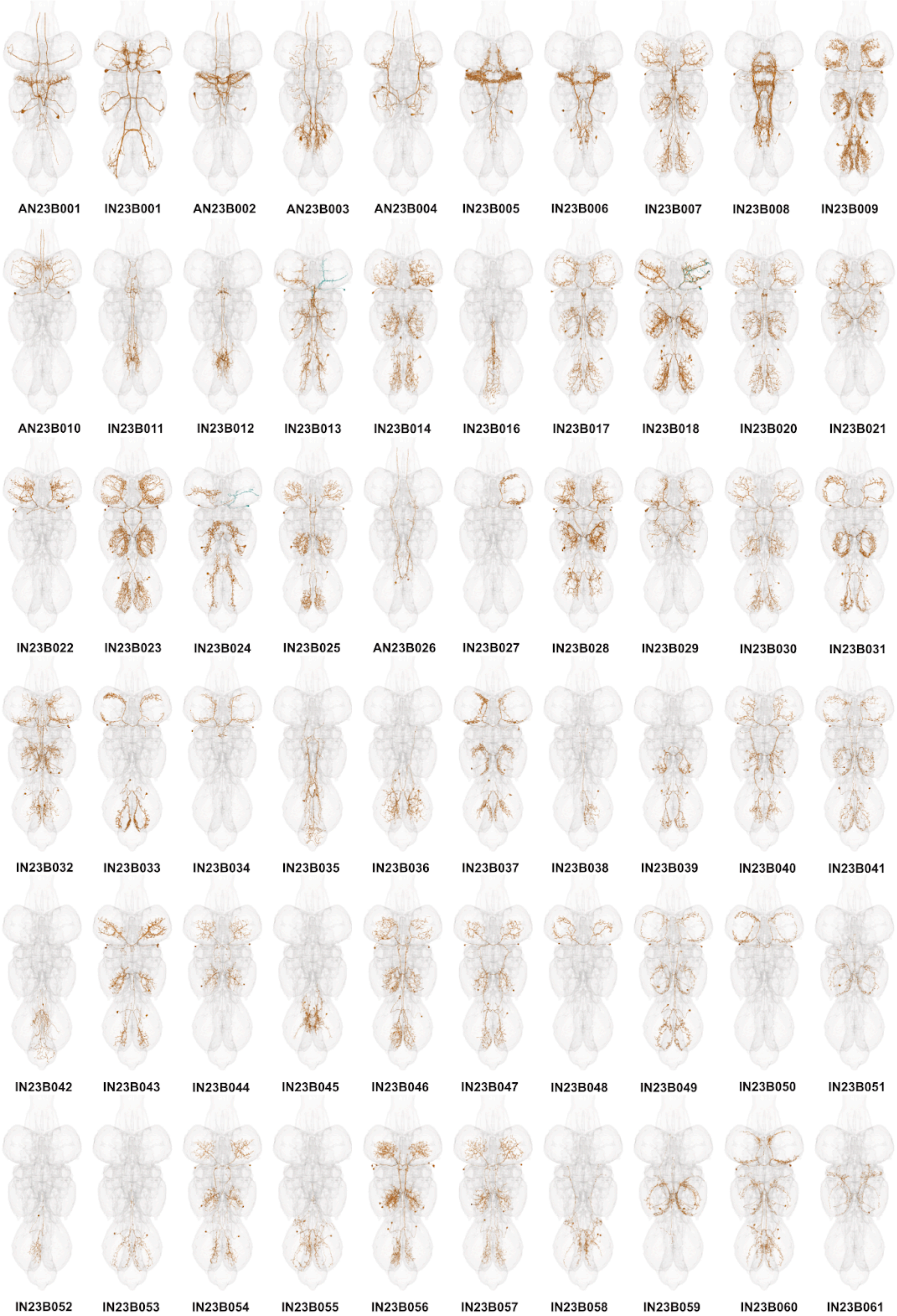

**Figure 46 - figure supplement 2. Systematic types of hemilineage 23B.** Systematic types have been arranged in numerical order, with neurons of the same type that belong to distinct classes (e.g., intrinsic neuron vs ascending neuron) plotted separately but placed adjacent to each other. Individual neuron meshes have been coloured based on predicted neurotransmitter: dark orange = acetylcholine, blue = gaba, marine = glutamate, dark purple = unknown.

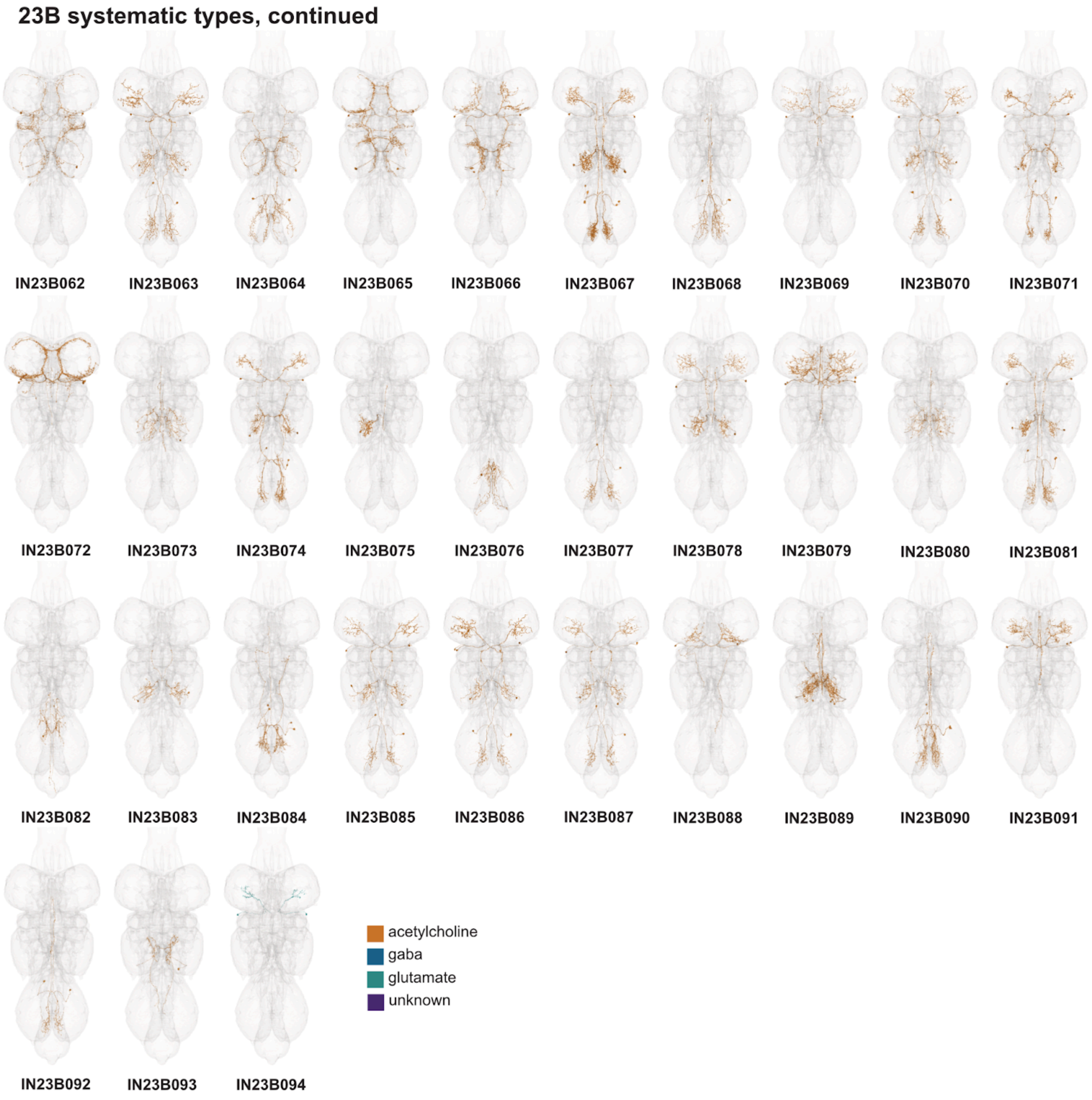

**Figure 46 - figure supplement 3. Systematic types of hemilineage 23B, continued.** Systematic types have been arranged in numerical order, with neurons of the same type that belong to distinct classes (e.g., intrinsic neuron vs ascending neuron) plotted separately but placed adjacent to each other. Individual neuron meshes have been coloured based on predicted neurotransmitter: dark orange = acetylcholine, blue = gaba, marine = glutamate, dark purple = unknown.

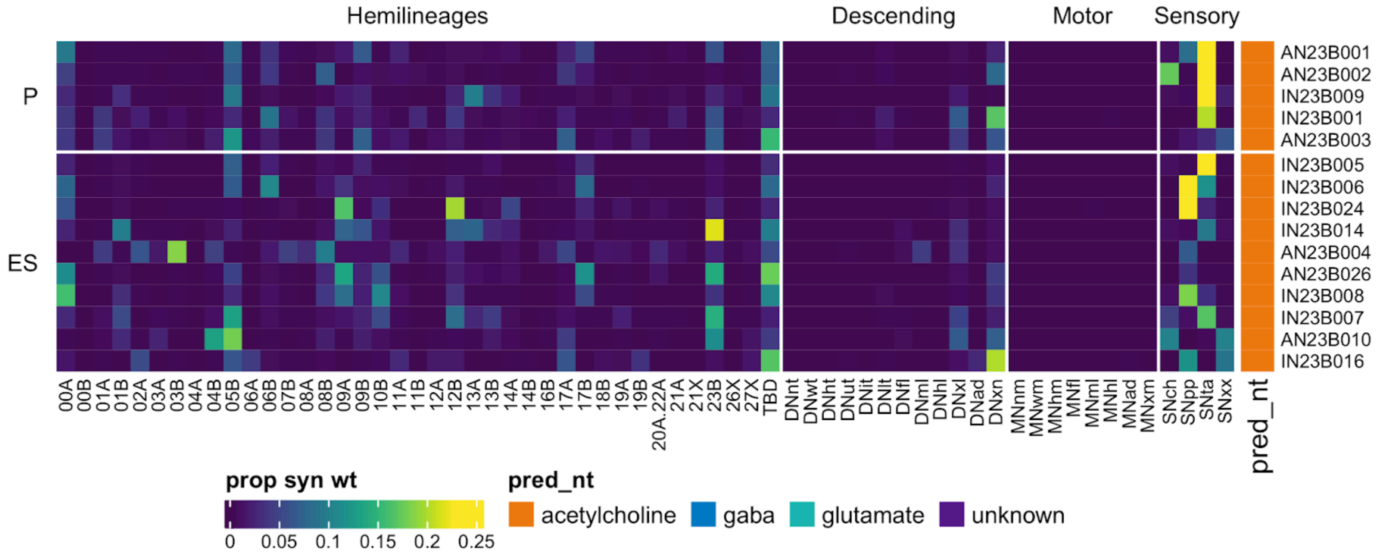

**Figure 46 - figure supplement 4. Connectivity to upstream partners by 23B primary and early secondary systematic types.** Proportions of synaptic weight to systematic types from upstream partners, normalised by row. 23B neurons have been clustered within each assigned birthtime window (P = primary, ES = early secondary, S = secondary) based on both upstream and downstream connectivity to hemilineages, descending neuron subclasses, motor neuron subclasses, and sensory neuron modalities. Annotation bar is coloured by the most common predicted neurotransmitter for the neurons of each type.

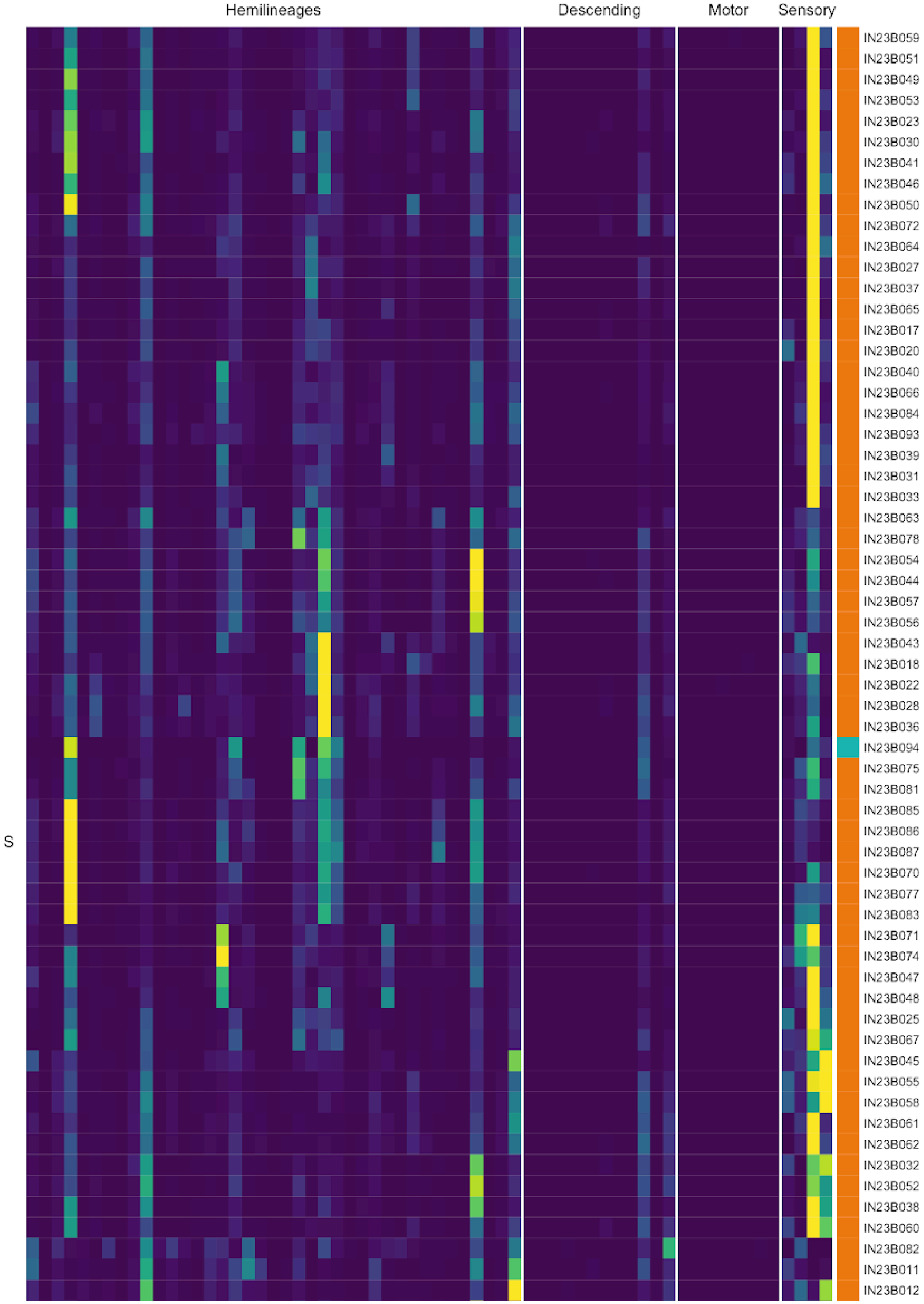
Hemilineages
Descending
Motor
Sensory
S
IN23B059
IN23B051
IN23B049
IN23B053
IN23B023
IN23B030
IN23B041
IN23B046
IN23B050
IN23B072
IN23B064
IN23B027
IN23B037
IN23B065
IN23B017
IN23B020
IN23B040
IN23B066
IN23B084
IN23B093
IN23B039
IN23B031
IN23B033
IN23B063
IN23B078
IN23B054
IN23B044
IN23B057
IN23B056
IN23B043
IN23B018
IN23B022
IN23B028
IN23B036
IN23B094
IN23B075
IN23B081
IN23B085
IN23B086
IN23B087
IN23B070
IN23B077
IN23B083
IN23B071
IN23B074
IN23B047
IN23B048
IN23B025
IN23B067
IN23B045
IN23B055
IN23B058
IN23B061
IN23B062
IN23B032
IN23B052
IN23B038
IN23B060
IN23B082
IN23B011
IN23B012

**Figure 46 - figure supplement 5. Connectivity to upstream partners by 23B secondary systematic types.** Proportions of synaptic weight to systematic types from upstream partners, normalised by row. 23B neurons have been clustered within each assigned birthtime window (P = primary, ES = early secondary, S = secondary) based on both upstream and downstream connectivity to hemilineages, descending neuron subclasses, motor neuron subclasses, and sensory neuron modalities. The annotation bar is coloured by the most common predicted neurotransmitter within each type.

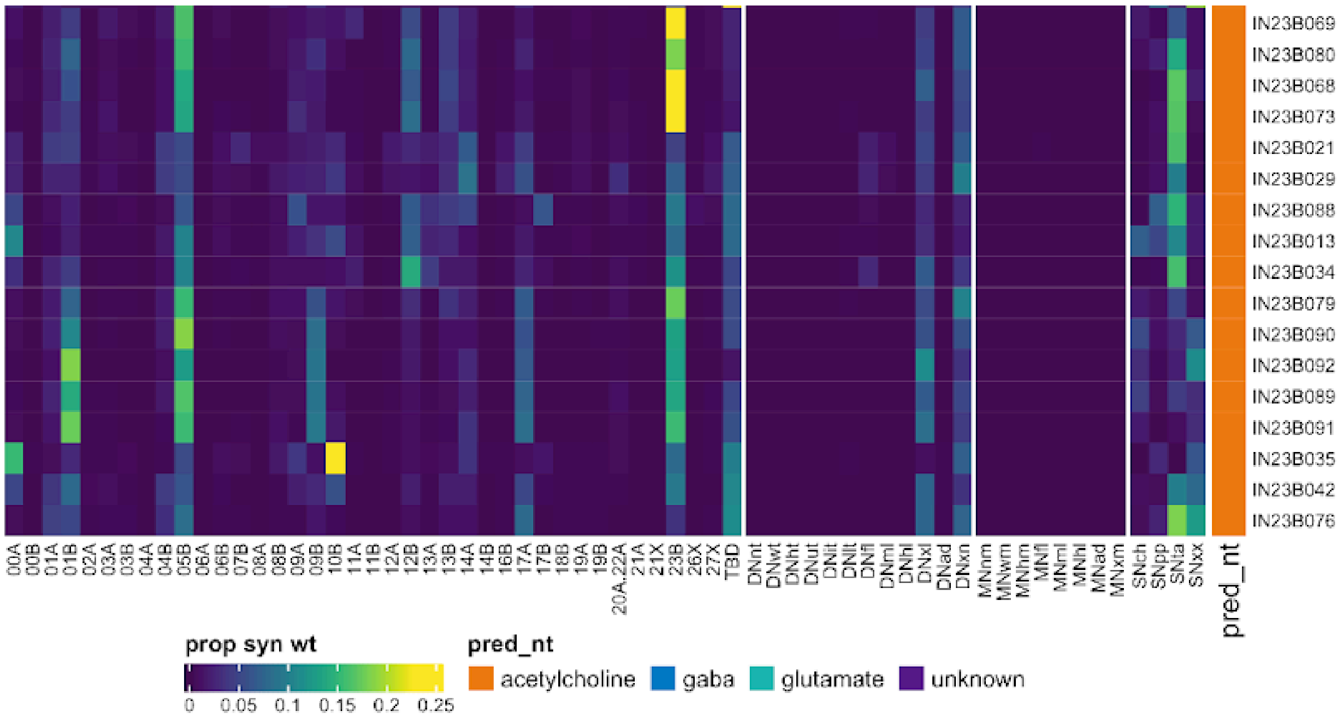

**Figure 46 - figure supplement 6.Connectivity to upstream partners by 23B secondary systematic types, continued.** Proportions of synaptic weight to systematic types from upstream partners, normalised by row. 23B neurons have been clustered within each assigned birthtime window (P = primary, ES = early secondary, S = secondary) based on both upstream and downstream connectivity to hemilineages, descending neuron subclasses, motor neuron subclasses, and sensory neuron modalities. The annotation bar is coloured by the most common predicted neurotransmitter within each type.

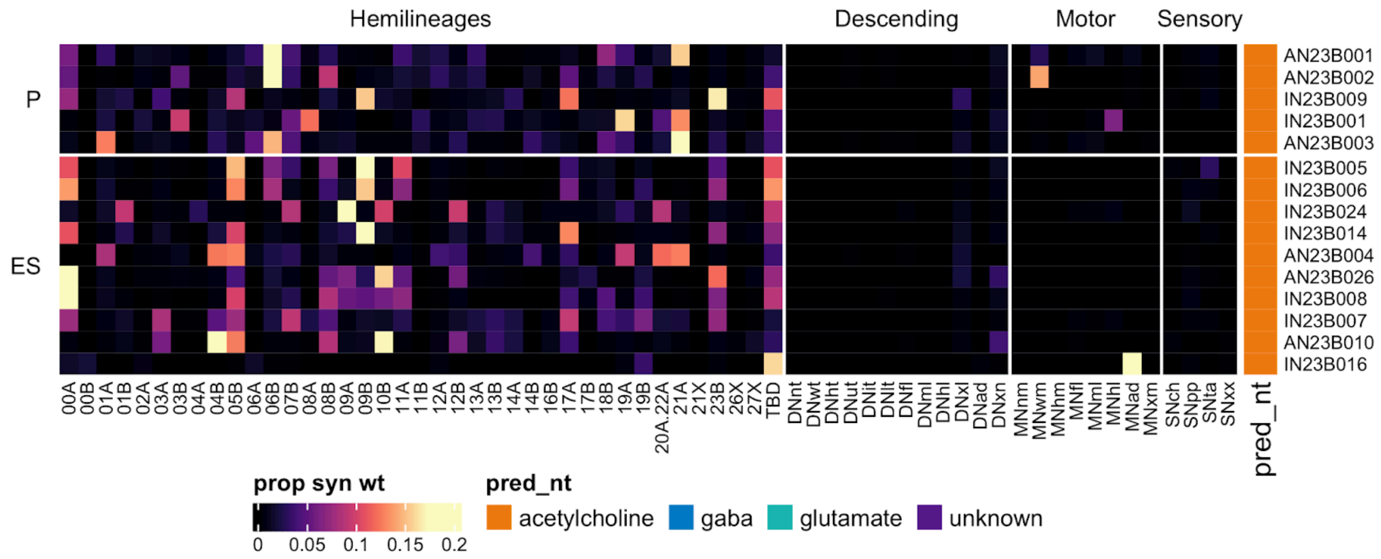

**Figure 46 - figure supplement 7. Connectivity to downstream partners by 23B primary and early secondary systematic types.** Proportions of synaptic weight from systematic types to downstream partners, normalised by row. 23B neurons have been clustered within each assigned birthtime window (P = primary, ES = early secondary, S = secondary) based on both upstream and downstream connectivity to hemilineages, descending neuron subclasses, motor neuron subclasses, and sensory neuron modalities. The annotation bar is coloured by the most common predicted neurotransmitter within each type.

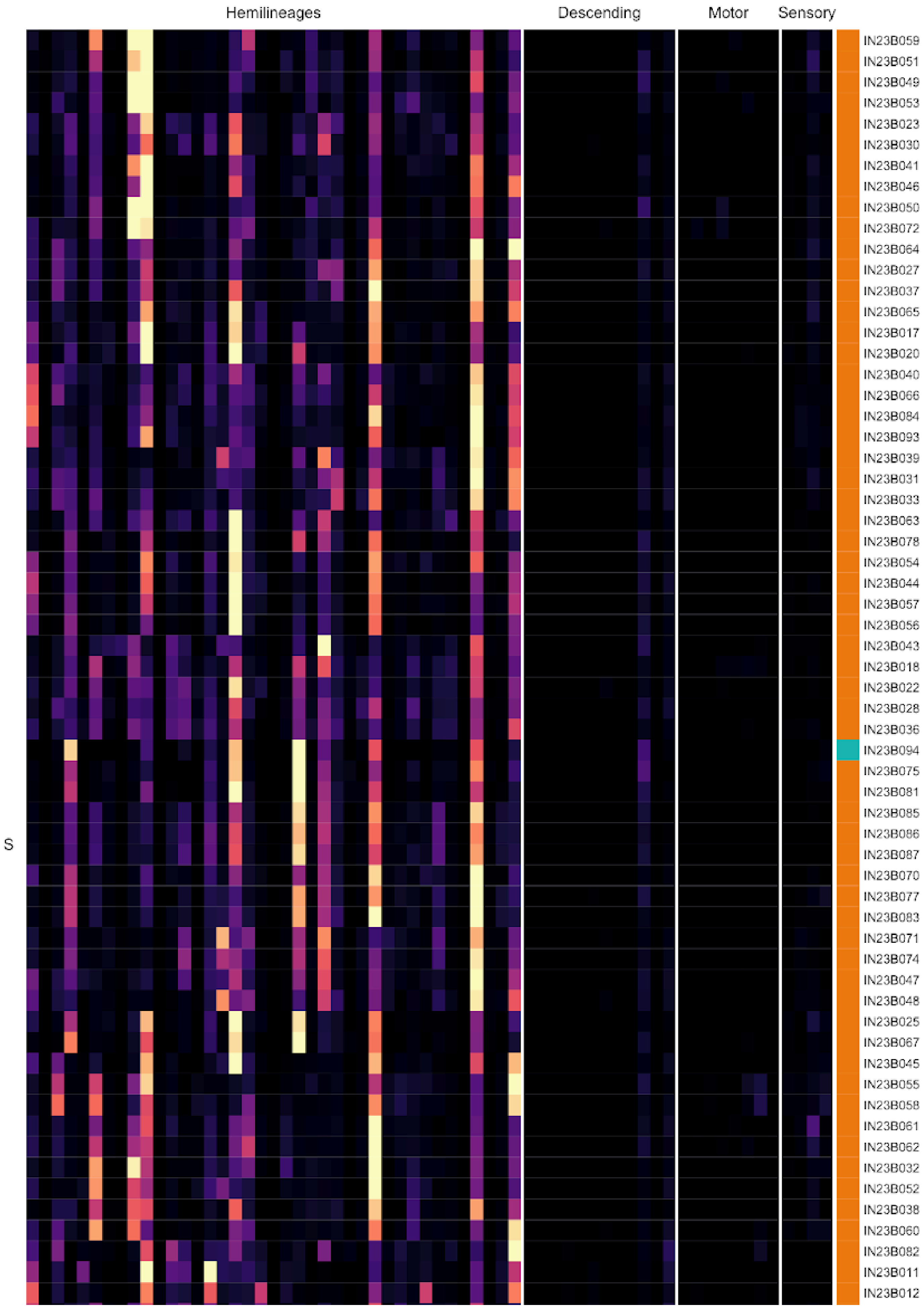
Hemilineages
Descending
Motor
Sensory
S
IN23B059
IN23B051
IN23B049
IN23B053
IN23B023
IN23B030
IN23B041
IN23B046
IN23B050
IN23B072
IN23B064
IN23B027
IN23B037
IN23B065
IN23B017
IN23B020
IN23B040
IN23B066
IN23B084
IN23B093
IN23B039
IN23B031
IN23B033
IN23B063
IN23B078
IN23B054
IN23B044
IN23B057
IN23B056
IN23B043
IN23B018
IN23B022
IN23B028
IN23B036
IN23B094
IN23B075
IN23B081
IN23B085
IN23B086
IN23B087
IN23B070
IN23B077
IN23B083
IN23B071
IN23B074
IN23B047
IN23B048
IN23B025
IN23B067
IN23B045
IN23B055
IN23B058
IN23B061
IN23B062
IN23B032
IN23B052
IN23B038
IN23B060
IN23B082
IN23B011
IN23B012

**Figure 46 - figure supplement 8. Connectivity to downstream partners by 23B secondary systematic types.** Proportions of synaptic weight from systematic types to downstream partners, normalised by row. 23B neurons have been clustered within each assigned birthtime window (P = primary, ES = early secondary, S = secondary) based on both upstream and downstream connectivity to hemilineages, descending neuron subclasses, motor neuron subclasses, and sensory neuron modalities. The annotation bar is coloured by the most common predicted neurotransmitter for the neurons of each type.

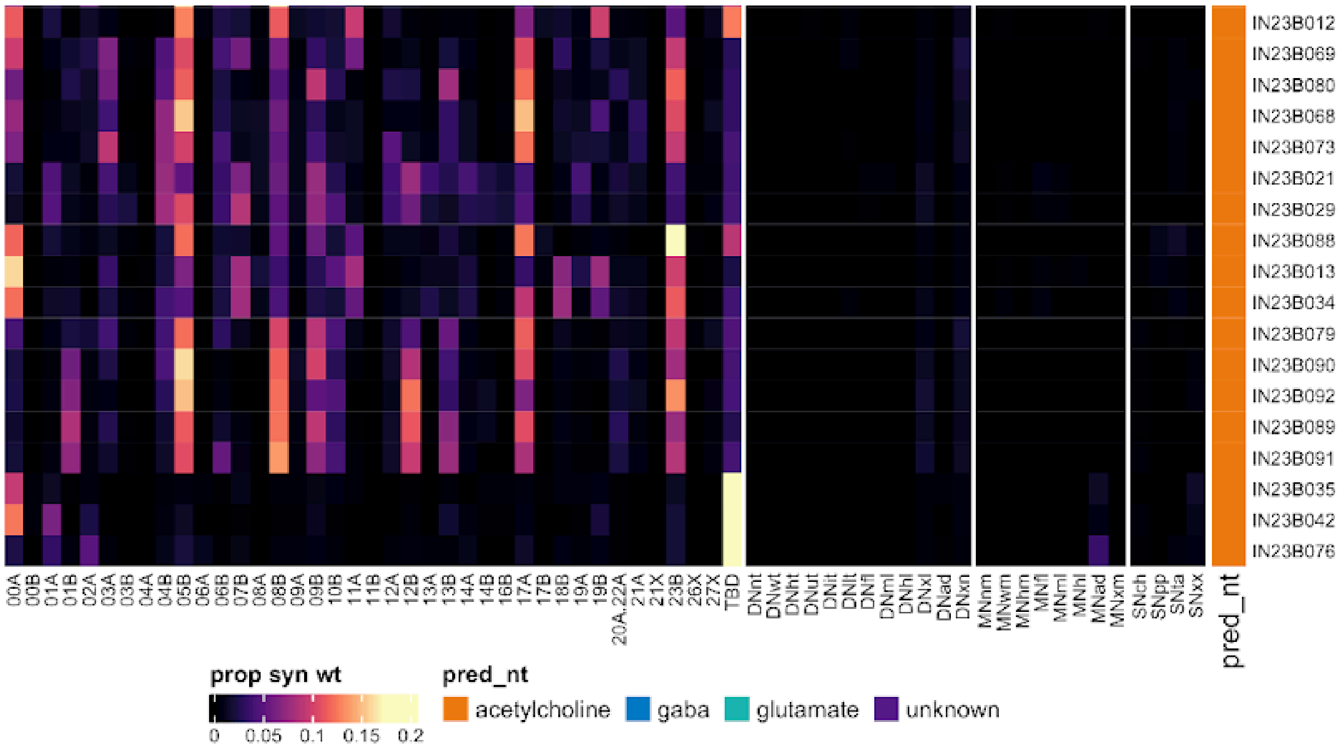

**Figure 46 - figure supplement 9. Connectivity to downstream partners by 23B secondary systematic types, continued.** Proportions of synaptic weight from systematic types to downstream partners, normalised by row. 23B neurons have been clustered within each assigned birthtime window (P = primary, ES = early secondary, S = secondary) based on both upstream and downstream connectivity to hemilineages, descending neuron subclasses, motor neuron subclasses, and sensory neuron modalities. The annotation bar is coloured by the most common predicted neurotransmitter for the neurons of each type.

Hemilineage 24B.25B

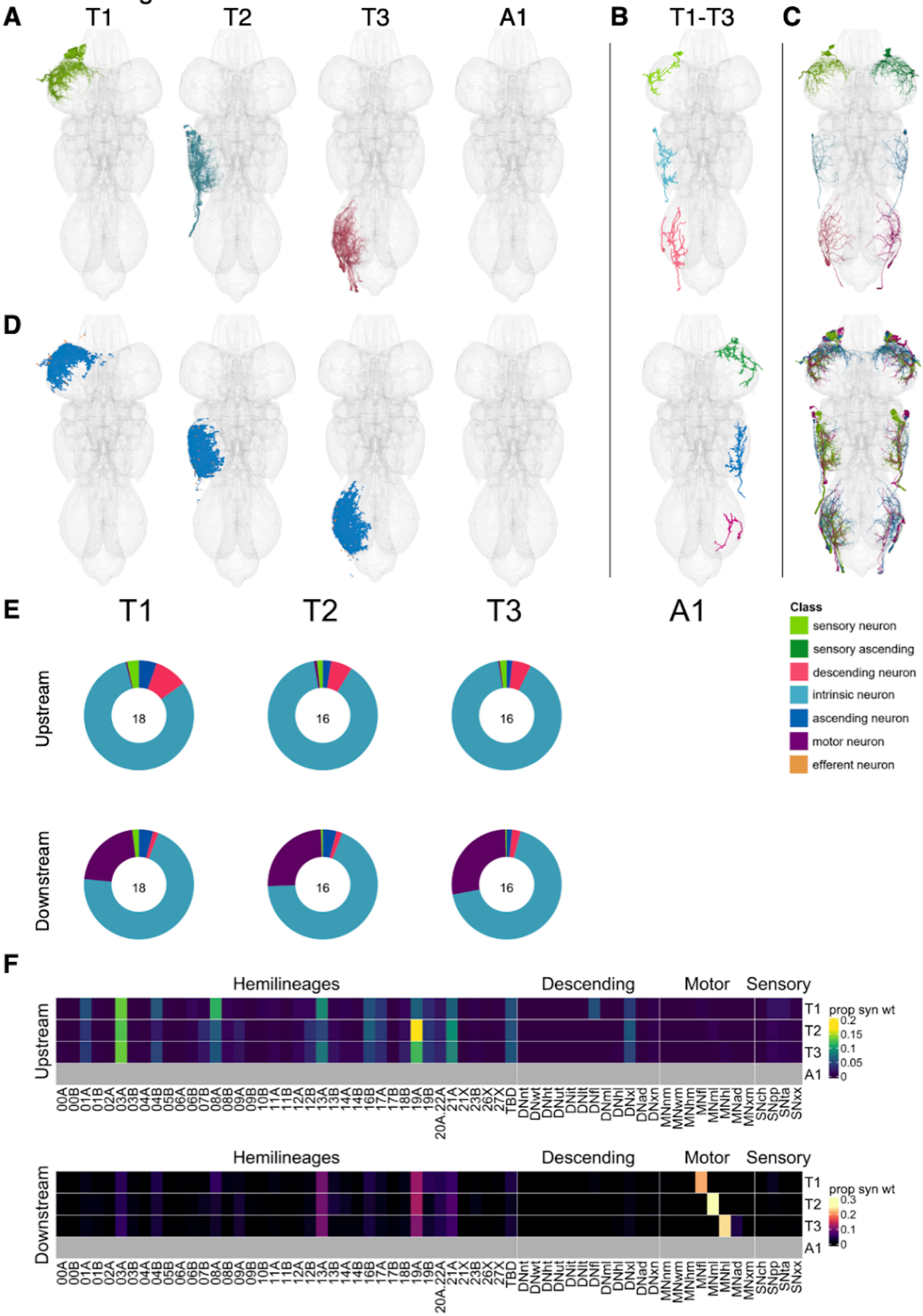

**Figure 47. Hemilineages 24B and 25B. A.** Meshes of all RHS secondary neurons plotted in neuromere-specific colours. **B.** "Representative" secondary neuron skeletons plotted in hemineuromere-specific colours. The skeleton with the top accumulated NBLAST score among all neurons from the hemilineage in a given hemineuromere was used. **C.** Neuron meshes of selected examples. Top: Fe reductor MN independent leg serial 11226 (T1-T3). Bottom: Serial populations exiting T1 via ProLN (blue), VProN (magenta), and ProAN (green) (T1-T3). **D.** Predicted synapses of RHS secondary neurons. Blue: postsynapses; dark orange: presynapses. **E.** Proportions of connections from secondary neurons to upstream or downstream partners, normalised by neuromere and coloured by broad class. Numbers of query neurons appear in the centre. **F.** Proportions of synaptic weight from secondary neurons originating in each neuromere to upstream or downstream partners, normalised by row.

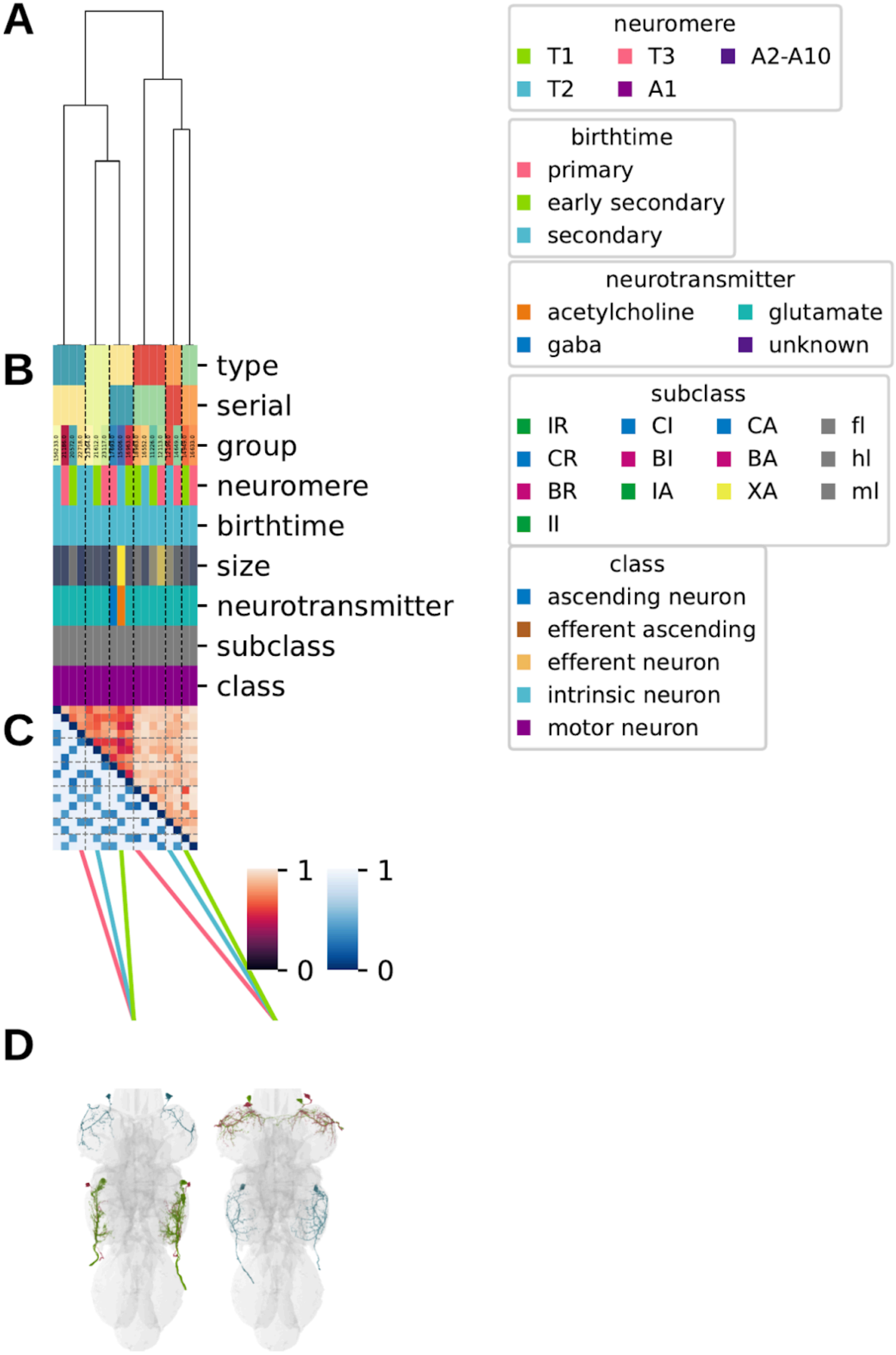
A
B
C
D
type
serial
group
neuromere
birthtime
size
neurotransmitter
subclass
class
1
0
1
0
neuromere
T1
T3
A2-A10
T2
A1
birthtime
primary
early secondary
secondary
neurotransmitter
acetylcholine
glutamate
gaba
unknown
subclass
IR
CI
CA
fl
CR
BI
BA
hl
BR
IA
XA
ml
II
class
ascending neuron
efferent ascending
efferent neuron
intrinsic neuron
motor neuron

**Figure 47 - figure supplement 1. Systematic typing of hemilineages 24B and 25B.** Types for motor neurons were assigned separately as outlined in our accompanying manuscript (Cheong et al., 2023). **A.** Hierarchical clustering dendrogram of hemilineage groups by laterally and serially aggregated connectivity cosine clustering. **B.** Categorical annotations of each hemilineage group, each column corresponding to the aligned leaf in A. Colours for cluster, serial set, and group are arbitrary for visualisation. Colours for neuromere, birthtime, neurotransmitter, subclass, and class are as in all other figures. **C.** Similarity distance heatmap for hemilineage. Cosine distance is in the upper triangle, while laterally symmetrised NBLAST distance is in the lower triangle. Systematic type names of some types are labelled. **D.** Morphologically representative groups from dendrogram subtrees. Each group, indicated by colour and line connecting to its column in B and C, is the most morphologically representative group (medoid of NBLAST distance) from a subtree of A. The subtrees (flat clusters) are equal height cuts of A determined to yield the number of groups per plot and plots in D.

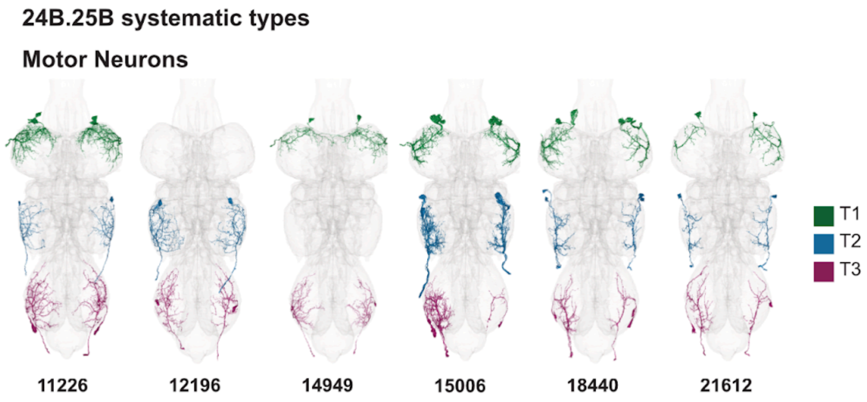

**Figure 47 - figure supplement 2. Systematic types of hemilineages 24B and 25B.** Motor neurons (typed separately in (Cheong et al., 2023)) have been plotted by serial set if identified in multiple neuromeres and by systematic type if not. Individual motor neuron meshes have been coloured based on soma neuromere: dark green = T1, blue = T2, magenta = T3.

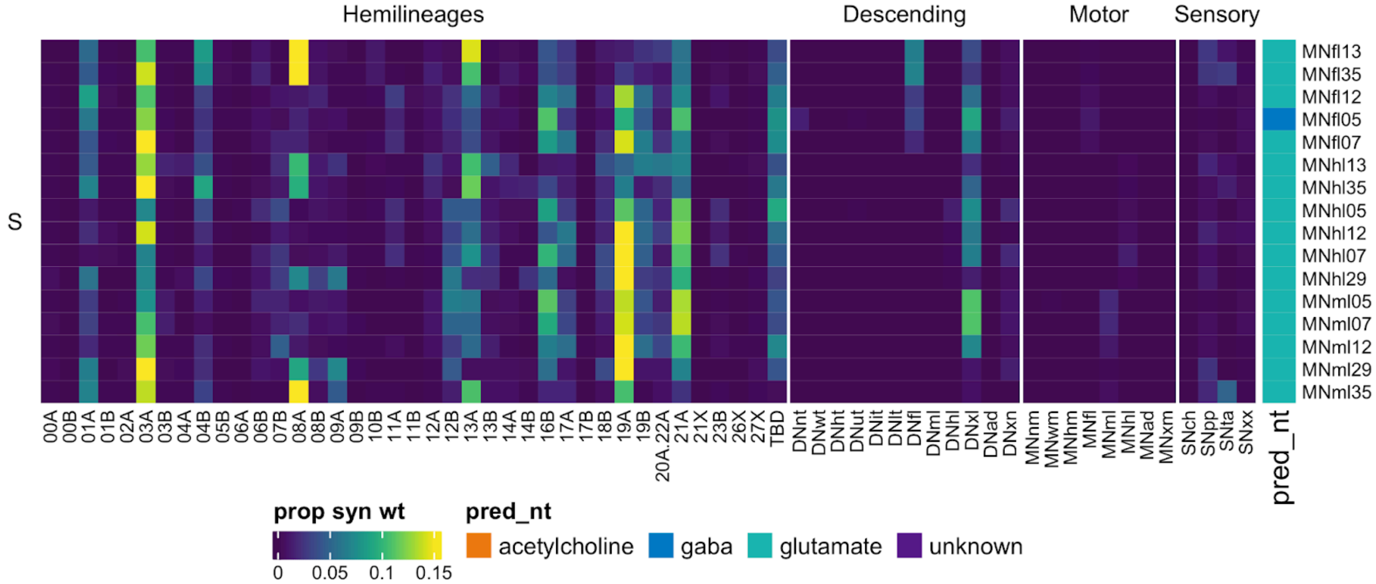

**Figure 47 - figure supplement 3. Connectivity to upstream partners by 24B.25B secondary systematic types.** Proportions of synaptic weight to systematic types from upstream partners, normalised by row. 24B.25B neurons have been clustered within each assigned birthtime window (P = primary, ES = early secondary, S = secondary) based on both upstream and downstream connectivity to hemilineages, descending neuron subclasses, motor neuron subclasses, and sensory neuron modalities. Annotation bar is coloured by the most common predicted neurotransmitter for the neurons of each type.

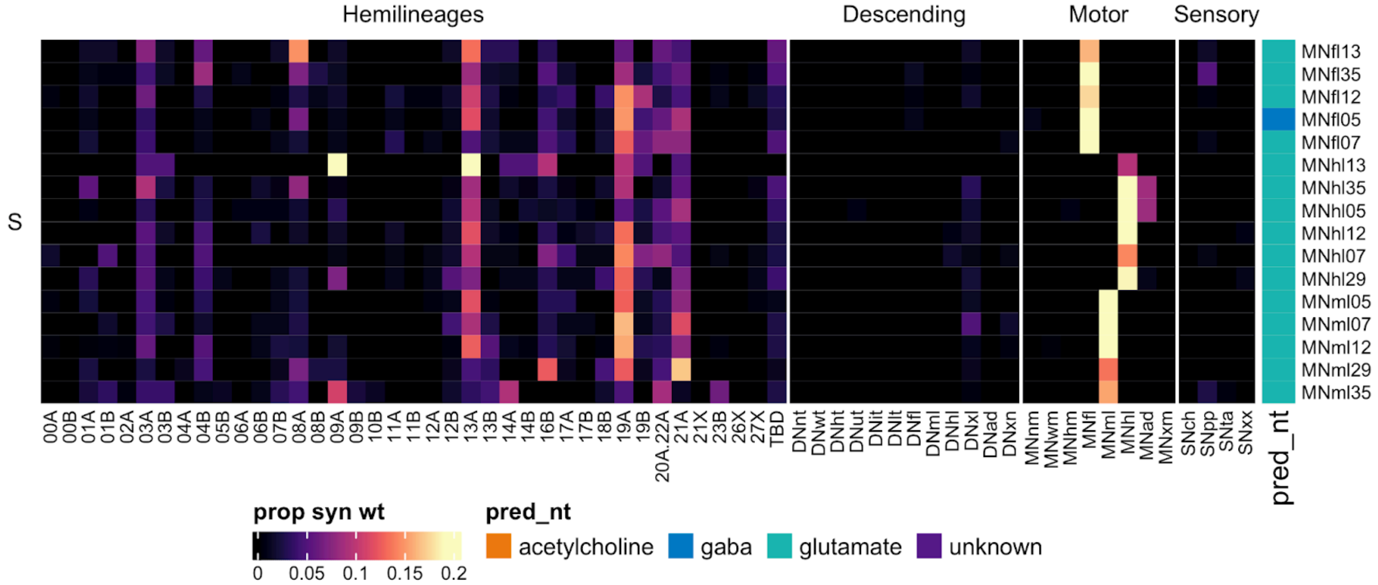

**Figure 47 - figure supplement 4. Connectivity to downstream partners by 24B.25B secondary systematic types.** Proportions of synaptic weight from systematic types to downstream partners, normalised by row. 24B.25B neurons have been clustered within each assigned birthtime window (P = primary, ES = early secondary, S = secondary) based on both upstream and downstream connectivity to hemilineages, descending neuron subclasses, motor neuron subclasses, and sensory neuron modalities. The annotation bar is coloured by the most common predicted neurotransmitter within each type.

#### “Primary” hemilineages

Several hemilineages do not include a significant secondary population, either because the parent neuroblast is eliminated after generating neurons in the embryo or because all/most of the secondary neurons die soon after birth (Truman et al., 2010). The morphologies of 00B, 04A, 14B, and 20B/21B/22B had been previously predicted and/or reported at light level. We tentatively assigned a neuroblast of origin to 5 populations of early-born neurons that are not associated with, or appear significantly distinct from, an identifiable secondary hemilineage (Figure 48). In some cases, light-level data regarding Notch expression was unavailable, so we could not confidently assign them to “A” vs “B” and instead opted for “X”. For reference, we plotted the postsynapses and presynapses for all primary neurons in T2 neuropils by annotated hemilineage (Figure 48 - figure supplement 1).

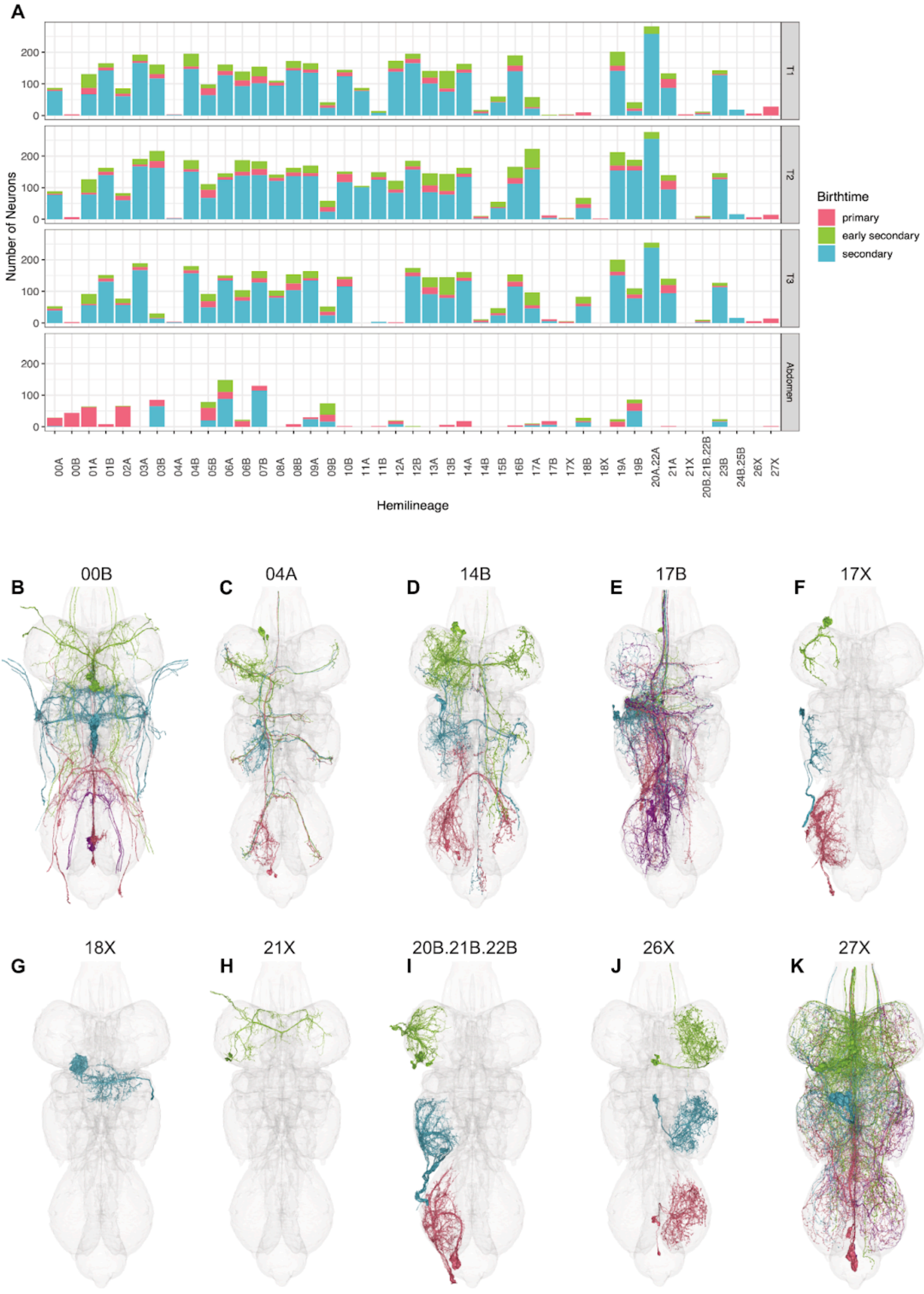
A
Number of Neurons
T1
T2
T3
Abdomen
Birthtime
primary
early secondary
secondary
Hemilineage
00A 00B 01A 01B 02A 03A 03B 04A 04B 05B 06A 06B 07B 08A 08B 09A 09B 10B 11A 11B 12A 12B 13A 13B 14A 14B 15B 16B 17A 17B 17X 18B 18X 19A 19B 20A.22A 21A 21X 20B.21B.22B 23B 24B.25B 26X 27X
B 00B
C 04A
D 14B
E 17B
F 17X
G 18X
H 21X
I 20B.21B.22B
J 26X
K 27X

**Figure 48. Hemilineages with few or no associated secondary neurons. A.** Estimated birthtime by hemilineage and soma neuromere (neurons of unknown hemilineage have been omitted and all abdominal neuromeres have been combined). **B.** Octopaminergic hemilineage 00B. **C.** Cholinergic hemilineage 04A. **D.** Gabaergic/glutamatergic hemilineage 14B. **E.** Gabaergic hemilineage "17B". **F.** Motor neurons from hemilineage "17X". **G.** Motor neurons from hemilineage "18X". **H.** Hemilineage "21X". **I.** Motor neurons from hemilineages 20B, 21B, or 22B. **J.** Gabergic hemilineage "26X". **K.** Putative neurosecretory hemilineage "27X". Neuron meshes are coloured by soma neuromere and side (green: T1 RHS, cyan: T2 RHS, pink: T3 RHS, purple: A1 RHS).

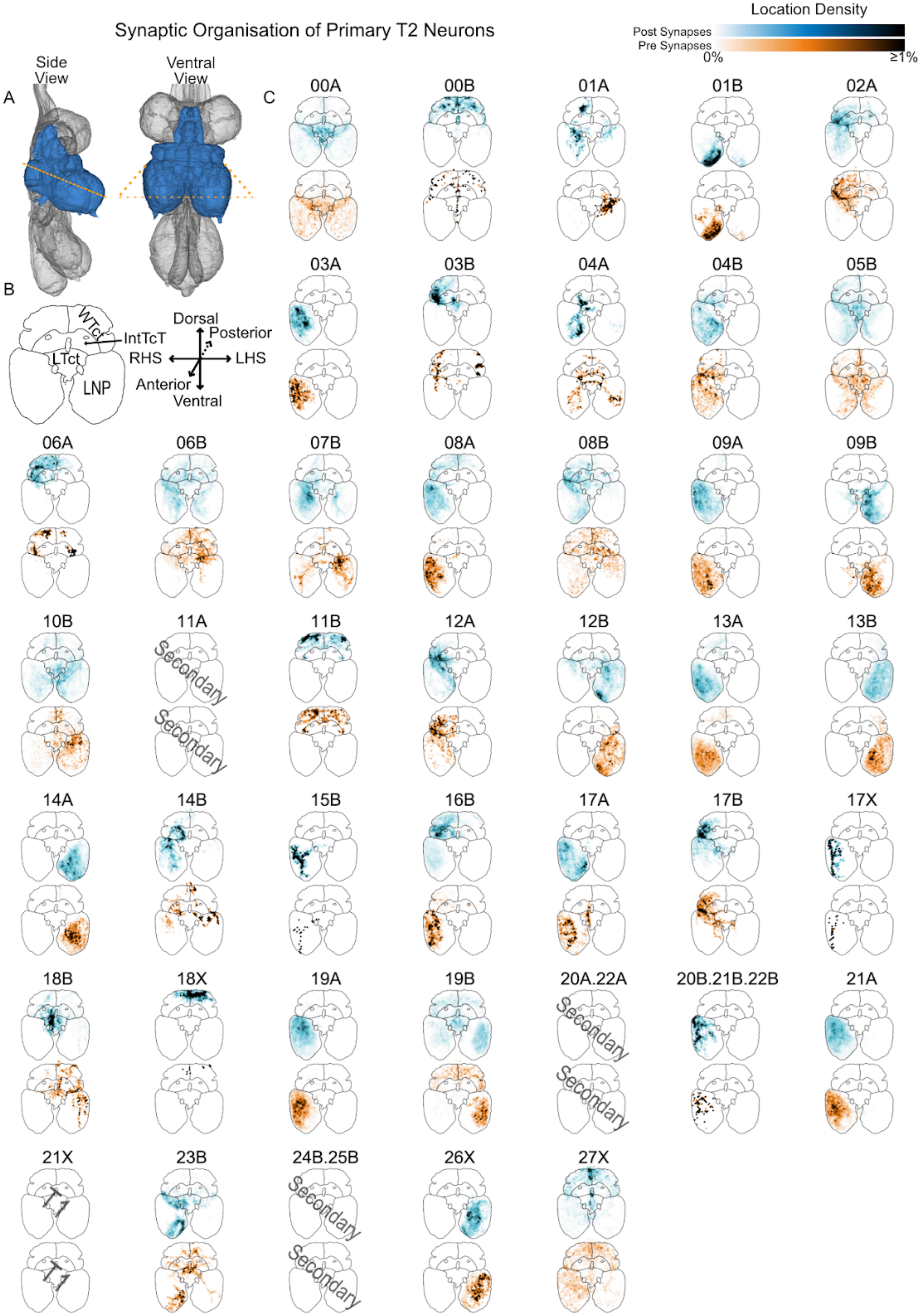

Synaptic Organisation of Primary T2 Neurons
Location Density
Post Synapses
Pre Synapses
0%
≥1%
Side View
Ventral View
A
B
C
WTct
IntTcT
LTct
LNP
Dorsal
Posterior
RHS
LHS
Anterior
Ventral
00A
00B
01A
01B
02A
03A
03B
04A
04B
05B
06A
06B
07B
08A
08B
09A
09B
10B
11A
11B
12A
12B
13A
13B
Secondary
14A
14B
15B
16B
17A
17B
17X
18B
18X
19A
19B
20A.22A
20B.21B.22B
21A
21X
23B
24B.25B
26X
27X
T1

**Figure 48 - figure supplement 1. Summary of the anatomical organisation of synapses for primary neurons in each hemilineage originating in T2 RHS. A.** Side and ventral views illustrating the neuropils surveyed: T2 Leg Neuropil (LNP), medial ventral association center (mVAC), ovoid (Ov), wing tectulum (WTct), lower tectulum (LTct), and Intermediate tectulum (IntTct). The orange dashed line shows the approximate location of the transverse section onto which their synapses have been projected. **B.** Template transverse section with major neuropils labelled and axes for orientation. **C.** Projected synapses for T2 RHS neurons of each hemilineage. Top (blue): postsynapses, connections to upstream neurons. Bottom (orange): presynapses, connections to downstream neurons. 21X is only observed in T1.

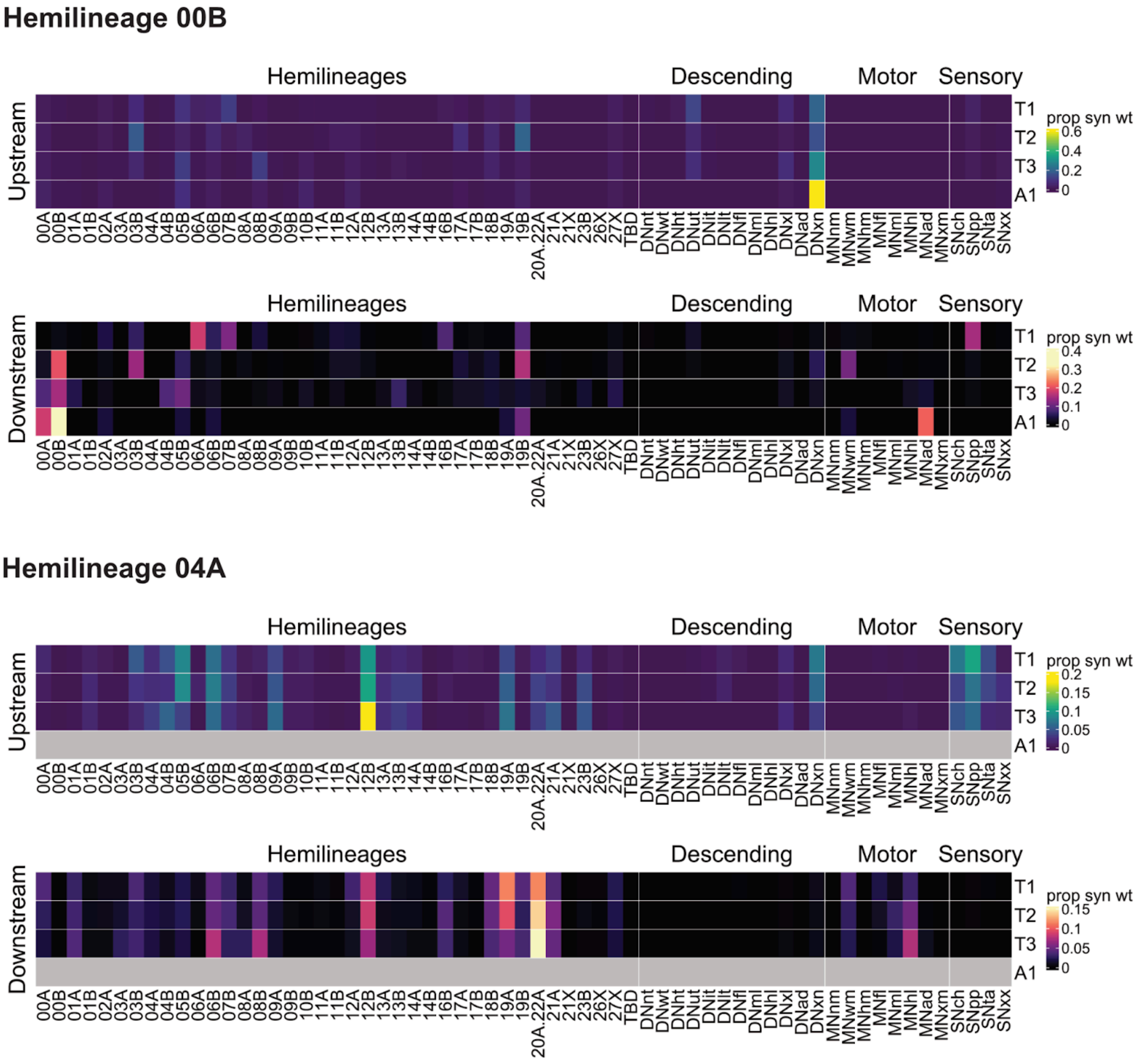

**Figure 48 - figure supplement 2. Connectivity heat maps for primary hemilineages 00B and 04A.** Ratios of connections from neurons originating in each neuromere to upstream or downstream partners, normalised by row.

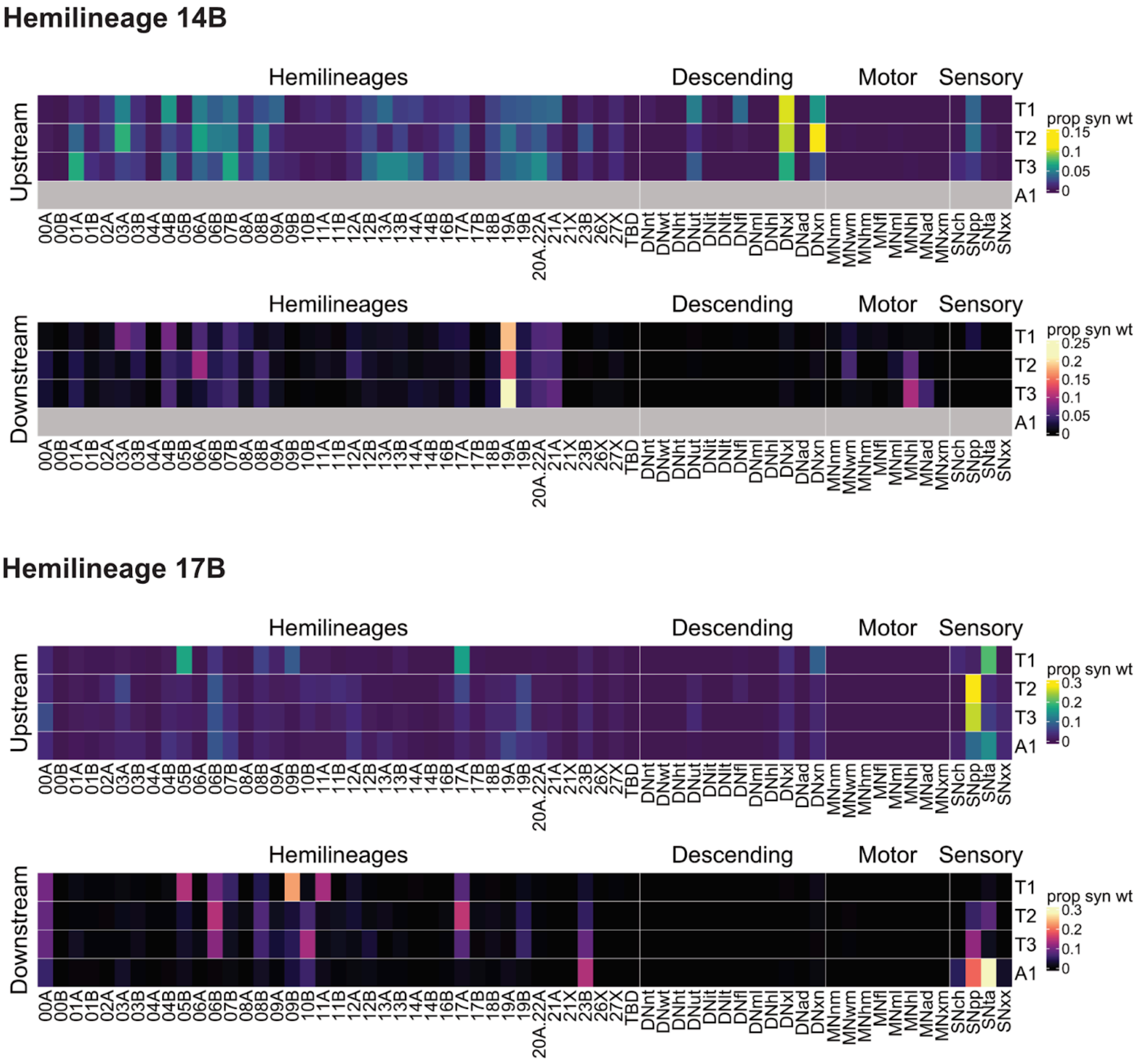

**Figure 48 - figure supplement 3. Connectivity heat maps for primary hemilineages 14B and 17B.** Ratios of connections from neurons originating in each neuromere to upstream or downstream partners, normalised by row.

#### Hemilineage 17X

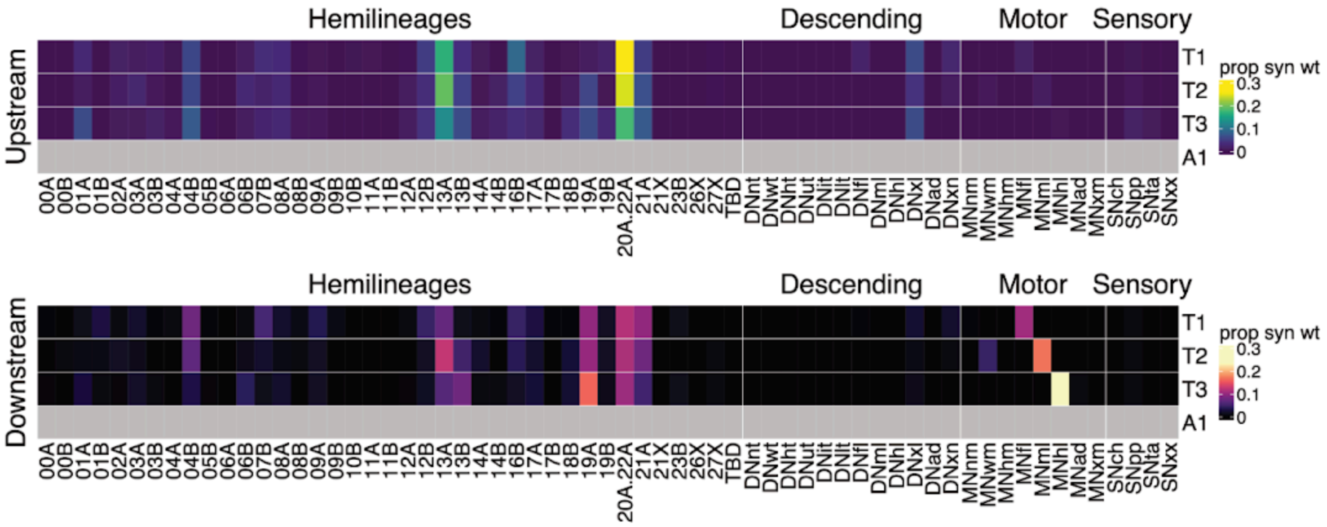

#### Hemilineage 18X

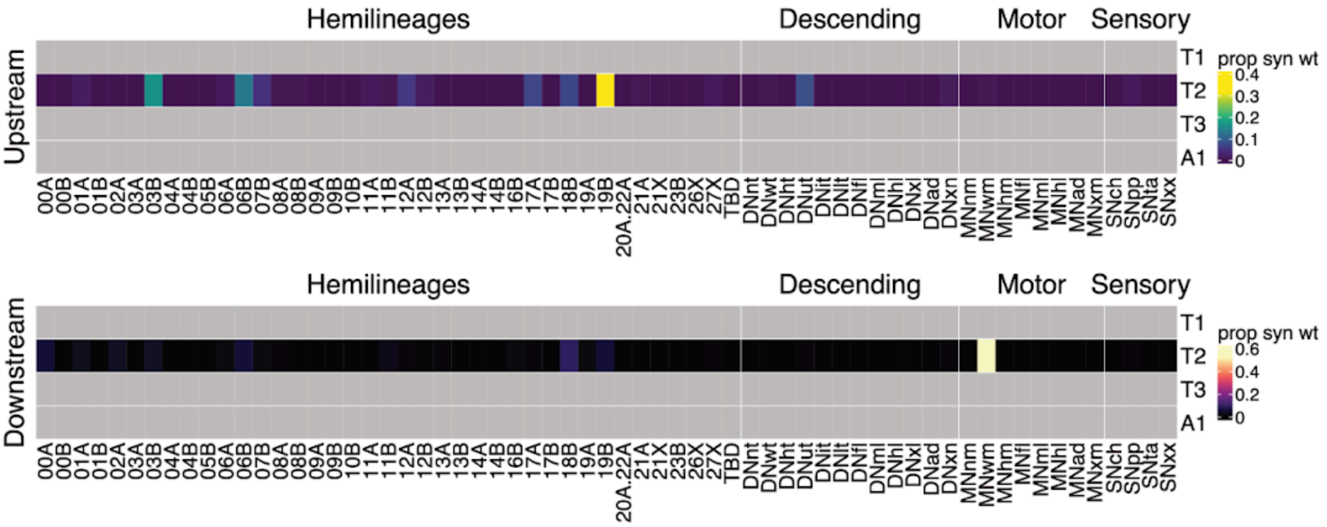

#### Hemilineage 20B.21B.22B

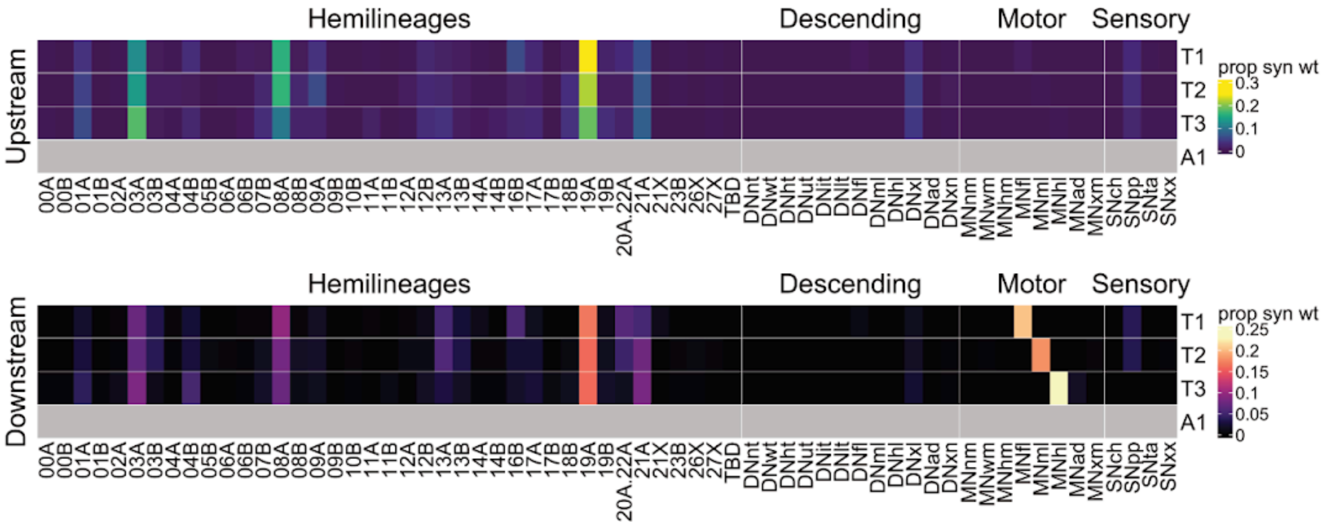

**Figure 48 - figure supplement 4. Connectivity heat maps for primary hemilineages 17X.18X and 20B.21B.22B.** Ratios of connections from neurons originating in each neuromere to upstream or downstream partners, normalised by row.

#### Hemilineage 21X

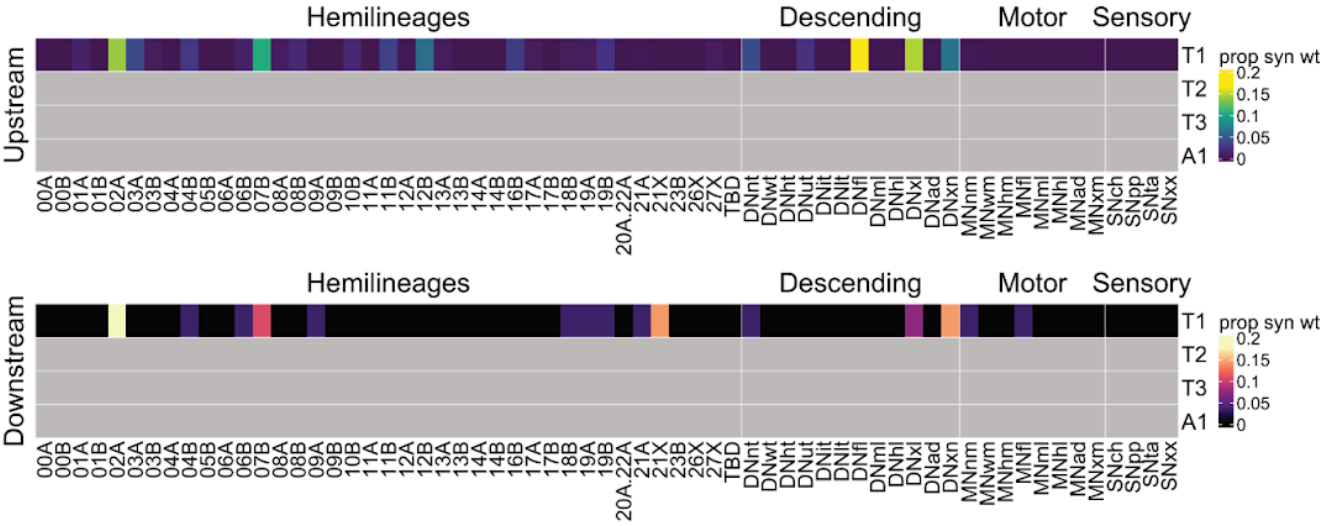

#### Hemilineage 26X

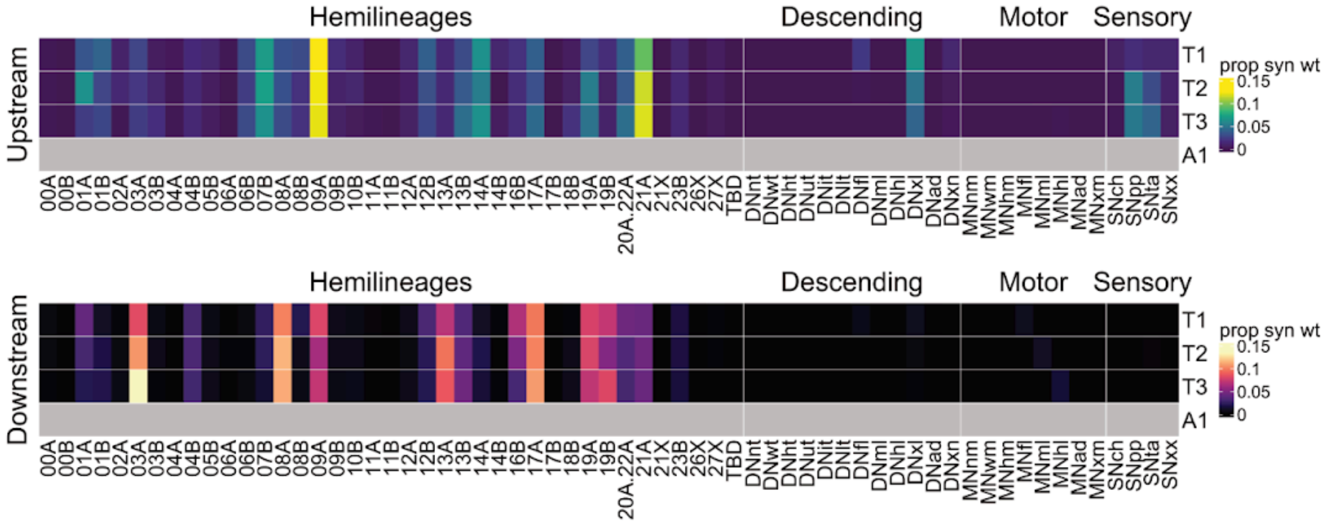

#### Hemilineage 27X

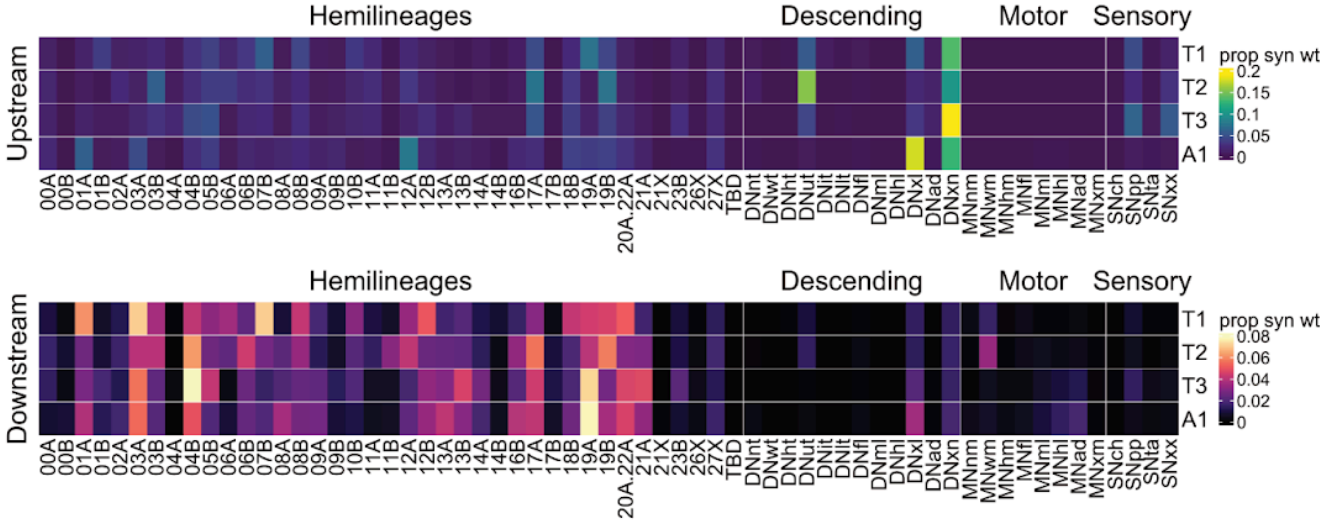

**Figure 48 - figure supplement 5. Connectivity heat maps for primary hemilineages 21X, 26X and 27X.** Ratios of connections from neurons originating in each neuromere to upstream or downstream partners, normalised by row.

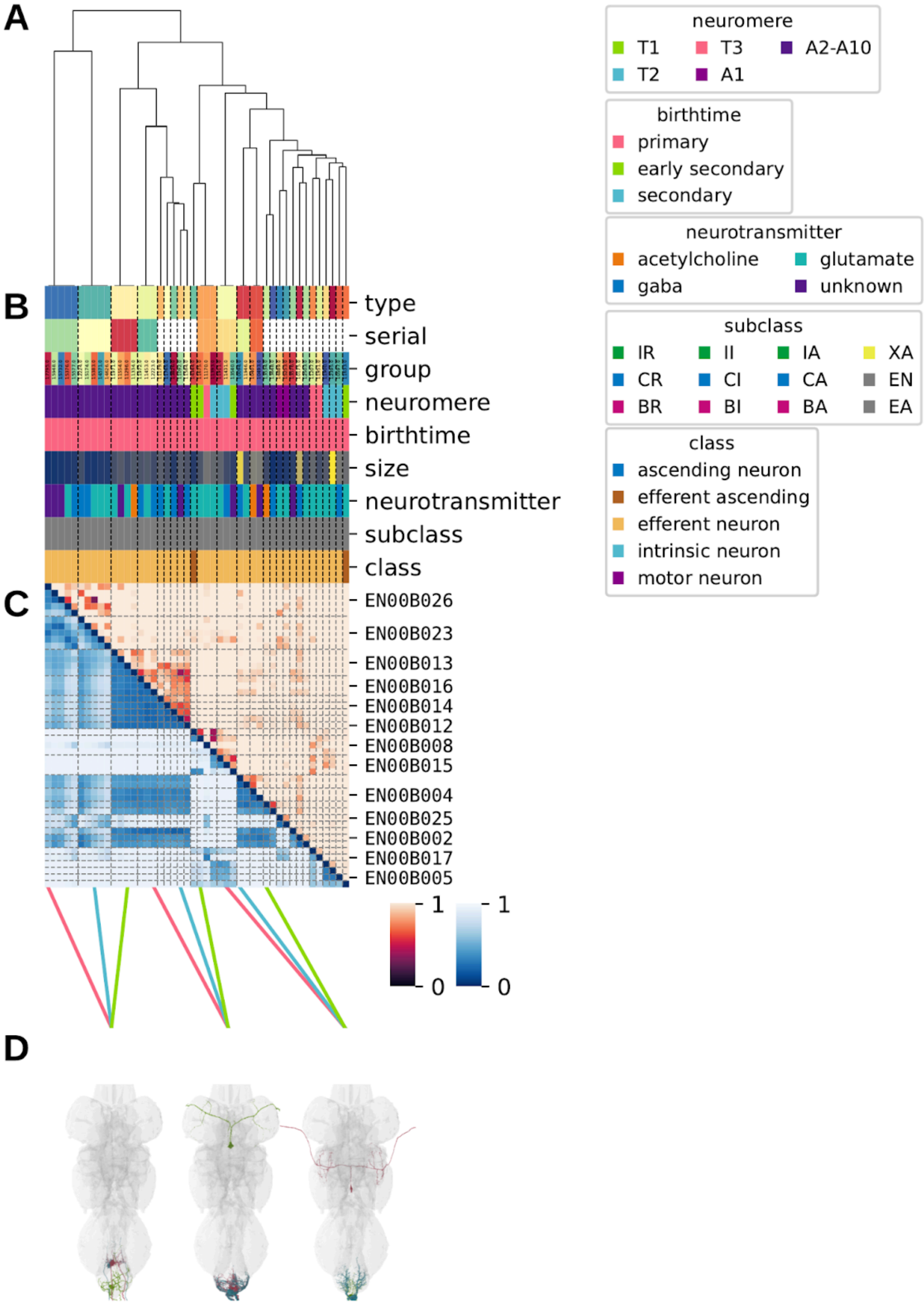

A
B
C
D
type
serial
group
neuromere
birthtime
size
neurotransmitter
subclass
class
EN00B026
EN00B023
EN00B013
EN00B016
EN00B014
EN00B012
EN00B008
EN00B015
EN00B004
EN00B025
EN00B002
EN00B017
EN00B005
1
0
1
0
neuromere
T1
T3
A2-A10
T2
A1
birthtime
primary
early secondary
secondary
neurotransmitter
acetylcholine
glutamate
gaba
unknown
subclass
IR
II
IA
XA
CR
CI
CA
EN
BR
BI
BA
EA
class
ascending neuron
efferent ascending
efferent neuron
intrinsic neuron
motor neuron

**Figure 48 - figure supplement 6. Systematic typing of hemilineage 00B. A.** Hierarchical clustering dendrogram of hemilineage groups by laterally and serially aggregated connectivity cosine clustering. **B.** Categorical annotations of each hemilineage group, each column corresponding to the aligned leaf in A. Colours for type, serial set, and group are arbitrary for visualisation. Colours for neuromere, birthtime, neurotransmitter, subclass, and class are as in all other figures. **C.** Similarity distance heatmap for hemilineage. Cosine distance is in the upper triangle, while laterally symmetrised NBLAST distance is in the lower triangle. Systematic type names of some types are labelled. **D.** Morphologically representative groups from dendrogram subtrees. Each group, indicated by colour and line connecting to its column in B and C, is the most morphologically representative group (medoid of NBLAST distance) from a subtree of A. The subtrees (flat clusters) are equal height cuts of A determined to yield the number of groups per plot and plots in D.

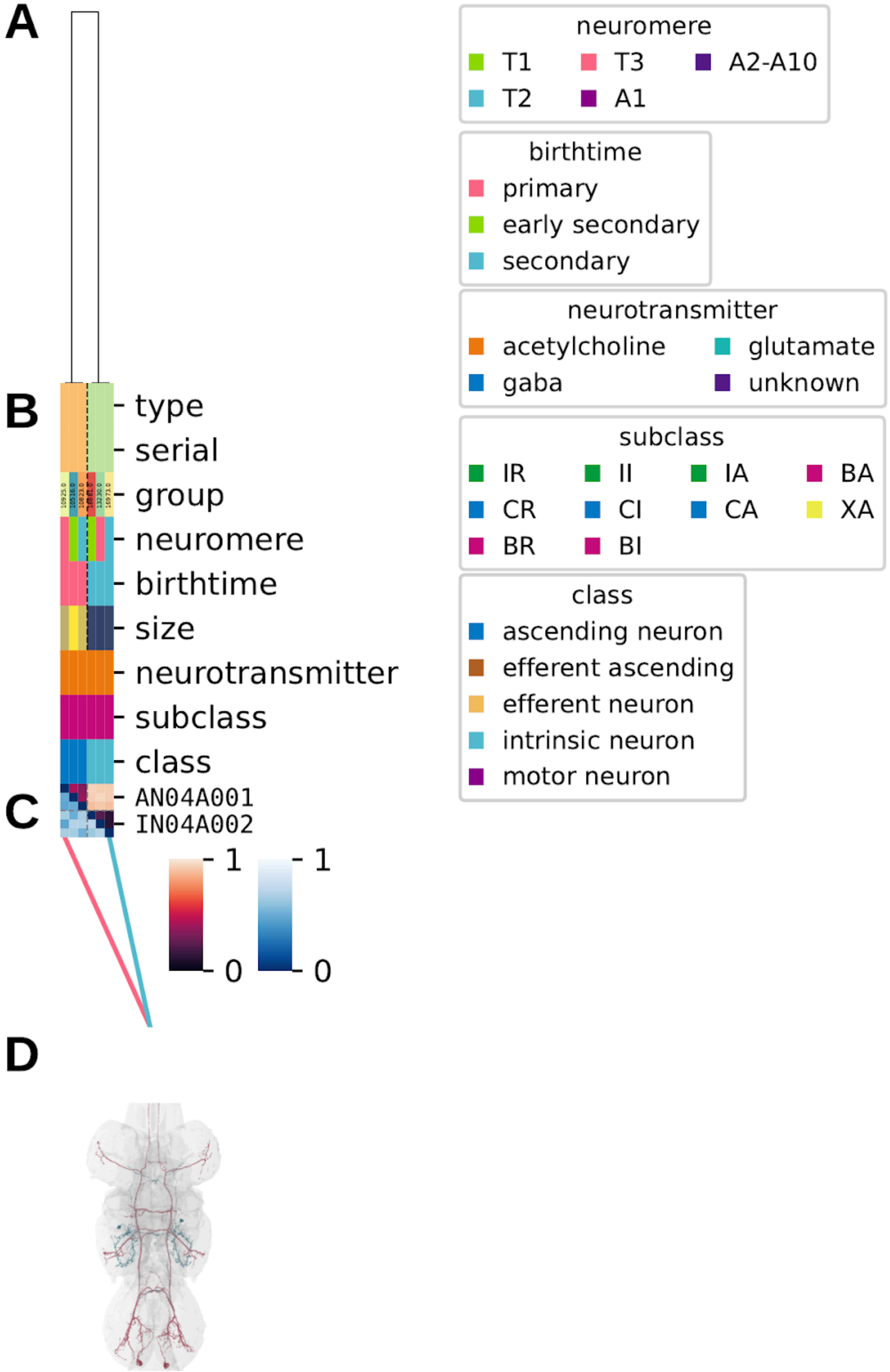

A
B
C
D
type
serial
group
neuromere
birthtime
size
neurotransmitter
subclass
class
AN04A001
IN04A002
1
0
1
0
neuromere
T1
T3
A2-A10
T2
A1
birthtime
primary
early secondary
secondary
neurotransmitter
acetylcholine
glutamate
gaba
unknown
subclass
IR
II
IA
BA
CR
CI
CA
XA
BR
BI
class
ascending neuron
efferent ascending
efferent neuron
intrinsic neuron
motor neuron

**Figure 48 - figure supplement 7. Systematic typing of hemilineage 04A. A.** Hierarchical clustering dendrogram of hemilineage groups by laterally and serially aggregated connectivity cosine clustering. **B.** Categorical annotations of each hemilineage group, each column corresponding to the aligned leaf in A. Colours for type, serial set, and group are arbitrary for visualisation. Colours for neuromere, birthtime, neurotransmitter, subclass, and class are as in all other figures. **C.** Similarity distance heatmap for hemilineage. Cosine distance is in the upper triangle, while laterally symmetrised NBLAST distance is in the lower triangle. Systematic type names of some types are labelled. **D.** Morphologically representative groups from dendrogram subtrees. Each group, indicated by colour and line connecting to its column in B and C, is the most morphologically representative group (medoid of NBLAST distance) from a subtree of A. The subtrees (flat clusters) are equal height cuts of A determined to yield the number of groups per plot and plots in D.

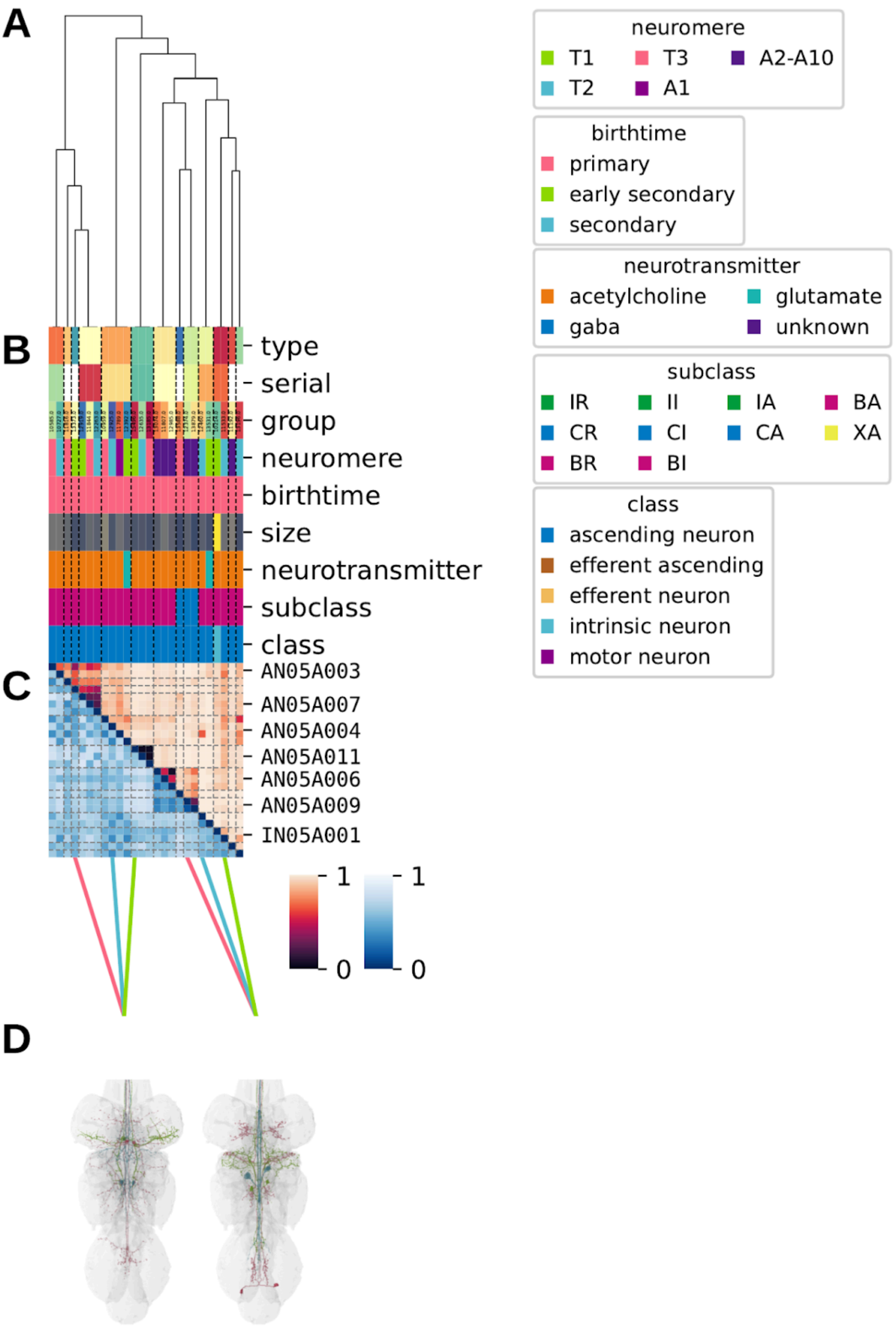
A
B
C
D
type
serial
group
neuromere
birthtime
size
neurotransmitter
subclass
class
AN05A003
AN05A007
AN05A004
AN05A011
AN05A006
AN05A009
IN05A001
1
0
1
0
neuromere
T1
T3
A2-A10
T2
A1
birthtime
primary
early secondary
secondary
neurotransmitter
acetylcholine
glutamate
gaba
unknown
subclass
IR
II
IA
BA
CR
CI
CA
XA
BR
BI
class
ascending neuron
efferent ascending
efferent neuron
intrinsic neuron
motor neuron

**Figure 48 - figure supplement 8. Systematic typing of hemilineage 05A. A.** Hierarchical clustering dendrogram of hemilineage groups by laterally and serially aggregated connectivity cosine clustering. **B.** Categorical annotations of each hemilineage group, each column corresponding to the aligned leaf in A. Colours for type, serial set, and group are arbitrary for visualisation. Colours for neuromere, birthtime, neurotransmitter, subclass, and class are as in all other figures. **C.** Similarity distance heatmap for hemilineage. Cosine distance is in the upper triangle, while laterally symmetrised NBLAST distance is in the lower triangle. Systematic type names of some types are labelled. **D.** Morphologically representative groups from dendrogram subtrees. Each group, indicated by colour and line connecting to its column in B and C, is the most morphologically representative group (medoid of NBLAST distance) from a subtree of A. The subtrees (flat clusters) are equal height cuts of A determined to yield the number of groups per plot and plots in D.

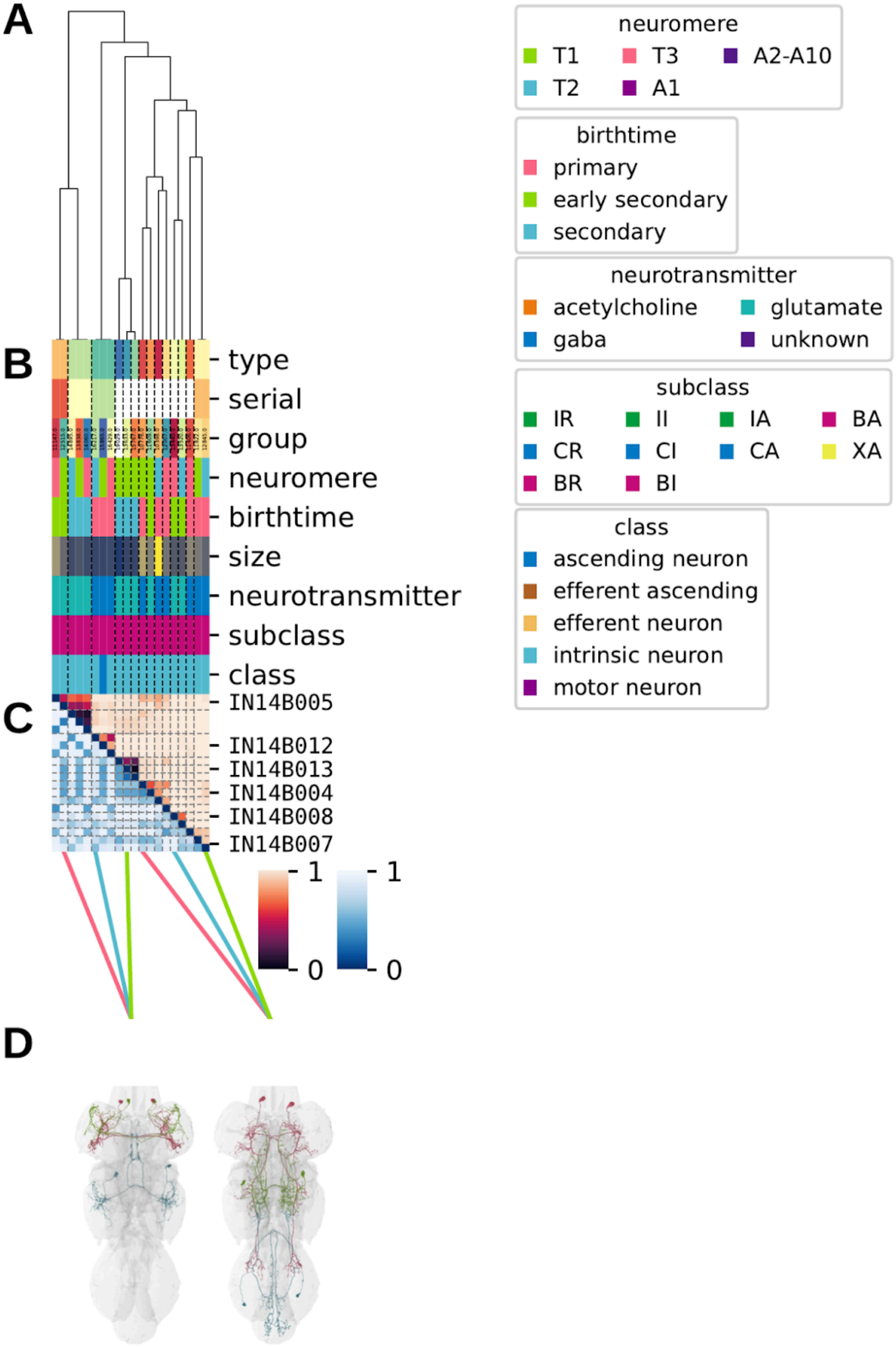

A
B
C
D
type
serial
group
neuromere
birthtime
size
neurotransmitter
subclass
class
IN14B005
IN14B012
IN14B013
IN14B004
IN14B008
IN14B007
1
0
1
0
neuromere
T1
T3
A2-A10
T2
A1
birthtime
primary
early secondary
secondary
neurotransmitter
acetylcholine
glutamate
gaba
unknown
subclass
IR
II
IA
BA
CR
CI
CA
XA
BR
BI
class
ascending neuron
efferent ascending
efferent neuron
intrinsic neuron
motor neuron

**Figure 48 - figure supplement 9. Systematic typing of hemilineage 14B. A.** Hierarchical clustering dendrogram of hemilineage groups by laterally and serially aggregated connectivity cosine clustering. **B.** Categorical annotations of each hemilineage group, each column corresponding to the aligned leaf in A. Colours for type, serial set, and group are arbitrary for visualisation. Colours for neuromere, birthtime, neurotransmitter, subclass, and class are as in all other figures. **C.** Similarity distance heatmap for hemilineage. Cosine distance is in the upper triangle, while laterally symmetrised NBLAST distance is in the lower triangle. Systematic type names of some types are labelled. **D.** Morphologically representative groups from dendrogram subtrees. Each group, indicated by colour and line connecting to its column in B and C, is the most morphologically representative group (medoid of NBLAST distance) from a subtree of A. The subtrees (flat clusters) are equal height cuts of A determined to yield the number of groups per plot and plots in D.

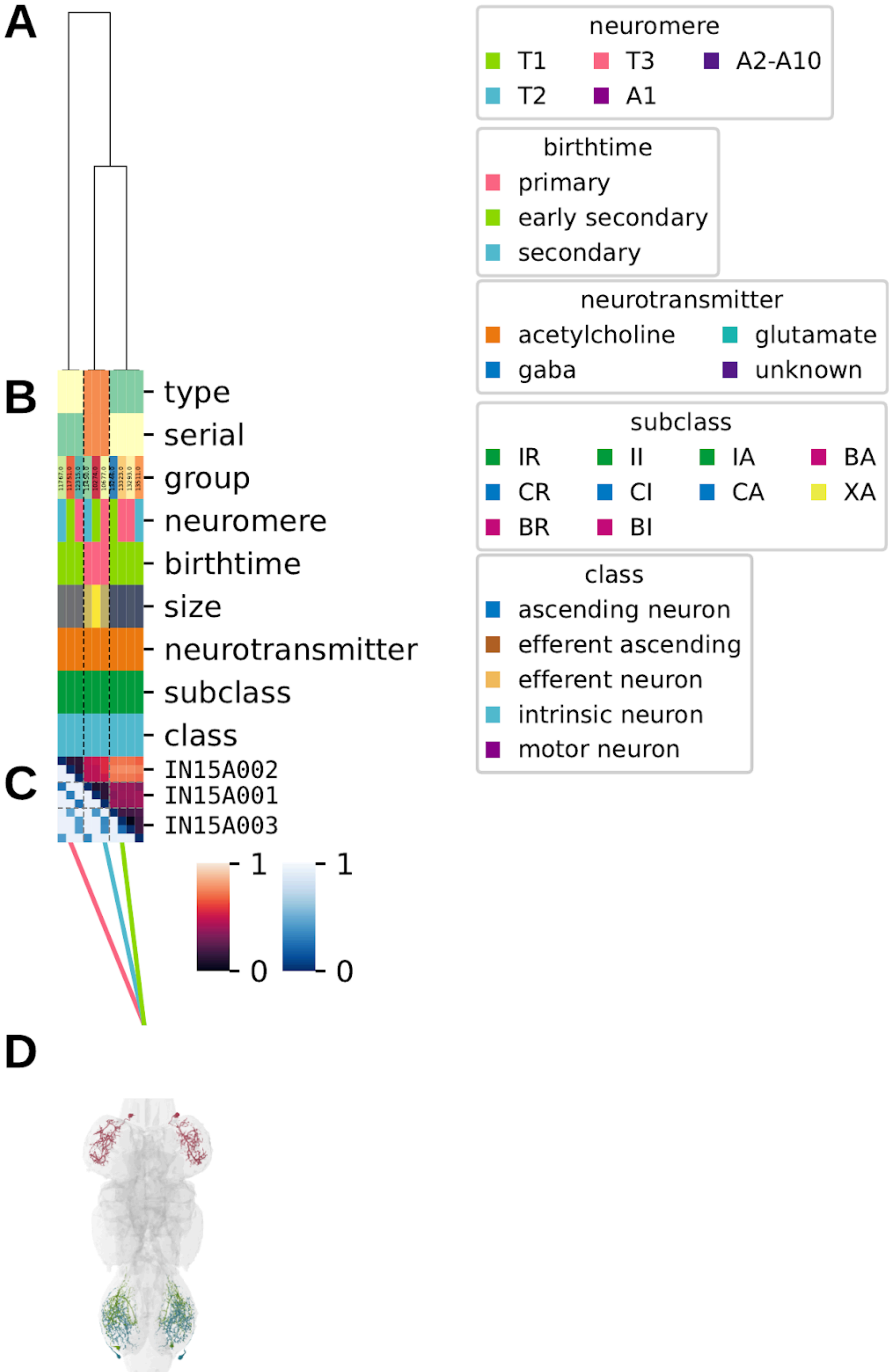
A
B
C
D
type
serial
group
neuromere
birthtime
size
neurotransmitter
subclass
class
IN15A002
IN15A001
IN15A003
1
0
1
0
neuromere
T1
T3
A2-A10
T2
A1
birthtime
primary
early secondary
secondary
neurotransmitter
acetylcholine
glutamate
gaba
unknown
subclass
IR
II
IA
BA
CR
CI
CA
XA
BR
BI
class
ascending neuron
efferent ascending
efferent neuron
intrinsic neuron
motor neuron

**Figure 48 - figure supplement 10. Systematic typing of hemilineage 15A. A.** Hierarchical clustering dendrogram of hemilineage groups by laterally and serially aggregated connectivity cosine clustering. **B.** Categorical annotations of each hemilineage group, each column corresponding to the aligned leaf in A. Colours for type, serial set, and group are arbitrary for visualisation. Colours for neuromere, birthtime, neurotransmitter, subclass, and class are as in all other figures. **C.** Similarity distance heatmap for hemilineage. Cosine distance is in the upper triangle, while laterally symmetrised NBLAST distance is in the lower triangle. Systematic type names of some types are labelled. **D.** Morphologically representative groups from dendrogram subtrees. Each group, indicated by colour and line connecting to its column in B and C, is the most morphologically representative group (medoid of NBLAST distance) from a subtree of A. The subtrees (flat clusters) are equal height cuts of A determined to yield the number of groups per plot and plots in D.

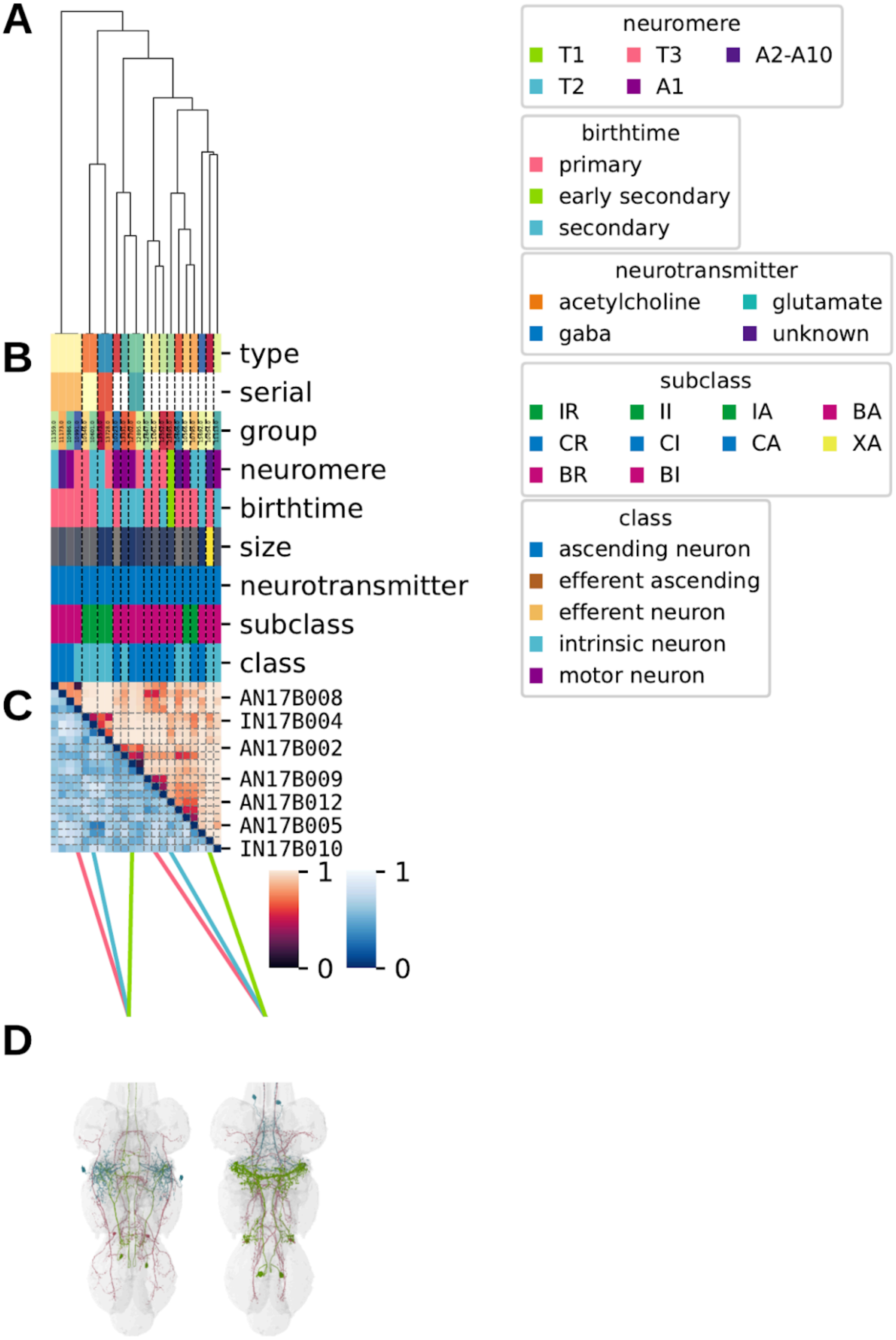
A
B
C
D
type
serial
group
neuromere
birthtime
size
neurotransmitter
subclass
class
AN17B008
IN17B004
AN17B002
AN17B009
AN17B012
AN17B005
IN17B010
1
0
1
0
neuromere
T1
T3
A2-A10
T2
A1
birthtime
primary
early secondary
secondary
neurotransmitter
acetylcholine
glutamate
gaba
unknown
subclass
IR
II
IA
BA
CR
CI
CA
XA
BR
BI
class
ascending neuron
efferent ascending
efferent neuron
intrinsic neuron
motor neuron

**Figure 48 - figure supplement 11. Systematic typing of hemilineage 17B. A.** Hierarchical clustering dendrogram of hemilineage groups by laterally and serially aggregated connectivity cosine clustering. **B.** Categorical annotations of each hemilineage group, each column corresponding to the aligned leaf in A. Colours for type, serial set, and group are arbitrary for visualisation. Colours for neuromere, birthtime, neurotransmitter, subclass, and class are as in all other figures. **C.** Similarity distance heatmap for hemilineage. Cosine distance is in the upper triangle, while laterally symmetrised NBLAST distance is in the lower triangle. Systematic type names of some types are labelled. **D.** Morphologically representative groups from dendrogram subtrees. Each group, indicated by colour and line connecting to its column in B and C, is the most morphologically representative group (medoid of NBLAST distance) from a subtree of A. The subtrees (flat clusters) are equal height cuts of A determined to yield the number of groups per plot and plots in D.

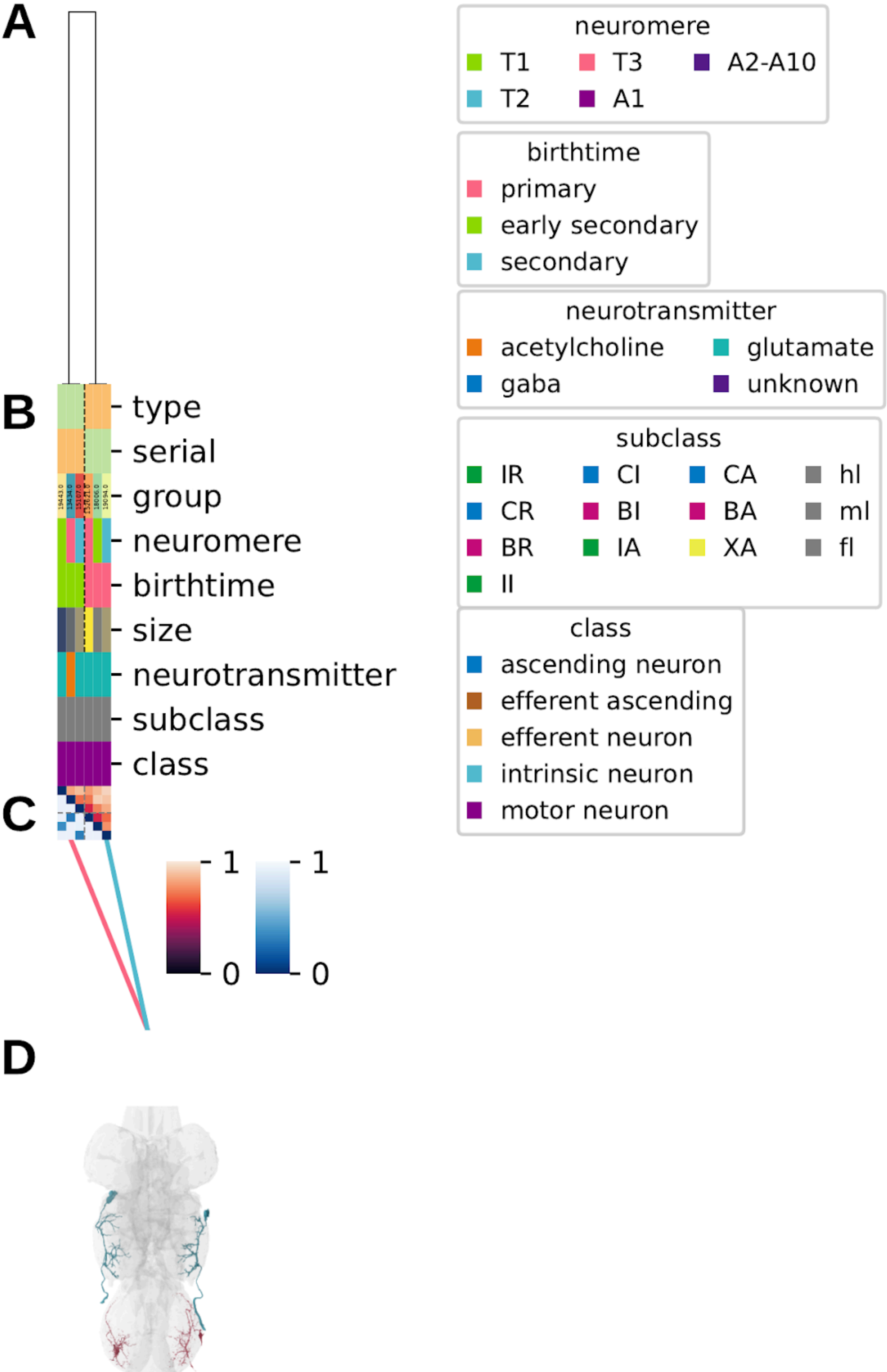

A
B
C
D
type
serial
group
neuromere
birthtime
size
neurotransmitter
subclass
class
1
0
1
0
neuromere
T1
T3
A2-A10
T2
A1
birthtime
primary
early secondary
secondary
neurotransmitter
acetylcholine
glutamate
gaba
unknown
subclass
IR
CI
CA
hl
CR
BI
BA
ml
BR
IA
XA
fl
II
class
ascending neuron
efferent ascending
efferent neuron
intrinsic neuron
motor neuron

**Figure 48 - figure supplement 12. Systematic typing of hemilineage 17X and 18X.** Types for motor neurons were assigned separately as outlined in our accompanying manuscript (Cheong et al., 2023). **A.** Hierarchical clustering dendrogram of hemilineage groups by laterally and serially aggregated connectivity cosine clustering. **B.** Categorical annotations of each hemilineage group, each column corresponding to the aligned leaf in A. Colours for cluster, serial set, and group are arbitrary for visualisation. Colours for neuromere, birthtime, neurotransmitter, subclass, and class are as in all other figures. **C.** Similarity distance heatmap for hemilineage. Cosine distance is in the upper triangle, while laterally symmetrised NBLAST distance is in the lower triangle. Systematic type names of some types are labelled. **D.** Morphologically representative groups from dendrogram subtrees. Each group, indicated by colour and line connecting to its column in B and C, is the most morphologically representative group (medoid of NBLAST distance) from a subtree of A. The subtrees (flat clusters) are equal height cuts of A determined to yield the number of groups per plot and plots in D.

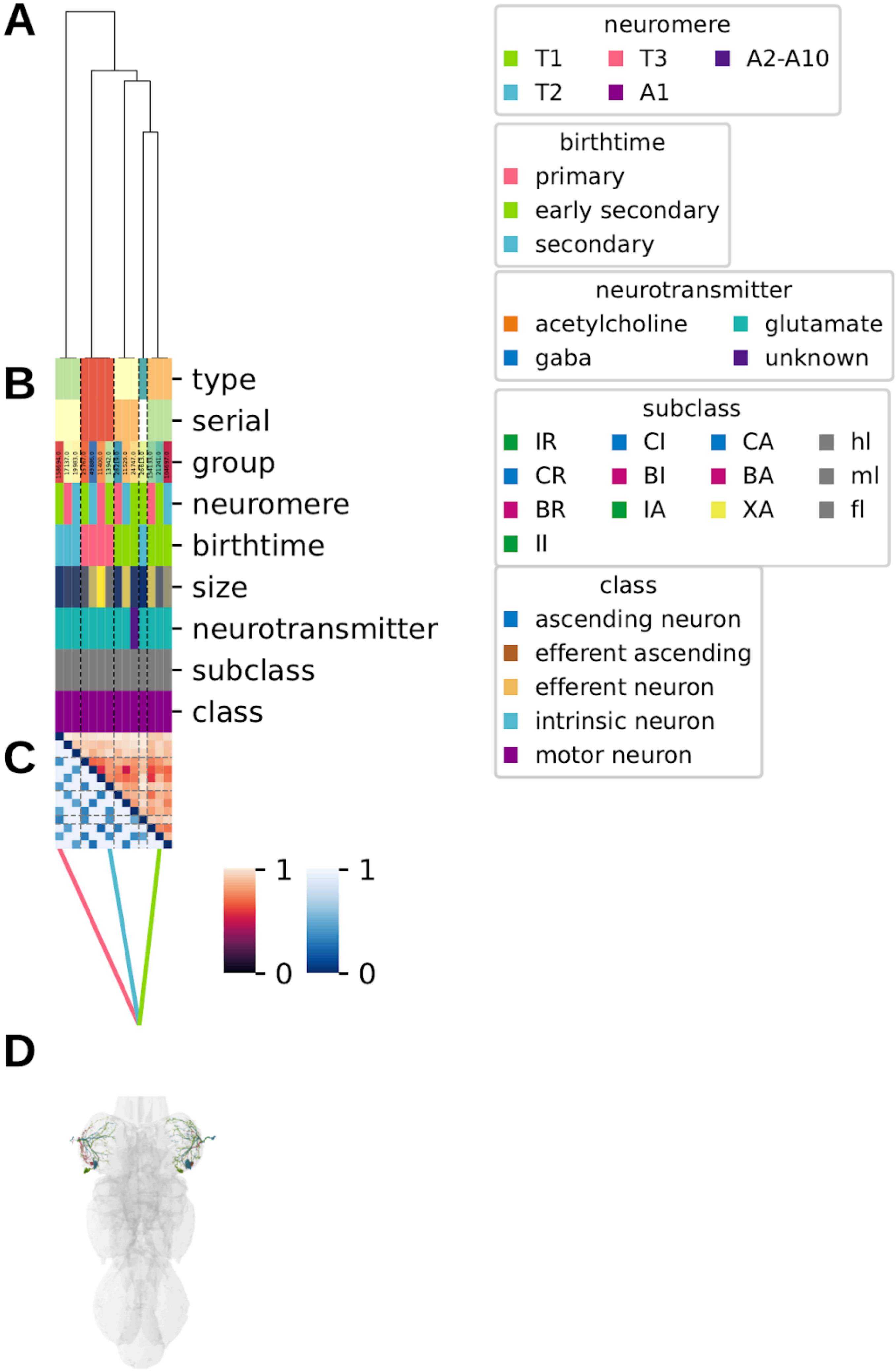
A
B
C
D
type
serial
group
neuromere
birthtime
size
neurotransmitter
subclass
class
1
0
1
0
neuromere
T1
T3
A2-A10
T2
A1
birthtime
primary
early secondary
secondary
neurotransmitter
acetylcholine
glutamate
gaba
unknown
subclass
IR
CI
CA
hl
CR
BI
BA
ml
BR
IA
XA
fl
II
class
ascending neuron
efferent ascending
efferent neuron
intrinsic neuron
motor neuron

**Figure 48 - figure supplement 13. Systematic typing of hemilineage 20B, 21B, and 22B.** Types for motor neurons were assigned separately as outlined in our accompanying manuscript (Cheong et al., 2023). **A.** Hierarchical clustering dendrogram of hemilineage groups by laterally and serially aggregated connectivity cosine clustering. **B.** Categorical annotations of each hemilineage group, each column corresponding to the aligned leaf in A. Colours for cluster, serial set, and group are arbitrary for visualisation. Colours for neuromere, birthtime, neurotransmitter, subclass, and class are as in all other figures. **C.** Similarity distance heatmap for hemilineage. Cosine distance is in the upper triangle, while laterally symmetrised NBLAST distance is in the lower triangle. Systematic type names of some types are labelled. **D.** Morphologically representative groups from dendrogram subtrees. Each group, indicated by colour and line connecting to its column in B and C, is the most morphologically representative group (medoid of NBLAST distance) from a subtree of A. The subtrees (flat clusters) are equal height cuts of A determined to yield the number of groups per plot and plots in D.

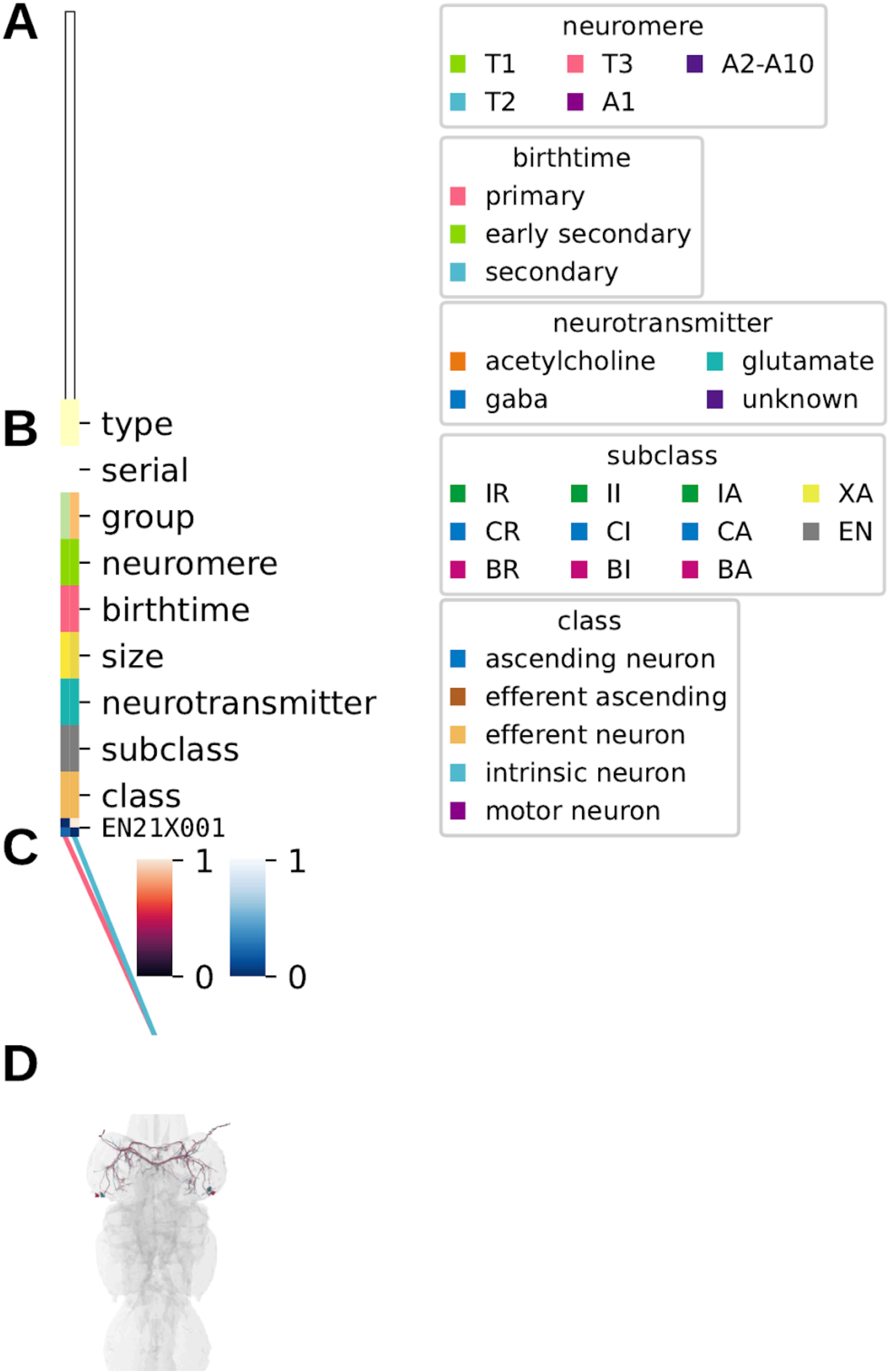
A
B
C
D
type
serial
group
neuromere
birthtime
size
neurotransmitter
subclass
class
EN21X001
1
0
1
0
neuromere
T1
T3
A2-A10
T2
A1
birthtime
primary
early secondary
secondary
neurotransmitter
acetylcholine
glutamate
gaba
unknown
subclass
IR
II
IA
XA
CR
CI
CA
EN
BR
BI
BA
class
ascending neuron
efferent ascending
efferent neuron
intrinsic neuron
motor neuron

**Figure 48 - figure supplement 14. Systematic typing of hemilineage 21X. A.** Hierarchical clustering dendrogram of hemilineage groups by laterally and serially aggregated connectivity cosine clustering. **B.** Categorical annotations of each hemilineage group, each column corresponding to the aligned leaf in A. Colours for type, serial set, and group are arbitrary for visualisation. Colours for neuromere, birthtime, neurotransmitter, subclass, and class are as in all other figures. **C.** Similarity distance heatmap for hemilineage. Cosine distance is in the upper triangle, while laterally symmetrised NBLAST distance is in the lower triangle. Systematic type names of some types are labelled. **D.** Morphologically representative groups from dendrogram subtrees. Each group, indicated by colour and line connecting to its column in B and C, is the most morphologically representative group (medoid of NBLAST distance) from a subtree of A. The subtrees (flat clusters) are equal height cuts of A determined to yield the number of groups per plot and plots in D.

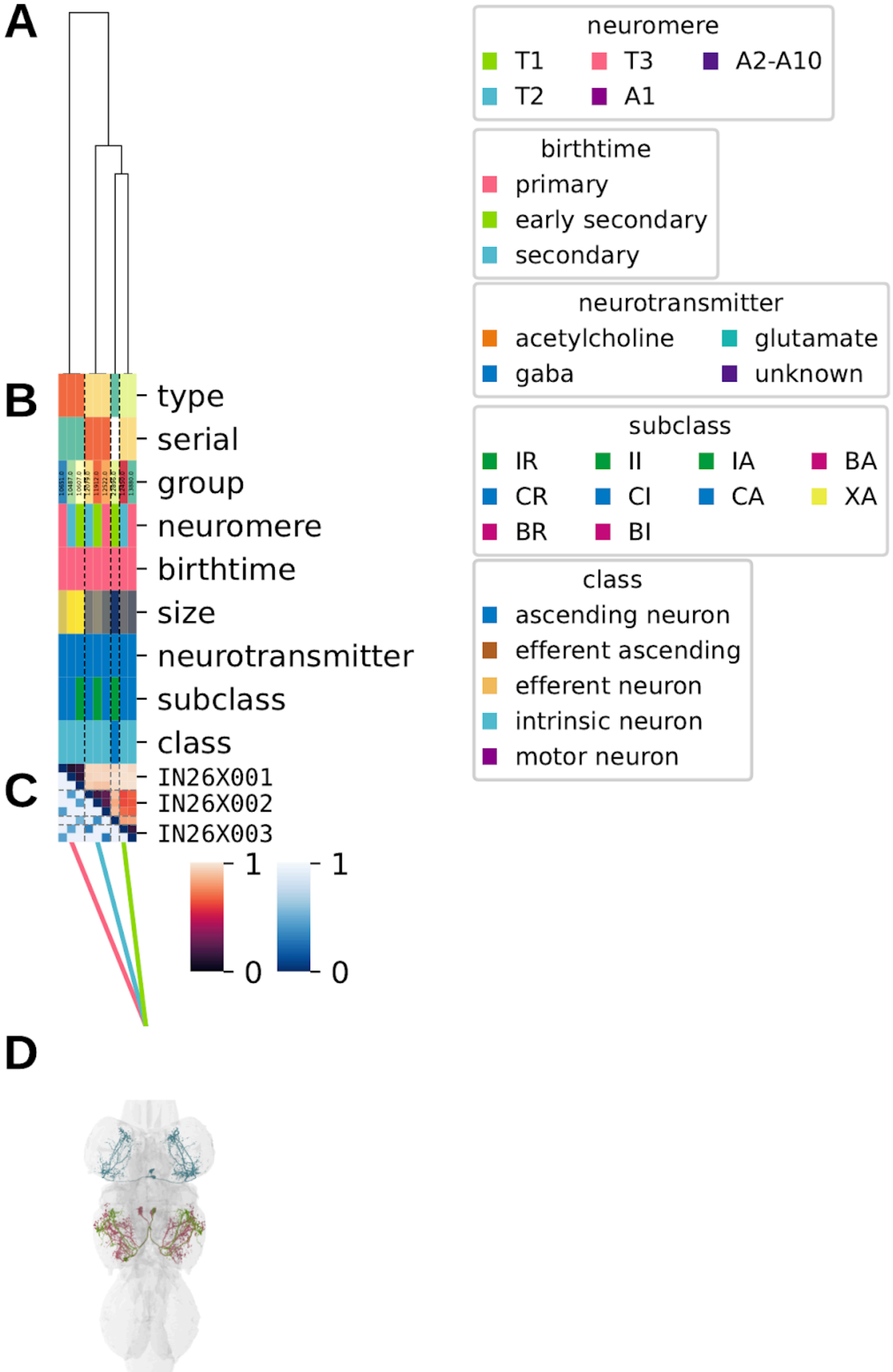
A
B
C
D
type
serial
group
neuromere
birthtime
size
neurotransmitter
subclass
class
IN26X001
IN26X002
IN26X003
1
0
1
0
neuromere
T1
T2
T3
A1
A2-A10
birthtime
primary
early secondary
secondary
neurotransmitter
acetylcholine
gaba
glutamate
unknown
subclass
IR
II
IA
BA
CR
CI
CA
XA
BR
BI
class
ascending neuron
efferent ascending
efferent neuron
intrinsic neuron
motor neuron

**Figure 48 - figure supplement 15. Systematic typing of hemilineage 26X. A.** Hierarchical clustering dendrogram of hemilineage groups by laterally and serially aggregated connectivity cosine clustering. **B.** Categorical annotations of each hemilineage group, each column corresponding to the aligned leaf in A. Colours for type, serial set, and group are arbitrary for visualisation. Colours for neuromere, birthtime, neurotransmitter, subclass, and class are as in all other figures. **C.** Similarity distance heatmap for hemilineage. Cosine distance is in the upper triangle, while laterally symmetrised NBLAST distance is in the lower triangle. Systematic type names of some types are labelled. **D.** Morphologically representative groups from dendrogram subtrees. Each group, indicated by colour and line connecting to its column in B and C, is the most morphologically representative group (medoid of NBLAST distance) from a subtree of A. The subtrees (flat clusters) are equal height cuts of A determined to yield the number of groups per plot and plots in D.

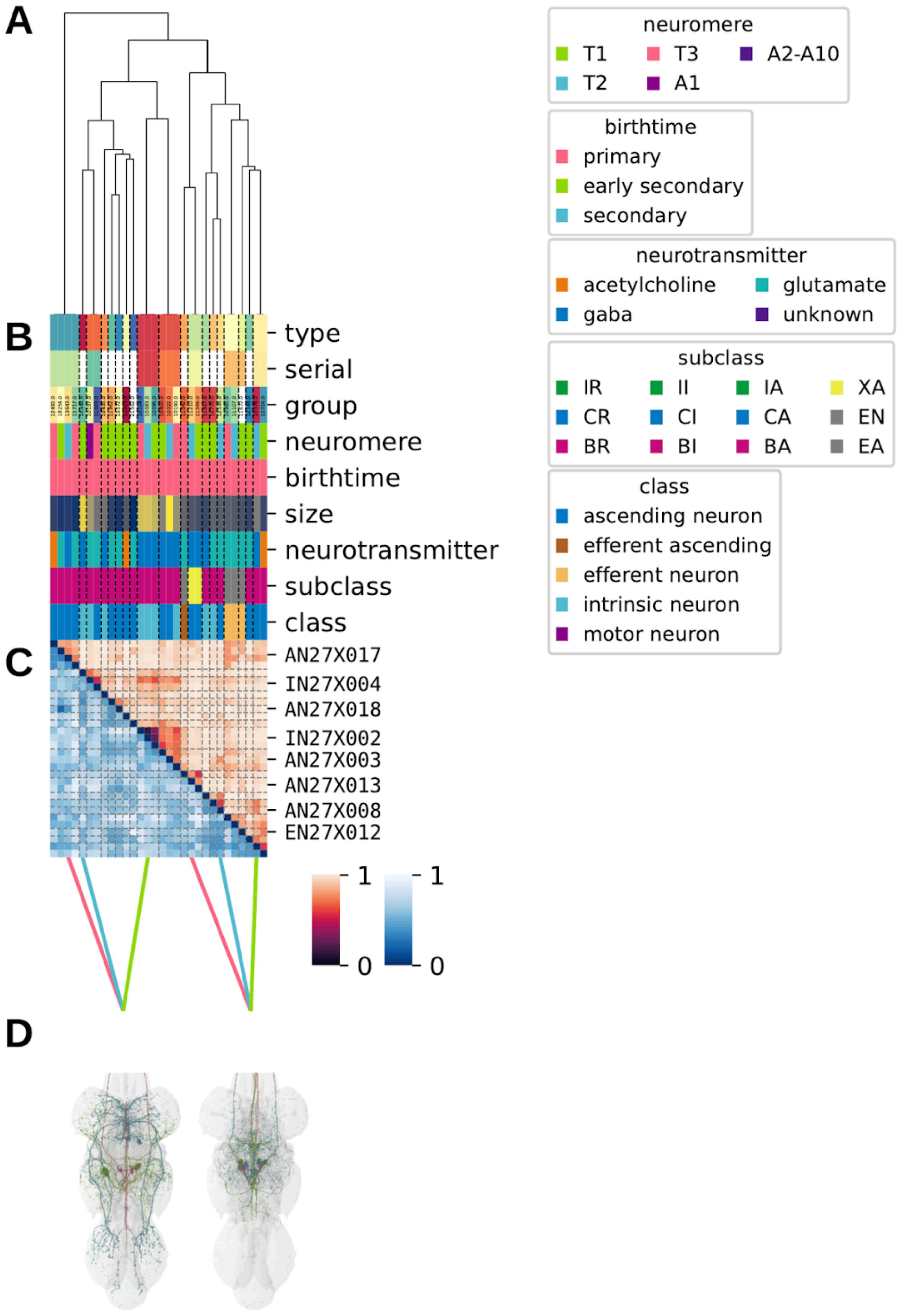
A
B
C
D
type
serial
group
neuromere
birthtime
size
neurotransmitter
subclass
class
AN27X017
IN27X004
AN27X018
IN27X002
AN27X003
AN27X013
AN27X008
EN27X012
1
0
1
0
neuromere
T1
T3
A2-A10
T2
A1
birthtime
primary
early secondary
secondary
neurotransmitter
acetylcholine
glutamate
gaba
unknown
subclass
IR
II
IA
XA
CR
CI
CA
EN
BR
BI
BA
EA
class
ascending neuron
efferent ascending
efferent neuron
intrinsic neuron
motor neuron

**Figure 48 - figure supplement 16. Systematic typing of hemilineage 27X. A.** Hierarchical clustering dendrogram of hemilineage groups by laterally and serially aggregated connectivity cosine clustering. **B.** Categorical annotations of each hemilineage group, each column corresponding to the aligned leaf in A. Colours for type, serial set, and group are arbitrary for visualisation. Colours for neuromere, birthtime, neurotransmitter, subclass, and class are as in all other figures. **C.** Similarity distance heatmap for hemilineage. Cosine distance is in the upper triangle, while laterally symmetrised NBLAST distance is in the lower triangle. Systematic type names of some types are labelled. **D.** Morphologically representative groups from dendrogram subtrees. Each group, indicated by colour and line connecting to its column in B and C, is the most morphologically representative group (medoid of NBLAST distance) from a subtree of A. The subtrees (flat clusters) are equal height cuts of A determined to yield the number of groups per plot and plots in D.

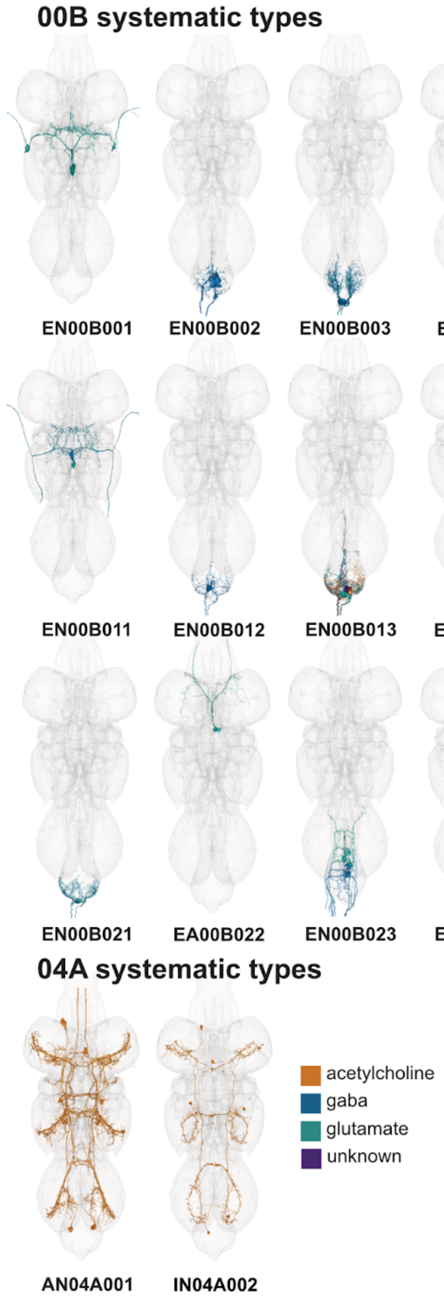

**Figure 48 - figure supplement 17. Systematic types of hemilineages 00B and 04A.** Systematic types have been arranged in numerical order, with neurons of the same type that belong to distinct classes (e.g., intrinsic neuron vs ascending neuron) plotted separately but placed adjacent to each other. Individual neuron meshes have been coloured based on their predicted neurotransmitters: dark orange = acetylcholine, blue = gaba, marine = glutamate, dark purple = unknown.

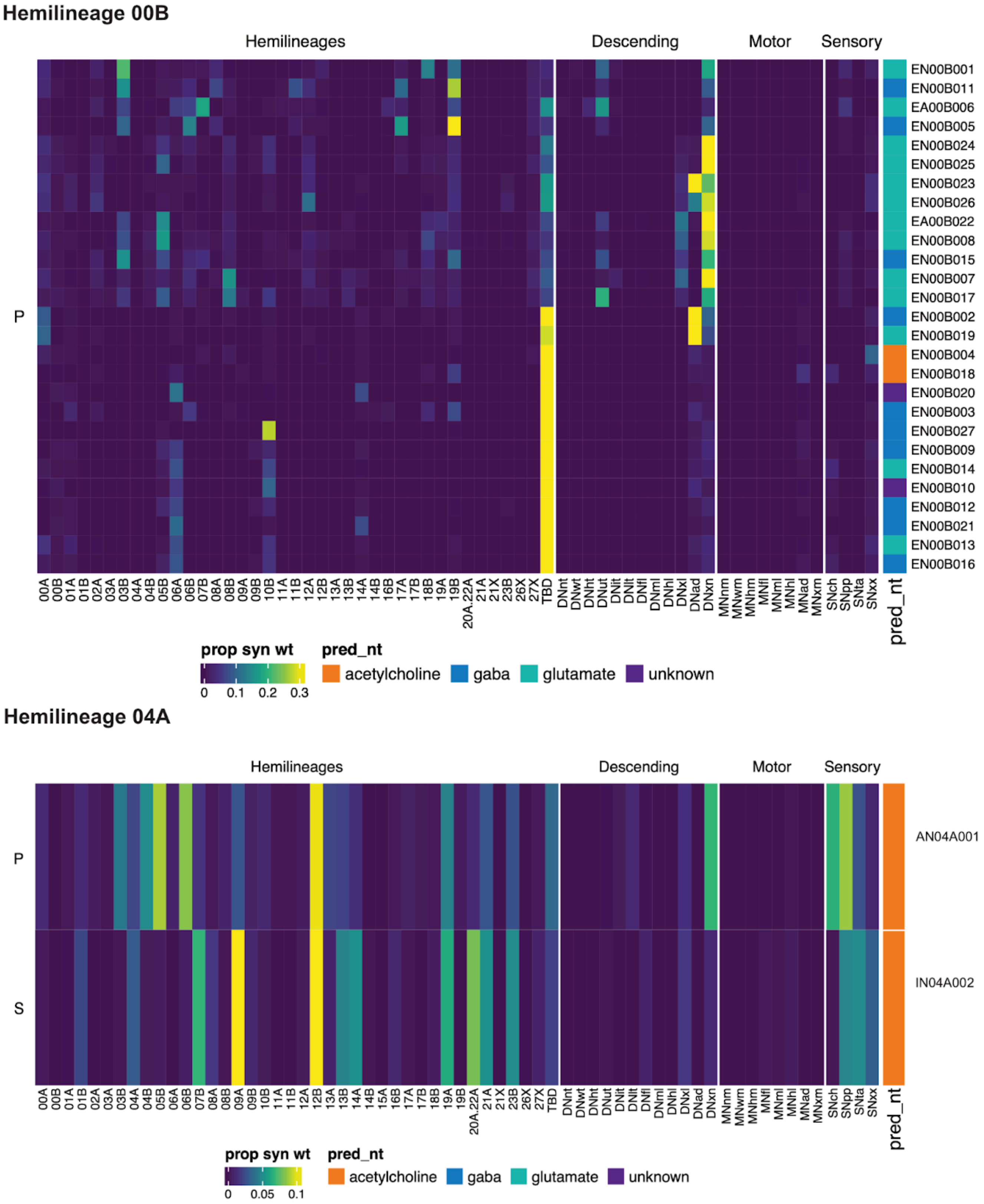

**Figure 48 - figure supplement 18. Connectivity to upstream partners by 00B and 04A systematic types.** Proportions of synaptic weight to systematic types from upstream partners, normalised by row. 00B and 04A neurons have been clustered within each assigned birthtime window (P = primary, ES = early secondary, S = secondary) based on both upstream and downstream connectivity to hemilineages, descending neuron subclasses, motor neuron subclasses, and sensory neuron modalities. Annotation bar is coloured by the most common predicted neurotransmitter for the neurons of each type.

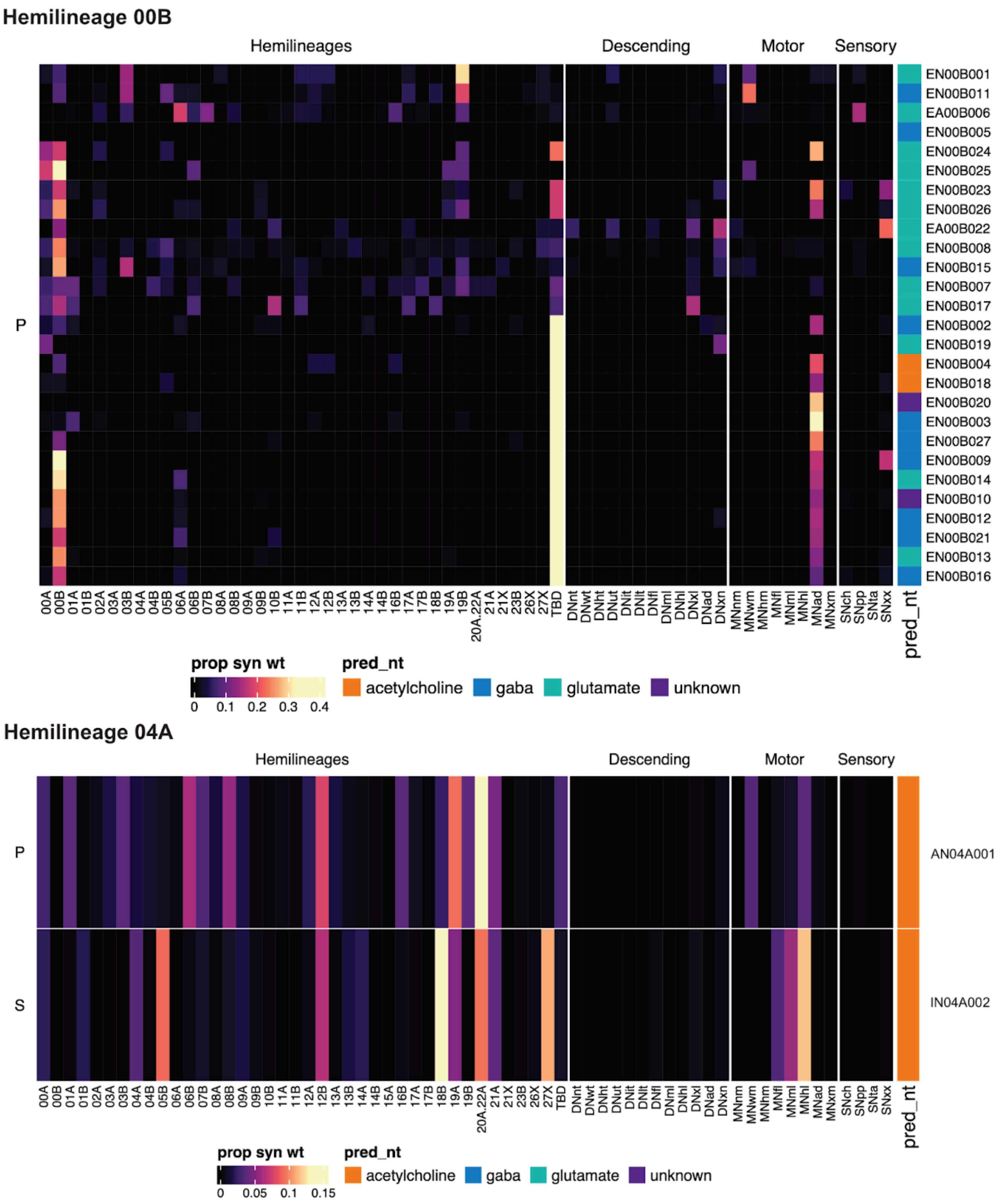

**Figure 48 - figure supplement 19. Connectivity to downstream partners by 00B and 04A systematic types.** Proportions of synaptic weight from systematic types to downstream partners, normalised by row. 00B and 04A neurons have been clustered within each assigned birthtime window (P = primary, ES = early secondary, S = secondary) based on both upstream and downstream connectivity to hemilineages, descending neuron subclasses, motor neuron subclasses, and sensory neuron modalities. The annotation bar is coloured by the most common predicted neurotransmitter within each type.

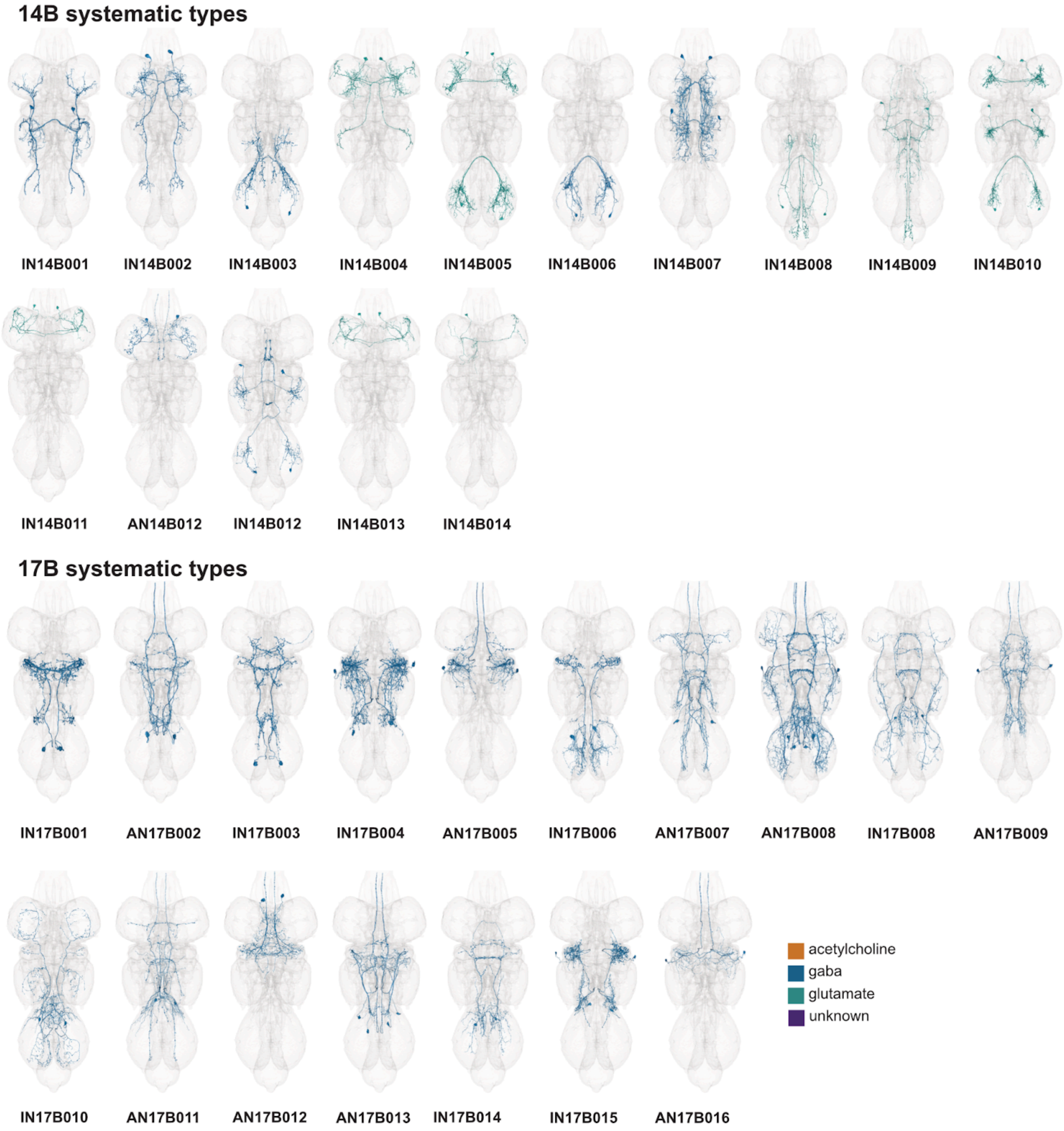

**Figure 48 - figure supplement 20. Systematic types of hemilineages 14B and 17B.** Systematic types have been arranged in numerical order, with neurons of the same type that belong to distinct classes (e.g., intrinsic neuron vs ascending neuron) plotted separately but placed adjacent to each other. Individual neuron meshes have been coloured based on their predicted neurotransmitters: dark orange = acetylcholine, blue = gaba, marine = glutamate, dark purple = unknown.

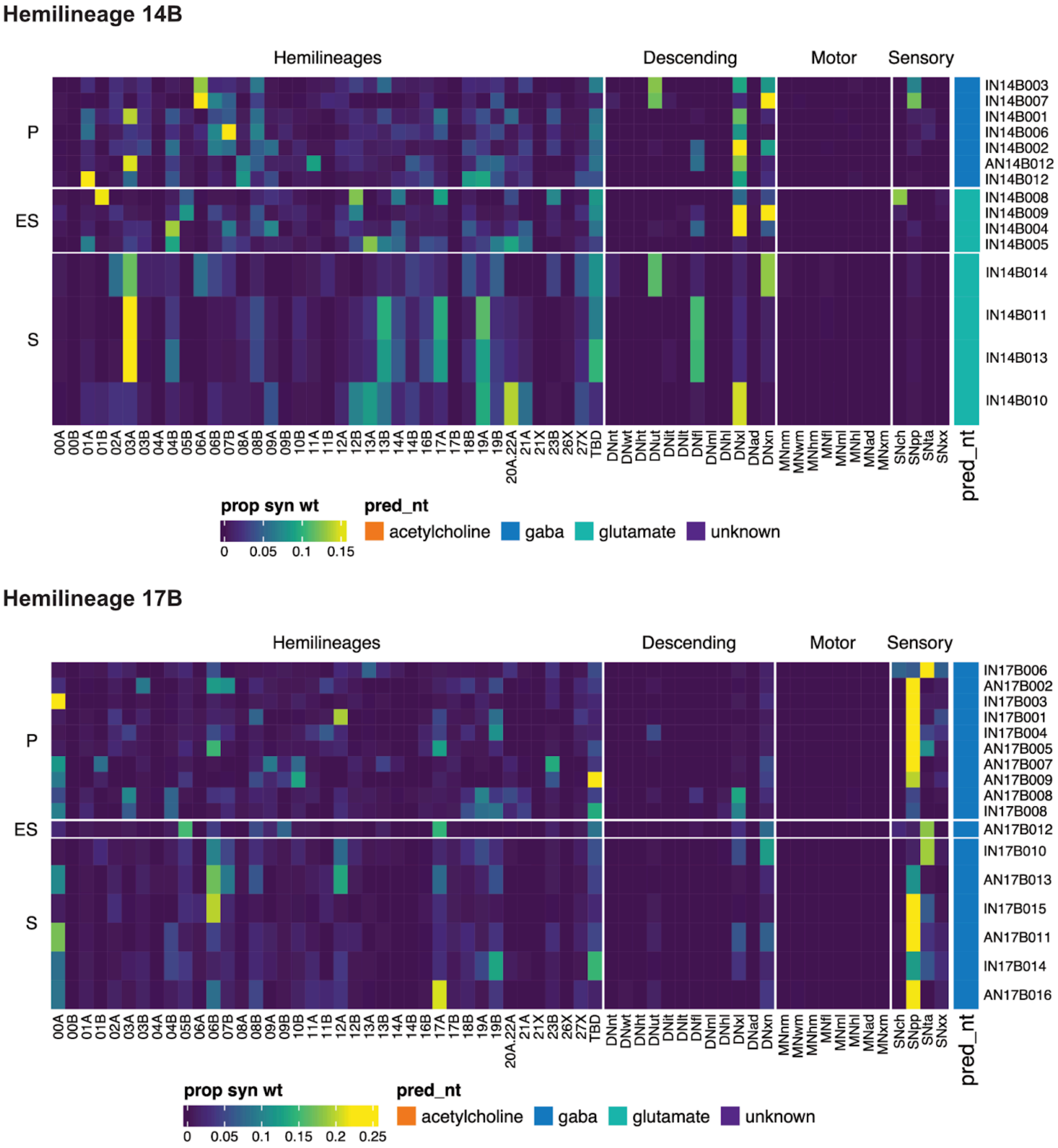

**Figure 48 - figure supplement 21. Connectivity to upstream partners by 14B and 17B systematic types.** Proportions of synaptic weight to systematic types from upstream partners, normalised by row. 14B and 17B have been clustered within each assigned birthtime window (P = primary, ES = early secondary, S = secondary) based on both upstream and downstream connectivity to hemilineages, descending neuron subclasses, motor neuron subclasses, and sensory neuron modalities. Annotation bar is coloured by the most common predicted neurotransmitter for the neurons of each type.

Hemilineage 14B

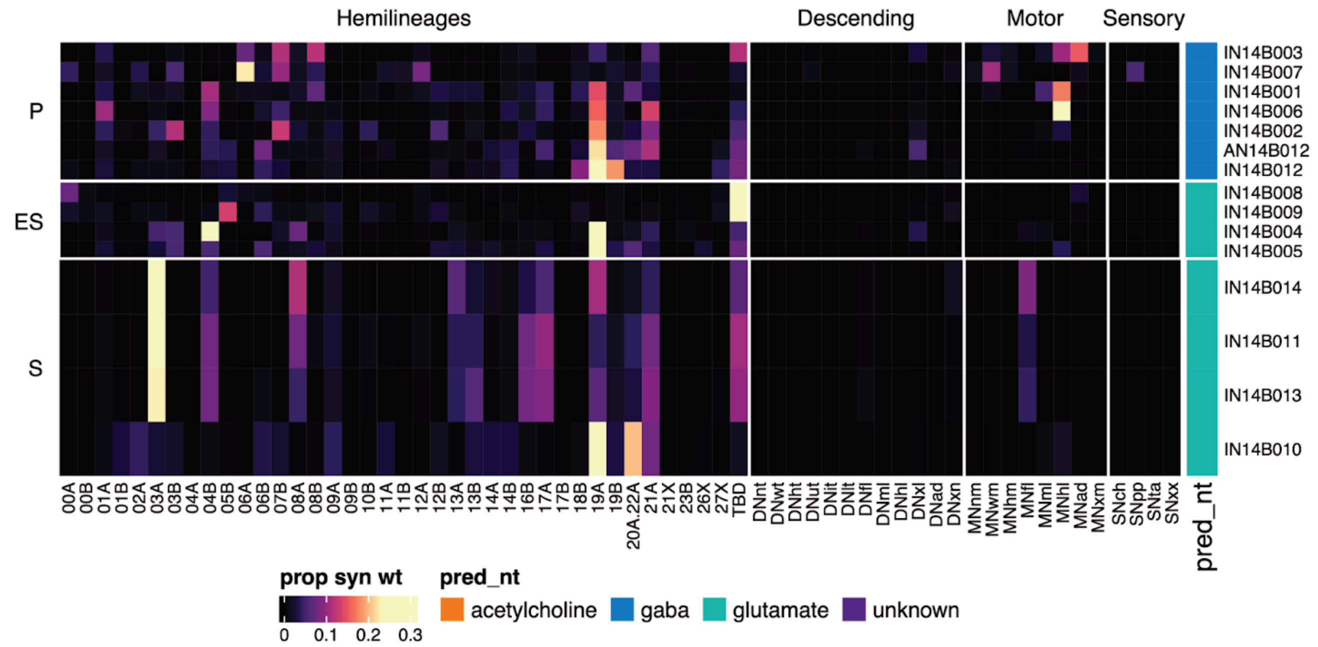

Hemilineage 17B

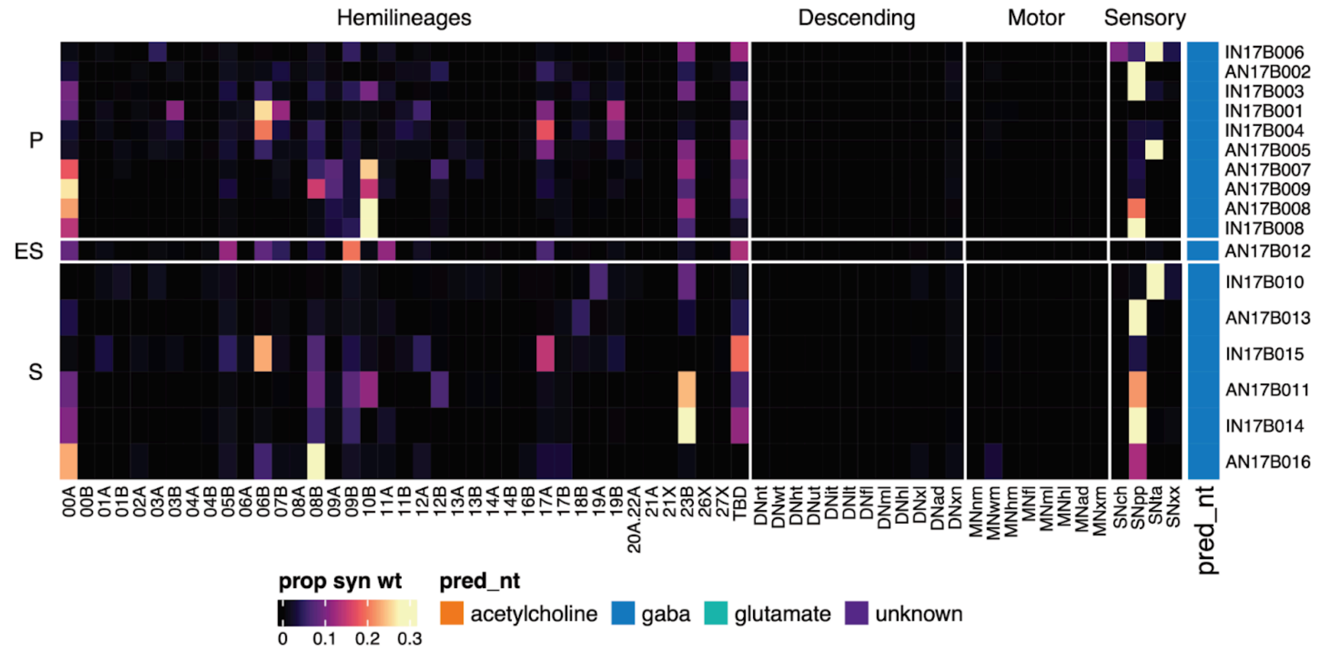

**Figure 48 - figure supplement 22. Connectivity to downstream partners by 14B and 17B systematic types.** Proportions of synaptic weight from systematic types to downstream partners, normalised by row. 14B and 17B neurons have been clustered within each assigned birthtime window (P = primary, ES = early secondary, S = secondary) based on both upstream and downstream connectivity to hemilineages, descending neuron subclasses, motor neuron subclasses, and sensory neuron modalities. The annotation bar is coloured by the most common predicted neurotransmitter within each type.

#### 17X systematic types

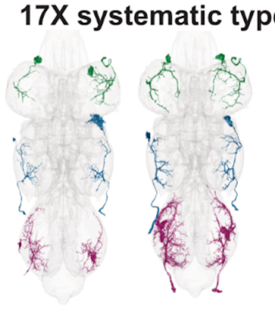

13434 18006

#### 18X systematic types

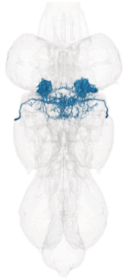

MNwm19

#### 20B.21B.22B systematic types

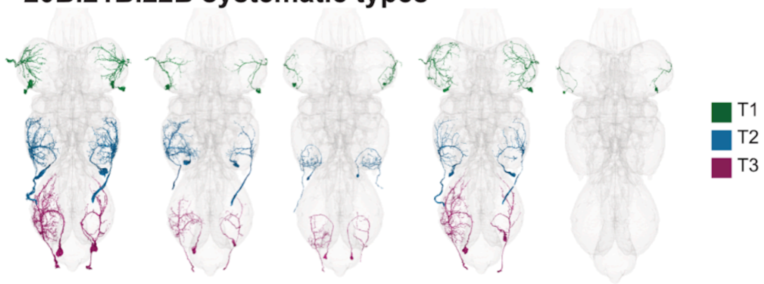

11400 11529 17137 21241 MNfl58

**Figure 48 - figure supplement 23. Systematic types of hemilineages 17X, 18X, and 20B.21B.22B.** Motor neurons (typed separately in (Cheong et al., 2023)) have been plotted by serial set if identified in multiple neuromeres and by systematic type if not. Individual motor neuron meshes have been coloured based on soma neuromere: dark green = T1, blue = T2, magenta = T3.

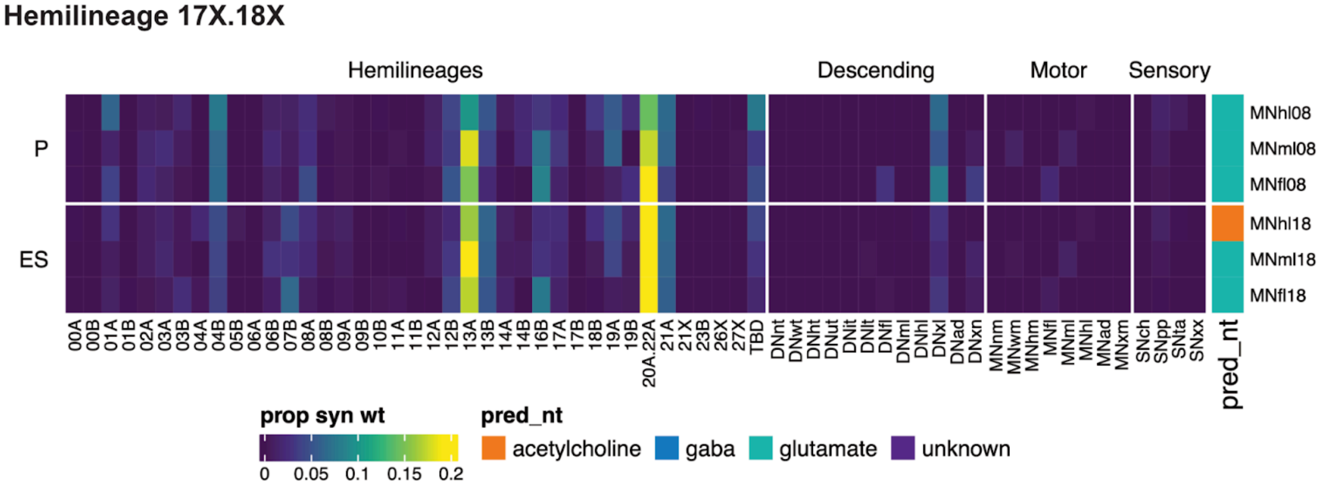

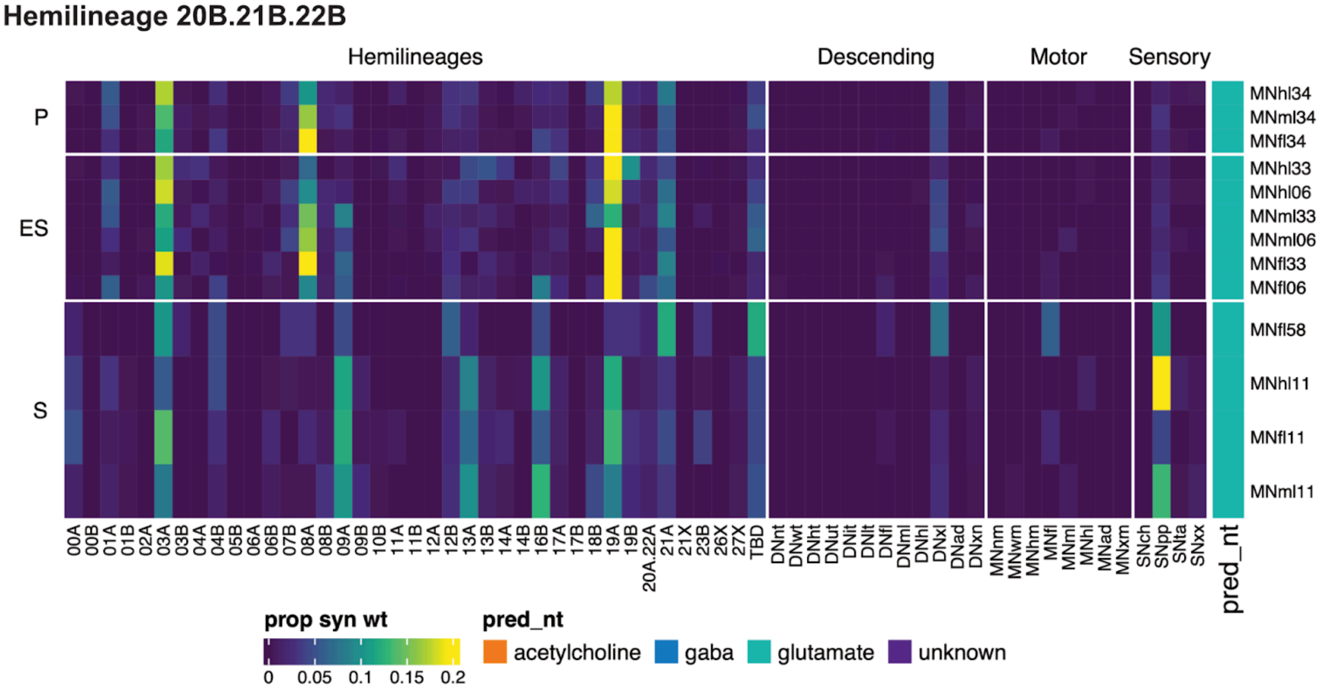

**Figure 48 - figure supplement 24. Connectivity to upstream partners by 17X, 18X, and 20B.21B.22B systematic types.** Proportions of synaptic weight to systematic types from upstream partners, normalised by row. 17X, 18X, and 20B.21B.22B neurons have been clustered within each assigned birthtime window (P = primary, ES = early secondary, S = secondary) based on both upstream and downstream connectivity to hemilineages, descending neuron subclasses, motor neuron subclasses, and sensory neuron modalities. Annotation bar is coloured by the most common predicted neurotransmitter for the neurons of each type.

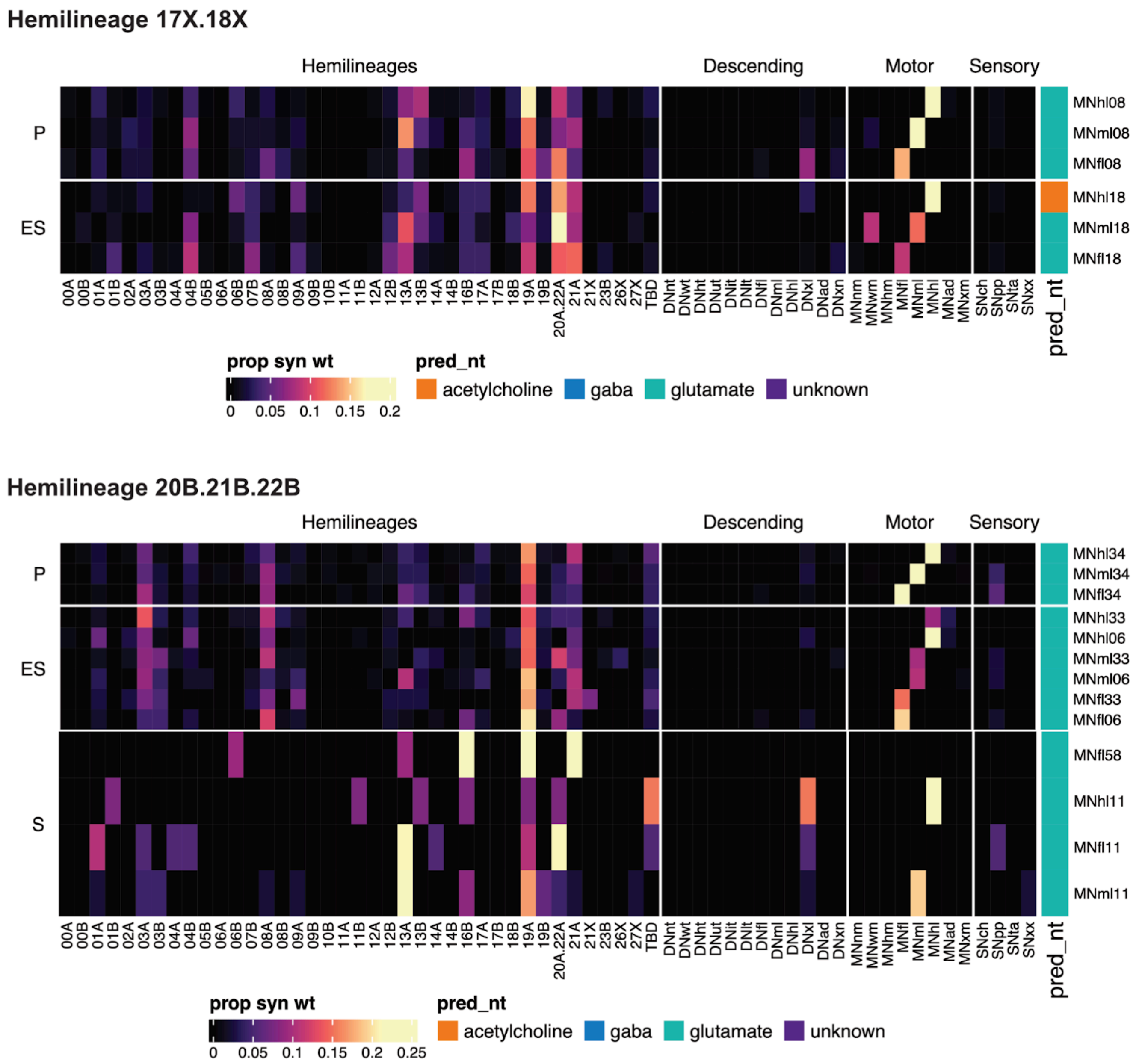

**Figure 48 - figure supplement 25. Connectivity to downstream partners by 17X, 18X, and 20B.21B.22B systematic types.** Proportions of synaptic weight from systematic types to downstream partners, normalised by row. 17X, 18X, and 20B.21B.22B neurons have been clustered within each assigned birthtime window (P = primary, ES = early secondary, S = secondary) based on both upstream and downstream connectivity to hemilineages, descending neuron subclasses, motor neuron subclasses, and sensory neuron modalities. The annotation bar is coloured by the most common predicted neurotransmitter within each type.

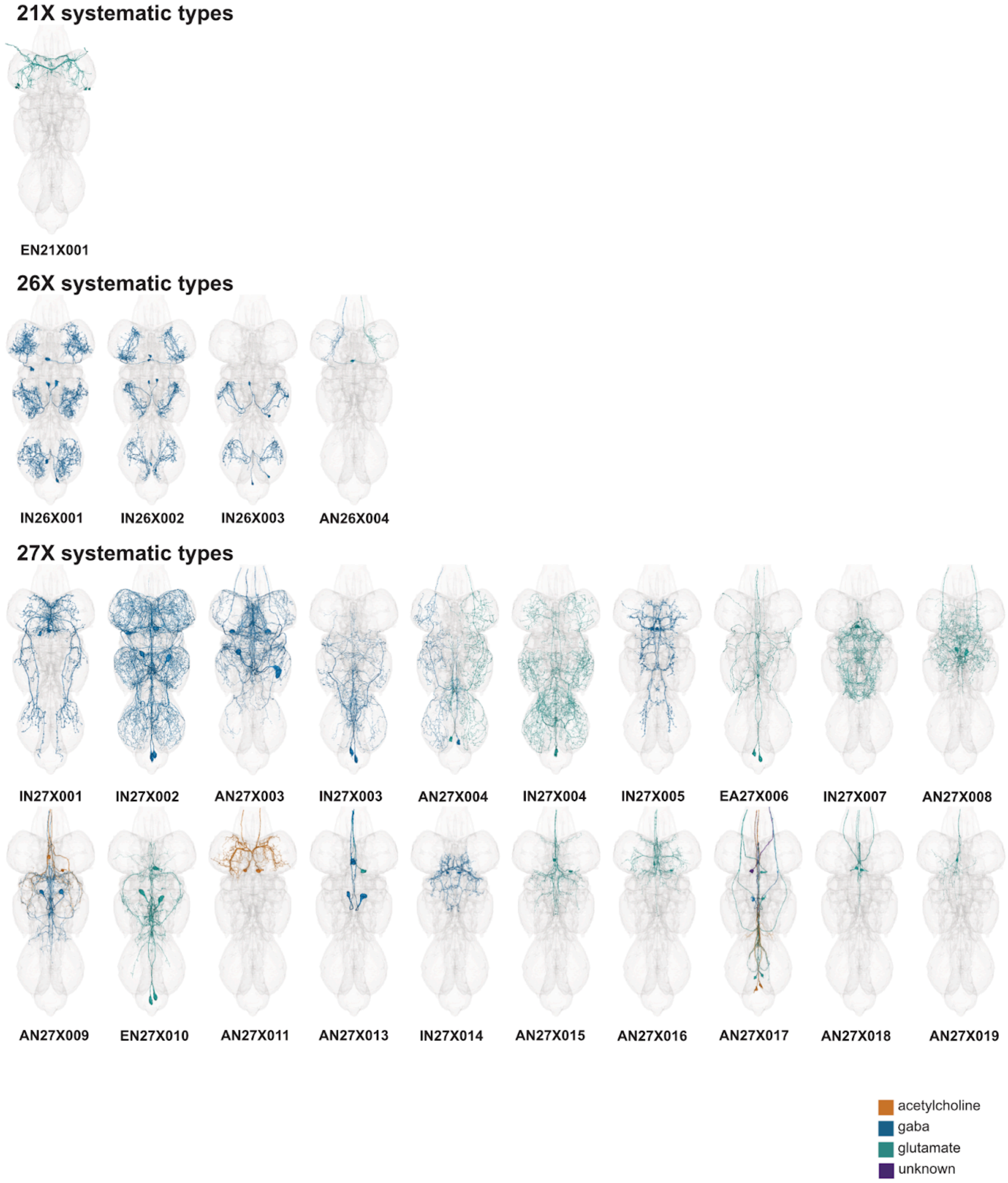

**Figure 48 - figure supplement 26. Systematic types of hemilineages 21X, 26X, and 27X.** Systematic types have been arranged in numerical order, with neurons of the same type that belong to distinct classes (e.g., intrinsic neuron vs ascending neuron) plotted separately but placed adjacent to each other. Individual neuron meshes have been coloured based on their predicted neurotransmitters: dark orange = acetylcholine, blue = gaba, marine = glutamate, dark purple = unknown.

**Hemilineage 26X**

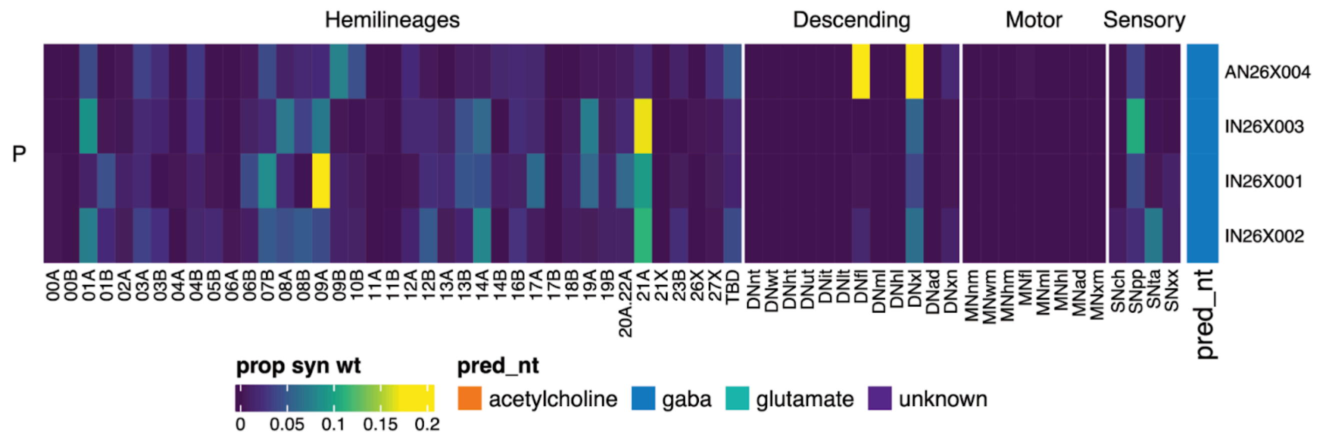

**Hemilineage 27X**

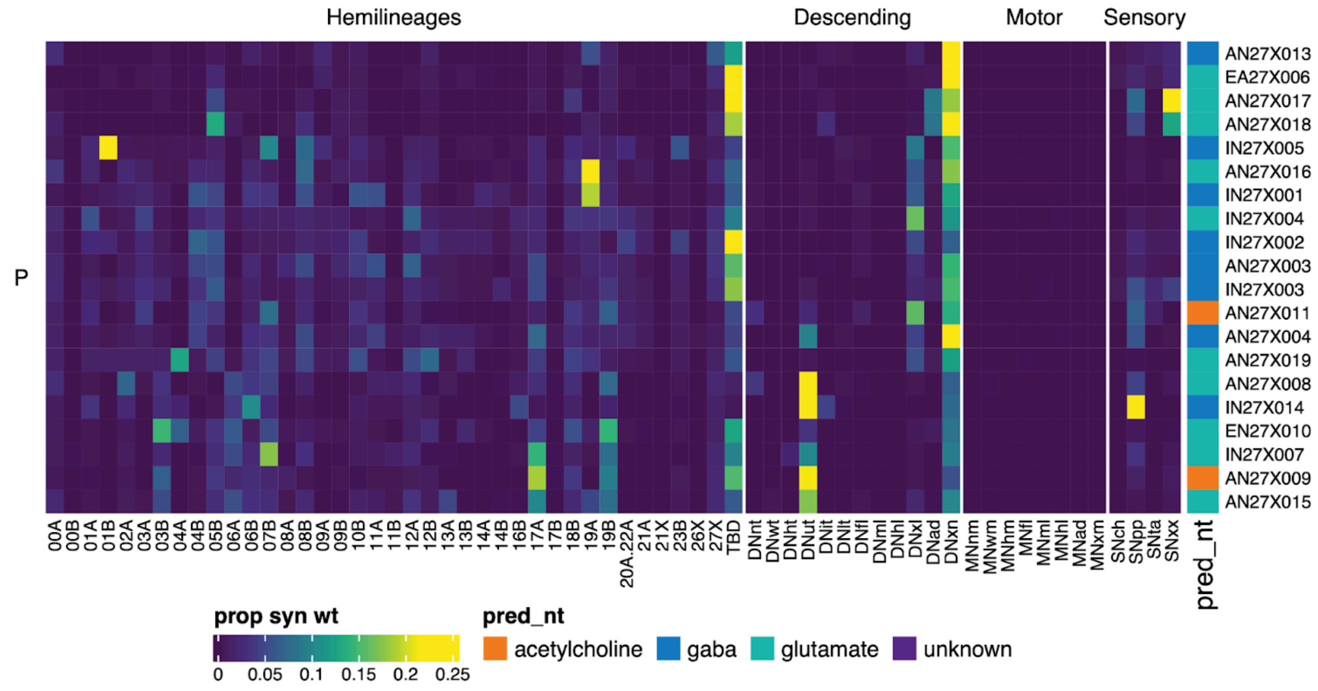

**Figure 48 - figure supplement 27. Connectivity to upstream partners by 26X and 27X systematic types.** Proportions of synaptic weight to systematic types from upstream partners, normalised by row. 26X and 27X neurons have been clustered within each assigned birthtime window (P = primary, ES = early secondary, S = secondary) based on both upstream and downstream connectivity to hemilineages, descending neuron subclasses, motor neuron subclasses, and sensory neuron modalities. Annotation bar is coloured by the most common predicted neurotransmitter for the neurons of each type.

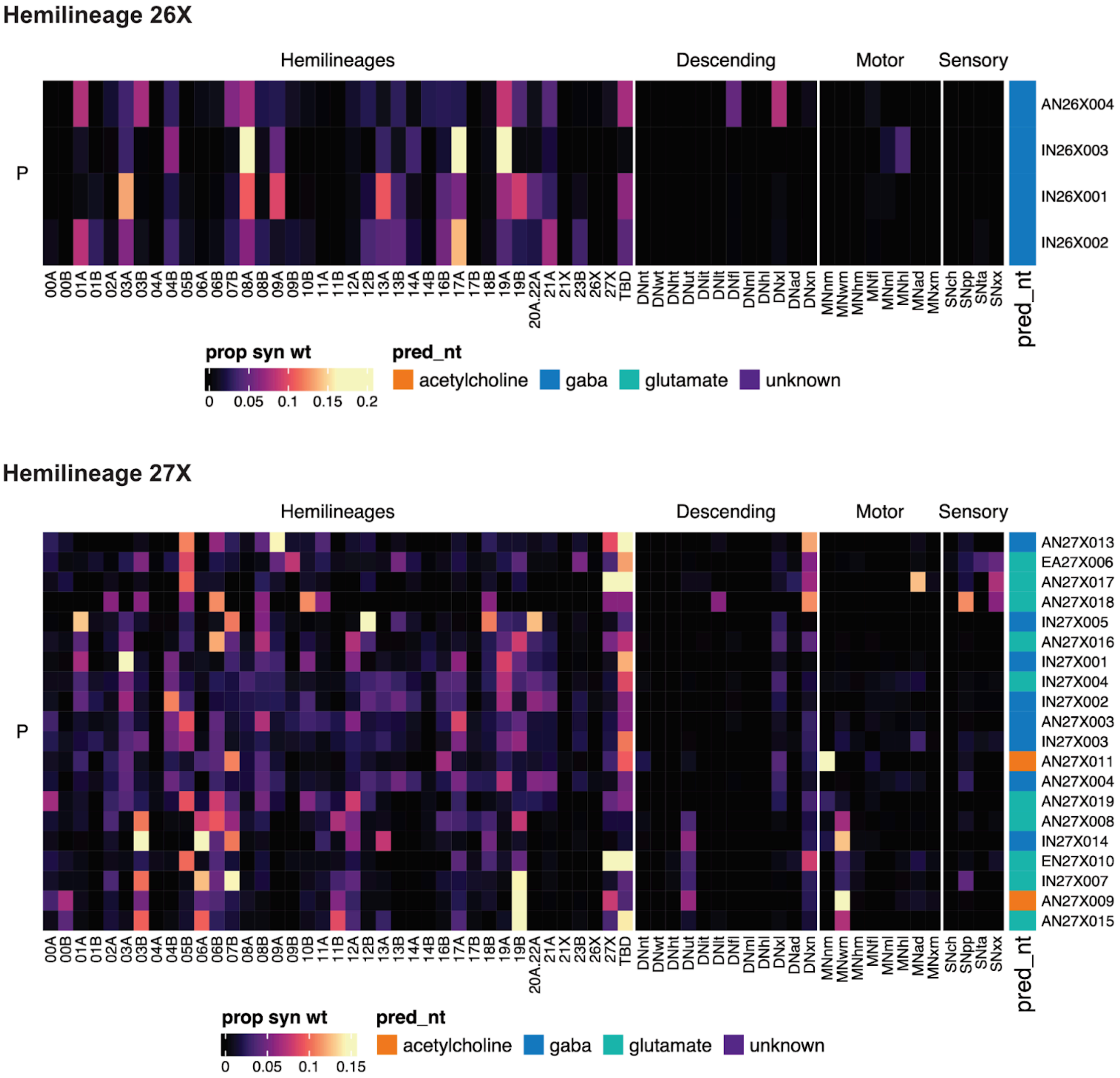

**Figure 48 - figure supplement 28. Connectivity to downstream partners by 26X and 27X systematic types.** Proportions of synaptic weight from systematic types to downstream partners, normalised by row. 26X and 27X neurons have been clustered within each assigned birthtime window (P = primary, ES = early secondary, S = secondary) based on both upstream and downstream connectivity to hemilineages, descending neuron subclasses, motor neuron subclasses, and sensory neuron modalities. The annotation bar is coloured by the most common predicted neurotransmitter within each type.

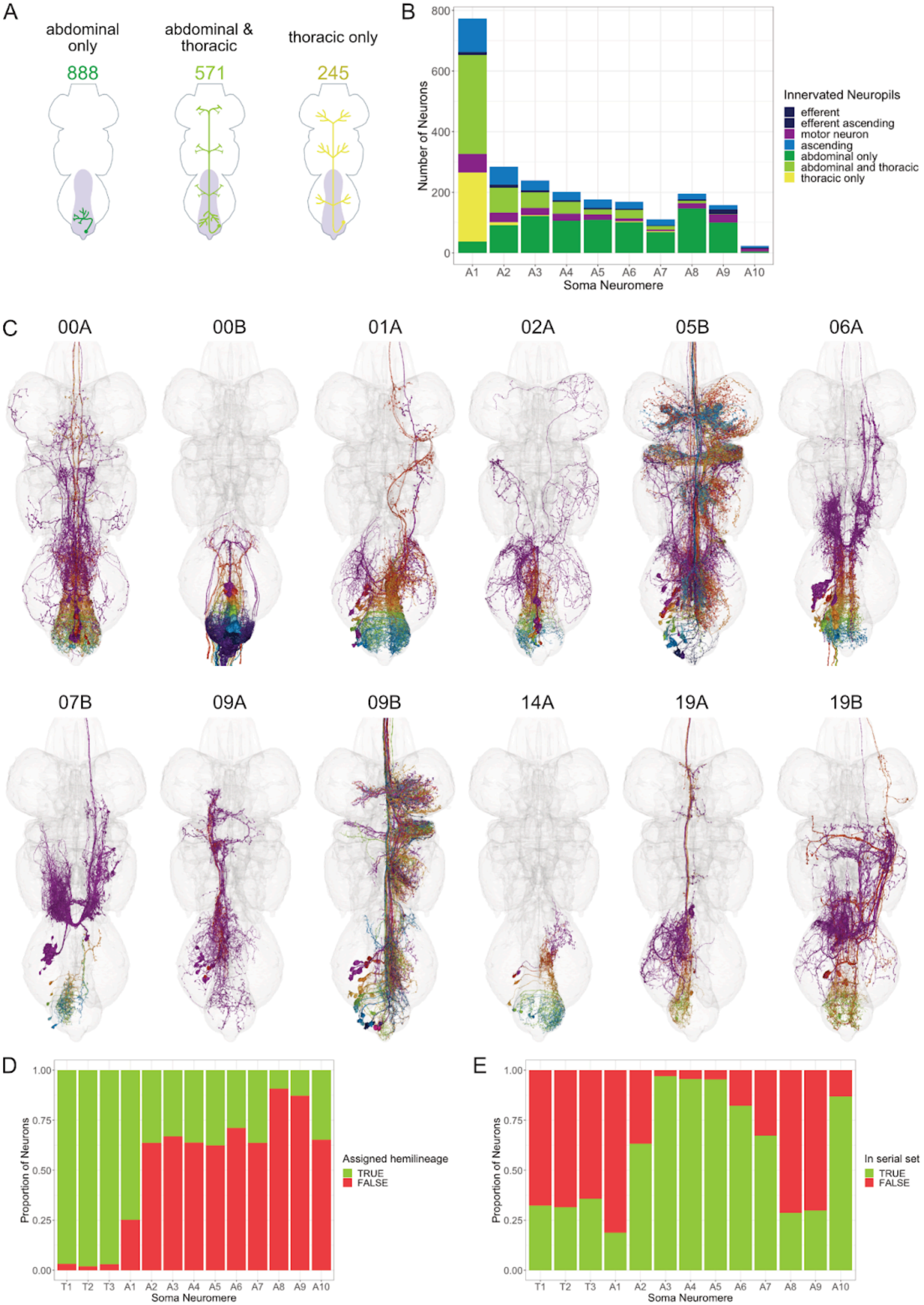
A
abdominal only
888
abdominal & thoracic
571
thoracic only
245
B
Number of Neurons
800
600
400
200
0
A1 A2 A3 A4 A5 A6 A7 A8 A9 A10
Soma Neuromere
Innervated Neuropils
efferent
efferent ascending
motor neuron
ascending
abdominal only
abdominal and thoracic
thoracic only
C
00A
00B
01A
02A
05B
06A
07B
09A
09B
14A
19A
19B
D
Proportion of Neurons
1.00
0.75
0.50
0.25
0.00
T1 T2 T3 A1 A2 A3 A4 A5 A6 A7 A8 A9 A10
Soma Neuromere
Assigned hemilineage
TRUE
FALSE
E
Proportion of Neurons
1.00
0.75
0.50
0.25
0.00
T1 T2 T3 A1 A2 A3 A4 A5 A6 A7 A8 A9 A10
Soma Neuromere
In serial set
TRUE
FALSE

**Figure 49. Organisation of abdominal ganglia. A.** Cartoon schematic depicting VNC innervation by neurons originating in abdominal neuromeres. Abdominal neurons have no more than 5 synapses outside of the ANm, abdominal & thoracic neurons have at least 5% of synapses in the ANm and more than 5 synapses outside of the ANm, and thoracic neurons have less than 5% of synapses in the ANm. **B.** Quantification of abdominal neuron VNC innervation categories by soma neuromere. **C.** Examples of hemilineages identified in abdominal neuromeres (midline or RHS soma side only). Neuron meshes are coloured by soma neuromere: purple = A1, red = A2, dark orange = A3, dark yellow = A4, green = A5, cyan = A6, blue = A7, navy = A8, dark purple = A9, magenta = A10. **D.** Proportion of neurons assigned to a hemilineage for each soma neuromere. **E.** Proportion of neurons assigned to a serial set for each soma neuromere.

A

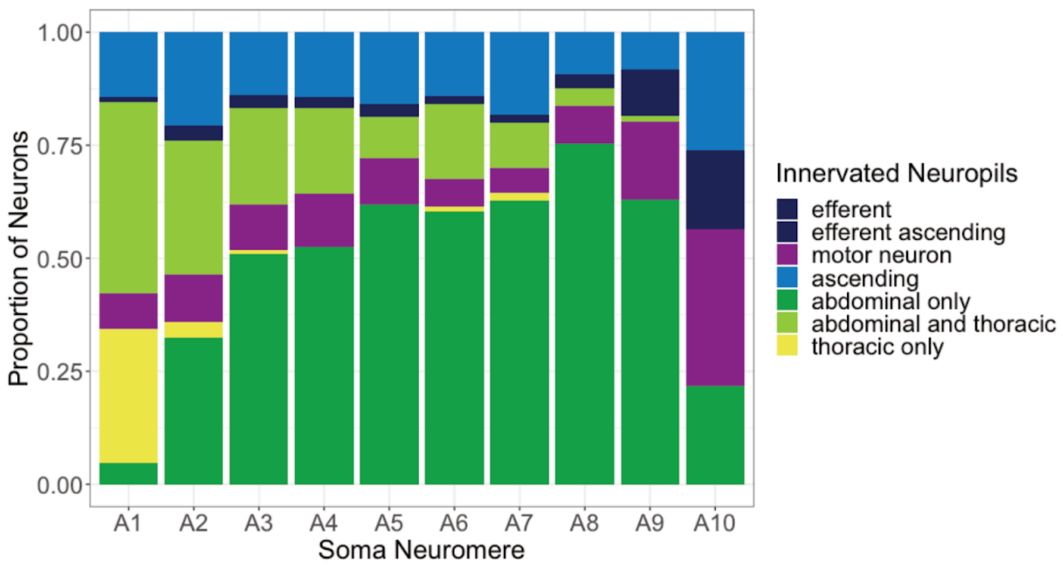

**Figure 49 - figure supplement 1.** Proportion of neurons originating in each abdominal neuromere that innervate the abdominal (dark green), thoracic (yellow), or abdominal and thoracic neuromeres (light green).

#### Hemilineage diversity

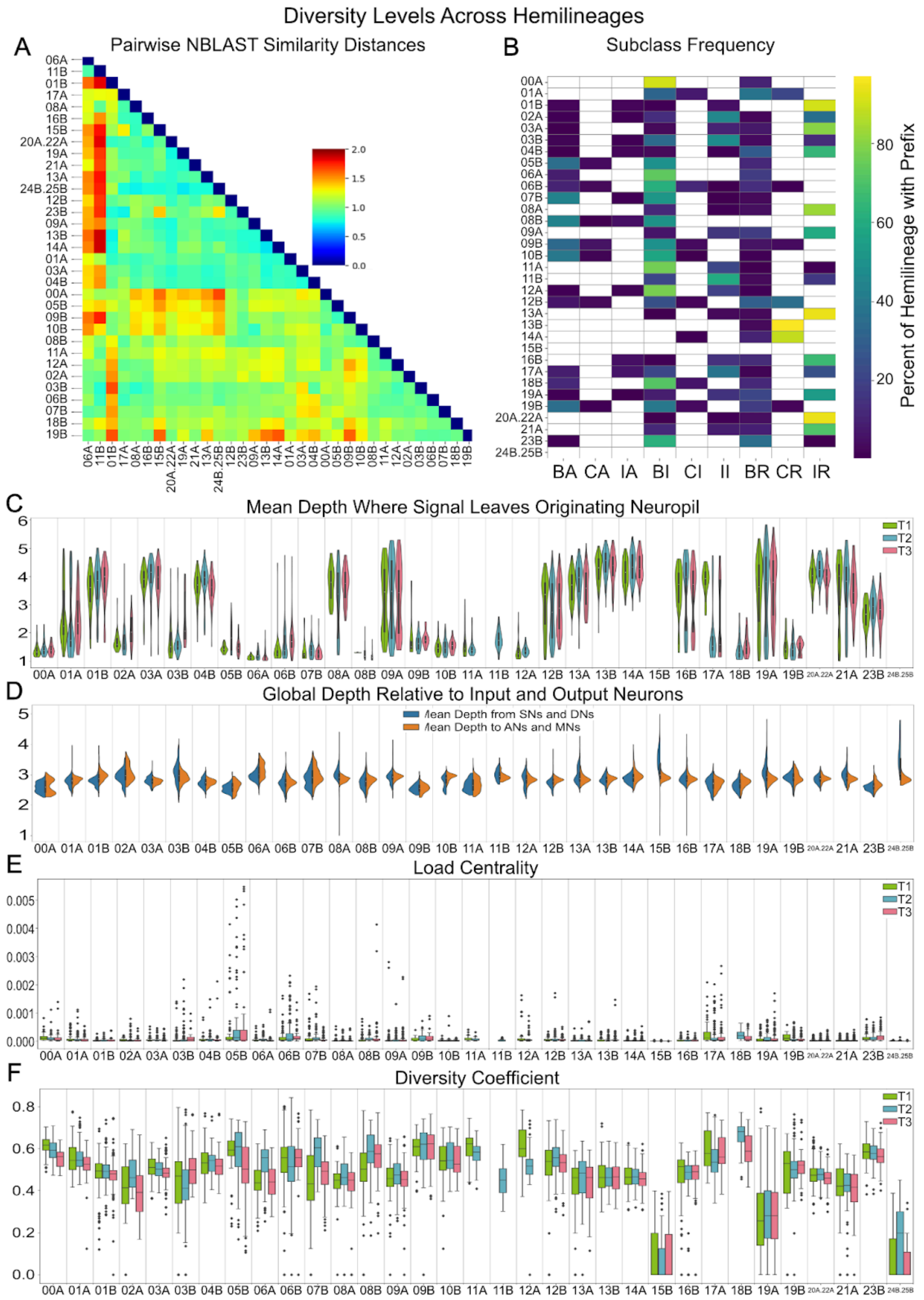
Diversity Levels Across Hemilineages
A
Pairwise NBLAST Similarity Distances
B
Subclass Frequency
Percent of Hemilineage with Prefix
BA CA IA BI CI II BR CR IR
C
Mean Depth Where Signal Leaves Originating Neuropil
T1
T2
T3
D
Global Depth Relative to Input and Output Neurons
Mean Depth from SNs and DNs
Mean Depth to ANs and MNs
E
Load Centrality
F
Diversity Coefficient
00A 01A 01B 02A 03A 03B 04B 05B 06A 06B 07B 08A 08B 09A 09B 10B 11A 11B 12A 12B 13A 13B 14A 15B 16B 17A 18B 19A 19B 20A.22A 21A 23B 24B.25B

**Figure 50. Diversity levels by hemilineage.** Data is shown for secondary neurons in T1, T2, and T3 with a hemilineage assignment. **A.** Morphologic similarity between hemilineages (only considering secondary neurons). NBLAST (Costa et al., 2016) score was calculated as the mean of all distance scores (from 0 to 2) between each neuron in the query target and its most similar match in the target hemilineage. **B.** Percentage of neurons in each hemilineage annotated with each respective subclass. White cell indicates no neurons annotated with respective prefix within hemilineage. Hemilineages 15B and and 24B.25B are motor neuron only hemilineages. **C.** Depth at which a signal leaves the originating neuropil, or else reaches a motor or ascending neuron. Higher values indicate a signal is likely to spend more time in its respective neuropil before exiting. **D.** Mean global shortest path distance to and from all output and input neurons, respectively. Calculated via a breadth-first search. **E.** Load centrality (Newman, 2001) distributions of all secondary neurons in each hemilineage, subdivided by neuromere. Load centrality is the fraction of all shortest paths between each pair of neurons that go through a given node. **F.** Diversity coefficient (inspired by (Eagle et al., 2010), except with weights to hemilineage-in-a-hemineuromere communities instead of individual nodes) distributions for all secondary neurons in each hemilineage, subdivided by neuromere. Diversity coefficient was calculated via BCT (Rubinov and Sporns, 2010). Data shown here were measured via iGraph (Csárdi and Nepusz, 2006) and NetworkX (Hagberg et al., 2008) graph implementations of the VNC.

Position and role in the VNC network also differ by birthtime. Despite there being far fewer primary neurons than secondary neurons, primary neurons are more centrally positioned within the VNC network (Figure 51A). Primary neurons are markedly more likely to be involved in the shortest path between pairs of neurons, an important measure of centrality called load centrality (Newman, 2001). Conversely, primary neurons are less insular than their secondary counterparts, connecting more abundantly and evenly with other hemilineages across the VNC (Figure 51A’). Their centrality in the network is reflected in their overrepresentation in the rich-club relative to their population (Figure 51B). In a network, a rich-club is a set of high-centrality nodes, here measured by load centrality, that are more strongly interconnected than other neurons (Colizza et al., 2006). Secondary neurons also tend to be further from input and output neurons (Figure 51C), and signals received by them take longer to leave their originating neuropil (Figure 51D). Overall, earlier born neurons seem to play more central and global roles than their later born counterparts.

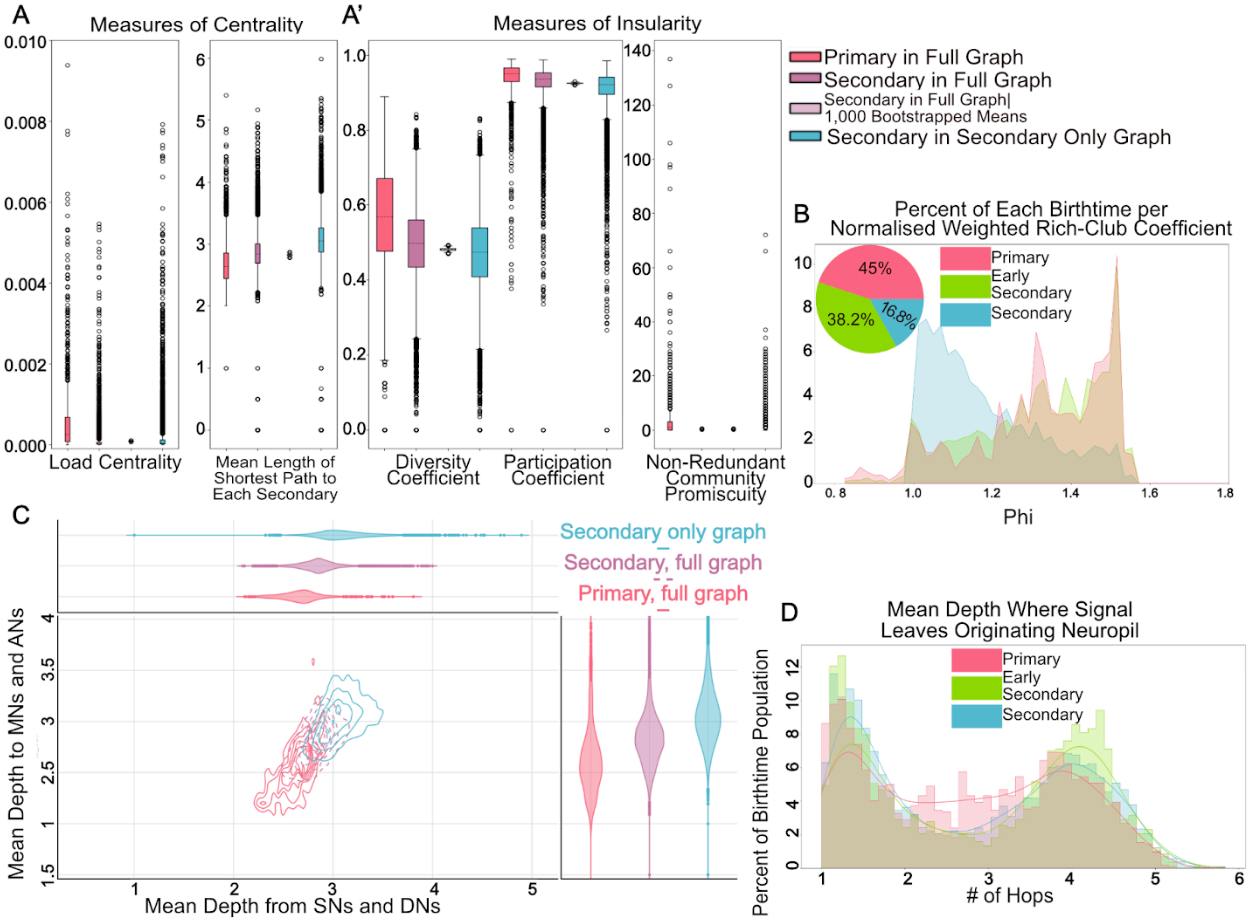

**Figure 51. Roles in the VNC network differ by birthtime. A-D**. Data is shown only for VNC-origin neurons in T1, T2 and T3. **A-A'.** Key measures of centrality and insularity within the network, compared between primary and secondary neurons in the full graph, as well as in a network excluding primary intrinsic neurons. One community is defined as all neurons of a hemilineage with a soma in a hemineuromere (e.g. all neurons identified as hemilineage 19A that have somas in the left side of T2 would be one community). **A.** Key measures of centrality. **A'.** Key measures of community insularity and diversity. Higher values suggest a higher variety of connection to communities. Non-redundant community promiscuity is a measure of how essential an individual neuron is in connecting its community to others in the graph. Primary neurons are uniquely responsible for connecting their respective communities to 1 other community (on median). **B.** Normalised weighted rich-club coefficient across birthtimes. Inset shows the percentage occurrence of the three birthtimes in the rich-club. **C.** Contour plot of depth from and to input neurons (all SNs and DNs) and output neurons (all MNs and ANs), respectively, calculated via a breadth-first search. **D.** Distribution of mean depth where a received signal leaves the neuropil at which it originated. Higher values indicate a signal is likely to spend more time in its respective neuropil before exiting. Data shown here were measured via iGraph (Csardi et al., 2006) and NetworkX (Hagberg et al., 2008) graph implementations of the VNC, based on either the full dataset or a subset of the dataset without primary intrinsic neurons (referred to as “secondary only graph” in the figure).

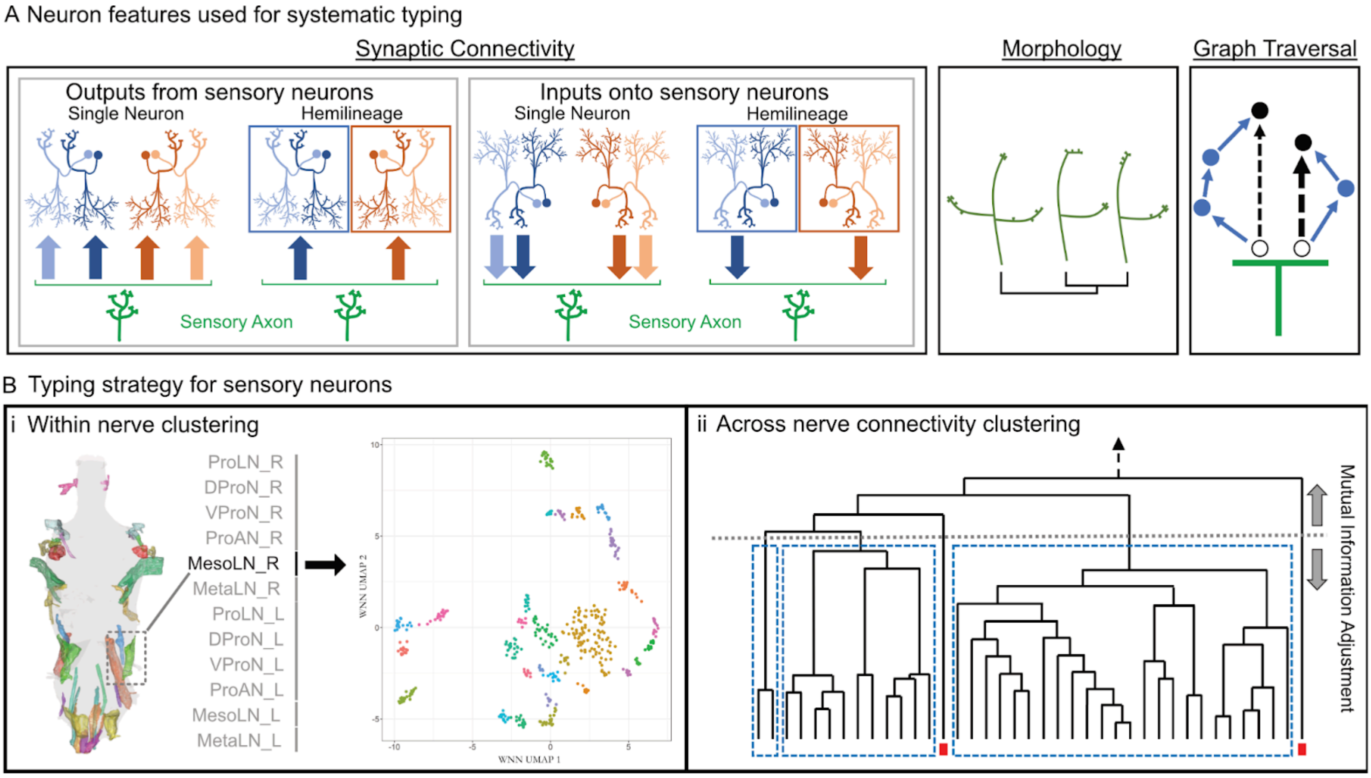

**Figure 52. Systematic typing of reconstructed sensory neurons. A.** Neuron features utilised for sensory neuron typing with a weighted nearest neighbour (WNN) approach included synaptic connectivity (synaptic outputs to individual neurons, synaptic outputs aggregated by hemilineage, synaptic inputs to individual neurons, synaptic inputs aggregated by hemilineage), morphology (inter-sensory neuron NBLAST scores), and graph measures (graph traversal distances). **B.** Depiction of typing process. **i.** Sensory neurons were initially clustered for each peripheral origin individually and individual nerve clusters were detected using HDBSCAN from a UMAP representation of the WNN output (Campello et al., 2013). **ii.** These clusterings were then utilised to inform a cut height of pan-region cosine clustering of synaptic connectivity onto serial neuron groupings.

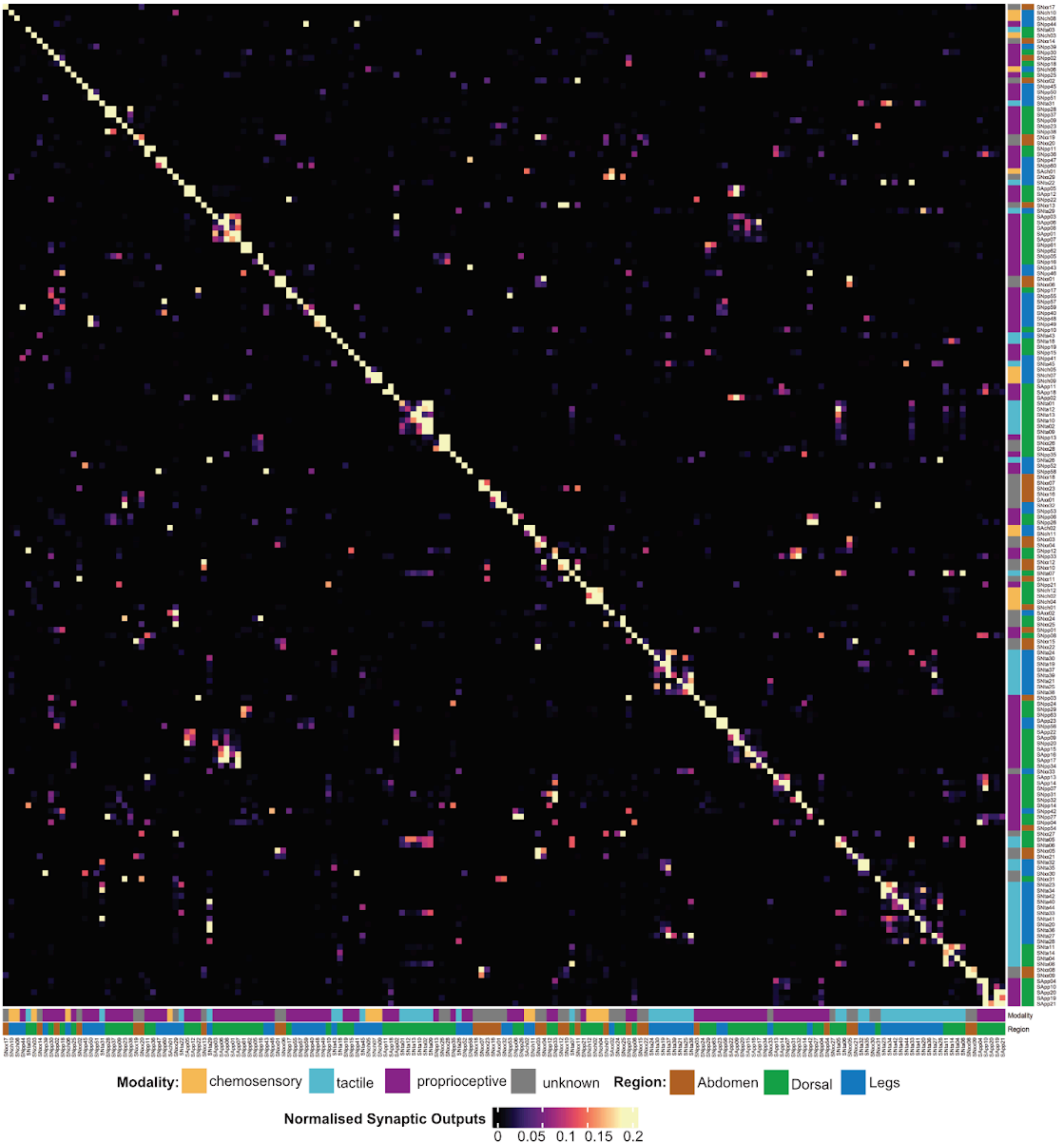

**Figure 52 - figure supplement 1. Sensory cell type interconnectivity.** Annotation bars indicate modality and peripheral region for each type. Rows are the upstream systematic type, with columns being the downstream systematic type.

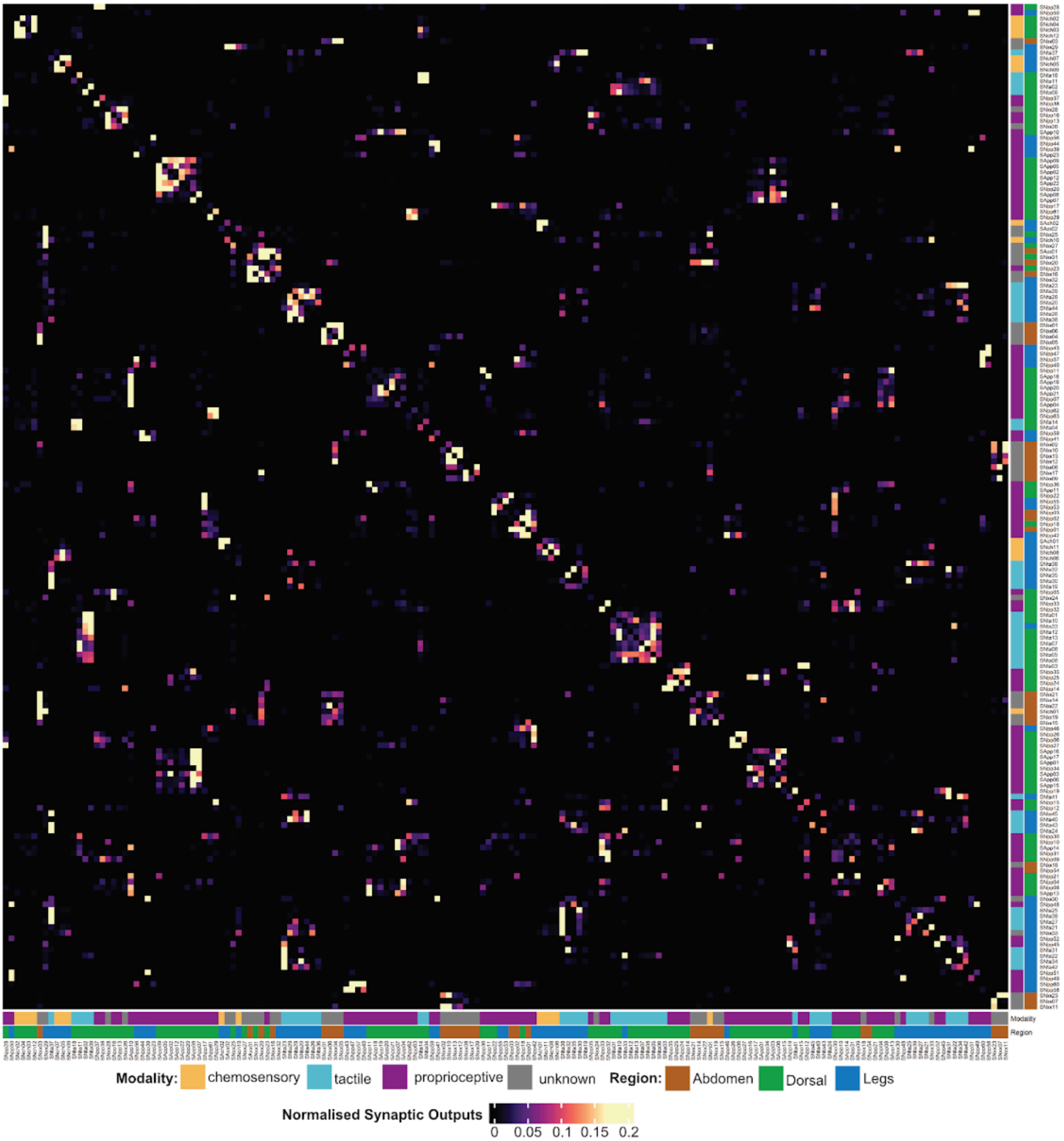

**Figure 52 - figure supplement 2. Sensory cell type interconnectivity, excluding intra-type connectivity.** Annotation bars indicate modality and peripheral region for each type. Rows are the upstream systematic type, with columns being the downstream systematic type.

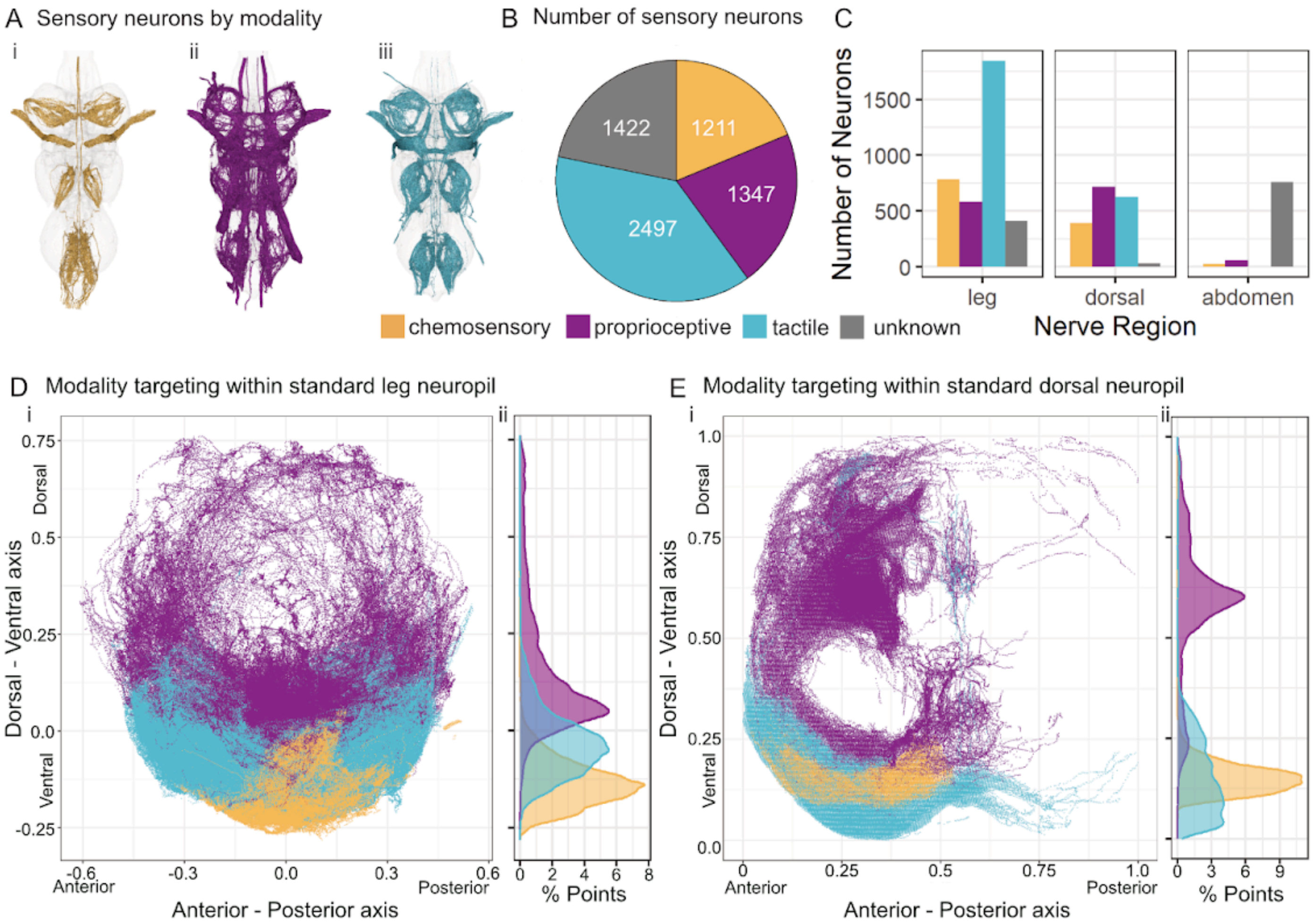

**Figure 53. Overview of sensory neurons by modality. A.** Plotted meshes of i. chemosensory neurons, ii. proprioceptive neurons, and iii. tactile neurons. **B.** Quantification of sensory neurons by assigned modality. C. Quantification of sensory neurons by modality, faceted by peripheral origin. **D.** i. Standard leg neuropil depiction, see Methods, of the regions innervated by sensory neurons of each modality. ii. Histogram showing the relative distribution of neuron innervation through the dorsal ventral axis. **E.** i. Standard dorsal neuropil depiction, see methods, of the regions innervated by sensory neurons of each modality. ii. Histogram showing the relative distribution of neuron innervation through the dorsal ventral axis.

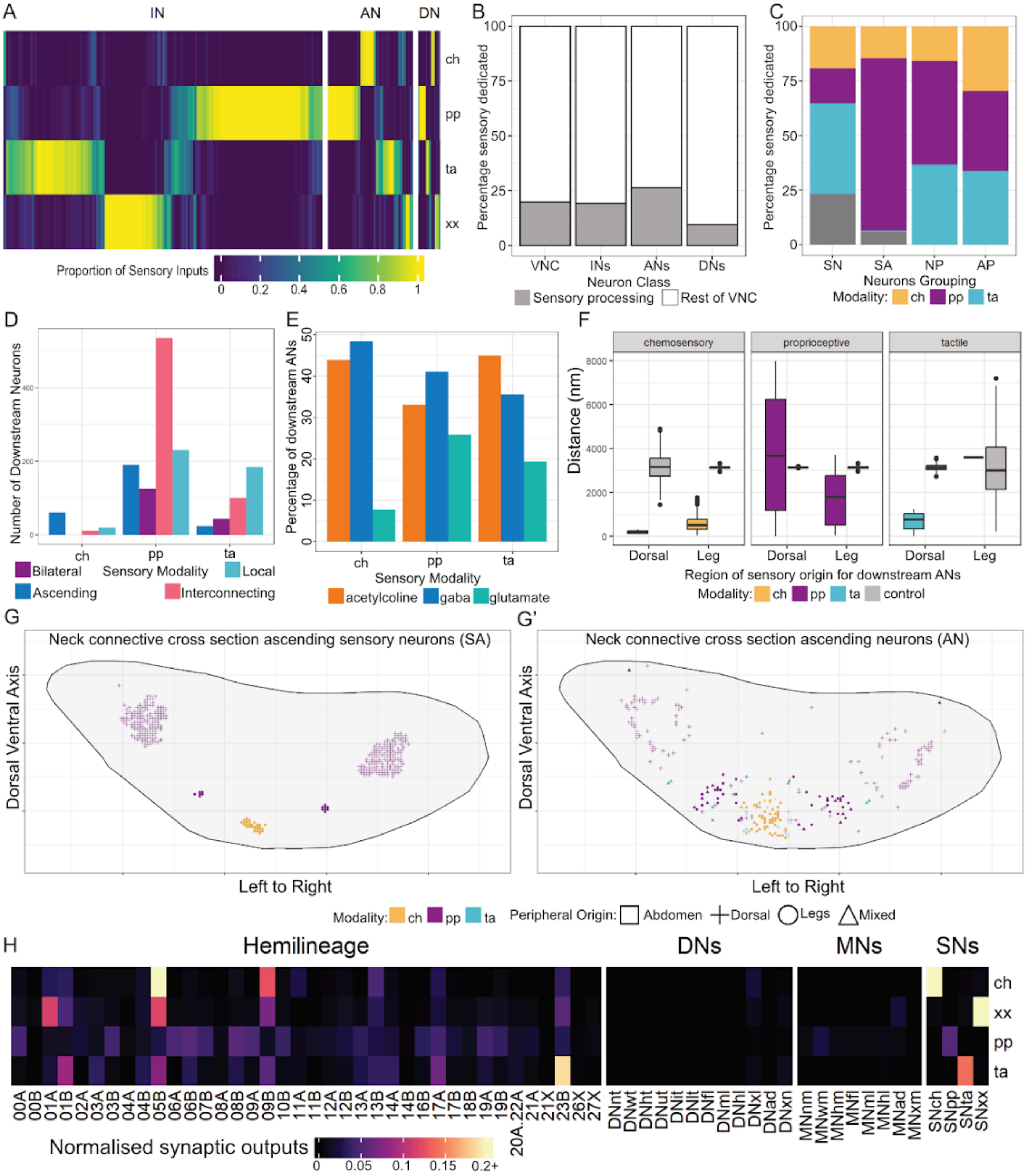

**Figure 54. Overview of sensory neuron targets by modality. A.** Proportion of inputs from sensory modalities to neurons implicated in direct sensory processing. **B.** Percentage of neurons from each class identified as directly sensory processing. VNC neurons = all VNC neurons (excluding sensory neurons), INs = intrinsic neurons, ANs = ascending neurons, and DNs = descending neurons. **C.** Proportion of neurons assigned to each modality for SN, sensory neurons, SA, sensory ascending neurons, NP, non ascending modality specific partners, AP, ascending modality specific partners. **D.** Number of downstream partners of each modality aggregated by expanded subclass: ascending (ANs), bilateral (connecting both hemispheres of the VNC), interconnecting (projecting across multiple neuromeres within a single hemisphere), and local (restricted to the hemi-neuromere associate to the entry nerve). **E.** Percentage breakdown for each modality of predicted neurotransmitter for ascending neurons implicated in direct sensory processing. **F.** Distance within a neck connective plane between neurons of the same region and modality vs.

a randomised control of all ascending neurons. **G.** Cross-section of the neck connective displaying the positioning of ascending sensory neurons. Coloured by modality. **G'.** Cross-section of the neck connective displaying the positioning of ascending partners of sensory neurons. Coloured by modality. **H.**Heatmap showing normalised sensory neuron modality connectivity to hemilineage, DN groups, MN groups, and SN modalities in the VNC. Connectivity to neurons of undetermined hemilineage (TBD) have been omitted.

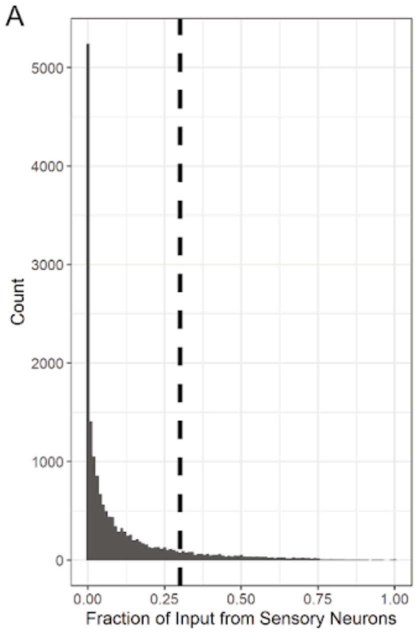

**Figure 54 - figure supplement 1. Sensory neuron targets.** Histogram of proportion of inputs to VNC neurons from sensory neurons. Dashed line represents the cutoff (0.30) used for classifying neurons as implicated in direct sensory processing.

Further breakdown of the downstream partner classes into coarse subclasses reveals additional diversity in downstream neuron targets (Figure 54D). The primary downstream subclass of proprioceptive neurons are neurons that interconnect thoracic neuromeres, suggesting a role in segmental coordination, but there are substantial bilateral and ascending populations as well. In contrast, chemosensory downstream targets are far more likely to be ascending, with few local or interconnecting examples, suggesting that the majority of chemosensory processing occurs in the brain. In contrast, tactile target neurons are typically local, with additional interconnecting and bilateral populations but with the lowest proportion of ascending partners, supporting a role in reflexive rather than directed behaviours.

Downstream ascending neurons of each modality are predicted to express roughly the same proportions of acetylcholine (excitatory) vs gaba (inhibitory), with slight biases towards gaba for chemosensory and proprioceptive and towards acetylcholine for tactile (Figure 53E). Chemosensory ascending neurons are notably less likely to express glutamate; however, the effects of glutamate depends on receptor expression in downstream partners, which cannot be determined here.

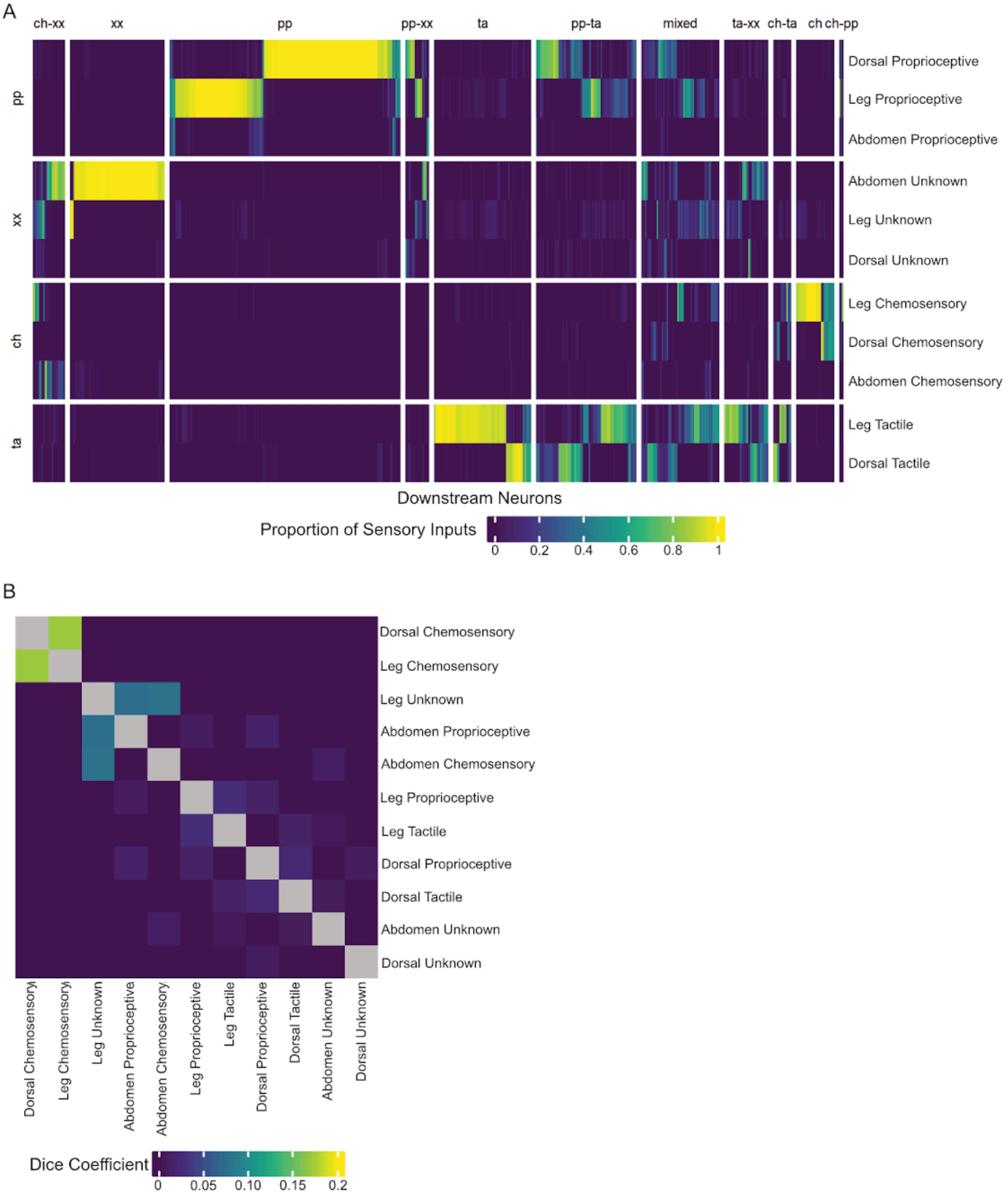

**Figure 54 - figure supplement 2. Sensory neuron targets by modality. A.** Heatmap showing relative specificity of sensory associated downstream neurons with each modality within each region, normalised by total inputs for each downstream neuron. **B.** Dice coefficient for downstream modality within each region.

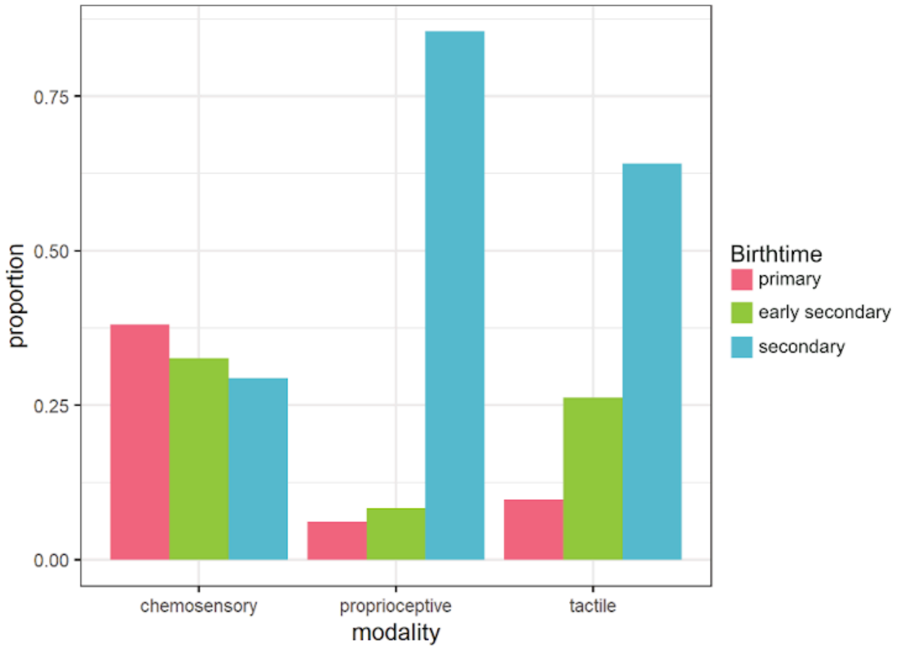

**Figure 54 - figure supplement 3. Proportion of downstream ascending neurons of each birthtime associated with each modality.** Pink = primary, green = early secondary and cyan = secondary.

Ascending sensory information appears to be organised by modality, with ascending sensory neurons of each modality retaining close association with the downstream ascending partners of that modality as they pass through the neck connective (Figure 54F). The pattern of organisation seen in the synapses of each modality in the leg neuropil (Figure 53D) appears to be largely maintained by ascending sensory and partner neurons' axons in the neck connective, with chemosensory axons near the ventral midline, tactile axons adjacent, and proprioceptive axons more dorsal (Figure 54G-G'). These ascending sensory and partner neurons' axons are also organised by peripheral origin, with neurons transmitting information from the leg and notum ascending in the ventromedial neck connective and neurons transmitting it from the neck, wing, and haltere ascending in progressively more dorsal positions (Figure 54 - figure supplement 4). It will be interesting in future to see whether and how the organisation of descending neurons reflects the sensory information they carry.

Finally, we see modality-specific patterns of connectivity when we break down second order neurons by hemilineage or subclass (Figure 54G). For example, chemosensory neurons mainly target hemilineages 05B and 09B, whilst tactile neurons target 01B, 05B, 09B, and especially 23B. Proprioceptive neurons exhibit a broader pattern of connectivity, possibly due to a higher number of discrete peripheral origins. Neurons of unknown modality have a distinct profile of hemilineage connectivity, likely because 1) few neurons in the abdomen (where the majority of unknown sensory neurons originate, Figure 53C) have been assigned to a hemilineage, 2) true connectivity is poorly represented due to incomplete reconstruction, and 3) multiple modalities are represented. Sensory neurons also exhibit a very high degree of intra-modality and low degree of inter-modality connectivity, presumably due to the aforementioned intratype connectivity, as was previously observed in the antennal lobe connectome (Marin et al., 2020).

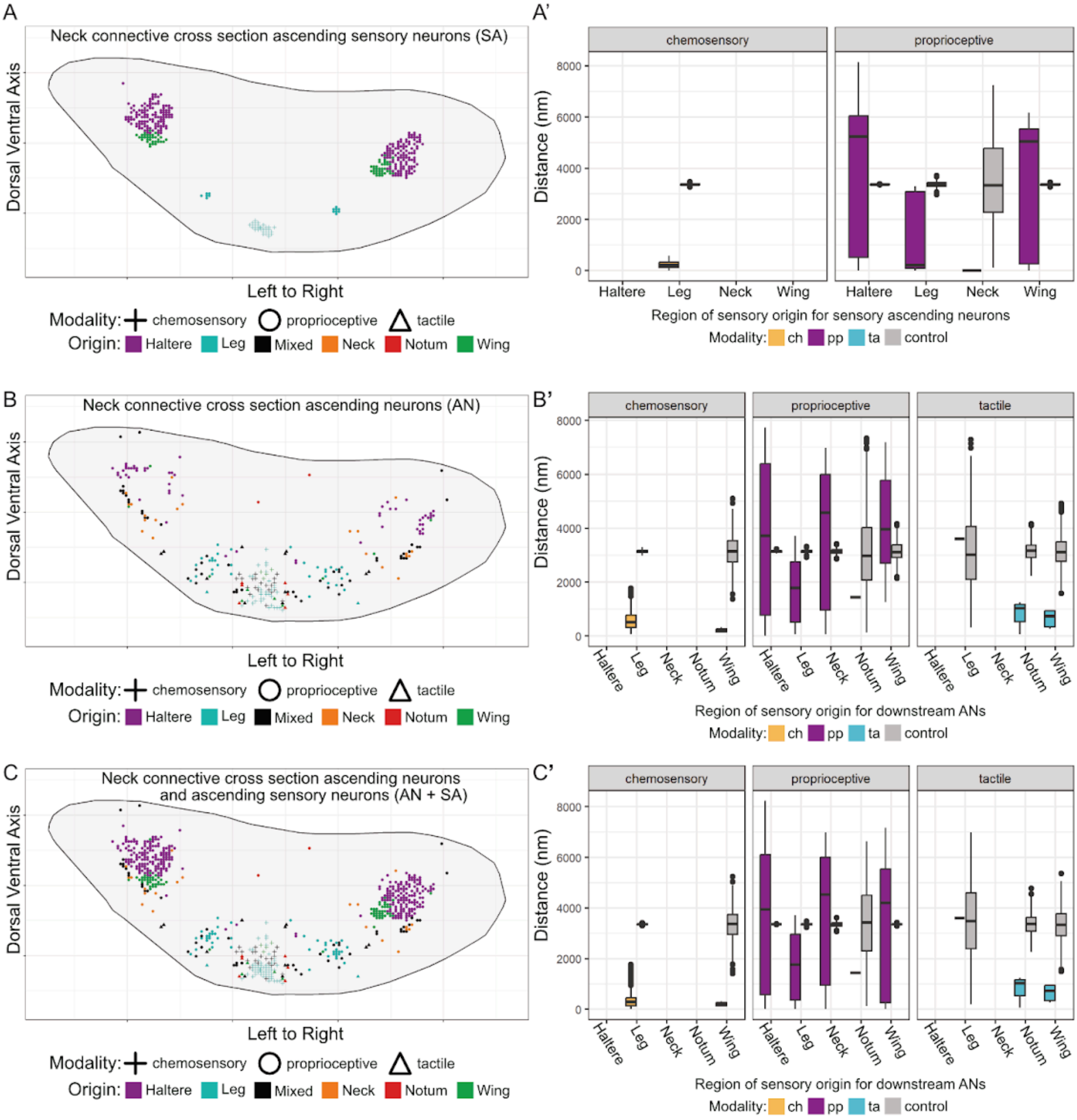

**Figure 54 - figure supplement 4. Targeting of sensory ascending neurons and ascending neurons within the neck connective. A.** Sensory ascending neuron targeting within the neck connective, coloured by region of peripheral origin. **A'.** Distance within the plane between neurons of the same region and modality vs. a randomised control of all sensory ascending neurons. **B.** Ascending neuron targeting within the neck connective, coloured by region of peripheral origin. **B'.** Distance within the plane between neurons of the same region and modality vs. a randomised control of all ascending neurons. **C.** Sensory ascending neuron and ascending neuron targeting within the neck connective, coloured by region of peripheral origin. **C'.** Distance within the plane between neurons of the same region and modality vs. a randomised control of all sensory ascending neurons and ascending neurons.

#### Leg Sensory Types

Sensory neurons from the front legs enter the prothoracic neuromere via four different nerves (ProLN, DProN, VProN, and ProAN), whilst those from the middle and hind legs enter the mesothoracic via the MesoLN and the metathoracic via the MetaLN, respectively (Figure 55A). As described above (Figure 52), neurons from each leg-associated nerve were clustered and then associated across leg nerves by connectivity to previously defined serial sets in order to define serial leg sensory types. Degradation and dark staining resulted in significant variance in the number and quality of sensory neurons recovered in each nerve: the fewest were recovered from the front legs, whilst in general more neurons were recovered on the right side than on the left (Figure 55B). Nevertheless, we successfully reconstructed the majority of leg sensory neurons, surpassing previous efforts (Phelps et al., 2021), and identified 63 distinct cell types (9 chemosensory, 22 proprioceptive, 27 tactile, and 5 of unknown modality). Here we highlight only a subset of these types, but reconstructions and connectivity heatmaps are provided for all (Figure 55 - figure supplement 1-4).

Broadly speaking, leg sensory neurons mainly target intrinsic neurons (~67%), followed by ascending neurons (~22%) and each other (~9%), with other classes receiving 1% or fewer of leg sensory outputs (Figure 55C). This is a significantly smaller proportion of ascending neuron targets than observed for dorsal sensory neurons, perhaps suggesting greater involvement in reflexive sensorimotor circuits that do not require communication with the brain. Leg sensory neurons of each modality tend to target neurons of specific hemilineages: chemosensory neurons mainly target 05B, tactile neurons mainly target 01B and 23B, and proprioceptive neurons have more diverse targets but show a preference for 09A (Figure 55E).

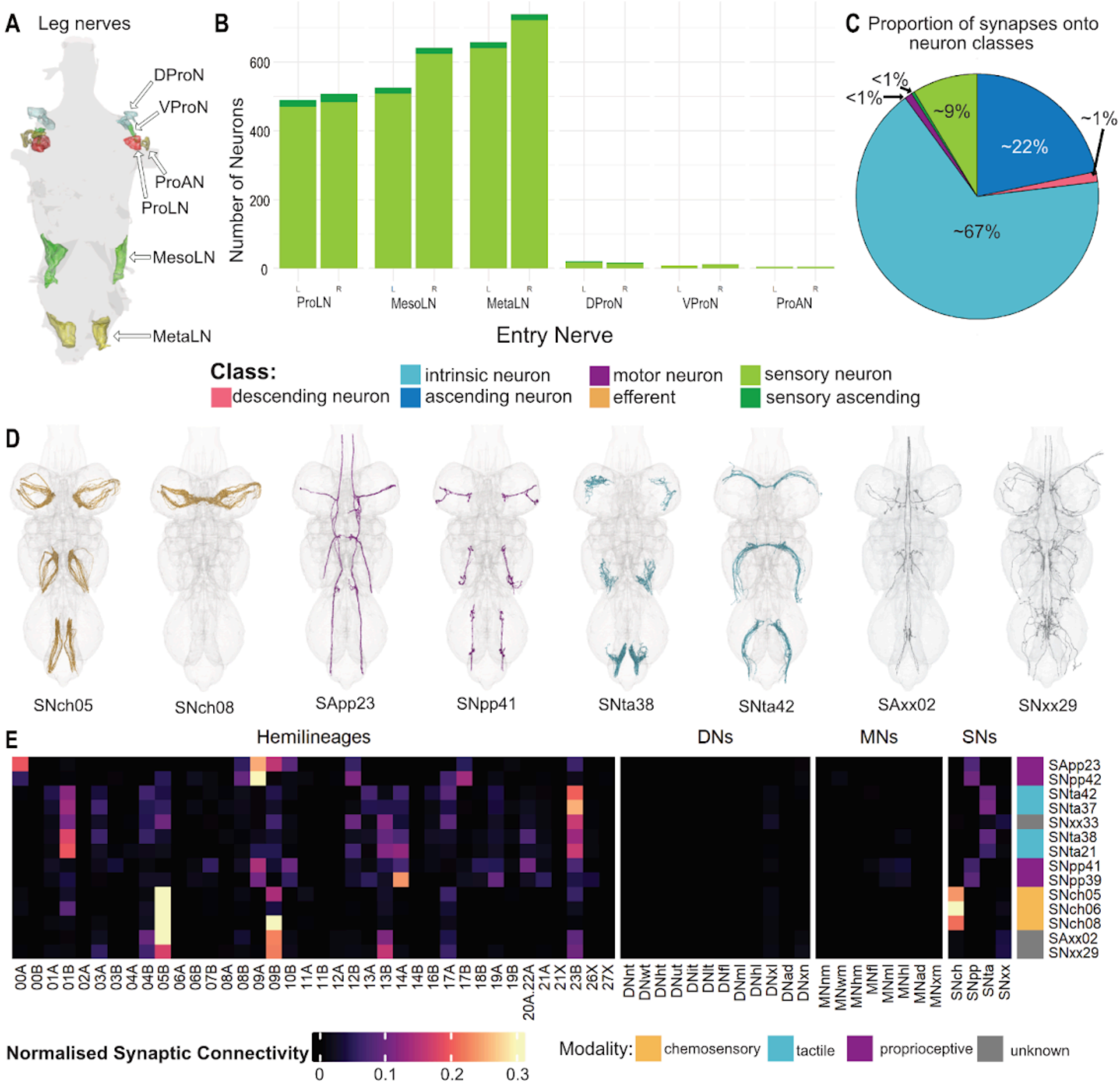

**Figure 55. Leg sensory neuron systematic types. A.** Anatomy of leg nerves (ventral view). **B.** Numbers of sensory neurons from each leg nerve. **C.** Proportions of downstream partners of leg sensory neurons, coloured by class. **D.** Plotted meshes of exemplar leg sensory types, coloured by assigned modality. **E.** Heatmap showing normalised sensory neuron cluster connectivity to hemilineages and other neuron subclasses in the VNC. Annotation bar displays annotated modality of component sensory neurons within the sensory neuron cluster.

#### Leg Sensory types

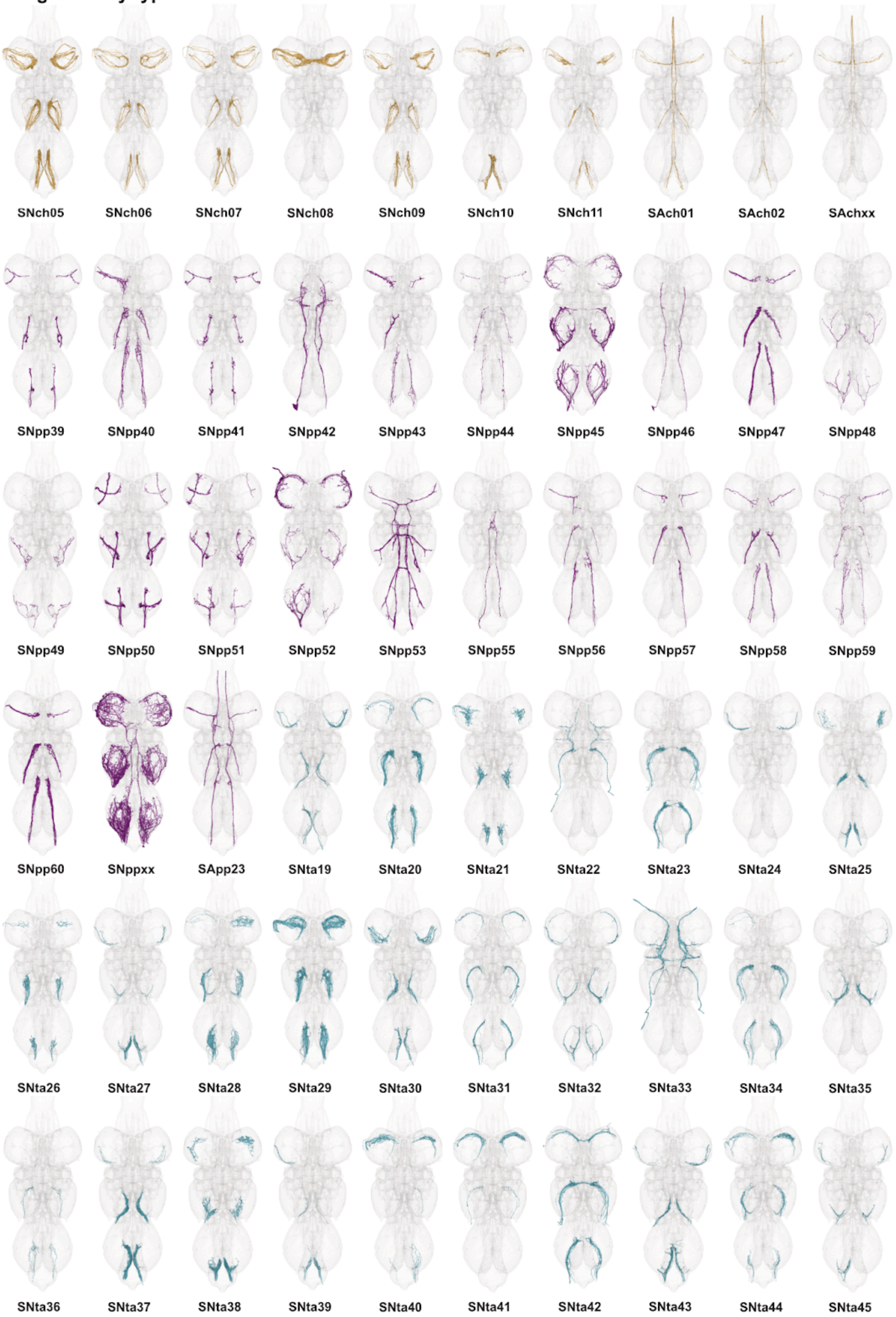

**Figure 55 - figure supplement 1. Leg sensory neuron types.** Plotted meshes of neurons that comprise each systematic type from leg nerves, coloured by modality: orange = chemosensory, purple = proprioceptive, cyan = tactile, grey = unknown.

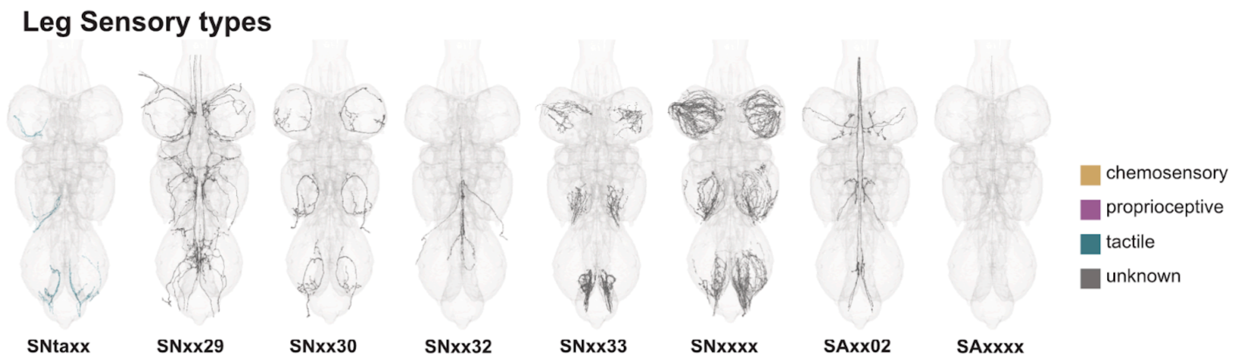

**Figure 55 - figure supplement 2. Leg sensory neuron types, continued.** Plotted meshes of neurons that comprise each systematic type from leg nerves, coloured by modality: orange = chemosensory, purple = proprioceptive, cyan = tactile, grey = unknown.

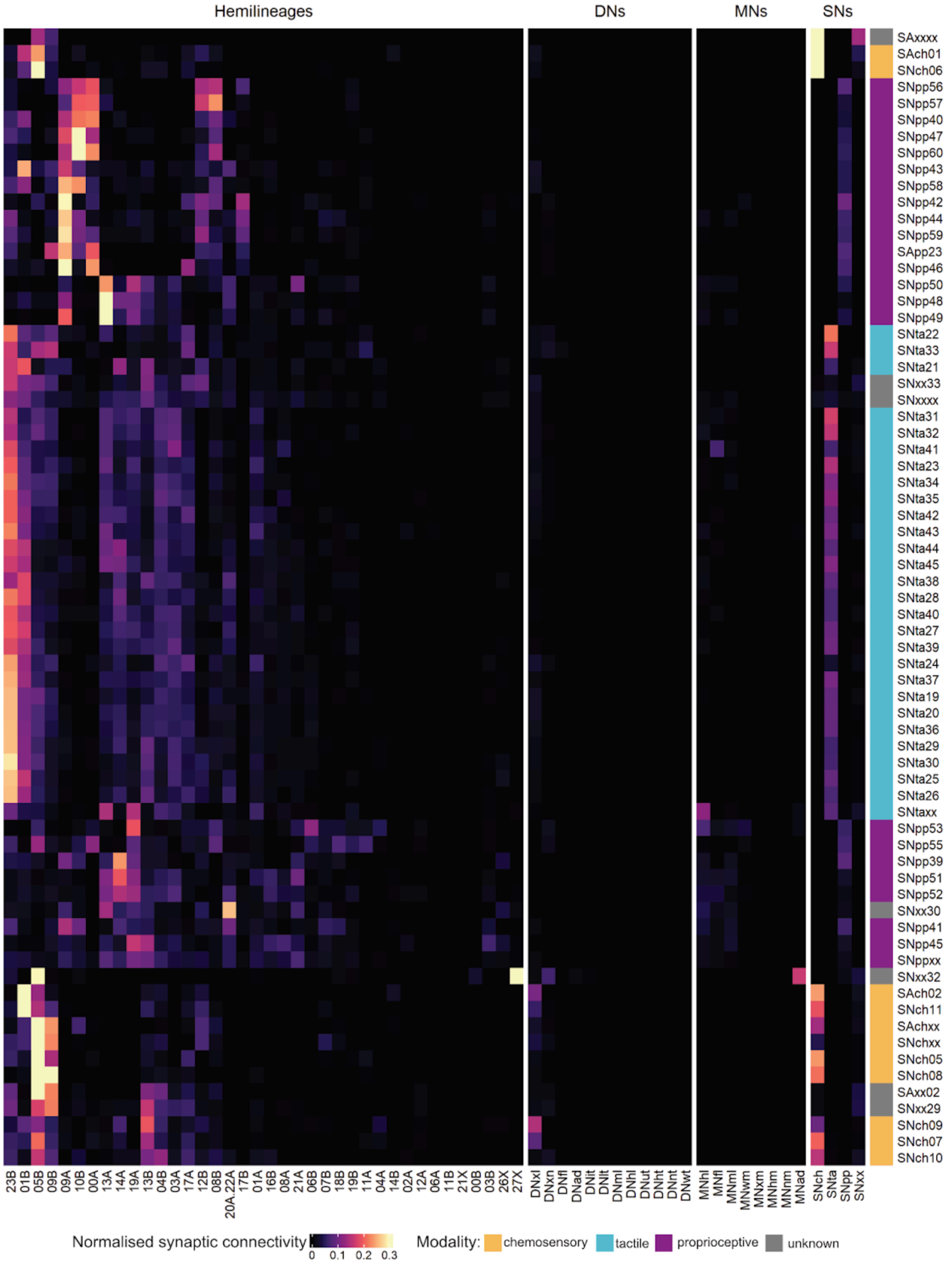

**Figure 55 - figure supplement 3. Leg sensory neuron type output connectivity by hemilineage.** Right annotation bar is coloured by modality: orange = chemosensory, purple = proprioceptive, cyan = tactile, grey = unknown. Rows and columns are ordered based on hierarchical clustering of data.

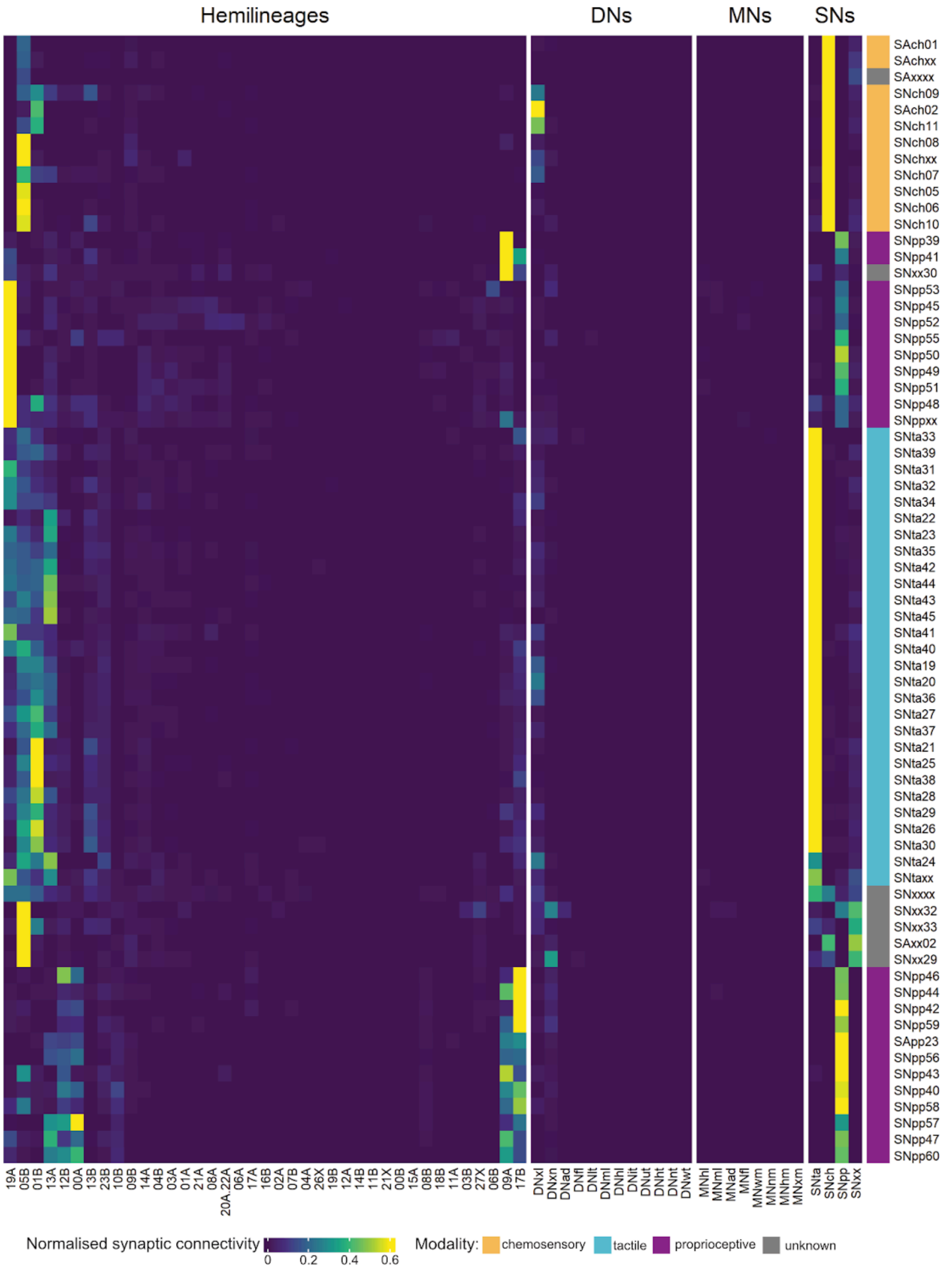

**Figure 55 - figure supplement 4. Leg sensory neuron type input connectivity by hemilineage.** Right annotation bar is coloured by modality: orange = chemosensory, purple = proprioceptive, cyan = tactile, grey = unknown. Rows and columns are ordered based on hierarchical clustering of data.

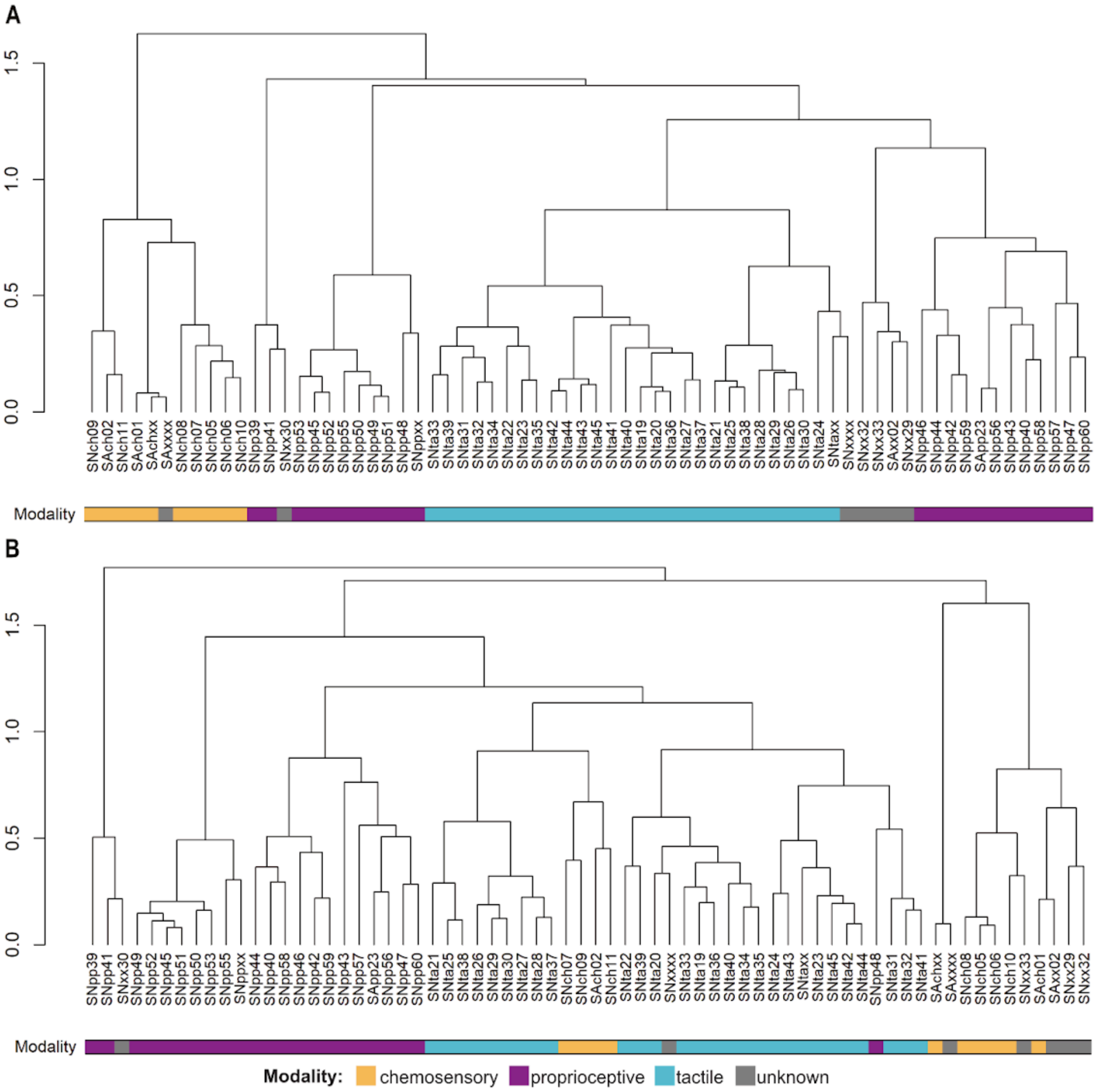

**Figure 55 - figure supplement 5. Leg sensory neuron type dendrograms showing combined inputs and outputs clustering. A.** Dendrogram showing combined clustering with intra-type connectivity included. **B.** Dendrogram showing combined clustering with intra-type connectivity excluded. Bottom bar is coloured by modality: orange = chemosensory, purple = proprioceptive, cyan = tactile, grey = unknown.

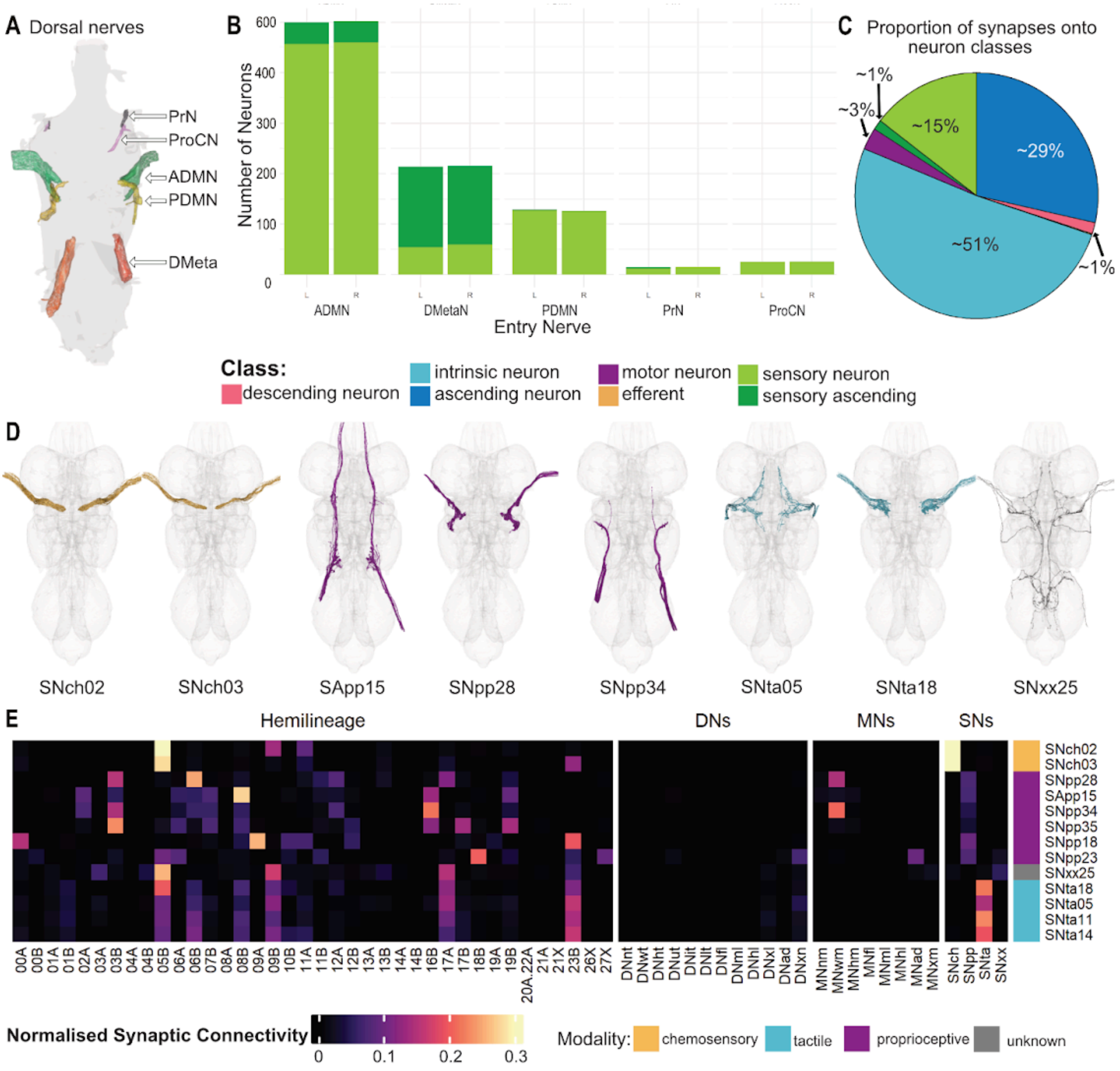

**Figure 56. Dorsal sensory neuron systematic types. A.** Anatomy of dorsal nerves (dorsal view). **B.** Numbers of sensory neurons from each dorsal nerve. **C.** Proportions of downstream partners of dorsal sensory neurons, coloured by class. **D.** Plotted meshes of exemplar dorsal types, coloured by assigned modality: orange = chemosensory, purple = proprioceptive, cyan = tactile, grey = unknown. **E.** Heatmap showing normalised sensory neuron cluster connectivity to hemilineages in the VNC. Annotation bar displays annotated modality of component sensory neurons within the sensory neuron cluster; orange = chemosensory, purple = proprioceptive, cyan = tactile, grey = unknown.

#### Dorsal Sensory types

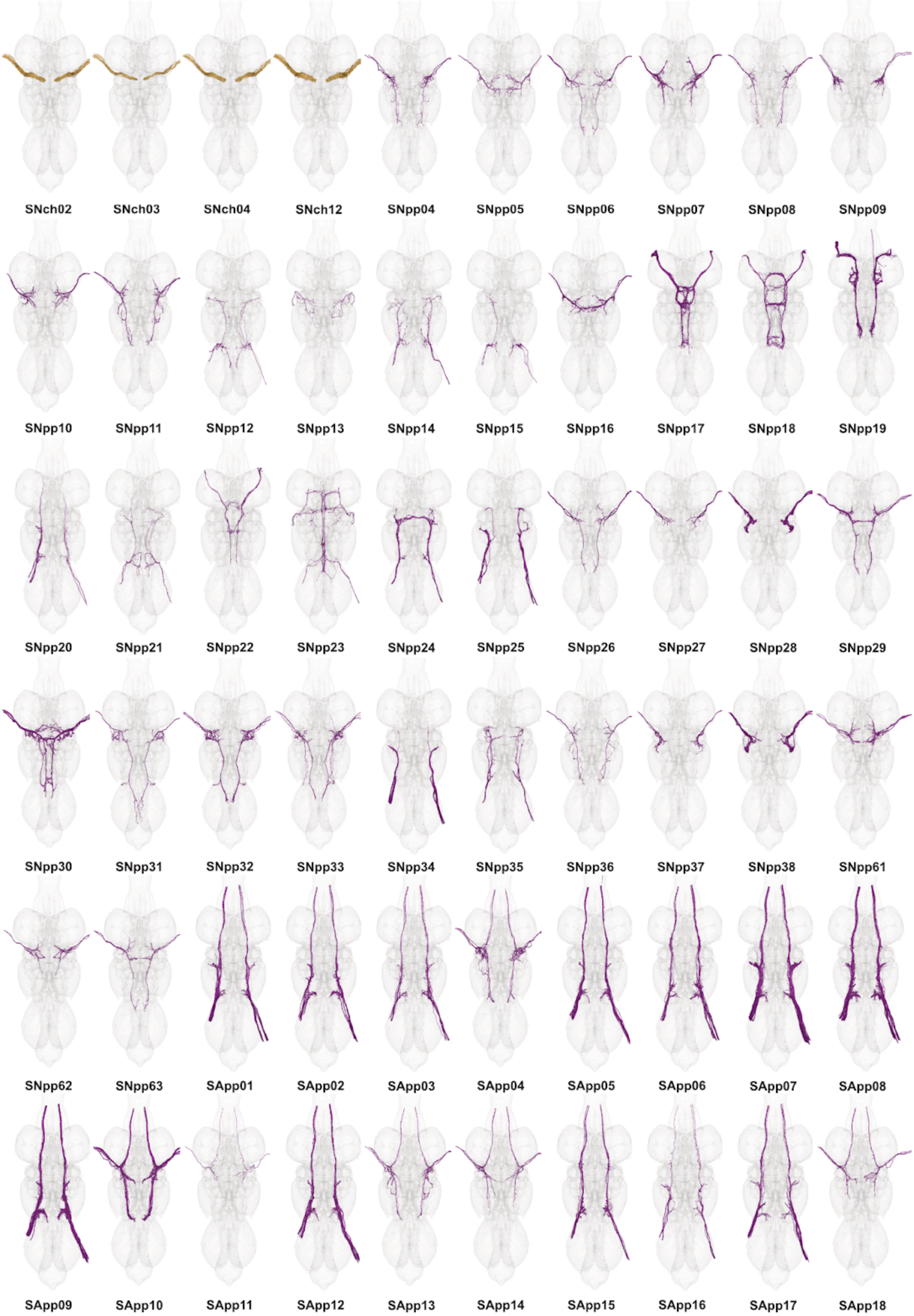

**Figure 56 - figure supplement 1. Dorsal sensory neuron types.** Plotted meshes of neurons that comprise each systematic type from dorsal nerves, coloured by modality: orange = chemosensory, purple = proprioceptive, cyan = tactile, grey = unknown.

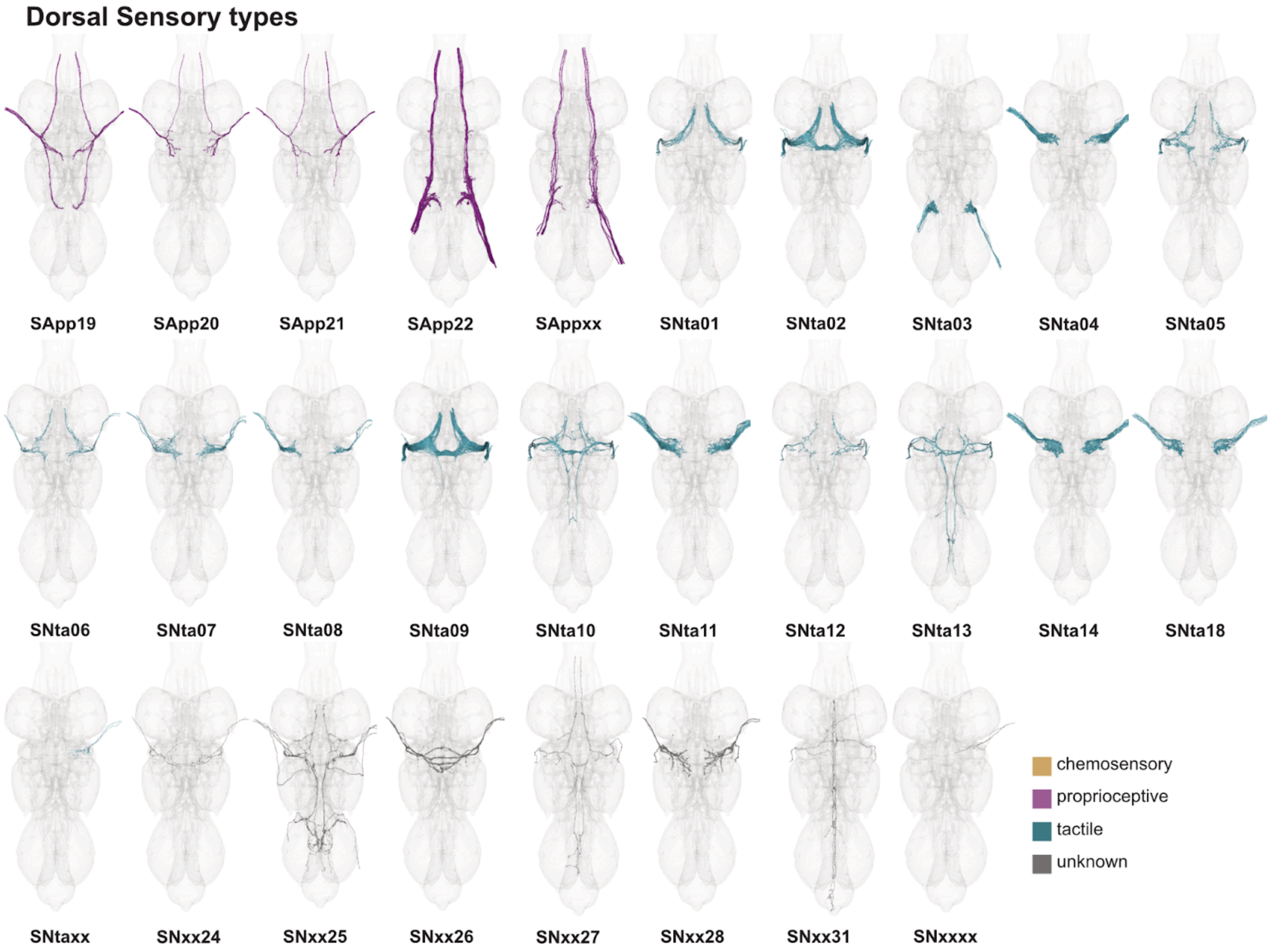

**Figure 56 - figure supplement 2.** Dorsal sensory neuron types, continued. Plotted meshes of neurons that comprise each systematic type from dorsal nerves, coloured by modality: orange = chemosensory, purple = proprioceptive, cyan = tactile, grey = unknown.

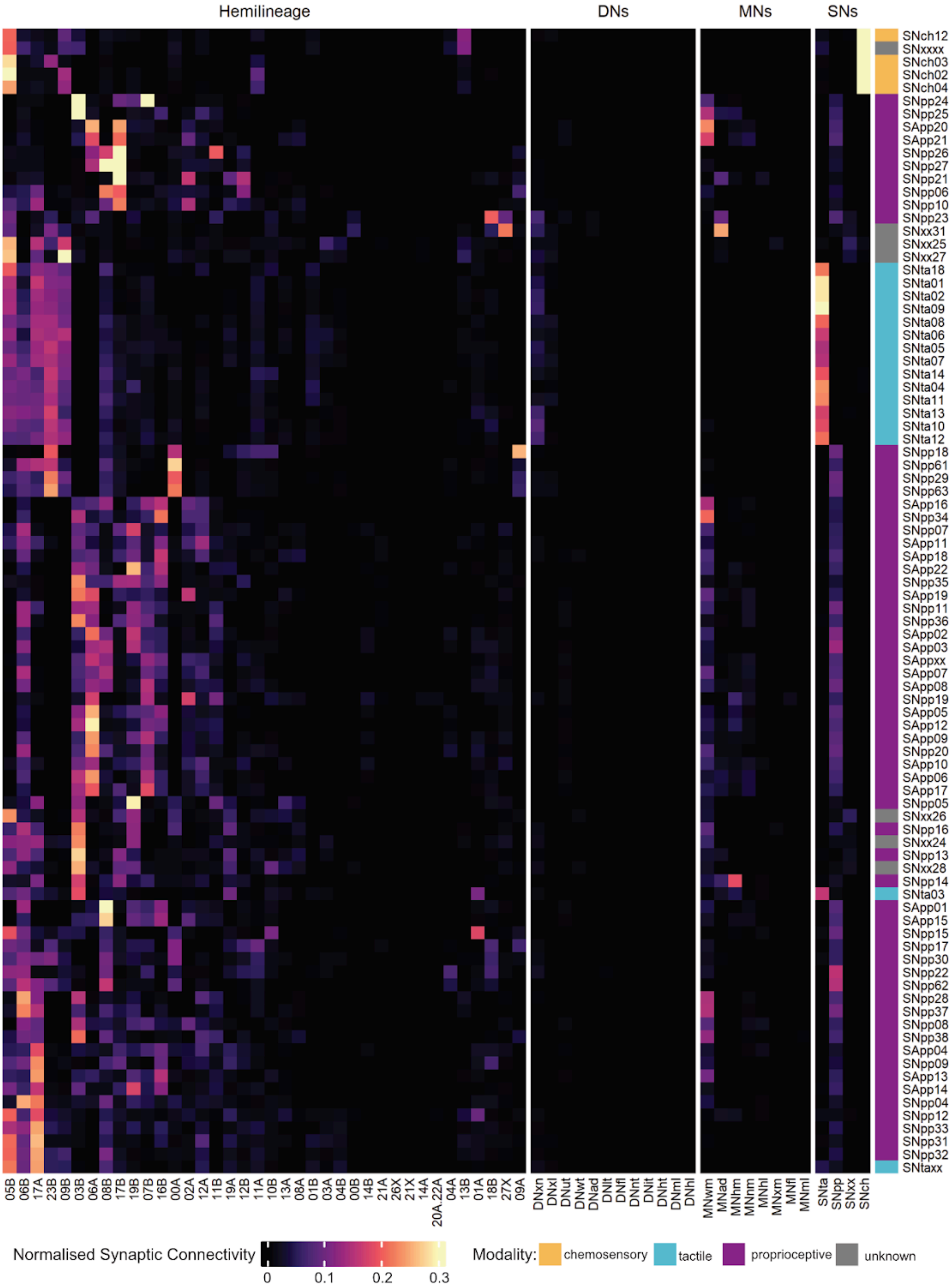

Hemilineage
DNs
MNs
SNs
SNch12 SNxxxx SNch03 SNch02 SNch04 SNpp24 SNpp25 SApp20 SApp21 SNpp26 SNpp27 SNpp21 SNpp06 SNpp10 SNpp23 SNxx31 SNxx25 SNxx27 SNta18 SNta01 SNta02 SNta09 SNta08 SNta06 SNta05 SNta07 SNta14 SNta04 SNta11 SNta13 SNta10 SNta12 SNpp18 SNpp61 SNpp29 SNpp63 SApp16 SNpp34 SNpp07 SApp11 SApp18 SApp22 SNpp35 SApp19 SNpp11 SNpp36 SApp02 SApp03 SAppxx SApp07 SApp08 SNpp19 SApp05 SApp12 SApp09 SNpp20 SApp10 SApp06 SApp17 SNpp05 SNxx26 SNpp16 SNxx24 SNpp13 SNxx28 SNpp14 SNta03 SApp01 SApp15 SNpp15 SNpp17 SNpp30 SNpp22 SNpp62 SNpp28 SNpp37 SNpp08 SNpp38 SApp04 SNpp09 SApp13 SApp14 SNpp04 SNpp12 SNpp33 SNpp31 SNpp32 SNtaxx
05B 06B 17A 23B 09B 03B 06A 08B 17B 19B 07B 16B 00A 02A 12A 11B 19A 12B 11A 10B 13A 08A 01B 03A 04B 00B 14B 21A 26X 21X 14A 20A.22A 04A 13B 01A 18B 27X 09A
DNxn DNxl DNut DNwt DNad DNlt DNfl DNnt DNit DNht DNml DNhl
MNwm MNad MNhm MNnm MNhl MNxm MNfl MNml
SNta SNpp SNxx SNch
Normalised Synaptic Connectivity
0 0.1 0.2 0.3
Modality: chemosensory tactile proprioceptive unknown

**Figure 56 - figure supplement 3.** Dorsal sensory neuron type output connectivity by hemilineage. Right annotation bar is coloured by modality: orange = chemosensory, purple = proprioceptive, cyan = tactile, grey = unknown. Rows and columns are ordered based on hierarchical clustering of data.

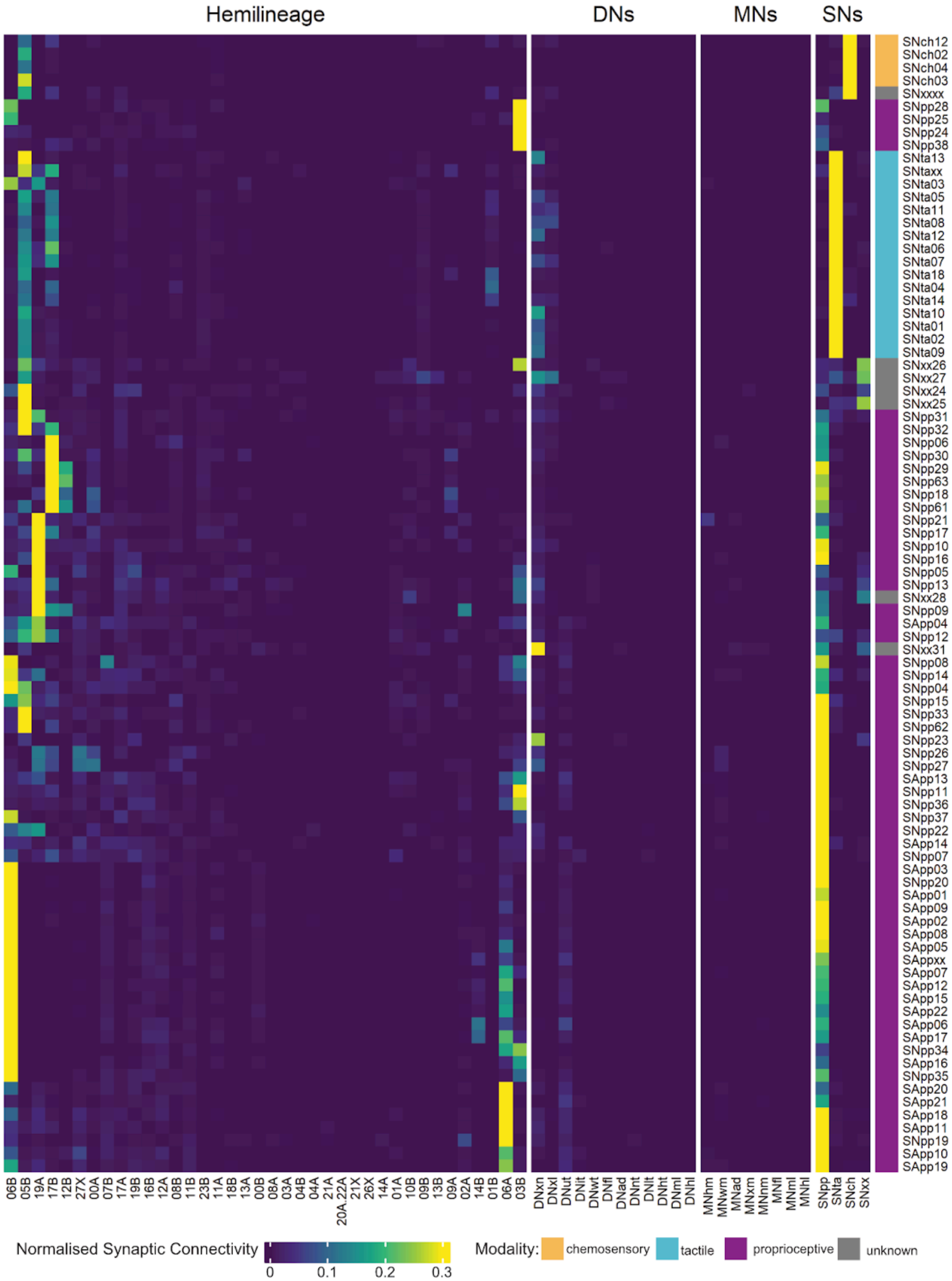
Hemilineage
DNs
MNs
SNs
SNch12
SNch02
SNch04
SNch03
SNxxxx
SNpp28
SNpp25
SNpp24
SNpp38
SNta13
SNtaxx
SNta03
SNta05
SNta11
SNta08
SNta12
SNta06
SNta07
SNta18
SNta04
SNta14
SNta10
SNta01
SNta02
SNta09
SNxx26
SNxx27
SNxx24
SNxx25
SNpp31
SNpp32
SNpp06
SNpp30
SNpp29
SNpp63
SNpp18
SNpp61
SNpp21
SNpp17
SNpp10
SNpp16
SNpp05
SNpp13
SNxx28
SNpp09
SApp04
SNpp12
SNxx31
SNpp08
SNpp14
SNpp04
SNpp15
SNpp33
SNpp62
SNpp23
SNpp26
SNpp27
SApp13
SNpp11
SNpp36
SNpp37
SNpp22
SApp14
SNpp07
SApp03
SNpp20
SApp01
SApp09
SApp02
SApp08
SApp05
SAppxx
SApp07
SApp12
SApp15
SApp22
SApp06
SApp17
SNpp34
SApp16
SNpp35
SApp20
SApp21
SApp18
SApp11
SNpp19
SApp10
SApp19
06B
05B
19A
17B
12B
27X
00A
07B
17A
19B
16B
12A
08B
11B
23B
11A
18B
13A
00B
08A
03A
04B
04A
21A
20A.22A
21X
26X
14A
01A
10B
09B
13B
09A
02A
14B
01B
06A
03B
DNxn
DNxl
DNut
DNit
DNwt
DNfl
DNad
DNnt
DNlt
DNht
DNml
DNhl
MNhm
MNwm
MNad
MNxm
MNnm
MNfl
MNml
MNhl
SNpp
SNta
SNch
SNxx
Normalised Synaptic Connectivity
0
0.1
0.2
0.3
Modality:
chemosensory
tactile
proprioceptive
unknown

**Figure 56 - figure supplement 4. Dorsal sensory neuron type input connectivity by hemilineage.** Right annotation bar is coloured by modality: orange = chemosensory, purple = proprioceptive, cyan = tactile, grey = unknown. Rows and columns are ordered based on hierarchical clustering of data.

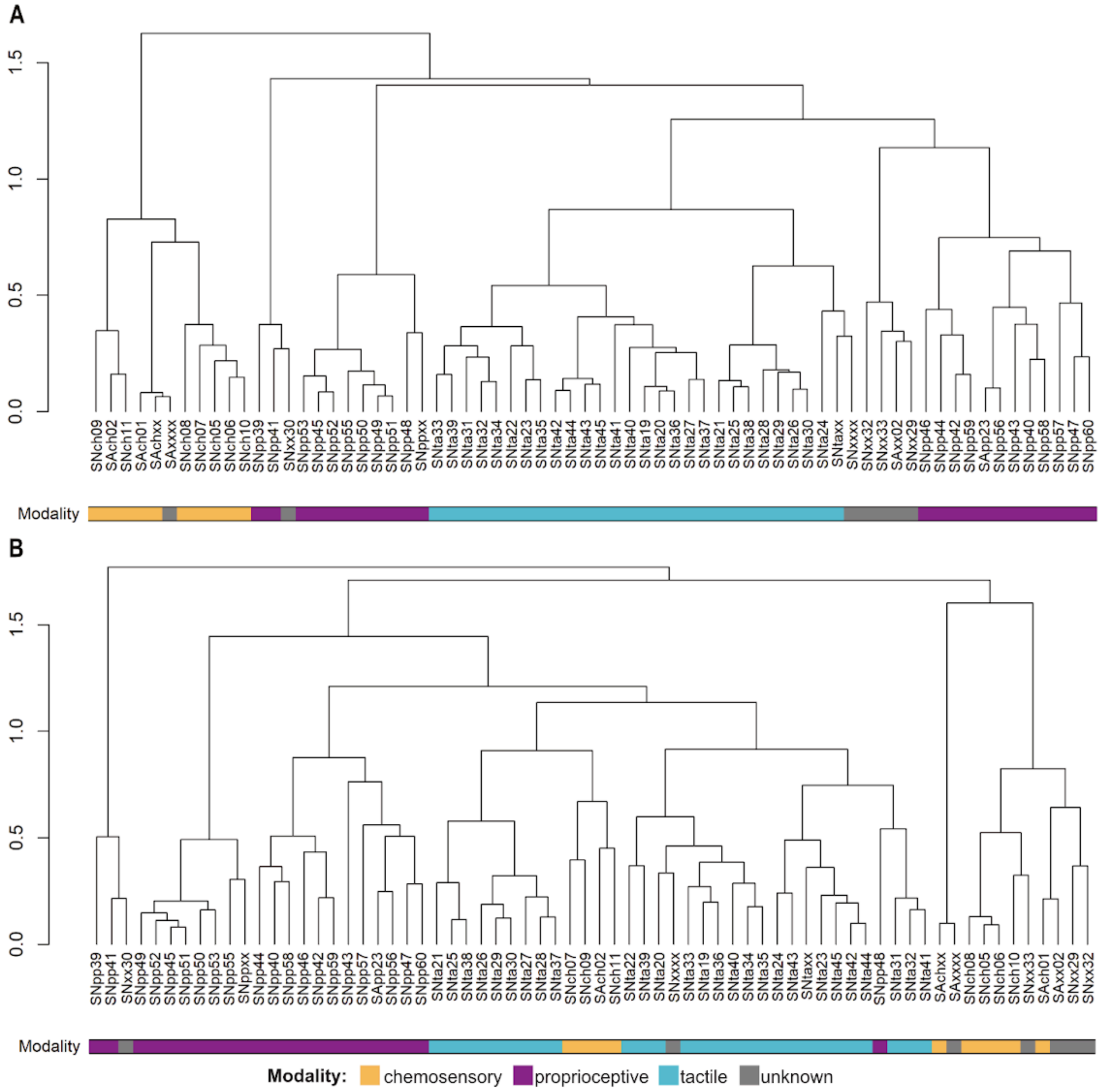

**Figure 56 - figure supplement 5. Dorsal sensory neuron type dendrograms showing combined inputs and outputs clustering. A.** Dendrogram showing combined clustering with intra-type connectivity included. **B.** Dendrogram showing combined clustering with intra-type connectivity excluded. Bottom bar is coloured by modality: orange = chemosensory, purple = proprioceptive, cyan = tactile, grey = unknown.

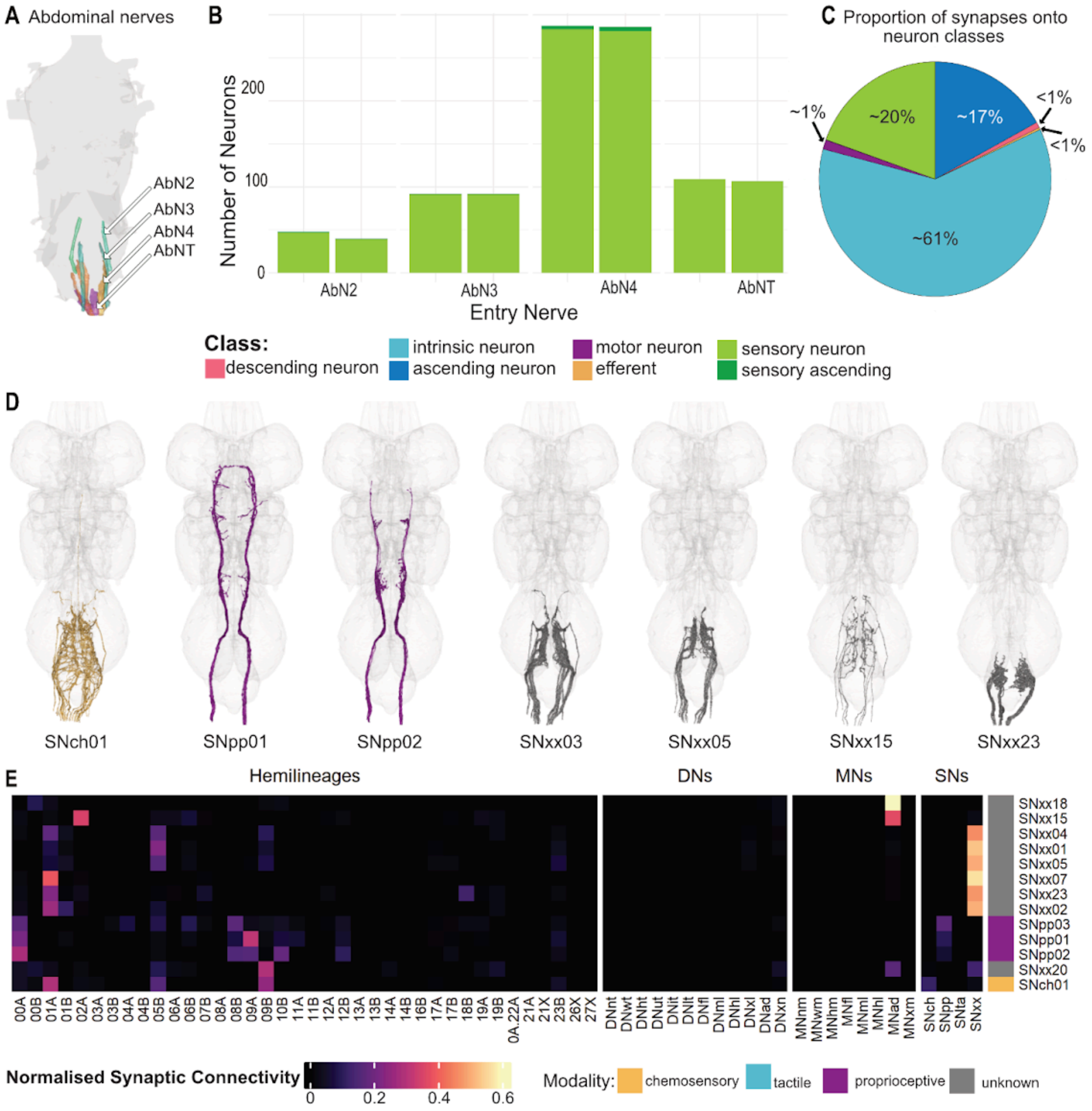

**Figure 57. Abdominal sensory neuron systematic types. A.** Anatomy of abdominal nerves (dorsal view). **B.** Numbers of sensory neurons from each abdominal nerve. **C.** Proportions of downstream partners of abdominal sensory neurons, coloured by class. **D.** Plotted meshes of exemplar abdominal types, coloured by assigned modality. **E.** Heatmap showing normalised sensory neuron cluster connectivity to hemilineages in the VNC. Annotation bar displays annotated modality of component sensory neurons within the sensory neuron cluster; orange = chemosensory, purple = proprioceptive, cyan = tactile, grey = unknown.

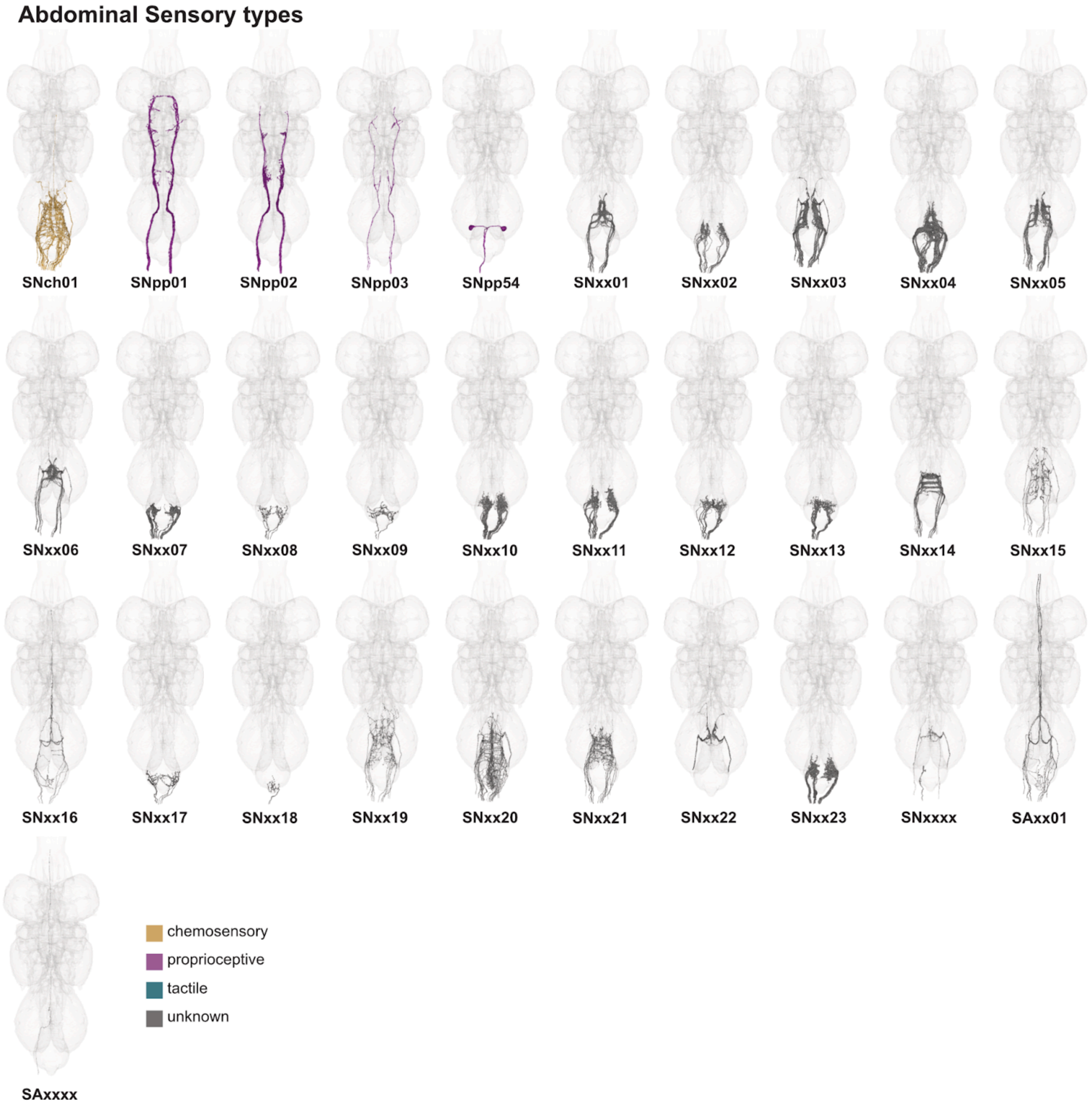

**Figure 57 - figure supplement 1. Abdominal sensory neuron types.** Plotted meshes of neurons that comprise each systematic type from abdominal nerves, coloured by modality: orange = chemosensory, purple = proprioceptive, cyan = tactile, grey = unknown.

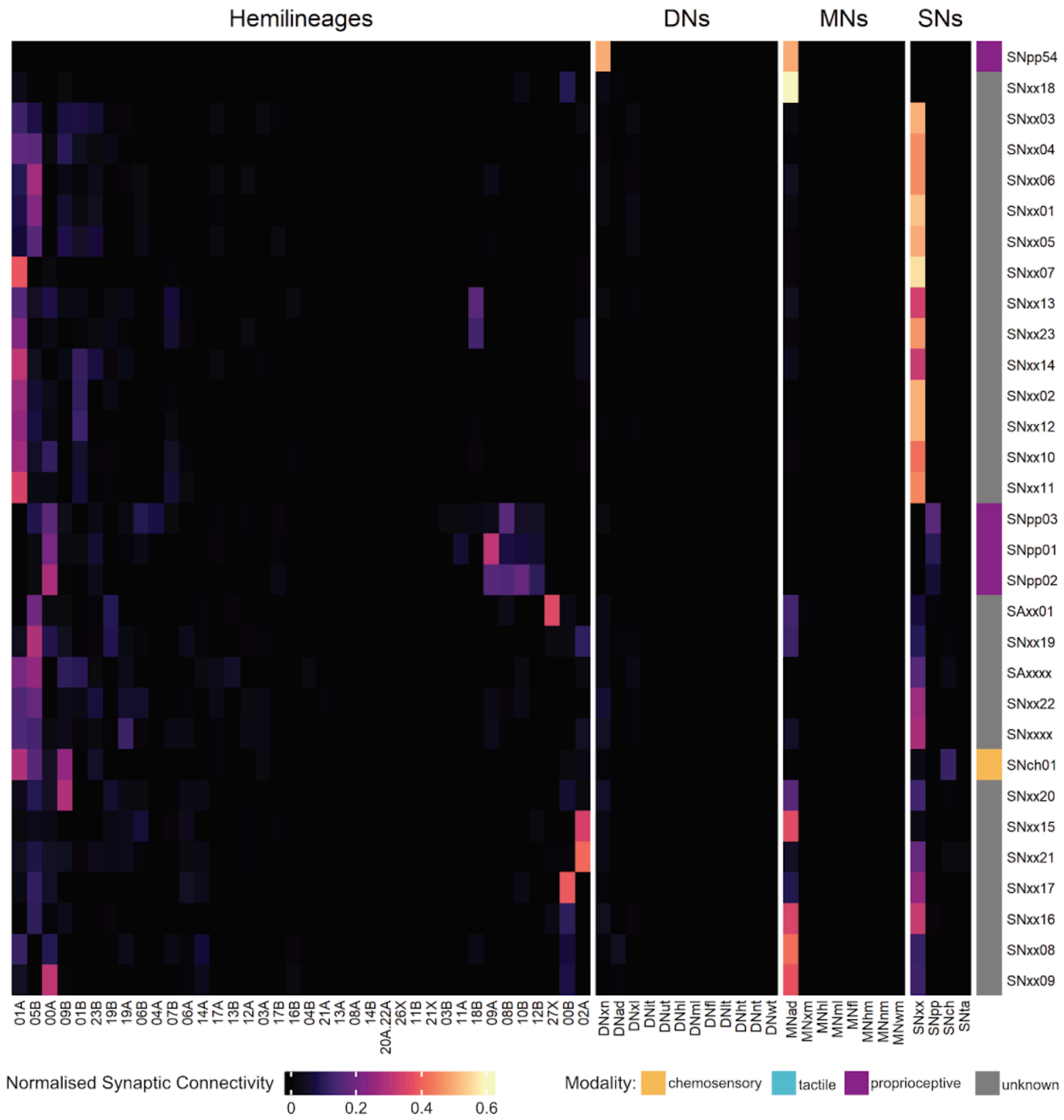

**Figure 57 - figure supplement 2. Abdominal sensory neuron type output connectivity by hemilineage.** Right annotation bar is coloured by modality: orange = chemosensory, purple = proprioceptive, cyan = tactile, grey = unknown. Rows and columns are ordered based on hierarchical clustering of data.

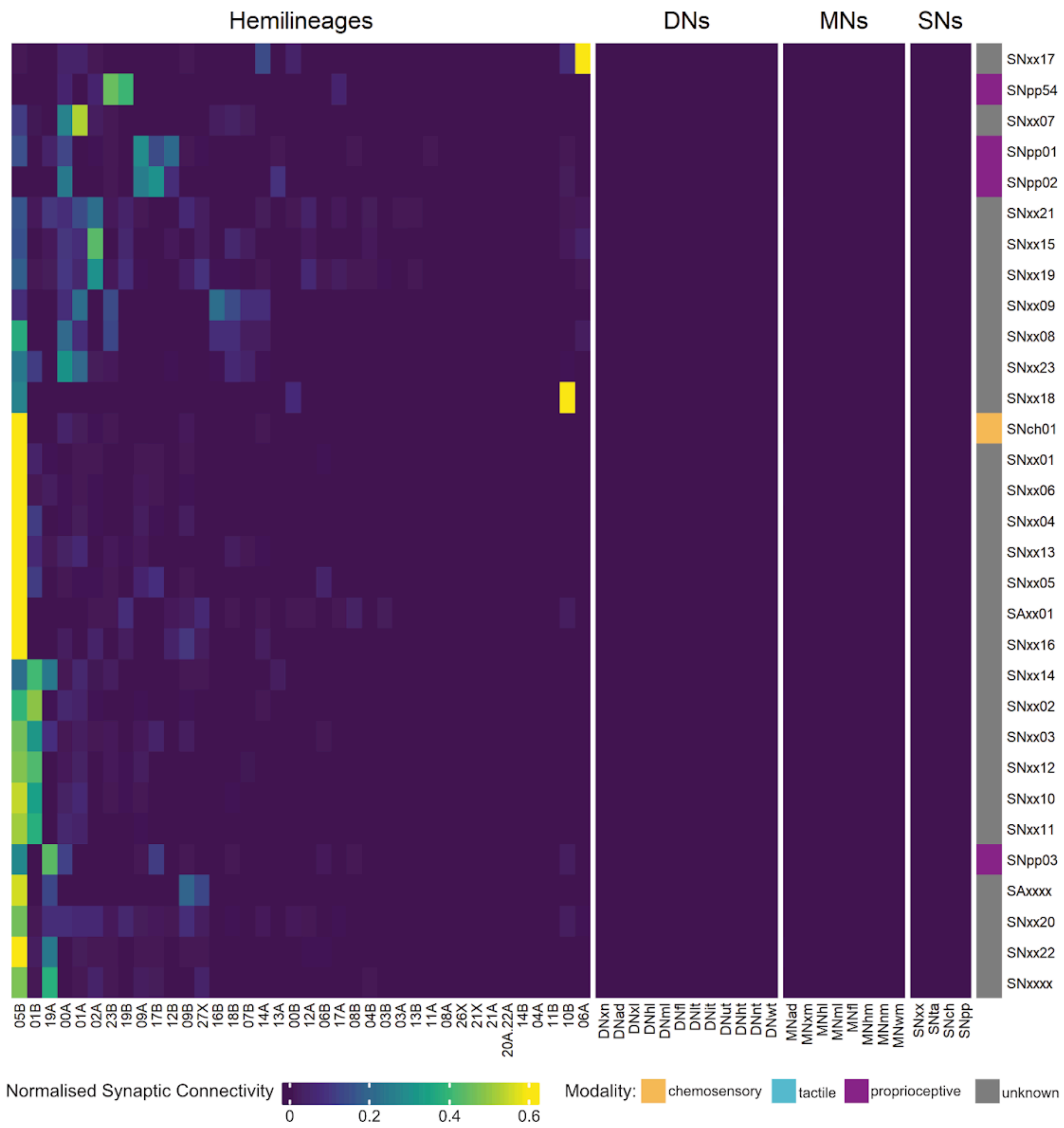

**Figure 57 - figure supplement 3. Abdominal sensory neuron type input connectivity by hemilineage.** Right annotation bar is coloured by modality: orange = chemosensory, purple = proprioceptive, cyan = tactile, grey = unknown. Rows and columns are ordered based on hierarchical clustering of data.

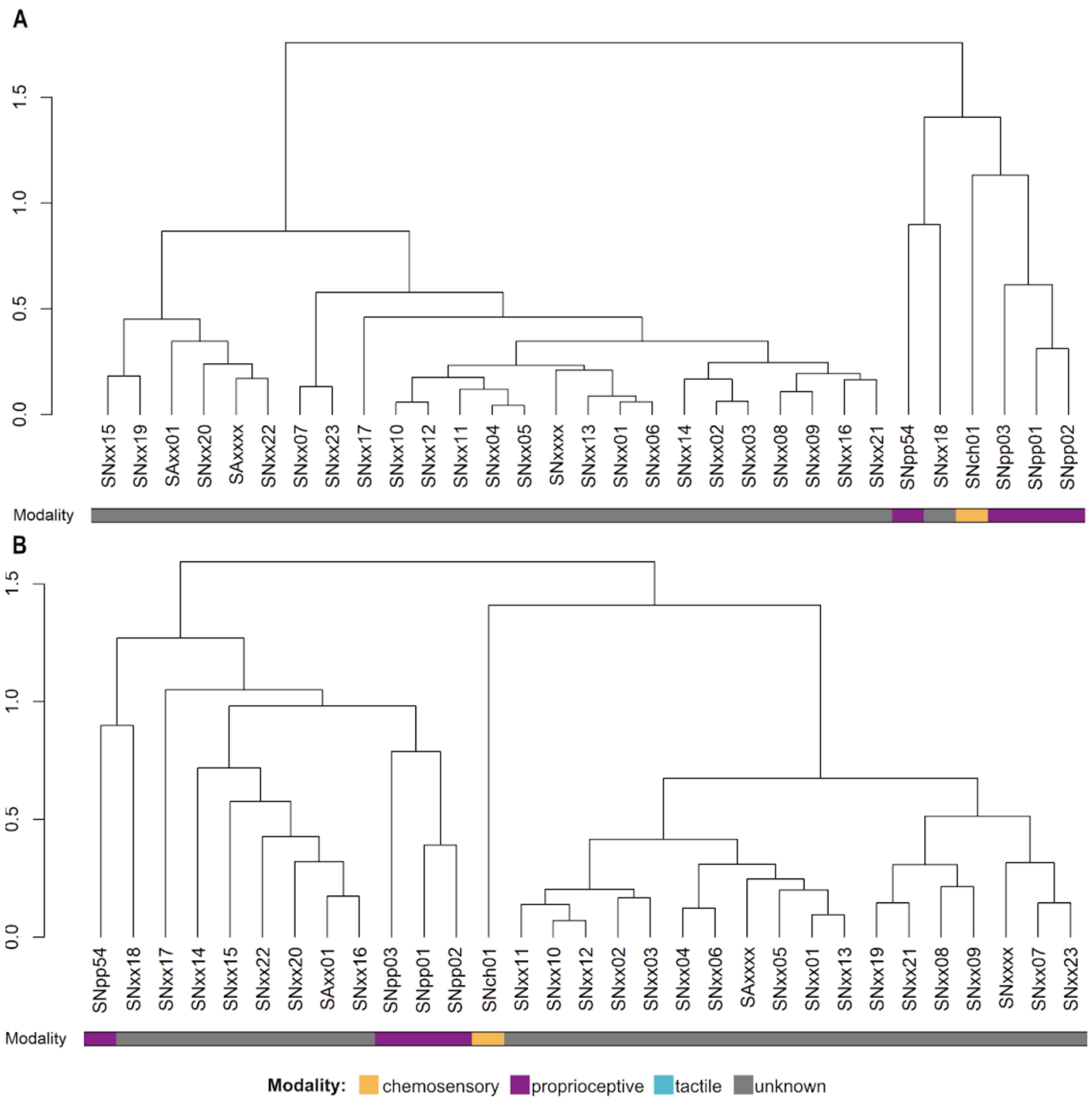

**Figure 57 - figure supplement 4. Abdominal sensory neuron type dendrograms showing combined inputs and outputs clustering. A.** Dendrogram showing combined clustering with intra-type connectivity included. **B.** Dendrogram showing combined clustering with intra-type connectivity excluded. Bottom bar is coloured by modality: orange = chemosensory, purple = proprioceptive, cyan = tactile, grey = unknown.

#### Inter-regional associations between sensory types

We suspected that there could be sensory types sharing similar functions across two or more of the three main regions (leg, dorsal, and abdominal). To address this, we used cosine clustering of outputs to serial neurons across the entire sensory neuron population (Figure 58 top, Figure 58 - figure supplement 1), rather than within each region as we had done originally to generate the types. We do not claim that the associations detected here represent genuine cross-regional types, although this may turn out to be true for at least a subset of cases; rather, we suggest that co-clustering implies functional association and may also indicate modality.

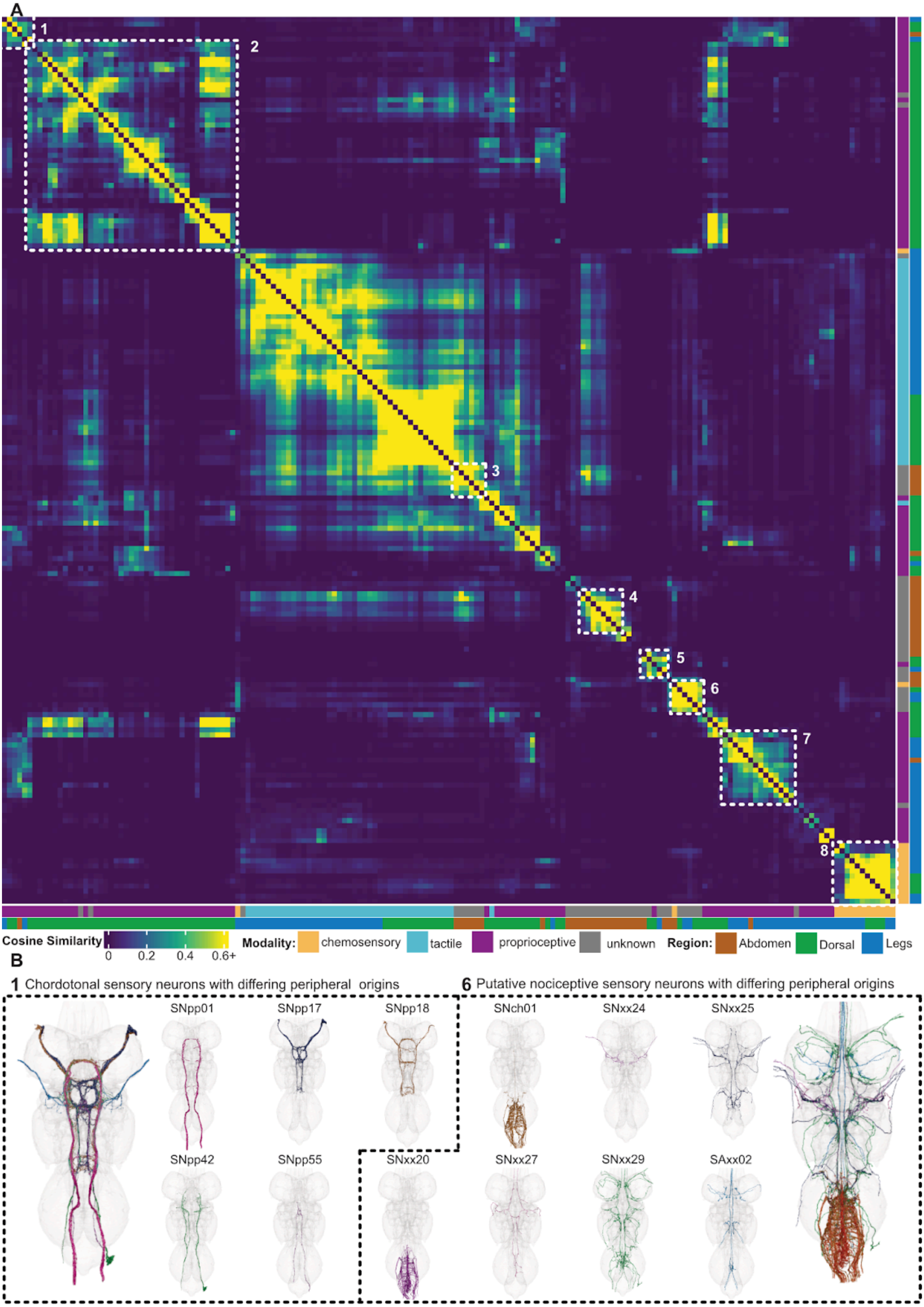
A
1
2
3
4
5
6
7
8
Cosine Similarity
0
0.2
0.4
0.6+
Modality:
chemosensory
tactile
proprioceptive
unknown
Region:
Abdomen
Dorsal
Legs
B
1 Chordotonal sensory neurons with differing peripheral origins
SNpp01
SNpp17
SNpp18
SNpp42
SNpp55
SNxx20
6 Putative nociceptive sensory neurons with differing peripheral origins
SNch01
SNxx24
SNxx25
SNxx27
SNxx29
SAxx02

**Figure 58. Inter-regional associations between sensory cell types. Top.** Heatmap showing cosine similarity between the outputs of sensory cell types on serial neuron groups. Coloured annotation bars indicate the assigned modality of the sensory cell type and its region of peripheral origin, inner and outer bars respectively. Heatmap row and column ordering generated through travelling sales problem optimisation. **Bottom.** Exemplar cell type clusters indicated by dashed line boxes 1 and 6 in heatmap.

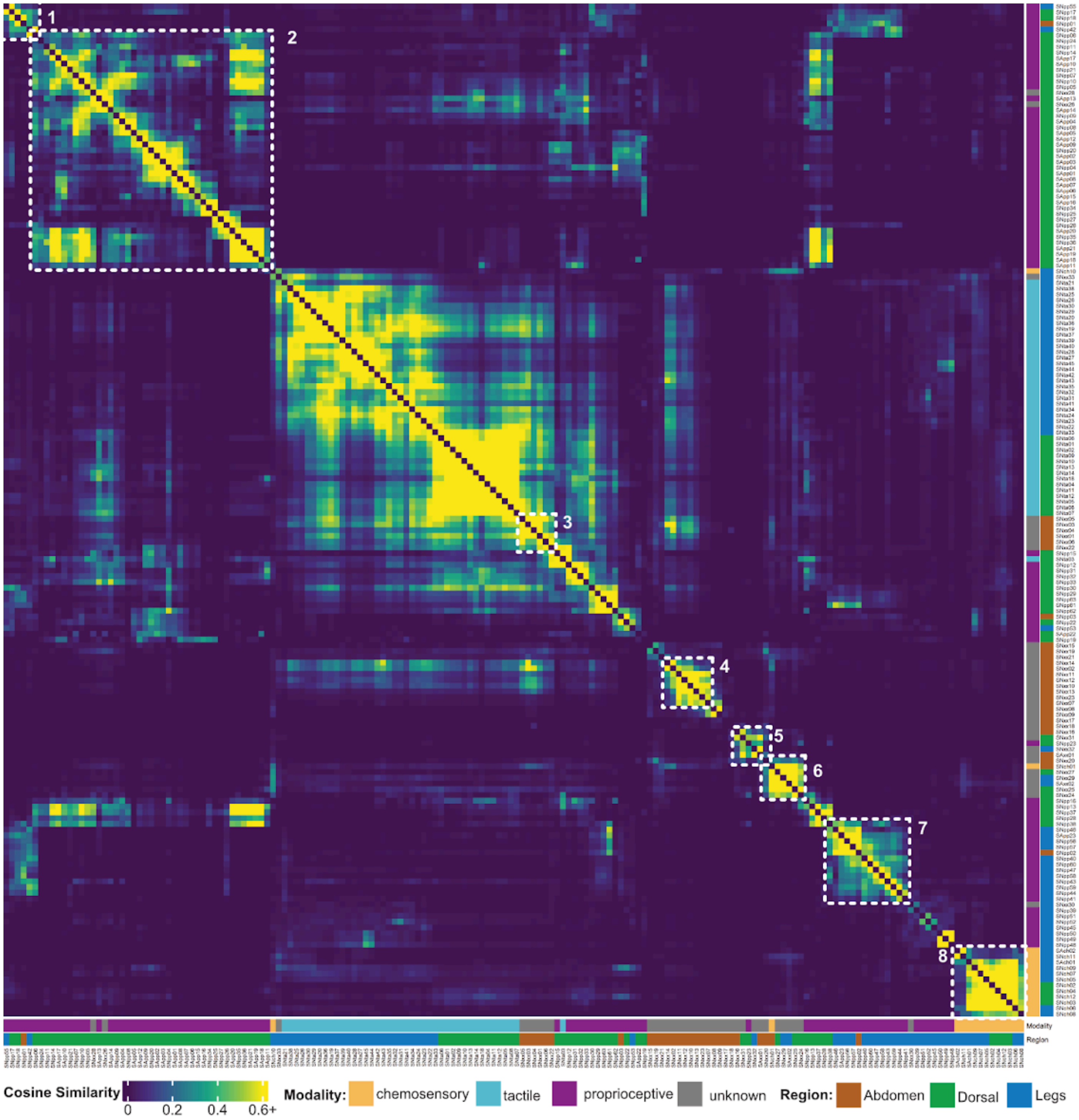

**Figure 58 - figure supplement 1. Cosine similarity between the outputs of sensory cell types (labelled) on serial neuron group.** Coloured annotation bars indicate the assigned modality of the sensory cell type and its region of peripheral origin, inner and outer bars respectively. Heatmap row and column ordering generated through travelling sales problem optimisation.

| Cluster | Systematic Type | Regional Origin | Sensory Modality |
|---|---|---|---|
| **1** | SNpp55 | Leg | proprioceptive |
| | SNpp17 | Dorsal | proprioceptive |
| | SNpp18 | Dorsal | proprioceptive |
| | SNpp01 | Abdomen | proprioceptive |
| | SNpp42 | Leg | proprioceptive |
| | SNpp06 | Dorsal | proprioceptive |
| **2** | SNpp06 | Dorsal | proprioceptive |
| | SNpp24 | Dorsal | proprioceptive |
| | SNpp11 | Dorsal | proprioceptive |
| | SNpp14 | Dorsal | proprioceptive |
| | SApp17 | Dorsal | proprioceptive |
| | SApp10 | Dorsal | proprioceptive |
| | SNpp21 | Dorsal | proprioceptive |
| | SNpp07 | Dorsal | proprioceptive |
| | SNpp10 | Dorsal | proprioceptive |
| | SNpp05 | Dorsal | proprioceptive |
| | SNxx26 | Dorsal | unknown |
| | SApp13 | Dorsal | proprioceptive |
| | SNxx26 | Dorsal | unknown |
| | SApp14 | Dorsal | proprioceptive |
| | SNpp09 | Dorsal | proprioceptive |
| | SApp04 | Dorsal | proprioceptive |
| | SNpp08 | Dorsal | proprioceptive |
| | SApp05 | Dorsal | proprioceptive |
| | SApp12 | Dorsal | proprioceptive |
| | SApp09 | Dorsal | proprioceptive |
| | SApp03 | Dorsal | proprioceptive |
| | SNpp04 | Dorsal | proprioceptive |
| | SApp01 | Dorsal | proprioceptive |
| | SApp08 | Dorsal | proprioceptive |
| | SApp07 | Dorsal | proprioceptive |
| | SApp06 | Dorsal | proprioceptive |
| | SApp15 | Dorsal | proprioceptive |
| | SApp16 | Dorsal | proprioceptive |
| | SNpp34 | Dorsal | proprioceptive |
| | SNpp25 | Dorsal | proprioceptive |
| | SNpp27 | Dorsal | proprioceptive |
| | SNpp26 | Dorsal | proprioceptive |
| | SApp20 | Dorsal | proprioceptive |
| | SNpp35 | Dorsal | proprioceptive |
| | SNpp36 | Dorsal | proprioceptive |
| | SApp21 | Dorsal | proprioceptive |
| | SApp19 | Dorsal | proprioceptive |
| | SApp18 | Dorsal | proprioceptive |
| | SApp11 | Dorsal | proprioceptive |
| 3 | SNxx05 | Abdomen | unknown |
| | SNxx03 | Abdomen | unknown |
| | SNxx04 | Abdomen | unknown |
| | SNxx01 | Abdomen | unknown |
| | SNxx06 | Abdomen | unknown |
| | SNxx22 | Abdomen | unknown |
| **4** | SNxx14 | Abdomen | unknown |
| | SNxx02 | Abdomen | unknown |
| | SNxx11 | Abdomen | unknown |
| | SNxx12 | Abdomen | unknown |
| | SNxx10 | Abdomen | unknown |
| | SNxx13 | Abdomen | unknown |
| | SNxx23 | Abdomen | unknown |
| | SNxx07 | Abdomen | unknown |
| **5** | SNxx16 | Abdomen | unknown |
| | SNxx31 | Dorsal | unknown |
| | SNpp23 | Dorsal | proprioceptive |
| | SNxx32 | Leg | unknown |
| | SAxx01 | Abdomen | unknown |
| | SNxx20 | Abdomen | unknown |
| **6** | SNxx20 | Abdomen | unknown |
| | SNch01 | Abdomen | chemosensory |
| | SNxx27 | Dorsal | unknown |
| | SNxx29 | Leg | unknown |
| | SAxx02 | Leg | unknown |
| | SNxx25 | Dorsal | unknown |
| | SNxx24 | Dorsal | unknown |
| **7** | SNpp38 | Dorsal | proprioceptive |
| | SNpp46 | Leg | proprioceptive |
| | SApp23 | Leg | proprioceptive |
| | SNpp56 | Leg | proprioceptive |
| | SNpp57 | Leg | proprioceptive |
| | SNpp02 | Abdomen | proprioceptive |
| | SNpp40 | Leg | proprioceptive |
| | SNpp60 | Leg | proprioceptive |
| | SNpp47 | Leg | proprioceptive |
| | SNpp58 | Leg | proprioceptive |
| | SNpp43 | Leg | proprioceptive |
| | SNpp59 | Leg | proprioceptive |
| | SNpp44 | Leg | proprioceptive |
| | SNpp41 | Leg | proprioceptive |
| **8** | SAch02 | Leg | chemosensory |
| | SNch11 | Leg | chemosensory |
| | SAch01 | Leg | chemosensory |
| | SNch09 | Leg | chemosensory |
| | SNch07 | Leg | chemosensory |
| | SNch05 | Leg | chemosensory |
| | SNch02 | Dorsal | chemosensory |
| | SNch04 | Dorsal | chemosensory |
| | SNch12 | Dorsal | chemosensory |
| | SNch03 | Dorsal | chemosensory |
| | SNch06 | Leg | chemosensory |
| | SNch08 | Leg | chemosensory |

**Figure 58 - table 1. Highlighted cosine similarity clusters from Figure 58A.** Cluster numbers, systematic types, regional origins, and annotated sensory modalities are provided.

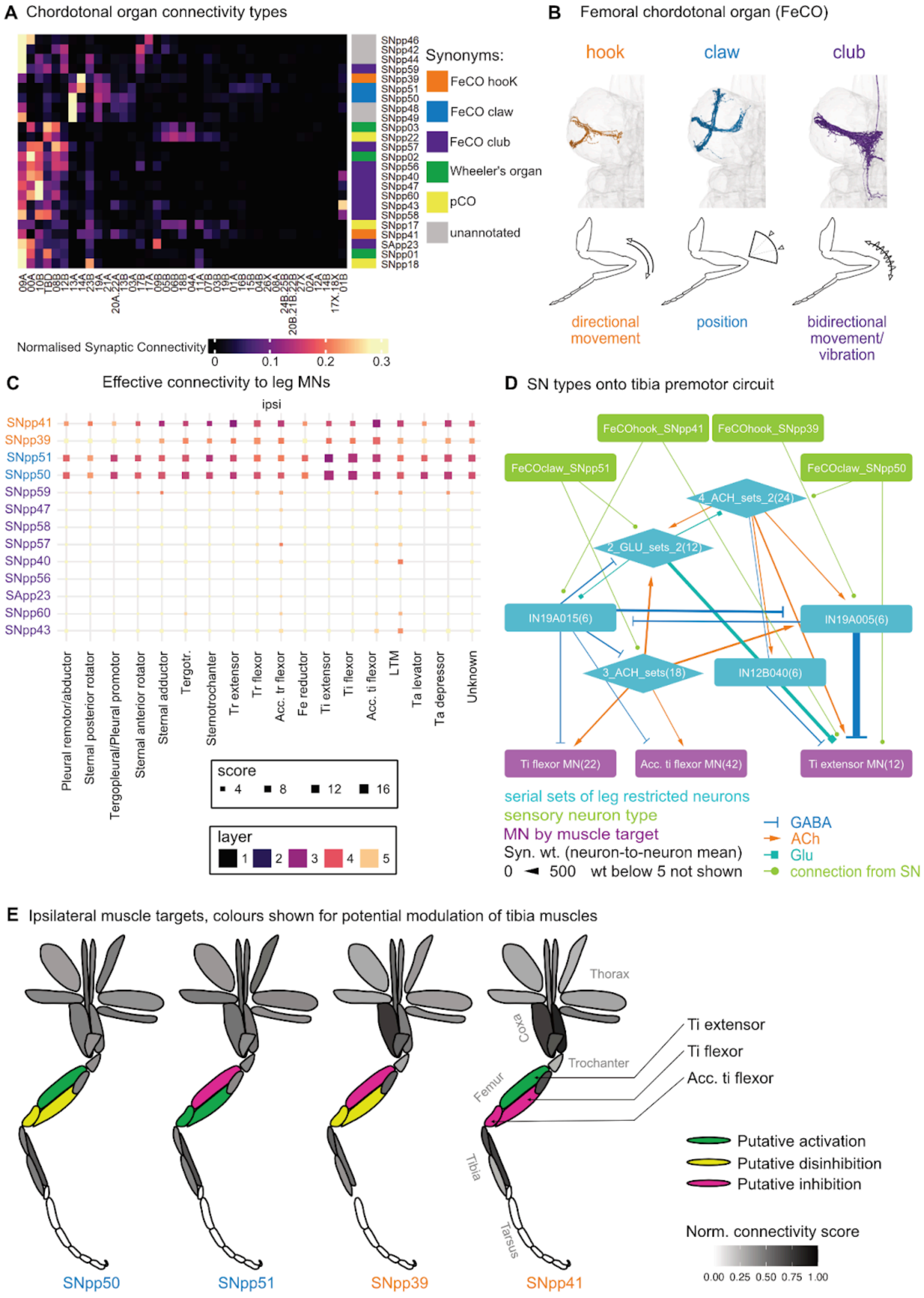

A Chordotonal organ connectivity types
SNpp46
SNpp42
SNpp44
SNpp59
SNpp39
SNpp51
SNpp50
SNpp48
SNpp49
SNpp03
SNpp22
SNpp57
SNpp02
SNpp56
SNpp40
SNpp47
SNpp60
SNpp43
SNpp58
SNpp17
SNpp41
SApp23
SNpp01
SNpp18
Synonyms:
FeCO hooK
FeCO claw
FeCO club
Wheeler's organ
pCO
unannotated
09A
00A
10B
TBD
08B
12B
13A
14A
23B
19A
21A
20A.22A
13B
03A
17B
17A
09B
05B
06B
18B
04A
11A
07B
03B
19B
01A
16B
15B
04B
26X
08A
24B.25B
20B.21B.22X
27X
02A
12A
14B
17X.18X
01B
Normalised Synaptic Connectivity
0
0.1
0.2
0.3
B Femoral chordotonal organ (FeCO)
hook
claw
club
directional movement
position
bidirectional movement/ vibration
C Effective connectivity to leg MNs
ipsi
SNpp41
SNpp39
SNpp51
SNpp50
SNpp59
SNpp47
SNpp58
SNpp57
SNpp40
SNpp56
SApp23
SNpp60
SNpp43
Pleural remotor/abductor
Sternal posterior rotator
Tergopleural/Pleural promotor
Sternal anterior rotator
Sternal adductor
Tergotr.
Sternotrochanter
Tr extensor
Tr flexor
Acc. tr flexor
Fe reductor
Ti extensor
Ti flexor
Acc. ti flexor
LTM
Ta levator
Ta depressor
Unknown
score
4
8
12
16
layer
1
2
3
4
5
D SN types onto tibia premotor circuit
FeCOhook_SNpp41
FeCOhook_SNpp39
FeCOclaw_SNpp51
FeCOclaw_SNpp50
4_ACH_sets_2(24)
2_GLU_sets_2(12)
IN19A015(6)
IN19A005(6)
3_ACH_sets(18)
IN12B040(6)
Ti flexor MN(22)
Acc. ti flexor MN(42)
Ti extensor MN(12)
serial sets of leg restricted neurons
sensory neuron type
MN by muscle target
Syn. wt. (neuron-to-neuron mean)
0
500
wt below 5 not shown
GABA
ACh
Glu
connection from SN
E Ipsilateral muscle targets, colours shown for potential modulation of tibia muscles
Thorax
Coxa
Trochanter
Femur
Tibia
Tarsus
Ti extensor
Ti flexor
Acc. ti flexor
Putative activation
Putative disinhibition
Putative inhibition
Norm. connectivity score
0.00
0.25
0.50
0.75
1.00
SNpp50
SNpp51
SNpp39
SNpp41

### FeCO leg motor circuits

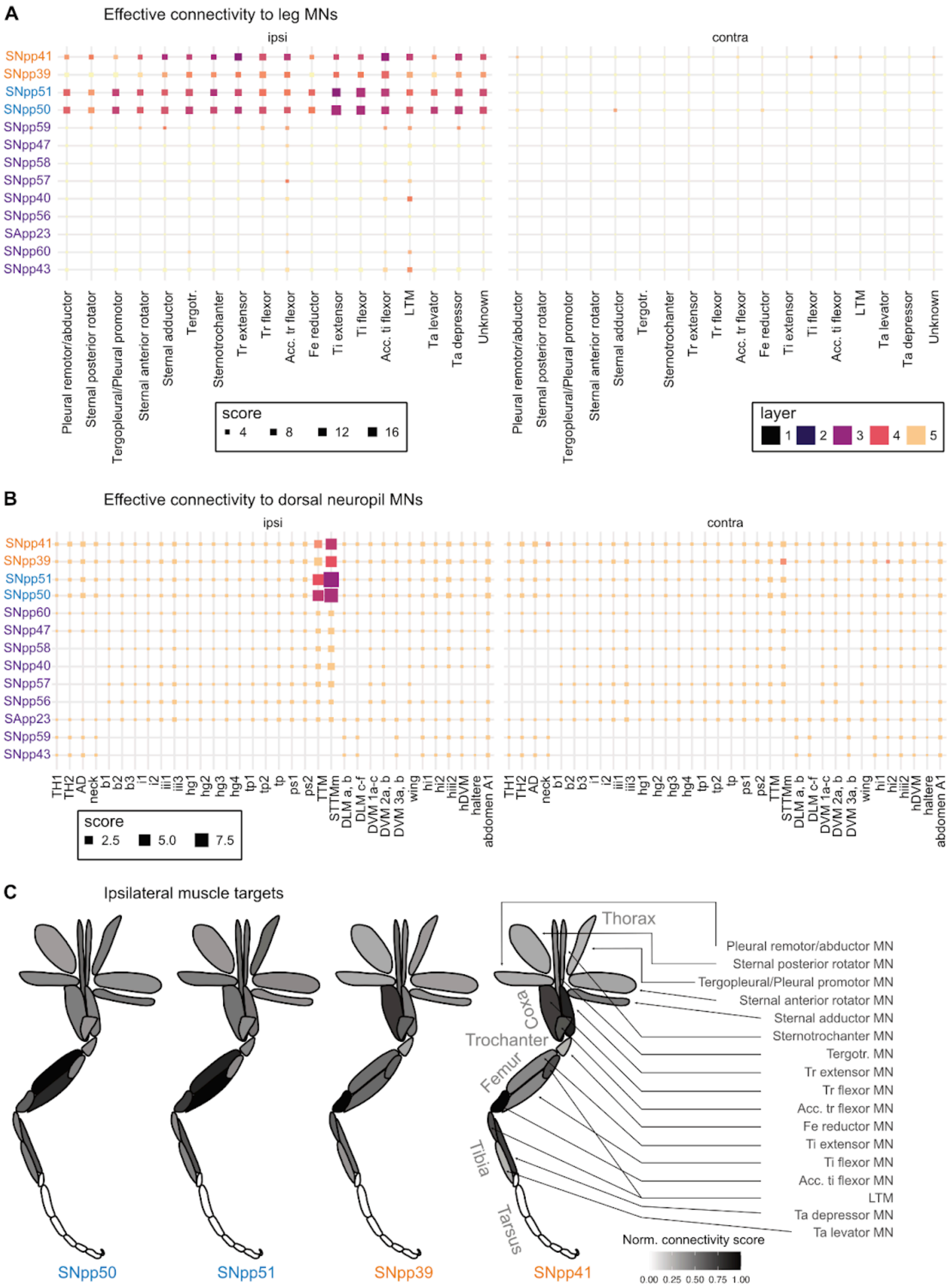

**Figure 59 - figure supplement 1. FeCO leg motor circuits. A.** Effective connectivity to leg motor neuron targets. **B.** Effective connectivity to dorsal motor neuron targets. **C.** Ipsilateral leg muscle targets for FeCO types.

#### Campaniform sensilla

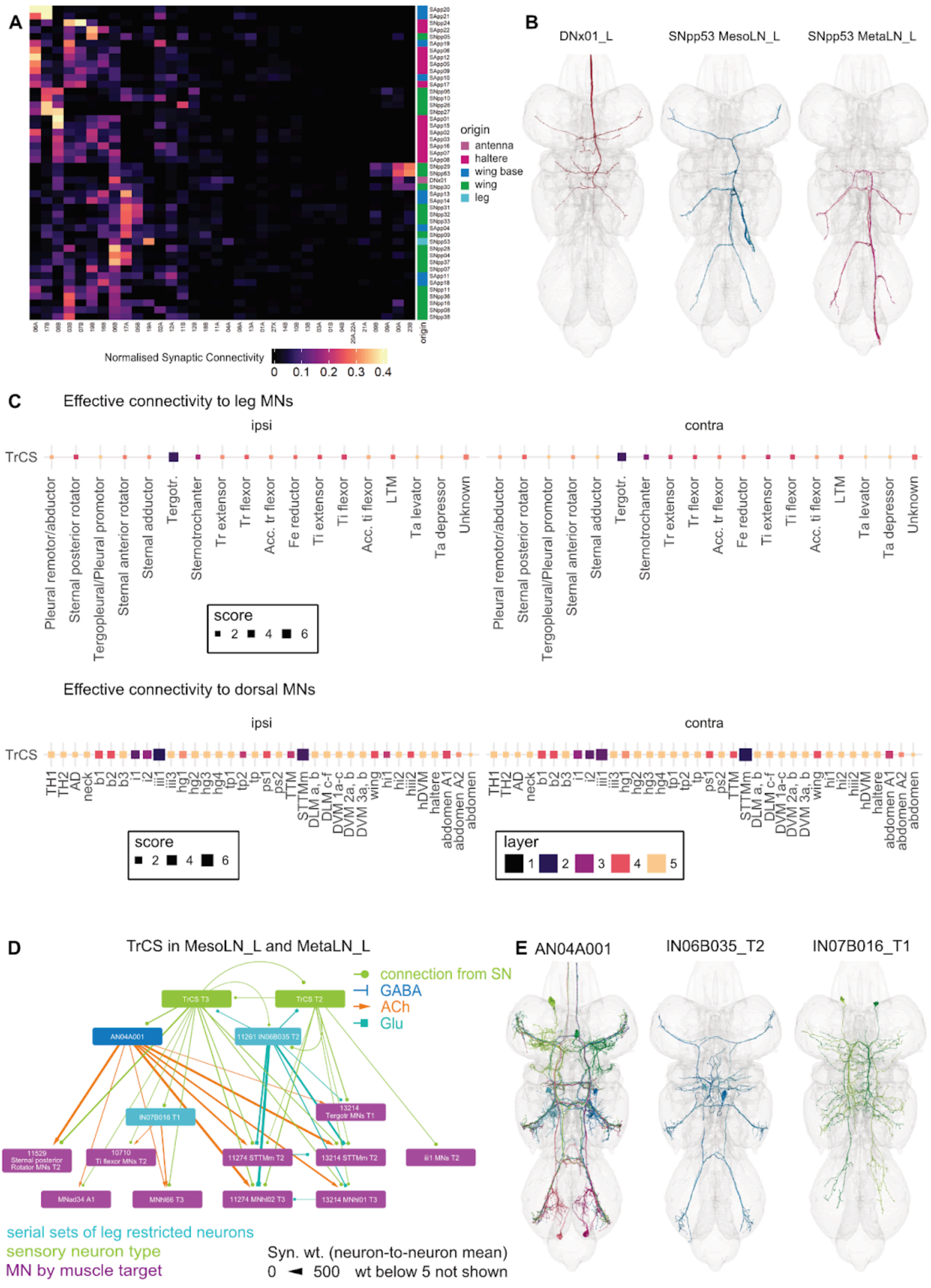

**Figure 60. Campaniform sensilla connectivity. A.** Campaniform sensilla types clustered by connectivity to hemilineages. **B.** Plotted meshes of bilateral TrCS from MesoLN_L vs MetaLN_L. **C.** Effective connectivity of TrCS to leg motor neuron targets vs dorsal motor neuron targets. **D.** Relationship of TrCS (MesoLN and MetaLN only) to standard leg premotor circuit. **E.** Plotted meshes of non-motor elements in the TrCS circuit shown in D.

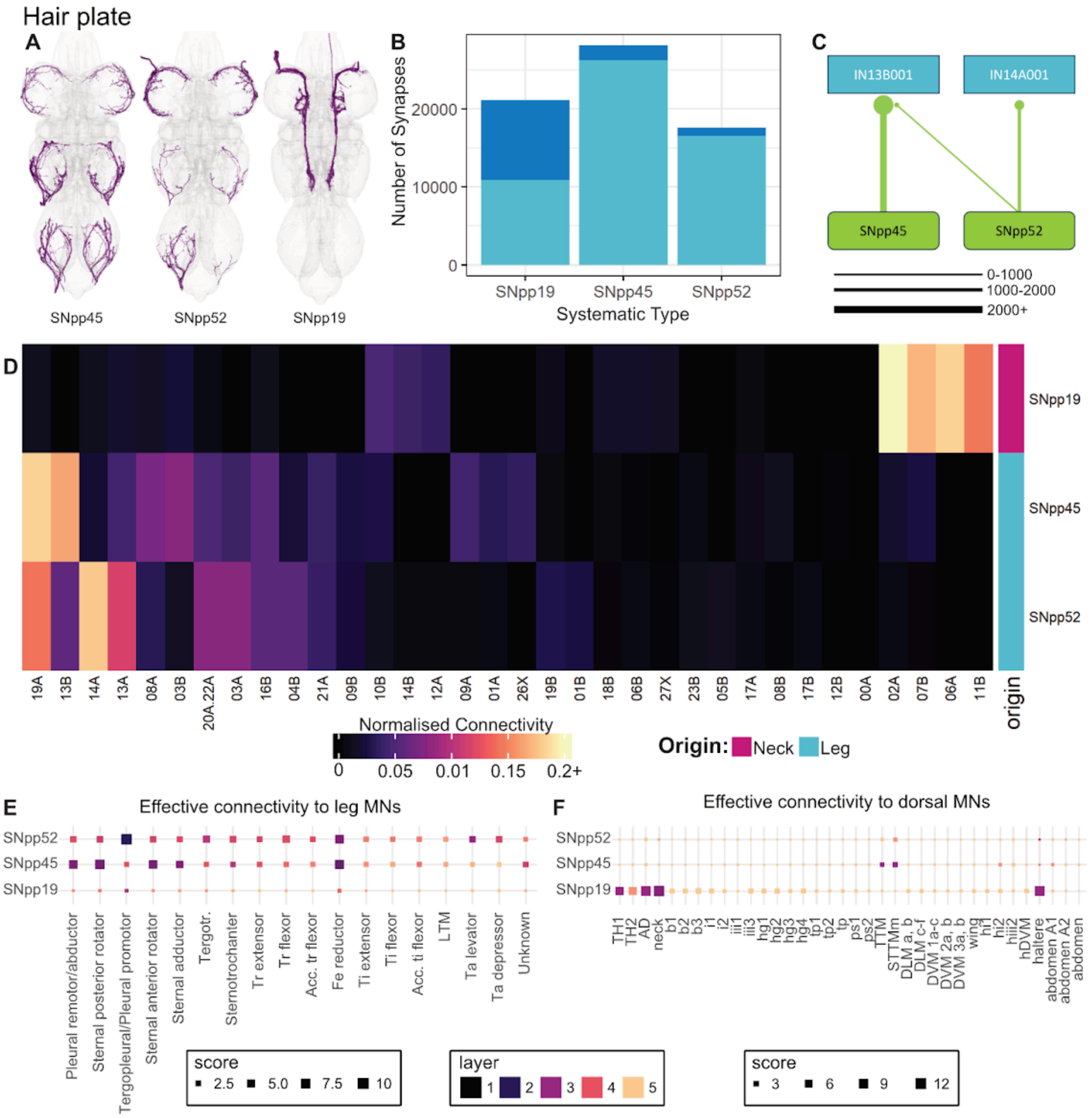

**Figure 61. Hair plate connectivity. A.** Plotted meshes of hair plate types. **B.** Number of synaptic outputs of hair plate types onto intrinsic neurons (cyan) and ascending neurons (blue). **C.** Diagram displaying the principal downstream connections of leg hairplate types. **D.** Hair plate types clustered by connectivity to hemilineages. **E.** Effective connectivity to ipsilateral leg motor neuron targets. **F.** Effective connectivity to ipsilateral dorsal motor neuron targets. Layer legend is shared between E and F.

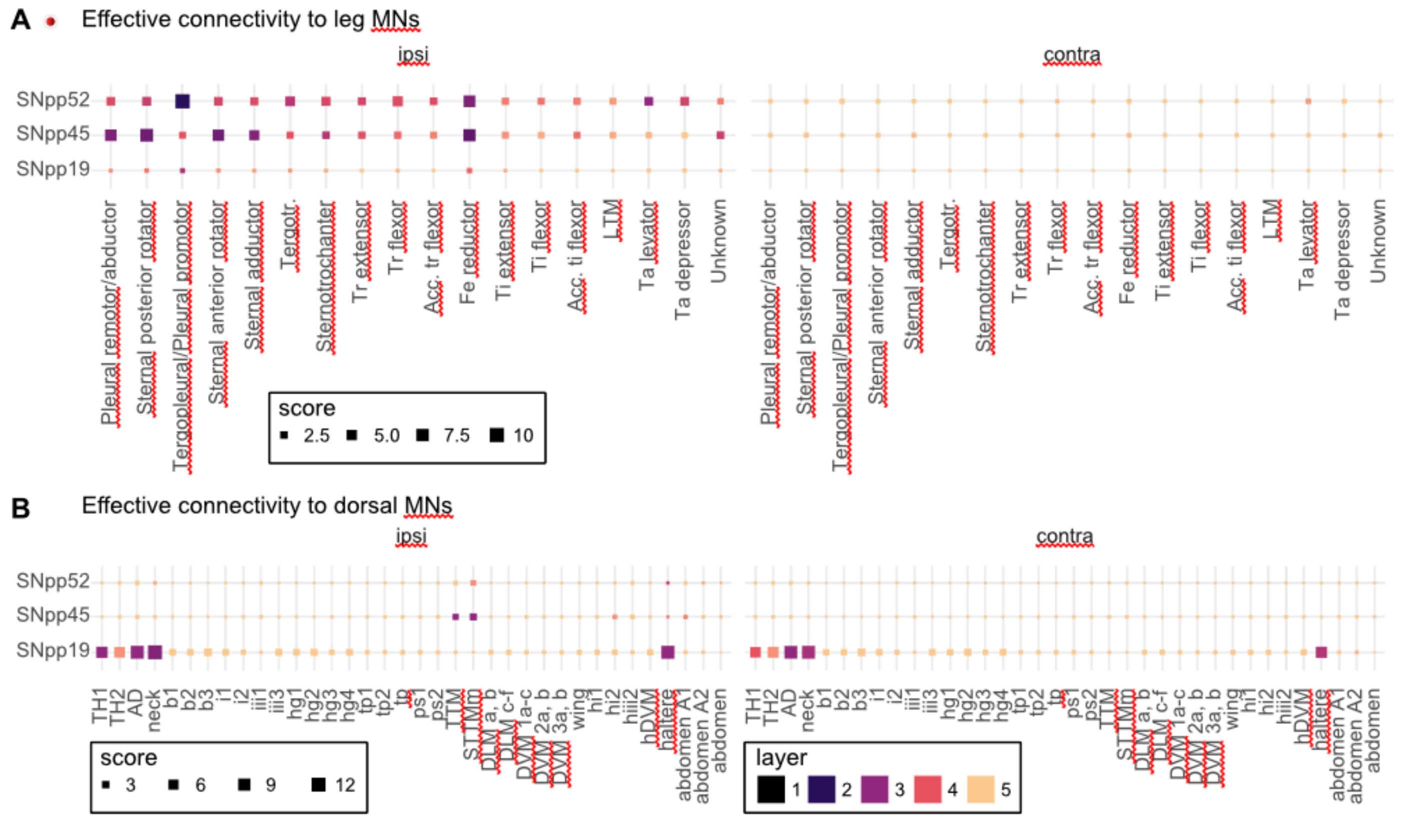

**Figure 61 - figure supplement 1. Hair plate effective connectivity. A.** Effective connectivity of selected hair plate types to ipsilateral and contralateral leg motor neurons. **B.** Effective connectivity of selected hair plate types to ipsilateral and contralateral dorsal motor neurons.

#### Tactile neuron types

Tactile hairs, referred to as "bristles" in the *Drosophila* literature, are the primary detectors of externally generated mechanical forces (reviewed in (Tuthill and Wilson, 2016a)). An adult fruit fly has several thousand bristles distributed over the cuticle: large macrochaetes in stereotyped positions and small microchaetes distributed with more variability but still in regularly spaced rows. Anatomical studies in various insect species have shown that bristle axons in the CNS are systematically organised depending on mechanoreceptor location, physiological tuning, and receptor type. For example, axon terminals of neurons from sensilla on distal leg segments terminate in the most central/ventral part of the leg neuropil, whilst those from proximal segments terminate in more peripheral rings (Tsubouchi et al., 2017).

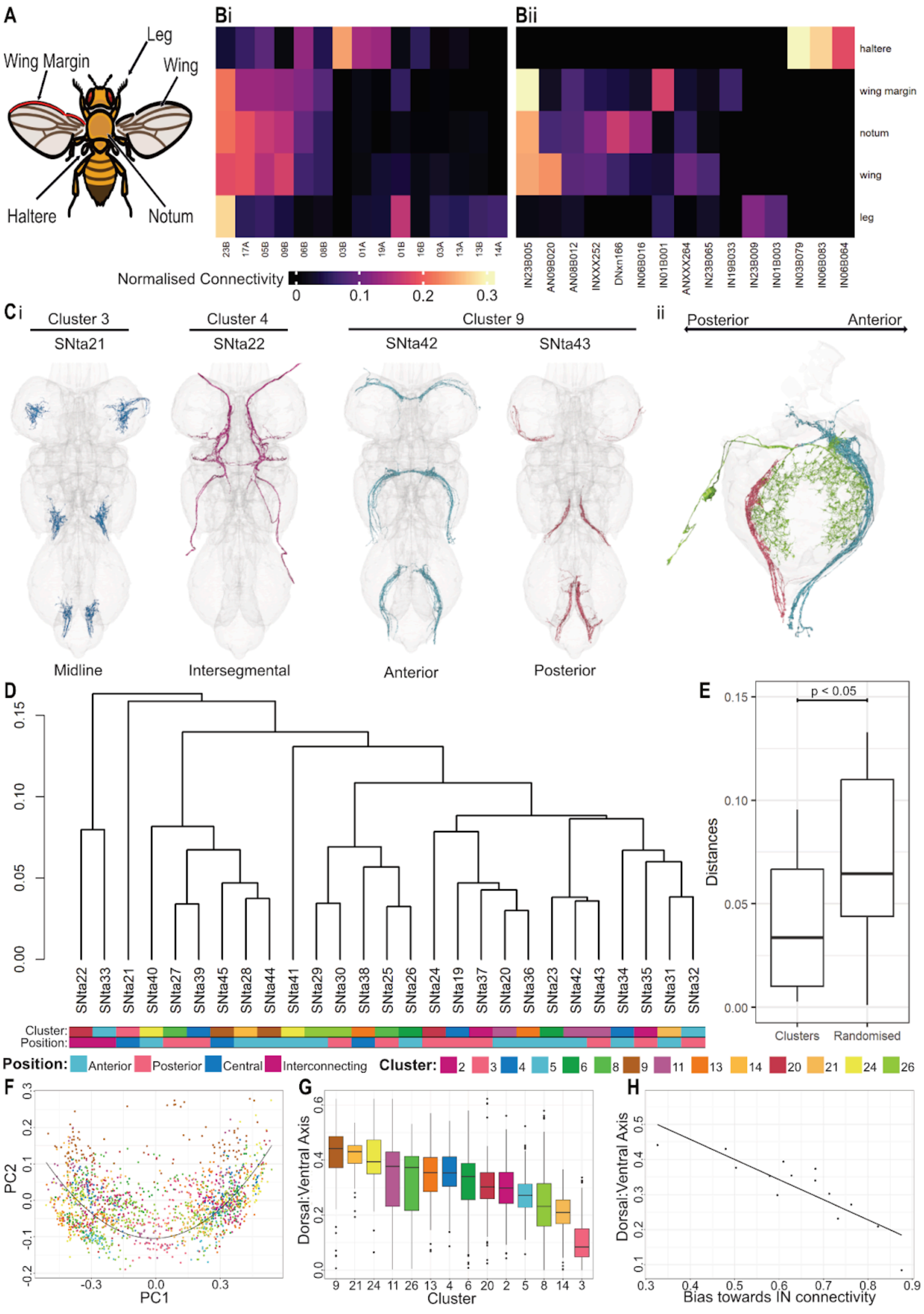
A
Wing Margin
Leg
Wing
Haltere
Notum
Bi
Bii
haltere
wing margin
notum
wing
leg
23B 17A 05B 09B 06B 08B 03B 01A 19A 01B 16B 03A 13A 13B 14A
IN23B005 AN09B020 AN08B012 INXXX252 DNxn166 IN06B016 IN01B001 ANXXX264 IN23B065 IN19B033 IN23B009 IN01B003 IN03B079 IN06B083 IN06B064
Normalised Connectivity
0 0.1 0.2 0.3
Ci
Cluster 3
SNta21
Cluster 4
SNta22
Cluster 9
SNta42
SNta43
ii
Posterior
Anterior
Midline
Intersegmental
Anterior
Posterior
D
E
p < 0.05
Distances
Clusters
Randomised
Cluster:
Position:
Position: Anterior Posterior Central Interconnecting
Cluster: 2 3 4 5 6 8 9 11 13 14 20 21 24 26
F
PC2
PC1
G
Dorsal:Ventral Axis
Cluster
9 21 24 11 26 13 4 6 20 2 5 8 14 3
H
Dorsal:Ventral Axis
Bias towards IN connectivity

**Figure 62. Tactile neuron types. A.** Principal regions of origin for tactile sensory neurons within the VNC. **B.** Heatmap showing downstream connectivity of tactile neurons to i. hemilineages, and ii. strongly associated downstream partner types, downstream partners shown receive a minimum of 5% of outputs from at least one regional group. **C.** i. Neuron plots showing type representatives from different positions and clusters. ii. Neuron plot showing overlap between SNta42 and SNta43 with a strong downstream partner (bodyid: 12405). **D.** Downstream connectivity of leg tactile sensory types. Row annotation displays relative position within the neuropil and the cluster from which these types were assigned (see methods). **E.** Quantification of intra-cluster distances from clustering outputs onto ascending and intrinsic neurons vs. a randomised control (p = 0.036). **F.** Relative positions of clusters within the standard leg neuropil. Colours are consistent to those indicated in C. **G.** Patterning of tactile clusters across the dorsal ventral axis. **H.** Relative bias towards intrinsic neuron connectivity for each cluster, broken down by birthtime.

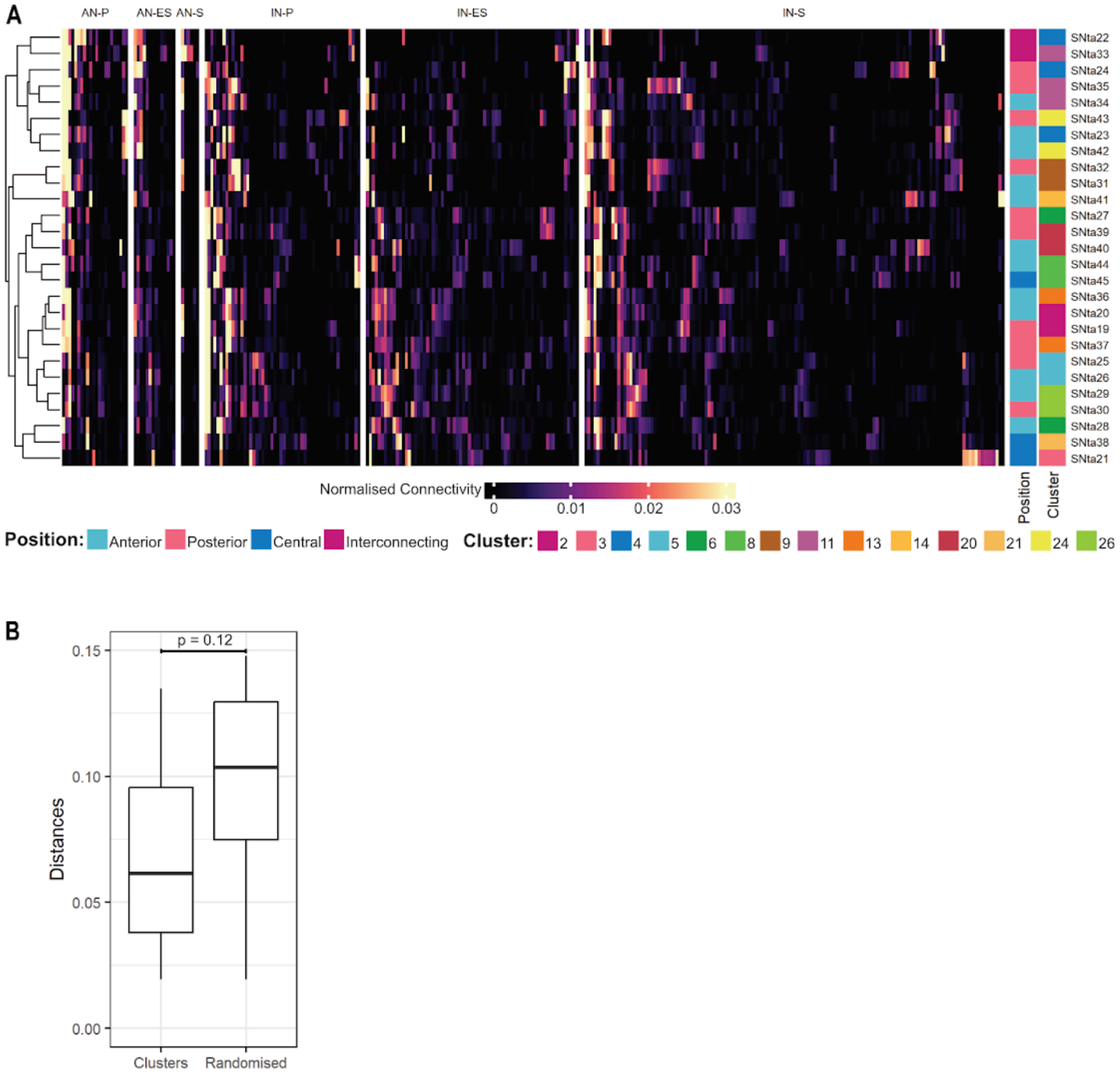

**Figure 62 - figure supplement 1. Tactile neuron types. A.** Connectivity clustering of tactile types across all birthtimes (AN-P = ascending neurons, primary; AN-ES = ascending neurons, early secondary; AN-S = ascending neurons, secondary; IN-P = intrinsic neurons, primary; IN-ES = intrinsic neurons, early secondary; IN-S = intrinsic neurons, secondary). **B.** Quantification of intra-cluster distances from clustering outputs onto ascending and intrinsic neurons across all birthtimes vs. a randomised control (p = 0.12).

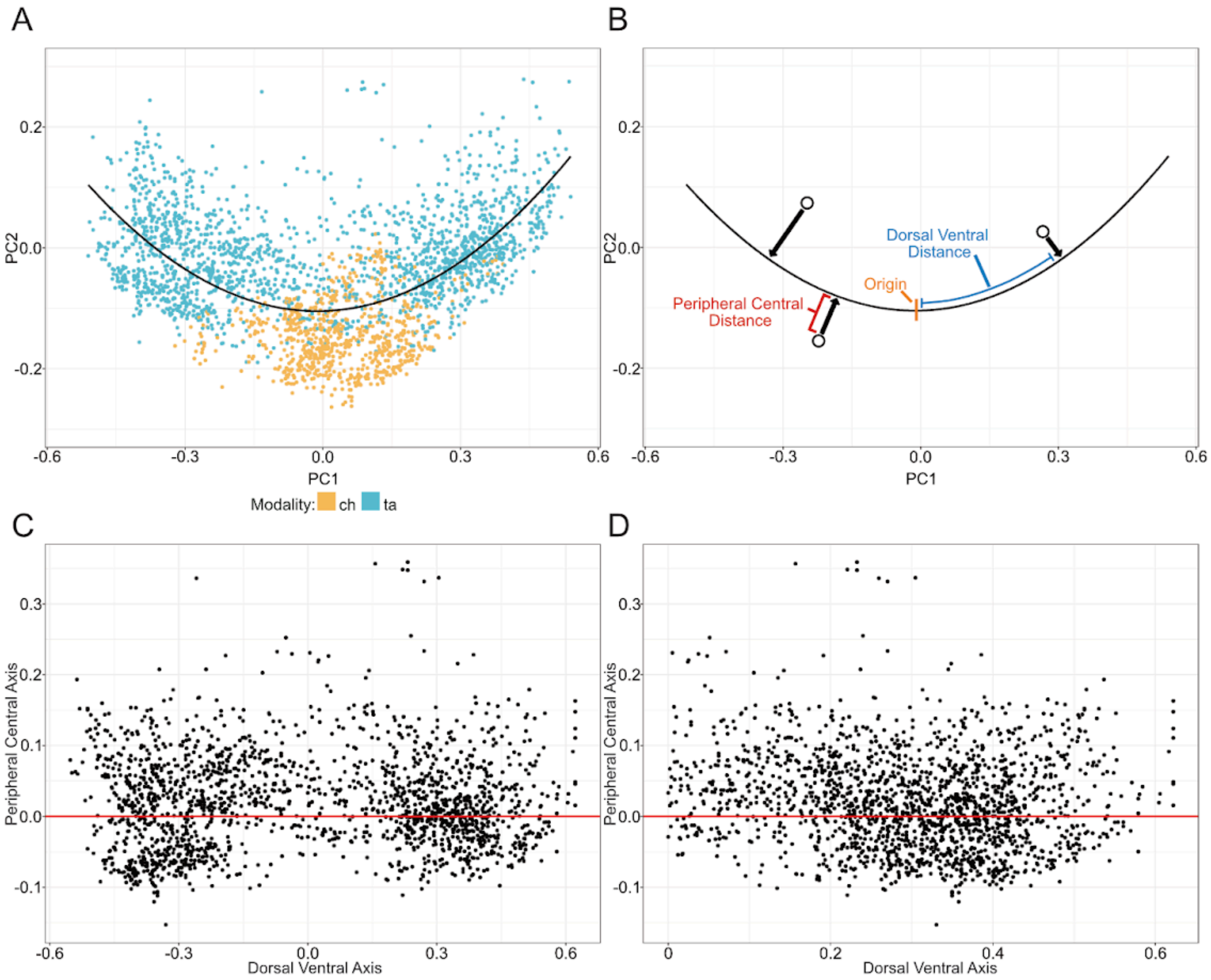

**Figure 62 - figure supplement 2. Adjusted dorsal-ventral axis methods. A.** Positioning within the standard leg neuropil of tactile (ta) and chemosensory (ch) leg-originating sensory neurons. Line (black) indicates the line of best fit through the data points. **B.** Demonstration of calculation of adjusted axes. Dorsal ventral distance is the distance along the line of best fit between the nearest point on the line for each neuron to the minimum point on the line (origin). The peripheral central distance is the distance from each datapoint to the line of best fit. **C** Replotted data points according to newly calculated axes. **D** Replotted data points showing the absolute value of the dorsal ventral axis used for data analysis.

#### Chemosensory neuron types

Besides the well-characterised chemosensory neurons on head structures that project to the antennal lobe and suboesophageal ganglion, flies possess chemosensory neurons on the tarsae and tibia of all legs and on the anterior wing margin (reviewed in (Stocker, 1994)). Taste neurons from the legs project to the ipsilateral VAC of the associated neuromere, with a small subset ascending directly to the brain, whereas those from the wing margin project to the ovoid. Expression patterns of all known gustatory receptor isoforms have elucidated the axon projection patterns of specific populations of leg and wing chemosensory neurons in the VNC (Kwon et al., 2014); however, in most cases these are too similar to distinguish from morphology alone.

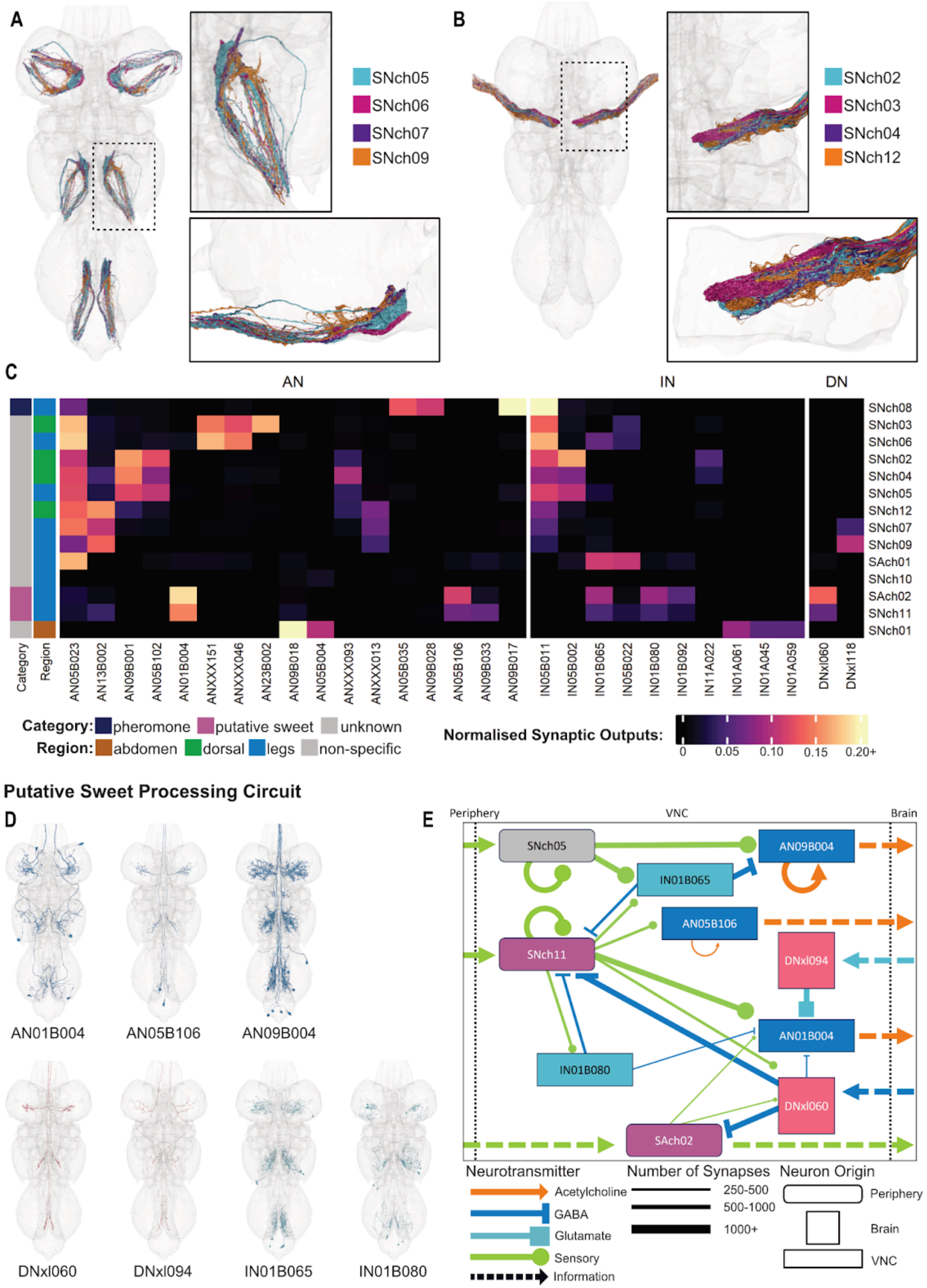
A
SNch05
SNch06
SNch07
SNch09
B
SNch02
SNch03
SNch04
SNch12
C
AN
IN
DN
SNch08
SNch03
SNch06
SNch02
SNch04
SNch05
SNch12
SNch07
SNch09
SAch01
SNch10
SAch02
SNch11
SNch01
Category
Region
AN05B023
AN13B002
AN09B001
AN05B102
AN01B004
ANXXX151
ANXXX046
AN23B002
AN09B018
AN05B004
ANXXX093
ANXXX013
AN05B035
AN09B028
AN05B106
AN09B033
AN09B017
IN05B011
IN05B002
IN01B065
IN05B022
IN01B080
IN01B092
IN11A022
IN01A061
IN01A045
IN01A059
DNxl060
DNxl118
Category: pheromone putative sweet unknown
Region: abdomen dorsal legs non-specific
Normalised Synaptic Outputs:
0
0.05
0.10
0.15
0.20+
Putative Sweet Processing Circuit
D
AN01B004
AN05B106
AN09B004
DNxl060
DNxl094
IN01B065
IN01B080
E
Periphery
VNC
Brain
SNch05
AN09B004
IN01B065
AN05B106
SNch11
DNxl094
AN01B004
IN01B080
DNxl060
SAch02
Neurotransmitter
Acetylcholine
GABA
Glutamate
Sensory
Information
Number of Synapses
250-500
500-1000
1000+
Neuron Origin
Periphery
Brain
VNC

**Figure 63. Chemosensory neurons and a putative sweet taste circuit. A.** Chemosensory neurons originating in the legs, coloured by type. i. All types plotted in symmetrised volume, ventral view. ii. Close-up of neuropil in dotted rectangle, ventral view. iii. Rotation of selected neuropil to reveal type-specific layers. **B.** Chemosensory neurons originating in the wings, coloured by type. i. All types plotted in symmetrised volume, ventral view. ii. Close-up of neuropil in dotted rectangle, ventral view. iii. Rotation of selected neuropil to reveal type-specific layers. **C.** Heatmap showing downstream connectivity to chemosensory implicated neuronal types. Row annotations indicate the region of origin and the tastant category assigned to the sensory neuron type. **D.** Plotted meshes of strongly connected downstream partners of putative sweet taste neurons. **E.** Putative sweet taste circuit with specific downstream partners.

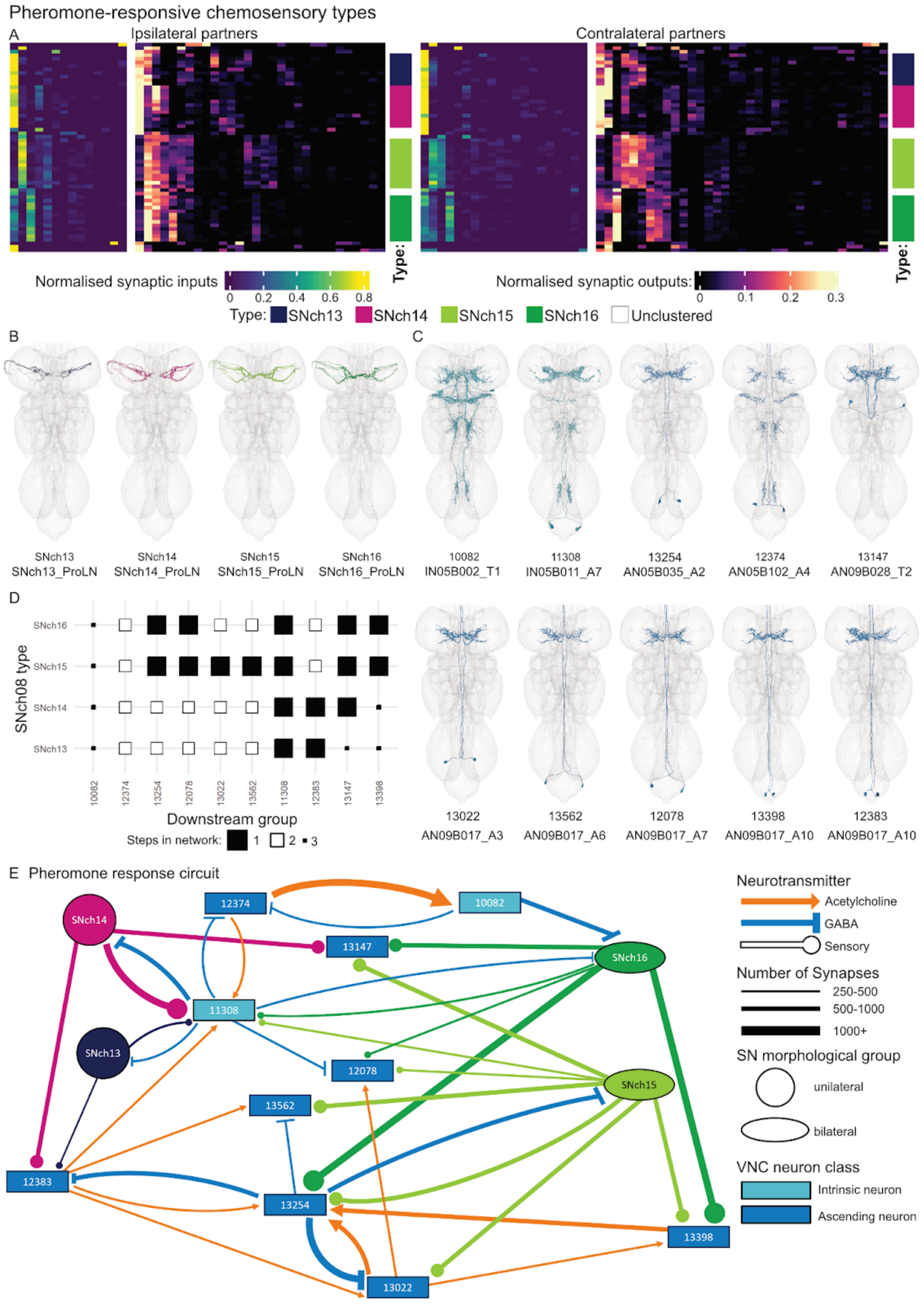

Pheromone-responsive chemosensory types
A
Ipsilateral partners
Contralateral partners
Normalised synaptic inputs
0 0.2 0.4 0.6 0.8
Type:
Normalised synaptic outputs:
0 0.1 0.2 0.3
Type:
Type: SNch13 SNch14 SNch15 SNch16 Unclustered
B
C
SNch13
SNch13_ProLN
SNch14
SNch14_ProLN
SNch15
SNch15_ProLN
SNch16
SNch16_ProLN
10082
IN05B002_T1
11308
IN05B011_A7
13254
AN05B035_A2
12374
AN05B102_A4
13147
AN09B028_T2
D
SNch16
SNch15
SNch14
SNch13
SNch08 type
10082 12374 13254 12078 13022 13562 11308 12383 13147 13398
Downstream group
Steps in network: 1 2 3
13022
AN09B017_A3
13562
AN09B017_A6
12078
AN09B017_A7
13398
AN09B017_A10
12383
AN09B017_A10
E Pheromone response circuit
SNch14
12374
10082
13147
SNch16
11308
SNch13
12078
SNch15
13562
12383
13254
13398
13022
Neurotransmitter
Acetylcholine
GABA
Sensory
Number of Synapses
250-500
500-1000
1000+
SN morphological group
unilateral
bilateral
VNC neuron class
Intrinsic neuron
Ascending neuron

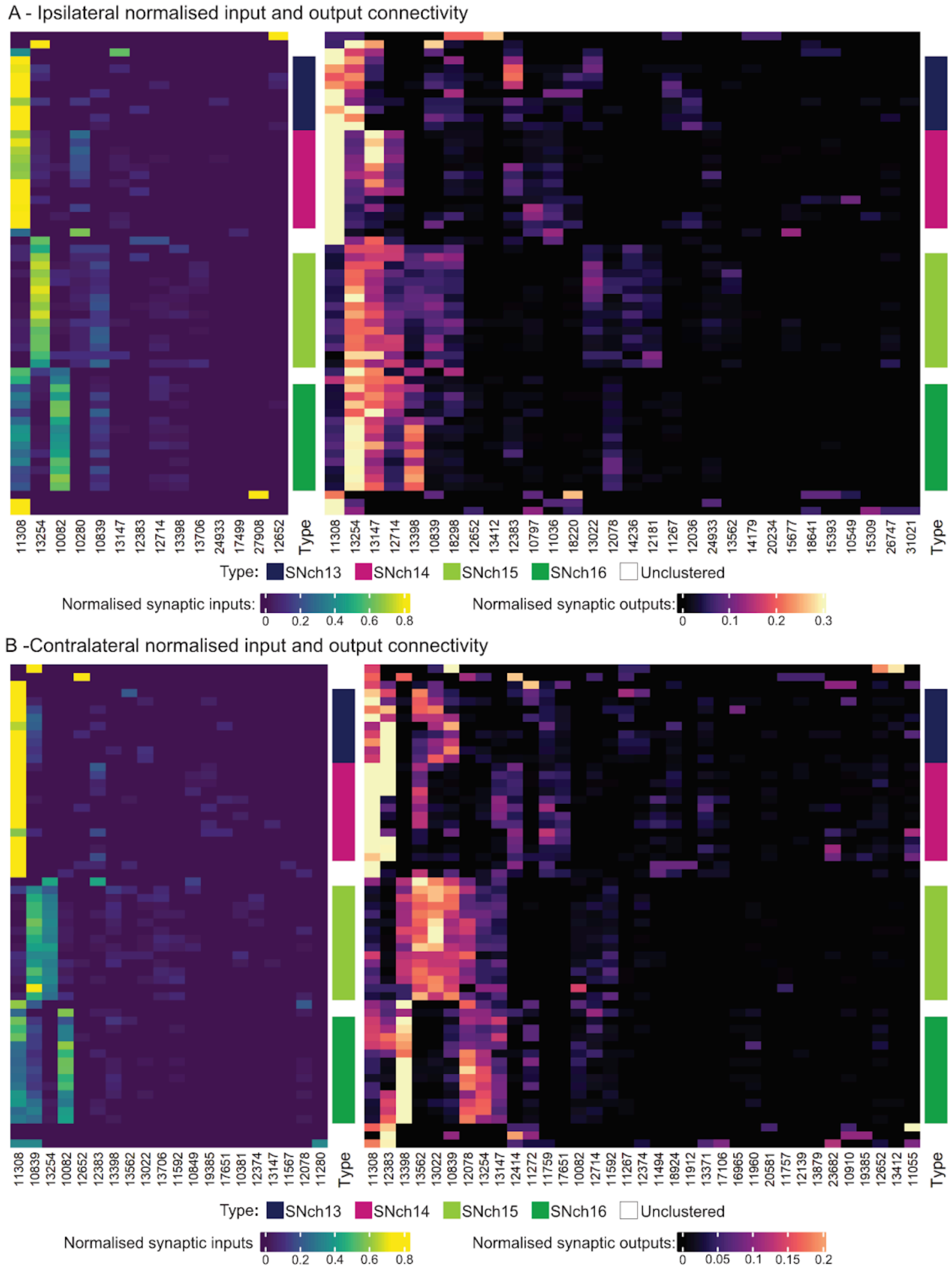

**Figure 64 - figure supplement 1. Pheromone response circuit. A.** Ipsilateral normalised input and output connectivity between four SNch08 subtypes and their strongest downstream targets. **B.** Contralateral normalised input and output connectivity between four SNch08 subtypes and their strongest downstream targets.

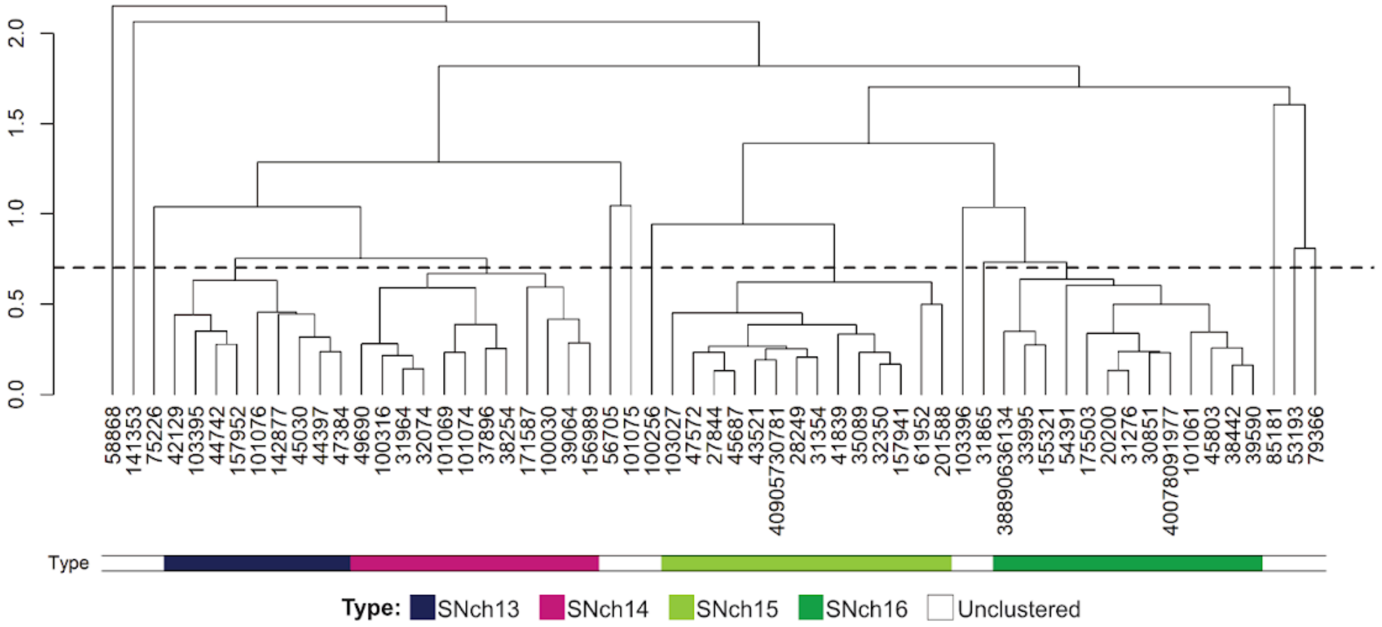

**Figure 64 - figure supplement 2. Pheromone response circuit.** Dendrogram used for hierarchical clustering of SNch08 neurons. The dashed line represents the cut height used to define the four main clusters/subtypes.

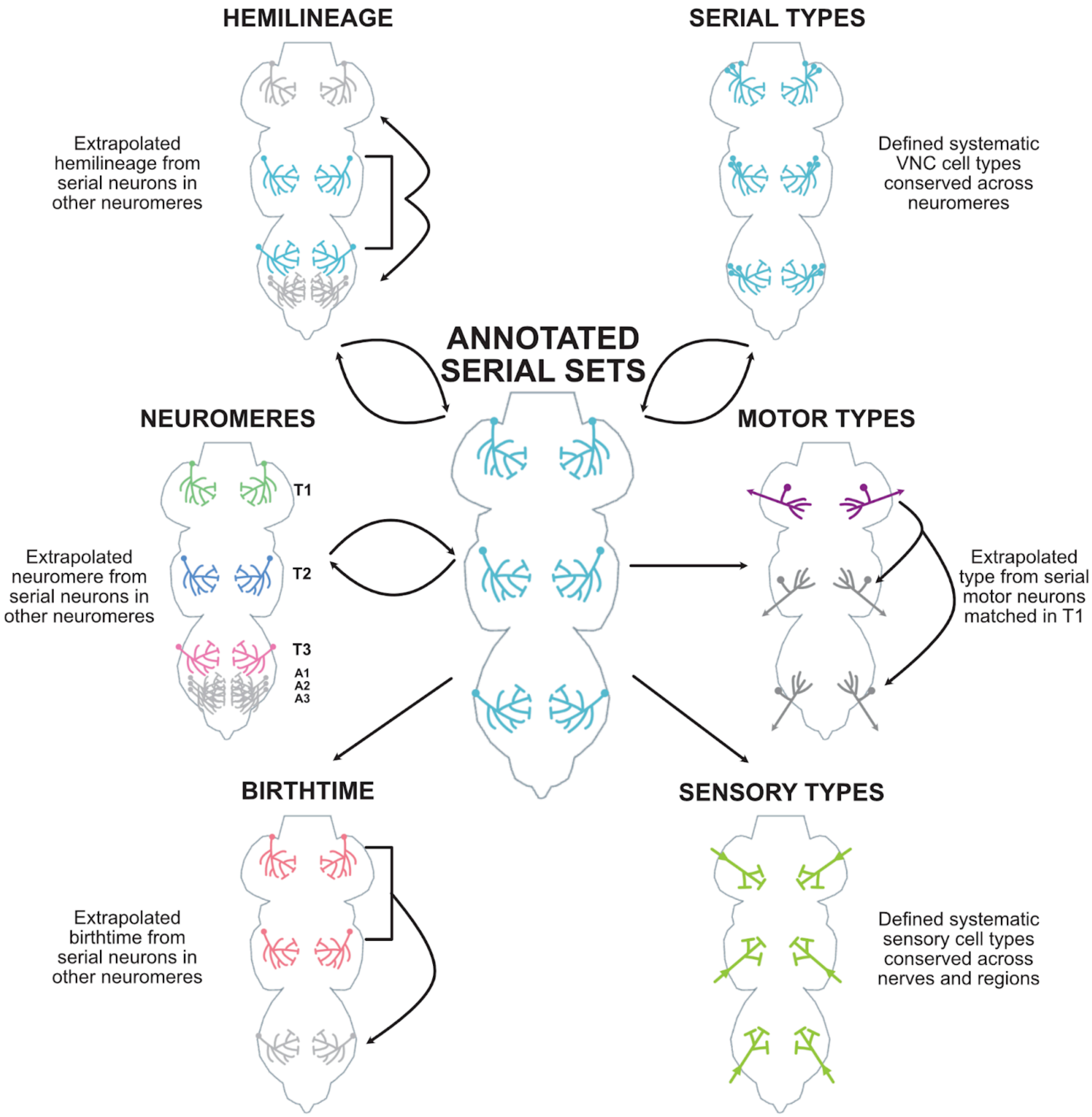

**Figure 65. Uses of serial homology for annotation and systematic cell typing in MANC.** The manual and programmatic identification of individual neurons that repeat in multiple neuromeres - referred to as serial sets (centre, cyan) - contributed to many aspects of the MANC annotation process: 1. Assignment of unknown neurons (grey) to the correct **hemilineage** based on serial homology to known neurons (cyan), especially in T1 or the abdomen. 2. Assignment of neurons to the correct left-right partner and/or soma **neuromere** in the abdomen (grey) based on serial homology to neurons in thoracic neuromeres (green = T1, blue = T2, magenta = T3). 3. Extrapolation of **birthtime** assignments in T1 and T2 (pink = primary) to inform those in T3 (grey), where somas are typically smaller. 4. Generation of systematic **serial types** (cyan) for neurons originating in the VNC. 5. Identification of leg **motor neuron types** in T2 and T3 (grey) based on serial homology to light level-matched cells in T1 (purple) (Cheong et al., 2023). 6. Generation of systematic **sensory cell types** (green) across nerves and peripheral regions.

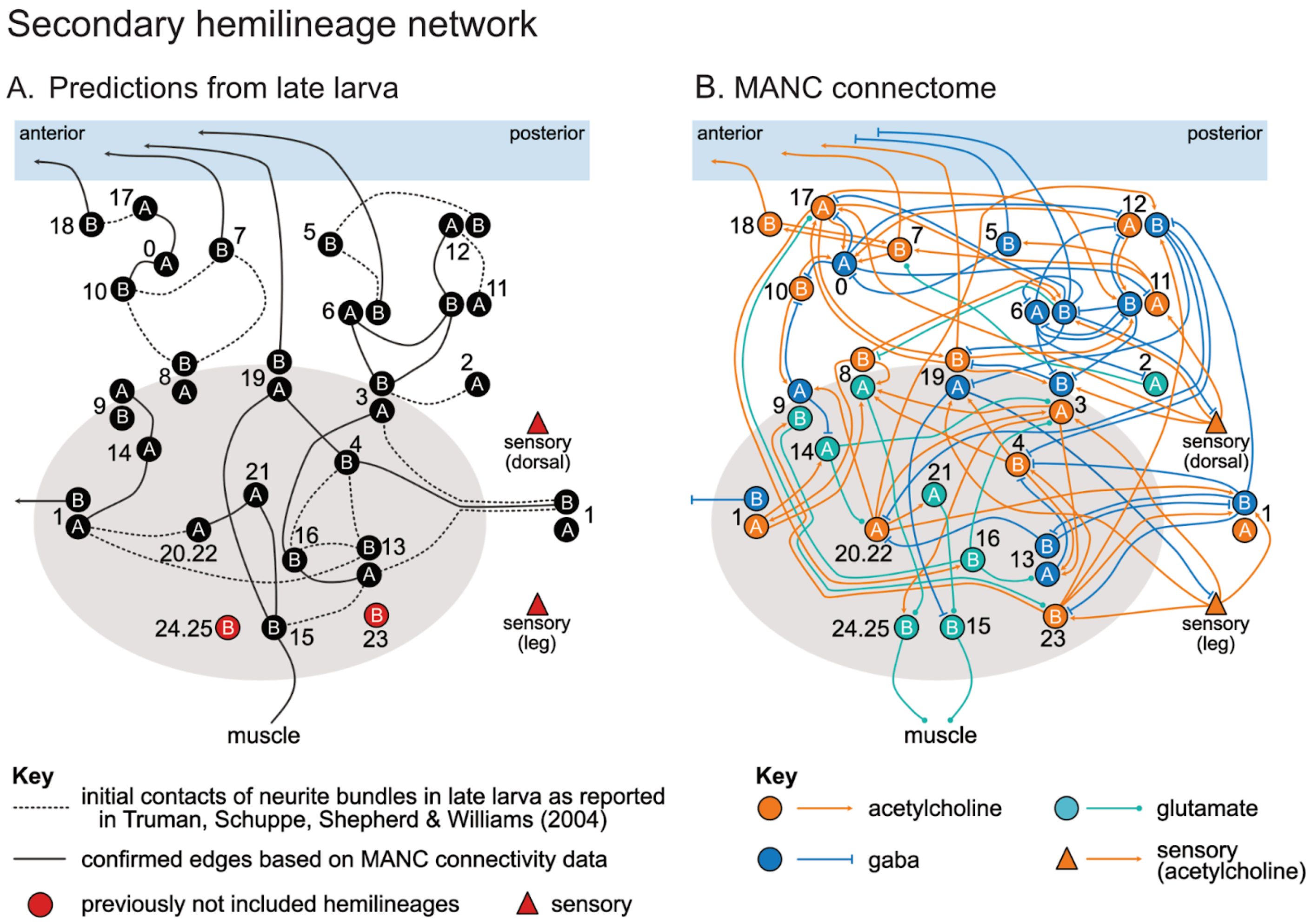

**Figure 66. Secondary hemilineage network.** Schematic summary of secondary hemilineage-to-hemilineage connectivity showing the top partners and consensus neurotransmitter prediction per hemilineage in the adult male T2 neuromere (B) compared against previously reported predictions from late larva (A). Within hemilineage connections are excluded from the schematic. **A.** The lines between hemilineages represent initial contacts of neurite bundles made in late larva as reported by Truman et. al., 2004. Bundles within the grey circle were hypothesised to terminate in the ventrolateral (leg) neuropil while those in blue stripe hypothesised to project in dorsal tracts. Solid black lines represent edges between hemilineages confirmed in the adult male based on a threshold of 5% of input or being in the top 4 partners. Thresholds were considered from the perspective of both hemilineage (i.e., hemilineage A receives at least 5% of its input from hemilineage B and hemilineage B provides at least 5% of its output to hemilinage A. **B.** All confirmed edges in (A) plus all newly reported hemilineage connections that are in the top 4 partners per hemilineage (considered from both hemilineage's perspective). Neurotransmitter identity is based on the upstream hemilinage's consensus neurotransmitter prediction.

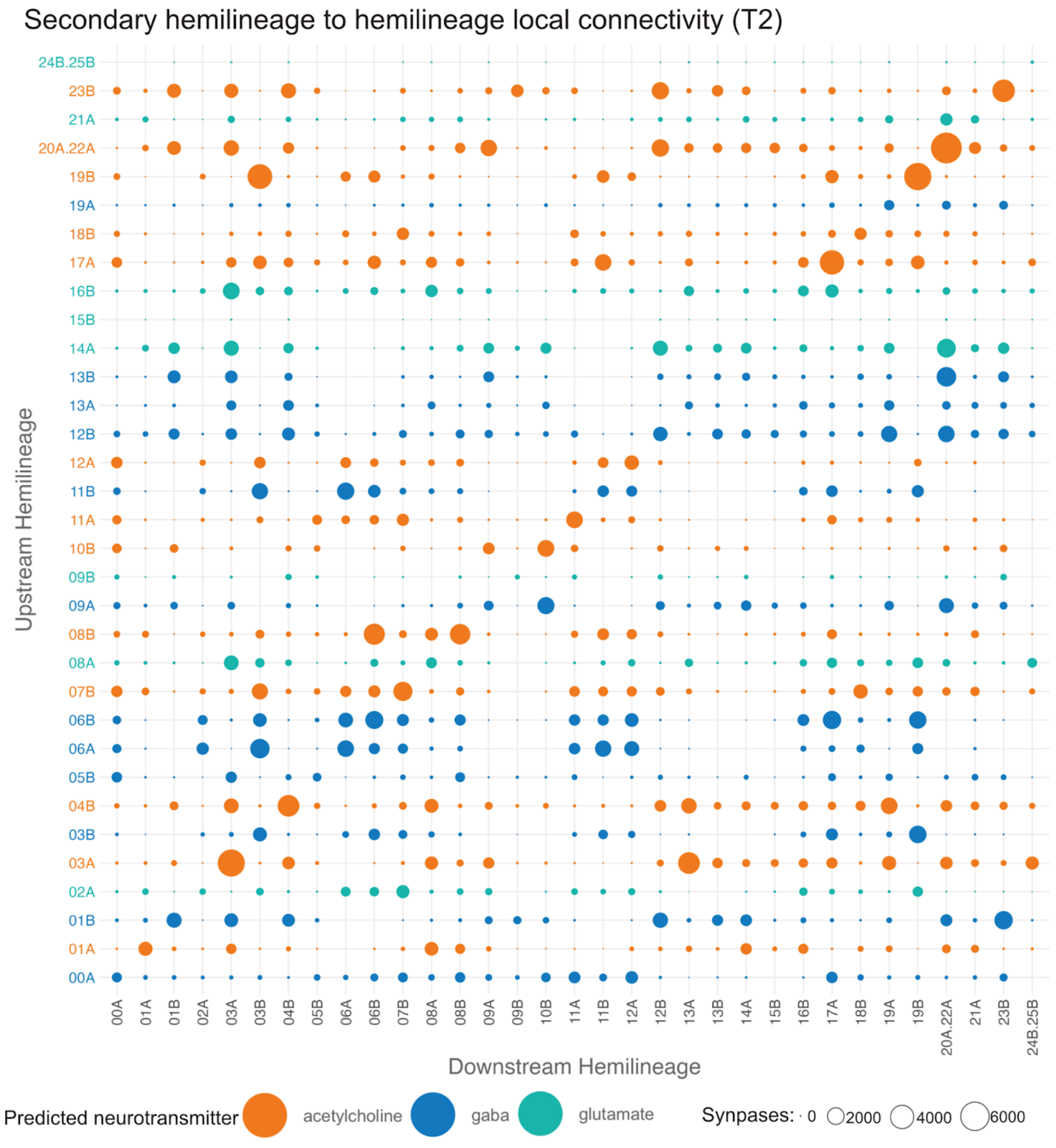

**Figure 66 - figure supplement 1. T2 secondary hemilineage to hemilineage connectivity.** Bubble plot showing secondary hemilineage to hemilineage connectivity restricted to T2. Colour of bubble is based on the consensus neurotransmitter prediction for the given upstream hemilineage. Size of each bubble is proportional to the synaptic weight between hemilineages.

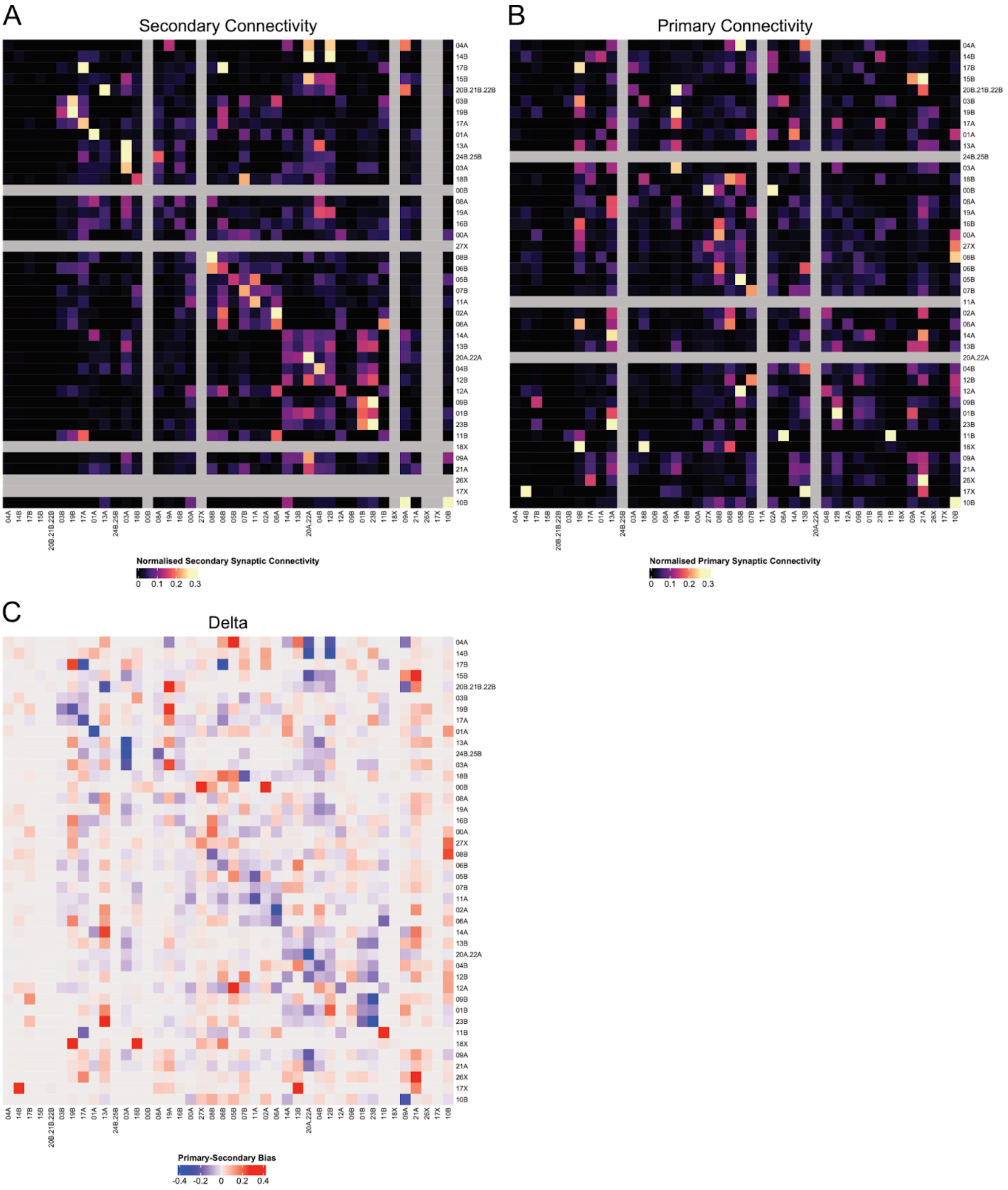

**Figure 66 - figure supplement 2. T2 hemilineage to hemilineage connectivity. A.** Clustering of secondary hemilineages by secondary hemilineage to secondary hemilineage synaptic connectivity (grey cells represent primary-only hemilineages). **B.** Heatmap of primary hemilineages by primary hemilineage to primary hemilineage synaptic connectivity (grey cells represent secondary-only hemilineages). Ordering enforced to be consistent with A. **C.** Bias in connectivity to hemilineages by birthtime, calculated by subtracting the normalised primary connectivity from the normalised secondary connectivity. In cases where a hemilineage is primary or secondary only, a normalised connectivity score of 0 is assigned to the respective matrix. Ordering enforced to be consistent with A.

#### Circuits for coordination of leg movements

*Drosophila* possess intricate leg circuits that enable them to navigate their environment with remarkable agility. While studies have successfully mapped the proprioceptive sensory input and motor output to individual leg neuropils (Azevedo et al., 2020; Brierley et al., 2012; Kuan et al., 2020; Tuthill and Wilson, 2016b), the INs that integrate these functions remain largely unknown. In our research, we have undertaken a significant effort to reconstruct and identify leg sensory neurons and (in a companion paper, (Cheong et al., 2023)) the leg motor neurons of the MANC dataset. Additionally, we have conducted a thorough annotation effort for the INs, encompassing cell typing, hemilineage annotation, birthtime, and innervation patterns, which enabled a comprehensive description of this neuron class.

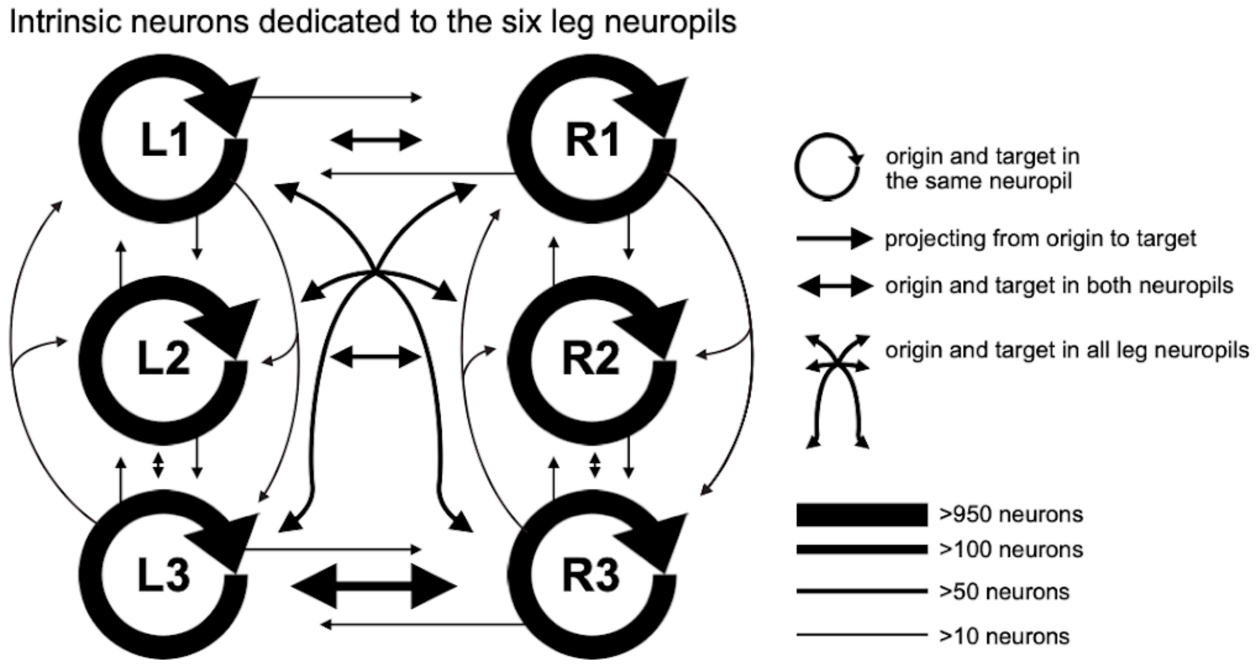

**Figure 67. Leg circuit schematic.** 6960 INs (>50%) have both origin and target specifically in the leg neuropil. Shown are the numbers of connections of these neurons, in the form of an arrow. Arrow is only shown if >10 neurons have the same origin and target. The great majority of intrinsic neurons of the leg have their origin and target within a single leg neuropil, over 950 neurons per leg neuropil. The number of neurons with the same leg connectivity pattern between legs ipsilaterally are all below 50. The most numerous bilateral connections between legs are in the T3 leg neuropils (>100).

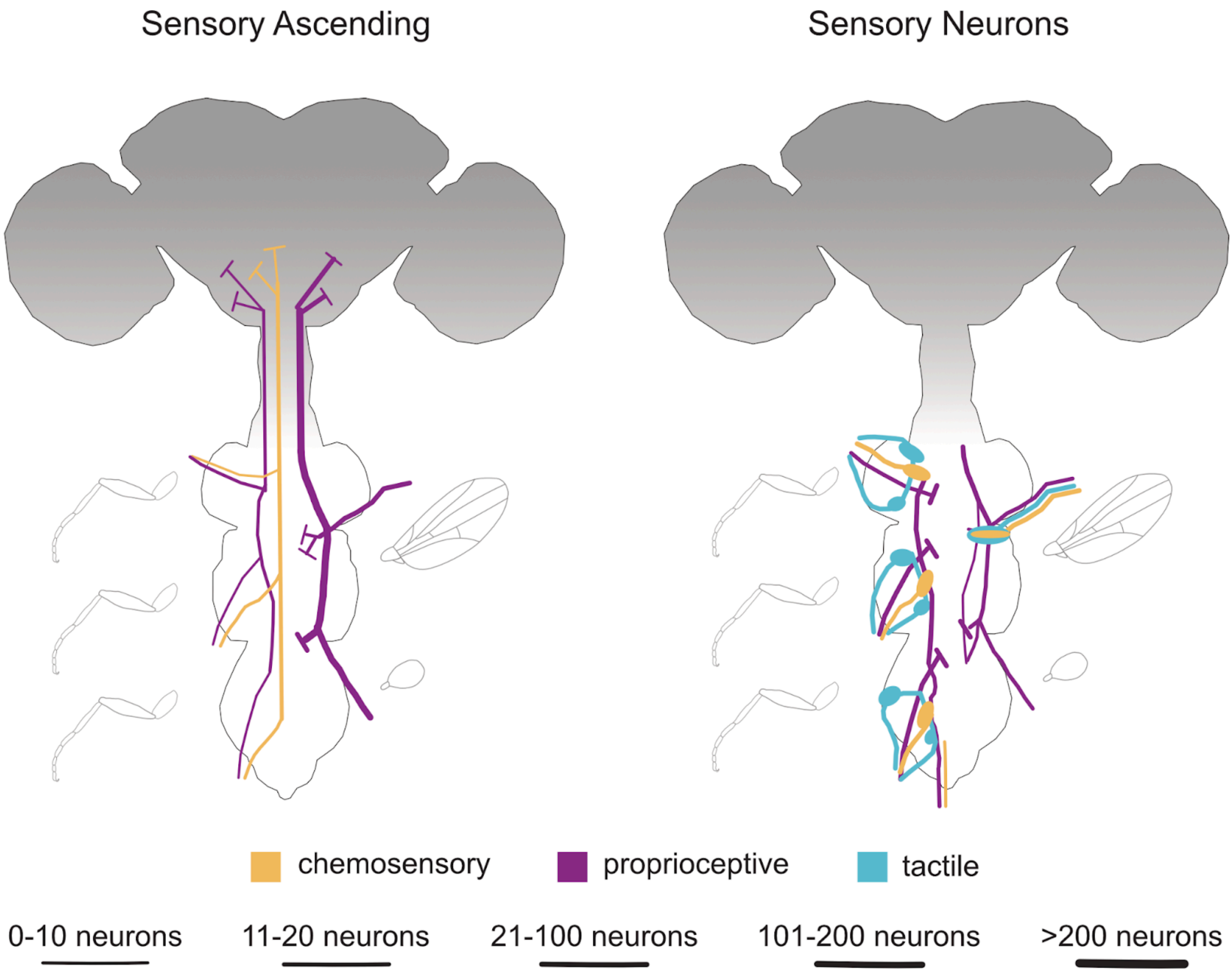

**Figure 68. Sensory flow schematic.** Global summary of MANC sensory neurons and their projections. Depicted origins are prothoracic leg (plus ventral prothorax), mesothoracic leg, metathoracic leg, wing (including wing margin), haltere, and abdomen. Line widths reflect higher neuron number when compared across sides and line colours reflect modality. Neurons of unknown modality have been omitted.
